## Supplemental Table 9 for "Region-specific reversal of epidermal planar polarity in the fancy *rosette* mouse"

| Chr | Position | SNP_rsID | N_miss | Allele_1 | Allele_0 | AF | Beta | S.E. | L_remle | L_mle | p_LRT | Outlier Beta | Gene name | Annotated feature | Impact |
|---|---|---|---|---|---|---|---|---|---|---|---|---|---|---|---|
| 15 | 55010273 | AX-168399809 | 2 | C | T | 0.036 | -6.30E-01 | 8.90E-02 | 1.07E+00 | 1.11E+00 | 1.70E-09 | Yes | Taf2 | downstream_gene_variant | MODIFIER |
| 15 | 26051946 | AX-170222819 | 0 | T | A | 0.026 | -7.41E-01 | 8.69E-02 | 2.91E-01 | 2.81E-01 | 2.52E-09 | Yes | Gm41279 | downstream_gene_variant | MODIFIER |
| 15 | 45318752 | AX-170223555 | 1 | T | C | 0.027 | -6.65E-01 | 8.91E-02 | 5.90E-01 | 5.94E-01 | 5.01E-09 | Yes | | intergenic_variant | MODIFIER |
| 15 | 56128978 | AX-97815157 | 0 | A | G | 0.035 | -6.12E-01 | 8.92E-02 | 1.04E+00 | 1.08E+00 | 5.02E-09 | Yes | | intergenic_variant | MODIFIER |
| 15 | 24625483 | AX-170011262 | 2 | G | A | 0.018 | -7.24E-01 | 7.89E-02 | 1.56E-01 | 1.32E-01 | 6.23E-09 | Yes | | intergenic_variant | MODIFIER |
| 15 | 38988917 | AX-170318380 | 0 | T | C | 0.026 | -6.83E-01 | 8.31E-02 | 3.07E-01 | 2.97E-01 | 7.78E-09 | Yes | Gm4740 | upstream_gene_variant | MODIFIER |
| 15 | 24042603 | AX-97789325 | 0 | T | A | 0.026 | -7.01E-01 | 9.06E-02 | 4.00E-01 | 3.95E-01 | 1.40E-08 | Yes | | intergenic_variant | MODIFIER |
| 15 | 24993289 | AX-170496603 | 0 | T | C | 0.053 | -3.50E-01 | 4.53E-02 | 4.00E-01 | 3.95E-01 | 1.40E-08 | Yes | Gm2824 | intron_variant,non_coding_transcript_variant | MODIFIER |
| 15 | 27477719 | AX-97789589 | 0 | A | T | 0.026 | -7.01E-01 | 9.06E-02 | 4.00E-01 | 3.95E-01 | 1.40E-08 | Yes | Ank | intron_variant | MODIFIER |
| 15 | 30616721 | AX-170248314 | 0 | G | A | 0.026 | -7.01E-01 | 9.06E-02 | 4.00E-01 | 3.95E-01 | 1.40E-08 | Yes | Ctnnd2 | intron_variant | MODIFIER |
| 15 | 34025585 | AX-170055212 | 0 | C | T | 0.053 | -3.50E-01 | 4.53E-02 | 4.00E-01 | 3.95E-01 | 1.40E-08 | Yes | Gm41290 | intron_variant,non_coding_transcript_variant | MODIFIER |
| 15 | 34571494 | AX-97828872 | 0 | A | G | 0.026 | -7.01E-01 | 9.06E-02 | 4.00E-01 | 3.95E-01 | 1.40E-08 | Yes | Nipal2 | downstream_gene_variant | MODIFIER |
| 15 | 34591700 | AX-97843120 | 0 | T | C | 0.053 | -3.50E-01 | 4.53E-02 | 4.00E-01 | 3.95E-01 | 1.40E-08 | Yes | Nipal2 | intron_variant | MODIFIER |
| 15 | 34776256 | AX-170378013 | 0 | A | C | 0.026 | -7.01E-01 | 9.06E-02 | 4.00E-01 | 3.95E-01 | 1.40E-08 | Yes | | intergenic_variant | MODIFIER |
| 15 | 31317841 | AX-170307968 | 1 | C | T | 0.027 | -6.86E-01 | 9.24E-02 | 4.97E-01 | 4.98E-01 | 1.58E-08 | Yes | Ankrd33b | intron_variant,NMD_transcript_variant | MODIFIER |
| 15 | 52517353 | AX-170055541 | 0 | T | A | 0.035 | -6.14E-01 | 9.52E-02 | 1.21E+00 | 1.25E+00 | 1.87E-08 | Yes | | intergenic_variant | MODIFIER |
| 15 | 53428203 | AX-170435385 | 0 | G | A | 0.035 | -6.14E-01 | 9.52E-02 | 1.21E+00 | 1.25E+00 | 1.87E-08 | Yes | | intergenic_variant | MODIFIER |
| 15 | 53510020 | AX-170055553 | 0 | C | G | 0.035 | -6.14E-01 | 9.52E-02 | 1.21E+00 | 1.25E+00 | 1.87E-08 | Yes | Samd12 | intron_variant | MODIFIER |
| 15 | 53554572 | AX-170189728 | 0 | T | A | 0.035 | -6.14E-01 | 9.52E-02 | 1.21E+00 | 1.25E+00 | 1.87E-08 | Yes | Samd12 | intron_variant | MODIFIER |
| 15 | 25612448 | AX-170335910 | 1 | T | A | 0.027 | -6.90E-01 | 9.08E-02 | 4.20E-01 | 4.18E-01 | 1.93E-08 | Yes | | intergenic_variant | MODIFIER |
| 15 | 38246668 | AX-170087823 | 0 | T | C | 0.026 | -6.23E-01 | 8.23E-02 | 4.05E-01 | 4.11E-01 | 2.83E-08 | Yes | Gm49313 | intron_variant,non_coding_transcript_variant | MODIFIER |
| 15 | 34750968 | AX-97813250 | 1 | C | T | 0.018 | -7.05E-01 | 8.28E-02 | 1.96E-01 | 1.69E-01 | 3.39E-08 | Yes | | intergenic_variant | MODIFIER |
| 15 | 25110296 | AX-170011276 | 1 | A | G | 0.054 | -3.51E-01 | 4.51E-02 | 3.29E-01 | 3.22E-01 | 3.80E-08 | Yes | Gm35996 | intron_variant,non_coding_transcript_variant | MODIFIER |
| 15 | 54872724 | AX-170469586 | 0 | A | G | 0.035 | -5.84E-01 | 8.88E-02 | 7.97E-01 | 8.16E-01 | 6.22E-08 | Yes | Enpp2 | intron_variant | MODIFIER |
| 15 | 54942659 | AX-97785366 | 0 | T | C | 0.035 | -5.84E-01 | 8.88E-02 | 7.97E-01 | 8.16E-01 | 6.22E-08 | Yes | Enpp2 | intron_variant | MODIFIER |
| 15 | 45342843 | AX-170513157 | 1 | A | G | 0.027 | -6.40E-01 | 9.17E-02 | 5.57E-01 | 5.49E-01 | 6.66E-08 | Yes | | intergenic_variant | MODIFIER |
| 15 | 44459323 | AX-170055414 | 0 | T | A | 0.035 | -6.19E-01 | 9.56E-02 | 8.27E-01 | 8.39E-01 | 8.44E-08 | Yes | Pkhd1l1 | intron_variant | MODIFIER |
| 15 | 44829884 | AX-97787793 | 0 | A | C | 0.07 | -3.09E-01 | 4.78E-02 | 8.27E-01 | 8.39E-01 | 8.44E-08 | Yes | A930017M01Rik | intron_variant,non_coding_transcript_variant | MODIFIER |
| 15 | 36450238 | AX-170513698 | 0 | G | A | 0.026 | -6.53E-01 | 8.67E-02 | 3.49E-01 | 3.31E-01 | 8.55E-08 | Yes | | regulatory_region_variant | MODIFIER |
| 15 | 36450304 | AX-170481529 | 0 | T | C | 0.026 | -6.53E-01 | 8.67E-02 | 3.49E-01 | 3.31E-01 | 8.55E-08 | Yes | | regulatory_region_variant | MODIFIER |
| 15 | 36732078 | AX-170337367 | 0 | T | C | 0.026 | -6.53E-01 | 8.67E-02 | 3.49E-01 | 3.31E-01 | 8.55E-08 | Yes | | regulatory_region_variant | MODIFIER |
| 15 | 36905695 | AX-97792421 | 0 | A | G | 0.026 | -6.53E-01 | 8.67E-02 | 3.49E-01 | 3.31E-01 | 8.55E-08 | Yes | Gm41300 | downstream_gene_variant | MODIFIER |
| 15 | 36974633 | AX-170207275 | 0 | G | T | 0.026 | -6.53E-01 | 8.67E-02 | 3.49E-01 | 3.31E-01 | 8.55E-08 | Yes | | regulatory_region_variant | MODIFIER |
| 15 | 36985047 | AX-97841191 | 0 | A | C | 0.026 | -6.53E-01 | 8.67E-02 | 3.49E-01 | 3.31E-01 | 8.55E-08 | Yes | | regulatory_region_variant | MODIFIER |
| 15 | 36990576 | AX-170244131 | 0 | C | T | 0.026 | -6.53E-01 | 8.67E-02 | 3.49E-01 | 3.31E-01 | 8.55E-08 | Yes | | regulatory_region_variant | MODIFIER |
| 15 | 37033266 | AX-97829429 | 0 | A | G | 0.026 | -6.53E-01 | 8.67E-02 | 3.49E-01 | 3.31E-01 | 8.55E-08 | Yes | Gm26766 | intron_variant,non_coding_transcript_variant | MODIFIER |
| 15 | 37034957 | AX-97793266 | 0 | C | T | 0.026 | -6.53E-01 | 8.67E-02 | 3.49E-01 | 3.31E-01 | 8.55E-08 | Yes | Gm26766 | intron_variant,non_coding_transcript_variant | MODIFIER |
| 15 | 37067942 | AX-170106256 | 0 | C | T | 0.026 | -6.53E-01 | 8.67E-02 | 3.49E-01 | 3.31E-01 | 8.55E-08 | Yes | | intergenic_variant | MODIFIER |
| 15 | 37081395 | AX-97829427 | 0 | T | C | 0.026 | -6.53E-01 | 8.67E-02 | 3.49E-01 | 3.31E-01 | 8.55E-08 | Yes | | intergenic_variant | MODIFIER |
| 15 | 37082297 | AX-170120293 | 0 | A | G | 0.026 | -6.53E-01 | 8.67E-02 | 3.49E-01 | 3.31E-01 | 8.55E-08 | Yes | | regulatory_region_variant | MODIFIER |
| 15 | 37100784 | AX-170413875 | 0 | T | C | 0.026 | -6.53E-01 | 8.67E-02 | 3.49E-01 | 3.31E-01 | 8.55E-08 | Yes | | regulatory_region_variant | MODIFIER |
| 15 | 37103393 | AX-170347564 | 0 | C | T | 0.026 | -6.53E-01 | 8.67E-02 | 3.49E-01 | 3.31E-01 | 8.55E-08 | Yes | | intergenic_variant | MODIFIER |
| 15 | 37110384 | AX-170327891 | 0 | A | G | 0.026 | -6.53E-01 | 8.67E-02 | 3.49E-01 | 3.31E-01 | 8.55E-08 | Yes | | intergenic_variant | MODIFIER |
| 15 | 37110736 | AX-170243781 | 0 | C | T | 0.026 | -6.53E-01 | 8.67E-02 | 3.49E-01 | 3.31E-01 | 8.55E-08 | Yes | | intergenic_variant | MODIFIER |
| 15 | 37111172 | AX-170526973 | 0 | A | G | 0.026 | -6.53E-01 | 8.67E-02 | 3.49E-01 | 3.31E-01 | 8.55E-08 | Yes | | intergenic_variant | MODIFIER |
| 15 | 37114617 | AX-170240410 | 0 | T | C | 0.026 | -6.53E-01 | 8.67E-02 | 3.49E-01 | 3.31E-01 | 8.55E-08 | Yes | | regulatory_region_variant | MODIFIER |
| 15 | 37124228 | AX-97837776 | 0 | T | C | 0.026 | -6.53E-01 | 8.67E-02 | 3.49E-01 | 3.31E-01 | 8.55E-08 | Yes | | regulatory_region_variant | MODIFIER |
| 15 | 37124635 | AX-170111721 | 0 | G | A | 0.026 | -6.53E-01 | 8.67E-02 | 3.49E-01 | 3.31E-01 | 8.55E-08 | Yes | | intergenic_variant | MODIFIER |
| 15 | 37127020 | AX-170294953 | 0 | G | A | 0.026 | -6.53E-01 | 8.67E-02 | 3.49E-01 | 3.31E-01 | 8.55E-08 | Yes | | intergenic_variant | MODIFIER |
| 15 | 37173935 | AX-170425278 | 0 | C | T | 0.026 | -6.53E-01 | 8.67E-02 | 3.49E-01 | 3.31E-01 | 8.55E-08 | Yes | Gm9509 | upstream_gene_variant | MODIFIER |
| 15 | 37183765 | AX-170273063 | 0 | G | A | 0.026 | -6.53E-01 | 8.67E-02 | 3.49E-01 | 3.31E-01 | 8.55E-08 | Yes | Gm8664 | upstream_gene_variant | MODIFIER |
| 15 | 37184109 | AX-170292403 | 0 | T | C | 0.026 | -6.53E-01 | 8.67E-02 | 3.49E-01 | 3.31E-01 | 8.55E-08 | Yes | Gm8664 | upstream_gene_variant | MODIFIER |
| 15 | 37230655 | AX-170269337 | 0 | G | A | 0.026 | -6.53E-01 | 8.67E-02 | 3.49E-01 | 3.31E-01 | 8.55E-08 | Yes | Grhl2 | upstream_gene_variant | MODIFIER |

| | | | | | | | | | | | | | | | |
|---|---|---|---|---|---|---|---|---|---|---|---|---|---|---|---|
| 15 | 37462413 | AX-170311270 | 0 | A | T | 0.026 | -6.53E-01 | 8.67E-02 | 3.49E-01 | 3.31E-01 | 8.55E-08 | Yes | Ncald | intron_variant | MODIFIER |
| 15 | 37534999 | AX-170011463 | 0 | T | C | 0.026 | -6.53E-01 | 8.67E-02 | 3.49E-01 | 3.31E-01 | 8.55E-08 | Yes | Ncald | intron_variant | MODIFIER |
| 15 | 37552122 | AX-97828974 | 0 | G | T | 0.026 | -6.53E-01 | 8.67E-02 | 3.49E-01 | 3.31E-01 | 8.55E-08 | Yes | Ncald | intron_variant | MODIFIER |
| 15 | 37564480 | AX-97832635 | 0 | T | C | 0.026 | -6.53E-01 | 8.67E-02 | 3.49E-01 | 3.31E-01 | 8.55E-08 | Yes | Ncald | intron_variant | MODIFIER |
| 15 | 37591823 | AX-170468258 | 0 | C | T | 0.026 | -6.53E-01 | 8.67E-02 | 3.49E-01 | 3.31E-01 | 8.55E-08 | Yes | Ncald | intron_variant | MODIFIER |
| 15 | 37595234 | AX-97831467 | 0 | G | A | 0.026 | -6.53E-01 | 8.67E-02 | 3.49E-01 | 3.31E-01 | 8.55E-08 | Yes | Ncald | intron_variant | MODIFIER |
| 15 | 37626238 | AX-170335148 | 0 | T | C | 0.026 | -6.53E-01 | 8.67E-02 | 3.49E-01 | 3.31E-01 | 8.55E-08 | Yes | Ncald | intron_variant | MODIFIER |
| 15 | 37633104 | AX-97820572 | 0 | A | G | 0.026 | -6.53E-01 | 8.67E-02 | 3.49E-01 | 3.31E-01 | 8.55E-08 | Yes | Ncald | intron_variant | MODIFIER |
| 15 | 37651780 | AX-170087821 | 0 | G | A | 0.026 | -6.53E-01 | 8.67E-02 | 3.49E-01 | 3.31E-01 | 8.55E-08 | Yes | Ncald | intron_variant | MODIFIER |
| 15 | 37656252 | AX-170285619 | 0 | T | C | 0.026 | -6.53E-01 | 8.67E-02 | 3.49E-01 | 3.31E-01 | 8.55E-08 | Yes | Ncald | intron_variant | MODIFIER |
| 15 | 37662412 | AX-170087822 | 0 | T | C | 0.026 | -6.53E-01 | 8.67E-02 | 3.49E-01 | 3.31E-01 | 8.55E-08 | Yes | Ncald | intron_variant | MODIFIER |
| 15 | 37719926 | AX-170202998 | 0 | A | G | 0.026 | -6.53E-01 | 8.67E-02 | 3.49E-01 | 3.31E-01 | 8.55E-08 | Yes | Ncald | intron_variant | MODIFIER |
| 15 | 37807633 | AX-170319416 | 0 | T | A | 0.026 | -6.53E-01 | 8.67E-02 | 3.49E-01 | 3.31E-01 | 8.55E-08 | Yes | | intergenic_variant | MODIFIER |
| 15 | 37976176 | AX-170011472 | 0 | T | C | 0.026 | -6.53E-01 | 8.67E-02 | 3.49E-01 | 3.31E-01 | 8.55E-08 | Yes | Ubr5 | synonymous_variant | LOW |
| 15 | 38228920 | AX-170105074 | 0 | C | A | 0.026 | -6.53E-01 | 8.67E-02 | 3.49E-01 | 3.31E-01 | 8.55E-08 | Yes | Odf1 | downstream_gene_variant | MODIFIER |
| 15 | 38230042 | AX-170362798 | 0 | C | T | 0.026 | -6.53E-01 | 8.67E-02 | 3.49E-01 | 3.31E-01 | 8.55E-08 | Yes | Odf1 | downstream_gene_variant | MODIFIER |
| 15 | 38279933 | AX-170055287 | 0 | A | G | 0.026 | -6.53E-01 | 8.67E-02 | 3.49E-01 | 3.31E-01 | 8.55E-08 | Yes | Gm49314 | upstream_gene_variant | MODIFIER |
| 15 | 38280296 | AX-170209007 | 0 | G | A | 0.026 | -6.53E-01 | 8.67E-02 | 3.49E-01 | 3.31E-01 | 8.55E-08 | Yes | Gm49314 | upstream_gene_variant | MODIFIER |
| 15 | 38325899 | AX-170393429 | 0 | C | T | 0.026 | -6.53E-01 | 8.67E-02 | 3.49E-01 | 3.31E-01 | 8.55E-08 | Yes | Gm35248 | intron_variant,non_coding_transcript_variant | MODIFIER |
| 15 | 38327290 | AX-170476936 | 0 | C | T | 0.026 | -6.53E-01 | 8.67E-02 | 3.49E-01 | 3.31E-01 | 8.55E-08 | Yes | Gm35248 | intron_variant,non_coding_transcript_variant | MODIFIER |
| 15 | 38358688 | AX-170456115 | 0 | A | C | 0.026 | -6.53E-01 | 8.67E-02 | 3.49E-01 | 3.31E-01 | 8.55E-08 | Yes | | intergenic_variant | MODIFIER |
| 15 | 38363048 | AX-170140252 | 0 | C | A | 0.026 | -6.53E-01 | 8.67E-02 | 3.49E-01 | 3.31E-01 | 8.55E-08 | Yes | | intergenic_variant | MODIFIER |
| 15 | 38887433 | AX-170145798 | 0 | G | A | 0.026 | -6.53E-01 | 8.67E-02 | 3.49E-01 | 3.31E-01 | 8.55E-08 | Yes | | intergenic_variant | MODIFIER |
| 15 | 38891367 | AX-170154359 | 0 | T | G | 0.026 | -6.53E-01 | 8.67E-02 | 3.49E-01 | 3.31E-01 | 8.55E-08 | Yes | | intergenic_variant | MODIFIER |
| 15 | 38895563 | AX-97841172 | 0 | G | A | 0.026 | -6.53E-01 | 8.67E-02 | 3.49E-01 | 3.31E-01 | 8.55E-08 | Yes | | intergenic_variant | MODIFIER |
| 15 | 38910152 | AX-170454581 | 0 | T | C | 0.026 | -6.53E-01 | 8.67E-02 | 3.49E-01 | 3.31E-01 | 8.55E-08 | Yes | | intergenic_variant | MODIFIER |
| 15 | 38910227 | AX-170290664 | 0 | A | G | 0.026 | -6.53E-01 | 8.67E-02 | 3.49E-01 | 3.31E-01 | 8.55E-08 | Yes | | intergenic_variant | MODIFIER |
| 15 | 38915970 | AX-170055291 | 0 | G | A | 0.026 | -6.53E-01 | 8.67E-02 | 3.49E-01 | 3.31E-01 | 8.55E-08 | Yes | | intergenic_variant | MODIFIER |
| 15 | 38929246 | AX-170227674 | 0 | G | T | 0.026 | -6.53E-01 | 8.67E-02 | 3.49E-01 | 3.31E-01 | 8.55E-08 | Yes | Baalc | upstream_gene_variant | MODIFIER |
| 15 | 38986550 | AX-170228289 | 0 | A | G | 0.026 | -6.53E-01 | 8.67E-02 | 3.49E-01 | 3.31E-01 | 8.55E-08 | Yes | Gm4740 | upstream_gene_variant | MODIFIER |
| 15 | 38991035 | AX-170503210 | 0 | C | T | 0.026 | -6.53E-01 | 8.67E-02 | 3.49E-01 | 3.31E-01 | 8.55E-08 | Yes | Gm4740 | non_coding_transcript_exon_variant | MODIFIER |
| 15 | 39026526 | AX-170196142 | 0 | T | C | 0.026 | -6.53E-01 | 8.67E-02 | 3.49E-01 | 3.31E-01 | 8.55E-08 | Yes | Fzd6 | intron_variant | MODIFIER |
| 15 | 39026877 | AX-170415579 | 0 | T | C | 0.026 | -6.53E-01 | 8.67E-02 | 3.49E-01 | 3.31E-01 | 8.55E-08 | Yes | Fzd6 | intron_variant | MODIFIER |
| 15 | 39083519 | AX-170272041 | 0 | A | G | 0.026 | -6.53E-01 | 8.67E-02 | 3.49E-01 | 3.31E-01 | 8.55E-08 | Yes | Cthrc1 | intron_variant | MODIFIER |
| 15 | 39125151 | AX-97820931 | 0 | A | C | 0.026 | -6.53E-01 | 8.67E-02 | 3.49E-01 | 3.31E-01 | 8.55E-08 | Yes | Dcaf13 | intron_variant | MODIFIER |
| 15 | 39603272 | AX-170263111 | 0 | A | G | 0.026 | -6.53E-01 | 8.67E-02 | 3.49E-01 | 3.31E-01 | 8.55E-08 | Yes | Rims2 | intron_variant | MODIFIER |
| 15 | 56680238 | AX-170140877 | 0 | G | C | 0.035 | -5.35E-01 | 8.40E-02 | 8.84E-01 | 9.20E-01 | 9.56E-08 | Yes | Has2 | intron_variant | MODIFIER |
| 15 | 41724822 | AX-170129058 | 1 | G | A | 0.045 | -5.79E-01 | 9.13E-02 | 8.85E-01 | 9.05E-01 | 1.12E-07 | Yes | Oxr1 | intron_variant | MODIFIER |
| 15 | 54574910 | AX-170055579 | 0 | T | A | 0.035 | -5.38E-01 | 8.42E-02 | 8.24E-01 | 8.53E-01 | 1.21E-07 | Yes | Mal2 | intron_variant | MODIFIER |
| 15 | 56683057 | AX-170420174 | 0 | T | A | 0.035 | -5.38E-01 | 8.42E-02 | 8.24E-01 | 8.53E-01 | 1.21E-07 | Yes | Has2 | intron_variant | MODIFIER |
| 15 | 29875341 | AX-170208249 | 0 | C | A | 0.026 | -6.73E-01 | 8.39E-02 | 2.29E-01 | 2.04E-01 | 1.30E-07 | Yes | | intergenic_variant | MODIFIER |
| 15 | 30336105 | AX-170055133 | 0 | T | A | 0.026 | -6.73E-01 | 8.39E-02 | 2.29E-01 | 2.04E-01 | 1.30E-07 | Yes | Ctnnd2 | intron_variant | MODIFIER |
| 15 | 36928815 | AX-170388369 | 1 | C | T | 0.027 | -6.43E-01 | 8.79E-02 | 3.74E-01 | 3.57E-01 | 1.52E-07 | Yes | Gm41300 | intron_variant,non_coding_transcript_variant | MODIFIER |
| 15 | 36922175 | AX-97782715 | 1 | T | C | 0.027 | -6.39E-01 | 8.77E-02 | 3.77E-01 | 3.61E-01 | 1.68E-07 | Yes | Gm41300 | intron_variant,non_coding_transcript_variant | MODIFIER |
| 15 | 53715791 | AX-97853398 | 0 | G | C | 0.035 | -5.19E-01 | 8.64E-02 | 1.14E+00 | 1.19E+00 | 1.69E-07 | Yes | Samd12 | intron_variant | MODIFIER |
| 15 | 25174176 | AX-170311354 | 0 | C | T | 0.026 | -7.00E-01 | 7.69E-02 | 9.52E-02 | 6.38E-02 | 2.02E-07 | Yes | 9230109A22Rik | intron_variant,non_coding_transcript_variant | MODIFIER |
| 15 | 26149773 | AX-97843559 | 0 | T | A | 0.026 | -7.00E-01 | 7.69E-02 | 9.52E-02 | 6.38E-02 | 2.02E-07 | Yes | | intergenic_variant | MODIFIER |
| 15 | 26441038 | AX-170332327 | 0 | G | A | 0.026 | -7.00E-01 | 7.69E-02 | 9.52E-02 | 6.38E-02 | 2.02E-07 | Yes | | intergenic_variant | MODIFIER |
| 15 | 36942427 | AX-170011450 | 1 | G | A | 0.027 | -6.47E-01 | 8.54E-02 | 2.96E-01 | 2.76E-01 | 2.02E-07 | Yes | Gm34590 | intron_variant,non_coding_transcript_variant | MODIFIER |
| 15 | 36361471 | AX-170249101 | 1 | C | T | 0.027 | -6.66E-01 | 8.51E-02 | 2.43E-01 | 2.21E-01 | 2.30E-07 | Yes | Gm33936 | intron_variant,non_coding_transcript_variant | MODIFIER |
| 15 | 25392213 | AX-170055049 | 1 | A | G | 0.027 | -6.70E-01 | 8.37E-02 | 2.07E-01 | 1.82E-01 | 2.66E-07 | Yes | Basp1 | intron_variant | MODIFIER |
| 15 | 25173337 | AX-170328839 | 0 | A | T | 0.026 | -6.98E-01 | 9.11E-02 | 2.67E-01 | 2.59E-01 | 2.66E-07 | Yes | 9230109A22Rik | intron_variant,non_coding_transcript_variant | MODIFIER |
| 15 | 37127934 | AX-170248451 | 0 | A | G | 0.026 | -5.89E-01 | 8.01E-02 | 3.20E-01 | 3.08E-01 | 3.49E-07 | Yes | | intergenic_variant | MODIFIER |
| 15 | 45934247 | AX-97829204 | 1 | C | A | 0.036 | -5.53E-01 | 8.80E-02 | 7.14E-01 | 7.31E-01 | 4.12E-07 | Yes | | intergenic_variant | MODIFIER |

| | | | | | | | | | | | | | | | |
|---|---|---|---|---|---|---|---|---|---|---|---|---|---|---|---|
| 15 | 53572842 | AX-170119417 | 0 | T | G | 0.035 | -5.68E-01 | 9.44E-02 | 8.64E-01 | 8.84E-01 | 5.16E-07 | Yes | Samd12 | intron_variant | MODIFIER |
| 15 | 37115680 | AX-170226877 | 0 | A | C | 0.026 | -6.16E-01 | 8.96E-02 | 4.22E-01 | 4.14E-01 | 5.72E-07 | Yes | | intergenic_variant | MODIFIER |
| 15 | 54568155 | AX-170468894 | 0 | G | A | 0.035 | -5.63E-01 | 9.30E-02 | 8.12E-01 | 8.22E-01 | 5.84E-07 | Yes | Mal2 | upstream_gene_variant | MODIFIER |
| 15 | 54580294 | AX-170055582 | 0 | C | A | 0.035 | -5.63E-01 | 9.30E-02 | 8.12E-01 | 8.22E-01 | 5.84E-07 | Yes | Mal2 | intron_variant | MODIFIER |
| 15 | 54588163 | AX-170363937 | 0 | T | C | 0.035 | -5.63E-01 | 9.30E-02 | 8.12E-01 | 8.22E-01 | 5.84E-07 | Yes | Mal2 | intron_variant | MODIFIER |
| 15 | 54619007 | AX-170273771 | 0 | G | A | 0.035 | -5.63E-01 | 9.30E-02 | 8.12E-01 | 8.22E-01 | 5.84E-07 | Yes | | intergenic_variant | MODIFIER |
| 15 | 54631856 | AX-170011728 | 0 | C | T | 0.035 | -5.63E-01 | 9.30E-02 | 8.12E-01 | 8.22E-01 | 5.84E-07 | Yes | | intergenic_variant | MODIFIER |
| 15 | 54646842 | AX-170378569 | 0 | T | A | 0.035 | -5.63E-01 | 9.30E-02 | 8.12E-01 | 8.22E-01 | 5.84E-07 | Yes | | intergenic_variant | MODIFIER |
| 15 | 54648982 | AX-170011734 | 0 | C | T | 0.035 | -5.63E-01 | 9.30E-02 | 8.12E-01 | 8.22E-01 | 5.84E-07 | Yes | | intergenic_variant | MODIFIER |
| 15 | 54661112 | AX-97828766 | 0 | T | C | 0.035 | -5.63E-01 | 9.30E-02 | 8.12E-01 | 8.22E-01 | 5.84E-07 | Yes | | intergenic_variant | MODIFIER |
| 15 | 54695307 | AX-170178523 | 0 | C | T | 0.035 | -5.63E-01 | 9.30E-02 | 8.12E-01 | 8.22E-01 | 5.84E-07 | Yes | | intergenic_variant | MODIFIER |
| 15 | 54695675 | AX-97812748 | 0 | G | A | 0.035 | -5.63E-01 | 9.30E-02 | 8.12E-01 | 8.22E-01 | 5.84E-07 | Yes | | intergenic_variant | MODIFIER |
| 15 | 54724262 | AX-170011731 | 0 | T | C | 0.035 | -5.63E-01 | 9.30E-02 | 8.12E-01 | 8.22E-01 | 5.84E-07 | Yes | | intergenic_variant | MODIFIER |
| 15 | 54736192 | AX-97837245 | 0 | A | G | 0.035 | -5.63E-01 | 9.30E-02 | 8.12E-01 | 8.22E-01 | 5.84E-07 | Yes | | intergenic_variant | MODIFIER |
| 15 | 54744436 | AX-170011729 | 0 | T | C | 0.035 | -5.63E-01 | 9.30E-02 | 8.12E-01 | 8.22E-01 | 5.84E-07 | Yes | Ccn3 | upstream_gene_variant | MODIFIER |
| 15 | 54761370 | AX-170011730 | 0 | T | C | 0.035 | -5.63E-01 | 9.30E-02 | 8.12E-01 | 8.22E-01 | 5.84E-07 | Yes | Gm7577 | downstream_gene_variant | MODIFIER |
| 15 | 54769124 | AX-170333599 | 0 | C | G | 0.035 | -5.63E-01 | 9.30E-02 | 8.12E-01 | 8.22E-01 | 5.84E-07 | Yes | Gm7577 | upstream_gene_variant | MODIFIER |
| 15 | 54772796 | AX-97786830 | 0 | G | T | 0.035 | -5.63E-01 | 9.30E-02 | 8.12E-01 | 8.22E-01 | 5.84E-07 | Yes | | intergenic_variant | MODIFIER |
| 15 | 54789738 | AX-170297203 | 0 | C | T | 0.035 | -5.63E-01 | 9.30E-02 | 8.12E-01 | 8.22E-01 | 5.84E-07 | Yes | | intergenic_variant | MODIFIER |
| 15 | 54793130 | AX-97789961 | 0 | A | G | 0.035 | -5.63E-01 | 9.30E-02 | 8.12E-01 | 8.22E-01 | 5.84E-07 | Yes | | intergenic_variant | MODIFIER |
| 15 | 54803989 | AX-170328039 | 0 | T | C | 0.035 | -5.63E-01 | 9.30E-02 | 8.12E-01 | 8.22E-01 | 5.84E-07 | Yes | | intergenic_variant | MODIFIER |
| 15 | 54810616 | AX-97845544 | 0 | C | A | 0.035 | -5.63E-01 | 9.30E-02 | 8.12E-01 | 8.22E-01 | 5.84E-07 | Yes | | regulatory_region_variant | MODIFIER |
| 15 | 54813244 | AX-170475172 | 0 | G | A | 0.035 | -5.63E-01 | 9.30E-02 | 8.12E-01 | 8.22E-01 | 5.84E-07 | Yes | | intergenic_variant | MODIFIER |
| 15 | 54817350 | AX-170127896 | 0 | G | A | 0.035 | -5.63E-01 | 9.30E-02 | 8.12E-01 | 8.22E-01 | 5.84E-07 | Yes | | intergenic_variant | MODIFIER |
| 15 | 54832217 | AX-170280307 | 0 | A | G | 0.035 | -5.63E-01 | 9.30E-02 | 8.12E-01 | 8.22E-01 | 5.84E-07 | Yes | | intergenic_variant | MODIFIER |
| 15 | 54838115 | AX-170289955 | 0 | G | A | 0.035 | -5.63E-01 | 9.30E-02 | 8.12E-01 | 8.22E-01 | 5.84E-07 | Yes | Enpp2 | downstream_gene_variant | MODIFIER |
| 15 | 54858291 | AX-170185933 | 0 | G | A | 0.035 | -5.63E-01 | 9.30E-02 | 8.12E-01 | 8.22E-01 | 5.84E-07 | Yes | Enpp2 | intron_variant | MODIFIER |
| 15 | 54881299 | AX-97798182 | 0 | T | C | 0.035 | -5.63E-01 | 9.30E-02 | 8.12E-01 | 8.22E-01 | 5.84E-07 | Yes | Enpp2 | intron_variant | MODIFIER |
| 15 | 54885770 | AX-97820782 | 0 | G | A | 0.035 | -5.63E-01 | 9.30E-02 | 8.12E-01 | 8.22E-01 | 5.84E-07 | Yes | Enpp2 | intron_variant | MODIFIER |
| 15 | 54909106 | AX-170240111 | 0 | G | A | 0.035 | -5.63E-01 | 9.30E-02 | 8.12E-01 | 8.22E-01 | 5.84E-07 | Yes | Enpp2 | intron_variant | MODIFIER |
| 15 | 54912060 | AX-170098369 | 0 | C | G | 0.035 | -5.63E-01 | 9.30E-02 | 8.12E-01 | 8.22E-01 | 5.84E-07 | Yes | Enpp2 | intron_variant | MODIFIER |
| 15 | 54912175 | AX-170494660 | 0 | G | A | 0.035 | -5.63E-01 | 9.30E-02 | 8.12E-01 | 8.22E-01 | 5.84E-07 | Yes | Enpp2 | intron_variant | MODIFIER |
| 15 | 54913824 | AX-97829744 | 0 | C | A | 0.035 | -5.63E-01 | 9.30E-02 | 8.12E-01 | 8.22E-01 | 5.84E-07 | Yes | Enpp2 | intron_variant | MODIFIER |
| 15 | 54932504 | AX-170150913 | 0 | G | A | 0.035 | -5.63E-01 | 9.30E-02 | 8.12E-01 | 8.22E-01 | 5.84E-07 | Yes | Enpp2 | intron_variant | MODIFIER |
| 15 | 54934421 | AX-170516732 | 0 | T | C | 0.035 | -5.63E-01 | 9.30E-02 | 8.12E-01 | 8.22E-01 | 5.84E-07 | Yes | Enpp2 | intron_variant | MODIFIER |
| 15 | 54936927 | AX-170186552 | 0 | G | A | 0.035 | -5.63E-01 | 9.30E-02 | 8.12E-01 | 8.22E-01 | 5.84E-07 | Yes | Enpp2 | intron_variant | MODIFIER |
| 15 | 54991244 | AX-97829842 | 0 | G | T | 0.035 | -5.63E-01 | 9.30E-02 | 8.12E-01 | 8.22E-01 | 5.84E-07 | Yes | 4930478E11Rik | upstream_gene_variant | MODIFIER |
| 15 | 54991325 | AX-170239257 | 0 | A | G | 0.035 | -5.63E-01 | 9.30E-02 | 8.12E-01 | 8.22E-01 | 5.84E-07 | Yes | 4930478E11Rik | upstream_gene_variant | MODIFIER |
| 15 | 55046147 | AX-97787224 | 0 | A | C | 0.035 | -5.63E-01 | 9.30E-02 | 8.12E-01 | 8.22E-01 | 5.84E-07 | Yes | Taf2 | intron_variant | MODIFIER |
| 15 | 55047215 | AX-170416122 | 0 | G | A | 0.035 | -5.63E-01 | 9.30E-02 | 8.12E-01 | 8.22E-01 | 5.84E-07 | Yes | Taf2 | intron_variant | MODIFIER |
| 15 | 55049192 | AX-170149614 | 0 | C | T | 0.035 | -5.63E-01 | 9.30E-02 | 8.12E-01 | 8.22E-01 | 5.84E-07 | Yes | Taf2 | intron_variant | MODIFIER |
| 15 | 55119027 | AX-97829843 | 0 | G | A | 0.035 | -5.63E-01 | 9.30E-02 | 8.12E-01 | 8.22E-01 | 5.84E-07 | Yes | Deptor | intron_variant | MODIFIER |
| 15 | 55124157 | AX-170329414 | 0 | G | A | 0.035 | -5.63E-01 | 9.30E-02 | 8.12E-01 | 8.22E-01 | 5.84E-07 | Yes | Deptor | intron_variant | MODIFIER |
| 15 | 55148866 | AX-170087868 | 0 | G | A | 0.035 | -5.63E-01 | 9.30E-02 | 8.12E-01 | 8.22E-01 | 5.84E-07 | Yes | Deptor | intron_variant | MODIFIER |
| 15 | 55169621 | AX-170011742 | 0 | C | T | 0.035 | -5.63E-01 | 9.30E-02 | 8.12E-01 | 8.22E-01 | 5.84E-07 | Yes | Deptor | intron_variant | MODIFIER |
| 15 | 55169874 | AX-170011736 | 0 | T | C | 0.035 | -5.63E-01 | 9.30E-02 | 8.12E-01 | 8.22E-01 | 5.84E-07 | Yes | Deptor | intron_variant | MODIFIER |
| 15 | 55184808 | AX-170055591 | 0 | T | A | 0.035 | -5.63E-01 | 9.30E-02 | 8.12E-01 | 8.22E-01 | 5.84E-07 | Yes | Deptor | intron_variant | MODIFIER |
| 15 | 55187980 | AX-97805283 | 0 | A | G | 0.035 | -5.63E-01 | 9.30E-02 | 8.12E-01 | 8.22E-01 | 5.84E-07 | Yes | Deptor | intron_variant | MODIFIER |
| 15 | 55194366 | AX-170113127 | 0 | A | C | 0.035 | -5.63E-01 | 9.30E-02 | 8.12E-01 | 8.22E-01 | 5.84E-07 | Yes | Deptor | intron_variant | MODIFIER |
| 15 | 55196400 | AX-170107064 | 0 | C | G | 0.035 | -5.63E-01 | 9.30E-02 | 8.12E-01 | 8.22E-01 | 5.84E-07 | Yes | Deptor | intron_variant | MODIFIER |
| 15 | 55222003 | AX-97789863 | 0 | A | G | 0.035 | -5.63E-01 | 9.30E-02 | 8.12E-01 | 8.22E-01 | 5.84E-07 | Yes | Deptor | intron_variant | MODIFIER |
| 15 | 55245631 | AX-97795794 | 0 | G | A | 0.035 | -5.63E-01 | 9.30E-02 | 8.12E-01 | 8.22E-01 | 5.84E-07 | Yes | Deptor | intron_variant | MODIFIER |
| 15 | 55256278 | AX-170420921 | 0 | A | G | 0.035 | -5.63E-01 | 9.30E-02 | 8.12E-01 | 8.22E-01 | 5.84E-07 | Yes | Deptor | downstream_gene_variant | MODIFIER |
| 15 | 55256808 | AX-170533096 | 0 | A | G | 0.035 | -5.63E-01 | 9.30E-02 | 8.12E-01 | 8.22E-01 | 5.84E-07 | Yes | Deptor | downstream_gene_variant | MODIFIER |

| | | | | | | | | | | | | | | | |
|---|---|---|---|---|---|---|---|---|---|---|---|---|---|---|---|
| 15 | 55268201 | AX-170011737 | 0 | C | A | 0.035 | -5.63E-01 | 9.30E-02 | 8.12E-01 | 8.22E-01 | 5.84E-07 | Yes | | regulatory_region_variant | MODIFIER |
| 15 | 55268247 | AX-170011740 | 0 | T | C | 0.035 | -5.63E-01 | 9.30E-02 | 8.12E-01 | 8.22E-01 | 5.84E-07 | Yes | | regulatory_region_variant | MODIFIER |
| 15 | 55268902 | AX-97796060 | 0 | C | T | 0.035 | -5.63E-01 | 9.30E-02 | 8.12E-01 | 8.22E-01 | 5.84E-07 | Yes | | regulatory_region_variant | MODIFIER |
| 15 | 55270441 | AX-170176930 | 0 | T | A | 0.035 | -5.63E-01 | 9.30E-02 | 8.12E-01 | 8.22E-01 | 5.84E-07 | Yes | | intergenic_variant | MODIFIER |
| 15 | 55435663 | AX-97784209 | 0 | T | C | 0.035 | -5.63E-01 | 9.30E-02 | 8.12E-01 | 8.22E-01 | 5.84E-07 | Yes | Col14a1 | intron_variant | MODIFIER |
| 15 | 55481623 | AX-170437893 | 0 | A | T | 0.035 | -5.63E-01 | 9.30E-02 | 8.12E-01 | 8.22E-01 | 5.84E-07 | Yes | Col14a1 | intron_variant | MODIFIER |
| 15 | 55482358 | AX-170117245 | 0 | T | C | 0.035 | -5.63E-01 | 9.30E-02 | 8.12E-01 | 8.22E-01 | 5.84E-07 | Yes | Col14a1 | intron_variant | MODIFIER |
| 15 | 55484077 | AX-170122840 | 0 | A | C | 0.035 | -5.63E-01 | 9.30E-02 | 8.12E-01 | 8.22E-01 | 5.84E-07 | Yes | Col14a1 | intron_variant | MODIFIER |
| 15 | 55494735 | AX-170113863 | 0 | T | C | 0.035 | -5.63E-01 | 9.30E-02 | 8.12E-01 | 8.22E-01 | 5.84E-07 | Yes | Col14a1 | intron_variant | MODIFIER |
| 15 | 56016364 | AX-170055608 | 0 | C | T | 0.035 | -5.63E-01 | 9.30E-02 | 8.12E-01 | 8.22E-01 | 5.84E-07 | Yes | | intergenic_variant | MODIFIER |
| 15 | 56020130 | AX-97807356 | 0 | A | G | 0.035 | -5.63E-01 | 9.30E-02 | 8.12E-01 | 8.22E-01 | 5.84E-07 | Yes | | intergenic_variant | MODIFIER |
| 15 | 56087720 | AX-97797926 | 0 | T | C | 0.035 | -5.63E-01 | 9.30E-02 | 8.12E-01 | 8.22E-01 | 5.84E-07 | Yes | | intergenic_variant | MODIFIER |
| 15 | 56157403 | AX-170448227 | 0 | C | T | 0.035 | -5.63E-01 | 9.30E-02 | 8.12E-01 | 8.22E-01 | 5.84E-07 | Yes | | intergenic_variant | MODIFIER |
| 15 | 56172033 | AX-97837736 | 0 | G | A | 0.035 | -5.63E-01 | 9.30E-02 | 8.12E-01 | 8.22E-01 | 5.84E-07 | Yes | Hba-ps3 | non_coding_transcript_exon_variant | MODIFIER |
| 15 | 56411448 | AX-170055613 | 0 | C | T | 0.035 | -5.63E-01 | 9.30E-02 | 8.12E-01 | 8.22E-01 | 5.84E-07 | Yes | Gm49213 | upstream_gene_variant | MODIFIER |
| 15 | 56411486 | AX-170513251 | 0 | C | T | 0.035 | -5.63E-01 | 9.30E-02 | 8.12E-01 | 8.22E-01 | 5.84E-07 | Yes | Gm49213 | upstream_gene_variant | MODIFIER |
| 15 | 56426333 | AX-97829380 | 0 | C | T | 0.035 | -5.63E-01 | 9.30E-02 | 8.12E-01 | 8.22E-01 | 5.84E-07 | Yes | | regulatory_region_variant | MODIFIER |
| 15 | 56428081 | AX-170310560 | 0 | G | T | 0.035 | -5.63E-01 | 9.30E-02 | 8.12E-01 | 8.22E-01 | 5.84E-07 | Yes | | intergenic_variant | MODIFIER |
| 15 | 56435628 | AX-97787651 | 0 | T | C | 0.035 | -5.63E-01 | 9.30E-02 | 8.12E-01 | 8.22E-01 | 5.84E-07 | Yes | | intergenic_variant | MODIFIER |
| 15 | 56469418 | AX-170519479 | 0 | G | T | 0.035 | -5.63E-01 | 9.30E-02 | 8.12E-01 | 8.22E-01 | 5.84E-07 | Yes | | intergenic_variant | MODIFIER |
| 15 | 56471660 | AX-170087871 | 0 | A | G | 0.035 | -5.63E-01 | 9.30E-02 | 8.12E-01 | 8.22E-01 | 5.84E-07 | Yes | | intergenic_variant | MODIFIER |
| 15 | 56515103 | AX-97829522 | 0 | T | C | 0.035 | -5.63E-01 | 9.30E-02 | 8.12E-01 | 8.22E-01 | 5.84E-07 | Yes | | intergenic_variant | MODIFIER |
| 15 | 56553845 | AX-170497046 | 0 | C | A | 0.035 | -5.63E-01 | 9.30E-02 | 8.12E-01 | 8.22E-01 | 5.84E-07 | Yes | | intergenic_variant | MODIFIER |
| 15 | 56554980 | AX-170324511 | 0 | T | C | 0.035 | -5.63E-01 | 9.30E-02 | 8.12E-01 | 8.22E-01 | 5.84E-07 | Yes | | intergenic_variant | MODIFIER |
| 15 | 56556074 | AX-170164411 | 0 | T | G | 0.035 | -5.63E-01 | 9.30E-02 | 8.12E-01 | 8.22E-01 | 5.84E-07 | Yes | | intergenic_variant | MODIFIER |
| 15 | 56559348 | AX-97851034 | 0 | A | G | 0.035 | -5.63E-01 | 9.30E-02 | 8.12E-01 | 8.22E-01 | 5.84E-07 | Yes | | intergenic_variant | MODIFIER |
| 15 | 56608715 | AX-170309840 | 0 | A | T | 0.035 | -5.63E-01 | 9.30E-02 | 8.12E-01 | 8.22E-01 | 5.84E-07 | Yes | | intergenic_variant | MODIFIER |
| 15 | 56616729 | AX-170278443 | 0 | C | T | 0.035 | -5.63E-01 | 9.30E-02 | 8.12E-01 | 8.22E-01 | 5.84E-07 | Yes | | regulatory_region_variant | MODIFIER |
| 15 | 56638583 | AX-170435581 | 0 | A | C | 0.035 | -5.63E-01 | 9.30E-02 | 8.12E-01 | 8.22E-01 | 5.84E-07 | Yes | | intergenic_variant | MODIFIER |
| 15 | 56650910 | AX-97847799 | 0 | A | C | 0.035 | -5.63E-01 | 9.30E-02 | 8.12E-01 | 8.22E-01 | 5.84E-07 | Yes | | intergenic_variant | MODIFIER |
| 15 | 56650978 | AX-170055615 | 0 | T | C | 0.035 | -5.63E-01 | 9.30E-02 | 8.12E-01 | 8.22E-01 | 5.84E-07 | Yes | | intergenic_variant | MODIFIER |
| 15 | 56655594 | AX-170055614 | 0 | G | A | 0.035 | -5.63E-01 | 9.30E-02 | 8.12E-01 | 8.22E-01 | 5.84E-07 | Yes | | intergenic_variant | MODIFIER |
| 15 | 56656285 | AX-170134058 | 0 | T | G | 0.035 | -5.63E-01 | 9.30E-02 | 8.12E-01 | 8.22E-01 | 5.84E-07 | Yes | | intergenic_variant | MODIFIER |
| 15 | 56700713 | AX-97851008 | 0 | A | G | 0.035 | -5.63E-01 | 9.30E-02 | 8.12E-01 | 8.22E-01 | 5.84E-07 | Yes | Has2os | intron_variant,non_coding_transcript_variant | MODIFIER |
| 15 | 56701266 | AX-170144447 | 0 | A | G | 0.035 | -5.63E-01 | 9.30E-02 | 8.12E-01 | 8.22E-01 | 5.84E-07 | Yes | Has2os | intron_variant,non_coding_transcript_variant | MODIFIER |
| 15 | 56701927 | AX-170456511 | 0 | T | A | 0.035 | -5.63E-01 | 9.30E-02 | 8.12E-01 | 8.22E-01 | 5.84E-07 | Yes | Has2os | intron_variant,non_coding_transcript_variant | MODIFIER |
| 15 | 56715066 | AX-170087872 | 0 | A | T | 0.035 | -5.63E-01 | 9.30E-02 | 8.12E-01 | 8.22E-01 | 5.84E-07 | Yes | Has2os | intron_variant,non_coding_transcript_variant | MODIFIER |
| 15 | 56727936 | AX-170534815 | 0 | G | C | 0.035 | -5.63E-01 | 9.30E-02 | 8.12E-01 | 8.22E-01 | 5.84E-07 | Yes | Has2os | intron_variant,non_coding_transcript_variant | MODIFIER |
| 15 | 56733820 | AX-170346662 | 0 | A | G | 0.035 | -5.63E-01 | 9.30E-02 | 8.12E-01 | 8.22E-01 | 5.84E-07 | Yes | Has2os | intron_variant,non_coding_transcript_variant | MODIFIER |
| 15 | 56738547 | AX-97829498 | 0 | A | G | 0.035 | -5.63E-01 | 9.30E-02 | 8.12E-01 | 8.22E-01 | 5.84E-07 | Yes | Has2os | intron_variant,non_coding_transcript_variant | MODIFIER |
| 15 | 56750293 | AX-170516196 | 0 | A | G | 0.035 | -5.63E-01 | 9.30E-02 | 8.12E-01 | 8.22E-01 | 5.84E-07 | Yes | Has2os | intron_variant,non_coding_transcript_variant | MODIFIER |
| 15 | 56796117 | AX-170287076 | 0 | G | A | 0.035 | -5.63E-01 | 9.30E-02 | 8.12E-01 | 8.22E-01 | 5.84E-07 | Yes | | intergenic_variant | MODIFIER |
| 15 | 56798499 | AX-168403535 | 0 | G | A | 0.035 | -5.63E-01 | 9.30E-02 | 8.12E-01 | 8.22E-01 | 5.84E-07 | Yes | | intergenic_variant | MODIFIER |
| 15 | 56816577 | AX-170232174 | 0 | G | A | 0.035 | -5.63E-01 | 9.30E-02 | 8.12E-01 | 8.22E-01 | 5.84E-07 | Yes | | intergenic_variant | MODIFIER |
| 15 | 56816793 | AX-170011768 | 0 | G | A | 0.035 | -5.63E-01 | 9.30E-02 | 8.12E-01 | 8.22E-01 | 5.84E-07 | Yes | | intergenic_variant | MODIFIER |
| 15 | 56816924 | AX-170011763 | 0 | C | T | 0.035 | -5.63E-01 | 9.30E-02 | 8.12E-01 | 8.22E-01 | 5.84E-07 | Yes | | intergenic_variant | MODIFIER |
| 15 | 56864840 | AX-170011761 | 0 | G | C | 0.035 | -5.63E-01 | 9.30E-02 | 8.12E-01 | 8.22E-01 | 5.84E-07 | Yes | | intergenic_variant | MODIFIER |
| 15 | 56864993 | AX-170170473 | 0 | G | C | 0.035 | -5.63E-01 | 9.30E-02 | 8.12E-01 | 8.22E-01 | 5.84E-07 | Yes | | intergenic_variant | MODIFIER |
| 15 | 56865678 | AX-170093262 | 0 | G | A | 0.035 | -5.63E-01 | 9.30E-02 | 8.12E-01 | 8.22E-01 | 5.84E-07 | Yes | | regulatory_region_variant | MODIFIER |
| 15 | 56872726 | AX-170011764 | 0 | T | A | 0.035 | -5.63E-01 | 9.30E-02 | 8.12E-01 | 8.22E-01 | 5.84E-07 | Yes | | intergenic_variant | MODIFIER |
| 15 | 56878202 | AX-170495346 | 0 | C | G | 0.035 | -5.63E-01 | 9.30E-02 | 8.12E-01 | 8.22E-01 | 5.84E-07 | Yes | Gm2582 | downstream_gene_variant | MODIFIER |
| 15 | 56898090 | AX-97829930 | 0 | T | C | 0.035 | -5.63E-01 | 9.30E-02 | 8.12E-01 | 8.22E-01 | 5.84E-07 | Yes | Gm2582 | intron_variant,non_coding_transcript_variant | MODIFIER |
| 15 | 56921270 | AX-97829922 | 0 | T | C | 0.035 | -5.63E-01 | 9.30E-02 | 8.12E-01 | 8.22E-01 | 5.84E-07 | Yes | Gm2582 | intron_variant,non_coding_transcript_variant | MODIFIER |
| 15 | 56924579 | AX-170402310 | 0 | A | G | 0.035 | -5.63E-01 | 9.30E-02 | 8.12E-01 | 8.22E-01 | 5.84E-07 | Yes | Gm2582 | intron_variant,non_coding_transcript_variant | MODIFIER |

| | | | | | | | | | | | | | | | |
|---|---|---|---|---|---|---|---|---|---|---|---|---|---|---|---|
| 15 | 56928213 | AX-170147498 | 0 | C | T | 0.035 | -5.63E-01 | 9.30E-02 | 8.12E-01 | 8.22E-01 | 5.84E-07 | Yes | Gm2582 | intron_variant,non_coding_transcript_variant | MODIFIER |
| 15 | 56951534 | AX-97824655 | 0 | A | G | 0.035 | -5.63E-01 | 9.30E-02 | 8.12E-01 | 8.22E-01 | 5.84E-07 | Yes | Gm2582 | intron_variant,non_coding_transcript_variant | MODIFIER |
| 15 | 56951736 | AX-170011760 | 0 | A | C | 0.035 | -5.63E-01 | 9.30E-02 | 8.12E-01 | 8.22E-01 | 5.84E-07 | Yes | Gm2582 | intron_variant,non_coding_transcript_variant | MODIFIER |
| 15 | 56971773 | AX-97818758 | 0 | C | T | 0.035 | -5.63E-01 | 9.30E-02 | 8.12E-01 | 8.22E-01 | 5.84E-07 | Yes | Gm2582 | intron_variant,non_coding_transcript_variant | MODIFIER |
| 15 | 56976643 | AX-170011766 | 0 | T | C | 0.035 | -5.63E-01 | 9.30E-02 | 8.12E-01 | 8.22E-01 | 5.84E-07 | Yes | Gm2582 | intron_variant,non_coding_transcript_variant | MODIFIER |
| 15 | 56977515 | AX-170271531 | 0 | T | C | 0.035 | -5.63E-01 | 9.30E-02 | 8.12E-01 | 8.22E-01 | 5.84E-07 | Yes | Gm2582 | intron_variant,non_coding_transcript_variant | MODIFIER |
| 15 | 56979214 | AX-170011762 | 0 | T | G | 0.035 | -5.63E-01 | 9.30E-02 | 8.12E-01 | 8.22E-01 | 5.84E-07 | Yes | Gm2582 | intron_variant,non_coding_transcript_variant | MODIFIER |
| 15 | 56980787 | AX-170303870 | 0 | T | C | 0.035 | -5.63E-01 | 9.30E-02 | 8.12E-01 | 8.22E-01 | 5.84E-07 | Yes | Gm2582 | intron_variant,non_coding_transcript_variant | MODIFIER |
| 15 | 57006051 | AX-97818146 | 0 | A | G | 0.035 | -5.63E-01 | 9.30E-02 | 8.12E-01 | 8.22E-01 | 5.84E-07 | Yes | Gm2582 | intron_variant,non_coding_transcript_variant | MODIFIER |
| 15 | 57013731 | AX-97800456 | 0 | A | C | 0.035 | -5.63E-01 | 9.30E-02 | 8.12E-01 | 8.22E-01 | 5.84E-07 | Yes | | intergenic_variant | MODIFIER |
| 15 | 57036867 | AX-170087873 | 0 | A | C | 0.035 | -5.63E-01 | 9.30E-02 | 8.12E-01 | 8.22E-01 | 5.84E-07 | Yes | | intergenic_variant | MODIFIER |
| 15 | 57037073 | AX-170011767 | 0 | A | G | 0.035 | -5.63E-01 | 9.30E-02 | 8.12E-01 | 8.22E-01 | 5.84E-07 | Yes | | intergenic_variant | MODIFIER |
| 15 | 57050331 | AX-170107438 | 0 | A | G | 0.035 | -5.63E-01 | 9.30E-02 | 8.12E-01 | 8.22E-01 | 5.84E-07 | Yes | | intergenic_variant | MODIFIER |
| 15 | 57075389 | AX-170208130 | 0 | G | C | 0.035 | -5.63E-01 | 9.30E-02 | 8.12E-01 | 8.22E-01 | 5.84E-07 | Yes | | intergenic_variant | MODIFIER |
| 15 | 57076007 | AX-170439781 | 0 | A | G | 0.035 | -5.63E-01 | 9.30E-02 | 8.12E-01 | 8.22E-01 | 5.84E-07 | Yes | | intergenic_variant | MODIFIER |
| 15 | 57103024 | AX-97846343 | 0 | A | G | 0.035 | -5.63E-01 | 9.30E-02 | 8.12E-01 | 8.22E-01 | 5.84E-07 | Yes | | intergenic_variant | MODIFIER |
| 15 | 57146843 | AX-97829947 | 0 | G | A | 0.035 | -5.63E-01 | 9.30E-02 | 8.12E-01 | 8.22E-01 | 5.84E-07 | Yes | | intergenic_variant | MODIFIER |
| 15 | 57169759 | AX-97829523 | 0 | A | G | 0.035 | -5.63E-01 | 9.30E-02 | 8.12E-01 | 8.22E-01 | 5.84E-07 | Yes | | intergenic_variant | MODIFIER |
| 15 | 57188538 | AX-170251680 | 0 | G | A | 0.035 | -5.63E-01 | 9.30E-02 | 8.12E-01 | 8.22E-01 | 5.84E-07 | Yes | | intergenic_variant | MODIFIER |
| 15 | 57188726 | AX-170437524 | 0 | T | C | 0.035 | -5.63E-01 | 9.30E-02 | 8.12E-01 | 8.22E-01 | 5.84E-07 | Yes | | intergenic_variant | MODIFIER |
| 15 | 57205045 | AX-170312036 | 0 | A | T | 0.035 | -5.63E-01 | 9.30E-02 | 8.12E-01 | 8.22E-01 | 5.84E-07 | Yes | | intergenic_variant | MODIFIER |
| 15 | 57205341 | AX-97829973 | 0 | T | G | 0.035 | -5.63E-01 | 9.30E-02 | 8.12E-01 | 8.22E-01 | 5.84E-07 | Yes | | intergenic_variant | MODIFIER |
| 15 | 57214150 | AX-170364610 | 0 | T | G | 0.035 | -5.63E-01 | 9.30E-02 | 8.12E-01 | 8.22E-01 | 5.84E-07 | Yes | | intergenic_variant | MODIFIER |
| 15 | 57214849 | AX-170456477 | 0 | T | C | 0.035 | -5.63E-01 | 9.30E-02 | 8.12E-01 | 8.22E-01 | 5.84E-07 | Yes | | intergenic_variant | MODIFIER |
| 15 | 57224172 | AX-170123682 | 0 | C | T | 0.035 | -5.63E-01 | 9.30E-02 | 8.12E-01 | 8.22E-01 | 5.84E-07 | Yes | | intergenic_variant | MODIFIER |
| 15 | 57224190 | AX-170506884 | 0 | A | C | 0.035 | -5.63E-01 | 9.30E-02 | 8.12E-01 | 8.22E-01 | 5.84E-07 | Yes | | intergenic_variant | MODIFIER |
| 15 | 57224234 | AX-170133505 | 0 | G | T | 0.035 | -5.63E-01 | 9.30E-02 | 8.12E-01 | 8.22E-01 | 5.84E-07 | Yes | | intergenic_variant | MODIFIER |
| 15 | 57268758 | AX-170518306 | 0 | A | T | 0.035 | -5.63E-01 | 9.30E-02 | 8.12E-01 | 8.22E-01 | 5.84E-07 | Yes | Slc22a22 | upstream_gene_variant | MODIFIER |
| 15 | 57288895 | AX-170211238 | 0 | A | G | 0.035 | -5.63E-01 | 9.30E-02 | 8.12E-01 | 8.22E-01 | 5.84E-07 | Yes | Slc22a22 | intron_variant | MODIFIER |
| 15 | 57289482 | AX-170147565 | 0 | T | C | 0.035 | -5.63E-01 | 9.30E-02 | 8.12E-01 | 8.22E-01 | 5.84E-07 | Yes | Slc22a22 | intron_variant | MODIFIER |
| 15 | 57299562 | AX-170220949 | 0 | C | T | 0.035 | -5.63E-01 | 9.30E-02 | 8.12E-01 | 8.22E-01 | 5.84E-07 | Yes | Slc22a22 | intron_variant | MODIFIER |
| 15 | 57299826 | AX-170168286 | 0 | T | C | 0.035 | -5.63E-01 | 9.30E-02 | 8.12E-01 | 8.22E-01 | 5.84E-07 | Yes | Slc22a22 | intron_variant | MODIFIER |
| 15 | 57303755 | AX-170430129 | 0 | A | G | 0.035 | -5.63E-01 | 9.30E-02 | 8.12E-01 | 8.22E-01 | 5.84E-07 | Yes | Slc22a22 | intron_variant | MODIFIER |
| 15 | 57316449 | AX-170055623 | 0 | C | A | 0.035 | -5.63E-01 | 9.30E-02 | 8.12E-01 | 8.22E-01 | 5.84E-07 | Yes | Slc22a22 | intron_variant | MODIFIER |
| 15 | 57321078 | AX-170055624 | 0 | A | G | 0.035 | -5.63E-01 | 9.30E-02 | 8.12E-01 | 8.22E-01 | 5.84E-07 | Yes | Slc22a22 | intron_variant | MODIFIER |
| 15 | 57326100 | AX-170411817 | 0 | A | G | 0.035 | -5.63E-01 | 9.30E-02 | 8.12E-01 | 8.22E-01 | 5.84E-07 | Yes | Slc22a22 | intron_variant | MODIFIER |
| 15 | 57335759 | AX-170474781 | 0 | T | A | 0.035 | -5.63E-01 | 9.30E-02 | 8.12E-01 | 8.22E-01 | 5.84E-07 | Yes | Slc22a22 | intron_variant | MODIFIER |
| 15 | 57349447 | AX-170223067 | 0 | C | T | 0.035 | -5.63E-01 | 9.30E-02 | 8.12E-01 | 8.22E-01 | 5.84E-07 | Yes | Slc22a22 | intron_variant | MODIFIER |
| 15 | 57368722 | AX-97809603 | 0 | C | T | 0.035 | -5.63E-01 | 9.30E-02 | 8.12E-01 | 8.22E-01 | 5.84E-07 | Yes | Slc22a22 | intron_variant | MODIFIER |
| 15 | 57380402 | AX-170256662 | 0 | G | A | 0.035 | -5.63E-01 | 9.30E-02 | 8.12E-01 | 8.22E-01 | 5.84E-07 | Yes | Slc22a22 | intron_variant | MODIFIER |
| 15 | 57380901 | AX-170188524 | 0 | C | A | 0.035 | -5.63E-01 | 9.30E-02 | 8.12E-01 | 8.22E-01 | 5.84E-07 | Yes | Slc22a22 | intron_variant | MODIFIER |
| 15 | 57399915 | AX-97814950 | 0 | T | C | 0.035 | -5.63E-01 | 9.30E-02 | 8.12E-01 | 8.22E-01 | 5.84E-07 | Yes | Slc22a22 | intron_variant | MODIFIER |
| 15 | 57400188 | AX-170533461 | 0 | G | A | 0.035 | -5.63E-01 | 9.30E-02 | 8.12E-01 | 8.22E-01 | 5.84E-07 | Yes | Slc22a22 | intron_variant | MODIFIER |
| 15 | 57402903 | AX-170011770 | 0 | G | A | 0.035 | -5.63E-01 | 9.30E-02 | 8.12E-01 | 8.22E-01 | 5.84E-07 | Yes | Slc22a22 | intron_variant | MODIFIER |
| 15 | 57428048 | AX-97829979 | 0 | G | A | 0.035 | -5.63E-01 | 9.30E-02 | 8.12E-01 | 8.22E-01 | 5.84E-07 | Yes | Slc22a22 | intron_variant | MODIFIER |
| 15 | 41734010 | AX-170326787 | 0 | C | A | 0.035 | -5.60E-01 | 8.63E-02 | 5.62E-01 | 5.73E-01 | 6.24E-07 | Yes | Oxr1 | intron_variant | MODIFIER |
| 15 | 54579849 | AX-170513693 | 1 | A | G | 0.036 | -5.57E-01 | 9.34E-02 | 8.73E-01 | 8.86E-01 | 6.28E-07 | Yes | Mal2 | intron_variant | MODIFIER |
| 15 | 54945374 | AX-170055588 | 1 | G | A | 0.036 | -5.57E-01 | 9.34E-02 | 8.73E-01 | 8.86E-01 | 6.28E-07 | Yes | Enpp2 | intron_variant | MODIFIER |
| 15 | 30570154 | AX-170330666 | 0 | G | A | 0.026 | -6.55E-01 | 8.92E-02 | 2.81E-01 | 2.64E-01 | 8.43E-07 | Yes | Ctnnd2 | intron_variant | MODIFIER |
| 15 | 24199688 | AX-170011252 | 0 | G | A | 0.026 | -6.59E-01 | 8.87E-02 | 2.63E-01 | 2.45E-01 | 8.88E-07 | Yes | Gm8408 | downstream_gene_variant | MODIFIER |
| 15 | 25604972 | AX-170011281 | 0 | A | T | 0.026 | -6.59E-01 | 8.87E-02 | 2.63E-01 | 2.45E-01 | 8.88E-07 | Yes | | intergenic_variant | MODIFIER |
| 15 | 27613324 | AX-97795131 | 0 | T | G | 0.026 | -6.59E-01 | 8.87E-02 | 2.63E-01 | 2.45E-01 | 8.88E-07 | Yes | Otulin | intron_variant | MODIFIER |
| 15 | 30463363 | AX-170291559 | 0 | C | A | 0.026 | -6.59E-01 | 8.87E-02 | 2.63E-01 | 2.45E-01 | 8.88E-07 | Yes | Ctnnd2 | intron_variant | MODIFIER |
| 15 | 55124009 | AX-170276769 | 1 | G | A | 0.036 | -5.51E-01 | 9.36E-02 | 8.75E-01 | 8.88E-01 | 8.89E-07 | Yes | Deptor | intron_variant | MODIFIER |

| | | | | | | | | | | | | | | | |
|---|---|---|---|---|---|---|---|---|---|---|---|---|---|---|---|
| 15 | 55169932 | AX-97793981 | 1 | T | C | 0.036 | -5.51E-01 | 9.36E-02 | 8.75E-01 | 8.88E-01 | 8.89E-07 | Yes | Deptor | intron_variant | MODIFIER |
| 15 | 56924772 | AX-170352458 | 1 | A | G | 0.036 | -5.51E-01 | 9.36E-02 | 8.75E-01 | 8.88E-01 | 8.89E-07 | Yes | Gm2582 | intron_variant,non_coding_transcript_variant | MODIFIER |
| 15 | 56894528 | AX-170106117 | 1 | C | G | 0.036 | -5.54E-01 | 9.30E-02 | 8.14E-01 | 8.25E-01 | 8.97E-07 | Yes | Gm2582 | intron_variant,non_coding_transcript_variant | MODIFIER |
| 15 | 23017725 | AX-97842048 | 0 | T | C | 0.026 | -6.48E-01 | 8.92E-02 | 2.95E-01 | 2.76E-01 | 9.03E-07 | Yes | Cdh18 | intron_variant | MODIFIER |
| 15 | 56072053 | AX-170312060 | 1 | C | G | 0.036 | -5.54E-01 | 9.32E-02 | 8.23E-01 | 8.33E-01 | 9.17E-07 | Yes | | intergenic_variant | MODIFIER |
| 15 | 56296404 | AX-97827108 | 1 | C | T | 0.036 | -5.54E-01 | 9.32E-02 | 8.23E-01 | 8.33E-01 | 9.17E-07 | Yes | | intergenic_variant | MODIFIER |
| 15 | 54631152 | AX-97829426 | 1 | C | A | 0.036 | -5.58E-01 | 9.50E-02 | 8.72E-01 | 8.83E-01 | 9.47E-07 | Yes | | intergenic_variant | MODIFIER |
| 15 | 56864962 | AX-170472987 | 0 | T | A | 0.035 | -4.93E-01 | 8.53E-02 | 9.44E-01 | 9.74E-01 | 9.71E-07 | Yes | | intergenic_variant | MODIFIER |
| 15 | 56976233 | AX-170128036 | 1 | C | G | 0.036 | -5.57E-01 | 9.38E-02 | 8.21E-01 | 8.31E-01 | 9.72E-07 | Yes | Gm2582 | intron_variant,non_coding_transcript_variant | MODIFIER |
| 15 | 23077036 | AX-170108159 | 1 | T | C | 0.018 | -6.07E-01 | 8.49E-02 | 3.18E-01 | 3.00E-01 | 9.91E-07 | Yes | Cdh18 | intron_variant,non_coding_transcript_variant | MODIFIER |
| 15 | 53511110 | AX-170451268 | 1 | G | C | 0.027 | -5.72E-01 | 9.74E-02 | 8.38E-01 | 8.48E-01 | 1.12E-06 | Yes | Samd12 | intron_variant | MODIFIER |
| 15 | 38280751 | AX-170494455 | 1 | C | A | 0.036 | -5.65E-01 | 9.21E-02 | 6.69E-01 | 6.72E-01 | 1.12E-06 | Yes | Gm49314 | upstream_gene_variant | MODIFIER |
| 15 | 23424640 | AX-97846558 | 1 | G | A | 0.027 | -6.42E-01 | 9.02E-02 | 3.18E-01 | 3.05E-01 | 1.13E-06 | Yes | Cdh18 | intron_variant,non_coding_transcript_variant | MODIFIER |
| 15 | 24083594 | AX-170011249 | 2 | T | C | 0.018 | -6.16E-01 | 8.77E-02 | 3.39E-01 | 3.27E-01 | 1.15E-06 | Yes | | intergenic_variant | MODIFIER |
| 15 | 37535141 | AX-170011461 | 0 | G | A | 0.035 | -5.72E-01 | 9.39E-02 | 6.85E-01 | 6.91E-01 | 1.22E-06 | Yes | Ncald | intron_variant | MODIFIER |
| 15 | 45271648 | AX-170327037 | 0 | A | G | 0.035 | -5.86E-01 | 9.56E-02 | 6.58E-01 | 6.52E-01 | 1.26E-06 | Yes | | intergenic_variant | MODIFIER |
| 15 | 45277030 | AX-170180236 | 0 | T | C | 0.035 | -5.86E-01 | 9.56E-02 | 6.58E-01 | 6.52E-01 | 1.26E-06 | Yes | | intergenic_variant | MODIFIER |
| 15 | 45277268 | AX-170460412 | 0 | G | A | 0.035 | -5.86E-01 | 9.56E-02 | 6.58E-01 | 6.52E-01 | 1.26E-06 | Yes | | intergenic_variant | MODIFIER |
| 15 | 45283952 | AX-170222065 | 0 | C | T | 0.035 | -5.86E-01 | 9.56E-02 | 6.58E-01 | 6.52E-01 | 1.26E-06 | Yes | | intergenic_variant | MODIFIER |
| 15 | 45288846 | AX-97820811 | 0 | C | A | 0.035 | -5.86E-01 | 9.56E-02 | 6.58E-01 | 6.52E-01 | 1.26E-06 | Yes | | intergenic_variant | MODIFIER |
| 15 | 45358172 | AX-170199707 | 0 | T | C | 0.035 | -5.86E-01 | 9.56E-02 | 6.58E-01 | 6.52E-01 | 1.26E-06 | Yes | | intergenic_variant | MODIFIER |
| 15 | 45399764 | AX-170369377 | 0 | A | G | 0.035 | -5.86E-01 | 9.56E-02 | 6.58E-01 | 6.52E-01 | 1.26E-06 | Yes | | intergenic_variant | MODIFIER |
| 15 | 45401367 | AX-97829373 | 0 | C | G | 0.035 | -5.86E-01 | 9.56E-02 | 6.58E-01 | 6.52E-01 | 1.26E-06 | Yes | | intergenic_variant | MODIFIER |
| 15 | 45420289 | AX-170011566 | 0 | C | T | 0.035 | -5.86E-01 | 9.56E-02 | 6.58E-01 | 6.52E-01 | 1.26E-06 | Yes | | intergenic_variant | MODIFIER |
| 15 | 45425459 | AX-170290667 | 0 | T | C | 0.035 | -5.86E-01 | 9.56E-02 | 6.58E-01 | 6.52E-01 | 1.26E-06 | Yes | | intergenic_variant | MODIFIER |
| 15 | 45456889 | AX-97835989 | 0 | A | G | 0.035 | -5.86E-01 | 9.56E-02 | 6.58E-01 | 6.52E-01 | 1.26E-06 | Yes | | intergenic_variant | MODIFIER |
| 15 | 45468135 | AX-170011569 | 0 | A | G | 0.035 | -5.86E-01 | 9.56E-02 | 6.58E-01 | 6.52E-01 | 1.26E-06 | Yes | | intergenic_variant | MODIFIER |
| 15 | 45472562 | AX-97780784 | 0 | C | T | 0.035 | -5.86E-01 | 9.56E-02 | 6.58E-01 | 6.52E-01 | 1.26E-06 | Yes | | intergenic_variant | MODIFIER |
| 15 | 45488885 | AX-97804481 | 0 | A | G | 0.035 | -5.86E-01 | 9.56E-02 | 6.58E-01 | 6.52E-01 | 1.26E-06 | Yes | | intergenic_variant | MODIFIER |
| 15 | 45488900 | AX-170318172 | 0 | A | G | 0.035 | -5.86E-01 | 9.56E-02 | 6.58E-01 | 6.52E-01 | 1.26E-06 | Yes | | intergenic_variant | MODIFIER |
| 15 | 45518701 | AX-170011567 | 0 | G | C | 0.035 | -5.86E-01 | 9.56E-02 | 6.58E-01 | 6.52E-01 | 1.26E-06 | Yes | | intergenic_variant | MODIFIER |
| 15 | 45632009 | AX-170441420 | 0 | C | A | 0.035 | -5.86E-01 | 9.56E-02 | 6.58E-01 | 6.52E-01 | 1.26E-06 | Yes | | intergenic_variant | MODIFIER |
| 15 | 45661576 | AX-170400812 | 0 | C | A | 0.035 | -5.86E-01 | 9.56E-02 | 6.58E-01 | 6.52E-01 | 1.26E-06 | Yes | | intergenic_variant | MODIFIER |
| 15 | 45663028 | AX-170463205 | 0 | C | T | 0.035 | -5.86E-01 | 9.56E-02 | 6.58E-01 | 6.52E-01 | 1.26E-06 | Yes | | intergenic_variant | MODIFIER |
| 15 | 45674132 | AX-170390688 | 0 | C | G | 0.035 | -5.86E-01 | 9.56E-02 | 6.58E-01 | 6.52E-01 | 1.26E-06 | Yes | | intergenic_variant | MODIFIER |
| 15 | 45745436 | AX-170153186 | 0 | C | T | 0.035 | -5.86E-01 | 9.56E-02 | 6.58E-01 | 6.52E-01 | 1.26E-06 | Yes | | intergenic_variant | MODIFIER |
| 15 | 45808077 | AX-170363653 | 0 | C | T | 0.035 | -5.86E-01 | 9.56E-02 | 6.58E-01 | 6.52E-01 | 1.26E-06 | Yes | | intergenic_variant | MODIFIER |
| 15 | 45826133 | AX-170534519 | 0 | G | T | 0.035 | -5.86E-01 | 9.56E-02 | 6.58E-01 | 6.52E-01 | 1.26E-06 | Yes | | intergenic_variant | MODIFIER |
| 15 | 45838023 | AX-170200198 | 0 | G | C | 0.035 | -5.86E-01 | 9.56E-02 | 6.58E-01 | 6.52E-01 | 1.26E-06 | Yes | | intergenic_variant | MODIFIER |
| 15 | 45916384 | AX-170275990 | 0 | A | G | 0.035 | -5.86E-01 | 9.56E-02 | 6.58E-01 | 6.52E-01 | 1.26E-06 | Yes | | intergenic_variant | MODIFIER |
| 15 | 45918994 | AX-170156160 | 0 | G | T | 0.035 | -5.86E-01 | 9.56E-02 | 6.58E-01 | 6.52E-01 | 1.26E-06 | Yes | | intergenic_variant | MODIFIER |
| 15 | 45919028 | AX-170476242 | 0 | C | T | 0.035 | -5.86E-01 | 9.56E-02 | 6.58E-01 | 6.52E-01 | 1.26E-06 | Yes | | intergenic_variant | MODIFIER |
| 15 | 45925062 | AX-170127813 | 0 | C | G | 0.035 | -5.86E-01 | 9.56E-02 | 6.58E-01 | 6.52E-01 | 1.26E-06 | Yes | | intergenic_variant | MODIFIER |
| 15 | 45925415 | AX-170311288 | 0 | G | A | 0.035 | -5.86E-01 | 9.56E-02 | 6.58E-01 | 6.52E-01 | 1.26E-06 | Yes | | intergenic_variant | MODIFIER |
| 15 | 45967541 | AX-170435876 | 0 | G | C | 0.035 | -5.86E-01 | 9.56E-02 | 6.58E-01 | 6.52E-01 | 1.26E-06 | Yes | Gm2203 | downstream_gene_variant | MODIFIER |
| 15 | 45982013 | AX-97836794 | 0 | T | C | 0.035 | -5.86E-01 | 9.56E-02 | 6.58E-01 | 6.52E-01 | 1.26E-06 | Yes | Gm5472 | upstream_gene_variant | MODIFIER |
| 15 | 45989736 | AX-97852165 | 0 | A | T | 0.035 | -5.86E-01 | 9.56E-02 | 6.58E-01 | 6.52E-01 | 1.26E-06 | Yes | | intergenic_variant | MODIFIER |
| 15 | 46009654 | AX-170055431 | 0 | A | G | 0.035 | -5.86E-01 | 9.56E-02 | 6.58E-01 | 6.52E-01 | 1.26E-06 | Yes | | intergenic_variant | MODIFIER |
| 15 | 46063785 | AX-97829374 | 0 | G | T | 0.035 | -5.86E-01 | 9.56E-02 | 6.58E-01 | 6.52E-01 | 1.26E-06 | Yes | | intergenic_variant | MODIFIER |
| 15 | 46104689 | AX-168404936 | 0 | C | T | 0.035 | -5.86E-01 | 9.56E-02 | 6.58E-01 | 6.52E-01 | 1.26E-06 | Yes | | intergenic_variant | MODIFIER |
| 15 | 46147892 | AX-170011582 | 0 | A | G | 0.035 | -5.86E-01 | 9.56E-02 | 6.58E-01 | 6.52E-01 | 1.26E-06 | Yes | Gm26063 | downstream_gene_variant | MODIFIER |
| 15 | 46227222 | AX-170011576 | 0 | T | C | 0.035 | -5.86E-01 | 9.56E-02 | 6.58E-01 | 6.52E-01 | 1.26E-06 | Yes | | intergenic_variant | MODIFIER |
| 15 | 46277484 | AX-97822582 | 0 | A | C | 0.035 | -5.86E-01 | 9.56E-02 | 6.58E-01 | 6.52E-01 | 1.26E-06 | Yes | | intergenic_variant | MODIFIER |
| 15 | 46286555 | AX-97838101 | 0 | G | A | 0.035 | -5.86E-01 | 9.56E-02 | 6.58E-01 | 6.52E-01 | 1.26E-06 | Yes | | intergenic_variant | MODIFIER |

| | | | | | | | | | | | | | | | |
|---|---|---|---|---|---|---|---|---|---|---|---|---|---|---|---|
| 15 | 30319525 | AX-170431172 | 0 | C | T | 0.026 | -6.58E-01 | 8.85E-02 | 2.48E-01 | 2.19E-01 | 1.28E-06 | Yes | Ctnnd2 | intron_variant | MODIFIER |
| 15 | 54904348 | AX-170496754 | 1 | C | T | 0.036 | -5.52E-01 | 9.33E-02 | 7.85E-01 | 7.94E-01 | 1.31E-06 | Yes | Enpp2 | intron_variant | MODIFIER |
| 15 | 26693363 | AX-170011300 | 0 | A | T | 0.026 | -6.60E-01 | 8.90E-02 | 2.50E-01 | 2.17E-01 | 1.32E-06 | Yes | Fbxl7 | intron_variant | MODIFIER |
| 15 | 45662125 | AX-97856784 | 1 | G | T | 0.036 | -5.79E-01 | 9.66E-02 | 7.24E-01 | 7.21E-01 | 1.38E-06 | Yes | | intergenic_variant | MODIFIER |
| 15 | 34058769 | AX-170359659 | 0 | A | G | 0.026 | -6.60E-01 | 8.73E-02 | 2.24E-01 | 1.93E-01 | 1.39E-06 | Yes | | intergenic_variant | MODIFIER |
| 15 | 45924894 | AX-170163428 | 1 | C | G | 0.036 | -5.80E-01 | 9.79E-02 | 7.60E-01 | 7.58E-01 | 1.47E-06 | Yes | | intergenic_variant | MODIFIER |
| 15 | 30597053 | AX-97781404 | 0 | A | G | 0.026 | -6.67E-01 | 8.65E-02 | 1.97E-01 | 1.62E-01 | 1.54E-06 | Yes | Ctnnd2 | intron_variant | MODIFIER |
| 15 | 34632379 | AX-170253824 | 0 | G | A | 0.026 | -6.67E-01 | 8.65E-02 | 1.97E-01 | 1.62E-01 | 1.54E-06 | Yes | Nipal2 | intron_variant | MODIFIER |
| 15 | 54831980 | AX-170165973 | 0 | A | T | 0.035 | -5.38E-01 | 9.50E-02 | 9.38E-01 | 9.55E-01 | 1.71E-06 | Yes | | intergenic_variant | MODIFIER |
| 15 | 54262866 | AX-170128292 | 0 | T | G | 0.035 | -5.43E-01 | 9.57E-02 | 9.25E-01 | 9.46E-01 | 1.72E-06 | Yes | Tnfrsf11b | intron_variant | MODIFIER |
| 15 | 53498485 | AX-170296783 | 0 | A | C | 0.035 | -5.05E-01 | 9.65E-02 | 1.54E+00 | 1.58E+00 | 1.77E-06 | Yes | Samd12 | intron_variant | MODIFIER |
| 15 | 52907759 | AX-170338585 | 0 | T | C | 0.035 | -4.96E-01 | 8.96E-02 | 1.03E+00 | 1.07E+00 | 1.96E-06 | Yes | | intergenic_variant | MODIFIER |
| 15 | 54176048 | AX-170011727 | 0 | A | G | 0.035 | -4.96E-01 | 8.96E-02 | 1.03E+00 | 1.07E+00 | 1.96E-06 | Yes | | intergenic_variant | MODIFIER |
| 15 | 56128030 | AX-170211069 | 0 | G | A | 0.044 | -5.11E-01 | 9.27E-02 | 1.04E+00 | 1.07E+00 | 2.01E-06 | Yes | | intergenic_variant | MODIFIER |
| 15 | 45420240 | AX-170011564 | 1 | T | C | 0.036 | -5.79E-01 | 9.71E-02 | 6.90E-01 | 6.86E-01 | 2.08E-06 | Yes | | intergenic_variant | MODIFIER |
| 15 | 42554087 | AX-170128336 | 0 | C | T | 0.035 | -5.67E-01 | 9.07E-02 | 5.43E-01 | 5.42E-01 | 2.13E-06 | Yes | Angpt1 | intron_variant | MODIFIER |
| 15 | 24091075 | AX-170466459 | 0 | A | T | 0.026 | -5.82E-01 | 8.90E-02 | 4.37E-01 | 4.45E-01 | 2.17E-06 | Yes | Gm48918 | downstream_gene_variant | MODIFIER |
| 15 | 45319809 | AX-170244166 | 1 | C | T | 0.036 | -5.78E-01 | 9.68E-02 | 6.80E-01 | 6.75E-01 | 2.19E-06 | Yes | | intergenic_variant | MODIFIER |
| 15 | 61575899 | AX-170135991 | 0 | A | T | 0.061 | -5.01E-01 | 9.39E-02 | 1.25E+00 | 1.29E+00 | 2.24E-06 | Yes | D030024E09Rik | intron_variant,non_coding_transcript_variant | MODIFIER |
| 15 | 45550532 | AX-170011565 | 1 | C | G | 0.036 | -5.78E-01 | 9.71E-02 | 6.81E-01 | 6.76E-01 | 2.31E-06 | Yes | | intergenic_variant | MODIFIER |
| 15 | 56796503 | AX-97793037 | 2 | G | A | 0.036 | -5.48E-01 | 9.41E-02 | 7.54E-01 | 7.61E-01 | 2.36E-06 | Yes | | intergenic_variant | MODIFIER |
| 15 | 24091788 | AX-97785231 | 0 | G | A | 0.026 | -5.88E-01 | 8.89E-02 | 4.06E-01 | 4.11E-01 | 2.45E-06 | Yes | Gm48918 | downstream_gene_variant | MODIFIER |
| 15 | 29715592 | AX-170011348 | 0 | G | C | 0.026 | -5.88E-01 | 8.89E-02 | 4.06E-01 | 4.11E-01 | 2.45E-06 | Yes | Mir3964 | upstream_gene_variant | MODIFIER |
| 15 | 30215258 | AX-97838776 | 0 | A | G | 0.026 | -5.88E-01 | 8.89E-02 | 4.06E-01 | 4.11E-01 | 2.45E-06 | Yes | Ctnnd2 | intron_variant | MODIFIER |
| 15 | 34776446 | AX-170011418 | 0 | C | T | 0.026 | -5.88E-01 | 8.89E-02 | 4.06E-01 | 4.11E-01 | 2.45E-06 | Yes | | intergenic_variant | MODIFIER |
| 15 | 51863584 | AX-97781287 | 0 | A | G | 0.035 | -4.99E-01 | 8.99E-02 | 9.58E-01 | 9.93E-01 | 2.53E-06 | Yes | Eif3h | intron_variant | MODIFIER |
| 15 | 54647310 | AX-97807625 | 0 | T | G | 0.035 | -5.06E-01 | 8.67E-02 | 7.30E-01 | 7.41E-01 | 2.61E-06 | Yes | | intergenic_variant | MODIFIER |
| 15 | 51785874 | AX-170138879 | 2 | C | A | 0.036 | -5.46E-01 | 9.45E-02 | 7.58E-01 | 7.66E-01 | 2.78E-06 | Yes | Eif3h | downstream_gene_variant | MODIFIER |
| 15 | 45471245 | AX-170314189 | 1 | A | T | 0.036 | -5.32E-01 | 9.36E-02 | 8.22E-01 | 8.42E-01 | 2.88E-06 | Yes | | intergenic_variant | MODIFIER |
| 15 | 44358250 | AX-170011547 | 0 | G | A | 0.035 | -5.88E-01 | 9.27E-02 | 4.76E-01 | 4.63E-01 | 3.10E-06 | Yes | | intergenic_variant | MODIFIER |
| 15 | 44229682 | AX-170233921 | 1 | G | A | 0.027 | -5.88E-01 | 9.54E-02 | 5.47E-01 | 5.40E-01 | 3.13E-06 | Yes | Trhr | intron_variant | MODIFIER |
| 15 | 45719627 | AX-170476734 | 0 | C | G | 0.035 | -5.18E-01 | 8.81E-02 | 6.82E-01 | 6.87E-01 | 3.21E-06 | Yes | | intergenic_variant | MODIFIER |
| 15 | 26385135 | AX-170499889 | 1 | G | A | 0.027 | -6.69E-01 | 8.48E-02 | 1.55E-01 | 1.17E-01 | 3.24E-06 | Yes | MAR11 | intron_variant | MODIFIER |
| 15 | 56684700 | AX-170357186 | 0 | T | C | 0.035 | -5.37E-01 | 9.49E-02 | 8.15E-01 | 8.27E-01 | 3.33E-06 | Yes | Has2 | intron_variant | MODIFIER |
| 15 | 41453529 | AX-170139833 | 0 | C | T | 0.035 | -5.90E-01 | 9.08E-02 | 4.22E-01 | 4.08E-01 | 3.36E-06 | Yes | Oxr1 | intron_variant | MODIFIER |
| 15 | 41454308 | AX-170131786 | 0 | G | A | 0.035 | -5.90E-01 | 9.08E-02 | 4.22E-01 | 4.08E-01 | 3.36E-06 | Yes | Oxr1 | intron_variant | MODIFIER |
| 15 | 41454938 | AX-170011498 | 0 | C | T | 0.035 | -5.90E-01 | 9.08E-02 | 4.22E-01 | 4.08E-01 | 3.36E-06 | Yes | Oxr1 | intron_variant | MODIFIER |
| 15 | 41460595 | AX-170258299 | 0 | T | A | 0.035 | -5.90E-01 | 9.08E-02 | 4.22E-01 | 4.08E-01 | 3.36E-06 | Yes | Oxr1 | intron_variant | MODIFIER |
| 15 | 41461077 | AX-170252479 | 0 | G | A | 0.035 | -5.90E-01 | 9.08E-02 | 4.22E-01 | 4.08E-01 | 3.36E-06 | Yes | Oxr1 | intron_variant | MODIFIER |
| 15 | 41484705 | AX-170412435 | 0 | T | C | 0.035 | -5.90E-01 | 9.08E-02 | 4.22E-01 | 4.08E-01 | 3.36E-06 | Yes | Oxr1 | intron_variant | MODIFIER |
| 15 | 41496556 | AX-170334050 | 0 | T | C | 0.035 | -5.90E-01 | 9.08E-02 | 4.22E-01 | 4.08E-01 | 3.36E-06 | Yes | Oxr1 | intron_variant | MODIFIER |
| 15 | 41501981 | AX-170314897 | 0 | T | C | 0.035 | -5.90E-01 | 9.08E-02 | 4.22E-01 | 4.08E-01 | 3.36E-06 | Yes | Oxr1 | intron_variant | MODIFIER |
| 15 | 41522834 | AX-168419981 | 0 | C | T | 0.035 | -5.90E-01 | 9.08E-02 | 4.22E-01 | 4.08E-01 | 3.36E-06 | Yes | Oxr1 | intron_variant | MODIFIER |
| 15 | 41617460 | AX-170131123 | 0 | C | A | 0.035 | -5.90E-01 | 9.08E-02 | 4.22E-01 | 4.08E-01 | 3.36E-06 | Yes | Oxr1 | intron_variant | MODIFIER |
| 15 | 41620463 | AX-170508931 | 0 | C | G | 0.035 | -5.90E-01 | 9.08E-02 | 4.22E-01 | 4.08E-01 | 3.36E-06 | Yes | Oxr1 | intron_variant | MODIFIER |
| 15 | 41625492 | AX-170200343 | 0 | G | T | 0.035 | -5.90E-01 | 9.08E-02 | 4.22E-01 | 4.08E-01 | 3.36E-06 | Yes | Oxr1 | intron_variant | MODIFIER |
| 15 | 41673024 | AX-170164962 | 0 | A | C | 0.035 | -5.90E-01 | 9.08E-02 | 4.22E-01 | 4.08E-01 | 3.36E-06 | Yes | Oxr1 | intron_variant | MODIFIER |
| 15 | 41681984 | AX-170055351 | 0 | C | A | 0.035 | -5.90E-01 | 9.08E-02 | 4.22E-01 | 4.08E-01 | 3.36E-06 | Yes | Oxr1 | intron_variant | MODIFIER |
| 15 | 41723600 | AX-170382387 | 0 | G | A | 0.035 | -5.90E-01 | 9.08E-02 | 4.22E-01 | 4.08E-01 | 3.36E-06 | Yes | Oxr1 | intron_variant | MODIFIER |
| 15 | 41727724 | AX-170483207 | 0 | C | T | 0.035 | -5.90E-01 | 9.08E-02 | 4.22E-01 | 4.08E-01 | 3.36E-06 | Yes | Oxr1 | intron_variant | MODIFIER |
| 15 | 41734172 | AX-170214367 | 0 | C | T | 0.035 | -5.90E-01 | 9.08E-02 | 4.22E-01 | 4.08E-01 | 3.36E-06 | Yes | Oxr1 | intron_variant | MODIFIER |
| 15 | 41764861 | AX-97843709 | 0 | A | G | 0.035 | -5.90E-01 | 9.08E-02 | 4.22E-01 | 4.08E-01 | 3.36E-06 | Yes | Oxr1 | intron_variant | MODIFIER |
| 15 | 52557533 | AX-170087861 | 1 | C | T | 0.036 | -5.36E-01 | 9.70E-02 | 9.21E-01 | 9.44E-01 | 3.41E-06 | Yes | | intergenic_variant | MODIFIER |
| 15 | 41672288 | AX-170087832 | 0 | C | A | 0.044 | -5.36E-01 | 9.32E-02 | 7.49E-01 | 7.63E-01 | 3.42E-06 | Yes | Oxr1 | intron_variant | MODIFIER |

| | | | | | | | | | | | | | | | |
|---|---|---|---|---|---|---|---|---|---|---|---|---|---|---|---|
| 15 | 29991037 | AX-170133327 | 0 | C | G | 0.035 | -5.93E-01 | 9.17E-02 | 4.26E-01 | 4.22E-01 | 3.52E-06 | Yes | | intergenic_variant | MODIFIER |
| 15 | 36917302 | AX-170011451 | 1 | C | T | 0.027 | -5.77E-01 | 9.71E-02 | 6.33E-01 | 6.34E-01 | 3.63E-06 | Yes | Gm41300 | intron_variant,non_coding_transcript_variant | MODIFIER |
| 15 | 22791872 | AX-170401086 | 1 | C | T | 0.027 | -5.80E-01 | 8.93E-02 | 4.14E-01 | 4.20E-01 | 3.78E-06 | Yes | Cdh18 | intron_variant | MODIFIER |
| 15 | 26596484 | AX-170195705 | 1 | T | C | 0.027 | -5.83E-01 | 9.01E-02 | 4.18E-01 | 4.23E-01 | 3.88E-06 | Yes | Fbxl7 | intron_variant | MODIFIER |
| 15 | 24676500 | AX-97828584 | 1 | G | A | 0.036 | -5.81E-01 | 9.31E-02 | 4.83E-01 | 4.78E-01 | 4.54E-06 | Yes | | intergenic_variant | MODIFIER |
| 13 | 93360417 | AX-170156281 | 1 | C | T | 0.045 | -4.74E-01 | 9.48E-02 | 1.54E+00 | 1.61E+00 | 4.71E-06 | Yes | Homer1 | intron_variant | MODIFIER |
| 15 | 53616927 | AX-170011720 | 0 | A | T | 0.035 | -4.63E-01 | 8.88E-02 | 1.16E+00 | 1.19E+00 | 4.87E-06 | Yes | Samd12 | intron_variant | MODIFIER |
| 15 | 26236580 | AX-170011290 | 1 | C | A | 0.036 | -5.77E-01 | 9.18E-02 | 4.63E-01 | 4.59E-01 | 4.97E-06 | Yes | | intergenic_variant | MODIFIER |
| 15 | 41628383 | AX-170285305 | 1 | A | T | 0.036 | -5.81E-01 | 9.14E-02 | 4.31E-01 | 4.19E-01 | 5.51E-06 | Yes | Oxr1 | intron_variant | MODIFIER |
| 15 | 41754371 | AX-170011516 | 1 | T | C | 0.036 | -5.75E-01 | 9.19E-02 | 4.62E-01 | 4.51E-01 | 5.72E-06 | Yes | Oxr1 | intron_variant | MODIFIER |
| 15 | 55006843 | AX-170152336 | 0 | G | A | 0.044 | -4.93E-01 | 9.25E-02 | 9.82E-01 | 1.01E+00 | 5.82E-06 | Yes | | intergenic_variant | MODIFIER |
| 15 | 45717036 | AX-168420024 | 0 | T | C | 0.035 | -5.51E-01 | 9.88E-02 | 7.84E-01 | 7.87E-01 | 5.83E-06 | Yes | | intergenic_variant | MODIFIER |
| 15 | 41458377 | AX-170298041 | 1 | T | C | 0.036 | -5.78E-01 | 9.10E-02 | 4.26E-01 | 4.14E-01 | 5.92E-06 | Yes | Oxr1 | intron_variant | MODIFIER |
| 15 | 43895779 | AX-97837321 | 0 | C | T | 0.035 | -5.06E-01 | 9.09E-02 | 7.75E-01 | 7.98E-01 | 6.42E-06 | Yes | | intergenic_variant | MODIFIER |
| 15 | 58485319 | AX-170463942 | 1 | A | T | 0.045 | -5.13E-01 | 9.48E-02 | 8.78E-01 | 8.88E-01 | 6.71E-06 | Yes | | intergenic_variant | MODIFIER |
| 15 | 52429669 | AX-170144602 | 0 | G | C | 0.035 | -4.85E-01 | 9.40E-02 | 1.13E+00 | 1.15E+00 | 6.87E-06 | Yes | | regulatory_region_variant | MODIFIER |
| 15 | 22115349 | AX-170337442 | 0 | C | T | 0.026 | -6.27E-01 | 9.15E-02 | 2.88E-01 | 2.61E-01 | 7.35E-06 | Yes | | intergenic_variant | MODIFIER |
| 15 | 22141684 | AX-170402140 | 0 | A | C | 0.026 | -6.27E-01 | 9.15E-02 | 2.88E-01 | 2.61E-01 | 7.35E-06 | Yes | | intergenic_variant | MODIFIER |
| 15 | 22144275 | AX-170188744 | 0 | A | G | 0.026 | -6.27E-01 | 9.15E-02 | 2.88E-01 | 2.61E-01 | 7.35E-06 | Yes | | intergenic_variant | MODIFIER |
| 15 | 22154772 | AX-170221118 | 0 | T | G | 0.026 | -6.27E-01 | 9.15E-02 | 2.88E-01 | 2.61E-01 | 7.35E-06 | Yes | | intergenic_variant | MODIFIER |
| 15 | 22280618 | AX-170418004 | 0 | A | G | 0.026 | -6.27E-01 | 9.15E-02 | 2.88E-01 | 2.61E-01 | 7.35E-06 | Yes | | intergenic_variant | MODIFIER |
| 15 | 22324894 | AX-170272979 | 0 | T | C | 0.026 | -6.27E-01 | 9.15E-02 | 2.88E-01 | 2.61E-01 | 7.35E-06 | Yes | | intergenic_variant | MODIFIER |
| 15 | 22326478 | AX-170222052 | 0 | C | T | 0.026 | -6.27E-01 | 9.15E-02 | 2.88E-01 | 2.61E-01 | 7.35E-06 | Yes | | intergenic_variant | MODIFIER |
| 15 | 22332259 | AX-170011211 | 0 | T | G | 0.026 | -6.27E-01 | 9.15E-02 | 2.88E-01 | 2.61E-01 | 7.35E-06 | Yes | | intergenic_variant | MODIFIER |
| 15 | 22418906 | AX-97828468 | 0 | G | A | 0.026 | -6.27E-01 | 9.15E-02 | 2.88E-01 | 2.61E-01 | 7.35E-06 | Yes | | intergenic_variant | MODIFIER |
| 15 | 22482996 | AX-97844673 | 0 | A | C | 0.026 | -6.27E-01 | 9.15E-02 | 2.88E-01 | 2.61E-01 | 7.35E-06 | Yes | | intergenic_variant | MODIFIER |
| 15 | 22496239 | AX-97830484 | 0 | T | C | 0.026 | -6.27E-01 | 9.15E-02 | 2.88E-01 | 2.61E-01 | 7.35E-06 | Yes | | intergenic_variant | MODIFIER |
| 15 | 22529285 | AX-97793528 | 0 | T | C | 0.026 | -6.27E-01 | 9.15E-02 | 2.88E-01 | 2.61E-01 | 7.35E-06 | Yes | | intergenic_variant | MODIFIER |
| 15 | 22569949 | AX-170364959 | 0 | T | C | 0.026 | -6.27E-01 | 9.15E-02 | 2.88E-01 | 2.61E-01 | 7.35E-06 | Yes | Cdh18 | intron_variant | MODIFIER |
| 15 | 22641512 | AX-170338452 | 0 | G | A | 0.026 | -6.27E-01 | 9.15E-02 | 2.88E-01 | 2.61E-01 | 7.35E-06 | Yes | Cdh18 | intron_variant | MODIFIER |
| 15 | 22651818 | AX-170526627 | 0 | C | T | 0.026 | -6.27E-01 | 9.15E-02 | 2.88E-01 | 2.61E-01 | 7.35E-06 | Yes | Cdh18 | intron_variant | MODIFIER |
| 15 | 22669354 | AX-170287954 | 0 | A | G | 0.026 | -6.27E-01 | 9.15E-02 | 2.88E-01 | 2.61E-01 | 7.35E-06 | Yes | Cdh18 | intron_variant | MODIFIER |
| 15 | 22723631 | AX-97791898 | 0 | A | G | 0.026 | -6.27E-01 | 9.15E-02 | 2.88E-01 | 2.61E-01 | 7.35E-06 | Yes | Cdh18 | intron_variant | MODIFIER |
| 15 | 22760571 | AX-170087794 | 0 | C | T | 0.026 | -6.27E-01 | 9.15E-02 | 2.88E-01 | 2.61E-01 | 7.35E-06 | Yes | Cdh18 | intron_variant | MODIFIER |
| 15 | 22791858 | AX-170199365 | 0 | T | G | 0.026 | -6.27E-01 | 9.15E-02 | 2.88E-01 | 2.61E-01 | 7.35E-06 | Yes | Cdh18 | intron_variant | MODIFIER |
| 15 | 22819788 | AX-97842850 | 0 | T | C | 0.026 | -6.27E-01 | 9.15E-02 | 2.88E-01 | 2.61E-01 | 7.35E-06 | Yes | Cdh18 | intron_variant | MODIFIER |
| 15 | 22841572 | AX-97808578 | 0 | G | A | 0.026 | -6.27E-01 | 9.15E-02 | 2.88E-01 | 2.61E-01 | 7.35E-06 | Yes | Cdh18 | intron_variant | MODIFIER |
| 15 | 22846203 | AX-170186004 | 0 | A | G | 0.026 | -6.27E-01 | 9.15E-02 | 2.88E-01 | 2.61E-01 | 7.35E-06 | Yes | Cdh18 | intron_variant | MODIFIER |
| 15 | 22917275 | AX-170230279 | 0 | T | C | 0.026 | -6.27E-01 | 9.15E-02 | 2.88E-01 | 2.61E-01 | 7.35E-06 | Yes | Cdh18 | intron_variant | MODIFIER |
| 15 | 22932895 | AX-170011227 | 0 | C | A | 0.026 | -6.27E-01 | 9.15E-02 | 2.88E-01 | 2.61E-01 | 7.35E-06 | Yes | Cdh18 | intron_variant | MODIFIER |
| 15 | 22951943 | AX-97857325 | 0 | T | C | 0.026 | -6.27E-01 | 9.15E-02 | 2.88E-01 | 2.61E-01 | 7.35E-06 | Yes | Cdh18 | intron_variant | MODIFIER |
| 15 | 22952130 | AX-170011222 | 0 | A | C | 0.026 | -6.27E-01 | 9.15E-02 | 2.88E-01 | 2.61E-01 | 7.35E-06 | Yes | Cdh18 | intron_variant | MODIFIER |
| 15 | 22988122 | AX-170011223 | 0 | A | G | 0.026 | -6.27E-01 | 9.15E-02 | 2.88E-01 | 2.61E-01 | 7.35E-06 | Yes | Cdh18 | intron_variant | MODIFIER |
| 15 | 23045508 | AX-170123195 | 0 | A | C | 0.026 | -6.27E-01 | 9.15E-02 | 2.88E-01 | 2.61E-01 | 7.35E-06 | Yes | Cdh18 | intron_variant,non_coding_transcript_variant | MODIFIER |
| 15 | 23059494 | AX-170256760 | 0 | T | C | 0.026 | -6.27E-01 | 9.15E-02 | 2.88E-01 | 2.61E-01 | 7.35E-06 | Yes | Cdh18 | intron_variant,non_coding_transcript_variant | MODIFIER |
| 15 | 23061037 | AX-170093183 | 0 | T | C | 0.026 | -6.27E-01 | 9.15E-02 | 2.88E-01 | 2.61E-01 | 7.35E-06 | Yes | Cdh18 | intron_variant,non_coding_transcript_variant | MODIFIER |
| 15 | 23168098 | AX-97785234 | 0 | T | G | 0.026 | -6.27E-01 | 9.15E-02 | 2.88E-01 | 2.61E-01 | 7.35E-06 | Yes | Cdh18 | intron_variant,non_coding_transcript_variant | MODIFIER |
| 15 | 23198920 | AX-97796860 | 0 | C | T | 0.026 | -6.27E-01 | 9.15E-02 | 2.88E-01 | 2.61E-01 | 7.35E-06 | Yes | Cdh18 | intron_variant,non_coding_transcript_variant | MODIFIER |
| 15 | 23222855 | AX-97789524 | 0 | T | C | 0.026 | -6.27E-01 | 9.15E-02 | 2.88E-01 | 2.61E-01 | 7.35E-06 | Yes | Cdh18 | intron_variant,non_coding_transcript_variant | MODIFIER |
| 15 | 23230569 | AX-170364301 | 0 | G | T | 0.026 | -6.27E-01 | 9.15E-02 | 2.88E-01 | 2.61E-01 | 7.35E-06 | Yes | Cdh18 | intron_variant,non_coding_transcript_variant | MODIFIER |
| 15 | 23236231 | AX-170241626 | 0 | T | A | 0.026 | -6.27E-01 | 9.15E-02 | 2.88E-01 | 2.61E-01 | 7.35E-06 | Yes | Cdh18 | intron_variant,non_coding_transcript_variant | MODIFIER |
| 15 | 23249904 | AX-170112904 | 0 | A | G | 0.026 | -6.27E-01 | 9.15E-02 | 2.88E-01 | 2.61E-01 | 7.35E-06 | Yes | Cdh18 | intron_variant,non_coding_transcript_variant | MODIFIER |
| 15 | 23277831 | AX-97783389 | 0 | A | G | 0.026 | -6.27E-01 | 9.15E-02 | 2.88E-01 | 2.61E-01 | 7.35E-06 | Yes | Cdh18 | intron_variant,non_coding_transcript_variant | MODIFIER |
| 15 | 23317541 | AX-170424244 | 0 | A | G | 0.026 | -6.27E-01 | 9.15E-02 | 2.88E-01 | 2.61E-01 | 7.35E-06 | Yes | Cdh18 | intron_variant,non_coding_transcript_variant | MODIFIER |

| | | | | | | | | | | | | | | | |
|---|---|---|---|---|---|---|---|---|---|---|---|---|---|---|---|
| 15 | 23332162 | AX-97841467 | 0 | G | A | 0.026 | -6.27E-01 | 9.15E-02 | 2.88E-01 | 2.61E-01 | 7.35E-06 | Yes | Cdh18 | intron_variant,non_coding_transcript_variant | MODIFIER |
| 15 | 23390494 | AX-170280358 | 0 | T | C | 0.026 | -6.27E-01 | 9.15E-02 | 2.88E-01 | 2.61E-01 | 7.35E-06 | Yes | Cdh18 | intron_variant,non_coding_transcript_variant | MODIFIER |
| 15 | 23514143 | AX-170011241 | 0 | C | T | 0.026 | -6.27E-01 | 9.15E-02 | 2.88E-01 | 2.61E-01 | 7.35E-06 | Yes | | intergenic_variant | MODIFIER |
| 15 | 23514245 | AX-170011238 | 0 | A | G | 0.026 | -6.27E-01 | 9.15E-02 | 2.88E-01 | 2.61E-01 | 7.35E-06 | Yes | | intergenic_variant | MODIFIER |
| 15 | 23514926 | AX-170126055 | 0 | G | A | 0.026 | -6.27E-01 | 9.15E-02 | 2.88E-01 | 2.61E-01 | 7.35E-06 | Yes | | intergenic_variant | MODIFIER |
| 15 | 23515036 | AX-97844782 | 0 | A | G | 0.026 | -6.27E-01 | 9.15E-02 | 2.88E-01 | 2.61E-01 | 7.35E-06 | Yes | | intergenic_variant | MODIFIER |
| 15 | 23555289 | AX-97846491 | 0 | A | C | 0.026 | -6.27E-01 | 9.15E-02 | 2.88E-01 | 2.61E-01 | 7.35E-06 | Yes | | intergenic_variant | MODIFIER |
| 15 | 23555438 | AX-170011237 | 0 | G | T | 0.026 | -6.27E-01 | 9.15E-02 | 2.88E-01 | 2.61E-01 | 7.35E-06 | Yes | | intergenic_variant | MODIFIER |
| 15 | 23606417 | AX-170111092 | 0 | T | A | 0.026 | -6.27E-01 | 9.15E-02 | 2.88E-01 | 2.61E-01 | 7.35E-06 | Yes | | intergenic_variant | MODIFIER |
| 15 | 23606523 | AX-170011242 | 0 | A | G | 0.026 | -6.27E-01 | 9.15E-02 | 2.88E-01 | 2.61E-01 | 7.35E-06 | Yes | | intergenic_variant | MODIFIER |
| 15 | 23624032 | AX-170011231 | 0 | T | C | 0.026 | -6.27E-01 | 9.15E-02 | 2.88E-01 | 2.61E-01 | 7.35E-06 | Yes | Gm48915 | non_coding_transcript_exon_variant | MODIFIER |
| 15 | 24014168 | AX-170011250 | 0 | T | C | 0.026 | -6.27E-01 | 9.15E-02 | 2.88E-01 | 2.61E-01 | 7.35E-06 | Yes | | intergenic_variant | MODIFIER |
| 15 | 24022548 | AX-97807100 | 0 | T | C | 0.026 | -6.27E-01 | 9.15E-02 | 2.88E-01 | 2.61E-01 | 7.35E-06 | Yes | | intergenic_variant | MODIFIER |
| 15 | 24059165 | AX-97829015 | 0 | T | G | 0.026 | -6.27E-01 | 9.15E-02 | 2.88E-01 | 2.61E-01 | 7.35E-06 | Yes | | intergenic_variant | MODIFIER |
| 15 | 24089073 | AX-170228325 | 0 | G | C | 0.026 | -6.27E-01 | 9.15E-02 | 2.88E-01 | 2.61E-01 | 7.35E-06 | Yes | | intergenic_variant | MODIFIER |
| 15 | 24100848 | AX-170011248 | 0 | A | G | 0.026 | -6.27E-01 | 9.15E-02 | 2.88E-01 | 2.61E-01 | 7.35E-06 | Yes | Gm48918 | upstream_gene_variant | MODIFIER |
| 15 | 24124060 | AX-170011253 | 0 | T | A | 0.026 | -6.27E-01 | 9.15E-02 | 2.88E-01 | 2.61E-01 | 7.35E-06 | Yes | | intergenic_variant | MODIFIER |
| 15 | 24129326 | AX-97857513 | 0 | C | T | 0.026 | -6.27E-01 | 9.15E-02 | 2.88E-01 | 2.61E-01 | 7.35E-06 | Yes | Gm48919 | downstream_gene_variant | MODIFIER |
| 15 | 24129578 | AX-170287238 | 0 | T | G | 0.026 | -6.27E-01 | 9.15E-02 | 2.88E-01 | 2.61E-01 | 7.35E-06 | Yes | Gm48919 | downstream_gene_variant | MODIFIER |
| 15 | 24129857 | AX-170011251 | 0 | A | G | 0.026 | -6.27E-01 | 9.15E-02 | 2.88E-01 | 2.61E-01 | 7.35E-06 | Yes | Gm48919 | non_coding_transcript_exon_variant | MODIFIER |
| 15 | 24140930 | AX-97853125 | 0 | A | G | 0.026 | -6.27E-01 | 9.15E-02 | 2.88E-01 | 2.61E-01 | 7.35E-06 | Yes | | intergenic_variant | MODIFIER |
| 15 | 24159614 | AX-170011245 | 0 | C | A | 0.026 | -6.27E-01 | 9.15E-02 | 2.88E-01 | 2.61E-01 | 7.35E-06 | Yes | | intergenic_variant | MODIFIER |
| 15 | 24163944 | AX-97842651 | 0 | A | G | 0.026 | -6.27E-01 | 9.15E-02 | 2.88E-01 | 2.61E-01 | 7.35E-06 | Yes | | intergenic_variant | MODIFIER |
| 15 | 24187791 | AX-170011258 | 0 | T | C | 0.026 | -6.27E-01 | 9.15E-02 | 2.88E-01 | 2.61E-01 | 7.35E-06 | Yes | | intergenic_variant | MODIFIER |
| 15 | 24615161 | AX-170011268 | 0 | G | A | 0.026 | -6.27E-01 | 9.15E-02 | 2.88E-01 | 2.61E-01 | 7.35E-06 | Yes | | intergenic_variant | MODIFIER |
| 15 | 24615894 | AX-170475401 | 0 | T | C | 0.026 | -6.27E-01 | 9.15E-02 | 2.88E-01 | 2.61E-01 | 7.35E-06 | Yes | | intergenic_variant | MODIFIER |
| 15 | 24625281 | AX-97799633 | 0 | G | A | 0.026 | -6.27E-01 | 9.15E-02 | 2.88E-01 | 2.61E-01 | 7.35E-06 | Yes | | intergenic_variant | MODIFIER |
| 15 | 24640270 | AX-170011265 | 0 | T | G | 0.026 | -6.27E-01 | 9.15E-02 | 2.88E-01 | 2.61E-01 | 7.35E-06 | Yes | | intergenic_variant | MODIFIER |
| 15 | 24676591 | AX-170011269 | 0 | C | G | 0.026 | -6.27E-01 | 9.15E-02 | 2.88E-01 | 2.61E-01 | 7.35E-06 | Yes | | intergenic_variant | MODIFIER |
| 15 | 24676673 | AX-170265301 | 0 | T | G | 0.026 | -6.27E-01 | 9.15E-02 | 2.88E-01 | 2.61E-01 | 7.35E-06 | Yes | | intergenic_variant | MODIFIER |
| 15 | 24677966 | AX-170114721 | 0 | C | A | 0.026 | -6.27E-01 | 9.15E-02 | 2.88E-01 | 2.61E-01 | 7.35E-06 | Yes | | intergenic_variant | MODIFIER |
| 15 | 24752183 | AX-97825661 | 0 | T | C | 0.026 | -6.27E-01 | 9.15E-02 | 2.88E-01 | 2.61E-01 | 7.35E-06 | Yes | | intergenic_variant | MODIFIER |
| 15 | 24752314 | AX-170212871 | 0 | T | C | 0.026 | -6.27E-01 | 9.15E-02 | 2.88E-01 | 2.61E-01 | 7.35E-06 | Yes | | intergenic_variant | MODIFIER |
| 15 | 24816864 | AX-97835688 | 0 | T | C | 0.026 | -6.27E-01 | 9.15E-02 | 2.88E-01 | 2.61E-01 | 7.35E-06 | Yes | Gm22810 | downstream_gene_variant | MODIFIER |
| 15 | 24820140 | AX-170413245 | 0 | T | C | 0.026 | -6.27E-01 | 9.15E-02 | 2.88E-01 | 2.61E-01 | 7.35E-06 | Yes | Gm22810 | downstream_gene_variant | MODIFIER |
| 15 | 24842778 | AX-97828585 | 0 | C | T | 0.026 | -6.27E-01 | 9.15E-02 | 2.88E-01 | 2.61E-01 | 7.35E-06 | Yes | | intergenic_variant | MODIFIER |
| 15 | 24867556 | AX-170165226 | 0 | T | A | 0.026 | -6.27E-01 | 9.15E-02 | 2.88E-01 | 2.61E-01 | 7.35E-06 | Yes | Gm35769 | intron_variant,non_coding_transcript_variant | MODIFIER |
| 15 | 24878007 | AX-170474428 | 0 | A | T | 0.026 | -6.27E-01 | 9.15E-02 | 2.88E-01 | 2.61E-01 | 7.35E-06 | Yes | Gm35769 | intron_variant,non_coding_transcript_variant | MODIFIER |
| 15 | 24880326 | AX-97821576 | 0 | C | A | 0.026 | -6.27E-01 | 9.15E-02 | 2.88E-01 | 2.61E-01 | 7.35E-06 | Yes | Gm35769 | intron_variant,non_coding_transcript_variant | MODIFIER |
| 15 | 24900104 | AX-170467995 | 0 | T | A | 0.026 | -6.27E-01 | 9.15E-02 | 2.88E-01 | 2.61E-01 | 7.35E-06 | Yes | Gm35769 | intron_variant,non_coding_transcript_variant | MODIFIER |
| 15 | 24917545 | AX-97780414 | 0 | T | C | 0.026 | -6.27E-01 | 9.15E-02 | 2.88E-01 | 2.61E-01 | 7.35E-06 | Yes | | intergenic_variant | MODIFIER |
| 15 | 24952896 | AX-97802837 | 0 | C | A | 0.026 | -6.27E-01 | 9.15E-02 | 2.88E-01 | 2.61E-01 | 7.35E-06 | Yes | | intergenic_variant | MODIFIER |
| 15 | 24964799 | AX-97852851 | 0 | G | T | 0.026 | -6.27E-01 | 9.15E-02 | 2.88E-01 | 2.61E-01 | 7.35E-06 | Yes | | intergenic_variant | MODIFIER |
| 15 | 24983878 | AX-170506984 | 0 | T | G | 0.026 | -6.27E-01 | 9.15E-02 | 2.88E-01 | 2.61E-01 | 7.35E-06 | Yes | Gm2824 | intron_variant,non_coding_transcript_variant | MODIFIER |
| 15 | 24988271 | AX-170438715 | 0 | G | A | 0.026 | -6.27E-01 | 9.15E-02 | 2.88E-01 | 2.61E-01 | 7.35E-06 | Yes | Gm2824 | intron_variant,non_coding_transcript_variant | MODIFIER |
| 15 | 25001391 | AX-170203857 | 0 | A | G | 0.026 | -6.27E-01 | 9.15E-02 | 2.88E-01 | 2.61E-01 | 7.35E-06 | Yes | Gm2824 | downstream_gene_variant | MODIFIER |
| 15 | 25002318 | AX-170448998 | 0 | G | T | 0.026 | -6.27E-01 | 9.15E-02 | 2.88E-01 | 2.61E-01 | 7.35E-06 | Yes | Gm2824 | downstream_gene_variant | MODIFIER |
| 15 | 25006227 | AX-97820421 | 0 | G | A | 0.026 | -6.27E-01 | 9.15E-02 | 2.88E-01 | 2.61E-01 | 7.35E-06 | Yes | Gm2824 | intron_variant,non_coding_transcript_variant | MODIFIER |
| 15 | 25007613 | AX-170240637 | 0 | T | C | 0.026 | -6.27E-01 | 9.15E-02 | 2.88E-01 | 2.61E-01 | 7.35E-06 | Yes | Gm2824 | intron_variant,non_coding_transcript_variant | MODIFIER |
| 15 | 25007720 | AX-170118865 | 0 | C | T | 0.026 | -6.27E-01 | 9.15E-02 | 2.88E-01 | 2.61E-01 | 7.35E-06 | Yes | Gm2824 | intron_variant,non_coding_transcript_variant | MODIFIER |
| 15 | 25033335 | AX-97855641 | 0 | G | C | 0.026 | -6.27E-01 | 9.15E-02 | 2.88E-01 | 2.61E-01 | 7.35E-06 | Yes | | intergenic_variant | MODIFIER |
| 15 | 25071513 | AX-170102553 | 0 | T | C | 0.026 | -6.27E-01 | 9.15E-02 | 2.88E-01 | 2.61E-01 | 7.35E-06 | Yes | | intergenic_variant | MODIFIER |
| 15 | 25093335 | AX-170011277 | 0 | A | G | 0.026 | -6.27E-01 | 9.15E-02 | 2.88E-01 | 2.61E-01 | 7.35E-06 | Yes | | intergenic_variant | MODIFIER |
| 15 | 25108148 | AX-170158821 | 0 | A | G | 0.026 | -6.27E-01 | 9.15E-02 | 2.88E-01 | 2.61E-01 | 7.35E-06 | Yes | Gm35996 | intron_variant,non_coding_transcript_variant | MODIFIER |

| | | | | | | | | | | | | | | | |
|---|---|---|---|---|---|---|---|---|---|---|---|---|---|---|---|
| 15 | 25128526 | AX-170011272 | 0 | G | C | 0.026 | -6.27E-01 | 9.15E-02 | 2.88E-01 | 2.61E-01 | 7.35E-06 | Yes | Gm35996 | upstream_gene_variant | MODIFIER |
| 15 | 25153422 | AX-170108179 | 0 | G | T | 0.026 | -6.27E-01 | 9.15E-02 | 2.88E-01 | 2.61E-01 | 7.35E-06 | Yes | 9230109A22Rik | upstream_gene_variant | MODIFIER |
| 15 | 25170881 | AX-170099391 | 0 | T | C | 0.026 | -6.27E-01 | 9.15E-02 | 2.88E-01 | 2.61E-01 | 7.35E-06 | Yes | 9230109A22Rik | intron_variant,non_coding_transcript_variant | MODIFIER |
| 15 | 25171174 | AX-97796107 | 0 | T | G | 0.026 | -6.27E-01 | 9.15E-02 | 2.88E-01 | 2.61E-01 | 7.35E-06 | Yes | 9230109A22Rik | intron_variant,non_coding_transcript_variant | MODIFIER |
| 15 | 25177550 | AX-170283649 | 0 | A | C | 0.026 | -6.27E-01 | 9.15E-02 | 2.88E-01 | 2.61E-01 | 7.35E-06 | Yes | 9230109A22Rik | intron_variant,non_coding_transcript_variant | MODIFIER |
| 15 | 25273746 | AX-97829100 | 0 | A | G | 0.026 | -6.27E-01 | 9.15E-02 | 2.88E-01 | 2.61E-01 | 7.35E-06 | Yes | | intergenic_variant | MODIFIER |
| 15 | 25288412 | AX-97827615 | 0 | T | C | 0.026 | -6.27E-01 | 9.15E-02 | 2.88E-01 | 2.61E-01 | 7.35E-06 | Yes | 4930445E18Rik | intron_variant,non_coding_transcript_variant | MODIFIER |
| 15 | 25313996 | AX-170207040 | 0 | G | A | 0.026 | -6.27E-01 | 9.15E-02 | 2.88E-01 | 2.61E-01 | 7.35E-06 | Yes | 4930445E18Rik | intron_variant,non_coding_transcript_variant | MODIFIER |
| 15 | 25322718 | AX-170156120 | 0 | T | C | 0.026 | -6.27E-01 | 9.15E-02 | 2.88E-01 | 2.61E-01 | 7.35E-06 | Yes | 4930445E18Rik | intron_variant,non_coding_transcript_variant | MODIFIER |
| 15 | 25371905 | AX-97828617 | 0 | T | C | 0.026 | -6.27E-01 | 9.15E-02 | 2.88E-01 | 2.61E-01 | 7.35E-06 | Yes | Basp1 | intron_variant | MODIFIER |
| 15 | 25390725 | AX-170396676 | 0 | G | A | 0.026 | -6.27E-01 | 9.15E-02 | 2.88E-01 | 2.61E-01 | 7.35E-06 | Yes | Basp1 | intron_variant | MODIFIER |
| 15 | 25395218 | AX-170378560 | 0 | C | T | 0.026 | -6.27E-01 | 9.15E-02 | 2.88E-01 | 2.61E-01 | 7.35E-06 | Yes | Basp1 | intron_variant | MODIFIER |
| 15 | 25395377 | AX-170206773 | 0 | G | T | 0.026 | -6.27E-01 | 9.15E-02 | 2.88E-01 | 2.61E-01 | 7.35E-06 | Yes | Basp1 | intron_variant | MODIFIER |
| 15 | 25395520 | AX-97833179 | 0 | C | A | 0.026 | -6.27E-01 | 9.15E-02 | 2.88E-01 | 2.61E-01 | 7.35E-06 | Yes | Basp1 | intron_variant | MODIFIER |
| 15 | 25404950 | AX-170535485 | 0 | A | G | 0.026 | -6.27E-01 | 9.15E-02 | 2.88E-01 | 2.61E-01 | 7.35E-06 | Yes | Basp1 | intron_variant | MODIFIER |
| 15 | 25407378 | AX-170252617 | 0 | A | G | 0.026 | -6.27E-01 | 9.15E-02 | 2.88E-01 | 2.61E-01 | 7.35E-06 | Yes | Basp1 | intron_variant | MODIFIER |
| 15 | 25450581 | AX-170146685 | 0 | G | T | 0.026 | -6.27E-01 | 9.15E-02 | 2.88E-01 | 2.61E-01 | 7.35E-06 | Yes | Gm48956 | upstream_gene_variant | MODIFIER |
| 15 | 25481108 | AX-170207250 | 0 | C | G | 0.026 | -6.27E-01 | 9.15E-02 | 2.88E-01 | 2.61E-01 | 7.35E-06 | Yes | | intergenic_variant | MODIFIER |
| 15 | 25495720 | AX-97824686 | 0 | A | G | 0.026 | -6.27E-01 | 9.15E-02 | 2.88E-01 | 2.61E-01 | 7.35E-06 | Yes | | regulatory_region_variant | MODIFIER |
| 15 | 25510204 | AX-170375182 | 0 | A | G | 0.026 | -6.27E-01 | 9.15E-02 | 2.88E-01 | 2.61E-01 | 7.35E-06 | Yes | | regulatory_region_variant | MODIFIER |
| 15 | 25532064 | AX-170189568 | 0 | C | T | 0.026 | -6.27E-01 | 9.15E-02 | 2.88E-01 | 2.61E-01 | 7.35E-06 | Yes | Gm36642 | intron_variant,non_coding_transcript_variant | MODIFIER |
| 15 | 25546915 | AX-170223902 | 0 | A | G | 0.026 | -6.27E-01 | 9.15E-02 | 2.88E-01 | 2.61E-01 | 7.35E-06 | Yes | Gm36642 | intron_variant,non_coding_transcript_variant | MODIFIER |
| 15 | 25554767 | AX-170011279 | 0 | T | C | 0.026 | -6.27E-01 | 9.15E-02 | 2.88E-01 | 2.61E-01 | 7.35E-06 | Yes | Gm36642 | intron_variant,non_coding_transcript_variant | MODIFIER |
| 15 | 25555471 | AX-170011280 | 0 | A | C | 0.026 | -6.27E-01 | 9.15E-02 | 2.88E-01 | 2.61E-01 | 7.35E-06 | Yes | Gm36642 | intron_variant,non_coding_transcript_variant | MODIFIER |
| 15 | 25586350 | AX-170420926 | 0 | C | A | 0.026 | -6.27E-01 | 9.15E-02 | 2.88E-01 | 2.61E-01 | 7.35E-06 | Yes | Gm36642 | intron_variant,non_coding_transcript_variant | MODIFIER |
| 15 | 25586632 | AX-97780751 | 0 | A | G | 0.026 | -6.27E-01 | 9.15E-02 | 2.88E-01 | 2.61E-01 | 7.35E-06 | Yes | Gm36642 | intron_variant,non_coding_transcript_variant | MODIFIER |
| 15 | 25605123 | AX-170011284 | 0 | A | G | 0.026 | -6.27E-01 | 9.15E-02 | 2.88E-01 | 2.61E-01 | 7.35E-06 | Yes | | intergenic_variant | MODIFIER |
| 15 | 25605286 | AX-170228594 | 0 | C | T | 0.026 | -6.27E-01 | 9.15E-02 | 2.88E-01 | 2.61E-01 | 7.35E-06 | Yes | | intergenic_variant | MODIFIER |
| 15 | 25709285 | AX-170055056 | 0 | A | G | 0.026 | -6.27E-01 | 9.15E-02 | 2.88E-01 | 2.61E-01 | 7.35E-06 | Yes | Myo10 | intron_variant | MODIFIER |
| 15 | 25719354 | AX-97822500 | 0 | A | G | 0.026 | -6.27E-01 | 9.15E-02 | 2.88E-01 | 2.61E-01 | 7.35E-06 | Yes | Myo10 | intron_variant | MODIFIER |
| 15 | 25719674 | AX-170497764 | 0 | G | A | 0.026 | -6.27E-01 | 9.15E-02 | 2.88E-01 | 2.61E-01 | 7.35E-06 | Yes | Myo10 | intron_variant | MODIFIER |
| 15 | 25726149 | AX-170308203 | 0 | A | C | 0.026 | -6.27E-01 | 9.15E-02 | 2.88E-01 | 2.61E-01 | 7.35E-06 | Yes | Myo10 | intron_variant | MODIFIER |
| 15 | 26047142 | AX-170380112 | 0 | C | T | 0.026 | -6.27E-01 | 9.15E-02 | 2.88E-01 | 2.61E-01 | 7.35E-06 | Yes | | intergenic_variant | MODIFIER |
| 15 | 26051233 | AX-97835316 | 0 | C | T | 0.026 | -6.27E-01 | 9.15E-02 | 2.88E-01 | 2.61E-01 | 7.35E-06 | Yes | Gm41279 | downstream_gene_variant | MODIFIER |
| 15 | 26051335 | AX-170229003 | 0 | A | G | 0.026 | -6.27E-01 | 9.15E-02 | 2.88E-01 | 2.61E-01 | 7.35E-06 | Yes | Gm41279 | downstream_gene_variant | MODIFIER |
| 15 | 26056990 | AX-170398131 | 0 | C | G | 0.026 | -6.27E-01 | 9.15E-02 | 2.88E-01 | 2.61E-01 | 7.35E-06 | Yes | Gm41279 | intron_variant,non_coding_transcript_variant | MODIFIER |
| 15 | 26057101 | AX-170055065 | 0 | A | G | 0.026 | -6.27E-01 | 9.15E-02 | 2.88E-01 | 2.61E-01 | 7.35E-06 | Yes | Gm41279 | intron_variant,non_coding_transcript_variant | MODIFIER |
| 15 | 26057700 | AX-170225425 | 0 | T | C | 0.026 | -6.27E-01 | 9.15E-02 | 2.88E-01 | 2.61E-01 | 7.35E-06 | Yes | Gm41279 | intron_variant,non_coding_transcript_variant | MODIFIER |
| 15 | 26067374 | AX-170011293 | 0 | T | C | 0.026 | -6.27E-01 | 9.15E-02 | 2.88E-01 | 2.61E-01 | 7.35E-06 | Yes | Gm41279 | upstream_gene_variant | MODIFIER |
| 15 | 26067456 | AX-170011291 | 0 | C | T | 0.026 | -6.27E-01 | 9.15E-02 | 2.88E-01 | 2.61E-01 | 7.35E-06 | Yes | | intergenic_variant | MODIFIER |
| 15 | 26099662 | AX-97845476 | 0 | T | C | 0.026 | -6.27E-01 | 9.15E-02 | 2.88E-01 | 2.61E-01 | 7.35E-06 | Yes | | intergenic_variant | MODIFIER |
| 15 | 26100231 | AX-170011289 | 0 | A | G | 0.026 | -6.27E-01 | 9.15E-02 | 2.88E-01 | 2.61E-01 | 7.35E-06 | Yes | | regulatory_region_variant | MODIFIER |
| 15 | 26129129 | AX-97820918 | 0 | C | T | 0.026 | -6.27E-01 | 9.15E-02 | 2.88E-01 | 2.61E-01 | 7.35E-06 | Yes | | intergenic_variant | MODIFIER |
| 15 | 26160100 | AX-170011287 | 0 | T | C | 0.026 | -6.27E-01 | 9.15E-02 | 2.88E-01 | 2.61E-01 | 7.35E-06 | Yes | | intergenic_variant | MODIFIER |
| 15 | 26181084 | AX-170144524 | 0 | G | T | 0.026 | -6.27E-01 | 9.15E-02 | 2.88E-01 | 2.61E-01 | 7.35E-06 | Yes | | intergenic_variant | MODIFIER |
| 15 | 26181729 | AX-170455907 | 0 | T | G | 0.026 | -6.27E-01 | 9.15E-02 | 2.88E-01 | 2.61E-01 | 7.35E-06 | Yes | | intergenic_variant | MODIFIER |
| 15 | 26226245 | AX-97829077 | 0 | C | T | 0.026 | -6.27E-01 | 9.15E-02 | 2.88E-01 | 2.61E-01 | 7.35E-06 | Yes | | intergenic_variant | MODIFIER |
| 15 | 26238374 | AX-170335269 | 0 | G | A | 0.026 | -6.27E-01 | 9.15E-02 | 2.88E-01 | 2.61E-01 | 7.35E-06 | Yes | | intergenic_variant | MODIFIER |
| 15 | 26238982 | AX-170259178 | 0 | C | T | 0.026 | -6.27E-01 | 9.15E-02 | 2.88E-01 | 2.61E-01 | 7.35E-06 | Yes | | intergenic_variant | MODIFIER |
| 15 | 26241685 | AX-97842461 | 0 | T | C | 0.026 | -6.27E-01 | 9.15E-02 | 2.88E-01 | 2.61E-01 | 7.35E-06 | Yes | | intergenic_variant | MODIFIER |
| 15 | 26271673 | AX-97841083 | 0 | T | C | 0.026 | -6.27E-01 | 9.15E-02 | 2.88E-01 | 2.61E-01 | 7.35E-06 | Yes | | intergenic_variant | MODIFIER |
| 15 | 26293454 | AX-170055067 | 0 | C | A | 0.026 | -6.27E-01 | 9.15E-02 | 2.88E-01 | 2.61E-01 | 7.35E-06 | Yes | | intergenic_variant | MODIFIER |
| 15 | 26293501 | AX-170055069 | 0 | T | C | 0.026 | -6.27E-01 | 9.15E-02 | 2.88E-01 | 2.61E-01 | 7.35E-06 | Yes | | intergenic_variant | MODIFIER |
| 15 | 26299169 | AX-97829079 | 0 | G | A | 0.026 | -6.27E-01 | 9.15E-02 | 2.88E-01 | 2.61E-01 | 7.35E-06 | Yes | | intergenic_variant | MODIFIER |

| | | | | | | | | | | | | | | | |
|---|---|---|---|---|---|---|---|---|---|---|---|---|---|---|---|
| 15 | 26299205 | AX-170011294 | 0 | C | T | 0.026 | -6.27E-01 | 9.15E-02 | 2.88E-01 | 2.61E-01 | 7.35E-06 | Yes | | intergenic_variant | MODIFIER |
| 15 | 26314335 | AX-170011285 | 0 | A | G | 0.026 | -6.27E-01 | 9.15E-02 | 2.88E-01 | 2.61E-01 | 7.35E-06 | Yes | MAR11 | downstream_gene_variant | MODIFIER |
| 15 | 26317423 | AX-170461841 | 0 | G | A | 0.026 | -6.27E-01 | 9.15E-02 | 2.88E-01 | 2.61E-01 | 7.35E-06 | Yes | MAR11 | intron_variant | MODIFIER |
| 15 | 26347824 | AX-170011292 | 0 | A | G | 0.026 | -6.27E-01 | 9.15E-02 | 2.88E-01 | 2.61E-01 | 7.35E-06 | Yes | MAR11 | intron_variant | MODIFIER |
| 15 | 26348847 | AX-170353500 | 0 | G | C | 0.026 | -6.27E-01 | 9.15E-02 | 2.88E-01 | 2.61E-01 | 7.35E-06 | Yes | MAR11 | intron_variant | MODIFIER |
| 15 | 26353386 | AX-97833712 | 0 | G | A | 0.026 | -6.27E-01 | 9.15E-02 | 2.88E-01 | 2.61E-01 | 7.35E-06 | Yes | MAR11 | intron_variant | MODIFIER |
| 15 | 26362950 | AX-97839885 | 0 | T | G | 0.026 | -6.27E-01 | 9.15E-02 | 2.88E-01 | 2.61E-01 | 7.35E-06 | Yes | MAR11 | intron_variant | MODIFIER |
| 15 | 26387507 | AX-170217128 | 0 | C | T | 0.026 | -6.27E-01 | 9.15E-02 | 2.88E-01 | 2.61E-01 | 7.35E-06 | Yes | MAR11 | intron_variant | MODIFIER |
| 15 | 26391693 | AX-97816440 | 0 | C | T | 0.026 | -6.27E-01 | 9.15E-02 | 2.88E-01 | 2.61E-01 | 7.35E-06 | Yes | MAR11 | intron_variant | MODIFIER |
| 15 | 26410605 | AX-97841598 | 0 | C | T | 0.026 | -6.27E-01 | 9.15E-02 | 2.88E-01 | 2.61E-01 | 7.35E-06 | Yes | MAR11 | downstream_gene_variant | MODIFIER |
| 15 | 26436040 | AX-170169048 | 0 | G | A | 0.026 | -6.27E-01 | 9.15E-02 | 2.88E-01 | 2.61E-01 | 7.35E-06 | Yes | | intergenic_variant | MODIFIER |
| 15 | 26436709 | AX-170168841 | 0 | T | A | 0.026 | -6.27E-01 | 9.15E-02 | 2.88E-01 | 2.61E-01 | 7.35E-06 | Yes | | intergenic_variant | MODIFIER |
| 15 | 26535837 | AX-97800437 | 0 | A | G | 0.026 | -6.27E-01 | 9.15E-02 | 2.88E-01 | 2.61E-01 | 7.35E-06 | Yes | Fbxl7 | downstream_gene_variant | MODIFIER |
| 15 | 26543736 | AX-170274678 | 0 | C | T | 0.026 | -6.27E-01 | 9.15E-02 | 2.88E-01 | 2.61E-01 | 7.35E-06 | Yes | Fbxl7 | missense_variant | MODERATE |
| 15 | 26588250 | AX-170322424 | 0 | A | G | 0.026 | -6.27E-01 | 9.15E-02 | 2.88E-01 | 2.61E-01 | 7.35E-06 | Yes | Fbxl7 | intron_variant | MODIFIER |
| 15 | 26588363 | AX-170195177 | 0 | A | G | 0.026 | -6.27E-01 | 9.15E-02 | 2.88E-01 | 2.61E-01 | 7.35E-06 | Yes | Fbxl7 | intron_variant | MODIFIER |
| 15 | 26588871 | AX-170011302 | 0 | T | G | 0.026 | -6.27E-01 | 9.15E-02 | 2.88E-01 | 2.61E-01 | 7.35E-06 | Yes | Fbxl7 | intron_variant | MODIFIER |
| 15 | 26593498 | AX-170011298 | 0 | A | G | 0.026 | -6.27E-01 | 9.15E-02 | 2.88E-01 | 2.61E-01 | 7.35E-06 | Yes | Fbxl7 | intron_variant | MODIFIER |
| 15 | 26594428 | AX-170011296 | 0 | G | A | 0.026 | -6.27E-01 | 9.15E-02 | 2.88E-01 | 2.61E-01 | 7.35E-06 | Yes | Fbxl7 | intron_variant | MODIFIER |
| 15 | 26596046 | AX-97815347 | 0 | C | A | 0.026 | -6.27E-01 | 9.15E-02 | 2.88E-01 | 2.61E-01 | 7.35E-06 | Yes | Fbxl7 | intron_variant | MODIFIER |
| 15 | 26598381 | AX-170225966 | 0 | G | A | 0.026 | -6.27E-01 | 9.15E-02 | 2.88E-01 | 2.61E-01 | 7.35E-06 | Yes | Fbxl7 | intron_variant | MODIFIER |
| 15 | 26618285 | AX-170354830 | 0 | T | C | 0.026 | -6.27E-01 | 9.15E-02 | 2.88E-01 | 2.61E-01 | 7.35E-06 | Yes | Fbxl7 | intron_variant | MODIFIER |
| 15 | 26622218 | AX-97794456 | 0 | G | A | 0.026 | -6.27E-01 | 9.15E-02 | 2.88E-01 | 2.61E-01 | 7.35E-06 | Yes | Fbxl7 | intron_variant | MODIFIER |
| 15 | 26622287 | AX-170011301 | 0 | T | C | 0.026 | -6.27E-01 | 9.15E-02 | 2.88E-01 | 2.61E-01 | 7.35E-06 | Yes | Fbxl7 | intron_variant | MODIFIER |
| 15 | 26624995 | AX-170460407 | 0 | G | T | 0.026 | -6.27E-01 | 9.15E-02 | 2.88E-01 | 2.61E-01 | 7.35E-06 | Yes | Fbxl7 | intron_variant | MODIFIER |
| 15 | 26627899 | AX-170011299 | 0 | T | C | 0.026 | -6.27E-01 | 9.15E-02 | 2.88E-01 | 2.61E-01 | 7.35E-06 | Yes | Fbxl7 | intron_variant | MODIFIER |
| 15 | 26685746 | AX-170011297 | 0 | C | T | 0.026 | -6.27E-01 | 9.15E-02 | 2.88E-01 | 2.61E-01 | 7.35E-06 | Yes | Fbxl7 | intron_variant | MODIFIER |
| 15 | 26713714 | AX-97829101 | 0 | A | G | 0.026 | -6.27E-01 | 9.15E-02 | 2.88E-01 | 2.61E-01 | 7.35E-06 | Yes | Fbxl7 | intron_variant | MODIFIER |
| 15 | 26719985 | AX-170372707 | 0 | T | G | 0.026 | -6.27E-01 | 9.15E-02 | 2.88E-01 | 2.61E-01 | 7.35E-06 | Yes | Fbxl7 | intron_variant | MODIFIER |
| 15 | 26767688 | AX-97849687 | 0 | G | A | 0.026 | -6.27E-01 | 9.15E-02 | 2.88E-01 | 2.61E-01 | 7.35E-06 | Yes | Fbxl7 | intron_variant | MODIFIER |
| 15 | 26824113 | AX-170259191 | 0 | A | T | 0.026 | -6.27E-01 | 9.15E-02 | 2.88E-01 | 2.61E-01 | 7.35E-06 | Yes | Fbxl7 | intron_variant | MODIFIER |
| 15 | 26827701 | AX-97782275 | 0 | T | C | 0.026 | -6.27E-01 | 9.15E-02 | 2.88E-01 | 2.61E-01 | 7.35E-06 | Yes | Fbxl7 | intron_variant | MODIFIER |
| 15 | 26858716 | AX-97829076 | 0 | T | C | 0.026 | -6.27E-01 | 9.15E-02 | 2.88E-01 | 2.61E-01 | 7.35E-06 | Yes | Fbxl7 | intron_variant | MODIFIER |
| 15 | 26864281 | AX-170309830 | 0 | C | G | 0.026 | -6.27E-01 | 9.15E-02 | 2.88E-01 | 2.61E-01 | 7.35E-06 | Yes | Fbxl7 | intron_variant | MODIFIER |
| 15 | 26916608 | AX-170137391 | 0 | A | G | 0.026 | -6.27E-01 | 9.15E-02 | 2.88E-01 | 2.61E-01 | 7.35E-06 | Yes | | intergenic_variant | MODIFIER |
| 15 | 27002478 | AX-170225756 | 0 | A | T | 0.026 | -6.27E-01 | 9.15E-02 | 2.88E-01 | 2.61E-01 | 7.35E-06 | Yes | | intergenic_variant | MODIFIER |
| 15 | 27003148 | AX-170488142 | 0 | A | C | 0.026 | -6.27E-01 | 9.15E-02 | 2.88E-01 | 2.61E-01 | 7.35E-06 | Yes | | intergenic_variant | MODIFIER |
| 15 | 27028789 | AX-97786924 | 0 | A | G | 0.026 | -6.27E-01 | 9.15E-02 | 2.88E-01 | 2.61E-01 | 7.35E-06 | Yes | Gm6576 | downstream_gene_variant | MODIFIER |
| 15 | 27033986 | AX-97852358 | 0 | G | T | 0.026 | -6.27E-01 | 9.15E-02 | 2.88E-01 | 2.61E-01 | 7.35E-06 | Yes | | intergenic_variant | MODIFIER |
| 15 | 27034663 | AX-170241074 | 0 | A | G | 0.026 | -6.27E-01 | 9.15E-02 | 2.88E-01 | 2.61E-01 | 7.35E-06 | Yes | | intergenic_variant | MODIFIER |
| 15 | 27042000 | AX-170055079 | 0 | T | C | 0.026 | -6.27E-01 | 9.15E-02 | 2.88E-01 | 2.61E-01 | 7.35E-06 | Yes | | intergenic_variant | MODIFIER |
| 15 | 27045506 | AX-170055084 | 0 | A | C | 0.026 | -6.27E-01 | 9.15E-02 | 2.88E-01 | 2.61E-01 | 7.35E-06 | Yes | | intergenic_variant | MODIFIER |
| 15 | 27045550 | AX-170055083 | 0 | C | T | 0.026 | -6.27E-01 | 9.15E-02 | 2.88E-01 | 2.61E-01 | 7.35E-06 | Yes | | intergenic_variant | MODIFIER |
| 15 | 27066188 | AX-97792509 | 0 | A | G | 0.026 | -6.27E-01 | 9.15E-02 | 2.88E-01 | 2.61E-01 | 7.35E-06 | Yes | Gm23903 | upstream_gene_variant | MODIFIER |
| 15 | 27076200 | AX-170011306 | 0 | G | C | 0.026 | -6.27E-01 | 9.15E-02 | 2.88E-01 | 2.61E-01 | 7.35E-06 | Yes | | intergenic_variant | MODIFIER |
| 15 | 27076351 | AX-170011304 | 0 | G | A | 0.026 | -6.27E-01 | 9.15E-02 | 2.88E-01 | 2.61E-01 | 7.35E-06 | Yes | | intergenic_variant | MODIFIER |
| 15 | 27089690 | AX-170400950 | 0 | T | A | 0.026 | -6.27E-01 | 9.15E-02 | 2.88E-01 | 2.61E-01 | 7.35E-06 | Yes | | intergenic_variant | MODIFIER |
| 15 | 27093971 | AX-170426889 | 0 | T | A | 0.026 | -6.27E-01 | 9.15E-02 | 2.88E-01 | 2.61E-01 | 7.35E-06 | Yes | | intergenic_variant | MODIFIER |
| 15 | 27096440 | AX-170011308 | 0 | C | G | 0.026 | -6.27E-01 | 9.15E-02 | 2.88E-01 | 2.61E-01 | 7.35E-06 | Yes | | intergenic_variant | MODIFIER |
| 15 | 27096574 | AX-97855835 | 0 | C | T | 0.026 | -6.27E-01 | 9.15E-02 | 2.88E-01 | 2.61E-01 | 7.35E-06 | Yes | | intergenic_variant | MODIFIER |
| 15 | 27120627 | AX-170087801 | 0 | A | G | 0.026 | -6.27E-01 | 9.15E-02 | 2.88E-01 | 2.61E-01 | 7.35E-06 | Yes | | intergenic_variant | MODIFIER |
| 15 | 27141869 | AX-97799126 | 0 | G | T | 0.026 | -6.27E-01 | 9.15E-02 | 2.88E-01 | 2.61E-01 | 7.35E-06 | Yes | | intergenic_variant | MODIFIER |
| 15 | 27167506 | AX-97828644 | 0 | A | C | 0.026 | -6.27E-01 | 9.15E-02 | 2.88E-01 | 2.61E-01 | 7.35E-06 | Yes | | intergenic_variant | MODIFIER |
| 15 | 27321140 | AX-170300871 | 0 | A | T | 0.026 | -6.27E-01 | 9.15E-02 | 2.88E-01 | 2.61E-01 | 7.35E-06 | Yes | | intergenic_variant | MODIFIER |

| | | | | | | | | | | | | | | | |
|---|---|---|---|---|---|---|---|---|---|---|---|---|---|---|---|
| 15 | 27327550 | AX-97817030 | 0 | C | T | 0.026 | -6.27E-01 | 9.15E-02 | 2.88E-01 | 2.61E-01 | 7.35E-06 | Yes | | intergenic_variant | MODIFIER |
| 15 | 27405249 | AX-170245871 | 0 | T | A | 0.026 | -6.27E-01 | 9.15E-02 | 2.88E-01 | 2.61E-01 | 7.35E-06 | Yes | | intergenic_variant | MODIFIER |
| 15 | 27405298 | AX-97789590 | 0 | T | C | 0.026 | -6.27E-01 | 9.15E-02 | 2.88E-01 | 2.61E-01 | 7.35E-06 | Yes | | intergenic_variant | MODIFIER |
| 15 | 27408300 | AX-170055089 | 0 | T | C | 0.026 | -6.27E-01 | 9.15E-02 | 2.88E-01 | 2.61E-01 | 7.35E-06 | Yes | | intergenic_variant | MODIFIER |
| 15 | 27411875 | AX-170325338 | 0 | G | A | 0.026 | -6.27E-01 | 9.15E-02 | 2.88E-01 | 2.61E-01 | 7.35E-06 | Yes | | intergenic_variant | MODIFIER |
| 15 | 27416628 | AX-97846817 | 0 | A | C | 0.026 | -6.27E-01 | 9.15E-02 | 2.88E-01 | 2.61E-01 | 7.35E-06 | Yes | | intergenic_variant | MODIFIER |
| 15 | 27426056 | AX-170166723 | 0 | T | C | 0.026 | -6.27E-01 | 9.15E-02 | 2.88E-01 | 2.61E-01 | 7.35E-06 | Yes | | intergenic_variant | MODIFIER |
| 15 | 27490288 | AX-170294295 | 0 | A | G | 0.026 | -6.27E-01 | 9.15E-02 | 2.88E-01 | 2.61E-01 | 7.35E-06 | Yes | Ank | intron_variant | MODIFIER |
| 15 | 27490357 | AX-170355476 | 0 | C | A | 0.026 | -6.27E-01 | 9.15E-02 | 2.88E-01 | 2.61E-01 | 7.35E-06 | Yes | Ank | intron_variant | MODIFIER |
| 15 | 27490781 | AX-170479036 | 0 | G | A | 0.026 | -6.27E-01 | 9.15E-02 | 2.88E-01 | 2.61E-01 | 7.35E-06 | Yes | Ank | intron_variant | MODIFIER |
| 15 | 27494214 | AX-170516702 | 0 | T | C | 0.026 | -6.27E-01 | 9.15E-02 | 2.88E-01 | 2.61E-01 | 7.35E-06 | Yes | Ank | intron_variant | MODIFIER |
| 15 | 27495411 | AX-97828643 | 0 | T | C | 0.026 | -6.27E-01 | 9.15E-02 | 2.88E-01 | 2.61E-01 | 7.35E-06 | Yes | Ank | intron_variant | MODIFIER |
| 15 | 27504324 | AX-170485446 | 0 | C | T | 0.026 | -6.27E-01 | 9.15E-02 | 2.88E-01 | 2.61E-01 | 7.35E-06 | Yes | Ank | intron_variant | MODIFIER |
| 15 | 27507388 | AX-170368128 | 0 | A | G | 0.026 | -6.27E-01 | 9.15E-02 | 2.88E-01 | 2.61E-01 | 7.35E-06 | Yes | Ank | intron_variant | MODIFIER |
| 15 | 27511498 | AX-170389104 | 0 | T | G | 0.026 | -6.27E-01 | 9.15E-02 | 2.88E-01 | 2.61E-01 | 7.35E-06 | Yes | Ank | intron_variant | MODIFIER |
| 15 | 27515292 | AX-170490531 | 0 | A | G | 0.026 | -6.27E-01 | 9.15E-02 | 2.88E-01 | 2.61E-01 | 7.35E-06 | Yes | Ank | intron_variant | MODIFIER |
| 15 | 27516291 | AX-97786158 | 0 | G | A | 0.026 | -6.27E-01 | 9.15E-02 | 2.88E-01 | 2.61E-01 | 7.35E-06 | Yes | Ank | intron_variant | MODIFIER |
| 15 | 27522358 | AX-170419294 | 0 | C | G | 0.026 | -6.27E-01 | 9.15E-02 | 2.88E-01 | 2.61E-01 | 7.35E-06 | Yes | Ank | intron_variant | MODIFIER |
| 15 | 27587088 | AX-170474157 | 0 | T | C | 0.026 | -6.27E-01 | 9.15E-02 | 2.88E-01 | 2.61E-01 | 7.35E-06 | Yes | Ank | intron_variant | MODIFIER |
| 15 | 27589494 | AX-170442506 | 0 | G | A | 0.026 | -6.27E-01 | 9.15E-02 | 2.88E-01 | 2.61E-01 | 7.35E-06 | Yes | Ank | intron_variant | MODIFIER |
| 15 | 27589622 | AX-170451016 | 0 | A | G | 0.026 | -6.27E-01 | 9.15E-02 | 2.88E-01 | 2.61E-01 | 7.35E-06 | Yes | Ank | intron_variant | MODIFIER |
| 15 | 27591527 | AX-97851403 | 0 | T | C | 0.026 | -6.27E-01 | 9.15E-02 | 2.88E-01 | 2.61E-01 | 7.35E-06 | Yes | Ank | synonymous_variant | LOW |
| 15 | 27606559 | AX-170011312 | 0 | A | G | 0.026 | -6.27E-01 | 9.15E-02 | 2.88E-01 | 2.61E-01 | 7.35E-06 | Yes | Otulin | intron_variant | MODIFIER |
| 15 | 27612852 | AX-170055090 | 0 | G | A | 0.026 | -6.27E-01 | 9.15E-02 | 2.88E-01 | 2.61E-01 | 7.35E-06 | Yes | Otulin | intron_variant | MODIFIER |
| 15 | 27612877 | AX-170143106 | 0 | T | G | 0.026 | -6.27E-01 | 9.15E-02 | 2.88E-01 | 2.61E-01 | 7.35E-06 | Yes | Otulin | intron_variant | MODIFIER |
| 15 | 27613058 | AX-170371715 | 0 | G | C | 0.026 | -6.27E-01 | 9.15E-02 | 2.88E-01 | 2.61E-01 | 7.35E-06 | Yes | Otulin | intron_variant | MODIFIER |
| 15 | 27698245 | AX-97828704 | 0 | A | T | 0.026 | -6.27E-01 | 9.15E-02 | 2.88E-01 | 2.61E-01 | 7.35E-06 | Yes | | intergenic_variant | MODIFIER |
| 15 | 28321743 | AX-97836292 | 0 | T | C | 0.026 | -6.27E-01 | 9.15E-02 | 2.88E-01 | 2.61E-01 | 7.35E-06 | Yes | Dnah5 | intron_variant | MODIFIER |
| 15 | 29572281 | AX-97829285 | 0 | C | A | 0.026 | -6.27E-01 | 9.15E-02 | 2.88E-01 | 2.61E-01 | 7.35E-06 | Yes | | intergenic_variant | MODIFIER |
| 15 | 29572850 | AX-170398607 | 0 | G | A | 0.026 | -6.27E-01 | 9.15E-02 | 2.88E-01 | 2.61E-01 | 7.35E-06 | Yes | | intergenic_variant | MODIFIER |
| 15 | 29574116 | AX-170248597 | 0 | A | G | 0.026 | -6.27E-01 | 9.15E-02 | 2.88E-01 | 2.61E-01 | 7.35E-06 | Yes | | intergenic_variant | MODIFIER |
| 15 | 29588019 | AX-170117036 | 0 | G | C | 0.026 | -6.27E-01 | 9.15E-02 | 2.88E-01 | 2.61E-01 | 7.35E-06 | Yes | | intergenic_variant | MODIFIER |
| 15 | 29616090 | AX-170093190 | 0 | C | T | 0.026 | -6.27E-01 | 9.15E-02 | 2.88E-01 | 2.61E-01 | 7.35E-06 | Yes | | intergenic_variant | MODIFIER |
| 15 | 29632422 | AX-170432190 | 0 | T | C | 0.026 | -6.27E-01 | 9.15E-02 | 2.88E-01 | 2.61E-01 | 7.35E-06 | Yes | | intergenic_variant | MODIFIER |
| 15 | 29634869 | AX-170346128 | 0 | A | G | 0.026 | -6.27E-01 | 9.15E-02 | 2.88E-01 | 2.61E-01 | 7.35E-06 | Yes | | intergenic_variant | MODIFIER |
| 15 | 29635358 | AX-97829284 | 0 | G | A | 0.026 | -6.27E-01 | 9.15E-02 | 2.88E-01 | 2.61E-01 | 7.35E-06 | Yes | | intergenic_variant | MODIFIER |
| 15 | 29689175 | AX-170055125 | 0 | G | A | 0.026 | -6.27E-01 | 9.15E-02 | 2.88E-01 | 2.61E-01 | 7.35E-06 | Yes | | intergenic_variant | MODIFIER |
| 15 | 29689249 | AX-170055122 | 0 | A | G | 0.026 | -6.27E-01 | 9.15E-02 | 2.88E-01 | 2.61E-01 | 7.35E-06 | Yes | | intergenic_variant | MODIFIER |
| 15 | 29715999 | AX-97835665 | 0 | A | G | 0.026 | -6.27E-01 | 9.15E-02 | 2.88E-01 | 2.61E-01 | 7.35E-06 | Yes | Mir3964 | upstream_gene_variant | MODIFIER |
| 15 | 29755802 | AX-170011355 | 0 | A | G | 0.026 | -6.27E-01 | 9.15E-02 | 2.88E-01 | 2.61E-01 | 7.35E-06 | Yes | | intergenic_variant | MODIFIER |
| 15 | 29756733 | AX-97829131 | 0 | G | A | 0.026 | -6.27E-01 | 9.15E-02 | 2.88E-01 | 2.61E-01 | 7.35E-06 | Yes | | intergenic_variant | MODIFIER |
| 15 | 29798700 | AX-170011349 | 0 | T | C | 0.026 | -6.27E-01 | 9.15E-02 | 2.88E-01 | 2.61E-01 | 7.35E-06 | Yes | | regulatory_region_variant | MODIFIER |
| 15 | 29831521 | AX-97848904 | 0 | G | C | 0.026 | -6.27E-01 | 9.15E-02 | 2.88E-01 | 2.61E-01 | 7.35E-06 | Yes | | intergenic_variant | MODIFIER |
| 15 | 29844515 | AX-170093191 | 0 | C | T | 0.026 | -6.27E-01 | 9.15E-02 | 2.88E-01 | 2.61E-01 | 7.35E-06 | Yes | | intergenic_variant | MODIFIER |
| 15 | 29859732 | AX-170416296 | 0 | C | T | 0.026 | -6.27E-01 | 9.15E-02 | 2.88E-01 | 2.61E-01 | 7.35E-06 | Yes | | intergenic_variant | MODIFIER |
| 15 | 29863633 | AX-170521866 | 0 | G | A | 0.026 | -6.27E-01 | 9.15E-02 | 2.88E-01 | 2.61E-01 | 7.35E-06 | Yes | | intergenic_variant | MODIFIER |
| 15 | 29863748 | AX-97785918 | 0 | A | C | 0.026 | -6.27E-01 | 9.15E-02 | 2.88E-01 | 2.61E-01 | 7.35E-06 | Yes | | intergenic_variant | MODIFIER |
| 15 | 29893263 | AX-170011357 | 0 | G | A | 0.026 | -6.27E-01 | 9.15E-02 | 2.88E-01 | 2.61E-01 | 7.35E-06 | Yes | Gm23033 | upstream_gene_variant | MODIFIER |
| 15 | 29957468 | AX-170313855 | 0 | G | A | 0.026 | -6.27E-01 | 9.15E-02 | 2.88E-01 | 2.61E-01 | 7.35E-06 | Yes | | intergenic_variant | MODIFIER |
| 15 | 29957498 | AX-170346728 | 0 | C | A | 0.026 | -6.27E-01 | 9.15E-02 | 2.88E-01 | 2.61E-01 | 7.35E-06 | Yes | | intergenic_variant | MODIFIER |
| 15 | 30004855 | AX-170166502 | 0 | T | C | 0.026 | -6.27E-01 | 9.15E-02 | 2.88E-01 | 2.61E-01 | 7.35E-06 | Yes | Gm23327 | upstream_gene_variant | MODIFIER |
| 15 | 30026234 | AX-170087805 | 0 | T | C | 0.026 | -6.27E-01 | 9.15E-02 | 2.88E-01 | 2.61E-01 | 7.35E-06 | Yes | | intergenic_variant | MODIFIER |
| 15 | 30033509 | AX-170462240 | 0 | C | A | 0.026 | -6.27E-01 | 9.15E-02 | 2.88E-01 | 2.61E-01 | 7.35E-06 | Yes | | intergenic_variant | MODIFIER |
| 15 | 30033969 | AX-170162479 | 0 | T | C | 0.026 | -6.27E-01 | 9.15E-02 | 2.88E-01 | 2.61E-01 | 7.35E-06 | Yes | | intergenic_variant | MODIFIER |

| | | | | | | | | | | | | | | | |
|---|---|---|---|---|---|---|---|---|---|---|---|---|---|---|---|
| 15 | 30056132 | AX-170055129 | 0 | C | T | 0.026 | -6.27E-01 | 9.15E-02 | 2.88E-01 | 2.61E-01 | 7.35E-06 | Yes | Gm26163 | upstream_gene_variant | MODIFIER |
| 15 | 30056178 | AX-97851324 | 0 | G | A | 0.026 | -6.27E-01 | 9.15E-02 | 2.88E-01 | 2.61E-01 | 7.35E-06 | Yes | Gm26163 | upstream_gene_variant | MODIFIER |
| 15 | 30134678 | AX-97857115 | 0 | A | G | 0.026 | -6.27E-01 | 9.15E-02 | 2.88E-01 | 2.61E-01 | 7.35E-06 | Yes | | intergenic_variant | MODIFIER |
| 15 | 30139203 | AX-170011361 | 0 | T | C | 0.026 | -6.27E-01 | 9.15E-02 | 2.88E-01 | 2.61E-01 | 7.35E-06 | Yes | | intergenic_variant | MODIFIER |
| 15 | 30185489 | AX-170515810 | 0 | C | T | 0.026 | -6.27E-01 | 9.15E-02 | 2.88E-01 | 2.61E-01 | 7.35E-06 | Yes | Ctnnd2 | intron_variant | MODIFIER |
| 15 | 30191170 | AX-170241011 | 0 | G | A | 0.026 | -6.27E-01 | 9.15E-02 | 2.88E-01 | 2.61E-01 | 7.35E-06 | Yes | Ctnnd2 | intron_variant | MODIFIER |
| 15 | 30191197 | AX-97784010 | 0 | T | C | 0.026 | -6.27E-01 | 9.15E-02 | 2.88E-01 | 2.61E-01 | 7.35E-06 | Yes | Ctnnd2 | intron_variant | MODIFIER |
| 15 | 30192616 | AX-170345616 | 0 | C | G | 0.026 | -6.27E-01 | 9.15E-02 | 2.88E-01 | 2.61E-01 | 7.35E-06 | Yes | Ctnnd2 | intron_variant | MODIFIER |
| 15 | 30202101 | AX-170011358 | 0 | C | A | 0.026 | -6.27E-01 | 9.15E-02 | 2.88E-01 | 2.61E-01 | 7.35E-06 | Yes | Ctnnd2 | intron_variant | MODIFIER |
| 15 | 30206182 | AX-170355992 | 0 | G | A | 0.026 | -6.27E-01 | 9.15E-02 | 2.88E-01 | 2.61E-01 | 7.35E-06 | Yes | Ctnnd2 | intron_variant | MODIFIER |
| 15 | 30207100 | AX-170344128 | 0 | C | A | 0.026 | -6.27E-01 | 9.15E-02 | 2.88E-01 | 2.61E-01 | 7.35E-06 | Yes | Ctnnd2 | intron_variant | MODIFIER |
| 15 | 30207177 | AX-170413130 | 0 | G | A | 0.026 | -6.27E-01 | 9.15E-02 | 2.88E-01 | 2.61E-01 | 7.35E-06 | Yes | Ctnnd2 | intron_variant | MODIFIER |
| 15 | 30207934 | AX-170424818 | 0 | T | C | 0.026 | -6.27E-01 | 9.15E-02 | 2.88E-01 | 2.61E-01 | 7.35E-06 | Yes | Ctnnd2 | intron_variant | MODIFIER |
| 15 | 30214385 | AX-170403815 | 0 | G | A | 0.026 | -6.27E-01 | 9.15E-02 | 2.88E-01 | 2.61E-01 | 7.35E-06 | Yes | Ctnnd2 | intron_variant | MODIFIER |
| 15 | 30233979 | AX-97782087 | 0 | T | C | 0.026 | -6.27E-01 | 9.15E-02 | 2.88E-01 | 2.61E-01 | 7.35E-06 | Yes | Ctnnd2 | intron_variant | MODIFIER |
| 15 | 30249808 | AX-170340901 | 0 | G | A | 0.026 | -6.27E-01 | 9.15E-02 | 2.88E-01 | 2.61E-01 | 7.35E-06 | Yes | Ctnnd2 | intron_variant | MODIFIER |
| 15 | 30259339 | AX-170129927 | 0 | G | T | 0.026 | -6.27E-01 | 9.15E-02 | 2.88E-01 | 2.61E-01 | 7.35E-06 | Yes | Ctnnd2 | intron_variant | MODIFIER |
| 15 | 30260640 | AX-170285381 | 0 | C | T | 0.026 | -6.27E-01 | 9.15E-02 | 2.88E-01 | 2.61E-01 | 7.35E-06 | Yes | Ctnnd2 | intron_variant | MODIFIER |
| 15 | 30260934 | AX-170241924 | 0 | T | C | 0.026 | -6.27E-01 | 9.15E-02 | 2.88E-01 | 2.61E-01 | 7.35E-06 | Yes | Ctnnd2 | intron_variant | MODIFIER |
| 15 | 30276182 | AX-97796944 | 0 | A | G | 0.026 | -6.27E-01 | 9.15E-02 | 2.88E-01 | 2.61E-01 | 7.35E-06 | Yes | Ctnnd2 | intron_variant | MODIFIER |
| 15 | 30277357 | AX-170370885 | 0 | G | C | 0.026 | -6.27E-01 | 9.15E-02 | 2.88E-01 | 2.61E-01 | 7.35E-06 | Yes | Ctnnd2 | intron_variant | MODIFIER |
| 15 | 30284540 | AX-170369028 | 0 | G | A | 0.026 | -6.27E-01 | 9.15E-02 | 2.88E-01 | 2.61E-01 | 7.35E-06 | Yes | Ctnnd2 | intron_variant | MODIFIER |
| 15 | 30284624 | AX-170437822 | 0 | A | G | 0.026 | -6.27E-01 | 9.15E-02 | 2.88E-01 | 2.61E-01 | 7.35E-06 | Yes | Ctnnd2 | intron_variant | MODIFIER |
| 15 | 30299790 | AX-97820467 | 0 | A | C | 0.026 | -6.27E-01 | 9.15E-02 | 2.88E-01 | 2.61E-01 | 7.35E-06 | Yes | Ctnnd2 | intron_variant | MODIFIER |
| 15 | 30300485 | AX-170287046 | 0 | A | G | 0.026 | -6.27E-01 | 9.15E-02 | 2.88E-01 | 2.61E-01 | 7.35E-06 | Yes | Ctnnd2 | intron_variant | MODIFIER |
| 15 | 30313364 | AX-170375196 | 0 | T | C | 0.026 | -6.27E-01 | 9.15E-02 | 2.88E-01 | 2.61E-01 | 7.35E-06 | Yes | Ctnnd2 | intron_variant | MODIFIER |
| 15 | 30318851 | AX-170272986 | 0 | T | C | 0.026 | -6.27E-01 | 9.15E-02 | 2.88E-01 | 2.61E-01 | 7.35E-06 | Yes | Ctnnd2 | intron_variant | MODIFIER |
| 15 | 30322280 | AX-170093192 | 0 | T | C | 0.026 | -6.27E-01 | 9.15E-02 | 2.88E-01 | 2.61E-01 | 7.35E-06 | Yes | Ctnnd2 | intron_variant | MODIFIER |
| 15 | 30323614 | AX-170232350 | 0 | A | G | 0.026 | -6.27E-01 | 9.15E-02 | 2.88E-01 | 2.61E-01 | 7.35E-06 | Yes | Ctnnd2 | intron_variant | MODIFIER |
| 15 | 30335628 | AX-97798582 | 0 | C | T | 0.026 | -6.27E-01 | 9.15E-02 | 2.88E-01 | 2.61E-01 | 7.35E-06 | Yes | Ctnnd2 | intron_variant | MODIFIER |
| 15 | 30336139 | AX-170055135 | 0 | C | G | 0.026 | -6.27E-01 | 9.15E-02 | 2.88E-01 | 2.61E-01 | 7.35E-06 | Yes | Ctnnd2 | intron_variant | MODIFIER |
| 15 | 30355399 | AX-170484343 | 0 | C | T | 0.026 | -6.27E-01 | 9.15E-02 | 2.88E-01 | 2.61E-01 | 7.35E-06 | Yes | Ctnnd2 | intron_variant | MODIFIER |
| 15 | 30380284 | AX-97826877 | 0 | C | T | 0.026 | -6.27E-01 | 9.15E-02 | 2.88E-01 | 2.61E-01 | 7.35E-06 | Yes | Ctnnd2 | intron_variant | MODIFIER |
| 15 | 30380308 | AX-170413350 | 0 | G | A | 0.026 | -6.27E-01 | 9.15E-02 | 2.88E-01 | 2.61E-01 | 7.35E-06 | Yes | Ctnnd2 | intron_variant | MODIFIER |
| 15 | 30389192 | AX-170531846 | 0 | G | A | 0.026 | -6.27E-01 | 9.15E-02 | 2.88E-01 | 2.61E-01 | 7.35E-06 | Yes | Ctnnd2 | intron_variant | MODIFIER |
| 15 | 30389261 | AX-97852213 | 0 | A | G | 0.026 | -6.27E-01 | 9.15E-02 | 2.88E-01 | 2.61E-01 | 7.35E-06 | Yes | Ctnnd2 | intron_variant | MODIFIER |
| 15 | 30407252 | AX-170418954 | 0 | G | C | 0.026 | -6.27E-01 | 9.15E-02 | 2.88E-01 | 2.61E-01 | 7.35E-06 | Yes | Ctnnd2 | intron_variant | MODIFIER |
| 15 | 30421505 | AX-97820473 | 0 | T | G | 0.026 | -6.27E-01 | 9.15E-02 | 2.88E-01 | 2.61E-01 | 7.35E-06 | Yes | Ctnnd2 | intron_variant | MODIFIER |
| 15 | 30421517 | AX-170163839 | 0 | A | C | 0.026 | -6.27E-01 | 9.15E-02 | 2.88E-01 | 2.61E-01 | 7.35E-06 | Yes | Ctnnd2 | intron_variant | MODIFIER |
| 15 | 30428061 | AX-170265248 | 0 | A | G | 0.026 | -6.27E-01 | 9.15E-02 | 2.88E-01 | 2.61E-01 | 7.35E-06 | Yes | Ctnnd2 | intron_variant | MODIFIER |
| 15 | 30430651 | AX-170362621 | 0 | A | G | 0.026 | -6.27E-01 | 9.15E-02 | 2.88E-01 | 2.61E-01 | 7.35E-06 | Yes | Ctnnd2 | intron_variant | MODIFIER |
| 15 | 30430756 | AX-170261113 | 0 | A | G | 0.026 | -6.27E-01 | 9.15E-02 | 2.88E-01 | 2.61E-01 | 7.35E-06 | Yes | Ctnnd2 | intron_variant | MODIFIER |
| 15 | 30433718 | AX-170434315 | 0 | T | A | 0.026 | -6.27E-01 | 9.15E-02 | 2.88E-01 | 2.61E-01 | 7.35E-06 | Yes | Ctnnd2 | intron_variant | MODIFIER |
| 15 | 30437309 | AX-170483847 | 0 | C | T | 0.026 | -6.27E-01 | 9.15E-02 | 2.88E-01 | 2.61E-01 | 7.35E-06 | Yes | Ctnnd2 | intron_variant | MODIFIER |
| 15 | 30451255 | AX-97843396 | 0 | C | T | 0.026 | -6.27E-01 | 9.15E-02 | 2.88E-01 | 2.61E-01 | 7.35E-06 | Yes | Ctnnd2 | intron_variant | MODIFIER |
| 15 | 30474767 | AX-97856601 | 0 | A | G | 0.026 | -6.27E-01 | 9.15E-02 | 2.88E-01 | 2.61E-01 | 7.35E-06 | Yes | Ctnnd2 | intron_variant | MODIFIER |
| 15 | 30475024 | AX-170145223 | 0 | C | T | 0.026 | -6.27E-01 | 9.15E-02 | 2.88E-01 | 2.61E-01 | 7.35E-06 | Yes | Ctnnd2 | intron_variant | MODIFIER |
| 15 | 30496497 | AX-170488412 | 0 | C | T | 0.026 | -6.27E-01 | 9.15E-02 | 2.88E-01 | 2.61E-01 | 7.35E-06 | Yes | Ctnnd2 | intron_variant | MODIFIER |
| 15 | 30501264 | AX-170196762 | 0 | A | C | 0.026 | -6.27E-01 | 9.15E-02 | 2.88E-01 | 2.61E-01 | 7.35E-06 | Yes | Ctnnd2 | intron_variant | MODIFIER |
| 15 | 30504983 | AX-170270474 | 0 | A | G | 0.026 | -6.27E-01 | 9.15E-02 | 2.88E-01 | 2.61E-01 | 7.35E-06 | Yes | Ctnnd2 | intron_variant | MODIFIER |
| 15 | 30560050 | AX-170440020 | 0 | T | C | 0.026 | -6.27E-01 | 9.15E-02 | 2.88E-01 | 2.61E-01 | 7.35E-06 | Yes | Ctnnd2 | intron_variant | MODIFIER |
| 15 | 30570127 | AX-170323413 | 0 | G | A | 0.026 | -6.27E-01 | 9.15E-02 | 2.88E-01 | 2.61E-01 | 7.35E-06 | Yes | Ctnnd2 | intron_variant | MODIFIER |
| 15 | 30571412 | AX-170457724 | 0 | A | G | 0.026 | -6.27E-01 | 9.15E-02 | 2.88E-01 | 2.61E-01 | 7.35E-06 | Yes | Ctnnd2 | intron_variant | MODIFIER |
| 15 | 30589196 | AX-170200036 | 0 | C | G | 0.026 | -6.27E-01 | 9.15E-02 | 2.88E-01 | 2.61E-01 | 7.35E-06 | Yes | Ctnnd2 | intron_variant | MODIFIER |

| | | | | | | | | | | | | | | | |
|---|---|---|---|---|---|---|---|---|---|---|---|---|---|---|---|
| 15 | 30591087 | AX-170238704 | 0 | A | G | 0.026 | -6.27E-01 | 9.15E-02 | 2.88E-01 | 2.61E-01 | 7.35E-06 | Yes | Ctnnd2 | intron_variant | MODIFIER |
| 15 | 30595676 | AX-170215418 | 0 | T | C | 0.026 | -6.27E-01 | 9.15E-02 | 2.88E-01 | 2.61E-01 | 7.35E-06 | Yes | Ctnnd2 | intron_variant | MODIFIER |
| 15 | 30603261 | AX-170347594 | 0 | C | A | 0.026 | -6.27E-01 | 9.15E-02 | 2.88E-01 | 2.61E-01 | 7.35E-06 | Yes | Ctnnd2 | intron_variant | MODIFIER |
| 15 | 30603381 | AX-170055141 | 0 | A | C | 0.026 | -6.27E-01 | 9.15E-02 | 2.88E-01 | 2.61E-01 | 7.35E-06 | Yes | Ctnnd2 | intron_variant | MODIFIER |
| 15 | 30615051 | AX-170254164 | 0 | G | A | 0.026 | -6.27E-01 | 9.15E-02 | 2.88E-01 | 2.61E-01 | 7.35E-06 | Yes | Ctnnd2 | intron_variant | MODIFIER |
| 15 | 30616850 | AX-170308755 | 0 | A | G | 0.026 | -6.27E-01 | 9.15E-02 | 2.88E-01 | 2.61E-01 | 7.35E-06 | Yes | Ctnnd2 | intron_variant | MODIFIER |
| 15 | 30628511 | AX-170462948 | 0 | T | C | 0.026 | -6.27E-01 | 9.15E-02 | 2.88E-01 | 2.61E-01 | 7.35E-06 | Yes | Ctnnd2 | intron_variant | MODIFIER |
| 15 | 30631812 | AX-170420273 | 0 | A | G | 0.026 | -6.27E-01 | 9.15E-02 | 2.88E-01 | 2.61E-01 | 7.35E-06 | Yes | Ctnnd2 | intron_variant | MODIFIER |
| 15 | 30637218 | AX-170444739 | 0 | A | C | 0.026 | -6.27E-01 | 9.15E-02 | 2.88E-01 | 2.61E-01 | 7.35E-06 | Yes | Ctnnd2 | intron_variant | MODIFIER |
| 15 | 30649565 | AX-170238268 | 0 | A | C | 0.026 | -6.27E-01 | 9.15E-02 | 2.88E-01 | 2.61E-01 | 7.35E-06 | Yes | Ctnnd2 | intron_variant | MODIFIER |
| 15 | 30653260 | AX-97791511 | 0 | C | T | 0.026 | -6.27E-01 | 9.15E-02 | 2.88E-01 | 2.61E-01 | 7.35E-06 | Yes | Ctnnd2 | intron_variant | MODIFIER |
| 15 | 30664979 | AX-170450881 | 0 | T | C | 0.026 | -6.27E-01 | 9.15E-02 | 2.88E-01 | 2.61E-01 | 7.35E-06 | Yes | Ctnnd2 | intron_variant | MODIFIER |
| 15 | 30666132 | AX-170223900 | 0 | G | A | 0.026 | -6.27E-01 | 9.15E-02 | 2.88E-01 | 2.61E-01 | 7.35E-06 | Yes | Ctnnd2 | intron_variant | MODIFIER |
| 15 | 30666228 | AX-170533023 | 0 | G | A | 0.026 | -6.27E-01 | 9.15E-02 | 2.88E-01 | 2.61E-01 | 7.35E-06 | Yes | Ctnnd2 | intron_variant | MODIFIER |
| 15 | 30669404 | AX-97820472 | 0 | C | A | 0.026 | -6.27E-01 | 9.15E-02 | 2.88E-01 | 2.61E-01 | 7.35E-06 | Yes | Ctnnd2 | splice_region_variant,intron_variant | LOW |
| 15 | 30673791 | AX-170011363 | 0 | A | G | 0.026 | -6.27E-01 | 9.15E-02 | 2.88E-01 | 2.61E-01 | 7.35E-06 | Yes | Ctnnd2 | intron_variant | MODIFIER |
| 15 | 30686346 | AX-170252475 | 0 | A | C | 0.026 | -6.27E-01 | 9.15E-02 | 2.88E-01 | 2.61E-01 | 7.35E-06 | Yes | Ctnnd2 | intron_variant | MODIFIER |
| 15 | 30692440 | AX-170297152 | 0 | G | A | 0.026 | -6.27E-01 | 9.15E-02 | 2.88E-01 | 2.61E-01 | 7.35E-06 | Yes | Ctnnd2 | intron_variant | MODIFIER |
| 15 | 30692591 | AX-170055144 | 0 | C | T | 0.026 | -6.27E-01 | 9.15E-02 | 2.88E-01 | 2.61E-01 | 7.35E-06 | Yes | Ctnnd2 | intron_variant | MODIFIER |
| 15 | 30693288 | AX-97810321 | 0 | C | A | 0.026 | -6.27E-01 | 9.15E-02 | 2.88E-01 | 2.61E-01 | 7.35E-06 | Yes | Ctnnd2 | intron_variant | MODIFIER |
| 15 | 30696768 | AX-170011362 | 0 | T | C | 0.026 | -6.27E-01 | 9.15E-02 | 2.88E-01 | 2.61E-01 | 7.35E-06 | Yes | Ctnnd2 | intron_variant | MODIFIER |
| 15 | 30712200 | AX-97816731 | 0 | G | A | 0.026 | -6.27E-01 | 9.15E-02 | 2.88E-01 | 2.61E-01 | 7.35E-06 | Yes | Ctnnd2 | intron_variant | MODIFIER |
| 15 | 30716346 | AX-170417662 | 0 | G | C | 0.026 | -6.27E-01 | 9.15E-02 | 2.88E-01 | 2.61E-01 | 7.35E-06 | Yes | Ctnnd2 | intron_variant | MODIFIER |
| 15 | 30718466 | AX-170177082 | 0 | G | C | 0.026 | -6.27E-01 | 9.15E-02 | 2.88E-01 | 2.61E-01 | 7.35E-06 | Yes | Ctnnd2 | intron_variant | MODIFIER |
| 15 | 30741877 | AX-170397319 | 0 | A | G | 0.026 | -6.27E-01 | 9.15E-02 | 2.88E-01 | 2.61E-01 | 7.35E-06 | Yes | Ctnnd2 | intron_variant | MODIFIER |
| 15 | 30757017 | AX-170480666 | 0 | G | T | 0.026 | -6.27E-01 | 9.15E-02 | 2.88E-01 | 2.61E-01 | 7.35E-06 | Yes | Ctnnd2 | intron_variant | MODIFIER |
| 15 | 30759078 | AX-170397403 | 0 | A | C | 0.026 | -6.27E-01 | 9.15E-02 | 2.88E-01 | 2.61E-01 | 7.35E-06 | Yes | Ctnnd2 | intron_variant | MODIFIER |
| 15 | 30759779 | AX-170327464 | 0 | G | A | 0.026 | -6.27E-01 | 9.15E-02 | 2.88E-01 | 2.61E-01 | 7.35E-06 | Yes | Ctnnd2 | intron_variant | MODIFIER |
| 15 | 30760759 | AX-170011365 | 0 | T | C | 0.026 | -6.27E-01 | 9.15E-02 | 2.88E-01 | 2.61E-01 | 7.35E-06 | Yes | Ctnnd2 | intron_variant | MODIFIER |
| 15 | 30761924 | AX-170418582 | 0 | G | C | 0.026 | -6.27E-01 | 9.15E-02 | 2.88E-01 | 2.61E-01 | 7.35E-06 | Yes | Ctnnd2 | intron_variant | MODIFIER |
| 15 | 30799482 | AX-170117697 | 0 | A | G | 0.026 | -6.27E-01 | 9.15E-02 | 2.88E-01 | 2.61E-01 | 7.35E-06 | Yes | Ctnnd2 | intron_variant | MODIFIER |
| 15 | 30817648 | AX-97854776 | 0 | C | T | 0.026 | -6.27E-01 | 9.15E-02 | 2.88E-01 | 2.61E-01 | 7.35E-06 | Yes | Ctnnd2 | intron_variant | MODIFIER |
| 15 | 30819363 | AX-170472866 | 0 | G | A | 0.026 | -6.27E-01 | 9.15E-02 | 2.88E-01 | 2.61E-01 | 7.35E-06 | Yes | Ctnnd2 | intron_variant | MODIFIER |
| 15 | 30820016 | AX-170349772 | 0 | G | A | 0.026 | -6.27E-01 | 9.15E-02 | 2.88E-01 | 2.61E-01 | 7.35E-06 | Yes | Ctnnd2 | intron_variant | MODIFIER |
| 15 | 30822265 | AX-170253684 | 0 | G | C | 0.026 | -6.27E-01 | 9.15E-02 | 2.88E-01 | 2.61E-01 | 7.35E-06 | Yes | Ctnnd2 | intron_variant | MODIFIER |
| 15 | 30822329 | AX-170483870 | 0 | C | T | 0.026 | -6.27E-01 | 9.15E-02 | 2.88E-01 | 2.61E-01 | 7.35E-06 | Yes | Ctnnd2 | intron_variant | MODIFIER |
| 15 | 30832197 | AX-170055148 | 0 | C | A | 0.026 | -6.27E-01 | 9.15E-02 | 2.88E-01 | 2.61E-01 | 7.35E-06 | Yes | Ctnnd2 | intron_variant | MODIFIER |
| 15 | 30841929 | AX-170087806 | 0 | T | C | 0.026 | -6.27E-01 | 9.15E-02 | 2.88E-01 | 2.61E-01 | 7.35E-06 | Yes | Ctnnd2 | intron_variant | MODIFIER |
| 15 | 30855771 | AX-170122800 | 0 | T | C | 0.026 | -6.27E-01 | 9.15E-02 | 2.88E-01 | 2.61E-01 | 7.35E-06 | Yes | Ctnnd2 | intron_variant | MODIFIER |
| 15 | 30863704 | AX-170496534 | 0 | G | T | 0.026 | -6.27E-01 | 9.15E-02 | 2.88E-01 | 2.61E-01 | 7.35E-06 | Yes | Ctnnd2 | intron_variant | MODIFIER |
| 15 | 30864681 | AX-97820490 | 0 | A | G | 0.026 | -6.27E-01 | 9.15E-02 | 2.88E-01 | 2.61E-01 | 7.35E-06 | Yes | Ctnnd2 | intron_variant | MODIFIER |
| 15 | 30869718 | AX-170259693 | 0 | T | C | 0.026 | -6.27E-01 | 9.15E-02 | 2.88E-01 | 2.61E-01 | 7.35E-06 | Yes | Ctnnd2 | intron_variant | MODIFIER |
| 15 | 30870690 | AX-170286043 | 0 | T | C | 0.026 | -6.27E-01 | 9.15E-02 | 2.88E-01 | 2.61E-01 | 7.35E-06 | Yes | Ctnnd2 | intron_variant | MODIFIER |
| 15 | 30880672 | AX-170179591 | 0 | T | C | 0.026 | -6.27E-01 | 9.15E-02 | 2.88E-01 | 2.61E-01 | 7.35E-06 | Yes | Ctnnd2 | intron_variant | MODIFIER |
| 15 | 30911984 | AX-170228824 | 0 | T | G | 0.026 | -6.27E-01 | 9.15E-02 | 2.88E-01 | 2.61E-01 | 7.35E-06 | Yes | Ctnnd2 | intron_variant | MODIFIER |
| 15 | 30912228 | AX-170300659 | 0 | G | A | 0.026 | -6.27E-01 | 9.15E-02 | 2.88E-01 | 2.61E-01 | 7.35E-06 | Yes | Ctnnd2 | intron_variant | MODIFIER |
| 15 | 30923374 | AX-170387738 | 0 | G | A | 0.026 | -6.27E-01 | 9.15E-02 | 2.88E-01 | 2.61E-01 | 7.35E-06 | Yes | Ctnnd2 | intron_variant | MODIFIER |
| 15 | 30923977 | AX-170093193 | 0 | C | T | 0.026 | -6.27E-01 | 9.15E-02 | 2.88E-01 | 2.61E-01 | 7.35E-06 | Yes | Ctnnd2 | intron_variant | MODIFIER |
| 15 | 30965192 | AX-97847955 | 0 | G | A | 0.026 | -6.27E-01 | 9.15E-02 | 2.88E-01 | 2.61E-01 | 7.35E-06 | Yes | Ctnnd2 | intron_variant | MODIFIER |
| 15 | 30977850 | AX-170315237 | 0 | T | A | 0.026 | -6.27E-01 | 9.15E-02 | 2.88E-01 | 2.61E-01 | 7.35E-06 | Yes | Ctnnd2 | intron_variant | MODIFIER |
| 15 | 30992547 | AX-170476639 | 0 | C | G | 0.026 | -6.27E-01 | 9.15E-02 | 2.88E-01 | 2.61E-01 | 7.35E-06 | Yes | Ctnnd2 | intron_variant | MODIFIER |
| 15 | 31009121 | AX-170329586 | 0 | C | T | 0.026 | -6.27E-01 | 9.15E-02 | 2.88E-01 | 2.61E-01 | 7.35E-06 | Yes | Ctnnd2 | synonymous_variant | LOW |
| 15 | 31009190 | AX-170147681 | 0 | G | A | 0.026 | -6.27E-01 | 9.15E-02 | 2.88E-01 | 2.61E-01 | 7.35E-06 | Yes | Ctnnd2 | intron_variant | MODIFIER |
| 15 | 31018593 | AX-97790539 | 0 | A | G | 0.026 | -6.27E-01 | 9.15E-02 | 2.88E-01 | 2.61E-01 | 7.35E-06 | Yes | Ctnnd2 | intron_variant | MODIFIER |

| | | | | | | | | | | | | | | | |
|---|---|---|---|---|---|---|---|---|---|---|---|---|---|---|---|
| 15 | 31024887 | AX-170273846 | 0 | T | C | 0.026 | -6.27E-01 | 9.15E-02 | 2.88E-01 | 2.61E-01 | 7.35E-06 | Yes | Ctnnd2 | intron_variant | MODIFIER |
| 15 | 31070508 | AX-97793190 | 0 | A | G | 0.026 | -6.27E-01 | 9.15E-02 | 2.88E-01 | 2.61E-01 | 7.35E-06 | Yes | | intergenic_variant | MODIFIER |
| 15 | 31085288 | AX-170249205 | 0 | G | A | 0.026 | -6.27E-01 | 9.15E-02 | 2.88E-01 | 2.61E-01 | 7.35E-06 | Yes | | intergenic_variant | MODIFIER |
| 15 | 31091699 | AX-170087808 | 0 | A | C | 0.026 | -6.27E-01 | 9.15E-02 | 2.88E-01 | 2.61E-01 | 7.35E-06 | Yes | | intergenic_variant | MODIFIER |
| 15 | 31093631 | AX-170055153 | 0 | G | A | 0.026 | -6.27E-01 | 9.15E-02 | 2.88E-01 | 2.61E-01 | 7.35E-06 | Yes | | intergenic_variant | MODIFIER |
| 15 | 31112501 | AX-170384926 | 0 | A | G | 0.026 | -6.27E-01 | 9.15E-02 | 2.88E-01 | 2.61E-01 | 7.35E-06 | Yes | | intergenic_variant | MODIFIER |
| 15 | 31119158 | AX-170011371 | 0 | A | G | 0.026 | -6.27E-01 | 9.15E-02 | 2.88E-01 | 2.61E-01 | 7.35E-06 | Yes | | intergenic_variant | MODIFIER |
| 15 | 31128976 | AX-170093194 | 0 | C | T | 0.026 | -6.27E-01 | 9.15E-02 | 2.88E-01 | 2.61E-01 | 7.35E-06 | Yes | | intergenic_variant | MODIFIER |
| 15 | 31129162 | AX-170346650 | 0 | A | G | 0.026 | -6.27E-01 | 9.15E-02 | 2.88E-01 | 2.61E-01 | 7.35E-06 | Yes | | intergenic_variant | MODIFIER |
| 15 | 31149667 | AX-97792163 | 0 | C | T | 0.026 | -6.27E-01 | 9.15E-02 | 2.88E-01 | 2.61E-01 | 7.35E-06 | Yes | | intergenic_variant | MODIFIER |
| 15 | 31151238 | AX-170417083 | 0 | C | A | 0.026 | -6.27E-01 | 9.15E-02 | 2.88E-01 | 2.61E-01 | 7.35E-06 | Yes | | regulatory_region_variant | MODIFIER |
| 15 | 31152860 | AX-170011369 | 0 | T | C | 0.026 | -6.27E-01 | 9.15E-02 | 2.88E-01 | 2.61E-01 | 7.35E-06 | Yes | | intergenic_variant | MODIFIER |
| 15 | 31167171 | AX-170160049 | 0 | T | A | 0.026 | -6.27E-01 | 9.15E-02 | 2.88E-01 | 2.61E-01 | 7.35E-06 | Yes | | intergenic_variant | MODIFIER |
| 15 | 31195088 | AX-97824541 | 0 | T | G | 0.026 | -6.27E-01 | 9.15E-02 | 2.88E-01 | 2.61E-01 | 7.35E-06 | Yes | Gm26416 | upstream_gene_variant | MODIFIER |
| 15 | 31198500 | AX-170354351 | 0 | G | T | 0.026 | -6.27E-01 | 9.15E-02 | 2.88E-01 | 2.61E-01 | 7.35E-06 | Yes | Gm26416 | upstream_gene_variant | MODIFIER |
| 15 | 31200088 | AX-170011373 | 0 | C | A | 0.026 | -6.27E-01 | 9.15E-02 | 2.88E-01 | 2.61E-01 | 7.35E-06 | Yes | | intergenic_variant | MODIFIER |
| 15 | 31200264 | AX-170011372 | 0 | A | G | 0.026 | -6.27E-01 | 9.15E-02 | 2.88E-01 | 2.61E-01 | 7.35E-06 | Yes | | intergenic_variant | MODIFIER |
| 15 | 31233181 | AX-170162140 | 0 | G | A | 0.026 | -6.27E-01 | 9.15E-02 | 2.88E-01 | 2.61E-01 | 7.35E-06 | Yes | Dap | intron_variant | MODIFIER |
| 15 | 31235742 | AX-170296011 | 0 | C | T | 0.026 | -6.27E-01 | 9.15E-02 | 2.88E-01 | 2.61E-01 | 7.35E-06 | Yes | Dap | intron_variant | MODIFIER |
| 15 | 31236389 | AX-170481694 | 0 | C | T | 0.026 | -6.27E-01 | 9.15E-02 | 2.88E-01 | 2.61E-01 | 7.35E-06 | Yes | Dap | intron_variant | MODIFIER |
| 15 | 31240684 | AX-170011370 | 0 | T | A | 0.026 | -6.27E-01 | 9.15E-02 | 2.88E-01 | 2.61E-01 | 7.35E-06 | Yes | Dap | intron_variant | MODIFIER |
| 15 | 31240918 | AX-170204494 | 0 | A | C | 0.026 | -6.27E-01 | 9.15E-02 | 2.88E-01 | 2.61E-01 | 7.35E-06 | Yes | Dap | intron_variant | MODIFIER |
| 15 | 31241603 | AX-170354673 | 0 | C | T | 0.026 | -6.27E-01 | 9.15E-02 | 2.88E-01 | 2.61E-01 | 7.35E-06 | Yes | Dap | intron_variant | MODIFIER |
| 15 | 31242652 | AX-170375170 | 0 | T | C | 0.026 | -6.27E-01 | 9.15E-02 | 2.88E-01 | 2.61E-01 | 7.35E-06 | Yes | Dap | intron_variant | MODIFIER |
| 15 | 31246335 | AX-97783319 | 0 | T | C | 0.026 | -6.27E-01 | 9.15E-02 | 2.88E-01 | 2.61E-01 | 7.35E-06 | Yes | Dap | intron_variant | MODIFIER |
| 15 | 31246978 | AX-170055156 | 0 | T | C | 0.026 | -6.27E-01 | 9.15E-02 | 2.88E-01 | 2.61E-01 | 7.35E-06 | Yes | Dap | intron_variant | MODIFIER |
| 15 | 31257242 | AX-170259963 | 0 | G | C | 0.026 | -6.27E-01 | 9.15E-02 | 2.88E-01 | 2.61E-01 | 7.35E-06 | Yes | Dap | intron_variant | MODIFIER |
| 15 | 31257605 | AX-170385123 | 0 | A | G | 0.026 | -6.27E-01 | 9.15E-02 | 2.88E-01 | 2.61E-01 | 7.35E-06 | Yes | Dap | intron_variant | MODIFIER |
| 15 | 31260792 | AX-170302249 | 0 | A | G | 0.026 | -6.27E-01 | 9.15E-02 | 2.88E-01 | 2.61E-01 | 7.35E-06 | Yes | Dap | intron_variant | MODIFIER |
| 15 | 31270054 | AX-170233197 | 0 | A | G | 0.026 | -6.27E-01 | 9.15E-02 | 2.88E-01 | 2.61E-01 | 7.35E-06 | Yes | Dap | intron_variant | MODIFIER |
| 15 | 31273455 | AX-170423142 | 0 | A | G | 0.026 | -6.27E-01 | 9.15E-02 | 2.88E-01 | 2.61E-01 | 7.35E-06 | Yes | Dap | 3_prime_UTR_variant | MODIFIER |
| 15 | 31273888 | AX-170301214 | 0 | A | G | 0.026 | -6.27E-01 | 9.15E-02 | 2.88E-01 | 2.61E-01 | 7.35E-06 | Yes | Dap | 3_prime_UTR_variant | MODIFIER |
| 15 | 31283304 | AX-97801409 | 0 | T | G | 0.026 | -6.27E-01 | 9.15E-02 | 2.88E-01 | 2.61E-01 | 7.35E-06 | Yes | Gm49296 | downstream_gene_variant | MODIFIER |
| 15 | 31304045 | AX-170428566 | 0 | T | C | 0.026 | -6.27E-01 | 9.15E-02 | 2.88E-01 | 2.61E-01 | 7.35E-06 | Yes | Ankrd33b | intron_variant,NMD_transcript_variant | MODIFIER |
| 15 | 31307789 | AX-97828853 | 0 | T | C | 0.026 | -6.27E-01 | 9.15E-02 | 2.88E-01 | 2.61E-01 | 7.35E-06 | Yes | Ankrd33b | intron_variant,NMD_transcript_variant | MODIFIER |
| 15 | 31311696 | AX-97802383 | 0 | A | G | 0.026 | -6.27E-01 | 9.15E-02 | 2.88E-01 | 2.61E-01 | 7.35E-06 | Yes | Ankrd33b | intron_variant,NMD_transcript_variant | MODIFIER |
| 15 | 31677181 | AX-170456757 | 0 | T | C | 0.026 | -6.27E-01 | 9.15E-02 | 2.88E-01 | 2.61E-01 | 7.35E-06 | Yes | | regulatory_region_variant | MODIFIER |
| 15 | 32333571 | AX-170055179 | 0 | A | G | 0.026 | -6.27E-01 | 9.15E-02 | 2.88E-01 | 2.61E-01 | 7.35E-06 | Yes | Sema5a | intron_variant | MODIFIER |
| 15 | 32353508 | AX-170435450 | 0 | A | T | 0.026 | -6.27E-01 | 9.15E-02 | 2.88E-01 | 2.61E-01 | 7.35E-06 | Yes | Sema5a | intron_variant | MODIFIER |
| 15 | 32397368 | AX-97817498 | 0 | A | G | 0.026 | -6.27E-01 | 9.15E-02 | 2.88E-01 | 2.61E-01 | 7.35E-06 | Yes | Sema5a | intron_variant | MODIFIER |
| 15 | 32402863 | AX-170322628 | 0 | G | A | 0.026 | -6.27E-01 | 9.15E-02 | 2.88E-01 | 2.61E-01 | 7.35E-06 | Yes | Sema5a | intron_variant | MODIFIER |
| 15 | 32439415 | AX-170204147 | 0 | A | G | 0.026 | -6.27E-01 | 9.15E-02 | 2.88E-01 | 2.61E-01 | 7.35E-06 | Yes | Sema5a | intron_variant | MODIFIER |
| 15 | 32443558 | AX-170272426 | 0 | G | A | 0.026 | -6.27E-01 | 9.15E-02 | 2.88E-01 | 2.61E-01 | 7.35E-06 | Yes | Sema5a | intron_variant | MODIFIER |
| 15 | 32446209 | AX-97790025 | 0 | C | T | 0.026 | -6.27E-01 | 9.15E-02 | 2.88E-01 | 2.61E-01 | 7.35E-06 | Yes | Sema5a | intron_variant | MODIFIER |
| 15 | 32450802 | AX-170258483 | 0 | G | A | 0.026 | -6.27E-01 | 9.15E-02 | 2.88E-01 | 2.61E-01 | 7.35E-06 | Yes | Sema5a | intron_variant | MODIFIER |
| 15 | 32522618 | AX-97829941 | 0 | T | C | 0.026 | -6.27E-01 | 9.15E-02 | 2.88E-01 | 2.61E-01 | 7.35E-06 | Yes | Sema5a | intron_variant | MODIFIER |
| 15 | 32531844 | AX-170264857 | 0 | A | G | 0.026 | -6.27E-01 | 9.15E-02 | 2.88E-01 | 2.61E-01 | 7.35E-06 | Yes | Sema5a | intron_variant | MODIFIER |
| 15 | 32543744 | AX-170478370 | 0 | A | G | 0.026 | -6.27E-01 | 9.15E-02 | 2.88E-01 | 2.61E-01 | 7.35E-06 | Yes | Sema5a | intron_variant | MODIFIER |
| 15 | 32543767 | AX-170522512 | 0 | A | G | 0.026 | -6.27E-01 | 9.15E-02 | 2.88E-01 | 2.61E-01 | 7.35E-06 | Yes | Sema5a | intron_variant | MODIFIER |
| 15 | 32557838 | AX-170276500 | 0 | G | A | 0.026 | -6.27E-01 | 9.15E-02 | 2.88E-01 | 2.61E-01 | 7.35E-06 | Yes | Sema5a | intron_variant | MODIFIER |
| 15 | 32616099 | AX-170285997 | 0 | G | A | 0.026 | -6.27E-01 | 9.15E-02 | 2.88E-01 | 2.61E-01 | 7.35E-06 | Yes | Sema5a | intron_variant | MODIFIER |
| 15 | 32617057 | AX-170055185 | 0 | G | T | 0.026 | -6.27E-01 | 9.15E-02 | 2.88E-01 | 2.61E-01 | 7.35E-06 | Yes | Sema5a | intron_variant | MODIFIER |
| 15 | 32618127 | AX-170335286 | 0 | A | G | 0.026 | -6.27E-01 | 9.15E-02 | 2.88E-01 | 2.61E-01 | 7.35E-06 | Yes | Sema5a | intron_variant | MODIFIER |
| 15 | 32618254 | AX-170055184 | 0 | C | T | 0.026 | -6.27E-01 | 9.15E-02 | 2.88E-01 | 2.61E-01 | 7.35E-06 | Yes | Sema5a | intron_variant | MODIFIER |

| | | | | | | | | | | | | | | | |
|---|---|---|---|---|---|---|---|---|---|---|---|---|---|---|---|
| 15 | 32621146 | AX-170339707 | 0 | G | A | 0.026 | -6.27E-01 | 9.15E-02 | 2.88E-01 | 2.61E-01 | 7.35E-06 | Yes | Sema5a | intron_variant | MODIFIER |
| 15 | 32621227 | AX-170352926 | 0 | G | A | 0.026 | -6.27E-01 | 9.15E-02 | 2.88E-01 | 2.61E-01 | 7.35E-06 | Yes | Sema5a | intron_variant | MODIFIER |
| 15 | 32623043 | AX-170161755 | 0 | C | T | 0.026 | -6.27E-01 | 9.15E-02 | 2.88E-01 | 2.61E-01 | 7.35E-06 | Yes | Sema5a | intron_variant | MODIFIER |
| 15 | 32623366 | AX-170011390 | 0 | T | C | 0.026 | -6.27E-01 | 9.15E-02 | 2.88E-01 | 2.61E-01 | 7.35E-06 | Yes | Sema5a | intron_variant | MODIFIER |
| 15 | 32922990 | AX-170168209 | 0 | G | A | 0.026 | -6.27E-01 | 9.15E-02 | 2.88E-01 | 2.61E-01 | 7.35E-06 | Yes | Sdc2 | intron_variant | MODIFIER |
| 15 | 32940151 | AX-170520117 | 0 | T | C | 0.026 | -6.27E-01 | 9.15E-02 | 2.88E-01 | 2.61E-01 | 7.35E-06 | Yes | Sdc2 | intron_variant | MODIFIER |
| 15 | 32946658 | AX-170093203 | 0 | T | C | 0.026 | -6.27E-01 | 9.15E-02 | 2.88E-01 | 2.61E-01 | 7.35E-06 | Yes | Sdc2 | intron_variant | MODIFIER |
| 15 | 32946721 | AX-170164725 | 0 | T | G | 0.026 | -6.27E-01 | 9.15E-02 | 2.88E-01 | 2.61E-01 | 7.35E-06 | Yes | Sdc2 | intron_variant | MODIFIER |
| 15 | 32947067 | AX-170123285 | 0 | T | C | 0.026 | -6.27E-01 | 9.15E-02 | 2.88E-01 | 2.61E-01 | 7.35E-06 | Yes | Sdc2 | intron_variant | MODIFIER |
| 15 | 32947187 | AX-170403664 | 0 | A | G | 0.026 | -6.27E-01 | 9.15E-02 | 2.88E-01 | 2.61E-01 | 7.35E-06 | Yes | Sdc2 | intron_variant | MODIFIER |
| 15 | 32965028 | AX-170301383 | 0 | C | T | 0.026 | -6.27E-01 | 9.15E-02 | 2.88E-01 | 2.61E-01 | 7.35E-06 | Yes | Sdc2 | intron_variant | MODIFIER |
| 15 | 32971626 | AX-170130425 | 0 | T | A | 0.026 | -6.27E-01 | 9.15E-02 | 2.88E-01 | 2.61E-01 | 7.35E-06 | Yes | Sdc2 | intron_variant | MODIFIER |
| 15 | 33303187 | AX-97799578 | 0 | A | G | 0.026 | -6.27E-01 | 9.15E-02 | 2.88E-01 | 2.61E-01 | 7.35E-06 | Yes | Cpq | intron_variant | MODIFIER |
| 15 | 33656435 | AX-97808750 | 0 | T | G | 0.026 | -6.27E-01 | 9.15E-02 | 2.88E-01 | 2.61E-01 | 7.35E-06 | Yes | 4930592A05Rik | upstream_gene_variant | MODIFIER |
| 15 | 33688099 | AX-97849298 | 0 | G | A | 0.026 | -6.27E-01 | 9.15E-02 | 2.88E-01 | 2.61E-01 | 7.35E-06 | Yes | Tspyl5 | upstream_gene_variant | MODIFIER |
| 15 | 33925932 | AX-97829347 | 0 | A | G | 0.026 | -6.27E-01 | 9.15E-02 | 2.88E-01 | 2.61E-01 | 7.35E-06 | Yes | | intergenic_variant | MODIFIER |
| 15 | 33950687 | AX-170170549 | 0 | G | A | 0.026 | -6.27E-01 | 9.15E-02 | 2.88E-01 | 2.61E-01 | 7.35E-06 | Yes | | intergenic_variant | MODIFIER |
| 15 | 33956550 | AX-170259744 | 0 | C | T | 0.026 | -6.27E-01 | 9.15E-02 | 2.88E-01 | 2.61E-01 | 7.35E-06 | Yes | | intergenic_variant | MODIFIER |
| 15 | 33963489 | AX-170341641 | 0 | G | A | 0.026 | -6.27E-01 | 9.15E-02 | 2.88E-01 | 2.61E-01 | 7.35E-06 | Yes | | intergenic_variant | MODIFIER |
| 15 | 33969904 | AX-97787354 | 0 | A | G | 0.026 | -6.27E-01 | 9.15E-02 | 2.88E-01 | 2.61E-01 | 7.35E-06 | Yes | | intergenic_variant | MODIFIER |
| 15 | 34025461 | AX-97843205 | 0 | C | A | 0.026 | -6.27E-01 | 9.15E-02 | 2.88E-01 | 2.61E-01 | 7.35E-06 | Yes | Gm41290 | intron_variant,non_coding_transcript_variant | MODIFIER |
| 15 | 34048446 | AX-170055213 | 0 | G | C | 0.026 | -6.27E-01 | 9.15E-02 | 2.88E-01 | 2.61E-01 | 7.35E-06 | Yes | Gm49091 | upstream_gene_variant | MODIFIER |
| 15 | 34048498 | AX-97789960 | 0 | A | G | 0.026 | -6.27E-01 | 9.15E-02 | 2.88E-01 | 2.61E-01 | 7.35E-06 | Yes | Gm49091 | upstream_gene_variant | MODIFIER |
| 15 | 34053361 | AX-170222489 | 0 | C | T | 0.026 | -6.27E-01 | 9.15E-02 | 2.88E-01 | 2.61E-01 | 7.35E-06 | Yes | Gm49091 | downstream_gene_variant | MODIFIER |
| 15 | 34053981 | AX-170425229 | 0 | T | C | 0.026 | -6.27E-01 | 9.15E-02 | 2.88E-01 | 2.61E-01 | 7.35E-06 | Yes | Gm49091 | downstream_gene_variant | MODIFIER |
| 15 | 34062900 | AX-170221075 | 0 | T | C | 0.026 | -6.27E-01 | 9.15E-02 | 2.88E-01 | 2.61E-01 | 7.35E-06 | Yes | | intergenic_variant | MODIFIER |
| 15 | 34079992 | AX-170485648 | 0 | G | A | 0.026 | -6.27E-01 | 9.15E-02 | 2.88E-01 | 2.61E-01 | 7.35E-06 | Yes | Mtdh | upstream_gene_variant | MODIFIER |
| 15 | 34082527 | AX-170055207 | 0 | C | A | 0.026 | -6.27E-01 | 9.15E-02 | 2.88E-01 | 2.61E-01 | 7.35E-06 | Yes | Mtdh | upstream_gene_variant | MODIFIER |
| 15 | 34097004 | AX-170529486 | 0 | C | T | 0.026 | -6.27E-01 | 9.15E-02 | 2.88E-01 | 2.61E-01 | 7.35E-06 | Yes | Mtdh | intron_variant | MODIFIER |
| 15 | 34116021 | AX-170404046 | 0 | T | C | 0.026 | -6.27E-01 | 9.15E-02 | 2.88E-01 | 2.61E-01 | 7.35E-06 | Yes | Mtdh | intron_variant | MODIFIER |
| 15 | 34143312 | AX-170190687 | 0 | C | A | 0.026 | -6.27E-01 | 9.15E-02 | 2.88E-01 | 2.61E-01 | 7.35E-06 | Yes | Mtdh | 3_prime_UTR_variant | MODIFIER |
| 15 | 34268895 | AX-170055215 | 0 | T | C | 0.026 | -6.27E-01 | 9.15E-02 | 2.88E-01 | 2.61E-01 | 7.35E-06 | Yes | Laptm4b | intron_variant | MODIFIER |
| 15 | 34287287 | AX-170011408 | 0 | A | G | 0.026 | -6.27E-01 | 9.15E-02 | 2.88E-01 | 2.61E-01 | 7.35E-06 | Yes | Laptm4b | downstream_gene_variant | MODIFIER |
| 15 | 34333293 | AX-97829346 | 0 | T | G | 0.026 | -6.27E-01 | 9.15E-02 | 2.88E-01 | 2.61E-01 | 7.35E-06 | Yes | Matn2 | intron_variant | MODIFIER |
| 15 | 34334501 | AX-170397617 | 0 | G | A | 0.026 | -6.27E-01 | 9.15E-02 | 2.88E-01 | 2.61E-01 | 7.35E-06 | Yes | Matn2 | intron_variant | MODIFIER |
| 15 | 34335100 | AX-170422896 | 0 | A | T | 0.026 | -6.27E-01 | 9.15E-02 | 2.88E-01 | 2.61E-01 | 7.35E-06 | Yes | Matn2 | intron_variant | MODIFIER |
| 15 | 34335712 | AX-170503659 | 0 | A | G | 0.026 | -6.27E-01 | 9.15E-02 | 2.88E-01 | 2.61E-01 | 7.35E-06 | Yes | Matn2 | intron_variant | MODIFIER |
| 15 | 34335937 | AX-97783168 | 0 | T | C | 0.026 | -6.27E-01 | 9.15E-02 | 2.88E-01 | 2.61E-01 | 7.35E-06 | Yes | Matn2 | intron_variant | MODIFIER |
| 15 | 34358765 | AX-170011407 | 0 | G | T | 0.026 | -6.27E-01 | 9.15E-02 | 2.88E-01 | 2.61E-01 | 7.35E-06 | Yes | Matn2 | intron_variant | MODIFIER |
| 15 | 34365439 | AX-97845842 | 0 | G | A | 0.026 | -6.27E-01 | 9.15E-02 | 2.88E-01 | 2.61E-01 | 7.35E-06 | Yes | Matn2 | intron_variant | MODIFIER |
| 15 | 34371361 | AX-170101079 | 0 | C | T | 0.026 | -6.27E-01 | 9.15E-02 | 2.88E-01 | 2.61E-01 | 7.35E-06 | Yes | Matn2 | intron_variant | MODIFIER |
| 15 | 34381102 | AX-170149217 | 0 | T | G | 0.026 | -6.27E-01 | 9.15E-02 | 2.88E-01 | 2.61E-01 | 7.35E-06 | Yes | Matn2 | intron_variant | MODIFIER |
| 15 | 34388402 | AX-170433933 | 0 | C | T | 0.026 | -6.27E-01 | 9.15E-02 | 2.88E-01 | 2.61E-01 | 7.35E-06 | Yes | Matn2 | intron_variant | MODIFIER |
| 15 | 34411573 | AX-170360935 | 0 | G | A | 0.026 | -6.27E-01 | 9.15E-02 | 2.88E-01 | 2.61E-01 | 7.35E-06 | Yes | Matn2 | intron_variant | MODIFIER |
| 15 | 34433315 | AX-97786042 | 0 | T | C | 0.026 | -6.27E-01 | 9.15E-02 | 2.88E-01 | 2.61E-01 | 7.35E-06 | Yes | Matn2 | intron_variant | MODIFIER |
| 15 | 34463865 | AX-170308193 | 0 | C | G | 0.026 | -6.27E-01 | 9.15E-02 | 2.88E-01 | 2.61E-01 | 7.35E-06 | Yes | Erich5 | intron_variant | MODIFIER |
| 15 | 34571516 | AX-170385749 | 0 | C | T | 0.026 | -6.27E-01 | 9.15E-02 | 2.88E-01 | 2.61E-01 | 7.35E-06 | Yes | Nipal2 | downstream_gene_variant | MODIFIER |
| 15 | 34571549 | AX-170330309 | 0 | T | C | 0.026 | -6.27E-01 | 9.15E-02 | 2.88E-01 | 2.61E-01 | 7.35E-06 | Yes | Nipal2 | downstream_gene_variant | MODIFIER |
| 15 | 34591630 | AX-170328259 | 0 | A | C | 0.026 | -6.27E-01 | 9.15E-02 | 2.88E-01 | 2.61E-01 | 7.35E-06 | Yes | Nipal2 | intron_variant | MODIFIER |
| 15 | 34606390 | AX-170240907 | 0 | C | A | 0.026 | -6.27E-01 | 9.15E-02 | 2.88E-01 | 2.61E-01 | 7.35E-06 | Yes | Nipal2 | intron_variant | MODIFIER |
| 15 | 34607066 | AX-170502350 | 0 | T | C | 0.026 | -6.27E-01 | 9.15E-02 | 2.88E-01 | 2.61E-01 | 7.35E-06 | Yes | Nipal2 | intron_variant | MODIFIER |
| 15 | 34647778 | AX-170093207 | 0 | C | T | 0.026 | -6.27E-01 | 9.15E-02 | 2.88E-01 | 2.61E-01 | 7.35E-06 | Yes | Nipal2 | intron_variant | MODIFIER |
| 15 | 34713566 | AX-170331756 | 0 | A | T | 0.026 | -6.27E-01 | 9.15E-02 | 2.88E-01 | 2.61E-01 | 7.35E-06 | Yes | | intergenic_variant | MODIFIER |
| 15 | 34726429 | AX-97856109 | 0 | A | C | 0.026 | -6.27E-01 | 9.15E-02 | 2.88E-01 | 2.61E-01 | 7.35E-06 | Yes | | intergenic_variant | MODIFIER |

| | | | | | | | | | | | | | | | |
|---|---|---|---|---|---|---|---|---|---|---|---|---|---|---|---|
| 15 | 34776117 | AX-97788748 | 0 | A | G | 0.026 | -6.27E-01 | 9.15E-02 | 2.88E-01 | 2.61E-01 | 7.35E-06 | Yes | | intergenic_variant | MODIFIER |
| 15 | 34776767 | AX-170139005 | 0 | G | T | 0.026 | -6.27E-01 | 9.15E-02 | 2.88E-01 | 2.61E-01 | 7.35E-06 | Yes | | intergenic_variant | MODIFIER |
| 15 | 34947395 | AX-97829232 | 0 | T | C | 0.026 | -6.27E-01 | 9.15E-02 | 2.88E-01 | 2.61E-01 | 7.35E-06 | Yes | Stk3 | intron_variant | MODIFIER |
| 15 | 35231334 | AX-97845279 | 0 | G | A | 0.026 | -6.27E-01 | 9.15E-02 | 2.88E-01 | 2.61E-01 | 7.35E-06 | Yes | Gm33497 | intron_variant,non_coding_transcript_variant | MODIFIER |
| 15 | 35416153 | AX-170152178 | 0 | A | G | 0.026 | -6.27E-01 | 9.15E-02 | 2.88E-01 | 2.61E-01 | 7.35E-06 | Yes | Vps13b | intron_variant | MODIFIER |
| 15 | 35477441 | AX-170251287 | 0 | C | T | 0.026 | -6.27E-01 | 9.15E-02 | 2.88E-01 | 2.61E-01 | 7.35E-06 | Yes | Vps13b | intron_variant | MODIFIER |
| 15 | 35547182 | AX-170522626 | 0 | A | T | 0.026 | -6.27E-01 | 9.15E-02 | 2.88E-01 | 2.61E-01 | 7.35E-06 | Yes | Vps13b | intron_variant | MODIFIER |
| 15 | 35827397 | AX-170011433 | 0 | C | T | 0.026 | -6.27E-01 | 9.15E-02 | 2.88E-01 | 2.61E-01 | 7.35E-06 | Yes | Vps13b | intron_variant | MODIFIER |
| 15 | 36354358 | AX-97806534 | 0 | G | A | 0.026 | -6.27E-01 | 9.15E-02 | 2.88E-01 | 2.61E-01 | 7.35E-06 | Yes | Gm33936 | intron_variant,non_coding_transcript_variant | MODIFIER |
| 15 | 36354460 | AX-170486303 | 0 | C | T | 0.026 | -6.27E-01 | 9.15E-02 | 2.88E-01 | 2.61E-01 | 7.35E-06 | Yes | Gm33936 | intron_variant,non_coding_transcript_variant | MODIFIER |
| 15 | 36354861 | AX-170055252 | 0 | T | C | 0.026 | -6.27E-01 | 9.15E-02 | 2.88E-01 | 2.61E-01 | 7.35E-06 | Yes | Gm33936 | intron_variant,non_coding_transcript_variant | MODIFIER |
| 15 | 36360692 | AX-170370818 | 0 | T | C | 0.026 | -6.27E-01 | 9.15E-02 | 2.88E-01 | 2.61E-01 | 7.35E-06 | Yes | Gm33936 | intron_variant,non_coding_transcript_variant | MODIFIER |
| 15 | 36402833 | AX-170501517 | 0 | T | C | 0.026 | -6.27E-01 | 9.15E-02 | 2.88E-01 | 2.61E-01 | 7.35E-06 | Yes | | intergenic_variant | MODIFIER |
| 15 | 36413405 | AX-170460250 | 0 | G | A | 0.026 | -6.27E-01 | 9.15E-02 | 2.88E-01 | 2.61E-01 | 7.35E-06 | Yes | | intergenic_variant | MODIFIER |
| 15 | 36421303 | AX-170232197 | 0 | G | A | 0.026 | -6.27E-01 | 9.15E-02 | 2.88E-01 | 2.61E-01 | 7.35E-06 | Yes | | regulatory_region_variant | MODIFIER |
| 15 | 36435821 | AX-170379650 | 0 | G | A | 0.026 | -6.27E-01 | 9.15E-02 | 2.88E-01 | 2.61E-01 | 7.35E-06 | Yes | Gm49224 | intron_variant,non_coding_transcript_variant | MODIFIER |
| 15 | 36444556 | AX-170182014 | 0 | A | G | 0.026 | -6.27E-01 | 9.15E-02 | 2.88E-01 | 2.61E-01 | 7.35E-06 | Yes | | intergenic_variant | MODIFIER |
| 15 | 36447224 | AX-170395537 | 0 | C | A | 0.026 | -6.27E-01 | 9.15E-02 | 2.88E-01 | 2.61E-01 | 7.35E-06 | Yes | | intergenic_variant | MODIFIER |
| 15 | 45674354 | AX-170143614 | 0 | G | C | 0.035 | -5.51E-01 | 9.90E-02 | 7.54E-01 | 7.57E-01 | 7.41E-06 | Yes | | intergenic_variant | MODIFIER |
| 15 | 45932908 | AX-170152797 | 0 | A | G | 0.035 | -5.50E-01 | 9.67E-02 | 6.79E-01 | 6.79E-01 | 7.57E-06 | Yes | | intergenic_variant | MODIFIER |
| 15 | 44832552 | AX-170417588 | 0 | A | G | 0.035 | -5.09E-01 | 9.10E-02 | 7.29E-01 | 7.47E-01 | 7.83E-06 | Yes | A930017M01Rik | intron_variant,non_coding_transcript_variant | MODIFIER |
| 15 | 56817609 | AX-97821909 | 0 | T | G | 0.044 | -4.92E-01 | 9.20E-02 | 9.01E-01 | 9.19E-01 | 7.91E-06 | Yes | | intergenic_variant | MODIFIER |
| 15 | 56428403 | AX-170329540 | 0 | A | G | 0.044 | -5.00E-01 | 9.44E-02 | 9.32E-01 | 9.52E-01 | 8.35E-06 | Yes | | intergenic_variant | MODIFIER |
| 15 | 57400211 | AX-170258144 | 0 | T | C | 0.044 | -5.00E-01 | 9.44E-02 | 9.32E-01 | 9.52E-01 | 8.35E-06 | Yes | Slc22a22 | intron_variant | MODIFIER |
| 15 | 54858317 | AX-170117206 | 0 | C | T | 0.035 | -5.44E-01 | 9.70E-02 | 7.08E-01 | 7.12E-01 | 8.59E-06 | Yes | Enpp2 | intron_variant | MODIFIER |
| 15 | 25514722 | AX-170425301 | 0 | C | G | 0.035 | -3.61E-01 | 5.89E-02 | 4.74E-01 | 4.72E-01 | 8.66E-06 | Yes | | intergenic_variant | MODIFIER |
| 15 | 56470419 | AX-170418013 | 2 | C | T | 0.045 | -4.44E-01 | 8.65E-02 | 1.08E+00 | 1.11E+00 | 8.98E-06 | Yes | | intergenic_variant | MODIFIER |
| 15 | 25001669 | AX-170120274 | 1 | G | C | 0.027 | -6.09E-01 | 9.44E-02 | 3.72E-01 | 3.52E-01 | 8.98E-06 | Yes | Gm2824 | downstream_gene_variant | MODIFIER |
| 15 | 34406970 | AX-170323507 | 1 | G | A | 0.027 | -6.09E-01 | 9.43E-02 | 3.68E-01 | 3.47E-01 | 9.25E-06 | Yes | Matn2 | intron_variant | MODIFIER |
| 15 | 45157955 | AX-170138511 | 0 | C | T | 0.035 | -5.59E-01 | 9.70E-02 | 6.18E-01 | 6.12E-01 | 9.31E-06 | Yes | | intergenic_variant | MODIFIER |
| 15 | 22523678 | AX-170011212 | 1 | C | G | 0.027 | -6.11E-01 | 9.30E-02 | 3.39E-01 | 3.17E-01 | 9.46E-06 | Yes | | intergenic_variant | MODIFIER |
| 15 | 57215655 | AX-97788690 | 2 | C | T | 0.045 | -4.79E-01 | 9.49E-02 | 1.16E+00 | 1.19E+00 | 9.79E-06 | Yes | | intergenic_variant | MODIFIER |
| 15 | 29887062 | AX-97840189 | 1 | T | C | 0.027 | -6.17E-01 | 9.25E-02 | 3.12E-01 | 2.86E-01 | 1.02E-05 | Yes | Gm23033 | downstream_gene_variant | MODIFIER |
| 15 | 32601430 | AX-97847693 | 1 | C | G | 0.027 | -6.17E-01 | 9.25E-02 | 3.12E-01 | 2.86E-01 | 1.02E-05 | Yes | Sema5a | intron_variant | MODIFIER |
| 15 | 58404546 | AX-170258498 | 0 | A | T | 0.053 | -4.55E-01 | 8.68E-02 | 9.31E-01 | 9.61E-01 | 1.04E-05 | Yes | Gm2675 | upstream_gene_variant | MODIFIER |
| 15 | 34397042 | AX-170246466 | 1 | G | A | 0.027 | -6.15E-01 | 9.20E-02 | 3.08E-01 | 2.85E-01 | 1.05E-05 | Yes | Matn2 | intron_variant | MODIFIER |
| 15 | 25158101 | AX-97826069 | 1 | G | A | 0.027 | -6.14E-01 | 9.31E-02 | 3.27E-01 | 3.03E-01 | 1.07E-05 | Yes | 9230109A22Rik | intron_variant,non_coding_transcript_variant | MODIFIER |
| 15 | 33950647 | AX-97829338 | 1 | C | T | 0.027 | -6.14E-01 | 9.31E-02 | 3.27E-01 | 3.03E-01 | 1.07E-05 | Yes | | intergenic_variant | MODIFIER |
| 15 | 53730936 | AX-170520271 | 0 | A | G | 0.044 | -4.45E-01 | 8.88E-02 | 1.16E+00 | 1.21E+00 | 1.09E-05 | Yes | Samd12 | intron_variant | MODIFIER |
| 15 | 55189047 | AX-170186301 | 0 | C | G | 0.044 | -4.96E-01 | 9.50E-02 | 9.47E-01 | 9.71E-01 | 1.09E-05 | Yes | Deptor | intron_variant | MODIFIER |
| 15 | 57303853 | AX-170087874 | 0 | C | T | 0.044 | -4.96E-01 | 9.50E-02 | 9.47E-01 | 9.71E-01 | 1.09E-05 | Yes | Slc22a22 | intron_variant | MODIFIER |
| 15 | 26057595 | AX-170473680 | 1 | G | T | 0.027 | -6.11E-01 | 9.40E-02 | 3.48E-01 | 3.24E-01 | 1.13E-05 | Yes | Gm41279 | intron_variant,non_coding_transcript_variant | MODIFIER |
| 15 | 25403913 | AX-170140091 | 1 | G | C | 0.027 | -6.11E-01 | 9.33E-02 | 3.34E-01 | 3.10E-01 | 1.15E-05 | Yes | Basp1 | intron_variant | MODIFIER |
| 15 | 26071295 | AX-97789358 | 1 | G | A | 0.027 | -6.11E-01 | 9.33E-02 | 3.34E-01 | 3.10E-01 | 1.15E-05 | Yes | | intergenic_variant | MODIFIER |
| 15 | 30277644 | AX-170187014 | 1 | C | T | 0.027 | -6.11E-01 | 9.33E-02 | 3.34E-01 | 3.10E-01 | 1.15E-05 | Yes | Ctnnd2 | intron_variant | MODIFIER |
| 15 | 22415067 | AX-170098859 | 1 | C | T | 0.027 | -6.15E-01 | 9.25E-02 | 3.11E-01 | 2.86E-01 | 1.15E-05 | Yes | | intergenic_variant | MODIFIER |
| 15 | 32960450 | AX-170460757 | 1 | A | G | 0.027 | -6.14E-01 | 9.47E-02 | 3.49E-01 | 3.26E-01 | 1.16E-05 | Yes | Sdc2 | intron_variant | MODIFIER |
| 15 | 32604998 | AX-170120993 | 1 | G | A | 0.027 | -6.11E-01 | 9.34E-02 | 3.35E-01 | 3.12E-01 | 1.17E-05 | Yes | Sema5a | intron_variant | MODIFIER |
| 15 | 56545162 | AX-170330735 | 1 | T | C | 0.036 | -4.90E-01 | 9.39E-02 | 9.33E-01 | 9.55E-01 | 1.18E-05 | Yes | | intergenic_variant | MODIFIER |
| 15 | 31219054 | AX-97838658 | 1 | A | G | 0.027 | -6.12E-01 | 9.32E-02 | 3.29E-01 | 3.07E-01 | 1.18E-05 | Yes | | intergenic_variant | MODIFIER |
| 15 | 31283150 | AX-170127590 | 1 | C | T | 0.027 | -6.12E-01 | 9.32E-02 | 3.29E-01 | 3.07E-01 | 1.18E-05 | Yes | Gm49296 | downstream_gene_variant | MODIFIER |
| 15 | 34711604 | AX-170214860 | 1 | A | C | 0.027 | -6.15E-01 | 9.29E-02 | 3.15E-01 | 2.89E-01 | 1.21E-05 | Yes | | intergenic_variant | MODIFIER |
| 15 | 41624422 | AX-170533377 | 0 | T | C | 0.035 | -4.98E-01 | 8.93E-02 | 6.78E-01 | 7.01E-01 | 1.22E-05 | Yes | Oxr1 | intron_variant | MODIFIER |
| 15 | 24017550 | AX-170055029 | 0 | C | T | 0.026 | -5.55E-01 | 9.30E-02 | 5.06E-01 | 5.17E-01 | 1.23E-05 | Yes | | intergenic_variant | MODIFIER |

| | | | | | | | | | | | | | | | |
|---|---|---|---|---|---|---|---|---|---|---|---|---|---|---|---|
| 15 | 56072016 | AX-170338316 | 0 | C | G | 0.035 | -5.05E-01 | 9.51E-02 | 8.47E-01 | 8.63E-01 | 1.25E-05 | Yes | | intergenic_variant | MODIFIER |
| 15 | 26755099 | AX-97828641 | 1 | G | A | 0.027 | -6.19E-01 | 9.15E-02 | 2.82E-01 | 2.55E-01 | 1.27E-05 | Yes | Fbxl7 | intron_variant | MODIFIER |
| 15 | 34185390 | AX-170186453 | 1 | G | T | 0.027 | -6.16E-01 | 9.31E-02 | 3.10E-01 | 2.84E-01 | 1.32E-05 | Yes | | intergenic_variant | MODIFIER |
| 15 | 22863109 | AX-170228172 | 1 | C | T | 0.027 | -6.17E-01 | 9.32E-02 | 3.11E-01 | 2.85E-01 | 1.32E-05 | Yes | Cdh18 | intron_variant | MODIFIER |
| 15 | 22650819 | AX-170492567 | 1 | C | A | 0.027 | -6.11E-01 | 9.28E-02 | 3.18E-01 | 2.92E-01 | 1.33E-05 | Yes | Cdh18 | intron_variant | MODIFIER |
| 15 | 26592540 | AX-170398026 | 1 | A | G | 0.027 | -6.11E-01 | 9.28E-02 | 3.18E-01 | 2.92E-01 | 1.33E-05 | Yes | Fbxl7 | intron_variant | MODIFIER |
| 15 | 29616009 | AX-170531205 | 1 | G | A | 0.027 | -6.11E-01 | 9.28E-02 | 3.18E-01 | 2.92E-01 | 1.33E-05 | Yes | | intergenic_variant | MODIFIER |
| 15 | 30433669 | AX-170378336 | 1 | T | C | 0.027 | -6.11E-01 | 9.28E-02 | 3.18E-01 | 2.92E-01 | 1.33E-05 | Yes | Ctnnd2 | intron_variant | MODIFIER |
| 15 | 30451162 | AX-170103692 | 1 | G | A | 0.027 | -6.11E-01 | 9.28E-02 | 3.18E-01 | 2.92E-01 | 1.33E-05 | Yes | Ctnnd2 | intron_variant | MODIFIER |
| 15 | 30800085 | AX-170221800 | 1 | G | A | 0.027 | -6.11E-01 | 9.28E-02 | 3.18E-01 | 2.92E-01 | 1.33E-05 | Yes | Ctnnd2 | intron_variant | MODIFIER |
| 15 | 30862900 | AX-170055149 | 1 | G | A | 0.027 | -6.11E-01 | 9.28E-02 | 3.18E-01 | 2.92E-01 | 1.33E-05 | Yes | Ctnnd2 | intron_variant | MODIFIER |
| 15 | 34709011 | AX-170517524 | 1 | C | A | 0.027 | -6.11E-01 | 9.28E-02 | 3.18E-01 | 2.92E-01 | 1.33E-05 | Yes | | intergenic_variant | MODIFIER |
| 15 | 29891824 | AX-170497799 | 1 | A | T | 0.027 | -6.19E-01 | 9.27E-02 | 2.98E-01 | 2.72E-01 | 1.33E-05 | Yes | Gm23033 | upstream_gene_variant | MODIFIER |
| 15 | 32620367 | AX-97809848 | 1 | G | A | 0.027 | -6.19E-01 | 9.32E-02 | 3.05E-01 | 2.77E-01 | 1.33E-05 | Yes | Sema5a | intron_variant | MODIFIER |
| 15 | 29693899 | AX-170100828 | 1 | A | G | 0.027 | -6.18E-01 | 9.18E-02 | 2.86E-01 | 2.59E-01 | 1.34E-05 | Yes | | intergenic_variant | MODIFIER |
| 15 | 34436368 | AX-97810979 | 2 | G | A | 0.027 | -5.95E-01 | 9.58E-02 | 4.13E-01 | 3.98E-01 | 1.39E-05 | Yes | Rpl30 | downstream_gene_variant | MODIFIER |
| 15 | 22641630 | AX-170418269 | 1 | A | T | 0.027 | -6.18E-01 | 9.20E-02 | 2.87E-01 | 2.59E-01 | 1.43E-05 | Yes | Cdh18 | intron_variant | MODIFIER |
| 15 | 44397482 | AX-170271729 | 1 | G | A | 0.036 | -5.00E-01 | 9.15E-02 | 7.21E-01 | 7.41E-01 | 1.43E-05 | Yes | Nudcd1 | intron_variant | MODIFIER |
| 15 | 51781285 | AX-170111939 | 0 | T | C | 0.035 | -5.18E-01 | 1.00E-01 | 9.28E-01 | 9.38E-01 | 1.43E-05 | Yes | | intergenic_variant | MODIFIER |
| 15 | 51785453 | AX-168408247 | 0 | T | C | 0.035 | -5.18E-01 | 1.00E-01 | 9.28E-01 | 9.38E-01 | 1.43E-05 | Yes | Eif3h | downstream_gene_variant | MODIFIER |
| 15 | 51786006 | AX-170273435 | 0 | T | G | 0.035 | -5.18E-01 | 1.00E-01 | 9.28E-01 | 9.38E-01 | 1.43E-05 | Yes | Eif3h | downstream_gene_variant | MODIFIER |
| 15 | 51798861 | AX-170313255 | 0 | T | C | 0.035 | -5.18E-01 | 1.00E-01 | 9.28E-01 | 9.38E-01 | 1.43E-05 | Yes | Eif3h | intron_variant | MODIFIER |
| 15 | 51802105 | AX-170222301 | 0 | A | G | 0.035 | -5.18E-01 | 1.00E-01 | 9.28E-01 | 9.38E-01 | 1.43E-05 | Yes | Eif3h | intron_variant | MODIFIER |
| 15 | 51813347 | AX-170534969 | 0 | G | A | 0.035 | -5.18E-01 | 1.00E-01 | 9.28E-01 | 9.38E-01 | 1.43E-05 | Yes | Eif3h | intron_variant | MODIFIER |
| 15 | 51816919 | AX-170087857 | 0 | T | C | 0.035 | -5.18E-01 | 1.00E-01 | 9.28E-01 | 9.38E-01 | 1.43E-05 | Yes | Eif3h | intron_variant | MODIFIER |
| 15 | 51857752 | AX-97843015 | 0 | A | G | 0.035 | -5.18E-01 | 1.00E-01 | 9.28E-01 | 9.38E-01 | 1.43E-05 | Yes | Eif3h | intron_variant | MODIFIER |
| 15 | 51936421 | AX-97782162 | 0 | G | A | 0.035 | -5.18E-01 | 1.00E-01 | 9.28E-01 | 9.38E-01 | 1.43E-05 | Yes | | intergenic_variant | MODIFIER |
| 15 | 51958646 | AX-170388805 | 0 | T | G | 0.035 | -5.18E-01 | 1.00E-01 | 9.28E-01 | 9.38E-01 | 1.43E-05 | Yes | Rad21 | downstream_gene_variant | MODIFIER |
| 15 | 51958764 | AX-170357907 | 0 | T | G | 0.035 | -5.18E-01 | 1.00E-01 | 9.28E-01 | 9.38E-01 | 1.43E-05 | Yes | Rad21 | downstream_gene_variant | MODIFIER |
| 15 | 51986941 | AX-97782400 | 0 | G | A | 0.035 | -5.18E-01 | 1.00E-01 | 9.28E-01 | 9.38E-01 | 1.43E-05 | Yes | Rad21 | intron_variant | MODIFIER |
| 15 | 52006915 | AX-170173623 | 0 | A | T | 0.035 | -5.18E-01 | 1.00E-01 | 9.28E-01 | 9.38E-01 | 1.43E-05 | Yes | | intergenic_variant | MODIFIER |
| 15 | 52008958 | AX-170313132 | 0 | T | C | 0.035 | -5.18E-01 | 1.00E-01 | 9.28E-01 | 9.38E-01 | 1.43E-05 | Yes | Gm48926 | upstream_gene_variant | MODIFIER |
| 15 | 52017575 | AX-170087858 | 0 | T | C | 0.035 | -5.18E-01 | 1.00E-01 | 9.28E-01 | 9.38E-01 | 1.43E-05 | Yes | Gm48926 | downstream_gene_variant | MODIFIER |
| 15 | 52019245 | AX-170011684 | 0 | T | C | 0.035 | -5.18E-01 | 1.00E-01 | 9.28E-01 | 9.38E-01 | 1.43E-05 | Yes | | intergenic_variant | MODIFIER |
| 15 | 52075489 | AX-97829451 | 0 | G | T | 0.035 | -5.18E-01 | 1.00E-01 | 9.28E-01 | 9.38E-01 | 1.43E-05 | Yes | | intergenic_variant | MODIFIER |
| 15 | 52075848 | AX-170011691 | 0 | T | C | 0.035 | -5.18E-01 | 1.00E-01 | 9.28E-01 | 9.38E-01 | 1.43E-05 | Yes | | intergenic_variant | MODIFIER |
| 15 | 52107963 | AX-170011687 | 0 | C | T | 0.035 | -5.18E-01 | 1.00E-01 | 9.28E-01 | 9.38E-01 | 1.43E-05 | Yes | | intergenic_variant | MODIFIER |
| 15 | 52108048 | AX-97829460 | 0 | C | A | 0.035 | -5.18E-01 | 1.00E-01 | 9.28E-01 | 9.38E-01 | 1.43E-05 | Yes | | intergenic_variant | MODIFIER |
| 15 | 52123264 | AX-170487078 | 0 | C | T | 0.035 | -5.18E-01 | 1.00E-01 | 9.28E-01 | 9.38E-01 | 1.43E-05 | Yes | | intergenic_variant | MODIFIER |
| 15 | 52132173 | AX-170011683 | 0 | C | T | 0.035 | -5.18E-01 | 1.00E-01 | 9.28E-01 | 9.38E-01 | 1.43E-05 | Yes | | intergenic_variant | MODIFIER |
| 15 | 52132927 | AX-170011682 | 0 | T | C | 0.035 | -5.18E-01 | 1.00E-01 | 9.28E-01 | 9.38E-01 | 1.43E-05 | Yes | | intergenic_variant | MODIFIER |
| 15 | 52360259 | AX-97797101 | 0 | T | C | 0.035 | -5.18E-01 | 1.00E-01 | 9.28E-01 | 9.38E-01 | 1.43E-05 | Yes | | intergenic_variant | MODIFIER |
| 15 | 52382771 | AX-170366490 | 0 | A | C | 0.035 | -5.18E-01 | 1.00E-01 | 9.28E-01 | 9.38E-01 | 1.43E-05 | Yes | | intergenic_variant | MODIFIER |
| 15 | 52384236 | AX-170127218 | 0 | C | T | 0.035 | -5.18E-01 | 1.00E-01 | 9.28E-01 | 9.38E-01 | 1.43E-05 | Yes | | regulatory_region_variant | MODIFIER |
| 15 | 52389953 | AX-170432659 | 0 | G | C | 0.035 | -5.18E-01 | 1.00E-01 | 9.28E-01 | 9.38E-01 | 1.43E-05 | Yes | | intergenic_variant | MODIFIER |
| 15 | 52395554 | AX-170251407 | 0 | C | T | 0.035 | -5.18E-01 | 1.00E-01 | 9.28E-01 | 9.38E-01 | 1.43E-05 | Yes | | intergenic_variant | MODIFIER |
| 15 | 52405449 | AX-97805865 | 0 | A | G | 0.035 | -5.18E-01 | 1.00E-01 | 9.28E-01 | 9.38E-01 | 1.43E-05 | Yes | | intergenic_variant | MODIFIER |
| 15 | 52405974 | AX-170450844 | 0 | C | A | 0.035 | -5.18E-01 | 1.00E-01 | 9.28E-01 | 9.38E-01 | 1.43E-05 | Yes | | intergenic_variant | MODIFIER |
| 15 | 52419477 | AX-170256659 | 0 | C | A | 0.035 | -5.18E-01 | 1.00E-01 | 9.28E-01 | 9.38E-01 | 1.43E-05 | Yes | | intergenic_variant | MODIFIER |
| 15 | 52426752 | AX-170482609 | 0 | T | C | 0.035 | -5.18E-01 | 1.00E-01 | 9.28E-01 | 9.38E-01 | 1.43E-05 | Yes | | intergenic_variant | MODIFIER |
| 15 | 52427173 | AX-97782844 | 0 | C | A | 0.035 | -5.18E-01 | 1.00E-01 | 9.28E-01 | 9.38E-01 | 1.43E-05 | Yes | | intergenic_variant | MODIFIER |
| 15 | 52427664 | AX-170162162 | 0 | T | C | 0.035 | -5.18E-01 | 1.00E-01 | 9.28E-01 | 9.38E-01 | 1.43E-05 | Yes | | intergenic_variant | MODIFIER |
| 15 | 52428340 | AX-170280257 | 0 | G | A | 0.035 | -5.18E-01 | 1.00E-01 | 9.28E-01 | 9.38E-01 | 1.43E-05 | Yes | | intergenic_variant | MODIFIER |
| 15 | 52431386 | AX-170265822 | 0 | C | T | 0.035 | -5.18E-01 | 1.00E-01 | 9.28E-01 | 9.38E-01 | 1.43E-05 | Yes | | intergenic_variant | MODIFIER |

| | | | | | | | | | | | | | | | |
|---|---|---|---|---|---|---|---|---|---|---|---|---|---|---|---|
| 15 | 52434023 | AX-97790644 | 0 | C | T | 0.035 | -5.18E-01 | 1.00E-01 | 9.28E-01 | 9.38E-01 | 1.43E-05 | Yes | | regulatory_region_variant | MODIFIER |
| 15 | 52445947 | AX-170407755 | 0 | G | A | 0.035 | -5.18E-01 | 1.00E-01 | 9.28E-01 | 9.38E-01 | 1.43E-05 | Yes | Gm41322 | intron_variant,non_coding_transcript_variant | MODIFIER |
| 15 | 52496572 | AX-97843075 | 0 | C | T | 0.035 | -5.18E-01 | 1.00E-01 | 9.28E-01 | 9.38E-01 | 1.43E-05 | Yes | Gm23545 | upstream_gene_variant | MODIFIER |
| 15 | 52509019 | AX-170126982 | 0 | T | A | 0.035 | -5.18E-01 | 1.00E-01 | 9.28E-01 | 9.38E-01 | 1.43E-05 | Yes | | regulatory_region_variant | MODIFIER |
| 15 | 52509206 | AX-170183504 | 0 | A | G | 0.035 | -5.18E-01 | 1.00E-01 | 9.28E-01 | 9.38E-01 | 1.43E-05 | Yes | | regulatory_region_variant | MODIFIER |
| 15 | 52515260 | AX-170325533 | 0 | A | G | 0.035 | -5.18E-01 | 1.00E-01 | 9.28E-01 | 9.38E-01 | 1.43E-05 | Yes | | intergenic_variant | MODIFIER |
| 15 | 52517035 | AX-170350729 | 0 | A | G | 0.035 | -5.18E-01 | 1.00E-01 | 9.28E-01 | 9.38E-01 | 1.43E-05 | Yes | | intergenic_variant | MODIFIER |
| 15 | 52517765 | AX-97783754 | 0 | T | C | 0.035 | -5.18E-01 | 1.00E-01 | 9.28E-01 | 9.38E-01 | 1.43E-05 | Yes | | intergenic_variant | MODIFIER |
| 15 | 52540158 | AX-170234995 | 0 | T | C | 0.035 | -5.18E-01 | 1.00E-01 | 9.28E-01 | 9.38E-01 | 1.43E-05 | Yes | | intergenic_variant | MODIFIER |
| 15 | 52588549 | AX-170011696 | 0 | A | G | 0.035 | -5.18E-01 | 1.00E-01 | 9.28E-01 | 9.38E-01 | 1.43E-05 | Yes | | intergenic_variant | MODIFIER |
| 15 | 52589179 | AX-170011697 | 0 | G | A | 0.035 | -5.18E-01 | 1.00E-01 | 9.28E-01 | 9.38E-01 | 1.43E-05 | Yes | | intergenic_variant | MODIFIER |
| 15 | 52591558 | AX-97829672 | 0 | A | C | 0.035 | -5.18E-01 | 1.00E-01 | 9.28E-01 | 9.38E-01 | 1.43E-05 | Yes | | intergenic_variant | MODIFIER |
| 15 | 52648501 | AX-97786417 | 0 | T | G | 0.035 | -5.18E-01 | 1.00E-01 | 9.28E-01 | 9.38E-01 | 1.43E-05 | Yes | | intergenic_variant | MODIFIER |
| 15 | 52655501 | AX-170011705 | 0 | C | T | 0.035 | -5.18E-01 | 1.00E-01 | 9.28E-01 | 9.38E-01 | 1.43E-05 | Yes | | intergenic_variant | MODIFIER |
| 15 | 52667966 | AX-170087862 | 0 | A | C | 0.035 | -5.18E-01 | 1.00E-01 | 9.28E-01 | 9.38E-01 | 1.43E-05 | Yes | | intergenic_variant | MODIFIER |
| 15 | 52693114 | AX-97848922 | 0 | G | C | 0.035 | -5.18E-01 | 1.00E-01 | 9.28E-01 | 9.38E-01 | 1.43E-05 | Yes | | intergenic_variant | MODIFIER |
| 15 | 52703916 | AX-170011706 | 0 | T | A | 0.035 | -5.18E-01 | 1.00E-01 | 9.28E-01 | 9.38E-01 | 1.43E-05 | Yes | | intergenic_variant | MODIFIER |
| 15 | 52756429 | AX-97829484 | 0 | T | A | 0.035 | -5.18E-01 | 1.00E-01 | 9.28E-01 | 9.38E-01 | 1.43E-05 | Yes | Gm34794 | intron_variant,non_coding_transcript_variant | MODIFIER |
| 15 | 52787332 | AX-170011698 | 0 | A | G | 0.035 | -5.18E-01 | 1.00E-01 | 9.28E-01 | 9.38E-01 | 1.43E-05 | Yes | Gm34794 | upstream_gene_variant | MODIFIER |
| 15 | 52787630 | AX-170011700 | 0 | A | G | 0.035 | -5.18E-01 | 1.00E-01 | 9.28E-01 | 9.38E-01 | 1.43E-05 | Yes | Gm34794 | upstream_gene_variant | MODIFIER |
| 15 | 52790826 | AX-170087863 | 0 | T | G | 0.035 | -5.18E-01 | 1.00E-01 | 9.28E-01 | 9.38E-01 | 1.43E-05 | Yes | Gm34794 | upstream_gene_variant | MODIFIER |
| 15 | 52802952 | AX-170185328 | 0 | A | G | 0.035 | -5.18E-01 | 1.00E-01 | 9.28E-01 | 9.38E-01 | 1.43E-05 | Yes | | intergenic_variant | MODIFIER |
| 15 | 52805558 | AX-170152843 | 0 | T | G | 0.035 | -5.18E-01 | 1.00E-01 | 9.28E-01 | 9.38E-01 | 1.43E-05 | Yes | | intergenic_variant | MODIFIER |
| 15 | 52810060 | AX-170011693 | 0 | G | A | 0.035 | -5.18E-01 | 1.00E-01 | 9.28E-01 | 9.38E-01 | 1.43E-05 | Yes | | intergenic_variant | MODIFIER |
| 15 | 52810087 | AX-170011694 | 0 | G | A | 0.035 | -5.18E-01 | 1.00E-01 | 9.28E-01 | 9.38E-01 | 1.43E-05 | Yes | | intergenic_variant | MODIFIER |
| 15 | 52810308 | AX-97843731 | 0 | A | G | 0.035 | -5.18E-01 | 1.00E-01 | 9.28E-01 | 9.38E-01 | 1.43E-05 | Yes | | intergenic_variant | MODIFIER |
| 15 | 52848609 | AX-170087864 | 0 | A | G | 0.035 | -5.18E-01 | 1.00E-01 | 9.28E-01 | 9.38E-01 | 1.43E-05 | Yes | | intergenic_variant | MODIFIER |
| 15 | 52858181 | AX-170011695 | 0 | T | G | 0.035 | -5.18E-01 | 1.00E-01 | 9.28E-01 | 9.38E-01 | 1.43E-05 | Yes | | intergenic_variant | MODIFIER |
| 15 | 52860650 | AX-97797643 | 0 | T | C | 0.035 | -5.18E-01 | 1.00E-01 | 9.28E-01 | 9.38E-01 | 1.43E-05 | Yes | | intergenic_variant | MODIFIER |
| 15 | 52904550 | AX-97829474 | 0 | G | A | 0.035 | -5.18E-01 | 1.00E-01 | 9.28E-01 | 9.38E-01 | 1.43E-05 | Yes | | intergenic_variant | MODIFIER |
| 15 | 52905270 | AX-170011702 | 0 | A | G | 0.035 | -5.18E-01 | 1.00E-01 | 9.28E-01 | 9.38E-01 | 1.43E-05 | Yes | | intergenic_variant | MODIFIER |
| 15 | 52908011 | AX-170268746 | 0 | A | G | 0.035 | -5.18E-01 | 1.00E-01 | 9.28E-01 | 9.38E-01 | 1.43E-05 | Yes | | intergenic_variant | MODIFIER |
| 15 | 52933509 | AX-170011699 | 0 | T | G | 0.035 | -5.18E-01 | 1.00E-01 | 9.28E-01 | 9.38E-01 | 1.43E-05 | Yes | | intergenic_variant | MODIFIER |
| 15 | 52941097 | AX-97829482 | 0 | T | G | 0.035 | -5.18E-01 | 1.00E-01 | 9.28E-01 | 9.38E-01 | 1.43E-05 | Yes | | intergenic_variant | MODIFIER |
| 15 | 52994255 | AX-170087865 | 0 | A | G | 0.035 | -5.18E-01 | 1.00E-01 | 9.28E-01 | 9.38E-01 | 1.43E-05 | Yes | | intergenic_variant | MODIFIER |
| 15 | 52994374 | AX-170269237 | 0 | T | A | 0.035 | -5.18E-01 | 1.00E-01 | 9.28E-01 | 9.38E-01 | 1.43E-05 | Yes | | intergenic_variant | MODIFIER |
| 15 | 53028570 | AX-170272811 | 0 | C | T | 0.035 | -5.18E-01 | 1.00E-01 | 9.28E-01 | 9.38E-01 | 1.43E-05 | Yes | | intergenic_variant | MODIFIER |
| 15 | 53031582 | AX-170434429 | 0 | C | T | 0.035 | -5.18E-01 | 1.00E-01 | 9.28E-01 | 9.38E-01 | 1.43E-05 | Yes | | intergenic_variant | MODIFIER |
| 15 | 53066321 | AX-97810030 | 0 | T | C | 0.035 | -5.18E-01 | 1.00E-01 | 9.28E-01 | 9.38E-01 | 1.43E-05 | Yes | Ext1 | 3_prime_UTR_variant | MODIFIER |
| 15 | 53069749 | AX-170329834 | 0 | A | T | 0.035 | -5.18E-01 | 1.00E-01 | 9.28E-01 | 9.38E-01 | 1.43E-05 | Yes | Ext1 | intron_variant | MODIFIER |
| 15 | 53109767 | AX-170503877 | 0 | C | G | 0.035 | -5.18E-01 | 1.00E-01 | 9.28E-01 | 9.38E-01 | 1.43E-05 | Yes | Ext1 | intron_variant | MODIFIER |
| 15 | 53120984 | AX-170011710 | 0 | A | G | 0.035 | -5.18E-01 | 1.00E-01 | 9.28E-01 | 9.38E-01 | 1.43E-05 | Yes | Ext1 | intron_variant | MODIFIER |
| 15 | 53136926 | AX-170011715 | 0 | G | C | 0.035 | -5.18E-01 | 1.00E-01 | 9.28E-01 | 9.38E-01 | 1.43E-05 | Yes | Ext1 | intron_variant | MODIFIER |
| 15 | 53137102 | AX-170011708 | 0 | G | A | 0.035 | -5.18E-01 | 1.00E-01 | 9.28E-01 | 9.38E-01 | 1.43E-05 | Yes | Ext1 | intron_variant | MODIFIER |
| 15 | 53142602 | AX-170465861 | 0 | C | T | 0.035 | -5.18E-01 | 1.00E-01 | 9.28E-01 | 9.38E-01 | 1.43E-05 | Yes | Ext1 | intron_variant | MODIFIER |
| 15 | 53142682 | AX-170102535 | 0 | A | G | 0.035 | -5.18E-01 | 1.00E-01 | 9.28E-01 | 9.38E-01 | 1.43E-05 | Yes | Ext1 | intron_variant | MODIFIER |
| 15 | 53143096 | AX-170093251 | 0 | G | A | 0.035 | -5.18E-01 | 1.00E-01 | 9.28E-01 | 9.38E-01 | 1.43E-05 | Yes | Ext1 | intron_variant | MODIFIER |
| 15 | 53149637 | AX-170266771 | 0 | A | G | 0.035 | -5.18E-01 | 1.00E-01 | 9.28E-01 | 9.38E-01 | 1.43E-05 | Yes | Ext1 | intron_variant | MODIFIER |
| 15 | 53174835 | AX-170011711 | 0 | A | G | 0.035 | -5.18E-01 | 1.00E-01 | 9.28E-01 | 9.38E-01 | 1.43E-05 | Yes | Ext1 | intron_variant | MODIFIER |
| 15 | 53184977 | AX-170011712 | 0 | T | C | 0.035 | -5.18E-01 | 1.00E-01 | 9.28E-01 | 9.38E-01 | 1.43E-05 | Yes | Ext1 | intron_variant | MODIFIER |
| 15 | 53185043 | AX-97849812 | 0 | T | C | 0.035 | -5.18E-01 | 1.00E-01 | 9.28E-01 | 9.38E-01 | 1.43E-05 | Yes | Ext1 | intron_variant | MODIFIER |
| 15 | 53233703 | AX-170011717 | 0 | A | G | 0.035 | -5.18E-01 | 1.00E-01 | 9.28E-01 | 9.38E-01 | 1.43E-05 | Yes | Ext1 | intron_variant | MODIFIER |
| 15 | 53242877 | AX-97831032 | 0 | A | G | 0.035 | -5.18E-01 | 1.00E-01 | 9.28E-01 | 9.38E-01 | 1.43E-05 | Yes | Ext1 | intron_variant | MODIFIER |
| 15 | 53246875 | AX-170011714 | 0 | C | G | 0.035 | -5.18E-01 | 1.00E-01 | 9.28E-01 | 9.38E-01 | 1.43E-05 | Yes | Ext1 | intron_variant | MODIFIER |

| | | | | | | | | | | | | | | | |
|---|---|---|---|---|---|---|---|---|---|---|---|---|---|---|---|
| 15 | 53252029 | AX-170055549 | 0 | G | A | 0.035 | -5.18E-01 | 1.00E-01 | 9.28E-01 | 9.38E-01 | 1.43E-05 | Yes | Ext1 | intron_variant | MODIFIER |
| 15 | 53281758 | AX-97854217 | 0 | A | G | 0.035 | -5.18E-01 | 1.00E-01 | 9.28E-01 | 9.38E-01 | 1.43E-05 | Yes | Ext1 | intron_variant | MODIFIER |
| 15 | 53285678 | AX-170011713 | 0 | A | G | 0.035 | -5.18E-01 | 1.00E-01 | 9.28E-01 | 9.38E-01 | 1.43E-05 | Yes | Ext1 | intron_variant | MODIFIER |
| 15 | 53289142 | AX-170341586 | 0 | T | C | 0.035 | -5.18E-01 | 1.00E-01 | 9.28E-01 | 9.38E-01 | 1.43E-05 | Yes | Ext1 | intron_variant | MODIFIER |
| 15 | 53301882 | AX-97799493 | 0 | A | G | 0.035 | -5.18E-01 | 1.00E-01 | 9.28E-01 | 9.38E-01 | 1.43E-05 | Yes | Ext1 | intron_variant | MODIFIER |
| 15 | 53302725 | AX-170011716 | 0 | G | C | 0.035 | -5.18E-01 | 1.00E-01 | 9.28E-01 | 9.38E-01 | 1.43E-05 | Yes | Ext1 | intron_variant | MODIFIER |
| 15 | 53305760 | AX-170423028 | 0 | G | A | 0.035 | -5.18E-01 | 1.00E-01 | 9.28E-01 | 9.38E-01 | 1.43E-05 | Yes | Ext1 | intron_variant | MODIFIER |
| 15 | 53342377 | AX-97838567 | 0 | A | G | 0.035 | -5.18E-01 | 1.00E-01 | 9.28E-01 | 9.38E-01 | 1.43E-05 | Yes | Ext1 | intron_variant | MODIFIER |
| 15 | 53342983 | AX-170145432 | 0 | G | A | 0.035 | -5.18E-01 | 1.00E-01 | 9.28E-01 | 9.38E-01 | 1.43E-05 | Yes | Ext1 | intron_variant | MODIFIER |
| 15 | 53362813 | AX-97785699 | 0 | T | C | 0.035 | -5.18E-01 | 1.00E-01 | 9.28E-01 | 9.38E-01 | 1.43E-05 | Yes | | intergenic_variant | MODIFIER |
| 15 | 53401612 | AX-97789712 | 0 | C | T | 0.035 | -5.18E-01 | 1.00E-01 | 9.28E-01 | 9.38E-01 | 1.43E-05 | Yes | | intergenic_variant | MODIFIER |
| 15 | 53401636 | AX-170274151 | 0 | G | A | 0.035 | -5.18E-01 | 1.00E-01 | 9.28E-01 | 9.38E-01 | 1.43E-05 | Yes | | intergenic_variant | MODIFIER |
| 15 | 53410053 | AX-170266211 | 0 | G | A | 0.035 | -5.18E-01 | 1.00E-01 | 9.28E-01 | 9.38E-01 | 1.43E-05 | Yes | | intergenic_variant | MODIFIER |
| 15 | 53443724 | AX-170222048 | 0 | T | C | 0.035 | -5.18E-01 | 1.00E-01 | 9.28E-01 | 9.38E-01 | 1.43E-05 | Yes | | intergenic_variant | MODIFIER |
| 15 | 53446094 | AX-97857058 | 0 | A | G | 0.035 | -5.18E-01 | 1.00E-01 | 9.28E-01 | 9.38E-01 | 1.43E-05 | Yes | | intergenic_variant | MODIFIER |
| 15 | 53448462 | AX-170446926 | 0 | T | C | 0.035 | -5.18E-01 | 1.00E-01 | 9.28E-01 | 9.38E-01 | 1.43E-05 | Yes | | intergenic_variant | MODIFIER |
| 15 | 53450585 | AX-168396056 | 0 | T | C | 0.035 | -5.18E-01 | 1.00E-01 | 9.28E-01 | 9.38E-01 | 1.43E-05 | Yes | Samd12 | downstream_gene_variant | MODIFIER |
| 15 | 53471379 | AX-97835690 | 0 | G | A | 0.035 | -5.18E-01 | 1.00E-01 | 9.28E-01 | 9.38E-01 | 1.43E-05 | Yes | Samd12 | intron_variant | MODIFIER |
| 15 | 53490203 | AX-170403506 | 0 | G | T | 0.035 | -5.18E-01 | 1.00E-01 | 9.28E-01 | 9.38E-01 | 1.43E-05 | Yes | Samd12 | intron_variant | MODIFIER |
| 15 | 53493364 | AX-170406455 | 0 | C | A | 0.035 | -5.18E-01 | 1.00E-01 | 9.28E-01 | 9.38E-01 | 1.43E-05 | Yes | Samd12 | intron_variant | MODIFIER |
| 15 | 53493403 | AX-170505035 | 0 | G | C | 0.035 | -5.18E-01 | 1.00E-01 | 9.28E-01 | 9.38E-01 | 1.43E-05 | Yes | Samd12 | intron_variant | MODIFIER |
| 15 | 53509754 | AX-170055552 | 0 | T | C | 0.035 | -5.18E-01 | 1.00E-01 | 9.28E-01 | 9.38E-01 | 1.43E-05 | Yes | Samd12 | intron_variant | MODIFIER |
| 15 | 53512887 | AX-170206127 | 0 | C | T | 0.035 | -5.18E-01 | 1.00E-01 | 9.28E-01 | 9.38E-01 | 1.43E-05 | Yes | Samd12 | intron_variant | MODIFIER |
| 15 | 53517390 | AX-170126132 | 0 | C | A | 0.035 | -5.18E-01 | 1.00E-01 | 9.28E-01 | 9.38E-01 | 1.43E-05 | Yes | Samd12 | intron_variant | MODIFIER |
| 15 | 53519163 | AX-97843035 | 0 | A | G | 0.035 | -5.18E-01 | 1.00E-01 | 9.28E-01 | 9.38E-01 | 1.43E-05 | Yes | Samd12 | intron_variant | MODIFIER |
| 15 | 53531691 | AX-170415214 | 0 | G | T | 0.035 | -5.18E-01 | 1.00E-01 | 9.28E-01 | 9.38E-01 | 1.43E-05 | Yes | Samd12 | intron_variant | MODIFIER |
| 15 | 53554549 | AX-170132073 | 0 | T | G | 0.035 | -5.18E-01 | 1.00E-01 | 9.28E-01 | 9.38E-01 | 1.43E-05 | Yes | Samd12 | intron_variant | MODIFIER |
| 15 | 53581745 | AX-97790870 | 0 | G | A | 0.035 | -5.18E-01 | 1.00E-01 | 9.28E-01 | 9.38E-01 | 1.43E-05 | Yes | Samd12 | intron_variant | MODIFIER |
| 15 | 53582316 | AX-170182095 | 0 | T | C | 0.035 | -5.18E-01 | 1.00E-01 | 9.28E-01 | 9.38E-01 | 1.43E-05 | Yes | Samd12 | intron_variant | MODIFIER |
| 15 | 53587888 | AX-170107189 | 0 | A | G | 0.035 | -5.18E-01 | 1.00E-01 | 9.28E-01 | 9.38E-01 | 1.43E-05 | Yes | Samd12 | intron_variant | MODIFIER |
| 15 | 53595380 | AX-170490977 | 0 | C | T | 0.035 | -5.18E-01 | 1.00E-01 | 9.28E-01 | 9.38E-01 | 1.43E-05 | Yes | Samd12 | intron_variant | MODIFIER |
| 15 | 53598261 | AX-170264778 | 0 | T | C | 0.035 | -5.18E-01 | 1.00E-01 | 9.28E-01 | 9.38E-01 | 1.43E-05 | Yes | Samd12 | intron_variant | MODIFIER |
| 15 | 53601977 | AX-170011718 | 0 | A | G | 0.035 | -5.18E-01 | 1.00E-01 | 9.28E-01 | 9.38E-01 | 1.43E-05 | Yes | Samd12 | intron_variant | MODIFIER |
| 15 | 53614096 | AX-170294242 | 0 | G | A | 0.035 | -5.18E-01 | 1.00E-01 | 9.28E-01 | 9.38E-01 | 1.43E-05 | Yes | Samd12 | intron_variant | MODIFIER |
| 15 | 53615834 | AX-170104891 | 0 | A | C | 0.035 | -5.18E-01 | 1.00E-01 | 9.28E-01 | 9.38E-01 | 1.43E-05 | Yes | Samd12 | intron_variant | MODIFIER |
| 15 | 53617003 | AX-97826594 | 0 | C | T | 0.035 | -5.18E-01 | 1.00E-01 | 9.28E-01 | 9.38E-01 | 1.43E-05 | Yes | Samd12 | intron_variant | MODIFIER |
| 15 | 53633724 | AX-170055556 | 0 | T | C | 0.035 | -5.18E-01 | 1.00E-01 | 9.28E-01 | 9.38E-01 | 1.43E-05 | Yes | Samd12 | intron_variant | MODIFIER |
| 15 | 53675801 | AX-170011721 | 0 | G | A | 0.035 | -5.18E-01 | 1.00E-01 | 9.28E-01 | 9.38E-01 | 1.43E-05 | Yes | Samd12 | intron_variant | MODIFIER |
| 15 | 53689116 | AX-170139984 | 0 | G | A | 0.035 | -5.18E-01 | 1.00E-01 | 9.28E-01 | 9.38E-01 | 1.43E-05 | Yes | Samd12 | intron_variant | MODIFIER |
| 15 | 53692954 | AX-170157134 | 0 | A | G | 0.035 | -5.18E-01 | 1.00E-01 | 9.28E-01 | 9.38E-01 | 1.43E-05 | Yes | Samd12 | intron_variant | MODIFIER |
| 15 | 53705215 | AX-170011722 | 0 | G | C | 0.035 | -5.18E-01 | 1.00E-01 | 9.28E-01 | 9.38E-01 | 1.43E-05 | Yes | Samd12 | intron_variant | MODIFIER |
| 15 | 53708369 | AX-170311157 | 0 | G | A | 0.035 | -5.18E-01 | 1.00E-01 | 9.28E-01 | 9.38E-01 | 1.43E-05 | Yes | Samd12 | intron_variant | MODIFIER |
| 15 | 53709982 | AX-170206216 | 0 | T | C | 0.035 | -5.18E-01 | 1.00E-01 | 9.28E-01 | 9.38E-01 | 1.43E-05 | Yes | Samd12 | intron_variant | MODIFIER |
| 15 | 53715538 | AX-170384000 | 0 | T | C | 0.035 | -5.18E-01 | 1.00E-01 | 9.28E-01 | 9.38E-01 | 1.43E-05 | Yes | Samd12 | intron_variant | MODIFIER |
| 15 | 53741722 | AX-170415975 | 0 | T | C | 0.035 | -5.18E-01 | 1.00E-01 | 9.28E-01 | 9.38E-01 | 1.43E-05 | Yes | Samd12 | intron_variant | MODIFIER |
| 15 | 53753201 | AX-97843809 | 0 | A | G | 0.035 | -5.18E-01 | 1.00E-01 | 9.28E-01 | 9.38E-01 | 1.43E-05 | Yes | Samd12 | intron_variant | MODIFIER |
| 15 | 53788935 | AX-170524839 | 0 | A | G | 0.035 | -5.18E-01 | 1.00E-01 | 9.28E-01 | 9.38E-01 | 1.43E-05 | Yes | Samd12 | intron_variant | MODIFIER |
| 15 | 53842851 | AX-170318374 | 0 | G | A | 0.035 | -5.18E-01 | 1.00E-01 | 9.28E-01 | 9.38E-01 | 1.43E-05 | Yes | Samd12 | intron_variant | MODIFIER |
| 15 | 53859945 | AX-170497745 | 0 | C | T | 0.035 | -5.18E-01 | 1.00E-01 | 9.28E-01 | 9.38E-01 | 1.43E-05 | Yes | Samd12 | intron_variant | MODIFIER |
| 15 | 53869844 | AX-170399174 | 0 | G | A | 0.035 | -5.18E-01 | 1.00E-01 | 9.28E-01 | 9.38E-01 | 1.43E-05 | Yes | Samd12 | intron_variant | MODIFIER |
| 15 | 53877630 | AX-170475866 | 0 | C | T | 0.035 | -5.18E-01 | 1.00E-01 | 9.28E-01 | 9.38E-01 | 1.43E-05 | Yes | Samd12 | intron_variant | MODIFIER |
| 15 | 53877806 | AX-170296442 | 0 | G | T | 0.035 | -5.18E-01 | 1.00E-01 | 9.28E-01 | 9.38E-01 | 1.43E-05 | Yes | Samd12 | intron_variant | MODIFIER |
| 15 | 53893394 | AX-170130522 | 0 | C | T | 0.035 | -5.18E-01 | 1.00E-01 | 9.28E-01 | 9.38E-01 | 1.43E-05 | Yes | Samd12 | intron_variant | MODIFIER |
| 15 | 53893756 | AX-170282196 | 0 | T | C | 0.035 | -5.18E-01 | 1.00E-01 | 9.28E-01 | 9.38E-01 | 1.43E-05 | Yes | Samd12 | intron_variant | MODIFIER |

| | | | | | | | | | | | | | | | |
|---|---|---|---|---|---|---|---|---|---|---|---|---|---|---|---|
| 15 | 53894889 | AX-170152306 | 0 | C | T | 0.035 | -5.18E-01 | 1.00E-01 | 9.28E-01 | 9.38E-01 | 1.43E-05 | Yes | Samd12 | intron_variant | MODIFIER |
| 15 | 53898207 | AX-170055564 | 0 | A | G | 0.035 | -5.18E-01 | 1.00E-01 | 9.28E-01 | 9.38E-01 | 1.43E-05 | Yes | Samd12 | intron_variant | MODIFIER |
| 15 | 53921606 | AX-170505093 | 0 | G | C | 0.035 | -5.18E-01 | 1.00E-01 | 9.28E-01 | 9.38E-01 | 1.43E-05 | Yes | | intergenic_variant | MODIFIER |
| 15 | 53961638 | AX-170426258 | 0 | G | A | 0.035 | -5.18E-01 | 1.00E-01 | 9.28E-01 | 9.38E-01 | 1.43E-05 | Yes | | intergenic_variant | MODIFIER |
| 15 | 53961740 | AX-170259380 | 0 | C | T | 0.035 | -5.18E-01 | 1.00E-01 | 9.28E-01 | 9.38E-01 | 1.43E-05 | Yes | | intergenic_variant | MODIFIER |
| 15 | 54002561 | AX-170318562 | 0 | G | A | 0.035 | -5.18E-01 | 1.00E-01 | 9.28E-01 | 9.38E-01 | 1.43E-05 | Yes | | intergenic_variant | MODIFIER |
| 15 | 54012536 | AX-170314538 | 0 | A | G | 0.035 | -5.18E-01 | 1.00E-01 | 9.28E-01 | 9.38E-01 | 1.43E-05 | Yes | | intergenic_variant | MODIFIER |
| 15 | 54012840 | AX-170501725 | 0 | A | G | 0.035 | -5.18E-01 | 1.00E-01 | 9.28E-01 | 9.38E-01 | 1.43E-05 | Yes | | intergenic_variant | MODIFIER |
| 15 | 54012913 | AX-170337918 | 0 | C | A | 0.035 | -5.18E-01 | 1.00E-01 | 9.28E-01 | 9.38E-01 | 1.43E-05 | Yes | | intergenic_variant | MODIFIER |
| 15 | 54015090 | AX-170102814 | 0 | T | G | 0.035 | -5.18E-01 | 1.00E-01 | 9.28E-01 | 9.38E-01 | 1.43E-05 | Yes | | intergenic_variant | MODIFIER |
| 15 | 54022839 | AX-97796086 | 0 | G | A | 0.035 | -5.18E-01 | 1.00E-01 | 9.28E-01 | 9.38E-01 | 1.43E-05 | Yes | | intergenic_variant | MODIFIER |
| 15 | 54035749 | AX-170154185 | 0 | G | A | 0.035 | -5.18E-01 | 1.00E-01 | 9.28E-01 | 9.38E-01 | 1.43E-05 | Yes | Gm26933 | non_coding_transcript_exon_variant | MODIFIER |
| 15 | 54110511 | AX-170267900 | 0 | A | G | 0.035 | -5.18E-01 | 1.00E-01 | 9.28E-01 | 9.38E-01 | 1.43E-05 | Yes | | intergenic_variant | MODIFIER |
| 15 | 54111303 | AX-170055567 | 0 | A | G | 0.035 | -5.18E-01 | 1.00E-01 | 9.28E-01 | 9.38E-01 | 1.43E-05 | Yes | | intergenic_variant | MODIFIER |
| 15 | 54114976 | AX-170055569 | 0 | A | T | 0.035 | -5.18E-01 | 1.00E-01 | 9.28E-01 | 9.38E-01 | 1.43E-05 | Yes | | intergenic_variant | MODIFIER |
| 15 | 54121224 | AX-170186762 | 0 | G | A | 0.035 | -5.18E-01 | 1.00E-01 | 9.28E-01 | 9.38E-01 | 1.43E-05 | Yes | | intergenic_variant | MODIFIER |
| 15 | 54129738 | AX-170425458 | 0 | A | G | 0.035 | -5.18E-01 | 1.00E-01 | 9.28E-01 | 9.38E-01 | 1.43E-05 | Yes | | intergenic_variant | MODIFIER |
| 15 | 54129886 | AX-170213694 | 0 | T | C | 0.035 | -5.18E-01 | 1.00E-01 | 9.28E-01 | 9.38E-01 | 1.43E-05 | Yes | | intergenic_variant | MODIFIER |
| 15 | 54139788 | AX-97848468 | 0 | G | T | 0.035 | -5.18E-01 | 1.00E-01 | 9.28E-01 | 9.38E-01 | 1.43E-05 | Yes | | intergenic_variant | MODIFIER |
| 15 | 54160532 | AX-170522395 | 0 | G | C | 0.035 | -5.18E-01 | 1.00E-01 | 9.28E-01 | 9.38E-01 | 1.43E-05 | Yes | | intergenic_variant | MODIFIER |
| 15 | 54161491 | AX-170011724 | 0 | T | A | 0.035 | -5.18E-01 | 1.00E-01 | 9.28E-01 | 9.38E-01 | 1.43E-05 | Yes | | intergenic_variant | MODIFIER |
| 15 | 54191318 | AX-170458874 | 0 | G | A | 0.035 | -5.18E-01 | 1.00E-01 | 9.28E-01 | 9.38E-01 | 1.43E-05 | Yes | | intergenic_variant | MODIFIER |
| 15 | 54216888 | AX-170533284 | 0 | C | G | 0.035 | -5.18E-01 | 1.00E-01 | 9.28E-01 | 9.38E-01 | 1.43E-05 | Yes | | intergenic_variant | MODIFIER |
| 15 | 54233482 | AX-170211019 | 0 | T | G | 0.035 | -5.18E-01 | 1.00E-01 | 9.28E-01 | 9.38E-01 | 1.43E-05 | Yes | | intergenic_variant | MODIFIER |
| 15 | 54263349 | AX-170420030 | 0 | C | G | 0.035 | -5.18E-01 | 1.00E-01 | 9.28E-01 | 9.38E-01 | 1.43E-05 | Yes | Tnfrsf11b | intron_variant | MODIFIER |
| 15 | 54267910 | AX-97810591 | 0 | T | C | 0.035 | -5.18E-01 | 1.00E-01 | 9.28E-01 | 9.38E-01 | 1.43E-05 | Yes | Tnfrsf11b | intron_variant | MODIFIER |
| 15 | 54283720 | AX-170055571 | 0 | T | G | 0.035 | -5.18E-01 | 1.00E-01 | 9.28E-01 | 9.38E-01 | 1.43E-05 | Yes | | intergenic_variant | MODIFIER |
| 15 | 54283867 | AX-170055572 | 0 | C | G | 0.035 | -5.18E-01 | 1.00E-01 | 9.28E-01 | 9.38E-01 | 1.43E-05 | Yes | | intergenic_variant | MODIFIER |
| 15 | 54303024 | AX-170153755 | 0 | A | C | 0.035 | -5.18E-01 | 1.00E-01 | 9.28E-01 | 9.38E-01 | 1.43E-05 | Yes | Gm7566 | downstream_gene_variant | MODIFIER |
| 15 | 54306653 | AX-170209810 | 0 | T | C | 0.035 | -5.18E-01 | 1.00E-01 | 9.28E-01 | 9.38E-01 | 1.43E-05 | Yes | Gm7566 | upstream_gene_variant | MODIFIER |
| 15 | 54307296 | AX-170093255 | 0 | A | G | 0.035 | -5.18E-01 | 1.00E-01 | 9.28E-01 | 9.38E-01 | 1.43E-05 | Yes | Gm5215 | downstream_gene_variant | MODIFIER |
| 15 | 54308411 | AX-170104837 | 0 | A | C | 0.035 | -5.18E-01 | 1.00E-01 | 9.28E-01 | 9.38E-01 | 1.43E-05 | Yes | Gm5215 | downstream_gene_variant | MODIFIER |
| 15 | 54318312 | AX-170284600 | 0 | A | C | 0.035 | -5.18E-01 | 1.00E-01 | 9.28E-01 | 9.38E-01 | 1.43E-05 | Yes | | intergenic_variant | MODIFIER |
| 15 | 54318526 | AX-97846827 | 0 | A | G | 0.035 | -5.18E-01 | 1.00E-01 | 9.28E-01 | 9.38E-01 | 1.43E-05 | Yes | | intergenic_variant | MODIFIER |
| 15 | 54336164 | AX-170087866 | 0 | A | G | 0.035 | -5.18E-01 | 1.00E-01 | 9.28E-01 | 9.38E-01 | 1.43E-05 | Yes | | intergenic_variant | MODIFIER |
| 15 | 54351889 | AX-170126444 | 0 | G | C | 0.035 | -5.18E-01 | 1.00E-01 | 9.28E-01 | 9.38E-01 | 1.43E-05 | Yes | | intergenic_variant | MODIFIER |
| 15 | 54359676 | AX-170478672 | 0 | T | C | 0.035 | -5.18E-01 | 1.00E-01 | 9.28E-01 | 9.38E-01 | 1.43E-05 | Yes | | intergenic_variant | MODIFIER |
| 15 | 54361215 | AX-170300621 | 0 | G | A | 0.035 | -5.18E-01 | 1.00E-01 | 9.28E-01 | 9.38E-01 | 1.43E-05 | Yes | | intergenic_variant | MODIFIER |
| 15 | 54367417 | AX-97789932 | 0 | T | C | 0.035 | -5.18E-01 | 1.00E-01 | 9.28E-01 | 9.38E-01 | 1.43E-05 | Yes | | intergenic_variant | MODIFIER |
| 15 | 54368059 | AX-170438729 | 0 | G | A | 0.035 | -5.18E-01 | 1.00E-01 | 9.28E-01 | 9.38E-01 | 1.43E-05 | Yes | | intergenic_variant | MODIFIER |
| 15 | 54368587 | AX-170458327 | 0 | G | C | 0.035 | -5.18E-01 | 1.00E-01 | 9.28E-01 | 9.38E-01 | 1.43E-05 | Yes | | intergenic_variant | MODIFIER |
| 15 | 54381229 | AX-170358585 | 0 | A | G | 0.035 | -5.18E-01 | 1.00E-01 | 9.28E-01 | 9.38E-01 | 1.43E-05 | Yes | | intergenic_variant | MODIFIER |
| 15 | 54387560 | AX-97846725 | 0 | T | C | 0.035 | -5.18E-01 | 1.00E-01 | 9.28E-01 | 9.38E-01 | 1.43E-05 | Yes | | intergenic_variant | MODIFIER |
| 15 | 54390732 | AX-170175185 | 0 | C | G | 0.035 | -5.18E-01 | 1.00E-01 | 9.28E-01 | 9.38E-01 | 1.43E-05 | Yes | | intergenic_variant | MODIFIER |
| 15 | 54408182 | AX-170237140 | 0 | G | A | 0.035 | -5.18E-01 | 1.00E-01 | 9.28E-01 | 9.38E-01 | 1.43E-05 | Yes | Colec10 | upstream_gene_variant | MODIFIER |
| 15 | 54412410 | AX-170366848 | 0 | A | G | 0.035 | -5.18E-01 | 1.00E-01 | 9.28E-01 | 9.38E-01 | 1.43E-05 | Yes | Colec10 | intron_variant | MODIFIER |
| 15 | 54417834 | AX-170055577 | 0 | G | A | 0.035 | -5.18E-01 | 1.00E-01 | 9.28E-01 | 9.38E-01 | 1.43E-05 | Yes | Colec10 | intron_variant | MODIFIER |
| 15 | 54428227 | AX-170055576 | 0 | C | T | 0.035 | -5.18E-01 | 1.00E-01 | 9.28E-01 | 9.38E-01 | 1.43E-05 | Yes | Colec10 | intron_variant | MODIFIER |
| 15 | 54428392 | AX-170251076 | 0 | A | G | 0.035 | -5.18E-01 | 1.00E-01 | 9.28E-01 | 9.38E-01 | 1.43E-05 | Yes | Colec10 | intron_variant | MODIFIER |
| 15 | 54430845 | AX-168403242 | 0 | C | A | 0.035 | -5.18E-01 | 1.00E-01 | 9.28E-01 | 9.38E-01 | 1.43E-05 | Yes | Colec10 | intron_variant | MODIFIER |
| 15 | 54430902 | AX-170035550 | 0 | C | T | 0.035 | -5.18E-01 | 1.00E-01 | 9.28E-01 | 9.38E-01 | 1.43E-05 | Yes | Colec10 | intron_variant | MODIFIER |
| 15 | 54431290 | AX-170155943 | 0 | A | G | 0.035 | -5.18E-01 | 1.00E-01 | 9.28E-01 | 9.38E-01 | 1.43E-05 | Yes | Colec10 | intron_variant | MODIFIER |
| 15 | 30832144 | AX-170206995 | 1 | G | A | 0.027 | -6.20E-01 | 9.17E-02 | 2.76E-01 | 2.47E-01 | 1.46E-05 | Yes | Ctnnd2 | intron_variant | MODIFIER |
| 15 | 43316586 | AX-97796106 | 0 | T | A | 0.044 | -4.75E-01 | 8.89E-02 | 7.94E-01 | 8.16E-01 | 1.49E-05 | Yes | | intergenic_variant | MODIFIER |

| | | | | | | | | | | | | | | | |
|---|---|---|---|---|---|---|---|---|---|---|---|---|---|---|---|
| 15 | 53616693 | AX-170011723 | 1 | G | C | 0.036 | -5.12E-01 | 1.00E-01 | 9.82E-01 | 9.96E-01 | 1.50E-05 | Yes | Samd12 | intron_variant | MODIFIER |
| 15 | 25336175 | AX-170430116 | 1 | C | T | 0.027 | -6.25E-01 | 9.21E-02 | 2.71E-01 | 2.40E-01 | 1.51E-05 | Yes | 4930445E18Rik | upstream_gene_variant | MODIFIER |
| 15 | 45486518 | AX-170011570 | 0 | C | G | 0.035 | -5.17E-01 | 9.46E-02 | 7.15E-01 | 7.31E-01 | 1.56E-05 | Yes | | intergenic_variant | MODIFIER |
| 15 | 30288512 | AX-170486699 | 1 | C | T | 0.027 | -6.25E-01 | 9.08E-02 | 2.50E-01 | 2.18E-01 | 1.57E-05 | Yes | Ctnnd2 | intron_variant | MODIFIER |
| 15 | 52728620 | AX-97829699 | 1 | G | C | 0.036 | -5.11E-01 | 1.00E-01 | 9.78E-01 | 9.90E-01 | 1.59E-05 | Yes | Med30 | intron_variant | MODIFIER |
| 15 | 54191048 | AX-170011725 | 1 | G | A | 0.036 | -5.10E-01 | 9.98E-02 | 9.61E-01 | 9.73E-01 | 1.61E-05 | Yes | | intergenic_variant | MODIFIER |
| 15 | 29587803 | AX-170303301 | 0 | T | C | 0.026 | -5.57E-01 | 9.01E-02 | 4.12E-01 | 3.97E-01 | 1.65E-05 | Yes | | intergenic_variant | MODIFIER |
| 15 | 26405247 | AX-170435601 | 1 | G | A | 0.027 | -6.29E-01 | 9.01E-02 | 2.32E-01 | 1.99E-01 | 1.66E-05 | Yes | MAR11 | intron_variant | MODIFIER |
| 15 | 22989959 | AX-170011226 | 2 | A | T | 0.027 | -5.97E-01 | 9.61E-02 | 4.00E-01 | 3.80E-01 | 1.67E-05 | Yes | Cdh18 | intron_variant | MODIFIER |
| 15 | 45274539 | AX-170093231 | 2 | A | G | 0.027 | -5.41E-01 | 9.60E-02 | 6.14E-01 | 6.14E-01 | 1.69E-05 | Yes | | intergenic_variant | MODIFIER |
| 15 | 54179916 | AX-97791257 | 1 | C | T | 0.036 | -5.07E-01 | 1.01E-01 | 9.98E-01 | 1.01E+00 | 1.90E-05 | Yes | | intergenic_variant | MODIFIER |
| 15 | 30301234 | AX-170390665 | 0 | A | G | 0.035 | -5.56E-01 | 8.98E-02 | 4.00E-01 | 3.87E-01 | 1.94E-05 | Yes | Ctnnd2 | intron_variant | MODIFIER |
| 15 | 24814954 | AX-170212396 | 0 | A | G | 0.026 | -5.64E-01 | 9.11E-02 | 3.93E-01 | 4.00E-01 | 2.06E-05 | Yes | | intergenic_variant | MODIFIER |
| 15 | 30344874 | AX-170196941 | 0 | A | G | 0.035 | -5.68E-01 | 9.33E-02 | 4.24E-01 | 4.12E-01 | 2.09E-05 | Yes | Ctnnd2 | intron_variant | MODIFIER |
| 15 | 38895404 | AX-170113116 | 0 | A | G | 0.044 | -5.07E-01 | 9.28E-02 | 6.77E-01 | 6.84E-01 | 2.09E-05 | Yes | | intergenic_variant | MODIFIER |
| 15 | 54417917 | AX-97807678 | 1 | A | G | 0.036 | -5.05E-01 | 1.00E-01 | 9.66E-01 | 9.80E-01 | 2.10E-05 | Yes | Colec10 | intron_variant | MODIFIER |
| 15 | 25397943 | AX-170163309 | 2 | G | A | 0.027 | -6.06E-01 | 9.29E-02 | 3.09E-01 | 2.85E-01 | 2.12E-05 | Yes | Basp1 | intron_variant | MODIFIER |
| 15 | 53498396 | AX-170257909 | 1 | C | T | 0.036 | -5.10E-01 | 1.00E-01 | 9.31E-01 | 9.41E-01 | 2.13E-05 | Yes | Samd12 | intron_variant | MODIFIER |
| 15 | 51781712 | AX-170130450 | 1 | A | T | 0.036 | -5.08E-01 | 1.01E-01 | 9.68E-01 | 9.80E-01 | 2.18E-05 | Yes | Eif3h | downstream_gene_variant | MODIFIER |
| 15 | 25117421 | AX-170011271 | 2 | C | A | 0.027 | -6.06E-01 | 9.32E-02 | 3.12E-01 | 2.88E-01 | 2.22E-05 | Yes | Gm35996 | intron_variant,non_coding_transcript_variant | MODIFIER |
| 15 | 43899252 | AX-170011546 | 0 | A | G | 0.035 | -4.86E-01 | 9.39E-02 | 8.37E-01 | 8.43E-01 | 2.38E-05 | Yes | | intergenic_variant | MODIFIER |
| 15 | 53675816 | AX-97829478 | 1 | C | T | 0.036 | -5.07E-01 | 1.01E-01 | 9.32E-01 | 9.43E-01 | 2.45E-05 | Yes | Samd12 | intron_variant | MODIFIER |
| 15 | 45718521 | AX-170399537 | 0 | G | A | 0.035 | -5.12E-01 | 9.98E-02 | 8.44E-01 | 8.56E-01 | 2.69E-05 | Yes | | intergenic_variant | MODIFIER |
| 15 | 51782756 | AX-170055531 | 1 | C | T | 0.036 | -5.10E-01 | 1.02E-01 | 9.30E-01 | 9.40E-01 | 2.79E-05 | Yes | Eif3h | downstream_gene_variant | MODIFIER |
| 15 | 52614510 | AX-170011703 | 1 | G | A | 0.036 | -5.05E-01 | 1.00E-01 | 8.96E-01 | 9.07E-01 | 2.86E-05 | Yes | | intergenic_variant | MODIFIER |
| 15 | 53550340 | AX-170261548 | 1 | C | T | 0.036 | -5.05E-01 | 1.00E-01 | 9.12E-01 | 9.22E-01 | 2.88E-05 | Yes | Samd12 | intron_variant | MODIFIER |
| 15 | 29834342 | AX-170011347 | 0 | A | C | 0.026 | -5.84E-01 | 9.82E-02 | 4.44E-01 | 4.31E-01 | 2.96E-05 | Yes | | intergenic_variant | MODIFIER |
| 15 | 31035477 | AX-170413296 | 0 | T | C | 0.026 | -5.84E-01 | 9.82E-02 | 4.44E-01 | 4.31E-01 | 2.96E-05 | Yes | 4930430F21Rik | upstream_gene_variant | MODIFIER |
| 15 | 32463349 | AX-170256922 | 0 | T | A | 0.026 | -5.84E-01 | 9.82E-02 | 4.44E-01 | 4.31E-01 | 2.96E-05 | Yes | Sema5a | intron_variant | MODIFIER |
| 15 | 25172828 | AX-170458501 | 2 | G | A | 0.027 | -6.21E-01 | 9.07E-02 | 2.34E-01 | 2.02E-01 | 2.98E-05 | Yes | 9230109A22Rik | intron_variant,non_coding_transcript_variant | MODIFIER |
| 15 | 52502949 | AX-170172047 | 1 | C | T | 0.027 | -5.03E-01 | 9.73E-02 | 7.97E-01 | 8.10E-01 | 3.00E-05 | Yes | Gm23267 | downstream_gene_variant | MODIFIER |
| 15 | 52857807 | AX-170011701 | 1 | G | T | 0.036 | -5.08E-01 | 1.01E-01 | 8.91E-01 | 8.99E-01 | 3.06E-05 | Yes | | intergenic_variant | MODIFIER |
| 15 | 30964465 | AX-170446710 | 0 | T | A | 0.026 | -5.72E-01 | 1.00E-01 | 5.19E-01 | 5.24E-01 | 3.44E-05 | Yes | Ctnnd2 | intron_variant | MODIFIER |
| 15 | 41510072 | AX-170170259 | 0 | G | A | 0.044 | -5.19E-01 | 9.26E-02 | 5.51E-01 | 5.48E-01 | 3.52E-05 | Yes | Oxr1 | intron_variant | MODIFIER |
| 15 | 27405270 | AX-170261973 | 0 | C | T | 0.035 | -5.32E-01 | 9.82E-02 | 6.28E-01 | 6.27E-01 | 3.74E-05 | Yes | | intergenic_variant | MODIFIER |
| 15 | 30863808 | AX-170055146 | 0 | C | T | 0.035 | -5.32E-01 | 9.82E-02 | 6.28E-01 | 6.27E-01 | 3.74E-05 | Yes | Ctnnd2 | intron_variant | MODIFIER |
| 15 | 30869888 | AX-170320638 | 0 | C | T | 0.035 | -5.32E-01 | 9.82E-02 | 6.28E-01 | 6.27E-01 | 3.74E-05 | Yes | Ctnnd2 | intron_variant | MODIFIER |
| 15 | 54159986 | AX-170011726 | 1 | G | T | 0.045 | -4.62E-01 | 9.89E-02 | 1.21E+00 | 1.23E+00 | 3.74E-05 | Yes | | intergenic_variant | MODIFIER |
| 15 | 23250016 | AX-97833833 | 2 | A | G | 0.027 | -5.28E-01 | 9.19E-02 | 4.91E-01 | 4.82E-01 | 3.86E-05 | Yes | Cdh18 | intron_variant,non_coding_transcript_variant | MODIFIER |
| 15 | 42787281 | AX-170011524 | 1 | A | T | 0.036 | -5.19E-01 | 1.00E-01 | 7.54E-01 | 7.55E-01 | 3.91E-05 | Yes | | intergenic_variant | MODIFIER |
| 15 | 26560297 | AX-170011303 | 0 | G | A | 0.035 | -5.36E-01 | 9.90E-02 | 6.27E-01 | 6.26E-01 | 3.95E-05 | Yes | Fbxl7 | intron_variant | MODIFIER |
| 15 | 26214336 | AX-170011286 | 0 | T | C | 0.026 | -5.94E-01 | 9.81E-02 | 3.91E-01 | 3.71E-01 | 4.03E-05 | Yes | | intergenic_variant | MODIFIER |
| 15 | 25035168 | AX-170011270 | 0 | G | A | 0.035 | -5.29E-01 | 9.83E-02 | 6.39E-01 | 6.38E-01 | 4.10E-05 | Yes | | intergenic_variant | MODIFIER |
| 15 | 43154685 | AX-170252752 | 2 | C | T | 0.036 | -5.15E-01 | 1.00E-01 | 7.80E-01 | 7.80E-01 | 4.11E-05 | Yes | Rspo2 | intron_variant | MODIFIER |
| 15 | 41778534 | AX-170011512 | 0 | G | A | 0.035 | -5.29E-01 | 9.88E-02 | 6.50E-01 | 6.45E-01 | 4.15E-05 | Yes | Oxr1 | intron_variant | MODIFIER |
| 15 | 41814974 | AX-170165810 | 0 | G | A | 0.035 | -5.29E-01 | 9.88E-02 | 6.50E-01 | 6.45E-01 | 4.15E-05 | Yes | Oxr1 | intron_variant | MODIFIER |
| 15 | 41839377 | AX-170011513 | 0 | G | A | 0.035 | -5.29E-01 | 9.88E-02 | 6.50E-01 | 6.45E-01 | 4.15E-05 | Yes | Oxr1 | intron_variant | MODIFIER |
| 15 | 41846839 | AX-170421503 | 0 | A | G | 0.035 | -5.29E-01 | 9.88E-02 | 6.50E-01 | 6.45E-01 | 4.15E-05 | Yes | Oxr1 | intron_variant | MODIFIER |
| 15 | 41861444 | AX-170323765 | 0 | A | C | 0.035 | -5.29E-01 | 9.88E-02 | 6.50E-01 | 6.45E-01 | 4.15E-05 | Yes | Oxr1 | downstream_gene_variant | MODIFIER |
| 15 | 41872025 | AX-170116333 | 0 | C | T | 0.035 | -5.29E-01 | 9.88E-02 | 6.50E-01 | 6.45E-01 | 4.15E-05 | Yes | Abra | upstream_gene_variant | MODIFIER |
| 15 | 41872548 | AX-170055355 | 0 | A | G | 0.035 | -5.29E-01 | 9.88E-02 | 6.50E-01 | 6.45E-01 | 4.15E-05 | Yes | Abra | upstream_gene_variant | MODIFIER |
| 15 | 41872946 | AX-170304742 | 0 | T | C | 0.035 | -5.29E-01 | 9.88E-02 | 6.50E-01 | 6.45E-01 | 4.15E-05 | Yes | Abra | upstream_gene_variant | MODIFIER |
| 15 | 41874322 | AX-170253662 | 0 | G | A | 0.035 | -5.29E-01 | 9.88E-02 | 6.50E-01 | 6.45E-01 | 4.15E-05 | Yes | Abra | upstream_gene_variant | MODIFIER |
| 15 | 41874376 | AX-170293114 | 0 | C | G | 0.035 | -5.29E-01 | 9.88E-02 | 6.50E-01 | 6.45E-01 | 4.15E-05 | Yes | Abra | upstream_gene_variant | MODIFIER |

| | | | | | | | | | | | | | | | |
|---|---|---|---|---|---|---|---|---|---|---|---|---|---|---|---|
| 15 | 41903394 | AX-170319490 | 0 | G | A | 0.035 | -5.29E-01 | 9.88E-02 | 6.50E-01 | 6.45E-01 | 4.15E-05 | Yes | | intergenic_variant | MODIFIER |
| 15 | 41920642 | AX-170011511 | 0 | C | T | 0.035 | -5.29E-01 | 9.88E-02 | 6.50E-01 | 6.45E-01 | 4.15E-05 | Yes | Gm5469 | upstream_gene_variant | MODIFIER |
| 15 | 41947897 | AX-170476080 | 0 | A | G | 0.035 | -5.29E-01 | 9.88E-02 | 6.50E-01 | 6.45E-01 | 4.15E-05 | Yes | Gm24751 | upstream_gene_variant | MODIFIER |
| 15 | 41963994 | AX-170394888 | 0 | A | G | 0.035 | -5.29E-01 | 9.88E-02 | 6.50E-01 | 6.45E-01 | 4.15E-05 | Yes | | intergenic_variant | MODIFIER |
| 15 | 41976469 | AX-97829502 | 0 | C | A | 0.035 | -5.29E-01 | 9.88E-02 | 6.50E-01 | 6.45E-01 | 4.15E-05 | Yes | | intergenic_variant | MODIFIER |
| 15 | 41976537 | AX-170376528 | 0 | T | C | 0.035 | -5.29E-01 | 9.88E-02 | 6.50E-01 | 6.45E-01 | 4.15E-05 | Yes | | intergenic_variant | MODIFIER |
| 15 | 41994109 | AX-170516084 | 0 | G | T | 0.035 | -5.29E-01 | 9.88E-02 | 6.50E-01 | 6.45E-01 | 4.15E-05 | Yes | | intergenic_variant | MODIFIER |
| 15 | 42031054 | AX-97805013 | 0 | A | C | 0.035 | -5.29E-01 | 9.88E-02 | 6.50E-01 | 6.45E-01 | 4.15E-05 | Yes | Gm5470 | downstream_gene_variant | MODIFIER |
| 15 | 42105905 | AX-170364661 | 0 | T | C | 0.035 | -5.29E-01 | 9.88E-02 | 6.50E-01 | 6.45E-01 | 4.15E-05 | Yes | | intergenic_variant | MODIFIER |
| 15 | 42189684 | AX-170102454 | 0 | C | T | 0.035 | -5.29E-01 | 9.88E-02 | 6.50E-01 | 6.45E-01 | 4.15E-05 | Yes | | intergenic_variant | MODIFIER |
| 15 | 42206586 | AX-170210550 | 0 | T | A | 0.035 | -5.29E-01 | 9.88E-02 | 6.50E-01 | 6.45E-01 | 4.15E-05 | Yes | | intergenic_variant | MODIFIER |
| 15 | 42246780 | AX-97829501 | 0 | G | A | 0.035 | -5.29E-01 | 9.88E-02 | 6.50E-01 | 6.45E-01 | 4.15E-05 | Yes | | intergenic_variant | MODIFIER |
| 15 | 42295010 | AX-170219992 | 0 | C | T | 0.035 | -5.29E-01 | 9.88E-02 | 6.50E-01 | 6.45E-01 | 4.15E-05 | Yes | | intergenic_variant | MODIFIER |
| 15 | 42295192 | AX-170286887 | 0 | A | G | 0.035 | -5.29E-01 | 9.88E-02 | 6.50E-01 | 6.45E-01 | 4.15E-05 | Yes | | intergenic_variant | MODIFIER |
| 15 | 42295511 | AX-97829181 | 0 | A | G | 0.035 | -5.29E-01 | 9.88E-02 | 6.50E-01 | 6.45E-01 | 4.15E-05 | Yes | | intergenic_variant | MODIFIER |
| 15 | 42299400 | AX-170213756 | 0 | G | A | 0.035 | -5.29E-01 | 9.88E-02 | 6.50E-01 | 6.45E-01 | 4.15E-05 | Yes | | intergenic_variant | MODIFIER |
| 15 | 42300559 | AX-170492529 | 0 | A | T | 0.035 | -5.29E-01 | 9.88E-02 | 6.50E-01 | 6.45E-01 | 4.15E-05 | Yes | | intergenic_variant | MODIFIER |
| 15 | 42300794 | AX-170516113 | 0 | G | T | 0.035 | -5.29E-01 | 9.88E-02 | 6.50E-01 | 6.45E-01 | 4.15E-05 | Yes | | intergenic_variant | MODIFIER |
| 15 | 42380132 | AX-170011519 | 0 | C | T | 0.035 | -5.29E-01 | 9.88E-02 | 6.50E-01 | 6.45E-01 | 4.15E-05 | Yes | | intergenic_variant | MODIFIER |
| 15 | 42411484 | AX-170346369 | 0 | G | A | 0.035 | -5.29E-01 | 9.88E-02 | 6.50E-01 | 6.45E-01 | 4.15E-05 | Yes | | intergenic_variant | MODIFIER |
| 15 | 42426344 | AX-170312602 | 0 | C | T | 0.035 | -5.29E-01 | 9.88E-02 | 6.50E-01 | 6.45E-01 | 4.15E-05 | Yes | Angpt1 | 3_prime_UTR_variant | MODIFIER |
| 15 | 42435213 | AX-170420434 | 0 | G | A | 0.035 | -5.29E-01 | 9.88E-02 | 6.50E-01 | 6.45E-01 | 4.15E-05 | Yes | Angpt1 | intron_variant | MODIFIER |
| 15 | 42435476 | AX-170146772 | 0 | C | T | 0.035 | -5.29E-01 | 9.88E-02 | 6.50E-01 | 6.45E-01 | 4.15E-05 | Yes | Angpt1 | intron_variant | MODIFIER |
| 15 | 42436000 | AX-97837239 | 0 | T | C | 0.035 | -5.29E-01 | 9.88E-02 | 6.50E-01 | 6.45E-01 | 4.15E-05 | Yes | Angpt1 | intron_variant | MODIFIER |
| 15 | 42471397 | AX-170226956 | 0 | C | A | 0.035 | -5.29E-01 | 9.88E-02 | 6.50E-01 | 6.45E-01 | 4.15E-05 | Yes | Angpt1 | intron_variant | MODIFIER |
| 15 | 42474832 | AX-170258645 | 0 | G | A | 0.035 | -5.29E-01 | 9.88E-02 | 6.50E-01 | 6.45E-01 | 4.15E-05 | Yes | Angpt1 | intron_variant | MODIFIER |
| 15 | 42475852 | AX-170219862 | 0 | T | C | 0.035 | -5.29E-01 | 9.88E-02 | 6.50E-01 | 6.45E-01 | 4.15E-05 | Yes | Angpt1 | intron_variant | MODIFIER |
| 15 | 42475965 | AX-170333778 | 0 | C | T | 0.035 | -5.29E-01 | 9.88E-02 | 6.50E-01 | 6.45E-01 | 4.15E-05 | Yes | Angpt1 | intron_variant | MODIFIER |
| 15 | 42505228 | AX-170230994 | 0 | A | C | 0.035 | -5.29E-01 | 9.88E-02 | 6.50E-01 | 6.45E-01 | 4.15E-05 | Yes | Angpt1 | intron_variant | MODIFIER |
| 15 | 42505259 | AX-170098156 | 0 | T | C | 0.035 | -5.29E-01 | 9.88E-02 | 6.50E-01 | 6.45E-01 | 4.15E-05 | Yes | Angpt1 | intron_variant | MODIFIER |
| 15 | 42550169 | AX-170284944 | 0 | T | A | 0.035 | -5.29E-01 | 9.88E-02 | 6.50E-01 | 6.45E-01 | 4.15E-05 | Yes | Angpt1 | intron_variant | MODIFIER |
| 15 | 42554402 | AX-170466507 | 0 | C | T | 0.035 | -5.29E-01 | 9.88E-02 | 6.50E-01 | 6.45E-01 | 4.15E-05 | Yes | Angpt1 | intron_variant | MODIFIER |
| 15 | 42615965 | AX-170432652 | 0 | A | G | 0.035 | -5.29E-01 | 9.88E-02 | 6.50E-01 | 6.45E-01 | 4.15E-05 | Yes | Angpt1 | intron_variant | MODIFIER |
| 15 | 42621582 | AX-170465810 | 0 | C | T | 0.035 | -5.29E-01 | 9.88E-02 | 6.50E-01 | 6.45E-01 | 4.15E-05 | Yes | Angpt1 | intron_variant | MODIFIER |
| 15 | 42624579 | AX-170473357 | 0 | G | A | 0.035 | -5.29E-01 | 9.88E-02 | 6.50E-01 | 6.45E-01 | 4.15E-05 | Yes | Angpt1 | intron_variant | MODIFIER |
| 15 | 42675393 | AX-97789556 | 0 | C | T | 0.035 | -5.29E-01 | 9.88E-02 | 6.50E-01 | 6.45E-01 | 4.15E-05 | Yes | Angpt1 | intron_variant | MODIFIER |
| 15 | 42690367 | AX-170227723 | 0 | A | C | 0.035 | -5.29E-01 | 9.88E-02 | 6.50E-01 | 6.45E-01 | 4.15E-05 | Yes | Gm17473 | intron_variant,non_coding_transcript_variant | MODIFIER |
| 15 | 42703500 | AX-170208004 | 0 | T | A | 0.035 | -5.29E-01 | 9.88E-02 | 6.50E-01 | 6.45E-01 | 4.15E-05 | Yes | Gm17473 | non_coding_transcript_exon_variant | MODIFIER |
| 15 | 42718309 | AX-170137171 | 0 | A | G | 0.035 | -5.29E-01 | 9.88E-02 | 6.50E-01 | 6.45E-01 | 4.15E-05 | Yes | | intergenic_variant | MODIFIER |
| 15 | 42885273 | AX-170011531 | 0 | G | C | 0.035 | -5.29E-01 | 9.88E-02 | 6.50E-01 | 6.45E-01 | 4.15E-05 | Yes | | intergenic_variant | MODIFIER |
| 15 | 43150784 | AX-97797125 | 0 | T | C | 0.035 | -5.29E-01 | 9.88E-02 | 6.50E-01 | 6.45E-01 | 4.15E-05 | Yes | Rspo2 | intron_variant | MODIFIER |
| 15 | 43202680 | AX-170301302 | 0 | G | A | 0.035 | -5.29E-01 | 9.88E-02 | 6.50E-01 | 6.45E-01 | 4.15E-05 | Yes | | regulatory_region_variant | MODIFIER |
| 15 | 43230340 | AX-97843063 | 0 | C | T | 0.035 | -5.29E-01 | 9.88E-02 | 6.50E-01 | 6.45E-01 | 4.15E-05 | Yes | | intergenic_variant | MODIFIER |
| 15 | 43246109 | AX-170267407 | 0 | A | T | 0.035 | -5.29E-01 | 9.88E-02 | 6.50E-01 | 6.45E-01 | 4.15E-05 | Yes | Eif3e | downstream_gene_variant | MODIFIER |
| 15 | 43254541 | AX-170108920 | 0 | C | T | 0.035 | -5.29E-01 | 9.88E-02 | 6.50E-01 | 6.45E-01 | 4.15E-05 | Yes | Eif3e | intron_variant | MODIFIER |
| 15 | 43268052 | AX-170431599 | 0 | C | T | 0.035 | -5.29E-01 | 9.88E-02 | 6.50E-01 | 6.45E-01 | 4.15E-05 | Yes | Eif3e | intron_variant | MODIFIER |
| 15 | 43272361 | AX-170087837 | 0 | A | C | 0.035 | -5.29E-01 | 9.88E-02 | 6.50E-01 | 6.45E-01 | 4.15E-05 | Yes | Eif3e | intron_variant | MODIFIER |
| 15 | 43354418 | AX-170011540 | 0 | G | C | 0.035 | -5.29E-01 | 9.88E-02 | 6.50E-01 | 6.45E-01 | 4.15E-05 | Yes | | intergenic_variant | MODIFIER |
| 15 | 43408252 | AX-170445629 | 0 | T | C | 0.035 | -5.29E-01 | 9.88E-02 | 6.50E-01 | 6.45E-01 | 4.15E-05 | Yes | | intergenic_variant | MODIFIER |
| 15 | 43415481 | AX-170011538 | 0 | G | A | 0.035 | -5.29E-01 | 9.88E-02 | 6.50E-01 | 6.45E-01 | 4.15E-05 | Yes | | intergenic_variant | MODIFIER |
| 15 | 43417250 | AX-170011539 | 0 | G | T | 0.035 | -5.29E-01 | 9.88E-02 | 6.50E-01 | 6.45E-01 | 4.15E-05 | Yes | | intergenic_variant | MODIFIER |
| 15 | 43494735 | AX-170199734 | 0 | G | T | 0.035 | -5.29E-01 | 9.88E-02 | 6.50E-01 | 6.45E-01 | 4.15E-05 | Yes | Emc2 | splice_region_variant,intron_variant | LOW |
| 15 | 43530171 | AX-170093228 | 0 | T | C | 0.035 | -5.29E-01 | 9.88E-02 | 6.50E-01 | 6.45E-01 | 4.15E-05 | Yes | Emc2 | downstream_gene_variant | MODIFIER |
| 15 | 43536309 | AX-170351212 | 0 | T | C | 0.035 | -5.29E-01 | 9.88E-02 | 6.50E-01 | 6.45E-01 | 4.15E-05 | Yes | | intergenic_variant | MODIFIER |

| | | | | | | | | | | | | | | | |
|---|---|---|---|---|---|---|---|---|---|---|---|---|---|---|---|
| 15 | 43581621 | AX-170251291 | 0 | C | G | 0.035 | -5.29E-01 | 9.88E-02 | 6.50E-01 | 6.45E-01 | 4.15E-05 | Yes | | intergenic_variant | MODIFIER |
| 15 | 43651467 | AX-170506315 | 0 | A | C | 0.035 | -5.29E-01 | 9.88E-02 | 6.50E-01 | 6.45E-01 | 4.15E-05 | Yes | | regulatory_region_variant | MODIFIER |
| 15 | 43664031 | AX-170499858 | 0 | G | A | 0.035 | -5.29E-01 | 9.88E-02 | 6.50E-01 | 6.45E-01 | 4.15E-05 | Yes | | intergenic_variant | MODIFIER |
| 15 | 43675701 | AX-170465209 | 0 | A | C | 0.035 | -5.29E-01 | 9.88E-02 | 6.50E-01 | 6.45E-01 | 4.15E-05 | Yes | | intergenic_variant | MODIFIER |
| 15 | 43697019 | AX-170252755 | 0 | T | G | 0.035 | -5.29E-01 | 9.88E-02 | 6.50E-01 | 6.45E-01 | 4.15E-05 | Yes | | intergenic_variant | MODIFIER |
| 15 | 43728974 | AX-170109875 | 0 | G | C | 0.035 | -5.29E-01 | 9.88E-02 | 6.50E-01 | 6.45E-01 | 4.15E-05 | Yes | | intergenic_variant | MODIFIER |
| 15 | 43731836 | AX-170327308 | 0 | T | C | 0.035 | -5.29E-01 | 9.88E-02 | 6.50E-01 | 6.45E-01 | 4.15E-05 | Yes | | intergenic_variant | MODIFIER |
| 15 | 43775396 | AX-170271893 | 0 | T | C | 0.035 | -5.29E-01 | 9.88E-02 | 6.50E-01 | 6.45E-01 | 4.15E-05 | Yes | | intergenic_variant | MODIFIER |
| 15 | 43808565 | AX-170525652 | 0 | T | G | 0.035 | -5.29E-01 | 9.88E-02 | 6.50E-01 | 6.45E-01 | 4.15E-05 | Yes | Gm2140 | non_coding_transcript_exon_variant | MODIFIER |
| 15 | 43848861 | AX-170055394 | 0 | C | T | 0.035 | -5.29E-01 | 9.88E-02 | 6.50E-01 | 6.45E-01 | 4.15E-05 | Yes | | intergenic_variant | MODIFIER |
| 15 | 43848999 | AX-97839707 | 0 | G | A | 0.035 | -5.29E-01 | 9.88E-02 | 6.50E-01 | 6.45E-01 | 4.15E-05 | Yes | | intergenic_variant | MODIFIER |
| 15 | 43851326 | AX-170255762 | 0 | C | T | 0.035 | -5.29E-01 | 9.88E-02 | 6.50E-01 | 6.45E-01 | 4.15E-05 | Yes | | intergenic_variant | MODIFIER |
| 15 | 43851831 | AX-170445595 | 0 | T | C | 0.035 | -5.29E-01 | 9.88E-02 | 6.50E-01 | 6.45E-01 | 4.15E-05 | Yes | | intergenic_variant | MODIFIER |
| 15 | 43853065 | AX-170011545 | 0 | G | A | 0.035 | -5.29E-01 | 9.88E-02 | 6.50E-01 | 6.45E-01 | 4.15E-05 | Yes | | intergenic_variant | MODIFIER |
| 15 | 43856587 | AX-170011541 | 0 | G | C | 0.035 | -5.29E-01 | 9.88E-02 | 6.50E-01 | 6.45E-01 | 4.15E-05 | Yes | | intergenic_variant | MODIFIER |
| 15 | 43890909 | AX-170011543 | 0 | G | T | 0.035 | -5.29E-01 | 9.88E-02 | 6.50E-01 | 6.45E-01 | 4.15E-05 | Yes | | intergenic_variant | MODIFIER |
| 15 | 43895069 | AX-170363152 | 0 | A | C | 0.035 | -5.29E-01 | 9.88E-02 | 6.50E-01 | 6.45E-01 | 4.15E-05 | Yes | | intergenic_variant | MODIFIER |
| 15 | 43914629 | AX-170534268 | 0 | T | C | 0.035 | -5.29E-01 | 9.88E-02 | 6.50E-01 | 6.45E-01 | 4.15E-05 | Yes | | intergenic_variant | MODIFIER |
| 15 | 43915045 | AX-170431848 | 0 | T | A | 0.035 | -5.29E-01 | 9.88E-02 | 6.50E-01 | 6.45E-01 | 4.15E-05 | Yes | | intergenic_variant | MODIFIER |
| 15 | 43915086 | AX-170370889 | 0 | A | G | 0.035 | -5.29E-01 | 9.88E-02 | 6.50E-01 | 6.45E-01 | 4.15E-05 | Yes | | intergenic_variant | MODIFIER |
| 15 | 43953318 | AX-170108958 | 0 | A | C | 0.035 | -5.29E-01 | 9.88E-02 | 6.50E-01 | 6.45E-01 | 4.15E-05 | Yes | | intergenic_variant | MODIFIER |
| 15 | 43953518 | AX-170486390 | 0 | A | T | 0.035 | -5.29E-01 | 9.88E-02 | 6.50E-01 | 6.45E-01 | 4.15E-05 | Yes | | intergenic_variant | MODIFIER |
| 15 | 43954175 | AX-170411757 | 0 | G | A | 0.035 | -5.29E-01 | 9.88E-02 | 6.50E-01 | 6.45E-01 | 4.15E-05 | Yes | | intergenic_variant | MODIFIER |
| 15 | 43975036 | AX-170087839 | 0 | A | C | 0.035 | -5.29E-01 | 9.88E-02 | 6.50E-01 | 6.45E-01 | 4.15E-05 | Yes | | intergenic_variant | MODIFIER |
| 15 | 43978320 | AX-170524715 | 0 | C | A | 0.035 | -5.29E-01 | 9.88E-02 | 6.50E-01 | 6.45E-01 | 4.15E-05 | Yes | | intergenic_variant | MODIFIER |
| 15 | 43982226 | AX-170472434 | 0 | G | A | 0.035 | -5.29E-01 | 9.88E-02 | 6.50E-01 | 6.45E-01 | 4.15E-05 | Yes | | intergenic_variant | MODIFIER |
| 15 | 44056764 | AX-170055397 | 0 | A | G | 0.035 | -5.29E-01 | 9.88E-02 | 6.50E-01 | 6.45E-01 | 4.15E-05 | Yes | | intergenic_variant | MODIFIER |
| 15 | 44056888 | AX-170055399 | 0 | G | A | 0.035 | -5.29E-01 | 9.88E-02 | 6.50E-01 | 6.45E-01 | 4.15E-05 | Yes | | intergenic_variant | MODIFIER |
| 15 | 44069080 | AX-170211037 | 0 | A | T | 0.035 | -5.29E-01 | 9.88E-02 | 6.50E-01 | 6.45E-01 | 4.15E-05 | Yes | | intergenic_variant | MODIFIER |
| 15 | 44071350 | AX-170358981 | 0 | C | T | 0.035 | -5.29E-01 | 9.88E-02 | 6.50E-01 | 6.45E-01 | 4.15E-05 | Yes | | intergenic_variant | MODIFIER |
| 15 | 44076966 | AX-170130226 | 0 | A | C | 0.035 | -5.29E-01 | 9.88E-02 | 6.50E-01 | 6.45E-01 | 4.15E-05 | Yes | | intergenic_variant | MODIFIER |
| 15 | 44097276 | AX-97786913 | 0 | C | T | 0.035 | -5.29E-01 | 9.88E-02 | 6.50E-01 | 6.45E-01 | 4.15E-05 | Yes | | intergenic_variant | MODIFIER |
| 15 | 44142065 | AX-97791359 | 0 | G | C | 0.035 | -5.29E-01 | 9.88E-02 | 6.50E-01 | 6.45E-01 | 4.15E-05 | Yes | | intergenic_variant | MODIFIER |
| 15 | 44181477 | AX-170115857 | 0 | G | A | 0.035 | -5.29E-01 | 9.88E-02 | 6.50E-01 | 6.45E-01 | 4.15E-05 | Yes | | intergenic_variant | MODIFIER |
| 15 | 44192099 | AX-170520729 | 0 | T | C | 0.035 | -5.29E-01 | 9.88E-02 | 6.50E-01 | 6.45E-01 | 4.15E-05 | Yes | Trhr | upstream_gene_variant | MODIFIER |
| 15 | 44194999 | AX-170456965 | 0 | G | A | 0.035 | -5.29E-01 | 9.88E-02 | 6.50E-01 | 6.45E-01 | 4.15E-05 | Yes | Trhr | upstream_gene_variant | MODIFIER |
| 15 | 44195817 | AX-170408592 | 0 | A | G | 0.035 | -5.29E-01 | 9.88E-02 | 6.50E-01 | 6.45E-01 | 4.15E-05 | Yes | Trhr | upstream_gene_variant | MODIFIER |
| 15 | 44197922 | AX-170321582 | 0 | A | G | 0.035 | -5.29E-01 | 9.88E-02 | 6.50E-01 | 6.45E-01 | 4.15E-05 | Yes | Trhr | intron_variant | MODIFIER |
| 15 | 44197997 | AX-170482479 | 0 | T | C | 0.035 | -5.29E-01 | 9.88E-02 | 6.50E-01 | 6.45E-01 | 4.15E-05 | Yes | Trhr | intron_variant | MODIFIER |
| 15 | 44226826 | AX-170224414 | 0 | C | G | 0.035 | -5.29E-01 | 9.88E-02 | 6.50E-01 | 6.45E-01 | 4.15E-05 | Yes | Trhr | intron_variant | MODIFIER |
| 15 | 44234785 | AX-170055408 | 0 | G | T | 0.035 | -5.29E-01 | 9.88E-02 | 6.50E-01 | 6.45E-01 | 4.15E-05 | Yes | Trhr | 3_prime_UTR_variant | MODIFIER |
| 15 | 44263358 | AX-170221098 | 0 | G | T | 0.035 | -5.29E-01 | 9.88E-02 | 6.50E-01 | 6.45E-01 | 4.15E-05 | Yes | | intergenic_variant | MODIFIER |
| 15 | 44304275 | AX-170132408 | 0 | T | C | 0.035 | -5.29E-01 | 9.88E-02 | 6.50E-01 | 6.45E-01 | 4.15E-05 | Yes | | intergenic_variant | MODIFIER |
| 15 | 44308111 | AX-170382484 | 0 | C | T | 0.035 | -5.29E-01 | 9.88E-02 | 6.50E-01 | 6.45E-01 | 4.15E-05 | Yes | | intergenic_variant | MODIFIER |
| 15 | 44322357 | AX-170424504 | 0 | A | C | 0.035 | -5.29E-01 | 9.88E-02 | 6.50E-01 | 6.45E-01 | 4.15E-05 | Yes | Gm19235 | upstream_gene_variant | MODIFIER |
| 15 | 44349384 | AX-170011553 | 0 | C | T | 0.035 | -5.29E-01 | 9.88E-02 | 6.50E-01 | 6.45E-01 | 4.15E-05 | Yes | Gm24903 | downstream_gene_variant | MODIFIER |
| 15 | 44351857 | AX-170011551 | 0 | A | G | 0.035 | -5.29E-01 | 9.88E-02 | 6.50E-01 | 6.45E-01 | 4.15E-05 | Yes | | intergenic_variant | MODIFIER |
| 15 | 44379851 | AX-170319644 | 0 | T | C | 0.035 | -5.29E-01 | 9.88E-02 | 6.50E-01 | 6.45E-01 | 4.15E-05 | Yes | Nudcd1 | intron_variant | MODIFIER |
| 15 | 44381084 | AX-170229123 | 0 | A | G | 0.035 | -5.29E-01 | 9.88E-02 | 6.50E-01 | 6.45E-01 | 4.15E-05 | Yes | Nudcd1 | intron_variant | MODIFIER |
| 15 | 44406303 | AX-170011552 | 0 | G | A | 0.035 | -5.29E-01 | 9.88E-02 | 6.50E-01 | 6.45E-01 | 4.15E-05 | Yes | Nudcd1 | intron_variant | MODIFIER |
| 15 | 44406367 | AX-170337979 | 0 | G | A | 0.035 | -5.29E-01 | 9.88E-02 | 6.50E-01 | 6.45E-01 | 4.15E-05 | Yes | Nudcd1 | intron_variant | MODIFIER |
| 15 | 44409860 | AX-170379729 | 0 | A | C | 0.035 | -5.29E-01 | 9.88E-02 | 6.50E-01 | 6.45E-01 | 4.15E-05 | Yes | Nudcd1 | intron_variant | MODIFIER |
| 15 | 44429881 | AX-170302064 | 0 | C | A | 0.035 | -5.29E-01 | 9.88E-02 | 6.50E-01 | 6.45E-01 | 4.15E-05 | Yes | Nudcd1 | upstream_gene_variant | MODIFIER |
| 15 | 44450479 | AX-170055410 | 0 | A | G | 0.035 | -5.29E-01 | 9.88E-02 | 6.50E-01 | 6.45E-01 | 4.15E-05 | Yes | | intergenic_variant | MODIFIER |

| | | | | | | | | | | | | | | | |
|---|---|---|---|---|---|---|---|---|---|---|---|---|---|---|---|
| 15 | 44460136 | AX-170201126 | 0 | C | T | 0.035 | -5.29E-01 | 9.88E-02 | 6.50E-01 | 6.45E-01 | 4.15E-05 | Yes | Pkhd1l1 | intron_variant | MODIFIER |
| 15 | 44460774 | AX-170301902 | 0 | C | T | 0.035 | -5.29E-01 | 9.88E-02 | 6.50E-01 | 6.45E-01 | 4.15E-05 | Yes | Pkhd1l1 | intron_variant | MODIFIER |
| 15 | 44463736 | AX-170527159 | 0 | A | G | 0.035 | -5.29E-01 | 9.88E-02 | 6.50E-01 | 6.45E-01 | 4.15E-05 | Yes | Pkhd1l1 | intron_variant | MODIFIER |
| 15 | 44463962 | AX-170356595 | 0 | A | G | 0.035 | -5.29E-01 | 9.88E-02 | 6.50E-01 | 6.45E-01 | 4.15E-05 | Yes | Pkhd1l1 | intron_variant | MODIFIER |
| 15 | 44494188 | AX-97799513 | 0 | T | C | 0.035 | -5.29E-01 | 9.88E-02 | 6.50E-01 | 6.45E-01 | 4.15E-05 | Yes | Pkhd1l1 | intron_variant | MODIFIER |
| 15 | 44511520 | AX-97796268 | 0 | C | T | 0.035 | -5.29E-01 | 9.88E-02 | 6.50E-01 | 6.45E-01 | 4.15E-05 | Yes | Pkhd1l1 | intron_variant | MODIFIER |
| 15 | 44550955 | AX-170487846 | 0 | A | G | 0.035 | -5.29E-01 | 9.88E-02 | 6.50E-01 | 6.45E-01 | 4.15E-05 | Yes | Pkhd1l1 | intron_variant | MODIFIER |
| 15 | 44558746 | AX-170464817 | 0 | C | A | 0.035 | -5.29E-01 | 9.88E-02 | 6.50E-01 | 6.45E-01 | 4.15E-05 | Yes | Pkhd1l1 | intron_variant | MODIFIER |
| 15 | 44565754 | AX-170259470 | 0 | G | A | 0.035 | -5.29E-01 | 9.88E-02 | 6.50E-01 | 6.45E-01 | 4.15E-05 | Yes | Pkhd1l1 | intron_variant | MODIFIER |
| 15 | 44575307 | AX-170263587 | 0 | A | G | 0.035 | -5.29E-01 | 9.88E-02 | 6.50E-01 | 6.45E-01 | 4.15E-05 | Yes | Pkhd1l1 | intron_variant | MODIFIER |
| 15 | 44577410 | AX-170309080 | 0 | C | A | 0.035 | -5.29E-01 | 9.88E-02 | 6.50E-01 | 6.45E-01 | 4.15E-05 | Yes | Pkhd1l1 | intron_variant | MODIFIER |
| 15 | 44590059 | AX-97829545 | 0 | C | T | 0.035 | -5.29E-01 | 9.88E-02 | 6.50E-01 | 6.45E-01 | 4.15E-05 | Yes | Pkhd1l1 | intron_variant | MODIFIER |
| 15 | 44609104 | AX-97795094 | 0 | T | C | 0.035 | -5.29E-01 | 9.88E-02 | 6.50E-01 | 6.45E-01 | 4.15E-05 | Yes | | intergenic_variant | MODIFIER |
| 15 | 44626246 | AX-170201650 | 0 | A | C | 0.035 | -5.29E-01 | 9.88E-02 | 6.50E-01 | 6.45E-01 | 4.15E-05 | Yes | Ebag9 | intron_variant | MODIFIER |
| 15 | 44636505 | AX-170502937 | 0 | G | C | 0.035 | -5.29E-01 | 9.88E-02 | 6.50E-01 | 6.45E-01 | 4.15E-05 | Yes | Ebag9 | intron_variant | MODIFIER |
| 15 | 44636594 | AX-97805933 | 0 | G | A | 0.035 | -5.29E-01 | 9.88E-02 | 6.50E-01 | 6.45E-01 | 4.15E-05 | Yes | Ebag9 | intron_variant | MODIFIER |
| 15 | 44676506 | AX-170055415 | 0 | G | A | 0.035 | -5.29E-01 | 9.88E-02 | 6.50E-01 | 6.45E-01 | 4.15E-05 | Yes | Sybu | intron_variant | MODIFIER |
| 15 | 44683690 | AX-170443757 | 0 | A | G | 0.035 | -5.29E-01 | 9.88E-02 | 6.50E-01 | 6.45E-01 | 4.15E-05 | Yes | Sybu | intron_variant | MODIFIER |
| 15 | 44683894 | AX-170307620 | 0 | C | T | 0.035 | -5.29E-01 | 9.88E-02 | 6.50E-01 | 6.45E-01 | 4.15E-05 | Yes | Sybu | intron_variant | MODIFIER |
| 15 | 44685141 | AX-97826812 | 0 | A | G | 0.035 | -5.29E-01 | 9.88E-02 | 6.50E-01 | 6.45E-01 | 4.15E-05 | Yes | Sybu | intron_variant | MODIFIER |
| 15 | 44686264 | AX-170398906 | 0 | T | C | 0.035 | -5.29E-01 | 9.88E-02 | 6.50E-01 | 6.45E-01 | 4.15E-05 | Yes | Sybu | intron_variant | MODIFIER |
| 15 | 44709966 | AX-170114052 | 0 | C | T | 0.035 | -5.29E-01 | 9.88E-02 | 6.50E-01 | 6.45E-01 | 4.15E-05 | Yes | Sybu | intron_variant | MODIFIER |
| 15 | 44711336 | AX-97799341 | 0 | C | T | 0.035 | -5.29E-01 | 9.88E-02 | 6.50E-01 | 6.45E-01 | 4.15E-05 | Yes | Sybu | intron_variant | MODIFIER |
| 15 | 44802809 | AX-170341166 | 0 | G | C | 0.035 | -5.29E-01 | 9.88E-02 | 6.50E-01 | 6.45E-01 | 4.15E-05 | Yes | A930017M01Rik | intron_variant,non_coding_transcript_variant | MODIFIER |
| 15 | 44802872 | AX-97787479 | 0 | C | T | 0.035 | -5.29E-01 | 9.88E-02 | 6.50E-01 | 6.45E-01 | 4.15E-05 | Yes | A930017M01Rik | intron_variant,non_coding_transcript_variant | MODIFIER |
| 15 | 44804829 | AX-170505020 | 0 | G | A | 0.035 | -5.29E-01 | 9.88E-02 | 6.50E-01 | 6.45E-01 | 4.15E-05 | Yes | A930017M01Rik | intron_variant,non_coding_transcript_variant | MODIFIER |
| 15 | 44818257 | AX-170201741 | 0 | T | C | 0.035 | -5.29E-01 | 9.88E-02 | 6.50E-01 | 6.45E-01 | 4.15E-05 | Yes | A930017M01Rik | intron_variant,non_coding_transcript_variant | MODIFIER |
| 15 | 44819200 | AX-170517487 | 0 | T | C | 0.035 | -5.29E-01 | 9.88E-02 | 6.50E-01 | 6.45E-01 | 4.15E-05 | Yes | A930017M01Rik | intron_variant,non_coding_transcript_variant | MODIFIER |
| 15 | 44822369 | AX-170437481 | 0 | G | C | 0.035 | -5.29E-01 | 9.88E-02 | 6.50E-01 | 6.45E-01 | 4.15E-05 | Yes | A930017M01Rik | intron_variant,non_coding_transcript_variant | MODIFIER |
| 15 | 44830290 | AX-170236132 | 0 | G | T | 0.035 | -5.29E-01 | 9.88E-02 | 6.50E-01 | 6.45E-01 | 4.15E-05 | Yes | A930017M01Rik | intron_variant,non_coding_transcript_variant | MODIFIER |
| 15 | 44850260 | AX-170397479 | 0 | A | G | 0.035 | -5.29E-01 | 9.88E-02 | 6.50E-01 | 6.45E-01 | 4.15E-05 | Yes | A930017M01Rik | downstream_gene_variant | MODIFIER |
| 15 | 44850392 | AX-97789588 | 0 | G | A | 0.035 | -5.29E-01 | 9.88E-02 | 6.50E-01 | 6.45E-01 | 4.15E-05 | Yes | A930017M01Rik | downstream_gene_variant | MODIFIER |
| 15 | 44873370 | AX-170055417 | 0 | C | G | 0.035 | -5.29E-01 | 9.88E-02 | 6.50E-01 | 6.45E-01 | 4.15E-05 | Yes | A930017M01Rik | intron_variant,non_coding_transcript_variant | MODIFIER |
| 15 | 44883195 | AX-170055419 | 0 | T | C | 0.035 | -5.29E-01 | 9.88E-02 | 6.50E-01 | 6.45E-01 | 4.15E-05 | Yes | A930017M01Rik | non_coding_transcript_exon_variant | MODIFIER |
| 15 | 44897681 | AX-170529376 | 0 | C | T | 0.035 | -5.29E-01 | 9.88E-02 | 6.50E-01 | 6.45E-01 | 4.15E-05 | Yes | | intergenic_variant | MODIFIER |
| 15 | 45021831 | AX-170011556 | 0 | G | A | 0.035 | -5.29E-01 | 9.88E-02 | 6.50E-01 | 6.45E-01 | 4.15E-05 | Yes | | intergenic_variant | MODIFIER |
| 15 | 45069690 | AX-170011563 | 0 | A | G | 0.035 | -5.29E-01 | 9.88E-02 | 6.50E-01 | 6.45E-01 | 4.15E-05 | Yes | | intergenic_variant | MODIFIER |
| 15 | 45071958 | AX-170011561 | 0 | G | T | 0.035 | -5.29E-01 | 9.88E-02 | 6.50E-01 | 6.45E-01 | 4.15E-05 | Yes | | intergenic_variant | MODIFIER |
| 15 | 45082430 | AX-170246912 | 0 | C | A | 0.035 | -5.29E-01 | 9.88E-02 | 6.50E-01 | 6.45E-01 | 4.15E-05 | Yes | | intergenic_variant | MODIFIER |
| 15 | 45160091 | AX-170055420 | 0 | G | T | 0.035 | -5.29E-01 | 9.88E-02 | 6.50E-01 | 6.45E-01 | 4.15E-05 | Yes | | intergenic_variant | MODIFIER |
| 15 | 45162843 | AX-170230662 | 0 | G | C | 0.035 | -5.29E-01 | 9.88E-02 | 6.50E-01 | 6.45E-01 | 4.15E-05 | Yes | | intergenic_variant | MODIFIER |
| 15 | 45166590 | AX-97846384 | 0 | G | T | 0.035 | -5.29E-01 | 9.88E-02 | 6.50E-01 | 6.45E-01 | 4.15E-05 | Yes | | intergenic_variant | MODIFIER |
| 15 | 45183968 | AX-97806595 | 0 | C | G | 0.035 | -5.29E-01 | 9.88E-02 | 6.50E-01 | 6.45E-01 | 4.15E-05 | Yes | | intergenic_variant | MODIFIER |
| 15 | 45184743 | AX-170339835 | 0 | T | C | 0.035 | -5.29E-01 | 9.88E-02 | 6.50E-01 | 6.45E-01 | 4.15E-05 | Yes | | intergenic_variant | MODIFIER |
| 15 | 45223649 | AX-170484601 | 0 | A | G | 0.035 | -5.29E-01 | 9.88E-02 | 6.50E-01 | 6.45E-01 | 4.15E-05 | Yes | | intergenic_variant | MODIFIER |
| 15 | 44003918 | AX-170334375 | 1 | T | C | 0.036 | -5.22E-01 | 9.96E-02 | 7.10E-01 | 7.08E-01 | 4.16E-05 | Yes | | intergenic_variant | MODIFIER |
| 15 | 42241459 | AX-170279610 | 1 | T | C | 0.036 | -5.23E-01 | 1.00E-01 | 7.15E-01 | 7.13E-01 | 4.20E-05 | Yes | | intergenic_variant | MODIFIER |
| 15 | 44305846 | AX-170482901 | 1 | G | A | 0.027 | -5.26E-01 | 1.01E-01 | 7.37E-01 | 7.44E-01 | 4.26E-05 | Yes | | intergenic_variant | MODIFIER |
| 15 | 35873305 | AX-97789713 | 0 | A | C | 0.026 | -5.83E-01 | 9.72E-02 | 4.02E-01 | 3.83E-01 | 4.35E-05 | Yes | Vps13b | intron_variant | MODIFIER |
| 15 | 24625951 | AX-170186775 | 0 | T | G | 0.026 | -5.91E-01 | 9.67E-02 | 3.70E-01 | 3.51E-01 | 4.49E-05 | Yes | | intergenic_variant | MODIFIER |
| 15 | 30568033 | AX-97849567 | 0 | A | G | 0.026 | -5.91E-01 | 9.67E-02 | 3.70E-01 | 3.51E-01 | 4.49E-05 | Yes | Ctnnd2 | intron_variant | MODIFIER |
| 15 | 44198833 | AX-170192328 | 1 | G | A | 0.036 | -5.21E-01 | 9.88E-02 | 6.74E-01 | 6.71E-01 | 4.72E-05 | Yes | Trhr | intron_variant | MODIFIER |
| 15 | 29612195 | AX-170128767 | 1 | C | G | 0.036 | -4.79E-01 | 9.11E-02 | 6.72E-01 | 7.00E-01 | 5.10E-05 | Yes | | intergenic_variant | MODIFIER |
| 15 | 25472405 | AX-97820424 | 0 | T | C | 0.026 | -5.87E-01 | 9.48E-02 | 3.40E-01 | 3.20E-01 | 5.19E-05 | Yes | | intergenic_variant | MODIFIER |

| | | | | | | | | | | | | | | | |
|---|---|---|---|---|---|---|---|---|---|---|---|---|---|---|---|
| 15 | 34194495 | AX-170122493 | 0 | T | C | 0.026 | -5.95E-01 | 9.53E-02 | 3.28E-01 | 3.06E-01 | 5.26E-05 | Yes | | intergenic_variant | MODIFIER |
| 15 | 44753990 | AX-97847471 | 1 | T | C | 0.036 | -5.21E-01 | 9.95E-02 | 6.82E-01 | 6.78E-01 | 5.30E-05 | Yes | Sybu | intron_variant | MODIFIER |
| 15 | 34279827 | AX-97783629 | 0 | A | C | 0.035 | -5.39E-01 | 9.44E-02 | 4.77E-01 | 4.70E-01 | 5.55E-05 | Yes | Laptm4b | intron_variant | MODIFIER |
| 15 | 58092036 | AX-97799063 | 0 | A | G | 0.053 | -4.64E-01 | 9.53E-02 | 8.96E-01 | 9.07E-01 | 5.90E-05 | Yes | Atad2 | downstream_gene_variant | MODIFIER |
| 15 | 58093306 | AX-170113401 | 0 | G | A | 0.053 | -4.64E-01 | 9.53E-02 | 8.96E-01 | 9.07E-01 | 5.90E-05 | Yes | Atad2 | downstream_gene_variant | MODIFIER |
| 15 | 58104929 | AX-170500068 | 0 | G | A | 0.053 | -4.64E-01 | 9.53E-02 | 8.96E-01 | 9.07E-01 | 5.90E-05 | Yes | Atad2 | intron_variant | MODIFIER |
| 15 | 58110721 | AX-170121777 | 0 | C | T | 0.053 | -4.64E-01 | 9.53E-02 | 8.96E-01 | 9.07E-01 | 5.90E-05 | Yes | Atad2 | intron_variant | MODIFIER |
| 15 | 58116939 | AX-97824853 | 0 | C | A | 0.053 | -4.64E-01 | 9.53E-02 | 8.96E-01 | 9.07E-01 | 5.90E-05 | Yes | Atad2 | intron_variant | MODIFIER |
| 15 | 58129631 | AX-170369604 | 0 | A | T | 0.053 | -4.64E-01 | 9.53E-02 | 8.96E-01 | 9.07E-01 | 5.90E-05 | Yes | Atad2 | intron_variant | MODIFIER |
| 15 | 58145003 | AX-170011785 | 0 | T | C | 0.053 | -4.64E-01 | 9.53E-02 | 8.96E-01 | 9.07E-01 | 5.90E-05 | Yes | Wdyhv1 | intron_variant | MODIFIER |
| 15 | 58146845 | AX-170447595 | 0 | C | G | 0.053 | -4.64E-01 | 9.53E-02 | 8.96E-01 | 9.07E-01 | 5.90E-05 | Yes | Wdyhv1 | intron_variant | MODIFIER |
| 15 | 58148041 | AX-170087875 | 0 | T | C | 0.053 | -4.64E-01 | 9.53E-02 | 8.96E-01 | 9.07E-01 | 5.90E-05 | Yes | Wdyhv1 | intron_variant | MODIFIER |
| 15 | 58162348 | AX-97821568 | 0 | C | T | 0.053 | -4.64E-01 | 9.53E-02 | 8.96E-01 | 9.07E-01 | 5.90E-05 | Yes | Wdyhv1 | downstream_gene_variant | MODIFIER |
| 15 | 58162518 | AX-170418421 | 0 | G | C | 0.053 | -4.64E-01 | 9.53E-02 | 8.96E-01 | 9.07E-01 | 5.90E-05 | Yes | Wdyhv1 | downstream_gene_variant | MODIFIER |
| 15 | 58308392 | AX-97797986 | 0 | T | C | 0.053 | -4.64E-01 | 9.53E-02 | 8.96E-01 | 9.07E-01 | 5.90E-05 | Yes | Gm15943 | intron_variant,non_coding_transcript_variant | MODIFIER |
| 15 | 58312462 | AX-170219687 | 0 | A | G | 0.053 | -4.64E-01 | 9.53E-02 | 8.96E-01 | 9.07E-01 | 5.90E-05 | Yes | Klhl38 | downstream_gene_variant | MODIFIER |
| 15 | 58313454 | AX-170177762 | 0 | C | A | 0.053 | -4.64E-01 | 9.53E-02 | 8.96E-01 | 9.07E-01 | 5.90E-05 | Yes | Klhl38 | downstream_gene_variant | MODIFIER |
| 15 | 58313972 | AX-170527388 | 0 | A | G | 0.053 | -4.64E-01 | 9.53E-02 | 8.96E-01 | 9.07E-01 | 5.90E-05 | Yes | Klhl38 | downstream_gene_variant | MODIFIER |
| 15 | 58313995 | AX-97803790 | 0 | G | A | 0.053 | -4.64E-01 | 9.53E-02 | 8.96E-01 | 9.07E-01 | 5.90E-05 | Yes | Klhl38 | downstream_gene_variant | MODIFIER |
| 15 | 58332419 | AX-170219293 | 0 | C | T | 0.053 | -4.64E-01 | 9.53E-02 | 8.96E-01 | 9.07E-01 | 5.90E-05 | Yes | | intergenic_variant | MODIFIER |
| 15 | 58334134 | AX-170200974 | 0 | T | G | 0.053 | -4.64E-01 | 9.53E-02 | 8.96E-01 | 9.07E-01 | 5.90E-05 | Yes | | intergenic_variant | MODIFIER |
| 15 | 58335883 | AX-170093263 | 0 | A | G | 0.053 | -4.64E-01 | 9.53E-02 | 8.96E-01 | 9.07E-01 | 5.90E-05 | Yes | | intergenic_variant | MODIFIER |
| 15 | 58337512 | AX-170446150 | 0 | T | C | 0.053 | -4.64E-01 | 9.53E-02 | 8.96E-01 | 9.07E-01 | 5.90E-05 | Yes | Anxa13 | downstream_gene_variant | MODIFIER |
| 15 | 58362226 | AX-170310526 | 0 | A | G | 0.053 | -4.64E-01 | 9.53E-02 | 8.96E-01 | 9.07E-01 | 5.90E-05 | Yes | Anxa13 | intron_variant,non_coding_transcript_variant | MODIFIER |
| 15 | 58372252 | AX-170222723 | 0 | C | A | 0.053 | -4.64E-01 | 9.53E-02 | 8.96E-01 | 9.07E-01 | 5.90E-05 | Yes | Anxa13 | intron_variant,non_coding_transcript_variant | MODIFIER |
| 15 | 58395954 | AX-97789204 | 0 | T | C | 0.053 | -4.64E-01 | 9.53E-02 | 8.96E-01 | 9.07E-01 | 5.90E-05 | Yes | | regulatory_region_variant | MODIFIER |
| 15 | 58396290 | AX-170055647 | 0 | G | A | 0.053 | -4.64E-01 | 9.53E-02 | 8.96E-01 | 9.07E-01 | 5.90E-05 | Yes | | intergenic_variant | MODIFIER |
| 15 | 58404893 | AX-170534846 | 0 | T | G | 0.053 | -4.64E-01 | 9.53E-02 | 8.96E-01 | 9.07E-01 | 5.90E-05 | Yes | Gm2675 | upstream_gene_variant | MODIFIER |
| 15 | 58410081 | AX-97845511 | 0 | G | A | 0.053 | -4.64E-01 | 9.53E-02 | 8.96E-01 | 9.07E-01 | 5.90E-05 | Yes | Gm2675 | intron_variant,non_coding_transcript_variant | MODIFIER |
| 15 | 58479245 | AX-170093264 | 0 | C | T | 0.053 | -4.64E-01 | 9.53E-02 | 8.96E-01 | 9.07E-01 | 5.90E-05 | Yes | | intergenic_variant | MODIFIER |
| 15 | 58488444 | AX-170467508 | 0 | A | C | 0.053 | -4.64E-01 | 9.53E-02 | 8.96E-01 | 9.07E-01 | 5.90E-05 | Yes | | intergenic_variant | MODIFIER |
| 15 | 58488946 | AX-170175639 | 0 | A | G | 0.053 | -4.64E-01 | 9.53E-02 | 8.96E-01 | 9.07E-01 | 5.90E-05 | Yes | | intergenic_variant | MODIFIER |
| 15 | 58490028 | AX-97827357 | 0 | G | A | 0.053 | -4.64E-01 | 9.53E-02 | 8.96E-01 | 9.07E-01 | 5.90E-05 | Yes | | intergenic_variant | MODIFIER |
| 15 | 58508202 | AX-170279072 | 0 | C | T | 0.053 | -4.64E-01 | 9.53E-02 | 8.96E-01 | 9.07E-01 | 5.90E-05 | Yes | Fer1l6 | upstream_gene_variant | MODIFIER |
| 15 | 58510653 | AX-170312785 | 0 | C | T | 0.053 | -4.64E-01 | 9.53E-02 | 8.96E-01 | 9.07E-01 | 5.90E-05 | Yes | Fer1l6 | intron_variant | MODIFIER |
| 15 | 58511889 | AX-170011788 | 0 | G | T | 0.053 | -4.64E-01 | 9.53E-02 | 8.96E-01 | 9.07E-01 | 5.90E-05 | Yes | Fer1l6 | intron_variant | MODIFIER |
| 15 | 58516092 | AX-97819363 | 0 | T | C | 0.053 | -4.64E-01 | 9.53E-02 | 8.96E-01 | 9.07E-01 | 5.90E-05 | Yes | Fer1l6 | intron_variant | MODIFIER |
| 15 | 58528728 | AX-170295931 | 0 | A | T | 0.053 | -4.64E-01 | 9.53E-02 | 8.96E-01 | 9.07E-01 | 5.90E-05 | Yes | Fer1l6 | intron_variant | MODIFIER |
| 15 | 58534541 | AX-170055649 | 0 | T | C | 0.053 | -4.64E-01 | 9.53E-02 | 8.96E-01 | 9.07E-01 | 5.90E-05 | Yes | Fer1l6 | intron_variant | MODIFIER |
| 15 | 58535848 | AX-170055648 | 0 | G | A | 0.053 | -4.64E-01 | 9.53E-02 | 8.96E-01 | 9.07E-01 | 5.90E-05 | Yes | Fer1l6 | intron_variant | MODIFIER |
| 15 | 58536087 | AX-170161202 | 0 | T | C | 0.053 | -4.64E-01 | 9.53E-02 | 8.96E-01 | 9.07E-01 | 5.90E-05 | Yes | Fer1l6 | intron_variant | MODIFIER |
| 15 | 58538517 | AX-170102602 | 0 | G | C | 0.053 | -4.64E-01 | 9.53E-02 | 8.96E-01 | 9.07E-01 | 5.90E-05 | Yes | Fer1l6 | intron_variant | MODIFIER |
| 15 | 58538615 | AX-170374474 | 0 | T | A | 0.053 | -4.64E-01 | 9.53E-02 | 8.96E-01 | 9.07E-01 | 5.90E-05 | Yes | Fer1l6 | intron_variant | MODIFIER |
| 15 | 58552491 | AX-97830043 | 0 | T | C | 0.053 | -4.64E-01 | 9.53E-02 | 8.96E-01 | 9.07E-01 | 5.90E-05 | Yes | Fer1l6 | intron_variant | MODIFIER |
| 15 | 58553065 | AX-170011792 | 0 | C | T | 0.053 | -4.64E-01 | 9.53E-02 | 8.96E-01 | 9.07E-01 | 5.90E-05 | Yes | Fer1l6 | intron_variant | MODIFIER |
| 15 | 58553602 | AX-170137237 | 0 | G | A | 0.053 | -4.64E-01 | 9.53E-02 | 8.96E-01 | 9.07E-01 | 5.90E-05 | Yes | Fer1l6 | intron_variant | MODIFIER |
| 15 | 58553619 | AX-170154394 | 0 | A | G | 0.053 | -4.64E-01 | 9.53E-02 | 8.96E-01 | 9.07E-01 | 5.90E-05 | Yes | Fer1l6 | intron_variant | MODIFIER |
| 15 | 58554177 | AX-170011790 | 0 | A | G | 0.053 | -4.64E-01 | 9.53E-02 | 8.96E-01 | 9.07E-01 | 5.90E-05 | Yes | Fer1l6 | intron_variant | MODIFIER |
| 15 | 58554467 | AX-170093265 | 0 | G | T | 0.053 | -4.64E-01 | 9.53E-02 | 8.96E-01 | 9.07E-01 | 5.90E-05 | Yes | Fer1l6 | intron_variant | MODIFIER |
| 15 | 60665939 | AX-170531239 | 0 | A | G | 0.061 | -4.12E-01 | 8.92E-02 | 1.14E+00 | 1.18E+00 | 5.91E-05 | Yes | Gm48946 | upstream_gene_variant | MODIFIER |
| 15 | 61248886 | AX-97820853 | 0 | G | A | 0.061 | -4.41E-01 | 9.49E-02 | 1.10E+00 | 1.13E+00 | 5.98E-05 | Yes | | intergenic_variant | MODIFIER |
| 15 | 30913202 | AX-170113000 | 0 | G | A | 0.035 | -5.82E-01 | 9.36E-02 | 3.28E-01 | 3.19E-01 | 6.00E-05 | Yes | Ctnnd2 | intron_variant | MODIFIER |
| 15 | 42162563 | AX-170197491 | 1 | C | A | 0.036 | -5.18E-01 | 9.97E-02 | 6.85E-01 | 6.82E-01 | 6.17E-05 | Yes | Gm49440 | upstream_gene_variant | MODIFIER |
| 15 | 43417393 | AX-170471387 | 1 | G | A | 0.036 | -5.20E-01 | 9.92E-02 | 6.60E-01 | 6.55E-01 | 6.18E-05 | Yes | | intergenic_variant | MODIFIER |

| | | | | | | | | | | | | | | | |
|---|---|---|---|---|---|---|---|---|---|---|---|---|---|---|---|
| 15 | 43604951 | AX-170312126 | 1 | A | G | 0.036 | -5.14E-01 | 9.95E-02 | 6.99E-01 | 6.98E-01 | 6.22E-05 | Yes | | intergenic_variant | MODIFIER |
| 15 | 43851199 | AX-170496538 | 1 | C | A | 0.036 | -5.14E-01 | 9.95E-02 | 6.99E-01 | 6.98E-01 | 6.22E-05 | Yes | | intergenic_variant | MODIFIER |
| 15 | 55141037 | AX-170297340 | 0 | G | A | 0.044 | -4.39E-01 | 9.46E-02 | 1.10E+00 | 1.14E+00 | 6.33E-05 | Yes | Deptor | intron_variant | MODIFIER |
| 15 | 45913882 | AX-170430367 | 0 | G | C | 0.035 | -5.14E-01 | 9.91E-02 | 6.80E-01 | 6.81E-01 | 6.62E-05 | Yes | | intergenic_variant | MODIFIER |
| 15 | 42672563 | AX-170143404 | 1 | G | T | 0.036 | -5.19E-01 | 1.00E-01 | 6.78E-01 | 6.75E-01 | 6.64E-05 | Yes | Angpt1 | intron_variant | MODIFIER |
| 15 | 57304020 | AX-170220079 | 1 | G | A | 0.054 | -4.23E-01 | 9.40E-02 | 1.25E+00 | 1.29E+00 | 6.69E-05 | Yes | Slc22a22 | intron_variant | MODIFIER |
| 15 | 25498872 | AX-170114929 | 0 | T | C | 0.053 | -3.10E-01 | 6.45E-02 | 9.26E-01 | 9.56E-01 | 6.72E-05 | Yes | | intergenic_variant | MODIFIER |
| 15 | 23638095 | AX-97828994 | 0 | C | A | 0.026 | -6.07E-01 | 9.23E-02 | 2.51E-01 | 2.21E-01 | 6.75E-05 | Yes | | intergenic_variant | MODIFIER |
| 15 | 31050143 | AX-97798436 | 0 | G | T | 0.026 | -6.07E-01 | 9.23E-02 | 2.51E-01 | 2.21E-01 | 6.75E-05 | Yes | | intergenic_variant | MODIFIER |
| 15 | 43864144 | AX-97792228 | 1 | G | A | 0.036 | -5.20E-01 | 9.90E-02 | 6.42E-01 | 6.38E-01 | 6.84E-05 | Yes | Tmem74 | downstream_gene_variant | MODIFIER |
| 15 | 44230575 | AX-170298249 | 1 | C | T | 0.036 | -5.20E-01 | 9.90E-02 | 6.42E-01 | 6.38E-01 | 6.84E-05 | Yes | Trhr | intron_variant | MODIFIER |
| 15 | 26897531 | AX-97829099 | 1 | A | G | 0.027 | -5.75E-01 | 9.75E-02 | 3.99E-01 | 3.83E-01 | 7.16E-05 | Yes | Fbxl7 | upstream_gene_variant | MODIFIER |
| 15 | 44621366 | AX-170132364 | 1 | G | A | 0.036 | -5.15E-01 | 9.92E-02 | 6.64E-01 | 6.62E-01 | 7.42E-05 | Yes | Ebag9 | intron_variant | MODIFIER |
| 15 | 44004170 | AX-170055400 | 1 | G | A | 0.036 | -5.19E-01 | 9.93E-02 | 6.46E-01 | 6.42E-01 | 7.47E-05 | Yes | | intergenic_variant | MODIFIER |
| 15 | 30205681 | AX-170192247 | 0 | C | T | 0.035 | -4.60E-01 | 9.07E-02 | 7.27E-01 | 7.61E-01 | 7.54E-05 | Yes | Ctnnd2 | intron_variant | MODIFIER |
| 15 | 29957535 | AX-97829161 | 2 | G | T | 0.018 | -5.82E-01 | 9.39E-02 | 3.20E-01 | 3.01E-01 | 7.73E-05 | Yes | | intergenic_variant | MODIFIER |
| 15 | 41953098 | AX-97829160 | 1 | A | G | 0.036 | -5.22E-01 | 9.93E-02 | 6.28E-01 | 6.23E-01 | 7.75E-05 | Yes | Gm24751 | downstream_gene_variant | MODIFIER |
| 15 | 42374886 | AX-97786072 | 1 | C | T | 0.036 | -5.20E-01 | 1.00E-01 | 6.48E-01 | 6.43E-01 | 8.42E-05 | Yes | | intergenic_variant | MODIFIER |
| 15 | 61041634 | AX-170404706 | 1 | T | A | 0.054 | -4.59E-01 | 9.73E-02 | 9.49E-01 | 9.60E-01 | 8.48E-05 | Yes | | intergenic_variant | MODIFIER |
| 15 | 31130158 | AX-170447699 | 0 | C | T | 0.035 | -5.25E-01 | 9.90E-02 | 5.95E-01 | 5.91E-01 | 8.55E-05 | Yes | | intergenic_variant | MODIFIER |
| 15 | 41871259 | AX-170011510 | 1 | G | A | 0.036 | -5.20E-01 | 9.93E-02 | 6.23E-01 | 6.18E-01 | 8.72E-05 | Yes | Abra | upstream_gene_variant | MODIFIER |
| 15 | 23334570 | AX-97789297 | 0 | A | T | 0.026 | -5.36E-01 | 1.01E-01 | 5.80E-01 | 5.76E-01 | 8.99E-05 | Yes | Cdh18 | intron_variant,non_coding_transcript_variant | MODIFIER |
| 13 | 11469881 | AX-170104874 | 1 | T | C | 0.375 | -2.52E-01 | 5.71E-02 | 6.17E+00 | 7.08E+00 | 9.56E-05 | Yes | | intergenic_variant | MODIFIER |
| 15 | 58485702 | AX-170508114 | 1 | T | A | 0.054 | -4.54E-01 | 9.60E-02 | 9.18E-01 | 9.31E-01 | 9.57E-05 | Yes | | intergenic_variant | MODIFIER |
| 15 | 26766164 | AX-170283721 | 1 | T | C | 0.036 | -5.14E-01 | 9.78E-02 | 6.00E-01 | 5.98E-01 | 9.90E-05 | Yes | Fbxl7 | intron_variant | MODIFIER |
| 15 | 22872808 | AX-170011225 | 0 | G | T | 0.035 | -5.32E-01 | 9.75E-02 | 5.08E-01 | 4.99E-01 | 1.13E-04 | Yes | Cdh18 | intron_variant | MODIFIER |
| 15 | 43405295 | AX-170397447 | 0 | T | C | 0.035 | -4.98E-01 | 1.01E-01 | 7.66E-01 | 7.69E-01 | 1.14E-04 | Yes | | intergenic_variant | MODIFIER |
| 15 | 53708894 | AX-170168684 | 0 | C | T | 0.044 | -4.44E-01 | 9.96E-02 | 1.13E+00 | 1.16E+00 | 1.14E-04 | Yes | Samd12 | intron_variant | MODIFIER |
| 15 | 45082136 | AX-170533218 | 0 | T | C | 0.035 | -4.31E-01 | 8.75E-02 | 7.48E-01 | 7.60E-01 | 1.15E-04 | Yes | | intergenic_variant | MODIFIER |
| 15 | 45922753 | AX-170270813 | 0 | A | G | 0.044 | -4.74E-01 | 1.05E-01 | 1.06E+00 | 1.07E+00 | 1.20E-04 | Yes | | intergenic_variant | MODIFIER |
| 15 | 52009262 | AX-170446887 | 0 | G | T | 0.044 | -4.54E-01 | 1.01E-01 | 1.07E+00 | 1.09E+00 | 1.25E-04 | Yes | Gm48926 | upstream_gene_variant | MODIFIER |
| 15 | 30176401 | AX-97789652 | 0 | G | A | 0.026 | -5.37E-01 | 9.74E-02 | 4.80E-01 | 4.76E-01 | 1.26E-04 | Yes | Ctnnd2 | intron_variant | MODIFIER |
| 15 | 42206147 | AX-170152420 | 2 | G | A | 0.036 | -5.09E-01 | 9.96E-02 | 6.37E-01 | 6.32E-01 | 1.28E-04 | Yes | Gm49441 | downstream_gene_variant | MODIFIER |
| 15 | 23605054 | AX-170515606 | 0 | G | T | 0.035 | -5.38E-01 | 9.52E-02 | 4.32E-01 | 4.21E-01 | 1.31E-04 | Yes | | intergenic_variant | MODIFIER |
| 15 | 25718808 | AX-170187290 | 0 | G | A | 0.035 | -5.38E-01 | 9.52E-02 | 4.32E-01 | 4.21E-01 | 1.31E-04 | Yes | Myo10 | intron_variant | MODIFIER |
| 15 | 25727670 | AX-170213499 | 0 | A | T | 0.035 | -5.38E-01 | 9.52E-02 | 4.32E-01 | 4.21E-01 | 1.31E-04 | Yes | Myo10 | intron_variant | MODIFIER |
| 15 | 30504825 | AX-97794608 | 0 | G | A | 0.035 | -5.38E-01 | 9.52E-02 | 4.32E-01 | 4.21E-01 | 1.31E-04 | Yes | Ctnnd2 | intron_variant | MODIFIER |
| 15 | 52008868 | AX-170177311 | 0 | C | T | 0.035 | -4.74E-01 | 1.01E-01 | 8.82E-01 | 8.96E-01 | 1.31E-04 | Yes | Gm48926 | upstream_gene_variant | MODIFIER |
| 15 | 53005724 | AX-170335356 | 0 | C | T | 0.035 | -4.74E-01 | 1.01E-01 | 8.82E-01 | 8.96E-01 | 1.31E-04 | Yes | | intergenic_variant | MODIFIER |
| 15 | 54429970 | AX-170326390 | 0 | G | C | 0.035 | -4.74E-01 | 1.01E-01 | 8.82E-01 | 8.96E-01 | 1.31E-04 | Yes | Colec10 | intron_variant | MODIFIER |
| 15 | 51779562 | AX-170135402 | 0 | A | G | 0.044 | -4.53E-01 | 1.00E-01 | 1.01E+00 | 1.03E+00 | 1.38E-04 | Yes | | intergenic_variant | MODIFIER |
| 15 | 58373308 | AX-170107790 | 2 | C | T | 0.055 | -4.45E-01 | 9.67E-02 | 9.45E-01 | 9.60E-01 | 1.40E-04 | Yes | Anxa13 | intron_variant,non_coding_transcript_variant | MODIFIER |
| 15 | 44004130 | AX-97848509 | 1 | T | G | 0.036 | -4.99E-01 | 1.05E-01 | 8.21E-01 | 8.25E-01 | 1.40E-04 | Yes | | intergenic_variant | MODIFIER |
| 15 | 44161838 | AX-170201011 | 0 | C | T | 0.035 | -4.98E-01 | 1.02E-01 | 7.45E-01 | 7.47E-01 | 1.49E-04 | Yes | | intergenic_variant | MODIFIER |
| 15 | 22819764 | AX-170319742 | 0 | C | T | 0.035 | -5.13E-01 | 9.64E-02 | 5.34E-01 | 5.33E-01 | 1.50E-04 | Yes | Cdh18 | intron_variant | MODIFIER |
| 15 | 30665929 | AX-170397189 | 0 | C | T | 0.035 | -5.13E-01 | 9.64E-02 | 5.34E-01 | 5.33E-01 | 1.50E-04 | Yes | Ctnnd2 | intron_variant | MODIFIER |
| 15 | 34056127 | AX-170255795 | 0 | C | A | 0.035 | -5.13E-01 | 9.64E-02 | 5.34E-01 | 5.33E-01 | 1.50E-04 | Yes | Gm49091 | downstream_gene_variant | MODIFIER |
| 15 | 34080514 | AX-170352614 | 0 | C | T | 0.035 | -5.45E-01 | 9.67E-02 | 4.23E-01 | 4.13E-01 | 1.61E-04 | Yes | Mtdh | upstream_gene_variant | MODIFIER |
| 15 | 53709217 | AX-170425921 | 0 | G | A | 0.044 | -4.50E-01 | 9.85E-02 | 9.36E-01 | 9.53E-01 | 1.66E-04 | Yes | Samd12 | intron_variant | MODIFIER |
| 15 | 32947619 | AX-170350618 | 0 | T | C | 0.026 | -5.53E-01 | 1.06E-01 | 5.70E-01 | 5.60E-01 | 1.68E-04 | Yes | Sdc2 | intron_variant | MODIFIER |
| 15 | 30479962 | AX-170055137 | 1 | G | A | 0.036 | -5.01E-01 | 9.73E-02 | 5.92E-01 | 5.96E-01 | 1.70E-04 | Yes | Ctnnd2 | intron_variant | MODIFIER |
| 15 | 30799427 | AX-97794026 | 0 | C | T | 0.035 | -5.35E-01 | 9.72E-02 | 4.57E-01 | 4.48E-01 | 1.75E-04 | Yes | Ctnnd2 | intron_variant | MODIFIER |
| 15 | 26760082 | AX-170514401 | 1 | G | A | 0.018 | -5.47E-01 | 1.02E-01 | 4.96E-01 | 4.90E-01 | 1.75E-04 | Yes | Fbxl7 | intron_variant | MODIFIER |
| 15 | 31085550 | AX-170380395 | 0 | C | T | 0.026 | -5.41E-01 | 9.69E-02 | 4.29E-01 | 4.19E-01 | 1.89E-04 | Yes | | intergenic_variant | MODIFIER |

| | | | | | | | | | | | | | | | |
|---|---|---|---|---|---|---|---|---|---|---|---|---|---|---|---|
| 15 | 29614898 | AX-170319511 | 0 | A | G | 0.026 | -5.27E-01 | 1.00E-01 | 5.26E-01 | 5.26E-01 | 2.00E-04 | Yes | | intergenic_variant | MODIFIER |
| 15 | 31292626 | AX-170231238 | 0 | G | A | 0.044 | -5.50E-01 | 8.91E-02 | 2.83E-01 | 2.68E-01 | 2.03E-04 | Yes | Ankrd33b | downstream_gene_variant | MODIFIER |
| 15 | 42625082 | AX-170283268 | 0 | G | A | 0.044 | -4.58E-01 | 9.95E-02 | 8.64E-01 | 8.72E-01 | 2.09E-04 | Yes | Angpt1 | intron_variant | MODIFIER |
| 15 | 33754298 | AX-97808697 | 0 | C | T | 0.035 | -5.44E-01 | 9.51E-02 | 3.80E-01 | 3.67E-01 | 2.16E-04 | Yes | Gm49088 | downstream_gene_variant | MODIFIER |
| 15 | 25215948 | AX-170461281 | 0 | A | G | 0.035 | -4.24E-01 | 9.36E-02 | 9.03E-01 | 9.30E-01 | 2.22E-04 | Yes | 9230109A22Rik | intron_variant,non_coding_transcript_variant | MODIFIER |
| 15 | 22923347 | AX-170197913 | 1 | C | T | 0.036 | -5.20E-01 | 9.88E-02 | 5.20E-01 | 5.15E-01 | 2.27E-04 | Yes | Cdh18 | intron_variant | MODIFIER |
| 15 | 22161184 | AX-97834518 | 0 | C | T | 0.035 | -5.42E-01 | 9.51E-02 | 3.82E-01 | 3.65E-01 | 2.32E-04 | Yes | | intergenic_variant | MODIFIER |
| 15 | 29615054 | AX-170209854 | 0 | G | A | 0.035 | -5.42E-01 | 9.51E-02 | 3.82E-01 | 3.65E-01 | 2.32E-04 | Yes | | intergenic_variant | MODIFIER |
| 15 | 62108336 | AX-170450103 | 0 | C | G | 0.07 | -3.99E-01 | 9.79E-02 | 1.40E+00 | 1.43E+00 | 2.35E-04 | Yes | Pvt1 | intron_variant,non_coding_transcript_variant | MODIFIER |
| 15 | 31119108 | AX-170379082 | 1 | G | A | 0.036 | -5.20E-01 | 1.00E-01 | 5.47E-01 | 5.42E-01 | 2.36E-04 | Yes | | intergenic_variant | MODIFIER |
| 15 | 46309979 | AX-97789802 | 1 | C | T | 0.054 | -4.23E-01 | 9.58E-02 | 9.82E-01 | 1.01E+00 | 2.40E-04 | Yes | | intergenic_variant | MODIFIER |
| 15 | 25206633 | AX-170185849 | 1 | C | T | 0.036 | -5.21E-01 | 9.89E-02 | 5.09E-01 | 5.02E-01 | 2.49E-04 | Yes | 9230109A22Rik | intron_variant,non_coding_transcript_variant | MODIFIER |
| 15 | 53595745 | AX-170184835 | 1 | A | T | 0.036 | -4.66E-01 | 1.00E-01 | 8.00E-01 | 8.12E-01 | 2.55E-04 | Yes | Samd12 | intron_variant | MODIFIER |
| 15 | 29693052 | AX-97820468 | 1 | A | C | 0.036 | -5.22E-01 | 9.77E-02 | 4.81E-01 | 4.76E-01 | 2.62E-04 | Yes | | intergenic_variant | MODIFIER |
| 15 | 44323947 | AX-170215769 | 0 | C | T | 0.044 | -4.57E-01 | 9.75E-02 | 7.70E-01 | 7.77E-01 | 2.72E-04 | Yes | Gm19235 | non_coding_transcript_exon_variant | MODIFIER |
| 15 | 56079001 | AX-170179342 | 0 | C | T | 0.044 | -4.25E-01 | 9.28E-02 | 8.27E-01 | 8.60E-01 | 2.79E-04 | Yes | | intergenic_variant | MODIFIER |
| 15 | 26261855 | AX-170011288 | 0 | C | T | 0.035 | -5.58E-01 | 9.64E-02 | 3.48E-01 | 3.25E-01 | 2.85E-04 | Yes | | intergenic_variant | MODIFIER |
| 15 | 54358689 | AX-170366444 | 0 | C | T | 0.026 | -4.40E-01 | 9.77E-02 | 8.79E-01 | 9.00E-01 | 2.86E-04 | Yes | | intergenic_variant | MODIFIER |
| 15 | 30770965 | AX-97802885 | 2 | G | A | 0.036 | -5.22E-01 | 9.70E-02 | 4.61E-01 | 4.50E-01 | 2.91E-04 | Yes | Ctnnd2 | intron_variant | MODIFIER |
| 15 | 44875677 | AX-170011560 | 0 | T | C | 0.044 | -4.85E-01 | 9.87E-02 | 6.37E-01 | 6.50E-01 | 2.93E-04 | Yes | A930017M01Rik | intron_variant,non_coding_transcript_variant | MODIFIER |
| 15 | 25210212 | AX-170055047 | 0 | A | T | 0.035 | -5.55E-01 | 9.18E-02 | 2.89E-01 | 2.64E-01 | 3.01E-04 | Yes | 9230109A22Rik | intron_variant,non_coding_transcript_variant | MODIFIER |
| 15 | 26658680 | AX-97836383 | 1 | G | T | 0.036 | -5.46E-01 | 9.20E-02 | 3.11E-01 | 2.93E-01 | 3.05E-04 | Yes | Fbxl7 | intron_variant | MODIFIER |
| 15 | 26989900 | AX-170229684 | 0 | G | A | 0.026 | -5.63E-01 | 9.73E-02 | 3.43E-01 | 3.14E-01 | 3.19E-04 | Yes | | intergenic_variant | MODIFIER |
| 15 | 29917347 | AX-97828698 | 0 | T | G | 0.026 | -5.63E-01 | 9.73E-02 | 3.43E-01 | 3.14E-01 | 3.19E-04 | Yes | | intergenic_variant | MODIFIER |
| 15 | 60648983 | AX-170087879 | 0 | A | G | 0.061 | -4.18E-01 | 9.91E-02 | 1.10E+00 | 1.11E+00 | 3.24E-04 | Yes | | intergenic_variant | MODIFIER |
| 15 | 60652755 | AX-170290272 | 0 | T | C | 0.061 | -4.18E-01 | 9.91E-02 | 1.10E+00 | 1.11E+00 | 3.24E-04 | Yes | | intergenic_variant | MODIFIER |
| 15 | 60652897 | AX-170216652 | 0 | G | C | 0.061 | -4.18E-01 | 9.91E-02 | 1.10E+00 | 1.11E+00 | 3.24E-04 | Yes | | intergenic_variant | MODIFIER |
| 15 | 60660661 | AX-170391374 | 0 | G | A | 0.061 | -4.18E-01 | 9.91E-02 | 1.10E+00 | 1.11E+00 | 3.24E-04 | Yes | | intergenic_variant | MODIFIER |
| 15 | 60665109 | AX-170160698 | 0 | G | A | 0.061 | -4.18E-01 | 9.91E-02 | 1.10E+00 | 1.11E+00 | 3.24E-04 | Yes | | intergenic_variant | MODIFIER |
| 15 | 60665223 | AX-97847362 | 0 | T | C | 0.061 | -4.18E-01 | 9.91E-02 | 1.10E+00 | 1.11E+00 | 3.24E-04 | Yes | | intergenic_variant | MODIFIER |
| 15 | 60665736 | AX-168385213 | 0 | C | T | 0.061 | -4.18E-01 | 9.91E-02 | 1.10E+00 | 1.11E+00 | 3.24E-04 | Yes | Gm48946 | upstream_gene_variant | MODIFIER |
| 15 | 60688384 | AX-170447752 | 0 | G | C | 0.061 | -4.18E-01 | 9.91E-02 | 1.10E+00 | 1.11E+00 | 3.24E-04 | Yes | | intergenic_variant | MODIFIER |
| 15 | 60727642 | AX-170055693 | 0 | C | T | 0.061 | -4.18E-01 | 9.91E-02 | 1.10E+00 | 1.11E+00 | 3.24E-04 | Yes | | intergenic_variant | MODIFIER |
| 15 | 60727967 | AX-170055691 | 0 | C | T | 0.061 | -4.18E-01 | 9.91E-02 | 1.10E+00 | 1.11E+00 | 3.24E-04 | Yes | | intergenic_variant | MODIFIER |
| 15 | 60733490 | AX-170220907 | 0 | C | T | 0.061 | -4.18E-01 | 9.91E-02 | 1.10E+00 | 1.11E+00 | 3.24E-04 | Yes | | intergenic_variant | MODIFIER |
| 15 | 60741210 | AX-170510104 | 0 | C | T | 0.061 | -4.18E-01 | 9.91E-02 | 1.10E+00 | 1.11E+00 | 3.24E-04 | Yes | Gm7086 | upstream_gene_variant | MODIFIER |
| 15 | 60743817 | AX-170342938 | 0 | G | C | 0.061 | -4.18E-01 | 9.91E-02 | 1.10E+00 | 1.11E+00 | 3.24E-04 | Yes | Gm7086 | intron_variant,non_coding_transcript_variant | MODIFIER |
| 15 | 60748588 | AX-170037297 | 0 | A | G | 0.061 | -4.18E-01 | 9.91E-02 | 1.10E+00 | 1.11E+00 | 3.24E-04 | Yes | Gm7086 | downstream_gene_variant | MODIFIER |
| 15 | 60750541 | AX-170011820 | 0 | G | A | 0.061 | -4.18E-01 | 9.91E-02 | 1.10E+00 | 1.11E+00 | 3.24E-04 | Yes | Gm7086 | downstream_gene_variant | MODIFIER |
| 15 | 60777811 | AX-97843281 | 0 | T | C | 0.061 | -4.18E-01 | 9.91E-02 | 1.10E+00 | 1.11E+00 | 3.24E-04 | Yes | | intergenic_variant | MODIFIER |
| 15 | 60787805 | AX-170226817 | 0 | G | T | 0.061 | -4.18E-01 | 9.91E-02 | 1.10E+00 | 1.11E+00 | 3.24E-04 | Yes | | intergenic_variant | MODIFIER |
| 15 | 60809062 | AX-97808561 | 0 | T | C | 0.061 | -4.18E-01 | 9.91E-02 | 1.10E+00 | 1.11E+00 | 3.24E-04 | Yes | | intergenic_variant | MODIFIER |
| 15 | 60809899 | AX-170448760 | 0 | T | C | 0.061 | -4.18E-01 | 9.91E-02 | 1.10E+00 | 1.11E+00 | 3.24E-04 | Yes | | intergenic_variant | MODIFIER |
| 15 | 60810156 | AX-170011822 | 0 | A | T | 0.061 | -4.18E-01 | 9.91E-02 | 1.10E+00 | 1.11E+00 | 3.24E-04 | Yes | | intergenic_variant | MODIFIER |
| 15 | 60811310 | AX-170513812 | 0 | G | A | 0.061 | -4.18E-01 | 9.91E-02 | 1.10E+00 | 1.11E+00 | 3.24E-04 | Yes | | intergenic_variant | MODIFIER |
| 15 | 60812787 | AX-170110667 | 0 | A | C | 0.061 | -4.18E-01 | 9.91E-02 | 1.10E+00 | 1.11E+00 | 3.24E-04 | Yes | | intergenic_variant | MODIFIER |
| 15 | 60814293 | AX-170190798 | 0 | C | T | 0.061 | -4.18E-01 | 9.91E-02 | 1.10E+00 | 1.11E+00 | 3.24E-04 | Yes | Fam84b | downstream_gene_variant | MODIFIER |
| 15 | 60814845 | AX-170055697 | 0 | C | G | 0.061 | -4.18E-01 | 9.91E-02 | 1.10E+00 | 1.11E+00 | 3.24E-04 | Yes | Fam84b | downstream_gene_variant | MODIFIER |
| 15 | 60815086 | AX-170011819 | 0 | C | T | 0.061 | -4.18E-01 | 9.91E-02 | 1.10E+00 | 1.11E+00 | 3.24E-04 | Yes | Fam84b | downstream_gene_variant | MODIFIER |
| 15 | 60817324 | AX-170011818 | 0 | A | T | 0.061 | -4.18E-01 | 9.91E-02 | 1.10E+00 | 1.11E+00 | 3.24E-04 | Yes | Fam84b | downstream_gene_variant | MODIFIER |
| 15 | 60830733 | AX-170011821 | 0 | A | G | 0.061 | -4.18E-01 | 9.91E-02 | 1.10E+00 | 1.11E+00 | 3.24E-04 | Yes | 9930014A18Rik | downstream_gene_variant | MODIFIER |
| 15 | 60851322 | AX-170393938 | 0 | C | T | 0.061 | -4.18E-01 | 9.91E-02 | 1.10E+00 | 1.11E+00 | 3.24E-04 | Yes | Fam84b | intron_variant,non_coding_transcript_variant | MODIFIER |
| 15 | 60865349 | AX-97826570 | 0 | A | G | 0.061 | -4.18E-01 | 9.91E-02 | 1.10E+00 | 1.11E+00 | 3.24E-04 | Yes | | intergenic_variant | MODIFIER |
| 15 | 60865380 | AX-170526555 | 0 | A | G | 0.061 | -4.18E-01 | 9.91E-02 | 1.10E+00 | 1.11E+00 | 3.24E-04 | Yes | | intergenic_variant | MODIFIER |

| | | | | | | | | | | | | | | | |
|---|---|---|---|---|---|---|---|---|---|---|---|---|---|---|---|
| 15 | 60900850 | AX-170185966 | 0 | C | A | 0.061 | -4.18E-01 | 9.91E-02 | 1.10E+00 | 1.11E+00 | 3.24E-04 | Yes | A1bg | intron_variant,non_coding_transcript_variant | MODIFIER |
| 15 | 60902462 | AX-97820990 | 0 | A | G | 0.061 | -4.18E-01 | 9.91E-02 | 1.10E+00 | 1.11E+00 | 3.24E-04 | Yes | A1bg | intron_variant,non_coding_transcript_variant | MODIFIER |
| 15 | 60913690 | AX-170331809 | 0 | T | A | 0.061 | -4.18E-01 | 9.91E-02 | 1.10E+00 | 1.11E+00 | 3.24E-04 | Yes | A1bg | downstream_gene_variant | MODIFIER |
| 15 | 60913773 | AX-170316848 | 0 | C | T | 0.061 | -4.18E-01 | 9.91E-02 | 1.10E+00 | 1.11E+00 | 3.24E-04 | Yes | A1bg | downstream_gene_variant | MODIFIER |
| 15 | 60916320 | AX-97812657 | 0 | G | T | 0.061 | -4.18E-01 | 9.91E-02 | 1.10E+00 | 1.11E+00 | 3.24E-04 | Yes | A1bg | downstream_gene_variant | MODIFIER |
| 15 | 60936745 | AX-170055698 | 0 | T | C | 0.061 | -4.18E-01 | 9.91E-02 | 1.10E+00 | 1.11E+00 | 3.24E-04 | Yes | Gm48951 | downstream_gene_variant | MODIFIER |
| 15 | 60948470 | AX-97832378 | 0 | G | T | 0.061 | -4.18E-01 | 9.91E-02 | 1.10E+00 | 1.11E+00 | 3.24E-04 | Yes | 4930402D18Rik | intron_variant,non_coding_transcript_variant | MODIFIER |
| 15 | 60948550 | AX-170472738 | 0 | T | G | 0.061 | -4.18E-01 | 9.91E-02 | 1.10E+00 | 1.11E+00 | 3.24E-04 | Yes | 4930402D18Rik | intron_variant,non_coding_transcript_variant | MODIFIER |
| 15 | 60967946 | AX-97844736 | 0 | G | A | 0.061 | -4.18E-01 | 9.91E-02 | 1.10E+00 | 1.11E+00 | 3.24E-04 | Yes | | regulatory_region_variant | MODIFIER |
| 15 | 61204721 | AX-170055705 | 0 | C | T | 0.061 | -4.18E-01 | 9.91E-02 | 1.10E+00 | 1.11E+00 | 3.24E-04 | Yes | Gm49497 | intron_variant,non_coding_transcript_variant | MODIFIER |
| 15 | 61208080 | AX-97789049 | 0 | A | G | 0.061 | -4.18E-01 | 9.91E-02 | 1.10E+00 | 1.11E+00 | 3.24E-04 | Yes | Gm49497 | intron_variant,non_coding_transcript_variant | MODIFIER |
| 15 | 61208275 | AX-170303516 | 0 | A | G | 0.061 | -4.18E-01 | 9.91E-02 | 1.10E+00 | 1.11E+00 | 3.24E-04 | Yes | Gm49497 | intron_variant,non_coding_transcript_variant | MODIFIER |
| 15 | 61214623 | AX-170498209 | 0 | G | C | 0.061 | -4.18E-01 | 9.91E-02 | 1.10E+00 | 1.11E+00 | 3.24E-04 | Yes | Gm49497 | intron_variant,non_coding_transcript_variant | MODIFIER |
| 15 | 61215142 | AX-170295513 | 0 | C | T | 0.061 | -4.18E-01 | 9.91E-02 | 1.10E+00 | 1.11E+00 | 3.24E-04 | Yes | Gm49497 | intron_variant,non_coding_transcript_variant | MODIFIER |
| 15 | 61218362 | AX-170189869 | 0 | A | T | 0.061 | -4.18E-01 | 9.91E-02 | 1.10E+00 | 1.11E+00 | 3.24E-04 | Yes | Gm49497 | intron_variant,non_coding_transcript_variant | MODIFIER |
| 15 | 61219342 | AX-170150015 | 0 | G | A | 0.061 | -4.18E-01 | 9.91E-02 | 1.10E+00 | 1.11E+00 | 3.24E-04 | Yes | Gm49497 | intron_variant,non_coding_transcript_variant | MODIFIER |
| 15 | 61219745 | AX-170123744 | 0 | G | T | 0.061 | -4.18E-01 | 9.91E-02 | 1.10E+00 | 1.11E+00 | 3.24E-04 | Yes | Gm49497 | intron_variant,non_coding_transcript_variant | MODIFIER |
| 15 | 61223514 | AX-97823745 | 0 | G | A | 0.061 | -4.18E-01 | 9.91E-02 | 1.10E+00 | 1.11E+00 | 3.24E-04 | Yes | Gm49497 | intron_variant,non_coding_transcript_variant | MODIFIER |
| 15 | 61227824 | AX-170276817 | 0 | G | T | 0.061 | -4.18E-01 | 9.91E-02 | 1.10E+00 | 1.11E+00 | 3.24E-04 | Yes | Gm49497 | intron_variant,non_coding_transcript_variant | MODIFIER |
| 15 | 61229196 | AX-170284926 | 0 | G | A | 0.061 | -4.18E-01 | 9.91E-02 | 1.10E+00 | 1.11E+00 | 3.24E-04 | Yes | Gm49497 | intron_variant,non_coding_transcript_variant | MODIFIER |
| 15 | 61239950 | AX-170510024 | 0 | A | T | 0.061 | -4.18E-01 | 9.91E-02 | 1.10E+00 | 1.11E+00 | 3.24E-04 | Yes | Gm49497 | non_coding_transcript_exon_variant | MODIFIER |
| 15 | 61247994 | AX-170399923 | 0 | A | G | 0.061 | -4.18E-01 | 9.91E-02 | 1.10E+00 | 1.11E+00 | 3.24E-04 | Yes | | intergenic_variant | MODIFIER |
| 15 | 61268874 | AX-97825998 | 0 | C | T | 0.061 | -4.18E-01 | 9.91E-02 | 1.10E+00 | 1.11E+00 | 3.24E-04 | Yes | | intergenic_variant | MODIFIER |
| 15 | 61298367 | AX-170452162 | 0 | T | C | 0.061 | -4.18E-01 | 9.91E-02 | 1.10E+00 | 1.11E+00 | 3.24E-04 | Yes | Gm29904 | intron_variant,non_coding_transcript_variant | MODIFIER |
| 15 | 61298743 | AX-170055706 | 0 | A | G | 0.061 | -4.18E-01 | 9.91E-02 | 1.10E+00 | 1.11E+00 | 3.24E-04 | Yes | Gm29904 | intron_variant,non_coding_transcript_variant | MODIFIER |
| 15 | 61300824 | AX-170011828 | 0 | A | G | 0.061 | -4.18E-01 | 9.91E-02 | 1.10E+00 | 1.11E+00 | 3.24E-04 | Yes | Gm29904 | intron_variant,non_coding_transcript_variant | MODIFIER |
| 15 | 61310202 | AX-170011825 | 0 | C | T | 0.061 | -4.18E-01 | 9.91E-02 | 1.10E+00 | 1.11E+00 | 3.24E-04 | Yes | Gm29904 | intron_variant,non_coding_transcript_variant | MODIFIER |
| 15 | 61310576 | AX-97805861 | 0 | A | G | 0.061 | -4.18E-01 | 9.91E-02 | 1.10E+00 | 1.11E+00 | 3.24E-04 | Yes | Gm29904 | intron_variant,non_coding_transcript_variant | MODIFIER |
| 15 | 61321694 | AX-170437633 | 0 | C | T | 0.061 | -4.18E-01 | 9.91E-02 | 1.10E+00 | 1.11E+00 | 3.24E-04 | Yes | Gm29904 | intron_variant,non_coding_transcript_variant | MODIFIER |
| 15 | 61324512 | AX-170317486 | 0 | G | T | 0.061 | -4.18E-01 | 9.91E-02 | 1.10E+00 | 1.11E+00 | 3.24E-04 | Yes | Gm29904 | intron_variant,non_coding_transcript_variant | MODIFIER |
| 15 | 61439765 | AX-170332749 | 0 | C | T | 0.061 | -4.18E-01 | 9.91E-02 | 1.10E+00 | 1.11E+00 | 3.24E-04 | Yes | | intergenic_variant | MODIFIER |
| 15 | 61440946 | AX-97837511 | 0 | A | G | 0.061 | -4.18E-01 | 9.91E-02 | 1.10E+00 | 1.11E+00 | 3.24E-04 | Yes | | intergenic_variant | MODIFIER |
| 15 | 61441452 | AX-168399894 | 0 | A | G | 0.061 | -4.18E-01 | 9.91E-02 | 1.10E+00 | 1.11E+00 | 3.24E-04 | Yes | | intergenic_variant | MODIFIER |
| 15 | 61441528 | AX-170353804 | 0 | A | G | 0.061 | -4.18E-01 | 9.91E-02 | 1.10E+00 | 1.11E+00 | 3.24E-04 | Yes | | intergenic_variant | MODIFIER |
| 15 | 61443782 | AX-170230614 | 0 | G | A | 0.061 | -4.18E-01 | 9.91E-02 | 1.10E+00 | 1.11E+00 | 3.24E-04 | Yes | | intergenic_variant | MODIFIER |
| 15 | 61466461 | AX-170093276 | 0 | A | G | 0.061 | -4.18E-01 | 9.91E-02 | 1.10E+00 | 1.11E+00 | 3.24E-04 | Yes | | intergenic_variant | MODIFIER |
| 15 | 61530102 | AX-170266662 | 0 | G | A | 0.061 | -4.18E-01 | 9.91E-02 | 1.10E+00 | 1.11E+00 | 3.24E-04 | Yes | Gm49498 | downstream_gene_variant | MODIFIER |
| 15 | 61531476 | AX-170470711 | 0 | G | A | 0.061 | -4.18E-01 | 9.91E-02 | 1.10E+00 | 1.11E+00 | 3.24E-04 | Yes | Gm49498 | downstream_gene_variant | MODIFIER |
| 15 | 61531876 | AX-170473061 | 0 | T | G | 0.061 | -4.18E-01 | 9.91E-02 | 1.10E+00 | 1.11E+00 | 3.24E-04 | Yes | Gm49498 | downstream_gene_variant | MODIFIER |
| 15 | 61556631 | AX-97790824 | 0 | T | G | 0.061 | -4.18E-01 | 9.91E-02 | 1.10E+00 | 1.11E+00 | 3.24E-04 | Yes | D030024E09Rik | intron_variant,non_coding_transcript_variant | MODIFIER |
| 15 | 61562018 | AX-97846808 | 0 | C | A | 0.061 | -4.18E-01 | 9.91E-02 | 1.10E+00 | 1.11E+00 | 3.24E-04 | Yes | D030024E09Rik | intron_variant,non_coding_transcript_variant | MODIFIER |
| 15 | 61609977 | AX-97834361 | 0 | C | T | 0.061 | -4.18E-01 | 9.91E-02 | 1.10E+00 | 1.11E+00 | 3.24E-04 | Yes | D030024E09Rik | intron_variant,non_coding_transcript_variant | MODIFIER |
| 15 | 61630132 | AX-170148633 | 0 | G | A | 0.061 | -4.18E-01 | 9.91E-02 | 1.10E+00 | 1.11E+00 | 3.24E-04 | Yes | D030024E09Rik | intron_variant,non_coding_transcript_variant | MODIFIER |
| 15 | 61693736 | AX-170474576 | 0 | A | G | 0.061 | -4.18E-01 | 9.91E-02 | 1.10E+00 | 1.11E+00 | 3.24E-04 | Yes | D030024E09Rik | intron_variant,non_coding_transcript_variant | MODIFIER |
| 15 | 61872742 | AX-170011832 | 0 | G | C | 0.061 | -4.18E-01 | 9.91E-02 | 1.10E+00 | 1.11E+00 | 3.24E-04 | Yes | | intergenic_variant | MODIFIER |
| 15 | 61918882 | AX-170280512 | 0 | A | G | 0.061 | -4.18E-01 | 9.91E-02 | 1.10E+00 | 1.11E+00 | 3.24E-04 | Yes | | intergenic_variant | MODIFIER |
| 15 | 62089406 | AX-170233868 | 0 | G | A | 0.061 | -4.18E-01 | 9.91E-02 | 1.10E+00 | 1.11E+00 | 3.24E-04 | Yes | Pvt1 | intron_variant,non_coding_transcript_variant | MODIFIER |
| 15 | 62095420 | AX-170288155 | 0 | T | C | 0.061 | -4.18E-01 | 9.91E-02 | 1.10E+00 | 1.11E+00 | 3.24E-04 | Yes | Pvt1 | intron_variant,non_coding_transcript_variant | MODIFIER |
| 15 | 62095466 | AX-170417905 | 0 | C | T | 0.061 | -4.18E-01 | 9.91E-02 | 1.10E+00 | 1.11E+00 | 3.24E-04 | Yes | Pvt1 | intron_variant,non_coding_transcript_variant | MODIFIER |
| 15 | 62101592 | AX-170311236 | 0 | A | C | 0.061 | -4.18E-01 | 9.91E-02 | 1.10E+00 | 1.11E+00 | 3.24E-04 | Yes | Pvt1 | intron_variant,non_coding_transcript_variant | MODIFIER |
| 15 | 62111945 | AX-170292685 | 0 | A | G | 0.061 | -4.18E-01 | 9.91E-02 | 1.10E+00 | 1.11E+00 | 3.24E-04 | Yes | Pvt1 | intron_variant,non_coding_transcript_variant | MODIFIER |
| 15 | 61294736 | AX-97782371 | 0 | C | A | 0.061 | -3.80E-01 | 9.10E-02 | 1.14E+00 | 1.17E+00 | 3.30E-04 | Yes | Gm29904 | intron_variant,non_coding_transcript_variant | MODIFIER |
| 15 | 52517698 | AX-170055543 | 1 | A | G | 0.036 | -4.22E-01 | 9.23E-02 | 8.10E-01 | 8.26E-01 | 3.35E-04 | Yes | | intergenic_variant | MODIFIER |
| 15 | 44667119 | AX-97789024 | 0 | G | C | 0.035 | -4.90E-01 | 1.02E-01 | 6.73E-01 | 6.72E-01 | 3.37E-04 | Yes | Sybu | downstream_gene_variant | MODIFIER |

| | | | | | | | | | | | | | | | |
|---|---|---|---|---|---|---|---|---|---|---|---|---|---|---|---|
| 15 | 42374766 | AX-170386161 | 0 | C | T | 0.053 | -4.03E-01 | 9.05E-02 | 8.84E-01 | 9.15E-01 | 3.40E-04 | Yes | | intergenic_variant | MODIFIER |
| 15 | 54350400 | AX-170472995 | 0 | A | G | 0.035 | -4.45E-01 | 1.04E-01 | 1.01E+00 | 1.03E+00 | 3.63E-04 | Yes | | intergenic_variant | MODIFIER |
| 15 | 42300483 | AX-170196441 | 0 | A | T | 0.035 | -4.75E-01 | 1.05E-01 | 8.15E-01 | 8.16E-01 | 3.68E-04 | Yes | | intergenic_variant | MODIFIER |
| 15 | 52348909 | AX-97805617 | 1 | A | G | 0.045 | -4.29E-01 | 9.94E-02 | 9.75E-01 | 9.96E-01 | 3.69E-04 | Yes | | intergenic_variant | MODIFIER |
| 15 | 29806746 | AX-170162800 | 1 | C | T | 0.036 | -4.84E-01 | 9.91E-02 | 6.23E-01 | 6.27E-01 | 3.78E-04 | Yes | | intergenic_variant | MODIFIER |
| 15 | 61298596 | AX-170099253 | 0 | C | G | 0.061 | -4.17E-01 | 9.97E-02 | 1.08E+00 | 1.11E+00 | 3.94E-04 | Yes | Gm29904 | intron_variant,non_coding_transcript_variant | MODIFIER |
| 15 | 42684570 | AX-170278467 | 0 | C | A | 0.044 | -4.55E-01 | 1.01E-01 | 8.20E-01 | 8.28E-01 | 4.13E-04 | Yes | Gm17473 | intron_variant,non_coding_transcript_variant | MODIFIER |
| 15 | 61143676 | AX-97819191 | 0 | C | T | 0.07 | -3.96E-01 | 9.65E-02 | 1.15E+00 | 1.17E+00 | 4.22E-04 | Yes | | intergenic_variant | MODIFIER |
| 15 | 60654873 | AX-170409480 | 1 | T | C | 0.062 | -4.06E-01 | 9.90E-02 | 1.15E+00 | 1.17E+00 | 4.30E-04 | Yes | | intergenic_variant | MODIFIER |
| 15 | 61555073 | AX-170055712 | 1 | G | T | 0.062 | -4.08E-01 | 9.86E-02 | 1.10E+00 | 1.12E+00 | 4.35E-04 | Yes | D030024E09Rik | intron_variant,non_coding_transcript_variant | MODIFIER |
| 13 | 12378431 | AX-170500006 | 1 | T | C | 0.179 | -4.29E-01 | 1.17E-01 | 3.86E+00 | 4.08E+00 | 4.43E-04 | Yes | Gm5445 | non_coding_transcript_exon_variant | MODIFIER |
| 15 | 51962618 | AX-97829322 | 2 | C | T | 0.036 | -4.27E-01 | 1.03E-01 | 1.10E+00 | 1.13E+00 | 4.45E-04 | Yes | Rad21 | 3_prime_UTR_variant | MODIFIER |
| 15 | 60918826 | AX-170483499 | 1 | C | A | 0.062 | -4.10E-01 | 1.01E-01 | 1.15E+00 | 1.17E+00 | 4.71E-04 | Yes | A1bg | intron_variant | MODIFIER |
| 15 | 44449865 | AX-97829549 | 1 | C | T | 0.036 | -4.88E-01 | 1.00E-01 | 5.97E-01 | 5.94E-01 | 4.85E-04 | Yes | | intergenic_variant | MODIFIER |
| 15 | 61470414 | AX-170506911 | 1 | G | T | 0.062 | -4.07E-01 | 9.97E-02 | 1.12E+00 | 1.14E+00 | 5.00E-04 | Yes | | intergenic_variant | MODIFIER |
| 15 | 60917889 | AX-170166702 | 1 | G | A | 0.062 | -4.09E-01 | 9.93E-02 | 1.08E+00 | 1.10E+00 | 5.02E-04 | Yes | A1bg | synonymous_variant | LOW |
| 15 | 22554403 | AX-170011219 | 0 | A | C | 0.272 | -2.80E-01 | 7.32E-02 | 1.44E+00 | 1.46E+00 | 5.26E-04 | Yes | Cdh18 | intron_variant | MODIFIER |
| 15 | 44857087 | AX-170191517 | 0 | C | A | 0.044 | -4.41E-01 | 9.87E-02 | 7.97E-01 | 8.11E-01 | 5.41E-04 | Yes | A930017M01Rik | non_coding_transcript_exon_variant | MODIFIER |
| 15 | 16433736 | AX-97828744 | 1 | G | A | 0.295 | -2.66E-01 | 7.32E-02 | 1.89E+00 | 1.93E+00 | 5.55E-04 | Yes | | intergenic_variant | MODIFIER |
| 15 | 17481218 | AX-170011141 | 0 | A | C | 0.281 | -2.66E-01 | 7.29E-02 | 1.83E+00 | 1.87E+00 | 5.56E-04 | Yes | | intergenic_variant | MODIFIER |
| 15 | 29587890 | AX-170087804 | 1 | G | A | 0.036 | -4.64E-01 | 9.25E-02 | 5.33E-01 | 5.34E-01 | 5.61E-04 | Yes | | intergenic_variant | MODIFIER |
| 15 | 58312372 | AX-170126649 | 0 | A | G | 0.053 | -4.32E-01 | 9.91E-02 | 8.62E-01 | 8.74E-01 | 5.65E-04 | Yes | Klhl38 | downstream_gene_variant | MODIFIER |
| 15 | 51856501 | AX-170284348 | 2 | G | C | 0.045 | -4.30E-01 | 1.03E-01 | 1.00E+00 | 1.02E+00 | 5.69E-04 | Yes | Eif3h | intron_variant | MODIFIER |
| 15 | 56700687 | AX-170429815 | 0 | A | T | 0.061 | -3.69E-01 | 8.87E-02 | 1.01E+00 | 1.06E+00 | 5.75E-04 | Yes | Has2os | intron_variant,non_coding_transcript_variant | MODIFIER |
| 15 | 31168592 | AX-97829386 | 1 | A | G | 0.036 | -4.59E-01 | 1.01E-01 | 7.30E-01 | 7.46E-01 | 5.93E-04 | Yes | | intergenic_variant | MODIFIER |
| 15 | 43650401 | AX-170055390 | 0 | A | G | 0.044 | -4.44E-01 | 9.88E-02 | 7.62E-01 | 7.70E-01 | 6.12E-04 | Yes | | intergenic_variant | MODIFIER |
| 15 | 30343327 | AX-170218844 | 1 | G | A | 0.036 | -5.63E-01 | 9.30E-02 | 2.59E-01 | 2.28E-01 | 6.15E-04 | Yes | Ctnnd2 | intron_variant | MODIFIER |
| 15 | 22923304 | AX-170312529 | 1 | G | T | 0.259 | -2.90E-01 | 7.59E-02 | 1.38E+00 | 1.39E+00 | 6.15E-04 | Yes | Cdh18 | intron_variant | MODIFIER |
| 15 | 22187859 | AX-170145898 | 0 | A | G | 0.263 | -2.89E-01 | 7.60E-02 | 1.39E+00 | 1.40E+00 | 6.41E-04 | Yes | | intergenic_variant | MODIFIER |
| 15 | 22278272 | AX-170506852 | 0 | C | T | 0.263 | -2.89E-01 | 7.60E-02 | 1.39E+00 | 1.40E+00 | 6.41E-04 | Yes | Gm27525 | downstream_gene_variant | MODIFIER |
| 15 | 22553812 | AX-97794075 | 0 | T | G | 0.263 | -2.89E-01 | 7.60E-02 | 1.39E+00 | 1.40E+00 | 6.41E-04 | Yes | Cdh18 | intron_variant | MODIFIER |
| 15 | 22562166 | AX-170011218 | 0 | G | A | 0.263 | -2.89E-01 | 7.60E-02 | 1.39E+00 | 1.40E+00 | 6.41E-04 | Yes | Cdh18 | intron_variant | MODIFIER |
| 15 | 22592640 | AX-97845676 | 0 | T | C | 0.263 | -2.89E-01 | 7.60E-02 | 1.39E+00 | 1.40E+00 | 6.41E-04 | Yes | Cdh18 | intron_variant | MODIFIER |
| 15 | 22691340 | AX-170435819 | 0 | A | T | 0.263 | -2.89E-01 | 7.60E-02 | 1.39E+00 | 1.40E+00 | 6.41E-04 | Yes | Cdh18 | intron_variant | MODIFIER |
| 15 | 22721596 | AX-170516851 | 0 | G | A | 0.263 | -2.89E-01 | 7.60E-02 | 1.39E+00 | 1.40E+00 | 6.41E-04 | Yes | Cdh18 | intron_variant | MODIFIER |
| 15 | 22821644 | AX-170267998 | 0 | C | G | 0.263 | -2.89E-01 | 7.60E-02 | 1.39E+00 | 1.40E+00 | 6.41E-04 | Yes | Cdh18 | intron_variant | MODIFIER |
| 15 | 22836513 | AX-170337352 | 0 | T | C | 0.263 | -2.89E-01 | 7.60E-02 | 1.39E+00 | 1.40E+00 | 6.41E-04 | Yes | Cdh18 | intron_variant | MODIFIER |
| 15 | 22859097 | AX-170284825 | 0 | A | C | 0.263 | -2.89E-01 | 7.60E-02 | 1.39E+00 | 1.40E+00 | 6.41E-04 | Yes | Cdh18 | intron_variant | MODIFIER |
| 15 | 23038156 | AX-97789296 | 0 | T | C | 0.263 | -2.89E-01 | 7.60E-02 | 1.39E+00 | 1.40E+00 | 6.41E-04 | Yes | Cdh18 | intron_variant,non_coding_transcript_variant | MODIFIER |
| 15 | 23038256 | AX-170360876 | 0 | G | A | 0.263 | -2.89E-01 | 7.60E-02 | 1.39E+00 | 1.40E+00 | 6.41E-04 | Yes | Cdh18 | intron_variant,non_coding_transcript_variant | MODIFIER |
| 15 | 42435950 | AX-170423051 | 0 | C | A | 0.035 | -5.12E-01 | 1.04E-01 | 5.42E-01 | 5.29E-01 | 6.60E-04 | Yes | Angpt1 | intron_variant | MODIFIER |
| 15 | 24017640 | AX-170055031 | 0 | T | A | 0.035 | -4.26E-01 | 9.44E-02 | 7.35E-01 | 7.55E-01 | 6.94E-04 | Yes | | intergenic_variant | MODIFIER |
| 3 | 116673670 | AX-170503851 | 0 | A | G | 0.193 | -3.96E-01 | 1.13E-01 | 3.53E+00 | 3.69E+00 | 7.16E-04 | Yes | Slc35a3 | synonymous_variant | LOW |
| 15 | 44853486 | AX-170455870 | 0 | C | T | 0.044 | -4.47E-01 | 9.87E-02 | 7.18E-01 | 7.24E-01 | 7.55E-04 | Yes | A930017M01Rik | intron_variant,non_coding_transcript_variant | MODIFIER |
| 15 | 61557472 | AX-170124686 | 0 | T | A | 0.07 | -3.76E-01 | 9.79E-02 | 1.27E+00 | 1.29E+00 | 7.60E-04 | Yes | D030024E09Rik | intron_variant,non_coding_transcript_variant | MODIFIER |
| 15 | 44773071 | AX-170087840 | 1 | C | A | 0.054 | -4.00E-01 | 9.63E-02 | 9.46E-01 | 9.66E-01 | 7.81E-04 | Yes | Sybu | intron_variant | MODIFIER |
| 15 | 22259388 | AX-170270141 | 0 | A | C | 0.263 | -2.85E-01 | 7.62E-02 | 1.39E+00 | 1.40E+00 | 8.00E-04 | Yes | | intergenic_variant | MODIFIER |
| 15 | 61444434 | AX-170167971 | 0 | A | G | 0.061 | -3.86E-01 | 1.01E-01 | 1.22E+00 | 1.25E+00 | 8.76E-04 | Yes | | intergenic_variant | MODIFIER |
| 15 | 22737729 | AX-170482037 | 1 | G | C | 0.268 | -2.79E-01 | 7.57E-02 | 1.43E+00 | 1.44E+00 | 8.84E-04 | Yes | Cdh18 | intron_variant | MODIFIER |
| 15 | 57349568 | AX-97797891 | 0 | A | G | 0.053 | -3.91E-01 | 9.89E-02 | 1.07E+00 | 1.10E+00 | 9.70E-04 | Yes | Slc22a22 | intron_variant | MODIFIER |
| 15 | 51786269 | AX-170127752 | 0 | C | T | 0.044 | -4.10E-01 | 1.04E-01 | 1.08E+00 | 1.11E+00 | 1.01E-03 | Yes | Eif3h | downstream_gene_variant | MODIFIER |
| 15 | 22539485 | AX-170254362 | 0 | T | A | 0.254 | -2.75E-01 | 7.45E-02 | 1.35E+00 | 1.37E+00 | 1.01E-03 | Yes | | intergenic_variant | MODIFIER |
| 15 | 25067425 | AX-97829075 | 0 | G | A | 0.044 | -4.63E-01 | 9.88E-02 | 5.99E-01 | 6.03E-01 | 1.08E-03 | Yes | | intergenic_variant | MODIFIER |
| 15 | 31004205 | AX-97816856 | 1 | C | T | 0.018 | -4.99E-01 | 9.80E-02 | 4.53E-01 | 4.52E-01 | 1.12E-03 | Yes | Ctnnd2 | intron_variant | MODIFIER |

| | | | | | | | | | | | | | | | |
|---|---|---|---|---|---|---|---|---|---|---|---|---|---|---|---|
| 15 | 20948053 | AX-97804006 | 1 | G | T | 0.062 | -3.98E-01 | 1.02E-01 | 1.07E+00 | 1.10E+00 | 1.16E-03 | Yes | Cdh12 | intron_variant,non_coding_transcript_variant | MODIFIER |
| 15 | 60808403 | AX-170532925 | 0 | C | T | 0.07 | -3.57E-01 | 9.62E-02 | 1.28E+00 | 1.31E+00 | 1.16E-03 | Yes | | intergenic_variant | MODIFIER |
| 15 | 54296657 | AX-170532343 | 0 | C | T | 0.053 | -3.88E-01 | 9.81E-02 | 1.02E+00 | 1.05E+00 | 1.19E-03 | Yes | | intergenic_variant | MODIFIER |
| 15 | 23312072 | AX-170420444 | 1 | C | T | 0.268 | -2.76E-01 | 7.62E-02 | 1.39E+00 | 1.40E+00 | 1.26E-03 | Yes | Cdh18 | intron_variant,non_coding_transcript_variant | MODIFIER |
| 15 | 21785682 | AX-97794581 | 0 | T | C | 0.061 | -3.34E-01 | 9.47E-02 | 1.50E+00 | 1.57E+00 | 1.34E-03 | Yes | | intergenic_variant | MODIFIER |
| 15 | 61014419 | AX-170087880 | 0 | G | T | 0.061 | -3.89E-01 | 1.03E-01 | 1.15E+00 | 1.17E+00 | 1.37E-03 | Yes | | intergenic_variant | MODIFIER |
| 15 | 62114570 | AX-97845755 | 0 | C | T | 0.061 | -3.58E-01 | 9.93E-02 | 1.34E+00 | 1.38E+00 | 1.43E-03 | Yes | Pvt1 | intron_variant,non_coding_transcript_variant | MODIFIER |
| 15 | 25634076 | AX-170355575 | 1 | T | G | 0.268 | -2.73E-01 | 7.59E-02 | 1.36E+00 | 1.37E+00 | 1.44E-03 | Yes | Myo10 | intron_variant | MODIFIER |
| 15 | 16217123 | AX-97787299 | 0 | T | C | 0.281 | -2.48E-01 | 7.33E-02 | 1.74E+00 | 1.77E+00 | 1.49E-03 | Yes | | intergenic_variant | MODIFIER |
| 15 | 44107066 | AX-170161234 | 2 | G | T | 0.045 | -4.32E-01 | 9.89E-02 | 7.09E-01 | 7.18E-01 | 1.51E-03 | Yes | | intergenic_variant | MODIFIER |
| 15 | 23215622 | AX-170433286 | 0 | T | C | 0.272 | -2.72E-01 | 7.64E-02 | 1.38E+00 | 1.39E+00 | 1.53E-03 | Yes | Cdh18 | intron_variant,non_coding_transcript_variant | MODIFIER |
| 15 | 23318774 | AX-170436642 | 0 | G | A | 0.272 | -2.72E-01 | 7.64E-02 | 1.38E+00 | 1.39E+00 | 1.53E-03 | Yes | Cdh18 | intron_variant,non_coding_transcript_variant | MODIFIER |
| 15 | 23589453 | AX-170011233 | 0 | G | T | 0.272 | -2.72E-01 | 7.64E-02 | 1.38E+00 | 1.39E+00 | 1.53E-03 | Yes | | intergenic_variant | MODIFIER |
| 15 | 23589477 | AX-97828992 | 0 | T | G | 0.272 | -2.72E-01 | 7.64E-02 | 1.38E+00 | 1.39E+00 | 1.53E-03 | Yes | | intergenic_variant | MODIFIER |
| 15 | 23664843 | AX-97829074 | 0 | G | T | 0.272 | -2.72E-01 | 7.64E-02 | 1.38E+00 | 1.39E+00 | 1.53E-03 | Yes | | intergenic_variant | MODIFIER |
| 15 | 23700139 | AX-97833946 | 0 | T | G | 0.272 | -2.72E-01 | 7.64E-02 | 1.38E+00 | 1.39E+00 | 1.53E-03 | Yes | | intergenic_variant | MODIFIER |
| 15 | 23764101 | AX-97829016 | 0 | C | T | 0.272 | -2.72E-01 | 7.64E-02 | 1.38E+00 | 1.39E+00 | 1.53E-03 | Yes | | intergenic_variant | MODIFIER |
| 15 | 23836511 | AX-170191437 | 0 | G | A | 0.272 | -2.72E-01 | 7.64E-02 | 1.38E+00 | 1.39E+00 | 1.53E-03 | Yes | | intergenic_variant | MODIFIER |
| 15 | 23839700 | AX-97828583 | 0 | A | G | 0.272 | -2.72E-01 | 7.64E-02 | 1.38E+00 | 1.39E+00 | 1.53E-03 | Yes | | intergenic_variant | MODIFIER |
| 15 | 23839986 | AX-170221131 | 0 | C | T | 0.272 | -2.72E-01 | 7.64E-02 | 1.38E+00 | 1.39E+00 | 1.53E-03 | Yes | | intergenic_variant | MODIFIER |
| 15 | 23859441 | AX-170291684 | 0 | G | T | 0.272 | -2.72E-01 | 7.64E-02 | 1.38E+00 | 1.39E+00 | 1.53E-03 | Yes | | intergenic_variant | MODIFIER |
| 15 | 23869253 | AX-97829018 | 0 | A | G | 0.272 | -2.72E-01 | 7.64E-02 | 1.38E+00 | 1.39E+00 | 1.53E-03 | Yes | | intergenic_variant | MODIFIER |
| 15 | 23902657 | AX-97847923 | 0 | C | T | 0.272 | -2.72E-01 | 7.64E-02 | 1.38E+00 | 1.39E+00 | 1.53E-03 | Yes | | intergenic_variant | MODIFIER |
| 15 | 23916247 | AX-170504371 | 0 | C | T | 0.272 | -2.72E-01 | 7.64E-02 | 1.38E+00 | 1.39E+00 | 1.53E-03 | Yes | | intergenic_variant | MODIFIER |
| 15 | 23920339 | AX-97821730 | 0 | A | G | 0.272 | -2.72E-01 | 7.64E-02 | 1.38E+00 | 1.39E+00 | 1.53E-03 | Yes | | intergenic_variant | MODIFIER |
| 15 | 23925648 | AX-170011255 | 0 | A | C | 0.272 | -2.72E-01 | 7.64E-02 | 1.38E+00 | 1.39E+00 | 1.53E-03 | Yes | | intergenic_variant | MODIFIER |
| 15 | 23965914 | AX-170011257 | 0 | C | G | 0.272 | -2.72E-01 | 7.64E-02 | 1.38E+00 | 1.39E+00 | 1.53E-03 | Yes | | intergenic_variant | MODIFIER |
| 15 | 23981092 | AX-97831848 | 0 | C | T | 0.272 | -2.72E-01 | 7.64E-02 | 1.38E+00 | 1.39E+00 | 1.53E-03 | Yes | | intergenic_variant | MODIFIER |
| 15 | 24153572 | AX-170011254 | 0 | A | C | 0.272 | -2.72E-01 | 7.64E-02 | 1.38E+00 | 1.39E+00 | 1.53E-03 | Yes | | intergenic_variant | MODIFIER |
| 15 | 24210360 | AX-170116703 | 0 | C | T | 0.272 | -2.72E-01 | 7.64E-02 | 1.38E+00 | 1.39E+00 | 1.53E-03 | Yes | Gm8408 | upstream_gene_variant | MODIFIER |
| 15 | 24257160 | AX-170471284 | 0 | C | A | 0.272 | -2.72E-01 | 7.64E-02 | 1.38E+00 | 1.39E+00 | 1.53E-03 | Yes | Gm49729 | intron_variant,non_coding_transcript_variant | MODIFIER |
| 15 | 24331298 | AX-97829045 | 0 | G | A | 0.272 | -2.72E-01 | 7.64E-02 | 1.38E+00 | 1.39E+00 | 1.53E-03 | Yes | Gm49729 | intron_variant,non_coding_transcript_variant | MODIFIER |
| 15 | 24354540 | AX-97856769 | 0 | G | A | 0.272 | -2.72E-01 | 7.64E-02 | 1.38E+00 | 1.39E+00 | 1.53E-03 | Yes | Gm36147 | upstream_gene_variant | MODIFIER |
| 15 | 24386245 | AX-97829914 | 0 | G | A | 0.272 | -2.72E-01 | 7.64E-02 | 1.38E+00 | 1.39E+00 | 1.53E-03 | Yes | | intergenic_variant | MODIFIER |
| 15 | 24935877 | AX-170108413 | 0 | A | G | 0.272 | -2.72E-01 | 7.64E-02 | 1.38E+00 | 1.39E+00 | 1.53E-03 | Yes | | intergenic_variant | MODIFIER |
| 15 | 25319824 | AX-170406756 | 0 | C | A | 0.272 | -2.72E-01 | 7.64E-02 | 1.38E+00 | 1.39E+00 | 1.53E-03 | Yes | 4930445E18Rik | intron_variant,non_coding_transcript_variant | MODIFIER |
| 15 | 25322025 | AX-97838055 | 0 | T | C | 0.272 | -2.72E-01 | 7.64E-02 | 1.38E+00 | 1.39E+00 | 1.53E-03 | Yes | 4930445E18Rik | intron_variant,non_coding_transcript_variant | MODIFIER |
| 15 | 25338750 | AX-170139784 | 0 | C | G | 0.272 | -2.72E-01 | 7.64E-02 | 1.38E+00 | 1.39E+00 | 1.53E-03 | Yes | | intergenic_variant | MODIFIER |
| 15 | 25344189 | AX-170448953 | 0 | G | A | 0.272 | -2.72E-01 | 7.64E-02 | 1.38E+00 | 1.39E+00 | 1.53E-03 | Yes | | intergenic_variant | MODIFIER |
| 15 | 25391610 | AX-170055048 | 0 | T | C | 0.272 | -2.72E-01 | 7.64E-02 | 1.38E+00 | 1.39E+00 | 1.53E-03 | Yes | Basp1 | intron_variant | MODIFIER |
| 15 | 25393779 | AX-170174605 | 0 | T | C | 0.272 | -2.72E-01 | 7.64E-02 | 1.38E+00 | 1.39E+00 | 1.53E-03 | Yes | Basp1 | intron_variant | MODIFIER |
| 15 | 25394750 | AX-170147697 | 0 | T | C | 0.272 | -2.72E-01 | 7.64E-02 | 1.38E+00 | 1.39E+00 | 1.53E-03 | Yes | Basp1 | intron_variant | MODIFIER |
| 15 | 25400007 | AX-170440930 | 0 | C | T | 0.272 | -2.72E-01 | 7.64E-02 | 1.38E+00 | 1.39E+00 | 1.53E-03 | Yes | Basp1 | intron_variant | MODIFIER |
| 15 | 25400515 | AX-170155148 | 0 | G | A | 0.272 | -2.72E-01 | 7.64E-02 | 1.38E+00 | 1.39E+00 | 1.53E-03 | Yes | Basp1 | intron_variant | MODIFIER |
| 15 | 25449925 | AX-170490511 | 0 | T | C | 0.272 | -2.72E-01 | 7.64E-02 | 1.38E+00 | 1.39E+00 | 1.53E-03 | Yes | Gm48956 | upstream_gene_variant | MODIFIER |
| 15 | 25464761 | AX-170496671 | 0 | T | G | 0.272 | -2.72E-01 | 7.64E-02 | 1.38E+00 | 1.39E+00 | 1.53E-03 | Yes | Gm48957 | downstream_gene_variant | MODIFIER |
| 15 | 25498070 | AX-170183083 | 0 | C | A | 0.272 | -2.72E-01 | 7.64E-02 | 1.38E+00 | 1.39E+00 | 1.53E-03 | Yes | | intergenic_variant | MODIFIER |
| 15 | 25498477 | AX-170276773 | 0 | A | C | 0.272 | -2.72E-01 | 7.64E-02 | 1.38E+00 | 1.39E+00 | 1.53E-03 | Yes | | intergenic_variant | MODIFIER |
| 15 | 25518691 | AX-170529324 | 0 | G | A | 0.272 | -2.72E-01 | 7.64E-02 | 1.38E+00 | 1.39E+00 | 1.53E-03 | Yes | | regulatory_region_variant | MODIFIER |
| 15 | 25539648 | AX-97806857 | 0 | A | G | 0.272 | -2.72E-01 | 7.64E-02 | 1.38E+00 | 1.39E+00 | 1.53E-03 | Yes | Gm36642 | intron_variant,non_coding_transcript_variant | MODIFIER |
| 15 | 25541699 | AX-170159255 | 0 | T | A | 0.272 | -2.72E-01 | 7.64E-02 | 1.38E+00 | 1.39E+00 | 1.53E-03 | Yes | Gm36642 | intron_variant,non_coding_transcript_variant | MODIFIER |
| 15 | 25581138 | AX-170440840 | 0 | C | T | 0.272 | -2.72E-01 | 7.64E-02 | 1.38E+00 | 1.39E+00 | 1.53E-03 | Yes | Gm36642 | intron_variant,non_coding_transcript_variant | MODIFIER |
| 15 | 25581322 | AX-170315510 | 0 | T | C | 0.272 | -2.72E-01 | 7.64E-02 | 1.38E+00 | 1.39E+00 | 1.53E-03 | Yes | Gm36642 | intron_variant,non_coding_transcript_variant | MODIFIER |
| 15 | 25636400 | AX-97828775 | 0 | A | G | 0.272 | -2.72E-01 | 7.64E-02 | 1.38E+00 | 1.39E+00 | 1.53E-03 | Yes | Myo10 | intron_variant | MODIFIER |

| | | | | | | | | | | | | | | | |
|---|---|---|---|---|---|---|---|---|---|---|---|---|---|---|---|
| 15 | 25650840 | AX-170370154 | 0 | A | T | 0.272 | -2.72E-01 | 7.64E-02 | 1.38E+00 | 1.39E+00 | 1.53E-03 | Yes | Myo10 | intron_variant | MODIFIER |
| 15 | 25681659 | AX-170195784 | 0 | C | T | 0.272 | -2.72E-01 | 7.64E-02 | 1.38E+00 | 1.39E+00 | 1.53E-03 | Yes | Myo10 | intron_variant | MODIFIER |
| 15 | 25709305 | AX-170055060 | 0 | C | T | 0.272 | -2.72E-01 | 7.64E-02 | 1.38E+00 | 1.39E+00 | 1.53E-03 | Yes | Myo10 | intron_variant | MODIFIER |
| 15 | 25711221 | AX-170479666 | 0 | C | T | 0.272 | -2.72E-01 | 7.64E-02 | 1.38E+00 | 1.39E+00 | 1.53E-03 | Yes | Myo10 | intron_variant | MODIFIER |
| 15 | 60649053 | AX-170397966 | 0 | C | G | 0.061 | -3.64E-01 | 1.00E-01 | 1.28E+00 | 1.31E+00 | 1.58E-03 | Yes | | intergenic_variant | MODIFIER |
| 15 | 20818544 | AX-97828898 | 2 | G | T | 0.064 | -3.69E-01 | 1.03E-01 | 1.32E+00 | 1.37E+00 | 1.61E-03 | Yes | Cdh12 | intron_variant,non_coding_transcript_variant | MODIFIER |
| 5 | 137656384 | AX-170272186 | 2 | A | G | 0.127 | -2.75E-01 | 8.60E-02 | 2.71E+00 | 2.83E+00 | 1.68E-03 | Yes | Agfg2 | intron_variant | MODIFIER |
| 15 | 61533947 | AX-170479147 | 0 | G | A | 0.07 | -3.61E-01 | 9.84E-02 | 1.21E+00 | 1.24E+00 | 1.69E-03 | Yes | Gm49498 | downstream_gene_variant | MODIFIER |
| 15 | 21633505 | AX-170101314 | 1 | C | G | 0.089 | -2.17E-01 | 6.19E-02 | 1.40E+00 | 1.45E+00 | 1.73E-03 | Yes | | intergenic_variant | MODIFIER |
| 15 | 25598562 | AX-170011283 | 0 | G | A | 0.044 | -4.24E-01 | 9.86E-02 | 7.22E-01 | 7.40E-01 | 1.76E-03 | Yes | | intergenic_variant | MODIFIER |
| 15 | 60919815 | AX-170055699 | 0 | C | T | 0.07 | -3.64E-01 | 9.74E-02 | 1.12E+00 | 1.14E+00 | 1.76E-03 | Yes | A1bg | synonymous_variant | LOW |
| 2 | 96919640 | AX-170502792 | 0 | G | A | 0.053 | -5.35E-01 | 1.63E-01 | 4.04E+00 | 4.36E+00 | 1.77E-03 | Yes | Lrrc4c | intron_variant | MODIFIER |
| 15 | 24258331 | AX-170357682 | 1 | A | C | 0.268 | -2.67E-01 | 7.63E-02 | 1.38E+00 | 1.39E+00 | 1.87E-03 | Yes | Gm49729 | intron_variant,non_coding_transcript_variant | MODIFIER |
| 13 | 10438176 | AX-97783303 | 0 | A | G | 0.368 | -1.90E-01 | 5.94E-02 | 3.60E+00 | 3.77E+00 | 2.04E-03 | Yes | Gm26861 | intron_variant,non_coding_transcript_variant | MODIFIER |
| 13 | 10740957 | AX-170007746 | 0 | C | G | 0.368 | -1.90E-01 | 5.94E-02 | 3.60E+00 | 3.77E+00 | 2.04E-03 | Yes | Gm26861 | downstream_gene_variant | MODIFIER |
| 3 | 19268471 | AX-170265390 | 0 | T | C | 0.132 | -2.81E-01 | 8.91E-02 | 3.27E+00 | 3.45E+00 | 2.06E-03 | Yes | Pde7a | upstream_gene_variant | MODIFIER |
| 15 | 23764417 | AX-170011244 | 1 | C | G | 0.259 | -2.42E-01 | 7.16E-02 | 1.52E+00 | 1.56E+00 | 2.08E-03 | Yes | | intergenic_variant | MODIFIER |
| 3 | 7239833 | AX-97854835 | 1 | C | T | 0.134 | -2.55E-01 | 8.18E-02 | 2.93E+00 | 3.08E+00 | 2.20E-03 | Yes | | intergenic_variant | MODIFIER |
| 13 | 8044918 | AX-170007707 | 1 | A | T | 0.188 | -3.73E-01 | 1.18E-01 | 3.67E+00 | 3.85E+00 | 2.25E-03 | Yes | | intergenic_variant | MODIFIER |
| 5 | 137873055 | AX-170026292 | 0 | A | C | 0.132 | -2.66E-01 | 8.60E-02 | 2.75E+00 | 2.88E+00 | 2.29E-03 | Yes | Pilrb2 | upstream_gene_variant | MODIFIER |
| 13 | 19530363 | AX-97835105 | 1 | T | C | 0.161 | -3.47E-01 | 1.12E-01 | 3.04E+00 | 3.20E+00 | 2.34E-03 | Yes | Gm18363 | downstream_gene_variant | MODIFIER |
| 15 | 23156843 | AX-170489231 | 0 | T | C | 0.263 | -2.56E-01 | 7.70E-02 | 1.55E+00 | 1.58E+00 | 2.45E-03 | Yes | Cdh18 | intron_variant,non_coding_transcript_variant | MODIFIER |
| 15 | 58362098 | AX-97851489 | 2 | G | A | 0.055 | -3.71E-01 | 9.83E-02 | 1.00E+00 | 1.03E+00 | 2.46E-03 | Yes | Anxa13 | intron_variant,non_coding_transcript_variant | MODIFIER |
| 15 | 24364599 | AX-97829044 | 0 | T | G | 0.272 | -2.57E-01 | 7.64E-02 | 1.43E+00 | 1.45E+00 | 2.65E-03 | Yes | Gm49729 | intron_variant,non_coding_transcript_variant | MODIFIER |
| 15 | 77624453 | AX-170406145 | 0 | C | G | 0.079 | -3.33E-01 | 1.03E-01 | 1.65E+00 | 1.71E+00 | 2.67E-03 | Yes | Gm8221 | intron_variant,non_coding_transcript_variant | MODIFIER |
| 15 | 25637640 | AX-170437020 | 1 | C | T | 0.277 | -2.60E-01 | 7.73E-02 | 1.42E+00 | 1.43E+00 | 2.69E-03 | Yes | Myo10 | intron_variant | MODIFIER |
| 5 | 53316706 | AX-170285645 | 0 | T | C | 0.491 | 2.96E-01 | 9.79E-02 | 2.87E+00 | 2.98E+00 | 2.84E-03 | Yes | | intergenic_variant | MODIFIER |
| 15 | 42684780 | AX-170234758 | 0 | A | C | 0.035 | -4.03E-01 | 1.02E-01 | 8.55E-01 | 8.73E-01 | 2.90E-03 | Yes | Gm17473 | intron_variant,non_coding_transcript_variant | MODIFIER |
| 15 | 23879450 | AX-170406198 | 2 | G | A | 0.282 | -2.57E-01 | 7.77E-02 | 1.46E+00 | 1.48E+00 | 3.02E-03 | Yes | | intergenic_variant | MODIFIER |
| 3 | 19201530 | AX-170115734 | 0 | A | C | 0.123 | -2.79E-01 | 9.15E-02 | 3.51E+00 | 3.72E+00 | 3.06E-03 | Yes | Mtfr1 | intron_variant,NMD_transcript_variant | MODIFIER |
| 15 | 43510394 | AX-170193225 | 0 | T | C | 0.035 | -3.79E-01 | 1.02E-01 | 1.01E+00 | 1.04E+00 | 3.17E-03 | Yes | Emc2 | intron_variant | MODIFIER |
| 15 | 21784589 | AX-170055002 | 1 | C | T | 0.062 | -3.54E-01 | 1.05E-01 | 1.33E+00 | 1.36E+00 | 3.29E-03 | Yes | | intergenic_variant | MODIFIER |
| 15 | 60693833 | AX-97821592 | 0 | C | T | 0.07 | -3.52E-01 | 9.96E-02 | 1.13E+00 | 1.15E+00 | 3.42E-03 | Yes | | regulatory_region_variant | MODIFIER |
| 15 | 27007597 | AX-97783354 | 0 | G | T | 0.044 | -4.32E-01 | 9.85E-02 | 6.07E-01 | 6.20E-01 | 3.42E-03 | Yes | | intergenic_variant | MODIFIER |
| 15 | 13766285 | AX-170112618 | 0 | T | A | 0.061 | -3.59E-01 | 1.05E-01 | 1.26E+00 | 1.29E+00 | 3.46E-03 | Yes | | intergenic_variant | MODIFIER |
| 15 | 15630671 | AX-170446239 | 0 | T | C | 0.061 | -3.59E-01 | 1.05E-01 | 1.26E+00 | 1.29E+00 | 3.46E-03 | Yes | | intergenic_variant | MODIFIER |
| 15 | 20882475 | AX-170428258 | 0 | T | C | 0.061 | -3.59E-01 | 1.05E-01 | 1.26E+00 | 1.29E+00 | 3.46E-03 | Yes | Cdh12 | intron_variant,non_coding_transcript_variant | MODIFIER |
| 15 | 20942228 | AX-170447277 | 0 | C | T | 0.061 | -3.59E-01 | 1.05E-01 | 1.26E+00 | 1.29E+00 | 3.46E-03 | Yes | Cdh12 | intron_variant,non_coding_transcript_variant | MODIFIER |
| 15 | 21154862 | AX-170011197 | 0 | C | A | 0.061 | -3.59E-01 | 1.05E-01 | 1.26E+00 | 1.29E+00 | 3.46E-03 | Yes | Cdh12 | intron_variant | MODIFIER |
| 15 | 21488358 | AX-170087791 | 0 | A | G | 0.061 | -3.59E-01 | 1.05E-01 | 1.26E+00 | 1.29E+00 | 3.46E-03 | Yes | Cdh12 | intron_variant | MODIFIER |
| 15 | 21589016 | AX-170087792 | 0 | T | C | 0.061 | -3.59E-01 | 1.05E-01 | 1.26E+00 | 1.29E+00 | 3.46E-03 | Yes | Cdh12 | 3_prime_UTR_variant | MODIFIER |
| 15 | 21596411 | AX-170323749 | 0 | G | T | 0.061 | -3.59E-01 | 1.05E-01 | 1.26E+00 | 1.29E+00 | 3.46E-03 | Yes | | intergenic_variant | MODIFIER |
| 15 | 21775306 | AX-170055003 | 0 | T | G | 0.061 | -3.59E-01 | 1.05E-01 | 1.26E+00 | 1.29E+00 | 3.46E-03 | Yes | | intergenic_variant | MODIFIER |
| 15 | 21775337 | AX-97790406 | 0 | T | C | 0.061 | -3.59E-01 | 1.05E-01 | 1.26E+00 | 1.29E+00 | 3.46E-03 | Yes | | intergenic_variant | MODIFIER |
| 15 | 21777898 | AX-170441513 | 0 | A | C | 0.061 | -3.59E-01 | 1.05E-01 | 1.26E+00 | 1.29E+00 | 3.46E-03 | Yes | | intergenic_variant | MODIFIER |
| 15 | 21800196 | AX-170011202 | 0 | C | T | 0.061 | -3.59E-01 | 1.05E-01 | 1.26E+00 | 1.29E+00 | 3.46E-03 | Yes | | intergenic_variant | MODIFIER |
| 15 | 21824462 | AX-170011206 | 0 | G | A | 0.061 | -3.59E-01 | 1.05E-01 | 1.26E+00 | 1.29E+00 | 3.46E-03 | Yes | | intergenic_variant | MODIFIER |
| 15 | 21824654 | AX-170011205 | 0 | A | G | 0.061 | -3.59E-01 | 1.05E-01 | 1.26E+00 | 1.29E+00 | 3.46E-03 | Yes | | intergenic_variant | MODIFIER |
| 15 | 21824710 | AX-170481191 | 0 | C | G | 0.061 | -3.59E-01 | 1.05E-01 | 1.26E+00 | 1.29E+00 | 3.46E-03 | Yes | | intergenic_variant | MODIFIER |
| 15 | 21825775 | AX-170011201 | 0 | G | A | 0.061 | -3.59E-01 | 1.05E-01 | 1.26E+00 | 1.29E+00 | 3.46E-03 | Yes | | intergenic_variant | MODIFIER |
| 15 | 21855737 | AX-170435016 | 0 | C | T | 0.061 | -3.59E-01 | 1.05E-01 | 1.26E+00 | 1.29E+00 | 3.46E-03 | Yes | | intergenic_variant | MODIFIER |
| 15 | 21898553 | AX-170254674 | 0 | T | C | 0.061 | -3.59E-01 | 1.05E-01 | 1.26E+00 | 1.29E+00 | 3.46E-03 | Yes | | intergenic_variant | MODIFIER |
| 15 | 21905080 | AX-97800045 | 0 | G | T | 0.061 | -3.59E-01 | 1.05E-01 | 1.26E+00 | 1.29E+00 | 3.46E-03 | Yes | | intergenic_variant | MODIFIER |
| 15 | 21925584 | AX-97796688 | 0 | C | T | 0.061 | -3.59E-01 | 1.05E-01 | 1.26E+00 | 1.29E+00 | 3.46E-03 | Yes | | intergenic_variant | MODIFIER |

| | | | | | | | | | | | | | | | |
|---|---|---|---|---|---|---|---|---|---|---|---|---|---|---|---|
| 15 | 21964397 | AX-97847553 | 0 | C | A | 0.061 | -3.59E-01 | 1.05E-01 | 1.26E+00 | 1.29E+00 | 3.46E-03 | Yes | Gm49187 | downstream_gene_variant | MODIFIER |
| 15 | 23834491 | AX-170170892 | 1 | C | T | 0.277 | -2.54E-01 | 7.60E-02 | 1.34E+00 | 1.36E+00 | 3.48E-03 | Yes | | intergenic_variant | MODIFIER |
| 15 | 22592518 | AX-170011210 | 0 | A | G | 0.272 | -2.45E-01 | 7.47E-02 | 1.43E+00 | 1.45E+00 | 3.59E-03 | Yes | Cdh18 | intron_variant | MODIFIER |
| 15 | 20948017 | AX-170490368 | 1 | C | A | 0.295 | -2.36E-01 | 7.66E-02 | 1.77E+00 | 1.79E+00 | 3.79E-03 | Yes | Cdh12 | intron_variant,non_coding_transcript_variant | MODIFIER |
| 15 | 21540574 | AX-170152404 | 0 | T | A | 0.307 | -2.35E-01 | 7.74E-02 | 1.86E+00 | 1.89E+00 | 3.88E-03 | Yes | Cdh12 | intron_variant | MODIFIER |
| 15 | 21886454 | AX-170276505 | 1 | C | T | 0.062 | -3.53E-01 | 1.06E-01 | 1.29E+00 | 1.32E+00 | 3.94E-03 | Yes | | intergenic_variant | MODIFIER |
| 15 | 24331005 | AX-170426828 | 1 | C | G | 0.277 | -2.51E-01 | 7.62E-02 | 1.36E+00 | 1.37E+00 | 3.97E-03 | Yes | Gm49729 | intron_variant,non_coding_transcript_variant | MODIFIER |
| 15 | 21479273 | AX-97828926 | 1 | T | C | 0.295 | -2.35E-01 | 7.70E-02 | 1.77E+00 | 1.79E+00 | 4.10E-03 | Yes | Cdh12 | intron_variant | MODIFIER |
| 15 | 17670242 | AX-170115293 | 1 | T | C | 0.295 | -2.22E-01 | 7.35E-02 | 1.85E+00 | 1.89E+00 | 4.21E-03 | Yes | | intergenic_variant | MODIFIER |
| 15 | 54306960 | AX-170218083 | 0 | T | C | 0.044 | -3.55E-01 | 1.05E-01 | 1.22E+00 | 1.25E+00 | 4.49E-03 | Yes | Gm7566 | upstream_gene_variant | MODIFIER |
| 15 | 14547561 | AX-97820986 | 0 | A | C | 0.298 | -2.33E-01 | 7.70E-02 | 1.77E+00 | 1.79E+00 | 4.52E-03 | Yes | | intergenic_variant | MODIFIER |
| 15 | 15699583 | AX-97821733 | 0 | T | A | 0.298 | -2.33E-01 | 7.70E-02 | 1.77E+00 | 1.79E+00 | 4.52E-03 | Yes | | intergenic_variant | MODIFIER |
| 15 | 15973864 | AX-170191277 | 0 | G | A | 0.298 | -2.33E-01 | 7.70E-02 | 1.77E+00 | 1.79E+00 | 4.52E-03 | Yes | | intergenic_variant | MODIFIER |
| 15 | 16081136 | AX-97828747 | 0 | C | A | 0.298 | -2.33E-01 | 7.70E-02 | 1.77E+00 | 1.79E+00 | 4.52E-03 | Yes | | intergenic_variant | MODIFIER |
| 15 | 16140738 | AX-170087778 | 0 | G | T | 0.298 | -2.33E-01 | 7.70E-02 | 1.77E+00 | 1.79E+00 | 4.52E-03 | Yes | | intergenic_variant | MODIFIER |
| 15 | 16273054 | AX-170011126 | 0 | T | C | 0.298 | -2.33E-01 | 7.70E-02 | 1.77E+00 | 1.79E+00 | 4.52E-03 | Yes | | intergenic_variant | MODIFIER |
| 15 | 16276366 | AX-97790669 | 0 | C | A | 0.298 | -2.33E-01 | 7.70E-02 | 1.77E+00 | 1.79E+00 | 4.52E-03 | Yes | | intergenic_variant | MODIFIER |
| 15 | 16324679 | AX-97796421 | 0 | T | C | 0.298 | -2.33E-01 | 7.70E-02 | 1.77E+00 | 1.79E+00 | 4.52E-03 | Yes | | intergenic_variant | MODIFIER |
| 15 | 16400479 | AX-97818473 | 0 | G | A | 0.298 | -2.33E-01 | 7.70E-02 | 1.77E+00 | 1.79E+00 | 4.52E-03 | Yes | | intergenic_variant | MODIFIER |
| 15 | 16400552 | AX-170011121 | 0 | G | A | 0.298 | -2.33E-01 | 7.70E-02 | 1.77E+00 | 1.79E+00 | 4.52E-03 | Yes | | intergenic_variant | MODIFIER |
| 15 | 16409750 | AX-170312509 | 0 | G | A | 0.298 | -2.33E-01 | 7.70E-02 | 1.77E+00 | 1.79E+00 | 4.52E-03 | Yes | | intergenic_variant | MODIFIER |
| 15 | 16489057 | AX-97789225 | 0 | A | G | 0.298 | -2.33E-01 | 7.70E-02 | 1.77E+00 | 1.79E+00 | 4.52E-03 | Yes | Gm41276 | non_coding_transcript_exon_variant | MODIFIER |
| 15 | 16498332 | AX-170011130 | 0 | A | G | 0.298 | -2.33E-01 | 7.70E-02 | 1.77E+00 | 1.79E+00 | 4.52E-03 | Yes | Gm49126 | intron_variant,non_coding_transcript_variant | MODIFIER |
| 15 | 16556760 | AX-170011128 | 0 | C | T | 0.298 | -2.33E-01 | 7.70E-02 | 1.77E+00 | 1.79E+00 | 4.52E-03 | Yes | | intergenic_variant | MODIFIER |
| 15 | 16575745 | AX-170011124 | 0 | T | A | 0.298 | -2.33E-01 | 7.70E-02 | 1.77E+00 | 1.79E+00 | 4.52E-03 | Yes | | intergenic_variant | MODIFIER |
| 15 | 17134910 | AX-170309307 | 0 | A | G | 0.298 | -2.33E-01 | 7.70E-02 | 1.77E+00 | 1.79E+00 | 4.52E-03 | Yes | | intergenic_variant | MODIFIER |
| 15 | 17202219 | AX-97853439 | 0 | A | C | 0.298 | -2.33E-01 | 7.70E-02 | 1.77E+00 | 1.79E+00 | 4.52E-03 | Yes | | intergenic_variant | MODIFIER |
| 15 | 17205950 | AX-170124249 | 0 | T | C | 0.298 | -2.33E-01 | 7.70E-02 | 1.77E+00 | 1.79E+00 | 4.52E-03 | Yes | | intergenic_variant | MODIFIER |
| 15 | 17315766 | AX-97803127 | 0 | A | G | 0.298 | -2.33E-01 | 7.70E-02 | 1.77E+00 | 1.79E+00 | 4.52E-03 | Yes | | intergenic_variant | MODIFIER |
| 15 | 17328268 | AX-170134231 | 0 | A | G | 0.298 | -2.33E-01 | 7.70E-02 | 1.77E+00 | 1.79E+00 | 4.52E-03 | Yes | | intergenic_variant | MODIFIER |
| 15 | 17328530 | AX-170141528 | 0 | G | A | 0.298 | -2.33E-01 | 7.70E-02 | 1.77E+00 | 1.79E+00 | 4.52E-03 | Yes | | intergenic_variant | MODIFIER |
| 15 | 17374855 | AX-97805578 | 0 | G | A | 0.298 | -2.33E-01 | 7.70E-02 | 1.77E+00 | 1.79E+00 | 4.52E-03 | Yes | | intergenic_variant | MODIFIER |
| 15 | 17396860 | AX-170087783 | 0 | A | G | 0.298 | -2.33E-01 | 7.70E-02 | 1.77E+00 | 1.79E+00 | 4.52E-03 | Yes | | intergenic_variant | MODIFIER |
| 15 | 17468297 | AX-170147634 | 0 | C | T | 0.298 | -2.33E-01 | 7.70E-02 | 1.77E+00 | 1.79E+00 | 4.52E-03 | Yes | | intergenic_variant | MODIFIER |
| 15 | 17468546 | AX-170396164 | 0 | A | T | 0.298 | -2.33E-01 | 7.70E-02 | 1.77E+00 | 1.79E+00 | 4.52E-03 | Yes | | intergenic_variant | MODIFIER |
| 15 | 17498111 | AX-170011145 | 0 | G | A | 0.298 | -2.33E-01 | 7.70E-02 | 1.77E+00 | 1.79E+00 | 4.52E-03 | Yes | | intergenic_variant | MODIFIER |
| 15 | 17602743 | AX-97828826 | 0 | G | A | 0.298 | -2.33E-01 | 7.70E-02 | 1.77E+00 | 1.79E+00 | 4.52E-03 | Yes | | intergenic_variant | MODIFIER |
| 15 | 17670114 | AX-170283656 | 0 | G | A | 0.298 | -2.33E-01 | 7.70E-02 | 1.77E+00 | 1.79E+00 | 4.52E-03 | Yes | | intergenic_variant | MODIFIER |
| 15 | 17689764 | AX-170166703 | 0 | G | T | 0.298 | -2.33E-01 | 7.70E-02 | 1.77E+00 | 1.79E+00 | 4.52E-03 | Yes | | intergenic_variant | MODIFIER |
| 15 | 17706259 | AX-170110182 | 0 | G | A | 0.298 | -2.33E-01 | 7.70E-02 | 1.77E+00 | 1.79E+00 | 4.52E-03 | Yes | | intergenic_variant | MODIFIER |
| 15 | 17712544 | AX-97819886 | 0 | A | C | 0.298 | -2.33E-01 | 7.70E-02 | 1.77E+00 | 1.79E+00 | 4.52E-03 | Yes | | intergenic_variant | MODIFIER |
| 15 | 17737829 | AX-170054917 | 0 | G | C | 0.298 | -2.33E-01 | 7.70E-02 | 1.77E+00 | 1.79E+00 | 4.52E-03 | Yes | | intergenic_variant | MODIFIER |
| 15 | 18721877 | AX-170087786 | 0 | T | C | 0.298 | -2.33E-01 | 7.70E-02 | 1.77E+00 | 1.79E+00 | 4.52E-03 | Yes | C030047K22Rik | intron_variant,non_coding_transcript_variant | MODIFIER |
| 15 | 20958905 | AX-170317416 | 0 | T | C | 0.298 | -2.33E-01 | 7.70E-02 | 1.77E+00 | 1.79E+00 | 4.52E-03 | Yes | Cdh12 | intron_variant,non_coding_transcript_variant | MODIFIER |
| 15 | 21136977 | AX-170175983 | 0 | T | A | 0.298 | -2.33E-01 | 7.70E-02 | 1.77E+00 | 1.79E+00 | 4.52E-03 | Yes | Cdh12 | intron_variant | MODIFIER |
| 15 | 21230531 | AX-170011199 | 0 | A | G | 0.298 | -2.33E-01 | 7.70E-02 | 1.77E+00 | 1.79E+00 | 4.52E-03 | Yes | Cdh12 | intron_variant | MODIFIER |
| 15 | 21236245 | AX-97805286 | 0 | C | T | 0.298 | -2.33E-01 | 7.70E-02 | 1.77E+00 | 1.79E+00 | 4.52E-03 | Yes | Cdh12 | intron_variant | MODIFIER |
| 15 | 21242900 | AX-170011196 | 0 | A | G | 0.298 | -2.33E-01 | 7.70E-02 | 1.77E+00 | 1.79E+00 | 4.52E-03 | Yes | Cdh12 | intron_variant | MODIFIER |
| 15 | 21383805 | AX-170534937 | 0 | G | A | 0.298 | -2.33E-01 | 7.70E-02 | 1.77E+00 | 1.79E+00 | 4.52E-03 | Yes | Cdh12 | intron_variant | MODIFIER |
| 15 | 21434833 | AX-170487403 | 0 | C | G | 0.298 | -2.33E-01 | 7.70E-02 | 1.77E+00 | 1.79E+00 | 4.52E-03 | Yes | Cdh12 | intron_variant | MODIFIER |
| 15 | 21457358 | AX-170054996 | 0 | C | T | 0.298 | -2.33E-01 | 7.70E-02 | 1.77E+00 | 1.79E+00 | 4.52E-03 | Yes | Cdh12 | intron_variant | MODIFIER |
| 15 | 21512958 | AX-170054993 | 0 | C | T | 0.298 | -2.33E-01 | 7.70E-02 | 1.77E+00 | 1.79E+00 | 4.52E-03 | Yes | Cdh12 | intron_variant | MODIFIER |
| 15 | 21515482 | AX-97801553 | 0 | C | T | 0.298 | -2.33E-01 | 7.70E-02 | 1.77E+00 | 1.79E+00 | 4.52E-03 | Yes | Cdh12 | intron_variant | MODIFIER |
| 15 | 21515642 | AX-170238093 | 0 | T | C | 0.298 | -2.33E-01 | 7.70E-02 | 1.77E+00 | 1.79E+00 | 4.52E-03 | Yes | Cdh12 | intron_variant | MODIFIER |

| | | | | | | | | | | | | | | | |
|---|---|---|---|---|---|---|---|---|---|---|---|---|---|---|---|
| 15 | 21535016 | AX-170389077 | 0 | A | C | 0.298 | -2.33E-01 | 7.70E-02 | 1.77E+00 | 1.79E+00 | 4.52E-03 | Yes | Cdh12 | intron_variant | MODIFIER |
| 15 | 21586962 | AX-170100550 | 0 | G | A | 0.298 | -2.33E-01 | 7.70E-02 | 1.77E+00 | 1.79E+00 | 4.52E-03 | Yes | Cdh12 | 3_prime_UTR_variant | MODIFIER |
| 15 | 21695800 | AX-97824007 | 0 | C | T | 0.298 | -2.33E-01 | 7.70E-02 | 1.77E+00 | 1.79E+00 | 4.52E-03 | Yes | | intergenic_variant | MODIFIER |
| 5 | 137615726 | AX-97795940 | 1 | T | C | 0.143 | -2.37E-01 | 8.36E-02 | 2.72E+00 | 2.86E+00 | 5.01E-03 | Yes | Pcolce | upstream_gene_variant | MODIFIER |
| 15 | 21540643 | AX-97794851 | 0 | T | C | 0.298 | -2.30E-01 | 7.71E-02 | 1.77E+00 | 1.79E+00 | 5.09E-03 | Yes | Cdh12 | intron_variant | MODIFIER |
| 13 | 4594361 | AX-170185428 | 0 | T | C | 0.404 | -1.69E-01 | 5.86E-02 | 3.54E+00 | 3.71E+00 | 5.12E-03 | no | Akr1e1 | intron_variant | MODIFIER |
| 15 | 17328701 | AX-170172799 | 0 | G | A | 0.298 | -2.30E-01 | 7.70E-02 | 1.77E+00 | 1.79E+00 | 5.14E-03 | Yes | | intergenic_variant | MODIFIER |
| 3 | 6412475 | AX-170522872 | 2 | A | T | 0.109 | -2.47E-01 | 8.81E-02 | 2.80E+00 | 2.92E+00 | 5.42E-03 | Yes | | intergenic_variant | MODIFIER |
| 15 | 21383891 | AX-170424741 | 1 | G | A | 0.304 | -2.26E-01 | 7.66E-02 | 1.81E+00 | 1.84E+00 | 5.47E-03 | Yes | Cdh12 | intron_variant | MODIFIER |
| 15 | 15913916 | AX-97828721 | 0 | T | A | 0.298 | -2.28E-01 | 7.70E-02 | 1.78E+00 | 1.80E+00 | 5.55E-03 | Yes | | intergenic_variant | MODIFIER |
| 15 | 21309389 | AX-170011200 | 2 | A | T | 0.3 | -2.26E-01 | 7.64E-02 | 1.76E+00 | 1.78E+00 | 5.60E-03 | Yes | Cdh12 | intron_variant | MODIFIER |
| 3 | 6631264 | AX-170019844 | 0 | T | C | 0.114 | -2.47E-01 | 8.80E-02 | 2.97E+00 | 3.12E+00 | 5.64E-03 | Yes | | intergenic_variant | MODIFIER |
| 15 | 21426926 | AX-97797323 | 1 | T | C | 0.304 | -2.26E-01 | 7.64E-02 | 1.76E+00 | 1.78E+00 | 5.72E-03 | Yes | Cdh12 | intron_variant | MODIFIER |
| 13 | 19477564 | AX-170007878 | 0 | G | A | 0.167 | -3.42E-01 | 1.21E-01 | 3.30E+00 | 3.45E+00 | 5.82E-03 | Yes | 9330199G10Rik | intron_variant,non_coding_transcript_variant | MODIFIER |
| 8 | 7551389 | AX-97845694 | 1 | A | G | 0.018 | -3.52E-01 | 1.25E-01 | 3.41E+00 | 3.68E+00 | 5.90E-03 | Yes | Gm30954 | intron_variant,non_coding_transcript_variant | MODIFIER |
| 15 | 17691492 | AX-170128122 | 1 | C | T | 0.295 | -2.26E-01 | 7.57E-02 | 1.65E+00 | 1.68E+00 | 5.92E-03 | Yes | | intergenic_variant | MODIFIER |
| 13 | 12258950 | AX-97796063 | 0 | A | G | 0.184 | -2.95E-01 | 1.07E-01 | 2.68E+00 | 2.79E+00 | 6.09E-03 | Yes | Mtr | upstream_gene_variant | MODIFIER |
| 5 | 129997721 | AX-170330741 | 0 | C | T | 0.149 | -2.32E-01 | 8.34E-02 | 2.33E+00 | 2.43E+00 | 6.25E-03 | Yes | Gusb | intron_variant | MODIFIER |
| 15 | 16381845 | AX-170011131 | 1 | C | T | 0.304 | -2.24E-01 | 7.67E-02 | 1.75E+00 | 1.77E+00 | 6.46E-03 | Yes | | intergenic_variant | MODIFIER |
| 14 | 100607110 | AX-170482745 | 2 | G | T | 0.036 | -4.09E-01 | 1.46E-01 | 3.63E+00 | 3.92E+00 | 6.68E-03 | Yes | | intergenic_variant | MODIFIER |
| 5 | 65772803 | AX-170466405 | 0 | T | G | 0.482 | 2.72E-01 | 9.94E-02 | 2.93E+00 | 3.03E+00 | 6.78E-03 | Yes | N4bp2 | intron_variant | MODIFIER |
| 13 | 5709469 | AX-170357746 | 0 | C | G | 0.202 | -3.08E-01 | 1.10E-01 | 3.54E+00 | 3.73E+00 | 6.82E-03 | Yes | Gm35330 | intron_variant,non_coding_transcript_variant | MODIFIER |
| 5 | 138005987 | AX-170241029 | 1 | T | G | 0.125 | -2.24E-01 | 8.10E-02 | 2.20E+00 | 2.30E+00 | 6.85E-03 | Yes | | intergenic_variant | MODIFIER |
| 15 | 11457226 | AX-170054807 | 0 | G | A | 0.298 | -2.28E-01 | 7.65E-02 | 1.56E+00 | 1.58E+00 | 6.98E-03 | Yes | | intergenic_variant | MODIFIER |
| 10 | 104572293 | AX-170502101 | 0 | C | A | 0.289 | -2.26E-01 | 8.21E-02 | 3.31E+00 | 3.44E+00 | 7.05E-03 | Yes | | intergenic_variant | MODIFIER |
| 15 | 21372040 | AX-170087790 | 1 | T | G | 0.304 | -2.21E-01 | 7.63E-02 | 1.72E+00 | 1.74E+00 | 7.18E-03 | Yes | Cdh12 | intron_variant | MODIFIER |
| 19 | 55398753 | AX-170520507 | 0 | T | C | 0.482 | 1.73E-01 | 6.39E-02 | 2.99E+00 | 3.08E+00 | 7.39E-03 | Yes | Vti1a | intron_variant | MODIFIER |
| 19 | 55331988 | AX-168415332 | 1 | C | G | 0.5 | 1.74E-01 | 6.42E-02 | 3.05E+00 | 3.15E+00 | 7.43E-03 | Yes | Vti1a | intron_variant | MODIFIER |
| 15 | 16366040 | AX-170011123 | 0 | A | T | 0.298 | -2.18E-01 | 7.77E-02 | 1.90E+00 | 1.93E+00 | 7.52E-03 | Yes | | intergenic_variant | MODIFIER |
| 15 | 21278338 | AX-97854162 | 0 | T | C | 0.298 | -2.18E-01 | 7.77E-02 | 1.90E+00 | 1.93E+00 | 7.52E-03 | Yes | Cdh12 | intron_variant | MODIFIER |
| 15 | 17468055 | AX-170087784 | 1 | T | G | 0.304 | -2.22E-01 | 7.82E-02 | 1.80E+00 | 1.83E+00 | 7.57E-03 | Yes | | intergenic_variant | MODIFIER |
| 15 | 21438843 | AX-97845955 | 1 | C | T | 0.304 | -2.22E-01 | 7.82E-02 | 1.80E+00 | 1.83E+00 | 7.57E-03 | Yes | Cdh12 | intron_variant | MODIFIER |
| 19 | 59408957 | AX-170361620 | 1 | T | A | 0.455 | -1.72E-01 | 6.33E-02 | 3.19E+00 | 3.30E+00 | 7.65E-03 | no | | intergenic_variant | MODIFIER |
| 19 | 56291093 | AX-170120117 | 1 | C | T | 0.455 | 1.73E-01 | 6.40E-02 | 3.12E+00 | 3.22E+00 | 7.67E-03 | Yes | Habp2 | intron_variant | MODIFIER |
| 5 | 62846346 | AX-170287223 | 1 | G | T | 0.473 | 2.61E-01 | 9.71E-02 | 2.86E+00 | 2.97E+00 | 7.79E-03 | Yes | Dthd1 | intron_variant | MODIFIER |
| 13 | 4581009 | AX-97841373 | 0 | G | T | 0.202 | -3.03E-01 | 1.11E-01 | 3.43E+00 | 3.60E+00 | 7.81E-03 | Yes | Akr1c21 | intron_variant | MODIFIER |
| 13 | 21439552 | AX-170188646 | 0 | T | C | 0.158 | -2.86E-01 | 1.06E-01 | 2.91E+00 | 3.04E+00 | 7.83E-03 | Yes | Zscan26 | downstream_gene_variant | MODIFIER |
| 15 | 23953913 | AX-97829017 | 1 | A | C | 0.277 | -2.30E-01 | 7.73E-02 | 1.51E+00 | 1.53E+00 | 7.89E-03 | Yes | | intergenic_variant | MODIFIER |
| 15 | 25107927 | AX-97802876 | 0 | C | T | 0.026 | -5.06E-01 | 9.86E-02 | 3.29E-01 | 3.09E-01 | 7.90E-03 | Yes | Gm35996 | intron_variant,non_coding_transcript_variant | MODIFIER |
| 16 | 66220962 | AX-170013282 | 1 | A | T | 0.518 | 2.07E-01 | 7.80E-02 | 2.68E+00 | 2.76E+00 | 8.40E-03 | Yes | | intergenic_variant | MODIFIER |
| 15 | 16401438 | AX-170224819 | 1 | C | T | 0.304 | -2.15E-01 | 7.63E-02 | 1.77E+00 | 1.80E+00 | 8.52E-03 | Yes | | intergenic_variant | MODIFIER |
| 15 | 17253817 | AX-170087782 | 1 | C | T | 0.304 | -2.15E-01 | 7.63E-02 | 1.77E+00 | 1.80E+00 | 8.52E-03 | Yes | | intergenic_variant | MODIFIER |
| 15 | 21457207 | AX-170460135 | 1 | A | G | 0.304 | -2.15E-01 | 7.63E-02 | 1.77E+00 | 1.80E+00 | 8.52E-03 | Yes | Cdh12 | intron_variant | MODIFIER |
| 15 | 17462782 | AX-170011140 | 0 | T | G | 0.298 | -2.18E-01 | 7.73E-02 | 1.74E+00 | 1.77E+00 | 8.77E-03 | Yes | | intergenic_variant | MODIFIER |
| 13 | 10137184 | AX-170326483 | 0 | A | T | 0.193 | -2.96E-01 | 1.12E-01 | 3.05E+00 | 3.18E+00 | 8.85E-03 | Yes | Chrm3 | intron_variant | MODIFIER |
| 19 | 53758821 | AX-168405769 | 1 | G | T | 0.509 | 1.72E-01 | 6.50E-02 | 3.03E+00 | 3.13E+00 | 8.86E-03 | Yes | Rbm20 | intron_variant | MODIFIER |
| 13 | 17080615 | AX-170007840 | 0 | G | A | 0.158 | -2.99E-01 | 1.13E-01 | 2.41E+00 | 2.49E+00 | 8.90E-03 | Yes | Sugct | intron_variant | MODIFIER |
| 16 | 63948705 | AX-97803933 | 0 | C | G | 0.518 | 2.26E-01 | 8.45E-02 | 3.42E+00 | 3.53E+00 | 8.90E-03 | Yes | | intergenic_variant | MODIFIER |
| 15 | 30865084 | AX-170214345 | 0 | G | T | 0.053 | -3.63E-01 | 1.02E-01 | 8.96E-01 | 9.25E-01 | 9.29E-03 | Yes | Ctnnd2 | intron_variant | MODIFIER |
| 15 | 43150517 | AX-170414590 | 0 | C | T | 0.053 | -3.71E-01 | 1.00E-01 | 8.08E-01 | 8.28E-01 | 9.39E-03 | Yes | Rspo2 | intron_variant | MODIFIER |
| 19 | 56200029 | AX-97788302 | 0 | C | A | 0.456 | 1.67E-01 | 6.34E-02 | 3.13E+00 | 3.23E+00 | 9.49E-03 | Yes | | regulatory_region_variant | MODIFIER |
| 18 | 69075883 | AX-170480308 | 0 | C | T | 0.167 | -3.01E-01 | 1.13E-01 | 3.53E+00 | 3.69E+00 | 9.53E-03 | Yes | | intergenic_variant | MODIFIER |
| 18 | 86119160 | AX-170016133 | 1 | C | T | 0.018 | -4.34E-01 | 1.67E-01 | 2.69E+00 | 2.86E+00 | 9.79E-03 | Yes | | intergenic_variant | MODIFIER |
| 5 | 137055253 | AX-170237520 | 0 | T | C | 0.132 | -2.12E-01 | 8.13E-02 | 2.45E+00 | 2.56E+00 | 9.81E-03 | Yes | | intergenic_variant | MODIFIER |

| | | | | | | | | | | | | | | | |
|---|---|---|---|---|---|---|---|---|---|---|---|---|---|---|---|
| 5 | 137757048 | AX-170261833 | 0 | T | C | 0.132 | -2.12E-01 | 8.13E-02 | 2.45E+00 | 2.56E+00 | 9.81E-03 | Yes | Tsc22d4 | upstream_gene_variant | MODIFIER |
| 19 | 58909525 | AX-170492546 | 1 | A | G | 0.464 | -1.67E-01 | 6.39E-02 | 3.18E+00 | 3.29E+00 | 9.83E-03 | no | Mir3086 | downstream_gene_variant | MODIFIER |
| 19 | 58760949 | AX-170281586 | 0 | C | T | 0.465 | -1.67E-01 | 6.39E-02 | 3.18E+00 | 3.28E+00 | 9.88E-03 | no | Pnliprp2 | intron_variant | MODIFIER |
| 19 | 58777222 | AX-170016931 | 0 | C | T | 0.465 | -1.67E-01 | 6.39E-02 | 3.18E+00 | 3.28E+00 | 9.88E-03 | no | Pnliprp2 | intron_variant | MODIFIER |
| 19 | 58804296 | AX-170146321 | 0 | A | G | 0.465 | -1.67E-01 | 6.39E-02 | 3.18E+00 | 3.28E+00 | 9.88E-03 | no | Hspa12a | intron_variant | MODIFIER |
| 19 | 58804317 | AX-170388766 | 0 | G | A | 0.465 | -1.67E-01 | 6.39E-02 | 3.18E+00 | 3.28E+00 | 9.88E-03 | no | Hspa12a | intron_variant | MODIFIER |
| 19 | 58818185 | AX-97854112 | 0 | T | C | 0.465 | -1.67E-01 | 6.39E-02 | 3.18E+00 | 3.28E+00 | 9.88E-03 | no | Hspa12a | intron_variant | MODIFIER |
| 19 | 58818396 | AX-170062824 | 0 | G | A | 0.465 | -1.67E-01 | 6.39E-02 | 3.18E+00 | 3.28E+00 | 9.88E-03 | no | Hspa12a | intron_variant | MODIFIER |
| 19 | 58829202 | AX-170473445 | 0 | T | C | 0.465 | -1.67E-01 | 6.39E-02 | 3.18E+00 | 3.28E+00 | 9.88E-03 | no | Hspa12a | intron_variant | MODIFIER |
| 19 | 58831262 | AX-170428162 | 0 | G | A | 0.465 | -1.67E-01 | 6.39E-02 | 3.18E+00 | 3.28E+00 | 9.88E-03 | no | Hspa12a | intron_variant | MODIFIER |
| 19 | 58844313 | AX-97856993 | 0 | G | A | 0.465 | -1.67E-01 | 6.39E-02 | 3.18E+00 | 3.28E+00 | 9.88E-03 | no | Hspa12a | intron_variant | MODIFIER |
| 19 | 58853135 | AX-170453291 | 0 | A | G | 0.465 | -1.67E-01 | 6.39E-02 | 3.18E+00 | 3.28E+00 | 9.88E-03 | no | Hspa12a | intron_variant | MODIFIER |
| 19 | 58853401 | AX-97781096 | 0 | T | C | 0.465 | -1.67E-01 | 6.39E-02 | 3.18E+00 | 3.28E+00 | 9.88E-03 | no | Hspa12a | intron_variant | MODIFIER |
| 19 | 58870736 | AX-170361090 | 0 | C | T | 0.465 | -1.67E-01 | 6.39E-02 | 3.18E+00 | 3.28E+00 | 9.88E-03 | no | Hspa12a | intron_variant | MODIFIER |
| 19 | 58885078 | AX-170467831 | 0 | A | C | 0.465 | -1.67E-01 | 6.39E-02 | 3.18E+00 | 3.28E+00 | 9.88E-03 | no | Hspa12a | intron_variant | MODIFIER |
| 19 | 58885631 | AX-170269458 | 0 | G | T | 0.465 | -1.67E-01 | 6.39E-02 | 3.18E+00 | 3.28E+00 | 9.88E-03 | no | Hspa12a | intron_variant | MODIFIER |
| 19 | 58893677 | AX-170352229 | 0 | T | C | 0.465 | -1.67E-01 | 6.39E-02 | 3.18E+00 | 3.28E+00 | 9.88E-03 | no | Hspa12a | intron_variant | MODIFIER |
| 19 | 58894807 | AX-170258707 | 0 | T | C | 0.465 | -1.67E-01 | 6.39E-02 | 3.18E+00 | 3.28E+00 | 9.88E-03 | no | Hspa12a | intron_variant | MODIFIER |
| 19 | 58912412 | AX-170372250 | 0 | G | A | 0.465 | -1.67E-01 | 6.39E-02 | 3.18E+00 | 3.28E+00 | 9.88E-03 | no | Mir3086 | upstream_gene_variant | MODIFIER |
| 19 | 58913720 | AX-170160283 | 0 | C | T | 0.465 | -1.67E-01 | 6.39E-02 | 3.18E+00 | 3.28E+00 | 9.88E-03 | no | Mir3086 | upstream_gene_variant | MODIFIER |
| 19 | 58935038 | AX-170312156 | 0 | A | G | 0.465 | -1.67E-01 | 6.39E-02 | 3.18E+00 | 3.28E+00 | 9.88E-03 | no | Hspa12a | intron_variant,non_coding_transcript_variant | MODIFIER |
| 19 | 58935561 | AX-170314276 | 0 | T | G | 0.465 | -1.67E-01 | 6.39E-02 | 3.18E+00 | 3.28E+00 | 9.88E-03 | no | Hspa12a | intron_variant,non_coding_transcript_variant | MODIFIER |
| 19 | 58938634 | AX-97836203 | 0 | C | T | 0.465 | -1.67E-01 | 6.39E-02 | 3.18E+00 | 3.28E+00 | 9.88E-03 | no | Eno4 | upstream_gene_variant | MODIFIER |
| 19 | 58938670 | AX-170261207 | 0 | T | C | 0.465 | -1.67E-01 | 6.39E-02 | 3.18E+00 | 3.28E+00 | 9.88E-03 | no | Eno4 | upstream_gene_variant | MODIFIER |
| 19 | 58958218 | AX-97855169 | 0 | A | G | 0.465 | -1.67E-01 | 6.39E-02 | 3.18E+00 | 3.28E+00 | 9.88E-03 | no | Eno4 | intron_variant | MODIFIER |
| 19 | 58973407 | AX-170395456 | 0 | C | T | 0.465 | -1.67E-01 | 6.39E-02 | 3.18E+00 | 3.28E+00 | 9.88E-03 | no | Shtn1 | downstream_gene_variant | MODIFIER |
| 19 | 58973524 | AX-170233142 | 0 | T | C | 0.465 | -1.67E-01 | 6.39E-02 | 3.18E+00 | 3.28E+00 | 9.88E-03 | no | Shtn1 | 3_prime_UTR_variant | MODIFIER |
| 19 | 58974307 | AX-170062827 | 0 | C | T | 0.465 | -1.67E-01 | 6.39E-02 | 3.18E+00 | 3.28E+00 | 9.88E-03 | no | Shtn1 | 3_prime_UTR_variant | MODIFIER |
| 19 | 58989732 | AX-168381864 | 0 | C | T | 0.465 | -1.67E-01 | 6.39E-02 | 3.18E+00 | 3.28E+00 | 9.88E-03 | no | Shtn1 | intron_variant | MODIFIER |
| 19 | 58996761 | AX-170290348 | 0 | G | A | 0.465 | -1.67E-01 | 6.39E-02 | 3.18E+00 | 3.28E+00 | 9.88E-03 | no | Shtn1 | intron_variant | MODIFIER |
| 19 | 58998836 | AX-170272373 | 0 | C | T | 0.465 | -1.67E-01 | 6.39E-02 | 3.18E+00 | 3.28E+00 | 9.88E-03 | no | Shtn1 | intron_variant | MODIFIER |
| 19 | 59176931 | AX-170529196 | 0 | G | A | 0.465 | -1.67E-01 | 6.39E-02 | 3.18E+00 | 3.28E+00 | 9.88E-03 | no | AC107710.1 | upstream_gene_variant | MODIFIER |
| 19 | 59203748 | AX-170102682 | 0 | T | C | 0.465 | -1.67E-01 | 6.39E-02 | 3.18E+00 | 3.28E+00 | 9.88E-03 | no | | intergenic_variant | MODIFIER |
| 19 | 59217800 | AX-97855691 | 0 | G | A | 0.465 | -1.67E-01 | 6.39E-02 | 3.18E+00 | 3.28E+00 | 9.88E-03 | no | Kcnk18 | upstream_gene_variant | MODIFIER |
| 19 | 59225957 | AX-170062830 | 0 | A | C | 0.465 | -1.67E-01 | 6.39E-02 | 3.18E+00 | 3.28E+00 | 9.88E-03 | no | Kcnk18 | intron_variant | MODIFIER |
| 19 | 59241076 | AX-97837114 | 0 | T | G | 0.465 | -1.67E-01 | 6.39E-02 | 3.18E+00 | 3.28E+00 | 9.88E-03 | no | Kcnk18 | downstream_gene_variant | MODIFIER |
| 19 | 59255991 | AX-170202951 | 0 | G | T | 0.465 | -1.67E-01 | 6.39E-02 | 3.18E+00 | 3.28E+00 | 9.88E-03 | no | Slc18a2 | upstream_gene_variant | MODIFIER |
| 19 | 59270619 | AX-170221767 | 0 | G | T | 0.465 | -1.67E-01 | 6.39E-02 | 3.18E+00 | 3.28E+00 | 9.88E-03 | no | Slc18a2 | intron_variant | MODIFIER |
| 19 | 59400918 | AX-170267689 | 0 | C | T | 0.465 | -1.67E-01 | 6.39E-02 | 3.18E+00 | 3.28E+00 | 9.88E-03 | no | Gm33756 | downstream_gene_variant | MODIFIER |
| 19 | 59404761 | AX-170162072 | 0 | A | G | 0.465 | -1.67E-01 | 6.39E-02 | 3.18E+00 | 3.28E+00 | 9.88E-03 | no | | regulatory_region_variant | MODIFIER |
| 19 | 59410769 | AX-170143633 | 0 | A | G | 0.465 | -1.67E-01 | 6.39E-02 | 3.18E+00 | 3.28E+00 | 9.88E-03 | no | | intergenic_variant | MODIFIER |
| 19 | 59424536 | AX-97805391 | 0 | G | A | 0.465 | -1.67E-01 | 6.39E-02 | 3.18E+00 | 3.28E+00 | 9.88E-03 | no | Emx2os | downstream_gene_variant | MODIFIER |
| 19 | 59424719 | AX-170175662 | 0 | T | C | 0.465 | -1.67E-01 | 6.39E-02 | 3.18E+00 | 3.28E+00 | 9.88E-03 | no | Emx2os | downstream_gene_variant | MODIFIER |
| 19 | 59429851 | AX-170288203 | 0 | T | C | 0.465 | -1.67E-01 | 6.39E-02 | 3.18E+00 | 3.28E+00 | 9.88E-03 | no | Emx2os | intron_variant,non_coding_transcript_variant | MODIFIER |
| 19 | 59438326 | AX-170481066 | 0 | T | C | 0.465 | -1.67E-01 | 6.39E-02 | 3.18E+00 | 3.28E+00 | 9.88E-03 | no | Emx2os | intron_variant,non_coding_transcript_variant | MODIFIER |
| 19 | 58921327 | AX-170331791 | 1 | A | G | 0.464 | -1.66E-01 | 6.38E-02 | 3.17E+00 | 3.27E+00 | 1.02E-02 | no | Hspa12a | intron_variant | MODIFIER |
| 15 | 21739832 | AX-170206915 | 1 | A | G | 0.304 | -2.06E-01 | 7.59E-02 | 1.88E+00 | 1.92E+00 | 1.02E-02 | Yes | | intergenic_variant | MODIFIER |
| 19 | 59016148 | AX-97809229 | 0 | C | T | 0.456 | -1.66E-01 | 6.37E-02 | 3.18E+00 | 3.28E+00 | 1.03E-02 | no | Shtn1 | intron_variant | MODIFIER |
| 19 | 59022702 | AX-170364030 | 0 | C | T | 0.456 | -1.66E-01 | 6.37E-02 | 3.18E+00 | 3.28E+00 | 1.03E-02 | no | Shtn1 | intron_variant | MODIFIER |
| 19 | 59259419 | AX-170155369 | 0 | C | T | 0.456 | -1.66E-01 | 6.37E-02 | 3.18E+00 | 3.28E+00 | 1.03E-02 | no | Slc18a2 | upstream_gene_variant | MODIFIER |
| 19 | 59269538 | AX-170094138 | 0 | G | A | 0.456 | -1.66E-01 | 6.37E-02 | 3.18E+00 | 3.28E+00 | 1.03E-02 | no | Slc18a2 | intron_variant | MODIFIER |
| 19 | 59273427 | AX-170529933 | 0 | G | A | 0.456 | -1.66E-01 | 6.37E-02 | 3.18E+00 | 3.28E+00 | 1.03E-02 | no | Slc18a2 | intron_variant | MODIFIER |
| 19 | 59273791 | AX-170481351 | 0 | G | A | 0.456 | -1.66E-01 | 6.37E-02 | 3.18E+00 | 3.28E+00 | 1.03E-02 | no | Slc18a2 | intron_variant | MODIFIER |
| 19 | 59278341 | AX-170088491 | 0 | G | A | 0.456 | -1.66E-01 | 6.37E-02 | 3.18E+00 | 3.28E+00 | 1.03E-02 | no | Slc18a2 | intron_variant | MODIFIER |

| | | | | | | | | | | | | | | | |
|---|---|---|---|---|---|---|---|---|---|---|---|---|---|---|---|
| 19 | 59281509 | AX-170135591 | 0 | A | C | 0.456 | -1.66E-01 | 6.37E-02 | 3.18E+00 | 3.28E+00 | 1.03E-02 | no | Slc18a2 | intron_variant | MODIFIER |
| 19 | 59284938 | AX-170016933 | 0 | G | A | 0.456 | -1.66E-01 | 6.37E-02 | 3.18E+00 | 3.28E+00 | 1.03E-02 | no | Slc18a2 | intron_variant | MODIFIER |
| 19 | 59290234 | AX-170016932 | 0 | G | A | 0.456 | -1.66E-01 | 6.37E-02 | 3.18E+00 | 3.28E+00 | 1.03E-02 | no | Slc18a2 | intron_variant | MODIFIER |
| 19 | 59323630 | AX-170016934 | 0 | A | G | 0.456 | -1.66E-01 | 6.37E-02 | 3.18E+00 | 3.28E+00 | 1.03E-02 | no | Rps12-ps3 | downstream_gene_variant | MODIFIER |
| 19 | 59332553 | AX-170448255 | 0 | G | A | 0.456 | -1.66E-01 | 6.37E-02 | 3.18E+00 | 3.28E+00 | 1.03E-02 | no | Pdzd8 | intron_variant | MODIFIER |
| 19 | 59335047 | AX-170446133 | 0 | T | A | 0.456 | -1.66E-01 | 6.37E-02 | 3.18E+00 | 3.28E+00 | 1.03E-02 | no | Pdzd8 | intron_variant | MODIFIER |
| 19 | 58846858 | AX-170226264 | 0 | T | C | 0.456 | -1.62E-01 | 6.24E-02 | 3.02E+00 | 3.11E+00 | 1.04E-02 | no | Hspa12a | intron_variant | MODIFIER |
| 15 | 77596320 | AX-170366438 | 0 | G | T | 0.079 | -2.95E-01 | 1.02E-01 | 1.49E+00 | 1.54E+00 | 1.04E-02 | Yes | Apol10b | upstream_gene_variant | MODIFIER |
| 19 | 59261958 | AX-170062833 | 1 | T | A | 0.455 | -1.65E-01 | 6.36E-02 | 3.17E+00 | 3.27E+00 | 1.04E-02 | no | Slc18a2 | intron_variant | MODIFIER |
| 8 | 60763681 | AX-170184104 | 0 | A | T | 0.14 | -3.48E-01 | 1.34E-01 | 3.05E+00 | 3.19E+00 | 1.04E-02 | Yes | | intergenic_variant | MODIFIER |
| 19 | 59342391 | AX-97851952 | 2 | A | G | 0.455 | -1.65E-01 | 6.37E-02 | 3.17E+00 | 3.27E+00 | 1.05E-02 | no | Pdzd8 | intron_variant | MODIFIER |
| 19 | 58462760 | AX-170286529 | 0 | A | G | 0.447 | 1.58E-01 | 6.20E-02 | 2.87E+00 | 2.96E+00 | 1.13E-02 | Yes | | intergenic_variant | MODIFIER |
| 15 | 16081596 | AX-168375752 | 0 | C | T | 0.298 | -2.04E-01 | 7.64E-02 | 1.90E+00 | 1.94E+00 | 1.13E-02 | Yes | | intergenic_variant | MODIFIER |
| 19 | 58588331 | AX-97808681 | 0 | G | A | 0.447 | 1.53E-01 | 6.00E-02 | 2.83E+00 | 2.92E+00 | 1.14E-02 | Yes | | intergenic_variant | MODIFIER |
| 3 | 6404465 | AX-170255202 | 0 | A | G | 0.132 | -2.29E-01 | 8.97E-02 | 2.87E+00 | 2.99E+00 | 1.14E-02 | Yes | | intergenic_variant | MODIFIER |
| 19 | 54414609 | AX-170016869 | 0 | C | T | 0.482 | 1.63E-01 | 6.41E-02 | 3.00E+00 | 3.09E+00 | 1.17E-02 | Yes | | intergenic_variant | MODIFIER |
| 19 | 54429631 | AX-170315790 | 0 | A | G | 0.482 | 1.63E-01 | 6.41E-02 | 3.00E+00 | 3.09E+00 | 1.17E-02 | Yes | | intergenic_variant | MODIFIER |
| 19 | 54536270 | AX-170530200 | 0 | T | C | 0.482 | 1.63E-01 | 6.41E-02 | 3.00E+00 | 3.09E+00 | 1.17E-02 | Yes | | intergenic_variant | MODIFIER |
| 19 | 54584574 | AX-170184055 | 0 | A | G | 0.482 | 1.63E-01 | 6.41E-02 | 3.00E+00 | 3.09E+00 | 1.17E-02 | Yes | | intergenic_variant | MODIFIER |
| 19 | 55016420 | AX-170016874 | 0 | A | G | 0.482 | 1.63E-01 | 6.41E-02 | 3.00E+00 | 3.09E+00 | 1.17E-02 | Yes | | intergenic_variant | MODIFIER |
| 19 | 55018467 | AX-170016875 | 0 | T | C | 0.482 | 1.63E-01 | 6.41E-02 | 3.00E+00 | 3.09E+00 | 1.17E-02 | Yes | | intergenic_variant | MODIFIER |
| 19 | 55025820 | AX-97837019 | 0 | A | G | 0.482 | 1.63E-01 | 6.41E-02 | 3.00E+00 | 3.09E+00 | 1.17E-02 | Yes | | intergenic_variant | MODIFIER |
| 19 | 55041524 | AX-170016877 | 0 | A | G | 0.482 | 1.63E-01 | 6.41E-02 | 3.00E+00 | 3.09E+00 | 1.17E-02 | Yes | | intergenic_variant | MODIFIER |
| 19 | 55047245 | AX-170247815 | 0 | G | C | 0.482 | 1.63E-01 | 6.41E-02 | 3.00E+00 | 3.09E+00 | 1.17E-02 | Yes | | intergenic_variant | MODIFIER |
| 19 | 55089100 | AX-170525132 | 0 | A | G | 0.482 | 1.63E-01 | 6.41E-02 | 3.00E+00 | 3.09E+00 | 1.17E-02 | Yes | Gpam | intron_variant | MODIFIER |
| 19 | 55161642 | AX-170270931 | 0 | C | G | 0.482 | 1.63E-01 | 6.41E-02 | 3.00E+00 | 3.09E+00 | 1.17E-02 | Yes | | intergenic_variant | MODIFIER |
| 19 | 55254661 | AX-170106404 | 0 | A | G | 0.482 | 1.63E-01 | 6.41E-02 | 3.00E+00 | 3.09E+00 | 1.17E-02 | Yes | Acsl5 | intron_variant | MODIFIER |
| 19 | 55262008 | AX-170206955 | 0 | G | C | 0.482 | 1.63E-01 | 6.41E-02 | 3.00E+00 | 3.09E+00 | 1.17E-02 | Yes | Acsl5 | intron_variant | MODIFIER |
| 19 | 55275469 | AX-170313928 | 0 | G | A | 0.482 | 1.63E-01 | 6.41E-02 | 3.00E+00 | 3.09E+00 | 1.17E-02 | Yes | Acsl5 | intron_variant | MODIFIER |
| 19 | 55276278 | AX-170170829 | 0 | C | G | 0.482 | 1.63E-01 | 6.41E-02 | 3.00E+00 | 3.09E+00 | 1.17E-02 | Yes | Acsl5 | intron_variant | MODIFIER |
| 19 | 55280015 | AX-170088486 | 0 | G | A | 0.482 | 1.63E-01 | 6.41E-02 | 3.00E+00 | 3.09E+00 | 1.17E-02 | Yes | Acsl5 | intron_variant | MODIFIER |
| 19 | 56190369 | AX-170270242 | 0 | G | A | 0.447 | 1.63E-01 | 6.40E-02 | 3.09E+00 | 3.19E+00 | 1.18E-02 | Yes | | regulatory_region_variant | MODIFIER |
| 19 | 56198118 | AX-170496198 | 0 | C | T | 0.447 | 1.63E-01 | 6.40E-02 | 3.09E+00 | 3.19E+00 | 1.18E-02 | Yes | | intergenic_variant | MODIFIER |
| 19 | 56202309 | AX-97813036 | 0 | G | A | 0.447 | 1.63E-01 | 6.40E-02 | 3.09E+00 | 3.19E+00 | 1.18E-02 | Yes | | intergenic_variant | MODIFIER |
| 19 | 56202576 | AX-170156294 | 0 | T | C | 0.447 | 1.63E-01 | 6.40E-02 | 3.09E+00 | 3.19E+00 | 1.18E-02 | Yes | | intergenic_variant | MODIFIER |
| 19 | 56213444 | AX-170260139 | 0 | T | A | 0.447 | 1.63E-01 | 6.40E-02 | 3.09E+00 | 3.19E+00 | 1.18E-02 | Yes | | intergenic_variant | MODIFIER |
| 19 | 56214199 | AX-170376676 | 0 | G | A | 0.447 | 1.63E-01 | 6.40E-02 | 3.09E+00 | 3.19E+00 | 1.18E-02 | Yes | | intergenic_variant | MODIFIER |
| 19 | 56215612 | AX-170062769 | 0 | G | A | 0.447 | 1.63E-01 | 6.40E-02 | 3.09E+00 | 3.19E+00 | 1.18E-02 | Yes | | regulatory_region_variant | MODIFIER |
| 19 | 56216807 | AX-170414332 | 0 | C | T | 0.447 | 1.63E-01 | 6.40E-02 | 3.09E+00 | 3.19E+00 | 1.18E-02 | Yes | | intergenic_variant | MODIFIER |
| 19 | 56225010 | AX-170527436 | 0 | G | C | 0.447 | 1.63E-01 | 6.40E-02 | 3.09E+00 | 3.19E+00 | 1.18E-02 | Yes | | regulatory_region_variant | MODIFIER |
| 19 | 56229329 | AX-170347781 | 0 | G | A | 0.447 | 1.63E-01 | 6.40E-02 | 3.09E+00 | 3.19E+00 | 1.18E-02 | Yes | | intergenic_variant | MODIFIER |
| 19 | 56229369 | AX-170140228 | 0 | G | A | 0.447 | 1.63E-01 | 6.40E-02 | 3.09E+00 | 3.19E+00 | 1.18E-02 | Yes | | intergenic_variant | MODIFIER |
| 19 | 56263303 | AX-97857175 | 0 | G | A | 0.447 | 1.63E-01 | 6.40E-02 | 3.09E+00 | 3.19E+00 | 1.18E-02 | Yes | | intergenic_variant | MODIFIER |
| 19 | 56270973 | AX-170399250 | 0 | T | C | 0.447 | 1.63E-01 | 6.40E-02 | 3.09E+00 | 3.19E+00 | 1.18E-02 | Yes | | intergenic_variant | MODIFIER |
| 19 | 56282782 | AX-170395254 | 0 | C | T | 0.447 | 1.63E-01 | 6.40E-02 | 3.09E+00 | 3.19E+00 | 1.18E-02 | Yes | Habp2 | upstream_gene_variant | MODIFIER |
| 19 | 56285726 | AX-170448848 | 0 | G | T | 0.447 | 1.63E-01 | 6.40E-02 | 3.09E+00 | 3.19E+00 | 1.18E-02 | Yes | Habp2 | upstream_gene_variant | MODIFIER |
| 19 | 56289845 | AX-170379336 | 0 | A | G | 0.447 | 1.63E-01 | 6.40E-02 | 3.09E+00 | 3.19E+00 | 1.18E-02 | Yes | Habp2 | intron_variant | MODIFIER |
| 19 | 56311235 | AX-170123950 | 0 | G | C | 0.447 | 1.63E-01 | 6.40E-02 | 3.09E+00 | 3.19E+00 | 1.18E-02 | Yes | Habp2 | intron_variant | MODIFIER |
| 19 | 56311277 | AX-170233088 | 0 | G | A | 0.447 | 1.63E-01 | 6.40E-02 | 3.09E+00 | 3.19E+00 | 1.18E-02 | Yes | Habp2 | intron_variant | MODIFIER |
| 19 | 56311316 | AX-170356001 | 0 | C | T | 0.447 | 1.63E-01 | 6.40E-02 | 3.09E+00 | 3.19E+00 | 1.18E-02 | Yes | Habp2 | intron_variant | MODIFIER |
| 19 | 56335269 | AX-170176400 | 0 | G | T | 0.447 | 1.63E-01 | 6.40E-02 | 3.09E+00 | 3.19E+00 | 1.18E-02 | Yes | Nrap | missense_variant | MODERATE |
| 19 | 56350054 | AX-170486280 | 0 | C | T | 0.447 | 1.63E-01 | 6.40E-02 | 3.09E+00 | 3.19E+00 | 1.18E-02 | Yes | Nrap | intron_variant | MODIFIER |
| 19 | 56354192 | AX-170131930 | 0 | G | A | 0.447 | 1.63E-01 | 6.40E-02 | 3.09E+00 | 3.19E+00 | 1.18E-02 | Yes | Nrap | synonymous_variant | LOW |
| 19 | 56355536 | AX-170519673 | 0 | G | A | 0.447 | 1.63E-01 | 6.40E-02 | 3.09E+00 | 3.19E+00 | 1.18E-02 | Yes | Nrap | synonymous_variant | LOW |

| | | | | | | | | | | | | | | | |
|---|---|---|---|---|---|---|---|---|---|---|---|---|---|---|---|
| 15 | 13097941 | AX-170436632 | 1 | T | G | 0.286 | -2.15E-01 | 7.75E-02 | 1.63E+00 | 1.65E+00 | 1.18E-02 | Yes | Cdh6 | intron_variant | MODIFIER |
| 8 | 65770954 | AX-170181635 | 0 | G | A | 0.14 | -3.48E-01 | 1.36E-01 | 3.29E+00 | 3.46E+00 | 1.18E-02 | Yes | MAR1 | intron_variant | MODIFIER |
| 15 | 21154668 | AX-170532593 | 0 | C | T | 0.298 | -2.06E-01 | 7.67E-02 | 1.81E+00 | 1.84E+00 | 1.19E-02 | Yes | Cdh12 | intron_variant | MODIFIER |
| 19 | 56350527 | AX-170062770 | 1 | A | G | 0.446 | 1.63E-01 | 6.41E-02 | 3.09E+00 | 3.19E+00 | 1.20E-02 | Yes | Nrap | intron_variant | MODIFIER |
| 18 | 68168663 | AX-170111303 | 0 | T | C | 0.351 | -1.56E-01 | 5.99E-02 | 3.69E+00 | 3.84E+00 | 1.22E-02 | no | Ldlrad4 | intron_variant | MODIFIER |
| 15 | 25476746 | AX-170463443 | 0 | T | C | 0.272 | -2.16E-01 | 7.45E-02 | 1.42E+00 | 1.44E+00 | 1.22E-02 | Yes | | intergenic_variant | MODIFIER |
| 19 | 54313637 | AX-97833864 | 0 | C | A | 0.491 | 1.61E-01 | 6.37E-02 | 3.00E+00 | 3.10E+00 | 1.24E-02 | Yes | | intergenic_variant | MODIFIER |
| 19 | 54316898 | AX-170016868 | 0 | C | T | 0.491 | 1.61E-01 | 6.37E-02 | 3.00E+00 | 3.10E+00 | 1.24E-02 | Yes | | intergenic_variant | MODIFIER |
| 19 | 54406694 | AX-97793509 | 0 | C | T | 0.491 | 1.61E-01 | 6.37E-02 | 3.00E+00 | 3.10E+00 | 1.24E-02 | Yes | | intergenic_variant | MODIFIER |
| 19 | 54407061 | AX-170016866 | 1 | T | A | 0.491 | 1.61E-01 | 6.37E-02 | 3.00E+00 | 3.10E+00 | 1.24E-02 | Yes | | intergenic_variant | MODIFIER |
| 19 | 54415584 | AX-170354074 | 0 | C | T | 0.491 | 1.61E-01 | 6.37E-02 | 3.00E+00 | 3.10E+00 | 1.24E-02 | Yes | | intergenic_variant | MODIFIER |
| 19 | 54429728 | AX-170435654 | 0 | C | T | 0.491 | 1.61E-01 | 6.37E-02 | 3.00E+00 | 3.10E+00 | 1.24E-02 | Yes | | intergenic_variant | MODIFIER |
| 19 | 54433350 | AX-170207720 | 0 | T | G | 0.491 | 1.61E-01 | 6.37E-02 | 3.00E+00 | 3.10E+00 | 1.24E-02 | Yes | | intergenic_variant | MODIFIER |
| 19 | 54447511 | AX-170445948 | 0 | G | A | 0.491 | 1.61E-01 | 6.37E-02 | 3.00E+00 | 3.10E+00 | 1.24E-02 | Yes | AC158127.1 | downstream_gene_variant | MODIFIER |
| 19 | 54544223 | AX-97836990 | 0 | G | A | 0.491 | 1.61E-01 | 6.37E-02 | 3.00E+00 | 3.10E+00 | 1.24E-02 | Yes | | intergenic_variant | MODIFIER |
| 19 | 54603722 | AX-170088484 | 0 | A | T | 0.491 | 1.61E-01 | 6.37E-02 | 3.00E+00 | 3.10E+00 | 1.24E-02 | Yes | | intergenic_variant | MODIFIER |
| 19 | 54669777 | AX-97794050 | 0 | A | G | 0.491 | 1.61E-01 | 6.37E-02 | 3.00E+00 | 3.10E+00 | 1.24E-02 | Yes | | intergenic_variant | MODIFIER |
| 19 | 54775646 | AX-170232055 | 0 | A | G | 0.491 | 1.61E-01 | 6.37E-02 | 3.00E+00 | 3.10E+00 | 1.24E-02 | Yes | | intergenic_variant | MODIFIER |
| 19 | 54819379 | AX-170380551 | 0 | A | G | 0.491 | 1.61E-01 | 6.37E-02 | 3.00E+00 | 3.10E+00 | 1.24E-02 | Yes | | intergenic_variant | MODIFIER |
| 19 | 54819609 | AX-97817063 | 0 | A | G | 0.491 | 1.61E-01 | 6.37E-02 | 3.00E+00 | 3.10E+00 | 1.24E-02 | Yes | | intergenic_variant | MODIFIER |
| 19 | 54828322 | AX-170180894 | 0 | T | C | 0.491 | 1.61E-01 | 6.37E-02 | 3.00E+00 | 3.10E+00 | 1.24E-02 | Yes | | intergenic_variant | MODIFIER |
| 19 | 54828360 | AX-170116992 | 0 | A | T | 0.491 | 1.61E-01 | 6.37E-02 | 3.00E+00 | 3.10E+00 | 1.24E-02 | Yes | | intergenic_variant | MODIFIER |
| 19 | 54833998 | AX-170336716 | 0 | G | A | 0.491 | 1.61E-01 | 6.37E-02 | 3.00E+00 | 3.10E+00 | 1.24E-02 | Yes | | intergenic_variant | MODIFIER |
| 19 | 54834144 | AX-170512646 | 0 | A | G | 0.491 | 1.61E-01 | 6.37E-02 | 3.00E+00 | 3.10E+00 | 1.24E-02 | Yes | | intergenic_variant | MODIFIER |
| 19 | 54838502 | AX-170113071 | 0 | C | T | 0.491 | 1.61E-01 | 6.37E-02 | 3.00E+00 | 3.10E+00 | 1.24E-02 | Yes | | intergenic_variant | MODIFIER |
| 19 | 54843571 | AX-170135182 | 0 | A | G | 0.491 | 1.61E-01 | 6.37E-02 | 3.00E+00 | 3.10E+00 | 1.24E-02 | Yes | | intergenic_variant | MODIFIER |
| 19 | 54847300 | AX-170294307 | 0 | A | G | 0.491 | 1.61E-01 | 6.37E-02 | 3.00E+00 | 3.10E+00 | 1.24E-02 | Yes | | intergenic_variant | MODIFIER |
| 19 | 55009064 | AX-168420030 | 0 | A | G | 0.491 | 1.61E-01 | 6.37E-02 | 3.00E+00 | 3.10E+00 | 1.24E-02 | Yes | | intergenic_variant | MODIFIER |
| 19 | 55283507 | AX-168383867 | 0 | T | C | 0.491 | 1.61E-01 | 6.37E-02 | 3.00E+00 | 3.10E+00 | 1.24E-02 | Yes | Acsl5 | intron_variant | MODIFIER |
| 17 | 71163791 | AX-170443875 | 1 | C | T | 0.045 | -3.71E-01 | 1.44E-01 | 3.54E+00 | 3.81E+00 | 1.25E-02 | Yes | Gm26561 | intron_variant,non_coding_transcript_variant | MODIFIER |
| 19 | 58565457 | AX-170062822 | 1 | A | T | 0.455 | 1.56E-01 | 6.19E-02 | 3.02E+00 | 3.12E+00 | 1.26E-02 | Yes | | intergenic_variant | MODIFIER |
| 5 | 138018106 | AX-170357637 | 0 | A | G | 0.132 | -2.07E-01 | 8.22E-02 | 2.47E+00 | 2.59E+00 | 1.28E-02 | Yes | Smok3a | upstream_gene_variant | MODIFIER |
| 19 | 58713434 | AX-170098969 | 0 | T | C | 0.447 | 1.58E-01 | 6.32E-02 | 3.00E+00 | 3.10E+00 | 1.30E-02 | Yes | 1810018F18Rik | intron_variant,non_coding_transcript_variant | MODIFIER |
| 5 | 63134659 | AX-170089501 | 0 | G | A | 0.491 | 2.31E-01 | 9.26E-02 | 2.79E+00 | 2.91E+00 | 1.31E-02 | Yes | | intergenic_variant | MODIFIER |
| 3 | 7043213 | AX-97829589 | 1 | T | C | 0.125 | -2.26E-01 | 9.07E-02 | 2.87E+00 | 2.98E+00 | 1.31E-02 | Yes | | intergenic_variant | MODIFIER |
| 15 | 22551715 | AX-170294673 | 0 | C | A | 0.272 | -2.29E-01 | 7.61E-02 | 1.25E+00 | 1.26E+00 | 1.33E-02 | Yes | Cdh18 | intron_variant | MODIFIER |
| 19 | 58767023 | AX-170252384 | 1 | C | T | 0.464 | -1.54E-01 | 6.15E-02 | 3.15E+00 | 3.26E+00 | 1.33E-02 | no | Pnliprp2 | intron_variant | MODIFIER |
| 18 | 61427083 | AX-170429810 | 0 | T | A | 0.175 | -2.92E-01 | 1.13E-01 | 3.79E+00 | 3.99E+00 | 1.35E-02 | Yes | | intergenic_variant | MODIFIER |
| 18 | 75226659 | AX-97835901 | 0 | C | A | 0.184 | -2.93E-01 | 1.15E-01 | 3.59E+00 | 3.76E+00 | 1.40E-02 | Yes | Dym | intron_variant | MODIFIER |
| 16 | 63304011 | AX-170343712 | 1 | C | T | 0.464 | -2.11E-01 | 8.53E-02 | 2.91E+00 | 2.99E+00 | 1.40E-02 | Yes | | intergenic_variant | MODIFIER |
| 13 | 16465490 | AX-170447612 | 0 | T | C | 0.184 | -2.85E-01 | 1.15E-01 | 2.70E+00 | 2.80E+00 | 1.41E-02 | Yes | | regulatory_region_variant | MODIFIER |
| 13 | 9824466 | AX-170273447 | 0 | T | C | 0.184 | -2.63E-01 | 1.06E-01 | 2.83E+00 | 2.96E+00 | 1.41E-02 | Yes | Gm47401 | downstream_gene_variant | MODIFIER |
| 15 | 71921937 | AX-170194879 | 0 | G | T | 0.079 | -2.59E-01 | 9.55E-02 | 1.63E+00 | 1.70E+00 | 1.41E-02 | Yes | Col22a1 | intron_variant | MODIFIER |
| 19 | 55011944 | AX-170477528 | 0 | T | C | 0.482 | 1.57E-01 | 6.35E-02 | 2.87E+00 | 2.96E+00 | 1.41E-02 | Yes | | intergenic_variant | MODIFIER |
| 19 | 54585398 | AX-97794049 | 0 | T | C | 0.491 | 1.57E-01 | 6.33E-02 | 3.07E+00 | 3.17E+00 | 1.42E-02 | Yes | | regulatory_region_variant | MODIFIER |
| 13 | 3762068 | AX-170287234 | 0 | T | C | 0.202 | -2.92E-01 | 1.17E-01 | 3.30E+00 | 3.44E+00 | 1.42E-02 | Yes | Gm35190 | downstream_gene_variant | MODIFIER |
| 13 | 4193855 | AX-170326581 | 0 | C | T | 0.202 | -2.92E-01 | 1.17E-01 | 3.30E+00 | 3.44E+00 | 1.42E-02 | Yes | Akr1c13 | intron_variant | MODIFIER |
| 13 | 5348397 | AX-170369975 | 0 | G | A | 0.202 | -2.92E-01 | 1.17E-01 | 3.30E+00 | 3.44E+00 | 1.42E-02 | Yes | | regulatory_region_variant | MODIFIER |
| 13 | 6601211 | AX-170304880 | 0 | T | A | 0.202 | -2.92E-01 | 1.17E-01 | 3.30E+00 | 3.44E+00 | 1.42E-02 | Yes | Pfkp | intron_variant | MODIFIER |
| 18 | 68947957 | AX-170326598 | 0 | C | T | 0.202 | -2.92E-01 | 1.17E-01 | 3.30E+00 | 3.44E+00 | 1.42E-02 | Yes | 4930546C10Rik | intron_variant,non_coding_transcript_variant | MODIFIER |
| 19 | 55608475 | AX-170421963 | 0 | A | G | 0.43 | 1.61E-01 | 6.50E-02 | 3.10E+00 | 3.20E+00 | 1.42E-02 | Yes | Vti1a | intron_variant | MODIFIER |
| 19 | 58896612 | AX-170165984 | 2 | T | C | 0.527 | 1.59E-01 | 6.43E-02 | 3.07E+00 | 3.16E+00 | 1.44E-02 | Yes | Hspa12a | intron_variant | MODIFIER |
| 5 | 138150597 | AX-170283625 | 0 | A | G | 0.132 | -2.12E-01 | 8.61E-02 | 2.43E+00 | 2.53E+00 | 1.46E-02 | Yes | Zfp113 | intron_variant | MODIFIER |
| 3 | 6318906 | AX-97848084 | 1 | T | C | 0.125 | -2.18E-01 | 8.90E-02 | 2.73E+00 | 2.84E+00 | 1.48E-02 | Yes | | intergenic_variant | MODIFIER |

| | | | | | | | | | | | | | | | |
|---|---|---|---|---|---|---|---|---|---|---|---|---|---|---|---|
| 15 | 22872901 | AX-170011229 | 0 | T | C | 0.281 | -2.16E-01 | 7.67E-02 | 1.43E+00 | 1.46E+00 | 1.53E-02 | Yes | Cdh18 | intron_variant | MODIFIER |
| 19 | 54427946 | AX-170121745 | 0 | G | A | 0.491 | 1.51E-01 | 6.18E-02 | 2.99E+00 | 3.09E+00 | 1.56E-02 | Yes | | regulatory_region_variant | MODIFIER |
| 19 | 54431479 | AX-170489980 | 2 | C | A | 0.5 | 1.50E-01 | 6.17E-02 | 2.89E+00 | 2.98E+00 | 1.56E-02 | Yes | | intergenic_variant | MODIFIER |
| 5 | 132135975 | AX-170451248 | 1 | C | T | 0.152 | -2.14E-01 | 8.76E-02 | 2.38E+00 | 2.48E+00 | 1.58E-02 | Yes | Auts2 | intron_variant,non_coding_transcript_variant | MODIFIER |
| 13 | 18783769 | AX-170354606 | 0 | T | A | 0.158 | -3.00E-01 | 1.24E-01 | 2.52E+00 | 2.59E+00 | 1.60E-02 | Yes | Vps41 | intron_variant | MODIFIER |
| 5 | 55672882 | AX-170396467 | 1 | A | G | 0.482 | 2.24E-01 | 9.25E-02 | 2.53E+00 | 2.61E+00 | 1.64E-02 | Yes | | intergenic_variant | MODIFIER |
| 2 | 143040111 | AX-97786921 | 0 | G | A | 0.088 | -2.55E-01 | 9.82E-02 | 1.75E+00 | 1.83E+00 | 1.64E-02 | Yes | | intergenic_variant | MODIFIER |
| 19 | 55409760 | AX-97837046 | 0 | G | A | 0.491 | 1.53E-01 | 6.35E-02 | 2.88E+00 | 2.97E+00 | 1.66E-02 | Yes | Vti1a | intron_variant | MODIFIER |
| 19 | 59309920 | AX-97843916 | 1 | A | C | 0.527 | 1.55E-01 | 6.44E-02 | 3.17E+00 | 3.28E+00 | 1.71E-02 | Yes | Pdzd8 | intron_variant | MODIFIER |
| 8 | 62795495 | AX-170102020 | 1 | A | T | 0.143 | -3.09E-01 | 1.28E-01 | 3.32E+00 | 3.51E+00 | 1.72E-02 | Yes | Gm45459 | intron_variant,non_coding_transcript_variant | MODIFIER |
| 19 | 57028792 | AX-170521578 | 0 | T | C | 0.447 | 1.55E-01 | 6.48E-02 | 2.91E+00 | 3.00E+00 | 1.75E-02 | Yes | Ablim1 | downstream_gene_variant | MODIFIER |
| 19 | 57050515 | AX-170444956 | 0 | C | T | 0.447 | 1.55E-01 | 6.48E-02 | 2.91E+00 | 3.00E+00 | 1.75E-02 | Yes | Ablim1 | intron_variant | MODIFIER |
| 19 | 57051543 | AX-170209207 | 0 | T | C | 0.447 | 1.55E-01 | 6.48E-02 | 2.91E+00 | 3.00E+00 | 1.75E-02 | Yes | Ablim1 | intron_variant | MODIFIER |
| 19 | 57051758 | AX-170401240 | 0 | T | C | 0.447 | 1.55E-01 | 6.48E-02 | 2.91E+00 | 3.00E+00 | 1.75E-02 | Yes | Ablim1 | intron_variant | MODIFIER |
| 19 | 57052251 | AX-170311869 | 0 | C | A | 0.447 | 1.55E-01 | 6.48E-02 | 2.91E+00 | 3.00E+00 | 1.75E-02 | Yes | Ablim1 | intron_variant | MODIFIER |
| 19 | 57054078 | AX-170368198 | 0 | A | G | 0.447 | 1.55E-01 | 6.48E-02 | 2.91E+00 | 3.00E+00 | 1.75E-02 | Yes | Ablim1 | intron_variant | MODIFIER |
| 9 | 97359335 | AX-170033078 | 1 | A | C | 0.125 | -2.65E-01 | 1.10E-01 | 3.19E+00 | 3.37E+00 | 1.78E-02 | Yes | Trim42 | intron_variant | MODIFIER |
| 8 | 63380637 | AX-170163019 | 1 | A | T | 0.143 | -3.14E-01 | 1.30E-01 | 3.31E+00 | 3.49E+00 | 1.80E-02 | Yes | | intergenic_variant | MODIFIER |
| 16 | 61403307 | AX-97828966 | 0 | A | C | 0.474 | -2.00E-01 | 8.43E-02 | 2.86E+00 | 2.93E+00 | 1.81E-02 | Yes | | intergenic_variant | MODIFIER |
| 16 | 61517799 | AX-170451540 | 0 | C | A | 0.474 | -2.00E-01 | 8.43E-02 | 2.86E+00 | 2.93E+00 | 1.81E-02 | Yes | | intergenic_variant | MODIFIER |
| 15 | 20958015 | AX-170371682 | 1 | A | C | 0.062 | -2.75E-01 | 1.04E-01 | 1.60E+00 | 1.66E+00 | 1.81E-02 | Yes | Cdh12 | intron_variant,non_coding_transcript_variant | MODIFIER |
| 16 | 65770212 | AX-170013278 | 0 | C | T | 0.509 | 1.91E-01 | 8.02E-02 | 2.89E+00 | 2.98E+00 | 1.81E-02 | Yes | | intergenic_variant | MODIFIER |
| 19 | 56670026 | AX-170016893 | 0 | C | G | 0.456 | 1.53E-01 | 6.46E-02 | 2.93E+00 | 3.01E+00 | 1.82E-02 | Yes | AC116849.1 | intron_variant,non_coding_transcript_variant | MODIFIER |
| 19 | 57038462 | AX-170440440 | 0 | G | A | 0.456 | 1.53E-01 | 6.46E-02 | 2.93E+00 | 3.01E+00 | 1.82E-02 | Yes | Ablim1 | intron_variant | MODIFIER |
| 19 | 57047911 | AX-170304036 | 0 | G | A | 0.456 | 1.53E-01 | 6.46E-02 | 2.93E+00 | 3.01E+00 | 1.82E-02 | Yes | Ablim1 | intron_variant | MODIFIER |
| 5 | 56806785 | AX-97799491 | 1 | G | A | 0.464 | 2.32E-01 | 9.74E-02 | 2.46E+00 | 2.52E+00 | 1.83E-02 | Yes | | intergenic_variant | MODIFIER |
| 19 | 57118807 | AX-170529724 | 0 | T | C | 0.456 | 1.49E-01 | 6.30E-02 | 2.80E+00 | 2.88E+00 | 1.84E-02 | Yes | Ablim1 | intron_variant | MODIFIER |
| 1 | 184223950 | AX-170043836 | 1 | A | G | 0.25 | 2.37E-01 | 1.00E-01 | 2.72E+00 | 2.83E+00 | 1.85E-02 | Yes | | intergenic_variant | MODIFIER |
| 3 | 7301279 | AX-97851813 | 0 | C | T | 0.14 | -2.07E-01 | 8.75E-02 | 2.85E+00 | 2.97E+00 | 1.85E-02 | Yes | | intergenic_variant | MODIFIER |
| 3 | 4365615 | AX-170289741 | 0 | T | C | 0.114 | -2.18E-01 | 9.25E-02 | 2.81E+00 | 2.93E+00 | 1.90E-02 | Yes | | intergenic_variant | MODIFIER |
| 3 | 6138225 | AX-170409668 | 0 | C | T | 0.114 | -2.18E-01 | 9.25E-02 | 2.81E+00 | 2.93E+00 | 1.90E-02 | Yes | Gm17934 | non_coding_transcript_exon_variant | MODIFIER |
| 3 | 6285719 | AX-170254192 | 0 | A | G | 0.114 | -2.18E-01 | 9.25E-02 | 2.81E+00 | 2.93E+00 | 1.90E-02 | Yes | | intergenic_variant | MODIFIER |
| 3 | 6292445 | AX-170065837 | 0 | T | G | 0.114 | -2.18E-01 | 9.25E-02 | 2.81E+00 | 2.93E+00 | 1.90E-02 | Yes | | intergenic_variant | MODIFIER |
| 3 | 6508179 | AX-170019838 | 0 | G | T | 0.114 | -2.18E-01 | 9.25E-02 | 2.81E+00 | 2.93E+00 | 1.90E-02 | Yes | | intergenic_variant | MODIFIER |
| 3 | 6720606 | AX-170019837 | 0 | A | C | 0.114 | -2.18E-01 | 9.25E-02 | 2.81E+00 | 2.93E+00 | 1.90E-02 | Yes | | intergenic_variant | MODIFIER |
| 3 | 6859137 | AX-170132308 | 0 | G | A | 0.114 | -2.18E-01 | 9.25E-02 | 2.81E+00 | 2.93E+00 | 1.90E-02 | Yes | | intergenic_variant | MODIFIER |
| 3 | 6859675 | AX-170118924 | 0 | G | C | 0.114 | -2.18E-01 | 9.25E-02 | 2.81E+00 | 2.93E+00 | 1.90E-02 | Yes | | regulatory_region_variant | MODIFIER |
| 3 | 6875075 | AX-170186596 | 0 | C | T | 0.114 | -2.18E-01 | 9.25E-02 | 2.81E+00 | 2.93E+00 | 1.90E-02 | Yes | | intergenic_variant | MODIFIER |
| 3 | 6907172 | AX-170200268 | 0 | G | T | 0.114 | -2.18E-01 | 9.25E-02 | 2.81E+00 | 2.93E+00 | 1.90E-02 | Yes | | intergenic_variant | MODIFIER |
| 3 | 6927153 | AX-97784736 | 0 | T | C | 0.114 | -2.18E-01 | 9.25E-02 | 2.81E+00 | 2.93E+00 | 1.90E-02 | Yes | | regulatory_region_variant | MODIFIER |
| 3 | 6956028 | AX-170370245 | 0 | A | G | 0.114 | -2.18E-01 | 9.25E-02 | 2.81E+00 | 2.93E+00 | 1.90E-02 | Yes | | intergenic_variant | MODIFIER |
| 3 | 6988052 | AX-97788487 | 0 | C | T | 0.114 | -2.18E-01 | 9.25E-02 | 2.81E+00 | 2.93E+00 | 1.90E-02 | Yes | | regulatory_region_variant | MODIFIER |
| 3 | 6996750 | AX-170525834 | 0 | C | T | 0.114 | -2.18E-01 | 9.25E-02 | 2.81E+00 | 2.93E+00 | 1.90E-02 | Yes | | intergenic_variant | MODIFIER |
| 3 | 6996794 | AX-170306726 | 0 | T | A | 0.114 | -2.18E-01 | 9.25E-02 | 2.81E+00 | 2.93E+00 | 1.90E-02 | Yes | | intergenic_variant | MODIFIER |
| 3 | 7241441 | AX-170065845 | 0 | C | A | 0.114 | -2.18E-01 | 9.25E-02 | 2.81E+00 | 2.93E+00 | 1.90E-02 | Yes | | intergenic_variant | MODIFIER |
| 3 | 7326375 | AX-170019849 | 0 | A | G | 0.114 | -2.18E-01 | 9.25E-02 | 2.81E+00 | 2.93E+00 | 1.90E-02 | Yes | | intergenic_variant | MODIFIER |
| 18 | 87647579 | AX-170472541 | 1 | G | A | 0.17 | -2.36E-01 | 1.00E-01 | 2.92E+00 | 3.05E+00 | 1.91E-02 | Yes | | intergenic_variant | MODIFIER |
| 13 | 17252518 | AX-170007848 | 0 | G | A | 0.175 | -3.01E-01 | 1.27E-01 | 2.40E+00 | 2.44E+00 | 1.94E-02 | Yes | Sugct | synonymous_variant | LOW |
| 13 | 8191768 | AX-170317917 | 0 | T | G | 0.184 | -2.52E-01 | 1.07E-01 | 2.66E+00 | 2.76E+00 | 1.95E-02 | Yes | | intergenic_variant | MODIFIER |
| 13 | 8248187 | AX-170121414 | 0 | T | C | 0.184 | -2.52E-01 | 1.07E-01 | 2.66E+00 | 2.76E+00 | 1.95E-02 | Yes | Adarb2 | intron_variant | MODIFIER |
| 13 | 10215471 | AX-170087363 | 0 | A | G | 0.184 | -2.52E-01 | 1.07E-01 | 2.66E+00 | 2.76E+00 | 1.95E-02 | Yes | Chrm3 | intron_variant | MODIFIER |
| 13 | 25026264 | AX-170498405 | 0 | A | T | 0.167 | -2.62E-01 | 1.12E-01 | 2.72E+00 | 2.83E+00 | 1.96E-02 | Yes | | intergenic_variant | MODIFIER |
| 15 | 18274177 | AX-168406861 | 0 | A | G | 0.307 | -1.96E-01 | 7.71E-02 | 1.71E+00 | 1.74E+00 | 1.98E-02 | Yes | 4921515E04Rik | intron_variant,non_coding_transcript_variant | MODIFIER |
| 16 | 66364725 | AX-170520326 | 2 | G | T | 0.527 | 1.83E-01 | 7.82E-02 | 2.66E+00 | 2.73E+00 | 2.01E-02 | Yes | | intergenic_variant | MODIFIER |

| | | | | | | | | | | | | | | | |
|---|---|---|---|---|---|---|---|---|---|---|---|---|---|---|---|
| 13 | 17022624 | AX-97831224 | 1 | C | T | 0.17 | -2.94E-01 | 1.26E-01 | 2.46E+00 | 2.51E+00 | 2.03E-02 | Yes | Sugct | intron_variant | MODIFIER |
| 19 | 52486705 | AX-170016826 | 0 | G | C | 0.491 | 1.47E-01 | 6.30E-02 | 2.94E+00 | 3.04E+00 | 2.05E-02 | Yes | | intergenic_variant | MODIFIER |
| 13 | 17555428 | AX-170153375 | 0 | T | G | 0.175 | -2.52E-01 | 1.08E-01 | 2.50E+00 | 2.59E+00 | 2.06E-02 | Yes | Sugct | intron_variant | MODIFIER |
| 19 | 55510371 | AX-170304068 | 0 | A | G | 0.439 | 1.52E-01 | 6.51E-02 | 2.99E+00 | 3.08E+00 | 2.06E-02 | Yes | Vti1a | intron_variant | MODIFIER |
| 19 | 55530562 | AX-168384071 | 0 | A | G | 0.439 | 1.52E-01 | 6.51E-02 | 2.99E+00 | 3.08E+00 | 2.06E-02 | Yes | Vti1a | intron_variant | MODIFIER |
| 19 | 55608544 | AX-170138893 | 0 | A | G | 0.439 | 1.52E-01 | 6.51E-02 | 2.99E+00 | 3.08E+00 | 2.06E-02 | Yes | Vti1a | intron_variant | MODIFIER |
| 19 | 55609156 | AX-170225819 | 0 | A | G | 0.439 | 1.52E-01 | 6.51E-02 | 2.99E+00 | 3.08E+00 | 2.06E-02 | Yes | Vti1a | intron_variant | MODIFIER |
| 19 | 55624303 | AX-97805702 | 0 | C | T | 0.439 | 1.52E-01 | 6.51E-02 | 2.99E+00 | 3.08E+00 | 2.06E-02 | Yes | Vti1a | 3_prime_UTR_variant | MODIFIER |
| 19 | 55624494 | AX-170521569 | 0 | G | T | 0.439 | 1.52E-01 | 6.51E-02 | 2.99E+00 | 3.08E+00 | 2.06E-02 | Yes | Vti1a | 3_prime_UTR_variant | MODIFIER |
| 19 | 55684911 | AX-170313462 | 0 | A | G | 0.439 | 1.52E-01 | 6.51E-02 | 2.99E+00 | 3.08E+00 | 2.06E-02 | Yes | 4930552P12Rik | intron_variant,non_coding_transcript_variant | MODIFIER |
| 19 | 55685493 | AX-97835053 | 0 | C | T | 0.439 | 1.52E-01 | 6.51E-02 | 2.99E+00 | 3.08E+00 | 2.06E-02 | Yes | 4930552P12Rik | intron_variant,non_coding_transcript_variant | MODIFIER |
| 19 | 55751636 | AX-170271153 | 0 | C | T | 0.439 | 1.52E-01 | 6.51E-02 | 2.99E+00 | 3.08E+00 | 2.06E-02 | Yes | Tcf7l2 | intron_variant | MODIFIER |
| 19 | 55804843 | AX-170433608 | 0 | C | T | 0.439 | 1.52E-01 | 6.51E-02 | 2.99E+00 | 3.08E+00 | 2.06E-02 | Yes | Tcf7l2 | intron_variant | MODIFIER |
| 19 | 55819058 | AX-170173769 | 0 | G | A | 0.439 | 1.52E-01 | 6.51E-02 | 2.99E+00 | 3.08E+00 | 2.06E-02 | Yes | Tcf7l2 | intron_variant | MODIFIER |
| 19 | 56000013 | AX-170149889 | 0 | A | G | 0.439 | 1.52E-01 | 6.51E-02 | 2.99E+00 | 3.08E+00 | 2.06E-02 | Yes | | regulatory_region_variant | MODIFIER |
| 19 | 56003059 | AX-170016892 | 0 | T | C | 0.439 | 1.52E-01 | 6.51E-02 | 2.99E+00 | 3.08E+00 | 2.06E-02 | Yes | | intergenic_variant | MODIFIER |
| 3 | 7111097 | AX-97797803 | 1 | G | A | 0.134 | -2.12E-01 | 9.08E-02 | 3.05E+00 | 3.18E+00 | 2.07E-02 | Yes | | intergenic_variant | MODIFIER |
| 19 | 58676933 | AX-170513446 | 0 | C | T | 0.447 | 1.48E-01 | 6.37E-02 | 2.97E+00 | 3.07E+00 | 2.07E-02 | Yes | Pnlip | intron_variant | MODIFIER |
| 19 | 58696883 | AX-170475858 | 0 | G | A | 0.447 | 1.48E-01 | 6.37E-02 | 2.97E+00 | 3.07E+00 | 2.07E-02 | Yes | 1810018F18Rik | upstream_gene_variant | MODIFIER |
| 19 | 58718532 | AX-170016926 | 0 | A | G | 0.447 | 1.48E-01 | 6.37E-02 | 2.97E+00 | 3.07E+00 | 2.07E-02 | Yes | 1810018F18Rik | downstream_gene_variant | MODIFIER |
| 19 | 58728639 | AX-170443404 | 0 | T | C | 0.447 | 1.48E-01 | 6.37E-02 | 2.97E+00 | 3.07E+00 | 2.07E-02 | Yes | Pnliprp1 | upstream_gene_variant | MODIFIER |
| 19 | 58733517 | AX-170217642 | 0 | A | G | 0.447 | 1.48E-01 | 6.37E-02 | 2.97E+00 | 3.07E+00 | 2.07E-02 | Yes | Pnliprp1 | intron_variant | MODIFIER |
| 19 | 58733539 | AX-97803212 | 0 | C | A | 0.447 | 1.48E-01 | 6.37E-02 | 2.97E+00 | 3.07E+00 | 2.07E-02 | Yes | Pnliprp1 | intron_variant | MODIFIER |
| 19 | 58734560 | AX-170420991 | 0 | T | G | 0.447 | 1.48E-01 | 6.37E-02 | 2.97E+00 | 3.07E+00 | 2.07E-02 | Yes | Pnliprp1 | intron_variant | MODIFIER |
| 19 | 58747774 | AX-170287593 | 0 | G | A | 0.447 | 1.48E-01 | 6.37E-02 | 2.97E+00 | 3.07E+00 | 2.07E-02 | Yes | Pnliprp1 | downstream_gene_variant | MODIFIER |
| 19 | 58749658 | AX-170364936 | 0 | C | T | 0.447 | 1.48E-01 | 6.37E-02 | 2.97E+00 | 3.07E+00 | 2.07E-02 | Yes | | intergenic_variant | MODIFIER |
| 19 | 58278708 | AX-97836197 | 0 | A | C | 0.447 | 1.44E-01 | 6.22E-02 | 2.84E+00 | 2.93E+00 | 2.08E-02 | Yes | Gfra1 | intron_variant | MODIFIER |
| 13 | 31914948 | AX-170321865 | 0 | A | T | 0.175 | -2.56E-01 | 1.09E-01 | 3.22E+00 | 3.38E+00 | 2.08E-02 | Yes | Gmds | intron_variant | MODIFIER |
| 13 | 7804389 | AX-97824692 | 1 | C | A | 0.188 | -2.71E-01 | 1.17E-01 | 2.86E+00 | 2.97E+00 | 2.09E-02 | Yes | Gm48253 | non_coding_transcript_exon_variant | MODIFIER |
| 3 | 5918058 | AX-170175542 | 0 | G | A | 0.132 | -2.10E-01 | 9.08E-02 | 2.83E+00 | 2.94E+00 | 2.10E-02 | Yes | | intergenic_variant | MODIFIER |
| 3 | 6166037 | AX-170483617 | 0 | T | C | 0.132 | -2.10E-01 | 9.08E-02 | 2.83E+00 | 2.94E+00 | 2.10E-02 | Yes | Gm6162 | non_coding_transcript_exon_variant | MODIFIER |
| 3 | 6166074 | AX-170156247 | 0 | G | A | 0.132 | -2.10E-01 | 9.08E-02 | 2.83E+00 | 2.94E+00 | 2.10E-02 | Yes | Gm6162 | non_coding_transcript_exon_variant | MODIFIER |
| 3 | 6166660 | AX-170397888 | 0 | A | G | 0.132 | -2.10E-01 | 9.08E-02 | 2.83E+00 | 2.94E+00 | 2.10E-02 | Yes | Gm6162 | non_coding_transcript_exon_variant | MODIFIER |
| 3 | 6203850 | AX-170456626 | 0 | A | G | 0.132 | -2.10E-01 | 9.08E-02 | 2.83E+00 | 2.94E+00 | 2.10E-02 | Yes | | intergenic_variant | MODIFIER |
| 3 | 6224015 | AX-170507779 | 0 | A | T | 0.132 | -2.10E-01 | 9.08E-02 | 2.83E+00 | 2.94E+00 | 2.10E-02 | Yes | | intergenic_variant | MODIFIER |
| 3 | 6242593 | AX-170487876 | 0 | C | G | 0.132 | -2.10E-01 | 9.08E-02 | 2.83E+00 | 2.94E+00 | 2.10E-02 | Yes | | intergenic_variant | MODIFIER |
| 3 | 6281761 | AX-170430335 | 0 | A | G | 0.132 | -2.10E-01 | 9.08E-02 | 2.83E+00 | 2.94E+00 | 2.10E-02 | Yes | | intergenic_variant | MODIFIER |
| 3 | 6281774 | AX-97782807 | 0 | C | T | 0.132 | -2.10E-01 | 9.08E-02 | 2.83E+00 | 2.94E+00 | 2.10E-02 | Yes | | intergenic_variant | MODIFIER |
| 3 | 6284976 | AX-170449415 | 0 | T | C | 0.132 | -2.10E-01 | 9.08E-02 | 2.83E+00 | 2.94E+00 | 2.10E-02 | Yes | | intergenic_variant | MODIFIER |
| 3 | 6291200 | AX-170203284 | 0 | A | T | 0.132 | -2.10E-01 | 9.08E-02 | 2.83E+00 | 2.94E+00 | 2.10E-02 | Yes | | intergenic_variant | MODIFIER |
| 3 | 6292253 | AX-97828338 | 0 | T | C | 0.132 | -2.10E-01 | 9.08E-02 | 2.83E+00 | 2.94E+00 | 2.10E-02 | Yes | | intergenic_variant | MODIFIER |
| 3 | 6292460 | AX-170065835 | 0 | T | G | 0.132 | -2.10E-01 | 9.08E-02 | 2.83E+00 | 2.94E+00 | 2.10E-02 | Yes | | intergenic_variant | MODIFIER |
| 3 | 6397188 | AX-97855584 | 0 | G | T | 0.132 | -2.10E-01 | 9.08E-02 | 2.83E+00 | 2.94E+00 | 2.10E-02 | Yes | | intergenic_variant | MODIFIER |
| 3 | 7022164 | AX-170508217 | 0 | T | C | 0.132 | -2.10E-01 | 9.08E-02 | 2.83E+00 | 2.94E+00 | 2.10E-02 | Yes | | intergenic_variant | MODIFIER |
| 3 | 7032864 | AX-170238553 | 0 | C | T | 0.132 | -2.10E-01 | 9.08E-02 | 2.83E+00 | 2.94E+00 | 2.10E-02 | Yes | | intergenic_variant | MODIFIER |
| 3 | 7035360 | AX-170508824 | 0 | A | G | 0.132 | -2.10E-01 | 9.08E-02 | 2.83E+00 | 2.94E+00 | 2.10E-02 | Yes | | intergenic_variant | MODIFIER |
| 3 | 7048328 | AX-170226418 | 0 | C | G | 0.132 | -2.10E-01 | 9.08E-02 | 2.83E+00 | 2.94E+00 | 2.10E-02 | Yes | | intergenic_variant | MODIFIER |
| 3 | 7058157 | AX-170179752 | 0 | C | T | 0.132 | -2.10E-01 | 9.08E-02 | 2.83E+00 | 2.94E+00 | 2.10E-02 | Yes | | intergenic_variant | MODIFIER |
| 3 | 7081168 | AX-97795733 | 0 | G | C | 0.132 | -2.10E-01 | 9.08E-02 | 2.83E+00 | 2.94E+00 | 2.10E-02 | Yes | | intergenic_variant | MODIFIER |
| 3 | 7105553 | AX-97796462 | 0 | A | G | 0.132 | -2.10E-01 | 9.08E-02 | 2.83E+00 | 2.94E+00 | 2.10E-02 | Yes | | intergenic_variant | MODIFIER |
| 3 | 7129705 | AX-170107492 | 0 | C | G | 0.132 | -2.10E-01 | 9.08E-02 | 2.83E+00 | 2.94E+00 | 2.10E-02 | Yes | | intergenic_variant | MODIFIER |
| 3 | 7167554 | AX-170019852 | 0 | A | C | 0.132 | -2.10E-01 | 9.08E-02 | 2.83E+00 | 2.94E+00 | 2.10E-02 | Yes | | intergenic_variant | MODIFIER |
| 3 | 7170072 | AX-97796166 | 0 | T | G | 0.132 | -2.10E-01 | 9.08E-02 | 2.83E+00 | 2.94E+00 | 2.10E-02 | Yes | | intergenic_variant | MODIFIER |
| 3 | 7176956 | AX-170019847 | 0 | A | G | 0.132 | -2.10E-01 | 9.08E-02 | 2.83E+00 | 2.94E+00 | 2.10E-02 | Yes | | intergenic_variant | MODIFIER |

| | | | | | | | | | | | | | | | |
|---|---|---|---|---|---|---|---|---|---|---|---|---|---|---|---|
| 3 | 7241031 | AX-170019854 | 0 | T | A | 0.132 | -2.10E-01 | 9.08E-02 | 2.83E+00 | 2.94E+00 | 2.10E-02 | Yes | | intergenic_variant | MODIFIER |
| 3 | 7245211 | AX-170065850 | 0 | C | T | 0.132 | -2.10E-01 | 9.08E-02 | 2.83E+00 | 2.94E+00 | 2.10E-02 | Yes | | regulatory_region_variant | MODIFIER |
| 3 | 7284716 | AX-170119413 | 0 | T | C | 0.132 | -2.10E-01 | 9.08E-02 | 2.83E+00 | 2.94E+00 | 2.10E-02 | Yes | | intergenic_variant | MODIFIER |
| 3 | 7318354 | AX-97818547 | 0 | C | T | 0.132 | -2.10E-01 | 9.08E-02 | 2.83E+00 | 2.94E+00 | 2.10E-02 | Yes | | intergenic_variant | MODIFIER |
| 16 | 61709499 | AX-97834676 | 0 | A | G | 0.465 | -1.94E-01 | 8.41E-02 | 2.74E+00 | 2.80E+00 | 2.13E-02 | Yes | | intergenic_variant | MODIFIER |
| 19 | 55448124 | AX-97805748 | 0 | T | C | 0.447 | 1.50E-01 | 6.49E-02 | 3.00E+00 | 3.10E+00 | 2.14E-02 | Yes | Vti1a | intron_variant | MODIFIER |
| 19 | 55452306 | AX-170016882 | 0 | C | G | 0.447 | 1.50E-01 | 6.49E-02 | 3.00E+00 | 3.10E+00 | 2.14E-02 | Yes | Vti1a | intron_variant | MODIFIER |
| 19 | 55486139 | AX-170016881 | 0 | A | G | 0.447 | 1.50E-01 | 6.49E-02 | 3.00E+00 | 3.10E+00 | 2.14E-02 | Yes | Vti1a | intron_variant | MODIFIER |
| 19 | 55486328 | AX-97835709 | 0 | C | T | 0.447 | 1.50E-01 | 6.49E-02 | 3.00E+00 | 3.10E+00 | 2.14E-02 | Yes | Vti1a | intron_variant | MODIFIER |
| 19 | 55491478 | AX-170261326 | 0 | G | A | 0.447 | 1.50E-01 | 6.49E-02 | 3.00E+00 | 3.10E+00 | 2.14E-02 | Yes | Vti1a | intron_variant | MODIFIER |
| 19 | 55509300 | AX-170509697 | 0 | C | G | 0.447 | 1.50E-01 | 6.49E-02 | 3.00E+00 | 3.10E+00 | 2.14E-02 | Yes | Vti1a | intron_variant | MODIFIER |
| 19 | 55580363 | AX-97804235 | 0 | A | G | 0.447 | 1.50E-01 | 6.49E-02 | 3.00E+00 | 3.10E+00 | 2.14E-02 | Yes | Vti1a | intron_variant | MODIFIER |
| 19 | 55609263 | AX-170099996 | 0 | G | A | 0.447 | 1.50E-01 | 6.49E-02 | 3.00E+00 | 3.10E+00 | 2.14E-02 | Yes | Vti1a | intron_variant | MODIFIER |
| 19 | 55609790 | AX-170459137 | 0 | G | A | 0.447 | 1.50E-01 | 6.49E-02 | 3.00E+00 | 3.10E+00 | 2.14E-02 | Yes | Vti1a | intron_variant | MODIFIER |
| 19 | 55613010 | AX-170104037 | 0 | C | A | 0.447 | 1.50E-01 | 6.49E-02 | 3.00E+00 | 3.10E+00 | 2.14E-02 | Yes | Vti1a | intron_variant | MODIFIER |
| 19 | 56002200 | AX-97836174 | 0 | A | T | 0.447 | 1.50E-01 | 6.49E-02 | 3.00E+00 | 3.10E+00 | 2.14E-02 | Yes | | intergenic_variant | MODIFIER |
| 19 | 56140572 | AX-97846268 | 0 | T | C | 0.447 | 1.50E-01 | 6.49E-02 | 3.00E+00 | 3.10E+00 | 2.14E-02 | Yes | AC161579.1 | non_coding_transcript_exon_variant | MODIFIER |
| 19 | 56165759 | AX-170235768 | 0 | C | T | 0.447 | 1.50E-01 | 6.49E-02 | 3.00E+00 | 3.10E+00 | 2.14E-02 | Yes | | intergenic_variant | MODIFIER |
| 19 | 56165791 | AX-170259765 | 0 | A | C | 0.447 | 1.50E-01 | 6.49E-02 | 3.00E+00 | 3.10E+00 | 2.14E-02 | Yes | | intergenic_variant | MODIFIER |
| 19 | 56172750 | AX-170480182 | 0 | C | T | 0.447 | 1.50E-01 | 6.49E-02 | 3.00E+00 | 3.10E+00 | 2.14E-02 | Yes | | intergenic_variant | MODIFIER |
| 19 | 56173106 | AX-170245537 | 0 | C | T | 0.447 | 1.50E-01 | 6.49E-02 | 3.00E+00 | 3.10E+00 | 2.14E-02 | Yes | | regulatory_region_variant | MODIFIER |
| 19 | 56175712 | AX-170331072 | 0 | C | T | 0.447 | 1.50E-01 | 6.49E-02 | 3.00E+00 | 3.10E+00 | 2.14E-02 | Yes | | intergenic_variant | MODIFIER |
| 19 | 56184378 | AX-170393789 | 0 | T | C | 0.447 | 1.50E-01 | 6.49E-02 | 3.00E+00 | 3.10E+00 | 2.14E-02 | Yes | | intergenic_variant | MODIFIER |
| 13 | 7334342 | AX-97822655 | 0 | T | C | 0.184 | -2.69E-01 | 1.16E-01 | 2.79E+00 | 2.89E+00 | 2.15E-02 | Yes | | intergenic_variant | MODIFIER |
| 13 | 8190622 | AX-170097805 | 0 | C | T | 0.184 | -2.69E-01 | 1.16E-01 | 2.79E+00 | 2.89E+00 | 2.15E-02 | Yes | | intergenic_variant | MODIFIER |
| 13 | 11147291 | AX-170180555 | 0 | C | A | 0.184 | -2.69E-01 | 1.16E-01 | 2.79E+00 | 2.89E+00 | 2.15E-02 | Yes | | intergenic_variant | MODIFIER |
| 5 | 61500746 | AX-170494565 | 0 | G | C | 0.482 | 2.26E-01 | 9.80E-02 | 2.64E+00 | 2.72E+00 | 2.15E-02 | Yes | | intergenic_variant | MODIFIER |
| 16 | 65337888 | AX-97845881 | 0 | C | A | 0.509 | 1.85E-01 | 8.02E-02 | 2.79E+00 | 2.87E+00 | 2.17E-02 | Yes | Gm49629 | upstream_gene_variant | MODIFIER |
| 19 | 58245196 | AX-97841542 | 0 | A | C | 0.447 | 1.43E-01 | 6.23E-02 | 2.84E+00 | 2.92E+00 | 2.18E-02 | no | Gfra1 | intron_variant | MODIFIER |
| 19 | 58260830 | AX-170181244 | 0 | G | A | 0.447 | 1.43E-01 | 6.23E-02 | 2.84E+00 | 2.92E+00 | 2.18E-02 | no | Gfra1 | intron_variant | MODIFIER |
| 19 | 58276379 | AX-170336726 | 0 | T | C | 0.447 | 1.43E-01 | 6.23E-02 | 2.84E+00 | 2.92E+00 | 2.18E-02 | no | Gfra1 | intron_variant | MODIFIER |
| 19 | 58282396 | AX-170207245 | 0 | A | G | 0.447 | 1.43E-01 | 6.23E-02 | 2.84E+00 | 2.92E+00 | 2.18E-02 | no | Gfra1 | intron_variant | MODIFIER |
| 19 | 58290497 | AX-170016916 | 0 | A | G | 0.447 | 1.43E-01 | 6.23E-02 | 2.84E+00 | 2.92E+00 | 2.18E-02 | no | Gfra1 | intron_variant | MODIFIER |
| 19 | 58304266 | AX-170322450 | 0 | T | C | 0.447 | 1.43E-01 | 6.23E-02 | 2.84E+00 | 2.92E+00 | 2.18E-02 | no | Gfra1 | intron_variant | MODIFIER |
| 19 | 58309834 | AX-170125957 | 0 | T | C | 0.447 | 1.43E-01 | 6.23E-02 | 2.84E+00 | 2.92E+00 | 2.18E-02 | no | Gfra1 | intron_variant | MODIFIER |
| 19 | 58322282 | AX-170477719 | 0 | A | C | 0.447 | 1.43E-01 | 6.23E-02 | 2.84E+00 | 2.92E+00 | 2.18E-02 | no | Gfra1 | intron_variant | MODIFIER |
| 19 | 58345157 | AX-97812861 | 0 | A | G | 0.447 | 1.43E-01 | 6.23E-02 | 2.84E+00 | 2.92E+00 | 2.18E-02 | no | Gfra1 | intron_variant | MODIFIER |
| 19 | 58387175 | AX-170184067 | 0 | T | A | 0.447 | 1.43E-01 | 6.23E-02 | 2.84E+00 | 2.92E+00 | 2.18E-02 | no | Gfra1 | intron_variant | MODIFIER |
| 19 | 58388599 | AX-170195971 | 0 | T | C | 0.447 | 1.43E-01 | 6.23E-02 | 2.84E+00 | 2.92E+00 | 2.18E-02 | no | Gfra1 | intron_variant | MODIFIER |
| 19 | 58395603 | AX-97830116 | 0 | A | G | 0.447 | 1.43E-01 | 6.23E-02 | 2.84E+00 | 2.92E+00 | 2.18E-02 | no | Gfra1 | intron_variant | MODIFIER |
| 19 | 58398498 | AX-170284363 | 0 | A | G | 0.447 | 1.43E-01 | 6.23E-02 | 2.84E+00 | 2.92E+00 | 2.18E-02 | no | Gfra1 | intron_variant | MODIFIER |
| 19 | 58426120 | AX-170037976 | 0 | G | A | 0.447 | 1.43E-01 | 6.23E-02 | 2.84E+00 | 2.92E+00 | 2.18E-02 | no | Gfra1 | intron_variant | MODIFIER |
| 19 | 58435440 | AX-170062815 | 0 | A | G | 0.447 | 1.43E-01 | 6.23E-02 | 2.84E+00 | 2.92E+00 | 2.18E-02 | no | Gfra1 | intron_variant | MODIFIER |
| 19 | 58450662 | AX-170459255 | 0 | T | G | 0.447 | 1.43E-01 | 6.23E-02 | 2.84E+00 | 2.92E+00 | 2.18E-02 | no | Gfra1 | intron_variant | MODIFIER |
| 19 | 58452024 | AX-170282563 | 0 | A | G | 0.447 | 1.43E-01 | 6.23E-02 | 2.84E+00 | 2.92E+00 | 2.18E-02 | no | Gfra1 | synonymous_variant | LOW |
| 19 | 58452494 | AX-170062818 | 0 | T | C | 0.447 | 1.43E-01 | 6.23E-02 | 2.84E+00 | 2.92E+00 | 2.18E-02 | no | Gfra1 | intron_variant | MODIFIER |
| 19 | 58460326 | AX-170300435 | 0 | C | T | 0.447 | 1.43E-01 | 6.23E-02 | 2.84E+00 | 2.92E+00 | 2.18E-02 | no | Gfra1 | upstream_gene_variant | MODIFIER |
| 19 | 58461602 | AX-170242226 | 0 | T | G | 0.447 | 1.43E-01 | 6.23E-02 | 2.84E+00 | 2.92E+00 | 2.18E-02 | no | | intergenic_variant | MODIFIER |
| 19 | 58470403 | AX-170213934 | 0 | C | T | 0.447 | 1.43E-01 | 6.23E-02 | 2.84E+00 | 2.92E+00 | 2.18E-02 | no | | intergenic_variant | MODIFIER |
| 19 | 58471069 | AX-170278353 | 0 | C | A | 0.447 | 1.43E-01 | 6.23E-02 | 2.84E+00 | 2.92E+00 | 2.18E-02 | no | | intergenic_variant | MODIFIER |
| 19 | 58471086 | AX-170191854 | 0 | C | A | 0.447 | 1.43E-01 | 6.23E-02 | 2.84E+00 | 2.92E+00 | 2.18E-02 | no | | intergenic_variant | MODIFIER |
| 19 | 58492300 | AX-170434126 | 0 | G | A | 0.447 | 1.43E-01 | 6.23E-02 | 2.84E+00 | 2.92E+00 | 2.18E-02 | no | AC102610.1 | downstream_gene_variant | MODIFIER |
| 19 | 58505580 | AX-170210262 | 0 | A | C | 0.447 | 1.43E-01 | 6.23E-02 | 2.84E+00 | 2.92E+00 | 2.18E-02 | no | | intergenic_variant | MODIFIER |
| 19 | 58505660 | AX-170409610 | 0 | A | C | 0.447 | 1.43E-01 | 6.23E-02 | 2.84E+00 | 2.92E+00 | 2.18E-02 | no | | intergenic_variant | MODIFIER |

| | | | | | | | | | | | | | | | |
|---|---|---|---|---|---|---|---|---|---|---|---|---|---|---|---|
| 19 | 58508120 | AX-170403504 | 0 | C | T | 0.447 | 1.43E-01 | 6.23E-02 | 2.84E+00 | 2.92E+00 | 2.18E-02 | no | Ccdc172 | upstream_gene_variant | MODIFIER |
| 19 | 58512674 | AX-170453960 | 0 | G | A | 0.447 | 1.43E-01 | 6.23E-02 | 2.84E+00 | 2.92E+00 | 2.18E-02 | no | Ccdc172 | missense_variant | MODERATE |
| 19 | 58512697 | AX-170241142 | 0 | T | A | 0.447 | 1.43E-01 | 6.23E-02 | 2.84E+00 | 2.92E+00 | 2.18E-02 | no | Ccdc172 | missense_variant | MODERATE |
| 19 | 58513481 | AX-170402615 | 0 | C | G | 0.447 | 1.43E-01 | 6.23E-02 | 2.84E+00 | 2.92E+00 | 2.18E-02 | no | Ccdc172 | intron_variant | MODIFIER |
| 19 | 58548436 | AX-170523569 | 0 | G | A | 0.447 | 1.43E-01 | 6.23E-02 | 2.84E+00 | 2.92E+00 | 2.18E-02 | no | Ccdc172 | intron_variant | MODIFIER |
| 19 | 58548887 | AX-170128948 | 0 | G | A | 0.447 | 1.43E-01 | 6.23E-02 | 2.84E+00 | 2.92E+00 | 2.18E-02 | no | Ccdc172 | intron_variant | MODIFIER |
| 19 | 58548929 | AX-170283697 | 0 | C | G | 0.447 | 1.43E-01 | 6.23E-02 | 2.84E+00 | 2.92E+00 | 2.18E-02 | no | Ccdc172 | intron_variant | MODIFIER |
| 19 | 58562168 | AX-170062820 | 0 | T | C | 0.447 | 1.43E-01 | 6.23E-02 | 2.84E+00 | 2.92E+00 | 2.18E-02 | no | | intergenic_variant | MODIFIER |
| 19 | 58567221 | AX-170405780 | 0 | T | C | 0.447 | 1.43E-01 | 6.23E-02 | 2.84E+00 | 2.92E+00 | 2.18E-02 | no | | intergenic_variant | MODIFIER |
| 19 | 58581260 | AX-170280545 | 0 | C | G | 0.447 | 1.43E-01 | 6.23E-02 | 2.84E+00 | 2.92E+00 | 2.18E-02 | no | | intergenic_variant | MODIFIER |
| 19 | 58581450 | AX-170290775 | 0 | C | T | 0.447 | 1.43E-01 | 6.23E-02 | 2.84E+00 | 2.92E+00 | 2.18E-02 | no | | intergenic_variant | MODIFIER |
| 19 | 58588346 | AX-170349961 | 0 | C | T | 0.447 | 1.43E-01 | 6.23E-02 | 2.84E+00 | 2.92E+00 | 2.18E-02 | no | | intergenic_variant | MODIFIER |
| 19 | 58620366 | AX-170318831 | 0 | G | T | 0.447 | 1.43E-01 | 6.23E-02 | 2.84E+00 | 2.92E+00 | 2.18E-02 | no | 1810007D17Rik | intron_variant,non_coding_transcript_variant | MODIFIER |
| 19 | 58659027 | AX-97854887 | 0 | C | T | 0.447 | 1.43E-01 | 6.23E-02 | 2.84E+00 | 2.92E+00 | 2.18E-02 | no | Pnlip | intron_variant,non_coding_transcript_variant | MODIFIER |
| 18 | 14859289 | AX-170060231 | 0 | G | T | 0.035 | 2.84E-01 | 1.23E-01 | 3.14E+00 | 3.39E+00 | 2.20E-02 | Yes | Taf4b | intron_variant | MODIFIER |
| 5 | 132142695 | AX-170435212 | 1 | A | T | 0.152 | -2.00E-01 | 8.63E-02 | 2.34E+00 | 2.43E+00 | 2.21E-02 | Yes | Auts2 | intron_variant,non_coding_transcript_variant | MODIFIER |
| 5 | 65153706 | AX-170072328 | 2 | T | C | 0.491 | 2.13E-01 | 9.30E-02 | 2.62E+00 | 2.72E+00 | 2.24E-02 | Yes | Klhl5 | intron_variant | MODIFIER |
| 13 | 17641858 | AX-170406096 | 1 | C | A | 0.179 | -2.46E-01 | 1.07E-01 | 2.53E+00 | 2.63E+00 | 2.25E-02 | Yes | Sugct | intron_variant | MODIFIER |
| 15 | 12429184 | AX-170517537 | 1 | G | A | 0.295 | -1.96E-01 | 7.72E-02 | 1.64E+00 | 1.67E+00 | 2.25E-02 | Yes | Pdzd2 | intron_variant | MODIFIER |
| 19 | 58257705 | AX-97801209 | 0 | A | C | 0.456 | 1.42E-01 | 6.20E-02 | 2.85E+00 | 2.94E+00 | 2.26E-02 | no | Gfra1 | intron_variant | MODIFIER |
| 19 | 58274954 | AX-170502886 | 0 | G | A | 0.456 | 1.42E-01 | 6.20E-02 | 2.85E+00 | 2.94E+00 | 2.26E-02 | no | Gfra1 | intron_variant | MODIFIER |
| 19 | 58287043 | AX-170037979 | 0 | G | A | 0.456 | 1.42E-01 | 6.20E-02 | 2.85E+00 | 2.94E+00 | 2.26E-02 | no | Gfra1 | intron_variant | MODIFIER |
| 19 | 58305297 | AX-170198098 | 0 | T | C | 0.456 | 1.42E-01 | 6.20E-02 | 2.85E+00 | 2.94E+00 | 2.26E-02 | no | Gfra1 | intron_variant | MODIFIER |
| 19 | 58314406 | AX-170211421 | 0 | G | A | 0.456 | 1.42E-01 | 6.20E-02 | 2.85E+00 | 2.94E+00 | 2.26E-02 | no | Gfra1 | intron_variant | MODIFIER |
| 19 | 58319929 | AX-170388371 | 0 | G | A | 0.456 | 1.42E-01 | 6.20E-02 | 2.85E+00 | 2.94E+00 | 2.26E-02 | no | Gfra1 | intron_variant | MODIFIER |
| 19 | 58326000 | AX-170330418 | 0 | C | T | 0.456 | 1.42E-01 | 6.20E-02 | 2.85E+00 | 2.94E+00 | 2.26E-02 | no | Gfra1 | intron_variant | MODIFIER |
| 19 | 58330924 | AX-170384746 | 0 | A | G | 0.456 | 1.42E-01 | 6.20E-02 | 2.85E+00 | 2.94E+00 | 2.26E-02 | no | Gfra1 | intron_variant | MODIFIER |
| 19 | 58332757 | AX-170062814 | 0 | G | A | 0.456 | 1.42E-01 | 6.20E-02 | 2.85E+00 | 2.94E+00 | 2.26E-02 | no | Gfra1 | intron_variant | MODIFIER |
| 19 | 58338020 | AX-170062811 | 0 | T | G | 0.456 | 1.42E-01 | 6.20E-02 | 2.85E+00 | 2.94E+00 | 2.26E-02 | no | Gfra1 | intron_variant | MODIFIER |
| 19 | 58375716 | AX-170494844 | 0 | A | C | 0.456 | 1.42E-01 | 6.20E-02 | 2.85E+00 | 2.94E+00 | 2.26E-02 | no | Gfra1 | intron_variant | MODIFIER |
| 19 | 58397233 | AX-170376601 | 0 | G | C | 0.456 | 1.42E-01 | 6.20E-02 | 2.85E+00 | 2.94E+00 | 2.26E-02 | no | Gfra1 | intron_variant | MODIFIER |
| 19 | 58405735 | AX-170225257 | 0 | A | G | 0.456 | 1.42E-01 | 6.20E-02 | 2.85E+00 | 2.94E+00 | 2.26E-02 | no | Gfra1 | intron_variant | MODIFIER |
| 19 | 58407979 | AX-170405677 | 0 | A | T | 0.456 | 1.42E-01 | 6.20E-02 | 2.85E+00 | 2.94E+00 | 2.26E-02 | no | Gfra1 | intron_variant | MODIFIER |
| 19 | 58409437 | AX-170192851 | 0 | A | C | 0.456 | 1.42E-01 | 6.20E-02 | 2.85E+00 | 2.94E+00 | 2.26E-02 | no | Gfra1 | intron_variant | MODIFIER |
| 19 | 58423356 | AX-170126091 | 0 | C | G | 0.456 | 1.42E-01 | 6.20E-02 | 2.85E+00 | 2.94E+00 | 2.26E-02 | no | Gfra1 | intron_variant | MODIFIER |
| 19 | 58565686 | AX-170062819 | 0 | A | C | 0.456 | 1.38E-01 | 6.06E-02 | 2.73E+00 | 2.81E+00 | 2.28E-02 | no | | intergenic_variant | MODIFIER |
| 9 | 90542910 | AX-170032959 | 0 | A | C | 0.132 | -2.66E-01 | 1.15E-01 | 3.26E+00 | 3.44E+00 | 2.29E-02 | Yes | | intergenic_variant | MODIFIER |
| 19 | 56160802 | AX-97810294 | 1 | T | C | 0.455 | 1.47E-01 | 6.43E-02 | 2.91E+00 | 3.00E+00 | 2.29E-02 | Yes | | intergenic_variant | MODIFIER |
| 16 | 55547868 | AX-170013102 | 0 | A | G | 0.535 | 1.90E-01 | 8.31E-02 | 2.80E+00 | 2.86E+00 | 2.31E-02 | Yes | | intergenic_variant | MODIFIER |
| 18 | 65202320 | AX-170307192 | 0 | A | C | 0.184 | -2.68E-01 | 1.15E-01 | 3.42E+00 | 3.57E+00 | 2.31E-02 | Yes | Nedd4l | intron_variant | MODIFIER |
| 4 | 100830103 | AX-170508700 | 0 | C | A | 0.228 | 2.61E-01 | 1.14E-01 | 2.86E+00 | 2.95E+00 | 2.31E-02 | Yes | Cachd1 | intron_variant | MODIFIER |
| 19 | 53488840 | AX-170181336 | 1 | A | G | 0.491 | 1.41E-01 | 6.21E-02 | 2.85E+00 | 2.95E+00 | 2.32E-02 | no | AC123702.1 | downstream_gene_variant | MODIFIER |
| 5 | 58085216 | AX-170100298 | 0 | A | G | 0.465 | 2.19E-01 | 9.63E-02 | 2.57E+00 | 2.65E+00 | 2.39E-02 | Yes | Pcdh7 | intron_variant | MODIFIER |
| 18 | 58105442 | AX-170465980 | 0 | T | C | 0.184 | -2.72E-01 | 1.17E-01 | 3.56E+00 | 3.71E+00 | 2.39E-02 | Yes | Fbn2 | intron_variant | MODIFIER |
| 18 | 64837532 | AX-170500958 | 0 | T | A | 0.184 | -2.72E-01 | 1.17E-01 | 3.56E+00 | 3.71E+00 | 2.39E-02 | Yes | | intergenic_variant | MODIFIER |
| 18 | 65839674 | AX-170216810 | 0 | G | A | 0.184 | -2.72E-01 | 1.17E-01 | 3.56E+00 | 3.71E+00 | 2.39E-02 | Yes | AC124578.3 | downstream_gene_variant | MODIFIER |
| 13 | 8087468 | AX-170394333 | 0 | C | T | 0.184 | -2.81E-01 | 1.24E-01 | 2.71E+00 | 2.77E+00 | 2.43E-02 | Yes | | intergenic_variant | MODIFIER |
| 13 | 9701100 | AX-170198844 | 0 | C | T | 0.184 | -2.81E-01 | 1.24E-01 | 2.71E+00 | 2.77E+00 | 2.43E-02 | Yes | Zmynd11 | intron_variant | MODIFIER |
| 19 | 56391717 | AX-170132323 | 0 | T | C | 0.447 | 1.50E-01 | 6.63E-02 | 2.97E+00 | 3.06E+00 | 2.44E-02 | Yes | Nrap | upstream_gene_variant | MODIFIER |
| 16 | 58030133 | AX-170326251 | 0 | G | T | 0.535 | 1.88E-01 | 8.34E-02 | 2.79E+00 | 2.84E+00 | 2.44E-02 | Yes | Gm49575 | intron_variant,non_coding_transcript_variant | MODIFIER |
| 5 | 137645683 | AX-170260767 | 0 | C | A | 0.132 | -2.05E-01 | 9.02E-02 | 2.37E+00 | 2.46E+00 | 2.46E-02 | Yes | Lrch4 | downstream_gene_variant | MODIFIER |
| 9 | 96758380 | AX-170307757 | 0 | T | C | 0.263 | -1.27E-01 | 5.60E-02 | 3.20E+00 | 3.36E+00 | 2.48E-02 | no | | intergenic_variant | MODIFIER |
| 15 | 11691681 | AX-170392739 | 1 | A | G | 0.286 | -1.99E-01 | 7.75E-02 | 1.53E+00 | 1.55E+00 | 2.49E-02 | Yes | | intergenic_variant | MODIFIER |
| 16 | 63844031 | AX-97797357 | 0 | G | A | 0.509 | 1.92E-01 | 8.50E-02 | 2.99E+00 | 3.07E+00 | 2.50E-02 | Yes | Epha3 | intron_variant | MODIFIER |

| | | | | | | | | | | | | | | | |
|---|---|---|---|---|---|---|---|---|---|---|---|---|---|---|---|
| 15 | 17327955 | AX-170230740 | 0 | G | A | 0.316 | -1.89E-01 | 7.67E-02 | 1.72E+00 | 1.75E+00 | 2.50E-02 | Yes | | intergenic_variant | MODIFIER |
| 19 | 57027997 | AX-170282904 | 0 | T | C | 0.447 | 1.45E-01 | 6.46E-02 | 2.97E+00 | 3.06E+00 | 2.52E-02 | Yes | Ablim1 | downstream_gene_variant | MODIFIER |
| 15 | 11381092 | AX-170363953 | 0 | G | C | 0.298 | -1.93E-01 | 7.75E-02 | 1.66E+00 | 1.68E+00 | 2.52E-02 | Yes | Tars | downstream_gene_variant | MODIFIER |
| 15 | 11389086 | AX-170300874 | 0 | G | A | 0.298 | -1.93E-01 | 7.75E-02 | 1.66E+00 | 1.68E+00 | 2.52E-02 | Yes | Tars | intron_variant | MODIFIER |
| 15 | 11438690 | AX-97827696 | 0 | C | A | 0.298 | -1.93E-01 | 7.75E-02 | 1.66E+00 | 1.68E+00 | 2.52E-02 | Yes | | intergenic_variant | MODIFIER |
| 15 | 11461696 | AX-170415787 | 0 | A | T | 0.298 | -1.93E-01 | 7.75E-02 | 1.66E+00 | 1.68E+00 | 2.52E-02 | Yes | | intergenic_variant | MODIFIER |
| 15 | 11462779 | AX-170138399 | 0 | A | G | 0.298 | -1.93E-01 | 7.75E-02 | 1.66E+00 | 1.68E+00 | 2.52E-02 | Yes | | intergenic_variant | MODIFIER |
| 15 | 11543288 | AX-97849135 | 0 | C | T | 0.298 | -1.93E-01 | 7.75E-02 | 1.66E+00 | 1.68E+00 | 2.52E-02 | Yes | | intergenic_variant | MODIFIER |
| 15 | 11570015 | AX-170116314 | 0 | G | C | 0.298 | -1.93E-01 | 7.75E-02 | 1.66E+00 | 1.68E+00 | 2.52E-02 | Yes | | intergenic_variant | MODIFIER |
| 15 | 11589496 | AX-170328942 | 0 | A | G | 0.298 | -1.93E-01 | 7.75E-02 | 1.66E+00 | 1.68E+00 | 2.52E-02 | Yes | | intergenic_variant | MODIFIER |
| 15 | 11617651 | AX-170108772 | 0 | A | C | 0.298 | -1.93E-01 | 7.75E-02 | 1.66E+00 | 1.68E+00 | 2.52E-02 | Yes | | intergenic_variant | MODIFIER |
| 15 | 11653701 | AX-97847524 | 0 | G | A | 0.298 | -1.93E-01 | 7.75E-02 | 1.66E+00 | 1.68E+00 | 2.52E-02 | Yes | | intergenic_variant | MODIFIER |
| 15 | 11670706 | AX-97828157 | 0 | G | A | 0.298 | -1.93E-01 | 7.75E-02 | 1.66E+00 | 1.68E+00 | 2.52E-02 | Yes | | intergenic_variant | MODIFIER |
| 15 | 11676789 | AX-170286351 | 0 | G | C | 0.298 | -1.93E-01 | 7.75E-02 | 1.66E+00 | 1.68E+00 | 2.52E-02 | Yes | | intergenic_variant | MODIFIER |
| 15 | 11694079 | AX-170440408 | 0 | G | A | 0.298 | -1.93E-01 | 7.75E-02 | 1.66E+00 | 1.68E+00 | 2.52E-02 | Yes | Gm41271 | downstream_gene_variant | MODIFIER |
| 15 | 11712887 | AX-170288010 | 0 | C | T | 0.298 | -1.93E-01 | 7.75E-02 | 1.66E+00 | 1.68E+00 | 2.52E-02 | Yes | | regulatory_region_variant | MODIFIER |
| 15 | 11714264 | AX-170236796 | 0 | T | C | 0.298 | -1.93E-01 | 7.75E-02 | 1.66E+00 | 1.68E+00 | 2.52E-02 | Yes | | intergenic_variant | MODIFIER |
| 15 | 11719995 | AX-170224335 | 0 | A | G | 0.298 | -1.93E-01 | 7.75E-02 | 1.66E+00 | 1.68E+00 | 2.52E-02 | Yes | | intergenic_variant | MODIFIER |
| 15 | 11721235 | AX-170343229 | 0 | T | C | 0.298 | -1.93E-01 | 7.75E-02 | 1.66E+00 | 1.68E+00 | 2.52E-02 | Yes | | intergenic_variant | MODIFIER |
| 15 | 11722768 | AX-97842189 | 0 | T | C | 0.298 | -1.93E-01 | 7.75E-02 | 1.66E+00 | 1.68E+00 | 2.52E-02 | Yes | | intergenic_variant | MODIFIER |
| 15 | 11960567 | AX-170011051 | 0 | G | A | 0.298 | -1.93E-01 | 7.75E-02 | 1.66E+00 | 1.68E+00 | 2.52E-02 | Yes | Gm24074 | upstream_gene_variant | MODIFIER |
| 15 | 11972637 | AX-97803981 | 0 | T | C | 0.298 | -1.93E-01 | 7.75E-02 | 1.66E+00 | 1.68E+00 | 2.52E-02 | Yes | 1700047G03Rik | downstream_gene_variant | MODIFIER |
| 15 | 11975517 | AX-170011046 | 0 | T | C | 0.298 | -1.93E-01 | 7.75E-02 | 1.66E+00 | 1.68E+00 | 2.52E-02 | Yes | | intergenic_variant | MODIFIER |
| 15 | 11981226 | AX-170233002 | 0 | T | C | 0.298 | -1.93E-01 | 7.75E-02 | 1.66E+00 | 1.68E+00 | 2.52E-02 | Yes | Sub1 | downstream_gene_variant | MODIFIER |
| 15 | 11986115 | AX-170011050 | 0 | G | A | 0.298 | -1.93E-01 | 7.75E-02 | 1.66E+00 | 1.68E+00 | 2.52E-02 | Yes | Sub1 | intron_variant | MODIFIER |
| 15 | 12003357 | AX-170178886 | 0 | A | C | 0.298 | -1.93E-01 | 7.75E-02 | 1.66E+00 | 1.68E+00 | 2.52E-02 | Yes | | intergenic_variant | MODIFIER |
| 15 | 12028828 | AX-97832052 | 0 | G | A | 0.298 | -1.93E-01 | 7.75E-02 | 1.66E+00 | 1.68E+00 | 2.52E-02 | Yes | Gm2559 | downstream_gene_variant | MODIFIER |
| 15 | 12051119 | AX-97825693 | 0 | A | G | 0.298 | -1.93E-01 | 7.75E-02 | 1.66E+00 | 1.68E+00 | 2.52E-02 | Yes | | intergenic_variant | MODIFIER |
| 15 | 12147304 | AX-170311319 | 0 | A | C | 0.298 | -1.93E-01 | 7.75E-02 | 1.66E+00 | 1.68E+00 | 2.52E-02 | Yes | Zfr | intron_variant | MODIFIER |
| 15 | 12227116 | AX-170329292 | 0 | T | C | 0.298 | -1.93E-01 | 7.75E-02 | 1.66E+00 | 1.68E+00 | 2.52E-02 | Yes | Mtmr12 | intron_variant | MODIFIER |
| 15 | 12227493 | AX-170190278 | 0 | T | A | 0.298 | -1.93E-01 | 7.75E-02 | 1.66E+00 | 1.68E+00 | 2.52E-02 | Yes | Mtmr12 | intron_variant | MODIFIER |
| 15 | 12231148 | AX-170246517 | 0 | T | C | 0.298 | -1.93E-01 | 7.75E-02 | 1.66E+00 | 1.68E+00 | 2.52E-02 | Yes | Mtmr12 | intron_variant | MODIFIER |
| 15 | 12231463 | AX-170395709 | 0 | G | T | 0.298 | -1.93E-01 | 7.75E-02 | 1.66E+00 | 1.68E+00 | 2.52E-02 | Yes | Mtmr12 | intron_variant | MODIFIER |
| 15 | 12231685 | AX-170309766 | 0 | A | G | 0.298 | -1.93E-01 | 7.75E-02 | 1.66E+00 | 1.68E+00 | 2.52E-02 | Yes | Mtmr12 | intron_variant | MODIFIER |
| 15 | 12264543 | AX-170351968 | 0 | A | T | 0.298 | -1.93E-01 | 7.75E-02 | 1.66E+00 | 1.68E+00 | 2.52E-02 | Yes | Mtmr12 | intron_variant | MODIFIER |
| 15 | 12270195 | AX-170466207 | 0 | T | G | 0.298 | -1.93E-01 | 7.75E-02 | 1.66E+00 | 1.68E+00 | 2.52E-02 | Yes | Mtmr12 | 3_prime_UTR_variant | MODIFIER |
| 15 | 12275332 | AX-170256435 | 0 | C | A | 0.298 | -1.93E-01 | 7.75E-02 | 1.66E+00 | 1.68E+00 | 2.52E-02 | Yes | Mtmr12 | downstream_gene_variant | MODIFIER |
| 15 | 12314801 | AX-97830876 | 0 | C | T | 0.298 | -1.93E-01 | 7.75E-02 | 1.66E+00 | 1.68E+00 | 2.52E-02 | Yes | | regulatory_region_variant | MODIFIER |
| 15 | 12346168 | AX-97841948 | 0 | C | T | 0.298 | -1.93E-01 | 7.75E-02 | 1.66E+00 | 1.68E+00 | 2.52E-02 | Yes | Golph3 | intron_variant | MODIFIER |
| 15 | 12368255 | AX-170054823 | 0 | T | C | 0.298 | -1.93E-01 | 7.75E-02 | 1.66E+00 | 1.68E+00 | 2.52E-02 | Yes | Pdzd2 | intron_variant | MODIFIER |
| 15 | 12370915 | AX-97815350 | 0 | C | T | 0.298 | -1.93E-01 | 7.75E-02 | 1.66E+00 | 1.68E+00 | 2.52E-02 | Yes | Pdzd2 | intron_variant | MODIFIER |
| 15 | 12389106 | AX-170093160 | 0 | G | A | 0.298 | -1.93E-01 | 7.75E-02 | 1.66E+00 | 1.68E+00 | 2.52E-02 | Yes | Pdzd2 | intron_variant | MODIFIER |
| 15 | 12429059 | AX-170283777 | 0 | T | C | 0.298 | -1.93E-01 | 7.75E-02 | 1.66E+00 | 1.68E+00 | 2.52E-02 | Yes | Pdzd2 | intron_variant | MODIFIER |
| 15 | 12429080 | AX-170275853 | 0 | A | G | 0.298 | -1.93E-01 | 7.75E-02 | 1.66E+00 | 1.68E+00 | 2.52E-02 | Yes | Pdzd2 | intron_variant | MODIFIER |
| 15 | 12464425 | AX-97833608 | 0 | T | C | 0.298 | -1.93E-01 | 7.75E-02 | 1.66E+00 | 1.68E+00 | 2.52E-02 | Yes | Pdzd2 | intron_variant | MODIFIER |
| 15 | 12558063 | AX-170405403 | 0 | A | G | 0.298 | -1.93E-01 | 7.75E-02 | 1.66E+00 | 1.68E+00 | 2.52E-02 | Yes | Pdzd2 | intron_variant | MODIFIER |
| 15 | 12613384 | AX-170419318 | 0 | A | G | 0.298 | -1.93E-01 | 7.75E-02 | 1.66E+00 | 1.68E+00 | 2.52E-02 | Yes | Pdzd2 | intron_variant,non_coding_transcript_variant | MODIFIER |
| 15 | 12615746 | AX-170370149 | 0 | G | A | 0.298 | -1.93E-01 | 7.75E-02 | 1.66E+00 | 1.68E+00 | 2.52E-02 | Yes | Pdzd2 | intron_variant,non_coding_transcript_variant | MODIFIER |
| 15 | 12628481 | AX-170087770 | 0 | G | A | 0.298 | -1.93E-01 | 7.75E-02 | 1.66E+00 | 1.68E+00 | 2.52E-02 | Yes | Pdzd2 | intron_variant,non_coding_transcript_variant | MODIFIER |
| 15 | 12631176 | AX-170288899 | 0 | C | T | 0.298 | -1.93E-01 | 7.75E-02 | 1.66E+00 | 1.68E+00 | 2.52E-02 | Yes | Pdzd2 | intron_variant,non_coding_transcript_variant | MODIFIER |
| 15 | 12631802 | AX-170221982 | 0 | T | G | 0.298 | -1.93E-01 | 7.75E-02 | 1.66E+00 | 1.68E+00 | 2.52E-02 | Yes | Pdzd2 | intron_variant,non_coding_transcript_variant | MODIFIER |
| 15 | 12636532 | AX-170358559 | 0 | T | C | 0.298 | -1.93E-01 | 7.75E-02 | 1.66E+00 | 1.68E+00 | 2.52E-02 | Yes | Pdzd2 | intron_variant,non_coding_transcript_variant | MODIFIER |
| 15 | 12636582 | AX-170444904 | 0 | C | T | 0.298 | -1.93E-01 | 7.75E-02 | 1.66E+00 | 1.68E+00 | 2.52E-02 | Yes | Pdzd2 | intron_variant,non_coding_transcript_variant | MODIFIER |
| 15 | 13109204 | AX-170345739 | 0 | T | C | 0.298 | -1.93E-01 | 7.75E-02 | 1.66E+00 | 1.68E+00 | 2.52E-02 | Yes | Cdh6 | intron_variant | MODIFIER |

| | | | | | | | | | | | | | | | |
|---|---|---|---|---|---|---|---|---|---|---|---|---|---|---|---|
| 15 | 13109393 | AX-97848135 | 0 | T | C | 0.298 | -1.93E-01 | 7.75E-02 | 1.66E+00 | 1.68E+00 | 2.52E-02 | Yes | Cdh6 | intron_variant | MODIFIER |
| 15 | 13122088 | AX-170440891 | 0 | A | G | 0.298 | -1.93E-01 | 7.75E-02 | 1.66E+00 | 1.68E+00 | 2.52E-02 | Yes | Cdh6 | intron_variant | MODIFIER |
| 15 | 13122388 | AX-170317913 | 0 | C | A | 0.298 | -1.93E-01 | 7.75E-02 | 1.66E+00 | 1.68E+00 | 2.52E-02 | Yes | Cdh6 | intron_variant | MODIFIER |
| 15 | 13122696 | AX-170054846 | 0 | C | A | 0.298 | -1.93E-01 | 7.75E-02 | 1.66E+00 | 1.68E+00 | 2.52E-02 | Yes | Cdh6 | intron_variant | MODIFIER |
| 15 | 13124052 | AX-170054847 | 0 | G | A | 0.298 | -1.93E-01 | 7.75E-02 | 1.66E+00 | 1.68E+00 | 2.52E-02 | Yes | Cdh6 | intron_variant | MODIFIER |
| 15 | 13133678 | AX-170118896 | 0 | C | T | 0.298 | -1.93E-01 | 7.75E-02 | 1.66E+00 | 1.68E+00 | 2.52E-02 | Yes | Cdh6 | intron_variant | MODIFIER |
| 15 | 17399950 | AX-170011144 | 0 | T | G | 0.298 | -1.93E-01 | 7.75E-02 | 1.66E+00 | 1.68E+00 | 2.52E-02 | Yes | | intergenic_variant | MODIFIER |
| 3 | 7080030 | AX-170430341 | 0 | C | A | 0.14 | -1.97E-01 | 8.80E-02 | 2.87E+00 | 2.99E+00 | 2.58E-02 | Yes | | intergenic_variant | MODIFIER |
| 5 | 52959921 | AX-170123552 | 1 | C | A | 0.5 | 2.25E-01 | 1.00E-01 | 2.40E+00 | 2.46E+00 | 2.59E-02 | Yes | Gm23532 | upstream_gene_variant | MODIFIER |
| 19 | 55812403 | AX-97855903 | 0 | C | T | 0.447 | 1.45E-01 | 6.51E-02 | 2.90E+00 | 2.99E+00 | 2.59E-02 | Yes | Tcf7l2 | intron_variant | MODIFIER |
| 5 | 52861581 | AX-170532591 | 0 | G | C | 0.482 | 2.19E-01 | 9.79E-02 | 2.54E+00 | 2.60E+00 | 2.60E-02 | Yes | Anapc4 | intron_variant | MODIFIER |
| 18 | 65871265 | AX-170111193 | 1 | C | G | 0.375 | -1.34E-01 | 5.82E-02 | 3.66E+00 | 3.83E+00 | 2.60E-02 | no | Grp | upstream_gene_variant | MODIFIER |
| 9 | 97210454 | AX-170311426 | 0 | A | T | 0.14 | -2.43E-01 | 1.08E-01 | 3.14E+00 | 3.30E+00 | 2.61E-02 | Yes | Gm5371 | non_coding_transcript_exon_variant | MODIFIER |
| 9 | 98341340 | AX-170491974 | 0 | G | C | 0.281 | -1.21E-01 | 5.41E-02 | 3.14E+00 | 3.30E+00 | 2.61E-02 | no | Nmnat3 | intron_variant | MODIFIER |
| 3 | 23033241 | AX-170126622 | 0 | C | T | 0.158 | -2.19E-01 | 9.80E-02 | 3.04E+00 | 3.18E+00 | 2.62E-02 | Yes | | intergenic_variant | MODIFIER |
| 3 | 24602324 | AX-170426278 | 0 | A | G | 0.316 | -1.10E-01 | 4.90E-02 | 3.04E+00 | 3.18E+00 | 2.62E-02 | no | Naaladl2 | intron_variant | MODIFIER |
| 19 | 53381790 | AX-97814370 | 0 | C | A | 0.491 | 1.45E-01 | 6.51E-02 | 2.89E+00 | 2.98E+00 | 2.64E-02 | Yes | Smndc1 | intron_variant | MODIFIER |
| 19 | 53387469 | AX-170232815 | 0 | T | C | 0.491 | 1.45E-01 | 6.51E-02 | 2.89E+00 | 2.98E+00 | 2.64E-02 | Yes | Smndc1 | intron_variant | MODIFIER |
| 19 | 53435151 | AX-170348793 | 0 | A | G | 0.491 | 1.45E-01 | 6.51E-02 | 2.89E+00 | 2.98E+00 | 2.64E-02 | Yes | | regulatory_region_variant | MODIFIER |
| 19 | 53435381 | AX-170279786 | 0 | C | G | 0.491 | 1.45E-01 | 6.51E-02 | 2.89E+00 | 2.98E+00 | 2.64E-02 | Yes | | regulatory_region_variant | MODIFIER |
| 19 | 53435647 | AX-170232994 | 0 | T | C | 0.491 | 1.45E-01 | 6.51E-02 | 2.89E+00 | 2.98E+00 | 2.64E-02 | Yes | | regulatory_region_variant | MODIFIER |
| 19 | 53437221 | AX-97799344 | 0 | G | T | 0.491 | 1.45E-01 | 6.51E-02 | 2.89E+00 | 2.98E+00 | 2.64E-02 | Yes | | regulatory_region_variant | MODIFIER |
| 19 | 53438119 | AX-170312348 | 0 | A | C | 0.491 | 1.45E-01 | 6.51E-02 | 2.89E+00 | 2.98E+00 | 2.64E-02 | Yes | | regulatory_region_variant | MODIFIER |
| 19 | 53445426 | AX-170376735 | 0 | A | G | 0.491 | 1.45E-01 | 6.51E-02 | 2.89E+00 | 2.98E+00 | 2.64E-02 | Yes | Mirt1 | intron_variant,non_coding_transcript_variant | MODIFIER |
| 19 | 53449115 | AX-170278609 | 0 | A | G | 0.491 | 1.45E-01 | 6.51E-02 | 2.89E+00 | 2.98E+00 | 2.64E-02 | Yes | Mirt1 | intron_variant,non_coding_transcript_variant | MODIFIER |
| 19 | 53456105 | AX-97815669 | 0 | A | G | 0.491 | 1.45E-01 | 6.51E-02 | 2.89E+00 | 2.98E+00 | 2.64E-02 | Yes | 4833407H14Rik | upstream_gene_variant | MODIFIER |
| 19 | 53472455 | AX-170398078 | 0 | G | T | 0.491 | 1.45E-01 | 6.51E-02 | 2.89E+00 | 2.98E+00 | 2.64E-02 | Yes | | intergenic_variant | MODIFIER |
| 19 | 53482751 | AX-170336303 | 0 | G | A | 0.491 | 1.45E-01 | 6.51E-02 | 2.89E+00 | 2.98E+00 | 2.64E-02 | Yes | AC123702.1 | upstream_gene_variant | MODIFIER |
| 19 | 53488905 | AX-170442641 | 0 | C | T | 0.491 | 1.45E-01 | 6.51E-02 | 2.89E+00 | 2.98E+00 | 2.64E-02 | Yes | AC123702.1 | downstream_gene_variant | MODIFIER |
| 19 | 53495790 | AX-170479842 | 0 | C | T | 0.491 | 1.45E-01 | 6.51E-02 | 2.89E+00 | 2.98E+00 | 2.64E-02 | Yes | | intergenic_variant | MODIFIER |
| 19 | 53499057 | AX-168414592 | 0 | T | C | 0.491 | 1.45E-01 | 6.51E-02 | 2.89E+00 | 2.98E+00 | 2.64E-02 | Yes | | intergenic_variant | MODIFIER |
| 19 | 53761523 | AX-97836089 | 0 | T | C | 0.491 | 1.45E-01 | 6.51E-02 | 2.89E+00 | 2.98E+00 | 2.64E-02 | Yes | Rbm20 | intron_variant | MODIFIER |
| 19 | 53771023 | AX-170282154 | 0 | A | G | 0.491 | 1.45E-01 | 6.51E-02 | 2.89E+00 | 2.98E+00 | 2.64E-02 | Yes | Gm16298 | upstream_gene_variant | MODIFIER |
| 19 | 53788952 | AX-97836112 | 0 | T | A | 0.491 | 1.45E-01 | 6.51E-02 | 2.89E+00 | 2.98E+00 | 2.64E-02 | Yes | Rbm20 | upstream_gene_variant | MODIFIER |
| 19 | 53834901 | AX-170362018 | 0 | T | C | 0.491 | 1.45E-01 | 6.51E-02 | 2.89E+00 | 2.98E+00 | 2.64E-02 | Yes | Rbm20 | intron_variant | MODIFIER |
| 19 | 53848814 | AX-170309438 | 0 | T | C | 0.491 | 1.45E-01 | 6.51E-02 | 2.89E+00 | 2.98E+00 | 2.64E-02 | Yes | Rbm20 | intron_variant,NMD_transcript_variant | MODIFIER |
| 19 | 53850853 | AX-170016857 | 0 | T | G | 0.491 | 1.45E-01 | 6.51E-02 | 2.89E+00 | 2.98E+00 | 2.64E-02 | Yes | Rbm20 | intron_variant,NMD_transcript_variant | MODIFIER |
| 19 | 53852665 | AX-170062722 | 0 | A | G | 0.491 | 1.45E-01 | 6.51E-02 | 2.89E+00 | 2.98E+00 | 2.64E-02 | Yes | Rbm20 | intron_variant,NMD_transcript_variant | MODIFIER |
| 19 | 53864807 | AX-97836114 | 0 | A | G | 0.491 | 1.45E-01 | 6.51E-02 | 2.89E+00 | 2.98E+00 | 2.64E-02 | Yes | Rbm20 | 3_prime_UTR_variant,NMD_transcript_variant | MODIFIER |
| 19 | 53875092 | AX-170062724 | 0 | C | G | 0.491 | 1.45E-01 | 6.51E-02 | 2.89E+00 | 2.98E+00 | 2.64E-02 | Yes | | regulatory_region_variant | MODIFIER |
| 19 | 53884740 | AX-170016859 | 0 | C | A | 0.491 | 1.45E-01 | 6.51E-02 | 2.89E+00 | 2.98E+00 | 2.64E-02 | Yes | AC117805.1 | downstream_gene_variant | MODIFIER |
| 19 | 53894398 | AX-170016856 | 0 | T | G | 0.491 | 1.45E-01 | 6.51E-02 | 2.89E+00 | 2.98E+00 | 2.64E-02 | Yes | Pdcd4 | intron_variant | MODIFIER |
| 19 | 54029760 | AX-170394176 | 0 | T | C | 0.491 | 1.45E-01 | 6.51E-02 | 2.89E+00 | 2.98E+00 | 2.64E-02 | Yes | Shoc2 | intron_variant | MODIFIER |
| 19 | 54039654 | AX-97793478 | 0 | G | A | 0.491 | 1.45E-01 | 6.51E-02 | 2.89E+00 | 2.98E+00 | 2.64E-02 | Yes | Adra2a | upstream_gene_variant | MODIFIER |
| 19 | 54116858 | AX-170516067 | 0 | T | G | 0.491 | 1.45E-01 | 6.51E-02 | 2.89E+00 | 2.98E+00 | 2.64E-02 | Yes | | intergenic_variant | MODIFIER |
| 19 | 54177085 | AX-170487251 | 0 | C | A | 0.491 | 1.45E-01 | 6.51E-02 | 2.89E+00 | 2.98E+00 | 2.64E-02 | Yes | | intergenic_variant | MODIFIER |
| 19 | 54261040 | AX-97836137 | 0 | T | C | 0.491 | 1.45E-01 | 6.51E-02 | 2.89E+00 | 2.98E+00 | 2.64E-02 | Yes | | intergenic_variant | MODIFIER |
| 3 | 7160260 | AX-170019853 | 1 | G | C | 0.134 | -2.03E-01 | 9.13E-02 | 2.79E+00 | 2.90E+00 | 2.65E-02 | Yes | | intergenic_variant | MODIFIER |
| 16 | 58246332 | AX-170110018 | 1 | A | G | 0.446 | -1.85E-01 | 8.30E-02 | 2.62E+00 | 2.67E+00 | 2.65E-02 | Yes | Gm49574 | intron_variant,non_coding_transcript_variant | MODIFIER |
| 6 | 134038851 | AX-97830016 | 0 | G | T | 0.044 | 3.32E-01 | 1.48E-01 | 3.26E+00 | 3.58E+00 | 2.66E-02 | Yes | Etv6 | intron_variant | MODIFIER |
| 19 | 54226036 | AX-170345985 | 0 | G | A | 0.491 | 1.47E-01 | 6.59E-02 | 2.99E+00 | 3.09E+00 | 2.69E-02 | Yes | | intergenic_variant | MODIFIER |
| 13 | 7450883 | AX-97826932 | 0 | C | T | 0.184 | -2.59E-01 | 1.17E-01 | 2.67E+00 | 2.75E+00 | 2.69E-02 | Yes | Gm36074 | intron_variant,non_coding_transcript_variant | MODIFIER |
| 13 | 7907879 | AX-170246394 | 0 | A | G | 0.184 | -2.59E-01 | 1.17E-01 | 2.67E+00 | 2.75E+00 | 2.69E-02 | Yes | | intergenic_variant | MODIFIER |
| 13 | 9667668 | AX-170238314 | 0 | A | G | 0.184 | -2.59E-01 | 1.17E-01 | 2.67E+00 | 2.75E+00 | 2.69E-02 | Yes | Dip2c | downstream_gene_variant | MODIFIER |

| | | | | | | | | | | | | | | | |
|---|---|---|---|---|---|---|---|---|---|---|---|---|---|---|---|
| 13 | 10886480 | AX-97844114 | 0 | A | G | 0.184 | -2.59E-01 | 1.17E-01 | 2.67E+00 | 2.75E+00 | 2.69E-02 | Yes | | intergenic_variant | MODIFIER |
| 13 | 11524312 | AX-170276806 | 0 | A | G | 0.184 | -2.59E-01 | 1.17E-01 | 2.67E+00 | 2.75E+00 | 2.69E-02 | Yes | Zp4-ps | intron_variant,non_coding_transcript_variant | MODIFIER |
| 5 | 129778875 | AX-170245538 | 0 | T | G | 0.149 | -1.84E-01 | 8.16E-02 | 2.23E+00 | 2.33E+00 | 2.70E-02 | Yes | Psph | intron_variant | MODIFIER |
| 5 | 129785059 | AX-170155481 | 0 | A | G | 0.149 | -1.84E-01 | 8.16E-02 | 2.23E+00 | 2.33E+00 | 2.70E-02 | Yes | Psph | intron_variant | MODIFIER |
| 13 | 18093301 | AX-97811091 | 0 | T | A | 0.149 | -2.62E-01 | 1.18E-01 | 2.56E+00 | 2.64E+00 | 2.77E-02 | Yes | 4930448F12Rik | intron_variant,non_coding_transcript_variant | MODIFIER |
| 5 | 53298828 | AX-170359063 | 1 | C | T | 0.5 | 2.21E-01 | 9.93E-02 | 2.42E+00 | 2.47E+00 | 2.78E-02 | Yes | | intergenic_variant | MODIFIER |
| 15 | 12001356 | AX-170415856 | 0 | T | C | 0.298 | -1.89E-01 | 7.76E-02 | 1.68E+00 | 1.70E+00 | 2.80E-02 | Yes | Sub1 | upstream_gene_variant | MODIFIER |
| 9 | 94262375 | AX-170170953 | 1 | C | T | 0.143 | -2.64E-01 | 1.18E-01 | 3.32E+00 | 3.48E+00 | 2.81E-02 | Yes | | intergenic_variant | MODIFIER |
| 5 | 53298394 | AX-170480539 | 2 | C | T | 0.5 | 2.20E-01 | 9.92E-02 | 2.42E+00 | 2.48E+00 | 2.81E-02 | Yes | | intergenic_variant | MODIFIER |
| 8 | 64860753 | AX-170361417 | 0 | G | A | 0.14 | -2.96E-01 | 1.33E-01 | 3.22E+00 | 3.38E+00 | 2.82E-02 | Yes | | intergenic_variant | MODIFIER |
| 5 | 128553122 | AX-170136553 | 0 | G | C | 0.149 | -1.89E-01 | 8.41E-02 | 2.21E+00 | 2.31E+00 | 2.83E-02 | Yes | | regulatory_region_variant | MODIFIER |
| 15 | 21848526 | AX-170508957 | 0 | G | T | 0.07 | -2.88E-01 | 1.07E-01 | 1.31E+00 | 1.35E+00 | 2.84E-02 | Yes | | intergenic_variant | MODIFIER |
| 19 | 53848667 | AX-170016860 | 0 | C | T | 0.482 | 1.45E-01 | 6.56E-02 | 2.97E+00 | 3.07E+00 | 2.84E-02 | Yes | Rbm20 | intron_variant,NMD_transcript_variant | MODIFIER |
| 5 | 61947925 | AX-97828364 | 0 | G | A | 0.465 | 2.17E-01 | 9.87E-02 | 2.55E+00 | 2.62E+00 | 2.85E-02 | Yes | | intergenic_variant | MODIFIER |
| 15 | 70867098 | AX-170011950 | 0 | C | T | 0.079 | -2.69E-01 | 1.07E-01 | 1.56E+00 | 1.60E+00 | 2.85E-02 | Yes | | intergenic_variant | MODIFIER |
| 19 | 58709621 | AX-170094136 | 1 | T | C | 0.438 | 1.43E-01 | 6.52E-02 | 3.00E+00 | 3.10E+00 | 2.86E-02 | no | 1810018F18Rik | intron_variant,non_coding_transcript_variant | MODIFIER |
| 19 | 53853238 | AX-170062725 | 1 | A | G | 0.482 | 1.40E-01 | 6.41E-02 | 2.91E+00 | 3.01E+00 | 2.91E-02 | no | Rbm20 | intron_variant,NMD_transcript_variant | MODIFIER |
| 16 | 56022551 | AX-170013113 | 0 | A | G | 0.456 | -1.79E-01 | 8.18E-02 | 2.71E+00 | 2.76E+00 | 2.93E-02 | Yes | Pcnp | intron_variant | MODIFIER |
| 3 | 6323214 | AX-170065836 | 1 | T | C | 0.134 | -1.98E-01 | 9.09E-02 | 2.76E+00 | 2.87E+00 | 2.93E-02 | Yes | | intergenic_variant | MODIFIER |
| 16 | 56585298 | AX-170013119 | 1 | C | T | 0.464 | -1.81E-01 | 8.28E-02 | 2.73E+00 | 2.79E+00 | 2.95E-02 | Yes | Abi3bp | intron_variant | MODIFIER |
| 15 | 11472546 | AX-170181903 | 1 | A | G | 0.304 | -1.85E-01 | 7.70E-02 | 1.71E+00 | 1.73E+00 | 2.95E-02 | Yes | | intergenic_variant | MODIFIER |
| 15 | 11538401 | AX-170335274 | 1 | G | A | 0.304 | -1.85E-01 | 7.70E-02 | 1.71E+00 | 1.73E+00 | 2.95E-02 | Yes | | intergenic_variant | MODIFIER |
| 13 | 19938932 | AX-170158573 | 0 | C | G | 0.167 | -2.35E-01 | 1.08E-01 | 2.73E+00 | 2.86E+00 | 2.96E-02 | Yes | | intergenic_variant | MODIFIER |
| 19 | 56214365 | AX-170062768 | 1 | G | C | 0.438 | 1.37E-01 | 6.29E-02 | 2.92E+00 | 3.01E+00 | 2.97E-02 | no | | intergenic_variant | MODIFIER |
| 13 | 16739390 | AX-170007834 | 1 | C | G | 0.17 | -2.80E-01 | 1.28E-01 | 2.45E+00 | 2.48E+00 | 2.97E-02 | Yes | | intergenic_variant | MODIFIER |
| 1 | 119386120 | AX-170311155 | 0 | C | T | 0.579 | 1.76E-01 | 8.10E-02 | 2.77E+00 | 2.85E+00 | 2.97E-02 | Yes | Gm29455 | downstream_gene_variant | MODIFIER |
| 5 | 136787879 | AX-97844614 | 0 | G | A | 0.149 | -1.67E-01 | 7.64E-02 | 2.45E+00 | 2.57E+00 | 2.99E-02 | no | Col26a1 | intron_variant | MODIFIER |
| 5 | 52656260 | AX-170277243 | 0 | A | C | 0.491 | 2.15E-01 | 9.83E-02 | 2.42E+00 | 2.47E+00 | 3.01E-02 | Yes | Sepsecs | missense_variant | MODERATE |
| 5 | 52688775 | AX-170380448 | 0 | G | A | 0.491 | 2.15E-01 | 9.83E-02 | 2.42E+00 | 2.47E+00 | 3.01E-02 | Yes | | regulatory_region_variant | MODIFIER |
| 5 | 64710391 | AX-170025160 | 1 | T | C | 0.482 | 2.11E-01 | 9.67E-02 | 2.47E+00 | 2.53E+00 | 3.01E-02 | Yes | Gm20033 | intron_variant,non_coding_transcript_variant | MODIFIER |
| 5 | 137108514 | AX-97800890 | 1 | T | C | 0.125 | -1.97E-01 | 8.99E-02 | 2.35E+00 | 2.43E+00 | 3.06E-02 | Yes | Trim56 | 3_prime_UTR_variant | MODIFIER |
| 5 | 134734487 | AX-170499426 | 1 | G | A | 0.152 | -1.74E-01 | 7.89E-02 | 2.26E+00 | 2.36E+00 | 3.08E-02 | Yes | Eln | intron_variant | MODIFIER |
| 15 | 11462139 | AX-170348603 | 1 | C | T | 0.304 | -1.81E-01 | 7.69E-02 | 1.76E+00 | 1.79E+00 | 3.09E-02 | Yes | | intergenic_variant | MODIFIER |
| 5 | 57562744 | AX-170241889 | 0 | G | A | 0.474 | 2.14E-01 | 9.71E-02 | 2.20E+00 | 2.26E+00 | 3.11E-02 | Yes | | intergenic_variant | MODIFIER |
| 15 | 12598679 | AX-170054831 | 0 | G | A | 0.307 | -1.82E-01 | 7.67E-02 | 1.72E+00 | 1.75E+00 | 3.11E-02 | Yes | Pdzd2 | intron_variant,non_coding_transcript_variant | MODIFIER |
| 5 | 137713227 | AX-170151728 | 1 | C | A | 0.125 | -1.96E-01 | 8.98E-02 | 2.34E+00 | 2.42E+00 | 3.11E-02 | Yes | | intergenic_variant | MODIFIER |
| 5 | 53712941 | AX-170269112 | 0 | C | T | 0.491 | 2.19E-01 | 1.01E-01 | 2.44E+00 | 2.49E+00 | 3.12E-02 | Yes | | intergenic_variant | MODIFIER |
| 13 | 12520300 | AX-170007771 | 0 | A | G | 0.184 | -2.66E-01 | 1.23E-01 | 2.53E+00 | 2.60E+00 | 3.12E-02 | Yes | Edaradd | 5_prime_UTR_variant | MODIFIER |
| 1 | 134661480 | AX-170395533 | 2 | C | A | 0.564 | 1.74E-01 | 8.07E-02 | 2.84E+00 | 2.94E+00 | 3.14E-02 | Yes | Syt2 | intron_variant | MODIFIER |
| 3 | 157115140 | AX-97855497 | 0 | A | G | 0.026 | -3.73E-01 | 1.73E-01 | 2.68E+00 | 2.82E+00 | 3.14E-02 | Yes | Negr1 | intron_variant | MODIFIER |
| 3 | 19238180 | AX-170486319 | 0 | A | G | 0.123 | -1.93E-01 | 8.96E-02 | 2.76E+00 | 2.89E+00 | 3.15E-02 | Yes | Pde7a | intron_variant | MODIFIER |
| 13 | 12345180 | AX-170477417 | 0 | A | G | 0.184 | -2.69E-01 | 1.25E-01 | 2.55E+00 | 2.60E+00 | 3.19E-02 | Yes | Actn2 | upstream_gene_variant | MODIFIER |
| 13 | 12360839 | AX-168394408 | 0 | T | C | 0.184 | -2.69E-01 | 1.25E-01 | 2.55E+00 | 2.60E+00 | 3.19E-02 | Yes | | intergenic_variant | MODIFIER |
| 13 | 12361500 | AX-170127803 | 0 | C | T | 0.184 | -2.69E-01 | 1.25E-01 | 2.55E+00 | 2.60E+00 | 3.19E-02 | Yes | | intergenic_variant | MODIFIER |
| 13 | 12361777 | AX-97824864 | 0 | C | T | 0.184 | -2.69E-01 | 1.25E-01 | 2.55E+00 | 2.60E+00 | 3.19E-02 | Yes | | intergenic_variant | MODIFIER |
| 13 | 12376588 | AX-170514560 | 0 | C | T | 0.184 | -2.69E-01 | 1.25E-01 | 2.55E+00 | 2.60E+00 | 3.19E-02 | Yes | Gm5445 | upstream_gene_variant | MODIFIER |
| 13 | 12380594 | AX-170277566 | 0 | A | G | 0.184 | -2.69E-01 | 1.25E-01 | 2.55E+00 | 2.60E+00 | 3.19E-02 | Yes | Gm5445 | downstream_gene_variant | MODIFIER |
| 13 | 12406786 | AX-97805977 | 0 | A | G | 0.184 | -2.69E-01 | 1.25E-01 | 2.55E+00 | 2.60E+00 | 3.19E-02 | Yes | Heatr1 | intron_variant | MODIFIER |
| 13 | 12413149 | AX-170232706 | 0 | C | T | 0.184 | -2.69E-01 | 1.25E-01 | 2.55E+00 | 2.60E+00 | 3.19E-02 | Yes | Heatr1 | intron_variant | MODIFIER |
| 13 | 12413268 | AX-170007770 | 0 | G | A | 0.184 | -2.69E-01 | 1.25E-01 | 2.55E+00 | 2.60E+00 | 3.19E-02 | Yes | Heatr1 | missense_variant | MODERATE |
| 13 | 12416064 | AX-170174691 | 0 | C | T | 0.184 | -2.69E-01 | 1.25E-01 | 2.55E+00 | 2.60E+00 | 3.19E-02 | Yes | Heatr1 | synonymous_variant | LOW |
| 13 | 12420318 | AX-170092662 | 0 | A | G | 0.184 | -2.69E-01 | 1.25E-01 | 2.55E+00 | 2.60E+00 | 3.19E-02 | Yes | Heatr1 | intron_variant | MODIFIER |
| 13 | 12427276 | AX-170007775 | 0 | G | A | 0.184 | -2.69E-01 | 1.25E-01 | 2.55E+00 | 2.60E+00 | 3.19E-02 | Yes | Heatr1 | intron_variant | MODIFIER |
| 13 | 12435920 | AX-170475979 | 0 | G | A | 0.184 | -2.69E-01 | 1.25E-01 | 2.55E+00 | 2.60E+00 | 3.19E-02 | Yes | Heatr1 | intron_variant | MODIFIER |

| | | | | | | | | | | | | | | | |
|---|---|---|---|---|---|---|---|---|---|---|---|---|---|---|---|
| 13 | 12454285 | AX-170141500 | 0 | G | A | 0.184 | -2.69E-01 | 1.25E-01 | 2.55E+00 | 2.60E+00 | 3.19E-02 | Yes | Lgals8 | intron_variant | MODIFIER |
| 13 | 12454674 | AX-170496255 | 0 | C | T | 0.184 | -2.69E-01 | 1.25E-01 | 2.55E+00 | 2.60E+00 | 3.19E-02 | Yes | Lgals8 | intron_variant | MODIFIER |
| 13 | 12455344 | AX-170250237 | 0 | C | G | 0.184 | -2.69E-01 | 1.25E-01 | 2.55E+00 | 2.60E+00 | 3.19E-02 | Yes | Lgals8 | intron_variant | MODIFIER |
| 13 | 12456830 | AX-170306483 | 0 | A | G | 0.184 | -2.69E-01 | 1.25E-01 | 2.55E+00 | 2.60E+00 | 3.19E-02 | Yes | Lgals8 | intron_variant | MODIFIER |
| 13 | 12507546 | AX-170524923 | 0 | A | G | 0.184 | -2.69E-01 | 1.25E-01 | 2.55E+00 | 2.60E+00 | 3.19E-02 | Yes | Edaradd | intron_variant | MODIFIER |
| 13 | 12507676 | AX-170146555 | 0 | C | A | 0.184 | -2.69E-01 | 1.25E-01 | 2.55E+00 | 2.60E+00 | 3.19E-02 | Yes | Edaradd | intron_variant | MODIFIER |
| 13 | 12507943 | AX-170104601 | 0 | A | T | 0.184 | -2.69E-01 | 1.25E-01 | 2.55E+00 | 2.60E+00 | 3.19E-02 | Yes | Edaradd | intron_variant | MODIFIER |
| 16 | 63598318 | AX-170013232 | 0 | G | A | 0.465 | -1.87E-01 | 8.70E-02 | 2.79E+00 | 2.85E+00 | 3.19E-02 | Yes | Epha3 | splice_region_variant,intron_variant | LOW |
| 8 | 33957554 | AX-170177438 | 0 | C | G | 0.14 | -2.65E-01 | 1.21E-01 | 3.33E+00 | 3.50E+00 | 3.19E-02 | Yes | | regulatory_region_variant | MODIFIER |
| 8 | 34309370 | AX-97804051 | 0 | G | A | 0.281 | -1.32E-01 | 6.06E-02 | 3.33E+00 | 3.50E+00 | 3.19E-02 | no | Gm6100 | upstream_gene_variant | MODIFIER |
| 8 | 34768947 | AX-170030497 | 0 | G | T | 0.281 | -1.32E-01 | 6.06E-02 | 3.33E+00 | 3.50E+00 | 3.19E-02 | no | | intergenic_variant | MODIFIER |
| 5 | 56807566 | AX-170152662 | 1 | T | C | 0.473 | 2.14E-01 | 9.90E-02 | 2.45E+00 | 2.52E+00 | 3.20E-02 | Yes | | intergenic_variant | MODIFIER |
| 16 | 63598192 | AX-170013234 | 1 | G | C | 0.464 | -1.87E-01 | 8.71E-02 | 2.79E+00 | 2.84E+00 | 3.21E-02 | Yes | Epha3 | missense_variant,splice_region_variant | MODERATE |
| 9 | 112552094 | AX-170152281 | 0 | G | A | 0.123 | -2.07E-01 | 9.37E-02 | 3.50E+00 | 3.77E+00 | 3.21E-02 | Yes | | intergenic_variant | MODIFIER |
| 16 | 63297266 | AX-170088044 | 1 | A | G | 0.464 | -1.87E-01 | 8.70E-02 | 2.79E+00 | 2.84E+00 | 3.22E-02 | Yes | | intergenic_variant | MODIFIER |
| 3 | 6405173 | AX-170327740 | 0 | C | T | 0.123 | -1.97E-01 | 9.21E-02 | 2.82E+00 | 2.94E+00 | 3.26E-02 | Yes | | intergenic_variant | MODIFIER |
| 19 | 60395119 | AX-170442559 | 0 | A | G | 0.526 | 1.35E-01 | 6.28E-02 | 2.97E+00 | 3.08E+00 | 3.26E-02 | no | | intergenic_variant | MODIFIER |
| 1 | 120559279 | AX-170141581 | 0 | C | T | 0.579 | 1.72E-01 | 8.06E-02 | 2.72E+00 | 2.80E+00 | 3.27E-02 | Yes | | intergenic_variant | MODIFIER |
| 16 | 63922118 | AX-170098436 | 1 | A | G | 0.518 | 1.75E-01 | 8.19E-02 | 2.89E+00 | 2.98E+00 | 3.27E-02 | Yes | | intergenic_variant | MODIFIER |
| 15 | 17136116 | AX-170515687 | 0 | G | A | 0.307 | -1.79E-01 | 7.72E-02 | 1.79E+00 | 1.82E+00 | 3.30E-02 | Yes | | regulatory_region_variant | MODIFIER |
| 3 | 6285268 | AX-170505261 | 1 | G | A | 0.116 | -1.99E-01 | 9.35E-02 | 2.75E+00 | 2.86E+00 | 3.30E-02 | Yes | | intergenic_variant | MODIFIER |
| 15 | 11453883 | AX-170011031 | 1 | T | A | 0.304 | -1.82E-01 | 7.75E-02 | 1.73E+00 | 1.75E+00 | 3.31E-02 | Yes | | intergenic_variant | MODIFIER |
| 19 | 60165005 | AX-170160267 | 0 | C | T | 0.456 | -1.34E-01 | 6.28E-02 | 3.00E+00 | 3.10E+00 | 3.33E-02 | no | E330013P04Rik | downstream_gene_variant | MODIFIER |
| 5 | 60086748 | AX-168413053 | 0 | C | T | 0.482 | 2.10E-01 | 9.85E-02 | 2.59E+00 | 2.66E+00 | 3.34E-02 | Yes | | intergenic_variant | MODIFIER |
| 5 | 60521720 | AX-170445766 | 0 | T | G | 0.482 | 2.10E-01 | 9.85E-02 | 2.59E+00 | 2.66E+00 | 3.34E-02 | Yes | | intergenic_variant | MODIFIER |
| 5 | 60689925 | AX-97813916 | 0 | T | C | 0.482 | 2.10E-01 | 9.85E-02 | 2.59E+00 | 2.66E+00 | 3.34E-02 | Yes | | intergenic_variant | MODIFIER |
| 5 | 55114980 | AX-170140990 | 0 | G | A | 0.491 | 2.06E-01 | 9.67E-02 | 2.49E+00 | 2.56E+00 | 3.37E-02 | Yes | | intergenic_variant | MODIFIER |
| 13 | 7184592 | AX-97826931 | 0 | A | G | 0.184 | -2.28E-01 | 1.07E-01 | 2.62E+00 | 2.72E+00 | 3.37E-02 | Yes | | intergenic_variant | MODIFIER |
| 16 | 58406111 | AX-170465093 | 0 | A | G | 0.439 | -1.72E-01 | 8.06E-02 | 2.54E+00 | 2.59E+00 | 3.38E-02 | no | Dcbld2 | upstream_gene_variant | MODIFIER |
| 19 | 51723003 | AX-170016821 | 1 | A | T | 0.491 | 1.37E-01 | 6.44E-02 | 2.97E+00 | 3.09E+00 | 3.38E-02 | no | | intergenic_variant | MODIFIER |
| 15 | 12648702 | AX-97786124 | 0 | T | C | 0.298 | -1.82E-01 | 7.78E-02 | 1.72E+00 | 1.75E+00 | 3.40E-02 | Yes | Pdzd2 | intron_variant,non_coding_transcript_variant | MODIFIER |
| 13 | 16338805 | AX-170257329 | 0 | G | A | 0.175 | -2.71E-01 | 1.27E-01 | 2.44E+00 | 2.48E+00 | 3.41E-02 | Yes | | intergenic_variant | MODIFIER |
| 13 | 16476006 | AX-170234915 | 0 | C | T | 0.175 | -2.71E-01 | 1.27E-01 | 2.44E+00 | 2.48E+00 | 3.41E-02 | Yes | | intergenic_variant | MODIFIER |
| 13 | 16636879 | AX-97822787 | 0 | A | G | 0.175 | -2.71E-01 | 1.27E-01 | 2.44E+00 | 2.48E+00 | 3.41E-02 | Yes | Gm7537 | downstream_gene_variant | MODIFIER |
| 13 | 16674290 | AX-170087377 | 0 | T | G | 0.175 | -2.71E-01 | 1.27E-01 | 2.44E+00 | 2.48E+00 | 3.41E-02 | Yes | | intergenic_variant | MODIFIER |
| 13 | 16725681 | AX-170007837 | 0 | C | T | 0.175 | -2.71E-01 | 1.27E-01 | 2.44E+00 | 2.48E+00 | 3.41E-02 | Yes | | intergenic_variant | MODIFIER |
| 13 | 16731829 | AX-97799498 | 0 | A | G | 0.175 | -2.71E-01 | 1.27E-01 | 2.44E+00 | 2.48E+00 | 3.41E-02 | Yes | | intergenic_variant | MODIFIER |
| 13 | 16732643 | AX-170007830 | 0 | G | C | 0.175 | -2.71E-01 | 1.27E-01 | 2.44E+00 | 2.48E+00 | 3.41E-02 | Yes | | intergenic_variant | MODIFIER |
| 13 | 16739324 | AX-170007828 | 0 | T | G | 0.175 | -2.71E-01 | 1.27E-01 | 2.44E+00 | 2.48E+00 | 3.41E-02 | Yes | | intergenic_variant | MODIFIER |
| 13 | 16739755 | AX-170087378 | 0 | C | T | 0.175 | -2.71E-01 | 1.27E-01 | 2.44E+00 | 2.48E+00 | 3.41E-02 | Yes | | intergenic_variant | MODIFIER |
| 13 | 16829983 | AX-97822920 | 0 | G | T | 0.175 | -2.71E-01 | 1.27E-01 | 2.44E+00 | 2.48E+00 | 3.41E-02 | Yes | | regulatory_region_variant | MODIFIER |
| 13 | 16833272 | AX-170225020 | 0 | T | C | 0.175 | -2.71E-01 | 1.27E-01 | 2.44E+00 | 2.48E+00 | 3.41E-02 | Yes | | intergenic_variant | MODIFIER |
| 13 | 16835108 | AX-170007824 | 0 | T | C | 0.175 | -2.71E-01 | 1.27E-01 | 2.44E+00 | 2.48E+00 | 3.41E-02 | Yes | | intergenic_variant | MODIFIER |
| 13 | 16838256 | AX-97822948 | 0 | G | A | 0.175 | -2.71E-01 | 1.27E-01 | 2.44E+00 | 2.48E+00 | 3.41E-02 | Yes | | intergenic_variant | MODIFIER |
| 13 | 16846245 | AX-170332093 | 0 | A | T | 0.175 | -2.71E-01 | 1.27E-01 | 2.44E+00 | 2.48E+00 | 3.41E-02 | Yes | | intergenic_variant | MODIFIER |
| 13 | 16851910 | AX-170195979 | 0 | A | G | 0.175 | -2.71E-01 | 1.27E-01 | 2.44E+00 | 2.48E+00 | 3.41E-02 | Yes | | intergenic_variant | MODIFIER |
| 13 | 16866131 | AX-170007836 | 0 | G | T | 0.175 | -2.71E-01 | 1.27E-01 | 2.44E+00 | 2.48E+00 | 3.41E-02 | Yes | Sugct | intron_variant | MODIFIER |
| 13 | 16903750 | AX-97850132 | 0 | G | A | 0.175 | -2.71E-01 | 1.27E-01 | 2.44E+00 | 2.48E+00 | 3.41E-02 | Yes | Sugct | intron_variant | MODIFIER |
| 13 | 16904484 | AX-170238392 | 0 | A | G | 0.175 | -2.71E-01 | 1.27E-01 | 2.44E+00 | 2.48E+00 | 3.41E-02 | Yes | Sugct | intron_variant | MODIFIER |
| 13 | 16907372 | AX-170007833 | 0 | C | T | 0.175 | -2.71E-01 | 1.27E-01 | 2.44E+00 | 2.48E+00 | 3.41E-02 | Yes | Sugct | intron_variant | MODIFIER |
| 13 | 16930674 | AX-97822837 | 0 | T | C | 0.175 | -2.71E-01 | 1.27E-01 | 2.44E+00 | 2.48E+00 | 3.41E-02 | Yes | Sugct | intron_variant | MODIFIER |
| 13 | 16935026 | AX-97814104 | 0 | G | A | 0.175 | -2.71E-01 | 1.27E-01 | 2.44E+00 | 2.48E+00 | 3.41E-02 | Yes | Sugct | intron_variant | MODIFIER |
| 13 | 16965961 | AX-170518591 | 0 | T | G | 0.175 | -2.71E-01 | 1.27E-01 | 2.44E+00 | 2.48E+00 | 3.41E-02 | Yes | Sugct | intron_variant | MODIFIER |
| 13 | 16967380 | AX-97806134 | 0 | A | C | 0.175 | -2.71E-01 | 1.27E-01 | 2.44E+00 | 2.48E+00 | 3.41E-02 | Yes | Sugct | intron_variant | MODIFIER |

| | | | | | | | | | | | | | | | |
|---|---|---|---|---|---|---|---|---|---|---|---|---|---|---|---|
| 13 | 16968391 | AX-170408218 | 0 | A | C | 0.175 | -2.71E-01 | 1.27E-01 | 2.44E+00 | 2.48E+00 | 3.41E-02 | Yes | Sugct | intron_variant | MODIFIER |
| 13 | 16972227 | AX-170135937 | 0 | G | A | 0.175 | -2.71E-01 | 1.27E-01 | 2.44E+00 | 2.48E+00 | 3.41E-02 | Yes | Sugct | intron_variant | MODIFIER |
| 13 | 16972473 | AX-170340173 | 0 | C | T | 0.175 | -2.71E-01 | 1.27E-01 | 2.44E+00 | 2.48E+00 | 3.41E-02 | Yes | Sugct | intron_variant | MODIFIER |
| 13 | 16972580 | AX-170146538 | 0 | A | G | 0.175 | -2.71E-01 | 1.27E-01 | 2.44E+00 | 2.48E+00 | 3.41E-02 | Yes | Sugct | intron_variant | MODIFIER |
| 13 | 16974782 | AX-170007823 | 0 | A | G | 0.175 | -2.71E-01 | 1.27E-01 | 2.44E+00 | 2.48E+00 | 3.41E-02 | Yes | Sugct | intron_variant | MODIFIER |
| 13 | 16975711 | AX-170380352 | 0 | T | A | 0.175 | -2.71E-01 | 1.27E-01 | 2.44E+00 | 2.48E+00 | 3.41E-02 | Yes | Sugct | intron_variant | MODIFIER |
| 13 | 16976108 | AX-170368346 | 0 | A | G | 0.175 | -2.71E-01 | 1.27E-01 | 2.44E+00 | 2.48E+00 | 3.41E-02 | Yes | Sugct | intron_variant | MODIFIER |
| 13 | 16976217 | AX-170481957 | 0 | A | G | 0.175 | -2.71E-01 | 1.27E-01 | 2.44E+00 | 2.48E+00 | 3.41E-02 | Yes | Sugct | synonymous_variant | LOW |
| 13 | 16981358 | AX-170509560 | 0 | G | A | 0.175 | -2.71E-01 | 1.27E-01 | 2.44E+00 | 2.48E+00 | 3.41E-02 | Yes | Sugct | intron_variant | MODIFIER |
| 13 | 16999339 | AX-170007827 | 0 | A | T | 0.175 | -2.71E-01 | 1.27E-01 | 2.44E+00 | 2.48E+00 | 3.41E-02 | Yes | Sugct | intron_variant | MODIFIER |
| 13 | 16999826 | AX-97786843 | 0 | A | G | 0.175 | -2.71E-01 | 1.27E-01 | 2.44E+00 | 2.48E+00 | 3.41E-02 | Yes | Sugct | intron_variant | MODIFIER |
| 13 | 17081332 | AX-170007844 | 0 | C | A | 0.175 | -2.71E-01 | 1.27E-01 | 2.44E+00 | 2.48E+00 | 3.41E-02 | Yes | Sugct | intron_variant | MODIFIER |
| 13 | 17083299 | AX-170431133 | 0 | C | T | 0.175 | -2.71E-01 | 1.27E-01 | 2.44E+00 | 2.48E+00 | 3.41E-02 | Yes | Sugct | intron_variant | MODIFIER |
| 13 | 17086709 | AX-170425241 | 0 | C | T | 0.175 | -2.71E-01 | 1.27E-01 | 2.44E+00 | 2.48E+00 | 3.41E-02 | Yes | Sugct | intron_variant | MODIFIER |
| 13 | 17098237 | AX-97856388 | 0 | A | C | 0.175 | -2.71E-01 | 1.27E-01 | 2.44E+00 | 2.48E+00 | 3.41E-02 | Yes | Sugct | intron_variant | MODIFIER |
| 13 | 17109356 | AX-170007845 | 0 | T | C | 0.175 | -2.71E-01 | 1.27E-01 | 2.44E+00 | 2.48E+00 | 3.41E-02 | Yes | Sugct | intron_variant | MODIFIER |
| 13 | 17114126 | AX-97793200 | 0 | T | C | 0.175 | -2.71E-01 | 1.27E-01 | 2.44E+00 | 2.48E+00 | 3.41E-02 | Yes | Sugct | intron_variant | MODIFIER |
| 13 | 17134696 | AX-97825013 | 0 | T | G | 0.175 | -2.71E-01 | 1.27E-01 | 2.44E+00 | 2.48E+00 | 3.41E-02 | Yes | Sugct | intron_variant | MODIFIER |
| 13 | 17159923 | AX-170210615 | 0 | T | C | 0.175 | -2.71E-01 | 1.27E-01 | 2.44E+00 | 2.48E+00 | 3.41E-02 | Yes | Sugct | intron_variant | MODIFIER |
| 13 | 17165926 | AX-170007843 | 0 | G | A | 0.175 | -2.71E-01 | 1.27E-01 | 2.44E+00 | 2.48E+00 | 3.41E-02 | Yes | Sugct | intron_variant | MODIFIER |
| 13 | 17168664 | AX-170444094 | 0 | G | C | 0.175 | -2.71E-01 | 1.27E-01 | 2.44E+00 | 2.48E+00 | 3.41E-02 | Yes | Sugct | intron_variant | MODIFIER |
| 13 | 17169160 | AX-97843943 | 0 | T | C | 0.175 | -2.71E-01 | 1.27E-01 | 2.44E+00 | 2.48E+00 | 3.41E-02 | Yes | Sugct | intron_variant | MODIFIER |
| 13 | 17175392 | AX-170199792 | 0 | G | A | 0.175 | -2.71E-01 | 1.27E-01 | 2.44E+00 | 2.48E+00 | 3.41E-02 | Yes | Sugct | intron_variant | MODIFIER |
| 13 | 17175897 | AX-170128797 | 0 | G | T | 0.175 | -2.71E-01 | 1.27E-01 | 2.44E+00 | 2.48E+00 | 3.41E-02 | Yes | Sugct | intron_variant | MODIFIER |
| 13 | 17176068 | AX-170195074 | 0 | C | T | 0.175 | -2.71E-01 | 1.27E-01 | 2.44E+00 | 2.48E+00 | 3.41E-02 | Yes | Sugct | intron_variant | MODIFIER |
| 13 | 17183519 | AX-170007841 | 0 | A | G | 0.175 | -2.71E-01 | 1.27E-01 | 2.44E+00 | 2.48E+00 | 3.41E-02 | Yes | Sugct | intron_variant | MODIFIER |
| 13 | 17187106 | AX-170447342 | 0 | T | C | 0.175 | -2.71E-01 | 1.27E-01 | 2.44E+00 | 2.48E+00 | 3.41E-02 | Yes | Sugct | intron_variant | MODIFIER |
| 13 | 17187133 | AX-170185404 | 0 | T | C | 0.175 | -2.71E-01 | 1.27E-01 | 2.44E+00 | 2.48E+00 | 3.41E-02 | Yes | Sugct | intron_variant | MODIFIER |
| 13 | 17194453 | AX-170521585 | 0 | G | C | 0.175 | -2.71E-01 | 1.27E-01 | 2.44E+00 | 2.48E+00 | 3.41E-02 | Yes | Sugct | intron_variant | MODIFIER |
| 13 | 17195032 | AX-97797525 | 0 | C | A | 0.175 | -2.71E-01 | 1.27E-01 | 2.44E+00 | 2.48E+00 | 3.41E-02 | Yes | Sugct | intron_variant | MODIFIER |
| 13 | 17224642 | AX-97784148 | 0 | T | G | 0.175 | -2.71E-01 | 1.27E-01 | 2.44E+00 | 2.48E+00 | 3.41E-02 | Yes | Sugct | intron_variant | MODIFIER |
| 13 | 17252740 | AX-170007842 | 0 | G | T | 0.175 | -2.71E-01 | 1.27E-01 | 2.44E+00 | 2.48E+00 | 3.41E-02 | Yes | Sugct | intron_variant | MODIFIER |
| 13 | 17256994 | AX-170296205 | 0 | T | C | 0.175 | -2.71E-01 | 1.27E-01 | 2.44E+00 | 2.48E+00 | 3.41E-02 | Yes | Sugct | intron_variant | MODIFIER |
| 13 | 17257169 | AX-170134828 | 0 | A | G | 0.175 | -2.71E-01 | 1.27E-01 | 2.44E+00 | 2.48E+00 | 3.41E-02 | Yes | Sugct | intron_variant | MODIFIER |
| 13 | 17291387 | AX-170156613 | 0 | C | T | 0.175 | -2.71E-01 | 1.27E-01 | 2.44E+00 | 2.48E+00 | 3.41E-02 | Yes | Sugct | intron_variant | MODIFIER |
| 13 | 17297249 | AX-97822921 | 0 | C | T | 0.175 | -2.71E-01 | 1.27E-01 | 2.44E+00 | 2.48E+00 | 3.41E-02 | Yes | Sugct | intron_variant | MODIFIER |
| 13 | 17298669 | AX-170007847 | 0 | G | A | 0.175 | -2.71E-01 | 1.27E-01 | 2.44E+00 | 2.48E+00 | 3.41E-02 | Yes | Sugct | intron_variant | MODIFIER |
| 13 | 17352049 | AX-170244786 | 0 | A | T | 0.175 | -2.71E-01 | 1.27E-01 | 2.44E+00 | 2.48E+00 | 3.41E-02 | Yes | Sugct | intron_variant | MODIFIER |
| 13 | 17390315 | AX-170456231 | 0 | G | A | 0.175 | -2.71E-01 | 1.27E-01 | 2.44E+00 | 2.48E+00 | 3.41E-02 | Yes | Sugct | intron_variant | MODIFIER |
| 13 | 17390557 | AX-170092672 | 0 | A | G | 0.175 | -2.71E-01 | 1.27E-01 | 2.44E+00 | 2.48E+00 | 3.41E-02 | Yes | Sugct | intron_variant | MODIFIER |
| 13 | 17391038 | AX-170249875 | 0 | T | C | 0.175 | -2.71E-01 | 1.27E-01 | 2.44E+00 | 2.48E+00 | 3.41E-02 | Yes | Sugct | intron_variant | MODIFIER |
| 13 | 17413052 | AX-170366982 | 0 | A | G | 0.175 | -2.71E-01 | 1.27E-01 | 2.44E+00 | 2.48E+00 | 3.41E-02 | Yes | Sugct | intron_variant | MODIFIER |
| 13 | 17413643 | AX-170283953 | 0 | C | A | 0.175 | -2.71E-01 | 1.27E-01 | 2.44E+00 | 2.48E+00 | 3.41E-02 | Yes | Sugct | intron_variant | MODIFIER |
| 13 | 17418157 | AX-97812436 | 0 | T | C | 0.175 | -2.71E-01 | 1.27E-01 | 2.44E+00 | 2.48E+00 | 3.41E-02 | Yes | Sugct | intron_variant | MODIFIER |
| 13 | 17421873 | AX-170361388 | 0 | G | A | 0.175 | -2.71E-01 | 1.27E-01 | 2.44E+00 | 2.48E+00 | 3.41E-02 | Yes | Sugct | intron_variant | MODIFIER |
| 15 | 71123225 | AX-170313651 | 0 | A | C | 0.096 | -2.21E-01 | 9.61E-02 | 1.79E+00 | 1.87E+00 | 3.46E-02 | Yes | | intergenic_variant | MODIFIER |
| 5 | 54429626 | AX-170400165 | 0 | A | G | 0.491 | 2.16E-01 | 1.01E-01 | 2.44E+00 | 2.49E+00 | 3.47E-02 | Yes | | intergenic_variant | MODIFIER |
| 19 | 58317213 | AX-170150806 | 0 | C | T | 0.447 | 1.29E-01 | 6.11E-02 | 2.81E+00 | 2.90E+00 | 3.47E-02 | no | Gfra1 | intron_variant | MODIFIER |
| 4 | 100750192 | AX-97846258 | 0 | G | T | 0.263 | 2.16E-01 | 1.02E-01 | 2.70E+00 | 2.79E+00 | 3.48E-02 | Yes | | intergenic_variant | MODIFIER |
| 5 | 65820029 | AX-97825292 | 0 | T | C | 0.491 | 2.04E-01 | 9.39E-02 | 2.17E+00 | 2.22E+00 | 3.49E-02 | Yes | N4bp2 | intron_variant | MODIFIER |
| 1 | 185350439 | AX-170043854 | 1 | T | C | 0.223 | 2.37E-01 | 1.13E-01 | 2.75E+00 | 2.86E+00 | 3.49E-02 | Yes | Bpnt1 | intron_variant | MODIFIER |
| 15 | 6814513 | AX-97783174 | 0 | G | A | 0.325 | -1.74E-01 | 7.50E-02 | 1.74E+00 | 1.78E+00 | 3.50E-02 | Yes | Osmr | 3_prime_UTR_variant | MODIFIER |
| 16 | 66745220 | AX-170013294 | 0 | G | A | 0.535 | 1.64E-01 | 7.78E-02 | 2.57E+00 | 2.64E+00 | 3.52E-02 | Yes | Cadm2 | intron_variant | MODIFIER |
| 15 | 64080909 | AX-170510175 | 1 | G | A | 0.33 | -1.59E-01 | 7.38E-02 | 2.18E+00 | 2.25E+00 | 3.56E-02 | no | | intergenic_variant | MODIFIER |

| | | | | | | | | | | | | | | | |
|---|---|---|---|---|---|---|---|---|---|---|---|---|---|---|---|
| 10 | 105867276 | AX-170152563 | 0 | C | T | 0.298 | -1.67E-01 | 7.94E-02 | 2.82E+00 | 2.92E+00 | 3.58E-02 | no | | regulatory_region_variant | MODIFIER |
| 10 | 104773613 | AX-170092081 | 1 | C | A | 0.286 | -1.76E-01 | 8.36E-02 | 2.85E+00 | 2.93E+00 | 3.59E-02 | Yes | | intergenic_variant | MODIFIER |
| 13 | 25143928 | AX-170273239 | 0 | A | C | 0.333 | -1.24E-01 | 5.88E-02 | 2.63E+00 | 2.72E+00 | 3.61E-02 | no | Dcdc2a | intron_variant | MODIFIER |
| 16 | 61681731 | AX-170446544 | 0 | A | C | 0.518 | 1.82E-01 | 8.67E-02 | 2.61E+00 | 2.66E+00 | 3.64E-02 | Yes | | intergenic_variant | MODIFIER |
| 14 | 110849098 | AX-170087723 | 0 | G | A | 0.421 | -1.11E-01 | 5.32E-02 | 2.86E+00 | 3.01E+00 | 3.64E-02 | no | | intergenic_variant | MODIFIER |
| 2 | 77380153 | AX-97838610 | 0 | G | A | 0.053 | -4.96E-01 | 2.25E-01 | 3.84E+00 | 4.07E+00 | 3.65E-02 | Yes | | intergenic_variant | MODIFIER |
| 2 | 92322023 | AX-170018332 | 0 | G | A | 0.053 | -4.96E-01 | 2.25E-01 | 3.84E+00 | 4.07E+00 | 3.65E-02 | Yes | Phf21a | intron_variant | MODIFIER |
| 2 | 96608828 | AX-97838138 | 0 | G | A | 0.053 | -4.96E-01 | 2.25E-01 | 3.84E+00 | 4.07E+00 | 3.65E-02 | Yes | Lrrc4c | intron_variant | MODIFIER |
| 5 | 24173157 | AX-97796203 | 0 | C | T | 0.053 | -4.96E-01 | 2.25E-01 | 3.84E+00 | 4.07E+00 | 3.65E-02 | Yes | Nupl2 | intron_variant | MODIFIER |
| 10 | 104825298 | AX-170433287 | 0 | G | A | 0.281 | -1.67E-01 | 8.00E-02 | 2.77E+00 | 2.87E+00 | 3.66E-02 | no | | intergenic_variant | MODIFIER |
| 18 | 84154317 | AX-170413197 | 0 | G | A | 0.158 | -2.17E-01 | 1.04E-01 | 2.74E+00 | 2.84E+00 | 3.66E-02 | Yes | Zfp407 | intron_variant | MODIFIER |
| 18 | 84405331 | AX-170016112 | 0 | T | C | 0.316 | -1.09E-01 | 5.20E-02 | 2.74E+00 | 2.84E+00 | 3.66E-02 | no | Zfp407 | intron_variant | MODIFIER |
| 18 | 84886138 | AX-170106961 | 0 | C | T | 0.158 | -2.17E-01 | 1.04E-01 | 2.74E+00 | 2.84E+00 | 3.66E-02 | Yes | | regulatory_region_variant | MODIFIER |
| 18 | 84942111 | AX-170215151 | 0 | T | A | 0.158 | -2.17E-01 | 1.04E-01 | 2.74E+00 | 2.84E+00 | 3.66E-02 | Yes | Timm21 | downstream_gene_variant | MODIFIER |
| 18 | 85391884 | AX-170423147 | 0 | T | A | 0.316 | -1.09E-01 | 5.20E-02 | 2.74E+00 | 2.84E+00 | 3.66E-02 | no | | intergenic_variant | MODIFIER |
| 13 | 10472003 | AX-170201092 | 1 | G | A | 0.188 | -2.57E-01 | 1.23E-01 | 2.64E+00 | 2.70E+00 | 3.67E-02 | Yes | Gm26861 | intron_variant,non_coding_transcript_variant | MODIFIER |
| 5 | 137919604 | AX-170251897 | 0 | G | A | 0.132 | -1.90E-01 | 8.96E-02 | 2.34E+00 | 2.42E+00 | 3.67E-02 | Yes | Cyp3a13 | intron_variant | MODIFIER |
| 13 | 17391925 | AX-170327726 | 1 | G | C | 0.17 | -2.68E-01 | 1.27E-01 | 2.47E+00 | 2.51E+00 | 3.67E-02 | Yes | Sugct | intron_variant | MODIFIER |
| 15 | 70435209 | AX-170386518 | 1 | A | G | 0.071 | -2.67E-01 | 1.08E-01 | 1.46E+00 | 1.50E+00 | 3.68E-02 | Yes | | intergenic_variant | MODIFIER |
| 1 | 181087652 | AX-170322290 | 2 | C | T | 0.218 | 2.33E-01 | 1.12E-01 | 2.72E+00 | 2.83E+00 | 3.70E-02 | Yes | Nvl | 3_prime_UTR_variant | MODIFIER |
| 13 | 10616104 | AX-170479836 | 1 | A | G | 0.179 | -2.39E-01 | 1.14E-01 | 2.52E+00 | 2.57E+00 | 3.71E-02 | Yes | Gm26861 | intron_variant,non_coding_transcript_variant | MODIFIER |
| 13 | 12455142 | AX-170259893 | 1 | G | A | 0.188 | -2.57E-01 | 1.23E-01 | 2.48E+00 | 2.53E+00 | 3.71E-02 | Yes | Lgals8 | intron_variant | MODIFIER |
| 9 | 98719421 | AX-170265214 | 0 | T | C | 0.132 | -2.31E-01 | 1.10E-01 | 3.15E+00 | 3.31E+00 | 3.73E-02 | Yes | | intergenic_variant | MODIFIER |
| 9 | 98736386 | AX-170196616 | 0 | T | A | 0.132 | -2.31E-01 | 1.10E-01 | 3.15E+00 | 3.31E+00 | 3.73E-02 | Yes | | intergenic_variant | MODIFIER |
| 13 | 20613233 | AX-170430465 | 1 | T | A | 0.17 | -2.47E-01 | 1.18E-01 | 2.55E+00 | 2.62E+00 | 3.74E-02 | Yes | Elmo1 | downstream_gene_variant | MODIFIER |
| 8 | 88406044 | AX-170232631 | 0 | A | T | 0.175 | -2.48E-01 | 1.17E-01 | 3.35E+00 | 3.53E+00 | 3.74E-02 | Yes | | intergenic_variant | MODIFIER |
| 1 | 193926814 | AX-170536089 | 0 | T | C | 0.281 | 1.78E-01 | 8.45E-02 | 2.39E+00 | 2.49E+00 | 3.74E-02 | Yes | | intergenic_variant | MODIFIER |
| 5 | 58332019 | AX-170412417 | 0 | C | G | 0.5 | 2.05E-01 | 9.80E-02 | 2.44E+00 | 2.51E+00 | 3.75E-02 | Yes | | intergenic_variant | MODIFIER |
| 16 | 56548057 | AX-170408998 | 0 | A | T | 0.465 | -1.72E-01 | 8.25E-02 | 2.71E+00 | 2.76E+00 | 3.76E-02 | no | Abi3bp | intron_variant | MODIFIER |
| 5 | 58150830 | AX-170072215 | 1 | G | A | 0.491 | 2.16E-01 | 1.04E-01 | 2.53E+00 | 2.58E+00 | 3.78E-02 | Yes | Gm42976 | upstream_gene_variant | MODIFIER |
| 5 | 55105180 | AX-170137024 | 1 | G | A | 0.491 | 2.06E-01 | 9.88E-02 | 2.50E+00 | 2.56E+00 | 3.79E-02 | Yes | | intergenic_variant | MODIFIER |
| 2 | 103418088 | AX-97803467 | 0 | G | T | 0.053 | -4.30E-01 | 2.05E-01 | 3.19E+00 | 3.36E+00 | 3.82E-02 | Yes | Elf5 | intron_variant,non_coding_transcript_variant | MODIFIER |
| 5 | 136785017 | AX-170265900 | 0 | A | G | 0.149 | -1.68E-01 | 8.05E-02 | 2.40E+00 | 2.50E+00 | 3.83E-02 | no | Col26a1 | intron_variant | MODIFIER |
| 18 | 88735904 | AX-170350532 | 0 | A | G | 0.158 | -1.76E-01 | 8.51E-02 | 2.65E+00 | 2.79E+00 | 3.83E-02 | Yes | Socs6 | intron_variant,non_coding_transcript_variant | MODIFIER |
| 4 | 77397402 | AX-170069742 | 0 | C | T | 0.333 | 2.26E-01 | 1.08E-01 | 3.14E+00 | 3.25E+00 | 3.83E-02 | Yes | Ptprd | intron_variant,non_coding_transcript_variant | MODIFIER |
| 16 | 58246171 | AX-170165281 | 1 | A | G | 0.446 | -1.79E-01 | 8.62E-02 | 2.69E+00 | 2.73E+00 | 3.85E-02 | Yes | Gm49574 | intron_variant,non_coding_transcript_variant | MODIFIER |
| 16 | 58423092 | AX-170480386 | 0 | C | T | 0.447 | -1.79E-01 | 8.62E-02 | 2.69E+00 | 2.73E+00 | 3.85E-02 | Yes | Dcbld2 | intron_variant | MODIFIER |
| 9 | 101672395 | AX-97848494 | 0 | T | G | 0.14 | -2.22E-01 | 1.06E-01 | 3.10E+00 | 3.25E+00 | 3.85E-02 | Yes | | intergenic_variant | MODIFIER |
| 5 | 137074984 | AX-97800889 | 1 | A | T | 0.134 | -1.87E-01 | 8.93E-02 | 2.39E+00 | 2.47E+00 | 3.86E-02 | Yes | Serpine1 | upstream_gene_variant | MODIFIER |
| 15 | 13097735 | AX-170425744 | 1 | G | A | 0.304 | -1.81E-01 | 7.90E-02 | 1.72E+00 | 1.75E+00 | 3.86E-02 | Yes | Cdh6 | intron_variant | MODIFIER |
| 19 | 58245263 | AX-170016918 | 0 | A | G | 0.439 | 1.34E-01 | 6.46E-02 | 2.80E+00 | 2.88E+00 | 3.88E-02 | no | Gfra1 | intron_variant | MODIFIER |
| 5 | 52941209 | AX-97784397 | 0 | A | G | 0.491 | 2.11E-01 | 1.02E-01 | 2.45E+00 | 2.50E+00 | 3.89E-02 | Yes | | intergenic_variant | MODIFIER |
| 5 | 52941530 | AX-170072121 | 0 | A | G | 0.491 | 2.11E-01 | 1.02E-01 | 2.45E+00 | 2.50E+00 | 3.89E-02 | Yes | | intergenic_variant | MODIFIER |
| 5 | 52941647 | AX-170072119 | 0 | G | A | 0.491 | 2.11E-01 | 1.02E-01 | 2.45E+00 | 2.50E+00 | 3.89E-02 | Yes | | intergenic_variant | MODIFIER |
| 5 | 52980164 | AX-170269353 | 0 | G | A | 0.491 | 2.11E-01 | 1.02E-01 | 2.45E+00 | 2.50E+00 | 3.89E-02 | Yes | 5033403H07Rik | intron_variant,non_coding_transcript_variant | MODIFIER |
| 5 | 52986182 | AX-97852929 | 0 | G | A | 0.491 | 2.11E-01 | 1.02E-01 | 2.45E+00 | 2.50E+00 | 3.89E-02 | Yes | 5033403H07Rik | downstream_gene_variant | MODIFIER |
| 5 | 53097596 | AX-170230992 | 0 | G | T | 0.491 | 2.11E-01 | 1.02E-01 | 2.45E+00 | 2.50E+00 | 3.89E-02 | Yes | | intergenic_variant | MODIFIER |
| 5 | 53231238 | AX-170228995 | 0 | G | T | 0.491 | 2.11E-01 | 1.02E-01 | 2.45E+00 | 2.50E+00 | 3.89E-02 | Yes | | intergenic_variant | MODIFIER |
| 5 | 53231297 | AX-97852466 | 0 | G | A | 0.491 | 2.11E-01 | 1.02E-01 | 2.45E+00 | 2.50E+00 | 3.89E-02 | Yes | | intergenic_variant | MODIFIER |
| 5 | 53231329 | AX-170146652 | 0 | A | G | 0.491 | 2.11E-01 | 1.02E-01 | 2.45E+00 | 2.50E+00 | 3.89E-02 | Yes | | intergenic_variant | MODIFIER |
| 5 | 53238805 | AX-170486158 | 0 | C | T | 0.491 | 2.11E-01 | 1.02E-01 | 2.45E+00 | 2.50E+00 | 3.89E-02 | Yes | | intergenic_variant | MODIFIER |
| 5 | 53241176 | AX-170271134 | 0 | T | G | 0.491 | 2.11E-01 | 1.02E-01 | 2.45E+00 | 2.50E+00 | 3.89E-02 | Yes | | regulatory_region_variant | MODIFIER |
| 5 | 53258052 | AX-170108345 | 0 | C | A | 0.491 | 2.11E-01 | 1.02E-01 | 2.45E+00 | 2.50E+00 | 3.89E-02 | Yes | | regulatory_region_variant | MODIFIER |
| 5 | 53280013 | AX-97810186 | 0 | C | T | 0.491 | 2.11E-01 | 1.02E-01 | 2.45E+00 | 2.50E+00 | 3.89E-02 | Yes | Smim20 | downstream_gene_variant | MODIFIER |

| | | | | | | | | | | | | | | | |
|---|---|---|---|---|---|---|---|---|---|---|---|---|---|---|---|
| 5 | 53287403 | AX-170106316 | 0 | G | A | 0.491 | 2.11E-01 | 1.02E-01 | 2.45E+00 | 2.50E+00 | 3.89E-02 | Yes | Gm45495 | downstream_gene_variant | MODIFIER |
| 5 | 53288541 | AX-170333255 | 0 | G | T | 0.491 | 2.11E-01 | 1.02E-01 | 2.45E+00 | 2.50E+00 | 3.89E-02 | Yes | | intergenic_variant | MODIFIER |
| 5 | 53291546 | AX-170111021 | 0 | G | A | 0.491 | 2.11E-01 | 1.02E-01 | 2.45E+00 | 2.50E+00 | 3.89E-02 | Yes | | intergenic_variant | MODIFIER |
| 5 | 53295568 | AX-170527713 | 0 | G | A | 0.491 | 2.11E-01 | 1.02E-01 | 2.45E+00 | 2.50E+00 | 3.89E-02 | Yes | | intergenic_variant | MODIFIER |
| 5 | 53297767 | AX-170126511 | 0 | C | G | 0.491 | 2.11E-01 | 1.02E-01 | 2.45E+00 | 2.50E+00 | 3.89E-02 | Yes | | intergenic_variant | MODIFIER |
| 5 | 53304131 | AX-97833383 | 0 | A | G | 0.491 | 2.11E-01 | 1.02E-01 | 2.45E+00 | 2.50E+00 | 3.89E-02 | Yes | | intergenic_variant | MODIFIER |
| 5 | 53320041 | AX-170371177 | 0 | T | A | 0.491 | 2.11E-01 | 1.02E-01 | 2.45E+00 | 2.50E+00 | 3.89E-02 | Yes | | intergenic_variant | MODIFIER |
| 5 | 53332547 | AX-97809936 | 0 | T | G | 0.491 | 2.11E-01 | 1.02E-01 | 2.45E+00 | 2.50E+00 | 3.89E-02 | Yes | | intergenic_variant | MODIFIER |
| 5 | 53370577 | AX-97833388 | 0 | A | G | 0.491 | 2.11E-01 | 1.02E-01 | 2.45E+00 | 2.50E+00 | 3.89E-02 | Yes | Gm42745 | intron_variant,non_coding_transcript_variant | MODIFIER |
| 5 | 53406787 | AX-97828844 | 0 | C | T | 0.491 | 2.11E-01 | 1.02E-01 | 2.45E+00 | 2.50E+00 | 3.89E-02 | Yes | | intergenic_variant | MODIFIER |
| 5 | 53409939 | AX-170380872 | 0 | A | G | 0.491 | 2.11E-01 | 1.02E-01 | 2.45E+00 | 2.50E+00 | 3.89E-02 | Yes | | regulatory_region_variant | MODIFIER |
| 5 | 53413020 | AX-170447115 | 0 | C | T | 0.491 | 2.11E-01 | 1.02E-01 | 2.45E+00 | 2.50E+00 | 3.89E-02 | Yes | | regulatory_region_variant | MODIFIER |
| 5 | 53425802 | AX-170450830 | 0 | T | C | 0.491 | 2.11E-01 | 1.02E-01 | 2.45E+00 | 2.50E+00 | 3.89E-02 | Yes | | intergenic_variant | MODIFIER |
| 5 | 53431827 | AX-170528407 | 0 | A | G | 0.491 | 2.11E-01 | 1.02E-01 | 2.45E+00 | 2.50E+00 | 3.89E-02 | Yes | | intergenic_variant | MODIFIER |
| 5 | 53432664 | AX-97850086 | 0 | T | C | 0.491 | 2.11E-01 | 1.02E-01 | 2.45E+00 | 2.50E+00 | 3.89E-02 | Yes | | intergenic_variant | MODIFIER |
| 5 | 53439191 | AX-170072131 | 0 | T | G | 0.491 | 2.11E-01 | 1.02E-01 | 2.45E+00 | 2.50E+00 | 3.89E-02 | Yes | | regulatory_region_variant | MODIFIER |
| 5 | 53449404 | AX-97810421 | 0 | A | G | 0.491 | 2.11E-01 | 1.02E-01 | 2.45E+00 | 2.50E+00 | 3.89E-02 | Yes | | intergenic_variant | MODIFIER |
| 5 | 53481941 | AX-170462803 | 0 | T | C | 0.491 | 2.11E-01 | 1.02E-01 | 2.45E+00 | 2.50E+00 | 3.89E-02 | Yes | Rbpj | intron_variant | MODIFIER |
| 5 | 53482084 | AX-97802403 | 0 | A | G | 0.491 | 2.11E-01 | 1.02E-01 | 2.45E+00 | 2.50E+00 | 3.89E-02 | Yes | Rbpj | intron_variant | MODIFIER |
| 5 | 53501460 | AX-170391192 | 0 | G | A | 0.491 | 2.11E-01 | 1.02E-01 | 2.45E+00 | 2.50E+00 | 3.89E-02 | Yes | Rbpj | intron_variant | MODIFIER |
| 5 | 53527683 | AX-170024962 | 0 | C | T | 0.491 | 2.11E-01 | 1.02E-01 | 2.45E+00 | 2.50E+00 | 3.89E-02 | Yes | Gm10441 | upstream_gene_variant | MODIFIER |
| 5 | 53535575 | AX-170024965 | 0 | C | T | 0.491 | 2.11E-01 | 1.02E-01 | 2.45E+00 | 2.50E+00 | 3.89E-02 | Yes | Rbpj | intron_variant | MODIFIER |
| 5 | 53546670 | AX-170102450 | 0 | A | G | 0.491 | 2.11E-01 | 1.02E-01 | 2.45E+00 | 2.50E+00 | 3.89E-02 | Yes | Rbpj | intron_variant | MODIFIER |
| 5 | 53551675 | AX-170178013 | 0 | A | G | 0.491 | 2.11E-01 | 1.02E-01 | 2.45E+00 | 2.50E+00 | 3.89E-02 | Yes | Rbpj | upstream_gene_variant | MODIFIER |
| 5 | 53552183 | AX-97794863 | 0 | A | C | 0.491 | 2.11E-01 | 1.02E-01 | 2.45E+00 | 2.50E+00 | 3.89E-02 | Yes | Rbpj | upstream_gene_variant | MODIFIER |
| 5 | 53569145 | AX-170024961 | 0 | G | A | 0.491 | 2.11E-01 | 1.02E-01 | 2.45E+00 | 2.50E+00 | 3.89E-02 | Yes | Rbpj | intron_variant | MODIFIER |
| 5 | 53643423 | AX-97839257 | 0 | T | C | 0.491 | 2.11E-01 | 1.02E-01 | 2.45E+00 | 2.50E+00 | 3.89E-02 | Yes | Rbpj | intron_variant | MODIFIER |
| 5 | 53691613 | AX-170072137 | 0 | C | T | 0.491 | 2.11E-01 | 1.02E-01 | 2.45E+00 | 2.50E+00 | 3.89E-02 | Yes | | intergenic_variant | MODIFIER |
| 5 | 53692560 | AX-97811109 | 0 | A | G | 0.491 | 2.11E-01 | 1.02E-01 | 2.45E+00 | 2.50E+00 | 3.89E-02 | Yes | | intergenic_variant | MODIFIER |
| 5 | 53698969 | AX-97852772 | 0 | T | C | 0.491 | 2.11E-01 | 1.02E-01 | 2.45E+00 | 2.50E+00 | 3.89E-02 | Yes | Cckar | 3_prime_UTR_variant | MODIFIER |
| 5 | 53699661 | AX-170024959 | 0 | G | T | 0.491 | 2.11E-01 | 1.02E-01 | 2.45E+00 | 2.50E+00 | 3.89E-02 | Yes | Cckar | 3_prime_UTR_variant | MODIFIER |
| 5 | 53699760 | AX-170024964 | 0 | G | T | 0.491 | 2.11E-01 | 1.02E-01 | 2.45E+00 | 2.50E+00 | 3.89E-02 | Yes | Cckar | missense_variant | MODERATE |
| 5 | 53701290 | AX-170072142 | 0 | A | G | 0.491 | 2.11E-01 | 1.02E-01 | 2.45E+00 | 2.50E+00 | 3.89E-02 | Yes | Cckar | synonymous_variant | LOW |
| 5 | 54234591 | AX-170024974 | 0 | A | C | 0.491 | 2.11E-01 | 1.02E-01 | 2.45E+00 | 2.50E+00 | 3.89E-02 | Yes | | intergenic_variant | MODIFIER |
| 5 | 54252584 | AX-97835843 | 0 | C | T | 0.491 | 2.11E-01 | 1.02E-01 | 2.45E+00 | 2.50E+00 | 3.89E-02 | Yes | | intergenic_variant | MODIFIER |
| 5 | 54261886 | AX-170024972 | 0 | G | A | 0.491 | 2.11E-01 | 1.02E-01 | 2.45E+00 | 2.50E+00 | 3.89E-02 | Yes | | intergenic_variant | MODIFIER |
| 5 | 54320031 | AX-170102804 | 0 | G | T | 0.491 | 2.11E-01 | 1.02E-01 | 2.45E+00 | 2.50E+00 | 3.89E-02 | Yes | | intergenic_variant | MODIFIER |
| 5 | 54325729 | AX-170198383 | 0 | C | T | 0.491 | 2.11E-01 | 1.02E-01 | 2.45E+00 | 2.50E+00 | 3.89E-02 | Yes | | intergenic_variant | MODIFIER |
| 5 | 54325825 | AX-170324579 | 0 | G | T | 0.491 | 2.11E-01 | 1.02E-01 | 2.45E+00 | 2.50E+00 | 3.89E-02 | Yes | | intergenic_variant | MODIFIER |
| 5 | 54327238 | AX-170132442 | 0 | A | G | 0.491 | 2.11E-01 | 1.02E-01 | 2.45E+00 | 2.50E+00 | 3.89E-02 | Yes | | intergenic_variant | MODIFIER |
| 5 | 54361656 | AX-170142233 | 0 | G | A | 0.491 | 2.11E-01 | 1.02E-01 | 2.45E+00 | 2.50E+00 | 3.89E-02 | Yes | Gm10440 | downstream_gene_variant | MODIFIER |
| 5 | 54366986 | AX-170296121 | 0 | T | C | 0.491 | 2.11E-01 | 1.02E-01 | 2.45E+00 | 2.50E+00 | 3.89E-02 | Yes | | intergenic_variant | MODIFIER |
| 5 | 54410516 | AX-97846344 | 0 | C | T | 0.491 | 2.11E-01 | 1.02E-01 | 2.45E+00 | 2.50E+00 | 3.89E-02 | Yes | | intergenic_variant | MODIFIER |
| 5 | 54411040 | AX-170236370 | 0 | C | G | 0.491 | 2.11E-01 | 1.02E-01 | 2.45E+00 | 2.50E+00 | 3.89E-02 | Yes | | intergenic_variant | MODIFIER |
| 5 | 54416224 | AX-170250762 | 0 | G | A | 0.491 | 2.11E-01 | 1.02E-01 | 2.45E+00 | 2.50E+00 | 3.89E-02 | Yes | | intergenic_variant | MODIFIER |
| 5 | 54429077 | AX-170362403 | 0 | A | C | 0.491 | 2.11E-01 | 1.02E-01 | 2.45E+00 | 2.50E+00 | 3.89E-02 | Yes | | intergenic_variant | MODIFIER |
| 5 | 54429568 | AX-170411711 | 0 | G | C | 0.491 | 2.11E-01 | 1.02E-01 | 2.45E+00 | 2.50E+00 | 3.89E-02 | Yes | | intergenic_variant | MODIFIER |
| 5 | 54436384 | AX-170440220 | 0 | C | T | 0.491 | 2.11E-01 | 1.02E-01 | 2.45E+00 | 2.50E+00 | 3.89E-02 | Yes | | intergenic_variant | MODIFIER |
| 5 | 54436845 | AX-170215255 | 0 | A | G | 0.491 | 2.11E-01 | 1.02E-01 | 2.45E+00 | 2.50E+00 | 3.89E-02 | Yes | | intergenic_variant | MODIFIER |
| 5 | 54436873 | AX-170170531 | 0 | C | G | 0.491 | 2.11E-01 | 1.02E-01 | 2.45E+00 | 2.50E+00 | 3.89E-02 | Yes | | intergenic_variant | MODIFIER |
| 5 | 54457248 | AX-97807229 | 0 | A | G | 0.491 | 2.11E-01 | 1.02E-01 | 2.45E+00 | 2.50E+00 | 3.89E-02 | Yes | | intergenic_variant | MODIFIER |
| 5 | 54462697 | AX-170528602 | 0 | G | A | 0.491 | 2.11E-01 | 1.02E-01 | 2.45E+00 | 2.50E+00 | 3.89E-02 | Yes | | intergenic_variant | MODIFIER |
| 5 | 54463618 | AX-170279653 | 0 | A | C | 0.491 | 2.11E-01 | 1.02E-01 | 2.45E+00 | 2.50E+00 | 3.89E-02 | Yes | | intergenic_variant | MODIFIER |
| 5 | 54534911 | AX-97799916 | 0 | C | G | 0.491 | 2.11E-01 | 1.02E-01 | 2.45E+00 | 2.50E+00 | 3.89E-02 | Yes | | intergenic_variant | MODIFIER |

| | | | | | | | | | | | | | | | |
|---|---|---|---|---|---|---|---|---|---|---|---|---|---|---|---|
| 5 | 54570224 | AX-170424560 | 0 | G | A | 0.491 | 2.11E-01 | 1.02E-01 | 2.45E+00 | 2.50E+00 | 3.89E-02 | Yes | Gm18451 | non_coding_transcript_exon_variant | MODIFIER |
| 5 | 54570405 | AX-170481922 | 0 | G | A | 0.491 | 2.11E-01 | 1.02E-01 | 2.45E+00 | 2.50E+00 | 3.89E-02 | Yes | Gm18451 | non_coding_transcript_exon_variant | MODIFIER |
| 5 | 54590017 | AX-170104117 | 0 | A | G | 0.491 | 2.11E-01 | 1.02E-01 | 2.45E+00 | 2.50E+00 | 3.89E-02 | Yes | | intergenic_variant | MODIFIER |
| 5 | 54913915 | AX-170305745 | 1 | G | A | 0.473 | 2.10E-01 | 1.01E-01 | 2.50E+00 | 2.56E+00 | 3.89E-02 | Yes | | intergenic_variant | MODIFIER |
| 5 | 62419837 | AX-170506198 | 1 | C | T | 0.473 | 2.10E-01 | 1.01E-01 | 2.50E+00 | 2.56E+00 | 3.89E-02 | Yes | | intergenic_variant | MODIFIER |
| 13 | 16103938 | AX-97822919 | 0 | T | C | 0.167 | -2.23E-01 | 1.05E-01 | 2.20E+00 | 2.26E+00 | 3.91E-02 | Yes | | intergenic_variant | MODIFIER |
| 13 | 16464910 | AX-170239190 | 1 | T | C | 0.179 | -2.58E-01 | 1.24E-01 | 2.46E+00 | 2.51E+00 | 3.93E-02 | Yes | | regulatory_region_variant | MODIFIER |
| 5 | 132611013 | AX-170231455 | 0 | G | A | 0.158 | -1.64E-01 | 7.87E-02 | 2.32E+00 | 2.43E+00 | 3.94E-02 | no | | intergenic_variant | MODIFIER |
| 8 | 56857308 | AX-97853192 | 1 | G | T | 0.152 | -2.85E-01 | 1.37E-01 | 3.08E+00 | 3.21E+00 | 3.94E-02 | Yes | Gm33501 | downstream_gene_variant | MODIFIER |
| 5 | 57145830 | AX-97815053 | 0 | A | G | 0.465 | 2.12E-01 | 1.02E-01 | 2.48E+00 | 2.54E+00 | 3.94E-02 | Yes | | intergenic_variant | MODIFIER |
| 4 | 100222479 | AX-170496846 | 0 | G | A | 0.228 | 2.36E-01 | 1.14E-01 | 2.76E+00 | 2.85E+00 | 3.95E-02 | Yes | Ror1 | intron_variant | MODIFIER |
| 13 | 19913929 | AX-170132357 | 0 | A | G | 0.167 | -2.25E-01 | 1.09E-01 | 2.59E+00 | 2.69E+00 | 3.97E-02 | Yes | | intergenic_variant | MODIFIER |
| 5 | 64219065 | AX-170025152 | 0 | G | A | 0.482 | 2.04E-01 | 9.92E-02 | 2.72E+00 | 2.80E+00 | 3.99E-02 | Yes | Tbc1d1 | intron_variant | MODIFIER |
| 16 | 54901091 | AX-170226573 | 0 | A | G | 0.526 | 1.70E-01 | 8.27E-02 | 2.73E+00 | 2.78E+00 | 4.01E-02 | Yes | | intergenic_variant | MODIFIER |
| 20 | 45447629 | AX-170084144 | 0 | T | C | 0.237 | 1.21E-01 | 5.89E-02 | 2.77E+00 | 2.92E+00 | 4.02E-02 | no | | intergenic_variant | MODIFIER |
| 15 | 12144050 | AX-170509688 | 0 | C | T | 0.307 | -1.71E-01 | 7.59E-02 | 1.75E+00 | 1.79E+00 | 4.03E-02 | no | Zfr | intron_variant | MODIFIER |
| 9 | 87510290 | AX-170350422 | 1 | C | T | 0.134 | -2.27E-01 | 1.10E-01 | 3.03E+00 | 3.20E+00 | 4.05E-02 | Yes | | intergenic_variant | MODIFIER |
| 16 | 61806240 | AX-170444767 | 1 | C | G | 0.464 | -1.75E-01 | 8.51E-02 | 2.55E+00 | 2.60E+00 | 4.07E-02 | Yes | | intergenic_variant | MODIFIER |
| 15 | 12429595 | AX-97857388 | 0 | G | A | 0.307 | -1.78E-01 | 7.70E-02 | 1.65E+00 | 1.68E+00 | 4.08E-02 | Yes | Pdzd2 | intron_variant | MODIFIER |
| 16 | 62116018 | AX-97790820 | 2 | C | T | 0.518 | 1.75E-01 | 8.58E-02 | 2.76E+00 | 2.82E+00 | 4.09E-02 | Yes | | intergenic_variant | MODIFIER |
| 5 | 24246722 | AX-170024538 | 0 | C | T | 0.053 | -4.24E-01 | 2.05E-01 | 3.21E+00 | 3.38E+00 | 4.10E-02 | Yes | Gm43827 | downstream_gene_variant | MODIFIER |
| 15 | 67851587 | AX-97844977 | 0 | G | A | 0.079 | -2.58E-01 | 1.08E-01 | 1.53E+00 | 1.58E+00 | 4.11E-02 | Yes | | intergenic_variant | MODIFIER |
| 9 | 90825832 | AX-170098483 | 0 | G | A | 0.123 | -2.41E-01 | 1.17E-01 | 3.11E+00 | 3.28E+00 | 4.13E-02 | Yes | Gm28703 | non_coding_transcript_exon_variant | MODIFIER |
| 8 | 86191963 | AX-170445971 | 0 | C | T | 0.202 | -2.64E-01 | 1.22E-01 | 3.81E+00 | 4.04E+00 | 4.14E-02 | Yes | | intergenic_variant | MODIFIER |
| 4 | 12555306 | AX-168414357 | 1 | C | T | 0.018 | -3.44E-01 | 1.67E-01 | 2.34E+00 | 2.47E+00 | 4.16E-02 | Yes | | intergenic_variant | MODIFIER |
| 4 | 82806957 | AX-170450684 | 1 | C | T | 0.357 | 2.30E-01 | 1.12E-01 | 3.04E+00 | 3.14E+00 | 4.16E-02 | Yes | Zdhhc21 | intron_variant | MODIFIER |
| 5 | 129601593 | AX-97851861 | 0 | A | G | 0.149 | -1.80E-01 | 8.60E-02 | 2.20E+00 | 2.28E+00 | 4.19E-02 | Yes | Mmp17 | intron_variant | MODIFIER |
| 5 | 111490704 | AX-170395681 | 0 | A | G | 0.447 | 2.02E-01 | 9.90E-02 | 3.05E+00 | 3.17E+00 | 4.20E-02 | Yes | | regulatory_region_variant | MODIFIER |
| 5 | 53487999 | AX-170157144 | 1 | T | C | 0.491 | 2.07E-01 | 1.01E-01 | 2.45E+00 | 2.50E+00 | 4.21E-02 | Yes | Rbpj | intron_variant | MODIFIER |
| 3 | 18816536 | AX-97781737 | 0 | T | C | 0.14 | -1.96E-01 | 9.64E-02 | 2.73E+00 | 2.83E+00 | 4.22E-02 | Yes | | intergenic_variant | MODIFIER |
| 4 | 106700130 | AX-97799070 | 1 | T | C | 0.241 | 2.20E-01 | 1.08E-01 | 2.72E+00 | 2.80E+00 | 4.22E-02 | Yes | Mroh7 | synonymous_variant | LOW |
| 13 | 11578417 | AX-170036856 | 2 | C | T | 0.191 | -2.43E-01 | 1.19E-01 | 2.52E+00 | 2.59E+00 | 4.23E-02 | Yes | Ryr2 | intron_variant | MODIFIER |
| 5 | 57880492 | AX-170203184 | 2 | C | A | 0.482 | 1.90E-01 | 9.36E-02 | 2.63E+00 | 2.72E+00 | 4.25E-02 | Yes | Pcdh7 | intron_variant | MODIFIER |
| 5 | 137005606 | AX-97815703 | 0 | A | C | 0.132 | -1.83E-01 | 8.94E-02 | 2.36E+00 | 2.45E+00 | 4.26E-02 | Yes | Gm26989 | downstream_gene_variant | MODIFIER |
| 5 | 137054375 | AX-97807146 | 0 | G | A | 0.132 | -1.83E-01 | 8.94E-02 | 2.36E+00 | 2.45E+00 | 4.26E-02 | Yes | | intergenic_variant | MODIFIER |
| 5 | 137125177 | AX-170026274 | 0 | T | C | 0.132 | -1.83E-01 | 8.94E-02 | 2.36E+00 | 2.45E+00 | 4.26E-02 | Yes | | intergenic_variant | MODIFIER |
| 5 | 137125241 | AX-97820848 | 0 | G | A | 0.132 | -1.83E-01 | 8.94E-02 | 2.36E+00 | 2.45E+00 | 4.26E-02 | Yes | | intergenic_variant | MODIFIER |
| 5 | 137280154 | AX-97827557 | 0 | A | C | 0.132 | -1.83E-01 | 8.94E-02 | 2.36E+00 | 2.45E+00 | 4.26E-02 | Yes | Gm42456 | intron_variant,non_coding_transcript_variant | MODIFIER |
| 5 | 137285891 | AX-170026283 | 0 | G | A | 0.132 | -1.83E-01 | 8.94E-02 | 2.36E+00 | 2.45E+00 | 4.26E-02 | Yes | Ache | upstream_gene_variant | MODIFIER |
| 5 | 137303779 | AX-97818030 | 0 | C | A | 0.132 | -1.83E-01 | 8.94E-02 | 2.36E+00 | 2.45E+00 | 4.26E-02 | Yes | Srrt | intron_variant | MODIFIER |
| 5 | 137321554 | AX-170142299 | 0 | T | C | 0.132 | -1.83E-01 | 8.94E-02 | 2.36E+00 | 2.45E+00 | 4.26E-02 | Yes | Slc12a9 | intron_variant | MODIFIER |
| 5 | 137328827 | AX-97844463 | 0 | T | C | 0.132 | -1.83E-01 | 8.94E-02 | 2.36E+00 | 2.45E+00 | 4.26E-02 | Yes | Slc12a9 | intron_variant | MODIFIER |
| 5 | 137359979 | AX-97803067 | 0 | C | T | 0.132 | -1.83E-01 | 8.94E-02 | 2.36E+00 | 2.45E+00 | 4.26E-02 | Yes | Ephb4 | intron_variant | MODIFIER |
| 5 | 137361246 | AX-170337713 | 0 | A | T | 0.132 | -1.83E-01 | 8.94E-02 | 2.36E+00 | 2.45E+00 | 4.26E-02 | Yes | Ephb4 | synonymous_variant | LOW |
| 5 | 137362343 | AX-170379699 | 0 | A | T | 0.132 | -1.83E-01 | 8.94E-02 | 2.36E+00 | 2.45E+00 | 4.26E-02 | Yes | Ephb4 | intron_variant | MODIFIER |
| 5 | 137373124 | AX-97839995 | 0 | A | G | 0.132 | -1.83E-01 | 8.94E-02 | 2.36E+00 | 2.45E+00 | 4.26E-02 | Yes | Ephb4 | intron_variant | MODIFIER |
| 5 | 137386223 | AX-170327386 | 0 | G | T | 0.132 | -1.83E-01 | 8.94E-02 | 2.36E+00 | 2.45E+00 | 4.26E-02 | Yes | Zan | intron_variant | MODIFIER |
| 5 | 137386485 | AX-170086123 | 0 | T | G | 0.132 | -1.83E-01 | 8.94E-02 | 2.36E+00 | 2.45E+00 | 4.26E-02 | Yes | Zan | missense_variant | MODERATE |
| 5 | 137390992 | AX-170210873 | 0 | C | T | 0.132 | -1.83E-01 | 8.94E-02 | 2.36E+00 | 2.45E+00 | 4.26E-02 | Yes | Zan | intron_variant | MODIFIER |
| 5 | 137391860 | AX-170401663 | 0 | G | A | 0.132 | -1.83E-01 | 8.94E-02 | 2.36E+00 | 2.45E+00 | 4.26E-02 | Yes | Zan | splice_region_variant,intron_variant | LOW |
| 5 | 137393013 | AX-170270795 | 0 | A | G | 0.132 | -1.83E-01 | 8.94E-02 | 2.36E+00 | 2.45E+00 | 4.26E-02 | Yes | Zan | synonymous_variant | LOW |
| 5 | 137467270 | AX-97820659 | 0 | C | T | 0.132 | -1.83E-01 | 8.94E-02 | 2.36E+00 | 2.45E+00 | 4.26E-02 | Yes | Zan | intron_variant | MODIFIER |
| 5 | 137472093 | AX-170039113 | 0 | A | G | 0.132 | -1.83E-01 | 8.94E-02 | 2.36E+00 | 2.45E+00 | 4.26E-02 | Yes | Zan | splice_region_variant,synonymous_variant | LOW |
| 5 | 137472985 | AX-170114280 | 0 | A | G | 0.132 | -1.83E-01 | 8.94E-02 | 2.36E+00 | 2.45E+00 | 4.26E-02 | Yes | Zan | intron_variant | MODIFIER |

| | | | | | | | | | | | | | | | |
|---|---|---|---|---|---|---|---|---|---|---|---|---|---|---|---|
| 5 | 137474613 | AX-170485495 | 0 | T | C | 0.132 | -1.83E-01 | 8.94E-02 | 2.36E+00 | 2.45E+00 | 4.26E-02 | Yes | Zan | intron_variant | MODIFIER |
| 5 | 137481552 | AX-170273098 | 0 | G | T | 0.132 | -1.83E-01 | 8.94E-02 | 2.36E+00 | 2.45E+00 | 4.26E-02 | Yes | Epo | downstream_gene_variant | MODIFIER |
| 5 | 137482991 | AX-170039114 | 0 | T | C | 0.132 | -1.83E-01 | 8.94E-02 | 2.36E+00 | 2.45E+00 | 4.26E-02 | Yes | Epo | downstream_gene_variant | MODIFIER |
| 5 | 137483059 | AX-170385995 | 0 | C | G | 0.132 | -1.83E-01 | 8.94E-02 | 2.36E+00 | 2.45E+00 | 4.26E-02 | Yes | Epo | 3_prime_UTR_variant | MODIFIER |
| 5 | 137483226 | AX-170405954 | 0 | C | T | 0.132 | -1.83E-01 | 8.94E-02 | 2.36E+00 | 2.45E+00 | 4.26E-02 | Yes | Epo | synonymous_variant | LOW |
| 5 | 137484639 | AX-170135955 | 0 | A | G | 0.132 | -1.83E-01 | 8.94E-02 | 2.36E+00 | 2.45E+00 | 4.26E-02 | Yes | Epo | intron_variant | MODIFIER |
| 5 | 137493677 | AX-170367574 | 0 | A | G | 0.132 | -1.83E-01 | 8.94E-02 | 2.36E+00 | 2.45E+00 | 4.26E-02 | Yes | Epo | intron_variant | MODIFIER |
| 5 | 137498666 | AX-170466529 | 0 | A | G | 0.132 | -1.83E-01 | 8.94E-02 | 2.36E+00 | 2.45E+00 | 4.26E-02 | Yes | Pop7 | downstream_gene_variant | MODIFIER |
| 5 | 137500541 | AX-170360802 | 0 | A | G | 0.132 | -1.83E-01 | 8.94E-02 | 2.36E+00 | 2.45E+00 | 4.26E-02 | Yes | Pop7 | downstream_gene_variant | MODIFIER |
| 5 | 137501143 | AX-170140170 | 0 | T | G | 0.132 | -1.83E-01 | 8.94E-02 | 2.36E+00 | 2.45E+00 | 4.26E-02 | Yes | Pop7 | downstream_gene_variant | MODIFIER |
| 5 | 137512768 | AX-97830647 | 0 | G | A | 0.132 | -1.83E-01 | 8.94E-02 | 2.36E+00 | 2.45E+00 | 4.26E-02 | Yes | Epo | intron_variant | MODIFIER |
| 5 | 137525528 | AX-170373152 | 0 | A | G | 0.132 | -1.83E-01 | 8.94E-02 | 2.36E+00 | 2.45E+00 | 4.26E-02 | Yes | Gnb2 | downstream_gene_variant | MODIFIER |
| 5 | 137543190 | AX-170288556 | 0 | A | G | 0.132 | -1.83E-01 | 8.94E-02 | 2.36E+00 | 2.45E+00 | 4.26E-02 | Yes | Gm17112 | downstream_gene_variant | MODIFIER |
| 5 | 137587089 | AX-97826466 | 0 | T | C | 0.132 | -1.83E-01 | 8.94E-02 | 2.36E+00 | 2.45E+00 | 4.26E-02 | Yes | Tfr2 | synonymous_variant | LOW |
| 5 | 137597503 | AX-170115867 | 0 | A | G | 0.132 | -1.83E-01 | 8.94E-02 | 2.36E+00 | 2.45E+00 | 4.26E-02 | Yes | Mospd3 | intron_variant | MODIFIER |
| 5 | 137603869 | AX-170289885 | 0 | A | T | 0.132 | -1.83E-01 | 8.94E-02 | 2.36E+00 | 2.45E+00 | 4.26E-02 | Yes | Pcolce | downstream_gene_variant | MODIFIER |
| 5 | 137616405 | AX-170103584 | 0 | T | A | 0.132 | -1.83E-01 | 8.94E-02 | 2.36E+00 | 2.45E+00 | 4.26E-02 | Yes | Pcolce | upstream_gene_variant | MODIFIER |
| 5 | 137623584 | AX-170491805 | 0 | A | G | 0.132 | -1.83E-01 | 8.94E-02 | 2.36E+00 | 2.45E+00 | 4.26E-02 | Yes | Fbxo24 | intron_variant | MODIFIER |
| 5 | 137632333 | AX-170167677 | 0 | C | A | 0.132 | -1.83E-01 | 8.94E-02 | 2.36E+00 | 2.45E+00 | 4.26E-02 | Yes | Lrch4 | intron_variant | MODIFIER |
| 5 | 137632834 | AX-97849747 | 0 | T | C | 0.132 | -1.83E-01 | 8.94E-02 | 2.36E+00 | 2.45E+00 | 4.26E-02 | Yes | Lrch4 | intron_variant | MODIFIER |
| 5 | 137641901 | AX-170240845 | 0 | C | G | 0.132 | -1.83E-01 | 8.94E-02 | 2.36E+00 | 2.45E+00 | 4.26E-02 | Yes | Lrch4 | downstream_gene_variant | MODIFIER |
| 5 | 137663645 | AX-170026291 | 0 | G | A | 0.132 | -1.83E-01 | 8.94E-02 | 2.36E+00 | 2.45E+00 | 4.26E-02 | Yes | Agfg2 | intron_variant | MODIFIER |
| 5 | 137685702 | AX-170348864 | 0 | C | G | 0.132 | -1.83E-01 | 8.94E-02 | 2.36E+00 | 2.45E+00 | 4.26E-02 | Yes | Agfg2 | upstream_gene_variant | MODIFIER |
| 5 | 137686203 | AX-170073653 | 0 | T | A | 0.132 | -1.83E-01 | 8.94E-02 | 2.36E+00 | 2.45E+00 | 4.26E-02 | Yes | Agfg2 | upstream_gene_variant | MODIFIER |
| 5 | 137687887 | AX-170233126 | 0 | C | T | 0.132 | -1.83E-01 | 8.94E-02 | 2.36E+00 | 2.45E+00 | 4.26E-02 | Yes | Agfg2 | upstream_gene_variant | MODIFIER |
| 5 | 137699983 | AX-170026286 | 0 | A | C | 0.132 | -1.83E-01 | 8.94E-02 | 2.36E+00 | 2.45E+00 | 4.26E-02 | Yes | | regulatory_region_variant | MODIFIER |
| 5 | 137713534 | AX-97797603 | 0 | C | T | 0.132 | -1.83E-01 | 8.94E-02 | 2.36E+00 | 2.45E+00 | 4.26E-02 | Yes | | regulatory_region_variant | MODIFIER |
| 5 | 137751273 | AX-170026289 | 0 | G | A | 0.132 | -1.83E-01 | 8.94E-02 | 2.36E+00 | 2.45E+00 | 4.26E-02 | Yes | Tsc22d4 | synonymous_variant | LOW |
| 5 | 137751285 | AX-170026284 | 0 | A | G | 0.132 | -1.83E-01 | 8.94E-02 | 2.36E+00 | 2.45E+00 | 4.26E-02 | Yes | Tsc22d4 | synonymous_variant | LOW |
| 5 | 137751383 | AX-97810770 | 0 | C | T | 0.132 | -1.83E-01 | 8.94E-02 | 2.36E+00 | 2.45E+00 | 4.26E-02 | Yes | Tsc22d4 | intron_variant | MODIFIER |
| 5 | 137752823 | AX-170104459 | 0 | A | G | 0.132 | -1.83E-01 | 8.94E-02 | 2.36E+00 | 2.45E+00 | 4.26E-02 | Yes | Tsc22d4 | intron_variant | MODIFIER |
| 5 | 137765211 | AX-170375155 | 0 | A | T | 0.132 | -1.83E-01 | 8.94E-02 | 2.36E+00 | 2.45E+00 | 4.26E-02 | Yes | Tsc22d4 | intron_variant | MODIFIER |
| 5 | 137773832 | AX-97845984 | 0 | G | T | 0.132 | -1.83E-01 | 8.94E-02 | 2.36E+00 | 2.45E+00 | 4.26E-02 | Yes | | intergenic_variant | MODIFIER |
| 5 | 137810689 | AX-170347953 | 0 | G | A | 0.132 | -1.83E-01 | 8.94E-02 | 2.36E+00 | 2.45E+00 | 4.26E-02 | Yes | Zcwpw1 | intron_variant | MODIFIER |
| 5 | 137811395 | AX-170298011 | 0 | A | G | 0.132 | -1.83E-01 | 8.94E-02 | 2.36E+00 | 2.45E+00 | 4.26E-02 | Yes | Zcwpw1 | intron_variant | MODIFIER |
| 5 | 137817662 | AX-97836820 | 0 | A | G | 0.132 | -1.83E-01 | 8.94E-02 | 2.36E+00 | 2.45E+00 | 4.26E-02 | Yes | Zcwpw1 | intron_variant | MODIFIER |
| 5 | 137829433 | AX-170313523 | 0 | T | C | 0.132 | -1.83E-01 | 8.94E-02 | 2.36E+00 | 2.45E+00 | 4.26E-02 | Yes | Pilra | intron_variant | MODIFIER |
| 5 | 137831277 | AX-97845401 | 0 | T | C | 0.132 | -1.83E-01 | 8.94E-02 | 2.36E+00 | 2.45E+00 | 4.26E-02 | Yes | Pilra | missense_variant | MODERATE |
| 5 | 137831399 | AX-170517641 | 0 | T | G | 0.132 | -1.83E-01 | 8.94E-02 | 2.36E+00 | 2.45E+00 | 4.26E-02 | Yes | Pilra | intron_variant | MODIFIER |
| 5 | 137875421 | AX-170106107 | 0 | A | G | 0.132 | -1.83E-01 | 8.94E-02 | 2.36E+00 | 2.45E+00 | 4.26E-02 | Yes | Pilrb2 | upstream_gene_variant | MODIFIER |
| 5 | 137875702 | AX-170026288 | 0 | T | C | 0.132 | -1.83E-01 | 8.94E-02 | 2.36E+00 | 2.45E+00 | 4.26E-02 | Yes | Pilrb2 | upstream_gene_variant | MODIFIER |
| 5 | 137877171 | AX-170124360 | 0 | T | C | 0.132 | -1.83E-01 | 8.94E-02 | 2.36E+00 | 2.45E+00 | 4.26E-02 | Yes | Gm43494 | upstream_gene_variant | MODIFIER |
| 5 | 137877923 | AX-170026285 | 0 | T | C | 0.132 | -1.83E-01 | 8.94E-02 | 2.36E+00 | 2.45E+00 | 4.26E-02 | Yes | Gm43494 | upstream_gene_variant | MODIFIER |
| 5 | 137878336 | AX-97798638 | 0 | A | G | 0.132 | -1.83E-01 | 8.94E-02 | 2.36E+00 | 2.45E+00 | 4.26E-02 | Yes | Gm43494 | upstream_gene_variant | MODIFIER |
| 5 | 137907551 | AX-170332161 | 0 | T | C | 0.132 | -1.83E-01 | 8.94E-02 | 2.36E+00 | 2.45E+00 | 4.26E-02 | Yes | Cyp3a13 | synonymous_variant | LOW |
| 5 | 137944616 | AX-97839772 | 0 | T | G | 0.132 | -1.83E-01 | 8.94E-02 | 2.36E+00 | 2.45E+00 | 4.26E-02 | Yes | | intergenic_variant | MODIFIER |
| 5 | 137947968 | AX-170201809 | 0 | T | G | 0.132 | -1.83E-01 | 8.94E-02 | 2.36E+00 | 2.45E+00 | 4.26E-02 | Yes | | intergenic_variant | MODIFIER |
| 5 | 137970903 | AX-97848851 | 0 | C | T | 0.132 | -1.83E-01 | 8.94E-02 | 2.36E+00 | 2.45E+00 | 4.26E-02 | Yes | | intergenic_variant | MODIFIER |
| 5 | 138009378 | AX-170097206 | 0 | A | G | 0.132 | -1.83E-01 | 8.94E-02 | 2.36E+00 | 2.45E+00 | 4.26E-02 | Yes | | intergenic_variant | MODIFIER |
| 5 | 138017714 | AX-170233129 | 0 | A | G | 0.132 | -1.83E-01 | 8.94E-02 | 2.36E+00 | 2.45E+00 | 4.26E-02 | Yes | Smok3a | upstream_gene_variant | MODIFIER |
| 5 | 138080437 | AX-170317193 | 0 | C | T | 0.132 | -1.83E-01 | 8.94E-02 | 2.36E+00 | 2.45E+00 | 4.26E-02 | Yes | Zkscan1 | upstream_gene_variant | MODIFIER |
| 5 | 138087558 | AX-97787173 | 0 | T | C | 0.132 | -1.83E-01 | 8.94E-02 | 2.36E+00 | 2.45E+00 | 4.26E-02 | Yes | Zkscan1 | intron_variant | MODIFIER |
| 5 | 138098266 | AX-170146263 | 0 | G | A | 0.132 | -1.83E-01 | 8.94E-02 | 2.36E+00 | 2.45E+00 | 4.26E-02 | Yes | Zkscan1 | intron_variant | MODIFIER |
| 5 | 138102165 | AX-170170703 | 0 | A | G | 0.132 | -1.83E-01 | 8.94E-02 | 2.36E+00 | 2.45E+00 | 4.26E-02 | Yes | Zkscan1 | 3_prime_UTR_variant | MODIFIER |

| | | | | | | | | | | | | | | | |
|---|---|---|---|---|---|---|---|---|---|---|---|---|---|---|---|
| 5 | 138104644 | AX-170259073 | 0 | T | G | 0.132 | -1.83E-01 | 8.94E-02 | 2.36E+00 | 2.45E+00 | 4.26E-02 | Yes | Zkscan1 | 3_prime_UTR_variant | MODIFIER |
| 5 | 138105069 | AX-170495622 | 0 | A | G | 0.132 | -1.83E-01 | 8.94E-02 | 2.36E+00 | 2.45E+00 | 4.26E-02 | Yes | Zkscan1 | 3_prime_UTR_variant | MODIFIER |
| 5 | 138120548 | AX-97794117 | 0 | G | A | 0.132 | -1.83E-01 | 8.94E-02 | 2.36E+00 | 2.45E+00 | 4.26E-02 | Yes | Zscan21 | intron_variant | MODIFIER |
| 5 | 138143763 | AX-170327397 | 0 | A | T | 0.132 | -1.83E-01 | 8.94E-02 | 2.36E+00 | 2.45E+00 | 4.26E-02 | Yes | Zfp113 | 3_prime_UTR_variant | MODIFIER |
| 5 | 138148723 | AX-170507260 | 0 | C | T | 0.132 | -1.83E-01 | 8.94E-02 | 2.36E+00 | 2.45E+00 | 4.26E-02 | Yes | Zfp113 | intron_variant | MODIFIER |
| 5 | 138152053 | AX-170089624 | 0 | A | G | 0.132 | -1.83E-01 | 8.94E-02 | 2.36E+00 | 2.45E+00 | 4.26E-02 | Yes | Zfp113 | intron_variant | MODIFIER |
| 5 | 138164451 | AX-170401989 | 0 | G | C | 0.132 | -1.83E-01 | 8.94E-02 | 2.36E+00 | 2.45E+00 | 4.26E-02 | Yes | Mcm7 | downstream_gene_variant | MODIFIER |
| 5 | 138171328 | AX-170073659 | 0 | G | A | 0.132 | -1.83E-01 | 8.94E-02 | 2.36E+00 | 2.45E+00 | 4.26E-02 | Yes | Mcm7 | intron_variant | MODIFIER |
| 5 | 138182264 | AX-97850082 | 0 | A | G | 0.132 | -1.83E-01 | 8.94E-02 | 2.36E+00 | 2.45E+00 | 4.26E-02 | Yes | Ap4m1 | downstream_gene_variant | MODIFIER |
| 5 | 138198503 | AX-170523187 | 0 | A | G | 0.132 | -1.83E-01 | 8.94E-02 | 2.36E+00 | 2.45E+00 | 4.26E-02 | Yes | Mblac1 | downstream_gene_variant | MODIFIER |
| 5 | 138223177 | AX-97785973 | 0 | C | A | 0.132 | -1.83E-01 | 8.94E-02 | 2.36E+00 | 2.45E+00 | 4.26E-02 | Yes | Nxpe5 | upstream_gene_variant | MODIFIER |
| 5 | 138228134 | AX-170198979 | 0 | G | C | 0.132 | -1.83E-01 | 8.94E-02 | 2.36E+00 | 2.45E+00 | 4.26E-02 | Yes | Nxpe5 | intron_variant | MODIFIER |
| 5 | 138228469 | AX-170291490 | 0 | T | C | 0.132 | -1.83E-01 | 8.94E-02 | 2.36E+00 | 2.45E+00 | 4.26E-02 | Yes | Nxpe5 | intron_variant | MODIFIER |
| 13 | 11586416 | AX-170342521 | 0 | T | C | 0.184 | -2.55E-01 | 1.25E-01 | 2.57E+00 | 2.62E+00 | 4.27E-02 | Yes | Ryr2 | intron_variant | MODIFIER |
| 13 | 21588709 | AX-170532753 | 1 | T | C | 0.17 | -2.16E-01 | 1.07E-01 | 2.66E+00 | 2.78E+00 | 4.28E-02 | Yes | | intergenic_variant | MODIFIER |
| 19 | 52800319 | AX-170016841 | 1 | G | T | 0.5 | 1.29E-01 | 6.39E-02 | 2.79E+00 | 2.88E+00 | 4.28E-02 | no | | intergenic_variant | MODIFIER |
| 5 | 64384481 | AX-170241754 | 0 | C | T | 0.491 | 1.90E-01 | 9.31E-02 | 2.44E+00 | 2.51E+00 | 4.28E-02 | Yes | Gm43837 | intron_variant,non_coding_transcript_variant | MODIFIER |
| 1 | 181010022 | AX-170157613 | 1 | A | G | 0.223 | 2.28E-01 | 1.13E-01 | 2.67E+00 | 2.77E+00 | 4.29E-02 | Yes | Ephx1 | intron_variant | MODIFIER |
| 10 | 103589319 | AX-97816672 | 0 | T | C | 0.289 | -1.68E-01 | 8.33E-02 | 2.80E+00 | 2.88E+00 | 4.33E-02 | no | | intergenic_variant | MODIFIER |
| 10 | 103592770 | AX-170247005 | 0 | T | C | 0.289 | -1.68E-01 | 8.33E-02 | 2.80E+00 | 2.88E+00 | 4.33E-02 | no | | intergenic_variant | MODIFIER |
| 10 | 103677829 | AX-170211509 | 0 | T | C | 0.289 | -1.68E-01 | 8.33E-02 | 2.80E+00 | 2.88E+00 | 4.33E-02 | no | | intergenic_variant | MODIFIER |
| 10 | 103760338 | AX-97816670 | 0 | A | G | 0.289 | -1.68E-01 | 8.33E-02 | 2.80E+00 | 2.88E+00 | 4.33E-02 | no | | intergenic_variant | MODIFIER |
| 10 | 103808703 | AX-170167363 | 0 | G | T | 0.289 | -1.68E-01 | 8.33E-02 | 2.80E+00 | 2.88E+00 | 4.33E-02 | no | | intergenic_variant | MODIFIER |
| 10 | 103820290 | AX-170319738 | 0 | G | A | 0.289 | -1.68E-01 | 8.33E-02 | 2.80E+00 | 2.88E+00 | 4.33E-02 | no | | intergenic_variant | MODIFIER |
| 10 | 104334027 | AX-97832979 | 0 | T | G | 0.289 | -1.68E-01 | 8.33E-02 | 2.80E+00 | 2.88E+00 | 4.33E-02 | no | | intergenic_variant | MODIFIER |
| 10 | 104408951 | AX-170533074 | 0 | A | G | 0.289 | -1.68E-01 | 8.33E-02 | 2.80E+00 | 2.88E+00 | 4.33E-02 | no | | intergenic_variant | MODIFIER |
| 10 | 104453874 | AX-170481117 | 0 | A | G | 0.289 | -1.68E-01 | 8.33E-02 | 2.80E+00 | 2.88E+00 | 4.33E-02 | no | | intergenic_variant | MODIFIER |
| 10 | 104479667 | AX-170315334 | 0 | A | G | 0.289 | -1.68E-01 | 8.33E-02 | 2.80E+00 | 2.88E+00 | 4.33E-02 | no | | intergenic_variant | MODIFIER |
| 10 | 104483447 | AX-97782243 | 0 | A | G | 0.289 | -1.68E-01 | 8.33E-02 | 2.80E+00 | 2.88E+00 | 4.33E-02 | no | | intergenic_variant | MODIFIER |
| 10 | 104515798 | AX-170003821 | 0 | C | G | 0.289 | -1.68E-01 | 8.33E-02 | 2.80E+00 | 2.88E+00 | 4.33E-02 | no | | regulatory_region_variant | MODIFIER |
| 10 | 104685227 | AX-170323842 | 0 | C | G | 0.289 | -1.68E-01 | 8.33E-02 | 2.80E+00 | 2.88E+00 | 4.33E-02 | no | | intergenic_variant | MODIFIER |
| 10 | 104755830 | AX-170392475 | 0 | G | C | 0.289 | -1.68E-01 | 8.33E-02 | 2.80E+00 | 2.88E+00 | 4.33E-02 | no | | intergenic_variant | MODIFIER |
| 10 | 104825135 | AX-170398284 | 0 | G | C | 0.289 | -1.68E-01 | 8.33E-02 | 2.80E+00 | 2.88E+00 | 4.33E-02 | no | | intergenic_variant | MODIFIER |
| 10 | 104825854 | AX-170092082 | 0 | G | A | 0.289 | -1.68E-01 | 8.33E-02 | 2.80E+00 | 2.88E+00 | 4.33E-02 | no | | intergenic_variant | MODIFIER |
| 10 | 104826723 | AX-170365445 | 0 | T | G | 0.289 | -1.68E-01 | 8.33E-02 | 2.80E+00 | 2.88E+00 | 4.33E-02 | no | | intergenic_variant | MODIFIER |
| 10 | 104866348 | AX-170131488 | 0 | T | C | 0.289 | -1.68E-01 | 8.33E-02 | 2.80E+00 | 2.88E+00 | 4.33E-02 | no | Gm25522 | upstream_gene_variant | MODIFIER |
| 10 | 104889489 | AX-170136889 | 0 | C | A | 0.289 | -1.68E-01 | 8.33E-02 | 2.80E+00 | 2.88E+00 | 4.33E-02 | no | | intergenic_variant | MODIFIER |
| 10 | 104924163 | AX-170523939 | 0 | C | T | 0.289 | -1.68E-01 | 8.33E-02 | 2.80E+00 | 2.88E+00 | 4.33E-02 | no | | intergenic_variant | MODIFIER |
| 10 | 104934422 | AX-170222141 | 0 | C | T | 0.289 | -1.68E-01 | 8.33E-02 | 2.80E+00 | 2.88E+00 | 4.33E-02 | no | | intergenic_variant | MODIFIER |
| 10 | 104936585 | AX-170434406 | 0 | T | G | 0.289 | -1.68E-01 | 8.33E-02 | 2.80E+00 | 2.88E+00 | 4.33E-02 | no | | intergenic_variant | MODIFIER |
| 10 | 104938049 | AX-170507630 | 0 | C | T | 0.289 | -1.68E-01 | 8.33E-02 | 2.80E+00 | 2.88E+00 | 4.33E-02 | no | | intergenic_variant | MODIFIER |
| 10 | 104938648 | AX-170479169 | 0 | A | G | 0.289 | -1.68E-01 | 8.33E-02 | 2.80E+00 | 2.88E+00 | 4.33E-02 | no | | intergenic_variant | MODIFIER |
| 10 | 104950108 | AX-170289966 | 0 | A | C | 0.289 | -1.68E-01 | 8.33E-02 | 2.80E+00 | 2.88E+00 | 4.33E-02 | no | | intergenic_variant | MODIFIER |
| 10 | 104962044 | AX-170045797 | 0 | A | G | 0.289 | -1.68E-01 | 8.33E-02 | 2.80E+00 | 2.88E+00 | 4.33E-02 | no | | intergenic_variant | MODIFIER |
| 10 | 104962110 | AX-170045790 | 0 | C | G | 0.289 | -1.68E-01 | 8.33E-02 | 2.80E+00 | 2.88E+00 | 4.33E-02 | no | | intergenic_variant | MODIFIER |
| 10 | 104978866 | AX-170327358 | 0 | T | G | 0.289 | -1.68E-01 | 8.33E-02 | 2.80E+00 | 2.88E+00 | 4.33E-02 | no | | intergenic_variant | MODIFIER |
| 10 | 105019950 | AX-170045791 | 0 | T | C | 0.289 | -1.68E-01 | 8.33E-02 | 2.80E+00 | 2.88E+00 | 4.33E-02 | no | | regulatory_region_variant | MODIFIER |
| 10 | 105022963 | AX-170045789 | 0 | G | A | 0.289 | -1.68E-01 | 8.33E-02 | 2.80E+00 | 2.88E+00 | 4.33E-02 | no | | intergenic_variant | MODIFIER |
| 10 | 105025300 | AX-170003829 | 0 | A | G | 0.289 | -1.68E-01 | 8.33E-02 | 2.80E+00 | 2.88E+00 | 4.33E-02 | no | | intergenic_variant | MODIFIER |
| 10 | 105064071 | AX-170480776 | 0 | T | C | 0.289 | -1.68E-01 | 8.33E-02 | 2.80E+00 | 2.88E+00 | 4.33E-02 | no | | intergenic_variant | MODIFIER |
| 10 | 105075380 | AX-97838057 | 0 | T | C | 0.289 | -1.68E-01 | 8.33E-02 | 2.80E+00 | 2.88E+00 | 4.33E-02 | no | | intergenic_variant | MODIFIER |
| 10 | 105090983 | AX-170423755 | 0 | G | A | 0.289 | -1.68E-01 | 8.33E-02 | 2.80E+00 | 2.88E+00 | 4.33E-02 | no | | intergenic_variant | MODIFIER |
| 10 | 105121152 | AX-170465089 | 0 | A | G | 0.289 | -1.68E-01 | 8.33E-02 | 2.80E+00 | 2.88E+00 | 4.33E-02 | no | | intergenic_variant | MODIFIER |
| 10 | 105546930 | AX-97842379 | 0 | A | G | 0.289 | -1.68E-01 | 8.33E-02 | 2.80E+00 | 2.88E+00 | 4.33E-02 | no | Tmtc2 | intron_variant | MODIFIER |

| | | | | | | | | | | | | | | | |
|---|---|---|---|---|---|---|---|---|---|---|---|---|---|---|---|
| 10 | 106030562 | AX-170003844 | 0 | T | C | 0.289 | -1.68E-01 | 8.33E-02 | 2.80E+00 | 2.88E+00 | 4.33E-02 | no | | intergenic_variant | MODIFIER |
| 10 | 106033069 | AX-170003842 | 0 | C | T | 0.289 | -1.68E-01 | 8.33E-02 | 2.80E+00 | 2.88E+00 | 4.33E-02 | no | | intergenic_variant | MODIFIER |
| 10 | 106036087 | AX-170003840 | 0 | T | C | 0.289 | -1.68E-01 | 8.33E-02 | 2.80E+00 | 2.88E+00 | 4.33E-02 | no | | intergenic_variant | MODIFIER |
| 13 | 3011969 | AX-170417293 | 1 | A | G | 0.196 | -2.17E-01 | 1.08E-01 | 2.86E+00 | 2.97E+00 | 4.33E-02 | Yes | Gm8598 | downstream_gene_variant | MODIFIER |
| 16 | 61863645 | AX-170013207 | 0 | T | A | 0.535 | 1.73E-01 | 8.52E-02 | 2.50E+00 | 2.55E+00 | 4.33E-02 | Yes | | intergenic_variant | MODIFIER |
| 16 | 67735147 | AX-170167140 | 2 | G | A | 0.5 | 1.51E-01 | 7.46E-02 | 2.63E+00 | 2.72E+00 | 4.34E-02 | Yes | Gm49648 | intron_variant,non_coding_transcript_variant | MODIFIER |
| 9 | 106018981 | AX-170365117 | 0 | G | A | 0.386 | 1.37E-01 | 6.80E-02 | 2.61E+00 | 2.76E+00 | 4.34E-02 | no | Col6a4 | intron_variant | MODIFIER |
| 11 | 91580841 | AX-170118246 | 1 | C | A | 0.027 | -4.28E-01 | 2.11E-01 | 2.98E+00 | 3.13E+00 | 4.34E-02 | Yes | Kif2b | upstream_gene_variant | MODIFIER |
| 16 | 56878076 | AX-170318521 | 0 | T | C | 0.456 | -1.66E-01 | 8.22E-02 | 2.66E+00 | 2.70E+00 | 4.35E-02 | no | Tmem45a | intron_variant | MODIFIER |
| 13 | 16980654 | AX-170309882 | 0 | A | G | 0.175 | -2.54E-01 | 1.25E-01 | 2.46E+00 | 2.51E+00 | 4.36E-02 | Yes | Sugct | intron_variant | MODIFIER |
| 19 | 52786478 | AX-170016836 | 1 | T | C | 0.5 | 1.27E-01 | 6.30E-02 | 2.77E+00 | 2.86E+00 | 4.36E-02 | no | | intergenic_variant | MODIFIER |
| 15 | 67846007 | AX-170011908 | 0 | A | G | 0.079 | -2.24E-01 | 9.80E-02 | 1.65E+00 | 1.71E+00 | 4.37E-02 | Yes | | intergenic_variant | MODIFIER |
| 13 | 38728738 | AX-170008142 | 0 | T | A | 0.175 | -2.08E-01 | 1.03E-01 | 2.84E+00 | 2.98E+00 | 4.38E-02 | Yes | Gm46392 | intron_variant,non_coding_transcript_variant | MODIFIER |
| 16 | 58538094 | AX-170488055 | 0 | G | A | 0.447 | -1.64E-01 | 8.15E-02 | 2.63E+00 | 2.69E+00 | 4.38E-02 | no | | intergenic_variant | MODIFIER |
| 13 | 25085702 | AX-170208777 | 2 | C | T | 0.173 | -2.27E-01 | 1.13E-01 | 2.62E+00 | 2.72E+00 | 4.38E-02 | Yes | Dcdc2a | intron_variant | MODIFIER |
| 5 | 52420189 | AX-170519240 | 0 | T | A | 0.482 | 2.12E-01 | 1.03E-01 | 2.24E+00 | 2.28E+00 | 4.39E-02 | Yes | Ccdc149 | intron_variant | MODIFIER |
| 13 | 11680934 | AX-170282992 | 1 | G | A | 0.188 | -2.33E-01 | 1.16E-01 | 2.76E+00 | 2.85E+00 | 4.40E-02 | Yes | Ryr2 | intron_variant | MODIFIER |
| 5 | 62029079 | AX-97819350 | 2 | C | A | 0.473 | 2.10E-01 | 1.04E-01 | 2.50E+00 | 2.56E+00 | 4.40E-02 | Yes | | intergenic_variant | MODIFIER |
| 4 | 84044305 | AX-170459534 | 1 | A | G | 0.42 | -1.66E-01 | 8.23E-02 | 2.69E+00 | 2.76E+00 | 4.42E-02 | no | 6030471H07Rik | intron_variant,non_coding_transcript_variant | MODIFIER |
| 3 | 19835733 | AX-170443613 | 1 | A | T | 0.125 | -2.04E-01 | 1.02E-01 | 2.77E+00 | 2.86E+00 | 4.43E-02 | Yes | | intergenic_variant | MODIFIER |
| 3 | 6986494 | AX-170160644 | 2 | T | A | 0.127 | -1.83E-01 | 9.11E-02 | 2.75E+00 | 2.87E+00 | 4.44E-02 | Yes | | intergenic_variant | MODIFIER |
| 15 | 11097126 | AX-170354046 | 0 | T | C | 0.061 | -2.69E-01 | 1.06E-01 | 1.33E+00 | 1.37E+00 | 4.44E-02 | Yes | Adamts12 | intron_variant | MODIFIER |
| 15 | 11447800 | AX-170100552 | 0 | C | T | 0.061 | -2.69E-01 | 1.06E-01 | 1.33E+00 | 1.37E+00 | 4.44E-02 | Yes | | intergenic_variant | MODIFIER |
| 1 | 110707932 | AX-170472103 | 0 | G | T | 0.044 | -2.66E-01 | 1.27E-01 | 3.67E+00 | 3.99E+00 | 4.44E-02 | Yes | | intergenic_variant | MODIFIER |
| 16 | 52425314 | AX-170288778 | 1 | T | C | 0.518 | 1.67E-01 | 8.33E-02 | 2.79E+00 | 2.85E+00 | 4.45E-02 | Yes | Alcam | intron_variant | MODIFIER |
| 16 | 63303895 | AX-170480237 | 1 | A | C | 0.527 | 1.68E-01 | 8.34E-02 | 2.60E+00 | 2.66E+00 | 4.45E-02 | Yes | | intergenic_variant | MODIFIER |
| 13 | 19126064 | AX-97839380 | 0 | T | G | 0.175 | -2.32E-01 | 1.16E-01 | 2.65E+00 | 2.74E+00 | 4.45E-02 | Yes | Amph | intron_variant | MODIFIER |
| 13 | 20662755 | AX-170421964 | 0 | G | C | 0.175 | -2.32E-01 | 1.16E-01 | 2.65E+00 | 2.74E+00 | 4.45E-02 | Yes | | intergenic_variant | MODIFIER |
| 5 | 137310667 | AX-170026277 | 1 | A | G | 0.125 | -1.85E-01 | 9.06E-02 | 2.28E+00 | 2.36E+00 | 4.48E-02 | Yes | Trip6 | synonymous_variant | LOW |
| 19 | 53513870 | AX-97820034 | 0 | G | A | 0.509 | 1.29E-01 | 6.46E-02 | 2.87E+00 | 2.98E+00 | 4.52E-02 | no | | intergenic_variant | MODIFIER |
| 5 | 137876732 | AX-170392168 | 0 | T | C | 0.132 | -1.73E-01 | 8.49E-02 | 2.31E+00 | 2.39E+00 | 4.52E-02 | no | Pilrb2 | upstream_gene_variant | MODIFIER |
| 5 | 137415201 | AX-170165429 | 1 | A | T | 0.134 | -1.79E-01 | 8.87E-02 | 2.38E+00 | 2.47E+00 | 4.53E-02 | Yes | Zan | intron_variant | MODIFIER |
| 5 | 137994556 | AX-97810242 | 1 | C | T | 0.134 | -1.79E-01 | 8.87E-02 | 2.38E+00 | 2.47E+00 | 4.53E-02 | Yes | Azgp1 | downstream_gene_variant | MODIFIER |
| 5 | 62408059 | AX-170291773 | 1 | G | A | 0.473 | 2.05E-01 | 1.02E-01 | 2.43E+00 | 2.49E+00 | 4.53E-02 | Yes | | intergenic_variant | MODIFIER |
| 5 | 47731771 | AX-170071997 | 1 | C | A | 0.509 | 1.94E-01 | 9.63E-02 | 2.49E+00 | 2.56E+00 | 4.54E-02 | Yes | | intergenic_variant | MODIFIER |
| 13 | 7322424 | AX-97793273 | 0 | C | T | 0.184 | -2.29E-01 | 1.14E-01 | 2.37E+00 | 2.42E+00 | 4.60E-02 | Yes | | intergenic_variant | MODIFIER |
| 9 | 106955420 | AX-170033227 | 0 | T | C | 0.377 | 1.34E-01 | 6.73E-02 | 2.66E+00 | 2.82E+00 | 4.62E-02 | no | Dock3 | intron_variant | MODIFIER |
| 9 | 107503557 | AX-170083392 | 0 | A | G | 0.377 | 1.34E-01 | 6.73E-02 | 2.66E+00 | 2.82E+00 | 4.62E-02 | no | Cacna2d2 | intron_variant | MODIFIER |
| 9 | 107516728 | AX-97813758 | 0 | T | C | 0.377 | 1.34E-01 | 6.73E-02 | 2.66E+00 | 2.82E+00 | 4.62E-02 | no | Cacna2d2 | intron_variant | MODIFIER |
| 9 | 107528320 | AX-170033240 | 0 | G | A | 0.377 | 1.34E-01 | 6.73E-02 | 2.66E+00 | 2.82E+00 | 4.62E-02 | no | Cacna2d2 | 3_prime_UTR_variant | MODIFIER |
| 9 | 107600732 | AX-170274664 | 0 | T | G | 0.377 | 1.34E-01 | 6.73E-02 | 2.66E+00 | 2.82E+00 | 4.62E-02 | no | Sema3b | intron_variant | MODIFIER |
| 10 | 104662325 | AX-170304567 | 0 | T | A | 0.298 | -1.65E-01 | 8.27E-02 | 2.82E+00 | 2.91E+00 | 4.63E-02 | no | | regulatory_region_variant | MODIFIER |
| 8 | 38029219 | AX-97833996 | 0 | T | C | 0.158 | 2.69E-01 | 1.29E-01 | 3.72E+00 | 3.90E+00 | 4.64E-02 | Yes | Sgcz | intron_variant | MODIFIER |
| 16 | 72688052 | AX-170280514 | 1 | T | A | 0.5 | 1.55E-01 | 7.78E-02 | 2.80E+00 | 2.89E+00 | 4.65E-02 | Yes | Robo1 | intron_variant | MODIFIER |
| 13 | 20906750 | AX-170104503 | 0 | T | C | 0.167 | -2.36E-01 | 1.19E-01 | 2.68E+00 | 2.77E+00 | 4.66E-02 | Yes | Aoah | intron_variant | MODIFIER |
| 1 | 185063173 | AX-170043852 | 0 | A | C | 0.228 | 2.32E-01 | 1.17E-01 | 2.71E+00 | 2.79E+00 | 4.66E-02 | Yes | | intergenic_variant | MODIFIER |
| 4 | 107548543 | AX-170473536 | 0 | G | A | 0.272 | 2.21E-01 | 1.11E-01 | 2.84E+00 | 2.92E+00 | 4.66E-02 | Yes | Glis1 | intron_variant | MODIFIER |
| 5 | 49704214 | AX-170425353 | 0 | G | A | 0.482 | 2.08E-01 | 1.04E-01 | 2.49E+00 | 2.54E+00 | 4.67E-02 | Yes | | intergenic_variant | MODIFIER |
| 16 | 65464751 | AX-97791139 | 2 | G | A | 0.527 | 1.62E-01 | 8.17E-02 | 2.67E+00 | 2.73E+00 | 4.67E-02 | Yes | | intergenic_variant | MODIFIER |
| 5 | 64338882 | AX-170179210 | 1 | C | T | 0.491 | 2.05E-01 | 1.03E-01 | 2.53E+00 | 2.58E+00 | 4.68E-02 | Yes | Tbc1d1 | intron_variant | MODIFIER |
| 15 | 67858745 | AX-170533485 | 0 | T | G | 0.079 | -2.28E-01 | 9.98E-02 | 1.60E+00 | 1.66E+00 | 4.69E-02 | Yes | | intergenic_variant | MODIFIER |
| 5 | 54410753 | AX-170335277 | 0 | G | A | 0.509 | 2.03E-01 | 1.02E-01 | 2.48E+00 | 2.53E+00 | 4.71E-02 | Yes | | intergenic_variant | MODIFIER |
| 19 | 51796698 | AX-168411131 | 0 | C | T | 0.482 | 1.23E-01 | 6.21E-02 | 2.90E+00 | 3.00E+00 | 4.71E-02 | no | Rpl31-ps18 | downstream_gene_variant | MODIFIER |
| 15 | 70561785 | AX-170055899 | 0 | G | A | 0.088 | -2.20E-01 | 9.85E-02 | 1.68E+00 | 1.75E+00 | 4.72E-02 | Yes | | intergenic_variant | MODIFIER |

| | | | | | | | | | | | | | | | |
|---|---|---|---|---|---|---|---|---|---|---|---|---|---|---|---|
| 15 | 3991628 | AX-97852939 | 0 | C | T | 0.14 | -1.14E-01 | 5.60E-02 | 2.23E+00 | 2.35E+00 | 4.73E-02 | no | AW549877 | intron_variant | MODIFIER |
| 13 | 19183627 | AX-170130284 | 1 | C | A | 0.161 | -2.35E-01 | 1.18E-01 | 2.60E+00 | 2.68E+00 | 4.75E-02 | Yes | Tcrg-V4 | upstream_gene_variant | MODIFIER |
| 14 | 110740464 | AX-170257789 | 0 | A | G | 0.421 | -1.06E-01 | 5.37E-02 | 2.85E+00 | 3.01E+00 | 4.75E-02 | no | | intergenic_variant | MODIFIER |
| 14 | 110740970 | AX-97846293 | 0 | G | A | 0.421 | -1.06E-01 | 5.37E-02 | 2.85E+00 | 3.01E+00 | 4.75E-02 | no | | intergenic_variant | MODIFIER |
| 14 | 110750267 | AX-170010688 | 0 | G | C | 0.421 | -1.06E-01 | 5.37E-02 | 2.85E+00 | 3.01E+00 | 4.75E-02 | no | Slitrk6 | synonymous_variant | LOW |
| 14 | 110779793 | AX-170010680 | 0 | G | C | 0.421 | -1.06E-01 | 5.37E-02 | 2.85E+00 | 3.01E+00 | 4.75E-02 | no | Gm9348 | non_coding_transcript_exon_variant | MODIFIER |
| 14 | 110849157 | AX-170345429 | 0 | T | C | 0.421 | -1.06E-01 | 5.37E-02 | 2.85E+00 | 3.01E+00 | 4.75E-02 | no | | intergenic_variant | MODIFIER |
| 14 | 110916051 | AX-97845222 | 0 | A | G | 0.421 | -1.06E-01 | 5.37E-02 | 2.85E+00 | 3.01E+00 | 4.75E-02 | no | | intergenic_variant | MODIFIER |
| 14 | 110971321 | AX-170131966 | 0 | T | C | 0.421 | -1.06E-01 | 5.37E-02 | 2.85E+00 | 3.01E+00 | 4.75E-02 | no | | intergenic_variant | MODIFIER |
| 14 | 110985286 | AX-170314636 | 0 | G | A | 0.421 | -1.06E-01 | 5.37E-02 | 2.85E+00 | 3.01E+00 | 4.75E-02 | no | | intergenic_variant | MODIFIER |
| 4 | 84639282 | AX-170264257 | 1 | A | T | 0.348 | 2.24E-01 | 1.11E-01 | 3.25E+00 | 3.37E+00 | 4.75E-02 | Yes | Bnc2 | intron_variant | MODIFIER |
| 8 | 65727400 | AX-170197782 | 1 | A | T | 0.134 | -2.60E-01 | 1.31E-01 | 2.91E+00 | 3.04E+00 | 4.76E-02 | Yes | MAR1 | intron_variant | MODIFIER |
| 5 | 135609266 | AX-97781441 | 0 | G | A | 0.149 | -1.72E-01 | 8.50E-02 | 2.23E+00 | 2.30E+00 | 4.78E-02 | no | Gm43123 | downstream_gene_variant | MODIFIER |
| 5 | 135645367 | AX-170073617 | 0 | A | G | 0.149 | -1.72E-01 | 8.50E-02 | 2.23E+00 | 2.30E+00 | 4.78E-02 | no | Rhbdd2 | 3_prime_UTR_variant | MODIFIER |
| 5 | 135645395 | AX-170242944 | 0 | G | A | 0.149 | -1.72E-01 | 8.50E-02 | 2.23E+00 | 2.30E+00 | 4.78E-02 | no | Rhbdd2 | 3_prime_UTR_variant | MODIFIER |
| 5 | 135646608 | AX-170219621 | 0 | G | A | 0.149 | -1.72E-01 | 8.50E-02 | 2.23E+00 | 2.30E+00 | 4.78E-02 | no | Rhbdd2 | downstream_gene_variant | MODIFIER |
| 19 | 59789655 | AX-170016940 | 1 | T | C | 0.464 | -1.26E-01 | 6.35E-02 | 2.88E+00 | 2.97E+00 | 4.78E-02 | no | | intergenic_variant | MODIFIER |
| 16 | 63731533 | AX-97781241 | 0 | G | T | 0.526 | 1.73E-01 | 8.75E-02 | 2.71E+00 | 2.75E+00 | 4.78E-02 | Yes | Epha3 | intron_variant | MODIFIER |
| 16 | 62784576 | AX-170088043 | 0 | T | C | 0.526 | 1.73E-01 | 8.76E-02 | 2.71E+00 | 2.75E+00 | 4.78E-02 | Yes | Nsun3 | intron_variant | MODIFIER |
| 5 | 129394082 | AX-170142508 | 0 | A | C | 0.149 | -1.76E-01 | 8.62E-02 | 2.16E+00 | 2.24E+00 | 4.79E-02 | Yes | | intergenic_variant | MODIFIER |
| 5 | 129447166 | AX-170171447 | 0 | C | T | 0.149 | -1.76E-01 | 8.62E-02 | 2.16E+00 | 2.24E+00 | 4.79E-02 | Yes | | intergenic_variant | MODIFIER |
| 16 | 63894734 | AX-97857686 | 0 | A | G | 0.518 | 1.66E-01 | 8.40E-02 | 2.81E+00 | 2.89E+00 | 4.80E-02 | Yes | | intergenic_variant | MODIFIER |
| 13 | 24661239 | AX-170432187 | 2 | T | G | 0.173 | -2.25E-01 | 1.13E-01 | 2.44E+00 | 2.52E+00 | 4.81E-02 | Yes | Ripor2 | intron_variant | MODIFIER |
| 13 | 16316718 | AX-97849854 | 0 | T | C | 0.167 | -2.46E-01 | 1.24E-01 | 2.44E+00 | 2.49E+00 | 4.83E-02 | Yes | | regulatory_region_variant | MODIFIER |
| 5 | 42411256 | AX-170117978 | 0 | C | G | 0.211 | -2.01E-01 | 1.01E-01 | 3.09E+00 | 3.24E+00 | 4.83E-02 | Yes | | intergenic_variant | MODIFIER |
| 13 | 33034503 | AX-170133784 | 0 | T | C | 0.175 | -2.30E-01 | 1.16E-01 | 2.76E+00 | 2.86E+00 | 4.83E-02 | Yes | Serpinb9b | intron_variant | MODIFIER |
| 1 | 52705746 | AX-170041230 | 0 | C | T | 0.105 | -3.74E-01 | 1.87E-01 | 3.26E+00 | 3.41E+00 | 4.83E-02 | Yes | Mfsd6 | intron_variant | MODIFIER |
| 3 | 19554757 | AX-170020062 | 0 | A | G | 0.123 | -1.91E-01 | 9.68E-02 | 2.74E+00 | 2.85E+00 | 4.84E-02 | Yes | Dnajc5b | intron_variant | MODIFIER |
| 16 | 58212604 | AX-97812659 | 2 | A | G | 0.436 | -1.64E-01 | 8.33E-02 | 2.64E+00 | 2.69E+00 | 4.84E-02 | no | | intergenic_variant | MODIFIER |
| 2 | 99295255 | AX-170018460 | 0 | G | A | 0.053 | -4.65E-01 | 2.25E-01 | 3.68E+00 | 3.89E+00 | 4.86E-02 | Yes | | intergenic_variant | MODIFIER |
| 16 | 60577088 | AX-170013183 | 0 | G | T | 0.526 | 1.72E-01 | 8.75E-02 | 2.71E+00 | 2.75E+00 | 4.88E-02 | Yes | Epha6 | intron_variant | MODIFIER |
| 16 | 60613670 | AX-97838818 | 0 | C | T | 0.526 | 1.72E-01 | 8.75E-02 | 2.71E+00 | 2.75E+00 | 4.88E-02 | Yes | Gm9017 | downstream_gene_variant | MODIFIER |
| 16 | 61481960 | AX-170266541 | 0 | G | C | 0.526 | 1.72E-01 | 8.75E-02 | 2.71E+00 | 2.75E+00 | 4.88E-02 | Yes | | intergenic_variant | MODIFIER |
| 16 | 61530493 | AX-170221857 | 0 | C | T | 0.526 | 1.72E-01 | 8.75E-02 | 2.71E+00 | 2.75E+00 | 4.88E-02 | Yes | | intergenic_variant | MODIFIER |
| 16 | 61548017 | AX-97823780 | 0 | T | C | 0.526 | 1.72E-01 | 8.75E-02 | 2.71E+00 | 2.75E+00 | 4.88E-02 | Yes | | intergenic_variant | MODIFIER |
| 16 | 61548198 | AX-170186581 | 0 | C | T | 0.526 | 1.72E-01 | 8.75E-02 | 2.71E+00 | 2.75E+00 | 4.88E-02 | Yes | | intergenic_variant | MODIFIER |
| 16 | 61557483 | AX-170088037 | 0 | A | G | 0.526 | 1.72E-01 | 8.75E-02 | 2.71E+00 | 2.75E+00 | 4.88E-02 | Yes | | intergenic_variant | MODIFIER |
| 16 | 61579551 | AX-170185122 | 0 | T | C | 0.526 | 1.72E-01 | 8.75E-02 | 2.71E+00 | 2.75E+00 | 4.88E-02 | Yes | | intergenic_variant | MODIFIER |
| 16 | 61598530 | AX-97832023 | 0 | A | G | 0.526 | 1.72E-01 | 8.75E-02 | 2.71E+00 | 2.75E+00 | 4.88E-02 | Yes | | intergenic_variant | MODIFIER |
| 16 | 61646242 | AX-170367331 | 0 | G | A | 0.526 | 1.72E-01 | 8.75E-02 | 2.71E+00 | 2.75E+00 | 4.88E-02 | Yes | | intergenic_variant | MODIFIER |
| 16 | 61650424 | AX-170313663 | 0 | G | A | 0.526 | 1.72E-01 | 8.75E-02 | 2.71E+00 | 2.75E+00 | 4.88E-02 | Yes | | intergenic_variant | MODIFIER |
| 16 | 61695315 | AX-170500764 | 0 | T | C | 0.526 | 1.72E-01 | 8.75E-02 | 2.71E+00 | 2.75E+00 | 4.88E-02 | Yes | | intergenic_variant | MODIFIER |
| 16 | 61749955 | AX-170088038 | 0 | A | G | 0.526 | 1.72E-01 | 8.75E-02 | 2.71E+00 | 2.75E+00 | 4.88E-02 | Yes | | intergenic_variant | MODIFIER |
| 16 | 61772079 | AX-170104871 | 0 | G | C | 0.526 | 1.72E-01 | 8.75E-02 | 2.71E+00 | 2.75E+00 | 4.88E-02 | Yes | | intergenic_variant | MODIFIER |
| 16 | 61785462 | AX-97852926 | 0 | G | A | 0.526 | 1.72E-01 | 8.75E-02 | 2.71E+00 | 2.75E+00 | 4.88E-02 | Yes | | intergenic_variant | MODIFIER |
| 16 | 61802790 | AX-170364072 | 0 | T | C | 0.526 | 1.72E-01 | 8.75E-02 | 2.71E+00 | 2.75E+00 | 4.88E-02 | Yes | | intergenic_variant | MODIFIER |
| 16 | 61825939 | AX-97831505 | 0 | A | G | 0.526 | 1.72E-01 | 8.75E-02 | 2.71E+00 | 2.75E+00 | 4.88E-02 | Yes | | intergenic_variant | MODIFIER |
| 16 | 61922293 | AX-170013203 | 0 | A | G | 0.526 | 1.72E-01 | 8.75E-02 | 2.71E+00 | 2.75E+00 | 4.88E-02 | Yes | | intergenic_variant | MODIFIER |
| 16 | 62037642 | AX-170013209 | 0 | A | G | 0.526 | 1.72E-01 | 8.75E-02 | 2.71E+00 | 2.75E+00 | 4.88E-02 | Yes | | intergenic_variant | MODIFIER |
| 16 | 62061542 | AX-170214140 | 0 | A | G | 0.526 | 1.72E-01 | 8.75E-02 | 2.71E+00 | 2.75E+00 | 4.88E-02 | Yes | | intergenic_variant | MODIFIER |
| 16 | 62063781 | AX-97832048 | 0 | A | G | 0.526 | 1.72E-01 | 8.75E-02 | 2.71E+00 | 2.75E+00 | 4.88E-02 | Yes | | intergenic_variant | MODIFIER |
| 16 | 62071780 | AX-170164215 | 0 | G | A | 0.526 | 1.72E-01 | 8.75E-02 | 2.71E+00 | 2.75E+00 | 4.88E-02 | Yes | | intergenic_variant | MODIFIER |
| 16 | 62084601 | AX-170013205 | 0 | G | A | 0.526 | 1.72E-01 | 8.75E-02 | 2.71E+00 | 2.75E+00 | 4.88E-02 | Yes | | intergenic_variant | MODIFIER |
| 16 | 62111954 | AX-170383144 | 0 | A | G | 0.526 | 1.72E-01 | 8.75E-02 | 2.71E+00 | 2.75E+00 | 4.88E-02 | Yes | | intergenic_variant | MODIFIER |

| | | | | | | | | | | | | | | | |
|---|---|---|---|---|---|---|---|---|---|---|---|---|---|---|---|
| 16 | 62139622 | AX-170124744 | 0 | C | A | 0.526 | 1.72E-01 | 8.75E-02 | 2.71E+00 | 2.75E+00 | 4.88E-02 | Yes | | intergenic_variant | MODIFIER |
| 16 | 62148045 | AX-170439475 | 0 | G | A | 0.526 | 1.72E-01 | 8.75E-02 | 2.71E+00 | 2.75E+00 | 4.88E-02 | Yes | | intergenic_variant | MODIFIER |
| 16 | 62161013 | AX-97832047 | 0 | A | T | 0.526 | 1.72E-01 | 8.75E-02 | 2.71E+00 | 2.75E+00 | 4.88E-02 | Yes | | intergenic_variant | MODIFIER |
| 16 | 62184133 | AX-170013198 | 0 | T | A | 0.526 | 1.72E-01 | 8.75E-02 | 2.71E+00 | 2.75E+00 | 4.88E-02 | Yes | | intergenic_variant | MODIFIER |
| 16 | 62184809 | AX-97823095 | 0 | G | C | 0.526 | 1.72E-01 | 8.75E-02 | 2.71E+00 | 2.75E+00 | 4.88E-02 | Yes | | intergenic_variant | MODIFIER |
| 16 | 62269706 | AX-170088040 | 0 | C | G | 0.526 | 1.72E-01 | 8.75E-02 | 2.71E+00 | 2.75E+00 | 4.88E-02 | Yes | | intergenic_variant | MODIFIER |
| 16 | 62295838 | AX-170290848 | 0 | C | T | 0.526 | 1.72E-01 | 8.75E-02 | 2.71E+00 | 2.75E+00 | 4.88E-02 | Yes | | intergenic_variant | MODIFIER |
| 16 | 62385811 | AX-97791111 | 0 | T | G | 0.526 | 1.72E-01 | 8.75E-02 | 2.71E+00 | 2.75E+00 | 4.88E-02 | Yes | | intergenic_variant | MODIFIER |
| 16 | 62458982 | AX-97843516 | 0 | G | A | 0.526 | 1.72E-01 | 8.75E-02 | 2.71E+00 | 2.75E+00 | 4.88E-02 | Yes | | intergenic_variant | MODIFIER |
| 16 | 62887542 | AX-97820550 | 0 | A | G | 0.526 | 1.72E-01 | 8.75E-02 | 2.71E+00 | 2.75E+00 | 4.88E-02 | Yes | Pros1 | intron_variant | MODIFIER |
| 16 | 63006265 | AX-170013220 | 0 | C | T | 0.526 | 1.72E-01 | 8.75E-02 | 2.71E+00 | 2.75E+00 | 4.88E-02 | Yes | Gm18870 | intron_variant,non_coding_transcript_variant | MODIFIER |
| 16 | 63010019 | AX-97838114 | 0 | C | T | 0.526 | 1.72E-01 | 8.75E-02 | 2.71E+00 | 2.75E+00 | 4.88E-02 | Yes | Gm18870 | intron_variant,non_coding_transcript_variant | MODIFIER |
| 16 | 63010380 | AX-170013224 | 0 | T | A | 0.526 | 1.72E-01 | 8.75E-02 | 2.71E+00 | 2.75E+00 | 4.88E-02 | Yes | Gm18870 | intron_variant,non_coding_transcript_variant | MODIFIER |
| 16 | 63029853 | AX-97802426 | 0 | C | T | 0.526 | 1.72E-01 | 8.75E-02 | 2.71E+00 | 2.75E+00 | 4.88E-02 | Yes | Gm18870 | intron_variant,non_coding_transcript_variant | MODIFIER |
| 16 | 63049809 | AX-170172115 | 0 | C | T | 0.526 | 1.72E-01 | 8.75E-02 | 2.71E+00 | 2.75E+00 | 4.88E-02 | Yes | Gm18870 | intron_variant,non_coding_transcript_variant | MODIFIER |
| 16 | 63049863 | AX-170277616 | 0 | A | G | 0.526 | 1.72E-01 | 8.75E-02 | 2.71E+00 | 2.75E+00 | 4.88E-02 | Yes | Gm18870 | intron_variant,non_coding_transcript_variant | MODIFIER |
| 16 | 63055690 | AX-170013225 | 0 | T | A | 0.526 | 1.72E-01 | 8.75E-02 | 2.71E+00 | 2.75E+00 | 4.88E-02 | Yes | Gm18870 | intron_variant,non_coding_transcript_variant | MODIFIER |
| 16 | 63079977 | AX-170013222 | 0 | C | T | 0.526 | 1.72E-01 | 8.75E-02 | 2.71E+00 | 2.75E+00 | 4.88E-02 | Yes | Gm18870 | intron_variant,non_coding_transcript_variant | MODIFIER |
| 16 | 63084367 | AX-97840253 | 0 | C | T | 0.526 | 1.72E-01 | 8.75E-02 | 2.71E+00 | 2.75E+00 | 4.88E-02 | Yes | Gm18870 | intron_variant,non_coding_transcript_variant | MODIFIER |
| 16 | 63090887 | AX-170013219 | 0 | G | C | 0.526 | 1.72E-01 | 8.75E-02 | 2.71E+00 | 2.75E+00 | 4.88E-02 | Yes | Gm18870 | intron_variant,non_coding_transcript_variant | MODIFIER |
| 16 | 63103180 | AX-170013226 | 0 | T | G | 0.526 | 1.72E-01 | 8.75E-02 | 2.71E+00 | 2.75E+00 | 4.88E-02 | Yes | Gm18870 | intron_variant,non_coding_transcript_variant | MODIFIER |
| 16 | 63123879 | AX-97856004 | 0 | A | G | 0.526 | 1.72E-01 | 8.75E-02 | 2.71E+00 | 2.75E+00 | 4.88E-02 | Yes | | intergenic_variant | MODIFIER |
| 16 | 63154899 | AX-170013223 | 0 | A | G | 0.526 | 1.72E-01 | 8.75E-02 | 2.71E+00 | 2.75E+00 | 4.88E-02 | Yes | | intergenic_variant | MODIFIER |
| 16 | 63155100 | AX-170013229 | 0 | G | A | 0.526 | 1.72E-01 | 8.75E-02 | 2.71E+00 | 2.75E+00 | 4.88E-02 | Yes | | intergenic_variant | MODIFIER |
| 16 | 63171137 | AX-97822114 | 0 | G | A | 0.526 | 1.72E-01 | 8.75E-02 | 2.71E+00 | 2.75E+00 | 4.88E-02 | Yes | | intergenic_variant | MODIFIER |
| 16 | 63236435 | AX-97800308 | 0 | G | A | 0.526 | 1.72E-01 | 8.75E-02 | 2.71E+00 | 2.75E+00 | 4.88E-02 | Yes | | intergenic_variant | MODIFIER |
| 16 | 63237440 | AX-170498032 | 0 | A | G | 0.526 | 1.72E-01 | 8.75E-02 | 2.71E+00 | 2.75E+00 | 4.88E-02 | Yes | | intergenic_variant | MODIFIER |
| 16 | 63263925 | AX-170236964 | 0 | A | G | 0.526 | 1.72E-01 | 8.75E-02 | 2.71E+00 | 2.75E+00 | 4.88E-02 | Yes | | intergenic_variant | MODIFIER |
| 16 | 63271749 | AX-97782442 | 0 | G | T | 0.526 | 1.72E-01 | 8.75E-02 | 2.71E+00 | 2.75E+00 | 4.88E-02 | Yes | | intergenic_variant | MODIFIER |
| 16 | 63305172 | AX-170113872 | 0 | C | G | 0.526 | 1.72E-01 | 8.75E-02 | 2.71E+00 | 2.75E+00 | 4.88E-02 | Yes | | intergenic_variant | MODIFIER |
| 16 | 63305438 | AX-170135587 | 0 | A | G | 0.526 | 1.72E-01 | 8.75E-02 | 2.71E+00 | 2.75E+00 | 4.88E-02 | Yes | | intergenic_variant | MODIFIER |
| 16 | 63313966 | AX-170447061 | 0 | A | G | 0.526 | 1.72E-01 | 8.75E-02 | 2.71E+00 | 2.75E+00 | 4.88E-02 | Yes | | intergenic_variant | MODIFIER |
| 16 | 63351304 | AX-97844466 | 0 | C | A | 0.526 | 1.72E-01 | 8.75E-02 | 2.71E+00 | 2.75E+00 | 4.88E-02 | Yes | | intergenic_variant | MODIFIER |
| 16 | 63351718 | AX-170166669 | 0 | A | G | 0.526 | 1.72E-01 | 8.75E-02 | 2.71E+00 | 2.75E+00 | 4.88E-02 | Yes | | regulatory_region_variant | MODIFIER |
| 16 | 63367847 | AX-170532745 | 0 | T | A | 0.526 | 1.72E-01 | 8.75E-02 | 2.71E+00 | 2.75E+00 | 4.88E-02 | Yes | | intergenic_variant | MODIFIER |
| 16 | 63393869 | AX-97835150 | 0 | G | C | 0.526 | 1.72E-01 | 8.75E-02 | 2.71E+00 | 2.75E+00 | 4.88E-02 | Yes | | intergenic_variant | MODIFIER |
| 16 | 63408931 | AX-97831546 | 0 | T | A | 0.526 | 1.72E-01 | 8.75E-02 | 2.71E+00 | 2.75E+00 | 4.88E-02 | Yes | | intergenic_variant | MODIFIER |
| 16 | 63464306 | AX-170179992 | 0 | T | C | 0.526 | 1.72E-01 | 8.75E-02 | 2.71E+00 | 2.75E+00 | 4.88E-02 | Yes | Gm49622 | downstream_gene_variant | MODIFIER |
| 16 | 63472325 | AX-170495769 | 0 | A | G | 0.526 | 1.72E-01 | 8.75E-02 | 2.71E+00 | 2.75E+00 | 4.88E-02 | Yes | Gm49622 | intron_variant,non_coding_transcript_variant | MODIFIER |
| 16 | 63473793 | AX-97782893 | 0 | G | A | 0.526 | 1.72E-01 | 8.75E-02 | 2.71E+00 | 2.75E+00 | 4.88E-02 | Yes | Gm49622 | intron_variant,non_coding_transcript_variant | MODIFIER |
| 16 | 63474341 | AX-170434475 | 0 | T | C | 0.526 | 1.72E-01 | 8.75E-02 | 2.71E+00 | 2.75E+00 | 4.88E-02 | Yes | Gm49622 | intron_variant,non_coding_transcript_variant | MODIFIER |
| 16 | 63489487 | AX-170108301 | 0 | T | C | 0.526 | 1.72E-01 | 8.75E-02 | 2.71E+00 | 2.75E+00 | 4.88E-02 | Yes | Gm49622 | intron_variant,non_coding_transcript_variant | MODIFIER |
| 16 | 63495313 | AX-170298695 | 0 | C | T | 0.526 | 1.72E-01 | 8.75E-02 | 2.71E+00 | 2.75E+00 | 4.88E-02 | Yes | Gm49622 | upstream_gene_variant | MODIFIER |
| 16 | 63495921 | AX-97831549 | 0 | A | C | 0.526 | 1.72E-01 | 8.75E-02 | 2.71E+00 | 2.75E+00 | 4.88E-02 | Yes | Gm49622 | upstream_gene_variant | MODIFIER |
| 16 | 63506511 | AX-170013233 | 0 | T | C | 0.526 | 1.72E-01 | 8.75E-02 | 2.71E+00 | 2.75E+00 | 4.88E-02 | Yes | | intergenic_variant | MODIFIER |
| 16 | 63520039 | AX-97822112 | 0 | T | G | 0.526 | 1.72E-01 | 8.75E-02 | 2.71E+00 | 2.75E+00 | 4.88E-02 | Yes | | intergenic_variant | MODIFIER |
| 16 | 63550965 | AX-170102515 | 0 | C | A | 0.526 | 1.72E-01 | 8.75E-02 | 2.71E+00 | 2.75E+00 | 4.88E-02 | Yes | Epha3 | intron_variant | MODIFIER |
| 16 | 63552689 | AX-97781485 | 0 | T | C | 0.526 | 1.72E-01 | 8.75E-02 | 2.71E+00 | 2.75E+00 | 4.88E-02 | Yes | Epha3 | intron_variant | MODIFIER |
| 16 | 63556735 | AX-170013235 | 0 | G | A | 0.526 | 1.72E-01 | 8.75E-02 | 2.71E+00 | 2.75E+00 | 4.88E-02 | Yes | Epha3 | intron_variant | MODIFIER |
| 16 | 63563079 | AX-170370905 | 0 | C | A | 0.526 | 1.72E-01 | 8.75E-02 | 2.71E+00 | 2.75E+00 | 4.88E-02 | Yes | Epha3 | intron_variant | MODIFIER |
| 16 | 63563211 | AX-97852498 | 0 | T | G | 0.526 | 1.72E-01 | 8.75E-02 | 2.71E+00 | 2.75E+00 | 4.88E-02 | Yes | Epha3 | intron_variant | MODIFIER |
| 16 | 63572590 | AX-170309368 | 0 | A | G | 0.526 | 1.72E-01 | 8.75E-02 | 2.71E+00 | 2.75E+00 | 4.88E-02 | Yes | Epha3 | intron_variant | MODIFIER |
| 16 | 63586434 | AX-170057635 | 0 | C | A | 0.526 | 1.72E-01 | 8.75E-02 | 2.71E+00 | 2.75E+00 | 4.88E-02 | Yes | Epha3 | intron_variant | MODIFIER |
| 16 | 63634569 | AX-170510686 | 0 | T | G | 0.526 | 1.72E-01 | 8.75E-02 | 2.71E+00 | 2.75E+00 | 4.88E-02 | Yes | Epha3 | intron_variant | MODIFIER |

| | | | | | | | | | | | | | | | |
|---|---|---|---|---|---|---|---|---|---|---|---|---|---|---|---|
| 16 | 63636827 | AX-170013236 | 0 | T | C | 0.526 | 1.72E-01 | 8.75E-02 | 2.71E+00 | 2.75E+00 | 4.88E-02 | Yes | Epha3 | intron_variant | MODIFIER |
| 16 | 63649540 | AX-170522566 | 0 | C | T | 0.526 | 1.72E-01 | 8.75E-02 | 2.71E+00 | 2.75E+00 | 4.88E-02 | Yes | Epha3 | intron_variant | MODIFIER |
| 16 | 63672376 | AX-97832050 | 0 | G | A | 0.526 | 1.72E-01 | 8.75E-02 | 2.71E+00 | 2.75E+00 | 4.88E-02 | Yes | Epha3 | intron_variant | MODIFIER |
| 16 | 63710413 | AX-97830801 | 0 | T | C | 0.526 | 1.72E-01 | 8.75E-02 | 2.71E+00 | 2.75E+00 | 4.88E-02 | Yes | Epha3 | intron_variant | MODIFIER |
| 1 | 119233607 | AX-170042460 | 0 | A | T | 0.184 | 2.42E-01 | 1.23E-01 | 2.55E+00 | 2.68E+00 | 4.89E-02 | Yes | | intergenic_variant | MODIFIER |
| 5 | 137659759 | AX-97856300 | 1 | A | G | 0.134 | -1.78E-01 | 8.93E-02 | 2.35E+00 | 2.44E+00 | 4.89E-02 | Yes | Agfg2 | intron_variant | MODIFIER |
| 19 | 52562613 | AX-170220435 | 1 | C | T | 0.5 | 1.26E-01 | 6.41E-02 | 2.72E+00 | 2.81E+00 | 4.91E-02 | no | | intergenic_variant | MODIFIER |
| 19 | 53010087 | AX-97836997 | 1 | T | A | 0.5 | 1.26E-01 | 6.41E-02 | 2.72E+00 | 2.81E+00 | 4.91E-02 | no | Xpnpep1 | downstream_gene_variant | MODIFIER |
| 18 | 76966764 | AX-170061377 | 1 | T | C | 0.429 | 1.69E-01 | 8.60E-02 | 2.66E+00 | 2.73E+00 | 4.93E-02 | Yes | Hdhd2 | intron_variant | MODIFIER |
| 2 | 72338167 | AX-97796889 | 0 | G | T | 0.044 | -3.96E-01 | 1.97E-01 | 3.37E+00 | 3.55E+00 | 4.93E-02 | Yes | Map3k20 | intron_variant | MODIFIER |
| 16 | 63637181 | AX-170280035 | 1 | G | A | 0.527 | 1.72E-01 | 8.75E-02 | 2.70E+00 | 2.75E+00 | 4.94E-02 | Yes | Epha3 | intron_variant | MODIFIER |
| 13 | 7733498 | AX-170087354 | 0 | G | A | 0.184 | -2.51E-01 | 1.28E-01 | 2.56E+00 | 2.60E+00 | 4.94E-02 | Yes | | intergenic_variant | MODIFIER |
| 6 | 103093037 | AX-170133801 | 0 | C | T | 0.079 | 2.96E-01 | 1.51E-01 | 2.96E+00 | 3.15E+00 | 4.96E-02 | Yes | | intergenic_variant | MODIFIER |
| 5 | 62258650 | AX-170329860 | 0 | G | A | 0.456 | 1.94E-01 | 9.82E-02 | 2.48E+00 | 2.54E+00 | 4.96E-02 | Yes | | intergenic_variant | MODIFIER |
| 5 | 61764571 | AX-170514956 | 0 | G | A | 0.482 | 1.97E-01 | 9.84E-02 | 2.25E+00 | 2.31E+00 | 4.98E-02 | Yes | | intergenic_variant | MODIFIER |
| 10 | 104675856 | AX-170133053 | 1 | G | A | 0.286 | -1.63E-01 | 8.34E-02 | 2.74E+00 | 2.82E+00 | 4.98E-02 | no | | intergenic_variant | MODIFIER |
| 5 | 55108976 | AX-170423807 | 0 | C | G | 0.482 | 2.02E-01 | 1.03E-01 | 2.52E+00 | 2.57E+00 | 4.99E-02 | Yes | | intergenic_variant | MODIFIER |
| 4 | 108054474 | AX-97799102 | 0 | G | A | 0.263 | 2.11E-01 | 1.08E-01 | 2.79E+00 | 2.88E+00 | 5.00E-02 | Yes | Scp2 | intron_variant | MODIFIER |
| 5 | 135273374 | AX-170428117 | 1 | T | C | 0.143 | -1.80E-01 | 8.98E-02 | 2.23E+00 | 2.30E+00 | 5.02E-02 | Yes | | intergenic_variant | MODIFIER |
| 10 | 104500044 | AX-170158123 | 0 | A | G | 0.281 | -1.59E-01 | 8.13E-02 | 2.79E+00 | 2.87E+00 | 5.05E-02 | no | | intergenic_variant | MODIFIER |
| 13 | 17179896 | AX-168373636 | 0 | G | A | 0.175 | -2.50E-01 | 1.27E-01 | 2.43E+00 | 2.47E+00 | 5.07E-02 | Yes | Sugct | intron_variant | MODIFIER |
| 13 | 16868976 | AX-170007825 | 1 | G | A | 0.179 | -2.49E-01 | 1.26E-01 | 2.42E+00 | 2.47E+00 | 5.07E-02 | Yes | Sugct | intron_variant | MODIFIER |
| 16 | 62303884 | AX-97823476 | 1 | A | G | 0.527 | 1.71E-01 | 8.76E-02 | 2.70E+00 | 2.74E+00 | 5.08E-02 | Yes | | intergenic_variant | MODIFIER |
| 3 | 6291364 | AX-170399831 | 0 | C | A | 0.14 | -1.75E-01 | 8.99E-02 | 2.71E+00 | 2.82E+00 | 5.09E-02 | Yes | | intergenic_variant | MODIFIER |
| 5 | 42993760 | AX-170024811 | 0 | T | A | 0.202 | -2.04E-01 | 1.04E-01 | 3.02E+00 | 3.15E+00 | 5.11E-02 | Yes | | intergenic_variant | MODIFIER |
| 3 | 76193619 | AX-97782367 | 0 | G | T | 0.053 | 2.53E-01 | 1.28E-01 | 3.22E+00 | 3.44E+00 | 5.15E-02 | Yes | Fstl5 | intron_variant | MODIFIER |
| 5 | 132676268 | AX-97796958 | 2 | A | G | 0.145 | -1.69E-01 | 8.44E-02 | 2.17E+00 | 2.24E+00 | 5.16E-02 | no | | intergenic_variant | MODIFIER |
| 5 | 52941503 | AX-170072118 | 1 | G | A | 0.491 | 1.85E-01 | 9.45E-02 | 2.46E+00 | 2.54E+00 | 5.19E-02 | Yes | | intergenic_variant | MODIFIER |
| 9 | 101205794 | AX-168377356 | 0 | T | C | 0.14 | -1.77E-01 | 9.14E-02 | 2.79E+00 | 2.93E+00 | 5.19E-02 | Yes | Ppp2r3a | intron_variant | MODIFIER |
| 18 | 52864219 | AX-170113312 | 0 | T | C | 0.193 | -2.09E-01 | 1.07E-01 | 2.99E+00 | 3.11E+00 | 5.21E-02 | Yes | Sncaip | intron_variant | MODIFIER |
| 3 | 17063539 | AX-170215400 | 0 | T | C | 0.123 | -1.94E-01 | 9.99E-02 | 2.70E+00 | 2.79E+00 | 5.23E-02 | Yes | | intergenic_variant | MODIFIER |
| 3 | 17078525 | AX-170358340 | 0 | A | C | 0.123 | -1.94E-01 | 9.99E-02 | 2.70E+00 | 2.79E+00 | 5.23E-02 | Yes | | intergenic_variant | MODIFIER |
| 3 | 17080695 | AX-170200466 | 0 | T | C | 0.123 | -1.94E-01 | 9.99E-02 | 2.70E+00 | 2.79E+00 | 5.23E-02 | Yes | | intergenic_variant | MODIFIER |
| 3 | 17088911 | AX-170412855 | 0 | A | G | 0.123 | -1.94E-01 | 9.99E-02 | 2.70E+00 | 2.79E+00 | 5.23E-02 | Yes | | intergenic_variant | MODIFIER |
| 5 | 60576031 | AX-170502761 | 0 | C | T | 0.491 | 1.95E-01 | 9.99E-02 | 2.46E+00 | 2.52E+00 | 5.24E-02 | Yes | Gm43390 | upstream_gene_variant | MODIFIER |
| 13 | 8986440 | AX-170450687 | 1 | A | G | 0.179 | -2.39E-01 | 1.23E-01 | 2.56E+00 | 2.61E+00 | 5.25E-02 | Yes | Gtpbp4 | intron_variant,NMD_transcript_variant | MODIFIER |
| 19 | 55510954 | AX-168395374 | 0 | A | G | 0.447 | 1.24E-01 | 6.43E-02 | 2.80E+00 | 2.88E+00 | 5.31E-02 | no | Vti1a | intron_variant | MODIFIER |
| 13 | 17672090 | AX-97827200 | 0 | G | A | 0.175 | -2.52E-01 | 1.29E-01 | 2.43E+00 | 2.46E+00 | 5.31E-02 | Yes | Sugct | intron_variant | MODIFIER |
| 19 | 55465987 | AX-170520390 | 0 | G | C | 0.447 | 1.26E-01 | 6.52E-02 | 2.89E+00 | 2.98E+00 | 5.31E-02 | no | Vti1a | intron_variant | MODIFIER |
| 16 | 65819377 | AX-97851921 | 1 | T | G | 0.509 | 1.60E-01 | 8.30E-02 | 2.68E+00 | 2.74E+00 | 5.31E-02 | Yes | Vgll3 | intron_variant | MODIFIER |
| 16 | 66220810 | AX-170013285 | 0 | C | A | 0.509 | 1.49E-01 | 7.69E-02 | 2.58E+00 | 2.64E+00 | 5.33E-02 | Yes | | intergenic_variant | MODIFIER |
| 16 | 66764405 | AX-97801103 | 0 | C | T | 0.526 | 1.53E-01 | 7.91E-02 | 2.53E+00 | 2.59E+00 | 5.34E-02 | Yes | Cadm2 | intron_variant | MODIFIER |
| 16 | 66776318 | AX-170013295 | 0 | C | T | 0.526 | 1.53E-01 | 7.91E-02 | 2.53E+00 | 2.59E+00 | 5.34E-02 | Yes | Cadm2 | intron_variant | MODIFIER |
| 16 | 66785262 | AX-97846907 | 0 | T | C | 0.526 | 1.53E-01 | 7.91E-02 | 2.53E+00 | 2.59E+00 | 5.34E-02 | Yes | Cadm2 | intron_variant | MODIFIER |
| 16 | 66794897 | AX-170490882 | 0 | A | C | 0.526 | 1.53E-01 | 7.91E-02 | 2.53E+00 | 2.59E+00 | 5.34E-02 | Yes | Cadm2 | intron_variant | MODIFIER |
| 16 | 66810074 | AX-97786091 | 0 | A | G | 0.526 | 1.53E-01 | 7.91E-02 | 2.53E+00 | 2.59E+00 | 5.34E-02 | Yes | Cadm2 | intron_variant | MODIFIER |
| 16 | 66815059 | AX-170013292 | 0 | T | C | 0.526 | 1.53E-01 | 7.91E-02 | 2.53E+00 | 2.59E+00 | 5.34E-02 | Yes | Cadm2 | intron_variant | MODIFIER |
| 16 | 66819934 | AX-170013296 | 0 | T | C | 0.526 | 1.53E-01 | 7.91E-02 | 2.53E+00 | 2.59E+00 | 5.34E-02 | Yes | Cadm2 | intron_variant | MODIFIER |
| 16 | 66827694 | AX-170160992 | 0 | T | G | 0.526 | 1.53E-01 | 7.91E-02 | 2.53E+00 | 2.59E+00 | 5.34E-02 | Yes | Cadm2 | intron_variant | MODIFIER |
| 1 | 181082307 | AX-170247689 | 0 | A | G | 0.246 | 2.02E-01 | 1.05E-01 | 2.60E+00 | 2.69E+00 | 5.35E-02 | Yes | Nvl | downstream_gene_variant | MODIFIER |
| 16 | 61708819 | AX-170425688 | 1 | C | T | 0.464 | -1.70E-01 | 8.81E-02 | 2.63E+00 | 2.67E+00 | 5.35E-02 | no | | intergenic_variant | MODIFIER |
| 16 | 70177743 | AX-170506958 | 0 | C | T | 0.491 | 1.46E-01 | 7.57E-02 | 2.75E+00 | 2.83E+00 | 5.35E-02 | Yes | | intergenic_variant | MODIFIER |
| 15 | 6499506 | AX-170526107 | 1 | A | G | 0.232 | -1.86E-01 | 9.65E-02 | 2.67E+00 | 2.77E+00 | 5.36E-02 | Yes | C9 | downstream_gene_variant | MODIFIER |
| 13 | 24422003 | AX-170117617 | 0 | A | T | 0.167 | -2.20E-01 | 1.13E-01 | 2.46E+00 | 2.54E+00 | 5.38E-02 | Yes | Cmah | intron_variant | MODIFIER |

| | | | | | | | | | | | | | | | |
|---|---|---|---|---|---|---|---|---|---|---|---|---|---|---|---|
| 13 | 6520213 | AX-97794600 | 0 | C | T | 0.202 | -2.05E-01 | 1.06E-01 | 2.83E+00 | 2.97E+00 | 5.39E-02 | Yes | | intergenic_variant | MODIFIER |
| 15 | 11575844 | AX-97783119 | 0 | C | T | 0.298 | -1.64E-01 | 7.64E-02 | 1.74E+00 | 1.78E+00 | 5.40E-02 | no | | intergenic_variant | MODIFIER |
| 16 | 62039355 | AX-170013201 | 1 | C | G | 0.464 | -1.64E-01 | 8.51E-02 | 2.58E+00 | 2.63E+00 | 5.41E-02 | no | | intergenic_variant | MODIFIER |
| 9 | 57780509 | AX-97802440 | 1 | G | T | 0.205 | -1.98E-01 | 1.01E-01 | 3.25E+00 | 3.43E+00 | 5.41E-02 | Yes | Gm17231 | intron_variant,non_coding_transcript_variant | MODIFIER |
| 16 | 63997866 | AX-97852995 | 1 | T | G | 0.5 | 1.67E-01 | 8.65E-02 | 2.88E+00 | 2.94E+00 | 5.41E-02 | Yes | | intergenic_variant | MODIFIER |
| 16 | 64066770 | AX-170013238 | 1 | T | A | 0.5 | 1.67E-01 | 8.65E-02 | 2.88E+00 | 2.94E+00 | 5.41E-02 | Yes | | intergenic_variant | MODIFIER |
| 16 | 63917865 | AX-170196381 | 0 | T | G | 0.5 | 1.67E-01 | 8.67E-02 | 2.87E+00 | 2.94E+00 | 5.42E-02 | Yes | | intergenic_variant | MODIFIER |
| 5 | 53661279 | AX-97825952 | 0 | G | A | 0.491 | 1.94E-01 | 1.01E-01 | 2.58E+00 | 2.65E+00 | 5.45E-02 | Yes | Rbpj | downstream_gene_variant | MODIFIER |
| 8 | 90844969 | AX-170524260 | 0 | G | C | 0.202 | -2.02E-01 | 1.03E-01 | 3.33E+00 | 3.51E+00 | 5.46E-02 | Yes | Chd9 | intron_variant | MODIFIER |
| 8 | 49410734 | AX-170193142 | 0 | A | T | 0.149 | -2.43E-01 | 1.26E-01 | 3.01E+00 | 3.14E+00 | 5.47E-02 | Yes | 4930555F03Rik | intron_variant,non_coding_transcript_variant | MODIFIER |
| 8 | 75384069 | AX-170351339 | 1 | T | C | 0.188 | -2.43E-01 | 1.23E-01 | 3.48E+00 | 3.68E+00 | 5.52E-02 | Yes | Umpk-ps | non_coding_transcript_exon_variant | MODIFIER |
| 8 | 73034289 | AX-170354153 | 0 | A | C | 0.184 | -2.48E-01 | 1.25E-01 | 3.49E+00 | 3.68E+00 | 5.53E-02 | Yes | Large1 | intron_variant | MODIFIER |
| 5 | 64429310 | AX-170280860 | 1 | T | C | 0.473 | 1.85E-01 | 9.56E-02 | 2.36E+00 | 2.43E+00 | 5.54E-02 | Yes | Gm31363 | upstream_gene_variant | MODIFIER |
| 5 | 64370219 | AX-170208701 | 0 | A | T | 0.491 | 1.87E-01 | 9.70E-02 | 2.46E+00 | 2.53E+00 | 5.54E-02 | Yes | | intergenic_variant | MODIFIER |
| 13 | 12432543 | AX-170007776 | 1 | C | T | 0.188 | -2.43E-01 | 1.27E-01 | 2.54E+00 | 2.60E+00 | 5.55E-02 | Yes | Heatr1 | intron_variant | MODIFIER |
| 16 | 66249904 | AX-97785782 | 0 | T | C | 0.518 | 1.54E-01 | 8.04E-02 | 2.55E+00 | 2.61E+00 | 5.57E-02 | Yes | Gm22448 | upstream_gene_variant | MODIFIER |
| 16 | 66260564 | AX-97832125 | 0 | G | T | 0.518 | 1.54E-01 | 8.04E-02 | 2.55E+00 | 2.61E+00 | 5.57E-02 | Yes | | intergenic_variant | MODIFIER |
| 16 | 66285287 | AX-170407295 | 0 | C | G | 0.518 | 1.54E-01 | 8.04E-02 | 2.55E+00 | 2.61E+00 | 5.57E-02 | Yes | | intergenic_variant | MODIFIER |
| 16 | 66285360 | AX-97828265 | 0 | T | C | 0.518 | 1.54E-01 | 8.04E-02 | 2.55E+00 | 2.61E+00 | 5.57E-02 | Yes | | intergenic_variant | MODIFIER |
| 16 | 66296558 | AX-170013288 | 0 | C | G | 0.518 | 1.54E-01 | 8.04E-02 | 2.55E+00 | 2.61E+00 | 5.57E-02 | Yes | | intergenic_variant | MODIFIER |
| 16 | 66337643 | AX-170013287 | 0 | T | C | 0.518 | 1.54E-01 | 8.04E-02 | 2.55E+00 | 2.61E+00 | 5.57E-02 | Yes | | intergenic_variant | MODIFIER |
| 16 | 66344162 | AX-170093510 | 0 | C | T | 0.518 | 1.54E-01 | 8.04E-02 | 2.55E+00 | 2.61E+00 | 5.57E-02 | Yes | | intergenic_variant | MODIFIER |
| 16 | 66361466 | AX-170507556 | 0 | G | T | 0.518 | 1.54E-01 | 8.04E-02 | 2.55E+00 | 2.61E+00 | 5.57E-02 | Yes | | intergenic_variant | MODIFIER |
| 16 | 66361654 | AX-170100411 | 0 | A | G | 0.518 | 1.54E-01 | 8.04E-02 | 2.55E+00 | 2.61E+00 | 5.57E-02 | Yes | | intergenic_variant | MODIFIER |
| 16 | 66372574 | AX-170438082 | 0 | A | G | 0.518 | 1.54E-01 | 8.04E-02 | 2.55E+00 | 2.61E+00 | 5.57E-02 | Yes | | intergenic_variant | MODIFIER |
| 16 | 66384921 | AX-170192725 | 0 | A | C | 0.518 | 1.54E-01 | 8.04E-02 | 2.55E+00 | 2.61E+00 | 5.57E-02 | Yes | | intergenic_variant | MODIFIER |
| 16 | 66394002 | AX-170508247 | 0 | G | A | 0.518 | 1.54E-01 | 8.04E-02 | 2.55E+00 | 2.61E+00 | 5.57E-02 | Yes | | intergenic_variant | MODIFIER |
| 10 | 104781689 | AX-170528029 | 0 | C | T | 0.298 | -1.47E-01 | 7.70E-02 | 2.62E+00 | 2.71E+00 | 5.59E-02 | no | | intergenic_variant | MODIFIER |
| 13 | 23165088 | AX-170472474 | 0 | G | A | 0.167 | -2.20E-01 | 1.15E-01 | 2.49E+00 | 2.57E+00 | 5.60E-02 | Yes | Vmn1r219 | downstream_gene_variant | MODIFIER |
| 9 | 86453517 | AX-170474780 | 1 | G | A | 0.116 | -2.23E-01 | 1.17E-01 | 2.60E+00 | 2.72E+00 | 5.60E-02 | Yes | Ube2cbp | intron_variant | MODIFIER |
| 18 | 77208524 | AX-170088378 | 0 | T | C | 0.158 | 1.23E-01 | 6.43E-02 | 2.86E+00 | 3.03E+00 | 5.62E-02 | no | St8sia5 | intron_variant | MODIFIER |
| 19 | 57049148 | AX-170094131 | 1 | T | C | 0.446 | 1.23E-01 | 6.44E-02 | 2.71E+00 | 2.78E+00 | 5.64E-02 | no | Ablim1 | intron_variant | MODIFIER |
| 5 | 56693450 | AX-170025009 | 1 | A | C | 0.464 | 1.99E-01 | 1.04E-01 | 2.48E+00 | 2.53E+00 | 5.64E-02 | Yes | | intergenic_variant | MODIFIER |
| 16 | 65300761 | AX-97806908 | 0 | G | A | 0.509 | 1.58E-01 | 8.30E-02 | 2.66E+00 | 2.72E+00 | 5.66E-02 | Yes | | intergenic_variant | MODIFIER |
| 16 | 65301073 | AX-170505303 | 0 | A | G | 0.509 | 1.58E-01 | 8.30E-02 | 2.66E+00 | 2.72E+00 | 5.66E-02 | Yes | | intergenic_variant | MODIFIER |
| 16 | 65323149 | AX-170013258 | 0 | A | C | 0.509 | 1.58E-01 | 8.30E-02 | 2.66E+00 | 2.72E+00 | 5.66E-02 | Yes | | intergenic_variant | MODIFIER |
| 16 | 65323437 | AX-170466103 | 0 | T | G | 0.509 | 1.58E-01 | 8.30E-02 | 2.66E+00 | 2.72E+00 | 5.66E-02 | Yes | | intergenic_variant | MODIFIER |
| 16 | 65336567 | AX-170417521 | 0 | A | G | 0.509 | 1.58E-01 | 8.30E-02 | 2.66E+00 | 2.72E+00 | 5.66E-02 | Yes | Gm49629 | upstream_gene_variant | MODIFIER |
| 16 | 65336855 | AX-170507019 | 0 | G | A | 0.509 | 1.58E-01 | 8.30E-02 | 2.66E+00 | 2.72E+00 | 5.66E-02 | Yes | Gm49629 | upstream_gene_variant | MODIFIER |
| 16 | 65385989 | AX-170057658 | 0 | C | G | 0.509 | 1.58E-01 | 8.30E-02 | 2.66E+00 | 2.72E+00 | 5.66E-02 | Yes | | intergenic_variant | MODIFIER |
| 16 | 65398145 | AX-170013256 | 0 | A | G | 0.509 | 1.58E-01 | 8.30E-02 | 2.66E+00 | 2.72E+00 | 5.66E-02 | Yes | | intergenic_variant | MODIFIER |
| 16 | 65405995 | AX-170467282 | 0 | G | T | 0.509 | 1.58E-01 | 8.30E-02 | 2.66E+00 | 2.72E+00 | 5.66E-02 | Yes | | intergenic_variant | MODIFIER |
| 16 | 65422211 | AX-170013266 | 0 | C | T | 0.509 | 1.58E-01 | 8.30E-02 | 2.66E+00 | 2.72E+00 | 5.66E-02 | Yes | | intergenic_variant | MODIFIER |
| 16 | 65424303 | AX-170207737 | 0 | A | G | 0.509 | 1.58E-01 | 8.30E-02 | 2.66E+00 | 2.72E+00 | 5.66E-02 | Yes | | intergenic_variant | MODIFIER |
| 16 | 65436432 | AX-170176511 | 0 | G | T | 0.509 | 1.58E-01 | 8.30E-02 | 2.66E+00 | 2.72E+00 | 5.66E-02 | Yes | Gm23659 | downstream_gene_variant | MODIFIER |
| 16 | 65443231 | AX-97814617 | 0 | C | T | 0.509 | 1.58E-01 | 8.30E-02 | 2.66E+00 | 2.72E+00 | 5.66E-02 | Yes | | intergenic_variant | MODIFIER |
| 16 | 65462270 | AX-170192521 | 0 | T | A | 0.509 | 1.58E-01 | 8.30E-02 | 2.66E+00 | 2.72E+00 | 5.66E-02 | Yes | | intergenic_variant | MODIFIER |
| 16 | 65486991 | AX-97789596 | 0 | C | A | 0.509 | 1.58E-01 | 8.30E-02 | 2.66E+00 | 2.72E+00 | 5.66E-02 | Yes | | intergenic_variant | MODIFIER |
| 16 | 65498341 | AX-170035567 | 0 | C | T | 0.509 | 1.58E-01 | 8.30E-02 | 2.66E+00 | 2.72E+00 | 5.66E-02 | Yes | | intergenic_variant | MODIFIER |
| 16 | 65499807 | AX-170013257 | 0 | C | G | 0.509 | 1.58E-01 | 8.30E-02 | 2.66E+00 | 2.72E+00 | 5.66E-02 | Yes | | intergenic_variant | MODIFIER |
| 16 | 65503327 | AX-170013263 | 0 | T | C | 0.509 | 1.58E-01 | 8.30E-02 | 2.66E+00 | 2.72E+00 | 5.66E-02 | Yes | | intergenic_variant | MODIFIER |
| 16 | 65578263 | AX-97802636 | 0 | G | A | 0.509 | 1.58E-01 | 8.30E-02 | 2.66E+00 | 2.72E+00 | 5.66E-02 | Yes | | intergenic_variant | MODIFIER |
| 16 | 65586200 | AX-97847988 | 0 | A | G | 0.509 | 1.58E-01 | 8.30E-02 | 2.66E+00 | 2.72E+00 | 5.66E-02 | Yes | | intergenic_variant | MODIFIER |
| 16 | 65588979 | AX-170418113 | 0 | A | G | 0.509 | 1.58E-01 | 8.30E-02 | 2.66E+00 | 2.72E+00 | 5.66E-02 | Yes | | intergenic_variant | MODIFIER |

| | | | | | | | | | | | | | | | |
|---|---|---|---|---|---|---|---|---|---|---|---|---|---|---|---|
| 16 | 65632892 | AX-170517171 | 0 | C | T | 0.509 | 1.58E-01 | 8.30E-02 | 2.66E+00 | 2.72E+00 | 5.66E-02 | Yes | | intergenic_variant | MODIFIER |
| 16 | 65661656 | AX-170159962 | 0 | G | A | 0.509 | 1.58E-01 | 8.30E-02 | 2.66E+00 | 2.72E+00 | 5.66E-02 | Yes | | intergenic_variant | MODIFIER |
| 16 | 65670432 | AX-97852265 | 0 | A | G | 0.509 | 1.58E-01 | 8.30E-02 | 2.66E+00 | 2.72E+00 | 5.66E-02 | Yes | | intergenic_variant | MODIFIER |
| 16 | 65707018 | AX-170088047 | 0 | G | T | 0.509 | 1.58E-01 | 8.30E-02 | 2.66E+00 | 2.72E+00 | 5.66E-02 | Yes | | intergenic_variant | MODIFIER |
| 16 | 65729392 | AX-170013280 | 0 | G | A | 0.509 | 1.58E-01 | 8.30E-02 | 2.66E+00 | 2.72E+00 | 5.66E-02 | Yes | | intergenic_variant | MODIFIER |
| 16 | 65729531 | AX-170406562 | 0 | A | G | 0.509 | 1.58E-01 | 8.30E-02 | 2.66E+00 | 2.72E+00 | 5.66E-02 | Yes | | regulatory_region_variant | MODIFIER |
| 16 | 65729779 | AX-170013279 | 0 | A | G | 0.509 | 1.58E-01 | 8.30E-02 | 2.66E+00 | 2.72E+00 | 5.66E-02 | Yes | | regulatory_region_variant | MODIFIER |
| 16 | 65819444 | AX-170119767 | 0 | C | T | 0.509 | 1.58E-01 | 8.30E-02 | 2.66E+00 | 2.72E+00 | 5.66E-02 | Yes | Vgll3 | intron_variant | MODIFIER |
| 16 | 65820343 | AX-170013273 | 0 | C | T | 0.509 | 1.58E-01 | 8.30E-02 | 2.66E+00 | 2.72E+00 | 5.66E-02 | Yes | Vgll3 | intron_variant | MODIFIER |
| 16 | 65829131 | AX-170399564 | 0 | G | A | 0.509 | 1.58E-01 | 8.30E-02 | 2.66E+00 | 2.72E+00 | 5.66E-02 | Yes | Vgll3 | intron_variant | MODIFIER |
| 16 | 65845186 | AX-170057667 | 0 | C | G | 0.509 | 1.58E-01 | 8.30E-02 | 2.66E+00 | 2.72E+00 | 5.66E-02 | Yes | Vgll3 | intron_variant | MODIFIER |
| 16 | 65845252 | AX-170057664 | 0 | T | A | 0.509 | 1.58E-01 | 8.30E-02 | 2.66E+00 | 2.72E+00 | 5.66E-02 | Yes | Vgll3 | intron_variant | MODIFIER |
| 16 | 65847028 | AX-170057665 | 0 | G | A | 0.509 | 1.58E-01 | 8.30E-02 | 2.66E+00 | 2.72E+00 | 5.66E-02 | Yes | Vgll3 | intron_variant | MODIFIER |
| 16 | 65865097 | AX-97801013 | 0 | C | A | 0.509 | 1.58E-01 | 8.30E-02 | 2.66E+00 | 2.72E+00 | 5.66E-02 | Yes | Vgll3 | 3_prime_UTR_variant | MODIFIER |
| 16 | 66021932 | AX-97833685 | 0 | A | G | 0.509 | 1.58E-01 | 8.30E-02 | 2.66E+00 | 2.72E+00 | 5.66E-02 | Yes | | intergenic_variant | MODIFIER |
| 16 | 66148762 | AX-97813804 | 0 | C | T | 0.509 | 1.58E-01 | 8.30E-02 | 2.66E+00 | 2.72E+00 | 5.66E-02 | Yes | | intergenic_variant | MODIFIER |
| 5 | 45752798 | AX-170514595 | 0 | C | G | 0.5 | 1.90E-01 | 9.82E-02 | 2.30E+00 | 2.37E+00 | 5.68E-02 | Yes | Lcorl | intron_variant | MODIFIER |
| 19 | 61167037 | AX-170340137 | 1 | C | T | 0.366 | -1.13E-01 | 5.89E-02 | 2.49E+00 | 2.59E+00 | 5.68E-02 | no | Gm22365 | upstream_gene_variant | MODIFIER |
| 3 | 38619911 | AX-170309849 | 0 | A | G | 0.211 | 1.80E-01 | 9.17E-02 | 3.48E+00 | 3.65E+00 | 5.69E-02 | Yes | Gm43538 | downstream_gene_variant | MODIFIER |
| 16 | 65735035 | AX-97832102 | 1 | C | T | 0.509 | 1.58E-01 | 8.31E-02 | 2.67E+00 | 2.73E+00 | 5.70E-02 | Yes | | intergenic_variant | MODIFIER |
| 16 | 58290395 | AX-170305769 | 0 | A | G | 0.456 | -1.65E-01 | 8.67E-02 | 2.62E+00 | 2.65E+00 | 5.71E-02 | no | | intergenic_variant | MODIFIER |
| 16 | 58330351 | AX-97831428 | 0 | T | C | 0.456 | -1.65E-01 | 8.68E-02 | 2.62E+00 | 2.65E+00 | 5.71E-02 | no | | intergenic_variant | MODIFIER |
| 13 | 20346221 | AX-170499495 | 0 | G | A | 0.175 | -2.31E-01 | 1.21E-01 | 2.53E+00 | 2.60E+00 | 5.72E-02 | Yes | Elmo1 | intron_variant | MODIFIER |
| 10 | 104592185 | AX-97805751 | 1 | A | T | 0.295 | -1.58E-01 | 8.31E-02 | 2.76E+00 | 2.84E+00 | 5.73E-02 | no | | regulatory_region_variant | MODIFIER |
| 15 | 67938529 | AX-170334072 | 2 | A | G | 0.082 | -2.32E-01 | 1.08E-01 | 1.72E+00 | 1.77E+00 | 5.73E-02 | Yes | Gm49409 | downstream_gene_variant | MODIFIER |
| 16 | 58136820 | AX-170013140 | 0 | T | C | 0.456 | -1.65E-01 | 8.68E-02 | 2.62E+00 | 2.65E+00 | 5.75E-02 | no | | regulatory_region_variant | MODIFIER |
| 16 | 58141048 | AX-170057537 | 0 | T | C | 0.456 | -1.65E-01 | 8.68E-02 | 2.62E+00 | 2.65E+00 | 5.75E-02 | no | | intergenic_variant | MODIFIER |
| 16 | 58150673 | AX-170013144 | 0 | T | G | 0.456 | -1.65E-01 | 8.68E-02 | 2.62E+00 | 2.65E+00 | 5.75E-02 | no | | intergenic_variant | MODIFIER |
| 16 | 58157733 | AX-170013145 | 0 | C | T | 0.456 | -1.65E-01 | 8.68E-02 | 2.62E+00 | 2.65E+00 | 5.75E-02 | no | | intergenic_variant | MODIFIER |
| 16 | 58157841 | AX-170155986 | 0 | C | T | 0.456 | -1.65E-01 | 8.68E-02 | 2.62E+00 | 2.65E+00 | 5.75E-02 | no | | intergenic_variant | MODIFIER |
| 16 | 58198001 | AX-97831452 | 0 | T | G | 0.456 | -1.65E-01 | 8.68E-02 | 2.62E+00 | 2.65E+00 | 5.75E-02 | no | | intergenic_variant | MODIFIER |
| 16 | 58209325 | AX-170220685 | 0 | T | C | 0.456 | -1.65E-01 | 8.68E-02 | 2.62E+00 | 2.65E+00 | 5.75E-02 | no | | intergenic_variant | MODIFIER |
| 16 | 58209510 | AX-170339013 | 0 | A | G | 0.456 | -1.65E-01 | 8.68E-02 | 2.62E+00 | 2.65E+00 | 5.75E-02 | no | | intergenic_variant | MODIFIER |
| 16 | 58230784 | AX-170526987 | 0 | A | G | 0.456 | -1.65E-01 | 8.68E-02 | 2.62E+00 | 2.65E+00 | 5.75E-02 | no | | intergenic_variant | MODIFIER |
| 16 | 58230929 | AX-170099125 | 0 | A | G | 0.456 | -1.65E-01 | 8.68E-02 | 2.62E+00 | 2.65E+00 | 5.75E-02 | no | | intergenic_variant | MODIFIER |
| 16 | 58236142 | AX-170249757 | 0 | T | G | 0.456 | -1.65E-01 | 8.68E-02 | 2.62E+00 | 2.65E+00 | 5.75E-02 | no | Gm49574 | upstream_gene_variant | MODIFIER |
| 16 | 58239097 | AX-170352126 | 0 | G | A | 0.456 | -1.65E-01 | 8.68E-02 | 2.62E+00 | 2.65E+00 | 5.75E-02 | no | Gm49574 | non_coding_transcript_exon_variant | MODIFIER |
| 16 | 58246229 | AX-170373726 | 0 | C | T | 0.456 | -1.65E-01 | 8.68E-02 | 2.62E+00 | 2.65E+00 | 5.75E-02 | no | Gm49574 | intron_variant,non_coding_transcript_variant | MODIFIER |
| 16 | 58258158 | AX-170138358 | 0 | T | C | 0.456 | -1.65E-01 | 8.68E-02 | 2.62E+00 | 2.65E+00 | 5.75E-02 | no | | intergenic_variant | MODIFIER |
| 16 | 58267831 | AX-97842043 | 0 | G | A | 0.456 | -1.65E-01 | 8.68E-02 | 2.62E+00 | 2.65E+00 | 5.75E-02 | no | | intergenic_variant | MODIFIER |
| 16 | 58288611 | AX-170399508 | 0 | G | A | 0.456 | -1.65E-01 | 8.68E-02 | 2.62E+00 | 2.65E+00 | 5.75E-02 | no | | intergenic_variant | MODIFIER |
| 16 | 58288706 | AX-97796067 | 0 | A | C | 0.456 | -1.65E-01 | 8.68E-02 | 2.62E+00 | 2.65E+00 | 5.75E-02 | no | | intergenic_variant | MODIFIER |
| 16 | 58304188 | AX-170223578 | 0 | T | C | 0.456 | -1.65E-01 | 8.68E-02 | 2.62E+00 | 2.65E+00 | 5.75E-02 | no | | intergenic_variant | MODIFIER |
| 16 | 58353729 | AX-170264422 | 0 | C | T | 0.456 | -1.65E-01 | 8.68E-02 | 2.62E+00 | 2.65E+00 | 5.75E-02 | no | | intergenic_variant | MODIFIER |
| 16 | 58357340 | AX-170369679 | 0 | A | G | 0.456 | -1.65E-01 | 8.68E-02 | 2.62E+00 | 2.65E+00 | 5.75E-02 | no | | intergenic_variant | MODIFIER |
| 16 | 58370402 | AX-170197187 | 0 | A | G | 0.456 | -1.65E-01 | 8.68E-02 | 2.62E+00 | 2.65E+00 | 5.75E-02 | no | | intergenic_variant | MODIFIER |
| 16 | 58377040 | AX-97832741 | 0 | A | G | 0.456 | -1.65E-01 | 8.68E-02 | 2.62E+00 | 2.65E+00 | 5.75E-02 | no | | intergenic_variant | MODIFIER |
| 16 | 58406093 | AX-170099553 | 0 | C | T | 0.456 | -1.65E-01 | 8.68E-02 | 2.62E+00 | 2.65E+00 | 5.75E-02 | no | Dcbld2 | upstream_gene_variant | MODIFIER |
| 16 | 58423859 | AX-170131382 | 0 | G | C | 0.456 | -1.65E-01 | 8.68E-02 | 2.62E+00 | 2.65E+00 | 5.75E-02 | no | Dcbld2 | intron_variant | MODIFIER |
| 16 | 58522140 | AX-170013149 | 0 | C | T | 0.456 | -1.65E-01 | 8.68E-02 | 2.62E+00 | 2.65E+00 | 5.75E-02 | no | St3gal6 | intron_variant | MODIFIER |
| 16 | 58538706 | AX-170013150 | 0 | C | A | 0.456 | -1.65E-01 | 8.68E-02 | 2.62E+00 | 2.65E+00 | 5.75E-02 | no | | intergenic_variant | MODIFIER |
| 16 | 58539148 | AX-170013146 | 0 | G | A | 0.456 | -1.65E-01 | 8.68E-02 | 2.62E+00 | 2.65E+00 | 5.75E-02 | no | | regulatory_region_variant | MODIFIER |
| 16 | 58543254 | AX-97829355 | 0 | T | C | 0.456 | -1.65E-01 | 8.68E-02 | 2.62E+00 | 2.65E+00 | 5.75E-02 | no | | regulatory_region_variant | MODIFIER |
| 16 | 58555882 | AX-97804999 | 0 | A | G | 0.456 | -1.65E-01 | 8.68E-02 | 2.62E+00 | 2.65E+00 | 5.75E-02 | no | Gm20942 | upstream_gene_variant | MODIFIER |

| | | | | | | | | | | | | | | | |
|---|---|---|---|---|---|---|---|---|---|---|---|---|---|---|---|
| 16 | 58583634 | AX-97842339 | 0 | T | C | 0.456 | -1.65E-01 | 8.68E-02 | 2.62E+00 | 2.65E+00 | 5.75E-02 | no | | intergenic_variant | MODIFIER |
| 16 | 58678865 | AX-170371907 | 0 | C | T | 0.456 | -1.65E-01 | 8.68E-02 | 2.62E+00 | 2.65E+00 | 5.75E-02 | no | Cpox | 3_prime_UTR_variant | MODIFIER |
| 16 | 58689731 | AX-168380613 | 0 | G | A | 0.456 | -1.65E-01 | 8.68E-02 | 2.62E+00 | 2.65E+00 | 5.75E-02 | no | Cpox | intron_variant,non_coding_transcript_variant | MODIFIER |
| 16 | 58701727 | AX-170446492 | 0 | A | G | 0.456 | -1.65E-01 | 8.68E-02 | 2.62E+00 | 2.65E+00 | 5.75E-02 | no | Cpox | intron_variant,non_coding_transcript_variant | MODIFIER |
| 16 | 58702705 | AX-170346708 | 0 | A | G | 0.456 | -1.65E-01 | 8.68E-02 | 2.62E+00 | 2.65E+00 | 5.75E-02 | no | Cpox | intron_variant,non_coding_transcript_variant | MODIFIER |
| 16 | 58702756 | AX-170169599 | 0 | A | G | 0.456 | -1.65E-01 | 8.68E-02 | 2.62E+00 | 2.65E+00 | 5.75E-02 | no | Cpox | intron_variant,non_coding_transcript_variant | MODIFIER |
| 16 | 58853327 | AX-170431834 | 0 | G | A | 0.456 | -1.65E-01 | 8.68E-02 | 2.62E+00 | 2.65E+00 | 5.75E-02 | no | | intergenic_variant | MODIFIER |
| 16 | 58918879 | AX-170144149 | 0 | T | C | 0.456 | -1.65E-01 | 8.68E-02 | 2.62E+00 | 2.65E+00 | 5.75E-02 | no | Olfr180 | upstream_gene_variant | MODIFIER |
| 16 | 59397226 | AX-170235695 | 0 | A | G | 0.456 | -1.65E-01 | 8.68E-02 | 2.62E+00 | 2.65E+00 | 5.75E-02 | no | | intergenic_variant | MODIFIER |
| 16 | 67696572 | AX-170249057 | 1 | C | T | 0.491 | 1.44E-01 | 7.58E-02 | 2.60E+00 | 2.68E+00 | 5.75E-02 | Yes | | intergenic_variant | MODIFIER |
| 13 | 16752438 | AX-170124195 | 0 | G | A | 0.184 | -2.30E-01 | 1.20E-01 | 2.36E+00 | 2.42E+00 | 5.77E-02 | Yes | | intergenic_variant | MODIFIER |
| 7 | 145310320 | AX-170343448 | 0 | T | C | 0.202 | -1.42E-01 | 7.51E-02 | 3.00E+00 | 3.17E+00 | 5.83E-02 | no | Mrgprf | downstream_gene_variant | MODIFIER |
| 5 | 44466312 | AX-97786606 | 2 | C | T | 0.209 | -1.94E-01 | 1.02E-01 | 2.95E+00 | 3.09E+00 | 5.83E-02 | Yes | | intergenic_variant | MODIFIER |
| 10 | 105078989 | AX-97824201 | 0 | G | A | 0.307 | -1.52E-01 | 8.03E-02 | 2.86E+00 | 2.97E+00 | 5.83E-02 | no | | intergenic_variant | MODIFIER |
| 15 | 20192902 | AX-170011184 | 2 | G | T | 0.227 | -1.83E-01 | 9.68E-02 | 2.85E+00 | 2.94E+00 | 5.84E-02 | Yes | | intergenic_variant | MODIFIER |
| 15 | 4269136 | AX-170054680 | 1 | A | T | 0.223 | -2.01E-01 | 1.07E-01 | 2.90E+00 | 2.98E+00 | 5.87E-02 | Yes | | intergenic_variant | MODIFIER |
| 5 | 54463935 | AX-170383754 | 0 | G | A | 0.491 | 1.88E-01 | 9.95E-02 | 2.61E+00 | 2.69E+00 | 5.87E-02 | Yes | | intergenic_variant | MODIFIER |
| 5 | 62474585 | AX-97807833 | 1 | A | C | 0.473 | 1.88E-01 | 9.85E-02 | 2.36E+00 | 2.42E+00 | 5.87E-02 | Yes | | intergenic_variant | MODIFIER |
| 3 | 19833411 | AX-97811959 | 0 | A | G | 0.123 | -1.93E-01 | 1.02E-01 | 2.70E+00 | 2.78E+00 | 5.88E-02 | Yes | | intergenic_variant | MODIFIER |
| 3 | 19835809 | AX-97830962 | 0 | A | G | 0.123 | -1.93E-01 | 1.02E-01 | 2.70E+00 | 2.78E+00 | 5.88E-02 | Yes | | intergenic_variant | MODIFIER |
| 3 | 19857565 | AX-170288958 | 0 | A | G | 0.123 | -1.93E-01 | 1.02E-01 | 2.70E+00 | 2.78E+00 | 5.88E-02 | Yes | | intergenic_variant | MODIFIER |
| 3 | 19860068 | AX-170094541 | 0 | G | A | 0.123 | -1.93E-01 | 1.02E-01 | 2.70E+00 | 2.78E+00 | 5.88E-02 | Yes | | intergenic_variant | MODIFIER |
| 3 | 19894310 | AX-97792742 | 0 | G | A | 0.123 | -1.93E-01 | 1.02E-01 | 2.70E+00 | 2.78E+00 | 5.88E-02 | Yes | 4632415L05Rik | upstream_gene_variant | MODIFIER |
| 3 | 19895350 | AX-170146967 | 0 | A | G | 0.123 | -1.93E-01 | 1.02E-01 | 2.70E+00 | 2.78E+00 | 5.88E-02 | Yes | 4632415L05Rik | non_coding_transcript_exon_variant | MODIFIER |
| 3 | 19895484 | AX-170118098 | 0 | A | G | 0.123 | -1.93E-01 | 1.02E-01 | 2.70E+00 | 2.78E+00 | 5.88E-02 | Yes | 4632415L05Rik | non_coding_transcript_exon_variant | MODIFIER |
| 8 | 32373334 | AX-170096217 | 2 | T | C | 0.136 | -2.23E-01 | 1.17E-01 | 3.15E+00 | 3.32E+00 | 5.88E-02 | Yes | Nrg1 | intron_variant | MODIFIER |
| 5 | 63830737 | AX-170086211 | 0 | C | T | 0.491 | 1.92E-01 | 1.01E-01 | 2.49E+00 | 2.55E+00 | 5.93E-02 | Yes | 0610040J01Rik | intron_variant | MODIFIER |
| 5 | 53239340 | AX-170171297 | 1 | T | G | 0.482 | 1.83E-01 | 9.54E-02 | 2.32E+00 | 2.38E+00 | 5.94E-02 | Yes | | intergenic_variant | MODIFIER |
| 16 | 58319057 | AX-170469587 | 1 | T | A | 0.464 | -1.58E-01 | 8.36E-02 | 2.57E+00 | 2.61E+00 | 5.97E-02 | no | Gm37825 | non_coding_transcript_exon_variant | MODIFIER |
| 18 | 76028611 | AX-170332661 | 0 | A | G | 0.167 | -1.96E-01 | 1.04E-01 | 2.86E+00 | 2.97E+00 | 5.98E-02 | Yes | Zbtb7c | intron_variant | MODIFIER |
| 5 | 52339409 | AX-170358504 | 0 | C | T | 0.5 | 1.95E-01 | 1.03E-01 | 2.47E+00 | 2.51E+00 | 5.98E-02 | Yes | Gm25601 | downstream_gene_variant | MODIFIER |
| 1 | 184386948 | AX-170398203 | 2 | C | T | 0.236 | 2.16E-01 | 1.15E-01 | 2.63E+00 | 2.70E+00 | 6.00E-02 | Yes | Gm37223 | intron_variant,non_coding_transcript_variant | MODIFIER |
| 4 | 103794094 | AX-170354063 | 0 | T | C | 0.246 | 1.98E-01 | 1.05E-01 | 2.86E+00 | 2.96E+00 | 6.00E-02 | Yes | Dab1 | intron_variant | MODIFIER |
| 5 | 57615776 | AX-97811240 | 0 | G | A | 0.474 | 1.83E-01 | 9.62E-02 | 2.34E+00 | 2.40E+00 | 6.02E-02 | Yes | 4932441J04Rik | intron_variant,non_coding_transcript_variant | MODIFIER |
| 5 | 64045964 | AX-170025148 | 1 | C | A | 0.491 | 1.93E-01 | 1.02E-01 | 2.49E+00 | 2.55E+00 | 6.04E-02 | Yes | 5830416I19Rik | intron_variant,non_coding_transcript_variant | MODIFIER |
| 13 | 8487844 | AX-170452240 | 0 | C | T | 0.184 | -2.25E-01 | 1.19E-01 | 2.41E+00 | 2.47E+00 | 6.05E-02 | Yes | Adarb2 | intron_variant | MODIFIER |
| 13 | 11418622 | AX-97830709 | 0 | T | C | 0.184 | -2.25E-01 | 1.19E-01 | 2.41E+00 | 2.47E+00 | 6.05E-02 | Yes | | intergenic_variant | MODIFIER |
| 1 | 135525020 | AX-170001771 | 0 | A | T | 0.544 | 1.54E-01 | 8.23E-02 | 2.79E+00 | 2.87E+00 | 6.05E-02 | Yes | Nav1 | intron_variant | MODIFIER |
| 10 | 105018451 | AX-97843027 | 1 | C | A | 0.295 | -1.57E-01 | 8.37E-02 | 2.70E+00 | 2.78E+00 | 6.05E-02 | no | | intergenic_variant | MODIFIER |
| 13 | 11311798 | AX-170050531 | 1 | G | T | 0.179 | -2.05E-01 | 1.09E-01 | 2.43E+00 | 2.50E+00 | 6.05E-02 | Yes | | intergenic_variant | MODIFIER |
| 5 | 56509474 | AX-170089482 | 0 | G | A | 0.465 | 1.94E-01 | 1.03E-01 | 2.55E+00 | 2.61E+00 | 6.06E-02 | Yes | | intergenic_variant | MODIFIER |
| 5 | 64987131 | AX-170494678 | 0 | G | A | 0.482 | 1.93E-01 | 1.02E-01 | 2.48E+00 | 2.53E+00 | 6.06E-02 | Yes | Fam114a1 | intron_variant | MODIFIER |
| 1 | 185767427 | AX-170219446 | 1 | A | G | 0.232 | 2.04E-01 | 1.09E-01 | 2.62E+00 | 2.69E+00 | 6.07E-02 | Yes | | intergenic_variant | MODIFIER |
| 15 | 77123393 | AX-170105141 | 1 | G | A | 0.08 | -2.29E-01 | 1.07E-01 | 1.68E+00 | 1.73E+00 | 6.09E-02 | Yes | Rbfox2 | intron_variant | MODIFIER |
| 5 | 135342311 | AX-170494704 | 2 | C | T | 0.145 | -1.73E-01 | 8.99E-02 | 2.21E+00 | 2.29E+00 | 6.09E-02 | no | Fkbp6 | intron_variant | MODIFIER |
| 13 | 18249320 | AX-170292918 | 1 | C | T | 0.152 | -2.33E-01 | 1.24E-01 | 2.46E+00 | 2.52E+00 | 6.10E-02 | Yes | Pou6f2 | intron_variant | MODIFIER |
| 15 | 19373752 | AX-170344195 | 0 | T | G | 0.228 | -2.03E-01 | 1.09E-01 | 2.79E+00 | 2.86E+00 | 6.12E-02 | Yes | Gm49162 | intron_variant,non_coding_transcript_variant | MODIFIER |
| 2 | 52443377 | AX-170321814 | 1 | C | T | 0.036 | -3.11E-01 | 1.67E-01 | 2.69E+00 | 2.81E+00 | 6.14E-02 | Yes | Cacnb4 | intron_variant | MODIFIER |
| 5 | 55014139 | AX-170072164 | 1 | A | C | 0.464 | 1.94E-01 | 1.03E-01 | 2.47E+00 | 2.52E+00 | 6.16E-02 | Yes | | intergenic_variant | MODIFIER |
| 15 | 12365586 | AX-170054822 | 0 | G | A | 0.298 | -1.59E-01 | 7.64E-02 | 1.76E+00 | 1.79E+00 | 6.17E-02 | no | Pdzd2 | intron_variant | MODIFIER |
| 13 | 19624856 | AX-170362654 | 0 | T | C | 0.167 | -2.33E-01 | 1.25E-01 | 2.56E+00 | 2.62E+00 | 6.17E-02 | Yes | Sfrp4 | intron_variant | MODIFIER |
| 16 | 56925156 | AX-170088030 | 0 | A | G | 0.465 | -1.52E-01 | 8.11E-02 | 2.58E+00 | 2.64E+00 | 6.18E-02 | no | Lnp1 | intron_variant | MODIFIER |
| 4 | 84294334 | AX-170533108 | 1 | G | A | 0.348 | 2.17E-01 | 1.15E-01 | 3.22E+00 | 3.33E+00 | 6.19E-02 | Yes | Bnc2 | intron_variant | MODIFIER |
| 1 | 43328108 | AX-170178081 | 0 | G | T | 0.096 | -3.38E-01 | 1.76E-01 | 3.40E+00 | 3.58E+00 | 6.19E-02 | Yes | | intergenic_variant | MODIFIER |

| | | | | | | | | | | | | | | | |
|---|---|---|---|---|---|---|---|---|---|---|---|---|---|---|---|
| 1 | 134529958 | AX-170154954 | 0 | C | T | 0.544 | 1.50E-01 | 8.05E-02 | 2.87E+00 | 2.96E+00 | 6.24E-02 | Yes | | intergenic_variant | MODIFIER |
| 19 | 60049924 | AX-170153934 | 0 | C | G | 0.526 | 1.17E-01 | 6.29E-02 | 2.77E+00 | 2.86E+00 | 6.26E-02 | no | | intergenic_variant | MODIFIER |
| 3 | 23416718 | AX-170318676 | 0 | G | A | 0.158 | -1.67E-01 | 9.02E-02 | 2.79E+00 | 2.93E+00 | 6.26E-02 | no | | intergenic_variant | MODIFIER |
| 3 | 23572826 | AX-97829702 | 0 | C | T | 0.158 | -1.67E-01 | 9.02E-02 | 2.79E+00 | 2.93E+00 | 6.26E-02 | no | | intergenic_variant | MODIFIER |
| 4 | 83820336 | AX-170525859 | 0 | C | T | 0.342 | 2.08E-01 | 1.12E-01 | 2.93E+00 | 3.01E+00 | 6.27E-02 | Yes | Ccdc171 | intron_variant | MODIFIER |
| 5 | 55576551 | AX-170429470 | 1 | A | G | 0.473 | 1.93E-01 | 1.03E-01 | 2.38E+00 | 2.43E+00 | 6.30E-02 | Yes | | intergenic_variant | MODIFIER |
| 5 | 58315676 | AX-170474944 | 1 | C | T | 0.473 | 1.93E-01 | 1.03E-01 | 2.38E+00 | 2.43E+00 | 6.30E-02 | Yes | | intergenic_variant | MODIFIER |
| 5 | 52686221 | AX-170437278 | 0 | C | T | 0.491 | 1.86E-01 | 9.85E-02 | 2.32E+00 | 2.37E+00 | 6.30E-02 | Yes | | intergenic_variant | MODIFIER |
| 5 | 58149030 | AX-170072214 | 1 | G | A | 0.464 | 1.93E-01 | 1.03E-01 | 2.46E+00 | 2.51E+00 | 6.32E-02 | Yes | Gm42976 | downstream_gene_variant | MODIFIER |
| 13 | 8827632 | AX-170213079 | 0 | G | A | 0.184 | -2.31E-01 | 1.24E-01 | 2.48E+00 | 2.53E+00 | 6.33E-02 | Yes | Wdr37 | intron_variant | MODIFIER |
| 13 | 9908660 | AX-170368756 | 0 | C | A | 0.184 | -2.31E-01 | 1.24E-01 | 2.48E+00 | 2.53E+00 | 6.33E-02 | Yes | Chrm3 | intron_variant | MODIFIER |
| 1 | 44440980 | AX-170254123 | 1 | A | C | 0.098 | -3.19E-01 | 1.70E-01 | 3.15E+00 | 3.30E+00 | 6.33E-02 | Yes | | intergenic_variant | MODIFIER |
| 5 | 135650925 | AX-97833286 | 1 | G | A | 0.152 | -1.61E-01 | 8.46E-02 | 2.21E+00 | 2.29E+00 | 6.34E-02 | no | Rhbdd2 | downstream_gene_variant | MODIFIER |
| 5 | 56451452 | AX-170121815 | 1 | C | T | 0.482 | 1.79E-01 | 9.43E-02 | 2.26E+00 | 2.33E+00 | 6.35E-02 | Yes | Gm42977 | intron_variant,non_coding_transcript_variant | MODIFIER |
| 5 | 135700695 | AX-170313955 | 0 | A | G | 0.149 | -1.59E-01 | 8.39E-02 | 2.27E+00 | 2.36E+00 | 6.35E-02 | no | Por | intron_variant | MODIFIER |
| 18 | 79965715 | AX-170016064 | 0 | G | A | 0.465 | 1.48E-01 | 7.95E-02 | 2.51E+00 | 2.59E+00 | 6.36E-02 | Yes | | intergenic_variant | MODIFIER |
| 3 | 38700121 | AX-170194596 | 0 | C | A | 0.211 | 1.64E-01 | 8.60E-02 | 3.38E+00 | 3.55E+00 | 6.37E-02 | Yes | | intergenic_variant | MODIFIER |
| 13 | 5790531 | AX-170488462 | 0 | A | G | 0.202 | -2.06E-01 | 1.11E-01 | 2.77E+00 | 2.86E+00 | 6.38E-02 | Yes | | intergenic_variant | MODIFIER |
| 5 | 58092909 | AX-170352999 | 0 | T | C | 0.456 | 1.86E-01 | 1.00E-01 | 2.49E+00 | 2.56E+00 | 6.39E-02 | Yes | Pcdh7 | intron_variant | MODIFIER |
| 9 | 50398383 | AX-170238385 | 0 | A | G | 0.211 | -1.90E-01 | 1.01E-01 | 3.28E+00 | 3.47E+00 | 6.40E-02 | Yes | | intergenic_variant | MODIFIER |
| 5 | 54970926 | AX-97799917 | 0 | C | T | 0.465 | 1.92E-01 | 1.03E-01 | 2.45E+00 | 2.50E+00 | 6.40E-02 | Yes | | intergenic_variant | MODIFIER |
| 5 | 54971311 | AX-170324125 | 0 | T | C | 0.465 | 1.92E-01 | 1.03E-01 | 2.45E+00 | 2.50E+00 | 6.40E-02 | Yes | | intergenic_variant | MODIFIER |
| 5 | 54971908 | AX-170160751 | 0 | T | C | 0.465 | 1.92E-01 | 1.03E-01 | 2.45E+00 | 2.50E+00 | 6.40E-02 | Yes | | intergenic_variant | MODIFIER |
| 5 | 55227125 | AX-170089481 | 0 | G | A | 0.465 | 1.92E-01 | 1.03E-01 | 2.45E+00 | 2.50E+00 | 6.40E-02 | Yes | | intergenic_variant | MODIFIER |
| 5 | 55284484 | AX-97783785 | 0 | T | C | 0.465 | 1.92E-01 | 1.03E-01 | 2.45E+00 | 2.50E+00 | 6.40E-02 | Yes | | intergenic_variant | MODIFIER |
| 5 | 55298570 | AX-170452693 | 0 | T | C | 0.465 | 1.92E-01 | 1.03E-01 | 2.45E+00 | 2.50E+00 | 6.40E-02 | Yes | Gm25513 | downstream_gene_variant | MODIFIER |
| 5 | 55302873 | AX-170024986 | 0 | A | G | 0.465 | 1.92E-01 | 1.03E-01 | 2.45E+00 | 2.50E+00 | 6.40E-02 | Yes | Gm25513 | downstream_gene_variant | MODIFIER |
| 5 | 55310595 | AX-170143454 | 0 | G | T | 0.465 | 1.92E-01 | 1.03E-01 | 2.45E+00 | 2.50E+00 | 6.40E-02 | Yes | | intergenic_variant | MODIFIER |
| 5 | 55312236 | AX-170242905 | 0 | T | C | 0.465 | 1.92E-01 | 1.03E-01 | 2.45E+00 | 2.50E+00 | 6.40E-02 | Yes | | intergenic_variant | MODIFIER |
| 5 | 55317247 | AX-170024990 | 0 | C | G | 0.465 | 1.92E-01 | 1.03E-01 | 2.45E+00 | 2.50E+00 | 6.40E-02 | Yes | | intergenic_variant | MODIFIER |
| 5 | 55346074 | AX-97851463 | 0 | T | C | 0.465 | 1.92E-01 | 1.03E-01 | 2.45E+00 | 2.50E+00 | 6.40E-02 | Yes | | intergenic_variant | MODIFIER |
| 5 | 55421164 | AX-97819499 | 0 | G | A | 0.465 | 1.92E-01 | 1.03E-01 | 2.45E+00 | 2.50E+00 | 6.40E-02 | Yes | | intergenic_variant | MODIFIER |
| 5 | 55452500 | AX-170024989 | 0 | A | T | 0.465 | 1.92E-01 | 1.03E-01 | 2.45E+00 | 2.50E+00 | 6.40E-02 | Yes | | intergenic_variant | MODIFIER |
| 5 | 55576882 | AX-170405153 | 0 | G | T | 0.465 | 1.92E-01 | 1.03E-01 | 2.45E+00 | 2.50E+00 | 6.40E-02 | Yes | | intergenic_variant | MODIFIER |
| 5 | 55660750 | AX-170327949 | 0 | T | C | 0.465 | 1.92E-01 | 1.03E-01 | 2.45E+00 | 2.50E+00 | 6.40E-02 | Yes | | intergenic_variant | MODIFIER |
| 5 | 55672975 | AX-170318081 | 0 | C | T | 0.465 | 1.92E-01 | 1.03E-01 | 2.45E+00 | 2.50E+00 | 6.40E-02 | Yes | | intergenic_variant | MODIFIER |
| 5 | 55692555 | AX-170211165 | 0 | T | G | 0.465 | 1.92E-01 | 1.03E-01 | 2.45E+00 | 2.50E+00 | 6.40E-02 | Yes | | intergenic_variant | MODIFIER |
| 5 | 55695975 | AX-170506182 | 0 | G | C | 0.465 | 1.92E-01 | 1.03E-01 | 2.45E+00 | 2.50E+00 | 6.40E-02 | Yes | | intergenic_variant | MODIFIER |
| 5 | 55714287 | AX-170135077 | 0 | T | C | 0.465 | 1.92E-01 | 1.03E-01 | 2.45E+00 | 2.50E+00 | 6.40E-02 | Yes | | intergenic_variant | MODIFIER |
| 5 | 55722720 | AX-170116420 | 0 | C | T | 0.465 | 1.92E-01 | 1.03E-01 | 2.45E+00 | 2.50E+00 | 6.40E-02 | Yes | Gm20223 | non_coding_transcript_exon_variant | MODIFIER |
| 5 | 55770260 | AX-170391449 | 0 | T | G | 0.465 | 1.92E-01 | 1.03E-01 | 2.45E+00 | 2.50E+00 | 6.40E-02 | Yes | | intergenic_variant | MODIFIER |
| 5 | 55771785 | AX-170409175 | 0 | G | T | 0.465 | 1.92E-01 | 1.03E-01 | 2.45E+00 | 2.50E+00 | 6.40E-02 | Yes | | intergenic_variant | MODIFIER |
| 5 | 55774109 | AX-170072175 | 0 | T | G | 0.465 | 1.92E-01 | 1.03E-01 | 2.45E+00 | 2.50E+00 | 6.40E-02 | Yes | | intergenic_variant | MODIFIER |
| 5 | 55813784 | AX-97832035 | 0 | T | C | 0.465 | 1.92E-01 | 1.03E-01 | 2.45E+00 | 2.50E+00 | 6.40E-02 | Yes | | intergenic_variant | MODIFIER |
| 5 | 55814002 | AX-170024997 | 0 | A | C | 0.465 | 1.92E-01 | 1.03E-01 | 2.45E+00 | 2.50E+00 | 6.40E-02 | Yes | | intergenic_variant | MODIFIER |
| 5 | 56078027 | AX-170520713 | 0 | C | T | 0.465 | 1.92E-01 | 1.03E-01 | 2.45E+00 | 2.50E+00 | 6.40E-02 | Yes | | intergenic_variant | MODIFIER |
| 5 | 56326454 | AX-97832113 | 0 | G | C | 0.465 | 1.92E-01 | 1.03E-01 | 2.45E+00 | 2.50E+00 | 6.40E-02 | Yes | | intergenic_variant | MODIFIER |
| 5 | 56366981 | AX-97807274 | 0 | A | G | 0.465 | 1.92E-01 | 1.03E-01 | 2.45E+00 | 2.50E+00 | 6.40E-02 | Yes | | intergenic_variant | MODIFIER |
| 5 | 56409124 | AX-97785640 | 0 | T | C | 0.465 | 1.92E-01 | 1.03E-01 | 2.45E+00 | 2.50E+00 | 6.40E-02 | Yes | | intergenic_variant | MODIFIER |
| 5 | 56415087 | AX-170025015 | 0 | A | G | 0.465 | 1.92E-01 | 1.03E-01 | 2.45E+00 | 2.50E+00 | 6.40E-02 | Yes | Gm42977 | intron_variant,non_coding_transcript_variant | MODIFIER |
| 5 | 56449111 | AX-170319011 | 0 | T | A | 0.465 | 1.92E-01 | 1.03E-01 | 2.45E+00 | 2.50E+00 | 6.40E-02 | Yes | Gm42977 | intron_variant,non_coding_transcript_variant | MODIFIER |
| 5 | 56509403 | AX-170025002 | 0 | C | T | 0.465 | 1.92E-01 | 1.03E-01 | 2.45E+00 | 2.50E+00 | 6.40E-02 | Yes | | intergenic_variant | MODIFIER |
| 5 | 56539813 | AX-170025001 | 0 | A | G | 0.465 | 1.92E-01 | 1.03E-01 | 2.45E+00 | 2.50E+00 | 6.40E-02 | Yes | | intergenic_variant | MODIFIER |
| 5 | 56617346 | AX-97794407 | 0 | C | A | 0.465 | 1.92E-01 | 1.03E-01 | 2.45E+00 | 2.50E+00 | 6.40E-02 | Yes | | intergenic_variant | MODIFIER |

| | | | | | | | | | | | | | | | |
|---|---|---|---|---|---|---|---|---|---|---|---|---|---|---|---|
| 5 | 56637941 | AX-97804447 | 0 | C | T | 0.465 | 1.92E-01 | 1.03E-01 | 2.45E+00 | 2.50E+00 | 6.40E-02 | Yes | | intergenic_variant | MODIFIER |
| 5 | 56649455 | AX-170025014 | 0 | A | G | 0.465 | 1.92E-01 | 1.03E-01 | 2.45E+00 | 2.50E+00 | 6.40E-02 | Yes | | intergenic_variant | MODIFIER |
| 5 | 56690171 | AX-170455039 | 0 | G | C | 0.465 | 1.92E-01 | 1.03E-01 | 2.45E+00 | 2.50E+00 | 6.40E-02 | Yes | | intergenic_variant | MODIFIER |
| 5 | 56721777 | AX-170025013 | 0 | A | G | 0.465 | 1.92E-01 | 1.03E-01 | 2.45E+00 | 2.50E+00 | 6.40E-02 | Yes | | intergenic_variant | MODIFIER |
| 5 | 56727705 | AX-170025005 | 0 | G | C | 0.465 | 1.92E-01 | 1.03E-01 | 2.45E+00 | 2.50E+00 | 6.40E-02 | Yes | | intergenic_variant | MODIFIER |
| 5 | 56738061 | AX-97803176 | 0 | T | G | 0.465 | 1.92E-01 | 1.03E-01 | 2.45E+00 | 2.50E+00 | 6.40E-02 | Yes | Gm23896 | downstream_gene_variant | MODIFIER |
| 5 | 56738134 | AX-170360879 | 0 | A | G | 0.465 | 1.92E-01 | 1.03E-01 | 2.45E+00 | 2.50E+00 | 6.40E-02 | Yes | Gm23896 | downstream_gene_variant | MODIFIER |
| 5 | 56797036 | AX-170101572 | 0 | T | C | 0.465 | 1.92E-01 | 1.03E-01 | 2.45E+00 | 2.50E+00 | 6.40E-02 | Yes | | intergenic_variant | MODIFIER |
| 5 | 56831672 | AX-97803207 | 0 | G | A | 0.465 | 1.92E-01 | 1.03E-01 | 2.45E+00 | 2.50E+00 | 6.40E-02 | Yes | | intergenic_variant | MODIFIER |
| 5 | 56838565 | AX-170160934 | 0 | C | T | 0.465 | 1.92E-01 | 1.03E-01 | 2.45E+00 | 2.50E+00 | 6.40E-02 | Yes | | intergenic_variant | MODIFIER |
| 5 | 56855826 | AX-97849408 | 0 | C | T | 0.465 | 1.92E-01 | 1.03E-01 | 2.45E+00 | 2.50E+00 | 6.40E-02 | Yes | | intergenic_variant | MODIFIER |
| 5 | 57082110 | AX-97799966 | 0 | T | G | 0.465 | 1.92E-01 | 1.03E-01 | 2.45E+00 | 2.50E+00 | 6.40E-02 | Yes | | intergenic_variant | MODIFIER |
| 5 | 57393122 | AX-170196095 | 0 | G | C | 0.465 | 1.92E-01 | 1.03E-01 | 2.45E+00 | 2.50E+00 | 6.40E-02 | Yes | | intergenic_variant | MODIFIER |
| 5 | 57394126 | AX-170465230 | 0 | G | C | 0.465 | 1.92E-01 | 1.03E-01 | 2.45E+00 | 2.50E+00 | 6.40E-02 | Yes | | intergenic_variant | MODIFIER |
| 5 | 57403943 | AX-170447114 | 0 | A | G | 0.465 | 1.92E-01 | 1.03E-01 | 2.45E+00 | 2.50E+00 | 6.40E-02 | Yes | | intergenic_variant | MODIFIER |
| 5 | 57410022 | AX-170072202 | 0 | T | G | 0.465 | 1.92E-01 | 1.03E-01 | 2.45E+00 | 2.50E+00 | 6.40E-02 | Yes | | intergenic_variant | MODIFIER |
| 5 | 57417477 | AX-170506208 | 0 | A | C | 0.465 | 1.92E-01 | 1.03E-01 | 2.45E+00 | 2.50E+00 | 6.40E-02 | Yes | | intergenic_variant | MODIFIER |
| 5 | 57457796 | AX-170072199 | 0 | C | G | 0.465 | 1.92E-01 | 1.03E-01 | 2.45E+00 | 2.50E+00 | 6.40E-02 | Yes | Gm43706 | downstream_gene_variant | MODIFIER |
| 5 | 57457846 | AX-170524092 | 0 | A | C | 0.465 | 1.92E-01 | 1.03E-01 | 2.45E+00 | 2.50E+00 | 6.40E-02 | Yes | Gm43706 | downstream_gene_variant | MODIFIER |
| 5 | 57487374 | AX-170366492 | 0 | A | G | 0.465 | 1.92E-01 | 1.03E-01 | 2.45E+00 | 2.50E+00 | 6.40E-02 | Yes | | intergenic_variant | MODIFIER |
| 5 | 57505366 | AX-170025028 | 0 | T | G | 0.465 | 1.92E-01 | 1.03E-01 | 2.45E+00 | 2.50E+00 | 6.40E-02 | Yes | | intergenic_variant | MODIFIER |
| 5 | 57505693 | AX-170025027 | 0 | C | G | 0.465 | 1.92E-01 | 1.03E-01 | 2.45E+00 | 2.50E+00 | 6.40E-02 | Yes | | intergenic_variant | MODIFIER |
| 5 | 57520100 | AX-170025030 | 0 | C | T | 0.465 | 1.92E-01 | 1.03E-01 | 2.45E+00 | 2.50E+00 | 6.40E-02 | Yes | | intergenic_variant | MODIFIER |
| 5 | 57529753 | AX-170417664 | 0 | G | A | 0.465 | 1.92E-01 | 1.03E-01 | 2.45E+00 | 2.50E+00 | 6.40E-02 | Yes | | intergenic_variant | MODIFIER |
| 5 | 57532417 | AX-170025031 | 0 | A | G | 0.465 | 1.92E-01 | 1.03E-01 | 2.45E+00 | 2.50E+00 | 6.40E-02 | Yes | | intergenic_variant | MODIFIER |
| 5 | 57737553 | AX-170127288 | 0 | C | G | 0.465 | 1.92E-01 | 1.03E-01 | 2.45E+00 | 2.50E+00 | 6.40E-02 | Yes | Pcdh7 | intron_variant | MODIFIER |
| 5 | 57743828 | AX-170298850 | 0 | G | A | 0.465 | 1.92E-01 | 1.03E-01 | 2.45E+00 | 2.50E+00 | 6.40E-02 | Yes | Pcdh7 | intron_variant | MODIFIER |
| 5 | 57755641 | AX-170232409 | 0 | A | G | 0.465 | 1.92E-01 | 1.03E-01 | 2.45E+00 | 2.50E+00 | 6.40E-02 | Yes | Pcdh7 | intron_variant | MODIFIER |
| 5 | 57761449 | AX-170320162 | 0 | A | G | 0.465 | 1.92E-01 | 1.03E-01 | 2.45E+00 | 2.50E+00 | 6.40E-02 | Yes | Pcdh7 | intron_variant | MODIFIER |
| 5 | 57885067 | AX-97842226 | 0 | C | A | 0.465 | 1.92E-01 | 1.03E-01 | 2.45E+00 | 2.50E+00 | 6.40E-02 | Yes | Pcdh7 | intron_variant | MODIFIER |
| 5 | 58085415 | AX-170177347 | 0 | A | C | 0.465 | 1.92E-01 | 1.03E-01 | 2.45E+00 | 2.50E+00 | 6.40E-02 | Yes | Pcdh7 | intron_variant | MODIFIER |
| 5 | 58085774 | AX-170223296 | 0 | T | A | 0.465 | 1.92E-01 | 1.03E-01 | 2.45E+00 | 2.50E+00 | 6.40E-02 | Yes | Pcdh7 | intron_variant | MODIFIER |
| 5 | 58086950 | AX-97849041 | 0 | T | C | 0.465 | 1.92E-01 | 1.03E-01 | 2.45E+00 | 2.50E+00 | 6.40E-02 | Yes | Pcdh7 | intron_variant | MODIFIER |
| 5 | 58151802 | AX-170072213 | 0 | A | G | 0.465 | 1.92E-01 | 1.03E-01 | 2.45E+00 | 2.50E+00 | 6.40E-02 | Yes | Gm42976 | upstream_gene_variant | MODIFIER |
| 5 | 58239428 | AX-170362608 | 0 | A | G | 0.465 | 1.92E-01 | 1.03E-01 | 2.45E+00 | 2.50E+00 | 6.40E-02 | Yes | | intergenic_variant | MODIFIER |
| 5 | 58253130 | AX-170139017 | 0 | C | T | 0.465 | 1.92E-01 | 1.03E-01 | 2.45E+00 | 2.50E+00 | 6.40E-02 | Yes | | intergenic_variant | MODIFIER |
| 5 | 58295186 | AX-170508442 | 0 | T | C | 0.465 | 1.92E-01 | 1.03E-01 | 2.45E+00 | 2.50E+00 | 6.40E-02 | Yes | | regulatory_region_variant | MODIFIER |
| 5 | 58341512 | AX-170414231 | 0 | A | T | 0.465 | 1.92E-01 | 1.03E-01 | 2.45E+00 | 2.50E+00 | 6.40E-02 | Yes | | intergenic_variant | MODIFIER |
| 5 | 59976051 | AX-97797118 | 0 | T | C | 0.465 | 1.92E-01 | 1.03E-01 | 2.45E+00 | 2.50E+00 | 6.40E-02 | Yes | 4933402J10Rik | intron_variant,non_coding_transcript_variant | MODIFIER |
| 5 | 60853648 | AX-97790681 | 0 | A | G | 0.465 | 1.92E-01 | 1.03E-01 | 2.45E+00 | 2.50E+00 | 6.40E-02 | Yes | | intergenic_variant | MODIFIER |
| 5 | 61423297 | AX-170173602 | 0 | A | T | 0.465 | 1.92E-01 | 1.03E-01 | 2.45E+00 | 2.50E+00 | 6.40E-02 | Yes | | intergenic_variant | MODIFIER |
| 5 | 61427003 | AX-97817003 | 0 | C | T | 0.465 | 1.92E-01 | 1.03E-01 | 2.45E+00 | 2.50E+00 | 6.40E-02 | Yes | | intergenic_variant | MODIFIER |
| 5 | 61438810 | AX-170257577 | 0 | G | A | 0.465 | 1.92E-01 | 1.03E-01 | 2.45E+00 | 2.50E+00 | 6.40E-02 | Yes | | intergenic_variant | MODIFIER |
| 5 | 61714920 | AX-97800047 | 0 | C | T | 0.465 | 1.92E-01 | 1.03E-01 | 2.45E+00 | 2.50E+00 | 6.40E-02 | Yes | Gm43384 | downstream_gene_variant | MODIFIER |
| 5 | 61731284 | AX-97854439 | 0 | T | C | 0.465 | 1.92E-01 | 1.03E-01 | 2.45E+00 | 2.50E+00 | 6.40E-02 | Yes | | intergenic_variant | MODIFIER |
| 5 | 61764697 | AX-170143142 | 0 | G | C | 0.465 | 1.92E-01 | 1.03E-01 | 2.45E+00 | 2.50E+00 | 6.40E-02 | Yes | | intergenic_variant | MODIFIER |
| 5 | 61820242 | AX-170131046 | 0 | A | T | 0.465 | 1.92E-01 | 1.03E-01 | 2.45E+00 | 2.50E+00 | 6.40E-02 | Yes | | intergenic_variant | MODIFIER |
| 5 | 61869286 | AX-97854176 | 0 | T | C | 0.465 | 1.92E-01 | 1.03E-01 | 2.45E+00 | 2.50E+00 | 6.40E-02 | Yes | | intergenic_variant | MODIFIER |
| 5 | 61874641 | AX-97831788 | 0 | C | G | 0.465 | 1.92E-01 | 1.03E-01 | 2.45E+00 | 2.50E+00 | 6.40E-02 | Yes | | intergenic_variant | MODIFIER |
| 5 | 61976903 | AX-170025115 | 0 | T | C | 0.465 | 1.92E-01 | 1.03E-01 | 2.45E+00 | 2.50E+00 | 6.40E-02 | Yes | | intergenic_variant | MODIFIER |
| 5 | 61986304 | AX-170389985 | 0 | G | T | 0.465 | 1.92E-01 | 1.03E-01 | 2.45E+00 | 2.50E+00 | 6.40E-02 | Yes | | intergenic_variant | MODIFIER |
| 5 | 61995968 | AX-170025113 | 0 | G | C | 0.465 | 1.92E-01 | 1.03E-01 | 2.45E+00 | 2.50E+00 | 6.40E-02 | Yes | | intergenic_variant | MODIFIER |
| 5 | 62060427 | AX-170331631 | 0 | T | C | 0.465 | 1.92E-01 | 1.03E-01 | 2.45E+00 | 2.50E+00 | 6.40E-02 | Yes | | intergenic_variant | MODIFIER |
| 5 | 62100728 | AX-170072276 | 0 | G | C | 0.465 | 1.92E-01 | 1.03E-01 | 2.45E+00 | 2.50E+00 | 6.40E-02 | Yes | | intergenic_variant | MODIFIER |

| | | | | | | | | | | | | | | | |
|---|---|---|---|---|---|---|---|---|---|---|---|---|---|---|---|
| 5 | 62204550 | AX-170329794 | 0 | C | T | 0.465 | 1.92E-01 | 1.03E-01 | 2.45E+00 | 2.50E+00 | 6.40E-02 | Yes | | intergenic_variant | MODIFIER |
| 5 | 62250890 | AX-170107832 | 0 | A | C | 0.465 | 1.92E-01 | 1.03E-01 | 2.45E+00 | 2.50E+00 | 6.40E-02 | Yes | | intergenic_variant | MODIFIER |
| 5 | 62250953 | AX-170089498 | 0 | T | C | 0.465 | 1.92E-01 | 1.03E-01 | 2.45E+00 | 2.50E+00 | 6.40E-02 | Yes | | intergenic_variant | MODIFIER |
| 5 | 62269704 | AX-170526125 | 0 | C | G | 0.465 | 1.92E-01 | 1.03E-01 | 2.45E+00 | 2.50E+00 | 6.40E-02 | Yes | | intergenic_variant | MODIFIER |
| 5 | 62300202 | AX-97834587 | 0 | A | G | 0.465 | 1.92E-01 | 1.03E-01 | 2.45E+00 | 2.50E+00 | 6.40E-02 | Yes | | intergenic_variant | MODIFIER |
| 5 | 62319536 | AX-170215350 | 0 | A | T | 0.465 | 1.92E-01 | 1.03E-01 | 2.45E+00 | 2.50E+00 | 6.40E-02 | Yes | | intergenic_variant | MODIFIER |
| 5 | 62322669 | AX-97849426 | 0 | T | C | 0.465 | 1.92E-01 | 1.03E-01 | 2.45E+00 | 2.50E+00 | 6.40E-02 | Yes | | intergenic_variant | MODIFIER |
| 5 | 62406208 | AX-97841116 | 0 | C | T | 0.465 | 1.92E-01 | 1.03E-01 | 2.45E+00 | 2.50E+00 | 6.40E-02 | Yes | | intergenic_variant | MODIFIER |
| 5 | 62419278 | AX-170295802 | 0 | G | A | 0.465 | 1.92E-01 | 1.03E-01 | 2.45E+00 | 2.50E+00 | 6.40E-02 | Yes | | intergenic_variant | MODIFIER |
| 5 | 62460890 | AX-97782109 | 0 | C | A | 0.465 | 1.92E-01 | 1.03E-01 | 2.45E+00 | 2.50E+00 | 6.40E-02 | Yes | | intergenic_variant | MODIFIER |
| 5 | 62507627 | AX-170025123 | 0 | C | G | 0.465 | 1.92E-01 | 1.03E-01 | 2.45E+00 | 2.50E+00 | 6.40E-02 | Yes | | intergenic_variant | MODIFIER |
| 5 | 62518828 | AX-97789139 | 0 | G | A | 0.465 | 1.92E-01 | 1.03E-01 | 2.45E+00 | 2.50E+00 | 6.40E-02 | Yes | Gm24079 | downstream_gene_variant | MODIFIER |
| 5 | 62540551 | AX-170025122 | 0 | T | G | 0.465 | 1.92E-01 | 1.03E-01 | 2.45E+00 | 2.50E+00 | 6.40E-02 | Yes | | intergenic_variant | MODIFIER |
| 5 | 62554877 | AX-97829966 | 0 | C | T | 0.465 | 1.92E-01 | 1.03E-01 | 2.45E+00 | 2.50E+00 | 6.40E-02 | Yes | | intergenic_variant | MODIFIER |
| 5 | 62560883 | AX-170025119 | 0 | C | T | 0.465 | 1.92E-01 | 1.03E-01 | 2.45E+00 | 2.50E+00 | 6.40E-02 | Yes | | intergenic_variant | MODIFIER |
| 5 | 62575703 | AX-97846985 | 0 | C | T | 0.465 | 1.92E-01 | 1.03E-01 | 2.45E+00 | 2.50E+00 | 6.40E-02 | Yes | | intergenic_variant | MODIFIER |
| 5 | 62770320 | AX-170089500 | 0 | T | C | 0.465 | 1.92E-01 | 1.03E-01 | 2.45E+00 | 2.50E+00 | 6.40E-02 | Yes | Arap2 | upstream_gene_variant | MODIFIER |
| 5 | 64168591 | AX-170292750 | 0 | G | A | 0.465 | 1.92E-01 | 1.03E-01 | 2.45E+00 | 2.50E+00 | 6.40E-02 | Yes | Tbc1d1 | intron_variant | MODIFIER |
| 5 | 64171066 | AX-170142555 | 0 | T | A | 0.465 | 1.92E-01 | 1.03E-01 | 2.45E+00 | 2.50E+00 | 6.40E-02 | Yes | Tbc1d1 | intron_variant | MODIFIER |
| 5 | 64178726 | AX-170189815 | 0 | T | C | 0.465 | 1.92E-01 | 1.03E-01 | 2.45E+00 | 2.50E+00 | 6.40E-02 | Yes | Tbc1d1 | intron_variant | MODIFIER |
| 5 | 65175721 | AX-170025168 | 0 | A | C | 0.465 | 1.92E-01 | 1.03E-01 | 2.45E+00 | 2.50E+00 | 6.40E-02 | Yes | | regulatory_region_variant | MODIFIER |
| 5 | 65257777 | AX-170025163 | 0 | C | T | 0.465 | 1.92E-01 | 1.03E-01 | 2.45E+00 | 2.50E+00 | 6.40E-02 | Yes | Wdr19 | synonymous_variant | LOW |
| 5 | 65279618 | AX-170266975 | 0 | C | T | 0.465 | 1.92E-01 | 1.03E-01 | 2.45E+00 | 2.50E+00 | 6.40E-02 | Yes | Rfc1 | missense_variant | MODERATE |
| 5 | 65358589 | AX-170393788 | 0 | C | T | 0.465 | 1.92E-01 | 1.03E-01 | 2.45E+00 | 2.50E+00 | 6.40E-02 | Yes | Klb | intron_variant | MODIFIER |
| 5 | 65367808 | AX-170124805 | 0 | G | T | 0.465 | 1.92E-01 | 1.03E-01 | 2.45E+00 | 2.50E+00 | 6.40E-02 | Yes | Klb | intron_variant | MODIFIER |
| 5 | 65370350 | AX-97816326 | 0 | T | G | 0.465 | 1.92E-01 | 1.03E-01 | 2.45E+00 | 2.50E+00 | 6.40E-02 | Yes | Klb | intron_variant | MODIFIER |
| 5 | 65371903 | AX-170535634 | 0 | G | C | 0.465 | 1.92E-01 | 1.03E-01 | 2.45E+00 | 2.50E+00 | 6.40E-02 | Yes | Klb | intron_variant | MODIFIER |
| 5 | 65375884 | AX-170249380 | 0 | T | C | 0.465 | 1.92E-01 | 1.03E-01 | 2.45E+00 | 2.50E+00 | 6.40E-02 | Yes | Klb | missense_variant | MODERATE |
| 5 | 65378396 | AX-170245890 | 0 | G | A | 0.465 | 1.92E-01 | 1.03E-01 | 2.45E+00 | 2.50E+00 | 6.40E-02 | Yes | Klb | intron_variant | MODIFIER |
| 5 | 65385332 | AX-170187205 | 0 | C | T | 0.465 | 1.92E-01 | 1.03E-01 | 2.45E+00 | 2.50E+00 | 6.40E-02 | Yes | Klb | downstream_gene_variant | MODIFIER |
| 5 | 65401676 | AX-170276464 | 0 | A | T | 0.465 | 1.92E-01 | 1.03E-01 | 2.45E+00 | 2.50E+00 | 6.40E-02 | Yes | Lias | intron_variant | MODIFIER |
| 5 | 65433756 | AX-170308320 | 0 | T | A | 0.465 | 1.92E-01 | 1.03E-01 | 2.45E+00 | 2.50E+00 | 6.40E-02 | Yes | Ugdh | intron_variant | MODIFIER |
| 5 | 65458151 | AX-170354187 | 0 | G | A | 0.465 | 1.92E-01 | 1.03E-01 | 2.45E+00 | 2.50E+00 | 6.40E-02 | Yes | Smim14 | intron_variant,NMD_transcript_variant | MODIFIER |
| 5 | 65460512 | AX-170497100 | 0 | A | G | 0.465 | 1.92E-01 | 1.03E-01 | 2.45E+00 | 2.50E+00 | 6.40E-02 | Yes | Smim14 | intron_variant,NMD_transcript_variant | MODIFIER |
| 5 | 65918412 | AX-170134432 | 0 | A | G | 0.465 | 1.92E-01 | 1.03E-01 | 2.45E+00 | 2.50E+00 | 6.40E-02 | Yes | Gm43769 | downstream_gene_variant | MODIFIER |
| 5 | 66050793 | AX-170444411 | 0 | T | C | 0.465 | 1.92E-01 | 1.03E-01 | 2.45E+00 | 2.50E+00 | 6.40E-02 | Yes | Rbm47 | intron_variant | MODIFIER |
| 5 | 66093548 | AX-170521769 | 0 | A | G | 0.465 | 1.92E-01 | 1.03E-01 | 2.45E+00 | 2.50E+00 | 6.40E-02 | Yes | Rbm47 | intron_variant | MODIFIER |
| 2 | 90538845 | AX-170513642 | 0 | C | T | 0.053 | -3.86E-01 | 2.07E-01 | 3.15E+00 | 3.31E+00 | 6.41E-02 | Yes | Ptprj | intron_variant | MODIFIER |
| 2 | 107615896 | AX-170435995 | 0 | G | A | 0.053 | -3.86E-01 | 2.07E-01 | 3.15E+00 | 3.31E+00 | 6.41E-02 | Yes | | intergenic_variant | MODIFIER |
| 8 | 61364395 | AX-170244376 | 0 | T | G | 0.14 | -2.22E-01 | 1.20E-01 | 2.51E+00 | 2.63E+00 | 6.44E-02 | Yes | Sh3rf1 | intron_variant | MODIFIER |
| 15 | 76445802 | AX-170209223 | 1 | A | C | 0.08 | -2.27E-01 | 1.07E-01 | 1.67E+00 | 1.72E+00 | 6.44E-02 | Yes | Mroh1 | intron_variant | MODIFIER |
| 15 | 77060613 | AX-170509485 | 1 | C | T | 0.08 | -2.27E-01 | 1.07E-01 | 1.67E+00 | 1.72E+00 | 6.44E-02 | Yes | Apol6 | downstream_gene_variant | MODIFIER |
| 5 | 55213342 | AX-170525379 | 1 | C | T | 0.464 | 1.91E-01 | 1.03E-01 | 2.44E+00 | 2.49E+00 | 6.45E-02 | Yes | | intergenic_variant | MODIFIER |
| 10 | 100263401 | AX-97800931 | 0 | G | T | 0.272 | -1.55E-01 | 8.41E-02 | 2.85E+00 | 2.94E+00 | 6.46E-02 | no | | intergenic_variant | MODIFIER |
| 10 | 100516213 | AX-170329148 | 0 | G | A | 0.272 | -1.55E-01 | 8.41E-02 | 2.85E+00 | 2.94E+00 | 6.46E-02 | no | Cep290 | synonymous_variant | LOW |
| 4 | 82905633 | AX-170253933 | 0 | G | C | 0.351 | 2.08E-01 | 1.12E-01 | 3.06E+00 | 3.16E+00 | 6.46E-02 | Yes | Frem1 | intron_variant | MODIFIER |
| 16 | 67720654 | AX-170327929 | 0 | C | T | 0.491 | 1.41E-01 | 7.65E-02 | 2.55E+00 | 2.62E+00 | 6.47E-02 | no | Gm49648 | intron_variant,non_coding_transcript_variant | MODIFIER |
| 5 | 57737222 | AX-170089485 | 0 | C | A | 0.474 | 1.84E-01 | 9.95E-02 | 2.53E+00 | 2.59E+00 | 6.48E-02 | Yes | Pcdh7 | intron_variant | MODIFIER |
| 5 | 49535738 | AX-170324210 | 0 | G | A | 0.482 | 1.81E-01 | 9.55E-02 | 2.22E+00 | 2.28E+00 | 6.48E-02 | Yes | | intergenic_variant | MODIFIER |
| 11 | 10002548 | AX-170169142 | 0 | G | A | 0.07 | 2.60E-01 | 1.41E-01 | 2.99E+00 | 3.18E+00 | 6.48E-02 | Yes | | intergenic_variant | MODIFIER |
| 15 | 71253553 | AX-170326860 | 1 | C | A | 0.08 | -2.29E-01 | 1.07E-01 | 1.64E+00 | 1.69E+00 | 6.48E-02 | Yes | | intergenic_variant | MODIFIER |
| 10 | 104959695 | AX-170045796 | 0 | A | G | 0.298 | -1.51E-01 | 8.21E-02 | 2.78E+00 | 2.86E+00 | 6.49E-02 | no | | intergenic_variant | MODIFIER |
| 16 | 63882634 | AX-170471474 | 1 | A | C | 0.509 | 1.61E-01 | 8.76E-02 | 2.81E+00 | 2.87E+00 | 6.51E-02 | Yes | | intergenic_variant | MODIFIER |
| 3 | 8007495 | AX-170144943 | 1 | T | C | 0.402 | -1.16E-01 | 6.22E-02 | 3.20E+00 | 3.41E+00 | 6.54E-02 | no | | regulatory_region_variant | MODIFIER |

| | | | | | | | | | | | | | | | |
|---|---|---|---|---|---|---|---|---|---|---|---|---|---|---|---|
| 16 | 63266746 | AX-170326335 | 0 | A | G | 0.518 | 1.58E-01 | 8.58E-02 | 2.72E+00 | 2.78E+00 | 6.54E-02 | Yes | | intergenic_variant | MODIFIER |
| 5 | 61660924 | AX-170273928 | 0 | C | T | 0.5 | 1.68E-01 | 9.05E-02 | 2.40E+00 | 2.49E+00 | 6.54E-02 | Yes | | intergenic_variant | MODIFIER |
| 19 | 54227984 | AX-168401735 | 0 | C | T | 0.491 | 1.19E-01 | 6.49E-02 | 2.71E+00 | 2.79E+00 | 6.55E-02 | no | | intergenic_variant | MODIFIER |
| 12 | 59780581 | AX-170049165 | 0 | G | A | 0.061 | -3.93E-01 | 2.13E-01 | 2.98E+00 | 3.11E+00 | 6.55E-02 | Yes | | intergenic_variant | MODIFIER |
| 5 | 132088002 | AX-170292987 | 0 | C | T | 0.149 | -1.69E-01 | 9.00E-02 | 2.27E+00 | 2.34E+00 | 6.56E-02 | no | Auts2 | intron_variant,non_coding_transcript_variant | MODIFIER |
| 5 | 132090003 | AX-170329658 | 0 | C | T | 0.149 | -1.69E-01 | 9.00E-02 | 2.27E+00 | 2.34E+00 | 6.56E-02 | no | Auts2 | intron_variant,non_coding_transcript_variant | MODIFIER |
| 5 | 132097362 | AX-170381508 | 0 | C | T | 0.149 | -1.69E-01 | 9.00E-02 | 2.27E+00 | 2.34E+00 | 6.56E-02 | no | Auts2 | intron_variant,non_coding_transcript_variant | MODIFIER |
| 5 | 132099512 | AX-170182136 | 0 | G | A | 0.149 | -1.69E-01 | 9.00E-02 | 2.27E+00 | 2.34E+00 | 6.56E-02 | no | Auts2 | intron_variant,non_coding_transcript_variant | MODIFIER |
| 5 | 132104344 | AX-170418624 | 0 | A | C | 0.149 | -1.69E-01 | 9.00E-02 | 2.27E+00 | 2.34E+00 | 6.56E-02 | no | Auts2 | intron_variant,non_coding_transcript_variant | MODIFIER |
| 5 | 132112255 | AX-170115809 | 0 | T | G | 0.149 | -1.69E-01 | 9.00E-02 | 2.27E+00 | 2.34E+00 | 6.56E-02 | no | Auts2 | intron_variant,non_coding_transcript_variant | MODIFIER |
| 5 | 132130116 | AX-97817245 | 0 | A | G | 0.149 | -1.69E-01 | 9.00E-02 | 2.27E+00 | 2.34E+00 | 6.56E-02 | no | Auts2 | intron_variant,non_coding_transcript_variant | MODIFIER |
| 5 | 132130540 | AX-170282909 | 0 | T | A | 0.149 | -1.69E-01 | 9.00E-02 | 2.27E+00 | 2.34E+00 | 6.56E-02 | no | Auts2 | intron_variant,non_coding_transcript_variant | MODIFIER |
| 5 | 132134444 | AX-170188641 | 0 | G | A | 0.149 | -1.69E-01 | 9.00E-02 | 2.27E+00 | 2.34E+00 | 6.56E-02 | no | Auts2 | intron_variant,non_coding_transcript_variant | MODIFIER |
| 5 | 132141215 | AX-170224102 | 0 | C | T | 0.149 | -1.69E-01 | 9.00E-02 | 2.27E+00 | 2.34E+00 | 6.56E-02 | no | Auts2 | intron_variant,non_coding_transcript_variant | MODIFIER |
| 5 | 132142150 | AX-170153348 | 0 | G | A | 0.149 | -1.69E-01 | 9.00E-02 | 2.27E+00 | 2.34E+00 | 6.56E-02 | no | Auts2 | intron_variant,non_coding_transcript_variant | MODIFIER |
| 5 | 132142225 | AX-170248995 | 0 | A | C | 0.149 | -1.69E-01 | 9.00E-02 | 2.27E+00 | 2.34E+00 | 6.56E-02 | no | Auts2 | intron_variant,non_coding_transcript_variant | MODIFIER |
| 5 | 132249515 | AX-170182651 | 0 | T | C | 0.149 | -1.69E-01 | 9.00E-02 | 2.27E+00 | 2.34E+00 | 6.56E-02 | no | Auts2 | intron_variant,non_coding_transcript_variant | MODIFIER |
| 16 | 63752888 | AX-170526392 | 0 | C | T | 0.509 | 1.61E-01 | 8.78E-02 | 2.81E+00 | 2.86E+00 | 6.57E-02 | Yes | Epha3 | intron_variant | MODIFIER |
| 16 | 63763545 | AX-97788937 | 0 | A | G | 0.509 | 1.61E-01 | 8.78E-02 | 2.81E+00 | 2.86E+00 | 6.57E-02 | Yes | Epha3 | intron_variant | MODIFIER |
| 16 | 63791849 | AX-170316889 | 0 | G | C | 0.509 | 1.61E-01 | 8.78E-02 | 2.81E+00 | 2.86E+00 | 6.57E-02 | Yes | Epha3 | intron_variant | MODIFIER |
| 16 | 63793814 | AX-97797497 | 0 | A | C | 0.509 | 1.61E-01 | 8.78E-02 | 2.81E+00 | 2.86E+00 | 6.57E-02 | Yes | Epha3 | intron_variant | MODIFIER |
| 16 | 63796707 | AX-170514662 | 0 | T | C | 0.509 | 1.61E-01 | 8.78E-02 | 2.81E+00 | 2.86E+00 | 6.57E-02 | Yes | Epha3 | intron_variant | MODIFIER |
| 16 | 63801914 | AX-170057639 | 0 | C | A | 0.509 | 1.61E-01 | 8.78E-02 | 2.81E+00 | 2.86E+00 | 6.57E-02 | Yes | Epha3 | intron_variant | MODIFIER |
| 16 | 63806983 | AX-170312215 | 0 | G | C | 0.509 | 1.61E-01 | 8.78E-02 | 2.81E+00 | 2.86E+00 | 6.57E-02 | Yes | Epha3 | intron_variant | MODIFIER |
| 16 | 63815028 | AX-97793600 | 0 | T | C | 0.509 | 1.61E-01 | 8.78E-02 | 2.81E+00 | 2.86E+00 | 6.57E-02 | Yes | Epha3 | intron_variant | MODIFIER |
| 16 | 63817762 | AX-170057641 | 0 | T | C | 0.509 | 1.61E-01 | 8.78E-02 | 2.81E+00 | 2.86E+00 | 6.57E-02 | Yes | Epha3 | intron_variant | MODIFIER |
| 16 | 63817778 | AX-170057640 | 0 | C | T | 0.509 | 1.61E-01 | 8.78E-02 | 2.81E+00 | 2.86E+00 | 6.57E-02 | Yes | Epha3 | intron_variant | MODIFIER |
| 16 | 63818074 | AX-170361798 | 0 | C | G | 0.509 | 1.61E-01 | 8.78E-02 | 2.81E+00 | 2.86E+00 | 6.57E-02 | Yes | Epha3 | intron_variant | MODIFIER |
| 16 | 63831025 | AX-170365703 | 0 | G | A | 0.509 | 1.61E-01 | 8.78E-02 | 2.81E+00 | 2.86E+00 | 6.57E-02 | Yes | Epha3 | intron_variant | MODIFIER |
| 16 | 63843374 | AX-170452224 | 0 | T | C | 0.509 | 1.61E-01 | 8.78E-02 | 2.81E+00 | 2.86E+00 | 6.57E-02 | Yes | Epha3 | intron_variant | MODIFIER |
| 16 | 63863131 | AX-170517052 | 0 | G | T | 0.509 | 1.61E-01 | 8.78E-02 | 2.81E+00 | 2.86E+00 | 6.57E-02 | Yes | Epha3 | intron_variant | MODIFIER |
| 16 | 63863172 | AX-97843794 | 0 | C | T | 0.509 | 1.61E-01 | 8.78E-02 | 2.81E+00 | 2.86E+00 | 6.57E-02 | Yes | Epha3 | intron_variant | MODIFIER |
| 16 | 63894937 | AX-170306622 | 0 | T | A | 0.509 | 1.61E-01 | 8.78E-02 | 2.81E+00 | 2.86E+00 | 6.57E-02 | Yes | | intergenic_variant | MODIFIER |
| 16 | 63907166 | AX-170304428 | 0 | T | G | 0.509 | 1.61E-01 | 8.78E-02 | 2.81E+00 | 2.86E+00 | 6.57E-02 | Yes | | intergenic_variant | MODIFIER |
| 16 | 63917440 | AX-170452761 | 0 | A | T | 0.509 | 1.61E-01 | 8.78E-02 | 2.81E+00 | 2.86E+00 | 6.57E-02 | Yes | | intergenic_variant | MODIFIER |
| 16 | 63918665 | AX-97836134 | 0 | T | C | 0.509 | 1.61E-01 | 8.78E-02 | 2.81E+00 | 2.86E+00 | 6.57E-02 | Yes | | intergenic_variant | MODIFIER |
| 16 | 63954714 | AX-170122035 | 0 | G | T | 0.509 | 1.61E-01 | 8.78E-02 | 2.81E+00 | 2.86E+00 | 6.57E-02 | Yes | Gm49627 | non_coding_transcript_exon_variant | MODIFIER |
| 16 | 63959174 | AX-170088045 | 0 | T | C | 0.509 | 1.61E-01 | 8.78E-02 | 2.81E+00 | 2.86E+00 | 6.57E-02 | Yes | Gm49627 | upstream_gene_variant | MODIFIER |
| 16 | 64018298 | AX-97832073 | 0 | C | A | 0.509 | 1.61E-01 | 8.78E-02 | 2.81E+00 | 2.86E+00 | 6.57E-02 | Yes | | intergenic_variant | MODIFIER |
| 16 | 64032812 | AX-170013244 | 0 | C | T | 0.509 | 1.61E-01 | 8.78E-02 | 2.81E+00 | 2.86E+00 | 6.57E-02 | Yes | | intergenic_variant | MODIFIER |
| 16 | 64034913 | AX-97844820 | 0 | G | A | 0.509 | 1.61E-01 | 8.78E-02 | 2.81E+00 | 2.86E+00 | 6.57E-02 | Yes | | intergenic_variant | MODIFIER |
| 16 | 64058770 | AX-170373717 | 0 | G | C | 0.509 | 1.61E-01 | 8.78E-02 | 2.81E+00 | 2.86E+00 | 6.57E-02 | Yes | | regulatory_region_variant | MODIFIER |
| 16 | 64058788 | AX-170118286 | 0 | G | A | 0.509 | 1.61E-01 | 8.78E-02 | 2.81E+00 | 2.86E+00 | 6.57E-02 | Yes | | regulatory_region_variant | MODIFIER |
| 16 | 64068150 | AX-170176381 | 0 | T | A | 0.509 | 1.61E-01 | 8.78E-02 | 2.81E+00 | 2.86E+00 | 6.57E-02 | Yes | | intergenic_variant | MODIFIER |
| 16 | 64068534 | AX-170515468 | 0 | T | C | 0.509 | 1.61E-01 | 8.78E-02 | 2.81E+00 | 2.86E+00 | 6.57E-02 | Yes | | intergenic_variant | MODIFIER |
| 16 | 64068677 | AX-97780796 | 0 | C | A | 0.509 | 1.61E-01 | 8.78E-02 | 2.81E+00 | 2.86E+00 | 6.57E-02 | Yes | | regulatory_region_variant | MODIFIER |
| 16 | 64078971 | AX-97831606 | 0 | C | T | 0.509 | 1.61E-01 | 8.78E-02 | 2.81E+00 | 2.86E+00 | 6.57E-02 | Yes | | intergenic_variant | MODIFIER |
| 16 | 54574635 | AX-170013089 | 2 | T | C | 0.455 | -1.47E-01 | 7.99E-02 | 2.58E+00 | 2.64E+00 | 6.58E-02 | no | | intergenic_variant | MODIFIER |
| 19 | 60210884 | AX-97824342 | 0 | A | G | 0.465 | -1.15E-01 | 6.26E-02 | 2.80E+00 | 2.89E+00 | 6.59E-02 | no | Fam204a | intron_variant | MODIFIER |
| 19 | 60278043 | AX-170448347 | 0 | G | A | 0.465 | -1.15E-01 | 6.26E-02 | 2.80E+00 | 2.89E+00 | 6.59E-02 | no | | intergenic_variant | MODIFIER |
| 5 | 64040771 | AX-170025145 | 1 | G | A | 0.491 | 1.90E-01 | 1.02E-01 | 2.41E+00 | 2.46E+00 | 6.60E-02 | Yes | 5830416I19Rik | upstream_gene_variant | MODIFIER |
| 4 | 98989024 | AX-170229781 | 1 | G | A | 0.241 | 2.12E-01 | 1.16E-01 | 2.83E+00 | 2.91E+00 | 6.62E-02 | Yes | Dock7 | synonymous_variant | LOW |
| 13 | 9546550 | AX-170321960 | 0 | T | C | 0.193 | -1.70E-01 | 9.28E-02 | 2.60E+00 | 2.73E+00 | 6.64E-02 | no | Dip2c | intron_variant | MODIFIER |
| 13 | 17076562 | AX-97834366 | 0 | A | G | 0.167 | -2.21E-01 | 1.18E-01 | 2.29E+00 | 2.35E+00 | 6.65E-02 | Yes | Sugct | intron_variant | MODIFIER |

| | | | | | | | | | | | | | | | |
|---|---|---|---|---|---|---|---|---|---|---|---|---|---|---|---|
| 16 | 58526965 | AX-170013147 | 0 | C | T | 0.535 | 1.53E-01 | 8.34E-02 | 2.61E+00 | 2.66E+00 | 6.65E-02 | Yes | St3gal6 | upstream_gene_variant | MODIFIER |
| 5 | 129993094 | AX-170299741 | 1 | A | T | 0.152 | -1.70E-01 | 8.87E-02 | 2.08E+00 | 2.15E+00 | 6.67E-02 | no | Gusb | intron_variant | MODIFIER |
| 18 | 74135414 | AX-170015988 | 0 | C | T | 0.175 | -2.14E-01 | 1.17E-01 | 2.83E+00 | 2.91E+00 | 6.67E-02 | Yes | | intergenic_variant | MODIFIER |
| 4 | 99237227 | AX-170421716 | 1 | G | A | 0.25 | 2.00E-01 | 1.09E-01 | 2.76E+00 | 2.85E+00 | 6.67E-02 | Yes | Atg4c | intron_variant | MODIFIER |
| 4 | 75957255 | AX-170349725 | 2 | C | T | 0.336 | 2.20E-01 | 1.19E-01 | 3.16E+00 | 3.25E+00 | 6.69E-02 | Yes | Ptprd | synonymous_variant | LOW |
| 8 | 31909748 | AX-170214377 | 0 | T | A | 0.14 | -2.21E-01 | 1.19E-01 | 3.21E+00 | 3.38E+00 | 6.70E-02 | Yes | Nrg1 | intron_variant | MODIFIER |
| 4 | 49276085 | AX-170492805 | 1 | C | T | 0.357 | -1.73E-01 | 9.28E-02 | 3.27E+00 | 3.42E+00 | 6.72E-02 | no | Plppr1 | intron_variant | MODIFIER |
| 5 | 66159658 | AX-170213697 | 1 | G | A | 0.491 | 1.71E-01 | 9.30E-02 | 2.45E+00 | 2.53E+00 | 6.72E-02 | Yes | Gm22929 | downstream_gene_variant | MODIFIER |
| 5 | 55473332 | AX-170024988 | 1 | C | G | 0.473 | 1.92E-01 | 1.03E-01 | 2.32E+00 | 2.37E+00 | 6.74E-02 | Yes | | intergenic_variant | MODIFIER |
| 10 | 105588086 | AX-97827426 | 0 | T | C | 0.289 | -1.55E-01 | 8.50E-02 | 2.75E+00 | 2.84E+00 | 6.74E-02 | no | Gm15663 | downstream_gene_variant | MODIFIER |
| 4 | 84445623 | AX-170314846 | 0 | A | T | 0.333 | 2.04E-01 | 1.12E-01 | 2.89E+00 | 2.96E+00 | 6.76E-02 | Yes | Bnc2 | intron_variant | MODIFIER |
| 16 | 63680101 | AX-97831574 | 1 | T | C | 0.518 | 1.61E-01 | 8.81E-02 | 2.66E+00 | 2.71E+00 | 6.76E-02 | Yes | Epha3 | intron_variant | MODIFIER |
| 15 | 67852850 | AX-170504937 | 0 | C | T | 0.079 | -2.28E-01 | 1.07E-01 | 1.64E+00 | 1.69E+00 | 6.77E-02 | Yes | | intergenic_variant | MODIFIER |
| 15 | 67859753 | AX-170205529 | 0 | C | G | 0.079 | -2.28E-01 | 1.07E-01 | 1.64E+00 | 1.69E+00 | 6.77E-02 | Yes | | intergenic_variant | MODIFIER |
| 15 | 67881737 | AX-170415861 | 0 | A | G | 0.079 | -2.28E-01 | 1.07E-01 | 1.64E+00 | 1.69E+00 | 6.77E-02 | Yes | | intergenic_variant | MODIFIER |
| 15 | 67883045 | AX-170011904 | 0 | C | T | 0.079 | -2.28E-01 | 1.07E-01 | 1.64E+00 | 1.69E+00 | 6.77E-02 | Yes | | intergenic_variant | MODIFIER |
| 15 | 67883077 | AX-170011909 | 0 | A | G | 0.079 | -2.28E-01 | 1.07E-01 | 1.64E+00 | 1.69E+00 | 6.77E-02 | Yes | | intergenic_variant | MODIFIER |
| 15 | 67922635 | AX-97846446 | 0 | A | G | 0.079 | -2.28E-01 | 1.07E-01 | 1.64E+00 | 1.69E+00 | 6.77E-02 | Yes | | intergenic_variant | MODIFIER |
| 15 | 67933188 | AX-170400228 | 0 | G | A | 0.079 | -2.28E-01 | 1.07E-01 | 1.64E+00 | 1.69E+00 | 6.77E-02 | Yes | | intergenic_variant | MODIFIER |
| 15 | 67936826 | AX-170190243 | 0 | A | G | 0.079 | -2.28E-01 | 1.07E-01 | 1.64E+00 | 1.69E+00 | 6.77E-02 | Yes | | intergenic_variant | MODIFIER |
| 15 | 67938416 | AX-170011903 | 0 | C | T | 0.079 | -2.28E-01 | 1.07E-01 | 1.64E+00 | 1.69E+00 | 6.77E-02 | Yes | Gm49409 | downstream_gene_variant | MODIFIER |
| 15 | 67938587 | AX-170179681 | 0 | T | G | 0.079 | -2.28E-01 | 1.07E-01 | 1.64E+00 | 1.69E+00 | 6.77E-02 | Yes | Gm49409 | downstream_gene_variant | MODIFIER |
| 15 | 67940479 | AX-170011911 | 0 | G | A | 0.079 | -2.28E-01 | 1.07E-01 | 1.64E+00 | 1.69E+00 | 6.77E-02 | Yes | Gm49409 | downstream_gene_variant | MODIFIER |
| 15 | 67944493 | AX-97800964 | 0 | T | C | 0.079 | -2.28E-01 | 1.07E-01 | 1.64E+00 | 1.69E+00 | 6.77E-02 | Yes | Gm49409 | upstream_gene_variant | MODIFIER |
| 15 | 67965135 | AX-170423148 | 0 | A | G | 0.079 | -2.28E-01 | 1.07E-01 | 1.64E+00 | 1.69E+00 | 6.77E-02 | Yes | | intergenic_variant | MODIFIER |
| 15 | 67965270 | AX-170404843 | 0 | G | A | 0.079 | -2.28E-01 | 1.07E-01 | 1.64E+00 | 1.69E+00 | 6.77E-02 | Yes | | intergenic_variant | MODIFIER |
| 15 | 67975238 | AX-97804330 | 0 | T | C | 0.079 | -2.28E-01 | 1.07E-01 | 1.64E+00 | 1.69E+00 | 6.77E-02 | Yes | | intergenic_variant | MODIFIER |
| 15 | 70389162 | AX-168412587 | 0 | A | G | 0.079 | -2.28E-01 | 1.07E-01 | 1.64E+00 | 1.69E+00 | 6.77E-02 | Yes | | intergenic_variant | MODIFIER |
| 15 | 70394922 | AX-97830355 | 0 | T | C | 0.079 | -2.28E-01 | 1.07E-01 | 1.64E+00 | 1.69E+00 | 6.77E-02 | Yes | | intergenic_variant | MODIFIER |
| 15 | 70402757 | AX-170246400 | 0 | C | T | 0.079 | -2.28E-01 | 1.07E-01 | 1.64E+00 | 1.69E+00 | 6.77E-02 | Yes | | intergenic_variant | MODIFIER |
| 15 | 70426020 | AX-170113083 | 0 | G | A | 0.079 | -2.28E-01 | 1.07E-01 | 1.64E+00 | 1.69E+00 | 6.77E-02 | Yes | | intergenic_variant | MODIFIER |
| 15 | 70455996 | AX-170368616 | 0 | G | A | 0.079 | -2.28E-01 | 1.07E-01 | 1.64E+00 | 1.69E+00 | 6.77E-02 | Yes | | intergenic_variant | MODIFIER |
| 15 | 70467403 | AX-170489035 | 0 | C | T | 0.079 | -2.28E-01 | 1.07E-01 | 1.64E+00 | 1.69E+00 | 6.77E-02 | Yes | | intergenic_variant | MODIFIER |
| 15 | 70467801 | AX-170206215 | 0 | C | T | 0.079 | -2.28E-01 | 1.07E-01 | 1.64E+00 | 1.69E+00 | 6.77E-02 | Yes | | intergenic_variant | MODIFIER |
| 15 | 70470290 | AX-170257889 | 0 | T | G | 0.079 | -2.28E-01 | 1.07E-01 | 1.64E+00 | 1.69E+00 | 6.77E-02 | Yes | | intergenic_variant | MODIFIER |
| 15 | 70493141 | AX-170364053 | 0 | C | A | 0.079 | -2.28E-01 | 1.07E-01 | 1.64E+00 | 1.69E+00 | 6.77E-02 | Yes | | intergenic_variant | MODIFIER |
| 15 | 70501126 | AX-170211771 | 0 | T | G | 0.079 | -2.28E-01 | 1.07E-01 | 1.64E+00 | 1.69E+00 | 6.77E-02 | Yes | | intergenic_variant | MODIFIER |
| 15 | 70545375 | AX-170193416 | 0 | C | T | 0.079 | -2.28E-01 | 1.07E-01 | 1.64E+00 | 1.69E+00 | 6.77E-02 | Yes | | intergenic_variant | MODIFIER |
| 15 | 70552673 | AX-170441793 | 0 | A | G | 0.079 | -2.28E-01 | 1.07E-01 | 1.64E+00 | 1.69E+00 | 6.77E-02 | Yes | | intergenic_variant | MODIFIER |
| 15 | 70562721 | AX-170407156 | 0 | T | C | 0.079 | -2.28E-01 | 1.07E-01 | 1.64E+00 | 1.69E+00 | 6.77E-02 | Yes | | intergenic_variant | MODIFIER |
| 15 | 70570405 | AX-170434830 | 0 | T | C | 0.079 | -2.28E-01 | 1.07E-01 | 1.64E+00 | 1.69E+00 | 6.77E-02 | Yes | | intergenic_variant | MODIFIER |
| 15 | 70593454 | AX-170128926 | 0 | G | A | 0.079 | -2.28E-01 | 1.07E-01 | 1.64E+00 | 1.69E+00 | 6.77E-02 | Yes | | intergenic_variant | MODIFIER |
| 15 | 70676868 | AX-170011952 | 0 | G | C | 0.079 | -2.28E-01 | 1.07E-01 | 1.64E+00 | 1.69E+00 | 6.77E-02 | Yes | | intergenic_variant | MODIFIER |
| 15 | 70676911 | AX-97830359 | 0 | T | G | 0.079 | -2.28E-01 | 1.07E-01 | 1.64E+00 | 1.69E+00 | 6.77E-02 | Yes | | intergenic_variant | MODIFIER |
| 15 | 70676956 | AX-170011945 | 0 | G | C | 0.079 | -2.28E-01 | 1.07E-01 | 1.64E+00 | 1.69E+00 | 6.77E-02 | Yes | | intergenic_variant | MODIFIER |
| 15 | 70717090 | AX-170011953 | 0 | C | G | 0.079 | -2.28E-01 | 1.07E-01 | 1.64E+00 | 1.69E+00 | 6.77E-02 | Yes | | intergenic_variant | MODIFIER |
| 15 | 70730550 | AX-97815706 | 0 | G | A | 0.079 | -2.28E-01 | 1.07E-01 | 1.64E+00 | 1.69E+00 | 6.77E-02 | Yes | | intergenic_variant | MODIFIER |
| 15 | 70732420 | AX-170011947 | 0 | G | C | 0.079 | -2.28E-01 | 1.07E-01 | 1.64E+00 | 1.69E+00 | 6.77E-02 | Yes | | intergenic_variant | MODIFIER |
| 15 | 70760667 | AX-97830357 | 0 | T | C | 0.079 | -2.28E-01 | 1.07E-01 | 1.64E+00 | 1.69E+00 | 6.77E-02 | Yes | Gm18155 | downstream_gene_variant | MODIFIER |
| 15 | 70773557 | AX-170514630 | 0 | T | C | 0.079 | -2.28E-01 | 1.07E-01 | 1.64E+00 | 1.69E+00 | 6.77E-02 | Yes | | intergenic_variant | MODIFIER |
| 15 | 70777379 | AX-170011944 | 0 | G | C | 0.079 | -2.28E-01 | 1.07E-01 | 1.64E+00 | 1.69E+00 | 6.77E-02 | Yes | | intergenic_variant | MODIFIER |
| 15 | 70807498 | AX-97835968 | 0 | T | C | 0.079 | -2.28E-01 | 1.07E-01 | 1.64E+00 | 1.69E+00 | 6.77E-02 | Yes | | intergenic_variant | MODIFIER |
| 15 | 70807846 | AX-170055903 | 0 | T | C | 0.079 | -2.28E-01 | 1.07E-01 | 1.64E+00 | 1.69E+00 | 6.77E-02 | Yes | | intergenic_variant | MODIFIER |
| 15 | 70826309 | AX-170011951 | 0 | A | C | 0.079 | -2.28E-01 | 1.07E-01 | 1.64E+00 | 1.69E+00 | 6.77E-02 | Yes | | intergenic_variant | MODIFIER |

| | | | | | | | | | | | | | | | |
|---|---|---|---|---|---|---|---|---|---|---|---|---|---|---|---|
| 15 | 70829919 | AX-170011949 | 0 | G | A | 0.079 | -2.28E-01 | 1.07E-01 | 1.64E+00 | 1.69E+00 | 6.77E-02 | Yes | | intergenic_variant | MODIFIER |
| 15 | 70840684 | AX-170157075 | 0 | C | T | 0.079 | -2.28E-01 | 1.07E-01 | 1.64E+00 | 1.69E+00 | 6.77E-02 | Yes | | intergenic_variant | MODIFIER |
| 15 | 70864754 | AX-170336751 | 0 | A | G | 0.079 | -2.28E-01 | 1.07E-01 | 1.64E+00 | 1.69E+00 | 6.77E-02 | Yes | | intergenic_variant | MODIFIER |
| 15 | 70864844 | AX-170126877 | 0 | A | T | 0.079 | -2.28E-01 | 1.07E-01 | 1.64E+00 | 1.69E+00 | 6.77E-02 | Yes | | intergenic_variant | MODIFIER |
| 15 | 70867131 | AX-170011954 | 0 | C | G | 0.079 | -2.28E-01 | 1.07E-01 | 1.64E+00 | 1.69E+00 | 6.77E-02 | Yes | | intergenic_variant | MODIFIER |
| 15 | 70882973 | AX-170011948 | 0 | T | C | 0.079 | -2.28E-01 | 1.07E-01 | 1.64E+00 | 1.69E+00 | 6.77E-02 | Yes | | intergenic_variant | MODIFIER |
| 15 | 70904510 | AX-170345476 | 0 | A | G | 0.079 | -2.28E-01 | 1.07E-01 | 1.64E+00 | 1.69E+00 | 6.77E-02 | Yes | | intergenic_variant | MODIFIER |
| 15 | 70904717 | AX-97820996 | 0 | C | T | 0.079 | -2.28E-01 | 1.07E-01 | 1.64E+00 | 1.69E+00 | 6.77E-02 | Yes | | intergenic_variant | MODIFIER |
| 15 | 70937296 | AX-170524325 | 0 | T | C | 0.079 | -2.28E-01 | 1.07E-01 | 1.64E+00 | 1.69E+00 | 6.77E-02 | Yes | | intergenic_variant | MODIFIER |
| 15 | 70945313 | AX-170473612 | 0 | A | C | 0.079 | -2.28E-01 | 1.07E-01 | 1.64E+00 | 1.69E+00 | 6.77E-02 | Yes | | intergenic_variant | MODIFIER |
| 15 | 70953118 | AX-97798212 | 0 | A | G | 0.079 | -2.28E-01 | 1.07E-01 | 1.64E+00 | 1.69E+00 | 6.77E-02 | Yes | | intergenic_variant | MODIFIER |
| 15 | 70988975 | AX-170314615 | 0 | C | A | 0.079 | -2.28E-01 | 1.07E-01 | 1.64E+00 | 1.69E+00 | 6.77E-02 | Yes | | regulatory_region_variant | MODIFIER |
| 15 | 70989080 | AX-170511511 | 0 | A | G | 0.079 | -2.28E-01 | 1.07E-01 | 1.64E+00 | 1.69E+00 | 6.77E-02 | Yes | | intergenic_variant | MODIFIER |
| 15 | 70993813 | AX-97829650 | 0 | C | T | 0.079 | -2.28E-01 | 1.07E-01 | 1.64E+00 | 1.69E+00 | 6.77E-02 | Yes | | intergenic_variant | MODIFIER |
| 15 | 71019664 | AX-170462322 | 0 | G | A | 0.079 | -2.28E-01 | 1.07E-01 | 1.64E+00 | 1.69E+00 | 6.77E-02 | Yes | | intergenic_variant | MODIFIER |
| 15 | 71032424 | AX-97846821 | 0 | T | C | 0.079 | -2.28E-01 | 1.07E-01 | 1.64E+00 | 1.69E+00 | 6.77E-02 | Yes | | intergenic_variant | MODIFIER |
| 15 | 71032967 | AX-170414253 | 0 | C | T | 0.079 | -2.28E-01 | 1.07E-01 | 1.64E+00 | 1.69E+00 | 6.77E-02 | Yes | | intergenic_variant | MODIFIER |
| 15 | 71070132 | AX-97789874 | 0 | C | T | 0.079 | -2.28E-01 | 1.07E-01 | 1.64E+00 | 1.69E+00 | 6.77E-02 | Yes | | intergenic_variant | MODIFIER |
| 15 | 71084051 | AX-97830358 | 0 | T | C | 0.079 | -2.28E-01 | 1.07E-01 | 1.64E+00 | 1.69E+00 | 6.77E-02 | Yes | | intergenic_variant | MODIFIER |
| 15 | 71113028 | AX-170184201 | 0 | G | A | 0.079 | -2.28E-01 | 1.07E-01 | 1.64E+00 | 1.69E+00 | 6.77E-02 | Yes | | intergenic_variant | MODIFIER |
| 15 | 71146580 | AX-170102808 | 0 | T | C | 0.079 | -2.28E-01 | 1.07E-01 | 1.64E+00 | 1.69E+00 | 6.77E-02 | Yes | | intergenic_variant | MODIFIER |
| 15 | 71157344 | AX-170142383 | 0 | A | C | 0.079 | -2.28E-01 | 1.07E-01 | 1.64E+00 | 1.69E+00 | 6.77E-02 | Yes | | intergenic_variant | MODIFIER |
| 15 | 71161011 | AX-168387969 | 0 | T | C | 0.079 | -2.28E-01 | 1.07E-01 | 1.64E+00 | 1.69E+00 | 6.77E-02 | Yes | | intergenic_variant | MODIFIER |
| 15 | 71161286 | AX-97843484 | 0 | T | C | 0.079 | -2.28E-01 | 1.07E-01 | 1.64E+00 | 1.69E+00 | 6.77E-02 | Yes | | intergenic_variant | MODIFIER |
| 15 | 71179429 | AX-170011964 | 0 | G | C | 0.079 | -2.28E-01 | 1.07E-01 | 1.64E+00 | 1.69E+00 | 6.77E-02 | Yes | | intergenic_variant | MODIFIER |
| 15 | 71193729 | AX-170011961 | 0 | G | A | 0.079 | -2.28E-01 | 1.07E-01 | 1.64E+00 | 1.69E+00 | 6.77E-02 | Yes | | intergenic_variant | MODIFIER |
| 15 | 71198147 | AX-170217497 | 0 | G | A | 0.079 | -2.28E-01 | 1.07E-01 | 1.64E+00 | 1.69E+00 | 6.77E-02 | Yes | | intergenic_variant | MODIFIER |
| 15 | 71203814 | AX-170499825 | 0 | T | C | 0.079 | -2.28E-01 | 1.07E-01 | 1.64E+00 | 1.69E+00 | 6.77E-02 | Yes | | intergenic_variant | MODIFIER |
| 15 | 71220988 | AX-170087901 | 0 | C | G | 0.079 | -2.28E-01 | 1.07E-01 | 1.64E+00 | 1.69E+00 | 6.77E-02 | Yes | | intergenic_variant | MODIFIER |
| 15 | 71242528 | AX-170011957 | 0 | G | A | 0.079 | -2.28E-01 | 1.07E-01 | 1.64E+00 | 1.69E+00 | 6.77E-02 | Yes | | intergenic_variant | MODIFIER |
| 15 | 71243451 | AX-97797173 | 0 | T | C | 0.079 | -2.28E-01 | 1.07E-01 | 1.64E+00 | 1.69E+00 | 6.77E-02 | Yes | | intergenic_variant | MODIFIER |
| 15 | 71344899 | AX-170210876 | 0 | T | C | 0.079 | -2.28E-01 | 1.07E-01 | 1.64E+00 | 1.69E+00 | 6.77E-02 | Yes | | intergenic_variant | MODIFIER |
| 15 | 71420018 | AX-170055910 | 0 | A | G | 0.079 | -2.28E-01 | 1.07E-01 | 1.64E+00 | 1.69E+00 | 6.77E-02 | Yes | | intergenic_variant | MODIFIER |
| 15 | 71436878 | AX-170249883 | 0 | T | C | 0.079 | -2.28E-01 | 1.07E-01 | 1.64E+00 | 1.69E+00 | 6.77E-02 | Yes | Fam135b | 3_prime_UTR_variant | MODIFIER |
| 15 | 71437855 | AX-170364432 | 0 | A | C | 0.079 | -2.28E-01 | 1.07E-01 | 1.64E+00 | 1.69E+00 | 6.77E-02 | Yes | Fam135b | 3_prime_UTR_variant | MODIFIER |
| 15 | 71441229 | AX-170249232 | 0 | A | G | 0.079 | -2.28E-01 | 1.07E-01 | 1.64E+00 | 1.69E+00 | 6.77E-02 | Yes | Fam135b | 3_prime_UTR_variant | MODIFIER |
| 15 | 71441518 | AX-170239887 | 0 | G | A | 0.079 | -2.28E-01 | 1.07E-01 | 1.64E+00 | 1.69E+00 | 6.77E-02 | Yes | Fam135b | 3_prime_UTR_variant | MODIFIER |
| 15 | 71595754 | AX-170192975 | 0 | A | G | 0.079 | -2.28E-01 | 1.07E-01 | 1.64E+00 | 1.69E+00 | 6.77E-02 | Yes | Fam135b | intron_variant | MODIFIER |
| 15 | 71625374 | AX-170252542 | 0 | A | G | 0.079 | -2.28E-01 | 1.07E-01 | 1.64E+00 | 1.69E+00 | 6.77E-02 | Yes | Fam135b | intron_variant | MODIFIER |
| 15 | 71626556 | AX-170239744 | 0 | A | G | 0.079 | -2.28E-01 | 1.07E-01 | 1.64E+00 | 1.69E+00 | 6.77E-02 | Yes | Fam135b | intron_variant | MODIFIER |
| 15 | 71632551 | AX-170222863 | 0 | T | C | 0.079 | -2.28E-01 | 1.07E-01 | 1.64E+00 | 1.69E+00 | 6.77E-02 | Yes | Fam135b | intron_variant | MODIFIER |
| 15 | 71666829 | AX-170467498 | 0 | A | G | 0.079 | -2.28E-01 | 1.07E-01 | 1.64E+00 | 1.69E+00 | 6.77E-02 | Yes | Fam135b | intron_variant | MODIFIER |
| 15 | 71674261 | AX-170087903 | 0 | C | A | 0.079 | -2.28E-01 | 1.07E-01 | 1.64E+00 | 1.69E+00 | 6.77E-02 | Yes | Fam135b | intron_variant | MODIFIER |
| 15 | 71734962 | AX-170342789 | 0 | T | C | 0.079 | -2.28E-01 | 1.07E-01 | 1.64E+00 | 1.69E+00 | 6.77E-02 | Yes | | intergenic_variant | MODIFIER |
| 15 | 71753200 | AX-97852679 | 0 | G | A | 0.079 | -2.28E-01 | 1.07E-01 | 1.64E+00 | 1.69E+00 | 6.77E-02 | Yes | | intergenic_variant | MODIFIER |
| 15 | 71759118 | AX-170108299 | 0 | C | G | 0.079 | -2.28E-01 | 1.07E-01 | 1.64E+00 | 1.69E+00 | 6.77E-02 | Yes | | intergenic_variant | MODIFIER |
| 15 | 71759337 | AX-97805157 | 0 | A | C | 0.079 | -2.28E-01 | 1.07E-01 | 1.64E+00 | 1.69E+00 | 6.77E-02 | Yes | | intergenic_variant | MODIFIER |
| 15 | 71773302 | AX-170011971 | 0 | C | T | 0.079 | -2.28E-01 | 1.07E-01 | 1.64E+00 | 1.69E+00 | 6.77E-02 | Yes | | intergenic_variant | MODIFIER |
| 15 | 71841199 | AX-170011969 | 0 | G | A | 0.079 | -2.28E-01 | 1.07E-01 | 1.64E+00 | 1.69E+00 | 6.77E-02 | Yes | Col22a1 | intron_variant | MODIFIER |
| 15 | 71841334 | AX-170269614 | 0 | G | A | 0.079 | -2.28E-01 | 1.07E-01 | 1.64E+00 | 1.69E+00 | 6.77E-02 | Yes | Col22a1 | splice_region_variant,intron_variant | LOW |
| 15 | 71851804 | AX-170278690 | 0 | G | A | 0.079 | -2.28E-01 | 1.07E-01 | 1.64E+00 | 1.69E+00 | 6.77E-02 | Yes | Col22a1 | intron_variant | MODIFIER |
| 15 | 71886869 | AX-170346746 | 0 | T | C | 0.079 | -2.28E-01 | 1.07E-01 | 1.64E+00 | 1.69E+00 | 6.77E-02 | Yes | Col22a1 | intron_variant | MODIFIER |
| 15 | 71887343 | AX-170512404 | 0 | T | A | 0.079 | -2.28E-01 | 1.07E-01 | 1.64E+00 | 1.69E+00 | 6.77E-02 | Yes | Col22a1 | intron_variant | MODIFIER |
| 15 | 71888082 | AX-170352629 | 0 | A | T | 0.079 | -2.28E-01 | 1.07E-01 | 1.64E+00 | 1.69E+00 | 6.77E-02 | Yes | Col22a1 | intron_variant | MODIFIER |

| | | | | | | | | | | | | | | | |
|---|---|---|---|---|---|---|---|---|---|---|---|---|---|---|---|
| 15 | 71905877 | AX-170227070 | 0 | A | C | 0.079 | -2.28E-01 | 1.07E-01 | 1.64E+00 | 1.69E+00 | 6.77E-02 | Yes | Col22a1 | intron_variant | MODIFIER |
| 15 | 71905916 | AX-170388833 | 0 | C | G | 0.079 | -2.28E-01 | 1.07E-01 | 1.64E+00 | 1.69E+00 | 6.77E-02 | Yes | Col22a1 | intron_variant | MODIFIER |
| 15 | 71919387 | AX-170055923 | 0 | C | T | 0.079 | -2.28E-01 | 1.07E-01 | 1.64E+00 | 1.69E+00 | 6.77E-02 | Yes | Col22a1 | intron_variant | MODIFIER |
| 15 | 71920089 | AX-170451953 | 0 | A | C | 0.079 | -2.28E-01 | 1.07E-01 | 1.64E+00 | 1.69E+00 | 6.77E-02 | Yes | Col22a1 | intron_variant | MODIFIER |
| 15 | 71922560 | AX-170143167 | 0 | T | C | 0.079 | -2.28E-01 | 1.07E-01 | 1.64E+00 | 1.69E+00 | 6.77E-02 | Yes | Col22a1 | intron_variant | MODIFIER |
| 15 | 71929964 | AX-170301094 | 0 | A | C | 0.079 | -2.28E-01 | 1.07E-01 | 1.64E+00 | 1.69E+00 | 6.77E-02 | Yes | Col22a1 | intron_variant | MODIFIER |
| 15 | 71938486 | AX-170285505 | 0 | G | A | 0.079 | -2.28E-01 | 1.07E-01 | 1.64E+00 | 1.69E+00 | 6.77E-02 | Yes | Col22a1 | intron_variant | MODIFIER |
| 15 | 71941494 | AX-170214115 | 0 | C | T | 0.079 | -2.28E-01 | 1.07E-01 | 1.64E+00 | 1.69E+00 | 6.77E-02 | Yes | Col22a1 | intron_variant | MODIFIER |
| 15 | 71941915 | AX-97816726 | 0 | G | T | 0.079 | -2.28E-01 | 1.07E-01 | 1.64E+00 | 1.69E+00 | 6.77E-02 | Yes | Col22a1 | intron_variant | MODIFIER |
| 15 | 71945754 | AX-170172205 | 0 | T | C | 0.079 | -2.28E-01 | 1.07E-01 | 1.64E+00 | 1.69E+00 | 6.77E-02 | Yes | Col22a1 | intron_variant | MODIFIER |
| 15 | 72032567 | AX-97856555 | 0 | A | G | 0.079 | -2.28E-01 | 1.07E-01 | 1.64E+00 | 1.69E+00 | 6.77E-02 | Yes | Col22a1 | intron_variant | MODIFIER |
| 15 | 72040484 | AX-170517613 | 0 | C | T | 0.079 | -2.28E-01 | 1.07E-01 | 1.64E+00 | 1.69E+00 | 6.77E-02 | Yes | | intergenic_variant | MODIFIER |
| 15 | 72040844 | AX-170326852 | 0 | C | A | 0.079 | -2.28E-01 | 1.07E-01 | 1.64E+00 | 1.69E+00 | 6.77E-02 | Yes | | intergenic_variant | MODIFIER |
| 15 | 72042624 | AX-170236447 | 0 | T | A | 0.079 | -2.28E-01 | 1.07E-01 | 1.64E+00 | 1.69E+00 | 6.77E-02 | Yes | | intergenic_variant | MODIFIER |
| 15 | 72053332 | AX-170197268 | 0 | T | C | 0.079 | -2.28E-01 | 1.07E-01 | 1.64E+00 | 1.69E+00 | 6.77E-02 | Yes | | intergenic_variant | MODIFIER |
| 15 | 72167692 | AX-97815455 | 0 | G | C | 0.079 | -2.28E-01 | 1.07E-01 | 1.64E+00 | 1.69E+00 | 6.77E-02 | Yes | Gm49457 | intron_variant,non_coding_transcript_variant | MODIFIER |
| 15 | 72226145 | AX-170055931 | 0 | T | G | 0.079 | -2.28E-01 | 1.07E-01 | 1.64E+00 | 1.69E+00 | 6.77E-02 | Yes | Gm49457 | intron_variant,non_coding_transcript_variant | MODIFIER |
| 15 | 72352625 | AX-170055933 | 0 | T | C | 0.079 | -2.28E-01 | 1.07E-01 | 1.64E+00 | 1.69E+00 | 6.77E-02 | Yes | | intergenic_variant | MODIFIER |
| 15 | 76472005 | AX-97829756 | 0 | C | A | 0.079 | -2.28E-01 | 1.07E-01 | 1.64E+00 | 1.69E+00 | 6.77E-02 | Yes | Bop1 | intron_variant | MODIFIER |
| 15 | 76502013 | AX-170474485 | 0 | A | G | 0.079 | -2.28E-01 | 1.07E-01 | 1.64E+00 | 1.69E+00 | 6.77E-02 | Yes | Dgat1 | downstream_gene_variant | MODIFIER |
| 15 | 76502794 | AX-170373900 | 0 | C | T | 0.079 | -2.28E-01 | 1.07E-01 | 1.64E+00 | 1.69E+00 | 6.77E-02 | Yes | Dgat1 | intron_variant | MODIFIER |
| 15 | 76512861 | AX-170510688 | 0 | C | T | 0.079 | -2.28E-01 | 1.07E-01 | 1.64E+00 | 1.69E+00 | 6.77E-02 | Yes | Dgat1 | upstream_gene_variant | MODIFIER |
| 15 | 76548385 | AX-170516985 | 0 | T | A | 0.079 | -2.28E-01 | 1.07E-01 | 1.64E+00 | 1.69E+00 | 6.77E-02 | Yes | Slc52a2 | downstream_gene_variant | MODIFIER |
| 15 | 76628915 | AX-97806573 | 0 | G | C | 0.079 | -2.28E-01 | 1.07E-01 | 1.64E+00 | 1.69E+00 | 6.77E-02 | Yes | Vps28 | upstream_gene_variant | MODIFIER |
| 15 | 76646783 | AX-97830454 | 0 | C | T | 0.079 | -2.28E-01 | 1.07E-01 | 1.64E+00 | 1.69E+00 | 6.77E-02 | Yes | Cyhr1 | intron_variant | MODIFIER |
| 15 | 76654047 | AX-170012028 | 0 | G | A | 0.079 | -2.28E-01 | 1.07E-01 | 1.64E+00 | 1.69E+00 | 6.77E-02 | Yes | Cyhr1 | downstream_gene_variant | MODIFIER |
| 15 | 76692569 | AX-97835138 | 0 | C | T | 0.079 | -2.28E-01 | 1.07E-01 | 1.64E+00 | 1.69E+00 | 6.77E-02 | Yes | Gpt | upstream_gene_variant | MODIFIER |
| 15 | 76699732 | AX-170531392 | 0 | T | C | 0.079 | -2.28E-01 | 1.07E-01 | 1.64E+00 | 1.69E+00 | 6.77E-02 | Yes | Mfsd3 | upstream_gene_variant | MODIFIER |
| 15 | 76707129 | AX-170012026 | 0 | G | T | 0.079 | -2.28E-01 | 1.07E-01 | 1.64E+00 | 1.69E+00 | 6.77E-02 | Yes | Mfsd3 | downstream_gene_variant | MODIFIER |
| 15 | 76715111 | AX-170012027 | 0 | A | G | 0.079 | -2.28E-01 | 1.07E-01 | 1.64E+00 | 1.69E+00 | 6.77E-02 | Yes | Lrrc14 | downstream_gene_variant | MODIFIER |
| 15 | 76715546 | AX-170093308 | 0 | A | G | 0.079 | -2.28E-01 | 1.07E-01 | 1.64E+00 | 1.69E+00 | 6.77E-02 | Yes | Lrrc14 | downstream_gene_variant | MODIFIER |
| 15 | 76730925 | AX-170012025 | 0 | C | T | 0.079 | -2.28E-01 | 1.07E-01 | 1.64E+00 | 1.69E+00 | 6.77E-02 | Yes | Arhgap39 | intron_variant | MODIFIER |
| 15 | 76751038 | AX-97831284 | 0 | C | T | 0.079 | -2.28E-01 | 1.07E-01 | 1.64E+00 | 1.69E+00 | 6.77E-02 | Yes | Arhgap39 | intron_variant | MODIFIER |
| 15 | 76759982 | AX-170322629 | 0 | C | G | 0.079 | -2.28E-01 | 1.07E-01 | 1.64E+00 | 1.69E+00 | 6.77E-02 | Yes | Arhgap39 | intron_variant | MODIFIER |
| 15 | 76760024 | AX-170197429 | 0 | A | G | 0.079 | -2.28E-01 | 1.07E-01 | 1.64E+00 | 1.69E+00 | 6.77E-02 | Yes | Arhgap39 | intron_variant | MODIFIER |
| 15 | 76760354 | AX-170331408 | 0 | A | G | 0.079 | -2.28E-01 | 1.07E-01 | 1.64E+00 | 1.69E+00 | 6.77E-02 | Yes | Arhgap39 | intron_variant | MODIFIER |
| 15 | 76766672 | AX-170249193 | 0 | G | T | 0.079 | -2.28E-01 | 1.07E-01 | 1.64E+00 | 1.69E+00 | 6.77E-02 | Yes | Arhgap39 | intron_variant | MODIFIER |
| 15 | 76768158 | AX-170117939 | 0 | A | C | 0.079 | -2.28E-01 | 1.07E-01 | 1.64E+00 | 1.69E+00 | 6.77E-02 | Yes | Arhgap39 | intron_variant | MODIFIER |
| 15 | 76773603 | AX-170524454 | 0 | T | C | 0.079 | -2.28E-01 | 1.07E-01 | 1.64E+00 | 1.69E+00 | 6.77E-02 | Yes | Arhgap39 | intron_variant | MODIFIER |
| 15 | 76784843 | AX-170295095 | 0 | C | A | 0.079 | -2.28E-01 | 1.07E-01 | 1.64E+00 | 1.69E+00 | 6.77E-02 | Yes | Arhgap39 | intron_variant | MODIFIER |
| 15 | 76784889 | AX-97803166 | 0 | G | A | 0.079 | -2.28E-01 | 1.07E-01 | 1.64E+00 | 1.69E+00 | 6.77E-02 | Yes | Arhgap39 | intron_variant | MODIFIER |
| 15 | 76784996 | AX-170242498 | 0 | T | A | 0.079 | -2.28E-01 | 1.07E-01 | 1.64E+00 | 1.69E+00 | 6.77E-02 | Yes | Arhgap39 | intron_variant | MODIFIER |
| 15 | 76873588 | AX-170374279 | 0 | A | C | 0.079 | -2.28E-01 | 1.07E-01 | 1.64E+00 | 1.69E+00 | 6.77E-02 | Yes | Zfp251 | upstream_gene_variant | MODIFIER |
| 15 | 76887668 | AX-170116501 | 0 | T | G | 0.079 | -2.28E-01 | 1.07E-01 | 1.64E+00 | 1.69E+00 | 6.77E-02 | Yes | Zfp7 | intron_variant | MODIFIER |
| 15 | 76888143 | AX-170270244 | 0 | T | G | 0.079 | -2.28E-01 | 1.07E-01 | 1.64E+00 | 1.69E+00 | 6.77E-02 | Yes | Zfp7 | intron_variant | MODIFIER |
| 15 | 76889580 | AX-170056030 | 0 | G | C | 0.079 | -2.28E-01 | 1.07E-01 | 1.64E+00 | 1.69E+00 | 6.77E-02 | Yes | Zfp7 | intron_variant | MODIFIER |
| 15 | 76905662 | AX-170462934 | 0 | C | T | 0.079 | -2.28E-01 | 1.07E-01 | 1.64E+00 | 1.69E+00 | 6.77E-02 | Yes | Rpl8 | intron_variant | MODIFIER |
| 15 | 76906520 | AX-170203632 | 0 | C | T | 0.079 | -2.28E-01 | 1.07E-01 | 1.64E+00 | 1.69E+00 | 6.77E-02 | Yes | Rpl8 | downstream_gene_variant | MODIFIER |
| 15 | 76987537 | AX-97829759 | 0 | G | A | 0.079 | -2.28E-01 | 1.07E-01 | 1.64E+00 | 1.69E+00 | 6.77E-02 | Yes | Gm35933 | upstream_gene_variant | MODIFIER |
| 15 | 76992942 | AX-170270635 | 0 | A | G | 0.079 | -2.28E-01 | 1.07E-01 | 1.64E+00 | 1.69E+00 | 6.77E-02 | Yes | Gm35933 | intron_variant,non_coding_transcript_variant | MODIFIER |
| 15 | 77054363 | AX-97789615 | 0 | T | C | 0.079 | -2.28E-01 | 1.07E-01 | 1.64E+00 | 1.69E+00 | 6.77E-02 | Yes | Apol6 | downstream_gene_variant | MODIFIER |
| 15 | 77054441 | AX-170139223 | 0 | A | G | 0.079 | -2.28E-01 | 1.07E-01 | 1.64E+00 | 1.69E+00 | 6.77E-02 | Yes | Apol6 | downstream_gene_variant | MODIFIER |
| 15 | 77057120 | AX-170319473 | 0 | C | T | 0.079 | -2.28E-01 | 1.07E-01 | 1.64E+00 | 1.69E+00 | 6.77E-02 | Yes | Apol6 | downstream_gene_variant | MODIFIER |
| 15 | 77060965 | AX-170465639 | 0 | T | A | 0.079 | -2.28E-01 | 1.07E-01 | 1.64E+00 | 1.69E+00 | 6.77E-02 | Yes | Apol6 | downstream_gene_variant | MODIFIER |

| | | | | | | | | | | | | | | | |
|---|---|---|---|---|---|---|---|---|---|---|---|---|---|---|---|
| 15 | 77062137 | AX-97807714 | 0 | A | G | 0.079 | -2.28E-01 | 1.07E-01 | 1.64E+00 | 1.69E+00 | 6.77E-02 | Yes | | intergenic_variant | MODIFIER |
| 15 | 77072624 | AX-170405953 | 0 | T | C | 0.079 | -2.28E-01 | 1.07E-01 | 1.64E+00 | 1.69E+00 | 6.77E-02 | Yes | Gm24753 | downstream_gene_variant | MODIFIER |
| 15 | 77091172 | AX-170087913 | 0 | G | A | 0.079 | -2.28E-01 | 1.07E-01 | 1.64E+00 | 1.69E+00 | 6.77E-02 | Yes | Rbfox2 | intron_variant | MODIFIER |
| 15 | 77094875 | AX-170056037 | 0 | C | A | 0.079 | -2.28E-01 | 1.07E-01 | 1.64E+00 | 1.69E+00 | 6.77E-02 | Yes | Rbfox2 | intron_variant | MODIFIER |
| 15 | 77103515 | AX-170056039 | 0 | T | A | 0.079 | -2.28E-01 | 1.07E-01 | 1.64E+00 | 1.69E+00 | 6.77E-02 | Yes | Rbfox2 | intron_variant | MODIFIER |
| 15 | 77114241 | AX-170468492 | 0 | T | C | 0.079 | -2.28E-01 | 1.07E-01 | 1.64E+00 | 1.69E+00 | 6.77E-02 | Yes | Rbfox2 | intron_variant | MODIFIER |
| 15 | 77115130 | AX-170311202 | 0 | T | C | 0.079 | -2.28E-01 | 1.07E-01 | 1.64E+00 | 1.69E+00 | 6.77E-02 | Yes | Rbfox2 | intron_variant | MODIFIER |
| 15 | 77147095 | AX-170160374 | 0 | G | A | 0.079 | -2.28E-01 | 1.07E-01 | 1.64E+00 | 1.69E+00 | 6.77E-02 | Yes | Rbfox2 | intron_variant | MODIFIER |
| 15 | 77207076 | AX-170012031 | 0 | C | T | 0.079 | -2.28E-01 | 1.07E-01 | 1.64E+00 | 1.69E+00 | 6.77E-02 | Yes | Rbfox2 | intron_variant | MODIFIER |
| 15 | 77224621 | AX-97829786 | 0 | C | T | 0.079 | -2.28E-01 | 1.07E-01 | 1.64E+00 | 1.69E+00 | 6.77E-02 | Yes | Rbfox2 | intron_variant | MODIFIER |
| 15 | 77246407 | AX-170093310 | 0 | G | A | 0.079 | -2.28E-01 | 1.07E-01 | 1.64E+00 | 1.69E+00 | 6.77E-02 | Yes | Rbfox2 | intron_variant | MODIFIER |
| 15 | 77298382 | AX-97830451 | 0 | C | T | 0.079 | -2.28E-01 | 1.07E-01 | 1.64E+00 | 1.69E+00 | 6.77E-02 | Yes | Rbfox2 | intron_variant | MODIFIER |
| 15 | 77437626 | AX-170012036 | 0 | T | C | 0.079 | -2.28E-01 | 1.07E-01 | 1.64E+00 | 1.69E+00 | 6.77E-02 | Yes | Apol7b | intron_variant | MODIFIER |
| 15 | 77445224 | AX-170126157 | 0 | T | C | 0.079 | -2.28E-01 | 1.07E-01 | 1.64E+00 | 1.69E+00 | 6.77E-02 | Yes | Apol7b | intron_variant | MODIFIER |
| 15 | 77531892 | AX-97782719 | 0 | T | C | 0.079 | -2.28E-01 | 1.07E-01 | 1.64E+00 | 1.69E+00 | 6.77E-02 | Yes | Apol7c | intron_variant | MODIFIER |
| 15 | 77541572 | AX-170164211 | 0 | A | C | 0.079 | -2.28E-01 | 1.07E-01 | 1.64E+00 | 1.69E+00 | 6.77E-02 | Yes | | intergenic_variant | MODIFIER |
| 15 | 77544014 | AX-170497378 | 0 | G | C | 0.079 | -2.28E-01 | 1.07E-01 | 1.64E+00 | 1.69E+00 | 6.77E-02 | Yes | | intergenic_variant | MODIFIER |
| 15 | 77564956 | AX-170087915 | 0 | C | T | 0.079 | -2.28E-01 | 1.07E-01 | 1.64E+00 | 1.69E+00 | 6.77E-02 | Yes | 1700025B11Rik | upstream_gene_variant | MODIFIER |
| 15 | 77567327 | AX-170402544 | 0 | C | T | 0.079 | -2.28E-01 | 1.07E-01 | 1.64E+00 | 1.69E+00 | 6.77E-02 | Yes | | intergenic_variant | MODIFIER |
| 15 | 77596181 | AX-97793822 | 0 | C | T | 0.079 | -2.28E-01 | 1.07E-01 | 1.64E+00 | 1.69E+00 | 6.77E-02 | Yes | Apol10b | upstream_gene_variant | MODIFIER |
| 15 | 77633593 | AX-97829787 | 0 | T | C | 0.079 | -2.28E-01 | 1.07E-01 | 1.64E+00 | 1.69E+00 | 6.77E-02 | Yes | Apol11b | downstream_gene_variant | MODIFIER |
| 15 | 77670151 | AX-97830306 | 0 | T | C | 0.079 | -2.28E-01 | 1.07E-01 | 1.64E+00 | 1.69E+00 | 6.77E-02 | Yes | Gm49407 | upstream_gene_variant | MODIFIER |
| 5 | 105608861 | AX-170163261 | 0 | A | T | 0.439 | 1.83E-01 | 1.00E-01 | 2.86E+00 | 2.96E+00 | 6.78E-02 | Yes | Lrrc8c | 3_prime_UTR_variant | MODIFIER |
| 5 | 65019083 | AX-170469167 | 0 | T | A | 0.482 | 1.89E-01 | 1.03E-01 | 2.48E+00 | 2.53E+00 | 6.80E-02 | Yes | Fam114a1 | intron_variant | MODIFIER |
| 5 | 14028382 | AX-170071351 | 1 | A | G | 0.321 | 1.56E-01 | 8.49E-02 | 3.11E+00 | 3.27E+00 | 6.80E-02 | Yes | Sema3e | intron_variant | MODIFIER |
| 5 | 135768357 | AX-97835473 | 1 | T | C | 0.161 | -1.45E-01 | 7.85E-02 | 2.36E+00 | 2.47E+00 | 6.80E-02 | no | Styxl1 | intron_variant | MODIFIER |
| 13 | 10550959 | AX-170335737 | 0 | G | T | 0.184 | -2.27E-01 | 1.24E-01 | 2.48E+00 | 2.53E+00 | 6.82E-02 | Yes | Gm26861 | intron_variant,non_coding_transcript_variant | MODIFIER |
| 13 | 11823018 | AX-170283725 | 0 | C | T | 0.184 | -2.27E-01 | 1.24E-01 | 2.48E+00 | 2.53E+00 | 6.82E-02 | Yes | Ryr2 | intron_variant | MODIFIER |
| 9 | 111364796 | AX-170228227 | 0 | T | C | 0.096 | -1.91E-01 | 1.05E-01 | 2.80E+00 | 2.94E+00 | 6.82E-02 | Yes | Trank1 | synonymous_variant | LOW |
| 19 | 52359370 | AX-170249028 | 0 | A | G | 0.482 | 1.16E-01 | 6.39E-02 | 2.71E+00 | 2.79E+00 | 6.82E-02 | no | | intergenic_variant | MODIFIER |
| 19 | 53111794 | AX-170294298 | 0 | C | T | 0.482 | 1.16E-01 | 6.39E-02 | 2.71E+00 | 2.79E+00 | 6.82E-02 | no | | regulatory_region_variant | MODIFIER |
| 19 | 53111888 | AX-170230167 | 0 | G | A | 0.482 | 1.16E-01 | 6.39E-02 | 2.71E+00 | 2.79E+00 | 6.82E-02 | no | | regulatory_region_variant | MODIFIER |
| 19 | 53112726 | AX-170103157 | 0 | C | A | 0.482 | 1.16E-01 | 6.39E-02 | 2.71E+00 | 2.79E+00 | 6.82E-02 | no | | regulatory_region_variant | MODIFIER |
| 19 | 52353874 | AX-170016831 | 1 | A | C | 0.482 | 1.17E-01 | 6.41E-02 | 2.74E+00 | 2.83E+00 | 6.82E-02 | no | | intergenic_variant | MODIFIER |
| 19 | 52728836 | AX-170421333 | 1 | A | G | 0.482 | 1.17E-01 | 6.41E-02 | 2.74E+00 | 2.83E+00 | 6.82E-02 | no | | intergenic_variant | MODIFIER |
| 5 | 57193219 | AX-168418383 | 1 | T | G | 0.455 | 1.79E-01 | 9.77E-02 | 2.50E+00 | 2.57E+00 | 6.85E-02 | Yes | | intergenic_variant | MODIFIER |
| 10 | 18189943 | AX-170511494 | 1 | T | A | 0.214 | 1.84E-01 | 9.67E-02 | 3.59E+00 | 3.77E+00 | 6.86E-02 | Yes | Ect2l | intron_variant | MODIFIER |
| 5 | 13880145 | AX-170270987 | 1 | G | T | 0.33 | 1.55E-01 | 8.49E-02 | 3.10E+00 | 3.26E+00 | 6.91E-02 | Yes | Dnd1-ps | non_coding_transcript_exon_variant | MODIFIER |
| 4 | 108869636 | AX-170070302 | 0 | G | A | 0.228 | 1.94E-01 | 1.07E-01 | 2.85E+00 | 2.98E+00 | 6.92E-02 | Yes | | intergenic_variant | MODIFIER |
| 1 | 185657950 | AX-97836243 | 0 | C | T | 0.237 | 1.98E-01 | 1.09E-01 | 2.53E+00 | 2.60E+00 | 6.94E-02 | Yes | | intergenic_variant | MODIFIER |
| 1 | 185669728 | AX-170407383 | 0 | C | T | 0.237 | 1.98E-01 | 1.09E-01 | 2.53E+00 | 2.60E+00 | 6.94E-02 | Yes | | regulatory_region_variant | MODIFIER |
| 1 | 185670006 | AX-170505257 | 0 | T | C | 0.237 | 1.98E-01 | 1.09E-01 | 2.53E+00 | 2.60E+00 | 6.94E-02 | Yes | | regulatory_region_variant | MODIFIER |
| 1 | 185670381 | AX-170156665 | 0 | T | C | 0.237 | 1.98E-01 | 1.09E-01 | 2.53E+00 | 2.60E+00 | 6.94E-02 | Yes | | regulatory_region_variant | MODIFIER |
| 1 | 185671169 | AX-170412335 | 0 | G | A | 0.237 | 1.98E-01 | 1.09E-01 | 2.53E+00 | 2.60E+00 | 6.94E-02 | Yes | | regulatory_region_variant | MODIFIER |
| 1 | 185674529 | AX-170181217 | 0 | T | C | 0.237 | 1.98E-01 | 1.09E-01 | 2.53E+00 | 2.60E+00 | 6.94E-02 | Yes | | intergenic_variant | MODIFIER |
| 1 | 185675771 | AX-170043863 | 0 | C | T | 0.237 | 1.98E-01 | 1.09E-01 | 2.53E+00 | 2.60E+00 | 6.94E-02 | Yes | | intergenic_variant | MODIFIER |
| 1 | 185677672 | AX-170325000 | 0 | T | C | 0.237 | 1.98E-01 | 1.09E-01 | 2.53E+00 | 2.60E+00 | 6.94E-02 | Yes | | intergenic_variant | MODIFIER |
| 1 | 185679032 | AX-170043866 | 0 | C | T | 0.237 | 1.98E-01 | 1.09E-01 | 2.53E+00 | 2.60E+00 | 6.94E-02 | Yes | | intergenic_variant | MODIFIER |
| 1 | 185687676 | AX-170090839 | 0 | C | T | 0.237 | 1.98E-01 | 1.09E-01 | 2.53E+00 | 2.60E+00 | 6.94E-02 | Yes | | intergenic_variant | MODIFIER |
| 1 | 185762630 | AX-170467552 | 0 | T | C | 0.237 | 1.98E-01 | 1.09E-01 | 2.53E+00 | 2.60E+00 | 6.94E-02 | Yes | Gm38093 | downstream_gene_variant | MODIFIER |
| 1 | 185767332 | AX-170151758 | 0 | G | C | 0.237 | 1.98E-01 | 1.09E-01 | 2.53E+00 | 2.60E+00 | 6.94E-02 | Yes | | intergenic_variant | MODIFIER |
| 1 | 185767821 | AX-170490816 | 0 | A | G | 0.237 | 1.98E-01 | 1.09E-01 | 2.53E+00 | 2.60E+00 | 6.94E-02 | Yes | | intergenic_variant | MODIFIER |
| 1 | 185769111 | AX-170156074 | 0 | C | T | 0.237 | 1.98E-01 | 1.09E-01 | 2.53E+00 | 2.60E+00 | 6.94E-02 | Yes | | intergenic_variant | MODIFIER |
| 1 | 185787602 | AX-97844005 | 0 | C | G | 0.237 | 1.98E-01 | 1.09E-01 | 2.53E+00 | 2.60E+00 | 6.94E-02 | Yes | | intergenic_variant | MODIFIER |

| | | | | | | | | | | | | | | | |
|---|---|---|---|---|---|---|---|---|---|---|---|---|---|---|---|
| 13 | 11868943 | AX-97783034 | 0 | T | C | 0.175 | -2.20E-01 | 1.21E-01 | 2.56E+00 | 2.61E+00 | 6.96E-02 | Yes | Ryr2 | intron_variant | MODIFIER |
| 6 | 135946555 | AX-170463493 | 0 | G | A | 0.579 | 1.59E-01 | 8.73E-02 | 2.45E+00 | 2.51E+00 | 6.96E-02 | Yes | Grin2b | intron_variant | MODIFIER |
| 6 | 136141088 | AX-170104043 | 0 | T | C | 0.579 | 1.59E-01 | 8.73E-02 | 2.45E+00 | 2.51E+00 | 6.96E-02 | Yes | Grin2b | intron_variant | MODIFIER |
| 6 | 137079191 | AX-170486801 | 0 | T | A | 0.579 | 1.59E-01 | 8.73E-02 | 2.45E+00 | 2.51E+00 | 6.96E-02 | Yes | Rerg | intron_variant | MODIFIER |
| 6 | 137561636 | AX-170228316 | 0 | A | G | 0.579 | 1.59E-01 | 8.73E-02 | 2.45E+00 | 2.51E+00 | 6.96E-02 | Yes | Eps8 | intron_variant | MODIFIER |
| 6 | 137586368 | AX-170208248 | 0 | T | A | 0.579 | 1.59E-01 | 8.73E-02 | 2.45E+00 | 2.51E+00 | 6.96E-02 | Yes | Eps8 | intron_variant | MODIFIER |
| 16 | 64068447 | AX-170219325 | 1 | T | A | 0.509 | 1.59E-01 | 8.77E-02 | 2.79E+00 | 2.85E+00 | 6.98E-02 | Yes | | intergenic_variant | MODIFIER |
| 5 | 56451231 | AX-97817199 | 1 | A | C | 0.473 | 1.84E-01 | 1.00E-01 | 2.35E+00 | 2.41E+00 | 6.98E-02 | Yes | Gm42977 | intron_variant,non_coding_transcript_variant | MODIFIER |
| 5 | 62554554 | AX-170025121 | 1 | C | T | 0.473 | 1.84E-01 | 1.00E-01 | 2.35E+00 | 2.41E+00 | 6.98E-02 | Yes | | intergenic_variant | MODIFIER |
| 19 | 51795407 | AX-170016819 | 0 | T | C | 0.482 | 1.17E-01 | 6.45E-02 | 2.82E+00 | 2.92E+00 | 6.98E-02 | no | Rpl31-ps18 | downstream_gene_variant | MODIFIER |
| 18 | 11192923 | AX-97834796 | 0 | A | C | 0.035 | -2.97E-01 | 1.64E-01 | 2.84E+00 | 2.99E+00 | 6.99E-02 | Yes | | intergenic_variant | MODIFIER |
| 18 | 75922793 | AX-170453470 | 1 | G | A | 0.446 | 1.54E-01 | 8.50E-02 | 2.56E+00 | 2.63E+00 | 7.00E-02 | Yes | Zbtb7c | intron_variant | MODIFIER |
| 13 | 3527712 | AX-97855047 | 0 | C | T | 0.202 | -1.94E-01 | 1.07E-01 | 2.71E+00 | 2.81E+00 | 7.02E-02 | Yes | | intergenic_variant | MODIFIER |
| 13 | 4548084 | AX-97822576 | 0 | T | A | 0.202 | -1.94E-01 | 1.07E-01 | 2.71E+00 | 2.81E+00 | 7.02E-02 | Yes | | intergenic_variant | MODIFIER |
| 5 | 66117011 | AX-170324630 | 1 | T | G | 0.482 | 1.85E-01 | 1.00E-01 | 2.28E+00 | 2.33E+00 | 7.03E-02 | Yes | Rbm47 | intron_variant | MODIFIER |
| 19 | 52173877 | AX-170470148 | 0 | T | G | 0.491 | 1.15E-01 | 6.36E-02 | 2.71E+00 | 2.80E+00 | 7.04E-02 | no | | intergenic_variant | MODIFIER |
| 19 | 52174066 | AX-170194260 | 0 | G | A | 0.491 | 1.15E-01 | 6.36E-02 | 2.71E+00 | 2.80E+00 | 7.04E-02 | no | | intergenic_variant | MODIFIER |
| 19 | 52202031 | AX-170126187 | 0 | C | G | 0.491 | 1.15E-01 | 6.36E-02 | 2.71E+00 | 2.80E+00 | 7.04E-02 | no | | intergenic_variant | MODIFIER |
| 19 | 52279478 | AX-170016834 | 0 | A | T | 0.491 | 1.15E-01 | 6.36E-02 | 2.71E+00 | 2.80E+00 | 7.04E-02 | no | | intergenic_variant | MODIFIER |
| 19 | 52285230 | AX-170016827 | 0 | G | A | 0.491 | 1.15E-01 | 6.36E-02 | 2.71E+00 | 2.80E+00 | 7.04E-02 | no | | intergenic_variant | MODIFIER |
| 19 | 52393555 | AX-170088479 | 0 | G | A | 0.491 | 1.15E-01 | 6.36E-02 | 2.71E+00 | 2.80E+00 | 7.04E-02 | no | | intergenic_variant | MODIFIER |
| 19 | 52479267 | AX-97852114 | 0 | C | T | 0.491 | 1.15E-01 | 6.36E-02 | 2.71E+00 | 2.80E+00 | 7.04E-02 | no | | intergenic_variant | MODIFIER |
| 19 | 52527951 | AX-170217801 | 0 | A | G | 0.491 | 1.15E-01 | 6.36E-02 | 2.71E+00 | 2.80E+00 | 7.04E-02 | no | | intergenic_variant | MODIFIER |
| 19 | 52562306 | AX-170237114 | 0 | A | C | 0.491 | 1.15E-01 | 6.36E-02 | 2.71E+00 | 2.80E+00 | 7.04E-02 | no | | intergenic_variant | MODIFIER |
| 19 | 52562381 | AX-170167305 | 0 | A | G | 0.491 | 1.15E-01 | 6.36E-02 | 2.71E+00 | 2.80E+00 | 7.04E-02 | no | | intergenic_variant | MODIFIER |
| 19 | 52562441 | AX-170427164 | 0 | T | C | 0.491 | 1.15E-01 | 6.36E-02 | 2.71E+00 | 2.80E+00 | 7.04E-02 | no | | intergenic_variant | MODIFIER |
| 19 | 52660061 | AX-97817449 | 0 | T | C | 0.491 | 1.15E-01 | 6.36E-02 | 2.71E+00 | 2.80E+00 | 7.04E-02 | no | | intergenic_variant | MODIFIER |
| 19 | 52684844 | AX-97793479 | 0 | G | A | 0.491 | 1.15E-01 | 6.36E-02 | 2.71E+00 | 2.80E+00 | 7.04E-02 | no | | intergenic_variant | MODIFIER |
| 19 | 52723102 | AX-170283856 | 0 | A | G | 0.491 | 1.15E-01 | 6.36E-02 | 2.71E+00 | 2.80E+00 | 7.04E-02 | no | | intergenic_variant | MODIFIER |
| 19 | 52723590 | AX-170016842 | 0 | A | G | 0.491 | 1.15E-01 | 6.36E-02 | 2.71E+00 | 2.80E+00 | 7.04E-02 | no | | intergenic_variant | MODIFIER |
| 19 | 52728743 | AX-170296829 | 0 | T | C | 0.491 | 1.15E-01 | 6.36E-02 | 2.71E+00 | 2.80E+00 | 7.04E-02 | no | | intergenic_variant | MODIFIER |
| 19 | 52730021 | AX-170357441 | 0 | C | A | 0.491 | 1.15E-01 | 6.36E-02 | 2.71E+00 | 2.80E+00 | 7.04E-02 | no | | intergenic_variant | MODIFIER |
| 19 | 52730672 | AX-170451470 | 0 | A | G | 0.491 | 1.15E-01 | 6.36E-02 | 2.71E+00 | 2.80E+00 | 7.04E-02 | no | | regulatory_region_variant | MODIFIER |
| 19 | 52730875 | AX-170347997 | 0 | G | T | 0.491 | 1.15E-01 | 6.36E-02 | 2.71E+00 | 2.80E+00 | 7.04E-02 | no | | regulatory_region_variant | MODIFIER |
| 19 | 52732201 | AX-170367905 | 0 | A | G | 0.491 | 1.15E-01 | 6.36E-02 | 2.71E+00 | 2.80E+00 | 7.04E-02 | no | | intergenic_variant | MODIFIER |
| 19 | 52738941 | AX-170503832 | 0 | T | C | 0.491 | 1.15E-01 | 6.36E-02 | 2.71E+00 | 2.80E+00 | 7.04E-02 | no | | intergenic_variant | MODIFIER |
| 19 | 52770709 | AX-170016837 | 0 | A | G | 0.491 | 1.15E-01 | 6.36E-02 | 2.71E+00 | 2.80E+00 | 7.04E-02 | no | | intergenic_variant | MODIFIER |
| 19 | 52772247 | AX-97794095 | 0 | A | C | 0.491 | 1.15E-01 | 6.36E-02 | 2.71E+00 | 2.80E+00 | 7.04E-02 | no | | intergenic_variant | MODIFIER |
| 19 | 52786451 | AX-170016844 | 0 | C | T | 0.491 | 1.15E-01 | 6.36E-02 | 2.71E+00 | 2.80E+00 | 7.04E-02 | no | | intergenic_variant | MODIFIER |
| 19 | 52863953 | AX-97836995 | 0 | G | A | 0.491 | 1.15E-01 | 6.36E-02 | 2.71E+00 | 2.80E+00 | 7.04E-02 | no | | intergenic_variant | MODIFIER |
| 19 | 52879560 | AX-97849890 | 0 | T | C | 0.491 | 1.15E-01 | 6.36E-02 | 2.71E+00 | 2.80E+00 | 7.04E-02 | no | | intergenic_variant | MODIFIER |
| 19 | 52880179 | AX-170016838 | 0 | G | A | 0.491 | 1.15E-01 | 6.36E-02 | 2.71E+00 | 2.80E+00 | 7.04E-02 | no | | intergenic_variant | MODIFIER |
| 19 | 52938774 | AX-170016840 | 0 | A | G | 0.491 | 1.15E-01 | 6.36E-02 | 2.71E+00 | 2.80E+00 | 7.04E-02 | no | Xpnpep1 | downstream_gene_variant | MODIFIER |
| 19 | 52942998 | AX-170344406 | 0 | C | T | 0.491 | 1.15E-01 | 6.36E-02 | 2.71E+00 | 2.80E+00 | 7.04E-02 | no | Xpnpep1 | downstream_gene_variant | MODIFIER |
| 19 | 52943043 | AX-170160948 | 0 | G | A | 0.491 | 1.15E-01 | 6.36E-02 | 2.71E+00 | 2.80E+00 | 7.04E-02 | no | Xpnpep1 | downstream_gene_variant | MODIFIER |
| 19 | 52943516 | AX-170477367 | 0 | T | A | 0.491 | 1.15E-01 | 6.36E-02 | 2.71E+00 | 2.80E+00 | 7.04E-02 | no | Xpnpep1 | downstream_gene_variant | MODIFIER |
| 19 | 52953351 | AX-170307290 | 0 | T | C | 0.491 | 1.15E-01 | 6.36E-02 | 2.71E+00 | 2.80E+00 | 7.04E-02 | no | Xpnpep1 | intron_variant,NMD_transcript_variant | MODIFIER |
| 19 | 52955246 | AX-170336666 | 0 | A | G | 0.491 | 1.15E-01 | 6.36E-02 | 2.71E+00 | 2.80E+00 | 7.04E-02 | no | Xpnpep1 | intron_variant,NMD_transcript_variant | MODIFIER |
| 19 | 52955378 | AX-170161315 | 0 | C | T | 0.491 | 1.15E-01 | 6.36E-02 | 2.71E+00 | 2.80E+00 | 7.04E-02 | no | Xpnpep1 | intron_variant,NMD_transcript_variant | MODIFIER |
| 19 | 52978385 | AX-170232682 | 0 | T | C | 0.491 | 1.15E-01 | 6.36E-02 | 2.71E+00 | 2.80E+00 | 7.04E-02 | no | Xpnpep1 | intron_variant,NMD_transcript_variant | MODIFIER |
| 19 | 52981793 | AX-170180413 | 0 | A | G | 0.491 | 1.15E-01 | 6.36E-02 | 2.71E+00 | 2.80E+00 | 7.04E-02 | no | Xpnpep1 | intron_variant,NMD_transcript_variant | MODIFIER |
| 19 | 52981953 | AX-170227818 | 0 | G | A | 0.491 | 1.15E-01 | 6.36E-02 | 2.71E+00 | 2.80E+00 | 7.04E-02 | no | Xpnpep1 | intron_variant,NMD_transcript_variant | MODIFIER |
| 19 | 52999502 | AX-170155235 | 0 | C | T | 0.491 | 1.15E-01 | 6.36E-02 | 2.71E+00 | 2.80E+00 | 7.04E-02 | no | Xpnpep1 | intron_variant,NMD_transcript_variant | MODIFIER |
| 19 | 52999774 | AX-97822827 | 0 | C | T | 0.491 | 1.15E-01 | 6.36E-02 | 2.71E+00 | 2.80E+00 | 7.04E-02 | no | Xpnpep1 | intron_variant,NMD_transcript_variant | MODIFIER |

| | | | | | | | | | | | | | | | |
|---|---|---|---|---|---|---|---|---|---|---|---|---|---|---|---|
| 19 | 52999880 | AX-170177465 | 0 | C | T | 0.491 | 1.15E-01 | 6.36E-02 | 2.71E+00 | 2.80E+00 | 7.04E-02 | no | Xpnpep1 | intron_variant,NMD_transcript_variant | MODIFIER |
| 19 | 53005000 | AX-170062701 | 0 | G | C | 0.491 | 1.15E-01 | 6.36E-02 | 2.71E+00 | 2.80E+00 | 7.04E-02 | no | Xpnpep1 | intron_variant,NMD_transcript_variant | MODIFIER |
| 19 | 53005559 | AX-170353201 | 0 | A | G | 0.491 | 1.15E-01 | 6.36E-02 | 2.71E+00 | 2.80E+00 | 7.04E-02 | no | Xpnpep1 | intron_variant,NMD_transcript_variant | MODIFIER |
| 19 | 53005898 | AX-170164987 | 0 | T | C | 0.491 | 1.15E-01 | 6.36E-02 | 2.71E+00 | 2.80E+00 | 7.04E-02 | no | Xpnpep1 | intron_variant,NMD_transcript_variant | MODIFIER |
| 17 | 69884363 | AX-170059575 | 1 | A | C | 0.054 | 2.75E-01 | 1.50E-01 | 3.24E+00 | 3.44E+00 | 7.05E-02 | Yes | | intergenic_variant | MODIFIER |
| 5 | 52329178 | AX-170203145 | 0 | C | T | 0.482 | 1.91E-01 | 1.05E-01 | 2.41E+00 | 2.45E+00 | 7.06E-02 | Yes | | intergenic_variant | MODIFIER |
| 5 | 52423941 | AX-170072100 | 0 | T | C | 0.482 | 1.91E-01 | 1.05E-01 | 2.41E+00 | 2.45E+00 | 7.06E-02 | Yes | Ccdc149 | intron_variant | MODIFIER |
| 5 | 52424577 | AX-170166192 | 0 | C | T | 0.482 | 1.91E-01 | 1.05E-01 | 2.41E+00 | 2.45E+00 | 7.06E-02 | Yes | Ccdc149 | intron_variant | MODIFIER |
| 5 | 52424875 | AX-170371322 | 0 | G | C | 0.482 | 1.91E-01 | 1.05E-01 | 2.41E+00 | 2.45E+00 | 7.06E-02 | Yes | Ccdc149 | intron_variant | MODIFIER |
| 5 | 52431070 | AX-170453023 | 0 | A | G | 0.482 | 1.91E-01 | 1.05E-01 | 2.41E+00 | 2.45E+00 | 7.06E-02 | Yes | Ccdc149 | intron_variant | MODIFIER |
| 5 | 52482096 | AX-170203661 | 0 | G | C | 0.482 | 1.91E-01 | 1.05E-01 | 2.41E+00 | 2.45E+00 | 7.06E-02 | Yes | | intergenic_variant | MODIFIER |
| 5 | 52492081 | AX-170347381 | 0 | A | G | 0.482 | 1.91E-01 | 1.05E-01 | 2.41E+00 | 2.45E+00 | 7.06E-02 | Yes | | intergenic_variant | MODIFIER |
| 5 | 52492593 | AX-170182365 | 0 | A | G | 0.482 | 1.91E-01 | 1.05E-01 | 2.41E+00 | 2.45E+00 | 7.06E-02 | Yes | | intergenic_variant | MODIFIER |
| 5 | 52504529 | AX-170336125 | 0 | G | A | 0.482 | 1.91E-01 | 1.05E-01 | 2.41E+00 | 2.45E+00 | 7.06E-02 | Yes | | intergenic_variant | MODIFIER |
| 5 | 52514948 | AX-97856618 | 0 | T | C | 0.482 | 1.91E-01 | 1.05E-01 | 2.41E+00 | 2.45E+00 | 7.06E-02 | Yes | | intergenic_variant | MODIFIER |
| 5 | 52527110 | AX-170089477 | 0 | G | A | 0.482 | 1.91E-01 | 1.05E-01 | 2.41E+00 | 2.45E+00 | 7.06E-02 | Yes | | intergenic_variant | MODIFIER |
| 5 | 52544409 | AX-170445601 | 0 | A | G | 0.482 | 1.91E-01 | 1.05E-01 | 2.41E+00 | 2.45E+00 | 7.06E-02 | Yes | Lgi2 | intron_variant | MODIFIER |
| 5 | 52550997 | AX-170271740 | 0 | A | G | 0.482 | 1.91E-01 | 1.05E-01 | 2.41E+00 | 2.45E+00 | 7.06E-02 | Yes | Lgi2 | intron_variant | MODIFIER |
| 5 | 52558112 | AX-170429168 | 0 | C | T | 0.482 | 1.91E-01 | 1.05E-01 | 2.41E+00 | 2.45E+00 | 7.06E-02 | Yes | Lgi2 | intron_variant | MODIFIER |
| 5 | 52563128 | AX-170250527 | 0 | C | G | 0.482 | 1.91E-01 | 1.05E-01 | 2.41E+00 | 2.45E+00 | 7.06E-02 | Yes | Lgi2 | intron_variant | MODIFIER |
| 5 | 52563446 | AX-170386308 | 0 | G | C | 0.482 | 1.91E-01 | 1.05E-01 | 2.41E+00 | 2.45E+00 | 7.06E-02 | Yes | Lgi2 | intron_variant | MODIFIER |
| 5 | 52590793 | AX-170351309 | 0 | T | C | 0.482 | 1.91E-01 | 1.05E-01 | 2.41E+00 | 2.45E+00 | 7.06E-02 | Yes | | intergenic_variant | MODIFIER |
| 5 | 52598709 | AX-170116552 | 0 | A | G | 0.482 | 1.91E-01 | 1.05E-01 | 2.41E+00 | 2.45E+00 | 7.06E-02 | Yes | | intergenic_variant | MODIFIER |
| 5 | 52610738 | AX-170072108 | 0 | G | A | 0.482 | 1.91E-01 | 1.05E-01 | 2.41E+00 | 2.45E+00 | 7.06E-02 | Yes | 8030423F21Rik | intron_variant,non_coding_transcript_variant | MODIFIER |
| 5 | 52613168 | AX-170268688 | 0 | C | G | 0.482 | 1.91E-01 | 1.05E-01 | 2.41E+00 | 2.45E+00 | 7.06E-02 | Yes | 8030423F21Rik | non_coding_transcript_exon_variant | MODIFIER |
| 5 | 52621753 | AX-170428787 | 0 | T | A | 0.482 | 1.91E-01 | 1.05E-01 | 2.41E+00 | 2.45E+00 | 7.06E-02 | Yes | 8030423F21Rik | upstream_gene_variant | MODIFIER |
| 5 | 52630224 | AX-170158240 | 0 | G | A | 0.482 | 1.91E-01 | 1.05E-01 | 2.41E+00 | 2.45E+00 | 7.06E-02 | Yes | | intergenic_variant | MODIFIER |
| 5 | 52632507 | AX-170388823 | 0 | G | A | 0.482 | 1.91E-01 | 1.05E-01 | 2.41E+00 | 2.45E+00 | 7.06E-02 | Yes | | intergenic_variant | MODIFIER |
| 5 | 52638482 | AX-170106927 | 0 | G | A | 0.482 | 1.91E-01 | 1.05E-01 | 2.41E+00 | 2.45E+00 | 7.06E-02 | Yes | Sepsecs | downstream_gene_variant | MODIFIER |
| 5 | 52647750 | AX-170411032 | 0 | G | A | 0.482 | 1.91E-01 | 1.05E-01 | 2.41E+00 | 2.45E+00 | 7.06E-02 | Yes | Sepsecs | intron_variant | MODIFIER |
| 5 | 52653717 | AX-170497041 | 0 | C | T | 0.482 | 1.91E-01 | 1.05E-01 | 2.41E+00 | 2.45E+00 | 7.06E-02 | Yes | Sepsecs | intron_variant | MODIFIER |
| 5 | 52657234 | AX-170418081 | 0 | A | G | 0.482 | 1.91E-01 | 1.05E-01 | 2.41E+00 | 2.45E+00 | 7.06E-02 | Yes | Sepsecs | intron_variant | MODIFIER |
| 5 | 52657635 | AX-170376960 | 0 | C | T | 0.482 | 1.91E-01 | 1.05E-01 | 2.41E+00 | 2.45E+00 | 7.06E-02 | Yes | Sepsecs | intron_variant | MODIFIER |
| 5 | 52658950 | AX-170099649 | 0 | C | T | 0.482 | 1.91E-01 | 1.05E-01 | 2.41E+00 | 2.45E+00 | 7.06E-02 | Yes | Sepsecs | intron_variant | MODIFIER |
| 5 | 52659745 | AX-170164512 | 0 | A | G | 0.482 | 1.91E-01 | 1.05E-01 | 2.41E+00 | 2.45E+00 | 7.06E-02 | Yes | Sepsecs | intron_variant | MODIFIER |
| 5 | 52670050 | AX-170494014 | 0 | C | T | 0.482 | 1.91E-01 | 1.05E-01 | 2.41E+00 | 2.45E+00 | 7.06E-02 | Yes | Sepsecs | upstream_gene_variant | MODIFIER |
| 5 | 52670133 | AX-170150747 | 0 | T | A | 0.482 | 1.91E-01 | 1.05E-01 | 2.41E+00 | 2.45E+00 | 7.06E-02 | Yes | Sepsecs | upstream_gene_variant | MODIFIER |
| 5 | 52672815 | AX-170484790 | 0 | C | T | 0.482 | 1.91E-01 | 1.05E-01 | 2.41E+00 | 2.45E+00 | 7.06E-02 | Yes | Sepsecs | upstream_gene_variant | MODIFIER |
| 5 | 52673630 | AX-170072111 | 0 | A | G | 0.482 | 1.91E-01 | 1.05E-01 | 2.41E+00 | 2.45E+00 | 7.06E-02 | Yes | Sepsecs | upstream_gene_variant | MODIFIER |
| 5 | 52674656 | AX-170072110 | 0 | T | C | 0.482 | 1.91E-01 | 1.05E-01 | 2.41E+00 | 2.45E+00 | 7.06E-02 | Yes | Sepsecs | upstream_gene_variant | MODIFIER |
| 5 | 52683327 | AX-97804641 | 0 | T | G | 0.482 | 1.91E-01 | 1.05E-01 | 2.41E+00 | 2.45E+00 | 7.06E-02 | Yes | | intergenic_variant | MODIFIER |
| 5 | 52699621 | AX-170355984 | 0 | G | T | 0.482 | 1.91E-01 | 1.05E-01 | 2.41E+00 | 2.45E+00 | 7.06E-02 | Yes | | intergenic_variant | MODIFIER |
| 5 | 52699752 | AX-170451488 | 0 | T | C | 0.482 | 1.91E-01 | 1.05E-01 | 2.41E+00 | 2.45E+00 | 7.06E-02 | Yes | | intergenic_variant | MODIFIER |
| 5 | 52705275 | AX-170520624 | 0 | C | T | 0.482 | 1.91E-01 | 1.05E-01 | 2.41E+00 | 2.45E+00 | 7.06E-02 | Yes | | intergenic_variant | MODIFIER |
| 5 | 52714189 | AX-97807982 | 0 | T | C | 0.482 | 1.91E-01 | 1.05E-01 | 2.41E+00 | 2.45E+00 | 7.06E-02 | Yes | | intergenic_variant | MODIFIER |
| 5 | 52720512 | AX-97802281 | 0 | C | T | 0.482 | 1.91E-01 | 1.05E-01 | 2.41E+00 | 2.45E+00 | 7.06E-02 | Yes | | intergenic_variant | MODIFIER |
| 5 | 52723558 | AX-170152676 | 0 | G | A | 0.482 | 1.91E-01 | 1.05E-01 | 2.41E+00 | 2.45E+00 | 7.06E-02 | Yes | | intergenic_variant | MODIFIER |
| 5 | 52744295 | AX-170425655 | 0 | G | T | 0.482 | 1.91E-01 | 1.05E-01 | 2.41E+00 | 2.45E+00 | 7.06E-02 | Yes | Pi4k2b | intron_variant | MODIFIER |
| 5 | 52756949 | AX-170219143 | 0 | C | T | 0.482 | 1.91E-01 | 1.05E-01 | 2.41E+00 | 2.45E+00 | 7.06E-02 | Yes | Pi4k2b | synonymous_variant | LOW |
| 5 | 52757119 | AX-170179522 | 0 | G | A | 0.482 | 1.91E-01 | 1.05E-01 | 2.41E+00 | 2.45E+00 | 7.06E-02 | Yes | Pi4k2b | intron_variant | MODIFIER |
| 5 | 52760643 | AX-97824467 | 0 | T | C | 0.482 | 1.91E-01 | 1.05E-01 | 2.41E+00 | 2.45E+00 | 7.06E-02 | Yes | Pi4k2b | intron_variant | MODIFIER |
| 5 | 52762829 | AX-170243176 | 0 | T | G | 0.482 | 1.91E-01 | 1.05E-01 | 2.41E+00 | 2.45E+00 | 7.06E-02 | Yes | Pi4k2b | intron_variant | MODIFIER |
| 5 | 52762866 | AX-170496308 | 0 | G | T | 0.482 | 1.91E-01 | 1.05E-01 | 2.41E+00 | 2.45E+00 | 7.06E-02 | Yes | Pi4k2b | intron_variant | MODIFIER |
| 5 | 52806638 | AX-170072114 | 0 | A | G | 0.482 | 1.91E-01 | 1.05E-01 | 2.41E+00 | 2.45E+00 | 7.06E-02 | Yes | Zcchc4 | intron_variant | MODIFIER |

| | | | | | | | | | | | | | | | |
|---|---|---|---|---|---|---|---|---|---|---|---|---|---|---|---|
| 5 | 52807367 | AX-97799202 | 0 | T | C | 0.482 | 1.91E-01 | 1.05E-01 | 2.41E+00 | 2.45E+00 | 7.06E-02 | Yes | Zcchc4 | intron_variant | MODIFIER |
| 5 | 52807877 | AX-170072116 | 0 | G | A | 0.482 | 1.91E-01 | 1.05E-01 | 2.41E+00 | 2.45E+00 | 7.06E-02 | Yes | Zcchc4 | intron_variant | MODIFIER |
| 5 | 52808529 | AX-170324498 | 0 | G | C | 0.482 | 1.91E-01 | 1.05E-01 | 2.41E+00 | 2.45E+00 | 7.06E-02 | Yes | Zcchc4 | intron_variant | MODIFIER |
| 5 | 52808780 | AX-170086209 | 0 | G | T | 0.482 | 1.91E-01 | 1.05E-01 | 2.41E+00 | 2.45E+00 | 7.06E-02 | Yes | Zcchc4 | intron_variant | MODIFIER |
| 5 | 52819077 | AX-170338950 | 0 | T | C | 0.482 | 1.91E-01 | 1.05E-01 | 2.41E+00 | 2.45E+00 | 7.06E-02 | Yes | Zcchc4 | intron_variant | MODIFIER |
| 5 | 52866027 | AX-170151841 | 0 | G | A | 0.482 | 1.91E-01 | 1.05E-01 | 2.41E+00 | 2.45E+00 | 7.06E-02 | Yes | Anapc4 | intron_variant | MODIFIER |
| 5 | 52870965 | AX-170412356 | 0 | G | A | 0.482 | 1.91E-01 | 1.05E-01 | 2.41E+00 | 2.45E+00 | 7.06E-02 | Yes | Anapc4 | downstream_gene_variant | MODIFIER |
| 5 | 52516769 | AX-170417357 | 1 | A | T | 0.473 | 1.91E-01 | 1.04E-01 | 2.36E+00 | 2.41E+00 | 7.06E-02 | Yes | | regulatory_region_variant | MODIFIER |
| 10 | 100251402 | AX-170092076 | 0 | C | T | 0.272 | -1.48E-01 | 8.24E-02 | 2.74E+00 | 2.83E+00 | 7.07E-02 | no | | intergenic_variant | MODIFIER |
| 4 | 99495911 | AX-170320440 | 1 | A | G | 0.223 | 2.18E-01 | 1.21E-01 | 2.76E+00 | 2.83E+00 | 7.07E-02 | Yes | | intergenic_variant | MODIFIER |
| 5 | 58231589 | AX-97797373 | 0 | A | G | 0.474 | 1.84E-01 | 1.01E-01 | 2.47E+00 | 2.52E+00 | 7.08E-02 | Yes | | intergenic_variant | MODIFIER |
| 3 | 6288722 | AX-170418156 | 0 | A | T | 0.132 | -1.64E-01 | 9.13E-02 | 2.67E+00 | 2.78E+00 | 7.10E-02 | no | | intergenic_variant | MODIFIER |
| 8 | 86517748 | AX-170173774 | 1 | C | A | 0.196 | -2.16E-01 | 1.18E-01 | 3.24E+00 | 3.39E+00 | 7.11E-02 | Yes | Abcc12 | intron_variant | MODIFIER |
| 5 | 56689877 | AX-170025006 | 0 | C | T | 0.465 | 1.79E-01 | 9.87E-02 | 2.48E+00 | 2.55E+00 | 7.11E-02 | Yes | | intergenic_variant | MODIFIER |
| 19 | 60336198 | AX-170251248 | 1 | C | G | 0.464 | -1.15E-01 | 6.42E-02 | 2.81E+00 | 2.89E+00 | 7.22E-02 | no | | intergenic_variant | MODIFIER |
| 4 | 64498932 | AX-170022966 | 1 | A | C | 0.33 | 2.00E-01 | 1.11E-01 | 3.07E+00 | 3.18E+00 | 7.23E-02 | Yes | | intergenic_variant | MODIFIER |
| 16 | 80726270 | AX-170483830 | 0 | T | G | 0.474 | 1.24E-01 | 6.93E-02 | 2.79E+00 | 2.89E+00 | 7.23E-02 | no | Gm19867 | upstream_gene_variant | MODIFIER |
| 5 | 54424123 | AX-170528128 | 1 | G | A | 0.482 | 1.76E-01 | 9.69E-02 | 2.34E+00 | 2.40E+00 | 7.27E-02 | Yes | | intergenic_variant | MODIFIER |
| 6 | 125873733 | AX-170220925 | 0 | T | A | 0.158 | -1.44E-01 | 7.94E-02 | 3.17E+00 | 3.40E+00 | 7.30E-02 | no | Ano2 | intron_variant,non_coding_transcript_variant | MODIFIER |
| 8 | 64389568 | AX-170236329 | 0 | A | C | 0.14 | -2.32E-01 | 1.30E-01 | 2.77E+00 | 2.88E+00 | 7.32E-02 | Yes | | intergenic_variant | MODIFIER |
| 5 | 52325404 | AX-170257684 | 1 | T | G | 0.482 | 1.89E-01 | 1.05E-01 | 2.43E+00 | 2.47E+00 | 7.32E-02 | Yes | | intergenic_variant | MODIFIER |
| 15 | 6900055 | AX-170437899 | 0 | T | A | 0.14 | -1.08E-01 | 5.81E-02 | 2.13E+00 | 2.23E+00 | 7.33E-02 | no | | intergenic_variant | MODIFIER |
| 18 | 58974457 | AX-170352775 | 2 | A | G | 0.255 | -1.21E-01 | 6.73E-02 | 2.97E+00 | 3.08E+00 | 7.33E-02 | no | Adamts19 | intron_variant | MODIFIER |
| 1 | 185658780 | AX-170215917 | 0 | C | T | 0.246 | 1.92E-01 | 1.07E-01 | 2.46E+00 | 2.53E+00 | 7.34E-02 | Yes | | intergenic_variant | MODIFIER |
| 5 | 44360332 | AX-170343286 | 1 | T | A | 0.214 | -1.77E-01 | 9.88E-02 | 2.63E+00 | 2.73E+00 | 7.34E-02 | Yes | Gm42698 | upstream_gene_variant | MODIFIER |
| 5 | 52941438 | AX-170072120 | 0 | G | A | 0.482 | 1.78E-01 | 9.88E-02 | 2.43E+00 | 2.49E+00 | 7.34E-02 | Yes | | intergenic_variant | MODIFIER |
| 2 | 101819716 | AX-170018498 | 0 | G | A | 0.053 | -3.31E-01 | 1.85E-01 | 2.95E+00 | 3.09E+00 | 7.35E-02 | Yes | Prr5l | intron_variant | MODIFIER |
| 16 | 56694243 | AX-97789999 | 2 | G | A | 0.455 | -1.51E-01 | 8.44E-02 | 2.58E+00 | 2.62E+00 | 7.35E-02 | no | Abi3bp | downstream_gene_variant | MODIFIER |
| 15 | 63698696 | AX-170055761 | 0 | G | T | 0.325 | -1.43E-01 | 7.66E-02 | 2.08E+00 | 2.14E+00 | 7.37E-02 | no | Gm41335 | intron_variant,non_coding_transcript_variant | MODIFIER |
| 15 | 63768823 | AX-170393542 | 0 | C | T | 0.325 | -1.43E-01 | 7.66E-02 | 2.08E+00 | 2.14E+00 | 7.37E-02 | no | | regulatory_region_variant | MODIFIER |
| 15 | 63769157 | AX-170334729 | 0 | G | T | 0.325 | -1.43E-01 | 7.66E-02 | 2.08E+00 | 2.14E+00 | 7.37E-02 | no | | regulatory_region_variant | MODIFIER |
| 15 | 63769282 | AX-170134729 | 0 | G | T | 0.325 | -1.43E-01 | 7.66E-02 | 2.08E+00 | 2.14E+00 | 7.37E-02 | no | | regulatory_region_variant | MODIFIER |
| 15 | 63774037 | AX-170177566 | 0 | G | T | 0.325 | -1.43E-01 | 7.66E-02 | 2.08E+00 | 2.14E+00 | 7.37E-02 | no | Gsdmc | downstream_gene_variant | MODIFIER |
| 15 | 63778740 | AX-170130750 | 0 | A | T | 0.325 | -1.43E-01 | 7.66E-02 | 2.08E+00 | 2.14E+00 | 7.37E-02 | no | Gsdmc | intron_variant | MODIFIER |
| 15 | 64047364 | AX-170246518 | 0 | C | T | 0.325 | -1.43E-01 | 7.66E-02 | 2.08E+00 | 2.14E+00 | 7.37E-02 | no | Fam49b | intron_variant | MODIFIER |
| 15 | 64048399 | AX-170488780 | 0 | A | G | 0.325 | -1.43E-01 | 7.66E-02 | 2.08E+00 | 2.14E+00 | 7.37E-02 | no | Fam49b | intron_variant | MODIFIER |
| 15 | 64048955 | AX-170144659 | 0 | G | C | 0.325 | -1.43E-01 | 7.66E-02 | 2.08E+00 | 2.14E+00 | 7.37E-02 | no | Fam49b | intron_variant | MODIFIER |
| 15 | 64049124 | AX-170176568 | 0 | G | A | 0.325 | -1.43E-01 | 7.66E-02 | 2.08E+00 | 2.14E+00 | 7.37E-02 | no | Fam49b | intron_variant | MODIFIER |
| 15 | 64067565 | AX-170255665 | 0 | A | G | 0.325 | -1.43E-01 | 7.66E-02 | 2.08E+00 | 2.14E+00 | 7.37E-02 | no | | regulatory_region_variant | MODIFIER |
| 15 | 64108795 | AX-170322120 | 0 | G | A | 0.325 | -1.43E-01 | 7.66E-02 | 2.08E+00 | 2.14E+00 | 7.37E-02 | no | Asap1 | intron_variant | MODIFIER |
| 15 | 64215725 | AX-170055768 | 0 | C | T | 0.325 | -1.43E-01 | 7.66E-02 | 2.08E+00 | 2.14E+00 | 7.37E-02 | no | Asap1 | intron_variant | MODIFIER |
| 15 | 64216225 | AX-170055767 | 0 | G | A | 0.325 | -1.43E-01 | 7.66E-02 | 2.08E+00 | 2.14E+00 | 7.37E-02 | no | Asap1 | intron_variant | MODIFIER |
| 15 | 64220767 | AX-170011860 | 0 | A | G | 0.325 | -1.43E-01 | 7.66E-02 | 2.08E+00 | 2.14E+00 | 7.37E-02 | no | Asap1 | intron_variant | MODIFIER |
| 15 | 64222989 | AX-170011858 | 0 | A | G | 0.325 | -1.43E-01 | 7.66E-02 | 2.08E+00 | 2.14E+00 | 7.37E-02 | no | Asap1 | intron_variant | MODIFIER |
| 15 | 64249062 | AX-97806914 | 0 | C | T | 0.325 | -1.43E-01 | 7.66E-02 | 2.08E+00 | 2.14E+00 | 7.37E-02 | no | Asap1 | intron_variant | MODIFIER |
| 15 | 64263973 | AX-170506508 | 0 | T | C | 0.325 | -1.43E-01 | 7.66E-02 | 2.08E+00 | 2.14E+00 | 7.37E-02 | no | Asap1 | intron_variant | MODIFIER |
| 16 | 63721664 | AX-170515430 | 0 | G | C | 0.474 | -1.52E-01 | 8.49E-02 | 2.63E+00 | 2.68E+00 | 7.38E-02 | no | Epha3 | intron_variant | MODIFIER |
| 3 | 7177081 | AX-170019850 | 0 | T | C | 0.132 | -1.64E-01 | 9.23E-02 | 2.73E+00 | 2.84E+00 | 7.39E-02 | no | | intergenic_variant | MODIFIER |
| 2 | 71842302 | AX-170018028 | 0 | C | T | 0.044 | -3.28E-01 | 1.84E-01 | 2.95E+00 | 3.10E+00 | 7.39E-02 | Yes | Itga6 | intron_variant | MODIFIER |
| 18 | 74370102 | AX-170205745 | 0 | T | C | 0.175 | -1.87E-01 | 1.05E-01 | 2.97E+00 | 3.12E+00 | 7.40E-02 | Yes | | intergenic_variant | MODIFIER |
| 16 | 65385788 | AX-97830843 | 1 | C | T | 0.5 | 1.42E-01 | 7.98E-02 | 2.67E+00 | 2.74E+00 | 7.41E-02 | no | | intergenic_variant | MODIFIER |
| 1 | 152497803 | AX-170043169 | 0 | A | T | 0.202 | 2.54E-01 | 1.41E-01 | 3.18E+00 | 3.26E+00 | 7.42E-02 | Yes | Colgalt2 | intron_variant | MODIFIER |
| 5 | 61503725 | AX-170476563 | 1 | T | A | 0.482 | 1.84E-01 | 1.03E-01 | 2.50E+00 | 2.55E+00 | 7.43E-02 | Yes | | intergenic_variant | MODIFIER |
| 5 | 64532332 | AX-170195619 | 1 | A | C | 0.482 | 1.84E-01 | 1.03E-01 | 2.50E+00 | 2.55E+00 | 7.43E-02 | Yes | | intergenic_variant | MODIFIER |

| | | | | | | | | | | | | | | | |
|---|---|---|---|---|---|---|---|---|---|---|---|---|---|---|---|
| 4 | 138911306 | AX-170283147 | 0 | A | T | 0.158 | 1.98E-01 | 1.11E-01 | 2.87E+00 | 2.95E+00 | 7.44E-02 | Yes | Otud3 | intron_variant | MODIFIER |
| 3 | 23282839 | AX-170189135 | 0 | A | T | 0.158 | -1.70E-01 | 9.55E-02 | 2.66E+00 | 2.77E+00 | 7.44E-02 | no | | intergenic_variant | MODIFIER |
| 13 | 22224201 | AX-170491037 | 0 | T | C | 0.158 | -2.12E-01 | 1.19E-01 | 2.55E+00 | 2.63E+00 | 7.44E-02 | Yes | Vmn1r193 | upstream_gene_variant | MODIFIER |
| 16 | 55548338 | AX-170190178 | 0 | T | C | 0.456 | -1.50E-01 | 8.44E-02 | 2.58E+00 | 2.62E+00 | 7.44E-02 | no | | intergenic_variant | MODIFIER |
| 8 | 24488174 | AX-170030363 | 2 | C | G | 0.327 | -1.16E-01 | 6.50E-02 | 2.58E+00 | 2.68E+00 | 7.48E-02 | no | | intergenic_variant | MODIFIER |
| 4 | 129611624 | AX-170023928 | 1 | A | G | 0.161 | 2.08E-01 | 1.17E-01 | 2.79E+00 | 2.85E+00 | 7.49E-02 | Yes | Iqcc | downstream_gene_variant | MODIFIER |
| 18 | 77211114 | AX-170061381 | 0 | G | T | 0.447 | 1.50E-01 | 8.43E-02 | 2.56E+00 | 2.63E+00 | 7.52E-02 | Yes | St8sia5 | intron_variant | MODIFIER |
| 5 | 60746823 | AX-170025094 | 1 | A | T | 0.482 | 1.84E-01 | 1.03E-01 | 2.49E+00 | 2.55E+00 | 7.52E-02 | Yes | | intergenic_variant | MODIFIER |
| 1 | 155718889 | AX-170167951 | 0 | T | C | 0.421 | -1.24E-01 | 6.29E-02 | 4.13E+00 | 4.46E+00 | 7.54E-02 | no | Lhx4 | intron_variant | MODIFIER |
| 5 | 132248136 | AX-170026193 | 1 | C | T | 0.152 | -1.63E-01 | 9.01E-02 | 2.28E+00 | 2.36E+00 | 7.55E-02 | no | Auts2 | intron_variant,non_coding_transcript_variant | MODIFIER |
| 1 | 45493701 | AX-170471186 | 0 | G | C | 0.096 | -3.07E-01 | 1.72E-01 | 3.10E+00 | 3.25E+00 | 7.55E-02 | Yes | Col5a2 | intron_variant | MODIFIER |
| 18 | 76071653 | AX-170245375 | 0 | C | G | 0.395 | -1.53E-01 | 8.63E-02 | 2.91E+00 | 3.02E+00 | 7.55E-02 | no | Zbtb7c | intron_variant | MODIFIER |
| 18 | 76333501 | AX-170513501 | 0 | C | T | 0.395 | -1.53E-01 | 8.63E-02 | 2.91E+00 | 3.02E+00 | 7.55E-02 | no | | intergenic_variant | MODIFIER |
| 16 | 66303598 | AX-170013284 | 1 | C | G | 0.509 | 1.38E-01 | 7.77E-02 | 2.55E+00 | 2.61E+00 | 7.58E-02 | no | | intergenic_variant | MODIFIER |
| 19 | 59475066 | AX-170148123 | 0 | T | A | 0.474 | -1.10E-01 | 6.19E-02 | 2.77E+00 | 2.86E+00 | 7.59E-02 | no | | intergenic_variant | MODIFIER |
| 5 | 55054026 | AX-97827221 | 0 | A | G | 0.482 | 1.84E-01 | 1.03E-01 | 2.48E+00 | 2.54E+00 | 7.60E-02 | Yes | | intergenic_variant | MODIFIER |
| 5 | 55091879 | AX-170135610 | 0 | G | C | 0.482 | 1.84E-01 | 1.03E-01 | 2.48E+00 | 2.54E+00 | 7.60E-02 | Yes | | intergenic_variant | MODIFIER |
| 5 | 55110548 | AX-170324038 | 0 | T | C | 0.482 | 1.84E-01 | 1.03E-01 | 2.48E+00 | 2.54E+00 | 7.60E-02 | Yes | | intergenic_variant | MODIFIER |
| 5 | 55112399 | AX-170214647 | 0 | T | C | 0.482 | 1.84E-01 | 1.03E-01 | 2.48E+00 | 2.54E+00 | 7.60E-02 | Yes | | intergenic_variant | MODIFIER |
| 5 | 55114939 | AX-170259241 | 0 | G | C | 0.482 | 1.84E-01 | 1.03E-01 | 2.48E+00 | 2.54E+00 | 7.60E-02 | Yes | | intergenic_variant | MODIFIER |
| 5 | 55186730 | AX-170530915 | 0 | G | A | 0.482 | 1.84E-01 | 1.03E-01 | 2.48E+00 | 2.54E+00 | 7.60E-02 | Yes | | intergenic_variant | MODIFIER |
| 5 | 55316749 | AX-170286313 | 0 | C | T | 0.482 | 1.84E-01 | 1.03E-01 | 2.48E+00 | 2.54E+00 | 7.60E-02 | Yes | | intergenic_variant | MODIFIER |
| 5 | 55360764 | AX-170265105 | 0 | A | G | 0.482 | 1.84E-01 | 1.03E-01 | 2.48E+00 | 2.54E+00 | 7.60E-02 | Yes | | intergenic_variant | MODIFIER |
| 5 | 55394603 | AX-170152637 | 0 | A | C | 0.482 | 1.84E-01 | 1.03E-01 | 2.48E+00 | 2.54E+00 | 7.60E-02 | Yes | | intergenic_variant | MODIFIER |
| 5 | 55411655 | AX-97793278 | 0 | C | A | 0.482 | 1.84E-01 | 1.03E-01 | 2.48E+00 | 2.54E+00 | 7.60E-02 | Yes | | intergenic_variant | MODIFIER |
| 5 | 56060645 | AX-170196743 | 0 | A | G | 0.482 | 1.84E-01 | 1.03E-01 | 2.48E+00 | 2.54E+00 | 7.60E-02 | Yes | | intergenic_variant | MODIFIER |
| 5 | 56184488 | AX-170424066 | 0 | G | A | 0.482 | 1.84E-01 | 1.03E-01 | 2.48E+00 | 2.54E+00 | 7.60E-02 | Yes | | intergenic_variant | MODIFIER |
| 5 | 56440446 | AX-170122167 | 0 | A | G | 0.482 | 1.84E-01 | 1.03E-01 | 2.48E+00 | 2.54E+00 | 7.60E-02 | Yes | Gm42977 | intron_variant,non_coding_transcript_variant | MODIFIER |
| 5 | 56776466 | AX-170203562 | 0 | C | A | 0.482 | 1.84E-01 | 1.03E-01 | 2.48E+00 | 2.54E+00 | 7.60E-02 | Yes | | intergenic_variant | MODIFIER |
| 5 | 57366136 | AX-170107486 | 0 | T | A | 0.482 | 1.84E-01 | 1.03E-01 | 2.48E+00 | 2.54E+00 | 7.60E-02 | Yes | | intergenic_variant | MODIFIER |
| 5 | 57885997 | AX-170415894 | 0 | C | T | 0.482 | 1.84E-01 | 1.03E-01 | 2.48E+00 | 2.54E+00 | 7.60E-02 | Yes | Pcdh7 | intron_variant | MODIFIER |
| 5 | 57962220 | AX-170469595 | 0 | C | T | 0.482 | 1.84E-01 | 1.03E-01 | 2.48E+00 | 2.54E+00 | 7.60E-02 | Yes | Pcdh7 | intron_variant | MODIFIER |
| 5 | 58098084 | AX-170226249 | 0 | A | G | 0.482 | 1.84E-01 | 1.03E-01 | 2.48E+00 | 2.54E+00 | 7.60E-02 | Yes | Pcdh7 | intron_variant | MODIFIER |
| 5 | 58154533 | AX-170406750 | 0 | T | C | 0.482 | 1.84E-01 | 1.03E-01 | 2.48E+00 | 2.54E+00 | 7.60E-02 | Yes | Gm42976 | upstream_gene_variant | MODIFIER |
| 5 | 58156490 | AX-97836411 | 0 | T | C | 0.482 | 1.84E-01 | 1.03E-01 | 2.48E+00 | 2.54E+00 | 7.60E-02 | Yes | | intergenic_variant | MODIFIER |
| 5 | 58161769 | AX-170255003 | 0 | T | C | 0.482 | 1.84E-01 | 1.03E-01 | 2.48E+00 | 2.54E+00 | 7.60E-02 | Yes | | intergenic_variant | MODIFIER |
| 5 | 58161873 | AX-170297187 | 0 | A | C | 0.482 | 1.84E-01 | 1.03E-01 | 2.48E+00 | 2.54E+00 | 7.60E-02 | Yes | | intergenic_variant | MODIFIER |
| 5 | 58289131 | AX-170380189 | 0 | G | A | 0.482 | 1.84E-01 | 1.03E-01 | 2.48E+00 | 2.54E+00 | 7.60E-02 | Yes | | intergenic_variant | MODIFIER |
| 5 | 58333845 | AX-170433384 | 0 | T | A | 0.482 | 1.84E-01 | 1.03E-01 | 2.48E+00 | 2.54E+00 | 7.60E-02 | Yes | | intergenic_variant | MODIFIER |
| 5 | 58370229 | AX-170160740 | 0 | G | A | 0.482 | 1.84E-01 | 1.03E-01 | 2.48E+00 | 2.54E+00 | 7.60E-02 | Yes | | intergenic_variant | MODIFIER |
| 5 | 60245146 | AX-97843195 | 0 | T | A | 0.482 | 1.84E-01 | 1.03E-01 | 2.48E+00 | 2.54E+00 | 7.60E-02 | Yes | | intergenic_variant | MODIFIER |
| 5 | 60293371 | AX-97798401 | 0 | C | T | 0.482 | 1.84E-01 | 1.03E-01 | 2.48E+00 | 2.54E+00 | 7.60E-02 | Yes | | intergenic_variant | MODIFIER |
| 5 | 60404181 | AX-97825091 | 0 | T | G | 0.482 | 1.84E-01 | 1.03E-01 | 2.48E+00 | 2.54E+00 | 7.60E-02 | Yes | | intergenic_variant | MODIFIER |
| 5 | 60420838 | AX-170521403 | 0 | G | A | 0.482 | 1.84E-01 | 1.03E-01 | 2.48E+00 | 2.54E+00 | 7.60E-02 | Yes | | intergenic_variant | MODIFIER |
| 5 | 60438697 | AX-170503654 | 0 | C | A | 0.482 | 1.84E-01 | 1.03E-01 | 2.48E+00 | 2.54E+00 | 7.60E-02 | Yes | | intergenic_variant | MODIFIER |
| 5 | 60490347 | AX-170197513 | 0 | G | C | 0.482 | 1.84E-01 | 1.03E-01 | 2.48E+00 | 2.54E+00 | 7.60E-02 | Yes | | intergenic_variant | MODIFIER |
| 5 | 60492553 | AX-97849545 | 0 | G | C | 0.482 | 1.84E-01 | 1.03E-01 | 2.48E+00 | 2.54E+00 | 7.60E-02 | Yes | | intergenic_variant | MODIFIER |
| 5 | 60562970 | AX-170110715 | 0 | A | G | 0.482 | 1.84E-01 | 1.03E-01 | 2.48E+00 | 2.54E+00 | 7.60E-02 | Yes | | intergenic_variant | MODIFIER |
| 5 | 60577890 | AX-97789527 | 0 | C | A | 0.482 | 1.84E-01 | 1.03E-01 | 2.48E+00 | 2.54E+00 | 7.60E-02 | Yes | Gm43390 | upstream_gene_variant | MODIFIER |
| 5 | 60589289 | AX-169999792 | 0 | A | G | 0.482 | 1.84E-01 | 1.03E-01 | 2.48E+00 | 2.54E+00 | 7.60E-02 | Yes | | intergenic_variant | MODIFIER |
| 5 | 60637824 | AX-170282254 | 0 | C | T | 0.482 | 1.84E-01 | 1.03E-01 | 2.48E+00 | 2.54E+00 | 7.60E-02 | Yes | | intergenic_variant | MODIFIER |
| 5 | 60695972 | AX-170120451 | 0 | G | A | 0.482 | 1.84E-01 | 1.03E-01 | 2.48E+00 | 2.54E+00 | 7.60E-02 | Yes | | intergenic_variant | MODIFIER |
| 5 | 60742599 | AX-170025087 | 0 | C | T | 0.482 | 1.84E-01 | 1.03E-01 | 2.48E+00 | 2.54E+00 | 7.60E-02 | Yes | | intergenic_variant | MODIFIER |
| 5 | 61446195 | AX-170431936 | 0 | C | T | 0.482 | 1.84E-01 | 1.03E-01 | 2.48E+00 | 2.54E+00 | 7.60E-02 | Yes | | intergenic_variant | MODIFIER |

| | | | | | | | | | | | | | | | |
|---|---|---|---|---|---|---|---|---|---|---|---|---|---|---|---|
| 5 | 61487050 | AX-170219404 | 0 | C | G | 0.482 | 1.84E-01 | 1.03E-01 | 2.48E+00 | 2.54E+00 | 7.60E-02 | Yes | | intergenic_variant | MODIFIER |
| 5 | 61491479 | AX-170089493 | 0 | T | C | 0.482 | 1.84E-01 | 1.03E-01 | 2.48E+00 | 2.54E+00 | 7.60E-02 | Yes | | intergenic_variant | MODIFIER |
| 5 | 61497560 | AX-168406659 | 0 | A | G | 0.482 | 1.84E-01 | 1.03E-01 | 2.48E+00 | 2.54E+00 | 7.60E-02 | Yes | | intergenic_variant | MODIFIER |
| 5 | 61525084 | AX-170448116 | 0 | A | T | 0.482 | 1.84E-01 | 1.03E-01 | 2.48E+00 | 2.54E+00 | 7.60E-02 | Yes | | intergenic_variant | MODIFIER |
| 5 | 61534678 | AX-97801184 | 0 | T | C | 0.482 | 1.84E-01 | 1.03E-01 | 2.48E+00 | 2.54E+00 | 7.60E-02 | Yes | | intergenic_variant | MODIFIER |
| 5 | 61614234 | AX-97793186 | 0 | C | A | 0.482 | 1.84E-01 | 1.03E-01 | 2.48E+00 | 2.54E+00 | 7.60E-02 | Yes | | intergenic_variant | MODIFIER |
| 5 | 61651209 | AX-97787957 | 0 | G | A | 0.482 | 1.84E-01 | 1.03E-01 | 2.48E+00 | 2.54E+00 | 7.60E-02 | Yes | | intergenic_variant | MODIFIER |
| 5 | 61676284 | AX-170226552 | 0 | C | G | 0.482 | 1.84E-01 | 1.03E-01 | 2.48E+00 | 2.54E+00 | 7.60E-02 | Yes | | intergenic_variant | MODIFIER |
| 5 | 61689420 | AX-170265121 | 0 | C | A | 0.482 | 1.84E-01 | 1.03E-01 | 2.48E+00 | 2.54E+00 | 7.60E-02 | Yes | | intergenic_variant | MODIFIER |
| 5 | 61717072 | AX-170225000 | 0 | C | T | 0.482 | 1.84E-01 | 1.03E-01 | 2.48E+00 | 2.54E+00 | 7.60E-02 | Yes | Gm43384 | non_coding_transcript_exon_variant | MODIFIER |
| 5 | 61813741 | AX-170522484 | 0 | A | G | 0.482 | 1.84E-01 | 1.03E-01 | 2.48E+00 | 2.54E+00 | 7.60E-02 | Yes | G6pd2 | downstream_gene_variant | MODIFIER |
| 5 | 62072901 | AX-170025116 | 0 | A | G | 0.482 | 1.84E-01 | 1.03E-01 | 2.48E+00 | 2.54E+00 | 7.60E-02 | Yes | | intergenic_variant | MODIFIER |
| 5 | 62688547 | AX-97813171 | 0 | C | A | 0.482 | 1.84E-01 | 1.03E-01 | 2.48E+00 | 2.54E+00 | 7.60E-02 | Yes | Arap2 | intron_variant | MODIFIER |
| 5 | 62811013 | AX-170151712 | 0 | C | T | 0.482 | 1.84E-01 | 1.03E-01 | 2.48E+00 | 2.54E+00 | 7.60E-02 | Yes | Dthd1 | upstream_gene_variant | MODIFIER |
| 5 | 62820195 | AX-170381360 | 0 | C | T | 0.482 | 1.84E-01 | 1.03E-01 | 2.48E+00 | 2.54E+00 | 7.60E-02 | Yes | Dthd1 | intron_variant | MODIFIER |
| 5 | 62850781 | AX-170215189 | 0 | T | C | 0.482 | 1.84E-01 | 1.03E-01 | 2.48E+00 | 2.54E+00 | 7.60E-02 | Yes | Dthd1 | intron_variant | MODIFIER |
| 5 | 62851500 | AX-170420235 | 0 | T | C | 0.482 | 1.84E-01 | 1.03E-01 | 2.48E+00 | 2.54E+00 | 7.60E-02 | Yes | Dthd1 | intron_variant | MODIFIER |
| 5 | 62854153 | AX-170429164 | 0 | C | T | 0.482 | 1.84E-01 | 1.03E-01 | 2.48E+00 | 2.54E+00 | 7.60E-02 | Yes | Dthd1 | intron_variant | MODIFIER |
| 5 | 62855676 | AX-170072287 | 0 | C | T | 0.482 | 1.84E-01 | 1.03E-01 | 2.48E+00 | 2.54E+00 | 7.60E-02 | Yes | Dthd1 | intron_variant | MODIFIER |
| 5 | 62858226 | AX-170222004 | 0 | T | G | 0.482 | 1.84E-01 | 1.03E-01 | 2.48E+00 | 2.54E+00 | 7.60E-02 | Yes | Dthd1 | intron_variant | MODIFIER |
| 5 | 62881095 | AX-170072286 | 0 | C | T | 0.482 | 1.84E-01 | 1.03E-01 | 2.48E+00 | 2.54E+00 | 7.60E-02 | Yes | Dthd1 | intron_variant | MODIFIER |
| 5 | 62889074 | AX-170298551 | 0 | C | G | 0.482 | 1.84E-01 | 1.03E-01 | 2.48E+00 | 2.54E+00 | 7.60E-02 | Yes | Dthd1 | downstream_gene_variant | MODIFIER |
| 5 | 62940575 | AX-170396550 | 0 | C | G | 0.482 | 1.84E-01 | 1.03E-01 | 2.48E+00 | 2.54E+00 | 7.60E-02 | Yes | | intergenic_variant | MODIFIER |
| 5 | 62951299 | AX-170506345 | 0 | G | A | 0.482 | 1.84E-01 | 1.03E-01 | 2.48E+00 | 2.54E+00 | 7.60E-02 | Yes | | intergenic_variant | MODIFIER |
| 5 | 62968014 | AX-168400132 | 0 | A | G | 0.482 | 1.84E-01 | 1.03E-01 | 2.48E+00 | 2.54E+00 | 7.60E-02 | Yes | | intergenic_variant | MODIFIER |
| 5 | 62976676 | AX-170176828 | 0 | G | A | 0.482 | 1.84E-01 | 1.03E-01 | 2.48E+00 | 2.54E+00 | 7.60E-02 | Yes | | intergenic_variant | MODIFIER |
| 5 | 63051165 | AX-170025132 | 0 | A | G | 0.482 | 1.84E-01 | 1.03E-01 | 2.48E+00 | 2.54E+00 | 7.60E-02 | Yes | | intergenic_variant | MODIFIER |
| 5 | 63100845 | AX-170518419 | 0 | G | A | 0.482 | 1.84E-01 | 1.03E-01 | 2.48E+00 | 2.54E+00 | 7.60E-02 | Yes | | intergenic_variant | MODIFIER |
| 5 | 63101849 | AX-170286316 | 0 | T | C | 0.482 | 1.84E-01 | 1.03E-01 | 2.48E+00 | 2.54E+00 | 7.60E-02 | Yes | | intergenic_variant | MODIFIER |
| 5 | 63131759 | AX-170308079 | 0 | C | G | 0.482 | 1.84E-01 | 1.03E-01 | 2.48E+00 | 2.54E+00 | 7.60E-02 | Yes | | intergenic_variant | MODIFIER |
| 5 | 63157781 | AX-97813975 | 0 | C | A | 0.482 | 1.84E-01 | 1.03E-01 | 2.48E+00 | 2.54E+00 | 7.60E-02 | Yes | | intergenic_variant | MODIFIER |
| 5 | 63158002 | AX-170172753 | 0 | G | A | 0.482 | 1.84E-01 | 1.03E-01 | 2.48E+00 | 2.54E+00 | 7.60E-02 | Yes | | intergenic_variant | MODIFIER |
| 5 | 63187214 | AX-170328475 | 0 | G | A | 0.482 | 1.84E-01 | 1.03E-01 | 2.48E+00 | 2.54E+00 | 7.60E-02 | Yes | | intergenic_variant | MODIFIER |
| 5 | 63190074 | AX-170502175 | 0 | C | G | 0.482 | 1.84E-01 | 1.03E-01 | 2.48E+00 | 2.54E+00 | 7.60E-02 | Yes | | intergenic_variant | MODIFIER |
| 5 | 63193597 | AX-170425038 | 0 | T | A | 0.482 | 1.84E-01 | 1.03E-01 | 2.48E+00 | 2.54E+00 | 7.60E-02 | Yes | | intergenic_variant | MODIFIER |
| 5 | 63203984 | AX-170169147 | 0 | A | C | 0.482 | 1.84E-01 | 1.03E-01 | 2.48E+00 | 2.54E+00 | 7.60E-02 | Yes | | intergenic_variant | MODIFIER |
| 5 | 63242659 | AX-170472846 | 0 | A | G | 0.482 | 1.84E-01 | 1.03E-01 | 2.48E+00 | 2.54E+00 | 7.60E-02 | Yes | | intergenic_variant | MODIFIER |
| 5 | 63243987 | AX-97797936 | 0 | T | C | 0.482 | 1.84E-01 | 1.03E-01 | 2.48E+00 | 2.54E+00 | 7.60E-02 | Yes | | intergenic_variant | MODIFIER |
| 5 | 63308269 | AX-170486854 | 0 | G | C | 0.482 | 1.84E-01 | 1.03E-01 | 2.48E+00 | 2.54E+00 | 7.60E-02 | Yes | | intergenic_variant | MODIFIER |
| 5 | 63428273 | AX-170089502 | 0 | A | G | 0.482 | 1.84E-01 | 1.03E-01 | 2.48E+00 | 2.54E+00 | 7.60E-02 | Yes | | intergenic_variant | MODIFIER |
| 5 | 63428483 | AX-170235731 | 0 | G | C | 0.482 | 1.84E-01 | 1.03E-01 | 2.48E+00 | 2.54E+00 | 7.60E-02 | Yes | | intergenic_variant | MODIFIER |
| 5 | 63428699 | AX-170393551 | 0 | A | C | 0.482 | 1.84E-01 | 1.03E-01 | 2.48E+00 | 2.54E+00 | 7.60E-02 | Yes | | intergenic_variant | MODIFIER |
| 5 | 63642390 | AX-170089503 | 0 | A | T | 0.482 | 1.84E-01 | 1.03E-01 | 2.48E+00 | 2.54E+00 | 7.60E-02 | Yes | Gm9954 | downstream_gene_variant | MODIFIER |
| 5 | 63784298 | AX-170228577 | 0 | G | A | 0.482 | 1.84E-01 | 1.03E-01 | 2.48E+00 | 2.54E+00 | 7.60E-02 | Yes | Nwd2 | intron_variant | MODIFIER |
| 5 | 63823013 | AX-170102524 | 0 | A | G | 0.482 | 1.84E-01 | 1.03E-01 | 2.48E+00 | 2.54E+00 | 7.60E-02 | Yes | 0610040J01Rik | intron_variant | MODIFIER |
| 5 | 63826537 | AX-170072304 | 0 | C | T | 0.482 | 1.84E-01 | 1.03E-01 | 2.48E+00 | 2.54E+00 | 7.60E-02 | Yes | 0610040J01Rik | intron_variant | MODIFIER |
| 5 | 63827324 | AX-170072303 | 0 | A | G | 0.482 | 1.84E-01 | 1.03E-01 | 2.48E+00 | 2.54E+00 | 7.60E-02 | Yes | 0610040J01Rik | intron_variant | MODIFIER |
| 5 | 63878826 | AX-97836144 | 0 | C | T | 0.482 | 1.84E-01 | 1.03E-01 | 2.48E+00 | 2.54E+00 | 7.60E-02 | Yes | 0610040J01Rik | intron_variant | MODIFIER |
| 5 | 63951665 | AX-170365291 | 0 | C | T | 0.482 | 1.84E-01 | 1.03E-01 | 2.48E+00 | 2.54E+00 | 7.60E-02 | Yes | Rell1 | intron_variant,NMD_transcript_variant | MODIFIER |
| 5 | 63958522 | AX-97847113 | 0 | C | T | 0.482 | 1.84E-01 | 1.03E-01 | 2.48E+00 | 2.54E+00 | 7.60E-02 | Yes | Rell1 | intron_variant,NMD_transcript_variant | MODIFIER |
| 5 | 63961842 | AX-170141203 | 0 | G | C | 0.482 | 1.84E-01 | 1.03E-01 | 2.48E+00 | 2.54E+00 | 7.60E-02 | Yes | Rell1 | intron_variant,NMD_transcript_variant | MODIFIER |
| 5 | 63998196 | AX-170383726 | 0 | C | T | 0.482 | 1.84E-01 | 1.03E-01 | 2.48E+00 | 2.54E+00 | 7.60E-02 | Yes | Gm25608 | downstream_gene_variant | MODIFIER |
| 5 | 64007764 | AX-97796456 | 0 | A | G | 0.482 | 1.84E-01 | 1.03E-01 | 2.48E+00 | 2.54E+00 | 7.60E-02 | Yes | | intergenic_variant | MODIFIER |
| 5 | 64022394 | AX-170268008 | 0 | T | C | 0.482 | 1.84E-01 | 1.03E-01 | 2.48E+00 | 2.54E+00 | 7.60E-02 | Yes | | intergenic_variant | MODIFIER |

| | | | | | | | | | | | | | | | |
|---|---|---|---|---|---|---|---|---|---|---|---|---|---|---|---|
| 5 | 64040707 | AX-170025144 | 0 | T | C | 0.482 | 1.84E-01 | 1.03E-01 | 2.48E+00 | 2.54E+00 | 7.60E-02 | Yes | 5830416I19Rik | upstream_gene_variant | MODIFIER |
| 5 | 64041701 | AX-170072308 | 0 | T | C | 0.482 | 1.84E-01 | 1.03E-01 | 2.48E+00 | 2.54E+00 | 7.60E-02 | Yes | 5830416I19Rik | upstream_gene_variant | MODIFIER |
| 5 | 64042164 | AX-170072309 | 0 | G | A | 0.482 | 1.84E-01 | 1.03E-01 | 2.48E+00 | 2.54E+00 | 7.60E-02 | Yes | 5830416I19Rik | upstream_gene_variant | MODIFIER |
| 5 | 64053352 | AX-170453830 | 0 | G | A | 0.482 | 1.84E-01 | 1.03E-01 | 2.48E+00 | 2.54E+00 | 7.60E-02 | Yes | 5830416I19Rik | downstream_gene_variant | MODIFIER |
| 5 | 64098407 | AX-170236366 | 0 | G | A | 0.482 | 1.84E-01 | 1.03E-01 | 2.48E+00 | 2.54E+00 | 7.60E-02 | Yes | Pgm2 | intron_variant | MODIFIER |
| 5 | 64126230 | AX-170025146 | 0 | G | T | 0.482 | 1.84E-01 | 1.03E-01 | 2.48E+00 | 2.54E+00 | 7.60E-02 | Yes | Pgm2 | intron_variant | MODIFIER |
| 5 | 64198348 | AX-170089504 | 0 | C | T | 0.482 | 1.84E-01 | 1.03E-01 | 2.48E+00 | 2.54E+00 | 7.60E-02 | Yes | Tbc1d1 | intron_variant | MODIFIER |
| 5 | 64211636 | AX-170311879 | 0 | T | C | 0.482 | 1.84E-01 | 1.03E-01 | 2.48E+00 | 2.54E+00 | 7.60E-02 | Yes | Tbc1d1 | intron_variant | MODIFIER |
| 5 | 64226112 | AX-170402592 | 0 | A | G | 0.482 | 1.84E-01 | 1.03E-01 | 2.48E+00 | 2.54E+00 | 7.60E-02 | Yes | Tbc1d1 | intron_variant | MODIFIER |
| 5 | 64229384 | AX-170089505 | 0 | A | G | 0.482 | 1.84E-01 | 1.03E-01 | 2.48E+00 | 2.54E+00 | 7.60E-02 | Yes | Tbc1d1 | intron_variant | MODIFIER |
| 5 | 64241256 | AX-170378145 | 0 | A | G | 0.482 | 1.84E-01 | 1.03E-01 | 2.48E+00 | 2.54E+00 | 7.60E-02 | Yes | Tbc1d1 | intron_variant | MODIFIER |
| 5 | 64254423 | AX-170095343 | 0 | G | A | 0.482 | 1.84E-01 | 1.03E-01 | 2.48E+00 | 2.54E+00 | 7.60E-02 | Yes | Tbc1d1 | intron_variant | MODIFIER |
| 5 | 64270527 | AX-170494151 | 0 | A | G | 0.482 | 1.84E-01 | 1.03E-01 | 2.48E+00 | 2.54E+00 | 7.60E-02 | Yes | Tbc1d1 | intron_variant | MODIFIER |
| 5 | 64285819 | AX-97841242 | 0 | G | A | 0.482 | 1.84E-01 | 1.03E-01 | 2.48E+00 | 2.54E+00 | 7.60E-02 | Yes | Tbc1d1 | intron_variant | MODIFIER |
| 5 | 64285868 | AX-170503231 | 0 | T | C | 0.482 | 1.84E-01 | 1.03E-01 | 2.48E+00 | 2.54E+00 | 7.60E-02 | Yes | Tbc1d1 | intron_variant | MODIFIER |
| 5 | 64314878 | AX-170105728 | 0 | G | A | 0.482 | 1.84E-01 | 1.03E-01 | 2.48E+00 | 2.54E+00 | 7.60E-02 | Yes | Tbc1d1 | intron_variant | MODIFIER |
| 5 | 64316073 | AX-170304093 | 0 | G | A | 0.482 | 1.84E-01 | 1.03E-01 | 2.48E+00 | 2.54E+00 | 7.60E-02 | Yes | Tbc1d1 | intron_variant | MODIFIER |
| 5 | 64317376 | AX-170161854 | 0 | A | G | 0.482 | 1.84E-01 | 1.03E-01 | 2.48E+00 | 2.54E+00 | 7.60E-02 | Yes | Tbc1d1 | intron_variant | MODIFIER |
| 5 | 64319889 | AX-170510327 | 0 | C | G | 0.482 | 1.84E-01 | 1.03E-01 | 2.48E+00 | 2.54E+00 | 7.60E-02 | Yes | Tbc1d1 | intron_variant | MODIFIER |
| 5 | 64333610 | AX-170144647 | 0 | G | A | 0.482 | 1.84E-01 | 1.03E-01 | 2.48E+00 | 2.54E+00 | 7.60E-02 | Yes | Tbc1d1 | intron_variant | MODIFIER |
| 5 | 64335579 | AX-170325297 | 0 | G | A | 0.482 | 1.84E-01 | 1.03E-01 | 2.48E+00 | 2.54E+00 | 7.60E-02 | Yes | Tbc1d1 | intron_variant | MODIFIER |
| 5 | 64336469 | AX-170387594 | 0 | G | C | 0.482 | 1.84E-01 | 1.03E-01 | 2.48E+00 | 2.54E+00 | 7.60E-02 | Yes | Tbc1d1 | intron_variant | MODIFIER |
| 5 | 64339408 | AX-170409569 | 0 | C | T | 0.482 | 1.84E-01 | 1.03E-01 | 2.48E+00 | 2.54E+00 | 7.60E-02 | Yes | Tbc1d1 | intron_variant | MODIFIER |
| 5 | 64393232 | AX-170174861 | 0 | G | A | 0.482 | 1.84E-01 | 1.03E-01 | 2.48E+00 | 2.54E+00 | 7.60E-02 | Yes | | regulatory_region_variant | MODIFIER |
| 5 | 64394390 | AX-170139365 | 0 | A | G | 0.482 | 1.84E-01 | 1.03E-01 | 2.48E+00 | 2.54E+00 | 7.60E-02 | Yes | | intergenic_variant | MODIFIER |
| 5 | 64396971 | AX-170148009 | 0 | A | G | 0.482 | 1.84E-01 | 1.03E-01 | 2.48E+00 | 2.54E+00 | 7.60E-02 | Yes | | intergenic_variant | MODIFIER |
| 5 | 64414703 | AX-170373462 | 0 | C | T | 0.482 | 1.84E-01 | 1.03E-01 | 2.48E+00 | 2.54E+00 | 7.60E-02 | Yes | | intergenic_variant | MODIFIER |
| 5 | 64430858 | AX-97800080 | 0 | G | A | 0.482 | 1.84E-01 | 1.03E-01 | 2.48E+00 | 2.54E+00 | 7.60E-02 | Yes | Gm31363 | upstream_gene_variant | MODIFIER |
| 5 | 64432947 | AX-170200320 | 0 | G | C | 0.482 | 1.84E-01 | 1.03E-01 | 2.48E+00 | 2.54E+00 | 7.60E-02 | Yes | Gm31363 | upstream_gene_variant | MODIFIER |
| 5 | 64438992 | AX-170097091 | 0 | C | A | 0.482 | 1.84E-01 | 1.03E-01 | 2.48E+00 | 2.54E+00 | 7.60E-02 | Yes | Gm31363 | intron_variant,non_coding_transcript_variant | MODIFIER |
| 5 | 64457105 | AX-170313590 | 0 | A | G | 0.482 | 1.84E-01 | 1.03E-01 | 2.48E+00 | 2.54E+00 | 7.60E-02 | Yes | 1700027F09Rik | intron_variant,non_coding_transcript_variant | MODIFIER |
| 5 | 64466706 | AX-97831123 | 0 | A | G | 0.482 | 1.84E-01 | 1.03E-01 | 2.48E+00 | 2.54E+00 | 7.60E-02 | Yes | 1700027F09Rik | intron_variant,non_coding_transcript_variant | MODIFIER |
| 5 | 64468632 | AX-170236267 | 0 | G | A | 0.482 | 1.84E-01 | 1.03E-01 | 2.48E+00 | 2.54E+00 | 7.60E-02 | Yes | 1700027F09Rik | upstream_gene_variant | MODIFIER |
| 5 | 64473355 | AX-170430781 | 0 | A | G | 0.482 | 1.84E-01 | 1.03E-01 | 2.48E+00 | 2.54E+00 | 7.60E-02 | Yes | C030018K13Rik | upstream_gene_variant | MODIFIER |
| 5 | 64473813 | AX-170393718 | 0 | G | A | 0.482 | 1.84E-01 | 1.03E-01 | 2.48E+00 | 2.54E+00 | 7.60E-02 | Yes | C030018K13Rik | upstream_gene_variant | MODIFIER |
| 5 | 64482935 | AX-170433465 | 0 | C | T | 0.482 | 1.84E-01 | 1.03E-01 | 2.48E+00 | 2.54E+00 | 7.60E-02 | Yes | C030018K13Rik | downstream_gene_variant | MODIFIER |
| 5 | 64483306 | AX-170263747 | 0 | C | T | 0.482 | 1.84E-01 | 1.03E-01 | 2.48E+00 | 2.54E+00 | 7.60E-02 | Yes | C030018K13Rik | downstream_gene_variant | MODIFIER |
| 5 | 64511739 | AX-97852500 | 0 | T | C | 0.482 | 1.84E-01 | 1.03E-01 | 2.48E+00 | 2.54E+00 | 7.60E-02 | Yes | | intergenic_variant | MODIFIER |
| 5 | 64538255 | AX-170511546 | 0 | A | T | 0.482 | 1.84E-01 | 1.03E-01 | 2.48E+00 | 2.54E+00 | 7.60E-02 | Yes | | regulatory_region_variant | MODIFIER |
| 5 | 64539433 | AX-170321851 | 0 | G | A | 0.482 | 1.84E-01 | 1.03E-01 | 2.48E+00 | 2.54E+00 | 7.60E-02 | Yes | | intergenic_variant | MODIFIER |
| 5 | 64557585 | AX-97831698 | 0 | A | G | 0.482 | 1.84E-01 | 1.03E-01 | 2.48E+00 | 2.54E+00 | 7.60E-02 | Yes | Gm43836 | upstream_gene_variant | MODIFIER |
| 5 | 64600625 | AX-170025157 | 0 | T | A | 0.482 | 1.84E-01 | 1.03E-01 | 2.48E+00 | 2.54E+00 | 7.60E-02 | Yes | Gm3716 | intron_variant,non_coding_transcript_variant | MODIFIER |
| 5 | 64603810 | AX-170025155 | 0 | C | T | 0.482 | 1.84E-01 | 1.03E-01 | 2.48E+00 | 2.54E+00 | 7.60E-02 | Yes | Gm3716 | upstream_gene_variant | MODIFIER |
| 5 | 64610652 | AX-170142225 | 0 | G | C | 0.482 | 1.84E-01 | 1.03E-01 | 2.48E+00 | 2.54E+00 | 7.60E-02 | Yes | Gm3716 | non_coding_transcript_exon_variant | MODIFIER |
| 5 | 64611914 | AX-170025159 | 0 | G | T | 0.482 | 1.84E-01 | 1.03E-01 | 2.48E+00 | 2.54E+00 | 7.60E-02 | Yes | Gm3716 | upstream_gene_variant | MODIFIER |
| 5 | 64615628 | AX-170025153 | 0 | A | G | 0.482 | 1.84E-01 | 1.03E-01 | 2.48E+00 | 2.54E+00 | 7.60E-02 | Yes | Gm3716 | upstream_gene_variant | MODIFIER |
| 5 | 64629765 | AX-97794012 | 0 | C | A | 0.482 | 1.84E-01 | 1.03E-01 | 2.48E+00 | 2.54E+00 | 7.60E-02 | Yes | | intergenic_variant | MODIFIER |
| 5 | 64662729 | AX-170025161 | 0 | T | C | 0.482 | 1.84E-01 | 1.03E-01 | 2.48E+00 | 2.54E+00 | 7.60E-02 | Yes | | regulatory_region_variant | MODIFIER |
| 5 | 64704434 | AX-170210260 | 0 | A | G | 0.482 | 1.84E-01 | 1.03E-01 | 2.48E+00 | 2.54E+00 | 7.60E-02 | Yes | Gm20033 | downstream_gene_variant | MODIFIER |
| 5 | 64706536 | AX-97797588 | 0 | G | A | 0.482 | 1.84E-01 | 1.03E-01 | 2.48E+00 | 2.54E+00 | 7.60E-02 | Yes | Gm20033 | downstream_gene_variant | MODIFIER |
| 5 | 64753258 | AX-170481804 | 0 | C | T | 0.482 | 1.84E-01 | 1.03E-01 | 2.48E+00 | 2.54E+00 | 7.60E-02 | Yes | Gm20033 | intron_variant,non_coding_transcript_variant | MODIFIER |
| 5 | 64757662 | AX-97804846 | 0 | C | T | 0.482 | 1.84E-01 | 1.03E-01 | 2.48E+00 | 2.54E+00 | 7.60E-02 | Yes | Gm20033 | intron_variant,non_coding_transcript_variant | MODIFIER |
| 5 | 64781126 | AX-170330126 | 0 | C | G | 0.482 | 1.84E-01 | 1.03E-01 | 2.48E+00 | 2.54E+00 | 7.60E-02 | Yes | Gm20033 | intron_variant,non_coding_transcript_variant | MODIFIER |
| 5 | 64814160 | AX-170146976 | 0 | C | T | 0.482 | 1.84E-01 | 1.03E-01 | 2.48E+00 | 2.54E+00 | 7.60E-02 | Yes | Klf3 | intron_variant | MODIFIER |

| | | | | | | | | | | | | | | | |
|---|---|---|---|---|---|---|---|---|---|---|---|---|---|---|---|
| 5 | 64814581 | AX-170286334 | 0 | A | G | 0.482 | 1.84E-01 | 1.03E-01 | 2.48E+00 | 2.54E+00 | 7.60E-02 | Yes | Klf3 | intron_variant | MODIFIER |
| 5 | 64815780 | AX-170126726 | 0 | T | C | 0.482 | 1.84E-01 | 1.03E-01 | 2.48E+00 | 2.54E+00 | 7.60E-02 | Yes | Klf3 | intron_variant | MODIFIER |
| 5 | 64896726 | AX-170130370 | 0 | T | G | 0.482 | 1.84E-01 | 1.03E-01 | 2.48E+00 | 2.54E+00 | 7.60E-02 | Yes | | regulatory_region_variant | MODIFIER |
| 5 | 64897866 | AX-97794299 | 0 | C | G | 0.482 | 1.84E-01 | 1.03E-01 | 2.48E+00 | 2.54E+00 | 7.60E-02 | Yes | | intergenic_variant | MODIFIER |
| 5 | 64926898 | AX-170383007 | 0 | G | A | 0.482 | 1.84E-01 | 1.03E-01 | 2.48E+00 | 2.54E+00 | 7.60E-02 | Yes | Tlr1 | missense_variant | MODERATE |
| 5 | 64932757 | AX-170247286 | 0 | C | T | 0.482 | 1.84E-01 | 1.03E-01 | 2.48E+00 | 2.54E+00 | 7.60E-02 | Yes | Tlr1 | 5_prime_UTR_variant | MODIFIER |
| 5 | 64983242 | AX-170477348 | 0 | A | G | 0.482 | 1.84E-01 | 1.03E-01 | 2.48E+00 | 2.54E+00 | 7.60E-02 | Yes | Fam114a1 | intron_variant | MODIFIER |
| 5 | 64983888 | AX-170235848 | 0 | C | T | 0.482 | 1.84E-01 | 1.03E-01 | 2.48E+00 | 2.54E+00 | 7.60E-02 | Yes | Fam114a1 | intron_variant | MODIFIER |
| 5 | 64986299 | AX-170468911 | 0 | A | C | 0.482 | 1.84E-01 | 1.03E-01 | 2.48E+00 | 2.54E+00 | 7.60E-02 | Yes | Fam114a1 | intron_variant | MODIFIER |
| 5 | 64988525 | AX-170299445 | 0 | A | G | 0.482 | 1.84E-01 | 1.03E-01 | 2.48E+00 | 2.54E+00 | 7.60E-02 | Yes | Fam114a1 | intron_variant | MODIFIER |
| 5 | 64989087 | AX-170535855 | 0 | C | T | 0.482 | 1.84E-01 | 1.03E-01 | 2.48E+00 | 2.54E+00 | 7.60E-02 | Yes | Fam114a1 | intron_variant | MODIFIER |
| 5 | 64989899 | AX-170204136 | 0 | G | T | 0.482 | 1.84E-01 | 1.03E-01 | 2.48E+00 | 2.54E+00 | 7.60E-02 | Yes | Fam114a1 | intron_variant | MODIFIER |
| 5 | 65005457 | AX-170158171 | 0 | C | T | 0.482 | 1.84E-01 | 1.03E-01 | 2.48E+00 | 2.54E+00 | 7.60E-02 | Yes | Fam114a1 | intron_variant | MODIFIER |
| 5 | 65006727 | AX-170072326 | 0 | C | T | 0.482 | 1.84E-01 | 1.03E-01 | 2.48E+00 | 2.54E+00 | 7.60E-02 | Yes | Fam114a1 | intron_variant | MODIFIER |
| 5 | 65007483 | AX-170470963 | 0 | C | G | 0.482 | 1.84E-01 | 1.03E-01 | 2.48E+00 | 2.54E+00 | 7.60E-02 | Yes | Fam114a1 | intron_variant | MODIFIER |
| 5 | 65018931 | AX-170297382 | 0 | C | G | 0.482 | 1.84E-01 | 1.03E-01 | 2.48E+00 | 2.54E+00 | 7.60E-02 | Yes | Fam114a1 | intron_variant | MODIFIER |
| 5 | 65033564 | AX-170404672 | 0 | C | T | 0.482 | 1.84E-01 | 1.03E-01 | 2.48E+00 | 2.54E+00 | 7.60E-02 | Yes | Fam114a1 | intron_variant | MODIFIER |
| 5 | 65115682 | AX-170025165 | 0 | T | C | 0.482 | 1.84E-01 | 1.03E-01 | 2.48E+00 | 2.54E+00 | 7.60E-02 | Yes | Klhl5 | intron_variant | MODIFIER |
| 5 | 65129502 | AX-170392960 | 0 | T | G | 0.482 | 1.84E-01 | 1.03E-01 | 2.48E+00 | 2.54E+00 | 7.60E-02 | Yes | Klhl5 | upstream_gene_variant | MODIFIER |
| 5 | 65142849 | AX-97796648 | 0 | A | T | 0.482 | 1.84E-01 | 1.03E-01 | 2.48E+00 | 2.54E+00 | 7.60E-02 | Yes | Klhl5 | intron_variant | MODIFIER |
| 5 | 65145001 | AX-170025162 | 0 | G | C | 0.482 | 1.84E-01 | 1.03E-01 | 2.48E+00 | 2.54E+00 | 7.60E-02 | Yes | Klhl5 | intron_variant | MODIFIER |
| 5 | 65145052 | AX-170292461 | 0 | A | G | 0.482 | 1.84E-01 | 1.03E-01 | 2.48E+00 | 2.54E+00 | 7.60E-02 | Yes | Klhl5 | intron_variant | MODIFIER |
| 5 | 65162988 | AX-170072327 | 0 | G | A | 0.482 | 1.84E-01 | 1.03E-01 | 2.48E+00 | 2.54E+00 | 7.60E-02 | Yes | Klhl5 | synonymous_variant | LOW |
| 5 | 65168050 | AX-97833633 | 0 | A | G | 0.482 | 1.84E-01 | 1.03E-01 | 2.48E+00 | 2.54E+00 | 7.60E-02 | Yes | Klhl5 | 3_prime_UTR_variant | MODIFIER |
| 5 | 65269319 | AX-170411041 | 0 | G | A | 0.482 | 1.84E-01 | 1.03E-01 | 2.48E+00 | 2.54E+00 | 7.60E-02 | Yes | Rfc1 | intron_variant | MODIFIER |
| 5 | 65312967 | AX-170355891 | 0 | A | T | 0.482 | 1.84E-01 | 1.03E-01 | 2.48E+00 | 2.54E+00 | 7.60E-02 | Yes | Rfc1 | missense_variant | MODERATE |
| 5 | 65703608 | AX-170025173 | 0 | C | A | 0.482 | 1.84E-01 | 1.03E-01 | 2.48E+00 | 2.54E+00 | 7.60E-02 | Yes | | regulatory_region_variant | MODIFIER |
| 5 | 65704193 | AX-97843238 | 0 | A | G | 0.482 | 1.84E-01 | 1.03E-01 | 2.48E+00 | 2.54E+00 | 7.60E-02 | Yes | | intergenic_variant | MODIFIER |
| 5 | 65718028 | AX-170334698 | 0 | G | A | 0.482 | 1.84E-01 | 1.03E-01 | 2.48E+00 | 2.54E+00 | 7.60E-02 | Yes | | regulatory_region_variant | MODIFIER |
| 5 | 65748178 | AX-97841160 | 0 | A | G | 0.482 | 1.84E-01 | 1.03E-01 | 2.48E+00 | 2.54E+00 | 7.60E-02 | Yes | | regulatory_region_variant | MODIFIER |
| 5 | 65770151 | AX-170153386 | 0 | A | T | 0.482 | 1.84E-01 | 1.03E-01 | 2.48E+00 | 2.54E+00 | 7.60E-02 | Yes | N4bp2 | intron_variant | MODIFIER |
| 5 | 65774187 | AX-170025177 | 0 | T | G | 0.482 | 1.84E-01 | 1.03E-01 | 2.48E+00 | 2.54E+00 | 7.60E-02 | Yes | N4bp2 | intron_variant | MODIFIER |
| 5 | 65775125 | AX-170025175 | 0 | C | G | 0.482 | 1.84E-01 | 1.03E-01 | 2.48E+00 | 2.54E+00 | 7.60E-02 | Yes | N4bp2 | intron_variant | MODIFIER |
| 5 | 65775605 | AX-170434990 | 0 | T | C | 0.482 | 1.84E-01 | 1.03E-01 | 2.48E+00 | 2.54E+00 | 7.60E-02 | Yes | N4bp2 | 5_prime_UTR_variant | MODIFIER |
| 5 | 65813842 | AX-170528644 | 0 | G | C | 0.482 | 1.84E-01 | 1.03E-01 | 2.48E+00 | 2.54E+00 | 7.60E-02 | Yes | N4bp2 | intron_variant | MODIFIER |
| 5 | 65988049 | AX-97811593 | 0 | G | A | 0.482 | 1.84E-01 | 1.03E-01 | 2.48E+00 | 2.54E+00 | 7.60E-02 | Yes | 9130230L23Rik | downstream_gene_variant | MODIFIER |
| 5 | 66050485 | AX-170366927 | 0 | A | C | 0.482 | 1.84E-01 | 1.03E-01 | 2.48E+00 | 2.54E+00 | 7.60E-02 | Yes | Rbm47 | intron_variant | MODIFIER |
| 5 | 66051292 | AX-170130094 | 0 | A | C | 0.482 | 1.84E-01 | 1.03E-01 | 2.48E+00 | 2.54E+00 | 7.60E-02 | Yes | Rbm47 | intron_variant | MODIFIER |
| 5 | 66057139 | AX-97842179 | 0 | C | T | 0.482 | 1.84E-01 | 1.03E-01 | 2.48E+00 | 2.54E+00 | 7.60E-02 | Yes | Rbm47 | intron_variant | MODIFIER |
| 5 | 66079230 | AX-97809329 | 0 | G | A | 0.482 | 1.84E-01 | 1.03E-01 | 2.48E+00 | 2.54E+00 | 7.60E-02 | Yes | Rbm47 | intron_variant | MODIFIER |
| 5 | 66100114 | AX-97833588 | 0 | A | G | 0.482 | 1.84E-01 | 1.03E-01 | 2.48E+00 | 2.54E+00 | 7.60E-02 | Yes | Rbm47 | intron_variant | MODIFIER |
| 5 | 66112975 | AX-170296784 | 0 | T | G | 0.482 | 1.84E-01 | 1.03E-01 | 2.48E+00 | 2.54E+00 | 7.60E-02 | Yes | Rbm47 | intron_variant | MODIFIER |
| 5 | 66118651 | AX-170409041 | 0 | A | G | 0.482 | 1.84E-01 | 1.03E-01 | 2.48E+00 | 2.54E+00 | 7.60E-02 | Yes | Rbm47 | intron_variant | MODIFIER |
| 5 | 66145330 | AX-170201755 | 0 | A | G | 0.482 | 1.84E-01 | 1.03E-01 | 2.48E+00 | 2.54E+00 | 7.60E-02 | Yes | Rbm47 | intron_variant | MODIFIER |
| 5 | 66158762 | AX-170229621 | 0 | C | T | 0.482 | 1.84E-01 | 1.03E-01 | 2.48E+00 | 2.54E+00 | 7.60E-02 | Yes | Gm26725 | intron_variant,non_coding_transcript_variant | MODIFIER |
| 5 | 66164281 | AX-170388528 | 0 | T | C | 0.482 | 1.84E-01 | 1.03E-01 | 2.48E+00 | 2.54E+00 | 7.60E-02 | Yes | Gm22929 | downstream_gene_variant | MODIFIER |
| 5 | 66180586 | AX-170283568 | 0 | T | C | 0.482 | 1.84E-01 | 1.03E-01 | 2.48E+00 | 2.54E+00 | 7.60E-02 | Yes | 1700126H18Rik | intron_variant,non_coding_transcript_variant | MODIFIER |
| 5 | 66180967 | AX-170369495 | 0 | A | G | 0.482 | 1.84E-01 | 1.03E-01 | 2.48E+00 | 2.54E+00 | 7.60E-02 | Yes | 1700126H18Rik | intron_variant,non_coding_transcript_variant | MODIFIER |
| 5 | 66219506 | AX-97832708 | 0 | A | G | 0.482 | 1.84E-01 | 1.03E-01 | 2.48E+00 | 2.54E+00 | 7.60E-02 | Yes | Gm3822 | downstream_gene_variant | MODIFIER |
| 5 | 66273536 | AX-170025179 | 0 | T | A | 0.482 | 1.84E-01 | 1.03E-01 | 2.48E+00 | 2.54E+00 | 7.60E-02 | Yes | Nsun7 | intron_variant | MODIFIER |
| 5 | 66298645 | AX-170025187 | 0 | T | C | 0.482 | 1.84E-01 | 1.03E-01 | 2.48E+00 | 2.54E+00 | 7.60E-02 | Yes | Nsun7 | downstream_gene_variant | MODIFIER |
| 5 | 66305694 | AX-170072348 | 0 | A | G | 0.482 | 1.84E-01 | 1.03E-01 | 2.48E+00 | 2.54E+00 | 7.60E-02 | Yes | Apbb2 | intron_variant | MODIFIER |
| 5 | 66311362 | AX-170072346 | 0 | T | C | 0.482 | 1.84E-01 | 1.03E-01 | 2.48E+00 | 2.54E+00 | 7.60E-02 | Yes | Apbb2 | intron_variant | MODIFIER |
| 16 | 65833263 | AX-170013272 | 2 | C | T | 0.518 | 1.55E-01 | 8.76E-02 | 2.72E+00 | 2.77E+00 | 7.60E-02 | Yes | Vgll3 | intron_variant | MODIFIER |

| | | | | | | | | | | | | | | | |
|---|---|---|---|---|---|---|---|---|---|---|---|---|---|---|---|
| 5 | 129846361 | AX-170443863 | 0 | A | C | 0.149 | -1.56E-01 | 8.46E-02 | 2.14E+00 | 2.22E+00 | 7.61E-02 | no | Cct6a | downstream_gene_variant | MODIFIER |
| 13 | 18163014 | AX-170355793 | 1 | T | C | 0.161 | -2.10E-01 | 1.18E-01 | 2.54E+00 | 2.60E+00 | 7.62E-02 | Yes | Pou6f2 | intron_variant | MODIFIER |
| 5 | 50636809 | AX-97789478 | 2 | A | G | 0.509 | 1.69E-01 | 9.49E-02 | 2.49E+00 | 2.57E+00 | 7.62E-02 | Yes | Gm43016 | intron_variant,non_coding_transcript_variant | MODIFIER |
| 4 | 86949274 | AX-170089306 | 0 | C | T | 0.351 | 1.87E-01 | 1.06E-01 | 2.88E+00 | 2.98E+00 | 7.62E-02 | Yes | | regulatory_region_variant | MODIFIER |
| 18 | 79771145 | AX-97780995 | 1 | G | A | 0.473 | 1.47E-01 | 8.23E-02 | 2.37E+00 | 2.44E+00 | 7.63E-02 | Yes | | intergenic_variant | MODIFIER |
| 10 | 92487706 | AX-170434685 | 0 | A | G | 0.307 | -1.40E-01 | 7.89E-02 | 2.61E+00 | 2.69E+00 | 7.63E-02 | no | Gm25884 | downstream_gene_variant | MODIFIER |
| 16 | 51960958 | AX-170129079 | 0 | T | C | 0.465 | -1.44E-01 | 8.16E-02 | 2.61E+00 | 2.66E+00 | 7.64E-02 | no | | intergenic_variant | MODIFIER |
| 16 | 53510809 | AX-170521167 | 0 | A | G | 0.465 | -1.44E-01 | 8.16E-02 | 2.61E+00 | 2.66E+00 | 7.64E-02 | no | | intergenic_variant | MODIFIER |
| 19 | 53107431 | AX-170343699 | 1 | T | C | 0.473 | 1.12E-01 | 6.36E-02 | 2.70E+00 | 2.78E+00 | 7.64E-02 | no | | regulatory_region_variant | MODIFIER |
| 16 | 70196677 | AX-170230114 | 0 | T | G | 0.491 | 1.31E-01 | 7.42E-02 | 2.67E+00 | 2.76E+00 | 7.65E-02 | no | | intergenic_variant | MODIFIER |
| 18 | 76217314 | AX-170451185 | 1 | G | C | 0.446 | 1.50E-01 | 8.44E-02 | 2.55E+00 | 2.62E+00 | 7.66E-02 | Yes | | intergenic_variant | MODIFIER |
| 5 | 148111818 | AX-170304548 | 1 | C | T | 0.054 | 3.06E-01 | 1.69E-01 | 3.33E+00 | 3.54E+00 | 7.66E-02 | Yes | Mtus2 | intron_variant | MODIFIER |
| 13 | 36359941 | AX-170452808 | 0 | T | A | 0.175 | -2.08E-01 | 1.18E-01 | 2.93E+00 | 3.05E+00 | 7.69E-02 | Yes | Fars2 | intron_variant | MODIFIER |
| 18 | 63348003 | AX-170382485 | 0 | G | A | 0.184 | -1.85E-01 | 1.04E-01 | 2.99E+00 | 3.13E+00 | 7.70E-02 | Yes | Piezo2 | intron_variant | MODIFIER |
| 18 | 63401830 | AX-170503774 | 0 | A | G | 0.184 | -1.85E-01 | 1.04E-01 | 2.99E+00 | 3.13E+00 | 7.70E-02 | Yes | Gm26972 | intron_variant,non_coding_transcript_variant | MODIFIER |
| 18 | 65098319 | AX-170390724 | 0 | G | A | 0.184 | -1.85E-01 | 1.04E-01 | 2.99E+00 | 3.13E+00 | 7.70E-02 | Yes | Nedd4l | intron_variant | MODIFIER |
| 16 | 52415715 | AX-170163369 | 1 | A | C | 0.464 | -1.44E-01 | 8.17E-02 | 2.60E+00 | 2.66E+00 | 7.72E-02 | no | Alcam | intron_variant | MODIFIER |
| 8 | 88390825 | AX-170031219 | 1 | C | A | 0.17 | -2.09E-01 | 1.18E-01 | 3.06E+00 | 3.19E+00 | 7.72E-02 | Yes | Gm45497 | upstream_gene_variant | MODIFIER |
| 5 | 132139064 | AX-170165377 | 1 | A | G | 0.152 | -1.62E-01 | 8.94E-02 | 2.25E+00 | 2.33E+00 | 7.74E-02 | no | Auts2 | intron_variant,non_coding_transcript_variant | MODIFIER |
| 4 | 47299414 | AX-170303492 | 1 | C | T | 0.366 | -1.64E-01 | 9.19E-02 | 3.13E+00 | 3.28E+00 | 7.74E-02 | no | Col15a1 | intron_variant | MODIFIER |
| 1 | 123552880 | AX-170173484 | 0 | A | G | 0.579 | 1.42E-01 | 8.02E-02 | 2.52E+00 | 2.60E+00 | 7.75E-02 | no | Dpp10 | intron_variant | MODIFIER |
| 16 | 65625273 | AX-97830840 | 0 | A | T | 0.518 | 1.42E-01 | 8.03E-02 | 2.57E+00 | 2.65E+00 | 7.80E-02 | no | | intergenic_variant | MODIFIER |
| 5 | 65772061 | AX-170122413 | 1 | T | A | 0.482 | 1.81E-01 | 1.03E-01 | 2.49E+00 | 2.55E+00 | 7.80E-02 | Yes | N4bp2 | intron_variant | MODIFIER |
| 5 | 42637460 | AX-170349147 | 1 | T | A | 0.196 | -1.82E-01 | 1.04E-01 | 2.85E+00 | 2.98E+00 | 7.81E-02 | Yes | | intergenic_variant | MODIFIER |
| 1 | 185677338 | AX-170485848 | 1 | T | A | 0.241 | 1.90E-01 | 1.08E-01 | 2.54E+00 | 2.61E+00 | 7.81E-02 | Yes | | regulatory_region_variant | MODIFIER |
| 13 | 20745054 | AX-170370781 | 1 | G | A | 0.179 | -2.13E-01 | 1.21E-01 | 2.49E+00 | 2.56E+00 | 7.81E-02 | Yes | Gm7614 | upstream_gene_variant | MODIFIER |
| 2 | 95169262 | AX-170307796 | 1 | G | T | 0.062 | -2.64E-01 | 1.51E-01 | 2.79E+00 | 2.94E+00 | 7.81E-02 | Yes | | intergenic_variant | MODIFIER |
| 3 | 19068916 | AX-97840512 | 1 | A | G | 0.116 | -1.74E-01 | 9.89E-02 | 2.64E+00 | 2.74E+00 | 7.82E-02 | Yes | | intergenic_variant | MODIFIER |
| 15 | 70469372 | AX-170296326 | 1 | G | C | 0.08 | -2.19E-01 | 1.06E-01 | 1.65E+00 | 1.70E+00 | 7.82E-02 | Yes | | intergenic_variant | MODIFIER |
| 5 | 52504631 | AX-170200805 | 0 | G | C | 0.482 | 1.82E-01 | 1.03E-01 | 2.44E+00 | 2.49E+00 | 7.83E-02 | Yes | | intergenic_variant | MODIFIER |
| 9 | 105734663 | AX-97852076 | 0 | G | A | 0.5 | 1.04E-01 | 5.90E-02 | 2.96E+00 | 3.13E+00 | 7.84E-02 | no | Col6a6 | intron_variant | MODIFIER |
| 9 | 107512026 | AX-170083396 | 0 | C | T | 0.5 | 1.04E-01 | 5.90E-02 | 2.96E+00 | 3.13E+00 | 7.84E-02 | no | Cacna2d2 | synonymous_variant | LOW |
| 9 | 117370411 | AX-170350744 | 0 | G | A | 0.123 | -1.65E-01 | 9.38E-02 | 2.95E+00 | 3.12E+00 | 7.85E-02 | no | Rbms3 | intron_variant | MODIFIER |
| 9 | 117447104 | AX-170123693 | 0 | A | T | 0.123 | -1.65E-01 | 9.38E-02 | 2.95E+00 | 3.12E+00 | 7.85E-02 | no | Rbms3 | intron_variant | MODIFIER |
| 15 | 77531695 | AX-170157357 | 1 | T | C | 0.08 | -2.17E-01 | 1.07E-01 | 1.71E+00 | 1.76E+00 | 7.85E-02 | Yes | Apol7c | intron_variant | MODIFIER |
| 16 | 65863230 | AX-170369885 | 1 | T | C | 0.5 | 1.40E-01 | 7.97E-02 | 2.64E+00 | 2.71E+00 | 7.85E-02 | no | Vgll3 | 3_prime_UTR_variant | MODIFIER |
| 18 | 77232151 | AX-170301188 | 1 | A | G | 0.446 | 1.50E-01 | 8.53E-02 | 2.55E+00 | 2.63E+00 | 7.87E-02 | Yes | St8sia5 | intron_variant | MODIFIER |
| 4 | 100882032 | AX-170150864 | 1 | A | G | 0.241 | 2.02E-01 | 1.15E-01 | 2.65E+00 | 2.71E+00 | 7.87E-02 | Yes | Cachd1 | intron_variant | MODIFIER |
| 8 | 84244817 | AX-170474885 | 0 | T | C | 0.193 | -2.17E-01 | 1.18E-01 | 3.58E+00 | 3.80E+00 | 7.87E-02 | Yes | D8Ertd738e | downstream_gene_variant | MODIFIER |
| 15 | 64263937 | AX-170386213 | 0 | C | T | 0.325 | -1.41E-01 | 7.64E-02 | 2.09E+00 | 2.15E+00 | 7.87E-02 | no | Asap1 | intron_variant | MODIFIER |
| 4 | 106888347 | AX-170230323 | 1 | G | A | 0.277 | 1.86E-01 | 1.06E-01 | 2.57E+00 | 2.64E+00 | 7.88E-02 | Yes | | intergenic_variant | MODIFIER |
| 1 | 122319992 | AX-170115455 | 1 | G | A | 0.58 | 1.48E-01 | 8.44E-02 | 2.59E+00 | 2.65E+00 | 7.89E-02 | Yes | | intergenic_variant | MODIFIER |
| 4 | 99307829 | AX-170297141 | 0 | G | A | 0.228 | 2.06E-01 | 1.17E-01 | 2.70E+00 | 2.77E+00 | 7.89E-02 | Yes | | intergenic_variant | MODIFIER |
| 4 | 99318330 | AX-170095069 | 0 | C | T | 0.228 | 2.06E-01 | 1.17E-01 | 2.70E+00 | 2.77E+00 | 7.89E-02 | Yes | | intergenic_variant | MODIFIER |
| 4 | 99322596 | AX-170201607 | 0 | C | T | 0.228 | 2.06E-01 | 1.17E-01 | 2.70E+00 | 2.77E+00 | 7.89E-02 | Yes | | intergenic_variant | MODIFIER |
| 4 | 99339999 | AX-170289919 | 0 | G | A | 0.228 | 2.06E-01 | 1.17E-01 | 2.70E+00 | 2.77E+00 | 7.89E-02 | Yes | | intergenic_variant | MODIFIER |
| 4 | 99340310 | AX-97818681 | 0 | T | C | 0.228 | 2.06E-01 | 1.17E-01 | 2.70E+00 | 2.77E+00 | 7.89E-02 | Yes | | intergenic_variant | MODIFIER |
| 4 | 99340348 | AX-170255831 | 0 | G | T | 0.228 | 2.06E-01 | 1.17E-01 | 2.70E+00 | 2.77E+00 | 7.89E-02 | Yes | | intergenic_variant | MODIFIER |
| 4 | 99363247 | AX-170429755 | 0 | C | T | 0.228 | 2.06E-01 | 1.17E-01 | 2.70E+00 | 2.77E+00 | 7.89E-02 | Yes | | intergenic_variant | MODIFIER |
| 4 | 99485759 | AX-97805491 | 0 | A | G | 0.228 | 2.06E-01 | 1.17E-01 | 2.70E+00 | 2.77E+00 | 7.89E-02 | Yes | | intergenic_variant | MODIFIER |
| 4 | 99496738 | AX-168388179 | 0 | T | C | 0.228 | 2.06E-01 | 1.17E-01 | 2.70E+00 | 2.77E+00 | 7.89E-02 | Yes | | intergenic_variant | MODIFIER |
| 4 | 99535165 | AX-97799996 | 0 | C | T | 0.228 | 2.06E-01 | 1.17E-01 | 2.70E+00 | 2.77E+00 | 7.89E-02 | Yes | Gm37117 | downstream_gene_variant | MODIFIER |
| 4 | 99547423 | AX-170479879 | 0 | C | T | 0.228 | 2.06E-01 | 1.17E-01 | 2.70E+00 | 2.77E+00 | 7.89E-02 | Yes | | intergenic_variant | MODIFIER |
| 4 | 99559643 | AX-170518728 | 0 | G | A | 0.228 | 2.06E-01 | 1.17E-01 | 2.70E+00 | 2.77E+00 | 7.89E-02 | Yes | | intergenic_variant | MODIFIER |

| | | | | | | | | | | | | | | | |
|---|---|---|---|---|---|---|---|---|---|---|---|---|---|---|---|
| 4 | 99681990 | AX-170406893 | 0 | T | C | 0.228 | 2.06E-01 | 1.17E-01 | 2.70E+00 | 2.77E+00 | 7.89E-02 | Yes | Gm12682 | intron_variant,non_coding_transcript_variant | MODIFIER |
| 4 | 100188360 | AX-170182155 | 0 | C | A | 0.228 | 2.06E-01 | 1.17E-01 | 2.70E+00 | 2.77E+00 | 7.89E-02 | Yes | Ror1 | intron_variant | MODIFIER |
| 4 | 100189445 | AX-170373380 | 0 | C | T | 0.228 | 2.06E-01 | 1.17E-01 | 2.70E+00 | 2.77E+00 | 7.89E-02 | Yes | Ror1 | intron_variant | MODIFIER |
| 4 | 100189463 | AX-170508684 | 0 | C | T | 0.228 | 2.06E-01 | 1.17E-01 | 2.70E+00 | 2.77E+00 | 7.89E-02 | Yes | Ror1 | intron_variant | MODIFIER |
| 4 | 100218126 | AX-170434549 | 0 | T | C | 0.228 | 2.06E-01 | 1.17E-01 | 2.70E+00 | 2.77E+00 | 7.89E-02 | Yes | Ror1 | intron_variant | MODIFIER |
| 4 | 100252161 | AX-170367741 | 0 | T | C | 0.228 | 2.06E-01 | 1.17E-01 | 2.70E+00 | 2.77E+00 | 7.89E-02 | Yes | Ror1 | intron_variant | MODIFIER |
| 4 | 100263250 | AX-170314418 | 0 | G | T | 0.228 | 2.06E-01 | 1.17E-01 | 2.70E+00 | 2.77E+00 | 7.89E-02 | Yes | Ror1 | intron_variant | MODIFIER |
| 4 | 100282789 | AX-170023512 | 0 | C | T | 0.228 | 2.06E-01 | 1.17E-01 | 2.70E+00 | 2.77E+00 | 7.89E-02 | Yes | Ror1 | intron_variant | MODIFIER |
| 4 | 100284725 | AX-170023511 | 0 | T | C | 0.228 | 2.06E-01 | 1.17E-01 | 2.70E+00 | 2.77E+00 | 7.89E-02 | Yes | Ror1 | intron_variant | MODIFIER |
| 4 | 100332046 | AX-170023515 | 0 | A | G | 0.228 | 2.06E-01 | 1.17E-01 | 2.70E+00 | 2.77E+00 | 7.89E-02 | Yes | Ror1 | intron_variant | MODIFIER |
| 4 | 100334662 | AX-170265609 | 0 | C | T | 0.228 | 2.06E-01 | 1.17E-01 | 2.70E+00 | 2.77E+00 | 7.89E-02 | Yes | Ror1 | intron_variant | MODIFIER |
| 4 | 100344901 | AX-170182467 | 0 | A | G | 0.228 | 2.06E-01 | 1.17E-01 | 2.70E+00 | 2.77E+00 | 7.89E-02 | Yes | Ror1 | intron_variant | MODIFIER |
| 4 | 100416056 | AX-170345291 | 0 | A | G | 0.228 | 2.06E-01 | 1.17E-01 | 2.70E+00 | 2.77E+00 | 7.89E-02 | Yes | Ror1 | intron_variant | MODIFIER |
| 4 | 100441171 | AX-170023510 | 0 | T | C | 0.228 | 2.06E-01 | 1.17E-01 | 2.70E+00 | 2.77E+00 | 7.89E-02 | Yes | Ror1 | synonymous_variant | LOW |
| 4 | 100442819 | AX-97806269 | 0 | A | G | 0.228 | 2.06E-01 | 1.17E-01 | 2.70E+00 | 2.77E+00 | 7.89E-02 | Yes | Ror1 | 3_prime_UTR_variant | MODIFIER |
| 4 | 100444464 | AX-170070152 | 0 | G | A | 0.228 | 2.06E-01 | 1.17E-01 | 2.70E+00 | 2.77E+00 | 7.89E-02 | Yes | Ror1 | 3_prime_UTR_variant | MODIFIER |
| 4 | 100824475 | AX-97792387 | 0 | T | C | 0.228 | 2.06E-01 | 1.17E-01 | 2.70E+00 | 2.77E+00 | 7.89E-02 | Yes | Cachd1 | intron_variant | MODIFIER |
| 4 | 100830523 | AX-170415186 | 0 | A | G | 0.228 | 2.06E-01 | 1.17E-01 | 2.70E+00 | 2.77E+00 | 7.89E-02 | Yes | Cachd1 | intron_variant | MODIFIER |
| 4 | 100830924 | AX-170023521 | 0 | T | C | 0.228 | 2.06E-01 | 1.17E-01 | 2.70E+00 | 2.77E+00 | 7.89E-02 | Yes | Cachd1 | intron_variant | MODIFIER |
| 4 | 100923226 | AX-97845226 | 0 | G | T | 0.228 | 2.06E-01 | 1.17E-01 | 2.70E+00 | 2.77E+00 | 7.89E-02 | Yes | Cachd1 | intron_variant | MODIFIER |
| 4 | 100923602 | AX-170070160 | 0 | A | G | 0.228 | 2.06E-01 | 1.17E-01 | 2.70E+00 | 2.77E+00 | 7.89E-02 | Yes | Cachd1 | intron_variant | MODIFIER |
| 4 | 100923739 | AX-170023524 | 0 | G | A | 0.228 | 2.06E-01 | 1.17E-01 | 2.70E+00 | 2.77E+00 | 7.89E-02 | Yes | Cachd1 | intron_variant | MODIFIER |
| 4 | 100931293 | AX-170023517 | 0 | G | A | 0.228 | 2.06E-01 | 1.17E-01 | 2.70E+00 | 2.77E+00 | 7.89E-02 | Yes | Cachd1 | intron_variant | MODIFIER |
| 4 | 100956217 | AX-170417308 | 0 | T | C | 0.228 | 2.06E-01 | 1.17E-01 | 2.70E+00 | 2.77E+00 | 7.89E-02 | Yes | Cachd1 | intron_variant | MODIFIER |
| 4 | 100957368 | AX-170279856 | 0 | A | G | 0.228 | 2.06E-01 | 1.17E-01 | 2.70E+00 | 2.77E+00 | 7.89E-02 | Yes | Cachd1 | intron_variant | MODIFIER |
| 4 | 100961889 | AX-170292382 | 0 | C | T | 0.228 | 2.06E-01 | 1.17E-01 | 2.70E+00 | 2.77E+00 | 7.89E-02 | Yes | Cachd1 | intron_variant | MODIFIER |
| 4 | 100962518 | AX-170363735 | 0 | A | G | 0.228 | 2.06E-01 | 1.17E-01 | 2.70E+00 | 2.77E+00 | 7.89E-02 | Yes | Cachd1 | intron_variant | MODIFIER |
| 4 | 100962564 | AX-97791120 | 0 | C | T | 0.228 | 2.06E-01 | 1.17E-01 | 2.70E+00 | 2.77E+00 | 7.89E-02 | Yes | Cachd1 | intron_variant | MODIFIER |
| 4 | 100978958 | AX-170516778 | 0 | C | A | 0.228 | 2.06E-01 | 1.17E-01 | 2.70E+00 | 2.77E+00 | 7.89E-02 | Yes | Cachd1 | intron_variant | MODIFIER |
| 4 | 100992710 | AX-170449884 | 0 | A | G | 0.228 | 2.06E-01 | 1.17E-01 | 2.70E+00 | 2.77E+00 | 7.89E-02 | Yes | Cachd1 | intron_variant | MODIFIER |
| 4 | 100994849 | AX-170295173 | 0 | G | C | 0.228 | 2.06E-01 | 1.17E-01 | 2.70E+00 | 2.77E+00 | 7.89E-02 | Yes | Cachd1 | intron_variant | MODIFIER |
| 4 | 100998547 | AX-170241327 | 0 | G | A | 0.228 | 2.06E-01 | 1.17E-01 | 2.70E+00 | 2.77E+00 | 7.89E-02 | Yes | Cachd1 | intron_variant | MODIFIER |
| 4 | 100999147 | AX-170291716 | 0 | A | C | 0.228 | 2.06E-01 | 1.17E-01 | 2.70E+00 | 2.77E+00 | 7.89E-02 | Yes | Cachd1 | intron_variant | MODIFIER |
| 4 | 101006862 | AX-97853349 | 0 | G | A | 0.228 | 2.06E-01 | 1.17E-01 | 2.70E+00 | 2.77E+00 | 7.89E-02 | Yes | Cachd1 | downstream_gene_variant | MODIFIER |
| 4 | 101037965 | AX-170220490 | 0 | G | A | 0.228 | 2.06E-01 | 1.17E-01 | 2.70E+00 | 2.77E+00 | 7.89E-02 | Yes | | intergenic_variant | MODIFIER |
| 4 | 101042539 | AX-170476054 | 0 | G | C | 0.228 | 2.06E-01 | 1.17E-01 | 2.70E+00 | 2.77E+00 | 7.89E-02 | Yes | | intergenic_variant | MODIFIER |
| 4 | 101045944 | AX-170195649 | 0 | T | C | 0.228 | 2.06E-01 | 1.17E-01 | 2.70E+00 | 2.77E+00 | 7.89E-02 | Yes | | intergenic_variant | MODIFIER |
| 4 | 101046121 | AX-170103538 | 0 | G | A | 0.228 | 2.06E-01 | 1.17E-01 | 2.70E+00 | 2.77E+00 | 7.89E-02 | Yes | | intergenic_variant | MODIFIER |
| 4 | 101046859 | AX-170270248 | 0 | A | G | 0.228 | 2.06E-01 | 1.17E-01 | 2.70E+00 | 2.77E+00 | 7.89E-02 | Yes | | intergenic_variant | MODIFIER |
| 4 | 101047910 | AX-170520286 | 0 | G | A | 0.228 | 2.06E-01 | 1.17E-01 | 2.70E+00 | 2.77E+00 | 7.89E-02 | Yes | | intergenic_variant | MODIFIER |
| 4 | 101048090 | AX-170467543 | 0 | A | G | 0.228 | 2.06E-01 | 1.17E-01 | 2.70E+00 | 2.77E+00 | 7.89E-02 | Yes | | intergenic_variant | MODIFIER |
| 4 | 101048790 | AX-170247156 | 0 | A | G | 0.228 | 2.06E-01 | 1.17E-01 | 2.70E+00 | 2.77E+00 | 7.89E-02 | Yes | | intergenic_variant | MODIFIER |
| 4 | 101049270 | AX-170382633 | 0 | A | G | 0.228 | 2.06E-01 | 1.17E-01 | 2.70E+00 | 2.77E+00 | 7.89E-02 | Yes | | intergenic_variant | MODIFIER |
| 4 | 101082610 | AX-97810657 | 0 | C | T | 0.228 | 2.06E-01 | 1.17E-01 | 2.70E+00 | 2.77E+00 | 7.89E-02 | Yes | Raver2 | intron_variant | MODIFIER |
| 5 | 27575302 | AX-170447741 | 0 | T | C | 0.053 | -3.70E-01 | 2.10E-01 | 3.06E+00 | 3.20E+00 | 7.90E-02 | Yes | Dpp6 | intron_variant | MODIFIER |
| 8 | 66167999 | AX-170121494 | 0 | A | G | 0.14 | -2.23E-01 | 1.27E-01 | 2.86E+00 | 2.98E+00 | 7.90E-02 | Yes | MAR1 | intron_variant | MODIFIER |
| 16 | 70608417 | AX-170318407 | 2 | C | T | 0.509 | 1.31E-01 | 7.43E-02 | 2.53E+00 | 2.60E+00 | 7.91E-02 | no | | intergenic_variant | MODIFIER |
| 3 | 23375435 | AX-170341942 | 0 | T | C | 0.158 | -1.59E-01 | 9.10E-02 | 2.68E+00 | 2.80E+00 | 7.91E-02 | no | Gm37492 | upstream_gene_variant | MODIFIER |
| 1 | 135389736 | AX-170359456 | 0 | A | T | 0.553 | 1.43E-01 | 8.20E-02 | 2.79E+00 | 2.88E+00 | 7.92E-02 | no | Ipo9 | intron_variant | MODIFIER |
| 13 | 18446982 | AX-170179670 | 0 | G | A | 0.158 | -2.25E-01 | 1.27E-01 | 2.38E+00 | 2.43E+00 | 7.92E-02 | Yes | | intergenic_variant | MODIFIER |
| 1 | 184307850 | AX-97835240 | 0 | T | C | 0.228 | 1.93E-01 | 1.10E-01 | 2.55E+00 | 2.63E+00 | 7.93E-02 | Yes | | intergenic_variant | MODIFIER |
| 6 | 15338475 | AX-97790815 | 1 | C | T | 0.036 | 2.78E-01 | 1.59E-01 | 2.65E+00 | 2.83E+00 | 7.93E-02 | Yes | Foxp2 | intron_variant | MODIFIER |
| 15 | 3461911 | AX-170155527 | 1 | G | A | 0.527 | 9.26E-02 | 5.29E-02 | 2.92E+00 | 3.01E+00 | 7.99E-02 | no | Ghr | intron_variant | MODIFIER |
| 13 | 10087909 | AX-170118774 | 1 | C | A | 0.179 | -2.17E-01 | 1.24E-01 | 2.54E+00 | 2.58E+00 | 8.01E-02 | Yes | Chrm3 | intron_variant | MODIFIER |

| | | | | | | | | | | | | | | | |
|---|---|---|---|---|---|---|---|---|---|---|---|---|---|---|---|
| 5 | 52942281 | AX-170140934 | 0 | G | A | 0.5 | 1.70E-01 | 9.61E-02 | 2.40E+00 | 2.48E+00 | 8.02E-02 | Yes | | intergenic_variant | MODIFIER |
| 19 | 59486460 | AX-170194327 | 1 | A | G | 0.464 | -1.10E-01 | 6.28E-02 | 2.77E+00 | 2.86E+00 | 8.02E-02 | no | | intergenic_variant | MODIFIER |
| 13 | 24541006 | AX-170260234 | 0 | A | G | 0.184 | -1.79E-01 | 1.01E-01 | 2.35E+00 | 2.44E+00 | 8.03E-02 | Yes | | intergenic_variant | MODIFIER |
| 5 | 64710415 | AX-170025156 | 1 | T | C | 0.473 | 1.71E-01 | 9.76E-02 | 2.53E+00 | 2.60E+00 | 8.04E-02 | Yes | Gm20033 | intron_variant,non_coding_transcript_variant | MODIFIER |
| 4 | 83188631 | AX-170244335 | 0 | T | C | 0.43 | -1.37E-01 | 7.88E-02 | 2.58E+00 | 2.66E+00 | 8.07E-02 | no | | intergenic_variant | MODIFIER |
| 15 | 5676295 | AX-170499058 | 0 | G | A | 0.228 | -1.71E-01 | 9.83E-02 | 2.76E+00 | 2.85E+00 | 8.12E-02 | no | | intergenic_variant | MODIFIER |
| 13 | 6954525 | AX-170050465 | 0 | G | A | 0.202 | -2.01E-01 | 1.15E-01 | 2.78E+00 | 2.88E+00 | 8.13E-02 | Yes | | intergenic_variant | MODIFIER |
| 4 | 83867156 | AX-170202708 | 1 | C | T | 0.438 | -1.41E-01 | 8.09E-02 | 2.62E+00 | 2.69E+00 | 8.15E-02 | no | Ccdc171 | downstream_gene_variant | MODIFIER |
| 4 | 78163315 | AX-97796387 | 0 | G | A | 0.316 | 1.90E-01 | 1.08E-01 | 3.09E+00 | 3.20E+00 | 8.15E-02 | Yes | Ptprd | intron_variant,non_coding_transcript_variant | MODIFIER |
| 16 | 63076245 | AX-97795535 | 0 | C | T | 0.456 | -1.40E-01 | 7.97E-02 | 2.45E+00 | 2.50E+00 | 8.16E-02 | no | Gm18870 | intron_variant,non_coding_transcript_variant | MODIFIER |
| 4 | 100945067 | AX-170095074 | 2 | T | C | 0.227 | 1.82E-01 | 1.05E-01 | 2.66E+00 | 2.75E+00 | 8.17E-02 | Yes | Cachd1 | intron_variant | MODIFIER |
| 4 | 100409299 | AX-97802163 | 0 | C | T | 0.43 | -1.32E-01 | 7.57E-02 | 2.55E+00 | 2.64E+00 | 8.17E-02 | no | Ror1 | intron_variant | MODIFIER |
| 5 | 80799936 | AX-170156578 | 0 | G | A | 0.439 | 1.75E-01 | 9.94E-02 | 3.15E+00 | 3.27E+00 | 8.18E-02 | Yes | | intergenic_variant | MODIFIER |
| 10 | 102399109 | AX-170003798 | 0 | A | T | 0.289 | -1.46E-01 | 8.40E-02 | 2.75E+00 | 2.82E+00 | 8.18E-02 | no | | intergenic_variant | MODIFIER |
| 10 | 102634290 | AX-170232721 | 0 | G | C | 0.289 | -1.46E-01 | 8.40E-02 | 2.75E+00 | 2.82E+00 | 8.18E-02 | no | | intergenic_variant | MODIFIER |
| 10 | 102734605 | AX-170112680 | 0 | A | G | 0.289 | -1.46E-01 | 8.40E-02 | 2.75E+00 | 2.82E+00 | 8.18E-02 | no | | intergenic_variant | MODIFIER |
| 10 | 102744203 | AX-170417822 | 0 | C | T | 0.289 | -1.46E-01 | 8.40E-02 | 2.75E+00 | 2.82E+00 | 8.18E-02 | no | | intergenic_variant | MODIFIER |
| 10 | 102838338 | AX-170254269 | 0 | G | C | 0.289 | -1.46E-01 | 8.40E-02 | 2.75E+00 | 2.82E+00 | 8.18E-02 | no | Gm5175 | non_coding_transcript_exon_variant | MODIFIER |
| 10 | 102892656 | AX-170141565 | 0 | C | A | 0.289 | -1.46E-01 | 8.40E-02 | 2.75E+00 | 2.82E+00 | 8.18E-02 | no | | intergenic_variant | MODIFIER |
| 10 | 102910315 | AX-97855481 | 0 | A | G | 0.289 | -1.46E-01 | 8.40E-02 | 2.75E+00 | 2.82E+00 | 8.18E-02 | no | | intergenic_variant | MODIFIER |
| 10 | 102910993 | AX-170003804 | 0 | G | A | 0.289 | -1.46E-01 | 8.40E-02 | 2.75E+00 | 2.82E+00 | 8.18E-02 | no | | intergenic_variant | MODIFIER |
| 10 | 102997014 | AX-97816598 | 0 | C | T | 0.289 | -1.46E-01 | 8.40E-02 | 2.75E+00 | 2.82E+00 | 8.18E-02 | no | Alx1 | downstream_gene_variant | MODIFIER |
| 10 | 103034514 | AX-170003803 | 0 | C | T | 0.289 | -1.46E-01 | 8.40E-02 | 2.75E+00 | 2.82E+00 | 8.18E-02 | no | Gm17028 | upstream_gene_variant | MODIFIER |
| 10 | 103254043 | AX-170477903 | 0 | G | A | 0.289 | -1.46E-01 | 8.40E-02 | 2.75E+00 | 2.82E+00 | 8.18E-02 | no | | regulatory_region_variant | MODIFIER |
| 10 | 103344534 | AX-170193433 | 0 | G | C | 0.289 | -1.46E-01 | 8.40E-02 | 2.75E+00 | 2.82E+00 | 8.18E-02 | no | | intergenic_variant | MODIFIER |
| 10 | 103366933 | AX-170408349 | 0 | C | A | 0.289 | -1.46E-01 | 8.40E-02 | 2.75E+00 | 2.82E+00 | 8.18E-02 | no | Slc6a15 | upstream_gene_variant | MODIFIER |
| 10 | 103367105 | AX-170501128 | 0 | T | G | 0.289 | -1.46E-01 | 8.40E-02 | 2.75E+00 | 2.82E+00 | 8.18E-02 | no | Slc6a15 | upstream_gene_variant | MODIFIER |
| 1 | 138762352 | AX-170227353 | 0 | C | A | 0.211 | 2.06E-01 | 1.19E-01 | 2.76E+00 | 2.84E+00 | 8.19E-02 | Yes | | intergenic_variant | MODIFIER |
| 15 | 3612399 | AX-170010914 | 1 | A | G | 0.33 | -1.32E-01 | 7.18E-02 | 2.06E+00 | 2.11E+00 | 8.20E-02 | no | | intergenic_variant | MODIFIER |
| 18 | 88519189 | AX-170155041 | 0 | G | A | 0.167 | -1.62E-01 | 9.36E-02 | 2.58E+00 | 2.70E+00 | 8.21E-02 | no | | intergenic_variant | MODIFIER |
| 13 | 21475767 | AX-170236062 | 0 | C | T | 0.167 | -2.25E-01 | 1.29E-01 | 2.43E+00 | 2.48E+00 | 8.21E-02 | Yes | Zkscan4 | upstream_gene_variant | MODIFIER |
| 13 | 8344669 | AX-170007704 | 1 | G | A | 0.188 | -2.14E-01 | 1.23E-01 | 2.52E+00 | 2.58E+00 | 8.22E-02 | Yes | Adarb2 | intron_variant | MODIFIER |
| 4 | 100967811 | AX-170256415 | 1 | C | A | 0.232 | 2.01E-01 | 1.16E-01 | 2.64E+00 | 2.70E+00 | 8.25E-02 | Yes | Cachd1 | intron_variant | MODIFIER |
| 8 | 21702830 | AX-170150920 | 0 | C | G | 0.167 | -1.76E-01 | 9.93E-02 | 3.35E+00 | 3.59E+00 | 8.26E-02 | Yes | AY761184 | intron_variant | MODIFIER |
| 4 | 98949576 | AX-170158864 | 0 | C | T | 0.237 | 2.02E-01 | 1.17E-01 | 2.72E+00 | 2.78E+00 | 8.26E-02 | Yes | Dock7 | intron_variant | MODIFIER |
| 4 | 98949627 | AX-170023486 | 0 | T | C | 0.237 | 2.02E-01 | 1.17E-01 | 2.72E+00 | 2.78E+00 | 8.26E-02 | Yes | Dock7 | intron_variant | MODIFIER |
| 4 | 98954282 | AX-170023484 | 0 | A | C | 0.237 | 2.02E-01 | 1.17E-01 | 2.72E+00 | 2.78E+00 | 8.26E-02 | Yes | Dock7 | intron_variant | MODIFIER |
| 4 | 98975789 | AX-170291866 | 0 | T | G | 0.237 | 2.02E-01 | 1.17E-01 | 2.72E+00 | 2.78E+00 | 8.26E-02 | Yes | Dock7 | intron_variant | MODIFIER |
| 4 | 98988261 | AX-170023483 | 0 | A | G | 0.237 | 2.02E-01 | 1.17E-01 | 2.72E+00 | 2.78E+00 | 8.26E-02 | Yes | Dock7 | intron_variant | MODIFIER |
| 4 | 98989298 | AX-97826444 | 0 | A | G | 0.237 | 2.02E-01 | 1.17E-01 | 2.72E+00 | 2.78E+00 | 8.26E-02 | Yes | Dock7 | synonymous_variant | LOW |
| 4 | 99032576 | AX-170448469 | 0 | T | A | 0.237 | 2.02E-01 | 1.17E-01 | 2.72E+00 | 2.78E+00 | 8.26E-02 | Yes | Angptl3 | intron_variant | MODIFIER |
| 4 | 99048594 | AX-170335591 | 0 | A | G | 0.237 | 2.02E-01 | 1.17E-01 | 2.72E+00 | 2.78E+00 | 8.26E-02 | Yes | Dock7 | intron_variant | MODIFIER |
| 4 | 99048978 | AX-97802752 | 0 | A | G | 0.237 | 2.02E-01 | 1.17E-01 | 2.72E+00 | 2.78E+00 | 8.26E-02 | Yes | Dock7 | intron_variant | MODIFIER |
| 4 | 99061484 | AX-170228145 | 0 | A | G | 0.237 | 2.02E-01 | 1.17E-01 | 2.72E+00 | 2.78E+00 | 8.26E-02 | Yes | Dock7 | intron_variant | MODIFIER |
| 4 | 99072339 | AX-97851106 | 0 | A | G | 0.237 | 2.02E-01 | 1.17E-01 | 2.72E+00 | 2.78E+00 | 8.26E-02 | Yes | Dock7 | intron_variant | MODIFIER |
| 4 | 99145268 | AX-170108631 | 0 | A | G | 0.237 | 2.02E-01 | 1.17E-01 | 2.72E+00 | 2.78E+00 | 8.26E-02 | Yes | Gm12681 | downstream_gene_variant | MODIFIER |
| 4 | 99198080 | AX-170314056 | 0 | C | T | 0.237 | 2.02E-01 | 1.17E-01 | 2.72E+00 | 2.78E+00 | 8.26E-02 | Yes | Atg4c | intron_variant | MODIFIER |
| 4 | 99209861 | AX-97842354 | 0 | T | G | 0.237 | 2.02E-01 | 1.17E-01 | 2.72E+00 | 2.78E+00 | 8.26E-02 | Yes | Atg4c | intron_variant | MODIFIER |
| 4 | 99232300 | AX-170023500 | 0 | C | T | 0.237 | 2.02E-01 | 1.17E-01 | 2.72E+00 | 2.78E+00 | 8.26E-02 | Yes | Atg4c | intron_variant | MODIFIER |
| 4 | 99237294 | AX-170372546 | 0 | T | C | 0.237 | 2.02E-01 | 1.17E-01 | 2.72E+00 | 2.78E+00 | 8.26E-02 | Yes | Atg4c | intron_variant | MODIFIER |
| 4 | 99264486 | AX-170023499 | 0 | A | G | 0.237 | 2.02E-01 | 1.17E-01 | 2.72E+00 | 2.78E+00 | 8.26E-02 | Yes | Atg4c | downstream_gene_variant | MODIFIER |
| 4 | 99274337 | AX-170070134 | 0 | C | T | 0.237 | 2.02E-01 | 1.17E-01 | 2.72E+00 | 2.78E+00 | 8.26E-02 | Yes | Gm10305 | non_coding_transcript_exon_variant | MODIFIER |
| 4 | 99274566 | AX-170070133 | 0 | T | C | 0.237 | 2.02E-01 | 1.17E-01 | 2.72E+00 | 2.78E+00 | 8.26E-02 | Yes | Gm10305 | non_coding_transcript_exon_variant | MODIFIER |
| 4 | 100216652 | AX-97818790 | 0 | G | A | 0.237 | 2.02E-01 | 1.17E-01 | 2.72E+00 | 2.78E+00 | 8.26E-02 | Yes | Ror1 | intron_variant | MODIFIER |

| | | | | | | | | | | | | | | | |
|---|---|---|---|---|---|---|---|---|---|---|---|---|---|---|---|
| 4 | 100486704 | AX-170247006 | 0 | A | G | 0.237 | 2.02E-01 | 1.17E-01 | 2.72E+00 | 2.78E+00 | 8.26E-02 | Yes | Ube2u | 5_prime_UTR_variant | MODIFIER |
| 4 | 100490366 | AX-170431972 | 0 | T | C | 0.237 | 2.02E-01 | 1.17E-01 | 2.72E+00 | 2.78E+00 | 8.26E-02 | Yes | Ube2u | intron_variant | MODIFIER |
| 4 | 100493960 | AX-170247446 | 0 | T | C | 0.237 | 2.02E-01 | 1.17E-01 | 2.72E+00 | 2.78E+00 | 8.26E-02 | Yes | Ube2u | intron_variant | MODIFIER |
| 4 | 100494195 | AX-170426726 | 0 | G | A | 0.237 | 2.02E-01 | 1.17E-01 | 2.72E+00 | 2.78E+00 | 8.26E-02 | Yes | Ube2u | intron_variant | MODIFIER |
| 4 | 100503768 | AX-170321902 | 0 | C | T | 0.237 | 2.02E-01 | 1.17E-01 | 2.72E+00 | 2.78E+00 | 8.26E-02 | Yes | Ube2u | intron_variant | MODIFIER |
| 4 | 100619197 | AX-170463585 | 0 | A | C | 0.237 | 2.02E-01 | 1.17E-01 | 2.72E+00 | 2.78E+00 | 8.26E-02 | Yes | | intergenic_variant | MODIFIER |
| 4 | 100644952 | AX-97790355 | 0 | A | G | 0.237 | 2.02E-01 | 1.17E-01 | 2.72E+00 | 2.78E+00 | 8.26E-02 | Yes | | intergenic_variant | MODIFIER |
| 4 | 100661010 | AX-170277726 | 0 | G | A | 0.237 | 2.02E-01 | 1.17E-01 | 2.72E+00 | 2.78E+00 | 8.26E-02 | Yes | | regulatory_region_variant | MODIFIER |
| 4 | 100672980 | AX-170070153 | 0 | T | G | 0.237 | 2.02E-01 | 1.17E-01 | 2.72E+00 | 2.78E+00 | 8.26E-02 | Yes | | intergenic_variant | MODIFIER |
| 4 | 100675055 | AX-170206027 | 0 | G | A | 0.237 | 2.02E-01 | 1.17E-01 | 2.72E+00 | 2.78E+00 | 8.26E-02 | Yes | | intergenic_variant | MODIFIER |
| 4 | 100717443 | AX-170103273 | 0 | T | C | 0.237 | 2.02E-01 | 1.17E-01 | 2.72E+00 | 2.78E+00 | 8.26E-02 | Yes | | intergenic_variant | MODIFIER |
| 4 | 100722500 | AX-170113641 | 0 | T | C | 0.237 | 2.02E-01 | 1.17E-01 | 2.72E+00 | 2.78E+00 | 8.26E-02 | Yes | | regulatory_region_variant | MODIFIER |
| 4 | 100727486 | AX-170185125 | 0 | T | A | 0.237 | 2.02E-01 | 1.17E-01 | 2.72E+00 | 2.78E+00 | 8.26E-02 | Yes | | intergenic_variant | MODIFIER |
| 4 | 100727538 | AX-170246037 | 0 | A | G | 0.237 | 2.02E-01 | 1.17E-01 | 2.72E+00 | 2.78E+00 | 8.26E-02 | Yes | | intergenic_variant | MODIFIER |
| 4 | 100727641 | AX-170100402 | 0 | T | G | 0.237 | 2.02E-01 | 1.17E-01 | 2.72E+00 | 2.78E+00 | 8.26E-02 | Yes | | intergenic_variant | MODIFIER |
| 4 | 100728377 | AX-170328056 | 0 | T | C | 0.237 | 2.02E-01 | 1.17E-01 | 2.72E+00 | 2.78E+00 | 8.26E-02 | Yes | | intergenic_variant | MODIFIER |
| 4 | 100744981 | AX-170453873 | 0 | A | T | 0.237 | 2.02E-01 | 1.17E-01 | 2.72E+00 | 2.78E+00 | 8.26E-02 | Yes | | intergenic_variant | MODIFIER |
| 4 | 100754496 | AX-97793369 | 0 | A | G | 0.237 | 2.02E-01 | 1.17E-01 | 2.72E+00 | 2.78E+00 | 8.26E-02 | Yes | | intergenic_variant | MODIFIER |
| 4 | 100852594 | AX-97793114 | 0 | A | G | 0.237 | 2.02E-01 | 1.17E-01 | 2.72E+00 | 2.78E+00 | 8.26E-02 | Yes | Cachd1 | intron_variant | MODIFIER |
| 4 | 100873183 | AX-170023522 | 0 | A | G | 0.237 | 2.02E-01 | 1.17E-01 | 2.72E+00 | 2.78E+00 | 8.26E-02 | Yes | Cachd1 | intron_variant | MODIFIER |
| 4 | 100879442 | AX-170200782 | 0 | T | C | 0.237 | 2.02E-01 | 1.17E-01 | 2.72E+00 | 2.78E+00 | 8.26E-02 | Yes | Cachd1 | intron_variant | MODIFIER |
| 4 | 100889246 | AX-97822934 | 0 | C | T | 0.237 | 2.02E-01 | 1.17E-01 | 2.72E+00 | 2.78E+00 | 8.26E-02 | Yes | Cachd1 | intron_variant | MODIFIER |
| 4 | 100894506 | AX-170023523 | 0 | C | T | 0.237 | 2.02E-01 | 1.17E-01 | 2.72E+00 | 2.78E+00 | 8.26E-02 | Yes | Cachd1 | intron_variant | MODIFIER |
| 4 | 100948617 | AX-168405670 | 0 | G | A | 0.237 | 2.02E-01 | 1.17E-01 | 2.72E+00 | 2.78E+00 | 8.26E-02 | Yes | Cachd1 | intron_variant | MODIFIER |
| 4 | 100955934 | AX-170364858 | 0 | C | T | 0.237 | 2.02E-01 | 1.17E-01 | 2.72E+00 | 2.78E+00 | 8.26E-02 | Yes | Cachd1 | intron_variant | MODIFIER |
| 5 | 136742999 | AX-170209366 | 0 | C | G | 0.158 | -1.42E-01 | 8.12E-02 | 2.40E+00 | 2.50E+00 | 8.27E-02 | no | Col26a1 | 3_prime_UTR_variant | MODIFIER |
| 5 | 9746133 | AX-97836340 | 0 | G | A | 0.325 | 1.48E-01 | 8.55E-02 | 2.91E+00 | 3.07E+00 | 8.28E-02 | Yes | | intergenic_variant | MODIFIER |
| 13 | 9702018 | AX-170007729 | 1 | A | G | 0.188 | -2.12E-01 | 1.22E-01 | 2.55E+00 | 2.60E+00 | 8.30E-02 | Yes | Zmynd11 | intron_variant | MODIFIER |
| 13 | 10188422 | AX-170485928 | 1 | G | A | 0.188 | -2.12E-01 | 1.22E-01 | 2.55E+00 | 2.60E+00 | 8.30E-02 | Yes | Chrm3 | intron_variant | MODIFIER |
| 3 | 7827499 | AX-170498087 | 2 | C | T | 0.391 | -1.19E-01 | 6.84E-02 | 3.04E+00 | 3.23E+00 | 8.31E-02 | no | | intergenic_variant | MODIFIER |
| 5 | 132104434 | AX-170335397 | 0 | C | T | 0.158 | -1.51E-01 | 8.58E-02 | 2.31E+00 | 2.40E+00 | 8.32E-02 | no | Auts2 | intron_variant,non_coding_transcript_variant | MODIFIER |
| 15 | 4015549 | AX-170010924 | 0 | C | T | 0.316 | -1.30E-01 | 7.03E-02 | 2.00E+00 | 2.06E+00 | 8.32E-02 | no | Gm2093 | non_coding_transcript_exon_variant | MODIFIER |
| 13 | 8502424 | AX-170129344 | 1 | C | T | 0.188 | -2.12E-01 | 1.23E-01 | 2.58E+00 | 2.64E+00 | 8.33E-02 | Yes | Adarb2 | intron_variant | MODIFIER |
| 16 | 29180671 | AX-170462177 | 1 | T | C | 0.509 | 1.36E-01 | 7.88E-02 | 2.76E+00 | 2.84E+00 | 8.34E-02 | no | | intergenic_variant | MODIFIER |
| 16 | 56647264 | AX-170057510 | 1 | A | G | 0.455 | -1.44E-01 | 8.29E-02 | 2.47E+00 | 2.51E+00 | 8.34E-02 | no | Abi3bp | intron_variant | MODIFIER |
| 5 | 66279120 | AX-97802538 | 1 | G | A | 0.482 | 1.81E-01 | 1.04E-01 | 2.48E+00 | 2.53E+00 | 8.34E-02 | Yes | Nsun7 | intron_variant | MODIFIER |
| 16 | 55273940 | AX-170133932 | 1 | A | G | 0.464 | -1.48E-01 | 8.57E-02 | 2.59E+00 | 2.63E+00 | 8.35E-02 | no | Zpld1 | intron_variant | MODIFIER |
| 4 | 106819459 | AX-170035693 | 1 | C | T | 0.241 | 1.98E-01 | 1.15E-01 | 2.64E+00 | 2.72E+00 | 8.35E-02 | Yes | | intergenic_variant | MODIFIER |
| 8 | 88421344 | AX-170424231 | 0 | G | A | 0.175 | -2.06E-01 | 1.19E-01 | 2.92E+00 | 3.04E+00 | 8.36E-02 | Yes | | intergenic_variant | MODIFIER |
| 15 | 76726957 | AX-170056028 | 1 | G | A | 0.08 | -2.08E-01 | 1.07E-01 | 1.79E+00 | 1.85E+00 | 8.36E-02 | Yes | Arhgap39 | intron_variant | MODIFIER |
| 5 | 55722939 | AX-170159254 | 0 | G | A | 0.465 | 1.75E-01 | 1.01E-01 | 2.48E+00 | 2.54E+00 | 8.37E-02 | Yes | Gm20223 | non_coding_transcript_exon_variant | MODIFIER |
| 15 | 12414646 | AX-170356113 | 0 | A | G | 0.298 | -1.55E-01 | 7.81E-02 | 1.72E+00 | 1.75E+00 | 8.37E-02 | no | Pdzd2 | intron_variant | MODIFIER |
| 5 | 64099519 | AX-170025149 | 1 | A | G | 0.482 | 1.80E-01 | 1.04E-01 | 2.50E+00 | 2.55E+00 | 8.38E-02 | Yes | Pgm2 | intron_variant | MODIFIER |
| 19 | 59678099 | AX-170299648 | 1 | T | A | 0.518 | 1.06E-01 | 6.15E-02 | 2.79E+00 | 2.89E+00 | 8.39E-02 | no | AC123621.1 | intron_variant,non_coding_transcript_variant | MODIFIER |
| 15 | 70414927 | AX-170176146 | 1 | C | T | 0.08 | -2.17E-01 | 1.08E-01 | 1.68E+00 | 1.74E+00 | 8.39E-02 | Yes | | intergenic_variant | MODIFIER |
| 13 | 10678661 | AX-97830338 | 0 | C | T | 0.193 | -2.02E-01 | 1.17E-01 | 2.55E+00 | 2.61E+00 | 8.40E-02 | Yes | Gm26861 | intron_variant,non_coding_transcript_variant | MODIFIER |
| 13 | 9639854 | AX-170504950 | 0 | G | A | 0.202 | -1.94E-01 | 1.12E-01 | 2.54E+00 | 2.61E+00 | 8.40E-02 | Yes | Dip2c | intron_variant | MODIFIER |
| 16 | 63368390 | AX-97800501 | 0 | C | T | 0.518 | 1.46E-01 | 8.49E-02 | 2.62E+00 | 2.68E+00 | 8.41E-02 | Yes | | intergenic_variant | MODIFIER |
| 4 | 84221912 | AX-170130266 | 1 | G | A | 0.339 | 2.02E-01 | 1.17E-01 | 2.98E+00 | 3.06E+00 | 8.42E-02 | Yes | | intergenic_variant | MODIFIER |
| 13 | 12383994 | AX-170425701 | 0 | G | A | 0.193 | -2.00E-01 | 1.15E-01 | 2.47E+00 | 2.54E+00 | 8.43E-02 | Yes | | intergenic_variant | MODIFIER |
| 5 | 63193650 | AX-170462769 | 1 | G | T | 0.491 | 1.74E-01 | 9.95E-02 | 2.39E+00 | 2.45E+00 | 8.43E-02 | Yes | | intergenic_variant | MODIFIER |
| 5 | 64414423 | AX-97850151 | 1 | G | A | 0.491 | 1.74E-01 | 9.95E-02 | 2.39E+00 | 2.45E+00 | 8.43E-02 | Yes | | intergenic_variant | MODIFIER |
| 5 | 66219476 | AX-170025184 | 1 | T | C | 0.491 | 1.74E-01 | 9.95E-02 | 2.39E+00 | 2.45E+00 | 8.43E-02 | Yes | Gm3822 | downstream_gene_variant | MODIFIER |
| 18 | 75922654 | AX-170424379 | 0 | A | G | 0.447 | 1.50E-01 | 8.66E-02 | 2.51E+00 | 2.57E+00 | 8.43E-02 | Yes | Zbtb7c | intron_variant | MODIFIER |

| | | | | | | | | | | | | | | | |
|---|---|---|---|---|---|---|---|---|---|---|---|---|---|---|---|
| 16 | 54804334 | AX-170150704 | 0 | A | C | 0.526 | 1.43E-01 | 8.27E-02 | 2.49E+00 | 2.53E+00 | 8.44E-02 | no | | intergenic_variant | MODIFIER |
| 7 | 123306553 | AX-170333070 | 1 | G | A | 0.411 | 1.36E-01 | 7.86E-02 | 3.05E+00 | 3.19E+00 | 8.47E-02 | no | Arhgap17 | intron_variant | MODIFIER |
| 19 | 61014587 | AX-170286108 | 0 | A | G | 0.377 | -1.03E-01 | 5.95E-02 | 2.46E+00 | 2.54E+00 | 8.48E-02 | no | Grk5 | intron_variant | MODIFIER |
| 19 | 61017482 | AX-170262040 | 0 | T | A | 0.377 | -1.03E-01 | 5.95E-02 | 2.46E+00 | 2.54E+00 | 8.48E-02 | no | Grk5 | intron_variant | MODIFIER |
| 19 | 61024100 | AX-170439655 | 0 | T | C | 0.377 | -1.03E-01 | 5.95E-02 | 2.46E+00 | 2.54E+00 | 8.48E-02 | no | Grk5 | intron_variant | MODIFIER |
| 19 | 61037774 | AX-170088495 | 0 | A | G | 0.377 | -1.03E-01 | 5.95E-02 | 2.46E+00 | 2.54E+00 | 8.48E-02 | no | Grk5 | intron_variant | MODIFIER |
| 19 | 61042912 | AX-170119394 | 0 | G | T | 0.377 | -1.03E-01 | 5.95E-02 | 2.46E+00 | 2.54E+00 | 8.48E-02 | no | Grk5 | intron_variant | MODIFIER |
| 19 | 61065893 | AX-97824572 | 0 | G | C | 0.377 | -1.03E-01 | 5.95E-02 | 2.46E+00 | 2.54E+00 | 8.48E-02 | no | Grk5 | intron_variant | MODIFIER |
| 19 | 61076510 | AX-170207556 | 0 | A | G | 0.377 | -1.03E-01 | 5.95E-02 | 2.46E+00 | 2.54E+00 | 8.48E-02 | no | Grk5 | intron_variant | MODIFIER |
| 19 | 61093456 | AX-170088496 | 0 | G | A | 0.377 | -1.03E-01 | 5.95E-02 | 2.46E+00 | 2.54E+00 | 8.48E-02 | no | Grk5 | 3_prime_UTR_variant | MODIFIER |
| 19 | 61094205 | AX-170062875 | 0 | C | T | 0.377 | -1.03E-01 | 5.95E-02 | 2.46E+00 | 2.54E+00 | 8.48E-02 | no | Grk5 | 3_prime_UTR_variant | MODIFIER |
| 19 | 61095832 | AX-170532424 | 0 | A | G | 0.377 | -1.03E-01 | 5.95E-02 | 2.46E+00 | 2.54E+00 | 8.48E-02 | no | Grk5 | 3_prime_UTR_variant | MODIFIER |
| 19 | 61166830 | AX-97836233 | 0 | G | A | 0.377 | -1.03E-01 | 5.95E-02 | 2.46E+00 | 2.54E+00 | 8.48E-02 | no | Gm22365 | upstream_gene_variant | MODIFIER |
| 19 | 61202790 | AX-170483942 | 0 | A | C | 0.377 | -1.03E-01 | 5.95E-02 | 2.46E+00 | 2.54E+00 | 8.48E-02 | no | | intergenic_variant | MODIFIER |
| 19 | 61208755 | AX-170102997 | 0 | C | T | 0.377 | -1.03E-01 | 5.95E-02 | 2.46E+00 | 2.54E+00 | 8.48E-02 | no | | intergenic_variant | MODIFIER |
| 19 | 61209233 | AX-170094144 | 0 | A | G | 0.377 | -1.03E-01 | 5.95E-02 | 2.46E+00 | 2.54E+00 | 8.48E-02 | no | | intergenic_variant | MODIFIER |
| 19 | 61256426 | AX-170295517 | 0 | T | C | 0.377 | -1.03E-01 | 5.95E-02 | 2.46E+00 | 2.54E+00 | 8.48E-02 | no | CR555305.1 | non_coding_transcript_exon_variant | MODIFIER |
| 19 | 61261300 | AX-97781726 | 0 | G | A | 0.377 | -1.03E-01 | 5.95E-02 | 2.46E+00 | 2.54E+00 | 8.48E-02 | no | CR555305.1 | downstream_gene_variant | MODIFIER |
| 5 | 62540446 | AX-97790541 | 0 | G | A | 0.465 | 1.77E-01 | 1.03E-01 | 2.57E+00 | 2.64E+00 | 8.53E-02 | Yes | | intergenic_variant | MODIFIER |
| 5 | 52328652 | AX-97794807 | 0 | C | A | 0.5 | 1.80E-01 | 1.04E-01 | 2.44E+00 | 2.49E+00 | 8.55E-02 | Yes | | intergenic_variant | MODIFIER |
| 5 | 52340286 | AX-170359994 | 0 | C | T | 0.5 | 1.80E-01 | 1.04E-01 | 2.44E+00 | 2.49E+00 | 8.55E-02 | Yes | Gm25601 | downstream_gene_variant | MODIFIER |
| 5 | 52419880 | AX-170408392 | 0 | A | G | 0.5 | 1.80E-01 | 1.04E-01 | 2.44E+00 | 2.49E+00 | 8.55E-02 | Yes | Ccdc149 | intron_variant | MODIFIER |
| 5 | 52420006 | AX-170272696 | 0 | G | A | 0.5 | 1.80E-01 | 1.04E-01 | 2.44E+00 | 2.49E+00 | 8.55E-02 | Yes | Ccdc149 | intron_variant | MODIFIER |
| 5 | 52424276 | AX-170154218 | 0 | C | T | 0.5 | 1.80E-01 | 1.04E-01 | 2.44E+00 | 2.49E+00 | 8.55E-02 | Yes | Ccdc149 | intron_variant | MODIFIER |
| 18 | 50101419 | AX-170060817 | 0 | A | T | 0.193 | -1.75E-01 | 1.02E-01 | 2.78E+00 | 2.89E+00 | 8.55E-02 | Yes | Tnfaip8 | intron_variant,non_coding_transcript_variant | MODIFIER |
| 2 | 169753570 | AX-170158255 | 0 | T | C | 0.175 | 2.65E-01 | 1.55E-01 | 2.85E+00 | 2.98E+00 | 8.56E-02 | Yes | Tshz2 | intron_variant | MODIFIER |
| 19 | 59622765 | AX-97795409 | 1 | A | G | 0.464 | -1.08E-01 | 6.30E-02 | 2.74E+00 | 2.82E+00 | 8.56E-02 | no | | intergenic_variant | MODIFIER |
| 13 | 11493297 | AX-170092660 | 1 | C | A | 0.188 | -2.14E-01 | 1.24E-01 | 2.57E+00 | 2.62E+00 | 8.56E-02 | Yes | | intergenic_variant | MODIFIER |
| 8 | 56821171 | AX-170449674 | 1 | G | A | 0.152 | -2.34E-01 | 1.36E-01 | 2.94E+00 | 3.05E+00 | 8.57E-02 | Yes | | intergenic_variant | MODIFIER |
| 10 | 102740663 | AX-170521294 | 1 | G | T | 0.295 | -1.40E-01 | 8.18E-02 | 2.73E+00 | 2.82E+00 | 8.59E-02 | no | | intergenic_variant | MODIFIER |
| 5 | 61524686 | AX-170316825 | 0 | T | C | 0.482 | 1.73E-01 | 1.00E-01 | 2.52E+00 | 2.58E+00 | 8.59E-02 | Yes | | intergenic_variant | MODIFIER |
| 5 | 128454226 | AX-170431747 | 0 | T | C | 0.149 | -1.45E-01 | 8.08E-02 | 2.11E+00 | 2.21E+00 | 8.60E-02 | no | Gm42500 | downstream_gene_variant | MODIFIER |
| 18 | 73741336 | AX-170015984 | 0 | A | G | 0.175 | -1.81E-01 | 1.06E-01 | 2.75E+00 | 2.86E+00 | 8.61E-02 | Yes | Elac1 | intron_variant | MODIFIER |
| 15 | 17712660 | AX-170495234 | 0 | G | A | 0.298 | -1.43E-01 | 7.56E-02 | 1.88E+00 | 1.91E+00 | 8.61E-02 | no | | intergenic_variant | MODIFIER |
| 18 | 76921561 | AX-170280854 | 1 | A | G | 0.446 | 1.50E-01 | 8.71E-02 | 2.47E+00 | 2.53E+00 | 8.62E-02 | Yes | | intergenic_variant | MODIFIER |
| 8 | 35648300 | AX-170097087 | 1 | A | G | 0.152 | -2.21E-01 | 1.29E-01 | 2.99E+00 | 3.11E+00 | 8.62E-02 | Yes | Mfhas1 | intron_variant | MODIFIER |
| 13 | 21141766 | AX-170007888 | 1 | A | C | 0.161 | -2.15E-01 | 1.25E-01 | 2.54E+00 | 2.61E+00 | 8.62E-02 | Yes | Olfr1368 | downstream_gene_variant | MODIFIER |
| 16 | 55529221 | AX-170407117 | 1 | A | G | 0.527 | 1.45E-01 | 8.47E-02 | 2.59E+00 | 2.62E+00 | 8.65E-02 | Yes | | intergenic_variant | MODIFIER |
| 16 | 56178715 | AX-170238075 | 1 | T | C | 0.527 | 1.45E-01 | 8.47E-02 | 2.59E+00 | 2.62E+00 | 8.65E-02 | Yes | Senp7 | intron_variant | MODIFIER |
| 16 | 57554304 | AX-170382728 | 1 | A | G | 0.527 | 1.45E-01 | 8.47E-02 | 2.59E+00 | 2.62E+00 | 8.65E-02 | Yes | Cmss1 | intron_variant | MODIFIER |
| 1 | 184916008 | AX-170464132 | 1 | A | G | 0.223 | 2.01E-01 | 1.18E-01 | 2.58E+00 | 2.66E+00 | 8.66E-02 | Yes | Mark1 | intron_variant | MODIFIER |
| 15 | 6919839 | AX-97808637 | 1 | C | T | 0.33 | -1.41E-01 | 7.51E-02 | 1.89E+00 | 1.93E+00 | 8.67E-02 | no | | intergenic_variant | MODIFIER |
| 4 | 106945536 | AX-170089331 | 1 | T | C | 0.259 | 1.93E-01 | 1.13E-01 | 2.71E+00 | 2.78E+00 | 8.69E-02 | Yes | Ssbp3 | intron_variant | MODIFIER |
| 5 | 132056349 | AX-97789077 | 2 | G | A | 0.155 | -1.57E-01 | 9.07E-02 | 2.34E+00 | 2.42E+00 | 8.69E-02 | no | Auts2 | intron_variant,non_coding_transcript_variant | MODIFIER |
| 5 | 61407696 | AX-170431464 | 1 | A | G | 0.455 | 1.78E-01 | 1.03E-01 | 2.35E+00 | 2.39E+00 | 8.72E-02 | Yes | | intergenic_variant | MODIFIER |
| 16 | 53868568 | AX-97819873 | 0 | A | G | 0.474 | -1.39E-01 | 8.15E-02 | 2.59E+00 | 2.64E+00 | 8.72E-02 | no | | intergenic_variant | MODIFIER |
| 5 | 58259169 | AX-97799962 | 1 | T | C | 0.482 | 1.70E-01 | 9.81E-02 | 2.37E+00 | 2.43E+00 | 8.74E-02 | Yes | | intergenic_variant | MODIFIER |
| 16 | 62110809 | AX-170139225 | 2 | A | T | 0.509 | 1.46E-01 | 8.55E-02 | 2.57E+00 | 2.62E+00 | 8.76E-02 | Yes | | intergenic_variant | MODIFIER |
| 16 | 57184663 | AX-170057521 | 0 | T | C | 0.535 | 1.46E-01 | 8.50E-02 | 2.47E+00 | 2.50E+00 | 8.78E-02 | Yes | Tbc1d23 | intron_variant | MODIFIER |
| 16 | 66392105 | AX-170408720 | 1 | T | G | 0.509 | 1.36E-01 | 7.96E-02 | 2.53E+00 | 2.59E+00 | 8.79E-02 | no | | intergenic_variant | MODIFIER |
| 4 | 100553803 | AX-170110207 | 0 | C | T | 0.237 | 1.96E-01 | 1.15E-01 | 2.77E+00 | 2.84E+00 | 8.79E-02 | Yes | Ube2u | downstream_gene_variant | MODIFIER |
| 15 | 77206811 | AX-170087914 | 1 | C | T | 0.08 | -2.14E-01 | 1.08E-01 | 1.69E+00 | 1.74E+00 | 8.81E-02 | Yes | Rbfox2 | intron_variant | MODIFIER |
| 13 | 10117113 | AX-97817088 | 0 | A | G | 0.193 | -1.84E-01 | 1.08E-01 | 2.55E+00 | 2.63E+00 | 8.82E-02 | Yes | Chrm3 | intron_variant | MODIFIER |
| 19 | 16008113 | AX-170099936 | 2 | G | T | 0.027 | -2.89E-01 | 1.71E-01 | 2.72E+00 | 2.89E+00 | 8.84E-02 | Yes | C130060C02Rik | intron_variant,non_coding_transcript_variant | MODIFIER |

| | | | | | | | | | | | | | | | |
|---|---|---|---|---|---|---|---|---|---|---|---|---|---|---|---|
| 5 | 135703419 | AX-97790275 | 0 | T | C | 0.149 | -1.45E-01 | 8.35E-02 | 2.30E+00 | 2.38E+00 | 8.84E-02 | no | Por | intron_variant | MODIFIER |
| 5 | 135769663 | AX-170188148 | 0 | T | C | 0.149 | -1.45E-01 | 8.35E-02 | 2.30E+00 | 2.38E+00 | 8.84E-02 | no | Styxl1 | intron_variant | MODIFIER |
| 5 | 135770489 | AX-170480732 | 0 | A | C | 0.149 | -1.45E-01 | 8.35E-02 | 2.30E+00 | 2.38E+00 | 8.84E-02 | no | Styxl1 | intron_variant | MODIFIER |
| 5 | 135789080 | AX-97809518 | 0 | G | T | 0.149 | -1.45E-01 | 8.35E-02 | 2.30E+00 | 2.38E+00 | 8.84E-02 | no | Mdh2 | intron_variant | MODIFIER |
| 5 | 135805049 | AX-97794448 | 0 | A | G | 0.149 | -1.45E-01 | 8.35E-02 | 2.30E+00 | 2.38E+00 | 8.84E-02 | no | Srrm3 | upstream_gene_variant | MODIFIER |
| 6 | 114486446 | AX-170295737 | 0 | G | T | 0.07 | -3.41E-01 | 2.00E-01 | 3.01E+00 | 3.16E+00 | 8.85E-02 | Yes | Hrh1 | downstream_gene_variant | MODIFIER |
| 10 | 18928042 | AX-170465487 | 2 | C | A | 0.2 | 1.68E-01 | 9.55E-02 | 3.40E+00 | 3.56E+00 | 8.85E-02 | Yes | | intergenic_variant | MODIFIER |
| 5 | 54569237 | AX-170072152 | 1 | A | C | 0.482 | 1.66E-01 | 9.63E-02 | 2.34E+00 | 2.41E+00 | 8.85E-02 | Yes | Gm18451 | upstream_gene_variant | MODIFIER |
| 4 | 54457382 | AX-170235382 | 0 | C | T | 0.368 | -1.74E-01 | 1.01E-01 | 3.13E+00 | 3.26E+00 | 8.87E-02 | Yes | | intergenic_variant | MODIFIER |
| 18 | 31676751 | AX-170317206 | 1 | T | A | 0.045 | -4.10E-01 | 2.37E-01 | 3.27E+00 | 3.45E+00 | 8.88E-02 | Yes | Sap130 | intron_variant | MODIFIER |
| 2 | 162133296 | AX-170298292 | 1 | T | A | 0.17 | 2.65E-01 | 1.56E-01 | 2.85E+00 | 2.99E+00 | 8.89E-02 | Yes | Ptprt | intron_variant | MODIFIER |
| 16 | 61775080 | AX-97841752 | 1 | C | T | 0.518 | 1.47E-01 | 8.67E-02 | 2.61E+00 | 2.65E+00 | 8.90E-02 | Yes | | intergenic_variant | MODIFIER |
| 5 | 52638909 | AX-170514758 | 1 | A | G | 0.482 | 1.81E-01 | 1.05E-01 | 2.39E+00 | 2.43E+00 | 8.91E-02 | Yes | Sepsecs | downstream_gene_variant | MODIFIER |
| 16 | 55287631 | AX-170530852 | 0 | C | T | 0.456 | -1.41E-01 | 8.23E-02 | 2.40E+00 | 2.44E+00 | 8.91E-02 | no | Zpld1 | upstream_gene_variant | MODIFIER |
| 3 | 31988331 | AX-170445590 | 0 | G | A | 0.158 | -1.74E-01 | 1.01E-01 | 3.22E+00 | 3.36E+00 | 8.91E-02 | Yes | Kcnmb2 | intron_variant | MODIFIER |
| 16 | 56607842 | AX-170435207 | 1 | C | T | 0.455 | -1.41E-01 | 8.24E-02 | 2.40E+00 | 2.44E+00 | 8.91E-02 | no | Abi3bp | intron_variant | MODIFIER |
| 5 | 112316747 | AX-97817884 | 0 | G | T | 0.447 | 1.68E-01 | 9.90E-02 | 2.71E+00 | 2.79E+00 | 8.91E-02 | Yes | Tpst2 | downstream_gene_variant | MODIFIER |
| 13 | 11643914 | AX-170098135 | 1 | G | A | 0.196 | -1.96E-01 | 1.15E-01 | 2.57E+00 | 2.64E+00 | 8.92E-02 | Yes | Ryr2 | intron_variant | MODIFIER |
| 9 | 91512619 | AX-170417806 | 0 | A | G | 0.132 | -1.75E-01 | 1.04E-01 | 2.73E+00 | 2.86E+00 | 8.92E-02 | Yes | Gm29478 | intron_variant,non_coding_transcript_variant | MODIFIER |
| 5 | 55734784 | AX-170419638 | 1 | A | G | 0.464 | 1.76E-01 | 1.03E-01 | 2.43E+00 | 2.48E+00 | 8.95E-02 | Yes | | intergenic_variant | MODIFIER |
| 3 | 24040984 | AX-170066105 | 1 | T | G | 0.161 | -1.54E-01 | 9.13E-02 | 2.73E+00 | 2.86E+00 | 8.96E-02 | no | Naaladl2 | intron_variant | MODIFIER |
| 16 | 63313466 | AX-97846945 | 0 | T | C | 0.518 | 1.47E-01 | 8.70E-02 | 2.60E+00 | 2.64E+00 | 8.96E-02 | Yes | | intergenic_variant | MODIFIER |
| 9 | 90689571 | AX-170102688 | 0 | T | A | 0.132 | -1.93E-01 | 1.14E-01 | 2.73E+00 | 2.84E+00 | 8.97E-02 | Yes | | intergenic_variant | MODIFIER |
| 9 | 97825313 | AX-170166068 | 0 | G | T | 0.14 | -1.66E-01 | 9.85E-02 | 2.68E+00 | 2.81E+00 | 8.97E-02 | no | Clstn2 | intron_variant | MODIFIER |
| 3 | 19134341 | AX-170020054 | 0 | A | G | 0.123 | -1.60E-01 | 9.45E-02 | 2.59E+00 | 2.68E+00 | 8.98E-02 | no | Armc1 | intron_variant | MODIFIER |
| 15 | 12226851 | AX-170290787 | 0 | A | C | 0.307 | -1.47E-01 | 7.56E-02 | 1.74E+00 | 1.77E+00 | 8.99E-02 | no | Mtmr12 | intron_variant | MODIFIER |
| 4 | 124650449 | AX-170376717 | 0 | A | G | 0.228 | 1.93E-01 | 1.14E-01 | 2.53E+00 | 2.60E+00 | 9.00E-02 | Yes | Gm2164 | intron_variant,non_coding_transcript_variant | MODIFIER |
| 5 | 57063776 | AX-170025021 | 1 | T | G | 0.482 | 1.75E-01 | 1.03E-01 | 2.45E+00 | 2.50E+00 | 9.00E-02 | Yes | | intergenic_variant | MODIFIER |
| 13 | 18439909 | AX-97810063 | 0 | T | C | 0.167 | -1.95E-01 | 1.14E-01 | 2.38E+00 | 2.46E+00 | 9.01E-02 | Yes | | intergenic_variant | MODIFIER |
| 5 | 50328220 | AX-170024908 | 0 | A | C | 0.491 | 1.73E-01 | 1.01E-01 | 2.40E+00 | 2.44E+00 | 9.01E-02 | Yes | Gm7205 | intron_variant,non_coding_transcript_variant | MODIFIER |
| 5 | 60538148 | AX-97799526 | 0 | G | A | 0.491 | 1.68E-01 | 9.76E-02 | 2.37E+00 | 2.43E+00 | 9.01E-02 | Yes | | intergenic_variant | MODIFIER |
| 5 | 66164925 | AX-170409042 | 0 | C | A | 0.491 | 1.68E-01 | 9.76E-02 | 2.37E+00 | 2.43E+00 | 9.01E-02 | Yes | Gm22929 | upstream_gene_variant | MODIFIER |
| 15 | 6250553 | AX-170303586 | 2 | C | T | 0.218 | -1.63E-01 | 9.64E-02 | 2.62E+00 | 2.69E+00 | 9.04E-02 | no | | intergenic_variant | MODIFIER |
| 6 | 147618503 | AX-170406248 | 0 | C | G | 0.044 | -2.93E-01 | 1.74E-01 | 2.91E+00 | 3.06E+00 | 9.05E-02 | Yes | Ccdc91 | intron_variant | MODIFIER |
| 5 | 136441796 | AX-170451864 | 0 | T | G | 0.14 | -1.44E-01 | 8.39E-02 | 2.36E+00 | 2.45E+00 | 9.05E-02 | no | Cux1 | intron_variant | MODIFIER |
| 2 | 72225385 | AX-170205095 | 0 | G | A | 0.044 | -3.11E-01 | 1.84E-01 | 2.86E+00 | 3.00E+00 | 9.07E-02 | Yes | Rapgef4 | intron_variant | MODIFIER |
| 1 | 124349965 | AX-170382532 | 0 | T | C | 0.561 | 1.41E-01 | 8.37E-02 | 2.64E+00 | 2.70E+00 | 9.07E-02 | no | | intergenic_variant | MODIFIER |
| 9 | 100948708 | AX-170033134 | 0 | G | A | 0.14 | -1.62E-01 | 9.63E-02 | 2.82E+00 | 2.97E+00 | 9.08E-02 | no | Stag1 | intron_variant | MODIFIER |
| 5 | 56689249 | AX-170089484 | 0 | T | G | 0.474 | 1.68E-01 | 9.81E-02 | 2.38E+00 | 2.43E+00 | 9.09E-02 | Yes | | intergenic_variant | MODIFIER |
| 10 | 101668697 | AX-97789421 | 0 | T | C | 0.272 | -1.42E-01 | 8.43E-02 | 2.75E+00 | 2.83E+00 | 9.09E-02 | no | | intergenic_variant | MODIFIER |
| 10 | 101670780 | AX-170308420 | 0 | G | T | 0.272 | -1.42E-01 | 8.43E-02 | 2.75E+00 | 2.83E+00 | 9.09E-02 | no | | intergenic_variant | MODIFIER |
| 10 | 101812900 | AX-170413790 | 0 | T | G | 0.272 | -1.42E-01 | 8.43E-02 | 2.75E+00 | 2.83E+00 | 9.09E-02 | no | Mgat4c | intron_variant | MODIFIER |
| 10 | 102152551 | AX-170143196 | 0 | T | C | 0.272 | -1.42E-01 | 8.43E-02 | 2.75E+00 | 2.83E+00 | 9.09E-02 | no | Mgat4c | intron_variant | MODIFIER |
| 10 | 102157654 | AX-170522226 | 0 | G | A | 0.272 | -1.42E-01 | 8.43E-02 | 2.75E+00 | 2.83E+00 | 9.09E-02 | no | Mgat4c | upstream_gene_variant | MODIFIER |
| 10 | 102164072 | AX-170045752 | 0 | C | T | 0.272 | -1.42E-01 | 8.43E-02 | 2.75E+00 | 2.83E+00 | 9.09E-02 | no | Mgat4c | intron_variant | MODIFIER |
| 10 | 102177694 | AX-170045753 | 0 | A | C | 0.272 | -1.42E-01 | 8.43E-02 | 2.75E+00 | 2.83E+00 | 9.09E-02 | no | Mgat4c | intron_variant | MODIFIER |
| 10 | 102203806 | AX-97850057 | 0 | A | G | 0.272 | -1.42E-01 | 8.43E-02 | 2.75E+00 | 2.83E+00 | 9.09E-02 | no | Mgat4c | intron_variant | MODIFIER |
| 10 | 102305584 | AX-170308767 | 0 | T | A | 0.272 | -1.42E-01 | 8.43E-02 | 2.75E+00 | 2.83E+00 | 9.09E-02 | no | Mgat4c | intron_variant | MODIFIER |
| 10 | 102305628 | AX-170204372 | 0 | C | T | 0.272 | -1.42E-01 | 8.43E-02 | 2.75E+00 | 2.83E+00 | 9.09E-02 | no | Mgat4c | intron_variant | MODIFIER |
| 10 | 102322304 | AX-170192093 | 0 | T | C | 0.272 | -1.42E-01 | 8.43E-02 | 2.75E+00 | 2.83E+00 | 9.09E-02 | no | Mgat4c | intron_variant | MODIFIER |
| 10 | 102366343 | AX-170086885 | 0 | G | A | 0.272 | -1.42E-01 | 8.43E-02 | 2.75E+00 | 2.83E+00 | 9.09E-02 | no | Mgat4c | intron_variant | MODIFIER |
| 10 | 102366539 | AX-170439056 | 0 | C | T | 0.272 | -1.42E-01 | 8.43E-02 | 2.75E+00 | 2.83E+00 | 9.09E-02 | no | Mgat4c | intron_variant | MODIFIER |
| 10 | 102366966 | AX-170139479 | 0 | C | T | 0.272 | -1.42E-01 | 8.43E-02 | 2.75E+00 | 2.83E+00 | 9.09E-02 | no | Mgat4c | intron_variant | MODIFIER |
| 10 | 102652406 | AX-170279717 | 0 | C | T | 0.272 | -1.42E-01 | 8.43E-02 | 2.75E+00 | 2.83E+00 | 9.09E-02 | no | | intergenic_variant | MODIFIER |

| | | | | | | | | | | | | | | | |
|---|---|---|---|---|---|---|---|---|---|---|---|---|---|---|---|
| 10 | 102667471 | AX-170113222 | 0 | G | A | 0.272 | -1.42E-01 | 8.43E-02 | 2.75E+00 | 2.83E+00 | 9.09E-02 | no | | regulatory_region_variant | MODIFIER |
| 10 | 102673186 | AX-170486597 | 0 | A | T | 0.272 | -1.42E-01 | 8.43E-02 | 2.75E+00 | 2.83E+00 | 9.09E-02 | no | | intergenic_variant | MODIFIER |
| 10 | 102703473 | AX-170133388 | 0 | C | T | 0.272 | -1.42E-01 | 8.43E-02 | 2.75E+00 | 2.83E+00 | 9.09E-02 | no | | intergenic_variant | MODIFIER |
| 10 | 102718660 | AX-97846358 | 0 | A | G | 0.272 | -1.42E-01 | 8.43E-02 | 2.75E+00 | 2.83E+00 | 9.09E-02 | no | | intergenic_variant | MODIFIER |
| 10 | 103042500 | AX-97816600 | 0 | C | T | 0.272 | -1.42E-01 | 8.43E-02 | 2.75E+00 | 2.83E+00 | 9.09E-02 | no | Lrriq1 | downstream_gene_variant | MODIFIER |
| 4 | 100979284 | AX-170395259 | 2 | T | A | 0.218 | 1.95E-01 | 1.16E-01 | 2.70E+00 | 2.78E+00 | 9.10E-02 | Yes | Cachd1 | intron_variant | MODIFIER |
| 5 | 62851158 | AX-170254355 | 0 | G | A | 0.482 | 1.75E-01 | 1.04E-01 | 2.59E+00 | 2.65E+00 | 9.10E-02 | Yes | Dthd1 | intron_variant | MODIFIER |
| 19 | 52068802 | AX-97852426 | 0 | C | T | 0.474 | 1.10E-01 | 6.52E-02 | 2.83E+00 | 2.93E+00 | 9.11E-02 | no | | intergenic_variant | MODIFIER |
| 18 | 88502907 | AX-170381412 | 0 | T | G | 0.158 | -1.58E-01 | 9.36E-02 | 2.59E+00 | 2.71E+00 | 9.13E-02 | no | | intergenic_variant | MODIFIER |
| 5 | 46511880 | AX-170449300 | 0 | C | T | 0.5 | 1.70E-01 | 1.00E-01 | 2.49E+00 | 2.55E+00 | 9.14E-02 | Yes | | intergenic_variant | MODIFIER |
| 13 | 18304369 | AX-97825287 | 1 | G | A | 0.179 | -2.13E-01 | 1.25E-01 | 2.40E+00 | 2.45E+00 | 9.15E-02 | Yes | Pou6f2 | intron_variant | MODIFIER |
| 16 | 56021240 | AX-97782780 | 2 | T | C | 0.527 | 1.43E-01 | 8.50E-02 | 2.57E+00 | 2.61E+00 | 9.15E-02 | no | Pcnp | intron_variant | MODIFIER |
| 13 | 24922092 | AX-170315257 | 1 | A | G | 0.179 | -1.66E-01 | 9.83E-02 | 2.49E+00 | 2.60E+00 | 9.16E-02 | no | Aldh5a1 | intron_variant | MODIFIER |
| 13 | 16473686 | AX-97837008 | 1 | C | T | 0.17 | -1.97E-01 | 1.16E-01 | 2.53E+00 | 2.61E+00 | 9.16E-02 | Yes | | intergenic_variant | MODIFIER |
| 1 | 124594438 | AX-170187616 | 1 | C | T | 0.58 | 1.34E-01 | 7.99E-02 | 2.67E+00 | 2.75E+00 | 9.18E-02 | no | Gm28928 | intron_variant,non_coding_transcript_variant | MODIFIER |
| 8 | 63397021 | AX-170521047 | 0 | C | T | 0.14 | -1.95E-01 | 1.16E-01 | 2.67E+00 | 2.81E+00 | 9.18E-02 | Yes | | intergenic_variant | MODIFIER |
| 16 | 73053516 | AX-170501943 | 2 | A | T | 0.491 | 1.27E-01 | 7.56E-02 | 2.66E+00 | 2.75E+00 | 9.18E-02 | no | | intergenic_variant | MODIFIER |
| 5 | 52424253 | AX-170412104 | 2 | G | A | 0.482 | 1.78E-01 | 1.05E-01 | 2.41E+00 | 2.45E+00 | 9.19E-02 | Yes | Ccdc149 | intron_variant | MODIFIER |
| 13 | 7994653 | AX-170497554 | 1 | G | T | 0.188 | -2.12E-01 | 1.26E-01 | 2.58E+00 | 2.63E+00 | 9.20E-02 | Yes | | intergenic_variant | MODIFIER |
| 4 | 125549318 | AX-170262867 | 1 | G | T | 0.214 | 1.76E-01 | 1.05E-01 | 2.69E+00 | 2.78E+00 | 9.22E-02 | Yes | Grik3 | intron_variant | MODIFIER |
| 10 | 106446839 | AX-97804888 | 1 | C | T | 0.295 | -1.43E-01 | 8.53E-02 | 2.68E+00 | 2.76E+00 | 9.24E-02 | no | Gm48380 | downstream_gene_variant | MODIFIER |
| 10 | 101848952 | AX-170111472 | 1 | C | T | 0.268 | -1.41E-01 | 8.41E-02 | 2.75E+00 | 2.83E+00 | 9.24E-02 | no | Mgat4c | intron_variant | MODIFIER |
| 2 | 169243666 | AX-170362657 | 1 | C | T | 0.17 | 2.59E-01 | 1.55E-01 | 2.83E+00 | 2.95E+00 | 9.24E-02 | Yes | | intergenic_variant | MODIFIER |
| 5 | 136676719 | AX-170133628 | 0 | C | T | 0.149 | -1.42E-01 | 8.31E-02 | 2.36E+00 | 2.45E+00 | 9.24E-02 | no | | intergenic_variant | MODIFIER |
| 5 | 136680628 | AX-97793003 | 0 | T | C | 0.149 | -1.42E-01 | 8.31E-02 | 2.36E+00 | 2.45E+00 | 9.24E-02 | no | | intergenic_variant | MODIFIER |
| 5 | 136695919 | AX-97793821 | 0 | G | A | 0.149 | -1.42E-01 | 8.31E-02 | 2.36E+00 | 2.45E+00 | 9.24E-02 | no | Myl10 | intron_variant | MODIFIER |
| 5 | 136747003 | AX-170407661 | 0 | A | G | 0.149 | -1.42E-01 | 8.31E-02 | 2.36E+00 | 2.45E+00 | 9.24E-02 | no | Col26a1 | intron_variant | MODIFIER |
| 5 | 136784587 | AX-170294569 | 0 | T | C | 0.149 | -1.42E-01 | 8.31E-02 | 2.36E+00 | 2.45E+00 | 9.24E-02 | no | Col26a1 | intron_variant | MODIFIER |
| 5 | 136819868 | AX-170424200 | 0 | T | C | 0.149 | -1.42E-01 | 8.31E-02 | 2.36E+00 | 2.45E+00 | 9.24E-02 | no | Col26a1 | intron_variant | MODIFIER |
| 5 | 136839581 | AX-97840340 | 0 | T | G | 0.149 | -1.42E-01 | 8.31E-02 | 2.36E+00 | 2.45E+00 | 9.24E-02 | no | Col26a1 | intron_variant | MODIFIER |
| 5 | 136846836 | AX-97822725 | 0 | A | G | 0.149 | -1.42E-01 | 8.31E-02 | 2.36E+00 | 2.45E+00 | 9.24E-02 | no | Col26a1 | intron_variant | MODIFIER |
| 5 | 136846885 | AX-170180123 | 0 | C | T | 0.149 | -1.42E-01 | 8.31E-02 | 2.36E+00 | 2.45E+00 | 9.24E-02 | no | Col26a1 | intron_variant | MODIFIER |
| 5 | 136856423 | AX-170173981 | 0 | T | A | 0.149 | -1.42E-01 | 8.31E-02 | 2.36E+00 | 2.45E+00 | 9.24E-02 | no | Col26a1 | intron_variant | MODIFIER |
| 5 | 136865106 | AX-170284089 | 0 | G | C | 0.149 | -1.42E-01 | 8.31E-02 | 2.36E+00 | 2.45E+00 | 9.24E-02 | no | Col26a1 | intron_variant | MODIFIER |
| 5 | 136870246 | AX-170227624 | 0 | T | G | 0.149 | -1.42E-01 | 8.31E-02 | 2.36E+00 | 2.45E+00 | 9.24E-02 | no | Col26a1 | intron_variant | MODIFIER |
| 5 | 136876906 | AX-170128732 | 0 | A | G | 0.149 | -1.42E-01 | 8.31E-02 | 2.36E+00 | 2.45E+00 | 9.24E-02 | no | Col26a1 | intron_variant | MODIFIER |
| 5 | 136877643 | AX-170527359 | 0 | T | C | 0.149 | -1.42E-01 | 8.31E-02 | 2.36E+00 | 2.45E+00 | 9.24E-02 | no | Col26a1 | intron_variant | MODIFIER |
| 5 | 136918116 | AX-97804310 | 0 | G | A | 0.149 | -1.42E-01 | 8.31E-02 | 2.36E+00 | 2.45E+00 | 9.24E-02 | no | Ift22 | downstream_gene_variant | MODIFIER |
| 5 | 136942518 | AX-170089623 | 0 | C | T | 0.149 | -1.42E-01 | 8.31E-02 | 2.36E+00 | 2.45E+00 | 9.24E-02 | no | | intergenic_variant | MODIFIER |
| 5 | 136950722 | AX-170312655 | 0 | T | C | 0.149 | -1.42E-01 | 8.31E-02 | 2.36E+00 | 2.45E+00 | 9.24E-02 | no | Fis1 | upstream_gene_variant | MODIFIER |
| 5 | 136957151 | AX-97820090 | 0 | C | A | 0.149 | -1.42E-01 | 8.31E-02 | 2.36E+00 | 2.45E+00 | 9.24E-02 | no | Fis1 | upstream_gene_variant | MODIFIER |
| 5 | 136980115 | AX-97805243 | 0 | T | C | 0.149 | -1.42E-01 | 8.31E-02 | 2.36E+00 | 2.45E+00 | 9.24E-02 | no | Cldn15 | downstream_gene_variant | MODIFIER |
| 5 | 136981407 | AX-170438330 | 0 | T | C | 0.149 | -1.42E-01 | 8.31E-02 | 2.36E+00 | 2.45E+00 | 9.24E-02 | no | Znhit1 | downstream_gene_variant | MODIFIER |
| 5 | 136998396 | AX-170486271 | 0 | C | T | 0.149 | -1.42E-01 | 8.31E-02 | 2.36E+00 | 2.45E+00 | 9.24E-02 | no | Plod3 | downstream_gene_variant | MODIFIER |
| 10 | 102184472 | AX-170525326 | 1 | A | G | 0.277 | -1.40E-01 | 8.38E-02 | 2.75E+00 | 2.84E+00 | 9.24E-02 | no | Mgat4c | intron_variant | MODIFIER |
| 13 | 7184305 | AX-170508554 | 0 | A | G | 0.184 | -2.11E-01 | 1.25E-01 | 2.53E+00 | 2.58E+00 | 9.25E-02 | Yes | | intergenic_variant | MODIFIER |
| 13 | 7184743 | AX-170492986 | 0 | G | C | 0.184 | -2.11E-01 | 1.25E-01 | 2.53E+00 | 2.58E+00 | 9.25E-02 | Yes | | intergenic_variant | MODIFIER |
| 13 | 7228838 | AX-97818747 | 0 | T | C | 0.184 | -2.11E-01 | 1.25E-01 | 2.53E+00 | 2.58E+00 | 9.25E-02 | Yes | | intergenic_variant | MODIFIER |
| 13 | 7228907 | AX-170429560 | 0 | C | T | 0.184 | -2.11E-01 | 1.25E-01 | 2.53E+00 | 2.58E+00 | 9.25E-02 | Yes | | intergenic_variant | MODIFIER |
| 13 | 7232700 | AX-170408196 | 0 | G | A | 0.184 | -2.11E-01 | 1.25E-01 | 2.53E+00 | 2.58E+00 | 9.25E-02 | Yes | | intergenic_variant | MODIFIER |
| 13 | 7236882 | AX-170370383 | 0 | A | T | 0.184 | -2.11E-01 | 1.25E-01 | 2.53E+00 | 2.58E+00 | 9.25E-02 | Yes | | intergenic_variant | MODIFIER |
| 13 | 7270684 | AX-97816430 | 0 | A | G | 0.184 | -2.11E-01 | 1.25E-01 | 2.53E+00 | 2.58E+00 | 9.25E-02 | Yes | | intergenic_variant | MODIFIER |
| 13 | 7288602 | AX-170521231 | 0 | T | C | 0.184 | -2.11E-01 | 1.25E-01 | 2.53E+00 | 2.58E+00 | 9.25E-02 | Yes | | intergenic_variant | MODIFIER |
| 13 | 7304657 | AX-170181180 | 0 | G | A | 0.184 | -2.11E-01 | 1.25E-01 | 2.53E+00 | 2.58E+00 | 9.25E-02 | Yes | Gm8725 | upstream_gene_variant | MODIFIER |

| | | | | | | | | | | | | | | | |
|---|---|---|---|---|---|---|---|---|---|---|---|---|---|---|---|
| 13 | 7313070 | AX-170512958 | 0 | A | G | 0.184 | -2.11E-01 | 1.25E-01 | 2.53E+00 | 2.58E+00 | 9.25E-02 | Yes | Gm8725 | downstream_gene_variant | MODIFIER |
| 13 | 7314751 | AX-170233853 | 0 | G | A | 0.184 | -2.11E-01 | 1.25E-01 | 2.53E+00 | 2.58E+00 | 9.25E-02 | Yes | | intergenic_variant | MODIFIER |
| 13 | 7320995 | AX-170513037 | 0 | C | A | 0.184 | -2.11E-01 | 1.25E-01 | 2.53E+00 | 2.58E+00 | 9.25E-02 | Yes | | intergenic_variant | MODIFIER |
| 13 | 7374927 | AX-97827140 | 0 | G | T | 0.184 | -2.11E-01 | 1.25E-01 | 2.53E+00 | 2.58E+00 | 9.25E-02 | Yes | | intergenic_variant | MODIFIER |
| 13 | 7375103 | AX-170131078 | 0 | C | T | 0.184 | -2.11E-01 | 1.25E-01 | 2.53E+00 | 2.58E+00 | 9.25E-02 | Yes | | regulatory_region_variant | MODIFIER |
| 13 | 7375174 | AX-170522799 | 0 | A | G | 0.184 | -2.11E-01 | 1.25E-01 | 2.53E+00 | 2.58E+00 | 9.25E-02 | Yes | | regulatory_region_variant | MODIFIER |
| 13 | 7379603 | AX-170007696 | 0 | G | A | 0.184 | -2.11E-01 | 1.25E-01 | 2.53E+00 | 2.58E+00 | 9.25E-02 | Yes | | intergenic_variant | MODIFIER |
| 13 | 7379843 | AX-170007691 | 0 | T | C | 0.184 | -2.11E-01 | 1.25E-01 | 2.53E+00 | 2.58E+00 | 9.25E-02 | Yes | | intergenic_variant | MODIFIER |
| 13 | 7384357 | AX-97857231 | 0 | G | T | 0.184 | -2.11E-01 | 1.25E-01 | 2.53E+00 | 2.58E+00 | 9.25E-02 | Yes | | intergenic_variant | MODIFIER |
| 13 | 7416946 | AX-170007688 | 0 | G | C | 0.184 | -2.11E-01 | 1.25E-01 | 2.53E+00 | 2.58E+00 | 9.25E-02 | Yes | | intergenic_variant | MODIFIER |
| 13 | 7417067 | AX-170007695 | 0 | A | G | 0.184 | -2.11E-01 | 1.25E-01 | 2.53E+00 | 2.58E+00 | 9.25E-02 | Yes | | intergenic_variant | MODIFIER |
| 13 | 7449443 | AX-170007693 | 0 | T | A | 0.184 | -2.11E-01 | 1.25E-01 | 2.53E+00 | 2.58E+00 | 9.25E-02 | Yes | Gm36074 | intron_variant,non_coding_transcript_variant | MODIFIER |
| 13 | 7450773 | AX-170007697 | 0 | G | T | 0.184 | -2.11E-01 | 1.25E-01 | 2.53E+00 | 2.58E+00 | 9.25E-02 | Yes | Gm36074 | intron_variant,non_coding_transcript_variant | MODIFIER |
| 13 | 7474670 | AX-97840145 | 0 | A | G | 0.184 | -2.11E-01 | 1.25E-01 | 2.53E+00 | 2.58E+00 | 9.25E-02 | Yes | | intergenic_variant | MODIFIER |
| 13 | 7490203 | AX-170484679 | 0 | G | T | 0.184 | -2.11E-01 | 1.25E-01 | 2.53E+00 | 2.58E+00 | 9.25E-02 | Yes | | intergenic_variant | MODIFIER |
| 13 | 7524956 | AX-170050474 | 0 | A | C | 0.184 | -2.11E-01 | 1.25E-01 | 2.53E+00 | 2.58E+00 | 9.25E-02 | Yes | | intergenic_variant | MODIFIER |
| 13 | 7530048 | AX-170050473 | 0 | C | T | 0.184 | -2.11E-01 | 1.25E-01 | 2.53E+00 | 2.58E+00 | 9.25E-02 | Yes | | intergenic_variant | MODIFIER |
| 13 | 7582185 | AX-170007694 | 0 | A | T | 0.184 | -2.11E-01 | 1.25E-01 | 2.53E+00 | 2.58E+00 | 9.25E-02 | Yes | | intergenic_variant | MODIFIER |
| 13 | 7582798 | AX-170007690 | 0 | A | G | 0.184 | -2.11E-01 | 1.25E-01 | 2.53E+00 | 2.58E+00 | 9.25E-02 | Yes | | intergenic_variant | MODIFIER |
| 13 | 7584897 | AX-97853340 | 0 | G | A | 0.184 | -2.11E-01 | 1.25E-01 | 2.53E+00 | 2.58E+00 | 9.25E-02 | Yes | | intergenic_variant | MODIFIER |
| 13 | 7586674 | AX-170168553 | 0 | G | A | 0.184 | -2.11E-01 | 1.25E-01 | 2.53E+00 | 2.58E+00 | 9.25E-02 | Yes | | intergenic_variant | MODIFIER |
| 13 | 7620490 | AX-170376455 | 0 | T | C | 0.184 | -2.11E-01 | 1.25E-01 | 2.53E+00 | 2.58E+00 | 9.25E-02 | Yes | | intergenic_variant | MODIFIER |
| 13 | 7653310 | AX-170325642 | 0 | G | C | 0.184 | -2.11E-01 | 1.25E-01 | 2.53E+00 | 2.58E+00 | 9.25E-02 | Yes | | intergenic_variant | MODIFIER |
| 13 | 7655732 | AX-170180394 | 0 | T | A | 0.184 | -2.11E-01 | 1.25E-01 | 2.53E+00 | 2.58E+00 | 9.25E-02 | Yes | | intergenic_variant | MODIFIER |
| 13 | 7660523 | AX-170455539 | 0 | G | T | 0.184 | -2.11E-01 | 1.25E-01 | 2.53E+00 | 2.58E+00 | 9.25E-02 | Yes | | intergenic_variant | MODIFIER |
| 13 | 7664882 | AX-97857638 | 0 | A | G | 0.184 | -2.11E-01 | 1.25E-01 | 2.53E+00 | 2.58E+00 | 9.25E-02 | Yes | | intergenic_variant | MODIFIER |
| 13 | 7665346 | AX-170150647 | 0 | C | G | 0.184 | -2.11E-01 | 1.25E-01 | 2.53E+00 | 2.58E+00 | 9.25E-02 | Yes | | intergenic_variant | MODIFIER |
| 13 | 7693698 | AX-170531177 | 0 | T | C | 0.184 | -2.11E-01 | 1.25E-01 | 2.53E+00 | 2.58E+00 | 9.25E-02 | Yes | | intergenic_variant | MODIFIER |
| 13 | 7699068 | AX-170114877 | 0 | C | G | 0.184 | -2.11E-01 | 1.25E-01 | 2.53E+00 | 2.58E+00 | 9.25E-02 | Yes | | intergenic_variant | MODIFIER |
| 13 | 7724991 | AX-170416714 | 0 | G | T | 0.184 | -2.11E-01 | 1.25E-01 | 2.53E+00 | 2.58E+00 | 9.25E-02 | Yes | | intergenic_variant | MODIFIER |
| 13 | 7725121 | AX-170101176 | 0 | T | C | 0.184 | -2.11E-01 | 1.25E-01 | 2.53E+00 | 2.58E+00 | 9.25E-02 | Yes | | intergenic_variant | MODIFIER |
| 13 | 7725499 | AX-170050482 | 0 | A | G | 0.184 | -2.11E-01 | 1.25E-01 | 2.53E+00 | 2.58E+00 | 9.25E-02 | Yes | | intergenic_variant | MODIFIER |
| 13 | 7726082 | AX-97830723 | 0 | C | A | 0.184 | -2.11E-01 | 1.25E-01 | 2.53E+00 | 2.58E+00 | 9.25E-02 | Yes | | intergenic_variant | MODIFIER |
| 13 | 7731981 | AX-170050479 | 0 | T | C | 0.184 | -2.11E-01 | 1.25E-01 | 2.53E+00 | 2.58E+00 | 9.25E-02 | Yes | | intergenic_variant | MODIFIER |
| 13 | 7733424 | AX-170186360 | 0 | G | T | 0.184 | -2.11E-01 | 1.25E-01 | 2.53E+00 | 2.58E+00 | 9.25E-02 | Yes | | intergenic_variant | MODIFIER |
| 13 | 7733932 | AX-170050476 | 0 | C | A | 0.184 | -2.11E-01 | 1.25E-01 | 2.53E+00 | 2.58E+00 | 9.25E-02 | Yes | | intergenic_variant | MODIFIER |
| 13 | 7770493 | AX-170050477 | 0 | C | G | 0.184 | -2.11E-01 | 1.25E-01 | 2.53E+00 | 2.58E+00 | 9.25E-02 | Yes | | intergenic_variant | MODIFIER |
| 13 | 7770582 | AX-170110663 | 0 | C | T | 0.184 | -2.11E-01 | 1.25E-01 | 2.53E+00 | 2.58E+00 | 9.25E-02 | Yes | | intergenic_variant | MODIFIER |
| 13 | 7770730 | AX-97798516 | 0 | T | C | 0.184 | -2.11E-01 | 1.25E-01 | 2.53E+00 | 2.58E+00 | 9.25E-02 | Yes | | intergenic_variant | MODIFIER |
| 13 | 7775367 | AX-170122127 | 0 | A | G | 0.184 | -2.11E-01 | 1.25E-01 | 2.53E+00 | 2.58E+00 | 9.25E-02 | Yes | | intergenic_variant | MODIFIER |
| 13 | 7775453 | AX-170441931 | 0 | T | C | 0.184 | -2.11E-01 | 1.25E-01 | 2.53E+00 | 2.58E+00 | 9.25E-02 | Yes | | intergenic_variant | MODIFIER |
| 13 | 7843321 | AX-170224528 | 0 | C | T | 0.184 | -2.11E-01 | 1.25E-01 | 2.53E+00 | 2.58E+00 | 9.25E-02 | Yes | | intergenic_variant | MODIFIER |
| 13 | 7880075 | AX-170396803 | 0 | T | G | 0.184 | -2.11E-01 | 1.25E-01 | 2.53E+00 | 2.58E+00 | 9.25E-02 | Yes | | intergenic_variant | MODIFIER |
| 13 | 7934597 | AX-97849646 | 0 | C | T | 0.184 | -2.11E-01 | 1.25E-01 | 2.53E+00 | 2.58E+00 | 9.25E-02 | Yes | | intergenic_variant | MODIFIER |
| 13 | 7937147 | AX-170382987 | 0 | T | C | 0.184 | -2.11E-01 | 1.25E-01 | 2.53E+00 | 2.58E+00 | 9.25E-02 | Yes | | intergenic_variant | MODIFIER |
| 13 | 7944911 | AX-170198038 | 0 | G | C | 0.184 | -2.11E-01 | 1.25E-01 | 2.53E+00 | 2.58E+00 | 9.25E-02 | Yes | | intergenic_variant | MODIFIER |
| 13 | 7944954 | AX-170506650 | 0 | G | A | 0.184 | -2.11E-01 | 1.25E-01 | 2.53E+00 | 2.58E+00 | 9.25E-02 | Yes | | intergenic_variant | MODIFIER |
| 13 | 7945000 | AX-170443755 | 0 | T | A | 0.184 | -2.11E-01 | 1.25E-01 | 2.53E+00 | 2.58E+00 | 9.25E-02 | Yes | | intergenic_variant | MODIFIER |
| 13 | 7956327 | AX-170339044 | 0 | G | A | 0.184 | -2.11E-01 | 1.25E-01 | 2.53E+00 | 2.58E+00 | 9.25E-02 | Yes | | intergenic_variant | MODIFIER |
| 13 | 7999300 | AX-97839286 | 0 | G | T | 0.184 | -2.11E-01 | 1.25E-01 | 2.53E+00 | 2.58E+00 | 9.25E-02 | Yes | | intergenic_variant | MODIFIER |
| 13 | 8028184 | AX-97786400 | 0 | G | C | 0.184 | -2.11E-01 | 1.25E-01 | 2.53E+00 | 2.58E+00 | 9.25E-02 | Yes | Gm9742 | intron_variant,non_coding_transcript_variant | MODIFIER |
| 13 | 8036042 | AX-170421858 | 0 | C | G | 0.184 | -2.11E-01 | 1.25E-01 | 2.53E+00 | 2.58E+00 | 9.25E-02 | Yes | Gm9742 | intron_variant,non_coding_transcript_variant | MODIFIER |
| 13 | 8044958 | AX-170087355 | 0 | T | G | 0.184 | -2.11E-01 | 1.25E-01 | 2.53E+00 | 2.58E+00 | 9.25E-02 | Yes | | intergenic_variant | MODIFIER |
| 13 | 8046660 | AX-170411234 | 0 | G | A | 0.184 | -2.11E-01 | 1.25E-01 | 2.53E+00 | 2.58E+00 | 9.25E-02 | Yes | | intergenic_variant | MODIFIER |

| | | | | | | | | | | | | | | | |
|---|---|---|---|---|---|---|---|---|---|---|---|---|---|---|---|
| 13 | 8064219 | AX-97822683 | 0 | A | G | 0.184 | -2.11E-01 | 1.25E-01 | 2.53E+00 | 2.58E+00 | 9.25E-02 | Yes | | intergenic_variant | MODIFIER |
| 13 | 8087684 | AX-170092650 | 0 | A | G | 0.184 | -2.11E-01 | 1.25E-01 | 2.53E+00 | 2.58E+00 | 9.25E-02 | Yes | | intergenic_variant | MODIFIER |
| 13 | 8089630 | AX-170007708 | 0 | C | T | 0.184 | -2.11E-01 | 1.25E-01 | 2.53E+00 | 2.58E+00 | 9.25E-02 | Yes | | intergenic_variant | MODIFIER |
| 13 | 8096113 | AX-170007706 | 0 | T | G | 0.184 | -2.11E-01 | 1.25E-01 | 2.53E+00 | 2.58E+00 | 9.25E-02 | Yes | | intergenic_variant | MODIFIER |
| 13 | 8119148 | AX-170087356 | 0 | G | T | 0.184 | -2.11E-01 | 1.25E-01 | 2.53E+00 | 2.58E+00 | 9.25E-02 | Yes | | intergenic_variant | MODIFIER |
| 13 | 8131848 | AX-170157053 | 0 | A | G | 0.184 | -2.11E-01 | 1.25E-01 | 2.53E+00 | 2.58E+00 | 9.25E-02 | Yes | Gm5191 | downstream_gene_variant | MODIFIER |
| 13 | 8140504 | AX-170007705 | 0 | C | T | 0.184 | -2.11E-01 | 1.25E-01 | 2.53E+00 | 2.58E+00 | 9.25E-02 | Yes | | intergenic_variant | MODIFIER |
| 13 | 8152219 | AX-170446677 | 0 | C | T | 0.184 | -2.11E-01 | 1.25E-01 | 2.53E+00 | 2.58E+00 | 9.25E-02 | Yes | | intergenic_variant | MODIFIER |
| 13 | 8152783 | AX-170349060 | 0 | G | C | 0.184 | -2.11E-01 | 1.25E-01 | 2.53E+00 | 2.58E+00 | 9.25E-02 | Yes | | intergenic_variant | MODIFIER |
| 13 | 8189815 | AX-170007709 | 0 | C | T | 0.184 | -2.11E-01 | 1.25E-01 | 2.53E+00 | 2.58E+00 | 9.25E-02 | Yes | | intergenic_variant | MODIFIER |
| 13 | 8190084 | AX-97835594 | 0 | A | G | 0.184 | -2.11E-01 | 1.25E-01 | 2.53E+00 | 2.58E+00 | 9.25E-02 | Yes | | intergenic_variant | MODIFIER |
| 13 | 8196670 | AX-170141499 | 0 | C | G | 0.184 | -2.11E-01 | 1.25E-01 | 2.53E+00 | 2.58E+00 | 9.25E-02 | Yes | Gm48257 | downstream_gene_variant | MODIFIER |
| 13 | 8211286 | AX-170137932 | 0 | G | A | 0.184 | -2.11E-01 | 1.25E-01 | 2.53E+00 | 2.58E+00 | 9.25E-02 | Yes | Adarb2 | intron_variant | MODIFIER |
| 13 | 8212199 | AX-170517752 | 0 | G | A | 0.184 | -2.11E-01 | 1.25E-01 | 2.53E+00 | 2.58E+00 | 9.25E-02 | Yes | Adarb2 | intron_variant | MODIFIER |
| 13 | 8212573 | AX-170457255 | 0 | A | G | 0.184 | -2.11E-01 | 1.25E-01 | 2.53E+00 | 2.58E+00 | 9.25E-02 | Yes | Adarb2 | intron_variant | MODIFIER |
| 13 | 8221134 | AX-97819218 | 0 | T | C | 0.184 | -2.11E-01 | 1.25E-01 | 2.53E+00 | 2.58E+00 | 9.25E-02 | Yes | Adarb2 | intron_variant | MODIFIER |
| 13 | 8223180 | AX-170050483 | 0 | C | T | 0.184 | -2.11E-01 | 1.25E-01 | 2.53E+00 | 2.58E+00 | 9.25E-02 | Yes | Adarb2 | intron_variant | MODIFIER |
| 13 | 8223481 | AX-170468648 | 0 | C | T | 0.184 | -2.11E-01 | 1.25E-01 | 2.53E+00 | 2.58E+00 | 9.25E-02 | Yes | Adarb2 | intron_variant | MODIFIER |
| 13 | 8226041 | AX-170398753 | 0 | T | C | 0.184 | -2.11E-01 | 1.25E-01 | 2.53E+00 | 2.58E+00 | 9.25E-02 | Yes | Adarb2 | intron_variant | MODIFIER |
| 13 | 8260159 | AX-170007710 | 0 | T | A | 0.184 | -2.11E-01 | 1.25E-01 | 2.53E+00 | 2.58E+00 | 9.25E-02 | Yes | Adarb2 | intron_variant | MODIFIER |
| 13 | 8260860 | AX-170087357 | 0 | G | T | 0.184 | -2.11E-01 | 1.25E-01 | 2.53E+00 | 2.58E+00 | 9.25E-02 | Yes | Adarb2 | intron_variant | MODIFIER |
| 13 | 8278497 | AX-170321810 | 0 | G | C | 0.184 | -2.11E-01 | 1.25E-01 | 2.53E+00 | 2.58E+00 | 9.25E-02 | Yes | Adarb2 | intron_variant | MODIFIER |
| 13 | 8281064 | AX-170405325 | 0 | C | T | 0.184 | -2.11E-01 | 1.25E-01 | 2.53E+00 | 2.58E+00 | 9.25E-02 | Yes | Adarb2 | intron_variant | MODIFIER |
| 13 | 8305868 | AX-170007699 | 0 | A | G | 0.184 | -2.11E-01 | 1.25E-01 | 2.53E+00 | 2.58E+00 | 9.25E-02 | Yes | Adarb2 | intron_variant | MODIFIER |
| 13 | 8311848 | AX-170356833 | 0 | T | C | 0.184 | -2.11E-01 | 1.25E-01 | 2.53E+00 | 2.58E+00 | 9.25E-02 | Yes | Adarb2 | intron_variant | MODIFIER |
| 13 | 8311986 | AX-170473953 | 0 | C | T | 0.184 | -2.11E-01 | 1.25E-01 | 2.53E+00 | 2.58E+00 | 9.25E-02 | Yes | Adarb2 | intron_variant | MODIFIER |
| 13 | 8317052 | AX-97831087 | 0 | G | A | 0.184 | -2.11E-01 | 1.25E-01 | 2.53E+00 | 2.58E+00 | 9.25E-02 | Yes | Adarb2 | intron_variant | MODIFIER |
| 13 | 8319014 | AX-170249505 | 0 | T | C | 0.184 | -2.11E-01 | 1.25E-01 | 2.53E+00 | 2.58E+00 | 9.25E-02 | Yes | Adarb2 | intron_variant | MODIFIER |
| 13 | 8322724 | AX-170395888 | 0 | C | T | 0.184 | -2.11E-01 | 1.25E-01 | 2.53E+00 | 2.58E+00 | 9.25E-02 | Yes | Adarb2 | intron_variant | MODIFIER |
| 13 | 8348523 | AX-97852906 | 0 | A | G | 0.184 | -2.11E-01 | 1.25E-01 | 2.53E+00 | 2.58E+00 | 9.25E-02 | Yes | Adarb2 | intron_variant | MODIFIER |
| 13 | 8348794 | AX-170302703 | 0 | G | A | 0.184 | -2.11E-01 | 1.25E-01 | 2.53E+00 | 2.58E+00 | 9.25E-02 | Yes | Adarb2 | intron_variant | MODIFIER |
| 13 | 8349207 | AX-170172872 | 0 | C | T | 0.184 | -2.11E-01 | 1.25E-01 | 2.53E+00 | 2.58E+00 | 9.25E-02 | Yes | Adarb2 | intron_variant | MODIFIER |
| 13 | 8354438 | AX-170050486 | 0 | C | T | 0.184 | -2.11E-01 | 1.25E-01 | 2.53E+00 | 2.58E+00 | 9.25E-02 | Yes | Adarb2 | intron_variant | MODIFIER |
| 13 | 8355364 | AX-170050485 | 0 | C | T | 0.184 | -2.11E-01 | 1.25E-01 | 2.53E+00 | 2.58E+00 | 9.25E-02 | Yes | Adarb2 | intron_variant | MODIFIER |
| 13 | 8379408 | AX-170087358 | 0 | T | C | 0.184 | -2.11E-01 | 1.25E-01 | 2.53E+00 | 2.58E+00 | 9.25E-02 | Yes | Adarb2 | intron_variant | MODIFIER |
| 13 | 8426629 | AX-97822704 | 0 | T | G | 0.184 | -2.11E-01 | 1.25E-01 | 2.53E+00 | 2.58E+00 | 9.25E-02 | Yes | Adarb2 | intron_variant | MODIFIER |
| 13 | 8426711 | AX-170425643 | 0 | T | C | 0.184 | -2.11E-01 | 1.25E-01 | 2.53E+00 | 2.58E+00 | 9.25E-02 | Yes | Adarb2 | intron_variant | MODIFIER |
| 13 | 8459340 | AX-170512379 | 0 | C | T | 0.184 | -2.11E-01 | 1.25E-01 | 2.53E+00 | 2.58E+00 | 9.25E-02 | Yes | Adarb2 | intron_variant | MODIFIER |
| 13 | 8485720 | AX-170125558 | 0 | C | G | 0.184 | -2.11E-01 | 1.25E-01 | 2.53E+00 | 2.58E+00 | 9.25E-02 | Yes | Adarb2 | intron_variant | MODIFIER |
| 13 | 8499114 | AX-170186687 | 0 | C | T | 0.184 | -2.11E-01 | 1.25E-01 | 2.53E+00 | 2.58E+00 | 9.25E-02 | Yes | Adarb2 | intron_variant | MODIFIER |
| 13 | 8523809 | AX-97809340 | 0 | G | A | 0.184 | -2.11E-01 | 1.25E-01 | 2.53E+00 | 2.58E+00 | 9.25E-02 | Yes | Adarb2 | intron_variant | MODIFIER |
| 13 | 8537163 | AX-170331022 | 0 | G | C | 0.184 | -2.11E-01 | 1.25E-01 | 2.53E+00 | 2.58E+00 | 9.25E-02 | Yes | Adarb2 | intron_variant | MODIFIER |
| 13 | 8539749 | AX-97853061 | 0 | A | G | 0.184 | -2.11E-01 | 1.25E-01 | 2.53E+00 | 2.58E+00 | 9.25E-02 | Yes | Adarb2 | intron_variant | MODIFIER |
| 13 | 8540197 | AX-170197128 | 0 | T | C | 0.184 | -2.11E-01 | 1.25E-01 | 2.53E+00 | 2.58E+00 | 9.25E-02 | Yes | Adarb2 | intron_variant | MODIFIER |
| 13 | 8569033 | AX-170007715 | 0 | C | T | 0.184 | -2.11E-01 | 1.25E-01 | 2.53E+00 | 2.58E+00 | 9.25E-02 | Yes | Adarb2 | intron_variant | MODIFIER |
| 13 | 8569178 | AX-97810075 | 0 | C | T | 0.184 | -2.11E-01 | 1.25E-01 | 2.53E+00 | 2.58E+00 | 9.25E-02 | Yes | Adarb2 | intron_variant | MODIFIER |
| 13 | 8585247 | AX-170240523 | 0 | A | G | 0.184 | -2.11E-01 | 1.25E-01 | 2.53E+00 | 2.58E+00 | 9.25E-02 | Yes | Adarb2 | intron_variant | MODIFIER |
| 13 | 8596712 | AX-97785818 | 0 | T | C | 0.184 | -2.11E-01 | 1.25E-01 | 2.53E+00 | 2.58E+00 | 9.25E-02 | Yes | Adarb2 | intron_variant | MODIFIER |
| 13 | 8600623 | AX-170418299 | 0 | G | A | 0.184 | -2.11E-01 | 1.25E-01 | 2.53E+00 | 2.58E+00 | 9.25E-02 | Yes | Adarb2 | intron_variant | MODIFIER |
| 13 | 8605348 | AX-170159577 | 0 | A | G | 0.184 | -2.11E-01 | 1.25E-01 | 2.53E+00 | 2.58E+00 | 9.25E-02 | Yes | Adarb2 | intron_variant | MODIFIER |
| 13 | 8607040 | AX-170285907 | 0 | G | C | 0.184 | -2.11E-01 | 1.25E-01 | 2.53E+00 | 2.58E+00 | 9.25E-02 | Yes | Adarb2 | intron_variant | MODIFIER |
| 13 | 8609415 | AX-170136736 | 0 | A | G | 0.184 | -2.11E-01 | 1.25E-01 | 2.53E+00 | 2.58E+00 | 9.25E-02 | Yes | Adarb2 | intron_variant | MODIFIER |
| 13 | 8616545 | AX-170306129 | 0 | G | A | 0.184 | -2.11E-01 | 1.25E-01 | 2.53E+00 | 2.58E+00 | 9.25E-02 | Yes | Adarb2 | intron_variant | MODIFIER |
| 13 | 8618531 | AX-97856393 | 0 | T | C | 0.184 | -2.11E-01 | 1.25E-01 | 2.53E+00 | 2.58E+00 | 9.25E-02 | Yes | Adarb2 | intron_variant | MODIFIER |

| | | | | | | | | | | | | | | | |
|---|---|---|---|---|---|---|---|---|---|---|---|---|---|---|---|
| 13 | 8622165 | AX-170007716 | 0 | A | C | 0.184 | -2.11E-01 | 1.25E-01 | 2.53E+00 | 2.58E+00 | 9.25E-02 | Yes | Adarb2 | intron_variant | MODIFIER |
| 13 | 8622363 | AX-170007717 | 0 | T | C | 0.184 | -2.11E-01 | 1.25E-01 | 2.53E+00 | 2.58E+00 | 9.25E-02 | Yes | Adarb2 | intron_variant | MODIFIER |
| 13 | 8622752 | AX-170007711 | 0 | C | T | 0.184 | -2.11E-01 | 1.25E-01 | 2.53E+00 | 2.58E+00 | 9.25E-02 | Yes | Adarb2 | intron_variant | MODIFIER |
| 13 | 8633753 | AX-97834230 | 0 | A | C | 0.184 | -2.11E-01 | 1.25E-01 | 2.53E+00 | 2.58E+00 | 9.25E-02 | Yes | Adarb2 | intron_variant | MODIFIER |
| 13 | 8633943 | AX-170050489 | 0 | G | A | 0.184 | -2.11E-01 | 1.25E-01 | 2.53E+00 | 2.58E+00 | 9.25E-02 | Yes | Adarb2 | intron_variant | MODIFIER |
| 13 | 8640989 | AX-170050488 | 0 | T | A | 0.184 | -2.11E-01 | 1.25E-01 | 2.53E+00 | 2.58E+00 | 9.25E-02 | Yes | Adarb2 | intron_variant | MODIFIER |
| 13 | 8662561 | AX-170007712 | 0 | A | G | 0.184 | -2.11E-01 | 1.25E-01 | 2.53E+00 | 2.58E+00 | 9.25E-02 | Yes | Adarb2 | intron_variant | MODIFIER |
| 13 | 8674844 | AX-170255562 | 0 | G | A | 0.184 | -2.11E-01 | 1.25E-01 | 2.53E+00 | 2.58E+00 | 9.25E-02 | Yes | Adarb2 | intron_variant | MODIFIER |
| 13 | 8679387 | AX-170136474 | 0 | G | A | 0.184 | -2.11E-01 | 1.25E-01 | 2.53E+00 | 2.58E+00 | 9.25E-02 | Yes | Adarb2 | intron_variant | MODIFIER |
| 13 | 8680852 | AX-170345981 | 0 | G | C | 0.184 | -2.11E-01 | 1.25E-01 | 2.53E+00 | 2.58E+00 | 9.25E-02 | Yes | Adarb2 | intron_variant | MODIFIER |
| 13 | 8694122 | AX-170092651 | 0 | G | A | 0.184 | -2.11E-01 | 1.25E-01 | 2.53E+00 | 2.58E+00 | 9.25E-02 | Yes | Adarb2 | intron_variant | MODIFIER |
| 13 | 8695463 | AX-170007718 | 0 | A | G | 0.184 | -2.11E-01 | 1.25E-01 | 2.53E+00 | 2.58E+00 | 9.25E-02 | Yes | Adarb2 | intron_variant | MODIFIER |
| 13 | 8696570 | AX-170036844 | 0 | C | A | 0.184 | -2.11E-01 | 1.25E-01 | 2.53E+00 | 2.58E+00 | 9.25E-02 | Yes | Adarb2 | intron_variant | MODIFIER |
| 13 | 8699521 | AX-170007714 | 0 | T | C | 0.184 | -2.11E-01 | 1.25E-01 | 2.53E+00 | 2.58E+00 | 9.25E-02 | Yes | Adarb2 | intron_variant | MODIFIER |
| 13 | 8714525 | AX-170092652 | 0 | C | T | 0.184 | -2.11E-01 | 1.25E-01 | 2.53E+00 | 2.58E+00 | 9.25E-02 | Yes | Adarb2 | intron_variant | MODIFIER |
| 13 | 8732759 | AX-97825967 | 0 | T | C | 0.184 | -2.11E-01 | 1.25E-01 | 2.53E+00 | 2.58E+00 | 9.25E-02 | Yes | Adarb2 | intron_variant | MODIFIER |
| 13 | 8736336 | AX-170304733 | 0 | G | A | 0.184 | -2.11E-01 | 1.25E-01 | 2.53E+00 | 2.58E+00 | 9.25E-02 | Yes | Adarb2 | intron_variant | MODIFIER |
| 13 | 8758400 | AX-170445160 | 0 | G | A | 0.184 | -2.11E-01 | 1.25E-01 | 2.53E+00 | 2.58E+00 | 9.25E-02 | Yes | Adarb2 | 3_prime_UTR_variant | MODIFIER |
| 13 | 8759411 | AX-170211356 | 0 | G | A | 0.184 | -2.11E-01 | 1.25E-01 | 2.53E+00 | 2.58E+00 | 9.25E-02 | Yes | Adarb2 | 3_prime_UTR_variant | MODIFIER |
| 13 | 8772412 | AX-170218425 | 0 | C | G | 0.184 | -2.11E-01 | 1.25E-01 | 2.53E+00 | 2.58E+00 | 9.25E-02 | Yes | Adarb2 | downstream_gene_variant | MODIFIER |
| 13 | 8772770 | AX-97812674 | 0 | T | C | 0.184 | -2.11E-01 | 1.25E-01 | 2.53E+00 | 2.58E+00 | 9.25E-02 | Yes | Adarb2 | downstream_gene_variant | MODIFIER |
| 13 | 8778674 | AX-170331785 | 0 | G | A | 0.184 | -2.11E-01 | 1.25E-01 | 2.53E+00 | 2.58E+00 | 9.25E-02 | Yes | | intergenic_variant | MODIFIER |
| 13 | 8785486 | AX-170087359 | 0 | A | G | 0.184 | -2.11E-01 | 1.25E-01 | 2.53E+00 | 2.58E+00 | 9.25E-02 | Yes | | regulatory_region_variant | MODIFIER |
| 13 | 8785516 | AX-170140684 | 0 | G | A | 0.184 | -2.11E-01 | 1.25E-01 | 2.53E+00 | 2.58E+00 | 9.25E-02 | Yes | | regulatory_region_variant | MODIFIER |
| 13 | 8822077 | AX-170493063 | 0 | A | T | 0.184 | -2.11E-01 | 1.25E-01 | 2.53E+00 | 2.58E+00 | 9.25E-02 | Yes | Wdr37 | intron_variant | MODIFIER |
| 13 | 8827247 | AX-97797452 | 0 | C | T | 0.184 | -2.11E-01 | 1.25E-01 | 2.53E+00 | 2.58E+00 | 9.25E-02 | Yes | Wdr37 | intron_variant | MODIFIER |
| 13 | 8831399 | AX-170329906 | 0 | T | C | 0.184 | -2.11E-01 | 1.25E-01 | 2.53E+00 | 2.58E+00 | 9.25E-02 | Yes | Wdr37 | intron_variant | MODIFIER |
| 13 | 8834867 | AX-170050490 | 0 | T | A | 0.184 | -2.11E-01 | 1.25E-01 | 2.53E+00 | 2.58E+00 | 9.25E-02 | Yes | Wdr37 | intron_variant | MODIFIER |
| 13 | 8847280 | AX-97816129 | 0 | G | A | 0.184 | -2.11E-01 | 1.25E-01 | 2.53E+00 | 2.58E+00 | 9.25E-02 | Yes | Wdr37 | intron_variant | MODIFIER |
| 13 | 8858014 | AX-170050496 | 0 | A | G | 0.184 | -2.11E-01 | 1.25E-01 | 2.53E+00 | 2.58E+00 | 9.25E-02 | Yes | Wdr37 | intron_variant | MODIFIER |
| 13 | 8858830 | AX-97815846 | 0 | C | T | 0.184 | -2.11E-01 | 1.25E-01 | 2.53E+00 | 2.58E+00 | 9.25E-02 | Yes | Wdr37 | intron_variant | MODIFIER |
| 13 | 8859017 | AX-170105786 | 0 | T | C | 0.184 | -2.11E-01 | 1.25E-01 | 2.53E+00 | 2.58E+00 | 9.25E-02 | Yes | Wdr37 | intron_variant | MODIFIER |
| 13 | 8888334 | AX-170504409 | 0 | G | T | 0.184 | -2.11E-01 | 1.25E-01 | 2.53E+00 | 2.58E+00 | 9.25E-02 | Yes | Idi1 | intron_variant | MODIFIER |
| 13 | 8908298 | AX-170464089 | 0 | A | C | 0.184 | -2.11E-01 | 1.25E-01 | 2.53E+00 | 2.58E+00 | 9.25E-02 | Yes | | intergenic_variant | MODIFIER |
| 13 | 8930136 | AX-170092654 | 0 | T | G | 0.184 | -2.11E-01 | 1.25E-01 | 2.53E+00 | 2.58E+00 | 9.25E-02 | Yes | | intergenic_variant | MODIFIER |
| 13 | 8930551 | AX-170036849 | 0 | C | T | 0.184 | -2.11E-01 | 1.25E-01 | 2.53E+00 | 2.58E+00 | 9.25E-02 | Yes | | intergenic_variant | MODIFIER |
| 13 | 8932398 | AX-97783477 | 0 | A | G | 0.184 | -2.11E-01 | 1.25E-01 | 2.53E+00 | 2.58E+00 | 9.25E-02 | Yes | | intergenic_variant | MODIFIER |
| 13 | 8940993 | AX-170283997 | 0 | C | A | 0.184 | -2.11E-01 | 1.25E-01 | 2.53E+00 | 2.58E+00 | 9.25E-02 | Yes | Gm9745 | downstream_gene_variant | MODIFIER |
| 13 | 8975268 | AX-170187908 | 0 | C | G | 0.184 | -2.11E-01 | 1.25E-01 | 2.53E+00 | 2.58E+00 | 9.25E-02 | Yes | Gtpbp4 | intron_variant,NMD_transcript_variant | MODIFIER |
| 13 | 9005122 | AX-170239317 | 0 | A | G | 0.184 | -2.11E-01 | 1.25E-01 | 2.53E+00 | 2.58E+00 | 9.25E-02 | Yes | Gm36264 | intron_variant,non_coding_transcript_variant | MODIFIER |
| 13 | 9023014 | AX-170087360 | 0 | A | T | 0.184 | -2.11E-01 | 1.25E-01 | 2.53E+00 | 2.58E+00 | 9.25E-02 | Yes | Gm36264 | intron_variant,non_coding_transcript_variant | MODIFIER |
| 13 | 9054177 | AX-97783259 | 0 | A | G | 0.184 | -2.11E-01 | 1.25E-01 | 2.53E+00 | 2.58E+00 | 9.25E-02 | Yes | Gm36264 | intron_variant,non_coding_transcript_variant | MODIFIER |
| 13 | 9057027 | AX-170522057 | 0 | T | C | 0.184 | -2.11E-01 | 1.25E-01 | 2.53E+00 | 2.58E+00 | 9.25E-02 | Yes | Gm36264 | intron_variant,non_coding_transcript_variant | MODIFIER |
| 13 | 9058848 | AX-170517655 | 0 | G | C | 0.184 | -2.11E-01 | 1.25E-01 | 2.53E+00 | 2.58E+00 | 9.25E-02 | Yes | Gm36264 | intron_variant,non_coding_transcript_variant | MODIFIER |
| 13 | 9058910 | AX-170147863 | 0 | T | C | 0.184 | -2.11E-01 | 1.25E-01 | 2.53E+00 | 2.58E+00 | 9.25E-02 | Yes | Gm36264 | intron_variant,non_coding_transcript_variant | MODIFIER |
| 13 | 9078344 | AX-170414296 | 0 | G | A | 0.184 | -2.11E-01 | 1.25E-01 | 2.53E+00 | 2.58E+00 | 9.25E-02 | Yes | Gm36264 | upstream_gene_variant | MODIFIER |
| 13 | 9097936 | AX-97844809 | 0 | T | C | 0.184 | -2.11E-01 | 1.25E-01 | 2.53E+00 | 2.58E+00 | 9.25E-02 | Yes | Larp4b | intron_variant | MODIFIER |
| 13 | 9124325 | AX-97793541 | 0 | A | G | 0.184 | -2.11E-01 | 1.25E-01 | 2.53E+00 | 2.58E+00 | 9.25E-02 | Yes | Larp4b | intron_variant | MODIFIER |
| 13 | 9127872 | AX-170050500 | 0 | A | G | 0.184 | -2.11E-01 | 1.25E-01 | 2.53E+00 | 2.58E+00 | 9.25E-02 | Yes | Larp4b | intron_variant | MODIFIER |
| 13 | 9143060 | AX-170212575 | 0 | C | T | 0.184 | -2.11E-01 | 1.25E-01 | 2.53E+00 | 2.58E+00 | 9.25E-02 | Yes | Larp4b | intron_variant | MODIFIER |
| 13 | 9143130 | AX-170470975 | 0 | G | A | 0.184 | -2.11E-01 | 1.25E-01 | 2.53E+00 | 2.58E+00 | 9.25E-02 | Yes | Larp4b | intron_variant | MODIFIER |
| 13 | 9143216 | AX-97826954 | 0 | G | A | 0.184 | -2.11E-01 | 1.25E-01 | 2.53E+00 | 2.58E+00 | 9.25E-02 | Yes | Larp4b | intron_variant | MODIFIER |
| 13 | 9151127 | AX-170050502 | 0 | T | C | 0.184 | -2.11E-01 | 1.25E-01 | 2.53E+00 | 2.58E+00 | 9.25E-02 | Yes | Larp4b | missense_variant | MODERATE |
| 13 | 9178999 | AX-97854642 | 0 | T | C | 0.184 | -2.11E-01 | 1.25E-01 | 2.53E+00 | 2.58E+00 | 9.25E-02 | Yes | Larp4b | downstream_gene_variant | MODIFIER |

| | | | | | | | | | | | | | | | |
|---|---|---|---|---|---|---|---|---|---|---|---|---|---|---|---|
| 13 | 9179060 | AX-170007721 | 0 | A | G | 0.184 | -2.11E-01 | 1.25E-01 | 2.53E+00 | 2.58E+00 | 9.25E-02 | Yes | Larp4b | downstream_gene_variant | MODIFIER |
| 13 | 9199583 | AX-170367738 | 0 | G | A | 0.184 | -2.11E-01 | 1.25E-01 | 2.53E+00 | 2.58E+00 | 9.25E-02 | Yes | | regulatory_region_variant | MODIFIER |
| 13 | 9199859 | AX-97830004 | 0 | A | G | 0.184 | -2.11E-01 | 1.25E-01 | 2.53E+00 | 2.58E+00 | 9.25E-02 | Yes | | intergenic_variant | MODIFIER |
| 13 | 9235238 | AX-170007723 | 0 | C | T | 0.184 | -2.11E-01 | 1.25E-01 | 2.53E+00 | 2.58E+00 | 9.25E-02 | Yes | | intergenic_variant | MODIFIER |
| 13 | 9239620 | AX-170007722 | 0 | G | C | 0.184 | -2.11E-01 | 1.25E-01 | 2.53E+00 | 2.58E+00 | 9.25E-02 | Yes | | intergenic_variant | MODIFIER |
| 13 | 9239651 | AX-170007720 | 0 | C | T | 0.184 | -2.11E-01 | 1.25E-01 | 2.53E+00 | 2.58E+00 | 9.25E-02 | Yes | | intergenic_variant | MODIFIER |
| 13 | 9240651 | AX-97817346 | 0 | A | G | 0.184 | -2.11E-01 | 1.25E-01 | 2.53E+00 | 2.58E+00 | 9.25E-02 | Yes | | intergenic_variant | MODIFIER |
| 13 | 9251398 | AX-170007719 | 0 | T | A | 0.184 | -2.11E-01 | 1.25E-01 | 2.53E+00 | 2.58E+00 | 9.25E-02 | Yes | | intergenic_variant | MODIFIER |
| 13 | 9251449 | AX-170524139 | 0 | C | T | 0.184 | -2.11E-01 | 1.25E-01 | 2.53E+00 | 2.58E+00 | 9.25E-02 | Yes | | intergenic_variant | MODIFIER |
| 13 | 9253365 | AX-170173552 | 0 | A | G | 0.184 | -2.11E-01 | 1.25E-01 | 2.53E+00 | 2.58E+00 | 9.25E-02 | Yes | | intergenic_variant | MODIFIER |
| 13 | 9262000 | AX-170158634 | 0 | G | A | 0.184 | -2.11E-01 | 1.25E-01 | 2.53E+00 | 2.58E+00 | 9.25E-02 | Yes | | intergenic_variant | MODIFIER |
| 13 | 9278692 | AX-170301281 | 0 | T | A | 0.184 | -2.11E-01 | 1.25E-01 | 2.53E+00 | 2.58E+00 | 9.25E-02 | Yes | Dip2c | intron_variant | MODIFIER |
| 13 | 9301494 | AX-170284666 | 0 | A | G | 0.184 | -2.11E-01 | 1.25E-01 | 2.53E+00 | 2.58E+00 | 9.25E-02 | Yes | Dip2c | intron_variant | MODIFIER |
| 13 | 9301795 | AX-170189602 | 0 | C | T | 0.184 | -2.11E-01 | 1.25E-01 | 2.53E+00 | 2.58E+00 | 9.25E-02 | Yes | Dip2c | intron_variant | MODIFIER |
| 13 | 9307067 | AX-170307777 | 0 | C | T | 0.184 | -2.11E-01 | 1.25E-01 | 2.53E+00 | 2.58E+00 | 9.25E-02 | Yes | Dip2c | intron_variant | MODIFIER |
| 13 | 9324995 | AX-97822735 | 0 | G | C | 0.184 | -2.11E-01 | 1.25E-01 | 2.53E+00 | 2.58E+00 | 9.25E-02 | Yes | Dip2c | intron_variant | MODIFIER |
| 13 | 9333037 | AX-170516079 | 0 | C | G | 0.184 | -2.11E-01 | 1.25E-01 | 2.53E+00 | 2.58E+00 | 9.25E-02 | Yes | Dip2c | intron_variant | MODIFIER |
| 13 | 9336501 | AX-170092656 | 0 | T | G | 0.184 | -2.11E-01 | 1.25E-01 | 2.53E+00 | 2.58E+00 | 9.25E-02 | Yes | Dip2c | intron_variant | MODIFIER |
| 13 | 9343115 | AX-170050506 | 0 | T | C | 0.184 | -2.11E-01 | 1.25E-01 | 2.53E+00 | 2.58E+00 | 9.25E-02 | Yes | Dip2c | intron_variant | MODIFIER |
| 13 | 9362805 | AX-97822734 | 0 | T | C | 0.184 | -2.11E-01 | 1.25E-01 | 2.53E+00 | 2.58E+00 | 9.25E-02 | Yes | Dip2c | intron_variant | MODIFIER |
| 13 | 9383032 | AX-170460723 | 0 | G | A | 0.184 | -2.11E-01 | 1.25E-01 | 2.53E+00 | 2.58E+00 | 9.25E-02 | Yes | Dip2c | intron_variant | MODIFIER |
| 13 | 9383675 | AX-170087361 | 0 | A | C | 0.184 | -2.11E-01 | 1.25E-01 | 2.53E+00 | 2.58E+00 | 9.25E-02 | Yes | Dip2c | intron_variant | MODIFIER |
| 13 | 9386705 | AX-170050504 | 0 | T | A | 0.184 | -2.11E-01 | 1.25E-01 | 2.53E+00 | 2.58E+00 | 9.25E-02 | Yes | Dip2c | intron_variant | MODIFIER |
| 13 | 9413998 | AX-170346692 | 0 | A | C | 0.184 | -2.11E-01 | 1.25E-01 | 2.53E+00 | 2.58E+00 | 9.25E-02 | Yes | Dip2c | intron_variant | MODIFIER |
| 13 | 9417738 | AX-97855697 | 0 | C | A | 0.184 | -2.11E-01 | 1.25E-01 | 2.53E+00 | 2.58E+00 | 9.25E-02 | Yes | Dip2c | intron_variant | MODIFIER |
| 13 | 9422639 | AX-170437859 | 0 | A | G | 0.184 | -2.11E-01 | 1.25E-01 | 2.53E+00 | 2.58E+00 | 9.25E-02 | Yes | Dip2c | intron_variant | MODIFIER |
| 13 | 9427317 | AX-170314007 | 0 | C | T | 0.184 | -2.11E-01 | 1.25E-01 | 2.53E+00 | 2.58E+00 | 9.25E-02 | Yes | Dip2c | intron_variant | MODIFIER |
| 13 | 9427350 | AX-170507392 | 0 | G | T | 0.184 | -2.11E-01 | 1.25E-01 | 2.53E+00 | 2.58E+00 | 9.25E-02 | Yes | Dip2c | intron_variant | MODIFIER |
| 13 | 9431423 | AX-170036850 | 0 | A | C | 0.184 | -2.11E-01 | 1.25E-01 | 2.53E+00 | 2.58E+00 | 9.25E-02 | Yes | Dip2c | intron_variant | MODIFIER |
| 13 | 9431892 | AX-170036843 | 0 | A | G | 0.184 | -2.11E-01 | 1.25E-01 | 2.53E+00 | 2.58E+00 | 9.25E-02 | Yes | Dip2c | intron_variant | MODIFIER |
| 13 | 9449152 | AX-97826955 | 0 | C | T | 0.184 | -2.11E-01 | 1.25E-01 | 2.53E+00 | 2.58E+00 | 9.25E-02 | Yes | Dip2c | intron_variant | MODIFIER |
| 13 | 9449414 | AX-170315914 | 0 | T | C | 0.184 | -2.11E-01 | 1.25E-01 | 2.53E+00 | 2.58E+00 | 9.25E-02 | Yes | Dip2c | intron_variant | MODIFIER |
| 13 | 9463927 | AX-170130355 | 0 | T | C | 0.184 | -2.11E-01 | 1.25E-01 | 2.53E+00 | 2.58E+00 | 9.25E-02 | Yes | Dip2c | intron_variant | MODIFIER |
| 13 | 9464814 | AX-170265705 | 0 | T | C | 0.184 | -2.11E-01 | 1.25E-01 | 2.53E+00 | 2.58E+00 | 9.25E-02 | Yes | Dip2c | intron_variant | MODIFIER |
| 13 | 9472608 | AX-97789117 | 0 | G | A | 0.184 | -2.11E-01 | 1.25E-01 | 2.53E+00 | 2.58E+00 | 9.25E-02 | Yes | Dip2c | intron_variant | MODIFIER |
| 13 | 9500651 | AX-97820815 | 0 | C | T | 0.184 | -2.11E-01 | 1.25E-01 | 2.53E+00 | 2.58E+00 | 9.25E-02 | Yes | Dip2c | intron_variant | MODIFIER |
| 13 | 9533626 | AX-170448863 | 0 | G | A | 0.184 | -2.11E-01 | 1.25E-01 | 2.53E+00 | 2.58E+00 | 9.25E-02 | Yes | Dip2c | intron_variant | MODIFIER |
| 13 | 9540681 | AX-97822230 | 0 | G | A | 0.184 | -2.11E-01 | 1.25E-01 | 2.53E+00 | 2.58E+00 | 9.25E-02 | Yes | Dip2c | intron_variant | MODIFIER |
| 13 | 9565873 | AX-170280641 | 0 | C | T | 0.184 | -2.11E-01 | 1.25E-01 | 2.53E+00 | 2.58E+00 | 9.25E-02 | Yes | Dip2c | intron_variant | MODIFIER |
| 13 | 9567274 | AX-170092657 | 0 | C | T | 0.184 | -2.11E-01 | 1.25E-01 | 2.53E+00 | 2.58E+00 | 9.25E-02 | Yes | Dip2c | intron_variant | MODIFIER |
| 13 | 9568771 | AX-170209383 | 0 | A | G | 0.184 | -2.11E-01 | 1.25E-01 | 2.53E+00 | 2.58E+00 | 9.25E-02 | Yes | Dip2c | intron_variant | MODIFIER |
| 13 | 9570824 | AX-170486705 | 0 | T | C | 0.184 | -2.11E-01 | 1.25E-01 | 2.53E+00 | 2.58E+00 | 9.25E-02 | Yes | Dip2c | intron_variant | MODIFIER |
| 13 | 9582082 | AX-170477049 | 0 | A | G | 0.184 | -2.11E-01 | 1.25E-01 | 2.53E+00 | 2.58E+00 | 9.25E-02 | Yes | Dip2c | intron_variant | MODIFIER |
| 13 | 9584077 | AX-97789005 | 0 | A | C | 0.184 | -2.11E-01 | 1.25E-01 | 2.53E+00 | 2.58E+00 | 9.25E-02 | Yes | Dip2c | intron_variant | MODIFIER |
| 13 | 9588514 | AX-170359614 | 0 | G | A | 0.184 | -2.11E-01 | 1.25E-01 | 2.53E+00 | 2.58E+00 | 9.25E-02 | Yes | Dip2c | intron_variant | MODIFIER |
| 13 | 9621165 | AX-170479883 | 0 | A | C | 0.184 | -2.11E-01 | 1.25E-01 | 2.53E+00 | 2.58E+00 | 9.25E-02 | Yes | Gm23496 | downstream_gene_variant | MODIFIER |
| 13 | 9623621 | AX-170184609 | 0 | T | C | 0.184 | -2.11E-01 | 1.25E-01 | 2.53E+00 | 2.58E+00 | 9.25E-02 | Yes | Gm23496 | non_coding_transcript_exon_variant | MODIFIER |
| 13 | 9647937 | AX-97815631 | 0 | T | C | 0.184 | -2.11E-01 | 1.25E-01 | 2.53E+00 | 2.58E+00 | 9.25E-02 | Yes | Dip2c | intron_variant | MODIFIER |
| 13 | 9648221 | AX-170007731 | 0 | G | A | 0.184 | -2.11E-01 | 1.25E-01 | 2.53E+00 | 2.58E+00 | 9.25E-02 | Yes | Dip2c | intron_variant | MODIFIER |
| 13 | 9648310 | AX-170007724 | 0 | G | A | 0.184 | -2.11E-01 | 1.25E-01 | 2.53E+00 | 2.58E+00 | 9.25E-02 | Yes | Dip2c | intron_variant | MODIFIER |
| 13 | 9648870 | AX-170459969 | 0 | G | T | 0.184 | -2.11E-01 | 1.25E-01 | 2.53E+00 | 2.58E+00 | 9.25E-02 | Yes | Dip2c | intron_variant | MODIFIER |
| 13 | 9692162 | AX-170111811 | 0 | T | C | 0.184 | -2.11E-01 | 1.25E-01 | 2.53E+00 | 2.58E+00 | 9.25E-02 | Yes | Zmynd11 | intron_variant | MODIFIER |
| 13 | 9700137 | AX-170007726 | 0 | G | A | 0.184 | -2.11E-01 | 1.25E-01 | 2.53E+00 | 2.58E+00 | 9.25E-02 | Yes | Zmynd11 | intron_variant | MODIFIER |
| 13 | 9721554 | AX-170479560 | 0 | C | T | 0.184 | -2.11E-01 | 1.25E-01 | 2.53E+00 | 2.58E+00 | 9.25E-02 | Yes | Zmynd11 | intron_variant | MODIFIER |

| | | | | | | | | | | | | | | | |
|---|---|---|---|---|---|---|---|---|---|---|---|---|---|---|---|
| 13 | 9728881 | AX-170007730 | 0 | C | T | 0.184 | -2.11E-01 | 1.25E-01 | 2.53E+00 | 2.58E+00 | 9.25E-02 | Yes | Zmynd11 | intron_variant | MODIFIER |
| 13 | 9732378 | AX-170007728 | 0 | T | C | 0.184 | -2.11E-01 | 1.25E-01 | 2.53E+00 | 2.58E+00 | 9.25E-02 | Yes | Zmynd11 | intron_variant | MODIFIER |
| 13 | 9753360 | AX-170474267 | 0 | T | C | 0.184 | -2.11E-01 | 1.25E-01 | 2.53E+00 | 2.58E+00 | 9.25E-02 | Yes | Zmynd11 | intron_variant | MODIFIER |
| 13 | 9777026 | AX-97822731 | 0 | G | A | 0.184 | -2.11E-01 | 1.25E-01 | 2.53E+00 | 2.58E+00 | 9.25E-02 | Yes | | intergenic_variant | MODIFIER |
| 13 | 9777745 | AX-170007725 | 0 | T | C | 0.184 | -2.11E-01 | 1.25E-01 | 2.53E+00 | 2.58E+00 | 9.25E-02 | Yes | | intergenic_variant | MODIFIER |
| 13 | 9785910 | AX-170435583 | 0 | T | C | 0.184 | -2.11E-01 | 1.25E-01 | 2.53E+00 | 2.58E+00 | 9.25E-02 | Yes | | intergenic_variant | MODIFIER |
| 13 | 9812035 | AX-170277337 | 0 | T | C | 0.184 | -2.11E-01 | 1.25E-01 | 2.53E+00 | 2.58E+00 | 9.25E-02 | Yes | | intergenic_variant | MODIFIER |
| 13 | 9821291 | AX-170502230 | 0 | C | G | 0.184 | -2.11E-01 | 1.25E-01 | 2.53E+00 | 2.58E+00 | 9.25E-02 | Yes | Gm47401 | downstream_gene_variant | MODIFIER |
| 13 | 9823288 | AX-97822761 | 0 | T | C | 0.184 | -2.11E-01 | 1.25E-01 | 2.53E+00 | 2.58E+00 | 9.25E-02 | Yes | Gm47401 | downstream_gene_variant | MODIFIER |
| 13 | 9844487 | AX-97788739 | 0 | G | A | 0.184 | -2.11E-01 | 1.25E-01 | 2.53E+00 | 2.58E+00 | 9.25E-02 | Yes | | intergenic_variant | MODIFIER |
| 13 | 9844667 | AX-170426418 | 0 | G | T | 0.184 | -2.11E-01 | 1.25E-01 | 2.53E+00 | 2.58E+00 | 9.25E-02 | Yes | | intergenic_variant | MODIFIER |
| 13 | 9847912 | AX-170259170 | 0 | G | A | 0.184 | -2.11E-01 | 1.25E-01 | 2.53E+00 | 2.58E+00 | 9.25E-02 | Yes | | intergenic_variant | MODIFIER |
| 13 | 9879540 | AX-170453738 | 0 | C | T | 0.184 | -2.11E-01 | 1.25E-01 | 2.53E+00 | 2.58E+00 | 9.25E-02 | Yes | Chrm3 | upstream_gene_variant | MODIFIER |
| 13 | 9880561 | AX-97822763 | 0 | A | G | 0.184 | -2.11E-01 | 1.25E-01 | 2.53E+00 | 2.58E+00 | 9.25E-02 | Yes | Chrm3 | upstream_gene_variant | MODIFIER |
| 13 | 9885912 | AX-170167840 | 0 | G | A | 0.184 | -2.11E-01 | 1.25E-01 | 2.53E+00 | 2.58E+00 | 9.25E-02 | Yes | Chrm3 | intron_variant | MODIFIER |
| 13 | 9896302 | AX-97822703 | 0 | A | C | 0.184 | -2.11E-01 | 1.25E-01 | 2.53E+00 | 2.58E+00 | 9.25E-02 | Yes | Chrm3 | intron_variant | MODIFIER |
| 13 | 9910028 | AX-170324130 | 0 | A | G | 0.184 | -2.11E-01 | 1.25E-01 | 2.53E+00 | 2.58E+00 | 9.25E-02 | Yes | Chrm3 | intron_variant | MODIFIER |
| 13 | 9910403 | AX-97808718 | 0 | T | C | 0.184 | -2.11E-01 | 1.25E-01 | 2.53E+00 | 2.58E+00 | 9.25E-02 | Yes | Chrm3 | intron_variant | MODIFIER |
| 13 | 9936122 | AX-170328987 | 0 | T | C | 0.184 | -2.11E-01 | 1.25E-01 | 2.53E+00 | 2.58E+00 | 9.25E-02 | Yes | Chrm3 | intron_variant | MODIFIER |
| 13 | 9938894 | AX-97841813 | 0 | T | C | 0.184 | -2.11E-01 | 1.25E-01 | 2.53E+00 | 2.58E+00 | 9.25E-02 | Yes | Chrm3 | intron_variant | MODIFIER |
| 13 | 9952643 | AX-170258349 | 0 | T | C | 0.184 | -2.11E-01 | 1.25E-01 | 2.53E+00 | 2.58E+00 | 9.25E-02 | Yes | Chrm3 | intron_variant | MODIFIER |
| 13 | 9953064 | AX-170200111 | 0 | T | A | 0.184 | -2.11E-01 | 1.25E-01 | 2.53E+00 | 2.58E+00 | 9.25E-02 | Yes | Chrm3 | intron_variant | MODIFIER |
| 13 | 9977417 | AX-97839427 | 0 | A | G | 0.184 | -2.11E-01 | 1.25E-01 | 2.53E+00 | 2.58E+00 | 9.25E-02 | Yes | Chrm3 | intron_variant | MODIFIER |
| 13 | 10009983 | AX-170092658 | 0 | A | G | 0.184 | -2.11E-01 | 1.25E-01 | 2.53E+00 | 2.58E+00 | 9.25E-02 | Yes | Chrm3 | intron_variant | MODIFIER |
| 13 | 10011093 | AX-170327927 | 0 | G | T | 0.184 | -2.11E-01 | 1.25E-01 | 2.53E+00 | 2.58E+00 | 9.25E-02 | Yes | Chrm3 | intron_variant | MODIFIER |
| 13 | 10013412 | AX-170050517 | 0 | G | A | 0.184 | -2.11E-01 | 1.25E-01 | 2.53E+00 | 2.58E+00 | 9.25E-02 | Yes | Chrm3 | intron_variant | MODIFIER |
| 13 | 10013762 | AX-170414673 | 0 | A | G | 0.184 | -2.11E-01 | 1.25E-01 | 2.53E+00 | 2.58E+00 | 9.25E-02 | Yes | Chrm3 | intron_variant | MODIFIER |
| 13 | 10045861 | AX-97810678 | 0 | C | T | 0.184 | -2.11E-01 | 1.25E-01 | 2.53E+00 | 2.58E+00 | 9.25E-02 | Yes | Chrm3 | intron_variant | MODIFIER |
| 13 | 10046159 | AX-170050516 | 0 | G | A | 0.184 | -2.11E-01 | 1.25E-01 | 2.53E+00 | 2.58E+00 | 9.25E-02 | Yes | Chrm3 | intron_variant | MODIFIER |
| 13 | 10047310 | AX-170103702 | 0 | C | T | 0.184 | -2.11E-01 | 1.25E-01 | 2.53E+00 | 2.58E+00 | 9.25E-02 | Yes | Chrm3 | intron_variant | MODIFIER |
| 13 | 10050338 | AX-170489425 | 0 | A | G | 0.184 | -2.11E-01 | 1.25E-01 | 2.53E+00 | 2.58E+00 | 9.25E-02 | Yes | Chrm3 | intron_variant | MODIFIER |
| 13 | 10066553 | AX-170156777 | 0 | T | C | 0.184 | -2.11E-01 | 1.25E-01 | 2.53E+00 | 2.58E+00 | 9.25E-02 | Yes | Chrm3 | intron_variant | MODIFIER |
| 13 | 10067281 | AX-97848221 | 0 | C | T | 0.184 | -2.11E-01 | 1.25E-01 | 2.53E+00 | 2.58E+00 | 9.25E-02 | Yes | Chrm3 | intron_variant | MODIFIER |
| 13 | 10069341 | AX-170405364 | 0 | C | G | 0.184 | -2.11E-01 | 1.25E-01 | 2.53E+00 | 2.58E+00 | 9.25E-02 | Yes | Chrm3 | intron_variant | MODIFIER |
| 13 | 10081957 | AX-170398848 | 0 | A | G | 0.184 | -2.11E-01 | 1.25E-01 | 2.53E+00 | 2.58E+00 | 9.25E-02 | Yes | Chrm3 | intron_variant | MODIFIER |
| 13 | 10086567 | AX-170423699 | 0 | A | G | 0.184 | -2.11E-01 | 1.25E-01 | 2.53E+00 | 2.58E+00 | 9.25E-02 | Yes | Chrm3 | intron_variant | MODIFIER |
| 13 | 10099446 | AX-170428204 | 0 | C | T | 0.184 | -2.11E-01 | 1.25E-01 | 2.53E+00 | 2.58E+00 | 9.25E-02 | Yes | Chrm3 | intron_variant | MODIFIER |
| 13 | 10133321 | AX-170425110 | 0 | G | A | 0.184 | -2.11E-01 | 1.25E-01 | 2.53E+00 | 2.58E+00 | 9.25E-02 | Yes | Chrm3 | intron_variant | MODIFIER |
| 13 | 10137156 | AX-170165897 | 0 | G | T | 0.184 | -2.11E-01 | 1.25E-01 | 2.53E+00 | 2.58E+00 | 9.25E-02 | Yes | Chrm3 | intron_variant | MODIFIER |
| 13 | 10139010 | AX-170087362 | 0 | G | T | 0.184 | -2.11E-01 | 1.25E-01 | 2.53E+00 | 2.58E+00 | 9.25E-02 | Yes | Chrm3 | intron_variant | MODIFIER |
| 13 | 10167917 | AX-170290498 | 0 | T | A | 0.184 | -2.11E-01 | 1.25E-01 | 2.53E+00 | 2.58E+00 | 9.25E-02 | Yes | Chrm3 | intron_variant | MODIFIER |
| 13 | 10188448 | AX-170007732 | 0 | G | T | 0.184 | -2.11E-01 | 1.25E-01 | 2.53E+00 | 2.58E+00 | 9.25E-02 | Yes | Chrm3 | intron_variant | MODIFIER |
| 13 | 10208750 | AX-170379126 | 0 | T | G | 0.184 | -2.11E-01 | 1.25E-01 | 2.53E+00 | 2.58E+00 | 9.25E-02 | Yes | Chrm3 | intron_variant | MODIFIER |
| 13 | 10214384 | AX-170317589 | 0 | A | C | 0.184 | -2.11E-01 | 1.25E-01 | 2.53E+00 | 2.58E+00 | 9.25E-02 | Yes | Chrm3 | intron_variant | MODIFIER |
| 13 | 10215403 | AX-170007734 | 0 | C | A | 0.184 | -2.11E-01 | 1.25E-01 | 2.53E+00 | 2.58E+00 | 9.25E-02 | Yes | Chrm3 | intron_variant | MODIFIER |
| 13 | 10216426 | AX-170007739 | 0 | T | C | 0.184 | -2.11E-01 | 1.25E-01 | 2.53E+00 | 2.58E+00 | 9.25E-02 | Yes | Chrm3 | intron_variant | MODIFIER |
| 13 | 10235907 | AX-97815791 | 0 | C | T | 0.184 | -2.11E-01 | 1.25E-01 | 2.53E+00 | 2.58E+00 | 9.25E-02 | Yes | Chrm3 | intron_variant | MODIFIER |
| 13 | 10266188 | AX-170146650 | 0 | C | T | 0.184 | -2.11E-01 | 1.25E-01 | 2.53E+00 | 2.58E+00 | 9.25E-02 | Yes | Chrm3 | intron_variant | MODIFIER |
| 13 | 10267618 | AX-170468961 | 0 | T | A | 0.184 | -2.11E-01 | 1.25E-01 | 2.53E+00 | 2.58E+00 | 9.25E-02 | Yes | Chrm3 | intron_variant | MODIFIER |
| 13 | 10270553 | AX-170007736 | 0 | T | C | 0.184 | -2.11E-01 | 1.25E-01 | 2.53E+00 | 2.58E+00 | 9.25E-02 | Yes | Chrm3 | intron_variant | MODIFIER |
| 13 | 10271219 | AX-170105112 | 0 | C | G | 0.184 | -2.11E-01 | 1.25E-01 | 2.53E+00 | 2.58E+00 | 9.25E-02 | Yes | Chrm3 | intron_variant | MODIFIER |
| 13 | 10282006 | AX-170239718 | 0 | T | C | 0.184 | -2.11E-01 | 1.25E-01 | 2.53E+00 | 2.58E+00 | 9.25E-02 | Yes | Chrm3 | intron_variant | MODIFIER |
| 13 | 10282245 | AX-170253542 | 0 | T | C | 0.184 | -2.11E-01 | 1.25E-01 | 2.53E+00 | 2.58E+00 | 9.25E-02 | Yes | Chrm3 | intron_variant | MODIFIER |
| 13 | 10282446 | AX-170502645 | 0 | A | C | 0.184 | -2.11E-01 | 1.25E-01 | 2.53E+00 | 2.58E+00 | 9.25E-02 | Yes | Chrm3 | intron_variant | MODIFIER |

| | | | | | | | | | | | | | | | |
|---|---|---|---|---|---|---|---|---|---|---|---|---|---|---|---|
| 13 | 10345501 | AX-170007733 | 0 | A | G | 0.184 | -2.11E-01 | 1.25E-01 | 2.53E+00 | 2.58E+00 | 9.25E-02 | Yes | Chrm3 | intron_variant | MODIFIER |
| 13 | 10350830 | AX-97822758 | 0 | T | C | 0.184 | -2.11E-01 | 1.25E-01 | 2.53E+00 | 2.58E+00 | 9.25E-02 | Yes | Chrm3 | intron_variant | MODIFIER |
| 13 | 10394007 | AX-170007738 | 0 | T | C | 0.184 | -2.11E-01 | 1.25E-01 | 2.53E+00 | 2.58E+00 | 9.25E-02 | Yes | Gm26861 | intron_variant,non_coding_transcript_variant | MODIFIER |
| 13 | 10394698 | AX-97822706 | 0 | G | A | 0.184 | -2.11E-01 | 1.25E-01 | 2.53E+00 | 2.58E+00 | 9.25E-02 | Yes | Gm26861 | intron_variant,non_coding_transcript_variant | MODIFIER |
| 13 | 10394785 | AX-170401254 | 0 | T | C | 0.184 | -2.11E-01 | 1.25E-01 | 2.53E+00 | 2.58E+00 | 9.25E-02 | Yes | Gm26861 | intron_variant,non_coding_transcript_variant | MODIFIER |
| 13 | 10410076 | AX-170223835 | 0 | C | T | 0.184 | -2.11E-01 | 1.25E-01 | 2.53E+00 | 2.58E+00 | 9.25E-02 | Yes | Gm26861 | intron_variant,non_coding_transcript_variant | MODIFIER |
| 13 | 10414913 | AX-170231184 | 0 | A | G | 0.184 | -2.11E-01 | 1.25E-01 | 2.53E+00 | 2.58E+00 | 9.25E-02 | Yes | Gm26861 | intron_variant,non_coding_transcript_variant | MODIFIER |
| 13 | 10421117 | AX-170087364 | 0 | T | G | 0.184 | -2.11E-01 | 1.25E-01 | 2.53E+00 | 2.58E+00 | 9.25E-02 | Yes | Gm26861 | intron_variant,non_coding_transcript_variant | MODIFIER |
| 13 | 10454884 | AX-170409004 | 0 | C | A | 0.184 | -2.11E-01 | 1.25E-01 | 2.53E+00 | 2.58E+00 | 9.25E-02 | Yes | Gm26861 | intron_variant,non_coding_transcript_variant | MODIFIER |
| 13 | 10454908 | AX-170141161 | 0 | A | G | 0.184 | -2.11E-01 | 1.25E-01 | 2.53E+00 | 2.58E+00 | 9.25E-02 | Yes | Gm26861 | intron_variant,non_coding_transcript_variant | MODIFIER |
| 13 | 10472112 | AX-97800922 | 0 | A | G | 0.184 | -2.11E-01 | 1.25E-01 | 2.53E+00 | 2.58E+00 | 9.25E-02 | Yes | Gm26861 | intron_variant,non_coding_transcript_variant | MODIFIER |
| 13 | 10477341 | AX-170106419 | 0 | A | C | 0.184 | -2.11E-01 | 1.25E-01 | 2.53E+00 | 2.58E+00 | 9.25E-02 | Yes | Gm26861 | intron_variant,non_coding_transcript_variant | MODIFIER |
| 13 | 10500237 | AX-170458112 | 0 | T | C | 0.184 | -2.11E-01 | 1.25E-01 | 2.53E+00 | 2.58E+00 | 9.25E-02 | Yes | Gm26861 | intron_variant,non_coding_transcript_variant | MODIFIER |
| 13 | 10544789 | AX-170277115 | 0 | G | A | 0.184 | -2.11E-01 | 1.25E-01 | 2.53E+00 | 2.58E+00 | 9.25E-02 | Yes | Gm26861 | intron_variant,non_coding_transcript_variant | MODIFIER |
| 13 | 10545336 | AX-170362651 | 0 | T | G | 0.184 | -2.11E-01 | 1.25E-01 | 2.53E+00 | 2.58E+00 | 9.25E-02 | Yes | Gm26861 | intron_variant,non_coding_transcript_variant | MODIFIER |
| 13 | 10545882 | AX-170372497 | 0 | A | C | 0.184 | -2.11E-01 | 1.25E-01 | 2.53E+00 | 2.58E+00 | 9.25E-02 | Yes | Gm26861 | intron_variant,non_coding_transcript_variant | MODIFIER |
| 13 | 10545967 | AX-170235969 | 0 | A | G | 0.184 | -2.11E-01 | 1.25E-01 | 2.53E+00 | 2.58E+00 | 9.25E-02 | Yes | Gm26861 | intron_variant,non_coding_transcript_variant | MODIFIER |
| 13 | 10556105 | AX-170121860 | 0 | A | G | 0.184 | -2.11E-01 | 1.25E-01 | 2.53E+00 | 2.58E+00 | 9.25E-02 | Yes | Gm26861 | intron_variant,non_coding_transcript_variant | MODIFIER |
| 13 | 10556185 | AX-170298843 | 0 | A | G | 0.184 | -2.11E-01 | 1.25E-01 | 2.53E+00 | 2.58E+00 | 9.25E-02 | Yes | Gm26861 | intron_variant,non_coding_transcript_variant | MODIFIER |
| 13 | 10591570 | AX-97833007 | 0 | A | G | 0.184 | -2.11E-01 | 1.25E-01 | 2.53E+00 | 2.58E+00 | 9.25E-02 | Yes | Gm23415 | downstream_gene_variant | MODIFIER |
| 13 | 10616928 | AX-170532972 | 0 | T | C | 0.184 | -2.11E-01 | 1.25E-01 | 2.53E+00 | 2.58E+00 | 9.25E-02 | Yes | Gm26861 | intron_variant,non_coding_transcript_variant | MODIFIER |
| 13 | 10621901 | AX-170484142 | 0 | G | T | 0.184 | -2.11E-01 | 1.25E-01 | 2.53E+00 | 2.58E+00 | 9.25E-02 | Yes | Gm26861 | intron_variant,non_coding_transcript_variant | MODIFIER |
| 13 | 10628688 | AX-170266425 | 0 | G | T | 0.184 | -2.11E-01 | 1.25E-01 | 2.53E+00 | 2.58E+00 | 9.25E-02 | Yes | Gm26861 | intron_variant,non_coding_transcript_variant | MODIFIER |
| 13 | 10630116 | AX-170050525 | 0 | C | T | 0.184 | -2.11E-01 | 1.25E-01 | 2.53E+00 | 2.58E+00 | 9.25E-02 | Yes | Gm26861 | intron_variant,non_coding_transcript_variant | MODIFIER |
| 13 | 10635717 | AX-170050527 | 0 | C | G | 0.184 | -2.11E-01 | 1.25E-01 | 2.53E+00 | 2.58E+00 | 9.25E-02 | Yes | Gm26861 | intron_variant,non_coding_transcript_variant | MODIFIER |
| 13 | 10643866 | AX-170379173 | 0 | T | C | 0.184 | -2.11E-01 | 1.25E-01 | 2.53E+00 | 2.58E+00 | 9.25E-02 | Yes | Gm26861 | intron_variant,non_coding_transcript_variant | MODIFIER |
| 13 | 10683952 | AX-170233587 | 0 | T | C | 0.184 | -2.11E-01 | 1.25E-01 | 2.53E+00 | 2.58E+00 | 9.25E-02 | Yes | Gm26861 | intron_variant,non_coding_transcript_variant | MODIFIER |
| 13 | 10692739 | AX-170385663 | 0 | G | T | 0.184 | -2.11E-01 | 1.25E-01 | 2.53E+00 | 2.58E+00 | 9.25E-02 | Yes | Gm26861 | intron_variant,non_coding_transcript_variant | MODIFIER |
| 13 | 10692898 | AX-170087365 | 0 | G | A | 0.184 | -2.11E-01 | 1.25E-01 | 2.53E+00 | 2.58E+00 | 9.25E-02 | Yes | Gm26861 | intron_variant,non_coding_transcript_variant | MODIFIER |
| 13 | 10717509 | AX-170136082 | 0 | G | C | 0.184 | -2.11E-01 | 1.25E-01 | 2.53E+00 | 2.58E+00 | 9.25E-02 | Yes | Gm26861 | intron_variant,non_coding_transcript_variant | MODIFIER |
| 13 | 10740903 | AX-170007752 | 0 | G | A | 0.184 | -2.11E-01 | 1.25E-01 | 2.53E+00 | 2.58E+00 | 9.25E-02 | Yes | Gm26861 | downstream_gene_variant | MODIFIER |
| 13 | 10741153 | AX-170087366 | 0 | G | A | 0.184 | -2.11E-01 | 1.25E-01 | 2.53E+00 | 2.58E+00 | 9.25E-02 | Yes | Gm26861 | downstream_gene_variant | MODIFIER |
| 13 | 10741881 | AX-170299034 | 0 | G | A | 0.184 | -2.11E-01 | 1.25E-01 | 2.53E+00 | 2.58E+00 | 9.25E-02 | Yes | Gm26861 | downstream_gene_variant | MODIFIER |
| 13 | 10766025 | AX-170007748 | 0 | C | T | 0.184 | -2.11E-01 | 1.25E-01 | 2.53E+00 | 2.58E+00 | 9.25E-02 | Yes | Gm47408 | intron_variant,non_coding_transcript_variant | MODIFIER |
| 13 | 10768560 | AX-170007751 | 0 | T | C | 0.184 | -2.11E-01 | 1.25E-01 | 2.53E+00 | 2.58E+00 | 9.25E-02 | Yes | Gm47408 | intron_variant,non_coding_transcript_variant | MODIFIER |
| 13 | 10819290 | AX-97794325 | 0 | G | A | 0.184 | -2.11E-01 | 1.25E-01 | 2.53E+00 | 2.58E+00 | 9.25E-02 | Yes | Gm47408 | intron_variant,non_coding_transcript_variant | MODIFIER |
| 13 | 10842095 | AX-170007750 | 0 | A | G | 0.184 | -2.11E-01 | 1.25E-01 | 2.53E+00 | 2.58E+00 | 9.25E-02 | Yes | | intergenic_variant | MODIFIER |
| 13 | 10842855 | AX-170087367 | 0 | G | A | 0.184 | -2.11E-01 | 1.25E-01 | 2.53E+00 | 2.58E+00 | 9.25E-02 | Yes | | intergenic_variant | MODIFIER |
| 13 | 10842927 | AX-170007749 | 0 | C | G | 0.184 | -2.11E-01 | 1.25E-01 | 2.53E+00 | 2.58E+00 | 9.25E-02 | Yes | | intergenic_variant | MODIFIER |
| 13 | 10886525 | AX-170007743 | 0 | A | C | 0.184 | -2.11E-01 | 1.25E-01 | 2.53E+00 | 2.58E+00 | 9.25E-02 | Yes | | intergenic_variant | MODIFIER |
| 13 | 10909558 | AX-97834494 | 0 | A | C | 0.184 | -2.11E-01 | 1.25E-01 | 2.53E+00 | 2.58E+00 | 9.25E-02 | Yes | | intergenic_variant | MODIFIER |
| 13 | 10941888 | AX-97821442 | 0 | C | T | 0.184 | -2.11E-01 | 1.25E-01 | 2.53E+00 | 2.58E+00 | 9.25E-02 | Yes | | intergenic_variant | MODIFIER |
| 13 | 10942473 | AX-170254704 | 0 | G | A | 0.184 | -2.11E-01 | 1.25E-01 | 2.53E+00 | 2.58E+00 | 9.25E-02 | Yes | | regulatory_region_variant | MODIFIER |
| 13 | 10968514 | AX-97807538 | 0 | T | G | 0.184 | -2.11E-01 | 1.25E-01 | 2.53E+00 | 2.58E+00 | 9.25E-02 | Yes | | intergenic_variant | MODIFIER |
| 13 | 10973881 | AX-170199719 | 0 | T | C | 0.184 | -2.11E-01 | 1.25E-01 | 2.53E+00 | 2.58E+00 | 9.25E-02 | Yes | Gm47486 | upstream_gene_variant | MODIFIER |
| 13 | 10980106 | AX-170209729 | 0 | T | C | 0.184 | -2.11E-01 | 1.25E-01 | 2.53E+00 | 2.58E+00 | 9.25E-02 | Yes | Gm47486 | downstream_gene_variant | MODIFIER |
| 13 | 10985401 | AX-170050528 | 0 | A | G | 0.184 | -2.11E-01 | 1.25E-01 | 2.53E+00 | 2.58E+00 | 9.25E-02 | Yes | | intergenic_variant | MODIFIER |
| 13 | 10987240 | AX-170163548 | 0 | T | C | 0.184 | -2.11E-01 | 1.25E-01 | 2.53E+00 | 2.58E+00 | 9.25E-02 | Yes | | intergenic_variant | MODIFIER |
| 13 | 11085148 | AX-170007745 | 0 | T | C | 0.184 | -2.11E-01 | 1.25E-01 | 2.53E+00 | 2.58E+00 | 9.25E-02 | Yes | | intergenic_variant | MODIFIER |
| 13 | 11101527 | AX-97815383 | 0 | A | G | 0.184 | -2.11E-01 | 1.25E-01 | 2.53E+00 | 2.58E+00 | 9.25E-02 | Yes | | intergenic_variant | MODIFIER |
| 13 | 11102874 | AX-170251393 | 0 | T | C | 0.184 | -2.11E-01 | 1.25E-01 | 2.53E+00 | 2.58E+00 | 9.25E-02 | Yes | | intergenic_variant | MODIFIER |
| 13 | 11117352 | AX-170228510 | 0 | T | C | 0.184 | -2.11E-01 | 1.25E-01 | 2.53E+00 | 2.58E+00 | 9.25E-02 | Yes | | intergenic_variant | MODIFIER |
| 13 | 11117384 | AX-97850590 | 0 | A | G | 0.184 | -2.11E-01 | 1.25E-01 | 2.53E+00 | 2.58E+00 | 9.25E-02 | Yes | | intergenic_variant | MODIFIER |
| 13 | 11117477 | AX-170468892 | 0 | C | G | 0.184 | -2.11E-01 | 1.25E-01 | 2.53E+00 | 2.58E+00 | 9.25E-02 | Yes | | intergenic_variant | MODIFIER |

| | | | | | | | | | | | | | | | |
|---|---|---|---|---|---|---|---|---|---|---|---|---|---|---|---|
| 13 | 11164133 | AX-97834785 | 0 | A | G | 0.184 | -2.11E-01 | 1.25E-01 | 2.53E+00 | 2.58E+00 | 9.25E-02 | Yes | | intergenic_variant | MODIFIER |
| 13 | 11176911 | AX-170175387 | 0 | G | A | 0.184 | -2.11E-01 | 1.25E-01 | 2.53E+00 | 2.58E+00 | 9.25E-02 | Yes | | intergenic_variant | MODIFIER |
| 13 | 11182719 | AX-170332325 | 0 | A | G | 0.184 | -2.11E-01 | 1.25E-01 | 2.53E+00 | 2.58E+00 | 9.25E-02 | Yes | | intergenic_variant | MODIFIER |
| 13 | 11184649 | AX-97838274 | 0 | T | G | 0.184 | -2.11E-01 | 1.25E-01 | 2.53E+00 | 2.58E+00 | 9.25E-02 | Yes | | intergenic_variant | MODIFIER |
| 13 | 11217603 | AX-97822813 | 0 | G | A | 0.184 | -2.11E-01 | 1.25E-01 | 2.53E+00 | 2.58E+00 | 9.25E-02 | Yes | | intergenic_variant | MODIFIER |
| 13 | 11219525 | AX-170303265 | 0 | G | A | 0.184 | -2.11E-01 | 1.25E-01 | 2.53E+00 | 2.58E+00 | 9.25E-02 | Yes | | intergenic_variant | MODIFIER |
| 13 | 11221964 | AX-170445265 | 0 | T | C | 0.184 | -2.11E-01 | 1.25E-01 | 2.53E+00 | 2.58E+00 | 9.25E-02 | Yes | | intergenic_variant | MODIFIER |
| 13 | 11224750 | AX-170422606 | 0 | T | C | 0.184 | -2.11E-01 | 1.25E-01 | 2.53E+00 | 2.58E+00 | 9.25E-02 | Yes | | intergenic_variant | MODIFIER |
| 13 | 11231033 | AX-170198335 | 0 | T | C | 0.184 | -2.11E-01 | 1.25E-01 | 2.53E+00 | 2.58E+00 | 9.25E-02 | Yes | | intergenic_variant | MODIFIER |
| 13 | 11239565 | AX-170178077 | 0 | A | T | 0.184 | -2.11E-01 | 1.25E-01 | 2.53E+00 | 2.58E+00 | 9.25E-02 | Yes | | intergenic_variant | MODIFIER |
| 13 | 11251361 | AX-170206975 | 0 | T | C | 0.184 | -2.11E-01 | 1.25E-01 | 2.53E+00 | 2.58E+00 | 9.25E-02 | Yes | | intergenic_variant | MODIFIER |
| 13 | 11252103 | AX-170193228 | 0 | T | C | 0.184 | -2.11E-01 | 1.25E-01 | 2.53E+00 | 2.58E+00 | 9.25E-02 | Yes | | intergenic_variant | MODIFIER |
| 13 | 11276029 | AX-97792528 | 0 | C | T | 0.184 | -2.11E-01 | 1.25E-01 | 2.53E+00 | 2.58E+00 | 9.25E-02 | Yes | | intergenic_variant | MODIFIER |
| 13 | 11280335 | AX-170369565 | 0 | T | C | 0.184 | -2.11E-01 | 1.25E-01 | 2.53E+00 | 2.58E+00 | 9.25E-02 | Yes | | intergenic_variant | MODIFIER |
| 13 | 11281200 | AX-170292444 | 0 | T | A | 0.184 | -2.11E-01 | 1.25E-01 | 2.53E+00 | 2.58E+00 | 9.25E-02 | Yes | | intergenic_variant | MODIFIER |
| 13 | 11281764 | AX-170138034 | 0 | T | C | 0.184 | -2.11E-01 | 1.25E-01 | 2.53E+00 | 2.58E+00 | 9.25E-02 | Yes | | intergenic_variant | MODIFIER |
| 13 | 11282559 | AX-170007755 | 0 | T | C | 0.184 | -2.11E-01 | 1.25E-01 | 2.53E+00 | 2.58E+00 | 9.25E-02 | Yes | | intergenic_variant | MODIFIER |
| 13 | 11309232 | AX-170050534 | 0 | A | G | 0.184 | -2.11E-01 | 1.25E-01 | 2.53E+00 | 2.58E+00 | 9.25E-02 | Yes | | intergenic_variant | MODIFIER |
| 13 | 11316743 | AX-97846842 | 0 | T | C | 0.184 | -2.11E-01 | 1.25E-01 | 2.53E+00 | 2.58E+00 | 9.25E-02 | Yes | | intergenic_variant | MODIFIER |
| 13 | 11332079 | AX-170386810 | 0 | T | C | 0.184 | -2.11E-01 | 1.25E-01 | 2.53E+00 | 2.58E+00 | 9.25E-02 | Yes | | regulatory_region_variant | MODIFIER |
| 13 | 11337865 | AX-170050532 | 0 | T | C | 0.184 | -2.11E-01 | 1.25E-01 | 2.53E+00 | 2.58E+00 | 9.25E-02 | Yes | | intergenic_variant | MODIFIER |
| 13 | 11338540 | AX-170092659 | 0 | G | A | 0.184 | -2.11E-01 | 1.25E-01 | 2.53E+00 | 2.58E+00 | 9.25E-02 | Yes | | regulatory_region_variant | MODIFIER |
| 13 | 11338557 | AX-170114590 | 0 | C | T | 0.184 | -2.11E-01 | 1.25E-01 | 2.53E+00 | 2.58E+00 | 9.25E-02 | Yes | | regulatory_region_variant | MODIFIER |
| 13 | 11392323 | AX-97782488 | 0 | T | C | 0.184 | -2.11E-01 | 1.25E-01 | 2.53E+00 | 2.58E+00 | 9.25E-02 | Yes | | intergenic_variant | MODIFIER |
| 13 | 11404756 | AX-170462259 | 0 | A | G | 0.184 | -2.11E-01 | 1.25E-01 | 2.53E+00 | 2.58E+00 | 9.25E-02 | Yes | | intergenic_variant | MODIFIER |
| 13 | 11406177 | AX-170411826 | 0 | T | C | 0.184 | -2.11E-01 | 1.25E-01 | 2.53E+00 | 2.58E+00 | 9.25E-02 | Yes | | intergenic_variant | MODIFIER |
| 13 | 11406964 | AX-170314118 | 0 | T | C | 0.184 | -2.11E-01 | 1.25E-01 | 2.53E+00 | 2.58E+00 | 9.25E-02 | Yes | | regulatory_region_variant | MODIFIER |
| 13 | 11410128 | AX-170109801 | 0 | T | C | 0.184 | -2.11E-01 | 1.25E-01 | 2.53E+00 | 2.58E+00 | 9.25E-02 | Yes | | intergenic_variant | MODIFIER |
| 13 | 11424112 | AX-170036855 | 0 | G | A | 0.184 | -2.11E-01 | 1.25E-01 | 2.53E+00 | 2.58E+00 | 9.25E-02 | Yes | Gm25496 | downstream_gene_variant | MODIFIER |
| 13 | 11431541 | AX-170263844 | 0 | G | A | 0.184 | -2.11E-01 | 1.25E-01 | 2.53E+00 | 2.58E+00 | 9.25E-02 | Yes | Gm25496 | upstream_gene_variant | MODIFIER |
| 13 | 11436814 | AX-170493423 | 0 | G | A | 0.184 | -2.11E-01 | 1.25E-01 | 2.53E+00 | 2.58E+00 | 9.25E-02 | Yes | | intergenic_variant | MODIFIER |
| 13 | 11443985 | AX-97822512 | 0 | C | T | 0.184 | -2.11E-01 | 1.25E-01 | 2.53E+00 | 2.58E+00 | 9.25E-02 | Yes | | intergenic_variant | MODIFIER |
| 13 | 11445807 | AX-170224315 | 0 | A | G | 0.184 | -2.11E-01 | 1.25E-01 | 2.53E+00 | 2.58E+00 | 9.25E-02 | Yes | | intergenic_variant | MODIFIER |
| 13 | 11468557 | AX-97814630 | 0 | C | T | 0.184 | -2.11E-01 | 1.25E-01 | 2.53E+00 | 2.58E+00 | 9.25E-02 | Yes | | intergenic_variant | MODIFIER |
| 13 | 11472191 | AX-170408601 | 0 | A | T | 0.184 | -2.11E-01 | 1.25E-01 | 2.53E+00 | 2.58E+00 | 9.25E-02 | Yes | | intergenic_variant | MODIFIER |
| 13 | 11490360 | AX-170190391 | 0 | C | T | 0.184 | -2.11E-01 | 1.25E-01 | 2.53E+00 | 2.58E+00 | 9.25E-02 | Yes | | intergenic_variant | MODIFIER |
| 13 | 11509054 | AX-170050541 | 0 | G | T | 0.184 | -2.11E-01 | 1.25E-01 | 2.53E+00 | 2.58E+00 | 9.25E-02 | Yes | | intergenic_variant | MODIFIER |
| 13 | 11514669 | AX-170050540 | 0 | A | C | 0.184 | -2.11E-01 | 1.25E-01 | 2.53E+00 | 2.58E+00 | 9.25E-02 | Yes | | intergenic_variant | MODIFIER |
| 13 | 11515035 | AX-170200971 | 0 | C | A | 0.184 | -2.11E-01 | 1.25E-01 | 2.53E+00 | 2.58E+00 | 9.25E-02 | Yes | | intergenic_variant | MODIFIER |
| 13 | 11523500 | AX-170050539 | 0 | A | G | 0.184 | -2.11E-01 | 1.25E-01 | 2.53E+00 | 2.58E+00 | 9.25E-02 | Yes | Zp4-ps | intron_variant,non_coding_transcript_variant | MODIFIER |
| 13 | 11568369 | AX-170154150 | 0 | G | A | 0.184 | -2.11E-01 | 1.25E-01 | 2.53E+00 | 2.58E+00 | 9.25E-02 | Yes | Ryr2 | intron_variant | MODIFIER |
| 13 | 11578226 | AX-97830859 | 0 | C | T | 0.184 | -2.11E-01 | 1.25E-01 | 2.53E+00 | 2.58E+00 | 9.25E-02 | Yes | Ryr2 | intron_variant | MODIFIER |
| 13 | 11584040 | AX-170171051 | 0 | A | G | 0.184 | -2.11E-01 | 1.25E-01 | 2.53E+00 | 2.58E+00 | 9.25E-02 | Yes | Ryr2 | intron_variant | MODIFIER |
| 13 | 11585467 | AX-170238175 | 0 | T | C | 0.184 | -2.11E-01 | 1.25E-01 | 2.53E+00 | 2.58E+00 | 9.25E-02 | Yes | Ryr2 | synonymous_variant | LOW |
| 13 | 11586511 | AX-170417102 | 0 | T | C | 0.184 | -2.11E-01 | 1.25E-01 | 2.53E+00 | 2.58E+00 | 9.25E-02 | Yes | Ryr2 | intron_variant | MODIFIER |
| 13 | 11598476 | AX-97830612 | 0 | G | T | 0.184 | -2.11E-01 | 1.25E-01 | 2.53E+00 | 2.58E+00 | 9.25E-02 | Yes | Ryr2 | intron_variant | MODIFIER |
| 13 | 11601738 | AX-170314262 | 0 | A | G | 0.184 | -2.11E-01 | 1.25E-01 | 2.53E+00 | 2.58E+00 | 9.25E-02 | Yes | Ryr2 | intron_variant | MODIFIER |
| 13 | 11605460 | AX-170504689 | 0 | A | G | 0.184 | -2.11E-01 | 1.25E-01 | 2.53E+00 | 2.58E+00 | 9.25E-02 | Yes | Ryr2 | intron_variant | MODIFIER |
| 13 | 11605729 | AX-170396971 | 0 | T | G | 0.184 | -2.11E-01 | 1.25E-01 | 2.53E+00 | 2.58E+00 | 9.25E-02 | Yes | Ryr2 | intron_variant | MODIFIER |
| 13 | 11607600 | AX-97818228 | 0 | G | A | 0.184 | -2.11E-01 | 1.25E-01 | 2.53E+00 | 2.58E+00 | 9.25E-02 | Yes | Ryr2 | intron_variant | MODIFIER |
| 13 | 11609792 | AX-170152665 | 0 | T | A | 0.184 | -2.11E-01 | 1.25E-01 | 2.53E+00 | 2.58E+00 | 9.25E-02 | Yes | Ryr2 | intron_variant | MODIFIER |
| 13 | 11644056 | AX-170087368 | 0 | A | G | 0.184 | -2.11E-01 | 1.25E-01 | 2.53E+00 | 2.58E+00 | 9.25E-02 | Yes | Ryr2 | intron_variant | MODIFIER |
| 13 | 11654560 | AX-170482308 | 0 | C | T | 0.184 | -2.11E-01 | 1.25E-01 | 2.53E+00 | 2.58E+00 | 9.25E-02 | Yes | Ryr2 | intron_variant | MODIFIER |
| 13 | 11665866 | AX-170276085 | 0 | G | A | 0.184 | -2.11E-01 | 1.25E-01 | 2.53E+00 | 2.58E+00 | 9.25E-02 | Yes | Ryr2 | intron_variant | MODIFIER |

| | | | | | | | | | | | | | | | |
|---|---|---|---|---|---|---|---|---|---|---|---|---|---|---|---|
| 13 | 11666088 | AX-170122925 | 0 | G | A | 0.184 | -2.11E-01 | 1.25E-01 | 2.53E+00 | 2.58E+00 | 9.25E-02 | Yes | Ryr2 | intron_variant | MODIFIER |
| 13 | 11668076 | AX-97806066 | 0 | A | G | 0.184 | -2.11E-01 | 1.25E-01 | 2.53E+00 | 2.58E+00 | 9.25E-02 | Yes | Ryr2 | intron_variant | MODIFIER |
| 13 | 11668705 | AX-170150167 | 0 | T | C | 0.184 | -2.11E-01 | 1.25E-01 | 2.53E+00 | 2.58E+00 | 9.25E-02 | Yes | Ryr2 | intron_variant | MODIFIER |
| 13 | 11676920 | AX-170167612 | 0 | C | G | 0.184 | -2.11E-01 | 1.25E-01 | 2.53E+00 | 2.58E+00 | 9.25E-02 | Yes | Ryr2 | intron_variant | MODIFIER |
| 13 | 11689833 | AX-170177966 | 0 | G | C | 0.184 | -2.11E-01 | 1.25E-01 | 2.53E+00 | 2.58E+00 | 9.25E-02 | Yes | Ryr2 | intron_variant | MODIFIER |
| 13 | 11689919 | AX-97831583 | 0 | G | A | 0.184 | -2.11E-01 | 1.25E-01 | 2.53E+00 | 2.58E+00 | 9.25E-02 | Yes | Ryr2 | intron_variant | MODIFIER |
| 13 | 11710814 | AX-97837190 | 0 | C | T | 0.184 | -2.11E-01 | 1.25E-01 | 2.53E+00 | 2.58E+00 | 9.25E-02 | Yes | Ryr2 | intron_variant | MODIFIER |
| 13 | 11716642 | AX-170433764 | 0 | C | G | 0.184 | -2.11E-01 | 1.25E-01 | 2.53E+00 | 2.58E+00 | 9.25E-02 | Yes | Ryr2 | intron_variant | MODIFIER |
| 13 | 11726592 | AX-170137345 | 0 | A | G | 0.184 | -2.11E-01 | 1.25E-01 | 2.53E+00 | 2.58E+00 | 9.25E-02 | Yes | Ryr2 | intron_variant | MODIFIER |
| 13 | 11738570 | AX-170182853 | 0 | T | C | 0.184 | -2.11E-01 | 1.25E-01 | 2.53E+00 | 2.58E+00 | 9.25E-02 | Yes | Ryr2 | intron_variant | MODIFIER |
| 13 | 11749540 | AX-97791730 | 0 | T | G | 0.184 | -2.11E-01 | 1.25E-01 | 2.53E+00 | 2.58E+00 | 9.25E-02 | Yes | Ryr2 | synonymous_variant | LOW |
| 13 | 11754573 | AX-170122464 | 0 | C | T | 0.184 | -2.11E-01 | 1.25E-01 | 2.53E+00 | 2.58E+00 | 9.25E-02 | Yes | Ryr2 | intron_variant | MODIFIER |
| 13 | 11769605 | AX-170488279 | 0 | G | A | 0.184 | -2.11E-01 | 1.25E-01 | 2.53E+00 | 2.58E+00 | 9.25E-02 | Yes | Ryr2 | intron_variant | MODIFIER |
| 13 | 11769626 | AX-170411063 | 0 | C | T | 0.184 | -2.11E-01 | 1.25E-01 | 2.53E+00 | 2.58E+00 | 9.25E-02 | Yes | Ryr2 | intron_variant | MODIFIER |
| 13 | 11770749 | AX-97831083 | 0 | A | G | 0.184 | -2.11E-01 | 1.25E-01 | 2.53E+00 | 2.58E+00 | 9.25E-02 | Yes | Ryr2 | intron_variant | MODIFIER |
| 13 | 11770938 | AX-170197572 | 0 | C | T | 0.184 | -2.11E-01 | 1.25E-01 | 2.53E+00 | 2.58E+00 | 9.25E-02 | Yes | Ryr2 | intron_variant | MODIFIER |
| 13 | 11773529 | AX-170372116 | 0 | G | A | 0.184 | -2.11E-01 | 1.25E-01 | 2.53E+00 | 2.58E+00 | 9.25E-02 | Yes | Ryr2 | intron_variant | MODIFIER |
| 13 | 11776330 | AX-170296750 | 0 | T | C | 0.184 | -2.11E-01 | 1.25E-01 | 2.53E+00 | 2.58E+00 | 9.25E-02 | Yes | Ryr2 | intron_variant | MODIFIER |
| 13 | 11778389 | AX-170261627 | 0 | G | A | 0.184 | -2.11E-01 | 1.25E-01 | 2.53E+00 | 2.58E+00 | 9.25E-02 | Yes | Ryr2 | intron_variant | MODIFIER |
| 13 | 11779598 | AX-170474852 | 0 | A | G | 0.184 | -2.11E-01 | 1.25E-01 | 2.53E+00 | 2.58E+00 | 9.25E-02 | Yes | Ryr2 | intron_variant | MODIFIER |
| 13 | 11785807 | AX-97852507 | 0 | C | T | 0.184 | -2.11E-01 | 1.25E-01 | 2.53E+00 | 2.58E+00 | 9.25E-02 | Yes | Ryr2 | intron_variant | MODIFIER |
| 13 | 11786416 | AX-170050548 | 0 | G | A | 0.184 | -2.11E-01 | 1.25E-01 | 2.53E+00 | 2.58E+00 | 9.25E-02 | Yes | Ryr2 | intron_variant | MODIFIER |
| 13 | 11802698 | AX-170346158 | 0 | A | T | 0.184 | -2.11E-01 | 1.25E-01 | 2.53E+00 | 2.58E+00 | 9.25E-02 | Yes | Ryr2 | intron_variant | MODIFIER |
| 13 | 11818464 | AX-170092661 | 0 | A | G | 0.184 | -2.11E-01 | 1.25E-01 | 2.53E+00 | 2.58E+00 | 9.25E-02 | Yes | Ryr2 | intron_variant | MODIFIER |
| 13 | 11823048 | AX-170325240 | 0 | T | G | 0.184 | -2.11E-01 | 1.25E-01 | 2.53E+00 | 2.58E+00 | 9.25E-02 | Yes | Ryr2 | intron_variant | MODIFIER |
| 13 | 11824216 | AX-170411013 | 0 | T | C | 0.184 | -2.11E-01 | 1.25E-01 | 2.53E+00 | 2.58E+00 | 9.25E-02 | Yes | Ryr2 | intron_variant | MODIFIER |
| 13 | 11827461 | AX-170404394 | 0 | A | G | 0.184 | -2.11E-01 | 1.25E-01 | 2.53E+00 | 2.58E+00 | 9.25E-02 | Yes | Ryr2 | intron_variant | MODIFIER |
| 13 | 11837595 | AX-170197378 | 0 | G | T | 0.184 | -2.11E-01 | 1.25E-01 | 2.53E+00 | 2.58E+00 | 9.25E-02 | Yes | Ryr2 | intron_variant | MODIFIER |
| 13 | 11842800 | AX-170103718 | 0 | G | A | 0.184 | -2.11E-01 | 1.25E-01 | 2.53E+00 | 2.58E+00 | 9.25E-02 | Yes | Ryr2 | intron_variant | MODIFIER |
| 13 | 11846940 | AX-97822815 | 0 | A | G | 0.184 | -2.11E-01 | 1.25E-01 | 2.53E+00 | 2.58E+00 | 9.25E-02 | Yes | Ryr2 | intron_variant | MODIFIER |
| 13 | 11850982 | AX-170162205 | 0 | T | C | 0.184 | -2.11E-01 | 1.25E-01 | 2.53E+00 | 2.58E+00 | 9.25E-02 | Yes | Ryr2 | intron_variant | MODIFIER |
| 13 | 11868847 | AX-170477768 | 0 | C | T | 0.184 | -2.11E-01 | 1.25E-01 | 2.53E+00 | 2.58E+00 | 9.25E-02 | Yes | Ryr2 | intron_variant | MODIFIER |
| 13 | 11872791 | AX-170499099 | 0 | G | A | 0.184 | -2.11E-01 | 1.25E-01 | 2.53E+00 | 2.58E+00 | 9.25E-02 | Yes | Ryr2 | intron_variant | MODIFIER |
| 13 | 11875430 | AX-170312393 | 0 | T | C | 0.184 | -2.11E-01 | 1.25E-01 | 2.53E+00 | 2.58E+00 | 9.25E-02 | Yes | Ryr2 | intron_variant | MODIFIER |
| 13 | 11876975 | AX-170219419 | 0 | G | A | 0.184 | -2.11E-01 | 1.25E-01 | 2.53E+00 | 2.58E+00 | 9.25E-02 | Yes | Ryr2 | intron_variant | MODIFIER |
| 13 | 11877094 | AX-170516956 | 0 | C | A | 0.184 | -2.11E-01 | 1.25E-01 | 2.53E+00 | 2.58E+00 | 9.25E-02 | Yes | Ryr2 | intron_variant | MODIFIER |
| 13 | 11886481 | AX-170007758 | 0 | A | G | 0.184 | -2.11E-01 | 1.25E-01 | 2.53E+00 | 2.58E+00 | 9.25E-02 | Yes | Ryr2 | intron_variant | MODIFIER |
| 13 | 11890814 | AX-97822659 | 0 | A | G | 0.184 | -2.11E-01 | 1.25E-01 | 2.53E+00 | 2.58E+00 | 9.25E-02 | Yes | Ryr2 | intron_variant | MODIFIER |
| 13 | 11924526 | AX-97801778 | 0 | C | T | 0.184 | -2.11E-01 | 1.25E-01 | 2.53E+00 | 2.58E+00 | 9.25E-02 | Yes | Ryr2 | intron_variant | MODIFIER |
| 13 | 11924579 | AX-170007766 | 0 | A | G | 0.184 | -2.11E-01 | 1.25E-01 | 2.53E+00 | 2.58E+00 | 9.25E-02 | Yes | Ryr2 | intron_variant | MODIFIER |
| 13 | 11955076 | AX-170432047 | 0 | C | T | 0.184 | -2.11E-01 | 1.25E-01 | 2.53E+00 | 2.58E+00 | 9.25E-02 | Yes | Ryr2 | intron_variant | MODIFIER |
| 13 | 11956249 | AX-170206641 | 0 | C | T | 0.184 | -2.11E-01 | 1.25E-01 | 2.53E+00 | 2.58E+00 | 9.25E-02 | Yes | Ryr2 | intron_variant | MODIFIER |
| 13 | 11960929 | AX-170007767 | 0 | G | A | 0.184 | -2.11E-01 | 1.25E-01 | 2.53E+00 | 2.58E+00 | 9.25E-02 | Yes | Ryr2 | intron_variant | MODIFIER |
| 13 | 11966124 | AX-170007763 | 0 | G | A | 0.184 | -2.11E-01 | 1.25E-01 | 2.53E+00 | 2.58E+00 | 9.25E-02 | Yes | Ryr2 | intron_variant | MODIFIER |
| 13 | 11972275 | AX-170216053 | 0 | C | T | 0.184 | -2.11E-01 | 1.25E-01 | 2.53E+00 | 2.58E+00 | 9.25E-02 | Yes | Ryr2 | intron_variant | MODIFIER |
| 13 | 11973246 | AX-170291728 | 0 | A | G | 0.184 | -2.11E-01 | 1.25E-01 | 2.53E+00 | 2.58E+00 | 9.25E-02 | Yes | Ryr2 | intron_variant | MODIFIER |
| 13 | 11979428 | AX-170202313 | 0 | T | G | 0.184 | -2.11E-01 | 1.25E-01 | 2.53E+00 | 2.58E+00 | 9.25E-02 | Yes | Ryr2 | intron_variant | MODIFIER |
| 13 | 11981143 | AX-170234281 | 0 | T | C | 0.184 | -2.11E-01 | 1.25E-01 | 2.53E+00 | 2.58E+00 | 9.25E-02 | Yes | Ryr2 | intron_variant | MODIFIER |
| 13 | 11982369 | AX-170263749 | 0 | T | G | 0.184 | -2.11E-01 | 1.25E-01 | 2.53E+00 | 2.58E+00 | 9.25E-02 | Yes | Ryr2 | intron_variant | MODIFIER |
| 13 | 11982407 | AX-97855577 | 0 | C | T | 0.184 | -2.11E-01 | 1.25E-01 | 2.53E+00 | 2.58E+00 | 9.25E-02 | Yes | Ryr2 | intron_variant | MODIFIER |
| 13 | 12013826 | AX-97783516 | 0 | C | T | 0.184 | -2.11E-01 | 1.25E-01 | 2.53E+00 | 2.58E+00 | 9.25E-02 | Yes | Ryr2 | intron_variant | MODIFIER |
| 13 | 12013967 | AX-170007759 | 0 | G | T | 0.184 | -2.11E-01 | 1.25E-01 | 2.53E+00 | 2.58E+00 | 9.25E-02 | Yes | Ryr2 | intron_variant | MODIFIER |
| 13 | 12024927 | AX-170388670 | 0 | A | G | 0.184 | -2.11E-01 | 1.25E-01 | 2.53E+00 | 2.58E+00 | 9.25E-02 | Yes | Ryr2 | intron_variant | MODIFIER |
| 13 | 12039966 | AX-170444287 | 0 | T | C | 0.184 | -2.11E-01 | 1.25E-01 | 2.53E+00 | 2.58E+00 | 9.25E-02 | Yes | Ryr2 | intron_variant | MODIFIER |

| | | | | | | | | | | | | | | | |
|---|---|---|---|---|---|---|---|---|---|---|---|---|---|---|---|
| 13 | 59892350 | AX-170008428 | 1 | G | A | 0.045 | 2.75E-01 | 1.62E-01 | 3.14E+00 | 3.35E+00 | 9.25E-02 | Yes | | regulatory_region_variant | MODIFIER |
| 5 | 52657655 | AX-170164419 | 1 | T | C | 0.473 | 1.67E-01 | 9.87E-02 | 2.47E+00 | 2.53E+00 | 9.27E-02 | Yes | Sepsecs | intron_variant | MODIFIER |
| 5 | 58334091 | AX-170247186 | 0 | G | T | 0.482 | 1.69E-01 | 1.01E-01 | 2.52E+00 | 2.59E+00 | 9.27E-02 | Yes | | intergenic_variant | MODIFIER |
| 15 | 19316835 | AX-97797517 | 0 | G | A | 0.237 | -1.76E-01 | 1.05E-01 | 2.79E+00 | 2.87E+00 | 9.29E-02 | Yes | Gm49162 | intron_variant,non_coding_transcript_variant | MODIFIER |
| 5 | 132660052 | AX-170212530 | 1 | A | G | 0.152 | -1.47E-01 | 8.54E-02 | 2.27E+00 | 2.35E+00 | 9.30E-02 | no | | intergenic_variant | MODIFIER |
| 16 | 55732483 | AX-170397345 | 1 | T | C | 0.527 | 1.42E-01 | 8.49E-02 | 2.57E+00 | 2.61E+00 | 9.30E-02 | no | | intergenic_variant | MODIFIER |
| 19 | 59771162 | AX-170322484 | 1 | T | C | 0.473 | -1.05E-01 | 6.30E-02 | 2.76E+00 | 2.84E+00 | 9.32E-02 | no | | intergenic_variant | MODIFIER |
| 15 | 71848011 | AX-97833568 | 1 | G | A | 0.08 | -2.11E-01 | 1.07E-01 | 1.68E+00 | 1.74E+00 | 9.33E-02 | Yes | Col22a1 | intron_variant | MODIFIER |
| 15 | 77647616 | AX-170178622 | 1 | G | A | 0.08 | -2.11E-01 | 1.07E-01 | 1.68E+00 | 1.74E+00 | 9.33E-02 | Yes | Apol11b | upstream_gene_variant | MODIFIER |
| 19 | 51725994 | AX-170016820 | 1 | G | C | 0.482 | 1.08E-01 | 6.49E-02 | 2.83E+00 | 2.93E+00 | 9.34E-02 | no | | intergenic_variant | MODIFIER |
| 1 | 185410091 | AX-170002472 | 0 | T | G | 0.246 | 1.90E-01 | 1.13E-01 | 2.46E+00 | 2.51E+00 | 9.35E-02 | Yes | Eprs | intron_variant | MODIFIER |
| 5 | 60242146 | AX-170487737 | 2 | C | T | 0.464 | 1.74E-01 | 1.03E-01 | 2.43E+00 | 2.48E+00 | 9.36E-02 | Yes | | intergenic_variant | MODIFIER |
| 19 | 50567384 | AX-170386168 | 0 | A | G | 0.482 | 1.08E-01 | 6.48E-02 | 2.83E+00 | 2.93E+00 | 9.36E-02 | no | Sorcs1 | intron_variant | MODIFIER |
| 19 | 51666672 | AX-97802103 | 0 | T | C | 0.482 | 1.08E-01 | 6.48E-02 | 2.83E+00 | 2.93E+00 | 9.36E-02 | no | | intergenic_variant | MODIFIER |
| 19 | 51668614 | AX-170422418 | 0 | T | C | 0.482 | 1.08E-01 | 6.48E-02 | 2.83E+00 | 2.93E+00 | 9.36E-02 | no | | intergenic_variant | MODIFIER |
| 19 | 51706628 | AX-97851285 | 0 | C | T | 0.482 | 1.08E-01 | 6.48E-02 | 2.83E+00 | 2.93E+00 | 9.36E-02 | no | | intergenic_variant | MODIFIER |
| 19 | 51749924 | AX-97794021 | 0 | G | A | 0.482 | 1.08E-01 | 6.48E-02 | 2.83E+00 | 2.93E+00 | 9.36E-02 | no | | intergenic_variant | MODIFIER |
| 19 | 51774323 | AX-97822770 | 0 | C | T | 0.482 | 1.08E-01 | 6.48E-02 | 2.83E+00 | 2.93E+00 | 9.36E-02 | no | | regulatory_region_variant | MODIFIER |
| 19 | 51792470 | AX-170062679 | 0 | T | G | 0.482 | 1.08E-01 | 6.48E-02 | 2.83E+00 | 2.93E+00 | 9.36E-02 | no | Rpl31-ps18 | non_coding_transcript_exon_variant | MODIFIER |
| 19 | 51793068 | AX-97836963 | 0 | T | C | 0.482 | 1.08E-01 | 6.48E-02 | 2.83E+00 | 2.93E+00 | 9.36E-02 | no | Rpl31-ps18 | downstream_gene_variant | MODIFIER |
| 19 | 51839087 | AX-97794353 | 0 | A | G | 0.482 | 1.08E-01 | 6.48E-02 | 2.83E+00 | 2.93E+00 | 9.36E-02 | no | | intergenic_variant | MODIFIER |
| 19 | 51994134 | AX-170192346 | 0 | C | T | 0.482 | 1.08E-01 | 6.48E-02 | 2.83E+00 | 2.93E+00 | 9.36E-02 | no | | intergenic_variant | MODIFIER |
| 19 | 52042693 | AX-170309961 | 0 | G | T | 0.482 | 1.08E-01 | 6.48E-02 | 2.83E+00 | 2.93E+00 | 9.36E-02 | no | | intergenic_variant | MODIFIER |
| 13 | 7314892 | AX-170170066 | 1 | T | A | 0.179 | -2.10E-01 | 1.25E-01 | 2.54E+00 | 2.58E+00 | 9.37E-02 | Yes | | intergenic_variant | MODIFIER |
| 16 | 65312978 | AX-97846923 | 1 | A | G | 0.509 | 1.36E-01 | 8.11E-02 | 2.56E+00 | 2.62E+00 | 9.39E-02 | no | | intergenic_variant | MODIFIER |
| 8 | 65384099 | AX-170300333 | 1 | C | T | 0.143 | -2.39E-01 | 1.43E-01 | 2.67E+00 | 2.75E+00 | 9.42E-02 | Yes | | intergenic_variant | MODIFIER |
| 1 | 135641763 | AX-170161539 | 0 | A | G | 0.544 | 1.32E-01 | 7.94E-02 | 2.77E+00 | 2.85E+00 | 9.42E-02 | no | Gm38399 | intron_variant,non_coding_transcript_variant | MODIFIER |
| 5 | 134396812 | AX-170026225 | 0 | T | C | 0.158 | -1.36E-01 | 7.99E-02 | 2.28E+00 | 2.38E+00 | 9.44E-02 | no | Gtf2ird1 | intron_variant | MODIFIER |
| 7 | 79658437 | AX-170170321 | 1 | C | T | 0.027 | -2.71E-01 | 1.63E-01 | 2.71E+00 | 2.90E+00 | 9.44E-02 | Yes | Ticrr | upstream_gene_variant | MODIFIER |
| 1 | 134612591 | AX-170184156 | 0 | C | T | 0.553 | 1.36E-01 | 8.15E-02 | 2.87E+00 | 2.97E+00 | 9.45E-02 | no | Kdm5b | intron_variant | MODIFIER |
| 5 | 132035558 | AX-168409758 | 1 | C | T | 0.161 | -1.52E-01 | 8.87E-02 | 2.27E+00 | 2.35E+00 | 9.45E-02 | no | Auts2 | intron_variant,non_coding_transcript_variant | MODIFIER |
| 10 | 101703962 | AX-170447846 | 1 | A | G | 0.277 | -1.39E-01 | 8.35E-02 | 2.76E+00 | 2.84E+00 | 9.46E-02 | no | Mgat4c | intron_variant | MODIFIER |
| 5 | 136855872 | AX-170254124 | 1 | C | G | 0.152 | -1.40E-01 | 8.28E-02 | 2.39E+00 | 2.48E+00 | 9.49E-02 | no | Col26a1 | intron_variant | MODIFIER |
| 5 | 18710388 | AX-97853022 | 0 | A | G | 0.333 | 1.42E-01 | 8.52E-02 | 2.97E+00 | 3.11E+00 | 9.49E-02 | no | | intergenic_variant | MODIFIER |
| 4 | 99152211 | AX-97835125 | 1 | T | G | 0.241 | 1.93E-01 | 1.16E-01 | 2.63E+00 | 2.69E+00 | 9.50E-02 | Yes | Gm12681 | intron_variant,non_coding_transcript_variant | MODIFIER |
| 1 | 184627594 | AX-170403812 | 1 | C | T | 0.241 | 1.85E-01 | 1.11E-01 | 2.54E+00 | 2.61E+00 | 9.51E-02 | Yes | Gm37800 | intron_variant,non_coding_transcript_variant | MODIFIER |
| 4 | 78897724 | AX-170126229 | 1 | G | A | 0.446 | -1.40E-01 | 8.41E-02 | 2.56E+00 | 2.63E+00 | 9.52E-02 | no | | intergenic_variant | MODIFIER |
| 15 | 76622959 | AX-170294368 | 2 | G | A | 0.082 | -2.08E-01 | 1.08E-01 | 1.73E+00 | 1.79E+00 | 9.52E-02 | Yes | Vps28 | intron_variant | MODIFIER |
| 5 | 62895779 | AX-97843930 | 1 | T | C | 0.482 | 1.74E-01 | 1.04E-01 | 2.46E+00 | 2.51E+00 | 9.53E-02 | Yes | | regulatory_region_variant | MODIFIER |
| 13 | 10269936 | AX-97820816 | 1 | C | T | 0.188 | -2.04E-01 | 1.22E-01 | 2.52E+00 | 2.56E+00 | 9.55E-02 | Yes | Chrm3 | intron_variant | MODIFIER |
| 5 | 135743745 | AX-97833710 | 1 | T | C | 0.152 | -1.41E-01 | 8.35E-02 | 2.33E+00 | 2.42E+00 | 9.55E-02 | no | Tmem120a | intron_variant | MODIFIER |
| 15 | 6169220 | AX-170316650 | 0 | G | A | 0.325 | -1.40E-01 | 7.59E-02 | 1.86E+00 | 1.90E+00 | 9.56E-02 | no | | intergenic_variant | MODIFIER |
| 15 | 6242763 | AX-170431213 | 0 | C | T | 0.325 | -1.40E-01 | 7.59E-02 | 1.86E+00 | 1.90E+00 | 9.56E-02 | no | | intergenic_variant | MODIFIER |
| 15 | 6329609 | AX-170249671 | 0 | C | A | 0.325 | -1.40E-01 | 7.59E-02 | 1.86E+00 | 1.90E+00 | 9.56E-02 | no | Dab2 | intron_variant | MODIFIER |
| 15 | 6332391 | AX-97785355 | 0 | A | G | 0.325 | -1.40E-01 | 7.59E-02 | 1.86E+00 | 1.90E+00 | 9.56E-02 | no | Dab2 | intron_variant | MODIFIER |
| 15 | 6335865 | AX-97826617 | 0 | C | T | 0.325 | -1.40E-01 | 7.59E-02 | 1.86E+00 | 1.90E+00 | 9.56E-02 | no | Dab2 | intron_variant | MODIFIER |
| 15 | 6773245 | AX-170244157 | 0 | A | G | 0.325 | -1.40E-01 | 7.59E-02 | 1.86E+00 | 1.90E+00 | 9.56E-02 | no | Rictor | intron_variant | MODIFIER |
| 15 | 6816948 | AX-170316652 | 0 | G | A | 0.325 | -1.40E-01 | 7.59E-02 | 1.86E+00 | 1.90E+00 | 9.56E-02 | no | Osmr | synonymous_variant | LOW |
| 15 | 6822314 | AX-170334847 | 0 | T | C | 0.325 | -1.40E-01 | 7.59E-02 | 1.86E+00 | 1.90E+00 | 9.56E-02 | no | Osmr | intron_variant | MODIFIER |
| 15 | 6844628 | AX-97820223 | 0 | C | T | 0.325 | -1.40E-01 | 7.59E-02 | 1.86E+00 | 1.90E+00 | 9.56E-02 | no | Osmr | intron_variant | MODIFIER |
| 15 | 6908409 | AX-170419530 | 0 | C | A | 0.325 | -1.40E-01 | 7.59E-02 | 1.86E+00 | 1.90E+00 | 9.56E-02 | no | | intergenic_variant | MODIFIER |
| 15 | 6914420 | AX-170106672 | 0 | G | A | 0.325 | -1.40E-01 | 7.59E-02 | 1.86E+00 | 1.90E+00 | 9.56E-02 | no | | intergenic_variant | MODIFIER |
| 10 | 102234073 | AX-170363969 | 2 | G | C | 0.273 | -1.38E-01 | 8.33E-02 | 2.69E+00 | 2.77E+00 | 9.56E-02 | no | Mgat4c | intron_variant | MODIFIER |
| 4 | 99145299 | AX-97855642 | 0 | C | T | 0.246 | 1.84E-01 | 1.11E-01 | 2.63E+00 | 2.70E+00 | 9.57E-02 | Yes | Gm12681 | downstream_gene_variant | MODIFIER |

| | | | | | | | | | | | | | | | |
|---|---|---|---|---|---|---|---|---|---|---|---|---|---|---|---|
| 9 | 107476599 | AX-170255700 | 0 | T | C | 0.5 | -9.99E-02 | 6.00E-02 | 2.98E+00 | 3.15E+00 | 9.57E-02 | no | Cacna2d2 | intron_variant | MODIFIER |
| 10 | 105045579 | AX-170500327 | 1 | T | C | 0.277 | -1.36E-01 | 8.15E-02 | 2.52E+00 | 2.60E+00 | 9.58E-02 | no | | intergenic_variant | MODIFIER |
| 5 | 52555406 | AX-97794812 | 0 | T | G | 0.482 | 1.69E-01 | 1.01E-01 | 2.44E+00 | 2.50E+00 | 9.59E-02 | Yes | Lgi2 | intron_variant | MODIFIER |
| 5 | 14230752 | AX-170420523 | 0 | C | T | 0.342 | 1.42E-01 | 8.52E-02 | 2.96E+00 | 3.10E+00 | 9.60E-02 | no | Sema3e | intron_variant | MODIFIER |
| 5 | 14324878 | AX-170531812 | 0 | A | G | 0.342 | 1.42E-01 | 8.52E-02 | 2.96E+00 | 3.10E+00 | 9.60E-02 | no | | intergenic_variant | MODIFIER |
| 4 | 103772586 | AX-170492177 | 0 | G | A | 0.246 | 1.85E-01 | 1.12E-01 | 2.61E+00 | 2.68E+00 | 9.61E-02 | Yes | Dab1 | intron_variant | MODIFIER |
| 19 | 59617526 | AX-170486763 | 1 | C | T | 0.473 | -1.04E-01 | 6.30E-02 | 2.75E+00 | 2.83E+00 | 9.63E-02 | no | | intergenic_variant | MODIFIER |
| 16 | 62288057 | AX-170013197 | 0 | G | A | 0.474 | -1.44E-01 | 8.69E-02 | 2.62E+00 | 2.66E+00 | 9.65E-02 | no | | intergenic_variant | MODIFIER |
| 13 | 10157989 | AX-170050521 | 1 | C | T | 0.188 | -2.05E-01 | 1.24E-01 | 2.54E+00 | 2.59E+00 | 9.66E-02 | Yes | Chrm3 | intron_variant | MODIFIER |
| 5 | 55302394 | AX-97832551 | 1 | C | T | 0.473 | 1.61E-01 | 9.62E-02 | 2.44E+00 | 2.51E+00 | 9.66E-02 | Yes | Gm25513 | downstream_gene_variant | MODIFIER |
| 10 | 70028175 | AX-170346340 | 1 | C | T | 0.411 | -1.26E-01 | 7.61E-02 | 2.85E+00 | 2.96E+00 | 9.66E-02 | no | Ank3 | downstream_gene_variant | MODIFIER |
| 8 | 60786324 | AX-97816946 | 1 | A | G | 0.143 | -2.28E-01 | 1.37E-01 | 2.55E+00 | 2.64E+00 | 9.69E-02 | Yes | | intergenic_variant | MODIFIER |
| 1 | 192266999 | AX-170423146 | 1 | C | A | 0.277 | 1.58E-01 | 9.39E-02 | 2.33E+00 | 2.40E+00 | 9.71E-02 | Yes | Kcnh1 | intron_variant | MODIFIER |
| 7 | 88212958 | AX-170463577 | 1 | A | G | 0.027 | -2.03E-01 | 1.23E-01 | 2.71E+00 | 2.91E+00 | 9.72E-02 | Yes | | intergenic_variant | MODIFIER |
| 13 | 17483093 | AX-170486507 | 0 | T | C | 0.175 | -2.12E-01 | 1.26E-01 | 2.40E+00 | 2.44E+00 | 9.73E-02 | Yes | Sugct | intron_variant | MODIFIER |
| 13 | 17493955 | AX-170181939 | 0 | A | T | 0.175 | -2.12E-01 | 1.26E-01 | 2.40E+00 | 2.44E+00 | 9.73E-02 | Yes | Sugct | intron_variant | MODIFIER |
| 13 | 17508990 | AX-170171254 | 0 | T | C | 0.175 | -2.12E-01 | 1.26E-01 | 2.40E+00 | 2.44E+00 | 9.73E-02 | Yes | Sugct | intron_variant | MODIFIER |
| 13 | 17512824 | AX-170143749 | 0 | T | A | 0.175 | -2.12E-01 | 1.26E-01 | 2.40E+00 | 2.44E+00 | 9.73E-02 | Yes | Sugct | intron_variant | MODIFIER |
| 13 | 17583395 | AX-170535917 | 0 | T | A | 0.175 | -2.12E-01 | 1.26E-01 | 2.40E+00 | 2.44E+00 | 9.73E-02 | Yes | Sugct | intron_variant | MODIFIER |
| 13 | 17606823 | AX-170102178 | 0 | A | G | 0.175 | -2.12E-01 | 1.26E-01 | 2.40E+00 | 2.44E+00 | 9.73E-02 | Yes | Sugct | intron_variant | MODIFIER |
| 13 | 17612312 | AX-170419847 | 0 | A | G | 0.175 | -2.12E-01 | 1.26E-01 | 2.40E+00 | 2.44E+00 | 9.73E-02 | Yes | Sugct | intron_variant | MODIFIER |
| 13 | 17642584 | AX-97795533 | 0 | A | G | 0.175 | -2.12E-01 | 1.26E-01 | 2.40E+00 | 2.44E+00 | 9.73E-02 | Yes | Sugct | intron_variant | MODIFIER |
| 13 | 17652740 | AX-170007858 | 0 | G | A | 0.175 | -2.12E-01 | 1.26E-01 | 2.40E+00 | 2.44E+00 | 9.73E-02 | Yes | Sugct | intron_variant | MODIFIER |
| 13 | 17696409 | AX-97822945 | 0 | G | T | 0.175 | -2.12E-01 | 1.26E-01 | 2.40E+00 | 2.44E+00 | 9.73E-02 | Yes | Mplkip | intron_variant | MODIFIER |
| 13 | 17702135 | AX-170007854 | 0 | T | G | 0.175 | -2.12E-01 | 1.26E-01 | 2.40E+00 | 2.44E+00 | 9.73E-02 | Yes | Mplkip | downstream_gene_variant | MODIFIER |
| 13 | 17746616 | AX-97841282 | 0 | T | C | 0.175 | -2.12E-01 | 1.26E-01 | 2.40E+00 | 2.44E+00 | 9.73E-02 | Yes | Cdk13 | intron_variant | MODIFIER |
| 13 | 17765180 | AX-170156546 | 0 | T | G | 0.175 | -2.12E-01 | 1.26E-01 | 2.40E+00 | 2.44E+00 | 9.73E-02 | Yes | Cdk13 | intron_variant | MODIFIER |
| 13 | 17778056 | AX-97829957 | 0 | T | G | 0.175 | -2.12E-01 | 1.26E-01 | 2.40E+00 | 2.44E+00 | 9.73E-02 | Yes | Cdk13 | intron_variant | MODIFIER |
| 13 | 17855135 | AX-170007859 | 0 | C | A | 0.175 | -2.12E-01 | 1.26E-01 | 2.40E+00 | 2.44E+00 | 9.73E-02 | Yes | | regulatory_region_variant | MODIFIER |
| 13 | 17855164 | AX-170007853 | 0 | C | T | 0.175 | -2.12E-01 | 1.26E-01 | 2.40E+00 | 2.44E+00 | 9.73E-02 | Yes | | regulatory_region_variant | MODIFIER |
| 13 | 18129278 | AX-170370370 | 0 | G | A | 0.175 | -2.12E-01 | 1.26E-01 | 2.40E+00 | 2.44E+00 | 9.73E-02 | Yes | Pou6f2 | intron_variant | MODIFIER |
| 13 | 18258686 | AX-97808731 | 0 | G | T | 0.175 | -2.12E-01 | 1.26E-01 | 2.40E+00 | 2.44E+00 | 9.73E-02 | Yes | Pou6f2 | intron_variant | MODIFIER |
| 13 | 18292269 | AX-170215703 | 0 | G | A | 0.175 | -2.12E-01 | 1.26E-01 | 2.40E+00 | 2.44E+00 | 9.73E-02 | Yes | Pou6f2 | intron_variant | MODIFIER |
| 13 | 18296794 | AX-170425042 | 0 | C | T | 0.175 | -2.12E-01 | 1.26E-01 | 2.40E+00 | 2.44E+00 | 9.73E-02 | Yes | Pou6f2 | intron_variant | MODIFIER |
| 13 | 18402648 | AX-170007867 | 0 | C | T | 0.175 | -2.12E-01 | 1.26E-01 | 2.40E+00 | 2.44E+00 | 9.73E-02 | Yes | Pou6f2 | upstream_gene_variant | MODIFIER |
| 13 | 18454170 | AX-170343640 | 0 | T | G | 0.175 | -2.12E-01 | 1.26E-01 | 2.40E+00 | 2.44E+00 | 9.73E-02 | Yes | | intergenic_variant | MODIFIER |
| 13 | 18462189 | AX-170455164 | 0 | G | A | 0.175 | -2.12E-01 | 1.26E-01 | 2.40E+00 | 2.44E+00 | 9.73E-02 | Yes | | intergenic_variant | MODIFIER |
| 15 | 76752814 | AX-170012024 | 0 | T | C | 0.088 | -1.92E-01 | 1.03E-01 | 1.82E+00 | 1.89E+00 | 9.74E-02 | Yes | Arhgap39 | intron_variant | MODIFIER |
| 19 | 60723747 | AX-170396222 | 1 | G | T | 0.455 | -1.04E-01 | 6.32E-02 | 2.77E+00 | 2.86E+00 | 9.74E-02 | no | | intergenic_variant | MODIFIER |
| 16 | 70205037 | AX-170137937 | 0 | A | G | 0.491 | 1.21E-01 | 7.31E-02 | 2.52E+00 | 2.60E+00 | 9.74E-02 | no | | intergenic_variant | MODIFIER |
| 13 | 5450052 | AX-170432677 | 0 | G | C | 0.202 | -1.91E-01 | 1.16E-01 | 2.69E+00 | 2.78E+00 | 9.75E-02 | Yes | Gm40660 | upstream_gene_variant | MODIFIER |
| 13 | 6055943 | AX-97827083 | 0 | A | G | 0.202 | -1.91E-01 | 1.16E-01 | 2.69E+00 | 2.78E+00 | 9.75E-02 | Yes | | intergenic_variant | MODIFIER |
| 13 | 6088458 | AX-170506912 | 0 | G | C | 0.202 | -1.91E-01 | 1.16E-01 | 2.69E+00 | 2.78E+00 | 9.75E-02 | Yes | | intergenic_variant | MODIFIER |
| 13 | 6505258 | AX-170036836 | 0 | A | G | 0.202 | -1.91E-01 | 1.16E-01 | 2.69E+00 | 2.78E+00 | 9.75E-02 | Yes | | intergenic_variant | MODIFIER |
| 19 | 59455875 | AX-170385279 | 0 | T | C | 0.474 | -1.04E-01 | 6.30E-02 | 2.75E+00 | 2.84E+00 | 9.77E-02 | no | Emx2 | upstream_gene_variant | MODIFIER |
| 19 | 59476042 | AX-170475289 | 0 | T | C | 0.474 | -1.04E-01 | 6.30E-02 | 2.75E+00 | 2.84E+00 | 9.77E-02 | no | | intergenic_variant | MODIFIER |
| 19 | 59482329 | AX-97830166 | 0 | A | G | 0.474 | -1.04E-01 | 6.30E-02 | 2.75E+00 | 2.84E+00 | 9.77E-02 | no | | intergenic_variant | MODIFIER |
| 19 | 59487308 | AX-170232563 | 0 | C | T | 0.474 | -1.04E-01 | 6.30E-02 | 2.75E+00 | 2.84E+00 | 9.77E-02 | no | | intergenic_variant | MODIFIER |
| 19 | 59487478 | AX-170420092 | 0 | A | G | 0.474 | -1.04E-01 | 6.30E-02 | 2.75E+00 | 2.84E+00 | 9.77E-02 | no | | intergenic_variant | MODIFIER |
| 19 | 59491942 | AX-170405869 | 0 | G | T | 0.474 | -1.04E-01 | 6.30E-02 | 2.75E+00 | 2.84E+00 | 9.77E-02 | no | 2700089I24Rik | upstream_gene_variant | MODIFIER |
| 19 | 59560409 | AX-170500833 | 0 | A | G | 0.474 | -1.04E-01 | 6.30E-02 | 2.75E+00 | 2.84E+00 | 9.77E-02 | no | 2700089I24Rik | downstream_gene_variant | MODIFIER |
| 19 | 59570744 | AX-170304556 | 0 | A | G | 0.474 | -1.04E-01 | 6.30E-02 | 2.75E+00 | 2.84E+00 | 9.77E-02 | no | | intergenic_variant | MODIFIER |
| 19 | 59570821 | AX-170393787 | 0 | A | G | 0.474 | -1.04E-01 | 6.30E-02 | 2.75E+00 | 2.84E+00 | 9.77E-02 | no | | intergenic_variant | MODIFIER |
| 19 | 59571628 | AX-170094139 | 0 | A | C | 0.474 | -1.04E-01 | 6.30E-02 | 2.75E+00 | 2.84E+00 | 9.77E-02 | no | | intergenic_variant | MODIFIER |

| | | | | | | | | | | | | | | | |
|---|---|---|---|---|---|---|---|---|---|---|---|---|---|---|---|
| 19 | 59595190 | AX-97812701 | 0 | C | G | 0.474 | -1.04E-01 | 6.30E-02 | 2.75E+00 | 2.84E+00 | 9.77E-02 | no | | intergenic_variant | MODIFIER |
| 19 | 59596866 | AX-170380488 | 0 | A | G | 0.474 | -1.04E-01 | 6.30E-02 | 2.75E+00 | 2.84E+00 | 9.77E-02 | no | | intergenic_variant | MODIFIER |
| 19 | 59602612 | AX-170062840 | 0 | A | G | 0.474 | -1.04E-01 | 6.30E-02 | 2.75E+00 | 2.84E+00 | 9.77E-02 | no | | intergenic_variant | MODIFIER |
| 19 | 59618996 | AX-170062839 | 0 | T | C | 0.474 | -1.04E-01 | 6.30E-02 | 2.75E+00 | 2.84E+00 | 9.77E-02 | no | | intergenic_variant | MODIFIER |
| 19 | 59665187 | AX-170138765 | 0 | T | C | 0.474 | -1.04E-01 | 6.30E-02 | 2.75E+00 | 2.84E+00 | 9.77E-02 | no | AC123621.1 | intron_variant,non_coding_transcript_variant | MODIFIER |
| 19 | 59688431 | AX-170290754 | 0 | C | T | 0.474 | -1.04E-01 | 6.30E-02 | 2.75E+00 | 2.84E+00 | 9.77E-02 | no | AC123621.2 | upstream_gene_variant | MODIFIER |
| 19 | 59724733 | AX-170118637 | 0 | A | C | 0.474 | -1.04E-01 | 6.30E-02 | 2.75E+00 | 2.84E+00 | 9.77E-02 | no | | intergenic_variant | MODIFIER |
| 19 | 59732623 | AX-170160822 | 0 | G | A | 0.474 | -1.04E-01 | 6.30E-02 | 2.75E+00 | 2.84E+00 | 9.77E-02 | no | | intergenic_variant | MODIFIER |
| 19 | 59733283 | AX-170125936 | 0 | G | C | 0.474 | -1.04E-01 | 6.30E-02 | 2.75E+00 | 2.84E+00 | 9.77E-02 | no | | intergenic_variant | MODIFIER |
| 19 | 59748042 | AX-170530418 | 0 | C | T | 0.474 | -1.04E-01 | 6.30E-02 | 2.75E+00 | 2.84E+00 | 9.77E-02 | no | | intergenic_variant | MODIFIER |
| 19 | 59757896 | AX-170447892 | 0 | A | G | 0.474 | -1.04E-01 | 6.30E-02 | 2.75E+00 | 2.84E+00 | 9.77E-02 | no | | intergenic_variant | MODIFIER |
| 19 | 59822841 | AX-170094141 | 0 | G | A | 0.474 | -1.04E-01 | 6.30E-02 | 2.75E+00 | 2.84E+00 | 9.77E-02 | no | | intergenic_variant | MODIFIER |
| 19 | 59839604 | AX-170016935 | 0 | C | G | 0.474 | -1.04E-01 | 6.30E-02 | 2.75E+00 | 2.84E+00 | 9.77E-02 | no | | intergenic_variant | MODIFIER |
| 19 | 59846661 | AX-170158419 | 0 | A | T | 0.474 | -1.04E-01 | 6.30E-02 | 2.75E+00 | 2.84E+00 | 9.77E-02 | no | | intergenic_variant | MODIFIER |
| 19 | 59858221 | AX-170433921 | 0 | T | C | 0.474 | -1.04E-01 | 6.30E-02 | 2.75E+00 | 2.84E+00 | 9.77E-02 | no | | intergenic_variant | MODIFIER |
| 19 | 59877986 | AX-170062845 | 0 | T | C | 0.474 | -1.04E-01 | 6.30E-02 | 2.75E+00 | 2.84E+00 | 9.77E-02 | no | | intergenic_variant | MODIFIER |
| 19 | 59878072 | AX-170016939 | 0 | C | T | 0.474 | -1.04E-01 | 6.30E-02 | 2.75E+00 | 2.84E+00 | 9.77E-02 | no | | intergenic_variant | MODIFIER |
| 19 | 59881668 | AX-170062847 | 0 | T | C | 0.474 | -1.04E-01 | 6.30E-02 | 2.75E+00 | 2.84E+00 | 9.77E-02 | no | | intergenic_variant | MODIFIER |
| 19 | 59881928 | AX-170062848 | 0 | T | C | 0.474 | -1.04E-01 | 6.30E-02 | 2.75E+00 | 2.84E+00 | 9.77E-02 | no | | intergenic_variant | MODIFIER |
| 19 | 59885962 | AX-97830182 | 0 | T | A | 0.474 | -1.04E-01 | 6.30E-02 | 2.75E+00 | 2.84E+00 | 9.77E-02 | no | | intergenic_variant | MODIFIER |
| 19 | 59901749 | AX-170433308 | 0 | A | G | 0.474 | -1.04E-01 | 6.30E-02 | 2.75E+00 | 2.84E+00 | 9.77E-02 | no | Rab11fip2 | downstream_gene_variant | MODIFIER |
| 19 | 59902462 | AX-170257233 | 0 | T | A | 0.474 | -1.04E-01 | 6.30E-02 | 2.75E+00 | 2.84E+00 | 9.77E-02 | no | Rab11fip2 | downstream_gene_variant | MODIFIER |
| 19 | 59902823 | AX-170138568 | 0 | C | G | 0.474 | -1.04E-01 | 6.30E-02 | 2.75E+00 | 2.84E+00 | 9.77E-02 | no | Rab11fip2 | downstream_gene_variant | MODIFIER |
| 19 | 59914733 | AX-170390960 | 0 | T | A | 0.474 | -1.04E-01 | 6.30E-02 | 2.75E+00 | 2.84E+00 | 9.77E-02 | no | Rab11fip2 | intron_variant | MODIFIER |
| 19 | 59918199 | AX-170382381 | 0 | C | T | 0.474 | -1.04E-01 | 6.30E-02 | 2.75E+00 | 2.84E+00 | 9.77E-02 | no | Rab11fip2 | intron_variant | MODIFIER |
| 19 | 59922095 | AX-170113034 | 0 | A | G | 0.474 | -1.04E-01 | 6.30E-02 | 2.75E+00 | 2.84E+00 | 9.77E-02 | no | Rab11fip2 | intron_variant | MODIFIER |
| 19 | 59936443 | AX-170094142 | 0 | G | A | 0.474 | -1.04E-01 | 6.30E-02 | 2.75E+00 | 2.84E+00 | 9.77E-02 | no | Rab11fip2 | intron_variant | MODIFIER |
| 19 | 59959859 | AX-97837146 | 0 | A | G | 0.474 | -1.04E-01 | 6.30E-02 | 2.75E+00 | 2.84E+00 | 9.77E-02 | no | | intergenic_variant | MODIFIER |
| 19 | 59966694 | AX-170431778 | 0 | A | G | 0.474 | -1.04E-01 | 6.30E-02 | 2.75E+00 | 2.84E+00 | 9.77E-02 | no | | intergenic_variant | MODIFIER |
| 19 | 59978868 | AX-170429784 | 0 | G | A | 0.474 | -1.04E-01 | 6.30E-02 | 2.75E+00 | 2.84E+00 | 9.77E-02 | no | | intergenic_variant | MODIFIER |
| 19 | 59994682 | AX-170165504 | 0 | C | A | 0.474 | -1.04E-01 | 6.30E-02 | 2.75E+00 | 2.84E+00 | 9.77E-02 | no | | intergenic_variant | MODIFIER |
| 19 | 60000265 | AX-170088492 | 0 | T | C | 0.474 | -1.04E-01 | 6.30E-02 | 2.75E+00 | 2.84E+00 | 9.77E-02 | no | | intergenic_variant | MODIFIER |
| 19 | 60006480 | AX-170513678 | 0 | G | C | 0.474 | -1.04E-01 | 6.30E-02 | 2.75E+00 | 2.84E+00 | 9.77E-02 | no | | intergenic_variant | MODIFIER |
| 19 | 60007179 | AX-170153053 | 0 | G | A | 0.474 | -1.04E-01 | 6.30E-02 | 2.75E+00 | 2.84E+00 | 9.77E-02 | no | | intergenic_variant | MODIFIER |
| 19 | 60015100 | AX-97834064 | 0 | G | A | 0.474 | -1.04E-01 | 6.30E-02 | 2.75E+00 | 2.84E+00 | 9.77E-02 | no | | regulatory_region_variant | MODIFIER |
| 19 | 60026030 | AX-97815539 | 0 | A | G | 0.474 | -1.04E-01 | 6.30E-02 | 2.75E+00 | 2.84E+00 | 9.77E-02 | no | | intergenic_variant | MODIFIER |
| 19 | 60057932 | AX-170196055 | 0 | T | C | 0.474 | -1.04E-01 | 6.30E-02 | 2.75E+00 | 2.84E+00 | 9.77E-02 | no | | intergenic_variant | MODIFIER |
| 19 | 60057993 | AX-170229714 | 0 | C | T | 0.474 | -1.04E-01 | 6.30E-02 | 2.75E+00 | 2.84E+00 | 9.77E-02 | no | | intergenic_variant | MODIFIER |
| 19 | 60058727 | AX-97821929 | 0 | A | C | 0.474 | -1.04E-01 | 6.30E-02 | 2.75E+00 | 2.84E+00 | 9.77E-02 | no | | intergenic_variant | MODIFIER |
| 19 | 60073497 | AX-170233820 | 0 | G | A | 0.474 | -1.04E-01 | 6.30E-02 | 2.75E+00 | 2.84E+00 | 9.77E-02 | no | | regulatory_region_variant | MODIFIER |
| 19 | 60075086 | AX-170162713 | 0 | C | A | 0.474 | -1.04E-01 | 6.30E-02 | 2.75E+00 | 2.84E+00 | 9.77E-02 | no | AC105061.2 | upstream_gene_variant | MODIFIER |
| 19 | 60075137 | AX-170399058 | 0 | A | G | 0.474 | -1.04E-01 | 6.30E-02 | 2.75E+00 | 2.84E+00 | 9.77E-02 | no | AC105061.2 | upstream_gene_variant | MODIFIER |
| 19 | 60075877 | AX-97789046 | 0 | C | T | 0.474 | -1.04E-01 | 6.30E-02 | 2.75E+00 | 2.84E+00 | 9.77E-02 | no | AC105061.2 | upstream_gene_variant | MODIFIER |
| 19 | 60076368 | AX-170237434 | 0 | A | G | 0.474 | -1.04E-01 | 6.30E-02 | 2.75E+00 | 2.84E+00 | 9.77E-02 | no | AC105061.2 | upstream_gene_variant | MODIFIER |
| 19 | 60108124 | AX-170236137 | 0 | A | G | 0.474 | -1.04E-01 | 6.30E-02 | 2.75E+00 | 2.84E+00 | 9.77E-02 | no | | intergenic_variant | MODIFIER |
| 19 | 60112788 | AX-170407929 | 0 | A | G | 0.474 | -1.04E-01 | 6.30E-02 | 2.75E+00 | 2.84E+00 | 9.77E-02 | no | | intergenic_variant | MODIFIER |
| 19 | 60112848 | AX-170088493 | 0 | G | A | 0.474 | -1.04E-01 | 6.30E-02 | 2.75E+00 | 2.84E+00 | 9.77E-02 | no | | intergenic_variant | MODIFIER |
| 19 | 60139122 | AX-97832506 | 0 | T | C | 0.474 | -1.04E-01 | 6.30E-02 | 2.75E+00 | 2.84E+00 | 9.77E-02 | no | | intergenic_variant | MODIFIER |
| 19 | 60152923 | AX-170176043 | 0 | A | G | 0.474 | -1.04E-01 | 6.30E-02 | 2.75E+00 | 2.84E+00 | 9.77E-02 | no | E330013P04Rik | intron_variant,non_coding_transcript_variant | MODIFIER |
| 19 | 60154032 | AX-170424738 | 0 | A | G | 0.474 | -1.04E-01 | 6.30E-02 | 2.75E+00 | 2.84E+00 | 9.77E-02 | no | E330013P04Rik | intron_variant,non_coding_transcript_variant | MODIFIER |
| 19 | 60167099 | AX-170487680 | 0 | T | G | 0.474 | -1.04E-01 | 6.30E-02 | 2.75E+00 | 2.84E+00 | 9.77E-02 | no | E330013P04Rik | downstream_gene_variant | MODIFIER |
| 19 | 60171299 | AX-170161790 | 0 | T | C | 0.474 | -1.04E-01 | 6.30E-02 | 2.75E+00 | 2.84E+00 | 9.77E-02 | no | | intergenic_variant | MODIFIER |
| 19 | 60218402 | AX-170062860 | 0 | T | C | 0.474 | -1.04E-01 | 6.30E-02 | 2.75E+00 | 2.84E+00 | 9.77E-02 | no | Fam204a | intron_variant | MODIFIER |
| 19 | 60223470 | AX-170062857 | 0 | A | G | 0.474 | -1.04E-01 | 6.30E-02 | 2.75E+00 | 2.84E+00 | 9.77E-02 | no | 4933412A08Rik | upstream_gene_variant | MODIFIER |

| | | | | | | | | | | | | | | | |
|---|---|---|---|---|---|---|---|---|---|---|---|---|---|---|---|
| 19 | 60229350 | AX-170256615 | 0 | A | G | 0.474 | -1.04E-01 | 6.30E-02 | 2.75E+00 | 2.84E+00 | 9.77E-02 | no | 4933412A08Rik | downstream_gene_variant | MODIFIER |
| 19 | 60243141 | AX-97855109 | 0 | G | A | 0.474 | -1.04E-01 | 6.30E-02 | 2.75E+00 | 2.84E+00 | 9.77E-02 | no | Gm28351 | intron_variant,non_coding_transcript_variant | MODIFIER |
| 19 | 60264336 | AX-97800002 | 0 | A | G | 0.474 | -1.04E-01 | 6.30E-02 | 2.75E+00 | 2.84E+00 | 9.77E-02 | no | | intergenic_variant | MODIFIER |
| 19 | 60293563 | AX-170143850 | 0 | G | C | 0.474 | -1.04E-01 | 6.30E-02 | 2.75E+00 | 2.84E+00 | 9.77E-02 | no | | intergenic_variant | MODIFIER |
| 19 | 60315846 | AX-170016947 | 0 | T | A | 0.474 | -1.04E-01 | 6.30E-02 | 2.75E+00 | 2.84E+00 | 9.77E-02 | no | | intergenic_variant | MODIFIER |
| 19 | 60341177 | AX-97808142 | 0 | A | G | 0.474 | -1.04E-01 | 6.30E-02 | 2.75E+00 | 2.84E+00 | 9.77E-02 | no | | intergenic_variant | MODIFIER |
| 19 | 60358302 | AX-170282761 | 0 | G | T | 0.474 | -1.04E-01 | 6.30E-02 | 2.75E+00 | 2.84E+00 | 9.77E-02 | no | | intergenic_variant | MODIFIER |
| 19 | 60362021 | AX-97795026 | 0 | T | C | 0.474 | -1.04E-01 | 6.30E-02 | 2.75E+00 | 2.84E+00 | 9.77E-02 | no | | intergenic_variant | MODIFIER |
| 19 | 60377542 | AX-170187877 | 0 | G | T | 0.474 | -1.04E-01 | 6.30E-02 | 2.75E+00 | 2.84E+00 | 9.77E-02 | no | | intergenic_variant | MODIFIER |
| 19 | 60379033 | AX-170016945 | 0 | G | A | 0.474 | -1.04E-01 | 6.30E-02 | 2.75E+00 | 2.84E+00 | 9.77E-02 | no | | intergenic_variant | MODIFIER |
| 19 | 60379277 | AX-170016948 | 0 | C | T | 0.474 | -1.04E-01 | 6.30E-02 | 2.75E+00 | 2.84E+00 | 9.77E-02 | no | | intergenic_variant | MODIFIER |
| 19 | 60389889 | AX-170016942 | 0 | G | A | 0.474 | -1.04E-01 | 6.30E-02 | 2.75E+00 | 2.84E+00 | 9.77E-02 | no | | intergenic_variant | MODIFIER |
| 19 | 60408369 | AX-170016946 | 0 | T | C | 0.474 | -1.04E-01 | 6.30E-02 | 2.75E+00 | 2.84E+00 | 9.77E-02 | no | | intergenic_variant | MODIFIER |
| 19 | 60440052 | AX-170512640 | 0 | G | A | 0.474 | -1.04E-01 | 6.30E-02 | 2.75E+00 | 2.84E+00 | 9.77E-02 | no | | intergenic_variant | MODIFIER |
| 19 | 60463783 | AX-97837699 | 0 | G | A | 0.474 | -1.04E-01 | 6.30E-02 | 2.75E+00 | 2.84E+00 | 9.77E-02 | no | Prlhr | downstream_gene_variant | MODIFIER |
| 13 | 21325851 | AX-170189658 | 0 | A | G | 0.167 | -2.06E-01 | 1.23E-01 | 2.39E+00 | 2.44E+00 | 9.78E-02 | Yes | | regulatory_region_variant | MODIFIER |
| 8 | 26097163 | AX-170030377 | 0 | A | C | 0.351 | -1.03E-01 | 6.28E-02 | 2.71E+00 | 2.82E+00 | 9.79E-02 | no | Hook3 | intron_variant | MODIFIER |
| 4 | 6417965 | AX-170445823 | 0 | C | T | 0.044 | 2.82E-01 | 1.70E-01 | 3.02E+00 | 3.23E+00 | 9.79E-02 | Yes | Nsmaf | intron_variant | MODIFIER |
| 4 | 106702352 | AX-170023606 | 0 | T | C | 0.237 | 1.94E-01 | 1.18E-01 | 2.63E+00 | 2.70E+00 | 9.79E-02 | Yes | Mroh7 | intron_variant | MODIFIER |
| 4 | 106707081 | AX-170394866 | 0 | C | G | 0.237 | 1.94E-01 | 1.18E-01 | 2.63E+00 | 2.70E+00 | 9.79E-02 | Yes | Mroh7 | intron_variant | MODIFIER |
| 4 | 106799146 | AX-170323580 | 0 | T | C | 0.237 | 1.94E-01 | 1.18E-01 | 2.63E+00 | 2.70E+00 | 9.79E-02 | Yes | Acot11 | intron_variant | MODIFIER |
| 4 | 106804490 | AX-170151412 | 0 | G | A | 0.237 | 1.94E-01 | 1.18E-01 | 2.63E+00 | 2.70E+00 | 9.79E-02 | Yes | Acot11 | intron_variant | MODIFIER |
| 4 | 106832717 | AX-97855491 | 0 | G | A | 0.237 | 1.94E-01 | 1.18E-01 | 2.63E+00 | 2.70E+00 | 9.79E-02 | Yes | | regulatory_region_variant | MODIFIER |
| 13 | 8000155 | AX-170101064 | 1 | A | G | 0.188 | -2.05E-01 | 1.24E-01 | 2.55E+00 | 2.61E+00 | 9.80E-02 | Yes | | intergenic_variant | MODIFIER |
| 13 | 8395903 | AX-97834287 | 1 | C | G | 0.188 | -2.05E-01 | 1.24E-01 | 2.55E+00 | 2.61E+00 | 9.80E-02 | Yes | Adarb2 | intron_variant | MODIFIER |
| 13 | 9287154 | AX-170218611 | 1 | G | T | 0.188 | -2.05E-01 | 1.24E-01 | 2.55E+00 | 2.61E+00 | 9.80E-02 | Yes | Dip2c | intron_variant | MODIFIER |
| 13 | 9670022 | AX-170289616 | 1 | A | G | 0.188 | -2.05E-01 | 1.24E-01 | 2.55E+00 | 2.61E+00 | 9.80E-02 | Yes | Dip2c | downstream_gene_variant | MODIFIER |
| 13 | 10064695 | AX-170481803 | 1 | G | A | 0.188 | -2.05E-01 | 1.24E-01 | 2.55E+00 | 2.61E+00 | 9.80E-02 | Yes | Chrm3 | intron_variant | MODIFIER |
| 13 | 10410304 | AX-170129484 | 1 | C | T | 0.188 | -2.05E-01 | 1.24E-01 | 2.55E+00 | 2.61E+00 | 9.80E-02 | Yes | Gm26861 | intron_variant,non_coding_transcript_variant | MODIFIER |
| 19 | 60165896 | AX-170296058 | 1 | G | A | 0.473 | -1.04E-01 | 6.29E-02 | 2.75E+00 | 2.83E+00 | 9.81E-02 | no | E330013P04Rik | downstream_gene_variant | MODIFIER |
| 5 | 58060106 | AX-97799968 | 2 | A | T | 0.455 | 1.65E-01 | 9.85E-02 | 2.36E+00 | 2.42E+00 | 9.81E-02 | Yes | Pcdh7 | intron_variant | MODIFIER |
| 18 | 86485688 | AX-97806983 | 0 | C | T | 0.158 | -1.62E-01 | 9.83E-02 | 2.54E+00 | 2.65E+00 | 9.82E-02 | no | Neto1 | intron_variant | MODIFIER |
| 13 | 9730489 | AX-170007727 | 1 | A | G | 0.188 | -2.04E-01 | 1.23E-01 | 2.56E+00 | 2.61E+00 | 9.82E-02 | Yes | Zmynd11 | intron_variant | MODIFIER |
| 13 | 10346560 | AX-170007740 | 1 | C | T | 0.188 | -2.04E-01 | 1.23E-01 | 2.56E+00 | 2.61E+00 | 9.82E-02 | Yes | Chrm3 | intron_variant | MODIFIER |
| 13 | 8436460 | AX-170190892 | 0 | T | G | 0.202 | -1.87E-01 | 1.13E-01 | 2.62E+00 | 2.67E+00 | 9.84E-02 | Yes | Adarb2 | intron_variant | MODIFIER |
| 3 | 38596049 | AX-97813473 | 0 | C | T | 0.202 | 1.56E-01 | 9.28E-02 | 3.20E+00 | 3.34E+00 | 9.85E-02 | Yes | | intergenic_variant | MODIFIER |
| 18 | 76203681 | AX-97812013 | 1 | G | C | 0.438 | 1.42E-01 | 8.60E-02 | 2.54E+00 | 2.60E+00 | 9.86E-02 | no | | intergenic_variant | MODIFIER |
| 5 | 66075903 | AX-170348544 | 0 | A | G | 0.482 | 1.66E-01 | 1.01E-01 | 2.51E+00 | 2.57E+00 | 9.86E-02 | Yes | Rbm47 | intron_variant | MODIFIER |
| 5 | 58106306 | AX-170179085 | 1 | C | T | 0.482 | 1.69E-01 | 1.02E-01 | 2.42E+00 | 2.48E+00 | 9.90E-02 | Yes | Pcdh7 | intron_variant | MODIFIER |
| 16 | 56060800 | AX-170483362 | 0 | C | T | 0.526 | 1.33E-01 | 8.08E-02 | 2.55E+00 | 2.59E+00 | 9.90E-02 | no | Senp7 | intron_variant,non_coding_transcript_variant | MODIFIER |
| 5 | 63386877 | AX-170404757 | 0 | C | A | 0.482 | 1.69E-01 | 1.03E-01 | 2.60E+00 | 2.67E+00 | 9.92E-02 | Yes | | intergenic_variant | MODIFIER |
| 4 | 68607406 | AX-170428998 | 0 | T | C | 0.342 | 1.81E-01 | 1.09E-01 | 3.06E+00 | 3.16E+00 | 9.92E-02 | Yes | | intergenic_variant | MODIFIER |
| 19 | 59903664 | AX-170479923 | 1 | A | T | 0.473 | -1.03E-01 | 6.29E-02 | 2.74E+00 | 2.83E+00 | 9.93E-02 | no | Rab11fip2 | downstream_gene_variant | MODIFIER |
| 14 | 122944755 | AX-170010880 | 0 | G | C | 0.307 | 1.29E-01 | 7.90E-02 | 2.73E+00 | 2.84E+00 | 9.96E-02 | no | Tmtc4 | synonymous_variant | LOW |
| 3 | 18625013 | AX-170486511 | 0 | T | G | 0.132 | -1.65E-01 | 1.01E-01 | 2.62E+00 | 2.71E+00 | 9.96E-02 | no | Gm42944 | non_coding_transcript_exon_variant | MODIFIER |
| 3 | 18634900 | AX-170228366 | 0 | G | A | 0.132 | -1.65E-01 | 1.01E-01 | 2.62E+00 | 2.71E+00 | 9.96E-02 | no | | intergenic_variant | MODIFIER |
| 7 | 99153838 | AX-170029611 | 1 | A | G | 0.375 | -1.68E-01 | 1.01E-01 | 3.05E+00 | 3.17E+00 | 9.98E-02 | no | Dgat2 | 3_prime_UTR_variant | MODIFIER |
| 13 | 19362745 | AX-170161125 | 0 | T | G | 0.184 | -1.85E-01 | 1.12E-01 | 2.50E+00 | 2.59E+00 | 9.98E-02 | Yes | Stard3nl | intron_variant | MODIFIER |
| 5 | 58259199 | AX-170112950 | 1 | C | G | 0.482 | 1.70E-01 | 1.03E-01 | 2.46E+00 | 2.51E+00 | 9.99E-02 | Yes | | intergenic_variant | MODIFIER |
| 13 | 10564279 | AX-170359088 | 1 | C | T | 0.188 | -2.03E-01 | 1.23E-01 | 2.55E+00 | 2.60E+00 | 1.00E-01 | Yes | Gm26861 | intron_variant,non_coding_transcript_variant | MODIFIER |
| 13 | 11432339 | AX-170386805 | 1 | C | T | 0.188 | -2.03E-01 | 1.23E-01 | 2.55E+00 | 2.60E+00 | 1.00E-01 | Yes | | regulatory_region_variant | MODIFIER |
| 12 | 108530962 | AX-170470888 | 0 | A | G | 0.132 | -2.85E-01 | 1.69E-01 | 3.34E+00 | 3.48E+00 | 1.00E-01 | Yes | Eml1 | intron_variant | MODIFIER |
| 10 | 104806876 | AX-170427524 | 0 | A | T | 0.307 | -1.32E-01 | 8.06E-02 | 2.78E+00 | 2.88E+00 | 1.00E-01 | no | | intergenic_variant | MODIFIER |
| 8 | 54135063 | AX-170351813 | 0 | T | A | 0.149 | -2.24E-01 | 1.37E-01 | 2.88E+00 | 2.99E+00 | 1.00E-01 | Yes | Vegfc | intron_variant | MODIFIER |

| | | | | | | | | | | | | | | | |
|---|---|---|---|---|---|---|---|---|---|---|---|---|---|---|---|
| 13 | 18399231 | AX-170227560 | 1 | G | A | 0.179 | -2.08E-01 | 1.26E-01 | 2.44E+00 | 2.49E+00 | 1.00E-01 | Yes | Pou6f2 | upstream_gene_variant | MODIFIER |
| 4 | 107784186 | AX-97854699 | 0 | C | T | 0.272 | 1.72E-01 | 1.05E-01 | 2.70E+00 | 2.78E+00 | 1.01E-01 | Yes | 4933424M12Rik | upstream_gene_variant | MODIFIER |
| 9 | 94554600 | AX-170033031 | 0 | T | C | 0.14 | -1.86E-01 | 1.14E-01 | 2.80E+00 | 2.91E+00 | 1.01E-01 | Yes | | intergenic_variant | MODIFIER |
| 18 | 76186867 | AX-170330496 | 0 | G | A | 0.439 | 1.42E-01 | 8.63E-02 | 2.54E+00 | 2.60E+00 | 1.01E-01 | no | | intergenic_variant | MODIFIER |
| 18 | 76188961 | AX-170061361 | 0 | T | G | 0.439 | 1.42E-01 | 8.63E-02 | 2.54E+00 | 2.60E+00 | 1.01E-01 | no | | intergenic_variant | MODIFIER |
| 18 | 76305461 | AX-170278389 | 0 | A | T | 0.439 | 1.42E-01 | 8.63E-02 | 2.54E+00 | 2.60E+00 | 1.01E-01 | no | Smad2 | downstream_gene_variant | MODIFIER |
| 18 | 76325399 | AX-97798124 | 0 | C | A | 0.439 | 1.42E-01 | 8.63E-02 | 2.54E+00 | 2.60E+00 | 1.01E-01 | no | AC164009.1 | downstream_gene_variant | MODIFIER |
| 18 | 76439827 | AX-97835932 | 0 | G | A | 0.439 | 1.42E-01 | 8.63E-02 | 2.54E+00 | 2.60E+00 | 1.01E-01 | no | | intergenic_variant | MODIFIER |
| 18 | 76626569 | AX-170093960 | 0 | T | C | 0.439 | 1.42E-01 | 8.63E-02 | 2.54E+00 | 2.60E+00 | 1.01E-01 | no | AC102081.1 | intron_variant,non_coding_transcript_variant | MODIFIER |
| 18 | 76627413 | AX-170360328 | 0 | A | G | 0.439 | 1.42E-01 | 8.63E-02 | 2.54E+00 | 2.60E+00 | 1.01E-01 | no | AC102081.1 | intron_variant,non_coding_transcript_variant | MODIFIER |
| 18 | 76628040 | AX-170179219 | 0 | T | C | 0.439 | 1.42E-01 | 8.63E-02 | 2.54E+00 | 2.60E+00 | 1.01E-01 | no | AC102081.1 | intron_variant,non_coding_transcript_variant | MODIFIER |
| 18 | 76628083 | AX-170146850 | 0 | C | T | 0.439 | 1.42E-01 | 8.63E-02 | 2.54E+00 | 2.60E+00 | 1.01E-01 | no | AC102081.1 | intron_variant,non_coding_transcript_variant | MODIFIER |
| 18 | 76644392 | AX-170439557 | 0 | C | T | 0.439 | 1.42E-01 | 8.63E-02 | 2.54E+00 | 2.60E+00 | 1.01E-01 | no | | intergenic_variant | MODIFIER |
| 18 | 76655344 | AX-97801657 | 0 | C | A | 0.439 | 1.42E-01 | 8.63E-02 | 2.54E+00 | 2.60E+00 | 1.01E-01 | no | | intergenic_variant | MODIFIER |
| 18 | 76662628 | AX-170248284 | 0 | T | C | 0.439 | 1.42E-01 | 8.63E-02 | 2.54E+00 | 2.60E+00 | 1.01E-01 | no | | intergenic_variant | MODIFIER |
| 18 | 76675853 | AX-170428202 | 0 | G | A | 0.439 | 1.42E-01 | 8.63E-02 | 2.54E+00 | 2.60E+00 | 1.01E-01 | no | | intergenic_variant | MODIFIER |
| 18 | 76675909 | AX-97790855 | 0 | T | C | 0.439 | 1.42E-01 | 8.63E-02 | 2.54E+00 | 2.60E+00 | 1.01E-01 | no | | intergenic_variant | MODIFIER |
| 18 | 76677667 | AX-170123588 | 0 | C | T | 0.439 | 1.42E-01 | 8.63E-02 | 2.54E+00 | 2.60E+00 | 1.01E-01 | no | | regulatory_region_variant | MODIFIER |
| 18 | 76684358 | AX-170383654 | 0 | T | C | 0.439 | 1.42E-01 | 8.63E-02 | 2.54E+00 | 2.60E+00 | 1.01E-01 | no | | intergenic_variant | MODIFIER |
| 18 | 76686429 | AX-170114865 | 0 | C | T | 0.439 | 1.42E-01 | 8.63E-02 | 2.54E+00 | 2.60E+00 | 1.01E-01 | no | | intergenic_variant | MODIFIER |
| 18 | 76716646 | AX-97841453 | 0 | C | T | 0.439 | 1.42E-01 | 8.63E-02 | 2.54E+00 | 2.60E+00 | 1.01E-01 | no | | intergenic_variant | MODIFIER |
| 18 | 76840370 | AX-170534518 | 0 | A | G | 0.439 | 1.42E-01 | 8.63E-02 | 2.54E+00 | 2.60E+00 | 1.01E-01 | no | | intergenic_variant | MODIFIER |
| 18 | 76846279 | AX-170121925 | 0 | A | C | 0.439 | 1.42E-01 | 8.63E-02 | 2.54E+00 | 2.60E+00 | 1.01E-01 | no | | intergenic_variant | MODIFIER |
| 18 | 76853051 | AX-97845628 | 0 | G | A | 0.439 | 1.42E-01 | 8.63E-02 | 2.54E+00 | 2.60E+00 | 1.01E-01 | no | Skor2 | upstream_gene_variant | MODIFIER |
| 18 | 76887877 | AX-170286175 | 0 | C | T | 0.439 | 1.42E-01 | 8.63E-02 | 2.54E+00 | 2.60E+00 | 1.01E-01 | no | Skor2 | intron_variant | MODIFIER |
| 18 | 76889219 | AX-170329784 | 0 | A | G | 0.439 | 1.42E-01 | 8.63E-02 | 2.54E+00 | 2.60E+00 | 1.01E-01 | no | Skor2 | intron_variant | MODIFIER |
| 18 | 76893942 | AX-170226746 | 0 | T | C | 0.439 | 1.42E-01 | 8.63E-02 | 2.54E+00 | 2.60E+00 | 1.01E-01 | no | Skor2 | intron_variant | MODIFIER |
| 18 | 76894113 | AX-170408872 | 0 | A | G | 0.439 | 1.42E-01 | 8.63E-02 | 2.54E+00 | 2.60E+00 | 1.01E-01 | no | Skor2 | synonymous_variant | LOW |
| 18 | 76894173 | AX-170251960 | 0 | C | T | 0.439 | 1.42E-01 | 8.63E-02 | 2.54E+00 | 2.60E+00 | 1.01E-01 | no | Skor2 | intron_variant | MODIFIER |
| 18 | 76963209 | AX-170061378 | 0 | G | A | 0.439 | 1.42E-01 | 8.63E-02 | 2.54E+00 | 2.60E+00 | 1.01E-01 | no | Hdhd2 | intron_variant | MODIFIER |
| 18 | 76988883 | AX-170016030 | 0 | C | T | 0.439 | 1.42E-01 | 8.63E-02 | 2.54E+00 | 2.60E+00 | 1.01E-01 | no | Katnal2 | downstream_gene_variant | MODIFIER |
| 18 | 76996663 | AX-170016028 | 0 | C | A | 0.439 | 1.42E-01 | 8.63E-02 | 2.54E+00 | 2.60E+00 | 1.01E-01 | no | Katnal2 | intron_variant | MODIFIER |
| 18 | 77007002 | AX-170016025 | 0 | T | C | 0.439 | 1.42E-01 | 8.63E-02 | 2.54E+00 | 2.60E+00 | 1.01E-01 | no | Katnal2 | intron_variant | MODIFIER |
| 18 | 77010595 | AX-170016031 | 0 | G | A | 0.439 | 1.42E-01 | 8.63E-02 | 2.54E+00 | 2.60E+00 | 1.01E-01 | no | Katnal2 | intron_variant | MODIFIER |
| 18 | 77048171 | AX-170016029 | 0 | T | A | 0.439 | 1.42E-01 | 8.63E-02 | 2.54E+00 | 2.60E+00 | 1.01E-01 | no | Katnal2 | upstream_gene_variant | MODIFIER |
| 18 | 77109350 | AX-170016027 | 0 | A | G | 0.439 | 1.42E-01 | 8.63E-02 | 2.54E+00 | 2.60E+00 | 1.01E-01 | no | Pias2 | intron_variant | MODIFIER |
| 18 | 77120912 | AX-170122239 | 0 | A | G | 0.439 | 1.42E-01 | 8.63E-02 | 2.54E+00 | 2.60E+00 | 1.01E-01 | no | Pias2 | intron_variant | MODIFIER |
| 18 | 77126776 | AX-170454373 | 0 | T | C | 0.439 | 1.42E-01 | 8.63E-02 | 2.54E+00 | 2.60E+00 | 1.01E-01 | no | Pias2 | intron_variant | MODIFIER |
| 18 | 77135637 | AX-170275773 | 0 | C | T | 0.439 | 1.42E-01 | 8.63E-02 | 2.54E+00 | 2.60E+00 | 1.01E-01 | no | Pias2 | intron_variant | MODIFIER |
| 18 | 77140617 | AX-170321561 | 0 | A | G | 0.439 | 1.42E-01 | 8.63E-02 | 2.54E+00 | 2.60E+00 | 1.01E-01 | no | Pias2 | intron_variant | MODIFIER |
| 18 | 77150728 | AX-170400266 | 0 | A | C | 0.439 | 1.42E-01 | 8.63E-02 | 2.54E+00 | 2.60E+00 | 1.01E-01 | no | Pias2 | downstream_gene_variant | MODIFIER |
| 18 | 77181512 | AX-170103774 | 0 | T | C | 0.439 | 1.42E-01 | 8.63E-02 | 2.54E+00 | 2.60E+00 | 1.01E-01 | no | St8sia5 | upstream_gene_variant | MODIFIER |
| 18 | 77194908 | AX-170098712 | 0 | T | A | 0.439 | 1.42E-01 | 8.63E-02 | 2.54E+00 | 2.60E+00 | 1.01E-01 | no | St8sia5 | intron_variant | MODIFIER |
| 18 | 77208008 | AX-170433540 | 0 | G | A | 0.439 | 1.42E-01 | 8.63E-02 | 2.54E+00 | 2.60E+00 | 1.01E-01 | no | St8sia5 | intron_variant | MODIFIER |
| 18 | 77215780 | AX-170323817 | 0 | A | T | 0.439 | 1.42E-01 | 8.63E-02 | 2.54E+00 | 2.60E+00 | 1.01E-01 | no | St8sia5 | intron_variant | MODIFIER |
| 18 | 77271975 | AX-97832891 | 0 | C | T | 0.439 | 1.42E-01 | 8.63E-02 | 2.54E+00 | 2.60E+00 | 1.01E-01 | no | | intergenic_variant | MODIFIER |
| 18 | 77295197 | AX-170088379 | 0 | T | C | 0.439 | 1.42E-01 | 8.63E-02 | 2.54E+00 | 2.60E+00 | 1.01E-01 | no | Loxhd1 | intron_variant | MODIFIER |
| 6 | 102502323 | AX-97786665 | 1 | C | T | 0.107 | 1.94E-01 | 1.18E-01 | 2.98E+00 | 3.19E+00 | 1.01E-01 | Yes | Cntn3 | intron_variant | MODIFIER |
| 13 | 19176799 | AX-170092679 | 0 | G | A | 0.175 | -2.08E-01 | 1.26E-01 | 2.48E+00 | 2.52E+00 | 1.01E-01 | Yes | Tcrg-V7 | upstream_gene_variant | MODIFIER |
| 4 | 84056656 | AX-170449705 | 1 | A | T | 0.312 | 1.78E-01 | 1.09E-01 | 2.85E+00 | 2.94E+00 | 1.01E-01 | Yes | | intergenic_variant | MODIFIER |
| 16 | 55699474 | AX-97848999 | 1 | A | T | 0.455 | -1.38E-01 | 8.41E-02 | 2.57E+00 | 2.61E+00 | 1.01E-01 | no | | regulatory_region_variant | MODIFIER |
| 1 | 121057563 | AX-170276787 | 2 | G | T | 0.6 | 1.30E-01 | 7.88E-02 | 2.47E+00 | 2.55E+00 | 1.01E-01 | no | | intergenic_variant | MODIFIER |
| 18 | 77981272 | AX-170501516 | 1 | C | A | 0.446 | 1.37E-01 | 8.37E-02 | 2.51E+00 | 2.58E+00 | 1.01E-01 | no | Epg5 | intron_variant | MODIFIER |
| 19 | 59538355 | AX-170366652 | 0 | A | T | 0.474 | -1.03E-01 | 6.30E-02 | 2.75E+00 | 2.83E+00 | 1.01E-01 | no | 2700089I24Rik | intron_variant,non_coding_transcript_variant | MODIFIER |

| | | | | | | | | | | | | | | | |
|---|---|---|---|---|---|---|---|---|---|---|---|---|---|---|---|
| 16 | 55265390 | AX-170057487 | 0 | A | C | 0.535 | 1.37E-01 | 8.29E-02 | 2.43E+00 | 2.47E+00 | 1.01E-01 | no | Zpld1 | intron_variant | MODIFIER |
| 16 | 55700104 | AX-170319839 | 0 | G | A | 0.535 | 1.37E-01 | 8.29E-02 | 2.43E+00 | 2.47E+00 | 1.01E-01 | no | | intergenic_variant | MODIFIER |
| 16 | 52300750 | AX-170519331 | 0 | A | C | 0.465 | -1.33E-01 | 8.09E-02 | 2.51E+00 | 2.56E+00 | 1.01E-01 | no | Alcam | intron_variant | MODIFIER |
| 6 | 104928386 | AX-170120440 | 0 | C | T | 0.07 | -3.55E-01 | 2.12E-01 | 3.30E+00 | 3.47E+00 | 1.01E-01 | Yes | | intergenic_variant | MODIFIER |
| 6 | 111494398 | AX-170230490 | 0 | G | A | 0.07 | -3.55E-01 | 2.12E-01 | 3.30E+00 | 3.47E+00 | 1.01E-01 | Yes | Grm7 | intron_variant | MODIFIER |
| 6 | 147302151 | AX-170307182 | 0 | C | G | 0.07 | -3.55E-01 | 2.12E-01 | 3.30E+00 | 3.47E+00 | 1.01E-01 | Yes | | regulatory_region_variant | MODIFIER |
| 5 | 66166700 | AX-170361309 | 1 | T | C | 0.473 | 1.63E-01 | 9.82E-02 | 2.39E+00 | 2.44E+00 | 1.01E-01 | Yes | Gm22929 | upstream_gene_variant | MODIFIER |
| 8 | 78130968 | AX-97791314 | 1 | T | C | 0.393 | -1.05E-01 | 6.32E-02 | 3.22E+00 | 3.36E+00 | 1.02E-01 | no | | intergenic_variant | MODIFIER |
| 5 | 52771680 | AX-170141061 | 1 | G | A | 0.473 | 1.71E-01 | 1.02E-01 | 2.32E+00 | 2.36E+00 | 1.02E-01 | Yes | Pi4k2b | downstream_gene_variant | MODIFIER |
| 18 | 62053827 | AX-170015838 | 0 | T | C | 0.184 | -1.70E-01 | 1.05E-01 | 2.84E+00 | 2.95E+00 | 1.02E-01 | no | | intergenic_variant | MODIFIER |
| 9 | 7935371 | AX-170166252 | 0 | C | T | 0.474 | 1.29E-01 | 7.85E-02 | 2.42E+00 | 2.52E+00 | 1.02E-01 | no | Yap1 | intron_variant | MODIFIER |
| 18 | 86188161 | AX-170161035 | 1 | T | C | 0.152 | -1.62E-01 | 9.95E-02 | 2.57E+00 | 2.68E+00 | 1.02E-01 | no | | intergenic_variant | MODIFIER |
| 7 | 20020858 | AX-170206517 | 2 | C | T | 0.427 | 1.46E-01 | 8.95E-02 | 2.62E+00 | 2.74E+00 | 1.02E-01 | Yes | Nlrp9b | intron_variant | MODIFIER |
| 18 | 79953783 | AX-170217879 | 0 | G | A | 0.465 | 1.35E-01 | 8.13E-02 | 2.34E+00 | 2.40E+00 | 1.02E-01 | no | | intergenic_variant | MODIFIER |
| 4 | 106761760 | AX-170023605 | 0 | C | T | 0.246 | 1.90E-01 | 1.17E-01 | 2.65E+00 | 2.71E+00 | 1.02E-01 | Yes | Acot11 | intron_variant | MODIFIER |
| 4 | 106769913 | AX-170380649 | 0 | A | G | 0.246 | 1.90E-01 | 1.17E-01 | 2.65E+00 | 2.71E+00 | 1.02E-01 | Yes | Acot11 | intron_variant | MODIFIER |
| 4 | 106769935 | AX-170463432 | 0 | C | T | 0.246 | 1.90E-01 | 1.17E-01 | 2.65E+00 | 2.71E+00 | 1.02E-01 | Yes | Acot11 | intron_variant | MODIFIER |
| 4 | 106770435 | AX-170070268 | 0 | T | C | 0.246 | 1.90E-01 | 1.17E-01 | 2.65E+00 | 2.71E+00 | 1.02E-01 | Yes | Acot11 | intron_variant | MODIFIER |
| 4 | 106771049 | AX-170070267 | 0 | A | G | 0.246 | 1.90E-01 | 1.17E-01 | 2.65E+00 | 2.71E+00 | 1.02E-01 | Yes | Acot11 | intron_variant | MODIFIER |
| 4 | 106792880 | AX-170070269 | 0 | T | C | 0.246 | 1.90E-01 | 1.17E-01 | 2.65E+00 | 2.71E+00 | 1.02E-01 | Yes | Acot11 | intron_variant | MODIFIER |
| 4 | 106793496 | AX-170304899 | 0 | C | T | 0.246 | 1.90E-01 | 1.17E-01 | 2.65E+00 | 2.71E+00 | 1.02E-01 | Yes | Acot11 | intron_variant | MODIFIER |
| 4 | 106799075 | AX-170528566 | 0 | T | A | 0.246 | 1.90E-01 | 1.17E-01 | 2.65E+00 | 2.71E+00 | 1.02E-01 | Yes | Acot11 | intron_variant | MODIFIER |
| 4 | 106819887 | AX-170139325 | 0 | G | A | 0.246 | 1.90E-01 | 1.17E-01 | 2.65E+00 | 2.71E+00 | 1.02E-01 | Yes | | intergenic_variant | MODIFIER |
| 4 | 125134425 | AX-170488844 | 0 | G | A | 0.246 | 1.73E-01 | 1.06E-01 | 2.65E+00 | 2.72E+00 | 1.02E-01 | Yes | | intergenic_variant | MODIFIER |
| 8 | 72763396 | AX-170350750 | 0 | A | G | 0.193 | -2.09E-01 | 1.26E-01 | 3.22E+00 | 3.36E+00 | 1.03E-01 | Yes | F2rl3 | 3_prime_UTR_variant | MODIFIER |
| 8 | 74298061 | AX-170472809 | 0 | G | T | 0.193 | -2.09E-01 | 1.26E-01 | 3.22E+00 | 3.36E+00 | 1.03E-01 | Yes | | intergenic_variant | MODIFIER |
| 8 | 74391739 | AX-170462124 | 0 | A | C | 0.386 | -1.05E-01 | 6.31E-02 | 3.22E+00 | 3.36E+00 | 1.03E-01 | no | | intergenic_variant | MODIFIER |
| 8 | 75357685 | AX-170511266 | 0 | T | C | 0.193 | -2.09E-01 | 1.26E-01 | 3.22E+00 | 3.36E+00 | 1.03E-01 | Yes | | intergenic_variant | MODIFIER |
| 8 | 75494001 | AX-170391447 | 0 | A | G | 0.386 | -1.05E-01 | 6.31E-02 | 3.22E+00 | 3.36E+00 | 1.03E-01 | no | Iqcm | intron_variant | MODIFIER |
| 8 | 77684670 | AX-170031068 | 0 | A | G | 0.386 | -1.05E-01 | 6.31E-02 | 3.22E+00 | 3.36E+00 | 1.03E-01 | no | Ednra | intron_variant | MODIFIER |
| 8 | 78106291 | AX-170031079 | 0 | T | C | 0.193 | -2.09E-01 | 1.26E-01 | 3.22E+00 | 3.36E+00 | 1.03E-01 | Yes | Gm45288 | upstream_gene_variant | MODIFIER |
| 13 | 8853215 | AX-170290126 | 1 | T | C | 0.188 | -2.02E-01 | 1.23E-01 | 2.49E+00 | 2.53E+00 | 1.03E-01 | Yes | Wdr37 | intron_variant | MODIFIER |
| 13 | 9482107 | AX-170219157 | 1 | A | G | 0.179 | -2.03E-01 | 1.24E-01 | 2.53E+00 | 2.58E+00 | 1.03E-01 | Yes | Dip2c | intron_variant | MODIFIER |
| 13 | 9782805 | AX-97826924 | 1 | G | A | 0.179 | -2.03E-01 | 1.24E-01 | 2.53E+00 | 2.58E+00 | 1.03E-01 | Yes | | intergenic_variant | MODIFIER |
| 18 | 76887265 | AX-170448840 | 1 | A | C | 0.446 | 1.40E-01 | 8.55E-02 | 2.48E+00 | 2.55E+00 | 1.03E-01 | no | Skor2 | intron_variant | MODIFIER |
| 15 | 67885987 | AX-170290074 | 0 | T | A | 0.088 | -1.86E-01 | 1.02E-01 | 1.82E+00 | 1.90E+00 | 1.03E-01 | Yes | | intergenic_variant | MODIFIER |
| 15 | 67886120 | AX-97820966 | 0 | G | A | 0.088 | -1.86E-01 | 1.02E-01 | 1.82E+00 | 1.90E+00 | 1.03E-01 | Yes | | intergenic_variant | MODIFIER |
| 16 | 56882641 | AX-170428358 | 0 | G | A | 0.465 | -1.39E-01 | 8.50E-02 | 2.53E+00 | 2.56E+00 | 1.03E-01 | no | Tmem45a | intron_variant | MODIFIER |
| 9 | 7974015 | AX-97813794 | 0 | T | C | 0.465 | 1.24E-01 | 7.57E-02 | 2.50E+00 | 2.60E+00 | 1.03E-01 | no | Yap1 | intron_variant | MODIFIER |
| 5 | 132029879 | AX-170227758 | 0 | T | C | 0.158 | -1.49E-01 | 8.90E-02 | 2.26E+00 | 2.33E+00 | 1.03E-01 | no | Auts2 | intron_variant,non_coding_transcript_variant | MODIFIER |
| 15 | 76906355 | AX-170284303 | 1 | G | A | 0.08 | -2.08E-01 | 1.09E-01 | 1.70E+00 | 1.76E+00 | 1.03E-01 | Yes | Rpl8 | downstream_gene_variant | MODIFIER |
| 15 | 77062538 | AX-170347668 | 1 | C | T | 0.08 | -2.08E-01 | 1.09E-01 | 1.70E+00 | 1.76E+00 | 1.03E-01 | Yes | | regulatory_region_variant | MODIFIER |
| 9 | 92757426 | AX-170254654 | 0 | T | C | 0.132 | -1.91E-01 | 1.18E-01 | 2.82E+00 | 2.93E+00 | 1.03E-01 | Yes | 4933400C23Rik | intron_variant,non_coding_transcript_variant | MODIFIER |
| 19 | 60501120 | AX-170325246 | 1 | C | T | 0.455 | -1.02E-01 | 6.31E-02 | 2.76E+00 | 2.85E+00 | 1.03E-01 | no | | intergenic_variant | MODIFIER |
| 5 | 132037347 | AX-170391014 | 0 | G | T | 0.149 | -1.49E-01 | 8.93E-02 | 2.25E+00 | 2.33E+00 | 1.03E-01 | no | Auts2 | intron_variant,non_coding_transcript_variant | MODIFIER |
| 5 | 132038558 | AX-170351183 | 0 | G | A | 0.149 | -1.49E-01 | 8.93E-02 | 2.25E+00 | 2.33E+00 | 1.03E-01 | no | Auts2 | intron_variant,non_coding_transcript_variant | MODIFIER |
| 5 | 132041492 | AX-170420909 | 0 | C | A | 0.149 | -1.49E-01 | 8.93E-02 | 2.25E+00 | 2.33E+00 | 1.03E-01 | no | Auts2 | intron_variant,non_coding_transcript_variant | MODIFIER |
| 16 | 58029573 | AX-170231843 | 1 | C | T | 0.464 | -1.39E-01 | 8.50E-02 | 2.53E+00 | 2.56E+00 | 1.03E-01 | no | Gm49575 | intron_variant,non_coding_transcript_variant | MODIFIER |
| 13 | 5891687 | AX-170277114 | 0 | C | G | 0.202 | -1.79E-01 | 1.10E-01 | 2.64E+00 | 2.71E+00 | 1.04E-01 | Yes | Gm47507 | intron_variant,non_coding_transcript_variant | MODIFIER |
| 19 | 60565637 | AX-170231336 | 1 | T | C | 0.455 | -1.02E-01 | 6.32E-02 | 2.75E+00 | 2.84E+00 | 1.04E-01 | no | Cacul1 | intron_variant | MODIFIER |
| 16 | 55611945 | AX-170151405 | 0 | T | C | 0.465 | -1.39E-01 | 8.50E-02 | 2.53E+00 | 2.56E+00 | 1.04E-01 | no | | intergenic_variant | MODIFIER |
| 1 | 181047231 | AX-170438930 | 1 | A | G | 0.241 | 1.82E-01 | 1.12E-01 | 2.46E+00 | 2.53E+00 | 1.04E-01 | Yes | 9130409I23Rik | upstream_gene_variant | MODIFIER |
| 4 | 116356613 | AX-97805679 | 0 | G | A | 0.228 | 1.80E-01 | 1.11E-01 | 2.74E+00 | 2.84E+00 | 1.04E-01 | Yes | Mast2 | intron_variant | MODIFIER |
| 18 | 77127544 | AX-170219860 | 1 | G | A | 0.446 | 1.39E-01 | 8.55E-02 | 2.48E+00 | 2.55E+00 | 1.04E-01 | no | Pias2 | intron_variant | MODIFIER |

| | | | | | | | | | | | | | | | |
|---|---|---|---|---|---|---|---|---|---|---|---|---|---|---|---|
| 13 | 17669189 | AX-170007856 | 1 | A | T | 0.179 | -2.04E-01 | 1.25E-01 | 2.43E+00 | 2.48E+00 | 1.04E-01 | Yes | Sugct | intron_variant | MODIFIER |
| 15 | 5811113 | AX-170226935 | 1 | A | G | 0.312 | -1.31E-01 | 7.43E-02 | 1.96E+00 | 2.02E+00 | 1.04E-01 | no | | intergenic_variant | MODIFIER |
| 5 | 128885754 | AX-170225511 | 1 | C | T | 0.152 | -1.52E-01 | 8.96E-02 | 2.16E+00 | 2.23E+00 | 1.04E-01 | no | Rimbp2 | intron_variant | MODIFIER |
| 8 | 55107040 | AX-97803258 | 0 | T | C | 0.149 | -1.95E-01 | 1.20E-01 | 2.96E+00 | 3.12E+00 | 1.04E-01 | Yes | | intergenic_variant | MODIFIER |
| 13 | 9887570 | AX-170442244 | 0 | C | A | 0.184 | -1.85E-01 | 1.13E-01 | 2.42E+00 | 2.48E+00 | 1.04E-01 | Yes | Chrm3 | intron_variant | MODIFIER |
| 10 | 101420652 | AX-170003781 | 0 | G | A | 0.272 | -1.37E-01 | 8.49E-02 | 2.61E+00 | 2.69E+00 | 1.04E-01 | no | | intergenic_variant | MODIFIER |
| 15 | 45488468 | AX-170128552 | 0 | A | C | 0.026 | -2.33E-01 | 1.06E-01 | 1.36E+00 | 1.43E+00 | 1.05E-01 | Yes | | intergenic_variant | MODIFIER |
| 1 | 185493986 | AX-170002474 | 0 | C | T | 0.228 | 1.58E-01 | 9.79E-02 | 2.61E+00 | 2.73E+00 | 1.05E-01 | Yes | 5033404E19Rik | upstream_gene_variant | MODIFIER |
| 16 | 55267430 | AX-170057486 | 0 | T | C | 0.465 | -1.38E-01 | 8.50E-02 | 2.53E+00 | 2.56E+00 | 1.05E-01 | no | Zpld1 | intron_variant | MODIFIER |
| 16 | 55267849 | AX-170250980 | 0 | G | C | 0.465 | -1.38E-01 | 8.50E-02 | 2.53E+00 | 2.56E+00 | 1.05E-01 | no | Zpld1 | intron_variant | MODIFIER |
| 16 | 55274075 | AX-170492333 | 0 | G | C | 0.465 | -1.38E-01 | 8.50E-02 | 2.53E+00 | 2.56E+00 | 1.05E-01 | no | Zpld1 | intron_variant | MODIFIER |
| 16 | 55321748 | AX-170120394 | 0 | C | T | 0.465 | -1.38E-01 | 8.50E-02 | 2.53E+00 | 2.56E+00 | 1.05E-01 | no | | regulatory_region_variant | MODIFIER |
| 16 | 55336859 | AX-170100421 | 0 | T | C | 0.465 | -1.38E-01 | 8.50E-02 | 2.53E+00 | 2.56E+00 | 1.05E-01 | no | | intergenic_variant | MODIFIER |
| 16 | 55348397 | AX-170417161 | 0 | A | C | 0.465 | -1.38E-01 | 8.50E-02 | 2.53E+00 | 2.56E+00 | 1.05E-01 | no | | intergenic_variant | MODIFIER |
| 16 | 55349500 | AX-170107475 | 0 | A | G | 0.465 | -1.38E-01 | 8.50E-02 | 2.53E+00 | 2.56E+00 | 1.05E-01 | no | | intergenic_variant | MODIFIER |
| 16 | 55361571 | AX-170461479 | 0 | G | A | 0.465 | -1.38E-01 | 8.50E-02 | 2.53E+00 | 2.56E+00 | 1.05E-01 | no | | intergenic_variant | MODIFIER |
| 16 | 55392460 | AX-170270916 | 0 | T | A | 0.465 | -1.38E-01 | 8.50E-02 | 2.53E+00 | 2.56E+00 | 1.05E-01 | no | | intergenic_variant | MODIFIER |
| 16 | 55416301 | AX-170296307 | 0 | G | A | 0.465 | -1.38E-01 | 8.50E-02 | 2.53E+00 | 2.56E+00 | 1.05E-01 | no | | intergenic_variant | MODIFIER |
| 16 | 55417203 | AX-97855506 | 0 | A | G | 0.465 | -1.38E-01 | 8.50E-02 | 2.53E+00 | 2.56E+00 | 1.05E-01 | no | | intergenic_variant | MODIFIER |
| 16 | 55451230 | AX-97801691 | 0 | A | C | 0.465 | -1.38E-01 | 8.50E-02 | 2.53E+00 | 2.56E+00 | 1.05E-01 | no | | intergenic_variant | MODIFIER |
| 16 | 55452865 | AX-170013107 | 0 | A | G | 0.465 | -1.38E-01 | 8.50E-02 | 2.53E+00 | 2.56E+00 | 1.05E-01 | no | | intergenic_variant | MODIFIER |
| 16 | 55524223 | AX-97822013 | 0 | G | A | 0.465 | -1.38E-01 | 8.50E-02 | 2.53E+00 | 2.56E+00 | 1.05E-01 | no | | intergenic_variant | MODIFIER |
| 16 | 55526517 | AX-170114336 | 0 | T | C | 0.465 | -1.38E-01 | 8.50E-02 | 2.53E+00 | 2.56E+00 | 1.05E-01 | no | | regulatory_region_variant | MODIFIER |
| 16 | 55527948 | AX-170444824 | 0 | T | C | 0.465 | -1.38E-01 | 8.50E-02 | 2.53E+00 | 2.56E+00 | 1.05E-01 | no | | intergenic_variant | MODIFIER |
| 16 | 55528910 | AX-170507404 | 0 | T | C | 0.465 | -1.38E-01 | 8.50E-02 | 2.53E+00 | 2.56E+00 | 1.05E-01 | no | | intergenic_variant | MODIFIER |
| 16 | 55530487 | AX-170357838 | 0 | A | G | 0.465 | -1.38E-01 | 8.50E-02 | 2.53E+00 | 2.56E+00 | 1.05E-01 | no | | intergenic_variant | MODIFIER |
| 16 | 55530535 | AX-170172437 | 0 | G | C | 0.465 | -1.38E-01 | 8.50E-02 | 2.53E+00 | 2.56E+00 | 1.05E-01 | no | | intergenic_variant | MODIFIER |
| 16 | 55572385 | AX-170356710 | 0 | T | G | 0.465 | -1.38E-01 | 8.50E-02 | 2.53E+00 | 2.56E+00 | 1.05E-01 | no | | intergenic_variant | MODIFIER |
| 16 | 55580582 | AX-97831907 | 0 | T | C | 0.465 | -1.38E-01 | 8.50E-02 | 2.53E+00 | 2.56E+00 | 1.05E-01 | no | | intergenic_variant | MODIFIER |
| 16 | 55603836 | AX-170235593 | 0 | A | G | 0.465 | -1.38E-01 | 8.50E-02 | 2.53E+00 | 2.56E+00 | 1.05E-01 | no | | intergenic_variant | MODIFIER |
| 16 | 55609645 | AX-170013105 | 0 | G | A | 0.465 | -1.38E-01 | 8.50E-02 | 2.53E+00 | 2.56E+00 | 1.05E-01 | no | | intergenic_variant | MODIFIER |
| 16 | 55611911 | AX-170013106 | 0 | C | G | 0.465 | -1.38E-01 | 8.50E-02 | 2.53E+00 | 2.56E+00 | 1.05E-01 | no | | intergenic_variant | MODIFIER |
| 16 | 55623334 | AX-170516453 | 0 | G | A | 0.465 | -1.38E-01 | 8.50E-02 | 2.53E+00 | 2.56E+00 | 1.05E-01 | no | | intergenic_variant | MODIFIER |
| 16 | 55671417 | AX-170159138 | 0 | T | C | 0.465 | -1.38E-01 | 8.50E-02 | 2.53E+00 | 2.56E+00 | 1.05E-01 | no | | intergenic_variant | MODIFIER |
| 16 | 55671467 | AX-170196374 | 0 | G | T | 0.465 | -1.38E-01 | 8.50E-02 | 2.53E+00 | 2.56E+00 | 1.05E-01 | no | | intergenic_variant | MODIFIER |
| 16 | 55671928 | AX-170375595 | 0 | T | C | 0.465 | -1.38E-01 | 8.50E-02 | 2.53E+00 | 2.56E+00 | 1.05E-01 | no | | intergenic_variant | MODIFIER |
| 16 | 55701887 | AX-170117880 | 0 | C | G | 0.465 | -1.38E-01 | 8.50E-02 | 2.53E+00 | 2.56E+00 | 1.05E-01 | no | | intergenic_variant | MODIFIER |
| 16 | 55715323 | AX-170224987 | 0 | C | T | 0.465 | -1.38E-01 | 8.50E-02 | 2.53E+00 | 2.56E+00 | 1.05E-01 | no | | intergenic_variant | MODIFIER |
| 16 | 55719459 | AX-170429057 | 0 | A | T | 0.465 | -1.38E-01 | 8.50E-02 | 2.53E+00 | 2.56E+00 | 1.05E-01 | no | | intergenic_variant | MODIFIER |
| 16 | 55732561 | AX-97831925 | 0 | A | G | 0.465 | -1.38E-01 | 8.50E-02 | 2.53E+00 | 2.56E+00 | 1.05E-01 | no | | intergenic_variant | MODIFIER |
| 16 | 55733535 | AX-170479186 | 0 | G | A | 0.465 | -1.38E-01 | 8.50E-02 | 2.53E+00 | 2.56E+00 | 1.05E-01 | no | | intergenic_variant | MODIFIER |
| 16 | 55733775 | AX-170057494 | 0 | G | A | 0.465 | -1.38E-01 | 8.50E-02 | 2.53E+00 | 2.56E+00 | 1.05E-01 | no | | regulatory_region_variant | MODIFIER |
| 16 | 55739611 | AX-170370304 | 0 | A | G | 0.465 | -1.38E-01 | 8.50E-02 | 2.53E+00 | 2.56E+00 | 1.05E-01 | no | Gm19771 | upstream_gene_variant | MODIFIER |
| 16 | 55781986 | AX-170269386 | 0 | A | G | 0.465 | -1.38E-01 | 8.50E-02 | 2.53E+00 | 2.56E+00 | 1.05E-01 | no | Gm26369 | upstream_gene_variant | MODIFIER |
| 16 | 55782015 | AX-170184037 | 0 | C | T | 0.465 | -1.38E-01 | 8.50E-02 | 2.53E+00 | 2.56E+00 | 1.05E-01 | no | Gm26369 | upstream_gene_variant | MODIFIER |
| 16 | 55798192 | AX-97814412 | 0 | C | A | 0.465 | -1.38E-01 | 8.50E-02 | 2.53E+00 | 2.56E+00 | 1.05E-01 | no | | intergenic_variant | MODIFIER |
| 16 | 55819815 | AX-170330280 | 0 | A | G | 0.465 | -1.38E-01 | 8.50E-02 | 2.53E+00 | 2.56E+00 | 1.05E-01 | no | Nfkbiz | intron_variant | MODIFIER |
| 16 | 55819857 | AX-170126935 | 0 | G | T | 0.465 | -1.38E-01 | 8.50E-02 | 2.53E+00 | 2.56E+00 | 1.05E-01 | no | Nfkbiz | intron_variant | MODIFIER |
| 16 | 55844692 | AX-170522439 | 0 | A | G | 0.465 | -1.38E-01 | 8.50E-02 | 2.53E+00 | 2.56E+00 | 1.05E-01 | no | Nxpe3 | synonymous_variant | LOW |
| 16 | 55845080 | AX-97824028 | 0 | T | C | 0.465 | -1.38E-01 | 8.50E-02 | 2.53E+00 | 2.56E+00 | 1.05E-01 | no | Nxpe3 | intron_variant | MODIFIER |
| 16 | 55869763 | AX-97782457 | 0 | C | T | 0.465 | -1.38E-01 | 8.50E-02 | 2.53E+00 | 2.56E+00 | 1.05E-01 | no | Nxpe3 | intron_variant | MODIFIER |
| 16 | 55872585 | AX-170200465 | 0 | G | A | 0.465 | -1.38E-01 | 8.50E-02 | 2.53E+00 | 2.56E+00 | 1.05E-01 | no | Nxpe3 | intron_variant | MODIFIER |
| 16 | 55874161 | AX-170411055 | 0 | A | G | 0.465 | -1.38E-01 | 8.50E-02 | 2.53E+00 | 2.56E+00 | 1.05E-01 | no | Nxpe3 | intron_variant | MODIFIER |
| 16 | 55874252 | AX-170532563 | 0 | A | C | 0.465 | -1.38E-01 | 8.50E-02 | 2.53E+00 | 2.56E+00 | 1.05E-01 | no | Nxpe3 | intron_variant | MODIFIER |

| | | | | | | | | | | | | | | | |
|---|---|---|---|---|---|---|---|---|---|---|---|---|---|---|---|
| 16 | 55884144 | AX-170526894 | 0 | C | G | 0.465 | -1.38E-01 | 8.50E-02 | 2.53E+00 | 2.56E+00 | 1.05E-01 | no | Nxpe3 | intron_variant | MODIFIER |
| 16 | 55918675 | AX-170444321 | 0 | G | A | 0.465 | -1.38E-01 | 8.50E-02 | 2.53E+00 | 2.56E+00 | 1.05E-01 | no | Cep97 | intron_variant | MODIFIER |
| 16 | 55919070 | AX-170155702 | 0 | C | T | 0.465 | -1.38E-01 | 8.50E-02 | 2.53E+00 | 2.56E+00 | 1.05E-01 | no | Cep97 | intron_variant | MODIFIER |
| 16 | 55948999 | AX-170013115 | 0 | C | A | 0.465 | -1.38E-01 | 8.50E-02 | 2.53E+00 | 2.56E+00 | 1.05E-01 | no | | regulatory_region_variant | MODIFIER |
| 16 | 55969663 | AX-170013112 | 0 | G | C | 0.465 | -1.38E-01 | 8.50E-02 | 2.53E+00 | 2.56E+00 | 1.05E-01 | no | Rpl24 | intron_variant | MODIFIER |
| 16 | 55973668 | AX-170013111 | 0 | C | G | 0.465 | -1.38E-01 | 8.50E-02 | 2.53E+00 | 2.56E+00 | 1.05E-01 | no | Rpl24 | downstream_gene_variant | MODIFIER |
| 16 | 55975813 | AX-170088028 | 0 | A | G | 0.465 | -1.38E-01 | 8.50E-02 | 2.53E+00 | 2.56E+00 | 1.05E-01 | no | Rpl24 | downstream_gene_variant | MODIFIER |
| 16 | 56026705 | AX-170309233 | 0 | A | G | 0.465 | -1.38E-01 | 8.50E-02 | 2.53E+00 | 2.56E+00 | 1.05E-01 | no | Pcnp | upstream_gene_variant | MODIFIER |
| 16 | 56027935 | AX-170455011 | 0 | T | A | 0.465 | -1.38E-01 | 8.50E-02 | 2.53E+00 | 2.56E+00 | 1.05E-01 | no | Trmt10c | downstream_gene_variant | MODIFIER |
| 16 | 56058509 | AX-170013114 | 0 | C | T | 0.465 | -1.38E-01 | 8.50E-02 | 2.53E+00 | 2.56E+00 | 1.05E-01 | no | Senp7 | intron_variant,non_coding_transcript_variant | MODIFIER |
| 16 | 56060828 | AX-170165221 | 0 | C | T | 0.465 | -1.38E-01 | 8.50E-02 | 2.53E+00 | 2.56E+00 | 1.05E-01 | no | Senp7 | intron_variant,non_coding_transcript_variant | MODIFIER |
| 16 | 56061732 | AX-97801938 | 0 | C | T | 0.465 | -1.38E-01 | 8.50E-02 | 2.53E+00 | 2.56E+00 | 1.05E-01 | no | Senp7 | intron_variant,non_coding_transcript_variant | MODIFIER |
| 16 | 56121856 | AX-170206243 | 0 | C | G | 0.465 | -1.38E-01 | 8.50E-02 | 2.53E+00 | 2.56E+00 | 1.05E-01 | no | Senp7 | intron_variant | MODIFIER |
| 16 | 56184212 | AX-97790721 | 0 | A | C | 0.465 | -1.38E-01 | 8.50E-02 | 2.53E+00 | 2.56E+00 | 1.05E-01 | no | Senp7 | intron_variant | MODIFIER |
| 16 | 56210836 | AX-170349665 | 0 | T | G | 0.465 | -1.38E-01 | 8.50E-02 | 2.53E+00 | 2.56E+00 | 1.05E-01 | no | Impg2 | intron_variant,non_coding_transcript_variant | MODIFIER |
| 16 | 56582893 | AX-170013118 | 0 | A | G | 0.465 | -1.38E-01 | 8.50E-02 | 2.53E+00 | 2.56E+00 | 1.05E-01 | no | Abi3bp | intron_variant | MODIFIER |
| 16 | 56584464 | AX-170057509 | 0 | A | G | 0.465 | -1.38E-01 | 8.50E-02 | 2.53E+00 | 2.56E+00 | 1.05E-01 | no | Abi3bp | intron_variant | MODIFIER |
| 16 | 56588165 | AX-97807404 | 0 | T | C | 0.465 | -1.38E-01 | 8.50E-02 | 2.53E+00 | 2.56E+00 | 1.05E-01 | no | Abi3bp | intron_variant | MODIFIER |
| 16 | 56591817 | AX-170333385 | 0 | A | G | 0.465 | -1.38E-01 | 8.50E-02 | 2.53E+00 | 2.56E+00 | 1.05E-01 | no | Abi3bp | intron_variant | MODIFIER |
| 16 | 56602883 | AX-170115025 | 0 | T | C | 0.465 | -1.38E-01 | 8.50E-02 | 2.53E+00 | 2.56E+00 | 1.05E-01 | no | Abi3bp | intron_variant | MODIFIER |
| 16 | 56604192 | AX-170263845 | 0 | T | C | 0.465 | -1.38E-01 | 8.50E-02 | 2.53E+00 | 2.56E+00 | 1.05E-01 | no | Abi3bp | intron_variant | MODIFIER |
| 16 | 56606787 | AX-170249178 | 0 | G | T | 0.465 | -1.38E-01 | 8.50E-02 | 2.53E+00 | 2.56E+00 | 1.05E-01 | no | Abi3bp | intron_variant | MODIFIER |
| 16 | 56613312 | AX-97835459 | 0 | A | G | 0.465 | -1.38E-01 | 8.50E-02 | 2.53E+00 | 2.56E+00 | 1.05E-01 | no | Abi3bp | intron_variant | MODIFIER |
| 16 | 56613562 | AX-170300000 | 0 | G | A | 0.465 | -1.38E-01 | 8.50E-02 | 2.53E+00 | 2.56E+00 | 1.05E-01 | no | Abi3bp | intron_variant | MODIFIER |
| 16 | 56616054 | AX-170151111 | 0 | C | T | 0.465 | -1.38E-01 | 8.50E-02 | 2.53E+00 | 2.56E+00 | 1.05E-01 | no | Abi3bp | intron_variant | MODIFIER |
| 16 | 56619718 | AX-170450887 | 0 | A | G | 0.465 | -1.38E-01 | 8.50E-02 | 2.53E+00 | 2.56E+00 | 1.05E-01 | no | Abi3bp | intron_variant | MODIFIER |
| 16 | 56632692 | AX-170180351 | 0 | A | G | 0.465 | -1.38E-01 | 8.50E-02 | 2.53E+00 | 2.56E+00 | 1.05E-01 | no | Abi3bp | intron_variant | MODIFIER |
| 16 | 56641715 | AX-170093497 | 0 | T | C | 0.465 | -1.38E-01 | 8.50E-02 | 2.53E+00 | 2.56E+00 | 1.05E-01 | no | Abi3bp | intron_variant | MODIFIER |
| 16 | 56654036 | AX-170321723 | 0 | G | A | 0.465 | -1.38E-01 | 8.50E-02 | 2.53E+00 | 2.56E+00 | 1.05E-01 | no | Abi3bp | synonymous_variant | LOW |
| 16 | 56670886 | AX-170375459 | 0 | T | C | 0.465 | -1.38E-01 | 8.50E-02 | 2.53E+00 | 2.56E+00 | 1.05E-01 | no | Abi3bp | intron_variant | MODIFIER |
| 16 | 56689676 | AX-170526066 | 0 | A | G | 0.465 | -1.38E-01 | 8.50E-02 | 2.53E+00 | 2.56E+00 | 1.05E-01 | no | Abi3bp | downstream_gene_variant | MODIFIER |
| 16 | 56722894 | AX-170182273 | 0 | T | C | 0.465 | -1.38E-01 | 8.50E-02 | 2.53E+00 | 2.56E+00 | 1.05E-01 | no | Adgrg7 | downstream_gene_variant | MODIFIER |
| 16 | 56729039 | AX-170139418 | 0 | G | C | 0.465 | -1.38E-01 | 8.50E-02 | 2.53E+00 | 2.56E+00 | 1.05E-01 | no | Adgrg7 | intron_variant | MODIFIER |
| 16 | 56748368 | AX-97814661 | 0 | T | G | 0.465 | -1.38E-01 | 8.50E-02 | 2.53E+00 | 2.56E+00 | 1.05E-01 | no | Adgrg7 | intron_variant | MODIFIER |
| 16 | 56753625 | AX-170163654 | 0 | A | G | 0.465 | -1.38E-01 | 8.50E-02 | 2.53E+00 | 2.56E+00 | 1.05E-01 | no | Adgrg7 | intron_variant | MODIFIER |
| 16 | 56844333 | AX-170402123 | 0 | T | A | 0.465 | -1.38E-01 | 8.50E-02 | 2.53E+00 | 2.56E+00 | 1.05E-01 | no | Tmem45a | intron_variant | MODIFIER |
| 16 | 56854305 | AX-170178998 | 0 | A | G | 0.465 | -1.38E-01 | 8.50E-02 | 2.53E+00 | 2.56E+00 | 1.05E-01 | no | Tmem45a | intron_variant | MODIFIER |
| 16 | 56868361 | AX-97788249 | 0 | A | G | 0.465 | -1.38E-01 | 8.50E-02 | 2.53E+00 | 2.56E+00 | 1.05E-01 | no | Tmem45a | intron_variant | MODIFIER |
| 16 | 56892898 | AX-170286472 | 0 | T | C | 0.465 | -1.38E-01 | 8.50E-02 | 2.53E+00 | 2.56E+00 | 1.05E-01 | no | Gm15839 | downstream_gene_variant | MODIFIER |
| 16 | 56915461 | AX-170159119 | 0 | G | T | 0.465 | -1.38E-01 | 8.50E-02 | 2.53E+00 | 2.56E+00 | 1.05E-01 | no | Lnp1 | intron_variant | MODIFIER |
| 16 | 56918653 | AX-170290806 | 0 | A | G | 0.465 | -1.38E-01 | 8.50E-02 | 2.53E+00 | 2.56E+00 | 1.05E-01 | no | Lnp1 | intron_variant | MODIFIER |
| 16 | 57019151 | AX-170013121 | 0 | G | A | 0.465 | -1.38E-01 | 8.50E-02 | 2.53E+00 | 2.56E+00 | 1.05E-01 | no | | intergenic_variant | MODIFIER |
| 16 | 57130568 | AX-97795780 | 0 | C | T | 0.465 | -1.38E-01 | 8.50E-02 | 2.53E+00 | 2.56E+00 | 1.05E-01 | no | Tomm70a | intron_variant | MODIFIER |
| 16 | 57134673 | AX-170013131 | 0 | C | A | 0.465 | -1.38E-01 | 8.50E-02 | 2.53E+00 | 2.56E+00 | 1.05E-01 | no | Tomm70a | synonymous_variant | LOW |
| 16 | 57141237 | AX-170013127 | 0 | G | C | 0.465 | -1.38E-01 | 8.50E-02 | 2.53E+00 | 2.56E+00 | 1.05E-01 | no | Tomm70a | intron_variant | MODIFIER |
| 16 | 57141289 | AX-170013128 | 0 | G | A | 0.465 | -1.38E-01 | 8.50E-02 | 2.53E+00 | 2.56E+00 | 1.05E-01 | no | Tomm70a | intron_variant | MODIFIER |
| 16 | 57169477 | AX-97811888 | 0 | G | A | 0.465 | -1.38E-01 | 8.50E-02 | 2.53E+00 | 2.56E+00 | 1.05E-01 | no | Tbc1d23 | 3_prime_UTR_variant | MODIFIER |
| 16 | 57184069 | AX-170219212 | 0 | C | T | 0.465 | -1.38E-01 | 8.50E-02 | 2.53E+00 | 2.56E+00 | 1.05E-01 | no | Tbc1d23 | intron_variant | MODIFIER |
| 16 | 57184245 | AX-170057520 | 0 | A | G | 0.465 | -1.38E-01 | 8.50E-02 | 2.53E+00 | 2.56E+00 | 1.05E-01 | no | Tbc1d23 | intron_variant | MODIFIER |
| 16 | 57184545 | AX-97791053 | 0 | T | C | 0.465 | -1.38E-01 | 8.50E-02 | 2.53E+00 | 2.56E+00 | 1.05E-01 | no | Tbc1d23 | intron_variant | MODIFIER |
| 16 | 57238428 | AX-97831380 | 0 | A | G | 0.465 | -1.38E-01 | 8.50E-02 | 2.53E+00 | 2.56E+00 | 1.05E-01 | no | | intergenic_variant | MODIFIER |
| 16 | 57303548 | AX-170282716 | 0 | T | G | 0.465 | -1.38E-01 | 8.50E-02 | 2.53E+00 | 2.56E+00 | 1.05E-01 | no | Cmss1 | intron_variant | MODIFIER |
| 16 | 57308052 | AX-170425570 | 0 | C | T | 0.465 | -1.38E-01 | 8.50E-02 | 2.53E+00 | 2.56E+00 | 1.05E-01 | no | Cmss1 | intron_variant | MODIFIER |
| 16 | 57308226 | AX-170493159 | 0 | C | G | 0.465 | -1.38E-01 | 8.50E-02 | 2.53E+00 | 2.56E+00 | 1.05E-01 | no | Cmss1 | intron_variant | MODIFIER |

| | | | | | | | | | | | | | | | |
|---|---|---|---|---|---|---|---|---|---|---|---|---|---|---|---|
| 16 | 57318699 | AX-170244373 | 0 | G | A | 0.465 | -1.38E-01 | 8.50E-02 | 2.53E+00 | 2.56E+00 | 1.05E-01 | no | Cmss1 | intron_variant | MODIFIER |
| 16 | 57320162 | AX-97801860 | 0 | C | T | 0.465 | -1.38E-01 | 8.50E-02 | 2.53E+00 | 2.56E+00 | 1.05E-01 | no | Cmss1 | intron_variant | MODIFIER |
| 16 | 57320245 | AX-170206405 | 0 | C | G | 0.465 | -1.38E-01 | 8.50E-02 | 2.53E+00 | 2.56E+00 | 1.05E-01 | no | Cmss1 | intron_variant | MODIFIER |
| 16 | 57336665 | AX-170057524 | 0 | G | A | 0.465 | -1.38E-01 | 8.50E-02 | 2.53E+00 | 2.56E+00 | 1.05E-01 | no | Cmss1 | intron_variant | MODIFIER |
| 16 | 57396708 | AX-170369913 | 0 | T | C | 0.465 | -1.38E-01 | 8.50E-02 | 2.53E+00 | 2.56E+00 | 1.05E-01 | no | Filip1l | intron_variant | MODIFIER |
| 16 | 57421945 | AX-170261640 | 0 | C | T | 0.465 | -1.38E-01 | 8.50E-02 | 2.53E+00 | 2.56E+00 | 1.05E-01 | no | Filip1l | intron_variant | MODIFIER |
| 16 | 57422460 | AX-170337177 | 0 | C | T | 0.465 | -1.38E-01 | 8.50E-02 | 2.53E+00 | 2.56E+00 | 1.05E-01 | no | Filip1l | intron_variant | MODIFIER |
| 16 | 57424058 | AX-97791054 | 0 | A | G | 0.465 | -1.38E-01 | 8.50E-02 | 2.53E+00 | 2.56E+00 | 1.05E-01 | no | Filip1l | intron_variant | MODIFIER |
| 16 | 57425389 | AX-170341603 | 0 | A | G | 0.465 | -1.38E-01 | 8.50E-02 | 2.53E+00 | 2.56E+00 | 1.05E-01 | no | Filip1l | intron_variant | MODIFIER |
| 16 | 57425457 | AX-170196244 | 0 | A | G | 0.465 | -1.38E-01 | 8.50E-02 | 2.53E+00 | 2.56E+00 | 1.05E-01 | no | Filip1l | intron_variant | MODIFIER |
| 16 | 57429453 | AX-170399275 | 0 | C | T | 0.465 | -1.38E-01 | 8.50E-02 | 2.53E+00 | 2.56E+00 | 1.05E-01 | no | Filip1l | intron_variant | MODIFIER |
| 16 | 57443240 | AX-170172708 | 0 | G | A | 0.465 | -1.38E-01 | 8.50E-02 | 2.53E+00 | 2.56E+00 | 1.05E-01 | no | Filip1l | intron_variant | MODIFIER |
| 16 | 57445664 | AX-170268402 | 0 | G | C | 0.465 | -1.38E-01 | 8.50E-02 | 2.53E+00 | 2.56E+00 | 1.05E-01 | no | Filip1l | intron_variant | MODIFIER |
| 16 | 57472761 | AX-97831381 | 0 | C | A | 0.465 | -1.38E-01 | 8.50E-02 | 2.53E+00 | 2.56E+00 | 1.05E-01 | no | Filip1l | intron_variant | MODIFIER |
| 16 | 57491003 | AX-170013137 | 0 | C | A | 0.465 | -1.38E-01 | 8.50E-02 | 2.53E+00 | 2.56E+00 | 1.05E-01 | no | Filip1l | intron_variant | MODIFIER |
| 16 | 57506304 | AX-170013132 | 0 | G | A | 0.465 | -1.38E-01 | 8.50E-02 | 2.53E+00 | 2.56E+00 | 1.05E-01 | no | Filip1l | intron_variant | MODIFIER |
| 16 | 57508897 | AX-170211135 | 0 | C | G | 0.465 | -1.38E-01 | 8.50E-02 | 2.53E+00 | 2.56E+00 | 1.05E-01 | no | Filip1l | intron_variant | MODIFIER |
| 16 | 57510685 | AX-170013134 | 0 | G | A | 0.465 | -1.38E-01 | 8.50E-02 | 2.53E+00 | 2.56E+00 | 1.05E-01 | no | Filip1l | intron_variant | MODIFIER |
| 16 | 57522847 | AX-170453572 | 0 | C | T | 0.465 | -1.38E-01 | 8.50E-02 | 2.53E+00 | 2.56E+00 | 1.05E-01 | no | Cmss1 | intron_variant | MODIFIER |
| 16 | 57524674 | AX-170013136 | 0 | A | G | 0.465 | -1.38E-01 | 8.50E-02 | 2.53E+00 | 2.56E+00 | 1.05E-01 | no | Cmss1 | intron_variant | MODIFIER |
| 16 | 57527575 | AX-170013135 | 0 | C | T | 0.465 | -1.38E-01 | 8.50E-02 | 2.53E+00 | 2.56E+00 | 1.05E-01 | no | Cmss1 | intron_variant | MODIFIER |
| 16 | 57538208 | AX-170200719 | 0 | G | C | 0.465 | -1.38E-01 | 8.50E-02 | 2.53E+00 | 2.56E+00 | 1.05E-01 | no | Cmss1 | intron_variant | MODIFIER |
| 16 | 57539424 | AX-170013138 | 0 | A | C | 0.465 | -1.38E-01 | 8.50E-02 | 2.53E+00 | 2.56E+00 | 1.05E-01 | no | Cmss1 | intron_variant | MODIFIER |
| 16 | 57556523 | AX-170493412 | 0 | A | G | 0.465 | -1.38E-01 | 8.50E-02 | 2.53E+00 | 2.56E+00 | 1.05E-01 | no | Cmss1 | intron_variant | MODIFIER |
| 16 | 57556547 | AX-170308839 | 0 | A | G | 0.465 | -1.38E-01 | 8.50E-02 | 2.53E+00 | 2.56E+00 | 1.05E-01 | no | Cmss1 | intron_variant | MODIFIER |
| 16 | 57654966 | AX-170520449 | 0 | G | A | 0.465 | -1.38E-01 | 8.50E-02 | 2.53E+00 | 2.56E+00 | 1.05E-01 | no | Col8a1 | intron_variant | MODIFIER |
| 16 | 57670988 | AX-170344860 | 0 | G | C | 0.465 | -1.38E-01 | 8.50E-02 | 2.53E+00 | 2.56E+00 | 1.05E-01 | no | Col8a1 | intron_variant | MODIFIER |
| 16 | 57738153 | AX-97791055 | 0 | G | T | 0.465 | -1.38E-01 | 8.50E-02 | 2.53E+00 | 2.56E+00 | 1.05E-01 | no | Col8a1 | intron_variant | MODIFIER |
| 16 | 57738274 | AX-170279173 | 0 | C | T | 0.465 | -1.38E-01 | 8.50E-02 | 2.53E+00 | 2.56E+00 | 1.05E-01 | no | Col8a1 | intron_variant | MODIFIER |
| 16 | 57759351 | AX-170386691 | 0 | T | C | 0.465 | -1.38E-01 | 8.50E-02 | 2.53E+00 | 2.56E+00 | 1.05E-01 | no | Col8a1 | upstream_gene_variant | MODIFIER |
| 16 | 57765821 | AX-97789694 | 0 | G | A | 0.465 | -1.38E-01 | 8.50E-02 | 2.53E+00 | 2.56E+00 | 1.05E-01 | no | Gm5674 | downstream_gene_variant | MODIFIER |
| 16 | 57960955 | AX-170433701 | 0 | C | T | 0.465 | -1.38E-01 | 8.50E-02 | 2.53E+00 | 2.56E+00 | 1.05E-01 | no | Gm49575 | intron_variant,non_coding_transcript_variant | MODIFIER |
| 16 | 57961297 | AX-170508849 | 0 | C | A | 0.465 | -1.38E-01 | 8.50E-02 | 2.53E+00 | 2.56E+00 | 1.05E-01 | no | Gm49575 | intron_variant,non_coding_transcript_variant | MODIFIER |
| 16 | 57961477 | AX-170093499 | 0 | C | A | 0.465 | -1.38E-01 | 8.50E-02 | 2.53E+00 | 2.56E+00 | 1.05E-01 | no | Gm49575 | intron_variant,non_coding_transcript_variant | MODIFIER |
| 16 | 57973075 | AX-170153009 | 0 | G | T | 0.465 | -1.38E-01 | 8.50E-02 | 2.53E+00 | 2.56E+00 | 1.05E-01 | no | Gm49575 | intron_variant,non_coding_transcript_variant | MODIFIER |
| 16 | 57982056 | AX-170344708 | 0 | C | A | 0.465 | -1.38E-01 | 8.50E-02 | 2.53E+00 | 2.56E+00 | 1.05E-01 | no | Gm49575 | intron_variant,non_coding_transcript_variant | MODIFIER |
| 16 | 57984983 | AX-97812468 | 0 | A | G | 0.465 | -1.38E-01 | 8.50E-02 | 2.53E+00 | 2.56E+00 | 1.05E-01 | no | Gm49575 | intron_variant,non_coding_transcript_variant | MODIFIER |
| 16 | 58019667 | AX-170498751 | 0 | C | T | 0.465 | -1.38E-01 | 8.50E-02 | 2.53E+00 | 2.56E+00 | 1.05E-01 | no | Gm49575 | intron_variant,non_coding_transcript_variant | MODIFIER |
| 16 | 58019964 | AX-170339604 | 0 | C | T | 0.465 | -1.38E-01 | 8.50E-02 | 2.53E+00 | 2.56E+00 | 1.05E-01 | no | Gm49575 | intron_variant,non_coding_transcript_variant | MODIFIER |
| 16 | 58029227 | AX-170395885 | 0 | C | A | 0.465 | -1.38E-01 | 8.50E-02 | 2.53E+00 | 2.56E+00 | 1.05E-01 | no | Gm49575 | non_coding_transcript_exon_variant | MODIFIER |
| 16 | 58029243 | AX-170279053 | 0 | A | G | 0.465 | -1.38E-01 | 8.50E-02 | 2.53E+00 | 2.56E+00 | 1.05E-01 | no | Gm49575 | non_coding_transcript_exon_variant | MODIFIER |
| 16 | 58029401 | AX-170497285 | 0 | T | A | 0.465 | -1.38E-01 | 8.50E-02 | 2.53E+00 | 2.56E+00 | 1.05E-01 | no | Gm49575 | intron_variant,non_coding_transcript_variant | MODIFIER |
| 16 | 58035027 | AX-170013143 | 0 | G | T | 0.465 | -1.38E-01 | 8.50E-02 | 2.53E+00 | 2.56E+00 | 1.05E-01 | no | Gm49575 | intron_variant,non_coding_transcript_variant | MODIFIER |
| 16 | 58036754 | AX-170398043 | 0 | A | G | 0.465 | -1.38E-01 | 8.50E-02 | 2.53E+00 | 2.56E+00 | 1.05E-01 | no | Gm49575 | intron_variant,non_coding_transcript_variant | MODIFIER |
| 16 | 58038910 | AX-170247545 | 0 | T | C | 0.465 | -1.38E-01 | 8.50E-02 | 2.53E+00 | 2.56E+00 | 1.05E-01 | no | Gm49575 | intron_variant,non_coding_transcript_variant | MODIFIER |
| 16 | 58040705 | AX-97831427 | 0 | C | T | 0.465 | -1.38E-01 | 8.50E-02 | 2.53E+00 | 2.56E+00 | 1.05E-01 | no | Gm49575 | intron_variant,non_coding_transcript_variant | MODIFIER |
| 16 | 55905873 | AX-170157295 | 1 | A | G | 0.536 | 1.38E-01 | 8.51E-02 | 2.53E+00 | 2.56E+00 | 1.05E-01 | no | Cep97 | splice_region_variant,intron_variant | LOW |
| 16 | 67686169 | AX-170523372 | 1 | A | T | 0.491 | 1.26E-01 | 7.79E-02 | 2.56E+00 | 2.62E+00 | 1.05E-01 | no | | intergenic_variant | MODIFIER |
| 5 | 14015315 | AX-170383751 | 1 | C | T | 0.348 | 1.37E-01 | 8.45E-02 | 2.94E+00 | 3.10E+00 | 1.05E-01 | no | | intergenic_variant | MODIFIER |
| 1 | 15433256 | AX-170325674 | 2 | T | G | 0.027 | 2.50E-01 | 1.55E-01 | 2.79E+00 | 2.99E+00 | 1.05E-01 | Yes | Kcnb2 | intron_variant | MODIFIER |
| 12 | 92124209 | AX-170135849 | 1 | A | T | 0.089 | 2.39E-01 | 1.47E-01 | 2.98E+00 | 3.16E+00 | 1.05E-01 | Yes | | intergenic_variant | MODIFIER |
| 4 | 100815442 | AX-170023526 | 1 | A | T | 0.232 | 1.88E-01 | 1.17E-01 | 2.70E+00 | 2.77E+00 | 1.05E-01 | Yes | Cachd1 | intron_variant | MODIFIER |
| 5 | 56539224 | AX-97843777 | 0 | G | A | 0.482 | 1.56E-01 | 9.50E-02 | 2.34E+00 | 2.42E+00 | 1.05E-01 | Yes | | intergenic_variant | MODIFIER |
| 4 | 77587250 | AX-97797557 | 1 | A | G | 0.33 | 1.69E-01 | 1.03E-01 | 3.13E+00 | 3.26E+00 | 1.05E-01 | Yes | Ptprd | intron_variant,non_coding_transcript_variant | MODIFIER |

| | | | | | | | | | | | | | | | |
|---|---|---|---|---|---|---|---|---|---|---|---|---|---|---|---|
| 15 | 67351821 | AX-97843937 | 0 | C | T | 0.333 | -1.32E-01 | 7.80E-02 | 2.15E+00 | 2.22E+00 | 1.05E-01 | no | 1700012I11Rik | intron_variant,non_coding_transcript_variant | MODIFIER |
| 5 | 54579891 | AX-170144366 | 0 | C | A | 0.491 | 1.64E-01 | 1.00E-01 | 2.38E+00 | 2.44E+00 | 1.06E-01 | Yes | Gm18451 | downstream_gene_variant | MODIFIER |
| 19 | 60278012 | AX-97805125 | 0 | C | T | 0.474 | -1.02E-01 | 6.31E-02 | 2.74E+00 | 2.82E+00 | 1.06E-01 | no | | intergenic_variant | MODIFIER |
| 19 | 59904810 | AX-170375063 | 0 | A | G | 0.456 | -1.02E-01 | 6.31E-02 | 2.76E+00 | 2.84E+00 | 1.06E-01 | no | Rab11fip2 | 3_prime_UTR_variant | MODIFIER |
| 19 | 60133792 | AX-170221030 | 0 | C | T | 0.456 | -1.02E-01 | 6.31E-02 | 2.76E+00 | 2.84E+00 | 1.06E-01 | no | | intergenic_variant | MODIFIER |
| 19 | 60150939 | AX-170245311 | 0 | A | G | 0.456 | -1.02E-01 | 6.31E-02 | 2.76E+00 | 2.84E+00 | 1.06E-01 | no | E330013P04Rik | intron_variant,non_coding_transcript_variant | MODIFIER |
| 19 | 60165408 | AX-170402879 | 0 | A | G | 0.456 | -1.02E-01 | 6.31E-02 | 2.76E+00 | 2.84E+00 | 1.06E-01 | no | E330013P04Rik | downstream_gene_variant | MODIFIER |
| 19 | 60165421 | AX-170126298 | 0 | A | G | 0.456 | -1.02E-01 | 6.31E-02 | 2.76E+00 | 2.84E+00 | 1.06E-01 | no | E330013P04Rik | downstream_gene_variant | MODIFIER |
| 19 | 60245298 | AX-168411612 | 0 | T | C | 0.456 | -1.02E-01 | 6.31E-02 | 2.76E+00 | 2.84E+00 | 1.06E-01 | no | Gm28351 | downstream_gene_variant | MODIFIER |
| 19 | 60488793 | AX-97852822 | 0 | G | A | 0.456 | -1.02E-01 | 6.31E-02 | 2.76E+00 | 2.84E+00 | 1.06E-01 | no | | intergenic_variant | MODIFIER |
| 19 | 60490049 | AX-170431170 | 0 | A | G | 0.456 | -1.02E-01 | 6.31E-02 | 2.76E+00 | 2.84E+00 | 1.06E-01 | no | | intergenic_variant | MODIFIER |
| 19 | 60501217 | AX-170491646 | 0 | T | A | 0.456 | -1.02E-01 | 6.31E-02 | 2.76E+00 | 2.84E+00 | 1.06E-01 | no | | intergenic_variant | MODIFIER |
| 19 | 60502734 | AX-170276345 | 0 | A | G | 0.456 | -1.02E-01 | 6.31E-02 | 2.76E+00 | 2.84E+00 | 1.06E-01 | no | | intergenic_variant | MODIFIER |
| 19 | 60505406 | AX-170062867 | 0 | T | C | 0.456 | -1.02E-01 | 6.31E-02 | 2.76E+00 | 2.84E+00 | 1.06E-01 | no | | intergenic_variant | MODIFIER |
| 19 | 60509149 | AX-170062865 | 0 | T | G | 0.456 | -1.02E-01 | 6.31E-02 | 2.76E+00 | 2.84E+00 | 1.06E-01 | no | | intergenic_variant | MODIFIER |
| 19 | 60511716 | AX-97821410 | 0 | G | T | 0.456 | -1.02E-01 | 6.31E-02 | 2.76E+00 | 2.84E+00 | 1.06E-01 | no | | intergenic_variant | MODIFIER |
| 19 | 60511837 | AX-170211250 | 0 | G | A | 0.456 | -1.02E-01 | 6.31E-02 | 2.76E+00 | 2.84E+00 | 1.06E-01 | no | | intergenic_variant | MODIFIER |
| 19 | 60515354 | AX-170178427 | 0 | T | C | 0.456 | -1.02E-01 | 6.31E-02 | 2.76E+00 | 2.84E+00 | 1.06E-01 | no | | intergenic_variant | MODIFIER |
| 19 | 60540452 | AX-170533425 | 0 | G | A | 0.456 | -1.02E-01 | 6.31E-02 | 2.76E+00 | 2.84E+00 | 1.06E-01 | no | Cacul1 | intron_variant | MODIFIER |
| 19 | 60569144 | AX-168380229 | 0 | G | T | 0.456 | -1.02E-01 | 6.31E-02 | 2.76E+00 | 2.84E+00 | 1.06E-01 | no | Cacul1 | intron_variant | MODIFIER |
| 19 | 60575617 | AX-170225013 | 0 | G | C | 0.456 | -1.02E-01 | 6.31E-02 | 2.76E+00 | 2.84E+00 | 1.06E-01 | no | Cacul1 | intron_variant | MODIFIER |
| 19 | 60575909 | AX-170339131 | 0 | G | A | 0.456 | -1.02E-01 | 6.31E-02 | 2.76E+00 | 2.84E+00 | 1.06E-01 | no | Cacul1 | intron_variant | MODIFIER |
| 19 | 60594903 | AX-170120661 | 0 | G | A | 0.456 | -1.02E-01 | 6.31E-02 | 2.76E+00 | 2.84E+00 | 1.06E-01 | no | | intergenic_variant | MODIFIER |
| 19 | 60596617 | AX-170094143 | 0 | C | T | 0.456 | -1.02E-01 | 6.31E-02 | 2.76E+00 | 2.84E+00 | 1.06E-01 | no | | intergenic_variant | MODIFIER |
| 19 | 60618880 | AX-97851377 | 0 | G | A | 0.456 | -1.02E-01 | 6.31E-02 | 2.76E+00 | 2.84E+00 | 1.06E-01 | no | AC108409.1 | intron_variant,non_coding_transcript_variant | MODIFIER |
| 19 | 60621050 | AX-170468157 | 0 | T | C | 0.456 | -1.02E-01 | 6.31E-02 | 2.76E+00 | 2.84E+00 | 1.06E-01 | no | AC108409.1 | intron_variant,non_coding_transcript_variant | MODIFIER |
| 19 | 60621105 | AX-170231071 | 0 | A | G | 0.456 | -1.02E-01 | 6.31E-02 | 2.76E+00 | 2.84E+00 | 1.06E-01 | no | AC108409.1 | intron_variant,non_coding_transcript_variant | MODIFIER |
| 19 | 60624792 | AX-170154216 | 0 | T | C | 0.456 | -1.02E-01 | 6.31E-02 | 2.76E+00 | 2.84E+00 | 1.06E-01 | no | AC108409.1 | intron_variant,non_coding_transcript_variant | MODIFIER |
| 19 | 60631488 | AX-97854086 | 0 | T | C | 0.456 | -1.02E-01 | 6.31E-02 | 2.76E+00 | 2.84E+00 | 1.06E-01 | no | | intergenic_variant | MODIFIER |
| 19 | 60650962 | AX-170225598 | 0 | A | C | 0.456 | -1.02E-01 | 6.31E-02 | 2.76E+00 | 2.84E+00 | 1.06E-01 | no | | intergenic_variant | MODIFIER |
| 19 | 60651097 | AX-170304109 | 0 | A | C | 0.456 | -1.02E-01 | 6.31E-02 | 2.76E+00 | 2.84E+00 | 1.06E-01 | no | | intergenic_variant | MODIFIER |
| 19 | 60663443 | AX-97851321 | 0 | T | C | 0.456 | -1.02E-01 | 6.31E-02 | 2.76E+00 | 2.84E+00 | 1.06E-01 | no | | intergenic_variant | MODIFIER |
| 19 | 60664062 | AX-170179708 | 0 | A | G | 0.456 | -1.02E-01 | 6.31E-02 | 2.76E+00 | 2.84E+00 | 1.06E-01 | no | | intergenic_variant | MODIFIER |
| 19 | 60666363 | AX-170386417 | 0 | T | C | 0.456 | -1.02E-01 | 6.31E-02 | 2.76E+00 | 2.84E+00 | 1.06E-01 | no | | intergenic_variant | MODIFIER |
| 19 | 60680623 | AX-170115306 | 0 | G | A | 0.456 | -1.02E-01 | 6.31E-02 | 2.76E+00 | 2.84E+00 | 1.06E-01 | no | | intergenic_variant | MODIFIER |
| 19 | 60680812 | AX-170133903 | 0 | A | G | 0.456 | -1.02E-01 | 6.31E-02 | 2.76E+00 | 2.84E+00 | 1.06E-01 | no | | intergenic_variant | MODIFIER |
| 19 | 60685507 | AX-170062870 | 0 | A | G | 0.456 | -1.02E-01 | 6.31E-02 | 2.76E+00 | 2.84E+00 | 1.06E-01 | no | | intergenic_variant | MODIFIER |
| 19 | 60686533 | AX-170088494 | 0 | A | G | 0.456 | -1.02E-01 | 6.31E-02 | 2.76E+00 | 2.84E+00 | 1.06E-01 | no | | intergenic_variant | MODIFIER |
| 19 | 60718393 | AX-170103291 | 0 | G | T | 0.456 | -1.02E-01 | 6.31E-02 | 2.76E+00 | 2.84E+00 | 1.06E-01 | no | | intergenic_variant | MODIFIER |
| 19 | 60723849 | AX-97837195 | 0 | T | C | 0.456 | -1.02E-01 | 6.31E-02 | 2.76E+00 | 2.84E+00 | 1.06E-01 | no | | intergenic_variant | MODIFIER |
| 19 | 60724211 | AX-170326225 | 0 | G | A | 0.456 | -1.02E-01 | 6.31E-02 | 2.76E+00 | 2.84E+00 | 1.06E-01 | no | | intergenic_variant | MODIFIER |
| 19 | 60741496 | AX-170346211 | 0 | C | T | 0.456 | -1.02E-01 | 6.31E-02 | 2.76E+00 | 2.84E+00 | 1.06E-01 | no | | intergenic_variant | MODIFIER |
| 19 | 60741694 | AX-97829029 | 0 | T | G | 0.456 | -1.02E-01 | 6.31E-02 | 2.76E+00 | 2.84E+00 | 1.06E-01 | no | | intergenic_variant | MODIFIER |
| 19 | 60745728 | AX-170368687 | 0 | T | C | 0.456 | -1.02E-01 | 6.31E-02 | 2.76E+00 | 2.84E+00 | 1.06E-01 | no | | intergenic_variant | MODIFIER |
| 19 | 60757991 | AX-170378737 | 0 | A | G | 0.456 | -1.02E-01 | 6.31E-02 | 2.76E+00 | 2.84E+00 | 1.06E-01 | no | Eif3a | downstream_gene_variant | MODIFIER |
| 19 | 60767535 | AX-97837193 | 0 | G | T | 0.456 | -1.02E-01 | 6.31E-02 | 2.76E+00 | 2.84E+00 | 1.06E-01 | no | Eif3a | intron_variant | MODIFIER |
| 19 | 60818718 | AX-170016953 | 0 | C | T | 0.456 | -1.02E-01 | 6.31E-02 | 2.76E+00 | 2.84E+00 | 1.06E-01 | no | Fam45a | intron_variant | MODIFIER |
| 19 | 60818903 | AX-170016956 | 0 | T | C | 0.456 | -1.02E-01 | 6.31E-02 | 2.76E+00 | 2.84E+00 | 1.06E-01 | no | Fam45a | intron_variant | MODIFIER |
| 19 | 60818995 | AX-170444108 | 0 | A | G | 0.456 | -1.02E-01 | 6.31E-02 | 2.76E+00 | 2.84E+00 | 1.06E-01 | no | Fam45a | intron_variant | MODIFIER |
| 19 | 60821198 | AX-170139340 | 0 | T | C | 0.456 | -1.02E-01 | 6.31E-02 | 2.76E+00 | 2.84E+00 | 1.06E-01 | no | Fam45a | intron_variant | MODIFIER |
| 19 | 60838768 | AX-170016955 | 0 | T | C | 0.456 | -1.02E-01 | 6.31E-02 | 2.76E+00 | 2.84E+00 | 1.06E-01 | no | Fam45a | downstream_gene_variant | MODIFIER |
| 19 | 60844749 | AX-170016954 | 0 | T | C | 0.456 | -1.02E-01 | 6.31E-02 | 2.76E+00 | 2.84E+00 | 1.06E-01 | no | Sfxn4 | intron_variant,NMD_transcript_variant | MODIFIER |
| 19 | 60846510 | AX-170478974 | 0 | A | T | 0.456 | -1.02E-01 | 6.31E-02 | 2.76E+00 | 2.84E+00 | 1.06E-01 | no | Sfxn4 | intron_variant,NMD_transcript_variant | MODIFIER |
| 19 | 60855639 | AX-97804555 | 0 | G | A | 0.456 | -1.02E-01 | 6.31E-02 | 2.76E+00 | 2.84E+00 | 1.06E-01 | no | Sfxn4 | intron_variant,NMD_transcript_variant | MODIFIER |

| | | | | | | | | | | | | | | | |
|---|---|---|---|---|---|---|---|---|---|---|---|---|---|---|---|
| 19 | 60900389 | AX-97836259 | 0 | G | A | 0.456 | -1.02E-01 | 6.31E-02 | 2.76E+00 | 2.84E+00 | 1.06E-01 | no | Grk5 | intron_variant | MODIFIER |
| 19 | 60901314 | AX-97793561 | 0 | A | C | 0.456 | -1.02E-01 | 6.31E-02 | 2.76E+00 | 2.84E+00 | 1.06E-01 | no | Grk5 | intron_variant | MODIFIER |
| 19 | 60917293 | AX-170016952 | 0 | T | C | 0.456 | -1.02E-01 | 6.31E-02 | 2.76E+00 | 2.84E+00 | 1.06E-01 | no | Grk5 | intron_variant | MODIFIER |
| 19 | 60935741 | AX-97794877 | 0 | C | G | 0.456 | -1.02E-01 | 6.31E-02 | 2.76E+00 | 2.84E+00 | 1.06E-01 | no | Grk5 | intron_variant | MODIFIER |
| 19 | 60952218 | AX-170016958 | 0 | G | A | 0.456 | -1.02E-01 | 6.31E-02 | 2.76E+00 | 2.84E+00 | 1.06E-01 | no | Grk5 | intron_variant | MODIFIER |
| 19 | 60958865 | AX-170133657 | 0 | C | A | 0.456 | -1.02E-01 | 6.31E-02 | 2.76E+00 | 2.84E+00 | 1.06E-01 | no | Grk5 | intron_variant | MODIFIER |
| 19 | 60973379 | AX-170186781 | 0 | C | T | 0.456 | -1.02E-01 | 6.31E-02 | 2.76E+00 | 2.84E+00 | 1.06E-01 | no | Grk5 | intron_variant | MODIFIER |
| 19 | 60978988 | AX-170215746 | 0 | G | A | 0.456 | -1.02E-01 | 6.31E-02 | 2.76E+00 | 2.84E+00 | 1.06E-01 | no | Grk5 | intron_variant | MODIFIER |
| 19 | 60979967 | AX-170508694 | 0 | T | G | 0.456 | -1.02E-01 | 6.31E-02 | 2.76E+00 | 2.84E+00 | 1.06E-01 | no | Grk5 | intron_variant | MODIFIER |
| 19 | 60980858 | AX-170197095 | 0 | C | T | 0.456 | -1.02E-01 | 6.31E-02 | 2.76E+00 | 2.84E+00 | 1.06E-01 | no | Grk5 | intron_variant | MODIFIER |
| 19 | 60990080 | AX-170102501 | 0 | A | C | 0.456 | -1.02E-01 | 6.31E-02 | 2.76E+00 | 2.84E+00 | 1.06E-01 | no | Grk5 | intron_variant | MODIFIER |
| 19 | 60991056 | AX-97790416 | 0 | C | T | 0.456 | -1.02E-01 | 6.31E-02 | 2.76E+00 | 2.84E+00 | 1.06E-01 | no | Grk5 | intron_variant | MODIFIER |
| 19 | 61001300 | AX-170100104 | 0 | A | C | 0.456 | -1.02E-01 | 6.31E-02 | 2.76E+00 | 2.84E+00 | 1.06E-01 | no | Grk5 | intron_variant | MODIFIER |
| 19 | 61001447 | AX-170467383 | 0 | A | C | 0.456 | -1.02E-01 | 6.31E-02 | 2.76E+00 | 2.84E+00 | 1.06E-01 | no | Grk5 | intron_variant | MODIFIER |
| 19 | 61004380 | AX-97836215 | 0 | G | A | 0.456 | -1.02E-01 | 6.31E-02 | 2.76E+00 | 2.84E+00 | 1.06E-01 | no | Grk5 | intron_variant | MODIFIER |
| 8 | 53337341 | AX-170030729 | 1 | A | T | 0.152 | -2.18E-01 | 1.35E-01 | 2.90E+00 | 3.01E+00 | 1.06E-01 | Yes | | intergenic_variant | MODIFIER |
| 16 | 56584863 | AX-170259588 | 1 | C | G | 0.464 | -1.38E-01 | 8.50E-02 | 2.53E+00 | 2.56E+00 | 1.06E-01 | no | Abi3bp | intron_variant | MODIFIER |
| 10 | 102668902 | AX-170512281 | 1 | G | A | 0.277 | -1.36E-01 | 8.43E-02 | 2.75E+00 | 2.83E+00 | 1.06E-01 | no | | intergenic_variant | MODIFIER |
| 4 | 109183866 | AX-170347952 | 1 | A | G | 0.259 | 1.82E-01 | 1.13E-01 | 2.67E+00 | 2.73E+00 | 1.06E-01 | Yes | Osbpl9 | intron_variant | MODIFIER |
| 3 | 18779184 | AX-97828606 | 1 | C | T | 0.143 | -1.63E-01 | 1.01E-01 | 2.64E+00 | 2.72E+00 | 1.06E-01 | no | | intergenic_variant | MODIFIER |
| 16 | 56887333 | AX-97788154 | 1 | T | C | 0.527 | 1.33E-01 | 8.19E-02 | 2.50E+00 | 2.54E+00 | 1.06E-01 | no | Tmem45a | upstream_gene_variant | MODIFIER |
| 10 | 101025064 | AX-170104024 | 1 | A | G | 0.286 | -1.37E-01 | 8.54E-02 | 2.67E+00 | 2.73E+00 | 1.06E-01 | no | | intergenic_variant | MODIFIER |
| 1 | 185424544 | AX-170099375 | 1 | T | C | 0.232 | 1.79E-01 | 1.11E-01 | 2.58E+00 | 2.66E+00 | 1.06E-01 | Yes | Eprs | intron_variant | MODIFIER |
| 6 | 111729257 | AX-170526451 | 0 | C | T | 0.061 | -2.20E-01 | 1.37E-01 | 2.84E+00 | 3.02E+00 | 1.06E-01 | Yes | | intergenic_variant | MODIFIER |
| 16 | 55498091 | AX-97836004 | 1 | T | C | 0.536 | 1.37E-01 | 8.50E-02 | 2.53E+00 | 2.56E+00 | 1.07E-01 | no | Mir5118 | downstream_gene_variant | MODIFIER |
| 16 | 72999979 | AX-170435673 | 1 | A | G | 0.5 | 1.23E-01 | 7.64E-02 | 2.64E+00 | 2.73E+00 | 1.07E-01 | no | Robo1 | intron_variant | MODIFIER |
| 19 | 60513684 | AX-170301546 | 1 | A | G | 0.455 | -1.01E-01 | 6.31E-02 | 2.76E+00 | 2.84E+00 | 1.07E-01 | no | | regulatory_region_variant | MODIFIER |
| 8 | 26138575 | AX-170390088 | 0 | G | T | 0.351 | -1.05E-01 | 6.55E-02 | 2.59E+00 | 2.68E+00 | 1.07E-01 | no | Rnf170 | intron_variant | MODIFIER |
| 16 | 57323477 | AX-170472265 | 1 | T | C | 0.464 | -1.37E-01 | 8.50E-02 | 2.53E+00 | 2.56E+00 | 1.07E-01 | no | Cmss1 | missense_variant | MODERATE |
| 5 | 50403740 | AX-170270680 | 0 | G | T | 0.509 | 1.63E-01 | 1.00E-01 | 2.43E+00 | 2.48E+00 | 1.07E-01 | Yes | | intergenic_variant | MODIFIER |
| 5 | 132653660 | AX-170311069 | 1 | A | C | 0.152 | -1.41E-01 | 8.51E-02 | 2.26E+00 | 2.34E+00 | 1.07E-01 | no | | regulatory_region_variant | MODIFIER |
| 4 | 83109048 | AX-170465787 | 1 | C | T | 0.339 | 1.84E-01 | 1.15E-01 | 2.89E+00 | 2.96E+00 | 1.07E-01 | Yes | | intergenic_variant | MODIFIER |
| 4 | 106603669 | AX-170381211 | 0 | C | T | 0.263 | 1.81E-01 | 1.13E-01 | 2.64E+00 | 2.71E+00 | 1.07E-01 | Yes | Lexm | intron_variant,NMD_transcript_variant | MODIFIER |
| 4 | 106604943 | AX-170250944 | 0 | G | A | 0.263 | 1.81E-01 | 1.13E-01 | 2.64E+00 | 2.71E+00 | 1.07E-01 | Yes | Lexm | intron_variant,NMD_transcript_variant | MODIFIER |
| 4 | 106605314 | AX-170156612 | 0 | A | G | 0.263 | 1.81E-01 | 1.13E-01 | 2.64E+00 | 2.71E+00 | 1.07E-01 | Yes | Lexm | intron_variant,NMD_transcript_variant | MODIFIER |
| 4 | 106864936 | AX-97809896 | 0 | C | T | 0.263 | 1.81E-01 | 1.13E-01 | 2.64E+00 | 2.71E+00 | 1.07E-01 | Yes | | regulatory_region_variant | MODIFIER |
| 4 | 106880703 | AX-170323237 | 0 | A | G | 0.263 | 1.81E-01 | 1.13E-01 | 2.64E+00 | 2.71E+00 | 1.07E-01 | Yes | | intergenic_variant | MODIFIER |
| 4 | 106888062 | AX-170250179 | 0 | T | C | 0.263 | 1.81E-01 | 1.13E-01 | 2.64E+00 | 2.71E+00 | 1.07E-01 | Yes | | intergenic_variant | MODIFIER |
| 4 | 106890099 | AX-170260654 | 0 | A | C | 0.263 | 1.81E-01 | 1.13E-01 | 2.64E+00 | 2.71E+00 | 1.07E-01 | Yes | | intergenic_variant | MODIFIER |
| 4 | 106890148 | AX-170113369 | 0 | T | A | 0.263 | 1.81E-01 | 1.13E-01 | 2.64E+00 | 2.71E+00 | 1.07E-01 | Yes | | intergenic_variant | MODIFIER |
| 4 | 106897702 | AX-170438752 | 0 | A | C | 0.263 | 1.81E-01 | 1.13E-01 | 2.64E+00 | 2.71E+00 | 1.07E-01 | Yes | | intergenic_variant | MODIFIER |
| 4 | 106917221 | AX-170147155 | 0 | T | C | 0.263 | 1.81E-01 | 1.13E-01 | 2.64E+00 | 2.71E+00 | 1.07E-01 | Yes | Ssbp3 | intron_variant | MODIFIER |
| 4 | 100321053 | AX-170023516 | 1 | C | T | 0.223 | 1.84E-01 | 1.15E-01 | 2.65E+00 | 2.73E+00 | 1.07E-01 | Yes | Ror1 | intron_variant | MODIFIER |
| 13 | 18572375 | AX-168415324 | 1 | C | T | 0.161 | -2.06E-01 | 1.27E-01 | 2.42E+00 | 2.46E+00 | 1.07E-01 | Yes | | intergenic_variant | MODIFIER |
| 13 | 19896188 | AX-170328384 | 0 | G | A | 0.167 | -2.11E-01 | 1.31E-01 | 2.50E+00 | 2.54E+00 | 1.08E-01 | Yes | | intergenic_variant | MODIFIER |
| 18 | 76189426 | AX-170061360 | 1 | T | C | 0.438 | 1.40E-01 | 8.71E-02 | 2.55E+00 | 2.60E+00 | 1.08E-01 | no | | intergenic_variant | MODIFIER |
| 18 | 80913584 | AX-97793449 | 1 | C | A | 0.205 | -1.07E-01 | 6.57E-02 | 2.41E+00 | 2.49E+00 | 1.08E-01 | no | Atp9b | intron_variant | MODIFIER |
| 4 | 104642053 | AX-170501963 | 1 | A | G | 0.241 | 1.83E-01 | 1.14E-01 | 2.75E+00 | 2.83E+00 | 1.08E-01 | Yes | Dab1 | intron_variant | MODIFIER |
| 4 | 103892947 | AX-97821005 | 0 | C | T | 0.246 | 1.75E-01 | 1.09E-01 | 2.74E+00 | 2.83E+00 | 1.08E-01 | Yes | Dab1 | intron_variant | MODIFIER |
| 1 | 184939009 | AX-170341936 | 1 | C | G | 0.241 | 1.79E-01 | 1.12E-01 | 2.55E+00 | 2.61E+00 | 1.08E-01 | Yes | Mark1 | intron_variant | MODIFIER |
| 10 | 100429816 | AX-97816577 | 0 | T | C | 0.272 | -1.34E-01 | 8.39E-02 | 2.61E+00 | 2.68E+00 | 1.08E-01 | no | Gm47673 | downstream_gene_variant | MODIFIER |
| 1 | 119370763 | AX-97808699 | 0 | C | T | 0.579 | 1.28E-01 | 8.01E-02 | 2.57E+00 | 2.63E+00 | 1.08E-01 | no | Gm29455 | intron_variant,non_coding_transcript_variant | MODIFIER |
| 5 | 62820109 | AX-170414610 | 0 | C | T | 0.482 | 1.59E-01 | 9.97E-02 | 2.64E+00 | 2.72E+00 | 1.09E-01 | Yes | Dthd1 | intron_variant | MODIFIER |
| 14 | 124361015 | AX-97805674 | 0 | G | T | 0.325 | 1.28E-01 | 8.02E-02 | 2.76E+00 | 2.88E+00 | 1.09E-01 | no | Fgf14 | intron_variant | MODIFIER |

| | | | | | | | | | | | | | | | |
|---|---|---|---|---|---|---|---|---|---|---|---|---|---|---|---|
| 19 | 60382830 | AX-97856463 | 0 | T | C | 0.474 | -1.01E-01 | 6.35E-02 | 2.82E+00 | 2.92E+00 | 1.09E-01 | no | | regulatory_region_variant | MODIFIER |
| 5 | 55382958 | AX-170024992 | 0 | G | A | 0.482 | 1.59E-01 | 9.88E-02 | 2.52E+00 | 2.58E+00 | 1.09E-01 | Yes | | intergenic_variant | MODIFIER |
| 5 | 64474131 | AX-170072315 | 0 | G | A | 0.482 | 1.59E-01 | 9.88E-02 | 2.52E+00 | 2.58E+00 | 1.09E-01 | Yes | C030018K13Rik | upstream_gene_variant | MODIFIER |
| 5 | 55452648 | AX-97805135 | 0 | T | C | 0.456 | 1.64E-01 | 1.01E-01 | 2.38E+00 | 2.43E+00 | 1.09E-01 | Yes | | intergenic_variant | MODIFIER |
| 5 | 55797688 | AX-170528740 | 0 | T | C | 0.456 | 1.64E-01 | 1.01E-01 | 2.38E+00 | 2.43E+00 | 1.09E-01 | Yes | Gm43705 | upstream_gene_variant | MODIFIER |
| 5 | 128543997 | AX-170479965 | 0 | T | C | 0.149 | -1.35E-01 | 8.18E-02 | 2.23E+00 | 2.34E+00 | 1.09E-01 | no | | intergenic_variant | MODIFIER |
| 1 | 134816399 | AX-170267791 | 1 | A | G | 0.562 | 1.30E-01 | 8.13E-02 | 2.67E+00 | 2.75E+00 | 1.09E-01 | no | Ppp1r12b | intron_variant | MODIFIER |
| 7 | 97689824 | AX-170145544 | 1 | G | A | 0.027 | -2.05E-01 | 1.29E-01 | 2.82E+00 | 3.02E+00 | 1.09E-01 | Yes | Rsf1 | 3_prime_UTR_variant | MODIFIER |
| 14 | 112419809 | AX-170352449 | 2 | C | T | 0.327 | 1.15E-01 | 7.24E-02 | 2.73E+00 | 2.85E+00 | 1.09E-01 | no | | intergenic_variant | MODIFIER |
| 19 | 59617968 | AX-170062842 | 1 | T | A | 0.518 | 1.00E-01 | 6.28E-02 | 2.73E+00 | 2.81E+00 | 1.09E-01 | no | | intergenic_variant | MODIFIER |
| 18 | 30780628 | AX-170093833 | 0 | T | A | 0.061 | -3.37E-01 | 2.11E-01 | 2.87E+00 | 2.98E+00 | 1.09E-01 | Yes | | intergenic_variant | MODIFIER |
| 5 | 130242098 | AX-170282785 | 1 | G | T | 0.152 | -1.48E-01 | 8.90E-02 | 2.17E+00 | 2.25E+00 | 1.09E-01 | no | Sbds | downstream_gene_variant | MODIFIER |
| 5 | 53487665 | AX-170421994 | 0 | C | T | 0.491 | 1.57E-01 | 9.69E-02 | 2.39E+00 | 2.46E+00 | 1.09E-01 | Yes | Rbpj | intron_variant | MODIFIER |
| 19 | 59757696 | AX-170496849 | 1 | A | G | 0.464 | -9.94E-02 | 6.24E-02 | 2.75E+00 | 2.84E+00 | 1.09E-01 | no | | intergenic_variant | MODIFIER |
| 15 | 72057016 | AX-170347323 | 0 | C | T | 0.079 | -1.96E-01 | 1.07E-01 | 1.77E+00 | 1.83E+00 | 1.09E-01 | Yes | | intergenic_variant | MODIFIER |
| 1 | 120208942 | AX-97854168 | 1 | G | A | 0.589 | 1.33E-01 | 8.27E-02 | 2.50E+00 | 2.56E+00 | 1.10E-01 | no | Steap3 | intron_variant | MODIFIER |
| 4 | 84607366 | AX-170164067 | 1 | C | T | 0.339 | 1.89E-01 | 1.18E-01 | 2.94E+00 | 3.01E+00 | 1.10E-01 | Yes | Bnc2 | intron_variant | MODIFIER |
| 1 | 144658288 | AX-170086635 | 0 | A | G | 0.219 | 1.90E-01 | 1.19E-01 | 2.85E+00 | 2.95E+00 | 1.10E-01 | Yes | | intergenic_variant | MODIFIER |
| 4 | 84577858 | AX-170331076 | 1 | G | A | 0.357 | 1.77E-01 | 1.11E-01 | 2.90E+00 | 2.99E+00 | 1.10E-01 | Yes | Bnc2 | intron_variant | MODIFIER |
| 4 | 104089462 | AX-170489382 | 1 | G | A | 0.241 | 1.85E-01 | 1.16E-01 | 2.67E+00 | 2.74E+00 | 1.10E-01 | Yes | Dab1 | intron_variant | MODIFIER |
| 9 | 67158125 | AX-170503394 | 0 | A | T | 0.193 | -1.93E-01 | 1.20E-01 | 3.11E+00 | 3.24E+00 | 1.10E-01 | Yes | Gm19299 | intron_variant,non_coding_transcript_variant | MODIFIER |
| 18 | 85833566 | AX-97796835 | 0 | T | C | 0.149 | -1.65E-01 | 1.04E-01 | 2.57E+00 | 2.68E+00 | 1.10E-01 | no | | intergenic_variant | MODIFIER |
| 4 | 82523394 | AX-97816738 | 0 | T | C | 0.342 | 1.89E-01 | 1.18E-01 | 2.95E+00 | 3.02E+00 | 1.10E-01 | Yes | Gm11266 | intron_variant,non_coding_transcript_variant | MODIFIER |
| 4 | 82579706 | AX-170089293 | 0 | G | A | 0.342 | 1.89E-01 | 1.18E-01 | 2.95E+00 | 3.02E+00 | 1.10E-01 | Yes | Nfib | intron_variant | MODIFIER |
| 4 | 82719540 | AX-97804075 | 0 | T | C | 0.342 | 1.89E-01 | 1.18E-01 | 2.95E+00 | 3.02E+00 | 1.10E-01 | Yes | | intergenic_variant | MODIFIER |
| 4 | 82725187 | AX-170296990 | 0 | T | C | 0.342 | 1.89E-01 | 1.18E-01 | 2.95E+00 | 3.02E+00 | 1.10E-01 | Yes | | intergenic_variant | MODIFIER |
| 4 | 82806226 | AX-170322801 | 0 | G | A | 0.342 | 1.89E-01 | 1.18E-01 | 2.95E+00 | 3.02E+00 | 1.10E-01 | Yes | Zdhhc21 | 3_prime_UTR_variant | MODIFIER |
| 4 | 82913961 | AX-170251914 | 0 | A | G | 0.342 | 1.89E-01 | 1.18E-01 | 2.95E+00 | 3.02E+00 | 1.10E-01 | Yes | Frem1 | splice_region_variant,intron_variant | LOW |
| 4 | 82918665 | AX-170229696 | 0 | C | A | 0.342 | 1.89E-01 | 1.18E-01 | 2.95E+00 | 3.02E+00 | 1.10E-01 | Yes | Frem1 | intron_variant | MODIFIER |
| 4 | 84109468 | AX-170515061 | 0 | G | A | 0.342 | 1.89E-01 | 1.18E-01 | 2.95E+00 | 3.02E+00 | 1.10E-01 | Yes | Gm12416 | downstream_gene_variant | MODIFIER |
| 4 | 84109983 | AX-170355547 | 0 | C | T | 0.342 | 1.89E-01 | 1.18E-01 | 2.95E+00 | 3.02E+00 | 1.10E-01 | Yes | Gm12416 | downstream_gene_variant | MODIFIER |
| 4 | 84112769 | AX-170156597 | 0 | G | A | 0.342 | 1.89E-01 | 1.18E-01 | 2.95E+00 | 3.02E+00 | 1.10E-01 | Yes | | intergenic_variant | MODIFIER |
| 4 | 84113211 | AX-97792786 | 0 | T | C | 0.342 | 1.89E-01 | 1.18E-01 | 2.95E+00 | 3.02E+00 | 1.10E-01 | Yes | | intergenic_variant | MODIFIER |
| 4 | 84113507 | AX-170506779 | 0 | A | G | 0.342 | 1.89E-01 | 1.18E-01 | 2.95E+00 | 3.02E+00 | 1.10E-01 | Yes | | intergenic_variant | MODIFIER |
| 4 | 84114668 | AX-170322263 | 0 | G | A | 0.342 | 1.89E-01 | 1.18E-01 | 2.95E+00 | 3.02E+00 | 1.10E-01 | Yes | | intergenic_variant | MODIFIER |
| 4 | 84115308 | AX-170023274 | 0 | C | T | 0.342 | 1.89E-01 | 1.18E-01 | 2.95E+00 | 3.02E+00 | 1.10E-01 | Yes | | intergenic_variant | MODIFIER |
| 4 | 84153999 | AX-170023276 | 0 | T | A | 0.342 | 1.89E-01 | 1.18E-01 | 2.95E+00 | 3.02E+00 | 1.10E-01 | Yes | | intergenic_variant | MODIFIER |
| 4 | 84156004 | AX-170023279 | 0 | A | G | 0.342 | 1.89E-01 | 1.18E-01 | 2.95E+00 | 3.02E+00 | 1.10E-01 | Yes | | intergenic_variant | MODIFIER |
| 4 | 84177034 | AX-170023275 | 0 | T | C | 0.342 | 1.89E-01 | 1.18E-01 | 2.95E+00 | 3.02E+00 | 1.10E-01 | Yes | | intergenic_variant | MODIFIER |
| 4 | 84263260 | AX-170237386 | 0 | T | C | 0.342 | 1.89E-01 | 1.18E-01 | 2.95E+00 | 3.02E+00 | 1.10E-01 | Yes | | regulatory_region_variant | MODIFIER |
| 4 | 84340371 | AX-170163985 | 0 | C | G | 0.342 | 1.89E-01 | 1.18E-01 | 2.95E+00 | 3.02E+00 | 1.10E-01 | Yes | Bnc2 | intron_variant | MODIFIER |
| 4 | 84638444 | AX-170523352 | 0 | G | A | 0.342 | 1.89E-01 | 1.18E-01 | 2.95E+00 | 3.02E+00 | 1.10E-01 | Yes | Bnc2 | intron_variant | MODIFIER |
| 4 | 84638525 | AX-170414341 | 0 | C | T | 0.342 | 1.89E-01 | 1.18E-01 | 2.95E+00 | 3.02E+00 | 1.10E-01 | Yes | Bnc2 | intron_variant | MODIFIER |
| 4 | 84667283 | AX-170363880 | 0 | C | T | 0.342 | 1.89E-01 | 1.18E-01 | 2.95E+00 | 3.02E+00 | 1.10E-01 | Yes | Bnc2 | intron_variant | MODIFIER |
| 4 | 84705503 | AX-170319071 | 0 | T | C | 0.342 | 1.89E-01 | 1.18E-01 | 2.95E+00 | 3.02E+00 | 1.10E-01 | Yes | | intergenic_variant | MODIFIER |
| 8 | 55103579 | AX-170195646 | 1 | C | G | 0.143 | -2.14E-01 | 1.34E-01 | 2.92E+00 | 3.04E+00 | 1.10E-01 | Yes | | intergenic_variant | MODIFIER |
| 1 | 181476767 | AX-170319343 | 1 | T | C | 0.232 | 1.77E-01 | 1.11E-01 | 2.53E+00 | 2.60E+00 | 1.10E-01 | Yes | | intergenic_variant | MODIFIER |
| 18 | 84121914 | AX-170316908 | 0 | A | G | 0.158 | -1.58E-01 | 9.94E-02 | 2.57E+00 | 2.66E+00 | 1.10E-01 | no | | regulatory_region_variant | MODIFIER |
| 6 | 104878787 | AX-170238836 | 0 | T | A | 0.053 | -2.48E-01 | 1.56E-01 | 2.60E+00 | 2.74E+00 | 1.10E-01 | Yes | | intergenic_variant | MODIFIER |
| 1 | 131072704 | AX-170042715 | 1 | C | T | 0.554 | 1.37E-01 | 8.62E-02 | 2.81E+00 | 2.89E+00 | 1.10E-01 | no | Mapkapk2 | intron_variant | MODIFIER |
| 3 | 31664544 | AX-97790220 | 0 | C | T | 0.158 | -1.61E-01 | 9.97E-02 | 3.18E+00 | 3.31E+00 | 1.11E-01 | no | Gm6558 | intron_variant,non_coding_transcript_variant | MODIFIER |
| 3 | 32344486 | AX-170020249 | 0 | C | T | 0.316 | -8.06E-02 | 4.99E-02 | 3.18E+00 | 3.31E+00 | 1.11E-01 | no | Zmat3 | intron_variant | MODIFIER |
| 4 | 86965076 | AX-97838314 | 2 | G | A | 0.345 | 1.74E-01 | 1.09E-01 | 2.84E+00 | 2.92E+00 | 1.11E-01 | Yes | | intergenic_variant | MODIFIER |
| 16 | 37847725 | AX-170161590 | 0 | T | C | 0.482 | 1.27E-01 | 7.98E-02 | 2.61E+00 | 2.66E+00 | 1.11E-01 | no | Gm36028 | downstream_gene_variant | MODIFIER |

| | | | | | | | | | | | | | | | |
|---|---|---|---|---|---|---|---|---|---|---|---|---|---|---|---|
| 16 | 58539074 | AX-170345363 | 0 | T | A | 0.456 | -1.38E-01 | 8.64E-02 | 2.55E+00 | 2.59E+00 | 1.11E-01 | no | | regulatory_region_variant | MODIFIER |
| 5 | 129001988 | AX-170367389 | 1 | G | A | 0.152 | -1.49E-01 | 8.93E-02 | 2.15E+00 | 2.22E+00 | 1.11E-01 | no | Stx2 | intron_variant | MODIFIER |
| 10 | 102274731 | AX-97816614 | 0 | T | C | 0.263 | -1.30E-01 | 8.20E-02 | 2.73E+00 | 2.82E+00 | 1.11E-01 | no | Mgat4c | intron_variant | MODIFIER |
| 1 | 135493184 | AX-170001766 | 1 | T | C | 0.527 | 1.29E-01 | 8.11E-02 | 2.75E+00 | 2.83E+00 | 1.11E-01 | no | Nav1 | intron_variant | MODIFIER |
| 5 | 8511862 | AX-170490130 | 1 | C | T | 0.036 | 1.99E-01 | 1.26E-01 | 2.91E+00 | 3.13E+00 | 1.11E-01 | Yes | Rundc3b | intron_variant | MODIFIER |
| 8 | 62731606 | AX-170506249 | 0 | T | G | 0.14 | -2.06E-01 | 1.30E-01 | 2.75E+00 | 2.87E+00 | 1.11E-01 | Yes | Gm45459 | intron_variant,non_coding_transcript_variant | MODIFIER |
| 16 | 40630413 | AX-97831026 | 1 | T | A | 0.5 | 1.22E-01 | 7.66E-02 | 2.51E+00 | 2.58E+00 | 1.11E-01 | no | Lsamp | intron_variant | MODIFIER |
| 1 | 127060586 | AX-170001647 | 0 | A | C | 0.544 | 1.33E-01 | 8.42E-02 | 2.73E+00 | 2.80E+00 | 1.12E-01 | no | | intergenic_variant | MODIFIER |
| 3 | 38498908 | AX-97814229 | 0 | C | T | 0.211 | 1.45E-01 | 9.01E-02 | 3.17E+00 | 3.30E+00 | 1.12E-01 | Yes | | intergenic_variant | MODIFIER |
| 16 | 63072380 | AX-170445281 | 1 | C | T | 0.518 | 1.36E-01 | 8.56E-02 | 2.64E+00 | 2.70E+00 | 1.12E-01 | no | Gm18870 | intron_variant,non_coding_transcript_variant | MODIFIER |
| 4 | 81938379 | AX-170332704 | 1 | G | A | 0.42 | -1.34E-01 | 8.46E-02 | 2.55E+00 | 2.61E+00 | 1.12E-01 | no | | intergenic_variant | MODIFIER |
| 12 | 91999125 | AX-170376629 | 0 | C | T | 0.079 | 2.70E-01 | 1.69E-01 | 3.04E+00 | 3.19E+00 | 1.12E-01 | Yes | | intergenic_variant | MODIFIER |
| 3 | 104251055 | AX-170235587 | 1 | G | A | 0.045 | 2.18E-01 | 1.38E-01 | 2.52E+00 | 2.66E+00 | 1.12E-01 | Yes | | intergenic_variant | MODIFIER |
| 19 | 60833742 | AX-170133537 | 1 | A | G | 0.464 | -1.01E-01 | 6.39E-02 | 2.81E+00 | 2.90E+00 | 1.12E-01 | no | Fam45a | intron_variant | MODIFIER |
| 18 | 80003867 | AX-170475737 | 1 | T | C | 0.473 | 1.27E-01 | 7.93E-02 | 2.44E+00 | 2.52E+00 | 1.12E-01 | no | | intergenic_variant | MODIFIER |
| 18 | 58227378 | AX-170353690 | 0 | T | C | 0.263 | -1.03E-01 | 6.53E-02 | 2.67E+00 | 2.77E+00 | 1.12E-01 | no | Gm25660 | downstream_gene_variant | MODIFIER |
| 10 | 101680788 | AX-170411804 | 0 | T | G | 0.263 | -1.32E-01 | 8.34E-02 | 2.59E+00 | 2.67E+00 | 1.12E-01 | no | Mgat4c | upstream_gene_variant | MODIFIER |
| 8 | 66022073 | AX-170324806 | 0 | A | G | 0.149 | -1.74E-01 | 1.10E-01 | 2.73E+00 | 2.88E+00 | 1.12E-01 | Yes | MAR1 | intron_variant | MODIFIER |
| 1 | 134191415 | AX-170234841 | 0 | G | T | 0.544 | 1.31E-01 | 8.30E-02 | 2.70E+00 | 2.77E+00 | 1.12E-01 | no | Mybph | upstream_gene_variant | MODIFIER |
| 13 | 7739815 | AX-170498803 | 0 | G | C | 0.167 | -1.97E-01 | 1.25E-01 | 2.55E+00 | 2.60E+00 | 1.13E-01 | Yes | | intergenic_variant | MODIFIER |
| 13 | 10361125 | AX-170282251 | 0 | T | G | 0.167 | -1.97E-01 | 1.25E-01 | 2.55E+00 | 2.60E+00 | 1.13E-01 | Yes | Gm26861 | intron_variant,non_coding_transcript_variant | MODIFIER |
| 3 | 23928990 | AX-170506167 | 0 | G | A | 0.158 | -1.53E-01 | 9.70E-02 | 2.60E+00 | 2.70E+00 | 1.13E-01 | no | Naaladl2 | intron_variant,non_coding_transcript_variant | MODIFIER |
| 5 | 49968615 | AX-170498718 | 0 | C | T | 0.482 | 1.62E-01 | 1.01E-01 | 2.40E+00 | 2.45E+00 | 1.13E-01 | Yes | Adgra3 | intron_variant | MODIFIER |
| 13 | 9618905 | AX-170288483 | 1 | A | G | 0.179 | -1.96E-01 | 1.23E-01 | 2.48E+00 | 2.52E+00 | 1.13E-01 | Yes | Gm23496 | downstream_gene_variant | MODIFIER |
| 13 | 11153704 | AX-170246880 | 1 | T | G | 0.179 | -1.96E-01 | 1.23E-01 | 2.48E+00 | 2.52E+00 | 1.13E-01 | Yes | | intergenic_variant | MODIFIER |
| 4 | 84037440 | AX-170515636 | 1 | A | G | 0.42 | -1.30E-01 | 8.19E-02 | 2.52E+00 | 2.58E+00 | 1.13E-01 | no | 6030471H07Rik | downstream_gene_variant | MODIFIER |
| 16 | 55715945 | AX-97810834 | 0 | G | A | 0.535 | 1.32E-01 | 8.25E-02 | 2.45E+00 | 2.49E+00 | 1.13E-01 | no | | intergenic_variant | MODIFIER |
| 15 | 76576282 | AX-170313516 | 0 | G | C | 0.079 | -1.91E-01 | 1.07E-01 | 1.85E+00 | 1.92E+00 | 1.13E-01 | Yes | Adck5 | upstream_gene_variant | MODIFIER |
| 16 | 56218345 | AX-170263647 | 1 | A | G | 0.527 | 1.32E-01 | 8.34E-02 | 2.53E+00 | 2.58E+00 | 1.13E-01 | no | Impg2 | downstream_gene_variant | MODIFIER |
| 6 | 136537598 | AX-170035741 | 2 | C | A | 0.564 | 1.25E-01 | 7.88E-02 | 2.52E+00 | 2.59E+00 | 1.13E-01 | no | Atf7ip | intron_variant | MODIFIER |
| 3 | 18559088 | AX-170020047 | 0 | A | G | 0.114 | -1.59E-01 | 1.01E-01 | 2.64E+00 | 2.73E+00 | 1.13E-01 | no | | intergenic_variant | MODIFIER |
| 3 | 18583337 | AX-97827827 | 0 | T | G | 0.114 | -1.59E-01 | 1.01E-01 | 2.64E+00 | 2.73E+00 | 1.13E-01 | no | | intergenic_variant | MODIFIER |
| 3 | 18627304 | AX-170318794 | 0 | G | C | 0.114 | -1.59E-01 | 1.01E-01 | 2.64E+00 | 2.73E+00 | 1.13E-01 | no | Gm42944 | downstream_gene_variant | MODIFIER |
| 3 | 18629504 | AX-170465523 | 0 | G | C | 0.114 | -1.59E-01 | 1.01E-01 | 2.64E+00 | 2.73E+00 | 1.13E-01 | no | Gm42944 | downstream_gene_variant | MODIFIER |
| 3 | 18630587 | AX-170211577 | 0 | C | T | 0.114 | -1.59E-01 | 1.01E-01 | 2.64E+00 | 2.73E+00 | 1.13E-01 | no | Gm42944 | downstream_gene_variant | MODIFIER |
| 3 | 18630762 | AX-170469248 | 0 | C | T | 0.114 | -1.59E-01 | 1.01E-01 | 2.64E+00 | 2.73E+00 | 1.13E-01 | no | Gm42944 | downstream_gene_variant | MODIFIER |
| 3 | 18652092 | AX-170020049 | 0 | A | G | 0.114 | -1.59E-01 | 1.01E-01 | 2.64E+00 | 2.73E+00 | 1.13E-01 | no | | intergenic_variant | MODIFIER |
| 3 | 18700130 | AX-97792782 | 0 | C | T | 0.114 | -1.59E-01 | 1.01E-01 | 2.64E+00 | 2.73E+00 | 1.13E-01 | no | | intergenic_variant | MODIFIER |
| 5 | 60490088 | AX-170122633 | 0 | C | T | 0.491 | 1.59E-01 | 1.00E-01 | 2.45E+00 | 2.52E+00 | 1.14E-01 | Yes | | intergenic_variant | MODIFIER |
| 5 | 129236134 | AX-170119762 | 1 | G | A | 0.152 | -1.49E-01 | 9.05E-02 | 2.17E+00 | 2.24E+00 | 1.14E-01 | no | | intergenic_variant | MODIFIER |
| 13 | 18250576 | AX-170087379 | 1 | G | A | 0.179 | -2.06E-01 | 1.29E-01 | 2.45E+00 | 2.48E+00 | 1.15E-01 | Yes | Pou6f2 | intron_variant | MODIFIER |
| 18 | 75134197 | AX-97820112 | 1 | C | T | 0.438 | 1.29E-01 | 8.20E-02 | 2.59E+00 | 2.67E+00 | 1.15E-01 | no | Dym | intron_variant | MODIFIER |
| 13 | 7379825 | AX-170007692 | 0 | C | T | 0.193 | -1.86E-01 | 1.18E-01 | 2.58E+00 | 2.64E+00 | 1.15E-01 | Yes | | intergenic_variant | MODIFIER |
| 13 | 10499231 | AX-97788183 | 0 | C | T | 0.193 | -1.86E-01 | 1.18E-01 | 2.58E+00 | 2.64E+00 | 1.15E-01 | Yes | Gm26861 | intron_variant,non_coding_transcript_variant | MODIFIER |
| 8 | 25176486 | AX-170329285 | 1 | A | G | 0.339 | -1.02E-01 | 6.52E-02 | 2.64E+00 | 2.74E+00 | 1.15E-01 | no | Tacc1 | intron_variant | MODIFIER |
| 1 | 122912050 | AX-170086599 | 1 | A | G | 0.571 | 1.33E-01 | 8.48E-02 | 2.65E+00 | 2.71E+00 | 1.15E-01 | no | | intergenic_variant | MODIFIER |
| 15 | 67040041 | AX-170122626 | 0 | C | A | 0.333 | -1.28E-01 | 7.81E-02 | 2.17E+00 | 2.23E+00 | 1.15E-01 | no | Gm31342 | upstream_gene_variant | MODIFIER |
| 4 | 81769059 | AX-170503803 | 0 | C | T | 0.421 | -1.33E-01 | 8.46E-02 | 2.55E+00 | 2.60E+00 | 1.15E-01 | no | | intergenic_variant | MODIFIER |
| 4 | 81792286 | AX-170247140 | 0 | T | C | 0.421 | -1.33E-01 | 8.46E-02 | 2.55E+00 | 2.60E+00 | 1.15E-01 | no | | intergenic_variant | MODIFIER |
| 4 | 81859857 | AX-97857660 | 0 | G | A | 0.421 | -1.33E-01 | 8.46E-02 | 2.55E+00 | 2.60E+00 | 1.15E-01 | no | | intergenic_variant | MODIFIER |
| 4 | 81933791 | AX-170130143 | 0 | G | A | 0.421 | -1.33E-01 | 8.46E-02 | 2.55E+00 | 2.60E+00 | 1.15E-01 | no | Gm12914 | upstream_gene_variant | MODIFIER |
| 4 | 82472234 | AX-170023251 | 0 | G | A | 0.421 | -1.33E-01 | 8.46E-02 | 2.55E+00 | 2.60E+00 | 1.15E-01 | no | Nfib | intron_variant | MODIFIER |
| 4 | 83150827 | AX-97801758 | 0 | C | T | 0.421 | -1.33E-01 | 8.46E-02 | 2.55E+00 | 2.60E+00 | 1.15E-01 | no | | intergenic_variant | MODIFIER |
| 8 | 68726311 | AX-170257065 | 0 | A | G | 0.184 | -1.82E-01 | 1.16E-01 | 2.74E+00 | 2.86E+00 | 1.15E-01 | Yes | Csgalnact1 | intron_variant | MODIFIER |

| | | | | | | | | | | | | | | | |
|---|---|---|---|---|---|---|---|---|---|---|---|---|---|---|---|
| 3 | 38872396 | AX-170468740 | 0 | A | G | 0.202 | 1.49E-01 | 9.32E-02 | 3.15E+00 | 3.28E+00 | 1.15E-01 | Yes | C230034O21Rik | downstream_gene_variant | MODIFIER |
| 3 | 24171642 | AX-170501668 | 0 | A | G | 0.158 | -1.53E-01 | 9.77E-02 | 2.66E+00 | 2.75E+00 | 1.15E-01 | no | Naaladl2 | missense_variant | MODERATE |
| 3 | 112324214 | AX-170094764 | 0 | A | C | 0.228 | -1.69E-01 | 1.08E-01 | 2.75E+00 | 2.82E+00 | 1.16E-01 | no | | intergenic_variant | MODIFIER |
| 15 | 3296016 | AX-170307213 | 0 | G | A | 0.544 | 8.43E-02 | 5.38E-02 | 2.75E+00 | 2.82E+00 | 1.16E-01 | no | Ccdc152 | intron_variant | MODIFIER |
| 15 | 5434139 | AX-170466960 | 0 | A | G | 0.228 | -1.69E-01 | 1.08E-01 | 2.75E+00 | 2.82E+00 | 1.16E-01 | no | | intergenic_variant | MODIFIER |
| 15 | 10624112 | AX-170267256 | 0 | A | G | 0.228 | -1.69E-01 | 1.08E-01 | 2.75E+00 | 2.82E+00 | 1.16E-01 | no | Rai14 | intron_variant | MODIFIER |
| 15 | 19421046 | AX-170499938 | 0 | T | A | 0.228 | -1.69E-01 | 1.08E-01 | 2.75E+00 | 2.82E+00 | 1.16E-01 | no | | intergenic_variant | MODIFIER |
| 15 | 20464394 | AX-170285974 | 0 | A | G | 0.228 | -1.69E-01 | 1.08E-01 | 2.75E+00 | 2.82E+00 | 1.16E-01 | no | Cdh12 | intron_variant,non_coding_transcript_variant | MODIFIER |
| 1 | 193952920 | AX-170091842 | 0 | A | G | 0.281 | 1.30E-01 | 8.19E-02 | 2.39E+00 | 2.48E+00 | 1.16E-01 | no | | intergenic_variant | MODIFIER |
| 4 | 81572016 | AX-170023237 | 0 | G | C | 0.333 | 1.82E-01 | 1.16E-01 | 2.93E+00 | 3.00E+00 | 1.16E-01 | Yes | | intergenic_variant | MODIFIER |
| 4 | 82345896 | AX-170222831 | 0 | A | G | 0.333 | 1.82E-01 | 1.16E-01 | 2.93E+00 | 3.00E+00 | 1.16E-01 | Yes | Nfib | intron_variant | MODIFIER |
| 4 | 82525936 | AX-170253762 | 0 | A | T | 0.333 | 1.82E-01 | 1.16E-01 | 2.93E+00 | 3.00E+00 | 1.16E-01 | Yes | Gm11266 | intron_variant,non_coding_transcript_variant | MODIFIER |
| 4 | 82528910 | AX-170422724 | 0 | G | A | 0.333 | 1.82E-01 | 1.16E-01 | 2.93E+00 | 3.00E+00 | 1.16E-01 | Yes | Gm11266 | intron_variant,non_coding_transcript_variant | MODIFIER |
| 4 | 82598224 | AX-170363754 | 0 | T | C | 0.333 | 1.82E-01 | 1.16E-01 | 2.93E+00 | 3.00E+00 | 1.16E-01 | Yes | Nfib | intron_variant | MODIFIER |
| 4 | 82604405 | AX-170097902 | 0 | G | A | 0.333 | 1.82E-01 | 1.16E-01 | 2.93E+00 | 3.00E+00 | 1.16E-01 | Yes | Nfib | intron_variant | MODIFIER |
| 4 | 82607007 | AX-97795686 | 0 | C | T | 0.333 | 1.82E-01 | 1.16E-01 | 2.93E+00 | 3.00E+00 | 1.16E-01 | Yes | Nfib | intron_variant | MODIFIER |
| 4 | 82668332 | AX-170069827 | 0 | C | T | 0.333 | 1.82E-01 | 1.16E-01 | 2.93E+00 | 3.00E+00 | 1.16E-01 | Yes | Nfib | intron_variant | MODIFIER |
| 4 | 82686826 | AX-170225641 | 0 | G | A | 0.333 | 1.82E-01 | 1.16E-01 | 2.93E+00 | 3.00E+00 | 1.16E-01 | Yes | Nfib | intron_variant | MODIFIER |
| 4 | 82769788 | AX-97820538 | 0 | G | A | 0.333 | 1.82E-01 | 1.16E-01 | 2.93E+00 | 3.00E+00 | 1.16E-01 | Yes | | intergenic_variant | MODIFIER |
| 4 | 82792960 | AX-170407177 | 0 | G | A | 0.333 | 1.82E-01 | 1.16E-01 | 2.93E+00 | 3.00E+00 | 1.16E-01 | Yes | | intergenic_variant | MODIFIER |
| 4 | 82806848 | AX-170522902 | 0 | G | A | 0.333 | 1.82E-01 | 1.16E-01 | 2.93E+00 | 3.00E+00 | 1.16E-01 | Yes | Zdhhc21 | intron_variant | MODIFIER |
| 4 | 82822144 | AX-170100209 | 0 | T | C | 0.333 | 1.82E-01 | 1.16E-01 | 2.93E+00 | 3.00E+00 | 1.16E-01 | Yes | Zdhhc21 | intron_variant | MODIFIER |
| 4 | 82837661 | AX-97785125 | 0 | A | C | 0.333 | 1.82E-01 | 1.16E-01 | 2.93E+00 | 3.00E+00 | 1.16E-01 | Yes | Zdhhc21 | intron_variant | MODIFIER |
| 4 | 83006747 | AX-170189023 | 0 | G | A | 0.333 | 1.82E-01 | 1.16E-01 | 2.93E+00 | 3.00E+00 | 1.16E-01 | Yes | Frem1 | intron_variant | MODIFIER |
| 4 | 83028204 | AX-170023261 | 0 | G | A | 0.333 | 1.82E-01 | 1.16E-01 | 2.93E+00 | 3.00E+00 | 1.16E-01 | Yes | Frem1 | intron_variant | MODIFIER |
| 4 | 83101634 | AX-97809527 | 0 | T | C | 0.333 | 1.82E-01 | 1.16E-01 | 2.93E+00 | 3.00E+00 | 1.16E-01 | Yes | | regulatory_region_variant | MODIFIER |
| 4 | 83108372 | AX-170023259 | 0 | C | T | 0.333 | 1.82E-01 | 1.16E-01 | 2.93E+00 | 3.00E+00 | 1.16E-01 | Yes | | regulatory_region_variant | MODIFIER |
| 4 | 83117223 | AX-170107093 | 0 | A | C | 0.333 | 1.82E-01 | 1.16E-01 | 2.93E+00 | 3.00E+00 | 1.16E-01 | Yes | Gm11248 | upstream_gene_variant | MODIFIER |
| 4 | 83147209 | AX-170023263 | 0 | G | A | 0.333 | 1.82E-01 | 1.16E-01 | 2.93E+00 | 3.00E+00 | 1.16E-01 | Yes | | intergenic_variant | MODIFIER |
| 4 | 83276069 | AX-170023258 | 0 | T | C | 0.333 | 1.82E-01 | 1.16E-01 | 2.93E+00 | 3.00E+00 | 1.16E-01 | Yes | Ttc39b | intron_variant | MODIFIER |
| 4 | 83342994 | AX-170495703 | 0 | C | A | 0.333 | 1.82E-01 | 1.16E-01 | 2.93E+00 | 3.00E+00 | 1.16E-01 | Yes | | regulatory_region_variant | MODIFIER |
| 4 | 83362473 | AX-170261086 | 0 | A | G | 0.333 | 1.82E-01 | 1.16E-01 | 2.93E+00 | 3.00E+00 | 1.16E-01 | Yes | | intergenic_variant | MODIFIER |
| 4 | 83376847 | AX-170412572 | 0 | C | A | 0.333 | 1.82E-01 | 1.16E-01 | 2.93E+00 | 3.00E+00 | 1.16E-01 | Yes | Gm11413 | downstream_gene_variant | MODIFIER |
| 4 | 83379315 | AX-170233948 | 0 | A | T | 0.333 | 1.82E-01 | 1.16E-01 | 2.93E+00 | 3.00E+00 | 1.16E-01 | Yes | Gm11184 | upstream_gene_variant | MODIFIER |
| 4 | 83396945 | AX-97813072 | 0 | A | G | 0.333 | 1.82E-01 | 1.16E-01 | 2.93E+00 | 3.00E+00 | 1.16E-01 | Yes | Gm11184 | downstream_gene_variant | MODIFIER |
| 4 | 83397148 | AX-170452235 | 0 | A | T | 0.333 | 1.82E-01 | 1.16E-01 | 2.93E+00 | 3.00E+00 | 1.16E-01 | Yes | Gm11184 | downstream_gene_variant | MODIFIER |
| 4 | 83422462 | AX-170348203 | 0 | A | C | 0.333 | 1.82E-01 | 1.16E-01 | 2.93E+00 | 3.00E+00 | 1.16E-01 | Yes | Snapc3 | intron_variant | MODIFIER |
| 4 | 83463029 | AX-170227088 | 0 | A | G | 0.333 | 1.82E-01 | 1.16E-01 | 2.93E+00 | 3.00E+00 | 1.16E-01 | Yes | Psip1 | intron_variant | MODIFIER |
| 4 | 83464062 | AX-170069841 | 0 | C | T | 0.333 | 1.82E-01 | 1.16E-01 | 2.93E+00 | 3.00E+00 | 1.16E-01 | Yes | Psip1 | intron_variant | MODIFIER |
| 4 | 83470040 | AX-170447855 | 0 | T | G | 0.333 | 1.82E-01 | 1.16E-01 | 2.93E+00 | 3.00E+00 | 1.16E-01 | Yes | Psip1 | intron_variant | MODIFIER |
| 4 | 83518426 | AX-170135882 | 0 | C | T | 0.333 | 1.82E-01 | 1.16E-01 | 2.93E+00 | 3.00E+00 | 1.16E-01 | Yes | | intergenic_variant | MODIFIER |
| 4 | 83657721 | AX-170464346 | 0 | A | C | 0.333 | 1.82E-01 | 1.16E-01 | 2.93E+00 | 3.00E+00 | 1.16E-01 | Yes | Ccdc171 | intron_variant | MODIFIER |
| 4 | 83832520 | AX-170112744 | 0 | A | G | 0.333 | 1.82E-01 | 1.16E-01 | 2.93E+00 | 3.00E+00 | 1.16E-01 | Yes | Ccdc171 | intron_variant | MODIFIER |
| 4 | 83899643 | AX-170410180 | 0 | A | C | 0.333 | 1.82E-01 | 1.16E-01 | 2.93E+00 | 3.00E+00 | 1.16E-01 | Yes | | intergenic_variant | MODIFIER |
| 4 | 84010431 | AX-170283789 | 0 | T | C | 0.333 | 1.82E-01 | 1.16E-01 | 2.93E+00 | 3.00E+00 | 1.16E-01 | Yes | | intergenic_variant | MODIFIER |
| 4 | 84059200 | AX-170241775 | 0 | A | T | 0.333 | 1.82E-01 | 1.16E-01 | 2.93E+00 | 3.00E+00 | 1.16E-01 | Yes | | intergenic_variant | MODIFIER |
| 4 | 84150727 | AX-170364992 | 0 | T | C | 0.333 | 1.82E-01 | 1.16E-01 | 2.93E+00 | 3.00E+00 | 1.16E-01 | Yes | | regulatory_region_variant | MODIFIER |
| 4 | 84175311 | AX-170233215 | 0 | C | T | 0.333 | 1.82E-01 | 1.16E-01 | 2.93E+00 | 3.00E+00 | 1.16E-01 | Yes | | intergenic_variant | MODIFIER |
| 4 | 84187879 | AX-170023280 | 0 | G | T | 0.333 | 1.82E-01 | 1.16E-01 | 2.93E+00 | 3.00E+00 | 1.16E-01 | Yes | | intergenic_variant | MODIFIER |
| 4 | 84190821 | AX-97807555 | 0 | A | G | 0.333 | 1.82E-01 | 1.16E-01 | 2.93E+00 | 3.00E+00 | 1.16E-01 | Yes | | intergenic_variant | MODIFIER |
| 4 | 84215359 | AX-170352728 | 0 | C | T | 0.333 | 1.82E-01 | 1.16E-01 | 2.93E+00 | 3.00E+00 | 1.16E-01 | Yes | | intergenic_variant | MODIFIER |
| 4 | 84218169 | AX-170277555 | 0 | G | A | 0.333 | 1.82E-01 | 1.16E-01 | 2.93E+00 | 3.00E+00 | 1.16E-01 | Yes | | intergenic_variant | MODIFIER |
| 4 | 84319935 | AX-170282527 | 0 | G | C | 0.333 | 1.82E-01 | 1.16E-01 | 2.93E+00 | 3.00E+00 | 1.16E-01 | Yes | Bnc2 | intron_variant | MODIFIER |
| 4 | 84320066 | AX-170514220 | 0 | A | G | 0.333 | 1.82E-01 | 1.16E-01 | 2.93E+00 | 3.00E+00 | 1.16E-01 | Yes | Bnc2 | intron_variant | MODIFIER |

| | | | | | | | | | | | | | | | |
|---|---|---|---|---|---|---|---|---|---|---|---|---|---|---|---|
| 4 | 84346929 | AX-170238847 | 0 | A | G | 0.333 | 1.82E-01 | 1.16E-01 | 2.93E+00 | 3.00E+00 | 1.16E-01 | Yes | Bnc2 | intron_variant | MODIFIER |
| 4 | 84351748 | AX-170342717 | 0 | C | T | 0.333 | 1.82E-01 | 1.16E-01 | 2.93E+00 | 3.00E+00 | 1.16E-01 | Yes | Bnc2 | intron_variant | MODIFIER |
| 4 | 84364265 | AX-170318510 | 0 | T | C | 0.333 | 1.82E-01 | 1.16E-01 | 2.93E+00 | 3.00E+00 | 1.16E-01 | Yes | Bnc2 | intron_variant | MODIFIER |
| 4 | 84379645 | AX-170317784 | 0 | C | T | 0.333 | 1.82E-01 | 1.16E-01 | 2.93E+00 | 3.00E+00 | 1.16E-01 | Yes | Bnc2 | intron_variant | MODIFIER |
| 4 | 84449895 | AX-170425493 | 0 | C | T | 0.333 | 1.82E-01 | 1.16E-01 | 2.93E+00 | 3.00E+00 | 1.16E-01 | Yes | Bnc2 | intron_variant | MODIFIER |
| 4 | 84528439 | AX-97848443 | 0 | T | G | 0.333 | 1.82E-01 | 1.16E-01 | 2.93E+00 | 3.00E+00 | 1.16E-01 | Yes | Bnc2 | intron_variant | MODIFIER |
| 4 | 84627295 | AX-170385363 | 0 | T | C | 0.333 | 1.82E-01 | 1.16E-01 | 2.93E+00 | 3.00E+00 | 1.16E-01 | Yes | Bnc2 | intron_variant | MODIFIER |
| 18 | 57899539 | AX-170140473 | 0 | T | G | 0.254 | -1.03E-01 | 6.59E-02 | 2.69E+00 | 2.79E+00 | 1.16E-01 | no | Slc12a2 | intron_variant | MODIFIER |
| 3 | 18803788 | AX-170519520 | 1 | C | T | 0.143 | -1.59E-01 | 1.01E-01 | 2.64E+00 | 2.72E+00 | 1.16E-01 | no | | intergenic_variant | MODIFIER |
| 9 | 102003364 | AX-170488924 | 0 | T | A | 0.14 | -1.51E-01 | 9.70E-02 | 2.70E+00 | 2.83E+00 | 1.16E-01 | no | Ephb1 | intron_variant | MODIFIER |
| 13 | 6152973 | AX-170452543 | 0 | G | C | 0.202 | -1.78E-01 | 1.14E-01 | 2.74E+00 | 2.84E+00 | 1.16E-01 | Yes | | intergenic_variant | MODIFIER |
| 13 | 19138778 | AX-170343302 | 1 | C | T | 0.17 | -1.92E-01 | 1.22E-01 | 2.46E+00 | 2.52E+00 | 1.16E-01 | Yes | Amph | intron_variant | MODIFIER |
| 4 | 54703651 | AX-170218825 | 0 | G | T | 0.368 | -1.60E-01 | 1.01E-01 | 3.05E+00 | 3.18E+00 | 1.16E-01 | no | | intergenic_variant | MODIFIER |
| 2 | 166947561 | AX-168391561 | 0 | G | A | 0.158 | 2.54E-01 | 1.62E-01 | 2.97E+00 | 3.11E+00 | 1.16E-01 | Yes | Cse1l | downstream_gene_variant | MODIFIER |
| 8 | 85217161 | AX-170520860 | 1 | G | C | 0.375 | -1.03E-01 | 6.37E-02 | 3.36E+00 | 3.53E+00 | 1.16E-01 | no | | intergenic_variant | MODIFIER |
| 4 | 106622341 | AX-170261272 | 0 | A | T | 0.272 | 1.73E-01 | 1.11E-01 | 2.66E+00 | 2.73E+00 | 1.16E-01 | Yes | Ttc22 | upstream_gene_variant | MODIFIER |
| 4 | 106926887 | AX-170070270 | 0 | C | T | 0.272 | 1.73E-01 | 1.11E-01 | 2.66E+00 | 2.73E+00 | 1.16E-01 | Yes | Ssbp3 | intron_variant | MODIFIER |
| 4 | 106969177 | AX-170449690 | 0 | G | A | 0.272 | 1.73E-01 | 1.11E-01 | 2.66E+00 | 2.73E+00 | 1.16E-01 | Yes | Ssbp3 | intron_variant | MODIFIER |
| 4 | 107013070 | AX-170519278 | 0 | A | T | 0.272 | 1.73E-01 | 1.11E-01 | 2.66E+00 | 2.73E+00 | 1.16E-01 | Yes | Ssbp3 | intron_variant | MODIFIER |
| 4 | 107393037 | AX-170385182 | 0 | G | A | 0.272 | 1.73E-01 | 1.11E-01 | 2.66E+00 | 2.73E+00 | 1.16E-01 | Yes | Ndc1 | intron_variant,NMD_transcript_variant | MODIFIER |
| 4 | 107393657 | AX-170244436 | 0 | C | T | 0.272 | 1.73E-01 | 1.11E-01 | 2.66E+00 | 2.73E+00 | 1.16E-01 | Yes | Ndc1 | intron_variant,NMD_transcript_variant | MODIFIER |
| 4 | 107411699 | AX-170136344 | 0 | G | A | 0.272 | 1.73E-01 | 1.11E-01 | 2.66E+00 | 2.73E+00 | 1.16E-01 | Yes | Ndc1 | intron_variant,NMD_transcript_variant | MODIFIER |
| 4 | 107412930 | AX-97793277 | 0 | G | A | 0.272 | 1.73E-01 | 1.11E-01 | 2.66E+00 | 2.73E+00 | 1.16E-01 | Yes | Ndc1 | intron_variant,NMD_transcript_variant | MODIFIER |
| 4 | 107417730 | AX-170449401 | 0 | A | T | 0.272 | 1.73E-01 | 1.11E-01 | 2.66E+00 | 2.73E+00 | 1.16E-01 | Yes | Ndc1 | downstream_gene_variant | MODIFIER |
| 4 | 107425256 | AX-170195400 | 0 | G | A | 0.272 | 1.73E-01 | 1.11E-01 | 2.66E+00 | 2.73E+00 | 1.16E-01 | Yes | | intergenic_variant | MODIFIER |
| 5 | 67324674 | AX-170314414 | 0 | C | T | 0.482 | 1.52E-01 | 9.69E-02 | 2.59E+00 | 2.68E+00 | 1.16E-01 | Yes | Slc30a9 | intron_variant | MODIFIER |
| 5 | 67327597 | AX-170464500 | 0 | C | T | 0.482 | 1.52E-01 | 9.69E-02 | 2.59E+00 | 2.68E+00 | 1.16E-01 | Yes | Slc30a9 | intron_variant | MODIFIER |
| 16 | 54648578 | AX-97856064 | 0 | T | C | 0.526 | 1.35E-01 | 8.63E-02 | 2.54E+00 | 2.57E+00 | 1.17E-01 | no | | intergenic_variant | MODIFIER |
| 16 | 54663184 | AX-170013084 | 0 | A | T | 0.526 | 1.35E-01 | 8.63E-02 | 2.54E+00 | 2.57E+00 | 1.17E-01 | no | | intergenic_variant | MODIFIER |
| 16 | 54663272 | AX-170093493 | 0 | T | C | 0.526 | 1.35E-01 | 8.63E-02 | 2.54E+00 | 2.57E+00 | 1.17E-01 | no | | intergenic_variant | MODIFIER |
| 16 | 54663536 | AX-170013082 | 0 | C | G | 0.526 | 1.35E-01 | 8.63E-02 | 2.54E+00 | 2.57E+00 | 1.17E-01 | no | | intergenic_variant | MODIFIER |
| 16 | 54668465 | AX-170218080 | 0 | C | G | 0.526 | 1.35E-01 | 8.63E-02 | 2.54E+00 | 2.57E+00 | 1.17E-01 | no | | intergenic_variant | MODIFIER |
| 16 | 54819744 | AX-97840355 | 0 | G | A | 0.526 | 1.35E-01 | 8.63E-02 | 2.54E+00 | 2.57E+00 | 1.17E-01 | no | | intergenic_variant | MODIFIER |
| 4 | 105505705 | AX-170035692 | 0 | T | C | 0.237 | 1.81E-01 | 1.16E-01 | 2.72E+00 | 2.79E+00 | 1.17E-01 | Yes | | intergenic_variant | MODIFIER |
| 4 | 105919263 | AX-97828985 | 0 | T | C | 0.237 | 1.81E-01 | 1.16E-01 | 2.72E+00 | 2.79E+00 | 1.17E-01 | Yes | | intergenic_variant | MODIFIER |
| 4 | 105923663 | AX-170373001 | 0 | T | C | 0.237 | 1.81E-01 | 1.16E-01 | 2.72E+00 | 2.79E+00 | 1.17E-01 | Yes | | intergenic_variant | MODIFIER |
| 4 | 105947089 | AX-170435285 | 0 | C | G | 0.237 | 1.81E-01 | 1.16E-01 | 2.72E+00 | 2.79E+00 | 1.17E-01 | Yes | | intergenic_variant | MODIFIER |
| 4 | 105978113 | AX-170368116 | 0 | T | G | 0.237 | 1.81E-01 | 1.16E-01 | 2.72E+00 | 2.79E+00 | 1.17E-01 | Yes | Gm24363 | upstream_gene_variant | MODIFIER |
| 4 | 105989343 | AX-170466205 | 0 | T | G | 0.237 | 1.81E-01 | 1.16E-01 | 2.72E+00 | 2.79E+00 | 1.17E-01 | Yes | | intergenic_variant | MODIFIER |
| 4 | 105992928 | AX-170229331 | 0 | G | A | 0.237 | 1.81E-01 | 1.16E-01 | 2.72E+00 | 2.79E+00 | 1.17E-01 | Yes | | regulatory_region_variant | MODIFIER |
| 4 | 105994128 | AX-170250236 | 0 | A | G | 0.237 | 1.81E-01 | 1.16E-01 | 2.72E+00 | 2.79E+00 | 1.17E-01 | Yes | | intergenic_variant | MODIFIER |
| 4 | 106008599 | AX-170373071 | 0 | A | G | 0.237 | 1.81E-01 | 1.16E-01 | 2.72E+00 | 2.79E+00 | 1.17E-01 | Yes | | intergenic_variant | MODIFIER |
| 16 | 29180049 | AX-170313452 | 0 | A | T | 0.5 | 1.26E-01 | 8.07E-02 | 2.71E+00 | 2.78E+00 | 1.17E-01 | no | | intergenic_variant | MODIFIER |
| 8 | 25049587 | AX-170233015 | 0 | A | T | 0.351 | -9.87E-02 | 6.33E-02 | 2.65E+00 | 2.75E+00 | 1.17E-01 | no | Plekha2 | intron_variant | MODIFIER |
| 13 | 18078498 | AX-97852225 | 0 | G | A | 0.158 | -2.00E-01 | 1.27E-01 | 2.43E+00 | 2.47E+00 | 1.17E-01 | Yes | | intergenic_variant | MODIFIER |
| 13 | 18082465 | AX-170139312 | 0 | G | C | 0.158 | -2.00E-01 | 1.27E-01 | 2.43E+00 | 2.47E+00 | 1.17E-01 | Yes | 4930448F12Rik | downstream_gene_variant | MODIFIER |
| 13 | 18137993 | AX-170528329 | 0 | C | G | 0.158 | -2.00E-01 | 1.27E-01 | 2.43E+00 | 2.47E+00 | 1.17E-01 | Yes | Pou6f2 | intron_variant | MODIFIER |
| 13 | 18138677 | AX-170280422 | 0 | T | C | 0.158 | -2.00E-01 | 1.27E-01 | 2.43E+00 | 2.47E+00 | 1.17E-01 | Yes | Pou6f2 | intron_variant | MODIFIER |
| 13 | 18150141 | AX-170237287 | 0 | T | A | 0.158 | -2.00E-01 | 1.27E-01 | 2.43E+00 | 2.47E+00 | 1.17E-01 | Yes | Pou6f2 | intron_variant | MODIFIER |
| 13 | 18150860 | AX-97808085 | 0 | C | T | 0.158 | -2.00E-01 | 1.27E-01 | 2.43E+00 | 2.47E+00 | 1.17E-01 | Yes | Pou6f2 | intron_variant | MODIFIER |
| 13 | 18153011 | AX-170489492 | 0 | A | G | 0.158 | -2.00E-01 | 1.27E-01 | 2.43E+00 | 2.47E+00 | 1.17E-01 | Yes | Pou6f2 | intron_variant | MODIFIER |
| 13 | 18154946 | AX-168390992 | 0 | T | C | 0.158 | -2.00E-01 | 1.27E-01 | 2.43E+00 | 2.47E+00 | 1.17E-01 | Yes | Pou6f2 | intron_variant | MODIFIER |
| 13 | 18162717 | AX-170423794 | 0 | A | C | 0.158 | -2.00E-01 | 1.27E-01 | 2.43E+00 | 2.47E+00 | 1.17E-01 | Yes | Pou6f2 | intron_variant | MODIFIER |
| 13 | 18162758 | AX-170487791 | 0 | C | T | 0.158 | -2.00E-01 | 1.27E-01 | 2.43E+00 | 2.47E+00 | 1.17E-01 | Yes | Pou6f2 | intron_variant | MODIFIER |

| | | | | | | | | | | | | | | | |
|---|---|---|---|---|---|---|---|---|---|---|---|---|---|---|---|
| 13 | 18168841 | AX-97789405 | 0 | T | C | 0.158 | -2.00E-01 | 1.27E-01 | 2.43E+00 | 2.47E+00 | 1.17E-01 | Yes | Pou6f2 | intron_variant | MODIFIER |
| 13 | 18230059 | AX-97844220 | 0 | A | G | 0.158 | -2.00E-01 | 1.27E-01 | 2.43E+00 | 2.47E+00 | 1.17E-01 | Yes | Pou6f2 | intron_variant | MODIFIER |
| 13 | 18252379 | AX-170489737 | 0 | T | C | 0.158 | -2.00E-01 | 1.27E-01 | 2.43E+00 | 2.47E+00 | 1.17E-01 | Yes | Pou6f2 | intron_variant | MODIFIER |
| 13 | 18271459 | AX-170163531 | 0 | T | G | 0.158 | -2.00E-01 | 1.27E-01 | 2.43E+00 | 2.47E+00 | 1.17E-01 | Yes | Pou6f2 | intron_variant | MODIFIER |
| 13 | 18297883 | AX-170284036 | 0 | T | G | 0.158 | -2.00E-01 | 1.27E-01 | 2.43E+00 | 2.47E+00 | 1.17E-01 | Yes | Pou6f2 | intron_variant | MODIFIER |
| 13 | 18315686 | AX-170175594 | 0 | A | G | 0.158 | -2.00E-01 | 1.27E-01 | 2.43E+00 | 2.47E+00 | 1.17E-01 | Yes | Pou6f2 | intron_variant | MODIFIER |
| 13 | 18325814 | AX-170129305 | 0 | G | A | 0.158 | -2.00E-01 | 1.27E-01 | 2.43E+00 | 2.47E+00 | 1.17E-01 | Yes | Pou6f2 | intron_variant | MODIFIER |
| 13 | 18339093 | AX-170143893 | 0 | A | C | 0.158 | -2.00E-01 | 1.27E-01 | 2.43E+00 | 2.47E+00 | 1.17E-01 | Yes | Pou6f2 | intron_variant | MODIFIER |
| 13 | 18363304 | AX-170433473 | 0 | G | A | 0.158 | -2.00E-01 | 1.27E-01 | 2.43E+00 | 2.47E+00 | 1.17E-01 | Yes | Pou6f2 | intron_variant | MODIFIER |
| 13 | 18379266 | AX-170218094 | 0 | C | A | 0.158 | -2.00E-01 | 1.27E-01 | 2.43E+00 | 2.47E+00 | 1.17E-01 | Yes | Pou6f2 | intron_variant | MODIFIER |
| 13 | 18381064 | AX-170209330 | 0 | A | G | 0.158 | -2.00E-01 | 1.27E-01 | 2.43E+00 | 2.47E+00 | 1.17E-01 | Yes | Pou6f2 | intron_variant | MODIFIER |
| 13 | 18398977 | AX-97852045 | 0 | C | T | 0.158 | -2.00E-01 | 1.27E-01 | 2.43E+00 | 2.47E+00 | 1.17E-01 | Yes | Pou6f2 | upstream_gene_variant | MODIFIER |
| 13 | 18403816 | AX-170152058 | 0 | A | G | 0.158 | -2.00E-01 | 1.27E-01 | 2.43E+00 | 2.47E+00 | 1.17E-01 | Yes | | intergenic_variant | MODIFIER |
| 13 | 18485752 | AX-97812639 | 0 | T | C | 0.158 | -2.00E-01 | 1.27E-01 | 2.43E+00 | 2.47E+00 | 1.17E-01 | Yes | | intergenic_variant | MODIFIER |
| 13 | 18489594 | AX-170401854 | 0 | A | G | 0.158 | -2.00E-01 | 1.27E-01 | 2.43E+00 | 2.47E+00 | 1.17E-01 | Yes | | intergenic_variant | MODIFIER |
| 13 | 18517901 | AX-170228267 | 0 | T | C | 0.158 | -2.00E-01 | 1.27E-01 | 2.43E+00 | 2.47E+00 | 1.17E-01 | Yes | | intergenic_variant | MODIFIER |
| 13 | 18518154 | AX-170170208 | 0 | G | A | 0.158 | -2.00E-01 | 1.27E-01 | 2.43E+00 | 2.47E+00 | 1.17E-01 | Yes | | intergenic_variant | MODIFIER |
| 13 | 18555898 | AX-170050648 | 0 | G | A | 0.158 | -2.00E-01 | 1.27E-01 | 2.43E+00 | 2.47E+00 | 1.17E-01 | Yes | | intergenic_variant | MODIFIER |
| 13 | 18568878 | AX-97822970 | 0 | C | A | 0.158 | -2.00E-01 | 1.27E-01 | 2.43E+00 | 2.47E+00 | 1.17E-01 | Yes | | intergenic_variant | MODIFIER |
| 13 | 18579545 | AX-170433189 | 0 | G | A | 0.158 | -2.00E-01 | 1.27E-01 | 2.43E+00 | 2.47E+00 | 1.17E-01 | Yes | | intergenic_variant | MODIFIER |
| 13 | 18592219 | AX-170407093 | 0 | A | G | 0.158 | -2.00E-01 | 1.27E-01 | 2.43E+00 | 2.47E+00 | 1.17E-01 | Yes | | intergenic_variant | MODIFIER |
| 13 | 18592516 | AX-170381823 | 0 | A | T | 0.158 | -2.00E-01 | 1.27E-01 | 2.43E+00 | 2.47E+00 | 1.17E-01 | Yes | | intergenic_variant | MODIFIER |
| 13 | 18593403 | AX-170127238 | 0 | G | A | 0.158 | -2.00E-01 | 1.27E-01 | 2.43E+00 | 2.47E+00 | 1.17E-01 | Yes | | intergenic_variant | MODIFIER |
| 13 | 18608601 | AX-168379755 | 0 | G | A | 0.158 | -2.00E-01 | 1.27E-01 | 2.43E+00 | 2.47E+00 | 1.17E-01 | Yes | | intergenic_variant | MODIFIER |
| 13 | 18623937 | AX-170410173 | 0 | C | T | 0.158 | -2.00E-01 | 1.27E-01 | 2.43E+00 | 2.47E+00 | 1.17E-01 | Yes | | intergenic_variant | MODIFIER |
| 13 | 18676438 | AX-170050652 | 0 | C | T | 0.158 | -2.00E-01 | 1.27E-01 | 2.43E+00 | 2.47E+00 | 1.17E-01 | Yes | | intergenic_variant | MODIFIER |
| 13 | 18676971 | AX-170108590 | 0 | C | T | 0.158 | -2.00E-01 | 1.27E-01 | 2.43E+00 | 2.47E+00 | 1.17E-01 | Yes | | intergenic_variant | MODIFIER |
| 13 | 18680545 | AX-170472633 | 0 | A | G | 0.158 | -2.00E-01 | 1.27E-01 | 2.43E+00 | 2.47E+00 | 1.17E-01 | Yes | | intergenic_variant | MODIFIER |
| 13 | 18704624 | AX-170345488 | 0 | G | A | 0.158 | -2.00E-01 | 1.27E-01 | 2.43E+00 | 2.47E+00 | 1.17E-01 | Yes | | intergenic_variant | MODIFIER |
| 13 | 18705160 | AX-170318096 | 0 | C | T | 0.158 | -2.00E-01 | 1.27E-01 | 2.43E+00 | 2.47E+00 | 1.17E-01 | Yes | | intergenic_variant | MODIFIER |
| 13 | 18715854 | AX-170381925 | 0 | A | G | 0.158 | -2.00E-01 | 1.27E-01 | 2.43E+00 | 2.47E+00 | 1.17E-01 | Yes | Vps41 | upstream_gene_variant | MODIFIER |
| 13 | 18739605 | AX-170347839 | 0 | A | G | 0.158 | -2.00E-01 | 1.27E-01 | 2.43E+00 | 2.47E+00 | 1.17E-01 | Yes | Vps41 | intron_variant | MODIFIER |
| 13 | 18741062 | AX-170182019 | 0 | G | C | 0.158 | -2.00E-01 | 1.27E-01 | 2.43E+00 | 2.47E+00 | 1.17E-01 | Yes | Vps41 | intron_variant | MODIFIER |
| 13 | 18741191 | AX-170288405 | 0 | T | C | 0.158 | -2.00E-01 | 1.27E-01 | 2.43E+00 | 2.47E+00 | 1.17E-01 | Yes | Vps41 | intron_variant | MODIFIER |
| 13 | 18745355 | AX-97848708 | 0 | A | C | 0.158 | -2.00E-01 | 1.27E-01 | 2.43E+00 | 2.47E+00 | 1.17E-01 | Yes | Vps41 | intron_variant | MODIFIER |
| 13 | 18761446 | AX-170303577 | 0 | G | A | 0.158 | -2.00E-01 | 1.27E-01 | 2.43E+00 | 2.47E+00 | 1.17E-01 | Yes | Vps41 | intron_variant | MODIFIER |
| 13 | 18763825 | AX-170334051 | 0 | C | T | 0.158 | -2.00E-01 | 1.27E-01 | 2.43E+00 | 2.47E+00 | 1.17E-01 | Yes | Vps41 | intron_variant | MODIFIER |
| 13 | 18763963 | AX-170351776 | 0 | G | A | 0.158 | -2.00E-01 | 1.27E-01 | 2.43E+00 | 2.47E+00 | 1.17E-01 | Yes | Vps41 | intron_variant | MODIFIER |
| 13 | 18767103 | AX-97781777 | 0 | T | C | 0.158 | -2.00E-01 | 1.27E-01 | 2.43E+00 | 2.47E+00 | 1.17E-01 | Yes | Vps41 | intron_variant | MODIFIER |
| 13 | 18770765 | AX-170197315 | 0 | G | C | 0.158 | -2.00E-01 | 1.27E-01 | 2.43E+00 | 2.47E+00 | 1.17E-01 | Yes | Vps41 | intron_variant | MODIFIER |
| 13 | 18786710 | AX-170441027 | 0 | A | G | 0.158 | -2.00E-01 | 1.27E-01 | 2.43E+00 | 2.47E+00 | 1.17E-01 | Yes | Vps41 | intron_variant | MODIFIER |
| 13 | 18801994 | AX-170427551 | 0 | C | T | 0.158 | -2.00E-01 | 1.27E-01 | 2.43E+00 | 2.47E+00 | 1.17E-01 | Yes | Vps41 | intron_variant | MODIFIER |
| 13 | 18818905 | AX-170254151 | 0 | A | G | 0.158 | -2.00E-01 | 1.27E-01 | 2.43E+00 | 2.47E+00 | 1.17E-01 | Yes | Vps41 | intron_variant | MODIFIER |
| 13 | 18827453 | AX-97852562 | 0 | C | T | 0.158 | -2.00E-01 | 1.27E-01 | 2.43E+00 | 2.47E+00 | 1.17E-01 | Yes | Vps41 | intron_variant | MODIFIER |
| 13 | 18839130 | AX-170493367 | 0 | G | A | 0.158 | -2.00E-01 | 1.27E-01 | 2.43E+00 | 2.47E+00 | 1.17E-01 | Yes | Vps41 | intron_variant | MODIFIER |
| 13 | 18840588 | AX-170139741 | 0 | C | G | 0.158 | -2.00E-01 | 1.27E-01 | 2.43E+00 | 2.47E+00 | 1.17E-01 | Yes | Vps41 | intron_variant | MODIFIER |
| 13 | 18842539 | AX-170406298 | 0 | A | G | 0.158 | -2.00E-01 | 1.27E-01 | 2.43E+00 | 2.47E+00 | 1.17E-01 | Yes | Vps41 | intron_variant | MODIFIER |
| 13 | 18847827 | AX-170389438 | 0 | C | G | 0.158 | -2.00E-01 | 1.27E-01 | 2.43E+00 | 2.47E+00 | 1.17E-01 | Yes | Vps41 | intron_variant | MODIFIER |
| 13 | 18851776 | AX-170007870 | 0 | C | T | 0.158 | -2.00E-01 | 1.27E-01 | 2.43E+00 | 2.47E+00 | 1.17E-01 | Yes | Vps41 | intron_variant | MODIFIER |
| 13 | 18854138 | AX-170007868 | 0 | C | T | 0.158 | -2.00E-01 | 1.27E-01 | 2.43E+00 | 2.47E+00 | 1.17E-01 | Yes | Vps41 | intron_variant | MODIFIER |
| 13 | 18862108 | AX-170233908 | 0 | C | A | 0.158 | -2.00E-01 | 1.27E-01 | 2.43E+00 | 2.47E+00 | 1.17E-01 | Yes | Vps41 | intron_variant | MODIFIER |
| 13 | 18862131 | AX-170252482 | 0 | G | A | 0.158 | -2.00E-01 | 1.27E-01 | 2.43E+00 | 2.47E+00 | 1.17E-01 | Yes | Vps41 | intron_variant | MODIFIER |
| 13 | 18864131 | AX-170007869 | 0 | G | A | 0.158 | -2.00E-01 | 1.27E-01 | 2.43E+00 | 2.47E+00 | 1.17E-01 | Yes | Vps41 | intron_variant | MODIFIER |
| 13 | 18987677 | AX-170050663 | 0 | T | C | 0.158 | -2.00E-01 | 1.27E-01 | 2.43E+00 | 2.47E+00 | 1.17E-01 | Yes | Amph | intron_variant | MODIFIER |

| | | | | | | | | | | | | | | | |
|---|---|---|---|---|---|---|---|---|---|---|---|---|---|---|---|
| 3 | 18709118 | AX-170136399 | 0 | C | T | 0.14 | -1.58E-01 | 1.01E-01 | 2.61E+00 | 2.69E+00 | 1.17E-01 | no | | intergenic_variant | MODIFIER |
| 3 | 18710067 | AX-170134365 | 0 | C | A | 0.14 | -1.58E-01 | 1.01E-01 | 2.61E+00 | 2.69E+00 | 1.17E-01 | no | | intergenic_variant | MODIFIER |
| 3 | 18710554 | AX-170020045 | 0 | C | A | 0.14 | -1.58E-01 | 1.01E-01 | 2.61E+00 | 2.69E+00 | 1.17E-01 | no | | intergenic_variant | MODIFIER |
| 3 | 18757994 | AX-170298623 | 0 | A | C | 0.14 | -1.58E-01 | 1.01E-01 | 2.61E+00 | 2.69E+00 | 1.17E-01 | no | Gm30341 | downstream_gene_variant | MODIFIER |
| 3 | 18758213 | AX-170394202 | 0 | A | G | 0.14 | -1.58E-01 | 1.01E-01 | 2.61E+00 | 2.69E+00 | 1.17E-01 | no | Gm30341 | downstream_gene_variant | MODIFIER |
| 3 | 18790304 | AX-170298113 | 0 | C | T | 0.14 | -1.58E-01 | 1.01E-01 | 2.61E+00 | 2.69E+00 | 1.17E-01 | no | | intergenic_variant | MODIFIER |
| 3 | 18807737 | AX-170088867 | 0 | G | T | 0.14 | -1.58E-01 | 1.01E-01 | 2.61E+00 | 2.69E+00 | 1.17E-01 | no | | intergenic_variant | MODIFIER |
| 3 | 18834019 | AX-170316863 | 0 | C | T | 0.14 | -1.58E-01 | 1.01E-01 | 2.61E+00 | 2.69E+00 | 1.17E-01 | no | | intergenic_variant | MODIFIER |
| 5 | 51339142 | AX-170024938 | 1 | A | G | 0.509 | 1.53E-01 | 9.67E-02 | 2.43E+00 | 2.51E+00 | 1.17E-01 | Yes | | intergenic_variant | MODIFIER |
| 13 | 20784988 | AX-170220165 | 0 | C | T | 0.175 | -1.95E-01 | 1.24E-01 | 2.42E+00 | 2.47E+00 | 1.17E-01 | Yes | | intergenic_variant | MODIFIER |
| 5 | 134365694 | AX-170026232 | 0 | G | A | 0.167 | -1.30E-01 | 8.01E-02 | 2.21E+00 | 2.31E+00 | 1.17E-01 | no | Gtf2ird1 | intron_variant | MODIFIER |
| 1 | 127904679 | AX-97844788 | 1 | T | C | 0.545 | 1.32E-01 | 8.45E-02 | 2.72E+00 | 2.79E+00 | 1.17E-01 | no | Rab3gap1 | intron_variant | MODIFIER |
| 6 | 112256091 | AX-170419005 | 0 | G | A | 0.053 | -3.52E-01 | 2.20E-01 | 3.30E+00 | 3.50E+00 | 1.17E-01 | Yes | | regulatory_region_variant | MODIFIER |
| 6 | 116943217 | AX-170181897 | 0 | A | G | 0.053 | -3.52E-01 | 2.20E-01 | 3.30E+00 | 3.50E+00 | 1.17E-01 | Yes | | intergenic_variant | MODIFIER |
| 9 | 86580533 | AX-170032891 | 0 | T | A | 0.132 | -1.95E-01 | 1.25E-01 | 2.86E+00 | 2.98E+00 | 1.17E-01 | Yes | Me1 | downstream_gene_variant | MODIFIER |
| 13 | 9422598 | AX-170505644 | 0 | C | G | 0.193 | -1.86E-01 | 1.19E-01 | 2.57E+00 | 2.62E+00 | 1.17E-01 | Yes | Dip2c | intron_variant | MODIFIER |
| 13 | 18162973 | AX-170203855 | 2 | A | T | 0.145 | -1.92E-01 | 1.21E-01 | 2.35E+00 | 2.41E+00 | 1.17E-01 | Yes | Pou6f2 | intron_variant | MODIFIER |
| 1 | 184595055 | AX-97840719 | 0 | T | G | 0.237 | 1.76E-01 | 1.12E-01 | 2.56E+00 | 2.62E+00 | 1.17E-01 | Yes | | regulatory_region_variant | MODIFIER |
| 1 | 184595189 | AX-170106362 | 0 | G | A | 0.237 | 1.76E-01 | 1.12E-01 | 2.56E+00 | 2.62E+00 | 1.17E-01 | Yes | | regulatory_region_variant | MODIFIER |
| 1 | 184609407 | AX-97845595 | 0 | T | C | 0.237 | 1.76E-01 | 1.12E-01 | 2.56E+00 | 2.62E+00 | 1.17E-01 | Yes | Gm37800 | downstream_gene_variant | MODIFIER |
| 1 | 184621030 | AX-170208682 | 0 | A | G | 0.237 | 1.76E-01 | 1.12E-01 | 2.56E+00 | 2.62E+00 | 1.17E-01 | Yes | Gm37800 | upstream_gene_variant | MODIFIER |
| 1 | 184630479 | AX-170358246 | 0 | A | G | 0.237 | 1.76E-01 | 1.12E-01 | 2.56E+00 | 2.62E+00 | 1.17E-01 | Yes | Gm37800 | upstream_gene_variant | MODIFIER |
| 1 | 184630550 | AX-170043844 | 0 | A | C | 0.237 | 1.76E-01 | 1.12E-01 | 2.56E+00 | 2.62E+00 | 1.17E-01 | Yes | Gm37800 | upstream_gene_variant | MODIFIER |
| 1 | 184640999 | AX-170189960 | 0 | T | C | 0.237 | 1.76E-01 | 1.12E-01 | 2.56E+00 | 2.62E+00 | 1.17E-01 | Yes | | intergenic_variant | MODIFIER |
| 1 | 184641457 | AX-168396661 | 0 | T | G | 0.237 | 1.76E-01 | 1.12E-01 | 2.56E+00 | 2.62E+00 | 1.17E-01 | Yes | | intergenic_variant | MODIFIER |
| 1 | 184641661 | AX-170125824 | 0 | G | C | 0.237 | 1.76E-01 | 1.12E-01 | 2.56E+00 | 2.62E+00 | 1.17E-01 | Yes | | intergenic_variant | MODIFIER |
| 1 | 184655267 | AX-170136328 | 0 | A | G | 0.237 | 1.76E-01 | 1.12E-01 | 2.56E+00 | 2.62E+00 | 1.17E-01 | Yes | | intergenic_variant | MODIFIER |
| 1 | 184667320 | AX-97791665 | 0 | C | A | 0.237 | 1.76E-01 | 1.12E-01 | 2.56E+00 | 2.62E+00 | 1.17E-01 | Yes | | intergenic_variant | MODIFIER |
| 1 | 184671096 | AX-170183366 | 0 | T | C | 0.237 | 1.76E-01 | 1.12E-01 | 2.56E+00 | 2.62E+00 | 1.17E-01 | Yes | | intergenic_variant | MODIFIER |
| 1 | 184677218 | AX-170446177 | 0 | A | G | 0.237 | 1.76E-01 | 1.12E-01 | 2.56E+00 | 2.62E+00 | 1.17E-01 | Yes | | intergenic_variant | MODIFIER |
| 1 | 184677235 | AX-97846605 | 0 | A | G | 0.237 | 1.76E-01 | 1.12E-01 | 2.56E+00 | 2.62E+00 | 1.17E-01 | Yes | | intergenic_variant | MODIFIER |
| 1 | 184707513 | AX-170456960 | 0 | G | A | 0.237 | 1.76E-01 | 1.12E-01 | 2.56E+00 | 2.62E+00 | 1.17E-01 | Yes | 4930488B22Rik | upstream_gene_variant | MODIFIER |
| 1 | 184707835 | AX-97796784 | 0 | C | T | 0.237 | 1.76E-01 | 1.12E-01 | 2.56E+00 | 2.62E+00 | 1.17E-01 | Yes | 4930488B22Rik | upstream_gene_variant | MODIFIER |
| 1 | 184746611 | AX-97798292 | 0 | C | T | 0.237 | 1.76E-01 | 1.12E-01 | 2.56E+00 | 2.62E+00 | 1.17E-01 | Yes | | intergenic_variant | MODIFIER |
| 1 | 184763199 | AX-170275354 | 0 | G | A | 0.237 | 1.76E-01 | 1.12E-01 | 2.56E+00 | 2.62E+00 | 1.17E-01 | Yes | | intergenic_variant | MODIFIER |
| 1 | 184792322 | AX-170383027 | 0 | T | C | 0.237 | 1.76E-01 | 1.12E-01 | 2.56E+00 | 2.62E+00 | 1.17E-01 | Yes | MAR1 | intron_variant | MODIFIER |
| 1 | 184793390 | AX-97815875 | 0 | A | G | 0.237 | 1.76E-01 | 1.12E-01 | 2.56E+00 | 2.62E+00 | 1.17E-01 | Yes | MAR1 | intron_variant | MODIFIER |
| 1 | 184803265 | AX-170002465 | 0 | C | T | 0.237 | 1.76E-01 | 1.12E-01 | 2.56E+00 | 2.62E+00 | 1.17E-01 | Yes | MAR1 | intron_variant | MODIFIER |
| 1 | 184914767 | AX-170139670 | 0 | A | G | 0.237 | 1.76E-01 | 1.12E-01 | 2.56E+00 | 2.62E+00 | 1.17E-01 | Yes | Mark1 | intron_variant | MODIFIER |
| 1 | 184916143 | AX-170442729 | 0 | A | G | 0.237 | 1.76E-01 | 1.12E-01 | 2.56E+00 | 2.62E+00 | 1.17E-01 | Yes | Mark1 | synonymous_variant | LOW |
| 1 | 184916725 | AX-170211408 | 0 | A | G | 0.237 | 1.76E-01 | 1.12E-01 | 2.56E+00 | 2.62E+00 | 1.17E-01 | Yes | Mark1 | intron_variant | MODIFIER |
| 1 | 185005736 | AX-170002469 | 0 | G | A | 0.237 | 1.76E-01 | 1.12E-01 | 2.56E+00 | 2.62E+00 | 1.17E-01 | Yes | | intergenic_variant | MODIFIER |
| 1 | 185017105 | AX-170277625 | 0 | C | T | 0.237 | 1.76E-01 | 1.12E-01 | 2.56E+00 | 2.62E+00 | 1.17E-01 | Yes | | intergenic_variant | MODIFIER |
| 1 | 185029040 | AX-170157033 | 0 | G | C | 0.237 | 1.76E-01 | 1.12E-01 | 2.56E+00 | 2.62E+00 | 1.17E-01 | Yes | | intergenic_variant | MODIFIER |
| 1 | 185031892 | AX-97830680 | 0 | A | G | 0.237 | 1.76E-01 | 1.12E-01 | 2.56E+00 | 2.62E+00 | 1.17E-01 | Yes | | regulatory_region_variant | MODIFIER |
| 1 | 185038099 | AX-170002460 | 0 | A | G | 0.237 | 1.76E-01 | 1.12E-01 | 2.56E+00 | 2.62E+00 | 1.17E-01 | Yes | | intergenic_variant | MODIFIER |
| 1 | 185038471 | AX-170002467 | 0 | G | T | 0.237 | 1.76E-01 | 1.12E-01 | 2.56E+00 | 2.62E+00 | 1.17E-01 | Yes | | intergenic_variant | MODIFIER |
| 1 | 185057226 | AX-170367272 | 0 | A | G | 0.237 | 1.76E-01 | 1.12E-01 | 2.56E+00 | 2.62E+00 | 1.17E-01 | Yes | | intergenic_variant | MODIFIER |
| 1 | 185061378 | AX-170043850 | 0 | A | T | 0.237 | 1.76E-01 | 1.12E-01 | 2.56E+00 | 2.62E+00 | 1.17E-01 | Yes | | intergenic_variant | MODIFIER |
| 1 | 185062906 | AX-97823686 | 0 | A | G | 0.237 | 1.76E-01 | 1.12E-01 | 2.56E+00 | 2.62E+00 | 1.17E-01 | Yes | | intergenic_variant | MODIFIER |
| 1 | 185065598 | AX-170441162 | 0 | A | G | 0.237 | 1.76E-01 | 1.12E-01 | 2.56E+00 | 2.62E+00 | 1.17E-01 | Yes | | intergenic_variant | MODIFIER |
| 1 | 185111100 | AX-170383317 | 0 | A | G | 0.237 | 1.76E-01 | 1.12E-01 | 2.56E+00 | 2.62E+00 | 1.17E-01 | Yes | | intergenic_variant | MODIFIER |
| 1 | 185111129 | AX-170454923 | 0 | A | G | 0.237 | 1.76E-01 | 1.12E-01 | 2.56E+00 | 2.62E+00 | 1.17E-01 | Yes | | intergenic_variant | MODIFIER |
| 1 | 185113770 | AX-170201640 | 0 | A | G | 0.237 | 1.76E-01 | 1.12E-01 | 2.56E+00 | 2.62E+00 | 1.17E-01 | Yes | | regulatory_region_variant | MODIFIER |

| | | | | | | | | | | | | | | | |
|---|---|---|---|---|---|---|---|---|---|---|---|---|---|---|---|
| 1 | 185114579 | AX-170191927 | 0 | T | C | 0.237 | 1.76E-01 | 1.12E-01 | 2.56E+00 | 2.62E+00 | 1.17E-01 | Yes | | intergenic_variant | MODIFIER |
| 1 | 185114607 | AX-170371562 | 0 | A | G | 0.237 | 1.76E-01 | 1.12E-01 | 2.56E+00 | 2.62E+00 | 1.17E-01 | Yes | | intergenic_variant | MODIFIER |
| 1 | 185117051 | AX-170464202 | 0 | T | C | 0.237 | 1.76E-01 | 1.12E-01 | 2.56E+00 | 2.62E+00 | 1.17E-01 | Yes | | intergenic_variant | MODIFIER |
| 1 | 185118176 | AX-170140825 | 0 | T | C | 0.237 | 1.76E-01 | 1.12E-01 | 2.56E+00 | 2.62E+00 | 1.17E-01 | Yes | | intergenic_variant | MODIFIER |
| 1 | 185121876 | AX-170226828 | 0 | C | T | 0.237 | 1.76E-01 | 1.12E-01 | 2.56E+00 | 2.62E+00 | 1.17E-01 | Yes | | intergenic_variant | MODIFIER |
| 1 | 185168164 | AX-170360859 | 0 | A | G | 0.237 | 1.76E-01 | 1.12E-01 | 2.56E+00 | 2.62E+00 | 1.17E-01 | Yes | | intergenic_variant | MODIFIER |
| 1 | 185201322 | AX-170509331 | 0 | G | A | 0.237 | 1.76E-01 | 1.12E-01 | 2.56E+00 | 2.62E+00 | 1.17E-01 | Yes | Rab3gap2 | upstream_gene_variant | MODIFIER |
| 1 | 185228140 | AX-170401686 | 0 | G | A | 0.237 | 1.76E-01 | 1.12E-01 | 2.56E+00 | 2.62E+00 | 1.17E-01 | Yes | Rab3gap2 | intron_variant | MODIFIER |
| 1 | 185266751 | AX-97794106 | 0 | C | T | 0.237 | 1.76E-01 | 1.12E-01 | 2.56E+00 | 2.62E+00 | 1.17E-01 | Yes | Rab3gap2 | intron_variant | MODIFIER |
| 1 | 185279947 | AX-170187370 | 0 | A | G | 0.237 | 1.76E-01 | 1.12E-01 | 2.56E+00 | 2.62E+00 | 1.17E-01 | Yes | Iars2 | downstream_gene_variant | MODIFIER |
| 1 | 185286937 | AX-170459258 | 0 | C | T | 0.237 | 1.76E-01 | 1.12E-01 | 2.56E+00 | 2.62E+00 | 1.17E-01 | Yes | Iars2 | 3_prime_UTR_variant | MODIFIER |
| 1 | 185308882 | AX-97787718 | 0 | T | C | 0.237 | 1.76E-01 | 1.12E-01 | 2.56E+00 | 2.62E+00 | 1.17E-01 | Yes | Iars2 | intron_variant | MODIFIER |
| 1 | 185316538 | AX-170287311 | 0 | A | G | 0.237 | 1.76E-01 | 1.12E-01 | 2.56E+00 | 2.62E+00 | 1.17E-01 | Yes | Iars2 | intron_variant | MODIFIER |
| 1 | 185323003 | AX-170099407 | 0 | C | T | 0.237 | 1.76E-01 | 1.12E-01 | 2.56E+00 | 2.62E+00 | 1.17E-01 | Yes | Iars2 | synonymous_variant | LOW |
| 1 | 185323153 | AX-97849364 | 0 | T | C | 0.237 | 1.76E-01 | 1.12E-01 | 2.56E+00 | 2.62E+00 | 1.17E-01 | Yes | Iars2 | intron_variant | MODIFIER |
| 1 | 185346747 | AX-170185934 | 0 | C | T | 0.237 | 1.76E-01 | 1.12E-01 | 2.56E+00 | 2.62E+00 | 1.17E-01 | Yes | Bpnt1 | intron_variant | MODIFIER |
| 1 | 185346778 | AX-170152147 | 0 | A | G | 0.237 | 1.76E-01 | 1.12E-01 | 2.56E+00 | 2.62E+00 | 1.17E-01 | Yes | Bpnt1 | intron_variant | MODIFIER |
| 1 | 185347011 | AX-170002478 | 0 | G | A | 0.237 | 1.76E-01 | 1.12E-01 | 2.56E+00 | 2.62E+00 | 1.17E-01 | Yes | Bpnt1 | intron_variant | MODIFIER |
| 1 | 185350175 | AX-170043858 | 0 | T | C | 0.237 | 1.76E-01 | 1.12E-01 | 2.56E+00 | 2.62E+00 | 1.17E-01 | Yes | Bpnt1 | intron_variant | MODIFIER |
| 1 | 185376818 | AX-97817975 | 0 | C | T | 0.237 | 1.76E-01 | 1.12E-01 | 2.56E+00 | 2.62E+00 | 1.17E-01 | Yes | Eprs | intron_variant | MODIFIER |
| 1 | 185402959 | AX-97814474 | 0 | T | G | 0.237 | 1.76E-01 | 1.12E-01 | 2.56E+00 | 2.62E+00 | 1.17E-01 | Yes | Eprs | intron_variant | MODIFIER |
| 1 | 185419958 | AX-170043855 | 0 | C | G | 0.237 | 1.76E-01 | 1.12E-01 | 2.56E+00 | 2.62E+00 | 1.17E-01 | Yes | Eprs | intron_variant | MODIFIER |
| 1 | 185426990 | AX-170002476 | 0 | G | T | 0.237 | 1.76E-01 | 1.12E-01 | 2.56E+00 | 2.62E+00 | 1.17E-01 | Yes | Eprs | intron_variant | MODIFIER |
| 1 | 185430354 | AX-170002471 | 0 | C | T | 0.237 | 1.76E-01 | 1.12E-01 | 2.56E+00 | 2.62E+00 | 1.17E-01 | Yes | Eprs | downstream_gene_variant | MODIFIER |
| 1 | 185437227 | AX-97789084 | 0 | T | A | 0.237 | 1.76E-01 | 1.12E-01 | 2.56E+00 | 2.62E+00 | 1.17E-01 | Yes | 9630028B13Rik | intron_variant,non_coding_transcript_variant | MODIFIER |
| 1 | 185454465 | AX-170002473 | 0 | T | C | 0.237 | 1.76E-01 | 1.12E-01 | 2.56E+00 | 2.62E+00 | 1.17E-01 | Yes | Slc30a10 | upstream_gene_variant | MODIFIER |
| 1 | 185470920 | AX-97794590 | 0 | C | T | 0.237 | 1.76E-01 | 1.12E-01 | 2.56E+00 | 2.62E+00 | 1.17E-01 | Yes | Slc30a10 | downstream_gene_variant | MODIFIER |
| 1 | 185473916 | AX-170301128 | 0 | C | T | 0.237 | 1.76E-01 | 1.12E-01 | 2.56E+00 | 2.62E+00 | 1.17E-01 | Yes | | regulatory_region_variant | MODIFIER |
| 1 | 185474544 | AX-97813742 | 0 | A | G | 0.237 | 1.76E-01 | 1.12E-01 | 2.56E+00 | 2.62E+00 | 1.17E-01 | Yes | | regulatory_region_variant | MODIFIER |
| 1 | 185476208 | AX-170002477 | 0 | C | T | 0.237 | 1.76E-01 | 1.12E-01 | 2.56E+00 | 2.62E+00 | 1.17E-01 | Yes | | intergenic_variant | MODIFIER |
| 1 | 185528488 | AX-170250659 | 0 | A | C | 0.237 | 1.76E-01 | 1.12E-01 | 2.56E+00 | 2.62E+00 | 1.17E-01 | Yes | | intergenic_variant | MODIFIER |
| 1 | 185535795 | AX-170152485 | 0 | T | C | 0.237 | 1.76E-01 | 1.12E-01 | 2.56E+00 | 2.62E+00 | 1.17E-01 | Yes | | intergenic_variant | MODIFIER |
| 1 | 185575241 | AX-170297163 | 0 | C | T | 0.237 | 1.76E-01 | 1.12E-01 | 2.56E+00 | 2.62E+00 | 1.17E-01 | Yes | | intergenic_variant | MODIFIER |
| 1 | 185594639 | AX-170191920 | 0 | T | C | 0.237 | 1.76E-01 | 1.12E-01 | 2.56E+00 | 2.62E+00 | 1.17E-01 | Yes | | intergenic_variant | MODIFIER |
| 1 | 185595385 | AX-170135200 | 0 | C | A | 0.237 | 1.76E-01 | 1.12E-01 | 2.56E+00 | 2.62E+00 | 1.17E-01 | Yes | | intergenic_variant | MODIFIER |
| 1 | 185596346 | AX-170440977 | 0 | G | A | 0.237 | 1.76E-01 | 1.12E-01 | 2.56E+00 | 2.62E+00 | 1.17E-01 | Yes | | intergenic_variant | MODIFIER |
| 1 | 185596405 | AX-170202878 | 0 | C | A | 0.237 | 1.76E-01 | 1.12E-01 | 2.56E+00 | 2.62E+00 | 1.17E-01 | Yes | | intergenic_variant | MODIFIER |
| 1 | 185596725 | AX-170181400 | 0 | A | G | 0.237 | 1.76E-01 | 1.12E-01 | 2.56E+00 | 2.62E+00 | 1.17E-01 | Yes | | intergenic_variant | MODIFIER |
| 1 | 185598933 | AX-170347858 | 0 | A | C | 0.237 | 1.76E-01 | 1.12E-01 | 2.56E+00 | 2.62E+00 | 1.17E-01 | Yes | | intergenic_variant | MODIFIER |
| 1 | 185601191 | AX-170246233 | 0 | C | T | 0.237 | 1.76E-01 | 1.12E-01 | 2.56E+00 | 2.62E+00 | 1.17E-01 | Yes | | intergenic_variant | MODIFIER |
| 1 | 185607931 | AX-170354254 | 0 | G | A | 0.237 | 1.76E-01 | 1.12E-01 | 2.56E+00 | 2.62E+00 | 1.17E-01 | Yes | | intergenic_variant | MODIFIER |
| 1 | 185608630 | AX-170148466 | 0 | T | C | 0.237 | 1.76E-01 | 1.12E-01 | 2.56E+00 | 2.62E+00 | 1.17E-01 | Yes | | regulatory_region_variant | MODIFIER |
| 1 | 185609072 | AX-97815275 | 0 | A | G | 0.237 | 1.76E-01 | 1.12E-01 | 2.56E+00 | 2.62E+00 | 1.17E-01 | Yes | | intergenic_variant | MODIFIER |
| 1 | 185619490 | AX-170376468 | 0 | G | A | 0.237 | 1.76E-01 | 1.12E-01 | 2.56E+00 | 2.62E+00 | 1.17E-01 | Yes | | intergenic_variant | MODIFIER |
| 1 | 168433554 | AX-97787313 | 1 | C | T | 0.223 | 2.18E-01 | 1.39E-01 | 2.89E+00 | 2.96E+00 | 1.18E-01 | Yes | Pbx1 | upstream_gene_variant | MODIFIER |
| 5 | 128862518 | AX-97823795 | 0 | A | G | 0.149 | -1.46E-01 | 8.94E-02 | 2.16E+00 | 2.23E+00 | 1.18E-01 | no | Rimbp2 | intron_variant | MODIFIER |
| 5 | 128876727 | AX-97822591 | 0 | C | T | 0.149 | -1.46E-01 | 8.94E-02 | 2.16E+00 | 2.23E+00 | 1.18E-01 | no | Rimbp2 | intron_variant | MODIFIER |
| 5 | 128897714 | AX-170130462 | 0 | G | A | 0.149 | -1.46E-01 | 8.94E-02 | 2.16E+00 | 2.23E+00 | 1.18E-01 | no | Rimbp2 | intron_variant | MODIFIER |
| 5 | 128897958 | AX-97798365 | 0 | T | C | 0.149 | -1.46E-01 | 8.94E-02 | 2.16E+00 | 2.23E+00 | 1.18E-01 | no | Rimbp2 | intron_variant | MODIFIER |
| 5 | 128906551 | AX-170026142 | 0 | A | G | 0.149 | -1.46E-01 | 8.94E-02 | 2.16E+00 | 2.23E+00 | 1.18E-01 | no | Rimbp2 | intron_variant | MODIFIER |
| 5 | 128942423 | AX-170204614 | 0 | G | A | 0.149 | -1.46E-01 | 8.94E-02 | 2.16E+00 | 2.23E+00 | 1.18E-01 | no | Rimbp2 | intron_variant | MODIFIER |
| 5 | 128943566 | AX-97838308 | 0 | T | C | 0.149 | -1.46E-01 | 8.94E-02 | 2.16E+00 | 2.23E+00 | 1.18E-01 | no | Rimbp2 | intron_variant | MODIFIER |
| 5 | 128956825 | AX-170306056 | 0 | A | G | 0.149 | -1.46E-01 | 8.94E-02 | 2.16E+00 | 2.23E+00 | 1.18E-01 | no | Rimbp2 | upstream_gene_variant | MODIFIER |
| 5 | 128963871 | AX-97809635 | 0 | C | T | 0.149 | -1.46E-01 | 8.94E-02 | 2.16E+00 | 2.23E+00 | 1.18E-01 | no | | intergenic_variant | MODIFIER |

| | | | | | | | | | | | | | | | |
|---|---|---|---|---|---|---|---|---|---|---|---|---|---|---|---|
| 5 | 128976540 | AX-170210110 | 0 | T | C | 0.149 | -1.46E-01 | 8.94E-02 | 2.16E+00 | 2.23E+00 | 1.18E-01 | no | | regulatory_region_variant | MODIFIER |
| 5 | 128983979 | AX-170330708 | 0 | T | C | 0.149 | -1.46E-01 | 8.94E-02 | 2.16E+00 | 2.23E+00 | 1.18E-01 | no | Stx2 | downstream_gene_variant | MODIFIER |
| 5 | 128986179 | AX-170296401 | 0 | C | A | 0.149 | -1.46E-01 | 8.94E-02 | 2.16E+00 | 2.23E+00 | 1.18E-01 | no | Stx2 | 3_prime_UTR_variant | MODIFIER |
| 5 | 128986309 | AX-170453911 | 0 | A | G | 0.149 | -1.46E-01 | 8.94E-02 | 2.16E+00 | 2.23E+00 | 1.18E-01 | no | Stx2 | 3_prime_UTR_variant | MODIFIER |
| 5 | 128988224 | AX-97821615 | 0 | A | G | 0.149 | -1.46E-01 | 8.94E-02 | 2.16E+00 | 2.23E+00 | 1.18E-01 | no | Stx2 | intron_variant | MODIFIER |
| 5 | 128988600 | AX-170454256 | 0 | G | A | 0.149 | -1.46E-01 | 8.94E-02 | 2.16E+00 | 2.23E+00 | 1.18E-01 | no | Stx2 | intron_variant | MODIFIER |
| 5 | 128997094 | AX-170170784 | 0 | T | C | 0.149 | -1.46E-01 | 8.94E-02 | 2.16E+00 | 2.23E+00 | 1.18E-01 | no | Stx2 | intron_variant | MODIFIER |
| 5 | 129003088 | AX-170522336 | 0 | C | T | 0.149 | -1.46E-01 | 8.94E-02 | 2.16E+00 | 2.23E+00 | 1.18E-01 | no | Stx2 | intron_variant | MODIFIER |
| 5 | 129005362 | AX-170402506 | 0 | G | A | 0.149 | -1.46E-01 | 8.94E-02 | 2.16E+00 | 2.23E+00 | 1.18E-01 | no | Stx2 | intron_variant | MODIFIER |
| 5 | 129013317 | AX-170154665 | 0 | T | G | 0.149 | -1.46E-01 | 8.94E-02 | 2.16E+00 | 2.23E+00 | 1.18E-01 | no | Stx2 | upstream_gene_variant | MODIFIER |
| 5 | 129032532 | AX-97834937 | 0 | C | T | 0.149 | -1.46E-01 | 8.94E-02 | 2.16E+00 | 2.23E+00 | 1.18E-01 | no | | intergenic_variant | MODIFIER |
| 5 | 129048655 | AX-170297683 | 0 | C | G | 0.149 | -1.46E-01 | 8.94E-02 | 2.16E+00 | 2.23E+00 | 1.18E-01 | no | | regulatory_region_variant | MODIFIER |
| 5 | 129049373 | AX-170416506 | 0 | C | T | 0.149 | -1.46E-01 | 8.94E-02 | 2.16E+00 | 2.23E+00 | 1.18E-01 | no | | regulatory_region_variant | MODIFIER |
| 5 | 129049479 | AX-170156884 | 0 | A | G | 0.149 | -1.46E-01 | 8.94E-02 | 2.16E+00 | 2.23E+00 | 1.18E-01 | no | | regulatory_region_variant | MODIFIER |
| 5 | 129061165 | AX-170265721 | 0 | A | G | 0.149 | -1.46E-01 | 8.94E-02 | 2.16E+00 | 2.23E+00 | 1.18E-01 | no | | intergenic_variant | MODIFIER |
| 5 | 129062905 | AX-97814897 | 0 | G | A | 0.149 | -1.46E-01 | 8.94E-02 | 2.16E+00 | 2.23E+00 | 1.18E-01 | no | | regulatory_region_variant | MODIFIER |
| 5 | 129066319 | AX-170181184 | 0 | T | C | 0.149 | -1.46E-01 | 8.94E-02 | 2.16E+00 | 2.23E+00 | 1.18E-01 | no | | regulatory_region_variant | MODIFIER |
| 5 | 129066451 | AX-170131523 | 0 | C | T | 0.149 | -1.46E-01 | 8.94E-02 | 2.16E+00 | 2.23E+00 | 1.18E-01 | no | | regulatory_region_variant | MODIFIER |
| 5 | 129072155 | AX-97813927 | 0 | G | A | 0.149 | -1.46E-01 | 8.94E-02 | 2.16E+00 | 2.23E+00 | 1.18E-01 | no | | regulatory_region_variant | MODIFIER |
| 5 | 129077230 | AX-170255402 | 0 | T | C | 0.149 | -1.46E-01 | 8.94E-02 | 2.16E+00 | 2.23E+00 | 1.18E-01 | no | | regulatory_region_variant | MODIFIER |
| 5 | 129122589 | AX-170134338 | 0 | C | T | 0.149 | -1.46E-01 | 8.94E-02 | 2.16E+00 | 2.23E+00 | 1.18E-01 | no | Adgrd1 | splice_region_variant,synonymous_variant | LOW |
| 5 | 129122677 | AX-170452696 | 0 | A | G | 0.149 | -1.46E-01 | 8.94E-02 | 2.16E+00 | 2.23E+00 | 1.18E-01 | no | Adgrd1 | intron_variant | MODIFIER |
| 5 | 129122725 | AX-170184467 | 0 | C | T | 0.149 | -1.46E-01 | 8.94E-02 | 2.16E+00 | 2.23E+00 | 1.18E-01 | no | Adgrd1 | intron_variant | MODIFIER |
| 5 | 129124684 | AX-170519678 | 0 | A | G | 0.149 | -1.46E-01 | 8.94E-02 | 2.16E+00 | 2.23E+00 | 1.18E-01 | no | Adgrd1 | intron_variant | MODIFIER |
| 5 | 129124849 | AX-97809299 | 0 | T | C | 0.149 | -1.46E-01 | 8.94E-02 | 2.16E+00 | 2.23E+00 | 1.18E-01 | no | Adgrd1 | intron_variant | MODIFIER |
| 5 | 129172043 | AX-97843958 | 0 | G | A | 0.149 | -1.46E-01 | 8.94E-02 | 2.16E+00 | 2.23E+00 | 1.18E-01 | no | Adgrd1 | intron_variant | MODIFIER |
| 5 | 129174457 | AX-170210743 | 0 | G | A | 0.149 | -1.46E-01 | 8.94E-02 | 2.16E+00 | 2.23E+00 | 1.18E-01 | no | Adgrd1 | intron_variant | MODIFIER |
| 5 | 129181294 | AX-170356826 | 0 | A | G | 0.149 | -1.46E-01 | 8.94E-02 | 2.16E+00 | 2.23E+00 | 1.18E-01 | no | Adgrd1 | intron_variant | MODIFIER |
| 5 | 129204842 | AX-170453749 | 0 | A | G | 0.149 | -1.46E-01 | 8.94E-02 | 2.16E+00 | 2.23E+00 | 1.18E-01 | no | Adgrd1 | downstream_gene_variant | MODIFIER |
| 5 | 129205876 | AX-97797818 | 0 | G | A | 0.149 | -1.46E-01 | 8.94E-02 | 2.16E+00 | 2.23E+00 | 1.18E-01 | no | Adgrd1 | downstream_gene_variant | MODIFIER |
| 5 | 129213209 | AX-170356749 | 0 | C | G | 0.149 | -1.46E-01 | 8.94E-02 | 2.16E+00 | 2.23E+00 | 1.18E-01 | no | | intergenic_variant | MODIFIER |
| 5 | 129230708 | AX-97814443 | 0 | A | G | 0.149 | -1.46E-01 | 8.94E-02 | 2.16E+00 | 2.23E+00 | 1.18E-01 | no | | intergenic_variant | MODIFIER |
| 5 | 129231342 | AX-170073494 | 0 | T | G | 0.149 | -1.46E-01 | 8.94E-02 | 2.16E+00 | 2.23E+00 | 1.18E-01 | no | | intergenic_variant | MODIFIER |
| 5 | 129232079 | AX-170178606 | 0 | T | C | 0.149 | -1.46E-01 | 8.94E-02 | 2.16E+00 | 2.23E+00 | 1.18E-01 | no | | intergenic_variant | MODIFIER |
| 5 | 129236809 | AX-170486648 | 0 | A | T | 0.149 | -1.46E-01 | 8.94E-02 | 2.16E+00 | 2.23E+00 | 1.18E-01 | no | | intergenic_variant | MODIFIER |
| 5 | 129256820 | AX-170095468 | 0 | T | C | 0.149 | -1.46E-01 | 8.94E-02 | 2.16E+00 | 2.23E+00 | 1.18E-01 | no | | intergenic_variant | MODIFIER |
| 5 | 129290166 | AX-170418938 | 0 | T | C | 0.149 | -1.46E-01 | 8.94E-02 | 2.16E+00 | 2.23E+00 | 1.18E-01 | no | | intergenic_variant | MODIFIER |
| 5 | 129291561 | AX-170285859 | 0 | T | G | 0.149 | -1.46E-01 | 8.94E-02 | 2.16E+00 | 2.23E+00 | 1.18E-01 | no | | intergenic_variant | MODIFIER |
| 5 | 129310181 | AX-170095469 | 0 | T | C | 0.149 | -1.46E-01 | 8.94E-02 | 2.16E+00 | 2.23E+00 | 1.18E-01 | no | | intergenic_variant | MODIFIER |
| 5 | 129310402 | AX-170379660 | 0 | G | T | 0.149 | -1.46E-01 | 8.94E-02 | 2.16E+00 | 2.23E+00 | 1.18E-01 | no | | intergenic_variant | MODIFIER |
| 5 | 129311194 | AX-170184224 | 0 | T | C | 0.149 | -1.46E-01 | 8.94E-02 | 2.16E+00 | 2.23E+00 | 1.18E-01 | no | | regulatory_region_variant | MODIFIER |
| 5 | 129312014 | AX-170461440 | 0 | A | C | 0.149 | -1.46E-01 | 8.94E-02 | 2.16E+00 | 2.23E+00 | 1.18E-01 | no | | intergenic_variant | MODIFIER |
| 5 | 129318039 | AX-170280627 | 0 | C | G | 0.149 | -1.46E-01 | 8.94E-02 | 2.16E+00 | 2.23E+00 | 1.18E-01 | no | | intergenic_variant | MODIFIER |
| 5 | 129318377 | AX-170337969 | 0 | A | G | 0.149 | -1.46E-01 | 8.94E-02 | 2.16E+00 | 2.23E+00 | 1.18E-01 | no | | intergenic_variant | MODIFIER |
| 5 | 129319100 | AX-170497802 | 0 | A | G | 0.149 | -1.46E-01 | 8.94E-02 | 2.16E+00 | 2.23E+00 | 1.18E-01 | no | | regulatory_region_variant | MODIFIER |
| 5 | 129319792 | AX-170518010 | 0 | G | A | 0.149 | -1.46E-01 | 8.94E-02 | 2.16E+00 | 2.23E+00 | 1.18E-01 | no | | regulatory_region_variant | MODIFIER |
| 5 | 129320003 | AX-97842315 | 0 | A | C | 0.149 | -1.46E-01 | 8.94E-02 | 2.16E+00 | 2.23E+00 | 1.18E-01 | no | | intergenic_variant | MODIFIER |
| 5 | 129321083 | AX-170225635 | 0 | C | T | 0.149 | -1.46E-01 | 8.94E-02 | 2.16E+00 | 2.23E+00 | 1.18E-01 | no | | intergenic_variant | MODIFIER |
| 5 | 129336952 | AX-170130473 | 0 | C | T | 0.149 | -1.46E-01 | 8.94E-02 | 2.16E+00 | 2.23E+00 | 1.18E-01 | no | | intergenic_variant | MODIFIER |
| 5 | 129343527 | AX-170026151 | 0 | T | C | 0.149 | -1.46E-01 | 8.94E-02 | 2.16E+00 | 2.23E+00 | 1.18E-01 | no | | intergenic_variant | MODIFIER |
| 5 | 129349604 | AX-170277171 | 0 | A | G | 0.149 | -1.46E-01 | 8.94E-02 | 2.16E+00 | 2.23E+00 | 1.18E-01 | no | | intergenic_variant | MODIFIER |
| 5 | 129350340 | AX-170026149 | 0 | T | C | 0.149 | -1.46E-01 | 8.94E-02 | 2.16E+00 | 2.23E+00 | 1.18E-01 | no | | intergenic_variant | MODIFIER |
| 5 | 129354259 | AX-97814319 | 0 | C | T | 0.149 | -1.46E-01 | 8.94E-02 | 2.16E+00 | 2.23E+00 | 1.18E-01 | no | | regulatory_region_variant | MODIFIER |
| 5 | 129357357 | AX-170196329 | 0 | C | T | 0.149 | -1.46E-01 | 8.94E-02 | 2.16E+00 | 2.23E+00 | 1.18E-01 | no | | intergenic_variant | MODIFIER |

| | | | | | | | | | | | | | | | |
|---|---|---|---|---|---|---|---|---|---|---|---|---|---|---|---|
| 5 | 129366247 | AX-170305940 | 0 | C | T | 0.149 | -1.46E-01 | 8.94E-02 | 2.16E+00 | 2.23E+00 | 1.18E-01 | no | Gm43001 | downstream_gene_variant | MODIFIER |
| 5 | 129367496 | AX-170476710 | 0 | G | A | 0.149 | -1.46E-01 | 8.94E-02 | 2.16E+00 | 2.23E+00 | 1.18E-01 | no | Gm43001 | downstream_gene_variant | MODIFIER |
| 5 | 129388901 | AX-97841604 | 0 | G | A | 0.149 | -1.46E-01 | 8.94E-02 | 2.16E+00 | 2.23E+00 | 1.18E-01 | no | | intergenic_variant | MODIFIER |
| 5 | 129389080 | AX-170026148 | 0 | T | C | 0.149 | -1.46E-01 | 8.94E-02 | 2.16E+00 | 2.23E+00 | 1.18E-01 | no | | intergenic_variant | MODIFIER |
| 5 | 129399107 | AX-170361621 | 0 | G | T | 0.149 | -1.46E-01 | 8.94E-02 | 2.16E+00 | 2.23E+00 | 1.18E-01 | no | | intergenic_variant | MODIFIER |
| 5 | 129400378 | AX-170423567 | 0 | T | C | 0.149 | -1.46E-01 | 8.94E-02 | 2.16E+00 | 2.23E+00 | 1.18E-01 | no | | intergenic_variant | MODIFIER |
| 5 | 129402126 | AX-170480036 | 0 | C | A | 0.149 | -1.46E-01 | 8.94E-02 | 2.16E+00 | 2.23E+00 | 1.18E-01 | no | | intergenic_variant | MODIFIER |
| 5 | 129404682 | AX-97844414 | 0 | C | T | 0.149 | -1.46E-01 | 8.94E-02 | 2.16E+00 | 2.23E+00 | 1.18E-01 | no | | intergenic_variant | MODIFIER |
| 5 | 129407940 | AX-170141082 | 0 | G | A | 0.149 | -1.46E-01 | 8.94E-02 | 2.16E+00 | 2.23E+00 | 1.18E-01 | no | | intergenic_variant | MODIFIER |
| 5 | 129443296 | AX-97849850 | 0 | C | T | 0.149 | -1.46E-01 | 8.94E-02 | 2.16E+00 | 2.23E+00 | 1.18E-01 | no | | intergenic_variant | MODIFIER |
| 5 | 129444630 | AX-170386444 | 0 | G | C | 0.149 | -1.46E-01 | 8.94E-02 | 2.16E+00 | 2.23E+00 | 1.18E-01 | no | | intergenic_variant | MODIFIER |
| 5 | 129449612 | AX-169999729 | 0 | T | C | 0.149 | -1.46E-01 | 8.94E-02 | 2.16E+00 | 2.23E+00 | 1.18E-01 | no | | intergenic_variant | MODIFIER |
| 5 | 129452839 | AX-170374046 | 0 | G | A | 0.149 | -1.46E-01 | 8.94E-02 | 2.16E+00 | 2.23E+00 | 1.18E-01 | no | | intergenic_variant | MODIFIER |
| 5 | 129453864 | AX-170279320 | 0 | A | T | 0.149 | -1.46E-01 | 8.94E-02 | 2.16E+00 | 2.23E+00 | 1.18E-01 | no | | intergenic_variant | MODIFIER |
| 5 | 129454007 | AX-170347231 | 0 | C | T | 0.149 | -1.46E-01 | 8.94E-02 | 2.16E+00 | 2.23E+00 | 1.18E-01 | no | | intergenic_variant | MODIFIER |
| 5 | 129456748 | AX-97793707 | 0 | T | C | 0.149 | -1.46E-01 | 8.94E-02 | 2.16E+00 | 2.23E+00 | 1.18E-01 | no | | intergenic_variant | MODIFIER |
| 5 | 129457684 | AX-170270329 | 0 | T | C | 0.149 | -1.46E-01 | 8.94E-02 | 2.16E+00 | 2.23E+00 | 1.18E-01 | no | | intergenic_variant | MODIFIER |
| 5 | 129457759 | AX-170268930 | 0 | C | A | 0.149 | -1.46E-01 | 8.94E-02 | 2.16E+00 | 2.23E+00 | 1.18E-01 | no | | intergenic_variant | MODIFIER |
| 5 | 129458441 | AX-170433321 | 0 | G | A | 0.149 | -1.46E-01 | 8.94E-02 | 2.16E+00 | 2.23E+00 | 1.18E-01 | no | | regulatory_region_variant | MODIFIER |
| 5 | 129519185 | AX-97843866 | 0 | A | G | 0.149 | -1.46E-01 | 8.94E-02 | 2.16E+00 | 2.23E+00 | 1.18E-01 | no | Sfswap | intron_variant | MODIFIER |
| 5 | 129535543 | AX-97807413 | 0 | G | A | 0.149 | -1.46E-01 | 8.94E-02 | 2.16E+00 | 2.23E+00 | 1.18E-01 | no | Sfswap | intron_variant | MODIFIER |
| 5 | 129537966 | AX-170183507 | 0 | A | G | 0.149 | -1.46E-01 | 8.94E-02 | 2.16E+00 | 2.23E+00 | 1.18E-01 | no | Sfswap | intron_variant | MODIFIER |
| 5 | 129556573 | AX-170456300 | 0 | G | C | 0.149 | -1.46E-01 | 8.94E-02 | 2.16E+00 | 2.23E+00 | 1.18E-01 | no | Sfswap | intron_variant | MODIFIER |
| 5 | 129650149 | AX-97840985 | 0 | T | A | 0.149 | -1.46E-01 | 8.94E-02 | 2.16E+00 | 2.23E+00 | 1.18E-01 | no | Zfp11 | downstream_gene_variant | MODIFIER |
| 5 | 129676360 | AX-170345692 | 0 | C | T | 0.149 | -1.46E-01 | 8.94E-02 | 2.16E+00 | 2.23E+00 | 1.18E-01 | no | | intergenic_variant | MODIFIER |
| 5 | 129730765 | AX-170533249 | 0 | A | G | 0.149 | -1.46E-01 | 8.94E-02 | 2.16E+00 | 2.23E+00 | 1.18E-01 | no | Nipsnap2 | intron_variant | MODIFIER |
| 5 | 129731412 | AX-97841791 | 0 | T | G | 0.149 | -1.46E-01 | 8.94E-02 | 2.16E+00 | 2.23E+00 | 1.18E-01 | no | Nipsnap2 | intron_variant | MODIFIER |
| 5 | 129732944 | AX-170105791 | 0 | G | A | 0.149 | -1.46E-01 | 8.94E-02 | 2.16E+00 | 2.23E+00 | 1.18E-01 | no | Nipsnap2 | intron_variant | MODIFIER |
| 5 | 129742165 | AX-170447350 | 0 | C | T | 0.149 | -1.46E-01 | 8.94E-02 | 2.16E+00 | 2.23E+00 | 1.18E-01 | no | Nipsnap2 | intron_variant | MODIFIER |
| 5 | 129744353 | AX-170073513 | 0 | C | T | 0.149 | -1.46E-01 | 8.94E-02 | 2.16E+00 | 2.23E+00 | 1.18E-01 | no | Nipsnap2 | intron_variant | MODIFIER |
| 5 | 129744591 | AX-170073512 | 0 | C | T | 0.149 | -1.46E-01 | 8.94E-02 | 2.16E+00 | 2.23E+00 | 1.18E-01 | no | Nipsnap2 | intron_variant | MODIFIER |
| 5 | 129746133 | AX-170426850 | 0 | C | T | 0.149 | -1.46E-01 | 8.94E-02 | 2.16E+00 | 2.23E+00 | 1.18E-01 | no | Nipsnap2 | intron_variant | MODIFIER |
| 5 | 129750327 | AX-170451709 | 0 | G | T | 0.149 | -1.46E-01 | 8.94E-02 | 2.16E+00 | 2.23E+00 | 1.18E-01 | no | Nipsnap2 | intron_variant | MODIFIER |
| 5 | 129781010 | AX-170501402 | 0 | G | A | 0.149 | -1.46E-01 | 8.94E-02 | 2.16E+00 | 2.23E+00 | 1.18E-01 | no | Psph | intron_variant | MODIFIER |
| 5 | 129781246 | AX-97805629 | 0 | A | C | 0.149 | -1.46E-01 | 8.94E-02 | 2.16E+00 | 2.23E+00 | 1.18E-01 | no | Psph | intron_variant | MODIFIER |
| 5 | 129785013 | AX-170109068 | 0 | A | G | 0.149 | -1.46E-01 | 8.94E-02 | 2.16E+00 | 2.23E+00 | 1.18E-01 | no | Psph | intron_variant | MODIFIER |
| 5 | 129790369 | AX-170316072 | 0 | G | A | 0.149 | -1.46E-01 | 8.94E-02 | 2.16E+00 | 2.23E+00 | 1.18E-01 | no | Psph | upstream_gene_variant | MODIFIER |
| 5 | 129873472 | AX-170422316 | 0 | G | A | 0.149 | -1.46E-01 | 8.94E-02 | 2.16E+00 | 2.23E+00 | 1.18E-01 | no | Phkg1 | intron_variant | MODIFIER |
| 5 | 129881072 | AX-97793654 | 0 | C | T | 0.149 | -1.46E-01 | 8.94E-02 | 2.16E+00 | 2.23E+00 | 1.18E-01 | no | Phkg1 | upstream_gene_variant | MODIFIER |
| 5 | 129888458 | AX-170474344 | 0 | G | C | 0.149 | -1.46E-01 | 8.94E-02 | 2.16E+00 | 2.23E+00 | 1.18E-01 | no | Chchd2 | upstream_gene_variant | MODIFIER |
| 5 | 129894096 | AX-170341652 | 0 | G | T | 0.149 | -1.46E-01 | 8.94E-02 | 2.16E+00 | 2.23E+00 | 1.18E-01 | no | Zbed5 | upstream_gene_variant | MODIFIER |
| 5 | 129922966 | AX-170410884 | 0 | G | A | 0.149 | -1.46E-01 | 8.94E-02 | 2.16E+00 | 2.23E+00 | 1.18E-01 | no | | intergenic_variant | MODIFIER |
| 5 | 129922989 | AX-170026160 | 0 | G | A | 0.149 | -1.46E-01 | 8.94E-02 | 2.16E+00 | 2.23E+00 | 1.18E-01 | no | | intergenic_variant | MODIFIER |
| 5 | 129925206 | AX-170403890 | 0 | T | C | 0.149 | -1.46E-01 | 8.94E-02 | 2.16E+00 | 2.23E+00 | 1.18E-01 | no | | regulatory_region_variant | MODIFIER |
| 5 | 129933274 | AX-97851690 | 0 | G | A | 0.149 | -1.46E-01 | 8.94E-02 | 2.16E+00 | 2.23E+00 | 1.18E-01 | no | | intergenic_variant | MODIFIER |
| 5 | 129964360 | AX-170026154 | 0 | C | G | 0.149 | -1.46E-01 | 8.94E-02 | 2.16E+00 | 2.23E+00 | 1.18E-01 | no | Vkorc1l1 | intron_variant | MODIFIER |
| 5 | 129965381 | AX-170089608 | 0 | C | T | 0.149 | -1.46E-01 | 8.94E-02 | 2.16E+00 | 2.23E+00 | 1.18E-01 | no | Vkorc1l1 | intron_variant | MODIFIER |
| 5 | 129974196 | AX-170026156 | 0 | T | A | 0.149 | -1.46E-01 | 8.94E-02 | 2.16E+00 | 2.23E+00 | 1.18E-01 | no | Vkorc1l1 | intron_variant | MODIFIER |
| 5 | 129999476 | AX-170351753 | 0 | T | C | 0.149 | -1.46E-01 | 8.94E-02 | 2.16E+00 | 2.23E+00 | 1.18E-01 | no | Gusb | synonymous_variant | LOW |
| 5 | 130010363 | AX-97799377 | 0 | C | T | 0.149 | -1.46E-01 | 8.94E-02 | 2.16E+00 | 2.23E+00 | 1.18E-01 | no | Asl | downstream_gene_variant | MODIFIER |
| 5 | 130010440 | AX-170501508 | 0 | G | A | 0.149 | -1.46E-01 | 8.94E-02 | 2.16E+00 | 2.23E+00 | 1.18E-01 | no | Asl | downstream_gene_variant | MODIFIER |
| 5 | 130020484 | AX-170026153 | 0 | A | G | 0.149 | -1.46E-01 | 8.94E-02 | 2.16E+00 | 2.23E+00 | 1.18E-01 | no | Asl | intron_variant | MODIFIER |
| 5 | 130051803 | AX-170026158 | 0 | C | T | 0.149 | -1.46E-01 | 8.94E-02 | 2.16E+00 | 2.23E+00 | 1.18E-01 | no | Crcp | intron_variant | MODIFIER |
| 5 | 130053075 | AX-170492902 | 0 | T | C | 0.149 | -1.46E-01 | 8.94E-02 | 2.16E+00 | 2.23E+00 | 1.18E-01 | no | Crcp | intron_variant | MODIFIER |

| | | | | | | | | | | | | | | | |
|---|---|---|---|---|---|---|---|---|---|---|---|---|---|---|---|
| 5 | 130061889 | AX-97857470 | 0 | T | C | 0.149 | -1.46E-01 | 8.94E-02 | 2.16E+00 | 2.23E+00 | 1.18E-01 | no | Crcp | downstream_gene_variant | MODIFIER |
| 5 | 130062850 | AX-170026159 | 0 | A | T | 0.149 | -1.46E-01 | 8.94E-02 | 2.16E+00 | 2.23E+00 | 1.18E-01 | no | Crcp | downstream_gene_variant | MODIFIER |
| 5 | 130103020 | AX-97805124 | 0 | T | C | 0.149 | -1.46E-01 | 8.94E-02 | 2.16E+00 | 2.23E+00 | 1.18E-01 | no | Tpst1 | intron_variant | MODIFIER |
| 5 | 130206646 | AX-97855038 | 0 | T | G | 0.149 | -1.46E-01 | 8.94E-02 | 2.16E+00 | 2.23E+00 | 1.18E-01 | no | Rabgef1 | intron_variant | MODIFIER |
| 5 | 130322740 | AX-97824197 | 0 | T | C | 0.149 | -1.46E-01 | 8.94E-02 | 2.16E+00 | 2.23E+00 | 1.18E-01 | no | Tyw1 | intron_variant | MODIFIER |
| 5 | 130348073 | AX-97800832 | 0 | T | C | 0.149 | -1.46E-01 | 8.94E-02 | 2.16E+00 | 2.23E+00 | 1.18E-01 | no | | intergenic_variant | MODIFIER |
| 10 | 106299448 | AX-170135806 | 0 | G | A | 0.289 | -1.34E-01 | 8.58E-02 | 2.68E+00 | 2.75E+00 | 1.18E-01 | no | | intergenic_variant | MODIFIER |
| 10 | 106367486 | AX-170036346 | 0 | C | T | 0.289 | -1.34E-01 | 8.58E-02 | 2.68E+00 | 2.75E+00 | 1.18E-01 | no | | intergenic_variant | MODIFIER |
| 10 | 106367788 | AX-97807823 | 0 | G | A | 0.289 | -1.34E-01 | 8.58E-02 | 2.68E+00 | 2.75E+00 | 1.18E-01 | no | | intergenic_variant | MODIFIER |
| 10 | 106419243 | AX-170524622 | 0 | G | A | 0.289 | -1.34E-01 | 8.58E-02 | 2.68E+00 | 2.75E+00 | 1.18E-01 | no | | intergenic_variant | MODIFIER |
| 10 | 106694956 | AX-170003848 | 0 | T | G | 0.289 | -1.34E-01 | 8.58E-02 | 2.68E+00 | 2.75E+00 | 1.18E-01 | no | Ppfia2 | intron_variant | MODIFIER |
| 10 | 106787355 | AX-170098051 | 0 | G | A | 0.289 | -1.34E-01 | 8.58E-02 | 2.68E+00 | 2.75E+00 | 1.18E-01 | no | Ppfia2 | intron_variant | MODIFIER |
| 10 | 106789209 | AX-170155451 | 0 | A | G | 0.289 | -1.34E-01 | 8.58E-02 | 2.68E+00 | 2.75E+00 | 1.18E-01 | no | Ppfia2 | intron_variant | MODIFIER |
| 10 | 106931316 | AX-170086893 | 0 | C | T | 0.289 | -1.34E-01 | 8.58E-02 | 2.68E+00 | 2.75E+00 | 1.18E-01 | no | Ppfia2 | 3_prime_UTR_variant | MODIFIER |
| 10 | 107436034 | AX-170086895 | 0 | G | A | 0.289 | -1.34E-01 | 8.58E-02 | 2.68E+00 | 2.75E+00 | 1.18E-01 | no | Gm38117 | downstream_gene_variant | MODIFIER |
| 10 | 107521802 | AX-170519728 | 0 | T | C | 0.289 | -1.34E-01 | 8.58E-02 | 2.68E+00 | 2.75E+00 | 1.18E-01 | no | Ptprq | intron_variant | MODIFIER |
| 1 | 194585860 | AX-170157645 | 1 | A | C | 0.277 | 1.47E-01 | 9.22E-02 | 2.34E+00 | 2.42E+00 | 1.18E-01 | Yes | | intergenic_variant | MODIFIER |
| 1 | 192239963 | AX-170481578 | 1 | C | T | 0.277 | 1.43E-01 | 9.06E-02 | 2.40E+00 | 2.49E+00 | 1.18E-01 | no | Kcnh1 | intron_variant | MODIFIER |
| 13 | 9075127 | AX-97857245 | 0 | C | T | 0.193 | -1.84E-01 | 1.18E-01 | 2.55E+00 | 2.62E+00 | 1.18E-01 | Yes | Gm36264 | intron_variant,non_coding_transcript_variant | MODIFIER |
| 13 | 10500187 | AX-170146290 | 0 | G | A | 0.193 | -1.84E-01 | 1.18E-01 | 2.55E+00 | 2.62E+00 | 1.18E-01 | Yes | Gm26861 | intron_variant,non_coding_transcript_variant | MODIFIER |
| 13 | 7515932 | AX-97799435 | 0 | G | A | 0.184 | -1.96E-01 | 1.26E-01 | 2.58E+00 | 2.63E+00 | 1.18E-01 | Yes | | intergenic_variant | MODIFIER |
| 5 | 9748446 | AX-170209900 | 0 | T | G | 0.316 | 1.40E-01 | 9.01E-02 | 2.85E+00 | 2.99E+00 | 1.18E-01 | no | | intergenic_variant | MODIFIER |
| 5 | 10117021 | AX-170071304 | 0 | G | T | 0.316 | 1.40E-01 | 9.01E-02 | 2.85E+00 | 2.99E+00 | 1.18E-01 | no | Gm5715 | non_coding_transcript_exon_variant | MODIFIER |
| 5 | 10117034 | AX-170071303 | 0 | C | T | 0.316 | 1.40E-01 | 9.01E-02 | 2.85E+00 | 2.99E+00 | 1.18E-01 | no | Gm5715 | non_coding_transcript_exon_variant | MODIFIER |
| 9 | 101618843 | AX-170033147 | 0 | C | T | 0.132 | -1.55E-01 | 9.96E-02 | 2.71E+00 | 2.85E+00 | 1.18E-01 | no | | intergenic_variant | MODIFIER |
| 13 | 20205439 | AX-170405605 | 1 | C | A | 0.179 | -1.92E-01 | 1.23E-01 | 2.53E+00 | 2.59E+00 | 1.18E-01 | Yes | Elmo1 | intron_variant | MODIFIER |
| 9 | 86307474 | AX-170334480 | 0 | T | C | 0.14 | -1.85E-01 | 1.19E-01 | 2.83E+00 | 2.94E+00 | 1.18E-01 | Yes | Ube2cbp | 3_prime_UTR_variant | MODIFIER |
| 9 | 88597743 | AX-170238108 | 0 | T | C | 0.14 | -1.85E-01 | 1.19E-01 | 2.83E+00 | 2.94E+00 | 1.18E-01 | Yes | Gm23017 | downstream_gene_variant | MODIFIER |
| 9 | 88736544 | AX-97825121 | 0 | C | A | 0.14 | -1.85E-01 | 1.19E-01 | 2.83E+00 | 2.94E+00 | 1.18E-01 | Yes | Bcl2a1d | upstream_gene_variant | MODIFIER |
| 18 | 74432566 | AX-170102603 | 0 | T | C | 0.193 | -1.74E-01 | 1.12E-01 | 2.71E+00 | 2.79E+00 | 1.18E-01 | Yes | | intergenic_variant | MODIFIER |
| 8 | 75380898 | AX-170514342 | 1 | T | A | 0.205 | -1.90E-01 | 1.20E-01 | 3.21E+00 | 3.38E+00 | 1.18E-01 | Yes | Umpk-ps | upstream_gene_variant | MODIFIER |
| 13 | 18307964 | AX-168378616 | 1 | C | T | 0.161 | -1.98E-01 | 1.26E-01 | 2.42E+00 | 2.47E+00 | 1.18E-01 | Yes | Pou6f2 | intron_variant | MODIFIER |
| 5 | 9682250 | AX-170357684 | 0 | T | C | 0.325 | 1.44E-01 | 9.30E-02 | 2.88E+00 | 3.02E+00 | 1.19E-01 | Yes | Grm3 | intron_variant | MODIFIER |
| 5 | 9707898 | AX-170071297 | 0 | T | C | 0.325 | 1.44E-01 | 9.30E-02 | 2.88E+00 | 3.02E+00 | 1.19E-01 | Yes | Grm3 | intron_variant | MODIFIER |
| 5 | 9733182 | AX-97789121 | 0 | A | G | 0.325 | 1.44E-01 | 9.30E-02 | 2.88E+00 | 3.02E+00 | 1.19E-01 | Yes | | intergenic_variant | MODIFIER |
| 5 | 9733432 | AX-170504075 | 0 | T | C | 0.325 | 1.44E-01 | 9.30E-02 | 2.88E+00 | 3.02E+00 | 1.19E-01 | Yes | | intergenic_variant | MODIFIER |
| 5 | 10125745 | AX-170301308 | 0 | G | A | 0.325 | 1.44E-01 | 9.30E-02 | 2.88E+00 | 3.02E+00 | 1.19E-01 | Yes | | intergenic_variant | MODIFIER |
| 1 | 180003917 | AX-97825311 | 1 | G | T | 0.241 | 1.76E-01 | 1.13E-01 | 2.63E+00 | 2.71E+00 | 1.19E-01 | Yes | Cdc42bpa | intron_variant | MODIFIER |
| 4 | 84113227 | AX-170208098 | 0 | G | C | 0.342 | 1.83E-01 | 1.17E-01 | 2.98E+00 | 3.06E+00 | 1.19E-01 | Yes | | intergenic_variant | MODIFIER |
| 16 | 63955116 | AX-170129111 | 1 | G | T | 0.5 | 1.34E-01 | 8.64E-02 | 2.67E+00 | 2.73E+00 | 1.19E-01 | no | Gm49627 | non_coding_transcript_exon_variant | MODIFIER |
| 13 | 8889683 | AX-170092653 | 0 | C | A | 0.184 | -1.91E-01 | 1.22E-01 | 2.51E+00 | 2.56E+00 | 1.19E-01 | Yes | Idi1 | intron_variant | MODIFIER |
| 4 | 76021994 | AX-170069714 | 0 | T | C | 0.325 | 1.83E-01 | 1.17E-01 | 2.99E+00 | 3.06E+00 | 1.19E-01 | Yes | Ptprd | intron_variant | MODIFIER |
| 4 | 78608800 | AX-170322562 | 0 | C | A | 0.325 | 1.83E-01 | 1.17E-01 | 2.99E+00 | 3.06E+00 | 1.19E-01 | Yes | | intergenic_variant | MODIFIER |
| 15 | 6578613 | AX-170421026 | 1 | C | T | 0.232 | -1.55E-01 | 1.00E-01 | 2.79E+00 | 2.87E+00 | 1.19E-01 | no | Fyb | upstream_gene_variant | MODIFIER |
| 8 | 47017011 | AX-170165532 | 0 | A | T | 0.149 | -2.12E-01 | 1.35E-01 | 3.08E+00 | 3.21E+00 | 1.19E-01 | Yes | Enpp6 | intron_variant | MODIFIER |
| 5 | 54995156 | AX-170401543 | 0 | G | A | 0.465 | 1.60E-01 | 1.02E-01 | 2.42E+00 | 2.48E+00 | 1.20E-01 | Yes | Gm43779 | upstream_gene_variant | MODIFIER |
| 15 | 4222214 | AX-170213890 | 0 | A | T | 0.228 | -1.55E-01 | 1.00E-01 | 2.77E+00 | 2.84E+00 | 1.20E-01 | no | | intergenic_variant | MODIFIER |
| 6 | 104054830 | AX-170237820 | 0 | T | A | 0.07 | 2.27E-01 | 1.47E-01 | 2.76E+00 | 2.93E+00 | 1.20E-01 | Yes | | intergenic_variant | MODIFIER |
| 2 | 113469974 | AX-170018719 | 0 | T | G | 0.053 | -3.46E-01 | 2.21E-01 | 3.07E+00 | 3.20E+00 | 1.20E-01 | Yes | Fmn1 | intron_variant | MODIFIER |
| 13 | 83983279 | AX-170462339 | 0 | T | C | 0.044 | -2.59E-01 | 1.68E-01 | 2.88E+00 | 3.03E+00 | 1.20E-01 | Yes | Gm4241 | intron_variant,non_coding_transcript_variant | MODIFIER |
| 4 | 155211632 | AX-170421987 | 0 | G | A | 0.132 | 2.21E-01 | 1.42E-01 | 2.95E+00 | 3.09E+00 | 1.20E-01 | Yes | Ski | intron_variant | MODIFIER |
| 5 | 34995326 | AX-170219899 | 2 | G | A | 0.345 | 1.53E-01 | 9.89E-02 | 2.84E+00 | 2.94E+00 | 1.20E-01 | Yes | Rgs12 | intron_variant | MODIFIER |
| 5 | 62932565 | AX-97849782 | 0 | G | A | 0.491 | 1.44E-01 | 9.11E-02 | 2.34E+00 | 2.44E+00 | 1.20E-01 | no | | intergenic_variant | MODIFIER |
| 3 | 18732021 | AX-170020048 | 0 | T | G | 0.14 | -1.50E-01 | 9.66E-02 | 2.49E+00 | 2.58E+00 | 1.20E-01 | no | | regulatory_region_variant | MODIFIER |

| | | | | | | | | | | | | | | | |
|---|---|---|---|---|---|---|---|---|---|---|---|---|---|---|---|
| 10 | 7005224 | AX-97810339 | 0 | T | A | 0.202 | -1.36E-01 | 8.80E-02 | 2.79E+00 | 2.93E+00 | 1.20E-01 | no | Oprm1 | intron_variant | MODIFIER |
| 8 | 34300518 | AX-170530341 | 0 | C | T | 0.14 | -1.68E-01 | 1.09E-01 | 2.77E+00 | 2.90E+00 | 1.20E-01 | no | Gm23177 | upstream_gene_variant | MODIFIER |
| 4 | 133708064 | AX-170387926 | 0 | T | C | 0.167 | 1.74E-01 | 1.13E-01 | 2.65E+00 | 2.71E+00 | 1.21E-01 | Yes | Arid1a | intron_variant | MODIFIER |
| 16 | 63758918 | AX-170479988 | 0 | C | T | 0.509 | 1.35E-01 | 8.75E-02 | 2.71E+00 | 2.77E+00 | 1.21E-01 | no | Epha3 | intron_variant | MODIFIER |
| 4 | 82368824 | AX-170360890 | 0 | G | A | 0.43 | -1.28E-01 | 8.29E-02 | 2.54E+00 | 2.60E+00 | 1.21E-01 | no | Nfib | intron_variant | MODIFIER |
| 4 | 82427156 | AX-170140761 | 0 | C | G | 0.43 | -1.28E-01 | 8.29E-02 | 2.54E+00 | 2.60E+00 | 1.21E-01 | no | Nfib | intron_variant | MODIFIER |
| 4 | 82431973 | AX-170191813 | 0 | G | A | 0.43 | -1.28E-01 | 8.29E-02 | 2.54E+00 | 2.60E+00 | 1.21E-01 | no | Nfib | intron_variant | MODIFIER |
| 4 | 82438150 | AX-170266156 | 0 | C | T | 0.43 | -1.28E-01 | 8.29E-02 | 2.54E+00 | 2.60E+00 | 1.21E-01 | no | Nfib | intron_variant | MODIFIER |
| 4 | 83355575 | AX-170480094 | 0 | A | C | 0.43 | -1.28E-01 | 8.29E-02 | 2.54E+00 | 2.60E+00 | 1.21E-01 | no | | intergenic_variant | MODIFIER |
| 4 | 83518735 | AX-97799866 | 0 | T | C | 0.43 | -1.28E-01 | 8.29E-02 | 2.54E+00 | 2.60E+00 | 1.21E-01 | no | | intergenic_variant | MODIFIER |
| 4 | 83627008 | AX-97844382 | 0 | T | C | 0.43 | -1.28E-01 | 8.29E-02 | 2.54E+00 | 2.60E+00 | 1.21E-01 | no | Ccdc171 | intron_variant | MODIFIER |
| 4 | 83824580 | AX-97805388 | 0 | A | C | 0.43 | -1.28E-01 | 8.29E-02 | 2.54E+00 | 2.60E+00 | 1.21E-01 | no | Ccdc171 | intron_variant | MODIFIER |
| 4 | 83847214 | AX-170441571 | 0 | A | G | 0.43 | -1.28E-01 | 8.29E-02 | 2.54E+00 | 2.60E+00 | 1.21E-01 | no | Ccdc171 | intron_variant | MODIFIER |
| 4 | 83852101 | AX-170122130 | 0 | G | A | 0.43 | -1.28E-01 | 8.29E-02 | 2.54E+00 | 2.60E+00 | 1.21E-01 | no | Ccdc171 | intron_variant | MODIFIER |
| 4 | 83857443 | AX-170184687 | 0 | G | A | 0.43 | -1.28E-01 | 8.29E-02 | 2.54E+00 | 2.60E+00 | 1.21E-01 | no | Ccdc171 | intron_variant | MODIFIER |
| 4 | 83868031 | AX-97852608 | 0 | T | G | 0.43 | -1.28E-01 | 8.29E-02 | 2.54E+00 | 2.60E+00 | 1.21E-01 | no | Ccdc171 | downstream_gene_variant | MODIFIER |
| 4 | 83868089 | AX-170139854 | 0 | T | C | 0.43 | -1.28E-01 | 8.29E-02 | 2.54E+00 | 2.60E+00 | 1.21E-01 | no | Ccdc171 | downstream_gene_variant | MODIFIER |
| 4 | 83917074 | AX-170487198 | 0 | A | G | 0.43 | -1.28E-01 | 8.29E-02 | 2.54E+00 | 2.60E+00 | 1.21E-01 | no | | intergenic_variant | MODIFIER |
| 4 | 83917436 | AX-97802228 | 0 | A | G | 0.43 | -1.28E-01 | 8.29E-02 | 2.54E+00 | 2.60E+00 | 1.21E-01 | no | | intergenic_variant | MODIFIER |
| 4 | 83926424 | AX-170307232 | 0 | A | G | 0.43 | -1.28E-01 | 8.29E-02 | 2.54E+00 | 2.60E+00 | 1.21E-01 | no | | regulatory_region_variant | MODIFIER |
| 4 | 83931660 | AX-170334241 | 0 | C | T | 0.43 | -1.28E-01 | 8.29E-02 | 2.54E+00 | 2.60E+00 | 1.21E-01 | no | | regulatory_region_variant | MODIFIER |
| 4 | 84007580 | AX-97823298 | 0 | G | A | 0.43 | -1.28E-01 | 8.29E-02 | 2.54E+00 | 2.60E+00 | 1.21E-01 | no | | intergenic_variant | MODIFIER |
| 4 | 84013649 | AX-170306910 | 0 | T | G | 0.43 | -1.28E-01 | 8.29E-02 | 2.54E+00 | 2.60E+00 | 1.21E-01 | no | | intergenic_variant | MODIFIER |
| 4 | 84026762 | AX-170366581 | 0 | A | G | 0.43 | -1.28E-01 | 8.29E-02 | 2.54E+00 | 2.60E+00 | 1.21E-01 | no | | intergenic_variant | MODIFIER |
| 4 | 84043134 | AX-170183039 | 0 | T | C | 0.43 | -1.28E-01 | 8.29E-02 | 2.54E+00 | 2.60E+00 | 1.21E-01 | no | 6030471H07Rik | intron_variant,non_coding_transcript_variant | MODIFIER |
| 16 | 51679814 | AX-170057429 | 1 | A | G | 0.464 | -1.30E-01 | 8.43E-02 | 2.56E+00 | 2.60E+00 | 1.21E-01 | no | | intergenic_variant | MODIFIER |
| 1 | 193341598 | AX-170365397 | 1 | T | A | 0.268 | 1.43E-01 | 9.10E-02 | 2.36E+00 | 2.44E+00 | 1.21E-01 | no | Lamb3 | intron_variant | MODIFIER |
| 15 | 64067253 | AX-170481127 | 0 | G | A | 0.325 | -1.23E-01 | 7.56E-02 | 2.15E+00 | 2.22E+00 | 1.22E-01 | no | | intergenic_variant | MODIFIER |
| 4 | 105012316 | AX-97831351 | 0 | A | G | 0.246 | 1.77E-01 | 1.15E-01 | 2.73E+00 | 2.81E+00 | 1.22E-01 | Yes | Fyb2 | intron_variant | MODIFIER |
| 4 | 105018432 | AX-170148560 | 0 | T | C | 0.246 | 1.77E-01 | 1.15E-01 | 2.73E+00 | 2.81E+00 | 1.22E-01 | Yes | Fyb2 | downstream_gene_variant | MODIFIER |
| 4 | 105022028 | AX-170300542 | 0 | T | C | 0.246 | 1.77E-01 | 1.15E-01 | 2.73E+00 | 2.81E+00 | 1.22E-01 | Yes | | intergenic_variant | MODIFIER |
| 4 | 105032884 | AX-97807850 | 0 | C | T | 0.246 | 1.77E-01 | 1.15E-01 | 2.73E+00 | 2.81E+00 | 1.22E-01 | Yes | Prkaa2 | 3_prime_UTR_variant | MODIFIER |
| 4 | 105094409 | AX-170439551 | 0 | G | A | 0.246 | 1.77E-01 | 1.15E-01 | 2.73E+00 | 2.81E+00 | 1.22E-01 | Yes | Prkaa2 | intron_variant | MODIFIER |
| 4 | 105950480 | AX-97801138 | 0 | C | T | 0.246 | 1.77E-01 | 1.15E-01 | 2.73E+00 | 2.81E+00 | 1.22E-01 | Yes | | intergenic_variant | MODIFIER |
| 15 | 67301765 | AX-170438264 | 1 | G | A | 0.321 | -1.19E-01 | 7.50E-02 | 2.26E+00 | 2.34E+00 | 1.22E-01 | no | 1700012I11Rik | intron_variant,non_coding_transcript_variant | MODIFIER |
| 13 | 17421559 | AX-170388418 | 0 | A | G | 0.184 | -1.80E-01 | 1.14E-01 | 2.37E+00 | 2.42E+00 | 1.22E-01 | Yes | Sugct | intron_variant | MODIFIER |
| 13 | 10544522 | AX-170323446 | 2 | G | A | 0.182 | -1.76E-01 | 1.14E-01 | 2.63E+00 | 2.72E+00 | 1.22E-01 | Yes | Gm26861 | intron_variant,non_coding_transcript_variant | MODIFIER |
| 18 | 45340419 | AX-97780608 | 0 | G | A | 0.061 | -3.25E-01 | 2.11E-01 | 2.84E+00 | 2.95E+00 | 1.22E-01 | Yes | Kcnn2 | intron_variant | MODIFIER |
| 6 | 101793783 | AX-170527707 | 0 | C | T | 0.026 | -2.19E-01 | 1.43E-01 | 2.91E+00 | 3.11E+00 | 1.22E-01 | Yes | Gm9871 | upstream_gene_variant | MODIFIER |
| 4 | 107833960 | AX-170374799 | 1 | C | T | 0.268 | 1.71E-01 | 1.11E-01 | 2.67E+00 | 2.74E+00 | 1.22E-01 | Yes | Lrp8os1 | upstream_gene_variant | MODIFIER |
| 2 | 16487470 | AX-97826533 | 1 | C | T | 0.027 | -2.05E-01 | 1.32E-01 | 2.45E+00 | 2.61E+00 | 1.22E-01 | Yes | Plxdc2 | intron_variant | MODIFIER |
| 5 | 64200968 | AX-170141451 | 1 | C | T | 0.491 | 1.55E-01 | 1.00E-01 | 2.56E+00 | 2.64E+00 | 1.23E-01 | Yes | Tbc1d1 | intron_variant | MODIFIER |
| 1 | 135725222 | AX-170473059 | 1 | T | C | 0.527 | 1.26E-01 | 8.22E-02 | 2.71E+00 | 2.78E+00 | 1.23E-01 | no | Csrp1 | upstream_gene_variant | MODIFIER |
| 16 | 35834111 | AX-170384904 | 2 | A | G | 0.5 | 1.16E-01 | 7.50E-02 | 2.56E+00 | 2.62E+00 | 1.23E-01 | no | Parp14 | 3_prime_UTR_variant | MODIFIER |
| 4 | 100232033 | AX-170424328 | 0 | T | C | 0.237 | 1.59E-01 | 1.03E-01 | 2.57E+00 | 2.65E+00 | 1.23E-01 | Yes | Ror1 | intron_variant | MODIFIER |
| 19 | 50579625 | AX-170016807 | 0 | T | C | 0.482 | 9.89E-02 | 6.46E-02 | 2.73E+00 | 2.82E+00 | 1.23E-01 | no | Sorcs1 | intron_variant | MODIFIER |
| 10 | 101122108 | AX-97819452 | 1 | G | A | 0.312 | -1.16E-01 | 7.50E-02 | 2.52E+00 | 2.62E+00 | 1.23E-01 | no | | intergenic_variant | MODIFIER |
| 19 | 59560365 | AX-170391094 | 1 | G | A | 0.518 | 9.70E-02 | 6.34E-02 | 2.70E+00 | 2.79E+00 | 1.24E-01 | no | 2700089I24Rik | downstream_gene_variant | MODIFIER |
| 5 | 43646156 | AX-170024819 | 0 | C | T | 0.184 | -1.50E-01 | 9.81E-02 | 2.79E+00 | 2.94E+00 | 1.24E-01 | no | | intergenic_variant | MODIFIER |
| 1 | 184512679 | AX-170086713 | 2 | A | G | 0.236 | 1.73E-01 | 1.12E-01 | 2.51E+00 | 2.58E+00 | 1.24E-01 | Yes | | intergenic_variant | MODIFIER |
| 5 | 48730715 | AX-170072019 | 1 | G | A | 0.491 | 1.59E-01 | 1.02E-01 | 2.33E+00 | 2.38E+00 | 1.24E-01 | Yes | Kcnip4 | intron_variant | MODIFIER |
| 5 | 50133243 | AX-97847769 | 1 | T | C | 0.491 | 1.59E-01 | 1.02E-01 | 2.33E+00 | 2.38E+00 | 1.24E-01 | Yes | | intergenic_variant | MODIFIER |
| 16 | 62103022 | AX-170526154 | 2 | G | A | 0.527 | 1.27E-01 | 8.25E-02 | 2.53E+00 | 2.58E+00 | 1.24E-01 | no | Gm49615 | intron_variant,non_coding_transcript_variant | MODIFIER |
| 18 | 75027239 | AX-170088376 | 0 | T | C | 0.175 | -1.77E-01 | 1.16E-01 | 2.73E+00 | 2.81E+00 | 1.24E-01 | Yes | Dym | intron_variant | MODIFIER |

| | | | | | | | | | | | | | | | |
|---|---|---|---|---|---|---|---|---|---|---|---|---|---|---|---|
| 13 | 25203111 | AX-170050781 | 0 | C | T | 0.167 | -1.66E-01 | 1.07E-01 | 2.39E+00 | 2.49E+00 | 1.24E-01 | no | Dcdc2a | intron_variant | MODIFIER |
| 1 | 189482874 | AX-97814553 | 0 | A | C | 0.272 | 1.47E-01 | 9.38E-02 | 2.32E+00 | 2.41E+00 | 1.25E-01 | Yes | | intergenic_variant | MODIFIER |
| 5 | 135374982 | AX-97793327 | 1 | G | C | 0.152 | -1.34E-01 | 8.52E-02 | 2.28E+00 | 2.36E+00 | 1.25E-01 | no | Nsun5 | missense_variant | MODERATE |
| 1 | 185141287 | AX-170217993 | 1 | A | G | 0.241 | 1.68E-01 | 1.10E-01 | 2.50E+00 | 2.57E+00 | 1.25E-01 | Yes | | intergenic_variant | MODIFIER |
| 1 | 134870153 | AX-170161986 | 1 | A | T | 0.536 | 1.21E-01 | 7.89E-02 | 2.56E+00 | 2.63E+00 | 1.25E-01 | no | Ppp1r12b | intron_variant | MODIFIER |
| 8 | 90042268 | AX-170235770 | 0 | A | G | 0.386 | -8.50E-02 | 5.56E-02 | 2.93E+00 | 3.05E+00 | 1.25E-01 | no | | intergenic_variant | MODIFIER |
| 1 | 166774664 | AX-170235475 | 0 | A | G | 0.219 | 2.11E-01 | 1.38E-01 | 2.78E+00 | 2.84E+00 | 1.26E-01 | Yes | Gm4954 | downstream_gene_variant | MODIFIER |
| 18 | 86574829 | AX-97851303 | 0 | A | G | 0.158 | -1.51E-01 | 9.89E-02 | 2.56E+00 | 2.66E+00 | 1.26E-01 | no | | intergenic_variant | MODIFIER |
| 5 | 129229975 | AX-170073495 | 0 | G | C | 0.158 | -1.41E-01 | 8.89E-02 | 2.20E+00 | 2.27E+00 | 1.26E-01 | no | | intergenic_variant | MODIFIER |
| 8 | 49363766 | AX-170030645 | 0 | T | A | 0.158 | -1.72E-01 | 1.13E-01 | 2.64E+00 | 2.78E+00 | 1.26E-01 | no | | intergenic_variant | MODIFIER |
| 5 | 62976367 | AX-170506570 | 0 | T | C | 0.474 | 1.56E-01 | 1.01E-01 | 2.41E+00 | 2.47E+00 | 1.26E-01 | Yes | | intergenic_variant | MODIFIER |
| 5 | 63055322 | AX-170072290 | 0 | C | A | 0.474 | 1.56E-01 | 1.01E-01 | 2.41E+00 | 2.47E+00 | 1.26E-01 | Yes | | intergenic_variant | MODIFIER |
| 5 | 63194160 | AX-170227981 | 0 | A | G | 0.474 | 1.56E-01 | 1.01E-01 | 2.41E+00 | 2.47E+00 | 1.26E-01 | Yes | | intergenic_variant | MODIFIER |
| 5 | 65019227 | AX-170254916 | 0 | A | T | 0.474 | 1.56E-01 | 1.01E-01 | 2.41E+00 | 2.47E+00 | 1.26E-01 | Yes | Fam114a1 | intron_variant | MODIFIER |
| 4 | 79357246 | AX-97798107 | 1 | A | G | 0.438 | -1.26E-01 | 8.25E-02 | 2.56E+00 | 2.63E+00 | 1.26E-01 | no | | intergenic_variant | MODIFIER |
| 16 | 58026878 | AX-97796639 | 1 | T | C | 0.455 | -1.25E-01 | 8.18E-02 | 2.50E+00 | 2.55E+00 | 1.26E-01 | no | Gm49575 | intron_variant,non_coding_transcript_variant | MODIFIER |
| 11 | 78689972 | AX-97815558 | 2 | G | A | 0.036 | -2.37E-01 | 1.56E-01 | 2.64E+00 | 2.78E+00 | 1.26E-01 | Yes | Nlk | intron_variant | MODIFIER |
| 5 | 62464436 | AX-168405189 | 1 | T | G | 0.464 | 1.48E-01 | 9.66E-02 | 2.46E+00 | 2.53E+00 | 1.26E-01 | Yes | | intergenic_variant | MODIFIER |
| 3 | 38610839 | AX-170296831 | 0 | G | A | 0.211 | 1.38E-01 | 8.93E-02 | 3.10E+00 | 3.23E+00 | 1.27E-01 | no | Gm43538 | upstream_gene_variant | MODIFIER |
| 4 | 106037438 | AX-170287505 | 0 | A | G | 0.263 | 1.69E-01 | 1.11E-01 | 2.72E+00 | 2.80E+00 | 1.27E-01 | Yes | | intergenic_variant | MODIFIER |
| 4 | 106184090 | AX-170456853 | 0 | T | G | 0.263 | 1.69E-01 | 1.11E-01 | 2.72E+00 | 2.80E+00 | 1.27E-01 | Yes | | intergenic_variant | MODIFIER |
| 4 | 72014200 | AX-170069654 | 0 | G | A | 0.456 | -1.29E-01 | 8.39E-02 | 2.45E+00 | 2.51E+00 | 1.27E-01 | no | | regulatory_region_variant | MODIFIER |
| 13 | 8759075 | AX-170232194 | 1 | G | A | 0.188 | -1.90E-01 | 1.24E-01 | 2.51E+00 | 2.56E+00 | 1.27E-01 | Yes | Adarb2 | 3_prime_UTR_variant | MODIFIER |
| 13 | 8908345 | AX-170444745 | 1 | G | A | 0.188 | -1.90E-01 | 1.24E-01 | 2.51E+00 | 2.56E+00 | 1.27E-01 | Yes | | intergenic_variant | MODIFIER |
| 16 | 55287819 | AX-170324618 | 2 | T | A | 0.455 | -1.25E-01 | 8.18E-02 | 2.50E+00 | 2.55E+00 | 1.27E-01 | no | Zpld1 | upstream_gene_variant | MODIFIER |
| 19 | 59687929 | AX-170179217 | 0 | G | A | 0.518 | 9.63E-02 | 6.35E-02 | 2.77E+00 | 2.86E+00 | 1.27E-01 | no | AC123621.2 | upstream_gene_variant | MODIFIER |
| 15 | 70398603 | AX-170154253 | 1 | T | C | 0.071 | -1.92E-01 | 1.06E-01 | 1.71E+00 | 1.78E+00 | 1.27E-01 | Yes | | intergenic_variant | MODIFIER |
| 19 | 59523182 | AX-170457069 | 2 | T | G | 0.482 | -9.62E-02 | 6.34E-02 | 2.70E+00 | 2.78E+00 | 1.27E-01 | no | 2700089I24Rik | intron_variant,non_coding_transcript_variant | MODIFIER |
| 16 | 9045857 | AX-97784489 | 0 | C | T | 0.044 | -2.75E-01 | 1.80E-01 | 3.03E+00 | 3.21E+00 | 1.27E-01 | Yes | Gm30567 | upstream_gene_variant | MODIFIER |
| 4 | 84577722 | AX-170245327 | 0 | A | C | 0.351 | 1.72E-01 | 1.13E-01 | 2.91E+00 | 2.99E+00 | 1.28E-01 | Yes | Bnc2 | intron_variant | MODIFIER |
| 1 | 191797120 | AX-170002578 | 1 | G | T | 0.277 | 1.42E-01 | 9.18E-02 | 2.36E+00 | 2.44E+00 | 1.28E-01 | no | | intergenic_variant | MODIFIER |
| 1 | 135813143 | AX-170399815 | 0 | C | T | 0.535 | 1.26E-01 | 8.31E-02 | 2.72E+00 | 2.79E+00 | 1.28E-01 | no | Tnni1 | downstream_gene_variant | MODIFIER |
| 6 | 104776183 | AX-170075759 | 0 | G | A | 0.07 | -3.07E-01 | 2.02E-01 | 2.97E+00 | 3.11E+00 | 1.28E-01 | Yes | Cntn6 | intron_variant | MODIFIER |
| 4 | 106214146 | AX-170023597 | 0 | A | G | 0.281 | 1.51E-01 | 1.00E-01 | 2.67E+00 | 2.76E+00 | 1.28E-01 | Yes | | intergenic_variant | MODIFIER |
| 5 | 10402730 | AX-170418226 | 0 | G | A | 0.342 | 1.26E-01 | 8.33E-02 | 2.74E+00 | 2.87E+00 | 1.28E-01 | no | | intergenic_variant | MODIFIER |
| 5 | 130041469 | AX-170341399 | 0 | G | A | 0.149 | -1.43E-01 | 9.07E-02 | 2.23E+00 | 2.30E+00 | 1.28E-01 | no | Crcp | intron_variant | MODIFIER |
| 8 | 75205390 | AX-170080443 | 1 | G | A | 0.196 | -1.92E-01 | 1.25E-01 | 3.15E+00 | 3.30E+00 | 1.28E-01 | Yes | Gm45903 | intron_variant,non_coding_transcript_variant | MODIFIER |
| 5 | 57507079 | AX-97852101 | 1 | G | T | 0.455 | 1.53E-01 | 9.92E-02 | 2.36E+00 | 2.42E+00 | 1.28E-01 | Yes | | intergenic_variant | MODIFIER |
| 1 | 168424870 | AX-170200331 | 1 | T | A | 0.232 | 1.68E-01 | 1.11E-01 | 2.77E+00 | 2.91E+00 | 1.29E-01 | Yes | Pbx1 | intron_variant | MODIFIER |
| 1 | 184023095 | AX-170382189 | 0 | A | G | 0.237 | 1.62E-01 | 1.06E-01 | 2.47E+00 | 2.55E+00 | 1.29E-01 | Yes | Dusp10 | intron_variant | MODIFIER |
| 4 | 108109993 | AX-170272026 | 1 | G | A | 0.25 | 1.75E-01 | 1.15E-01 | 2.62E+00 | 2.68E+00 | 1.29E-01 | Yes | Scp2 | intron_variant | MODIFIER |
| 5 | 128271181 | AX-170165056 | 0 | G | A | 0.149 | -1.30E-01 | 8.22E-02 | 2.19E+00 | 2.29E+00 | 1.29E-01 | no | Tmem132d | intron_variant | MODIFIER |
| 1 | 151012562 | AX-170246278 | 1 | A | G | 0.205 | 1.63E-01 | 1.08E-01 | 2.65E+00 | 2.77E+00 | 1.29E-01 | Yes | | intergenic_variant | MODIFIER |
| 12 | 107375783 | AX-170308972 | 0 | G | T | 0.123 | -2.60E-01 | 1.66E-01 | 3.32E+00 | 3.46E+00 | 1.29E-01 | Yes | | intergenic_variant | MODIFIER |
| 3 | 19340564 | AX-170115893 | 1 | C | T | 0.125 | -1.52E-01 | 1.01E-01 | 2.62E+00 | 2.71E+00 | 1.29E-01 | no | | intergenic_variant | MODIFIER |
| 16 | 56785200 | AX-170231431 | 1 | A | T | 0.464 | -1.25E-01 | 8.28E-02 | 2.59E+00 | 2.64E+00 | 1.29E-01 | no | Adgrg7 | intron_variant | MODIFIER |
| 15 | 3705702 | AX-170255238 | 0 | C | T | 0.088 | -1.49E-01 | 8.96E-02 | 1.95E+00 | 2.05E+00 | 1.29E-01 | no | | intergenic_variant | MODIFIER |
| 15 | 3828827 | AX-97844094 | 0 | A | G | 0.088 | -1.49E-01 | 8.96E-02 | 1.95E+00 | 2.05E+00 | 1.29E-01 | no | | intergenic_variant | MODIFIER |
| 5 | 130047789 | AX-170026157 | 0 | G | T | 0.149 | -1.38E-01 | 8.78E-02 | 2.22E+00 | 2.30E+00 | 1.29E-01 | no | Crcp | intron_variant | MODIFIER |
| 1 | 54769885 | AX-170109636 | 0 | G | C | 0.123 | 2.22E-01 | 1.47E-01 | 2.94E+00 | 3.15E+00 | 1.30E-01 | Yes | Ankrd44 | intron_variant | MODIFIER |
| 1 | 127440742 | AX-97813204 | 1 | G | A | 0.562 | 1.21E-01 | 8.06E-02 | 2.66E+00 | 2.74E+00 | 1.30E-01 | no | Mgat5 | intron_variant | MODIFIER |
| 1 | 134999538 | AX-170001762 | 0 | A | G | 0.544 | 1.25E-01 | 8.32E-02 | 2.72E+00 | 2.79E+00 | 1.30E-01 | no | Lgr6 | intron_variant | MODIFIER |
| 13 | 8305983 | AX-97822630 | 1 | T | C | 0.188 | -1.86E-01 | 1.23E-01 | 2.52E+00 | 2.57E+00 | 1.30E-01 | Yes | Adarb2 | intron_variant | MODIFIER |
| 5 | 129893384 | AX-170276927 | 0 | G | A | 0.158 | -1.37E-01 | 8.71E-02 | 2.20E+00 | 2.28E+00 | 1.30E-01 | no | Zbed5 | upstream_gene_variant | MODIFIER |

| | | | | | | | | | | | | | | | |
|---|---|---|---|---|---|---|---|---|---|---|---|---|---|---|---|
| 13 | 18627979 | AX-170152578 | 1 | T | C | 0.161 | -1.91E-01 | 1.25E-01 | 2.43E+00 | 2.48E+00 | 1.30E-01 | Yes | | intergenic_variant | MODIFIER |
| 4 | 64058400 | AX-170022957 | 0 | G | A | 0.342 | 1.64E-01 | 1.08E-01 | 3.08E+00 | 3.18E+00 | 1.30E-01 | Yes | 8030451A03Rik | intron_variant,non_coding_transcript_variant | MODIFIER |
| 13 | 9622799 | AX-170459131 | 1 | G | C | 0.188 | -1.87E-01 | 1.24E-01 | 2.55E+00 | 2.61E+00 | 1.30E-01 | Yes | Gm23496 | downstream_gene_variant | MODIFIER |
| 4 | 85097405 | AX-97798823 | 0 | C | T | 0.351 | 1.66E-01 | 1.10E-01 | 2.95E+00 | 3.05E+00 | 1.30E-01 | Yes | Cntln | intron_variant | MODIFIER |
| 4 | 76101935 | AX-170095019 | 0 | T | A | 0.316 | 1.75E-01 | 1.16E-01 | 2.91E+00 | 2.97E+00 | 1.30E-01 | Yes | Ptprd | intron_variant | MODIFIER |
| 13 | 18375667 | AX-170087380 | 0 | G | T | 0.158 | -1.81E-01 | 1.20E-01 | 2.56E+00 | 2.65E+00 | 1.30E-01 | Yes | Pou6f2 | intron_variant | MODIFIER |
| 1 | 184824653 | AX-170399834 | 0 | G | A | 0.246 | 1.60E-01 | 1.06E-01 | 2.48E+00 | 2.57E+00 | 1.30E-01 | Yes | MAR2 | intron_variant | MODIFIER |
| 4 | 108628660 | AX-170217158 | 0 | T | C | 0.228 | 1.79E-01 | 1.19E-01 | 2.60E+00 | 2.66E+00 | 1.31E-01 | Yes | Cc2d1b | synonymous_variant | LOW |
| 4 | 108629166 | AX-170362613 | 0 | A | C | 0.228 | 1.79E-01 | 1.19E-01 | 2.60E+00 | 2.66E+00 | 1.31E-01 | Yes | Cc2d1b | intron_variant | MODIFIER |
| 4 | 108632997 | AX-170089339 | 0 | A | G | 0.228 | 1.79E-01 | 1.19E-01 | 2.60E+00 | 2.66E+00 | 1.31E-01 | Yes | Cc2d1b | intron_variant | MODIFIER |
| 4 | 108674720 | AX-170232393 | 0 | T | G | 0.228 | 1.79E-01 | 1.19E-01 | 2.60E+00 | 2.66E+00 | 1.31E-01 | Yes | Zfyve9 | intron_variant | MODIFIER |
| 4 | 108694942 | AX-97819899 | 0 | G | A | 0.228 | 1.79E-01 | 1.19E-01 | 2.60E+00 | 2.66E+00 | 1.31E-01 | Yes | Zfyve9 | intron_variant | MODIFIER |
| 4 | 108710295 | AX-170089340 | 0 | T | C | 0.228 | 1.79E-01 | 1.19E-01 | 2.60E+00 | 2.66E+00 | 1.31E-01 | Yes | Zfyve9 | intron_variant | MODIFIER |
| 4 | 108729615 | AX-170132020 | 0 | T | C | 0.228 | 1.79E-01 | 1.19E-01 | 2.60E+00 | 2.66E+00 | 1.31E-01 | Yes | Zfyve9 | intron_variant | MODIFIER |
| 4 | 108738297 | AX-97852307 | 0 | G | A | 0.228 | 1.79E-01 | 1.19E-01 | 2.60E+00 | 2.66E+00 | 1.31E-01 | Yes | Zfyve9 | intron_variant | MODIFIER |
| 4 | 108746213 | AX-170143550 | 0 | G | A | 0.228 | 1.79E-01 | 1.19E-01 | 2.60E+00 | 2.66E+00 | 1.31E-01 | Yes | Zfyve9 | intron_variant | MODIFIER |
| 4 | 108761322 | AX-170035689 | 0 | T | C | 0.228 | 1.79E-01 | 1.19E-01 | 2.60E+00 | 2.66E+00 | 1.31E-01 | Yes | Zfyve9 | intron_variant | MODIFIER |
| 4 | 108762775 | AX-170338994 | 0 | T | C | 0.228 | 1.79E-01 | 1.19E-01 | 2.60E+00 | 2.66E+00 | 1.31E-01 | Yes | Zfyve9 | intron_variant | MODIFIER |
| 4 | 108762905 | AX-97791135 | 0 | C | T | 0.228 | 1.79E-01 | 1.19E-01 | 2.60E+00 | 2.66E+00 | 1.31E-01 | Yes | Zfyve9 | intron_variant | MODIFIER |
| 4 | 108763062 | AX-170198681 | 0 | T | G | 0.228 | 1.79E-01 | 1.19E-01 | 2.60E+00 | 2.66E+00 | 1.31E-01 | Yes | Zfyve9 | intron_variant | MODIFIER |
| 4 | 108841443 | AX-97798452 | 0 | A | C | 0.228 | 1.79E-01 | 1.19E-01 | 2.60E+00 | 2.66E+00 | 1.31E-01 | Yes | Txndc12 | intron_variant | MODIFIER |
| 4 | 108846124 | AX-170023655 | 0 | T | C | 0.228 | 1.79E-01 | 1.19E-01 | 2.60E+00 | 2.66E+00 | 1.31E-01 | Yes | Txndc12 | intron_variant | MODIFIER |
| 4 | 108876538 | AX-170023651 | 0 | T | C | 0.228 | 1.79E-01 | 1.19E-01 | 2.60E+00 | 2.66E+00 | 1.31E-01 | Yes | Rab3b | upstream_gene_variant | MODIFIER |
| 4 | 108915120 | AX-170023648 | 0 | T | C | 0.228 | 1.79E-01 | 1.19E-01 | 2.60E+00 | 2.66E+00 | 1.31E-01 | Yes | Rab3b | intron_variant | MODIFIER |
| 4 | 108976161 | AX-170276152 | 0 | C | T | 0.228 | 1.79E-01 | 1.19E-01 | 2.60E+00 | 2.66E+00 | 1.31E-01 | Yes | 8030443G20Rik | upstream_gene_variant | MODIFIER |
| 4 | 109077305 | AX-168414081 | 0 | A | G | 0.228 | 1.79E-01 | 1.19E-01 | 2.60E+00 | 2.66E+00 | 1.31E-01 | Yes | Osbpl9 | intron_variant | MODIFIER |
| 4 | 109087811 | AX-97807689 | 0 | T | C | 0.228 | 1.79E-01 | 1.19E-01 | 2.60E+00 | 2.66E+00 | 1.31E-01 | Yes | Osbpl9 | intron_variant | MODIFIER |
| 4 | 109874495 | AX-170023663 | 0 | A | G | 0.228 | 1.79E-01 | 1.19E-01 | 2.60E+00 | 2.66E+00 | 1.31E-01 | Yes | Faf1 | intron_variant | MODIFIER |
| 5 | 65359515 | AX-170343779 | 0 | G | A | 0.465 | 1.53E-01 | 1.00E-01 | 2.42E+00 | 2.48E+00 | 1.31E-01 | Yes | Klb | intron_variant | MODIFIER |
| 5 | 132549984 | AX-170519724 | 0 | C | T | 0.158 | -1.32E-01 | 8.51E-02 | 2.29E+00 | 2.37E+00 | 1.31E-01 | no | Gm28616 | downstream_gene_variant | MODIFIER |
| 5 | 132565737 | AX-170520868 | 0 | C | A | 0.158 | -1.32E-01 | 8.51E-02 | 2.29E+00 | 2.37E+00 | 1.31E-01 | no | | intergenic_variant | MODIFIER |
| 5 | 132584971 | AX-170493853 | 0 | A | G | 0.158 | -1.32E-01 | 8.51E-02 | 2.29E+00 | 2.37E+00 | 1.31E-01 | no | | intergenic_variant | MODIFIER |
| 5 | 132585770 | AX-170221587 | 0 | A | G | 0.158 | -1.32E-01 | 8.51E-02 | 2.29E+00 | 2.37E+00 | 1.31E-01 | no | | intergenic_variant | MODIFIER |
| 5 | 132608252 | AX-97815825 | 0 | G | A | 0.158 | -1.32E-01 | 8.51E-02 | 2.29E+00 | 2.37E+00 | 1.31E-01 | no | | intergenic_variant | MODIFIER |
| 5 | 132608887 | AX-170278506 | 0 | T | C | 0.158 | -1.32E-01 | 8.51E-02 | 2.29E+00 | 2.37E+00 | 1.31E-01 | no | | intergenic_variant | MODIFIER |
| 5 | 132608910 | AX-170534896 | 0 | G | C | 0.158 | -1.32E-01 | 8.51E-02 | 2.29E+00 | 2.37E+00 | 1.31E-01 | no | | intergenic_variant | MODIFIER |
| 5 | 132609818 | AX-170336259 | 0 | G | A | 0.158 | -1.32E-01 | 8.51E-02 | 2.29E+00 | 2.37E+00 | 1.31E-01 | no | | intergenic_variant | MODIFIER |
| 5 | 132611209 | AX-170360696 | 0 | T | A | 0.158 | -1.32E-01 | 8.51E-02 | 2.29E+00 | 2.37E+00 | 1.31E-01 | no | | intergenic_variant | MODIFIER |
| 5 | 132615984 | AX-170298059 | 0 | C | T | 0.158 | -1.32E-01 | 8.51E-02 | 2.29E+00 | 2.37E+00 | 1.31E-01 | no | | intergenic_variant | MODIFIER |
| 5 | 132653723 | AX-170445732 | 0 | A | G | 0.158 | -1.32E-01 | 8.51E-02 | 2.29E+00 | 2.37E+00 | 1.31E-01 | no | | regulatory_region_variant | MODIFIER |
| 5 | 132657600 | AX-170406505 | 0 | C | T | 0.158 | -1.32E-01 | 8.51E-02 | 2.29E+00 | 2.37E+00 | 1.31E-01 | no | | intergenic_variant | MODIFIER |
| 5 | 132694231 | AX-170121818 | 0 | T | C | 0.158 | -1.32E-01 | 8.51E-02 | 2.29E+00 | 2.37E+00 | 1.31E-01 | no | | intergenic_variant | MODIFIER |
| 5 | 132763286 | AX-97800573 | 0 | G | A | 0.158 | -1.32E-01 | 8.51E-02 | 2.29E+00 | 2.37E+00 | 1.31E-01 | no | | intergenic_variant | MODIFIER |
| 5 | 133062784 | AX-170448172 | 0 | C | T | 0.158 | -1.32E-01 | 8.51E-02 | 2.29E+00 | 2.37E+00 | 1.31E-01 | no | | intergenic_variant | MODIFIER |
| 4 | 82307240 | AX-170397251 | 1 | A | C | 0.348 | 1.77E-01 | 1.18E-01 | 2.86E+00 | 2.93E+00 | 1.31E-01 | Yes | Nfib | intron_variant | MODIFIER |
| 13 | 10970944 | AX-170324855 | 0 | T | C | 0.184 | -1.87E-01 | 1.24E-01 | 2.51E+00 | 2.55E+00 | 1.31E-01 | Yes | Gm47486 | upstream_gene_variant | MODIFIER |
| 18 | 53228014 | AX-170392127 | 0 | T | C | 0.053 | 3.39E-01 | 2.25E-01 | 3.01E+00 | 3.18E+00 | 1.31E-01 | Yes | | intergenic_variant | MODIFIER |
| 13 | 8540534 | AX-170098174 | 1 | T | G | 0.188 | -1.85E-01 | 1.22E-01 | 2.49E+00 | 2.54E+00 | 1.31E-01 | Yes | Adarb2 | intron_variant | MODIFIER |
| 13 | 9117493 | AX-170414790 | 1 | C | T | 0.188 | -1.85E-01 | 1.22E-01 | 2.49E+00 | 2.54E+00 | 1.31E-01 | Yes | Larp4b | intron_variant | MODIFIER |
| 8 | 65498434 | AX-170163554 | 1 | G | T | 0.152 | -1.94E-01 | 1.30E-01 | 2.65E+00 | 2.75E+00 | 1.31E-01 | Yes | | intergenic_variant | MODIFIER |
| 5 | 129721122 | AX-170166572 | 0 | A | G | 0.158 | -1.43E-01 | 9.03E-02 | 2.15E+00 | 2.22E+00 | 1.31E-01 | no | Mrps17 | downstream_gene_variant | MODIFIER |
| 1 | 181757603 | AX-170356005 | 0 | T | C | 0.237 | 1.64E-01 | 1.08E-01 | 2.43E+00 | 2.51E+00 | 1.31E-01 | Yes | Dnah14 | downstream_gene_variant | MODIFIER |
| 16 | 67080475 | AX-170103662 | 0 | G | A | 0.482 | 1.17E-01 | 7.76E-02 | 2.53E+00 | 2.59E+00 | 1.32E-01 | no | Cadm2 | intron_variant | MODIFIER |
| 16 | 67296741 | AX-170013305 | 0 | A | G | 0.482 | 1.17E-01 | 7.76E-02 | 2.53E+00 | 2.59E+00 | 1.32E-01 | no | Cadm2 | intron_variant | MODIFIER |

| | | | | | | | | | | | | | | | |
|---|---|---|---|---|---|---|---|---|---|---|---|---|---|---|---|
| 16 | 67337996 | AX-97782352 | 0 | C | T | 0.482 | 1.17E-01 | 7.76E-02 | 2.53E+00 | 2.59E+00 | 1.32E-01 | no | Cadm2 | intron_variant | MODIFIER |
| 16 | 67350807 | AX-170125765 | 0 | C | T | 0.482 | 1.17E-01 | 7.76E-02 | 2.53E+00 | 2.59E+00 | 1.32E-01 | no | Cadm2 | intron_variant | MODIFIER |
| 16 | 67381326 | AX-170503977 | 0 | G | T | 0.482 | 1.17E-01 | 7.76E-02 | 2.53E+00 | 2.59E+00 | 1.32E-01 | no | Cadm2 | intron_variant | MODIFIER |
| 16 | 67382229 | AX-170159109 | 0 | C | G | 0.482 | 1.17E-01 | 7.76E-02 | 2.53E+00 | 2.59E+00 | 1.32E-01 | no | Cadm2 | intron_variant | MODIFIER |
| 16 | 67422678 | AX-170482525 | 0 | A | G | 0.482 | 1.17E-01 | 7.76E-02 | 2.53E+00 | 2.59E+00 | 1.32E-01 | no | Cadm2 | intron_variant | MODIFIER |
| 16 | 67433553 | AX-170057691 | 0 | A | G | 0.482 | 1.17E-01 | 7.76E-02 | 2.53E+00 | 2.59E+00 | 1.32E-01 | no | Cadm2 | intron_variant | MODIFIER |
| 16 | 67589257 | AX-170473363 | 0 | C | T | 0.482 | 1.17E-01 | 7.76E-02 | 2.53E+00 | 2.59E+00 | 1.32E-01 | no | Cadm2 | intron_variant | MODIFIER |
| 16 | 67589332 | AX-170519404 | 0 | C | A | 0.482 | 1.17E-01 | 7.76E-02 | 2.53E+00 | 2.59E+00 | 1.32E-01 | no | Cadm2 | intron_variant | MODIFIER |
| 16 | 67658831 | AX-170369421 | 0 | C | A | 0.482 | 1.17E-01 | 7.76E-02 | 2.53E+00 | 2.59E+00 | 1.32E-01 | no | | intergenic_variant | MODIFIER |
| 16 | 67662860 | AX-170351024 | 0 | A | G | 0.482 | 1.17E-01 | 7.76E-02 | 2.53E+00 | 2.59E+00 | 1.32E-01 | no | | intergenic_variant | MODIFIER |
| 16 | 67665034 | AX-170088050 | 0 | A | C | 0.482 | 1.17E-01 | 7.76E-02 | 2.53E+00 | 2.59E+00 | 1.32E-01 | no | | intergenic_variant | MODIFIER |
| 16 | 67696822 | AX-170455912 | 0 | C | G | 0.482 | 1.17E-01 | 7.76E-02 | 2.53E+00 | 2.59E+00 | 1.32E-01 | no | Gm49648 | downstream_gene_variant | MODIFIER |
| 16 | 67724879 | AX-170102046 | 0 | A | G | 0.482 | 1.17E-01 | 7.76E-02 | 2.53E+00 | 2.59E+00 | 1.32E-01 | no | Gm49648 | intron_variant,non_coding_transcript_variant | MODIFIER |
| 16 | 67724902 | AX-170451349 | 0 | G | A | 0.482 | 1.17E-01 | 7.76E-02 | 2.53E+00 | 2.59E+00 | 1.32E-01 | no | Gm49648 | intron_variant,non_coding_transcript_variant | MODIFIER |
| 16 | 67736750 | AX-170392300 | 0 | A | T | 0.482 | 1.17E-01 | 7.76E-02 | 2.53E+00 | 2.59E+00 | 1.32E-01 | no | Gm49648 | intron_variant,non_coding_transcript_variant | MODIFIER |
| 1 | 134926691 | AX-170458077 | 1 | G | A | 0.545 | 1.25E-01 | 8.31E-02 | 2.73E+00 | 2.80E+00 | 1.32E-01 | no | Ppp1r12b | intron_variant | MODIFIER |
| 1 | 53034648 | AX-170336534 | 0 | G | A | 0.105 | -2.97E-01 | 1.92E-01 | 3.28E+00 | 3.41E+00 | 1.32E-01 | Yes | | intergenic_variant | MODIFIER |
| 1 | 54229978 | AX-97844815 | 0 | C | T | 0.105 | -2.97E-01 | 1.92E-01 | 3.28E+00 | 3.41E+00 | 1.32E-01 | Yes | | intergenic_variant | MODIFIER |
| 1 | 194526059 | AX-170148008 | 1 | A | T | 0.277 | 1.40E-01 | 9.16E-02 | 2.35E+00 | 2.43E+00 | 1.32E-01 | no | | intergenic_variant | MODIFIER |
| 1 | 16057508 | AX-170115937 | 0 | A | G | 0.377 | 1.17E-01 | 7.77E-02 | 2.59E+00 | 2.66E+00 | 1.32E-01 | no | Gm7634 | downstream_gene_variant | MODIFIER |
| 13 | 7545679 | AX-97812378 | 1 | C | T | 0.196 | -1.80E-01 | 1.19E-01 | 2.52E+00 | 2.57E+00 | 1.32E-01 | Yes | | intergenic_variant | MODIFIER |
| 13 | 11971913 | AX-97822811 | 0 | T | C | 0.184 | -1.88E-01 | 1.25E-01 | 2.53E+00 | 2.58E+00 | 1.32E-01 | Yes | Ryr2 | intron_variant | MODIFIER |
| 1 | 191824113 | AX-170002580 | 1 | C | T | 0.277 | 1.30E-01 | 8.49E-02 | 2.33E+00 | 2.42E+00 | 1.32E-01 | no | Nek2 | intron_variant | MODIFIER |
| 12 | 69637458 | AX-170421688 | 0 | G | A | 0.061 | -3.18E-01 | 2.12E-01 | 2.84E+00 | 2.95E+00 | 1.32E-01 | Yes | Sos2 | intron_variant | MODIFIER |
| 4 | 98956064 | AX-97819264 | 2 | G | A | 0.236 | 1.61E-01 | 1.08E-01 | 2.75E+00 | 2.85E+00 | 1.32E-01 | Yes | Dock7 | intron_variant | MODIFIER |
| 19 | 60857703 | AX-170270054 | 1 | T | C | 0.464 | -9.51E-02 | 6.36E-02 | 2.71E+00 | 2.80E+00 | 1.32E-01 | no | Sfxn4 | intron_variant,NMD_transcript_variant | MODIFIER |
| 1 | 135048487 | AX-97790858 | 1 | G | A | 0.545 | 1.24E-01 | 8.32E-02 | 2.73E+00 | 2.80E+00 | 1.32E-01 | no | Lgr6 | intron_variant | MODIFIER |
| 5 | 13880011 | AX-170288574 | 1 | A | G | 0.348 | 1.26E-01 | 8.41E-02 | 2.86E+00 | 3.01E+00 | 1.32E-01 | no | Dnd1-ps | non_coding_transcript_exon_variant | MODIFIER |
| 5 | 137096372 | AX-168411289 | 0 | T | G | 0.14 | -1.28E-01 | 8.48E-02 | 2.46E+00 | 2.56E+00 | 1.32E-01 | no | | regulatory_region_variant | MODIFIER |
| 4 | 108675456 | AX-170420614 | 1 | G | A | 0.259 | 1.70E-01 | 1.13E-01 | 2.57E+00 | 2.63E+00 | 1.32E-01 | Yes | Zfyve9 | intron_variant | MODIFIER |
| 12 | 90170069 | AX-170007229 | 0 | A | G | 0.105 | 2.37E-01 | 1.58E-01 | 2.89E+00 | 3.01E+00 | 1.32E-01 | Yes | Nrxn3 | intron_variant | MODIFIER |
| 4 | 124334352 | AX-97805841 | 0 | A | G | 0.228 | 1.60E-01 | 1.06E-01 | 2.60E+00 | 2.68E+00 | 1.32E-01 | Yes | 4933435F18Rik | upstream_gene_variant | MODIFIER |
| 4 | 124650074 | AX-170254780 | 0 | C | G | 0.228 | 1.60E-01 | 1.06E-01 | 2.60E+00 | 2.68E+00 | 1.32E-01 | Yes | Gm2164 | intron_variant,non_coding_transcript_variant | MODIFIER |
| 4 | 125548823 | AX-170187741 | 0 | G | T | 0.228 | 1.60E-01 | 1.06E-01 | 2.60E+00 | 2.68E+00 | 1.32E-01 | Yes | Grik3 | intron_variant | MODIFIER |
| 3 | 8268639 | AX-170329224 | 0 | C | T | 0.412 | -9.39E-02 | 6.26E-02 | 2.95E+00 | 3.12E+00 | 1.33E-01 | no | | intergenic_variant | MODIFIER |
| 3 | 18863522 | AX-170421465 | 0 | G | C | 0.123 | -1.52E-01 | 1.01E-01 | 2.63E+00 | 2.71E+00 | 1.33E-01 | no | | intergenic_variant | MODIFIER |
| 3 | 18868279 | AX-170086160 | 0 | C | T | 0.123 | -1.52E-01 | 1.01E-01 | 2.63E+00 | 2.71E+00 | 1.33E-01 | no | | intergenic_variant | MODIFIER |
| 3 | 18877414 | AX-170335996 | 0 | A | G | 0.123 | -1.52E-01 | 1.01E-01 | 2.63E+00 | 2.71E+00 | 1.33E-01 | no | | regulatory_region_variant | MODIFIER |
| 3 | 18890999 | AX-170473384 | 0 | T | C | 0.123 | -1.52E-01 | 1.01E-01 | 2.63E+00 | 2.71E+00 | 1.33E-01 | no | | intergenic_variant | MODIFIER |
| 3 | 18892719 | AX-97807257 | 0 | A | G | 0.123 | -1.52E-01 | 1.01E-01 | 2.63E+00 | 2.71E+00 | 1.33E-01 | no | | intergenic_variant | MODIFIER |
| 3 | 18995711 | AX-170020050 | 0 | T | C | 0.123 | -1.52E-01 | 1.01E-01 | 2.63E+00 | 2.71E+00 | 1.33E-01 | no | | intergenic_variant | MODIFIER |
| 3 | 19007187 | AX-170020058 | 0 | T | C | 0.123 | -1.52E-01 | 1.01E-01 | 2.63E+00 | 2.71E+00 | 1.33E-01 | no | | intergenic_variant | MODIFIER |
| 3 | 19015019 | AX-97829791 | 0 | G | A | 0.123 | -1.52E-01 | 1.01E-01 | 2.63E+00 | 2.71E+00 | 1.33E-01 | no | | intergenic_variant | MODIFIER |
| 3 | 19025824 | AX-170222289 | 0 | C | G | 0.123 | -1.52E-01 | 1.01E-01 | 2.63E+00 | 2.71E+00 | 1.33E-01 | no | | intergenic_variant | MODIFIER |
| 3 | 19105156 | AX-97844450 | 0 | A | C | 0.123 | -1.52E-01 | 1.01E-01 | 2.63E+00 | 2.71E+00 | 1.33E-01 | no | | intergenic_variant | MODIFIER |
| 3 | 19105963 | AX-170020059 | 0 | A | T | 0.123 | -1.52E-01 | 1.01E-01 | 2.63E+00 | 2.71E+00 | 1.33E-01 | no | | intergenic_variant | MODIFIER |
| 3 | 19127486 | AX-97796310 | 0 | G | A | 0.123 | -1.52E-01 | 1.01E-01 | 2.63E+00 | 2.71E+00 | 1.33E-01 | no | Armc1 | downstream_gene_variant | MODIFIER |
| 3 | 19128792 | AX-170212814 | 0 | C | G | 0.123 | -1.52E-01 | 1.01E-01 | 2.63E+00 | 2.71E+00 | 1.33E-01 | no | Armc1 | downstream_gene_variant | MODIFIER |
| 3 | 19131409 | AX-170182340 | 0 | T | C | 0.123 | -1.52E-01 | 1.01E-01 | 2.63E+00 | 2.71E+00 | 1.33E-01 | no | Armc1 | 3_prime_UTR_variant | MODIFIER |
| 3 | 19131838 | AX-170506415 | 0 | A | C | 0.123 | -1.52E-01 | 1.01E-01 | 2.63E+00 | 2.71E+00 | 1.33E-01 | no | Armc1 | 3_prime_UTR_variant | MODIFIER |
| 3 | 19134113 | AX-170244022 | 0 | G | A | 0.123 | -1.52E-01 | 1.01E-01 | 2.63E+00 | 2.71E+00 | 1.33E-01 | no | Armc1 | synonymous_variant | LOW |
| 3 | 19134355 | AX-97802480 | 0 | T | C | 0.123 | -1.52E-01 | 1.01E-01 | 2.63E+00 | 2.71E+00 | 1.33E-01 | no | Armc1 | intron_variant | MODIFIER |
| 3 | 19137650 | AX-170020052 | 0 | G | A | 0.123 | -1.52E-01 | 1.01E-01 | 2.63E+00 | 2.71E+00 | 1.33E-01 | no | Armc1 | intron_variant | MODIFIER |
| 3 | 19137874 | AX-170020053 | 0 | C | T | 0.123 | -1.52E-01 | 1.01E-01 | 2.63E+00 | 2.71E+00 | 1.33E-01 | no | Armc1 | intron_variant | MODIFIER |

| | | | | | | | | | | | | | | | |
|---|---|---|---|---|---|---|---|---|---|---|---|---|---|---|---|
| 3 | 19194737 | AX-170020055 | 0 | G | C | 0.123 | -1.52E-01 | 1.01E-01 | 2.63E+00 | 2.71E+00 | 1.33E-01 | no | Mtfr1 | intron_variant,NMD_transcript_variant | MODIFIER |
| 3 | 19195009 | AX-97811648 | 0 | T | G | 0.123 | -1.52E-01 | 1.01E-01 | 2.63E+00 | 2.71E+00 | 1.33E-01 | no | Mtfr1 | intron_variant,NMD_transcript_variant | MODIFIER |
| 3 | 19201103 | AX-170316436 | 0 | A | G | 0.123 | -1.52E-01 | 1.01E-01 | 2.63E+00 | 2.71E+00 | 1.33E-01 | no | Mtfr1 | intron_variant,NMD_transcript_variant | MODIFIER |
| 3 | 19202074 | AX-170363512 | 0 | A | G | 0.123 | -1.52E-01 | 1.01E-01 | 2.63E+00 | 2.71E+00 | 1.33E-01 | no | Mtfr1 | intron_variant,NMD_transcript_variant | MODIFIER |
| 3 | 19203065 | AX-170393722 | 0 | C | T | 0.123 | -1.52E-01 | 1.01E-01 | 2.63E+00 | 2.71E+00 | 1.33E-01 | no | Mtfr1 | intron_variant,NMD_transcript_variant | MODIFIER |
| 3 | 19203447 | AX-170020051 | 0 | A | G | 0.123 | -1.52E-01 | 1.01E-01 | 2.63E+00 | 2.71E+00 | 1.33E-01 | no | Mtfr1 | intron_variant,NMD_transcript_variant | MODIFIER |
| 3 | 19211071 | AX-170020057 | 0 | T | C | 0.123 | -1.52E-01 | 1.01E-01 | 2.63E+00 | 2.71E+00 | 1.33E-01 | no | Mtfr1 | intron_variant,NMD_transcript_variant | MODIFIER |
| 3 | 19216041 | AX-170524274 | 0 | G | C | 0.123 | -1.52E-01 | 1.01E-01 | 2.63E+00 | 2.71E+00 | 1.33E-01 | no | Mtfr1 | intron_variant,NMD_transcript_variant | MODIFIER |
| 3 | 19226062 | AX-170020056 | 0 | T | C | 0.123 | -1.52E-01 | 1.01E-01 | 2.63E+00 | 2.71E+00 | 1.33E-01 | no | Pde7a | 3_prime_UTR_variant | MODIFIER |
| 3 | 19248216 | AX-170492274 | 0 | A | G | 0.123 | -1.52E-01 | 1.01E-01 | 2.63E+00 | 2.71E+00 | 1.33E-01 | no | Pde7a | intron_variant | MODIFIER |
| 3 | 19255857 | AX-170270344 | 0 | C | T | 0.123 | -1.52E-01 | 1.01E-01 | 2.63E+00 | 2.71E+00 | 1.33E-01 | no | Pde7a | intron_variant | MODIFIER |
| 3 | 19298078 | AX-97796307 | 0 | C | T | 0.123 | -1.52E-01 | 1.01E-01 | 2.63E+00 | 2.71E+00 | 1.33E-01 | no | Pde7a | intron_variant | MODIFIER |
| 3 | 19349465 | AX-170503618 | 0 | C | G | 0.123 | -1.52E-01 | 1.01E-01 | 2.63E+00 | 2.71E+00 | 1.33E-01 | no | | intergenic_variant | MODIFIER |
| 3 | 19350879 | AX-170320699 | 0 | C | G | 0.123 | -1.52E-01 | 1.01E-01 | 2.63E+00 | 2.71E+00 | 1.33E-01 | no | | intergenic_variant | MODIFIER |
| 3 | 19381616 | AX-170114968 | 0 | C | T | 0.123 | -1.52E-01 | 1.01E-01 | 2.63E+00 | 2.71E+00 | 1.33E-01 | no | | intergenic_variant | MODIFIER |
| 3 | 19382735 | AX-97811337 | 0 | T | C | 0.123 | -1.52E-01 | 1.01E-01 | 2.63E+00 | 2.71E+00 | 1.33E-01 | no | | intergenic_variant | MODIFIER |
| 3 | 19382896 | AX-170106317 | 0 | G | A | 0.123 | -1.52E-01 | 1.01E-01 | 2.63E+00 | 2.71E+00 | 1.33E-01 | no | | intergenic_variant | MODIFIER |
| 3 | 19383193 | AX-170145531 | 0 | A | C | 0.123 | -1.52E-01 | 1.01E-01 | 2.63E+00 | 2.71E+00 | 1.33E-01 | no | | intergenic_variant | MODIFIER |
| 3 | 19390562 | AX-170167275 | 0 | A | G | 0.123 | -1.52E-01 | 1.01E-01 | 2.63E+00 | 2.71E+00 | 1.33E-01 | no | | intergenic_variant | MODIFIER |
| 3 | 19401200 | AX-170506326 | 0 | G | T | 0.123 | -1.52E-01 | 1.01E-01 | 2.63E+00 | 2.71E+00 | 1.33E-01 | no | | intergenic_variant | MODIFIER |
| 3 | 19429795 | AX-97849141 | 0 | T | C | 0.123 | -1.52E-01 | 1.01E-01 | 2.63E+00 | 2.71E+00 | 1.33E-01 | no | | intergenic_variant | MODIFIER |
| 3 | 19454919 | AX-170066023 | 0 | C | T | 0.123 | -1.52E-01 | 1.01E-01 | 2.63E+00 | 2.71E+00 | 1.33E-01 | no | | intergenic_variant | MODIFIER |
| 3 | 19498226 | AX-170245594 | 0 | A | T | 0.123 | -1.52E-01 | 1.01E-01 | 2.63E+00 | 2.71E+00 | 1.33E-01 | no | | intergenic_variant | MODIFIER |
| 3 | 19509322 | AX-170020070 | 0 | A | G | 0.123 | -1.52E-01 | 1.01E-01 | 2.63E+00 | 2.71E+00 | 1.33E-01 | no | Dnajc5b | intron_variant | MODIFIER |
| 3 | 19525662 | AX-97827564 | 0 | T | G | 0.123 | -1.52E-01 | 1.01E-01 | 2.63E+00 | 2.71E+00 | 1.33E-01 | no | Dnajc5b | intron_variant | MODIFIER |
| 3 | 19527971 | AX-170493949 | 0 | G | A | 0.123 | -1.52E-01 | 1.01E-01 | 2.63E+00 | 2.71E+00 | 1.33E-01 | no | Dnajc5b | intron_variant | MODIFIER |
| 3 | 19554651 | AX-170020061 | 0 | C | T | 0.123 | -1.52E-01 | 1.01E-01 | 2.63E+00 | 2.71E+00 | 1.33E-01 | no | Dnajc5b | intron_variant | MODIFIER |
| 3 | 19557000 | AX-170020071 | 0 | G | A | 0.123 | -1.52E-01 | 1.01E-01 | 2.63E+00 | 2.71E+00 | 1.33E-01 | no | Dnajc5b | intron_variant | MODIFIER |
| 3 | 19573725 | AX-97844208 | 0 | C | T | 0.123 | -1.52E-01 | 1.01E-01 | 2.63E+00 | 2.71E+00 | 1.33E-01 | no | Dnajc5b | intron_variant | MODIFIER |
| 3 | 19687833 | AX-170152332 | 0 | G | A | 0.123 | -1.52E-01 | 1.01E-01 | 2.63E+00 | 2.71E+00 | 1.33E-01 | no | Trim55 | intron_variant | MODIFIER |
| 3 | 19688049 | AX-170160622 | 0 | T | C | 0.123 | -1.52E-01 | 1.01E-01 | 2.63E+00 | 2.71E+00 | 1.33E-01 | no | Trim55 | intron_variant | MODIFIER |
| 13 | 9472718 | AX-170323002 | 0 | T | C | 0.193 | -1.62E-01 | 1.08E-01 | 2.55E+00 | 2.62E+00 | 1.33E-01 | no | Dip2c | intron_variant | MODIFIER |
| 1 | 135418071 | AX-170042825 | 1 | A | G | 0.536 | 1.20E-01 | 8.05E-02 | 2.70E+00 | 2.77E+00 | 1.33E-01 | no | Ipo9 | intron_variant | MODIFIER |
| 15 | 4567259 | AX-170295577 | 0 | A | G | 0.237 | -1.56E-01 | 1.04E-01 | 2.67E+00 | 2.72E+00 | 1.33E-01 | no | Plcxd3 | intron_variant | MODIFIER |
| 5 | 64985453 | AX-97804337 | 0 | C | T | 0.474 | 1.45E-01 | 9.65E-02 | 2.52E+00 | 2.60E+00 | 1.33E-01 | Yes | Fam114a1 | intron_variant | MODIFIER |
| 1 | 133929944 | AX-170091699 | 0 | G | A | 0.535 | 1.24E-01 | 8.30E-02 | 2.72E+00 | 2.79E+00 | 1.33E-01 | no | | intergenic_variant | MODIFIER |
| 1 | 133967204 | AX-170460327 | 0 | G | A | 0.535 | 1.24E-01 | 8.30E-02 | 2.72E+00 | 2.79E+00 | 1.33E-01 | no | Gm1627 | intron_variant,non_coding_transcript_variant | MODIFIER |
| 1 | 133968232 | AX-170264814 | 0 | T | G | 0.535 | 1.24E-01 | 8.30E-02 | 2.72E+00 | 2.79E+00 | 1.33E-01 | no | Gm1627 | intron_variant,non_coding_transcript_variant | MODIFIER |
| 1 | 133970511 | AX-170001756 | 0 | C | T | 0.535 | 1.24E-01 | 8.30E-02 | 2.72E+00 | 2.79E+00 | 1.33E-01 | no | Gm1627 | non_coding_transcript_exon_variant | MODIFIER |
| 1 | 133977596 | AX-170368442 | 0 | G | T | 0.535 | 1.24E-01 | 8.30E-02 | 2.72E+00 | 2.79E+00 | 1.33E-01 | no | | regulatory_region_variant | MODIFIER |
| 1 | 134027247 | AX-170347664 | 0 | G | A | 0.535 | 1.24E-01 | 8.30E-02 | 2.72E+00 | 2.79E+00 | 1.33E-01 | no | | intergenic_variant | MODIFIER |
| 1 | 134031098 | AX-170435807 | 0 | C | A | 0.535 | 1.24E-01 | 8.30E-02 | 2.72E+00 | 2.79E+00 | 1.33E-01 | no | Gm18970 | downstream_gene_variant | MODIFIER |
| 1 | 134033083 | AX-170296709 | 0 | T | C | 0.535 | 1.24E-01 | 8.30E-02 | 2.72E+00 | 2.79E+00 | 1.33E-01 | no | Fmod | upstream_gene_variant | MODIFIER |
| 1 | 134035465 | AX-97853644 | 0 | C | T | 0.535 | 1.24E-01 | 8.30E-02 | 2.72E+00 | 2.79E+00 | 1.33E-01 | no | Fmod | upstream_gene_variant | MODIFIER |
| 1 | 134035865 | AX-170327552 | 0 | A | G | 0.535 | 1.24E-01 | 8.30E-02 | 2.72E+00 | 2.79E+00 | 1.33E-01 | no | Fmod | upstream_gene_variant | MODIFIER |
| 1 | 134037261 | AX-170165096 | 0 | C | T | 0.535 | 1.24E-01 | 8.30E-02 | 2.72E+00 | 2.79E+00 | 1.33E-01 | no | Fmod | 5_prime_UTR_variant | MODIFIER |
| 1 | 134038412 | AX-170480321 | 0 | C | T | 0.535 | 1.24E-01 | 8.30E-02 | 2.72E+00 | 2.79E+00 | 1.33E-01 | no | Fmod | intron_variant | MODIFIER |
| 1 | 134079725 | AX-170243843 | 0 | C | G | 0.535 | 1.24E-01 | 8.30E-02 | 2.72E+00 | 2.79E+00 | 1.33E-01 | no | Btg2 | upstream_gene_variant | MODIFIER |
| 1 | 134204381 | AX-170142736 | 0 | A | G | 0.535 | 1.24E-01 | 8.30E-02 | 2.72E+00 | 2.79E+00 | 1.33E-01 | no | Adora1 | intron_variant | MODIFIER |
| 1 | 134214371 | AX-170186010 | 0 | C | G | 0.535 | 1.24E-01 | 8.30E-02 | 2.72E+00 | 2.79E+00 | 1.33E-01 | no | Adora1 | intron_variant | MODIFIER |
| 1 | 134215971 | AX-170513108 | 0 | T | C | 0.535 | 1.24E-01 | 8.30E-02 | 2.72E+00 | 2.79E+00 | 1.33E-01 | no | Adora1 | intron_variant | MODIFIER |
| 1 | 134379245 | AX-97781701 | 0 | T | G | 0.535 | 1.24E-01 | 8.30E-02 | 2.72E+00 | 2.79E+00 | 1.33E-01 | no | | intergenic_variant | MODIFIER |
| 1 | 134515359 | AX-170349693 | 0 | T | C | 0.535 | 1.24E-01 | 8.30E-02 | 2.72E+00 | 2.79E+00 | 1.33E-01 | no | Gm15454 | upstream_gene_variant | MODIFIER |
| 1 | 134520934 | AX-97792746 | 0 | C | T | 0.535 | 1.24E-01 | 8.30E-02 | 2.72E+00 | 2.79E+00 | 1.33E-01 | no | Gm15454 | downstream_gene_variant | MODIFIER |

| | | | | | | | | | | | | | | | |
|---|---|---|---|---|---|---|---|---|---|---|---|---|---|---|---|
| 1 | 134523821 | AX-97856248 | 0 | T | C | 0.535 | 1.24E-01 | 8.30E-02 | 2.72E+00 | 2.79E+00 | 1.33E-01 | no | Gm29376 | intron_variant,non_coding_transcript_variant | MODIFIER |
| 1 | 134530222 | AX-170173644 | 0 | G | A | 0.535 | 1.24E-01 | 8.30E-02 | 2.72E+00 | 2.79E+00 | 1.33E-01 | no | | regulatory_region_variant | MODIFIER |
| 1 | 135170649 | AX-97811594 | 0 | T | C | 0.535 | 1.24E-01 | 8.30E-02 | 2.72E+00 | 2.79E+00 | 1.33E-01 | no | Gpr37l1 | upstream_gene_variant | MODIFIER |
| 1 | 135525096 | AX-170001772 | 0 | T | C | 0.535 | 1.24E-01 | 8.30E-02 | 2.72E+00 | 2.79E+00 | 1.33E-01 | no | Nav1 | intron_variant | MODIFIER |
| 1 | 135533281 | AX-97813039 | 0 | A | G | 0.535 | 1.24E-01 | 8.30E-02 | 2.72E+00 | 2.79E+00 | 1.33E-01 | no | Nav1 | intron_variant | MODIFIER |
| 1 | 135594126 | AX-170001767 | 0 | T | C | 0.535 | 1.24E-01 | 8.30E-02 | 2.72E+00 | 2.79E+00 | 1.33E-01 | no | Gm4793 | intron_variant,non_coding_transcript_variant | MODIFIER |
| 1 | 135594178 | AX-170001770 | 0 | T | G | 0.535 | 1.24E-01 | 8.30E-02 | 2.72E+00 | 2.79E+00 | 1.33E-01 | no | Gm4793 | intron_variant,non_coding_transcript_variant | MODIFIER |
| 1 | 135598344 | AX-170428238 | 0 | G | A | 0.535 | 1.24E-01 | 8.30E-02 | 2.72E+00 | 2.79E+00 | 1.33E-01 | no | Gm4793 | intron_variant,non_coding_transcript_variant | MODIFIER |
| 1 | 135607473 | AX-97814378 | 0 | T | C | 0.535 | 1.24E-01 | 8.30E-02 | 2.72E+00 | 2.79E+00 | 1.33E-01 | no | Gm38399 | non_coding_transcript_exon_variant | MODIFIER |
| 1 | 135626669 | AX-170122112 | 0 | A | G | 0.535 | 1.24E-01 | 8.30E-02 | 2.72E+00 | 2.79E+00 | 1.33E-01 | no | Gm38399 | intron_variant,non_coding_transcript_variant | MODIFIER |
| 1 | 135631338 | AX-170456968 | 0 | T | C | 0.535 | 1.24E-01 | 8.30E-02 | 2.72E+00 | 2.79E+00 | 1.33E-01 | no | Gm38399 | intron_variant,non_coding_transcript_variant | MODIFIER |
| 1 | 135645057 | AX-170298123 | 0 | T | C | 0.535 | 1.24E-01 | 8.30E-02 | 2.72E+00 | 2.79E+00 | 1.33E-01 | no | Gm38399 | intron_variant,non_coding_transcript_variant | MODIFIER |
| 1 | 135658579 | AX-170237988 | 0 | A | G | 0.535 | 1.24E-01 | 8.30E-02 | 2.72E+00 | 2.79E+00 | 1.33E-01 | no | Gm38399 | intron_variant,non_coding_transcript_variant | MODIFIER |
| 1 | 135744820 | AX-170311508 | 0 | T | C | 0.535 | 1.24E-01 | 8.30E-02 | 2.72E+00 | 2.79E+00 | 1.33E-01 | no | Csrp1 | intron_variant | MODIFIER |
| 1 | 135765556 | AX-170256197 | 0 | A | T | 0.535 | 1.24E-01 | 8.30E-02 | 2.72E+00 | 2.79E+00 | 1.33E-01 | no | Phlda3 | upstream_gene_variant | MODIFIER |
| 1 | 135776509 | AX-170397287 | 0 | A | G | 0.535 | 1.24E-01 | 8.30E-02 | 2.72E+00 | 2.79E+00 | 1.33E-01 | no | Tnni1 | upstream_gene_variant | MODIFIER |
| 1 | 135780116 | AX-170220991 | 0 | T | C | 0.535 | 1.24E-01 | 8.30E-02 | 2.72E+00 | 2.79E+00 | 1.33E-01 | no | Tnni1 | upstream_gene_variant | MODIFIER |
| 1 | 135781752 | AX-97810550 | 0 | T | C | 0.535 | 1.24E-01 | 8.30E-02 | 2.72E+00 | 2.79E+00 | 1.33E-01 | no | Tnni1 | upstream_gene_variant | MODIFIER |
| 1 | 135789137 | AX-170284871 | 0 | A | G | 0.535 | 1.24E-01 | 8.30E-02 | 2.72E+00 | 2.79E+00 | 1.33E-01 | no | Tnni1 | intron_variant | MODIFIER |
| 1 | 135789165 | AX-170171100 | 0 | T | G | 0.535 | 1.24E-01 | 8.30E-02 | 2.72E+00 | 2.79E+00 | 1.33E-01 | no | Tnni1 | intron_variant | MODIFIER |
| 1 | 135796127 | AX-170423943 | 0 | A | G | 0.535 | 1.24E-01 | 8.30E-02 | 2.72E+00 | 2.79E+00 | 1.33E-01 | no | Tnni1 | intron_variant | MODIFIER |
| 1 | 135799623 | AX-170309721 | 0 | A | G | 0.535 | 1.24E-01 | 8.30E-02 | 2.72E+00 | 2.79E+00 | 1.33E-01 | no | Tnni1 | intron_variant | MODIFIER |
| 1 | 135809273 | AX-170372556 | 0 | A | T | 0.535 | 1.24E-01 | 8.30E-02 | 2.72E+00 | 2.79E+00 | 1.33E-01 | no | Tnni1 | intron_variant | MODIFIER |
| 4 | 84271717 | AX-97836449 | 1 | G | A | 0.339 | 1.76E-01 | 1.17E-01 | 2.87E+00 | 2.94E+00 | 1.33E-01 | Yes | Bnc2 | downstream_gene_variant | MODIFIER |
| 5 | 134295595 | AX-170118920 | 0 | A | G | 0.149 | -1.33E-01 | 8.58E-02 | 2.26E+00 | 2.34E+00 | 1.33E-01 | no | Gtf2i | missense_variant | MODERATE |
| 5 | 135113372 | AX-97831813 | 0 | G | A | 0.149 | -1.33E-01 | 8.58E-02 | 2.26E+00 | 2.34E+00 | 1.33E-01 | no | Mlxipl | intron_variant | MODIFIER |
| 5 | 135203924 | AX-170216107 | 0 | A | G | 0.149 | -1.33E-01 | 8.58E-02 | 2.26E+00 | 2.34E+00 | 1.33E-01 | no | Baz1b | intron_variant | MODIFIER |
| 5 | 135435102 | AX-97806483 | 0 | A | G | 0.149 | -1.33E-01 | 8.58E-02 | 2.26E+00 | 2.34E+00 | 1.33E-01 | no | Hip1 | missense_variant | MODERATE |
| 5 | 135567899 | AX-97814689 | 0 | T | C | 0.149 | -1.33E-01 | 8.58E-02 | 2.26E+00 | 2.34E+00 | 1.33E-01 | no | Ccl24 | downstream_gene_variant | MODIFIER |
| 2 | 123729809 | AX-170018905 | 0 | G | A | 0.184 | 1.85E-01 | 1.24E-01 | 2.62E+00 | 2.75E+00 | 1.33E-01 | Yes | | intergenic_variant | MODIFIER |
| 13 | 8280296 | AX-170112233 | 0 | G | A | 0.193 | -1.80E-01 | 1.20E-01 | 2.52E+00 | 2.58E+00 | 1.33E-01 | Yes | Adarb2 | intron_variant | MODIFIER |
| 1 | 133816455 | AX-170345494 | 0 | A | G | 0.544 | 1.24E-01 | 8.33E-02 | 2.73E+00 | 2.80E+00 | 1.33E-01 | no | | regulatory_region_variant | MODIFIER |
| 1 | 133816519 | AX-170466191 | 0 | A | G | 0.544 | 1.24E-01 | 8.33E-02 | 2.73E+00 | 2.80E+00 | 1.33E-01 | no | | regulatory_region_variant | MODIFIER |
| 1 | 133955409 | AX-97808383 | 0 | G | A | 0.544 | 1.24E-01 | 8.33E-02 | 2.73E+00 | 2.80E+00 | 1.33E-01 | no | | regulatory_region_variant | MODIFIER |
| 1 | 133972959 | AX-170001754 | 0 | A | G | 0.544 | 1.24E-01 | 8.33E-02 | 2.73E+00 | 2.80E+00 | 1.33E-01 | no | Gm1627 | downstream_gene_variant | MODIFIER |
| 1 | 133975569 | AX-97843284 | 0 | A | G | 0.544 | 1.24E-01 | 8.33E-02 | 2.73E+00 | 2.80E+00 | 1.33E-01 | no | Gm1627 | downstream_gene_variant | MODIFIER |
| 1 | 134188907 | AX-170309205 | 0 | A | G | 0.544 | 1.24E-01 | 8.33E-02 | 2.73E+00 | 2.80E+00 | 1.33E-01 | no | Mybph | upstream_gene_variant | MODIFIER |
| 1 | 134190049 | AX-170412119 | 0 | A | G | 0.544 | 1.24E-01 | 8.33E-02 | 2.73E+00 | 2.80E+00 | 1.33E-01 | no | Mybph | upstream_gene_variant | MODIFIER |
| 1 | 134207811 | AX-170306597 | 0 | G | A | 0.544 | 1.24E-01 | 8.33E-02 | 2.73E+00 | 2.80E+00 | 1.33E-01 | no | Adora1 | intron_variant | MODIFIER |
| 1 | 134214658 | AX-170279926 | 0 | C | T | 0.544 | 1.24E-01 | 8.33E-02 | 2.73E+00 | 2.80E+00 | 1.33E-01 | no | Adora1 | intron_variant | MODIFIER |
| 1 | 134451215 | AX-170372555 | 0 | C | T | 0.544 | 1.24E-01 | 8.33E-02 | 2.73E+00 | 2.80E+00 | 1.33E-01 | no | Klhl12 | upstream_gene_variant | MODIFIER |
| 1 | 134458816 | AX-170219864 | 0 | T | C | 0.544 | 1.24E-01 | 8.33E-02 | 2.73E+00 | 2.80E+00 | 1.33E-01 | no | Klhl12 | intron_variant | MODIFIER |
| 1 | 134464139 | AX-170210458 | 0 | A | G | 0.544 | 1.24E-01 | 8.33E-02 | 2.73E+00 | 2.80E+00 | 1.33E-01 | no | Klhl12 | intron_variant | MODIFIER |
| 1 | 134472588 | AX-170194276 | 0 | C | T | 0.544 | 1.24E-01 | 8.33E-02 | 2.73E+00 | 2.80E+00 | 1.33E-01 | no | Klhl12 | intron_variant | MODIFIER |
| 1 | 134474898 | AX-170160823 | 0 | C | T | 0.544 | 1.24E-01 | 8.33E-02 | 2.73E+00 | 2.80E+00 | 1.33E-01 | no | Klhl12 | intron_variant | MODIFIER |
| 1 | 134484611 | AX-170463093 | 0 | T | A | 0.544 | 1.24E-01 | 8.33E-02 | 2.73E+00 | 2.80E+00 | 1.33E-01 | no | Klhl12 | intron_variant | MODIFIER |
| 1 | 134495313 | AX-170335632 | 0 | T | C | 0.544 | 1.24E-01 | 8.33E-02 | 2.73E+00 | 2.80E+00 | 1.33E-01 | no | Klhl12 | downstream_gene_variant | MODIFIER |
| 1 | 134507315 | AX-170343812 | 0 | A | G | 0.544 | 1.24E-01 | 8.33E-02 | 2.73E+00 | 2.80E+00 | 1.33E-01 | no | Rabif | 3_prime_UTR_variant | MODIFIER |
| 1 | 134517457 | AX-170288204 | 0 | C | T | 0.544 | 1.24E-01 | 8.33E-02 | 2.73E+00 | 2.80E+00 | 1.33E-01 | no | Gm15454 | upstream_gene_variant | MODIFIER |
| 1 | 134552878 | AX-97823581 | 0 | G | A | 0.544 | 1.24E-01 | 8.33E-02 | 2.73E+00 | 2.80E+00 | 1.33E-01 | no | Mgat4e | upstream_gene_variant | MODIFIER |
| 1 | 134554142 | AX-170122631 | 0 | A | T | 0.544 | 1.24E-01 | 8.33E-02 | 2.73E+00 | 2.80E+00 | 1.33E-01 | no | Mgat4e | upstream_gene_variant | MODIFIER |
| 1 | 134570662 | AX-170293634 | 0 | A | T | 0.544 | 1.24E-01 | 8.33E-02 | 2.73E+00 | 2.80E+00 | 1.33E-01 | no | Kdm5b | intron_variant | MODIFIER |
| 1 | 134590127 | AX-170143237 | 0 | C | A | 0.544 | 1.24E-01 | 8.33E-02 | 2.73E+00 | 2.80E+00 | 1.33E-01 | no | Kdm5b | intron_variant | MODIFIER |
| 1 | 134599089 | AX-170305607 | 0 | C | A | 0.544 | 1.24E-01 | 8.33E-02 | 2.73E+00 | 2.80E+00 | 1.33E-01 | no | Kdm5b | synonymous_variant | LOW |

| | | | | | | | | | | | | | | | |
|---|---|---|---|---|---|---|---|---|---|---|---|---|---|---|---|
| 1 | 134604909 | AX-170138655 | 0 | C | T | 0.544 | 1.24E-01 | 8.33E-02 | 2.73E+00 | 2.80E+00 | 1.33E-01 | no | Kdm5b | intron_variant | MODIFIER |
| 1 | 134613797 | AX-170146485 | 0 | C | T | 0.544 | 1.24E-01 | 8.33E-02 | 2.73E+00 | 2.80E+00 | 1.33E-01 | no | Kdm5b | intron_variant | MODIFIER |
| 1 | 134641914 | AX-170177030 | 0 | T | C | 0.544 | 1.24E-01 | 8.33E-02 | 2.73E+00 | 2.80E+00 | 1.33E-01 | no | Gm26783 | downstream_gene_variant | MODIFIER |
| 1 | 134653305 | AX-170500505 | 0 | C | A | 0.544 | 1.24E-01 | 8.33E-02 | 2.73E+00 | 2.80E+00 | 1.33E-01 | no | Syt2 | intron_variant | MODIFIER |
| 1 | 134653566 | AX-170086172 | 0 | T | G | 0.544 | 1.24E-01 | 8.33E-02 | 2.73E+00 | 2.80E+00 | 1.33E-01 | no | Syt2 | intron_variant | MODIFIER |
| 1 | 134661058 | AX-170406878 | 0 | G | A | 0.544 | 1.24E-01 | 8.33E-02 | 2.73E+00 | 2.80E+00 | 1.33E-01 | no | Syt2 | intron_variant | MODIFIER |
| 1 | 134661345 | AX-170131451 | 0 | A | G | 0.544 | 1.24E-01 | 8.33E-02 | 2.73E+00 | 2.80E+00 | 1.33E-01 | no | Syt2 | intron_variant | MODIFIER |
| 1 | 134672611 | AX-97815988 | 0 | G | A | 0.544 | 1.24E-01 | 8.33E-02 | 2.73E+00 | 2.80E+00 | 1.33E-01 | no | Syt2 | intron_variant | MODIFIER |
| 1 | 134673045 | AX-170042805 | 0 | G | C | 0.544 | 1.24E-01 | 8.33E-02 | 2.73E+00 | 2.80E+00 | 1.33E-01 | no | Syt2 | intron_variant | MODIFIER |
| 1 | 134677554 | AX-170042802 | 0 | A | G | 0.544 | 1.24E-01 | 8.33E-02 | 2.73E+00 | 2.80E+00 | 1.33E-01 | no | Syt2 | intron_variant | MODIFIER |
| 1 | 134697953 | AX-170348450 | 0 | C | T | 0.544 | 1.24E-01 | 8.33E-02 | 2.73E+00 | 2.80E+00 | 1.33E-01 | no | Syt2 | intron_variant | MODIFIER |
| 1 | 134698553 | AX-170491458 | 0 | T | C | 0.544 | 1.24E-01 | 8.33E-02 | 2.73E+00 | 2.80E+00 | 1.33E-01 | no | Syt2 | intron_variant | MODIFIER |
| 1 | 134698913 | AX-170330220 | 0 | C | A | 0.544 | 1.24E-01 | 8.33E-02 | 2.73E+00 | 2.80E+00 | 1.33E-01 | no | Syt2 | intron_variant | MODIFIER |
| 1 | 134704323 | AX-170260183 | 0 | T | C | 0.544 | 1.24E-01 | 8.33E-02 | 2.73E+00 | 2.80E+00 | 1.33E-01 | no | Syt2 | upstream_gene_variant | MODIFIER |
| 1 | 134704725 | AX-170258871 | 0 | C | T | 0.544 | 1.24E-01 | 8.33E-02 | 2.73E+00 | 2.80E+00 | 1.33E-01 | no | Syt2 | upstream_gene_variant | MODIFIER |
| 1 | 134704890 | AX-170322588 | 0 | C | G | 0.544 | 1.24E-01 | 8.33E-02 | 2.73E+00 | 2.80E+00 | 1.33E-01 | no | Syt2 | upstream_gene_variant | MODIFIER |
| 1 | 134714303 | AX-97855511 | 0 | C | T | 0.544 | 1.24E-01 | 8.33E-02 | 2.73E+00 | 2.80E+00 | 1.33E-01 | no | Syt2 | intron_variant | MODIFIER |
| 1 | 134743154 | AX-170452087 | 0 | G | T | 0.544 | 1.24E-01 | 8.33E-02 | 2.73E+00 | 2.80E+00 | 1.33E-01 | no | Syt2 | intron_variant | MODIFIER |
| 1 | 134747924 | AX-170285568 | 0 | T | A | 0.544 | 1.24E-01 | 8.33E-02 | 2.73E+00 | 2.80E+00 | 1.33E-01 | no | Syt2 | 3_prime_UTR_variant | MODIFIER |
| 1 | 134753304 | AX-170483105 | 0 | C | T | 0.544 | 1.24E-01 | 8.33E-02 | 2.73E+00 | 2.80E+00 | 1.33E-01 | no | Ppp1r12b | downstream_gene_variant | MODIFIER |
| 1 | 134767929 | AX-170236347 | 0 | C | T | 0.544 | 1.24E-01 | 8.33E-02 | 2.73E+00 | 2.80E+00 | 1.33E-01 | no | Ppp1r12b | intron_variant | MODIFIER |
| 1 | 134779627 | AX-170322435 | 0 | C | G | 0.544 | 1.24E-01 | 8.33E-02 | 2.73E+00 | 2.80E+00 | 1.33E-01 | no | Ppp1r12b | intron_variant | MODIFIER |
| 1 | 134796864 | AX-170474162 | 0 | C | T | 0.544 | 1.24E-01 | 8.33E-02 | 2.73E+00 | 2.80E+00 | 1.33E-01 | no | Ppp1r12b | intron_variant | MODIFIER |
| 1 | 134810521 | AX-170401563 | 0 | G | A | 0.544 | 1.24E-01 | 8.33E-02 | 2.73E+00 | 2.80E+00 | 1.33E-01 | no | Ppp1r12b | intron_variant | MODIFIER |
| 1 | 134813566 | AX-170097418 | 0 | A | G | 0.544 | 1.24E-01 | 8.33E-02 | 2.73E+00 | 2.80E+00 | 1.33E-01 | no | Ppp1r12b | intron_variant | MODIFIER |
| 1 | 134816466 | AX-170524648 | 0 | T | C | 0.544 | 1.24E-01 | 8.33E-02 | 2.73E+00 | 2.80E+00 | 1.33E-01 | no | Ppp1r12b | intron_variant | MODIFIER |
| 1 | 134825367 | AX-170172662 | 0 | G | A | 0.544 | 1.24E-01 | 8.33E-02 | 2.73E+00 | 2.80E+00 | 1.33E-01 | no | Ppp1r12b | intron_variant | MODIFIER |
| 1 | 134837806 | AX-170339527 | 0 | C | T | 0.544 | 1.24E-01 | 8.33E-02 | 2.73E+00 | 2.80E+00 | 1.33E-01 | no | Ppp1r12b | intron_variant | MODIFIER |
| 1 | 134846443 | AX-170161365 | 0 | G | A | 0.544 | 1.24E-01 | 8.33E-02 | 2.73E+00 | 2.80E+00 | 1.33E-01 | no | Ppp1r12b | intron_variant | MODIFIER |
| 1 | 134847799 | AX-170516833 | 0 | G | A | 0.544 | 1.24E-01 | 8.33E-02 | 2.73E+00 | 2.80E+00 | 1.33E-01 | no | Ppp1r12b | intron_variant | MODIFIER |
| 1 | 134853630 | AX-170042813 | 0 | T | G | 0.544 | 1.24E-01 | 8.33E-02 | 2.73E+00 | 2.80E+00 | 1.33E-01 | no | Ppp1r12b | intron_variant | MODIFIER |
| 1 | 134873145 | AX-170042808 | 0 | C | T | 0.544 | 1.24E-01 | 8.33E-02 | 2.73E+00 | 2.80E+00 | 1.33E-01 | no | Ppp1r12b | intron_variant | MODIFIER |
| 1 | 134896129 | AX-170292067 | 0 | C | G | 0.544 | 1.24E-01 | 8.33E-02 | 2.73E+00 | 2.80E+00 | 1.33E-01 | no | Ppp1r12b | intron_variant | MODIFIER |
| 1 | 134900322 | AX-97815827 | 0 | A | G | 0.544 | 1.24E-01 | 8.33E-02 | 2.73E+00 | 2.80E+00 | 1.33E-01 | no | Ppp1r12b | intron_variant | MODIFIER |
| 1 | 134929949 | AX-170297546 | 0 | C | T | 0.544 | 1.24E-01 | 8.33E-02 | 2.73E+00 | 2.80E+00 | 1.33E-01 | no | Ppp1r12b | intron_variant | MODIFIER |
| 1 | 134933543 | AX-170001761 | 0 | C | T | 0.544 | 1.24E-01 | 8.33E-02 | 2.73E+00 | 2.80E+00 | 1.33E-01 | no | Ppp1r12b | intron_variant | MODIFIER |
| 1 | 134935493 | AX-170374581 | 0 | G | A | 0.544 | 1.24E-01 | 8.33E-02 | 2.73E+00 | 2.80E+00 | 1.33E-01 | no | Ppp1r12b | intron_variant | MODIFIER |
| 1 | 134973611 | AX-170001760 | 0 | A | G | 0.544 | 1.24E-01 | 8.33E-02 | 2.73E+00 | 2.80E+00 | 1.33E-01 | no | Ube2t | intron_variant | MODIFIER |
| 1 | 134973762 | AX-170001764 | 0 | T | C | 0.544 | 1.24E-01 | 8.33E-02 | 2.73E+00 | 2.80E+00 | 1.33E-01 | no | Ube2t | intron_variant | MODIFIER |
| 1 | 134978049 | AX-97824834 | 0 | T | G | 0.544 | 1.24E-01 | 8.33E-02 | 2.73E+00 | 2.80E+00 | 1.33E-01 | no | Ube2t | downstream_gene_variant | MODIFIER |
| 1 | 135002606 | AX-170001765 | 0 | G | A | 0.544 | 1.24E-01 | 8.33E-02 | 2.73E+00 | 2.80E+00 | 1.33E-01 | no | Lgr6 | intron_variant | MODIFIER |
| 1 | 135002945 | AX-170385074 | 0 | G | C | 0.544 | 1.24E-01 | 8.33E-02 | 2.73E+00 | 2.80E+00 | 1.33E-01 | no | Lgr6 | intron_variant | MODIFIER |
| 1 | 135014783 | AX-97787512 | 0 | C | T | 0.544 | 1.24E-01 | 8.33E-02 | 2.73E+00 | 2.80E+00 | 1.33E-01 | no | Lgr6 | intron_variant | MODIFIER |
| 1 | 135040343 | AX-170183350 | 0 | G | C | 0.544 | 1.24E-01 | 8.33E-02 | 2.73E+00 | 2.80E+00 | 1.33E-01 | no | Lgr6 | intron_variant | MODIFIER |
| 1 | 135041701 | AX-170448659 | 0 | A | T | 0.544 | 1.24E-01 | 8.33E-02 | 2.73E+00 | 2.80E+00 | 1.33E-01 | no | Lgr6 | intron_variant | MODIFIER |
| 1 | 135047993 | AX-170489731 | 0 | C | T | 0.544 | 1.24E-01 | 8.33E-02 | 2.73E+00 | 2.80E+00 | 1.33E-01 | no | Lgr6 | intron_variant | MODIFIER |
| 1 | 135059438 | AX-170425196 | 0 | C | T | 0.544 | 1.24E-01 | 8.33E-02 | 2.73E+00 | 2.80E+00 | 1.33E-01 | no | Lgr6 | intron_variant | MODIFIER |
| 1 | 135110118 | AX-170142476 | 0 | A | T | 0.544 | 1.24E-01 | 8.33E-02 | 2.73E+00 | 2.80E+00 | 1.33E-01 | no | Lgr6 | upstream_gene_variant | MODIFIER |
| 1 | 135124768 | AX-170200843 | 0 | T | C | 0.544 | 1.24E-01 | 8.33E-02 | 2.73E+00 | 2.80E+00 | 1.33E-01 | no | Ptprv | downstream_gene_variant | MODIFIER |
| 1 | 135126298 | AX-170335082 | 0 | T | C | 0.544 | 1.24E-01 | 8.33E-02 | 2.73E+00 | 2.80E+00 | 1.33E-01 | no | Ptprv | downstream_gene_variant | MODIFIER |
| 1 | 135157252 | AX-170344078 | 0 | G | A | 0.544 | 1.24E-01 | 8.33E-02 | 2.73E+00 | 2.80E+00 | 1.33E-01 | no | Gpr37l1 | downstream_gene_variant | MODIFIER |
| 1 | 135157652 | AX-170297230 | 0 | T | C | 0.544 | 1.24E-01 | 8.33E-02 | 2.73E+00 | 2.80E+00 | 1.33E-01 | no | Gpr37l1 | downstream_gene_variant | MODIFIER |
| 1 | 135159274 | AX-170042816 | 0 | A | G | 0.544 | 1.24E-01 | 8.33E-02 | 2.73E+00 | 2.80E+00 | 1.33E-01 | no | Gpr37l1 | downstream_gene_variant | MODIFIER |
| 1 | 135160325 | AX-170042818 | 0 | T | G | 0.544 | 1.24E-01 | 8.33E-02 | 2.73E+00 | 2.80E+00 | 1.33E-01 | no | Gpr37l1 | 3_prime_UTR_variant | MODIFIER |

| | | | | | | | | | | | | | | | |
|---|---|---|---|---|---|---|---|---|---|---|---|---|---|---|---|
| 1 | 135164694 | AX-170164679 | 0 | G | A | 0.544 | 1.24E-01 | 8.33E-02 | 2.73E+00 | 2.80E+00 | 1.33E-01 | no | Gpr37l1 | intron_variant | MODIFIER |
| 1 | 135166606 | AX-170533553 | 0 | G | A | 0.544 | 1.24E-01 | 8.33E-02 | 2.73E+00 | 2.80E+00 | 1.33E-01 | no | Gpr37l1 | intron_variant | MODIFIER |
| 1 | 135170264 | AX-170531234 | 0 | A | G | 0.544 | 1.24E-01 | 8.33E-02 | 2.73E+00 | 2.80E+00 | 1.33E-01 | no | Gpr37l1 | upstream_gene_variant | MODIFIER |
| 1 | 135173222 | AX-170345231 | 0 | G | A | 0.544 | 1.24E-01 | 8.33E-02 | 2.73E+00 | 2.80E+00 | 1.33E-01 | no | | intergenic_variant | MODIFIER |
| 1 | 135174658 | AX-170451974 | 0 | A | C | 0.544 | 1.24E-01 | 8.33E-02 | 2.73E+00 | 2.80E+00 | 1.33E-01 | no | | intergenic_variant | MODIFIER |
| 1 | 135193805 | AX-170534264 | 0 | A | C | 0.544 | 1.24E-01 | 8.33E-02 | 2.73E+00 | 2.80E+00 | 1.33E-01 | no | | intergenic_variant | MODIFIER |
| 1 | 135211177 | AX-170271826 | 0 | C | T | 0.544 | 1.24E-01 | 8.33E-02 | 2.73E+00 | 2.80E+00 | 1.33E-01 | no | | intergenic_variant | MODIFIER |
| 1 | 135239907 | AX-170091702 | 0 | A | C | 0.544 | 1.24E-01 | 8.33E-02 | 2.73E+00 | 2.80E+00 | 1.33E-01 | no | | intergenic_variant | MODIFIER |
| 1 | 135241151 | AX-170312383 | 0 | G | A | 0.544 | 1.24E-01 | 8.33E-02 | 2.73E+00 | 2.80E+00 | 1.33E-01 | no | | intergenic_variant | MODIFIER |
| 1 | 135246458 | AX-170287976 | 0 | G | A | 0.544 | 1.24E-01 | 8.33E-02 | 2.73E+00 | 2.80E+00 | 1.33E-01 | no | Gm26642 | upstream_gene_variant | MODIFIER |
| 1 | 135247511 | AX-170163746 | 0 | C | A | 0.544 | 1.24E-01 | 8.33E-02 | 2.73E+00 | 2.80E+00 | 1.33E-01 | no | Gm26642 | upstream_gene_variant | MODIFIER |
| 1 | 135248473 | AX-170306291 | 0 | C | T | 0.544 | 1.24E-01 | 8.33E-02 | 2.73E+00 | 2.80E+00 | 1.33E-01 | no | Gm26642 | upstream_gene_variant | MODIFIER |
| 1 | 135251294 | AX-97824836 | 0 | T | C | 0.544 | 1.24E-01 | 8.33E-02 | 2.73E+00 | 2.80E+00 | 1.33E-01 | no | Elf3 | downstream_gene_variant | MODIFIER |
| 1 | 135268315 | AX-170470358 | 0 | A | G | 0.544 | 1.24E-01 | 8.33E-02 | 2.73E+00 | 2.80E+00 | 1.33E-01 | no | Rnpep | intron_variant | MODIFIER |
| 1 | 135268417 | AX-170122724 | 0 | A | G | 0.544 | 1.24E-01 | 8.33E-02 | 2.73E+00 | 2.80E+00 | 1.33E-01 | no | Rnpep | intron_variant | MODIFIER |
| 1 | 135271653 | AX-170509858 | 0 | G | A | 0.544 | 1.24E-01 | 8.33E-02 | 2.73E+00 | 2.80E+00 | 1.33E-01 | no | Rnpep | synonymous_variant | LOW |
| 1 | 135283198 | AX-170163265 | 0 | T | A | 0.544 | 1.24E-01 | 8.33E-02 | 2.73E+00 | 2.80E+00 | 1.33E-01 | no | Rnpep | intron_variant | MODIFIER |
| 1 | 135312467 | AX-170042821 | 0 | C | T | 0.544 | 1.24E-01 | 8.33E-02 | 2.73E+00 | 2.80E+00 | 1.33E-01 | no | Timm17a | intron_variant | MODIFIER |
| 1 | 135328566 | AX-170423014 | 0 | A | C | 0.544 | 1.24E-01 | 8.33E-02 | 2.73E+00 | 2.80E+00 | 1.33E-01 | no | Lmod1 | intron_variant | MODIFIER |
| 1 | 135328669 | AX-170488363 | 0 | C | T | 0.544 | 1.24E-01 | 8.33E-02 | 2.73E+00 | 2.80E+00 | 1.33E-01 | no | Lmod1 | intron_variant | MODIFIER |
| 1 | 135331215 | AX-170161653 | 0 | C | T | 0.544 | 1.24E-01 | 8.33E-02 | 2.73E+00 | 2.80E+00 | 1.33E-01 | no | Lmod1 | intron_variant | MODIFIER |
| 1 | 135331263 | AX-170352248 | 0 | C | G | 0.544 | 1.24E-01 | 8.33E-02 | 2.73E+00 | 2.80E+00 | 1.33E-01 | no | Lmod1 | intron_variant | MODIFIER |
| 1 | 135331547 | AX-170485939 | 0 | T | C | 0.544 | 1.24E-01 | 8.33E-02 | 2.73E+00 | 2.80E+00 | 1.33E-01 | no | Lmod1 | intron_variant | MODIFIER |
| 1 | 135331902 | AX-170139531 | 0 | T | G | 0.544 | 1.24E-01 | 8.33E-02 | 2.73E+00 | 2.80E+00 | 1.33E-01 | no | Lmod1 | intron_variant | MODIFIER |
| 1 | 135334159 | AX-170267447 | 0 | G | A | 0.544 | 1.24E-01 | 8.33E-02 | 2.73E+00 | 2.80E+00 | 1.33E-01 | no | Lmod1 | intron_variant | MODIFIER |
| 1 | 135348333 | AX-170154260 | 0 | T | C | 0.544 | 1.24E-01 | 8.33E-02 | 2.73E+00 | 2.80E+00 | 1.33E-01 | no | Lmod1 | intron_variant | MODIFIER |
| 1 | 135355655 | AX-170246216 | 0 | T | C | 0.544 | 1.24E-01 | 8.33E-02 | 2.73E+00 | 2.80E+00 | 1.33E-01 | no | Lmod1 | intron_variant | MODIFIER |
| 1 | 135355810 | AX-170484098 | 0 | C | T | 0.544 | 1.24E-01 | 8.33E-02 | 2.73E+00 | 2.80E+00 | 1.33E-01 | no | Lmod1 | intron_variant | MODIFIER |
| 1 | 135359908 | AX-170159596 | 0 | A | G | 0.544 | 1.24E-01 | 8.33E-02 | 2.73E+00 | 2.80E+00 | 1.33E-01 | no | Lmod1 | intron_variant | MODIFIER |
| 1 | 135363879 | AX-170383907 | 0 | C | T | 0.544 | 1.24E-01 | 8.33E-02 | 2.73E+00 | 2.80E+00 | 1.33E-01 | no | Lmod1 | synonymous_variant | LOW |
| 1 | 135366343 | AX-170223914 | 0 | A | G | 0.544 | 1.24E-01 | 8.33E-02 | 2.73E+00 | 2.80E+00 | 1.33E-01 | no | Shisa4 | downstream_gene_variant | MODIFIER |
| 1 | 135366664 | AX-170476850 | 0 | A | G | 0.544 | 1.24E-01 | 8.33E-02 | 2.73E+00 | 2.80E+00 | 1.33E-01 | no | Shisa4 | downstream_gene_variant | MODIFIER |
| 1 | 135366924 | AX-170510242 | 0 | C | G | 0.544 | 1.24E-01 | 8.33E-02 | 2.73E+00 | 2.80E+00 | 1.33E-01 | no | Shisa4 | downstream_gene_variant | MODIFIER |
| 1 | 135389717 | AX-97830461 | 0 | G | A | 0.544 | 1.24E-01 | 8.33E-02 | 2.73E+00 | 2.80E+00 | 1.33E-01 | no | Ipo9 | intron_variant | MODIFIER |
| 1 | 135396319 | AX-97782302 | 0 | T | C | 0.544 | 1.24E-01 | 8.33E-02 | 2.73E+00 | 2.80E+00 | 1.33E-01 | no | Ipo9 | intron_variant | MODIFIER |
| 1 | 135403866 | AX-170324248 | 0 | C | T | 0.544 | 1.24E-01 | 8.33E-02 | 2.73E+00 | 2.80E+00 | 1.33E-01 | no | Ipo9 | intron_variant | MODIFIER |
| 1 | 135404021 | AX-170525018 | 0 | C | T | 0.544 | 1.24E-01 | 8.33E-02 | 2.73E+00 | 2.80E+00 | 1.33E-01 | no | Ipo9 | intron_variant | MODIFIER |
| 1 | 135406094 | AX-170215502 | 0 | C | T | 0.544 | 1.24E-01 | 8.33E-02 | 2.73E+00 | 2.80E+00 | 1.33E-01 | no | Ipo9 | intron_variant | MODIFIER |
| 1 | 135409276 | AX-170151963 | 0 | G | A | 0.544 | 1.24E-01 | 8.33E-02 | 2.73E+00 | 2.80E+00 | 1.33E-01 | no | Ipo9 | intron_variant | MODIFIER |
| 1 | 135409603 | AX-170042824 | 0 | G | A | 0.544 | 1.24E-01 | 8.33E-02 | 2.73E+00 | 2.80E+00 | 1.33E-01 | no | Ipo9 | intron_variant | MODIFIER |
| 1 | 135422249 | AX-170476687 | 0 | C | T | 0.544 | 1.24E-01 | 8.33E-02 | 2.73E+00 | 2.80E+00 | 1.33E-01 | no | Ipo9 | intron_variant | MODIFIER |
| 1 | 135427544 | AX-170331412 | 0 | T | G | 0.544 | 1.24E-01 | 8.33E-02 | 2.73E+00 | 2.80E+00 | 1.33E-01 | no | Ipo9 | intron_variant | MODIFIER |
| 1 | 135427634 | AX-170121026 | 0 | A | G | 0.544 | 1.24E-01 | 8.33E-02 | 2.73E+00 | 2.80E+00 | 1.33E-01 | no | Ipo9 | intron_variant | MODIFIER |
| 1 | 135450848 | AX-170265575 | 0 | T | C | 0.544 | 1.24E-01 | 8.33E-02 | 2.73E+00 | 2.80E+00 | 1.33E-01 | no | Nav1 | synonymous_variant | LOW |
| 1 | 135469058 | AX-170187154 | 0 | G | A | 0.544 | 1.24E-01 | 8.33E-02 | 2.73E+00 | 2.80E+00 | 1.33E-01 | no | Nav1 | intron_variant | MODIFIER |
| 1 | 135470238 | AX-170323160 | 0 | G | A | 0.544 | 1.24E-01 | 8.33E-02 | 2.73E+00 | 2.80E+00 | 1.33E-01 | no | Nav1 | intron_variant | MODIFIER |
| 1 | 135473866 | AX-170350233 | 0 | G | A | 0.544 | 1.24E-01 | 8.33E-02 | 2.73E+00 | 2.80E+00 | 1.33E-01 | no | Nav1 | intron_variant | MODIFIER |
| 1 | 135474748 | AX-170105946 | 0 | T | C | 0.544 | 1.24E-01 | 8.33E-02 | 2.73E+00 | 2.80E+00 | 1.33E-01 | no | Nav1 | intron_variant | MODIFIER |
| 1 | 135486593 | AX-170001768 | 0 | C | T | 0.544 | 1.24E-01 | 8.33E-02 | 2.73E+00 | 2.80E+00 | 1.33E-01 | no | Nav1 | intron_variant | MODIFIER |
| 1 | 135492384 | AX-170171983 | 0 | A | G | 0.544 | 1.24E-01 | 8.33E-02 | 2.73E+00 | 2.80E+00 | 1.33E-01 | no | Nav1 | intron_variant | MODIFIER |
| 1 | 135499263 | AX-170237033 | 0 | A | G | 0.544 | 1.24E-01 | 8.33E-02 | 2.73E+00 | 2.80E+00 | 1.33E-01 | no | Nav1 | intron_variant | MODIFIER |
| 1 | 135520925 | AX-170327154 | 0 | G | A | 0.544 | 1.24E-01 | 8.33E-02 | 2.73E+00 | 2.80E+00 | 1.33E-01 | no | Nav1 | intron_variant | MODIFIER |
| 1 | 135560686 | AX-170042827 | 0 | T | C | 0.544 | 1.24E-01 | 8.33E-02 | 2.73E+00 | 2.80E+00 | 1.33E-01 | no | Nav1 | intron_variant | MODIFIER |
| 1 | 135595968 | AX-170366676 | 0 | A | T | 0.544 | 1.24E-01 | 8.33E-02 | 2.73E+00 | 2.80E+00 | 1.33E-01 | no | Gm4793 | intron_variant,non_coding_transcript_variant | MODIFIER |

| | | | | | | | | | | | | | | | |
|---|---|---|---|---|---|---|---|---|---|---|---|---|---|---|---|
| 1 | 135625228 | AX-170415805 | 0 | A | G | 0.544 | 1.24E-01 | 8.33E-02 | 2.73E+00 | 2.80E+00 | 1.33E-01 | no | Gm38399 | intron_variant,non_coding_transcript_variant | MODIFIER |
| 1 | 135643049 | AX-170526271 | 0 | C | T | 0.544 | 1.24E-01 | 8.33E-02 | 2.73E+00 | 2.80E+00 | 1.33E-01 | no | Gm38399 | intron_variant,non_coding_transcript_variant | MODIFIER |
| 1 | 135644122 | AX-170149227 | 0 | C | T | 0.544 | 1.24E-01 | 8.33E-02 | 2.73E+00 | 2.80E+00 | 1.33E-01 | no | Gm38399 | intron_variant,non_coding_transcript_variant | MODIFIER |
| 1 | 135644565 | AX-170120694 | 0 | C | T | 0.544 | 1.24E-01 | 8.33E-02 | 2.73E+00 | 2.80E+00 | 1.33E-01 | no | Gm38399 | intron_variant,non_coding_transcript_variant | MODIFIER |
| 1 | 135658178 | AX-170374567 | 0 | G | A | 0.544 | 1.24E-01 | 8.33E-02 | 2.73E+00 | 2.80E+00 | 1.33E-01 | no | Gm38399 | intron_variant,non_coding_transcript_variant | MODIFIER |
| 1 | 135658387 | AX-170518243 | 0 | T | G | 0.544 | 1.24E-01 | 8.33E-02 | 2.73E+00 | 2.80E+00 | 1.33E-01 | no | Gm38399 | intron_variant,non_coding_transcript_variant | MODIFIER |
| 1 | 135658472 | AX-170180741 | 0 | A | G | 0.544 | 1.24E-01 | 8.33E-02 | 2.73E+00 | 2.80E+00 | 1.33E-01 | no | Gm38399 | intron_variant,non_coding_transcript_variant | MODIFIER |
| 1 | 135662856 | AX-170142230 | 0 | C | T | 0.544 | 1.24E-01 | 8.33E-02 | 2.73E+00 | 2.80E+00 | 1.33E-01 | no | Gm38399 | intron_variant,non_coding_transcript_variant | MODIFIER |
| 1 | 135680930 | AX-170405824 | 0 | G | A | 0.544 | 1.24E-01 | 8.33E-02 | 2.73E+00 | 2.80E+00 | 1.33E-01 | no | Gm38399 | intron_variant,non_coding_transcript_variant | MODIFIER |
| 1 | 135689183 | AX-170188346 | 0 | A | G | 0.544 | 1.24E-01 | 8.33E-02 | 2.73E+00 | 2.80E+00 | 1.33E-01 | no | Gm38399 | upstream_gene_variant | MODIFIER |
| 1 | 135689500 | AX-170414111 | 0 | G | C | 0.544 | 1.24E-01 | 8.33E-02 | 2.73E+00 | 2.80E+00 | 1.33E-01 | no | Gm38399 | upstream_gene_variant | MODIFIER |
| 1 | 135702750 | AX-170042830 | 0 | G | T | 0.544 | 1.24E-01 | 8.33E-02 | 2.73E+00 | 2.80E+00 | 1.33E-01 | no | | intergenic_variant | MODIFIER |
| 1 | 135705141 | AX-170042832 | 0 | G | A | 0.544 | 1.24E-01 | 8.33E-02 | 2.73E+00 | 2.80E+00 | 1.33E-01 | no | | intergenic_variant | MODIFIER |
| 1 | 135705677 | AX-170042831 | 0 | A | G | 0.544 | 1.24E-01 | 8.33E-02 | 2.73E+00 | 2.80E+00 | 1.33E-01 | no | | intergenic_variant | MODIFIER |
| 1 | 135733511 | AX-170157983 | 0 | A | G | 0.544 | 1.24E-01 | 8.33E-02 | 2.73E+00 | 2.80E+00 | 1.33E-01 | no | Csrp1 | intron_variant | MODIFIER |
| 1 | 135749383 | AX-170418591 | 0 | A | G | 0.544 | 1.24E-01 | 8.33E-02 | 2.73E+00 | 2.80E+00 | 1.33E-01 | no | Csrp1 | intron_variant | MODIFIER |
| 1 | 135749860 | AX-170518032 | 0 | A | C | 0.544 | 1.24E-01 | 8.33E-02 | 2.73E+00 | 2.80E+00 | 1.33E-01 | no | Csrp1 | intron_variant | MODIFIER |
| 1 | 135755770 | AX-170254665 | 0 | G | A | 0.544 | 1.24E-01 | 8.33E-02 | 2.73E+00 | 2.80E+00 | 1.33E-01 | no | Csrp1 | downstream_gene_variant | MODIFIER |
| 1 | 135756829 | AX-170172742 | 0 | T | C | 0.544 | 1.24E-01 | 8.33E-02 | 2.73E+00 | 2.80E+00 | 1.33E-01 | no | Csrp1 | downstream_gene_variant | MODIFIER |
| 1 | 135790003 | AX-170290327 | 0 | G | A | 0.544 | 1.24E-01 | 8.33E-02 | 2.73E+00 | 2.80E+00 | 1.33E-01 | no | Tnni1 | intron_variant | MODIFIER |
| 1 | 135800221 | AX-170106102 | 0 | C | T | 0.544 | 1.24E-01 | 8.33E-02 | 2.73E+00 | 2.80E+00 | 1.33E-01 | no | Tnni1 | upstream_gene_variant | MODIFIER |
| 1 | 135809890 | AX-170350457 | 0 | A | G | 0.544 | 1.24E-01 | 8.33E-02 | 2.73E+00 | 2.80E+00 | 1.33E-01 | no | Tnni1 | 3_prime_UTR_variant | MODIFIER |
| 1 | 135810402 | AX-170360055 | 0 | C | G | 0.544 | 1.24E-01 | 8.33E-02 | 2.73E+00 | 2.80E+00 | 1.33E-01 | no | Tnni1 | downstream_gene_variant | MODIFIER |
| 13 | 11360063 | AX-97844683 | 1 | A | C | 0.188 | -1.83E-01 | 1.22E-01 | 2.52E+00 | 2.57E+00 | 1.33E-01 | Yes | | intergenic_variant | MODIFIER |
| 8 | 66521309 | AX-170080299 | 0 | T | C | 0.14 | -1.97E-01 | 1.32E-01 | 2.66E+00 | 2.77E+00 | 1.33E-01 | Yes | Tktl2 | downstream_gene_variant | MODIFIER |
| 15 | 3140615 | AX-170428150 | 2 | A | C | 0.236 | -1.55E-01 | 1.04E-01 | 2.74E+00 | 2.82E+00 | 1.33E-01 | no | Gm49231 | intron_variant,non_coding_transcript_variant | MODIFIER |
| 1 | 115286057 | AX-170528964 | 0 | T | G | 0.175 | -2.38E-01 | 1.52E-01 | 3.48E+00 | 3.72E+00 | 1.34E-01 | Yes | | intergenic_variant | MODIFIER |
| 1 | 135167755 | AX-170260115 | 1 | A | G | 0.545 | 1.24E-01 | 8.32E-02 | 2.73E+00 | 2.80E+00 | 1.34E-01 | no | Gpr37l1 | upstream_gene_variant | MODIFIER |
| 5 | 134261898 | AX-170176506 | 0 | A | G | 0.158 | -1.32E-01 | 8.53E-02 | 2.26E+00 | 2.34E+00 | 1.34E-01 | no | Gtf2i | synonymous_variant | LOW |
| 5 | 134285927 | AX-170447984 | 0 | C | T | 0.158 | -1.32E-01 | 8.53E-02 | 2.26E+00 | 2.34E+00 | 1.34E-01 | no | Gtf2i | intron_variant | MODIFIER |
| 5 | 134289399 | AX-170355372 | 0 | G | A | 0.158 | -1.32E-01 | 8.53E-02 | 2.26E+00 | 2.34E+00 | 1.34E-01 | no | Gtf2i | synonymous_variant | LOW |
| 5 | 134324269 | AX-97800627 | 0 | C | T | 0.158 | -1.32E-01 | 8.53E-02 | 2.26E+00 | 2.34E+00 | 1.34E-01 | no | Mir3965 | upstream_gene_variant | MODIFIER |
| 5 | 134381113 | AX-97808113 | 0 | C | T | 0.158 | -1.32E-01 | 8.53E-02 | 2.26E+00 | 2.34E+00 | 1.34E-01 | no | Gtf2ird1 | intron_variant | MODIFIER |
| 5 | 134404954 | AX-170508573 | 0 | G | A | 0.158 | -1.32E-01 | 8.53E-02 | 2.26E+00 | 2.34E+00 | 1.34E-01 | no | Gtf2ird1 | synonymous_variant | LOW |
| 5 | 134572980 | AX-97814626 | 0 | A | G | 0.158 | -1.32E-01 | 8.53E-02 | 2.26E+00 | 2.34E+00 | 1.34E-01 | no | Mir7228 | downstream_gene_variant | MODIFIER |
| 5 | 134598127 | AX-97797662 | 0 | T | C | 0.158 | -1.32E-01 | 8.53E-02 | 2.26E+00 | 2.34E+00 | 1.34E-01 | no | Rfc2 | 3_prime_UTR_variant | MODIFIER |
| 5 | 134619152 | AX-170144087 | 0 | T | G | 0.158 | -1.32E-01 | 8.53E-02 | 2.26E+00 | 2.34E+00 | 1.34E-01 | no | Eif4h | downstream_gene_variant | MODIFIER |
| 5 | 134668287 | AX-170026230 | 0 | T | C | 0.158 | -1.32E-01 | 8.53E-02 | 2.26E+00 | 2.34E+00 | 1.34E-01 | no | Limk1 | intron_variant | MODIFIER |
| 5 | 134668435 | AX-170478086 | 0 | T | C | 0.158 | -1.32E-01 | 8.53E-02 | 2.26E+00 | 2.34E+00 | 1.34E-01 | no | Limk1 | intron_variant | MODIFIER |
| 5 | 134668890 | AX-170348200 | 0 | C | T | 0.158 | -1.32E-01 | 8.53E-02 | 2.26E+00 | 2.34E+00 | 1.34E-01 | no | Limk1 | intron_variant | MODIFIER |
| 5 | 134680906 | AX-170488455 | 0 | A | C | 0.158 | -1.32E-01 | 8.53E-02 | 2.26E+00 | 2.34E+00 | 1.34E-01 | no | Limk1 | intron_variant | MODIFIER |
| 5 | 134684473 | AX-170315214 | 0 | G | A | 0.158 | -1.32E-01 | 8.53E-02 | 2.26E+00 | 2.34E+00 | 1.34E-01 | no | Limk1 | intron_variant | MODIFIER |
| 5 | 134685869 | AX-170026226 | 0 | T | G | 0.158 | -1.32E-01 | 8.53E-02 | 2.26E+00 | 2.34E+00 | 1.34E-01 | no | Limk1 | intron_variant | MODIFIER |
| 5 | 134686233 | AX-170295117 | 0 | T | C | 0.158 | -1.32E-01 | 8.53E-02 | 2.26E+00 | 2.34E+00 | 1.34E-01 | no | Limk1 | intron_variant | MODIFIER |
| 5 | 134687380 | AX-170026221 | 0 | T | C | 0.158 | -1.32E-01 | 8.53E-02 | 2.26E+00 | 2.34E+00 | 1.34E-01 | no | Limk1 | intron_variant | MODIFIER |
| 5 | 134734452 | AX-97799514 | 0 | T | C | 0.158 | -1.32E-01 | 8.53E-02 | 2.26E+00 | 2.34E+00 | 1.34E-01 | no | Eln | intron_variant | MODIFIER |
| 5 | 134736980 | AX-170429499 | 0 | G | C | 0.158 | -1.32E-01 | 8.53E-02 | 2.26E+00 | 2.34E+00 | 1.34E-01 | no | Eln | intron_variant | MODIFIER |
| 5 | 134856114 | AX-97823388 | 0 | T | C | 0.158 | -1.32E-01 | 8.53E-02 | 2.26E+00 | 2.34E+00 | 1.34E-01 | no | | intergenic_variant | MODIFIER |
| 5 | 135067625 | AX-97806444 | 0 | T | C | 0.158 | -1.32E-01 | 8.53E-02 | 2.26E+00 | 2.34E+00 | 1.34E-01 | no | Dnajc30 | downstream_gene_variant | MODIFIER |
| 5 | 135134926 | AX-170073610 | 0 | A | G | 0.158 | -1.32E-01 | 8.53E-02 | 2.26E+00 | 2.34E+00 | 1.34E-01 | no | Mlxipl | intron_variant | MODIFIER |
| 5 | 135232030 | AX-170112720 | 0 | T | C | 0.158 | -1.32E-01 | 8.53E-02 | 2.26E+00 | 2.34E+00 | 1.34E-01 | no | Baz1b | intron_variant | MODIFIER |
| 5 | 135342438 | AX-170364448 | 0 | C | T | 0.158 | -1.32E-01 | 8.53E-02 | 2.26E+00 | 2.34E+00 | 1.34E-01 | no | Fkbp6 | intron_variant | MODIFIER |
| 5 | 135350009 | AX-170356323 | 0 | T | G | 0.158 | -1.32E-01 | 8.53E-02 | 2.26E+00 | 2.34E+00 | 1.34E-01 | no | Fkbp6 | 5_prime_UTR_variant | MODIFIER |
| 5 | 135353802 | AX-170293217 | 0 | T | G | 0.158 | -1.32E-01 | 8.53E-02 | 2.26E+00 | 2.34E+00 | 1.34E-01 | no | Fkbp6 | upstream_gene_variant | MODIFIER |

| | | | | | | | | | | | | | | | |
|---|---|---|---|---|---|---|---|---|---|---|---|---|---|---|---|
| 5 | 135399328 | AX-170254752 | 0 | A | G | 0.158 | -1.32E-01 | 8.53E-02 | 2.26E+00 | 2.34E+00 | 1.34E-01 | no | Pom121 | upstream_gene_variant | MODIFIER |
| 5 | 135405161 | AX-97811162 | 0 | T | A | 0.158 | -1.32E-01 | 8.53E-02 | 2.26E+00 | 2.34E+00 | 1.34E-01 | no | Hip1 | downstream_gene_variant | MODIFIER |
| 5 | 135441628 | AX-170296468 | 0 | A | G | 0.158 | -1.32E-01 | 8.53E-02 | 2.26E+00 | 2.34E+00 | 1.34E-01 | no | Hip1 | intron_variant | MODIFIER |
| 5 | 135442435 | AX-170416950 | 0 | G | A | 0.158 | -1.32E-01 | 8.53E-02 | 2.26E+00 | 2.34E+00 | 1.34E-01 | no | Hip1 | intron_variant | MODIFIER |
| 5 | 135485068 | AX-170026247 | 0 | C | T | 0.158 | -1.32E-01 | 8.53E-02 | 2.26E+00 | 2.34E+00 | 1.34E-01 | no | Hip1 | intron_variant | MODIFIER |
| 13 | 18602734 | AX-170477954 | 1 | T | A | 0.161 | -1.96E-01 | 1.30E-01 | 2.47E+00 | 2.51E+00 | 1.34E-01 | Yes | | intergenic_variant | MODIFIER |
| 1 | 192400091 | AX-97838172 | 2 | C | T | 0.273 | 1.39E-01 | 9.01E-02 | 2.25E+00 | 2.34E+00 | 1.34E-01 | no | Kcnh1 | intron_variant | MODIFIER |
| 5 | 10634756 | AX-170156124 | 0 | G | A | 0.351 | 1.19E-01 | 7.96E-02 | 2.65E+00 | 2.79E+00 | 1.34E-01 | no | | intergenic_variant | MODIFIER |
| 10 | 69919936 | AX-168404519 | 0 | T | C | 0.412 | -1.17E-01 | 7.87E-02 | 2.73E+00 | 2.82E+00 | 1.34E-01 | no | Ank3 | intron_variant | MODIFIER |
| 1 | 96425584 | AX-170320028 | 1 | C | T | 0.161 | 2.21E-01 | 1.49E-01 | 2.69E+00 | 2.82E+00 | 1.34E-01 | Yes | | intergenic_variant | MODIFIER |
| 6 | 139755436 | AX-97781909 | 0 | G | A | 0.07 | 2.58E-01 | 1.73E-01 | 2.89E+00 | 3.06E+00 | 1.34E-01 | Yes | Pik3c2g | intron_variant | MODIFIER |
| 5 | 66484100 | AX-97800128 | 1 | C | T | 0.491 | 1.53E-01 | 1.02E-01 | 2.55E+00 | 2.62E+00 | 1.34E-01 | Yes | Apbb2 | intron_variant | MODIFIER |
| 18 | 79879105 | AX-97792933 | 0 | T | C | 0.474 | 1.18E-01 | 7.80E-02 | 2.41E+00 | 2.50E+00 | 1.34E-01 | no | Gm26977 | upstream_gene_variant | MODIFIER |
| 13 | 18715910 | AX-170456214 | 0 | C | T | 0.149 | -1.74E-01 | 1.16E-01 | 2.48E+00 | 2.56E+00 | 1.34E-01 | Yes | Vps41 | upstream_gene_variant | MODIFIER |
| 15 | 43080848 | AX-170121660 | 1 | T | C | 0.036 | -2.66E-01 | 1.15E-01 | 1.17E+00 | 1.23E+00 | 1.35E-01 | Yes | Rspo2 | intron_variant | MODIFIER |
| 5 | 130339629 | AX-170195072 | 1 | G | A | 0.152 | -1.40E-01 | 8.91E-02 | 2.16E+00 | 2.23E+00 | 1.35E-01 | no | Tyw1 | intron_variant | MODIFIER |
| 6 | 137054729 | AX-97802385 | 1 | T | C | 0.58 | 1.23E-01 | 8.19E-02 | 2.47E+00 | 2.54E+00 | 1.35E-01 | no | Rerg | downstream_gene_variant | MODIFIER |
| 3 | 19573632 | AX-170066027 | 2 | C | T | 0.127 | -1.50E-01 | 1.01E-01 | 2.66E+00 | 2.75E+00 | 1.35E-01 | no | Dnajc5b | intron_variant | MODIFIER |
| 10 | 102159806 | AX-170504343 | 0 | C | A | 0.272 | -1.28E-01 | 8.64E-02 | 2.74E+00 | 2.82E+00 | 1.35E-01 | no | Mgat4c | intron_variant | MODIFIER |
| 5 | 11949662 | AX-170071317 | 1 | C | T | 0.357 | 1.22E-01 | 8.19E-02 | 2.68E+00 | 2.82E+00 | 1.35E-01 | no | | intergenic_variant | MODIFIER |
| 5 | 85661315 | AX-170207767 | 0 | A | C | 0.43 | 1.62E-01 | 1.06E-01 | 3.28E+00 | 3.40E+00 | 1.35E-01 | Yes | | intergenic_variant | MODIFIER |
| 1 | 135509520 | AX-170238166 | 1 | G | A | 0.545 | 1.24E-01 | 8.33E-02 | 2.72E+00 | 2.79E+00 | 1.35E-01 | no | Nav1 | intron_variant | MODIFIER |
| 18 | 63164233 | AX-170390721 | 0 | T | C | 0.404 | 1.33E-01 | 8.90E-02 | 2.51E+00 | 2.57E+00 | 1.35E-01 | no | Piezo2 | intron_variant | MODIFIER |
| 13 | 11330945 | AX-170050536 | 0 | C | T | 0.184 | -1.90E-01 | 1.27E-01 | 2.52E+00 | 2.56E+00 | 1.35E-01 | Yes | | intergenic_variant | MODIFIER |
| 13 | 8498763 | AX-97797861 | 1 | C | T | 0.188 | -1.86E-01 | 1.25E-01 | 2.53E+00 | 2.58E+00 | 1.35E-01 | Yes | Adarb2 | intron_variant | MODIFIER |
| 14 | 111886928 | AX-170010708 | 1 | T | G | 0.321 | 1.02E-01 | 6.89E-02 | 2.67E+00 | 2.79E+00 | 1.35E-01 | no | | intergenic_variant | MODIFIER |
| 16 | 71297542 | AX-170013355 | 0 | A | C | 0.482 | 1.17E-01 | 7.84E-02 | 2.60E+00 | 2.66E+00 | 1.35E-01 | no | | intergenic_variant | MODIFIER |
| 4 | 103288614 | AX-97815358 | 0 | T | G | 0.228 | 1.74E-01 | 1.18E-01 | 2.69E+00 | 2.76E+00 | 1.36E-01 | Yes | 4921539E11Rik | intron_variant | MODIFIER |
| 4 | 103301971 | AX-170339817 | 0 | T | G | 0.228 | 1.74E-01 | 1.18E-01 | 2.69E+00 | 2.76E+00 | 1.36E-01 | Yes | 4930456L15Rik | intron_variant,non_coding_transcript_variant | MODIFIER |
| 4 | 103303080 | AX-170453657 | 0 | C | T | 0.228 | 1.74E-01 | 1.18E-01 | 2.69E+00 | 2.76E+00 | 1.36E-01 | Yes | 4930456L15Rik | intron_variant,non_coding_transcript_variant | MODIFIER |
| 4 | 103311971 | AX-170086007 | 0 | A | G | 0.228 | 1.74E-01 | 1.18E-01 | 2.69E+00 | 2.76E+00 | 1.36E-01 | Yes | Oma1 | upstream_gene_variant | MODIFIER |
| 4 | 103313735 | AX-170419602 | 0 | A | C | 0.228 | 1.74E-01 | 1.18E-01 | 2.69E+00 | 2.76E+00 | 1.36E-01 | Yes | Oma1 | upstream_gene_variant | MODIFIER |
| 4 | 103314261 | AX-170441085 | 0 | G | C | 0.228 | 1.74E-01 | 1.18E-01 | 2.69E+00 | 2.76E+00 | 1.36E-01 | Yes | Oma1 | intron_variant | MODIFIER |
| 4 | 103314413 | AX-170275820 | 0 | C | G | 0.228 | 1.74E-01 | 1.18E-01 | 2.69E+00 | 2.76E+00 | 1.36E-01 | Yes | Oma1 | intron_variant | MODIFIER |
| 4 | 103481922 | AX-97825728 | 0 | T | C | 0.228 | 1.74E-01 | 1.18E-01 | 2.69E+00 | 2.76E+00 | 1.36E-01 | Yes | Gm12718 | intron_variant,non_coding_transcript_variant | MODIFIER |
| 4 | 103554075 | AX-170395244 | 0 | T | C | 0.228 | 1.74E-01 | 1.18E-01 | 2.69E+00 | 2.76E+00 | 1.36E-01 | Yes | Gm12720 | upstream_gene_variant | MODIFIER |
| 4 | 103615369 | AX-170120456 | 0 | G | T | 0.228 | 1.74E-01 | 1.18E-01 | 2.69E+00 | 2.76E+00 | 1.36E-01 | Yes | Dab1 | upstream_gene_variant | MODIFIER |
| 4 | 104072544 | AX-170400674 | 0 | T | A | 0.228 | 1.74E-01 | 1.18E-01 | 2.69E+00 | 2.76E+00 | 1.36E-01 | Yes | Dab1 | intron_variant | MODIFIER |
| 4 | 104080275 | AX-170533263 | 0 | T | G | 0.228 | 1.74E-01 | 1.18E-01 | 2.69E+00 | 2.76E+00 | 1.36E-01 | Yes | Dab1 | intron_variant | MODIFIER |
| 4 | 104102438 | AX-170035694 | 0 | T | C | 0.228 | 1.74E-01 | 1.18E-01 | 2.69E+00 | 2.76E+00 | 1.36E-01 | Yes | Dab1 | intron_variant | MODIFIER |
| 4 | 104355190 | AX-170274903 | 0 | C | T | 0.228 | 1.74E-01 | 1.18E-01 | 2.69E+00 | 2.76E+00 | 1.36E-01 | Yes | Dab1 | intron_variant | MODIFIER |
| 4 | 104585337 | AX-170380214 | 0 | G | A | 0.228 | 1.74E-01 | 1.18E-01 | 2.69E+00 | 2.76E+00 | 1.36E-01 | Yes | Dab1 | intron_variant | MODIFIER |
| 1 | 182468450 | AX-170124121 | 1 | A | T | 0.232 | 1.68E-01 | 1.13E-01 | 2.52E+00 | 2.59E+00 | 1.36E-01 | Yes | Capn2 | intron_variant | MODIFIER |
| 5 | 132127384 | AX-170481845 | 0 | T | C | 0.149 | -1.35E-01 | 8.84E-02 | 2.29E+00 | 2.37E+00 | 1.36E-01 | no | Auts2 | intron_variant,non_coding_transcript_variant | MODIFIER |
| 10 | 96662886 | AX-97816504 | 0 | T | A | 0.281 | -1.28E-01 | 8.59E-02 | 2.57E+00 | 2.63E+00 | 1.36E-01 | no | | intergenic_variant | MODIFIER |
| 10 | 96749466 | AX-170466638 | 0 | T | C | 0.281 | -1.28E-01 | 8.59E-02 | 2.57E+00 | 2.63E+00 | 1.36E-01 | no | | intergenic_variant | MODIFIER |
| 10 | 96873111 | AX-170530562 | 0 | C | T | 0.281 | -1.28E-01 | 8.59E-02 | 2.57E+00 | 2.63E+00 | 1.36E-01 | no | Gm48521 | intron_variant,non_coding_transcript_variant | MODIFIER |
| 7 | 99663437 | AX-170242842 | 0 | C | T | 0.377 | -1.54E-01 | 1.03E-01 | 2.95E+00 | 3.05E+00 | 1.36E-01 | no | Slco2b1 | intron_variant | MODIFIER |
| 19 | 60919344 | AX-170016957 | 0 | A | T | 0.465 | -9.45E-02 | 6.38E-02 | 2.70E+00 | 2.80E+00 | 1.36E-01 | no | Grk5 | intron_variant | MODIFIER |
| 5 | 58316021 | AX-170434984 | 1 | C | T | 0.473 | 1.49E-01 | 9.83E-02 | 2.35E+00 | 2.41E+00 | 1.36E-01 | Yes | | intergenic_variant | MODIFIER |
| 9 | 77956800 | AX-170279175 | 0 | A | G | 0.395 | -1.00E-01 | 6.61E-02 | 3.19E+00 | 3.29E+00 | 1.36E-01 | no | Elovl5 | intron_variant | MODIFIER |
| 1 | 191697388 | AX-97828413 | 1 | A | G | 0.286 | 1.33E-01 | 8.84E-02 | 2.40E+00 | 2.49E+00 | 1.36E-01 | no | | regulatory_region_variant | MODIFIER |
| 4 | 105965136 | AX-170136739 | 0 | C | T | 0.272 | 1.62E-01 | 1.09E-01 | 2.73E+00 | 2.81E+00 | 1.36E-01 | Yes | | intergenic_variant | MODIFIER |
| 4 | 105965404 | AX-170215010 | 0 | G | A | 0.272 | 1.62E-01 | 1.09E-01 | 2.73E+00 | 2.81E+00 | 1.36E-01 | Yes | | intergenic_variant | MODIFIER |

| | | | | | | | | | | | | | | | |
|---|---|---|---|---|---|---|---|---|---|---|---|---|---|---|---|
| 4 | 105966291 | AX-170115753 | 0 | A | G | 0.272 | 1.62E-01 | 1.09E-01 | 2.73E+00 | 2.81E+00 | 1.36E-01 | Yes | | intergenic_variant | MODIFIER |
| 4 | 105966481 | AX-170329012 | 0 | G | A | 0.272 | 1.62E-01 | 1.09E-01 | 2.73E+00 | 2.81E+00 | 1.36E-01 | Yes | | intergenic_variant | MODIFIER |
| 4 | 105966506 | AX-170070247 | 0 | G | T | 0.272 | 1.62E-01 | 1.09E-01 | 2.73E+00 | 2.81E+00 | 1.36E-01 | Yes | | intergenic_variant | MODIFIER |
| 4 | 105975542 | AX-170117800 | 0 | C | T | 0.272 | 1.62E-01 | 1.09E-01 | 2.73E+00 | 2.81E+00 | 1.36E-01 | Yes | Gm24363 | downstream_gene_variant | MODIFIER |
| 4 | 106165636 | AX-170248790 | 0 | A | G | 0.272 | 1.62E-01 | 1.09E-01 | 2.73E+00 | 2.81E+00 | 1.36E-01 | Yes | | intergenic_variant | MODIFIER |
| 4 | 106174242 | AX-170275309 | 0 | G | A | 0.272 | 1.62E-01 | 1.09E-01 | 2.73E+00 | 2.81E+00 | 1.36E-01 | Yes | | intergenic_variant | MODIFIER |
| 4 | 79913057 | AX-170089288 | 0 | T | C | 0.447 | -1.26E-01 | 8.43E-02 | 2.47E+00 | 2.53E+00 | 1.36E-01 | no | | intergenic_variant | MODIFIER |
| 6 | 103440329 | AX-170439105 | 1 | C | G | 0.045 | 2.89E-01 | 1.95E-01 | 2.97E+00 | 3.17E+00 | 1.36E-01 | Yes | | intergenic_variant | MODIFIER |
| 19 | 60511645 | AX-170062864 | 0 | A | G | 0.465 | -9.41E-02 | 6.36E-02 | 2.78E+00 | 2.87E+00 | 1.36E-01 | no | | intergenic_variant | MODIFIER |
| 18 | 78309447 | AX-170292495 | 0 | G | A | 0.447 | 1.23E-01 | 8.26E-02 | 2.52E+00 | 2.60E+00 | 1.37E-01 | no | Slc14a2 | intron_variant | MODIFIER |
| 19 | 60545311 | AX-170309013 | 1 | A | G | 0.464 | -9.38E-02 | 6.35E-02 | 2.71E+00 | 2.79E+00 | 1.37E-01 | no | Cacul1 | intron_variant | MODIFIER |
| 1 | 124738731 | AX-170458775 | 0 | G | A | 0.57 | 1.19E-01 | 8.07E-02 | 2.67E+00 | 2.74E+00 | 1.37E-01 | no | Gm28928 | intron_variant,non_coding_transcript_variant | MODIFIER |
| 5 | 52105697 | AX-97818369 | 1 | G | A | 0.491 | 1.58E-01 | 1.05E-01 | 2.34E+00 | 2.38E+00 | 1.37E-01 | Yes | | regulatory_region_variant | MODIFIER |
| 13 | 7649864 | AX-170204644 | 0 | C | T | 0.184 | -1.76E-01 | 1.19E-01 | 2.58E+00 | 2.64E+00 | 1.38E-01 | Yes | | intergenic_variant | MODIFIER |
| 4 | 107483583 | AX-170478930 | 0 | C | T | 0.254 | 1.70E-01 | 1.15E-01 | 2.63E+00 | 2.69E+00 | 1.38E-01 | Yes | Glis1 | intron_variant | MODIFIER |
| 4 | 107533472 | AX-170245964 | 0 | T | C | 0.254 | 1.70E-01 | 1.15E-01 | 2.63E+00 | 2.69E+00 | 1.38E-01 | Yes | Glis1 | intron_variant | MODIFIER |
| 4 | 107548685 | AX-170212964 | 0 | G | A | 0.254 | 1.70E-01 | 1.15E-01 | 2.63E+00 | 2.69E+00 | 1.38E-01 | Yes | Glis1 | intron_variant | MODIFIER |
| 4 | 107582555 | AX-170300190 | 0 | G | A | 0.254 | 1.70E-01 | 1.15E-01 | 2.63E+00 | 2.69E+00 | 1.38E-01 | Yes | Glis1 | intron_variant | MODIFIER |
| 4 | 107582751 | AX-170336889 | 0 | A | G | 0.254 | 1.70E-01 | 1.15E-01 | 2.63E+00 | 2.69E+00 | 1.38E-01 | Yes | Glis1 | intron_variant | MODIFIER |
| 4 | 107620247 | AX-170427887 | 0 | T | C | 0.254 | 1.70E-01 | 1.15E-01 | 2.63E+00 | 2.69E+00 | 1.38E-01 | Yes | Glis1 | intron_variant | MODIFIER |
| 4 | 107691660 | AX-170203573 | 0 | A | G | 0.254 | 1.70E-01 | 1.15E-01 | 2.63E+00 | 2.69E+00 | 1.38E-01 | Yes | Gm12899 | downstream_gene_variant | MODIFIER |
| 4 | 107803002 | AX-170402665 | 0 | A | G | 0.254 | 1.70E-01 | 1.15E-01 | 2.63E+00 | 2.69E+00 | 1.38E-01 | Yes | Lrp8 | intron_variant | MODIFIER |
| 4 | 108028123 | AX-97821456 | 0 | T | C | 0.254 | 1.70E-01 | 1.15E-01 | 2.63E+00 | 2.69E+00 | 1.38E-01 | Yes | Podn | intron_variant | MODIFIER |
| 4 | 108095568 | AX-97799105 | 0 | G | A | 0.254 | 1.70E-01 | 1.15E-01 | 2.63E+00 | 2.69E+00 | 1.38E-01 | Yes | Scp2 | intron_variant | MODIFIER |
| 4 | 108110949 | AX-170186096 | 0 | T | C | 0.254 | 1.70E-01 | 1.15E-01 | 2.63E+00 | 2.69E+00 | 1.38E-01 | Yes | Scp2 | intron_variant | MODIFIER |
| 4 | 108112953 | AX-97807150 | 0 | G | T | 0.254 | 1.70E-01 | 1.15E-01 | 2.63E+00 | 2.69E+00 | 1.38E-01 | Yes | Scp2 | intron_variant | MODIFIER |
| 4 | 108117788 | AX-170409315 | 0 | A | C | 0.254 | 1.70E-01 | 1.15E-01 | 2.63E+00 | 2.69E+00 | 1.38E-01 | Yes | Scp2 | intron_variant | MODIFIER |
| 4 | 108120789 | AX-170419875 | 0 | G | C | 0.254 | 1.70E-01 | 1.15E-01 | 2.63E+00 | 2.69E+00 | 1.38E-01 | Yes | Scp2 | upstream_gene_variant | MODIFIER |
| 4 | 108121672 | AX-170238357 | 0 | C | G | 0.254 | 1.70E-01 | 1.15E-01 | 2.63E+00 | 2.69E+00 | 1.38E-01 | Yes | Scp2 | upstream_gene_variant | MODIFIER |
| 4 | 108328486 | AX-97798444 | 0 | A | G | 0.254 | 1.70E-01 | 1.15E-01 | 2.63E+00 | 2.69E+00 | 1.38E-01 | Yes | Coa7 | intron_variant,non_coding_transcript_variant | MODIFIER |
| 4 | 108880014 | AX-170023650 | 0 | C | T | 0.254 | 1.70E-01 | 1.15E-01 | 2.63E+00 | 2.69E+00 | 1.38E-01 | Yes | Rab3b | intron_variant | MODIFIER |
| 4 | 108891242 | AX-170023654 | 0 | G | T | 0.254 | 1.70E-01 | 1.15E-01 | 2.63E+00 | 2.69E+00 | 1.38E-01 | Yes | Rab3b | intron_variant | MODIFIER |
| 4 | 108930999 | AX-170242009 | 0 | A | C | 0.254 | 1.70E-01 | 1.15E-01 | 2.63E+00 | 2.69E+00 | 1.38E-01 | Yes | Rab3b | intron_variant | MODIFIER |
| 4 | 108943193 | AX-97788949 | 0 | A | G | 0.254 | 1.70E-01 | 1.15E-01 | 2.63E+00 | 2.69E+00 | 1.38E-01 | Yes | Rab3b | downstream_gene_variant | MODIFIER |
| 4 | 108943935 | AX-170496171 | 0 | C | G | 0.254 | 1.70E-01 | 1.15E-01 | 2.63E+00 | 2.69E+00 | 1.38E-01 | Yes | Rab3b | downstream_gene_variant | MODIFIER |
| 4 | 109169766 | AX-97798442 | 0 | C | A | 0.254 | 1.70E-01 | 1.15E-01 | 2.63E+00 | 2.69E+00 | 1.38E-01 | Yes | Osbpl9 | intron_variant | MODIFIER |
| 4 | 109176380 | AX-170261357 | 0 | C | A | 0.254 | 1.70E-01 | 1.15E-01 | 2.63E+00 | 2.69E+00 | 1.38E-01 | Yes | Osbpl9 | intron_variant | MODIFIER |
| 4 | 109185508 | AX-170507136 | 0 | C | T | 0.254 | 1.70E-01 | 1.15E-01 | 2.63E+00 | 2.69E+00 | 1.38E-01 | Yes | Osbpl9 | intron_variant | MODIFIER |
| 4 | 109196471 | AX-97782502 | 0 | A | G | 0.254 | 1.70E-01 | 1.15E-01 | 2.63E+00 | 2.69E+00 | 1.38E-01 | Yes | Osbpl9 | intron_variant | MODIFIER |
| 4 | 109204065 | AX-97824732 | 0 | A | G | 0.254 | 1.70E-01 | 1.15E-01 | 2.63E+00 | 2.69E+00 | 1.38E-01 | Yes | Osbpl9 | upstream_gene_variant | MODIFIER |
| 4 | 105112663 | AX-170172389 | 0 | G | C | 0.254 | 1.70E-01 | 1.16E-01 | 2.75E+00 | 2.82E+00 | 1.38E-01 | Yes | Prkaa2 | upstream_gene_variant | MODIFIER |
| 16 | 35626858 | AX-170328613 | 1 | C | T | 0.518 | 1.18E-01 | 7.94E-02 | 2.54E+00 | 2.59E+00 | 1.38E-01 | no | Sema5b | intron_variant | MODIFIER |
| 1 | 189484547 | AX-170294039 | 0 | T | A | 0.263 | 1.43E-01 | 9.52E-02 | 2.41E+00 | 2.50E+00 | 1.38E-01 | no | | intergenic_variant | MODIFIER |
| 3 | 38699857 | AX-97841883 | 1 | G | A | 0.214 | 1.34E-01 | 8.90E-02 | 3.16E+00 | 3.31E+00 | 1.38E-01 | no | | intergenic_variant | MODIFIER |
| 13 | 10012828 | AX-170171075 | 2 | G | T | 0.191 | -1.79E-01 | 1.21E-01 | 2.52E+00 | 2.57E+00 | 1.38E-01 | Yes | Chrm3 | intron_variant | MODIFIER |
| 1 | 135246179 | AX-170253005 | 1 | C | T | 0.545 | 1.23E-01 | 8.33E-02 | 2.72E+00 | 2.79E+00 | 1.39E-01 | no | Gm26642 | upstream_gene_variant | MODIFIER |
| 19 | 59649731 | AX-97836192 | 0 | T | C | 0.518 | 9.20E-02 | 6.25E-02 | 2.70E+00 | 2.78E+00 | 1.39E-01 | no | | intergenic_variant | MODIFIER |
| 19 | 58644085 | AX-170384218 | 0 | A | G | 0.447 | 9.26E-02 | 6.27E-02 | 2.57E+00 | 2.66E+00 | 1.39E-01 | no | | intergenic_variant | MODIFIER |
| 13 | 18562177 | AX-170050649 | 0 | A | C | 0.158 | -1.86E-01 | 1.25E-01 | 2.45E+00 | 2.51E+00 | 1.39E-01 | Yes | | intergenic_variant | MODIFIER |
| 9 | 92733355 | AX-170033007 | 0 | C | T | 0.132 | -1.88E-01 | 1.28E-01 | 2.80E+00 | 2.89E+00 | 1.39E-01 | Yes | 4933400C23Rik | intron_variant,non_coding_transcript_variant | MODIFIER |
| 8 | 86983173 | AX-170533680 | 0 | T | A | 0.211 | -1.74E-01 | 1.16E-01 | 3.18E+00 | 3.34E+00 | 1.39E-01 | Yes | | intergenic_variant | MODIFIER |
| 19 | 59617919 | AX-170152282 | 1 | C | G | 0.518 | 9.20E-02 | 6.26E-02 | 2.70E+00 | 2.78E+00 | 1.39E-01 | no | | intergenic_variant | MODIFIER |
| 3 | 7516496 | AX-170365573 | 0 | C | T | 0.404 | -9.25E-02 | 6.30E-02 | 2.82E+00 | 2.98E+00 | 1.39E-01 | no | Zc2hc1a | synonymous_variant | LOW |
| 18 | 68952153 | AX-97842631 | 0 | C | T | 0.175 | -1.78E-01 | 1.21E-01 | 2.75E+00 | 2.81E+00 | 1.39E-01 | Yes | 4930546C10Rik | upstream_gene_variant | MODIFIER |

| | | | | | | | | | | | | | | | |
|---|---|---|---|---|---|---|---|---|---|---|---|---|---|---|---|
| 15 | 6349485 | AX-170195376 | 0 | C | T | 0.219 | -1.54E-01 | 1.05E-01 | 2.58E+00 | 2.64E+00 | 1.40E-01 | no | Dab2 | intron_variant | MODIFIER |
| 4 | 103334832 | AX-170201743 | 0 | G | A | 0.237 | 1.72E-01 | 1.17E-01 | 2.71E+00 | 2.78E+00 | 1.40E-01 | Yes | Oma1 | intron_variant | MODIFIER |
| 4 | 103347509 | AX-170326233 | 0 | T | A | 0.237 | 1.72E-01 | 1.17E-01 | 2.71E+00 | 2.78E+00 | 1.40E-01 | Yes | Oma1 | intron_variant | MODIFIER |
| 4 | 103347558 | AX-97854875 | 0 | A | G | 0.237 | 1.72E-01 | 1.17E-01 | 2.71E+00 | 2.78E+00 | 1.40E-01 | Yes | Oma1 | intron_variant | MODIFIER |
| 4 | 103635688 | AX-97816106 | 0 | C | A | 0.237 | 1.72E-01 | 1.17E-01 | 2.71E+00 | 2.78E+00 | 1.40E-01 | Yes | Dab1 | intron_variant | MODIFIER |
| 4 | 103648113 | AX-170181547 | 0 | A | C | 0.237 | 1.72E-01 | 1.17E-01 | 2.71E+00 | 2.78E+00 | 1.40E-01 | Yes | Dab1 | intron_variant | MODIFIER |
| 4 | 103649060 | AX-170482323 | 0 | C | A | 0.237 | 1.72E-01 | 1.17E-01 | 2.71E+00 | 2.78E+00 | 1.40E-01 | Yes | Dab1 | intron_variant | MODIFIER |
| 4 | 103649561 | AX-170387543 | 0 | A | G | 0.237 | 1.72E-01 | 1.17E-01 | 2.71E+00 | 2.78E+00 | 1.40E-01 | Yes | Dab1 | intron_variant | MODIFIER |
| 4 | 103661921 | AX-170108733 | 0 | C | T | 0.237 | 1.72E-01 | 1.17E-01 | 2.71E+00 | 2.78E+00 | 1.40E-01 | Yes | Dab1 | intron_variant | MODIFIER |
| 4 | 103678269 | AX-97804517 | 0 | G | C | 0.237 | 1.72E-01 | 1.17E-01 | 2.71E+00 | 2.78E+00 | 1.40E-01 | Yes | Dab1 | intron_variant | MODIFIER |
| 4 | 103707881 | AX-170375295 | 0 | T | C | 0.237 | 1.72E-01 | 1.17E-01 | 2.71E+00 | 2.78E+00 | 1.40E-01 | Yes | Dab1 | intron_variant | MODIFIER |
| 4 | 103716174 | AX-170355166 | 0 | G | A | 0.237 | 1.72E-01 | 1.17E-01 | 2.71E+00 | 2.78E+00 | 1.40E-01 | Yes | Dab1 | intron_variant | MODIFIER |
| 4 | 103724356 | AX-170197382 | 0 | T | C | 0.237 | 1.72E-01 | 1.17E-01 | 2.71E+00 | 2.78E+00 | 1.40E-01 | Yes | Dab1 | intron_variant | MODIFIER |
| 4 | 103777322 | AX-170397698 | 0 | C | G | 0.237 | 1.72E-01 | 1.17E-01 | 2.71E+00 | 2.78E+00 | 1.40E-01 | Yes | Dab1 | intron_variant | MODIFIER |
| 4 | 103780739 | AX-170301129 | 0 | A | T | 0.237 | 1.72E-01 | 1.17E-01 | 2.71E+00 | 2.78E+00 | 1.40E-01 | Yes | Dab1 | intron_variant | MODIFIER |
| 4 | 103780767 | AX-170239967 | 0 | A | C | 0.237 | 1.72E-01 | 1.17E-01 | 2.71E+00 | 2.78E+00 | 1.40E-01 | Yes | Dab1 | intron_variant | MODIFIER |
| 4 | 103785441 | AX-170177409 | 0 | C | G | 0.237 | 1.72E-01 | 1.17E-01 | 2.71E+00 | 2.78E+00 | 1.40E-01 | Yes | Dab1 | intron_variant | MODIFIER |
| 4 | 103786620 | AX-170300753 | 0 | T | A | 0.237 | 1.72E-01 | 1.17E-01 | 2.71E+00 | 2.78E+00 | 1.40E-01 | Yes | Dab1 | intron_variant | MODIFIER |
| 4 | 103786671 | AX-97820987 | 0 | T | C | 0.237 | 1.72E-01 | 1.17E-01 | 2.71E+00 | 2.78E+00 | 1.40E-01 | Yes | Dab1 | intron_variant | MODIFIER |
| 4 | 103787950 | AX-170247795 | 0 | C | T | 0.237 | 1.72E-01 | 1.17E-01 | 2.71E+00 | 2.78E+00 | 1.40E-01 | Yes | Dab1 | intron_variant | MODIFIER |
| 4 | 103791881 | AX-170443541 | 0 | C | G | 0.237 | 1.72E-01 | 1.17E-01 | 2.71E+00 | 2.78E+00 | 1.40E-01 | Yes | Dab1 | intron_variant | MODIFIER |
| 4 | 103792168 | AX-170388613 | 0 | T | C | 0.237 | 1.72E-01 | 1.17E-01 | 2.71E+00 | 2.78E+00 | 1.40E-01 | Yes | Dab1 | intron_variant | MODIFIER |
| 4 | 103792412 | AX-170270481 | 0 | T | A | 0.237 | 1.72E-01 | 1.17E-01 | 2.71E+00 | 2.78E+00 | 1.40E-01 | Yes | Dab1 | intron_variant | MODIFIER |
| 4 | 103793344 | AX-170393516 | 0 | C | T | 0.237 | 1.72E-01 | 1.17E-01 | 2.71E+00 | 2.78E+00 | 1.40E-01 | Yes | Dab1 | intron_variant | MODIFIER |
| 4 | 103793565 | AX-170519106 | 0 | C | G | 0.237 | 1.72E-01 | 1.17E-01 | 2.71E+00 | 2.78E+00 | 1.40E-01 | Yes | Dab1 | intron_variant | MODIFIER |
| 4 | 103802307 | AX-170123259 | 0 | A | C | 0.237 | 1.72E-01 | 1.17E-01 | 2.71E+00 | 2.78E+00 | 1.40E-01 | Yes | Dab1 | intron_variant | MODIFIER |
| 4 | 103802370 | AX-170218228 | 0 | C | T | 0.237 | 1.72E-01 | 1.17E-01 | 2.71E+00 | 2.78E+00 | 1.40E-01 | Yes | Dab1 | intron_variant | MODIFIER |
| 4 | 103804551 | AX-170151151 | 0 | G | C | 0.237 | 1.72E-01 | 1.17E-01 | 2.71E+00 | 2.78E+00 | 1.40E-01 | Yes | Dab1 | intron_variant | MODIFIER |
| 4 | 103817023 | AX-170343995 | 0 | G | A | 0.237 | 1.72E-01 | 1.17E-01 | 2.71E+00 | 2.78E+00 | 1.40E-01 | Yes | Dab1 | intron_variant | MODIFIER |
| 4 | 103817132 | AX-170070203 | 0 | T | G | 0.237 | 1.72E-01 | 1.17E-01 | 2.71E+00 | 2.78E+00 | 1.40E-01 | Yes | Dab1 | intron_variant | MODIFIER |
| 4 | 103823655 | AX-97803686 | 0 | C | T | 0.237 | 1.72E-01 | 1.17E-01 | 2.71E+00 | 2.78E+00 | 1.40E-01 | Yes | Dab1 | intron_variant | MODIFIER |
| 4 | 103855781 | AX-170106781 | 0 | C | T | 0.237 | 1.72E-01 | 1.17E-01 | 2.71E+00 | 2.78E+00 | 1.40E-01 | Yes | Dab1 | intron_variant | MODIFIER |
| 4 | 103866854 | AX-170427770 | 0 | T | C | 0.237 | 1.72E-01 | 1.17E-01 | 2.71E+00 | 2.78E+00 | 1.40E-01 | Yes | Dab1 | intron_variant | MODIFIER |
| 4 | 103873428 | AX-168381388 | 0 | A | G | 0.237 | 1.72E-01 | 1.17E-01 | 2.71E+00 | 2.78E+00 | 1.40E-01 | Yes | Dab1 | intron_variant | MODIFIER |
| 4 | 103875085 | AX-170426699 | 0 | T | C | 0.237 | 1.72E-01 | 1.17E-01 | 2.71E+00 | 2.78E+00 | 1.40E-01 | Yes | Dab1 | intron_variant | MODIFIER |
| 4 | 103893304 | AX-170227020 | 0 | A | G | 0.237 | 1.72E-01 | 1.17E-01 | 2.71E+00 | 2.78E+00 | 1.40E-01 | Yes | Dab1 | intron_variant | MODIFIER |
| 4 | 103900945 | AX-170415836 | 0 | C | T | 0.237 | 1.72E-01 | 1.17E-01 | 2.71E+00 | 2.78E+00 | 1.40E-01 | Yes | Dab1 | intron_variant | MODIFIER |
| 4 | 103919150 | AX-170266734 | 0 | T | A | 0.237 | 1.72E-01 | 1.17E-01 | 2.71E+00 | 2.78E+00 | 1.40E-01 | Yes | Dab1 | intron_variant | MODIFIER |
| 4 | 103926078 | AX-170023578 | 0 | A | C | 0.237 | 1.72E-01 | 1.17E-01 | 2.71E+00 | 2.78E+00 | 1.40E-01 | Yes | Dab1 | intron_variant | MODIFIER |
| 4 | 103932308 | AX-170401991 | 0 | C | T | 0.237 | 1.72E-01 | 1.17E-01 | 2.71E+00 | 2.78E+00 | 1.40E-01 | Yes | Dab1 | intron_variant | MODIFIER |
| 4 | 103933027 | AX-170165962 | 0 | G | T | 0.237 | 1.72E-01 | 1.17E-01 | 2.71E+00 | 2.78E+00 | 1.40E-01 | Yes | Dab1 | intron_variant | MODIFIER |
| 4 | 103962063 | AX-170023575 | 0 | G | T | 0.237 | 1.72E-01 | 1.17E-01 | 2.71E+00 | 2.78E+00 | 1.40E-01 | Yes | Dab1 | intron_variant | MODIFIER |
| 4 | 103965387 | AX-170070206 | 0 | A | C | 0.237 | 1.72E-01 | 1.17E-01 | 2.71E+00 | 2.78E+00 | 1.40E-01 | Yes | Dab1 | intron_variant | MODIFIER |
| 4 | 103965901 | AX-170095081 | 0 | A | G | 0.237 | 1.72E-01 | 1.17E-01 | 2.71E+00 | 2.78E+00 | 1.40E-01 | Yes | Dab1 | intron_variant | MODIFIER |
| 4 | 104045768 | AX-97819483 | 0 | T | G | 0.237 | 1.72E-01 | 1.17E-01 | 2.71E+00 | 2.78E+00 | 1.40E-01 | Yes | Dab1 | intron_variant | MODIFIER |
| 4 | 104084549 | AX-170379818 | 0 | A | T | 0.237 | 1.72E-01 | 1.17E-01 | 2.71E+00 | 2.78E+00 | 1.40E-01 | Yes | Dab1 | intron_variant | MODIFIER |
| 4 | 104084646 | AX-170277958 | 0 | C | T | 0.237 | 1.72E-01 | 1.17E-01 | 2.71E+00 | 2.78E+00 | 1.40E-01 | Yes | Dab1 | intron_variant | MODIFIER |
| 4 | 104313333 | AX-97828295 | 0 | A | G | 0.237 | 1.72E-01 | 1.17E-01 | 2.71E+00 | 2.78E+00 | 1.40E-01 | Yes | Dab1 | intron_variant | MODIFIER |
| 4 | 104642009 | AX-97834857 | 0 | A | C | 0.237 | 1.72E-01 | 1.17E-01 | 2.71E+00 | 2.78E+00 | 1.40E-01 | Yes | Dab1 | intron_variant | MODIFIER |
| 4 | 104855389 | AX-170492762 | 0 | C | T | 0.237 | 1.72E-01 | 1.17E-01 | 2.71E+00 | 2.78E+00 | 1.40E-01 | Yes | C8a | intron_variant | MODIFIER |
| 4 | 104923403 | AX-170120632 | 0 | A | G | 0.237 | 1.72E-01 | 1.17E-01 | 2.71E+00 | 2.78E+00 | 1.40E-01 | Yes | Fyb2 | intron_variant | MODIFIER |
| 4 | 104923689 | AX-170104367 | 0 | A | G | 0.237 | 1.72E-01 | 1.17E-01 | 2.71E+00 | 2.78E+00 | 1.40E-01 | Yes | Fyb2 | intron_variant | MODIFIER |
| 4 | 104926060 | AX-170473218 | 0 | A | G | 0.237 | 1.72E-01 | 1.17E-01 | 2.71E+00 | 2.78E+00 | 1.40E-01 | Yes | Fyb2 | intron_variant | MODIFIER |
| 4 | 104929033 | AX-170253305 | 0 | A | C | 0.237 | 1.72E-01 | 1.17E-01 | 2.71E+00 | 2.78E+00 | 1.40E-01 | Yes | Fyb2 | intron_variant | MODIFIER |

| | | | | | | | | | | | | | | | |
|---|---|---|---|---|---|---|---|---|---|---|---|---|---|---|---|
| 4 | 104933263 | AX-170403306 | 0 | C | T | 0.237 | 1.72E-01 | 1.17E-01 | 2.71E+00 | 2.78E+00 | 1.40E-01 | Yes | Fyb2 | intron_variant | MODIFIER |
| 4 | 104940227 | AX-97844207 | 0 | C | T | 0.237 | 1.72E-01 | 1.17E-01 | 2.71E+00 | 2.78E+00 | 1.40E-01 | Yes | Fyb2 | intron_variant | MODIFIER |
| 4 | 104948175 | AX-170247750 | 0 | A | C | 0.237 | 1.72E-01 | 1.17E-01 | 2.71E+00 | 2.78E+00 | 1.40E-01 | Yes | Fyb2 | intron_variant | MODIFIER |
| 4 | 104952988 | AX-170329859 | 0 | T | A | 0.237 | 1.72E-01 | 1.17E-01 | 2.71E+00 | 2.78E+00 | 1.40E-01 | Yes | Fyb2 | intron_variant | MODIFIER |
| 4 | 104955811 | AX-170462071 | 0 | A | G | 0.237 | 1.72E-01 | 1.17E-01 | 2.71E+00 | 2.78E+00 | 1.40E-01 | Yes | Fyb2 | intron_variant | MODIFIER |
| 4 | 104969773 | AX-170289166 | 0 | C | T | 0.237 | 1.72E-01 | 1.17E-01 | 2.71E+00 | 2.78E+00 | 1.40E-01 | Yes | Fyb2 | intron_variant | MODIFIER |
| 4 | 104972707 | AX-170421866 | 0 | G | A | 0.237 | 1.72E-01 | 1.17E-01 | 2.71E+00 | 2.78E+00 | 1.40E-01 | Yes | Fyb2 | intron_variant | MODIFIER |
| 4 | 104973051 | AX-97819240 | 0 | G | A | 0.237 | 1.72E-01 | 1.17E-01 | 2.71E+00 | 2.78E+00 | 1.40E-01 | Yes | Fyb2 | intron_variant | MODIFIER |
| 4 | 104973627 | AX-170234374 | 0 | T | C | 0.237 | 1.72E-01 | 1.17E-01 | 2.71E+00 | 2.78E+00 | 1.40E-01 | Yes | Fyb2 | intron_variant | MODIFIER |
| 4 | 104991020 | AX-170349081 | 0 | G | C | 0.237 | 1.72E-01 | 1.17E-01 | 2.71E+00 | 2.78E+00 | 1.40E-01 | Yes | Fyb2 | intron_variant | MODIFIER |
| 4 | 104993743 | AX-170196228 | 0 | G | A | 0.237 | 1.72E-01 | 1.17E-01 | 2.71E+00 | 2.78E+00 | 1.40E-01 | Yes | Fyb2 | intron_variant | MODIFIER |
| 4 | 104993967 | AX-97804029 | 0 | T | C | 0.237 | 1.72E-01 | 1.17E-01 | 2.71E+00 | 2.78E+00 | 1.40E-01 | Yes | Fyb2 | intron_variant | MODIFIER |
| 4 | 104994165 | AX-170298184 | 0 | C | T | 0.237 | 1.72E-01 | 1.17E-01 | 2.71E+00 | 2.78E+00 | 1.40E-01 | Yes | Fyb2 | intron_variant | MODIFIER |
| 4 | 104996167 | AX-170252092 | 0 | T | C | 0.237 | 1.72E-01 | 1.17E-01 | 2.71E+00 | 2.78E+00 | 1.40E-01 | Yes | Fyb2 | intron_variant | MODIFIER |
| 14 | 123580987 | AX-170010895 | 0 | A | G | 0.333 | 1.11E-01 | 7.59E-02 | 2.72E+00 | 2.85E+00 | 1.40E-01 | no | Nalcn | intron_variant | MODIFIER |
| 5 | 54343020 | AX-170491204 | 0 | G | T | 0.491 | 1.42E-01 | 9.61E-02 | 2.51E+00 | 2.59E+00 | 1.40E-01 | no | | intergenic_variant | MODIFIER |
| 1 | 180404263 | AX-170280089 | 0 | G | A | 0.237 | 1.64E-01 | 1.11E-01 | 2.51E+00 | 2.57E+00 | 1.40E-01 | Yes | Itpkb | intron_variant | MODIFIER |
| 1 | 180434953 | AX-97793421 | 0 | G | T | 0.237 | 1.64E-01 | 1.11E-01 | 2.51E+00 | 2.57E+00 | 1.40E-01 | Yes | Stum | 3_prime_UTR_variant | MODIFIER |
| 1 | 180440616 | AX-170281536 | 0 | A | G | 0.237 | 1.64E-01 | 1.11E-01 | 2.51E+00 | 2.57E+00 | 1.40E-01 | Yes | Stum | intron_variant | MODIFIER |
| 1 | 180470532 | AX-170355748 | 0 | G | A | 0.237 | 1.64E-01 | 1.11E-01 | 2.51E+00 | 2.57E+00 | 1.40E-01 | Yes | Stum | intron_variant | MODIFIER |
| 1 | 180470785 | AX-170241853 | 0 | G | A | 0.237 | 1.64E-01 | 1.11E-01 | 2.51E+00 | 2.57E+00 | 1.40E-01 | Yes | Stum | intron_variant | MODIFIER |
| 1 | 180739006 | AX-97819709 | 0 | C | T | 0.237 | 1.64E-01 | 1.11E-01 | 2.51E+00 | 2.57E+00 | 1.40E-01 | Yes | Acbd3 | intron_variant | MODIFIER |
| 1 | 180801706 | AX-170372831 | 0 | T | A | 0.237 | 1.64E-01 | 1.11E-01 | 2.51E+00 | 2.57E+00 | 1.40E-01 | Yes | H3f3a | downstream_gene_variant | MODIFIER |
| 1 | 180911045 | AX-97827888 | 0 | C | T | 0.237 | 1.64E-01 | 1.11E-01 | 2.51E+00 | 2.57E+00 | 1.40E-01 | Yes | Pycr2 | downstream_gene_variant | MODIFIER |
| 1 | 180915472 | AX-170189476 | 0 | A | T | 0.237 | 1.64E-01 | 1.11E-01 | 2.51E+00 | 2.57E+00 | 1.40E-01 | Yes | | intergenic_variant | MODIFIER |
| 1 | 180925141 | AX-170375319 | 0 | C | G | 0.237 | 1.64E-01 | 1.11E-01 | 2.51E+00 | 2.57E+00 | 1.40E-01 | Yes | | intergenic_variant | MODIFIER |
| 1 | 180945296 | AX-170293936 | 0 | G | T | 0.237 | 1.64E-01 | 1.11E-01 | 2.51E+00 | 2.57E+00 | 1.40E-01 | Yes | Tmem63a | intron_variant | MODIFIER |
| 1 | 180946146 | AX-170513997 | 0 | T | C | 0.237 | 1.64E-01 | 1.11E-01 | 2.51E+00 | 2.57E+00 | 1.40E-01 | Yes | Tmem63a | intron_variant | MODIFIER |
| 1 | 180949688 | AX-170323079 | 0 | A | G | 0.237 | 1.64E-01 | 1.11E-01 | 2.51E+00 | 2.57E+00 | 1.40E-01 | Yes | Tmem63a | intron_variant | MODIFIER |
| 1 | 180956235 | AX-170240666 | 0 | C | T | 0.237 | 1.64E-01 | 1.11E-01 | 2.51E+00 | 2.57E+00 | 1.40E-01 | Yes | Tmem63a | intron_variant | MODIFIER |
| 1 | 180960724 | AX-97855663 | 0 | C | T | 0.237 | 1.64E-01 | 1.11E-01 | 2.51E+00 | 2.57E+00 | 1.40E-01 | Yes | Tmem63a | intron_variant | MODIFIER |
| 1 | 180961026 | AX-170232641 | 0 | C | T | 0.237 | 1.64E-01 | 1.11E-01 | 2.51E+00 | 2.57E+00 | 1.40E-01 | Yes | Tmem63a | intron_variant | MODIFIER |
| 1 | 180961377 | AX-170449143 | 0 | G | A | 0.237 | 1.64E-01 | 1.11E-01 | 2.51E+00 | 2.57E+00 | 1.40E-01 | Yes | Tmem63a | intron_variant | MODIFIER |
| 1 | 180982979 | AX-170333405 | 0 | C | G | 0.237 | 1.64E-01 | 1.11E-01 | 2.51E+00 | 2.57E+00 | 1.40E-01 | Yes | Ephx1 | intron_variant | MODIFIER |
| 1 | 181094354 | AX-170142642 | 0 | G | A | 0.237 | 1.64E-01 | 1.11E-01 | 2.51E+00 | 2.57E+00 | 1.40E-01 | Yes | Nvl | intron_variant | MODIFIER |
| 1 | 181095289 | AX-170043778 | 0 | T | G | 0.237 | 1.64E-01 | 1.11E-01 | 2.51E+00 | 2.57E+00 | 1.40E-01 | Yes | Nvl | intron_variant | MODIFIER |
| 1 | 181102408 | AX-170002424 | 0 | C | G | 0.237 | 1.64E-01 | 1.11E-01 | 2.51E+00 | 2.57E+00 | 1.40E-01 | Yes | Nvl | intron_variant | MODIFIER |
| 1 | 181102987 | AX-170002420 | 0 | T | C | 0.237 | 1.64E-01 | 1.11E-01 | 2.51E+00 | 2.57E+00 | 1.40E-01 | Yes | Nvl | intron_variant | MODIFIER |
| 1 | 181114227 | AX-170043779 | 0 | G | A | 0.237 | 1.64E-01 | 1.11E-01 | 2.51E+00 | 2.57E+00 | 1.40E-01 | Yes | Nvl | synonymous_variant | LOW |
| 1 | 181129982 | AX-97787834 | 0 | T | C | 0.237 | 1.64E-01 | 1.11E-01 | 2.51E+00 | 2.57E+00 | 1.40E-01 | Yes | Nvl | intron_variant | MODIFIER |
| 1 | 181168361 | AX-170091809 | 0 | G | A | 0.237 | 1.64E-01 | 1.11E-01 | 2.51E+00 | 2.57E+00 | 1.40E-01 | Yes | Cnih4 | downstream_gene_variant | MODIFIER |
| 1 | 181168393 | AX-170321854 | 0 | C | T | 0.237 | 1.64E-01 | 1.11E-01 | 2.51E+00 | 2.57E+00 | 1.40E-01 | Yes | Cnih4 | downstream_gene_variant | MODIFIER |
| 1 | 181168618 | AX-170355307 | 0 | C | T | 0.237 | 1.64E-01 | 1.11E-01 | 2.51E+00 | 2.57E+00 | 1.40E-01 | Yes | Cnih4 | downstream_gene_variant | MODIFIER |
| 1 | 181393308 | AX-170086708 | 0 | T | G | 0.237 | 1.64E-01 | 1.11E-01 | 2.51E+00 | 2.57E+00 | 1.40E-01 | Yes | Cnih3 | intron_variant | MODIFIER |
| 1 | 181406628 | AX-170122674 | 0 | C | G | 0.237 | 1.64E-01 | 1.11E-01 | 2.51E+00 | 2.57E+00 | 1.40E-01 | Yes | Cnih3 | intron_variant | MODIFIER |
| 1 | 181410728 | AX-97816547 | 0 | T | C | 0.237 | 1.64E-01 | 1.11E-01 | 2.51E+00 | 2.57E+00 | 1.40E-01 | Yes | Cnih3 | intron_variant | MODIFIER |
| 1 | 181429602 | AX-170264262 | 0 | G | C | 0.237 | 1.64E-01 | 1.11E-01 | 2.51E+00 | 2.57E+00 | 1.40E-01 | Yes | Cnih3 | intron_variant | MODIFIER |
| 9 | 117117901 | AX-170307670 | 0 | T | A | 0.123 | -1.26E-01 | 8.64E-02 | 2.79E+00 | 2.96E+00 | 1.40E-01 | no | Rbms3 | intron_variant | MODIFIER |
| 9 | 117563424 | AX-170033372 | 0 | A | T | 0.123 | -1.26E-01 | 8.64E-02 | 2.79E+00 | 2.96E+00 | 1.40E-01 | no | Rbms3 | intron_variant | MODIFIER |
| 9 | 117829439 | AX-170508597 | 0 | G | A | 0.123 | -1.26E-01 | 8.64E-02 | 2.79E+00 | 2.96E+00 | 1.40E-01 | no | Gm45881 | intron_variant,non_coding_transcript_variant | MODIFIER |
| 5 | 67677887 | AX-170375301 | 0 | C | T | 0.482 | 1.48E-01 | 1.01E-01 | 2.55E+00 | 2.62E+00 | 1.40E-01 | Yes | Atp8a1 | intron_variant | MODIFIER |
| 1 | 183946193 | AX-170322849 | 1 | T | C | 0.232 | 1.68E-01 | 1.14E-01 | 2.52E+00 | 2.59E+00 | 1.41E-01 | Yes | | intergenic_variant | MODIFIER |
| 1 | 184707293 | AX-170385661 | 1 | A | T | 0.232 | 1.68E-01 | 1.14E-01 | 2.52E+00 | 2.59E+00 | 1.41E-01 | Yes | | intergenic_variant | MODIFIER |
| 15 | 66444792 | AX-97794639 | 0 | T | C | 0.325 | -1.11E-01 | 7.15E-02 | 2.13E+00 | 2.21E+00 | 1.41E-01 | no | Lrrc6 | intron_variant | MODIFIER |

| | | | | | | | | | | | | | | | |
|---|---|---|---|---|---|---|---|---|---|---|---|---|---|---|---|
| 18 | 76325632 | AX-170319746 | 0 | A | T | 0.439 | 1.21E-01 | 8.24E-02 | 2.63E+00 | 2.71E+00 | 1.41E-01 | no | AC164009.1 | downstream_gene_variant | MODIFIER |
| 5 | 67646942 | AX-170128148 | 0 | T | C | 0.456 | 1.49E-01 | 9.96E-02 | 2.36E+00 | 2.43E+00 | 1.41E-01 | Yes | Atp8a1 | intron_variant | MODIFIER |
| 4 | 84361540 | AX-170097583 | 1 | T | C | 0.321 | 1.53E-01 | 1.05E-01 | 2.75E+00 | 2.85E+00 | 1.41E-01 | Yes | Bnc2 | intron_variant | MODIFIER |
| 5 | 129412357 | AX-170420284 | 1 | C | T | 0.152 | -1.39E-01 | 9.06E-02 | 2.20E+00 | 2.27E+00 | 1.41E-01 | no | | regulatory_region_variant | MODIFIER |
| 4 | 85475947 | AX-170069873 | 1 | T | C | 0.357 | 1.48E-01 | 1.00E-01 | 2.94E+00 | 3.05E+00 | 1.41E-01 | Yes | Sh3gl2 | intron_variant | MODIFIER |
| 5 | 57520009 | AX-170025026 | 0 | T | C | 0.456 | 1.46E-01 | 9.74E-02 | 2.35E+00 | 2.42E+00 | 1.41E-01 | Yes | | intergenic_variant | MODIFIER |
| 4 | 63314890 | AX-97792114 | 1 | G | T | 0.348 | 1.71E-01 | 1.16E-01 | 3.00E+00 | 3.08E+00 | 1.41E-01 | Yes | Col27a1 | intron_variant | MODIFIER |
| 13 | 5499259 | AX-170087351 | 0 | A | C | 0.202 | -1.85E-01 | 1.26E-01 | 2.63E+00 | 2.69E+00 | 1.41E-01 | Yes | Gm40660 | intron_variant,non_coding_transcript_variant | MODIFIER |
| 2 | 71771472 | AX-97838541 | 0 | G | A | 0.044 | -2.84E-01 | 1.94E-01 | 2.71E+00 | 2.81E+00 | 1.41E-01 | Yes | Itga6 | intron_variant,non_coding_transcript_variant | MODIFIER |
| 8 | 16544371 | AX-97780351 | 0 | A | T | 0.228 | -1.06E-01 | 7.27E-02 | 2.67E+00 | 2.77E+00 | 1.41E-01 | no | Csmd1 | intron_variant | MODIFIER |
| 5 | 64932949 | AX-170459888 | 0 | C | A | 0.482 | 1.51E-01 | 1.02E-01 | 2.46E+00 | 2.51E+00 | 1.42E-01 | Yes | Tlr1 | upstream_gene_variant | MODIFIER |
| 15 | 25596346 | AX-170265488 | 1 | T | A | 0.232 | -1.51E-01 | 1.03E-01 | 2.72E+00 | 2.80E+00 | 1.42E-01 | no | Gm36642 | upstream_gene_variant | MODIFIER |
| 4 | 107720113 | AX-97837387 | 1 | G | C | 0.259 | 1.65E-01 | 1.13E-01 | 2.70E+00 | 2.77E+00 | 1.42E-01 | Yes | 1700047F07Rik | downstream_gene_variant | MODIFIER |
| 1 | 135125054 | AX-170280240 | 1 | A | T | 0.545 | 1.22E-01 | 8.32E-02 | 2.71E+00 | 2.78E+00 | 1.42E-01 | no | Ptprv | downstream_gene_variant | MODIFIER |
| 18 | 77231472 | AX-170093962 | 0 | C | G | 0.439 | 1.26E-01 | 8.60E-02 | 2.51E+00 | 2.57E+00 | 1.42E-01 | no | St8sia5 | intron_variant | MODIFIER |
| 3 | 22974569 | AX-170066082 | 0 | T | C | 0.167 | -1.39E-01 | 9.52E-02 | 2.63E+00 | 2.73E+00 | 1.42E-01 | no | | intergenic_variant | MODIFIER |
| 4 | 109175823 | AX-170166886 | 1 | G | C | 0.25 | 1.66E-01 | 1.14E-01 | 2.61E+00 | 2.68E+00 | 1.42E-01 | Yes | Osbpl9 | intron_variant | MODIFIER |
| 18 | 76073509 | AX-170379415 | 1 | A | T | 0.402 | -1.24E-01 | 8.50E-02 | 2.84E+00 | 2.94E+00 | 1.43E-01 | no | Zbtb7c | intron_variant | MODIFIER |
| 18 | 68769651 | AX-170451276 | 0 | G | A | 0.175 | -1.73E-01 | 1.19E-01 | 2.68E+00 | 2.74E+00 | 1.43E-01 | Yes | | intergenic_variant | MODIFIER |
| 5 | 66349451 | AX-170221106 | 0 | C | A | 0.482 | 1.49E-01 | 1.02E-01 | 2.50E+00 | 2.56E+00 | 1.43E-01 | Yes | Apbb2 | intron_variant | MODIFIER |
| 5 | 66377765 | AX-170025181 | 0 | C | T | 0.482 | 1.49E-01 | 1.02E-01 | 2.50E+00 | 2.56E+00 | 1.43E-01 | Yes | Apbb2 | intron_variant | MODIFIER |
| 5 | 66384354 | AX-97844171 | 0 | C | T | 0.482 | 1.49E-01 | 1.02E-01 | 2.50E+00 | 2.56E+00 | 1.43E-01 | Yes | Apbb2 | intron_variant | MODIFIER |
| 5 | 66385031 | AX-170025182 | 0 | A | G | 0.482 | 1.49E-01 | 1.02E-01 | 2.50E+00 | 2.56E+00 | 1.43E-01 | Yes | Apbb2 | intron_variant | MODIFIER |
| 9 | 59332214 | AX-170131577 | 0 | C | T | 0.211 | -1.76E-01 | 1.19E-01 | 3.12E+00 | 3.24E+00 | 1.43E-01 | Yes | Bbs4 | intron_variant | MODIFIER |
| 1 | 135481252 | AX-170428892 | 1 | T | A | 0.527 | 1.19E-01 | 8.14E-02 | 2.65E+00 | 2.72E+00 | 1.43E-01 | no | Nav1 | intron_variant | MODIFIER |
| 3 | 38641145 | AX-170256621 | 1 | C | T | 0.205 | 1.37E-01 | 9.29E-02 | 3.11E+00 | 3.24E+00 | 1.43E-01 | no | | intergenic_variant | MODIFIER |
| 16 | 51980309 | AX-170111616 | 0 | T | A | 0.465 | -1.21E-01 | 8.29E-02 | 2.55E+00 | 2.59E+00 | 1.43E-01 | no | Gm29686 | downstream_gene_variant | MODIFIER |
| 5 | 134650833 | AX-97807602 | 1 | T | C | 0.161 | -1.28E-01 | 8.54E-02 | 2.30E+00 | 2.38E+00 | 1.43E-01 | no | | regulatory_region_variant | MODIFIER |
| 1 | 166156147 | AX-170267530 | 0 | T | C | 0.237 | 1.85E-01 | 1.27E-01 | 2.79E+00 | 2.86E+00 | 1.43E-01 | Yes | Gpa33 | downstream_gene_variant | MODIFIER |
| 16 | 51651523 | AX-170013038 | 1 | A | G | 0.464 | -1.21E-01 | 8.29E-02 | 2.55E+00 | 2.59E+00 | 1.43E-01 | no | | intergenic_variant | MODIFIER |
| 16 | 53871768 | AX-170281235 | 1 | C | T | 0.464 | -1.21E-01 | 8.29E-02 | 2.55E+00 | 2.59E+00 | 1.43E-01 | no | | intergenic_variant | MODIFIER |
| 4 | 104952484 | AX-170320101 | 1 | C | T | 0.241 | 1.68E-01 | 1.16E-01 | 2.66E+00 | 2.73E+00 | 1.43E-01 | Yes | Fyb2 | intron_variant | MODIFIER |
| 4 | 68734101 | AX-170089265 | 0 | A | G | 0.342 | 1.67E-01 | 1.14E-01 | 2.99E+00 | 3.06E+00 | 1.44E-01 | Yes | | intergenic_variant | MODIFIER |
| 5 | 50856172 | AX-170024927 | 1 | C | T | 0.509 | 1.50E-01 | 1.01E-01 | 2.37E+00 | 2.42E+00 | 1.44E-01 | Yes | | intergenic_variant | MODIFIER |
| 5 | 51357965 | AX-170283282 | 1 | T | A | 0.509 | 1.50E-01 | 1.01E-01 | 2.37E+00 | 2.42E+00 | 1.44E-01 | Yes | | intergenic_variant | MODIFIER |
| 1 | 135804531 | AX-170283338 | 0 | G | A | 0.553 | 1.18E-01 | 8.14E-02 | 2.69E+00 | 2.78E+00 | 1.44E-01 | no | Tnni1 | upstream_gene_variant | MODIFIER |
| 6 | 127087927 | AX-97834226 | 1 | G | C | 0.545 | 1.17E-01 | 8.06E-02 | 2.58E+00 | 2.66E+00 | 1.44E-01 | no | Tigar | missense_variant | MODERATE |
| 13 | 9282400 | AX-170036851 | 2 | C | T | 0.191 | -1.69E-01 | 1.16E-01 | 2.50E+00 | 2.57E+00 | 1.44E-01 | no | Dip2c | intron_variant | MODIFIER |
| 5 | 129601490 | AX-170527502 | 0 | A | G | 0.158 | -1.34E-01 | 8.87E-02 | 2.25E+00 | 2.33E+00 | 1.44E-01 | no | Mmp17 | intron_variant | MODIFIER |
| 1 | 131707464 | AX-170492995 | 1 | G | A | 0.545 | 1.19E-01 | 8.22E-02 | 2.76E+00 | 2.84E+00 | 1.44E-01 | no | Rab7b | intron_variant | MODIFIER |
| 16 | 63171116 | AX-170057630 | 0 | T | C | 0.474 | -1.28E-01 | 8.76E-02 | 2.59E+00 | 2.63E+00 | 1.44E-01 | no | | intergenic_variant | MODIFIER |
| 9 | 76792259 | AX-170082861 | 0 | T | C | 0.193 | -1.54E-01 | 1.06E-01 | 2.95E+00 | 3.09E+00 | 1.44E-01 | no | | intergenic_variant | MODIFIER |
| 16 | 40453423 | AX-170411220 | 0 | G | T | 0.5 | 1.18E-01 | 8.09E-02 | 2.61E+00 | 2.67E+00 | 1.44E-01 | no | Lsamp | intron_variant | MODIFIER |
| 5 | 4364133 | AX-97809264 | 0 | G | A | 0.43 | -9.63E-02 | 6.60E-02 | 2.51E+00 | 2.62E+00 | 1.45E-01 | no | | intergenic_variant | MODIFIER |
| 13 | 18370017 | AX-170050646 | 0 | G | A | 0.167 | -1.75E-01 | 1.19E-01 | 2.46E+00 | 2.53E+00 | 1.45E-01 | Yes | Pou6f2 | intron_variant | MODIFIER |
| 5 | 64378200 | AX-97797217 | 1 | T | C | 0.473 | 1.46E-01 | 9.94E-02 | 2.40E+00 | 2.46E+00 | 1.45E-01 | Yes | | intergenic_variant | MODIFIER |
| 13 | 18418772 | AX-97788370 | 1 | G | A | 0.179 | -1.82E-01 | 1.23E-01 | 2.40E+00 | 2.44E+00 | 1.46E-01 | Yes | | intergenic_variant | MODIFIER |
| 1 | 191812874 | AX-170399187 | 0 | G | A | 0.281 | 1.37E-01 | 9.11E-02 | 2.26E+00 | 2.34E+00 | 1.46E-01 | no | | regulatory_region_variant | MODIFIER |
| 3 | 7909562 | AX-170287416 | 0 | G | A | 0.412 | -8.85E-02 | 6.12E-02 | 2.86E+00 | 3.02E+00 | 1.46E-01 | no | | intergenic_variant | MODIFIER |
| 5 | 106311967 | AX-97853706 | 0 | T | C | 0.456 | 1.61E-01 | 1.11E-01 | 2.81E+00 | 2.87E+00 | 1.46E-01 | Yes | | intergenic_variant | MODIFIER |
| 5 | 113081588 | AX-170114744 | 0 | A | G | 0.456 | 1.61E-01 | 1.11E-01 | 2.81E+00 | 2.87E+00 | 1.46E-01 | Yes | 2900026A02Rik | downstream_gene_variant | MODIFIER |
| 1 | 135804988 | AX-97818425 | 1 | A | G | 0.527 | 1.17E-01 | 8.11E-02 | 2.73E+00 | 2.81E+00 | 1.46E-01 | no | Tnni1 | upstream_gene_variant | MODIFIER |
| 4 | 63270895 | AX-170239552 | 0 | C | T | 0.342 | 1.59E-01 | 1.10E-01 | 2.87E+00 | 2.95E+00 | 1.46E-01 | Yes | Col27a1 | intron_variant | MODIFIER |
| 18 | 58351823 | AX-170060966 | 0 | A | G | 0.316 | -9.41E-02 | 6.45E-02 | 3.05E+00 | 3.17E+00 | 1.46E-01 | no | | intergenic_variant | MODIFIER |

| | | | | | | | | | | | | | | | |
|---|---|---|---|---|---|---|---|---|---|---|---|---|---|---|---|
| 18 | 79308325 | AX-170016057 | 0 | A | C | 0.456 | 1.26E-01 | 8.40E-02 | 2.27E+00 | 2.34E+00 | 1.46E-01 | no | AC140980.1 | upstream_gene_variant | MODIFIER |
| 5 | 56210813 | AX-170281495 | 0 | G | A | 0.491 | 1.48E-01 | 1.00E-01 | 2.39E+00 | 2.45E+00 | 1.46E-01 | Yes | | intergenic_variant | MODIFIER |
| 4 | 81661683 | AX-170253379 | 0 | C | T | 0.421 | -1.19E-01 | 8.21E-02 | 2.58E+00 | 2.64E+00 | 1.46E-01 | no | | intergenic_variant | MODIFIER |
| 4 | 54221451 | AX-97790972 | 0 | G | A | 0.368 | -1.33E-01 | 9.22E-02 | 2.83E+00 | 2.94E+00 | 1.47E-01 | no | | intergenic_variant | MODIFIER |
| 19 | 61093750 | AX-170535732 | 0 | C | T | 0.377 | -8.62E-02 | 5.89E-02 | 2.42E+00 | 2.51E+00 | 1.47E-01 | no | Grk5 | 3_prime_UTR_variant | MODIFIER |
| 9 | 61195926 | AX-170306300 | 0 | T | A | 0.193 | -1.95E-01 | 1.32E-01 | 3.19E+00 | 3.29E+00 | 1.47E-01 | Yes | | intergenic_variant | MODIFIER |
| 9 | 64338713 | AX-170232038 | 0 | A | G | 0.193 | -1.95E-01 | 1.32E-01 | 3.19E+00 | 3.29E+00 | 1.47E-01 | Yes | Dis3l | intron_variant | MODIFIER |
| 9 | 69714855 | AX-170189625 | 0 | T | A | 0.193 | -1.95E-01 | 1.32E-01 | 3.19E+00 | 3.29E+00 | 1.47E-01 | Yes | | regulatory_region_variant | MODIFIER |
| 9 | 78783870 | AX-170341544 | 0 | A | G | 0.386 | -9.76E-02 | 6.61E-02 | 3.19E+00 | 3.29E+00 | 1.47E-01 | no | | regulatory_region_variant | MODIFIER |
| 11 | 22452761 | AX-170130769 | 0 | G | A | 0.044 | -2.72E-01 | 1.89E-01 | 2.78E+00 | 2.94E+00 | 1.47E-01 | Yes | | intergenic_variant | MODIFIER |
| 19 | 52586866 | AX-97850728 | 0 | T | C | 0.491 | 9.14E-02 | 6.33E-02 | 2.60E+00 | 2.69E+00 | 1.47E-01 | no | | intergenic_variant | MODIFIER |
| 19 | 52851646 | AX-170016843 | 0 | A | G | 0.491 | 9.14E-02 | 6.33E-02 | 2.60E+00 | 2.69E+00 | 1.47E-01 | no | | intergenic_variant | MODIFIER |
| 19 | 53149636 | AX-97845878 | 0 | A | G | 0.491 | 9.14E-02 | 6.33E-02 | 2.60E+00 | 2.69E+00 | 1.47E-01 | no | Add3 | intron_variant | MODIFIER |
| 4 | 107491701 | AX-170070280 | 0 | C | G | 0.263 | 1.64E-01 | 1.13E-01 | 2.65E+00 | 2.70E+00 | 1.47E-01 | Yes | Glis1 | intron_variant | MODIFIER |
| 4 | 107501413 | AX-170337162 | 0 | G | A | 0.263 | 1.64E-01 | 1.13E-01 | 2.65E+00 | 2.70E+00 | 1.47E-01 | Yes | Glis1 | intron_variant | MODIFIER |
| 4 | 107503156 | AX-170449121 | 0 | T | C | 0.263 | 1.64E-01 | 1.13E-01 | 2.65E+00 | 2.70E+00 | 1.47E-01 | Yes | Glis1 | intron_variant | MODIFIER |
| 4 | 107503224 | AX-170485847 | 0 | G | A | 0.263 | 1.64E-01 | 1.13E-01 | 2.65E+00 | 2.70E+00 | 1.47E-01 | Yes | Glis1 | intron_variant | MODIFIER |
| 4 | 107505332 | AX-170379871 | 0 | G | A | 0.263 | 1.64E-01 | 1.13E-01 | 2.65E+00 | 2.70E+00 | 1.47E-01 | Yes | Glis1 | intron_variant | MODIFIER |
| 4 | 107536269 | AX-170353332 | 0 | T | C | 0.263 | 1.64E-01 | 1.13E-01 | 2.65E+00 | 2.70E+00 | 1.47E-01 | Yes | Glis1 | intron_variant | MODIFIER |
| 4 | 107538470 | AX-97837465 | 0 | G | A | 0.263 | 1.64E-01 | 1.13E-01 | 2.65E+00 | 2.70E+00 | 1.47E-01 | Yes | Glis1 | intron_variant | MODIFIER |
| 4 | 107549760 | AX-170156360 | 0 | A | G | 0.263 | 1.64E-01 | 1.13E-01 | 2.65E+00 | 2.70E+00 | 1.47E-01 | Yes | Glis1 | intron_variant | MODIFIER |
| 4 | 107550480 | AX-170408066 | 0 | A | G | 0.263 | 1.64E-01 | 1.13E-01 | 2.65E+00 | 2.70E+00 | 1.47E-01 | Yes | Glis1 | intron_variant | MODIFIER |
| 4 | 107558419 | AX-170228007 | 0 | A | C | 0.263 | 1.64E-01 | 1.13E-01 | 2.65E+00 | 2.70E+00 | 1.47E-01 | Yes | Glis1 | intron_variant | MODIFIER |
| 4 | 107559463 | AX-170373075 | 0 | C | T | 0.263 | 1.64E-01 | 1.13E-01 | 2.65E+00 | 2.70E+00 | 1.47E-01 | Yes | Glis1 | intron_variant | MODIFIER |
| 4 | 107563444 | AX-170346501 | 0 | C | G | 0.263 | 1.64E-01 | 1.13E-01 | 2.65E+00 | 2.70E+00 | 1.47E-01 | Yes | Glis1 | intron_variant | MODIFIER |
| 4 | 107566476 | AX-170366497 | 0 | G | A | 0.263 | 1.64E-01 | 1.13E-01 | 2.65E+00 | 2.70E+00 | 1.47E-01 | Yes | Glis1 | intron_variant | MODIFIER |
| 4 | 107581319 | AX-170475304 | 0 | A | G | 0.263 | 1.64E-01 | 1.13E-01 | 2.65E+00 | 2.70E+00 | 1.47E-01 | Yes | Glis1 | intron_variant | MODIFIER |
| 4 | 107632435 | AX-170262374 | 0 | C | T | 0.263 | 1.64E-01 | 1.13E-01 | 2.65E+00 | 2.70E+00 | 1.47E-01 | Yes | Glis1 | splice_region_variant,intron_variant | LOW |
| 4 | 107637715 | AX-170379091 | 0 | C | T | 0.263 | 1.64E-01 | 1.13E-01 | 2.65E+00 | 2.70E+00 | 1.47E-01 | Yes | Glis1 | downstream_gene_variant | MODIFIER |
| 4 | 107679950 | AX-97793605 | 0 | C | T | 0.263 | 1.64E-01 | 1.13E-01 | 2.65E+00 | 2.70E+00 | 1.47E-01 | Yes | Dmrtb1 | intron_variant | MODIFIER |
| 4 | 107725119 | AX-170357634 | 0 | C | G | 0.263 | 1.64E-01 | 1.13E-01 | 2.65E+00 | 2.70E+00 | 1.47E-01 | Yes | 1700047F07Rik | intron_variant,non_coding_transcript_variant | MODIFIER |
| 4 | 107735217 | AX-170023631 | 0 | C | G | 0.263 | 1.64E-01 | 1.13E-01 | 2.65E+00 | 2.70E+00 | 1.47E-01 | Yes | 1700047F07Rik | intron_variant,non_coding_transcript_variant | MODIFIER |
| 4 | 107735531 | AX-97788490 | 0 | C | A | 0.263 | 1.64E-01 | 1.13E-01 | 2.65E+00 | 2.70E+00 | 1.47E-01 | Yes | 1700047F07Rik | intron_variant,non_coding_transcript_variant | MODIFIER |
| 4 | 107746835 | AX-170213113 | 0 | G | A | 0.263 | 1.64E-01 | 1.13E-01 | 2.65E+00 | 2.70E+00 | 1.47E-01 | Yes | 1700047F07Rik | intron_variant,non_coding_transcript_variant | MODIFIER |
| 4 | 107761835 | AX-170023633 | 0 | A | G | 0.263 | 1.64E-01 | 1.13E-01 | 2.65E+00 | 2.70E+00 | 1.47E-01 | Yes | 1700047F07Rik | downstream_gene_variant | MODIFIER |
| 4 | 107771355 | AX-170478498 | 0 | C | T | 0.263 | 1.64E-01 | 1.13E-01 | 2.65E+00 | 2.70E+00 | 1.47E-01 | Yes | Gm12907 | downstream_gene_variant | MODIFIER |
| 4 | 107773860 | AX-170402629 | 0 | A | T | 0.263 | 1.64E-01 | 1.13E-01 | 2.65E+00 | 2.70E+00 | 1.47E-01 | Yes | Gm12907 | downstream_gene_variant | MODIFIER |
| 4 | 107784256 | AX-170070288 | 0 | T | C | 0.263 | 1.64E-01 | 1.13E-01 | 2.65E+00 | 2.70E+00 | 1.47E-01 | Yes | 4933424M12Rik | upstream_gene_variant | MODIFIER |
| 4 | 107831358 | AX-170023627 | 0 | G | A | 0.263 | 1.64E-01 | 1.13E-01 | 2.65E+00 | 2.70E+00 | 1.47E-01 | Yes | Lrp8os1 | upstream_gene_variant | MODIFIER |
| 4 | 107834688 | AX-170507324 | 0 | G | T | 0.263 | 1.64E-01 | 1.13E-01 | 2.65E+00 | 2.70E+00 | 1.47E-01 | Yes | Lrp8os1 | upstream_gene_variant | MODIFIER |
| 4 | 107997371 | AX-97823397 | 0 | A | G | 0.263 | 1.64E-01 | 1.13E-01 | 2.65E+00 | 2.70E+00 | 1.47E-01 | Yes | Slc1a7 | intron_variant | MODIFIER |
| 4 | 108254521 | AX-170192530 | 0 | T | C | 0.263 | 1.64E-01 | 1.13E-01 | 2.65E+00 | 2.70E+00 | 1.47E-01 | Yes | Zyg11b | intron_variant | MODIFIER |
| 4 | 108265837 | AX-170223387 | 0 | A | G | 0.263 | 1.64E-01 | 1.13E-01 | 2.65E+00 | 2.70E+00 | 1.47E-01 | Yes | Zyg11b | missense_variant | MODERATE |
| 4 | 108488314 | AX-170023634 | 0 | T | C | 0.263 | 1.64E-01 | 1.13E-01 | 2.65E+00 | 2.70E+00 | 1.47E-01 | Yes | Tut4 | intron_variant | MODIFIER |
| 4 | 108603272 | AX-170380937 | 0 | A | C | 0.263 | 1.64E-01 | 1.13E-01 | 2.65E+00 | 2.70E+00 | 1.47E-01 | Yes | Orc1 | intron_variant | MODIFIER |
| 4 | 108614058 | AX-170070298 | 0 | C | T | 0.263 | 1.64E-01 | 1.13E-01 | 2.65E+00 | 2.70E+00 | 1.47E-01 | Yes | Orc1 | intron_variant | MODIFIER |
| 4 | 108903436 | AX-170293801 | 0 | A | T | 0.263 | 1.64E-01 | 1.13E-01 | 2.65E+00 | 2.70E+00 | 1.47E-01 | Yes | Rab3b | intron_variant | MODIFIER |
| 4 | 108907150 | AX-170416005 | 0 | G | A | 0.263 | 1.64E-01 | 1.13E-01 | 2.65E+00 | 2.70E+00 | 1.47E-01 | Yes | Rab3b | intron_variant | MODIFIER |
| 4 | 108914226 | AX-170177378 | 0 | T | G | 0.263 | 1.64E-01 | 1.13E-01 | 2.65E+00 | 2.70E+00 | 1.47E-01 | Yes | Rab3b | intron_variant | MODIFIER |
| 4 | 108923636 | AX-170343132 | 0 | C | T | 0.263 | 1.64E-01 | 1.13E-01 | 2.65E+00 | 2.70E+00 | 1.47E-01 | Yes | Rab3b | intron_variant | MODIFIER |
| 4 | 108960023 | AX-170070303 | 0 | G | A | 0.263 | 1.64E-01 | 1.13E-01 | 2.65E+00 | 2.70E+00 | 1.47E-01 | Yes | 8030443G20Rik | non_coding_transcript_exon_variant | MODIFIER |
| 4 | 108960050 | AX-170095092 | 0 | G | A | 0.263 | 1.64E-01 | 1.13E-01 | 2.65E+00 | 2.70E+00 | 1.47E-01 | Yes | 8030443G20Rik | non_coding_transcript_exon_variant | MODIFIER |
| 4 | 109090279 | AX-170391826 | 0 | C | T | 0.263 | 1.64E-01 | 1.13E-01 | 2.65E+00 | 2.70E+00 | 1.47E-01 | Yes | Osbpl9 | intron_variant | MODIFIER |
| 1 | 135748196 | AX-170254404 | 1 | T | C | 0.536 | 1.17E-01 | 8.11E-02 | 2.74E+00 | 2.81E+00 | 1.47E-01 | no | Csrp1 | intron_variant | MODIFIER |

| | | | | | | | | | | | | | | | |
|---|---|---|---|---|---|---|---|---|---|---|---|---|---|---|---|
| 16 | 65697529 | AX-97833336 | 1 | A | G | 0.5 | 1.22E-01 | 8.38E-02 | 2.51E+00 | 2.56E+00 | 1.47E-01 | no | | intergenic_variant | MODIFIER |
| 1 | 181548704 | AX-170274537 | 1 | A | G | 0.232 | 1.62E-01 | 1.12E-01 | 2.48E+00 | 2.54E+00 | 1.48E-01 | Yes | | intergenic_variant | MODIFIER |
| 4 | 84344782 | AX-97783980 | 1 | T | C | 0.33 | 1.65E-01 | 1.14E-01 | 2.85E+00 | 2.92E+00 | 1.48E-01 | Yes | Bnc2 | intron_variant | MODIFIER |
| 12 | 104927288 | AX-170007453 | 0 | G | A | 0.132 | -2.39E-01 | 1.66E-01 | 2.85E+00 | 2.95E+00 | 1.48E-01 | Yes | Syne3 | downstream_gene_variant | MODIFIER |
| 18 | 59293271 | AX-170118074 | 0 | T | A | 0.184 | -1.64E-01 | 1.14E-01 | 2.82E+00 | 2.90E+00 | 1.48E-01 | no | Chsy3 | intron_variant | MODIFIER |
| 18 | 61504029 | AX-170139704 | 0 | T | C | 0.184 | -1.64E-01 | 1.14E-01 | 2.82E+00 | 2.90E+00 | 1.48E-01 | no | Arhgef37 | intron_variant | MODIFIER |
| 5 | 64488870 | AX-170114484 | 2 | T | G | 0.473 | 1.46E-01 | 9.97E-02 | 2.40E+00 | 2.46E+00 | 1.48E-01 | Yes | C030018K13Rik | downstream_gene_variant | MODIFIER |
| 11 | 30169042 | AX-170400444 | 0 | T | C | 0.088 | 2.38E-01 | 1.65E-01 | 2.83E+00 | 2.95E+00 | 1.48E-01 | Yes | Sptbn1 | intron_variant | MODIFIER |
| 4 | 95901731 | AX-170292295 | 0 | T | C | 0.246 | 1.64E-01 | 1.13E-01 | 2.49E+00 | 2.55E+00 | 1.48E-01 | Yes | Fggy | intron_variant | MODIFIER |
| 5 | 129189817 | AX-170515608 | 1 | C | A | 0.152 | -1.35E-01 | 8.91E-02 | 2.16E+00 | 2.24E+00 | 1.48E-01 | no | Adgrd1 | intron_variant | MODIFIER |
| 18 | 72331037 | AX-97835816 | 0 | G | A | 0.184 | -1.45E-01 | 1.01E-01 | 2.85E+00 | 2.98E+00 | 1.48E-01 | no | Dcc | intron_variant | MODIFIER |
| 15 | 71438716 | AX-170429006 | 0 | C | A | 0.088 | -1.68E-01 | 1.05E-01 | 1.97E+00 | 2.05E+00 | 1.48E-01 | no | Fam135b | 3_prime_UTR_variant | MODIFIER |
| 9 | 97317143 | AX-170477083 | 0 | A | G | 0.14 | -1.53E-01 | 1.06E-01 | 2.56E+00 | 2.64E+00 | 1.49E-01 | no | | intergenic_variant | MODIFIER |
| 18 | 69707185 | AX-168387931 | 1 | G | A | 0.268 | -1.34E-01 | 9.29E-02 | 2.91E+00 | 3.03E+00 | 1.49E-01 | no | | intergenic_variant | MODIFIER |
| 5 | 51452281 | AX-170509277 | 1 | G | A | 0.491 | 1.53E-01 | 1.04E-01 | 2.38E+00 | 2.42E+00 | 1.49E-01 | Yes | Ppargc1a | downstream_gene_variant | MODIFIER |
| 3 | 7376018 | AX-170019851 | 0 | T | G | 0.386 | -9.14E-02 | 6.37E-02 | 2.91E+00 | 3.08E+00 | 1.50E-01 | no | Pkia | intron_variant | MODIFIER |
| 3 | 7385070 | AX-170019855 | 0 | G | A | 0.386 | -9.14E-02 | 6.37E-02 | 2.91E+00 | 3.08E+00 | 1.50E-01 | no | Pkia | intron_variant | MODIFIER |
| 3 | 7385289 | AX-170019846 | 0 | G | A | 0.386 | -9.14E-02 | 6.37E-02 | 2.91E+00 | 3.08E+00 | 1.50E-01 | no | Pkia | intron_variant | MODIFIER |
| 3 | 7468198 | AX-97798055 | 0 | T | G | 0.386 | -9.14E-02 | 6.37E-02 | 2.91E+00 | 3.08E+00 | 1.50E-01 | no | | intergenic_variant | MODIFIER |
| 3 | 7514467 | AX-170065855 | 0 | T | A | 0.386 | -9.14E-02 | 6.37E-02 | 2.91E+00 | 3.08E+00 | 1.50E-01 | no | Zc2hc1a | intron_variant | MODIFIER |
| 3 | 7522214 | AX-97792304 | 0 | G | A | 0.386 | -9.14E-02 | 6.37E-02 | 2.91E+00 | 3.08E+00 | 1.50E-01 | no | Zc2hc1a | intron_variant | MODIFIER |
| 3 | 7530827 | AX-170150335 | 0 | A | G | 0.386 | -9.14E-02 | 6.37E-02 | 2.91E+00 | 3.08E+00 | 1.50E-01 | no | Zc2hc1a | intron_variant | MODIFIER |
| 3 | 7539486 | AX-170201308 | 0 | G | C | 0.386 | -9.14E-02 | 6.37E-02 | 2.91E+00 | 3.08E+00 | 1.50E-01 | no | Zc2hc1a | intron_variant | MODIFIER |
| 3 | 7551616 | AX-170370653 | 0 | T | C | 0.386 | -9.14E-02 | 6.37E-02 | 2.91E+00 | 3.08E+00 | 1.50E-01 | no | Zc2hc1a | 3_prime_UTR_variant | MODIFIER |
| 3 | 7572141 | AX-170065854 | 0 | A | T | 0.386 | -9.14E-02 | 6.37E-02 | 2.91E+00 | 3.08E+00 | 1.50E-01 | no | Il7 | 3_prime_UTR_variant | MODIFIER |
| 3 | 7575007 | AX-170116702 | 0 | A | T | 0.386 | -9.14E-02 | 6.37E-02 | 2.91E+00 | 3.08E+00 | 1.50E-01 | no | Il7 | intron_variant | MODIFIER |
| 3 | 7587608 | AX-97795660 | 0 | G | A | 0.386 | -9.14E-02 | 6.37E-02 | 2.91E+00 | 3.08E+00 | 1.50E-01 | no | Il7 | intron_variant | MODIFIER |
| 3 | 7612801 | AX-170185628 | 0 | A | G | 0.386 | -9.14E-02 | 6.37E-02 | 2.91E+00 | 3.08E+00 | 1.50E-01 | no | Gm16685 | non_coding_transcript_exon_variant | MODIFIER |
| 3 | 7614643 | AX-170461266 | 0 | T | C | 0.386 | -9.14E-02 | 6.37E-02 | 2.91E+00 | 3.08E+00 | 1.50E-01 | no | Gm16685 | intron_variant,non_coding_transcript_variant | MODIFIER |
| 3 | 7719774 | AX-170409815 | 0 | C | T | 0.386 | -9.14E-02 | 6.37E-02 | 2.91E+00 | 3.08E+00 | 1.50E-01 | no | | intergenic_variant | MODIFIER |
| 3 | 7827070 | AX-170437465 | 0 | A | C | 0.386 | -9.14E-02 | 6.37E-02 | 2.91E+00 | 3.08E+00 | 1.50E-01 | no | | intergenic_variant | MODIFIER |
| 3 | 7874393 | AX-170019862 | 0 | A | G | 0.386 | -9.14E-02 | 6.37E-02 | 2.91E+00 | 3.08E+00 | 1.50E-01 | no | | intergenic_variant | MODIFIER |
| 3 | 7954276 | AX-170159711 | 0 | C | T | 0.386 | -9.14E-02 | 6.37E-02 | 2.91E+00 | 3.08E+00 | 1.50E-01 | no | 1700010I02Rik | intron_variant,non_coding_transcript_variant | MODIFIER |
| 3 | 8023301 | AX-170315387 | 0 | C | T | 0.386 | -9.14E-02 | 6.37E-02 | 2.91E+00 | 3.08E+00 | 1.50E-01 | no | | intergenic_variant | MODIFIER |
| 3 | 8271807 | AX-170186670 | 0 | G | A | 0.386 | -9.14E-02 | 6.37E-02 | 2.91E+00 | 3.08E+00 | 1.50E-01 | no | | intergenic_variant | MODIFIER |
| 3 | 8554492 | AX-170175428 | 0 | T | G | 0.386 | -9.14E-02 | 6.37E-02 | 2.91E+00 | 3.08E+00 | 1.50E-01 | no | Stmn2 | intron_variant | MODIFIER |
| 5 | 52577913 | AX-170455379 | 1 | A | T | 0.473 | 1.44E-01 | 9.82E-02 | 2.34E+00 | 2.40E+00 | 1.50E-01 | Yes | Gm5866 | upstream_gene_variant | MODIFIER |
| 3 | 26341604 | AX-170224958 | 0 | T | C | 0.149 | -1.33E-01 | 9.30E-02 | 2.66E+00 | 2.77E+00 | 1.50E-01 | no | | regulatory_region_variant | MODIFIER |
| 13 | 24964973 | AX-170460684 | 0 | G | A | 0.158 | -1.70E-01 | 1.17E-01 | 2.41E+00 | 2.48E+00 | 1.50E-01 | no | Gpld1 | intron_variant | MODIFIER |
| 1 | 192197605 | AX-170470731 | 1 | G | T | 0.268 | 1.34E-01 | 9.17E-02 | 2.35E+00 | 2.43E+00 | 1.50E-01 | no | Kcnh1 | intron_variant | MODIFIER |
| 4 | 84757128 | AX-170023283 | 1 | G | A | 0.348 | 1.64E-01 | 1.14E-01 | 2.93E+00 | 3.01E+00 | 1.51E-01 | Yes | | intergenic_variant | MODIFIER |
| 18 | 77208079 | AX-170384578 | 1 | C | T | 0.446 | 1.23E-01 | 8.60E-02 | 2.55E+00 | 2.62E+00 | 1.51E-01 | no | St8sia5 | intron_variant | MODIFIER |
| 4 | 107764208 | AX-97848746 | 1 | A | G | 0.259 | 1.61E-01 | 1.13E-01 | 2.66E+00 | 2.72E+00 | 1.51E-01 | Yes | Gm12907 | upstream_gene_variant | MODIFIER |
| 1 | 123734767 | AX-170292143 | 1 | G | A | 0.562 | 1.15E-01 | 8.05E-02 | 2.63E+00 | 2.69E+00 | 1.51E-01 | no | Dpp10 | intron_variant | MODIFIER |
| 5 | 129878347 | AX-170162687 | 0 | A | C | 0.158 | -1.33E-01 | 8.76E-02 | 2.14E+00 | 2.21E+00 | 1.51E-01 | no | Phkg1 | intron_variant | MODIFIER |
| 1 | 127154464 | AX-97839816 | 1 | C | A | 0.562 | 1.22E-01 | 8.55E-02 | 2.72E+00 | 2.80E+00 | 1.51E-01 | no | | intergenic_variant | MODIFIER |
| 5 | 60400323 | AX-170349797 | 1 | G | A | 0.464 | 1.43E-01 | 9.98E-02 | 2.51E+00 | 2.58E+00 | 1.51E-01 | no | | intergenic_variant | MODIFIER |
| 1 | 126657091 | AX-97814691 | 0 | G | A | 0.57 | 1.15E-01 | 8.07E-02 | 2.63E+00 | 2.69E+00 | 1.51E-01 | no | Nckap5 | intron_variant | MODIFIER |
| 1 | 193601723 | AX-170484483 | 1 | T | C | 0.277 | 1.36E-01 | 9.22E-02 | 2.30E+00 | 2.38E+00 | 1.51E-01 | no | | intergenic_variant | MODIFIER |
| 4 | 48154729 | AX-170497168 | 1 | G | A | 0.357 | -1.35E-01 | 9.44E-02 | 2.96E+00 | 3.08E+00 | 1.51E-01 | no | Stx17 | intron_variant | MODIFIER |
| 13 | 17390127 | AX-170350765 | 0 | T | A | 0.184 | -1.77E-01 | 1.20E-01 | 2.29E+00 | 2.34E+00 | 1.51E-01 | Yes | Sugct | intron_variant | MODIFIER |
| 14 | 124331816 | AX-170054655 | 0 | A | G | 0.333 | 1.09E-01 | 7.63E-02 | 2.73E+00 | 2.86E+00 | 1.52E-01 | no | Fgf14 | intron_variant | MODIFIER |
| 1 | 75585891 | AX-170225003 | 0 | C | T | 0.132 | -2.29E-01 | 1.61E-01 | 2.80E+00 | 2.92E+00 | 1.52E-01 | Yes | | intergenic_variant | MODIFIER |
| 18 | 79719039 | AX-170198550 | 1 | A | C | 0.464 | 1.19E-01 | 8.25E-02 | 2.41E+00 | 2.48E+00 | 1.52E-01 | no | | intergenic_variant | MODIFIER |

| | | | | | | | | | | | | | | | |
|---|---|---|---|---|---|---|---|---|---|---|---|---|---|---|---|
| 4 | 79075052 | AX-97850346 | 0 | C | T | 0.439 | -1.18E-01 | 8.23E-02 | 2.56E+00 | 2.63E+00 | 1.52E-01 | no | | intergenic_variant | MODIFIER |
| 5 | 67974947 | AX-170186387 | 0 | A | C | 0.474 | 1.42E-01 | 9.89E-02 | 2.48E+00 | 2.55E+00 | 1.52E-01 | no | Gm5108 | intron_variant | MODIFIER |
| 1 | 170767851 | AX-170104415 | 2 | G | A | 0.209 | 1.82E-01 | 1.28E-01 | 2.91E+00 | 3.02E+00 | 1.52E-01 | Yes | Atf6 | intron_variant | MODIFIER |
| 18 | 79551436 | AX-170061432 | 0 | T | G | 0.465 | 1.21E-01 | 8.33E-02 | 2.40E+00 | 2.47E+00 | 1.52E-01 | no | Gm25824 | downstream_gene_variant | MODIFIER |
| 18 | 79556406 | AX-170061431 | 0 | T | C | 0.465 | 1.21E-01 | 8.33E-02 | 2.40E+00 | 2.47E+00 | 1.52E-01 | no | Gm25824 | upstream_gene_variant | MODIFIER |
| 18 | 79559198 | AX-170257026 | 0 | C | G | 0.465 | 1.21E-01 | 8.33E-02 | 2.40E+00 | 2.47E+00 | 1.52E-01 | no | | intergenic_variant | MODIFIER |
| 18 | 79566446 | AX-170161217 | 0 | C | G | 0.465 | 1.21E-01 | 8.33E-02 | 2.40E+00 | 2.47E+00 | 1.52E-01 | no | | intergenic_variant | MODIFIER |
| 18 | 79636258 | AX-170534978 | 0 | T | C | 0.465 | 1.21E-01 | 8.33E-02 | 2.40E+00 | 2.47E+00 | 1.52E-01 | no | | intergenic_variant | MODIFIER |
| 18 | 79670442 | AX-170126297 | 0 | C | T | 0.465 | 1.21E-01 | 8.33E-02 | 2.40E+00 | 2.47E+00 | 1.52E-01 | no | AC122051.2 | downstream_gene_variant | MODIFIER |
| 18 | 79672663 | AX-170262019 | 0 | C | T | 0.465 | 1.21E-01 | 8.33E-02 | 2.40E+00 | 2.47E+00 | 1.52E-01 | no | AC122051.2 | downstream_gene_variant | MODIFIER |
| 18 | 79681039 | AX-170349430 | 0 | C | T | 0.465 | 1.21E-01 | 8.33E-02 | 2.40E+00 | 2.47E+00 | 1.52E-01 | no | | intergenic_variant | MODIFIER |
| 18 | 79712877 | AX-97848997 | 0 | A | G | 0.465 | 1.21E-01 | 8.33E-02 | 2.40E+00 | 2.47E+00 | 1.52E-01 | no | | intergenic_variant | MODIFIER |
| 18 | 79718873 | AX-97845044 | 0 | G | A | 0.465 | 1.21E-01 | 8.33E-02 | 2.40E+00 | 2.47E+00 | 1.52E-01 | no | | intergenic_variant | MODIFIER |
| 18 | 79765768 | AX-170159622 | 0 | A | G | 0.465 | 1.21E-01 | 8.33E-02 | 2.40E+00 | 2.47E+00 | 1.52E-01 | no | | intergenic_variant | MODIFIER |
| 18 | 79770894 | AX-170131273 | 0 | G | A | 0.465 | 1.21E-01 | 8.33E-02 | 2.40E+00 | 2.47E+00 | 1.52E-01 | no | | intergenic_variant | MODIFIER |
| 18 | 79798589 | AX-170419442 | 0 | T | A | 0.465 | 1.21E-01 | 8.33E-02 | 2.40E+00 | 2.47E+00 | 1.52E-01 | no | | intergenic_variant | MODIFIER |
| 18 | 79799895 | AX-170088383 | 0 | A | G | 0.465 | 1.21E-01 | 8.33E-02 | 2.40E+00 | 2.47E+00 | 1.52E-01 | no | | intergenic_variant | MODIFIER |
| 18 | 79805031 | AX-170016067 | 0 | G | T | 0.465 | 1.21E-01 | 8.33E-02 | 2.40E+00 | 2.47E+00 | 1.52E-01 | no | | intergenic_variant | MODIFIER |
| 18 | 79845005 | AX-170061442 | 0 | G | A | 0.465 | 1.21E-01 | 8.33E-02 | 2.40E+00 | 2.47E+00 | 1.52E-01 | no | | intergenic_variant | MODIFIER |
| 18 | 79845807 | AX-170088384 | 0 | G | A | 0.465 | 1.21E-01 | 8.33E-02 | 2.40E+00 | 2.47E+00 | 1.52E-01 | no | | intergenic_variant | MODIFIER |
| 18 | 79867874 | AX-170016065 | 0 | C | G | 0.465 | 1.21E-01 | 8.33E-02 | 2.40E+00 | 2.47E+00 | 1.52E-01 | no | | intergenic_variant | MODIFIER |
| 18 | 79879204 | AX-170016070 | 0 | T | C | 0.465 | 1.21E-01 | 8.33E-02 | 2.40E+00 | 2.47E+00 | 1.52E-01 | no | Gm26977 | upstream_gene_variant | MODIFIER |
| 18 | 79887907 | AX-170016068 | 0 | A | G | 0.465 | 1.21E-01 | 8.33E-02 | 2.40E+00 | 2.47E+00 | 1.52E-01 | no | | intergenic_variant | MODIFIER |
| 18 | 79936476 | AX-97786973 | 0 | T | C | 0.465 | 1.21E-01 | 8.33E-02 | 2.40E+00 | 2.47E+00 | 1.52E-01 | no | AC139296.1 | downstream_gene_variant | MODIFIER |
| 18 | 79950355 | AX-97802582 | 0 | G | A | 0.465 | 1.21E-01 | 8.33E-02 | 2.40E+00 | 2.47E+00 | 1.52E-01 | no | | regulatory_region_variant | MODIFIER |
| 18 | 79965147 | AX-170016072 | 0 | A | G | 0.465 | 1.21E-01 | 8.33E-02 | 2.40E+00 | 2.47E+00 | 1.52E-01 | no | | intergenic_variant | MODIFIER |
| 18 | 79998321 | AX-97799538 | 0 | T | C | 0.465 | 1.21E-01 | 8.33E-02 | 2.40E+00 | 2.47E+00 | 1.52E-01 | no | | intergenic_variant | MODIFIER |
| 18 | 80006869 | AX-170436782 | 0 | A | G | 0.465 | 1.21E-01 | 8.33E-02 | 2.40E+00 | 2.47E+00 | 1.52E-01 | no | | intergenic_variant | MODIFIER |
| 18 | 80007627 | AX-170404195 | 0 | A | G | 0.465 | 1.21E-01 | 8.33E-02 | 2.40E+00 | 2.47E+00 | 1.52E-01 | no | AC127331.1 | upstream_gene_variant | MODIFIER |
| 18 | 80060524 | AX-170321187 | 0 | T | C | 0.465 | 1.21E-01 | 8.33E-02 | 2.40E+00 | 2.47E+00 | 1.52E-01 | no | Pard6g | intron_variant | MODIFIER |
| 18 | 80063542 | AX-170453198 | 0 | T | C | 0.465 | 1.21E-01 | 8.33E-02 | 2.40E+00 | 2.47E+00 | 1.52E-01 | no | Pard6g | intron_variant | MODIFIER |
| 18 | 80073421 | AX-170468370 | 0 | T | C | 0.465 | 1.21E-01 | 8.33E-02 | 2.40E+00 | 2.47E+00 | 1.52E-01 | no | Pard6g | intron_variant | MODIFIER |
| 18 | 80073795 | AX-97846669 | 0 | T | G | 0.465 | 1.21E-01 | 8.33E-02 | 2.40E+00 | 2.47E+00 | 1.52E-01 | no | Pard6g | intron_variant | MODIFIER |
| 18 | 80083245 | AX-170354201 | 0 | T | C | 0.465 | 1.21E-01 | 8.33E-02 | 2.40E+00 | 2.47E+00 | 1.52E-01 | no | Pard6g | intron_variant | MODIFIER |
| 18 | 80380286 | AX-170166451 | 0 | T | C | 0.465 | 1.21E-01 | 8.33E-02 | 2.40E+00 | 2.47E+00 | 1.52E-01 | no | | intergenic_variant | MODIFIER |
| 18 | 80380828 | AX-170479031 | 0 | A | C | 0.465 | 1.21E-01 | 8.33E-02 | 2.40E+00 | 2.47E+00 | 1.52E-01 | no | | intergenic_variant | MODIFIER |
| 18 | 80445704 | AX-170276755 | 0 | A | C | 0.465 | 1.21E-01 | 8.33E-02 | 2.40E+00 | 2.47E+00 | 1.52E-01 | no | Ctdp1 | intron_variant | MODIFIER |
| 18 | 80543069 | AX-170098070 | 0 | C | T | 0.465 | 1.21E-01 | 8.33E-02 | 2.40E+00 | 2.47E+00 | 1.52E-01 | no | AC127256.2 | intron_variant,non_coding_transcript_variant | MODIFIER |
| 18 | 80543401 | AX-170016074 | 0 | A | G | 0.465 | 1.21E-01 | 8.33E-02 | 2.40E+00 | 2.47E+00 | 1.52E-01 | no | AC127256.2 | non_coding_transcript_exon_variant | MODIFIER |
| 18 | 80553141 | AX-97847119 | 0 | A | G | 0.465 | 1.21E-01 | 8.33E-02 | 2.40E+00 | 2.47E+00 | 1.52E-01 | no | | intergenic_variant | MODIFIER |
| 18 | 80576683 | AX-97811885 | 0 | T | G | 0.465 | 1.21E-01 | 8.33E-02 | 2.40E+00 | 2.47E+00 | 1.52E-01 | no | AC149054.1 | intron_variant,non_coding_transcript_variant | MODIFIER |
| 18 | 80594163 | AX-97789604 | 0 | T | C | 0.465 | 1.21E-01 | 8.33E-02 | 2.40E+00 | 2.47E+00 | 1.52E-01 | no | AC149054.1 | intron_variant,non_coding_transcript_variant | MODIFIER |
| 18 | 80599921 | AX-170189619 | 0 | C | A | 0.465 | 1.21E-01 | 8.33E-02 | 2.40E+00 | 2.47E+00 | 1.52E-01 | no | AC149054.1 | intron_variant,non_coding_transcript_variant | MODIFIER |
| 18 | 80614953 | AX-170016080 | 0 | C | T | 0.465 | 1.21E-01 | 8.33E-02 | 2.40E+00 | 2.47E+00 | 1.52E-01 | no | Nfatc1 | intron_variant | MODIFIER |
| 8 | 25538213 | AX-170030374 | 1 | T | C | 0.339 | -9.22E-02 | 6.44E-02 | 2.53E+00 | 2.62E+00 | 1.52E-01 | no | Fgfr1 | intron_variant | MODIFIER |
| 10 | 102744247 | AX-170457175 | 0 | T | A | 0.289 | -1.20E-01 | 8.44E-02 | 2.68E+00 | 2.76E+00 | 1.52E-01 | no | | intergenic_variant | MODIFIER |
| 18 | 61809419 | AX-170371625 | 0 | C | T | 0.404 | 1.28E-01 | 8.91E-02 | 2.46E+00 | 2.52E+00 | 1.52E-01 | no | Ablim3 | intron_variant | MODIFIER |
| 8 | 90513298 | AX-170031246 | 1 | T | C | 0.188 | -1.27E-01 | 8.91E-02 | 2.62E+00 | 2.75E+00 | 1.52E-01 | no | | regulatory_region_variant | MODIFIER |
| 4 | 85614919 | AX-170150207 | 1 | T | A | 0.339 | 1.53E-01 | 1.07E-01 | 3.04E+00 | 3.16E+00 | 1.52E-01 | Yes | Sh3gl2 | intron_variant | MODIFIER |
| 4 | 100826566 | AX-170351371 | 2 | A | G | 0.427 | -1.09E-01 | 7.62E-02 | 2.58E+00 | 2.66E+00 | 1.53E-01 | no | Cachd1 | intron_variant | MODIFIER |
| 3 | 38234211 | AX-170134829 | 0 | C | G | 0.211 | 1.25E-01 | 8.74E-02 | 3.01E+00 | 3.12E+00 | 1.53E-01 | no | | regulatory_region_variant | MODIFIER |
| 1 | 135004921 | AX-170199684 | 1 | C | T | 0.536 | 1.17E-01 | 8.23E-02 | 2.72E+00 | 2.79E+00 | 1.53E-01 | no | Lgr6 | intron_variant | MODIFIER |
| 5 | 13977457 | AX-170404886 | 1 | A | G | 0.339 | 1.19E-01 | 8.40E-02 | 2.84E+00 | 2.98E+00 | 1.53E-01 | no | | intergenic_variant | MODIFIER |
| 5 | 10397123 | AX-97788336 | 0 | T | G | 0.342 | 1.17E-01 | 8.22E-02 | 2.71E+00 | 2.85E+00 | 1.53E-01 | no | | intergenic_variant | MODIFIER |

| | | | | | | | | | | | | | | | |
|---|---|---|---|---|---|---|---|---|---|---|---|---|---|---|---|
| 5 | 10400298 | AX-170458592 | 0 | T | A | 0.342 | 1.17E-01 | 8.22E-02 | 2.71E+00 | 2.85E+00 | 1.53E-01 | no | | intergenic_variant | MODIFIER |
| 5 | 10416664 | AX-170024403 | 0 | G | A | 0.342 | 1.17E-01 | 8.22E-02 | 2.71E+00 | 2.85E+00 | 1.53E-01 | no | | intergenic_variant | MODIFIER |
| 5 | 10418144 | AX-170140318 | 0 | A | G | 0.342 | 1.17E-01 | 8.22E-02 | 2.71E+00 | 2.85E+00 | 1.53E-01 | no | | intergenic_variant | MODIFIER |
| 5 | 10493650 | AX-97828758 | 0 | T | C | 0.342 | 1.17E-01 | 8.22E-02 | 2.71E+00 | 2.85E+00 | 1.53E-01 | no | | intergenic_variant | MODIFIER |
| 5 | 10633672 | AX-97816382 | 0 | C | T | 0.342 | 1.17E-01 | 8.22E-02 | 2.71E+00 | 2.85E+00 | 1.53E-01 | no | | intergenic_variant | MODIFIER |
| 5 | 11948420 | AX-170472404 | 0 | A | G | 0.342 | 1.17E-01 | 8.22E-02 | 2.71E+00 | 2.85E+00 | 1.53E-01 | no | | intergenic_variant | MODIFIER |
| 5 | 36913715 | AX-170147245 | 1 | A | G | 0.33 | 1.60E-01 | 1.12E-01 | 3.03E+00 | 3.15E+00 | 1.53E-01 | Yes | Ppp2r2c | intron_variant | MODIFIER |
| 3 | 7422362 | AX-170499111 | 0 | G | T | 0.404 | -8.95E-02 | 6.29E-02 | 2.91E+00 | 3.07E+00 | 1.53E-01 | no | Pkia | intron_variant | MODIFIER |
| 3 | 7506129 | AX-97815155 | 0 | T | C | 0.404 | -8.95E-02 | 6.29E-02 | 2.91E+00 | 3.07E+00 | 1.53E-01 | no | Zc2hc1a | intron_variant | MODIFIER |
| 3 | 7571198 | AX-170158153 | 0 | T | G | 0.404 | -8.95E-02 | 6.29E-02 | 2.91E+00 | 3.07E+00 | 1.53E-01 | no | Il7 | downstream_gene_variant | MODIFIER |
| 3 | 7571387 | AX-170436163 | 0 | C | G | 0.404 | -8.95E-02 | 6.29E-02 | 2.91E+00 | 3.07E+00 | 1.53E-01 | no | Il7 | downstream_gene_variant | MODIFIER |
| 3 | 7575079 | AX-170526734 | 0 | A | G | 0.404 | -8.95E-02 | 6.29E-02 | 2.91E+00 | 3.07E+00 | 1.53E-01 | no | Il7 | intron_variant | MODIFIER |
| 3 | 7618664 | AX-97827940 | 0 | C | T | 0.404 | -8.95E-02 | 6.29E-02 | 2.91E+00 | 3.07E+00 | 1.53E-01 | no | Gm16685 | intron_variant,non_coding_transcript_variant | MODIFIER |
| 3 | 7630063 | AX-170339270 | 0 | C | T | 0.404 | -8.95E-02 | 6.29E-02 | 2.91E+00 | 3.07E+00 | 1.53E-01 | no | Gm16685 | intron_variant,non_coding_transcript_variant | MODIFIER |
| 3 | 7679213 | AX-170493962 | 0 | T | C | 0.404 | -8.95E-02 | 6.29E-02 | 2.91E+00 | 3.07E+00 | 1.53E-01 | no | Gm16685 | intron_variant,non_coding_transcript_variant | MODIFIER |
| 3 | 7679245 | AX-170514177 | 0 | G | A | 0.404 | -8.95E-02 | 6.29E-02 | 2.91E+00 | 3.07E+00 | 1.53E-01 | no | Gm16685 | intron_variant,non_coding_transcript_variant | MODIFIER |
| 3 | 7679863 | AX-170454817 | 0 | G | T | 0.404 | -8.95E-02 | 6.29E-02 | 2.91E+00 | 3.07E+00 | 1.53E-01 | no | Gm16685 | intron_variant,non_coding_transcript_variant | MODIFIER |
| 3 | 7685093 | AX-170493157 | 0 | T | A | 0.404 | -8.95E-02 | 6.29E-02 | 2.91E+00 | 3.07E+00 | 1.53E-01 | no | Gm16685 | intron_variant,non_coding_transcript_variant | MODIFIER |
| 3 | 7698167 | AX-97787286 | 0 | A | G | 0.404 | -8.95E-02 | 6.29E-02 | 2.91E+00 | 3.07E+00 | 1.53E-01 | no | | intergenic_variant | MODIFIER |
| 3 | 7746965 | AX-97856849 | 0 | T | G | 0.404 | -8.95E-02 | 6.29E-02 | 2.91E+00 | 3.07E+00 | 1.53E-01 | no | | intergenic_variant | MODIFIER |
| 3 | 7767213 | AX-170094514 | 0 | T | C | 0.404 | -8.95E-02 | 6.29E-02 | 2.91E+00 | 3.07E+00 | 1.53E-01 | no | | intergenic_variant | MODIFIER |
| 3 | 7873707 | AX-170019861 | 0 | A | G | 0.404 | -8.95E-02 | 6.29E-02 | 2.91E+00 | 3.07E+00 | 1.53E-01 | no | | intergenic_variant | MODIFIER |
| 3 | 7874045 | AX-170246194 | 0 | C | T | 0.404 | -8.95E-02 | 6.29E-02 | 2.91E+00 | 3.07E+00 | 1.53E-01 | no | | intergenic_variant | MODIFIER |
| 3 | 7874206 | AX-170094515 | 0 | C | T | 0.404 | -8.95E-02 | 6.29E-02 | 2.91E+00 | 3.07E+00 | 1.53E-01 | no | | intergenic_variant | MODIFIER |
| 3 | 7874698 | AX-170127408 | 0 | C | T | 0.404 | -8.95E-02 | 6.29E-02 | 2.91E+00 | 3.07E+00 | 1.53E-01 | no | | intergenic_variant | MODIFIER |
| 3 | 7875073 | AX-170226871 | 0 | A | C | 0.404 | -8.95E-02 | 6.29E-02 | 2.91E+00 | 3.07E+00 | 1.53E-01 | no | | intergenic_variant | MODIFIER |
| 3 | 7908291 | AX-97837929 | 0 | C | T | 0.404 | -8.95E-02 | 6.29E-02 | 2.91E+00 | 3.07E+00 | 1.53E-01 | no | | intergenic_variant | MODIFIER |
| 3 | 7910290 | AX-170186925 | 0 | G | C | 0.404 | -8.95E-02 | 6.29E-02 | 2.91E+00 | 3.07E+00 | 1.53E-01 | no | | intergenic_variant | MODIFIER |
| 3 | 7917857 | AX-170019864 | 0 | G | A | 0.404 | -8.95E-02 | 6.29E-02 | 2.91E+00 | 3.07E+00 | 1.53E-01 | no | | intergenic_variant | MODIFIER |
| 3 | 7924880 | AX-170329155 | 0 | A | G | 0.404 | -8.95E-02 | 6.29E-02 | 2.91E+00 | 3.07E+00 | 1.53E-01 | no | 1700010I02Rik | downstream_gene_variant | MODIFIER |
| 3 | 7981851 | AX-97846534 | 0 | G | A | 0.404 | -8.95E-02 | 6.29E-02 | 2.91E+00 | 3.07E+00 | 1.53E-01 | no | | intergenic_variant | MODIFIER |
| 3 | 7986227 | AX-97786352 | 0 | A | G | 0.404 | -8.95E-02 | 6.29E-02 | 2.91E+00 | 3.07E+00 | 1.53E-01 | no | | intergenic_variant | MODIFIER |
| 3 | 8014270 | AX-97846960 | 0 | C | T | 0.404 | -8.95E-02 | 6.29E-02 | 2.91E+00 | 3.07E+00 | 1.53E-01 | no | | intergenic_variant | MODIFIER |
| 4 | 107954423 | AX-170023630 | 1 | G | A | 0.268 | 1.59E-01 | 1.11E-01 | 2.60E+00 | 2.66E+00 | 1.53E-01 | Yes | 4930407G08Rik | intron_variant,non_coding_transcript_variant | MODIFIER |
| 1 | 13636526 | AX-170286530 | 1 | A | G | 0.357 | 1.06E-01 | 7.45E-02 | 2.61E+00 | 2.69E+00 | 1.53E-01 | no | Lactb2 | intron_variant | MODIFIER |
| 3 | 7798154 | AX-170019863 | 2 | A | T | 0.373 | -9.37E-02 | 6.60E-02 | 2.87E+00 | 3.03E+00 | 1.53E-01 | no | | intergenic_variant | MODIFIER |
| 4 | 99901052 | AX-170228463 | 0 | T | C | 0.421 | -1.11E-01 | 7.83E-02 | 2.58E+00 | 2.66E+00 | 1.53E-01 | no | Efcab7 | intron_variant | MODIFIER |
| 4 | 100553654 | AX-170205965 | 0 | A | G | 0.421 | -1.11E-01 | 7.83E-02 | 2.58E+00 | 2.66E+00 | 1.53E-01 | no | Ube2u | downstream_gene_variant | MODIFIER |
| 4 | 107409300 | AX-170330677 | 1 | T | C | 0.268 | 1.57E-01 | 1.10E-01 | 2.61E+00 | 2.68E+00 | 1.54E-01 | Yes | Ndc1 | intron_variant,NMD_transcript_variant | MODIFIER |
| 1 | 100646229 | AX-170329495 | 0 | C | T | 0.167 | 2.09E-01 | 1.48E-01 | 2.64E+00 | 2.76E+00 | 1.54E-01 | Yes | Gm18185 | downstream_gene_variant | MODIFIER |
| 8 | 61086371 | AX-170311880 | 1 | G | A | 0.143 | -1.94E-01 | 1.34E-01 | 2.39E+00 | 2.47E+00 | 1.54E-01 | Yes | Nek1 | intron_variant | MODIFIER |
| 3 | 8014511 | AX-170106918 | 0 | C | T | 0.412 | -8.90E-02 | 6.26E-02 | 2.94E+00 | 3.10E+00 | 1.54E-01 | no | | intergenic_variant | MODIFIER |
| 12 | 116677303 | AX-170300210 | 1 | G | A | 0.125 | -2.20E-01 | 1.55E-01 | 2.91E+00 | 3.06E+00 | 1.54E-01 | Yes | Ptprn2 | intron_variant | MODIFIER |
| 3 | 38876277 | AX-170359271 | 1 | A | G | 0.205 | 1.34E-01 | 9.33E-02 | 3.07E+00 | 3.19E+00 | 1.54E-01 | no | C230034O21Rik | intron_variant,non_coding_transcript_variant | MODIFIER |
| 5 | 56654595 | AX-97795719 | 0 | C | T | 0.465 | 1.47E-01 | 1.02E-01 | 2.41E+00 | 2.46E+00 | 1.54E-01 | Yes | | intergenic_variant | MODIFIER |
| 4 | 19562501 | AX-97855546 | 1 | G | A | 0.054 | -2.05E-01 | 1.45E-01 | 2.90E+00 | 3.07E+00 | 1.54E-01 | Yes | Cpne3 | intron_variant | MODIFIER |
| 9 | 88701290 | AX-170032933 | 1 | A | G | 0.143 | -1.69E-01 | 1.19E-01 | 2.78E+00 | 2.88E+00 | 1.54E-01 | no | Mthfsl | intron_variant | MODIFIER |
| 1 | 122929271 | AX-170522784 | 0 | T | G | 0.561 | 1.15E-01 | 8.07E-02 | 2.62E+00 | 2.69E+00 | 1.54E-01 | no | | intergenic_variant | MODIFIER |
| 3 | 26667433 | AX-170257832 | 1 | G | A | 0.134 | -1.46E-01 | 1.03E-01 | 2.67E+00 | 2.76E+00 | 1.54E-01 | no | Spata16 | synonymous_variant | LOW |
| 7 | 98065098 | AX-170473281 | 0 | G | T | 0.132 | -2.51E-01 | 1.75E-01 | 3.08E+00 | 3.23E+00 | 1.54E-01 | Yes | Myo7a | intron_variant | MODIFIER |
| 5 | 129005652 | AX-97809548 | 0 | G | A | 0.158 | -1.27E-01 | 8.66E-02 | 2.26E+00 | 2.35E+00 | 1.54E-01 | no | Stx2 | intron_variant | MODIFIER |
| 1 | 20442651 | AX-170100063 | 0 | G | A | 0.404 | 1.13E-01 | 7.98E-02 | 2.66E+00 | 2.73E+00 | 1.54E-01 | no | Pkhd1 | intron_variant | MODIFIER |
| 5 | 49601133 | AX-170072036 | 0 | A | T | 0.482 | 1.52E-01 | 1.02E-01 | 2.19E+00 | 2.23E+00 | 1.54E-01 | Yes | | intergenic_variant | MODIFIER |
| 16 | 71088973 | AX-170452560 | 0 | A | T | 0.491 | 1.08E-01 | 7.54E-02 | 2.44E+00 | 2.51E+00 | 1.54E-01 | no | | intergenic_variant | MODIFIER |

| | | | | | | | | | | | | | | | |
|---|---|---|---|---|---|---|---|---|---|---|---|---|---|---|---|
| 16 | 71957862 | AX-170500511 | 0 | T | C | 0.491 | 1.08E-01 | 7.54E-02 | 2.44E+00 | 2.51E+00 | 1.54E-01 | no | | intergenic_variant | MODIFIER |
| 5 | 13468933 | AX-170365383 | 0 | G | A | 0.342 | 1.19E-01 | 8.42E-02 | 2.84E+00 | 2.98E+00 | 1.54E-01 | no | Sema3a | intron_variant | MODIFIER |
| 5 | 13470565 | AX-170264357 | 0 | C | T | 0.342 | 1.19E-01 | 8.42E-02 | 2.84E+00 | 2.98E+00 | 1.54E-01 | no | Sema3a | intron_variant | MODIFIER |
| 5 | 13668898 | AX-170170754 | 0 | T | C | 0.342 | 1.19E-01 | 8.42E-02 | 2.84E+00 | 2.98E+00 | 1.54E-01 | no | | intergenic_variant | MODIFIER |
| 5 | 13685831 | AX-170214684 | 0 | G | C | 0.342 | 1.19E-01 | 8.42E-02 | 2.84E+00 | 2.98E+00 | 1.54E-01 | no | | intergenic_variant | MODIFIER |
| 5 | 13686400 | AX-170497824 | 0 | A | G | 0.342 | 1.19E-01 | 8.42E-02 | 2.84E+00 | 2.98E+00 | 1.54E-01 | no | | intergenic_variant | MODIFIER |
| 5 | 13690140 | AX-170024432 | 0 | A | C | 0.342 | 1.19E-01 | 8.42E-02 | 2.84E+00 | 2.98E+00 | 1.54E-01 | no | | intergenic_variant | MODIFIER |
| 5 | 13709339 | AX-97798674 | 0 | T | G | 0.342 | 1.19E-01 | 8.42E-02 | 2.84E+00 | 2.98E+00 | 1.54E-01 | no | | intergenic_variant | MODIFIER |
| 5 | 13711740 | AX-170486744 | 0 | G | T | 0.342 | 1.19E-01 | 8.42E-02 | 2.84E+00 | 2.98E+00 | 1.54E-01 | no | | intergenic_variant | MODIFIER |
| 5 | 13731612 | AX-97823953 | 0 | A | C | 0.342 | 1.19E-01 | 8.42E-02 | 2.84E+00 | 2.98E+00 | 1.54E-01 | no | | intergenic_variant | MODIFIER |
| 5 | 13735873 | AX-170210699 | 0 | A | G | 0.342 | 1.19E-01 | 8.42E-02 | 2.84E+00 | 2.98E+00 | 1.54E-01 | no | | intergenic_variant | MODIFIER |
| 5 | 13740407 | AX-170071344 | 0 | C | G | 0.342 | 1.19E-01 | 8.42E-02 | 2.84E+00 | 2.98E+00 | 1.54E-01 | no | | intergenic_variant | MODIFIER |
| 5 | 13756688 | AX-97837010 | 0 | A | G | 0.342 | 1.19E-01 | 8.42E-02 | 2.84E+00 | 2.98E+00 | 1.54E-01 | no | 4930568A13Rik | intron_variant,non_coding_transcript_variant | MODIFIER |
| 5 | 13818988 | AX-170095225 | 0 | G | C | 0.342 | 1.19E-01 | 8.42E-02 | 2.84E+00 | 2.98E+00 | 1.54E-01 | no | Gm6650 | downstream_gene_variant | MODIFIER |
| 5 | 13822885 | AX-170292456 | 0 | C | T | 0.342 | 1.19E-01 | 8.42E-02 | 2.84E+00 | 2.98E+00 | 1.54E-01 | no | | intergenic_variant | MODIFIER |
| 5 | 13823359 | AX-170178073 | 0 | C | T | 0.342 | 1.19E-01 | 8.42E-02 | 2.84E+00 | 2.98E+00 | 1.54E-01 | no | | intergenic_variant | MODIFIER |
| 5 | 13826079 | AX-170353768 | 0 | G | A | 0.342 | 1.19E-01 | 8.42E-02 | 2.84E+00 | 2.98E+00 | 1.54E-01 | no | | intergenic_variant | MODIFIER |
| 5 | 13879429 | AX-97799035 | 0 | A | G | 0.342 | 1.19E-01 | 8.42E-02 | 2.84E+00 | 2.98E+00 | 1.54E-01 | no | Dnd1-ps | non_coding_transcript_exon_variant | MODIFIER |
| 5 | 13977433 | AX-170207306 | 0 | A | C | 0.342 | 1.19E-01 | 8.42E-02 | 2.84E+00 | 2.98E+00 | 1.54E-01 | no | | intergenic_variant | MODIFIER |
| 5 | 13985871 | AX-170217851 | 0 | A | G | 0.342 | 1.19E-01 | 8.42E-02 | 2.84E+00 | 2.98E+00 | 1.54E-01 | no | | intergenic_variant | MODIFIER |
| 5 | 14015082 | AX-170220457 | 0 | G | A | 0.342 | 1.19E-01 | 8.42E-02 | 2.84E+00 | 2.98E+00 | 1.54E-01 | no | | intergenic_variant | MODIFIER |
| 5 | 14020569 | AX-170243560 | 0 | A | G | 0.342 | 1.19E-01 | 8.42E-02 | 2.84E+00 | 2.98E+00 | 1.54E-01 | no | Sema3e | upstream_gene_variant | MODIFIER |
| 5 | 14021387 | AX-170071350 | 0 | C | T | 0.342 | 1.19E-01 | 8.42E-02 | 2.84E+00 | 2.98E+00 | 1.54E-01 | no | Sema3e | upstream_gene_variant | MODIFIER |
| 5 | 14021503 | AX-170071348 | 0 | T | C | 0.342 | 1.19E-01 | 8.42E-02 | 2.84E+00 | 2.98E+00 | 1.54E-01 | no | Sema3e | upstream_gene_variant | MODIFIER |
| 5 | 14022063 | AX-97803204 | 0 | G | A | 0.342 | 1.19E-01 | 8.42E-02 | 2.84E+00 | 2.98E+00 | 1.54E-01 | no | Sema3e | upstream_gene_variant | MODIFIER |
| 5 | 14060220 | AX-97844777 | 0 | C | T | 0.342 | 1.19E-01 | 8.42E-02 | 2.84E+00 | 2.98E+00 | 1.54E-01 | no | Sema3e | intron_variant | MODIFIER |
| 8 | 22433881 | AX-170030342 | 0 | T | C | 0.167 | -1.60E-01 | 1.12E-01 | 3.06E+00 | 3.21E+00 | 1.54E-01 | no | Mrps31 | downstream_gene_variant | MODIFIER |
| 8 | 61440618 | AX-170267127 | 1 | G | T | 0.134 | -2.11E-01 | 1.47E-01 | 2.46E+00 | 2.53E+00 | 1.54E-01 | Yes | | intergenic_variant | MODIFIER |
| 16 | 70832144 | AX-170013350 | 0 | T | C | 0.491 | 1.09E-01 | 7.62E-02 | 2.49E+00 | 2.56E+00 | 1.54E-01 | no | | intergenic_variant | MODIFIER |
| 16 | 71391801 | AX-170101847 | 0 | T | C | 0.491 | 1.09E-01 | 7.62E-02 | 2.49E+00 | 2.56E+00 | 1.54E-01 | no | | regulatory_region_variant | MODIFIER |
| 1 | 134219678 | AX-170347474 | 0 | T | C | 0.526 | 1.12E-01 | 7.90E-02 | 2.73E+00 | 2.81E+00 | 1.54E-01 | no | Adora1 | intron_variant | MODIFIER |
| 16 | 72063946 | AX-170237049 | 1 | A | T | 0.491 | 1.09E-01 | 7.63E-02 | 2.50E+00 | 2.57E+00 | 1.55E-01 | no | | intergenic_variant | MODIFIER |
| 1 | 134900912 | AX-170042812 | 0 | G | A | 0.535 | 1.12E-01 | 7.92E-02 | 2.74E+00 | 2.82E+00 | 1.55E-01 | no | Ppp1r12b | intron_variant | MODIFIER |
| 16 | 70211883 | AX-97830791 | 0 | T | C | 0.491 | 1.09E-01 | 7.58E-02 | 2.45E+00 | 2.52E+00 | 1.55E-01 | no | | intergenic_variant | MODIFIER |
| 5 | 55390843 | AX-97797512 | 0 | T | C | 0.482 | 1.43E-01 | 9.96E-02 | 2.45E+00 | 2.51E+00 | 1.55E-01 | no | | intergenic_variant | MODIFIER |
| 1 | 124842452 | AX-170212672 | 0 | G | A | 0.579 | 1.18E-01 | 8.34E-02 | 2.57E+00 | 2.63E+00 | 1.55E-01 | no | Gm28928 | intron_variant,non_coding_transcript_variant | MODIFIER |
| 16 | 52605800 | AX-170013050 | 0 | G | C | 0.518 | 1.16E-01 | 8.05E-02 | 2.39E+00 | 2.43E+00 | 1.55E-01 | no | | intergenic_variant | MODIFIER |
| 4 | 54266449 | AX-170198347 | 0 | T | C | 0.368 | -1.47E-01 | 1.03E-01 | 2.97E+00 | 3.08E+00 | 1.55E-01 | no | | intergenic_variant | MODIFIER |
| 3 | 8284105 | AX-170019867 | 0 | G | A | 0.386 | -9.00E-02 | 6.36E-02 | 2.91E+00 | 3.07E+00 | 1.55E-01 | no | | intergenic_variant | MODIFIER |
| 4 | 70244456 | AX-97801126 | 1 | A | G | 0.438 | -1.15E-01 | 8.06E-02 | 2.48E+00 | 2.55E+00 | 1.55E-01 | no | Cdk5rap2 | intron_variant,NMD_transcript_variant | MODIFIER |
| 6 | 114386592 | AX-170255198 | 0 | C | T | 0.061 | 2.97E-01 | 2.10E-01 | 2.88E+00 | 3.02E+00 | 1.55E-01 | Yes | | intergenic_variant | MODIFIER |
| 18 | 86964057 | AX-170365869 | 0 | G | C | 0.175 | -1.52E-01 | 1.07E-01 | 2.60E+00 | 2.69E+00 | 1.56E-01 | no | | intergenic_variant | MODIFIER |
| 16 | 39589993 | AX-170363810 | 0 | A | G | 0.5 | 1.13E-01 | 7.97E-02 | 2.61E+00 | 2.67E+00 | 1.56E-01 | no | | intergenic_variant | MODIFIER |
| 4 | 80791921 | AX-170430249 | 0 | A | G | 0.333 | 1.62E-01 | 1.15E-01 | 2.83E+00 | 2.90E+00 | 1.56E-01 | Yes | | intergenic_variant | MODIFIER |
| 1 | 193376229 | AX-170494924 | 1 | A | G | 0.268 | 1.32E-01 | 9.12E-02 | 2.36E+00 | 2.44E+00 | 1.56E-01 | no | | intergenic_variant | MODIFIER |
| 15 | 83520715 | AX-97805447 | 0 | G | A | 0.096 | -1.78E-01 | 9.99E-02 | 1.63E+00 | 1.71E+00 | 1.56E-01 | Yes | Gm24575 | downstream_gene_variant | MODIFIER |
| 6 | 138855396 | AX-170257973 | 0 | C | T | 0.07 | -2.87E-01 | 2.03E-01 | 2.89E+00 | 3.02E+00 | 1.56E-01 | Yes | Gm9038 | downstream_gene_variant | MODIFIER |
| 16 | 33108967 | AX-170164873 | 0 | G | A | 0.474 | 1.08E-01 | 7.62E-02 | 2.62E+00 | 2.69E+00 | 1.56E-01 | no | Lmln | intron_variant | MODIFIER |
| 4 | 107558297 | AX-170265616 | 1 | G | A | 0.268 | 1.61E-01 | 1.14E-01 | 2.62E+00 | 2.67E+00 | 1.56E-01 | Yes | Glis1 | intron_variant | MODIFIER |
| 15 | 5754697 | AX-97851430 | 1 | G | A | 0.339 | -1.13E-01 | 7.23E-02 | 1.97E+00 | 2.04E+00 | 1.56E-01 | no | | intergenic_variant | MODIFIER |
| 15 | 71433360 | AX-168400162 | 0 | T | C | 0.079 | -1.95E-01 | 1.12E-01 | 1.69E+00 | 1.75E+00 | 1.56E-01 | Yes | Fam135b | 3_prime_UTR_variant | MODIFIER |
| 5 | 52601977 | AX-170320020 | 0 | C | A | 0.482 | 1.48E-01 | 1.03E-01 | 2.40E+00 | 2.45E+00 | 1.56E-01 | Yes | | intergenic_variant | MODIFIER |
| 6 | 49050200 | AX-170353881 | 0 | G | A | 0.053 | 2.01E-01 | 1.43E-01 | 2.90E+00 | 3.08E+00 | 1.56E-01 | Yes | Gpnmb | intron_variant | MODIFIER |
| 9 | 5855920 | AX-170522343 | 1 | C | T | 0.473 | 1.11E-01 | 7.80E-02 | 2.48E+00 | 2.57E+00 | 1.57E-01 | no | | intergenic_variant | MODIFIER |

| | | | | | | | | | | | | | | | |
|---|---|---|---|---|---|---|---|---|---|---|---|---|---|---|---|
| 10 | 71127193 | AX-170276862 | 0 | T | C | 0.412 | -1.11E-01 | 7.92E-02 | 2.75E+00 | 2.84E+00 | 1.57E-01 | no | Bicc1 | intron_variant | MODIFIER |
| 5 | 11441409 | AX-170213149 | 0 | C | G | 0.351 | 1.18E-01 | 8.41E-02 | 2.73E+00 | 2.86E+00 | 1.57E-01 | no | | intergenic_variant | MODIFIER |
| 1 | 123790817 | AX-170308873 | 0 | G | A | 0.316 | 7.93E-02 | 5.62E-02 | 2.57E+00 | 2.69E+00 | 1.57E-01 | no | Dpp10 | intron_variant | MODIFIER |
| 8 | 93531499 | AX-170260885 | 0 | T | A | 0.211 | -1.62E-01 | 1.13E-01 | 3.11E+00 | 3.24E+00 | 1.57E-01 | no | Ces5a | intron_variant | MODIFIER |
| 8 | 93700532 | AX-170204966 | 0 | A | T | 0.211 | -1.62E-01 | 1.13E-01 | 3.11E+00 | 3.24E+00 | 1.57E-01 | no | | intergenic_variant | MODIFIER |
| 8 | 93736930 | AX-170247458 | 0 | A | T | 0.211 | -1.62E-01 | 1.13E-01 | 3.11E+00 | 3.24E+00 | 1.57E-01 | no | | intergenic_variant | MODIFIER |
| 8 | 93935283 | AX-170235659 | 0 | T | G | 0.211 | -1.62E-01 | 1.13E-01 | 3.11E+00 | 3.24E+00 | 1.57E-01 | no | Gnao1 | intron_variant | MODIFIER |
| 8 | 94844852 | AX-170189001 | 0 | A | T | 0.211 | -1.62E-01 | 1.13E-01 | 3.11E+00 | 3.24E+00 | 1.57E-01 | no | Coq9 | synonymous_variant | LOW |
| 3 | 151588169 | AX-97851115 | 0 | T | A | 0.333 | 1.10E-01 | 7.76E-02 | 2.96E+00 | 3.07E+00 | 1.57E-01 | no | | intergenic_variant | MODIFIER |
| 6 | 135809945 | AX-170213727 | 0 | G | A | 0.579 | 1.21E-01 | 8.52E-02 | 2.45E+00 | 2.51E+00 | 1.57E-01 | no | Grin2b | intron_variant | MODIFIER |
| 9 | 70902075 | AX-170250748 | 1 | T | C | 0.045 | 1.60E-01 | 1.14E-01 | 2.75E+00 | 2.95E+00 | 1.57E-01 | Yes | Lipc | intron_variant | MODIFIER |
| 14 | 117227157 | AX-97789671 | 0 | G | A | 0.325 | 1.02E-01 | 7.28E-02 | 2.73E+00 | 2.85E+00 | 1.57E-01 | no | Gpc6 | intron_variant | MODIFIER |
| 5 | 129291582 | AX-170073498 | 2 | C | T | 0.164 | -1.22E-01 | 8.30E-02 | 2.24E+00 | 2.33E+00 | 1.57E-01 | no | | intergenic_variant | MODIFIER |
| 1 | 55746935 | AX-170256921 | 0 | A | G | 0.105 | -2.60E-01 | 1.84E-01 | 2.98E+00 | 3.11E+00 | 1.57E-01 | Yes | Plcl1 | intron_variant | MODIFIER |
| 4 | 86192495 | AX-170403119 | 1 | G | C | 0.348 | 1.57E-01 | 1.12E-01 | 2.71E+00 | 2.78E+00 | 1.57E-01 | Yes | Adamtsl1 | intron_variant | MODIFIER |
| 1 | 181528298 | AX-170107758 | 1 | G | A | 0.232 | 1.59E-01 | 1.12E-01 | 2.46E+00 | 2.53E+00 | 1.58E-01 | Yes | | intergenic_variant | MODIFIER |
| 1 | 191716695 | AX-170426846 | 1 | C | A | 0.286 | 1.31E-01 | 8.99E-02 | 2.25E+00 | 2.33E+00 | 1.58E-01 | no | Lpgat1 | upstream_gene_variant | MODIFIER |
| 3 | 7930433 | AX-97781595 | 0 | A | C | 0.404 | -9.06E-02 | 6.43E-02 | 2.96E+00 | 3.12E+00 | 1.58E-01 | no | 1700010I02Rik | intron_variant,non_coding_transcript_variant | MODIFIER |
| 15 | 67303107 | AX-170011892 | 0 | A | G | 0.333 | -1.09E-01 | 7.42E-02 | 2.24E+00 | 2.31E+00 | 1.58E-01 | no | 1700012I11Rik | intron_variant,non_coding_transcript_variant | MODIFIER |
| 2 | 97191562 | AX-97798933 | 0 | T | C | 0.053 | -2.94E-01 | 2.10E-01 | 2.76E+00 | 2.86E+00 | 1.58E-01 | Yes | Lrrc4c | intron_variant | MODIFIER |
| 2 | 102474449 | AX-170371154 | 0 | C | A | 0.053 | -2.94E-01 | 2.10E-01 | 2.76E+00 | 2.86E+00 | 1.58E-01 | Yes | | intergenic_variant | MODIFIER |
| 9 | 9265824 | AX-170365136 | 1 | A | G | 0.464 | 1.09E-01 | 7.69E-02 | 2.49E+00 | 2.60E+00 | 1.58E-01 | no | Gm16833 | downstream_gene_variant | MODIFIER |
| 7 | 23340472 | AX-170076835 | 1 | C | T | 0.42 | 1.23E-01 | 8.75E-02 | 2.62E+00 | 2.75E+00 | 1.58E-01 | no | Nlrp4e | missense_variant | MODERATE |
| 18 | 29795424 | AX-170088313 | 1 | A | G | 0.054 | 3.29E-01 | 2.33E-01 | 2.95E+00 | 3.10E+00 | 1.58E-01 | Yes | | intergenic_variant | MODIFIER |
| 8 | 94271122 | AX-170371221 | 1 | G | A | 0.214 | -1.61E-01 | 1.13E-01 | 3.14E+00 | 3.27E+00 | 1.58E-01 | no | Nup93 | intron_variant | MODIFIER |
| 15 | 3295724 | AX-170054663 | 2 | A | T | 0.245 | -1.44E-01 | 1.02E-01 | 2.68E+00 | 2.75E+00 | 1.58E-01 | no | Ccdc152 | intron_variant | MODIFIER |
| 16 | 57654892 | AX-170503576 | 1 | T | C | 0.527 | 1.18E-01 | 8.37E-02 | 2.51E+00 | 2.55E+00 | 1.58E-01 | no | Col8a1 | intron_variant | MODIFIER |
| 1 | 168233689 | AX-170174099 | 0 | G | A | 0.228 | 1.85E-01 | 1.32E-01 | 2.81E+00 | 2.87E+00 | 1.58E-01 | Yes | Pbx1 | intron_variant | MODIFIER |
| 5 | 57691113 | AX-170072204 | 1 | A | T | 0.482 | 1.41E-01 | 9.97E-02 | 2.52E+00 | 2.59E+00 | 1.58E-01 | no | 4932441J04Rik | intron_variant,non_coding_transcript_variant | MODIFIER |
| 4 | 96854621 | AX-170483805 | 0 | G | C | 0.246 | 1.51E-01 | 1.08E-01 | 2.64E+00 | 2.72E+00 | 1.59E-01 | Yes | | intergenic_variant | MODIFIER |
| 4 | 104073776 | AX-170509594 | 0 | A | T | 0.246 | 1.63E-01 | 1.17E-01 | 2.72E+00 | 2.79E+00 | 1.60E-01 | Yes | Dab1 | intron_variant | MODIFIER |
| 4 | 64664126 | AX-97797968 | 0 | T | G | 0.333 | 1.54E-01 | 1.10E-01 | 2.74E+00 | 2.81E+00 | 1.60E-01 | Yes | | intergenic_variant | MODIFIER |
| 4 | 69032377 | AX-170515748 | 0 | A | G | 0.342 | 1.63E-01 | 1.17E-01 | 2.90E+00 | 2.96E+00 | 1.60E-01 | Yes | | intergenic_variant | MODIFIER |
| 5 | 130048965 | AX-170331667 | 0 | T | C | 0.158 | -1.23E-01 | 8.43E-02 | 2.23E+00 | 2.31E+00 | 1.60E-01 | no | Crcp | synonymous_variant | LOW |
| 1 | 166261046 | AX-170183988 | 1 | G | A | 0.232 | 1.92E-01 | 1.37E-01 | 2.82E+00 | 2.88E+00 | 1.60E-01 | Yes | Ildr2 | intron_variant | MODIFIER |
| 4 | 139523566 | AX-170205907 | 0 | C | T | 0.184 | 1.45E-01 | 1.04E-01 | 2.71E+00 | 2.78E+00 | 1.60E-01 | Yes | | intergenic_variant | MODIFIER |
| 4 | 63330068 | AX-97853685 | 0 | A | G | 0.342 | 1.62E-01 | 1.16E-01 | 2.92E+00 | 2.98E+00 | 1.60E-01 | Yes | Col27a1 | intron_variant | MODIFIER |
| 13 | 18859812 | AX-97793554 | 0 | G | A | 0.167 | -1.70E-01 | 1.20E-01 | 2.44E+00 | 2.50E+00 | 1.60E-01 | no | Vps41 | intron_variant | MODIFIER |
| 16 | 67696665 | AX-170302750 | 0 | T | C | 0.491 | 1.08E-01 | 7.68E-02 | 2.49E+00 | 2.55E+00 | 1.60E-01 | no | | intergenic_variant | MODIFIER |
| 7 | 71060454 | AX-97803187 | 0 | T | C | 0.36 | -1.38E-01 | 9.81E-02 | 3.01E+00 | 3.12E+00 | 1.60E-01 | no | Gm42397 | intron_variant,non_coding_transcript_variant | MODIFIER |
| 4 | 133589274 | AX-170326964 | 1 | G | T | 0.152 | 1.60E-01 | 1.15E-01 | 2.65E+00 | 2.71E+00 | 1.61E-01 | Yes | Gpn2 | intron_variant | MODIFIER |
| 3 | 38802777 | AX-97845144 | 0 | T | C | 0.202 | 1.27E-01 | 9.09E-02 | 2.90E+00 | 3.02E+00 | 1.61E-01 | no | | intergenic_variant | MODIFIER |
| 3 | 38266065 | AX-170066405 | 0 | A | T | 0.211 | 1.26E-01 | 9.01E-02 | 2.93E+00 | 3.04E+00 | 1.61E-01 | no | Gm25574 | downstream_gene_variant | MODIFIER |
| 3 | 38076158 | AX-170155103 | 0 | T | C | 0.202 | 1.32E-01 | 9.33E-02 | 3.07E+00 | 3.19E+00 | 1.62E-01 | no | | intergenic_variant | MODIFIER |
| 3 | 38093580 | AX-97831019 | 0 | G | A | 0.202 | 1.32E-01 | 9.33E-02 | 3.07E+00 | 3.19E+00 | 1.62E-01 | no | Gm43821 | downstream_gene_variant | MODIFIER |
| 3 | 38093705 | AX-170161316 | 0 | A | G | 0.202 | 1.32E-01 | 9.33E-02 | 3.07E+00 | 3.19E+00 | 1.62E-01 | no | Gm43821 | downstream_gene_variant | MODIFIER |
| 3 | 38108867 | AX-170088910 | 0 | C | T | 0.202 | 1.32E-01 | 9.33E-02 | 3.07E+00 | 3.19E+00 | 1.62E-01 | no | Rpl21-ps10 | upstream_gene_variant | MODIFIER |
| 3 | 38116003 | AX-170223686 | 0 | A | G | 0.202 | 1.32E-01 | 9.33E-02 | 3.07E+00 | 3.19E+00 | 1.62E-01 | no | Gm43821 | intron_variant,non_coding_transcript_variant | MODIFIER |
| 3 | 38117840 | AX-170203296 | 0 | G | A | 0.202 | 1.32E-01 | 9.33E-02 | 3.07E+00 | 3.19E+00 | 1.62E-01 | no | Gm43821 | intron_variant,non_coding_transcript_variant | MODIFIER |
| 3 | 38118619 | AX-170245269 | 0 | C | T | 0.202 | 1.32E-01 | 9.33E-02 | 3.07E+00 | 3.19E+00 | 1.62E-01 | no | Gm43821 | intron_variant,non_coding_transcript_variant | MODIFIER |
| 3 | 38134797 | AX-170503145 | 0 | T | C | 0.202 | 1.32E-01 | 9.33E-02 | 3.07E+00 | 3.19E+00 | 1.62E-01 | no | Gm43821 | upstream_gene_variant | MODIFIER |
| 3 | 38145355 | AX-97821602 | 0 | C | T | 0.202 | 1.32E-01 | 9.33E-02 | 3.07E+00 | 3.19E+00 | 1.62E-01 | no | | regulatory_region_variant | MODIFIER |
| 3 | 38150286 | AX-170324132 | 0 | C | T | 0.202 | 1.32E-01 | 9.33E-02 | 3.07E+00 | 3.19E+00 | 1.62E-01 | no | | intergenic_variant | MODIFIER |
| 3 | 38186812 | AX-170364027 | 0 | G | T | 0.202 | 1.32E-01 | 9.33E-02 | 3.07E+00 | 3.19E+00 | 1.62E-01 | no | | intergenic_variant | MODIFIER |

| | | | | | | | | | | | | | | | |
|---|---|---|---|---|---|---|---|---|---|---|---|---|---|---|---|
| 3 | 38194185 | AX-170302380 | 0 | G | A | 0.202 | 1.32E-01 | 9.33E-02 | 3.07E+00 | 3.19E+00 | 1.62E-01 | no | | regulatory_region_variant | MODIFIER |
| 3 | 38194505 | AX-170429303 | 0 | T | C | 0.202 | 1.32E-01 | 9.33E-02 | 3.07E+00 | 3.19E+00 | 1.62E-01 | no | | intergenic_variant | MODIFIER |
| 3 | 38234568 | AX-170020324 | 0 | A | C | 0.202 | 1.32E-01 | 9.33E-02 | 3.07E+00 | 3.19E+00 | 1.62E-01 | no | | regulatory_region_variant | MODIFIER |
| 3 | 38249225 | AX-97798118 | 0 | T | G | 0.202 | 1.32E-01 | 9.33E-02 | 3.07E+00 | 3.19E+00 | 1.62E-01 | no | | regulatory_region_variant | MODIFIER |
| 3 | 38264330 | AX-97851423 | 0 | G | A | 0.202 | 1.32E-01 | 9.33E-02 | 3.07E+00 | 3.19E+00 | 1.62E-01 | no | Gm25574 | downstream_gene_variant | MODIFIER |
| 3 | 38277847 | AX-170020326 | 0 | T | C | 0.202 | 1.32E-01 | 9.33E-02 | 3.07E+00 | 3.19E+00 | 1.62E-01 | no | | intergenic_variant | MODIFIER |
| 3 | 38278048 | AX-170020325 | 0 | C | T | 0.202 | 1.32E-01 | 9.33E-02 | 3.07E+00 | 3.19E+00 | 1.62E-01 | no | | intergenic_variant | MODIFIER |
| 3 | 38314221 | AX-170443064 | 0 | G | T | 0.202 | 1.32E-01 | 9.33E-02 | 3.07E+00 | 3.19E+00 | 1.62E-01 | no | | intergenic_variant | MODIFIER |
| 3 | 38316974 | AX-170333092 | 0 | C | G | 0.202 | 1.32E-01 | 9.33E-02 | 3.07E+00 | 3.19E+00 | 1.62E-01 | no | | intergenic_variant | MODIFIER |
| 3 | 38334436 | AX-170442667 | 0 | C | A | 0.202 | 1.32E-01 | 9.33E-02 | 3.07E+00 | 3.19E+00 | 1.62E-01 | no | | intergenic_variant | MODIFIER |
| 3 | 38340455 | AX-170270682 | 0 | A | G | 0.202 | 1.32E-01 | 9.33E-02 | 3.07E+00 | 3.19E+00 | 1.62E-01 | no | | intergenic_variant | MODIFIER |
| 3 | 38394666 | AX-170197959 | 0 | C | T | 0.202 | 1.32E-01 | 9.33E-02 | 3.07E+00 | 3.19E+00 | 1.62E-01 | no | | intergenic_variant | MODIFIER |
| 3 | 38399256 | AX-170362440 | 0 | G | A | 0.202 | 1.32E-01 | 9.33E-02 | 3.07E+00 | 3.19E+00 | 1.62E-01 | no | | intergenic_variant | MODIFIER |
| 3 | 38399360 | AX-170193390 | 0 | T | C | 0.202 | 1.32E-01 | 9.33E-02 | 3.07E+00 | 3.19E+00 | 1.62E-01 | no | | intergenic_variant | MODIFIER |
| 3 | 38425116 | AX-170418447 | 0 | A | G | 0.202 | 1.32E-01 | 9.33E-02 | 3.07E+00 | 3.19E+00 | 1.62E-01 | no | Gm36823 | downstream_gene_variant | MODIFIER |
| 3 | 38425256 | AX-170143667 | 0 | T | C | 0.202 | 1.32E-01 | 9.33E-02 | 3.07E+00 | 3.19E+00 | 1.62E-01 | no | Gm36823 | downstream_gene_variant | MODIFIER |
| 3 | 38446027 | AX-170363192 | 0 | G | C | 0.202 | 1.32E-01 | 9.33E-02 | 3.07E+00 | 3.19E+00 | 1.62E-01 | no | Ankrd50 | downstream_gene_variant | MODIFIER |
| 3 | 38452302 | AX-170520781 | 0 | T | C | 0.202 | 1.32E-01 | 9.33E-02 | 3.07E+00 | 3.19E+00 | 1.62E-01 | no | Ankrd50 | 3_prime_UTR_variant | MODIFIER |
| 3 | 38459572 | AX-170312408 | 0 | C | T | 0.202 | 1.32E-01 | 9.33E-02 | 3.07E+00 | 3.19E+00 | 1.62E-01 | no | Ankrd50 | upstream_gene_variant | MODIFIER |
| 3 | 38471854 | AX-97848246 | 0 | C | A | 0.202 | 1.32E-01 | 9.33E-02 | 3.07E+00 | 3.19E+00 | 1.62E-01 | no | Ankrd50 | intron_variant | MODIFIER |
| 3 | 38471938 | AX-170066411 | 0 | T | A | 0.202 | 1.32E-01 | 9.33E-02 | 3.07E+00 | 3.19E+00 | 1.62E-01 | no | Ankrd50 | intron_variant | MODIFIER |
| 3 | 38482576 | AX-170225480 | 0 | T | C | 0.202 | 1.32E-01 | 9.33E-02 | 3.07E+00 | 3.19E+00 | 1.62E-01 | no | Ankrd50 | intron_variant | MODIFIER |
| 3 | 38490085 | AX-170162046 | 0 | T | C | 0.202 | 1.32E-01 | 9.33E-02 | 3.07E+00 | 3.19E+00 | 1.62E-01 | no | | intergenic_variant | MODIFIER |
| 3 | 38515987 | AX-170212745 | 0 | C | T | 0.202 | 1.32E-01 | 9.33E-02 | 3.07E+00 | 3.19E+00 | 1.62E-01 | no | | intergenic_variant | MODIFIER |
| 3 | 38527799 | AX-170504394 | 0 | G | A | 0.202 | 1.32E-01 | 9.33E-02 | 3.07E+00 | 3.19E+00 | 1.62E-01 | no | | intergenic_variant | MODIFIER |
| 3 | 38532894 | AX-170374608 | 0 | A | G | 0.202 | 1.32E-01 | 9.33E-02 | 3.07E+00 | 3.19E+00 | 1.62E-01 | no | | intergenic_variant | MODIFIER |
| 3 | 38538556 | AX-170459517 | 0 | C | T | 0.202 | 1.32E-01 | 9.33E-02 | 3.07E+00 | 3.19E+00 | 1.62E-01 | no | | intergenic_variant | MODIFIER |
| 3 | 38539335 | AX-170249676 | 0 | A | T | 0.202 | 1.32E-01 | 9.33E-02 | 3.07E+00 | 3.19E+00 | 1.62E-01 | no | | intergenic_variant | MODIFIER |
| 3 | 38550892 | AX-170134978 | 0 | G | A | 0.202 | 1.32E-01 | 9.33E-02 | 3.07E+00 | 3.19E+00 | 1.62E-01 | no | | intergenic_variant | MODIFIER |
| 3 | 38569217 | AX-97795031 | 0 | G | A | 0.202 | 1.32E-01 | 9.33E-02 | 3.07E+00 | 3.19E+00 | 1.62E-01 | no | | regulatory_region_variant | MODIFIER |
| 3 | 38570952 | AX-170275451 | 0 | T | C | 0.202 | 1.32E-01 | 9.33E-02 | 3.07E+00 | 3.19E+00 | 1.62E-01 | no | | intergenic_variant | MODIFIER |
| 3 | 38572661 | AX-170423811 | 0 | A | G | 0.202 | 1.32E-01 | 9.33E-02 | 3.07E+00 | 3.19E+00 | 1.62E-01 | no | | intergenic_variant | MODIFIER |
| 3 | 38595784 | AX-170220450 | 0 | G | T | 0.202 | 1.32E-01 | 9.33E-02 | 3.07E+00 | 3.19E+00 | 1.62E-01 | no | | intergenic_variant | MODIFIER |
| 3 | 38624057 | AX-170367339 | 0 | C | T | 0.202 | 1.32E-01 | 9.33E-02 | 3.07E+00 | 3.19E+00 | 1.62E-01 | no | | intergenic_variant | MODIFIER |
| 3 | 38625556 | AX-97828275 | 0 | C | T | 0.202 | 1.32E-01 | 9.33E-02 | 3.07E+00 | 3.19E+00 | 1.62E-01 | no | | regulatory_region_variant | MODIFIER |
| 3 | 38636823 | AX-170141136 | 0 | C | T | 0.202 | 1.32E-01 | 9.33E-02 | 3.07E+00 | 3.19E+00 | 1.62E-01 | no | | intergenic_variant | MODIFIER |
| 3 | 38637677 | AX-170246788 | 0 | C | T | 0.202 | 1.32E-01 | 9.33E-02 | 3.07E+00 | 3.19E+00 | 1.62E-01 | no | | intergenic_variant | MODIFIER |
| 3 | 38645386 | AX-170302619 | 0 | A | G | 0.202 | 1.32E-01 | 9.33E-02 | 3.07E+00 | 3.19E+00 | 1.62E-01 | no | | intergenic_variant | MODIFIER |
| 3 | 38663448 | AX-170408343 | 0 | T | C | 0.202 | 1.32E-01 | 9.33E-02 | 3.07E+00 | 3.19E+00 | 1.62E-01 | no | | regulatory_region_variant | MODIFIER |
| 3 | 38671794 | AX-170469718 | 0 | A | G | 0.202 | 1.32E-01 | 9.33E-02 | 3.07E+00 | 3.19E+00 | 1.62E-01 | no | | intergenic_variant | MODIFIER |
| 3 | 38672913 | AX-170135391 | 0 | C | T | 0.202 | 1.32E-01 | 9.33E-02 | 3.07E+00 | 3.19E+00 | 1.62E-01 | no | | intergenic_variant | MODIFIER |
| 3 | 38683429 | AX-97828746 | 0 | T | C | 0.202 | 1.32E-01 | 9.33E-02 | 3.07E+00 | 3.19E+00 | 1.62E-01 | no | | intergenic_variant | MODIFIER |
| 3 | 38705782 | AX-170085994 | 0 | A | G | 0.202 | 1.32E-01 | 9.33E-02 | 3.07E+00 | 3.19E+00 | 1.62E-01 | no | | intergenic_variant | MODIFIER |
| 3 | 38713031 | AX-170296871 | 0 | T | C | 0.202 | 1.32E-01 | 9.33E-02 | 3.07E+00 | 3.19E+00 | 1.62E-01 | no | | regulatory_region_variant | MODIFIER |
| 3 | 38713188 | AX-97826420 | 0 | A | G | 0.202 | 1.32E-01 | 9.33E-02 | 3.07E+00 | 3.19E+00 | 1.62E-01 | no | | regulatory_region_variant | MODIFIER |
| 3 | 38732857 | AX-170244639 | 0 | A | G | 0.202 | 1.32E-01 | 9.33E-02 | 3.07E+00 | 3.19E+00 | 1.62E-01 | no | | intergenic_variant | MODIFIER |
| 3 | 38738851 | AX-170517490 | 0 | G | A | 0.202 | 1.32E-01 | 9.33E-02 | 3.07E+00 | 3.19E+00 | 1.62E-01 | no | | intergenic_variant | MODIFIER |
| 3 | 38744064 | AX-170020328 | 0 | G | A | 0.202 | 1.32E-01 | 9.33E-02 | 3.07E+00 | 3.19E+00 | 1.62E-01 | no | | intergenic_variant | MODIFIER |
| 3 | 38802689 | AX-170020331 | 0 | C | T | 0.202 | 1.32E-01 | 9.33E-02 | 3.07E+00 | 3.19E+00 | 1.62E-01 | no | | intergenic_variant | MODIFIER |
| 3 | 38829136 | AX-170020327 | 0 | A | G | 0.202 | 1.32E-01 | 9.33E-02 | 3.07E+00 | 3.19E+00 | 1.62E-01 | no | | intergenic_variant | MODIFIER |
| 3 | 38830005 | AX-170088911 | 0 | A | G | 0.202 | 1.32E-01 | 9.33E-02 | 3.07E+00 | 3.19E+00 | 1.62E-01 | no | | intergenic_variant | MODIFIER |
| 3 | 38830132 | AX-170325981 | 0 | C | T | 0.202 | 1.32E-01 | 9.33E-02 | 3.07E+00 | 3.19E+00 | 1.62E-01 | no | | intergenic_variant | MODIFIER |
| 3 | 38842214 | AX-170487858 | 0 | G | A | 0.202 | 1.32E-01 | 9.33E-02 | 3.07E+00 | 3.19E+00 | 1.62E-01 | no | | intergenic_variant | MODIFIER |
| 3 | 38878925 | AX-170335206 | 0 | C | G | 0.202 | 1.32E-01 | 9.33E-02 | 3.07E+00 | 3.19E+00 | 1.62E-01 | no | C230034O21Rik | intron_variant,non_coding_transcript_variant | MODIFIER |

| | | | | | | | | | | | | | | | |
|---|---|---|---|---|---|---|---|---|---|---|---|---|---|---|---|
| 3 | 38884980 | AX-170509079 | 0 | T | C | 0.202 | 1.32E-01 | 9.33E-02 | 3.07E+00 | 3.19E+00 | 1.62E-01 | no | Fat4 | upstream_gene_variant | MODIFIER |
| 3 | 38904450 | AX-170438551 | 0 | A | G | 0.202 | 1.32E-01 | 9.33E-02 | 3.07E+00 | 3.19E+00 | 1.62E-01 | no | Fat4 | intron_variant | MODIFIER |
| 3 | 38905064 | AX-170448096 | 0 | A | G | 0.202 | 1.32E-01 | 9.33E-02 | 3.07E+00 | 3.19E+00 | 1.62E-01 | no | Fat4 | intron_variant | MODIFIER |
| 3 | 38905613 | AX-97794647 | 0 | A | G | 0.202 | 1.32E-01 | 9.33E-02 | 3.07E+00 | 3.19E+00 | 1.62E-01 | no | Fat4 | intron_variant | MODIFIER |
| 3 | 38905714 | AX-170142000 | 0 | G | C | 0.202 | 1.32E-01 | 9.33E-02 | 3.07E+00 | 3.19E+00 | 1.62E-01 | no | Fat4 | intron_variant | MODIFIER |
| 3 | 38912764 | AX-97836452 | 0 | G | A | 0.202 | 1.32E-01 | 9.33E-02 | 3.07E+00 | 3.19E+00 | 1.62E-01 | no | Fat4 | intron_variant | MODIFIER |
| 3 | 38919124 | AX-170276903 | 0 | G | A | 0.202 | 1.32E-01 | 9.33E-02 | 3.07E+00 | 3.19E+00 | 1.62E-01 | no | Fat4 | intron_variant | MODIFIER |
| 3 | 38919533 | AX-170167070 | 0 | C | T | 0.202 | 1.32E-01 | 9.33E-02 | 3.07E+00 | 3.19E+00 | 1.62E-01 | no | Fat4 | intron_variant | MODIFIER |
| 3 | 38942100 | AX-170066425 | 0 | T | C | 0.202 | 1.32E-01 | 9.33E-02 | 3.07E+00 | 3.19E+00 | 1.62E-01 | no | Fat4 | intron_variant | MODIFIER |
| 3 | 38946514 | AX-97842989 | 0 | G | A | 0.202 | 1.32E-01 | 9.33E-02 | 3.07E+00 | 3.19E+00 | 1.62E-01 | no | Fat4 | intron_variant | MODIFIER |
| 3 | 38946863 | AX-170378723 | 0 | C | T | 0.202 | 1.32E-01 | 9.33E-02 | 3.07E+00 | 3.19E+00 | 1.62E-01 | no | Fat4 | intron_variant | MODIFIER |
| 3 | 38948226 | AX-170487202 | 0 | A | C | 0.202 | 1.32E-01 | 9.33E-02 | 3.07E+00 | 3.19E+00 | 1.62E-01 | no | Fat4 | intron_variant | MODIFIER |
| 3 | 19131319 | AX-170339305 | 1 | C | T | 0.134 | -1.36E-01 | 9.73E-02 | 2.60E+00 | 2.69E+00 | 1.62E-01 | no | Armc1 | downstream_gene_variant | MODIFIER |
| 9 | 15745488 | AX-170349680 | 1 | T | A | 0.482 | 1.14E-01 | 8.17E-02 | 2.61E+00 | 2.70E+00 | 1.62E-01 | no | Slc36a4 | downstream_gene_variant | MODIFIER |
| 18 | 80396259 | AX-170312162 | 0 | C | T | 0.456 | 1.09E-01 | 7.74E-02 | 2.46E+00 | 2.55E+00 | 1.62E-01 | no | | intergenic_variant | MODIFIER |
| 13 | 6741474 | AX-170448207 | 0 | A | G | 0.202 | -1.57E-01 | 1.12E-01 | 2.58E+00 | 2.65E+00 | 1.62E-01 | no | | intergenic_variant | MODIFIER |
| 7 | 36993651 | AX-170263134 | 0 | C | T | 0.351 | 1.18E-01 | 8.48E-02 | 2.57E+00 | 2.68E+00 | 1.62E-01 | no | | intergenic_variant | MODIFIER |
| 20 | 48645661 | AX-170499925 | 1 | A | C | 0.241 | 8.14E-02 | 5.86E-02 | 2.62E+00 | 2.74E+00 | 1.62E-01 | no | Slc25a14 | intron_variant | MODIFIER |
| 16 | 52726933 | AX-170383674 | 0 | A | G | 0.526 | 1.18E-01 | 8.42E-02 | 2.47E+00 | 2.51E+00 | 1.62E-01 | no | | intergenic_variant | MODIFIER |
| 16 | 52730639 | AX-170487523 | 0 | T | C | 0.526 | 1.18E-01 | 8.42E-02 | 2.47E+00 | 2.51E+00 | 1.62E-01 | no | | intergenic_variant | MODIFIER |
| 4 | 64322563 | AX-97856135 | 1 | G | A | 0.321 | 1.67E-01 | 1.20E-01 | 2.93E+00 | 2.99E+00 | 1.62E-01 | Yes | | intergenic_variant | MODIFIER |
| 9 | 52662830 | AX-170082457 | 0 | G | A | 0.561 | 7.79E-02 | 5.56E-02 | 3.02E+00 | 3.14E+00 | 1.63E-01 | no | | intergenic_variant | MODIFIER |
| 18 | 80175891 | AX-170173148 | 0 | G | A | 0.386 | -1.16E-01 | 8.38E-02 | 2.78E+00 | 2.89E+00 | 1.63E-01 | no | | intergenic_variant | MODIFIER |
| 4 | 100568199 | AX-170396095 | 0 | G | A | 0.237 | 1.62E-01 | 1.17E-01 | 2.61E+00 | 2.67E+00 | 1.63E-01 | Yes | Ccdc50-ps | upstream_gene_variant | MODIFIER |
| 9 | 41818503 | AX-170120248 | 0 | G | A | 0.035 | 2.08E-01 | 1.51E-01 | 2.75E+00 | 2.93E+00 | 1.63E-01 | Yes | Gm35940 | intron_variant,non_coding_transcript_variant | MODIFIER |
| 4 | 124650554 | AX-170301528 | 0 | G | A | 0.246 | 1.45E-01 | 1.04E-01 | 2.62E+00 | 2.70E+00 | 1.63E-01 | Yes | Gm2164 | intron_variant,non_coding_transcript_variant | MODIFIER |
| 1 | 63324408 | AX-97789952 | 0 | C | T | 0.132 | -2.60E-01 | 1.87E-01 | 2.96E+00 | 3.09E+00 | 1.63E-01 | Yes | | intergenic_variant | MODIFIER |
| 4 | 100349423 | AX-97793897 | 0 | G | C | 0.43 | -1.06E-01 | 7.63E-02 | 2.57E+00 | 2.65E+00 | 1.63E-01 | no | Ror1 | intron_variant | MODIFIER |
| 4 | 100363426 | AX-97814029 | 0 | T | G | 0.43 | -1.06E-01 | 7.63E-02 | 2.57E+00 | 2.65E+00 | 1.63E-01 | no | Ror1 | intron_variant | MODIFIER |
| 4 | 100363831 | AX-170023513 | 0 | T | A | 0.43 | -1.06E-01 | 7.63E-02 | 2.57E+00 | 2.65E+00 | 1.63E-01 | no | Ror1 | intron_variant | MODIFIER |
| 4 | 100419556 | AX-170309185 | 0 | C | T | 0.43 | -1.06E-01 | 7.63E-02 | 2.57E+00 | 2.65E+00 | 1.63E-01 | no | Ror1 | intron_variant | MODIFIER |
| 4 | 100429808 | AX-170023508 | 0 | T | C | 0.43 | -1.06E-01 | 7.63E-02 | 2.57E+00 | 2.65E+00 | 1.63E-01 | no | Ror1 | intron_variant | MODIFIER |
| 4 | 100792137 | AX-170023520 | 0 | A | G | 0.43 | -1.06E-01 | 7.63E-02 | 2.57E+00 | 2.65E+00 | 1.63E-01 | no | Cachd1 | intron_variant | MODIFIER |
| 4 | 100793182 | AX-170121139 | 0 | T | C | 0.43 | -1.06E-01 | 7.63E-02 | 2.57E+00 | 2.65E+00 | 1.63E-01 | no | Cachd1 | intron_variant | MODIFIER |
| 4 | 100794725 | AX-170255018 | 0 | T | C | 0.43 | -1.06E-01 | 7.63E-02 | 2.57E+00 | 2.65E+00 | 1.63E-01 | no | Cachd1 | intron_variant | MODIFIER |
| 4 | 100796429 | AX-170023519 | 0 | G | T | 0.43 | -1.06E-01 | 7.63E-02 | 2.57E+00 | 2.65E+00 | 1.63E-01 | no | Cachd1 | intron_variant | MODIFIER |
| 4 | 100824018 | AX-170132606 | 0 | T | C | 0.43 | -1.06E-01 | 7.63E-02 | 2.57E+00 | 2.65E+00 | 1.63E-01 | no | Cachd1 | intron_variant | MODIFIER |
| 4 | 100824720 | AX-170310094 | 0 | A | C | 0.43 | -1.06E-01 | 7.63E-02 | 2.57E+00 | 2.65E+00 | 1.63E-01 | no | Cachd1 | intron_variant | MODIFIER |
| 1 | 181477907 | AX-170383039 | 1 | T | C | 0.223 | 1.56E-01 | 1.12E-01 | 2.52E+00 | 2.59E+00 | 1.63E-01 | Yes | | intergenic_variant | MODIFIER |
| 13 | 10581907 | AX-97791473 | 0 | A | G | 0.193 | -1.62E-01 | 1.16E-01 | 2.50E+00 | 2.55E+00 | 1.63E-01 | no | Gm26861 | intron_variant,non_coding_transcript_variant | MODIFIER |
| 18 | 76714565 | AX-170294957 | 1 | A | G | 0.429 | 1.13E-01 | 8.13E-02 | 2.57E+00 | 2.66E+00 | 1.63E-01 | no | | intergenic_variant | MODIFIER |
| 8 | 66042773 | AX-170454843 | 1 | A | G | 0.152 | -1.56E-01 | 1.13E-01 | 2.66E+00 | 2.80E+00 | 1.63E-01 | no | MAR1 | intron_variant | MODIFIER |
| 5 | 54186745 | AX-97835848 | 1 | A | C | 0.482 | 1.37E-01 | 9.58E-02 | 2.30E+00 | 2.37E+00 | 1.63E-01 | no | | intergenic_variant | MODIFIER |
| 1 | 65933237 | AX-170000699 | 1 | G | T | 0.161 | 2.14E-01 | 1.55E-01 | 2.75E+00 | 2.89E+00 | 1.63E-01 | Yes | Crygf | downstream_gene_variant | MODIFIER |
| 5 | 103967539 | AX-170295715 | 0 | T | A | 0.439 | 1.51E-01 | 1.09E-01 | 2.78E+00 | 2.84E+00 | 1.64E-01 | Yes | Hsd17b13 | intron_variant | MODIFIER |
| 16 | 80472889 | AX-97810159 | 0 | G | T | 0.474 | 1.03E-01 | 7.45E-02 | 2.67E+00 | 2.75E+00 | 1.64E-01 | no | | intergenic_variant | MODIFIER |
| 8 | 72712538 | AX-170408483 | 0 | A | T | 0.211 | -1.41E-01 | 1.02E-01 | 2.91E+00 | 3.09E+00 | 1.64E-01 | no | Nwd1 | 3_prime_UTR_variant | MODIFIER |
| 1 | 153019440 | AX-170315025 | 1 | A | G | 0.062 | -1.33E-01 | 9.68E-02 | 2.82E+00 | 3.03E+00 | 1.64E-01 | no | Nmnat2 | intron_variant | MODIFIER |
| 15 | 66596227 | AX-170055819 | 1 | C | A | 0.321 | -1.10E-01 | 7.56E-02 | 2.22E+00 | 2.28E+00 | 1.64E-01 | no | Phf20l1 | intron_variant | MODIFIER |
| 1 | 93623370 | AX-170310386 | 2 | C | T | 0.127 | 1.93E-01 | 1.40E-01 | 2.65E+00 | 2.78E+00 | 1.65E-01 | Yes | Stk25 | intron_variant | MODIFIER |
| 4 | 76532762 | AX-170519119 | 0 | G | T | 0.325 | 1.55E-01 | 1.13E-01 | 2.75E+00 | 2.83E+00 | 1.65E-01 | Yes | Ptprd | intron_variant,non_coding_transcript_variant | MODIFIER |
| 14 | 124243226 | AX-97802259 | 1 | T | C | 0.304 | 1.04E-01 | 7.56E-02 | 2.68E+00 | 2.79E+00 | 1.65E-01 | no | Fgf14 | intron_variant | MODIFIER |
| 10 | 100061982 | AX-170275243 | 0 | G | A | 0.272 | -1.15E-01 | 8.34E-02 | 2.63E+00 | 2.70E+00 | 1.65E-01 | no | Kitl | intron_variant | MODIFIER |
| 3 | 38443317 | AX-170149343 | 1 | C | T | 0.223 | 1.24E-01 | 8.89E-02 | 3.00E+00 | 3.13E+00 | 1.65E-01 | no | Gm7815 | upstream_gene_variant | MODIFIER |

| | | | | | | | | | | | | | | | |
|---|---|---|---|---|---|---|---|---|---|---|---|---|---|---|---|
| 5 | 14009301 | AX-97833859 | 0 | C | T | 0.333 | 1.14E-01 | 8.26E-02 | 2.86E+00 | 3.01E+00 | 1.65E-01 | no | | intergenic_variant | MODIFIER |
| 5 | 60277164 | AX-170025077 | 1 | G | A | 0.473 | 1.42E-01 | 1.01E-01 | 2.39E+00 | 2.45E+00 | 1.65E-01 | no | | intergenic_variant | MODIFIER |
| 10 | 23704435 | AX-170496300 | 0 | A | G | 0.184 | 1.32E-01 | 9.54E-02 | 2.99E+00 | 3.12E+00 | 1.65E-01 | no | | intergenic_variant | MODIFIER |
| 13 | 5710854 | AX-170007672 | 0 | C | G | 0.193 | -1.60E-01 | 1.16E-01 | 2.64E+00 | 2.72E+00 | 1.65E-01 | no | Gm35330 | intron_variant,non_coding_transcript_variant | MODIFIER |
| 1 | 123054153 | AX-170042540 | 1 | G | A | 0.571 | 1.15E-01 | 8.34E-02 | 2.59E+00 | 2.66E+00 | 1.65E-01 | no | | intergenic_variant | MODIFIER |
| 13 | 19114677 | AX-170394186 | 0 | C | T | 0.175 | -1.73E-01 | 1.24E-01 | 2.47E+00 | 2.52E+00 | 1.65E-01 | no | Amph | intron_variant | MODIFIER |
| 13 | 19123607 | AX-170387607 | 0 | T | G | 0.175 | -1.73E-01 | 1.24E-01 | 2.47E+00 | 2.52E+00 | 1.65E-01 | no | Amph | intron_variant | MODIFIER |
| 13 | 19125379 | AX-170523324 | 0 | G | A | 0.175 | -1.73E-01 | 1.24E-01 | 2.47E+00 | 2.52E+00 | 1.65E-01 | no | Amph | intron_variant | MODIFIER |
| 13 | 19125818 | AX-170335790 | 0 | G | A | 0.175 | -1.73E-01 | 1.24E-01 | 2.47E+00 | 2.52E+00 | 1.65E-01 | no | Amph | intron_variant | MODIFIER |
| 13 | 19138849 | AX-97840838 | 0 | A | G | 0.175 | -1.73E-01 | 1.24E-01 | 2.47E+00 | 2.52E+00 | 1.65E-01 | no | Amph | intron_variant | MODIFIER |
| 13 | 19149822 | AX-170050666 | 0 | C | T | 0.175 | -1.73E-01 | 1.24E-01 | 2.47E+00 | 2.52E+00 | 1.65E-01 | no | Amph | synonymous_variant | LOW |
| 13 | 19153177 | AX-170481160 | 0 | C | G | 0.175 | -1.73E-01 | 1.24E-01 | 2.47E+00 | 2.52E+00 | 1.65E-01 | no | Amph | downstream_gene_variant | MODIFIER |
| 13 | 19154290 | AX-170184560 | 0 | T | A | 0.175 | -1.73E-01 | 1.24E-01 | 2.47E+00 | 2.52E+00 | 1.65E-01 | no | Amph | downstream_gene_variant | MODIFIER |
| 13 | 19154940 | AX-170438755 | 0 | C | G | 0.175 | -1.73E-01 | 1.24E-01 | 2.47E+00 | 2.52E+00 | 1.65E-01 | no | Amph | downstream_gene_variant | MODIFIER |
| 13 | 19158000 | AX-170411407 | 0 | A | G | 0.175 | -1.73E-01 | 1.24E-01 | 2.47E+00 | 2.52E+00 | 1.65E-01 | no | Gm31887 | downstream_gene_variant | MODIFIER |
| 13 | 19183355 | AX-97850003 | 0 | C | T | 0.175 | -1.73E-01 | 1.24E-01 | 2.47E+00 | 2.52E+00 | 1.65E-01 | no | Tcrg-V7 | downstream_gene_variant | MODIFIER |
| 13 | 19360609 | AX-170213842 | 0 | G | A | 0.175 | -1.73E-01 | 1.24E-01 | 2.47E+00 | 2.52E+00 | 1.65E-01 | no | Stard3nl | intron_variant | MODIFIER |
| 13 | 19362579 | AX-170165094 | 0 | T | G | 0.175 | -1.73E-01 | 1.24E-01 | 2.47E+00 | 2.52E+00 | 1.65E-01 | no | Stard3nl | intron_variant | MODIFIER |
| 13 | 19362956 | AX-170152411 | 0 | G | A | 0.175 | -1.73E-01 | 1.24E-01 | 2.47E+00 | 2.52E+00 | 1.65E-01 | no | Stard3nl | intron_variant | MODIFIER |
| 13 | 19827014 | AX-170108260 | 0 | G | C | 0.175 | -1.73E-01 | 1.24E-01 | 2.47E+00 | 2.52E+00 | 1.65E-01 | no | Gpr141 | upstream_gene_variant | MODIFIER |
| 13 | 19902079 | AX-170104531 | 0 | G | A | 0.175 | -1.73E-01 | 1.24E-01 | 2.47E+00 | 2.52E+00 | 1.65E-01 | no | | intergenic_variant | MODIFIER |
| 13 | 19902644 | AX-97844542 | 0 | T | G | 0.175 | -1.73E-01 | 1.24E-01 | 2.47E+00 | 2.52E+00 | 1.65E-01 | no | | intergenic_variant | MODIFIER |
| 13 | 19983356 | AX-170050689 | 0 | T | C | 0.175 | -1.73E-01 | 1.24E-01 | 2.47E+00 | 2.52E+00 | 1.65E-01 | no | | intergenic_variant | MODIFIER |
| 13 | 19984106 | AX-170007881 | 0 | C | T | 0.175 | -1.73E-01 | 1.24E-01 | 2.47E+00 | 2.52E+00 | 1.65E-01 | no | | intergenic_variant | MODIFIER |
| 13 | 20097045 | AX-170229421 | 0 | C | G | 0.175 | -1.73E-01 | 1.24E-01 | 2.47E+00 | 2.52E+00 | 1.65E-01 | no | Elmo1 | intron_variant | MODIFIER |
| 13 | 20111249 | AX-170050690 | 0 | A | G | 0.175 | -1.73E-01 | 1.24E-01 | 2.47E+00 | 2.52E+00 | 1.65E-01 | no | Elmo1 | intron_variant | MODIFIER |
| 13 | 20146126 | AX-170193218 | 0 | A | G | 0.175 | -1.73E-01 | 1.24E-01 | 2.47E+00 | 2.52E+00 | 1.65E-01 | no | Elmo1 | intron_variant | MODIFIER |
| 13 | 20146238 | AX-170534916 | 0 | C | T | 0.175 | -1.73E-01 | 1.24E-01 | 2.47E+00 | 2.52E+00 | 1.65E-01 | no | Elmo1 | intron_variant | MODIFIER |
| 13 | 20146439 | AX-170353356 | 0 | T | C | 0.175 | -1.73E-01 | 1.24E-01 | 2.47E+00 | 2.52E+00 | 1.65E-01 | no | Elmo1 | intron_variant | MODIFIER |
| 13 | 20175301 | AX-170360599 | 0 | A | G | 0.175 | -1.73E-01 | 1.24E-01 | 2.47E+00 | 2.52E+00 | 1.65E-01 | no | Elmo1 | intron_variant | MODIFIER |
| 13 | 20175773 | AX-97812498 | 0 | T | C | 0.175 | -1.73E-01 | 1.24E-01 | 2.47E+00 | 2.52E+00 | 1.65E-01 | no | Elmo1 | intron_variant | MODIFIER |
| 13 | 20176239 | AX-170050694 | 0 | A | G | 0.175 | -1.73E-01 | 1.24E-01 | 2.47E+00 | 2.52E+00 | 1.65E-01 | no | Elmo1 | intron_variant | MODIFIER |
| 13 | 20177276 | AX-170307274 | 0 | A | T | 0.175 | -1.73E-01 | 1.24E-01 | 2.47E+00 | 2.52E+00 | 1.65E-01 | no | Elmo1 | intron_variant | MODIFIER |
| 13 | 20178809 | AX-170134494 | 0 | A | C | 0.175 | -1.73E-01 | 1.24E-01 | 2.47E+00 | 2.52E+00 | 1.65E-01 | no | Elmo1 | intron_variant | MODIFIER |
| 13 | 20186083 | AX-170472676 | 0 | C | T | 0.175 | -1.73E-01 | 1.24E-01 | 2.47E+00 | 2.52E+00 | 1.65E-01 | no | Elmo1 | intron_variant | MODIFIER |
| 13 | 20189181 | AX-170330201 | 0 | T | C | 0.175 | -1.73E-01 | 1.24E-01 | 2.47E+00 | 2.52E+00 | 1.65E-01 | no | Elmo1 | intron_variant | MODIFIER |
| 13 | 20201243 | AX-170474194 | 0 | T | G | 0.175 | -1.73E-01 | 1.24E-01 | 2.47E+00 | 2.52E+00 | 1.65E-01 | no | Elmo1 | intron_variant | MODIFIER |
| 13 | 20211248 | AX-170449620 | 0 | T | G | 0.175 | -1.73E-01 | 1.24E-01 | 2.47E+00 | 2.52E+00 | 1.65E-01 | no | Elmo1 | intron_variant | MODIFIER |
| 13 | 20246823 | AX-170186297 | 0 | G | A | 0.175 | -1.73E-01 | 1.24E-01 | 2.47E+00 | 2.52E+00 | 1.65E-01 | no | Elmo1 | intron_variant | MODIFIER |
| 13 | 20249595 | AX-170509443 | 0 | T | C | 0.175 | -1.73E-01 | 1.24E-01 | 2.47E+00 | 2.52E+00 | 1.65E-01 | no | Elmo1 | intron_variant | MODIFIER |
| 13 | 20251144 | AX-170358391 | 0 | T | C | 0.175 | -1.73E-01 | 1.24E-01 | 2.47E+00 | 2.52E+00 | 1.65E-01 | no | Elmo1 | synonymous_variant | LOW |
| 13 | 20251448 | AX-170368844 | 0 | C | G | 0.175 | -1.73E-01 | 1.24E-01 | 2.47E+00 | 2.52E+00 | 1.65E-01 | no | Elmo1 | intron_variant | MODIFIER |
| 13 | 20270139 | AX-170497700 | 0 | C | T | 0.175 | -1.73E-01 | 1.24E-01 | 2.47E+00 | 2.52E+00 | 1.65E-01 | no | Elmo1 | intron_variant | MODIFIER |
| 13 | 20270397 | AX-170328454 | 0 | C | T | 0.175 | -1.73E-01 | 1.24E-01 | 2.47E+00 | 2.52E+00 | 1.65E-01 | no | Elmo1 | intron_variant | MODIFIER |
| 13 | 20271827 | AX-170050696 | 0 | T | G | 0.175 | -1.73E-01 | 1.24E-01 | 2.47E+00 | 2.52E+00 | 1.65E-01 | no | Elmo1 | intron_variant | MODIFIER |
| 13 | 20283531 | AX-170425256 | 0 | A | G | 0.175 | -1.73E-01 | 1.24E-01 | 2.47E+00 | 2.52E+00 | 1.65E-01 | no | Elmo1 | intron_variant | MODIFIER |
| 13 | 20283716 | AX-170035524 | 0 | T | C | 0.175 | -1.73E-01 | 1.24E-01 | 2.47E+00 | 2.52E+00 | 1.65E-01 | no | Elmo1 | intron_variant | MODIFIER |
| 13 | 20286966 | AX-170141838 | 0 | T | G | 0.175 | -1.73E-01 | 1.24E-01 | 2.47E+00 | 2.52E+00 | 1.65E-01 | no | Elmo1 | intron_variant | MODIFIER |
| 13 | 20301917 | AX-97808362 | 0 | C | A | 0.175 | -1.73E-01 | 1.24E-01 | 2.47E+00 | 2.52E+00 | 1.65E-01 | no | Elmo1 | intron_variant | MODIFIER |
| 13 | 20312500 | AX-170248018 | 0 | G | A | 0.175 | -1.73E-01 | 1.24E-01 | 2.47E+00 | 2.52E+00 | 1.65E-01 | no | Elmo1 | intron_variant | MODIFIER |
| 13 | 20315428 | AX-170453909 | 0 | G | A | 0.175 | -1.73E-01 | 1.24E-01 | 2.47E+00 | 2.52E+00 | 1.65E-01 | no | Elmo1 | intron_variant | MODIFIER |
| 13 | 20322074 | AX-170227329 | 0 | T | C | 0.175 | -1.73E-01 | 1.24E-01 | 2.47E+00 | 2.52E+00 | 1.65E-01 | no | Elmo1 | intron_variant | MODIFIER |
| 13 | 20322495 | AX-170471637 | 0 | G | T | 0.175 | -1.73E-01 | 1.24E-01 | 2.47E+00 | 2.52E+00 | 1.65E-01 | no | Elmo1 | intron_variant | MODIFIER |
| 13 | 20335379 | AX-170404723 | 0 | C | T | 0.175 | -1.73E-01 | 1.24E-01 | 2.47E+00 | 2.52E+00 | 1.65E-01 | no | Elmo1 | intron_variant | MODIFIER |

| | | | | | | | | | | | | | | | |
|---|---|---|---|---|---|---|---|---|---|---|---|---|---|---|---|
| 13 | 20342541 | AX-170491516 | 0 | G | A | 0.175 | -1.73E-01 | 1.24E-01 | 2.47E+00 | 2.52E+00 | 1.65E-01 | no | Elmo1 | intron_variant | MODIFIER |
| 13 | 20348391 | AX-170366460 | 0 | G | T | 0.175 | -1.73E-01 | 1.24E-01 | 2.47E+00 | 2.52E+00 | 1.65E-01 | no | Elmo1 | intron_variant | MODIFIER |
| 13 | 20348705 | AX-170328221 | 0 | G | A | 0.175 | -1.73E-01 | 1.24E-01 | 2.47E+00 | 2.52E+00 | 1.65E-01 | no | Elmo1 | intron_variant | MODIFIER |
| 13 | 20364137 | AX-97830242 | 0 | A | C | 0.175 | -1.73E-01 | 1.24E-01 | 2.47E+00 | 2.52E+00 | 1.65E-01 | no | Elmo1 | intron_variant | MODIFIER |
| 13 | 20365143 | AX-170452375 | 0 | C | A | 0.175 | -1.73E-01 | 1.24E-01 | 2.47E+00 | 2.52E+00 | 1.65E-01 | no | Elmo1 | intron_variant | MODIFIER |
| 13 | 20365366 | AX-170476028 | 0 | C | T | 0.175 | -1.73E-01 | 1.24E-01 | 2.47E+00 | 2.52E+00 | 1.65E-01 | no | Elmo1 | intron_variant | MODIFIER |
| 13 | 20374082 | AX-170243957 | 0 | A | C | 0.175 | -1.73E-01 | 1.24E-01 | 2.47E+00 | 2.52E+00 | 1.65E-01 | no | Elmo1 | intron_variant | MODIFIER |
| 13 | 20453161 | AX-170204501 | 0 | T | C | 0.175 | -1.73E-01 | 1.24E-01 | 2.47E+00 | 2.52E+00 | 1.65E-01 | no | Elmo1 | intron_variant | MODIFIER |
| 13 | 20482029 | AX-170481610 | 0 | A | G | 0.175 | -1.73E-01 | 1.24E-01 | 2.47E+00 | 2.52E+00 | 1.65E-01 | no | Elmo1 | intron_variant | MODIFIER |
| 13 | 20493356 | AX-170343910 | 0 | A | T | 0.175 | -1.73E-01 | 1.24E-01 | 2.47E+00 | 2.52E+00 | 1.65E-01 | no | Elmo1 | intron_variant | MODIFIER |
| 13 | 20497559 | AX-170194588 | 0 | T | C | 0.175 | -1.73E-01 | 1.24E-01 | 2.47E+00 | 2.52E+00 | 1.65E-01 | no | Elmo1 | intron_variant | MODIFIER |
| 13 | 20505481 | AX-170163776 | 0 | C | T | 0.175 | -1.73E-01 | 1.24E-01 | 2.47E+00 | 2.52E+00 | 1.65E-01 | no | Elmo1 | intron_variant | MODIFIER |
| 13 | 20539530 | AX-170172089 | 0 | G | T | 0.175 | -1.73E-01 | 1.24E-01 | 2.47E+00 | 2.52E+00 | 1.65E-01 | no | Elmo1 | intron_variant | MODIFIER |
| 13 | 20548169 | AX-170374269 | 0 | C | T | 0.175 | -1.73E-01 | 1.24E-01 | 2.47E+00 | 2.52E+00 | 1.65E-01 | no | Elmo1 | intron_variant | MODIFIER |
| 13 | 20570254 | AX-170412149 | 0 | T | A | 0.175 | -1.73E-01 | 1.24E-01 | 2.47E+00 | 2.52E+00 | 1.65E-01 | no | Elmo1 | intron_variant | MODIFIER |
| 13 | 20572718 | AX-170401338 | 0 | T | C | 0.175 | -1.73E-01 | 1.24E-01 | 2.47E+00 | 2.52E+00 | 1.65E-01 | no | Elmo1 | intron_variant | MODIFIER |
| 13 | 20572861 | AX-170007886 | 0 | G | A | 0.175 | -1.73E-01 | 1.24E-01 | 2.47E+00 | 2.52E+00 | 1.65E-01 | no | Elmo1 | intron_variant | MODIFIER |
| 13 | 20574340 | AX-170007887 | 0 | C | G | 0.175 | -1.73E-01 | 1.24E-01 | 2.47E+00 | 2.52E+00 | 1.65E-01 | no | Elmo1 | intron_variant | MODIFIER |
| 13 | 20576915 | AX-170167050 | 0 | T | A | 0.175 | -1.73E-01 | 1.24E-01 | 2.47E+00 | 2.52E+00 | 1.65E-01 | no | Elmo1 | intron_variant | MODIFIER |
| 13 | 20591404 | AX-97840999 | 0 | G | A | 0.175 | -1.73E-01 | 1.24E-01 | 2.47E+00 | 2.52E+00 | 1.65E-01 | no | Elmo1 | intron_variant | MODIFIER |
| 13 | 20603927 | AX-170149431 | 0 | T | G | 0.175 | -1.73E-01 | 1.24E-01 | 2.47E+00 | 2.52E+00 | 1.65E-01 | no | Elmo1 | intron_variant | MODIFIER |
| 13 | 20606674 | AX-170236578 | 0 | T | C | 0.175 | -1.73E-01 | 1.24E-01 | 2.47E+00 | 2.52E+00 | 1.65E-01 | no | Elmo1 | 3_prime_UTR_variant | MODIFIER |
| 13 | 20607257 | AX-170382981 | 0 | G | A | 0.175 | -1.73E-01 | 1.24E-01 | 2.47E+00 | 2.52E+00 | 1.65E-01 | no | Elmo1 | 3_prime_UTR_variant | MODIFIER |
| 13 | 20618964 | AX-170307696 | 0 | G | C | 0.175 | -1.73E-01 | 1.24E-01 | 2.47E+00 | 2.52E+00 | 1.65E-01 | no | | intergenic_variant | MODIFIER |
| 13 | 20619016 | AX-170447677 | 0 | A | G | 0.175 | -1.73E-01 | 1.24E-01 | 2.47E+00 | 2.52E+00 | 1.65E-01 | no | | intergenic_variant | MODIFIER |
| 13 | 20627062 | AX-170479980 | 0 | C | T | 0.175 | -1.73E-01 | 1.24E-01 | 2.47E+00 | 2.52E+00 | 1.65E-01 | no | | intergenic_variant | MODIFIER |
| 13 | 20646676 | AX-170529748 | 0 | A | G | 0.175 | -1.73E-01 | 1.24E-01 | 2.47E+00 | 2.52E+00 | 1.65E-01 | no | Gm47721 | non_coding_transcript_exon_variant | MODIFIER |
| 13 | 20648331 | AX-170120877 | 0 | G | C | 0.175 | -1.73E-01 | 1.24E-01 | 2.47E+00 | 2.52E+00 | 1.65E-01 | no | Gm47721 | downstream_gene_variant | MODIFIER |
| 13 | 20648398 | AX-170234412 | 0 | T | G | 0.175 | -1.73E-01 | 1.24E-01 | 2.47E+00 | 2.52E+00 | 1.65E-01 | no | Gm47721 | downstream_gene_variant | MODIFIER |
| 13 | 20648560 | AX-170177291 | 0 | A | G | 0.175 | -1.73E-01 | 1.24E-01 | 2.47E+00 | 2.52E+00 | 1.65E-01 | no | Gm47721 | downstream_gene_variant | MODIFIER |
| 13 | 20660366 | AX-97836267 | 0 | G | A | 0.175 | -1.73E-01 | 1.24E-01 | 2.47E+00 | 2.52E+00 | 1.65E-01 | no | | intergenic_variant | MODIFIER |
| 13 | 20725859 | AX-170035526 | 0 | A | G | 0.175 | -1.73E-01 | 1.24E-01 | 2.47E+00 | 2.52E+00 | 1.65E-01 | no | | intergenic_variant | MODIFIER |
| 13 | 20734445 | AX-170131789 | 0 | C | T | 0.175 | -1.73E-01 | 1.24E-01 | 2.47E+00 | 2.52E+00 | 1.65E-01 | no | | intergenic_variant | MODIFIER |
| 13 | 20772824 | AX-170133869 | 0 | T | A | 0.175 | -1.73E-01 | 1.24E-01 | 2.47E+00 | 2.52E+00 | 1.65E-01 | no | | intergenic_variant | MODIFIER |
| 13 | 20772911 | AX-170435866 | 0 | A | G | 0.175 | -1.73E-01 | 1.24E-01 | 2.47E+00 | 2.52E+00 | 1.65E-01 | no | | intergenic_variant | MODIFIER |
| 13 | 20813699 | AX-170185573 | 0 | A | G | 0.175 | -1.73E-01 | 1.24E-01 | 2.47E+00 | 2.52E+00 | 1.65E-01 | no | Aoah | intron_variant | MODIFIER |
| 13 | 20855214 | AX-170171531 | 0 | A | G | 0.175 | -1.73E-01 | 1.24E-01 | 2.47E+00 | 2.52E+00 | 1.65E-01 | no | Aoah | intron_variant | MODIFIER |
| 4 | 134091257 | AX-170302332 | 0 | T | C | 0.167 | 1.49E-01 | 1.08E-01 | 2.57E+00 | 2.63E+00 | 1.66E-01 | Yes | Cd52 | downstream_gene_variant | MODIFIER |
| 5 | 88202568 | AX-170295246 | 0 | C | G | 0.43 | 1.51E-01 | 1.08E-01 | 3.09E+00 | 3.18E+00 | 1.66E-01 | Yes | Gm42621 | downstream_gene_variant | MODIFIER |
| 5 | 52812801 | AX-170534557 | 1 | C | A | 0.5 | 1.37E-01 | 9.56E-02 | 2.26E+00 | 2.33E+00 | 1.66E-01 | no | Zcchc4 | intron_variant | MODIFIER |
| 4 | 107840270 | AX-170023623 | 1 | C | T | 0.268 | 1.54E-01 | 1.12E-01 | 2.62E+00 | 2.68E+00 | 1.66E-01 | Yes | Lrp8 | intron_variant | MODIFIER |
| 3 | 19459872 | AX-97826776 | 1 | C | T | 0.125 | -1.39E-01 | 1.01E-01 | 2.60E+00 | 2.68E+00 | 1.66E-01 | no | | intergenic_variant | MODIFIER |
| 15 | 76498342 | AX-170056025 | 0 | A | G | 0.088 | -1.56E-01 | 9.86E-02 | 1.88E+00 | 1.95E+00 | 1.66E-01 | no | Dgat1 | downstream_gene_variant | MODIFIER |
| 13 | 18338962 | AX-170450597 | 1 | G | A | 0.161 | -1.76E-01 | 1.26E-01 | 2.39E+00 | 2.44E+00 | 1.66E-01 | Yes | Pou6f2 | intron_variant | MODIFIER |
| 16 | 55371162 | AX-170419615 | 1 | T | C | 0.527 | 1.18E-01 | 8.41E-02 | 2.42E+00 | 2.46E+00 | 1.66E-01 | no | | intergenic_variant | MODIFIER |
| 3 | 7630146 | AX-170102625 | 0 | G | A | 0.404 | -8.68E-02 | 6.30E-02 | 2.90E+00 | 3.05E+00 | 1.66E-01 | no | Gm16685 | intron_variant,non_coding_transcript_variant | MODIFIER |
| 16 | 32648323 | AX-170468311 | 1 | G | A | 0.509 | 1.09E-01 | 7.93E-02 | 2.62E+00 | 2.69E+00 | 1.66E-01 | no | Tnk2 | intron_variant | MODIFIER |
| 5 | 114599325 | AX-170275120 | 0 | G | A | 0.465 | 1.41E-01 | 1.03E-01 | 2.75E+00 | 2.82E+00 | 1.67E-01 | no | Fam222a | intron_variant | MODIFIER |
| 16 | 40904660 | AX-170012886 | 0 | A | T | 0.491 | 1.12E-01 | 8.15E-02 | 2.60E+00 | 2.65E+00 | 1.67E-01 | no | Lsamp | intron_variant | MODIFIER |
| 16 | 40916480 | AX-170012889 | 0 | C | T | 0.491 | 1.12E-01 | 8.15E-02 | 2.60E+00 | 2.65E+00 | 1.67E-01 | no | Lsamp | intron_variant | MODIFIER |
| 1 | 191659792 | AX-97791543 | 0 | T | C | 0.272 | 1.29E-01 | 9.18E-02 | 2.35E+00 | 2.43E+00 | 1.67E-01 | no | | intergenic_variant | MODIFIER |
| 1 | 191669088 | AX-170043980 | 0 | G | C | 0.272 | 1.29E-01 | 9.18E-02 | 2.35E+00 | 2.43E+00 | 1.67E-01 | no | | intergenic_variant | MODIFIER |
| 1 | 191673478 | AX-170002581 | 0 | G | T | 0.272 | 1.29E-01 | 9.18E-02 | 2.35E+00 | 2.43E+00 | 1.67E-01 | no | | intergenic_variant | MODIFIER |
| 1 | 191805058 | AX-170295230 | 0 | C | T | 0.272 | 1.29E-01 | 9.18E-02 | 2.35E+00 | 2.43E+00 | 1.67E-01 | no | | intergenic_variant | MODIFIER |

| | | | | | | | | | | | | | | | |
|---|---|---|---|---|---|---|---|---|---|---|---|---|---|---|---|
| 1 | 191808638 | AX-97832868 | 0 | A | G | 0.272 | 1.29E-01 | 9.18E-02 | 2.35E+00 | 2.43E+00 | 1.67E-01 | no | | regulatory_region_variant | MODIFIER |
| 1 | 191822255 | AX-170158914 | 0 | A | C | 0.272 | 1.29E-01 | 9.18E-02 | 2.35E+00 | 2.43E+00 | 1.67E-01 | no | Nek2 | missense_variant | MODERATE |
| 1 | 191826427 | AX-170002574 | 0 | G | A | 0.272 | 1.29E-01 | 9.18E-02 | 2.35E+00 | 2.43E+00 | 1.67E-01 | no | Nek2 | intron_variant | MODIFIER |
| 1 | 191836666 | AX-170505015 | 0 | A | C | 0.272 | 1.29E-01 | 9.18E-02 | 2.35E+00 | 2.43E+00 | 1.67E-01 | no | Nek2 | downstream_gene_variant | MODIFIER |
| 1 | 191836870 | AX-170377616 | 0 | T | C | 0.272 | 1.29E-01 | 9.18E-02 | 2.35E+00 | 2.43E+00 | 1.67E-01 | no | Nek2 | downstream_gene_variant | MODIFIER |
| 1 | 191837227 | AX-170294282 | 0 | A | C | 0.272 | 1.29E-01 | 9.18E-02 | 2.35E+00 | 2.43E+00 | 1.67E-01 | no | Nek2 | downstream_gene_variant | MODIFIER |
| 1 | 191865869 | AX-170528790 | 0 | G | T | 0.272 | 1.29E-01 | 9.18E-02 | 2.35E+00 | 2.43E+00 | 1.67E-01 | no | | intergenic_variant | MODIFIER |
| 1 | 191885346 | AX-97795511 | 0 | G | A | 0.272 | 1.29E-01 | 9.18E-02 | 2.35E+00 | 2.43E+00 | 1.67E-01 | no | 1700034H15Rik | downstream_gene_variant | MODIFIER |
| 1 | 191885518 | AX-170002575 | 0 | A | G | 0.272 | 1.29E-01 | 9.18E-02 | 2.35E+00 | 2.43E+00 | 1.67E-01 | no | 1700034H15Rik | downstream_gene_variant | MODIFIER |
| 1 | 191886210 | AX-170043983 | 0 | A | G | 0.272 | 1.29E-01 | 9.18E-02 | 2.35E+00 | 2.43E+00 | 1.67E-01 | no | 1700034H15Rik | downstream_gene_variant | MODIFIER |
| 1 | 191890897 | AX-170043984 | 0 | G | A | 0.272 | 1.29E-01 | 9.18E-02 | 2.35E+00 | 2.43E+00 | 1.67E-01 | no | 1700034H15Rik | downstream_gene_variant | MODIFIER |
| 1 | 191965516 | AX-170480247 | 0 | T | C | 0.272 | 1.29E-01 | 9.18E-02 | 2.35E+00 | 2.43E+00 | 1.67E-01 | no | | regulatory_region_variant | MODIFIER |
| 1 | 191967696 | AX-170368168 | 0 | A | G | 0.272 | 1.29E-01 | 9.18E-02 | 2.35E+00 | 2.43E+00 | 1.67E-01 | no | | intergenic_variant | MODIFIER |
| 1 | 192091867 | AX-97814657 | 0 | G | A | 0.272 | 1.29E-01 | 9.18E-02 | 2.35E+00 | 2.43E+00 | 1.67E-01 | no | Traf5 | intron_variant | MODIFIER |
| 1 | 192105460 | AX-170086724 | 0 | C | T | 0.272 | 1.29E-01 | 9.18E-02 | 2.35E+00 | 2.43E+00 | 1.67E-01 | no | Rcor3 | intron_variant | MODIFIER |
| 1 | 192141559 | AX-170411594 | 0 | A | G | 0.272 | 1.29E-01 | 9.18E-02 | 2.35E+00 | 2.43E+00 | 1.67E-01 | no | Rcor3 | upstream_gene_variant | MODIFIER |
| 1 | 192168780 | AX-170246633 | 0 | G | T | 0.272 | 1.29E-01 | 9.18E-02 | 2.35E+00 | 2.43E+00 | 1.67E-01 | no | | intergenic_variant | MODIFIER |
| 1 | 192171580 | AX-170365107 | 0 | T | C | 0.272 | 1.29E-01 | 9.18E-02 | 2.35E+00 | 2.43E+00 | 1.67E-01 | no | | intergenic_variant | MODIFIER |
| 1 | 192172948 | AX-170532524 | 0 | G | A | 0.272 | 1.29E-01 | 9.18E-02 | 2.35E+00 | 2.43E+00 | 1.67E-01 | no | | intergenic_variant | MODIFIER |
| 1 | 192208679 | AX-97826670 | 0 | G | T | 0.272 | 1.29E-01 | 9.18E-02 | 2.35E+00 | 2.43E+00 | 1.67E-01 | no | Kcnh1 | intron_variant | MODIFIER |
| 1 | 192208782 | AX-170043988 | 0 | A | G | 0.272 | 1.29E-01 | 9.18E-02 | 2.35E+00 | 2.43E+00 | 1.67E-01 | no | Kcnh1 | intron_variant | MODIFIER |
| 1 | 192210960 | AX-170043986 | 0 | T | C | 0.272 | 1.29E-01 | 9.18E-02 | 2.35E+00 | 2.43E+00 | 1.67E-01 | no | Kcnh1 | intron_variant | MODIFIER |
| 1 | 192211139 | AX-170524823 | 0 | A | G | 0.272 | 1.29E-01 | 9.18E-02 | 2.35E+00 | 2.43E+00 | 1.67E-01 | no | Kcnh1 | intron_variant | MODIFIER |
| 1 | 192214672 | AX-170349044 | 0 | A | G | 0.272 | 1.29E-01 | 9.18E-02 | 2.35E+00 | 2.43E+00 | 1.67E-01 | no | Kcnh1 | intron_variant | MODIFIER |
| 1 | 192227354 | AX-170505883 | 0 | A | G | 0.272 | 1.29E-01 | 9.18E-02 | 2.35E+00 | 2.43E+00 | 1.67E-01 | no | Kcnh1 | intron_variant | MODIFIER |
| 1 | 192234929 | AX-170322061 | 0 | T | C | 0.272 | 1.29E-01 | 9.18E-02 | 2.35E+00 | 2.43E+00 | 1.67E-01 | no | Kcnh1 | intron_variant | MODIFIER |
| 1 | 192235413 | AX-170415185 | 0 | T | C | 0.272 | 1.29E-01 | 9.18E-02 | 2.35E+00 | 2.43E+00 | 1.67E-01 | no | Kcnh1 | intron_variant | MODIFIER |
| 1 | 192237163 | AX-170216982 | 0 | T | C | 0.272 | 1.29E-01 | 9.18E-02 | 2.35E+00 | 2.43E+00 | 1.67E-01 | no | Kcnh1 | intron_variant | MODIFIER |
| 1 | 192238816 | AX-170401222 | 0 | T | C | 0.272 | 1.29E-01 | 9.18E-02 | 2.35E+00 | 2.43E+00 | 1.67E-01 | no | Kcnh1 | synonymous_variant | LOW |
| 1 | 192240115 | AX-170177512 | 0 | T | C | 0.272 | 1.29E-01 | 9.18E-02 | 2.35E+00 | 2.43E+00 | 1.67E-01 | no | Kcnh1 | intron_variant | MODIFIER |
| 1 | 192242308 | AX-170495270 | 0 | A | T | 0.272 | 1.29E-01 | 9.18E-02 | 2.35E+00 | 2.43E+00 | 1.67E-01 | no | Kcnh1 | intron_variant | MODIFIER |
| 1 | 192248133 | AX-170002583 | 0 | T | C | 0.272 | 1.29E-01 | 9.18E-02 | 2.35E+00 | 2.43E+00 | 1.67E-01 | no | Kcnh1 | intron_variant | MODIFIER |
| 1 | 192251210 | AX-170002587 | 0 | C | G | 0.272 | 1.29E-01 | 9.18E-02 | 2.35E+00 | 2.43E+00 | 1.67E-01 | no | Kcnh1 | intron_variant | MODIFIER |
| 1 | 192265245 | AX-170520922 | 0 | C | T | 0.272 | 1.29E-01 | 9.18E-02 | 2.35E+00 | 2.43E+00 | 1.67E-01 | no | Kcnh1 | intron_variant | MODIFIER |
| 1 | 192266974 | AX-170333684 | 0 | C | T | 0.272 | 1.29E-01 | 9.18E-02 | 2.35E+00 | 2.43E+00 | 1.67E-01 | no | Kcnh1 | intron_variant | MODIFIER |
| 1 | 192268028 | AX-170043991 | 0 | A | C | 0.272 | 1.29E-01 | 9.18E-02 | 2.35E+00 | 2.43E+00 | 1.67E-01 | no | Kcnh1 | intron_variant | MODIFIER |
| 1 | 192273752 | AX-170002586 | 0 | C | T | 0.272 | 1.29E-01 | 9.18E-02 | 2.35E+00 | 2.43E+00 | 1.67E-01 | no | Kcnh1 | intron_variant | MODIFIER |
| 1 | 192281713 | AX-170002584 | 0 | G | T | 0.272 | 1.29E-01 | 9.18E-02 | 2.35E+00 | 2.43E+00 | 1.67E-01 | no | Kcnh1 | intron_variant | MODIFIER |
| 1 | 192343890 | AX-170460621 | 0 | A | G | 0.272 | 1.29E-01 | 9.18E-02 | 2.35E+00 | 2.43E+00 | 1.67E-01 | no | Kcnh1 | intron_variant | MODIFIER |
| 1 | 192400302 | AX-170392076 | 0 | C | T | 0.272 | 1.29E-01 | 9.18E-02 | 2.35E+00 | 2.43E+00 | 1.67E-01 | no | Kcnh1 | intron_variant | MODIFIER |
| 1 | 192413462 | AX-170491129 | 0 | T | C | 0.272 | 1.29E-01 | 9.18E-02 | 2.35E+00 | 2.43E+00 | 1.67E-01 | no | Kcnh1 | intron_variant | MODIFIER |
| 1 | 192430302 | AX-170104745 | 0 | T | C | 0.272 | 1.29E-01 | 9.18E-02 | 2.35E+00 | 2.43E+00 | 1.67E-01 | no | Kcnh1 | intron_variant | MODIFIER |
| 1 | 192442075 | AX-97783549 | 0 | A | G | 0.272 | 1.29E-01 | 9.18E-02 | 2.35E+00 | 2.43E+00 | 1.67E-01 | no | Kcnh1 | intron_variant | MODIFIER |
| 1 | 192443379 | AX-170400854 | 0 | A | G | 0.272 | 1.29E-01 | 9.18E-02 | 2.35E+00 | 2.43E+00 | 1.67E-01 | no | Kcnh1 | intron_variant | MODIFIER |
| 1 | 192490237 | AX-170242823 | 0 | T | C | 0.272 | 1.29E-01 | 9.18E-02 | 2.35E+00 | 2.43E+00 | 1.67E-01 | no | Kcnh1 | intron_variant | MODIFIER |
| 1 | 192536425 | AX-170086725 | 0 | T | C | 0.272 | 1.29E-01 | 9.18E-02 | 2.35E+00 | 2.43E+00 | 1.67E-01 | no | Hhat | intron_variant | MODIFIER |
| 1 | 192549408 | AX-170217912 | 0 | T | C | 0.272 | 1.29E-01 | 9.18E-02 | 2.35E+00 | 2.43E+00 | 1.67E-01 | no | Hhat | intron_variant | MODIFIER |
| 1 | 192558198 | AX-170379487 | 0 | G | A | 0.272 | 1.29E-01 | 9.18E-02 | 2.35E+00 | 2.43E+00 | 1.67E-01 | no | Hhat | intron_variant | MODIFIER |
| 1 | 192595729 | AX-169999714 | 0 | A | G | 0.272 | 1.29E-01 | 9.18E-02 | 2.35E+00 | 2.43E+00 | 1.67E-01 | no | Hhat | intron_variant | MODIFIER |
| 1 | 192595751 | AX-170086173 | 0 | T | C | 0.272 | 1.29E-01 | 9.18E-02 | 2.35E+00 | 2.43E+00 | 1.67E-01 | no | Hhat | intron_variant | MODIFIER |
| 1 | 192600709 | AX-170419827 | 0 | A | G | 0.272 | 1.29E-01 | 9.18E-02 | 2.35E+00 | 2.43E+00 | 1.67E-01 | no | Hhat | intron_variant | MODIFIER |
| 1 | 192601016 | AX-170344051 | 0 | T | A | 0.272 | 1.29E-01 | 9.18E-02 | 2.35E+00 | 2.43E+00 | 1.67E-01 | no | Hhat | intron_variant | MODIFIER |
| 1 | 192636100 | AX-97850952 | 0 | T | G | 0.272 | 1.29E-01 | 9.18E-02 | 2.35E+00 | 2.43E+00 | 1.67E-01 | no | Hhat | intron_variant | MODIFIER |
| 1 | 192723000 | AX-97803950 | 0 | T | C | 0.272 | 1.29E-01 | 9.18E-02 | 2.35E+00 | 2.43E+00 | 1.67E-01 | no | Hhat | intron_variant | MODIFIER |

| | | | | | | | | | | | | | | | |
|---|---|---|---|---|---|---|---|---|---|---|---|---|---|---|---|
| 1 | 192757780 | AX-97802018 | 0 | G | A | 0.272 | 1.29E-01 | 9.18E-02 | 2.35E+00 | 2.43E+00 | 1.67E-01 | no | Hhat | intron_variant | MODIFIER |
| 1 | 192785134 | AX-97792156 | 0 | C | T | 0.272 | 1.29E-01 | 9.18E-02 | 2.35E+00 | 2.43E+00 | 1.67E-01 | no | | regulatory_region_variant | MODIFIER |
| 1 | 192812079 | AX-97831983 | 0 | C | T | 0.272 | 1.29E-01 | 9.18E-02 | 2.35E+00 | 2.43E+00 | 1.67E-01 | no | | regulatory_region_variant | MODIFIER |
| 1 | 192820483 | AX-170130506 | 0 | G | C | 0.272 | 1.29E-01 | 9.18E-02 | 2.35E+00 | 2.43E+00 | 1.67E-01 | no | Gm38360 | downstream_gene_variant | MODIFIER |
| 1 | 192836951 | AX-97798297 | 0 | A | G | 0.272 | 1.29E-01 | 9.18E-02 | 2.35E+00 | 2.43E+00 | 1.67E-01 | no | A730013G03Rik | upstream_gene_variant | MODIFIER |
| 1 | 192837711 | AX-170002591 | 0 | G | A | 0.272 | 1.29E-01 | 9.18E-02 | 2.35E+00 | 2.43E+00 | 1.67E-01 | no | A730013G03Rik | upstream_gene_variant | MODIFIER |
| 1 | 192845072 | AX-170002593 | 0 | C | A | 0.272 | 1.29E-01 | 9.18E-02 | 2.35E+00 | 2.43E+00 | 1.67E-01 | no | Sertad4 | 3_prime_UTR_variant | MODIFIER |
| 1 | 192869205 | AX-97794650 | 0 | G | A | 0.272 | 1.29E-01 | 9.18E-02 | 2.35E+00 | 2.43E+00 | 1.67E-01 | no | | intergenic_variant | MODIFIER |
| 1 | 192876732 | AX-170002598 | 0 | A | C | 0.272 | 1.29E-01 | 9.18E-02 | 2.35E+00 | 2.43E+00 | 1.67E-01 | no | | intergenic_variant | MODIFIER |
| 1 | 192883545 | AX-170002596 | 0 | A | G | 0.272 | 1.29E-01 | 9.18E-02 | 2.35E+00 | 2.43E+00 | 1.67E-01 | no | | intergenic_variant | MODIFIER |
| 1 | 192909094 | AX-97818623 | 0 | T | C | 0.272 | 1.29E-01 | 9.18E-02 | 2.35E+00 | 2.43E+00 | 1.67E-01 | no | Syt14 | intron_variant | MODIFIER |
| 1 | 192967389 | AX-170002590 | 0 | A | G | 0.272 | 1.29E-01 | 9.18E-02 | 2.35E+00 | 2.43E+00 | 1.67E-01 | no | Syt14 | intron_variant | MODIFIER |
| 1 | 193058287 | AX-170271492 | 0 | T | A | 0.272 | 1.29E-01 | 9.18E-02 | 2.35E+00 | 2.43E+00 | 1.67E-01 | no | | intergenic_variant | MODIFIER |
| 1 | 193079781 | AX-170346111 | 0 | A | C | 0.272 | 1.29E-01 | 9.18E-02 | 2.35E+00 | 2.43E+00 | 1.67E-01 | no | | intergenic_variant | MODIFIER |
| 1 | 193079974 | AX-170255949 | 0 | A | G | 0.272 | 1.29E-01 | 9.18E-02 | 2.35E+00 | 2.43E+00 | 1.67E-01 | no | | intergenic_variant | MODIFIER |
| 1 | 193111218 | AX-170252876 | 0 | T | C | 0.272 | 1.29E-01 | 9.18E-02 | 2.35E+00 | 2.43E+00 | 1.67E-01 | no | Diexf | intron_variant | MODIFIER |
| 1 | 193111508 | AX-170458391 | 0 | A | T | 0.272 | 1.29E-01 | 9.18E-02 | 2.35E+00 | 2.43E+00 | 1.67E-01 | no | Diexf | intron_variant | MODIFIER |
| 1 | 193112330 | AX-170280281 | 0 | T | C | 0.272 | 1.29E-01 | 9.18E-02 | 2.35E+00 | 2.43E+00 | 1.67E-01 | no | Diexf | intron_variant | MODIFIER |
| 1 | 193155878 | AX-97829489 | 0 | T | C | 0.272 | 1.29E-01 | 9.18E-02 | 2.35E+00 | 2.43E+00 | 1.67E-01 | no | Irf6 | intron_variant | MODIFIER |
| 1 | 193164315 | AX-170443529 | 0 | C | T | 0.272 | 1.29E-01 | 9.18E-02 | 2.35E+00 | 2.43E+00 | 1.67E-01 | no | Irf6 | intron_variant | MODIFIER |
| 1 | 193184690 | AX-170123697 | 0 | G | A | 0.272 | 1.29E-01 | 9.18E-02 | 2.35E+00 | 2.43E+00 | 1.67E-01 | no | Traf3ip3 | intron_variant | MODIFIER |
| 1 | 193191965 | AX-97824979 | 0 | G | A | 0.272 | 1.29E-01 | 9.18E-02 | 2.35E+00 | 2.43E+00 | 1.67E-01 | no | Traf3ip3 | intron_variant | MODIFIER |
| 1 | 193195642 | AX-170252254 | 0 | T | C | 0.272 | 1.29E-01 | 9.18E-02 | 2.35E+00 | 2.43E+00 | 1.67E-01 | no | Traf3ip3 | intron_variant | MODIFIER |
| 1 | 193219228 | AX-170044008 | 0 | C | T | 0.272 | 1.29E-01 | 9.18E-02 | 2.35E+00 | 2.43E+00 | 1.67E-01 | no | Hsd11b1 | downstream_gene_variant | MODIFIER |
| 1 | 193241778 | AX-97843920 | 0 | A | G | 0.272 | 1.29E-01 | 9.18E-02 | 2.35E+00 | 2.43E+00 | 1.67E-01 | no | Hsd11b1 | intron_variant | MODIFIER |
| 1 | 193250497 | AX-170450177 | 0 | A | G | 0.272 | 1.29E-01 | 9.18E-02 | 2.35E+00 | 2.43E+00 | 1.67E-01 | no | Hsd11b1 | intron_variant | MODIFIER |
| 1 | 193327455 | AX-170525406 | 0 | G | A | 0.272 | 1.29E-01 | 9.18E-02 | 2.35E+00 | 2.43E+00 | 1.67E-01 | no | Lamb3 | intron_variant | MODIFIER |
| 1 | 193336432 | AX-170491783 | 0 | A | C | 0.272 | 1.29E-01 | 9.18E-02 | 2.35E+00 | 2.43E+00 | 1.67E-01 | no | Lamb3 | intron_variant | MODIFIER |
| 1 | 193344054 | AX-170165128 | 0 | C | T | 0.272 | 1.29E-01 | 9.18E-02 | 2.35E+00 | 2.43E+00 | 1.67E-01 | no | Lamb3 | downstream_gene_variant | MODIFIER |
| 1 | 193355376 | AX-170289681 | 0 | T | C | 0.272 | 1.29E-01 | 9.18E-02 | 2.35E+00 | 2.43E+00 | 1.67E-01 | no | Camk1g | intron_variant | MODIFIER |
| 1 | 193357732 | AX-170394273 | 0 | A | G | 0.272 | 1.29E-01 | 9.18E-02 | 2.35E+00 | 2.43E+00 | 1.67E-01 | no | Camk1g | intron_variant | MODIFIER |
| 1 | 193357843 | AX-170002602 | 0 | A | C | 0.272 | 1.29E-01 | 9.18E-02 | 2.35E+00 | 2.43E+00 | 1.67E-01 | no | Camk1g | intron_variant | MODIFIER |
| 1 | 193357872 | AX-170002601 | 0 | T | C | 0.272 | 1.29E-01 | 9.18E-02 | 2.35E+00 | 2.43E+00 | 1.67E-01 | no | Camk1g | intron_variant | MODIFIER |
| 1 | 193358450 | AX-97841050 | 0 | A | C | 0.272 | 1.29E-01 | 9.18E-02 | 2.35E+00 | 2.43E+00 | 1.67E-01 | no | Camk1g | intron_variant | MODIFIER |
| 1 | 193360352 | AX-170176340 | 0 | C | T | 0.272 | 1.29E-01 | 9.18E-02 | 2.35E+00 | 2.43E+00 | 1.67E-01 | no | Camk1g | intron_variant | MODIFIER |
| 1 | 193389795 | AX-170379526 | 0 | T | C | 0.272 | 1.29E-01 | 9.18E-02 | 2.35E+00 | 2.43E+00 | 1.67E-01 | no | | regulatory_region_variant | MODIFIER |
| 1 | 193392005 | AX-170044013 | 0 | A | G | 0.272 | 1.29E-01 | 9.18E-02 | 2.35E+00 | 2.43E+00 | 1.67E-01 | no | | intergenic_variant | MODIFIER |
| 1 | 193507393 | AX-170148676 | 0 | A | G | 0.272 | 1.29E-01 | 9.18E-02 | 2.35E+00 | 2.43E+00 | 1.67E-01 | no | Mir205 | downstream_gene_variant | MODIFIER |
| 1 | 193509145 | AX-170269157 | 0 | C | A | 0.272 | 1.29E-01 | 9.18E-02 | 2.35E+00 | 2.43E+00 | 1.67E-01 | no | Mir205 | upstream_gene_variant | MODIFIER |
| 1 | 193510506 | AX-170266284 | 0 | A | C | 0.272 | 1.29E-01 | 9.18E-02 | 2.35E+00 | 2.43E+00 | 1.67E-01 | no | Mir205 | upstream_gene_variant | MODIFIER |
| 1 | 193511713 | AX-170338327 | 0 | C | T | 0.272 | 1.29E-01 | 9.18E-02 | 2.35E+00 | 2.43E+00 | 1.67E-01 | no | Mir205 | upstream_gene_variant | MODIFIER |
| 1 | 193515361 | AX-170501210 | 0 | T | G | 0.272 | 1.29E-01 | 9.18E-02 | 2.35E+00 | 2.43E+00 | 1.67E-01 | no | | intergenic_variant | MODIFIER |
| 1 | 193570694 | AX-170044015 | 0 | G | A | 0.272 | 1.29E-01 | 9.18E-02 | 2.35E+00 | 2.43E+00 | 1.67E-01 | no | | intergenic_variant | MODIFIER |
| 1 | 193586992 | AX-97824974 | 0 | G | A | 0.272 | 1.29E-01 | 9.18E-02 | 2.35E+00 | 2.43E+00 | 1.67E-01 | no | | regulatory_region_variant | MODIFIER |
| 1 | 193599016 | AX-170097036 | 0 | G | A | 0.272 | 1.29E-01 | 9.18E-02 | 2.35E+00 | 2.43E+00 | 1.67E-01 | no | | intergenic_variant | MODIFIER |
| 1 | 193611597 | AX-97800590 | 0 | A | C | 0.272 | 1.29E-01 | 9.18E-02 | 2.35E+00 | 2.43E+00 | 1.67E-01 | no | | regulatory_region_variant | MODIFIER |
| 1 | 193611923 | AX-170144691 | 0 | T | C | 0.272 | 1.29E-01 | 9.18E-02 | 2.35E+00 | 2.43E+00 | 1.67E-01 | no | | intergenic_variant | MODIFIER |
| 1 | 193647812 | AX-170172443 | 0 | T | C | 0.272 | 1.29E-01 | 9.18E-02 | 2.35E+00 | 2.43E+00 | 1.67E-01 | no | | regulatory_region_variant | MODIFIER |
| 1 | 193650073 | AX-170262264 | 0 | T | C | 0.272 | 1.29E-01 | 9.18E-02 | 2.35E+00 | 2.43E+00 | 1.67E-01 | no | | intergenic_variant | MODIFIER |
| 1 | 193653244 | AX-170413826 | 0 | T | C | 0.272 | 1.29E-01 | 9.18E-02 | 2.35E+00 | 2.43E+00 | 1.67E-01 | no | | intergenic_variant | MODIFIER |
| 1 | 193664463 | AX-170512887 | 0 | A | G | 0.272 | 1.29E-01 | 9.18E-02 | 2.35E+00 | 2.43E+00 | 1.67E-01 | no | | regulatory_region_variant | MODIFIER |
| 1 | 193673297 | AX-170122917 | 0 | A | G | 0.272 | 1.29E-01 | 9.18E-02 | 2.35E+00 | 2.43E+00 | 1.67E-01 | no | | regulatory_region_variant | MODIFIER |
| 1 | 193700986 | AX-170258959 | 0 | G | A | 0.272 | 1.29E-01 | 9.18E-02 | 2.35E+00 | 2.43E+00 | 1.67E-01 | no | | intergenic_variant | MODIFIER |
| 1 | 193703565 | AX-170343835 | 0 | C | G | 0.272 | 1.29E-01 | 9.18E-02 | 2.35E+00 | 2.43E+00 | 1.67E-01 | no | | regulatory_region_variant | MODIFIER |

| | | | | | | | | | | | | | | |
|---|---|---|---|---|---|---|---|---|---|---|---|---|---|---|
| 1 | 193706138 | AX-170525374 | 0 | T | C | 0.272 | 1.29E-01 | 9.18E-02 | 2.35E+00 | 2.43E+00 | 1.67E-01 | no | intergenic_variant | MODIFIER |
| 1 | 193706438 | AX-170371837 | 0 | C | A | 0.272 | 1.29E-01 | 9.18E-02 | 2.35E+00 | 2.43E+00 | 1.67E-01 | no | intergenic_variant | MODIFIER |
| 1 | 193709507 | AX-170468692 | 0 | A | C | 0.272 | 1.29E-01 | 9.18E-02 | 2.35E+00 | 2.43E+00 | 1.67E-01 | no | intergenic_variant | MODIFIER |
| 1 | 193709621 | AX-170442561 | 0 | A | G | 0.272 | 1.29E-01 | 9.18E-02 | 2.35E+00 | 2.43E+00 | 1.67E-01 | no | intergenic_variant | MODIFIER |
| 1 | 193739085 | AX-170168633 | 0 | C | T | 0.272 | 1.29E-01 | 9.18E-02 | 2.35E+00 | 2.43E+00 | 1.67E-01 | no | regulatory_region_variant | MODIFIER |
| 1 | 193760613 | AX-170278220 | 0 | A | G | 0.272 | 1.29E-01 | 9.18E-02 | 2.35E+00 | 2.43E+00 | 1.67E-01 | no | intergenic_variant | MODIFIER |
| 1 | 193760946 | AX-97847191 | 0 | A | G | 0.272 | 1.29E-01 | 9.18E-02 | 2.35E+00 | 2.43E+00 | 1.67E-01 | no | intergenic_variant | MODIFIER |
| 1 | 193768884 | AX-170214820 | 0 | C | T | 0.272 | 1.29E-01 | 9.18E-02 | 2.35E+00 | 2.43E+00 | 1.67E-01 | no | intergenic_variant | MODIFIER |
| 1 | 193769085 | AX-170437907 | 0 | A | C | 0.272 | 1.29E-01 | 9.18E-02 | 2.35E+00 | 2.43E+00 | 1.67E-01 | no | intergenic_variant | MODIFIER |
| 1 | 193843543 | AX-170478677 | 0 | G | T | 0.272 | 1.29E-01 | 9.18E-02 | 2.35E+00 | 2.43E+00 | 1.67E-01 | no | intergenic_variant | MODIFIER |
| 1 | 193921799 | AX-170002610 | 0 | C | G | 0.272 | 1.29E-01 | 9.18E-02 | 2.35E+00 | 2.43E+00 | 1.67E-01 | no | intergenic_variant | MODIFIER |
| 1 | 193922578 | AX-170159737 | 0 | C | G | 0.272 | 1.29E-01 | 9.18E-02 | 2.35E+00 | 2.43E+00 | 1.67E-01 | no | intergenic_variant | MODIFIER |
| 1 | 193933722 | AX-170413138 | 0 | T | C | 0.272 | 1.29E-01 | 9.18E-02 | 2.35E+00 | 2.43E+00 | 1.67E-01 | no | intergenic_variant | MODIFIER |
| 1 | 193942383 | AX-170324790 | 0 | A | G | 0.272 | 1.29E-01 | 9.18E-02 | 2.35E+00 | 2.43E+00 | 1.67E-01 | no | intergenic_variant | MODIFIER |
| 1 | 193945981 | AX-97838200 | 0 | C | T | 0.272 | 1.29E-01 | 9.18E-02 | 2.35E+00 | 2.43E+00 | 1.67E-01 | no | intergenic_variant | MODIFIER |
| 1 | 193946700 | AX-170309812 | 0 | G | A | 0.272 | 1.29E-01 | 9.18E-02 | 2.35E+00 | 2.43E+00 | 1.67E-01 | no | intergenic_variant | MODIFIER |
| 1 | 193952036 | AX-170218087 | 0 | T | C | 0.272 | 1.29E-01 | 9.18E-02 | 2.35E+00 | 2.43E+00 | 1.67E-01 | no | regulatory_region_variant | MODIFIER |
| 1 | 193952793 | AX-170458750 | 0 | A | G | 0.272 | 1.29E-01 | 9.18E-02 | 2.35E+00 | 2.43E+00 | 1.67E-01 | no | intergenic_variant | MODIFIER |
| 1 | 193960095 | AX-170132605 | 0 | T | C | 0.272 | 1.29E-01 | 9.18E-02 | 2.35E+00 | 2.43E+00 | 1.67E-01 | no | intergenic_variant | MODIFIER |
| 1 | 193960323 | AX-170127997 | 0 | A | G | 0.272 | 1.29E-01 | 9.18E-02 | 2.35E+00 | 2.43E+00 | 1.67E-01 | no | intergenic_variant | MODIFIER |
| 1 | 193965539 | AX-170300829 | 0 | C | G | 0.272 | 1.29E-01 | 9.18E-02 | 2.35E+00 | 2.43E+00 | 1.67E-01 | no | intergenic_variant | MODIFIER |
| 1 | 193965566 | AX-170143006 | 0 | T | C | 0.272 | 1.29E-01 | 9.18E-02 | 2.35E+00 | 2.43E+00 | 1.67E-01 | no | intergenic_variant | MODIFIER |
| 1 | 194004335 | AX-170225677 | 0 | T | C | 0.272 | 1.29E-01 | 9.18E-02 | 2.35E+00 | 2.43E+00 | 1.67E-01 | no | intergenic_variant | MODIFIER |
| 1 | 194038901 | AX-170240450 | 0 | T | G | 0.272 | 1.29E-01 | 9.18E-02 | 2.35E+00 | 2.43E+00 | 1.67E-01 | no | intergenic_variant | MODIFIER |
| 1 | 194042074 | AX-170284905 | 0 | G | A | 0.272 | 1.29E-01 | 9.18E-02 | 2.35E+00 | 2.43E+00 | 1.67E-01 | no | regulatory_region_variant | MODIFIER |
| 1 | 194331412 | AX-170238158 | 0 | C | T | 0.272 | 1.29E-01 | 9.18E-02 | 2.35E+00 | 2.43E+00 | 1.67E-01 | no | intergenic_variant | MODIFIER |
| 1 | 194335959 | AX-170002612 | 0 | C | T | 0.272 | 1.29E-01 | 9.18E-02 | 2.35E+00 | 2.43E+00 | 1.67E-01 | no | regulatory_region_variant | MODIFIER |
| 1 | 194342767 | AX-170044033 | 0 | A | G | 0.272 | 1.29E-01 | 9.18E-02 | 2.35E+00 | 2.43E+00 | 1.67E-01 | no | intergenic_variant | MODIFIER |
| 1 | 194350543 | AX-170044030 | 0 | C | T | 0.272 | 1.29E-01 | 9.18E-02 | 2.35E+00 | 2.43E+00 | 1.67E-01 | no | intergenic_variant | MODIFIER |
| 1 | 194350570 | AX-170481888 | 0 | C | T | 0.272 | 1.29E-01 | 9.18E-02 | 2.35E+00 | 2.43E+00 | 1.67E-01 | no | intergenic_variant | MODIFIER |
| 1 | 194359431 | AX-97827689 | 0 | G | T | 0.272 | 1.29E-01 | 9.18E-02 | 2.35E+00 | 2.43E+00 | 1.67E-01 | no | intergenic_variant | MODIFIER |
| 1 | 194360549 | AX-170002615 | 0 | T | A | 0.272 | 1.29E-01 | 9.18E-02 | 2.35E+00 | 2.43E+00 | 1.67E-01 | no | intergenic_variant | MODIFIER |
| 1 | 194386872 | AX-97796432 | 0 | C | A | 0.272 | 1.29E-01 | 9.18E-02 | 2.35E+00 | 2.43E+00 | 1.67E-01 | no | intergenic_variant | MODIFIER |
| 1 | 194404067 | AX-170002611 | 0 | T | C | 0.272 | 1.29E-01 | 9.18E-02 | 2.35E+00 | 2.43E+00 | 1.67E-01 | no | intergenic_variant | MODIFIER |
| 1 | 194404235 | AX-97790908 | 0 | C | T | 0.272 | 1.29E-01 | 9.18E-02 | 2.35E+00 | 2.43E+00 | 1.67E-01 | no | intergenic_variant | MODIFIER |
| 1 | 194424978 | AX-170316124 | 0 | T | C | 0.272 | 1.29E-01 | 9.18E-02 | 2.35E+00 | 2.43E+00 | 1.67E-01 | no | regulatory_region_variant | MODIFIER |
| 1 | 194439762 | AX-170333574 | 0 | G | A | 0.272 | 1.29E-01 | 9.18E-02 | 2.35E+00 | 2.43E+00 | 1.67E-01 | no | intergenic_variant | MODIFIER |
| 1 | 194440114 | AX-97787205 | 0 | A | G | 0.272 | 1.29E-01 | 9.18E-02 | 2.35E+00 | 2.43E+00 | 1.67E-01 | no | intergenic_variant | MODIFIER |
| 1 | 194443938 | AX-170273848 | 0 | T | G | 0.272 | 1.29E-01 | 9.18E-02 | 2.35E+00 | 2.43E+00 | 1.67E-01 | no | intergenic_variant | MODIFIER |
| 1 | 194443968 | AX-170327655 | 0 | A | G | 0.272 | 1.29E-01 | 9.18E-02 | 2.35E+00 | 2.43E+00 | 1.67E-01 | no | intergenic_variant | MODIFIER |
| 1 | 194453754 | AX-170308339 | 0 | G | A | 0.272 | 1.29E-01 | 9.18E-02 | 2.35E+00 | 2.43E+00 | 1.67E-01 | no | intergenic_variant | MODIFIER |
| 1 | 194456078 | AX-170508271 | 0 | C | T | 0.272 | 1.29E-01 | 9.18E-02 | 2.35E+00 | 2.43E+00 | 1.67E-01 | no | intergenic_variant | MODIFIER |
| 1 | 194457538 | AX-170492027 | 0 | T | G | 0.272 | 1.29E-01 | 9.18E-02 | 2.35E+00 | 2.43E+00 | 1.67E-01 | no | intergenic_variant | MODIFIER |
| 1 | 194475554 | AX-170224250 | 0 | A | G | 0.272 | 1.29E-01 | 9.18E-02 | 2.35E+00 | 2.43E+00 | 1.67E-01 | no | intergenic_variant | MODIFIER |
| 1 | 194479472 | AX-170137071 | 0 | C | A | 0.272 | 1.29E-01 | 9.18E-02 | 2.35E+00 | 2.43E+00 | 1.67E-01 | no | intergenic_variant | MODIFIER |
| 1 | 194482311 | AX-170295474 | 0 | T | C | 0.272 | 1.29E-01 | 9.18E-02 | 2.35E+00 | 2.43E+00 | 1.67E-01 | no | regulatory_region_variant | MODIFIER |
| 1 | 194482351 | AX-170157103 | 0 | A | G | 0.272 | 1.29E-01 | 9.18E-02 | 2.35E+00 | 2.43E+00 | 1.67E-01 | no | regulatory_region_variant | MODIFIER |
| 1 | 194482694 | AX-170044035 | 0 | A | G | 0.272 | 1.29E-01 | 9.18E-02 | 2.35E+00 | 2.43E+00 | 1.67E-01 | no | regulatory_region_variant | MODIFIER |
| 1 | 194484064 | AX-170044034 | 0 | G | A | 0.272 | 1.29E-01 | 9.18E-02 | 2.35E+00 | 2.43E+00 | 1.67E-01 | no | intergenic_variant | MODIFIER |
| 1 | 194485106 | AX-170416531 | 0 | C | T | 0.272 | 1.29E-01 | 9.18E-02 | 2.35E+00 | 2.43E+00 | 1.67E-01 | no | intergenic_variant | MODIFIER |
| 1 | 194488569 | AX-170254689 | 0 | T | C | 0.272 | 1.29E-01 | 9.18E-02 | 2.35E+00 | 2.43E+00 | 1.67E-01 | no | intergenic_variant | MODIFIER |
| 1 | 194503384 | AX-170145061 | 0 | C | G | 0.272 | 1.29E-01 | 9.18E-02 | 2.35E+00 | 2.43E+00 | 1.67E-01 | no | intergenic_variant | MODIFIER |
| 1 | 194503955 | AX-170468286 | 0 | C | T | 0.272 | 1.29E-01 | 9.18E-02 | 2.35E+00 | 2.43E+00 | 1.67E-01 | no | intergenic_variant | MODIFIER |
| 1 | 194504299 | AX-170475900 | 0 | A | G | 0.272 | 1.29E-01 | 9.18E-02 | 2.35E+00 | 2.43E+00 | 1.67E-01 | no | regulatory_region_variant | MODIFIER |

| | | | | | | | | | | | | | | | |
|---|---|---|---|---|---|---|---|---|---|---|---|---|---|---|---|
| 1 | 194528939 | AX-170398661 | 0 | G | A | 0.272 | 1.29E-01 | 9.18E-02 | 2.35E+00 | 2.43E+00 | 1.67E-01 | no | | intergenic_variant | MODIFIER |
| 1 | 194541745 | AX-170158751 | 0 | C | T | 0.272 | 1.29E-01 | 9.18E-02 | 2.35E+00 | 2.43E+00 | 1.67E-01 | no | | intergenic_variant | MODIFIER |
| 1 | 194578092 | AX-170190915 | 0 | G | A | 0.272 | 1.29E-01 | 9.18E-02 | 2.35E+00 | 2.43E+00 | 1.67E-01 | no | | intergenic_variant | MODIFIER |
| 1 | 194587761 | AX-170140362 | 0 | A | G | 0.272 | 1.29E-01 | 9.18E-02 | 2.35E+00 | 2.43E+00 | 1.67E-01 | no | | regulatory_region_variant | MODIFIER |
| 1 | 194631263 | AX-170257903 | 0 | T | C | 0.272 | 1.29E-01 | 9.18E-02 | 2.35E+00 | 2.43E+00 | 1.67E-01 | no | Plxna2 | intron_variant | MODIFIER |
| 1 | 194632509 | AX-170463428 | 0 | T | C | 0.272 | 1.29E-01 | 9.18E-02 | 2.35E+00 | 2.43E+00 | 1.67E-01 | no | Plxna2 | intron_variant | MODIFIER |
| 1 | 194633479 | AX-170044038 | 0 | C | G | 0.272 | 1.29E-01 | 9.18E-02 | 2.35E+00 | 2.43E+00 | 1.67E-01 | no | Plxna2 | intron_variant | MODIFIER |
| 1 | 194641426 | AX-170044039 | 0 | C | A | 0.272 | 1.29E-01 | 9.18E-02 | 2.35E+00 | 2.43E+00 | 1.67E-01 | no | Plxna2 | intron_variant | MODIFIER |
| 1 | 194759288 | AX-170327453 | 0 | C | A | 0.272 | 1.29E-01 | 9.18E-02 | 2.35E+00 | 2.43E+00 | 1.67E-01 | no | Plxna2 | intron_variant | MODIFIER |
| 1 | 194786367 | AX-170495912 | 0 | A | G | 0.272 | 1.29E-01 | 9.18E-02 | 2.35E+00 | 2.43E+00 | 1.67E-01 | no | Plxna2 | intron_variant | MODIFIER |
| 1 | 194786400 | AX-97795904 | 0 | A | C | 0.272 | 1.29E-01 | 9.18E-02 | 2.35E+00 | 2.43E+00 | 1.67E-01 | no | Plxna2 | intron_variant | MODIFIER |
| 1 | 194805857 | AX-97847581 | 0 | T | C | 0.272 | 1.29E-01 | 9.18E-02 | 2.35E+00 | 2.43E+00 | 1.67E-01 | no | Plxna2 | intron_variant | MODIFIER |
| 1 | 194845984 | AX-170250015 | 0 | T | C | 0.272 | 1.29E-01 | 9.18E-02 | 2.35E+00 | 2.43E+00 | 1.67E-01 | no | | intergenic_variant | MODIFIER |
| 1 | 194881722 | AX-170002617 | 0 | C | T | 0.272 | 1.29E-01 | 9.18E-02 | 2.35E+00 | 2.43E+00 | 1.67E-01 | no | | intergenic_variant | MODIFIER |
| 1 | 194884893 | AX-170366818 | 0 | A | C | 0.272 | 1.29E-01 | 9.18E-02 | 2.35E+00 | 2.43E+00 | 1.67E-01 | no | | intergenic_variant | MODIFIER |
| 1 | 194885110 | AX-170149953 | 0 | C | G | 0.272 | 1.29E-01 | 9.18E-02 | 2.35E+00 | 2.43E+00 | 1.67E-01 | no | | regulatory_region_variant | MODIFIER |
| 1 | 194886567 | AX-97831545 | 0 | G | T | 0.272 | 1.29E-01 | 9.18E-02 | 2.35E+00 | 2.43E+00 | 1.67E-01 | no | | regulatory_region_variant | MODIFIER |
| 1 | 194889500 | AX-170279667 | 0 | T | G | 0.272 | 1.29E-01 | 9.18E-02 | 2.35E+00 | 2.43E+00 | 1.67E-01 | no | | intergenic_variant | MODIFIER |
| 1 | 194895568 | AX-170429476 | 0 | T | G | 0.272 | 1.29E-01 | 9.18E-02 | 2.35E+00 | 2.43E+00 | 1.67E-01 | no | | intergenic_variant | MODIFIER |
| 1 | 194924887 | AX-97815528 | 0 | G | A | 0.272 | 1.29E-01 | 9.18E-02 | 2.35E+00 | 2.43E+00 | 1.67E-01 | no | | intergenic_variant | MODIFIER |
| 1 | 194925250 | AX-170002621 | 0 | T | C | 0.272 | 1.29E-01 | 9.18E-02 | 2.35E+00 | 2.43E+00 | 1.67E-01 | no | | regulatory_region_variant | MODIFIER |
| 1 | 194925324 | AX-170002619 | 0 | A | G | 0.272 | 1.29E-01 | 9.18E-02 | 2.35E+00 | 2.43E+00 | 1.67E-01 | no | | regulatory_region_variant | MODIFIER |
| 1 | 194925812 | AX-170523508 | 0 | A | G | 0.272 | 1.29E-01 | 9.18E-02 | 2.35E+00 | 2.43E+00 | 1.67E-01 | no | | regulatory_region_variant | MODIFIER |
| 1 | 194940300 | AX-170279902 | 0 | A | G | 0.272 | 1.29E-01 | 9.18E-02 | 2.35E+00 | 2.43E+00 | 1.67E-01 | no | Cd34 | intron_variant | MODIFIER |
| 1 | 194960643 | AX-97790167 | 0 | A | C | 0.272 | 1.29E-01 | 9.18E-02 | 2.35E+00 | 2.43E+00 | 1.67E-01 | no | Cd34 | 3_prime_UTR_variant | MODIFIER |
| 10 | 107810834 | AX-170003862 | 0 | C | A | 0.316 | -1.14E-01 | 8.30E-02 | 2.81E+00 | 2.91E+00 | 1.67E-01 | no | Otogl | intron_variant | MODIFIER |
| 16 | 29194103 | AX-170408155 | 0 | T | C | 0.5 | 1.08E-01 | 7.90E-02 | 2.66E+00 | 2.72E+00 | 1.67E-01 | no | | regulatory_region_variant | MODIFIER |
| 5 | 3300304 | AX-97789601 | 0 | G | T | 0.43 | -9.17E-02 | 6.64E-02 | 2.51E+00 | 2.62E+00 | 1.67E-01 | no | | intergenic_variant | MODIFIER |
| 9 | 9563982 | AX-170400123 | 1 | T | C | 0.464 | 1.13E-01 | 8.17E-02 | 2.47E+00 | 2.57E+00 | 1.67E-01 | no | Gm47334 | upstream_gene_variant | MODIFIER |
| 13 | 9847573 | AX-170224138 | 1 | C | G | 0.188 | -1.71E-01 | 1.23E-01 | 2.46E+00 | 2.51E+00 | 1.67E-01 | no | | intergenic_variant | MODIFIER |
| 5 | 112949108 | AX-170233231 | 0 | A | G | 0.456 | 1.50E-01 | 1.09E-01 | 2.89E+00 | 2.97E+00 | 1.67E-01 | Yes | Grk3 | intron_variant | MODIFIER |
| 4 | 86575528 | AX-170239409 | 1 | C | T | 0.339 | 1.49E-01 | 1.09E-01 | 2.70E+00 | 2.78E+00 | 1.67E-01 | Yes | Haus6 | downstream_gene_variant | MODIFIER |
| 18 | 76692800 | AX-97810542 | 0 | G | A | 0.439 | 1.17E-01 | 8.50E-02 | 2.51E+00 | 2.57E+00 | 1.68E-01 | no | | intergenic_variant | MODIFIER |
| 18 | 76924945 | AX-170287143 | 0 | A | G | 0.439 | 1.17E-01 | 8.50E-02 | 2.51E+00 | 2.57E+00 | 1.68E-01 | no | | intergenic_variant | MODIFIER |
| 1 | 163132699 | AX-170035632 | 0 | A | T | 0.211 | 1.98E-01 | 1.44E-01 | 2.85E+00 | 2.90E+00 | 1.68E-01 | Yes | | intergenic_variant | MODIFIER |
| 15 | 6003774 | AX-170054705 | 1 | A | G | 0.321 | -1.12E-01 | 7.42E-02 | 2.01E+00 | 2.06E+00 | 1.68E-01 | no | | intergenic_variant | MODIFIER |
| 10 | 97649553 | AX-170003717 | 1 | C | T | 0.286 | -1.18E-01 | 8.57E-02 | 2.56E+00 | 2.62E+00 | 1.68E-01 | no | Epyc | intron_variant | MODIFIER |
| 8 | 128112465 | AX-170505066 | 0 | T | C | 0.053 | 2.41E-01 | 1.75E-01 | 2.94E+00 | 3.10E+00 | 1.68E-01 | Yes | | regulatory_region_variant | MODIFIER |
| 5 | 129212879 | AX-170459880 | 1 | C | A | 0.152 | -1.29E-01 | 8.88E-02 | 2.17E+00 | 2.25E+00 | 1.68E-01 | no | | intergenic_variant | MODIFIER |
| 5 | 129405759 | AX-170026150 | 1 | G | C | 0.152 | -1.29E-01 | 8.88E-02 | 2.17E+00 | 2.25E+00 | 1.68E-01 | no | | regulatory_region_variant | MODIFIER |
| 15 | 6171648 | AX-170137989 | 1 | C | T | 0.339 | -1.18E-01 | 7.62E-02 | 1.94E+00 | 1.99E+00 | 1.68E-01 | no | | intergenic_variant | MODIFIER |
| 4 | 80835402 | AX-170213152 | 1 | A | G | 0.438 | -1.17E-01 | 8.46E-02 | 2.53E+00 | 2.59E+00 | 1.68E-01 | no | Tyrp1 | missense_variant | MODERATE |
| 14 | 123321744 | AX-97855474 | 1 | C | A | 0.33 | 1.08E-01 | 7.89E-02 | 2.64E+00 | 2.75E+00 | 1.68E-01 | no | Nalcn | intron_variant | MODIFIER |
| 13 | 25325558 | AX-170007948 | 0 | T | C | 0.167 | -1.58E-01 | 1.13E-01 | 2.37E+00 | 2.42E+00 | 1.69E-01 | no | | intergenic_variant | MODIFIER |
| 3 | 38280105 | AX-170020323 | 1 | G | C | 0.205 | 1.27E-01 | 9.22E-02 | 3.02E+00 | 3.14E+00 | 1.69E-01 | no | | intergenic_variant | MODIFIER |
| 13 | 6890657 | AX-170382049 | 0 | C | T | 0.202 | -1.73E-01 | 1.26E-01 | 2.59E+00 | 2.65E+00 | 1.69E-01 | Yes | | intergenic_variant | MODIFIER |
| 15 | 3531273 | AX-97841011 | 0 | C | T | 0.325 | -1.12E-01 | 7.41E-02 | 2.01E+00 | 2.06E+00 | 1.69E-01 | no | Ghr | intron_variant | MODIFIER |
| 15 | 3569753 | AX-170010921 | 0 | C | T | 0.325 | -1.12E-01 | 7.41E-02 | 2.01E+00 | 2.06E+00 | 1.69E-01 | no | Ghr | intron_variant | MODIFIER |
| 15 | 3569912 | AX-97852821 | 0 | G | A | 0.325 | -1.12E-01 | 7.41E-02 | 2.01E+00 | 2.06E+00 | 1.69E-01 | no | Ghr | intron_variant | MODIFIER |
| 15 | 3593700 | AX-170145418 | 0 | G | C | 0.325 | -1.12E-01 | 7.41E-02 | 2.01E+00 | 2.06E+00 | 1.69E-01 | no | | intergenic_variant | MODIFIER |
| 15 | 3607412 | AX-97821097 | 0 | A | G | 0.325 | -1.12E-01 | 7.41E-02 | 2.01E+00 | 2.06E+00 | 1.69E-01 | no | Gm22031 | upstream_gene_variant | MODIFIER |
| 15 | 3635698 | AX-170010917 | 0 | G | A | 0.325 | -1.12E-01 | 7.41E-02 | 2.01E+00 | 2.06E+00 | 1.69E-01 | no | | intergenic_variant | MODIFIER |
| 15 | 3642628 | AX-97820859 | 0 | A | G | 0.325 | -1.12E-01 | 7.41E-02 | 2.01E+00 | 2.06E+00 | 1.69E-01 | no | | intergenic_variant | MODIFIER |
| 15 | 3698947 | AX-170249067 | 0 | G | T | 0.325 | -1.12E-01 | 7.41E-02 | 2.01E+00 | 2.06E+00 | 1.69E-01 | no | | intergenic_variant | MODIFIER |

| | | | | | | | | | | | | | | | |
|---|---|---|---|---|---|---|---|---|---|---|---|---|---|---|---|
| 15 | 3704504 | AX-170122352 | 0 | T | C | 0.325 | -1.12E-01 | 7.41E-02 | 2.01E+00 | 2.06E+00 | 1.69E-01 | no | | intergenic_variant | MODIFIER |
| 15 | 3734438 | AX-170516604 | 0 | A | C | 0.325 | -1.12E-01 | 7.41E-02 | 2.01E+00 | 2.06E+00 | 1.69E-01 | no | | intergenic_variant | MODIFIER |
| 15 | 3735082 | AX-97837902 | 0 | C | T | 0.325 | -1.12E-01 | 7.41E-02 | 2.01E+00 | 2.06E+00 | 1.69E-01 | no | | intergenic_variant | MODIFIER |
| 15 | 3759526 | AX-170217731 | 0 | A | G | 0.325 | -1.12E-01 | 7.41E-02 | 2.01E+00 | 2.06E+00 | 1.69E-01 | no | | intergenic_variant | MODIFIER |
| 15 | 3760278 | AX-170233835 | 0 | C | T | 0.325 | -1.12E-01 | 7.41E-02 | 2.01E+00 | 2.06E+00 | 1.69E-01 | no | | intergenic_variant | MODIFIER |
| 15 | 3826295 | AX-170360192 | 0 | C | T | 0.325 | -1.12E-01 | 7.41E-02 | 2.01E+00 | 2.06E+00 | 1.69E-01 | no | | intergenic_variant | MODIFIER |
| 15 | 3989309 | AX-170168513 | 0 | A | T | 0.325 | -1.12E-01 | 7.41E-02 | 2.01E+00 | 2.06E+00 | 1.69E-01 | no | AW549877 | intron_variant | MODIFIER |
| 15 | 4008233 | AX-170298089 | 0 | G | A | 0.325 | -1.12E-01 | 7.41E-02 | 2.01E+00 | 2.06E+00 | 1.69E-01 | no | Gm2093 | intron_variant,non_coding_transcript_variant | MODIFIER |
| 15 | 4093188 | AX-97828273 | 0 | G | C | 0.325 | -1.12E-01 | 7.41E-02 | 2.01E+00 | 2.06E+00 | 1.69E-01 | no | Oxct1 | intron_variant | MODIFIER |
| 15 | 4918498 | AX-170151951 | 0 | T | C | 0.325 | -1.12E-01 | 7.41E-02 | 2.01E+00 | 2.06E+00 | 1.69E-01 | no | Mroh2b | intron_variant | MODIFIER |
| 15 | 4938423 | AX-97828022 | 0 | G | T | 0.325 | -1.12E-01 | 7.41E-02 | 2.01E+00 | 2.06E+00 | 1.69E-01 | no | Mroh2b | missense_variant | MODERATE |
| 15 | 5188035 | AX-97828302 | 0 | T | C | 0.325 | -1.12E-01 | 7.41E-02 | 2.01E+00 | 2.06E+00 | 1.69E-01 | no | Ttc33 | intron_variant | MODIFIER |
| 15 | 5243509 | AX-97784961 | 0 | T | G | 0.325 | -1.12E-01 | 7.41E-02 | 2.01E+00 | 2.06E+00 | 1.69E-01 | no | Ptger4 | intron_variant | MODIFIER |
| 15 | 5914165 | AX-97800927 | 0 | G | A | 0.325 | -1.12E-01 | 7.41E-02 | 2.01E+00 | 2.06E+00 | 1.69E-01 | no | | intergenic_variant | MODIFIER |
| 15 | 5922425 | AX-170010951 | 0 | A | C | 0.325 | -1.12E-01 | 7.41E-02 | 2.01E+00 | 2.06E+00 | 1.69E-01 | no | | intergenic_variant | MODIFIER |
| 15 | 5940909 | AX-97848422 | 0 | C | T | 0.325 | -1.12E-01 | 7.41E-02 | 2.01E+00 | 2.06E+00 | 1.69E-01 | no | | regulatory_region_variant | MODIFIER |
| 15 | 5942477 | AX-170113272 | 0 | G | A | 0.325 | -1.12E-01 | 7.41E-02 | 2.01E+00 | 2.06E+00 | 1.69E-01 | no | | regulatory_region_variant | MODIFIER |
| 15 | 5967661 | AX-170215241 | 0 | A | G | 0.325 | -1.12E-01 | 7.41E-02 | 2.01E+00 | 2.06E+00 | 1.69E-01 | no | | intergenic_variant | MODIFIER |
| 15 | 6002296 | AX-170175440 | 0 | C | T | 0.325 | -1.12E-01 | 7.41E-02 | 2.01E+00 | 2.06E+00 | 1.69E-01 | no | | intergenic_variant | MODIFIER |
| 15 | 6023161 | AX-97843007 | 0 | G | T | 0.325 | -1.12E-01 | 7.41E-02 | 2.01E+00 | 2.06E+00 | 1.69E-01 | no | | intergenic_variant | MODIFIER |
| 15 | 6081228 | AX-97828332 | 0 | G | A | 0.325 | -1.12E-01 | 7.41E-02 | 2.01E+00 | 2.06E+00 | 1.69E-01 | no | | regulatory_region_variant | MODIFIER |
| 15 | 6089078 | AX-170100822 | 0 | C | T | 0.325 | -1.12E-01 | 7.41E-02 | 2.01E+00 | 2.06E+00 | 1.69E-01 | no | | intergenic_variant | MODIFIER |
| 3 | 36786672 | AX-97838937 | 0 | G | A | 0.202 | 1.25E-01 | 9.09E-02 | 3.01E+00 | 3.13E+00 | 1.69E-01 | no | | regulatory_region_variant | MODIFIER |
| 15 | 7085846 | AX-170429900 | 0 | C | T | 0.228 | -1.37E-01 | 1.01E-01 | 2.78E+00 | 2.88E+00 | 1.69E-01 | no | Lifr | upstream_gene_variant | MODIFIER |
| 16 | 58196587 | AX-170205455 | 0 | A | T | 0.456 | -1.18E-01 | 8.55E-02 | 2.54E+00 | 2.59E+00 | 1.69E-01 | no | | intergenic_variant | MODIFIER |
| 4 | 128910669 | AX-170374704 | 1 | T | A | 0.161 | 1.58E-01 | 1.15E-01 | 2.52E+00 | 2.57E+00 | 1.69E-01 | Yes | Trim62 | 3_prime_UTR_variant | MODIFIER |
| 16 | 73048606 | AX-170238362 | 1 | G | A | 0.5 | 1.06E-01 | 7.72E-02 | 2.59E+00 | 2.66E+00 | 1.69E-01 | no | Robo1 | downstream_gene_variant | MODIFIER |
| 1 | 66298800 | AX-97822205 | 0 | C | T | 0.114 | -2.66E-01 | 1.90E-01 | 3.21E+00 | 3.35E+00 | 1.70E-01 | Yes | Map2 | intron_variant | MODIFIER |
| 1 | 68927644 | AX-170195286 | 0 | G | A | 0.114 | -2.66E-01 | 1.90E-01 | 3.21E+00 | 3.35E+00 | 1.70E-01 | Yes | Erbb4 | intron_variant | MODIFIER |
| 4 | 63687274 | AX-170466334 | 0 | A | G | 0.342 | 1.60E-01 | 1.17E-01 | 2.88E+00 | 2.94E+00 | 1.70E-01 | Yes | | intergenic_variant | MODIFIER |
| 4 | 120806658 | AX-97796071 | 0 | A | C | 0.211 | 1.53E-01 | 1.11E-01 | 2.48E+00 | 2.55E+00 | 1.70E-01 | Yes | Nfyc | intron_variant | MODIFIER |
| 5 | 89813454 | AX-170480584 | 1 | A | G | 0.438 | 1.53E-01 | 1.10E-01 | 3.13E+00 | 3.23E+00 | 1.70E-01 | Yes | Adamts3 | intron_variant | MODIFIER |
| 14 | 122276804 | AX-170476151 | 0 | G | T | 0.298 | 1.08E-01 | 7.91E-02 | 2.67E+00 | 2.78E+00 | 1.70E-01 | no | Clybl | intron_variant | MODIFIER |
| 14 | 122322450 | AX-97782401 | 0 | A | C | 0.298 | 1.08E-01 | 7.91E-02 | 2.67E+00 | 2.78E+00 | 1.70E-01 | no | Clybl | intron_variant | MODIFIER |
| 14 | 122469694 | AX-170010871 | 0 | T | C | 0.298 | 1.08E-01 | 7.91E-02 | 2.67E+00 | 2.78E+00 | 1.70E-01 | no | Zic5 | upstream_gene_variant | MODIFIER |
| 14 | 122470233 | AX-97828854 | 0 | T | C | 0.298 | 1.08E-01 | 7.91E-02 | 2.67E+00 | 2.78E+00 | 1.70E-01 | no | Zic5 | upstream_gene_variant | MODIFIER |
| 14 | 122512210 | AX-97856854 | 0 | G | A | 0.298 | 1.08E-01 | 7.91E-02 | 2.67E+00 | 2.78E+00 | 1.70E-01 | no | Gm18143 | downstream_gene_variant | MODIFIER |
| 14 | 122513569 | AX-170054637 | 0 | T | C | 0.298 | 1.08E-01 | 7.91E-02 | 2.67E+00 | 2.78E+00 | 1.70E-01 | no | Gm18143 | non_coding_transcript_exon_variant | MODIFIER |
| 14 | 122882246 | AX-170093134 | 0 | G | A | 0.298 | 1.08E-01 | 7.91E-02 | 2.67E+00 | 2.78E+00 | 1.70E-01 | no | Pcca | intron_variant | MODIFIER |
| 14 | 122944791 | AX-170010876 | 0 | T | C | 0.298 | 1.08E-01 | 7.91E-02 | 2.67E+00 | 2.78E+00 | 1.70E-01 | no | Tmtc4 | synonymous_variant | LOW |
| 14 | 122962725 | AX-97820817 | 0 | A | G | 0.298 | 1.08E-01 | 7.91E-02 | 2.67E+00 | 2.78E+00 | 1.70E-01 | no | Tmtc4 | intron_variant | MODIFIER |
| 14 | 122987003 | AX-170010877 | 0 | C | T | 0.298 | 1.08E-01 | 7.91E-02 | 2.67E+00 | 2.78E+00 | 1.70E-01 | no | Tmtc4 | upstream_gene_variant | MODIFIER |
| 14 | 122988269 | AX-170010878 | 0 | T | C | 0.298 | 1.08E-01 | 7.91E-02 | 2.67E+00 | 2.78E+00 | 1.70E-01 | no | Tmtc4 | upstream_gene_variant | MODIFIER |
| 14 | 122990812 | AX-97828182 | 0 | G | A | 0.298 | 1.08E-01 | 7.91E-02 | 2.67E+00 | 2.78E+00 | 1.70E-01 | no | | regulatory_region_variant | MODIFIER |
| 14 | 123008537 | AX-170295611 | 0 | C | T | 0.298 | 1.08E-01 | 7.91E-02 | 2.67E+00 | 2.78E+00 | 1.70E-01 | no | | regulatory_region_variant | MODIFIER |
| 14 | 123024025 | AX-170093135 | 0 | T | G | 0.298 | 1.08E-01 | 7.91E-02 | 2.67E+00 | 2.78E+00 | 1.70E-01 | no | | intergenic_variant | MODIFIER |
| 14 | 123037757 | AX-170010875 | 0 | G | A | 0.298 | 1.08E-01 | 7.91E-02 | 2.67E+00 | 2.78E+00 | 1.70E-01 | no | | intergenic_variant | MODIFIER |
| 14 | 123043789 | AX-97798795 | 0 | C | A | 0.298 | 1.08E-01 | 7.91E-02 | 2.67E+00 | 2.78E+00 | 1.70E-01 | no | AC158982.1 | intron_variant,non_coding_transcript_variant | MODIFIER |
| 14 | 123046673 | AX-170010882 | 0 | T | G | 0.298 | 1.08E-01 | 7.91E-02 | 2.67E+00 | 2.78E+00 | 1.70E-01 | no | AC158982.1 | intron_variant,non_coding_transcript_variant | MODIFIER |
| 14 | 123075619 | AX-170010872 | 0 | G | C | 0.298 | 1.08E-01 | 7.91E-02 | 2.67E+00 | 2.78E+00 | 1.70E-01 | no | AC158982.1 | intron_variant,non_coding_transcript_variant | MODIFIER |
| 14 | 123096707 | AX-170449427 | 0 | C | T | 0.298 | 1.08E-01 | 7.91E-02 | 2.67E+00 | 2.78E+00 | 1.70E-01 | no | AC158982.1 | intron_variant,non_coding_transcript_variant | MODIFIER |
| 14 | 123116943 | AX-170514994 | 0 | A | G | 0.298 | 1.08E-01 | 7.91E-02 | 2.67E+00 | 2.78E+00 | 1.70E-01 | no | AC158982.1 | intron_variant,non_coding_transcript_variant | MODIFIER |
| 14 | 123134383 | AX-170202532 | 0 | G | A | 0.298 | 1.08E-01 | 7.91E-02 | 2.67E+00 | 2.78E+00 | 1.70E-01 | no | AC158982.1 | intron_variant,non_coding_transcript_variant | MODIFIER |
| 14 | 123136189 | AX-170514920 | 0 | G | A | 0.298 | 1.08E-01 | 7.91E-02 | 2.67E+00 | 2.78E+00 | 1.70E-01 | no | AC158982.1 | intron_variant,non_coding_transcript_variant | MODIFIER |

| | | | | | | | | | | | | | | | |
|---|---|---|---|---|---|---|---|---|---|---|---|---|---|---|---|
| 14 | 123141855 | AX-170399179 | 0 | A | G | 0.298 | 1.08E-01 | 7.91E-02 | 2.67E+00 | 2.78E+00 | 1.70E-01 | no | AC158982.1 | intron_variant,non_coding_transcript_variant | MODIFIER |
| 14 | 123158669 | AX-170473031 | 0 | C | T | 0.298 | 1.08E-01 | 7.91E-02 | 2.67E+00 | 2.78E+00 | 1.70E-01 | no | AC158982.1 | intron_variant,non_coding_transcript_variant | MODIFIER |
| 14 | 123175148 | AX-97828179 | 0 | A | G | 0.298 | 1.08E-01 | 7.91E-02 | 2.67E+00 | 2.78E+00 | 1.70E-01 | no | AC158982.1 | intron_variant,non_coding_transcript_variant | MODIFIER |
| 14 | 123178989 | AX-170054645 | 0 | G | A | 0.298 | 1.08E-01 | 7.91E-02 | 2.67E+00 | 2.78E+00 | 1.70E-01 | no | AC158982.1 | intron_variant,non_coding_transcript_variant | MODIFIER |
| 14 | 123179203 | AX-170054643 | 0 | T | C | 0.298 | 1.08E-01 | 7.91E-02 | 2.67E+00 | 2.78E+00 | 1.70E-01 | no | AC158982.1 | intron_variant,non_coding_transcript_variant | MODIFIER |
| 14 | 123184797 | AX-170054644 | 0 | C | T | 0.298 | 1.08E-01 | 7.91E-02 | 2.67E+00 | 2.78E+00 | 1.70E-01 | no | AC158982.1 | intron_variant,non_coding_transcript_variant | MODIFIER |
| 14 | 123220154 | AX-97855879 | 0 | C | T | 0.298 | 1.08E-01 | 7.91E-02 | 2.67E+00 | 2.78E+00 | 1.70E-01 | no | AC158982.1 | intron_variant,non_coding_transcript_variant | MODIFIER |
| 14 | 123232580 | AX-170194793 | 0 | C | T | 0.298 | 1.08E-01 | 7.91E-02 | 2.67E+00 | 2.78E+00 | 1.70E-01 | no | AC158982.1 | intron_variant,non_coding_transcript_variant | MODIFIER |
| 14 | 123236963 | AX-170187799 | 0 | G | A | 0.298 | 1.08E-01 | 7.91E-02 | 2.67E+00 | 2.78E+00 | 1.70E-01 | no | AC158982.1 | intron_variant,non_coding_transcript_variant | MODIFIER |
| 14 | 123239982 | AX-170209249 | 0 | T | C | 0.298 | 1.08E-01 | 7.91E-02 | 2.67E+00 | 2.78E+00 | 1.70E-01 | no | AC158982.1 | intron_variant,non_coding_transcript_variant | MODIFIER |
| 14 | 123242990 | AX-97828181 | 0 | A | G | 0.298 | 1.08E-01 | 7.91E-02 | 2.67E+00 | 2.78E+00 | 1.70E-01 | no | AC158982.1 | intron_variant,non_coding_transcript_variant | MODIFIER |
| 14 | 123260340 | AX-170362970 | 0 | A | G | 0.298 | 1.08E-01 | 7.91E-02 | 2.67E+00 | 2.78E+00 | 1.70E-01 | no | AC158982.1 | intron_variant,non_coding_transcript_variant | MODIFIER |
| 14 | 123276172 | AX-97799657 | 0 | G | A | 0.298 | 1.08E-01 | 7.91E-02 | 2.67E+00 | 2.78E+00 | 1.70E-01 | no | Nalcn | downstream_gene_variant | MODIFIER |
| 14 | 123276544 | AX-170343423 | 0 | A | G | 0.298 | 1.08E-01 | 7.91E-02 | 2.67E+00 | 2.78E+00 | 1.70E-01 | no | Nalcn | downstream_gene_variant | MODIFIER |
| 14 | 123289908 | AX-170330542 | 0 | T | C | 0.298 | 1.08E-01 | 7.91E-02 | 2.67E+00 | 2.78E+00 | 1.70E-01 | no | Nalcn | intron_variant | MODIFIER |
| 14 | 123290626 | AX-170454147 | 0 | C | T | 0.298 | 1.08E-01 | 7.91E-02 | 2.67E+00 | 2.78E+00 | 1.70E-01 | no | Nalcn | intron_variant | MODIFIER |
| 14 | 123297958 | AX-170417711 | 0 | T | C | 0.298 | 1.08E-01 | 7.91E-02 | 2.67E+00 | 2.78E+00 | 1.70E-01 | no | Nalcn | intron_variant | MODIFIER |
| 14 | 123309569 | AX-170512304 | 0 | C | T | 0.298 | 1.08E-01 | 7.91E-02 | 2.67E+00 | 2.78E+00 | 1.70E-01 | no | Nalcn | intron_variant | MODIFIER |
| 14 | 123839377 | AX-168382502 | 0 | C | A | 0.298 | 1.08E-01 | 7.91E-02 | 2.67E+00 | 2.78E+00 | 1.70E-01 | no | Itgbl1 | intron_variant | MODIFIER |
| 14 | 124238816 | AX-170261436 | 0 | T | C | 0.298 | 1.08E-01 | 7.91E-02 | 2.67E+00 | 2.78E+00 | 1.70E-01 | no | Fgf14 | intron_variant | MODIFIER |
| 14 | 124245636 | AX-170170810 | 0 | T | A | 0.298 | 1.08E-01 | 7.91E-02 | 2.67E+00 | 2.78E+00 | 1.70E-01 | no | Fgf14 | intron_variant | MODIFIER |
| 14 | 124247006 | AX-170515907 | 0 | C | A | 0.298 | 1.08E-01 | 7.91E-02 | 2.67E+00 | 2.78E+00 | 1.70E-01 | no | Fgf14 | intron_variant | MODIFIER |
| 14 | 124247552 | AX-170455686 | 0 | A | G | 0.298 | 1.08E-01 | 7.91E-02 | 2.67E+00 | 2.78E+00 | 1.70E-01 | no | Fgf14 | intron_variant | MODIFIER |
| 14 | 124257732 | AX-170284781 | 0 | G | T | 0.298 | 1.08E-01 | 7.91E-02 | 2.67E+00 | 2.78E+00 | 1.70E-01 | no | Fgf14 | intron_variant | MODIFIER |
| 14 | 124275004 | AX-170393360 | 0 | G | A | 0.298 | 1.08E-01 | 7.91E-02 | 2.67E+00 | 2.78E+00 | 1.70E-01 | no | Fgf14 | intron_variant | MODIFIER |
| 14 | 124441001 | AX-97855980 | 0 | T | C | 0.298 | 1.08E-01 | 7.91E-02 | 2.67E+00 | 2.78E+00 | 1.70E-01 | no | Fgf14 | intron_variant | MODIFIER |
| 14 | 124462781 | AX-170231656 | 0 | C | A | 0.298 | 1.08E-01 | 7.91E-02 | 2.67E+00 | 2.78E+00 | 1.70E-01 | no | Fgf14 | intron_variant | MODIFIER |
| 14 | 124465521 | AX-97787637 | 0 | A | G | 0.298 | 1.08E-01 | 7.91E-02 | 2.67E+00 | 2.78E+00 | 1.70E-01 | no | Fgf14 | intron_variant | MODIFIER |
| 14 | 124552855 | AX-97828230 | 0 | T | G | 0.298 | 1.08E-01 | 7.91E-02 | 2.67E+00 | 2.78E+00 | 1.70E-01 | no | Fgf14 | intron_variant | MODIFIER |
| 18 | 78775042 | AX-170405162 | 0 | A | G | 0.447 | 1.20E-01 | 8.66E-02 | 2.43E+00 | 2.50E+00 | 1.70E-01 | no | Setbp1 | intron_variant | MODIFIER |
| 18 | 78776148 | AX-170016055 | 0 | T | C | 0.447 | 1.20E-01 | 8.66E-02 | 2.43E+00 | 2.50E+00 | 1.70E-01 | no | Setbp1 | intron_variant | MODIFIER |
| 14 | 122944280 | AX-97846482 | 1 | A | G | 0.295 | 1.08E-01 | 7.91E-02 | 2.69E+00 | 2.80E+00 | 1.70E-01 | no | Tmtc4 | intron_variant | MODIFIER |
| 9 | 9342457 | AX-97839270 | 0 | T | C | 0.465 | 1.12E-01 | 8.15E-02 | 2.46E+00 | 2.56E+00 | 1.70E-01 | no | | intergenic_variant | MODIFIER |
| 4 | 48886872 | AX-170401536 | 0 | G | A | 0.36 | -1.34E-01 | 9.78E-02 | 2.95E+00 | 3.05E+00 | 1.70E-01 | no | | intergenic_variant | MODIFIER |
| 7 | 24309888 | AX-170363707 | 0 | G | A | 0.412 | 1.15E-01 | 8.46E-02 | 2.60E+00 | 2.72E+00 | 1.70E-01 | no | Zfp94 | intron_variant | MODIFIER |
| 4 | 100793814 | AX-170125863 | 0 | G | A | 0.439 | -1.03E-01 | 7.54E-02 | 2.60E+00 | 2.69E+00 | 1.70E-01 | no | Cachd1 | intron_variant | MODIFIER |
| 16 | 79214875 | AX-97812393 | 0 | T | G | 0.482 | 9.38E-02 | 6.88E-02 | 2.64E+00 | 2.72E+00 | 1.70E-01 | no | | intergenic_variant | MODIFIER |
| 18 | 80489553 | AX-170297626 | 0 | T | C | 0.228 | -8.55E-02 | 6.29E-02 | 2.70E+00 | 2.82E+00 | 1.70E-01 | no | | intergenic_variant | MODIFIER |
| 4 | 124371752 | AX-97811523 | 1 | T | C | 0.223 | 1.56E-01 | 1.14E-01 | 2.57E+00 | 2.64E+00 | 1.70E-01 | Yes | | intergenic_variant | MODIFIER |
| 4 | 124706287 | AX-170070554 | 1 | T | C | 0.223 | 1.56E-01 | 1.14E-01 | 2.57E+00 | 2.64E+00 | 1.70E-01 | Yes | Fhl3 | intron_variant | MODIFIER |
| 4 | 125134268 | AX-97816265 | 1 | T | G | 0.223 | 1.56E-01 | 1.14E-01 | 2.57E+00 | 2.64E+00 | 1.70E-01 | Yes | | intergenic_variant | MODIFIER |
| 9 | 52654484 | AX-97823711 | 0 | G | T | 0.561 | 7.82E-02 | 5.63E-02 | 3.12E+00 | 3.25E+00 | 1.70E-01 | no | | intergenic_variant | MODIFIER |
| 16 | 50830877 | AX-170436208 | 1 | A | G | 0.473 | -1.11E-01 | 8.10E-02 | 2.50E+00 | 2.55E+00 | 1.70E-01 | no | | intergenic_variant | MODIFIER |
| 1 | 165495044 | AX-168406487 | 1 | A | G | 0.223 | 1.80E-01 | 1.32E-01 | 2.84E+00 | 2.92E+00 | 1.70E-01 | Yes | Adcy10 | intron_variant | MODIFIER |
| 15 | 84242973 | AX-170392055 | 0 | T | C | 0.105 | -1.49E-01 | 9.44E-02 | 1.86E+00 | 1.95E+00 | 1.71E-01 | no | Parvb | intron_variant | MODIFIER |
| 1 | 134517231 | AX-170316334 | 0 | T | C | 0.526 | 1.12E-01 | 8.23E-02 | 2.71E+00 | 2.79E+00 | 1.71E-01 | no | Gm15454 | upstream_gene_variant | MODIFIER |
| 12 | 89355353 | AX-170049733 | 0 | G | T | 0.088 | 2.18E-01 | 1.59E-01 | 2.96E+00 | 3.09E+00 | 1.71E-01 | Yes | Nrxn3 | intron_variant | MODIFIER |
| 5 | 48520218 | AX-170024881 | 1 | T | C | 0.5 | 1.45E-01 | 1.02E-01 | 2.25E+00 | 2.29E+00 | 1.71E-01 | Yes | Kcnip4 | intron_variant | MODIFIER |
| 4 | 64818900 | AX-97849795 | 0 | C | T | 0.333 | 1.51E-01 | 1.11E-01 | 2.88E+00 | 2.96E+00 | 1.71E-01 | Yes | | intergenic_variant | MODIFIER |
| 4 | 68768660 | AX-170023036 | 0 | G | A | 0.333 | 1.51E-01 | 1.11E-01 | 2.88E+00 | 2.96E+00 | 1.71E-01 | Yes | Brinp1 | intron_variant | MODIFIER |
| 13 | 20288722 | AX-170493905 | 2 | C | A | 0.182 | -1.69E-01 | 1.24E-01 | 2.53E+00 | 2.59E+00 | 1.71E-01 | no | Elmo1 | intron_variant | MODIFIER |
| 16 | 52794730 | AX-170415849 | 0 | C | G | 0.465 | -1.14E-01 | 8.33E-02 | 2.54E+00 | 2.57E+00 | 1.71E-01 | no | 4930404A05Rik | downstream_gene_variant | MODIFIER |
| 9 | 118156912 | AX-170142443 | 0 | T | C | 0.123 | -1.18E-01 | 8.71E-02 | 2.69E+00 | 2.85E+00 | 1.72E-01 | no | Gm17399 | intron_variant,non_coding_transcript_variant | MODIFIER |
| 9 | 114243111 | AX-170083518 | 0 | C | T | 0.132 | -1.37E-01 | 1.01E-01 | 2.95E+00 | 3.10E+00 | 1.72E-01 | no | 4921528I07Rik | intron_variant,non_coding_transcript_variant | MODIFIER |

| | | | | | | | | | | | | | | | |
|---|---|---|---|---|---|---|---|---|---|---|---|---|---|---|---|
| 14 | 113394157 | AX-170420288 | 1 | G | A | 0.036 | 1.83E-01 | 1.35E-01 | 2.58E+00 | 2.76E+00 | 1.72E-01 | Yes | | intergenic_variant | MODIFIER |
| 10 | 70347638 | AX-97781370 | 1 | G | T | 0.402 | -1.05E-01 | 7.73E-02 | 2.70E+00 | 2.79E+00 | 1.72E-01 | no | | intergenic_variant | MODIFIER |
| 5 | 63779123 | AX-170379766 | 1 | G | A | 0.491 | 1.30E-01 | 9.50E-02 | 2.48E+00 | 2.57E+00 | 1.72E-01 | no | Nwd2 | intron_variant | MODIFIER |
| 16 | 54556464 | AX-97839236 | 0 | A | C | 0.526 | 1.12E-01 | 8.10E-02 | 2.40E+00 | 2.45E+00 | 1.72E-01 | no | | intergenic_variant | MODIFIER |
| 16 | 53181115 | AX-170013061 | 0 | T | C | 0.526 | 1.12E-01 | 8.12E-02 | 2.41E+00 | 2.45E+00 | 1.72E-01 | no | | intergenic_variant | MODIFIER |
| 16 | 70694896 | AX-170433662 | 1 | T | C | 0.491 | 1.05E-01 | 7.68E-02 | 2.56E+00 | 2.63E+00 | 1.72E-01 | no | | intergenic_variant | MODIFIER |
| 16 | 52089155 | AX-97831785 | 1 | T | C | 0.464 | -1.11E-01 | 8.10E-02 | 2.49E+00 | 2.54E+00 | 1.73E-01 | no | Cblb | intron_variant | MODIFIER |
| 4 | 124062276 | AX-170264915 | 0 | G | C | 0.219 | 1.54E-01 | 1.13E-01 | 2.56E+00 | 2.63E+00 | 1.73E-01 | Yes | | intergenic_variant | MODIFIER |
| 4 | 124063874 | AX-170147662 | 0 | G | A | 0.219 | 1.54E-01 | 1.13E-01 | 2.56E+00 | 2.63E+00 | 1.73E-01 | Yes | | intergenic_variant | MODIFIER |
| 4 | 124091920 | AX-170376217 | 0 | G | A | 0.219 | 1.54E-01 | 1.13E-01 | 2.56E+00 | 2.63E+00 | 1.73E-01 | Yes | | intergenic_variant | MODIFIER |
| 4 | 124119565 | AX-170147001 | 0 | C | T | 0.219 | 1.54E-01 | 1.13E-01 | 2.56E+00 | 2.63E+00 | 1.73E-01 | Yes | | intergenic_variant | MODIFIER |
| 4 | 124128404 | AX-170378475 | 0 | T | C | 0.219 | 1.54E-01 | 1.13E-01 | 2.56E+00 | 2.63E+00 | 1.73E-01 | Yes | | intergenic_variant | MODIFIER |
| 4 | 124129294 | AX-170137453 | 0 | A | T | 0.219 | 1.54E-01 | 1.13E-01 | 2.56E+00 | 2.63E+00 | 1.73E-01 | Yes | | intergenic_variant | MODIFIER |
| 4 | 124144272 | AX-97822747 | 0 | G | T | 0.219 | 1.54E-01 | 1.13E-01 | 2.56E+00 | 2.63E+00 | 1.73E-01 | Yes | | intergenic_variant | MODIFIER |
| 4 | 124144606 | AX-170070541 | 0 | A | G | 0.219 | 1.54E-01 | 1.13E-01 | 2.56E+00 | 2.63E+00 | 1.73E-01 | Yes | | intergenic_variant | MODIFIER |
| 4 | 124159185 | AX-170070540 | 0 | A | G | 0.219 | 1.54E-01 | 1.13E-01 | 2.56E+00 | 2.63E+00 | 1.73E-01 | Yes | | intergenic_variant | MODIFIER |
| 4 | 124160475 | AX-170257728 | 0 | C | T | 0.219 | 1.54E-01 | 1.13E-01 | 2.56E+00 | 2.63E+00 | 1.73E-01 | Yes | | intergenic_variant | MODIFIER |
| 4 | 124163720 | AX-170462249 | 0 | C | T | 0.219 | 1.54E-01 | 1.13E-01 | 2.56E+00 | 2.63E+00 | 1.73E-01 | Yes | | intergenic_variant | MODIFIER |
| 4 | 124165951 | AX-170095132 | 0 | G | C | 0.219 | 1.54E-01 | 1.13E-01 | 2.56E+00 | 2.63E+00 | 1.73E-01 | Yes | | intergenic_variant | MODIFIER |
| 4 | 124190437 | AX-170184006 | 0 | A | G | 0.219 | 1.54E-01 | 1.13E-01 | 2.56E+00 | 2.63E+00 | 1.73E-01 | Yes | | intergenic_variant | MODIFIER |
| 4 | 124195953 | AX-170343070 | 0 | G | A | 0.219 | 1.54E-01 | 1.13E-01 | 2.56E+00 | 2.63E+00 | 1.73E-01 | Yes | | intergenic_variant | MODIFIER |
| 4 | 124210083 | AX-170442528 | 0 | T | C | 0.219 | 1.54E-01 | 1.13E-01 | 2.56E+00 | 2.63E+00 | 1.73E-01 | Yes | | intergenic_variant | MODIFIER |
| 4 | 124212569 | AX-170235733 | 0 | T | C | 0.219 | 1.54E-01 | 1.13E-01 | 2.56E+00 | 2.63E+00 | 1.73E-01 | Yes | | intergenic_variant | MODIFIER |
| 4 | 124235830 | AX-170350877 | 0 | C | A | 0.219 | 1.54E-01 | 1.13E-01 | 2.56E+00 | 2.63E+00 | 1.73E-01 | Yes | | intergenic_variant | MODIFIER |
| 4 | 124252491 | AX-170301003 | 0 | A | G | 0.219 | 1.54E-01 | 1.13E-01 | 2.56E+00 | 2.63E+00 | 1.73E-01 | Yes | | intergenic_variant | MODIFIER |
| 4 | 124268784 | AX-97826028 | 0 | T | C | 0.219 | 1.54E-01 | 1.13E-01 | 2.56E+00 | 2.63E+00 | 1.73E-01 | Yes | | intergenic_variant | MODIFIER |
| 4 | 124270218 | AX-170449353 | 0 | T | C | 0.219 | 1.54E-01 | 1.13E-01 | 2.56E+00 | 2.63E+00 | 1.73E-01 | Yes | | intergenic_variant | MODIFIER |
| 4 | 124270983 | AX-170207594 | 0 | G | A | 0.219 | 1.54E-01 | 1.13E-01 | 2.56E+00 | 2.63E+00 | 1.73E-01 | Yes | | intergenic_variant | MODIFIER |
| 4 | 124292503 | AX-170070544 | 0 | T | C | 0.219 | 1.54E-01 | 1.13E-01 | 2.56E+00 | 2.63E+00 | 1.73E-01 | Yes | Gm12916 | downstream_gene_variant | MODIFIER |
| 4 | 124292516 | AX-97783166 | 0 | G | A | 0.219 | 1.54E-01 | 1.13E-01 | 2.56E+00 | 2.63E+00 | 1.73E-01 | Yes | Gm12916 | downstream_gene_variant | MODIFIER |
| 4 | 124301453 | AX-170172360 | 0 | T | C | 0.219 | 1.54E-01 | 1.13E-01 | 2.56E+00 | 2.63E+00 | 1.73E-01 | Yes | Gm12916 | upstream_gene_variant | MODIFIER |
| 4 | 124314030 | AX-170231473 | 0 | T | G | 0.219 | 1.54E-01 | 1.13E-01 | 2.56E+00 | 2.63E+00 | 1.73E-01 | Yes | 1700021L23Rik | upstream_gene_variant | MODIFIER |
| 4 | 124317608 | AX-170130740 | 0 | G | C | 0.219 | 1.54E-01 | 1.13E-01 | 2.56E+00 | 2.63E+00 | 1.73E-01 | Yes | | intergenic_variant | MODIFIER |
| 4 | 124328939 | AX-170378743 | 0 | C | A | 0.219 | 1.54E-01 | 1.13E-01 | 2.56E+00 | 2.63E+00 | 1.73E-01 | Yes | 4933435F18Rik | downstream_gene_variant | MODIFIER |
| 4 | 124332383 | AX-170173177 | 0 | G | A | 0.219 | 1.54E-01 | 1.13E-01 | 2.56E+00 | 2.63E+00 | 1.73E-01 | Yes | 4933435F18Rik | non_coding_transcript_exon_variant | MODIFIER |
| 4 | 124338202 | AX-170349597 | 0 | A | G | 0.219 | 1.54E-01 | 1.13E-01 | 2.56E+00 | 2.63E+00 | 1.73E-01 | Yes | Gm22983 | upstream_gene_variant | MODIFIER |
| 4 | 124385897 | AX-170070547 | 0 | A | G | 0.219 | 1.54E-01 | 1.13E-01 | 2.56E+00 | 2.63E+00 | 1.73E-01 | Yes | | regulatory_region_variant | MODIFIER |
| 4 | 124402665 | AX-170161379 | 0 | A | G | 0.219 | 1.54E-01 | 1.13E-01 | 2.56E+00 | 2.63E+00 | 1.73E-01 | Yes | | intergenic_variant | MODIFIER |
| 4 | 124415341 | AX-170169422 | 0 | C | T | 0.219 | 1.54E-01 | 1.13E-01 | 2.56E+00 | 2.63E+00 | 1.73E-01 | Yes | 1700057H15Rik | intron_variant,non_coding_transcript_variant | MODIFIER |
| 4 | 124426887 | AX-170475778 | 0 | T | C | 0.219 | 1.54E-01 | 1.13E-01 | 2.56E+00 | 2.63E+00 | 1.73E-01 | Yes | 1700057H15Rik | intron_variant,non_coding_transcript_variant | MODIFIER |
| 4 | 124487453 | AX-170023870 | 0 | A | C | 0.219 | 1.54E-01 | 1.13E-01 | 2.56E+00 | 2.63E+00 | 1.73E-01 | Yes | 1700057H15Rik | upstream_gene_variant | MODIFIER |
| 4 | 124506870 | AX-170023865 | 0 | T | C | 0.219 | 1.54E-01 | 1.13E-01 | 2.56E+00 | 2.63E+00 | 1.73E-01 | Yes | | intergenic_variant | MODIFIER |
| 4 | 124516705 | AX-170445338 | 0 | A | G | 0.219 | 1.54E-01 | 1.13E-01 | 2.56E+00 | 2.63E+00 | 1.73E-01 | Yes | | intergenic_variant | MODIFIER |
| 4 | 124537679 | AX-170338244 | 0 | T | C | 0.219 | 1.54E-01 | 1.13E-01 | 2.56E+00 | 2.63E+00 | 1.73E-01 | Yes | Gm2164 | intron_variant,non_coding_transcript_variant | MODIFIER |
| 4 | 124544108 | AX-97839769 | 0 | T | C | 0.219 | 1.54E-01 | 1.13E-01 | 2.56E+00 | 2.63E+00 | 1.73E-01 | Yes | Gm2164 | intron_variant,non_coding_transcript_variant | MODIFIER |
| 4 | 124549170 | AX-170334717 | 0 | A | G | 0.219 | 1.54E-01 | 1.13E-01 | 2.56E+00 | 2.63E+00 | 1.73E-01 | Yes | Gm2164 | intron_variant,non_coding_transcript_variant | MODIFIER |
| 4 | 124553779 | AX-170144846 | 0 | T | G | 0.219 | 1.54E-01 | 1.13E-01 | 2.56E+00 | 2.63E+00 | 1.73E-01 | Yes | Gm2164 | intron_variant,non_coding_transcript_variant | MODIFIER |
| 4 | 124557375 | AX-170516676 | 0 | T | C | 0.219 | 1.54E-01 | 1.13E-01 | 2.56E+00 | 2.63E+00 | 1.73E-01 | Yes | Gm2164 | intron_variant,non_coding_transcript_variant | MODIFIER |
| 4 | 124592291 | AX-97808056 | 0 | G | A | 0.219 | 1.54E-01 | 1.13E-01 | 2.56E+00 | 2.63E+00 | 1.73E-01 | Yes | Gm2164 | intron_variant,non_coding_transcript_variant | MODIFIER |
| 4 | 124608883 | AX-97839367 | 0 | A | C | 0.219 | 1.54E-01 | 1.13E-01 | 2.56E+00 | 2.63E+00 | 1.73E-01 | Yes | 3100002H09Rik | downstream_gene_variant | MODIFIER |
| 4 | 124630491 | AX-170335000 | 0 | A | G | 0.219 | 1.54E-01 | 1.13E-01 | 2.56E+00 | 2.63E+00 | 1.73E-01 | Yes | Gm2164 | intron_variant,non_coding_transcript_variant | MODIFIER |
| 4 | 124633838 | AX-170023869 | 0 | G | C | 0.219 | 1.54E-01 | 1.13E-01 | 2.56E+00 | 2.63E+00 | 1.73E-01 | Yes | Gm2164 | intron_variant,non_coding_transcript_variant | MODIFIER |
| 4 | 124635719 | AX-170334584 | 0 | A | C | 0.219 | 1.54E-01 | 1.13E-01 | 2.56E+00 | 2.63E+00 | 1.73E-01 | Yes | Gm2164 | intron_variant,non_coding_transcript_variant | MODIFIER |
| 4 | 124693777 | AX-170115175 | 0 | A | G | 0.219 | 1.54E-01 | 1.13E-01 | 2.56E+00 | 2.63E+00 | 1.73E-01 | Yes | Utp11 | upstream_gene_variant | MODIFIER |

| | | | | | | | | | | | | | | | |
|---|---|---|---|---|---|---|---|---|---|---|---|---|---|---|---|
| 4 | 124718116 | AX-170455305 | 0 | A | G | 0.219 | 1.54E-01 | 1.13E-01 | 2.56E+00 | 2.63E+00 | 1.73E-01 | Yes | Sf3a3 | intron_variant | MODIFIER |
| 4 | 125068158 | AX-170023876 | 0 | A | G | 0.219 | 1.54E-01 | 1.13E-01 | 2.56E+00 | 2.63E+00 | 1.73E-01 | Yes | Dnali1 | upstream_gene_variant | MODIFIER |
| 4 | 125223118 | AX-170371704 | 0 | T | C | 0.219 | 1.54E-01 | 1.13E-01 | 2.56E+00 | 2.63E+00 | 1.73E-01 | Yes | | intergenic_variant | MODIFIER |
| 4 | 125223198 | AX-170350265 | 0 | A | G | 0.219 | 1.54E-01 | 1.13E-01 | 2.56E+00 | 2.63E+00 | 1.73E-01 | Yes | | intergenic_variant | MODIFIER |
| 4 | 125223488 | AX-170348181 | 0 | T | C | 0.219 | 1.54E-01 | 1.13E-01 | 2.56E+00 | 2.63E+00 | 1.73E-01 | Yes | | intergenic_variant | MODIFIER |
| 4 | 125231551 | AX-170155468 | 0 | T | G | 0.219 | 1.54E-01 | 1.13E-01 | 2.56E+00 | 2.63E+00 | 1.73E-01 | Yes | 1700041M05Rik | intron_variant,non_coding_transcript_variant | MODIFIER |
| 4 | 125261799 | AX-170226823 | 0 | G | T | 0.219 | 1.54E-01 | 1.13E-01 | 2.56E+00 | 2.63E+00 | 1.73E-01 | Yes | | intergenic_variant | MODIFIER |
| 4 | 125316816 | AX-170203428 | 0 | C | G | 0.219 | 1.54E-01 | 1.13E-01 | 2.56E+00 | 2.63E+00 | 1.73E-01 | Yes | | intergenic_variant | MODIFIER |
| 4 | 125318613 | AX-170230665 | 0 | G | A | 0.219 | 1.54E-01 | 1.13E-01 | 2.56E+00 | 2.63E+00 | 1.73E-01 | Yes | | intergenic_variant | MODIFIER |
| 4 | 125318809 | AX-170365568 | 0 | A | G | 0.219 | 1.54E-01 | 1.13E-01 | 2.56E+00 | 2.63E+00 | 1.73E-01 | Yes | | regulatory_region_variant | MODIFIER |
| 4 | 125319261 | AX-170405568 | 0 | C | G | 0.219 | 1.54E-01 | 1.13E-01 | 2.56E+00 | 2.63E+00 | 1.73E-01 | Yes | | regulatory_region_variant | MODIFIER |
| 4 | 125360674 | AX-170128042 | 0 | G | A | 0.219 | 1.54E-01 | 1.13E-01 | 2.56E+00 | 2.63E+00 | 1.73E-01 | Yes | | intergenic_variant | MODIFIER |
| 4 | 125524998 | AX-170449692 | 0 | A | C | 0.219 | 1.54E-01 | 1.13E-01 | 2.56E+00 | 2.63E+00 | 1.73E-01 | Yes | Grik3 | intron_variant | MODIFIER |
| 4 | 125541656 | AX-170415002 | 0 | T | C | 0.219 | 1.54E-01 | 1.13E-01 | 2.56E+00 | 2.63E+00 | 1.73E-01 | Yes | Grik3 | intron_variant | MODIFIER |
| 4 | 125552480 | AX-170444837 | 0 | T | C | 0.219 | 1.54E-01 | 1.13E-01 | 2.56E+00 | 2.63E+00 | 1.73E-01 | Yes | Grik3 | intron_variant | MODIFIER |
| 4 | 125620837 | AX-170532708 | 0 | T | C | 0.219 | 1.54E-01 | 1.13E-01 | 2.56E+00 | 2.63E+00 | 1.73E-01 | Yes | Grik3 | intron_variant | MODIFIER |
| 4 | 125621738 | AX-170200567 | 0 | C | G | 0.219 | 1.54E-01 | 1.13E-01 | 2.56E+00 | 2.63E+00 | 1.73E-01 | Yes | Grik3 | intron_variant | MODIFIER |
| 4 | 125626701 | AX-170329836 | 0 | C | G | 0.219 | 1.54E-01 | 1.13E-01 | 2.56E+00 | 2.63E+00 | 1.73E-01 | Yes | Grik3 | intron_variant | MODIFIER |
| 4 | 125626786 | AX-170472969 | 0 | A | G | 0.219 | 1.54E-01 | 1.13E-01 | 2.56E+00 | 2.63E+00 | 1.73E-01 | Yes | Grik3 | intron_variant | MODIFIER |
| 4 | 125630111 | AX-170301204 | 0 | A | G | 0.219 | 1.54E-01 | 1.13E-01 | 2.56E+00 | 2.63E+00 | 1.73E-01 | Yes | Grik3 | intron_variant | MODIFIER |
| 4 | 125630258 | AX-170370130 | 0 | T | G | 0.219 | 1.54E-01 | 1.13E-01 | 2.56E+00 | 2.63E+00 | 1.73E-01 | Yes | Grik3 | intron_variant | MODIFIER |
| 4 | 125630767 | AX-170535908 | 0 | A | G | 0.219 | 1.54E-01 | 1.13E-01 | 2.56E+00 | 2.63E+00 | 1.73E-01 | Yes | Grik3 | intron_variant | MODIFIER |
| 4 | 125633759 | AX-170023881 | 0 | C | T | 0.219 | 1.54E-01 | 1.13E-01 | 2.56E+00 | 2.63E+00 | 1.73E-01 | Yes | Grik3 | intron_variant | MODIFIER |
| 4 | 125664901 | AX-170303991 | 0 | C | T | 0.219 | 1.54E-01 | 1.13E-01 | 2.56E+00 | 2.63E+00 | 1.73E-01 | Yes | Grik3 | intron_variant | MODIFIER |
| 1 | 180960132 | AX-170331385 | 0 | A | T | 0.237 | 1.40E-01 | 1.03E-01 | 2.51E+00 | 2.60E+00 | 1.73E-01 | no | Tmem63a | intron_variant | MODIFIER |
| 16 | 80843129 | AX-170433394 | 1 | A | T | 0.464 | 9.75E-02 | 7.20E-02 | 2.69E+00 | 2.79E+00 | 1.73E-01 | no | | intergenic_variant | MODIFIER |
| 16 | 29477269 | AX-170531493 | 2 | C | T | 0.518 | 1.06E-01 | 7.75E-02 | 2.55E+00 | 2.62E+00 | 1.73E-01 | no | Atp13a4 | intron_variant | MODIFIER |
| 1 | 168178594 | AX-170103783 | 0 | C | T | 0.228 | 1.80E-01 | 1.33E-01 | 2.80E+00 | 2.87E+00 | 1.73E-01 | Yes | Pbx1 | intron_variant | MODIFIER |
| 18 | 80620957 | AX-170224937 | 0 | A | C | 0.149 | -1.47E-01 | 1.07E-01 | 2.44E+00 | 2.50E+00 | 1.73E-01 | no | Nfatc1 | intron_variant | MODIFIER |
| 5 | 50038264 | AX-170072046 | 1 | T | C | 0.491 | 1.45E-01 | 1.04E-01 | 2.33E+00 | 2.37E+00 | 1.73E-01 | Yes | Adgra3 | intron_variant | MODIFIER |
| 4 | 54703614 | AX-170256528 | 0 | C | T | 0.368 | -1.35E-01 | 9.93E-02 | 2.88E+00 | 2.99E+00 | 1.73E-01 | no | | intergenic_variant | MODIFIER |
| 9 | 8023158 | AX-97803983 | 0 | G | A | 0.456 | 1.11E-01 | 8.10E-02 | 2.48E+00 | 2.57E+00 | 1.73E-01 | no | Cfap300 | intron_variant | MODIFIER |
| 7 | 38212282 | AX-170126133 | 0 | C | A | 0.351 | 1.21E-01 | 8.93E-02 | 2.66E+00 | 2.75E+00 | 1.73E-01 | no | Plekhf1 | downstream_gene_variant | MODIFIER |
| 5 | 49846397 | AX-170309706 | 2 | C | T | 0.518 | 1.40E-01 | 1.01E-01 | 2.34E+00 | 2.39E+00 | 1.73E-01 | no | | intergenic_variant | MODIFIER |
| 1 | 180437653 | AX-170431853 | 0 | G | A | 0.246 | 1.47E-01 | 1.08E-01 | 2.54E+00 | 2.61E+00 | 1.74E-01 | Yes | Stum | 3_prime_UTR_variant | MODIFIER |
| 6 | 127934832 | AX-170155848 | 0 | C | T | 0.544 | 1.13E-01 | 8.29E-02 | 2.51E+00 | 2.58E+00 | 1.74E-01 | no | Tspan11 | intron_variant | MODIFIER |
| 5 | 55299975 | AX-170024987 | 0 | C | G | 0.456 | 1.38E-01 | 9.97E-02 | 2.37E+00 | 2.43E+00 | 1.74E-01 | no | Gm25513 | downstream_gene_variant | MODIFIER |
| 3 | 66417293 | AX-170404301 | 1 | G | A | 0.045 | 1.73E-01 | 1.29E-01 | 2.76E+00 | 2.96E+00 | 1.74E-01 | Yes | | intergenic_variant | MODIFIER |
| 16 | 35713367 | AX-170194699 | 1 | C | T | 0.5 | 1.07E-01 | 7.84E-02 | 2.52E+00 | 2.57E+00 | 1.74E-01 | no | Dirc2 | intron_variant | MODIFIER |
| 5 | 135476072 | AX-170281237 | 2 | G | C | 0.164 | -1.20E-01 | 8.56E-02 | 2.30E+00 | 2.38E+00 | 1.74E-01 | no | Hip1 | intron_variant | MODIFIER |
| 8 | 128250138 | AX-170472805 | 0 | C | T | 0.061 | 2.42E-01 | 1.78E-01 | 2.97E+00 | 3.12E+00 | 1.74E-01 | Yes | | intergenic_variant | MODIFIER |
| 16 | 69885400 | AX-170035568 | 0 | C | T | 0.491 | 1.04E-01 | 7.69E-02 | 2.56E+00 | 2.62E+00 | 1.74E-01 | no | | intergenic_variant | MODIFIER |
| 16 | 70168394 | AX-170166560 | 0 | T | C | 0.491 | 1.04E-01 | 7.69E-02 | 2.56E+00 | 2.62E+00 | 1.74E-01 | no | | intergenic_variant | MODIFIER |
| 16 | 70168522 | AX-170275935 | 0 | G | A | 0.491 | 1.04E-01 | 7.69E-02 | 2.56E+00 | 2.62E+00 | 1.74E-01 | no | | intergenic_variant | MODIFIER |
| 16 | 70169444 | AX-170148880 | 0 | G | A | 0.491 | 1.04E-01 | 7.69E-02 | 2.56E+00 | 2.62E+00 | 1.74E-01 | no | | intergenic_variant | MODIFIER |
| 16 | 70177014 | AX-170414229 | 0 | C | T | 0.491 | 1.04E-01 | 7.69E-02 | 2.56E+00 | 2.62E+00 | 1.74E-01 | no | | intergenic_variant | MODIFIER |
| 16 | 70183362 | AX-170174250 | 0 | C | T | 0.491 | 1.04E-01 | 7.69E-02 | 2.56E+00 | 2.62E+00 | 1.74E-01 | no | | intergenic_variant | MODIFIER |
| 16 | 70183424 | AX-170272037 | 0 | G | A | 0.491 | 1.04E-01 | 7.69E-02 | 2.56E+00 | 2.62E+00 | 1.74E-01 | no | | intergenic_variant | MODIFIER |
| 16 | 70183818 | AX-97843624 | 0 | T | C | 0.491 | 1.04E-01 | 7.69E-02 | 2.56E+00 | 2.62E+00 | 1.74E-01 | no | | intergenic_variant | MODIFIER |
| 16 | 70194133 | AX-170482762 | 0 | G | T | 0.491 | 1.04E-01 | 7.69E-02 | 2.56E+00 | 2.62E+00 | 1.74E-01 | no | Gm49656 | downstream_gene_variant | MODIFIER |
| 16 | 70194759 | AX-170265543 | 0 | C | G | 0.491 | 1.04E-01 | 7.69E-02 | 2.56E+00 | 2.62E+00 | 1.74E-01 | no | Gm49656 | downstream_gene_variant | MODIFIER |
| 16 | 70194970 | AX-170221297 | 0 | A | G | 0.491 | 1.04E-01 | 7.69E-02 | 2.56E+00 | 2.62E+00 | 1.74E-01 | no | Gm49656 | downstream_gene_variant | MODIFIER |
| 16 | 70203484 | AX-170013341 | 0 | G | C | 0.491 | 1.04E-01 | 7.69E-02 | 2.56E+00 | 2.62E+00 | 1.74E-01 | no | | intergenic_variant | MODIFIER |
| 16 | 70204637 | AX-170013343 | 0 | T | C | 0.491 | 1.04E-01 | 7.69E-02 | 2.56E+00 | 2.62E+00 | 1.74E-01 | no | | intergenic_variant | MODIFIER |

| | | | | | | | | | | | | | | | |
|---|---|---|---|---|---|---|---|---|---|---|---|---|---|---|---|
| 16 | 70232954 | AX-170260628 | 0 | C | G | 0.491 | 1.04E-01 | 7.69E-02 | 2.56E+00 | 2.62E+00 | 1.74E-01 | no | | regulatory_region_variant | MODIFIER |
| 16 | 70238093 | AX-170057727 | 0 | A | G | 0.491 | 1.04E-01 | 7.69E-02 | 2.56E+00 | 2.62E+00 | 1.74E-01 | no | | intergenic_variant | MODIFIER |
| 16 | 70238169 | AX-170093516 | 0 | T | C | 0.491 | 1.04E-01 | 7.69E-02 | 2.56E+00 | 2.62E+00 | 1.74E-01 | no | | intergenic_variant | MODIFIER |
| 16 | 70241422 | AX-170013340 | 0 | A | G | 0.491 | 1.04E-01 | 7.69E-02 | 2.56E+00 | 2.62E+00 | 1.74E-01 | no | | intergenic_variant | MODIFIER |
| 16 | 70266937 | AX-97794159 | 0 | A | C | 0.491 | 1.04E-01 | 7.69E-02 | 2.56E+00 | 2.62E+00 | 1.74E-01 | no | | intergenic_variant | MODIFIER |
| 16 | 70279328 | AX-170425583 | 0 | G | C | 0.491 | 1.04E-01 | 7.69E-02 | 2.56E+00 | 2.62E+00 | 1.74E-01 | no | | intergenic_variant | MODIFIER |
| 16 | 70307923 | AX-170013344 | 0 | A | G | 0.491 | 1.04E-01 | 7.69E-02 | 2.56E+00 | 2.62E+00 | 1.74E-01 | no | | intergenic_variant | MODIFIER |
| 16 | 70331279 | AX-170315965 | 0 | C | T | 0.491 | 1.04E-01 | 7.69E-02 | 2.56E+00 | 2.62E+00 | 1.74E-01 | no | Gbe1 | intron_variant | MODIFIER |
| 16 | 70348311 | AX-170103941 | 0 | G | A | 0.491 | 1.04E-01 | 7.69E-02 | 2.56E+00 | 2.62E+00 | 1.74E-01 | no | Gbe1 | intron_variant | MODIFIER |
| 16 | 70355028 | AX-170445144 | 0 | C | T | 0.491 | 1.04E-01 | 7.69E-02 | 2.56E+00 | 2.62E+00 | 1.74E-01 | no | Gbe1 | intron_variant | MODIFIER |
| 16 | 70388987 | AX-170190634 | 0 | A | G | 0.491 | 1.04E-01 | 7.69E-02 | 2.56E+00 | 2.62E+00 | 1.74E-01 | no | Gbe1 | intron_variant | MODIFIER |
| 16 | 70407237 | AX-170131736 | 0 | C | T | 0.491 | 1.04E-01 | 7.69E-02 | 2.56E+00 | 2.62E+00 | 1.74E-01 | no | Gbe1 | intron_variant | MODIFIER |
| 16 | 70420536 | AX-170135887 | 0 | T | G | 0.491 | 1.04E-01 | 7.69E-02 | 2.56E+00 | 2.62E+00 | 1.74E-01 | no | Gbe1 | intron_variant | MODIFIER |
| 16 | 70424611 | AX-170306400 | 0 | G | T | 0.491 | 1.04E-01 | 7.69E-02 | 2.56E+00 | 2.62E+00 | 1.74E-01 | no | Gbe1 | intron_variant | MODIFIER |
| 16 | 70424629 | AX-170387702 | 0 | G | A | 0.491 | 1.04E-01 | 7.69E-02 | 2.56E+00 | 2.62E+00 | 1.74E-01 | no | Gbe1 | intron_variant | MODIFIER |
| 16 | 70433488 | AX-170358897 | 0 | T | G | 0.491 | 1.04E-01 | 7.69E-02 | 2.56E+00 | 2.62E+00 | 1.74E-01 | no | Gbe1 | intron_variant | MODIFIER |
| 16 | 70465013 | AX-170143841 | 0 | A | G | 0.491 | 1.04E-01 | 7.69E-02 | 2.56E+00 | 2.62E+00 | 1.74E-01 | no | Gbe1 | intron_variant | MODIFIER |
| 16 | 70469423 | AX-170149720 | 0 | A | G | 0.491 | 1.04E-01 | 7.69E-02 | 2.56E+00 | 2.62E+00 | 1.74E-01 | no | Gbe1 | intron_variant | MODIFIER |
| 16 | 70495772 | AX-170166400 | 0 | T | C | 0.491 | 1.04E-01 | 7.69E-02 | 2.56E+00 | 2.62E+00 | 1.74E-01 | no | Gbe1 | intron_variant | MODIFIER |
| 16 | 70511359 | AX-170381473 | 0 | G | A | 0.491 | 1.04E-01 | 7.69E-02 | 2.56E+00 | 2.62E+00 | 1.74E-01 | no | Gbe1 | intron_variant | MODIFIER |
| 16 | 70522962 | AX-170349492 | 0 | C | T | 0.491 | 1.04E-01 | 7.69E-02 | 2.56E+00 | 2.62E+00 | 1.74E-01 | no | Gbe1 | intron_variant | MODIFIER |
| 16 | 70523865 | AX-170408428 | 0 | A | C | 0.491 | 1.04E-01 | 7.69E-02 | 2.56E+00 | 2.62E+00 | 1.74E-01 | no | Gbe1 | intron_variant | MODIFIER |
| 16 | 70523988 | AX-170108749 | 0 | T | C | 0.491 | 1.04E-01 | 7.69E-02 | 2.56E+00 | 2.62E+00 | 1.74E-01 | no | Gbe1 | intron_variant | MODIFIER |
| 16 | 70526254 | AX-170194438 | 0 | G | A | 0.491 | 1.04E-01 | 7.69E-02 | 2.56E+00 | 2.62E+00 | 1.74E-01 | no | Gbe1 | intron_variant | MODIFIER |
| 16 | 70530043 | AX-170126518 | 0 | T | C | 0.491 | 1.04E-01 | 7.69E-02 | 2.56E+00 | 2.62E+00 | 1.74E-01 | no | Gbe1 | intron_variant | MODIFIER |
| 16 | 70531810 | AX-170328944 | 0 | C | G | 0.491 | 1.04E-01 | 7.69E-02 | 2.56E+00 | 2.62E+00 | 1.74E-01 | no | Gbe1 | intron_variant | MODIFIER |
| 16 | 70533197 | AX-170191485 | 0 | T | G | 0.491 | 1.04E-01 | 7.69E-02 | 2.56E+00 | 2.62E+00 | 1.74E-01 | no | Gbe1 | intron_variant | MODIFIER |
| 16 | 70536580 | AX-170181223 | 0 | G | T | 0.491 | 1.04E-01 | 7.69E-02 | 2.56E+00 | 2.62E+00 | 1.74E-01 | no | Gbe1 | intron_variant | MODIFIER |
| 16 | 70550773 | AX-170281020 | 0 | G | T | 0.491 | 1.04E-01 | 7.69E-02 | 2.56E+00 | 2.62E+00 | 1.74E-01 | no | Gbe1 | intron_variant | MODIFIER |
| 16 | 70560078 | AX-170486383 | 0 | A | T | 0.491 | 1.04E-01 | 7.69E-02 | 2.56E+00 | 2.62E+00 | 1.74E-01 | no | Gbe1 | intron_variant | MODIFIER |
| 16 | 70560095 | AX-97831675 | 0 | A | G | 0.491 | 1.04E-01 | 7.69E-02 | 2.56E+00 | 2.62E+00 | 1.74E-01 | no | Gbe1 | intron_variant | MODIFIER |
| 16 | 70575245 | AX-170411684 | 0 | T | C | 0.491 | 1.04E-01 | 7.69E-02 | 2.56E+00 | 2.62E+00 | 1.74E-01 | no | | intergenic_variant | MODIFIER |
| 16 | 70600857 | AX-170176019 | 0 | A | G | 0.491 | 1.04E-01 | 7.69E-02 | 2.56E+00 | 2.62E+00 | 1.74E-01 | no | | intergenic_variant | MODIFIER |
| 16 | 70602639 | AX-170168103 | 0 | T | G | 0.491 | 1.04E-01 | 7.69E-02 | 2.56E+00 | 2.62E+00 | 1.74E-01 | no | | intergenic_variant | MODIFIER |
| 16 | 70606987 | AX-170228038 | 0 | A | C | 0.491 | 1.04E-01 | 7.69E-02 | 2.56E+00 | 2.62E+00 | 1.74E-01 | no | | intergenic_variant | MODIFIER |
| 16 | 70608358 | AX-170468013 | 0 | G | A | 0.491 | 1.04E-01 | 7.69E-02 | 2.56E+00 | 2.62E+00 | 1.74E-01 | no | | intergenic_variant | MODIFIER |
| 16 | 70608500 | AX-170297860 | 0 | T | C | 0.491 | 1.04E-01 | 7.69E-02 | 2.56E+00 | 2.62E+00 | 1.74E-01 | no | | intergenic_variant | MODIFIER |
| 16 | 70655412 | AX-170284566 | 0 | A | G | 0.491 | 1.04E-01 | 7.69E-02 | 2.56E+00 | 2.62E+00 | 1.74E-01 | no | | intergenic_variant | MODIFIER |
| 16 | 70655441 | AX-170291363 | 0 | G | A | 0.491 | 1.04E-01 | 7.69E-02 | 2.56E+00 | 2.62E+00 | 1.74E-01 | no | | intergenic_variant | MODIFIER |
| 16 | 70655492 | AX-97797181 | 0 | A | C | 0.491 | 1.04E-01 | 7.69E-02 | 2.56E+00 | 2.62E+00 | 1.74E-01 | no | | intergenic_variant | MODIFIER |
| 16 | 70741788 | AX-97851529 | 0 | G | A | 0.491 | 1.04E-01 | 7.69E-02 | 2.56E+00 | 2.62E+00 | 1.74E-01 | no | | intergenic_variant | MODIFIER |
| 16 | 70785875 | AX-170013345 | 0 | G | A | 0.491 | 1.04E-01 | 7.69E-02 | 2.56E+00 | 2.62E+00 | 1.74E-01 | no | | intergenic_variant | MODIFIER |
| 16 | 70816675 | AX-170013349 | 0 | A | C | 0.491 | 1.04E-01 | 7.69E-02 | 2.56E+00 | 2.62E+00 | 1.74E-01 | no | | intergenic_variant | MODIFIER |
| 16 | 70817005 | AX-97856821 | 0 | C | A | 0.491 | 1.04E-01 | 7.69E-02 | 2.56E+00 | 2.62E+00 | 1.74E-01 | no | | intergenic_variant | MODIFIER |
| 16 | 70853853 | AX-170013347 | 0 | C | A | 0.491 | 1.04E-01 | 7.69E-02 | 2.56E+00 | 2.62E+00 | 1.74E-01 | no | | intergenic_variant | MODIFIER |
| 16 | 70935475 | AX-170309410 | 0 | T | C | 0.491 | 1.04E-01 | 7.69E-02 | 2.56E+00 | 2.62E+00 | 1.74E-01 | no | | intergenic_variant | MODIFIER |
| 16 | 70938946 | AX-170435257 | 0 | G | A | 0.491 | 1.04E-01 | 7.69E-02 | 2.56E+00 | 2.62E+00 | 1.74E-01 | no | | intergenic_variant | MODIFIER |
| 16 | 70951869 | AX-170403605 | 0 | C | T | 0.491 | 1.04E-01 | 7.69E-02 | 2.56E+00 | 2.62E+00 | 1.74E-01 | no | Gm29823 | upstream_gene_variant | MODIFIER |
| 16 | 70987285 | AX-97832256 | 0 | G | C | 0.491 | 1.04E-01 | 7.69E-02 | 2.56E+00 | 2.62E+00 | 1.74E-01 | no | Gm29823 | intron_variant,non_coding_transcript_variant | MODIFIER |
| 16 | 70993800 | AX-170186033 | 0 | A | G | 0.491 | 1.04E-01 | 7.69E-02 | 2.56E+00 | 2.62E+00 | 1.74E-01 | no | Gm29823 | intron_variant,non_coding_transcript_variant | MODIFIER |
| 16 | 71008977 | AX-97832283 | 0 | T | C | 0.491 | 1.04E-01 | 7.69E-02 | 2.56E+00 | 2.62E+00 | 1.74E-01 | no | Gm29823 | intron_variant,non_coding_transcript_variant | MODIFIER |
| 16 | 71061002 | AX-170481199 | 0 | A | G | 0.491 | 1.04E-01 | 7.69E-02 | 2.56E+00 | 2.62E+00 | 1.74E-01 | no | | intergenic_variant | MODIFIER |
| 16 | 71090631 | AX-170531482 | 0 | A | G | 0.491 | 1.04E-01 | 7.69E-02 | 2.56E+00 | 2.62E+00 | 1.74E-01 | no | | intergenic_variant | MODIFIER |
| 16 | 71095342 | AX-170506286 | 0 | G | C | 0.491 | 1.04E-01 | 7.69E-02 | 2.56E+00 | 2.62E+00 | 1.74E-01 | no | | intergenic_variant | MODIFIER |

| | | | | | | | | | | | | | | | |
|---|---|---|---|---|---|---|---|---|---|---|---|---|---|---|---|
| 16 | 71135157 | AX-170512808 | 0 | C | G | 0.491 | 1.04E-01 | 7.69E-02 | 2.56E+00 | 2.62E+00 | 1.74E-01 | no | 4931420L22Rik | splice_region_variant,intron_variant,non_coding_transcript_variant | LOW |
| 16 | 71135677 | AX-170448168 | 0 | G | T | 0.491 | 1.04E-01 | 7.69E-02 | 2.56E+00 | 2.62E+00 | 1.74E-01 | no | 4931420L22Rik | intron_variant,non_coding_transcript_variant | MODIFIER |
| 16 | 71220135 | AX-170415929 | 0 | C | A | 0.491 | 1.04E-01 | 7.69E-02 | 2.56E+00 | 2.62E+00 | 1.74E-01 | no | Gm49666 | upstream_gene_variant | MODIFIER |
| 16 | 71255231 | AX-170013357 | 0 | G | C | 0.491 | 1.04E-01 | 7.69E-02 | 2.56E+00 | 2.62E+00 | 1.74E-01 | no | 4930567J20Rik | intron_variant,non_coding_transcript_variant | MODIFIER |
| 16 | 71255528 | AX-97832285 | 0 | A | G | 0.491 | 1.04E-01 | 7.69E-02 | 2.56E+00 | 2.62E+00 | 1.74E-01 | no | 4930567J20Rik | intron_variant,non_coding_transcript_variant | MODIFIER |
| 16 | 71268195 | AX-170013353 | 0 | A | C | 0.491 | 1.04E-01 | 7.69E-02 | 2.56E+00 | 2.62E+00 | 1.74E-01 | no | 4930567J20Rik | downstream_gene_variant | MODIFIER |
| 16 | 71271665 | AX-170191659 | 0 | A | C | 0.491 | 1.04E-01 | 7.69E-02 | 2.56E+00 | 2.62E+00 | 1.74E-01 | no | 4930567J20Rik | downstream_gene_variant | MODIFIER |
| 16 | 71295728 | AX-97850699 | 0 | T | C | 0.491 | 1.04E-01 | 7.69E-02 | 2.56E+00 | 2.62E+00 | 1.74E-01 | no | | intergenic_variant | MODIFIER |
| 16 | 71376192 | AX-170325420 | 0 | C | T | 0.491 | 1.04E-01 | 7.69E-02 | 2.56E+00 | 2.62E+00 | 1.74E-01 | no | | intergenic_variant | MODIFIER |
| 16 | 71533449 | AX-170124870 | 0 | C | T | 0.491 | 1.04E-01 | 7.69E-02 | 2.56E+00 | 2.62E+00 | 1.74E-01 | no | | intergenic_variant | MODIFIER |
| 16 | 71769505 | AX-170013358 | 0 | A | G | 0.491 | 1.04E-01 | 7.69E-02 | 2.56E+00 | 2.62E+00 | 1.74E-01 | no | | intergenic_variant | MODIFIER |
| 16 | 71808739 | AX-170013359 | 0 | A | G | 0.491 | 1.04E-01 | 7.69E-02 | 2.56E+00 | 2.62E+00 | 1.74E-01 | no | | intergenic_variant | MODIFIER |
| 16 | 71935686 | AX-170104561 | 0 | C | T | 0.491 | 1.04E-01 | 7.69E-02 | 2.56E+00 | 2.62E+00 | 1.74E-01 | no | | intergenic_variant | MODIFIER |
| 16 | 71945305 | AX-170142726 | 0 | C | T | 0.491 | 1.04E-01 | 7.69E-02 | 2.56E+00 | 2.62E+00 | 1.74E-01 | no | | intergenic_variant | MODIFIER |
| 16 | 71959389 | AX-170254324 | 0 | C | T | 0.491 | 1.04E-01 | 7.69E-02 | 2.56E+00 | 2.62E+00 | 1.74E-01 | no | | intergenic_variant | MODIFIER |
| 16 | 71959779 | AX-170256630 | 0 | G | C | 0.491 | 1.04E-01 | 7.69E-02 | 2.56E+00 | 2.62E+00 | 1.74E-01 | no | | intergenic_variant | MODIFIER |
| 16 | 71960240 | AX-97812394 | 0 | T | C | 0.491 | 1.04E-01 | 7.69E-02 | 2.56E+00 | 2.62E+00 | 1.74E-01 | no | | intergenic_variant | MODIFIER |
| 16 | 72067783 | AX-170364954 | 0 | A | G | 0.491 | 1.04E-01 | 7.69E-02 | 2.56E+00 | 2.62E+00 | 1.74E-01 | no | | intergenic_variant | MODIFIER |
| 16 | 72067858 | AX-170396816 | 0 | C | T | 0.491 | 1.04E-01 | 7.69E-02 | 2.56E+00 | 2.62E+00 | 1.74E-01 | no | | intergenic_variant | MODIFIER |
| 16 | 72067917 | AX-170330257 | 0 | C | G | 0.491 | 1.04E-01 | 7.69E-02 | 2.56E+00 | 2.62E+00 | 1.74E-01 | no | | intergenic_variant | MODIFIER |
| 16 | 72085101 | AX-97822215 | 0 | A | G | 0.491 | 1.04E-01 | 7.69E-02 | 2.56E+00 | 2.62E+00 | 1.74E-01 | no | | intergenic_variant | MODIFIER |
| 16 | 72099135 | AX-170357679 | 0 | T | G | 0.491 | 1.04E-01 | 7.69E-02 | 2.56E+00 | 2.62E+00 | 1.74E-01 | no | | intergenic_variant | MODIFIER |
| 16 | 72099359 | AX-170456383 | 0 | C | A | 0.491 | 1.04E-01 | 7.69E-02 | 2.56E+00 | 2.62E+00 | 1.74E-01 | no | | intergenic_variant | MODIFIER |
| 16 | 72103219 | AX-170291063 | 0 | T | C | 0.491 | 1.04E-01 | 7.69E-02 | 2.56E+00 | 2.62E+00 | 1.74E-01 | no | | intergenic_variant | MODIFIER |
| 16 | 72103363 | AX-170116859 | 0 | C | G | 0.491 | 1.04E-01 | 7.69E-02 | 2.56E+00 | 2.62E+00 | 1.74E-01 | no | | intergenic_variant | MODIFIER |
| 16 | 72105025 | AX-97798533 | 0 | C | A | 0.491 | 1.04E-01 | 7.69E-02 | 2.56E+00 | 2.62E+00 | 1.74E-01 | no | | intergenic_variant | MODIFIER |
| 16 | 72105652 | AX-170202305 | 0 | C | A | 0.491 | 1.04E-01 | 7.69E-02 | 2.56E+00 | 2.62E+00 | 1.74E-01 | no | | intergenic_variant | MODIFIER |
| 16 | 72118383 | AX-170172652 | 0 | C | T | 0.491 | 1.04E-01 | 7.69E-02 | 2.56E+00 | 2.62E+00 | 1.74E-01 | no | | intergenic_variant | MODIFIER |
| 16 | 72135582 | AX-170314143 | 0 | T | C | 0.491 | 1.04E-01 | 7.69E-02 | 2.56E+00 | 2.62E+00 | 1.74E-01 | no | | intergenic_variant | MODIFIER |
| 16 | 72143416 | AX-170171061 | 0 | G | A | 0.491 | 1.04E-01 | 7.69E-02 | 2.56E+00 | 2.62E+00 | 1.74E-01 | no | | regulatory_region_variant | MODIFIER |
| 1 | 192540517 | AX-170414967 | 0 | C | T | 0.281 | 1.25E-01 | 8.95E-02 | 2.30E+00 | 2.38E+00 | 1.74E-01 | no | Hhat | intron_variant | MODIFIER |
| 1 | 194628785 | AX-170044042 | 0 | G | T | 0.281 | 1.25E-01 | 8.95E-02 | 2.30E+00 | 2.38E+00 | 1.74E-01 | no | Plxna2 | intron_variant | MODIFIER |
| 8 | 84838777 | AX-170080633 | 1 | T | C | 0.196 | -1.76E-01 | 1.27E-01 | 3.24E+00 | 3.38E+00 | 1.74E-01 | Yes | Rad23a | intron_variant | MODIFIER |
| 10 | 104746185 | AX-170428185 | 0 | T | C | 0.289 | -1.15E-01 | 8.53E-02 | 2.64E+00 | 2.71E+00 | 1.75E-01 | no | | intergenic_variant | MODIFIER |
| 20 | 46658012 | AX-170084155 | 0 | A | G | 0.254 | 7.57E-02 | 5.62E-02 | 2.62E+00 | 2.75E+00 | 1.75E-01 | no | Gm29242 | intron_variant,non_coding_transcript_variant | MODIFIER |
| 15 | 19069275 | AX-170011169 | 2 | A | C | 0.236 | -1.41E-01 | 1.04E-01 | 2.72E+00 | 2.79E+00 | 1.75E-01 | no | Gm8328 | upstream_gene_variant | MODIFIER |
| 15 | 6401276 | AX-170146323 | 0 | G | A | 0.237 | -1.40E-01 | 1.04E-01 | 2.65E+00 | 2.73E+00 | 1.75E-01 | no | Dab2 | intron_variant | MODIFIER |
| 5 | 10785521 | AX-170106878 | 0 | G | A | 0.351 | 1.15E-01 | 8.55E-02 | 2.72E+00 | 2.85E+00 | 1.75E-01 | no | | intergenic_variant | MODIFIER |
| 10 | 19062476 | AX-170363043 | 0 | G | A | 0.193 | 1.40E-01 | 1.01E-01 | 3.24E+00 | 3.37E+00 | 1.75E-01 | no | | regulatory_region_variant | MODIFIER |
| 3 | 78759578 | AX-170168588 | 0 | C | T | 0.044 | -2.06E-01 | 1.54E-01 | 2.78E+00 | 2.94E+00 | 1.75E-01 | Yes | Gm18952 | downstream_gene_variant | MODIFIER |
| 1 | 186263107 | AX-170369428 | 1 | A | T | 0.232 | 1.49E-01 | 1.08E-01 | 2.38E+00 | 2.45E+00 | 1.75E-01 | Yes | | regulatory_region_variant | MODIFIER |
| 1 | 155804817 | AX-170259368 | 0 | T | C | 0.439 | -9.77E-02 | 6.70E-02 | 3.61E+00 | 3.80E+00 | 1.76E-01 | no | Qsox1 | intron_variant | MODIFIER |
| 15 | 71920272 | AX-170093300 | 1 | T | C | 0.089 | -1.74E-01 | 1.10E-01 | 1.84E+00 | 1.91E+00 | 1.76E-01 | Yes | Col22a1 | intron_variant | MODIFIER |
| 11 | 47897814 | AX-97805798 | 0 | G | A | 0.088 | -2.11E-01 | 1.57E-01 | 2.69E+00 | 2.81E+00 | 1.76E-01 | Yes | Sgcd | intron_variant,non_coding_transcript_variant | MODIFIER |
| 9 | 101437135 | AX-170487134 | 1 | C | G | 0.134 | -1.46E-01 | 1.08E-01 | 2.68E+00 | 2.79E+00 | 1.76E-01 | no | Gm29385 | upstream_gene_variant | MODIFIER |
| 1 | 133997267 | AX-97813298 | 0 | G | A | 0.526 | 1.13E-01 | 8.41E-02 | 2.70E+00 | 2.77E+00 | 1.76E-01 | no | | intergenic_variant | MODIFIER |
| 12 | 118435224 | AX-170173979 | 1 | C | T | 0.125 | -2.36E-01 | 1.71E-01 | 3.17E+00 | 3.31E+00 | 1.76E-01 | Yes | | intergenic_variant | MODIFIER |
| 4 | 103812297 | AX-170070201 | 0 | C | T | 0.246 | 1.44E-01 | 1.07E-01 | 2.61E+00 | 2.68E+00 | 1.76E-01 | no | Dab1 | intron_variant | MODIFIER |
| 3 | 7384409 | AX-170250087 | 0 | G | A | 0.395 | -8.53E-02 | 6.37E-02 | 2.76E+00 | 2.91E+00 | 1.76E-01 | no | Pkia | intron_variant | MODIFIER |
| 10 | 101057213 | AX-170439763 | 0 | G | A | 0.298 | -1.12E-01 | 8.36E-02 | 2.64E+00 | 2.70E+00 | 1.76E-01 | no | | intergenic_variant | MODIFIER |
| 18 | 62050002 | AX-97834956 | 0 | G | A | 0.184 | -1.49E-01 | 1.11E-01 | 2.70E+00 | 2.76E+00 | 1.76E-01 | no | | regulatory_region_variant | MODIFIER |
| 13 | 30682656 | AX-170474143 | 0 | T | C | 0.175 | -1.42E-01 | 1.06E-01 | 2.69E+00 | 2.81E+00 | 1.77E-01 | no | Dusp22 | intron_variant | MODIFIER |
| 13 | 33044130 | AX-170008047 | 0 | T | A | 0.175 | -1.42E-01 | 1.06E-01 | 2.69E+00 | 2.81E+00 | 1.77E-01 | no | Serpinb9b | downstream_gene_variant | MODIFIER |
| 4 | 120700613 | AX-170179719 | 0 | G | A | 0.211 | 1.58E-01 | 1.18E-01 | 2.57E+00 | 2.63E+00 | 1.77E-01 | Yes | Kcnq4 | intron_variant | MODIFIER |

| | | | | | | | | | | | | | | | |
|---|---|---|---|---|---|---|---|---|---|---|---|---|---|---|---|
| 4 | 120715355 | AX-97799509 | 0 | G | A | 0.211 | 1.58E-01 | 1.18E-01 | 2.57E+00 | 2.63E+00 | 1.77E-01 | Yes | Kcnq4 | synonymous_variant | LOW |
| 4 | 120715650 | AX-170070489 | 0 | T | C | 0.211 | 1.58E-01 | 1.18E-01 | 2.57E+00 | 2.63E+00 | 1.77E-01 | Yes | Kcnq4 | intron_variant | MODIFIER |
| 4 | 120792967 | AX-170181592 | 0 | C | T | 0.211 | 1.58E-01 | 1.18E-01 | 2.57E+00 | 2.63E+00 | 1.77E-01 | Yes | Nfyc | intron_variant | MODIFIER |
| 4 | 120823162 | AX-170470304 | 0 | A | C | 0.211 | 1.58E-01 | 1.18E-01 | 2.57E+00 | 2.63E+00 | 1.77E-01 | Yes | Nfyc | intron_variant | MODIFIER |
| 4 | 120930701 | AX-170023837 | 0 | T | C | 0.211 | 1.58E-01 | 1.18E-01 | 2.57E+00 | 2.63E+00 | 1.77E-01 | Yes | Zfp69 | synonymous_variant | LOW |
| 4 | 120944415 | AX-97853267 | 0 | A | G | 0.211 | 1.58E-01 | 1.18E-01 | 2.57E+00 | 2.63E+00 | 1.77E-01 | Yes | Zfp69 | intron_variant | MODIFIER |
| 4 | 120999327 | AX-170343845 | 0 | C | T | 0.211 | 1.58E-01 | 1.18E-01 | 2.57E+00 | 2.63E+00 | 1.77E-01 | Yes | Smap2 | intron_variant | MODIFIER |
| 4 | 120999576 | AX-170126483 | 0 | G | T | 0.211 | 1.58E-01 | 1.18E-01 | 2.57E+00 | 2.63E+00 | 1.77E-01 | Yes | Smap2 | intron_variant | MODIFIER |
| 4 | 121000856 | AX-170351455 | 0 | T | C | 0.211 | 1.58E-01 | 1.18E-01 | 2.57E+00 | 2.63E+00 | 1.77E-01 | Yes | Smap2 | intron_variant | MODIFIER |
| 4 | 121001095 | AX-170253599 | 0 | G | A | 0.211 | 1.58E-01 | 1.18E-01 | 2.57E+00 | 2.63E+00 | 1.77E-01 | Yes | Smap2 | intron_variant | MODIFIER |
| 4 | 121006873 | AX-170070498 | 0 | T | C | 0.211 | 1.58E-01 | 1.18E-01 | 2.57E+00 | 2.63E+00 | 1.77E-01 | Yes | Smap2 | intron_variant | MODIFIER |
| 4 | 121019230 | AX-170023841 | 0 | T | C | 0.211 | 1.58E-01 | 1.18E-01 | 2.57E+00 | 2.63E+00 | 1.77E-01 | Yes | Smap2 | upstream_gene_variant | MODIFIER |
| 4 | 121019393 | AX-170421532 | 0 | G | A | 0.211 | 1.58E-01 | 1.18E-01 | 2.57E+00 | 2.63E+00 | 1.77E-01 | Yes | Smap2 | upstream_gene_variant | MODIFIER |
| 4 | 121019433 | AX-170103223 | 0 | A | T | 0.211 | 1.58E-01 | 1.18E-01 | 2.57E+00 | 2.63E+00 | 1.77E-01 | Yes | Smap2 | upstream_gene_variant | MODIFIER |
| 4 | 121023912 | AX-170376094 | 0 | C | T | 0.211 | 1.58E-01 | 1.18E-01 | 2.57E+00 | 2.63E+00 | 1.77E-01 | Yes | | regulatory_region_variant | MODIFIER |
| 18 | 59545118 | AX-170500572 | 2 | A | C | 0.418 | 1.15E-01 | 8.38E-02 | 2.40E+00 | 2.48E+00 | 1.77E-01 | no | | intergenic_variant | MODIFIER |
| 10 | 70265548 | AX-170045167 | 0 | A | G | 0.412 | -1.06E-01 | 7.90E-02 | 2.72E+00 | 2.80E+00 | 1.77E-01 | no | Slc16a9 | intron_variant | MODIFIER |
| 13 | 19364559 | AX-170288883 | 0 | G | A | 0.167 | -1.73E-01 | 1.27E-01 | 2.49E+00 | 2.53E+00 | 1.77E-01 | no | Stard3nl | intron_variant | MODIFIER |
| 13 | 19370083 | AX-170239517 | 0 | G | C | 0.167 | -1.73E-01 | 1.27E-01 | 2.49E+00 | 2.53E+00 | 1.77E-01 | no | Stard3nl | intron_variant | MODIFIER |
| 13 | 19371085 | AX-170137874 | 0 | T | C | 0.167 | -1.73E-01 | 1.27E-01 | 2.49E+00 | 2.53E+00 | 1.77E-01 | no | Stard3nl | intron_variant | MODIFIER |
| 13 | 19372996 | AX-170527518 | 0 | T | G | 0.167 | -1.73E-01 | 1.27E-01 | 2.49E+00 | 2.53E+00 | 1.77E-01 | no | Stard3nl | intron_variant | MODIFIER |
| 13 | 19376089 | AX-170258724 | 0 | C | G | 0.167 | -1.73E-01 | 1.27E-01 | 2.49E+00 | 2.53E+00 | 1.77E-01 | no | Stard3nl | intron_variant | MODIFIER |
| 13 | 19477857 | AX-170050673 | 0 | A | G | 0.167 | -1.73E-01 | 1.27E-01 | 2.49E+00 | 2.53E+00 | 1.77E-01 | no | 9330199G10Rik | intron_variant,non_coding_transcript_variant | MODIFIER |
| 13 | 19477902 | AX-170050672 | 0 | T | C | 0.167 | -1.73E-01 | 1.27E-01 | 2.49E+00 | 2.53E+00 | 1.77E-01 | no | 9330199G10Rik | intron_variant,non_coding_transcript_variant | MODIFIER |
| 13 | 19481173 | AX-170007877 | 0 | G | A | 0.167 | -1.73E-01 | 1.27E-01 | 2.49E+00 | 2.53E+00 | 1.77E-01 | no | 9330199G10Rik | intron_variant,non_coding_transcript_variant | MODIFIER |
| 13 | 19481350 | AX-170378432 | 0 | G | T | 0.167 | -1.73E-01 | 1.27E-01 | 2.49E+00 | 2.53E+00 | 1.77E-01 | no | 9330199G10Rik | intron_variant,non_coding_transcript_variant | MODIFIER |
| 13 | 19493791 | AX-170360221 | 0 | A | G | 0.167 | -1.73E-01 | 1.27E-01 | 2.49E+00 | 2.53E+00 | 1.77E-01 | no | 9330199G10Rik | intron_variant,non_coding_transcript_variant | MODIFIER |
| 13 | 19499138 | AX-97834568 | 0 | G | A | 0.167 | -1.73E-01 | 1.27E-01 | 2.49E+00 | 2.53E+00 | 1.77E-01 | no | 9330199G10Rik | intron_variant,non_coding_transcript_variant | MODIFIER |
| 13 | 19530940 | AX-170007880 | 0 | A | G | 0.167 | -1.73E-01 | 1.27E-01 | 2.49E+00 | 2.53E+00 | 1.77E-01 | no | Gm18363 | downstream_gene_variant | MODIFIER |
| 13 | 19556363 | AX-97834931 | 0 | A | G | 0.167 | -1.73E-01 | 1.27E-01 | 2.49E+00 | 2.53E+00 | 1.77E-01 | no | | intergenic_variant | MODIFIER |
| 13 | 19558275 | AX-168375954 | 0 | C | G | 0.167 | -1.73E-01 | 1.27E-01 | 2.49E+00 | 2.53E+00 | 1.77E-01 | no | | intergenic_variant | MODIFIER |
| 13 | 19580508 | AX-97800551 | 0 | G | A | 0.167 | -1.73E-01 | 1.27E-01 | 2.49E+00 | 2.53E+00 | 1.77E-01 | no | | intergenic_variant | MODIFIER |
| 13 | 19607458 | AX-170226762 | 0 | A | G | 0.167 | -1.73E-01 | 1.27E-01 | 2.49E+00 | 2.53E+00 | 1.77E-01 | no | Epdr1 | intron_variant | MODIFIER |
| 13 | 19618223 | AX-170357699 | 0 | A | G | 0.167 | -1.73E-01 | 1.27E-01 | 2.49E+00 | 2.53E+00 | 1.77E-01 | no | Sfrp4 | upstream_gene_variant | MODIFIER |
| 13 | 19625352 | AX-170453371 | 0 | A | G | 0.167 | -1.73E-01 | 1.27E-01 | 2.49E+00 | 2.53E+00 | 1.77E-01 | no | Sfrp4 | intron_variant | MODIFIER |
| 13 | 19626000 | AX-170413551 | 0 | G | T | 0.167 | -1.73E-01 | 1.27E-01 | 2.49E+00 | 2.53E+00 | 1.77E-01 | no | Sfrp4 | intron_variant | MODIFIER |
| 13 | 19626529 | AX-170203170 | 0 | C | G | 0.167 | -1.73E-01 | 1.27E-01 | 2.49E+00 | 2.53E+00 | 1.77E-01 | no | Sfrp4 | intron_variant | MODIFIER |
| 13 | 19633514 | AX-97791936 | 0 | C | T | 0.167 | -1.73E-01 | 1.27E-01 | 2.49E+00 | 2.53E+00 | 1.77E-01 | no | Sfrp4 | downstream_gene_variant | MODIFIER |
| 13 | 19633577 | AX-170518118 | 0 | A | G | 0.167 | -1.73E-01 | 1.27E-01 | 2.49E+00 | 2.53E+00 | 1.77E-01 | no | Sfrp4 | downstream_gene_variant | MODIFIER |
| 13 | 19658174 | AX-170200746 | 0 | T | C | 0.167 | -1.73E-01 | 1.27E-01 | 2.49E+00 | 2.53E+00 | 1.77E-01 | no | Nme8 | intron_variant | MODIFIER |
| 13 | 19658269 | AX-170282631 | 0 | G | A | 0.167 | -1.73E-01 | 1.27E-01 | 2.49E+00 | 2.53E+00 | 1.77E-01 | no | Nme8 | intron_variant | MODIFIER |
| 13 | 19660228 | AX-170124090 | 0 | A | C | 0.167 | -1.73E-01 | 1.27E-01 | 2.49E+00 | 2.53E+00 | 1.77E-01 | no | Nme8 | intron_variant | MODIFIER |
| 13 | 19660246 | AX-170479546 | 0 | T | C | 0.167 | -1.73E-01 | 1.27E-01 | 2.49E+00 | 2.53E+00 | 1.77E-01 | no | Nme8 | intron_variant | MODIFIER |
| 13 | 19902153 | AX-170517589 | 0 | A | G | 0.167 | -1.73E-01 | 1.27E-01 | 2.49E+00 | 2.53E+00 | 1.77E-01 | no | | intergenic_variant | MODIFIER |
| 13 | 19902262 | AX-170132030 | 0 | T | C | 0.167 | -1.73E-01 | 1.27E-01 | 2.49E+00 | 2.53E+00 | 1.77E-01 | no | | intergenic_variant | MODIFIER |
| 13 | 19913471 | AX-170104556 | 0 | G | A | 0.167 | -1.73E-01 | 1.27E-01 | 2.49E+00 | 2.53E+00 | 1.77E-01 | no | | intergenic_variant | MODIFIER |
| 13 | 19913583 | AX-170490245 | 0 | C | T | 0.167 | -1.73E-01 | 1.27E-01 | 2.49E+00 | 2.53E+00 | 1.77E-01 | no | | intergenic_variant | MODIFIER |
| 13 | 19922670 | AX-97807558 | 0 | A | C | 0.167 | -1.73E-01 | 1.27E-01 | 2.49E+00 | 2.53E+00 | 1.77E-01 | no | | intergenic_variant | MODIFIER |
| 13 | 19953494 | AX-97780405 | 0 | C | A | 0.167 | -1.73E-01 | 1.27E-01 | 2.49E+00 | 2.53E+00 | 1.77E-01 | no | | intergenic_variant | MODIFIER |
| 13 | 20031192 | AX-170411644 | 0 | G | A | 0.167 | -1.73E-01 | 1.27E-01 | 2.49E+00 | 2.53E+00 | 1.77E-01 | no | | intergenic_variant | MODIFIER |
| 13 | 20045103 | AX-170007883 | 0 | G | A | 0.167 | -1.73E-01 | 1.27E-01 | 2.49E+00 | 2.53E+00 | 1.77E-01 | no | | intergenic_variant | MODIFIER |
| 13 | 20060822 | AX-170409408 | 0 | T | C | 0.167 | -1.73E-01 | 1.27E-01 | 2.49E+00 | 2.53E+00 | 1.77E-01 | no | | intergenic_variant | MODIFIER |
| 13 | 20069741 | AX-170320253 | 0 | G | A | 0.167 | -1.73E-01 | 1.27E-01 | 2.49E+00 | 2.53E+00 | 1.77E-01 | no | | intergenic_variant | MODIFIER |
| 13 | 20077916 | AX-170110113 | 0 | C | T | 0.167 | -1.73E-01 | 1.27E-01 | 2.49E+00 | 2.53E+00 | 1.77E-01 | no | | intergenic_variant | MODIFIER |

| | | | | | | | | | | | | | | | |
|---|---|---|---|---|---|---|---|---|---|---|---|---|---|---|---|
| 13 | 20094382 | AX-170302620 | 0 | C | T | 0.167 | -1.73E-01 | 1.27E-01 | 2.49E+00 | 2.53E+00 | 1.77E-01 | no | Elmo1 | intron_variant | MODIFIER |
| 13 | 20898486 | AX-170234975 | 0 | T | A | 0.167 | -1.73E-01 | 1.27E-01 | 2.49E+00 | 2.53E+00 | 1.77E-01 | no | Aoah | intron_variant | MODIFIER |
| 13 | 20900461 | AX-170247718 | 0 | C | T | 0.167 | -1.73E-01 | 1.27E-01 | 2.49E+00 | 2.53E+00 | 1.77E-01 | no | Aoah | intron_variant | MODIFIER |
| 13 | 20904049 | AX-170390274 | 0 | T | C | 0.167 | -1.73E-01 | 1.27E-01 | 2.49E+00 | 2.53E+00 | 1.77E-01 | no | Aoah | intron_variant | MODIFIER |
| 13 | 20907017 | AX-170297046 | 0 | A | C | 0.167 | -1.73E-01 | 1.27E-01 | 2.49E+00 | 2.53E+00 | 1.77E-01 | no | Aoah | intron_variant | MODIFIER |
| 13 | 20913252 | AX-170443253 | 0 | G | A | 0.167 | -1.73E-01 | 1.27E-01 | 2.49E+00 | 2.53E+00 | 1.77E-01 | no | Aoah | intron_variant | MODIFIER |
| 13 | 20917703 | AX-170520845 | 0 | T | G | 0.167 | -1.73E-01 | 1.27E-01 | 2.49E+00 | 2.53E+00 | 1.77E-01 | no | Aoah | intron_variant | MODIFIER |
| 13 | 20940383 | AX-170307895 | 0 | A | G | 0.167 | -1.73E-01 | 1.27E-01 | 2.49E+00 | 2.53E+00 | 1.77E-01 | no | Aoah | intron_variant | MODIFIER |
| 13 | 20948142 | AX-170272381 | 0 | T | C | 0.167 | -1.73E-01 | 1.27E-01 | 2.49E+00 | 2.53E+00 | 1.77E-01 | no | Aoah | intron_variant | MODIFIER |
| 13 | 20958044 | AX-170299588 | 0 | T | C | 0.167 | -1.73E-01 | 1.27E-01 | 2.49E+00 | 2.53E+00 | 1.77E-01 | no | Aoah | intron_variant | MODIFIER |
| 13 | 20982379 | AX-170312363 | 0 | A | G | 0.167 | -1.73E-01 | 1.27E-01 | 2.49E+00 | 2.53E+00 | 1.77E-01 | no | Aoah | intron_variant | MODIFIER |
| 13 | 20997006 | AX-170419181 | 0 | G | A | 0.167 | -1.73E-01 | 1.27E-01 | 2.49E+00 | 2.53E+00 | 1.77E-01 | no | Aoah | intron_variant | MODIFIER |
| 13 | 20997054 | AX-97822944 | 0 | A | C | 0.167 | -1.73E-01 | 1.27E-01 | 2.49E+00 | 2.53E+00 | 1.77E-01 | no | Aoah | intron_variant | MODIFIER |
| 13 | 20997241 | AX-170368951 | 0 | A | G | 0.167 | -1.73E-01 | 1.27E-01 | 2.49E+00 | 2.53E+00 | 1.77E-01 | no | Aoah | intron_variant | MODIFIER |
| 13 | 20997374 | AX-170447881 | 0 | G | A | 0.167 | -1.73E-01 | 1.27E-01 | 2.49E+00 | 2.53E+00 | 1.77E-01 | no | Aoah | intron_variant | MODIFIER |
| 13 | 21012708 | AX-170312624 | 0 | C | T | 0.167 | -1.73E-01 | 1.27E-01 | 2.49E+00 | 2.53E+00 | 1.77E-01 | no | Aoah | intron_variant | MODIFIER |
| 13 | 21073323 | AX-170149703 | 0 | T | C | 0.167 | -1.73E-01 | 1.27E-01 | 2.49E+00 | 2.53E+00 | 1.77E-01 | no | Olfr1370 | upstream_gene_variant | MODIFIER |
| 13 | 21073778 | AX-170375364 | 0 | G | A | 0.167 | -1.73E-01 | 1.27E-01 | 2.49E+00 | 2.53E+00 | 1.77E-01 | no | Olfr1370 | upstream_gene_variant | MODIFIER |
| 13 | 21089576 | AX-170092683 | 0 | A | G | 0.167 | -1.73E-01 | 1.27E-01 | 2.49E+00 | 2.53E+00 | 1.77E-01 | no | | intergenic_variant | MODIFIER |
| 13 | 21132052 | AX-170404050 | 0 | A | G | 0.167 | -1.73E-01 | 1.27E-01 | 2.49E+00 | 2.53E+00 | 1.77E-01 | no | Olfr263 | upstream_gene_variant | MODIFIER |
| 13 | 21191046 | AX-97822946 | 0 | A | G | 0.167 | -1.73E-01 | 1.27E-01 | 2.49E+00 | 2.53E+00 | 1.77E-01 | no | Trim27 | intron_variant | MODIFIER |
| 13 | 21195220 | AX-170007891 | 0 | C | T | 0.167 | -1.73E-01 | 1.27E-01 | 2.49E+00 | 2.53E+00 | 1.77E-01 | no | Trim27 | downstream_gene_variant | MODIFIER |
| 13 | 21203227 | AX-170007892 | 0 | G | A | 0.167 | -1.73E-01 | 1.27E-01 | 2.49E+00 | 2.53E+00 | 1.77E-01 | no | | intergenic_variant | MODIFIER |
| 13 | 21222788 | AX-170258909 | 0 | C | T | 0.167 | -1.73E-01 | 1.27E-01 | 2.49E+00 | 2.53E+00 | 1.77E-01 | no | | regulatory_region_variant | MODIFIER |
| 13 | 21246628 | AX-170500431 | 0 | G | A | 0.167 | -1.73E-01 | 1.27E-01 | 2.49E+00 | 2.53E+00 | 1.77E-01 | no | | intergenic_variant | MODIFIER |
| 13 | 21266397 | AX-170122048 | 0 | G | T | 0.167 | -1.73E-01 | 1.27E-01 | 2.49E+00 | 2.53E+00 | 1.77E-01 | no | | regulatory_region_variant | MODIFIER |
| 13 | 21288200 | AX-170285653 | 0 | A | C | 0.167 | -1.73E-01 | 1.27E-01 | 2.49E+00 | 2.53E+00 | 1.77E-01 | no | Gpx5 | intron_variant | MODIFIER |
| 13 | 21291191 | AX-170288663 | 0 | A | C | 0.167 | -1.73E-01 | 1.27E-01 | 2.49E+00 | 2.53E+00 | 1.77E-01 | no | Gpx5 | intron_variant | MODIFIER |
| 13 | 21291450 | AX-170327149 | 0 | A | G | 0.167 | -1.73E-01 | 1.27E-01 | 2.49E+00 | 2.53E+00 | 1.77E-01 | no | Gpx5 | synonymous_variant | LOW |
| 13 | 21294807 | AX-170402997 | 0 | G | T | 0.167 | -1.73E-01 | 1.27E-01 | 2.49E+00 | 2.53E+00 | 1.77E-01 | no | Gpx5 | upstream_gene_variant | MODIFIER |
| 13 | 21295461 | AX-170227554 | 0 | G | A | 0.167 | -1.73E-01 | 1.27E-01 | 2.49E+00 | 2.53E+00 | 1.77E-01 | no | Gpx5 | upstream_gene_variant | MODIFIER |
| 13 | 21298355 | AX-170179194 | 0 | A | T | 0.167 | -1.73E-01 | 1.27E-01 | 2.49E+00 | 2.53E+00 | 1.77E-01 | no | Gm11270 | upstream_gene_variant | MODIFIER |
| 13 | 21326283 | AX-170523716 | 0 | A | G | 0.167 | -1.73E-01 | 1.27E-01 | 2.49E+00 | 2.53E+00 | 1.77E-01 | no | | regulatory_region_variant | MODIFIER |
| 13 | 21336986 | AX-170146636 | 0 | C | G | 0.167 | -1.73E-01 | 1.27E-01 | 2.49E+00 | 2.53E+00 | 1.77E-01 | no | | intergenic_variant | MODIFIER |
| 13 | 21345484 | AX-170306089 | 0 | A | G | 0.167 | -1.73E-01 | 1.27E-01 | 2.49E+00 | 2.53E+00 | 1.77E-01 | no | Olfr1367 | upstream_gene_variant | MODIFIER |
| 13 | 21358042 | AX-170323924 | 0 | A | G | 0.167 | -1.73E-01 | 1.27E-01 | 2.49E+00 | 2.53E+00 | 1.77E-01 | no | Zscan12 | upstream_gene_variant | MODIFIER |
| 13 | 21358295 | AX-170398863 | 0 | A | T | 0.167 | -1.73E-01 | 1.27E-01 | 2.49E+00 | 2.53E+00 | 1.77E-01 | no | Zscan12 | upstream_gene_variant | MODIFIER |
| 13 | 21370751 | AX-97852628 | 0 | A | G | 0.167 | -1.73E-01 | 1.27E-01 | 2.49E+00 | 2.53E+00 | 1.77E-01 | no | Zscan12 | downstream_gene_variant | MODIFIER |
| 13 | 21382903 | AX-170281666 | 0 | C | G | 0.167 | -1.73E-01 | 1.27E-01 | 2.49E+00 | 2.53E+00 | 1.77E-01 | no | Zkscan3 | downstream_gene_variant | MODIFIER |
| 13 | 21394573 | AX-170186389 | 0 | C | T | 0.167 | -1.73E-01 | 1.27E-01 | 2.49E+00 | 2.53E+00 | 1.77E-01 | no | Zkscan3 | missense_variant | MODERATE |
| 13 | 21403793 | AX-97816747 | 0 | C | G | 0.167 | -1.73E-01 | 1.27E-01 | 2.49E+00 | 2.53E+00 | 1.77E-01 | no | Zkscan3 | upstream_gene_variant | MODIFIER |
| 13 | 21404743 | AX-170141731 | 0 | G | A | 0.167 | -1.73E-01 | 1.27E-01 | 2.49E+00 | 2.53E+00 | 1.77E-01 | no | Zkscan3 | upstream_gene_variant | MODIFIER |
| 13 | 21416826 | AX-170102643 | 0 | A | G | 0.167 | -1.73E-01 | 1.27E-01 | 2.49E+00 | 2.53E+00 | 1.77E-01 | no | Pgbd1 | downstream_gene_variant | MODIFIER |
| 13 | 21431109 | AX-170445089 | 0 | T | G | 0.167 | -1.73E-01 | 1.27E-01 | 2.49E+00 | 2.53E+00 | 1.77E-01 | no | Pgbd1 | intron_variant | MODIFIER |
| 13 | 21433570 | AX-170167769 | 0 | A | C | 0.167 | -1.73E-01 | 1.27E-01 | 2.49E+00 | 2.53E+00 | 1.77E-01 | no | Pgbd1 | intron_variant | MODIFIER |
| 13 | 21464509 | AX-97830205 | 0 | T | C | 0.167 | -1.73E-01 | 1.27E-01 | 2.49E+00 | 2.53E+00 | 1.77E-01 | no | Nkapl | downstream_gene_variant | MODIFIER |
| 13 | 21478406 | AX-170197706 | 0 | G | T | 0.167 | -1.73E-01 | 1.27E-01 | 2.49E+00 | 2.53E+00 | 1.77E-01 | no | Zkscan4 | upstream_gene_variant | MODIFIER |
| 13 | 21480777 | AX-170218355 | 0 | C | A | 0.167 | -1.73E-01 | 1.27E-01 | 2.49E+00 | 2.53E+00 | 1.77E-01 | no | Zkscan4 | intron_variant | MODIFIER |
| 13 | 21497433 | AX-170379213 | 0 | G | A | 0.167 | -1.73E-01 | 1.27E-01 | 2.49E+00 | 2.53E+00 | 1.77E-01 | no | Gm11273 | downstream_gene_variant | MODIFIER |
| 13 | 21525937 | AX-170035529 | 0 | A | G | 0.167 | -1.73E-01 | 1.27E-01 | 2.49E+00 | 2.53E+00 | 1.77E-01 | no | Zkscan8 | intron_variant | MODIFIER |
| 13 | 21537934 | AX-170035525 | 0 | G | C | 0.167 | -1.73E-01 | 1.27E-01 | 2.49E+00 | 2.53E+00 | 1.77E-01 | no | Olfr1366 | missense_variant | MODERATE |
| 13 | 21547736 | AX-170364939 | 0 | A | G | 0.167 | -1.73E-01 | 1.27E-01 | 2.49E+00 | 2.53E+00 | 1.77E-01 | no | Olfr1366 | upstream_gene_variant | MODIFIER |
| 13 | 21551395 | AX-170254891 | 0 | G | T | 0.167 | -1.73E-01 | 1.27E-01 | 2.49E+00 | 2.53E+00 | 1.77E-01 | no | Olfr1535 | downstream_gene_variant | MODIFIER |
| 13 | 21551484 | AX-97810703 | 0 | C | G | 0.167 | -1.73E-01 | 1.27E-01 | 2.49E+00 | 2.53E+00 | 1.77E-01 | no | Olfr1535 | downstream_gene_variant | MODIFIER |

| | | | | | | | | | | | | | | | |
|---|---|---|---|---|---|---|---|---|---|---|---|---|---|---|---|
| 13 | 21559866 | AX-170262673 | 0 | A | G | 0.167 | -1.73E-01 | 1.27E-01 | 2.49E+00 | 2.53E+00 | 1.77E-01 | no | Olfr1535 | upstream_gene_variant | MODIFIER |
| 13 | 21574439 | AX-170307491 | 0 | T | C | 0.167 | -1.73E-01 | 1.27E-01 | 2.49E+00 | 2.53E+00 | 1.77E-01 | no | Olfr1364 | missense_variant | MODERATE |
| 13 | 21588736 | AX-170406014 | 0 | A | G | 0.167 | -1.73E-01 | 1.27E-01 | 2.49E+00 | 2.53E+00 | 1.77E-01 | no | | intergenic_variant | MODIFIER |
| 13 | 21596351 | AX-170221561 | 0 | T | G | 0.167 | -1.73E-01 | 1.27E-01 | 2.49E+00 | 2.53E+00 | 1.77E-01 | no | | intergenic_variant | MODIFIER |
| 13 | 21614278 | AX-170217855 | 0 | G | A | 0.167 | -1.73E-01 | 1.27E-01 | 2.49E+00 | 2.53E+00 | 1.77E-01 | no | Olfr1362 | upstream_gene_variant | MODIFIER |
| 13 | 21642485 | AX-170132537 | 0 | A | C | 0.167 | -1.73E-01 | 1.27E-01 | 2.49E+00 | 2.53E+00 | 1.77E-01 | no | Olfr11 | upstream_gene_variant | MODIFIER |
| 13 | 21658075 | AX-97786617 | 0 | T | C | 0.167 | -1.73E-01 | 1.27E-01 | 2.49E+00 | 2.53E+00 | 1.77E-01 | no | Olfr1361 | downstream_gene_variant | MODIFIER |
| 13 | 21659298 | AX-170007894 | 0 | C | T | 0.167 | -1.73E-01 | 1.27E-01 | 2.49E+00 | 2.53E+00 | 1.77E-01 | no | Olfr1361 | synonymous_variant | LOW |
| 15 | 66186851 | AX-170522285 | 0 | C | A | 0.325 | -1.07E-01 | 7.55E-02 | 2.21E+00 | 2.27E+00 | 1.77E-01 | no | Kcnq3 | intron_variant | MODIFIER |
| 15 | 66225793 | AX-170324862 | 0 | T | C | 0.325 | -1.07E-01 | 7.55E-02 | 2.21E+00 | 2.27E+00 | 1.77E-01 | no | Kcnq3 | intron_variant | MODIFIER |
| 15 | 66226269 | AX-170397878 | 0 | A | G | 0.325 | -1.07E-01 | 7.55E-02 | 2.21E+00 | 2.27E+00 | 1.77E-01 | no | Kcnq3 | intron_variant | MODIFIER |
| 15 | 66228655 | AX-170011883 | 0 | C | A | 0.325 | -1.07E-01 | 7.55E-02 | 2.21E+00 | 2.27E+00 | 1.77E-01 | no | Kcnq3 | intron_variant | MODIFIER |
| 15 | 66235957 | AX-97816036 | 0 | T | C | 0.325 | -1.07E-01 | 7.55E-02 | 2.21E+00 | 2.27E+00 | 1.77E-01 | no | Kcnq3 | intron_variant | MODIFIER |
| 15 | 66244636 | AX-170011882 | 0 | A | G | 0.325 | -1.07E-01 | 7.55E-02 | 2.21E+00 | 2.27E+00 | 1.77E-01 | no | Kcnq3 | intron_variant | MODIFIER |
| 15 | 66268315 | AX-170347368 | 0 | C | T | 0.325 | -1.07E-01 | 7.55E-02 | 2.21E+00 | 2.27E+00 | 1.77E-01 | no | Kcnq3 | intron_variant | MODIFIER |
| 15 | 66268462 | AX-97788515 | 0 | A | C | 0.325 | -1.07E-01 | 7.55E-02 | 2.21E+00 | 2.27E+00 | 1.77E-01 | no | Kcnq3 | intron_variant | MODIFIER |
| 15 | 66279138 | AX-170235783 | 0 | T | G | 0.325 | -1.07E-01 | 7.55E-02 | 2.21E+00 | 2.27E+00 | 1.77E-01 | no | Kcnq3 | intron_variant | MODIFIER |
| 15 | 66279213 | AX-170113597 | 0 | C | T | 0.325 | -1.07E-01 | 7.55E-02 | 2.21E+00 | 2.27E+00 | 1.77E-01 | no | Kcnq3 | intron_variant | MODIFIER |
| 15 | 66291922 | AX-97786894 | 0 | G | C | 0.325 | -1.07E-01 | 7.55E-02 | 2.21E+00 | 2.27E+00 | 1.77E-01 | no | Gm27242 | downstream_gene_variant | MODIFIER |
| 15 | 66291974 | AX-170432727 | 0 | T | G | 0.325 | -1.07E-01 | 7.55E-02 | 2.21E+00 | 2.27E+00 | 1.77E-01 | no | Gm27242 | downstream_gene_variant | MODIFIER |
| 15 | 66295453 | AX-170486025 | 0 | G | C | 0.325 | -1.07E-01 | 7.55E-02 | 2.21E+00 | 2.27E+00 | 1.77E-01 | no | | intergenic_variant | MODIFIER |
| 15 | 66296015 | AX-170351614 | 0 | C | T | 0.325 | -1.07E-01 | 7.55E-02 | 2.21E+00 | 2.27E+00 | 1.77E-01 | no | | intergenic_variant | MODIFIER |
| 15 | 66371152 | AX-170288706 | 0 | A | C | 0.325 | -1.07E-01 | 7.55E-02 | 2.21E+00 | 2.27E+00 | 1.77E-01 | no | | intergenic_variant | MODIFIER |
| 15 | 66372938 | AX-170131871 | 0 | T | C | 0.325 | -1.07E-01 | 7.55E-02 | 2.21E+00 | 2.27E+00 | 1.77E-01 | no | | intergenic_variant | MODIFIER |
| 15 | 66375768 | AX-170435327 | 0 | C | G | 0.325 | -1.07E-01 | 7.55E-02 | 2.21E+00 | 2.27E+00 | 1.77E-01 | no | Lrrc6 | downstream_gene_variant | MODIFIER |
| 15 | 66443577 | AX-170147478 | 0 | C | T | 0.325 | -1.07E-01 | 7.55E-02 | 2.21E+00 | 2.27E+00 | 1.77E-01 | no | Lrrc6 | intron_variant | MODIFIER |
| 15 | 66444163 | AX-170443118 | 0 | C | T | 0.325 | -1.07E-01 | 7.55E-02 | 2.21E+00 | 2.27E+00 | 1.77E-01 | no | Lrrc6 | intron_variant | MODIFIER |
| 15 | 66464707 | AX-170279693 | 0 | G | A | 0.325 | -1.07E-01 | 7.55E-02 | 2.21E+00 | 2.27E+00 | 1.77E-01 | no | Lrrc6 | intron_variant | MODIFIER |
| 15 | 66507711 | AX-170508198 | 0 | T | C | 0.325 | -1.07E-01 | 7.55E-02 | 2.21E+00 | 2.27E+00 | 1.77E-01 | no | | intergenic_variant | MODIFIER |
| 15 | 66517107 | AX-97802105 | 0 | G | A | 0.325 | -1.07E-01 | 7.55E-02 | 2.21E+00 | 2.27E+00 | 1.77E-01 | no | | intergenic_variant | MODIFIER |
| 15 | 66549102 | AX-170495690 | 0 | T | C | 0.325 | -1.07E-01 | 7.55E-02 | 2.21E+00 | 2.27E+00 | 1.77E-01 | no | Tmem71 | intron_variant | MODIFIER |
| 15 | 66667765 | AX-170501177 | 0 | T | A | 0.325 | -1.07E-01 | 7.55E-02 | 2.21E+00 | 2.27E+00 | 1.77E-01 | no | Tg | upstream_gene_variant | MODIFIER |
| 15 | 66669014 | AX-170122994 | 0 | A | G | 0.325 | -1.07E-01 | 7.55E-02 | 2.21E+00 | 2.27E+00 | 1.77E-01 | no | Tg | upstream_gene_variant | MODIFIER |
| 15 | 66676342 | AX-97855243 | 0 | C | A | 0.325 | -1.07E-01 | 7.55E-02 | 2.21E+00 | 2.27E+00 | 1.77E-01 | no | Tg | intron_variant | MODIFIER |
| 4 | 55193257 | AX-170094979 | 0 | G | A | 0.368 | -1.38E-01 | 1.02E-01 | 2.94E+00 | 3.05E+00 | 1.77E-01 | no | Gm12514 | intron_variant,non_coding_transcript_variant | MODIFIER |
| 16 | 79449476 | AX-170129042 | 1 | C | T | 0.473 | 1.00E-01 | 7.48E-02 | 2.66E+00 | 2.74E+00 | 1.77E-01 | no | | intergenic_variant | MODIFIER |
| 20 | 46723746 | AX-170084161 | 1 | C | T | 0.241 | 7.86E-02 | 5.87E-02 | 2.60E+00 | 2.73E+00 | 1.77E-01 | no | Gm29242 | intron_variant,non_coding_transcript_variant | MODIFIER |
| 9 | 97958726 | AX-170462681 | 0 | C | T | 0.132 | -1.60E-01 | 1.20E-01 | 2.68E+00 | 2.76E+00 | 1.78E-01 | no | Clstn2 | intron_variant | MODIFIER |
| 9 | 101031586 | AX-170378131 | 0 | C | T | 0.132 | -1.46E-01 | 1.09E-01 | 2.69E+00 | 2.80E+00 | 1.78E-01 | no | Pccb | intron_variant | MODIFIER |
| 1 | 120859175 | AX-170383208 | 1 | A | G | 0.571 | 1.14E-01 | 8.44E-02 | 2.54E+00 | 2.60E+00 | 1.78E-01 | no | Gm29347 | downstream_gene_variant | MODIFIER |
| 1 | 121215035 | AX-170109795 | 1 | C | A | 0.571 | 1.14E-01 | 8.44E-02 | 2.54E+00 | 2.60E+00 | 1.78E-01 | no | | intergenic_variant | MODIFIER |
| 11 | 60776467 | AX-97846932 | 0 | G | T | 0.044 | -2.16E-01 | 1.62E-01 | 2.79E+00 | 2.93E+00 | 1.78E-01 | Yes | Top3a | intron_variant | MODIFIER |
| 13 | 9952710 | AX-170254015 | 1 | T | A | 0.188 | -1.70E-01 | 1.25E-01 | 2.45E+00 | 2.49E+00 | 1.78E-01 | no | Chrm3 | intron_variant | MODIFIER |
| 16 | 71729118 | AX-170406519 | 1 | T | C | 0.482 | 9.77E-02 | 7.30E-02 | 2.62E+00 | 2.72E+00 | 1.79E-01 | no | | intergenic_variant | MODIFIER |
| 2 | 117454342 | AX-170094382 | 0 | C | T | 0.14 | 2.05E-01 | 1.53E-01 | 2.91E+00 | 3.04E+00 | 1.79E-01 | Yes | | intergenic_variant | MODIFIER |
| 16 | 31582892 | AX-170012758 | 0 | T | C | 0.5 | 1.05E-01 | 7.87E-02 | 2.62E+00 | 2.68E+00 | 1.79E-01 | no | | intergenic_variant | MODIFIER |
| 13 | 5891763 | AX-170180787 | 0 | T | C | 0.211 | -1.34E-01 | 1.00E-01 | 2.63E+00 | 2.75E+00 | 1.79E-01 | no | Gm47507 | intron_variant,non_coding_transcript_variant | MODIFIER |
| 1 | 192239981 | AX-170245504 | 1 | C | T | 0.286 | 1.23E-01 | 8.99E-02 | 2.39E+00 | 2.48E+00 | 1.79E-01 | no | Kcnh1 | intron_variant | MODIFIER |
| 8 | 16540026 | AX-170408191 | 0 | T | G | 0.228 | -9.81E-02 | 7.32E-02 | 2.56E+00 | 2.66E+00 | 1.79E-01 | no | Csmd1 | intron_variant | MODIFIER |
| 1 | 121185645 | AX-170476247 | 0 | T | C | 0.57 | 1.13E-01 | 8.44E-02 | 2.54E+00 | 2.60E+00 | 1.79E-01 | no | | intergenic_variant | MODIFIER |
| 4 | 54638725 | AX-170370921 | 1 | C | T | 0.366 | -1.38E-01 | 1.02E-01 | 3.00E+00 | 3.11E+00 | 1.79E-01 | no | | intergenic_variant | MODIFIER |
| 4 | 54666233 | AX-170142095 | 1 | A | G | 0.366 | -1.38E-01 | 1.02E-01 | 3.00E+00 | 3.11E+00 | 1.79E-01 | no | Gm12477 | downstream_gene_variant | MODIFIER |
| 16 | 70598976 | AX-97853660 | 0 | A | G | 0.482 | 1.06E-01 | 7.94E-02 | 2.63E+00 | 2.70E+00 | 1.79E-01 | no | | intergenic_variant | MODIFIER |
| 5 | 46737281 | AX-170100834 | 0 | C | A | 0.5 | 1.33E-01 | 9.83E-02 | 2.43E+00 | 2.49E+00 | 1.79E-01 | no | | intergenic_variant | MODIFIER |

| | | | | | | | | | | | | | | | |
|---|---|---|---|---|---|---|---|---|---|---|---|---|---|---|---|
| 3 | 20271574 | AX-97784943 | 0 | T | C | 0.132 | -1.36E-01 | 1.02E-01 | 2.54E+00 | 2.61E+00 | 1.79E-01 | no | Cpb1 | intron_variant | MODIFIER |
| 8 | 72837978 | AX-170501570 | 1 | A | T | 0.188 | -1.38E-01 | 1.04E-01 | 2.83E+00 | 2.98E+00 | 1.79E-01 | no | Large1 | intron_variant | MODIFIER |
| 13 | 9582410 | AX-170349649 | 0 | T | A | 0.184 | -1.71E-01 | 1.27E-01 | 2.50E+00 | 2.55E+00 | 1.80E-01 | no | Dip2c | intron_variant | MODIFIER |
| 16 | 79301392 | AX-170093533 | 0 | C | T | 0.474 | 9.65E-02 | 7.23E-02 | 2.65E+00 | 2.73E+00 | 1.80E-01 | no | | intergenic_variant | MODIFIER |
| 1 | 166535105 | AX-97852985 | 0 | G | A | 0.237 | 1.73E-01 | 1.30E-01 | 2.65E+00 | 2.71E+00 | 1.80E-01 | Yes | | intergenic_variant | MODIFIER |
| 13 | 20644382 | AX-97846884 | 1 | T | C | 0.17 | -1.66E-01 | 1.22E-01 | 2.40E+00 | 2.45E+00 | 1.80E-01 | no | Gm47721 | upstream_gene_variant | MODIFIER |
| 1 | 181448885 | AX-170427367 | 0 | A | G | 0.228 | 1.52E-01 | 1.13E-01 | 2.49E+00 | 2.56E+00 | 1.80E-01 | Yes | Cnih3 | intron_variant | MODIFIER |
| 1 | 181452705 | AX-170290034 | 0 | A | G | 0.228 | 1.52E-01 | 1.13E-01 | 2.49E+00 | 2.56E+00 | 1.80E-01 | Yes | Cnih3 | intron_variant | MODIFIER |
| 1 | 181454418 | AX-170224831 | 0 | T | C | 0.228 | 1.52E-01 | 1.13E-01 | 2.49E+00 | 2.56E+00 | 1.80E-01 | Yes | Cnih3 | intron_variant | MODIFIER |
| 1 | 181466571 | AX-170106277 | 0 | T | C | 0.228 | 1.52E-01 | 1.13E-01 | 2.49E+00 | 2.56E+00 | 1.80E-01 | Yes | | regulatory_region_variant | MODIFIER |
| 1 | 181492791 | AX-97795185 | 0 | A | G | 0.228 | 1.52E-01 | 1.13E-01 | 2.49E+00 | 2.56E+00 | 1.80E-01 | Yes | | intergenic_variant | MODIFIER |
| 1 | 181521282 | AX-170406825 | 0 | A | G | 0.228 | 1.52E-01 | 1.13E-01 | 2.49E+00 | 2.56E+00 | 1.80E-01 | Yes | | regulatory_region_variant | MODIFIER |
| 1 | 181523886 | AX-170197277 | 0 | G | A | 0.228 | 1.52E-01 | 1.13E-01 | 2.49E+00 | 2.56E+00 | 1.80E-01 | Yes | | regulatory_region_variant | MODIFIER |
| 1 | 181528723 | AX-170527243 | 0 | A | G | 0.228 | 1.52E-01 | 1.13E-01 | 2.49E+00 | 2.56E+00 | 1.80E-01 | Yes | | regulatory_region_variant | MODIFIER |
| 1 | 181530284 | AX-170332638 | 0 | C | G | 0.228 | 1.52E-01 | 1.13E-01 | 2.49E+00 | 2.56E+00 | 1.80E-01 | Yes | | intergenic_variant | MODIFIER |
| 1 | 181530374 | AX-170532134 | 0 | A | G | 0.228 | 1.52E-01 | 1.13E-01 | 2.49E+00 | 2.56E+00 | 1.80E-01 | Yes | | intergenic_variant | MODIFIER |
| 1 | 181643209 | AX-170002427 | 0 | A | G | 0.228 | 1.52E-01 | 1.13E-01 | 2.49E+00 | 2.56E+00 | 1.80E-01 | Yes | Dnah14 | intron_variant | MODIFIER |
| 1 | 181995691 | AX-170375567 | 0 | G | C | 0.228 | 1.52E-01 | 1.13E-01 | 2.49E+00 | 2.56E+00 | 1.80E-01 | Yes | Enah | intron_variant | MODIFIER |
| 1 | 182072041 | AX-170525824 | 0 | C | T | 0.228 | 1.52E-01 | 1.13E-01 | 2.49E+00 | 2.56E+00 | 1.80E-01 | Yes | Gm37177 | downstream_gene_variant | MODIFIER |
| 1 | 182072111 | AX-170144972 | 0 | G | A | 0.228 | 1.52E-01 | 1.13E-01 | 2.49E+00 | 2.56E+00 | 1.80E-01 | Yes | Gm37177 | downstream_gene_variant | MODIFIER |
| 1 | 182090516 | AX-170002434 | 0 | C | T | 0.228 | 1.52E-01 | 1.13E-01 | 2.49E+00 | 2.56E+00 | 1.80E-01 | Yes | | regulatory_region_variant | MODIFIER |
| 1 | 182209434 | AX-170043801 | 0 | G | A | 0.228 | 1.52E-01 | 1.13E-01 | 2.49E+00 | 2.56E+00 | 1.80E-01 | Yes | | intergenic_variant | MODIFIER |
| 1 | 182272823 | AX-170086710 | 0 | A | G | 0.228 | 1.52E-01 | 1.13E-01 | 2.49E+00 | 2.56E+00 | 1.80E-01 | Yes | Degs1 | downstream_gene_variant | MODIFIER |
| 1 | 182389035 | AX-97856948 | 0 | A | C | 0.228 | 1.52E-01 | 1.13E-01 | 2.49E+00 | 2.56E+00 | 1.80E-01 | Yes | Gm5706 | downstream_gene_variant | MODIFIER |
| 1 | 182390307 | AX-170205496 | 0 | T | C | 0.228 | 1.52E-01 | 1.13E-01 | 2.49E+00 | 2.56E+00 | 1.80E-01 | Yes | Gm5706 | downstream_gene_variant | MODIFIER |
| 1 | 182390972 | AX-170502847 | 0 | A | G | 0.228 | 1.52E-01 | 1.13E-01 | 2.49E+00 | 2.56E+00 | 1.80E-01 | Yes | Gm5706 | downstream_gene_variant | MODIFIER |
| 1 | 182400076 | AX-97795089 | 0 | C | T | 0.228 | 1.52E-01 | 1.13E-01 | 2.49E+00 | 2.56E+00 | 1.80E-01 | Yes | | regulatory_region_variant | MODIFIER |
| 1 | 182417378 | AX-170295627 | 0 | C | T | 0.228 | 1.52E-01 | 1.13E-01 | 2.49E+00 | 2.56E+00 | 1.80E-01 | Yes | Trp53bp2 | intron_variant | MODIFIER |
| 1 | 182433670 | AX-170487499 | 0 | C | G | 0.228 | 1.52E-01 | 1.13E-01 | 2.49E+00 | 2.56E+00 | 1.80E-01 | Yes | Trp53bp2 | intron_variant | MODIFIER |
| 1 | 182438329 | AX-170421045 | 0 | A | G | 0.228 | 1.52E-01 | 1.13E-01 | 2.49E+00 | 2.56E+00 | 1.80E-01 | Yes | Trp53bp2 | intron_variant | MODIFIER |
| 1 | 182438450 | AX-170385738 | 0 | T | C | 0.228 | 1.52E-01 | 1.13E-01 | 2.49E+00 | 2.56E+00 | 1.80E-01 | Yes | Trp53bp2 | intron_variant | MODIFIER |
| 1 | 182438487 | AX-170264254 | 0 | A | C | 0.228 | 1.52E-01 | 1.13E-01 | 2.49E+00 | 2.56E+00 | 1.80E-01 | Yes | Trp53bp2 | intron_variant | MODIFIER |
| 1 | 182441856 | AX-170298822 | 0 | A | G | 0.228 | 1.52E-01 | 1.13E-01 | 2.49E+00 | 2.56E+00 | 1.80E-01 | Yes | Trp53bp2 | intron_variant | MODIFIER |
| 1 | 182465743 | AX-170136402 | 0 | C | G | 0.228 | 1.52E-01 | 1.13E-01 | 2.49E+00 | 2.56E+00 | 1.80E-01 | Yes | Capn2 | downstream_gene_variant | MODIFIER |
| 1 | 182471985 | AX-170043810 | 0 | A | T | 0.228 | 1.52E-01 | 1.13E-01 | 2.49E+00 | 2.56E+00 | 1.80E-01 | Yes | Capn2 | intron_variant | MODIFIER |
| 1 | 182479212 | AX-170521333 | 0 | C | T | 0.228 | 1.52E-01 | 1.13E-01 | 2.49E+00 | 2.56E+00 | 1.80E-01 | Yes | Capn2 | intron_variant | MODIFIER |
| 1 | 182479264 | AX-170231480 | 0 | A | G | 0.228 | 1.52E-01 | 1.13E-01 | 2.49E+00 | 2.56E+00 | 1.80E-01 | Yes | Capn2 | intron_variant | MODIFIER |
| 1 | 182493249 | AX-170370940 | 0 | C | G | 0.228 | 1.52E-01 | 1.13E-01 | 2.49E+00 | 2.56E+00 | 1.80E-01 | Yes | Capn2 | intron_variant | MODIFIER |
| 1 | 183712408 | AX-170161542 | 0 | A | G | 0.228 | 1.52E-01 | 1.13E-01 | 2.49E+00 | 2.56E+00 | 1.80E-01 | Yes | | intergenic_variant | MODIFIER |
| 1 | 183752663 | AX-170334950 | 0 | G | A | 0.228 | 1.52E-01 | 1.13E-01 | 2.49E+00 | 2.56E+00 | 1.80E-01 | Yes | Gm34068 | intron_variant,non_coding_transcript_variant | MODIFIER |
| 1 | 183753311 | AX-170207307 | 0 | G | A | 0.228 | 1.52E-01 | 1.13E-01 | 2.49E+00 | 2.56E+00 | 1.80E-01 | Yes | Gm34068 | intron_variant,non_coding_transcript_variant | MODIFIER |
| 1 | 183755224 | AX-170362738 | 0 | G | A | 0.228 | 1.52E-01 | 1.13E-01 | 2.49E+00 | 2.56E+00 | 1.80E-01 | Yes | Gm34068 | intron_variant,non_coding_transcript_variant | MODIFIER |
| 1 | 183762695 | AX-170002446 | 0 | A | G | 0.228 | 1.52E-01 | 1.13E-01 | 2.49E+00 | 2.56E+00 | 1.80E-01 | Yes | Gm34068 | intron_variant,non_coding_transcript_variant | MODIFIER |
| 1 | 183763138 | AX-97857417 | 0 | G | A | 0.228 | 1.52E-01 | 1.13E-01 | 2.49E+00 | 2.56E+00 | 1.80E-01 | Yes | Gm34068 | intron_variant,non_coding_transcript_variant | MODIFIER |
| 1 | 183796636 | AX-170510510 | 0 | C | T | 0.228 | 1.52E-01 | 1.13E-01 | 2.49E+00 | 2.56E+00 | 1.80E-01 | Yes | Gm34068 | intron_variant,non_coding_transcript_variant | MODIFIER |
| 1 | 183910140 | AX-170128118 | 0 | T | G | 0.228 | 1.52E-01 | 1.13E-01 | 2.49E+00 | 2.56E+00 | 1.80E-01 | Yes | | intergenic_variant | MODIFIER |
| 1 | 183917088 | AX-97815404 | 0 | C | T | 0.228 | 1.52E-01 | 1.13E-01 | 2.49E+00 | 2.56E+00 | 1.80E-01 | Yes | Gm38332 | upstream_gene_variant | MODIFIER |
| 1 | 183920455 | AX-170244135 | 0 | C | T | 0.228 | 1.52E-01 | 1.13E-01 | 2.49E+00 | 2.56E+00 | 1.80E-01 | Yes | Gm38332 | downstream_gene_variant | MODIFIER |
| 1 | 183943893 | AX-170291757 | 0 | C | T | 0.228 | 1.52E-01 | 1.13E-01 | 2.49E+00 | 2.56E+00 | 1.80E-01 | Yes | | intergenic_variant | MODIFIER |
| 1 | 183944299 | AX-170483491 | 0 | G | A | 0.228 | 1.52E-01 | 1.13E-01 | 2.49E+00 | 2.56E+00 | 1.80E-01 | Yes | | intergenic_variant | MODIFIER |
| 1 | 183945269 | AX-170206464 | 0 | C | T | 0.228 | 1.52E-01 | 1.13E-01 | 2.49E+00 | 2.56E+00 | 1.80E-01 | Yes | | intergenic_variant | MODIFIER |
| 1 | 183947093 | AX-170510813 | 0 | A | C | 0.228 | 1.52E-01 | 1.13E-01 | 2.49E+00 | 2.56E+00 | 1.80E-01 | Yes | | regulatory_region_variant | MODIFIER |
| 1 | 183952760 | AX-170336133 | 0 | A | C | 0.228 | 1.52E-01 | 1.13E-01 | 2.49E+00 | 2.56E+00 | 1.80E-01 | Yes | | intergenic_variant | MODIFIER |
| 1 | 183952947 | AX-170338293 | 0 | C | T | 0.228 | 1.52E-01 | 1.13E-01 | 2.49E+00 | 2.56E+00 | 1.80E-01 | Yes | | intergenic_variant | MODIFIER |

| | | | | | | | | | | | | | | | |
|---|---|---|---|---|---|---|---|---|---|---|---|---|---|---|---|
| 1 | 183953079 | AX-170493198 | 0 | G | A | 0.228 | 1.52E-01 | 1.13E-01 | 2.49E+00 | 2.56E+00 | 1.80E-01 | Yes | | intergenic_variant | MODIFIER |
| 1 | 183954493 | AX-170132173 | 0 | C | T | 0.228 | 1.52E-01 | 1.13E-01 | 2.49E+00 | 2.56E+00 | 1.80E-01 | Yes | | intergenic_variant | MODIFIER |
| 1 | 183954628 | AX-170228842 | 0 | T | C | 0.228 | 1.52E-01 | 1.13E-01 | 2.49E+00 | 2.56E+00 | 1.80E-01 | Yes | | regulatory_region_variant | MODIFIER |
| 1 | 183956818 | AX-170171376 | 0 | A | G | 0.228 | 1.52E-01 | 1.13E-01 | 2.49E+00 | 2.56E+00 | 1.80E-01 | Yes | | intergenic_variant | MODIFIER |
| 1 | 183963804 | AX-170234908 | 0 | G | T | 0.228 | 1.52E-01 | 1.13E-01 | 2.49E+00 | 2.56E+00 | 1.80E-01 | Yes | | intergenic_variant | MODIFIER |
| 1 | 183963860 | AX-170427315 | 0 | C | T | 0.228 | 1.52E-01 | 1.13E-01 | 2.49E+00 | 2.56E+00 | 1.80E-01 | Yes | | intergenic_variant | MODIFIER |
| 1 | 183966400 | AX-97794887 | 0 | A | C | 0.228 | 1.52E-01 | 1.13E-01 | 2.49E+00 | 2.56E+00 | 1.80E-01 | Yes | | intergenic_variant | MODIFIER |
| 1 | 183966670 | AX-170446301 | 0 | A | G | 0.228 | 1.52E-01 | 1.13E-01 | 2.49E+00 | 2.56E+00 | 1.80E-01 | Yes | | intergenic_variant | MODIFIER |
| 1 | 183979479 | AX-170407409 | 0 | T | C | 0.228 | 1.52E-01 | 1.13E-01 | 2.49E+00 | 2.56E+00 | 1.80E-01 | Yes | | intergenic_variant | MODIFIER |
| 1 | 183986980 | AX-170481113 | 0 | A | C | 0.228 | 1.52E-01 | 1.13E-01 | 2.49E+00 | 2.56E+00 | 1.80E-01 | Yes | | intergenic_variant | MODIFIER |
| 1 | 183987251 | AX-170498072 | 0 | C | A | 0.228 | 1.52E-01 | 1.13E-01 | 2.49E+00 | 2.56E+00 | 1.80E-01 | Yes | | intergenic_variant | MODIFIER |
| 1 | 183987302 | AX-170133254 | 0 | T | G | 0.228 | 1.52E-01 | 1.13E-01 | 2.49E+00 | 2.56E+00 | 1.80E-01 | Yes | | intergenic_variant | MODIFIER |
| 1 | 183989700 | AX-170177764 | 0 | T | C | 0.228 | 1.52E-01 | 1.13E-01 | 2.49E+00 | 2.56E+00 | 1.80E-01 | Yes | | intergenic_variant | MODIFIER |
| 1 | 183998947 | AX-170399400 | 0 | A | G | 0.228 | 1.52E-01 | 1.13E-01 | 2.49E+00 | 2.56E+00 | 1.80E-01 | Yes | | intergenic_variant | MODIFIER |
| 1 | 183999429 | AX-170503051 | 0 | C | A | 0.228 | 1.52E-01 | 1.13E-01 | 2.49E+00 | 2.56E+00 | 1.80E-01 | Yes | | intergenic_variant | MODIFIER |
| 1 | 183999529 | AX-170444123 | 0 | A | G | 0.228 | 1.52E-01 | 1.13E-01 | 2.49E+00 | 2.56E+00 | 1.80E-01 | Yes | | intergenic_variant | MODIFIER |
| 1 | 184004907 | AX-170043827 | 0 | G | A | 0.228 | 1.52E-01 | 1.13E-01 | 2.49E+00 | 2.56E+00 | 1.80E-01 | Yes | | intergenic_variant | MODIFIER |
| 1 | 184005385 | AX-170317049 | 0 | T | A | 0.228 | 1.52E-01 | 1.13E-01 | 2.49E+00 | 2.56E+00 | 1.80E-01 | Yes | | regulatory_region_variant | MODIFIER |
| 1 | 184020913 | AX-170043832 | 0 | T | C | 0.228 | 1.52E-01 | 1.13E-01 | 2.49E+00 | 2.56E+00 | 1.80E-01 | Yes | Dusp10 | intron_variant | MODIFIER |
| 1 | 184045721 | AX-170375707 | 0 | T | C | 0.228 | 1.52E-01 | 1.13E-01 | 2.49E+00 | 2.56E+00 | 1.80E-01 | Yes | Dusp10 | intron_variant | MODIFIER |
| 1 | 184113664 | AX-170285990 | 0 | C | A | 0.228 | 1.52E-01 | 1.13E-01 | 2.49E+00 | 2.56E+00 | 1.80E-01 | Yes | | intergenic_variant | MODIFIER |
| 1 | 184114781 | AX-170411989 | 0 | T | C | 0.228 | 1.52E-01 | 1.13E-01 | 2.49E+00 | 2.56E+00 | 1.80E-01 | Yes | | intergenic_variant | MODIFIER |
| 1 | 184128967 | AX-97836947 | 0 | C | T | 0.228 | 1.52E-01 | 1.13E-01 | 2.49E+00 | 2.56E+00 | 1.80E-01 | Yes | | intergenic_variant | MODIFIER |
| 1 | 184160910 | AX-97814451 | 0 | C | T | 0.228 | 1.52E-01 | 1.13E-01 | 2.49E+00 | 2.56E+00 | 1.80E-01 | Yes | | intergenic_variant | MODIFIER |
| 1 | 184161243 | AX-170281510 | 0 | T | G | 0.228 | 1.52E-01 | 1.13E-01 | 2.49E+00 | 2.56E+00 | 1.80E-01 | Yes | | intergenic_variant | MODIFIER |
| 1 | 184177126 | AX-170141259 | 0 | G | C | 0.228 | 1.52E-01 | 1.13E-01 | 2.49E+00 | 2.56E+00 | 1.80E-01 | Yes | | intergenic_variant | MODIFIER |
| 1 | 184215922 | AX-170043837 | 0 | C | A | 0.228 | 1.52E-01 | 1.13E-01 | 2.49E+00 | 2.56E+00 | 1.80E-01 | Yes | | intergenic_variant | MODIFIER |
| 1 | 184238263 | AX-170002451 | 0 | A | G | 0.228 | 1.52E-01 | 1.13E-01 | 2.49E+00 | 2.56E+00 | 1.80E-01 | Yes | | intergenic_variant | MODIFIER |
| 1 | 184238279 | AX-170002452 | 0 | T | C | 0.228 | 1.52E-01 | 1.13E-01 | 2.49E+00 | 2.56E+00 | 1.80E-01 | Yes | | intergenic_variant | MODIFIER |
| 1 | 184254725 | AX-97823974 | 0 | T | C | 0.228 | 1.52E-01 | 1.13E-01 | 2.49E+00 | 2.56E+00 | 1.80E-01 | Yes | | intergenic_variant | MODIFIER |
| 1 | 184254850 | AX-170216106 | 0 | T | G | 0.228 | 1.52E-01 | 1.13E-01 | 2.49E+00 | 2.56E+00 | 1.80E-01 | Yes | | intergenic_variant | MODIFIER |
| 1 | 184279231 | AX-170002458 | 0 | G | A | 0.228 | 1.52E-01 | 1.13E-01 | 2.49E+00 | 2.56E+00 | 1.80E-01 | Yes | | intergenic_variant | MODIFIER |
| 1 | 184286617 | AX-170002457 | 0 | A | G | 0.228 | 1.52E-01 | 1.13E-01 | 2.49E+00 | 2.56E+00 | 1.80E-01 | Yes | | intergenic_variant | MODIFIER |
| 1 | 184293801 | AX-97844420 | 0 | T | C | 0.228 | 1.52E-01 | 1.13E-01 | 2.49E+00 | 2.56E+00 | 1.80E-01 | Yes | | intergenic_variant | MODIFIER |
| 1 | 184305220 | AX-170002454 | 0 | A | G | 0.228 | 1.52E-01 | 1.13E-01 | 2.49E+00 | 2.56E+00 | 1.80E-01 | Yes | | intergenic_variant | MODIFIER |
| 1 | 184305551 | AX-170106633 | 0 | A | G | 0.228 | 1.52E-01 | 1.13E-01 | 2.49E+00 | 2.56E+00 | 1.80E-01 | Yes | | intergenic_variant | MODIFIER |
| 1 | 184307777 | AX-170002455 | 0 | A | G | 0.228 | 1.52E-01 | 1.13E-01 | 2.49E+00 | 2.56E+00 | 1.80E-01 | Yes | | intergenic_variant | MODIFIER |
| 1 | 184310053 | AX-170002453 | 0 | A | G | 0.228 | 1.52E-01 | 1.13E-01 | 2.49E+00 | 2.56E+00 | 1.80E-01 | Yes | | intergenic_variant | MODIFIER |
| 1 | 184318170 | AX-170173667 | 0 | A | C | 0.228 | 1.52E-01 | 1.13E-01 | 2.49E+00 | 2.56E+00 | 1.80E-01 | Yes | | intergenic_variant | MODIFIER |
| 1 | 184323105 | AX-170201774 | 0 | T | A | 0.228 | 1.52E-01 | 1.13E-01 | 2.49E+00 | 2.56E+00 | 1.80E-01 | Yes | | intergenic_variant | MODIFIER |
| 1 | 184343006 | AX-97794378 | 0 | G | A | 0.228 | 1.52E-01 | 1.13E-01 | 2.49E+00 | 2.56E+00 | 1.80E-01 | Yes | | intergenic_variant | MODIFIER |
| 1 | 184384540 | AX-170239816 | 0 | A | C | 0.228 | 1.52E-01 | 1.13E-01 | 2.49E+00 | 2.56E+00 | 1.80E-01 | Yes | Gm37223 | intron_variant,non_coding_transcript_variant | MODIFIER |
| 1 | 184385408 | AX-170427021 | 0 | G | A | 0.228 | 1.52E-01 | 1.13E-01 | 2.49E+00 | 2.56E+00 | 1.80E-01 | Yes | Gm37223 | intron_variant,non_coding_transcript_variant | MODIFIER |
| 1 | 184387847 | AX-170242558 | 0 | T | C | 0.228 | 1.52E-01 | 1.13E-01 | 2.49E+00 | 2.56E+00 | 1.80E-01 | Yes | Gm37223 | intron_variant,non_coding_transcript_variant | MODIFIER |
| 1 | 184393467 | AX-97814472 | 0 | A | G | 0.228 | 1.52E-01 | 1.13E-01 | 2.49E+00 | 2.56E+00 | 1.80E-01 | Yes | Gm37223 | intron_variant,non_coding_transcript_variant | MODIFIER |
| 1 | 184435975 | AX-97839837 | 0 | C | A | 0.228 | 1.52E-01 | 1.13E-01 | 2.49E+00 | 2.56E+00 | 1.80E-01 | Yes | | intergenic_variant | MODIFIER |
| 1 | 184445172 | AX-170035637 | 0 | G | A | 0.228 | 1.52E-01 | 1.13E-01 | 2.49E+00 | 2.56E+00 | 1.80E-01 | Yes | 2900092O11Rik | downstream_gene_variant | MODIFIER |
| 1 | 184469452 | AX-97833460 | 0 | G | A | 0.228 | 1.52E-01 | 1.13E-01 | 2.49E+00 | 2.56E+00 | 1.80E-01 | Yes | | intergenic_variant | MODIFIER |
| 1 | 184478133 | AX-170283698 | 0 | A | G | 0.228 | 1.52E-01 | 1.13E-01 | 2.49E+00 | 2.56E+00 | 1.80E-01 | Yes | | regulatory_region_variant | MODIFIER |
| 1 | 184478657 | AX-170186318 | 0 | T | C | 0.228 | 1.52E-01 | 1.13E-01 | 2.49E+00 | 2.56E+00 | 1.80E-01 | Yes | | intergenic_variant | MODIFIER |
| 1 | 184480198 | AX-170380558 | 0 | C | G | 0.228 | 1.52E-01 | 1.13E-01 | 2.49E+00 | 2.56E+00 | 1.80E-01 | Yes | | intergenic_variant | MODIFIER |
| 1 | 184484299 | AX-170043841 | 0 | A | C | 0.228 | 1.52E-01 | 1.13E-01 | 2.49E+00 | 2.56E+00 | 1.80E-01 | Yes | | intergenic_variant | MODIFIER |
| 1 | 184536003 | AX-97814250 | 0 | T | C | 0.228 | 1.52E-01 | 1.13E-01 | 2.49E+00 | 2.56E+00 | 1.80E-01 | Yes | 1700112H15Rik | intron_variant,non_coding_transcript_variant | MODIFIER |
| 2 | 143661793 | AX-170065122 | 0 | G | A | 0.14 | -2.60E-01 | 1.87E-01 | 3.32E+00 | 3.45E+00 | 1.80E-01 | Yes | Pcsk2 | intron_variant | MODIFIER |

| | | | | | | | | | | | | | | | |
|---|---|---|---|---|---|---|---|---|---|---|---|---|---|---|---|
| 5 | 64100051 | AX-97822231 | 0 | G | A | 0.482 | 1.37E-01 | 1.02E-01 | 2.44E+00 | 2.50E+00 | 1.80E-01 | no | Pgm2 | intron_variant | MODIFIER |
| 15 | 76501099 | AX-170056023 | 0 | G | A | 0.088 | -1.65E-01 | 1.05E-01 | 1.83E+00 | 1.90E+00 | 1.80E-01 | no | Dgat1 | downstream_gene_variant | MODIFIER |
| 13 | 7613562 | AX-97781037 | 0 | G | T | 0.193 | -1.60E-01 | 1.19E-01 | 2.49E+00 | 2.55E+00 | 1.80E-01 | no | | intergenic_variant | MODIFIER |
| 13 | 9848555 | AX-170469490 | 0 | C | T | 0.193 | -1.60E-01 | 1.19E-01 | 2.49E+00 | 2.55E+00 | 1.80E-01 | no | | intergenic_variant | MODIFIER |
| 13 | 10260139 | AX-170235506 | 0 | T | C | 0.193 | -1.60E-01 | 1.19E-01 | 2.49E+00 | 2.55E+00 | 1.80E-01 | no | Chrm3 | intron_variant | MODIFIER |
| 5 | 44058855 | AX-170161486 | 0 | A | T | 0.202 | -1.36E-01 | 1.03E-01 | 2.72E+00 | 2.83E+00 | 1.81E-01 | no | Prom1 | intron_variant | MODIFIER |
| 18 | 71923972 | AX-170326605 | 0 | A | G | 0.184 | -1.47E-01 | 1.11E-01 | 2.80E+00 | 2.89E+00 | 1.81E-01 | no | Dcc | intron_variant | MODIFIER |
| 18 | 72022581 | AX-170015952 | 0 | T | A | 0.184 | -1.47E-01 | 1.11E-01 | 2.80E+00 | 2.89E+00 | 1.81E-01 | no | Dcc | intron_variant | MODIFIER |
| 1 | 193297247 | AX-170310878 | 1 | C | T | 0.277 | 1.25E-01 | 9.12E-02 | 2.34E+00 | 2.41E+00 | 1.81E-01 | no | Lamb3 | upstream_gene_variant | MODIFIER |
| 19 | 59732418 | AX-170348949 | 1 | T | G | 0.5 | 8.46E-02 | 6.36E-02 | 2.63E+00 | 2.72E+00 | 1.81E-01 | no | | intergenic_variant | MODIFIER |
| 16 | 40628095 | AX-97843526 | 1 | G | A | 0.5 | 1.10E-01 | 8.22E-02 | 2.55E+00 | 2.60E+00 | 1.81E-01 | no | Lsamp | intron_variant | MODIFIER |
| 3 | 36973623 | AX-170066367 | 0 | A | T | 0.202 | 1.15E-01 | 8.59E-02 | 2.95E+00 | 3.07E+00 | 1.81E-01 | no | 4932438A13Rik | intron_variant | MODIFIER |
| 8 | 25435666 | AX-170521536 | 0 | A | T | 0.368 | -8.65E-02 | 6.51E-02 | 2.64E+00 | 2.74E+00 | 1.81E-01 | no | Gm45237 | upstream_gene_variant | MODIFIER |
| 18 | 48604507 | AX-170126416 | 0 | G | A | 0.044 | -2.56E-01 | 1.93E-01 | 2.58E+00 | 2.70E+00 | 1.81E-01 | Yes | | intergenic_variant | MODIFIER |
| 4 | 87177147 | AX-170158451 | 1 | G | A | 0.348 | 1.37E-01 | 1.04E-01 | 2.78E+00 | 2.89E+00 | 1.81E-01 | no | Slc24a2 | intron_variant | MODIFIER |
| 7 | 144898237 | AX-170428746 | 0 | T | A | 0.202 | -9.78E-02 | 7.39E-02 | 2.69E+00 | 2.83E+00 | 1.82E-01 | no | Fgf15 | intron_variant | MODIFIER |
| 4 | 76437569 | AX-168420001 | 1 | T | C | 0.321 | 1.55E-01 | 1.16E-01 | 2.83E+00 | 2.89E+00 | 1.82E-01 | Yes | Ptprd | intron_variant,non_coding_transcript_variant | MODIFIER |
| 1 | 139521091 | AX-170332830 | 1 | G | T | 0.205 | 1.67E-01 | 1.26E-01 | 2.73E+00 | 2.80E+00 | 1.82E-01 | Yes | F13b | intron_variant | MODIFIER |
| 1 | 162871730 | AX-170142638 | 0 | A | T | 0.219 | 1.87E-01 | 1.41E-01 | 2.84E+00 | 2.90E+00 | 1.82E-01 | Yes | Fmo2 | downstream_gene_variant | MODIFIER |
| 1 | 164287541 | AX-168391671 | 0 | G | C | 0.219 | 1.87E-01 | 1.41E-01 | 2.84E+00 | 2.90E+00 | 1.82E-01 | Yes | Blzf1 | downstream_gene_variant | MODIFIER |
| 1 | 164356422 | AX-170219739 | 0 | G | A | 0.219 | 1.87E-01 | 1.41E-01 | 2.84E+00 | 2.90E+00 | 1.82E-01 | Yes | Nme7 | intron_variant | MODIFIER |
| 1 | 164545091 | AX-170357471 | 0 | T | C | 0.219 | 1.87E-01 | 1.41E-01 | 2.84E+00 | 2.90E+00 | 1.82E-01 | Yes | | intergenic_variant | MODIFIER |
| 1 | 164660550 | AX-97813984 | 0 | C | A | 0.219 | 1.87E-01 | 1.41E-01 | 2.84E+00 | 2.90E+00 | 1.82E-01 | Yes | Gm37754 | downstream_gene_variant | MODIFIER |
| 1 | 164754109 | AX-170446587 | 0 | A | G | 0.219 | 1.87E-01 | 1.41E-01 | 2.84E+00 | 2.90E+00 | 1.82E-01 | Yes | | intergenic_variant | MODIFIER |
| 1 | 164760786 | AX-170155423 | 0 | T | C | 0.219 | 1.87E-01 | 1.41E-01 | 2.84E+00 | 2.90E+00 | 1.82E-01 | Yes | | intergenic_variant | MODIFIER |
| 1 | 164762832 | AX-97808128 | 0 | G | A | 0.219 | 1.87E-01 | 1.41E-01 | 2.84E+00 | 2.90E+00 | 1.82E-01 | Yes | | intergenic_variant | MODIFIER |
| 1 | 164768253 | AX-170379100 | 0 | A | T | 0.219 | 1.87E-01 | 1.41E-01 | 2.84E+00 | 2.90E+00 | 1.82E-01 | Yes | | intergenic_variant | MODIFIER |
| 1 | 164771396 | AX-97813988 | 0 | C | T | 0.219 | 1.87E-01 | 1.41E-01 | 2.84E+00 | 2.90E+00 | 1.82E-01 | Yes | | regulatory_region_variant | MODIFIER |
| 1 | 164774762 | AX-170464374 | 0 | A | G | 0.219 | 1.87E-01 | 1.41E-01 | 2.84E+00 | 2.90E+00 | 1.82E-01 | Yes | | intergenic_variant | MODIFIER |
| 1 | 164821431 | AX-97838396 | 0 | T | C | 0.219 | 1.87E-01 | 1.41E-01 | 2.84E+00 | 2.90E+00 | 1.82E-01 | Yes | Dpt | intron_variant | MODIFIER |
| 1 | 164847814 | AX-170086674 | 0 | A | G | 0.219 | 1.87E-01 | 1.41E-01 | 2.84E+00 | 2.90E+00 | 1.82E-01 | Yes | Gm32569 | intron_variant,non_coding_transcript_variant | MODIFIER |
| 18 | 62503566 | AX-170061055 | 0 | C | T | 0.184 | -1.60E-01 | 1.21E-01 | 2.80E+00 | 2.86E+00 | 1.82E-01 | no | Fbxo38 | downstream_gene_variant | MODIFIER |
| 18 | 62955697 | AX-170497092 | 0 | G | A | 0.184 | -1.60E-01 | 1.21E-01 | 2.80E+00 | 2.86E+00 | 1.82E-01 | no | Apcdd1 | downstream_gene_variant | MODIFIER |
| 4 | 83688040 | AX-97802143 | 1 | G | A | 0.321 | 1.46E-01 | 1.10E-01 | 2.65E+00 | 2.74E+00 | 1.82E-01 | Yes | Ccdc171 | intron_variant | MODIFIER |
| 16 | 70936528 | AX-97850149 | 1 | G | A | 0.491 | 1.02E-01 | 7.67E-02 | 2.55E+00 | 2.62E+00 | 1.82E-01 | no | | intergenic_variant | MODIFIER |
| 15 | 11108244 | AX-170494430 | 0 | G | C | 0.237 | -1.41E-01 | 1.06E-01 | 2.66E+00 | 2.72E+00 | 1.82E-01 | no | Adamts12 | intron_variant | MODIFIER |
| 4 | 84195922 | AX-170023278 | 0 | T | A | 0.333 | 1.52E-01 | 1.14E-01 | 2.83E+00 | 2.90E+00 | 1.82E-01 | Yes | | intergenic_variant | MODIFIER |
| 5 | 49968122 | AX-170228270 | 0 | C | G | 0.5 | 1.41E-01 | 1.01E-01 | 2.23E+00 | 2.28E+00 | 1.83E-01 | no | Adgra3 | intron_variant | MODIFIER |
| 20 | 48501227 | AX-170395301 | 1 | T | C | 0.241 | 7.82E-02 | 5.90E-02 | 2.59E+00 | 2.72E+00 | 1.83E-01 | no | Aifm1 | intron_variant | MODIFIER |
| 3 | 22827171 | AX-168387518 | 1 | T | C | 0.152 | -1.32E-01 | 9.97E-02 | 2.58E+00 | 2.66E+00 | 1.83E-01 | no | | intergenic_variant | MODIFIER |
| 10 | 5967871 | AX-170099449 | 0 | C | T | 0.202 | -1.26E-01 | 9.55E-02 | 2.88E+00 | 3.03E+00 | 1.83E-01 | no | | regulatory_region_variant | MODIFIER |
| 15 | 6346481 | AX-170010959 | 0 | G | A | 0.325 | -1.09E-01 | 7.53E-02 | 2.07E+00 | 2.14E+00 | 1.83E-01 | no | Dab2 | intron_variant | MODIFIER |
| 16 | 78406190 | AX-170526896 | 0 | A | G | 0.456 | 9.98E-02 | 7.53E-02 | 2.61E+00 | 2.69E+00 | 1.83E-01 | no | | intergenic_variant | MODIFIER |
| 1 | 168762673 | AX-170321748 | 0 | A | G | 0.219 | 1.85E-01 | 1.40E-01 | 2.74E+00 | 2.79E+00 | 1.83E-01 | Yes | | intergenic_variant | MODIFIER |
| 5 | 129676412 | AX-97781133 | 1 | A | G | 0.152 | -1.27E-01 | 9.04E-02 | 2.17E+00 | 2.24E+00 | 1.83E-01 | no | | intergenic_variant | MODIFIER |
| 4 | 78638958 | AX-97855138 | 1 | G | T | 0.321 | 1.51E-01 | 1.14E-01 | 2.78E+00 | 2.85E+00 | 1.83E-01 | Yes | | intergenic_variant | MODIFIER |
| 9 | 67906849 | AX-170438116 | 1 | C | T | 0.196 | -1.40E-01 | 1.06E-01 | 2.73E+00 | 2.84E+00 | 1.83E-01 | no | Vps13c | intron_variant | MODIFIER |
| 16 | 71066950 | AX-170125930 | 2 | T | A | 0.491 | 1.02E-01 | 7.68E-02 | 2.55E+00 | 2.62E+00 | 1.83E-01 | no | | intergenic_variant | MODIFIER |
| 16 | 53778496 | AX-97831805 | 0 | T | C | 0.518 | 1.09E-01 | 8.20E-02 | 2.50E+00 | 2.54E+00 | 1.83E-01 | no | | intergenic_variant | MODIFIER |
| 4 | 47031882 | AX-170198839 | 0 | A | T | 0.351 | -1.24E-01 | 9.31E-02 | 2.95E+00 | 3.08E+00 | 1.83E-01 | no | Anks6 | intron_variant | MODIFIER |
| 7 | 37035780 | AX-97844175 | 1 | G | A | 0.357 | 1.17E-01 | 8.84E-02 | 2.59E+00 | 2.68E+00 | 1.83E-01 | no | | intergenic_variant | MODIFIER |
| 1 | 133991622 | AX-170356772 | 1 | T | C | 0.527 | 1.11E-01 | 8.38E-02 | 2.69E+00 | 2.76E+00 | 1.83E-01 | no | | regulatory_region_variant | MODIFIER |
| 1 | 134670008 | AX-97814101 | 1 | C | T | 0.527 | 1.11E-01 | 8.38E-02 | 2.69E+00 | 2.76E+00 | 1.83E-01 | no | Syt2 | intron_variant | MODIFIER |
| 1 | 135750966 | AX-170164751 | 1 | T | G | 0.527 | 1.11E-01 | 8.38E-02 | 2.69E+00 | 2.76E+00 | 1.83E-01 | no | Csrp1 | intron_variant | MODIFIER |

| | | | | | | | | | | | | | | | |
|---|---|---|---|---|---|---|---|---|---|---|---|---|---|---|---|
| 5 | 67981051 | AX-170190987 | 1 | C | G | 0.482 | 1.33E-01 | 1.00E-01 | 2.55E+00 | 2.62E+00 | 1.84E-01 | no | Gm5108 | downstream_gene_variant | MODIFIER |
| 14 | 64913322 | AX-97815801 | 0 | C | T | 0.044 | -2.44E-01 | 1.85E-01 | 2.65E+00 | 2.76E+00 | 1.84E-01 | Yes | Hmbox1 | intron_variant | MODIFIER |
| 15 | 6249290 | AX-97798113 | 1 | T | G | 0.08 | -1.49E-01 | 9.47E-02 | 1.82E+00 | 1.90E+00 | 1.84E-01 | no | | intergenic_variant | MODIFIER |
| 4 | 127716968 | AX-170219778 | 0 | G | A | 0.14 | 1.53E-01 | 1.15E-01 | 2.60E+00 | 2.67E+00 | 1.84E-01 | Yes | | intergenic_variant | MODIFIER |
| 10 | 99789236 | AX-170086881 | 0 | G | A | 0.272 | -1.12E-01 | 8.51E-02 | 2.63E+00 | 2.69E+00 | 1.84E-01 | no | Gm35206 | intron_variant,non_coding_transcript_variant | MODIFIER |
| 4 | 76909740 | AX-97812594 | 0 | T | C | 0.333 | 1.51E-01 | 1.14E-01 | 2.90E+00 | 2.98E+00 | 1.84E-01 | Yes | Ptprd | intron_variant,non_coding_transcript_variant | MODIFIER |
| 12 | 109837860 | AX-170178610 | 0 | C | T | 0.132 | -2.05E-01 | 1.55E-01 | 2.88E+00 | 3.00E+00 | 1.84E-01 | Yes | | intergenic_variant | MODIFIER |
| 1 | 69283850 | AX-170311231 | 0 | G | A | 0.132 | 2.35E-01 | 1.78E-01 | 2.85E+00 | 3.00E+00 | 1.84E-01 | Yes | | intergenic_variant | MODIFIER |
| 18 | 57044418 | AX-170307629 | 0 | A | G | 0.193 | -1.35E-01 | 1.02E-01 | 2.60E+00 | 2.70E+00 | 1.85E-01 | no | | intergenic_variant | MODIFIER |
| 19 | 59649834 | AX-170257598 | 0 | T | G | 0.518 | 8.30E-02 | 6.29E-02 | 2.63E+00 | 2.72E+00 | 1.85E-01 | no | | intergenic_variant | MODIFIER |
| 3 | 7680041 | AX-97824522 | 1 | T | C | 0.393 | -8.57E-02 | 6.51E-02 | 2.86E+00 | 3.02E+00 | 1.85E-01 | no | Gm16685 | intron_variant,non_coding_transcript_variant | MODIFIER |
| 5 | 13731590 | AX-170338764 | 1 | T | C | 0.339 | 1.11E-01 | 8.42E-02 | 2.83E+00 | 2.97E+00 | 1.85E-01 | no | | intergenic_variant | MODIFIER |
| 20 | 42622568 | AX-170356357 | 0 | A | G | 0.237 | 7.74E-02 | 5.88E-02 | 2.61E+00 | 2.73E+00 | 1.85E-01 | no | Tenm1 | intron_variant | MODIFIER |
| 20 | 42630871 | AX-170224357 | 0 | T | C | 0.237 | 7.74E-02 | 5.88E-02 | 2.61E+00 | 2.73E+00 | 1.85E-01 | no | Tenm1 | intron_variant | MODIFIER |
| 20 | 42917290 | AX-170352803 | 0 | C | T | 0.237 | 7.74E-02 | 5.88E-02 | 2.61E+00 | 2.73E+00 | 1.85E-01 | no | Tenm1 | intron_variant | MODIFIER |
| 20 | 42958968 | AX-170086387 | 0 | C | T | 0.237 | 7.74E-02 | 5.88E-02 | 2.61E+00 | 2.73E+00 | 1.85E-01 | no | Tenm1 | intron_variant | MODIFIER |
| 20 | 44242946 | AX-170169509 | 0 | T | A | 0.237 | 7.74E-02 | 5.88E-02 | 2.61E+00 | 2.73E+00 | 1.85E-01 | no | | intergenic_variant | MODIFIER |
| 20 | 44467217 | AX-170084131 | 0 | C | T | 0.237 | 7.74E-02 | 5.88E-02 | 2.61E+00 | 2.73E+00 | 1.85E-01 | no | Itpa-ps3 | non_coding_transcript_exon_variant | MODIFIER |
| 20 | 45222962 | AX-97796507 | 0 | T | A | 0.237 | 7.74E-02 | 5.88E-02 | 2.61E+00 | 2.73E+00 | 1.85E-01 | no | | intergenic_variant | MODIFIER |
| 20 | 45300809 | AX-170425335 | 0 | G | A | 0.237 | 7.74E-02 | 5.88E-02 | 2.61E+00 | 2.73E+00 | 1.85E-01 | no | | intergenic_variant | MODIFIER |
| 20 | 45464635 | AX-170084139 | 0 | C | A | 0.237 | 7.74E-02 | 5.88E-02 | 2.61E+00 | 2.73E+00 | 1.85E-01 | no | | intergenic_variant | MODIFIER |
| 20 | 45549084 | AX-170248664 | 0 | T | A | 0.237 | 7.74E-02 | 5.88E-02 | 2.61E+00 | 2.73E+00 | 1.85E-01 | no | Gm5386 | non_coding_transcript_exon_variant | MODIFIER |
| 20 | 45712797 | AX-170098725 | 0 | A | G | 0.237 | 7.74E-02 | 5.88E-02 | 2.61E+00 | 2.73E+00 | 1.85E-01 | no | | intergenic_variant | MODIFIER |
| 20 | 46116214 | AX-170206891 | 0 | T | C | 0.237 | 7.74E-02 | 5.88E-02 | 2.61E+00 | 2.73E+00 | 1.85E-01 | no | | intergenic_variant | MODIFIER |
| 20 | 46117980 | AX-170084149 | 0 | A | G | 0.237 | 7.74E-02 | 5.88E-02 | 2.61E+00 | 2.73E+00 | 1.85E-01 | no | | intergenic_variant | MODIFIER |
| 20 | 46131536 | AX-97800847 | 0 | G | A | 0.237 | 7.74E-02 | 5.88E-02 | 2.61E+00 | 2.73E+00 | 1.85E-01 | no | Gm14653 | downstream_gene_variant | MODIFIER |
| 20 | 46175374 | AX-170033774 | 0 | G | A | 0.237 | 7.74E-02 | 5.88E-02 | 2.61E+00 | 2.73E+00 | 1.85E-01 | no | | intergenic_variant | MODIFIER |
| 20 | 46248726 | AX-170084151 | 0 | G | A | 0.237 | 7.74E-02 | 5.88E-02 | 2.61E+00 | 2.73E+00 | 1.85E-01 | no | | intergenic_variant | MODIFIER |
| 20 | 46251765 | AX-170235409 | 0 | A | G | 0.237 | 7.74E-02 | 5.88E-02 | 2.61E+00 | 2.73E+00 | 1.85E-01 | no | | intergenic_variant | MODIFIER |
| 20 | 46426501 | AX-170139274 | 0 | T | A | 0.237 | 7.74E-02 | 5.88E-02 | 2.61E+00 | 2.73E+00 | 1.85E-01 | no | | intergenic_variant | MODIFIER |
| 20 | 46431134 | AX-170513499 | 0 | C | T | 0.237 | 7.74E-02 | 5.88E-02 | 2.61E+00 | 2.73E+00 | 1.85E-01 | no | | intergenic_variant | MODIFIER |
| 20 | 46539664 | AX-170275169 | 0 | C | T | 0.237 | 7.74E-02 | 5.88E-02 | 2.61E+00 | 2.73E+00 | 1.85E-01 | no | Gm7722 | upstream_gene_variant | MODIFIER |
| 20 | 46604204 | AX-170084156 | 0 | C | T | 0.237 | 7.74E-02 | 5.88E-02 | 2.61E+00 | 2.73E+00 | 1.85E-01 | no | Gm29242 | intron_variant,non_coding_transcript_variant | MODIFIER |
| 20 | 46722012 | AX-170084154 | 0 | A | C | 0.237 | 7.74E-02 | 5.88E-02 | 2.61E+00 | 2.73E+00 | 1.85E-01 | no | Gm29242 | intron_variant,non_coding_transcript_variant | MODIFIER |
| 20 | 46735156 | AX-170241503 | 0 | G | C | 0.237 | 7.74E-02 | 5.88E-02 | 2.61E+00 | 2.73E+00 | 1.85E-01 | no | Gm29242 | intron_variant,non_coding_transcript_variant | MODIFIER |
| 20 | 46735375 | AX-170465820 | 0 | A | G | 0.237 | 7.74E-02 | 5.88E-02 | 2.61E+00 | 2.73E+00 | 1.85E-01 | no | Gm29242 | intron_variant,non_coding_transcript_variant | MODIFIER |
| 20 | 46739774 | AX-97819876 | 0 | A | G | 0.237 | 7.74E-02 | 5.88E-02 | 2.61E+00 | 2.73E+00 | 1.85E-01 | no | Gm29242 | intron_variant,non_coding_transcript_variant | MODIFIER |
| 20 | 47432035 | AX-170227933 | 0 | T | C | 0.237 | 7.74E-02 | 5.88E-02 | 2.61E+00 | 2.73E+00 | 1.85E-01 | no | | intergenic_variant | MODIFIER |
| 20 | 47658221 | AX-170084171 | 0 | C | A | 0.237 | 7.74E-02 | 5.88E-02 | 2.61E+00 | 2.73E+00 | 1.85E-01 | no | | intergenic_variant | MODIFIER |
| 20 | 47718215 | AX-97783504 | 0 | C | T | 0.237 | 7.74E-02 | 5.88E-02 | 2.61E+00 | 2.73E+00 | 1.85E-01 | no | | intergenic_variant | MODIFIER |
| 20 | 47955155 | AX-97844598 | 0 | A | G | 0.237 | 7.74E-02 | 5.88E-02 | 2.61E+00 | 2.73E+00 | 1.85E-01 | no | Ocrl | intron_variant | MODIFIER |
| 20 | 48543343 | AX-170102264 | 0 | T | C | 0.237 | 7.74E-02 | 5.88E-02 | 2.61E+00 | 2.73E+00 | 1.85E-01 | no | Zfp280c | intron_variant | MODIFIER |
| 20 | 48546001 | AX-170418747 | 0 | T | G | 0.237 | 7.74E-02 | 5.88E-02 | 2.61E+00 | 2.73E+00 | 1.85E-01 | no | Zfp280c | intron_variant | MODIFIER |
| 20 | 48644603 | AX-97852651 | 0 | C | T | 0.237 | 7.74E-02 | 5.88E-02 | 2.61E+00 | 2.73E+00 | 1.85E-01 | no | Slc25a14 | intron_variant | MODIFIER |
| 18 | 67482593 | AX-170491412 | 0 | C | T | 0.281 | -1.21E-01 | 9.13E-02 | 2.93E+00 | 3.05E+00 | 1.85E-01 | no | Prelid3a | downstream_gene_variant | MODIFIER |
| 20 | 46297296 | AX-97821161 | 1 | A | T | 0.241 | 7.74E-02 | 5.87E-02 | 2.59E+00 | 2.72E+00 | 1.85E-01 | no | 4930515L19Rik | intron_variant,non_coding_transcript_variant | MODIFIER |
| 9 | 7661100 | AX-170151571 | 2 | T | A | 0.455 | 1.05E-01 | 7.94E-02 | 2.50E+00 | 2.60E+00 | 1.85E-01 | no | Mmp20 | intron_variant | MODIFIER |
| 10 | 101194477 | AX-170285086 | 1 | A | C | 0.295 | -1.12E-01 | 8.54E-02 | 2.64E+00 | 2.70E+00 | 1.85E-01 | no | | intergenic_variant | MODIFIER |
| 16 | 55636806 | AX-170093495 | 0 | C | T | 0.465 | -1.13E-01 | 8.46E-02 | 2.50E+00 | 2.53E+00 | 1.85E-01 | no | | regulatory_region_variant | MODIFIER |
| 14 | 122283750 | AX-170272562 | 0 | G | A | 0.325 | 1.05E-01 | 7.96E-02 | 2.69E+00 | 2.80E+00 | 1.85E-01 | no | Clybl | intron_variant | MODIFIER |
| 14 | 122286460 | AX-170108242 | 0 | C | G | 0.325 | 1.05E-01 | 7.96E-02 | 2.69E+00 | 2.80E+00 | 1.85E-01 | no | Clybl | intron_variant | MODIFIER |
| 14 | 122315628 | AX-170106268 | 0 | T | C | 0.325 | 1.05E-01 | 7.96E-02 | 2.69E+00 | 2.80E+00 | 1.85E-01 | no | Clybl | intron_variant | MODIFIER |
| 14 | 122323716 | AX-170222237 | 0 | C | T | 0.325 | 1.05E-01 | 7.96E-02 | 2.69E+00 | 2.80E+00 | 1.85E-01 | no | Clybl | intron_variant | MODIFIER |
| 14 | 122327504 | AX-170353666 | 0 | G | A | 0.325 | 1.05E-01 | 7.96E-02 | 2.69E+00 | 2.80E+00 | 1.85E-01 | no | Clybl | intron_variant | MODIFIER |

| | | | | | | | | | | | | | | | |
|---|---|---|---|---|---|---|---|---|---|---|---|---|---|---|---|
| 14 | 122349333 | AX-97830862 | 0 | A | G | 0.325 | 1.05E-01 | 7.96E-02 | 2.69E+00 | 2.80E+00 | 1.85E-01 | no | Clybl | intron_variant | MODIFIER |
| 14 | 122416661 | AX-170010868 | 0 | T | A | 0.325 | 1.05E-01 | 7.96E-02 | 2.69E+00 | 2.80E+00 | 1.85E-01 | no | | regulatory_region_variant | MODIFIER |
| 14 | 122416691 | AX-170010870 | 0 | A | T | 0.325 | 1.05E-01 | 7.96E-02 | 2.69E+00 | 2.80E+00 | 1.85E-01 | no | | regulatory_region_variant | MODIFIER |
| 14 | 122417705 | AX-97827841 | 0 | G | T | 0.325 | 1.05E-01 | 7.96E-02 | 2.69E+00 | 2.80E+00 | 1.85E-01 | no | | intergenic_variant | MODIFIER |
| 14 | 122441146 | AX-97827845 | 0 | A | G | 0.325 | 1.05E-01 | 7.96E-02 | 2.69E+00 | 2.80E+00 | 1.85E-01 | no | Gm5089 | intron_variant | MODIFIER |
| 14 | 123339511 | AX-97846805 | 0 | C | T | 0.325 | 1.05E-01 | 7.96E-02 | 2.69E+00 | 2.80E+00 | 1.85E-01 | no | Nalcn | intron_variant | MODIFIER |
| 14 | 123371456 | AX-170414473 | 0 | A | G | 0.325 | 1.05E-01 | 7.96E-02 | 2.69E+00 | 2.80E+00 | 1.85E-01 | no | Nalcn | intron_variant | MODIFIER |
| 14 | 123374343 | AX-170292538 | 0 | A | G | 0.325 | 1.05E-01 | 7.96E-02 | 2.69E+00 | 2.80E+00 | 1.85E-01 | no | Nalcn | intron_variant | MODIFIER |
| 14 | 123402250 | AX-170087749 | 0 | C | A | 0.325 | 1.05E-01 | 7.96E-02 | 2.69E+00 | 2.80E+00 | 1.85E-01 | no | Nalcn | intron_variant | MODIFIER |
| 14 | 123410278 | AX-97791503 | 0 | A | G | 0.325 | 1.05E-01 | 7.96E-02 | 2.69E+00 | 2.80E+00 | 1.85E-01 | no | Nalcn | intron_variant | MODIFIER |
| 14 | 123445934 | AX-97854866 | 0 | G | A | 0.325 | 1.05E-01 | 7.96E-02 | 2.69E+00 | 2.80E+00 | 1.85E-01 | no | Nalcn | intron_variant | MODIFIER |
| 14 | 123463332 | AX-97828178 | 0 | A | G | 0.325 | 1.05E-01 | 7.96E-02 | 2.69E+00 | 2.80E+00 | 1.85E-01 | no | Nalcn | intron_variant | MODIFIER |
| 14 | 123651983 | AX-170010886 | 0 | A | C | 0.325 | 1.05E-01 | 7.96E-02 | 2.69E+00 | 2.80E+00 | 1.85E-01 | no | | intergenic_variant | MODIFIER |
| 14 | 123661159 | AX-97830694 | 0 | T | C | 0.325 | 1.05E-01 | 7.96E-02 | 2.69E+00 | 2.80E+00 | 1.85E-01 | no | Itgbl1 | intron_variant | MODIFIER |
| 14 | 123839169 | AX-168419360 | 0 | G | A | 0.325 | 1.05E-01 | 7.96E-02 | 2.69E+00 | 2.80E+00 | 1.85E-01 | no | Itgbl1 | intron_variant | MODIFIER |
| 14 | 123843234 | AX-97828227 | 0 | T | C | 0.325 | 1.05E-01 | 7.96E-02 | 2.69E+00 | 2.80E+00 | 1.85E-01 | no | Itgbl1 | intron_variant | MODIFIER |
| 14 | 123973953 | AX-97821312 | 0 | C | T | 0.325 | 1.05E-01 | 7.96E-02 | 2.69E+00 | 2.80E+00 | 1.85E-01 | no | Fgf14 | downstream_gene_variant | MODIFIER |
| 14 | 124145336 | AX-97827973 | 0 | G | A | 0.325 | 1.05E-01 | 7.96E-02 | 2.69E+00 | 2.80E+00 | 1.85E-01 | no | Fgf14 | intron_variant | MODIFIER |
| 14 | 124179071 | AX-97828229 | 0 | G | A | 0.325 | 1.05E-01 | 7.96E-02 | 2.69E+00 | 2.80E+00 | 1.85E-01 | no | Fgf14 | intron_variant | MODIFIER |
| 14 | 124306812 | AX-97782000 | 0 | G | T | 0.325 | 1.05E-01 | 7.96E-02 | 2.69E+00 | 2.80E+00 | 1.85E-01 | no | Fgf14 | intron_variant | MODIFIER |
| 14 | 124315887 | AX-97787169 | 0 | C | T | 0.325 | 1.05E-01 | 7.96E-02 | 2.69E+00 | 2.80E+00 | 1.85E-01 | no | Fgf14 | intron_variant | MODIFIER |
| 14 | 124436772 | AX-170274067 | 0 | T | A | 0.325 | 1.05E-01 | 7.96E-02 | 2.69E+00 | 2.80E+00 | 1.85E-01 | no | Fgf14 | intron_variant | MODIFIER |
| 14 | 124669981 | AX-170453687 | 0 | C | T | 0.325 | 1.05E-01 | 7.96E-02 | 2.69E+00 | 2.80E+00 | 1.85E-01 | no | Fgf14 | intron_variant | MODIFIER |
| 18 | 58041367 | AX-170495684 | 1 | G | A | 0.339 | -8.25E-02 | 6.21E-02 | 3.03E+00 | 3.16E+00 | 1.86E-01 | no | Fbn2 | intron_variant | MODIFIER |
| 5 | 64489776 | AX-168409870 | 0 | G | A | 0.482 | 1.30E-01 | 9.78E-02 | 2.45E+00 | 2.51E+00 | 1.86E-01 | no | | regulatory_region_variant | MODIFIER |
| 15 | 72053157 | AX-97811941 | 1 | G | A | 0.08 | -1.69E-01 | 1.08E-01 | 1.82E+00 | 1.89E+00 | 1.86E-01 | no | | intergenic_variant | MODIFIER |
| 18 | 58678570 | AX-170015793 | 1 | A | G | 0.33 | -8.63E-02 | 6.51E-02 | 3.00E+00 | 3.12E+00 | 1.86E-01 | no | Isoc1 | 3_prime_UTR_variant | MODIFIER |
| 16 | 51537306 | AX-170510071 | 0 | A | G | 0.482 | -1.07E-01 | 8.09E-02 | 2.50E+00 | 2.55E+00 | 1.86E-01 | no | | intergenic_variant | MODIFIER |
| 1 | 194631451 | AX-170413767 | 0 | G | T | 0.272 | 1.22E-01 | 9.11E-02 | 2.37E+00 | 2.46E+00 | 1.86E-01 | no | Plxna2 | intron_variant | MODIFIER |
| 12 | 106834575 | AX-97781682 | 0 | G | A | 0.123 | -2.08E-01 | 1.58E-01 | 2.96E+00 | 3.08E+00 | 1.86E-01 | Yes | 1700013N06Rik | upstream_gene_variant | MODIFIER |
| 4 | 88201441 | AX-170153591 | 0 | C | T | 0.412 | -1.11E-01 | 8.28E-02 | 2.37E+00 | 2.42E+00 | 1.86E-01 | no | Focad | intron_variant | MODIFIER |
| 4 | 88440405 | AX-170069930 | 0 | T | C | 0.412 | -1.11E-01 | 8.28E-02 | 2.37E+00 | 2.42E+00 | 1.86E-01 | no | Hacd4 | upstream_gene_variant | MODIFIER |
| 5 | 134683251 | AX-97810253 | 1 | G | A | 0.152 | -1.16E-01 | 8.46E-02 | 2.26E+00 | 2.34E+00 | 1.87E-01 | no | Limk1 | intron_variant | MODIFIER |
| 18 | 58536574 | AX-170112518 | 0 | G | T | 0.333 | -8.60E-02 | 6.50E-02 | 3.01E+00 | 3.13E+00 | 1.87E-01 | no | | intergenic_variant | MODIFIER |
| 16 | 71041902 | AX-170136625 | 1 | A | C | 0.482 | 9.84E-02 | 7.47E-02 | 2.56E+00 | 2.64E+00 | 1.87E-01 | no | | intergenic_variant | MODIFIER |
| 13 | 9501272 | AX-170238805 | 1 | C | T | 0.196 | -1.52E-01 | 1.14E-01 | 2.46E+00 | 2.51E+00 | 1.87E-01 | no | Dip2c | intron_variant | MODIFIER |
| 9 | 98068866 | AX-170257822 | 1 | C | T | 0.152 | -1.40E-01 | 1.07E-01 | 2.69E+00 | 2.81E+00 | 1.87E-01 | no | | intergenic_variant | MODIFIER |
| 4 | 82757725 | AX-170442071 | 1 | A | T | 0.33 | 1.46E-01 | 1.11E-01 | 2.71E+00 | 2.79E+00 | 1.87E-01 | Yes | | regulatory_region_variant | MODIFIER |
| 16 | 70232908 | AX-170275015 | 1 | C | A | 0.491 | 1.01E-01 | 7.67E-02 | 2.55E+00 | 2.61E+00 | 1.88E-01 | no | | regulatory_region_variant | MODIFIER |
| 7 | 12832654 | AX-97844842 | 0 | C | A | 0.36 | 9.42E-02 | 7.19E-02 | 2.60E+00 | 2.73E+00 | 1.88E-01 | no | Zfp110 | upstream_gene_variant | MODIFIER |
| 16 | 52648384 | AX-170239393 | 1 | A | G | 0.473 | -1.11E-01 | 8.38E-02 | 2.52E+00 | 2.56E+00 | 1.88E-01 | no | | intergenic_variant | MODIFIER |
| 9 | 9130686 | AX-97832693 | 0 | A | G | 0.474 | 1.08E-01 | 8.16E-02 | 2.44E+00 | 2.53E+00 | 1.88E-01 | no | Arhgap42 | intron_variant | MODIFIER |
| 9 | 9257602 | AX-170096460 | 0 | T | C | 0.474 | 1.08E-01 | 8.16E-02 | 2.44E+00 | 2.53E+00 | 1.88E-01 | no | Gm16833 | upstream_gene_variant | MODIFIER |
| 9 | 9260436 | AX-170337420 | 0 | G | A | 0.474 | 1.08E-01 | 8.16E-02 | 2.44E+00 | 2.53E+00 | 1.88E-01 | no | Gm16833 | upstream_gene_variant | MODIFIER |
| 9 | 9261295 | AX-170199414 | 0 | C | T | 0.474 | 1.08E-01 | 8.16E-02 | 2.44E+00 | 2.53E+00 | 1.88E-01 | no | Gm16833 | downstream_gene_variant | MODIFIER |
| 9 | 9261989 | AX-170443108 | 0 | A | G | 0.474 | 1.08E-01 | 8.16E-02 | 2.44E+00 | 2.53E+00 | 1.88E-01 | no | Gm16833 | downstream_gene_variant | MODIFIER |
| 9 | 9264165 | AX-170318790 | 0 | T | C | 0.474 | 1.08E-01 | 8.16E-02 | 2.44E+00 | 2.53E+00 | 1.88E-01 | no | Gm16833 | downstream_gene_variant | MODIFIER |
| 9 | 9275950 | AX-97815097 | 0 | G | T | 0.474 | 1.08E-01 | 8.16E-02 | 2.44E+00 | 2.53E+00 | 1.88E-01 | no | | intergenic_variant | MODIFIER |
| 9 | 9288645 | AX-170268050 | 0 | G | T | 0.474 | 1.08E-01 | 8.16E-02 | 2.44E+00 | 2.53E+00 | 1.88E-01 | no | | intergenic_variant | MODIFIER |
| 9 | 9289911 | AX-170460874 | 0 | T | C | 0.474 | 1.08E-01 | 8.16E-02 | 2.44E+00 | 2.53E+00 | 1.88E-01 | no | | intergenic_variant | MODIFIER |
| 9 | 9290093 | AX-97830597 | 0 | G | A | 0.474 | 1.08E-01 | 8.16E-02 | 2.44E+00 | 2.53E+00 | 1.88E-01 | no | | intergenic_variant | MODIFIER |
| 9 | 9290484 | AX-170115866 | 0 | A | G | 0.474 | 1.08E-01 | 8.16E-02 | 2.44E+00 | 2.53E+00 | 1.88E-01 | no | | intergenic_variant | MODIFIER |
| 9 | 9292434 | AX-170360016 | 0 | G | A | 0.474 | 1.08E-01 | 8.16E-02 | 2.44E+00 | 2.53E+00 | 1.88E-01 | no | | intergenic_variant | MODIFIER |
| 9 | 9303220 | AX-170134352 | 0 | G | T | 0.474 | 1.08E-01 | 8.16E-02 | 2.44E+00 | 2.53E+00 | 1.88E-01 | no | Gm22815 | downstream_gene_variant | MODIFIER |

| | | | | | | | | | | | | | | | |
|---|---|---|---|---|---|---|---|---|---|---|---|---|---|---|---|
| 9 | 9365178 | AX-170228555 | 0 | A | G | 0.474 | 1.08E-01 | 8.16E-02 | 2.44E+00 | 2.53E+00 | 1.88E-01 | no | Gm46102 | intron_variant,non_coding_transcript_variant | MODIFIER |
| 9 | 9370343 | AX-97784318 | 0 | T | C | 0.474 | 1.08E-01 | 8.16E-02 | 2.44E+00 | 2.53E+00 | 1.88E-01 | no | Gm46102 | intron_variant,non_coding_transcript_variant | MODIFIER |
| 9 | 9395866 | AX-170270670 | 0 | C | T | 0.474 | 1.08E-01 | 8.16E-02 | 2.44E+00 | 2.53E+00 | 1.88E-01 | no | Gm46102 | intron_variant,non_coding_transcript_variant | MODIFIER |
| 9 | 9404627 | AX-170323215 | 0 | T | C | 0.474 | 1.08E-01 | 8.16E-02 | 2.44E+00 | 2.53E+00 | 1.88E-01 | no | Gm46102 | intron_variant,non_coding_transcript_variant | MODIFIER |
| 9 | 9505233 | AX-97797117 | 0 | T | C | 0.474 | 1.08E-01 | 8.16E-02 | 2.44E+00 | 2.53E+00 | 1.88E-01 | no | Gm46102 | intron_variant,non_coding_transcript_variant | MODIFIER |
| 9 | 9565623 | AX-97849578 | 0 | T | C | 0.474 | 1.08E-01 | 8.16E-02 | 2.44E+00 | 2.53E+00 | 1.88E-01 | no | Gm47334 | intron_variant,non_coding_transcript_variant | MODIFIER |
| 9 | 9593270 | AX-170096462 | 0 | G | A | 0.474 | 1.08E-01 | 8.16E-02 | 2.44E+00 | 2.53E+00 | 1.88E-01 | no | | intergenic_variant | MODIFIER |
| 9 | 9630553 | AX-97813167 | 0 | A | G | 0.474 | 1.08E-01 | 8.16E-02 | 2.44E+00 | 2.53E+00 | 1.88E-01 | no | | intergenic_variant | MODIFIER |
| 16 | 80665946 | AX-170175349 | 0 | T | A | 0.456 | 9.78E-02 | 7.48E-02 | 2.63E+00 | 2.71E+00 | 1.88E-01 | no | | intergenic_variant | MODIFIER |
| 7 | 79672889 | AX-170127932 | 0 | C | A | 0.114 | -2.33E-01 | 1.75E-01 | 3.08E+00 | 3.24E+00 | 1.88E-01 | Yes | Ticrr | intron_variant | MODIFIER |
| 18 | 74203404 | AX-170362360 | 0 | G | A | 0.175 | -1.60E-01 | 1.22E-01 | 2.68E+00 | 2.74E+00 | 1.88E-01 | no | Ska1 | intron_variant | MODIFIER |
| 4 | 108905734 | AX-170375904 | 1 | G | A | 0.259 | 1.46E-01 | 1.12E-01 | 2.63E+00 | 2.69E+00 | 1.88E-01 | Yes | Rab3b | intron_variant | MODIFIER |
| 1 | 124745450 | AX-170100106 | 1 | C | T | 0.58 | 1.10E-01 | 8.36E-02 | 2.58E+00 | 2.65E+00 | 1.88E-01 | no | Gm28928 | intron_variant,non_coding_transcript_variant | MODIFIER |
| 13 | 10871207 | AX-170090922 | 0 | G | A | 0.193 | -1.54E-01 | 1.16E-01 | 2.46E+00 | 2.52E+00 | 1.88E-01 | no | | intergenic_variant | MODIFIER |
| 1 | 184005350 | AX-168380110 | 1 | C | T | 0.232 | 1.46E-01 | 1.10E-01 | 2.45E+00 | 2.52E+00 | 1.88E-01 | Yes | | regulatory_region_variant | MODIFIER |
| 13 | 10181272 | AX-170461257 | 0 | C | T | 0.193 | -1.56E-01 | 1.19E-01 | 2.56E+00 | 2.63E+00 | 1.88E-01 | no | Chrm3 | intron_variant | MODIFIER |
| 4 | 132707202 | AX-170302019 | 0 | T | C | 0.158 | 1.51E-01 | 1.15E-01 | 2.57E+00 | 2.62E+00 | 1.88E-01 | Yes | Eya3 | intron_variant | MODIFIER |
| 19 | 50895890 | AX-170288269 | 1 | T | C | 0.482 | 8.37E-02 | 6.42E-02 | 2.70E+00 | 2.80E+00 | 1.88E-01 | no | | intergenic_variant | MODIFIER |
| 16 | 53539759 | AX-97790695 | 0 | G | A | 0.474 | -1.10E-01 | 8.37E-02 | 2.52E+00 | 2.56E+00 | 1.88E-01 | no | | intergenic_variant | MODIFIER |
| 16 | 52651947 | AX-170494604 | 0 | C | A | 0.474 | -1.10E-01 | 8.37E-02 | 2.52E+00 | 2.56E+00 | 1.89E-01 | no | | intergenic_variant | MODIFIER |
| 13 | 9452608 | AX-170363929 | 0 | T | C | 0.193 | -1.53E-01 | 1.16E-01 | 2.50E+00 | 2.57E+00 | 1.89E-01 | no | Dip2c | intron_variant | MODIFIER |
| 4 | 70143948 | AX-97798562 | 0 | G | A | 0.456 | -1.06E-01 | 8.09E-02 | 2.52E+00 | 2.57E+00 | 1.89E-01 | no | | intergenic_variant | MODIFIER |
| 10 | 71268185 | AX-170505577 | 0 | T | C | 0.404 | -9.88E-02 | 7.58E-02 | 2.69E+00 | 2.78E+00 | 1.89E-01 | no | Ube2d1 | intron_variant | MODIFIER |
| 16 | 50413770 | AX-97831758 | 0 | A | G | 0.482 | -1.07E-01 | 8.10E-02 | 2.50E+00 | 2.55E+00 | 1.89E-01 | no | Bbx | intron_variant | MODIFIER |
| 16 | 53635220 | AX-170093490 | 1 | G | A | 0.473 | -1.10E-01 | 8.38E-02 | 2.52E+00 | 2.56E+00 | 1.89E-01 | no | | intergenic_variant | MODIFIER |
| 1 | 166574363 | AX-97855100 | 0 | G | A | 0.237 | 1.68E-01 | 1.29E-01 | 2.79E+00 | 2.86E+00 | 1.89E-01 | Yes | | intergenic_variant | MODIFIER |
| 15 | 10505814 | AX-170253398 | 0 | T | G | 0.237 | -1.38E-01 | 1.05E-01 | 2.65E+00 | 2.70E+00 | 1.89E-01 | no | Ttc23l | intron_variant | MODIFIER |
| 13 | 7107066 | AX-170444188 | 1 | G | A | 0.196 | -1.63E-01 | 1.25E-01 | 2.62E+00 | 2.67E+00 | 1.89E-01 | no | | intergenic_variant | MODIFIER |
| 10 | 14129987 | AX-170002777 | 0 | A | T | 0.202 | -1.36E-01 | 1.04E-01 | 2.80E+00 | 2.90E+00 | 1.89E-01 | no | Hivep2 | synonymous_variant | LOW |
| 13 | 4738013 | AX-170257793 | 1 | T | C | 0.205 | -1.51E-01 | 1.16E-01 | 2.59E+00 | 2.68E+00 | 1.89E-01 | no | Gm40658 | intron_variant,non_coding_transcript_variant | MODIFIER |
| 5 | 62934927 | AX-170204440 | 1 | T | C | 0.473 | 1.27E-01 | 9.55E-02 | 2.38E+00 | 2.45E+00 | 1.90E-01 | no | | intergenic_variant | MODIFIER |
| 14 | 123290265 | AX-170296909 | 1 | T | G | 0.295 | 1.04E-01 | 7.96E-02 | 2.65E+00 | 2.76E+00 | 1.90E-01 | no | Nalcn | intron_variant | MODIFIER |
| 4 | 124076107 | AX-170360747 | 0 | A | G | 0.237 | 1.39E-01 | 1.06E-01 | 2.56E+00 | 2.63E+00 | 1.90E-01 | no | | intergenic_variant | MODIFIER |
| 6 | 44592155 | AX-170027112 | 1 | C | T | 0.045 | 2.11E-01 | 1.61E-01 | 3.03E+00 | 3.26E+00 | 1.90E-01 | Yes | | intergenic_variant | MODIFIER |
| 3 | 38740591 | AX-170020330 | 0 | G | A | 0.202 | 1.23E-01 | 9.34E-02 | 3.06E+00 | 3.18E+00 | 1.90E-01 | no | | intergenic_variant | MODIFIER |
| 3 | 38878001 | AX-170240179 | 0 | G | A | 0.202 | 1.23E-01 | 9.34E-02 | 3.06E+00 | 3.18E+00 | 1.90E-01 | no | C230034O21Rik | intron_variant,non_coding_transcript_variant | MODIFIER |
| 9 | 84214168 | AX-170377595 | 0 | T | G | 0.219 | 1.40E-01 | 1.07E-01 | 3.04E+00 | 3.17E+00 | 1.90E-01 | no | | intergenic_variant | MODIFIER |
| 18 | 53333758 | AX-170397405 | 0 | A | T | 0.237 | -1.29E-01 | 9.93E-02 | 2.66E+00 | 2.76E+00 | 1.90E-01 | no | Snx24 | intron_variant | MODIFIER |
| 18 | 74653605 | AX-170404911 | 1 | G | A | 0.438 | 1.12E-01 | 8.51E-02 | 2.49E+00 | 2.55E+00 | 1.90E-01 | no | Myo5b | intron_variant | MODIFIER |
| 16 | 51695743 | AX-97802155 | 0 | G | A | 0.474 | -1.10E-01 | 8.37E-02 | 2.52E+00 | 2.56E+00 | 1.91E-01 | no | | intergenic_variant | MODIFIER |
| 16 | 51701547 | AX-170013033 | 0 | A | G | 0.474 | -1.10E-01 | 8.37E-02 | 2.52E+00 | 2.56E+00 | 1.91E-01 | no | | intergenic_variant | MODIFIER |
| 16 | 51702223 | AX-170013043 | 0 | C | A | 0.474 | -1.10E-01 | 8.37E-02 | 2.52E+00 | 2.56E+00 | 1.91E-01 | no | | intergenic_variant | MODIFIER |
| 16 | 51704743 | AX-170093484 | 0 | T | C | 0.474 | -1.10E-01 | 8.37E-02 | 2.52E+00 | 2.56E+00 | 1.91E-01 | no | | intergenic_variant | MODIFIER |
| 16 | 51723021 | AX-170013042 | 0 | G | C | 0.474 | -1.10E-01 | 8.37E-02 | 2.52E+00 | 2.56E+00 | 1.91E-01 | no | | regulatory_region_variant | MODIFIER |
| 16 | 51785717 | AX-170465839 | 0 | A | G | 0.474 | -1.10E-01 | 8.37E-02 | 2.52E+00 | 2.56E+00 | 1.91E-01 | no | | intergenic_variant | MODIFIER |
| 16 | 51786456 | AX-170482419 | 0 | A | G | 0.474 | -1.10E-01 | 8.37E-02 | 2.52E+00 | 2.56E+00 | 1.91E-01 | no | | intergenic_variant | MODIFIER |
| 16 | 51804539 | AX-170494092 | 0 | C | T | 0.474 | -1.10E-01 | 8.37E-02 | 2.52E+00 | 2.56E+00 | 1.91E-01 | no | | intergenic_variant | MODIFIER |
| 16 | 51806697 | AX-170331346 | 0 | G | C | 0.474 | -1.10E-01 | 8.37E-02 | 2.52E+00 | 2.56E+00 | 1.91E-01 | no | | intergenic_variant | MODIFIER |
| 16 | 51807671 | AX-170470770 | 0 | G | A | 0.474 | -1.10E-01 | 8.37E-02 | 2.52E+00 | 2.56E+00 | 1.91E-01 | no | | regulatory_region_variant | MODIFIER |
| 16 | 51925439 | AX-170408706 | 0 | A | G | 0.474 | -1.10E-01 | 8.37E-02 | 2.52E+00 | 2.56E+00 | 1.91E-01 | no | | intergenic_variant | MODIFIER |
| 16 | 51963558 | AX-97798388 | 0 | C | T | 0.474 | -1.10E-01 | 8.37E-02 | 2.52E+00 | 2.56E+00 | 1.91E-01 | no | | intergenic_variant | MODIFIER |
| 16 | 52001383 | AX-170200096 | 0 | C | T | 0.474 | -1.10E-01 | 8.37E-02 | 2.52E+00 | 2.56E+00 | 1.91E-01 | no | Gm29686 | intron_variant,non_coding_transcript_variant | MODIFIER |
| 16 | 52024770 | AX-170526455 | 0 | C | T | 0.474 | -1.10E-01 | 8.37E-02 | 2.52E+00 | 2.56E+00 | 1.91E-01 | no | | intergenic_variant | MODIFIER |
| 16 | 52064539 | AX-170013045 | 0 | G | A | 0.474 | -1.10E-01 | 8.37E-02 | 2.52E+00 | 2.56E+00 | 1.91E-01 | no | Cblb | intron_variant | MODIFIER |

| | | | | | | | | | | | | | | | |
|---|---|---|---|---|---|---|---|---|---|---|---|---|---|---|---|
| 16 | 52074967 | AX-97836958 | 0 | C | T | 0.474 | -1.10E-01 | 8.37E-02 | 2.52E+00 | 2.56E+00 | 1.91E-01 | no | Cblb | intron_variant | MODIFIER |
| 16 | 52126329 | AX-170196026 | 0 | A | G | 0.526 | 1.10E-01 | 8.37E-02 | 2.52E+00 | 2.56E+00 | 1.91E-01 | no | Cblb | intron_variant | MODIFIER |
| 16 | 52126503 | AX-170221397 | 0 | G | C | 0.474 | -1.10E-01 | 8.37E-02 | 2.52E+00 | 2.56E+00 | 1.91E-01 | no | Cblb | intron_variant | MODIFIER |
| 16 | 52130239 | AX-170219044 | 0 | G | A | 0.474 | -1.10E-01 | 8.37E-02 | 2.52E+00 | 2.56E+00 | 1.91E-01 | no | Cblb | intron_variant | MODIFIER |
| 16 | 52134994 | AX-170374016 | 0 | T | G | 0.474 | -1.10E-01 | 8.37E-02 | 2.52E+00 | 2.56E+00 | 1.91E-01 | no | Cblb | intron_variant | MODIFIER |
| 16 | 52138541 | AX-170489024 | 0 | A | G | 0.474 | -1.10E-01 | 8.37E-02 | 2.52E+00 | 2.56E+00 | 1.91E-01 | no | Cblb | intron_variant | MODIFIER |
| 16 | 52146393 | AX-170454139 | 0 | T | G | 0.474 | -1.10E-01 | 8.37E-02 | 2.52E+00 | 2.56E+00 | 1.91E-01 | no | Cblb | intron_variant | MODIFIER |
| 16 | 52150528 | AX-170176724 | 0 | T | C | 0.474 | -1.10E-01 | 8.37E-02 | 2.52E+00 | 2.56E+00 | 1.91E-01 | no | Cblb | intron_variant | MODIFIER |
| 16 | 52152579 | AX-170492734 | 0 | C | T | 0.474 | -1.10E-01 | 8.37E-02 | 2.52E+00 | 2.56E+00 | 1.91E-01 | no | Cblb | synonymous_variant | LOW |
| 16 | 52156861 | AX-170315059 | 0 | G | A | 0.474 | -1.10E-01 | 8.37E-02 | 2.52E+00 | 2.56E+00 | 1.91E-01 | no | Cblb | intron_variant | MODIFIER |
| 16 | 52175172 | AX-97831237 | 0 | C | T | 0.474 | -1.10E-01 | 8.37E-02 | 2.52E+00 | 2.56E+00 | 1.91E-01 | no | Cblb | intron_variant | MODIFIER |
| 16 | 52177829 | AX-170227649 | 0 | A | T | 0.474 | -1.10E-01 | 8.37E-02 | 2.52E+00 | 2.56E+00 | 1.91E-01 | no | Cblb | intron_variant | MODIFIER |
| 16 | 52187286 | AX-170386552 | 0 | T | C | 0.474 | -1.10E-01 | 8.37E-02 | 2.52E+00 | 2.56E+00 | 1.91E-01 | no | Cblb | intron_variant | MODIFIER |
| 16 | 52335360 | AX-170142060 | 0 | G | A | 0.474 | -1.10E-01 | 8.37E-02 | 2.52E+00 | 2.56E+00 | 1.91E-01 | no | Alcam | intron_variant | MODIFIER |
| 16 | 52339533 | AX-170215504 | 0 | A | G | 0.474 | -1.10E-01 | 8.37E-02 | 2.52E+00 | 2.56E+00 | 1.91E-01 | no | Alcam | intron_variant | MODIFIER |
| 16 | 52341228 | AX-170403326 | 0 | G | A | 0.474 | -1.10E-01 | 8.37E-02 | 2.52E+00 | 2.56E+00 | 1.91E-01 | no | Alcam | intron_variant | MODIFIER |
| 16 | 52359913 | AX-170160015 | 0 | G | A | 0.474 | -1.10E-01 | 8.37E-02 | 2.52E+00 | 2.56E+00 | 1.91E-01 | no | Alcam | intron_variant | MODIFIER |
| 16 | 52374919 | AX-97854095 | 0 | T | G | 0.474 | -1.10E-01 | 8.37E-02 | 2.52E+00 | 2.56E+00 | 1.91E-01 | no | Alcam | intron_variant | MODIFIER |
| 16 | 52380467 | AX-170183069 | 0 | T | G | 0.474 | -1.10E-01 | 8.37E-02 | 2.52E+00 | 2.56E+00 | 1.91E-01 | no | Alcam | intron_variant | MODIFIER |
| 16 | 52383220 | AX-170430910 | 0 | C | T | 0.474 | -1.10E-01 | 8.37E-02 | 2.52E+00 | 2.56E+00 | 1.91E-01 | no | Alcam | intron_variant | MODIFIER |
| 16 | 52383584 | AX-170197581 | 0 | T | C | 0.474 | -1.10E-01 | 8.37E-02 | 2.52E+00 | 2.56E+00 | 1.91E-01 | no | Alcam | intron_variant | MODIFIER |
| 16 | 52415548 | AX-170093486 | 0 | C | T | 0.474 | -1.10E-01 | 8.37E-02 | 2.52E+00 | 2.56E+00 | 1.91E-01 | no | Alcam | intron_variant | MODIFIER |
| 16 | 52433242 | AX-170164332 | 0 | A | C | 0.474 | -1.10E-01 | 8.37E-02 | 2.52E+00 | 2.56E+00 | 1.91E-01 | no | Alcam | intron_variant | MODIFIER |
| 16 | 52453605 | AX-170170157 | 0 | T | C | 0.474 | -1.10E-01 | 8.37E-02 | 2.52E+00 | 2.56E+00 | 1.91E-01 | no | Alcam | 5_prime_UTR_variant | MODIFIER |
| 16 | 52464255 | AX-170320613 | 0 | C | A | 0.474 | -1.10E-01 | 8.37E-02 | 2.52E+00 | 2.56E+00 | 1.91E-01 | no | | intergenic_variant | MODIFIER |
| 16 | 52483271 | AX-170194448 | 0 | C | T | 0.474 | -1.10E-01 | 8.37E-02 | 2.52E+00 | 2.56E+00 | 1.91E-01 | no | | intergenic_variant | MODIFIER |
| 16 | 52607331 | AX-170180468 | 0 | A | G | 0.474 | -1.10E-01 | 8.37E-02 | 2.52E+00 | 2.56E+00 | 1.91E-01 | no | | intergenic_variant | MODIFIER |
| 16 | 52609260 | AX-170013052 | 0 | G | A | 0.474 | -1.10E-01 | 8.37E-02 | 2.52E+00 | 2.56E+00 | 1.91E-01 | no | | intergenic_variant | MODIFIER |
| 16 | 52617974 | AX-97813190 | 0 | A | G | 0.474 | -1.10E-01 | 8.37E-02 | 2.52E+00 | 2.56E+00 | 1.91E-01 | no | | intergenic_variant | MODIFIER |
| 16 | 52654207 | AX-170093487 | 0 | C | G | 0.474 | -1.10E-01 | 8.37E-02 | 2.52E+00 | 2.56E+00 | 1.91E-01 | no | | intergenic_variant | MODIFIER |
| 16 | 52667585 | AX-170107903 | 0 | T | C | 0.474 | -1.10E-01 | 8.37E-02 | 2.52E+00 | 2.56E+00 | 1.91E-01 | no | | intergenic_variant | MODIFIER |
| 16 | 52679485 | AX-170454572 | 0 | G | A | 0.474 | -1.10E-01 | 8.37E-02 | 2.52E+00 | 2.56E+00 | 1.91E-01 | no | | intergenic_variant | MODIFIER |
| 16 | 52720585 | AX-170459264 | 0 | C | G | 0.474 | -1.10E-01 | 8.37E-02 | 2.52E+00 | 2.56E+00 | 1.91E-01 | no | | intergenic_variant | MODIFIER |
| 16 | 52726769 | AX-170535400 | 0 | C | T | 0.474 | -1.10E-01 | 8.37E-02 | 2.52E+00 | 2.56E+00 | 1.91E-01 | no | | intergenic_variant | MODIFIER |
| 16 | 52727111 | AX-170057442 | 0 | A | C | 0.474 | -1.10E-01 | 8.37E-02 | 2.52E+00 | 2.56E+00 | 1.91E-01 | no | | regulatory_region_variant | MODIFIER |
| 16 | 52729398 | AX-170057444 | 0 | T | C | 0.474 | -1.10E-01 | 8.37E-02 | 2.52E+00 | 2.56E+00 | 1.91E-01 | no | | intergenic_variant | MODIFIER |
| 16 | 52730882 | AX-170498100 | 0 | T | C | 0.474 | -1.10E-01 | 8.37E-02 | 2.52E+00 | 2.56E+00 | 1.91E-01 | no | | intergenic_variant | MODIFIER |
| 16 | 52756285 | AX-170524146 | 0 | A | C | 0.474 | -1.10E-01 | 8.37E-02 | 2.52E+00 | 2.56E+00 | 1.91E-01 | no | | intergenic_variant | MODIFIER |
| 16 | 52823217 | AX-170175782 | 0 | C | T | 0.474 | -1.10E-01 | 8.37E-02 | 2.52E+00 | 2.56E+00 | 1.91E-01 | no | | intergenic_variant | MODIFIER |
| 16 | 52850490 | AX-170226813 | 0 | T | C | 0.474 | -1.10E-01 | 8.37E-02 | 2.52E+00 | 2.56E+00 | 1.91E-01 | no | 4930405D01Rik | intron_variant,non_coding_transcript_variant | MODIFIER |
| 16 | 52850577 | AX-170262936 | 0 | T | A | 0.474 | -1.10E-01 | 8.37E-02 | 2.52E+00 | 2.56E+00 | 1.91E-01 | no | 4930405D01Rik | intron_variant,non_coding_transcript_variant | MODIFIER |
| 16 | 52872264 | AX-170234426 | 0 | G | A | 0.474 | -1.10E-01 | 8.37E-02 | 2.52E+00 | 2.56E+00 | 1.91E-01 | no | 4930405D01Rik | intron_variant,non_coding_transcript_variant | MODIFIER |
| 16 | 52880854 | AX-97831265 | 0 | T | C | 0.474 | -1.10E-01 | 8.37E-02 | 2.52E+00 | 2.56E+00 | 1.91E-01 | no | 4930405D01Rik | downstream_gene_variant | MODIFIER |
| 16 | 52881191 | AX-170239891 | 0 | C | T | 0.474 | -1.10E-01 | 8.37E-02 | 2.52E+00 | 2.56E+00 | 1.91E-01 | no | 4930405D01Rik | downstream_gene_variant | MODIFIER |
| 16 | 52978258 | AX-170183191 | 0 | C | T | 0.474 | -1.10E-01 | 8.37E-02 | 2.52E+00 | 2.56E+00 | 1.91E-01 | no | | intergenic_variant | MODIFIER |
| 16 | 53022182 | AX-170288604 | 0 | T | A | 0.474 | -1.10E-01 | 8.37E-02 | 2.52E+00 | 2.56E+00 | 1.91E-01 | no | | intergenic_variant | MODIFIER |
| 16 | 53060957 | AX-170242948 | 0 | T | C | 0.474 | -1.10E-01 | 8.37E-02 | 2.52E+00 | 2.56E+00 | 1.91E-01 | no | | intergenic_variant | MODIFIER |
| 16 | 53078059 | AX-170530438 | 0 | A | T | 0.474 | -1.10E-01 | 8.37E-02 | 2.52E+00 | 2.56E+00 | 1.91E-01 | no | | intergenic_variant | MODIFIER |
| 16 | 53231727 | AX-170057453 | 0 | A | G | 0.474 | -1.10E-01 | 8.37E-02 | 2.52E+00 | 2.56E+00 | 1.91E-01 | no | | intergenic_variant | MODIFIER |
| 16 | 53460734 | AX-170517221 | 0 | C | T | 0.474 | -1.10E-01 | 8.37E-02 | 2.52E+00 | 2.56E+00 | 1.91E-01 | no | | intergenic_variant | MODIFIER |
| 16 | 53489107 | AX-170295658 | 0 | T | A | 0.474 | -1.10E-01 | 8.37E-02 | 2.52E+00 | 2.56E+00 | 1.91E-01 | no | | intergenic_variant | MODIFIER |
| 16 | 53516444 | AX-97808940 | 0 | G | A | 0.474 | -1.10E-01 | 8.37E-02 | 2.52E+00 | 2.56E+00 | 1.91E-01 | no | | intergenic_variant | MODIFIER |
| 16 | 53517088 | AX-170484615 | 0 | T | C | 0.526 | 1.10E-01 | 8.37E-02 | 2.52E+00 | 2.56E+00 | 1.91E-01 | no | | intergenic_variant | MODIFIER |
| 16 | 53539680 | AX-170097530 | 0 | G | C | 0.474 | -1.10E-01 | 8.37E-02 | 2.52E+00 | 2.56E+00 | 1.91E-01 | no | | intergenic_variant | MODIFIER |

| | | | | | | | | | | | | | | | |
|---|---|---|---|---|---|---|---|---|---|---|---|---|---|---|---|
| 16 | 53565947 | AX-170202344 | 0 | A | G | 0.474 | -1.10E-01 | 8.37E-02 | 2.52E+00 | 2.56E+00 | 1.91E-01 | no | | intergenic_variant | MODIFIER |
| 16 | 53617069 | AX-170483242 | 0 | T | A | 0.474 | -1.10E-01 | 8.37E-02 | 2.52E+00 | 2.56E+00 | 1.91E-01 | no | Gm49638 | downstream_gene_variant | MODIFIER |
| 16 | 53617149 | AX-170312048 | 0 | C | A | 0.474 | -1.10E-01 | 8.37E-02 | 2.52E+00 | 2.56E+00 | 1.91E-01 | no | Gm49638 | downstream_gene_variant | MODIFIER |
| 16 | 53617197 | AX-97855647 | 0 | A | G | 0.474 | -1.10E-01 | 8.37E-02 | 2.52E+00 | 2.56E+00 | 1.91E-01 | no | Gm49638 | downstream_gene_variant | MODIFIER |
| 16 | 53635250 | AX-170136543 | 0 | T | G | 0.474 | -1.10E-01 | 8.37E-02 | 2.52E+00 | 2.56E+00 | 1.91E-01 | no | | intergenic_variant | MODIFIER |
| 16 | 53656681 | AX-97804328 | 0 | G | T | 0.474 | -1.10E-01 | 8.37E-02 | 2.52E+00 | 2.56E+00 | 1.91E-01 | no | | intergenic_variant | MODIFIER |
| 16 | 53704459 | AX-170296723 | 0 | T | C | 0.474 | -1.10E-01 | 8.37E-02 | 2.52E+00 | 2.56E+00 | 1.91E-01 | no | | intergenic_variant | MODIFIER |
| 16 | 53742620 | AX-170013072 | 0 | C | T | 0.474 | -1.10E-01 | 8.37E-02 | 2.52E+00 | 2.56E+00 | 1.91E-01 | no | | intergenic_variant | MODIFIER |
| 16 | 53746138 | AX-170088021 | 0 | G | A | 0.474 | -1.10E-01 | 8.37E-02 | 2.52E+00 | 2.56E+00 | 1.91E-01 | no | | intergenic_variant | MODIFIER |
| 16 | 53778707 | AX-170013063 | 0 | A | G | 0.474 | -1.10E-01 | 8.37E-02 | 2.52E+00 | 2.56E+00 | 1.91E-01 | no | | intergenic_variant | MODIFIER |
| 16 | 53822539 | AX-97831802 | 0 | G | T | 0.526 | 1.10E-01 | 8.37E-02 | 2.52E+00 | 2.56E+00 | 1.91E-01 | no | | intergenic_variant | MODIFIER |
| 16 | 53858825 | AX-170013067 | 0 | G | A | 0.474 | -1.10E-01 | 8.37E-02 | 2.52E+00 | 2.56E+00 | 1.91E-01 | no | | intergenic_variant | MODIFIER |
| 16 | 53868358 | AX-170013069 | 0 | A | G | 0.474 | -1.10E-01 | 8.37E-02 | 2.52E+00 | 2.56E+00 | 1.91E-01 | no | | intergenic_variant | MODIFIER |
| 16 | 53871933 | AX-170115573 | 0 | C | G | 0.474 | -1.10E-01 | 8.37E-02 | 2.52E+00 | 2.56E+00 | 1.91E-01 | no | | intergenic_variant | MODIFIER |
| 16 | 53873192 | AX-170442201 | 0 | G | A | 0.474 | -1.10E-01 | 8.37E-02 | 2.52E+00 | 2.56E+00 | 1.91E-01 | no | | intergenic_variant | MODIFIER |
| 16 | 53897606 | AX-170088022 | 0 | G | C | 0.474 | -1.10E-01 | 8.37E-02 | 2.52E+00 | 2.56E+00 | 1.91E-01 | no | | intergenic_variant | MODIFIER |
| 16 | 53958303 | AX-170315014 | 0 | C | T | 0.474 | -1.10E-01 | 8.37E-02 | 2.52E+00 | 2.56E+00 | 1.91E-01 | no | | intergenic_variant | MODIFIER |
| 16 | 54032884 | AX-97831830 | 0 | G | A | 0.474 | -1.10E-01 | 8.37E-02 | 2.52E+00 | 2.56E+00 | 1.91E-01 | no | | intergenic_variant | MODIFIER |
| 16 | 54061518 | AX-170088024 | 0 | G | A | 0.474 | -1.10E-01 | 8.37E-02 | 2.52E+00 | 2.56E+00 | 1.91E-01 | no | | intergenic_variant | MODIFIER |
| 16 | 54098247 | AX-170093492 | 0 | G | A | 0.474 | -1.10E-01 | 8.37E-02 | 2.52E+00 | 2.56E+00 | 1.91E-01 | no | | intergenic_variant | MODIFIER |
| 16 | 54350967 | AX-170483931 | 0 | C | T | 0.474 | -1.10E-01 | 8.37E-02 | 2.52E+00 | 2.56E+00 | 1.91E-01 | no | Gm25853 | upstream_gene_variant | MODIFIER |
| 16 | 54411427 | AX-97851187 | 0 | C | A | 0.474 | -1.10E-01 | 8.37E-02 | 2.52E+00 | 2.56E+00 | 1.91E-01 | no | | intergenic_variant | MODIFIER |
| 16 | 54458825 | AX-170013081 | 0 | G | T | 0.474 | -1.10E-01 | 8.37E-02 | 2.52E+00 | 2.56E+00 | 1.91E-01 | no | | intergenic_variant | MODIFIER |
| 16 | 54485450 | AX-97820187 | 0 | A | T | 0.474 | -1.10E-01 | 8.37E-02 | 2.52E+00 | 2.56E+00 | 1.91E-01 | no | | intergenic_variant | MODIFIER |
| 16 | 54508760 | AX-170013079 | 0 | C | A | 0.474 | -1.10E-01 | 8.37E-02 | 2.52E+00 | 2.56E+00 | 1.91E-01 | no | | intergenic_variant | MODIFIER |
| 16 | 54547478 | AX-170200007 | 0 | G | A | 0.474 | -1.10E-01 | 8.37E-02 | 2.52E+00 | 2.56E+00 | 1.91E-01 | no | | intergenic_variant | MODIFIER |
| 16 | 54610494 | AX-168376009 | 0 | G | A | 0.474 | -1.10E-01 | 8.37E-02 | 2.52E+00 | 2.56E+00 | 1.91E-01 | no | | intergenic_variant | MODIFIER |
| 16 | 54610567 | AX-97831859 | 0 | C | T | 0.474 | -1.10E-01 | 8.37E-02 | 2.52E+00 | 2.56E+00 | 1.91E-01 | no | | intergenic_variant | MODIFIER |
| 15 | 5870846 | AX-97855118 | 0 | A | G | 0.325 | -1.06E-01 | 7.45E-02 | 2.09E+00 | 2.15E+00 | 1.91E-01 | no | Gm8043 | downstream_gene_variant | MODIFIER |
| 19 | 50737093 | AX-170214130 | 0 | C | T | 0.491 | 8.41E-02 | 6.49E-02 | 2.74E+00 | 2.84E+00 | 1.91E-01 | no | Gm26629 | intron_variant,non_coding_transcript_variant | MODIFIER |
| 4 | 81082704 | AX-97809995 | 0 | T | C | 0.439 | -1.11E-01 | 8.52E-02 | 2.54E+00 | 2.59E+00 | 1.91E-01 | no | | regulatory_region_variant | MODIFIER |
| 4 | 81093763 | AX-170464793 | 0 | C | A | 0.439 | -1.11E-01 | 8.52E-02 | 2.54E+00 | 2.59E+00 | 1.91E-01 | no | | intergenic_variant | MODIFIER |
| 4 | 81108597 | AX-170416124 | 0 | G | A | 0.439 | -1.11E-01 | 8.52E-02 | 2.54E+00 | 2.59E+00 | 1.91E-01 | no | | intergenic_variant | MODIFIER |
| 4 | 81381967 | AX-170095030 | 0 | A | G | 0.439 | -1.11E-01 | 8.52E-02 | 2.54E+00 | 2.59E+00 | 1.91E-01 | no | Mpdz | intron_variant | MODIFIER |
| 1 | 149412015 | AX-170145919 | 1 | A | G | 0.196 | 1.50E-01 | 1.15E-01 | 2.64E+00 | 2.73E+00 | 1.92E-01 | Yes | Gm29398 | downstream_gene_variant | MODIFIER |
| 5 | 64430761 | AX-170102933 | 0 | C | T | 0.482 | 1.38E-01 | 1.04E-01 | 2.42E+00 | 2.46E+00 | 1.92E-01 | no | Gm31363 | upstream_gene_variant | MODIFIER |
| 12 | 58206555 | AX-170107028 | 0 | C | T | 0.07 | 2.29E-01 | 1.76E-01 | 2.96E+00 | 3.13E+00 | 1.92E-01 | Yes | | intergenic_variant | MODIFIER |
| 1 | 127654643 | AX-170152530 | 0 | G | A | 0.553 | 1.08E-01 | 8.33E-02 | 2.63E+00 | 2.70E+00 | 1.92E-01 | no | Tmem163 | intron_variant,NMD_transcript_variant | MODIFIER |
| 9 | 8247355 | AX-170031768 | 1 | C | T | 0.464 | 1.06E-01 | 8.06E-02 | 2.46E+00 | 2.55E+00 | 1.92E-01 | no | Mir1899 | downstream_gene_variant | MODIFIER |
| 4 | 139197867 | AX-170179502 | 1 | C | T | 0.17 | 1.45E-01 | 1.12E-01 | 2.62E+00 | 2.68E+00 | 1.92E-01 | Yes | Capzb | intron_variant | MODIFIER |
| 4 | 132943072 | AX-170140357 | 1 | T | C | 0.161 | 1.48E-01 | 1.13E-01 | 2.56E+00 | 2.61E+00 | 1.92E-01 | Yes | | regulatory_region_variant | MODIFIER |
| 15 | 4702992 | AX-170343319 | 1 | G | A | 0.232 | -1.36E-01 | 1.05E-01 | 2.70E+00 | 2.75E+00 | 1.92E-01 | no | | intergenic_variant | MODIFIER |
| 16 | 46250559 | AX-170230453 | 0 | A | G | 0.465 | -1.04E-01 | 8.00E-02 | 2.56E+00 | 2.61E+00 | 1.92E-01 | no | | intergenic_variant | MODIFIER |
| 4 | 139548692 | AX-170275252 | 0 | A | G | 0.167 | 1.36E-01 | 1.05E-01 | 2.72E+00 | 2.81E+00 | 1.92E-01 | no | Iffo2 | intron_variant | MODIFIER |
| 10 | 17789411 | AX-170044232 | 1 | A | C | 0.214 | 1.33E-01 | 9.99E-02 | 3.20E+00 | 3.34E+00 | 1.92E-01 | no | Gm20655 | downstream_gene_variant | MODIFIER |
| 4 | 88204785 | AX-97817729 | 0 | A | G | 0.421 | -1.08E-01 | 8.14E-02 | 2.38E+00 | 2.43E+00 | 1.92E-01 | no | Focad | intron_variant | MODIFIER |
| 4 | 124687297 | AX-170089364 | 1 | G | A | 0.241 | 1.38E-01 | 1.06E-01 | 2.55E+00 | 2.62E+00 | 1.92E-01 | no | Utp11 | intron_variant | MODIFIER |
| 4 | 71854414 | AX-170411047 | 0 | T | C | 0.439 | -1.07E-01 | 8.25E-02 | 2.56E+00 | 2.63E+00 | 1.92E-01 | no | | intergenic_variant | MODIFIER |
| 3 | 14868564 | AX-170523030 | 2 | C | T | 0.436 | -8.36E-02 | 6.43E-02 | 2.97E+00 | 3.14E+00 | 1.93E-01 | no | Car3 | intron_variant | MODIFIER |
| 5 | 52070582 | AX-170167784 | 2 | C | T | 0.491 | 1.39E-01 | 1.05E-01 | 2.37E+00 | 2.41E+00 | 1.93E-01 | no | | intergenic_variant | MODIFIER |
| 1 | 193183764 | AX-170188719 | 0 | C | T | 0.281 | 1.19E-01 | 8.91E-02 | 2.33E+00 | 2.42E+00 | 1.93E-01 | no | Traf3ip3 | intron_variant | MODIFIER |
| 1 | 184242546 | AX-97785690 | 0 | G | T | 0.237 | 1.43E-01 | 1.09E-01 | 2.45E+00 | 2.51E+00 | 1.93E-01 | no | | intergenic_variant | MODIFIER |
| 5 | 49593073 | AX-97814962 | 1 | C | A | 0.482 | 1.41E-01 | 1.06E-01 | 2.37E+00 | 2.41E+00 | 1.93E-01 | no | | intergenic_variant | MODIFIER |
| 13 | 20188891 | AX-170422576 | 1 | C | T | 0.179 | -1.64E-01 | 1.24E-01 | 2.44E+00 | 2.50E+00 | 1.93E-01 | no | Elmo1 | intron_variant | MODIFIER |

| | | | | | | | | | | | | | | | |
|---|---|---|---|---|---|---|---|---|---|---|---|---|---|---|---|
| 8 | 34298365 | AX-170486410 | 0 | T | C | 0.132 | -1.61E-01 | 1.24E-01 | 2.62E+00 | 2.71E+00 | 1.93E-01 | no | Gm23177 | upstream_gene_variant | MODIFIER |
| 1 | 172154578 | AX-170043593 | 1 | G | A | 0.214 | 1.75E-01 | 1.35E-01 | 2.66E+00 | 2.71E+00 | 1.93E-01 | Yes | Dcaf8 | intron_variant | MODIFIER |
| 16 | 70522395 | AX-170107849 | 0 | T | C | 0.482 | 9.61E-02 | 7.41E-02 | 2.58E+00 | 2.65E+00 | 1.93E-01 | no | Gbe1 | intron_variant | MODIFIER |
| 1 | 193947210 | AX-170320223 | 0 | G | A | 0.281 | 1.18E-01 | 8.88E-02 | 2.35E+00 | 2.44E+00 | 1.93E-01 | no | | intergenic_variant | MODIFIER |
| 5 | 50271598 | AX-170209964 | 0 | A | G | 0.482 | 1.40E-01 | 1.05E-01 | 2.35E+00 | 2.39E+00 | 1.93E-01 | no | Gm7205 | intron_variant,non_coding_transcript_variant | MODIFIER |
| 14 | 123293669 | AX-97810956 | 1 | T | C | 0.295 | 1.02E-01 | 7.88E-02 | 2.67E+00 | 2.78E+00 | 1.93E-01 | no | Nalcn | intron_variant | MODIFIER |
| 16 | 52353632 | AX-170057436 | 1 | G | A | 0.473 | -1.09E-01 | 8.37E-02 | 2.52E+00 | 2.56E+00 | 1.93E-01 | no | Alcam | intron_variant | MODIFIER |
| 15 | 3612040 | AX-170323569 | 0 | A | G | 0.325 | -1.06E-01 | 7.39E-02 | 2.02E+00 | 2.08E+00 | 1.93E-01 | no | | intergenic_variant | MODIFIER |
| 4 | 124307257 | AX-170070543 | 1 | T | C | 0.223 | 1.46E-01 | 1.12E-01 | 2.50E+00 | 2.57E+00 | 1.93E-01 | Yes | 1700021L23Rik | intron_variant,non_coding_transcript_variant | MODIFIER |
| 9 | 4453819 | AX-97787168 | 0 | C | T | 0.465 | 1.05E-01 | 8.03E-02 | 2.45E+00 | 2.54E+00 | 1.94E-01 | no | Gria4 | intron_variant | MODIFIER |
| 9 | 4503307 | AX-97857161 | 0 | G | A | 0.465 | 1.05E-01 | 8.03E-02 | 2.45E+00 | 2.54E+00 | 1.94E-01 | no | Gria4 | intron_variant | MODIFIER |
| 9 | 4518335 | AX-170081555 | 0 | C | T | 0.465 | 1.05E-01 | 8.03E-02 | 2.45E+00 | 2.54E+00 | 1.94E-01 | no | Gria4 | intron_variant | MODIFIER |
| 9 | 4518367 | AX-170081560 | 0 | C | T | 0.465 | 1.05E-01 | 8.03E-02 | 2.45E+00 | 2.54E+00 | 1.94E-01 | no | Gria4 | intron_variant | MODIFIER |
| 9 | 4536196 | AX-170483660 | 0 | A | G | 0.465 | 1.05E-01 | 8.03E-02 | 2.45E+00 | 2.54E+00 | 1.94E-01 | no | Gria4 | intron_variant | MODIFIER |
| 9 | 4555908 | AX-170239058 | 0 | G | T | 0.465 | 1.05E-01 | 8.03E-02 | 2.45E+00 | 2.54E+00 | 1.94E-01 | no | Gria4 | intron_variant | MODIFIER |
| 9 | 4556056 | AX-170525816 | 0 | C | T | 0.465 | 1.05E-01 | 8.03E-02 | 2.45E+00 | 2.54E+00 | 1.94E-01 | no | Gria4 | intron_variant | MODIFIER |
| 9 | 4562123 | AX-170372641 | 0 | T | C | 0.465 | 1.05E-01 | 8.03E-02 | 2.45E+00 | 2.54E+00 | 1.94E-01 | no | Gria4 | intron_variant | MODIFIER |
| 9 | 4575322 | AX-97783079 | 0 | C | T | 0.465 | 1.05E-01 | 8.03E-02 | 2.45E+00 | 2.54E+00 | 1.94E-01 | no | Gria4 | intron_variant | MODIFIER |
| 9 | 4761699 | AX-97843527 | 0 | G | A | 0.465 | 1.05E-01 | 8.03E-02 | 2.45E+00 | 2.54E+00 | 1.94E-01 | no | Gria4 | intron_variant | MODIFIER |
| 9 | 4798232 | AX-168397188 | 0 | C | T | 0.465 | 1.05E-01 | 8.03E-02 | 2.45E+00 | 2.54E+00 | 1.94E-01 | no | Gria4 | upstream_gene_variant | MODIFIER |
| 9 | 6178277 | AX-168387017 | 0 | A | C | 0.465 | 1.05E-01 | 8.03E-02 | 2.45E+00 | 2.54E+00 | 1.94E-01 | no | Pdgfd | intron_variant | MODIFIER |
| 9 | 6254580 | AX-97805281 | 0 | C | T | 0.465 | 1.05E-01 | 8.03E-02 | 2.45E+00 | 2.54E+00 | 1.94E-01 | no | Pdgfd | intron_variant | MODIFIER |
| 9 | 6346281 | AX-97786426 | 0 | C | T | 0.465 | 1.05E-01 | 8.03E-02 | 2.45E+00 | 2.54E+00 | 1.94E-01 | no | Pdgfd | intron_variant | MODIFIER |
| 9 | 6995886 | AX-168402421 | 0 | T | G | 0.465 | 1.05E-01 | 8.03E-02 | 2.45E+00 | 2.54E+00 | 1.94E-01 | no | Dync2h1 | intron_variant | MODIFIER |
| 9 | 7367150 | AX-168408687 | 0 | T | C | 0.465 | 1.05E-01 | 8.03E-02 | 2.45E+00 | 2.54E+00 | 1.94E-01 | no | Mmp1b | downstream_gene_variant | MODIFIER |
| 9 | 7609172 | AX-170118316 | 0 | G | T | 0.465 | 1.05E-01 | 8.03E-02 | 2.45E+00 | 2.54E+00 | 1.94E-01 | no | | intergenic_variant | MODIFIER |
| 9 | 7617993 | AX-170081612 | 0 | A | G | 0.465 | 1.05E-01 | 8.03E-02 | 2.45E+00 | 2.54E+00 | 1.94E-01 | no | | intergenic_variant | MODIFIER |
| 9 | 7618061 | AX-170096457 | 0 | A | G | 0.465 | 1.05E-01 | 8.03E-02 | 2.45E+00 | 2.54E+00 | 1.94E-01 | no | | intergenic_variant | MODIFIER |
| 9 | 7620884 | AX-170081611 | 0 | C | T | 0.465 | 1.05E-01 | 8.03E-02 | 2.45E+00 | 2.54E+00 | 1.94E-01 | no | | intergenic_variant | MODIFIER |
| 9 | 7621310 | AX-170081610 | 0 | T | C | 0.465 | 1.05E-01 | 8.03E-02 | 2.45E+00 | 2.54E+00 | 1.94E-01 | no | | intergenic_variant | MODIFIER |
| 9 | 7628700 | AX-170340190 | 0 | T | C | 0.465 | 1.05E-01 | 8.03E-02 | 2.45E+00 | 2.54E+00 | 1.94E-01 | no | Mmp20 | intron_variant | MODIFIER |
| 9 | 7643347 | AX-97841640 | 0 | A | G | 0.465 | 1.05E-01 | 8.03E-02 | 2.45E+00 | 2.54E+00 | 1.94E-01 | no | Mmp20 | intron_variant | MODIFIER |
| 9 | 7644583 | AX-168409351 | 0 | T | C | 0.465 | 1.05E-01 | 8.03E-02 | 2.45E+00 | 2.54E+00 | 1.94E-01 | no | Mmp20 | intron_variant | MODIFIER |
| 9 | 7644976 | AX-170289487 | 0 | A | C | 0.465 | 1.05E-01 | 8.03E-02 | 2.45E+00 | 2.54E+00 | 1.94E-01 | no | Mmp20 | intron_variant | MODIFIER |
| 9 | 7655369 | AX-170468729 | 0 | T | C | 0.465 | 1.05E-01 | 8.03E-02 | 2.45E+00 | 2.54E+00 | 1.94E-01 | no | Mmp20 | intron_variant | MODIFIER |
| 9 | 7657574 | AX-170201394 | 0 | C | T | 0.465 | 1.05E-01 | 8.03E-02 | 2.45E+00 | 2.54E+00 | 1.94E-01 | no | Mmp20 | intron_variant | MODIFIER |
| 9 | 7662377 | AX-170365088 | 0 | A | G | 0.465 | 1.05E-01 | 8.03E-02 | 2.45E+00 | 2.54E+00 | 1.94E-01 | no | Mmp20 | intron_variant | MODIFIER |
| 9 | 7662433 | AX-170443023 | 0 | T | C | 0.465 | 1.05E-01 | 8.03E-02 | 2.45E+00 | 2.54E+00 | 1.94E-01 | no | Mmp20 | intron_variant | MODIFIER |
| 9 | 7682009 | AX-170174851 | 0 | T | C | 0.465 | 1.05E-01 | 8.03E-02 | 2.45E+00 | 2.54E+00 | 1.94E-01 | no | AV064505 | intron_variant,non_coding_transcript_variant | MODIFIER |
| 9 | 7683293 | AX-170241166 | 0 | T | A | 0.465 | 1.05E-01 | 8.03E-02 | 2.45E+00 | 2.54E+00 | 1.94E-01 | no | AV064505 | intron_variant,non_coding_transcript_variant | MODIFIER |
| 9 | 7692262 | AX-97839842 | 0 | A | G | 0.465 | 1.05E-01 | 8.03E-02 | 2.45E+00 | 2.54E+00 | 1.94E-01 | no | Mmp7 | intron_variant | MODIFIER |
| 9 | 7966565 | AX-170123125 | 0 | T | C | 0.465 | 1.05E-01 | 8.03E-02 | 2.45E+00 | 2.54E+00 | 1.94E-01 | no | Yap1 | intron_variant | MODIFIER |
| 9 | 7966587 | AX-170081618 | 0 | T | C | 0.465 | 1.05E-01 | 8.03E-02 | 2.45E+00 | 2.54E+00 | 1.94E-01 | no | Yap1 | intron_variant | MODIFIER |
| 9 | 7966669 | AX-170081619 | 0 | G | A | 0.465 | 1.05E-01 | 8.03E-02 | 2.45E+00 | 2.54E+00 | 1.94E-01 | no | Yap1 | intron_variant | MODIFIER |
| 9 | 7975515 | AX-170118219 | 0 | A | G | 0.465 | 1.05E-01 | 8.03E-02 | 2.45E+00 | 2.54E+00 | 1.94E-01 | no | Yap1 | intron_variant | MODIFIER |
| 9 | 7990499 | AX-170177757 | 0 | A | G | 0.465 | 1.05E-01 | 8.03E-02 | 2.45E+00 | 2.54E+00 | 1.94E-01 | no | Yap1 | intron_variant | MODIFIER |
| 9 | 8007284 | AX-170222606 | 0 | G | C | 0.465 | 1.05E-01 | 8.03E-02 | 2.45E+00 | 2.54E+00 | 1.94E-01 | no | Yap1 | upstream_gene_variant | MODIFIER |
| 9 | 8041511 | AX-97834497 | 0 | T | A | 0.465 | 1.05E-01 | 8.03E-02 | 2.45E+00 | 2.54E+00 | 1.94E-01 | no | Cfap300 | intron_variant | MODIFIER |
| 9 | 8055696 | AX-170144997 | 0 | A | G | 0.465 | 1.05E-01 | 8.03E-02 | 2.45E+00 | 2.54E+00 | 1.94E-01 | no | | intergenic_variant | MODIFIER |
| 9 | 8071101 | AX-170408870 | 0 | G | C | 0.465 | 1.05E-01 | 8.03E-02 | 2.45E+00 | 2.54E+00 | 1.94E-01 | no | | intergenic_variant | MODIFIER |
| 9 | 8098944 | AX-170448524 | 0 | C | T | 0.465 | 1.05E-01 | 8.03E-02 | 2.45E+00 | 2.54E+00 | 1.94E-01 | no | Cep126 | intron_variant | MODIFIER |
| 9 | 8099904 | AX-170463767 | 0 | C | T | 0.465 | 1.05E-01 | 8.03E-02 | 2.45E+00 | 2.54E+00 | 1.94E-01 | no | Cep126 | synonymous_variant | LOW |
| 9 | 8108472 | AX-97853369 | 0 | C | T | 0.465 | 1.05E-01 | 8.03E-02 | 2.45E+00 | 2.54E+00 | 1.94E-01 | no | Cep126 | intron_variant | MODIFIER |
| 9 | 8122612 | AX-170193017 | 0 | G | A | 0.465 | 1.05E-01 | 8.03E-02 | 2.45E+00 | 2.54E+00 | 1.94E-01 | no | Cep126 | intron_variant | MODIFIER |

| | | | | | | | | | | | | | | | |
|---|---|---|---|---|---|---|---|---|---|---|---|---|---|---|---|
| 9 | 8157066 | AX-97803108 | 0 | G | A | 0.465 | 1.05E-01 | 8.03E-02 | 2.45E+00 | 2.54E+00 | 1.94E-01 | no | Gm19324 | intron_variant,non_coding_transcript_variant | MODIFIER |
| 9 | 8177361 | AX-170464507 | 0 | C | G | 0.465 | 1.05E-01 | 8.03E-02 | 2.45E+00 | 2.54E+00 | 1.94E-01 | no | Gm19324 | intron_variant,non_coding_transcript_variant | MODIFIER |
| 9 | 8177593 | AX-170227222 | 0 | A | G | 0.465 | 1.05E-01 | 8.03E-02 | 2.45E+00 | 2.54E+00 | 1.94E-01 | no | Gm19324 | intron_variant,non_coding_transcript_variant | MODIFIER |
| 9 | 8185141 | AX-170443671 | 0 | A | G | 0.465 | 1.05E-01 | 8.03E-02 | 2.45E+00 | 2.54E+00 | 1.94E-01 | no | Gm19324 | downstream_gene_variant | MODIFIER |
| 9 | 8256031 | AX-170031762 | 0 | C | T | 0.465 | 1.05E-01 | 8.03E-02 | 2.45E+00 | 2.54E+00 | 1.94E-01 | no | | intergenic_variant | MODIFIER |
| 9 | 8326413 | AX-170031764 | 0 | T | C | 0.465 | 1.05E-01 | 8.03E-02 | 2.45E+00 | 2.54E+00 | 1.94E-01 | no | | intergenic_variant | MODIFIER |
| 9 | 8329759 | AX-97816709 | 0 | T | C | 0.465 | 1.05E-01 | 8.03E-02 | 2.45E+00 | 2.54E+00 | 1.94E-01 | no | | intergenic_variant | MODIFIER |
| 9 | 8389654 | AX-170031766 | 0 | T | C | 0.465 | 1.05E-01 | 8.03E-02 | 2.45E+00 | 2.54E+00 | 1.94E-01 | no | | intergenic_variant | MODIFIER |
| 9 | 8402208 | AX-170347330 | 0 | G | C | 0.465 | 1.05E-01 | 8.03E-02 | 2.45E+00 | 2.54E+00 | 1.94E-01 | no | | intergenic_variant | MODIFIER |
| 9 | 8402332 | AX-170377873 | 0 | T | G | 0.465 | 1.05E-01 | 8.03E-02 | 2.45E+00 | 2.54E+00 | 1.94E-01 | no | | intergenic_variant | MODIFIER |
| 9 | 8426126 | AX-97784345 | 0 | C | A | 0.465 | 1.05E-01 | 8.03E-02 | 2.45E+00 | 2.54E+00 | 1.94E-01 | no | | intergenic_variant | MODIFIER |
| 9 | 8497595 | AX-170283603 | 0 | A | C | 0.465 | 1.05E-01 | 8.03E-02 | 2.45E+00 | 2.54E+00 | 1.94E-01 | no | | intergenic_variant | MODIFIER |
| 9 | 8551835 | AX-97832843 | 0 | G | T | 0.465 | 1.05E-01 | 8.03E-02 | 2.45E+00 | 2.54E+00 | 1.94E-01 | no | Trpc6 | intron_variant | MODIFIER |
| 9 | 8554868 | AX-170186707 | 0 | G | T | 0.465 | 1.05E-01 | 8.03E-02 | 2.45E+00 | 2.54E+00 | 1.94E-01 | no | Trpc6 | intron_variant | MODIFIER |
| 9 | 8637372 | AX-170256849 | 0 | G | A | 0.465 | 1.05E-01 | 8.03E-02 | 2.45E+00 | 2.54E+00 | 1.94E-01 | no | Trpc6 | intron_variant | MODIFIER |
| 9 | 8817019 | AX-170429131 | 0 | A | G | 0.465 | 1.05E-01 | 8.03E-02 | 2.45E+00 | 2.54E+00 | 1.94E-01 | no | | intergenic_variant | MODIFIER |
| 9 | 8822633 | AX-170090242 | 0 | A | G | 0.465 | 1.05E-01 | 8.03E-02 | 2.45E+00 | 2.54E+00 | 1.94E-01 | no | | intergenic_variant | MODIFIER |
| 9 | 8828445 | AX-170140351 | 0 | T | C | 0.465 | 1.05E-01 | 8.03E-02 | 2.45E+00 | 2.54E+00 | 1.94E-01 | no | | intergenic_variant | MODIFIER |
| 9 | 8833074 | AX-170104924 | 0 | A | G | 0.465 | 1.05E-01 | 8.03E-02 | 2.45E+00 | 2.54E+00 | 1.94E-01 | no | | intergenic_variant | MODIFIER |
| 9 | 8853059 | AX-97786243 | 0 | C | T | 0.465 | 1.05E-01 | 8.03E-02 | 2.45E+00 | 2.54E+00 | 1.94E-01 | no | | intergenic_variant | MODIFIER |
| 9 | 8887410 | AX-170397997 | 0 | A | G | 0.465 | 1.05E-01 | 8.03E-02 | 2.45E+00 | 2.54E+00 | 1.94E-01 | no | | intergenic_variant | MODIFIER |
| 9 | 8896863 | AX-97824631 | 0 | T | C | 0.465 | 1.05E-01 | 8.03E-02 | 2.45E+00 | 2.54E+00 | 1.94E-01 | no | Pgr | upstream_gene_variant | MODIFIER |
| 9 | 8897003 | AX-170031777 | 0 | C | T | 0.465 | 1.05E-01 | 8.03E-02 | 2.45E+00 | 2.54E+00 | 1.94E-01 | no | Pgr | upstream_gene_variant | MODIFIER |
| 9 | 8897206 | AX-170446353 | 0 | A | G | 0.465 | 1.05E-01 | 8.03E-02 | 2.45E+00 | 2.54E+00 | 1.94E-01 | no | Pgr | upstream_gene_variant | MODIFIER |
| 9 | 8966972 | AX-97790031 | 0 | C | T | 0.465 | 1.05E-01 | 8.03E-02 | 2.45E+00 | 2.54E+00 | 1.94E-01 | no | Pgr | downstream_gene_variant | MODIFIER |
| 9 | 9003424 | AX-170325805 | 0 | G | A | 0.465 | 1.05E-01 | 8.03E-02 | 2.45E+00 | 2.54E+00 | 1.94E-01 | no | Arhgap42 | intron_variant | MODIFIER |
| 9 | 9004631 | AX-170161703 | 0 | A | T | 0.465 | 1.05E-01 | 8.03E-02 | 2.45E+00 | 2.54E+00 | 1.94E-01 | no | Arhgap42 | intron_variant | MODIFIER |
| 9 | 9005335 | AX-170304239 | 0 | G | A | 0.465 | 1.05E-01 | 8.03E-02 | 2.45E+00 | 2.54E+00 | 1.94E-01 | no | Arhgap42 | intron_variant | MODIFIER |
| 9 | 9010547 | AX-97787033 | 0 | T | C | 0.465 | 1.05E-01 | 8.03E-02 | 2.45E+00 | 2.54E+00 | 1.94E-01 | no | Arhgap42 | intron_variant | MODIFIER |
| 9 | 9011234 | AX-170031773 | 0 | A | G | 0.465 | 1.05E-01 | 8.03E-02 | 2.45E+00 | 2.54E+00 | 1.94E-01 | no | Arhgap42 | intron_variant | MODIFIER |
| 9 | 9013247 | AX-170031770 | 0 | T | C | 0.465 | 1.05E-01 | 8.03E-02 | 2.45E+00 | 2.54E+00 | 1.94E-01 | no | Arhgap42 | intron_variant | MODIFIER |
| 9 | 9046349 | AX-170031775 | 0 | A | G | 0.465 | 1.05E-01 | 8.03E-02 | 2.45E+00 | 2.54E+00 | 1.94E-01 | no | Arhgap42 | intron_variant | MODIFIER |
| 9 | 9046846 | AX-170031774 | 0 | A | G | 0.465 | 1.05E-01 | 8.03E-02 | 2.45E+00 | 2.54E+00 | 1.94E-01 | no | Arhgap42 | intron_variant | MODIFIER |
| 9 | 9082262 | AX-170211640 | 0 | A | G | 0.465 | 1.05E-01 | 8.03E-02 | 2.45E+00 | 2.54E+00 | 1.94E-01 | no | Arhgap42 | intron_variant | MODIFIER |
| 9 | 9082438 | AX-97845281 | 0 | G | A | 0.465 | 1.05E-01 | 8.03E-02 | 2.45E+00 | 2.54E+00 | 1.94E-01 | no | Arhgap42 | intron_variant | MODIFIER |
| 9 | 9082485 | AX-170479492 | 0 | C | G | 0.465 | 1.05E-01 | 8.03E-02 | 2.45E+00 | 2.54E+00 | 1.94E-01 | no | Arhgap42 | intron_variant | MODIFIER |
| 9 | 9114972 | AX-170342273 | 0 | T | C | 0.465 | 1.05E-01 | 8.03E-02 | 2.45E+00 | 2.54E+00 | 1.94E-01 | no | Arhgap42 | intron_variant | MODIFIER |
| 9 | 9121703 | AX-170188097 | 0 | C | T | 0.465 | 1.05E-01 | 8.03E-02 | 2.45E+00 | 2.54E+00 | 1.94E-01 | no | Arhgap42 | intron_variant | MODIFIER |
| 9 | 9121753 | AX-170201581 | 0 | C | T | 0.465 | 1.05E-01 | 8.03E-02 | 2.45E+00 | 2.54E+00 | 1.94E-01 | no | Arhgap42 | intron_variant | MODIFIER |
| 9 | 9126553 | AX-170204920 | 0 | G | A | 0.465 | 1.05E-01 | 8.03E-02 | 2.45E+00 | 2.54E+00 | 1.94E-01 | no | Arhgap42 | intron_variant | MODIFIER |
| 9 | 9126857 | AX-170117488 | 0 | C | T | 0.465 | 1.05E-01 | 8.03E-02 | 2.45E+00 | 2.54E+00 | 1.94E-01 | no | Arhgap42 | intron_variant | MODIFIER |
| 9 | 9136824 | AX-170263050 | 0 | G | T | 0.465 | 1.05E-01 | 8.03E-02 | 2.45E+00 | 2.54E+00 | 1.94E-01 | no | Arhgap42 | intron_variant | MODIFIER |
| 9 | 9138768 | AX-170238562 | 0 | A | C | 0.465 | 1.05E-01 | 8.03E-02 | 2.45E+00 | 2.54E+00 | 1.94E-01 | no | Arhgap42 | intron_variant | MODIFIER |
| 9 | 9141014 | AX-97837763 | 0 | A | C | 0.465 | 1.05E-01 | 8.03E-02 | 2.45E+00 | 2.54E+00 | 1.94E-01 | no | Arhgap42 | intron_variant | MODIFIER |
| 8 | 86025514 | AX-170429632 | 1 | T | G | 0.188 | -1.47E-01 | 1.14E-01 | 2.78E+00 | 2.90E+00 | 1.94E-01 | no | Phkb | intron_variant | MODIFIER |
| 4 | 124221983 | AX-97784335 | 0 | A | G | 0.211 | 1.40E-01 | 1.08E-01 | 2.58E+00 | 2.67E+00 | 1.94E-01 | no | | intergenic_variant | MODIFIER |
| 13 | 23075554 | AX-170050746 | 0 | A | G | 0.167 | -1.40E-01 | 1.06E-01 | 2.39E+00 | 2.48E+00 | 1.94E-01 | no | Vmn1r215 | upstream_gene_variant | MODIFIER |
| 13 | 23622105 | AX-170269606 | 0 | A | G | 0.167 | -1.40E-01 | 1.06E-01 | 2.39E+00 | 2.48E+00 | 1.94E-01 | no | Hist1h2be | upstream_gene_variant | MODIFIER |
| 5 | 93732629 | AX-170025637 | 1 | G | T | 0.446 | 1.63E-01 | 1.26E-01 | 2.56E+00 | 2.72E+00 | 1.94E-01 | Yes | | intergenic_variant | MODIFIER |
| 10 | 19256597 | AX-97814100 | 0 | C | T | 0.184 | 1.26E-01 | 9.48E-02 | 3.22E+00 | 3.38E+00 | 1.94E-01 | no | | intergenic_variant | MODIFIER |
| 1 | 170201745 | AX-170141316 | 0 | C | G | 0.211 | -1.72E-01 | 1.25E-01 | 3.46E+00 | 3.65E+00 | 1.94E-01 | no | Uhmk1 | intron_variant | MODIFIER |
| 1 | 138418104 | AX-97846931 | 0 | A | G | 0.211 | 1.51E-01 | 1.17E-01 | 2.66E+00 | 2.74E+00 | 1.94E-01 | Yes | | intergenic_variant | MODIFIER |
| 10 | 10634595 | AX-97799384 | 0 | T | C | 0.228 | -1.24E-01 | 9.62E-02 | 2.89E+00 | 3.02E+00 | 1.95E-01 | no | | intergenic_variant | MODIFIER |
| 13 | 5445877 | AX-170499356 | 1 | A | T | 0.188 | -1.54E-01 | 1.20E-01 | 2.69E+00 | 2.76E+00 | 1.95E-01 | no | | intergenic_variant | MODIFIER |

| | | | | | | | | | | | | | | | |
|---|---|---|---|---|---|---|---|---|---|---|---|---|---|---|---|
| 1 | 67462169 | AX-97847439 | 0 | G | T | 0.114 | -2.37E-01 | 1.83E-01 | 2.97E+00 | 3.10E+00 | 1.95E-01 | Yes | | intergenic_variant | MODIFIER |
| 1 | 132398487 | AX-97849122 | 1 | A | G | 0.536 | 1.05E-01 | 8.19E-02 | 2.70E+00 | 2.79E+00 | 1.95E-01 | no | | intergenic_variant | MODIFIER |
| 9 | 7663306 | AX-97850847 | 1 | A | G | 0.464 | 1.05E-01 | 8.02E-02 | 2.46E+00 | 2.55E+00 | 1.95E-01 | no | Mmp20 | intron_variant | MODIFIER |
| 9 | 8071863 | AX-97851502 | 1 | G | A | 0.464 | 1.05E-01 | 8.02E-02 | 2.46E+00 | 2.55E+00 | 1.95E-01 | no | Cep126 | downstream_gene_variant | MODIFIER |
| 16 | 50399187 | AX-97806722 | 0 | A | G | 0.518 | 1.06E-01 | 8.07E-02 | 2.45E+00 | 2.48E+00 | 1.95E-01 | no | Bbx | intron_variant | MODIFIER |
| 7 | 100973897 | AX-97803456 | 0 | T | C | 0.351 | -1.25E-01 | 9.67E-02 | 2.92E+00 | 3.03E+00 | 1.95E-01 | no | P2ry6 | intron_variant,non_coding_transcript_variant | MODIFIER |
| 4 | 101044067 | AX-97831493 | 1 | T | C | 0.223 | 1.36E-01 | 1.05E-01 | 2.62E+00 | 2.72E+00 | 1.95E-01 | no | | intergenic_variant | MODIFIER |
| 1 | 173999694 | AX-97850803 | 0 | C | T | 0.228 | -1.37E-01 | 1.02E-01 | 3.29E+00 | 3.49E+00 | 1.95E-01 | no | Gm18445 | intron_variant,non_coding_transcript_variant | MODIFIER |
| 11 | 13118721 | AX-170410646 | 0 | T | C | 0.044 | 2.38E-01 | 1.85E-01 | 2.89E+00 | 3.08E+00 | 1.95E-01 | Yes | | intergenic_variant | MODIFIER |
| 18 | 78576087 | AX-170256519 | 0 | A | C | 0.158 | -1.40E-01 | 1.09E-01 | 2.57E+00 | 2.65E+00 | 1.95E-01 | no | Slc14a2 | intron_variant | MODIFIER |
| 13 | 7582451 | AX-170007689 | 1 | C | A | 0.196 | -1.49E-01 | 1.15E-01 | 2.49E+00 | 2.55E+00 | 1.96E-01 | no | | intergenic_variant | MODIFIER |
| 4 | 136581396 | AX-170434143 | 0 | C | T | 0.167 | 1.47E-01 | 1.14E-01 | 2.59E+00 | 2.64E+00 | 1.96E-01 | Yes | Kdm1a | intron_variant | MODIFIER |
| 8 | 32721162 | AX-170199339 | 0 | A | T | 0.14 | -1.37E-01 | 1.07E-01 | 2.76E+00 | 2.90E+00 | 1.96E-01 | no | Nrg1 | intron_variant | MODIFIER |
| 10 | 93987371 | AX-170410420 | 1 | C | T | 0.295 | -1.09E-01 | 8.44E-02 | 2.55E+00 | 2.61E+00 | 1.96E-01 | no | Vezt | intron_variant | MODIFIER |
| 1 | 193673725 | AX-170140519 | 0 | T | A | 0.272 | 1.23E-01 | 9.36E-02 | 2.37E+00 | 2.45E+00 | 1.96E-01 | no | | regulatory_region_variant | MODIFIER |
| 8 | 29427720 | AX-170485681 | 0 | T | C | 0.158 | -1.55E-01 | 1.19E-01 | 2.99E+00 | 3.12E+00 | 1.96E-01 | no | | intergenic_variant | MODIFIER |
| 4 | 54230435 | AX-170368396 | 1 | T | C | 0.357 | -1.26E-01 | 9.79E-02 | 2.95E+00 | 3.07E+00 | 1.96E-01 | no | | intergenic_variant | MODIFIER |
| 16 | 79078081 | AX-170291675 | 0 | A | G | 0.465 | 9.54E-02 | 7.43E-02 | 2.64E+00 | 2.72E+00 | 1.96E-01 | no | Tmprss15 | intron_variant | MODIFIER |
| 16 | 79230912 | AX-97796837 | 0 | A | G | 0.465 | 9.54E-02 | 7.43E-02 | 2.64E+00 | 2.72E+00 | 1.96E-01 | no | | intergenic_variant | MODIFIER |
| 1 | 139446585 | AX-97813380 | 0 | T | C | 0.202 | 1.69E-01 | 1.32E-01 | 2.68E+00 | 2.73E+00 | 1.96E-01 | Yes | Zbtb41 | intron_variant | MODIFIER |
| 3 | 19674052 | AX-170256834 | 0 | T | G | 0.123 | -1.31E-01 | 1.02E-01 | 2.61E+00 | 2.69E+00 | 1.96E-01 | no | Trim55 | intron_variant | MODIFIER |
| 5 | 46812613 | AX-170024852 | 1 | T | A | 0.5 | 1.29E-01 | 9.84E-02 | 2.43E+00 | 2.49E+00 | 1.96E-01 | no | | intergenic_variant | MODIFIER |
| 16 | 62081070 | AX-170013199 | 0 | T | C | 0.526 | 1.08E-01 | 8.29E-02 | 2.46E+00 | 2.52E+00 | 1.97E-01 | no | | intergenic_variant | MODIFIER |
| 1 | 124369486 | AX-170306978 | 1 | G | A | 0.571 | 1.04E-01 | 8.03E-02 | 2.51E+00 | 2.58E+00 | 1.97E-01 | no | | intergenic_variant | MODIFIER |
| 4 | 124494735 | AX-170070551 | 0 | C | T | 0.246 | 1.32E-01 | 1.02E-01 | 2.49E+00 | 2.56E+00 | 1.97E-01 | no | | intergenic_variant | MODIFIER |
| 5 | 45745128 | AX-170137955 | 1 | T | G | 0.518 | 1.28E-01 | 9.60E-02 | 2.28E+00 | 2.34E+00 | 1.97E-01 | no | Lcorl | intron_variant | MODIFIER |
| 1 | 179915665 | AX-170251936 | 0 | T | A | 0.246 | 1.36E-01 | 1.06E-01 | 2.61E+00 | 2.70E+00 | 1.97E-01 | no | | intergenic_variant | MODIFIER |
| 15 | 6171785 | AX-97817163 | 1 | A | C | 0.125 | -9.39E-02 | 6.11E-02 | 1.82E+00 | 1.91E+00 | 1.97E-01 | no | | intergenic_variant | MODIFIER |
| 13 | 23353621 | AX-170530921 | 0 | A | G | 0.167 | -1.37E-01 | 1.05E-01 | 2.41E+00 | 2.51E+00 | 1.97E-01 | no | Zfp322a | 3_prime_UTR_variant | MODIFIER |
| 20 | 45066963 | AX-170288994 | 1 | T | C | 0.241 | 7.56E-02 | 5.90E-02 | 2.59E+00 | 2.72E+00 | 1.97E-01 | no | | intergenic_variant | MODIFIER |
| 15 | 66198165 | AX-170011880 | 0 | C | T | 0.333 | -1.02E-01 | 7.56E-02 | 2.21E+00 | 2.27E+00 | 1.97E-01 | no | Kcnq3 | intron_variant | MODIFIER |
| 8 | 24871958 | AX-170479721 | 1 | A | G | 0.348 | -8.29E-02 | 6.45E-02 | 2.59E+00 | 2.68E+00 | 1.97E-01 | no | Adam32 | intron_variant | MODIFIER |
| 10 | 100282965 | AX-170332565 | 1 | A | T | 0.268 | -1.10E-01 | 8.57E-02 | 2.64E+00 | 2.70E+00 | 1.97E-01 | no | | intergenic_variant | MODIFIER |
| 5 | 59731800 | AX-170025063 | 0 | C | G | 0.465 | 1.31E-01 | 1.00E-01 | 2.41E+00 | 2.47E+00 | 1.98E-01 | no | | intergenic_variant | MODIFIER |
| 4 | 133866519 | AX-170373399 | 0 | T | C | 0.167 | 1.45E-01 | 1.13E-01 | 2.60E+00 | 2.66E+00 | 1.98E-01 | Yes | Rps6ka1 | intron_variant | MODIFIER |
| 13 | 117494695 | AX-97835911 | 2 | T | C | 0.045 | 1.75E-01 | 1.37E-01 | 2.89E+00 | 3.10E+00 | 1.98E-01 | Yes | | intergenic_variant | MODIFIER |
| 3 | 32120417 | AX-97833790 | 0 | T | C | 0.158 | -1.16E-01 | 9.06E-02 | 2.91E+00 | 3.05E+00 | 1.98E-01 | no | Kcnmb2 | intron_variant | MODIFIER |
| 9 | 8307298 | AX-170031767 | 0 | C | T | 0.491 | 1.08E-01 | 8.33E-02 | 2.45E+00 | 2.55E+00 | 1.98E-01 | no | | intergenic_variant | MODIFIER |
| 4 | 81020718 | AX-97798766 | 0 | G | A | 0.447 | -1.07E-01 | 8.35E-02 | 2.54E+00 | 2.59E+00 | 1.98E-01 | no | | intergenic_variant | MODIFIER |
| 4 | 81039316 | AX-170509189 | 0 | T | A | 0.447 | -1.07E-01 | 8.35E-02 | 2.54E+00 | 2.59E+00 | 1.98E-01 | no | | intergenic_variant | MODIFIER |
| 4 | 81040460 | AX-170309505 | 0 | T | C | 0.447 | -1.07E-01 | 8.35E-02 | 2.54E+00 | 2.59E+00 | 1.98E-01 | no | | intergenic_variant | MODIFIER |
| 4 | 81040806 | AX-170399185 | 0 | C | T | 0.447 | -1.07E-01 | 8.35E-02 | 2.54E+00 | 2.59E+00 | 1.98E-01 | no | | intergenic_variant | MODIFIER |
| 4 | 81040880 | AX-170069800 | 0 | T | C | 0.447 | -1.07E-01 | 8.35E-02 | 2.54E+00 | 2.59E+00 | 1.98E-01 | no | | intergenic_variant | MODIFIER |
| 4 | 81040943 | AX-170069802 | 0 | C | T | 0.447 | -1.07E-01 | 8.35E-02 | 2.54E+00 | 2.59E+00 | 1.98E-01 | no | | intergenic_variant | MODIFIER |
| 4 | 81076245 | AX-97848162 | 0 | G | A | 0.447 | -1.07E-01 | 8.35E-02 | 2.54E+00 | 2.59E+00 | 1.98E-01 | no | | intergenic_variant | MODIFIER |
| 4 | 81420906 | AX-170133732 | 0 | G | C | 0.447 | -1.07E-01 | 8.35E-02 | 2.54E+00 | 2.59E+00 | 1.98E-01 | no | Mpdz | intron_variant | MODIFIER |
| 4 | 81432497 | AX-170393871 | 0 | C | T | 0.447 | -1.07E-01 | 8.35E-02 | 2.54E+00 | 2.59E+00 | 1.98E-01 | no | Mpdz | intron_variant | MODIFIER |
| 4 | 86162433 | AX-170319087 | 0 | G | T | 0.342 | 1.43E-01 | 1.12E-01 | 2.72E+00 | 2.79E+00 | 1.98E-01 | no | Adamtsl1 | intron_variant | MODIFIER |
| 4 | 86169450 | AX-170308619 | 0 | G | C | 0.342 | 1.43E-01 | 1.12E-01 | 2.72E+00 | 2.79E+00 | 1.98E-01 | no | Adamtsl1 | intron_variant | MODIFIER |
| 4 | 86176671 | AX-170095040 | 0 | G | T | 0.342 | 1.43E-01 | 1.12E-01 | 2.72E+00 | 2.79E+00 | 1.98E-01 | no | Adamtsl1 | intron_variant | MODIFIER |
| 4 | 86181651 | AX-170160482 | 0 | A | C | 0.342 | 1.43E-01 | 1.12E-01 | 2.72E+00 | 2.79E+00 | 1.98E-01 | no | Adamtsl1 | intron_variant | MODIFIER |
| 4 | 86182843 | AX-170517090 | 0 | T | C | 0.342 | 1.43E-01 | 1.12E-01 | 2.72E+00 | 2.79E+00 | 1.98E-01 | no | Adamtsl1 | intron_variant | MODIFIER |
| 4 | 86183937 | AX-170268439 | 0 | G | T | 0.342 | 1.43E-01 | 1.12E-01 | 2.72E+00 | 2.79E+00 | 1.98E-01 | no | Adamtsl1 | intron_variant | MODIFIER |
| 4 | 86192645 | AX-97838584 | 0 | C | T | 0.342 | 1.43E-01 | 1.12E-01 | 2.72E+00 | 2.79E+00 | 1.98E-01 | no | Adamtsl1 | intron_variant | MODIFIER |

| | | | | | | | | | | | | | | | |
|---|---|---|---|---|---|---|---|---|---|---|---|---|---|---|---|
| 4 | 86192926 | AX-170532505 | 0 | C | T | 0.342 | 1.43E-01 | 1.12E-01 | 2.72E+00 | 2.79E+00 | 1.98E-01 | no | Adamtsl1 | intron_variant | MODIFIER |
| 4 | 86218977 | AX-170023312 | 0 | G | C | 0.342 | 1.43E-01 | 1.12E-01 | 2.72E+00 | 2.79E+00 | 1.98E-01 | no | Adamtsl1 | intron_variant | MODIFIER |
| 4 | 86221235 | AX-170023316 | 0 | A | G | 0.342 | 1.43E-01 | 1.12E-01 | 2.72E+00 | 2.79E+00 | 1.98E-01 | no | Adamtsl1 | intron_variant | MODIFIER |
| 4 | 86252599 | AX-170023310 | 0 | A | G | 0.342 | 1.43E-01 | 1.12E-01 | 2.72E+00 | 2.79E+00 | 1.98E-01 | no | Adamtsl1 | intron_variant | MODIFIER |
| 4 | 86281257 | AX-97790016 | 0 | T | G | 0.342 | 1.43E-01 | 1.12E-01 | 2.72E+00 | 2.79E+00 | 1.98E-01 | no | Adamtsl1 | intron_variant,non_coding_transcript_variant | MODIFIER |
| 4 | 86281428 | AX-170023311 | 0 | A | T | 0.342 | 1.43E-01 | 1.12E-01 | 2.72E+00 | 2.79E+00 | 1.98E-01 | no | Adamtsl1 | intron_variant,non_coding_transcript_variant | MODIFIER |
| 4 | 86301668 | AX-170023314 | 0 | C | T | 0.342 | 1.43E-01 | 1.12E-01 | 2.72E+00 | 2.79E+00 | 1.98E-01 | no | Adamtsl1 | intron_variant,non_coding_transcript_variant | MODIFIER |
| 4 | 86303421 | AX-170253452 | 0 | A | G | 0.342 | 1.43E-01 | 1.12E-01 | 2.72E+00 | 2.79E+00 | 1.98E-01 | no | Adamtsl1 | intron_variant,non_coding_transcript_variant | MODIFIER |
| 4 | 86306835 | AX-170023313 | 0 | G | A | 0.342 | 1.43E-01 | 1.12E-01 | 2.72E+00 | 2.79E+00 | 1.98E-01 | no | Adamtsl1 | intron_variant,non_coding_transcript_variant | MODIFIER |
| 4 | 86318490 | AX-170137548 | 0 | G | A | 0.342 | 1.43E-01 | 1.12E-01 | 2.72E+00 | 2.79E+00 | 1.98E-01 | no | Adamtsl1 | intron_variant,non_coding_transcript_variant | MODIFIER |
| 4 | 86324973 | AX-170366085 | 0 | A | G | 0.342 | 1.43E-01 | 1.12E-01 | 2.72E+00 | 2.79E+00 | 1.98E-01 | no | Adamtsl1 | intron_variant,non_coding_transcript_variant | MODIFIER |
| 4 | 86345356 | AX-170023307 | 0 | T | C | 0.342 | 1.43E-01 | 1.12E-01 | 2.72E+00 | 2.79E+00 | 1.98E-01 | no | Adamtsl1 | intron_variant,non_coding_transcript_variant | MODIFIER |
| 4 | 86346927 | AX-97818277 | 0 | C | T | 0.342 | 1.43E-01 | 1.12E-01 | 2.72E+00 | 2.79E+00 | 1.98E-01 | no | Adamtsl1 | intron_variant,non_coding_transcript_variant | MODIFIER |
| 4 | 86386256 | AX-97837623 | 0 | G | A | 0.342 | 1.43E-01 | 1.12E-01 | 2.72E+00 | 2.79E+00 | 1.98E-01 | no | Adamtsl1 | intron_variant,non_coding_transcript_variant | MODIFIER |
| 4 | 86390271 | AX-170458000 | 0 | T | C | 0.342 | 1.43E-01 | 1.12E-01 | 2.72E+00 | 2.79E+00 | 1.98E-01 | no | Adamtsl1 | intron_variant,non_coding_transcript_variant | MODIFIER |
| 4 | 86401290 | AX-170511529 | 0 | G | A | 0.342 | 1.43E-01 | 1.12E-01 | 2.72E+00 | 2.79E+00 | 1.98E-01 | no | Adamtsl1 | intron_variant,non_coding_transcript_variant | MODIFIER |
| 4 | 86450721 | AX-97808640 | 0 | G | A | 0.342 | 1.43E-01 | 1.12E-01 | 2.72E+00 | 2.79E+00 | 1.98E-01 | no | Saxo1 | intron_variant | MODIFIER |
| 4 | 86451739 | AX-170485538 | 0 | T | C | 0.342 | 1.43E-01 | 1.12E-01 | 2.72E+00 | 2.79E+00 | 1.98E-01 | no | Saxo1 | intron_variant | MODIFIER |
| 4 | 86466460 | AX-97823540 | 0 | A | G | 0.342 | 1.43E-01 | 1.12E-01 | 2.72E+00 | 2.79E+00 | 1.98E-01 | no | Saxo1 | intron_variant | MODIFIER |
| 4 | 86480133 | AX-170278513 | 0 | A | G | 0.342 | 1.43E-01 | 1.12E-01 | 2.72E+00 | 2.79E+00 | 1.98E-01 | no | Saxo1 | intron_variant | MODIFIER |
| 4 | 86488575 | AX-170461938 | 0 | G | A | 0.342 | 1.43E-01 | 1.12E-01 | 2.72E+00 | 2.79E+00 | 1.98E-01 | no | Saxo1 | intron_variant | MODIFIER |
| 4 | 86489414 | AX-97793725 | 0 | A | G | 0.342 | 1.43E-01 | 1.12E-01 | 2.72E+00 | 2.79E+00 | 1.98E-01 | no | Saxo1 | intron_variant | MODIFIER |
| 4 | 86496114 | AX-170183133 | 0 | T | C | 0.342 | 1.43E-01 | 1.12E-01 | 2.72E+00 | 2.79E+00 | 1.98E-01 | no | Saxo1 | intron_variant | MODIFIER |
| 4 | 86506638 | AX-170273744 | 0 | A | G | 0.342 | 1.43E-01 | 1.12E-01 | 2.72E+00 | 2.79E+00 | 1.98E-01 | no | Saxo1 | intron_variant | MODIFIER |
| 4 | 86518508 | AX-97794158 | 0 | A | G | 0.342 | 1.43E-01 | 1.12E-01 | 2.72E+00 | 2.79E+00 | 1.98E-01 | no | Saxo1 | intron_variant | MODIFIER |
| 4 | 86762058 | AX-170023323 | 0 | C | G | 0.342 | 1.43E-01 | 1.12E-01 | 2.72E+00 | 2.79E+00 | 1.98E-01 | no | Dennd4c | intron_variant | MODIFIER |
| 4 | 86777353 | AX-170203538 | 0 | T | C | 0.342 | 1.43E-01 | 1.12E-01 | 2.72E+00 | 2.79E+00 | 1.98E-01 | no | Dennd4c | intron_variant | MODIFIER |
| 4 | 86778040 | AX-97782035 | 0 | T | C | 0.342 | 1.43E-01 | 1.12E-01 | 2.72E+00 | 2.79E+00 | 1.98E-01 | no | Dennd4c | intron_variant | MODIFIER |
| 4 | 86802119 | AX-170250041 | 0 | C | T | 0.342 | 1.43E-01 | 1.12E-01 | 2.72E+00 | 2.79E+00 | 1.98E-01 | no | Dennd4c | intron_variant | MODIFIER |
| 4 | 86802389 | AX-170023329 | 0 | T | C | 0.342 | 1.43E-01 | 1.12E-01 | 2.72E+00 | 2.79E+00 | 1.98E-01 | no | Dennd4c | intron_variant | MODIFIER |
| 4 | 86822102 | AX-97823541 | 0 | G | A | 0.342 | 1.43E-01 | 1.12E-01 | 2.72E+00 | 2.79E+00 | 1.98E-01 | no | Dennd4c | intron_variant | MODIFIER |
| 4 | 86827247 | AX-170023327 | 0 | G | A | 0.342 | 1.43E-01 | 1.12E-01 | 2.72E+00 | 2.79E+00 | 1.98E-01 | no | Dennd4c | intron_variant | MODIFIER |
| 4 | 86834497 | AX-97794005 | 0 | G | A | 0.342 | 1.43E-01 | 1.12E-01 | 2.72E+00 | 2.79E+00 | 1.98E-01 | no | Dennd4c | downstream_gene_variant | MODIFIER |
| 4 | 86878534 | AX-170023328 | 0 | T | C | 0.342 | 1.43E-01 | 1.12E-01 | 2.72E+00 | 2.79E+00 | 1.98E-01 | no | Acer2 | intron_variant | MODIFIER |
| 4 | 86887466 | AX-97801018 | 0 | G | A | 0.342 | 1.43E-01 | 1.12E-01 | 2.72E+00 | 2.79E+00 | 1.98E-01 | no | Acer2 | intron_variant | MODIFIER |
| 4 | 86901817 | AX-170023322 | 0 | T | G | 0.342 | 1.43E-01 | 1.12E-01 | 2.72E+00 | 2.79E+00 | 1.98E-01 | no | Acer2 | intron_variant | MODIFIER |
| 4 | 86957783 | AX-170023321 | 0 | C | T | 0.342 | 1.43E-01 | 1.12E-01 | 2.72E+00 | 2.79E+00 | 1.98E-01 | no | | regulatory_region_variant | MODIFIER |
| 4 | 86987405 | AX-97788979 | 0 | T | C | 0.342 | 1.43E-01 | 1.12E-01 | 2.72E+00 | 2.79E+00 | 1.98E-01 | no | Slc24a2 | downstream_gene_variant | MODIFIER |
| 4 | 86997221 | AX-170023324 | 0 | G | A | 0.342 | 1.43E-01 | 1.12E-01 | 2.72E+00 | 2.79E+00 | 1.98E-01 | no | Slc24a2 | intron_variant | MODIFIER |
| 4 | 87002322 | AX-170315780 | 0 | A | C | 0.342 | 1.43E-01 | 1.12E-01 | 2.72E+00 | 2.79E+00 | 1.98E-01 | no | Slc24a2 | intron_variant | MODIFIER |
| 4 | 87023277 | AX-170135539 | 0 | T | C | 0.342 | 1.43E-01 | 1.12E-01 | 2.72E+00 | 2.79E+00 | 1.98E-01 | no | Slc24a2 | intron_variant | MODIFIER |
| 4 | 87025331 | AX-170454013 | 0 | C | T | 0.342 | 1.43E-01 | 1.12E-01 | 2.72E+00 | 2.79E+00 | 1.98E-01 | no | Slc24a2 | intron_variant | MODIFIER |
| 4 | 87027285 | AX-170162295 | 0 | A | G | 0.342 | 1.43E-01 | 1.12E-01 | 2.72E+00 | 2.79E+00 | 1.98E-01 | no | Slc24a2 | intron_variant | MODIFIER |
| 4 | 87168857 | AX-97849880 | 0 | C | T | 0.342 | 1.43E-01 | 1.12E-01 | 2.72E+00 | 2.79E+00 | 1.98E-01 | no | Slc24a2 | intron_variant | MODIFIER |
| 4 | 87168912 | AX-170267648 | 0 | C | T | 0.342 | 1.43E-01 | 1.12E-01 | 2.72E+00 | 2.79E+00 | 1.98E-01 | no | Slc24a2 | intron_variant | MODIFIER |
| 4 | 87177247 | AX-170069905 | 0 | T | C | 0.342 | 1.43E-01 | 1.12E-01 | 2.72E+00 | 2.79E+00 | 1.98E-01 | no | Slc24a2 | intron_variant | MODIFIER |
| 4 | 87179720 | AX-97797292 | 0 | T | C | 0.342 | 1.43E-01 | 1.12E-01 | 2.72E+00 | 2.79E+00 | 1.98E-01 | no | Slc24a2 | intron_variant | MODIFIER |
| 4 | 87287665 | AX-97797793 | 0 | G | A | 0.342 | 1.43E-01 | 1.12E-01 | 2.72E+00 | 2.79E+00 | 1.98E-01 | no | | intergenic_variant | MODIFIER |
| 4 | 87326605 | AX-97815812 | 0 | C | T | 0.342 | 1.43E-01 | 1.12E-01 | 2.72E+00 | 2.79E+00 | 1.98E-01 | no | | intergenic_variant | MODIFIER |
| 4 | 87775303 | AX-170503674 | 0 | G | A | 0.342 | 1.43E-01 | 1.12E-01 | 2.72E+00 | 2.79E+00 | 1.98E-01 | no | Mllt3 | intron_variant | MODIFIER |
| 4 | 87788950 | AX-170104184 | 0 | C | T | 0.342 | 1.43E-01 | 1.12E-01 | 2.72E+00 | 2.79E+00 | 1.98E-01 | no | Mllt3 | intron_variant | MODIFIER |
| 4 | 87789089 | AX-170522320 | 0 | T | G | 0.342 | 1.43E-01 | 1.12E-01 | 2.72E+00 | 2.79E+00 | 1.98E-01 | no | Mllt3 | intron_variant | MODIFIER |
| 4 | 87797512 | AX-170312587 | 0 | G | A | 0.342 | 1.43E-01 | 1.12E-01 | 2.72E+00 | 2.79E+00 | 1.98E-01 | no | Mllt3 | intron_variant | MODIFIER |
| 6 | 127788621 | AX-170244532 | 0 | A | T | 0.535 | 1.03E-01 | 8.02E-02 | 2.50E+00 | 2.57E+00 | 1.98E-01 | no | Mir7233 | upstream_gene_variant | MODIFIER |

| | | | | | | | | | | | | | | | |
|---|---|---|---|---|---|---|---|---|---|---|---|---|---|---|---|
| 10 | 13834962 | AX-170489843 | 0 | A | T | 0.193 | -1.37E-01 | 1.07E-01 | 2.81E+00 | 2.91E+00 | 1.99E-01 | no | Aig1 | intron_variant | MODIFIER |
| 8 | 68730935 | AX-170149605 | 0 | A | C | 0.184 | -1.44E-01 | 1.13E-01 | 2.77E+00 | 2.90E+00 | 1.99E-01 | no | Csgalnact1 | intron_variant | MODIFIER |
| 13 | 21028870 | AX-170050715 | 1 | A | G | 0.17 | -1.63E-01 | 1.26E-01 | 2.47E+00 | 2.52E+00 | 1.99E-01 | no | Aoah | downstream_gene_variant | MODIFIER |
| 18 | 57186776 | AX-170515232 | 0 | T | A | 0.193 | -1.23E-01 | 9.64E-02 | 2.75E+00 | 2.88E+00 | 1.99E-01 | no | Megf10 | intron_variant | MODIFIER |
| 1 | 127226571 | AX-170390029 | 0 | A | G | 0.561 | 1.07E-01 | 8.41E-02 | 2.67E+00 | 2.74E+00 | 1.99E-01 | no | Mgat5 | intron_variant | MODIFIER |
| 9 | 105641593 | AX-170207159 | 0 | T | C | 0.123 | -1.45E-01 | 1.14E-01 | 2.69E+00 | 2.78E+00 | 1.99E-01 | no | Pik3r4 | upstream_gene_variant | MODIFIER |
| 1 | 139675084 | AX-170327020 | 0 | T | A | 0.202 | 1.59E-01 | 1.25E-01 | 2.71E+00 | 2.79E+00 | 1.99E-01 | Yes | Gm26271 | downstream_gene_variant | MODIFIER |
| 3 | 22846689 | AX-170094549 | 0 | C | G | 0.158 | -1.33E-01 | 1.04E-01 | 2.59E+00 | 2.66E+00 | 1.99E-01 | no | | intergenic_variant | MODIFIER |
| 7 | 144423988 | AX-97782605 | 0 | A | G | 0.211 | -9.46E-02 | 7.45E-02 | 2.81E+00 | 2.97E+00 | 1.99E-01 | no | Shank2 | downstream_gene_variant | MODIFIER |
| 18 | 86238828 | AX-170528959 | 0 | T | C | 0.167 | -1.19E-01 | 9.27E-02 | 2.51E+00 | 2.64E+00 | 2.00E-01 | no | AC132351.1 | downstream_gene_variant | MODIFIER |
| 3 | 37927225 | AX-170161964 | 0 | C | A | 0.202 | 1.16E-01 | 9.04E-02 | 3.00E+00 | 3.13E+00 | 2.00E-01 | no | Gm20755 | intron_variant,non_coding_transcript_variant | MODIFIER |
| 1 | 191701210 | AX-170294755 | 0 | C | T | 0.272 | 1.20E-01 | 9.21E-02 | 2.40E+00 | 2.48E+00 | 2.00E-01 | no | | intergenic_variant | MODIFIER |
| 16 | 71959698 | AX-170351100 | 1 | T | A | 0.5 | 9.92E-02 | 7.69E-02 | 2.47E+00 | 2.54E+00 | 2.00E-01 | no | | intergenic_variant | MODIFIER |
| 6 | 130997660 | AX-97781998 | 1 | C | G | 0.562 | 1.08E-01 | 8.38E-02 | 2.53E+00 | 2.59E+00 | 2.00E-01 | no | Gm44133 | intron_variant,non_coding_transcript_variant | MODIFIER |
| 9 | 6359914 | AX-170144700 | 0 | C | T | 0.482 | 1.07E-01 | 8.25E-02 | 2.46E+00 | 2.55E+00 | 2.00E-01 | no | Pdgfd | synonymous_variant | LOW |
| 9 | 6361572 | AX-170455391 | 0 | C | T | 0.482 | 1.07E-01 | 8.25E-02 | 2.46E+00 | 2.55E+00 | 2.00E-01 | no | Pdgfd | intron_variant | MODIFIER |
| 9 | 6362053 | AX-170487064 | 0 | A | G | 0.482 | 1.07E-01 | 8.25E-02 | 2.46E+00 | 2.55E+00 | 2.00E-01 | no | Pdgfd | intron_variant | MODIFIER |
| 9 | 6400400 | AX-97847493 | 0 | T | C | 0.482 | 1.07E-01 | 8.25E-02 | 2.46E+00 | 2.55E+00 | 2.00E-01 | no | | intergenic_variant | MODIFIER |
| 9 | 6411782 | AX-170497991 | 0 | C | T | 0.482 | 1.07E-01 | 8.25E-02 | 2.46E+00 | 2.55E+00 | 2.00E-01 | no | | intergenic_variant | MODIFIER |
| 9 | 6414579 | AX-97805455 | 0 | C | T | 0.482 | 1.07E-01 | 8.25E-02 | 2.46E+00 | 2.55E+00 | 2.00E-01 | no | | intergenic_variant | MODIFIER |
| 9 | 6455720 | AX-170425593 | 0 | C | A | 0.482 | 1.07E-01 | 8.25E-02 | 2.46E+00 | 2.55E+00 | 2.00E-01 | no | | intergenic_variant | MODIFIER |
| 9 | 6483766 | AX-170031747 | 0 | C | T | 0.482 | 1.07E-01 | 8.25E-02 | 2.46E+00 | 2.55E+00 | 2.00E-01 | no | | intergenic_variant | MODIFIER |
| 9 | 6519359 | AX-170031740 | 0 | T | C | 0.482 | 1.07E-01 | 8.25E-02 | 2.46E+00 | 2.55E+00 | 2.00E-01 | no | | intergenic_variant | MODIFIER |
| 9 | 6562302 | AX-170031741 | 0 | C | T | 0.482 | 1.07E-01 | 8.25E-02 | 2.46E+00 | 2.55E+00 | 2.00E-01 | no | | intergenic_variant | MODIFIER |
| 9 | 6562488 | AX-170031745 | 0 | A | G | 0.482 | 1.07E-01 | 8.25E-02 | 2.46E+00 | 2.55E+00 | 2.00E-01 | no | | intergenic_variant | MODIFIER |
| 9 | 6562680 | AX-170031744 | 0 | A | C | 0.482 | 1.07E-01 | 8.25E-02 | 2.46E+00 | 2.55E+00 | 2.00E-01 | no | | intergenic_variant | MODIFIER |
| 9 | 6562852 | AX-170482109 | 0 | C | G | 0.482 | 1.07E-01 | 8.25E-02 | 2.46E+00 | 2.55E+00 | 2.00E-01 | no | | intergenic_variant | MODIFIER |
| 9 | 7000856 | AX-168404074 | 0 | A | G | 0.482 | 1.07E-01 | 8.25E-02 | 2.46E+00 | 2.55E+00 | 2.00E-01 | no | Dync2h1 | intron_variant | MODIFIER |
| 2 | 62712493 | AX-170206209 | 0 | G | A | 0.053 | -1.48E-01 | 1.16E-01 | 2.57E+00 | 2.75E+00 | 2.00E-01 | no | Kcnh7 | intron_variant | MODIFIER |
| 8 | 34639055 | AX-170384030 | 0 | C | T | 0.149 | -1.47E-01 | 1.16E-01 | 2.75E+00 | 2.87E+00 | 2.00E-01 | no | B930018H19Rik | intron_variant,non_coding_transcript_variant | MODIFIER |
| 4 | 137405161 | AX-170286949 | 2 | C | G | 0.173 | 1.44E-01 | 1.13E-01 | 2.69E+00 | 2.76E+00 | 2.00E-01 | Yes | Cela3a | intron_variant | MODIFIER |
| 20 | 38899709 | AX-170502191 | 0 | C | G | 0.228 | 7.45E-02 | 5.87E-02 | 2.65E+00 | 2.78E+00 | 2.00E-01 | no | 6030498E09Rik | intron_variant | MODIFIER |
| 1 | 121708276 | AX-97851113 | 1 | G | A | 0.58 | 1.06E-01 | 8.18E-02 | 2.45E+00 | 2.51E+00 | 2.00E-01 | no | Gm37920 | downstream_gene_variant | MODIFIER |
| 13 | 7654231 | AX-97814968 | 0 | G | A | 0.193 | -1.52E-01 | 1.18E-01 | 2.46E+00 | 2.52E+00 | 2.00E-01 | no | | intergenic_variant | MODIFIER |
| 13 | 9168668 | AX-170370857 | 0 | C | T | 0.193 | -1.52E-01 | 1.18E-01 | 2.46E+00 | 2.52E+00 | 2.00E-01 | no | Larp4b | intron_variant | MODIFIER |
| 13 | 9501407 | AX-170291428 | 0 | C | A | 0.193 | -1.52E-01 | 1.18E-01 | 2.46E+00 | 2.52E+00 | 2.00E-01 | no | Dip2c | intron_variant | MODIFIER |
| 16 | 63800267 | AX-170493919 | 1 | G | A | 0.491 | 1.08E-01 | 8.53E-02 | 2.69E+00 | 2.76E+00 | 2.00E-01 | no | Epha3 | intron_variant | MODIFIER |
| 5 | 63668062 | AX-170329296 | 1 | C | T | 0.473 | 1.26E-01 | 9.85E-02 | 2.59E+00 | 2.67E+00 | 2.00E-01 | no | Nwd2 | intron_variant | MODIFIER |
| 2 | 139015174 | AX-170208921 | 0 | C | A | 0.149 | 2.25E-01 | 1.77E-01 | 2.87E+00 | 2.97E+00 | 2.01E-01 | Yes | | intergenic_variant | MODIFIER |
| 8 | 16581128 | AX-170096179 | 1 | C | T | 0.232 | -9.70E-02 | 7.61E-02 | 2.58E+00 | 2.68E+00 | 2.01E-01 | no | Csmd1 | intron_variant | MODIFIER |
| 4 | 128910853 | AX-170136593 | 0 | C | A | 0.158 | 1.49E-01 | 1.16E-01 | 2.53E+00 | 2.57E+00 | 2.01E-01 | Yes | Trim62 | 3_prime_UTR_variant | MODIFIER |
| 4 | 128913305 | AX-170023926 | 0 | T | C | 0.158 | 1.49E-01 | 1.16E-01 | 2.53E+00 | 2.57E+00 | 2.01E-01 | Yes | Trim62 | downstream_gene_variant | MODIFIER |
| 4 | 128915275 | AX-170089368 | 0 | A | G | 0.158 | 1.49E-01 | 1.16E-01 | 2.53E+00 | 2.57E+00 | 2.01E-01 | Yes | Trim62 | downstream_gene_variant | MODIFIER |
| 4 | 128917867 | AX-170023923 | 0 | A | G | 0.158 | 1.49E-01 | 1.16E-01 | 2.53E+00 | 2.57E+00 | 2.01E-01 | Yes | | regulatory_region_variant | MODIFIER |
| 12 | 71383688 | AX-170490754 | 0 | A | G | 0.079 | -1.82E-01 | 1.44E-01 | 2.67E+00 | 2.81E+00 | 2.01E-01 | Yes | 1700083H02Rik | intron_variant,non_coding_transcript_variant | MODIFIER |
| 16 | 50314759 | AX-170088014 | 0 | C | T | 0.482 | -1.02E-01 | 7.96E-02 | 2.58E+00 | 2.63E+00 | 2.01E-01 | no | Bbx | intron_variant | MODIFIER |
| 7 | 37870729 | AX-170028850 | 1 | T | C | 0.348 | 1.22E-01 | 9.62E-02 | 2.62E+00 | 2.71E+00 | 2.01E-01 | no | | regulatory_region_variant | MODIFIER |
| 3 | 97933447 | AX-170067460 | 1 | A | C | 0.018 | -1.72E-01 | 1.34E-01 | 2.47E+00 | 2.62E+00 | 2.01E-01 | no | Gm5544 | intron_variant,non_coding_transcript_variant | MODIFIER |
| 4 | 88913374 | AX-170378745 | 0 | A | T | 0.421 | -1.02E-01 | 7.89E-02 | 2.40E+00 | 2.46E+00 | 2.01E-01 | no | Mir31 | upstream_gene_variant | MODIFIER |
| 1 | 150316664 | AX-170346003 | 1 | G | A | 0.214 | 1.48E-01 | 1.17E-01 | 2.71E+00 | 2.80E+00 | 2.01E-01 | Yes | Pdc | upstream_gene_variant | MODIFIER |
| 10 | 72383235 | AX-97815909 | 0 | G | A | 0.412 | -1.01E-01 | 7.95E-02 | 2.67E+00 | 2.74E+00 | 2.01E-01 | no | | intergenic_variant | MODIFIER |
| 16 | 50307797 | AX-170013005 | 1 | G | A | 0.482 | -1.01E-01 | 7.97E-02 | 2.58E+00 | 2.63E+00 | 2.02E-01 | no | Bbx | intron_variant | MODIFIER |
| 5 | 50004564 | AX-170393733 | 0 | G | C | 0.5 | 1.36E-01 | 1.05E-01 | 2.39E+00 | 2.43E+00 | 2.02E-01 | no | Adgra3 | intron_variant | MODIFIER |
| 15 | 72054235 | AX-170468513 | 0 | G | A | 0.105 | -1.35E-01 | 9.49E-02 | 2.01E+00 | 2.11E+00 | 2.02E-01 | no | | intergenic_variant | MODIFIER |

| | | | | | | | | | | | | | | | |
|---|---|---|---|---|---|---|---|---|---|---|---|---|---|---|---|
| 15 | 68560169 | AX-170182351 | 1 | T | C | 0.348 | -9.75E-02 | 7.40E-02 | 2.29E+00 | 2.38E+00 | 2.02E-01 | no | | intergenic_variant | MODIFIER |
| 4 | 124049032 | AX-170378489 | 0 | C | T | 0.237 | 1.36E-01 | 1.07E-01 | 2.60E+00 | 2.67E+00 | 2.02E-01 | no | | regulatory_region_variant | MODIFIER |
| 4 | 124051761 | AX-170378807 | 0 | C | T | 0.237 | 1.36E-01 | 1.07E-01 | 2.60E+00 | 2.67E+00 | 2.02E-01 | no | | intergenic_variant | MODIFIER |
| 4 | 124055410 | AX-170341073 | 0 | G | A | 0.237 | 1.36E-01 | 1.07E-01 | 2.60E+00 | 2.67E+00 | 2.02E-01 | no | | intergenic_variant | MODIFIER |
| 4 | 124065371 | AX-170346177 | 0 | G | A | 0.237 | 1.36E-01 | 1.07E-01 | 2.60E+00 | 2.67E+00 | 2.02E-01 | no | | intergenic_variant | MODIFIER |
| 4 | 124071866 | AX-170242506 | 0 | A | G | 0.237 | 1.36E-01 | 1.07E-01 | 2.60E+00 | 2.67E+00 | 2.02E-01 | no | | intergenic_variant | MODIFIER |
| 4 | 124281735 | AX-170406593 | 0 | G | A | 0.237 | 1.36E-01 | 1.07E-01 | 2.60E+00 | 2.67E+00 | 2.02E-01 | no | Gm37667 | upstream_gene_variant | MODIFIER |
| 4 | 124308002 | AX-170301144 | 0 | C | T | 0.237 | 1.36E-01 | 1.07E-01 | 2.60E+00 | 2.67E+00 | 2.02E-01 | no | 1700021L23Rik | intron_variant,non_coding_transcript_variant | MODIFIER |
| 4 | 124371993 | AX-170321411 | 0 | G | A | 0.237 | 1.36E-01 | 1.07E-01 | 2.60E+00 | 2.67E+00 | 2.02E-01 | no | | intergenic_variant | MODIFIER |
| 4 | 124377197 | AX-170099184 | 0 | C | A | 0.237 | 1.36E-01 | 1.07E-01 | 2.60E+00 | 2.67E+00 | 2.02E-01 | no | | intergenic_variant | MODIFIER |
| 4 | 124380648 | AX-170238960 | 0 | A | G | 0.237 | 1.36E-01 | 1.07E-01 | 2.60E+00 | 2.67E+00 | 2.02E-01 | no | | intergenic_variant | MODIFIER |
| 4 | 124380758 | AX-170070549 | 0 | T | G | 0.237 | 1.36E-01 | 1.07E-01 | 2.60E+00 | 2.67E+00 | 2.02E-01 | no | | intergenic_variant | MODIFIER |
| 4 | 124391363 | AX-97853218 | 0 | G | A | 0.237 | 1.36E-01 | 1.07E-01 | 2.60E+00 | 2.67E+00 | 2.02E-01 | no | | intergenic_variant | MODIFIER |
| 4 | 124398909 | AX-170475689 | 0 | C | T | 0.237 | 1.36E-01 | 1.07E-01 | 2.60E+00 | 2.67E+00 | 2.02E-01 | no | | regulatory_region_variant | MODIFIER |
| 4 | 124399042 | AX-170105949 | 0 | G | A | 0.237 | 1.36E-01 | 1.07E-01 | 2.60E+00 | 2.67E+00 | 2.02E-01 | no | | regulatory_region_variant | MODIFIER |
| 4 | 124407081 | AX-170217970 | 0 | G | A | 0.237 | 1.36E-01 | 1.07E-01 | 2.60E+00 | 2.67E+00 | 2.02E-01 | no | | intergenic_variant | MODIFIER |
| 4 | 124415260 | AX-170524305 | 0 | C | T | 0.237 | 1.36E-01 | 1.07E-01 | 2.60E+00 | 2.67E+00 | 2.02E-01 | no | 1700057H15Rik | intron_variant,non_coding_transcript_variant | MODIFIER |
| 4 | 124453987 | AX-170023868 | 0 | T | C | 0.237 | 1.36E-01 | 1.07E-01 | 2.60E+00 | 2.67E+00 | 2.02E-01 | no | 1700057H15Rik | upstream_gene_variant | MODIFIER |
| 4 | 124487567 | AX-170236349 | 0 | G | A | 0.237 | 1.36E-01 | 1.07E-01 | 2.60E+00 | 2.67E+00 | 2.02E-01 | no | 1700057H15Rik | upstream_gene_variant | MODIFIER |
| 4 | 124513838 | AX-170210256 | 0 | T | C | 0.237 | 1.36E-01 | 1.07E-01 | 2.60E+00 | 2.67E+00 | 2.02E-01 | no | | regulatory_region_variant | MODIFIER |
| 4 | 124517045 | AX-170463167 | 0 | C | G | 0.237 | 1.36E-01 | 1.07E-01 | 2.60E+00 | 2.67E+00 | 2.02E-01 | no | | intergenic_variant | MODIFIER |
| 4 | 124530432 | AX-170209057 | 0 | G | A | 0.237 | 1.36E-01 | 1.07E-01 | 2.60E+00 | 2.67E+00 | 2.02E-01 | no | Gm2164 | intron_variant,non_coding_transcript_variant | MODIFIER |
| 4 | 124537773 | AX-170311538 | 0 | G | A | 0.237 | 1.36E-01 | 1.07E-01 | 2.60E+00 | 2.67E+00 | 2.02E-01 | no | Gm2164 | intron_variant,non_coding_transcript_variant | MODIFIER |
| 4 | 124538676 | AX-170346070 | 0 | C | G | 0.237 | 1.36E-01 | 1.07E-01 | 2.60E+00 | 2.67E+00 | 2.02E-01 | no | Gm2164 | intron_variant,non_coding_transcript_variant | MODIFIER |
| 4 | 124545366 | AX-170023867 | 0 | G | C | 0.237 | 1.36E-01 | 1.07E-01 | 2.60E+00 | 2.67E+00 | 2.02E-01 | no | Gm2164 | intron_variant,non_coding_transcript_variant | MODIFIER |
| 4 | 124607683 | AX-170381034 | 0 | C | T | 0.237 | 1.36E-01 | 1.07E-01 | 2.60E+00 | 2.67E+00 | 2.02E-01 | no | 3100002H09Rik | downstream_gene_variant | MODIFIER |
| 4 | 124634462 | AX-170023873 | 0 | A | T | 0.237 | 1.36E-01 | 1.07E-01 | 2.60E+00 | 2.67E+00 | 2.02E-01 | no | Gm2164 | intron_variant,non_coding_transcript_variant | MODIFIER |
| 4 | 124643055 | AX-170241944 | 0 | A | C | 0.237 | 1.36E-01 | 1.07E-01 | 2.60E+00 | 2.67E+00 | 2.02E-01 | no | Gm2164 | intron_variant,non_coding_transcript_variant | MODIFIER |
| 4 | 124649619 | AX-170140984 | 0 | G | A | 0.237 | 1.36E-01 | 1.07E-01 | 2.60E+00 | 2.67E+00 | 2.02E-01 | no | Gm2164 | intron_variant,non_coding_transcript_variant | MODIFIER |
| 4 | 124680159 | AX-170267012 | 0 | G | A | 0.237 | 1.36E-01 | 1.07E-01 | 2.60E+00 | 2.67E+00 | 2.02E-01 | no | Utp11 | intron_variant | MODIFIER |
| 4 | 124693674 | AX-170364323 | 0 | G | A | 0.237 | 1.36E-01 | 1.07E-01 | 2.60E+00 | 2.67E+00 | 2.02E-01 | no | Utp11 | upstream_gene_variant | MODIFIER |
| 4 | 124733669 | AX-170482512 | 0 | A | G | 0.237 | 1.36E-01 | 1.07E-01 | 2.60E+00 | 2.67E+00 | 2.02E-01 | no | Sf3a3 | downstream_gene_variant | MODIFIER |
| 4 | 124757683 | AX-170275372 | 0 | T | C | 0.237 | 1.36E-01 | 1.07E-01 | 2.60E+00 | 2.67E+00 | 2.02E-01 | no | Inpp5b | intron_variant | MODIFIER |
| 4 | 124888014 | AX-170199852 | 0 | T | A | 0.237 | 1.36E-01 | 1.07E-01 | 2.60E+00 | 2.67E+00 | 2.02E-01 | no | Epha10 | downstream_gene_variant | MODIFIER |
| 4 | 124914331 | AX-97857074 | 0 | G | T | 0.237 | 1.36E-01 | 1.07E-01 | 2.60E+00 | 2.67E+00 | 2.02E-01 | no | Cdca8 | downstream_gene_variant | MODIFIER |
| 4 | 124921700 | AX-170228333 | 0 | C | T | 0.237 | 1.36E-01 | 1.07E-01 | 2.60E+00 | 2.67E+00 | 2.02E-01 | no | Cdca8 | intron_variant | MODIFIER |
| 4 | 125198655 | AX-170314365 | 0 | G | A | 0.237 | 1.36E-01 | 1.07E-01 | 2.60E+00 | 2.67E+00 | 2.02E-01 | no | | intergenic_variant | MODIFIER |
| 4 | 125199946 | AX-170491295 | 0 | A | C | 0.237 | 1.36E-01 | 1.07E-01 | 2.60E+00 | 2.67E+00 | 2.02E-01 | no | | intergenic_variant | MODIFIER |
| 4 | 125206579 | AX-97794036 | 0 | G | A | 0.237 | 1.36E-01 | 1.07E-01 | 2.60E+00 | 2.67E+00 | 2.02E-01 | no | | intergenic_variant | MODIFIER |
| 4 | 125236064 | AX-170391975 | 0 | T | C | 0.237 | 1.36E-01 | 1.07E-01 | 2.60E+00 | 2.67E+00 | 2.02E-01 | no | 1700041M05Rik | intron_variant,non_coding_transcript_variant | MODIFIER |
| 4 | 125236716 | AX-97818211 | 0 | C | T | 0.237 | 1.36E-01 | 1.07E-01 | 2.60E+00 | 2.67E+00 | 2.02E-01 | no | 1700041M05Rik | intron_variant,non_coding_transcript_variant | MODIFIER |
| 4 | 125302635 | AX-170478634 | 0 | G | A | 0.237 | 1.36E-01 | 1.07E-01 | 2.60E+00 | 2.67E+00 | 2.02E-01 | no | | intergenic_variant | MODIFIER |
| 4 | 125312826 | AX-170523921 | 0 | C | T | 0.237 | 1.36E-01 | 1.07E-01 | 2.60E+00 | 2.67E+00 | 2.02E-01 | no | | intergenic_variant | MODIFIER |
| 4 | 125361609 | AX-170516503 | 0 | T | C | 0.237 | 1.36E-01 | 1.07E-01 | 2.60E+00 | 2.67E+00 | 2.02E-01 | no | | intergenic_variant | MODIFIER |
| 4 | 125363610 | AX-170268013 | 0 | G | A | 0.237 | 1.36E-01 | 1.07E-01 | 2.60E+00 | 2.67E+00 | 2.02E-01 | no | | intergenic_variant | MODIFIER |
| 4 | 125364753 | AX-170380710 | 0 | C | T | 0.237 | 1.36E-01 | 1.07E-01 | 2.60E+00 | 2.67E+00 | 2.02E-01 | no | | intergenic_variant | MODIFIER |
| 4 | 125470300 | AX-170476893 | 0 | C | G | 0.237 | 1.36E-01 | 1.07E-01 | 2.60E+00 | 2.67E+00 | 2.02E-01 | no | | intergenic_variant | MODIFIER |
| 4 | 125470333 | AX-170473242 | 0 | T | C | 0.237 | 1.36E-01 | 1.07E-01 | 2.60E+00 | 2.67E+00 | 2.02E-01 | no | | intergenic_variant | MODIFIER |
| 4 | 125470385 | AX-170222752 | 0 | T | C | 0.237 | 1.36E-01 | 1.07E-01 | 2.60E+00 | 2.67E+00 | 2.02E-01 | no | | intergenic_variant | MODIFIER |
| 4 | 125531486 | AX-170517434 | 0 | G | A | 0.237 | 1.36E-01 | 1.07E-01 | 2.60E+00 | 2.67E+00 | 2.02E-01 | no | Grik3 | intron_variant | MODIFIER |
| 4 | 125544025 | AX-170400368 | 0 | C | T | 0.237 | 1.36E-01 | 1.07E-01 | 2.60E+00 | 2.67E+00 | 2.02E-01 | no | Grik3 | intron_variant | MODIFIER |
| 4 | 125551664 | AX-170070575 | 0 | G | T | 0.237 | 1.36E-01 | 1.07E-01 | 2.60E+00 | 2.67E+00 | 2.02E-01 | no | Grik3 | intron_variant | MODIFIER |
| 4 | 125557662 | AX-170296827 | 0 | A | G | 0.237 | 1.36E-01 | 1.07E-01 | 2.60E+00 | 2.67E+00 | 2.02E-01 | no | Grik3 | intron_variant | MODIFIER |
| 4 | 125561631 | AX-170150178 | 0 | T | C | 0.237 | 1.36E-01 | 1.07E-01 | 2.60E+00 | 2.67E+00 | 2.02E-01 | no | Grik3 | intron_variant | MODIFIER |

| | | | | | | | | | | | | | | | |
|---|---|---|---|---|---|---|---|---|---|---|---|---|---|---|---|
| 4 | 125562850 | AX-97848731 | 0 | G | T | 0.237 | 1.36E-01 | 1.07E-01 | 2.60E+00 | 2.67E+00 | 2.02E-01 | no | Grik3 | intron_variant | MODIFIER |
| 4 | 125596356 | AX-170374171 | 0 | G | T | 0.237 | 1.36E-01 | 1.07E-01 | 2.60E+00 | 2.67E+00 | 2.02E-01 | no | Grik3 | intron_variant | MODIFIER |
| 4 | 125599891 | AX-170318901 | 0 | G | T | 0.237 | 1.36E-01 | 1.07E-01 | 2.60E+00 | 2.67E+00 | 2.02E-01 | no | Grik3 | intron_variant | MODIFIER |
| 11 | 11557231 | AX-170004286 | 0 | C | T | 0.044 | -2.39E-01 | 1.90E-01 | 2.76E+00 | 2.92E+00 | 2.02E-01 | Yes | Gm11998 | upstream_gene_variant | MODIFIER |
| 9 | 17118184 | AX-97824323 | 1 | C | T | 0.491 | 1.10E-01 | 8.65E-02 | 2.58E+00 | 2.66E+00 | 2.02E-01 | no | | intergenic_variant | MODIFIER |
| 16 | 39973285 | AX-170132803 | 0 | A | T | 0.5 | 1.04E-01 | 8.13E-02 | 2.49E+00 | 2.54E+00 | 2.02E-01 | no | | intergenic_variant | MODIFIER |
| 20 | 46055652 | AX-97854623 | 0 | G | A | 0.237 | 7.44E-02 | 5.89E-02 | 2.67E+00 | 2.80E+00 | 2.02E-01 | no | | intergenic_variant | MODIFIER |
| 4 | 107639324 | AX-97792821 | 0 | A | G | 0.263 | 1.42E-01 | 1.12E-01 | 2.65E+00 | 2.71E+00 | 2.02E-01 | no | Glis1 | downstream_gene_variant | MODIFIER |
| 2 | 169451668 | AX-170535030 | 1 | A | T | 0.17 | 1.94E-01 | 1.54E-01 | 2.68E+00 | 2.82E+00 | 2.02E-01 | Yes | | intergenic_variant | MODIFIER |
| 15 | 25490336 | AX-170535100 | 0 | G | A | 0.237 | -1.33E-01 | 1.05E-01 | 2.68E+00 | 2.75E+00 | 2.02E-01 | no | | intergenic_variant | MODIFIER |
| 18 | 75946394 | AX-170500998 | 1 | T | A | 0.402 | -1.10E-01 | 8.73E-02 | 2.77E+00 | 2.85E+00 | 2.02E-01 | no | Zbtb7c | intron_variant | MODIFIER |
| 18 | 76341914 | AX-170061363 | 1 | T | C | 0.402 | -1.10E-01 | 8.73E-02 | 2.77E+00 | 2.85E+00 | 2.02E-01 | no | | intergenic_variant | MODIFIER |
| 8 | 25464202 | AX-170079552 | 1 | A | C | 0.357 | -8.21E-02 | 6.47E-02 | 2.58E+00 | 2.67E+00 | 2.03E-01 | no | Gm31045 | intron_variant,non_coding_transcript_variant | MODIFIER |
| 9 | 17002002 | AX-170031857 | 2 | T | A | 0.491 | 1.11E-01 | 8.77E-02 | 2.58E+00 | 2.65E+00 | 2.03E-01 | no | | intergenic_variant | MODIFIER |
| 16 | 81079196 | AX-97825008 | 1 | T | C | 0.455 | 9.27E-02 | 7.32E-02 | 2.63E+00 | 2.72E+00 | 2.03E-01 | no | | intergenic_variant | MODIFIER |
| 9 | 8830335 | AX-170189725 | 0 | T | C | 0.456 | 9.90E-02 | 7.71E-02 | 2.44E+00 | 2.53E+00 | 2.03E-01 | no | | intergenic_variant | MODIFIER |
| 8 | 34635303 | AX-170319818 | 0 | C | T | 0.14 | -1.61E-01 | 1.26E-01 | 2.55E+00 | 2.62E+00 | 2.03E-01 | no | B930018H19Rik | intron_variant,non_coding_transcript_variant | MODIFIER |
| 13 | 38773557 | AX-170098161 | 1 | C | T | 0.17 | -1.29E-01 | 1.01E-01 | 2.45E+00 | 2.57E+00 | 2.03E-01 | no | Gm46392 | intron_variant,non_coding_transcript_variant | MODIFIER |
| 10 | 67749693 | AX-97798672 | 0 | A | G | 0.412 | -9.90E-02 | 7.85E-02 | 2.71E+00 | 2.80E+00 | 2.04E-01 | no | | intergenic_variant | MODIFIER |
| 9 | 9091122 | AX-97837758 | 1 | A | G | 0.473 | 1.02E-01 | 8.00E-02 | 2.46E+00 | 2.56E+00 | 2.04E-01 | no | Arhgap42 | intron_variant | MODIFIER |
| 4 | 54496601 | AX-97780625 | 0 | T | C | 0.368 | -1.23E-01 | 9.70E-02 | 2.93E+00 | 3.05E+00 | 2.04E-01 | no | | intergenic_variant | MODIFIER |
| 15 | 5595013 | AX-97828024 | 2 | T | G | 0.236 | -1.32E-01 | 1.05E-01 | 2.69E+00 | 2.75E+00 | 2.04E-01 | no | 5430437J10Rik | downstream_gene_variant | MODIFIER |
| 20 | 45251751 | AX-170033767 | 2 | A | G | 0.227 | 7.44E-02 | 5.90E-02 | 2.63E+00 | 2.76E+00 | 2.04E-01 | no | | intergenic_variant | MODIFIER |
| 14 | 112118707 | AX-170093117 | 1 | T | A | 0.312 | 9.04E-02 | 7.15E-02 | 2.58E+00 | 2.69E+00 | 2.04E-01 | no | | intergenic_variant | MODIFIER |
| 15 | 66873122 | AX-170224027 | 0 | A | C | 0.351 | -9.29E-02 | 7.17E-02 | 2.37E+00 | 2.44E+00 | 2.04E-01 | no | | intergenic_variant | MODIFIER |
| 4 | 100581716 | AX-170335510 | 0 | C | A | 0.237 | 1.45E-01 | 1.14E-01 | 2.58E+00 | 2.65E+00 | 2.04E-01 | Yes | Gm12700 | non_coding_transcript_exon_variant | MODIFIER |
| 15 | 5939353 | AX-170227271 | 0 | G | A | 0.325 | -1.02E-01 | 7.45E-02 | 2.11E+00 | 2.17E+00 | 2.04E-01 | no | | intergenic_variant | MODIFIER |
| 13 | 11487479 | AX-170349018 | 0 | G | A | 0.184 | -1.59E-01 | 1.25E-01 | 2.54E+00 | 2.59E+00 | 2.04E-01 | no | | intergenic_variant | MODIFIER |
| 5 | 129356854 | AX-170474429 | 0 | G | A | 0.158 | -1.17E-01 | 8.69E-02 | 2.19E+00 | 2.26E+00 | 2.05E-01 | no | | intergenic_variant | MODIFIER |
| 18 | 74370187 | AX-170218263 | 1 | A | G | 0.17 | -1.54E-01 | 1.23E-01 | 2.68E+00 | 2.74E+00 | 2.05E-01 | no | | intergenic_variant | MODIFIER |
| 10 | 100555437 | AX-97825483 | 1 | C | A | 0.277 | -1.08E-01 | 8.55E-02 | 2.64E+00 | 2.71E+00 | 2.05E-01 | no | Cep290 | intron_variant | MODIFIER |
| 13 | 5622181 | AX-170007674 | 1 | A | C | 0.196 | -1.45E-01 | 1.14E-01 | 2.52E+00 | 2.58E+00 | 2.05E-01 | no | Gm35330 | intron_variant,non_coding_transcript_variant | MODIFIER |
| 1 | 171425719 | AX-170343824 | 2 | T | C | 0.218 | 1.67E-01 | 1.32E-01 | 2.61E+00 | 2.67E+00 | 2.05E-01 | Yes | Gm26641 | intron_variant,non_coding_transcript_variant | MODIFIER |
| 2 | 101135043 | AX-170263852 | 1 | A | G | 0.054 | -2.72E-01 | 2.16E-01 | 2.70E+00 | 2.79E+00 | 2.05E-01 | Yes | | intergenic_variant | MODIFIER |
| 9 | 8500104 | AX-170117725 | 1 | C | A | 0.464 | 1.02E-01 | 8.00E-02 | 2.44E+00 | 2.53E+00 | 2.05E-01 | no | | intergenic_variant | MODIFIER |
| 4 | 86195157 | AX-170309571 | 0 | T | C | 0.333 | 1.38E-01 | 1.10E-01 | 2.71E+00 | 2.78E+00 | 2.05E-01 | no | Adamtsl1 | intron_variant | MODIFIER |
| 4 | 86195499 | AX-170512218 | 0 | A | G | 0.333 | 1.38E-01 | 1.10E-01 | 2.71E+00 | 2.78E+00 | 2.05E-01 | no | Adamtsl1 | intron_variant | MODIFIER |
| 4 | 86201020 | AX-170366606 | 0 | C | T | 0.333 | 1.38E-01 | 1.10E-01 | 2.71E+00 | 2.78E+00 | 2.05E-01 | no | Adamtsl1 | intron_variant | MODIFIER |
| 4 | 86201525 | AX-170138654 | 0 | C | A | 0.333 | 1.38E-01 | 1.10E-01 | 2.71E+00 | 2.78E+00 | 2.05E-01 | no | Adamtsl1 | intron_variant | MODIFIER |
| 4 | 86204780 | AX-170408419 | 0 | T | C | 0.333 | 1.38E-01 | 1.10E-01 | 2.71E+00 | 2.78E+00 | 2.05E-01 | no | Adamtsl1 | intron_variant | MODIFIER |
| 4 | 86328817 | AX-170390212 | 0 | T | C | 0.333 | 1.38E-01 | 1.10E-01 | 2.71E+00 | 2.78E+00 | 2.05E-01 | no | Adamtsl1 | intron_variant,non_coding_transcript_variant | MODIFIER |
| 4 | 86366918 | AX-170459868 | 0 | G | A | 0.333 | 1.38E-01 | 1.10E-01 | 2.71E+00 | 2.78E+00 | 2.05E-01 | no | Adamtsl1 | intron_variant,non_coding_transcript_variant | MODIFIER |
| 4 | 86370016 | AX-97816084 | 0 | T | A | 0.333 | 1.38E-01 | 1.10E-01 | 2.71E+00 | 2.78E+00 | 2.05E-01 | no | Adamtsl1 | intron_variant,non_coding_transcript_variant | MODIFIER |
| 4 | 86370680 | AX-170393151 | 0 | C | T | 0.333 | 1.38E-01 | 1.10E-01 | 2.71E+00 | 2.78E+00 | 2.05E-01 | no | Adamtsl1 | intron_variant,non_coding_transcript_variant | MODIFIER |
| 4 | 86423848 | AX-170432773 | 0 | G | C | 0.333 | 1.38E-01 | 1.10E-01 | 2.71E+00 | 2.78E+00 | 2.05E-01 | no | Adamtsl1 | intron_variant,non_coding_transcript_variant | MODIFIER |
| 4 | 86560972 | AX-170332577 | 0 | A | G | 0.333 | 1.38E-01 | 1.10E-01 | 2.71E+00 | 2.78E+00 | 2.05E-01 | no | Saxo1 | upstream_gene_variant | MODIFIER |
| 4 | 86595736 | AX-170412545 | 0 | A | C | 0.333 | 1.38E-01 | 1.10E-01 | 2.71E+00 | 2.78E+00 | 2.05E-01 | no | Haus6 | intron_variant | MODIFIER |
| 4 | 86888623 | AX-170214053 | 0 | G | A | 0.333 | 1.38E-01 | 1.10E-01 | 2.71E+00 | 2.78E+00 | 2.05E-01 | no | Acer2 | intron_variant | MODIFIER |
| 4 | 86892371 | AX-170461590 | 0 | G | A | 0.333 | 1.38E-01 | 1.10E-01 | 2.71E+00 | 2.78E+00 | 2.05E-01 | no | Acer2 | intron_variant | MODIFIER |
| 4 | 87106198 | AX-170172072 | 0 | T | C | 0.333 | 1.38E-01 | 1.10E-01 | 2.71E+00 | 2.78E+00 | 2.05E-01 | no | Slc24a2 | intron_variant | MODIFIER |
| 4 | 87117402 | AX-170154804 | 0 | A | T | 0.333 | 1.38E-01 | 1.10E-01 | 2.71E+00 | 2.78E+00 | 2.05E-01 | no | Slc24a2 | intron_variant | MODIFIER |
| 4 | 87143055 | AX-97804032 | 0 | T | C | 0.333 | 1.38E-01 | 1.10E-01 | 2.71E+00 | 2.78E+00 | 2.05E-01 | no | Slc24a2 | intron_variant | MODIFIER |
| 4 | 87143478 | AX-170438740 | 0 | C | G | 0.333 | 1.38E-01 | 1.10E-01 | 2.71E+00 | 2.78E+00 | 2.05E-01 | no | Slc24a2 | intron_variant | MODIFIER |
| 4 | 87401438 | AX-170069910 | 0 | G | T | 0.333 | 1.38E-01 | 1.10E-01 | 2.71E+00 | 2.78E+00 | 2.05E-01 | no | | regulatory_region_variant | MODIFIER |

| | | | | | | | | | | | | | | | |
|---|---|---|---|---|---|---|---|---|---|---|---|---|---|---|---|
| 4 | 87648465 | AX-170343559 | 0 | A | G | 0.333 | 1.38E-01 | 1.10E-01 | 2.71E+00 | 2.78E+00 | 2.05E-01 | no | | regulatory_region_variant | MODIFIER |
| 4 | 87673412 | AX-170518232 | 0 | C | T | 0.333 | 1.38E-01 | 1.10E-01 | 2.71E+00 | 2.78E+00 | 2.05E-01 | no | | intergenic_variant | MODIFIER |
| 4 | 87673571 | AX-170434363 | 0 | C | T | 0.333 | 1.38E-01 | 1.10E-01 | 2.71E+00 | 2.78E+00 | 2.05E-01 | no | | intergenic_variant | MODIFIER |
| 4 | 87748076 | AX-170418563 | 0 | C | A | 0.333 | 1.38E-01 | 1.10E-01 | 2.71E+00 | 2.78E+00 | 2.05E-01 | no | | intergenic_variant | MODIFIER |
| 4 | 87839347 | AX-170238195 | 0 | C | T | 0.333 | 1.38E-01 | 1.10E-01 | 2.71E+00 | 2.78E+00 | 2.05E-01 | no | Mllt3 | intron_variant | MODIFIER |
| 12 | 64027169 | AX-170168688 | 1 | A | T | 0.054 | 2.93E-01 | 2.33E-01 | 2.88E+00 | 3.03E+00 | 2.05E-01 | Yes | | intergenic_variant | MODIFIER |
| 6 | 126507993 | AX-97822412 | 1 | A | G | 0.562 | 1.04E-01 | 8.24E-02 | 2.53E+00 | 2.60E+00 | 2.06E-01 | no | | intergenic_variant | MODIFIER |
| 6 | 129381949 | AX-170315243 | 1 | G | A | 0.562 | 1.04E-01 | 8.24E-02 | 2.53E+00 | 2.60E+00 | 2.06E-01 | no | Clec12b | intron_variant | MODIFIER |
| 1 | 16239039 | AX-170299971 | 0 | T | C | 0.377 | 9.46E-02 | 7.49E-02 | 2.55E+00 | 2.63E+00 | 2.06E-01 | no | Stau2 | intron_variant | MODIFIER |
| 4 | 124629965 | AX-97838185 | 2 | G | T | 0.236 | 1.34E-01 | 1.06E-01 | 2.62E+00 | 2.69E+00 | 2.06E-01 | no | Gm2164 | intron_variant,non_coding_transcript_variant | MODIFIER |
| 16 | 62691269 | AX-170198633 | 0 | T | C | 0.474 | -1.09E-01 | 8.57E-02 | 2.51E+00 | 2.55E+00 | 2.06E-01 | no | | intergenic_variant | MODIFIER |
| 10 | 93776314 | AX-170240180 | 0 | A | G | 0.281 | -1.11E-01 | 8.79E-02 | 2.54E+00 | 2.59E+00 | 2.06E-01 | no | Gm3571 | upstream_gene_variant | MODIFIER |
| 11 | 8075332 | AX-170397252 | 0 | T | C | 0.053 | 2.31E-01 | 1.85E-01 | 2.82E+00 | 2.99E+00 | 2.06E-01 | Yes | | intergenic_variant | MODIFIER |
| 10 | 99868476 | AX-170254684 | 0 | T | C | 0.289 | -1.07E-01 | 8.53E-02 | 2.61E+00 | 2.67E+00 | 2.06E-01 | no | Gm47579 | upstream_gene_variant | MODIFIER |
| 10 | 99868542 | AX-170214284 | 0 | C | A | 0.289 | -1.07E-01 | 8.53E-02 | 2.61E+00 | 2.67E+00 | 2.06E-01 | no | Gm47579 | upstream_gene_variant | MODIFIER |
| 10 | 99897341 | AX-170163630 | 0 | G | A | 0.289 | -1.07E-01 | 8.53E-02 | 2.61E+00 | 2.67E+00 | 2.06E-01 | no | | intergenic_variant | MODIFIER |
| 10 | 99901914 | AX-170253145 | 0 | A | G | 0.289 | -1.07E-01 | 8.53E-02 | 2.61E+00 | 2.67E+00 | 2.06E-01 | no | | intergenic_variant | MODIFIER |
| 10 | 99907156 | AX-168409232 | 0 | T | G | 0.289 | -1.07E-01 | 8.53E-02 | 2.61E+00 | 2.67E+00 | 2.06E-01 | no | | intergenic_variant | MODIFIER |
| 10 | 99948872 | AX-170390997 | 0 | C | T | 0.289 | -1.07E-01 | 8.53E-02 | 2.61E+00 | 2.67E+00 | 2.06E-01 | no | | regulatory_region_variant | MODIFIER |
| 10 | 99950036 | AX-170373084 | 0 | C | T | 0.289 | -1.07E-01 | 8.53E-02 | 2.61E+00 | 2.67E+00 | 2.06E-01 | no | | regulatory_region_variant | MODIFIER |
| 10 | 100667217 | AX-170424320 | 0 | C | T | 0.289 | -1.07E-01 | 8.53E-02 | 2.61E+00 | 2.67E+00 | 2.06E-01 | no | | intergenic_variant | MODIFIER |
| 10 | 100667337 | AX-170158297 | 0 | G | A | 0.289 | -1.07E-01 | 8.53E-02 | 2.61E+00 | 2.67E+00 | 2.06E-01 | no | | intergenic_variant | MODIFIER |
| 10 | 100681948 | AX-170092077 | 0 | A | G | 0.289 | -1.07E-01 | 8.53E-02 | 2.61E+00 | 2.67E+00 | 2.06E-01 | no | | intergenic_variant | MODIFIER |
| 10 | 100704930 | AX-170449862 | 0 | A | G | 0.289 | -1.07E-01 | 8.53E-02 | 2.61E+00 | 2.67E+00 | 2.06E-01 | no | Gm35722 | downstream_gene_variant | MODIFIER |
| 10 | 100708148 | AX-170086883 | 0 | T | G | 0.289 | -1.07E-01 | 8.53E-02 | 2.61E+00 | 2.67E+00 | 2.06E-01 | no | Gm35722 | non_coding_transcript_exon_variant | MODIFIER |
| 10 | 100708431 | AX-170097432 | 0 | C | T | 0.289 | -1.07E-01 | 8.53E-02 | 2.61E+00 | 2.67E+00 | 2.06E-01 | no | Gm35722 | non_coding_transcript_exon_variant | MODIFIER |
| 10 | 100777698 | AX-170155357 | 0 | G | C | 0.289 | -1.07E-01 | 8.53E-02 | 2.61E+00 | 2.67E+00 | 2.06E-01 | no | | intergenic_variant | MODIFIER |
| 10 | 100778473 | AX-170003767 | 0 | G | T | 0.289 | -1.07E-01 | 8.53E-02 | 2.61E+00 | 2.67E+00 | 2.06E-01 | no | | intergenic_variant | MODIFIER |
| 10 | 100778818 | AX-97816555 | 0 | A | G | 0.289 | -1.07E-01 | 8.53E-02 | 2.61E+00 | 2.67E+00 | 2.06E-01 | no | | intergenic_variant | MODIFIER |
| 10 | 100819046 | AX-170291656 | 0 | A | C | 0.289 | -1.07E-01 | 8.53E-02 | 2.61E+00 | 2.67E+00 | 2.06E-01 | no | | intergenic_variant | MODIFIER |
| 10 | 100916579 | AX-170003763 | 0 | C | T | 0.289 | -1.07E-01 | 8.53E-02 | 2.61E+00 | 2.67E+00 | 2.06E-01 | no | | intergenic_variant | MODIFIER |
| 10 | 100916929 | AX-97798833 | 0 | G | A | 0.289 | -1.07E-01 | 8.53E-02 | 2.61E+00 | 2.67E+00 | 2.06E-01 | no | | intergenic_variant | MODIFIER |
| 10 | 100948584 | AX-97816578 | 0 | G | A | 0.289 | -1.07E-01 | 8.53E-02 | 2.61E+00 | 2.67E+00 | 2.06E-01 | no | | intergenic_variant | MODIFIER |
| 10 | 100952900 | AX-170045737 | 0 | C | T | 0.289 | -1.07E-01 | 8.53E-02 | 2.61E+00 | 2.67E+00 | 2.06E-01 | no | | intergenic_variant | MODIFIER |
| 10 | 100970120 | AX-170198457 | 0 | C | A | 0.289 | -1.07E-01 | 8.53E-02 | 2.61E+00 | 2.67E+00 | 2.06E-01 | no | | intergenic_variant | MODIFIER |
| 10 | 100992329 | AX-97796797 | 0 | C | T | 0.289 | -1.07E-01 | 8.53E-02 | 2.61E+00 | 2.67E+00 | 2.06E-01 | no | | intergenic_variant | MODIFIER |
| 10 | 100995368 | AX-170003768 | 0 | A | G | 0.289 | -1.07E-01 | 8.53E-02 | 2.61E+00 | 2.67E+00 | 2.06E-01 | no | | intergenic_variant | MODIFIER |
| 10 | 101000810 | AX-170003770 | 0 | G | A | 0.289 | -1.07E-01 | 8.53E-02 | 2.61E+00 | 2.67E+00 | 2.06E-01 | no | | intergenic_variant | MODIFIER |
| 10 | 101003990 | AX-170045739 | 0 | A | G | 0.289 | -1.07E-01 | 8.53E-02 | 2.61E+00 | 2.67E+00 | 2.06E-01 | no | | intergenic_variant | MODIFIER |
| 10 | 101018151 | AX-170092078 | 0 | T | C | 0.289 | -1.07E-01 | 8.53E-02 | 2.61E+00 | 2.67E+00 | 2.06E-01 | no | | intergenic_variant | MODIFIER |
| 10 | 101019265 | AX-170045740 | 0 | G | A | 0.289 | -1.07E-01 | 8.53E-02 | 2.61E+00 | 2.67E+00 | 2.06E-01 | no | | intergenic_variant | MODIFIER |
| 10 | 101025165 | AX-170403713 | 0 | T | G | 0.289 | -1.07E-01 | 8.53E-02 | 2.61E+00 | 2.67E+00 | 2.06E-01 | no | | intergenic_variant | MODIFIER |
| 10 | 101059824 | AX-170295460 | 0 | A | G | 0.289 | -1.07E-01 | 8.53E-02 | 2.61E+00 | 2.67E+00 | 2.06E-01 | no | | intergenic_variant | MODIFIER |
| 10 | 101061272 | AX-97834303 | 0 | A | C | 0.289 | -1.07E-01 | 8.53E-02 | 2.61E+00 | 2.67E+00 | 2.06E-01 | no | | intergenic_variant | MODIFIER |
| 10 | 101074954 | AX-170129813 | 0 | G | T | 0.289 | -1.07E-01 | 8.53E-02 | 2.61E+00 | 2.67E+00 | 2.06E-01 | no | | intergenic_variant | MODIFIER |
| 10 | 101081199 | AX-97833320 | 0 | T | A | 0.289 | -1.07E-01 | 8.53E-02 | 2.61E+00 | 2.67E+00 | 2.06E-01 | no | | intergenic_variant | MODIFIER |
| 10 | 101089135 | AX-170514696 | 0 | A | G | 0.289 | -1.07E-01 | 8.53E-02 | 2.61E+00 | 2.67E+00 | 2.06E-01 | no | | intergenic_variant | MODIFIER |
| 10 | 101126208 | AX-170190195 | 0 | A | G | 0.289 | -1.07E-01 | 8.53E-02 | 2.61E+00 | 2.67E+00 | 2.06E-01 | no | | intergenic_variant | MODIFIER |
| 10 | 101145198 | AX-170422425 | 0 | C | T | 0.289 | -1.07E-01 | 8.53E-02 | 2.61E+00 | 2.67E+00 | 2.06E-01 | no | | intergenic_variant | MODIFIER |
| 5 | 67390979 | AX-170072366 | 0 | A | G | 0.465 | 1.28E-01 | 1.01E-01 | 2.50E+00 | 2.57E+00 | 2.06E-01 | no | Bend4 | downstream_gene_variant | MODIFIER |
| 5 | 67392155 | AX-170072364 | 0 | A | G | 0.465 | 1.28E-01 | 1.01E-01 | 2.50E+00 | 2.57E+00 | 2.06E-01 | no | Bend4 | 3_prime_UTR_variant | MODIFIER |
| 5 | 67392643 | AX-170072368 | 0 | C | T | 0.465 | 1.28E-01 | 1.01E-01 | 2.50E+00 | 2.57E+00 | 2.06E-01 | no | Bend4 | 3_prime_UTR_variant | MODIFIER |
| 5 | 67401582 | AX-170318364 | 0 | A | G | 0.465 | 1.28E-01 | 1.01E-01 | 2.50E+00 | 2.57E+00 | 2.06E-01 | no | Bend4 | intron_variant | MODIFIER |
| 18 | 59445535 | AX-170144951 | 1 | G | A | 0.402 | 1.20E-01 | 9.40E-02 | 2.44E+00 | 2.49E+00 | 2.07E-01 | no | Chsy3 | intron_variant,NMD_transcript_variant | MODIFIER |

| | | | | | | | | | | | | | | | |
|---|---|---|---|---|---|---|---|---|---|---|---|---|---|---|---|
| 18 | 69725066 | AX-170439916 | 0 | T | G | 0.263 | -1.15E-01 | 9.23E-02 | 2.76E+00 | 2.86E+00 | 2.07E-01 | no | | intergenic_variant | MODIFIER |
| 18 | 63069941 | AX-170210129 | 0 | A | G | 0.404 | 1.20E-01 | 9.38E-02 | 2.43E+00 | 2.47E+00 | 2.07E-01 | no | Piezo2 | intron_variant | MODIFIER |
| 4 | 81414198 | AX-170260782 | 0 | C | T | 0.456 | -1.08E-01 | 8.63E-02 | 2.61E+00 | 2.66E+00 | 2.07E-01 | no | Mpdz | intron_variant | MODIFIER |
| 13 | 10909732 | AX-170221716 | 0 | A | G | 0.193 | -1.51E-01 | 1.19E-01 | 2.43E+00 | 2.49E+00 | 2.07E-01 | no | | intergenic_variant | MODIFIER |
| 1 | 173330386 | AX-170229062 | 0 | C | T | 0.228 | -1.55E-01 | 1.13E-01 | 3.62E+00 | 3.86E+00 | 2.07E-01 | no | Ackr1 | downstream_gene_variant | MODIFIER |
| 18 | 72275034 | AX-97818140 | 1 | T | G | 0.179 | -1.44E-01 | 1.15E-01 | 2.74E+00 | 2.82E+00 | 2.07E-01 | no | Dcc | intron_variant | MODIFIER |
| 4 | 85722267 | AX-170023306 | 0 | T | A | 0.342 | 1.37E-01 | 1.09E-01 | 2.81E+00 | 2.89E+00 | 2.07E-01 | no | Adamtsl1 | intron_variant,non_coding_transcript_variant | MODIFIER |
| 9 | 97204469 | AX-168373343 | 0 | C | T | 0.132 | -1.39E-01 | 1.10E-01 | 2.61E+00 | 2.70E+00 | 2.07E-01 | no | Gm5371 | downstream_gene_variant | MODIFIER |
| 18 | 78707298 | AX-170016053 | 0 | A | G | 0.456 | 1.10E-01 | 8.57E-02 | 2.41E+00 | 2.48E+00 | 2.07E-01 | no | | intergenic_variant | MODIFIER |
| 18 | 78711027 | AX-170526432 | 0 | T | C | 0.456 | 1.10E-01 | 8.57E-02 | 2.41E+00 | 2.48E+00 | 2.07E-01 | no | | intergenic_variant | MODIFIER |
| 18 | 78747359 | AX-170016049 | 0 | C | A | 0.456 | 1.10E-01 | 8.57E-02 | 2.41E+00 | 2.48E+00 | 2.07E-01 | no | Setbp1 | downstream_gene_variant | MODIFIER |
| 18 | 78761171 | AX-97835558 | 0 | G | T | 0.456 | 1.10E-01 | 8.57E-02 | 2.41E+00 | 2.48E+00 | 2.07E-01 | no | Setbp1 | intron_variant | MODIFIER |
| 18 | 78771178 | AX-170456021 | 0 | A | G | 0.456 | 1.10E-01 | 8.57E-02 | 2.41E+00 | 2.48E+00 | 2.07E-01 | no | Setbp1 | intron_variant | MODIFIER |
| 18 | 78775697 | AX-170102212 | 0 | G | A | 0.456 | 1.10E-01 | 8.57E-02 | 2.41E+00 | 2.48E+00 | 2.07E-01 | no | Setbp1 | intron_variant | MODIFIER |
| 18 | 78775956 | AX-97816929 | 0 | T | C | 0.456 | 1.10E-01 | 8.57E-02 | 2.41E+00 | 2.48E+00 | 2.07E-01 | no | Setbp1 | intron_variant | MODIFIER |
| 18 | 78941101 | AX-170143130 | 0 | G | T | 0.456 | 1.10E-01 | 8.57E-02 | 2.41E+00 | 2.48E+00 | 2.07E-01 | no | Setbp1 | intron_variant | MODIFIER |
| 18 | 78946112 | AX-170489000 | 0 | T | C | 0.456 | 1.10E-01 | 8.57E-02 | 2.41E+00 | 2.48E+00 | 2.07E-01 | no | Setbp1 | intron_variant | MODIFIER |
| 18 | 78947814 | AX-170492775 | 0 | A | G | 0.456 | 1.10E-01 | 8.57E-02 | 2.41E+00 | 2.48E+00 | 2.07E-01 | no | Setbp1 | intron_variant | MODIFIER |
| 18 | 78948144 | AX-170167475 | 0 | T | C | 0.456 | 1.10E-01 | 8.57E-02 | 2.41E+00 | 2.48E+00 | 2.07E-01 | no | Setbp1 | intron_variant | MODIFIER |
| 18 | 78951352 | AX-97853621 | 0 | C | T | 0.456 | 1.10E-01 | 8.57E-02 | 2.41E+00 | 2.48E+00 | 2.07E-01 | no | Setbp1 | intron_variant | MODIFIER |
| 18 | 78967978 | AX-170459495 | 0 | G | A | 0.456 | 1.10E-01 | 8.57E-02 | 2.41E+00 | 2.48E+00 | 2.07E-01 | no | Setbp1 | intron_variant | MODIFIER |
| 18 | 78971979 | AX-170132406 | 0 | T | C | 0.456 | 1.10E-01 | 8.57E-02 | 2.41E+00 | 2.48E+00 | 2.07E-01 | no | Setbp1 | intron_variant | MODIFIER |
| 18 | 78975836 | AX-97819645 | 0 | G | A | 0.456 | 1.10E-01 | 8.57E-02 | 2.41E+00 | 2.48E+00 | 2.07E-01 | no | Setbp1 | intron_variant | MODIFIER |
| 18 | 78976172 | AX-170509869 | 0 | T | C | 0.456 | 1.10E-01 | 8.57E-02 | 2.41E+00 | 2.48E+00 | 2.07E-01 | no | Setbp1 | intron_variant | MODIFIER |
| 18 | 78976250 | AX-170440730 | 0 | C | T | 0.456 | 1.10E-01 | 8.57E-02 | 2.41E+00 | 2.48E+00 | 2.07E-01 | no | Setbp1 | intron_variant | MODIFIER |
| 18 | 79030076 | AX-97793591 | 0 | A | G | 0.456 | 1.10E-01 | 8.57E-02 | 2.41E+00 | 2.48E+00 | 2.07E-01 | no | Setbp1 | intron_variant | MODIFIER |
| 18 | 79074114 | AX-97831832 | 0 | G | A | 0.456 | 1.10E-01 | 8.57E-02 | 2.41E+00 | 2.48E+00 | 2.07E-01 | no | Setbp1 | intron_variant | MODIFIER |
| 18 | 79090542 | AX-170234300 | 0 | G | A | 0.456 | 1.10E-01 | 8.57E-02 | 2.41E+00 | 2.48E+00 | 2.07E-01 | no | Setbp1 | intron_variant | MODIFIER |
| 18 | 79095042 | AX-97849988 | 0 | A | G | 0.456 | 1.10E-01 | 8.57E-02 | 2.41E+00 | 2.48E+00 | 2.07E-01 | no | Setbp1 | intron_variant | MODIFIER |
| 18 | 79099942 | AX-170159604 | 0 | C | T | 0.456 | 1.10E-01 | 8.57E-02 | 2.41E+00 | 2.48E+00 | 2.07E-01 | no | Setbp1 | intron_variant | MODIFIER |
| 18 | 79119266 | AX-170381600 | 0 | C | T | 0.456 | 1.10E-01 | 8.57E-02 | 2.41E+00 | 2.48E+00 | 2.07E-01 | no | | intergenic_variant | MODIFIER |
| 18 | 79149310 | AX-97789215 | 0 | A | G | 0.456 | 1.10E-01 | 8.57E-02 | 2.41E+00 | 2.48E+00 | 2.07E-01 | no | | intergenic_variant | MODIFIER |
| 18 | 79190940 | AX-170278186 | 0 | C | T | 0.456 | 1.10E-01 | 8.57E-02 | 2.41E+00 | 2.48E+00 | 2.07E-01 | no | | intergenic_variant | MODIFIER |
| 18 | 79296583 | AX-170016059 | 0 | T | C | 0.456 | 1.10E-01 | 8.57E-02 | 2.41E+00 | 2.48E+00 | 2.07E-01 | no | | intergenic_variant | MODIFIER |
| 18 | 79300299 | AX-170217908 | 0 | A | G | 0.456 | 1.10E-01 | 8.57E-02 | 2.41E+00 | 2.48E+00 | 2.07E-01 | no | AC140980.1 | downstream_gene_variant | MODIFIER |
| 18 | 79307988 | AX-170016063 | 0 | C | T | 0.456 | 1.10E-01 | 8.57E-02 | 2.41E+00 | 2.48E+00 | 2.07E-01 | no | AC140980.1 | upstream_gene_variant | MODIFIER |
| 18 | 79308935 | AX-170285160 | 0 | A | G | 0.456 | 1.10E-01 | 8.57E-02 | 2.41E+00 | 2.48E+00 | 2.07E-01 | no | AC140980.1 | upstream_gene_variant | MODIFIER |
| 18 | 79373122 | AX-170399758 | 0 | A | G | 0.456 | 1.10E-01 | 8.57E-02 | 2.41E+00 | 2.48E+00 | 2.07E-01 | no | | regulatory_region_variant | MODIFIER |
| 18 | 79389210 | AX-97788084 | 0 | A | C | 0.456 | 1.10E-01 | 8.57E-02 | 2.41E+00 | 2.48E+00 | 2.07E-01 | no | | intergenic_variant | MODIFIER |
| 18 | 79419474 | AX-170491819 | 0 | C | T | 0.456 | 1.10E-01 | 8.57E-02 | 2.41E+00 | 2.48E+00 | 2.07E-01 | no | | intergenic_variant | MODIFIER |
| 18 | 79419768 | AX-170370309 | 0 | A | G | 0.456 | 1.10E-01 | 8.57E-02 | 2.41E+00 | 2.48E+00 | 2.07E-01 | no | | intergenic_variant | MODIFIER |
| 18 | 79419791 | AX-170397926 | 0 | T | G | 0.456 | 1.10E-01 | 8.57E-02 | 2.41E+00 | 2.48E+00 | 2.07E-01 | no | | intergenic_variant | MODIFIER |
| 18 | 79427562 | AX-170183925 | 0 | A | G | 0.456 | 1.10E-01 | 8.57E-02 | 2.41E+00 | 2.48E+00 | 2.07E-01 | no | | intergenic_variant | MODIFIER |
| 18 | 79434816 | AX-97813498 | 0 | C | T | 0.456 | 1.10E-01 | 8.57E-02 | 2.41E+00 | 2.48E+00 | 2.07E-01 | no | | intergenic_variant | MODIFIER |
| 18 | 79461907 | AX-170223171 | 0 | A | G | 0.456 | 1.10E-01 | 8.57E-02 | 2.41E+00 | 2.48E+00 | 2.07E-01 | no | | intergenic_variant | MODIFIER |
| 18 | 79464928 | AX-170440631 | 0 | A | G | 0.456 | 1.10E-01 | 8.57E-02 | 2.41E+00 | 2.48E+00 | 2.07E-01 | no | | intergenic_variant | MODIFIER |
| 18 | 79487963 | AX-170037857 | 0 | G | A | 0.456 | 1.10E-01 | 8.57E-02 | 2.41E+00 | 2.48E+00 | 2.07E-01 | no | | intergenic_variant | MODIFIER |
| 18 | 79487998 | AX-170182466 | 0 | C | T | 0.456 | 1.10E-01 | 8.57E-02 | 2.41E+00 | 2.48E+00 | 2.07E-01 | no | | intergenic_variant | MODIFIER |
| 2 | 102314368 | AX-170386847 | 1 | G | A | 0.054 | -2.70E-01 | 2.16E-01 | 2.83E+00 | 2.94E+00 | 2.08E-01 | Yes | Trim44 | intron_variant | MODIFIER |
| 1 | 14579022 | AX-169999986 | 0 | T | C | 0.377 | 9.93E-02 | 7.92E-02 | 2.62E+00 | 2.67E+00 | 2.08E-01 | no | | intergenic_variant | MODIFIER |
| 9 | 92324854 | AX-170090377 | 0 | G | A | 0.123 | -1.59E-01 | 1.27E-01 | 2.67E+00 | 2.76E+00 | 2.08E-01 | no | 1700057G04Rik | intron_variant | MODIFIER |
| 16 | 72555890 | AX-170477245 | 0 | A | G | 0.491 | 9.61E-02 | 7.58E-02 | 2.47E+00 | 2.54E+00 | 2.08E-01 | no | Robo1 | intron_variant | MODIFIER |
| 5 | 42810447 | AX-170527286 | 0 | C | G | 0.202 | -1.28E-01 | 1.03E-01 | 2.66E+00 | 2.76E+00 | 2.08E-01 | no | | intergenic_variant | MODIFIER |
| 7 | 99612318 | AX-170371548 | 1 | T | C | 0.375 | -1.32E-01 | 1.05E-01 | 2.89E+00 | 2.98E+00 | 2.08E-01 | no | | intergenic_variant | MODIFIER |

| | | | | | | | | | | | | | | | |
|---|---|---|---|---|---|---|---|---|---|---|---|---|---|---|---|
| 4 | 85395210 | AX-170185976 | 1 | G | A | 0.339 | 1.44E-01 | 1.15E-01 | 2.85E+00 | 2.92E+00 | 2.08E-01 | Yes | Sh3gl2 | intron_variant | MODIFIER |
| 4 | 63411825 | AX-170022946 | 1 | A | G | 0.348 | 1.45E-01 | 1.16E-01 | 2.85E+00 | 2.91E+00 | 2.08E-01 | Yes | Whrn | downstream_gene_variant | MODIFIER |
| 16 | 53030628 | AX-170419332 | 1 | A | T | 0.464 | -1.01E-01 | 7.99E-02 | 2.46E+00 | 2.51E+00 | 2.08E-01 | no | | intergenic_variant | MODIFIER |
| 18 | 59379228 | AX-170126120 | 0 | A | T | 0.404 | 1.21E-01 | 9.27E-02 | 2.28E+00 | 2.32E+00 | 2.08E-01 | no | Chsy3 | intron_variant | MODIFIER |
| 18 | 59758051 | AX-170138448 | 0 | T | A | 0.404 | 1.21E-01 | 9.27E-02 | 2.28E+00 | 2.32E+00 | 2.08E-01 | no | | intergenic_variant | MODIFIER |
| 4 | 84757392 | AX-170337598 | 0 | A | G | 0.342 | 1.44E-01 | 1.15E-01 | 2.85E+00 | 2.92E+00 | 2.08E-01 | Yes | | intergenic_variant | MODIFIER |
| 4 | 84785654 | AX-170530943 | 0 | C | T | 0.342 | 1.44E-01 | 1.15E-01 | 2.85E+00 | 2.92E+00 | 2.08E-01 | Yes | | intergenic_variant | MODIFIER |
| 4 | 84956583 | AX-170223808 | 0 | T | G | 0.342 | 1.44E-01 | 1.15E-01 | 2.85E+00 | 2.92E+00 | 2.08E-01 | Yes | Cntln | intron_variant | MODIFIER |
| 4 | 84960466 | AX-97831113 | 0 | G | A | 0.342 | 1.44E-01 | 1.15E-01 | 2.85E+00 | 2.92E+00 | 2.08E-01 | Yes | Cntln | intron_variant | MODIFIER |
| 4 | 84962993 | AX-170405106 | 0 | T | C | 0.342 | 1.44E-01 | 1.15E-01 | 2.85E+00 | 2.92E+00 | 2.08E-01 | Yes | Cntln | intron_variant | MODIFIER |
| 4 | 84988280 | AX-170234379 | 0 | G | A | 0.342 | 1.44E-01 | 1.15E-01 | 2.85E+00 | 2.92E+00 | 2.08E-01 | Yes | Cntln | intron_variant | MODIFIER |
| 4 | 85026574 | AX-170484149 | 0 | G | A | 0.342 | 1.44E-01 | 1.15E-01 | 2.85E+00 | 2.92E+00 | 2.08E-01 | Yes | Cntln | intron_variant | MODIFIER |
| 4 | 85056300 | AX-170413750 | 0 | A | T | 0.342 | 1.44E-01 | 1.15E-01 | 2.85E+00 | 2.92E+00 | 2.08E-01 | Yes | Cntln | intron_variant | MODIFIER |
| 4 | 85070668 | AX-170069868 | 0 | C | T | 0.342 | 1.44E-01 | 1.15E-01 | 2.85E+00 | 2.92E+00 | 2.08E-01 | Yes | Cntln | intron_variant | MODIFIER |
| 4 | 85070709 | AX-170431113 | 0 | G | C | 0.342 | 1.44E-01 | 1.15E-01 | 2.85E+00 | 2.92E+00 | 2.08E-01 | Yes | Cntln | intron_variant | MODIFIER |
| 4 | 85071491 | AX-97798196 | 0 | A | C | 0.342 | 1.44E-01 | 1.15E-01 | 2.85E+00 | 2.92E+00 | 2.08E-01 | Yes | Cntln | intron_variant | MODIFIER |
| 4 | 85077102 | AX-170110410 | 0 | C | T | 0.342 | 1.44E-01 | 1.15E-01 | 2.85E+00 | 2.92E+00 | 2.08E-01 | Yes | Cntln | intron_variant | MODIFIER |
| 4 | 85090675 | AX-170220906 | 0 | C | G | 0.342 | 1.44E-01 | 1.15E-01 | 2.85E+00 | 2.92E+00 | 2.08E-01 | Yes | Cntln | intron_variant | MODIFIER |
| 4 | 85109351 | AX-170471793 | 0 | G | A | 0.342 | 1.44E-01 | 1.15E-01 | 2.85E+00 | 2.92E+00 | 2.08E-01 | Yes | Cntln | intron_variant | MODIFIER |
| 4 | 85122337 | AX-170271171 | 0 | G | A | 0.342 | 1.44E-01 | 1.15E-01 | 2.85E+00 | 2.92E+00 | 2.08E-01 | Yes | Cntln | intron_variant | MODIFIER |
| 4 | 85128163 | AX-170118091 | 0 | C | T | 0.342 | 1.44E-01 | 1.15E-01 | 2.85E+00 | 2.92E+00 | 2.08E-01 | Yes | Cntln | intron_variant | MODIFIER |
| 4 | 85188055 | AX-170367579 | 0 | A | G | 0.342 | 1.44E-01 | 1.15E-01 | 2.85E+00 | 2.92E+00 | 2.08E-01 | Yes | | intergenic_variant | MODIFIER |
| 4 | 85188318 | AX-97855420 | 0 | C | T | 0.342 | 1.44E-01 | 1.15E-01 | 2.85E+00 | 2.92E+00 | 2.08E-01 | Yes | | intergenic_variant | MODIFIER |
| 4 | 85216579 | AX-170527645 | 0 | G | T | 0.342 | 1.44E-01 | 1.15E-01 | 2.85E+00 | 2.92E+00 | 2.08E-01 | Yes | Sh3gl2 | intron_variant | MODIFIER |
| 4 | 85232803 | AX-170023296 | 0 | G | A | 0.342 | 1.44E-01 | 1.15E-01 | 2.85E+00 | 2.92E+00 | 2.08E-01 | Yes | Sh3gl2 | intron_variant | MODIFIER |
| 4 | 85262791 | AX-97827902 | 0 | G | A | 0.342 | 1.44E-01 | 1.15E-01 | 2.85E+00 | 2.92E+00 | 2.08E-01 | Yes | Sh3gl2 | intron_variant | MODIFIER |
| 4 | 85292693 | AX-170023292 | 0 | G | A | 0.342 | 1.44E-01 | 1.15E-01 | 2.85E+00 | 2.92E+00 | 2.08E-01 | Yes | Sh3gl2 | intron_variant | MODIFIER |
| 4 | 85333545 | AX-170413852 | 0 | A | T | 0.342 | 1.44E-01 | 1.15E-01 | 2.85E+00 | 2.92E+00 | 2.08E-01 | Yes | Sh3gl2 | intron_variant | MODIFIER |
| 4 | 85334480 | AX-97825515 | 0 | G | T | 0.342 | 1.44E-01 | 1.15E-01 | 2.85E+00 | 2.92E+00 | 2.08E-01 | Yes | Sh3gl2 | intron_variant | MODIFIER |
| 4 | 85382168 | AX-170163910 | 0 | G | A | 0.342 | 1.44E-01 | 1.15E-01 | 2.85E+00 | 2.92E+00 | 2.08E-01 | Yes | Sh3gl2 | intron_variant | MODIFIER |
| 4 | 85389590 | AX-97798195 | 0 | G | A | 0.342 | 1.44E-01 | 1.15E-01 | 2.85E+00 | 2.92E+00 | 2.08E-01 | Yes | Sh3gl2 | downstream_gene_variant | MODIFIER |
| 4 | 85391451 | AX-170271096 | 0 | T | G | 0.342 | 1.44E-01 | 1.15E-01 | 2.85E+00 | 2.92E+00 | 2.08E-01 | Yes | Sh3gl2 | downstream_gene_variant | MODIFIER |
| 4 | 85394862 | AX-170475518 | 0 | G | A | 0.342 | 1.44E-01 | 1.15E-01 | 2.85E+00 | 2.92E+00 | 2.08E-01 | Yes | Sh3gl2 | intron_variant | MODIFIER |
| 4 | 85395151 | AX-170197945 | 0 | A | G | 0.342 | 1.44E-01 | 1.15E-01 | 2.85E+00 | 2.92E+00 | 2.08E-01 | Yes | Sh3gl2 | intron_variant | MODIFIER |
| 4 | 85396244 | AX-170397582 | 0 | T | C | 0.342 | 1.44E-01 | 1.15E-01 | 2.85E+00 | 2.92E+00 | 2.08E-01 | Yes | Sh3gl2 | intron_variant | MODIFIER |
| 4 | 85416987 | AX-170217494 | 0 | C | T | 0.342 | 1.44E-01 | 1.15E-01 | 2.85E+00 | 2.92E+00 | 2.08E-01 | Yes | Sh3gl2 | intron_variant | MODIFIER |
| 4 | 85419318 | AX-170315635 | 0 | C | T | 0.342 | 1.44E-01 | 1.15E-01 | 2.85E+00 | 2.92E+00 | 2.08E-01 | Yes | Sh3gl2 | intron_variant | MODIFIER |
| 4 | 85433519 | AX-170256460 | 0 | G | A | 0.342 | 1.44E-01 | 1.15E-01 | 2.85E+00 | 2.92E+00 | 2.08E-01 | Yes | Sh3gl2 | intron_variant | MODIFIER |
| 4 | 85434914 | AX-170286923 | 0 | T | C | 0.342 | 1.44E-01 | 1.15E-01 | 2.85E+00 | 2.92E+00 | 2.08E-01 | Yes | Sh3gl2 | intron_variant | MODIFIER |
| 4 | 85448627 | AX-170195811 | 0 | A | G | 0.342 | 1.44E-01 | 1.15E-01 | 2.85E+00 | 2.92E+00 | 2.08E-01 | Yes | Sh3gl2 | intron_variant | MODIFIER |
| 4 | 85455543 | AX-170173137 | 0 | G | T | 0.342 | 1.44E-01 | 1.15E-01 | 2.85E+00 | 2.92E+00 | 2.08E-01 | Yes | Sh3gl2 | intron_variant | MODIFIER |
| 4 | 85468195 | AX-170069872 | 0 | A | G | 0.342 | 1.44E-01 | 1.15E-01 | 2.85E+00 | 2.92E+00 | 2.08E-01 | Yes | Sh3gl2 | intron_variant | MODIFIER |
| 4 | 85472475 | AX-170069876 | 0 | C | T | 0.342 | 1.44E-01 | 1.15E-01 | 2.85E+00 | 2.92E+00 | 2.08E-01 | Yes | Sh3gl2 | intron_variant | MODIFIER |
| 4 | 85613370 | AX-168409455 | 0 | A | G | 0.342 | 1.44E-01 | 1.15E-01 | 2.85E+00 | 2.92E+00 | 2.08E-01 | Yes | Sh3gl2 | intron_variant | MODIFIER |
| 4 | 85772453 | AX-170023302 | 0 | A | G | 0.342 | 1.44E-01 | 1.15E-01 | 2.85E+00 | 2.92E+00 | 2.08E-01 | Yes | Adamtsl1 | intron_variant,non_coding_transcript_variant | MODIFIER |
| 4 | 85781536 | AX-170023301 | 0 | A | G | 0.342 | 1.44E-01 | 1.15E-01 | 2.85E+00 | 2.92E+00 | 2.08E-01 | Yes | Adamtsl1 | intron_variant,non_coding_transcript_variant | MODIFIER |
| 4 | 85812075 | AX-170496544 | 0 | A | G | 0.342 | 1.44E-01 | 1.15E-01 | 2.85E+00 | 2.92E+00 | 2.08E-01 | Yes | Adamtsl1 | intron_variant | MODIFIER |
| 4 | 85812392 | AX-170299154 | 0 | G | A | 0.342 | 1.44E-01 | 1.15E-01 | 2.85E+00 | 2.92E+00 | 2.08E-01 | Yes | Adamtsl1 | intron_variant | MODIFIER |
| 4 | 85812505 | AX-170213163 | 0 | A | G | 0.342 | 1.44E-01 | 1.15E-01 | 2.85E+00 | 2.92E+00 | 2.08E-01 | Yes | Adamtsl1 | intron_variant | MODIFIER |
| 4 | 85814657 | AX-170095038 | 0 | C | T | 0.342 | 1.44E-01 | 1.15E-01 | 2.85E+00 | 2.92E+00 | 2.08E-01 | Yes | Adamtsl1 | intron_variant | MODIFIER |
| 4 | 85817429 | AX-168409755 | 0 | T | G | 0.342 | 1.44E-01 | 1.15E-01 | 2.85E+00 | 2.92E+00 | 2.08E-01 | Yes | Adamtsl1 | intron_variant | MODIFIER |
| 4 | 85821007 | AX-170023305 | 0 | C | G | 0.342 | 1.44E-01 | 1.15E-01 | 2.85E+00 | 2.92E+00 | 2.08E-01 | Yes | Adamtsl1 | intron_variant | MODIFIER |
| 4 | 85821163 | AX-170069885 | 0 | T | C | 0.342 | 1.44E-01 | 1.15E-01 | 2.85E+00 | 2.92E+00 | 2.08E-01 | Yes | Adamtsl1 | intron_variant | MODIFIER |
| 4 | 85836892 | AX-168375671 | 0 | C | T | 0.342 | 1.44E-01 | 1.15E-01 | 2.85E+00 | 2.92E+00 | 2.08E-01 | Yes | Adamtsl1 | intron_variant | MODIFIER |

| | | | | | | | | | | | | | | | |
|---|---|---|---|---|---|---|---|---|---|---|---|---|---|---|---|
| 4 | 85842226 | AX-170436008 | 0 | T | C | 0.342 | 1.44E-01 | 1.15E-01 | 2.85E+00 | 2.92E+00 | 2.08E-01 | Yes | Adamtsl1 | intron_variant | MODIFIER |
| 4 | 85945880 | AX-170444285 | 0 | T | C | 0.342 | 1.44E-01 | 1.15E-01 | 2.85E+00 | 2.92E+00 | 2.08E-01 | Yes | Adamtsl1 | intron_variant | MODIFIER |
| 4 | 85952117 | AX-170320604 | 0 | G | A | 0.342 | 1.44E-01 | 1.15E-01 | 2.85E+00 | 2.92E+00 | 2.08E-01 | Yes | Adamtsl1 | intron_variant | MODIFIER |
| 4 | 85982640 | AX-170187769 | 0 | A | G | 0.342 | 1.44E-01 | 1.15E-01 | 2.85E+00 | 2.92E+00 | 2.08E-01 | Yes | Adamtsl1 | intron_variant | MODIFIER |
| 4 | 86026252 | AX-170379675 | 0 | A | G | 0.342 | 1.44E-01 | 1.15E-01 | 2.85E+00 | 2.92E+00 | 2.08E-01 | Yes | Adamtsl1 | intron_variant | MODIFIER |
| 4 | 86026312 | AX-170397146 | 0 | T | G | 0.342 | 1.44E-01 | 1.15E-01 | 2.85E+00 | 2.92E+00 | 2.08E-01 | Yes | Adamtsl1 | intron_variant | MODIFIER |
| 4 | 86066064 | AX-170089303 | 0 | T | G | 0.342 | 1.44E-01 | 1.15E-01 | 2.85E+00 | 2.92E+00 | 2.08E-01 | Yes | Adamtsl1 | intron_variant | MODIFIER |
| 4 | 75266860 | AX-97823741 | 1 | G | A | 0.321 | 1.48E-01 | 1.19E-01 | 2.86E+00 | 2.92E+00 | 2.08E-01 | Yes | | regulatory_region_variant | MODIFIER |
| 6 | 147977394 | AX-170475493 | 0 | T | A | 0.044 | -2.29E-01 | 1.84E-01 | 2.75E+00 | 2.86E+00 | 2.09E-01 | Yes | | intergenic_variant | MODIFIER |
| 1 | 171401623 | AX-170043577 | 1 | C | T | 0.214 | 1.69E-01 | 1.35E-01 | 2.63E+00 | 2.68E+00 | 2.09E-01 | Yes | Arhgap30 | intron_variant | MODIFIER |
| 4 | 125264776 | AX-170355785 | 1 | T | A | 0.232 | 1.32E-01 | 1.06E-01 | 2.61E+00 | 2.68E+00 | 2.09E-01 | no | | intergenic_variant | MODIFIER |
| 9 | 90724145 | AX-170101978 | 0 | G | A | 0.14 | -1.48E-01 | 1.18E-01 | 2.73E+00 | 2.83E+00 | 2.09E-01 | no | | intergenic_variant | MODIFIER |
| 4 | 104968338 | AX-170317171 | 0 | G | A | 0.237 | 1.45E-01 | 1.16E-01 | 2.65E+00 | 2.73E+00 | 2.09E-01 | Yes | Fyb2 | intron_variant | MODIFIER |
| 9 | 9261076 | AX-170135569 | 1 | A | G | 0.473 | 1.03E-01 | 8.14E-02 | 2.44E+00 | 2.53E+00 | 2.09E-01 | no | Gm16833 | downstream_gene_variant | MODIFIER |
| 8 | 122919021 | AX-170081406 | 0 | T | C | 0.035 | -2.53E-01 | 2.02E-01 | 2.61E+00 | 2.70E+00 | 2.09E-01 | Yes | Ankrd11 | upstream_gene_variant | MODIFIER |
| 3 | 20458264 | AX-170298145 | 0 | G | T | 0.132 | -1.28E-01 | 1.02E-01 | 2.53E+00 | 2.61E+00 | 2.09E-01 | no | | intergenic_variant | MODIFIER |
| 3 | 20492440 | AX-97847087 | 0 | C | T | 0.132 | -1.28E-01 | 1.02E-01 | 2.53E+00 | 2.61E+00 | 2.09E-01 | no | | intergenic_variant | MODIFIER |
| 8 | 24984544 | AX-170174971 | 1 | T | A | 0.357 | -8.06E-02 | 6.45E-02 | 2.58E+00 | 2.67E+00 | 2.10E-01 | no | Adam9 | intron_variant | MODIFIER |
| 5 | 129213781 | AX-170384640 | 0 | G | T | 0.158 | -1.09E-01 | 8.25E-02 | 2.22E+00 | 2.30E+00 | 2.10E-01 | no | | intergenic_variant | MODIFIER |
| 18 | 80626740 | AX-97796304 | 0 | C | T | 0.386 | -1.05E-01 | 8.47E-02 | 2.73E+00 | 2.83E+00 | 2.10E-01 | no | Nfatc1 | intron_variant | MODIFIER |
| 8 | 25338050 | AX-170099584 | 1 | T | C | 0.348 | -8.07E-02 | 6.46E-02 | 2.58E+00 | 2.67E+00 | 2.10E-01 | no | | intergenic_variant | MODIFIER |
| 1 | 148313882 | AX-170373111 | 0 | C | G | 0.202 | -1.61E-01 | 1.25E-01 | 3.23E+00 | 3.40E+00 | 2.10E-01 | no | | intergenic_variant | MODIFIER |
| 8 | 24294904 | AX-170079530 | 0 | A | G | 0.351 | -8.07E-02 | 6.47E-02 | 2.58E+00 | 2.67E+00 | 2.10E-01 | no | | regulatory_region_variant | MODIFIER |
| 8 | 24295901 | AX-97814276 | 0 | A | G | 0.351 | -8.07E-02 | 6.47E-02 | 2.58E+00 | 2.67E+00 | 2.10E-01 | no | | intergenic_variant | MODIFIER |
| 8 | 24298417 | AX-170520223 | 0 | C | T | 0.351 | -8.07E-02 | 6.47E-02 | 2.58E+00 | 2.67E+00 | 2.10E-01 | no | | intergenic_variant | MODIFIER |
| 8 | 24305481 | AX-170079532 | 0 | G | T | 0.351 | -8.07E-02 | 6.47E-02 | 2.58E+00 | 2.67E+00 | 2.10E-01 | no | | intergenic_variant | MODIFIER |
| 8 | 24379427 | AX-170175439 | 0 | G | A | 0.351 | -8.07E-02 | 6.47E-02 | 2.58E+00 | 2.67E+00 | 2.10E-01 | no | | intergenic_variant | MODIFIER |
| 8 | 24382366 | AX-170411911 | 0 | G | A | 0.351 | -8.07E-02 | 6.47E-02 | 2.58E+00 | 2.67E+00 | 2.10E-01 | no | | intergenic_variant | MODIFIER |
| 8 | 24452120 | AX-170200526 | 0 | G | A | 0.351 | -8.07E-02 | 6.47E-02 | 2.58E+00 | 2.67E+00 | 2.10E-01 | no | Gm44620 | downstream_gene_variant | MODIFIER |
| 8 | 24452248 | AX-170079536 | 0 | G | A | 0.351 | -8.07E-02 | 6.47E-02 | 2.58E+00 | 2.67E+00 | 2.10E-01 | no | Gm44620 | downstream_gene_variant | MODIFIER |
| 8 | 24452337 | AX-170079537 | 0 | A | G | 0.351 | -8.07E-02 | 6.47E-02 | 2.58E+00 | 2.67E+00 | 2.10E-01 | no | Gm44620 | downstream_gene_variant | MODIFIER |
| 8 | 24453317 | AX-170484046 | 0 | G | A | 0.351 | -8.07E-02 | 6.47E-02 | 2.58E+00 | 2.67E+00 | 2.10E-01 | no | Gm44620 | downstream_gene_variant | MODIFIER |
| 8 | 24453500 | AX-170131779 | 0 | G | A | 0.351 | -8.07E-02 | 6.47E-02 | 2.58E+00 | 2.67E+00 | 2.10E-01 | no | Gm44620 | downstream_gene_variant | MODIFIER |
| 8 | 24470949 | AX-170159661 | 0 | C | T | 0.351 | -8.07E-02 | 6.47E-02 | 2.58E+00 | 2.67E+00 | 2.10E-01 | no | | intergenic_variant | MODIFIER |
| 8 | 24490191 | AX-170271620 | 0 | C | T | 0.351 | -8.07E-02 | 6.47E-02 | 2.58E+00 | 2.67E+00 | 2.10E-01 | no | | regulatory_region_variant | MODIFIER |
| 8 | 24502335 | AX-170030365 | 0 | A | G | 0.351 | -8.07E-02 | 6.47E-02 | 2.58E+00 | 2.67E+00 | 2.10E-01 | no | 9130214F15Rik | upstream_gene_variant | MODIFIER |
| 8 | 24509157 | AX-170229735 | 0 | T | C | 0.351 | -8.07E-02 | 6.47E-02 | 2.58E+00 | 2.67E+00 | 2.10E-01 | no | 9130214F15Rik | downstream_gene_variant | MODIFIER |
| 8 | 24512630 | AX-170269000 | 0 | T | C | 0.351 | -8.07E-02 | 6.47E-02 | 2.58E+00 | 2.67E+00 | 2.10E-01 | no | 9130214F15Rik | downstream_gene_variant | MODIFIER |
| 8 | 24512754 | AX-170173370 | 0 | C | A | 0.351 | -8.07E-02 | 6.47E-02 | 2.58E+00 | 2.67E+00 | 2.10E-01 | no | 9130214F15Rik | downstream_gene_variant | MODIFIER |
| 8 | 24512774 | AX-170342507 | 0 | C | T | 0.351 | -8.07E-02 | 6.47E-02 | 2.58E+00 | 2.67E+00 | 2.10E-01 | no | 9130214F15Rik | downstream_gene_variant | MODIFIER |
| 8 | 24555808 | AX-170401688 | 0 | C | T | 0.351 | -8.07E-02 | 6.47E-02 | 2.58E+00 | 2.67E+00 | 2.10E-01 | no | Ido2 | intron_variant | MODIFIER |
| 8 | 24631150 | AX-170103640 | 0 | G | T | 0.351 | -8.07E-02 | 6.47E-02 | 2.58E+00 | 2.67E+00 | 2.10E-01 | no | Adam18 | intron_variant | MODIFIER |
| 8 | 24650066 | AX-170079539 | 0 | T | C | 0.351 | -8.07E-02 | 6.47E-02 | 2.58E+00 | 2.67E+00 | 2.10E-01 | no | Adam18 | intron_variant | MODIFIER |
| 8 | 24650367 | AX-170079542 | 0 | G | A | 0.351 | -8.07E-02 | 6.47E-02 | 2.58E+00 | 2.67E+00 | 2.10E-01 | no | Adam18 | intron_variant | MODIFIER |
| 8 | 24652789 | AX-170256307 | 0 | T | C | 0.351 | -8.07E-02 | 6.47E-02 | 2.58E+00 | 2.67E+00 | 2.10E-01 | no | Adam18 | intron_variant | MODIFIER |
| 8 | 24675965 | AX-170402967 | 0 | T | C | 0.351 | -8.07E-02 | 6.47E-02 | 2.58E+00 | 2.67E+00 | 2.10E-01 | no | Adam18 | upstream_gene_variant | MODIFIER |
| 8 | 24745207 | AX-170106336 | 0 | C | T | 0.351 | -8.07E-02 | 6.47E-02 | 2.58E+00 | 2.67E+00 | 2.10E-01 | no | Adam5 | intron_variant | MODIFIER |
| 8 | 24864244 | AX-170216878 | 0 | A | G | 0.351 | -8.07E-02 | 6.47E-02 | 2.58E+00 | 2.67E+00 | 2.10E-01 | no | Adam32 | intron_variant | MODIFIER |
| 8 | 24881093 | AX-170129693 | 0 | C | T | 0.351 | -8.07E-02 | 6.47E-02 | 2.58E+00 | 2.67E+00 | 2.10E-01 | no | Adam32 | intron_variant | MODIFIER |
| 8 | 24881143 | AX-170490159 | 0 | C | T | 0.351 | -8.07E-02 | 6.47E-02 | 2.58E+00 | 2.67E+00 | 2.10E-01 | no | Adam32 | intron_variant | MODIFIER |
| 8 | 24904178 | AX-170111411 | 0 | G | A | 0.351 | -8.07E-02 | 6.47E-02 | 2.58E+00 | 2.67E+00 | 2.10E-01 | no | Adam32 | intron_variant | MODIFIER |
| 8 | 24907439 | AX-170079545 | 0 | C | A | 0.351 | -8.07E-02 | 6.47E-02 | 2.58E+00 | 2.67E+00 | 2.10E-01 | no | Adam32 | intron_variant | MODIFIER |
| 8 | 24945538 | AX-170299874 | 0 | G | A | 0.351 | -8.07E-02 | 6.47E-02 | 2.58E+00 | 2.67E+00 | 2.10E-01 | no | Adam9 | downstream_gene_variant | MODIFIER |
| 8 | 24946617 | AX-170288487 | 0 | T | A | 0.351 | -8.07E-02 | 6.47E-02 | 2.58E+00 | 2.67E+00 | 2.10E-01 | no | Adam9 | downstream_gene_variant | MODIFIER |

| | | | | | | | | | | | | | | | |
|---|---|---|---|---|---|---|---|---|---|---|---|---|---|---|---|
| 8 | 24965627 | AX-170402653 | 0 | T | C | 0.351 | -8.07E-02 | 6.47E-02 | 2.58E+00 | 2.67E+00 | 2.10E-01 | no | Adam9 | intron_variant | MODIFIER |
| 8 | 24977492 | AX-170232066 | 0 | G | A | 0.351 | -8.07E-02 | 6.47E-02 | 2.58E+00 | 2.67E+00 | 2.10E-01 | no | Adam9 | intron_variant | MODIFIER |
| 8 | 24989776 | AX-170385138 | 0 | T | C | 0.351 | -8.07E-02 | 6.47E-02 | 2.58E+00 | 2.67E+00 | 2.10E-01 | no | Adam9 | intron_variant | MODIFIER |
| 8 | 25116367 | AX-170030367 | 0 | G | C | 0.351 | -8.07E-02 | 6.47E-02 | 2.58E+00 | 2.67E+00 | 2.10E-01 | no | | intergenic_variant | MODIFIER |
| 8 | 25145355 | AX-97842539 | 0 | G | A | 0.351 | -8.07E-02 | 6.47E-02 | 2.58E+00 | 2.67E+00 | 2.10E-01 | no | | regulatory_region_variant | MODIFIER |
| 8 | 25156568 | AX-170079550 | 0 | A | G | 0.351 | -8.07E-02 | 6.47E-02 | 2.58E+00 | 2.67E+00 | 2.10E-01 | no | Tacc1 | 3_prime_UTR_variant | MODIFIER |
| 8 | 25156625 | AX-170079547 | 0 | A | G | 0.351 | -8.07E-02 | 6.47E-02 | 2.58E+00 | 2.67E+00 | 2.10E-01 | no | Tacc1 | 3_prime_UTR_variant | MODIFIER |
| 8 | 25160940 | AX-170079549 | 0 | T | C | 0.351 | -8.07E-02 | 6.47E-02 | 2.58E+00 | 2.67E+00 | 2.10E-01 | no | Tacc1 | intron_variant | MODIFIER |
| 8 | 25175179 | AX-170516125 | 0 | C | T | 0.351 | -8.07E-02 | 6.47E-02 | 2.58E+00 | 2.67E+00 | 2.10E-01 | no | Tacc1 | synonymous_variant | LOW |
| 8 | 25189831 | AX-170129840 | 0 | A | G | 0.351 | -8.07E-02 | 6.47E-02 | 2.58E+00 | 2.67E+00 | 2.10E-01 | no | Tacc1 | intron_variant | MODIFIER |
| 8 | 25213315 | AX-97810498 | 0 | C | T | 0.351 | -8.07E-02 | 6.47E-02 | 2.58E+00 | 2.67E+00 | 2.10E-01 | no | Tacc1 | intron_variant,NMD_transcript_variant | MODIFIER |
| 8 | 25236478 | AX-170191906 | 0 | T | G | 0.351 | -8.07E-02 | 6.47E-02 | 2.58E+00 | 2.67E+00 | 2.10E-01 | no | Tacc1 | intron_variant,NMD_transcript_variant | MODIFIER |
| 8 | 25248366 | AX-170178750 | 0 | C | T | 0.351 | -8.07E-02 | 6.47E-02 | 2.58E+00 | 2.67E+00 | 2.10E-01 | no | Tacc1 | intron_variant,NMD_transcript_variant | MODIFIER |
| 8 | 25252912 | AX-170246044 | 0 | G | A | 0.351 | -8.07E-02 | 6.47E-02 | 2.58E+00 | 2.67E+00 | 2.10E-01 | no | Tacc1 | intron_variant,NMD_transcript_variant | MODIFIER |
| 8 | 25253038 | AX-170205313 | 0 | C | A | 0.351 | -8.07E-02 | 6.47E-02 | 2.58E+00 | 2.67E+00 | 2.10E-01 | no | Tacc1 | intron_variant,NMD_transcript_variant | MODIFIER |
| 8 | 25277999 | AX-170115040 | 0 | G | A | 0.351 | -8.07E-02 | 6.47E-02 | 2.58E+00 | 2.67E+00 | 2.10E-01 | no | | intergenic_variant | MODIFIER |
| 8 | 25316089 | AX-170252060 | 0 | A | G | 0.351 | -8.07E-02 | 6.47E-02 | 2.58E+00 | 2.67E+00 | 2.10E-01 | no | | intergenic_variant | MODIFIER |
| 8 | 25322294 | AX-170171970 | 0 | T | C | 0.351 | -8.07E-02 | 6.47E-02 | 2.58E+00 | 2.67E+00 | 2.10E-01 | no | | intergenic_variant | MODIFIER |
| 8 | 25341188 | AX-170514054 | 0 | G | A | 0.351 | -8.07E-02 | 6.47E-02 | 2.58E+00 | 2.67E+00 | 2.10E-01 | no | | intergenic_variant | MODIFIER |
| 8 | 25354671 | AX-170462527 | 0 | G | C | 0.351 | -8.07E-02 | 6.47E-02 | 2.58E+00 | 2.67E+00 | 2.10E-01 | no | | regulatory_region_variant | MODIFIER |
| 8 | 25376298 | AX-170461298 | 0 | C | G | 0.351 | -8.07E-02 | 6.47E-02 | 2.58E+00 | 2.67E+00 | 2.10E-01 | no | | regulatory_region_variant | MODIFIER |
| 8 | 25379138 | AX-170270274 | 0 | A | G | 0.351 | -8.07E-02 | 6.47E-02 | 2.58E+00 | 2.67E+00 | 2.10E-01 | no | | intergenic_variant | MODIFIER |
| 8 | 25382251 | AX-97801847 | 0 | C | A | 0.351 | -8.07E-02 | 6.47E-02 | 2.58E+00 | 2.67E+00 | 2.10E-01 | no | | intergenic_variant | MODIFIER |
| 8 | 25385651 | AX-170166625 | 0 | A | G | 0.351 | -8.07E-02 | 6.47E-02 | 2.58E+00 | 2.67E+00 | 2.10E-01 | no | | regulatory_region_variant | MODIFIER |
| 8 | 25388364 | AX-170419811 | 0 | G | T | 0.351 | -8.07E-02 | 6.47E-02 | 2.58E+00 | 2.67E+00 | 2.10E-01 | no | | intergenic_variant | MODIFIER |
| 8 | 25400209 | AX-170272469 | 0 | A | G | 0.351 | -8.07E-02 | 6.47E-02 | 2.58E+00 | 2.67E+00 | 2.10E-01 | no | Gm39147 | intron_variant,non_coding_transcript_variant | MODIFIER |
| 8 | 25400261 | AX-170329277 | 0 | C | T | 0.351 | -8.07E-02 | 6.47E-02 | 2.58E+00 | 2.67E+00 | 2.10E-01 | no | Gm39147 | intron_variant,non_coding_transcript_variant | MODIFIER |
| 8 | 25437760 | AX-170301388 | 0 | C | T | 0.351 | -8.07E-02 | 6.47E-02 | 2.58E+00 | 2.67E+00 | 2.10E-01 | no | Gm45237 | upstream_gene_variant | MODIFIER |
| 8 | 25441963 | AX-170079555 | 0 | A | G | 0.351 | -8.07E-02 | 6.47E-02 | 2.58E+00 | 2.67E+00 | 2.10E-01 | no | Gm45238 | downstream_gene_variant | MODIFIER |
| 8 | 25451632 | AX-170079554 | 0 | T | C | 0.351 | -8.07E-02 | 6.47E-02 | 2.58E+00 | 2.67E+00 | 2.10E-01 | no | Gm10689 | upstream_gene_variant | MODIFIER |
| 8 | 25487761 | AX-170525970 | 0 | C | G | 0.351 | -8.07E-02 | 6.47E-02 | 2.58E+00 | 2.67E+00 | 2.10E-01 | no | Gm31045 | upstream_gene_variant | MODIFIER |
| 8 | 25488107 | AX-170151351 | 0 | A | G | 0.351 | -8.07E-02 | 6.47E-02 | 2.58E+00 | 2.67E+00 | 2.10E-01 | no | Gm31045 | upstream_gene_variant | MODIFIER |
| 8 | 25500823 | AX-170114792 | 0 | C | A | 0.351 | -8.07E-02 | 6.47E-02 | 2.58E+00 | 2.67E+00 | 2.10E-01 | no | | intergenic_variant | MODIFIER |
| 8 | 25505038 | AX-170119502 | 0 | A | G | 0.351 | -8.07E-02 | 6.47E-02 | 2.58E+00 | 2.67E+00 | 2.10E-01 | no | | intergenic_variant | MODIFIER |
| 8 | 25513377 | AX-170195758 | 0 | G | T | 0.351 | -8.07E-02 | 6.47E-02 | 2.58E+00 | 2.67E+00 | 2.10E-01 | no | Fgfr1 | upstream_gene_variant | MODIFIER |
| 8 | 25515994 | AX-170488764 | 0 | G | A | 0.351 | -8.07E-02 | 6.47E-02 | 2.58E+00 | 2.67E+00 | 2.10E-01 | no | Fgfr1 | upstream_gene_variant | MODIFIER |
| 8 | 25523467 | AX-170491391 | 0 | C | T | 0.351 | -8.07E-02 | 6.47E-02 | 2.58E+00 | 2.67E+00 | 2.10E-01 | no | Fgfr1 | intron_variant | MODIFIER |
| 8 | 25527844 | AX-170378847 | 0 | G | A | 0.351 | -8.07E-02 | 6.47E-02 | 2.58E+00 | 2.67E+00 | 2.10E-01 | no | Fgfr1 | intron_variant | MODIFIER |
| 8 | 25532982 | AX-170308723 | 0 | T | C | 0.351 | -8.07E-02 | 6.47E-02 | 2.58E+00 | 2.67E+00 | 2.10E-01 | no | Fgfr1 | intron_variant | MODIFIER |
| 8 | 25538586 | AX-170167446 | 0 | C | T | 0.351 | -8.07E-02 | 6.47E-02 | 2.58E+00 | 2.67E+00 | 2.10E-01 | no | Fgfr1 | intron_variant | MODIFIER |
| 8 | 25552747 | AX-170030373 | 0 | G | A | 0.351 | -8.07E-02 | 6.47E-02 | 2.58E+00 | 2.67E+00 | 2.10E-01 | no | Fgfr1 | intron_variant | MODIFIER |
| 8 | 25580044 | AX-170286422 | 0 | A | G | 0.351 | -8.07E-02 | 6.47E-02 | 2.58E+00 | 2.67E+00 | 2.10E-01 | no | Letm2 | intron_variant | MODIFIER |
| 8 | 25592335 | AX-170407806 | 0 | C | G | 0.351 | -8.07E-02 | 6.47E-02 | 2.58E+00 | 2.67E+00 | 2.10E-01 | no | D830025C05Rik | upstream_gene_variant | MODIFIER |
| 8 | 25592369 | AX-170359787 | 0 | T | C | 0.351 | -8.07E-02 | 6.47E-02 | 2.58E+00 | 2.67E+00 | 2.10E-01 | no | D830025C05Rik | upstream_gene_variant | MODIFIER |
| 8 | 25592771 | AX-170030372 | 0 | T | C | 0.351 | -8.07E-02 | 6.47E-02 | 2.58E+00 | 2.67E+00 | 2.10E-01 | no | D830025C05Rik | upstream_gene_variant | MODIFIER |
| 8 | 25632546 | AX-170479226 | 0 | C | T | 0.351 | -8.07E-02 | 6.47E-02 | 2.58E+00 | 2.67E+00 | 2.10E-01 | no | Nsd3 | intron_variant | MODIFIER |
| 8 | 25637305 | AX-170178187 | 0 | G | A | 0.351 | -8.07E-02 | 6.47E-02 | 2.58E+00 | 2.67E+00 | 2.10E-01 | no | Nsd3 | upstream_gene_variant | MODIFIER |
| 8 | 25654783 | AX-170116422 | 0 | T | C | 0.351 | -8.07E-02 | 6.47E-02 | 2.58E+00 | 2.67E+00 | 2.10E-01 | no | Nsd3 | intron_variant | MODIFIER |
| 8 | 25669821 | AX-170404196 | 0 | G | C | 0.351 | -8.07E-02 | 6.47E-02 | 2.58E+00 | 2.67E+00 | 2.10E-01 | no | Nsd3 | intron_variant | MODIFIER |
| 8 | 25686956 | AX-170522251 | 0 | A | G | 0.351 | -8.07E-02 | 6.47E-02 | 2.58E+00 | 2.67E+00 | 2.10E-01 | no | Nsd3 | intron_variant | MODIFIER |
| 8 | 25698285 | AX-170288902 | 0 | T | G | 0.351 | -8.07E-02 | 6.47E-02 | 2.58E+00 | 2.67E+00 | 2.10E-01 | no | Nsd3 | intron_variant | MODIFIER |
| 8 | 25702751 | AX-170079557 | 0 | A | C | 0.351 | -8.07E-02 | 6.47E-02 | 2.58E+00 | 2.67E+00 | 2.10E-01 | no | Nsd3 | intron_variant | MODIFIER |
| 8 | 25732522 | AX-170079559 | 0 | G | A | 0.351 | -8.07E-02 | 6.47E-02 | 2.58E+00 | 2.67E+00 | 2.10E-01 | no | Ddhd2 | intron_variant | MODIFIER |
| 8 | 25750435 | AX-170299059 | 0 | G | A | 0.351 | -8.07E-02 | 6.47E-02 | 2.58E+00 | 2.67E+00 | 2.10E-01 | no | Ddhd2 | intron_variant | MODIFIER |

| | | | | | | | | | | | | | | | |
|---|---|---|---|---|---|---|---|---|---|---|---|---|---|---|---|
| 8 | 25757451 | AX-170163743 | 0 | C | G | 0.351 | -8.07E-02 | 6.47E-02 | 2.58E+00 | 2.67E+00 | 2.10E-01 | no | Ddhd2 | upstream_gene_variant | MODIFIER |
| 8 | 25891099 | AX-170408368 | 0 | G | T | 0.351 | -8.07E-02 | 6.47E-02 | 2.58E+00 | 2.67E+00 | 2.10E-01 | no | Kcnu1 | intron_variant | MODIFIER |
| 8 | 25894967 | AX-170349138 | 0 | T | A | 0.351 | -8.07E-02 | 6.47E-02 | 2.58E+00 | 2.67E+00 | 2.10E-01 | no | Kcnu1 | intron_variant | MODIFIER |
| 8 | 25937851 | AX-170290523 | 0 | T | C | 0.351 | -8.07E-02 | 6.47E-02 | 2.58E+00 | 2.67E+00 | 2.10E-01 | no | Kcnu1 | 3_prime_UTR_variant | MODIFIER |
| 8 | 25944336 | AX-170420990 | 0 | C | T | 0.351 | -8.07E-02 | 6.47E-02 | 2.58E+00 | 2.67E+00 | 2.10E-01 | no | Hgsnat | downstream_gene_variant | MODIFIER |
| 8 | 25966769 | AX-170107876 | 0 | C | T | 0.351 | -8.07E-02 | 6.47E-02 | 2.58E+00 | 2.67E+00 | 2.10E-01 | no | Hgsnat | intron_variant | MODIFIER |
| 8 | 25968367 | AX-97822864 | 0 | G | A | 0.351 | -8.07E-02 | 6.47E-02 | 2.58E+00 | 2.67E+00 | 2.10E-01 | no | Hgsnat | synonymous_variant | LOW |
| 8 | 25982869 | AX-170475907 | 0 | C | T | 0.351 | -8.07E-02 | 6.47E-02 | 2.58E+00 | 2.67E+00 | 2.10E-01 | no | Pomk | 3_prime_UTR_variant | MODIFIER |
| 8 | 26024382 | AX-170410464 | 0 | T | C | 0.351 | -8.07E-02 | 6.47E-02 | 2.58E+00 | 2.67E+00 | 2.10E-01 | no | Hook3 | 3_prime_UTR_variant | MODIFIER |
| 8 | 26024808 | AX-170419564 | 0 | C | T | 0.351 | -8.07E-02 | 6.47E-02 | 2.58E+00 | 2.67E+00 | 2.10E-01 | no | Hook3 | 3_prime_UTR_variant | MODIFIER |
| 8 | 26032557 | AX-170163608 | 0 | G | A | 0.351 | -8.07E-02 | 6.47E-02 | 2.58E+00 | 2.67E+00 | 2.10E-01 | no | Hook3 | intron_variant | MODIFIER |
| 8 | 26041690 | AX-97804549 | 0 | T | C | 0.351 | -8.07E-02 | 6.47E-02 | 2.58E+00 | 2.67E+00 | 2.10E-01 | no | Hook3 | intron_variant | MODIFIER |
| 8 | 26041737 | AX-170318545 | 0 | G | A | 0.351 | -8.07E-02 | 6.47E-02 | 2.58E+00 | 2.67E+00 | 2.10E-01 | no | Hook3 | intron_variant | MODIFIER |
| 8 | 26041905 | AX-170266922 | 0 | T | C | 0.351 | -8.07E-02 | 6.47E-02 | 2.58E+00 | 2.67E+00 | 2.10E-01 | no | Hook3 | intron_variant | MODIFIER |
| 8 | 26055909 | AX-170430610 | 0 | G | T | 0.351 | -8.07E-02 | 6.47E-02 | 2.58E+00 | 2.67E+00 | 2.10E-01 | no | Hook3 | intron_variant | MODIFIER |
| 8 | 26070985 | AX-170030376 | 0 | T | C | 0.351 | -8.07E-02 | 6.47E-02 | 2.58E+00 | 2.67E+00 | 2.10E-01 | no | Hook3 | intron_variant | MODIFIER |
| 8 | 26075661 | AX-170372596 | 0 | C | T | 0.351 | -8.07E-02 | 6.47E-02 | 2.58E+00 | 2.67E+00 | 2.10E-01 | no | Hook3 | intron_variant | MODIFIER |
| 8 | 26091993 | AX-170335983 | 0 | T | C | 0.351 | -8.07E-02 | 6.47E-02 | 2.58E+00 | 2.67E+00 | 2.10E-01 | no | Hook3 | intron_variant | MODIFIER |
| 8 | 26093772 | AX-170123185 | 0 | C | A | 0.351 | -8.07E-02 | 6.47E-02 | 2.58E+00 | 2.67E+00 | 2.10E-01 | no | Hook3 | intron_variant | MODIFIER |
| 8 | 26099928 | AX-170499517 | 0 | C | T | 0.351 | -8.07E-02 | 6.47E-02 | 2.58E+00 | 2.67E+00 | 2.10E-01 | no | Hook3 | intron_variant | MODIFIER |
| 8 | 26108706 | AX-170492890 | 0 | G | T | 0.351 | -8.07E-02 | 6.47E-02 | 2.58E+00 | 2.67E+00 | 2.10E-01 | no | Hook3 | intron_variant | MODIFIER |
| 8 | 26108798 | AX-170198145 | 0 | C | G | 0.351 | -8.07E-02 | 6.47E-02 | 2.58E+00 | 2.67E+00 | 2.10E-01 | no | Hook3 | intron_variant | MODIFIER |
| 8 | 26238389 | AX-170335112 | 0 | C | T | 0.351 | -8.07E-02 | 6.47E-02 | 2.58E+00 | 2.67E+00 | 2.10E-01 | no | Gm22227 | downstream_gene_variant | MODIFIER |
| 8 | 26317047 | AX-170284298 | 0 | T | C | 0.351 | -8.07E-02 | 6.47E-02 | 2.58E+00 | 2.67E+00 | 2.10E-01 | no | Gm31784 | upstream_gene_variant | MODIFIER |
| 8 | 26338967 | AX-170079572 | 0 | C | T | 0.351 | -8.07E-02 | 6.47E-02 | 2.58E+00 | 2.67E+00 | 2.10E-01 | no | Gm45307 | intron_variant,non_coding_transcript_variant | MODIFIER |
| 8 | 26339371 | AX-170079570 | 0 | A | G | 0.351 | -8.07E-02 | 6.47E-02 | 2.58E+00 | 2.67E+00 | 2.10E-01 | no | Gm45307 | intron_variant,non_coding_transcript_variant | MODIFIER |
| 8 | 26345451 | AX-170079569 | 0 | T | G | 0.351 | -8.07E-02 | 6.47E-02 | 2.58E+00 | 2.67E+00 | 2.10E-01 | no | Gm45307 | intron_variant,non_coding_transcript_variant | MODIFIER |
| 8 | 26345924 | AX-170144007 | 0 | G | A | 0.351 | -8.07E-02 | 6.47E-02 | 2.58E+00 | 2.67E+00 | 2.10E-01 | no | Gm45307 | intron_variant,non_coding_transcript_variant | MODIFIER |
| 8 | 26353267 | AX-170365271 | 0 | A | T | 0.351 | -8.07E-02 | 6.47E-02 | 2.58E+00 | 2.67E+00 | 2.10E-01 | no | Gm45307 | intron_variant,non_coding_transcript_variant | MODIFIER |
| 8 | 26382741 | AX-170470189 | 0 | G | A | 0.351 | -8.07E-02 | 6.47E-02 | 2.58E+00 | 2.67E+00 | 2.10E-01 | no | | intergenic_variant | MODIFIER |
| 8 | 26383513 | AX-170348107 | 0 | G | A | 0.351 | -8.07E-02 | 6.47E-02 | 2.58E+00 | 2.67E+00 | 2.10E-01 | no | | intergenic_variant | MODIFIER |
| 8 | 26430853 | AX-170373406 | 0 | A | G | 0.351 | -8.07E-02 | 6.47E-02 | 2.58E+00 | 2.67E+00 | 2.10E-01 | no | Gm31898 | non_coding_transcript_exon_variant | MODIFIER |
| 8 | 26431350 | AX-170238447 | 0 | T | A | 0.351 | -8.07E-02 | 6.47E-02 | 2.58E+00 | 2.67E+00 | 2.10E-01 | no | Gm31898 | intron_variant,non_coding_transcript_variant | MODIFIER |
| 8 | 26432539 | AX-170194140 | 0 | T | C | 0.351 | -8.07E-02 | 6.47E-02 | 2.58E+00 | 2.67E+00 | 2.10E-01 | no | Gm31898 | intron_variant,non_coding_transcript_variant | MODIFIER |
| 8 | 26432751 | AX-170282959 | 0 | C | A | 0.351 | -8.07E-02 | 6.47E-02 | 2.58E+00 | 2.67E+00 | 2.10E-01 | no | Gm31898 | intron_variant,non_coding_transcript_variant | MODIFIER |
| 8 | 26441371 | AX-97821477 | 0 | C | T | 0.351 | -8.07E-02 | 6.47E-02 | 2.58E+00 | 2.67E+00 | 2.10E-01 | no | Gm45580 | downstream_gene_variant | MODIFIER |
| 8 | 26442133 | AX-170458986 | 0 | G | A | 0.351 | -8.07E-02 | 6.47E-02 | 2.58E+00 | 2.67E+00 | 2.10E-01 | no | Gm45580 | downstream_gene_variant | MODIFIER |
| 8 | 26473574 | AX-170263276 | 0 | C | T | 0.351 | -8.07E-02 | 6.47E-02 | 2.58E+00 | 2.67E+00 | 2.10E-01 | no | Gm31983 | intron_variant,non_coding_transcript_variant | MODIFIER |
| 8 | 26482319 | AX-97801483 | 0 | G | A | 0.351 | -8.07E-02 | 6.47E-02 | 2.58E+00 | 2.67E+00 | 2.10E-01 | no | Gm31983 | intron_variant,non_coding_transcript_variant | MODIFIER |
| 8 | 26488662 | AX-170212520 | 0 | C | T | 0.351 | -8.07E-02 | 6.47E-02 | 2.58E+00 | 2.67E+00 | 2.10E-01 | no | Gm31983 | upstream_gene_variant | MODIFIER |
| 4 | 86190713 | AX-170220657 | 0 | T | C | 0.342 | 1.36E-01 | 1.09E-01 | 2.63E+00 | 2.70E+00 | 2.10E-01 | no | Adamtsl1 | intron_variant | MODIFIER |
| 3 | 151849432 | AX-170361698 | 1 | A | G | 0.339 | 9.05E-02 | 7.25E-02 | 2.92E+00 | 3.03E+00 | 2.10E-01 | no | | intergenic_variant | MODIFIER |
| 4 | 77328040 | AX-170162184 | 0 | C | T | 0.316 | 1.42E-01 | 1.14E-01 | 2.87E+00 | 2.94E+00 | 2.10E-01 | no | Ptprd | intron_variant,non_coding_transcript_variant | MODIFIER |
| 16 | 72188420 | AX-170388984 | 0 | T | C | 0.491 | 9.58E-02 | 7.63E-02 | 2.51E+00 | 2.58E+00 | 2.10E-01 | no | | intergenic_variant | MODIFIER |
| 3 | 152975919 | AX-97821365 | 1 | C | A | 0.357 | 8.59E-02 | 6.85E-02 | 3.00E+00 | 3.12E+00 | 2.11E-01 | no | St6galnac5 | intron_variant | MODIFIER |
| 5 | 56517726 | AX-170403620 | 0 | G | A | 0.465 | 1.30E-01 | 1.02E-01 | 2.39E+00 | 2.45E+00 | 2.11E-01 | no | | intergenic_variant | MODIFIER |
| 18 | 72399376 | AX-170015966 | 0 | G | T | 0.184 | -1.38E-01 | 1.11E-01 | 2.72E+00 | 2.81E+00 | 2.11E-01 | no | | intergenic_variant | MODIFIER |
| 18 | 3012495 | AX-170060038 | 0 | T | C | 0.412 | 1.31E-01 | 1.06E-01 | 2.59E+00 | 2.72E+00 | 2.11E-01 | no | AC125218.3 | upstream_gene_variant | MODIFIER |
| 5 | 48793900 | AX-170108185 | 0 | T | C | 0.465 | 1.27E-01 | 1.00E-01 | 2.41E+00 | 2.45E+00 | 2.11E-01 | no | Kcnip4 | intron_variant | MODIFIER |
| 19 | 60069442 | AX-170514445 | 0 | A | C | 0.509 | 8.03E-02 | 6.47E-02 | 2.66E+00 | 2.74E+00 | 2.11E-01 | no | | intergenic_variant | MODIFIER |
| 18 | 79115764 | AX-170417302 | 1 | C | T | 0.464 | 1.08E-01 | 8.50E-02 | 2.37E+00 | 2.44E+00 | 2.11E-01 | no | | intergenic_variant | MODIFIER |
| 18 | 63468479 | AX-170396822 | 0 | C | G | 0.184 | -1.43E-01 | 1.15E-01 | 2.68E+00 | 2.75E+00 | 2.11E-01 | no | | intergenic_variant | MODIFIER |
| 16 | 35792089 | AX-97804566 | 0 | A | G | 0.482 | 9.58E-02 | 7.66E-02 | 2.53E+00 | 2.58E+00 | 2.11E-01 | no | Hspbap1 | intron_variant,NMD_transcript_variant | MODIFIER |
| 13 | 20146576 | AX-170323852 | 1 | C | A | 0.179 | -1.56E-01 | 1.24E-01 | 2.47E+00 | 2.52E+00 | 2.11E-01 | no | Elmo1 | intron_variant | MODIFIER |

| | | | | | | | | | | | | | | |
|---|---|---|---|---|---|---|---|---|---|---|---|---|---|---|
| 13 | 20179306 | AX-170278591 | 1 | G | T | 0.179 | -1.56E-01 | 1.24E-01 | 2.47E+00 | 2.52E+00 | 2.11E-01 | no | Elmo1 | intron_variant | MODIFIER |
| 4 | 107512994 | AX-97839699 | 1 | T | C | 0.268 | 1.20E-01 | 9.70E-02 | 2.61E+00 | 2.70E+00 | 2.11E-01 | no | Glis1 | intron_variant | MODIFIER |
| 2 | 42889675 | AX-170125702 | 1 | T | C | 0.036 | -2.13E-01 | 1.71E-01 | 2.54E+00 | 2.65E+00 | 2.12E-01 | Yes | | intergenic_variant | MODIFIER |
| 8 | 83450489 | AX-170464314 | 0 | T | C | 0.211 | -1.22E-01 | 9.83E-02 | 2.90E+00 | 3.06E+00 | 2.12E-01 | no | Scoc | intron_variant | MODIFIER |
| 4 | 85155422 | AX-170381700 | 1 | A | T | 0.348 | 1.40E-01 | 1.13E-01 | 2.85E+00 | 2.93E+00 | 2.12E-01 | no | | intergenic_variant | MODIFIER |
| 13 | 71518426 | AX-170310257 | 1 | A | T | 0.062 | 1.71E-01 | 1.39E-01 | 2.87E+00 | 3.04E+00 | 2.12E-01 | Yes | 1700112M02Rik | intron_variant,non_coding_transcript_variant | MODIFIER |
| 4 | 120670253 | AX-97793289 | 0 | G | A | 0.228 | 1.37E-01 | 1.10E-01 | 2.61E+00 | 2.67E+00 | 2.12E-01 | no | Cited4 | downstream_gene_variant | MODIFIER |
| 4 | 120672395 | AX-170120174 | 0 | C | G | 0.228 | 1.37E-01 | 1.10E-01 | 2.61E+00 | 2.67E+00 | 2.12E-01 | no | Cited4 | downstream_gene_variant | MODIFIER |
| 4 | 120672883 | AX-170262825 | 0 | C | G | 0.228 | 1.37E-01 | 1.10E-01 | 2.61E+00 | 2.67E+00 | 2.12E-01 | no | | regulatory_region_variant | MODIFIER |
| 4 | 120693128 | AX-170095126 | 0 | G | A | 0.228 | 1.37E-01 | 1.10E-01 | 2.61E+00 | 2.67E+00 | 2.12E-01 | no | Kcnq4 | downstream_gene_variant | MODIFIER |
| 4 | 120702858 | AX-170375717 | 0 | A | G | 0.228 | 1.37E-01 | 1.10E-01 | 2.61E+00 | 2.67E+00 | 2.12E-01 | no | Kcnq4 | intron_variant | MODIFIER |
| 4 | 120709727 | AX-170162901 | 0 | T | C | 0.228 | 1.37E-01 | 1.10E-01 | 2.61E+00 | 2.67E+00 | 2.12E-01 | no | Kcnq4 | intron_variant | MODIFIER |
| 4 | 120725165 | AX-170070490 | 0 | A | G | 0.228 | 1.37E-01 | 1.10E-01 | 2.61E+00 | 2.67E+00 | 2.12E-01 | no | Kcnq4 | intron_variant | MODIFIER |
| 4 | 120727885 | AX-170174418 | 0 | A | G | 0.228 | 1.37E-01 | 1.10E-01 | 2.61E+00 | 2.67E+00 | 2.12E-01 | no | Kcnq4 | intron_variant | MODIFIER |
| 4 | 120758549 | AX-170252033 | 0 | G | T | 0.228 | 1.37E-01 | 1.10E-01 | 2.61E+00 | 2.67E+00 | 2.12E-01 | no | Nfyc | intron_variant | MODIFIER |
| 4 | 120759769 | AX-170310581 | 0 | C | A | 0.228 | 1.37E-01 | 1.10E-01 | 2.61E+00 | 2.67E+00 | 2.12E-01 | no | Nfyc | intron_variant | MODIFIER |
| 4 | 120760053 | AX-170166879 | 0 | G | C | 0.228 | 1.37E-01 | 1.10E-01 | 2.61E+00 | 2.67E+00 | 2.12E-01 | no | Nfyc | intron_variant | MODIFIER |
| 4 | 120761555 | AX-170124694 | 0 | T | G | 0.228 | 1.37E-01 | 1.10E-01 | 2.61E+00 | 2.67E+00 | 2.12E-01 | no | Nfyc | intron_variant | MODIFIER |
| 4 | 120800884 | AX-170238506 | 0 | C | T | 0.228 | 1.37E-01 | 1.10E-01 | 2.61E+00 | 2.67E+00 | 2.12E-01 | no | Nfyc | intron_variant | MODIFIER |
| 4 | 120807849 | AX-170338887 | 0 | A | G | 0.228 | 1.37E-01 | 1.10E-01 | 2.61E+00 | 2.67E+00 | 2.12E-01 | no | Nfyc | intron_variant | MODIFIER |
| 4 | 120812397 | AX-170401149 | 0 | T | C | 0.228 | 1.37E-01 | 1.10E-01 | 2.61E+00 | 2.67E+00 | 2.12E-01 | no | Nfyc | intron_variant | MODIFIER |
| 4 | 120840355 | AX-170329676 | 0 | C | G | 0.228 | 1.37E-01 | 1.10E-01 | 2.61E+00 | 2.67E+00 | 2.12E-01 | no | Gm23936 | downstream_gene_variant | MODIFIER |
| 4 | 120840795 | AX-97785799 | 0 | C | T | 0.228 | 1.37E-01 | 1.10E-01 | 2.61E+00 | 2.67E+00 | 2.12E-01 | no | Gm23936 | downstream_gene_variant | MODIFIER |
| 1 | 185907577 | AX-170276596 | 0 | T | C | 0.228 | 1.37E-01 | 1.09E-01 | 2.40E+00 | 2.47E+00 | 2.13E-01 | no | | intergenic_variant | MODIFIER |
| 1 | 185931736 | AX-170374105 | 0 | G | A | 0.228 | 1.37E-01 | 1.09E-01 | 2.40E+00 | 2.47E+00 | 2.13E-01 | no | | intergenic_variant | MODIFIER |
| 1 | 185966205 | AX-170205205 | 0 | T | C | 0.228 | 1.37E-01 | 1.09E-01 | 2.40E+00 | 2.47E+00 | 2.13E-01 | no | | intergenic_variant | MODIFIER |
| 1 | 185972512 | AX-170137212 | 0 | C | T | 0.228 | 1.37E-01 | 1.09E-01 | 2.40E+00 | 2.47E+00 | 2.13E-01 | no | | intergenic_variant | MODIFIER |
| 1 | 185972680 | AX-170532331 | 0 | C | T | 0.228 | 1.37E-01 | 1.09E-01 | 2.40E+00 | 2.47E+00 | 2.13E-01 | no | | intergenic_variant | MODIFIER |
| 1 | 185972705 | AX-170313658 | 0 | C | T | 0.228 | 1.37E-01 | 1.09E-01 | 2.40E+00 | 2.47E+00 | 2.13E-01 | no | | intergenic_variant | MODIFIER |
| 1 | 185972906 | AX-170525765 | 0 | T | A | 0.228 | 1.37E-01 | 1.09E-01 | 2.40E+00 | 2.47E+00 | 2.13E-01 | no | | intergenic_variant | MODIFIER |
| 1 | 186137910 | AX-97824926 | 0 | G | A | 0.228 | 1.37E-01 | 1.09E-01 | 2.40E+00 | 2.47E+00 | 2.13E-01 | no | | regulatory_region_variant | MODIFIER |
| 1 | 186138396 | AX-170281503 | 0 | T | A | 0.228 | 1.37E-01 | 1.09E-01 | 2.40E+00 | 2.47E+00 | 2.13E-01 | no | | intergenic_variant | MODIFIER |
| 1 | 186139418 | AX-170443762 | 0 | C | A | 0.228 | 1.37E-01 | 1.09E-01 | 2.40E+00 | 2.47E+00 | 2.13E-01 | no | | intergenic_variant | MODIFIER |
| 1 | 186141712 | AX-170273539 | 0 | A | G | 0.228 | 1.37E-01 | 1.09E-01 | 2.40E+00 | 2.47E+00 | 2.13E-01 | no | | intergenic_variant | MODIFIER |
| 1 | 186146821 | AX-170512053 | 0 | G | A | 0.228 | 1.37E-01 | 1.09E-01 | 2.40E+00 | 2.47E+00 | 2.13E-01 | no | | intergenic_variant | MODIFIER |
| 1 | 186159757 | AX-170250631 | 0 | C | G | 0.228 | 1.37E-01 | 1.09E-01 | 2.40E+00 | 2.47E+00 | 2.13E-01 | no | | intergenic_variant | MODIFIER |
| 1 | 186160144 | AX-170414582 | 0 | T | G | 0.228 | 1.37E-01 | 1.09E-01 | 2.40E+00 | 2.47E+00 | 2.13E-01 | no | | intergenic_variant | MODIFIER |
| 1 | 186160286 | AX-97788883 | 0 | C | T | 0.228 | 1.37E-01 | 1.09E-01 | 2.40E+00 | 2.47E+00 | 2.13E-01 | no | | intergenic_variant | MODIFIER |
| 1 | 186168567 | AX-170461674 | 0 | A | G | 0.228 | 1.37E-01 | 1.09E-01 | 2.40E+00 | 2.47E+00 | 2.13E-01 | no | | intergenic_variant | MODIFIER |
| 1 | 186168587 | AX-170242016 | 0 | T | C | 0.228 | 1.37E-01 | 1.09E-01 | 2.40E+00 | 2.47E+00 | 2.13E-01 | no | | intergenic_variant | MODIFIER |
| 1 | 186202778 | AX-170383815 | 0 | A | G | 0.228 | 1.37E-01 | 1.09E-01 | 2.40E+00 | 2.47E+00 | 2.13E-01 | no | Gm37134 | intron_variant,non_coding_transcript_variant | MODIFIER |
| 1 | 186205432 | AX-170161360 | 0 | C | T | 0.228 | 1.37E-01 | 1.09E-01 | 2.40E+00 | 2.47E+00 | 2.13E-01 | no | Gm37134 | intron_variant,non_coding_transcript_variant | MODIFIER |
| 1 | 186233779 | AX-170114211 | 0 | A | C | 0.228 | 1.37E-01 | 1.09E-01 | 2.40E+00 | 2.47E+00 | 2.13E-01 | no | | intergenic_variant | MODIFIER |
| 1 | 186234612 | AX-170389260 | 0 | C | A | 0.228 | 1.37E-01 | 1.09E-01 | 2.40E+00 | 2.47E+00 | 2.13E-01 | no | | regulatory_region_variant | MODIFIER |
| 1 | 186234711 | AX-170330314 | 0 | G | A | 0.228 | 1.37E-01 | 1.09E-01 | 2.40E+00 | 2.47E+00 | 2.13E-01 | no | | regulatory_region_variant | MODIFIER |
| 1 | 186236586 | AX-170386012 | 0 | A | T | 0.228 | 1.37E-01 | 1.09E-01 | 2.40E+00 | 2.47E+00 | 2.13E-01 | no | | regulatory_region_variant | MODIFIER |
| 1 | 186236967 | AX-97855104 | 0 | A | G | 0.228 | 1.37E-01 | 1.09E-01 | 2.40E+00 | 2.47E+00 | 2.13E-01 | no | | regulatory_region_variant | MODIFIER |
| 1 | 186254300 | AX-170491864 | 0 | T | C | 0.228 | 1.37E-01 | 1.09E-01 | 2.40E+00 | 2.47E+00 | 2.13E-01 | no | | regulatory_region_variant | MODIFIER |
| 1 | 186261888 | AX-170102555 | 0 | G | A | 0.228 | 1.37E-01 | 1.09E-01 | 2.40E+00 | 2.47E+00 | 2.13E-01 | no | | regulatory_region_variant | MODIFIER |
| 1 | 186263217 | AX-170342645 | 0 | A | G | 0.228 | 1.37E-01 | 1.09E-01 | 2.40E+00 | 2.47E+00 | 2.13E-01 | no | | regulatory_region_variant | MODIFIER |
| 1 | 186271959 | AX-170318761 | 0 | G | A | 0.228 | 1.37E-01 | 1.09E-01 | 2.40E+00 | 2.47E+00 | 2.13E-01 | no | | intergenic_variant | MODIFIER |
| 1 | 186585743 | AX-170002497 | 0 | T | G | 0.228 | 1.37E-01 | 1.09E-01 | 2.40E+00 | 2.47E+00 | 2.13E-01 | no | | regulatory_region_variant | MODIFIER |
| 1 | 186670040 | AX-170356872 | 0 | A | G | 0.228 | 1.37E-01 | 1.09E-01 | 2.40E+00 | 2.47E+00 | 2.13E-01 | no | Tgfb2 | intron_variant | MODIFIER |
| 1 | 186672937 | AX-170363887 | 0 | G | A | 0.228 | 1.37E-01 | 1.09E-01 | 2.40E+00 | 2.47E+00 | 2.13E-01 | no | Tgfb2 | intron_variant | MODIFIER |

| | | | | | | | | | | | | | | | |
|---|---|---|---|---|---|---|---|---|---|---|---|---|---|---|---|
| 1 | 186690942 | AX-170420828 | 0 | A | G | 0.228 | 1.37E-01 | 1.09E-01 | 2.40E+00 | 2.47E+00 | 2.13E-01 | no | Tgfb2 | intron_variant | MODIFIER |
| 1 | 186694193 | AX-170142460 | 0 | T | C | 0.228 | 1.37E-01 | 1.09E-01 | 2.40E+00 | 2.47E+00 | 2.13E-01 | no | Tgfb2 | intron_variant | MODIFIER |
| 1 | 186894572 | AX-170248428 | 0 | A | C | 0.228 | 1.37E-01 | 1.09E-01 | 2.40E+00 | 2.47E+00 | 2.13E-01 | no | | intergenic_variant | MODIFIER |
| 1 | 186894983 | AX-170043883 | 0 | A | G | 0.228 | 1.37E-01 | 1.09E-01 | 2.40E+00 | 2.47E+00 | 2.13E-01 | no | | intergenic_variant | MODIFIER |
| 3 | 36548983 | AX-170520222 | 0 | C | T | 0.211 | 1.13E-01 | 9.10E-02 | 2.93E+00 | 3.05E+00 | 2.13E-01 | no | Exosc9 | upstream_gene_variant | MODIFIER |
| 7 | 11991442 | AX-97797848 | 1 | T | C | 0.357 | 9.91E-02 | 8.01E-02 | 2.62E+00 | 2.74E+00 | 2.13E-01 | no | Vmn1r-ps59 | upstream_gene_variant | MODIFIER |
| 9 | 8181205 | AX-97790869 | 1 | C | T | 0.464 | 1.01E-01 | 8.07E-02 | 2.45E+00 | 2.53E+00 | 2.13E-01 | no | Gm19324 | intron_variant,non_coding_transcript_variant | MODIFIER |
| 1 | 132268461 | AX-170187449 | 1 | G | A | 0.545 | 1.04E-01 | 8.42E-02 | 2.62E+00 | 2.69E+00 | 2.13E-01 | no | | intergenic_variant | MODIFIER |
| 16 | 72563280 | AX-170306059 | 1 | C | G | 0.5 | 9.59E-02 | 7.69E-02 | 2.51E+00 | 2.58E+00 | 2.13E-01 | no | Robo1 | intron_variant | MODIFIER |
| 4 | 131598608 | AX-170023954 | 0 | T | A | 0.167 | 1.39E-01 | 1.12E-01 | 2.60E+00 | 2.66E+00 | 2.13E-01 | no | | intergenic_variant | MODIFIER |
| 4 | 136988467 | AX-170291758 | 0 | C | A | 0.167 | 1.39E-01 | 1.12E-01 | 2.60E+00 | 2.66E+00 | 2.13E-01 | no | Zbtb40 | intron_variant | MODIFIER |
| 8 | 26095652 | AX-170500479 | 0 | T | A | 0.351 | -7.99E-02 | 6.45E-02 | 2.59E+00 | 2.68E+00 | 2.13E-01 | no | Hook3 | intron_variant | MODIFIER |
| 2 | 124667439 | AX-170199758 | 0 | A | G | 0.14 | -2.23E-01 | 1.78E-01 | 3.05E+00 | 3.16E+00 | 2.13E-01 | Yes | Sema6d | downstream_gene_variant | MODIFIER |
| 2 | 156584210 | AX-97795399 | 0 | C | T | 0.14 | -2.23E-01 | 1.78E-01 | 3.05E+00 | 3.16E+00 | 2.13E-01 | Yes | | regulatory_region_variant | MODIFIER |
| 8 | 25632626 | AX-170425035 | 2 | G | A | 0.345 | -7.97E-02 | 6.40E-02 | 2.51E+00 | 2.61E+00 | 2.14E-01 | no | Nsd3 | intron_variant | MODIFIER |
| 5 | 103549729 | AX-97852811 | 1 | G | A | 0.036 | 1.34E-01 | 1.10E-01 | 2.81E+00 | 3.02E+00 | 2.14E-01 | no | Ptpn13 | intron_variant | MODIFIER |
| 4 | 103928331 | AX-170281579 | 0 | G | A | 0.219 | 1.24E-01 | 1.01E-01 | 2.65E+00 | 2.77E+00 | 2.14E-01 | no | Dab1 | intron_variant | MODIFIER |
| 4 | 69985574 | AX-170191949 | 1 | G | A | 0.312 | 1.44E-01 | 1.16E-01 | 2.85E+00 | 2.91E+00 | 2.14E-01 | no | | intergenic_variant | MODIFIER |
| 4 | 125612235 | AX-170023885 | 0 | C | T | 0.228 | 1.35E-01 | 1.09E-01 | 2.49E+00 | 2.56E+00 | 2.14E-01 | no | Grik3 | intron_variant | MODIFIER |
| 8 | 25316453 | AX-170187518 | 0 | A | T | 0.351 | -7.94E-02 | 6.42E-02 | 2.61E+00 | 2.70E+00 | 2.14E-01 | no | | intergenic_variant | MODIFIER |
| 14 | 112170018 | AX-97827873 | 1 | C | T | 0.321 | 8.98E-02 | 7.27E-02 | 2.61E+00 | 2.71E+00 | 2.14E-01 | no | | intergenic_variant | MODIFIER |
| 1 | 121605203 | AX-170395433 | 1 | A | G | 0.58 | 1.04E-01 | 8.33E-02 | 2.50E+00 | 2.56E+00 | 2.14E-01 | no | | regulatory_region_variant | MODIFIER |
| 5 | 60521745 | AX-170218812 | 0 | T | C | 0.474 | 1.25E-01 | 9.89E-02 | 2.37E+00 | 2.44E+00 | 2.14E-01 | no | | intergenic_variant | MODIFIER |
| 5 | 63827213 | AX-97834313 | 0 | A | G | 0.474 | 1.25E-01 | 9.89E-02 | 2.37E+00 | 2.44E+00 | 2.14E-01 | no | 0610040J01Rik | intron_variant | MODIFIER |
| 4 | 137439988 | AX-170070824 | 0 | T | C | 0.175 | 1.32E-01 | 1.07E-01 | 2.62E+00 | 2.69E+00 | 2.14E-01 | no | Gm13010 | upstream_gene_variant | MODIFIER |
| 2 | 123714541 | AX-170503680 | 1 | G | A | 0.161 | 1.83E-01 | 1.49E-01 | 2.68E+00 | 2.81E+00 | 2.15E-01 | Yes | | intergenic_variant | MODIFIER |
| 4 | 86545703 | AX-97805319 | 1 | A | G | 0.33 | 1.28E-01 | 1.04E-01 | 2.63E+00 | 2.72E+00 | 2.15E-01 | no | Saxo1 | intron_variant | MODIFIER |
| 5 | 67501453 | AX-170322943 | 0 | T | C | 0.456 | 1.25E-01 | 1.00E-01 | 2.49E+00 | 2.56E+00 | 2.15E-01 | no | | intergenic_variant | MODIFIER |
| 3 | 28845650 | AX-170238585 | 1 | A | T | 0.161 | -1.12E-01 | 9.06E-02 | 2.88E+00 | 3.01E+00 | 2.15E-01 | no | | intergenic_variant | MODIFIER |
| 7 | 93689391 | AX-170078096 | 0 | G | A | 0.105 | -1.92E-01 | 1.56E-01 | 2.81E+00 | 2.97E+00 | 2.15E-01 | Yes | | intergenic_variant | MODIFIER |
| 1 | 126607364 | AX-170271466 | 1 | C | T | 0.571 | 1.04E-01 | 8.33E-02 | 2.50E+00 | 2.56E+00 | 2.15E-01 | no | Nckap5 | intron_variant | MODIFIER |
| 1 | 134604646 | AX-170153032 | 0 | A | G | 0.526 | 1.00E-01 | 8.15E-02 | 2.64E+00 | 2.71E+00 | 2.15E-01 | no | Kdm5b | intron_variant | MODIFIER |
| 1 | 120362932 | AX-170366872 | 1 | A | G | 0.58 | 1.04E-01 | 8.34E-02 | 2.51E+00 | 2.56E+00 | 2.15E-01 | no | | intergenic_variant | MODIFIER |
| 4 | 95998552 | AX-170431351 | 1 | C | T | 0.259 | 1.25E-01 | 9.97E-02 | 2.43E+00 | 2.51E+00 | 2.15E-01 | no | Hook1 | intron_variant | MODIFIER |
| 5 | 67403195 | AX-170228212 | 1 | C | T | 0.473 | 1.23E-01 | 9.85E-02 | 2.44E+00 | 2.52E+00 | 2.16E-01 | no | Bend4 | intron_variant | MODIFIER |
| 4 | 83333145 | AX-170138407 | 0 | C | A | 0.333 | 1.39E-01 | 1.13E-01 | 2.84E+00 | 2.91E+00 | 2.16E-01 | no | | intergenic_variant | MODIFIER |
| 4 | 84775717 | AX-170223020 | 0 | A | G | 0.333 | 1.39E-01 | 1.13E-01 | 2.84E+00 | 2.91E+00 | 2.16E-01 | no | | intergenic_variant | MODIFIER |
| 4 | 84833855 | AX-170300791 | 0 | G | A | 0.333 | 1.39E-01 | 1.13E-01 | 2.84E+00 | 2.91E+00 | 2.16E-01 | no | | intergenic_variant | MODIFIER |
| 4 | 84835172 | AX-170318913 | 0 | T | A | 0.333 | 1.39E-01 | 1.13E-01 | 2.84E+00 | 2.91E+00 | 2.16E-01 | no | | intergenic_variant | MODIFIER |
| 4 | 84859153 | AX-97802952 | 0 | C | T | 0.333 | 1.39E-01 | 1.13E-01 | 2.84E+00 | 2.91E+00 | 2.16E-01 | no | | intergenic_variant | MODIFIER |
| 4 | 84881557 | AX-97849446 | 0 | G | A | 0.333 | 1.39E-01 | 1.13E-01 | 2.84E+00 | 2.91E+00 | 2.16E-01 | no | Cntln | upstream_gene_variant | MODIFIER |
| 4 | 84887775 | AX-170333384 | 0 | C | G | 0.333 | 1.39E-01 | 1.13E-01 | 2.84E+00 | 2.91E+00 | 2.16E-01 | no | Cntln | intron_variant | MODIFIER |
| 4 | 84897442 | AX-170400829 | 0 | A | G | 0.333 | 1.39E-01 | 1.13E-01 | 2.84E+00 | 2.91E+00 | 2.16E-01 | no | Cntln | intron_variant | MODIFIER |
| 4 | 85272228 | AX-170150035 | 0 | T | G | 0.333 | 1.39E-01 | 1.13E-01 | 2.84E+00 | 2.91E+00 | 2.16E-01 | no | Sh3gl2 | intron_variant | MODIFIER |
| 4 | 85272603 | AX-170432999 | 0 | T | C | 0.333 | 1.39E-01 | 1.13E-01 | 2.84E+00 | 2.91E+00 | 2.16E-01 | no | Sh3gl2 | intron_variant | MODIFIER |
| 4 | 85273414 | AX-170417488 | 0 | A | G | 0.333 | 1.39E-01 | 1.13E-01 | 2.84E+00 | 2.91E+00 | 2.16E-01 | no | Sh3gl2 | intron_variant | MODIFIER |
| 4 | 85293912 | AX-97842821 | 0 | T | C | 0.333 | 1.39E-01 | 1.13E-01 | 2.84E+00 | 2.91E+00 | 2.16E-01 | no | Sh3gl2 | intron_variant | MODIFIER |
| 4 | 85296155 | AX-170510951 | 0 | T | C | 0.333 | 1.39E-01 | 1.13E-01 | 2.84E+00 | 2.91E+00 | 2.16E-01 | no | Sh3gl2 | intron_variant | MODIFIER |
| 4 | 85296193 | AX-170408190 | 0 | G | C | 0.333 | 1.39E-01 | 1.13E-01 | 2.84E+00 | 2.91E+00 | 2.16E-01 | no | Sh3gl2 | intron_variant | MODIFIER |
| 4 | 85300261 | AX-170324641 | 0 | T | C | 0.333 | 1.39E-01 | 1.13E-01 | 2.84E+00 | 2.91E+00 | 2.16E-01 | no | Sh3gl2 | intron_variant | MODIFIER |
| 4 | 85584154 | AX-97784125 | 0 | T | G | 0.333 | 1.39E-01 | 1.13E-01 | 2.84E+00 | 2.91E+00 | 2.16E-01 | no | Sh3gl2 | intron_variant | MODIFIER |
| 4 | 85590241 | AX-170324651 | 0 | G | C | 0.333 | 1.39E-01 | 1.13E-01 | 2.84E+00 | 2.91E+00 | 2.16E-01 | no | Sh3gl2 | intron_variant | MODIFIER |
| 4 | 85628003 | AX-170342682 | 0 | C | T | 0.333 | 1.39E-01 | 1.13E-01 | 2.84E+00 | 2.91E+00 | 2.16E-01 | no | Sh3gl2 | intron_variant | MODIFIER |
| 4 | 85638402 | AX-170089300 | 0 | C | A | 0.333 | 1.39E-01 | 1.13E-01 | 2.84E+00 | 2.91E+00 | 2.16E-01 | no | Sh3gl2 | 3_prime_UTR_variant | MODIFIER |

| | | | | | | | | | | | | | | | |
|---|---|---|---|---|---|---|---|---|---|---|---|---|---|---|---|
| 4 | 85678552 | AX-170186476 | 0 | G | A | 0.333 | 1.39E-01 | 1.13E-01 | 2.84E+00 | 2.91E+00 | 2.16E-01 | no | Adamtsl1 | intron_variant,non_coding_transcript_variant | MODIFIER |
| 4 | 85684089 | AX-170457660 | 0 | G | A | 0.333 | 1.39E-01 | 1.13E-01 | 2.84E+00 | 2.91E+00 | 2.16E-01 | no | Adamtsl1 | intron_variant,non_coding_transcript_variant | MODIFIER |
| 4 | 85705407 | AX-170089301 | 0 | T | C | 0.333 | 1.39E-01 | 1.13E-01 | 2.84E+00 | 2.91E+00 | 2.16E-01 | no | Adamtsl1 | intron_variant,non_coding_transcript_variant | MODIFIER |
| 4 | 85728212 | AX-170023300 | 0 | A | C | 0.333 | 1.39E-01 | 1.13E-01 | 2.84E+00 | 2.91E+00 | 2.16E-01 | no | Adamtsl1 | intron_variant,non_coding_transcript_variant | MODIFIER |
| 4 | 85736063 | AX-170395689 | 0 | G | C | 0.333 | 1.39E-01 | 1.13E-01 | 2.84E+00 | 2.91E+00 | 2.16E-01 | no | Adamtsl1 | intron_variant,non_coding_transcript_variant | MODIFIER |
| 4 | 85738608 | AX-170101927 | 0 | C | G | 0.333 | 1.39E-01 | 1.13E-01 | 2.84E+00 | 2.91E+00 | 2.16E-01 | no | Adamtsl1 | intron_variant,non_coding_transcript_variant | MODIFIER |
| 4 | 85739670 | AX-97821025 | 0 | C | T | 0.333 | 1.39E-01 | 1.13E-01 | 2.84E+00 | 2.91E+00 | 2.16E-01 | no | Adamtsl1 | intron_variant,non_coding_transcript_variant | MODIFIER |
| 4 | 85836861 | AX-97785898 | 0 | A | G | 0.333 | 1.39E-01 | 1.13E-01 | 2.84E+00 | 2.91E+00 | 2.16E-01 | no | Adamtsl1 | intron_variant | MODIFIER |
| 4 | 86022798 | AX-170342472 | 0 | G | A | 0.333 | 1.39E-01 | 1.13E-01 | 2.84E+00 | 2.91E+00 | 2.16E-01 | no | Adamtsl1 | intron_variant | MODIFIER |
| 4 | 86023475 | AX-170503928 | 0 | C | T | 0.333 | 1.39E-01 | 1.13E-01 | 2.84E+00 | 2.91E+00 | 2.16E-01 | no | Adamtsl1 | intron_variant | MODIFIER |
| 12 | 107825093 | AX-170408548 | 0 | C | T | 0.123 | -1.86E-01 | 1.49E-01 | 3.07E+00 | 3.23E+00 | 2.16E-01 | Yes | | intergenic_variant | MODIFIER |
| 18 | 87917927 | AX-170409595 | 0 | A | C | 0.158 | -1.32E-01 | 1.06E-01 | 2.47E+00 | 2.56E+00 | 2.16E-01 | no | | intergenic_variant | MODIFIER |
| 2 | 105070720 | AX-170150036 | 0 | C | T | 0.07 | 1.86E-01 | 1.51E-01 | 2.58E+00 | 2.72E+00 | 2.16E-01 | Yes | | intergenic_variant | MODIFIER |
| 3 | 104228177 | AX-170021298 | 0 | T | C | 0.035 | -2.12E-01 | 1.72E-01 | 2.55E+00 | 2.65E+00 | 2.16E-01 | Yes | | regulatory_region_variant | MODIFIER |
| 9 | 9082114 | AX-170375706 | 1 | A | G | 0.455 | 9.65E-02 | 7.76E-02 | 2.48E+00 | 2.58E+00 | 2.16E-01 | no | Arhgap42 | intron_variant | MODIFIER |
| 4 | 64608110 | AX-170022965 | 1 | T | A | 0.304 | 1.32E-01 | 1.08E-01 | 2.62E+00 | 2.71E+00 | 2.16E-01 | no | | intergenic_variant | MODIFIER |
| 10 | 100068581 | AX-170309910 | 1 | T | C | 0.277 | -1.05E-01 | 8.52E-02 | 2.67E+00 | 2.74E+00 | 2.16E-01 | no | Kitl | intron_variant | MODIFIER |
| 16 | 19021073 | AX-170468394 | 0 | G | A | 0.14 | 1.15E-01 | 9.41E-02 | 2.82E+00 | 2.96E+00 | 2.16E-01 | no | Gm18747 | upstream_gene_variant | MODIFIER |
| 18 | 85563891 | AX-170016126 | 1 | G | A | 0.143 | -1.34E-01 | 1.08E-01 | 2.52E+00 | 2.62E+00 | 2.16E-01 | no | | intergenic_variant | MODIFIER |
| 7 | 124153349 | AX-170471432 | 1 | C | T | 0.42 | 1.01E-01 | 8.23E-02 | 2.89E+00 | 3.00E+00 | 2.16E-01 | no | Hs3st4 | intron_variant | MODIFIER |
| 13 | 7972731 | AX-170339443 | 2 | A | G | 0.209 | -1.37E-01 | 1.11E-01 | 2.57E+00 | 2.66E+00 | 2.16E-01 | no | | intergenic_variant | MODIFIER |
| 7 | 99781899 | AX-97784889 | 1 | A | G | 0.375 | -1.30E-01 | 1.06E-01 | 2.85E+00 | 2.94E+00 | 2.16E-01 | no | F730035P03Rik | upstream_gene_variant | MODIFIER |
| 9 | 8036391 | AX-170345522 | 1 | C | T | 0.464 | 9.97E-02 | 8.00E-02 | 2.45E+00 | 2.53E+00 | 2.16E-01 | no | Cfap300 | intron_variant | MODIFIER |
| 9 | 6521818 | AX-170031748 | 0 | A | G | 0.474 | 1.00E-01 | 8.06E-02 | 2.48E+00 | 2.58E+00 | 2.16E-01 | no | | intergenic_variant | MODIFIER |
| 7 | 38313212 | AX-170269511 | 0 | G | C | 0.351 | 1.17E-01 | 9.47E-02 | 2.57E+00 | 2.65E+00 | 2.16E-01 | no | Gm9220 | intron_variant,non_coding_transcript_variant | MODIFIER |
| 1 | 119279024 | AX-170373460 | 0 | A | G | 0.57 | 1.03E-01 | 8.32E-02 | 2.50E+00 | 2.56E+00 | 2.17E-01 | no | | regulatory_region_variant | MODIFIER |
| 1 | 119302025 | AX-170390850 | 0 | T | C | 0.57 | 1.03E-01 | 8.32E-02 | 2.50E+00 | 2.56E+00 | 2.17E-01 | no | Gm29456 | downstream_gene_variant | MODIFIER |
| 1 | 119309223 | AX-170189623 | 0 | A | G | 0.57 | 1.03E-01 | 8.32E-02 | 2.50E+00 | 2.56E+00 | 2.17E-01 | no | Gm29456 | intron_variant,non_coding_transcript_variant | MODIFIER |
| 1 | 119354289 | AX-170207359 | 0 | C | T | 0.57 | 1.03E-01 | 8.32E-02 | 2.50E+00 | 2.56E+00 | 2.17E-01 | no | Gm29455 | intron_variant,non_coding_transcript_variant | MODIFIER |
| 1 | 119442713 | AX-170204203 | 0 | A | G | 0.57 | 1.03E-01 | 8.32E-02 | 2.50E+00 | 2.56E+00 | 2.17E-01 | no | | intergenic_variant | MODIFIER |
| 1 | 119442950 | AX-170281749 | 0 | C | T | 0.57 | 1.03E-01 | 8.32E-02 | 2.50E+00 | 2.56E+00 | 2.17E-01 | no | | regulatory_region_variant | MODIFIER |
| 1 | 119844930 | AX-170001540 | 0 | C | T | 0.57 | 1.03E-01 | 8.32E-02 | 2.50E+00 | 2.56E+00 | 2.17E-01 | no | | intergenic_variant | MODIFIER |
| 1 | 119912357 | AX-170143579 | 0 | A | T | 0.57 | 1.03E-01 | 8.32E-02 | 2.50E+00 | 2.56E+00 | 2.17E-01 | no | Tmem177 | intron_variant | MODIFIER |
| 1 | 119928950 | AX-170423891 | 0 | A | G | 0.57 | 1.03E-01 | 8.32E-02 | 2.50E+00 | 2.56E+00 | 2.17E-01 | no | Cfap221 | intron_variant | MODIFIER |
| 1 | 120192015 | AX-170372258 | 0 | T | C | 0.57 | 1.03E-01 | 8.32E-02 | 2.50E+00 | 2.56E+00 | 2.17E-01 | no | 3110009E18Rik | downstream_gene_variant | MODIFIER |
| 1 | 120193033 | AX-170315159 | 0 | C | T | 0.57 | 1.03E-01 | 8.32E-02 | 2.50E+00 | 2.56E+00 | 2.17E-01 | no | 3110009E18Rik | downstream_gene_variant | MODIFIER |
| 1 | 120401478 | AX-170001553 | 0 | C | T | 0.57 | 1.03E-01 | 8.32E-02 | 2.50E+00 | 2.56E+00 | 2.17E-01 | no | | intergenic_variant | MODIFIER |
| 1 | 120434953 | AX-170396191 | 0 | A | G | 0.57 | 1.03E-01 | 8.32E-02 | 2.50E+00 | 2.56E+00 | 2.17E-01 | no | 1700012E03Rik | upstream_gene_variant | MODIFIER |
| 1 | 120435180 | AX-170481515 | 0 | G | A | 0.57 | 1.03E-01 | 8.32E-02 | 2.50E+00 | 2.56E+00 | 2.17E-01 | no | 1700012E03Rik | upstream_gene_variant | MODIFIER |
| 1 | 120440192 | AX-170319044 | 0 | T | C | 0.57 | 1.03E-01 | 8.32E-02 | 2.50E+00 | 2.56E+00 | 2.17E-01 | no | 1700012E03Rik | downstream_gene_variant | MODIFIER |
| 1 | 120454767 | AX-170107722 | 0 | A | G | 0.57 | 1.03E-01 | 8.32E-02 | 2.50E+00 | 2.56E+00 | 2.17E-01 | no | | intergenic_variant | MODIFIER |
| 1 | 120460802 | AX-170383228 | 0 | A | G | 0.57 | 1.03E-01 | 8.32E-02 | 2.50E+00 | 2.56E+00 | 2.17E-01 | no | | intergenic_variant | MODIFIER |
| 1 | 120464490 | AX-170475534 | 0 | G | T | 0.57 | 1.03E-01 | 8.32E-02 | 2.50E+00 | 2.56E+00 | 2.17E-01 | no | | intergenic_variant | MODIFIER |
| 1 | 120467777 | AX-170042484 | 0 | G | A | 0.57 | 1.03E-01 | 8.32E-02 | 2.50E+00 | 2.56E+00 | 2.17E-01 | no | | intergenic_variant | MODIFIER |
| 1 | 120558673 | AX-170241949 | 0 | T | A | 0.57 | 1.03E-01 | 8.32E-02 | 2.50E+00 | 2.56E+00 | 2.17E-01 | no | | intergenic_variant | MODIFIER |
| 1 | 120559054 | AX-170178131 | 0 | C | G | 0.57 | 1.03E-01 | 8.32E-02 | 2.50E+00 | 2.56E+00 | 2.17E-01 | no | | intergenic_variant | MODIFIER |
| 1 | 120559240 | AX-170441329 | 0 | G | A | 0.57 | 1.03E-01 | 8.32E-02 | 2.50E+00 | 2.56E+00 | 2.17E-01 | no | | intergenic_variant | MODIFIER |
| 1 | 120560726 | AX-170320395 | 0 | C | T | 0.57 | 1.03E-01 | 8.32E-02 | 2.50E+00 | 2.56E+00 | 2.17E-01 | no | | intergenic_variant | MODIFIER |
| 1 | 120561150 | AX-97794319 | 0 | T | C | 0.57 | 1.03E-01 | 8.32E-02 | 2.50E+00 | 2.56E+00 | 2.17E-01 | no | | intergenic_variant | MODIFIER |
| 1 | 120561191 | AX-170490616 | 0 | G | A | 0.57 | 1.03E-01 | 8.32E-02 | 2.50E+00 | 2.56E+00 | 2.17E-01 | no | | intergenic_variant | MODIFIER |
| 1 | 120569812 | AX-170447150 | 0 | G | A | 0.57 | 1.03E-01 | 8.32E-02 | 2.50E+00 | 2.56E+00 | 2.17E-01 | no | | intergenic_variant | MODIFIER |
| 1 | 120579332 | AX-97855802 | 0 | C | A | 0.57 | 1.03E-01 | 8.32E-02 | 2.50E+00 | 2.56E+00 | 2.17E-01 | no | | intergenic_variant | MODIFIER |
| 1 | 120624392 | AX-170135362 | 0 | C | G | 0.57 | 1.03E-01 | 8.32E-02 | 2.50E+00 | 2.56E+00 | 2.17E-01 | no | 2610027F03Rik | intron_variant,non_coding_transcript_variant | MODIFIER |
| 1 | 120673400 | AX-170133600 | 0 | G | A | 0.57 | 1.03E-01 | 8.32E-02 | 2.50E+00 | 2.56E+00 | 2.17E-01 | no | 2610027F03Rik | intron_variant,non_coding_transcript_variant | MODIFIER |

| | | | | | | | | | | | | | | | |
|---|---|---|---|---|---|---|---|---|---|---|---|---|---|---|---|
| 1 | 120679667 | AX-170532411 | 0 | A | G | 0.57 | 1.03E-01 | 8.32E-02 | 2.50E+00 | 2.56E+00 | 2.17E-01 | no | 2610027F03Rik | intron_variant,non_coding_transcript_variant | MODIFIER |
| 1 | 120682915 | AX-170367558 | 0 | A | G | 0.57 | 1.03E-01 | 8.32E-02 | 2.50E+00 | 2.56E+00 | 2.17E-01 | no | 2610027F03Rik | downstream_gene_variant | MODIFIER |
| 1 | 120754377 | AX-170446697 | 0 | G | A | 0.57 | 1.03E-01 | 8.32E-02 | 2.50E+00 | 2.56E+00 | 2.17E-01 | no | Gm29346 | upstream_gene_variant | MODIFIER |
| 1 | 121393974 | AX-170390115 | 0 | A | G | 0.57 | 1.03E-01 | 8.32E-02 | 2.50E+00 | 2.56E+00 | 2.17E-01 | no | | intergenic_variant | MODIFIER |
| 1 | 121608677 | AX-170485768 | 0 | A | G | 0.57 | 1.03E-01 | 8.32E-02 | 2.50E+00 | 2.56E+00 | 2.17E-01 | no | | intergenic_variant | MODIFIER |
| 1 | 121620104 | AX-170440605 | 0 | C | T | 0.57 | 1.03E-01 | 8.32E-02 | 2.50E+00 | 2.56E+00 | 2.17E-01 | no | | intergenic_variant | MODIFIER |
| 1 | 121620460 | AX-170494588 | 0 | G | A | 0.57 | 1.03E-01 | 8.32E-02 | 2.50E+00 | 2.56E+00 | 2.17E-01 | no | | intergenic_variant | MODIFIER |
| 1 | 121620477 | AX-170242557 | 0 | C | G | 0.57 | 1.03E-01 | 8.32E-02 | 2.50E+00 | 2.56E+00 | 2.17E-01 | no | | intergenic_variant | MODIFIER |
| 1 | 121644370 | AX-170378140 | 0 | A | G | 0.57 | 1.03E-01 | 8.32E-02 | 2.50E+00 | 2.56E+00 | 2.17E-01 | no | Gm28591 | downstream_gene_variant | MODIFIER |
| 1 | 121670809 | AX-170360525 | 0 | G | A | 0.57 | 1.03E-01 | 8.32E-02 | 2.50E+00 | 2.56E+00 | 2.17E-01 | no | | intergenic_variant | MODIFIER |
| 1 | 121694884 | AX-170119694 | 0 | A | G | 0.57 | 1.03E-01 | 8.32E-02 | 2.50E+00 | 2.56E+00 | 2.17E-01 | no | | intergenic_variant | MODIFIER |
| 1 | 121734777 | AX-170396434 | 0 | G | A | 0.57 | 1.03E-01 | 8.32E-02 | 2.50E+00 | 2.56E+00 | 2.17E-01 | no | | regulatory_region_variant | MODIFIER |
| 1 | 121809696 | AX-97824706 | 0 | T | C | 0.57 | 1.03E-01 | 8.32E-02 | 2.50E+00 | 2.56E+00 | 2.17E-01 | no | | intergenic_variant | MODIFIER |
| 1 | 121834378 | AX-170274025 | 0 | C | T | 0.57 | 1.03E-01 | 8.32E-02 | 2.50E+00 | 2.56E+00 | 2.17E-01 | no | | intergenic_variant | MODIFIER |
| 1 | 121907695 | AX-97830708 | 0 | C | A | 0.57 | 1.03E-01 | 8.32E-02 | 2.50E+00 | 2.56E+00 | 2.17E-01 | no | | intergenic_variant | MODIFIER |
| 1 | 121907909 | AX-170140099 | 0 | C | G | 0.57 | 1.03E-01 | 8.32E-02 | 2.50E+00 | 2.56E+00 | 2.17E-01 | no | | intergenic_variant | MODIFIER |
| 1 | 121963865 | AX-97827018 | 0 | A | G | 0.57 | 1.03E-01 | 8.32E-02 | 2.50E+00 | 2.56E+00 | 2.17E-01 | no | | intergenic_variant | MODIFIER |
| 1 | 122280217 | AX-97836943 | 0 | C | T | 0.57 | 1.03E-01 | 8.32E-02 | 2.50E+00 | 2.56E+00 | 2.17E-01 | no | | intergenic_variant | MODIFIER |
| 1 | 122296698 | AX-170042520 | 0 | G | A | 0.57 | 1.03E-01 | 8.32E-02 | 2.50E+00 | 2.56E+00 | 2.17E-01 | no | | intergenic_variant | MODIFIER |
| 1 | 122297222 | AX-170350830 | 0 | T | C | 0.57 | 1.03E-01 | 8.32E-02 | 2.50E+00 | 2.56E+00 | 2.17E-01 | no | | intergenic_variant | MODIFIER |
| 1 | 122299475 | AX-170101384 | 0 | G | T | 0.57 | 1.03E-01 | 8.32E-02 | 2.50E+00 | 2.56E+00 | 2.17E-01 | no | | intergenic_variant | MODIFIER |
| 1 | 122305268 | AX-97800082 | 0 | G | A | 0.57 | 1.03E-01 | 8.32E-02 | 2.50E+00 | 2.56E+00 | 2.17E-01 | no | | intergenic_variant | MODIFIER |
| 1 | 122322237 | AX-170427501 | 0 | C | T | 0.57 | 1.03E-01 | 8.32E-02 | 2.50E+00 | 2.56E+00 | 2.17E-01 | no | | intergenic_variant | MODIFIER |
| 13 | 18370301 | AX-170480268 | 0 | A | G | 0.167 | -1.43E-01 | 1.14E-01 | 2.42E+00 | 2.48E+00 | 2.17E-01 | no | Pou6f2 | intron_variant | MODIFIER |
| 1 | 119309525 | AX-97813086 | 1 | A | C | 0.571 | 1.03E-01 | 8.32E-02 | 2.50E+00 | 2.56E+00 | 2.17E-01 | no | Gm29456 | intron_variant,non_coding_transcript_variant | MODIFIER |
| 1 | 119309250 | AX-170001539 | 1 | G | A | 0.571 | 1.03E-01 | 8.32E-02 | 2.50E+00 | 2.56E+00 | 2.17E-01 | no | Gm29456 | intron_variant,non_coding_transcript_variant | MODIFIER |
| 4 | 90580465 | AX-97849745 | 1 | C | A | 0.25 | 1.35E-01 | 1.08E-01 | 2.45E+00 | 2.50E+00 | 2.17E-01 | no | | intergenic_variant | MODIFIER |
| 13 | 40191122 | AX-97805279 | 0 | A | G | 0.167 | -1.26E-01 | 1.01E-01 | 2.40E+00 | 2.51E+00 | 2.17E-01 | no | Ofcc1 | intron_variant | MODIFIER |
| 3 | 14897270 | AX-170483344 | 1 | G | A | 0.42 | -8.02E-02 | 6.54E-02 | 2.89E+00 | 3.05E+00 | 2.17E-01 | no | Car2 | intron_variant | MODIFIER |
| 5 | 66504304 | AX-170250845 | 0 | T | C | 0.474 | 1.23E-01 | 9.95E-02 | 2.52E+00 | 2.58E+00 | 2.17E-01 | no | Apbb2 | intron_variant | MODIFIER |
| 18 | 79465371 | AX-170452617 | 1 | A | G | 0.455 | 1.08E-01 | 8.64E-02 | 2.41E+00 | 2.47E+00 | 2.18E-01 | no | | intergenic_variant | MODIFIER |
| 5 | 66534121 | AX-170350150 | 1 | A | G | 0.473 | 1.22E-01 | 9.85E-02 | 2.51E+00 | 2.58E+00 | 2.18E-01 | no | Apbb2 | intron_variant | MODIFIER |
| 18 | 68021207 | AX-170318774 | 1 | G | T | 0.17 | -1.48E-01 | 1.21E-01 | 2.65E+00 | 2.70E+00 | 2.18E-01 | no | Ldlrad4 | intron_variant | MODIFIER |
| 16 | 29275301 | AX-170233206 | 2 | G | A | 0.518 | 9.85E-02 | 8.00E-02 | 2.55E+00 | 2.61E+00 | 2.18E-01 | no | Atp13a5 | intron_variant | MODIFIER |
| 4 | 72376492 | AX-170330940 | 1 | C | T | 0.321 | 1.47E-01 | 1.20E-01 | 2.83E+00 | 2.89E+00 | 2.18E-01 | Yes | | intergenic_variant | MODIFIER |
| 13 | 21249701 | AX-97830287 | 0 | C | T | 0.175 | -1.50E-01 | 1.20E-01 | 2.44E+00 | 2.50E+00 | 2.18E-01 | no | | regulatory_region_variant | MODIFIER |
| 8 | 106740106 | AX-170090202 | 0 | G | A | 0.219 | 1.30E-01 | 1.06E-01 | 2.89E+00 | 2.99E+00 | 2.18E-01 | no | Tango6 | intron_variant | MODIFIER |
| 4 | 124953454 | AX-170258493 | 0 | C | T | 0.237 | 1.33E-01 | 1.08E-01 | 2.61E+00 | 2.68E+00 | 2.18E-01 | no | Gm12930 | downstream_gene_variant | MODIFIER |
| 4 | 85112423 | AX-170411189 | 1 | G | A | 0.339 | 1.42E-01 | 1.16E-01 | 2.89E+00 | 2.96E+00 | 2.18E-01 | no | Cntln | intron_variant | MODIFIER |
| 13 | 33043528 | AX-97798701 | 0 | A | G | 0.175 | -1.30E-01 | 1.07E-01 | 2.60E+00 | 2.71E+00 | 2.18E-01 | no | Serpinb9b | downstream_gene_variant | MODIFIER |
| 13 | 35928341 | AX-170008113 | 0 | T | C | 0.175 | -1.30E-01 | 1.07E-01 | 2.60E+00 | 2.71E+00 | 2.18E-01 | no | | intergenic_variant | MODIFIER |
| 4 | 64271874 | AX-97801397 | 0 | C | T | 0.325 | 1.47E-01 | 1.20E-01 | 2.84E+00 | 2.89E+00 | 2.18E-01 | Yes | Gm11217 | intron_variant,non_coding_transcript_variant | MODIFIER |
| 4 | 64276301 | AX-170022971 | 0 | A | C | 0.325 | 1.47E-01 | 1.20E-01 | 2.84E+00 | 2.89E+00 | 2.18E-01 | Yes | Gm11217 | intron_variant,non_coding_transcript_variant | MODIFIER |
| 4 | 64276362 | AX-170022962 | 0 | T | C | 0.325 | 1.47E-01 | 1.20E-01 | 2.84E+00 | 2.89E+00 | 2.18E-01 | Yes | Gm11217 | intron_variant,non_coding_transcript_variant | MODIFIER |
| 4 | 64285779 | AX-170207238 | 0 | T | G | 0.325 | 1.47E-01 | 1.20E-01 | 2.84E+00 | 2.89E+00 | 2.18E-01 | Yes | | intergenic_variant | MODIFIER |
| 4 | 64285838 | AX-97845820 | 0 | C | T | 0.325 | 1.47E-01 | 1.20E-01 | 2.84E+00 | 2.89E+00 | 2.18E-01 | Yes | | intergenic_variant | MODIFIER |
| 4 | 64302141 | AX-170120125 | 0 | A | G | 0.325 | 1.47E-01 | 1.20E-01 | 2.84E+00 | 2.89E+00 | 2.18E-01 | Yes | | intergenic_variant | MODIFIER |
| 4 | 64345935 | AX-97817623 | 0 | T | C | 0.325 | 1.47E-01 | 1.20E-01 | 2.84E+00 | 2.89E+00 | 2.18E-01 | Yes | | intergenic_variant | MODIFIER |
| 4 | 64348209 | AX-170495126 | 0 | T | C | 0.325 | 1.47E-01 | 1.20E-01 | 2.84E+00 | 2.89E+00 | 2.18E-01 | Yes | | intergenic_variant | MODIFIER |
| 4 | 64374269 | AX-170022969 | 0 | G | A | 0.325 | 1.47E-01 | 1.20E-01 | 2.84E+00 | 2.89E+00 | 2.18E-01 | Yes | | intergenic_variant | MODIFIER |
| 4 | 64390967 | AX-170022963 | 0 | C | G | 0.325 | 1.47E-01 | 1.20E-01 | 2.84E+00 | 2.89E+00 | 2.18E-01 | Yes | | intergenic_variant | MODIFIER |
| 4 | 64397358 | AX-97798399 | 0 | G | T | 0.325 | 1.47E-01 | 1.20E-01 | 2.84E+00 | 2.89E+00 | 2.18E-01 | Yes | | intergenic_variant | MODIFIER |
| 4 | 64403273 | AX-97829093 | 0 | T | G | 0.325 | 1.47E-01 | 1.20E-01 | 2.84E+00 | 2.89E+00 | 2.18E-01 | Yes | | intergenic_variant | MODIFIER |
| 4 | 64441920 | AX-97797965 | 0 | A | G | 0.325 | 1.47E-01 | 1.20E-01 | 2.84E+00 | 2.89E+00 | 2.18E-01 | Yes | Gm23950 | upstream_gene_variant | MODIFIER |

| | | | | | | | | | | | | | | | |
|---|---|---|---|---|---|---|---|---|---|---|---|---|---|---|---|
| 4 | 64472616 | AX-97807334 | 0 | A | C | 0.325 | 1.47E-01 | 1.20E-01 | 2.84E+00 | 2.89E+00 | 2.18E-01 | Yes | | intergenic_variant | MODIFIER |
| 4 | 64500581 | AX-97855904 | 0 | A | G | 0.325 | 1.47E-01 | 1.20E-01 | 2.84E+00 | 2.89E+00 | 2.18E-01 | Yes | | intergenic_variant | MODIFIER |
| 4 | 64527218 | AX-170022975 | 0 | A | G | 0.325 | 1.47E-01 | 1.20E-01 | 2.84E+00 | 2.89E+00 | 2.18E-01 | Yes | | intergenic_variant | MODIFIER |
| 4 | 64535013 | AX-170022970 | 0 | T | C | 0.325 | 1.47E-01 | 1.20E-01 | 2.84E+00 | 2.89E+00 | 2.18E-01 | Yes | | intergenic_variant | MODIFIER |
| 4 | 64535131 | AX-97841348 | 0 | C | T | 0.325 | 1.47E-01 | 1.20E-01 | 2.84E+00 | 2.89E+00 | 2.18E-01 | Yes | | intergenic_variant | MODIFIER |
| 4 | 64558449 | AX-170251685 | 0 | A | G | 0.325 | 1.47E-01 | 1.20E-01 | 2.84E+00 | 2.89E+00 | 2.18E-01 | Yes | | intergenic_variant | MODIFIER |
| 4 | 64565364 | AX-170113941 | 0 | T | C | 0.325 | 1.47E-01 | 1.20E-01 | 2.84E+00 | 2.89E+00 | 2.18E-01 | Yes | | intergenic_variant | MODIFIER |
| 4 | 64575144 | AX-170022974 | 0 | C | T | 0.325 | 1.47E-01 | 1.20E-01 | 2.84E+00 | 2.89E+00 | 2.18E-01 | Yes | | intergenic_variant | MODIFIER |
| 4 | 64581257 | AX-170022973 | 0 | T | C | 0.325 | 1.47E-01 | 1.20E-01 | 2.84E+00 | 2.89E+00 | 2.18E-01 | Yes | | intergenic_variant | MODIFIER |
| 4 | 64594207 | AX-97807848 | 0 | A | G | 0.325 | 1.47E-01 | 1.20E-01 | 2.84E+00 | 2.89E+00 | 2.18E-01 | Yes | | intergenic_variant | MODIFIER |
| 4 | 64601692 | AX-170244914 | 0 | T | C | 0.325 | 1.47E-01 | 1.20E-01 | 2.84E+00 | 2.89E+00 | 2.18E-01 | Yes | | intergenic_variant | MODIFIER |
| 4 | 64609384 | AX-97804376 | 0 | T | G | 0.325 | 1.47E-01 | 1.20E-01 | 2.84E+00 | 2.89E+00 | 2.18E-01 | Yes | | intergenic_variant | MODIFIER |
| 4 | 64648707 | AX-97814980 | 0 | T | C | 0.325 | 1.47E-01 | 1.20E-01 | 2.84E+00 | 2.89E+00 | 2.18E-01 | Yes | | intergenic_variant | MODIFIER |
| 4 | 64665754 | AX-170183581 | 0 | C | A | 0.325 | 1.47E-01 | 1.20E-01 | 2.84E+00 | 2.89E+00 | 2.18E-01 | Yes | | intergenic_variant | MODIFIER |
| 4 | 64668959 | AX-170143914 | 0 | G | T | 0.325 | 1.47E-01 | 1.20E-01 | 2.84E+00 | 2.89E+00 | 2.18E-01 | Yes | | intergenic_variant | MODIFIER |
| 4 | 64671038 | AX-170325281 | 0 | G | A | 0.325 | 1.47E-01 | 1.20E-01 | 2.84E+00 | 2.89E+00 | 2.18E-01 | Yes | | intergenic_variant | MODIFIER |
| 4 | 64671215 | AX-170347896 | 0 | C | G | 0.325 | 1.47E-01 | 1.20E-01 | 2.84E+00 | 2.89E+00 | 2.18E-01 | Yes | | intergenic_variant | MODIFIER |
| 4 | 64683706 | AX-170331166 | 0 | C | T | 0.325 | 1.47E-01 | 1.20E-01 | 2.84E+00 | 2.89E+00 | 2.18E-01 | Yes | | intergenic_variant | MODIFIER |
| 4 | 64683852 | AX-97818851 | 0 | T | C | 0.325 | 1.47E-01 | 1.20E-01 | 2.84E+00 | 2.89E+00 | 2.18E-01 | Yes | | intergenic_variant | MODIFIER |
| 4 | 64722939 | AX-170329090 | 0 | G | A | 0.325 | 1.47E-01 | 1.20E-01 | 2.84E+00 | 2.89E+00 | 2.18E-01 | Yes | | intergenic_variant | MODIFIER |
| 4 | 64741664 | AX-170454344 | 0 | A | G | 0.325 | 1.47E-01 | 1.20E-01 | 2.84E+00 | 2.89E+00 | 2.18E-01 | Yes | | intergenic_variant | MODIFIER |
| 4 | 64756925 | AX-170132047 | 0 | G | A | 0.325 | 1.47E-01 | 1.20E-01 | 2.84E+00 | 2.89E+00 | 2.18E-01 | Yes | | intergenic_variant | MODIFIER |
| 4 | 64842216 | AX-97780269 | 0 | A | C | 0.325 | 1.47E-01 | 1.20E-01 | 2.84E+00 | 2.89E+00 | 2.18E-01 | Yes | | intergenic_variant | MODIFIER |
| 4 | 64853134 | AX-97790004 | 0 | G | A | 0.325 | 1.47E-01 | 1.20E-01 | 2.84E+00 | 2.89E+00 | 2.18E-01 | Yes | Gm11218 | downstream_gene_variant | MODIFIER |
| 4 | 64854423 | AX-170022979 | 0 | T | A | 0.325 | 1.47E-01 | 1.20E-01 | 2.84E+00 | 2.89E+00 | 2.18E-01 | Yes | Gm11218 | downstream_gene_variant | MODIFIER |
| 4 | 65132348 | AX-170089257 | 0 | T | C | 0.325 | 1.47E-01 | 1.20E-01 | 2.84E+00 | 2.89E+00 | 2.18E-01 | Yes | Pappa | intron_variant | MODIFIER |
| 4 | 68598371 | AX-170527028 | 0 | A | G | 0.325 | 1.47E-01 | 1.20E-01 | 2.84E+00 | 2.89E+00 | 2.18E-01 | Yes | | intergenic_variant | MODIFIER |
| 4 | 68633777 | AX-170242960 | 0 | A | G | 0.325 | 1.47E-01 | 1.20E-01 | 2.84E+00 | 2.89E+00 | 2.18E-01 | Yes | | intergenic_variant | MODIFIER |
| 4 | 68643252 | AX-170315551 | 0 | T | C | 0.325 | 1.47E-01 | 1.20E-01 | 2.84E+00 | 2.89E+00 | 2.18E-01 | Yes | | intergenic_variant | MODIFIER |
| 4 | 68675177 | AX-170420175 | 0 | T | C | 0.325 | 1.47E-01 | 1.20E-01 | 2.84E+00 | 2.89E+00 | 2.18E-01 | Yes | | intergenic_variant | MODIFIER |
| 4 | 68736450 | AX-170023031 | 0 | A | G | 0.325 | 1.47E-01 | 1.20E-01 | 2.84E+00 | 2.89E+00 | 2.18E-01 | Yes | | intergenic_variant | MODIFIER |
| 4 | 68768942 | AX-97791060 | 0 | G | T | 0.325 | 1.47E-01 | 1.20E-01 | 2.84E+00 | 2.89E+00 | 2.18E-01 | Yes | Brinp1 | intron_variant | MODIFIER |
| 4 | 68772269 | AX-170233810 | 0 | C | T | 0.325 | 1.47E-01 | 1.20E-01 | 2.84E+00 | 2.89E+00 | 2.18E-01 | Yes | Brinp1 | intron_variant | MODIFIER |
| 4 | 69025600 | AX-170139441 | 0 | A | G | 0.325 | 1.47E-01 | 1.20E-01 | 2.84E+00 | 2.89E+00 | 2.18E-01 | Yes | | intergenic_variant | MODIFIER |
| 4 | 69038470 | AX-170443788 | 0 | C | G | 0.325 | 1.47E-01 | 1.20E-01 | 2.84E+00 | 2.89E+00 | 2.18E-01 | Yes | | intergenic_variant | MODIFIER |
| 4 | 69042541 | AX-170396762 | 0 | C | T | 0.325 | 1.47E-01 | 1.20E-01 | 2.84E+00 | 2.89E+00 | 2.18E-01 | Yes | | intergenic_variant | MODIFIER |
| 4 | 69053950 | AX-170276203 | 0 | C | T | 0.325 | 1.47E-01 | 1.20E-01 | 2.84E+00 | 2.89E+00 | 2.18E-01 | Yes | | intergenic_variant | MODIFIER |
| 4 | 69078584 | AX-170133539 | 0 | G | A | 0.325 | 1.47E-01 | 1.20E-01 | 2.84E+00 | 2.89E+00 | 2.18E-01 | Yes | | intergenic_variant | MODIFIER |
| 4 | 69085300 | AX-170510673 | 0 | C | A | 0.325 | 1.47E-01 | 1.20E-01 | 2.84E+00 | 2.89E+00 | 2.18E-01 | Yes | | intergenic_variant | MODIFIER |
| 4 | 69215284 | AX-170023042 | 0 | T | G | 0.325 | 1.47E-01 | 1.20E-01 | 2.84E+00 | 2.89E+00 | 2.18E-01 | Yes | | intergenic_variant | MODIFIER |
| 4 | 69291807 | AX-170384169 | 0 | C | T | 0.325 | 1.47E-01 | 1.20E-01 | 2.84E+00 | 2.89E+00 | 2.18E-01 | Yes | Gm11404 | non_coding_transcript_exon_variant | MODIFIER |
| 4 | 69496836 | AX-170517917 | 0 | A | G | 0.325 | 1.47E-01 | 1.20E-01 | 2.84E+00 | 2.89E+00 | 2.18E-01 | Yes | | intergenic_variant | MODIFIER |
| 4 | 69533513 | AX-170069606 | 0 | A | G | 0.325 | 1.47E-01 | 1.20E-01 | 2.84E+00 | 2.89E+00 | 2.18E-01 | Yes | | intergenic_variant | MODIFIER |
| 4 | 69536558 | AX-170108411 | 0 | C | A | 0.325 | 1.47E-01 | 1.20E-01 | 2.84E+00 | 2.89E+00 | 2.18E-01 | Yes | | intergenic_variant | MODIFIER |
| 4 | 69603500 | AX-170238747 | 0 | T | C | 0.325 | 1.47E-01 | 1.20E-01 | 2.84E+00 | 2.89E+00 | 2.18E-01 | Yes | | intergenic_variant | MODIFIER |
| 4 | 69614564 | AX-170281505 | 0 | T | A | 0.325 | 1.47E-01 | 1.20E-01 | 2.84E+00 | 2.89E+00 | 2.18E-01 | Yes | | intergenic_variant | MODIFIER |
| 4 | 69615227 | AX-170023044 | 0 | A | G | 0.325 | 1.47E-01 | 1.20E-01 | 2.84E+00 | 2.89E+00 | 2.18E-01 | Yes | | intergenic_variant | MODIFIER |
| 4 | 69644093 | AX-170257203 | 0 | T | C | 0.325 | 1.47E-01 | 1.20E-01 | 2.84E+00 | 2.89E+00 | 2.18E-01 | Yes | | intergenic_variant | MODIFIER |
| 4 | 69682119 | AX-170504003 | 0 | A | C | 0.325 | 1.47E-01 | 1.20E-01 | 2.84E+00 | 2.89E+00 | 2.18E-01 | Yes | | intergenic_variant | MODIFIER |
| 4 | 69692306 | AX-170023049 | 0 | T | C | 0.325 | 1.47E-01 | 1.20E-01 | 2.84E+00 | 2.89E+00 | 2.18E-01 | Yes | Gm11221 | non_coding_transcript_exon_variant | MODIFIER |
| 4 | 69751998 | AX-170383111 | 0 | T | C | 0.325 | 1.47E-01 | 1.20E-01 | 2.84E+00 | 2.89E+00 | 2.18E-01 | Yes | | intergenic_variant | MODIFIER |
| 4 | 69758400 | AX-170100959 | 0 | C | A | 0.325 | 1.47E-01 | 1.20E-01 | 2.84E+00 | 2.89E+00 | 2.18E-01 | Yes | | intergenic_variant | MODIFIER |
| 4 | 69811646 | AX-170128128 | 0 | T | C | 0.325 | 1.47E-01 | 1.20E-01 | 2.84E+00 | 2.89E+00 | 2.18E-01 | Yes | | intergenic_variant | MODIFIER |
| 4 | 69821409 | AX-170205810 | 0 | A | G | 0.325 | 1.47E-01 | 1.20E-01 | 2.84E+00 | 2.89E+00 | 2.18E-01 | Yes | | intergenic_variant | MODIFIER |

| | | | | | | | | | | | | | | | |
|---|---|---|---|---|---|---|---|---|---|---|---|---|---|---|---|
| 4 | 69830948 | AX-170130183 | 0 | G | A | 0.325 | 1.47E-01 | 1.20E-01 | 2.84E+00 | 2.89E+00 | 2.18E-01 | Yes | | intergenic_variant | MODIFIER |
| 4 | 69839115 | AX-97790895 | 0 | T | C | 0.325 | 1.47E-01 | 1.20E-01 | 2.84E+00 | 2.89E+00 | 2.18E-01 | Yes | | intergenic_variant | MODIFIER |
| 4 | 69868913 | AX-97841689 | 0 | T | C | 0.325 | 1.47E-01 | 1.20E-01 | 2.84E+00 | 2.89E+00 | 2.18E-01 | Yes | | intergenic_variant | MODIFIER |
| 4 | 69869046 | AX-170359092 | 0 | G | A | 0.325 | 1.47E-01 | 1.20E-01 | 2.84E+00 | 2.89E+00 | 2.18E-01 | Yes | | intergenic_variant | MODIFIER |
| 4 | 69885839 | AX-97807767 | 0 | G | T | 0.325 | 1.47E-01 | 1.20E-01 | 2.84E+00 | 2.89E+00 | 2.18E-01 | Yes | | intergenic_variant | MODIFIER |
| 4 | 69904468 | AX-170069615 | 0 | A | C | 0.325 | 1.47E-01 | 1.20E-01 | 2.84E+00 | 2.89E+00 | 2.18E-01 | Yes | | intergenic_variant | MODIFIER |
| 4 | 69920674 | AX-170367765 | 0 | G | C | 0.325 | 1.47E-01 | 1.20E-01 | 2.84E+00 | 2.89E+00 | 2.18E-01 | Yes | Gm11223 | downstream_gene_variant | MODIFIER |
| 4 | 69920849 | AX-170115054 | 0 | A | G | 0.325 | 1.47E-01 | 1.20E-01 | 2.84E+00 | 2.89E+00 | 2.18E-01 | Yes | Gm11223 | downstream_gene_variant | MODIFIER |
| 4 | 69932417 | AX-170343849 | 0 | A | T | 0.325 | 1.47E-01 | 1.20E-01 | 2.84E+00 | 2.89E+00 | 2.18E-01 | Yes | | intergenic_variant | MODIFIER |
| 4 | 69932621 | AX-170521502 | 0 | G | C | 0.325 | 1.47E-01 | 1.20E-01 | 2.84E+00 | 2.89E+00 | 2.18E-01 | Yes | | intergenic_variant | MODIFIER |
| 4 | 70032904 | AX-170069620 | 0 | T | A | 0.325 | 1.47E-01 | 1.20E-01 | 2.84E+00 | 2.89E+00 | 2.18E-01 | Yes | | intergenic_variant | MODIFIER |
| 4 | 71989279 | AX-170480043 | 0 | G | T | 0.325 | 1.47E-01 | 1.20E-01 | 2.84E+00 | 2.89E+00 | 2.18E-01 | Yes | Gm11234 | intron_variant,non_coding_transcript_variant | MODIFIER |
| 4 | 71999397 | AX-170439405 | 0 | C | T | 0.325 | 1.47E-01 | 1.20E-01 | 2.84E+00 | 2.89E+00 | 2.18E-01 | Yes | Gm11234 | non_coding_transcript_exon_variant | MODIFIER |
| 4 | 72003025 | AX-170249771 | 0 | G | A | 0.325 | 1.47E-01 | 1.20E-01 | 2.84E+00 | 2.89E+00 | 2.18E-01 | Yes | Gm11234 | downstream_gene_variant | MODIFIER |
| 4 | 72061271 | AX-170509650 | 0 | A | C | 0.325 | 1.47E-01 | 1.20E-01 | 2.84E+00 | 2.89E+00 | 2.18E-01 | Yes | | regulatory_region_variant | MODIFIER |
| 4 | 72101405 | AX-170095009 | 0 | C | T | 0.325 | 1.47E-01 | 1.20E-01 | 2.84E+00 | 2.89E+00 | 2.18E-01 | Yes | | intergenic_variant | MODIFIER |
| 4 | 72107657 | AX-97798486 | 0 | A | G | 0.325 | 1.47E-01 | 1.20E-01 | 2.84E+00 | 2.89E+00 | 2.18E-01 | Yes | | intergenic_variant | MODIFIER |
| 4 | 72134929 | AX-170254846 | 0 | T | A | 0.325 | 1.47E-01 | 1.20E-01 | 2.84E+00 | 2.89E+00 | 2.18E-01 | Yes | Tle1 | intron_variant | MODIFIER |
| 4 | 72136330 | AX-170324949 | 0 | C | A | 0.325 | 1.47E-01 | 1.20E-01 | 2.84E+00 | 2.89E+00 | 2.18E-01 | Yes | Tle1 | intron_variant | MODIFIER |
| 4 | 72242626 | AX-170448553 | 0 | T | C | 0.325 | 1.47E-01 | 1.20E-01 | 2.84E+00 | 2.89E+00 | 2.18E-01 | Yes | | intergenic_variant | MODIFIER |
| 4 | 72252527 | AX-170447694 | 0 | G | A | 0.325 | 1.47E-01 | 1.20E-01 | 2.84E+00 | 2.89E+00 | 2.18E-01 | Yes | | intergenic_variant | MODIFIER |
| 4 | 72252542 | AX-97784217 | 0 | C | T | 0.325 | 1.47E-01 | 1.20E-01 | 2.84E+00 | 2.89E+00 | 2.18E-01 | Yes | | intergenic_variant | MODIFIER |
| 4 | 72317221 | AX-170023089 | 0 | C | T | 0.325 | 1.47E-01 | 1.20E-01 | 2.84E+00 | 2.89E+00 | 2.18E-01 | Yes | | intergenic_variant | MODIFIER |
| 4 | 72353383 | AX-170176478 | 0 | T | G | 0.325 | 1.47E-01 | 1.20E-01 | 2.84E+00 | 2.89E+00 | 2.18E-01 | Yes | | intergenic_variant | MODIFIER |
| 4 | 72383142 | AX-170023092 | 0 | G | A | 0.325 | 1.47E-01 | 1.20E-01 | 2.84E+00 | 2.89E+00 | 2.18E-01 | Yes | | regulatory_region_variant | MODIFIER |
| 4 | 72426548 | AX-170500865 | 0 | A | G | 0.325 | 1.47E-01 | 1.20E-01 | 2.84E+00 | 2.89E+00 | 2.18E-01 | Yes | | intergenic_variant | MODIFIER |
| 4 | 74065137 | AX-170181891 | 0 | A | G | 0.325 | 1.47E-01 | 1.20E-01 | 2.84E+00 | 2.89E+00 | 2.18E-01 | Yes | Frmd3 | intron_variant | MODIFIER |
| 4 | 74083846 | AX-170069690 | 0 | A | G | 0.325 | 1.47E-01 | 1.20E-01 | 2.84E+00 | 2.89E+00 | 2.18E-01 | Yes | Frmd3 | intron_variant | MODIFIER |
| 4 | 74741112 | AX-170264854 | 0 | G | T | 0.325 | 1.47E-01 | 1.20E-01 | 2.84E+00 | 2.89E+00 | 2.18E-01 | Yes | | intergenic_variant | MODIFIER |
| 4 | 75120766 | AX-170511912 | 0 | A | T | 0.325 | 1.47E-01 | 1.20E-01 | 2.84E+00 | 2.89E+00 | 2.18E-01 | Yes | | intergenic_variant | MODIFIER |
| 4 | 75128878 | AX-170023145 | 0 | A | G | 0.325 | 1.47E-01 | 1.20E-01 | 2.84E+00 | 2.89E+00 | 2.18E-01 | Yes | Gm11257 | non_coding_transcript_exon_variant | MODIFIER |
| 4 | 75142239 | AX-170197550 | 0 | C | G | 0.325 | 1.47E-01 | 1.20E-01 | 2.84E+00 | 2.89E+00 | 2.18E-01 | Yes | | intergenic_variant | MODIFIER |
| 4 | 75148789 | AX-97836117 | 0 | A | G | 0.325 | 1.47E-01 | 1.20E-01 | 2.84E+00 | 2.89E+00 | 2.18E-01 | Yes | Gm12912 | non_coding_transcript_exon_variant | MODIFIER |
| 4 | 75234132 | AX-170155700 | 0 | T | C | 0.325 | 1.47E-01 | 1.20E-01 | 2.84E+00 | 2.89E+00 | 2.18E-01 | Yes | | intergenic_variant | MODIFIER |
| 4 | 75235405 | AX-170302007 | 0 | C | T | 0.325 | 1.47E-01 | 1.20E-01 | 2.84E+00 | 2.89E+00 | 2.18E-01 | Yes | | intergenic_variant | MODIFIER |
| 4 | 75271502 | AX-170527522 | 0 | T | G | 0.325 | 1.47E-01 | 1.20E-01 | 2.84E+00 | 2.89E+00 | 2.18E-01 | Yes | | intergenic_variant | MODIFIER |
| 4 | 75271604 | AX-170456061 | 0 | C | T | 0.325 | 1.47E-01 | 1.20E-01 | 2.84E+00 | 2.89E+00 | 2.18E-01 | Yes | | intergenic_variant | MODIFIER |
| 4 | 75381459 | AX-170439182 | 0 | C | T | 0.325 | 1.47E-01 | 1.20E-01 | 2.84E+00 | 2.89E+00 | 2.18E-01 | Yes | Gm11259 | non_coding_transcript_exon_variant | MODIFIER |
| 4 | 75389322 | AX-170256243 | 0 | G | C | 0.325 | 1.47E-01 | 1.20E-01 | 2.84E+00 | 2.89E+00 | 2.18E-01 | Yes | | intergenic_variant | MODIFIER |
| 4 | 75439868 | AX-170426550 | 0 | T | C | 0.325 | 1.47E-01 | 1.20E-01 | 2.84E+00 | 2.89E+00 | 2.18E-01 | Yes | | intergenic_variant | MODIFIER |
| 4 | 75467138 | AX-170533734 | 0 | G | A | 0.325 | 1.47E-01 | 1.20E-01 | 2.84E+00 | 2.89E+00 | 2.18E-01 | Yes | | regulatory_region_variant | MODIFIER |
| 4 | 75515399 | AX-97784244 | 0 | G | A | 0.325 | 1.47E-01 | 1.20E-01 | 2.84E+00 | 2.89E+00 | 2.18E-01 | Yes | | intergenic_variant | MODIFIER |
| 4 | 75634984 | AX-170023150 | 0 | T | C | 0.325 | 1.47E-01 | 1.20E-01 | 2.84E+00 | 2.89E+00 | 2.18E-01 | Yes | | intergenic_variant | MODIFIER |
| 4 | 75715812 | AX-170069708 | 0 | C | T | 0.325 | 1.47E-01 | 1.20E-01 | 2.84E+00 | 2.89E+00 | 2.18E-01 | Yes | | intergenic_variant | MODIFIER |
| 4 | 75908630 | AX-170191725 | 0 | C | A | 0.325 | 1.47E-01 | 1.20E-01 | 2.84E+00 | 2.89E+00 | 2.18E-01 | Yes | | intergenic_variant | MODIFIER |
| 4 | 75957526 | AX-170516210 | 0 | A | G | 0.325 | 1.47E-01 | 1.20E-01 | 2.84E+00 | 2.89E+00 | 2.18E-01 | Yes | Ptprd | intron_variant | MODIFIER |
| 4 | 76015998 | AX-97797472 | 0 | G | A | 0.325 | 1.47E-01 | 1.20E-01 | 2.84E+00 | 2.89E+00 | 2.18E-01 | Yes | Ptprd | intron_variant | MODIFIER |
| 4 | 76092411 | AX-170434969 | 0 | C | T | 0.325 | 1.47E-01 | 1.20E-01 | 2.84E+00 | 2.89E+00 | 2.18E-01 | Yes | Ptprd | intron_variant | MODIFIER |
| 4 | 76268990 | AX-170498988 | 0 | C | A | 0.325 | 1.47E-01 | 1.20E-01 | 2.84E+00 | 2.89E+00 | 2.18E-01 | Yes | Ptprd | intron_variant | MODIFIER |
| 4 | 76428537 | AX-170459470 | 0 | T | C | 0.325 | 1.47E-01 | 1.20E-01 | 2.84E+00 | 2.89E+00 | 2.18E-01 | Yes | Ptprd | intron_variant,non_coding_transcript_variant | MODIFIER |
| 4 | 76430078 | AX-170205385 | 0 | G | T | 0.325 | 1.47E-01 | 1.20E-01 | 2.84E+00 | 2.89E+00 | 2.18E-01 | Yes | Ptprd | intron_variant,non_coding_transcript_variant | MODIFIER |
| 4 | 76435397 | AX-170224212 | 0 | T | C | 0.325 | 1.47E-01 | 1.20E-01 | 2.84E+00 | 2.89E+00 | 2.18E-01 | Yes | Ptprd | intron_variant,non_coding_transcript_variant | MODIFIER |
| 4 | 76501609 | AX-170455877 | 0 | T | A | 0.325 | 1.47E-01 | 1.20E-01 | 2.84E+00 | 2.89E+00 | 2.18E-01 | Yes | Ptprd | intron_variant,non_coding_transcript_variant | MODIFIER |
| 4 | 76508560 | AX-170397825 | 0 | T | C | 0.325 | 1.47E-01 | 1.20E-01 | 2.84E+00 | 2.89E+00 | 2.18E-01 | Yes | Ptprd | intron_variant,non_coding_transcript_variant | MODIFIER |

| | | | | | | | | | | | | | | | |
|---|---|---|---|---|---|---|---|---|---|---|---|---|---|---|---|
| 4 | 76574209 | AX-170069730 | 0 | C | T | 0.325 | 1.47E-01 | 1.20E-01 | 2.84E+00 | 2.89E+00 | 2.18E-01 | Yes | Ptprd | intron_variant,non_coding_transcript_variant | MODIFIER |
| 4 | 76607196 | AX-170069726 | 0 | C | T | 0.325 | 1.47E-01 | 1.20E-01 | 2.84E+00 | 2.89E+00 | 2.18E-01 | Yes | Ptprd | intron_variant,non_coding_transcript_variant | MODIFIER |
| 4 | 76619116 | AX-170128932 | 0 | G | A | 0.325 | 1.47E-01 | 1.20E-01 | 2.84E+00 | 2.89E+00 | 2.18E-01 | Yes | Ptprd | intron_variant,non_coding_transcript_variant | MODIFIER |
| 4 | 76631259 | AX-170023163 | 0 | G | A | 0.325 | 1.47E-01 | 1.20E-01 | 2.84E+00 | 2.89E+00 | 2.18E-01 | Yes | Ptprd | intron_variant,non_coding_transcript_variant | MODIFIER |
| 4 | 76850161 | AX-170305894 | 0 | C | T | 0.325 | 1.47E-01 | 1.20E-01 | 2.84E+00 | 2.89E+00 | 2.18E-01 | Yes | Ptprd | intron_variant,non_coding_transcript_variant | MODIFIER |
| 4 | 76851744 | AX-170069735 | 0 | C | T | 0.325 | 1.47E-01 | 1.20E-01 | 2.84E+00 | 2.89E+00 | 2.18E-01 | Yes | Ptprd | intron_variant,non_coding_transcript_variant | MODIFIER |
| 4 | 76856088 | AX-170287160 | 0 | A | G | 0.325 | 1.47E-01 | 1.20E-01 | 2.84E+00 | 2.89E+00 | 2.18E-01 | Yes | Ptprd | intron_variant,non_coding_transcript_variant | MODIFIER |
| 4 | 76910208 | AX-170187398 | 0 | T | C | 0.325 | 1.47E-01 | 1.20E-01 | 2.84E+00 | 2.89E+00 | 2.18E-01 | Yes | Ptprd | intron_variant,non_coding_transcript_variant | MODIFIER |
| 4 | 76911284 | AX-170415109 | 0 | G | A | 0.325 | 1.47E-01 | 1.20E-01 | 2.84E+00 | 2.89E+00 | 2.18E-01 | Yes | Ptprd | intron_variant,non_coding_transcript_variant | MODIFIER |
| 4 | 77257916 | AX-170320418 | 0 | G | A | 0.325 | 1.47E-01 | 1.20E-01 | 2.84E+00 | 2.89E+00 | 2.18E-01 | Yes | Ptprd | intron_variant,non_coding_transcript_variant | MODIFIER |
| 4 | 77258171 | AX-170331328 | 0 | C | T | 0.325 | 1.47E-01 | 1.20E-01 | 2.84E+00 | 2.89E+00 | 2.18E-01 | Yes | Ptprd | intron_variant,non_coding_transcript_variant | MODIFIER |
| 4 | 77316975 | AX-170350128 | 0 | C | T | 0.325 | 1.47E-01 | 1.20E-01 | 2.84E+00 | 2.89E+00 | 2.18E-01 | Yes | Ptprd | intron_variant,non_coding_transcript_variant | MODIFIER |
| 4 | 77333541 | AX-170207831 | 0 | T | G | 0.325 | 1.47E-01 | 1.20E-01 | 2.84E+00 | 2.89E+00 | 2.18E-01 | Yes | Ptprd | intron_variant,non_coding_transcript_variant | MODIFIER |
| 4 | 77356409 | AX-170338982 | 0 | C | T | 0.325 | 1.47E-01 | 1.20E-01 | 2.84E+00 | 2.89E+00 | 2.18E-01 | Yes | Ptprd | intron_variant,non_coding_transcript_variant | MODIFIER |
| 4 | 77379974 | AX-170069740 | 0 | A | G | 0.325 | 1.47E-01 | 1.20E-01 | 2.84E+00 | 2.89E+00 | 2.18E-01 | Yes | Ptprd | intron_variant,non_coding_transcript_variant | MODIFIER |
| 4 | 77382264 | AX-170328008 | 0 | G | A | 0.325 | 1.47E-01 | 1.20E-01 | 2.84E+00 | 2.89E+00 | 2.18E-01 | Yes | Ptprd | intron_variant,non_coding_transcript_variant | MODIFIER |
| 4 | 77398737 | AX-170069744 | 0 | G | T | 0.325 | 1.47E-01 | 1.20E-01 | 2.84E+00 | 2.89E+00 | 2.18E-01 | Yes | Ptprd | intron_variant,non_coding_transcript_variant | MODIFIER |
| 4 | 77474689 | AX-97790924 | 0 | C | T | 0.325 | 1.47E-01 | 1.20E-01 | 2.84E+00 | 2.89E+00 | 2.18E-01 | Yes | Ptprd | intron_variant,non_coding_transcript_variant | MODIFIER |
| 4 | 77528661 | AX-170089284 | 0 | A | G | 0.325 | 1.47E-01 | 1.20E-01 | 2.84E+00 | 2.89E+00 | 2.18E-01 | Yes | Ptprd | intron_variant,non_coding_transcript_variant | MODIFIER |
| 4 | 77530622 | AX-170123403 | 0 | C | T | 0.325 | 1.47E-01 | 1.20E-01 | 2.84E+00 | 2.89E+00 | 2.18E-01 | Yes | Ptprd | intron_variant,non_coding_transcript_variant | MODIFIER |
| 4 | 77585645 | AX-170425299 | 0 | G | A | 0.325 | 1.47E-01 | 1.20E-01 | 2.84E+00 | 2.89E+00 | 2.18E-01 | Yes | Ptprd | intron_variant,non_coding_transcript_variant | MODIFIER |
| 4 | 77616077 | AX-170129687 | 0 | C | A | 0.325 | 1.47E-01 | 1.20E-01 | 2.84E+00 | 2.89E+00 | 2.18E-01 | Yes | Ptprd | intron_variant,non_coding_transcript_variant | MODIFIER |
| 4 | 77650472 | AX-170149196 | 0 | T | C | 0.325 | 1.47E-01 | 1.20E-01 | 2.84E+00 | 2.89E+00 | 2.18E-01 | Yes | Ptprd | intron_variant,non_coding_transcript_variant | MODIFIER |
| 4 | 77683957 | AX-170281098 | 0 | C | T | 0.325 | 1.47E-01 | 1.20E-01 | 2.84E+00 | 2.89E+00 | 2.18E-01 | Yes | Ptprd | intron_variant,non_coding_transcript_variant | MODIFIER |
| 4 | 77702398 | AX-170532218 | 0 | G | C | 0.325 | 1.47E-01 | 1.20E-01 | 2.84E+00 | 2.89E+00 | 2.18E-01 | Yes | Ptprd | intron_variant,non_coding_transcript_variant | MODIFIER |
| 4 | 77702421 | AX-170265200 | 0 | C | G | 0.325 | 1.47E-01 | 1.20E-01 | 2.84E+00 | 2.89E+00 | 2.18E-01 | Yes | Ptprd | intron_variant,non_coding_transcript_variant | MODIFIER |
| 4 | 77704793 | AX-170523102 | 0 | T | C | 0.325 | 1.47E-01 | 1.20E-01 | 2.84E+00 | 2.89E+00 | 2.18E-01 | Yes | Ptprd | intron_variant,non_coding_transcript_variant | MODIFIER |
| 4 | 78142585 | AX-170069763 | 0 | C | A | 0.325 | 1.47E-01 | 1.20E-01 | 2.84E+00 | 2.89E+00 | 2.18E-01 | Yes | Ptprd | intron_variant,non_coding_transcript_variant | MODIFIER |
| 4 | 78232474 | AX-170185754 | 0 | G | A | 0.325 | 1.47E-01 | 1.20E-01 | 2.84E+00 | 2.89E+00 | 2.18E-01 | Yes | Gm11260 | upstream_gene_variant | MODIFIER |
| 4 | 78430967 | AX-170529474 | 0 | C | T | 0.325 | 1.47E-01 | 1.20E-01 | 2.84E+00 | 2.89E+00 | 2.18E-01 | Yes | | intergenic_variant | MODIFIER |
| 4 | 78431414 | AX-170523083 | 0 | G | A | 0.325 | 1.47E-01 | 1.20E-01 | 2.84E+00 | 2.89E+00 | 2.18E-01 | Yes | | intergenic_variant | MODIFIER |
| 4 | 78431494 | AX-170444207 | 0 | T | C | 0.325 | 1.47E-01 | 1.20E-01 | 2.84E+00 | 2.89E+00 | 2.18E-01 | Yes | | intergenic_variant | MODIFIER |
| 4 | 80810586 | AX-97855252 | 0 | A | C | 0.325 | 1.47E-01 | 1.20E-01 | 2.84E+00 | 2.89E+00 | 2.18E-01 | Yes | | intergenic_variant | MODIFIER |
| 1 | 171944748 | AX-170043585 | 0 | C | G | 0.211 | 1.67E-01 | 1.36E-01 | 2.60E+00 | 2.65E+00 | 2.18E-01 | Yes | Slamf6 | downstream_gene_variant | MODIFIER |
| 4 | 69957358 | AX-170528694 | 0 | G | A | 0.316 | 1.42E-01 | 1.16E-01 | 2.78E+00 | 2.85E+00 | 2.18E-01 | no | Gm11224 | upstream_gene_variant | MODIFIER |
| 1 | 121155223 | AX-170505634 | 1 | T | A | 0.58 | 1.03E-01 | 8.33E-02 | 2.50E+00 | 2.56E+00 | 2.18E-01 | no | Celrr | intron_variant,non_coding_transcript_variant | MODIFIER |
| 1 | 119372458 | AX-170534314 | 0 | G | A | 0.579 | 1.03E-01 | 8.34E-02 | 2.50E+00 | 2.56E+00 | 2.18E-01 | no | Gm29455 | intron_variant,non_coding_transcript_variant | MODIFIER |
| 1 | 119374220 | AX-170042467 | 0 | G | A | 0.579 | 1.03E-01 | 8.34E-02 | 2.50E+00 | 2.56E+00 | 2.18E-01 | no | Gm29455 | intron_variant,non_coding_transcript_variant | MODIFIER |
| 1 | 119379258 | AX-170042468 | 0 | C | A | 0.579 | 1.03E-01 | 8.34E-02 | 2.50E+00 | 2.56E+00 | 2.18E-01 | no | Gm29455 | intron_variant,non_coding_transcript_variant | MODIFIER |
| 1 | 119437551 | AX-170375500 | 0 | C | A | 0.579 | 1.03E-01 | 8.34E-02 | 2.50E+00 | 2.56E+00 | 2.18E-01 | no | | intergenic_variant | MODIFIER |
| 1 | 119437887 | AX-170238419 | 0 | C | T | 0.579 | 1.03E-01 | 8.34E-02 | 2.50E+00 | 2.56E+00 | 2.18E-01 | no | | intergenic_variant | MODIFIER |
| 1 | 120021076 | AX-170042476 | 0 | A | G | 0.579 | 1.03E-01 | 8.34E-02 | 2.50E+00 | 2.56E+00 | 2.18E-01 | no | Sctr | intron_variant | MODIFIER |
| 1 | 120024288 | AX-170042475 | 0 | A | G | 0.579 | 1.03E-01 | 8.34E-02 | 2.50E+00 | 2.56E+00 | 2.18E-01 | no | Sctr | intron_variant | MODIFIER |
| 1 | 120075853 | AX-170371586 | 0 | C | A | 0.579 | 1.03E-01 | 8.34E-02 | 2.50E+00 | 2.56E+00 | 2.18E-01 | no | Tmem37 | upstream_gene_variant | MODIFIER |
| 1 | 120108735 | AX-97826202 | 0 | T | C | 0.579 | 1.03E-01 | 8.34E-02 | 2.50E+00 | 2.56E+00 | 2.18E-01 | no | Dbi | downstream_gene_variant | MODIFIER |
| 1 | 120112911 | AX-170360985 | 0 | A | G | 0.579 | 1.03E-01 | 8.34E-02 | 2.50E+00 | 2.56E+00 | 2.18E-01 | no | Dbi | downstream_gene_variant | MODIFIER |
| 1 | 120116189 | AX-170245380 | 0 | A | T | 0.579 | 1.03E-01 | 8.34E-02 | 2.50E+00 | 2.56E+00 | 2.18E-01 | no | Dbi | intron_variant | MODIFIER |
| 1 | 120135044 | AX-170346933 | 0 | A | G | 0.579 | 1.03E-01 | 8.34E-02 | 2.50E+00 | 2.56E+00 | 2.18E-01 | no | 3110009E18Rik | intron_variant | MODIFIER |
| 1 | 120135263 | AX-170440259 | 0 | G | C | 0.579 | 1.03E-01 | 8.34E-02 | 2.50E+00 | 2.56E+00 | 2.18E-01 | no | 3110009E18Rik | intron_variant | MODIFIER |
| 1 | 120135978 | AX-170419216 | 0 | C | G | 0.579 | 1.03E-01 | 8.34E-02 | 2.50E+00 | 2.56E+00 | 2.18E-01 | no | 3110009E18Rik | intron_variant | MODIFIER |
| 1 | 120136281 | AX-170263529 | 0 | T | C | 0.579 | 1.03E-01 | 8.34E-02 | 2.50E+00 | 2.56E+00 | 2.18E-01 | no | 3110009E18Rik | intron_variant | MODIFIER |
| 1 | 120152551 | AX-170412751 | 0 | T | A | 0.579 | 1.03E-01 | 8.34E-02 | 2.50E+00 | 2.56E+00 | 2.18E-01 | no | 3110009E18Rik | intron_variant | MODIFIER |
| 1 | 120233750 | AX-97839423 | 0 | T | C | 0.579 | 1.03E-01 | 8.34E-02 | 2.50E+00 | 2.56E+00 | 2.18E-01 | no | Steap3 | intron_variant | MODIFIER |
| 1 | 120233925 | AX-170042477 | 0 | C | T | 0.579 | 1.03E-01 | 8.34E-02 | 2.50E+00 | 2.56E+00 | 2.18E-01 | no | Steap3 | intron_variant | MODIFIER |

| | | | | | | | | | | | | | | | |
|---|---|---|---|---|---|---|---|---|---|---|---|---|---|---|---|
| 1 | 120267181 | AX-170443421 | 0 | T | G | 0.579 | 1.03E-01 | 8.34E-02 | 2.50E+00 | 2.56E+00 | 2.18E-01 | no | Steap3 | intron_variant | MODIFIER |
| 1 | 120312411 | AX-170001555 | 0 | A | C | 0.579 | 1.03E-01 | 8.34E-02 | 2.50E+00 | 2.56E+00 | 2.18E-01 | no | | intergenic_variant | MODIFIER |
| 1 | 120342761 | AX-170001550 | 0 | T | C | 0.579 | 1.03E-01 | 8.34E-02 | 2.50E+00 | 2.56E+00 | 2.18E-01 | no | C1ql2 | 3_prime_UTR_variant | MODIFIER |
| 1 | 120352712 | AX-97813113 | 0 | A | C | 0.579 | 1.03E-01 | 8.34E-02 | 2.50E+00 | 2.56E+00 | 2.18E-01 | no | | intergenic_variant | MODIFIER |
| 1 | 120363785 | AX-170476797 | 0 | C | T | 0.579 | 1.03E-01 | 8.34E-02 | 2.50E+00 | 2.56E+00 | 2.18E-01 | no | | intergenic_variant | MODIFIER |
| 1 | 120365718 | AX-170001548 | 0 | G | A | 0.579 | 1.03E-01 | 8.34E-02 | 2.50E+00 | 2.56E+00 | 2.18E-01 | no | | intergenic_variant | MODIFIER |
| 1 | 120367114 | AX-170443534 | 0 | G | C | 0.579 | 1.03E-01 | 8.34E-02 | 2.50E+00 | 2.56E+00 | 2.18E-01 | no | | intergenic_variant | MODIFIER |
| 1 | 120390750 | AX-170419572 | 0 | T | C | 0.579 | 1.03E-01 | 8.34E-02 | 2.50E+00 | 2.56E+00 | 2.18E-01 | no | | regulatory_region_variant | MODIFIER |
| 1 | 120395474 | AX-97845429 | 0 | T | C | 0.579 | 1.03E-01 | 8.34E-02 | 2.50E+00 | 2.56E+00 | 2.18E-01 | no | | intergenic_variant | MODIFIER |
| 1 | 120599453 | AX-170255895 | 0 | A | C | 0.579 | 1.03E-01 | 8.34E-02 | 2.50E+00 | 2.56E+00 | 2.18E-01 | no | En1 | upstream_gene_variant | MODIFIER |
| 1 | 120600410 | AX-170348594 | 0 | T | C | 0.579 | 1.03E-01 | 8.34E-02 | 2.50E+00 | 2.56E+00 | 2.18E-01 | no | En1 | upstream_gene_variant | MODIFIER |
| 1 | 120605054 | AX-170201618 | 0 | C | A | 0.579 | 1.03E-01 | 8.34E-02 | 2.50E+00 | 2.56E+00 | 2.18E-01 | no | En1 | intron_variant | MODIFIER |
| 1 | 120616286 | AX-170228453 | 0 | G | A | 0.579 | 1.03E-01 | 8.34E-02 | 2.50E+00 | 2.56E+00 | 2.18E-01 | no | | intergenic_variant | MODIFIER |
| 1 | 120616662 | AX-170042486 | 0 | G | T | 0.579 | 1.03E-01 | 8.34E-02 | 2.50E+00 | 2.56E+00 | 2.18E-01 | no | | intergenic_variant | MODIFIER |
| 1 | 120622608 | AX-170277541 | 0 | T | G | 0.579 | 1.03E-01 | 8.34E-02 | 2.50E+00 | 2.56E+00 | 2.18E-01 | no | 2610027F03Rik | intron_variant,non_coding_transcript_variant | MODIFIER |
| 1 | 120664615 | AX-170358398 | 0 | C | T | 0.579 | 1.03E-01 | 8.34E-02 | 2.50E+00 | 2.56E+00 | 2.18E-01 | no | 2610027F03Rik | intron_variant,non_coding_transcript_variant | MODIFIER |
| 1 | 120668419 | AX-170513881 | 0 | C | T | 0.579 | 1.03E-01 | 8.34E-02 | 2.50E+00 | 2.56E+00 | 2.18E-01 | no | 2610027F03Rik | intron_variant,non_coding_transcript_variant | MODIFIER |
| 1 | 120682961 | AX-170141035 | 0 | C | T | 0.579 | 1.03E-01 | 8.34E-02 | 2.50E+00 | 2.56E+00 | 2.18E-01 | no | 2610027F03Rik | downstream_gene_variant | MODIFIER |
| 1 | 120786052 | AX-170001560 | 0 | G | A | 0.579 | 1.03E-01 | 8.34E-02 | 2.50E+00 | 2.56E+00 | 2.18E-01 | no | | intergenic_variant | MODIFIER |
| 1 | 120799795 | AX-170406586 | 0 | G | T | 0.579 | 1.03E-01 | 8.34E-02 | 2.50E+00 | 2.56E+00 | 2.18E-01 | no | | intergenic_variant | MODIFIER |
| 1 | 120804021 | AX-170514427 | 0 | T | G | 0.579 | 1.03E-01 | 8.34E-02 | 2.50E+00 | 2.56E+00 | 2.18E-01 | no | | intergenic_variant | MODIFIER |
| 1 | 120812763 | AX-170397003 | 0 | G | A | 0.579 | 1.03E-01 | 8.34E-02 | 2.50E+00 | 2.56E+00 | 2.18E-01 | no | Gm29345 | upstream_gene_variant | MODIFIER |
| 1 | 120814047 | AX-170383453 | 0 | G | A | 0.579 | 1.03E-01 | 8.34E-02 | 2.50E+00 | 2.56E+00 | 2.18E-01 | no | Gm29345 | upstream_gene_variant | MODIFIER |
| 1 | 120814107 | AX-170185763 | 0 | T | C | 0.579 | 1.03E-01 | 8.34E-02 | 2.50E+00 | 2.56E+00 | 2.18E-01 | no | Gm29345 | upstream_gene_variant | MODIFIER |
| 1 | 120828699 | AX-170001556 | 0 | C | T | 0.579 | 1.03E-01 | 8.34E-02 | 2.50E+00 | 2.56E+00 | 2.18E-01 | no | Gm29344 | upstream_gene_variant | MODIFIER |
| 1 | 120829177 | AX-170001559 | 0 | T | A | 0.579 | 1.03E-01 | 8.34E-02 | 2.50E+00 | 2.56E+00 | 2.18E-01 | no | Gm29344 | upstream_gene_variant | MODIFIER |
| 1 | 120835662 | AX-97811539 | 0 | T | C | 0.579 | 1.03E-01 | 8.34E-02 | 2.50E+00 | 2.56E+00 | 2.18E-01 | no | Gm29345 | intron_variant,non_coding_transcript_variant | MODIFIER |
| 1 | 120840853 | AX-170284508 | 0 | G | T | 0.579 | 1.03E-01 | 8.34E-02 | 2.50E+00 | 2.56E+00 | 2.18E-01 | no | Gm29345 | downstream_gene_variant | MODIFIER |
| 1 | 120853658 | AX-170487393 | 0 | A | T | 0.579 | 1.03E-01 | 8.34E-02 | 2.50E+00 | 2.56E+00 | 2.18E-01 | no | Gm29347 | upstream_gene_variant | MODIFIER |
| 1 | 120857963 | AX-170091661 | 0 | G | A | 0.579 | 1.03E-01 | 8.34E-02 | 2.50E+00 | 2.56E+00 | 2.18E-01 | no | Gm29347 | non_coding_transcript_exon_variant | MODIFIER |
| 1 | 120859401 | AX-170195384 | 0 | A | G | 0.579 | 1.03E-01 | 8.34E-02 | 2.50E+00 | 2.56E+00 | 2.18E-01 | no | Gm29347 | downstream_gene_variant | MODIFIER |
| 1 | 120860732 | AX-170042494 | 0 | C | T | 0.579 | 1.03E-01 | 8.34E-02 | 2.50E+00 | 2.56E+00 | 2.18E-01 | no | Gm29347 | downstream_gene_variant | MODIFIER |
| 1 | 120872307 | AX-170001557 | 0 | A | G | 0.579 | 1.03E-01 | 8.34E-02 | 2.50E+00 | 2.56E+00 | 2.18E-01 | no | Gm37043 | downstream_gene_variant | MODIFIER |
| 1 | 120873791 | AX-97824715 | 0 | T | C | 0.579 | 1.03E-01 | 8.34E-02 | 2.50E+00 | 2.56E+00 | 2.18E-01 | no | Gm37043 | downstream_gene_variant | MODIFIER |
| 1 | 120877368 | AX-170528521 | 0 | C | T | 0.579 | 1.03E-01 | 8.34E-02 | 2.50E+00 | 2.56E+00 | 2.18E-01 | no | | intergenic_variant | MODIFIER |
| 1 | 120883383 | AX-170399493 | 0 | C | T | 0.579 | 1.03E-01 | 8.34E-02 | 2.50E+00 | 2.56E+00 | 2.18E-01 | no | | regulatory_region_variant | MODIFIER |
| 1 | 120908326 | AX-170514464 | 0 | G | T | 0.579 | 1.03E-01 | 8.34E-02 | 2.50E+00 | 2.56E+00 | 2.18E-01 | no | | intergenic_variant | MODIFIER |
| 1 | 120915834 | AX-170192279 | 0 | G | T | 0.579 | 1.03E-01 | 8.34E-02 | 2.50E+00 | 2.56E+00 | 2.18E-01 | no | | intergenic_variant | MODIFIER |
| 1 | 120916657 | AX-170479973 | 0 | T | C | 0.579 | 1.03E-01 | 8.34E-02 | 2.50E+00 | 2.56E+00 | 2.18E-01 | no | | intergenic_variant | MODIFIER |
| 1 | 120942292 | AX-97827916 | 0 | T | C | 0.579 | 1.03E-01 | 8.34E-02 | 2.50E+00 | 2.56E+00 | 2.18E-01 | no | | intergenic_variant | MODIFIER |
| 1 | 120943576 | AX-170526884 | 0 | A | C | 0.579 | 1.03E-01 | 8.34E-02 | 2.50E+00 | 2.56E+00 | 2.18E-01 | no | | intergenic_variant | MODIFIER |
| 1 | 120949306 | AX-170350239 | 0 | A | G | 0.579 | 1.03E-01 | 8.34E-02 | 2.50E+00 | 2.56E+00 | 2.18E-01 | no | | intergenic_variant | MODIFIER |
| 1 | 120950178 | AX-170507387 | 0 | A | G | 0.579 | 1.03E-01 | 8.34E-02 | 2.50E+00 | 2.56E+00 | 2.18E-01 | no | | intergenic_variant | MODIFIER |
| 1 | 120951331 | AX-97833679 | 0 | A | C | 0.579 | 1.03E-01 | 8.34E-02 | 2.50E+00 | 2.56E+00 | 2.18E-01 | no | | intergenic_variant | MODIFIER |
| 1 | 120951479 | AX-170234844 | 0 | A | G | 0.579 | 1.03E-01 | 8.34E-02 | 2.50E+00 | 2.56E+00 | 2.18E-01 | no | | intergenic_variant | MODIFIER |
| 1 | 120966634 | AX-170405531 | 0 | A | G | 0.579 | 1.03E-01 | 8.34E-02 | 2.50E+00 | 2.56E+00 | 2.18E-01 | no | | intergenic_variant | MODIFIER |
| 1 | 120968649 | AX-170494684 | 0 | T | C | 0.579 | 1.03E-01 | 8.34E-02 | 2.50E+00 | 2.56E+00 | 2.18E-01 | no | | intergenic_variant | MODIFIER |
| 1 | 120970310 | AX-170283154 | 0 | G | C | 0.579 | 1.03E-01 | 8.34E-02 | 2.50E+00 | 2.56E+00 | 2.18E-01 | no | | intergenic_variant | MODIFIER |
| 1 | 120983663 | AX-97780244 | 0 | A | C | 0.579 | 1.03E-01 | 8.34E-02 | 2.50E+00 | 2.56E+00 | 2.18E-01 | no | | intergenic_variant | MODIFIER |
| 1 | 120991684 | AX-170331949 | 0 | T | C | 0.579 | 1.03E-01 | 8.34E-02 | 2.50E+00 | 2.56E+00 | 2.18E-01 | no | | intergenic_variant | MODIFIER |
| 1 | 120992228 | AX-170395883 | 0 | T | C | 0.579 | 1.03E-01 | 8.34E-02 | 2.50E+00 | 2.56E+00 | 2.18E-01 | no | | intergenic_variant | MODIFIER |
| 1 | 121015022 | AX-170091662 | 0 | C | A | 0.579 | 1.03E-01 | 8.34E-02 | 2.50E+00 | 2.56E+00 | 2.18E-01 | no | | intergenic_variant | MODIFIER |
| 1 | 121015332 | AX-170455858 | 0 | G | A | 0.579 | 1.03E-01 | 8.34E-02 | 2.50E+00 | 2.56E+00 | 2.18E-01 | no | | intergenic_variant | MODIFIER |
| 1 | 121016565 | AX-170383926 | 0 | A | G | 0.579 | 1.03E-01 | 8.34E-02 | 2.50E+00 | 2.56E+00 | 2.18E-01 | no | | intergenic_variant | MODIFIER |

| | | | | | | | | | | | | | | | |
|---|---|---|---|---|---|---|---|---|---|---|---|---|---|---|---|
| 1 | 121027728 | AX-170454621 | 0 | A | C | 0.579 | 1.03E-01 | 8.34E-02 | 2.50E+00 | 2.56E+00 | 2.18E-01 | no | | intergenic_variant | MODIFIER |
| 1 | 121034280 | AX-170086595 | 0 | G | A | 0.579 | 1.03E-01 | 8.34E-02 | 2.50E+00 | 2.56E+00 | 2.18E-01 | no | | intergenic_variant | MODIFIER |
| 1 | 121034382 | AX-170297558 | 0 | A | G | 0.579 | 1.03E-01 | 8.34E-02 | 2.50E+00 | 2.56E+00 | 2.18E-01 | no | | intergenic_variant | MODIFIER |
| 1 | 121056689 | AX-170514138 | 0 | A | C | 0.579 | 1.03E-01 | 8.34E-02 | 2.50E+00 | 2.56E+00 | 2.18E-01 | no | | intergenic_variant | MODIFIER |
| 1 | 121056979 | AX-170419395 | 0 | A | C | 0.579 | 1.03E-01 | 8.34E-02 | 2.50E+00 | 2.56E+00 | 2.18E-01 | no | | intergenic_variant | MODIFIER |
| 1 | 121057672 | AX-170091663 | 0 | A | G | 0.579 | 1.03E-01 | 8.34E-02 | 2.50E+00 | 2.56E+00 | 2.18E-01 | no | | intergenic_variant | MODIFIER |
| 1 | 121060847 | AX-170236537 | 0 | T | C | 0.579 | 1.03E-01 | 8.34E-02 | 2.50E+00 | 2.56E+00 | 2.18E-01 | no | | intergenic_variant | MODIFIER |
| 1 | 121070250 | AX-170237796 | 0 | C | T | 0.579 | 1.03E-01 | 8.34E-02 | 2.50E+00 | 2.56E+00 | 2.18E-01 | no | | intergenic_variant | MODIFIER |
| 1 | 121085889 | AX-170207539 | 0 | G | A | 0.579 | 1.03E-01 | 8.34E-02 | 2.50E+00 | 2.56E+00 | 2.18E-01 | no | Celrr | upstream_gene_variant | MODIFIER |
| 1 | 121099362 | AX-170184653 | 0 | T | C | 0.579 | 1.03E-01 | 8.34E-02 | 2.50E+00 | 2.56E+00 | 2.18E-01 | no | Celrr | intron_variant,non_coding_transcript_variant | MODIFIER |
| 1 | 121113498 | AX-170516999 | 0 | T | A | 0.579 | 1.03E-01 | 8.34E-02 | 2.50E+00 | 2.56E+00 | 2.18E-01 | no | Celrr | intron_variant,non_coding_transcript_variant | MODIFIER |
| 1 | 121132705 | AX-170488084 | 0 | G | A | 0.579 | 1.03E-01 | 8.34E-02 | 2.50E+00 | 2.56E+00 | 2.18E-01 | no | Celrr | intron_variant,non_coding_transcript_variant | MODIFIER |
| 1 | 121138998 | AX-170344963 | 0 | A | G | 0.579 | 1.03E-01 | 8.34E-02 | 2.50E+00 | 2.56E+00 | 2.18E-01 | no | Celrr | intron_variant,non_coding_transcript_variant | MODIFIER |
| 1 | 121152175 | AX-170155308 | 0 | G | C | 0.579 | 1.03E-01 | 8.34E-02 | 2.50E+00 | 2.56E+00 | 2.18E-01 | no | Celrr | intron_variant,non_coding_transcript_variant | MODIFIER |
| 1 | 121153711 | AX-170517898 | 0 | C | G | 0.579 | 1.03E-01 | 8.34E-02 | 2.50E+00 | 2.56E+00 | 2.18E-01 | no | Celrr | intron_variant,non_coding_transcript_variant | MODIFIER |
| 1 | 121156360 | AX-170480944 | 0 | G | A | 0.579 | 1.03E-01 | 8.34E-02 | 2.50E+00 | 2.56E+00 | 2.18E-01 | no | Celrr | intron_variant,non_coding_transcript_variant | MODIFIER |
| 1 | 121157957 | AX-170356791 | 0 | T | G | 0.579 | 1.03E-01 | 8.34E-02 | 2.50E+00 | 2.56E+00 | 2.18E-01 | no | Celrr | intron_variant,non_coding_transcript_variant | MODIFIER |
| 1 | 121167407 | AX-170348925 | 0 | A | G | 0.579 | 1.03E-01 | 8.34E-02 | 2.50E+00 | 2.56E+00 | 2.18E-01 | no | Celrr | intron_variant,non_coding_transcript_variant | MODIFIER |
| 1 | 121169283 | AX-170337985 | 0 | T | C | 0.579 | 1.03E-01 | 8.34E-02 | 2.50E+00 | 2.56E+00 | 2.18E-01 | no | Celrr | intron_variant,non_coding_transcript_variant | MODIFIER |
| 1 | 121182299 | AX-170226690 | 0 | A | G | 0.579 | 1.03E-01 | 8.34E-02 | 2.50E+00 | 2.56E+00 | 2.18E-01 | no | Celrr | downstream_gene_variant | MODIFIER |
| 1 | 121190053 | AX-97830418 | 0 | A | G | 0.579 | 1.03E-01 | 8.34E-02 | 2.50E+00 | 2.56E+00 | 2.18E-01 | no | | intergenic_variant | MODIFIER |
| 1 | 121201118 | AX-170389313 | 0 | T | C | 0.579 | 1.03E-01 | 8.34E-02 | 2.50E+00 | 2.56E+00 | 2.18E-01 | no | | intergenic_variant | MODIFIER |
| 1 | 121212098 | AX-97839719 | 0 | G | A | 0.579 | 1.03E-01 | 8.34E-02 | 2.50E+00 | 2.56E+00 | 2.18E-01 | no | | intergenic_variant | MODIFIER |
| 1 | 121212169 | AX-170175889 | 0 | T | C | 0.579 | 1.03E-01 | 8.34E-02 | 2.50E+00 | 2.56E+00 | 2.18E-01 | no | | intergenic_variant | MODIFIER |
| 1 | 121232048 | AX-170509723 | 0 | A | G | 0.579 | 1.03E-01 | 8.34E-02 | 2.50E+00 | 2.56E+00 | 2.18E-01 | no | Gm29359 | downstream_gene_variant | MODIFIER |
| 1 | 121237321 | AX-97786385 | 0 | A | G | 0.579 | 1.03E-01 | 8.34E-02 | 2.50E+00 | 2.56E+00 | 2.18E-01 | no | | intergenic_variant | MODIFIER |
| 1 | 121247051 | AX-170228976 | 0 | A | G | 0.579 | 1.03E-01 | 8.34E-02 | 2.50E+00 | 2.56E+00 | 2.18E-01 | no | | regulatory_region_variant | MODIFIER |
| 1 | 121250698 | AX-170213366 | 0 | G | A | 0.579 | 1.03E-01 | 8.34E-02 | 2.50E+00 | 2.56E+00 | 2.18E-01 | no | | intergenic_variant | MODIFIER |
| 1 | 121253822 | AX-170281886 | 0 | A | G | 0.579 | 1.03E-01 | 8.34E-02 | 2.50E+00 | 2.56E+00 | 2.18E-01 | no | | regulatory_region_variant | MODIFIER |
| 1 | 121304811 | AX-170234045 | 0 | A | T | 0.579 | 1.03E-01 | 8.34E-02 | 2.50E+00 | 2.56E+00 | 2.18E-01 | no | Insig2 | 3_prime_UTR_variant | MODIFIER |
| 1 | 121315223 | AX-170177061 | 0 | G | T | 0.579 | 1.03E-01 | 8.34E-02 | 2.50E+00 | 2.56E+00 | 2.18E-01 | no | Insig2 | intron_variant | MODIFIER |
| 1 | 121316084 | AX-170239667 | 0 | C | T | 0.579 | 1.03E-01 | 8.34E-02 | 2.50E+00 | 2.56E+00 | 2.18E-01 | no | Insig2 | intron_variant | MODIFIER |
| 1 | 121316411 | AX-170196779 | 0 | T | C | 0.579 | 1.03E-01 | 8.34E-02 | 2.50E+00 | 2.56E+00 | 2.18E-01 | no | Insig2 | intron_variant | MODIFIER |
| 1 | 121318092 | AX-97834012 | 0 | A | G | 0.579 | 1.03E-01 | 8.34E-02 | 2.50E+00 | 2.56E+00 | 2.18E-01 | no | Insig2 | intron_variant | MODIFIER |
| 1 | 121323027 | AX-170421086 | 0 | C | T | 0.579 | 1.03E-01 | 8.34E-02 | 2.50E+00 | 2.56E+00 | 2.18E-01 | no | Insig2 | intron_variant | MODIFIER |
| 1 | 121325145 | AX-170482889 | 0 | C | T | 0.579 | 1.03E-01 | 8.34E-02 | 2.50E+00 | 2.56E+00 | 2.18E-01 | no | Insig2 | intron_variant | MODIFIER |
| 1 | 121325585 | AX-170097868 | 0 | A | G | 0.579 | 1.03E-01 | 8.34E-02 | 2.50E+00 | 2.56E+00 | 2.18E-01 | no | Insig2 | intron_variant | MODIFIER |
| 1 | 121325652 | AX-170213648 | 0 | A | G | 0.579 | 1.03E-01 | 8.34E-02 | 2.50E+00 | 2.56E+00 | 2.18E-01 | no | Insig2 | intron_variant | MODIFIER |
| 1 | 121325774 | AX-170162122 | 0 | A | G | 0.579 | 1.03E-01 | 8.34E-02 | 2.50E+00 | 2.56E+00 | 2.18E-01 | no | Insig2 | intron_variant | MODIFIER |
| 1 | 121326016 | AX-170429226 | 0 | G | A | 0.579 | 1.03E-01 | 8.34E-02 | 2.50E+00 | 2.56E+00 | 2.18E-01 | no | Insig2 | intron_variant | MODIFIER |
| 1 | 121552302 | AX-170138485 | 0 | G | C | 0.579 | 1.03E-01 | 8.34E-02 | 2.50E+00 | 2.56E+00 | 2.18E-01 | no | Ddx18 | downstream_gene_variant | MODIFIER |
| 1 | 121556906 | AX-170370443 | 0 | C | T | 0.579 | 1.03E-01 | 8.34E-02 | 2.50E+00 | 2.56E+00 | 2.18E-01 | no | Ddx18 | intron_variant | MODIFIER |
| 1 | 121594689 | AX-170263005 | 0 | T | C | 0.579 | 1.03E-01 | 8.34E-02 | 2.50E+00 | 2.56E+00 | 2.18E-01 | no | | intergenic_variant | MODIFIER |
| 1 | 121602373 | AX-170371990 | 0 | G | A | 0.579 | 1.03E-01 | 8.34E-02 | 2.50E+00 | 2.56E+00 | 2.18E-01 | no | | intergenic_variant | MODIFIER |
| 1 | 121605408 | AX-170414110 | 0 | G | T | 0.579 | 1.03E-01 | 8.34E-02 | 2.50E+00 | 2.56E+00 | 2.18E-01 | no | | regulatory_region_variant | MODIFIER |
| 1 | 121605634 | AX-170424542 | 0 | A | C | 0.579 | 1.03E-01 | 8.34E-02 | 2.50E+00 | 2.56E+00 | 2.18E-01 | no | | regulatory_region_variant | MODIFIER |
| 1 | 121651390 | AX-170486178 | 0 | G | C | 0.579 | 1.03E-01 | 8.34E-02 | 2.50E+00 | 2.56E+00 | 2.18E-01 | no | | intergenic_variant | MODIFIER |
| 1 | 121753454 | AX-170116629 | 0 | T | C | 0.579 | 1.03E-01 | 8.34E-02 | 2.50E+00 | 2.56E+00 | 2.18E-01 | no | | intergenic_variant | MODIFIER |
| 1 | 121796741 | AX-170176975 | 0 | G | A | 0.579 | 1.03E-01 | 8.34E-02 | 2.50E+00 | 2.56E+00 | 2.18E-01 | no | | intergenic_variant | MODIFIER |
| 1 | 121809674 | AX-170192532 | 0 | C | T | 0.579 | 1.03E-01 | 8.34E-02 | 2.50E+00 | 2.56E+00 | 2.18E-01 | no | | intergenic_variant | MODIFIER |
| 1 | 121811319 | AX-170319799 | 0 | C | G | 0.579 | 1.03E-01 | 8.34E-02 | 2.50E+00 | 2.56E+00 | 2.18E-01 | no | | intergenic_variant | MODIFIER |
| 1 | 121811597 | AX-170262488 | 0 | A | G | 0.579 | 1.03E-01 | 8.34E-02 | 2.50E+00 | 2.56E+00 | 2.18E-01 | no | | intergenic_variant | MODIFIER |
| 1 | 121818420 | AX-170321362 | 0 | A | G | 0.579 | 1.03E-01 | 8.34E-02 | 2.50E+00 | 2.56E+00 | 2.18E-01 | no | | intergenic_variant | MODIFIER |
| 1 | 121835269 | AX-170350717 | 0 | G | T | 0.579 | 1.03E-01 | 8.34E-02 | 2.50E+00 | 2.56E+00 | 2.18E-01 | no | | intergenic_variant | MODIFIER |

| | | | | | | | | | | | | | | | |
|---|---|---|---|---|---|---|---|---|---|---|---|---|---|---|---|
| 1 | 121848362 | AX-170347633 | 0 | A | C | 0.579 | 1.03E-01 | 8.34E-02 | 2.50E+00 | 2.56E+00 | 2.18E-01 | no | | intergenic_variant | MODIFIER |
| 1 | 121848529 | AX-170263877 | 0 | G | C | 0.579 | 1.03E-01 | 8.34E-02 | 2.50E+00 | 2.56E+00 | 2.18E-01 | no | | intergenic_variant | MODIFIER |
| 1 | 121848839 | AX-170137573 | 0 | T | C | 0.579 | 1.03E-01 | 8.34E-02 | 2.50E+00 | 2.56E+00 | 2.18E-01 | no | | intergenic_variant | MODIFIER |
| 1 | 121888712 | AX-170001565 | 0 | G | A | 0.579 | 1.03E-01 | 8.34E-02 | 2.50E+00 | 2.56E+00 | 2.18E-01 | no | | intergenic_variant | MODIFIER |
| 1 | 121954790 | AX-170379063 | 0 | T | G | 0.579 | 1.03E-01 | 8.34E-02 | 2.50E+00 | 2.56E+00 | 2.18E-01 | no | | intergenic_variant | MODIFIER |
| 1 | 121956714 | AX-170496044 | 0 | T | G | 0.579 | 1.03E-01 | 8.34E-02 | 2.50E+00 | 2.56E+00 | 2.18E-01 | no | | intergenic_variant | MODIFIER |
| 1 | 121964645 | AX-170409309 | 0 | G | A | 0.579 | 1.03E-01 | 8.34E-02 | 2.50E+00 | 2.56E+00 | 2.18E-01 | no | Gm15394 | downstream_gene_variant | MODIFIER |
| 1 | 122261369 | AX-170279368 | 0 | A | G | 0.579 | 1.03E-01 | 8.34E-02 | 2.50E+00 | 2.56E+00 | 2.18E-01 | no | | intergenic_variant | MODIFIER |
| 1 | 122274991 | AX-170424345 | 0 | T | A | 0.579 | 1.03E-01 | 8.34E-02 | 2.50E+00 | 2.56E+00 | 2.18E-01 | no | | intergenic_variant | MODIFIER |
| 1 | 122298228 | AX-170345094 | 0 | G | A | 0.579 | 1.03E-01 | 8.34E-02 | 2.50E+00 | 2.56E+00 | 2.18E-01 | no | | intergenic_variant | MODIFIER |
| 4 | 77317941 | AX-170325618 | 1 | C | T | 0.321 | 1.47E-01 | 1.20E-01 | 2.86E+00 | 2.92E+00 | 2.18E-01 | Yes | Ptprd | intron_variant,non_coding_transcript_variant | MODIFIER |
| 16 | 72613715 | AX-170509377 | 0 | A | C | 0.482 | 9.08E-02 | 7.42E-02 | 2.59E+00 | 2.67E+00 | 2.19E-01 | no | Robo1 | intron_variant | MODIFIER |
| 18 | 79377710 | AX-170318482 | 0 | G | A | 0.465 | 1.04E-01 | 8.30E-02 | 2.38E+00 | 2.44E+00 | 2.19E-01 | no | | intergenic_variant | MODIFIER |
| 20 | 46720035 | AX-170084152 | 0 | C | T | 0.254 | 7.11E-02 | 5.81E-02 | 2.56E+00 | 2.68E+00 | 2.19E-01 | no | Gm29242 | intron_variant,non_coding_transcript_variant | MODIFIER |
| 8 | 36823086 | AX-170309483 | 0 | T | A | 0.149 | -1.69E-01 | 1.37E-01 | 2.94E+00 | 3.04E+00 | 2.19E-01 | no | Dlc1 | intron_variant | MODIFIER |
| 8 | 37114440 | AX-170310088 | 0 | A | T | 0.149 | -1.69E-01 | 1.37E-01 | 2.94E+00 | 3.04E+00 | 2.19E-01 | no | Gm45524 | intron_variant,non_coding_transcript_variant | MODIFIER |
| 8 | 37441325 | AX-170030518 | 0 | A | G | 0.149 | -1.69E-01 | 1.37E-01 | 2.94E+00 | 3.04E+00 | 2.19E-01 | no | | intergenic_variant | MODIFIER |
| 8 | 38935071 | AX-170079822 | 0 | G | A | 0.298 | -8.44E-02 | 6.87E-02 | 2.94E+00 | 3.04E+00 | 2.19E-01 | no | | intergenic_variant | MODIFIER |
| 8 | 41097752 | AX-170496602 | 0 | C | T | 0.149 | -1.69E-01 | 1.37E-01 | 2.94E+00 | 3.04E+00 | 2.19E-01 | no | Mtus1 | intron_variant | MODIFIER |
| 8 | 42068369 | AX-170161026 | 0 | A | T | 0.149 | -1.69E-01 | 1.37E-01 | 2.94E+00 | 3.04E+00 | 2.19E-01 | no | | intergenic_variant | MODIFIER |
| 8 | 42190912 | AX-170242419 | 0 | G | A | 0.149 | -1.69E-01 | 1.37E-01 | 2.94E+00 | 3.04E+00 | 2.19E-01 | no | | intergenic_variant | MODIFIER |
| 16 | 51704507 | AX-170506734 | 1 | C | T | 0.464 | -9.93E-02 | 8.05E-02 | 2.51E+00 | 2.56E+00 | 2.19E-01 | no | | intergenic_variant | MODIFIER |
| 16 | 40578526 | AX-170166828 | 0 | A | T | 0.5 | 1.02E-01 | 8.31E-02 | 2.58E+00 | 2.62E+00 | 2.19E-01 | no | Lsamp | intron_variant | MODIFIER |
| 16 | 73040988 | AX-170367679 | 0 | C | T | 0.491 | 9.46E-02 | 7.72E-02 | 2.56E+00 | 2.63E+00 | 2.19E-01 | no | Robo1 | intron_variant | MODIFIER |
| 3 | 26285534 | AX-170146350 | 1 | A | G | 0.125 | -1.28E-01 | 1.04E-01 | 2.54E+00 | 2.62E+00 | 2.19E-01 | no | Nlgn1 | intron_variant,non_coding_transcript_variant | MODIFIER |
| 5 | 84056051 | AX-170139027 | 2 | A | C | 0.436 | 1.36E-01 | 1.09E-01 | 3.10E+00 | 3.20E+00 | 2.19E-01 | no | Epha5 | 3_prime_UTR_variant | MODIFIER |
| 1 | 171536047 | AX-170086685 | 0 | G | C | 0.211 | 1.67E-01 | 1.36E-01 | 2.59E+00 | 2.63E+00 | 2.19E-01 | Yes | Itln1 | upstream_gene_variant | MODIFIER |
| 16 | 51702856 | AX-170156591 | 0 | A | G | 0.465 | -9.91E-02 | 8.05E-02 | 2.51E+00 | 2.56E+00 | 2.19E-01 | no | | intergenic_variant | MODIFIER |
| 1 | 120991776 | AX-170451520 | 1 | T | G | 0.571 | 9.94E-02 | 8.12E-02 | 2.58E+00 | 2.66E+00 | 2.19E-01 | no | | intergenic_variant | MODIFIER |
| 19 | 50613126 | AX-97809179 | 0 | T | G | 0.482 | 7.78E-02 | 6.39E-02 | 2.67E+00 | 2.76E+00 | 2.20E-01 | no | Sorcs1 | intron_variant | MODIFIER |
| 12 | 104175018 | AX-170279352 | 0 | A | C | 0.132 | -2.03E-01 | 1.67E-01 | 2.80E+00 | 2.89E+00 | 2.20E-01 | Yes | Serpina3d-ps | intron_variant,non_coding_transcript_variant | MODIFIER |
| 4 | 128701040 | AX-170070649 | 0 | G | A | 0.167 | 1.29E-01 | 1.06E-01 | 2.70E+00 | 2.77E+00 | 2.20E-01 | no | Phc2 | intron_variant | MODIFIER |
| 8 | 24525128 | AX-170261747 | 1 | C | T | 0.348 | -7.89E-02 | 6.45E-02 | 2.58E+00 | 2.67E+00 | 2.20E-01 | no | | regulatory_region_variant | MODIFIER |
| 4 | 116596402 | AX-170501741 | 0 | T | C | 0.219 | 1.29E-01 | 1.06E-01 | 2.65E+00 | 2.74E+00 | 2.20E-01 | no | Nasp | downstream_gene_variant | MODIFIER |
| 1 | 134970908 | AX-170106512 | 1 | T | C | 0.536 | 9.81E-02 | 8.05E-02 | 2.64E+00 | 2.71E+00 | 2.20E-01 | no | Ube2t | intron_variant | MODIFIER |
| 1 | 135450659 | AX-97836448 | 1 | T | G | 0.536 | 9.81E-02 | 8.05E-02 | 2.64E+00 | 2.71E+00 | 2.20E-01 | no | Nav1 | intron_variant | MODIFIER |
| 5 | 103342874 | AX-170523539 | 1 | G | A | 0.455 | 1.35E-01 | 1.11E-01 | 2.69E+00 | 2.74E+00 | 2.20E-01 | no | Gm42619 | upstream_gene_variant | MODIFIER |
| 1 | 120692225 | AX-170492692 | 1 | G | A | 0.58 | 1.03E-01 | 8.33E-02 | 2.50E+00 | 2.56E+00 | 2.20E-01 | no | 2610027F03Rik | downstream_gene_variant | MODIFIER |
| 5 | 104088225 | AX-97835855 | 0 | A | G | 0.439 | 1.33E-01 | 1.09E-01 | 2.59E+00 | 2.65E+00 | 2.20E-01 | no | Sparcl1 | intron_variant | MODIFIER |
| 5 | 105268387 | AX-170409257 | 0 | C | T | 0.439 | 1.33E-01 | 1.09E-01 | 2.59E+00 | 2.65E+00 | 2.20E-01 | no | Gm43302 | intron_variant | MODIFIER |
| 5 | 107017911 | AX-170103004 | 0 | T | A | 0.439 | 1.33E-01 | 1.09E-01 | 2.59E+00 | 2.65E+00 | 2.20E-01 | no | | regulatory_region_variant | MODIFIER |
| 5 | 45559158 | AX-170117363 | 0 | G | A | 0.509 | 1.23E-01 | 9.80E-02 | 2.36E+00 | 2.41E+00 | 2.20E-01 | no | Fam184b | intron_variant | MODIFIER |
| 4 | 134684980 | AX-97819990 | 0 | A | G | 0.158 | 1.41E-01 | 1.16E-01 | 2.60E+00 | 2.68E+00 | 2.20E-01 | no | Man1c1 | intron_variant | MODIFIER |
| 4 | 100822809 | AX-170127505 | 1 | A | G | 0.429 | -9.43E-02 | 7.70E-02 | 2.55E+00 | 2.63E+00 | 2.20E-01 | no | Cachd1 | intron_variant | MODIFIER |
| 4 | 55467454 | AX-170217034 | 1 | G | T | 0.366 | -1.24E-01 | 1.02E-01 | 2.90E+00 | 3.00E+00 | 2.20E-01 | no | Gm22150 | downstream_gene_variant | MODIFIER |
| 12 | 73026785 | AX-170344696 | 0 | G | A | 0.07 | -2.11E-01 | 1.73E-01 | 2.89E+00 | 3.05E+00 | 2.20E-01 | Yes | | intergenic_variant | MODIFIER |
| 10 | 72390277 | AX-170219140 | 2 | T | G | 0.391 | -9.36E-02 | 7.69E-02 | 2.63E+00 | 2.72E+00 | 2.21E-01 | no | | intergenic_variant | MODIFIER |
| 1 | 128682616 | AX-170042664 | 1 | G | A | 0.554 | 1.04E-01 | 8.56E-02 | 2.62E+00 | 2.68E+00 | 2.21E-01 | no | | regulatory_region_variant | MODIFIER |
| 10 | 99874238 | AX-170445809 | 1 | C | T | 0.295 | -1.04E-01 | 8.54E-02 | 2.64E+00 | 2.70E+00 | 2.21E-01 | no | Gm47578 | intron_variant,non_coding_transcript_variant | MODIFIER |
| 10 | 100952390 | AX-170003771 | 1 | G | A | 0.295 | -1.04E-01 | 8.54E-02 | 2.64E+00 | 2.70E+00 | 2.21E-01 | no | | intergenic_variant | MODIFIER |
| 15 | 19275840 | AX-170054948 | 0 | C | A | 0.237 | -1.22E-01 | 1.00E-01 | 2.58E+00 | 2.64E+00 | 2.21E-01 | no | | intergenic_variant | MODIFIER |
| 5 | 43615684 | AX-170024822 | 0 | A | G | 0.184 | -1.28E-01 | 1.05E-01 | 2.67E+00 | 2.77E+00 | 2.21E-01 | no | C1qtnf7 | synonymous_variant | LOW |
| 5 | 44400083 | AX-170128160 | 0 | T | G | 0.184 | -1.28E-01 | 1.05E-01 | 2.67E+00 | 2.77E+00 | 2.21E-01 | no | | intergenic_variant | MODIFIER |
| 5 | 49924049 | AX-97850690 | 1 | A | C | 0.473 | 1.31E-01 | 1.04E-01 | 2.33E+00 | 2.38E+00 | 2.21E-01 | no | | intergenic_variant | MODIFIER |

| | | | | | | | | | | | | | | | |
|---|---|---|---|---|---|---|---|---|---|---|---|---|---|---|---|
| 4 | 86184024 | AX-170375568 | 0 | C | T | 0.342 | 1.37E-01 | 1.13E-01 | 2.73E+00 | 2.80E+00 | 2.21E-01 | no | Adamtsl1 | intron_variant | MODIFIER |
| 8 | 25671463 | AX-170271436 | 0 | C | T | 0.351 | -7.90E-02 | 6.49E-02 | 2.58E+00 | 2.67E+00 | 2.21E-01 | no | Nsd3 | intron_variant | MODIFIER |
| 1 | 146614606 | AX-97851915 | 0 | A | G | 0.202 | 1.67E-01 | 1.37E-01 | 2.77E+00 | 2.83E+00 | 2.21E-01 | Yes | Brinp3 | intron_variant | MODIFIER |
| 16 | 35662190 | AX-97831510 | 0 | C | T | 0.491 | 9.74E-02 | 7.94E-02 | 2.51E+00 | 2.55E+00 | 2.21E-01 | no | Sema5b | intron_variant | MODIFIER |
| 5 | 81300566 | AX-97804489 | 1 | G | T | 0.455 | 1.22E-01 | 1.00E-01 | 2.80E+00 | 2.88E+00 | 2.21E-01 | no | Adgrl3 | intron_variant | MODIFIER |
| 1 | 82197231 | AX-170186616 | 0 | A | G | 0.167 | 2.09E-01 | 1.72E-01 | 2.71E+00 | 2.81E+00 | 2.22E-01 | Yes | | intergenic_variant | MODIFIER |
| 1 | 99252940 | AX-97824636 | 0 | A | G | 0.167 | 2.09E-01 | 1.72E-01 | 2.71E+00 | 2.81E+00 | 2.22E-01 | Yes | | intergenic_variant | MODIFIER |
| 1 | 171684529 | AX-170043579 | 1 | C | T | 0.223 | 1.57E-01 | 1.28E-01 | 2.50E+00 | 2.56E+00 | 2.22E-01 | Yes | Cd48 | intron_variant | MODIFIER |
| 3 | 14642090 | AX-97815430 | 0 | G | A | 0.421 | -7.94E-02 | 6.55E-02 | 2.89E+00 | 3.04E+00 | 2.22E-01 | no | Car13 | intron_variant | MODIFIER |
| 3 | 14806332 | AX-170019984 | 0 | T | C | 0.421 | -7.94E-02 | 6.55E-02 | 2.89E+00 | 3.04E+00 | 2.22E-01 | no | Car1 | intron_variant | MODIFIER |
| 3 | 14819202 | AX-97818565 | 0 | G | T | 0.421 | -7.94E-02 | 6.55E-02 | 2.89E+00 | 3.04E+00 | 2.22E-01 | no | | intergenic_variant | MODIFIER |
| 3 | 14829975 | AX-170326291 | 0 | A | C | 0.421 | -7.94E-02 | 6.55E-02 | 2.89E+00 | 3.04E+00 | 2.22E-01 | no | | intergenic_variant | MODIFIER |
| 3 | 14830000 | AX-170361818 | 0 | T | C | 0.421 | -7.94E-02 | 6.55E-02 | 2.89E+00 | 3.04E+00 | 2.22E-01 | no | | intergenic_variant | MODIFIER |
| 3 | 14838849 | AX-170395128 | 0 | G | A | 0.421 | -7.94E-02 | 6.55E-02 | 2.89E+00 | 3.04E+00 | 2.22E-01 | no | | intergenic_variant | MODIFIER |
| 3 | 14841429 | AX-97793044 | 0 | G | A | 0.421 | -7.94E-02 | 6.55E-02 | 2.89E+00 | 3.04E+00 | 2.22E-01 | no | | intergenic_variant | MODIFIER |
| 3 | 14858724 | AX-97827516 | 0 | A | G | 0.421 | -7.94E-02 | 6.55E-02 | 2.89E+00 | 3.04E+00 | 2.22E-01 | no | Car3 | upstream_gene_variant | MODIFIER |
| 3 | 14863892 | AX-170103727 | 0 | G | A | 0.421 | -7.94E-02 | 6.55E-02 | 2.89E+00 | 3.04E+00 | 2.22E-01 | no | Car3 | intron_variant | MODIFIER |
| 3 | 14867043 | AX-170155625 | 0 | C | T | 0.421 | -7.94E-02 | 6.55E-02 | 2.89E+00 | 3.04E+00 | 2.22E-01 | no | Car3 | intron_variant | MODIFIER |
| 3 | 14869372 | AX-170431475 | 0 | A | G | 0.421 | -7.94E-02 | 6.55E-02 | 2.89E+00 | 3.04E+00 | 2.22E-01 | no | Car3 | intron_variant | MODIFIER |
| 3 | 14872889 | AX-170435213 | 0 | G | A | 0.421 | -7.94E-02 | 6.55E-02 | 2.89E+00 | 3.04E+00 | 2.22E-01 | no | Car3 | downstream_gene_variant | MODIFIER |
| 3 | 14876548 | AX-170136674 | 0 | A | C | 0.421 | -7.94E-02 | 6.55E-02 | 2.89E+00 | 3.04E+00 | 2.22E-01 | no | Car3 | downstream_gene_variant | MODIFIER |
| 3 | 14877740 | AX-170503315 | 0 | C | T | 0.421 | -7.94E-02 | 6.55E-02 | 2.89E+00 | 3.04E+00 | 2.22E-01 | no | | regulatory_region_variant | MODIFIER |
| 3 | 14879132 | AX-170403618 | 0 | C | A | 0.421 | -7.94E-02 | 6.55E-02 | 2.89E+00 | 3.04E+00 | 2.22E-01 | no | | regulatory_region_variant | MODIFIER |
| 3 | 14881149 | AX-170396099 | 0 | A | G | 0.421 | -7.94E-02 | 6.55E-02 | 2.89E+00 | 3.04E+00 | 2.22E-01 | no | | regulatory_region_variant | MODIFIER |
| 3 | 14891806 | AX-170100385 | 0 | C | A | 0.421 | -7.94E-02 | 6.55E-02 | 2.89E+00 | 3.04E+00 | 2.22E-01 | no | Car2 | intron_variant | MODIFIER |
| 3 | 14892055 | AX-170256611 | 0 | C | G | 0.421 | -7.94E-02 | 6.55E-02 | 2.89E+00 | 3.04E+00 | 2.22E-01 | no | Car2 | intron_variant | MODIFIER |
| 13 | 19162031 | AX-170355143 | 0 | A | G | 0.175 | -1.51E-01 | 1.23E-01 | 2.47E+00 | 2.52E+00 | 2.22E-01 | no | Gm31887 | downstream_gene_variant | MODIFIER |
| 5 | 134618177 | AX-170026224 | 0 | G | A | 0.167 | -1.05E-01 | 8.29E-02 | 2.27E+00 | 2.36E+00 | 2.22E-01 | no | Eif4h | downstream_gene_variant | MODIFIER |
| 1 | 120093406 | AX-170520440 | 2 | T | C | 0.582 | 1.02E-01 | 8.34E-02 | 2.50E+00 | 2.56E+00 | 2.22E-01 | no | | regulatory_region_variant | MODIFIER |
| 16 | 45344145 | AX-170290913 | 0 | T | C | 0.518 | 9.82E-02 | 8.08E-02 | 2.61E+00 | 2.66E+00 | 2.22E-01 | no | | intergenic_variant | MODIFIER |
| 16 | 45865987 | AX-170139131 | 0 | T | C | 0.518 | 9.82E-02 | 8.08E-02 | 2.61E+00 | 2.66E+00 | 2.22E-01 | no | Phldb2 | intron_variant | MODIFIER |
| 13 | 6603683 | AX-170207407 | 1 | T | C | 0.205 | -1.24E-01 | 1.03E-01 | 2.68E+00 | 2.80E+00 | 2.22E-01 | no | Pfkp | intron_variant | MODIFIER |
| 4 | 107499345 | AX-170444686 | 0 | G | C | 0.263 | 1.36E-01 | 1.12E-01 | 2.60E+00 | 2.67E+00 | 2.22E-01 | no | Glis1 | intron_variant | MODIFIER |
| 4 | 107811790 | AX-97787635 | 0 | C | T | 0.263 | 1.36E-01 | 1.12E-01 | 2.60E+00 | 2.67E+00 | 2.22E-01 | no | Lrp8 | intron_variant | MODIFIER |
| 16 | 53633203 | AX-170434358 | 0 | C | T | 0.518 | 9.97E-02 | 8.19E-02 | 2.57E+00 | 2.62E+00 | 2.22E-01 | no | | intergenic_variant | MODIFIER |
| 1 | 70192168 | AX-170135225 | 1 | A | G | 0.143 | -2.38E-01 | 1.96E-01 | 2.89E+00 | 2.99E+00 | 2.22E-01 | Yes | Spag16 | intron_variant | MODIFIER |
| 14 | 113562354 | AX-170460110 | 0 | G | C | 0.325 | 8.56E-02 | 7.07E-02 | 2.62E+00 | 2.73E+00 | 2.22E-01 | no | | intergenic_variant | MODIFIER |
| 1 | 185597335 | AX-170525860 | 2 | G | A | 0.236 | 1.33E-01 | 1.08E-01 | 2.46E+00 | 2.53E+00 | 2.22E-01 | no | | intergenic_variant | MODIFIER |
| 18 | 79686411 | AX-170263378 | 0 | A | G | 0.447 | 1.01E-01 | 8.14E-02 | 2.40E+00 | 2.48E+00 | 2.23E-01 | no | | intergenic_variant | MODIFIER |
| 6 | 136083088 | AX-170179645 | 0 | C | T | 0.579 | 1.02E-01 | 8.38E-02 | 2.52E+00 | 2.60E+00 | 2.23E-01 | no | Grin2b | intron_variant | MODIFIER |
| 7 | 94197019 | AX-97798779 | 1 | A | T | 0.125 | 1.57E-01 | 1.30E-01 | 2.62E+00 | 2.79E+00 | 2.23E-01 | Yes | Gm5899 | upstream_gene_variant | MODIFIER |
| 9 | 91313396 | AX-170032977 | 0 | G | A | 0.123 | -1.51E-01 | 1.24E-01 | 2.57E+00 | 2.66E+00 | 2.23E-01 | no | | intergenic_variant | MODIFIER |
| 6 | 147401618 | AX-170491502 | 0 | G | A | 0.044 | -1.92E-01 | 1.57E-01 | 2.48E+00 | 2.59E+00 | 2.23E-01 | Yes | | intergenic_variant | MODIFIER |
| 4 | 64398999 | AX-170022964 | 1 | A | C | 0.321 | 1.44E-01 | 1.19E-01 | 2.84E+00 | 2.91E+00 | 2.23E-01 | no | | intergenic_variant | MODIFIER |
| 3 | 14869348 | AX-170527460 | 1 | A | G | 0.429 | -8.09E-02 | 6.68E-02 | 2.92E+00 | 3.08E+00 | 2.23E-01 | no | Car3 | intron_variant | MODIFIER |
| 4 | 137157744 | AX-170497165 | 2 | G | A | 0.2 | 1.20E-01 | 9.94E-02 | 2.63E+00 | 2.71E+00 | 2.23E-01 | no | Gm13003 | downstream_gene_variant | MODIFIER |
| 1 | 191444520 | AX-170461867 | 1 | G | T | 0.268 | 1.20E-01 | 9.58E-02 | 2.32E+00 | 2.40E+00 | 2.24E-01 | no | Gm32200 | intron_variant,non_coding_transcript_variant | MODIFIER |
| 13 | 20449313 | AX-170128601 | 0 | A | G | 0.184 | -1.42E-01 | 1.15E-01 | 2.43E+00 | 2.51E+00 | 2.24E-01 | no | Elmo1 | intron_variant | MODIFIER |
| 8 | 26346479 | AX-170343066 | 1 | G | A | 0.357 | -7.83E-02 | 6.46E-02 | 2.57E+00 | 2.67E+00 | 2.24E-01 | no | Gm45307 | intron_variant,non_coding_transcript_variant | MODIFIER |
| 18 | 79121222 | AX-170254713 | 1 | T | C | 0.455 | 1.06E-01 | 8.57E-02 | 2.42E+00 | 2.49E+00 | 2.24E-01 | no | | intergenic_variant | MODIFIER |
| 4 | 70211267 | AX-170023055 | 0 | T | C | 0.447 | -9.77E-02 | 8.05E-02 | 2.56E+00 | 2.64E+00 | 2.24E-01 | no | | intergenic_variant | MODIFIER |
| 9 | 92569387 | AX-170515445 | 0 | G | A | 0.14 | -1.45E-01 | 1.20E-01 | 2.65E+00 | 2.73E+00 | 2.24E-01 | no | Plod2 | intron_variant | MODIFIER |
| 1 | 44204843 | AX-170375309 | 0 | C | T | 0.096 | -2.08E-01 | 1.72E-01 | 2.79E+00 | 2.91E+00 | 2.24E-01 | Yes | Mettl21e | 3_prime_UTR_variant | MODIFIER |
| 1 | 179865000 | AX-97824929 | 0 | C | T | 0.237 | 1.38E-01 | 1.14E-01 | 2.56E+00 | 2.63E+00 | 2.24E-01 | no | | regulatory_region_variant | MODIFIER |

| | | | | | | | | | | | | | | | |
|---|---|---|---|---|---|---|---|---|---|---|---|---|---|---|---|
| 1 | 179865143 | AX-170275882 | 0 | G | A | 0.237 | 1.38E-01 | 1.14E-01 | 2.56E+00 | 2.63E+00 | 2.24E-01 | no | | regulatory_region_variant | MODIFIER |
| 1 | 179912991 | AX-97855690 | 0 | T | C | 0.237 | 1.38E-01 | 1.14E-01 | 2.56E+00 | 2.63E+00 | 2.24E-01 | no | | intergenic_variant | MODIFIER |
| 1 | 179998036 | AX-170108389 | 0 | G | A | 0.237 | 1.38E-01 | 1.14E-01 | 2.56E+00 | 2.63E+00 | 2.24E-01 | no | Cdc42bpa | intron_variant | MODIFIER |
| 1 | 179998403 | AX-170152468 | 0 | C | T | 0.237 | 1.38E-01 | 1.14E-01 | 2.56E+00 | 2.63E+00 | 2.24E-01 | no | Cdc42bpa | intron_variant | MODIFIER |
| 1 | 180003110 | AX-170445193 | 0 | G | A | 0.237 | 1.38E-01 | 1.14E-01 | 2.56E+00 | 2.63E+00 | 2.24E-01 | no | Cdc42bpa | intron_variant | MODIFIER |
| 1 | 180003215 | AX-170434294 | 0 | C | T | 0.237 | 1.38E-01 | 1.14E-01 | 2.56E+00 | 2.63E+00 | 2.24E-01 | no | Cdc42bpa | intron_variant | MODIFIER |
| 1 | 180004136 | AX-170359241 | 0 | A | G | 0.237 | 1.38E-01 | 1.14E-01 | 2.56E+00 | 2.63E+00 | 2.24E-01 | no | Cdc42bpa | intron_variant | MODIFIER |
| 1 | 180005861 | AX-170488504 | 0 | C | G | 0.237 | 1.38E-01 | 1.14E-01 | 2.56E+00 | 2.63E+00 | 2.24E-01 | no | Cdc42bpa | intron_variant | MODIFIER |
| 1 | 180045066 | AX-170091805 | 0 | C | A | 0.237 | 1.38E-01 | 1.14E-01 | 2.56E+00 | 2.63E+00 | 2.24E-01 | no | Cdc42bpa | intron_variant | MODIFIER |
| 1 | 180045256 | AX-170452151 | 0 | G | A | 0.237 | 1.38E-01 | 1.14E-01 | 2.56E+00 | 2.63E+00 | 2.24E-01 | no | Cdc42bpa | intron_variant | MODIFIER |
| 1 | 180073572 | AX-170002404 | 0 | C | T | 0.237 | 1.38E-01 | 1.14E-01 | 2.56E+00 | 2.63E+00 | 2.24E-01 | no | Cdc42bpa | intron_variant | MODIFIER |
| 1 | 180088755 | AX-170252960 | 0 | C | T | 0.237 | 1.38E-01 | 1.14E-01 | 2.56E+00 | 2.63E+00 | 2.24E-01 | no | Cdc42bpa | intron_variant | MODIFIER |
| 1 | 180089039 | AX-170502165 | 0 | T | C | 0.237 | 1.38E-01 | 1.14E-01 | 2.56E+00 | 2.63E+00 | 2.24E-01 | no | Cdc42bpa | intron_variant | MODIFIER |
| 1 | 180089052 | AX-170043751 | 0 | A | C | 0.237 | 1.38E-01 | 1.14E-01 | 2.56E+00 | 2.63E+00 | 2.24E-01 | no | Cdc42bpa | intron_variant | MODIFIER |
| 1 | 180095409 | AX-97820568 | 0 | A | G | 0.237 | 1.38E-01 | 1.14E-01 | 2.56E+00 | 2.63E+00 | 2.24E-01 | no | Cdc42bpa | intron_variant | MODIFIER |
| 1 | 180107925 | AX-97793634 | 0 | A | G | 0.237 | 1.38E-01 | 1.14E-01 | 2.56E+00 | 2.63E+00 | 2.24E-01 | no | Cdc42bpa | intron_variant | MODIFIER |
| 1 | 180152071 | AX-170172179 | 0 | A | G | 0.237 | 1.38E-01 | 1.14E-01 | 2.56E+00 | 2.63E+00 | 2.24E-01 | no | Cdc42bpa | intron_variant | MODIFIER |
| 1 | 180166238 | AX-170248210 | 0 | T | C | 0.237 | 1.38E-01 | 1.14E-01 | 2.56E+00 | 2.63E+00 | 2.24E-01 | no | Coq8a | 3_prime_UTR_variant | MODIFIER |
| 1 | 180166416 | AX-97813845 | 0 | G | T | 0.237 | 1.38E-01 | 1.14E-01 | 2.56E+00 | 2.63E+00 | 2.24E-01 | no | Coq8a | 3_prime_UTR_variant | MODIFIER |
| 1 | 180168281 | AX-170388476 | 0 | C | T | 0.237 | 1.38E-01 | 1.14E-01 | 2.56E+00 | 2.63E+00 | 2.24E-01 | no | Coq8a | intron_variant | MODIFIER |
| 1 | 180171972 | AX-170303880 | 0 | A | G | 0.237 | 1.38E-01 | 1.14E-01 | 2.56E+00 | 2.63E+00 | 2.24E-01 | no | Coq8a | intron_variant | MODIFIER |
| 1 | 180172932 | AX-170353592 | 0 | G | A | 0.237 | 1.38E-01 | 1.14E-01 | 2.56E+00 | 2.63E+00 | 2.24E-01 | no | Coq8a | intron_variant | MODIFIER |
| 1 | 180178867 | AX-170237173 | 0 | C | T | 0.237 | 1.38E-01 | 1.14E-01 | 2.56E+00 | 2.63E+00 | 2.24E-01 | no | Coq8a | intron_variant | MODIFIER |
| 1 | 180183312 | AX-170002407 | 0 | C | T | 0.237 | 1.38E-01 | 1.14E-01 | 2.56E+00 | 2.63E+00 | 2.24E-01 | no | Coq8a | intron_variant | MODIFIER |
| 1 | 180188668 | AX-97782153 | 0 | T | G | 0.237 | 1.38E-01 | 1.14E-01 | 2.56E+00 | 2.63E+00 | 2.24E-01 | no | Coq8a | intron_variant | MODIFIER |
| 1 | 180193714 | AX-170136925 | 0 | C | T | 0.237 | 1.38E-01 | 1.14E-01 | 2.56E+00 | 2.63E+00 | 2.24E-01 | no | Coq8a | intron_variant | MODIFIER |
| 1 | 180202036 | AX-170333849 | 0 | G | A | 0.237 | 1.38E-01 | 1.14E-01 | 2.56E+00 | 2.63E+00 | 2.24E-01 | no | Coq8a | upstream_gene_variant | MODIFIER |
| 1 | 180202229 | AX-170371651 | 0 | A | C | 0.237 | 1.38E-01 | 1.14E-01 | 2.56E+00 | 2.63E+00 | 2.24E-01 | no | Coq8a | upstream_gene_variant | MODIFIER |
| 1 | 180207994 | AX-97847706 | 0 | T | C | 0.237 | 1.38E-01 | 1.14E-01 | 2.56E+00 | 2.63E+00 | 2.24E-01 | no | Gm37336 | intron_variant,non_coding_transcript_variant | MODIFIER |
| 1 | 180233041 | AX-170245731 | 0 | A | G | 0.237 | 1.38E-01 | 1.14E-01 | 2.56E+00 | 2.63E+00 | 2.24E-01 | no | Psen2 | intron_variant | MODIFIER |
| 1 | 180233288 | AX-97832889 | 0 | T | C | 0.237 | 1.38E-01 | 1.14E-01 | 2.56E+00 | 2.63E+00 | 2.24E-01 | no | Psen2 | intron_variant | MODIFIER |
| 1 | 180308989 | AX-97854391 | 0 | T | C | 0.237 | 1.38E-01 | 1.14E-01 | 2.56E+00 | 2.63E+00 | 2.24E-01 | no | | regulatory_region_variant | MODIFIER |
| 1 | 180310393 | AX-170469700 | 0 | A | G | 0.237 | 1.38E-01 | 1.14E-01 | 2.56E+00 | 2.63E+00 | 2.24E-01 | no | | intergenic_variant | MODIFIER |
| 1 | 180314338 | AX-170373184 | 0 | C | G | 0.237 | 1.38E-01 | 1.14E-01 | 2.56E+00 | 2.63E+00 | 2.24E-01 | no | | regulatory_region_variant | MODIFIER |
| 1 | 180347767 | AX-170447420 | 0 | T | C | 0.237 | 1.38E-01 | 1.14E-01 | 2.56E+00 | 2.63E+00 | 2.24E-01 | no | Itpkb | intron_variant | MODIFIER |
| 14 | 107115683 | AX-170522475 | 1 | C | T | 0.036 | -2.46E-01 | 2.04E-01 | 2.69E+00 | 2.79E+00 | 2.25E-01 | Yes | | intergenic_variant | MODIFIER |
| 4 | 85223409 | AX-170023295 | 1 | C | T | 0.339 | 1.38E-01 | 1.15E-01 | 2.87E+00 | 2.95E+00 | 2.25E-01 | no | Sh3gl2 | intron_variant | MODIFIER |
| 4 | 154873395 | AX-170468711 | 0 | A | T | 0.114 | 1.87E-01 | 1.55E-01 | 2.85E+00 | 2.98E+00 | 2.25E-01 | Yes | Mmel1 | intron_variant | MODIFIER |
| 5 | 52059585 | AX-170072092 | 0 | G | A | 0.474 | 1.26E-01 | 1.01E-01 | 2.36E+00 | 2.40E+00 | 2.25E-01 | no | | intergenic_variant | MODIFIER |
| 4 | 80070916 | AX-97798131 | 0 | C | T | 0.439 | -9.92E-02 | 8.17E-02 | 2.54E+00 | 2.60E+00 | 2.25E-01 | no | | intergenic_variant | MODIFIER |
| 15 | 81780062 | AX-170243939 | 0 | G | C | 0.553 | 9.50E-02 | 7.83E-02 | 2.54E+00 | 2.62E+00 | 2.25E-01 | no | Zc3h7b | intron_variant | MODIFIER |
| 4 | 133004499 | AX-170500793 | 2 | C | T | 0.164 | 1.36E-01 | 1.12E-01 | 2.49E+00 | 2.55E+00 | 2.25E-01 | no | Fgr | downstream_gene_variant | MODIFIER |
| 3 | 23626049 | AX-170088876 | 0 | T | C | 0.167 | -1.16E-01 | 9.66E-02 | 2.66E+00 | 2.74E+00 | 2.25E-01 | no | | intergenic_variant | MODIFIER |
| 10 | 99628783 | AX-170231463 | 0 | G | A | 0.272 | -1.03E-01 | 8.56E-02 | 2.62E+00 | 2.68E+00 | 2.25E-01 | no | Gm20110 | intron_variant,non_coding_transcript_variant | MODIFIER |
| 10 | 99629179 | AX-170199898 | 0 | G | T | 0.272 | -1.03E-01 | 8.56E-02 | 2.62E+00 | 2.68E+00 | 2.25E-01 | no | Gm20110 | intron_variant,non_coding_transcript_variant | MODIFIER |
| 10 | 99629207 | AX-170486245 | 0 | T | C | 0.272 | -1.03E-01 | 8.56E-02 | 2.62E+00 | 2.68E+00 | 2.25E-01 | no | Gm20110 | intron_variant,non_coding_transcript_variant | MODIFIER |
| 10 | 99647015 | AX-170092073 | 0 | G | T | 0.272 | -1.03E-01 | 8.56E-02 | 2.62E+00 | 2.68E+00 | 2.25E-01 | no | Gm20110 | intron_variant,non_coding_transcript_variant | MODIFIER |
| 10 | 99656978 | AX-170045718 | 0 | G | A | 0.272 | -1.03E-01 | 8.56E-02 | 2.62E+00 | 2.68E+00 | 2.25E-01 | no | Gm20110 | non_coding_transcript_exon_variant | MODIFIER |
| 10 | 99660738 | AX-97853220 | 0 | C | T | 0.272 | -1.03E-01 | 8.56E-02 | 2.62E+00 | 2.68E+00 | 2.25E-01 | no | Gm20110 | downstream_gene_variant | MODIFIER |
| 10 | 99675708 | AX-170003751 | 0 | G | A | 0.272 | -1.03E-01 | 8.56E-02 | 2.62E+00 | 2.68E+00 | 2.25E-01 | no | Gm35101 | downstream_gene_variant | MODIFIER |
| 10 | 99676217 | AX-170003747 | 0 | A | G | 0.272 | -1.03E-01 | 8.56E-02 | 2.62E+00 | 2.68E+00 | 2.25E-01 | no | Gm35101 | downstream_gene_variant | MODIFIER |
| 10 | 99692346 | AX-170086880 | 0 | T | C | 0.272 | -1.03E-01 | 8.56E-02 | 2.62E+00 | 2.68E+00 | 2.25E-01 | no | | intergenic_variant | MODIFIER |
| 10 | 99694086 | AX-170003746 | 0 | C | G | 0.272 | -1.03E-01 | 8.56E-02 | 2.62E+00 | 2.68E+00 | 2.25E-01 | no | | intergenic_variant | MODIFIER |
| 10 | 99742473 | AX-170003750 | 0 | T | C | 0.272 | -1.03E-01 | 8.56E-02 | 2.62E+00 | 2.68E+00 | 2.25E-01 | no | Gm47578 | intron_variant,non_coding_transcript_variant | MODIFIER |

| | | | | | | | | | | | | | | | |
|---|---|---|---|---|---|---|---|---|---|---|---|---|---|---|---|
| 10 | 99742785 | AX-170003748 | 0 | A | C | 0.272 | -1.03E-01 | 8.56E-02 | 2.62E+00 | 2.68E+00 | 2.25E-01 | no | Gm47578 | intron_variant,non_coding_transcript_variant | MODIFIER |
| 10 | 99742852 | AX-170003752 | 0 | A | T | 0.272 | -1.03E-01 | 8.56E-02 | 2.62E+00 | 2.68E+00 | 2.25E-01 | no | Gm47578 | intron_variant,non_coding_transcript_variant | MODIFIER |
| 10 | 99789217 | AX-170003749 | 0 | G | A | 0.272 | -1.03E-01 | 8.56E-02 | 2.62E+00 | 2.68E+00 | 2.25E-01 | no | Gm35206 | intron_variant,non_coding_transcript_variant | MODIFIER |
| 10 | 99796005 | AX-170517964 | 0 | C | G | 0.272 | -1.03E-01 | 8.56E-02 | 2.62E+00 | 2.68E+00 | 2.25E-01 | no | Gm35206 | intron_variant,non_coding_transcript_variant | MODIFIER |
| 10 | 99830136 | AX-97834993 | 0 | A | C | 0.272 | -1.03E-01 | 8.56E-02 | 2.62E+00 | 2.68E+00 | 2.25E-01 | no | Gm35206 | non_coding_transcript_exon_variant | MODIFIER |
| 10 | 99834576 | AX-170377464 | 0 | A | G | 0.272 | -1.03E-01 | 8.56E-02 | 2.62E+00 | 2.68E+00 | 2.25E-01 | no | Gm35206 | downstream_gene_variant | MODIFIER |
| 10 | 99840327 | AX-170251475 | 0 | T | C | 0.272 | -1.03E-01 | 8.56E-02 | 2.62E+00 | 2.68E+00 | 2.25E-01 | no | Gm47578 | intron_variant,non_coding_transcript_variant | MODIFIER |
| 10 | 99860082 | AX-170477432 | 0 | A | G | 0.272 | -1.03E-01 | 8.56E-02 | 2.62E+00 | 2.68E+00 | 2.25E-01 | no | Gm47578 | intron_variant,non_coding_transcript_variant | MODIFIER |
| 10 | 99860303 | AX-170107855 | 0 | T | G | 0.272 | -1.03E-01 | 8.56E-02 | 2.62E+00 | 2.68E+00 | 2.25E-01 | no | Gm47578 | intron_variant,non_coding_transcript_variant | MODIFIER |
| 10 | 99860743 | AX-170241003 | 0 | C | A | 0.272 | -1.03E-01 | 8.56E-02 | 2.62E+00 | 2.68E+00 | 2.25E-01 | no | Gm47579 | downstream_gene_variant | MODIFIER |
| 10 | 99895829 | AX-170318838 | 0 | T | C | 0.272 | -1.03E-01 | 8.56E-02 | 2.62E+00 | 2.68E+00 | 2.25E-01 | no | Gm47578 | downstream_gene_variant | MODIFIER |
| 10 | 99902871 | AX-170045722 | 0 | C | T | 0.272 | -1.03E-01 | 8.56E-02 | 2.62E+00 | 2.68E+00 | 2.25E-01 | no | | regulatory_region_variant | MODIFIER |
| 10 | 99903422 | AX-170045723 | 0 | A | G | 0.272 | -1.03E-01 | 8.56E-02 | 2.62E+00 | 2.68E+00 | 2.25E-01 | no | | intergenic_variant | MODIFIER |
| 10 | 99989282 | AX-170191962 | 0 | C | T | 0.272 | -1.03E-01 | 8.56E-02 | 2.62E+00 | 2.68E+00 | 2.25E-01 | no | Gm17870 | non_coding_transcript_exon_variant | MODIFIER |
| 10 | 99989569 | AX-170227059 | 0 | T | A | 0.272 | -1.03E-01 | 8.56E-02 | 2.62E+00 | 2.68E+00 | 2.25E-01 | no | Gm17870 | non_coding_transcript_exon_variant | MODIFIER |
| 10 | 99990198 | AX-97847285 | 0 | G | A | 0.272 | -1.03E-01 | 8.56E-02 | 2.62E+00 | 2.68E+00 | 2.25E-01 | no | Gm17870 | downstream_gene_variant | MODIFIER |
| 10 | 99992665 | AX-170206424 | 0 | T | C | 0.272 | -1.03E-01 | 8.56E-02 | 2.62E+00 | 2.68E+00 | 2.25E-01 | no | Gm17870 | downstream_gene_variant | MODIFIER |
| 10 | 99992712 | AX-170146528 | 0 | T | C | 0.272 | -1.03E-01 | 8.56E-02 | 2.62E+00 | 2.68E+00 | 2.25E-01 | no | Gm17870 | downstream_gene_variant | MODIFIER |
| 10 | 99994200 | AX-170397158 | 0 | T | C | 0.272 | -1.03E-01 | 8.56E-02 | 2.62E+00 | 2.68E+00 | 2.25E-01 | no | Gm17870 | downstream_gene_variant | MODIFIER |
| 10 | 99995221 | AX-170489939 | 0 | G | A | 0.272 | -1.03E-01 | 8.56E-02 | 2.62E+00 | 2.68E+00 | 2.25E-01 | no | | intergenic_variant | MODIFIER |
| 10 | 100017863 | AX-170427133 | 0 | A | G | 0.272 | -1.03E-01 | 8.56E-02 | 2.62E+00 | 2.68E+00 | 2.25E-01 | no | Kitl | intron_variant | MODIFIER |
| 10 | 100021431 | AX-170273855 | 0 | G | A | 0.272 | -1.03E-01 | 8.56E-02 | 2.62E+00 | 2.68E+00 | 2.25E-01 | no | Kitl | intron_variant | MODIFIER |
| 10 | 100021602 | AX-97787212 | 0 | G | A | 0.272 | -1.03E-01 | 8.56E-02 | 2.62E+00 | 2.68E+00 | 2.25E-01 | no | Kitl | intron_variant | MODIFIER |
| 10 | 100068385 | AX-170423169 | 0 | A | G | 0.272 | -1.03E-01 | 8.56E-02 | 2.62E+00 | 2.68E+00 | 2.25E-01 | no | Kitl | intron_variant | MODIFIER |
| 10 | 100068478 | AX-170462740 | 0 | T | G | 0.272 | -1.03E-01 | 8.56E-02 | 2.62E+00 | 2.68E+00 | 2.25E-01 | no | Kitl | intron_variant | MODIFIER |
| 10 | 100068557 | AX-97834664 | 0 | G | A | 0.272 | -1.03E-01 | 8.56E-02 | 2.62E+00 | 2.68E+00 | 2.25E-01 | no | Kitl | intron_variant | MODIFIER |
| 10 | 100081341 | AX-170155039 | 0 | T | G | 0.272 | -1.03E-01 | 8.56E-02 | 2.62E+00 | 2.68E+00 | 2.25E-01 | no | Kitl | intron_variant | MODIFIER |
| 10 | 100085352 | AX-170374453 | 0 | A | G | 0.272 | -1.03E-01 | 8.56E-02 | 2.62E+00 | 2.68E+00 | 2.25E-01 | no | Kitl | intron_variant | MODIFIER |
| 10 | 100098410 | AX-170322074 | 0 | G | A | 0.272 | -1.03E-01 | 8.56E-02 | 2.62E+00 | 2.68E+00 | 2.25E-01 | no | Kitl | downstream_gene_variant | MODIFIER |
| 10 | 100115558 | AX-170124111 | 0 | A | C | 0.272 | -1.03E-01 | 8.56E-02 | 2.62E+00 | 2.68E+00 | 2.25E-01 | no | | intergenic_variant | MODIFIER |
| 10 | 100116413 | AX-170257094 | 0 | C | T | 0.272 | -1.03E-01 | 8.56E-02 | 2.62E+00 | 2.68E+00 | 2.25E-01 | no | | intergenic_variant | MODIFIER |
| 10 | 100121120 | AX-170045728 | 0 | T | C | 0.272 | -1.03E-01 | 8.56E-02 | 2.62E+00 | 2.68E+00 | 2.25E-01 | no | | intergenic_variant | MODIFIER |
| 10 | 100131463 | AX-170422152 | 0 | A | T | 0.272 | -1.03E-01 | 8.56E-02 | 2.62E+00 | 2.68E+00 | 2.25E-01 | no | Gm35533 | intron_variant,non_coding_transcript_variant | MODIFIER |
| 10 | 100135605 | AX-170500378 | 0 | A | G | 0.272 | -1.03E-01 | 8.56E-02 | 2.62E+00 | 2.68E+00 | 2.25E-01 | no | Gm22918 | upstream_gene_variant | MODIFIER |
| 10 | 100139256 | AX-97853099 | 0 | G | A | 0.272 | -1.03E-01 | 8.56E-02 | 2.62E+00 | 2.68E+00 | 2.25E-01 | no | Gm25287 | upstream_gene_variant | MODIFIER |
| 10 | 100157296 | AX-170429359 | 0 | T | C | 0.272 | -1.03E-01 | 8.56E-02 | 2.62E+00 | 2.68E+00 | 2.25E-01 | no | | intergenic_variant | MODIFIER |
| 10 | 100157931 | AX-170288402 | 0 | A | G | 0.272 | -1.03E-01 | 8.56E-02 | 2.62E+00 | 2.68E+00 | 2.25E-01 | no | | intergenic_variant | MODIFIER |
| 10 | 100158600 | AX-170086882 | 0 | T | C | 0.272 | -1.03E-01 | 8.56E-02 | 2.62E+00 | 2.68E+00 | 2.25E-01 | no | | intergenic_variant | MODIFIER |
| 10 | 100158878 | AX-170258261 | 0 | C | T | 0.272 | -1.03E-01 | 8.56E-02 | 2.62E+00 | 2.68E+00 | 2.25E-01 | no | | intergenic_variant | MODIFIER |
| 10 | 100186157 | AX-170524884 | 0 | G | A | 0.272 | -1.03E-01 | 8.56E-02 | 2.62E+00 | 2.68E+00 | 2.25E-01 | no | | intergenic_variant | MODIFIER |
| 10 | 100186800 | AX-170461679 | 0 | C | A | 0.272 | -1.03E-01 | 8.56E-02 | 2.62E+00 | 2.68E+00 | 2.25E-01 | no | | intergenic_variant | MODIFIER |
| 10 | 100212327 | AX-97816469 | 0 | C | T | 0.272 | -1.03E-01 | 8.56E-02 | 2.62E+00 | 2.68E+00 | 2.25E-01 | no | | intergenic_variant | MODIFIER |
| 10 | 100222124 | AX-170409016 | 0 | G | C | 0.272 | -1.03E-01 | 8.56E-02 | 2.62E+00 | 2.68E+00 | 2.25E-01 | no | | intergenic_variant | MODIFIER |
| 10 | 100226521 | AX-170003761 | 0 | C | G | 0.272 | -1.03E-01 | 8.56E-02 | 2.62E+00 | 2.68E+00 | 2.25E-01 | no | | intergenic_variant | MODIFIER |
| 10 | 100262338 | AX-170003755 | 0 | G | A | 0.272 | -1.03E-01 | 8.56E-02 | 2.62E+00 | 2.68E+00 | 2.25E-01 | no | | intergenic_variant | MODIFIER |
| 10 | 100270949 | AX-170003760 | 0 | C | T | 0.272 | -1.03E-01 | 8.56E-02 | 2.62E+00 | 2.68E+00 | 2.25E-01 | no | | intergenic_variant | MODIFIER |
| 10 | 100293630 | AX-97856196 | 0 | G | A | 0.272 | -1.03E-01 | 8.56E-02 | 2.62E+00 | 2.68E+00 | 2.25E-01 | no | Gm8689 | downstream_gene_variant | MODIFIER |
| 10 | 100293924 | AX-170460941 | 0 | C | A | 0.272 | -1.03E-01 | 8.56E-02 | 2.62E+00 | 2.68E+00 | 2.25E-01 | no | Gm8689 | downstream_gene_variant | MODIFIER |
| 10 | 100405450 | AX-97830575 | 0 | G | A | 0.272 | -1.03E-01 | 8.56E-02 | 2.62E+00 | 2.68E+00 | 2.25E-01 | no | | intergenic_variant | MODIFIER |
| 10 | 100408198 | AX-170003754 | 0 | A | T | 0.272 | -1.03E-01 | 8.56E-02 | 2.62E+00 | 2.68E+00 | 2.25E-01 | no | | intergenic_variant | MODIFIER |
| 10 | 100431273 | AX-170003758 | 0 | C | T | 0.272 | -1.03E-01 | 8.56E-02 | 2.62E+00 | 2.68E+00 | 2.25E-01 | no | Gm47673 | downstream_gene_variant | MODIFIER |
| 10 | 100444356 | AX-170504976 | 0 | G | C | 0.272 | -1.03E-01 | 8.56E-02 | 2.62E+00 | 2.68E+00 | 2.25E-01 | no | Tmtc3 | 3_prime_UTR_variant | MODIFIER |
| 10 | 100464540 | AX-97816847 | 0 | T | G | 0.272 | -1.03E-01 | 8.56E-02 | 2.62E+00 | 2.68E+00 | 2.25E-01 | no | Tmtc3 | intron_variant | MODIFIER |
| 10 | 100464608 | AX-170502013 | 0 | C | G | 0.272 | -1.03E-01 | 8.56E-02 | 2.62E+00 | 2.68E+00 | 2.25E-01 | no | Tmtc3 | intron_variant | MODIFIER |

| | | | | | | | | | | | | | | | |
|---|---|---|---|---|---|---|---|---|---|---|---|---|---|---|---|
| 10 | 100527465 | AX-170285162 | 0 | T | A | 0.272 | -1.03E-01 | 8.56E-02 | 2.62E+00 | 2.68E+00 | 2.25E-01 | no | Cep290 | intron_variant | MODIFIER |
| 10 | 100541761 | AX-97807987 | 0 | C | A | 0.272 | -1.03E-01 | 8.56E-02 | 2.62E+00 | 2.68E+00 | 2.25E-01 | no | Cep290 | missense_variant | MODERATE |
| 10 | 100555344 | AX-170252612 | 0 | C | T | 0.272 | -1.03E-01 | 8.56E-02 | 2.62E+00 | 2.68E+00 | 2.25E-01 | no | Cep290 | intron_variant | MODIFIER |
| 10 | 100555524 | AX-170442437 | 0 | C | T | 0.272 | -1.03E-01 | 8.56E-02 | 2.62E+00 | 2.68E+00 | 2.25E-01 | no | Cep290 | intron_variant | MODIFIER |
| 10 | 100568680 | AX-170045730 | 0 | A | G | 0.272 | -1.03E-01 | 8.56E-02 | 2.62E+00 | 2.68E+00 | 2.25E-01 | no | 4930430F08Rik | downstream_gene_variant | MODIFIER |
| 10 | 100588705 | AX-170357563 | 0 | T | C | 0.272 | -1.03E-01 | 8.56E-02 | 2.62E+00 | 2.68E+00 | 2.25E-01 | no | 1700017N19Rik | upstream_gene_variant | MODIFIER |
| 10 | 100589222 | AX-97816418 | 0 | T | C | 0.272 | -1.03E-01 | 8.56E-02 | 2.62E+00 | 2.68E+00 | 2.25E-01 | no | 1700017N19Rik | upstream_gene_variant | MODIFIER |
| 10 | 100589309 | AX-170187961 | 0 | T | C | 0.272 | -1.03E-01 | 8.56E-02 | 2.62E+00 | 2.68E+00 | 2.25E-01 | no | 1700017N19Rik | upstream_gene_variant | MODIFIER |
| 10 | 100603950 | AX-170443340 | 0 | G | A | 0.272 | -1.03E-01 | 8.56E-02 | 2.62E+00 | 2.68E+00 | 2.25E-01 | no | 1700017N19Rik | intron_variant | MODIFIER |
| 10 | 100606411 | AX-170492207 | 0 | A | G | 0.272 | -1.03E-01 | 8.56E-02 | 2.62E+00 | 2.68E+00 | 2.25E-01 | no | 1700017N19Rik | intron_variant | MODIFIER |
| 10 | 100608355 | AX-170149047 | 0 | T | C | 0.272 | -1.03E-01 | 8.56E-02 | 2.62E+00 | 2.68E+00 | 2.25E-01 | no | 1700017N19Rik | intron_variant | MODIFIER |
| 10 | 100622327 | AX-170472384 | 0 | T | C | 0.272 | -1.03E-01 | 8.56E-02 | 2.62E+00 | 2.68E+00 | 2.25E-01 | no | 1700017N19Rik | downstream_gene_variant | MODIFIER |
| 10 | 100626439 | AX-97801083 | 0 | T | C | 0.272 | -1.03E-01 | 8.56E-02 | 2.62E+00 | 2.68E+00 | 2.25E-01 | no | | intergenic_variant | MODIFIER |
| 10 | 100626527 | AX-170255303 | 0 | A | G | 0.272 | -1.03E-01 | 8.56E-02 | 2.62E+00 | 2.68E+00 | 2.25E-01 | no | | intergenic_variant | MODIFIER |
| 10 | 100626541 | AX-170275398 | 0 | C | T | 0.272 | -1.03E-01 | 8.56E-02 | 2.62E+00 | 2.68E+00 | 2.25E-01 | no | | intergenic_variant | MODIFIER |
| 10 | 100626643 | AX-170256648 | 0 | T | C | 0.272 | -1.03E-01 | 8.56E-02 | 2.62E+00 | 2.68E+00 | 2.25E-01 | no | | intergenic_variant | MODIFIER |
| 10 | 101387032 | AX-170003779 | 0 | T | A | 0.272 | -1.03E-01 | 8.56E-02 | 2.62E+00 | 2.68E+00 | 2.25E-01 | no | | intergenic_variant | MODIFIER |
| 4 | 69998508 | AX-168378674 | 0 | G | A | 0.316 | 1.41E-01 | 1.17E-01 | 2.82E+00 | 2.88E+00 | 2.26E-01 | no | | intergenic_variant | MODIFIER |
| 4 | 71616842 | AX-97782721 | 0 | T | G | 0.316 | 1.41E-01 | 1.17E-01 | 2.82E+00 | 2.88E+00 | 2.26E-01 | no | | intergenic_variant | MODIFIER |
| 4 | 71654960 | AX-170241810 | 0 | G | T | 0.316 | 1.41E-01 | 1.17E-01 | 2.82E+00 | 2.88E+00 | 2.26E-01 | no | | intergenic_variant | MODIFIER |
| 4 | 71757522 | AX-97807224 | 0 | T | G | 0.316 | 1.41E-01 | 1.17E-01 | 2.82E+00 | 2.88E+00 | 2.26E-01 | no | Gm11232 | intron_variant | MODIFIER |
| 4 | 72083918 | AX-170485372 | 0 | A | G | 0.316 | 1.41E-01 | 1.17E-01 | 2.82E+00 | 2.88E+00 | 2.26E-01 | no | | intergenic_variant | MODIFIER |
| 4 | 72087609 | AX-170211918 | 0 | T | C | 0.316 | 1.41E-01 | 1.17E-01 | 2.82E+00 | 2.88E+00 | 2.26E-01 | no | | regulatory_region_variant | MODIFIER |
| 4 | 72128445 | AX-170132939 | 0 | G | A | 0.316 | 1.41E-01 | 1.17E-01 | 2.82E+00 | 2.88E+00 | 2.26E-01 | no | Tle1 | intron_variant | MODIFIER |
| 4 | 72135658 | AX-170261480 | 0 | T | A | 0.316 | 1.41E-01 | 1.17E-01 | 2.82E+00 | 2.88E+00 | 2.26E-01 | no | Tle1 | intron_variant | MODIFIER |
| 4 | 72251227 | AX-170132241 | 0 | T | C | 0.316 | 1.41E-01 | 1.17E-01 | 2.82E+00 | 2.88E+00 | 2.26E-01 | no | | intergenic_variant | MODIFIER |
| 4 | 72902664 | AX-170339820 | 0 | G | T | 0.316 | 1.41E-01 | 1.17E-01 | 2.82E+00 | 2.88E+00 | 2.26E-01 | no | Gm25653 | downstream_gene_variant | MODIFIER |
| 4 | 73363291 | AX-170069680 | 0 | T | G | 0.316 | 1.41E-01 | 1.17E-01 | 2.82E+00 | 2.88E+00 | 2.26E-01 | no | | intergenic_variant | MODIFIER |
| 4 | 74106796 | AX-170069689 | 0 | T | G | 0.316 | 1.41E-01 | 1.17E-01 | 2.82E+00 | 2.88E+00 | 2.26E-01 | no | Frmd3 | intron_variant | MODIFIER |
| 4 | 74763448 | AX-170375919 | 0 | A | G | 0.316 | 1.41E-01 | 1.17E-01 | 2.82E+00 | 2.88E+00 | 2.26E-01 | no | | intergenic_variant | MODIFIER |
| 4 | 74993451 | AX-170437547 | 0 | A | G | 0.316 | 1.41E-01 | 1.17E-01 | 2.82E+00 | 2.88E+00 | 2.26E-01 | no | | intergenic_variant | MODIFIER |
| 4 | 75121111 | AX-170023146 | 0 | T | C | 0.316 | 1.41E-01 | 1.17E-01 | 2.82E+00 | 2.88E+00 | 2.26E-01 | no | | intergenic_variant | MODIFIER |
| 4 | 75234396 | AX-170333015 | 0 | T | G | 0.316 | 1.41E-01 | 1.17E-01 | 2.82E+00 | 2.88E+00 | 2.26E-01 | no | | intergenic_variant | MODIFIER |
| 4 | 75239698 | AX-170407803 | 0 | C | T | 0.316 | 1.41E-01 | 1.17E-01 | 2.82E+00 | 2.88E+00 | 2.26E-01 | no | | intergenic_variant | MODIFIER |
| 4 | 75256084 | AX-170218420 | 0 | A | G | 0.316 | 1.41E-01 | 1.17E-01 | 2.82E+00 | 2.88E+00 | 2.26E-01 | no | | intergenic_variant | MODIFIER |
| 4 | 75384038 | AX-170300820 | 0 | T | C | 0.316 | 1.41E-01 | 1.17E-01 | 2.82E+00 | 2.88E+00 | 2.26E-01 | no | Gm11259 | downstream_gene_variant | MODIFIER |
| 4 | 75457431 | AX-170387574 | 0 | G | A | 0.316 | 1.41E-01 | 1.17E-01 | 2.82E+00 | 2.88E+00 | 2.26E-01 | no | | intergenic_variant | MODIFIER |
| 4 | 75514837 | AX-170107266 | 0 | A | T | 0.316 | 1.41E-01 | 1.17E-01 | 2.82E+00 | 2.88E+00 | 2.26E-01 | no | | intergenic_variant | MODIFIER |
| 4 | 75646491 | AX-170183748 | 0 | T | G | 0.316 | 1.41E-01 | 1.17E-01 | 2.82E+00 | 2.88E+00 | 2.26E-01 | no | | intergenic_variant | MODIFIER |
| 4 | 75747024 | AX-170023156 | 0 | A | C | 0.316 | 1.41E-01 | 1.17E-01 | 2.82E+00 | 2.88E+00 | 2.26E-01 | no | | intergenic_variant | MODIFIER |
| 4 | 75787766 | AX-170123475 | 0 | G | A | 0.316 | 1.41E-01 | 1.17E-01 | 2.82E+00 | 2.88E+00 | 2.26E-01 | no | | regulatory_region_variant | MODIFIER |
| 4 | 75855535 | AX-170390139 | 0 | A | G | 0.316 | 1.41E-01 | 1.17E-01 | 2.82E+00 | 2.88E+00 | 2.26E-01 | no | Gm11256 | downstream_gene_variant | MODIFIER |
| 4 | 75956564 | AX-170351432 | 0 | A | G | 0.316 | 1.41E-01 | 1.17E-01 | 2.82E+00 | 2.88E+00 | 2.26E-01 | no | Ptprd | intron_variant | MODIFIER |
| 4 | 76009390 | AX-170069711 | 0 | G | T | 0.316 | 1.41E-01 | 1.17E-01 | 2.82E+00 | 2.88E+00 | 2.26E-01 | no | Ptprd | intron_variant | MODIFIER |
| 4 | 76022146 | AX-170069713 | 0 | C | T | 0.316 | 1.41E-01 | 1.17E-01 | 2.82E+00 | 2.88E+00 | 2.26E-01 | no | Ptprd | intron_variant | MODIFIER |
| 4 | 76081565 | AX-170104419 | 0 | C | T | 0.316 | 1.41E-01 | 1.17E-01 | 2.82E+00 | 2.88E+00 | 2.26E-01 | no | Ptprd | intron_variant | MODIFIER |
| 4 | 76083344 | AX-170333271 | 0 | T | C | 0.316 | 1.41E-01 | 1.17E-01 | 2.82E+00 | 2.88E+00 | 2.26E-01 | no | Ptprd | intron_variant | MODIFIER |
| 4 | 76165561 | AX-97787217 | 0 | G | A | 0.316 | 1.41E-01 | 1.17E-01 | 2.82E+00 | 2.88E+00 | 2.26E-01 | no | Ptprd | intron_variant | MODIFIER |
| 4 | 76218363 | AX-170168833 | 0 | T | C | 0.316 | 1.41E-01 | 1.17E-01 | 2.82E+00 | 2.88E+00 | 2.26E-01 | no | Ptprd | intron_variant | MODIFIER |
| 4 | 76267313 | AX-97855255 | 0 | C | A | 0.316 | 1.41E-01 | 1.17E-01 | 2.82E+00 | 2.88E+00 | 2.26E-01 | no | Ptprd | intron_variant | MODIFIER |
| 4 | 76304828 | AX-170348457 | 0 | G | A | 0.316 | 1.41E-01 | 1.17E-01 | 2.82E+00 | 2.88E+00 | 2.26E-01 | no | Ptprd | intron_variant | MODIFIER |
| 4 | 76415687 | AX-97806254 | 0 | G | A | 0.316 | 1.41E-01 | 1.17E-01 | 2.82E+00 | 2.88E+00 | 2.26E-01 | no | Ptprd | intron_variant,non_coding_transcript_variant | MODIFIER |
| 4 | 76525698 | AX-97850868 | 0 | C | T | 0.316 | 1.41E-01 | 1.17E-01 | 2.82E+00 | 2.88E+00 | 2.26E-01 | no | Ptprd | intron_variant,non_coding_transcript_variant | MODIFIER |
| 4 | 76533129 | AX-170251272 | 0 | G | A | 0.316 | 1.41E-01 | 1.17E-01 | 2.82E+00 | 2.88E+00 | 2.26E-01 | no | Ptprd | intron_variant,non_coding_transcript_variant | MODIFIER |

| | | | | | | | | | | | | | | | |
|---|---|---|---|---|---|---|---|---|---|---|---|---|---|---|---|
| 4 | 76562981 | AX-170135900 | 0 | A | G | 0.316 | 1.41E-01 | 1.17E-01 | 2.82E+00 | 2.88E+00 | 2.26E-01 | no | Ptprd | intron_variant,non_coding_transcript_variant | MODIFIER |
| 4 | 76609230 | AX-170069731 | 0 | T | C | 0.316 | 1.41E-01 | 1.17E-01 | 2.82E+00 | 2.88E+00 | 2.26E-01 | no | Ptprd | intron_variant,non_coding_transcript_variant | MODIFIER |
| 4 | 76609588 | AX-170095022 | 0 | C | T | 0.316 | 1.41E-01 | 1.17E-01 | 2.82E+00 | 2.88E+00 | 2.26E-01 | no | Ptprd | intron_variant,non_coding_transcript_variant | MODIFIER |
| 4 | 76616036 | AX-170431092 | 0 | G | C | 0.316 | 1.41E-01 | 1.17E-01 | 2.82E+00 | 2.88E+00 | 2.26E-01 | no | Ptprd | intron_variant,non_coding_transcript_variant | MODIFIER |
| 4 | 76632109 | AX-97830000 | 0 | T | C | 0.316 | 1.41E-01 | 1.17E-01 | 2.82E+00 | 2.88E+00 | 2.26E-01 | no | Ptprd | intron_variant,non_coding_transcript_variant | MODIFIER |
| 4 | 76632679 | AX-170310590 | 0 | A | G | 0.316 | 1.41E-01 | 1.17E-01 | 2.82E+00 | 2.88E+00 | 2.26E-01 | no | Ptprd | intron_variant,non_coding_transcript_variant | MODIFIER |
| 4 | 76855927 | AX-170069732 | 0 | A | G | 0.316 | 1.41E-01 | 1.17E-01 | 2.82E+00 | 2.88E+00 | 2.26E-01 | no | Ptprd | intron_variant,non_coding_transcript_variant | MODIFIER |
| 4 | 76895569 | AX-170493165 | 0 | C | T | 0.316 | 1.41E-01 | 1.17E-01 | 2.82E+00 | 2.88E+00 | 2.26E-01 | no | Ptprd | intron_variant,non_coding_transcript_variant | MODIFIER |
| 4 | 76958885 | AX-170496405 | 0 | A | G | 0.316 | 1.41E-01 | 1.17E-01 | 2.82E+00 | 2.88E+00 | 2.26E-01 | no | Ptprd | intron_variant,non_coding_transcript_variant | MODIFIER |
| 4 | 77344791 | AX-170095023 | 0 | A | C | 0.316 | 1.41E-01 | 1.17E-01 | 2.82E+00 | 2.88E+00 | 2.26E-01 | no | Ptprd | intron_variant,non_coding_transcript_variant | MODIFIER |
| 4 | 77370222 | AX-170069741 | 0 | G | A | 0.316 | 1.41E-01 | 1.17E-01 | 2.82E+00 | 2.88E+00 | 2.26E-01 | no | Ptprd | intron_variant,non_coding_transcript_variant | MODIFIER |
| 4 | 77476846 | AX-170351576 | 0 | A | G | 0.316 | 1.41E-01 | 1.17E-01 | 2.82E+00 | 2.88E+00 | 2.26E-01 | no | Ptprd | intron_variant,non_coding_transcript_variant | MODIFIER |
| 4 | 77493956 | AX-170314584 | 0 | G | A | 0.316 | 1.41E-01 | 1.17E-01 | 2.82E+00 | 2.88E+00 | 2.26E-01 | no | Ptprd | intron_variant,non_coding_transcript_variant | MODIFIER |
| 4 | 77498797 | AX-168406839 | 0 | A | G | 0.316 | 1.41E-01 | 1.17E-01 | 2.82E+00 | 2.88E+00 | 2.26E-01 | no | Ptprd | intron_variant,non_coding_transcript_variant | MODIFIER |
| 4 | 77530244 | AX-170413196 | 0 | A | G | 0.316 | 1.41E-01 | 1.17E-01 | 2.82E+00 | 2.88E+00 | 2.26E-01 | no | Ptprd | intron_variant,non_coding_transcript_variant | MODIFIER |
| 4 | 77551747 | AX-170069748 | 0 | A | T | 0.316 | 1.41E-01 | 1.17E-01 | 2.82E+00 | 2.88E+00 | 2.26E-01 | no | Ptprd | intron_variant,non_coding_transcript_variant | MODIFIER |
| 4 | 77555206 | AX-170069747 | 0 | T | C | 0.316 | 1.41E-01 | 1.17E-01 | 2.82E+00 | 2.88E+00 | 2.26E-01 | no | Ptprd | intron_variant,non_coding_transcript_variant | MODIFIER |
| 4 | 77566142 | AX-97854254 | 0 | G | A | 0.316 | 1.41E-01 | 1.17E-01 | 2.82E+00 | 2.88E+00 | 2.26E-01 | no | Ptprd | intron_variant,non_coding_transcript_variant | MODIFIER |
| 4 | 77566209 | AX-170069746 | 0 | C | T | 0.316 | 1.41E-01 | 1.17E-01 | 2.82E+00 | 2.88E+00 | 2.26E-01 | no | Ptprd | intron_variant,non_coding_transcript_variant | MODIFIER |
| 4 | 77632839 | AX-170528165 | 0 | C | T | 0.316 | 1.41E-01 | 1.17E-01 | 2.82E+00 | 2.88E+00 | 2.26E-01 | no | Ptprd | intron_variant,non_coding_transcript_variant | MODIFIER |
| 4 | 77637108 | AX-170450615 | 0 | T | G | 0.316 | 1.41E-01 | 1.17E-01 | 2.82E+00 | 2.88E+00 | 2.26E-01 | no | Ptprd | intron_variant,non_coding_transcript_variant | MODIFIER |
| 4 | 78275265 | AX-170431300 | 0 | C | G | 0.316 | 1.41E-01 | 1.17E-01 | 2.82E+00 | 2.88E+00 | 2.26E-01 | no | | intergenic_variant | MODIFIER |
| 4 | 78282881 | AX-170465992 | 0 | A | T | 0.316 | 1.41E-01 | 1.17E-01 | 2.82E+00 | 2.88E+00 | 2.26E-01 | no | | intergenic_variant | MODIFIER |
| 4 | 78285286 | AX-170466991 | 0 | T | C | 0.316 | 1.41E-01 | 1.17E-01 | 2.82E+00 | 2.88E+00 | 2.26E-01 | no | | intergenic_variant | MODIFIER |
| 4 | 78339006 | AX-170492460 | 0 | A | C | 0.316 | 1.41E-01 | 1.17E-01 | 2.82E+00 | 2.88E+00 | 2.26E-01 | no | | intergenic_variant | MODIFIER |
| 4 | 78431246 | AX-170220984 | 0 | C | A | 0.316 | 1.41E-01 | 1.17E-01 | 2.82E+00 | 2.88E+00 | 2.26E-01 | no | | intergenic_variant | MODIFIER |
| 4 | 78599524 | AX-170502691 | 0 | A | G | 0.316 | 1.41E-01 | 1.17E-01 | 2.82E+00 | 2.88E+00 | 2.26E-01 | no | | intergenic_variant | MODIFIER |
| 4 | 78663271 | AX-170069778 | 0 | A | G | 0.316 | 1.41E-01 | 1.17E-01 | 2.82E+00 | 2.88E+00 | 2.26E-01 | no | Gm11261 | upstream_gene_variant | MODIFIER |
| 4 | 78663553 | AX-170069777 | 0 | G | C | 0.316 | 1.41E-01 | 1.17E-01 | 2.82E+00 | 2.88E+00 | 2.26E-01 | no | Gm11261 | upstream_gene_variant | MODIFIER |
| 4 | 78663638 | AX-170069775 | 0 | C | T | 0.316 | 1.41E-01 | 1.17E-01 | 2.82E+00 | 2.88E+00 | 2.26E-01 | no | Gm11261 | upstream_gene_variant | MODIFIER |
| 4 | 78687049 | AX-170437357 | 0 | G | A | 0.316 | 1.41E-01 | 1.17E-01 | 2.82E+00 | 2.88E+00 | 2.26E-01 | no | | intergenic_variant | MODIFIER |
| 4 | 78697331 | AX-170335756 | 0 | C | G | 0.316 | 1.41E-01 | 1.17E-01 | 2.82E+00 | 2.88E+00 | 2.26E-01 | no | Gm11262 | upstream_gene_variant | MODIFIER |
| 4 | 78709088 | AX-170342874 | 0 | A | C | 0.316 | 1.41E-01 | 1.17E-01 | 2.82E+00 | 2.88E+00 | 2.26E-01 | no | | intergenic_variant | MODIFIER |
| 4 | 78720349 | AX-170023184 | 0 | G | A | 0.316 | 1.41E-01 | 1.17E-01 | 2.82E+00 | 2.88E+00 | 2.26E-01 | no | | intergenic_variant | MODIFIER |
| 4 | 78744268 | AX-170023186 | 0 | C | T | 0.316 | 1.41E-01 | 1.17E-01 | 2.82E+00 | 2.88E+00 | 2.26E-01 | no | | intergenic_variant | MODIFIER |
| 4 | 79142512 | AX-170423054 | 0 | C | T | 0.316 | 1.41E-01 | 1.17E-01 | 2.82E+00 | 2.88E+00 | 2.26E-01 | no | | intergenic_variant | MODIFIER |
| 4 | 79169277 | AX-170326259 | 0 | A | G | 0.316 | 1.41E-01 | 1.17E-01 | 2.82E+00 | 2.88E+00 | 2.26E-01 | no | | intergenic_variant | MODIFIER |
| 4 | 79191445 | AX-170231239 | 0 | G | T | 0.316 | 1.41E-01 | 1.17E-01 | 2.82E+00 | 2.88E+00 | 2.26E-01 | no | | intergenic_variant | MODIFIER |
| 4 | 79221257 | AX-170200988 | 0 | A | G | 0.316 | 1.41E-01 | 1.17E-01 | 2.82E+00 | 2.88E+00 | 2.26E-01 | no | | intergenic_variant | MODIFIER |
| 4 | 79272080 | AX-170492413 | 0 | A | C | 0.316 | 1.41E-01 | 1.17E-01 | 2.82E+00 | 2.88E+00 | 2.26E-01 | no | | intergenic_variant | MODIFIER |
| 4 | 79285290 | AX-170265935 | 0 | C | G | 0.316 | 1.41E-01 | 1.17E-01 | 2.82E+00 | 2.88E+00 | 2.26E-01 | no | | intergenic_variant | MODIFIER |
| 4 | 80839410 | AX-170509892 | 0 | G | T | 0.316 | 1.41E-01 | 1.17E-01 | 2.82E+00 | 2.88E+00 | 2.26E-01 | no | Tyrp1 | intron_variant | MODIFIER |
| 4 | 81044183 | AX-170259988 | 0 | T | G | 0.316 | 1.41E-01 | 1.17E-01 | 2.82E+00 | 2.88E+00 | 2.26E-01 | no | | intergenic_variant | MODIFIER |
| 4 | 81136553 | AX-97842863 | 0 | A | G | 0.316 | 1.41E-01 | 1.17E-01 | 2.82E+00 | 2.88E+00 | 2.26E-01 | no | | intergenic_variant | MODIFIER |
| 4 | 81158158 | AX-170212837 | 0 | T | G | 0.316 | 1.41E-01 | 1.17E-01 | 2.82E+00 | 2.88E+00 | 2.26E-01 | no | | intergenic_variant | MODIFIER |
| 8 | 25982763 | AX-170153847 | 1 | G | A | 0.357 | -7.80E-02 | 6.48E-02 | 2.62E+00 | 2.71E+00 | 2.26E-01 | no | Pomk | 3_prime_UTR_variant | MODIFIER |
| 1 | 151363745 | AX-170218559 | 2 | A | G | 0.218 | 1.63E-01 | 1.35E-01 | 2.80E+00 | 2.86E+00 | 2.26E-01 | Yes | Ivns1abp | 3_prime_UTR_variant | MODIFIER |
| 10 | 71136633 | AX-170331467 | 1 | A | C | 0.402 | -9.79E-02 | 8.13E-02 | 2.62E+00 | 2.69E+00 | 2.26E-01 | no | Bicc1 | intron_variant | MODIFIER |
| 10 | 72281573 | AX-170192657 | 1 | A | G | 0.402 | -9.79E-02 | 8.13E-02 | 2.62E+00 | 2.69E+00 | 2.26E-01 | no | | intergenic_variant | MODIFIER |
| 4 | 54421620 | AX-170331326 | 0 | C | T | 0.368 | -1.18E-01 | 9.82E-02 | 2.80E+00 | 2.91E+00 | 2.26E-01 | no | | intergenic_variant | MODIFIER |
| 1 | 191681847 | AX-97852458 | 0 | C | T | 0.272 | 1.14E-01 | 9.19E-02 | 2.36E+00 | 2.44E+00 | 2.26E-01 | no | | regulatory_region_variant | MODIFIER |
| 1 | 194525235 | AX-170120147 | 0 | G | A | 0.272 | 1.14E-01 | 9.19E-02 | 2.36E+00 | 2.44E+00 | 2.26E-01 | no | | intergenic_variant | MODIFIER |
| 4 | 137758906 | AX-170321695 | 1 | C | T | 0.161 | 1.33E-01 | 1.11E-01 | 2.66E+00 | 2.73E+00 | 2.26E-01 | no | Alpl | intron_variant | MODIFIER |
| 8 | 68132811 | AX-170413423 | 0 | A | G | 0.149 | -1.56E-01 | 1.30E-01 | 2.59E+00 | 2.69E+00 | 2.26E-01 | no | Psd3 | intron_variant | MODIFIER |

| | | | | | | | | | | | | | | | |
|---|---|---|---|---|---|---|---|---|---|---|---|---|---|---|---|
| 1 | 54699197 | AX-170200376 | 0 | C | T | 0.114 | 1.98E-01 | 1.65E-01 | 2.84E+00 | 2.99E+00 | 2.27E-01 | Yes | Ankrd44 | intron_variant | MODIFIER |
| 4 | 86571485 | AX-97854501 | 0 | C | A | 0.342 | 1.32E-01 | 1.10E-01 | 2.72E+00 | 2.80E+00 | 2.27E-01 | no | Rraga | upstream_gene_variant | MODIFIER |
| 4 | 87401989 | AX-170023337 | 0 | G | A | 0.342 | 1.32E-01 | 1.10E-01 | 2.72E+00 | 2.80E+00 | 2.27E-01 | no | | regulatory_region_variant | MODIFIER |
| 5 | 46969263 | AX-170071985 | 1 | A | G | 0.491 | 1.22E-01 | 9.90E-02 | 2.41E+00 | 2.47E+00 | 2.27E-01 | no | | intergenic_variant | MODIFIER |
| 16 | 70531727 | AX-170099980 | 0 | G | A | 0.491 | 9.33E-02 | 7.74E-02 | 2.57E+00 | 2.64E+00 | 2.27E-01 | no | Gbe1 | intron_variant | MODIFIER |
| 5 | 51948459 | AX-97802908 | 1 | C | T | 0.482 | 1.31E-01 | 1.06E-01 | 2.37E+00 | 2.41E+00 | 2.27E-01 | no | | intergenic_variant | MODIFIER |
| 8 | 65528812 | AX-97842122 | 1 | G | A | 0.134 | -1.69E-01 | 1.40E-01 | 2.57E+00 | 2.65E+00 | 2.27E-01 | no | | intergenic_variant | MODIFIER |
| 15 | 3588844 | AX-170010919 | 1 | G | A | 0.33 | -9.81E-02 | 7.35E-02 | 2.07E+00 | 2.13E+00 | 2.27E-01 | no | | intergenic_variant | MODIFIER |
| 1 | 102067751 | AX-170287555 | 0 | C | A | 0.14 | 1.62E-01 | 1.35E-01 | 2.67E+00 | 2.81E+00 | 2.27E-01 | Yes | | intergenic_variant | MODIFIER |
| 1 | 171582972 | AX-170120658 | 1 | C | T | 0.214 | 1.62E-01 | 1.35E-01 | 2.64E+00 | 2.69E+00 | 2.27E-01 | Yes | Cd244a | 3_prime_UTR_variant | MODIFIER |
| 7 | 145161211 | AX-170079135 | 0 | G | A | 0.211 | -8.89E-02 | 7.45E-02 | 2.72E+00 | 2.86E+00 | 2.27E-01 | no | Gm45181 | upstream_gene_variant | MODIFIER |
| 13 | 9732251 | AX-170185380 | 2 | T | A | 0.191 | -1.49E-01 | 1.22E-01 | 2.45E+00 | 2.50E+00 | 2.27E-01 | no | Zmynd11 | intron_variant | MODIFIER |
| 18 | 80845904 | AX-170344985 | 1 | T | C | 0.143 | -1.41E-01 | 1.14E-01 | 2.38E+00 | 2.45E+00 | 2.27E-01 | no | Atp9b | intron_variant | MODIFIER |
| 5 | 66891167 | AX-97785656 | 1 | A | C | 0.482 | 1.22E-01 | 1.01E-01 | 2.54E+00 | 2.61E+00 | 2.28E-01 | no | Limch1 | intron_variant | MODIFIER |
| 1 | 171589271 | AX-170309382 | 1 | C | T | 0.214 | 1.62E-01 | 1.35E-01 | 2.66E+00 | 2.71E+00 | 2.28E-01 | Yes | Ly9 | intron_variant | MODIFIER |
| 19 | 50533972 | AX-97794022 | 0 | G | A | 0.491 | 7.53E-02 | 6.30E-02 | 2.71E+00 | 2.81E+00 | 2.28E-01 | no | Sorcs1 | intron_variant | MODIFIER |
| 9 | 86366922 | AX-170083024 | 0 | C | T | 0.14 | -1.26E-01 | 1.06E-01 | 2.67E+00 | 2.80E+00 | 2.28E-01 | no | Ube2cbp | intron_variant | MODIFIER |
| 4 | 54479714 | AX-170252208 | 1 | G | A | 0.366 | -1.22E-01 | 1.02E-01 | 2.89E+00 | 2.99E+00 | 2.28E-01 | no | | intergenic_variant | MODIFIER |
| 15 | 66902047 | AX-170311784 | 1 | C | T | 0.33 | -9.64E-02 | 7.67E-02 | 2.27E+00 | 2.34E+00 | 2.28E-01 | no | Ccn4 | intron_variant | MODIFIER |
| 5 | 97870909 | AX-170415531 | 0 | A | C | 0.439 | 1.28E-01 | 1.07E-01 | 2.94E+00 | 3.02E+00 | 2.28E-01 | no | | intergenic_variant | MODIFIER |
| 13 | 5660913 | AX-170087352 | 0 | C | T | 0.211 | -1.41E-01 | 1.17E-01 | 2.61E+00 | 2.68E+00 | 2.28E-01 | no | Gm35330 | intron_variant,non_coding_transcript_variant | MODIFIER |
| 8 | 26300269 | AX-170312619 | 0 | G | A | 0.36 | -7.74E-02 | 6.47E-02 | 2.62E+00 | 2.72E+00 | 2.28E-01 | no | Gm31784 | intron_variant,non_coding_transcript_variant | MODIFIER |
| 5 | 49095741 | AX-170324507 | 1 | C | T | 0.482 | 1.30E-01 | 1.06E-01 | 2.36E+00 | 2.40E+00 | 2.28E-01 | no | Kcnip4 | intron_variant | MODIFIER |
| 12 | 73106092 | AX-170533053 | 0 | T | C | 0.061 | -2.59E-01 | 2.17E-01 | 2.64E+00 | 2.72E+00 | 2.28E-01 | Yes | Six4 | intron_variant | MODIFIER |
| 18 | 39698692 | AX-97793395 | 0 | C | T | 0.061 | -2.59E-01 | 2.17E-01 | 2.64E+00 | 2.72E+00 | 2.28E-01 | Yes | AC124518.1 | intron_variant,non_coding_transcript_variant | MODIFIER |
| 18 | 43332053 | AX-97856005 | 0 | G | A | 0.061 | -2.59E-01 | 2.17E-01 | 2.64E+00 | 2.72E+00 | 2.28E-01 | Yes | Dpysl3 | intron_variant | MODIFIER |
| 15 | 67651860 | AX-97789851 | 1 | G | A | 0.33 | -9.65E-02 | 7.68E-02 | 2.27E+00 | 2.34E+00 | 2.28E-01 | no | | intergenic_variant | MODIFIER |
| 5 | 48621038 | AX-170024878 | 0 | G | A | 0.482 | 1.31E-01 | 1.06E-01 | 2.37E+00 | 2.40E+00 | 2.29E-01 | no | Kcnip4 | intron_variant | MODIFIER |
| 5 | 48649072 | AX-170024887 | 0 | A | G | 0.482 | 1.31E-01 | 1.06E-01 | 2.37E+00 | 2.40E+00 | 2.29E-01 | no | Kcnip4 | intron_variant | MODIFIER |
| 5 | 48678721 | AX-170131499 | 0 | A | T | 0.482 | 1.31E-01 | 1.06E-01 | 2.37E+00 | 2.40E+00 | 2.29E-01 | no | Kcnip4 | intron_variant | MODIFIER |
| 5 | 48736462 | AX-170424930 | 0 | T | C | 0.482 | 1.31E-01 | 1.06E-01 | 2.37E+00 | 2.40E+00 | 2.29E-01 | no | Kcnip4 | intron_variant | MODIFIER |
| 5 | 48746174 | AX-97794113 | 0 | C | A | 0.482 | 1.31E-01 | 1.06E-01 | 2.37E+00 | 2.40E+00 | 2.29E-01 | no | Kcnip4 | intron_variant | MODIFIER |
| 5 | 49302184 | AX-170259187 | 0 | A | G | 0.482 | 1.31E-01 | 1.06E-01 | 2.37E+00 | 2.40E+00 | 2.29E-01 | no | Kcnip4 | intron_variant | MODIFIER |
| 5 | 49308122 | AX-170136894 | 0 | T | C | 0.482 | 1.31E-01 | 1.06E-01 | 2.37E+00 | 2.40E+00 | 2.29E-01 | no | Kcnip4 | intron_variant | MODIFIER |
| 5 | 49309087 | AX-97832416 | 0 | A | C | 0.482 | 1.31E-01 | 1.06E-01 | 2.37E+00 | 2.40E+00 | 2.29E-01 | no | Kcnip4 | intron_variant | MODIFIER |
| 5 | 49318358 | AX-170109650 | 0 | A | G | 0.482 | 1.31E-01 | 1.06E-01 | 2.37E+00 | 2.40E+00 | 2.29E-01 | no | Kcnip4 | intron_variant | MODIFIER |
| 5 | 49338748 | AX-170112894 | 0 | G | A | 0.482 | 1.31E-01 | 1.06E-01 | 2.37E+00 | 2.40E+00 | 2.29E-01 | no | Kcnip4 | intron_variant | MODIFIER |
| 5 | 49342986 | AX-170072032 | 0 | A | G | 0.482 | 1.31E-01 | 1.06E-01 | 2.37E+00 | 2.40E+00 | 2.29E-01 | no | Kcnip4 | intron_variant | MODIFIER |
| 5 | 49346495 | AX-170504102 | 0 | C | T | 0.482 | 1.31E-01 | 1.06E-01 | 2.37E+00 | 2.40E+00 | 2.29E-01 | no | Kcnip4 | intron_variant | MODIFIER |
| 5 | 49348187 | AX-170402537 | 0 | C | T | 0.482 | 1.31E-01 | 1.06E-01 | 2.37E+00 | 2.40E+00 | 2.29E-01 | no | Kcnip4 | intron_variant | MODIFIER |
| 5 | 49348608 | AX-170221609 | 0 | C | T | 0.482 | 1.31E-01 | 1.06E-01 | 2.37E+00 | 2.40E+00 | 2.29E-01 | no | Kcnip4 | intron_variant | MODIFIER |
| 5 | 49350922 | AX-97798514 | 0 | A | C | 0.482 | 1.31E-01 | 1.06E-01 | 2.37E+00 | 2.40E+00 | 2.29E-01 | no | Kcnip4 | intron_variant | MODIFIER |
| 5 | 49387168 | AX-170109703 | 0 | C | A | 0.482 | 1.31E-01 | 1.06E-01 | 2.37E+00 | 2.40E+00 | 2.29E-01 | no | Kcnip4 | intron_variant | MODIFIER |
| 5 | 49389863 | AX-170125088 | 0 | C | T | 0.482 | 1.31E-01 | 1.06E-01 | 2.37E+00 | 2.40E+00 | 2.29E-01 | no | Kcnip4 | intron_variant | MODIFIER |
| 5 | 49399412 | AX-170484215 | 0 | T | C | 0.482 | 1.31E-01 | 1.06E-01 | 2.37E+00 | 2.40E+00 | 2.29E-01 | no | Kcnip4 | intron_variant | MODIFIER |
| 5 | 49428628 | AX-170446131 | 0 | C | T | 0.482 | 1.31E-01 | 1.06E-01 | 2.37E+00 | 2.40E+00 | 2.29E-01 | no | Kcnip4 | intron_variant | MODIFIER |
| 5 | 49443537 | AX-170072034 | 0 | C | T | 0.482 | 1.31E-01 | 1.06E-01 | 2.37E+00 | 2.40E+00 | 2.29E-01 | no | Kcnip4 | intron_variant | MODIFIER |
| 5 | 49444914 | AX-170375740 | 0 | A | C | 0.482 | 1.31E-01 | 1.06E-01 | 2.37E+00 | 2.40E+00 | 2.29E-01 | no | Kcnip4 | intron_variant | MODIFIER |
| 5 | 49449930 | AX-170458637 | 0 | A | C | 0.482 | 1.31E-01 | 1.06E-01 | 2.37E+00 | 2.40E+00 | 2.29E-01 | no | Kcnip4 | intron_variant | MODIFIER |
| 5 | 49450109 | AX-170176515 | 0 | T | G | 0.482 | 1.31E-01 | 1.06E-01 | 2.37E+00 | 2.40E+00 | 2.29E-01 | no | Kcnip4 | intron_variant | MODIFIER |
| 5 | 49460128 | AX-170306248 | 0 | C | G | 0.482 | 1.31E-01 | 1.06E-01 | 2.37E+00 | 2.40E+00 | 2.29E-01 | no | Kcnip4 | intron_variant | MODIFIER |
| 5 | 49466516 | AX-170214911 | 0 | G | C | 0.482 | 1.31E-01 | 1.06E-01 | 2.37E+00 | 2.40E+00 | 2.29E-01 | no | Kcnip4 | intron_variant | MODIFIER |
| 5 | 49537881 | AX-170223726 | 0 | T | C | 0.482 | 1.31E-01 | 1.06E-01 | 2.37E+00 | 2.40E+00 | 2.29E-01 | no | | intergenic_variant | MODIFIER |
| 5 | 49555191 | AX-170398200 | 0 | A | G | 0.482 | 1.31E-01 | 1.06E-01 | 2.37E+00 | 2.40E+00 | 2.29E-01 | no | Gm5555 | downstream_gene_variant | MODIFIER |

| | | | | | | | | | | | | | | | |
|---|---|---|---|---|---|---|---|---|---|---|---|---|---|---|---|
| 5 | 49555604 | AX-170170106 | 0 | G | T | 0.482 | 1.31E-01 | 1.06E-01 | 2.37E+00 | 2.40E+00 | 2.29E-01 | no | Gm5555 | downstream_gene_variant | MODIFIER |
| 5 | 49581275 | AX-170024900 | 0 | C | G | 0.482 | 1.31E-01 | 1.06E-01 | 2.37E+00 | 2.40E+00 | 2.29E-01 | no | | intergenic_variant | MODIFIER |
| 5 | 49581380 | AX-170024896 | 0 | T | A | 0.482 | 1.31E-01 | 1.06E-01 | 2.37E+00 | 2.40E+00 | 2.29E-01 | no | | intergenic_variant | MODIFIER |
| 5 | 49652110 | AX-170072035 | 0 | G | A | 0.482 | 1.31E-01 | 1.06E-01 | 2.37E+00 | 2.40E+00 | 2.29E-01 | no | | intergenic_variant | MODIFIER |
| 5 | 49653036 | AX-170365661 | 0 | C | G | 0.482 | 1.31E-01 | 1.06E-01 | 2.37E+00 | 2.40E+00 | 2.29E-01 | no | | intergenic_variant | MODIFIER |
| 5 | 49657575 | AX-170196416 | 0 | T | G | 0.482 | 1.31E-01 | 1.06E-01 | 2.37E+00 | 2.40E+00 | 2.29E-01 | no | | intergenic_variant | MODIFIER |
| 5 | 49660025 | AX-170253031 | 0 | A | C | 0.482 | 1.31E-01 | 1.06E-01 | 2.37E+00 | 2.40E+00 | 2.29E-01 | no | | intergenic_variant | MODIFIER |
| 5 | 49666660 | AX-170225092 | 0 | T | C | 0.482 | 1.31E-01 | 1.06E-01 | 2.37E+00 | 2.40E+00 | 2.29E-01 | no | | intergenic_variant | MODIFIER |
| 5 | 49703889 | AX-170317787 | 0 | T | A | 0.482 | 1.31E-01 | 1.06E-01 | 2.37E+00 | 2.40E+00 | 2.29E-01 | no | | intergenic_variant | MODIFIER |
| 5 | 49707402 | AX-170336190 | 0 | G | A | 0.482 | 1.31E-01 | 1.06E-01 | 2.37E+00 | 2.40E+00 | 2.29E-01 | no | | intergenic_variant | MODIFIER |
| 5 | 49718218 | AX-170239662 | 0 | T | C | 0.482 | 1.31E-01 | 1.06E-01 | 2.37E+00 | 2.40E+00 | 2.29E-01 | no | | intergenic_variant | MODIFIER |
| 5 | 49767981 | AX-170149537 | 0 | T | C | 0.482 | 1.31E-01 | 1.06E-01 | 2.37E+00 | 2.40E+00 | 2.29E-01 | no | | intergenic_variant | MODIFIER |
| 5 | 49779457 | AX-170072041 | 0 | A | G | 0.482 | 1.31E-01 | 1.06E-01 | 2.37E+00 | 2.40E+00 | 2.29E-01 | no | | intergenic_variant | MODIFIER |
| 5 | 49784083 | AX-97783265 | 0 | G | A | 0.482 | 1.31E-01 | 1.06E-01 | 2.37E+00 | 2.40E+00 | 2.29E-01 | no | | intergenic_variant | MODIFIER |
| 5 | 49808498 | AX-170072038 | 0 | A | G | 0.482 | 1.31E-01 | 1.06E-01 | 2.37E+00 | 2.40E+00 | 2.29E-01 | no | | intergenic_variant | MODIFIER |
| 5 | 49809548 | AX-97813675 | 0 | A | G | 0.482 | 1.31E-01 | 1.06E-01 | 2.37E+00 | 2.40E+00 | 2.29E-01 | no | | intergenic_variant | MODIFIER |
| 5 | 49853281 | AX-170167972 | 0 | C | T | 0.482 | 1.31E-01 | 1.06E-01 | 2.37E+00 | 2.40E+00 | 2.29E-01 | no | | intergenic_variant | MODIFIER |
| 5 | 49873969 | AX-170426912 | 0 | T | C | 0.482 | 1.31E-01 | 1.06E-01 | 2.37E+00 | 2.40E+00 | 2.29E-01 | no | | intergenic_variant | MODIFIER |
| 5 | 49892728 | AX-170156552 | 0 | C | G | 0.482 | 1.31E-01 | 1.06E-01 | 2.37E+00 | 2.40E+00 | 2.29E-01 | no | | intergenic_variant | MODIFIER |
| 5 | 49900272 | AX-170236736 | 0 | C | G | 0.482 | 1.31E-01 | 1.06E-01 | 2.37E+00 | 2.40E+00 | 2.29E-01 | no | Gm7988 | non_coding_transcript_exon_variant | MODIFIER |
| 5 | 49917363 | AX-170095315 | 0 | T | C | 0.482 | 1.31E-01 | 1.06E-01 | 2.37E+00 | 2.40E+00 | 2.29E-01 | no | | intergenic_variant | MODIFIER |
| 5 | 49960915 | AX-170276594 | 0 | G | T | 0.482 | 1.31E-01 | 1.06E-01 | 2.37E+00 | 2.40E+00 | 2.29E-01 | no | Adgra3 | missense_variant | MODERATE |
| 5 | 49984203 | AX-170290835 | 0 | C | G | 0.482 | 1.31E-01 | 1.06E-01 | 2.37E+00 | 2.40E+00 | 2.29E-01 | no | Adgra3 | intron_variant | MODIFIER |
| 5 | 49991593 | AX-170500113 | 0 | C | T | 0.482 | 1.31E-01 | 1.06E-01 | 2.37E+00 | 2.40E+00 | 2.29E-01 | no | Adgra3 | intron_variant | MODIFIER |
| 5 | 50024688 | AX-170072043 | 0 | T | C | 0.482 | 1.31E-01 | 1.06E-01 | 2.37E+00 | 2.40E+00 | 2.29E-01 | no | Adgra3 | intron_variant | MODIFIER |
| 5 | 50031838 | AX-97787977 | 0 | C | T | 0.482 | 1.31E-01 | 1.06E-01 | 2.37E+00 | 2.40E+00 | 2.29E-01 | no | Adgra3 | intron_variant | MODIFIER |
| 5 | 50036783 | AX-170072045 | 0 | C | T | 0.482 | 1.31E-01 | 1.06E-01 | 2.37E+00 | 2.40E+00 | 2.29E-01 | no | Adgra3 | intron_variant | MODIFIER |
| 5 | 50048320 | AX-170072047 | 0 | T | C | 0.482 | 1.31E-01 | 1.06E-01 | 2.37E+00 | 2.40E+00 | 2.29E-01 | no | Adgra3 | intron_variant | MODIFIER |
| 5 | 50062159 | AX-170429652 | 0 | T | C | 0.482 | 1.31E-01 | 1.06E-01 | 2.37E+00 | 2.40E+00 | 2.29E-01 | no | Adgra3 | upstream_gene_variant | MODIFIER |
| 5 | 50064919 | AX-170176144 | 0 | A | C | 0.482 | 1.31E-01 | 1.06E-01 | 2.37E+00 | 2.40E+00 | 2.29E-01 | no | Gm42460 | downstream_gene_variant | MODIFIER |
| 5 | 50066301 | AX-170531505 | 0 | G | A | 0.482 | 1.31E-01 | 1.06E-01 | 2.37E+00 | 2.40E+00 | 2.29E-01 | no | Gm42460 | downstream_gene_variant | MODIFIER |
| 5 | 50068490 | AX-170424286 | 0 | G | A | 0.482 | 1.31E-01 | 1.06E-01 | 2.37E+00 | 2.40E+00 | 2.29E-01 | no | | intergenic_variant | MODIFIER |
| 5 | 50069079 | AX-170327863 | 0 | A | G | 0.482 | 1.31E-01 | 1.06E-01 | 2.37E+00 | 2.40E+00 | 2.29E-01 | no | | intergenic_variant | MODIFIER |
| 5 | 50137468 | AX-170126743 | 0 | C | A | 0.482 | 1.31E-01 | 1.06E-01 | 2.37E+00 | 2.40E+00 | 2.29E-01 | no | | intergenic_variant | MODIFIER |
| 5 | 50140346 | AX-170401234 | 0 | G | A | 0.482 | 1.31E-01 | 1.06E-01 | 2.37E+00 | 2.40E+00 | 2.29E-01 | no | | intergenic_variant | MODIFIER |
| 5 | 50140478 | AX-170507193 | 0 | G | A | 0.482 | 1.31E-01 | 1.06E-01 | 2.37E+00 | 2.40E+00 | 2.29E-01 | no | | intergenic_variant | MODIFIER |
| 5 | 50143522 | AX-168381613 | 0 | C | T | 0.482 | 1.31E-01 | 1.06E-01 | 2.37E+00 | 2.40E+00 | 2.29E-01 | no | | intergenic_variant | MODIFIER |
| 5 | 50162512 | AX-97853557 | 0 | G | A | 0.482 | 1.31E-01 | 1.06E-01 | 2.37E+00 | 2.40E+00 | 2.29E-01 | no | | intergenic_variant | MODIFIER |
| 5 | 50163760 | AX-170144711 | 0 | T | C | 0.482 | 1.31E-01 | 1.06E-01 | 2.37E+00 | 2.40E+00 | 2.29E-01 | no | | intergenic_variant | MODIFIER |
| 5 | 50201745 | AX-170024906 | 0 | G | A | 0.482 | 1.31E-01 | 1.06E-01 | 2.37E+00 | 2.40E+00 | 2.29E-01 | no | | intergenic_variant | MODIFIER |
| 5 | 50202412 | AX-170072050 | 0 | T | G | 0.482 | 1.31E-01 | 1.06E-01 | 2.37E+00 | 2.40E+00 | 2.29E-01 | no | | regulatory_region_variant | MODIFIER |
| 5 | 50254076 | AX-168384772 | 0 | G | T | 0.482 | 1.31E-01 | 1.06E-01 | 2.37E+00 | 2.40E+00 | 2.29E-01 | no | Mir8117 | downstream_gene_variant | MODIFIER |
| 5 | 50259280 | AX-170349089 | 0 | T | G | 0.482 | 1.31E-01 | 1.06E-01 | 2.37E+00 | 2.40E+00 | 2.29E-01 | no | Gm7205 | intron_variant,non_coding_transcript_variant | MODIFIER |
| 5 | 50259295 | AX-170147898 | 0 | G | A | 0.482 | 1.31E-01 | 1.06E-01 | 2.37E+00 | 2.40E+00 | 2.29E-01 | no | Gm7205 | intron_variant,non_coding_transcript_variant | MODIFIER |
| 5 | 50284704 | AX-170532786 | 0 | A | C | 0.482 | 1.31E-01 | 1.06E-01 | 2.37E+00 | 2.40E+00 | 2.29E-01 | no | Gm7205 | intron_variant,non_coding_transcript_variant | MODIFIER |
| 5 | 50298687 | AX-170024904 | 0 | T | C | 0.482 | 1.31E-01 | 1.06E-01 | 2.37E+00 | 2.40E+00 | 2.29E-01 | no | Gm7205 | non_coding_transcript_exon_variant | MODIFIER |
| 5 | 50319799 | AX-170500618 | 0 | G | A | 0.482 | 1.31E-01 | 1.06E-01 | 2.37E+00 | 2.40E+00 | 2.29E-01 | no | Gm7205 | intron_variant,non_coding_transcript_variant | MODIFIER |
| 5 | 50328252 | AX-170024907 | 0 | G | A | 0.482 | 1.31E-01 | 1.06E-01 | 2.37E+00 | 2.40E+00 | 2.29E-01 | no | Gm7205 | intron_variant,non_coding_transcript_variant | MODIFIER |
| 5 | 50328643 | AX-170340820 | 0 | G | A | 0.482 | 1.31E-01 | 1.06E-01 | 2.37E+00 | 2.40E+00 | 2.29E-01 | no | Gm7205 | intron_variant,non_coding_transcript_variant | MODIFIER |
| 5 | 50344292 | AX-170305679 | 0 | T | A | 0.482 | 1.31E-01 | 1.06E-01 | 2.37E+00 | 2.40E+00 | 2.29E-01 | no | Gm7205 | intron_variant,non_coding_transcript_variant | MODIFIER |
| 5 | 50382583 | AX-97799886 | 0 | G | A | 0.482 | 1.31E-01 | 1.06E-01 | 2.37E+00 | 2.40E+00 | 2.29E-01 | no | Gm7205 | intron_variant,non_coding_transcript_variant | MODIFIER |
| 5 | 50401500 | AX-170107938 | 0 | C | A | 0.482 | 1.31E-01 | 1.06E-01 | 2.37E+00 | 2.40E+00 | 2.29E-01 | no | | intergenic_variant | MODIFIER |
| 5 | 50403820 | AX-170505029 | 0 | A | G | 0.482 | 1.31E-01 | 1.06E-01 | 2.37E+00 | 2.40E+00 | 2.29E-01 | no | | intergenic_variant | MODIFIER |
| 5 | 50403902 | AX-170271193 | 0 | A | C | 0.482 | 1.31E-01 | 1.06E-01 | 2.37E+00 | 2.40E+00 | 2.29E-01 | no | | intergenic_variant | MODIFIER |

| | | | | | | | | | | | | | | | |
|---|---|---|---|---|---|---|---|---|---|---|---|---|---|---|---|
| 5 | 50404099 | AX-170483691 | 0 | C | T | 0.482 | 1.31E-01 | 1.06E-01 | 2.37E+00 | 2.40E+00 | 2.29E-01 | no | | intergenic_variant | MODIFIER |
| 5 | 50407041 | AX-170166844 | 0 | T | C | 0.482 | 1.31E-01 | 1.06E-01 | 2.37E+00 | 2.40E+00 | 2.29E-01 | no | | intergenic_variant | MODIFIER |
| 5 | 50408321 | AX-170387885 | 0 | A | G | 0.482 | 1.31E-01 | 1.06E-01 | 2.37E+00 | 2.40E+00 | 2.29E-01 | no | | intergenic_variant | MODIFIER |
| 5 | 50409741 | AX-170436552 | 0 | A | T | 0.482 | 1.31E-01 | 1.06E-01 | 2.37E+00 | 2.40E+00 | 2.29E-01 | no | | intergenic_variant | MODIFIER |
| 5 | 50409996 | AX-170448834 | 0 | G | A | 0.482 | 1.31E-01 | 1.06E-01 | 2.37E+00 | 2.40E+00 | 2.29E-01 | no | | intergenic_variant | MODIFIER |
| 5 | 50410040 | AX-170471785 | 0 | C | T | 0.482 | 1.31E-01 | 1.06E-01 | 2.37E+00 | 2.40E+00 | 2.29E-01 | no | | intergenic_variant | MODIFIER |
| 5 | 50419743 | AX-168399574 | 0 | C | T | 0.482 | 1.31E-01 | 1.06E-01 | 2.37E+00 | 2.40E+00 | 2.29E-01 | no | | intergenic_variant | MODIFIER |
| 5 | 50421641 | AX-168384804 | 0 | G | T | 0.482 | 1.31E-01 | 1.06E-01 | 2.37E+00 | 2.40E+00 | 2.29E-01 | no | | intergenic_variant | MODIFIER |
| 5 | 50422071 | AX-97803489 | 0 | C | A | 0.482 | 1.31E-01 | 1.06E-01 | 2.37E+00 | 2.40E+00 | 2.29E-01 | no | | intergenic_variant | MODIFIER |
| 5 | 50476358 | AX-170072053 | 0 | C | A | 0.482 | 1.31E-01 | 1.06E-01 | 2.37E+00 | 2.40E+00 | 2.29E-01 | no | Gm36814 | intron_variant,non_coding_transcript_variant | MODIFIER |
| 5 | 50516565 | AX-170469021 | 0 | T | C | 0.482 | 1.31E-01 | 1.06E-01 | 2.37E+00 | 2.40E+00 | 2.29E-01 | no | Gm36814 | intron_variant,non_coding_transcript_variant | MODIFIER |
| 5 | 50517532 | AX-170117637 | 0 | T | C | 0.482 | 1.31E-01 | 1.06E-01 | 2.37E+00 | 2.40E+00 | 2.29E-01 | no | Gm36814 | intron_variant,non_coding_transcript_variant | MODIFIER |
| 5 | 50518508 | AX-170118898 | 0 | G | C | 0.482 | 1.31E-01 | 1.06E-01 | 2.37E+00 | 2.40E+00 | 2.29E-01 | no | Gm36814 | intron_variant,non_coding_transcript_variant | MODIFIER |
| 5 | 50530639 | AX-170285093 | 0 | T | G | 0.482 | 1.31E-01 | 1.06E-01 | 2.37E+00 | 2.40E+00 | 2.29E-01 | no | Gm36814 | intron_variant,non_coding_transcript_variant | MODIFIER |
| 5 | 50543867 | AX-170426330 | 0 | T | C | 0.482 | 1.31E-01 | 1.06E-01 | 2.37E+00 | 2.40E+00 | 2.29E-01 | no | Gm43016 | intron_variant,non_coding_transcript_variant | MODIFIER |
| 5 | 50551085 | AX-170443318 | 0 | A | G | 0.482 | 1.31E-01 | 1.06E-01 | 2.37E+00 | 2.40E+00 | 2.29E-01 | no | Gm43016 | intron_variant,non_coding_transcript_variant | MODIFIER |
| 5 | 50553144 | AX-170438929 | 0 | C | A | 0.482 | 1.31E-01 | 1.06E-01 | 2.37E+00 | 2.40E+00 | 2.29E-01 | no | Gm43016 | intron_variant,non_coding_transcript_variant | MODIFIER |
| 5 | 50571591 | AX-170189464 | 0 | T | A | 0.482 | 1.31E-01 | 1.06E-01 | 2.37E+00 | 2.40E+00 | 2.29E-01 | no | Gm43016 | intron_variant,non_coding_transcript_variant | MODIFIER |
| 5 | 50588869 | AX-170117712 | 0 | T | C | 0.482 | 1.31E-01 | 1.06E-01 | 2.37E+00 | 2.40E+00 | 2.29E-01 | no | Gm43016 | intron_variant,non_coding_transcript_variant | MODIFIER |
| 5 | 50592984 | AX-170024925 | 0 | A | C | 0.482 | 1.31E-01 | 1.06E-01 | 2.37E+00 | 2.40E+00 | 2.29E-01 | no | Gm43016 | intron_variant,non_coding_transcript_variant | MODIFIER |
| 5 | 50612999 | AX-97799884 | 0 | A | G | 0.482 | 1.31E-01 | 1.06E-01 | 2.37E+00 | 2.40E+00 | 2.29E-01 | no | Gm43016 | intron_variant,non_coding_transcript_variant | MODIFIER |
| 5 | 50641792 | AX-170024928 | 0 | A | C | 0.482 | 1.31E-01 | 1.06E-01 | 2.37E+00 | 2.40E+00 | 2.29E-01 | no | Gm43016 | intron_variant,non_coding_transcript_variant | MODIFIER |
| 5 | 50680022 | AX-97805978 | 0 | T | C | 0.482 | 1.31E-01 | 1.06E-01 | 2.37E+00 | 2.40E+00 | 2.29E-01 | no | Gm43016 | intron_variant,non_coding_transcript_variant | MODIFIER |
| 5 | 50730019 | AX-97801508 | 0 | G | T | 0.482 | 1.31E-01 | 1.06E-01 | 2.37E+00 | 2.40E+00 | 2.29E-01 | no | | intergenic_variant | MODIFIER |
| 5 | 50743688 | AX-170110370 | 0 | A | C | 0.482 | 1.31E-01 | 1.06E-01 | 2.37E+00 | 2.40E+00 | 2.29E-01 | no | | intergenic_variant | MODIFIER |
| 5 | 50744633 | AX-170491178 | 0 | T | C | 0.482 | 1.31E-01 | 1.06E-01 | 2.37E+00 | 2.40E+00 | 2.29E-01 | no | | intergenic_variant | MODIFIER |
| 5 | 50747635 | AX-170313596 | 0 | A | G | 0.482 | 1.31E-01 | 1.06E-01 | 2.37E+00 | 2.40E+00 | 2.29E-01 | no | | intergenic_variant | MODIFIER |
| 5 | 50749973 | AX-168394885 | 0 | G | A | 0.482 | 1.31E-01 | 1.06E-01 | 2.37E+00 | 2.40E+00 | 2.29E-01 | no | | intergenic_variant | MODIFIER |
| 5 | 50754238 | AX-170341855 | 0 | G | A | 0.482 | 1.31E-01 | 1.06E-01 | 2.37E+00 | 2.40E+00 | 2.29E-01 | no | | intergenic_variant | MODIFIER |
| 5 | 50759154 | AX-170427071 | 0 | C | T | 0.482 | 1.31E-01 | 1.06E-01 | 2.37E+00 | 2.40E+00 | 2.29E-01 | no | | intergenic_variant | MODIFIER |
| 5 | 50767291 | AX-170475481 | 0 | A | G | 0.482 | 1.31E-01 | 1.06E-01 | 2.37E+00 | 2.40E+00 | 2.29E-01 | no | | intergenic_variant | MODIFIER |
| 5 | 50780729 | AX-170489498 | 0 | C | G | 0.482 | 1.31E-01 | 1.06E-01 | 2.37E+00 | 2.40E+00 | 2.29E-01 | no | | intergenic_variant | MODIFIER |
| 5 | 50796045 | AX-170024919 | 0 | T | C | 0.482 | 1.31E-01 | 1.06E-01 | 2.37E+00 | 2.40E+00 | 2.29E-01 | no | | intergenic_variant | MODIFIER |
| 5 | 50832471 | AX-97792820 | 0 | C | T | 0.482 | 1.31E-01 | 1.06E-01 | 2.37E+00 | 2.40E+00 | 2.29E-01 | no | | intergenic_variant | MODIFIER |
| 5 | 50848238 | AX-170165942 | 0 | G | A | 0.482 | 1.31E-01 | 1.06E-01 | 2.37E+00 | 2.40E+00 | 2.29E-01 | no | | intergenic_variant | MODIFIER |
| 5 | 50856970 | AX-170072064 | 0 | G | C | 0.482 | 1.31E-01 | 1.06E-01 | 2.37E+00 | 2.40E+00 | 2.29E-01 | no | | intergenic_variant | MODIFIER |
| 5 | 50857496 | AX-170196278 | 0 | A | G | 0.482 | 1.31E-01 | 1.06E-01 | 2.37E+00 | 2.40E+00 | 2.29E-01 | no | | intergenic_variant | MODIFIER |
| 5 | 50883470 | AX-170089474 | 0 | T | C | 0.482 | 1.31E-01 | 1.06E-01 | 2.37E+00 | 2.40E+00 | 2.29E-01 | no | | intergenic_variant | MODIFIER |
| 5 | 50911532 | AX-170454510 | 0 | T | C | 0.482 | 1.31E-01 | 1.06E-01 | 2.37E+00 | 2.40E+00 | 2.29E-01 | no | | intergenic_variant | MODIFIER |
| 5 | 50949693 | AX-170201413 | 0 | G | A | 0.482 | 1.31E-01 | 1.06E-01 | 2.37E+00 | 2.40E+00 | 2.29E-01 | no | | intergenic_variant | MODIFIER |
| 5 | 50949991 | AX-170095318 | 0 | A | C | 0.482 | 1.31E-01 | 1.06E-01 | 2.37E+00 | 2.40E+00 | 2.29E-01 | no | | intergenic_variant | MODIFIER |
| 5 | 50967917 | AX-170462090 | 0 | A | T | 0.482 | 1.31E-01 | 1.06E-01 | 2.37E+00 | 2.40E+00 | 2.29E-01 | no | | intergenic_variant | MODIFIER |
| 5 | 51039403 | AX-170320703 | 0 | A | G | 0.482 | 1.31E-01 | 1.06E-01 | 2.37E+00 | 2.40E+00 | 2.29E-01 | no | | intergenic_variant | MODIFIER |
| 5 | 51041346 | AX-170121890 | 0 | A | G | 0.482 | 1.31E-01 | 1.06E-01 | 2.37E+00 | 2.40E+00 | 2.29E-01 | no | | regulatory_region_variant | MODIFIER |
| 5 | 51042252 | AX-170272102 | 0 | A | G | 0.482 | 1.31E-01 | 1.06E-01 | 2.37E+00 | 2.40E+00 | 2.29E-01 | no | | intergenic_variant | MODIFIER |
| 5 | 51042694 | AX-170154950 | 0 | G | A | 0.482 | 1.31E-01 | 1.06E-01 | 2.37E+00 | 2.40E+00 | 2.29E-01 | no | | intergenic_variant | MODIFIER |
| 5 | 51051030 | AX-170506044 | 0 | C | A | 0.482 | 1.31E-01 | 1.06E-01 | 2.37E+00 | 2.40E+00 | 2.29E-01 | no | Gm40319 | upstream_gene_variant | MODIFIER |
| 5 | 51205971 | AX-170228562 | 0 | C | A | 0.482 | 1.31E-01 | 1.06E-01 | 2.37E+00 | 2.40E+00 | 2.29E-01 | no | | intergenic_variant | MODIFIER |
| 5 | 51945913 | AX-170465531 | 0 | T | C | 0.482 | 1.31E-01 | 1.06E-01 | 2.37E+00 | 2.40E+00 | 2.29E-01 | no | | intergenic_variant | MODIFIER |
| 5 | 51945928 | AX-170159433 | 0 | C | A | 0.482 | 1.31E-01 | 1.06E-01 | 2.37E+00 | 2.40E+00 | 2.29E-01 | no | | intergenic_variant | MODIFIER |
| 5 | 51950709 | AX-170180600 | 0 | A | C | 0.482 | 1.31E-01 | 1.06E-01 | 2.37E+00 | 2.40E+00 | 2.29E-01 | no | | intergenic_variant | MODIFIER |
| 5 | 51973401 | AX-170382363 | 0 | A | G | 0.482 | 1.31E-01 | 1.06E-01 | 2.37E+00 | 2.40E+00 | 2.29E-01 | no | | intergenic_variant | MODIFIER |
| 5 | 52002687 | AX-170127089 | 0 | T | C | 0.482 | 1.31E-01 | 1.06E-01 | 2.37E+00 | 2.40E+00 | 2.29E-01 | no | | intergenic_variant | MODIFIER |
| 5 | 52009002 | AX-170299346 | 0 | C | T | 0.482 | 1.31E-01 | 1.06E-01 | 2.37E+00 | 2.40E+00 | 2.29E-01 | no | | intergenic_variant | MODIFIER |

| | | | | | | | | | | | | | | | |
|---|---|---|---|---|---|---|---|---|---|---|---|---|---|---|---|
| 5 | 52034639 | AX-170095322 | 0 | T | G | 0.482 | 1.31E-01 | 1.06E-01 | 2.37E+00 | 2.40E+00 | 2.29E-01 | no | | intergenic_variant | MODIFIER |
| 5 | 52040882 | AX-170236246 | 0 | C | T | 0.482 | 1.31E-01 | 1.06E-01 | 2.37E+00 | 2.40E+00 | 2.29E-01 | no | | regulatory_region_variant | MODIFIER |
| 5 | 52046785 | AX-170385275 | 0 | T | C | 0.482 | 1.31E-01 | 1.06E-01 | 2.37E+00 | 2.40E+00 | 2.29E-01 | no | | regulatory_region_variant | MODIFIER |
| 5 | 52049057 | AX-170320393 | 0 | G | A | 0.482 | 1.31E-01 | 1.06E-01 | 2.37E+00 | 2.40E+00 | 2.29E-01 | no | | intergenic_variant | MODIFIER |
| 5 | 52049417 | AX-170105393 | 0 | T | C | 0.482 | 1.31E-01 | 1.06E-01 | 2.37E+00 | 2.40E+00 | 2.29E-01 | no | | intergenic_variant | MODIFIER |
| 5 | 52055418 | AX-170197809 | 0 | C | T | 0.482 | 1.31E-01 | 1.06E-01 | 2.37E+00 | 2.40E+00 | 2.29E-01 | no | | intergenic_variant | MODIFIER |
| 5 | 52081527 | AX-170293877 | 0 | C | T | 0.482 | 1.31E-01 | 1.06E-01 | 2.37E+00 | 2.40E+00 | 2.29E-01 | no | Gm43174 | non_coding_transcript_exon_variant | MODIFIER |
| 5 | 52085874 | AX-170188631 | 0 | G | A | 0.482 | 1.31E-01 | 1.06E-01 | 2.37E+00 | 2.40E+00 | 2.29E-01 | no | Gm43174 | upstream_gene_variant | MODIFIER |
| 5 | 52086246 | AX-170140729 | 0 | G | A | 0.482 | 1.31E-01 | 1.06E-01 | 2.37E+00 | 2.40E+00 | 2.29E-01 | no | Gm43174 | upstream_gene_variant | MODIFIER |
| 5 | 52086435 | AX-170115729 | 0 | C | T | 0.482 | 1.31E-01 | 1.06E-01 | 2.37E+00 | 2.40E+00 | 2.29E-01 | no | Gm43174 | upstream_gene_variant | MODIFIER |
| 5 | 52087436 | AX-170386004 | 0 | T | G | 0.482 | 1.31E-01 | 1.06E-01 | 2.37E+00 | 2.40E+00 | 2.29E-01 | no | | intergenic_variant | MODIFIER |
| 5 | 52124623 | AX-97802973 | 0 | T | C | 0.482 | 1.31E-01 | 1.06E-01 | 2.37E+00 | 2.40E+00 | 2.29E-01 | no | | intergenic_variant | MODIFIER |
| 5 | 52140126 | AX-170404230 | 0 | A | G | 0.482 | 1.31E-01 | 1.06E-01 | 2.37E+00 | 2.40E+00 | 2.29E-01 | no | Mir6417 | upstream_gene_variant | MODIFIER |
| 5 | 52171758 | AX-170196124 | 0 | G | A | 0.482 | 1.31E-01 | 1.06E-01 | 2.37E+00 | 2.40E+00 | 2.29E-01 | no | Dhx15 | intron_variant | MODIFIER |
| 5 | 52205145 | AX-170532102 | 0 | A | G | 0.482 | 1.31E-01 | 1.06E-01 | 2.37E+00 | 2.40E+00 | 2.29E-01 | no | C130083M11Rik | intron_variant,non_coding_transcript_variant | MODIFIER |
| 5 | 52213988 | AX-170512061 | 0 | T | C | 0.482 | 1.31E-01 | 1.06E-01 | 2.37E+00 | 2.40E+00 | 2.29E-01 | no | C130083M11Rik | intron_variant,non_coding_transcript_variant | MODIFIER |
| 5 | 52214257 | AX-170097098 | 0 | G | T | 0.482 | 1.31E-01 | 1.06E-01 | 2.37E+00 | 2.40E+00 | 2.29E-01 | no | C130083M11Rik | intron_variant,non_coding_transcript_variant | MODIFIER |
| 5 | 52214530 | AX-170355477 | 0 | G | A | 0.482 | 1.31E-01 | 1.06E-01 | 2.37E+00 | 2.40E+00 | 2.29E-01 | no | C130083M11Rik | intron_variant,non_coding_transcript_variant | MODIFIER |
| 5 | 52267888 | AX-97850256 | 0 | A | G | 0.482 | 1.31E-01 | 1.06E-01 | 2.37E+00 | 2.40E+00 | 2.29E-01 | no | 9230114K14Rik | intron_variant,non_coding_transcript_variant | MODIFIER |
| 5 | 52281940 | AX-170112519 | 0 | G | A | 0.482 | 1.31E-01 | 1.06E-01 | 2.37E+00 | 2.40E+00 | 2.29E-01 | no | 9230114K14Rik | downstream_gene_variant | MODIFIER |
| 5 | 52282741 | AX-170242662 | 0 | T | C | 0.482 | 1.31E-01 | 1.06E-01 | 2.37E+00 | 2.40E+00 | 2.29E-01 | no | 9230114K14Rik | downstream_gene_variant | MODIFIER |
| 3 | 145218240 | AX-97796616 | 0 | T | C | 0.325 | -8.25E-02 | 6.83E-02 | 3.02E+00 | 3.16E+00 | 2.29E-01 | no | | intergenic_variant | MODIFIER |
| 9 | 101182346 | AX-170358564 | 0 | A | G | 0.132 | -1.37E-01 | 1.15E-01 | 2.66E+00 | 2.75E+00 | 2.29E-01 | no | Ppp2r3a | intron_variant | MODIFIER |
| 8 | 24489835 | AX-170406775 | 1 | C | T | 0.357 | -7.65E-02 | 6.39E-02 | 2.58E+00 | 2.68E+00 | 2.29E-01 | no | | regulatory_region_variant | MODIFIER |
| 18 | 68209244 | AX-170294401 | 1 | T | A | 0.277 | -1.10E-01 | 9.23E-02 | 2.77E+00 | 2.87E+00 | 2.29E-01 | no | Ldlrad4 | intron_variant | MODIFIER |
| 16 | 50055830 | AX-170238472 | 0 | A | C | 0.482 | -9.74E-02 | 8.11E-02 | 2.58E+00 | 2.62E+00 | 2.29E-01 | no | | intergenic_variant | MODIFIER |
| 4 | 47029781 | AX-170293193 | 0 | A | G | 0.36 | -1.15E-01 | 9.60E-02 | 2.88E+00 | 2.98E+00 | 2.29E-01 | no | Anks6 | intron_variant | MODIFIER |
| 1 | 172823941 | AX-170201185 | 1 | G | A | 0.214 | 1.57E-01 | 1.31E-01 | 2.60E+00 | 2.66E+00 | 2.29E-01 | Yes | Crp | intron_variant,non_coding_transcript_variant | MODIFIER |
| 1 | 173299438 | AX-170002304 | 1 | C | T | 0.214 | 1.57E-01 | 1.31E-01 | 2.60E+00 | 2.66E+00 | 2.29E-01 | Yes | | intergenic_variant | MODIFIER |
| 4 | 75844400 | AX-170311895 | 0 | T | G | 0.325 | 1.30E-01 | 1.09E-01 | 2.72E+00 | 2.78E+00 | 2.29E-01 | no | | intergenic_variant | MODIFIER |
| 1 | 187964021 | AX-170417550 | 1 | G | A | 0.241 | 1.21E-01 | 9.64E-02 | 2.27E+00 | 2.36E+00 | 2.29E-01 | no | Esrrg | intron_variant | MODIFIER |
| 5 | 44378900 | AX-170071936 | 1 | G | A | 0.188 | -1.27E-01 | 1.07E-01 | 2.70E+00 | 2.81E+00 | 2.29E-01 | no | | intergenic_variant | MODIFIER |
| 13 | 8736238 | AX-170405489 | 0 | T | C | 0.193 | -1.49E-01 | 1.24E-01 | 2.54E+00 | 2.60E+00 | 2.29E-01 | no | Adarb2 | intron_variant | MODIFIER |
| 8 | 66048126 | AX-170251579 | 0 | A | G | 0.14 | -1.74E-01 | 1.45E-01 | 2.55E+00 | 2.62E+00 | 2.29E-01 | Yes | MAR1 | intron_variant | MODIFIER |
| 7 | 70913129 | AX-170477478 | 1 | C | A | 0.375 | -1.24E-01 | 1.03E-01 | 2.95E+00 | 3.05E+00 | 2.29E-01 | no | Gm42397 | intron_variant,non_coding_transcript_variant | MODIFIER |
| 10 | 101816460 | AX-170196989 | 1 | A | G | 0.277 | -9.79E-02 | 8.19E-02 | 2.59E+00 | 2.67E+00 | 2.30E-01 | no | Mgat4c | intron_variant | MODIFIER |
| 16 | 44301638 | AX-170147145 | 1 | G | A | 0.518 | 9.66E-02 | 8.08E-02 | 2.60E+00 | 2.66E+00 | 2.30E-01 | no | Sidt1 | intron_variant | MODIFIER |
| 4 | 129741737 | AX-170384611 | 1 | T | C | 0.152 | 1.40E-01 | 1.17E-01 | 2.62E+00 | 2.67E+00 | 2.30E-01 | no | Khdrbs1 | intron_variant | MODIFIER |
| 18 | 53502142 | AX-170426365 | 1 | T | A | 0.241 | -1.19E-01 | 1.00E-01 | 2.62E+00 | 2.71E+00 | 2.30E-01 | no | Prdm6 | intron_variant | MODIFIER |
| 1 | 181167906 | AX-170002421 | 1 | G | C | 0.232 | 1.37E-01 | 1.13E-01 | 2.46E+00 | 2.52E+00 | 2.30E-01 | no | Cnih4 | downstream_gene_variant | MODIFIER |
| 15 | 22136810 | AX-170116178 | 0 | A | G | 0.237 | -1.18E-01 | 9.90E-02 | 2.60E+00 | 2.67E+00 | 2.30E-01 | no | | intergenic_variant | MODIFIER |
| 2 | 120214150 | AX-170097719 | 0 | T | C | 0.149 | 2.16E-01 | 1.81E-01 | 2.83E+00 | 2.93E+00 | 2.30E-01 | Yes | Pla2g4e | intron_variant | MODIFIER |
| 2 | 121317292 | AX-170355607 | 0 | T | C | 0.149 | 2.16E-01 | 1.81E-01 | 2.83E+00 | 2.93E+00 | 2.30E-01 | Yes | Ppip5k1 | intron_variant | MODIFIER |
| 18 | 76347671 | AX-170061365 | 1 | T | C | 0.393 | -1.04E-01 | 8.77E-02 | 2.72E+00 | 2.80E+00 | 2.30E-01 | no | | intergenic_variant | MODIFIER |
| 7 | 38380007 | AX-170077108 | 1 | A | G | 0.304 | 1.17E-01 | 9.76E-02 | 2.54E+00 | 2.64E+00 | 2.30E-01 | no | Gm22638 | upstream_gene_variant | MODIFIER |
| 8 | 52408088 | AX-170030705 | 0 | A | T | 0.158 | -1.34E-01 | 1.13E-01 | 2.70E+00 | 2.83E+00 | 2.30E-01 | no | 1190028D05Rik | downstream_gene_variant | MODIFIER |
| 4 | 137912982 | AX-170158995 | 1 | G | A | 0.17 | 1.37E-01 | 1.15E-01 | 2.65E+00 | 2.70E+00 | 2.31E-01 | no | Ece1 | intron_variant | MODIFIER |
| 10 | 100098469 | AX-170513420 | 1 | T | A | 0.268 | -9.90E-02 | 8.24E-02 | 2.51E+00 | 2.59E+00 | 2.31E-01 | no | Kitl | downstream_gene_variant | MODIFIER |
| 7 | 100886076 | AX-170078241 | 0 | A | G | 0.36 | -1.20E-01 | 1.01E-01 | 2.85E+00 | 2.95E+00 | 2.31E-01 | no | Arhgef17 | intron_variant | MODIFIER |
| 5 | 45016427 | AX-170533297 | 0 | C | T | 0.202 | -1.20E-01 | 1.01E-01 | 2.63E+00 | 2.71E+00 | 2.31E-01 | no | D5Ertd615e | intron_variant,non_coding_transcript_variant | MODIFIER |
| 16 | 18864892 | AX-170122653 | 2 | G | A | 0.127 | 1.14E-01 | 9.58E-02 | 2.91E+00 | 3.05E+00 | 2.31E-01 | no | Gm28539 | intron_variant,NMD_transcript_variant | MODIFIER |
| 18 | 75922207 | AX-170356032 | 0 | G | A | 0.395 | -1.04E-01 | 8.76E-02 | 2.73E+00 | 2.81E+00 | 2.31E-01 | no | Zbtb7c | intron_variant | MODIFIER |
| 18 | 75930793 | AX-170016013 | 0 | G | T | 0.395 | -1.04E-01 | 8.76E-02 | 2.73E+00 | 2.81E+00 | 2.31E-01 | no | Zbtb7c | intron_variant | MODIFIER |
| 18 | 75930997 | AX-170016016 | 0 | G | C | 0.395 | -1.04E-01 | 8.76E-02 | 2.73E+00 | 2.81E+00 | 2.31E-01 | no | Zbtb7c | intron_variant | MODIFIER |

| | | | | | | | | | | | | | | | |
|---|---|---|---|---|---|---|---|---|---|---|---|---|---|---|---|
| 18 | 75976339 | AX-170016018 | 0 | G | C | 0.395 | -1.04E-01 | 8.76E-02 | 2.73E+00 | 2.81E+00 | 2.31E-01 | no | Zbtb7c | intron_variant | MODIFIER |
| 18 | 75992340 | AX-97807261 | 0 | G | A | 0.395 | -1.04E-01 | 8.76E-02 | 2.73E+00 | 2.81E+00 | 2.31E-01 | no | Zbtb7c | intron_variant | MODIFIER |
| 18 | 76019882 | AX-170016014 | 0 | G | A | 0.395 | -1.04E-01 | 8.76E-02 | 2.73E+00 | 2.81E+00 | 2.31E-01 | no | Zbtb7c | intron_variant | MODIFIER |
| 18 | 76021492 | AX-170016012 | 0 | T | C | 0.395 | -1.04E-01 | 8.76E-02 | 2.73E+00 | 2.81E+00 | 2.31E-01 | no | Zbtb7c | intron_variant | MODIFIER |
| 18 | 76034999 | AX-170194810 | 0 | A | G | 0.395 | -1.04E-01 | 8.76E-02 | 2.73E+00 | 2.81E+00 | 2.31E-01 | no | Zbtb7c | intron_variant | MODIFIER |
| 18 | 76046330 | AX-170149034 | 0 | A | G | 0.395 | -1.04E-01 | 8.76E-02 | 2.73E+00 | 2.81E+00 | 2.31E-01 | no | Zbtb7c | intron_variant | MODIFIER |
| 18 | 76046710 | AX-170353771 | 0 | T | C | 0.395 | -1.04E-01 | 8.76E-02 | 2.73E+00 | 2.81E+00 | 2.31E-01 | no | Zbtb7c | intron_variant | MODIFIER |
| 18 | 76152743 | AX-170386339 | 0 | G | A | 0.395 | -1.04E-01 | 8.76E-02 | 2.73E+00 | 2.81E+00 | 2.31E-01 | no | Zbtb7c | downstream_gene_variant | MODIFIER |
| 18 | 76163047 | AX-170410856 | 0 | C | T | 0.395 | -1.04E-01 | 8.76E-02 | 2.73E+00 | 2.81E+00 | 2.31E-01 | no | | intergenic_variant | MODIFIER |
| 18 | 76163401 | AX-170116012 | 0 | G | A | 0.395 | -1.04E-01 | 8.76E-02 | 2.73E+00 | 2.81E+00 | 2.31E-01 | no | | intergenic_variant | MODIFIER |
| 18 | 76173304 | AX-170103521 | 0 | G | C | 0.395 | -1.04E-01 | 8.76E-02 | 2.73E+00 | 2.81E+00 | 2.31E-01 | no | Mir6358 | downstream_gene_variant | MODIFIER |
| 18 | 76209102 | AX-170329137 | 0 | T | C | 0.395 | -1.04E-01 | 8.76E-02 | 2.73E+00 | 2.81E+00 | 2.31E-01 | no | | intergenic_variant | MODIFIER |
| 18 | 76214882 | AX-170388545 | 0 | G | A | 0.395 | -1.04E-01 | 8.76E-02 | 2.73E+00 | 2.81E+00 | 2.31E-01 | no | | intergenic_variant | MODIFIER |
| 18 | 76253846 | AX-170219094 | 0 | G | T | 0.395 | -1.04E-01 | 8.76E-02 | 2.73E+00 | 2.81E+00 | 2.31E-01 | no | Smad2 | intron_variant | MODIFIER |
| 18 | 76263395 | AX-170472243 | 0 | G | A | 0.395 | -1.04E-01 | 8.76E-02 | 2.73E+00 | 2.81E+00 | 2.31E-01 | no | Smad2 | intron_variant | MODIFIER |
| 18 | 76263409 | AX-170132585 | 0 | G | A | 0.395 | -1.04E-01 | 8.76E-02 | 2.73E+00 | 2.81E+00 | 2.31E-01 | no | Smad2 | intron_variant | MODIFIER |
| 18 | 76264277 | AX-170109779 | 0 | G | A | 0.395 | -1.04E-01 | 8.76E-02 | 2.73E+00 | 2.81E+00 | 2.31E-01 | no | Smad2 | intron_variant | MODIFIER |
| 18 | 76279945 | AX-170091046 | 0 | C | T | 0.395 | -1.04E-01 | 8.76E-02 | 2.73E+00 | 2.81E+00 | 2.31E-01 | no | Smad2 | intron_variant | MODIFIER |
| 18 | 76307208 | AX-170272540 | 0 | C | T | 0.395 | -1.04E-01 | 8.76E-02 | 2.73E+00 | 2.81E+00 | 2.31E-01 | no | Smad2 | downstream_gene_variant | MODIFIER |
| 18 | 76325234 | AX-170265024 | 0 | G | A | 0.395 | -1.04E-01 | 8.76E-02 | 2.73E+00 | 2.81E+00 | 2.31E-01 | no | AC164009.1 | downstream_gene_variant | MODIFIER |
| 18 | 76325360 | AX-170403526 | 0 | T | C | 0.395 | -1.04E-01 | 8.76E-02 | 2.73E+00 | 2.81E+00 | 2.31E-01 | no | AC164009.1 | downstream_gene_variant | MODIFIER |
| 18 | 76332984 | AX-170016019 | 0 | T | C | 0.395 | -1.04E-01 | 8.76E-02 | 2.73E+00 | 2.81E+00 | 2.31E-01 | no | | intergenic_variant | MODIFIER |
| 18 | 76333272 | AX-170016022 | 0 | G | A | 0.395 | -1.04E-01 | 8.76E-02 | 2.73E+00 | 2.81E+00 | 2.31E-01 | no | | intergenic_variant | MODIFIER |
| 18 | 76341047 | AX-97795246 | 0 | T | C | 0.395 | -1.04E-01 | 8.76E-02 | 2.73E+00 | 2.81E+00 | 2.31E-01 | no | | intergenic_variant | MODIFIER |
| 18 | 76348169 | AX-170268200 | 0 | A | T | 0.395 | -1.04E-01 | 8.76E-02 | 2.73E+00 | 2.81E+00 | 2.31E-01 | no | | intergenic_variant | MODIFIER |
| 18 | 76360408 | AX-170016024 | 0 | A | C | 0.395 | -1.04E-01 | 8.76E-02 | 2.73E+00 | 2.81E+00 | 2.31E-01 | no | | intergenic_variant | MODIFIER |
| 18 | 76393125 | AX-97835191 | 0 | C | T | 0.395 | -1.04E-01 | 8.76E-02 | 2.73E+00 | 2.81E+00 | 2.31E-01 | no | | regulatory_region_variant | MODIFIER |
| 18 | 76395254 | AX-170228758 | 0 | G | A | 0.395 | -1.04E-01 | 8.76E-02 | 2.73E+00 | 2.81E+00 | 2.31E-01 | no | | intergenic_variant | MODIFIER |
| 18 | 76896742 | AX-97792532 | 0 | T | C | 0.395 | -1.04E-01 | 8.76E-02 | 2.73E+00 | 2.81E+00 | 2.31E-01 | no | Skor2 | intron_variant | MODIFIER |
| 18 | 76933431 | AX-170468036 | 0 | A | G | 0.395 | -1.04E-01 | 8.76E-02 | 2.73E+00 | 2.81E+00 | 2.31E-01 | no | Ier3ip1 | intron_variant | MODIFIER |
| 18 | 76967317 | AX-170313359 | 0 | A | G | 0.395 | -1.04E-01 | 8.76E-02 | 2.73E+00 | 2.81E+00 | 2.31E-01 | no | Hdhd2 | intron_variant | MODIFIER |
| 18 | 76968652 | AX-170316331 | 0 | G | A | 0.395 | -1.04E-01 | 8.76E-02 | 2.73E+00 | 2.81E+00 | 2.31E-01 | no | Hdhd2 | intron_variant | MODIFIER |
| 18 | 76969248 | AX-170294491 | 0 | G | A | 0.395 | -1.04E-01 | 8.76E-02 | 2.73E+00 | 2.81E+00 | 2.31E-01 | no | Hdhd2 | intron_variant | MODIFIER |
| 18 | 77165123 | AX-97782554 | 0 | C | T | 0.395 | -1.04E-01 | 8.76E-02 | 2.73E+00 | 2.81E+00 | 2.31E-01 | no | | intergenic_variant | MODIFIER |
| 15 | 10548521 | AX-170143517 | 0 | C | A | 0.237 | -1.26E-01 | 1.06E-01 | 2.64E+00 | 2.69E+00 | 2.31E-01 | no | Ttc23l | intron_variant | MODIFIER |
| 15 | 11334208 | AX-170011033 | 0 | C | T | 0.237 | -1.26E-01 | 1.06E-01 | 2.64E+00 | 2.69E+00 | 2.31E-01 | no | Adamts12 | intron_variant | MODIFIER |
| 15 | 22551761 | AX-170342795 | 0 | C | T | 0.237 | -1.26E-01 | 1.06E-01 | 2.64E+00 | 2.69E+00 | 2.31E-01 | no | Cdh18 | intron_variant | MODIFIER |
| 3 | 10761628 | AX-170406259 | 1 | C | T | 0.402 | -7.62E-02 | 6.44E-02 | 2.81E+00 | 2.97E+00 | 2.31E-01 | no | | intergenic_variant | MODIFIER |
| 4 | 69195196 | AX-170069601 | 0 | G | A | 0.325 | 1.42E-01 | 1.20E-01 | 2.87E+00 | 2.93E+00 | 2.31E-01 | no | | intergenic_variant | MODIFIER |
| 15 | 3618048 | AX-170087751 | 0 | A | C | 0.333 | -9.79E-02 | 7.41E-02 | 2.07E+00 | 2.13E+00 | 2.31E-01 | no | | intergenic_variant | MODIFIER |
| 7 | 100912860 | AX-170078242 | 1 | G | A | 0.375 | -1.27E-01 | 1.07E-01 | 2.85E+00 | 2.94E+00 | 2.31E-01 | no | Arhgef17 | intron_variant | MODIFIER |
| 7 | 101653717 | AX-170112718 | 1 | A | G | 0.375 | -1.27E-01 | 1.07E-01 | 2.85E+00 | 2.94E+00 | 2.31E-01 | no | | regulatory_region_variant | MODIFIER |
| 16 | 36611781 | AX-170391825 | 1 | C | T | 0.491 | 9.55E-02 | 7.95E-02 | 2.51E+00 | 2.56E+00 | 2.32E-01 | no | Cd86 | intron_variant | MODIFIER |
| 4 | 86788532 | AX-170419606 | 1 | T | A | 0.339 | 1.35E-01 | 1.14E-01 | 2.74E+00 | 2.81E+00 | 2.32E-01 | no | Dennd4c | intron_variant | MODIFIER |
| 5 | 104981384 | AX-170142580 | 0 | A | G | 0.447 | 1.37E-01 | 1.16E-01 | 2.75E+00 | 2.80E+00 | 2.32E-01 | no | Abcg3 | intron_variant | MODIFIER |
| 5 | 66504403 | AX-170358238 | 0 | A | G | 0.482 | 1.21E-01 | 1.01E-01 | 2.53E+00 | 2.60E+00 | 2.32E-01 | no | Apbb2 | intron_variant | MODIFIER |
| 5 | 66509339 | AX-97801797 | 0 | G | A | 0.482 | 1.21E-01 | 1.01E-01 | 2.53E+00 | 2.60E+00 | 2.32E-01 | no | Apbb2 | intron_variant | MODIFIER |
| 5 | 66527144 | AX-170238389 | 0 | C | A | 0.482 | 1.21E-01 | 1.01E-01 | 2.53E+00 | 2.60E+00 | 2.32E-01 | no | Apbb2 | intron_variant | MODIFIER |
| 5 | 66533251 | AX-97832710 | 0 | A | G | 0.482 | 1.21E-01 | 1.01E-01 | 2.53E+00 | 2.60E+00 | 2.32E-01 | no | Apbb2 | intron_variant | MODIFIER |
| 5 | 66636089 | AX-97812410 | 0 | A | G | 0.482 | 1.21E-01 | 1.01E-01 | 2.53E+00 | 2.60E+00 | 2.32E-01 | no | Uchl1os | intron_variant,non_coding_transcript_variant | MODIFIER |
| 5 | 66639869 | AX-170225987 | 0 | G | A | 0.482 | 1.21E-01 | 1.01E-01 | 2.53E+00 | 2.60E+00 | 2.32E-01 | no | Uchl1os | intron_variant,non_coding_transcript_variant | MODIFIER |
| 5 | 66658528 | AX-97807960 | 0 | A | G | 0.482 | 1.21E-01 | 1.01E-01 | 2.53E+00 | 2.60E+00 | 2.32E-01 | no | Uchl1os | intron_variant,non_coding_transcript_variant | MODIFIER |
| 5 | 66668772 | AX-170469143 | 0 | G | A | 0.482 | 1.21E-01 | 1.01E-01 | 2.53E+00 | 2.60E+00 | 2.32E-01 | no | Uchl1os | intron_variant,non_coding_transcript_variant | MODIFIER |
| 5 | 66668856 | AX-170523180 | 0 | G | A | 0.482 | 1.21E-01 | 1.01E-01 | 2.53E+00 | 2.60E+00 | 2.32E-01 | no | Uchl1os | intron_variant,non_coding_transcript_variant | MODIFIER |

| | | | | | | | | | | | | | | | |
|---|---|---|---|---|---|---|---|---|---|---|---|---|---|---|---|
| 5 | 66675350 | AX-97809422 | 0 | A | G | 0.482 | 1.21E-01 | 1.01E-01 | 2.53E+00 | 2.60E+00 | 2.32E-01 | no | Uchl1 | upstream_gene_variant | MODIFIER |
| 5 | 66681000 | AX-170360210 | 0 | C | T | 0.482 | 1.21E-01 | 1.01E-01 | 2.53E+00 | 2.60E+00 | 2.32E-01 | no | Uchl1 | intron_variant | MODIFIER |
| 5 | 66681133 | AX-170113039 | 0 | C | T | 0.482 | 1.21E-01 | 1.01E-01 | 2.53E+00 | 2.60E+00 | 2.32E-01 | no | Uchl1 | intron_variant | MODIFIER |
| 5 | 66683622 | AX-170072352 | 0 | A | G | 0.482 | 1.21E-01 | 1.01E-01 | 2.53E+00 | 2.60E+00 | 2.32E-01 | no | Uchl1 | intron_variant | MODIFIER |
| 5 | 66684447 | AX-170333474 | 0 | T | C | 0.482 | 1.21E-01 | 1.01E-01 | 2.53E+00 | 2.60E+00 | 2.32E-01 | no | Uchl1 | intron_variant | MODIFIER |
| 5 | 66689289 | AX-170213704 | 0 | G | A | 0.482 | 1.21E-01 | 1.01E-01 | 2.53E+00 | 2.60E+00 | 2.32E-01 | no | Uchl1 | downstream_gene_variant | MODIFIER |
| 5 | 66731947 | AX-97832706 | 0 | G | A | 0.482 | 1.21E-01 | 1.01E-01 | 2.53E+00 | 2.60E+00 | 2.32E-01 | no | | regulatory_region_variant | MODIFIER |
| 5 | 66764209 | AX-170442374 | 0 | A | G | 0.482 | 1.21E-01 | 1.01E-01 | 2.53E+00 | 2.60E+00 | 2.32E-01 | no | Limch1 | intron_variant | MODIFIER |
| 5 | 66778943 | AX-170025190 | 0 | A | C | 0.482 | 1.21E-01 | 1.01E-01 | 2.53E+00 | 2.60E+00 | 2.32E-01 | no | Limch1 | intron_variant | MODIFIER |
| 5 | 66786797 | AX-170095346 | 0 | T | C | 0.482 | 1.21E-01 | 1.01E-01 | 2.53E+00 | 2.60E+00 | 2.32E-01 | no | Limch1 | intron_variant | MODIFIER |
| 5 | 66881129 | AX-170115673 | 0 | C | T | 0.482 | 1.21E-01 | 1.01E-01 | 2.53E+00 | 2.60E+00 | 2.32E-01 | no | Limch1 | intron_variant | MODIFIER |
| 5 | 66891718 | AX-170486045 | 0 | G | C | 0.482 | 1.21E-01 | 1.01E-01 | 2.53E+00 | 2.60E+00 | 2.32E-01 | no | Limch1 | intron_variant | MODIFIER |
| 5 | 66894904 | AX-170072357 | 0 | A | G | 0.482 | 1.21E-01 | 1.01E-01 | 2.53E+00 | 2.60E+00 | 2.32E-01 | no | Limch1 | intron_variant | MODIFIER |
| 5 | 66899128 | AX-97831876 | 0 | C | A | 0.482 | 1.21E-01 | 1.01E-01 | 2.53E+00 | 2.60E+00 | 2.32E-01 | no | Limch1 | intron_variant | MODIFIER |
| 5 | 66921126 | AX-170488626 | 0 | G | A | 0.482 | 1.21E-01 | 1.01E-01 | 2.53E+00 | 2.60E+00 | 2.32E-01 | no | Limch1 | intron_variant | MODIFIER |
| 5 | 67024805 | AX-170098864 | 0 | T | C | 0.482 | 1.21E-01 | 1.01E-01 | 2.53E+00 | 2.60E+00 | 2.32E-01 | no | Limch1 | intron_variant | MODIFIER |
| 5 | 67055307 | AX-170393604 | 0 | T | C | 0.482 | 1.21E-01 | 1.01E-01 | 2.53E+00 | 2.60E+00 | 2.32E-01 | no | Limch1 | 3_prime_UTR_variant | MODIFIER |
| 5 | 67055909 | AX-97807457 | 0 | C | T | 0.482 | 1.21E-01 | 1.01E-01 | 2.53E+00 | 2.60E+00 | 2.32E-01 | no | Limch1 | 3_prime_UTR_variant | MODIFIER |
| 5 | 67061273 | AX-170425448 | 0 | G | A | 0.482 | 1.21E-01 | 1.01E-01 | 2.53E+00 | 2.60E+00 | 2.32E-01 | no | Limch1 | downstream_gene_variant | MODIFIER |
| 5 | 67063121 | AX-170430437 | 0 | T | C | 0.482 | 1.21E-01 | 1.01E-01 | 2.53E+00 | 2.60E+00 | 2.32E-01 | no | | intergenic_variant | MODIFIER |
| 5 | 67078749 | AX-170248398 | 0 | A | C | 0.482 | 1.21E-01 | 1.01E-01 | 2.53E+00 | 2.60E+00 | 2.32E-01 | no | | intergenic_variant | MODIFIER |
| 5 | 67078770 | AX-170184483 | 0 | C | A | 0.482 | 1.21E-01 | 1.01E-01 | 2.53E+00 | 2.60E+00 | 2.32E-01 | no | | intergenic_variant | MODIFIER |
| 5 | 67079177 | AX-97845212 | 0 | C | T | 0.482 | 1.21E-01 | 1.01E-01 | 2.53E+00 | 2.60E+00 | 2.32E-01 | no | | intergenic_variant | MODIFIER |
| 5 | 67099963 | AX-97841003 | 0 | C | T | 0.482 | 1.21E-01 | 1.01E-01 | 2.53E+00 | 2.60E+00 | 2.32E-01 | no | Phox2b | upstream_gene_variant | MODIFIER |
| 5 | 67100235 | AX-170510831 | 0 | A | G | 0.482 | 1.21E-01 | 1.01E-01 | 2.53E+00 | 2.60E+00 | 2.32E-01 | no | Phox2b | upstream_gene_variant | MODIFIER |
| 5 | 67211706 | AX-97814393 | 0 | G | A | 0.482 | 1.21E-01 | 1.01E-01 | 2.53E+00 | 2.60E+00 | 2.32E-01 | no | | intergenic_variant | MODIFIER |
| 5 | 67259015 | AX-170421790 | 0 | C | T | 0.482 | 1.21E-01 | 1.01E-01 | 2.53E+00 | 2.60E+00 | 2.32E-01 | no | Tmem33 | upstream_gene_variant | MODIFIER |
| 5 | 67266848 | AX-170447625 | 0 | C | T | 0.482 | 1.21E-01 | 1.01E-01 | 2.53E+00 | 2.60E+00 | 2.32E-01 | no | Tmem33 | intron_variant | MODIFIER |
| 5 | 67299563 | AX-170480816 | 0 | C | T | 0.482 | 1.21E-01 | 1.01E-01 | 2.53E+00 | 2.60E+00 | 2.32E-01 | no | | regulatory_region_variant | MODIFIER |
| 5 | 67330585 | AX-170151189 | 0 | G | A | 0.482 | 1.21E-01 | 1.01E-01 | 2.53E+00 | 2.60E+00 | 2.32E-01 | no | Slc30a9 | intron_variant | MODIFIER |
| 5 | 67331780 | AX-170508363 | 0 | A | C | 0.482 | 1.21E-01 | 1.01E-01 | 2.53E+00 | 2.60E+00 | 2.32E-01 | no | Slc30a9 | intron_variant | MODIFIER |
| 5 | 67345401 | AX-170356552 | 0 | A | G | 0.482 | 1.21E-01 | 1.01E-01 | 2.53E+00 | 2.60E+00 | 2.32E-01 | no | Slc30a9 | intron_variant | MODIFIER |
| 5 | 67347975 | AX-170025197 | 0 | A | G | 0.482 | 1.21E-01 | 1.01E-01 | 2.53E+00 | 2.60E+00 | 2.32E-01 | no | Slc30a9 | synonymous_variant | LOW |
| 5 | 67348023 | AX-170025196 | 0 | T | C | 0.482 | 1.21E-01 | 1.01E-01 | 2.53E+00 | 2.60E+00 | 2.32E-01 | no | Slc30a9 | synonymous_variant | LOW |
| 5 | 67349359 | AX-170173295 | 0 | G | A | 0.482 | 1.21E-01 | 1.01E-01 | 2.53E+00 | 2.60E+00 | 2.32E-01 | no | Slc30a9 | intron_variant | MODIFIER |
| 5 | 67361021 | AX-170089508 | 0 | T | C | 0.482 | 1.21E-01 | 1.01E-01 | 2.53E+00 | 2.60E+00 | 2.32E-01 | no | Slc30a9 | downstream_gene_variant | MODIFIER |
| 5 | 67390960 | AX-170072365 | 0 | C | G | 0.482 | 1.21E-01 | 1.01E-01 | 2.53E+00 | 2.60E+00 | 2.32E-01 | no | Bend4 | downstream_gene_variant | MODIFIER |
| 18 | 84063887 | AX-170061537 | 1 | C | T | 0.161 | -1.28E-01 | 1.06E-01 | 2.49E+00 | 2.58E+00 | 2.32E-01 | no | Tshz1 | intron_variant | MODIFIER |
| 12 | 88773373 | AX-97833284 | 0 | G | T | 0.044 | -2.38E-01 | 2.01E-01 | 2.62E+00 | 2.72E+00 | 2.32E-01 | Yes | Nrxn3 | intron_variant,non_coding_transcript_variant | MODIFIER |
| 16 | 72248328 | AX-170121099 | 1 | A | C | 0.491 | 9.22E-02 | 7.74E-02 | 2.57E+00 | 2.64E+00 | 2.32E-01 | no | | intergenic_variant | MODIFIER |
| 1 | 134191987 | AX-170157215 | 1 | T | C | 0.536 | 9.90E-02 | 8.35E-02 | 2.65E+00 | 2.72E+00 | 2.32E-01 | no | Mybph | upstream_gene_variant | MODIFIER |
| 19 | 60596003 | AX-170403142 | 0 | A | G | 0.456 | -7.70E-02 | 6.50E-02 | 2.67E+00 | 2.75E+00 | 2.32E-01 | no | | intergenic_variant | MODIFIER |
| 5 | 67462321 | AX-170521859 | 0 | T | C | 0.456 | 1.20E-01 | 1.00E-01 | 2.50E+00 | 2.57E+00 | 2.33E-01 | no | C330024D21Rik | upstream_gene_variant | MODIFIER |
| 5 | 67468888 | AX-170248204 | 0 | C | A | 0.456 | 1.20E-01 | 1.00E-01 | 2.50E+00 | 2.57E+00 | 2.33E-01 | no | C330024D21Rik | non_coding_transcript_exon_variant | MODIFIER |
| 5 | 67498430 | AX-170282264 | 0 | G | A | 0.456 | 1.20E-01 | 1.00E-01 | 2.50E+00 | 2.57E+00 | 2.33E-01 | no | | intergenic_variant | MODIFIER |
| 5 | 67499379 | AX-170489032 | 0 | T | C | 0.456 | 1.20E-01 | 1.00E-01 | 2.50E+00 | 2.57E+00 | 2.33E-01 | no | | intergenic_variant | MODIFIER |
| 5 | 67545596 | AX-170072370 | 0 | T | C | 0.456 | 1.20E-01 | 1.00E-01 | 2.50E+00 | 2.57E+00 | 2.33E-01 | no | | intergenic_variant | MODIFIER |
| 5 | 67550501 | AX-170072372 | 0 | C | T | 0.456 | 1.20E-01 | 1.00E-01 | 2.50E+00 | 2.57E+00 | 2.33E-01 | no | | regulatory_region_variant | MODIFIER |
| 5 | 67564476 | AX-170072369 | 0 | T | C | 0.456 | 1.20E-01 | 1.00E-01 | 2.50E+00 | 2.57E+00 | 2.33E-01 | no | | intergenic_variant | MODIFIER |
| 5 | 67632536 | AX-170202243 | 0 | T | G | 0.456 | 1.20E-01 | 1.00E-01 | 2.50E+00 | 2.57E+00 | 2.33E-01 | no | Atp8a1 | intron_variant | MODIFIER |
| 5 | 67660341 | AX-170095348 | 0 | A | G | 0.456 | 1.20E-01 | 1.00E-01 | 2.50E+00 | 2.57E+00 | 2.33E-01 | no | Atp8a1 | intron_variant | MODIFIER |
| 5 | 67665100 | AX-170515436 | 0 | T | C | 0.456 | 1.20E-01 | 1.00E-01 | 2.50E+00 | 2.57E+00 | 2.33E-01 | no | Atp8a1 | intron_variant | MODIFIER |
| 5 | 67682967 | AX-170480485 | 0 | A | G | 0.456 | 1.20E-01 | 1.00E-01 | 2.50E+00 | 2.57E+00 | 2.33E-01 | no | Atp8a1 | intron_variant | MODIFIER |
| 6 | 134247868 | AX-170230189 | 0 | G | T | 0.553 | 1.00E-01 | 8.36E-02 | 2.47E+00 | 2.54E+00 | 2.33E-01 | no | Etv6 | intron_variant | MODIFIER |

| | | | | | | | | | | | | | | | |
|---|---|---|---|---|---|---|---|---|---|---|---|---|---|---|---|
| 13 | 20343189 | AX-170129593 | 1 | C | G | 0.179 | -1.46E-01 | 1.21E-01 | 2.44E+00 | 2.50E+00 | 2.33E-01 | no | Elmo1 | intron_variant | MODIFIER |
| 13 | 19369548 | AX-170450726 | 1 | G | T | 0.17 | -1.51E-01 | 1.27E-01 | 2.51E+00 | 2.57E+00 | 2.33E-01 | no | Stard3nl | intron_variant | MODIFIER |
| 16 | 40218911 | AX-170420049 | 1 | T | G | 0.5 | 9.72E-02 | 8.16E-02 | 2.57E+00 | 2.61E+00 | 2.33E-01 | no | Lsamp | intron_variant | MODIFIER |
| 3 | 23907639 | AX-170020131 | 0 | G | A | 0.158 | -1.21E-01 | 1.01E-01 | 2.47E+00 | 2.54E+00 | 2.33E-01 | no | Naaladl2 | intron_variant,non_coding_transcript_variant | MODIFIER |
| 16 | 19501462 | AX-170037391 | 1 | A | G | 0.143 | 1.11E-01 | 9.38E-02 | 2.85E+00 | 3.00E+00 | 2.33E-01 | no | | intergenic_variant | MODIFIER |
| 13 | 10272109 | AX-170414004 | 1 | G | T | 0.196 | -1.46E-01 | 1.23E-01 | 2.55E+00 | 2.61E+00 | 2.33E-01 | no | Chrm3 | intron_variant | MODIFIER |
| 4 | 77000378 | AX-170476775 | 0 | T | G | 0.333 | 1.38E-01 | 1.17E-01 | 2.76E+00 | 2.82E+00 | 2.33E-01 | no | Ptprd | intron_variant,non_coding_transcript_variant | MODIFIER |
| 9 | 106400622 | AX-97813722 | 1 | T | C | 0.25 | -6.79E-02 | 5.74E-02 | 2.71E+00 | 2.80E+00 | 2.33E-01 | no | | regulatory_region_variant | MODIFIER |
| 16 | 52224285 | AX-97856907 | 0 | T | G | 0.518 | 9.46E-02 | 7.95E-02 | 2.56E+00 | 2.62E+00 | 2.33E-01 | no | | intergenic_variant | MODIFIER |
| 16 | 72164748 | AX-97797670 | 0 | C | T | 0.491 | 9.18E-02 | 7.72E-02 | 2.57E+00 | 2.64E+00 | 2.33E-01 | no | | intergenic_variant | MODIFIER |
| 16 | 72166242 | AX-170057766 | 0 | T | C | 0.491 | 9.18E-02 | 7.72E-02 | 2.57E+00 | 2.64E+00 | 2.33E-01 | no | | intergenic_variant | MODIFIER |
| 16 | 72177293 | AX-170168379 | 0 | A | C | 0.491 | 9.18E-02 | 7.72E-02 | 2.57E+00 | 2.64E+00 | 2.33E-01 | no | | intergenic_variant | MODIFIER |
| 16 | 72188750 | AX-170155693 | 0 | A | G | 0.491 | 9.18E-02 | 7.72E-02 | 2.57E+00 | 2.64E+00 | 2.33E-01 | no | | intergenic_variant | MODIFIER |
| 16 | 72188994 | AX-170135285 | 0 | T | C | 0.491 | 9.18E-02 | 7.72E-02 | 2.57E+00 | 2.64E+00 | 2.33E-01 | no | | intergenic_variant | MODIFIER |
| 16 | 72189329 | AX-97818419 | 0 | T | C | 0.491 | 9.18E-02 | 7.72E-02 | 2.57E+00 | 2.64E+00 | 2.33E-01 | no | | intergenic_variant | MODIFIER |
| 16 | 72221253 | AX-170294395 | 0 | A | G | 0.491 | 9.18E-02 | 7.72E-02 | 2.57E+00 | 2.64E+00 | 2.33E-01 | no | | intergenic_variant | MODIFIER |
| 16 | 72222365 | AX-97820564 | 0 | T | C | 0.491 | 9.18E-02 | 7.72E-02 | 2.57E+00 | 2.64E+00 | 2.33E-01 | no | 8030451O07Rik | downstream_gene_variant | MODIFIER |
| 16 | 72557631 | AX-170477440 | 0 | T | A | 0.491 | 9.18E-02 | 7.72E-02 | 2.57E+00 | 2.64E+00 | 2.33E-01 | no | Robo1 | intron_variant | MODIFIER |
| 16 | 72563186 | AX-170528464 | 0 | G | A | 0.491 | 9.18E-02 | 7.72E-02 | 2.57E+00 | 2.64E+00 | 2.33E-01 | no | Robo1 | intron_variant | MODIFIER |
| 16 | 72607689 | AX-170157824 | 0 | T | A | 0.491 | 9.18E-02 | 7.72E-02 | 2.57E+00 | 2.64E+00 | 2.33E-01 | no | Robo1 | intron_variant | MODIFIER |
| 16 | 72615421 | AX-170253140 | 0 | A | T | 0.491 | 9.18E-02 | 7.72E-02 | 2.57E+00 | 2.64E+00 | 2.33E-01 | no | Robo1 | intron_variant | MODIFIER |
| 16 | 72631595 | AX-170057779 | 0 | A | G | 0.491 | 9.18E-02 | 7.72E-02 | 2.57E+00 | 2.64E+00 | 2.33E-01 | no | Robo1 | intron_variant | MODIFIER |
| 16 | 72641577 | AX-170057778 | 0 | A | G | 0.491 | 9.18E-02 | 7.72E-02 | 2.57E+00 | 2.64E+00 | 2.33E-01 | no | Robo1 | intron_variant | MODIFIER |
| 16 | 72690576 | AX-170416768 | 0 | G | A | 0.491 | 9.18E-02 | 7.72E-02 | 2.57E+00 | 2.64E+00 | 2.33E-01 | no | Robo1 | intron_variant | MODIFIER |
| 16 | 72886154 | AX-170249302 | 0 | C | T | 0.491 | 9.18E-02 | 7.72E-02 | 2.57E+00 | 2.64E+00 | 2.33E-01 | no | Robo1 | intron_variant | MODIFIER |
| 16 | 72909640 | AX-97832310 | 0 | A | T | 0.491 | 9.18E-02 | 7.72E-02 | 2.57E+00 | 2.64E+00 | 2.33E-01 | no | Robo1 | intron_variant | MODIFIER |
| 16 | 73025449 | AX-170181828 | 0 | A | G | 0.491 | 9.18E-02 | 7.72E-02 | 2.57E+00 | 2.64E+00 | 2.33E-01 | no | Robo1 | intron_variant | MODIFIER |
| 16 | 73026705 | AX-170296802 | 0 | G | T | 0.491 | 9.18E-02 | 7.72E-02 | 2.57E+00 | 2.64E+00 | 2.33E-01 | no | Robo1 | intron_variant | MODIFIER |
| 16 | 73036959 | AX-97808958 | 0 | A | G | 0.491 | 9.18E-02 | 7.72E-02 | 2.57E+00 | 2.64E+00 | 2.33E-01 | no | Robo1 | intron_variant | MODIFIER |
| 16 | 73095527 | AX-170348623 | 0 | T | C | 0.491 | 9.18E-02 | 7.72E-02 | 2.57E+00 | 2.64E+00 | 2.33E-01 | no | 4930500H12Rik | upstream_gene_variant | MODIFIER |
| 12 | 38056493 | AX-170517716 | 2 | T | C | 0.018 | -1.60E-01 | 1.35E-01 | 2.58E+00 | 2.75E+00 | 2.34E-01 | no | Dgkb | intron_variant | MODIFIER |
| 9 | 4531563 | AX-170435204 | 0 | C | A | 0.474 | 9.57E-02 | 7.92E-02 | 2.41E+00 | 2.50E+00 | 2.34E-01 | no | Gria4 | intron_variant | MODIFIER |
| 16 | 35603364 | AX-97810796 | 0 | A | G | 0.491 | 9.49E-02 | 7.94E-02 | 2.51E+00 | 2.56E+00 | 2.34E-01 | no | Sema5b | intron_variant | MODIFIER |
| 16 | 35662121 | AX-170519991 | 0 | G | A | 0.491 | 9.49E-02 | 7.94E-02 | 2.51E+00 | 2.56E+00 | 2.34E-01 | no | Sema5b | intron_variant | MODIFIER |
| 16 | 35664625 | AX-170242649 | 0 | A | T | 0.491 | 9.49E-02 | 7.94E-02 | 2.51E+00 | 2.56E+00 | 2.34E-01 | no | Sema5b | downstream_gene_variant | MODIFIER |
| 16 | 35680189 | AX-170150614 | 0 | G | T | 0.491 | 9.49E-02 | 7.94E-02 | 2.51E+00 | 2.56E+00 | 2.34E-01 | no | | intergenic_variant | MODIFIER |
| 16 | 35692276 | AX-170057124 | 0 | A | G | 0.491 | 9.49E-02 | 7.94E-02 | 2.51E+00 | 2.56E+00 | 2.34E-01 | no | Dirc2 | downstream_gene_variant | MODIFIER |
| 16 | 35693661 | AX-170057125 | 0 | A | G | 0.491 | 9.49E-02 | 7.94E-02 | 2.51E+00 | 2.56E+00 | 2.34E-01 | no | Dirc2 | downstream_gene_variant | MODIFIER |
| 16 | 35694403 | AX-170312543 | 0 | C | A | 0.491 | 9.49E-02 | 7.94E-02 | 2.51E+00 | 2.56E+00 | 2.34E-01 | no | Dirc2 | 3_prime_UTR_variant | MODIFIER |
| 16 | 35696757 | AX-170361263 | 0 | T | C | 0.491 | 9.49E-02 | 7.94E-02 | 2.51E+00 | 2.56E+00 | 2.34E-01 | no | Dirc2 | 3_prime_UTR_variant | MODIFIER |
| 16 | 35698132 | AX-170121796 | 0 | T | C | 0.491 | 9.49E-02 | 7.94E-02 | 2.51E+00 | 2.56E+00 | 2.34E-01 | no | Dirc2 | intron_variant | MODIFIER |
| 16 | 35791902 | AX-170345701 | 0 | T | C | 0.491 | 9.49E-02 | 7.94E-02 | 2.51E+00 | 2.56E+00 | 2.34E-01 | no | Hspbap1 | intron_variant,NMD_transcript_variant | MODIFIER |
| 16 | 35826745 | AX-170057129 | 0 | A | G | 0.491 | 9.49E-02 | 7.94E-02 | 2.51E+00 | 2.56E+00 | 2.34E-01 | no | Hspbap1 | 3_prime_UTR_variant,NMD_transcript_variant | MODIFIER |
| 16 | 35833995 | AX-97819602 | 0 | T | C | 0.491 | 9.49E-02 | 7.94E-02 | 2.51E+00 | 2.56E+00 | 2.34E-01 | no | Parp14 | 3_prime_UTR_variant | MODIFIER |
| 16 | 35856486 | AX-170012822 | 0 | G | A | 0.491 | 9.49E-02 | 7.94E-02 | 2.51E+00 | 2.56E+00 | 2.34E-01 | no | Parp14 | synonymous_variant | LOW |
| 16 | 35862787 | AX-170012824 | 0 | C | T | 0.491 | 9.49E-02 | 7.94E-02 | 2.51E+00 | 2.56E+00 | 2.34E-01 | no | Parp14 | intron_variant | MODIFIER |
| 16 | 35873076 | AX-97812034 | 0 | G | A | 0.491 | 9.49E-02 | 7.94E-02 | 2.51E+00 | 2.56E+00 | 2.34E-01 | no | Parp14 | upstream_gene_variant | MODIFIER |
| 16 | 35873119 | AX-170386635 | 0 | T | G | 0.491 | 9.49E-02 | 7.94E-02 | 2.51E+00 | 2.56E+00 | 2.34E-01 | no | Parp14 | upstream_gene_variant | MODIFIER |
| 16 | 35884434 | AX-170121289 | 0 | G | A | 0.491 | 9.49E-02 | 7.94E-02 | 2.51E+00 | 2.56E+00 | 2.34E-01 | no | | intergenic_variant | MODIFIER |
| 16 | 35884630 | AX-97851333 | 0 | G | A | 0.491 | 9.49E-02 | 7.94E-02 | 2.51E+00 | 2.56E+00 | 2.34E-01 | no | | intergenic_variant | MODIFIER |
| 16 | 35886727 | AX-170223383 | 0 | C | T | 0.491 | 9.49E-02 | 7.94E-02 | 2.51E+00 | 2.56E+00 | 2.34E-01 | no | Gm41467 | upstream_gene_variant | MODIFIER |
| 16 | 36026794 | AX-97850311 | 0 | G | A | 0.491 | 9.49E-02 | 7.94E-02 | 2.51E+00 | 2.56E+00 | 2.34E-01 | no | Kpna1 | intron_variant | MODIFIER |
| 16 | 36506777 | AX-97793755 | 0 | C | T | 0.491 | 9.49E-02 | 7.94E-02 | 2.51E+00 | 2.56E+00 | 2.34E-01 | no | Casr | intron_variant | MODIFIER |
| 16 | 36614139 | AX-170120880 | 0 | G | A | 0.491 | 9.49E-02 | 7.94E-02 | 2.51E+00 | 2.56E+00 | 2.34E-01 | no | Cd86 | intron_variant | MODIFIER |

| | | | | | | | | | | | | | | |
|---|---|---|---|---|---|---|---|---|---|---|---|---|---|---|
| 16 | 36618252 | AX-97811146 | 0 | G | T | 0.491 | 9.49E-02 | 7.94E-02 | 2.51E+00 | 2.56E+00 | 2.34E-01 | no | Cd86 | intron_variant | MODIFIER |
| 16 | 36622467 | AX-170468018 | 0 | G | T | 0.491 | 9.49E-02 | 7.94E-02 | 2.51E+00 | 2.56E+00 | 2.34E-01 | no | Cd86 | intron_variant | MODIFIER |
| 16 | 36626467 | AX-170499800 | 0 | A | G | 0.491 | 9.49E-02 | 7.94E-02 | 2.51E+00 | 2.56E+00 | 2.34E-01 | no | Cd86 | intron_variant | MODIFIER |
| 13 | 6591099 | AX-170312827 | 0 | T | A | 0.202 | -1.32E-01 | 1.10E-01 | 2.48E+00 | 2.55E+00 | 2.34E-01 | no | Pfkp | intron_variant | MODIFIER |
| 16 | 29259665 | AX-170182437 | 0 | G | A | 0.5 | 9.76E-02 | 8.21E-02 | 2.56E+00 | 2.61E+00 | 2.34E-01 | no | Atp13a5 | intron_variant | MODIFIER |
| 3 | 23624688 | AX-170162944 | 2 | T | A | 0.164 | -1.19E-01 | 1.01E-01 | 2.68E+00 | 2.77E+00 | 2.34E-01 | no | | intergenic_variant | MODIFIER |
| 5 | 49858088 | AX-170095314 | 2 | A | G | 0.491 | 1.27E-01 | 1.03E-01 | 2.32E+00 | 2.36E+00 | 2.34E-01 | no | | intergenic_variant | MODIFIER |
| 1 | 126605809 | AX-170086170 | 1 | G | A | 0.562 | 9.05E-02 | 7.57E-02 | 2.49E+00 | 2.56E+00 | 2.35E-01 | no | Nckap5 | intron_variant | MODIFIER |
| 8 | 55660290 | AX-170250642 | 0 | T | G | 0.149 | -1.40E-01 | 1.19E-01 | 2.65E+00 | 2.77E+00 | 2.35E-01 | no | Gm18320 | downstream_gene_variant | MODIFIER |
| 16 | 70334463 | AX-97831679 | 0 | A | G | 0.491 | 9.31E-02 | 7.83E-02 | 2.55E+00 | 2.61E+00 | 2.35E-01 | no | Gbe1 | intron_variant | MODIFIER |
| 7 | 101515623 | AX-170169344 | 0 | A | G | 0.386 | -1.17E-01 | 9.96E-02 | 2.82E+00 | 2.92E+00 | 2.35E-01 | no | Gm45837 | downstream_gene_variant | MODIFIER |
| 12 | 70193335 | AX-170092513 | 0 | G | A | 0.07 | 2.63E-01 | 2.21E-01 | 3.00E+00 | 3.12E+00 | 2.35E-01 | Yes | Pygl | intron_variant | MODIFIER |
| 9 | 56004396 | AX-170163643 | 1 | G | A | 0.205 | -1.21E-01 | 1.02E-01 | 3.01E+00 | 3.18E+00 | 2.35E-01 | no | 4930563M21Rik | intron_variant | MODIFIER |
| 4 | 127300686 | AX-170113727 | 1 | G | A | 0.152 | 1.40E-01 | 1.19E-01 | 2.61E+00 | 2.68E+00 | 2.35E-01 | no | Gm12948 | upstream_gene_variant | MODIFIER |
| 4 | 54194913 | AX-170267231 | 0 | A | G | 0.368 | -1.21E-01 | 1.03E-01 | 2.88E+00 | 2.99E+00 | 2.35E-01 | no | | intergenic_variant | MODIFIER |
| 4 | 54195482 | AX-170285968 | 0 | T | C | 0.368 | -1.21E-01 | 1.03E-01 | 2.88E+00 | 2.99E+00 | 2.35E-01 | no | | intergenic_variant | MODIFIER |
| 4 | 54195768 | AX-97854717 | 0 | G | A | 0.368 | -1.21E-01 | 1.03E-01 | 2.88E+00 | 2.99E+00 | 2.35E-01 | no | | intergenic_variant | MODIFIER |
| 4 | 54230055 | AX-170324436 | 0 | G | A | 0.368 | -1.21E-01 | 1.03E-01 | 2.88E+00 | 2.99E+00 | 2.35E-01 | no | | intergenic_variant | MODIFIER |
| 4 | 54235125 | AX-170423549 | 0 | A | G | 0.368 | -1.21E-01 | 1.03E-01 | 2.88E+00 | 2.99E+00 | 2.35E-01 | no | Gm12469 | downstream_gene_variant | MODIFIER |
| 4 | 54266740 | AX-170257019 | 0 | A | G | 0.368 | -1.21E-01 | 1.03E-01 | 2.88E+00 | 2.99E+00 | 2.35E-01 | no | | intergenic_variant | MODIFIER |
| 4 | 54296866 | AX-170304219 | 0 | C | G | 0.368 | -1.21E-01 | 1.03E-01 | 2.88E+00 | 2.99E+00 | 2.35E-01 | no | | intergenic_variant | MODIFIER |
| 4 | 54327407 | AX-170318933 | 0 | G | C | 0.368 | -1.21E-01 | 1.03E-01 | 2.88E+00 | 2.99E+00 | 2.35E-01 | no | | intergenic_variant | MODIFIER |
| 4 | 54327883 | AX-170512628 | 0 | T | C | 0.368 | -1.21E-01 | 1.03E-01 | 2.88E+00 | 2.99E+00 | 2.35E-01 | no | | intergenic_variant | MODIFIER |
| 4 | 54328718 | AX-170350866 | 0 | C | T | 0.368 | -1.21E-01 | 1.03E-01 | 2.88E+00 | 2.99E+00 | 2.35E-01 | no | | regulatory_region_variant | MODIFIER |
| 4 | 54328818 | AX-170466187 | 0 | A | C | 0.368 | -1.21E-01 | 1.03E-01 | 2.88E+00 | 2.99E+00 | 2.35E-01 | no | | regulatory_region_variant | MODIFIER |
| 4 | 54330045 | AX-170513582 | 0 | A | G | 0.368 | -1.21E-01 | 1.03E-01 | 2.88E+00 | 2.99E+00 | 2.35E-01 | no | | regulatory_region_variant | MODIFIER |
| 4 | 54330818 | AX-170462533 | 0 | C | G | 0.368 | -1.21E-01 | 1.03E-01 | 2.88E+00 | 2.99E+00 | 2.35E-01 | no | | regulatory_region_variant | MODIFIER |
| 4 | 54334097 | AX-170389980 | 0 | T | C | 0.368 | -1.21E-01 | 1.03E-01 | 2.88E+00 | 2.99E+00 | 2.35E-01 | no | | regulatory_region_variant | MODIFIER |
| 4 | 54341940 | AX-170386507 | 0 | G | A | 0.368 | -1.21E-01 | 1.03E-01 | 2.88E+00 | 2.99E+00 | 2.35E-01 | no | | intergenic_variant | MODIFIER |
| 4 | 54360664 | AX-97823284 | 0 | T | C | 0.368 | -1.21E-01 | 1.03E-01 | 2.88E+00 | 2.99E+00 | 2.35E-01 | no | | intergenic_variant | MODIFIER |
| 4 | 54386877 | AX-170069389 | 0 | C | T | 0.368 | -1.21E-01 | 1.03E-01 | 2.88E+00 | 2.99E+00 | 2.35E-01 | no | | intergenic_variant | MODIFIER |
| 4 | 54388112 | AX-170069388 | 0 | C | T | 0.368 | -1.21E-01 | 1.03E-01 | 2.88E+00 | 2.99E+00 | 2.35E-01 | no | | regulatory_region_variant | MODIFIER |
| 4 | 54476380 | AX-97786945 | 0 | C | G | 0.368 | -1.21E-01 | 1.03E-01 | 2.88E+00 | 2.99E+00 | 2.35E-01 | no | | intergenic_variant | MODIFIER |
| 4 | 54476909 | AX-170328346 | 0 | T | C | 0.368 | -1.21E-01 | 1.03E-01 | 2.88E+00 | 2.99E+00 | 2.35E-01 | no | | intergenic_variant | MODIFIER |
| 4 | 54477379 | AX-170458135 | 0 | G | A | 0.368 | -1.21E-01 | 1.03E-01 | 2.88E+00 | 2.99E+00 | 2.35E-01 | no | | intergenic_variant | MODIFIER |
| 4 | 54479802 | AX-170238835 | 0 | A | T | 0.368 | -1.21E-01 | 1.03E-01 | 2.88E+00 | 2.99E+00 | 2.35E-01 | no | | intergenic_variant | MODIFIER |
| 4 | 54480415 | AX-170370372 | 0 | A | G | 0.368 | -1.21E-01 | 1.03E-01 | 2.88E+00 | 2.99E+00 | 2.35E-01 | no | | intergenic_variant | MODIFIER |
| 4 | 54497176 | AX-170335858 | 0 | C | T | 0.368 | -1.21E-01 | 1.03E-01 | 2.88E+00 | 2.99E+00 | 2.35E-01 | no | | intergenic_variant | MODIFIER |
| 4 | 54524818 | AX-170300450 | 0 | G | A | 0.368 | -1.21E-01 | 1.03E-01 | 2.88E+00 | 2.99E+00 | 2.35E-01 | no | | intergenic_variant | MODIFIER |
| 4 | 54525342 | AX-170410346 | 0 | G | A | 0.368 | -1.21E-01 | 1.03E-01 | 2.88E+00 | 2.99E+00 | 2.35E-01 | no | | regulatory_region_variant | MODIFIER |
| 4 | 54527873 | AX-170308793 | 0 | A | G | 0.368 | -1.21E-01 | 1.03E-01 | 2.88E+00 | 2.99E+00 | 2.35E-01 | no | | regulatory_region_variant | MODIFIER |
| 4 | 54531881 | AX-170477044 | 0 | C | G | 0.368 | -1.21E-01 | 1.03E-01 | 2.88E+00 | 2.99E+00 | 2.35E-01 | no | | regulatory_region_variant | MODIFIER |
| 4 | 54532118 | AX-97788555 | 0 | T | G | 0.368 | -1.21E-01 | 1.03E-01 | 2.88E+00 | 2.99E+00 | 2.35E-01 | no | | regulatory_region_variant | MODIFIER |
| 4 | 54533828 | AX-170407065 | 0 | G | T | 0.368 | -1.21E-01 | 1.03E-01 | 2.88E+00 | 2.99E+00 | 2.35E-01 | no | | regulatory_region_variant | MODIFIER |
| 4 | 54563216 | AX-170022822 | 0 | A | G | 0.368 | -1.21E-01 | 1.03E-01 | 2.88E+00 | 2.99E+00 | 2.35E-01 | no | Tpt1-ps2 | upstream_gene_variant | MODIFIER |
| 4 | 54566308 | AX-170222879 | 0 | G | T | 0.368 | -1.21E-01 | 1.03E-01 | 2.88E+00 | 2.99E+00 | 2.35E-01 | no | Tpt1-ps2 | downstream_gene_variant | MODIFIER |
| 4 | 54594705 | AX-170069397 | 0 | G | C | 0.368 | -1.21E-01 | 1.03E-01 | 2.88E+00 | 2.99E+00 | 2.35E-01 | no | | intergenic_variant | MODIFIER |
| 4 | 54601163 | AX-170022824 | 0 | G | T | 0.368 | -1.21E-01 | 1.03E-01 | 2.88E+00 | 2.99E+00 | 2.35E-01 | no | | intergenic_variant | MODIFIER |
| 4 | 54609608 | AX-170022826 | 0 | G | A | 0.368 | -1.21E-01 | 1.03E-01 | 2.88E+00 | 2.99E+00 | 2.35E-01 | no | | intergenic_variant | MODIFIER |
| 4 | 54610487 | AX-170432444 | 0 | C | A | 0.368 | -1.21E-01 | 1.03E-01 | 2.88E+00 | 2.99E+00 | 2.35E-01 | no | | intergenic_variant | MODIFIER |
| 4 | 54614662 | AX-170451114 | 0 | G | C | 0.368 | -1.21E-01 | 1.03E-01 | 2.88E+00 | 2.99E+00 | 2.35E-01 | no | | intergenic_variant | MODIFIER |
| 4 | 54614881 | AX-170022823 | 0 | A | G | 0.368 | -1.21E-01 | 1.03E-01 | 2.88E+00 | 2.99E+00 | 2.35E-01 | no | | intergenic_variant | MODIFIER |
| 4 | 54636774 | AX-170426896 | 0 | C | A | 0.368 | -1.21E-01 | 1.03E-01 | 2.88E+00 | 2.99E+00 | 2.35E-01 | no | | intergenic_variant | MODIFIER |
| 4 | 54639730 | AX-97831947 | 0 | T | C | 0.368 | -1.21E-01 | 1.03E-01 | 2.88E+00 | 2.99E+00 | 2.35E-01 | no | | intergenic_variant | MODIFIER |

| | | | | | | | | | | | | | | | |
|---|---|---|---|---|---|---|---|---|---|---|---|---|---|---|---|
| 4 | 54649039 | AX-170297526 | 0 | C | G | 0.368 | -1.21E-01 | 1.03E-01 | 2.88E+00 | 2.99E+00 | 2.35E-01 | no | Gm12478 | intron_variant,non_coding_transcript_variant | MODIFIER |
| 4 | 54664352 | AX-170470668 | 0 | C | T | 0.368 | -1.21E-01 | 1.03E-01 | 2.88E+00 | 2.99E+00 | 2.35E-01 | no | Gm12478 | intron_variant,non_coding_transcript_variant | MODIFIER |
| 4 | 54689414 | AX-170161229 | 0 | T | C | 0.368 | -1.21E-01 | 1.03E-01 | 2.88E+00 | 2.99E+00 | 2.35E-01 | no | Gm12478 | non_coding_transcript_exon_variant | MODIFIER |
| 4 | 54705301 | AX-170478129 | 0 | C | T | 0.368 | -1.21E-01 | 1.03E-01 | 2.88E+00 | 2.99E+00 | 2.35E-01 | no | | intergenic_variant | MODIFIER |
| 4 | 54723259 | AX-97802344 | 0 | C | A | 0.368 | -1.21E-01 | 1.03E-01 | 2.88E+00 | 2.99E+00 | 2.35E-01 | no | | intergenic_variant | MODIFIER |
| 4 | 54723498 | AX-170490898 | 0 | G | C | 0.368 | -1.21E-01 | 1.03E-01 | 2.88E+00 | 2.99E+00 | 2.35E-01 | no | | intergenic_variant | MODIFIER |
| 4 | 54729469 | AX-170425661 | 0 | C | G | 0.368 | -1.21E-01 | 1.03E-01 | 2.88E+00 | 2.99E+00 | 2.35E-01 | no | | intergenic_variant | MODIFIER |
| 4 | 54732329 | AX-170493684 | 0 | T | C | 0.368 | -1.21E-01 | 1.03E-01 | 2.88E+00 | 2.99E+00 | 2.35E-01 | no | | intergenic_variant | MODIFIER |
| 4 | 54736054 | AX-170069398 | 0 | T | G | 0.368 | -1.21E-01 | 1.03E-01 | 2.88E+00 | 2.99E+00 | 2.35E-01 | no | | intergenic_variant | MODIFIER |
| 4 | 54736315 | AX-170069399 | 0 | G | T | 0.368 | -1.21E-01 | 1.03E-01 | 2.88E+00 | 2.99E+00 | 2.35E-01 | no | | intergenic_variant | MODIFIER |
| 4 | 54736335 | AX-170236296 | 0 | A | T | 0.368 | -1.21E-01 | 1.03E-01 | 2.88E+00 | 2.99E+00 | 2.35E-01 | no | | intergenic_variant | MODIFIER |
| 4 | 54747930 | AX-170109840 | 0 | G | A | 0.368 | -1.21E-01 | 1.03E-01 | 2.88E+00 | 2.99E+00 | 2.35E-01 | no | | intergenic_variant | MODIFIER |
| 4 | 54748150 | AX-170417630 | 0 | A | G | 0.368 | -1.21E-01 | 1.03E-01 | 2.88E+00 | 2.99E+00 | 2.35E-01 | no | | intergenic_variant | MODIFIER |
| 4 | 54753846 | AX-170415680 | 0 | T | G | 0.368 | -1.21E-01 | 1.03E-01 | 2.88E+00 | 2.99E+00 | 2.35E-01 | no | | intergenic_variant | MODIFIER |
| 4 | 54761085 | AX-170450113 | 0 | A | G | 0.368 | -1.21E-01 | 1.03E-01 | 2.88E+00 | 2.99E+00 | 2.35E-01 | no | | intergenic_variant | MODIFIER |
| 4 | 54761352 | AX-170492880 | 0 | T | C | 0.368 | -1.21E-01 | 1.03E-01 | 2.88E+00 | 2.99E+00 | 2.35E-01 | no | | intergenic_variant | MODIFIER |
| 4 | 54762589 | AX-170128839 | 0 | G | A | 0.368 | -1.21E-01 | 1.03E-01 | 2.88E+00 | 2.99E+00 | 2.35E-01 | no | | intergenic_variant | MODIFIER |
| 4 | 54763398 | AX-170353817 | 0 | G | C | 0.368 | -1.21E-01 | 1.03E-01 | 2.88E+00 | 2.99E+00 | 2.35E-01 | no | | intergenic_variant | MODIFIER |
| 4 | 54764579 | AX-170335394 | 0 | T | C | 0.368 | -1.21E-01 | 1.03E-01 | 2.88E+00 | 2.99E+00 | 2.35E-01 | no | | intergenic_variant | MODIFIER |
| 4 | 54946798 | AX-170506607 | 0 | G | A | 0.368 | -1.21E-01 | 1.03E-01 | 2.88E+00 | 2.99E+00 | 2.35E-01 | no | Zfp462 | upstream_gene_variant | MODIFIER |
| 4 | 54951001 | AX-170207843 | 0 | G | A | 0.368 | -1.21E-01 | 1.03E-01 | 2.88E+00 | 2.99E+00 | 2.35E-01 | no | Zfp462 | intron_variant | MODIFIER |
| 4 | 54958395 | AX-170483366 | 0 | C | T | 0.368 | -1.21E-01 | 1.03E-01 | 2.88E+00 | 2.99E+00 | 2.35E-01 | no | Zfp462 | intron_variant | MODIFIER |
| 4 | 54958560 | AX-170361420 | 0 | C | T | 0.368 | -1.21E-01 | 1.03E-01 | 2.88E+00 | 2.99E+00 | 2.35E-01 | no | Zfp462 | intron_variant | MODIFIER |
| 4 | 54959336 | AX-170094978 | 0 | T | C | 0.368 | -1.21E-01 | 1.03E-01 | 2.88E+00 | 2.99E+00 | 2.35E-01 | no | Zfp462 | intron_variant | MODIFIER |
| 4 | 54959443 | AX-170398297 | 0 | G | A | 0.368 | -1.21E-01 | 1.03E-01 | 2.88E+00 | 2.99E+00 | 2.35E-01 | no | Zfp462 | intron_variant | MODIFIER |
| 4 | 54986461 | AX-97783860 | 0 | A | G | 0.368 | -1.21E-01 | 1.03E-01 | 2.88E+00 | 2.99E+00 | 2.35E-01 | no | Zfp462 | intron_variant | MODIFIER |
| 4 | 54989937 | AX-170249833 | 0 | A | G | 0.368 | -1.21E-01 | 1.03E-01 | 2.88E+00 | 2.99E+00 | 2.35E-01 | no | Zfp462 | intron_variant | MODIFIER |
| 4 | 55005775 | AX-170298326 | 0 | A | G | 0.368 | -1.21E-01 | 1.03E-01 | 2.88E+00 | 2.99E+00 | 2.35E-01 | no | Zfp462 | intron_variant | MODIFIER |
| 4 | 55018970 | AX-170171800 | 0 | C | T | 0.368 | -1.21E-01 | 1.03E-01 | 2.88E+00 | 2.99E+00 | 2.35E-01 | no | Zfp462 | intron_variant | MODIFIER |
| 4 | 55023881 | AX-170022827 | 0 | C | T | 0.368 | -1.21E-01 | 1.03E-01 | 2.88E+00 | 2.99E+00 | 2.35E-01 | no | Zfp462 | intron_variant | MODIFIER |
| 4 | 55024737 | AX-170256980 | 0 | A | G | 0.368 | -1.21E-01 | 1.03E-01 | 2.88E+00 | 2.99E+00 | 2.35E-01 | no | Zfp462 | intron_variant | MODIFIER |
| 4 | 55053652 | AX-170405349 | 0 | T | C | 0.368 | -1.21E-01 | 1.03E-01 | 2.88E+00 | 2.99E+00 | 2.35E-01 | no | Zfp462 | intron_variant | MODIFIER |
| 4 | 55054249 | AX-170447155 | 0 | C | G | 0.368 | -1.21E-01 | 1.03E-01 | 2.88E+00 | 2.99E+00 | 2.35E-01 | no | Zfp462 | intron_variant | MODIFIER |
| 4 | 55056645 | AX-170132703 | 0 | T | G | 0.368 | -1.21E-01 | 1.03E-01 | 2.88E+00 | 2.99E+00 | 2.35E-01 | no | Zfp462 | intron_variant | MODIFIER |
| 4 | 55060244 | AX-170480667 | 0 | G | A | 0.368 | -1.21E-01 | 1.03E-01 | 2.88E+00 | 2.99E+00 | 2.35E-01 | no | Zfp462 | splice_region_variant,intron_variant | LOW |
| 4 | 55070004 | AX-170294498 | 0 | T | A | 0.368 | -1.21E-01 | 1.03E-01 | 2.88E+00 | 2.99E+00 | 2.35E-01 | no | Zfp462 | intron_variant | MODIFIER |
| 4 | 55083613 | AX-170069410 | 0 | G | A | 0.368 | -1.21E-01 | 1.03E-01 | 2.88E+00 | 2.99E+00 | 2.35E-01 | no | Zfp462 | downstream_gene_variant | MODIFIER |
| 4 | 55084229 | AX-170069405 | 0 | G | A | 0.368 | -1.21E-01 | 1.03E-01 | 2.88E+00 | 2.99E+00 | 2.35E-01 | no | Zfp462 | downstream_gene_variant | MODIFIER |
| 4 | 55116700 | AX-170069404 | 0 | C | T | 0.368 | -1.21E-01 | 1.03E-01 | 2.88E+00 | 2.99E+00 | 2.35E-01 | no | Gm12515 | non_coding_transcript_exon_variant | MODIFIER |
| 4 | 55123042 | AX-170292689 | 0 | A | G | 0.368 | -1.21E-01 | 1.03E-01 | 2.88E+00 | 2.99E+00 | 2.35E-01 | no | | intergenic_variant | MODIFIER |
| 4 | 55143146 | AX-170111132 | 0 | T | C | 0.368 | -1.21E-01 | 1.03E-01 | 2.88E+00 | 2.99E+00 | 2.35E-01 | no | Gm12514 | intron_variant,non_coding_transcript_variant | MODIFIER |
| 4 | 55176548 | AX-170473737 | 0 | T | C | 0.368 | -1.21E-01 | 1.03E-01 | 2.88E+00 | 2.99E+00 | 2.35E-01 | no | Gm12514 | intron_variant,non_coding_transcript_variant | MODIFIER |
| 4 | 55179204 | AX-170523243 | 0 | C | T | 0.368 | -1.21E-01 | 1.03E-01 | 2.88E+00 | 2.99E+00 | 2.35E-01 | no | Gm12514 | intron_variant,non_coding_transcript_variant | MODIFIER |
| 4 | 55194908 | AX-170212632 | 0 | A | G | 0.368 | -1.21E-01 | 1.03E-01 | 2.88E+00 | 2.99E+00 | 2.35E-01 | no | Gm12514 | intron_variant,non_coding_transcript_variant | MODIFIER |
| 4 | 55223766 | AX-97826120 | 0 | T | C | 0.368 | -1.21E-01 | 1.03E-01 | 2.88E+00 | 2.99E+00 | 2.35E-01 | no | Gm12514 | intron_variant,non_coding_transcript_variant | MODIFIER |
| 4 | 55243950 | AX-170218509 | 0 | A | G | 0.368 | -1.21E-01 | 1.03E-01 | 2.88E+00 | 2.99E+00 | 2.35E-01 | no | Gm12514 | intron_variant,non_coding_transcript_variant | MODIFIER |
| 4 | 55253093 | AX-170112511 | 0 | G | A | 0.368 | -1.21E-01 | 1.03E-01 | 2.88E+00 | 2.99E+00 | 2.35E-01 | no | Gm12508 | upstream_gene_variant | MODIFIER |
| 4 | 55275504 | AX-170496085 | 0 | C | T | 0.368 | -1.21E-01 | 1.03E-01 | 2.88E+00 | 2.99E+00 | 2.35E-01 | no | Gm12514 | intron_variant,non_coding_transcript_variant | MODIFIER |
| 4 | 55293447 | AX-170238858 | 0 | T | C | 0.368 | -1.21E-01 | 1.03E-01 | 2.88E+00 | 2.99E+00 | 2.35E-01 | no | | regulatory_region_variant | MODIFIER |
| 4 | 55326751 | AX-170522410 | 0 | C | T | 0.368 | -1.21E-01 | 1.03E-01 | 2.88E+00 | 2.99E+00 | 2.35E-01 | no | | intergenic_variant | MODIFIER |
| 4 | 55326769 | AX-170285665 | 0 | C | T | 0.368 | -1.21E-01 | 1.03E-01 | 2.88E+00 | 2.99E+00 | 2.35E-01 | no | | intergenic_variant | MODIFIER |
| 4 | 55361041 | AX-170146796 | 0 | C | G | 0.368 | -1.21E-01 | 1.03E-01 | 2.88E+00 | 2.99E+00 | 2.35E-01 | no | Rad23b | intron_variant | MODIFIER |
| 4 | 55394534 | AX-170245544 | 0 | A | G | 0.368 | -1.21E-01 | 1.03E-01 | 2.88E+00 | 2.99E+00 | 2.35E-01 | no | Rad23b | downstream_gene_variant | MODIFIER |
| 4 | 55398106 | AX-170069414 | 0 | A | G | 0.368 | -1.21E-01 | 1.03E-01 | 2.88E+00 | 2.99E+00 | 2.35E-01 | no | | regulatory_region_variant | MODIFIER |

| | | | | | | | | | | | | | | | |
|---|---|---|---|---|---|---|---|---|---|---|---|---|---|---|---|
| 4 | 55407307 | AX-170069413 | 0 | C | T | 0.368 | -1.21E-01 | 1.03E-01 | 2.88E+00 | 2.99E+00 | 2.35E-01 | no | Gm12513 | downstream_gene_variant | MODIFIER |
| 4 | 55433548 | AX-170435222 | 0 | G | C | 0.368 | -1.21E-01 | 1.03E-01 | 2.88E+00 | 2.99E+00 | 2.35E-01 | no | | intergenic_variant | MODIFIER |
| 4 | 55444241 | AX-170239763 | 0 | C | T | 0.368 | -1.21E-01 | 1.03E-01 | 2.88E+00 | 2.99E+00 | 2.35E-01 | no | | intergenic_variant | MODIFIER |
| 4 | 55450857 | AX-170069411 | 0 | A | G | 0.368 | -1.21E-01 | 1.03E-01 | 2.88E+00 | 2.99E+00 | 2.35E-01 | no | Gm12510 | non_coding_transcript_exon_variant | MODIFIER |
| 4 | 55470290 | AX-170069415 | 0 | A | C | 0.368 | -1.21E-01 | 1.03E-01 | 2.88E+00 | 2.99E+00 | 2.35E-01 | no | Gm22150 | upstream_gene_variant | MODIFIER |
| 4 | 55471054 | AX-170375695 | 0 | C | T | 0.368 | -1.21E-01 | 1.03E-01 | 2.88E+00 | 2.99E+00 | 2.35E-01 | no | Gm22150 | upstream_gene_variant | MODIFIER |
| 4 | 55513949 | AX-170246984 | 0 | A | G | 0.368 | -1.21E-01 | 1.03E-01 | 2.88E+00 | 2.99E+00 | 2.35E-01 | no | | regulatory_region_variant | MODIFIER |
| 4 | 55552343 | AX-170409490 | 0 | G | A | 0.368 | -1.21E-01 | 1.03E-01 | 2.88E+00 | 2.99E+00 | 2.35E-01 | no | | intergenic_variant | MODIFIER |
| 4 | 55552368 | AX-170477091 | 0 | T | G | 0.368 | -1.21E-01 | 1.03E-01 | 2.88E+00 | 2.99E+00 | 2.35E-01 | no | | intergenic_variant | MODIFIER |
| 4 | 55570483 | AX-170266093 | 0 | C | T | 0.368 | -1.21E-01 | 1.03E-01 | 2.88E+00 | 2.99E+00 | 2.35E-01 | no | Gm12511 | intron_variant,non_coding_transcript_variant | MODIFIER |
| 4 | 55576848 | AX-170407963 | 0 | G | A | 0.368 | -1.21E-01 | 1.03E-01 | 2.88E+00 | 2.99E+00 | 2.35E-01 | no | Gm12511 | intron_variant,non_coding_transcript_variant | MODIFIER |
| 4 | 55577554 | AX-97844636 | 0 | A | G | 0.368 | -1.21E-01 | 1.03E-01 | 2.88E+00 | 2.99E+00 | 2.35E-01 | no | Gm12511 | intron_variant,non_coding_transcript_variant | MODIFIER |
| 4 | 55585814 | AX-170475170 | 0 | T | C | 0.368 | -1.21E-01 | 1.03E-01 | 2.88E+00 | 2.99E+00 | 2.35E-01 | no | Gm12511 | intron_variant,non_coding_transcript_variant | MODIFIER |
| 4 | 55586148 | AX-170490684 | 0 | C | T | 0.368 | -1.21E-01 | 1.03E-01 | 2.88E+00 | 2.99E+00 | 2.35E-01 | no | Gm12511 | intron_variant,non_coding_transcript_variant | MODIFIER |
| 4 | 55601247 | AX-170465859 | 0 | C | T | 0.368 | -1.21E-01 | 1.03E-01 | 2.88E+00 | 2.99E+00 | 2.35E-01 | no | Gm12506 | upstream_gene_variant | MODIFIER |
| 4 | 55601322 | AX-170523535 | 0 | T | C | 0.368 | -1.21E-01 | 1.03E-01 | 2.88E+00 | 2.99E+00 | 2.35E-01 | no | Gm12506 | upstream_gene_variant | MODIFIER |
| 4 | 55601374 | AX-170116482 | 0 | T | C | 0.368 | -1.21E-01 | 1.03E-01 | 2.88E+00 | 2.99E+00 | 2.35E-01 | no | Gm12506 | upstream_gene_variant | MODIFIER |
| 4 | 55602372 | AX-170466585 | 0 | C | T | 0.368 | -1.21E-01 | 1.03E-01 | 2.88E+00 | 2.99E+00 | 2.35E-01 | no | Gm12506 | upstream_gene_variant | MODIFIER |
| 4 | 55606526 | AX-170300430 | 0 | G | T | 0.368 | -1.21E-01 | 1.03E-01 | 2.88E+00 | 2.99E+00 | 2.35E-01 | no | Gm12506 | intron_variant,non_coding_transcript_variant | MODIFIER |
| 4 | 64704208 | AX-170111460 | 1 | G | A | 0.33 | 1.40E-01 | 1.19E-01 | 2.82E+00 | 2.88E+00 | 2.35E-01 | no | | intergenic_variant | MODIFIER |
| 4 | 48659845 | AX-97816027 | 1 | T | C | 0.348 | -1.12E-01 | 9.47E-02 | 2.94E+00 | 3.06E+00 | 2.35E-01 | no | Tmeff1 | intron_variant | MODIFIER |
| 7 | 38542119 | AX-170168886 | 1 | G | T | 0.357 | 1.12E-01 | 9.47E-02 | 2.59E+00 | 2.66E+00 | 2.35E-01 | no | | intergenic_variant | MODIFIER |
| 1 | 152655671 | AX-170136755 | 0 | T | C | 0.193 | -1.45E-01 | 1.19E-01 | 3.22E+00 | 3.40E+00 | 2.36E-01 | no | Rgl1 | intron_variant | MODIFIER |
| 9 | 52804205 | AX-170032363 | 1 | G | A | 0.223 | -1.33E-01 | 1.12E-01 | 2.99E+00 | 3.11E+00 | 2.36E-01 | no | | intergenic_variant | MODIFIER |
| 8 | 91161068 | AX-170187638 | 1 | C | G | 0.188 | -1.16E-01 | 9.87E-02 | 2.69E+00 | 2.81E+00 | 2.36E-01 | no | Aktip | intron_variant | MODIFIER |
| 10 | 100001629 | AX-170389536 | 1 | A | G | 0.277 | -1.02E-01 | 8.65E-02 | 2.63E+00 | 2.70E+00 | 2.36E-01 | no | | regulatory_region_variant | MODIFIER |
| 2 | 147285938 | AX-97786808 | 1 | G | A | 0.143 | -2.26E-01 | 1.85E-01 | 3.22E+00 | 3.34E+00 | 2.36E-01 | Yes | Gm14115 | upstream_gene_variant | MODIFIER |
| 4 | 82657791 | AX-97834255 | 1 | G | T | 0.339 | 1.27E-01 | 1.08E-01 | 2.81E+00 | 2.89E+00 | 2.36E-01 | no | Nfib | intron_variant | MODIFIER |
| 5 | 52141159 | AX-170333208 | 1 | G | A | 0.491 | 1.26E-01 | 1.03E-01 | 2.31E+00 | 2.36E+00 | 2.36E-01 | no | Gm43176 | non_coding_transcript_exon_variant | MODIFIER |
| 16 | 78532014 | AX-170128504 | 1 | T | G | 0.455 | 9.04E-02 | 7.67E-02 | 2.59E+00 | 2.66E+00 | 2.36E-01 | no | | regulatory_region_variant | MODIFIER |
| 5 | 66744857 | AX-170025194 | 0 | A | T | 0.491 | 1.19E-01 | 9.98E-02 | 2.48E+00 | 2.55E+00 | 2.37E-01 | no | Limch1 | upstream_gene_variant | MODIFIER |
| 18 | 79207569 | AX-170417723 | 1 | G | A | 0.464 | 1.05E-01 | 8.68E-02 | 2.39E+00 | 2.46E+00 | 2.37E-01 | no | AC167812.1 | intron_variant,non_coding_transcript_variant | MODIFIER |
| 3 | 151251457 | AX-170533823 | 2 | G | A | 0.364 | 8.53E-02 | 7.23E-02 | 2.93E+00 | 3.04E+00 | 2.37E-01 | no | | intergenic_variant | MODIFIER |
| 15 | 66698612 | AX-97788924 | 0 | G | A | 0.333 | -9.47E-02 | 7.68E-02 | 2.28E+00 | 2.35E+00 | 2.37E-01 | no | Tg | synonymous_variant | LOW |
| 15 | 66701217 | AX-170294210 | 0 | A | G | 0.333 | -9.47E-02 | 7.68E-02 | 2.28E+00 | 2.35E+00 | 2.37E-01 | no | Tg | intron_variant | MODIFIER |
| 15 | 66718157 | AX-97840474 | 0 | G | T | 0.333 | -9.47E-02 | 7.68E-02 | 2.28E+00 | 2.35E+00 | 2.37E-01 | no | Tg | intron_variant | MODIFIER |
| 15 | 66738251 | AX-170512647 | 0 | T | C | 0.333 | -9.47E-02 | 7.68E-02 | 2.28E+00 | 2.35E+00 | 2.37E-01 | no | Tg | intron_variant | MODIFIER |
| 15 | 66741751 | AX-170216261 | 0 | T | C | 0.333 | -9.47E-02 | 7.68E-02 | 2.28E+00 | 2.35E+00 | 2.37E-01 | no | Tg | intron_variant | MODIFIER |
| 15 | 66744228 | AX-170364725 | 0 | T | C | 0.333 | -9.47E-02 | 7.68E-02 | 2.28E+00 | 2.35E+00 | 2.37E-01 | no | Tg | intron_variant | MODIFIER |
| 15 | 66772326 | AX-170087891 | 0 | T | C | 0.333 | -9.47E-02 | 7.68E-02 | 2.28E+00 | 2.35E+00 | 2.37E-01 | no | Tg | intron_variant | MODIFIER |
| 15 | 66804175 | AX-170138342 | 0 | A | G | 0.333 | -9.47E-02 | 7.68E-02 | 2.28E+00 | 2.35E+00 | 2.37E-01 | no | Tg | intron_variant | MODIFIER |
| 15 | 66823860 | AX-170423911 | 0 | T | C | 0.333 | -9.47E-02 | 7.68E-02 | 2.28E+00 | 2.35E+00 | 2.37E-01 | no | Tg | intron_variant | MODIFIER |
| 15 | 66848959 | AX-170431207 | 0 | G | A | 0.333 | -9.47E-02 | 7.68E-02 | 2.28E+00 | 2.35E+00 | 2.37E-01 | no | Tg | intron_variant | MODIFIER |
| 15 | 66862943 | AX-170183495 | 0 | C | A | 0.333 | -9.47E-02 | 7.68E-02 | 2.28E+00 | 2.35E+00 | 2.37E-01 | no | | intergenic_variant | MODIFIER |
| 15 | 66869583 | AX-170011887 | 0 | A | G | 0.333 | -9.47E-02 | 7.68E-02 | 2.28E+00 | 2.35E+00 | 2.37E-01 | no | | intergenic_variant | MODIFIER |
| 15 | 66886562 | AX-170011886 | 0 | G | A | 0.333 | -9.47E-02 | 7.68E-02 | 2.28E+00 | 2.35E+00 | 2.37E-01 | no | Ccn4 | upstream_gene_variant | MODIFIER |
| 15 | 66903288 | AX-170516791 | 0 | G | A | 0.333 | -9.47E-02 | 7.68E-02 | 2.28E+00 | 2.35E+00 | 2.37E-01 | no | Ccn4 | intron_variant | MODIFIER |
| 15 | 66904336 | AX-170011888 | 0 | C | G | 0.333 | -9.47E-02 | 7.68E-02 | 2.28E+00 | 2.35E+00 | 2.37E-01 | no | Ccn4 | intron_variant | MODIFIER |
| 15 | 66945933 | AX-97829550 | 0 | T | C | 0.333 | -9.47E-02 | 7.68E-02 | 2.28E+00 | 2.35E+00 | 2.37E-01 | no | Ndrg1 | intron_variant | MODIFIER |
| 15 | 66960621 | AX-97851779 | 0 | G | A | 0.333 | -9.47E-02 | 7.68E-02 | 2.28E+00 | 2.35E+00 | 2.37E-01 | no | Ndrg1 | intron_variant | MODIFIER |
| 15 | 66972194 | AX-170287727 | 0 | G | A | 0.333 | -9.47E-02 | 7.68E-02 | 2.28E+00 | 2.35E+00 | 2.37E-01 | no | Ndrg1 | upstream_gene_variant | MODIFIER |
| 15 | 66972511 | AX-170160812 | 0 | G | C | 0.333 | -9.47E-02 | 7.68E-02 | 2.28E+00 | 2.35E+00 | 2.37E-01 | no | Ndrg1 | upstream_gene_variant | MODIFIER |
| 15 | 67025763 | AX-170166532 | 0 | C | T | 0.333 | -9.47E-02 | 7.68E-02 | 2.28E+00 | 2.35E+00 | 2.37E-01 | no | | intergenic_variant | MODIFIER |
| 15 | 67035035 | AX-170243968 | 0 | C | T | 0.333 | -9.47E-02 | 7.68E-02 | 2.28E+00 | 2.35E+00 | 2.37E-01 | no | | intergenic_variant | MODIFIER |

| | | | | | | | | | | | | | | | |
|---|---|---|---|---|---|---|---|---|---|---|---|---|---|---|---|
| 15 | 67042524 | AX-170258240 | 0 | T | G | 0.333 | -9.47E-02 | 7.68E-02 | 2.28E+00 | 2.35E+00 | 2.37E-01 | no | Gm31342 | downstream_gene_variant | MODIFIER |
| 15 | 67048030 | AX-170494940 | 0 | T | G | 0.333 | -9.47E-02 | 7.68E-02 | 2.28E+00 | 2.35E+00 | 2.37E-01 | no | | regulatory_region_variant | MODIFIER |
| 15 | 67193743 | AX-97823056 | 0 | G | C | 0.333 | -9.47E-02 | 7.68E-02 | 2.28E+00 | 2.35E+00 | 2.37E-01 | no | | intergenic_variant | MODIFIER |
| 15 | 67301622 | AX-170087893 | 0 | G | A | 0.333 | -9.47E-02 | 7.68E-02 | 2.28E+00 | 2.35E+00 | 2.37E-01 | no | 1700012I11Rik | intron_variant,non_coding_transcript_variant | MODIFIER |
| 15 | 67324413 | AX-97854430 | 0 | G | C | 0.333 | -9.47E-02 | 7.68E-02 | 2.28E+00 | 2.35E+00 | 2.37E-01 | no | 1700012I11Rik | intron_variant,non_coding_transcript_variant | MODIFIER |
| 15 | 67333200 | AX-170011902 | 0 | C | T | 0.333 | -9.47E-02 | 7.68E-02 | 2.28E+00 | 2.35E+00 | 2.37E-01 | no | 1700012I11Rik | intron_variant,non_coding_transcript_variant | MODIFIER |
| 15 | 67352568 | AX-170011896 | 0 | G | T | 0.333 | -9.47E-02 | 7.68E-02 | 2.28E+00 | 2.35E+00 | 2.37E-01 | no | 1700012I11Rik | intron_variant,non_coding_transcript_variant | MODIFIER |
| 15 | 67354009 | AX-170011901 | 0 | G | A | 0.333 | -9.47E-02 | 7.68E-02 | 2.28E+00 | 2.35E+00 | 2.37E-01 | no | 1700012I11Rik | intron_variant,non_coding_transcript_variant | MODIFIER |
| 15 | 67356343 | AX-170472903 | 0 | G | A | 0.333 | -9.47E-02 | 7.68E-02 | 2.28E+00 | 2.35E+00 | 2.37E-01 | no | 1700012I11Rik | intron_variant,non_coding_transcript_variant | MODIFIER |
| 15 | 67357392 | AX-170446705 | 0 | T | C | 0.333 | -9.47E-02 | 7.68E-02 | 2.28E+00 | 2.35E+00 | 2.37E-01 | no | 1700012I11Rik | intron_variant,non_coding_transcript_variant | MODIFIER |
| 15 | 67357443 | AX-170470097 | 0 | A | G | 0.333 | -9.47E-02 | 7.68E-02 | 2.28E+00 | 2.35E+00 | 2.37E-01 | no | 1700012I11Rik | intron_variant,non_coding_transcript_variant | MODIFIER |
| 15 | 67357536 | AX-170437597 | 0 | G | A | 0.333 | -9.47E-02 | 7.68E-02 | 2.28E+00 | 2.35E+00 | 2.37E-01 | no | 1700012I11Rik | intron_variant,non_coding_transcript_variant | MODIFIER |
| 15 | 67357938 | AX-170371994 | 0 | A | C | 0.333 | -9.47E-02 | 7.68E-02 | 2.28E+00 | 2.35E+00 | 2.37E-01 | no | 1700012I11Rik | intron_variant,non_coding_transcript_variant | MODIFIER |
| 15 | 67358036 | AX-170444588 | 0 | G | A | 0.333 | -9.47E-02 | 7.68E-02 | 2.28E+00 | 2.35E+00 | 2.37E-01 | no | 1700012I11Rik | intron_variant,non_coding_transcript_variant | MODIFIER |
| 15 | 67373563 | AX-97855323 | 0 | G | A | 0.333 | -9.47E-02 | 7.68E-02 | 2.28E+00 | 2.35E+00 | 2.37E-01 | no | 1700012I11Rik | intron_variant,non_coding_transcript_variant | MODIFIER |
| 15 | 67373704 | AX-170011897 | 0 | C | T | 0.333 | -9.47E-02 | 7.68E-02 | 2.28E+00 | 2.35E+00 | 2.37E-01 | no | 1700012I11Rik | intron_variant,non_coding_transcript_variant | MODIFIER |
| 15 | 67388799 | AX-170227754 | 0 | G | A | 0.333 | -9.47E-02 | 7.68E-02 | 2.28E+00 | 2.35E+00 | 2.37E-01 | no | | regulatory_region_variant | MODIFIER |
| 15 | 67391672 | AX-170093289 | 0 | T | C | 0.333 | -9.47E-02 | 7.68E-02 | 2.28E+00 | 2.35E+00 | 2.37E-01 | no | | intergenic_variant | MODIFIER |
| 15 | 67403552 | AX-170011895 | 0 | C | G | 0.333 | -9.47E-02 | 7.68E-02 | 2.28E+00 | 2.35E+00 | 2.37E-01 | no | | intergenic_variant | MODIFIER |
| 15 | 67413531 | AX-170414380 | 0 | T | C | 0.333 | -9.47E-02 | 7.68E-02 | 2.28E+00 | 2.35E+00 | 2.37E-01 | no | | intergenic_variant | MODIFIER |
| 15 | 67413646 | AX-170119065 | 0 | G | A | 0.333 | -9.47E-02 | 7.68E-02 | 2.28E+00 | 2.35E+00 | 2.37E-01 | no | | intergenic_variant | MODIFIER |
| 15 | 67415155 | AX-170326369 | 0 | T | G | 0.333 | -9.47E-02 | 7.68E-02 | 2.28E+00 | 2.35E+00 | 2.37E-01 | no | | intergenic_variant | MODIFIER |
| 15 | 67425822 | AX-97835323 | 0 | C | T | 0.333 | -9.47E-02 | 7.68E-02 | 2.28E+00 | 2.35E+00 | 2.37E-01 | no | | intergenic_variant | MODIFIER |
| 15 | 67441842 | AX-170011893 | 0 | A | G | 0.333 | -9.47E-02 | 7.68E-02 | 2.28E+00 | 2.35E+00 | 2.37E-01 | no | | intergenic_variant | MODIFIER |
| 15 | 67441938 | AX-97791895 | 0 | G | A | 0.333 | -9.47E-02 | 7.68E-02 | 2.28E+00 | 2.35E+00 | 2.37E-01 | no | | intergenic_variant | MODIFIER |
| 15 | 67548272 | AX-97849912 | 0 | A | G | 0.333 | -9.47E-02 | 7.68E-02 | 2.28E+00 | 2.35E+00 | 2.37E-01 | no | | intergenic_variant | MODIFIER |
| 15 | 67554053 | AX-170097993 | 0 | G | A | 0.333 | -9.47E-02 | 7.68E-02 | 2.28E+00 | 2.35E+00 | 2.37E-01 | no | | intergenic_variant | MODIFIER |
| 15 | 67685733 | AX-97799266 | 0 | T | C | 0.333 | -9.47E-02 | 7.68E-02 | 2.28E+00 | 2.35E+00 | 2.37E-01 | no | | intergenic_variant | MODIFIER |
| 15 | 67712942 | AX-170143357 | 0 | C | T | 0.333 | -9.47E-02 | 7.68E-02 | 2.28E+00 | 2.35E+00 | 2.37E-01 | no | | intergenic_variant | MODIFIER |
| 15 | 67721439 | AX-97846903 | 0 | T | C | 0.333 | -9.47E-02 | 7.68E-02 | 2.28E+00 | 2.35E+00 | 2.37E-01 | no | | intergenic_variant | MODIFIER |
| 15 | 67730646 | AX-170345150 | 0 | T | C | 0.333 | -9.47E-02 | 7.68E-02 | 2.28E+00 | 2.35E+00 | 2.37E-01 | no | | intergenic_variant | MODIFIER |
| 15 | 67771884 | AX-97797746 | 0 | C | T | 0.333 | -9.47E-02 | 7.68E-02 | 2.28E+00 | 2.35E+00 | 2.37E-01 | no | | intergenic_variant | MODIFIER |
| 4 | 55109107 | AX-170185557 | 1 | G | A | 0.366 | -1.20E-01 | 1.03E-01 | 2.86E+00 | 2.96E+00 | 2.37E-01 | no | | intergenic_variant | MODIFIER |
| 2 | 118560352 | AX-170522341 | 1 | C | A | 0.143 | 1.95E-01 | 1.66E-01 | 2.87E+00 | 2.98E+00 | 2.37E-01 | Yes | | intergenic_variant | MODIFIER |
| 9 | 17642843 | AX-170237177 | 1 | C | T | 0.482 | 9.53E-02 | 8.12E-02 | 2.63E+00 | 2.74E+00 | 2.37E-01 | no | | intergenic_variant | MODIFIER |
| 7 | 99173487 | AX-170202480 | 0 | T | C | 0.377 | -1.24E-01 | 1.06E-01 | 2.86E+00 | 2.95E+00 | 2.37E-01 | no | Dgat2 | intron_variant | MODIFIER |
| 7 | 99181520 | AX-170271330 | 0 | C | T | 0.377 | -1.24E-01 | 1.06E-01 | 2.86E+00 | 2.95E+00 | 2.37E-01 | no | Dgat2 | intron_variant | MODIFIER |
| 7 | 99289971 | AX-97800428 | 0 | T | C | 0.377 | -1.24E-01 | 1.06E-01 | 2.86E+00 | 2.95E+00 | 2.37E-01 | no | Map6 | intron_variant | MODIFIER |
| 7 | 99302296 | AX-97831233 | 0 | G | T | 0.377 | -1.24E-01 | 1.06E-01 | 2.86E+00 | 2.95E+00 | 2.37E-01 | no | Map6 | intron_variant | MODIFIER |
| 7 | 99315461 | AX-170369761 | 0 | A | G | 0.377 | -1.24E-01 | 1.06E-01 | 2.86E+00 | 2.95E+00 | 2.37E-01 | no | Map6 | intron_variant | MODIFIER |
| 7 | 99320744 | AX-170356460 | 0 | C | G | 0.377 | -1.24E-01 | 1.06E-01 | 2.86E+00 | 2.95E+00 | 2.37E-01 | no | Map6 | intron_variant | MODIFIER |
| 7 | 99335480 | AX-170427863 | 0 | C | T | 0.377 | -1.24E-01 | 1.06E-01 | 2.86E+00 | 2.95E+00 | 2.37E-01 | no | Map6 | intron_variant | MODIFIER |
| 7 | 99385094 | AX-170427436 | 0 | A | G | 0.377 | -1.24E-01 | 1.06E-01 | 2.86E+00 | 2.95E+00 | 2.37E-01 | no | Gdpd5 | intron_variant | MODIFIER |
| 7 | 99484623 | AX-170233273 | 0 | C | T | 0.377 | -1.24E-01 | 1.06E-01 | 2.86E+00 | 2.95E+00 | 2.37E-01 | no | Rps3 | upstream_gene_variant | MODIFIER |
| 7 | 99485366 | AX-170319466 | 0 | T | G | 0.377 | -1.24E-01 | 1.06E-01 | 2.86E+00 | 2.95E+00 | 2.37E-01 | no | Rps3 | upstream_gene_variant | MODIFIER |
| 7 | 99496728 | AX-170469436 | 0 | T | C | 0.377 | -1.24E-01 | 1.06E-01 | 2.86E+00 | 2.95E+00 | 2.37E-01 | no | n-R5s156 | upstream_gene_variant | MODIFIER |
| 7 | 99499496 | AX-170172757 | 0 | T | C | 0.377 | -1.24E-01 | 1.06E-01 | 2.86E+00 | 2.95E+00 | 2.37E-01 | no | n-R5s156 | downstream_gene_variant | MODIFIER |
| 7 | 99499825 | AX-170288345 | 0 | C | T | 0.377 | -1.24E-01 | 1.06E-01 | 2.86E+00 | 2.95E+00 | 2.37E-01 | no | n-R5s156 | downstream_gene_variant | MODIFIER |
| 7 | 99521474 | AX-170512857 | 0 | C | A | 0.377 | -1.24E-01 | 1.06E-01 | 2.86E+00 | 2.95E+00 | 2.37E-01 | no | | intergenic_variant | MODIFIER |
| 7 | 99588744 | AX-170297993 | 0 | C | T | 0.377 | -1.24E-01 | 1.06E-01 | 2.86E+00 | 2.95E+00 | 2.37E-01 | no | Arrb1 | intron_variant | MODIFIER |
| 7 | 99593780 | AX-170029619 | 0 | A | T | 0.377 | -1.24E-01 | 1.06E-01 | 2.86E+00 | 2.95E+00 | 2.37E-01 | no | Arrb1 | intron_variant | MODIFIER |
| 7 | 99600717 | AX-97850938 | 0 | G | A | 0.377 | -1.24E-01 | 1.06E-01 | 2.86E+00 | 2.95E+00 | 2.37E-01 | no | Arrb1 | intron_variant | MODIFIER |
| 7 | 99705825 | AX-170522328 | 0 | G | A | 0.377 | -1.24E-01 | 1.06E-01 | 2.86E+00 | 2.95E+00 | 2.37E-01 | no | Slco2b1 | intron_variant | MODIFIER |
| 7 | 99719836 | AX-170029618 | 0 | C | T | 0.377 | -1.24E-01 | 1.06E-01 | 2.86E+00 | 2.95E+00 | 2.37E-01 | no | | intergenic_variant | MODIFIER |

| | | | | | | | | | | | | | | | |
|---|---|---|---|---|---|---|---|---|---|---|---|---|---|---|---|
| 7 | 99734242 | AX-170179996 | 0 | C | T | 0.377 | -1.24E-01 | 1.06E-01 | 2.86E+00 | 2.95E+00 | 2.37E-01 | no | Olfr520 | upstream_gene_variant | MODIFIER |
| 7 | 99800669 | AX-170168813 | 0 | A | G | 0.377 | -1.24E-01 | 1.06E-01 | 2.86E+00 | 2.95E+00 | 2.37E-01 | no | | intergenic_variant | MODIFIER |
| 7 | 99828606 | AX-97786846 | 0 | G | A | 0.377 | -1.24E-01 | 1.06E-01 | 2.86E+00 | 2.95E+00 | 2.37E-01 | no | Neu3 | upstream_gene_variant | MODIFIER |
| 7 | 99837738 | AX-170193287 | 0 | T | G | 0.377 | -1.24E-01 | 1.06E-01 | 2.86E+00 | 2.95E+00 | 2.37E-01 | no | Spcs2 | 3_prime_UTR_variant | MODIFIER |
| 7 | 99853503 | AX-170483270 | 0 | C | T | 0.377 | -1.24E-01 | 1.06E-01 | 2.86E+00 | 2.95E+00 | 2.37E-01 | no | Spcs2 | intron_variant | MODIFIER |
| 7 | 99911491 | AX-170380041 | 0 | T | C | 0.377 | -1.24E-01 | 1.06E-01 | 2.86E+00 | 2.95E+00 | 2.37E-01 | no | Xrra1 | intron_variant | MODIFIER |
| 7 | 99912190 | AX-97783395 | 0 | C | T | 0.377 | -1.24E-01 | 1.06E-01 | 2.86E+00 | 2.95E+00 | 2.37E-01 | no | Xrra1 | intron_variant | MODIFIER |
| 7 | 99990608 | AX-97802863 | 0 | A | G | 0.377 | -1.24E-01 | 1.06E-01 | 2.86E+00 | 2.95E+00 | 2.37E-01 | no | | regulatory_region_variant | MODIFIER |
| 7 | 99991099 | AX-170148578 | 0 | A | C | 0.377 | -1.24E-01 | 1.06E-01 | 2.86E+00 | 2.95E+00 | 2.37E-01 | no | | regulatory_region_variant | MODIFIER |
| 7 | 100019677 | AX-97824154 | 0 | C | T | 0.377 | -1.24E-01 | 1.06E-01 | 2.86E+00 | 2.95E+00 | 2.37E-01 | no | Chrdl2 | intron_variant | MODIFIER |
| 7 | 100024839 | AX-170174607 | 0 | C | T | 0.377 | -1.24E-01 | 1.06E-01 | 2.86E+00 | 2.95E+00 | 2.37E-01 | no | Chrdl2 | intron_variant | MODIFIER |
| 7 | 100028749 | AX-170530567 | 0 | T | C | 0.377 | -1.24E-01 | 1.06E-01 | 2.86E+00 | 2.95E+00 | 2.37E-01 | no | Chrdl2 | synonymous_variant | LOW |
| 7 | 100046386 | AX-170376370 | 0 | C | T | 0.377 | -1.24E-01 | 1.06E-01 | 2.86E+00 | 2.95E+00 | 2.37E-01 | no | | intergenic_variant | MODIFIER |
| 7 | 100049637 | AX-170507976 | 0 | G | A | 0.377 | -1.24E-01 | 1.06E-01 | 2.86E+00 | 2.95E+00 | 2.37E-01 | no | | intergenic_variant | MODIFIER |
| 7 | 100056618 | AX-170177961 | 0 | T | C | 0.377 | -1.24E-01 | 1.06E-01 | 2.86E+00 | 2.95E+00 | 2.37E-01 | no | | intergenic_variant | MODIFIER |
| 7 | 100059833 | AX-170167643 | 0 | T | C | 0.377 | -1.24E-01 | 1.06E-01 | 2.86E+00 | 2.95E+00 | 2.37E-01 | no | | intergenic_variant | MODIFIER |
| 7 | 100063095 | AX-170254270 | 0 | G | A | 0.377 | -1.24E-01 | 1.06E-01 | 2.86E+00 | 2.95E+00 | 2.37E-01 | no | | intergenic_variant | MODIFIER |
| 7 | 100067147 | AX-170336743 | 0 | C | T | 0.377 | -1.24E-01 | 1.06E-01 | 2.86E+00 | 2.95E+00 | 2.37E-01 | no | | intergenic_variant | MODIFIER |
| 7 | 100086546 | AX-170365292 | 0 | A | G | 0.377 | -1.24E-01 | 1.06E-01 | 2.86E+00 | 2.95E+00 | 2.37E-01 | no | Pold3 | intron_variant | MODIFIER |
| 7 | 100106034 | AX-170423652 | 0 | A | G | 0.377 | -1.24E-01 | 1.06E-01 | 2.86E+00 | 2.95E+00 | 2.37E-01 | no | Pold3 | intron_variant | MODIFIER |
| 7 | 100111873 | AX-170096042 | 0 | G | A | 0.377 | -1.24E-01 | 1.06E-01 | 2.86E+00 | 2.95E+00 | 2.37E-01 | no | Pold3 | intron_variant | MODIFIER |
| 7 | 100112081 | AX-170269900 | 0 | T | C | 0.377 | -1.24E-01 | 1.06E-01 | 2.86E+00 | 2.95E+00 | 2.37E-01 | no | Pold3 | intron_variant | MODIFIER |
| 7 | 100184106 | AX-170473343 | 0 | G | A | 0.377 | -1.24E-01 | 1.06E-01 | 2.86E+00 | 2.95E+00 | 2.37E-01 | no | Kcne3 | intron_variant | MODIFIER |
| 7 | 100184554 | AX-170138205 | 0 | G | A | 0.377 | -1.24E-01 | 1.06E-01 | 2.86E+00 | 2.95E+00 | 2.37E-01 | no | Kcne3 | 3_prime_UTR_variant | MODIFIER |
| 7 | 100184930 | AX-170499072 | 0 | T | C | 0.377 | -1.24E-01 | 1.06E-01 | 2.86E+00 | 2.95E+00 | 2.37E-01 | no | Kcne3 | downstream_gene_variant | MODIFIER |
| 7 | 100189730 | AX-170337410 | 0 | A | G | 0.377 | -1.24E-01 | 1.06E-01 | 2.86E+00 | 2.95E+00 | 2.37E-01 | no | Kcne3 | downstream_gene_variant | MODIFIER |
| 7 | 100205876 | AX-170393446 | 0 | A | G | 0.377 | -1.24E-01 | 1.06E-01 | 2.86E+00 | 2.95E+00 | 2.37E-01 | no | Gm34821 | intron_variant,non_coding_transcript_variant | MODIFIER |
| 7 | 100697888 | AX-170345276 | 0 | G | A | 0.377 | -1.24E-01 | 1.06E-01 | 2.86E+00 | 2.95E+00 | 2.37E-01 | no | | intergenic_variant | MODIFIER |
| 7 | 100704456 | AX-97793148 | 0 | T | C | 0.377 | -1.24E-01 | 1.06E-01 | 2.86E+00 | 2.95E+00 | 2.37E-01 | no | Fam168a | upstream_gene_variant | MODIFIER |
| 7 | 100743096 | AX-170264044 | 0 | C | G | 0.377 | -1.24E-01 | 1.06E-01 | 2.86E+00 | 2.95E+00 | 2.37E-01 | no | Fam168a | intron_variant | MODIFIER |
| 7 | 100883152 | AX-97850717 | 0 | C | A | 0.377 | -1.24E-01 | 1.06E-01 | 2.86E+00 | 2.95E+00 | 2.37E-01 | no | Arhgef17 | synonymous_variant | LOW |
| 7 | 100934731 | AX-97856246 | 0 | C | T | 0.377 | -1.24E-01 | 1.06E-01 | 2.86E+00 | 2.95E+00 | 2.37E-01 | no | P2ry6 | downstream_gene_variant | MODIFIER |
| 7 | 100934793 | AX-170513261 | 0 | G | A | 0.377 | -1.24E-01 | 1.06E-01 | 2.86E+00 | 2.95E+00 | 2.37E-01 | no | P2ry6 | downstream_gene_variant | MODIFIER |
| 7 | 101002824 | AX-170202981 | 0 | T | G | 0.377 | -1.24E-01 | 1.06E-01 | 2.86E+00 | 2.95E+00 | 2.37E-01 | no | P2ry2 | intron_variant | MODIFIER |
| 7 | 101005420 | AX-97836644 | 0 | T | C | 0.377 | -1.24E-01 | 1.06E-01 | 2.86E+00 | 2.95E+00 | 2.37E-01 | no | P2ry2 | intron_variant | MODIFIER |
| 7 | 101066108 | AX-170269483 | 0 | C | T | 0.377 | -1.24E-01 | 1.06E-01 | 2.86E+00 | 2.95E+00 | 2.37E-01 | no | Gm5735 | upstream_gene_variant | MODIFIER |
| 7 | 101066637 | AX-170460690 | 0 | G | A | 0.377 | -1.24E-01 | 1.06E-01 | 2.86E+00 | 2.95E+00 | 2.37E-01 | no | | intergenic_variant | MODIFIER |
| 7 | 101099707 | AX-170303837 | 0 | A | C | 0.377 | -1.24E-01 | 1.06E-01 | 2.86E+00 | 2.95E+00 | 2.37E-01 | no | Fchsd2 | intron_variant | MODIFIER |
| 7 | 101191225 | AX-170399481 | 0 | T | C | 0.377 | -1.24E-01 | 1.06E-01 | 2.86E+00 | 2.95E+00 | 2.37E-01 | no | Fchsd2 | intron_variant | MODIFIER |
| 7 | 101310970 | AX-170373678 | 0 | T | C | 0.377 | -1.24E-01 | 1.06E-01 | 2.86E+00 | 2.95E+00 | 2.37E-01 | no | | regulatory_region_variant | MODIFIER |
| 7 | 101356451 | AX-170145493 | 0 | T | C | 0.377 | -1.24E-01 | 1.06E-01 | 2.86E+00 | 2.95E+00 | 2.37E-01 | no | Arap1 | intron_variant | MODIFIER |
| 7 | 101362337 | AX-170415811 | 0 | G | A | 0.377 | -1.24E-01 | 1.06E-01 | 2.86E+00 | 2.95E+00 | 2.37E-01 | no | Arap1 | intron_variant | MODIFIER |
| 7 | 101365346 | AX-170078252 | 0 | C | T | 0.377 | -1.24E-01 | 1.06E-01 | 2.86E+00 | 2.95E+00 | 2.37E-01 | no | Arap1 | intron_variant | MODIFIER |
| 7 | 101370661 | AX-170354421 | 0 | T | C | 0.377 | -1.24E-01 | 1.06E-01 | 2.86E+00 | 2.95E+00 | 2.37E-01 | no | Arap1 | upstream_gene_variant | MODIFIER |
| 7 | 101375644 | AX-170289062 | 0 | A | G | 0.377 | -1.24E-01 | 1.06E-01 | 2.86E+00 | 2.95E+00 | 2.37E-01 | no | Arap1 | upstream_gene_variant | MODIFIER |
| 7 | 101410366 | AX-170321257 | 0 | T | C | 0.377 | -1.24E-01 | 1.06E-01 | 2.86E+00 | 2.95E+00 | 2.37E-01 | no | Gm45837 | upstream_gene_variant | MODIFIER |
| 7 | 101444378 | AX-170209536 | 0 | G | A | 0.377 | -1.24E-01 | 1.06E-01 | 2.86E+00 | 2.95E+00 | 2.37E-01 | no | Gm45837 | intron_variant | MODIFIER |
| 7 | 101444433 | AX-170177493 | 0 | A | G | 0.377 | -1.24E-01 | 1.06E-01 | 2.86E+00 | 2.95E+00 | 2.37E-01 | no | Gm45837 | intron_variant | MODIFIER |
| 7 | 101459744 | AX-170195866 | 0 | T | A | 0.377 | -1.24E-01 | 1.06E-01 | 2.86E+00 | 2.95E+00 | 2.37E-01 | no | Gm45837 | intron_variant | MODIFIER |
| 7 | 101463662 | AX-170172269 | 0 | T | C | 0.377 | -1.24E-01 | 1.06E-01 | 2.86E+00 | 2.95E+00 | 2.37E-01 | no | Gm45837 | intron_variant | MODIFIER |
| 7 | 101482009 | AX-170294226 | 0 | G | A | 0.377 | -1.24E-01 | 1.06E-01 | 2.86E+00 | 2.95E+00 | 2.37E-01 | no | Gm45837 | intron_variant | MODIFIER |
| 7 | 101482804 | AX-170469456 | 0 | A | G | 0.377 | -1.24E-01 | 1.06E-01 | 2.86E+00 | 2.95E+00 | 2.37E-01 | no | Gm45837 | intron_variant | MODIFIER |
| 7 | 101484577 | AX-170332373 | 0 | T | C | 0.377 | -1.24E-01 | 1.06E-01 | 2.86E+00 | 2.95E+00 | 2.37E-01 | no | Gm45837 | synonymous_variant | LOW |
| 7 | 101492946 | AX-170432248 | 0 | G | C | 0.377 | -1.24E-01 | 1.06E-01 | 2.86E+00 | 2.95E+00 | 2.37E-01 | no | Gm45837 | intron_variant | MODIFIER |

| | | | | | | | | | | | | | | | |
|---|---|---|---|---|---|---|---|---|---|---|---|---|---|---|---|
| 7 | 101505599 | AX-170432752 | 0 | G | A | 0.377 | -1.24E-01 | 1.06E-01 | 2.86E+00 | 2.95E+00 | 2.37E-01 | no | Gm45837 | intron_variant | MODIFIER |
| 7 | 101514025 | AX-97816278 | 0 | C | T | 0.377 | -1.24E-01 | 1.06E-01 | 2.86E+00 | 2.95E+00 | 2.37E-01 | no | Gm45837 | downstream_gene_variant | MODIFIER |
| 7 | 101524001 | AX-170276639 | 0 | T | G | 0.377 | -1.24E-01 | 1.06E-01 | 2.86E+00 | 2.95E+00 | 2.37E-01 | no | Gm35363 | downstream_gene_variant | MODIFIER |
| 7 | 101525317 | AX-170103496 | 0 | C | A | 0.377 | -1.24E-01 | 1.06E-01 | 2.86E+00 | 2.95E+00 | 2.37E-01 | no | Gm35363 | downstream_gene_variant | MODIFIER |
| 7 | 101528932 | AX-170280917 | 0 | T | C | 0.377 | -1.24E-01 | 1.06E-01 | 2.86E+00 | 2.95E+00 | 2.37E-01 | no | Gm35363 | intron_variant,non_coding_transcript_variant | MODIFIER |
| 7 | 101574486 | AX-97850786 | 0 | T | C | 0.377 | -1.24E-01 | 1.06E-01 | 2.86E+00 | 2.95E+00 | 2.37E-01 | no | Art2b | downstream_gene_variant | MODIFIER |
| 7 | 101637178 | AX-170402535 | 0 | A | G | 0.377 | -1.24E-01 | 1.06E-01 | 2.86E+00 | 2.95E+00 | 2.37E-01 | no | | intergenic_variant | MODIFIER |
| 1 | 171983630 | AX-170473545 | 1 | C | T | 0.214 | 1.58E-01 | 1.34E-01 | 2.57E+00 | 2.62E+00 | 2.37E-01 | Yes | | intergenic_variant | MODIFIER |
| 2 | 127226512 | AX-170473335 | 0 | G | A | 0.158 | 1.76E-01 | 1.51E-01 | 2.69E+00 | 2.82E+00 | 2.37E-01 | Yes | Snrnp200 | intron_variant | MODIFIER |
| 13 | 5821103 | AX-170510640 | 1 | G | A | 0.205 | -1.44E-01 | 1.22E-01 | 2.61E+00 | 2.67E+00 | 2.37E-01 | no | Gm26043 | downstream_gene_variant | MODIFIER |
| 13 | 5822800 | AX-170533688 | 1 | C | G | 0.205 | -1.44E-01 | 1.22E-01 | 2.61E+00 | 2.67E+00 | 2.37E-01 | no | Gm26043 | upstream_gene_variant | MODIFIER |
| 4 | 97659456 | AX-170310741 | 1 | T | C | 0.241 | 1.36E-01 | 1.15E-01 | 2.62E+00 | 2.68E+00 | 2.37E-01 | no | E130114P18Rik | intron_variant | MODIFIER |
| 1 | 120156081 | AX-97796515 | 0 | A | G | 0.579 | 9.83E-02 | 8.32E-02 | 2.55E+00 | 2.61E+00 | 2.37E-01 | no | 3110009E18Rik | intron_variant | MODIFIER |
| 3 | 10693383 | AX-170019910 | 1 | G | A | 0.402 | -7.60E-02 | 6.52E-02 | 2.72E+00 | 2.87E+00 | 2.37E-01 | no | | intergenic_variant | MODIFIER |
| 1 | 166796560 | AX-170462119 | 0 | G | A | 0.211 | 1.56E-01 | 1.34E-01 | 2.75E+00 | 2.83E+00 | 2.38E-01 | Yes | Gm37904 | downstream_gene_variant | MODIFIER |
| 18 | 74417106 | AX-170503831 | 1 | A | G | 0.17 | -1.44E-01 | 1.22E-01 | 2.66E+00 | 2.71E+00 | 2.38E-01 | no | AC147992.2 | upstream_gene_variant | MODIFIER |
| 11 | 32790215 | AX-170124130 | 0 | T | C | 0.088 | 1.93E-01 | 1.65E-01 | 2.64E+00 | 2.75E+00 | 2.38E-01 | Yes | 4930555O08Rik | intron_variant,non_coding_transcript_variant | MODIFIER |
| 2 | 119087843 | AX-170018822 | 0 | A | G | 0.14 | -2.17E-01 | 1.83E-01 | 2.99E+00 | 3.09E+00 | 2.38E-01 | Yes | Knl1 | intron_variant | MODIFIER |
| 2 | 138321019 | AX-97801908 | 0 | G | A | 0.14 | -2.17E-01 | 1.83E-01 | 2.99E+00 | 3.09E+00 | 2.38E-01 | Yes | Btbd3 | intron_variant,non_coding_transcript_variant | MODIFIER |
| 2 | 156513453 | AX-97813704 | 0 | C | T | 0.14 | -2.17E-01 | 1.83E-01 | 2.99E+00 | 3.09E+00 | 2.38E-01 | Yes | Epb41l1 | intron_variant | MODIFIER |
| 2 | 156539143 | AX-170426416 | 0 | G | A | 0.14 | -2.17E-01 | 1.83E-01 | 2.99E+00 | 3.09E+00 | 2.38E-01 | Yes | Epb41l1 | intron_variant | MODIFIER |
| 7 | 30918118 | AX-170242585 | 1 | T | C | 0.045 | 1.70E-01 | 1.42E-01 | 2.37E+00 | 2.54E+00 | 2.38E-01 | Yes | Mag | upstream_gene_variant | MODIFIER |
| 5 | 83628884 | AX-170156691 | 2 | A | T | 0.455 | 1.28E-01 | 1.09E-01 | 2.96E+00 | 3.04E+00 | 2.38E-01 | no | | intergenic_variant | MODIFIER |
| 18 | 69798057 | AX-170249118 | 0 | C | A | 0.263 | -1.13E-01 | 9.68E-02 | 2.79E+00 | 2.88E+00 | 2.38E-01 | no | | intergenic_variant | MODIFIER |
| 8 | 64357441 | AX-170266756 | 1 | G | A | 0.143 | -1.51E-01 | 1.29E-01 | 2.69E+00 | 2.81E+00 | 2.38E-01 | no | | intergenic_variant | MODIFIER |
| 4 | 109009510 | AX-97825337 | 0 | A | G | 0.228 | 1.39E-01 | 1.18E-01 | 2.57E+00 | 2.64E+00 | 2.38E-01 | no | Nrd1 | intron_variant | MODIFIER |
| 15 | 3693878 | AX-170340593 | 1 | A | G | 0.232 | -1.26E-01 | 1.08E-01 | 2.66E+00 | 2.71E+00 | 2.38E-01 | no | | intergenic_variant | MODIFIER |
| 15 | 5012298 | AX-170313035 | 1 | T | C | 0.232 | -1.26E-01 | 1.08E-01 | 2.66E+00 | 2.71E+00 | 2.38E-01 | no | C7 | intron_variant | MODIFIER |
| 15 | 23067925 | AX-170511417 | 1 | C | T | 0.232 | -1.26E-01 | 1.08E-01 | 2.66E+00 | 2.71E+00 | 2.38E-01 | no | Cdh18 | intron_variant,non_coding_transcript_variant | MODIFIER |
| 3 | 38585201 | AX-170458531 | 0 | G | T | 0.202 | 1.10E-01 | 9.34E-02 | 2.99E+00 | 3.10E+00 | 2.38E-01 | no | | intergenic_variant | MODIFIER |
| 16 | 38008448 | AX-170331146 | 1 | T | C | 0.509 | 9.41E-02 | 7.99E-02 | 2.56E+00 | 2.62E+00 | 2.38E-01 | no | Gpr156 | downstream_gene_variant | MODIFIER |
| 7 | 37837632 | AX-170077103 | 1 | T | C | 0.348 | 1.12E-01 | 9.58E-02 | 2.61E+00 | 2.69E+00 | 2.38E-01 | no | | regulatory_region_variant | MODIFIER |
| 4 | 129664484 | AX-97836621 | 1 | A | G | 0.161 | 1.39E-01 | 1.18E-01 | 2.59E+00 | 2.63E+00 | 2.39E-01 | no | Kpna6 | upstream_gene_variant | MODIFIER |
| 18 | 86640615 | AX-170458671 | 1 | T | A | 0.152 | -1.28E-01 | 1.08E-01 | 2.48E+00 | 2.56E+00 | 2.39E-01 | no | | intergenic_variant | MODIFIER |
| 16 | 70290209 | AX-97829120 | 1 | A | G | 0.482 | 9.04E-02 | 7.68E-02 | 2.56E+00 | 2.62E+00 | 2.39E-01 | no | Gm49657 | downstream_gene_variant | MODIFIER |
| 16 | 71960216 | AX-170418471 | 1 | C | T | 0.482 | 9.04E-02 | 7.68E-02 | 2.56E+00 | 2.62E+00 | 2.39E-01 | no | | intergenic_variant | MODIFIER |
| 5 | 84170252 | AX-170237531 | 1 | C | T | 0.446 | 1.25E-01 | 1.06E-01 | 3.01E+00 | 3.09E+00 | 2.39E-01 | no | Epha5 | intron_variant | MODIFIER |
| 8 | 86199674 | AX-97815573 | 0 | C | T | 0.202 | -1.42E-01 | 1.21E-01 | 2.91E+00 | 3.02E+00 | 2.39E-01 | no | | intergenic_variant | MODIFIER |
| 17 | 89389746 | AX-170402889 | 1 | T | G | 0.027 | -1.58E-01 | 1.33E-01 | 2.45E+00 | 2.60E+00 | 2.39E-01 | no | CT025565.1 | downstream_gene_variant | MODIFIER |
| 16 | 30031705 | AX-97830788 | 1 | A | G | 0.5 | 8.93E-02 | 7.62E-02 | 2.59E+00 | 2.65E+00 | 2.39E-01 | no | 9030404E10Rik | intron_variant,non_coding_transcript_variant | MODIFIER |
| 7 | 4364776 | AX-170235880 | 1 | A | C | 0.17 | 9.13E-02 | 7.79E-02 | 2.93E+00 | 3.09E+00 | 2.39E-01 | no | Gp6 | downstream_gene_variant | MODIFIER |
| 13 | 6898116 | AX-170289856 | 1 | T | A | 0.214 | -1.37E-01 | 1.17E-01 | 2.58E+00 | 2.66E+00 | 2.39E-01 | no | | intergenic_variant | MODIFIER |
| 18 | 79984081 | AX-170373028 | 1 | G | C | 0.473 | 9.91E-02 | 8.22E-02 | 2.37E+00 | 2.44E+00 | 2.39E-01 | no | | intergenic_variant | MODIFIER |
| 14 | 118999893 | AX-170249974 | 1 | G | T | 0.348 | 8.59E-02 | 7.36E-02 | 2.63E+00 | 2.74E+00 | 2.39E-01 | no | Uggt2 | intron_variant,non_coding_transcript_variant | MODIFIER |
| 18 | 76307186 | AX-170146744 | 1 | C | T | 0.393 | -1.02E-01 | 8.71E-02 | 2.72E+00 | 2.79E+00 | 2.40E-01 | no | Smad2 | downstream_gene_variant | MODIFIER |
| 5 | 83007150 | AX-170104204 | 0 | T | G | 0.43 | 1.17E-01 | 1.00E-01 | 2.77E+00 | 2.84E+00 | 2.40E-01 | no | | intergenic_variant | MODIFIER |
| 8 | 74375346 | AX-170080430 | 0 | G | A | 0.193 | -1.31E-01 | 1.12E-01 | 2.84E+00 | 2.98E+00 | 2.40E-01 | no | | intergenic_variant | MODIFIER |
| 8 | 76393147 | AX-97851704 | 0 | C | T | 0.193 | -1.31E-01 | 1.12E-01 | 2.84E+00 | 2.98E+00 | 2.40E-01 | no | Gm45740 | intron_variant,non_coding_transcript_variant | MODIFIER |
| 16 | 18355031 | AX-97808529 | 0 | C | T | 0.14 | 1.08E-01 | 9.24E-02 | 2.88E+00 | 3.04E+00 | 2.40E-01 | no | Arvcf | intron_variant | MODIFIER |
| 4 | 116163763 | AX-97789110 | 0 | A | G | 0.219 | 1.38E-01 | 1.18E-01 | 2.62E+00 | 2.68E+00 | 2.40E-01 | no | Tspan1 | synonymous_variant | LOW |
| 4 | 139855154 | AX-170070863 | 1 | A | T | 0.152 | 1.35E-01 | 1.16E-01 | 2.66E+00 | 2.71E+00 | 2.40E-01 | no | | intergenic_variant | MODIFIER |
| 4 | 140180978 | AX-170387097 | 1 | C | T | 0.152 | 1.35E-01 | 1.16E-01 | 2.66E+00 | 2.71E+00 | 2.40E-01 | no | Igsf21 | intron_variant | MODIFIER |
| 4 | 140777497 | AX-170070881 | 2 | C | T | 0.182 | 1.28E-01 | 1.09E-01 | 2.58E+00 | 2.64E+00 | 2.40E-01 | no | Padi4 | upstream_gene_variant | MODIFIER |
| 16 | 54550697 | AX-170505387 | 1 | A | G | 0.464 | -9.67E-02 | 8.22E-02 | 2.52E+00 | 2.57E+00 | 2.40E-01 | no | | intergenic_variant | MODIFIER |

| | | | | | | | | | | | | | | | |
|---|---|---|---|---|---|---|---|---|---|---|---|---|---|---|---|
| 5 | 96873136 | AX-170260209 | 0 | C | A | 0.439 | 1.22E-01 | 1.04E-01 | 2.86E+00 | 2.94E+00 | 2.40E-01 | no | | intergenic_variant | MODIFIER |
| 18 | 78300222 | AX-170162536 | 0 | A | G | 0.447 | 1.00E-01 | 8.48E-02 | 2.48E+00 | 2.55E+00 | 2.40E-01 | no | Slc14a2 | intron_variant | MODIFIER |
| 18 | 78300745 | AX-170459119 | 0 | A | G | 0.447 | 1.00E-01 | 8.48E-02 | 2.48E+00 | 2.55E+00 | 2.40E-01 | no | Slc14a2 | intron_variant | MODIFIER |
| 18 | 78320192 | AX-170510616 | 0 | C | T | 0.447 | 1.00E-01 | 8.48E-02 | 2.48E+00 | 2.55E+00 | 2.40E-01 | no | Slc14a2 | intron_variant | MODIFIER |
| 18 | 78323862 | AX-170204394 | 0 | G | A | 0.447 | 1.00E-01 | 8.48E-02 | 2.48E+00 | 2.55E+00 | 2.40E-01 | no | Slc14a2 | intron_variant | MODIFIER |
| 18 | 78324653 | AX-170217572 | 0 | A | G | 0.447 | 1.00E-01 | 8.48E-02 | 2.48E+00 | 2.55E+00 | 2.40E-01 | no | Slc14a2 | intron_variant | MODIFIER |
| 18 | 78325367 | AX-97802305 | 0 | T | C | 0.447 | 1.00E-01 | 8.48E-02 | 2.48E+00 | 2.55E+00 | 2.40E-01 | no | Slc14a2 | intron_variant | MODIFIER |
| 18 | 78330746 | AX-170313407 | 0 | T | G | 0.447 | 1.00E-01 | 8.48E-02 | 2.48E+00 | 2.55E+00 | 2.40E-01 | no | Slc14a2 | intron_variant | MODIFIER |
| 18 | 78349098 | AX-170375324 | 0 | C | T | 0.447 | 1.00E-01 | 8.48E-02 | 2.48E+00 | 2.55E+00 | 2.40E-01 | no | Gm6133 | upstream_gene_variant | MODIFIER |
| 18 | 78363409 | AX-170061407 | 0 | A | G | 0.447 | 1.00E-01 | 8.48E-02 | 2.48E+00 | 2.55E+00 | 2.40E-01 | no | Slc14a2 | intron_variant | MODIFIER |
| 18 | 78363909 | AX-170257199 | 0 | C | T | 0.447 | 1.00E-01 | 8.48E-02 | 2.48E+00 | 2.55E+00 | 2.40E-01 | no | Slc14a2 | intron_variant | MODIFIER |
| 16 | 38025948 | AX-170419371 | 0 | T | C | 0.482 | 9.52E-02 | 8.03E-02 | 2.45E+00 | 2.49E+00 | 2.41E-01 | no | | intergenic_variant | MODIFIER |
| 11 | 29669151 | AX-170304765 | 0 | C | T | 0.079 | 1.91E-01 | 1.65E-01 | 2.73E+00 | 2.85E+00 | 2.41E-01 | Yes | | intergenic_variant | MODIFIER |
| 16 | 45223188 | AX-170411207 | 2 | A | G | 0.464 | -9.31E-02 | 7.93E-02 | 2.52E+00 | 2.58E+00 | 2.41E-01 | no | Btla | upstream_gene_variant | MODIFIER |
| 7 | 36386929 | AX-97854518 | 0 | A | G | 0.316 | 1.00E-01 | 8.59E-02 | 2.59E+00 | 2.68E+00 | 2.41E-01 | no | | regulatory_region_variant | MODIFIER |
| 13 | 17624943 | AX-97826959 | 1 | G | T | 0.179 | -1.50E-01 | 1.26E-01 | 2.42E+00 | 2.47E+00 | 2.41E-01 | no | Sugct | intron_variant | MODIFIER |
| 4 | 64059196 | AX-97797294 | 0 | T | C | 0.333 | 1.34E-01 | 1.15E-01 | 2.82E+00 | 2.88E+00 | 2.41E-01 | no | 8030451A03Rik | intron_variant,non_coding_transcript_variant | MODIFIER |
| 4 | 54542008 | AX-170515318 | 0 | T | C | 0.36 | -1.19E-01 | 1.02E-01 | 2.89E+00 | 2.99E+00 | 2.41E-01 | no | | intergenic_variant | MODIFIER |
| 4 | 55129059 | AX-170069407 | 0 | T | C | 0.36 | -1.19E-01 | 1.02E-01 | 2.89E+00 | 2.99E+00 | 2.41E-01 | no | Gm12514 | downstream_gene_variant | MODIFIER |
| 4 | 77013991 | AX-170211990 | 0 | A | C | 0.325 | 1.41E-01 | 1.21E-01 | 2.84E+00 | 2.90E+00 | 2.41E-01 | no | Ptprd | intron_variant,non_coding_transcript_variant | MODIFIER |
| 9 | 94284292 | AX-170508074 | 0 | G | A | 0.14 | -1.38E-01 | 1.19E-01 | 2.60E+00 | 2.68E+00 | 2.41E-01 | no | Gm24200 | upstream_gene_variant | MODIFIER |
| 8 | 17319460 | AX-97782271 | 0 | G | T | 0.228 | -8.89E-02 | 7.61E-02 | 2.57E+00 | 2.66E+00 | 2.41E-01 | no | Csmd1 | intron_variant | MODIFIER |
| 8 | 61204859 | AX-170446299 | 1 | G | A | 0.143 | -1.78E-01 | 1.50E-01 | 2.46E+00 | 2.51E+00 | 2.41E-01 | Yes | | intergenic_variant | MODIFIER |
| 14 | 106667466 | AX-168419956 | 0 | C | T | 0.044 | -1.87E-01 | 1.61E-01 | 2.84E+00 | 3.00E+00 | 2.42E-01 | Yes | | intergenic_variant | MODIFIER |
| 4 | 127158422 | AX-170023902 | 0 | C | G | 0.184 | 1.29E-01 | 1.10E-01 | 2.56E+00 | 2.64E+00 | 2.42E-01 | no | | intergenic_variant | MODIFIER |
| 5 | 49387213 | AX-97808946 | 1 | A | C | 0.491 | 1.29E-01 | 1.05E-01 | 2.28E+00 | 2.32E+00 | 2.42E-01 | no | Kcnip4 | intron_variant | MODIFIER |
| 3 | 31983911 | AX-170263909 | 0 | T | C | 0.158 | -1.06E-01 | 9.12E-02 | 2.81E+00 | 2.93E+00 | 2.42E-01 | no | Kcnmb2 | intron_variant | MODIFIER |
| 16 | 72615706 | AX-170294135 | 1 | T | G | 0.491 | 9.01E-02 | 7.71E-02 | 2.56E+00 | 2.63E+00 | 2.42E-01 | no | Robo1 | intron_variant | MODIFIER |
| 18 | 60356216 | AX-170291803 | 1 | C | T | 0.411 | 1.11E-01 | 9.33E-02 | 2.43E+00 | 2.48E+00 | 2.42E-01 | no | AC132320.1 | upstream_gene_variant | MODIFIER |
| 13 | 19631716 | AX-170328163 | 1 | C | T | 0.17 | -1.39E-01 | 1.18E-01 | 2.48E+00 | 2.56E+00 | 2.42E-01 | no | Sfrp4 | intron_variant | MODIFIER |
| 16 | 77874173 | AX-170307033 | 0 | A | G | 0.465 | 9.09E-02 | 7.76E-02 | 2.52E+00 | 2.59E+00 | 2.42E-01 | no | | intergenic_variant | MODIFIER |
| 1 | 175549598 | AX-170114130 | 2 | T | C | 0.245 | 1.29E-01 | 1.10E-01 | 2.52E+00 | 2.59E+00 | 2.42E-01 | no | | intergenic_variant | MODIFIER |
| 4 | 78604587 | AX-170319388 | 1 | A | C | 0.312 | 1.36E-01 | 1.17E-01 | 2.84E+00 | 2.90E+00 | 2.42E-01 | no | | intergenic_variant | MODIFIER |
| 9 | 68502483 | AX-97843136 | 0 | G | C | 0.193 | -1.46E-01 | 1.26E-01 | 2.79E+00 | 2.86E+00 | 2.43E-01 | no | 4930502A04Rik | intron_variant,non_coding_transcript_variant | MODIFIER |
| 9 | 7235684 | AX-97828323 | 1 | A | T | 0.455 | 9.32E-02 | 7.89E-02 | 2.44E+00 | 2.53E+00 | 2.43E-01 | no | Gm47741 | downstream_gene_variant | MODIFIER |
| 16 | 51224932 | AX-170370405 | 0 | G | C | 0.518 | 1.00E-01 | 8.52E-02 | 2.47E+00 | 2.50E+00 | 2.43E-01 | no | | intergenic_variant | MODIFIER |
| 16 | 52871724 | AX-97849391 | 2 | T | G | 0.5 | 9.37E-02 | 8.03E-02 | 2.55E+00 | 2.61E+00 | 2.43E-01 | no | 4930405D01Rik | intron_variant,non_coding_transcript_variant | MODIFIER |
| 16 | 37861253 | AX-170438924 | 1 | T | A | 0.473 | 9.31E-02 | 8.00E-02 | 2.59E+00 | 2.65E+00 | 2.43E-01 | no | Gm36028 | upstream_gene_variant | MODIFIER |
| 1 | 50339898 | AX-170086466 | 0 | T | C | 0.114 | 2.06E-01 | 1.78E-01 | 2.85E+00 | 2.99E+00 | 2.43E-01 | Yes | | intergenic_variant | MODIFIER |
| 12 | 17654388 | AX-170487025 | 1 | G | A | 0.045 | -2.04E-01 | 1.76E-01 | 2.68E+00 | 2.78E+00 | 2.43E-01 | Yes | | intergenic_variant | MODIFIER |
| 5 | 89781558 | AX-170220458 | 2 | T | A | 0.436 | 1.19E-01 | 1.02E-01 | 2.94E+00 | 3.03E+00 | 2.43E-01 | no | Adamts3 | intron_variant | MODIFIER |
| 16 | 72641869 | AX-170468748 | 2 | G | T | 0.491 | 8.97E-02 | 7.70E-02 | 2.56E+00 | 2.64E+00 | 2.43E-01 | no | Robo1 | intron_variant | MODIFIER |
| 9 | 73139562 | AX-170032698 | 0 | C | T | 0.211 | -1.48E-01 | 1.28E-01 | 2.70E+00 | 2.76E+00 | 2.43E-01 | no | Gm38215 | downstream_gene_variant | MODIFIER |
| 5 | 49424629 | AX-170514735 | 1 | A | G | 0.482 | 1.27E-01 | 1.06E-01 | 2.38E+00 | 2.41E+00 | 2.44E-01 | no | Kcnip4 | intron_variant | MODIFIER |
| 5 | 48675594 | AX-170434659 | 0 | C | T | 0.491 | 1.23E-01 | 1.01E-01 | 2.30E+00 | 2.35E+00 | 2.44E-01 | no | Kcnip4 | intron_variant | MODIFIER |
| 5 | 50087965 | AX-170362512 | 0 | C | T | 0.491 | 1.23E-01 | 1.01E-01 | 2.30E+00 | 2.35E+00 | 2.44E-01 | no | | intergenic_variant | MODIFIER |
| 10 | 14746376 | AX-170044192 | 0 | T | C | 0.202 | -1.19E-01 | 1.03E-01 | 2.75E+00 | 2.85E+00 | 2.44E-01 | no | Gm8355 | non_coding_transcript_exon_variant | MODIFIER |
| 18 | 59561082 | AX-170178188 | 0 | T | G | 0.395 | 1.04E-01 | 8.78E-02 | 2.41E+00 | 2.48E+00 | 2.44E-01 | no | | intergenic_variant | MODIFIER |
| 6 | 135433444 | AX-97800619 | 1 | A | G | 0.554 | 1.03E-01 | 8.70E-02 | 2.43E+00 | 2.48E+00 | 2.44E-01 | no | | intergenic_variant | MODIFIER |
| 5 | 104806578 | AX-170127584 | 0 | C | G | 0.351 | 1.15E-01 | 9.91E-02 | 2.57E+00 | 2.72E+00 | 2.44E-01 | no | | intergenic_variant | MODIFIER |
| 7 | 100112189 | AX-170078223 | 0 | C | T | 0.377 | -1.20E-01 | 1.04E-01 | 2.81E+00 | 2.90E+00 | 2.44E-01 | no | Pold3 | intron_variant | MODIFIER |
| 5 | 50883512 | AX-170024922 | 0 | C | G | 0.509 | 1.23E-01 | 1.03E-01 | 2.38E+00 | 2.42E+00 | 2.44E-01 | no | | intergenic_variant | MODIFIER |
| 6 | 136025526 | AX-97815127 | 0 | C | T | 0.588 | 1.01E-01 | 8.42E-02 | 2.37E+00 | 2.42E+00 | 2.44E-01 | no | Grin2b | intron_variant | MODIFIER |
| 20 | 151670995 | AX-170344428 | 0 | A | T | 0.035 | -1.40E-01 | 1.22E-01 | 2.74E+00 | 2.93E+00 | 2.44E-01 | no | | intergenic_variant | MODIFIER |

| | | | | | | | | | | | | | | | |
|---|---|---|---|---|---|---|---|---|---|---|---|---|---|---|---|
| 2 | 165172188 | AX-170188241 | 1 | C | T | 0.17 | 1.93E-01 | 1.67E-01 | 2.64E+00 | 2.75E+00 | 2.44E-01 | Yes | Cdh22 | intron_variant | MODIFIER |
| 18 | 74592616 | AX-170461326 | 1 | A | G | 0.42 | 1.01E-01 | 8.69E-02 | 2.54E+00 | 2.60E+00 | 2.44E-01 | no | Myo5b | intron_variant | MODIFIER |
| 13 | 108449675 | AX-170507927 | 0 | G | A | 0.044 | 1.88E-01 | 1.63E-01 | 2.89E+00 | 3.08E+00 | 2.45E-01 | Yes | Pde4d | upstream_gene_variant | MODIFIER |
| 5 | 104813915 | AX-170089575 | 0 | T | G | 0.456 | 1.29E-01 | 1.11E-01 | 2.61E+00 | 2.67E+00 | 2.45E-01 | no | Zfp951 | downstream_gene_variant | MODIFIER |
| 5 | 108653923 | AX-170329783 | 0 | T | C | 0.456 | 1.29E-01 | 1.11E-01 | 2.61E+00 | 2.67E+00 | 2.45E-01 | no | Dgkq | intron_variant | MODIFIER |
| 1 | 166718607 | AX-170086677 | 0 | A | G | 0.219 | 1.64E-01 | 1.42E-01 | 2.74E+00 | 2.79E+00 | 2.45E-01 | Yes | | intergenic_variant | MODIFIER |
| 1 | 166720232 | AX-170495933 | 0 | T | C | 0.219 | 1.64E-01 | 1.42E-01 | 2.74E+00 | 2.79E+00 | 2.45E-01 | Yes | | intergenic_variant | MODIFIER |
| 1 | 166731672 | AX-97782315 | 0 | A | T | 0.219 | 1.64E-01 | 1.42E-01 | 2.74E+00 | 2.79E+00 | 2.45E-01 | Yes | | intergenic_variant | MODIFIER |
| 1 | 166758300 | AX-170284342 | 0 | G | A | 0.219 | 1.64E-01 | 1.42E-01 | 2.74E+00 | 2.79E+00 | 2.45E-01 | Yes | Gm4954 | upstream_gene_variant | MODIFIER |
| 1 | 166760862 | AX-170043446 | 0 | C | T | 0.219 | 1.64E-01 | 1.42E-01 | 2.74E+00 | 2.79E+00 | 2.45E-01 | Yes | Gm4954 | upstream_gene_variant | MODIFIER |
| 1 | 166784017 | AX-170405629 | 0 | G | T | 0.219 | 1.64E-01 | 1.42E-01 | 2.74E+00 | 2.79E+00 | 2.45E-01 | Yes | | intergenic_variant | MODIFIER |
| 1 | 166799244 | AX-170307116 | 0 | G | A | 0.219 | 1.64E-01 | 1.42E-01 | 2.74E+00 | 2.79E+00 | 2.45E-01 | Yes | Gm37904 | upstream_gene_variant | MODIFIER |
| 1 | 166815837 | AX-97814084 | 0 | A | G | 0.219 | 1.64E-01 | 1.42E-01 | 2.74E+00 | 2.79E+00 | 2.45E-01 | Yes | | intergenic_variant | MODIFIER |
| 1 | 166844412 | AX-170424845 | 0 | A | G | 0.219 | 1.64E-01 | 1.42E-01 | 2.74E+00 | 2.79E+00 | 2.45E-01 | Yes | | intergenic_variant | MODIFIER |
| 1 | 166844702 | AX-97811975 | 0 | C | T | 0.219 | 1.64E-01 | 1.42E-01 | 2.74E+00 | 2.79E+00 | 2.45E-01 | Yes | | intergenic_variant | MODIFIER |
| 1 | 166851464 | AX-170429779 | 0 | T | C | 0.219 | 1.64E-01 | 1.42E-01 | 2.74E+00 | 2.79E+00 | 2.45E-01 | Yes | | intergenic_variant | MODIFIER |
| 1 | 166856844 | AX-170406490 | 0 | T | A | 0.219 | 1.64E-01 | 1.42E-01 | 2.74E+00 | 2.79E+00 | 2.45E-01 | Yes | | intergenic_variant | MODIFIER |
| 1 | 166881762 | AX-97811483 | 0 | A | G | 0.219 | 1.64E-01 | 1.42E-01 | 2.74E+00 | 2.79E+00 | 2.45E-01 | Yes | | intergenic_variant | MODIFIER |
| 1 | 166888560 | AX-170336307 | 0 | C | T | 0.219 | 1.64E-01 | 1.42E-01 | 2.74E+00 | 2.79E+00 | 2.45E-01 | Yes | | intergenic_variant | MODIFIER |
| 1 | 166901861 | AX-170303283 | 0 | G | A | 0.219 | 1.64E-01 | 1.42E-01 | 2.74E+00 | 2.79E+00 | 2.45E-01 | Yes | | intergenic_variant | MODIFIER |
| 1 | 166903449 | AX-170481702 | 0 | G | A | 0.219 | 1.64E-01 | 1.42E-01 | 2.74E+00 | 2.79E+00 | 2.45E-01 | Yes | | intergenic_variant | MODIFIER |
| 1 | 166908302 | AX-170091781 | 0 | C | T | 0.219 | 1.64E-01 | 1.42E-01 | 2.74E+00 | 2.79E+00 | 2.45E-01 | Yes | | intergenic_variant | MODIFIER |
| 1 | 166943679 | AX-170166909 | 0 | C | T | 0.219 | 1.64E-01 | 1.42E-01 | 2.74E+00 | 2.79E+00 | 2.45E-01 | Yes | Gm6286 | downstream_gene_variant | MODIFIER |
| 1 | 167025321 | AX-170002235 | 0 | A | G | 0.219 | 1.64E-01 | 1.42E-01 | 2.74E+00 | 2.79E+00 | 2.45E-01 | Yes | Fam78b | intron_variant,NMD_transcript_variant | MODIFIER |
| 1 | 167089912 | AX-97812911 | 0 | T | C | 0.219 | 1.64E-01 | 1.42E-01 | 2.74E+00 | 2.79E+00 | 2.45E-01 | Yes | Fam78b | downstream_gene_variant | MODIFIER |
| 1 | 167097423 | AX-97821149 | 0 | G | A | 0.219 | 1.64E-01 | 1.42E-01 | 2.74E+00 | 2.79E+00 | 2.45E-01 | Yes | | intergenic_variant | MODIFIER |
| 1 | 167098200 | AX-170207923 | 0 | T | G | 0.219 | 1.64E-01 | 1.42E-01 | 2.74E+00 | 2.79E+00 | 2.45E-01 | Yes | | intergenic_variant | MODIFIER |
| 1 | 167098229 | AX-170308842 | 0 | C | T | 0.219 | 1.64E-01 | 1.42E-01 | 2.74E+00 | 2.79E+00 | 2.45E-01 | Yes | | intergenic_variant | MODIFIER |
| 1 | 167100204 | AX-170451659 | 0 | G | A | 0.219 | 1.64E-01 | 1.42E-01 | 2.74E+00 | 2.79E+00 | 2.45E-01 | Yes | | intergenic_variant | MODIFIER |
| 1 | 167101182 | AX-170310702 | 0 | G | T | 0.219 | 1.64E-01 | 1.42E-01 | 2.74E+00 | 2.79E+00 | 2.45E-01 | Yes | | intergenic_variant | MODIFIER |
| 1 | 167101572 | AX-170503911 | 0 | A | G | 0.219 | 1.64E-01 | 1.42E-01 | 2.74E+00 | 2.79E+00 | 2.45E-01 | Yes | | intergenic_variant | MODIFIER |
| 1 | 167120735 | AX-170175568 | 0 | T | C | 0.219 | 1.64E-01 | 1.42E-01 | 2.74E+00 | 2.79E+00 | 2.45E-01 | Yes | | intergenic_variant | MODIFIER |
| 1 | 167148678 | AX-97833873 | 0 | C | A | 0.219 | 1.64E-01 | 1.42E-01 | 2.74E+00 | 2.79E+00 | 2.45E-01 | Yes | | intergenic_variant | MODIFIER |
| 1 | 167229370 | AX-170276621 | 0 | C | T | 0.219 | 1.64E-01 | 1.42E-01 | 2.74E+00 | 2.79E+00 | 2.45E-01 | Yes | Uck2 | intron_variant | MODIFIER |
| 1 | 167287392 | AX-170247552 | 0 | G | T | 0.219 | 1.64E-01 | 1.42E-01 | 2.74E+00 | 2.79E+00 | 2.45E-01 | Yes | Uck2 | upstream_gene_variant | MODIFIER |
| 1 | 167293937 | AX-97850179 | 0 | T | G | 0.219 | 1.64E-01 | 1.42E-01 | 2.74E+00 | 2.79E+00 | 2.45E-01 | Yes | Mir6347 | downstream_gene_variant | MODIFIER |
| 1 | 167434191 | AX-170489343 | 0 | A | G | 0.219 | 1.64E-01 | 1.42E-01 | 2.74E+00 | 2.79E+00 | 2.45E-01 | Yes | | intergenic_variant | MODIFIER |
| 1 | 167461767 | AX-168408468 | 0 | T | C | 0.219 | 1.64E-01 | 1.42E-01 | 2.74E+00 | 2.79E+00 | 2.45E-01 | Yes | Lrrc52 | intron_variant | MODIFIER |
| 1 | 167463627 | AX-170376355 | 0 | G | A | 0.219 | 1.64E-01 | 1.42E-01 | 2.74E+00 | 2.79E+00 | 2.45E-01 | Yes | Lrrc52 | intron_variant | MODIFIER |
| 1 | 167466155 | AX-170189747 | 0 | A | G | 0.219 | 1.64E-01 | 1.42E-01 | 2.74E+00 | 2.79E+00 | 2.45E-01 | Yes | Lrrc52 | missense_variant | MODERATE |
| 1 | 167503808 | AX-170255537 | 0 | G | A | 0.219 | 1.64E-01 | 1.42E-01 | 2.74E+00 | 2.79E+00 | 2.45E-01 | Yes | | intergenic_variant | MODIFIER |
| 1 | 167509126 | AX-97810383 | 0 | A | G | 0.219 | 1.64E-01 | 1.42E-01 | 2.74E+00 | 2.79E+00 | 2.45E-01 | Yes | | intergenic_variant | MODIFIER |
| 1 | 167509808 | AX-170285357 | 0 | T | C | 0.219 | 1.64E-01 | 1.42E-01 | 2.74E+00 | 2.79E+00 | 2.45E-01 | Yes | | intergenic_variant | MODIFIER |
| 1 | 167512126 | AX-170292132 | 0 | T | C | 0.219 | 1.64E-01 | 1.42E-01 | 2.74E+00 | 2.79E+00 | 2.45E-01 | Yes | | intergenic_variant | MODIFIER |
| 1 | 167518690 | AX-170253401 | 0 | T | C | 0.219 | 1.64E-01 | 1.42E-01 | 2.74E+00 | 2.79E+00 | 2.45E-01 | Yes | | intergenic_variant | MODIFIER |
| 1 | 167518830 | AX-170528812 | 0 | C | T | 0.219 | 1.64E-01 | 1.42E-01 | 2.74E+00 | 2.79E+00 | 2.45E-01 | Yes | | intergenic_variant | MODIFIER |
| 1 | 167519374 | AX-170414311 | 0 | T | A | 0.219 | 1.64E-01 | 1.42E-01 | 2.74E+00 | 2.79E+00 | 2.45E-01 | Yes | | intergenic_variant | MODIFIER |
| 1 | 167519635 | AX-170496180 | 0 | C | A | 0.219 | 1.64E-01 | 1.42E-01 | 2.74E+00 | 2.79E+00 | 2.45E-01 | Yes | | intergenic_variant | MODIFIER |
| 1 | 167523139 | AX-170002240 | 0 | A | T | 0.219 | 1.64E-01 | 1.42E-01 | 2.74E+00 | 2.79E+00 | 2.45E-01 | Yes | | intergenic_variant | MODIFIER |
| 1 | 167524305 | AX-97811667 | 0 | C | T | 0.219 | 1.64E-01 | 1.42E-01 | 2.74E+00 | 2.79E+00 | 2.45E-01 | Yes | | intergenic_variant | MODIFIER |
| 1 | 167531938 | AX-170002245 | 0 | A | G | 0.219 | 1.64E-01 | 1.42E-01 | 2.74E+00 | 2.79E+00 | 2.45E-01 | Yes | | intergenic_variant | MODIFIER |
| 1 | 167553209 | AX-170284131 | 0 | T | G | 0.219 | 1.64E-01 | 1.42E-01 | 2.74E+00 | 2.79E+00 | 2.45E-01 | Yes | | intergenic_variant | MODIFIER |
| 1 | 167553763 | AX-170002242 | 0 | A | G | 0.219 | 1.64E-01 | 1.42E-01 | 2.74E+00 | 2.79E+00 | 2.45E-01 | Yes | | regulatory_region_variant | MODIFIER |
| 1 | 167555279 | AX-97787524 | 0 | A | C | 0.219 | 1.64E-01 | 1.42E-01 | 2.74E+00 | 2.79E+00 | 2.45E-01 | Yes | | intergenic_variant | MODIFIER |

| | | | | | | | | | | | | | | | |
|---|---|---|---|---|---|---|---|---|---|---|---|---|---|---|---|
| 1 | 167560015 | AX-170123903 | 0 | A | C | 0.219 | 1.64E-01 | 1.42E-01 | 2.74E+00 | 2.79E+00 | 2.45E-01 | Yes | | intergenic_variant | MODIFIER |
| 1 | 168101879 | AX-168418817 | 0 | T | A | 0.219 | 1.64E-01 | 1.42E-01 | 2.74E+00 | 2.79E+00 | 2.45E-01 | Yes | Gm20711 | intron_variant,non_coding_transcript_variant | MODIFIER |
| 1 | 168133507 | AX-170133282 | 0 | G | A | 0.219 | 1.64E-01 | 1.42E-01 | 2.74E+00 | 2.79E+00 | 2.45E-01 | Yes | Pbx1 | intron_variant | MODIFIER |
| 1 | 168159865 | AX-170343654 | 0 | G | A | 0.219 | 1.64E-01 | 1.42E-01 | 2.74E+00 | 2.79E+00 | 2.45E-01 | Yes | Pbx1 | intron_variant | MODIFIER |
| 1 | 168162610 | AX-170091783 | 0 | A | G | 0.219 | 1.64E-01 | 1.42E-01 | 2.74E+00 | 2.79E+00 | 2.45E-01 | Yes | Pbx1 | intron_variant | MODIFIER |
| 1 | 168165690 | AX-170172614 | 0 | T | A | 0.219 | 1.64E-01 | 1.42E-01 | 2.74E+00 | 2.79E+00 | 2.45E-01 | Yes | Pbx1 | intron_variant | MODIFIER |
| 1 | 168174447 | AX-170375625 | 0 | G | A | 0.219 | 1.64E-01 | 1.42E-01 | 2.74E+00 | 2.79E+00 | 2.45E-01 | Yes | Pbx1 | intron_variant | MODIFIER |
| 1 | 168181150 | AX-170505521 | 0 | T | C | 0.219 | 1.64E-01 | 1.42E-01 | 2.74E+00 | 2.79E+00 | 2.45E-01 | Yes | Pbx1 | intron_variant | MODIFIER |
| 1 | 168187409 | AX-170491526 | 0 | C | T | 0.219 | 1.64E-01 | 1.42E-01 | 2.74E+00 | 2.79E+00 | 2.45E-01 | Yes | Pbx1 | intron_variant | MODIFIER |
| 1 | 168189503 | AX-170043478 | 0 | A | T | 0.219 | 1.64E-01 | 1.42E-01 | 2.74E+00 | 2.79E+00 | 2.45E-01 | Yes | Pbx1 | intron_variant | MODIFIER |
| 1 | 168194251 | AX-170043477 | 0 | A | T | 0.219 | 1.64E-01 | 1.42E-01 | 2.74E+00 | 2.79E+00 | 2.45E-01 | Yes | Pbx1 | intron_variant | MODIFIER |
| 1 | 168194737 | AX-170043479 | 0 | C | G | 0.219 | 1.64E-01 | 1.42E-01 | 2.74E+00 | 2.79E+00 | 2.45E-01 | Yes | Pbx1 | intron_variant | MODIFIER |
| 1 | 168195611 | AX-170378651 | 0 | T | C | 0.219 | 1.64E-01 | 1.42E-01 | 2.74E+00 | 2.79E+00 | 2.45E-01 | Yes | Pbx1 | intron_variant | MODIFIER |
| 1 | 168207693 | AX-170228852 | 0 | A | G | 0.219 | 1.64E-01 | 1.42E-01 | 2.74E+00 | 2.79E+00 | 2.45E-01 | Yes | Pbx1 | intron_variant | MODIFIER |
| 1 | 168208459 | AX-170462512 | 0 | C | T | 0.219 | 1.64E-01 | 1.42E-01 | 2.74E+00 | 2.79E+00 | 2.45E-01 | Yes | Pbx1 | intron_variant | MODIFIER |
| 1 | 168225823 | AX-170260298 | 0 | A | T | 0.219 | 1.64E-01 | 1.42E-01 | 2.74E+00 | 2.79E+00 | 2.45E-01 | Yes | Pbx1 | intron_variant | MODIFIER |
| 1 | 168232715 | AX-97856914 | 0 | C | T | 0.219 | 1.64E-01 | 1.42E-01 | 2.74E+00 | 2.79E+00 | 2.45E-01 | Yes | Pbx1 | intron_variant | MODIFIER |
| 1 | 168238000 | AX-170433723 | 0 | T | C | 0.219 | 1.64E-01 | 1.42E-01 | 2.74E+00 | 2.79E+00 | 2.45E-01 | Yes | Pbx1 | intron_variant | MODIFIER |
| 1 | 168247908 | AX-170131815 | 0 | A | C | 0.219 | 1.64E-01 | 1.42E-01 | 2.74E+00 | 2.79E+00 | 2.45E-01 | Yes | Pbx1 | intron_variant | MODIFIER |
| 1 | 168251792 | AX-170132697 | 0 | G | C | 0.219 | 1.64E-01 | 1.42E-01 | 2.74E+00 | 2.79E+00 | 2.45E-01 | Yes | Pbx1 | intron_variant | MODIFIER |
| 1 | 168271256 | AX-170150214 | 0 | C | G | 0.219 | 1.64E-01 | 1.42E-01 | 2.74E+00 | 2.79E+00 | 2.45E-01 | Yes | Pbx1 | intron_variant | MODIFIER |
| 1 | 168274338 | AX-170170604 | 0 | T | C | 0.219 | 1.64E-01 | 1.42E-01 | 2.74E+00 | 2.79E+00 | 2.45E-01 | Yes | Pbx1 | intron_variant | MODIFIER |
| 1 | 168274701 | AX-170326215 | 0 | C | A | 0.219 | 1.64E-01 | 1.42E-01 | 2.74E+00 | 2.79E+00 | 2.45E-01 | Yes | Pbx1 | intron_variant | MODIFIER |
| 1 | 168275128 | AX-170499785 | 0 | A | G | 0.219 | 1.64E-01 | 1.42E-01 | 2.74E+00 | 2.79E+00 | 2.45E-01 | Yes | Pbx1 | intron_variant | MODIFIER |
| 1 | 168275373 | AX-170201290 | 0 | G | A | 0.219 | 1.64E-01 | 1.42E-01 | 2.74E+00 | 2.79E+00 | 2.45E-01 | Yes | Pbx1 | intron_variant | MODIFIER |
| 1 | 168276427 | AX-170290718 | 0 | G | C | 0.219 | 1.64E-01 | 1.42E-01 | 2.74E+00 | 2.79E+00 | 2.45E-01 | Yes | Pbx1 | intron_variant | MODIFIER |
| 1 | 168279295 | AX-170427714 | 0 | G | A | 0.219 | 1.64E-01 | 1.42E-01 | 2.74E+00 | 2.79E+00 | 2.45E-01 | Yes | Pbx1 | intron_variant | MODIFIER |
| 1 | 168305435 | AX-170179623 | 0 | T | C | 0.219 | 1.64E-01 | 1.42E-01 | 2.74E+00 | 2.79E+00 | 2.45E-01 | Yes | Pbx1 | intron_variant | MODIFIER |
| 1 | 168310599 | AX-170134114 | 0 | T | C | 0.219 | 1.64E-01 | 1.42E-01 | 2.74E+00 | 2.79E+00 | 2.45E-01 | Yes | Pbx1 | intron_variant | MODIFIER |
| 1 | 168334424 | AX-170430838 | 0 | A | C | 0.219 | 1.64E-01 | 1.42E-01 | 2.74E+00 | 2.79E+00 | 2.45E-01 | Yes | Pbx1 | intron_variant | MODIFIER |
| 1 | 168342982 | AX-97808628 | 0 | G | A | 0.219 | 1.64E-01 | 1.42E-01 | 2.74E+00 | 2.79E+00 | 2.45E-01 | Yes | Pbx1 | intron_variant | MODIFIER |
| 1 | 168343016 | AX-170180282 | 0 | C | A | 0.219 | 1.64E-01 | 1.42E-01 | 2.74E+00 | 2.79E+00 | 2.45E-01 | Yes | Pbx1 | intron_variant | MODIFIER |
| 1 | 168343252 | AX-170228478 | 0 | A | G | 0.219 | 1.64E-01 | 1.42E-01 | 2.74E+00 | 2.79E+00 | 2.45E-01 | Yes | Pbx1 | intron_variant | MODIFIER |
| 1 | 168350212 | AX-170273822 | 0 | A | G | 0.219 | 1.64E-01 | 1.42E-01 | 2.74E+00 | 2.79E+00 | 2.45E-01 | Yes | Pbx1 | intron_variant | MODIFIER |
| 1 | 168354936 | AX-170364064 | 0 | A | G | 0.219 | 1.64E-01 | 1.42E-01 | 2.74E+00 | 2.79E+00 | 2.45E-01 | Yes | Pbx1 | intron_variant | MODIFIER |
| 1 | 168368991 | AX-170415618 | 0 | T | C | 0.219 | 1.64E-01 | 1.42E-01 | 2.74E+00 | 2.79E+00 | 2.45E-01 | Yes | Pbx1 | intron_variant | MODIFIER |
| 1 | 168374470 | AX-170289840 | 0 | T | C | 0.219 | 1.64E-01 | 1.42E-01 | 2.74E+00 | 2.79E+00 | 2.45E-01 | Yes | Pbx1 | intron_variant | MODIFIER |
| 1 | 168382492 | AX-170399774 | 0 | G | A | 0.219 | 1.64E-01 | 1.42E-01 | 2.74E+00 | 2.79E+00 | 2.45E-01 | Yes | Pbx1 | intron_variant | MODIFIER |
| 1 | 168386547 | AX-97812944 | 0 | T | G | 0.219 | 1.64E-01 | 1.42E-01 | 2.74E+00 | 2.79E+00 | 2.45E-01 | Yes | Pbx1 | intron_variant | MODIFIER |
| 1 | 168386740 | AX-170297555 | 0 | C | T | 0.219 | 1.64E-01 | 1.42E-01 | 2.74E+00 | 2.79E+00 | 2.45E-01 | Yes | Pbx1 | intron_variant | MODIFIER |
| 1 | 168414905 | AX-170043484 | 0 | A | G | 0.219 | 1.64E-01 | 1.42E-01 | 2.74E+00 | 2.79E+00 | 2.45E-01 | Yes | Pbx1 | intron_variant | MODIFIER |
| 1 | 168470588 | AX-97824389 | 0 | C | A | 0.219 | 1.64E-01 | 1.42E-01 | 2.74E+00 | 2.79E+00 | 2.45E-01 | Yes | | intergenic_variant | MODIFIER |
| 1 | 168476227 | AX-170086681 | 0 | C | T | 0.219 | 1.64E-01 | 1.42E-01 | 2.74E+00 | 2.79E+00 | 2.45E-01 | Yes | | regulatory_region_variant | MODIFIER |
| 1 | 168477306 | AX-170182070 | 0 | T | C | 0.219 | 1.64E-01 | 1.42E-01 | 2.74E+00 | 2.79E+00 | 2.45E-01 | Yes | | regulatory_region_variant | MODIFIER |
| 1 | 168478139 | AX-170480342 | 0 | A | C | 0.219 | 1.64E-01 | 1.42E-01 | 2.74E+00 | 2.79E+00 | 2.45E-01 | Yes | | intergenic_variant | MODIFIER |
| 1 | 168486445 | AX-170463148 | 0 | C | G | 0.219 | 1.64E-01 | 1.42E-01 | 2.74E+00 | 2.79E+00 | 2.45E-01 | Yes | Mir6348 | upstream_gene_variant | MODIFIER |
| 1 | 168490346 | AX-170496237 | 0 | T | C | 0.219 | 1.64E-01 | 1.42E-01 | 2.74E+00 | 2.79E+00 | 2.45E-01 | Yes | Mir6348 | upstream_gene_variant | MODIFIER |
| 1 | 168502007 | AX-170153253 | 0 | C | T | 0.219 | 1.64E-01 | 1.42E-01 | 2.74E+00 | 2.79E+00 | 2.45E-01 | Yes | | regulatory_region_variant | MODIFIER |
| 1 | 168508049 | AX-170002253 | 0 | C | T | 0.219 | 1.64E-01 | 1.42E-01 | 2.74E+00 | 2.79E+00 | 2.45E-01 | Yes | | intergenic_variant | MODIFIER |
| 1 | 168517484 | AX-170173540 | 0 | A | G | 0.219 | 1.64E-01 | 1.42E-01 | 2.74E+00 | 2.79E+00 | 2.45E-01 | Yes | | intergenic_variant | MODIFIER |
| 1 | 168531961 | AX-170490486 | 0 | A | G | 0.219 | 1.64E-01 | 1.42E-01 | 2.74E+00 | 2.79E+00 | 2.45E-01 | Yes | | intergenic_variant | MODIFIER |
| 1 | 168533128 | AX-170294529 | 0 | A | G | 0.219 | 1.64E-01 | 1.42E-01 | 2.74E+00 | 2.79E+00 | 2.45E-01 | Yes | | intergenic_variant | MODIFIER |
| 1 | 168533143 | AX-170294902 | 0 | C | G | 0.219 | 1.64E-01 | 1.42E-01 | 2.74E+00 | 2.79E+00 | 2.45E-01 | Yes | | intergenic_variant | MODIFIER |
| 1 | 168533188 | AX-170002255 | 0 | C | T | 0.219 | 1.64E-01 | 1.42E-01 | 2.74E+00 | 2.79E+00 | 2.45E-01 | Yes | | intergenic_variant | MODIFIER |

| | | | | | | | | | | | | | | | |
|---|---|---|---|---|---|---|---|---|---|---|---|---|---|---|---|
| 1 | 168534634 | AX-170002254 | 0 | A | G | 0.219 | 1.64E-01 | 1.42E-01 | 2.74E+00 | 2.79E+00 | 2.45E-01 | Yes | | intergenic_variant | MODIFIER |
| 1 | 168538956 | AX-170462038 | 0 | G | A | 0.219 | 1.64E-01 | 1.42E-01 | 2.74E+00 | 2.79E+00 | 2.45E-01 | Yes | | intergenic_variant | MODIFIER |
| 1 | 168549684 | AX-97854161 | 0 | T | C | 0.219 | 1.64E-01 | 1.42E-01 | 2.74E+00 | 2.79E+00 | 2.45E-01 | Yes | | regulatory_region_variant | MODIFIER |
| 1 | 168560846 | AX-170213784 | 0 | A | G | 0.219 | 1.64E-01 | 1.42E-01 | 2.74E+00 | 2.79E+00 | 2.45E-01 | Yes | | intergenic_variant | MODIFIER |
| 1 | 168584195 | AX-170459930 | 0 | A | C | 0.219 | 1.64E-01 | 1.42E-01 | 2.74E+00 | 2.79E+00 | 2.45E-01 | Yes | | intergenic_variant | MODIFIER |
| 1 | 168584233 | AX-170207300 | 0 | A | T | 0.219 | 1.64E-01 | 1.42E-01 | 2.74E+00 | 2.79E+00 | 2.45E-01 | Yes | | intergenic_variant | MODIFIER |
| 1 | 168585227 | AX-170273119 | 0 | A | G | 0.219 | 1.64E-01 | 1.42E-01 | 2.74E+00 | 2.79E+00 | 2.45E-01 | Yes | | intergenic_variant | MODIFIER |
| 1 | 168595066 | AX-97812943 | 0 | C | T | 0.219 | 1.64E-01 | 1.42E-01 | 2.74E+00 | 2.79E+00 | 2.45E-01 | Yes | | intergenic_variant | MODIFIER |
| 1 | 168632843 | AX-170308083 | 0 | T | G | 0.219 | 1.64E-01 | 1.42E-01 | 2.74E+00 | 2.79E+00 | 2.45E-01 | Yes | | intergenic_variant | MODIFIER |
| 1 | 168667648 | AX-170504772 | 0 | G | A | 0.219 | 1.64E-01 | 1.42E-01 | 2.74E+00 | 2.79E+00 | 2.45E-01 | Yes | | intergenic_variant | MODIFIER |
| 1 | 168693396 | AX-170251563 | 0 | T | C | 0.219 | 1.64E-01 | 1.42E-01 | 2.74E+00 | 2.79E+00 | 2.45E-01 | Yes | | intergenic_variant | MODIFIER |
| 1 | 168709701 | AX-170043489 | 0 | T | C | 0.219 | 1.64E-01 | 1.42E-01 | 2.74E+00 | 2.79E+00 | 2.45E-01 | Yes | | regulatory_region_variant | MODIFIER |
| 1 | 168709916 | AX-170412485 | 0 | A | G | 0.219 | 1.64E-01 | 1.42E-01 | 2.74E+00 | 2.79E+00 | 2.45E-01 | Yes | | regulatory_region_variant | MODIFIER |
| 1 | 168746790 | AX-170043491 | 0 | T | C | 0.219 | 1.64E-01 | 1.42E-01 | 2.74E+00 | 2.79E+00 | 2.45E-01 | Yes | | intergenic_variant | MODIFIER |
| 1 | 168760605 | AX-170263752 | 0 | A | C | 0.219 | 1.64E-01 | 1.42E-01 | 2.74E+00 | 2.79E+00 | 2.45E-01 | Yes | | intergenic_variant | MODIFIER |
| 1 | 168760729 | AX-170369715 | 0 | C | T | 0.219 | 1.64E-01 | 1.42E-01 | 2.74E+00 | 2.79E+00 | 2.45E-01 | Yes | | intergenic_variant | MODIFIER |
| 1 | 168763592 | AX-97812588 | 0 | A | G | 0.219 | 1.64E-01 | 1.42E-01 | 2.74E+00 | 2.79E+00 | 2.45E-01 | Yes | | intergenic_variant | MODIFIER |
| 1 | 168772405 | AX-170242057 | 0 | C | T | 0.219 | 1.64E-01 | 1.42E-01 | 2.74E+00 | 2.79E+00 | 2.45E-01 | Yes | | intergenic_variant | MODIFIER |
| 1 | 168776892 | AX-170375163 | 0 | G | T | 0.219 | 1.64E-01 | 1.42E-01 | 2.74E+00 | 2.79E+00 | 2.45E-01 | Yes | | intergenic_variant | MODIFIER |
| 1 | 168801029 | AX-170508951 | 0 | T | C | 0.219 | 1.64E-01 | 1.42E-01 | 2.74E+00 | 2.79E+00 | 2.45E-01 | Yes | | intergenic_variant | MODIFIER |
| 1 | 168834600 | AX-170482837 | 0 | T | C | 0.219 | 1.64E-01 | 1.42E-01 | 2.74E+00 | 2.79E+00 | 2.45E-01 | Yes | | intergenic_variant | MODIFIER |
| 1 | 168844255 | AX-170208195 | 0 | G | C | 0.219 | 1.64E-01 | 1.42E-01 | 2.74E+00 | 2.79E+00 | 2.45E-01 | Yes | | intergenic_variant | MODIFIER |
| 1 | 168847586 | AX-170160602 | 0 | A | C | 0.219 | 1.64E-01 | 1.42E-01 | 2.74E+00 | 2.79E+00 | 2.45E-01 | Yes | | intergenic_variant | MODIFIER |
| 1 | 168867509 | AX-170285585 | 0 | G | A | 0.219 | 1.64E-01 | 1.42E-01 | 2.74E+00 | 2.79E+00 | 2.45E-01 | Yes | | intergenic_variant | MODIFIER |
| 1 | 168869089 | AX-97787120 | 0 | T | C | 0.219 | 1.64E-01 | 1.42E-01 | 2.74E+00 | 2.79E+00 | 2.45E-01 | Yes | | intergenic_variant | MODIFIER |
| 1 | 168891839 | AX-97780829 | 0 | C | T | 0.219 | 1.64E-01 | 1.42E-01 | 2.74E+00 | 2.79E+00 | 2.45E-01 | Yes | | intergenic_variant | MODIFIER |
| 1 | 168892201 | AX-170311635 | 0 | T | C | 0.219 | 1.64E-01 | 1.42E-01 | 2.74E+00 | 2.79E+00 | 2.45E-01 | Yes | | intergenic_variant | MODIFIER |
| 1 | 168921625 | AX-170133040 | 0 | A | G | 0.219 | 1.64E-01 | 1.42E-01 | 2.74E+00 | 2.79E+00 | 2.45E-01 | Yes | | intergenic_variant | MODIFIER |
| 1 | 168924713 | AX-170135014 | 0 | G | A | 0.219 | 1.64E-01 | 1.42E-01 | 2.74E+00 | 2.79E+00 | 2.45E-01 | Yes | | regulatory_region_variant | MODIFIER |
| 1 | 168945762 | AX-170513553 | 0 | A | T | 0.219 | 1.64E-01 | 1.42E-01 | 2.74E+00 | 2.79E+00 | 2.45E-01 | Yes | | intergenic_variant | MODIFIER |
| 1 | 168946077 | AX-170142948 | 0 | C | T | 0.219 | 1.64E-01 | 1.42E-01 | 2.74E+00 | 2.79E+00 | 2.45E-01 | Yes | | intergenic_variant | MODIFIER |
| 1 | 168970973 | AX-170272277 | 0 | A | G | 0.219 | 1.64E-01 | 1.42E-01 | 2.74E+00 | 2.79E+00 | 2.45E-01 | Yes | | intergenic_variant | MODIFIER |
| 1 | 168990358 | AX-170500232 | 0 | G | C | 0.219 | 1.64E-01 | 1.42E-01 | 2.74E+00 | 2.79E+00 | 2.45E-01 | Yes | | intergenic_variant | MODIFIER |
| 1 | 168999603 | AX-170134917 | 0 | T | G | 0.219 | 1.64E-01 | 1.42E-01 | 2.74E+00 | 2.79E+00 | 2.45E-01 | Yes | | intergenic_variant | MODIFIER |
| 1 | 169015691 | AX-170521218 | 0 | C | T | 0.219 | 1.64E-01 | 1.42E-01 | 2.74E+00 | 2.79E+00 | 2.45E-01 | Yes | Mir6354 | downstream_gene_variant | MODIFIER |
| 1 | 169017764 | AX-170415962 | 0 | A | G | 0.219 | 1.64E-01 | 1.42E-01 | 2.74E+00 | 2.79E+00 | 2.45E-01 | Yes | Mir6354 | downstream_gene_variant | MODIFIER |
| 1 | 169019070 | AX-170104538 | 0 | A | G | 0.219 | 1.64E-01 | 1.42E-01 | 2.74E+00 | 2.79E+00 | 2.45E-01 | Yes | Mir6354 | non_coding_transcript_exon_variant | MODIFIER |
| 1 | 169019782 | AX-170440675 | 0 | G | A | 0.219 | 1.64E-01 | 1.42E-01 | 2.74E+00 | 2.79E+00 | 2.45E-01 | Yes | Mir6354 | upstream_gene_variant | MODIFIER |
| 1 | 169020028 | AX-170375593 | 0 | C | G | 0.219 | 1.64E-01 | 1.42E-01 | 2.74E+00 | 2.79E+00 | 2.45E-01 | Yes | Mir6354 | upstream_gene_variant | MODIFIER |
| 1 | 169034432 | AX-170002258 | 0 | C | T | 0.219 | 1.64E-01 | 1.42E-01 | 2.74E+00 | 2.79E+00 | 2.45E-01 | Yes | | intergenic_variant | MODIFIER |
| 1 | 169041068 | AX-97850662 | 0 | T | C | 0.219 | 1.64E-01 | 1.42E-01 | 2.74E+00 | 2.79E+00 | 2.45E-01 | Yes | | intergenic_variant | MODIFIER |
| 1 | 169042682 | AX-170233929 | 0 | A | G | 0.219 | 1.64E-01 | 1.42E-01 | 2.74E+00 | 2.79E+00 | 2.45E-01 | Yes | | intergenic_variant | MODIFIER |
| 1 | 169044470 | AX-170164304 | 0 | A | G | 0.219 | 1.64E-01 | 1.42E-01 | 2.74E+00 | 2.79E+00 | 2.45E-01 | Yes | | intergenic_variant | MODIFIER |
| 1 | 169045177 | AX-170474797 | 0 | G | A | 0.219 | 1.64E-01 | 1.42E-01 | 2.74E+00 | 2.79E+00 | 2.45E-01 | Yes | | intergenic_variant | MODIFIER |
| 1 | 169049981 | AX-170191107 | 0 | G | C | 0.219 | 1.64E-01 | 1.42E-01 | 2.74E+00 | 2.79E+00 | 2.45E-01 | Yes | | intergenic_variant | MODIFIER |
| 1 | 169051931 | AX-170296290 | 0 | T | C | 0.219 | 1.64E-01 | 1.42E-01 | 2.74E+00 | 2.79E+00 | 2.45E-01 | Yes | | regulatory_region_variant | MODIFIER |
| 1 | 169059941 | AX-170002259 | 0 | G | A | 0.219 | 1.64E-01 | 1.42E-01 | 2.74E+00 | 2.79E+00 | 2.45E-01 | Yes | | intergenic_variant | MODIFIER |
| 1 | 169060872 | AX-170002262 | 0 | G | C | 0.219 | 1.64E-01 | 1.42E-01 | 2.74E+00 | 2.79E+00 | 2.45E-01 | Yes | | intergenic_variant | MODIFIER |
| 10 | 103746978 | AX-170193471 | 1 | G | A | 0.277 | -1.01E-01 | 8.71E-02 | 2.52E+00 | 2.64E+00 | 2.45E-01 | no | | intergenic_variant | MODIFIER |
| 2 | 163943063 | AX-170065505 | 0 | C | T | 0.167 | 1.90E-01 | 1.65E-01 | 2.81E+00 | 2.94E+00 | 2.45E-01 | Yes | | intergenic_variant | MODIFIER |
| 10 | 92748719 | AX-97816347 | 0 | T | C | 0.298 | -9.62E-02 | 8.26E-02 | 2.51E+00 | 2.57E+00 | 2.46E-01 | no | | intergenic_variant | MODIFIER |
| 1 | 123638561 | AX-97826309 | 0 | A | C | 0.553 | 9.79E-02 | 8.47E-02 | 2.59E+00 | 2.65E+00 | 2.46E-01 | no | Dpp10 | intron_variant | MODIFIER |
| 1 | 123893608 | AX-97781663 | 0 | C | G | 0.553 | 9.79E-02 | 8.47E-02 | 2.59E+00 | 2.65E+00 | 2.46E-01 | no | Dpp10 | intron_variant | MODIFIER |
| 4 | 134432256 | AX-170470592 | 2 | G | T | 0.155 | 1.33E-01 | 1.15E-01 | 2.57E+00 | 2.63E+00 | 2.46E-01 | no | Pafah2 | downstream_gene_variant | MODIFIER |

| | | | | | | | | | | | | | | | |
|---|---|---|---|---|---|---|---|---|---|---|---|---|---|---|---|
| 13 | 3789174 | AX-170268517 | 0 | T | A | 0.202 | -1.36E-01 | 1.16E-01 | 2.47E+00 | 2.53E+00 | 2.46E-01 | no | | intergenic_variant | MODIFIER |
| 18 | 79868341 | AX-170016071 | 1 | G | A | 0.464 | 9.68E-02 | 8.20E-02 | 2.41E+00 | 2.49E+00 | 2.46E-01 | no | | intergenic_variant | MODIFIER |
| 13 | 8662638 | AX-170007713 | 0 | C | T | 0.184 | -1.38E-01 | 1.18E-01 | 2.49E+00 | 2.55E+00 | 2.46E-01 | no | Adarb2 | intron_variant | MODIFIER |
| 1 | 139020884 | AX-170394796 | 1 | T | C | 0.214 | 1.43E-01 | 1.24E-01 | 2.70E+00 | 2.77E+00 | 2.46E-01 | no | Dennd1b | intron_variant | MODIFIER |
| 18 | 78710730 | AX-170235752 | 1 | C | T | 0.455 | 1.00E-01 | 8.52E-02 | 2.41E+00 | 2.48E+00 | 2.47E-01 | no | | intergenic_variant | MODIFIER |
| 16 | 51394013 | AX-170115970 | 0 | A | T | 0.474 | -9.00E-02 | 7.71E-02 | 2.48E+00 | 2.53E+00 | 2.47E-01 | no | | intergenic_variant | MODIFIER |
| 1 | 145753800 | AX-97780362 | 1 | T | G | 0.214 | 1.59E-01 | 1.39E-01 | 2.79E+00 | 2.84E+00 | 2.47E-01 | Yes | | intergenic_variant | MODIFIER |
| 13 | 5954671 | AX-170339238 | 1 | G | A | 0.205 | -1.43E-01 | 1.24E-01 | 2.62E+00 | 2.68E+00 | 2.47E-01 | no | Gm46401 | intron_variant,non_coding_transcript_variant | MODIFIER |
| 14 | 112173383 | AX-170498380 | 1 | T | A | 0.312 | 8.38E-02 | 7.29E-02 | 2.62E+00 | 2.72E+00 | 2.47E-01 | no | | intergenic_variant | MODIFIER |
| 6 | 136679552 | AX-170319053 | 2 | T | G | 0.582 | 1.02E-01 | 8.59E-02 | 2.37E+00 | 2.43E+00 | 2.47E-01 | no | | intergenic_variant | MODIFIER |
| 5 | 80379215 | AX-170497760 | 1 | G | C | 0.446 | 1.26E-01 | 1.10E-01 | 2.86E+00 | 2.93E+00 | 2.47E-01 | no | | intergenic_variant | MODIFIER |
| 5 | 81932148 | AX-170160056 | 1 | T | C | 0.438 | 1.10E-01 | 9.61E-02 | 2.88E+00 | 2.98E+00 | 2.47E-01 | no | | intergenic_variant | MODIFIER |
| 10 | 6925274 | AX-170044055 | 0 | C | A | 0.228 | -1.09E-01 | 9.54E-02 | 2.80E+00 | 2.93E+00 | 2.47E-01 | no | Oprm1 | intron_variant | MODIFIER |
| 7 | 12951464 | AX-170254755 | 1 | A | G | 0.348 | 9.41E-02 | 8.17E-02 | 2.57E+00 | 2.69E+00 | 2.48E-01 | no | 1810019N24Rik | intron_variant,non_coding_transcript_variant | MODIFIER |
| 9 | 102036254 | AX-170347500 | 1 | C | T | 0.143 | -1.30E-01 | 1.13E-01 | 2.66E+00 | 2.75E+00 | 2.48E-01 | no | Ephb1 | intron_variant | MODIFIER |
| 15 | 66862821 | AX-170394757 | 0 | G | A | 0.333 | -9.24E-02 | 7.66E-02 | 2.28E+00 | 2.35E+00 | 2.48E-01 | no | | intergenic_variant | MODIFIER |
| 9 | 6483842 | AX-170237977 | 1 | A | G | 0.473 | 9.35E-02 | 8.03E-02 | 2.47E+00 | 2.57E+00 | 2.48E-01 | no | | intergenic_variant | MODIFIER |
| 9 | 92260259 | AX-170455510 | 0 | C | T | 0.132 | -1.42E-01 | 1.23E-01 | 2.59E+00 | 2.67E+00 | 2.48E-01 | no | Plscr1 | intron_variant | MODIFIER |
| 18 | 79119305 | AX-170435867 | 0 | G | A | 0.465 | 1.02E-01 | 8.65E-02 | 2.39E+00 | 2.45E+00 | 2.48E-01 | no | | intergenic_variant | MODIFIER |
| 5 | 50990556 | AX-97791833 | 0 | G | A | 0.491 | 1.24E-01 | 1.05E-01 | 2.37E+00 | 2.42E+00 | 2.48E-01 | no | | intergenic_variant | MODIFIER |
| 10 | 18409803 | AX-170002846 | 0 | G | A | 0.202 | 1.20E-01 | 1.03E-01 | 3.09E+00 | 3.20E+00 | 2.48E-01 | no | Nhsl1 | intron_variant | MODIFIER |
| 7 | 18746679 | AX-170086029 | 1 | C | A | 0.375 | 9.37E-02 | 8.18E-02 | 2.64E+00 | 2.78E+00 | 2.48E-01 | no | Gm24464 | upstream_gene_variant | MODIFIER |
| 2 | 123537631 | AX-170214083 | 1 | C | T | 0.134 | 1.86E-01 | 1.63E-01 | 2.78E+00 | 2.91E+00 | 2.48E-01 | Yes | | intergenic_variant | MODIFIER |
| 18 | 59452657 | AX-170431606 | 1 | G | A | 0.411 | 1.11E-01 | 9.46E-02 | 2.43E+00 | 2.48E+00 | 2.48E-01 | no | Chsy3 | intron_variant,NMD_transcript_variant | MODIFIER |
| 18 | 60268161 | AX-170529094 | 1 | A | G | 0.411 | 1.11E-01 | 9.46E-02 | 2.43E+00 | 2.48E+00 | 2.48E-01 | no | Gm4841 | downstream_gene_variant | MODIFIER |
| 13 | 11672796 | AX-170126993 | 1 | T | C | 0.188 | -1.31E-01 | 1.13E-01 | 2.51E+00 | 2.57E+00 | 2.48E-01 | no | Ryr2 | intron_variant | MODIFIER |
| 4 | 75451268 | AX-170236013 | 1 | G | A | 0.304 | 1.22E-01 | 1.07E-01 | 2.71E+00 | 2.80E+00 | 2.48E-01 | no | | intergenic_variant | MODIFIER |
| 16 | 40457724 | AX-97785788 | 2 | T | G | 0.491 | 9.33E-02 | 8.03E-02 | 2.49E+00 | 2.53E+00 | 2.48E-01 | no | Lsamp | intron_variant | MODIFIER |
| 5 | 112052416 | AX-170455269 | 0 | A | G | 0.447 | 1.25E-01 | 1.09E-01 | 2.61E+00 | 2.66E+00 | 2.48E-01 | no | | intergenic_variant | MODIFIER |
| 1 | 181451384 | AX-170290602 | 0 | A | C | 0.246 | 1.11E-01 | 9.51E-02 | 2.45E+00 | 2.53E+00 | 2.48E-01 | no | Cnih3 | intron_variant | MODIFIER |
| 4 | 64766145 | AX-97846799 | 1 | C | T | 0.33 | 1.39E-01 | 1.22E-01 | 2.79E+00 | 2.85E+00 | 2.48E-01 | no | | intergenic_variant | MODIFIER |
| 4 | 91354676 | AX-97843257 | 1 | A | C | 0.42 | -9.59E-02 | 8.21E-02 | 2.45E+00 | 2.50E+00 | 2.48E-01 | no | Elavl2 | intron_variant | MODIFIER |
| 8 | 75593584 | AX-97837017 | 1 | A | T | 0.188 | -1.42E-01 | 1.24E-01 | 2.80E+00 | 2.89E+00 | 2.48E-01 | no | Iqcm | intron_variant | MODIFIER |
| 4 | 155184573 | AX-170071136 | 0 | A | G | 0.114 | 1.63E-01 | 1.43E-01 | 2.68E+00 | 2.81E+00 | 2.48E-01 | Yes | Ski | intron_variant | MODIFIER |
| 4 | 120703865 | AX-170240386 | 0 | C | T | 0.211 | 1.18E-01 | 1.03E-01 | 2.59E+00 | 2.67E+00 | 2.48E-01 | no | Kcnq4 | intron_variant | MODIFIER |
| 15 | 83385691 | AX-170341870 | 1 | G | A | 0.098 | -1.41E-01 | 1.02E-01 | 1.89E+00 | 1.97E+00 | 2.48E-01 | no | Pacsin2 | intron_variant | MODIFIER |
| 7 | 145114249 | AX-170269916 | 0 | C | T | 0.202 | -8.44E-02 | 7.39E-02 | 2.65E+00 | 2.79E+00 | 2.48E-01 | no | | regulatory_region_variant | MODIFIER |
| 4 | 72517511 | AX-170420279 | 1 | A | C | 0.33 | 1.37E-01 | 1.20E-01 | 2.78E+00 | 2.84E+00 | 2.49E-01 | no | | intergenic_variant | MODIFIER |
| 5 | 114366744 | AX-170487825 | 0 | C | T | 0.456 | 1.24E-01 | 1.09E-01 | 2.78E+00 | 2.85E+00 | 2.49E-01 | no | Kctd10 | intron_variant | MODIFIER |
| 4 | 136524094 | AX-170298741 | 1 | A | G | 0.17 | 1.22E-01 | 1.06E-01 | 2.49E+00 | 2.55E+00 | 2.49E-01 | no | Luzp1 | intron_variant | MODIFIER |
| 5 | 37147456 | AX-170024698 | 1 | T | A | 0.348 | 1.27E-01 | 1.10E-01 | 2.91E+00 | 2.98E+00 | 2.49E-01 | no | Jakmip1 | intron_variant | MODIFIER |
| 1 | 139600790 | AX-170516280 | 0 | A | G | 0.202 | 1.47E-01 | 1.29E-01 | 2.71E+00 | 2.77E+00 | 2.49E-01 | Yes | Cfhr3 | intron_variant,non_coding_transcript_variant | MODIFIER |
| 1 | 139628559 | AX-170250869 | 0 | G | A | 0.202 | 1.47E-01 | 1.29E-01 | 2.71E+00 | 2.77E+00 | 2.49E-01 | Yes | Cfhr3 | intron_variant,non_coding_transcript_variant | MODIFIER |
| 1 | 139631406 | AX-170282701 | 0 | C | A | 0.202 | 1.47E-01 | 1.29E-01 | 2.71E+00 | 2.77E+00 | 2.49E-01 | Yes | Cfhr3 | intron_variant,non_coding_transcript_variant | MODIFIER |
| 1 | 139790289 | AX-97819902 | 0 | G | C | 0.202 | 1.47E-01 | 1.29E-01 | 2.71E+00 | 2.77E+00 | 2.49E-01 | Yes | Gm25619 | downstream_gene_variant | MODIFIER |
| 1 | 139807571 | AX-97846018 | 0 | C | T | 0.202 | 1.47E-01 | 1.29E-01 | 2.71E+00 | 2.77E+00 | 2.49E-01 | Yes | Cfhr2 | downstream_gene_variant | MODIFIER |
| 1 | 139861998 | AX-170230208 | 0 | T | C | 0.202 | 1.47E-01 | 1.29E-01 | 2.71E+00 | 2.77E+00 | 2.49E-01 | Yes | Cfhr2 | upstream_gene_variant | MODIFIER |
| 1 | 139863064 | AX-170042910 | 0 | C | T | 0.202 | 1.47E-01 | 1.29E-01 | 2.71E+00 | 2.77E+00 | 2.49E-01 | Yes | Cfhr2 | upstream_gene_variant | MODIFIER |
| 1 | 140175668 | AX-97824833 | 0 | T | C | 0.202 | 1.47E-01 | 1.29E-01 | 2.71E+00 | 2.77E+00 | 2.49E-01 | Yes | Cfh | intron_variant | MODIFIER |
| 1 | 140306487 | AX-97840619 | 0 | T | C | 0.202 | 1.47E-01 | 1.29E-01 | 2.71E+00 | 2.77E+00 | 2.49E-01 | Yes | Kcnt2 | intron_variant | MODIFIER |
| 1 | 140341383 | AX-97836055 | 0 | T | C | 0.202 | 1.47E-01 | 1.29E-01 | 2.71E+00 | 2.77E+00 | 2.49E-01 | Yes | Kcnt2 | intron_variant | MODIFIER |
| 1 | 140354075 | AX-170226613 | 0 | A | G | 0.202 | 1.47E-01 | 1.29E-01 | 2.71E+00 | 2.77E+00 | 2.49E-01 | Yes | Kcnt2 | intron_variant | MODIFIER |
| 4 | 69930500 | AX-170231708 | 0 | C | T | 0.325 | 1.34E-01 | 1.17E-01 | 2.83E+00 | 2.89E+00 | 2.49E-01 | no | Gm11223 | upstream_gene_variant | MODIFIER |
| 4 | 69054467 | AX-97785217 | 1 | A | G | 0.321 | 1.37E-01 | 1.20E-01 | 2.80E+00 | 2.85E+00 | 2.49E-01 | no | | intergenic_variant | MODIFIER |

| | | | | | | | | | | | | | | | |
|---|---|---|---|---|---|---|---|---|---|---|---|---|---|---|---|
| 18 | 73266806 | AX-170093947 | 1 | T | C | 0.188 | -1.37E-01 | 1.20E-01 | 2.63E+00 | 2.68E+00 | 2.49E-01 | no | | intergenic_variant | MODIFIER |
| 3 | 78812210 | AX-170436376 | 1 | C | T | 0.054 | -1.52E-01 | 1.33E-01 | 2.61E+00 | 2.75E+00 | 2.49E-01 | no | | intergenic_variant | MODIFIER |
| 5 | 51104409 | AX-97796965 | 0 | C | T | 0.474 | 1.21E-01 | 1.03E-01 | 2.42E+00 | 2.47E+00 | 2.49E-01 | no | | intergenic_variant | MODIFIER |
| 16 | 77973629 | AX-170265169 | 1 | G | A | 0.464 | 8.69E-02 | 7.56E-02 | 2.55E+00 | 2.63E+00 | 2.50E-01 | no | | intergenic_variant | MODIFIER |
| 18 | 59657537 | AX-170383078 | 1 | G | C | 0.411 | 1.10E-01 | 9.37E-02 | 2.40E+00 | 2.45E+00 | 2.50E-01 | no | | intergenic_variant | MODIFIER |
| 11 | 51255056 | AX-170388298 | 0 | C | A | 0.07 | -1.91E-01 | 1.67E-01 | 2.61E+00 | 2.70E+00 | 2.50E-01 | Yes | BC049762 | intron_variant | MODIFIER |
| 6 | 125243259 | AX-97820102 | 0 | C | A | 0.184 | -9.25E-02 | 8.13E-02 | 2.82E+00 | 2.97E+00 | 2.50E-01 | no | Vamp1 | downstream_gene_variant | MODIFIER |
| 5 | 82071257 | AX-170025419 | 0 | G | C | 0.421 | 1.17E-01 | 1.02E-01 | 2.82E+00 | 2.89E+00 | 2.50E-01 | no | | intergenic_variant | MODIFIER |
| 4 | 64704107 | AX-170209572 | 1 | G | T | 0.321 | 1.36E-01 | 1.19E-01 | 2.80E+00 | 2.86E+00 | 2.50E-01 | no | | intergenic_variant | MODIFIER |
| 5 | 50084187 | AX-170182522 | 0 | C | T | 0.491 | 1.26E-01 | 1.04E-01 | 2.26E+00 | 2.31E+00 | 2.50E-01 | no | | intergenic_variant | MODIFIER |
| 5 | 52070498 | AX-97782653 | 0 | G | A | 0.491 | 1.26E-01 | 1.04E-01 | 2.26E+00 | 2.31E+00 | 2.50E-01 | no | | intergenic_variant | MODIFIER |
| 1 | 124691052 | AX-170411795 | 0 | C | T | 0.57 | 9.27E-02 | 8.10E-02 | 2.60E+00 | 2.67E+00 | 2.50E-01 | no | Gm28928 | intron_variant,non_coding_transcript_variant | MODIFIER |
| 10 | 92527241 | AX-97844753 | 0 | C | T | 0.289 | -9.65E-02 | 8.42E-02 | 2.57E+00 | 2.65E+00 | 2.50E-01 | no | | intergenic_variant | MODIFIER |
| 4 | 77595767 | AX-170292916 | 1 | A | G | 0.321 | 1.25E-01 | 1.09E-01 | 2.75E+00 | 2.82E+00 | 2.50E-01 | no | Ptprd | intron_variant,non_coding_transcript_variant | MODIFIER |
| 18 | 85259677 | AX-170469199 | 0 | T | A | 0.175 | -1.07E-01 | 9.18E-02 | 2.45E+00 | 2.56E+00 | 2.50E-01 | no | | intergenic_variant | MODIFIER |
| 4 | 85837284 | AX-170522281 | 0 | C | G | 0.351 | 1.20E-01 | 1.05E-01 | 2.73E+00 | 2.81E+00 | 2.51E-01 | no | Adamtsl1 | intron_variant | MODIFIER |
| 10 | 102274944 | AX-170206012 | 0 | A | G | 0.272 | -9.19E-02 | 8.07E-02 | 2.65E+00 | 2.74E+00 | 2.51E-01 | no | Mgat4c | intron_variant | MODIFIER |
| 18 | 52844180 | AX-170349485 | 1 | T | C | 0.205 | -1.24E-01 | 1.08E-01 | 2.62E+00 | 2.70E+00 | 2.51E-01 | no | Sncaip | intron_variant | MODIFIER |
| 13 | 3633710 | AX-170408485 | 1 | C | T | 0.196 | -1.42E-01 | 1.24E-01 | 2.54E+00 | 2.60E+00 | 2.51E-01 | no | Asb13 | upstream_gene_variant | MODIFIER |
| 4 | 143710593 | AX-170089392 | 0 | T | C | 0.088 | 1.92E-01 | 1.69E-01 | 2.75E+00 | 2.87E+00 | 2.51E-01 | Yes | | intergenic_variant | MODIFIER |
| 3 | 20985438 | AX-170066049 | 1 | G | A | 0.134 | -1.16E-01 | 1.01E-01 | 2.57E+00 | 2.66E+00 | 2.52E-01 | no | | regulatory_region_variant | MODIFIER |
| 1 | 123429998 | AX-170468041 | 1 | A | G | 0.554 | 9.38E-02 | 8.20E-02 | 2.57E+00 | 2.63E+00 | 2.52E-01 | no | Dpp10 | intron_variant | MODIFIER |
| 1 | 123719697 | AX-170525782 | 1 | T | C | 0.554 | 9.38E-02 | 8.20E-02 | 2.57E+00 | 2.63E+00 | 2.52E-01 | no | Dpp10 | intron_variant | MODIFIER |
| 1 | 183882671 | AX-170104801 | 0 | C | T | 0.237 | 1.21E-01 | 1.04E-01 | 2.46E+00 | 2.55E+00 | 2.52E-01 | no | | intergenic_variant | MODIFIER |
| 5 | 111013661 | AX-170364179 | 0 | G | C | 0.465 | 1.24E-01 | 1.09E-01 | 2.76E+00 | 2.81E+00 | 2.52E-01 | no | Ttc28 | intron_variant | MODIFIER |
| 5 | 50459506 | AX-170436027 | 1 | G | A | 0.491 | 1.16E-01 | 1.00E-01 | 2.44E+00 | 2.50E+00 | 2.52E-01 | no | Gm36814 | intron_variant,non_coding_transcript_variant | MODIFIER |
| 16 | 19307797 | AX-97830684 | 0 | A | G | 0.14 | 1.05E-01 | 9.21E-02 | 2.87E+00 | 3.02E+00 | 2.52E-01 | no | Gm22158 | non_coding_transcript_exon_variant | MODIFIER |
| 5 | 68171260 | AX-170453382 | 1 | T | C | 0.473 | 1.14E-01 | 9.98E-02 | 2.53E+00 | 2.60E+00 | 2.52E-01 | no | Grxcr1 | downstream_gene_variant | MODIFIER |
| 1 | 126662642 | AX-170294349 | 1 | C | A | 0.562 | 9.41E-02 | 8.24E-02 | 2.57E+00 | 2.64E+00 | 2.52E-01 | no | Nckap5 | intron_variant | MODIFIER |
| 5 | 69157080 | AX-170241913 | 2 | C | G | 0.482 | 1.01E-01 | 8.95E-02 | 2.78E+00 | 2.92E+00 | 2.52E-01 | no | Kctd8 | intron_variant | MODIFIER |
| 6 | 129384315 | AX-170305068 | 1 | A | G | 0.554 | 9.25E-02 | 8.01E-02 | 2.46E+00 | 2.54E+00 | 2.52E-01 | no | Clec12b | intron_variant | MODIFIER |
| 3 | 151816942 | AX-97797701 | 0 | A | G | 0.342 | 7.99E-02 | 7.04E-02 | 2.83E+00 | 2.93E+00 | 2.52E-01 | no | Ptgfr | intron_variant | MODIFIER |
| 15 | 6632851 | AX-170349116 | 0 | A | G | 0.237 | -1.20E-01 | 1.06E-01 | 2.62E+00 | 2.68E+00 | 2.52E-01 | no | Fyb | intron_variant | MODIFIER |
| 15 | 4629586 | AX-170407179 | 1 | A | G | 0.232 | -1.20E-01 | 1.06E-01 | 2.66E+00 | 2.71E+00 | 2.53E-01 | no | | intergenic_variant | MODIFIER |
| 15 | 11266045 | AX-170472756 | 1 | A | T | 0.232 | -1.20E-01 | 1.06E-01 | 2.66E+00 | 2.71E+00 | 2.53E-01 | no | Adamts12 | intron_variant | MODIFIER |
| 9 | 117758899 | AX-170379639 | 0 | T | G | 0.123 | -1.05E-01 | 9.30E-02 | 2.66E+00 | 2.80E+00 | 2.53E-01 | no | Gm45881 | intron_variant,non_coding_transcript_variant | MODIFIER |
| 4 | 154754224 | AX-170283099 | 0 | A | G | 0.105 | 1.77E-01 | 1.56E-01 | 2.70E+00 | 2.81E+00 | 2.53E-01 | Yes | | intergenic_variant | MODIFIER |
| 1 | 176114412 | AX-97799949 | 0 | G | A | 0.237 | 1.17E-01 | 1.03E-01 | 2.55E+00 | 2.65E+00 | 2.53E-01 | no | Pld5 | intron_variant | MODIFIER |
| 14 | 124691843 | AX-170193720 | 1 | A | G | 0.321 | 8.58E-02 | 7.56E-02 | 2.61E+00 | 2.73E+00 | 2.53E-01 | no | | intergenic_variant | MODIFIER |
| 16 | 35636514 | AX-97790828 | 0 | A | C | 0.509 | 8.56E-02 | 7.41E-02 | 2.46E+00 | 2.51E+00 | 2.53E-01 | no | Sema5b | intron_variant | MODIFIER |
| 4 | 124706165 | AX-170379329 | 1 | C | A | 0.232 | 1.20E-01 | 1.06E-01 | 2.59E+00 | 2.66E+00 | 2.53E-01 | no | Fhl3 | intron_variant | MODIFIER |
| 16 | 51648797 | AX-170391214 | 0 | A | G | 0.518 | 9.04E-02 | 7.83E-02 | 2.46E+00 | 2.51E+00 | 2.53E-01 | no | | intergenic_variant | MODIFIER |
| 16 | 51223879 | AX-170369981 | 1 | T | C | 0.473 | -9.56E-02 | 8.31E-02 | 2.49E+00 | 2.53E+00 | 2.53E-01 | no | | intergenic_variant | MODIFIER |
| 5 | 129779048 | AX-170364755 | 1 | C | T | 0.161 | -1.04E-01 | 8.41E-02 | 2.15E+00 | 2.24E+00 | 2.53E-01 | no | Psph | 5_prime_UTR_variant | MODIFIER |
| 1 | 124749878 | AX-170309611 | 0 | C | T | 0.561 | 9.41E-02 | 8.26E-02 | 2.57E+00 | 2.63E+00 | 2.53E-01 | no | Gm28928 | intron_variant,non_coding_transcript_variant | MODIFIER |
| 18 | 78365800 | AX-170340378 | 1 | T | G | 0.446 | 9.77E-02 | 8.50E-02 | 2.49E+00 | 2.56E+00 | 2.53E-01 | no | Slc14a2 | intron_variant | MODIFIER |
| 9 | 94494246 | AX-170356217 | 1 | G | A | 0.143 | -1.37E-01 | 1.21E-01 | 2.71E+00 | 2.79E+00 | 2.53E-01 | no | | intergenic_variant | MODIFIER |
| 13 | 6682155 | AX-97822515 | 0 | G | T | 0.202 | -1.41E-01 | 1.23E-01 | 2.52E+00 | 2.57E+00 | 2.53E-01 | no | | intergenic_variant | MODIFIER |
| 1 | 138481493 | AX-170134410 | 1 | G | A | 0.027 | 1.55E-01 | 1.38E-01 | 2.76E+00 | 2.98E+00 | 2.53E-01 | Yes | Nek7 | downstream_gene_variant | MODIFIER |
| 4 | 77546032 | AX-170509768 | 1 | T | G | 0.312 | 1.34E-01 | 1.18E-01 | 2.82E+00 | 2.88E+00 | 2.53E-01 | no | Ptprd | intron_variant,non_coding_transcript_variant | MODIFIER |
| 16 | 35512898 | AX-170526265 | 0 | G | A | 0.509 | 9.13E-02 | 7.98E-02 | 2.53E+00 | 2.57E+00 | 2.54E-01 | no | 1600019K03Rik | upstream_gene_variant | MODIFIER |
| 16 | 35513720 | AX-170057120 | 0 | C | A | 0.509 | 9.13E-02 | 7.98E-02 | 2.53E+00 | 2.57E+00 | 2.54E-01 | no | 1600019K03Rik | upstream_gene_variant | MODIFIER |
| 16 | 35515629 | AX-97838613 | 0 | A | G | 0.509 | 9.13E-02 | 7.98E-02 | 2.53E+00 | 2.57E+00 | 2.54E-01 | no | | intergenic_variant | MODIFIER |
| 16 | 35536285 | AX-97831512 | 0 | T | C | 0.509 | 9.13E-02 | 7.98E-02 | 2.53E+00 | 2.57E+00 | 2.54E-01 | no | Sema5b | upstream_gene_variant | MODIFIER |

| | | | | | | | | | | | | | | | |
|---|---|---|---|---|---|---|---|---|---|---|---|---|---|---|---|
| 16 | 35549774 | AX-170057123 | 0 | C | T | 0.509 | 9.13E-02 | 7.98E-02 | 2.53E+00 | 2.57E+00 | 2.54E-01 | no | Sema5b | intron_variant | MODIFIER |
| 16 | 35550505 | AX-170307020 | 0 | T | C | 0.509 | 9.13E-02 | 7.98E-02 | 2.53E+00 | 2.57E+00 | 2.54E-01 | no | Sema5b | intron_variant | MODIFIER |
| 16 | 35556972 | AX-97819672 | 0 | G | A | 0.509 | 9.13E-02 | 7.98E-02 | 2.53E+00 | 2.57E+00 | 2.54E-01 | no | Sema5b | intron_variant | MODIFIER |
| 16 | 35558444 | AX-170395325 | 0 | C | T | 0.509 | 9.13E-02 | 7.98E-02 | 2.53E+00 | 2.57E+00 | 2.54E-01 | no | Sema5b | intron_variant | MODIFIER |
| 16 | 35590534 | AX-170123987 | 0 | T | G | 0.509 | 9.13E-02 | 7.98E-02 | 2.53E+00 | 2.57E+00 | 2.54E-01 | no | Sema5b | intron_variant | MODIFIER |
| 16 | 35598396 | AX-170035562 | 0 | C | T | 0.509 | 9.13E-02 | 7.98E-02 | 2.53E+00 | 2.57E+00 | 2.54E-01 | no | Sema5b | intron_variant | MODIFIER |
| 16 | 35629222 | AX-170439364 | 0 | C | T | 0.509 | 9.13E-02 | 7.98E-02 | 2.53E+00 | 2.57E+00 | 2.54E-01 | no | Sema5b | intron_variant | MODIFIER |
| 16 | 35702172 | AX-170353871 | 0 | C | G | 0.509 | 9.13E-02 | 7.98E-02 | 2.53E+00 | 2.57E+00 | 2.54E-01 | no | Dirc2 | intron_variant | MODIFIER |
| 16 | 35745405 | AX-170459277 | 0 | C | T | 0.509 | 9.13E-02 | 7.98E-02 | 2.53E+00 | 2.57E+00 | 2.54E-01 | no | Dirc2 | intron_variant | MODIFIER |
| 16 | 35798350 | AX-170035561 | 0 | G | A | 0.509 | 9.13E-02 | 7.98E-02 | 2.53E+00 | 2.57E+00 | 2.54E-01 | no | Hspbap1 | intron_variant,NMD_transcript_variant | MODIFIER |
| 16 | 35823316 | AX-170012826 | 0 | G | A | 0.509 | 9.13E-02 | 7.98E-02 | 2.53E+00 | 2.57E+00 | 2.54E-01 | no | Hspbap1 | intron_variant,NMD_transcript_variant | MODIFIER |
| 16 | 35824331 | AX-170012823 | 0 | A | C | 0.509 | 9.13E-02 | 7.98E-02 | 2.53E+00 | 2.57E+00 | 2.54E-01 | no | Hspbap1 | intron_variant,NMD_transcript_variant | MODIFIER |
| 16 | 35826635 | AX-170057127 | 0 | A | G | 0.509 | 9.13E-02 | 7.98E-02 | 2.53E+00 | 2.57E+00 | 2.54E-01 | no | Hspbap1 | 3_prime_UTR_variant,NMD_transcript_variant | MODIFIER |
| 16 | 35826683 | AX-170057128 | 0 | C | T | 0.509 | 9.13E-02 | 7.98E-02 | 2.53E+00 | 2.57E+00 | 2.54E-01 | no | Hspbap1 | 3_prime_UTR_variant,NMD_transcript_variant | MODIFIER |
| 16 | 35826815 | AX-170057130 | 0 | G | A | 0.509 | 9.13E-02 | 7.98E-02 | 2.53E+00 | 2.57E+00 | 2.54E-01 | no | Hspbap1 | 3_prime_UTR_variant,NMD_transcript_variant | MODIFIER |
| 16 | 35829094 | AX-170236401 | 0 | T | A | 0.509 | 9.13E-02 | 7.98E-02 | 2.53E+00 | 2.57E+00 | 2.54E-01 | no | Hspbap1 | downstream_gene_variant | MODIFIER |
| 16 | 35833181 | AX-170261275 | 0 | G | A | 0.509 | 9.13E-02 | 7.98E-02 | 2.53E+00 | 2.57E+00 | 2.54E-01 | no | Hspbap1 | downstream_gene_variant | MODIFIER |
| 3 | 23694923 | AX-170206068 | 2 | C | G | 0.164 | -1.10E-01 | 9.66E-02 | 2.54E+00 | 2.63E+00 | 2.54E-01 | no | | intergenic_variant | MODIFIER |
| 1 | 167958425 | AX-170531131 | 1 | G | C | 0.232 | 1.52E-01 | 1.34E-01 | 2.74E+00 | 2.80E+00 | 2.54E-01 | Yes | | intergenic_variant | MODIFIER |
| 7 | 36544977 | AX-170077067 | 0 | A | T | 0.36 | 9.76E-02 | 8.60E-02 | 2.60E+00 | 2.69E+00 | 2.54E-01 | no | | intergenic_variant | MODIFIER |
| 4 | 95902582 | AX-170242803 | 0 | A | G | 0.263 | 1.24E-01 | 1.08E-01 | 2.50E+00 | 2.56E+00 | 2.54E-01 | no | Fggy | intron_variant | MODIFIER |
| 4 | 86179990 | AX-170169237 | 0 | A | G | 0.325 | 1.22E-01 | 1.07E-01 | 2.58E+00 | 2.67E+00 | 2.54E-01 | no | Adamtsl1 | intron_variant | MODIFIER |
| 18 | 30718912 | AX-170325764 | 1 | G | T | 0.062 | -2.44E-01 | 2.15E-01 | 2.62E+00 | 2.71E+00 | 2.54E-01 | Yes | | intergenic_variant | MODIFIER |
| 18 | 74387397 | AX-170368955 | 1 | T | G | 0.17 | -1.38E-01 | 1.22E-01 | 2.61E+00 | 2.66E+00 | 2.54E-01 | no | | intergenic_variant | MODIFIER |
| 4 | 64745265 | AX-97825042 | 0 | C | A | 0.325 | 1.32E-01 | 1.17E-01 | 2.86E+00 | 2.93E+00 | 2.54E-01 | no | | intergenic_variant | MODIFIER |
| 4 | 132584577 | AX-170354875 | 1 | A | C | 0.179 | 1.25E-01 | 1.10E-01 | 2.62E+00 | 2.67E+00 | 2.54E-01 | no | Ptafr | downstream_gene_variant | MODIFIER |
| 16 | 40238181 | AX-170156696 | 1 | G | C | 0.5 | 9.33E-02 | 8.19E-02 | 2.56E+00 | 2.61E+00 | 2.54E-01 | no | Lsamp | intron_variant | MODIFIER |
| 12 | 68793123 | AX-170415853 | 0 | A | G | 0.061 | 2.43E-01 | 2.14E-01 | 2.93E+00 | 3.07E+00 | 2.54E-01 | Yes | | intergenic_variant | MODIFIER |
| 12 | 70390825 | AX-170434183 | 0 | T | C | 0.061 | 2.43E-01 | 2.14E-01 | 2.93E+00 | 3.07E+00 | 2.54E-01 | Yes | | intergenic_variant | MODIFIER |
| 12 | 72303493 | AX-170232824 | 0 | A | G | 0.061 | 2.43E-01 | 2.14E-01 | 2.93E+00 | 3.07E+00 | 2.54E-01 | Yes | Rtn1 | intron_variant | MODIFIER |
| 18 | 37363371 | AX-170161048 | 0 | T | C | 0.061 | 2.43E-01 | 2.14E-01 | 2.93E+00 | 3.07E+00 | 2.54E-01 | Yes | Pcdhb8 | downstream_gene_variant | MODIFIER |
| 18 | 45036611 | AX-170157087 | 0 | G | A | 0.061 | 2.43E-01 | 2.14E-01 | 2.93E+00 | 3.07E+00 | 2.54E-01 | Yes | | intergenic_variant | MODIFIER |
| 4 | 136281092 | AX-170115805 | 1 | C | G | 0.17 | 1.31E-01 | 1.15E-01 | 2.63E+00 | 2.68E+00 | 2.55E-01 | no | Zfp46 | upstream_gene_variant | MODIFIER |
| 4 | 103634906 | AX-170517800 | 0 | C | A | 0.237 | 1.32E-01 | 1.17E-01 | 2.66E+00 | 2.73E+00 | 2.55E-01 | no | Dab1 | intron_variant | MODIFIER |
| 10 | 20817211 | AX-97803061 | 0 | G | T | 0.044 | 2.06E-01 | 1.83E-01 | 2.75E+00 | 2.91E+00 | 2.55E-01 | Yes | Gm48651 | intron_variant,non_coding_transcript_variant | MODIFIER |
| 6 | 137032648 | AX-170271166 | 0 | T | C | 0.588 | 1.01E-01 | 8.78E-02 | 2.43E+00 | 2.48E+00 | 2.55E-01 | no | | intergenic_variant | MODIFIER |
| 6 | 78797184 | AX-170243194 | 1 | T | C | 0.027 | -1.63E-01 | 1.46E-01 | 2.67E+00 | 2.84E+00 | 2.55E-01 | no | | regulatory_region_variant | MODIFIER |
| 18 | 88315714 | AX-170141023 | 1 | T | G | 0.152 | -1.21E-01 | 1.06E-01 | 2.50E+00 | 2.58E+00 | 2.55E-01 | no | | intergenic_variant | MODIFIER |
| 4 | 77460965 | AX-170263408 | 0 | T | C | 0.316 | 1.27E-01 | 1.12E-01 | 2.84E+00 | 2.90E+00 | 2.55E-01 | no | Ptprd | intron_variant,non_coding_transcript_variant | MODIFIER |
| 6 | 88577009 | AX-97811623 | 0 | G | A | 0.035 | -1.89E-01 | 1.65E-01 | 2.48E+00 | 2.59E+00 | 2.56E-01 | Yes | Kbtbd12 | intron_variant | MODIFIER |
| 6 | 43996759 | AX-170027110 | 0 | T | C | 0.035 | -1.99E-01 | 1.75E-01 | 2.51E+00 | 2.61E+00 | 2.56E-01 | Yes | | intergenic_variant | MODIFIER |
| 4 | 85735590 | AX-170134691 | 1 | T | C | 0.348 | 1.23E-01 | 1.09E-01 | 2.67E+00 | 2.75E+00 | 2.56E-01 | no | Adamtsl1 | intron_variant,non_coding_transcript_variant | MODIFIER |
| 3 | 23295186 | AX-170499249 | 1 | A | T | 0.161 | -1.16E-01 | 1.03E-01 | 2.60E+00 | 2.67E+00 | 2.56E-01 | no | | intergenic_variant | MODIFIER |
| 18 | 77233690 | AX-170345275 | 0 | A | G | 0.43 | 1.00E-01 | 8.75E-02 | 2.46E+00 | 2.51E+00 | 2.56E-01 | no | St8sia5 | intron_variant | MODIFIER |
| 8 | 86020968 | AX-97847424 | 0 | C | T | 0.202 | -1.39E-01 | 1.23E-01 | 2.78E+00 | 2.87E+00 | 2.56E-01 | no | Phkb | intron_variant | MODIFIER |
| 1 | 168208117 | AX-170527773 | 1 | C | T | 0.223 | 1.56E-01 | 1.38E-01 | 2.68E+00 | 2.74E+00 | 2.57E-01 | Yes | Pbx1 | intron_variant | MODIFIER |
| 3 | 20686635 | AX-170189748 | 0 | G | T | 0.14 | -1.17E-01 | 1.03E-01 | 2.57E+00 | 2.65E+00 | 2.57E-01 | no | | intergenic_variant | MODIFIER |
| 4 | 133583489 | AX-170374225 | 1 | T | A | 0.161 | 1.33E-01 | 1.18E-01 | 2.57E+00 | 2.61E+00 | 2.57E-01 | no | Gpn2 | upstream_gene_variant | MODIFIER |
| 4 | 137020599 | AX-170466770 | 1 | A | T | 0.161 | 1.35E-01 | 1.20E-01 | 2.58E+00 | 2.62E+00 | 2.57E-01 | no | Zbtb40 | intron_variant | MODIFIER |
| 5 | 81827503 | AX-97800795 | 0 | T | G | 0.447 | 1.14E-01 | 1.02E-01 | 2.79E+00 | 2.86E+00 | 2.57E-01 | no | Adgrl3 | downstream_gene_variant | MODIFIER |
| 10 | 9283778 | AX-170334299 | 1 | T | A | 0.589 | 5.64E-02 | 5.03E-02 | 2.72E+00 | 2.82E+00 | 2.57E-01 | no | | intergenic_variant | MODIFIER |
| 7 | 99661819 | AX-170196108 | 0 | T | C | 0.36 | -1.15E-01 | 1.03E-01 | 2.84E+00 | 2.93E+00 | 2.57E-01 | no | Slco2b1 | intron_variant | MODIFIER |
| 1 | 193509213 | AX-97810768 | 0 | T | C | 0.281 | 1.00E-01 | 8.58E-02 | 2.34E+00 | 2.42E+00 | 2.57E-01 | no | Mir205 | upstream_gene_variant | MODIFIER |
| 13 | 3967327 | AX-170336804 | 2 | A | T | 0.209 | -1.37E-01 | 1.22E-01 | 2.64E+00 | 2.70E+00 | 2.57E-01 | no | | intergenic_variant | MODIFIER |

| | | | | | | | | | | | | | | | |
|---|---|---|---|---|---|---|---|---|---|---|---|---|---|---|---|
| 15 | 6792181 | AX-97795123 | 1 | A | G | 0.089 | -1.32E-01 | 9.85E-02 | 1.93E+00 | 2.02E+00 | 2.57E-01 | no | Rictor | intron_variant | MODIFIER |
| 16 | 15522869 | AX-170315585 | 0 | C | G | 0.14 | 1.09E-01 | 9.67E-02 | 2.88E+00 | 3.02E+00 | 2.57E-01 | no | | intergenic_variant | MODIFIER |
| 6 | 130826386 | AX-170412219 | 1 | A | G | 0.545 | 9.21E-02 | 8.14E-02 | 2.54E+00 | 2.62E+00 | 2.58E-01 | no | Gm44118 | downstream_gene_variant | MODIFIER |
| 1 | 190328911 | AX-170334582 | 0 | G | A | 0.254 | 1.12E-01 | 9.63E-02 | 2.37E+00 | 2.45E+00 | 2.58E-01 | no | | intergenic_variant | MODIFIER |
| 2 | 162967074 | AX-170179038 | 0 | T | C | 0.167 | 2.01E-01 | 1.79E-01 | 2.83E+00 | 2.94E+00 | 2.58E-01 | Yes | L3mbtl1 | synonymous_variant | LOW |
| 6 | 104509964 | AX-170305375 | 0 | G | A | 0.07 | -2.34E-01 | 2.09E-01 | 2.70E+00 | 2.79E+00 | 2.58E-01 | Yes | Cntn6 | intron_variant | MODIFIER |
| 4 | 155015338 | AX-170410045 | 1 | C | T | 0.134 | 1.59E-01 | 1.42E-01 | 2.73E+00 | 2.86E+00 | 2.58E-01 | Yes | Plch2 | upstream_gene_variant | MODIFIER |
| 4 | 127192710 | AX-170510822 | 0 | G | A | 0.149 | 1.34E-01 | 1.19E-01 | 2.60E+00 | 2.67E+00 | 2.58E-01 | no | Dlgap3 | intron_variant | MODIFIER |
| 4 | 127196323 | AX-170459113 | 0 | C | T | 0.149 | 1.34E-01 | 1.19E-01 | 2.60E+00 | 2.67E+00 | 2.58E-01 | no | Dlgap3 | intron_variant | MODIFIER |
| 4 | 127253387 | AX-170277812 | 0 | C | T | 0.149 | 1.34E-01 | 1.19E-01 | 2.60E+00 | 2.67E+00 | 2.58E-01 | no | | intergenic_variant | MODIFIER |
| 4 | 127279503 | AX-170070610 | 0 | G | A | 0.149 | 1.34E-01 | 1.19E-01 | 2.60E+00 | 2.67E+00 | 2.58E-01 | no | | regulatory_region_variant | MODIFIER |
| 4 | 127280105 | AX-170229391 | 0 | G | A | 0.149 | 1.34E-01 | 1.19E-01 | 2.60E+00 | 2.67E+00 | 2.58E-01 | no | Gm22221 | downstream_gene_variant | MODIFIER |
| 4 | 127285214 | AX-170507902 | 0 | T | C | 0.149 | 1.34E-01 | 1.19E-01 | 2.60E+00 | 2.67E+00 | 2.58E-01 | no | Gm22221 | upstream_gene_variant | MODIFIER |
| 4 | 127290112 | AX-170277607 | 0 | T | C | 0.149 | 1.34E-01 | 1.19E-01 | 2.60E+00 | 2.67E+00 | 2.58E-01 | no | | intergenic_variant | MODIFIER |
| 4 | 127291260 | AX-170137730 | 0 | A | G | 0.149 | 1.34E-01 | 1.19E-01 | 2.60E+00 | 2.67E+00 | 2.58E-01 | no | | intergenic_variant | MODIFIER |
| 4 | 127292397 | AX-170480527 | 0 | G | A | 0.149 | 1.34E-01 | 1.19E-01 | 2.60E+00 | 2.67E+00 | 2.58E-01 | no | Gm12948 | downstream_gene_variant | MODIFIER |
| 4 | 127292859 | AX-170283511 | 0 | G | T | 0.149 | 1.34E-01 | 1.19E-01 | 2.60E+00 | 2.67E+00 | 2.58E-01 | no | Gm12948 | downstream_gene_variant | MODIFIER |
| 4 | 127297374 | AX-170390813 | 0 | T | C | 0.149 | 1.34E-01 | 1.19E-01 | 2.60E+00 | 2.67E+00 | 2.58E-01 | no | Gm12948 | non_coding_transcript_exon_variant | MODIFIER |
| 4 | 127297394 | AX-170373267 | 0 | C | T | 0.149 | 1.34E-01 | 1.19E-01 | 2.60E+00 | 2.67E+00 | 2.58E-01 | no | Gm12948 | non_coding_transcript_exon_variant | MODIFIER |
| 4 | 127297520 | AX-170023901 | 0 | T | C | 0.149 | 1.34E-01 | 1.19E-01 | 2.60E+00 | 2.67E+00 | 2.58E-01 | no | Gm12948 | upstream_gene_variant | MODIFIER |
| 4 | 127336771 | AX-170137892 | 0 | A | G | 0.149 | 1.34E-01 | 1.19E-01 | 2.60E+00 | 2.67E+00 | 2.58E-01 | no | | intergenic_variant | MODIFIER |
| 4 | 127346350 | AX-170213593 | 0 | C | T | 0.149 | 1.34E-01 | 1.19E-01 | 2.60E+00 | 2.67E+00 | 2.58E-01 | no | Gjb4 | downstream_gene_variant | MODIFIER |
| 4 | 127346499 | AX-97832519 | 0 | A | G | 0.149 | 1.34E-01 | 1.19E-01 | 2.60E+00 | 2.67E+00 | 2.58E-01 | no | Gjb4 | downstream_gene_variant | MODIFIER |
| 4 | 127348665 | AX-170392339 | 0 | G | A | 0.149 | 1.34E-01 | 1.19E-01 | 2.60E+00 | 2.67E+00 | 2.58E-01 | no | Gjb4 | downstream_gene_variant | MODIFIER |
| 4 | 127351302 | AX-170239602 | 0 | T | A | 0.149 | 1.34E-01 | 1.19E-01 | 2.60E+00 | 2.67E+00 | 2.58E-01 | no | Gjb5 | downstream_gene_variant | MODIFIER |
| 4 | 127356955 | AX-170070612 | 0 | G | T | 0.149 | 1.34E-01 | 1.19E-01 | 2.60E+00 | 2.67E+00 | 2.58E-01 | no | Gjb5 | intron_variant | MODIFIER |
| 4 | 127360080 | AX-170070614 | 0 | T | C | 0.149 | 1.34E-01 | 1.19E-01 | 2.60E+00 | 2.67E+00 | 2.58E-01 | no | Gjb5 | upstream_gene_variant | MODIFIER |
| 4 | 127360256 | AX-170070615 | 0 | T | C | 0.149 | 1.34E-01 | 1.19E-01 | 2.60E+00 | 2.67E+00 | 2.58E-01 | no | Gjb5 | upstream_gene_variant | MODIFIER |
| 4 | 127366292 | AX-170070613 | 0 | T | C | 0.149 | 1.34E-01 | 1.19E-01 | 2.60E+00 | 2.67E+00 | 2.58E-01 | no | | intergenic_variant | MODIFIER |
| 4 | 127368831 | AX-170211334 | 0 | G | C | 0.149 | 1.34E-01 | 1.19E-01 | 2.60E+00 | 2.67E+00 | 2.58E-01 | no | | regulatory_region_variant | MODIFIER |
| 4 | 127368932 | AX-170393806 | 0 | T | C | 0.149 | 1.34E-01 | 1.19E-01 | 2.60E+00 | 2.67E+00 | 2.58E-01 | no | | regulatory_region_variant | MODIFIER |
| 4 | 127372702 | AX-170385541 | 0 | A | G | 0.149 | 1.34E-01 | 1.19E-01 | 2.60E+00 | 2.67E+00 | 2.58E-01 | no | | intergenic_variant | MODIFIER |
| 4 | 127391216 | AX-170193480 | 0 | A | G | 0.149 | 1.34E-01 | 1.19E-01 | 2.60E+00 | 2.67E+00 | 2.58E-01 | no | | intergenic_variant | MODIFIER |
| 4 | 127399095 | AX-170269881 | 0 | G | A | 0.149 | 1.34E-01 | 1.19E-01 | 2.60E+00 | 2.67E+00 | 2.58E-01 | no | | intergenic_variant | MODIFIER |
| 4 | 127407210 | AX-170095143 | 0 | C | T | 0.149 | 1.34E-01 | 1.19E-01 | 2.60E+00 | 2.67E+00 | 2.58E-01 | no | Gm12947 | downstream_gene_variant | MODIFIER |
| 4 | 127416830 | AX-170367938 | 0 | A | G | 0.149 | 1.34E-01 | 1.19E-01 | 2.60E+00 | 2.67E+00 | 2.58E-01 | no | | regulatory_region_variant | MODIFIER |
| 4 | 127423861 | AX-170469766 | 0 | G | A | 0.149 | 1.34E-01 | 1.19E-01 | 2.60E+00 | 2.67E+00 | 2.58E-01 | no | | intergenic_variant | MODIFIER |
| 4 | 127424023 | AX-170220628 | 0 | G | A | 0.149 | 1.34E-01 | 1.19E-01 | 2.60E+00 | 2.67E+00 | 2.58E-01 | no | | regulatory_region_variant | MODIFIER |
| 4 | 127428834 | AX-170297921 | 0 | C | T | 0.149 | 1.34E-01 | 1.19E-01 | 2.60E+00 | 2.67E+00 | 2.58E-01 | no | | intergenic_variant | MODIFIER |
| 4 | 127442962 | AX-170251300 | 0 | A | G | 0.149 | 1.34E-01 | 1.19E-01 | 2.60E+00 | 2.67E+00 | 2.58E-01 | no | | intergenic_variant | MODIFIER |
| 4 | 127447944 | AX-170101805 | 0 | G | T | 0.149 | 1.34E-01 | 1.19E-01 | 2.60E+00 | 2.67E+00 | 2.58E-01 | no | | intergenic_variant | MODIFIER |
| 4 | 133131597 | AX-170023982 | 0 | A | T | 0.175 | 1.17E-01 | 1.04E-01 | 2.62E+00 | 2.69E+00 | 2.58E-01 | no | Wasf2 | intron_variant | MODIFIER |
| 4 | 95868441 | AX-170319526 | 1 | C | T | 0.259 | 1.22E-01 | 1.07E-01 | 2.49E+00 | 2.56E+00 | 2.58E-01 | no | Fggy | intron_variant | MODIFIER |
| 10 | 100818588 | AX-97801389 | 1 | C | A | 0.295 | -9.59E-02 | 8.51E-02 | 2.58E+00 | 2.64E+00 | 2.58E-01 | no | | intergenic_variant | MODIFIER |
| 6 | 125972595 | AX-170394519 | 0 | A | C | 0.184 | -8.49E-02 | 7.60E-02 | 2.79E+00 | 2.96E+00 | 2.58E-01 | no | Ano2 | intron_variant,NMD_transcript_variant | MODIFIER |
| 9 | 118714002 | AX-170033384 | 0 | C | T | 0.123 | -9.54E-02 | 8.54E-02 | 2.65E+00 | 2.80E+00 | 2.58E-01 | no | Itga9 | intron_variant | MODIFIER |
| 16 | 19239400 | AX-170250047 | 0 | A | G | 0.132 | 1.07E-01 | 9.55E-02 | 2.83E+00 | 2.98E+00 | 2.59E-01 | no | Iglv3 | downstream_gene_variant | MODIFIER |
| 4 | 68799222 | AX-170069591 | 0 | C | A | 0.333 | 1.31E-01 | 1.16E-01 | 2.82E+00 | 2.88E+00 | 2.59E-01 | no | Brinp1 | intron_variant | MODIFIER |
| 6 | 129641419 | AX-97837709 | 1 | G | A | 0.562 | 9.22E-02 | 8.06E-02 | 2.44E+00 | 2.51E+00 | 2.59E-01 | no | Klrc3 | missense_variant | MODERATE |
| 13 | 20177219 | AX-170134816 | 0 | A | C | 0.175 | -1.37E-01 | 1.20E-01 | 2.46E+00 | 2.52E+00 | 2.59E-01 | no | Elmo1 | intron_variant | MODIFIER |
| 8 | 25750672 | AX-170239410 | 1 | C | T | 0.357 | -7.26E-02 | 6.44E-02 | 2.55E+00 | 2.64E+00 | 2.59E-01 | no | Ddhd2 | intron_variant | MODIFIER |
| 1 | 75642143 | AX-170103843 | 0 | A | T | 0.14 | 2.10E-01 | 1.88E-01 | 2.80E+00 | 2.92E+00 | 2.59E-01 | Yes | Gm5257 | downstream_gene_variant | MODIFIER |
| 1 | 76454556 | AX-170118815 | 0 | A | G | 0.14 | 2.10E-01 | 1.88E-01 | 2.80E+00 | 2.92E+00 | 2.59E-01 | Yes | Gm29069 | intron_variant,non_coding_transcript_variant | MODIFIER |
| 13 | 11083368 | AX-170345208 | 0 | T | C | 0.193 | -1.29E-01 | 1.13E-01 | 2.49E+00 | 2.55E+00 | 2.59E-01 | no | | intergenic_variant | MODIFIER |

| | | | | | | | | | | | | | | | |
|---|---|---|---|---|---|---|---|---|---|---|---|---|---|---|---|
| 16 | 19210174 | AX-170530983 | 1 | C | T | 0.143 | 1.08E-01 | 9.64E-02 | 2.86E+00 | 3.00E+00 | 2.59E-01 | no | Gm43388 | downstream_gene_variant | MODIFIER |
| 8 | 37788793 | AX-170219489 | 1 | T | C | 0.152 | -1.52E-01 | 1.35E-01 | 2.88E+00 | 2.98E+00 | 2.59E-01 | no | Gm16205 | upstream_gene_variant | MODIFIER |
| 6 | 136834235 | AX-170477753 | 0 | A | G | 0.57 | 1.01E-01 | 8.79E-02 | 2.44E+00 | 2.49E+00 | 2.59E-01 | no | BC049715 | intron_variant | MODIFIER |
| 4 | 80926102 | AX-170287028 | 2 | T | C | 0.455 | -9.56E-02 | 8.44E-02 | 2.51E+00 | 2.56E+00 | 2.59E-01 | no | Lurap1l | intron_variant | MODIFIER |
| 16 | 77067780 | AX-170013428 | 0 | A | G | 0.456 | 8.33E-02 | 7.43E-02 | 2.60E+00 | 2.68E+00 | 2.59E-01 | no | Usp25 | intron_variant | MODIFIER |
| 7 | 39690897 | AX-170028871 | 0 | G | A | 0.351 | 1.04E-01 | 9.29E-02 | 2.58E+00 | 2.66E+00 | 2.59E-01 | no | | intergenic_variant | MODIFIER |
| 4 | 63026019 | AX-170022936 | 0 | T | C | 0.342 | 1.31E-01 | 1.17E-01 | 2.84E+00 | 2.89E+00 | 2.60E-01 | no | Zfp618 | intron_variant | MODIFIER |
| 4 | 63028263 | AX-97822345 | 0 | T | C | 0.342 | 1.31E-01 | 1.17E-01 | 2.84E+00 | 2.89E+00 | 2.60E-01 | no | Zfp618 | intron_variant | MODIFIER |
| 4 | 63069254 | AX-97839971 | 0 | T | C | 0.342 | 1.31E-01 | 1.17E-01 | 2.84E+00 | 2.89E+00 | 2.60E-01 | no | Zfp618 | intron_variant | MODIFIER |
| 4 | 63069840 | AX-170319939 | 0 | G | C | 0.342 | 1.31E-01 | 1.17E-01 | 2.84E+00 | 2.89E+00 | 2.60E-01 | no | Zfp618 | intron_variant | MODIFIER |
| 4 | 63076950 | AX-168386561 | 0 | T | C | 0.342 | 1.31E-01 | 1.17E-01 | 2.84E+00 | 2.89E+00 | 2.60E-01 | no | Zfp618 | intron_variant | MODIFIER |
| 4 | 63084121 | AX-170153082 | 0 | C | T | 0.342 | 1.31E-01 | 1.17E-01 | 2.84E+00 | 2.89E+00 | 2.60E-01 | no | Zfp618 | intron_variant | MODIFIER |
| 4 | 63087880 | AX-170128214 | 0 | C | T | 0.342 | 1.31E-01 | 1.17E-01 | 2.84E+00 | 2.89E+00 | 2.60E-01 | no | Zfp618 | intron_variant | MODIFIER |
| 4 | 63087914 | AX-97815238 | 0 | G | A | 0.342 | 1.31E-01 | 1.17E-01 | 2.84E+00 | 2.89E+00 | 2.60E-01 | no | Zfp618 | intron_variant | MODIFIER |
| 4 | 63100906 | AX-170254568 | 0 | C | T | 0.342 | 1.31E-01 | 1.17E-01 | 2.84E+00 | 2.89E+00 | 2.60E-01 | no | Zfp618 | intron_variant | MODIFIER |
| 4 | 63105593 | AX-170385006 | 0 | A | G | 0.342 | 1.31E-01 | 1.17E-01 | 2.84E+00 | 2.89E+00 | 2.60E-01 | no | Zfp618 | intron_variant | MODIFIER |
| 4 | 63258995 | AX-170089254 | 0 | T | C | 0.342 | 1.31E-01 | 1.17E-01 | 2.84E+00 | 2.89E+00 | 2.60E-01 | no | Col27a1 | intron_variant | MODIFIER |
| 4 | 63269943 | AX-170022941 | 0 | C | T | 0.342 | 1.31E-01 | 1.17E-01 | 2.84E+00 | 2.89E+00 | 2.60E-01 | no | Col27a1 | intron_variant | MODIFIER |
| 4 | 63272102 | AX-170266301 | 0 | G | A | 0.342 | 1.31E-01 | 1.17E-01 | 2.84E+00 | 2.89E+00 | 2.60E-01 | no | Col27a1 | intron_variant | MODIFIER |
| 4 | 63273175 | AX-170381143 | 0 | C | G | 0.342 | 1.31E-01 | 1.17E-01 | 2.84E+00 | 2.89E+00 | 2.60E-01 | no | Col27a1 | intron_variant | MODIFIER |
| 4 | 63278326 | AX-170123670 | 0 | T | C | 0.342 | 1.31E-01 | 1.17E-01 | 2.84E+00 | 2.89E+00 | 2.60E-01 | no | Col27a1 | intron_variant | MODIFIER |
| 4 | 63280683 | AX-170268340 | 0 | A | G | 0.342 | 1.31E-01 | 1.17E-01 | 2.84E+00 | 2.89E+00 | 2.60E-01 | no | Col27a1 | intron_variant | MODIFIER |
| 4 | 63289996 | AX-170022940 | 0 | G | C | 0.342 | 1.31E-01 | 1.17E-01 | 2.84E+00 | 2.89E+00 | 2.60E-01 | no | Col27a1 | intron_variant | MODIFIER |
| 4 | 63290409 | AX-97842183 | 0 | A | G | 0.342 | 1.31E-01 | 1.17E-01 | 2.84E+00 | 2.89E+00 | 2.60E-01 | no | Col27a1 | intron_variant | MODIFIER |
| 4 | 63305495 | AX-170022939 | 0 | G | A | 0.342 | 1.31E-01 | 1.17E-01 | 2.84E+00 | 2.89E+00 | 2.60E-01 | no | Col27a1 | intron_variant | MODIFIER |
| 4 | 63306898 | AX-170183636 | 0 | G | T | 0.342 | 1.31E-01 | 1.17E-01 | 2.84E+00 | 2.89E+00 | 2.60E-01 | no | Col27a1 | intron_variant | MODIFIER |
| 4 | 63375796 | AX-97815720 | 0 | T | C | 0.342 | 1.31E-01 | 1.17E-01 | 2.84E+00 | 2.89E+00 | 2.60E-01 | no | Akna | intron_variant | MODIFIER |
| 4 | 63381447 | AX-170089255 | 0 | T | G | 0.342 | 1.31E-01 | 1.17E-01 | 2.84E+00 | 2.89E+00 | 2.60E-01 | no | Akna | intron_variant | MODIFIER |
| 4 | 63412261 | AX-97828658 | 0 | A | G | 0.342 | 1.31E-01 | 1.17E-01 | 2.84E+00 | 2.89E+00 | 2.60E-01 | no | Whrn | downstream_gene_variant | MODIFIER |
| 4 | 63413276 | AX-170022945 | 0 | A | G | 0.342 | 1.31E-01 | 1.17E-01 | 2.84E+00 | 2.89E+00 | 2.60E-01 | no | Whrn | downstream_gene_variant | MODIFIER |
| 4 | 63476985 | AX-170022937 | 0 | G | A | 0.342 | 1.31E-01 | 1.17E-01 | 2.84E+00 | 2.89E+00 | 2.60E-01 | no | Whrn | intron_variant | MODIFIER |
| 4 | 63477955 | AX-170089256 | 0 | A | G | 0.342 | 1.31E-01 | 1.17E-01 | 2.84E+00 | 2.89E+00 | 2.60E-01 | no | Whrn | intron_variant | MODIFIER |
| 4 | 63504119 | AX-170278231 | 0 | G | T | 0.342 | 1.31E-01 | 1.17E-01 | 2.84E+00 | 2.89E+00 | 2.60E-01 | no | | intergenic_variant | MODIFIER |
| 4 | 63506608 | AX-170022947 | 0 | G | A | 0.342 | 1.31E-01 | 1.17E-01 | 2.84E+00 | 2.89E+00 | 2.60E-01 | no | | intergenic_variant | MODIFIER |
| 4 | 63508879 | AX-97820670 | 0 | G | A | 0.342 | 1.31E-01 | 1.17E-01 | 2.84E+00 | 2.89E+00 | 2.60E-01 | no | | regulatory_region_variant | MODIFIER |
| 4 | 63511867 | AX-170454125 | 0 | A | G | 0.342 | 1.31E-01 | 1.17E-01 | 2.84E+00 | 2.89E+00 | 2.60E-01 | no | | intergenic_variant | MODIFIER |
| 4 | 63520351 | AX-170022938 | 0 | G | A | 0.342 | 1.31E-01 | 1.17E-01 | 2.84E+00 | 2.89E+00 | 2.60E-01 | no | | intergenic_variant | MODIFIER |
| 4 | 63520904 | AX-170241630 | 0 | C | T | 0.342 | 1.31E-01 | 1.17E-01 | 2.84E+00 | 2.89E+00 | 2.60E-01 | no | | intergenic_variant | MODIFIER |
| 4 | 63525689 | AX-170501260 | 0 | A | C | 0.342 | 1.31E-01 | 1.17E-01 | 2.84E+00 | 2.89E+00 | 2.60E-01 | no | | intergenic_variant | MODIFIER |
| 4 | 63533537 | AX-170158696 | 0 | T | C | 0.342 | 1.31E-01 | 1.17E-01 | 2.84E+00 | 2.89E+00 | 2.60E-01 | no | | intergenic_variant | MODIFIER |
| 4 | 63534928 | AX-97820768 | 0 | G | A | 0.342 | 1.31E-01 | 1.17E-01 | 2.84E+00 | 2.89E+00 | 2.60E-01 | no | | intergenic_variant | MODIFIER |
| 4 | 63535775 | AX-168412231 | 0 | C | T | 0.342 | 1.31E-01 | 1.17E-01 | 2.84E+00 | 2.89E+00 | 2.60E-01 | no | | intergenic_variant | MODIFIER |
| 4 | 63568455 | AX-170094988 | 0 | A | G | 0.342 | 1.31E-01 | 1.17E-01 | 2.84E+00 | 2.89E+00 | 2.60E-01 | no | Tmem268 | splice_region_variant,intron_variant | LOW |
| 4 | 63606714 | AX-170310339 | 0 | G | A | 0.342 | 1.31E-01 | 1.17E-01 | 2.84E+00 | 2.89E+00 | 2.60E-01 | no | Tex48 | downstream_gene_variant | MODIFIER |
| 4 | 63617484 | AX-97818907 | 0 | A | G | 0.342 | 1.31E-01 | 1.17E-01 | 2.84E+00 | 2.89E+00 | 2.60E-01 | no | Tex48 | intron_variant | MODIFIER |
| 4 | 63621068 | AX-170477570 | 0 | A | G | 0.342 | 1.31E-01 | 1.17E-01 | 2.84E+00 | 2.89E+00 | 2.60E-01 | no | Tex48 | intron_variant | MODIFIER |
| 4 | 63623104 | AX-170393104 | 0 | C | T | 0.342 | 1.31E-01 | 1.17E-01 | 2.84E+00 | 2.89E+00 | 2.60E-01 | no | Tex48 | upstream_gene_variant | MODIFIER |
| 4 | 63635498 | AX-97852601 | 0 | G | A | 0.342 | 1.31E-01 | 1.17E-01 | 2.84E+00 | 2.89E+00 | 2.60E-01 | no | Gm24234 | upstream_gene_variant | MODIFIER |
| 4 | 63635937 | AX-170445247 | 0 | A | T | 0.342 | 1.31E-01 | 1.17E-01 | 2.84E+00 | 2.89E+00 | 2.60E-01 | no | Gm24234 | upstream_gene_variant | MODIFIER |
| 4 | 63669806 | AX-97842857 | 0 | T | C | 0.342 | 1.31E-01 | 1.17E-01 | 2.84E+00 | 2.89E+00 | 2.60E-01 | no | | intergenic_variant | MODIFIER |
| 4 | 63677074 | AX-170343837 | 0 | T | C | 0.342 | 1.31E-01 | 1.17E-01 | 2.84E+00 | 2.89E+00 | 2.60E-01 | no | | intergenic_variant | MODIFIER |
| 4 | 63683412 | AX-170488367 | 0 | G | A | 0.342 | 1.31E-01 | 1.17E-01 | 2.84E+00 | 2.89E+00 | 2.60E-01 | no | | intergenic_variant | MODIFIER |
| 4 | 63724497 | AX-97786997 | 0 | C | T | 0.342 | 1.31E-01 | 1.17E-01 | 2.84E+00 | 2.89E+00 | 2.60E-01 | no | Tnfsf15 | downstream_gene_variant | MODIFIER |
| 4 | 63726539 | AX-170288496 | 0 | T | C | 0.342 | 1.31E-01 | 1.17E-01 | 2.84E+00 | 2.89E+00 | 2.60E-01 | no | Tnfsf15 | downstream_gene_variant | MODIFIER |

| | | | | | | | | | | | | | | | |
|---|---|---|---|---|---|---|---|---|---|---|---|---|---|---|---|
| 4 | 63743081 | AX-170370357 | 0 | T | C | 0.342 | 1.31E-01 | 1.17E-01 | 2.84E+00 | 2.89E+00 | 2.60E-01 | no | Tnfsf15 | intron_variant | MODIFIER |
| 4 | 63752857 | AX-97805794 | 0 | G | A | 0.342 | 1.31E-01 | 1.17E-01 | 2.84E+00 | 2.89E+00 | 2.60E-01 | no | | regulatory_region_variant | MODIFIER |
| 4 | 63754324 | AX-170022960 | 0 | T | A | 0.342 | 1.31E-01 | 1.17E-01 | 2.84E+00 | 2.89E+00 | 2.60E-01 | no | | intergenic_variant | MODIFIER |
| 4 | 63784225 | AX-97841022 | 0 | C | T | 0.342 | 1.31E-01 | 1.17E-01 | 2.84E+00 | 2.89E+00 | 2.60E-01 | no | | intergenic_variant | MODIFIER |
| 4 | 63810261 | AX-97808607 | 0 | T | C | 0.342 | 1.31E-01 | 1.17E-01 | 2.84E+00 | 2.89E+00 | 2.60E-01 | no | | intergenic_variant | MODIFIER |
| 4 | 63824669 | AX-170022956 | 0 | A | G | 0.342 | 1.31E-01 | 1.17E-01 | 2.84E+00 | 2.89E+00 | 2.60E-01 | no | | intergenic_variant | MODIFIER |
| 4 | 63930350 | AX-97837595 | 0 | A | G | 0.342 | 1.31E-01 | 1.17E-01 | 2.84E+00 | 2.89E+00 | 2.60E-01 | no | | intergenic_variant | MODIFIER |
| 4 | 64003932 | AX-97812593 | 0 | A | C | 0.342 | 1.31E-01 | 1.17E-01 | 2.84E+00 | 2.89E+00 | 2.60E-01 | no | Tnc | intron_variant | MODIFIER |
| 4 | 64025423 | AX-170022955 | 0 | T | C | 0.342 | 1.31E-01 | 1.17E-01 | 2.84E+00 | 2.89E+00 | 2.60E-01 | no | Tnc | upstream_gene_variant | MODIFIER |
| 4 | 64979273 | AX-170122533 | 0 | G | A | 0.342 | 1.31E-01 | 1.17E-01 | 2.84E+00 | 2.89E+00 | 2.60E-01 | no | | intergenic_variant | MODIFIER |
| 4 | 68598421 | AX-97817671 | 0 | G | T | 0.342 | 1.31E-01 | 1.17E-01 | 2.84E+00 | 2.89E+00 | 2.60E-01 | no | | intergenic_variant | MODIFIER |
| 4 | 68613460 | AX-170023033 | 0 | G | C | 0.342 | 1.31E-01 | 1.17E-01 | 2.84E+00 | 2.89E+00 | 2.60E-01 | no | | intergenic_variant | MODIFIER |
| 4 | 68634564 | AX-97788888 | 0 | G | A | 0.342 | 1.31E-01 | 1.17E-01 | 2.84E+00 | 2.89E+00 | 2.60E-01 | no | | intergenic_variant | MODIFIER |
| 4 | 68717551 | AX-97814607 | 0 | A | G | 0.342 | 1.31E-01 | 1.17E-01 | 2.84E+00 | 2.89E+00 | 2.60E-01 | no | | intergenic_variant | MODIFIER |
| 4 | 68778146 | AX-170191666 | 0 | T | A | 0.342 | 1.31E-01 | 1.17E-01 | 2.84E+00 | 2.89E+00 | 2.60E-01 | no | Brinp1 | intron_variant | MODIFIER |
| 4 | 68952986 | AX-170069593 | 0 | C | A | 0.342 | 1.31E-01 | 1.17E-01 | 2.84E+00 | 2.89E+00 | 2.60E-01 | no | Brinp1 | intron_variant | MODIFIER |
| 4 | 68990606 | AX-170319808 | 0 | T | C | 0.342 | 1.31E-01 | 1.17E-01 | 2.84E+00 | 2.89E+00 | 2.60E-01 | no | | intergenic_variant | MODIFIER |
| 4 | 69028401 | AX-170527684 | 0 | C | T | 0.342 | 1.31E-01 | 1.17E-01 | 2.84E+00 | 2.89E+00 | 2.60E-01 | no | | intergenic_variant | MODIFIER |
| 4 | 69084436 | AX-170458118 | 0 | C | A | 0.342 | 1.31E-01 | 1.17E-01 | 2.84E+00 | 2.89E+00 | 2.60E-01 | no | | intergenic_variant | MODIFIER |
| 4 | 69117356 | AX-170405981 | 0 | A | G | 0.342 | 1.31E-01 | 1.17E-01 | 2.84E+00 | 2.89E+00 | 2.60E-01 | no | | intergenic_variant | MODIFIER |
| 4 | 69157248 | AX-170069599 | 0 | G | T | 0.342 | 1.31E-01 | 1.17E-01 | 2.84E+00 | 2.89E+00 | 2.60E-01 | no | | intergenic_variant | MODIFIER |
| 4 | 69215314 | AX-170023038 | 0 | C | T | 0.342 | 1.31E-01 | 1.17E-01 | 2.84E+00 | 2.89E+00 | 2.60E-01 | no | | intergenic_variant | MODIFIER |
| 4 | 69248705 | AX-170023037 | 0 | G | A | 0.342 | 1.31E-01 | 1.17E-01 | 2.84E+00 | 2.89E+00 | 2.60E-01 | no | | intergenic_variant | MODIFIER |
| 4 | 69252769 | AX-170252702 | 0 | G | A | 0.342 | 1.31E-01 | 1.17E-01 | 2.84E+00 | 2.89E+00 | 2.60E-01 | no | | intergenic_variant | MODIFIER |
| 4 | 69252820 | AX-170339925 | 0 | C | T | 0.342 | 1.31E-01 | 1.17E-01 | 2.84E+00 | 2.89E+00 | 2.60E-01 | no | | intergenic_variant | MODIFIER |
| 4 | 69358927 | AX-170358701 | 0 | A | C | 0.342 | 1.31E-01 | 1.17E-01 | 2.84E+00 | 2.89E+00 | 2.60E-01 | no | | intergenic_variant | MODIFIER |
| 4 | 69493445 | AX-170208222 | 0 | T | C | 0.342 | 1.31E-01 | 1.17E-01 | 2.84E+00 | 2.89E+00 | 2.60E-01 | no | | intergenic_variant | MODIFIER |
| 4 | 69528530 | AX-170069604 | 0 | T | C | 0.342 | 1.31E-01 | 1.17E-01 | 2.84E+00 | 2.89E+00 | 2.60E-01 | no | | intergenic_variant | MODIFIER |
| 4 | 69531922 | AX-170069607 | 0 | C | T | 0.342 | 1.31E-01 | 1.17E-01 | 2.84E+00 | 2.89E+00 | 2.60E-01 | no | | intergenic_variant | MODIFIER |
| 4 | 69533541 | AX-97810446 | 0 | A | G | 0.342 | 1.31E-01 | 1.17E-01 | 2.84E+00 | 2.89E+00 | 2.60E-01 | no | | intergenic_variant | MODIFIER |
| 4 | 69534527 | AX-170352413 | 0 | T | C | 0.342 | 1.31E-01 | 1.17E-01 | 2.84E+00 | 2.89E+00 | 2.60E-01 | no | | intergenic_variant | MODIFIER |
| 4 | 69534755 | AX-170535224 | 0 | T | C | 0.342 | 1.31E-01 | 1.17E-01 | 2.84E+00 | 2.89E+00 | 2.60E-01 | no | | intergenic_variant | MODIFIER |
| 4 | 69589486 | AX-97823763 | 0 | T | G | 0.342 | 1.31E-01 | 1.17E-01 | 2.84E+00 | 2.89E+00 | 2.60E-01 | no | | intergenic_variant | MODIFIER |
| 4 | 69593177 | AX-170164240 | 0 | G | A | 0.342 | 1.31E-01 | 1.17E-01 | 2.84E+00 | 2.89E+00 | 2.60E-01 | no | | intergenic_variant | MODIFIER |
| 16 | 29534295 | AX-170441857 | 1 | C | A | 0.509 | 8.96E-02 | 7.97E-02 | 2.58E+00 | 2.63E+00 | 2.60E-01 | no | Atp13a4 | intron_variant | MODIFIER |
| 4 | 64819038 | AX-170022980 | 1 | A | T | 0.304 | 1.28E-01 | 1.15E-01 | 2.71E+00 | 2.78E+00 | 2.60E-01 | no | | intergenic_variant | MODIFIER |
| 7 | 4211398 | AX-170521286 | 0 | G | T | 0.175 | 8.76E-02 | 7.83E-02 | 2.88E+00 | 3.03E+00 | 2.60E-01 | no | Gm15926 | downstream_gene_variant | MODIFIER |
| 7 | 4215253 | AX-170163202 | 0 | C | T | 0.175 | 8.76E-02 | 7.83E-02 | 2.88E+00 | 3.03E+00 | 2.60E-01 | no | | intergenic_variant | MODIFIER |
| 7 | 4215498 | AX-170266943 | 0 | C | A | 0.175 | 8.76E-02 | 7.83E-02 | 2.88E+00 | 3.03E+00 | 2.60E-01 | no | | intergenic_variant | MODIFIER |
| 7 | 4235946 | AX-170182188 | 0 | T | C | 0.175 | 8.76E-02 | 7.83E-02 | 2.88E+00 | 3.03E+00 | 2.60E-01 | no | Lilra5 | upstream_gene_variant | MODIFIER |
| 7 | 4338073 | AX-170403950 | 0 | C | T | 0.175 | 8.76E-02 | 7.83E-02 | 2.88E+00 | 3.03E+00 | 2.60E-01 | no | Ncr1 | intron_variant | MODIFIER |
| 7 | 4350873 | AX-170137898 | 0 | T | C | 0.175 | 8.76E-02 | 7.83E-02 | 2.88E+00 | 3.03E+00 | 2.60E-01 | no | | intergenic_variant | MODIFIER |
| 7 | 4373428 | AX-170188866 | 0 | C | A | 0.175 | 8.76E-02 | 7.83E-02 | 2.88E+00 | 3.03E+00 | 2.60E-01 | no | Gp6 | intron_variant | MODIFIER |
| 9 | 112575344 | AX-170147718 | 0 | T | C | 0.123 | -1.21E-01 | 1.08E-01 | 2.82E+00 | 2.93E+00 | 2.60E-01 | no | | intergenic_variant | MODIFIER |
| 3 | 26192364 | AX-170138006 | 0 | T | C | 0.132 | -1.18E-01 | 1.05E-01 | 2.58E+00 | 2.66E+00 | 2.60E-01 | no | Nlgn1 | intron_variant,non_coding_transcript_variant | MODIFIER |
| 3 | 26192622 | AX-97846299 | 0 | A | G | 0.132 | -1.18E-01 | 1.05E-01 | 2.58E+00 | 2.66E+00 | 2.60E-01 | no | Nlgn1 | intron_variant,non_coding_transcript_variant | MODIFIER |
| 3 | 26199828 | AX-170161641 | 0 | C | T | 0.132 | -1.18E-01 | 1.05E-01 | 2.58E+00 | 2.66E+00 | 2.60E-01 | no | Nlgn1 | intron_variant,non_coding_transcript_variant | MODIFIER |
| 3 | 26221408 | AX-170181931 | 0 | A | G | 0.132 | -1.18E-01 | 1.05E-01 | 2.58E+00 | 2.66E+00 | 2.60E-01 | no | Nlgn1 | intron_variant,non_coding_transcript_variant | MODIFIER |
| 3 | 26230830 | AX-170020159 | 0 | C | T | 0.132 | -1.18E-01 | 1.05E-01 | 2.58E+00 | 2.66E+00 | 2.60E-01 | no | Nlgn1 | intron_variant,non_coding_transcript_variant | MODIFIER |
| 3 | 26280422 | AX-170020164 | 0 | C | G | 0.132 | -1.18E-01 | 1.05E-01 | 2.58E+00 | 2.66E+00 | 2.60E-01 | no | Nlgn1 | intron_variant,non_coding_transcript_variant | MODIFIER |
| 3 | 26288089 | AX-170444073 | 0 | C | T | 0.132 | -1.18E-01 | 1.05E-01 | 2.58E+00 | 2.66E+00 | 2.60E-01 | no | Nlgn1 | intron_variant,non_coding_transcript_variant | MODIFIER |
| 3 | 26288137 | AX-170410938 | 0 | G | C | 0.132 | -1.18E-01 | 1.05E-01 | 2.58E+00 | 2.66E+00 | 2.60E-01 | no | Nlgn1 | intron_variant,non_coding_transcript_variant | MODIFIER |
| 3 | 26288492 | AX-170383980 | 0 | T | C | 0.132 | -1.18E-01 | 1.05E-01 | 2.58E+00 | 2.66E+00 | 2.60E-01 | no | Nlgn1 | intron_variant,non_coding_transcript_variant | MODIFIER |
| 3 | 26296626 | AX-170094555 | 0 | C | T | 0.132 | -1.18E-01 | 1.05E-01 | 2.58E+00 | 2.66E+00 | 2.60E-01 | no | Nlgn1 | intron_variant,non_coding_transcript_variant | MODIFIER |

| | | | | | | | | | | | | | | | |
|---|---|---|---|---|---|---|---|---|---|---|---|---|---|---|---|
| 3 | 26308402 | AX-170191391 | 0 | A | G | 0.132 | -1.18E-01 | 1.05E-01 | 2.58E+00 | 2.66E+00 | 2.60E-01 | no | Nlgn1 | intron_variant,non_coding_transcript_variant | MODIFIER |
| 3 | 26308536 | AX-170470369 | 0 | T | C | 0.132 | -1.18E-01 | 1.05E-01 | 2.58E+00 | 2.66E+00 | 2.60E-01 | no | Nlgn1 | intron_variant,non_coding_transcript_variant | MODIFIER |
| 3 | 26318859 | AX-170088888 | 0 | T | C | 0.132 | -1.18E-01 | 1.05E-01 | 2.58E+00 | 2.66E+00 | 2.60E-01 | no | Nlgn1 | intron_variant,non_coding_transcript_variant | MODIFIER |
| 3 | 26341446 | AX-170225649 | 0 | C | T | 0.132 | -1.18E-01 | 1.05E-01 | 2.58E+00 | 2.66E+00 | 2.60E-01 | no | | regulatory_region_variant | MODIFIER |
| 3 | 26341481 | AX-170284366 | 0 | G | A | 0.132 | -1.18E-01 | 1.05E-01 | 2.58E+00 | 2.66E+00 | 2.60E-01 | no | | regulatory_region_variant | MODIFIER |
| 3 | 26355113 | AX-170279904 | 0 | C | T | 0.132 | -1.18E-01 | 1.05E-01 | 2.58E+00 | 2.66E+00 | 2.60E-01 | no | | regulatory_region_variant | MODIFIER |
| 3 | 26361614 | AX-170240020 | 0 | T | C | 0.132 | -1.18E-01 | 1.05E-01 | 2.58E+00 | 2.66E+00 | 2.60E-01 | no | | intergenic_variant | MODIFIER |
| 3 | 26379437 | AX-170392883 | 0 | A | C | 0.132 | -1.18E-01 | 1.05E-01 | 2.58E+00 | 2.66E+00 | 2.60E-01 | no | Gm4855 | upstream_gene_variant | MODIFIER |
| 3 | 26382693 | AX-97828183 | 0 | T | C | 0.132 | -1.18E-01 | 1.05E-01 | 2.58E+00 | 2.66E+00 | 2.60E-01 | no | Gm4855 | downstream_gene_variant | MODIFIER |
| 3 | 26388135 | AX-97796604 | 0 | A | C | 0.132 | -1.18E-01 | 1.05E-01 | 2.58E+00 | 2.66E+00 | 2.60E-01 | no | | intergenic_variant | MODIFIER |
| 3 | 26433924 | AX-170146971 | 0 | T | C | 0.132 | -1.18E-01 | 1.05E-01 | 2.58E+00 | 2.66E+00 | 2.60E-01 | no | | intergenic_variant | MODIFIER |
| 3 | 26443064 | AX-170270426 | 0 | G | T | 0.132 | -1.18E-01 | 1.05E-01 | 2.58E+00 | 2.66E+00 | 2.60E-01 | no | | regulatory_region_variant | MODIFIER |
| 3 | 26643055 | AX-170066156 | 0 | C | T | 0.132 | -1.18E-01 | 1.05E-01 | 2.58E+00 | 2.66E+00 | 2.60E-01 | no | Spata16 | intron_variant | MODIFIER |
| 3 | 26647608 | AX-170405369 | 0 | A | G | 0.132 | -1.18E-01 | 1.05E-01 | 2.58E+00 | 2.66E+00 | 2.60E-01 | no | Spata16 | intron_variant | MODIFIER |
| 3 | 26669424 | AX-170174116 | 0 | C | T | 0.132 | -1.18E-01 | 1.05E-01 | 2.58E+00 | 2.66E+00 | 2.60E-01 | no | Spata16 | intron_variant | MODIFIER |
| 3 | 26743742 | AX-170371574 | 0 | T | C | 0.132 | -1.18E-01 | 1.05E-01 | 2.58E+00 | 2.66E+00 | 2.60E-01 | no | Spata16 | intron_variant | MODIFIER |
| 3 | 26756272 | AX-170502852 | 0 | G | C | 0.132 | -1.18E-01 | 1.05E-01 | 2.58E+00 | 2.66E+00 | 2.60E-01 | no | Spata16 | intron_variant | MODIFIER |
| 3 | 26773837 | AX-170020171 | 0 | C | A | 0.132 | -1.18E-01 | 1.05E-01 | 2.58E+00 | 2.66E+00 | 2.60E-01 | no | Spata16 | intron_variant | MODIFIER |
| 3 | 26776604 | AX-170020167 | 0 | G | T | 0.132 | -1.18E-01 | 1.05E-01 | 2.58E+00 | 2.66E+00 | 2.60E-01 | no | Spata16 | intron_variant | MODIFIER |
| 3 | 26781051 | AX-170315313 | 0 | T | C | 0.132 | -1.18E-01 | 1.05E-01 | 2.58E+00 | 2.66E+00 | 2.60E-01 | no | Spata16 | intron_variant | MODIFIER |
| 3 | 26793411 | AX-97807459 | 0 | A | G | 0.132 | -1.18E-01 | 1.05E-01 | 2.58E+00 | 2.66E+00 | 2.60E-01 | no | Spata16 | intron_variant | MODIFIER |
| 3 | 26818164 | AX-170315462 | 0 | C | T | 0.132 | -1.18E-01 | 1.05E-01 | 2.58E+00 | 2.66E+00 | 2.60E-01 | no | Spata16 | intron_variant | MODIFIER |
| 3 | 26818237 | AX-170094557 | 0 | G | A | 0.132 | -1.18E-01 | 1.05E-01 | 2.58E+00 | 2.66E+00 | 2.60E-01 | no | Spata16 | intron_variant | MODIFIER |
| 3 | 26831317 | AX-170020172 | 0 | C | T | 0.132 | -1.18E-01 | 1.05E-01 | 2.58E+00 | 2.66E+00 | 2.60E-01 | no | Spata16 | intron_variant | MODIFIER |
| 3 | 26831566 | AX-170499485 | 0 | T | C | 0.132 | -1.18E-01 | 1.05E-01 | 2.58E+00 | 2.66E+00 | 2.60E-01 | no | Spata16 | intron_variant | MODIFIER |
| 3 | 26841231 | AX-170104376 | 0 | C | G | 0.132 | -1.18E-01 | 1.05E-01 | 2.58E+00 | 2.66E+00 | 2.60E-01 | no | Spata16 | intron_variant | MODIFIER |
| 3 | 26872316 | AX-170066162 | 0 | G | A | 0.132 | -1.18E-01 | 1.05E-01 | 2.58E+00 | 2.66E+00 | 2.60E-01 | no | Spata16 | intron_variant | MODIFIER |
| 3 | 26878451 | AX-170066161 | 0 | G | C | 0.132 | -1.18E-01 | 1.05E-01 | 2.58E+00 | 2.66E+00 | 2.60E-01 | no | Spata16 | intron_variant | MODIFIER |
| 3 | 26878589 | AX-170066163 | 0 | G | A | 0.132 | -1.18E-01 | 1.05E-01 | 2.58E+00 | 2.66E+00 | 2.60E-01 | no | Spata16 | synonymous_variant | LOW |
| 3 | 26924535 | AX-97789163 | 0 | C | T | 0.132 | -1.18E-01 | 1.05E-01 | 2.58E+00 | 2.66E+00 | 2.60E-01 | no | Spata16 | intron_variant | MODIFIER |
| 3 | 26983274 | AX-170212532 | 0 | A | G | 0.132 | -1.18E-01 | 1.05E-01 | 2.58E+00 | 2.66E+00 | 2.60E-01 | no | Spata16 | downstream_gene_variant | MODIFIER |
| 3 | 26983471 | AX-170251727 | 0 | T | C | 0.132 | -1.18E-01 | 1.05E-01 | 2.58E+00 | 2.66E+00 | 2.60E-01 | no | Spata16 | downstream_gene_variant | MODIFIER |
| 3 | 26984659 | AX-170282015 | 0 | A | G | 0.132 | -1.18E-01 | 1.05E-01 | 2.58E+00 | 2.66E+00 | 2.60E-01 | no | Spata16 | downstream_gene_variant | MODIFIER |
| 3 | 26995411 | AX-170020165 | 0 | T | G | 0.132 | -1.18E-01 | 1.05E-01 | 2.58E+00 | 2.66E+00 | 2.60E-01 | no | | intergenic_variant | MODIFIER |
| 3 | 27047986 | AX-170252047 | 0 | T | C | 0.132 | -1.18E-01 | 1.05E-01 | 2.58E+00 | 2.66E+00 | 2.60E-01 | no | Gm7558 | upstream_gene_variant | MODIFIER |
| 3 | 27056448 | AX-97846784 | 0 | G | T | 0.132 | -1.18E-01 | 1.05E-01 | 2.58E+00 | 2.66E+00 | 2.60E-01 | no | | intergenic_variant | MODIFIER |
| 3 | 27192039 | AX-170314302 | 0 | T | C | 0.132 | -1.18E-01 | 1.05E-01 | 2.58E+00 | 2.66E+00 | 2.60E-01 | no | Nceh1 | intron_variant | MODIFIER |
| 1 | 138057904 | AX-97855538 | 0 | G | A | 0.211 | 1.42E-01 | 1.27E-01 | 2.71E+00 | 2.78E+00 | 2.60E-01 | no | Ptprc | downstream_gene_variant | MODIFIER |
| 1 | 138064955 | AX-170260766 | 0 | A | G | 0.211 | 1.42E-01 | 1.27E-01 | 2.71E+00 | 2.78E+00 | 2.60E-01 | no | Ptprc | intron_variant,non_coding_transcript_variant | MODIFIER |
| 1 | 138108870 | AX-170042879 | 0 | C | G | 0.211 | 1.42E-01 | 1.27E-01 | 2.71E+00 | 2.78E+00 | 2.60E-01 | no | Ptprc | intron_variant,non_coding_transcript_variant | MODIFIER |
| 1 | 138111059 | AX-170042881 | 0 | A | G | 0.211 | 1.42E-01 | 1.27E-01 | 2.71E+00 | 2.78E+00 | 2.60E-01 | no | Ptprc | intron_variant,non_coding_transcript_variant | MODIFIER |
| 1 | 138115368 | AX-170416434 | 0 | C | T | 0.211 | 1.42E-01 | 1.27E-01 | 2.71E+00 | 2.78E+00 | 2.60E-01 | no | Ptprc | intron_variant,non_coding_transcript_variant | MODIFIER |
| 1 | 138141443 | AX-170129664 | 0 | T | C | 0.211 | 1.42E-01 | 1.27E-01 | 2.71E+00 | 2.78E+00 | 2.60E-01 | no | Ptprc | intron_variant,non_coding_transcript_variant | MODIFIER |
| 1 | 138141510 | AX-97786328 | 0 | C | T | 0.211 | 1.42E-01 | 1.27E-01 | 2.71E+00 | 2.78E+00 | 2.60E-01 | no | Ptprc | intron_variant,non_coding_transcript_variant | MODIFIER |
| 1 | 138141883 | AX-170472143 | 0 | T | C | 0.211 | 1.42E-01 | 1.27E-01 | 2.71E+00 | 2.78E+00 | 2.60E-01 | no | Ptprc | intron_variant,non_coding_transcript_variant | MODIFIER |
| 1 | 138158457 | AX-170480318 | 0 | A | G | 0.211 | 1.42E-01 | 1.27E-01 | 2.71E+00 | 2.78E+00 | 2.60E-01 | no | Ptprc | intron_variant,non_coding_transcript_variant | MODIFIER |
| 1 | 138159265 | AX-170430445 | 0 | A | G | 0.211 | 1.42E-01 | 1.27E-01 | 2.71E+00 | 2.78E+00 | 2.60E-01 | no | Ptprc | intron_variant,non_coding_transcript_variant | MODIFIER |
| 1 | 138159821 | AX-170001810 | 0 | T | C | 0.211 | 1.42E-01 | 1.27E-01 | 2.71E+00 | 2.78E+00 | 2.60E-01 | no | Ptprc | intron_variant,non_coding_transcript_variant | MODIFIER |
| 1 | 138163461 | AX-170451726 | 0 | G | T | 0.211 | 1.42E-01 | 1.27E-01 | 2.71E+00 | 2.78E+00 | 2.60E-01 | no | Ptprc | intron_variant,non_coding_transcript_variant | MODIFIER |
| 1 | 138500735 | AX-97781848 | 0 | C | T | 0.211 | 1.42E-01 | 1.27E-01 | 2.71E+00 | 2.78E+00 | 2.60E-01 | no | Nek7 | intron_variant | MODIFIER |
| 1 | 138602464 | AX-97800340 | 0 | A | G | 0.211 | 1.42E-01 | 1.27E-01 | 2.71E+00 | 2.78E+00 | 2.60E-01 | no | Nek7 | intron_variant | MODIFIER |
| 1 | 138632715 | AX-170086626 | 0 | A | G | 0.211 | 1.42E-01 | 1.27E-01 | 2.71E+00 | 2.78E+00 | 2.60E-01 | no | | intergenic_variant | MODIFIER |
| 1 | 138660724 | AX-170430222 | 0 | A | G | 0.211 | 1.42E-01 | 1.27E-01 | 2.71E+00 | 2.78E+00 | 2.60E-01 | no | Gm5833 | intron_variant,non_coding_transcript_variant | MODIFIER |
| 1 | 138666268 | AX-170325482 | 0 | G | A | 0.211 | 1.42E-01 | 1.27E-01 | 2.71E+00 | 2.78E+00 | 2.60E-01 | no | Gm5833 | intron_variant,non_coding_transcript_variant | MODIFIER |

| | | | | | | | | | | | | | | | |
|---|---|---|---|---|---|---|---|---|---|---|---|---|---|---|---|
| 1 | 138666439 | AX-170380821 | 0 | G | C | 0.211 | 1.42E-01 | 1.27E-01 | 2.71E+00 | 2.78E+00 | 2.60E-01 | no | Gm5833 | downstream_gene_variant | MODIFIER |
| 1 | 138666662 | AX-170371507 | 0 | G | A | 0.211 | 1.42E-01 | 1.27E-01 | 2.71E+00 | 2.78E+00 | 2.60E-01 | no | Gm5833 | downstream_gene_variant | MODIFIER |
| 1 | 138690744 | AX-170512399 | 0 | T | A | 0.211 | 1.42E-01 | 1.27E-01 | 2.71E+00 | 2.78E+00 | 2.60E-01 | no | Gm8790 | upstream_gene_variant | MODIFIER |
| 1 | 138696025 | AX-97830316 | 0 | G | A | 0.211 | 1.42E-01 | 1.27E-01 | 2.71E+00 | 2.78E+00 | 2.60E-01 | no | | intergenic_variant | MODIFIER |
| 1 | 138738131 | AX-170091709 | 0 | A | G | 0.211 | 1.42E-01 | 1.27E-01 | 2.71E+00 | 2.78E+00 | 2.60E-01 | no | | intergenic_variant | MODIFIER |
| 1 | 138798328 | AX-170513554 | 0 | C | T | 0.211 | 1.42E-01 | 1.27E-01 | 2.71E+00 | 2.78E+00 | 2.60E-01 | no | | intergenic_variant | MODIFIER |
| 1 | 138798497 | AX-170171656 | 0 | T | C | 0.211 | 1.42E-01 | 1.27E-01 | 2.71E+00 | 2.78E+00 | 2.60E-01 | no | | intergenic_variant | MODIFIER |
| 1 | 138802795 | AX-170001815 | 0 | G | A | 0.211 | 1.42E-01 | 1.27E-01 | 2.71E+00 | 2.78E+00 | 2.60E-01 | no | | intergenic_variant | MODIFIER |
| 1 | 138802812 | AX-170001813 | 0 | A | G | 0.211 | 1.42E-01 | 1.27E-01 | 2.71E+00 | 2.78E+00 | 2.60E-01 | no | | intergenic_variant | MODIFIER |
| 1 | 138802947 | AX-170203338 | 0 | T | A | 0.211 | 1.42E-01 | 1.27E-01 | 2.71E+00 | 2.78E+00 | 2.60E-01 | no | | intergenic_variant | MODIFIER |
| 1 | 138841388 | AX-97816453 | 0 | A | C | 0.211 | 1.42E-01 | 1.27E-01 | 2.71E+00 | 2.78E+00 | 2.60E-01 | no | Lhx9 | intron_variant | MODIFIER |
| 1 | 138846628 | AX-97813427 | 0 | T | C | 0.211 | 1.42E-01 | 1.27E-01 | 2.71E+00 | 2.78E+00 | 2.60E-01 | no | Lhx9 | upstream_gene_variant | MODIFIER |
| 1 | 139005207 | AX-170500855 | 0 | G | C | 0.211 | 1.42E-01 | 1.27E-01 | 2.71E+00 | 2.78E+00 | 2.60E-01 | no | Dennd1b | intron_variant | MODIFIER |
| 1 | 139028757 | AX-170149164 | 0 | T | C | 0.211 | 1.42E-01 | 1.27E-01 | 2.71E+00 | 2.78E+00 | 2.60E-01 | no | Dennd1b | intron_variant | MODIFIER |
| 1 | 139042753 | AX-170387533 | 0 | A | G | 0.211 | 1.42E-01 | 1.27E-01 | 2.71E+00 | 2.78E+00 | 2.60E-01 | no | Dennd1b | intron_variant | MODIFIER |
| 1 | 139048063 | AX-170391872 | 0 | A | C | 0.211 | 1.42E-01 | 1.27E-01 | 2.71E+00 | 2.78E+00 | 2.60E-01 | no | Dennd1b | intron_variant | MODIFIER |
| 1 | 139048104 | AX-170187237 | 0 | G | C | 0.211 | 1.42E-01 | 1.27E-01 | 2.71E+00 | 2.78E+00 | 2.60E-01 | no | Dennd1b | intron_variant | MODIFIER |
| 1 | 139048189 | AX-170153198 | 0 | G | T | 0.211 | 1.42E-01 | 1.27E-01 | 2.71E+00 | 2.78E+00 | 2.60E-01 | no | Dennd1b | intron_variant | MODIFIER |
| 1 | 139050419 | AX-170280544 | 0 | G | T | 0.211 | 1.42E-01 | 1.27E-01 | 2.71E+00 | 2.78E+00 | 2.60E-01 | no | Dennd1b | intron_variant | MODIFIER |
| 1 | 139057940 | AX-170488145 | 0 | C | A | 0.211 | 1.42E-01 | 1.27E-01 | 2.71E+00 | 2.78E+00 | 2.60E-01 | no | Dennd1b | intron_variant | MODIFIER |
| 1 | 139067645 | AX-170430712 | 0 | C | T | 0.211 | 1.42E-01 | 1.27E-01 | 2.71E+00 | 2.78E+00 | 2.60E-01 | no | Dennd1b | intron_variant | MODIFIER |
| 1 | 139077329 | AX-170151646 | 0 | A | G | 0.211 | 1.42E-01 | 1.27E-01 | 2.71E+00 | 2.78E+00 | 2.60E-01 | no | Dennd1b | intron_variant | MODIFIER |
| 1 | 139077684 | AX-97813424 | 0 | A | G | 0.211 | 1.42E-01 | 1.27E-01 | 2.71E+00 | 2.78E+00 | 2.60E-01 | no | Dennd1b | intron_variant | MODIFIER |
| 1 | 139097135 | AX-97781013 | 0 | T | C | 0.211 | 1.42E-01 | 1.27E-01 | 2.71E+00 | 2.78E+00 | 2.60E-01 | no | Dennd1b | intron_variant | MODIFIER |
| 1 | 139230029 | AX-170086627 | 0 | A | G | 0.211 | 1.42E-01 | 1.27E-01 | 2.71E+00 | 2.78E+00 | 2.60E-01 | no | Crb1 | intron_variant | MODIFIER |
| 1 | 139237779 | AX-170318636 | 0 | A | G | 0.211 | 1.42E-01 | 1.27E-01 | 2.71E+00 | 2.78E+00 | 2.60E-01 | no | Crb1 | intron_variant | MODIFIER |
| 1 | 140483128 | AX-97842875 | 0 | A | G | 0.211 | 1.42E-01 | 1.27E-01 | 2.71E+00 | 2.78E+00 | 2.60E-01 | no | Kcnt2 | intron_variant | MODIFIER |
| 1 | 140530476 | AX-97808868 | 0 | C | T | 0.211 | 1.42E-01 | 1.27E-01 | 2.71E+00 | 2.78E+00 | 2.60E-01 | no | Kcnt2 | intron_variant | MODIFIER |
| 1 | 140532066 | AX-170295369 | 0 | T | C | 0.211 | 1.42E-01 | 1.27E-01 | 2.71E+00 | 2.78E+00 | 2.60E-01 | no | Kcnt2 | intron_variant | MODIFIER |
| 1 | 140553347 | AX-170001839 | 0 | C | T | 0.211 | 1.42E-01 | 1.27E-01 | 2.71E+00 | 2.78E+00 | 2.60E-01 | no | Kcnt2 | intron_variant | MODIFIER |
| 1 | 140621266 | AX-170001840 | 0 | A | G | 0.211 | 1.42E-01 | 1.27E-01 | 2.71E+00 | 2.78E+00 | 2.60E-01 | no | 4930590L20Rik | intron_variant,non_coding_transcript_variant | MODIFIER |
| 16 | 75697482 | AX-170013404 | 0 | T | C | 0.491 | 8.44E-02 | 7.45E-02 | 2.49E+00 | 2.57E+00 | 2.60E-01 | no | Gm30162 | intron_variant,non_coding_transcript_variant | MODIFIER |
| 1 | 176986960 | AX-170534062 | 1 | A | C | 0.232 | 1.29E-01 | 1.15E-01 | 2.64E+00 | 2.72E+00 | 2.60E-01 | no | Sdccag8 | intron_variant | MODIFIER |
| 5 | 51950236 | AX-170199099 | 1 | A | T | 0.473 | 1.15E-01 | 1.01E-01 | 2.43E+00 | 2.48E+00 | 2.60E-01 | no | | intergenic_variant | MODIFIER |
| 15 | 19336932 | AX-97807241 | 2 | C | T | 0.245 | -1.19E-01 | 1.06E-01 | 2.68E+00 | 2.74E+00 | 2.60E-01 | no | Gm49162 | intron_variant,non_coding_transcript_variant | MODIFIER |
| 9 | 98188410 | AX-170287264 | 1 | C | T | 0.143 | -1.28E-01 | 1.15E-01 | 2.59E+00 | 2.66E+00 | 2.60E-01 | no | 2610303G11Rik | upstream_gene_variant | MODIFIER |
| 16 | 57208355 | AX-97831423 | 0 | A | G | 0.465 | -9.51E-02 | 8.41E-02 | 2.51E+00 | 2.55E+00 | 2.61E-01 | no | Tbc1d23 | intron_variant | MODIFIER |
| 8 | 55106769 | AX-170453079 | 0 | C | A | 0.158 | -1.27E-01 | 1.14E-01 | 2.68E+00 | 2.82E+00 | 2.61E-01 | no | | intergenic_variant | MODIFIER |
| 1 | 20311074 | AX-170268434 | 0 | C | T | 0.404 | 9.12E-02 | 8.13E-02 | 2.58E+00 | 2.64E+00 | 2.61E-01 | no | Pkhd1 | intron_variant | MODIFIER |
| 1 | 21756553 | AX-170451814 | 0 | A | T | 0.404 | 9.12E-02 | 8.13E-02 | 2.58E+00 | 2.64E+00 | 2.61E-01 | no | Kcnq5 | intron_variant | MODIFIER |
| 15 | 3205655 | AX-170429922 | 0 | G | A | 0.246 | -1.16E-01 | 1.04E-01 | 2.66E+00 | 2.72E+00 | 2.61E-01 | no | | intergenic_variant | MODIFIER |
| 4 | 86117051 | AX-97827043 | 1 | A | C | 0.33 | 1.24E-01 | 1.12E-01 | 2.78E+00 | 2.85E+00 | 2.61E-01 | no | Adamtsl1 | intron_variant | MODIFIER |
| 4 | 95753851 | AX-170070062 | 0 | G | A | 0.254 | 1.28E-01 | 1.12E-01 | 2.40E+00 | 2.45E+00 | 2.61E-01 | no | Fggy | intron_variant | MODIFIER |
| 4 | 95761919 | AX-170401910 | 0 | C | T | 0.254 | 1.28E-01 | 1.12E-01 | 2.40E+00 | 2.45E+00 | 2.61E-01 | no | Fggy | intron_variant | MODIFIER |
| 4 | 95762460 | AX-170089320 | 0 | A | G | 0.254 | 1.28E-01 | 1.12E-01 | 2.40E+00 | 2.45E+00 | 2.61E-01 | no | Fggy | intron_variant | MODIFIER |
| 13 | 18758115 | AX-170165381 | 1 | A | G | 0.17 | -1.29E-01 | 1.14E-01 | 2.50E+00 | 2.58E+00 | 2.62E-01 | no | Vps41 | intron_variant | MODIFIER |
| 16 | 80212951 | AX-170443396 | 0 | G | A | 0.474 | 8.27E-02 | 7.41E-02 | 2.61E+00 | 2.68E+00 | 2.62E-01 | no | | intergenic_variant | MODIFIER |
| 7 | 72826074 | AX-170127163 | 0 | C | G | 0.368 | -1.13E-01 | 1.02E-01 | 2.80E+00 | 2.89E+00 | 2.62E-01 | no | | intergenic_variant | MODIFIER |
| 5 | 66723677 | AX-170495204 | 1 | T | C | 0.482 | 1.13E-01 | 1.01E-01 | 2.54E+00 | 2.60E+00 | 2.62E-01 | no | Gm43280 | downstream_gene_variant | MODIFIER |
| 1 | 149022803 | AX-170223646 | 2 | A | T | 0.209 | 1.53E-01 | 1.37E-01 | 2.72E+00 | 2.78E+00 | 2.62E-01 | Yes | Gm15443 | downstream_gene_variant | MODIFIER |
| 8 | 38262341 | AX-170377787 | 1 | T | A | 0.161 | -1.45E-01 | 1.29E-01 | 2.92E+00 | 3.03E+00 | 2.62E-01 | no | Sgcz | intron_variant | MODIFIER |
| 4 | 101525424 | AX-170392286 | 0 | G | A | 0.368 | 9.95E-02 | 8.95E-02 | 2.65E+00 | 2.76E+00 | 2.62E-01 | no | Dnajc6 | intron_variant | MODIFIER |
| 1 | 165183225 | AX-170002198 | 1 | G | A | 0.241 | 1.26E-01 | 1.13E-01 | 2.91E+00 | 3.06E+00 | 2.62E-01 | no | Sft2d2 | intron_variant | MODIFIER |
| 18 | 53947271 | AX-170490626 | 0 | C | T | 0.053 | 2.45E-01 | 2.21E-01 | 2.71E+00 | 2.85E+00 | 2.62E-01 | Yes | Csnk1g3 | intron_variant | MODIFIER |

| | | | | | | | | | | | | | | | |
|---|---|---|---|---|---|---|---|---|---|---|---|---|---|---|---|
| 13 | 6056002 | AX-170229907 | 1 | G | T | 0.205 | -1.31E-01 | 1.17E-01 | 2.63E+00 | 2.70E+00 | 2.62E-01 | no | | intergenic_variant | MODIFIER |
| 15 | 3614154 | AX-170251194 | 2 | C | T | 0.245 | -1.15E-01 | 1.04E-01 | 2.66E+00 | 2.72E+00 | 2.62E-01 | no | | intergenic_variant | MODIFIER |
| 18 | 80493728 | AX-170061454 | 1 | T | G | 0.143 | -1.33E-01 | 1.18E-01 | 2.50E+00 | 2.56E+00 | 2.62E-01 | no | | intergenic_variant | MODIFIER |
| 1 | 96387855 | AX-97812419 | 2 | C | T | 0.155 | 1.97E-01 | 1.77E-01 | 2.74E+00 | 2.85E+00 | 2.62E-01 | Yes | | intergenic_variant | MODIFIER |
| 18 | 79978186 | AX-97803746 | 0 | A | G | 0.447 | 9.34E-02 | 8.21E-02 | 2.43E+00 | 2.51E+00 | 2.62E-01 | no | | intergenic_variant | MODIFIER |
| 16 | 86229383 | AX-97832626 | 0 | T | C | 0.447 | 8.63E-02 | 7.77E-02 | 2.78E+00 | 2.86E+00 | 2.62E-01 | no | | intergenic_variant | MODIFIER |
| 5 | 67539560 | AX-170102339 | 0 | A | C | 0.474 | 1.12E-01 | 1.00E-01 | 2.53E+00 | 2.59E+00 | 2.63E-01 | no | | intergenic_variant | MODIFIER |
| 5 | 67590533 | AX-170281571 | 0 | A | G | 0.474 | 1.12E-01 | 1.00E-01 | 2.53E+00 | 2.59E+00 | 2.63E-01 | no | | intergenic_variant | MODIFIER |
| 5 | 67590895 | AX-170169975 | 0 | T | C | 0.474 | 1.12E-01 | 1.00E-01 | 2.53E+00 | 2.59E+00 | 2.63E-01 | no | | intergenic_variant | MODIFIER |
| 5 | 67682741 | AX-170152694 | 0 | C | T | 0.474 | 1.12E-01 | 1.00E-01 | 2.53E+00 | 2.59E+00 | 2.63E-01 | no | Atp8a1 | intron_variant | MODIFIER |
| 5 | 67961831 | AX-170496684 | 0 | C | T | 0.474 | 1.12E-01 | 1.00E-01 | 2.53E+00 | 2.59E+00 | 2.63E-01 | no | Gm5108 | intron_variant | MODIFIER |
| 5 | 67961967 | AX-170181177 | 0 | G | A | 0.474 | 1.12E-01 | 1.00E-01 | 2.53E+00 | 2.59E+00 | 2.63E-01 | no | Gm5108 | intron_variant | MODIFIER |
| 5 | 67962306 | AX-170210044 | 0 | G | A | 0.474 | 1.12E-01 | 1.00E-01 | 2.53E+00 | 2.59E+00 | 2.63E-01 | no | Gm5108 | intron_variant | MODIFIER |
| 5 | 67965789 | AX-170160731 | 0 | T | C | 0.474 | 1.12E-01 | 1.00E-01 | 2.53E+00 | 2.59E+00 | 2.63E-01 | no | Gm5108 | intron_variant | MODIFIER |
| 5 | 67974783 | AX-97794439 | 0 | A | C | 0.474 | 1.12E-01 | 1.00E-01 | 2.53E+00 | 2.59E+00 | 2.63E-01 | no | Gm5108 | intron_variant | MODIFIER |
| 5 | 67980651 | AX-170525683 | 0 | A | G | 0.474 | 1.12E-01 | 1.00E-01 | 2.53E+00 | 2.59E+00 | 2.63E-01 | no | Gm5108 | downstream_gene_variant | MODIFIER |
| 5 | 67983314 | AX-170474811 | 0 | G | T | 0.474 | 1.12E-01 | 1.00E-01 | 2.53E+00 | 2.59E+00 | 2.63E-01 | no | Gm43027 | non_coding_transcript_exon_variant | MODIFIER |
| 5 | 68007838 | AX-170236540 | 0 | T | C | 0.474 | 1.12E-01 | 1.00E-01 | 2.53E+00 | 2.59E+00 | 2.63E-01 | no | | intergenic_variant | MODIFIER |
| 5 | 68033763 | AX-170155461 | 0 | C | T | 0.474 | 1.12E-01 | 1.00E-01 | 2.53E+00 | 2.59E+00 | 2.63E-01 | no | Grxcr1 | intron_variant | MODIFIER |
| 5 | 68061745 | AX-170497758 | 0 | T | A | 0.474 | 1.12E-01 | 1.00E-01 | 2.53E+00 | 2.59E+00 | 2.63E-01 | no | Grxcr1 | intron_variant | MODIFIER |
| 5 | 68083120 | AX-170313835 | 0 | G | A | 0.474 | 1.12E-01 | 1.00E-01 | 2.53E+00 | 2.59E+00 | 2.63E-01 | no | Grxcr1 | intron_variant | MODIFIER |
| 5 | 68084809 | AX-170253648 | 0 | T | C | 0.474 | 1.12E-01 | 1.00E-01 | 2.53E+00 | 2.59E+00 | 2.63E-01 | no | Grxcr1 | intron_variant | MODIFIER |
| 5 | 68088235 | AX-97856814 | 0 | T | G | 0.474 | 1.12E-01 | 1.00E-01 | 2.53E+00 | 2.59E+00 | 2.63E-01 | no | Grxcr1 | intron_variant | MODIFIER |
| 5 | 68089323 | AX-170072378 | 0 | T | C | 0.474 | 1.12E-01 | 1.00E-01 | 2.53E+00 | 2.59E+00 | 2.63E-01 | no | Grxcr1 | intron_variant | MODIFIER |
| 5 | 68089724 | AX-170072380 | 0 | A | C | 0.474 | 1.12E-01 | 1.00E-01 | 2.53E+00 | 2.59E+00 | 2.63E-01 | no | Grxcr1 | intron_variant | MODIFIER |
| 5 | 68110635 | AX-170206592 | 0 | G | A | 0.474 | 1.12E-01 | 1.00E-01 | 2.53E+00 | 2.59E+00 | 2.63E-01 | no | Grxcr1 | intron_variant | MODIFIER |
| 5 | 68119490 | AX-170297958 | 0 | C | A | 0.474 | 1.12E-01 | 1.00E-01 | 2.53E+00 | 2.59E+00 | 2.63E-01 | no | Grxcr1 | intron_variant | MODIFIER |
| 5 | 68263525 | AX-170214265 | 0 | G | T | 0.474 | 1.12E-01 | 1.00E-01 | 2.53E+00 | 2.59E+00 | 2.63E-01 | no | | intergenic_variant | MODIFIER |
| 5 | 68294939 | AX-170228956 | 0 | A | G | 0.474 | 1.12E-01 | 1.00E-01 | 2.53E+00 | 2.59E+00 | 2.63E-01 | no | | intergenic_variant | MODIFIER |
| 5 | 68295327 | AX-170168241 | 0 | G | A | 0.474 | 1.12E-01 | 1.00E-01 | 2.53E+00 | 2.59E+00 | 2.63E-01 | no | | intergenic_variant | MODIFIER |
| 5 | 68365994 | AX-170535061 | 0 | G | T | 0.474 | 1.12E-01 | 1.00E-01 | 2.53E+00 | 2.59E+00 | 2.63E-01 | no | | intergenic_variant | MODIFIER |
| 5 | 68366174 | AX-170072388 | 0 | T | C | 0.474 | 1.12E-01 | 1.00E-01 | 2.53E+00 | 2.59E+00 | 2.63E-01 | no | | intergenic_variant | MODIFIER |
| 5 | 68366565 | AX-97795757 | 0 | C | T | 0.474 | 1.12E-01 | 1.00E-01 | 2.53E+00 | 2.59E+00 | 2.63E-01 | no | | intergenic_variant | MODIFIER |
| 8 | 49712690 | AX-170448728 | 0 | A | T | 0.158 | -1.35E-01 | 1.22E-01 | 2.80E+00 | 2.92E+00 | 2.63E-01 | no | | intergenic_variant | MODIFIER |
| 13 | 19913782 | AX-170290170 | 0 | A | G | 0.167 | -1.44E-01 | 1.28E-01 | 2.49E+00 | 2.54E+00 | 2.63E-01 | no | | intergenic_variant | MODIFIER |
| 5 | 48054718 | AX-170284859 | 0 | C | T | 0.5 | 1.20E-01 | 1.05E-01 | 2.40E+00 | 2.44E+00 | 2.63E-01 | no | Slit2 | intron_variant,NMD_transcript_variant | MODIFIER |
| 5 | 48506243 | AX-170509574 | 0 | T | C | 0.5 | 1.20E-01 | 1.05E-01 | 2.40E+00 | 2.44E+00 | 2.63E-01 | no | Kcnip4 | intron_variant | MODIFIER |
| 5 | 48506277 | AX-170024885 | 0 | A | G | 0.5 | 1.20E-01 | 1.05E-01 | 2.40E+00 | 2.44E+00 | 2.63E-01 | no | Kcnip4 | intron_variant | MODIFIER |
| 5 | 48507467 | AX-170363063 | 0 | T | C | 0.5 | 1.20E-01 | 1.05E-01 | 2.40E+00 | 2.44E+00 | 2.63E-01 | no | Kcnip4 | intron_variant | MODIFIER |
| 5 | 48513238 | AX-170336547 | 0 | C | G | 0.5 | 1.20E-01 | 1.05E-01 | 2.40E+00 | 2.44E+00 | 2.63E-01 | no | Kcnip4 | intron_variant | MODIFIER |
| 5 | 48520747 | AX-170395601 | 0 | A | G | 0.5 | 1.20E-01 | 1.05E-01 | 2.40E+00 | 2.44E+00 | 2.63E-01 | no | Kcnip4 | intron_variant | MODIFIER |
| 5 | 48534542 | AX-170024882 | 0 | C | T | 0.5 | 1.20E-01 | 1.05E-01 | 2.40E+00 | 2.44E+00 | 2.63E-01 | no | Kcnip4 | intron_variant | MODIFIER |
| 5 | 48542243 | AX-170228389 | 0 | G | A | 0.5 | 1.20E-01 | 1.05E-01 | 2.40E+00 | 2.44E+00 | 2.63E-01 | no | Kcnip4 | intron_variant | MODIFIER |
| 5 | 48543002 | AX-170024879 | 0 | A | G | 0.5 | 1.20E-01 | 1.05E-01 | 2.40E+00 | 2.44E+00 | 2.63E-01 | no | Kcnip4 | intron_variant | MODIFIER |
| 5 | 48559443 | AX-170129758 | 0 | C | G | 0.5 | 1.20E-01 | 1.05E-01 | 2.40E+00 | 2.44E+00 | 2.63E-01 | no | Kcnip4 | intron_variant | MODIFIER |
| 5 | 48559983 | AX-170135021 | 0 | C | T | 0.5 | 1.20E-01 | 1.05E-01 | 2.40E+00 | 2.44E+00 | 2.63E-01 | no | Kcnip4 | intron_variant | MODIFIER |
| 5 | 48566630 | AX-97818136 | 0 | A | G | 0.5 | 1.20E-01 | 1.05E-01 | 2.40E+00 | 2.44E+00 | 2.63E-01 | no | Kcnip4 | intron_variant | MODIFIER |
| 5 | 48568510 | AX-170024884 | 0 | C | A | 0.5 | 1.20E-01 | 1.05E-01 | 2.40E+00 | 2.44E+00 | 2.63E-01 | no | Kcnip4 | intron_variant | MODIFIER |
| 5 | 48627101 | AX-170497142 | 0 | C | T | 0.5 | 1.20E-01 | 1.05E-01 | 2.40E+00 | 2.44E+00 | 2.63E-01 | no | Kcnip4 | intron_variant | MODIFIER |
| 5 | 48627300 | AX-170521414 | 0 | C | T | 0.5 | 1.20E-01 | 1.05E-01 | 2.40E+00 | 2.44E+00 | 2.63E-01 | no | Kcnip4 | intron_variant | MODIFIER |
| 5 | 48628418 | AX-170024886 | 0 | A | T | 0.5 | 1.20E-01 | 1.05E-01 | 2.40E+00 | 2.44E+00 | 2.63E-01 | no | Kcnip4 | intron_variant | MODIFIER |
| 5 | 48632640 | AX-170100382 | 0 | A | G | 0.5 | 1.20E-01 | 1.05E-01 | 2.40E+00 | 2.44E+00 | 2.63E-01 | no | Kcnip4 | intron_variant | MODIFIER |
| 5 | 48634881 | AX-170072014 | 0 | C | G | 0.5 | 1.20E-01 | 1.05E-01 | 2.40E+00 | 2.44E+00 | 2.63E-01 | no | Kcnip4 | intron_variant | MODIFIER |
| 5 | 48651168 | AX-97805759 | 0 | C | T | 0.5 | 1.20E-01 | 1.05E-01 | 2.40E+00 | 2.44E+00 | 2.63E-01 | no | Kcnip4 | intron_variant | MODIFIER |

| | | | | | | | | | | | | | | | |
|---|---|---|---|---|---|---|---|---|---|---|---|---|---|---|---|
| 5 | 48683931 | AX-170490231 | 0 | G | A | 0.5 | 1.20E-01 | 1.05E-01 | 2.40E+00 | 2.44E+00 | 2.63E-01 | no | Kcnip4 | intron_variant | MODIFIER |
| 5 | 48687686 | AX-170296566 | 0 | G | A | 0.5 | 1.20E-01 | 1.05E-01 | 2.40E+00 | 2.44E+00 | 2.63E-01 | no | Kcnip4 | intron_variant | MODIFIER |
| 5 | 48719222 | AX-170430683 | 0 | C | A | 0.5 | 1.20E-01 | 1.05E-01 | 2.40E+00 | 2.44E+00 | 2.63E-01 | no | Kcnip4 | intron_variant | MODIFIER |
| 5 | 48776408 | AX-97831504 | 0 | C | T | 0.5 | 1.20E-01 | 1.05E-01 | 2.40E+00 | 2.44E+00 | 2.63E-01 | no | Kcnip4 | intron_variant | MODIFIER |
| 5 | 48777183 | AX-170217347 | 0 | A | C | 0.5 | 1.20E-01 | 1.05E-01 | 2.40E+00 | 2.44E+00 | 2.63E-01 | no | Kcnip4 | intron_variant | MODIFIER |
| 5 | 48789512 | AX-170535499 | 0 | G | A | 0.5 | 1.20E-01 | 1.05E-01 | 2.40E+00 | 2.44E+00 | 2.63E-01 | no | Kcnip4 | intron_variant | MODIFIER |
| 5 | 48789889 | AX-170315829 | 0 | C | A | 0.5 | 1.20E-01 | 1.05E-01 | 2.40E+00 | 2.44E+00 | 2.63E-01 | no | Kcnip4 | intron_variant | MODIFIER |
| 5 | 48829351 | AX-97792972 | 0 | C | T | 0.5 | 1.20E-01 | 1.05E-01 | 2.40E+00 | 2.44E+00 | 2.63E-01 | no | Kcnip4 | intron_variant | MODIFIER |
| 5 | 48854881 | AX-170100810 | 0 | C | G | 0.5 | 1.20E-01 | 1.05E-01 | 2.40E+00 | 2.44E+00 | 2.63E-01 | no | Kcnip4 | intron_variant | MODIFIER |
| 5 | 48879227 | AX-97817161 | 0 | C | T | 0.5 | 1.20E-01 | 1.05E-01 | 2.40E+00 | 2.44E+00 | 2.63E-01 | no | Kcnip4 | intron_variant | MODIFIER |
| 5 | 48879613 | AX-170243390 | 0 | A | C | 0.5 | 1.20E-01 | 1.05E-01 | 2.40E+00 | 2.44E+00 | 2.63E-01 | no | Kcnip4 | intron_variant | MODIFIER |
| 5 | 48908487 | AX-168402753 | 0 | C | T | 0.5 | 1.20E-01 | 1.05E-01 | 2.40E+00 | 2.44E+00 | 2.63E-01 | no | Kcnip4 | intron_variant | MODIFIER |
| 5 | 49346840 | AX-170322149 | 0 | T | G | 0.5 | 1.20E-01 | 1.05E-01 | 2.40E+00 | 2.44E+00 | 2.63E-01 | no | Kcnip4 | intron_variant | MODIFIER |
| 5 | 49347029 | AX-170108490 | 0 | G | A | 0.5 | 1.20E-01 | 1.05E-01 | 2.40E+00 | 2.44E+00 | 2.63E-01 | no | Kcnip4 | intron_variant | MODIFIER |
| 5 | 49369985 | AX-170407458 | 0 | T | C | 0.5 | 1.20E-01 | 1.05E-01 | 2.40E+00 | 2.44E+00 | 2.63E-01 | no | Kcnip4 | intron_variant | MODIFIER |
| 5 | 49374659 | AX-170467380 | 0 | A | G | 0.5 | 1.20E-01 | 1.05E-01 | 2.40E+00 | 2.44E+00 | 2.63E-01 | no | Kcnip4 | intron_variant | MODIFIER |
| 5 | 49374848 | AX-170531953 | 0 | A | G | 0.5 | 1.20E-01 | 1.05E-01 | 2.40E+00 | 2.44E+00 | 2.63E-01 | no | Kcnip4 | intron_variant | MODIFIER |
| 5 | 49375797 | AX-170242710 | 0 | G | A | 0.5 | 1.20E-01 | 1.05E-01 | 2.40E+00 | 2.44E+00 | 2.63E-01 | no | Kcnip4 | intron_variant | MODIFIER |
| 5 | 49427619 | AX-170190150 | 0 | C | T | 0.5 | 1.20E-01 | 1.05E-01 | 2.40E+00 | 2.44E+00 | 2.63E-01 | no | Kcnip4 | intron_variant | MODIFIER |
| 5 | 49428175 | AX-170440721 | 0 | G | A | 0.5 | 1.20E-01 | 1.05E-01 | 2.40E+00 | 2.44E+00 | 2.63E-01 | no | Kcnip4 | intron_variant | MODIFIER |
| 5 | 49440616 | AX-170492463 | 0 | A | T | 0.5 | 1.20E-01 | 1.05E-01 | 2.40E+00 | 2.44E+00 | 2.63E-01 | no | Kcnip4 | intron_variant | MODIFIER |
| 5 | 49476134 | AX-170224139 | 0 | A | G | 0.5 | 1.20E-01 | 1.05E-01 | 2.40E+00 | 2.44E+00 | 2.63E-01 | no | Kcnip4 | intron_variant | MODIFIER |
| 5 | 49483583 | AX-170180049 | 0 | T | C | 0.5 | 1.20E-01 | 1.05E-01 | 2.40E+00 | 2.44E+00 | 2.63E-01 | no | Kcnip4 | intron_variant | MODIFIER |
| 5 | 49546424 | AX-170293575 | 0 | C | T | 0.5 | 1.20E-01 | 1.05E-01 | 2.40E+00 | 2.44E+00 | 2.63E-01 | no | | intergenic_variant | MODIFIER |
| 5 | 49546804 | AX-170424878 | 0 | A | G | 0.5 | 1.20E-01 | 1.05E-01 | 2.40E+00 | 2.44E+00 | 2.63E-01 | no | | intergenic_variant | MODIFIER |
| 5 | 49546836 | AX-170295103 | 0 | A | C | 0.5 | 1.20E-01 | 1.05E-01 | 2.40E+00 | 2.44E+00 | 2.63E-01 | no | | intergenic_variant | MODIFIER |
| 5 | 49589452 | AX-170159346 | 0 | G | A | 0.5 | 1.20E-01 | 1.05E-01 | 2.40E+00 | 2.44E+00 | 2.63E-01 | no | | intergenic_variant | MODIFIER |
| 5 | 49594275 | AX-170248512 | 0 | T | C | 0.5 | 1.20E-01 | 1.05E-01 | 2.40E+00 | 2.44E+00 | 2.63E-01 | no | | intergenic_variant | MODIFIER |
| 5 | 49665119 | AX-170265544 | 0 | T | C | 0.5 | 1.20E-01 | 1.05E-01 | 2.40E+00 | 2.44E+00 | 2.63E-01 | no | | intergenic_variant | MODIFIER |
| 5 | 49666353 | AX-170024898 | 0 | C | A | 0.5 | 1.20E-01 | 1.05E-01 | 2.40E+00 | 2.44E+00 | 2.63E-01 | no | | intergenic_variant | MODIFIER |
| 5 | 49714809 | AX-170109800 | 0 | A | G | 0.5 | 1.20E-01 | 1.05E-01 | 2.40E+00 | 2.44E+00 | 2.63E-01 | no | | intergenic_variant | MODIFIER |
| 5 | 49760476 | AX-97782790 | 0 | C | T | 0.5 | 1.20E-01 | 1.05E-01 | 2.40E+00 | 2.44E+00 | 2.63E-01 | no | | intergenic_variant | MODIFIER |
| 5 | 49761671 | AX-170024895 | 0 | G | A | 0.5 | 1.20E-01 | 1.05E-01 | 2.40E+00 | 2.44E+00 | 2.63E-01 | no | | intergenic_variant | MODIFIER |
| 5 | 49960166 | AX-170124779 | 0 | T | G | 0.5 | 1.20E-01 | 1.05E-01 | 2.40E+00 | 2.44E+00 | 2.63E-01 | no | Adgra3 | 3_prime_UTR_variant | MODIFIER |
| 5 | 50527461 | AX-97836873 | 0 | C | G | 0.5 | 1.20E-01 | 1.05E-01 | 2.40E+00 | 2.44E+00 | 2.63E-01 | no | Gm36814 | non_coding_transcript_exon_variant | MODIFIER |
| 5 | 50573005 | AX-170531613 | 0 | G | A | 0.5 | 1.20E-01 | 1.05E-01 | 2.40E+00 | 2.44E+00 | 2.63E-01 | no | Gm43016 | intron_variant,non_coding_transcript_variant | MODIFIER |
| 5 | 50574682 | AX-170463901 | 0 | C | T | 0.5 | 1.20E-01 | 1.05E-01 | 2.40E+00 | 2.44E+00 | 2.63E-01 | no | Gm43016 | intron_variant,non_coding_transcript_variant | MODIFIER |
| 5 | 50658860 | AX-97854392 | 0 | A | C | 0.5 | 1.20E-01 | 1.05E-01 | 2.40E+00 | 2.44E+00 | 2.63E-01 | no | Gm43016 | intron_variant,non_coding_transcript_variant | MODIFIER |
| 5 | 50703046 | AX-168408191 | 0 | C | T | 0.5 | 1.20E-01 | 1.05E-01 | 2.40E+00 | 2.44E+00 | 2.63E-01 | no | | intergenic_variant | MODIFIER |
| 5 | 50724051 | AX-170024920 | 0 | C | T | 0.5 | 1.20E-01 | 1.05E-01 | 2.40E+00 | 2.44E+00 | 2.63E-01 | no | | intergenic_variant | MODIFIER |
| 5 | 50736310 | AX-168399164 | 0 | G | A | 0.5 | 1.20E-01 | 1.05E-01 | 2.40E+00 | 2.44E+00 | 2.63E-01 | no | | intergenic_variant | MODIFIER |
| 5 | 50744601 | AX-170452780 | 0 | T | C | 0.5 | 1.20E-01 | 1.05E-01 | 2.40E+00 | 2.44E+00 | 2.63E-01 | no | | intergenic_variant | MODIFIER |
| 5 | 50754015 | AX-97808196 | 0 | G | T | 0.5 | 1.20E-01 | 1.05E-01 | 2.40E+00 | 2.44E+00 | 2.63E-01 | no | | intergenic_variant | MODIFIER |
| 5 | 50788269 | AX-97800046 | 0 | T | C | 0.5 | 1.20E-01 | 1.05E-01 | 2.40E+00 | 2.44E+00 | 2.63E-01 | no | | intergenic_variant | MODIFIER |
| 5 | 50848323 | AX-170208830 | 0 | G | A | 0.5 | 1.20E-01 | 1.05E-01 | 2.40E+00 | 2.44E+00 | 2.63E-01 | no | | intergenic_variant | MODIFIER |
| 5 | 50856340 | AX-170024926 | 0 | G | T | 0.5 | 1.20E-01 | 1.05E-01 | 2.40E+00 | 2.44E+00 | 2.63E-01 | no | | intergenic_variant | MODIFIER |
| 5 | 50856701 | AX-97803350 | 0 | G | A | 0.5 | 1.20E-01 | 1.05E-01 | 2.40E+00 | 2.44E+00 | 2.63E-01 | no | | intergenic_variant | MODIFIER |
| 5 | 50922498 | AX-170024929 | 0 | G | A | 0.5 | 1.20E-01 | 1.05E-01 | 2.40E+00 | 2.44E+00 | 2.63E-01 | no | Gm43015 | downstream_gene_variant | MODIFIER |
| 5 | 50924699 | AX-97781055 | 0 | G | A | 0.5 | 1.20E-01 | 1.05E-01 | 2.40E+00 | 2.44E+00 | 2.63E-01 | no | Gm43015 | downstream_gene_variant | MODIFIER |
| 5 | 51051382 | AX-170315064 | 0 | T | C | 0.5 | 1.20E-01 | 1.05E-01 | 2.40E+00 | 2.44E+00 | 2.63E-01 | no | Gm40319 | upstream_gene_variant | MODIFIER |
| 5 | 51074612 | AX-170313231 | 0 | T | C | 0.5 | 1.20E-01 | 1.05E-01 | 2.40E+00 | 2.44E+00 | 2.63E-01 | no | | intergenic_variant | MODIFIER |
| 5 | 51078359 | AX-97806564 | 0 | C | A | 0.5 | 1.20E-01 | 1.05E-01 | 2.40E+00 | 2.44E+00 | 2.63E-01 | no | | intergenic_variant | MODIFIER |
| 5 | 51088592 | AX-170072070 | 0 | T | G | 0.5 | 1.20E-01 | 1.05E-01 | 2.40E+00 | 2.44E+00 | 2.63E-01 | no | | intergenic_variant | MODIFIER |
| 5 | 51109438 | AX-170459226 | 0 | C | T | 0.5 | 1.20E-01 | 1.05E-01 | 2.40E+00 | 2.44E+00 | 2.63E-01 | no | | intergenic_variant | MODIFIER |

| | | | | | | | | | | | | | | | |
|---|---|---|---|---|---|---|---|---|---|---|---|---|---|---|---|
| 5 | 51110034 | AX-170228645 | 0 | G | T | 0.5 | 1.20E-01 | 1.05E-01 | 2.40E+00 | 2.44E+00 | 2.63E-01 | no | | intergenic_variant | MODIFIER |
| 5 | 51152839 | AX-170411153 | 0 | T | A | 0.5 | 1.20E-01 | 1.05E-01 | 2.40E+00 | 2.44E+00 | 2.63E-01 | no | | intergenic_variant | MODIFIER |
| 5 | 51184132 | AX-170278455 | 0 | C | T | 0.5 | 1.20E-01 | 1.05E-01 | 2.40E+00 | 2.44E+00 | 2.63E-01 | no | | intergenic_variant | MODIFIER |
| 5 | 51184312 | AX-97786637 | 0 | C | A | 0.5 | 1.20E-01 | 1.05E-01 | 2.40E+00 | 2.44E+00 | 2.63E-01 | no | | intergenic_variant | MODIFIER |
| 5 | 51194678 | AX-97804041 | 0 | C | T | 0.5 | 1.20E-01 | 1.05E-01 | 2.40E+00 | 2.44E+00 | 2.63E-01 | no | | intergenic_variant | MODIFIER |
| 5 | 51258847 | AX-170024933 | 0 | A | G | 0.5 | 1.20E-01 | 1.05E-01 | 2.40E+00 | 2.44E+00 | 2.63E-01 | no | | intergenic_variant | MODIFIER |
| 5 | 51278584 | AX-97799885 | 0 | C | T | 0.5 | 1.20E-01 | 1.05E-01 | 2.40E+00 | 2.44E+00 | 2.63E-01 | no | | intergenic_variant | MODIFIER |
| 5 | 51295426 | AX-97848108 | 0 | T | C | 0.5 | 1.20E-01 | 1.05E-01 | 2.40E+00 | 2.44E+00 | 2.63E-01 | no | | intergenic_variant | MODIFIER |
| 5 | 51337107 | AX-170282899 | 0 | T | C | 0.5 | 1.20E-01 | 1.05E-01 | 2.40E+00 | 2.44E+00 | 2.63E-01 | no | | intergenic_variant | MODIFIER |
| 5 | 51339242 | AX-97848367 | 0 | G | A | 0.5 | 1.20E-01 | 1.05E-01 | 2.40E+00 | 2.44E+00 | 2.63E-01 | no | | intergenic_variant | MODIFIER |
| 5 | 51343664 | AX-170331427 | 0 | A | G | 0.5 | 1.20E-01 | 1.05E-01 | 2.40E+00 | 2.44E+00 | 2.63E-01 | no | | intergenic_variant | MODIFIER |
| 5 | 51350126 | AX-97783824 | 0 | T | C | 0.5 | 1.20E-01 | 1.05E-01 | 2.40E+00 | 2.44E+00 | 2.63E-01 | no | | intergenic_variant | MODIFIER |
| 5 | 51350356 | AX-170522041 | 0 | A | G | 0.5 | 1.20E-01 | 1.05E-01 | 2.40E+00 | 2.44E+00 | 2.63E-01 | no | | intergenic_variant | MODIFIER |
| 5 | 51350487 | AX-170435297 | 0 | A | G | 0.5 | 1.20E-01 | 1.05E-01 | 2.40E+00 | 2.44E+00 | 2.63E-01 | no | | intergenic_variant | MODIFIER |
| 5 | 51392984 | AX-170024936 | 0 | T | G | 0.5 | 1.20E-01 | 1.05E-01 | 2.40E+00 | 2.44E+00 | 2.63E-01 | no | | intergenic_variant | MODIFIER |
| 5 | 51452349 | AX-170161237 | 0 | C | T | 0.5 | 1.20E-01 | 1.05E-01 | 2.40E+00 | 2.44E+00 | 2.63E-01 | no | Ppargc1a | downstream_gene_variant | MODIFIER |
| 5 | 51492570 | AX-170257140 | 0 | A | G | 0.5 | 1.20E-01 | 1.05E-01 | 2.40E+00 | 2.44E+00 | 2.63E-01 | no | Ppargc1a | intron_variant,NMD_transcript_variant | MODIFIER |
| 5 | 51500042 | AX-170511926 | 0 | T | C | 0.5 | 1.20E-01 | 1.05E-01 | 2.40E+00 | 2.44E+00 | 2.63E-01 | no | Ppargc1a | intron_variant,NMD_transcript_variant | MODIFIER |
| 5 | 51516432 | AX-97837199 | 0 | T | G | 0.5 | 1.20E-01 | 1.05E-01 | 2.40E+00 | 2.44E+00 | 2.63E-01 | no | Ppargc1a | intron_variant,NMD_transcript_variant | MODIFIER |
| 5 | 51544446 | AX-170072078 | 0 | C | T | 0.5 | 1.20E-01 | 1.05E-01 | 2.40E+00 | 2.44E+00 | 2.63E-01 | no | Ppargc1a | intron_variant,NMD_transcript_variant | MODIFIER |
| 5 | 51545847 | AX-170072080 | 0 | T | G | 0.5 | 1.20E-01 | 1.05E-01 | 2.40E+00 | 2.44E+00 | 2.63E-01 | no | Ppargc1a | intron_variant,NMD_transcript_variant | MODIFIER |
| 5 | 51545903 | AX-170329117 | 0 | C | T | 0.5 | 1.20E-01 | 1.05E-01 | 2.40E+00 | 2.44E+00 | 2.63E-01 | no | Ppargc1a | intron_variant,NMD_transcript_variant | MODIFIER |
| 5 | 51546378 | AX-170072079 | 0 | C | T | 0.5 | 1.20E-01 | 1.05E-01 | 2.40E+00 | 2.44E+00 | 2.63E-01 | no | Ppargc1a | intron_variant,NMD_transcript_variant | MODIFIER |
| 5 | 51546936 | AX-170409465 | 0 | T | C | 0.5 | 1.20E-01 | 1.05E-01 | 2.40E+00 | 2.44E+00 | 2.63E-01 | no | Ppargc1a | intron_variant,NMD_transcript_variant | MODIFIER |
| 5 | 51552376 | AX-170292996 | 0 | T | C | 0.5 | 1.20E-01 | 1.05E-01 | 2.40E+00 | 2.44E+00 | 2.63E-01 | no | Ppargc1a | intron_variant,NMD_transcript_variant | MODIFIER |
| 5 | 51561076 | AX-170455740 | 0 | T | C | 0.5 | 1.20E-01 | 1.05E-01 | 2.40E+00 | 2.44E+00 | 2.63E-01 | no | Ppargc1a | intron_variant | MODIFIER |
| 5 | 51620241 | AX-97834579 | 0 | A | G | 0.5 | 1.20E-01 | 1.05E-01 | 2.40E+00 | 2.44E+00 | 2.63E-01 | no | | intergenic_variant | MODIFIER |
| 5 | 51626106 | AX-170024931 | 0 | G | C | 0.5 | 1.20E-01 | 1.05E-01 | 2.40E+00 | 2.44E+00 | 2.63E-01 | no | | intergenic_variant | MODIFIER |
| 5 | 51661283 | AX-97816381 | 0 | A | G | 0.5 | 1.20E-01 | 1.05E-01 | 2.40E+00 | 2.44E+00 | 2.63E-01 | no | | intergenic_variant | MODIFIER |
| 5 | 51801513 | AX-170072087 | 0 | C | T | 0.5 | 1.20E-01 | 1.05E-01 | 2.40E+00 | 2.44E+00 | 2.63E-01 | no | | intergenic_variant | MODIFIER |
| 5 | 51808992 | AX-97809162 | 0 | G | T | 0.5 | 1.20E-01 | 1.05E-01 | 2.40E+00 | 2.44E+00 | 2.63E-01 | no | | intergenic_variant | MODIFIER |
| 5 | 51825372 | AX-170273274 | 0 | C | T | 0.5 | 1.20E-01 | 1.05E-01 | 2.40E+00 | 2.44E+00 | 2.63E-01 | no | | intergenic_variant | MODIFIER |
| 5 | 51838354 | AX-170024943 | 0 | G | A | 0.5 | 1.20E-01 | 1.05E-01 | 2.40E+00 | 2.44E+00 | 2.63E-01 | no | | intergenic_variant | MODIFIER |
| 5 | 51839139 | AX-170110186 | 0 | T | C | 0.5 | 1.20E-01 | 1.05E-01 | 2.40E+00 | 2.44E+00 | 2.63E-01 | no | | regulatory_region_variant | MODIFIER |
| 5 | 51839655 | AX-170186569 | 0 | G | A | 0.5 | 1.20E-01 | 1.05E-01 | 2.40E+00 | 2.44E+00 | 2.63E-01 | no | | regulatory_region_variant | MODIFIER |
| 5 | 51839717 | AX-170518214 | 0 | G | A | 0.5 | 1.20E-01 | 1.05E-01 | 2.40E+00 | 2.44E+00 | 2.63E-01 | no | | regulatory_region_variant | MODIFIER |
| 5 | 51840662 | AX-170456683 | 0 | T | C | 0.5 | 1.20E-01 | 1.05E-01 | 2.40E+00 | 2.44E+00 | 2.63E-01 | no | | regulatory_region_variant | MODIFIER |
| 5 | 51858981 | AX-170377244 | 0 | A | G | 0.5 | 1.20E-01 | 1.05E-01 | 2.40E+00 | 2.44E+00 | 2.63E-01 | no | Gm43606 | upstream_gene_variant | MODIFIER |
| 5 | 51859418 | AX-170024944 | 0 | C | T | 0.5 | 1.20E-01 | 1.05E-01 | 2.40E+00 | 2.44E+00 | 2.63E-01 | no | Gm43606 | upstream_gene_variant | MODIFIER |
| 5 | 51862368 | AX-170198084 | 0 | A | G | 0.5 | 1.20E-01 | 1.05E-01 | 2.40E+00 | 2.44E+00 | 2.63E-01 | no | Gm43606 | upstream_gene_variant | MODIFIER |
| 5 | 52041513 | AX-170194286 | 0 | A | G | 0.5 | 1.20E-01 | 1.05E-01 | 2.40E+00 | 2.44E+00 | 2.63E-01 | no | | regulatory_region_variant | MODIFIER |
| 5 | 52072645 | AX-170299851 | 0 | C | T | 0.5 | 1.20E-01 | 1.05E-01 | 2.40E+00 | 2.44E+00 | 2.63E-01 | no | | intergenic_variant | MODIFIER |
| 5 | 52202398 | AX-170100475 | 0 | A | G | 0.5 | 1.20E-01 | 1.05E-01 | 2.40E+00 | 2.44E+00 | 2.63E-01 | no | 9230114K14Rik | downstream_gene_variant | MODIFIER |
| 5 | 52207754 | AX-170203343 | 0 | G | A | 0.5 | 1.20E-01 | 1.05E-01 | 2.40E+00 | 2.44E+00 | 2.63E-01 | no | C130083M11Rik | intron_variant,non_coding_transcript_variant | MODIFIER |
| 5 | 52259954 | AX-170494171 | 0 | C | T | 0.5 | 1.20E-01 | 1.05E-01 | 2.40E+00 | 2.44E+00 | 2.63E-01 | no | 9230114K14Rik | intron_variant,non_coding_transcript_variant | MODIFIER |
| 5 | 52278738 | AX-97843964 | 0 | T | C | 0.5 | 1.20E-01 | 1.05E-01 | 2.40E+00 | 2.44E+00 | 2.63E-01 | no | 9230114K14Rik | intron_variant,non_coding_transcript_variant | MODIFIER |
| 5 | 52279367 | AX-170072096 | 0 | G | A | 0.5 | 1.20E-01 | 1.05E-01 | 2.40E+00 | 2.44E+00 | 2.63E-01 | no | 9230114K14Rik | downstream_gene_variant | MODIFIER |
| 5 | 52280851 | AX-170187778 | 0 | G | A | 0.5 | 1.20E-01 | 1.05E-01 | 2.40E+00 | 2.44E+00 | 2.63E-01 | no | 9230114K14Rik | downstream_gene_variant | MODIFIER |
| 5 | 52281652 | AX-170487295 | 0 | C | T | 0.5 | 1.20E-01 | 1.05E-01 | 2.40E+00 | 2.44E+00 | 2.63E-01 | no | 9230114K14Rik | downstream_gene_variant | MODIFIER |
| 4 | 97832222 | AX-97780592 | 0 | C | T | 0.246 | 1.20E-01 | 1.08E-01 | 2.60E+00 | 2.68E+00 | 2.63E-01 | no | Nfia | intron_variant | MODIFIER |
| 5 | 52034858 | AX-170133519 | 0 | T | A | 0.482 | 1.18E-01 | 1.00E-01 | 2.28E+00 | 2.34E+00 | 2.63E-01 | no | | intergenic_variant | MODIFIER |
| 12 | 71474024 | AX-170347297 | 0 | A | G | 0.07 | 2.47E-01 | 2.21E-01 | 2.95E+00 | 3.08E+00 | 2.63E-01 | Yes | Gm40438 | intron_variant,non_coding_transcript_variant | MODIFIER |
| 4 | 88175386 | AX-170089307 | 0 | A | G | 0.246 | 1.28E-01 | 1.12E-01 | 2.44E+00 | 2.49E+00 | 2.63E-01 | no | Focad | intron_variant | MODIFIER |
| 4 | 88175425 | AX-170301058 | 0 | G | A | 0.246 | 1.28E-01 | 1.12E-01 | 2.44E+00 | 2.49E+00 | 2.63E-01 | no | Focad | intron_variant | MODIFIER |

| | | | | | | | | | | | | | | | |
|---|---|---|---|---|---|---|---|---|---|---|---|---|---|---|---|
| 4 | 88337740 | AX-97798251 | 0 | T | G | 0.246 | 1.28E-01 | 1.12E-01 | 2.44E+00 | 2.49E+00 | 2.63E-01 | no | Focad | intron_variant | MODIFIER |
| 4 | 88369107 | AX-170023355 | 0 | C | G | 0.246 | 1.28E-01 | 1.12E-01 | 2.44E+00 | 2.49E+00 | 2.63E-01 | no | Focad | intron_variant | MODIFIER |
| 4 | 88391678 | AX-170023349 | 0 | A | G | 0.246 | 1.28E-01 | 1.12E-01 | 2.44E+00 | 2.49E+00 | 2.63E-01 | no | Focad | intron_variant | MODIFIER |
| 4 | 88824224 | AX-170241801 | 0 | T | C | 0.246 | 1.28E-01 | 1.12E-01 | 2.44E+00 | 2.49E+00 | 2.63E-01 | no | Ifna6 | upstream_gene_variant | MODIFIER |
| 4 | 88841837 | AX-170305589 | 0 | T | C | 0.246 | 1.28E-01 | 1.12E-01 | 2.44E+00 | 2.49E+00 | 2.63E-01 | no | Ifna4 | upstream_gene_variant | MODIFIER |
| 4 | 88858696 | AX-97819912 | 0 | C | A | 0.246 | 1.28E-01 | 1.12E-01 | 2.44E+00 | 2.49E+00 | 2.63E-01 | no | | intergenic_variant | MODIFIER |
| 4 | 89014514 | AX-170319704 | 0 | G | A | 0.246 | 1.28E-01 | 1.12E-01 | 2.44E+00 | 2.49E+00 | 2.63E-01 | no | AL805899.2 | intron_variant,non_coding_transcript_variant | MODIFIER |
| 4 | 89023452 | AX-170141990 | 0 | T | G | 0.246 | 1.28E-01 | 1.12E-01 | 2.44E+00 | 2.49E+00 | 2.63E-01 | no | AL805899.2 | intron_variant,non_coding_transcript_variant | MODIFIER |
| 4 | 89024735 | AX-97786977 | 0 | G | A | 0.246 | 1.28E-01 | 1.12E-01 | 2.44E+00 | 2.49E+00 | 2.63E-01 | no | AL805899.2 | intron_variant,non_coding_transcript_variant | MODIFIER |
| 4 | 89038468 | AX-170023363 | 0 | T | A | 0.246 | 1.28E-01 | 1.12E-01 | 2.44E+00 | 2.49E+00 | 2.63E-01 | no | AL805899.2 | intron_variant,non_coding_transcript_variant | MODIFIER |
| 4 | 89246433 | AX-170330240 | 0 | A | G | 0.246 | 1.28E-01 | 1.12E-01 | 2.44E+00 | 2.49E+00 | 2.63E-01 | no | Gm12606 | intron_variant,non_coding_transcript_variant | MODIFIER |
| 4 | 89367130 | AX-170141743 | 0 | A | G | 0.246 | 1.28E-01 | 1.12E-01 | 2.44E+00 | 2.49E+00 | 2.63E-01 | no | | regulatory_region_variant | MODIFIER |
| 4 | 89373697 | AX-170240349 | 0 | A | G | 0.246 | 1.28E-01 | 1.12E-01 | 2.44E+00 | 2.49E+00 | 2.63E-01 | no | | regulatory_region_variant | MODIFIER |
| 4 | 89645326 | AX-170023365 | 0 | T | C | 0.246 | 1.28E-01 | 1.12E-01 | 2.44E+00 | 2.49E+00 | 2.63E-01 | no | | intergenic_variant | MODIFIER |
| 4 | 90239108 | AX-97855029 | 0 | G | A | 0.246 | 1.28E-01 | 1.12E-01 | 2.44E+00 | 2.49E+00 | 2.63E-01 | no | | intergenic_variant | MODIFIER |
| 4 | 90450282 | AX-170152836 | 0 | T | G | 0.246 | 1.28E-01 | 1.12E-01 | 2.44E+00 | 2.49E+00 | 2.63E-01 | no | Gm12635 | upstream_gene_variant | MODIFIER |
| 4 | 90731873 | AX-170023385 | 0 | T | C | 0.246 | 1.28E-01 | 1.12E-01 | 2.44E+00 | 2.49E+00 | 2.63E-01 | no | | intergenic_variant | MODIFIER |
| 4 | 90926763 | AX-170494706 | 0 | C | T | 0.246 | 1.28E-01 | 1.12E-01 | 2.44E+00 | 2.49E+00 | 2.63E-01 | no | | intergenic_variant | MODIFIER |
| 4 | 91029311 | AX-170090781 | 0 | A | T | 0.246 | 1.28E-01 | 1.12E-01 | 2.44E+00 | 2.49E+00 | 2.63E-01 | no | | intergenic_variant | MODIFIER |
| 8 | 50429493 | AX-170096264 | 2 | G | A | 0.027 | 1.79E-01 | 1.62E-01 | 2.62E+00 | 2.80E+00 | 2.63E-01 | Yes | Gm45455 | intron_variant,non_coding_transcript_variant | MODIFIER |
| 2 | 137900371 | AX-170019157 | 1 | C | T | 0.152 | 2.07E-01 | 1.87E-01 | 2.86E+00 | 2.95E+00 | 2.63E-01 | Yes | | intergenic_variant | MODIFIER |
| 7 | 144070274 | AX-170235234 | 1 | G | A | 0.196 | -8.38E-02 | 7.58E-02 | 2.68E+00 | 2.81E+00 | 2.63E-01 | no | Shank2 | intron_variant | MODIFIER |
| 16 | 39177400 | AX-170516230 | 1 | T | C | 0.473 | 8.87E-02 | 7.96E-02 | 2.60E+00 | 2.65E+00 | 2.64E-01 | no | | intergenic_variant | MODIFIER |
| 4 | 127228726 | AX-170462109 | 0 | C | A | 0.149 | 1.33E-01 | 1.20E-01 | 2.64E+00 | 2.71E+00 | 2.64E-01 | no | Dlgap3 | intron_variant | MODIFIER |
| 10 | 94233699 | AX-170003682 | 0 | G | A | 0.289 | -9.63E-02 | 8.59E-02 | 2.51E+00 | 2.57E+00 | 2.64E-01 | no | | intergenic_variant | MODIFIER |
| 1 | 181451370 | AX-97824628 | 2 | G | A | 0.227 | 1.27E-01 | 1.12E-01 | 2.49E+00 | 2.55E+00 | 2.64E-01 | no | Cnih3 | intron_variant | MODIFIER |
| 12 | 69400695 | AX-170310435 | 0 | C | T | 0.061 | -2.47E-01 | 2.22E-01 | 2.60E+00 | 2.68E+00 | 2.64E-01 | Yes | | regulatory_region_variant | MODIFIER |
| 1 | 76013512 | AX-170149007 | 0 | G | A | 0.14 | 2.02E-01 | 1.82E-01 | 2.79E+00 | 2.90E+00 | 2.64E-01 | Yes | Gm29065 | downstream_gene_variant | MODIFIER |
| 1 | 164960959 | AX-97801177 | 0 | A | G | 0.228 | 1.53E-01 | 1.38E-01 | 2.74E+00 | 2.79E+00 | 2.64E-01 | Yes | 1700029M03Rik | non_coding_transcript_exon_variant | MODIFIER |
| 1 | 164962999 | AX-170444464 | 0 | C | T | 0.228 | 1.53E-01 | 1.38E-01 | 2.74E+00 | 2.79E+00 | 2.64E-01 | Yes | 1700029M03Rik | downstream_gene_variant | MODIFIER |
| 1 | 164963623 | AX-170002197 | 0 | C | T | 0.228 | 1.53E-01 | 1.38E-01 | 2.74E+00 | 2.79E+00 | 2.64E-01 | Yes | 1700029M03Rik | downstream_gene_variant | MODIFIER |
| 1 | 164977037 | AX-97818529 | 0 | G | A | 0.228 | 1.53E-01 | 1.38E-01 | 2.74E+00 | 2.79E+00 | 2.64E-01 | Yes | | intergenic_variant | MODIFIER |
| 1 | 165008080 | AX-97814062 | 0 | A | G | 0.228 | 1.53E-01 | 1.38E-01 | 2.74E+00 | 2.79E+00 | 2.64E-01 | Yes | | intergenic_variant | MODIFIER |
| 1 | 165057225 | AX-170002191 | 0 | T | C | 0.228 | 1.53E-01 | 1.38E-01 | 2.74E+00 | 2.79E+00 | 2.64E-01 | Yes | | intergenic_variant | MODIFIER |
| 1 | 165068158 | AX-97831495 | 0 | T | C | 0.228 | 1.53E-01 | 1.38E-01 | 2.74E+00 | 2.79E+00 | 2.64E-01 | Yes | 4930568G15Rik | upstream_gene_variant | MODIFIER |
| 1 | 165139662 | AX-170002194 | 0 | G | A | 0.228 | 1.53E-01 | 1.38E-01 | 2.74E+00 | 2.79E+00 | 2.64E-01 | Yes | Tbx19 | intron_variant | MODIFIER |
| 1 | 165140064 | AX-97799354 | 0 | C | T | 0.228 | 1.53E-01 | 1.38E-01 | 2.74E+00 | 2.79E+00 | 2.64E-01 | Yes | Tbx19 | intron_variant | MODIFIER |
| 1 | 165183286 | AX-97814085 | 0 | A | C | 0.228 | 1.53E-01 | 1.38E-01 | 2.74E+00 | 2.79E+00 | 2.64E-01 | Yes | Sft2d2 | intron_variant | MODIFIER |
| 1 | 165190843 | AX-170173330 | 0 | G | C | 0.228 | 1.53E-01 | 1.38E-01 | 2.74E+00 | 2.79E+00 | 2.64E-01 | Yes | Sft2d2 | intron_variant | MODIFIER |
| 1 | 165214293 | AX-97827552 | 0 | C | T | 0.228 | 1.53E-01 | 1.38E-01 | 2.74E+00 | 2.79E+00 | 2.64E-01 | Yes | Tiprl | 3_prime_UTR_variant | MODIFIER |
| 1 | 165215931 | AX-170130456 | 0 | G | A | 0.228 | 1.53E-01 | 1.38E-01 | 2.74E+00 | 2.79E+00 | 2.64E-01 | Yes | Tiprl | intron_variant | MODIFIER |
| 1 | 165225741 | AX-97850413 | 0 | G | A | 0.228 | 1.53E-01 | 1.38E-01 | 2.74E+00 | 2.79E+00 | 2.64E-01 | Yes | Tiprl | intron_variant | MODIFIER |
| 1 | 165267159 | AX-170002193 | 0 | C | T | 0.228 | 1.53E-01 | 1.38E-01 | 2.74E+00 | 2.79E+00 | 2.64E-01 | Yes | | intergenic_variant | MODIFIER |
| 1 | 165273810 | AX-97841720 | 0 | C | T | 0.228 | 1.53E-01 | 1.38E-01 | 2.74E+00 | 2.79E+00 | 2.64E-01 | Yes | | intergenic_variant | MODIFIER |
| 1 | 165307276 | AX-170240124 | 0 | G | A | 0.228 | 1.53E-01 | 1.38E-01 | 2.74E+00 | 2.79E+00 | 2.64E-01 | Yes | Gpr161 | intron_variant | MODIFIER |
| 1 | 165308235 | AX-97815613 | 0 | A | G | 0.228 | 1.53E-01 | 1.38E-01 | 2.74E+00 | 2.79E+00 | 2.64E-01 | Yes | Gpr161 | intron_variant | MODIFIER |
| 1 | 165308844 | AX-170002192 | 0 | G | T | 0.228 | 1.53E-01 | 1.38E-01 | 2.74E+00 | 2.79E+00 | 2.64E-01 | Yes | Gpr161 | intron_variant | MODIFIER |
| 1 | 165308941 | AX-170149342 | 0 | C | A | 0.228 | 1.53E-01 | 1.38E-01 | 2.74E+00 | 2.79E+00 | 2.64E-01 | Yes | Gpr161 | intron_variant | MODIFIER |
| 1 | 165308966 | AX-170502966 | 0 | C | G | 0.228 | 1.53E-01 | 1.38E-01 | 2.74E+00 | 2.79E+00 | 2.64E-01 | Yes | Gpr161 | intron_variant | MODIFIER |
| 1 | 165394763 | AX-97804200 | 0 | T | C | 0.228 | 1.53E-01 | 1.38E-01 | 2.74E+00 | 2.79E+00 | 2.64E-01 | Yes | Dcaf6 | intron_variant | MODIFIER |
| 1 | 165539068 | AX-97849332 | 0 | C | T | 0.228 | 1.53E-01 | 1.38E-01 | 2.74E+00 | 2.79E+00 | 2.64E-01 | Yes | Adcy10 | intron_variant | MODIFIER |
| 1 | 165541618 | AX-170002210 | 0 | G | A | 0.228 | 1.53E-01 | 1.38E-01 | 2.74E+00 | 2.79E+00 | 2.64E-01 | Yes | Adcy10 | synonymous_variant | LOW |
| 1 | 165585443 | AX-97814108 | 0 | C | A | 0.228 | 1.53E-01 | 1.38E-01 | 2.74E+00 | 2.79E+00 | 2.64E-01 | Yes | Gm38210 | non_coding_transcript_exon_variant | MODIFIER |
| 1 | 165589668 | AX-170002216 | 0 | A | G | 0.228 | 1.53E-01 | 1.38E-01 | 2.74E+00 | 2.79E+00 | 2.64E-01 | Yes | Mpzl1 | downstream_gene_variant | MODIFIER |

| | | | | | | | | | | | | | | | |
|---|---|---|---|---|---|---|---|---|---|---|---|---|---|---|---|
| 1 | 165639088 | AX-97849087 | 0 | A | T | 0.228 | 1.53E-01 | 1.38E-01 | 2.74E+00 | 2.79E+00 | 2.64E-01 | Yes | Mpzl1 | upstream_gene_variant | MODIFIER |
| 1 | 165650766 | AX-97831333 | 0 | T | C | 0.228 | 1.53E-01 | 1.38E-01 | 2.74E+00 | 2.79E+00 | 2.64E-01 | Yes | Rcsd1 | intron_variant | MODIFIER |
| 1 | 165652880 | AX-170174349 | 0 | G | C | 0.228 | 1.53E-01 | 1.38E-01 | 2.74E+00 | 2.79E+00 | 2.64E-01 | Yes | Rcsd1 | intron_variant | MODIFIER |
| 1 | 165679627 | AX-97855201 | 0 | A | G | 0.228 | 1.53E-01 | 1.38E-01 | 2.74E+00 | 2.79E+00 | 2.64E-01 | Yes | Rcsd1 | intron_variant | MODIFIER |
| 1 | 165773348 | AX-170307839 | 0 | A | G | 0.228 | 1.53E-01 | 1.38E-01 | 2.74E+00 | 2.79E+00 | 2.64E-01 | Yes | Creg1 | intron_variant | MODIFIER |
| 1 | 165776182 | AX-97814137 | 0 | G | A | 0.228 | 1.53E-01 | 1.38E-01 | 2.74E+00 | 2.79E+00 | 2.64E-01 | Yes | Creg1 | downstream_gene_variant | MODIFIER |
| 1 | 165794428 | AX-170298330 | 0 | A | G | 0.228 | 1.53E-01 | 1.38E-01 | 2.74E+00 | 2.79E+00 | 2.64E-01 | Yes | Cd247 | intron_variant | MODIFIER |
| 1 | 165808478 | AX-97791106 | 0 | C | T | 0.228 | 1.53E-01 | 1.38E-01 | 2.74E+00 | 2.79E+00 | 2.64E-01 | Yes | Cd247 | intron_variant | MODIFIER |
| 1 | 165817515 | AX-170281953 | 0 | A | G | 0.228 | 1.53E-01 | 1.38E-01 | 2.74E+00 | 2.79E+00 | 2.64E-01 | Yes | Cd247 | intron_variant | MODIFIER |
| 1 | 165821317 | AX-170396236 | 0 | C | T | 0.228 | 1.53E-01 | 1.38E-01 | 2.74E+00 | 2.79E+00 | 2.64E-01 | Yes | Cd247 | intron_variant | MODIFIER |
| 1 | 165824313 | AX-170043424 | 0 | G | A | 0.228 | 1.53E-01 | 1.38E-01 | 2.74E+00 | 2.79E+00 | 2.64E-01 | Yes | Cd247 | intron_variant | MODIFIER |
| 1 | 165825838 | AX-170442224 | 0 | A | G | 0.228 | 1.53E-01 | 1.38E-01 | 2.74E+00 | 2.79E+00 | 2.64E-01 | Yes | Cd247 | intron_variant | MODIFIER |
| 1 | 165826067 | AX-97854672 | 0 | G | A | 0.228 | 1.53E-01 | 1.38E-01 | 2.74E+00 | 2.79E+00 | 2.64E-01 | Yes | Cd247 | intron_variant | MODIFIER |
| 1 | 165826132 | AX-170100003 | 0 | A | G | 0.228 | 1.53E-01 | 1.38E-01 | 2.74E+00 | 2.79E+00 | 2.64E-01 | Yes | Cd247 | intron_variant | MODIFIER |
| 1 | 165993513 | AX-97826650 | 0 | T | C | 0.228 | 1.53E-01 | 1.38E-01 | 2.74E+00 | 2.79E+00 | 2.64E-01 | Yes | Pou2f1 | intron_variant | MODIFIER |
| 1 | 166164492 | AX-97850309 | 0 | G | A | 0.228 | 1.53E-01 | 1.38E-01 | 2.74E+00 | 2.79E+00 | 2.64E-01 | Yes | Gpa33 | intron_variant | MODIFIER |
| 1 | 166167830 | AX-170246977 | 0 | G | A | 0.228 | 1.53E-01 | 1.38E-01 | 2.74E+00 | 2.79E+00 | 2.64E-01 | Yes | Gpa33 | downstream_gene_variant | MODIFIER |
| 1 | 166171228 | AX-170451049 | 0 | G | C | 0.228 | 1.53E-01 | 1.38E-01 | 2.74E+00 | 2.79E+00 | 2.64E-01 | Yes | Gpa33 | downstream_gene_variant | MODIFIER |
| 1 | 166173512 | AX-97792840 | 0 | A | G | 0.228 | 1.53E-01 | 1.38E-01 | 2.74E+00 | 2.79E+00 | 2.64E-01 | Yes | | intergenic_variant | MODIFIER |
| 1 | 166231877 | AX-170298706 | 0 | C | T | 0.228 | 1.53E-01 | 1.38E-01 | 2.74E+00 | 2.79E+00 | 2.64E-01 | Yes | Mael | intron_variant | MODIFIER |
| 1 | 166258837 | AX-97830573 | 0 | G | A | 0.228 | 1.53E-01 | 1.38E-01 | 2.74E+00 | 2.79E+00 | 2.64E-01 | Yes | Ildr2 | intron_variant | MODIFIER |
| 1 | 166260063 | AX-170227529 | 0 | T | C | 0.228 | 1.53E-01 | 1.38E-01 | 2.74E+00 | 2.79E+00 | 2.64E-01 | Yes | Ildr2 | intron_variant | MODIFIER |
| 1 | 166260971 | AX-170043435 | 0 | G | A | 0.228 | 1.53E-01 | 1.38E-01 | 2.74E+00 | 2.79E+00 | 2.64E-01 | Yes | Ildr2 | intron_variant | MODIFIER |
| 1 | 166264882 | AX-170288111 | 0 | T | C | 0.228 | 1.53E-01 | 1.38E-01 | 2.74E+00 | 2.79E+00 | 2.64E-01 | Yes | Ildr2 | intron_variant | MODIFIER |
| 1 | 166270358 | AX-170043437 | 0 | C | T | 0.228 | 1.53E-01 | 1.38E-01 | 2.74E+00 | 2.79E+00 | 2.64E-01 | Yes | Ildr2 | intron_variant | MODIFIER |
| 1 | 166283524 | AX-97831653 | 0 | T | C | 0.228 | 1.53E-01 | 1.38E-01 | 2.74E+00 | 2.79E+00 | 2.64E-01 | Yes | Ildr2 | intron_variant | MODIFIER |
| 1 | 166286241 | AX-170125792 | 0 | T | C | 0.228 | 1.53E-01 | 1.38E-01 | 2.74E+00 | 2.79E+00 | 2.64E-01 | Yes | Ildr2 | intron_variant | MODIFIER |
| 1 | 166292472 | AX-170290872 | 0 | A | T | 0.228 | 1.53E-01 | 1.38E-01 | 2.74E+00 | 2.79E+00 | 2.64E-01 | Yes | Ildr2 | intron_variant | MODIFIER |
| 1 | 166294598 | AX-170043438 | 0 | G | A | 0.228 | 1.53E-01 | 1.38E-01 | 2.74E+00 | 2.79E+00 | 2.64E-01 | Yes | Ildr2 | synonymous_variant | LOW |
| 1 | 166344021 | AX-97787394 | 0 | T | C | 0.228 | 1.53E-01 | 1.38E-01 | 2.74E+00 | 2.79E+00 | 2.64E-01 | Yes | | intergenic_variant | MODIFIER |
| 1 | 166396114 | AX-97814059 | 0 | T | C | 0.228 | 1.53E-01 | 1.38E-01 | 2.74E+00 | 2.79E+00 | 2.64E-01 | Yes | Tada1 | downstream_gene_variant | MODIFIER |
| 1 | 166479050 | AX-97851562 | 0 | T | C | 0.228 | 1.53E-01 | 1.38E-01 | 2.74E+00 | 2.79E+00 | 2.64E-01 | Yes | Gm4846 | downstream_gene_variant | MODIFIER |
| 1 | 166500851 | AX-170002223 | 0 | T | A | 0.228 | 1.53E-01 | 1.38E-01 | 2.74E+00 | 2.79E+00 | 2.64E-01 | Yes | Gm4846 | upstream_gene_variant | MODIFIER |
| 1 | 166516355 | AX-170086676 | 0 | A | C | 0.228 | 1.53E-01 | 1.38E-01 | 2.74E+00 | 2.79E+00 | 2.64E-01 | Yes | | intergenic_variant | MODIFIER |
| 1 | 166575001 | AX-170187440 | 0 | A | T | 0.228 | 1.53E-01 | 1.38E-01 | 2.74E+00 | 2.79E+00 | 2.64E-01 | Yes | | intergenic_variant | MODIFIER |
| 1 | 166575483 | AX-170264633 | 0 | A | G | 0.228 | 1.53E-01 | 1.38E-01 | 2.74E+00 | 2.79E+00 | 2.64E-01 | Yes | | intergenic_variant | MODIFIER |
| 1 | 166585502 | AX-170429726 | 0 | C | T | 0.228 | 1.53E-01 | 1.38E-01 | 2.74E+00 | 2.79E+00 | 2.64E-01 | Yes | | intergenic_variant | MODIFIER |
| 1 | 166627913 | AX-170176976 | 0 | A | G | 0.228 | 1.53E-01 | 1.38E-01 | 2.74E+00 | 2.79E+00 | 2.64E-01 | Yes | Gm4847 | downstream_gene_variant | MODIFIER |
| 1 | 166628307 | AX-170043442 | 0 | C | G | 0.228 | 1.53E-01 | 1.38E-01 | 2.74E+00 | 2.79E+00 | 2.64E-01 | Yes | Gm4847 | downstream_gene_variant | MODIFIER |
| 1 | 166634558 | AX-168399872 | 0 | C | T | 0.228 | 1.53E-01 | 1.38E-01 | 2.74E+00 | 2.79E+00 | 2.64E-01 | Yes | Gm4847 | intron_variant | MODIFIER |
| 1 | 166679584 | AX-97820461 | 0 | G | A | 0.228 | 1.53E-01 | 1.38E-01 | 2.74E+00 | 2.79E+00 | 2.64E-01 | Yes | Fmo9 | intron_variant | MODIFIER |
| 1 | 167228659 | AX-170375253 | 0 | C | T | 0.228 | 1.53E-01 | 1.38E-01 | 2.74E+00 | 2.79E+00 | 2.64E-01 | Yes | Uck2 | intron_variant | MODIFIER |
| 1 | 167355904 | AX-170497611 | 0 | T | C | 0.228 | 1.53E-01 | 1.38E-01 | 2.74E+00 | 2.79E+00 | 2.64E-01 | Yes | Aldh9a1 | intron_variant | MODIFIER |
| 1 | 167356731 | AX-97817507 | 0 | G | A | 0.228 | 1.53E-01 | 1.38E-01 | 2.74E+00 | 2.79E+00 | 2.64E-01 | Yes | Aldh9a1 | splice_region_variant,intron_variant | LOW |
| 1 | 167461540 | AX-170237925 | 0 | G | C | 0.228 | 1.53E-01 | 1.38E-01 | 2.74E+00 | 2.79E+00 | 2.64E-01 | Yes | Lrrc52 | intron_variant | MODIFIER |
| 1 | 167634344 | AX-97809702 | 0 | T | C | 0.228 | 1.53E-01 | 1.38E-01 | 2.74E+00 | 2.79E+00 | 2.64E-01 | Yes | Rxrg | intron_variant | MODIFIER |
| 1 | 167804528 | AX-97788751 | 0 | C | T | 0.228 | 1.53E-01 | 1.38E-01 | 2.74E+00 | 2.79E+00 | 2.64E-01 | Yes | Lmx1a | intron_variant | MODIFIER |
| 1 | 167818327 | AX-170378991 | 0 | G | A | 0.228 | 1.53E-01 | 1.38E-01 | 2.74E+00 | 2.79E+00 | 2.64E-01 | Yes | Lmx1a | intron_variant | MODIFIER |
| 1 | 167818596 | AX-170043464 | 0 | C | T | 0.228 | 1.53E-01 | 1.38E-01 | 2.74E+00 | 2.79E+00 | 2.64E-01 | Yes | Lmx1a | intron_variant | MODIFIER |
| 1 | 167820385 | AX-170397877 | 0 | G | T | 0.228 | 1.53E-01 | 1.38E-01 | 2.74E+00 | 2.79E+00 | 2.64E-01 | Yes | Lmx1a | intron_variant | MODIFIER |
| 1 | 167821263 | AX-170218568 | 0 | T | C | 0.228 | 1.53E-01 | 1.38E-01 | 2.74E+00 | 2.79E+00 | 2.64E-01 | Yes | Lmx1a | intron_variant | MODIFIER |
| 1 | 167821953 | AX-170163080 | 0 | A | G | 0.228 | 1.53E-01 | 1.38E-01 | 2.74E+00 | 2.79E+00 | 2.64E-01 | Yes | Lmx1a | intron_variant | MODIFIER |
| 1 | 167821996 | AX-170357622 | 0 | C | T | 0.228 | 1.53E-01 | 1.38E-01 | 2.74E+00 | 2.79E+00 | 2.64E-01 | Yes | Lmx1a | intron_variant | MODIFIER |
| 1 | 167822070 | AX-170156627 | 0 | G | A | 0.228 | 1.53E-01 | 1.38E-01 | 2.74E+00 | 2.79E+00 | 2.64E-01 | Yes | Lmx1a | intron_variant | MODIFIER |

| | | | | | | | | | | | | | | | |
|---|---|---|---|---|---|---|---|---|---|---|---|---|---|---|---|
| 1 | 167823578 | AX-170294874 | 0 | G | A | 0.228 | 1.53E-01 | 1.38E-01 | 2.74E+00 | 2.79E+00 | 2.64E-01 | Yes | Lmx1a | intron_variant | MODIFIER |
| 1 | 167831699 | AX-170195851 | 0 | A | G | 0.228 | 1.53E-01 | 1.38E-01 | 2.74E+00 | 2.79E+00 | 2.64E-01 | Yes | Lmx1a | intron_variant | MODIFIER |
| 1 | 167831908 | AX-170226948 | 0 | C | G | 0.228 | 1.53E-01 | 1.38E-01 | 2.74E+00 | 2.79E+00 | 2.64E-01 | Yes | Lmx1a | intron_variant | MODIFIER |
| 1 | 167833654 | AX-97817285 | 0 | G | A | 0.228 | 1.53E-01 | 1.38E-01 | 2.74E+00 | 2.79E+00 | 2.64E-01 | Yes | Lmx1a | intron_variant | MODIFIER |
| 1 | 167840571 | AX-170264708 | 0 | C | A | 0.228 | 1.53E-01 | 1.38E-01 | 2.74E+00 | 2.79E+00 | 2.64E-01 | Yes | Lmx1a | intron_variant | MODIFIER |
| 1 | 167861390 | AX-97814188 | 0 | T | C | 0.228 | 1.53E-01 | 1.38E-01 | 2.74E+00 | 2.79E+00 | 2.64E-01 | Yes | | intergenic_variant | MODIFIER |
| 1 | 167887228 | AX-170392070 | 0 | G | A | 0.228 | 1.53E-01 | 1.38E-01 | 2.74E+00 | 2.79E+00 | 2.64E-01 | Yes | | intergenic_variant | MODIFIER |
| 1 | 167887252 | AX-170369152 | 0 | G | A | 0.228 | 1.53E-01 | 1.38E-01 | 2.74E+00 | 2.79E+00 | 2.64E-01 | Yes | | intergenic_variant | MODIFIER |
| 1 | 167889855 | AX-170145093 | 0 | G | A | 0.228 | 1.53E-01 | 1.38E-01 | 2.74E+00 | 2.79E+00 | 2.64E-01 | Yes | | intergenic_variant | MODIFIER |
| 1 | 167892574 | AX-170303145 | 0 | G | A | 0.228 | 1.53E-01 | 1.38E-01 | 2.74E+00 | 2.79E+00 | 2.64E-01 | Yes | | regulatory_region_variant | MODIFIER |
| 1 | 167895597 | AX-170441841 | 0 | T | C | 0.228 | 1.53E-01 | 1.38E-01 | 2.74E+00 | 2.79E+00 | 2.64E-01 | Yes | | intergenic_variant | MODIFIER |
| 1 | 167902155 | AX-170043466 | 0 | C | T | 0.228 | 1.53E-01 | 1.38E-01 | 2.74E+00 | 2.79E+00 | 2.64E-01 | Yes | | intergenic_variant | MODIFIER |
| 1 | 167911669 | AX-170185297 | 0 | A | G | 0.228 | 1.53E-01 | 1.38E-01 | 2.74E+00 | 2.79E+00 | 2.64E-01 | Yes | | intergenic_variant | MODIFIER |
| 1 | 167921049 | AX-170043468 | 0 | C | T | 0.228 | 1.53E-01 | 1.38E-01 | 2.74E+00 | 2.79E+00 | 2.64E-01 | Yes | | intergenic_variant | MODIFIER |
| 1 | 167929996 | AX-170313267 | 0 | G | A | 0.228 | 1.53E-01 | 1.38E-01 | 2.74E+00 | 2.79E+00 | 2.64E-01 | Yes | | regulatory_region_variant | MODIFIER |
| 1 | 167943921 | AX-170291092 | 0 | C | G | 0.228 | 1.53E-01 | 1.38E-01 | 2.74E+00 | 2.79E+00 | 2.64E-01 | Yes | | intergenic_variant | MODIFIER |
| 1 | 167944126 | AX-170376887 | 0 | T | C | 0.228 | 1.53E-01 | 1.38E-01 | 2.74E+00 | 2.79E+00 | 2.64E-01 | Yes | | intergenic_variant | MODIFIER |
| 1 | 167956311 | AX-170355263 | 0 | T | C | 0.228 | 1.53E-01 | 1.38E-01 | 2.74E+00 | 2.79E+00 | 2.64E-01 | Yes | | intergenic_variant | MODIFIER |
| 1 | 167977703 | AX-170348576 | 0 | G | C | 0.228 | 1.53E-01 | 1.38E-01 | 2.74E+00 | 2.79E+00 | 2.64E-01 | Yes | | intergenic_variant | MODIFIER |
| 1 | 168114181 | AX-170183387 | 0 | A | G | 0.228 | 1.53E-01 | 1.38E-01 | 2.74E+00 | 2.79E+00 | 2.64E-01 | Yes | Gm20711 | intron_variant,non_coding_transcript_variant | MODIFIER |
| 1 | 168119695 | AX-170146985 | 0 | C | G | 0.228 | 1.53E-01 | 1.38E-01 | 2.74E+00 | 2.79E+00 | 2.64E-01 | Yes | Gm20711 | intron_variant,non_coding_transcript_variant | MODIFIER |
| 1 | 168131456 | AX-168379992 | 0 | T | C | 0.228 | 1.53E-01 | 1.38E-01 | 2.74E+00 | 2.79E+00 | 2.64E-01 | Yes | Pbx1 | intron_variant | MODIFIER |
| 16 | 53722677 | AX-170013074 | 0 | T | G | 0.474 | -9.34E-02 | 8.34E-02 | 2.53E+00 | 2.57E+00 | 2.64E-01 | no | | intergenic_variant | MODIFIER |
| 4 | 74863317 | AX-170023128 | 0 | A | T | 0.456 | -9.35E-02 | 8.40E-02 | 2.57E+00 | 2.64E+00 | 2.64E-01 | no | | intergenic_variant | MODIFIER |
| 18 | 58266586 | AX-170285791 | 2 | G | A | 0.355 | -6.57E-02 | 5.91E-02 | 2.89E+00 | 3.01E+00 | 2.64E-01 | no | | regulatory_region_variant | MODIFIER |
| 4 | 129924468 | AX-170402380 | 0 | T | G | 0.167 | 1.18E-01 | 1.05E-01 | 2.50E+00 | 2.56E+00 | 2.64E-01 | no | Spocd1 | upstream_gene_variant | MODIFIER |
| 4 | 130057366 | AX-170070674 | 0 | G | A | 0.167 | 1.18E-01 | 1.05E-01 | 2.50E+00 | 2.56E+00 | 2.64E-01 | no | Col16a1 | intron_variant | MODIFIER |
| 3 | 10469653 | AX-170329088 | 0 | A | C | 0.395 | -7.12E-02 | 6.46E-02 | 2.78E+00 | 2.94E+00 | 2.64E-01 | no | | intergenic_variant | MODIFIER |
| 3 | 10523284 | AX-97842255 | 0 | C | T | 0.395 | -7.12E-02 | 6.46E-02 | 2.78E+00 | 2.94E+00 | 2.64E-01 | no | | intergenic_variant | MODIFIER |
| 3 | 10554386 | AX-97789735 | 0 | T | G | 0.395 | -7.12E-02 | 6.46E-02 | 2.78E+00 | 2.94E+00 | 2.64E-01 | no | | intergenic_variant | MODIFIER |
| 3 | 10660182 | AX-97802723 | 0 | T | C | 0.395 | -7.12E-02 | 6.46E-02 | 2.78E+00 | 2.94E+00 | 2.64E-01 | no | | intergenic_variant | MODIFIER |
| 3 | 10692820 | AX-170390093 | 0 | C | T | 0.395 | -7.12E-02 | 6.46E-02 | 2.78E+00 | 2.94E+00 | 2.64E-01 | no | | intergenic_variant | MODIFIER |
| 3 | 10693043 | AX-170494555 | 0 | T | A | 0.395 | -7.12E-02 | 6.46E-02 | 2.78E+00 | 2.94E+00 | 2.64E-01 | no | | intergenic_variant | MODIFIER |
| 3 | 10696615 | AX-170254055 | 0 | A | G | 0.395 | -7.12E-02 | 6.46E-02 | 2.78E+00 | 2.94E+00 | 2.64E-01 | no | | intergenic_variant | MODIFIER |
| 3 | 10696687 | AX-97827490 | 0 | A | G | 0.395 | -7.12E-02 | 6.46E-02 | 2.78E+00 | 2.94E+00 | 2.64E-01 | no | | intergenic_variant | MODIFIER |
| 3 | 10700543 | AX-170019908 | 0 | G | C | 0.395 | -7.12E-02 | 6.46E-02 | 2.78E+00 | 2.94E+00 | 2.64E-01 | no | | intergenic_variant | MODIFIER |
| 3 | 10784200 | AX-170019911 | 0 | C | T | 0.395 | -7.12E-02 | 6.46E-02 | 2.78E+00 | 2.94E+00 | 2.64E-01 | no | | intergenic_variant | MODIFIER |
| 6 | 136243565 | AX-170383183 | 0 | T | C | 0.561 | 1.00E-01 | 8.79E-02 | 2.40E+00 | 2.45E+00 | 2.64E-01 | no | | intergenic_variant | MODIFIER |
| 1 | 176982608 | AX-170325091 | 1 | C | A | 0.232 | 1.29E-01 | 1.16E-01 | 2.58E+00 | 2.65E+00 | 2.64E-01 | no | Sdccag8 | intron_variant | MODIFIER |
| 3 | 20875937 | AX-170469628 | 1 | C | T | 0.143 | -1.07E-01 | 9.66E-02 | 2.58E+00 | 2.66E+00 | 2.64E-01 | no | | intergenic_variant | MODIFIER |
| 14 | 117740209 | AX-97850935 | 1 | C | A | 0.357 | 8.02E-02 | 7.26E-02 | 2.65E+00 | 2.76E+00 | 2.64E-01 | no | Gpc6 | intron_variant | MODIFIER |
| 4 | 64346651 | AX-170099984 | 2 | A | G | 0.327 | 1.21E-01 | 1.09E-01 | 2.78E+00 | 2.87E+00 | 2.65E-01 | no | | regulatory_region_variant | MODIFIER |
| 4 | 51487114 | AX-170022789 | 1 | T | G | 0.357 | -1.06E-01 | 9.59E-02 | 2.88E+00 | 2.99E+00 | 2.65E-01 | no | | intergenic_variant | MODIFIER |
| 8 | 97515782 | AX-170383461 | 0 | T | C | 0.237 | 1.08E-01 | 9.79E-02 | 2.87E+00 | 2.98E+00 | 2.65E-01 | no | | intergenic_variant | MODIFIER |
| 1 | 23742982 | AX-170526697 | 0 | T | C | 0.421 | 9.27E-02 | 8.37E-02 | 2.62E+00 | 2.70E+00 | 2.65E-01 | no | | intergenic_variant | MODIFIER |
| 11 | 49255339 | AX-170206808 | 0 | A | G | 0.088 | -1.72E-01 | 1.56E-01 | 2.66E+00 | 2.77E+00 | 2.65E-01 | no | Mgat1 | intron_variant | MODIFIER |
| 10 | 23690230 | AX-170101381 | 0 | A | G | 0.184 | 1.12E-01 | 1.00E-01 | 2.93E+00 | 3.04E+00 | 2.65E-01 | no | | intergenic_variant | MODIFIER |
| 10 | 23690646 | AX-170186588 | 0 | G | A | 0.184 | 1.12E-01 | 1.00E-01 | 2.93E+00 | 3.04E+00 | 2.65E-01 | no | | intergenic_variant | MODIFIER |
| 16 | 53557187 | AX-170057462 | 0 | T | C | 0.518 | 9.23E-02 | 8.25E-02 | 2.52E+00 | 2.56E+00 | 2.65E-01 | no | | intergenic_variant | MODIFIER |
| 4 | 73942635 | AX-170211081 | 1 | G | A | 0.438 | -9.20E-02 | 8.21E-02 | 2.51E+00 | 2.56E+00 | 2.65E-01 | no | 2310002L09Rik | intron_variant | MODIFIER |
| 1 | 146500586 | AX-170497982 | 0 | A | G | 0.211 | 1.18E-01 | 1.05E-01 | 2.48E+00 | 2.60E+00 | 2.65E-01 | no | Brinp3 | intron_variant | MODIFIER |
| 3 | 10475653 | AX-170189452 | 0 | T | A | 0.404 | -7.34E-02 | 6.67E-02 | 2.71E+00 | 2.86E+00 | 2.65E-01 | no | | intergenic_variant | MODIFIER |
| 4 | 88916243 | AX-170411746 | 0 | A | C | 0.421 | -9.27E-02 | 8.21E-02 | 2.45E+00 | 2.50E+00 | 2.65E-01 | no | AL805899.1 | intron_variant,non_coding_transcript_variant | MODIFIER |
| 4 | 88921879 | AX-170198875 | 0 | T | C | 0.421 | -9.27E-02 | 8.21E-02 | 2.45E+00 | 2.50E+00 | 2.65E-01 | no | AL805899.1 | intron_variant,non_coding_transcript_variant | MODIFIER |

| | | | | | | | | | | | | | | | |
|---|---|---|---|---|---|---|---|---|---|---|---|---|---|---|---|
| 4 | 89539341 | AX-97808553 | 0 | G | A | 0.421 | -9.27E-02 | 8.21E-02 | 2.45E+00 | 2.50E+00 | 2.65E-01 | no | | intergenic_variant | MODIFIER |
| 4 | 89539883 | AX-170276461 | 0 | C | T | 0.421 | -9.27E-02 | 8.21E-02 | 2.45E+00 | 2.50E+00 | 2.65E-01 | no | | intergenic_variant | MODIFIER |
| 4 | 89547928 | AX-170511668 | 0 | C | A | 0.421 | -9.27E-02 | 8.21E-02 | 2.45E+00 | 2.50E+00 | 2.65E-01 | no | | intergenic_variant | MODIFIER |
| 4 | 90575396 | AX-97828817 | 0 | T | C | 0.421 | -9.27E-02 | 8.21E-02 | 2.45E+00 | 2.50E+00 | 2.65E-01 | no | | intergenic_variant | MODIFIER |
| 4 | 90622445 | AX-97832726 | 0 | A | G | 0.421 | -9.27E-02 | 8.21E-02 | 2.45E+00 | 2.50E+00 | 2.65E-01 | no | | intergenic_variant | MODIFIER |
| 4 | 90628983 | AX-170069970 | 0 | T | A | 0.421 | -9.27E-02 | 8.21E-02 | 2.45E+00 | 2.50E+00 | 2.65E-01 | no | | intergenic_variant | MODIFIER |
| 4 | 90643581 | AX-170378197 | 0 | T | C | 0.421 | -9.27E-02 | 8.21E-02 | 2.45E+00 | 2.50E+00 | 2.65E-01 | no | | intergenic_variant | MODIFIER |
| 4 | 90644122 | AX-170023384 | 0 | A | G | 0.421 | -9.27E-02 | 8.21E-02 | 2.45E+00 | 2.50E+00 | 2.65E-01 | no | | intergenic_variant | MODIFIER |
| 4 | 90685700 | AX-170531004 | 0 | T | C | 0.421 | -9.27E-02 | 8.21E-02 | 2.45E+00 | 2.50E+00 | 2.65E-01 | no | | intergenic_variant | MODIFIER |
| 4 | 90690518 | AX-97804322 | 0 | G | A | 0.421 | -9.27E-02 | 8.21E-02 | 2.45E+00 | 2.50E+00 | 2.65E-01 | no | | intergenic_variant | MODIFIER |
| 4 | 91312379 | AX-170391101 | 0 | G | T | 0.421 | -9.27E-02 | 8.21E-02 | 2.45E+00 | 2.50E+00 | 2.65E-01 | no | Elavl2 | intron_variant | MODIFIER |
| 4 | 91315667 | AX-170500269 | 0 | A | G | 0.421 | -9.27E-02 | 8.21E-02 | 2.45E+00 | 2.50E+00 | 2.65E-01 | no | Elavl2 | intron_variant | MODIFIER |
| 4 | 91315894 | AX-170277962 | 0 | A | G | 0.421 | -9.27E-02 | 8.21E-02 | 2.45E+00 | 2.50E+00 | 2.65E-01 | no | Elavl2 | intron_variant | MODIFIER |
| 4 | 91316814 | AX-170528260 | 0 | G | A | 0.421 | -9.27E-02 | 8.21E-02 | 2.45E+00 | 2.50E+00 | 2.65E-01 | no | Elavl2 | intron_variant | MODIFIER |
| 4 | 91322304 | AX-170323731 | 0 | C | A | 0.421 | -9.27E-02 | 8.21E-02 | 2.45E+00 | 2.50E+00 | 2.65E-01 | no | Elavl2 | intron_variant | MODIFIER |
| 4 | 91350256 | AX-97810390 | 0 | A | G | 0.421 | -9.27E-02 | 8.21E-02 | 2.45E+00 | 2.50E+00 | 2.65E-01 | no | Elavl2 | intron_variant | MODIFIER |
| 4 | 91354460 | AX-170392551 | 0 | A | G | 0.421 | -9.27E-02 | 8.21E-02 | 2.45E+00 | 2.50E+00 | 2.65E-01 | no | Elavl2 | intron_variant | MODIFIER |
| 4 | 91355325 | AX-170069979 | 0 | C | T | 0.421 | -9.27E-02 | 8.21E-02 | 2.45E+00 | 2.50E+00 | 2.65E-01 | no | Elavl2 | intron_variant | MODIFIER |
| 4 | 137460988 | AX-170323807 | 0 | G | A | 0.167 | 1.26E-01 | 1.14E-01 | 2.58E+00 | 2.63E+00 | 2.65E-01 | no | | intergenic_variant | MODIFIER |
| 1 | 166942307 | AX-170002236 | 1 | T | C | 0.223 | 1.54E-01 | 1.39E-01 | 2.73E+00 | 2.78E+00 | 2.65E-01 | Yes | Gm6286 | downstream_gene_variant | MODIFIER |
| 4 | 148179124 | AX-170070984 | 2 | A | C | 0.1 | 1.83E-01 | 1.66E-01 | 2.77E+00 | 2.89E+00 | 2.66E-01 | Yes | | intergenic_variant | MODIFIER |
| 8 | 65539109 | AX-170420546 | 1 | C | T | 0.134 | -1.56E-01 | 1.41E-01 | 2.59E+00 | 2.67E+00 | 2.66E-01 | no | | intergenic_variant | MODIFIER |
| 18 | 85270533 | AX-170135427 | 1 | A | C | 0.152 | -1.22E-01 | 1.08E-01 | 2.45E+00 | 2.54E+00 | 2.66E-01 | no | | intergenic_variant | MODIFIER |
| 13 | 3051683 | AX-170301514 | 0 | C | T | 0.202 | -1.38E-01 | 1.25E-01 | 2.60E+00 | 2.65E+00 | 2.66E-01 | no | | intergenic_variant | MODIFIER |
| 13 | 3258245 | AX-97822490 | 0 | C | G | 0.202 | -1.38E-01 | 1.25E-01 | 2.60E+00 | 2.65E+00 | 2.66E-01 | no | | intergenic_variant | MODIFIER |
| 13 | 3301109 | AX-97822493 | 0 | C | T | 0.202 | -1.38E-01 | 1.25E-01 | 2.60E+00 | 2.65E+00 | 2.66E-01 | no | Gm9616 | downstream_gene_variant | MODIFIER |
| 13 | 3344521 | AX-170526997 | 0 | A | G | 0.202 | -1.38E-01 | 1.25E-01 | 2.60E+00 | 2.65E+00 | 2.66E-01 | no | Gm48530 | intron_variant,non_coding_transcript_variant | MODIFIER |
| 13 | 3375559 | AX-170188105 | 0 | A | G | 0.202 | -1.38E-01 | 1.25E-01 | 2.60E+00 | 2.65E+00 | 2.66E-01 | no | | intergenic_variant | MODIFIER |
| 13 | 3522235 | AX-170520676 | 0 | A | G | 0.202 | -1.38E-01 | 1.25E-01 | 2.60E+00 | 2.65E+00 | 2.66E-01 | no | | intergenic_variant | MODIFIER |
| 13 | 3533681 | AX-97822608 | 0 | A | G | 0.202 | -1.38E-01 | 1.25E-01 | 2.60E+00 | 2.65E+00 | 2.66E-01 | no | Gdi2 | upstream_gene_variant | MODIFIER |
| 13 | 3564202 | AX-170007658 | 0 | T | C | 0.202 | -1.38E-01 | 1.25E-01 | 2.60E+00 | 2.65E+00 | 2.66E-01 | no | Gdi2 | intron_variant | MODIFIER |
| 13 | 3567905 | AX-97822491 | 0 | T | C | 0.202 | -1.38E-01 | 1.25E-01 | 2.60E+00 | 2.65E+00 | 2.66E-01 | no | Gdi2 | downstream_gene_variant | MODIFIER |
| 13 | 3573252 | AX-170007657 | 0 | C | G | 0.202 | -1.38E-01 | 1.25E-01 | 2.60E+00 | 2.65E+00 | 2.66E-01 | no | Fam208b | intron_variant | MODIFIER |
| 13 | 3633285 | AX-170007652 | 0 | A | G | 0.202 | -1.38E-01 | 1.25E-01 | 2.60E+00 | 2.65E+00 | 2.66E-01 | no | Asb13 | upstream_gene_variant | MODIFIER |
| 13 | 3640164 | AX-170487067 | 0 | G | C | 0.202 | -1.38E-01 | 1.25E-01 | 2.60E+00 | 2.65E+00 | 2.66E-01 | no | Asb13 | intron_variant | MODIFIER |
| 13 | 3657344 | AX-170151065 | 0 | C | T | 0.202 | -1.38E-01 | 1.25E-01 | 2.60E+00 | 2.65E+00 | 2.66E-01 | no | Asb13 | downstream_gene_variant | MODIFIER |
| 13 | 3659287 | AX-170501998 | 0 | C | A | 0.202 | -1.38E-01 | 1.25E-01 | 2.60E+00 | 2.65E+00 | 2.66E-01 | no | | regulatory_region_variant | MODIFIER |
| 13 | 3681548 | AX-170294588 | 0 | C | T | 0.202 | -1.38E-01 | 1.25E-01 | 2.60E+00 | 2.65E+00 | 2.66E-01 | no | | intergenic_variant | MODIFIER |
| 13 | 3695121 | AX-170200987 | 0 | G | A | 0.202 | -1.38E-01 | 1.25E-01 | 2.60E+00 | 2.65E+00 | 2.66E-01 | no | | regulatory_region_variant | MODIFIER |
| 13 | 3700668 | AX-170289927 | 0 | C | T | 0.202 | -1.38E-01 | 1.25E-01 | 2.60E+00 | 2.65E+00 | 2.66E-01 | no | | regulatory_region_variant | MODIFIER |
| 13 | 3709494 | AX-170221253 | 0 | C | T | 0.202 | -1.38E-01 | 1.25E-01 | 2.60E+00 | 2.65E+00 | 2.66E-01 | no | | regulatory_region_variant | MODIFIER |
| 13 | 3726165 | AX-170151048 | 0 | C | G | 0.202 | -1.38E-01 | 1.25E-01 | 2.60E+00 | 2.65E+00 | 2.66E-01 | no | | regulatory_region_variant | MODIFIER |
| 13 | 3750784 | AX-170137589 | 0 | A | T | 0.202 | -1.38E-01 | 1.25E-01 | 2.60E+00 | 2.65E+00 | 2.66E-01 | no | | regulatory_region_variant | MODIFIER |
| 13 | 3752572 | AX-170488751 | 0 | A | G | 0.202 | -1.38E-01 | 1.25E-01 | 2.60E+00 | 2.65E+00 | 2.66E-01 | no | | intergenic_variant | MODIFIER |
| 13 | 3788175 | AX-170286579 | 0 | A | G | 0.202 | -1.38E-01 | 1.25E-01 | 2.60E+00 | 2.65E+00 | 2.66E-01 | no | | intergenic_variant | MODIFIER |
| 13 | 3817935 | AX-170449192 | 0 | T | C | 0.202 | -1.38E-01 | 1.25E-01 | 2.60E+00 | 2.65E+00 | 2.66E-01 | no | | intergenic_variant | MODIFIER |
| 13 | 3818615 | AX-170301747 | 0 | T | G | 0.202 | -1.38E-01 | 1.25E-01 | 2.60E+00 | 2.65E+00 | 2.66E-01 | no | | intergenic_variant | MODIFIER |
| 13 | 3831960 | AX-170414683 | 0 | A | G | 0.202 | -1.38E-01 | 1.25E-01 | 2.60E+00 | 2.65E+00 | 2.66E-01 | no | Gm40655 | intron_variant,non_coding_transcript_variant | MODIFIER |
| 13 | 3832117 | AX-170174644 | 0 | A | G | 0.202 | -1.38E-01 | 1.25E-01 | 2.60E+00 | 2.65E+00 | 2.66E-01 | no | Gm40655 | intron_variant,non_coding_transcript_variant | MODIFIER |
| 13 | 3837986 | AX-170386375 | 0 | G | C | 0.202 | -1.38E-01 | 1.25E-01 | 2.60E+00 | 2.65E+00 | 2.66E-01 | no | Calm4 | missense_variant | MODERATE |
| 13 | 3899384 | AX-97822518 | 0 | A | G | 0.202 | -1.38E-01 | 1.25E-01 | 2.60E+00 | 2.65E+00 | 2.66E-01 | no | Net1 | intron_variant | MODIFIER |
| 13 | 3900443 | AX-170050400 | 0 | C | T | 0.202 | -1.38E-01 | 1.25E-01 | 2.60E+00 | 2.65E+00 | 2.66E-01 | no | Net1 | intron_variant | MODIFIER |
| 13 | 3905163 | AX-170506234 | 0 | T | C | 0.202 | -1.38E-01 | 1.25E-01 | 2.60E+00 | 2.65E+00 | 2.66E-01 | no | Net1 | intron_variant | MODIFIER |
| 13 | 3912986 | AX-97804609 | 0 | C | A | 0.202 | -1.38E-01 | 1.25E-01 | 2.60E+00 | 2.65E+00 | 2.66E-01 | no | Net1 | intron_variant | MODIFIER |

| | | | | | | | | | | | | | | | |
|---|---|---|---|---|---|---|---|---|---|---|---|---|---|---|---|
| 13 | 3916599 | AX-170050402 | 0 | T | G | 0.202 | -1.38E-01 | 1.25E-01 | 2.60E+00 | 2.65E+00 | 2.66E-01 | no | Net1 | intron_variant | MODIFIER |
| 13 | 3921782 | AX-170050403 | 0 | T | C | 0.202 | -1.38E-01 | 1.25E-01 | 2.60E+00 | 2.65E+00 | 2.66E-01 | no | Tubal3 | upstream_gene_variant | MODIFIER |
| 13 | 3930073 | AX-170419949 | 0 | T | C | 0.202 | -1.38E-01 | 1.25E-01 | 2.60E+00 | 2.65E+00 | 2.66E-01 | no | Tubal3 | intron_variant | MODIFIER |
| 13 | 3931232 | AX-170326121 | 0 | A | G | 0.202 | -1.38E-01 | 1.25E-01 | 2.60E+00 | 2.65E+00 | 2.66E-01 | no | Tubal3 | intron_variant | MODIFIER |
| 13 | 3941678 | AX-97797275 | 0 | T | C | 0.202 | -1.38E-01 | 1.25E-01 | 2.60E+00 | 2.65E+00 | 2.66E-01 | no | Ucn3 | intron_variant | MODIFIER |
| 13 | 3952493 | AX-170192636 | 0 | A | G | 0.202 | -1.38E-01 | 1.25E-01 | 2.60E+00 | 2.65E+00 | 2.66E-01 | no | Gm22460 | downstream_gene_variant | MODIFIER |
| 13 | 3952682 | AX-170102788 | 0 | G | A | 0.202 | -1.38E-01 | 1.25E-01 | 2.60E+00 | 2.65E+00 | 2.66E-01 | no | Gm22460 | downstream_gene_variant | MODIFIER |
| 13 | 3992002 | AX-170342962 | 0 | C | A | 0.202 | -1.38E-01 | 1.25E-01 | 2.60E+00 | 2.65E+00 | 2.66E-01 | no | | intergenic_variant | MODIFIER |
| 13 | 4029227 | AX-170335884 | 0 | A | G | 0.202 | -1.38E-01 | 1.25E-01 | 2.60E+00 | 2.65E+00 | 2.66E-01 | no | Gm47853 | downstream_gene_variant | MODIFIER |
| 13 | 4035430 | AX-170460534 | 0 | A | G | 0.202 | -1.38E-01 | 1.25E-01 | 2.60E+00 | 2.65E+00 | 2.66E-01 | no | Gm18857 | intron_variant,non_coding_transcript_variant | MODIFIER |
| 13 | 4037990 | AX-170219216 | 0 | T | C | 0.202 | -1.38E-01 | 1.25E-01 | 2.60E+00 | 2.65E+00 | 2.66E-01 | no | Gm18857 | non_coding_transcript_exon_variant | MODIFIER |
| 13 | 4045177 | AX-170298612 | 0 | T | C | 0.202 | -1.38E-01 | 1.25E-01 | 2.60E+00 | 2.65E+00 | 2.66E-01 | no | Akr1c14 | upstream_gene_variant | MODIFIER |
| 13 | 4051243 | AX-97822606 | 0 | G | T | 0.202 | -1.38E-01 | 1.25E-01 | 2.60E+00 | 2.65E+00 | 2.66E-01 | no | Akr1c14 | 5_prime_UTR_variant | MODIFIER |
| 13 | 4063012 | AX-170183344 | 0 | G | A | 0.202 | -1.38E-01 | 1.25E-01 | 2.60E+00 | 2.65E+00 | 2.66E-01 | no | Akr1c14 | intron_variant | MODIFIER |
| 13 | 4079567 | AX-170363942 | 0 | G | C | 0.202 | -1.38E-01 | 1.25E-01 | 2.60E+00 | 2.65E+00 | 2.66E-01 | no | Akr1c14 | intron_variant | MODIFIER |
| 13 | 4086256 | AX-170402316 | 0 | C | A | 0.202 | -1.38E-01 | 1.25E-01 | 2.60E+00 | 2.65E+00 | 2.66E-01 | no | Akr1c14 | intron_variant | MODIFIER |
| 13 | 4103132 | AX-170409587 | 0 | G | A | 0.202 | -1.38E-01 | 1.25E-01 | 2.60E+00 | 2.65E+00 | 2.66E-01 | no | Gm47905 | downstream_gene_variant | MODIFIER |
| 13 | 4115676 | AX-170529549 | 0 | G | T | 0.202 | -1.38E-01 | 1.25E-01 | 2.60E+00 | 2.65E+00 | 2.66E-01 | no | | intergenic_variant | MODIFIER |
| 13 | 4119125 | AX-170532121 | 0 | C | A | 0.202 | -1.38E-01 | 1.25E-01 | 2.60E+00 | 2.65E+00 | 2.66E-01 | no | | regulatory_region_variant | MODIFIER |
| 13 | 4137477 | AX-170007663 | 0 | A | G | 0.202 | -1.38E-01 | 1.25E-01 | 2.60E+00 | 2.65E+00 | 2.66E-01 | no | Akr1c18 | intron_variant | MODIFIER |
| 13 | 4160084 | AX-170050405 | 0 | C | T | 0.202 | -1.38E-01 | 1.25E-01 | 2.60E+00 | 2.65E+00 | 2.66E-01 | no | | intergenic_variant | MODIFIER |
| 13 | 4166121 | AX-170224142 | 0 | T | G | 0.202 | -1.38E-01 | 1.25E-01 | 2.60E+00 | 2.65E+00 | 2.66E-01 | no | Gm47925 | upstream_gene_variant | MODIFIER |
| 13 | 4170689 | AX-170050404 | 0 | T | C | 0.202 | -1.38E-01 | 1.25E-01 | 2.60E+00 | 2.65E+00 | 2.66E-01 | no | Gm47925 | intron_variant,non_coding_transcript_variant | MODIFIER |
| 13 | 4171120 | AX-170007662 | 0 | T | C | 0.202 | -1.38E-01 | 1.25E-01 | 2.60E+00 | 2.65E+00 | 2.66E-01 | no | Gm47925 | intron_variant,non_coding_transcript_variant | MODIFIER |
| 13 | 4176610 | AX-170217473 | 0 | G | C | 0.202 | -1.38E-01 | 1.25E-01 | 2.60E+00 | 2.65E+00 | 2.66E-01 | no | Gm47925 | downstream_gene_variant | MODIFIER |
| 13 | 4178940 | AX-97821290 | 0 | A | C | 0.202 | -1.38E-01 | 1.25E-01 | 2.60E+00 | 2.65E+00 | 2.66E-01 | no | | intergenic_variant | MODIFIER |
| 13 | 4196661 | AX-97833466 | 0 | G | T | 0.202 | -1.38E-01 | 1.25E-01 | 2.60E+00 | 2.65E+00 | 2.66E-01 | no | Akr1c13 | intron_variant | MODIFIER |
| 13 | 4237692 | AX-170252257 | 0 | A | G | 0.202 | -1.38E-01 | 1.25E-01 | 2.60E+00 | 2.65E+00 | 2.66E-01 | no | Akr1c19 | intron_variant | MODIFIER |
| 13 | 4238671 | AX-170112484 | 0 | T | G | 0.202 | -1.38E-01 | 1.25E-01 | 2.60E+00 | 2.65E+00 | 2.66E-01 | no | Akr1c19 | intron_variant | MODIFIER |
| 13 | 4238773 | AX-170362205 | 0 | G | A | 0.202 | -1.38E-01 | 1.25E-01 | 2.60E+00 | 2.65E+00 | 2.66E-01 | no | Akr1c19 | intron_variant | MODIFIER |
| 13 | 4248745 | AX-170464121 | 0 | A | G | 0.202 | -1.38E-01 | 1.25E-01 | 2.60E+00 | 2.65E+00 | 2.66E-01 | no | Akr1c19 | downstream_gene_variant | MODIFIER |
| 13 | 4279899 | AX-97826903 | 0 | C | G | 0.202 | -1.38E-01 | 1.25E-01 | 2.60E+00 | 2.65E+00 | 2.66E-01 | no | Akr1c12 | upstream_gene_variant | MODIFIER |
| 13 | 4291910 | AX-170315747 | 0 | T | C | 0.202 | -1.38E-01 | 1.25E-01 | 2.60E+00 | 2.65E+00 | 2.66E-01 | no | | intergenic_variant | MODIFIER |
| 13 | 4298122 | AX-170302567 | 0 | C | T | 0.202 | -1.38E-01 | 1.25E-01 | 2.60E+00 | 2.65E+00 | 2.66E-01 | no | | intergenic_variant | MODIFIER |
| 13 | 4313995 | AX-170198158 | 0 | C | T | 0.202 | -1.38E-01 | 1.25E-01 | 2.60E+00 | 2.65E+00 | 2.66E-01 | no | | intergenic_variant | MODIFIER |
| 13 | 4359678 | AX-170227083 | 0 | A | G | 0.202 | -1.38E-01 | 1.25E-01 | 2.60E+00 | 2.65E+00 | 2.66E-01 | no | Gm9973 | intron_variant,non_coding_transcript_variant | MODIFIER |
| 13 | 4427384 | AX-170509692 | 0 | G | C | 0.202 | -1.38E-01 | 1.25E-01 | 2.60E+00 | 2.65E+00 | 2.66E-01 | no | | intergenic_variant | MODIFIER |
| 13 | 4427437 | AX-170434800 | 0 | C | T | 0.202 | -1.38E-01 | 1.25E-01 | 2.60E+00 | 2.65E+00 | 2.66E-01 | no | | intergenic_variant | MODIFIER |
| 13 | 4434266 | AX-170509642 | 0 | C | A | 0.202 | -1.38E-01 | 1.25E-01 | 2.60E+00 | 2.65E+00 | 2.66E-01 | no | Akr1c6 | upstream_gene_variant | MODIFIER |
| 13 | 4448197 | AX-170050410 | 0 | C | G | 0.202 | -1.38E-01 | 1.25E-01 | 2.60E+00 | 2.65E+00 | 2.66E-01 | no | Akr1c6 | intron_variant | MODIFIER |
| 13 | 4465085 | AX-170098291 | 0 | T | C | 0.202 | -1.38E-01 | 1.25E-01 | 2.60E+00 | 2.65E+00 | 2.66E-01 | no | Gm48010 | downstream_gene_variant | MODIFIER |
| 13 | 4465115 | AX-170427125 | 0 | C | T | 0.202 | -1.38E-01 | 1.25E-01 | 2.60E+00 | 2.65E+00 | 2.66E-01 | no | Gm48010 | downstream_gene_variant | MODIFIER |
| 13 | 4466555 | AX-170050413 | 0 | A | G | 0.202 | -1.38E-01 | 1.25E-01 | 2.60E+00 | 2.65E+00 | 2.66E-01 | no | Gm48010 | non_coding_transcript_exon_variant | MODIFIER |
| 13 | 4467154 | AX-97825130 | 0 | C | T | 0.202 | -1.38E-01 | 1.25E-01 | 2.60E+00 | 2.65E+00 | 2.66E-01 | no | Gm48010 | upstream_gene_variant | MODIFIER |
| 13 | 4468860 | AX-170467623 | 0 | G | C | 0.202 | -1.38E-01 | 1.25E-01 | 2.60E+00 | 2.65E+00 | 2.66E-01 | no | Gm48010 | upstream_gene_variant | MODIFIER |
| 13 | 4469885 | AX-170498227 | 0 | T | A | 0.202 | -1.38E-01 | 1.25E-01 | 2.60E+00 | 2.65E+00 | 2.66E-01 | no | Gm48010 | upstream_gene_variant | MODIFIER |
| 13 | 4506405 | AX-170249296 | 0 | T | A | 0.202 | -1.38E-01 | 1.25E-01 | 2.60E+00 | 2.65E+00 | 2.66E-01 | no | Akr1c20 | downstream_gene_variant | MODIFIER |
| 13 | 4580138 | AX-170215535 | 0 | C | T | 0.202 | -1.38E-01 | 1.25E-01 | 2.60E+00 | 2.65E+00 | 2.66E-01 | no | Akr1c21 | intron_variant | MODIFIER |
| 13 | 4584988 | AX-170342421 | 0 | C | T | 0.202 | -1.38E-01 | 1.25E-01 | 2.60E+00 | 2.65E+00 | 2.66E-01 | no | Akr1c21 | intron_variant | MODIFIER |
| 13 | 4598791 | AX-170204280 | 0 | C | T | 0.202 | -1.38E-01 | 1.25E-01 | 2.60E+00 | 2.65E+00 | 2.66E-01 | no | Akr1e1 | synonymous_variant | LOW |
| 13 | 4644196 | AX-170007668 | 0 | T | C | 0.202 | -1.38E-01 | 1.25E-01 | 2.60E+00 | 2.65E+00 | 2.66E-01 | no | Gm40658 | intron_variant,non_coding_transcript_variant | MODIFIER |
| 13 | 4644551 | AX-170007667 | 0 | C | G | 0.202 | -1.38E-01 | 1.25E-01 | 2.60E+00 | 2.65E+00 | 2.66E-01 | no | Gm40658 | intron_variant,non_coding_transcript_variant | MODIFIER |
| 13 | 4672902 | AX-170126755 | 0 | G | A | 0.202 | -1.38E-01 | 1.25E-01 | 2.60E+00 | 2.65E+00 | 2.66E-01 | no | Gm40658 | intron_variant,non_coding_transcript_variant | MODIFIER |
| 13 | 4690486 | AX-170279832 | 0 | G | T | 0.202 | -1.38E-01 | 1.25E-01 | 2.60E+00 | 2.65E+00 | 2.66E-01 | no | Gm40658 | intron_variant,non_coding_transcript_variant | MODIFIER |

| | | | | | | | | | | | | | | | |
|---|---|---|---|---|---|---|---|---|---|---|---|---|---|---|---|
| 13 | 4709740 | AX-170050420 | 0 | A | G | 0.202 | -1.38E-01 | 1.25E-01 | 2.60E+00 | 2.65E+00 | 2.66E-01 | no | Gm40658 | intron_variant,non_coding_transcript_variant | MODIFIER |
| 13 | 4710299 | AX-170050415 | 0 | G | C | 0.202 | -1.38E-01 | 1.25E-01 | 2.60E+00 | 2.65E+00 | 2.66E-01 | no | Gm40658 | intron_variant,non_coding_transcript_variant | MODIFIER |
| 13 | 4745345 | AX-170050421 | 0 | A | G | 0.202 | -1.38E-01 | 1.25E-01 | 2.60E+00 | 2.65E+00 | 2.66E-01 | no | Gm40658 | intron_variant,non_coding_transcript_variant | MODIFIER |
| 13 | 4773064 | AX-170342997 | 0 | G | A | 0.202 | -1.38E-01 | 1.25E-01 | 2.60E+00 | 2.65E+00 | 2.66E-01 | no | Gm5444 | intron_variant,non_coding_transcript_variant | MODIFIER |
| 13 | 4807138 | AX-170271844 | 0 | C | T | 0.202 | -1.38E-01 | 1.25E-01 | 2.60E+00 | 2.65E+00 | 2.66E-01 | no | Gm5444 | intron_variant,non_coding_transcript_variant | MODIFIER |
| 13 | 4807295 | AX-170352969 | 0 | A | G | 0.202 | -1.38E-01 | 1.25E-01 | 2.60E+00 | 2.65E+00 | 2.66E-01 | no | Gm5444 | intron_variant,non_coding_transcript_variant | MODIFIER |
| 13 | 4833516 | AX-170445871 | 0 | T | C | 0.202 | -1.38E-01 | 1.25E-01 | 2.60E+00 | 2.65E+00 | 2.66E-01 | no | Gm5444 | splice_region_variant,intron_variant,non_coding_transcript_variant | LOW |
| 13 | 4882624 | AX-170036834 | 0 | C | G | 0.202 | -1.38E-01 | 1.25E-01 | 2.60E+00 | 2.65E+00 | 2.66E-01 | no | | intergenic_variant | MODIFIER |
| 13 | 4897382 | AX-170520244 | 0 | G | T | 0.202 | -1.38E-01 | 1.25E-01 | 2.60E+00 | 2.65E+00 | 2.66E-01 | no | | intergenic_variant | MODIFIER |
| 13 | 4916709 | AX-170261083 | 0 | A | G | 0.202 | -1.38E-01 | 1.25E-01 | 2.60E+00 | 2.65E+00 | 2.66E-01 | no | Gm47450 | intron_variant,non_coding_transcript_variant | MODIFIER |
| 13 | 4920649 | AX-97780558 | 0 | C | A | 0.202 | -1.38E-01 | 1.25E-01 | 2.60E+00 | 2.65E+00 | 2.66E-01 | no | Gm47450 | intron_variant,non_coding_transcript_variant | MODIFIER |
| 13 | 4922138 | AX-170217759 | 0 | C | G | 0.202 | -1.38E-01 | 1.25E-01 | 2.60E+00 | 2.65E+00 | 2.66E-01 | no | Gm47450 | intron_variant,non_coding_transcript_variant | MODIFIER |
| 13 | 4970017 | AX-170092638 | 0 | G | A | 0.202 | -1.38E-01 | 1.25E-01 | 2.60E+00 | 2.65E+00 | 2.66E-01 | no | | intergenic_variant | MODIFIER |
| 13 | 4976856 | AX-170346073 | 0 | T | G | 0.202 | -1.38E-01 | 1.25E-01 | 2.60E+00 | 2.65E+00 | 2.66E-01 | no | | intergenic_variant | MODIFIER |
| 13 | 4999471 | AX-170213918 | 0 | A | T | 0.202 | -1.38E-01 | 1.25E-01 | 2.60E+00 | 2.65E+00 | 2.66E-01 | no | | intergenic_variant | MODIFIER |
| 13 | 5000637 | AX-170092639 | 0 | A | G | 0.202 | -1.38E-01 | 1.25E-01 | 2.60E+00 | 2.65E+00 | 2.66E-01 | no | | intergenic_variant | MODIFIER |
| 13 | 5024211 | AX-170414125 | 0 | T | C | 0.202 | -1.38E-01 | 1.25E-01 | 2.60E+00 | 2.65E+00 | 2.66E-01 | no | | intergenic_variant | MODIFIER |
| 13 | 5050705 | AX-97790800 | 0 | A | G | 0.202 | -1.38E-01 | 1.25E-01 | 2.60E+00 | 2.65E+00 | 2.66E-01 | no | | intergenic_variant | MODIFIER |
| 13 | 5054244 | AX-170050423 | 0 | G | A | 0.202 | -1.38E-01 | 1.25E-01 | 2.60E+00 | 2.65E+00 | 2.66E-01 | no | | intergenic_variant | MODIFIER |
| 13 | 5055420 | AX-170050424 | 0 | T | C | 0.202 | -1.38E-01 | 1.25E-01 | 2.60E+00 | 2.65E+00 | 2.66E-01 | no | | intergenic_variant | MODIFIER |
| 13 | 5055478 | AX-170050426 | 0 | A | G | 0.202 | -1.38E-01 | 1.25E-01 | 2.60E+00 | 2.65E+00 | 2.66E-01 | no | | intergenic_variant | MODIFIER |
| 13 | 5061687 | AX-97801506 | 0 | T | C | 0.202 | -1.38E-01 | 1.25E-01 | 2.60E+00 | 2.65E+00 | 2.66E-01 | no | | intergenic_variant | MODIFIER |
| 13 | 5076312 | AX-170036831 | 0 | G | C | 0.202 | -1.38E-01 | 1.25E-01 | 2.60E+00 | 2.65E+00 | 2.66E-01 | no | | intergenic_variant | MODIFIER |
| 13 | 5101291 | AX-170143388 | 0 | G | T | 0.202 | -1.38E-01 | 1.25E-01 | 2.60E+00 | 2.65E+00 | 2.66E-01 | no | | intergenic_variant | MODIFIER |
| 13 | 5101316 | AX-170482511 | 0 | G | T | 0.202 | -1.38E-01 | 1.25E-01 | 2.60E+00 | 2.65E+00 | 2.66E-01 | no | | intergenic_variant | MODIFIER |
| 13 | 5106286 | AX-170269987 | 0 | A | G | 0.202 | -1.38E-01 | 1.25E-01 | 2.60E+00 | 2.65E+00 | 2.66E-01 | no | | intergenic_variant | MODIFIER |
| 13 | 5121565 | AX-170415860 | 0 | A | G | 0.202 | -1.38E-01 | 1.25E-01 | 2.60E+00 | 2.65E+00 | 2.66E-01 | no | | intergenic_variant | MODIFIER |
| 13 | 5155760 | AX-170007671 | 0 | A | G | 0.202 | -1.38E-01 | 1.25E-01 | 2.60E+00 | 2.65E+00 | 2.66E-01 | no | | intergenic_variant | MODIFIER |
| 13 | 5166193 | AX-170087350 | 0 | G | A | 0.202 | -1.38E-01 | 1.25E-01 | 2.60E+00 | 2.65E+00 | 2.66E-01 | no | | intergenic_variant | MODIFIER |
| 13 | 5188631 | AX-170436873 | 0 | A | G | 0.202 | -1.38E-01 | 1.25E-01 | 2.60E+00 | 2.65E+00 | 2.66E-01 | no | Gm24790 | downstream_gene_variant | MODIFIER |
| 13 | 5188660 | AX-170377901 | 0 | G | T | 0.202 | -1.38E-01 | 1.25E-01 | 2.60E+00 | 2.65E+00 | 2.66E-01 | no | Gm24790 | downstream_gene_variant | MODIFIER |
| 13 | 5234032 | AX-170265397 | 0 | C | T | 0.202 | -1.38E-01 | 1.25E-01 | 2.60E+00 | 2.65E+00 | 2.66E-01 | no | | intergenic_variant | MODIFIER |
| 13 | 5240854 | AX-97851480 | 0 | G | T | 0.202 | -1.38E-01 | 1.25E-01 | 2.60E+00 | 2.65E+00 | 2.66E-01 | no | | intergenic_variant | MODIFIER |
| 13 | 5243317 | AX-170217239 | 0 | C | A | 0.202 | -1.38E-01 | 1.25E-01 | 2.60E+00 | 2.65E+00 | 2.66E-01 | no | | intergenic_variant | MODIFIER |
| 13 | 5250336 | AX-170322381 | 0 | G | A | 0.202 | -1.38E-01 | 1.25E-01 | 2.60E+00 | 2.65E+00 | 2.66E-01 | no | | intergenic_variant | MODIFIER |
| 13 | 5259696 | AX-170259419 | 0 | A | G | 0.202 | -1.38E-01 | 1.25E-01 | 2.60E+00 | 2.65E+00 | 2.66E-01 | no | | intergenic_variant | MODIFIER |
| 13 | 5259845 | AX-170101190 | 0 | A | G | 0.202 | -1.38E-01 | 1.25E-01 | 2.60E+00 | 2.65E+00 | 2.66E-01 | no | | intergenic_variant | MODIFIER |
| 13 | 5288055 | AX-170384991 | 0 | T | C | 0.202 | -1.38E-01 | 1.25E-01 | 2.60E+00 | 2.65E+00 | 2.66E-01 | no | | intergenic_variant | MODIFIER |
| 13 | 5288280 | AX-170194033 | 0 | A | G | 0.202 | -1.38E-01 | 1.25E-01 | 2.60E+00 | 2.65E+00 | 2.66E-01 | no | | intergenic_variant | MODIFIER |
| 13 | 5291146 | AX-170131213 | 0 | T | C | 0.202 | -1.38E-01 | 1.25E-01 | 2.60E+00 | 2.65E+00 | 2.66E-01 | no | | intergenic_variant | MODIFIER |
| 13 | 5292689 | AX-170525895 | 0 | G | A | 0.202 | -1.38E-01 | 1.25E-01 | 2.60E+00 | 2.65E+00 | 2.66E-01 | no | | intergenic_variant | MODIFIER |
| 13 | 5298040 | AX-170090920 | 0 | T | C | 0.202 | -1.38E-01 | 1.25E-01 | 2.60E+00 | 2.65E+00 | 2.66E-01 | no | | intergenic_variant | MODIFIER |
| 13 | 5300858 | AX-170476731 | 0 | T | C | 0.202 | -1.38E-01 | 1.25E-01 | 2.60E+00 | 2.65E+00 | 2.66E-01 | no | | intergenic_variant | MODIFIER |
| 13 | 5318371 | AX-170519118 | 0 | A | G | 0.202 | -1.38E-01 | 1.25E-01 | 2.60E+00 | 2.65E+00 | 2.66E-01 | no | | intergenic_variant | MODIFIER |
| 13 | 5338277 | AX-170505903 | 0 | A | C | 0.202 | -1.38E-01 | 1.25E-01 | 2.60E+00 | 2.65E+00 | 2.66E-01 | no | | intergenic_variant | MODIFIER |
| 13 | 5405707 | AX-170135651 | 0 | G | A | 0.202 | -1.38E-01 | 1.25E-01 | 2.60E+00 | 2.65E+00 | 2.66E-01 | no | | intergenic_variant | MODIFIER |
| 13 | 5418954 | AX-97798711 | 0 | C | A | 0.202 | -1.38E-01 | 1.25E-01 | 2.60E+00 | 2.65E+00 | 2.66E-01 | no | | intergenic_variant | MODIFIER |
| 13 | 5439798 | AX-170418564 | 0 | T | G | 0.202 | -1.38E-01 | 1.25E-01 | 2.60E+00 | 2.65E+00 | 2.66E-01 | no | | intergenic_variant | MODIFIER |
| 13 | 5477187 | AX-97845623 | 0 | G | A | 0.202 | -1.38E-01 | 1.25E-01 | 2.60E+00 | 2.65E+00 | 2.66E-01 | no | Gm40660 | intron_variant,non_coding_transcript_variant | MODIFIER |
| 13 | 5492132 | AX-170283702 | 0 | T | A | 0.202 | -1.38E-01 | 1.25E-01 | 2.60E+00 | 2.65E+00 | 2.66E-01 | no | Gm40660 | intron_variant,non_coding_transcript_variant | MODIFIER |
| 13 | 5496940 | AX-170399598 | 0 | T | A | 0.202 | -1.38E-01 | 1.25E-01 | 2.60E+00 | 2.65E+00 | 2.66E-01 | no | Gm40660 | intron_variant,non_coding_transcript_variant | MODIFIER |
| 13 | 5497954 | AX-170115589 | 0 | A | G | 0.202 | -1.38E-01 | 1.25E-01 | 2.60E+00 | 2.65E+00 | 2.66E-01 | no | Gm40660 | intron_variant,non_coding_transcript_variant | MODIFIER |
| 13 | 5499339 | AX-170489872 | 0 | C | A | 0.202 | -1.38E-01 | 1.25E-01 | 2.60E+00 | 2.65E+00 | 2.66E-01 | no | Gm40660 | intron_variant,non_coding_transcript_variant | MODIFIER |
| 13 | 5508433 | AX-97806578 | 0 | G | A | 0.202 | -1.38E-01 | 1.25E-01 | 2.60E+00 | 2.65E+00 | 2.66E-01 | no | Gm40660 | intron_variant,non_coding_transcript_variant | MODIFIER |

| | | | | | | | | | | | | | | | |
|---|---|---|---|---|---|---|---|---|---|---|---|---|---|---|---|
| 13 | 5509118 | AX-170350151 | 0 | T | C | 0.202 | -1.38E-01 | 1.25E-01 | 2.60E+00 | 2.65E+00 | 2.66E-01 | no | Gm40660 | downstream_gene_variant | MODIFIER |
| 13 | 5532432 | AX-170092640 | 0 | G | T | 0.202 | -1.38E-01 | 1.25E-01 | 2.60E+00 | 2.65E+00 | 2.66E-01 | no | Gm40660 | intron_variant,non_coding_transcript_variant | MODIFIER |
| 13 | 5542423 | AX-170216168 | 0 | C | G | 0.202 | -1.38E-01 | 1.25E-01 | 2.60E+00 | 2.65E+00 | 2.66E-01 | no | Gm35330 | intron_variant,non_coding_transcript_variant | MODIFIER |
| 13 | 5544002 | AX-170404810 | 0 | G | T | 0.202 | -1.38E-01 | 1.25E-01 | 2.60E+00 | 2.65E+00 | 2.66E-01 | no | Gm35330 | intron_variant,non_coding_transcript_variant | MODIFIER |
| 13 | 5555709 | AX-170472339 | 0 | A | T | 0.202 | -1.38E-01 | 1.25E-01 | 2.60E+00 | 2.65E+00 | 2.66E-01 | no | Gm35330 | upstream_gene_variant | MODIFIER |
| 13 | 5556830 | AX-170181610 | 0 | G | A | 0.202 | -1.38E-01 | 1.25E-01 | 2.60E+00 | 2.65E+00 | 2.66E-01 | no | Gm35330 | intron_variant,non_coding_transcript_variant | MODIFIER |
| 13 | 5557414 | AX-170531960 | 0 | C | A | 0.202 | -1.38E-01 | 1.25E-01 | 2.60E+00 | 2.65E+00 | 2.66E-01 | no | Gm35330 | intron_variant,non_coding_transcript_variant | MODIFIER |
| 13 | 5576516 | AX-170339348 | 0 | C | G | 0.202 | -1.38E-01 | 1.25E-01 | 2.60E+00 | 2.65E+00 | 2.66E-01 | no | Gm35330 | intron_variant,non_coding_transcript_variant | MODIFIER |
| 13 | 5614100 | AX-170135352 | 0 | G | A | 0.202 | -1.38E-01 | 1.25E-01 | 2.60E+00 | 2.65E+00 | 2.66E-01 | no | Gm35330 | intron_variant,non_coding_transcript_variant | MODIFIER |
| 13 | 5614236 | AX-170507418 | 0 | C | T | 0.202 | -1.38E-01 | 1.25E-01 | 2.60E+00 | 2.65E+00 | 2.66E-01 | no | Gm35330 | intron_variant,non_coding_transcript_variant | MODIFIER |
| 13 | 5614444 | AX-170223546 | 0 | A | G | 0.202 | -1.38E-01 | 1.25E-01 | 2.60E+00 | 2.65E+00 | 2.66E-01 | no | Gm35330 | intron_variant,non_coding_transcript_variant | MODIFIER |
| 13 | 5641707 | AX-170291638 | 0 | T | G | 0.202 | -1.38E-01 | 1.25E-01 | 2.60E+00 | 2.65E+00 | 2.66E-01 | no | Gm35330 | upstream_gene_variant | MODIFIER |
| 13 | 5661006 | AX-170007673 | 0 | G | A | 0.202 | -1.38E-01 | 1.25E-01 | 2.60E+00 | 2.65E+00 | 2.66E-01 | no | Gm35330 | intron_variant,non_coding_transcript_variant | MODIFIER |
| 13 | 5673570 | AX-170317952 | 0 | C | T | 0.202 | -1.38E-01 | 1.25E-01 | 2.60E+00 | 2.65E+00 | 2.66E-01 | no | Gm35330 | intron_variant,non_coding_transcript_variant | MODIFIER |
| 13 | 5675675 | AX-170453628 | 0 | C | A | 0.202 | -1.38E-01 | 1.25E-01 | 2.60E+00 | 2.65E+00 | 2.66E-01 | no | Gm35330 | intron_variant,non_coding_transcript_variant | MODIFIER |
| 13 | 5708958 | AX-97788346 | 0 | G | A | 0.202 | -1.38E-01 | 1.25E-01 | 2.60E+00 | 2.65E+00 | 2.66E-01 | no | Gm35330 | intron_variant,non_coding_transcript_variant | MODIFIER |
| 13 | 5709041 | AX-170007676 | 0 | A | G | 0.202 | -1.38E-01 | 1.25E-01 | 2.60E+00 | 2.65E+00 | 2.66E-01 | no | Gm35330 | intron_variant,non_coding_transcript_variant | MODIFIER |
| 13 | 5717358 | AX-170297560 | 0 | C | T | 0.202 | -1.38E-01 | 1.25E-01 | 2.60E+00 | 2.65E+00 | 2.66E-01 | no | Gm35330 | intron_variant,non_coding_transcript_variant | MODIFIER |
| 13 | 5720823 | AX-170234439 | 0 | T | C | 0.202 | -1.38E-01 | 1.25E-01 | 2.60E+00 | 2.65E+00 | 2.66E-01 | no | Gm35330 | intron_variant,non_coding_transcript_variant | MODIFIER |
| 13 | 5740880 | AX-170486836 | 0 | G | T | 0.202 | -1.38E-01 | 1.25E-01 | 2.60E+00 | 2.65E+00 | 2.66E-01 | no | | intergenic_variant | MODIFIER |
| 13 | 5741429 | AX-170421275 | 0 | G | T | 0.202 | -1.38E-01 | 1.25E-01 | 2.60E+00 | 2.65E+00 | 2.66E-01 | no | | intergenic_variant | MODIFIER |
| 13 | 5744538 | AX-170188805 | 0 | C | T | 0.202 | -1.38E-01 | 1.25E-01 | 2.60E+00 | 2.65E+00 | 2.66E-01 | no | | intergenic_variant | MODIFIER |
| 13 | 5744941 | AX-170529923 | 0 | C | T | 0.202 | -1.38E-01 | 1.25E-01 | 2.60E+00 | 2.65E+00 | 2.66E-01 | no | | intergenic_variant | MODIFIER |
| 13 | 5771311 | AX-170153687 | 0 | G | A | 0.202 | -1.38E-01 | 1.25E-01 | 2.60E+00 | 2.65E+00 | 2.66E-01 | no | | intergenic_variant | MODIFIER |
| 13 | 5771471 | AX-170120858 | 0 | A | C | 0.202 | -1.38E-01 | 1.25E-01 | 2.60E+00 | 2.65E+00 | 2.66E-01 | no | | intergenic_variant | MODIFIER |
| 13 | 5790563 | AX-170201940 | 0 | T | G | 0.202 | -1.38E-01 | 1.25E-01 | 2.60E+00 | 2.65E+00 | 2.66E-01 | no | | intergenic_variant | MODIFIER |
| 13 | 5790962 | AX-170460786 | 0 | T | C | 0.202 | -1.38E-01 | 1.25E-01 | 2.60E+00 | 2.65E+00 | 2.66E-01 | no | | intergenic_variant | MODIFIER |
| 13 | 5800562 | AX-170139134 | 0 | C | G | 0.202 | -1.38E-01 | 1.25E-01 | 2.60E+00 | 2.65E+00 | 2.66E-01 | no | | intergenic_variant | MODIFIER |
| 13 | 5813761 | AX-170110493 | 0 | A | G | 0.202 | -1.38E-01 | 1.25E-01 | 2.60E+00 | 2.65E+00 | 2.66E-01 | no | | regulatory_region_variant | MODIFIER |
| 13 | 5815551 | AX-170122523 | 0 | A | G | 0.202 | -1.38E-01 | 1.25E-01 | 2.60E+00 | 2.65E+00 | 2.66E-01 | no | | intergenic_variant | MODIFIER |
| 13 | 5817180 | AX-97822626 | 0 | A | G | 0.202 | -1.38E-01 | 1.25E-01 | 2.60E+00 | 2.65E+00 | 2.66E-01 | no | Gm26043 | downstream_gene_variant | MODIFIER |
| 13 | 5823227 | AX-170113479 | 0 | T | A | 0.202 | -1.38E-01 | 1.25E-01 | 2.60E+00 | 2.65E+00 | 2.66E-01 | no | Gm26043 | upstream_gene_variant | MODIFIER |
| 13 | 5823366 | AX-170452290 | 0 | A | G | 0.202 | -1.38E-01 | 1.25E-01 | 2.60E+00 | 2.65E+00 | 2.66E-01 | no | Gm26043 | upstream_gene_variant | MODIFIER |
| 13 | 5831643 | AX-170164916 | 0 | T | C | 0.202 | -1.38E-01 | 1.25E-01 | 2.60E+00 | 2.65E+00 | 2.66E-01 | no | | intergenic_variant | MODIFIER |
| 13 | 5837160 | AX-170451252 | 0 | G | A | 0.202 | -1.38E-01 | 1.25E-01 | 2.60E+00 | 2.65E+00 | 2.66E-01 | no | | intergenic_variant | MODIFIER |
| 13 | 5866618 | AX-170501589 | 0 | A | G | 0.202 | -1.38E-01 | 1.25E-01 | 2.60E+00 | 2.65E+00 | 2.66E-01 | no | Klf6 | intron_variant | MODIFIER |
| 13 | 5866727 | AX-97794896 | 0 | C | T | 0.202 | -1.38E-01 | 1.25E-01 | 2.60E+00 | 2.65E+00 | 2.66E-01 | no | Klf6 | synonymous_variant | LOW |
| 13 | 5881450 | AX-170171454 | 0 | G | A | 0.202 | -1.38E-01 | 1.25E-01 | 2.60E+00 | 2.65E+00 | 2.66E-01 | no | | intergenic_variant | MODIFIER |
| 13 | 5900610 | AX-170087353 | 0 | T | C | 0.202 | -1.38E-01 | 1.25E-01 | 2.60E+00 | 2.65E+00 | 2.66E-01 | no | Gm47507 | intron_variant,non_coding_transcript_variant | MODIFIER |
| 13 | 5901943 | AX-170263556 | 0 | T | C | 0.202 | -1.38E-01 | 1.25E-01 | 2.60E+00 | 2.65E+00 | 2.66E-01 | no | Gm47507 | intron_variant,non_coding_transcript_variant | MODIFIER |
| 13 | 5932567 | AX-170450173 | 0 | T | A | 0.202 | -1.38E-01 | 1.25E-01 | 2.60E+00 | 2.65E+00 | 2.66E-01 | no | Gm47507 | downstream_gene_variant | MODIFIER |
| 13 | 5940016 | AX-97823265 | 0 | G | A | 0.202 | -1.38E-01 | 1.25E-01 | 2.60E+00 | 2.65E+00 | 2.66E-01 | no | Gm46401 | intron_variant,non_coding_transcript_variant | MODIFIER |
| 13 | 5957550 | AX-170319357 | 0 | A | G | 0.202 | -1.38E-01 | 1.25E-01 | 2.60E+00 | 2.65E+00 | 2.66E-01 | no | Gm46401 | intron_variant,non_coding_transcript_variant | MODIFIER |
| 13 | 5977706 | AX-97820948 | 0 | A | G | 0.202 | -1.38E-01 | 1.25E-01 | 2.60E+00 | 2.65E+00 | 2.66E-01 | no | Gm46401 | upstream_gene_variant | MODIFIER |
| 13 | 5977858 | AX-170461765 | 0 | C | T | 0.202 | -1.38E-01 | 1.25E-01 | 2.60E+00 | 2.65E+00 | 2.66E-01 | no | Gm46401 | non_coding_transcript_exon_variant | MODIFIER |
| 13 | 5979069 | AX-170342159 | 0 | A | G | 0.202 | -1.38E-01 | 1.25E-01 | 2.60E+00 | 2.65E+00 | 2.66E-01 | no | Gm46401 | intron_variant,non_coding_transcript_variant | MODIFIER |
| 13 | 5979999 | AX-170217202 | 0 | A | G | 0.202 | -1.38E-01 | 1.25E-01 | 2.60E+00 | 2.65E+00 | 2.66E-01 | no | Gm46401 | intron_variant,non_coding_transcript_variant | MODIFIER |
| 13 | 5984274 | AX-170343119 | 0 | G | A | 0.202 | -1.38E-01 | 1.25E-01 | 2.60E+00 | 2.65E+00 | 2.66E-01 | no | Gm46401 | intron_variant,non_coding_transcript_variant | MODIFIER |
| 13 | 5990993 | AX-97787329 | 0 | T | C | 0.202 | -1.38E-01 | 1.25E-01 | 2.60E+00 | 2.65E+00 | 2.66E-01 | no | Gm46401 | intron_variant,non_coding_transcript_variant | MODIFIER |
| 13 | 6025544 | AX-170143934 | 0 | G | A | 0.202 | -1.38E-01 | 1.25E-01 | 2.60E+00 | 2.65E+00 | 2.66E-01 | no | Gm47514 | intron_variant,non_coding_transcript_variant | MODIFIER |
| 13 | 6026231 | AX-170345814 | 0 | C | T | 0.202 | -1.38E-01 | 1.25E-01 | 2.60E+00 | 2.65E+00 | 2.66E-01 | no | Gm47514 | non_coding_transcript_exon_variant | MODIFIER |
| 13 | 6057829 | AX-170435010 | 0 | A | C | 0.202 | -1.38E-01 | 1.25E-01 | 2.60E+00 | 2.65E+00 | 2.66E-01 | no | | intergenic_variant | MODIFIER |
| 13 | 6058774 | AX-170230223 | 0 | C | T | 0.202 | -1.38E-01 | 1.25E-01 | 2.60E+00 | 2.65E+00 | 2.66E-01 | no | | intergenic_variant | MODIFIER |
| 13 | 6065519 | AX-170339263 | 0 | C | T | 0.202 | -1.38E-01 | 1.25E-01 | 2.60E+00 | 2.65E+00 | 2.66E-01 | no | | intergenic_variant | MODIFIER |

| | | | | | | | | | | | | | | | |
|---|---|---|---|---|---|---|---|---|---|---|---|---|---|---|---|
| 13 | 6065540 | AX-170426586 | 0 | C | T | 0.202 | -1.38E-01 | 1.25E-01 | 2.60E+00 | 2.65E+00 | 2.66E-01 | no | | intergenic_variant | MODIFIER |
| 13 | 6079244 | AX-170512292 | 0 | T | C | 0.202 | -1.38E-01 | 1.25E-01 | 2.60E+00 | 2.65E+00 | 2.66E-01 | no | | intergenic_variant | MODIFIER |
| 13 | 6087220 | AX-170206983 | 0 | G | A | 0.202 | -1.38E-01 | 1.25E-01 | 2.60E+00 | 2.65E+00 | 2.66E-01 | no | | intergenic_variant | MODIFIER |
| 13 | 6099461 | AX-170328207 | 0 | C | G | 0.202 | -1.38E-01 | 1.25E-01 | 2.60E+00 | 2.65E+00 | 2.66E-01 | no | | intergenic_variant | MODIFIER |
| 13 | 6101759 | AX-170375796 | 0 | T | C | 0.202 | -1.38E-01 | 1.25E-01 | 2.60E+00 | 2.65E+00 | 2.66E-01 | no | | intergenic_variant | MODIFIER |
| 13 | 6101917 | AX-170335946 | 0 | G | A | 0.202 | -1.38E-01 | 1.25E-01 | 2.60E+00 | 2.65E+00 | 2.66E-01 | no | | intergenic_variant | MODIFIER |
| 13 | 6131634 | AX-170240700 | 0 | G | A | 0.202 | -1.38E-01 | 1.25E-01 | 2.60E+00 | 2.65E+00 | 2.66E-01 | no | | intergenic_variant | MODIFIER |
| 13 | 6137335 | AX-170306077 | 0 | G | A | 0.202 | -1.38E-01 | 1.25E-01 | 2.60E+00 | 2.65E+00 | 2.66E-01 | no | | regulatory_region_variant | MODIFIER |
| 13 | 6153710 | AX-170327531 | 0 | G | C | 0.202 | -1.38E-01 | 1.25E-01 | 2.60E+00 | 2.65E+00 | 2.66E-01 | no | | intergenic_variant | MODIFIER |
| 13 | 6160789 | AX-97792158 | 0 | C | A | 0.202 | -1.38E-01 | 1.25E-01 | 2.60E+00 | 2.65E+00 | 2.66E-01 | no | | intergenic_variant | MODIFIER |
| 13 | 6176079 | AX-170498812 | 0 | G | A | 0.202 | -1.38E-01 | 1.25E-01 | 2.60E+00 | 2.65E+00 | 2.66E-01 | no | | intergenic_variant | MODIFIER |
| 13 | 6179184 | AX-170529962 | 0 | C | A | 0.202 | -1.38E-01 | 1.25E-01 | 2.60E+00 | 2.65E+00 | 2.66E-01 | no | | intergenic_variant | MODIFIER |
| 13 | 6181945 | AX-170188885 | 0 | T | C | 0.202 | -1.38E-01 | 1.25E-01 | 2.60E+00 | 2.65E+00 | 2.66E-01 | no | | intergenic_variant | MODIFIER |
| 13 | 6182572 | AX-170407421 | 0 | G | A | 0.202 | -1.38E-01 | 1.25E-01 | 2.60E+00 | 2.65E+00 | 2.66E-01 | no | | intergenic_variant | MODIFIER |
| 13 | 6198178 | AX-170500330 | 0 | T | C | 0.202 | -1.38E-01 | 1.25E-01 | 2.60E+00 | 2.65E+00 | 2.66E-01 | no | | intergenic_variant | MODIFIER |
| 13 | 6202955 | AX-170007677 | 0 | T | C | 0.202 | -1.38E-01 | 1.25E-01 | 2.60E+00 | 2.65E+00 | 2.66E-01 | no | | intergenic_variant | MODIFIER |
| 13 | 6204037 | AX-170226206 | 0 | T | C | 0.202 | -1.38E-01 | 1.25E-01 | 2.60E+00 | 2.65E+00 | 2.66E-01 | no | | intergenic_variant | MODIFIER |
| 13 | 6225589 | AX-170347046 | 0 | G | A | 0.202 | -1.38E-01 | 1.25E-01 | 2.60E+00 | 2.65E+00 | 2.66E-01 | no | | intergenic_variant | MODIFIER |
| 13 | 6229579 | AX-170007678 | 0 | C | T | 0.202 | -1.38E-01 | 1.25E-01 | 2.60E+00 | 2.65E+00 | 2.66E-01 | no | | intergenic_variant | MODIFIER |
| 13 | 6235569 | AX-170306519 | 0 | A | G | 0.202 | -1.38E-01 | 1.25E-01 | 2.60E+00 | 2.65E+00 | 2.66E-01 | no | | intergenic_variant | MODIFIER |
| 13 | 6239496 | AX-97785254 | 0 | A | G | 0.202 | -1.38E-01 | 1.25E-01 | 2.60E+00 | 2.65E+00 | 2.66E-01 | no | | intergenic_variant | MODIFIER |
| 13 | 6240479 | AX-170223945 | 0 | A | G | 0.202 | -1.38E-01 | 1.25E-01 | 2.60E+00 | 2.65E+00 | 2.66E-01 | no | | intergenic_variant | MODIFIER |
| 13 | 6241025 | AX-170107912 | 0 | T | A | 0.202 | -1.38E-01 | 1.25E-01 | 2.60E+00 | 2.65E+00 | 2.66E-01 | no | | regulatory_region_variant | MODIFIER |
| 13 | 6257029 | AX-170007679 | 0 | T | C | 0.202 | -1.38E-01 | 1.25E-01 | 2.60E+00 | 2.65E+00 | 2.66E-01 | no | | intergenic_variant | MODIFIER |
| 13 | 6306681 | AX-170188419 | 0 | G | C | 0.202 | -1.38E-01 | 1.25E-01 | 2.60E+00 | 2.65E+00 | 2.66E-01 | no | | intergenic_variant | MODIFIER |
| 13 | 6307621 | AX-97781296 | 0 | G | T | 0.202 | -1.38E-01 | 1.25E-01 | 2.60E+00 | 2.65E+00 | 2.66E-01 | no | | regulatory_region_variant | MODIFIER |
| 13 | 6340184 | AX-170513839 | 0 | G | A | 0.202 | -1.38E-01 | 1.25E-01 | 2.60E+00 | 2.65E+00 | 2.66E-01 | no | | intergenic_variant | MODIFIER |
| 13 | 6340213 | AX-170121687 | 0 | A | G | 0.202 | -1.38E-01 | 1.25E-01 | 2.60E+00 | 2.65E+00 | 2.66E-01 | no | | intergenic_variant | MODIFIER |
| 13 | 6342163 | AX-170050446 | 0 | T | C | 0.202 | -1.38E-01 | 1.25E-01 | 2.60E+00 | 2.65E+00 | 2.66E-01 | no | | intergenic_variant | MODIFIER |
| 13 | 6365078 | AX-170092642 | 0 | G | A | 0.202 | -1.38E-01 | 1.25E-01 | 2.60E+00 | 2.65E+00 | 2.66E-01 | no | | intergenic_variant | MODIFIER |
| 13 | 6374588 | AX-170347424 | 0 | T | G | 0.202 | -1.38E-01 | 1.25E-01 | 2.60E+00 | 2.65E+00 | 2.66E-01 | no | | intergenic_variant | MODIFIER |
| 13 | 6377493 | AX-170492275 | 0 | A | G | 0.202 | -1.38E-01 | 1.25E-01 | 2.60E+00 | 2.65E+00 | 2.66E-01 | no | | intergenic_variant | MODIFIER |
| 13 | 6386862 | AX-170316725 | 0 | T | C | 0.202 | -1.38E-01 | 1.25E-01 | 2.60E+00 | 2.65E+00 | 2.66E-01 | no | | intergenic_variant | MODIFIER |
| 13 | 6416532 | AX-170249943 | 0 | G | A | 0.202 | -1.38E-01 | 1.25E-01 | 2.60E+00 | 2.65E+00 | 2.66E-01 | no | | intergenic_variant | MODIFIER |
| 13 | 6432649 | AX-170213004 | 0 | G | A | 0.202 | -1.38E-01 | 1.25E-01 | 2.60E+00 | 2.65E+00 | 2.66E-01 | no | | intergenic_variant | MODIFIER |
| 13 | 6433242 | AX-170126075 | 0 | A | G | 0.202 | -1.38E-01 | 1.25E-01 | 2.60E+00 | 2.65E+00 | 2.66E-01 | no | | intergenic_variant | MODIFIER |
| 13 | 6434115 | AX-170439467 | 0 | T | C | 0.202 | -1.38E-01 | 1.25E-01 | 2.60E+00 | 2.65E+00 | 2.66E-01 | no | | intergenic_variant | MODIFIER |
| 13 | 6457593 | AX-170355568 | 0 | A | G | 0.202 | -1.38E-01 | 1.25E-01 | 2.60E+00 | 2.65E+00 | 2.66E-01 | no | | intergenic_variant | MODIFIER |
| 13 | 6461127 | AX-170129743 | 0 | G | A | 0.202 | -1.38E-01 | 1.25E-01 | 2.60E+00 | 2.65E+00 | 2.66E-01 | no | | intergenic_variant | MODIFIER |
| 13 | 6470576 | AX-170050452 | 0 | G | A | 0.202 | -1.38E-01 | 1.25E-01 | 2.60E+00 | 2.65E+00 | 2.66E-01 | no | | regulatory_region_variant | MODIFIER |
| 13 | 6505122 | AX-170036842 | 0 | T | C | 0.202 | -1.38E-01 | 1.25E-01 | 2.60E+00 | 2.65E+00 | 2.66E-01 | no | | intergenic_variant | MODIFIER |
| 13 | 6505404 | AX-170036841 | 0 | T | C | 0.202 | -1.38E-01 | 1.25E-01 | 2.60E+00 | 2.65E+00 | 2.66E-01 | no | | regulatory_region_variant | MODIFIER |
| 13 | 6507773 | AX-170168389 | 0 | A | T | 0.202 | -1.38E-01 | 1.25E-01 | 2.60E+00 | 2.65E+00 | 2.66E-01 | no | | intergenic_variant | MODIFIER |
| 13 | 6552621 | AX-97854680 | 0 | A | T | 0.202 | -1.38E-01 | 1.25E-01 | 2.60E+00 | 2.65E+00 | 2.66E-01 | no | Pitrm1 | intron_variant | MODIFIER |
| 13 | 6560538 | AX-170395114 | 0 | A | G | 0.202 | -1.38E-01 | 1.25E-01 | 2.60E+00 | 2.65E+00 | 2.66E-01 | no | Pitrm1 | intron_variant | MODIFIER |
| 13 | 6564365 | AX-170329900 | 0 | C | T | 0.202 | -1.38E-01 | 1.25E-01 | 2.60E+00 | 2.65E+00 | 2.66E-01 | no | Pitrm1 | intron_variant | MODIFIER |
| 13 | 6564680 | AX-170104136 | 0 | T | C | 0.202 | -1.38E-01 | 1.25E-01 | 2.60E+00 | 2.65E+00 | 2.66E-01 | no | Pitrm1 | intron_variant | MODIFIER |
| 13 | 6566908 | AX-170404256 | 0 | G | A | 0.202 | -1.38E-01 | 1.25E-01 | 2.60E+00 | 2.65E+00 | 2.66E-01 | no | Pitrm1 | intron_variant | MODIFIER |
| 13 | 6567622 | AX-97847264 | 0 | G | A | 0.202 | -1.38E-01 | 1.25E-01 | 2.60E+00 | 2.65E+00 | 2.66E-01 | no | Pitrm1 | intron_variant | MODIFIER |
| 13 | 6579264 | AX-170106276 | 0 | A | G | 0.202 | -1.38E-01 | 1.25E-01 | 2.60E+00 | 2.65E+00 | 2.66E-01 | no | Pitrm1 | intron_variant | MODIFIER |
| 13 | 6581470 | AX-170398220 | 0 | T | C | 0.202 | -1.38E-01 | 1.25E-01 | 2.60E+00 | 2.65E+00 | 2.66E-01 | no | Pitrm1 | downstream_gene_variant | MODIFIER |
| 13 | 6581505 | AX-170445083 | 0 | A | G | 0.202 | -1.38E-01 | 1.25E-01 | 2.60E+00 | 2.65E+00 | 2.66E-01 | no | Pitrm1 | downstream_gene_variant | MODIFIER |
| 13 | 6590670 | AX-170286380 | 0 | G | A | 0.202 | -1.38E-01 | 1.25E-01 | 2.60E+00 | 2.65E+00 | 2.66E-01 | no | Pfkp | intron_variant | MODIFIER |
| 13 | 6600671 | AX-170294260 | 0 | C | T | 0.202 | -1.38E-01 | 1.25E-01 | 2.60E+00 | 2.65E+00 | 2.66E-01 | no | Pfkp | splice_region_variant,intron_variant | LOW |

| | | | | | | | | | | | | | | | |
|---|---|---|---|---|---|---|---|---|---|---|---|---|---|---|---|
| 13 | 6601243 | AX-170204509 | 0 | T | C | 0.202 | -1.38E-01 | 1.25E-01 | 2.60E+00 | 2.65E+00 | 2.66E-01 | no | Pfkp | intron_variant | MODIFIER |
| 13 | 6614837 | AX-170122788 | 0 | A | G | 0.202 | -1.38E-01 | 1.25E-01 | 2.60E+00 | 2.65E+00 | 2.66E-01 | no | Pfkp | intron_variant | MODIFIER |
| 13 | 6615249 | AX-170346139 | 0 | G | A | 0.202 | -1.38E-01 | 1.25E-01 | 2.60E+00 | 2.65E+00 | 2.66E-01 | no | Pfkp | intron_variant | MODIFIER |
| 13 | 6620596 | AX-170423664 | 0 | A | G | 0.202 | -1.38E-01 | 1.25E-01 | 2.60E+00 | 2.65E+00 | 2.66E-01 | no | Pfkp | intron_variant | MODIFIER |
| 13 | 6624734 | AX-170050458 | 0 | T | C | 0.202 | -1.38E-01 | 1.25E-01 | 2.60E+00 | 2.65E+00 | 2.66E-01 | no | Pfkp | intron_variant | MODIFIER |
| 13 | 6626261 | AX-170050455 | 0 | C | T | 0.202 | -1.38E-01 | 1.25E-01 | 2.60E+00 | 2.65E+00 | 2.66E-01 | no | Pfkp | intron_variant | MODIFIER |
| 13 | 6638917 | AX-170221483 | 0 | G | A | 0.202 | -1.38E-01 | 1.25E-01 | 2.60E+00 | 2.65E+00 | 2.66E-01 | no | Pfkp | intron_variant | MODIFIER |
| 13 | 6639213 | AX-97801239 | 0 | C | T | 0.202 | -1.38E-01 | 1.25E-01 | 2.60E+00 | 2.65E+00 | 2.66E-01 | no | Pfkp | intron_variant | MODIFIER |
| 13 | 6641077 | AX-170148693 | 0 | T | C | 0.202 | -1.38E-01 | 1.25E-01 | 2.60E+00 | 2.65E+00 | 2.66E-01 | no | Pfkp | intron_variant | MODIFIER |
| 13 | 6645666 | AX-170421437 | 0 | C | T | 0.202 | -1.38E-01 | 1.25E-01 | 2.60E+00 | 2.65E+00 | 2.66E-01 | no | Pfkp | intron_variant | MODIFIER |
| 13 | 6660231 | AX-170452985 | 0 | A | C | 0.202 | -1.38E-01 | 1.25E-01 | 2.60E+00 | 2.65E+00 | 2.66E-01 | no | Gm10029 | downstream_gene_variant | MODIFIER |
| 13 | 6666822 | AX-97824224 | 0 | A | G | 0.202 | -1.38E-01 | 1.25E-01 | 2.60E+00 | 2.65E+00 | 2.66E-01 | no | Gm10029 | upstream_gene_variant | MODIFIER |
| 13 | 6666900 | AX-170302180 | 0 | C | G | 0.202 | -1.38E-01 | 1.25E-01 | 2.60E+00 | 2.65E+00 | 2.66E-01 | no | Gm10029 | upstream_gene_variant | MODIFIER |
| 13 | 6676655 | AX-170182924 | 0 | T | C | 0.202 | -1.38E-01 | 1.25E-01 | 2.60E+00 | 2.65E+00 | 2.66E-01 | no | Gm48238 | downstream_gene_variant | MODIFIER |
| 13 | 6719209 | AX-170335831 | 0 | T | G | 0.202 | -1.38E-01 | 1.25E-01 | 2.60E+00 | 2.65E+00 | 2.66E-01 | no | | intergenic_variant | MODIFIER |
| 13 | 6720892 | AX-170154860 | 0 | T | C | 0.202 | -1.38E-01 | 1.25E-01 | 2.60E+00 | 2.65E+00 | 2.66E-01 | no | | intergenic_variant | MODIFIER |
| 13 | 6739369 | AX-170340043 | 0 | G | A | 0.202 | -1.38E-01 | 1.25E-01 | 2.60E+00 | 2.65E+00 | 2.66E-01 | no | | intergenic_variant | MODIFIER |
| 13 | 6742568 | AX-170500794 | 0 | G | A | 0.202 | -1.38E-01 | 1.25E-01 | 2.60E+00 | 2.65E+00 | 2.66E-01 | no | | intergenic_variant | MODIFIER |
| 13 | 6748780 | AX-170483693 | 0 | G | T | 0.202 | -1.38E-01 | 1.25E-01 | 2.60E+00 | 2.65E+00 | 2.66E-01 | no | | intergenic_variant | MODIFIER |
| 13 | 6748812 | AX-97829417 | 0 | C | T | 0.202 | -1.38E-01 | 1.25E-01 | 2.60E+00 | 2.65E+00 | 2.66E-01 | no | | intergenic_variant | MODIFIER |
| 13 | 6768710 | AX-170007686 | 0 | T | C | 0.202 | -1.38E-01 | 1.25E-01 | 2.60E+00 | 2.65E+00 | 2.66E-01 | no | | intergenic_variant | MODIFIER |
| 13 | 6769090 | AX-170092646 | 0 | T | C | 0.202 | -1.38E-01 | 1.25E-01 | 2.60E+00 | 2.65E+00 | 2.66E-01 | no | | intergenic_variant | MODIFIER |
| 13 | 6772798 | AX-170036839 | 0 | A | G | 0.202 | -1.38E-01 | 1.25E-01 | 2.60E+00 | 2.65E+00 | 2.66E-01 | no | | intergenic_variant | MODIFIER |
| 13 | 6772877 | AX-170036837 | 0 | T | G | 0.202 | -1.38E-01 | 1.25E-01 | 2.60E+00 | 2.65E+00 | 2.66E-01 | no | | intergenic_variant | MODIFIER |
| 13 | 6803388 | AX-170116093 | 0 | G | A | 0.202 | -1.38E-01 | 1.25E-01 | 2.60E+00 | 2.65E+00 | 2.66E-01 | no | | intergenic_variant | MODIFIER |
| 13 | 6837789 | AX-170007685 | 0 | G | C | 0.202 | -1.38E-01 | 1.25E-01 | 2.60E+00 | 2.65E+00 | 2.66E-01 | no | | intergenic_variant | MODIFIER |
| 13 | 6851468 | AX-170187002 | 0 | A | G | 0.202 | -1.38E-01 | 1.25E-01 | 2.60E+00 | 2.65E+00 | 2.66E-01 | no | | intergenic_variant | MODIFIER |
| 13 | 6852626 | AX-170007683 | 0 | C | T | 0.202 | -1.38E-01 | 1.25E-01 | 2.60E+00 | 2.65E+00 | 2.66E-01 | no | | intergenic_variant | MODIFIER |
| 13 | 6852667 | AX-170292024 | 0 | A | C | 0.202 | -1.38E-01 | 1.25E-01 | 2.60E+00 | 2.65E+00 | 2.66E-01 | no | | intergenic_variant | MODIFIER |
| 13 | 6873875 | AX-170007684 | 0 | T | C | 0.202 | -1.38E-01 | 1.25E-01 | 2.60E+00 | 2.65E+00 | 2.66E-01 | no | | intergenic_variant | MODIFIER |
| 13 | 6874004 | AX-97822578 | 0 | T | C | 0.202 | -1.38E-01 | 1.25E-01 | 2.60E+00 | 2.65E+00 | 2.66E-01 | no | | intergenic_variant | MODIFIER |
| 13 | 6952785 | AX-170309545 | 0 | A | C | 0.202 | -1.38E-01 | 1.25E-01 | 2.60E+00 | 2.65E+00 | 2.66E-01 | no | | intergenic_variant | MODIFIER |
| 13 | 6956342 | AX-170242760 | 0 | A | T | 0.202 | -1.38E-01 | 1.25E-01 | 2.60E+00 | 2.65E+00 | 2.66E-01 | no | | intergenic_variant | MODIFIER |
| 13 | 6956435 | AX-97788530 | 0 | T | C | 0.202 | -1.38E-01 | 1.25E-01 | 2.60E+00 | 2.65E+00 | 2.66E-01 | no | | intergenic_variant | MODIFIER |
| 13 | 6964669 | AX-170482161 | 0 | G | C | 0.202 | -1.38E-01 | 1.25E-01 | 2.60E+00 | 2.65E+00 | 2.66E-01 | no | | intergenic_variant | MODIFIER |
| 13 | 6964731 | AX-170493072 | 0 | G | A | 0.202 | -1.38E-01 | 1.25E-01 | 2.60E+00 | 2.65E+00 | 2.66E-01 | no | | intergenic_variant | MODIFIER |
| 13 | 6979117 | AX-97845130 | 0 | A | G | 0.202 | -1.38E-01 | 1.25E-01 | 2.60E+00 | 2.65E+00 | 2.66E-01 | no | | intergenic_variant | MODIFIER |
| 13 | 6979173 | AX-170036840 | 0 | T | C | 0.202 | -1.38E-01 | 1.25E-01 | 2.60E+00 | 2.65E+00 | 2.66E-01 | no | | intergenic_variant | MODIFIER |
| 13 | 6983267 | AX-97786078 | 0 | C | T | 0.202 | -1.38E-01 | 1.25E-01 | 2.60E+00 | 2.65E+00 | 2.66E-01 | no | | intergenic_variant | MODIFIER |
| 13 | 6983327 | AX-170372400 | 0 | G | A | 0.202 | -1.38E-01 | 1.25E-01 | 2.60E+00 | 2.65E+00 | 2.66E-01 | no | | intergenic_variant | MODIFIER |
| 13 | 7023585 | AX-170253630 | 0 | G | A | 0.202 | -1.38E-01 | 1.25E-01 | 2.60E+00 | 2.65E+00 | 2.66E-01 | no | Gm40662 | intron_variant,non_coding_transcript_variant | MODIFIER |
| 13 | 7024004 | AX-170136616 | 0 | A | G | 0.202 | -1.38E-01 | 1.25E-01 | 2.60E+00 | 2.65E+00 | 2.66E-01 | no | Gm40662 | intron_variant,non_coding_transcript_variant | MODIFIER |
| 13 | 7028836 | AX-97818450 | 0 | T | C | 0.202 | -1.38E-01 | 1.25E-01 | 2.60E+00 | 2.65E+00 | 2.66E-01 | no | Gm40662 | intron_variant,non_coding_transcript_variant | MODIFIER |
| 13 | 7062416 | AX-97822660 | 0 | C | T | 0.202 | -1.38E-01 | 1.25E-01 | 2.60E+00 | 2.65E+00 | 2.66E-01 | no | Gm40662 | intron_variant,non_coding_transcript_variant | MODIFIER |
| 13 | 7088545 | AX-170430874 | 0 | T | C | 0.202 | -1.38E-01 | 1.25E-01 | 2.60E+00 | 2.65E+00 | 2.66E-01 | no | | intergenic_variant | MODIFIER |
| 13 | 7100640 | AX-97822574 | 0 | G | T | 0.202 | -1.38E-01 | 1.25E-01 | 2.60E+00 | 2.65E+00 | 2.66E-01 | no | | intergenic_variant | MODIFIER |
| 13 | 7101265 | AX-170105047 | 0 | G | C | 0.202 | -1.38E-01 | 1.25E-01 | 2.60E+00 | 2.65E+00 | 2.66E-01 | no | | intergenic_variant | MODIFIER |
| 13 | 7103669 | AX-170314173 | 0 | G | A | 0.202 | -1.38E-01 | 1.25E-01 | 2.60E+00 | 2.65E+00 | 2.66E-01 | no | | intergenic_variant | MODIFIER |
| 13 | 7149885 | AX-170107729 | 0 | G | A | 0.202 | -1.38E-01 | 1.25E-01 | 2.60E+00 | 2.65E+00 | 2.66E-01 | no | Gm40663 | downstream_gene_variant | MODIFIER |
| 13 | 7149961 | AX-170209662 | 0 | T | A | 0.202 | -1.38E-01 | 1.25E-01 | 2.60E+00 | 2.65E+00 | 2.66E-01 | no | Gm40663 | downstream_gene_variant | MODIFIER |
| 13 | 7150617 | AX-170347201 | 0 | T | C | 0.202 | -1.38E-01 | 1.25E-01 | 2.60E+00 | 2.65E+00 | 2.66E-01 | no | Gm40663 | downstream_gene_variant | MODIFIER |
| 13 | 7157247 | AX-170436826 | 0 | A | G | 0.202 | -1.38E-01 | 1.25E-01 | 2.60E+00 | 2.65E+00 | 2.66E-01 | no | Gm40663 | intron_variant,non_coding_transcript_variant | MODIFIER |
| 13 | 7158063 | AX-170136764 | 0 | T | C | 0.202 | -1.38E-01 | 1.25E-01 | 2.60E+00 | 2.65E+00 | 2.66E-01 | no | Gm40663 | intron_variant,non_coding_transcript_variant | MODIFIER |
| 13 | 7158959 | AX-170390561 | 0 | C | A | 0.202 | -1.38E-01 | 1.25E-01 | 2.60E+00 | 2.65E+00 | 2.66E-01 | no | Gm40663 | intron_variant,non_coding_transcript_variant | MODIFIER |

| | | | | | | | | | | | | | | | |
|---|---|---|---|---|---|---|---|---|---|---|---|---|---|---|---|
| 13 | 7159082 | AX-170092648 | 0 | A | G | 0.202 | -1.38E-01 | 1.25E-01 | 2.60E+00 | 2.65E+00 | 2.66E-01 | no | Gm40663 | intron_variant,non_coding_transcript_variant | MODIFIER |
| 13 | 7163940 | AX-170104482 | 0 | C | T | 0.202 | -1.38E-01 | 1.25E-01 | 2.60E+00 | 2.65E+00 | 2.66E-01 | no | Gm40663 | intron_variant,non_coding_transcript_variant | MODIFIER |
| 3 | 26670884 | AX-97839799 | 0 | T | C | 0.132 | -1.07E-01 | 9.64E-02 | 2.52E+00 | 2.61E+00 | 2.66E-01 | no | Spata16 | intron_variant | MODIFIER |
| 3 | 149619673 | AX-170341511 | 0 | T | C | 0.36 | 7.68E-02 | 6.96E-02 | 2.85E+00 | 2.95E+00 | 2.66E-01 | no | Gm31121 | intron_variant,non_coding_transcript_variant | MODIFIER |
| 16 | 70407435 | AX-97815124 | 2 | A | G | 0.473 | 8.26E-02 | 7.22E-02 | 2.35E+00 | 2.43E+00 | 2.67E-01 | no | Gbe1 | intron_variant | MODIFIER |
| 5 | 60752065 | AX-170025089 | 1 | T | C | 0.473 | 1.10E-01 | 9.67E-02 | 2.37E+00 | 2.44E+00 | 2.67E-01 | no | | intergenic_variant | MODIFIER |
| 7 | 37817125 | AX-170243397 | 1 | A | G | 0.33 | 1.03E-01 | 9.33E-02 | 2.60E+00 | 2.69E+00 | 2.67E-01 | no | | intergenic_variant | MODIFIER |
| 3 | 9618387 | AX-170262903 | 0 | T | C | 0.412 | -7.00E-02 | 6.38E-02 | 2.78E+00 | 2.93E+00 | 2.67E-01 | no | | intergenic_variant | MODIFIER |
| 3 | 9619325 | AX-170319695 | 0 | G | A | 0.412 | -7.00E-02 | 6.38E-02 | 2.78E+00 | 2.93E+00 | 2.67E-01 | no | | intergenic_variant | MODIFIER |
| 3 | 9620098 | AX-170310013 | 0 | T | G | 0.412 | -7.00E-02 | 6.38E-02 | 2.78E+00 | 2.93E+00 | 2.67E-01 | no | | intergenic_variant | MODIFIER |
| 3 | 9620640 | AX-170065889 | 0 | C | T | 0.412 | -7.00E-02 | 6.38E-02 | 2.78E+00 | 2.93E+00 | 2.67E-01 | no | | intergenic_variant | MODIFIER |
| 3 | 9623494 | AX-170065891 | 0 | T | C | 0.412 | -7.00E-02 | 6.38E-02 | 2.78E+00 | 2.93E+00 | 2.67E-01 | no | | intergenic_variant | MODIFIER |
| 3 | 9623828 | AX-170309298 | 0 | C | T | 0.412 | -7.00E-02 | 6.38E-02 | 2.78E+00 | 2.93E+00 | 2.67E-01 | no | | intergenic_variant | MODIFIER |
| 3 | 9624200 | AX-170164562 | 0 | C | T | 0.412 | -7.00E-02 | 6.38E-02 | 2.78E+00 | 2.93E+00 | 2.67E-01 | no | | intergenic_variant | MODIFIER |
| 12 | 92124180 | AX-170315920 | 0 | A | T | 0.088 | 1.68E-01 | 1.53E-01 | 2.76E+00 | 2.89E+00 | 2.67E-01 | Yes | | intergenic_variant | MODIFIER |
| 8 | 42913168 | AX-170079907 | 1 | G | A | 0.161 | 1.51E-01 | 1.34E-01 | 3.07E+00 | 3.17E+00 | 2.67E-01 | Yes | | intergenic_variant | MODIFIER |
| 18 | 76189915 | AX-170493148 | 0 | T | C | 0.395 | -9.39E-02 | 8.54E-02 | 2.78E+00 | 2.88E+00 | 2.67E-01 | no | | intergenic_variant | MODIFIER |
| 10 | 28064417 | AX-97811576 | 1 | T | C | 0.196 | 1.04E-01 | 9.49E-02 | 2.87E+00 | 3.00E+00 | 2.67E-01 | no | Gm18507 | downstream_gene_variant | MODIFIER |
| 1 | 139233027 | AX-170042898 | 1 | C | T | 0.214 | 1.37E-01 | 1.24E-01 | 2.67E+00 | 2.74E+00 | 2.67E-01 | no | Crb1 | intron_variant | MODIFIER |
| 7 | 100493132 | AX-97833334 | 0 | A | G | 0.36 | -1.10E-01 | 1.00E-01 | 2.75E+00 | 2.84E+00 | 2.67E-01 | no | Ucp2 | upstream_gene_variant | MODIFIER |
| 5 | 128834914 | AX-170310960 | 2 | G | A | 0.155 | -1.05E-01 | 8.74E-02 | 2.17E+00 | 2.26E+00 | 2.67E-01 | no | Rimbp2 | intron_variant | MODIFIER |
| 1 | 171751194 | AX-170126049 | 0 | T | C | 0.219 | 1.37E-01 | 1.23E-01 | 2.50E+00 | 2.56E+00 | 2.67E-01 | no | | intergenic_variant | MODIFIER |
| 4 | 136970892 | AX-170287899 | 1 | C | T | 0.161 | 1.30E-01 | 1.18E-01 | 2.58E+00 | 2.63E+00 | 2.67E-01 | no | | regulatory_region_variant | MODIFIER |
| 8 | 61523198 | AX-170332125 | 0 | T | A | 0.149 | -1.68E-01 | 1.50E-01 | 2.45E+00 | 2.51E+00 | 2.67E-01 | no | Palld | intron_variant | MODIFIER |
| 5 | 43335655 | AX-170413178 | 0 | G | T | 0.184 | -1.21E-01 | 1.10E-01 | 2.65E+00 | 2.74E+00 | 2.67E-01 | no | | intergenic_variant | MODIFIER |
| 4 | 138315956 | AX-170465144 | 0 | G | A | 0.175 | 1.19E-01 | 1.08E-01 | 2.64E+00 | 2.70E+00 | 2.68E-01 | no | Pink1 | intron_variant | MODIFIER |
| 13 | 8226800 | AX-170317307 | 1 | G | A | 0.196 | -1.31E-01 | 1.16E-01 | 2.42E+00 | 2.48E+00 | 2.68E-01 | no | Adarb2 | intron_variant | MODIFIER |
| 5 | 89779536 | AX-170414492 | 0 | T | C | 0.43 | 1.19E-01 | 1.08E-01 | 2.82E+00 | 2.89E+00 | 2.68E-01 | no | Adamts3 | intron_variant | MODIFIER |
| 4 | 54328062 | AX-170488766 | 1 | C | T | 0.375 | -1.13E-01 | 1.02E-01 | 2.81E+00 | 2.90E+00 | 2.68E-01 | no | | regulatory_region_variant | MODIFIER |
| 4 | 88392281 | AX-170338662 | 0 | A | G | 0.412 | -9.34E-02 | 8.27E-02 | 2.41E+00 | 2.47E+00 | 2.68E-01 | no | Focad | intron_variant | MODIFIER |
| 18 | 72983000 | AX-170500308 | 0 | T | C | 0.184 | -1.34E-01 | 1.22E-01 | 2.68E+00 | 2.73E+00 | 2.68E-01 | no | | intergenic_variant | MODIFIER |
| 7 | 70851782 | AX-170422164 | 0 | C | A | 0.377 | -1.13E-01 | 1.02E-01 | 2.92E+00 | 3.03E+00 | 2.68E-01 | no | | intergenic_variant | MODIFIER |
| 4 | 138106003 | AX-170024065 | 1 | G | T | 0.17 | 1.30E-01 | 1.18E-01 | 2.63E+00 | 2.68E+00 | 2.68E-01 | no | Eif4g3 | intron_variant | MODIFIER |
| 8 | 21533779 | AX-170416879 | 0 | A | C | 0.175 | -1.12E-01 | 1.02E-01 | 2.93E+00 | 3.09E+00 | 2.68E-01 | no | Defa32 | downstream_gene_variant | MODIFIER |
| 1 | 23701849 | AX-97807618 | 1 | C | T | 0.446 | 8.97E-02 | 8.17E-02 | 2.66E+00 | 2.74E+00 | 2.68E-01 | no | | intergenic_variant | MODIFIER |
| 12 | 82175602 | AX-170135786 | 0 | A | T | 0.053 | 2.18E-01 | 1.99E-01 | 2.78E+00 | 2.91E+00 | 2.68E-01 | Yes | Sipa1l1 | intron_variant | MODIFIER |
| 6 | 135167877 | AX-170028311 | 1 | C | G | 0.562 | 9.59E-02 | 8.38E-02 | 2.34E+00 | 2.40E+00 | 2.68E-01 | no | Hebp1 | intron_variant | MODIFIER |
| 1 | 148920718 | AX-170001962 | 1 | A | G | 0.205 | 1.54E-01 | 1.41E-01 | 2.75E+00 | 2.80E+00 | 2.68E-01 | Yes | | intergenic_variant | MODIFIER |
| 1 | 149029008 | AX-170508797 | 1 | A | G | 0.205 | 1.54E-01 | 1.41E-01 | 2.75E+00 | 2.80E+00 | 2.68E-01 | Yes | Gm15443 | upstream_gene_variant | MODIFIER |
| 1 | 152844544 | AX-170533942 | 1 | T | A | 0.205 | 1.54E-01 | 1.41E-01 | 2.75E+00 | 2.80E+00 | 2.68E-01 | Yes | Smg7 | intron_variant | MODIFIER |
| 7 | 72349187 | AX-170029274 | 1 | A | T | 0.366 | -1.12E-01 | 1.02E-01 | 2.86E+00 | 2.95E+00 | 2.68E-01 | no | | intergenic_variant | MODIFIER |
| 13 | 5957466 | AX-170328928 | 1 | C | T | 0.205 | -1.34E-01 | 1.22E-01 | 2.58E+00 | 2.64E+00 | 2.69E-01 | no | Gm46401 | intron_variant,non_coding_transcript_variant | MODIFIER |
| 18 | 75174909 | AX-170016001 | 0 | T | A | 0.43 | 9.59E-02 | 8.62E-02 | 2.50E+00 | 2.56E+00 | 2.69E-01 | no | Dym | intron_variant | MODIFIER |
| 8 | 86434374 | AX-170426583 | 1 | G | A | 0.214 | -1.25E-01 | 1.14E-01 | 2.89E+00 | 3.01E+00 | 2.69E-01 | no | | intergenic_variant | MODIFIER |
| 4 | 140325736 | AX-170024113 | 1 | G | A | 0.17 | 1.21E-01 | 1.10E-01 | 2.58E+00 | 2.65E+00 | 2.69E-01 | no | Gm9867 | upstream_gene_variant | MODIFIER |
| 4 | 73169125 | AX-170119531 | 1 | T | C | 0.446 | -9.40E-02 | 8.49E-02 | 2.52E+00 | 2.57E+00 | 2.69E-01 | no | | intergenic_variant | MODIFIER |
| 1 | 139006248 | AX-97824462 | 1 | G | A | 0.214 | 1.40E-01 | 1.28E-01 | 2.72E+00 | 2.78E+00 | 2.70E-01 | no | Dennd1b | intron_variant | MODIFIER |
| 1 | 136002034 | AX-170166013 | 2 | C | T | 0.218 | 1.26E-01 | 1.15E-01 | 2.72E+00 | 2.81E+00 | 2.70E-01 | no | Igfn1 | intron_variant | MODIFIER |
| 4 | 140870691 | AX-170024123 | 1 | C | T | 0.17 | 1.27E-01 | 1.16E-01 | 2.63E+00 | 2.68E+00 | 2.70E-01 | no | 4930515B02Rik | upstream_gene_variant | MODIFIER |
| 19 | 50666807 | AX-170175415 | 1 | G | C | 0.482 | 7.03E-02 | 6.43E-02 | 2.69E+00 | 2.78E+00 | 2.70E-01 | no | Sorcs1 | intron_variant | MODIFIER |
| 3 | 22766774 | AX-97854463 | 0 | A | C | 0.158 | -1.13E-01 | 1.03E-01 | 2.58E+00 | 2.65E+00 | 2.70E-01 | no | | intergenic_variant | MODIFIER |
| 3 | 22789481 | AX-170020116 | 0 | G | A | 0.158 | -1.13E-01 | 1.03E-01 | 2.58E+00 | 2.65E+00 | 2.70E-01 | no | | intergenic_variant | MODIFIER |
| 3 | 22789502 | AX-97833170 | 0 | T | C | 0.158 | -1.13E-01 | 1.03E-01 | 2.58E+00 | 2.65E+00 | 2.70E-01 | no | | intergenic_variant | MODIFIER |
| 3 | 22826621 | AX-170476018 | 0 | G | A | 0.158 | -1.13E-01 | 1.03E-01 | 2.58E+00 | 2.65E+00 | 2.70E-01 | no | | intergenic_variant | MODIFIER |

| | | | | | | | | | | | | | | | |
|---|---|---|---|---|---|---|---|---|---|---|---|---|---|---|---|
| 3 | 22863704 | AX-168391098 | 0 | G | A | 0.158 | -1.13E-01 | 1.03E-01 | 2.58E+00 | 2.65E+00 | 2.70E-01 | no | | intergenic_variant | MODIFIER |
| 3 | 22931143 | AX-170428131 | 0 | G | A | 0.158 | -1.13E-01 | 1.03E-01 | 2.58E+00 | 2.65E+00 | 2.70E-01 | no | | intergenic_variant | MODIFIER |
| 3 | 22948453 | AX-97833165 | 0 | C | T | 0.158 | -1.13E-01 | 1.03E-01 | 2.58E+00 | 2.65E+00 | 2.70E-01 | no | | intergenic_variant | MODIFIER |
| 3 | 22968237 | AX-97798296 | 0 | T | G | 0.158 | -1.13E-01 | 1.03E-01 | 2.58E+00 | 2.65E+00 | 2.70E-01 | no | | intergenic_variant | MODIFIER |
| 3 | 22992289 | AX-170418071 | 0 | G | A | 0.158 | -1.13E-01 | 1.03E-01 | 2.58E+00 | 2.65E+00 | 2.70E-01 | no | | intergenic_variant | MODIFIER |
| 3 | 23032848 | AX-170158670 | 0 | A | G | 0.158 | -1.13E-01 | 1.03E-01 | 2.58E+00 | 2.65E+00 | 2.70E-01 | no | | intergenic_variant | MODIFIER |
| 3 | 23282700 | AX-170219646 | 0 | T | C | 0.158 | -1.13E-01 | 1.03E-01 | 2.58E+00 | 2.65E+00 | 2.70E-01 | no | | intergenic_variant | MODIFIER |
| 3 | 23283294 | AX-170493955 | 0 | T | C | 0.158 | -1.13E-01 | 1.03E-01 | 2.58E+00 | 2.65E+00 | 2.70E-01 | no | | intergenic_variant | MODIFIER |
| 3 | 23300548 | AX-170527559 | 0 | T | C | 0.158 | -1.13E-01 | 1.03E-01 | 2.58E+00 | 2.65E+00 | 2.70E-01 | no | | intergenic_variant | MODIFIER |
| 3 | 23372725 | AX-97836945 | 0 | T | G | 0.158 | -1.13E-01 | 1.03E-01 | 2.58E+00 | 2.65E+00 | 2.70E-01 | no | Gm37492 | downstream_gene_variant | MODIFIER |
| 3 | 23394154 | AX-170066088 | 0 | G | A | 0.158 | -1.13E-01 | 1.03E-01 | 2.58E+00 | 2.65E+00 | 2.70E-01 | no | | intergenic_variant | MODIFIER |
| 3 | 23394538 | AX-170066090 | 0 | T | C | 0.158 | -1.13E-01 | 1.03E-01 | 2.58E+00 | 2.65E+00 | 2.70E-01 | no | | intergenic_variant | MODIFIER |
| 3 | 23406546 | AX-170066091 | 0 | G | A | 0.158 | -1.13E-01 | 1.03E-01 | 2.58E+00 | 2.65E+00 | 2.70E-01 | no | | intergenic_variant | MODIFIER |
| 3 | 23427627 | AX-170373060 | 0 | A | C | 0.158 | -1.13E-01 | 1.03E-01 | 2.58E+00 | 2.65E+00 | 2.70E-01 | no | | intergenic_variant | MODIFIER |
| 3 | 23427889 | AX-170198338 | 0 | A | G | 0.158 | -1.13E-01 | 1.03E-01 | 2.58E+00 | 2.65E+00 | 2.70E-01 | no | | intergenic_variant | MODIFIER |
| 3 | 23428936 | AX-170301759 | 0 | C | A | 0.158 | -1.13E-01 | 1.03E-01 | 2.58E+00 | 2.65E+00 | 2.70E-01 | no | | intergenic_variant | MODIFIER |
| 3 | 23434701 | AX-170328248 | 0 | G | A | 0.158 | -1.13E-01 | 1.03E-01 | 2.58E+00 | 2.65E+00 | 2.70E-01 | no | | intergenic_variant | MODIFIER |
| 3 | 23454548 | AX-170094550 | 0 | C | T | 0.158 | -1.13E-01 | 1.03E-01 | 2.58E+00 | 2.65E+00 | 2.70E-01 | no | | intergenic_variant | MODIFIER |
| 3 | 23460857 | AX-170533886 | 0 | C | T | 0.158 | -1.13E-01 | 1.03E-01 | 2.58E+00 | 2.65E+00 | 2.70E-01 | no | 4930412E21Rik | upstream_gene_variant | MODIFIER |
| 3 | 23473325 | AX-170440842 | 0 | C | G | 0.158 | -1.13E-01 | 1.03E-01 | 2.58E+00 | 2.65E+00 | 2.70E-01 | no | 4930412E21Rik | intron_variant,non_coding_transcript_variant | MODIFIER |
| 3 | 23510955 | AX-170272941 | 0 | C | T | 0.158 | -1.13E-01 | 1.03E-01 | 2.58E+00 | 2.65E+00 | 2.70E-01 | no | 4930412E21Rik | intron_variant,non_coding_transcript_variant | MODIFIER |
| 3 | 23518221 | AX-170302611 | 0 | C | A | 0.158 | -1.13E-01 | 1.03E-01 | 2.58E+00 | 2.65E+00 | 2.70E-01 | no | 4930412E21Rik | intron_variant,non_coding_transcript_variant | MODIFIER |
| 3 | 23542510 | AX-170294605 | 0 | T | C | 0.158 | -1.13E-01 | 1.03E-01 | 2.58E+00 | 2.65E+00 | 2.70E-01 | no | | intergenic_variant | MODIFIER |
| 3 | 23544545 | AX-97833941 | 0 | T | C | 0.158 | -1.13E-01 | 1.03E-01 | 2.58E+00 | 2.65E+00 | 2.70E-01 | no | | intergenic_variant | MODIFIER |
| 3 | 23592516 | AX-170066096 | 0 | T | C | 0.158 | -1.13E-01 | 1.03E-01 | 2.58E+00 | 2.65E+00 | 2.70E-01 | no | | regulatory_region_variant | MODIFIER |
| 3 | 23663758 | AX-97837062 | 0 | T | C | 0.158 | -1.13E-01 | 1.03E-01 | 2.58E+00 | 2.65E+00 | 2.70E-01 | no | | intergenic_variant | MODIFIER |
| 3 | 23663881 | AX-170266887 | 0 | C | T | 0.158 | -1.13E-01 | 1.03E-01 | 2.58E+00 | 2.65E+00 | 2.70E-01 | no | | intergenic_variant | MODIFIER |
| 3 | 23708708 | AX-170526370 | 0 | C | T | 0.158 | -1.13E-01 | 1.03E-01 | 2.58E+00 | 2.65E+00 | 2.70E-01 | no | | intergenic_variant | MODIFIER |
| 3 | 23749753 | AX-170526626 | 0 | T | C | 0.158 | -1.13E-01 | 1.03E-01 | 2.58E+00 | 2.65E+00 | 2.70E-01 | no | | intergenic_variant | MODIFIER |
| 3 | 23749826 | AX-170490272 | 0 | T | C | 0.158 | -1.13E-01 | 1.03E-01 | 2.58E+00 | 2.65E+00 | 2.70E-01 | no | | intergenic_variant | MODIFIER |
| 3 | 23750973 | AX-97795472 | 0 | T | C | 0.158 | -1.13E-01 | 1.03E-01 | 2.58E+00 | 2.65E+00 | 2.70E-01 | no | | intergenic_variant | MODIFIER |
| 3 | 23778679 | AX-97853529 | 0 | T | C | 0.158 | -1.13E-01 | 1.03E-01 | 2.58E+00 | 2.65E+00 | 2.70E-01 | no | | intergenic_variant | MODIFIER |
| 3 | 23778706 | AX-170531242 | 0 | A | T | 0.158 | -1.13E-01 | 1.03E-01 | 2.58E+00 | 2.65E+00 | 2.70E-01 | no | | intergenic_variant | MODIFIER |
| 3 | 23782162 | AX-170488545 | 0 | G | T | 0.158 | -1.13E-01 | 1.03E-01 | 2.58E+00 | 2.65E+00 | 2.70E-01 | no | | intergenic_variant | MODIFIER |
| 3 | 23837880 | AX-170066101 | 0 | T | C | 0.158 | -1.13E-01 | 1.03E-01 | 2.58E+00 | 2.65E+00 | 2.70E-01 | no | Naaladl2 | intron_variant,non_coding_transcript_variant | MODIFIER |
| 3 | 23837983 | AX-170066098 | 0 | G | C | 0.158 | -1.13E-01 | 1.03E-01 | 2.58E+00 | 2.65E+00 | 2.70E-01 | no | Naaladl2 | intron_variant,non_coding_transcript_variant | MODIFIER |
| 3 | 23848266 | AX-170138303 | 0 | T | C | 0.158 | -1.13E-01 | 1.03E-01 | 2.58E+00 | 2.65E+00 | 2.70E-01 | no | Naaladl2 | intron_variant,non_coding_transcript_variant | MODIFIER |
| 3 | 23848447 | AX-170231888 | 0 | A | G | 0.158 | -1.13E-01 | 1.03E-01 | 2.58E+00 | 2.65E+00 | 2.70E-01 | no | Naaladl2 | intron_variant,non_coding_transcript_variant | MODIFIER |
| 3 | 23865185 | AX-170504398 | 0 | A | C | 0.158 | -1.13E-01 | 1.03E-01 | 2.58E+00 | 2.65E+00 | 2.70E-01 | no | Naaladl2 | intron_variant,non_coding_transcript_variant | MODIFIER |
| 3 | 23866948 | AX-170020130 | 0 | C | A | 0.158 | -1.13E-01 | 1.03E-01 | 2.58E+00 | 2.65E+00 | 2.70E-01 | no | Naaladl2 | intron_variant,non_coding_transcript_variant | MODIFIER |
| 3 | 23869060 | AX-170020126 | 0 | A | G | 0.158 | -1.13E-01 | 1.03E-01 | 2.58E+00 | 2.65E+00 | 2.70E-01 | no | Naaladl2 | intron_variant,non_coding_transcript_variant | MODIFIER |
| 3 | 23909215 | AX-170020127 | 0 | T | C | 0.158 | -1.13E-01 | 1.03E-01 | 2.58E+00 | 2.65E+00 | 2.70E-01 | no | Naaladl2 | intron_variant,non_coding_transcript_variant | MODIFIER |
| 3 | 23919627 | AX-170428144 | 0 | G | C | 0.158 | -1.13E-01 | 1.03E-01 | 2.58E+00 | 2.65E+00 | 2.70E-01 | no | Naaladl2 | intron_variant,non_coding_transcript_variant | MODIFIER |
| 3 | 23920004 | AX-170232486 | 0 | A | C | 0.158 | -1.13E-01 | 1.03E-01 | 2.58E+00 | 2.65E+00 | 2.70E-01 | no | Naaladl2 | intron_variant,non_coding_transcript_variant | MODIFIER |
| 3 | 23945544 | AX-170020124 | 0 | T | C | 0.158 | -1.13E-01 | 1.03E-01 | 2.58E+00 | 2.65E+00 | 2.70E-01 | no | Naaladl2 | intron_variant | MODIFIER |
| 3 | 24008627 | AX-168377252 | 0 | G | A | 0.158 | -1.13E-01 | 1.03E-01 | 2.58E+00 | 2.65E+00 | 2.70E-01 | no | Naaladl2 | intron_variant | MODIFIER |
| 3 | 24044031 | AX-170391939 | 0 | A | G | 0.158 | -1.13E-01 | 1.03E-01 | 2.58E+00 | 2.65E+00 | 2.70E-01 | no | Naaladl2 | intron_variant | MODIFIER |
| 3 | 24100465 | AX-170221691 | 0 | C | G | 0.158 | -1.13E-01 | 1.03E-01 | 2.58E+00 | 2.65E+00 | 2.70E-01 | no | Naaladl2 | intron_variant | MODIFIER |
| 3 | 24107912 | AX-170501730 | 0 | C | T | 0.158 | -1.13E-01 | 1.03E-01 | 2.58E+00 | 2.65E+00 | 2.70E-01 | no | Naaladl2 | intron_variant | MODIFIER |
| 3 | 24131215 | AX-170349343 | 0 | A | G | 0.158 | -1.13E-01 | 1.03E-01 | 2.58E+00 | 2.65E+00 | 2.70E-01 | no | Naaladl2 | intron_variant | MODIFIER |
| 3 | 24150862 | AX-170264165 | 0 | C | T | 0.158 | -1.13E-01 | 1.03E-01 | 2.58E+00 | 2.65E+00 | 2.70E-01 | no | Naaladl2 | intron_variant | MODIFIER |
| 3 | 24172720 | AX-97785042 | 0 | C | T | 0.158 | -1.13E-01 | 1.03E-01 | 2.58E+00 | 2.65E+00 | 2.70E-01 | no | Naaladl2 | intron_variant | MODIFIER |
| 3 | 24172994 | AX-170434007 | 0 | T | G | 0.158 | -1.13E-01 | 1.03E-01 | 2.58E+00 | 2.65E+00 | 2.70E-01 | no | Naaladl2 | intron_variant | MODIFIER |
| 3 | 24176617 | AX-170198358 | 0 | C | G | 0.158 | -1.13E-01 | 1.03E-01 | 2.58E+00 | 2.65E+00 | 2.70E-01 | no | Naaladl2 | intron_variant | MODIFIER |

| | | | | | | | | | | | | | | | |
|---|---|---|---|---|---|---|---|---|---|---|---|---|---|---|---|
| 3 | 24216516 | AX-170266895 | 0 | C | A | 0.158 | -1.13E-01 | 1.03E-01 | 2.58E+00 | 2.65E+00 | 2.70E-01 | no | Naaladl2 | intron_variant | MODIFIER |
| 3 | 24420587 | AX-97792507 | 0 | A | C | 0.158 | -1.13E-01 | 1.03E-01 | 2.58E+00 | 2.65E+00 | 2.70E-01 | no | Naaladl2 | intron_variant | MODIFIER |
| 3 | 24438462 | AX-170424488 | 0 | C | T | 0.158 | -1.13E-01 | 1.03E-01 | 2.58E+00 | 2.65E+00 | 2.70E-01 | no | Naaladl2 | intron_variant | MODIFIER |
| 3 | 24441387 | AX-170419174 | 0 | C | T | 0.158 | -1.13E-01 | 1.03E-01 | 2.58E+00 | 2.65E+00 | 2.70E-01 | no | Naaladl2 | intron_variant | MODIFIER |
| 3 | 24493059 | AX-170391590 | 0 | G | T | 0.158 | -1.13E-01 | 1.03E-01 | 2.58E+00 | 2.65E+00 | 2.70E-01 | no | Naaladl2 | intron_variant | MODIFIER |
| 3 | 24508036 | AX-170020134 | 0 | C | T | 0.158 | -1.13E-01 | 1.03E-01 | 2.58E+00 | 2.65E+00 | 2.70E-01 | no | Naaladl2 | intron_variant | MODIFIER |
| 3 | 24516626 | AX-97840769 | 0 | G | A | 0.158 | -1.13E-01 | 1.03E-01 | 2.58E+00 | 2.65E+00 | 2.70E-01 | no | Naaladl2 | intron_variant | MODIFIER |
| 3 | 24602378 | AX-170364344 | 0 | G | C | 0.158 | -1.13E-01 | 1.03E-01 | 2.58E+00 | 2.65E+00 | 2.70E-01 | no | Naaladl2 | intron_variant | MODIFIER |
| 3 | 24604457 | AX-97792044 | 0 | T | C | 0.158 | -1.13E-01 | 1.03E-01 | 2.58E+00 | 2.65E+00 | 2.70E-01 | no | Naaladl2 | intron_variant | MODIFIER |
| 3 | 24629793 | AX-97843242 | 0 | T | G | 0.158 | -1.13E-01 | 1.03E-01 | 2.58E+00 | 2.65E+00 | 2.70E-01 | no | Naaladl2 | intron_variant | MODIFIER |
| 3 | 24638234 | AX-170242244 | 0 | G | A | 0.158 | -1.13E-01 | 1.03E-01 | 2.58E+00 | 2.65E+00 | 2.70E-01 | no | Naaladl2 | intron_variant | MODIFIER |
| 3 | 24667736 | AX-97814544 | 0 | C | T | 0.158 | -1.13E-01 | 1.03E-01 | 2.58E+00 | 2.65E+00 | 2.70E-01 | no | Naaladl2 | intron_variant | MODIFIER |
| 6 | 137333835 | AX-97808501 | 1 | C | T | 0.589 | 9.57E-02 | 8.50E-02 | 2.40E+00 | 2.47E+00 | 2.70E-01 | no | Ptpro | intron_variant | MODIFIER |
| 7 | 73882737 | AX-170226223 | 0 | C | G | 0.36 | -1.05E-01 | 9.53E-02 | 2.91E+00 | 3.02E+00 | 2.70E-01 | no | | intergenic_variant | MODIFIER |
| 6 | 138413449 | AX-170498716 | 0 | C | G | 0.561 | 9.27E-02 | 8.26E-02 | 2.42E+00 | 2.49E+00 | 2.70E-01 | no | Lmo3 | intron_variant,NMD_transcript_variant | MODIFIER |
| 9 | 19176904 | AX-97805179 | 1 | T | C | 0.473 | 9.91E-02 | 9.00E-02 | 2.56E+00 | 2.62E+00 | 2.70E-01 | no | Olfr839-ps1 | 5_prime_UTR_variant | MODIFIER |
| 1 | 168162752 | AX-170432996 | 0 | T | A | 0.228 | 1.36E-01 | 1.24E-01 | 2.60E+00 | 2.66E+00 | 2.70E-01 | no | Pbx1 | intron_variant | MODIFIER |
| 8 | 24852623 | AX-170419057 | 2 | G | A | 0.364 | -6.95E-02 | 6.33E-02 | 2.57E+00 | 2.66E+00 | 2.70E-01 | no | Adam32 | intron_variant | MODIFIER |
| 19 | 60536514 | AX-170201872 | 0 | C | G | 0.447 | -6.64E-02 | 6.05E-02 | 2.58E+00 | 2.67E+00 | 2.70E-01 | no | Cacul1 | intron_variant | MODIFIER |
| 4 | 129835144 | AX-170089370 | 0 | T | A | 0.167 | 1.17E-01 | 1.06E-01 | 2.54E+00 | 2.60E+00 | 2.70E-01 | no | Ptp4a2 | intron_variant | MODIFIER |
| 6 | 136884220 | AX-170028335 | 1 | C | T | 0.571 | 9.70E-02 | 8.70E-02 | 2.45E+00 | 2.51E+00 | 2.70E-01 | no | | intergenic_variant | MODIFIER |
| 6 | 137478893 | AX-170028342 | 1 | C | T | 0.571 | 9.70E-02 | 8.70E-02 | 2.45E+00 | 2.51E+00 | 2.70E-01 | no | Eps8 | 3_prime_UTR_variant | MODIFIER |
| 16 | 38422189 | AX-170177994 | 0 | G | C | 0.491 | 8.95E-02 | 8.14E-02 | 2.58E+00 | 2.63E+00 | 2.71E-01 | no | Pla1a | intron_variant | MODIFIER |
| 16 | 39432102 | AX-170252700 | 0 | G | A | 0.491 | 8.95E-02 | 8.14E-02 | 2.58E+00 | 2.63E+00 | 2.71E-01 | no | | intergenic_variant | MODIFIER |
| 1 | 121955595 | AX-170466741 | 0 | T | C | 0.57 | 9.17E-02 | 8.30E-02 | 2.52E+00 | 2.58E+00 | 2.71E-01 | no | | intergenic_variant | MODIFIER |
| 15 | 66883777 | AX-170279226 | 0 | A | G | 0.333 | -8.85E-02 | 7.83E-02 | 2.38E+00 | 2.45E+00 | 2.71E-01 | no | | regulatory_region_variant | MODIFIER |
| 15 | 67048848 | AX-170373537 | 0 | A | T | 0.333 | -8.85E-02 | 7.83E-02 | 2.38E+00 | 2.45E+00 | 2.71E-01 | no | | regulatory_region_variant | MODIFIER |
| 18 | 50171262 | AX-97836797 | 0 | G | T | 0.184 | -1.04E-01 | 9.45E-02 | 2.55E+00 | 2.65E+00 | 2.71E-01 | no | Hsd17b4 | intron_variant | MODIFIER |
| 19 | 50926919 | AX-170244982 | 1 | T | A | 0.482 | 7.02E-02 | 6.43E-02 | 2.68E+00 | 2.77E+00 | 2.71E-01 | no | | intergenic_variant | MODIFIER |
| 15 | 3572369 | AX-170193079 | 0 | C | T | 0.246 | -1.13E-01 | 1.03E-01 | 2.67E+00 | 2.73E+00 | 2.71E-01 | no | Ghr | intron_variant | MODIFIER |
| 15 | 4370283 | AX-97852123 | 0 | C | T | 0.246 | -1.13E-01 | 1.03E-01 | 2.67E+00 | 2.73E+00 | 2.71E-01 | no | | intergenic_variant | MODIFIER |
| 14 | 111943566 | AX-170010702 | 2 | G | T | 0.327 | 8.03E-02 | 7.33E-02 | 2.59E+00 | 2.69E+00 | 2.71E-01 | no | | intergenic_variant | MODIFIER |
| 9 | 22498551 | AX-170090262 | 0 | C | A | 0.474 | 9.58E-02 | 8.67E-02 | 2.51E+00 | 2.59E+00 | 2.71E-01 | no | Bbs9 | intron_variant | MODIFIER |
| 8 | 48929092 | AX-170030638 | 0 | T | C | 0.167 | 1.35E-01 | 1.22E-01 | 3.05E+00 | 3.20E+00 | 2.71E-01 | no | | intergenic_variant | MODIFIER |
| 1 | 42001396 | AX-170180150 | 1 | T | A | 0.411 | -7.79E-02 | 7.14E-02 | 2.66E+00 | 2.77E+00 | 2.71E-01 | no | | intergenic_variant | MODIFIER |
| 1 | 140382211 | AX-170191735 | 1 | A | C | 0.196 | 1.36E-01 | 1.24E-01 | 2.70E+00 | 2.78E+00 | 2.71E-01 | no | Kcnt2 | intron_variant | MODIFIER |
| 9 | 8458880 | AX-170284020 | 0 | G | T | 0.465 | 8.97E-02 | 8.06E-02 | 2.46E+00 | 2.55E+00 | 2.71E-01 | no | | intergenic_variant | MODIFIER |
| 4 | 89025501 | AX-170203204 | 0 | C | T | 0.43 | -8.99E-02 | 8.07E-02 | 2.45E+00 | 2.51E+00 | 2.71E-01 | no | AL805899.2 | intron_variant,non_coding_transcript_variant | MODIFIER |
| 4 | 89025529 | AX-170300678 | 0 | C | T | 0.43 | -8.99E-02 | 8.07E-02 | 2.45E+00 | 2.51E+00 | 2.71E-01 | no | AL805899.2 | intron_variant,non_coding_transcript_variant | MODIFIER |
| 4 | 89036387 | AX-170305661 | 0 | G | A | 0.43 | -8.99E-02 | 8.07E-02 | 2.45E+00 | 2.51E+00 | 2.71E-01 | no | AL805899.2 | intron_variant,non_coding_transcript_variant | MODIFIER |
| 4 | 89038689 | AX-97785066 | 0 | A | C | 0.43 | -8.99E-02 | 8.07E-02 | 2.45E+00 | 2.51E+00 | 2.71E-01 | no | AL805899.2 | intron_variant,non_coding_transcript_variant | MODIFIER |
| 4 | 89038715 | AX-170489043 | 0 | C | T | 0.43 | -8.99E-02 | 8.07E-02 | 2.45E+00 | 2.51E+00 | 2.71E-01 | no | AL805899.2 | intron_variant,non_coding_transcript_variant | MODIFIER |
| 4 | 89038743 | AX-170023364 | 0 | G | A | 0.43 | -8.99E-02 | 8.07E-02 | 2.45E+00 | 2.51E+00 | 2.71E-01 | no | AL805899.2 | intron_variant,non_coding_transcript_variant | MODIFIER |
| 4 | 89040126 | AX-170305166 | 0 | A | G | 0.43 | -8.99E-02 | 8.07E-02 | 2.45E+00 | 2.51E+00 | 2.71E-01 | no | AL805899.2 | upstream_gene_variant | MODIFIER |
| 4 | 89306404 | AX-97795034 | 0 | T | C | 0.43 | -8.99E-02 | 8.07E-02 | 2.45E+00 | 2.51E+00 | 2.71E-01 | no | Cdkn2b | 3_prime_UTR_variant | MODIFIER |
| 4 | 89306692 | AX-170475339 | 0 | G | A | 0.43 | -8.99E-02 | 8.07E-02 | 2.45E+00 | 2.51E+00 | 2.71E-01 | no | Cdkn2b | 3_prime_UTR_variant | MODIFIER |
| 4 | 89384316 | AX-170525718 | 0 | G | A | 0.43 | -8.99E-02 | 8.07E-02 | 2.45E+00 | 2.51E+00 | 2.71E-01 | no | Gm12609 | intron_variant,non_coding_transcript_variant | MODIFIER |
| 4 | 89489986 | AX-170201207 | 0 | C | A | 0.43 | -8.99E-02 | 8.07E-02 | 2.45E+00 | 2.51E+00 | 2.71E-01 | no | | intergenic_variant | MODIFIER |
| 4 | 89491246 | AX-170069949 | 0 | A | G | 0.43 | -8.99E-02 | 8.07E-02 | 2.45E+00 | 2.51E+00 | 2.71E-01 | no | | intergenic_variant | MODIFIER |
| 4 | 89501219 | AX-97848070 | 0 | C | T | 0.43 | -8.99E-02 | 8.07E-02 | 2.45E+00 | 2.51E+00 | 2.71E-01 | no | | intergenic_variant | MODIFIER |
| 4 | 89508483 | AX-170023367 | 0 | G | A | 0.43 | -8.99E-02 | 8.07E-02 | 2.45E+00 | 2.51E+00 | 2.71E-01 | no | | regulatory_region_variant | MODIFIER |
| 4 | 89508814 | AX-170535227 | 0 | G | T | 0.43 | -8.99E-02 | 8.07E-02 | 2.45E+00 | 2.51E+00 | 2.71E-01 | no | | regulatory_region_variant | MODIFIER |
| 4 | 89550176 | AX-170111807 | 0 | T | C | 0.43 | -8.99E-02 | 8.07E-02 | 2.45E+00 | 2.51E+00 | 2.71E-01 | no | | intergenic_variant | MODIFIER |
| 4 | 89577169 | AX-170426450 | 0 | A | G | 0.43 | -8.99E-02 | 8.07E-02 | 2.45E+00 | 2.51E+00 | 2.71E-01 | no | | intergenic_variant | MODIFIER |

| | | | | | | | | | | | | | | | |
|---|---|---|---|---|---|---|---|---|---|---|---|---|---|---|---|
| 4 | 89585952 | AX-170207904 | 0 | A | G | 0.43 | -8.99E-02 | 8.07E-02 | 2.45E+00 | 2.51E+00 | 2.71E-01 | no | | intergenic_variant | MODIFIER |
| 4 | 89590206 | AX-97839003 | 0 | A | G | 0.43 | -8.99E-02 | 8.07E-02 | 2.45E+00 | 2.51E+00 | 2.71E-01 | no | | intergenic_variant | MODIFIER |
| 4 | 89591349 | AX-170299382 | 0 | C | T | 0.43 | -8.99E-02 | 8.07E-02 | 2.45E+00 | 2.51E+00 | 2.71E-01 | no | | intergenic_variant | MODIFIER |
| 4 | 89628613 | AX-170515169 | 0 | A | G | 0.43 | -8.99E-02 | 8.07E-02 | 2.45E+00 | 2.51E+00 | 2.71E-01 | no | | intergenic_variant | MODIFIER |
| 4 | 89629002 | AX-170479338 | 0 | T | C | 0.43 | -8.99E-02 | 8.07E-02 | 2.45E+00 | 2.51E+00 | 2.71E-01 | no | | intergenic_variant | MODIFIER |
| 4 | 89629039 | AX-170168659 | 0 | A | G | 0.43 | -8.99E-02 | 8.07E-02 | 2.45E+00 | 2.51E+00 | 2.71E-01 | no | | intergenic_variant | MODIFIER |
| 4 | 89658068 | AX-170146297 | 0 | C | T | 0.43 | -8.99E-02 | 8.07E-02 | 2.45E+00 | 2.51E+00 | 2.71E-01 | no | | intergenic_variant | MODIFIER |
| 4 | 89658589 | AX-170422624 | 0 | T | G | 0.43 | -8.99E-02 | 8.07E-02 | 2.45E+00 | 2.51E+00 | 2.71E-01 | no | | intergenic_variant | MODIFIER |
| 4 | 89660538 | AX-170386751 | 0 | C | T | 0.43 | -8.99E-02 | 8.07E-02 | 2.45E+00 | 2.51E+00 | 2.71E-01 | no | | intergenic_variant | MODIFIER |
| 4 | 80144039 | AX-97838904 | 1 | C | A | 0.446 | -9.38E-02 | 8.50E-02 | 2.52E+00 | 2.57E+00 | 2.71E-01 | no | | intergenic_variant | MODIFIER |
| 15 | 6168047 | AX-170204268 | 0 | A | C | 0.088 | -1.30E-01 | 9.90E-02 | 1.92E+00 | 2.01E+00 | 2.71E-01 | no | | intergenic_variant | MODIFIER |
| 15 | 6171864 | AX-170241375 | 0 | A | G | 0.088 | -1.30E-01 | 9.90E-02 | 1.92E+00 | 2.01E+00 | 2.71E-01 | no | | intergenic_variant | MODIFIER |
| 15 | 6311659 | AX-170010957 | 0 | C | T | 0.088 | -1.30E-01 | 9.90E-02 | 1.92E+00 | 2.01E+00 | 2.71E-01 | no | Dab2 | intron_variant | MODIFIER |
| 15 | 6320780 | AX-170010958 | 0 | T | C | 0.088 | -1.30E-01 | 9.90E-02 | 1.92E+00 | 2.01E+00 | 2.71E-01 | no | Dab2 | intron_variant | MODIFIER |
| 15 | 6323227 | AX-170010956 | 0 | A | G | 0.088 | -1.30E-01 | 9.90E-02 | 1.92E+00 | 2.01E+00 | 2.71E-01 | no | Dab2 | intron_variant | MODIFIER |
| 15 | 6346916 | AX-170390067 | 0 | C | T | 0.088 | -1.30E-01 | 9.90E-02 | 1.92E+00 | 2.01E+00 | 2.71E-01 | no | Dab2 | intron_variant | MODIFIER |
| 15 | 6767531 | AX-97840491 | 0 | G | A | 0.088 | -1.30E-01 | 9.90E-02 | 1.92E+00 | 2.01E+00 | 2.71E-01 | no | Rictor | intron_variant | MODIFIER |
| 15 | 6843820 | AX-170054721 | 0 | G | A | 0.088 | -1.30E-01 | 9.90E-02 | 1.92E+00 | 2.01E+00 | 2.71E-01 | no | Osmr | intron_variant | MODIFIER |
| 15 | 6886965 | AX-170010961 | 0 | C | G | 0.088 | -1.30E-01 | 9.90E-02 | 1.92E+00 | 2.01E+00 | 2.71E-01 | no | | intergenic_variant | MODIFIER |
| 15 | 6900878 | AX-97840765 | 0 | T | C | 0.088 | -1.30E-01 | 9.90E-02 | 1.92E+00 | 2.01E+00 | 2.71E-01 | no | | intergenic_variant | MODIFIER |
| 15 | 6914326 | AX-170010962 | 0 | T | C | 0.088 | -1.30E-01 | 9.90E-02 | 1.92E+00 | 2.01E+00 | 2.71E-01 | no | | intergenic_variant | MODIFIER |
| 15 | 6920940 | AX-170215214 | 0 | C | G | 0.088 | -1.30E-01 | 9.90E-02 | 1.92E+00 | 2.01E+00 | 2.71E-01 | no | | intergenic_variant | MODIFIER |
| 15 | 7167980 | AX-170144970 | 0 | A | G | 0.088 | -1.30E-01 | 9.90E-02 | 1.92E+00 | 2.01E+00 | 2.71E-01 | no | Lifr | intron_variant | MODIFIER |
| 6 | 144996145 | AX-170130415 | 2 | C | A | 0.1 | 1.72E-01 | 1.58E-01 | 2.69E+00 | 2.81E+00 | 2.71E-01 | Yes | Bcat1 | 3_prime_UTR_variant | MODIFIER |
| 16 | 19060357 | AX-170441526 | 0 | C | G | 0.132 | 1.02E-01 | 9.38E-02 | 2.80E+00 | 2.94E+00 | 2.71E-01 | no | 2010309G21Rik | intron_variant,non_coding_transcript_variant | MODIFIER |
| 4 | 95895955 | AX-170023454 | 0 | C | T | 0.263 | 1.24E-01 | 1.10E-01 | 2.41E+00 | 2.46E+00 | 2.72E-01 | no | Fggy | intron_variant | MODIFIER |
| 8 | 68837361 | AX-170104861 | 1 | G | T | 0.188 | -1.28E-01 | 1.18E-01 | 2.77E+00 | 2.88E+00 | 2.72E-01 | no | | intergenic_variant | MODIFIER |
| 7 | 15024643 | AX-170085866 | 0 | T | G | 0.325 | 9.26E-02 | 8.48E-02 | 2.58E+00 | 2.72E+00 | 2.72E-01 | no | Obox4-ps9 | intron_variant,non_coding_transcript_variant | MODIFIER |
| 18 | 80778223 | AX-97807110 | 0 | T | C | 0.544 | 8.00E-02 | 7.19E-02 | 2.44E+00 | 2.52E+00 | 2.72E-01 | no | Atp9b | intron_variant | MODIFIER |
| 16 | 80712125 | AX-170261009 | 1 | A | G | 0.464 | 8.12E-02 | 7.41E-02 | 2.58E+00 | 2.65E+00 | 2.72E-01 | no | Gm49578 | downstream_gene_variant | MODIFIER |
| 9 | 92413805 | AX-170534731 | 0 | G | A | 0.132 | -1.33E-01 | 1.22E-01 | 2.61E+00 | 2.70E+00 | 2.72E-01 | no | | regulatory_region_variant | MODIFIER |
| 19 | 50486828 | AX-97834757 | 0 | T | C | 0.482 | 7.00E-02 | 6.44E-02 | 2.68E+00 | 2.77E+00 | 2.72E-01 | no | Sorcs1 | intron_variant | MODIFIER |
| 19 | 50497512 | AX-170194211 | 0 | G | A | 0.482 | 7.00E-02 | 6.44E-02 | 2.68E+00 | 2.77E+00 | 2.72E-01 | no | Sorcs1 | intron_variant | MODIFIER |
| 19 | 50497579 | AX-170379132 | 0 | A | C | 0.482 | 7.00E-02 | 6.44E-02 | 2.68E+00 | 2.77E+00 | 2.72E-01 | no | Sorcs1 | intron_variant | MODIFIER |
| 19 | 50532043 | AX-170402021 | 0 | A | G | 0.482 | 7.00E-02 | 6.44E-02 | 2.68E+00 | 2.77E+00 | 2.72E-01 | no | Sorcs1 | intron_variant | MODIFIER |
| 19 | 50533676 | AX-170308816 | 0 | C | G | 0.482 | 7.00E-02 | 6.44E-02 | 2.68E+00 | 2.77E+00 | 2.72E-01 | no | Sorcs1 | intron_variant | MODIFIER |
| 19 | 50559155 | AX-170110276 | 0 | A | G | 0.482 | 7.00E-02 | 6.44E-02 | 2.68E+00 | 2.77E+00 | 2.72E-01 | no | Sorcs1 | intron_variant | MODIFIER |
| 19 | 50567521 | AX-170401799 | 0 | T | C | 0.482 | 7.00E-02 | 6.44E-02 | 2.68E+00 | 2.77E+00 | 2.72E-01 | no | Sorcs1 | intron_variant | MODIFIER |
| 19 | 50572917 | AX-170088478 | 0 | T | C | 0.482 | 7.00E-02 | 6.44E-02 | 2.68E+00 | 2.77E+00 | 2.72E-01 | no | Sorcs1 | intron_variant | MODIFIER |
| 19 | 50581209 | AX-170320224 | 0 | G | A | 0.482 | 7.00E-02 | 6.44E-02 | 2.68E+00 | 2.77E+00 | 2.72E-01 | no | Sorcs1 | intron_variant | MODIFIER |
| 19 | 50582933 | AX-170236243 | 0 | C | T | 0.482 | 7.00E-02 | 6.44E-02 | 2.68E+00 | 2.77E+00 | 2.72E-01 | no | Sorcs1 | intron_variant | MODIFIER |
| 19 | 50583534 | AX-170016806 | 0 | T | C | 0.482 | 7.00E-02 | 6.44E-02 | 2.68E+00 | 2.77E+00 | 2.72E-01 | no | Sorcs1 | intron_variant | MODIFIER |
| 19 | 50591416 | AX-170243305 | 0 | C | G | 0.482 | 7.00E-02 | 6.44E-02 | 2.68E+00 | 2.77E+00 | 2.72E-01 | no | Sorcs1 | intron_variant | MODIFIER |
| 19 | 50608936 | AX-170016801 | 0 | T | C | 0.482 | 7.00E-02 | 6.44E-02 | 2.68E+00 | 2.77E+00 | 2.72E-01 | no | Sorcs1 | intron_variant | MODIFIER |
| 19 | 50613755 | AX-170518947 | 0 | C | A | 0.482 | 7.00E-02 | 6.44E-02 | 2.68E+00 | 2.77E+00 | 2.72E-01 | no | Sorcs1 | intron_variant | MODIFIER |
| 19 | 50617776 | AX-170436871 | 0 | G | A | 0.482 | 7.00E-02 | 6.44E-02 | 2.68E+00 | 2.77E+00 | 2.72E-01 | no | Sorcs1 | intron_variant | MODIFIER |
| 19 | 50618424 | AX-170328063 | 0 | A | G | 0.482 | 7.00E-02 | 6.44E-02 | 2.68E+00 | 2.77E+00 | 2.72E-01 | no | Sorcs1 | intron_variant | MODIFIER |
| 19 | 50629648 | AX-170016805 | 0 | G | A | 0.482 | 7.00E-02 | 6.44E-02 | 2.68E+00 | 2.77E+00 | 2.72E-01 | no | Sorcs1 | intron_variant | MODIFIER |
| 19 | 50639262 | AX-170329487 | 0 | A | C | 0.482 | 7.00E-02 | 6.44E-02 | 2.68E+00 | 2.77E+00 | 2.72E-01 | no | Sorcs1 | intron_variant | MODIFIER |
| 19 | 50640180 | AX-170512002 | 0 | C | T | 0.482 | 7.00E-02 | 6.44E-02 | 2.68E+00 | 2.77E+00 | 2.72E-01 | no | Sorcs1 | intron_variant | MODIFIER |
| 19 | 50659927 | AX-97836060 | 0 | G | A | 0.482 | 7.00E-02 | 6.44E-02 | 2.68E+00 | 2.77E+00 | 2.72E-01 | no | Sorcs1 | intron_variant | MODIFIER |
| 19 | 50667231 | AX-170296425 | 0 | C | G | 0.482 | 7.00E-02 | 6.44E-02 | 2.68E+00 | 2.77E+00 | 2.72E-01 | no | Sorcs1 | intron_variant | MODIFIER |
| 19 | 50671593 | AX-170125224 | 0 | C | G | 0.482 | 7.00E-02 | 6.44E-02 | 2.68E+00 | 2.77E+00 | 2.72E-01 | no | Sorcs1 | intron_variant | MODIFIER |
| 19 | 50695118 | AX-170016803 | 0 | T | C | 0.482 | 7.00E-02 | 6.44E-02 | 2.68E+00 | 2.77E+00 | 2.72E-01 | no | Gm26629 | intron_variant,non_coding_transcript_variant | MODIFIER |

| | | | | | | | | | | | | | | | |
|---|---|---|---|---|---|---|---|---|---|---|---|---|---|---|---|
| 19 | 50712903 | AX-170099251 | 0 | T | C | 0.482 | 7.00E-02 | 6.44E-02 | 2.68E+00 | 2.77E+00 | 2.72E-01 | no | Gm26629 | intron_variant,non_coding_transcript_variant | MODIFIER |
| 19 | 50722724 | AX-170141194 | 0 | G | T | 0.482 | 7.00E-02 | 6.44E-02 | 2.68E+00 | 2.77E+00 | 2.72E-01 | no | Gm26629 | intron_variant,non_coding_transcript_variant | MODIFIER |
| 19 | 50722797 | AX-170170056 | 0 | A | G | 0.482 | 7.00E-02 | 6.44E-02 | 2.68E+00 | 2.77E+00 | 2.72E-01 | no | Gm26629 | intron_variant,non_coding_transcript_variant | MODIFIER |
| 19 | 50758527 | AX-170062656 | 0 | C | T | 0.482 | 7.00E-02 | 6.44E-02 | 2.68E+00 | 2.77E+00 | 2.72E-01 | no | Gm26629 | intron_variant,non_coding_transcript_variant | MODIFIER |
| 19 | 50763490 | AX-97848870 | 0 | A | G | 0.482 | 7.00E-02 | 6.44E-02 | 2.68E+00 | 2.77E+00 | 2.72E-01 | no | Gm26629 | intron_variant,non_coding_transcript_variant | MODIFIER |
| 19 | 50792417 | AX-170162051 | 0 | C | G | 0.482 | 7.00E-02 | 6.44E-02 | 2.68E+00 | 2.77E+00 | 2.72E-01 | no | | intergenic_variant | MODIFIER |
| 19 | 50792831 | AX-170526705 | 0 | T | C | 0.482 | 7.00E-02 | 6.44E-02 | 2.68E+00 | 2.77E+00 | 2.72E-01 | no | | intergenic_variant | MODIFIER |
| 19 | 50792900 | AX-170215654 | 0 | T | A | 0.482 | 7.00E-02 | 6.44E-02 | 2.68E+00 | 2.77E+00 | 2.72E-01 | no | | intergenic_variant | MODIFIER |
| 19 | 50793476 | AX-170242991 | 0 | T | A | 0.482 | 7.00E-02 | 6.44E-02 | 2.68E+00 | 2.77E+00 | 2.72E-01 | no | | intergenic_variant | MODIFIER |
| 19 | 50794845 | AX-170377207 | 0 | C | A | 0.482 | 7.00E-02 | 6.44E-02 | 2.68E+00 | 2.77E+00 | 2.72E-01 | no | | intergenic_variant | MODIFIER |
| 19 | 50795715 | AX-170503290 | 0 | T | G | 0.482 | 7.00E-02 | 6.44E-02 | 2.68E+00 | 2.77E+00 | 2.72E-01 | no | | intergenic_variant | MODIFIER |
| 19 | 50797525 | AX-97818981 | 0 | C | A | 0.482 | 7.00E-02 | 6.44E-02 | 2.68E+00 | 2.77E+00 | 2.72E-01 | no | | intergenic_variant | MODIFIER |
| 19 | 50810851 | AX-97837690 | 0 | G | T | 0.482 | 7.00E-02 | 6.44E-02 | 2.68E+00 | 2.77E+00 | 2.72E-01 | no | Gm25659 | upstream_gene_variant | MODIFIER |
| 19 | 50836765 | AX-170408471 | 0 | C | T | 0.482 | 7.00E-02 | 6.44E-02 | 2.68E+00 | 2.77E+00 | 2.72E-01 | no | | intergenic_variant | MODIFIER |
| 19 | 50836913 | AX-170153646 | 0 | T | C | 0.482 | 7.00E-02 | 6.44E-02 | 2.68E+00 | 2.77E+00 | 2.72E-01 | no | | intergenic_variant | MODIFIER |
| 19 | 50837233 | AX-97786960 | 0 | C | T | 0.482 | 7.00E-02 | 6.44E-02 | 2.68E+00 | 2.77E+00 | 2.72E-01 | no | | intergenic_variant | MODIFIER |
| 19 | 50837624 | AX-170104219 | 0 | C | G | 0.482 | 7.00E-02 | 6.44E-02 | 2.68E+00 | 2.77E+00 | 2.72E-01 | no | | intergenic_variant | MODIFIER |
| 19 | 50875840 | AX-170435381 | 0 | C | G | 0.482 | 7.00E-02 | 6.44E-02 | 2.68E+00 | 2.77E+00 | 2.72E-01 | no | | intergenic_variant | MODIFIER |
| 19 | 50893813 | AX-170094115 | 0 | A | G | 0.482 | 7.00E-02 | 6.44E-02 | 2.68E+00 | 2.77E+00 | 2.72E-01 | no | | intergenic_variant | MODIFIER |
| 19 | 50894791 | AX-168379130 | 0 | T | C | 0.482 | 7.00E-02 | 6.44E-02 | 2.68E+00 | 2.77E+00 | 2.72E-01 | no | | intergenic_variant | MODIFIER |
| 19 | 50895002 | AX-170218448 | 0 | T | C | 0.482 | 7.00E-02 | 6.44E-02 | 2.68E+00 | 2.77E+00 | 2.72E-01 | no | | intergenic_variant | MODIFIER |
| 19 | 50895315 | AX-170478693 | 0 | C | T | 0.482 | 7.00E-02 | 6.44E-02 | 2.68E+00 | 2.77E+00 | 2.72E-01 | no | | intergenic_variant | MODIFIER |
| 19 | 50943655 | AX-170062659 | 0 | A | T | 0.482 | 7.00E-02 | 6.44E-02 | 2.68E+00 | 2.77E+00 | 2.72E-01 | no | | intergenic_variant | MODIFIER |
| 19 | 51166199 | AX-170016816 | 0 | T | C | 0.482 | 7.00E-02 | 6.44E-02 | 2.68E+00 | 2.77E+00 | 2.72E-01 | no | | intergenic_variant | MODIFIER |
| 19 | 51166266 | AX-170016813 | 0 | G | A | 0.482 | 7.00E-02 | 6.44E-02 | 2.68E+00 | 2.77E+00 | 2.72E-01 | no | | intergenic_variant | MODIFIER |
| 19 | 51521192 | AX-168385876 | 0 | A | C | 0.482 | 7.00E-02 | 6.44E-02 | 2.68E+00 | 2.77E+00 | 2.72E-01 | no | | intergenic_variant | MODIFIER |
| 19 | 51536816 | AX-97841907 | 0 | A | G | 0.482 | 7.00E-02 | 6.44E-02 | 2.68E+00 | 2.77E+00 | 2.72E-01 | no | | intergenic_variant | MODIFIER |
| 19 | 51542641 | AX-170239183 | 0 | G | C | 0.482 | 7.00E-02 | 6.44E-02 | 2.68E+00 | 2.77E+00 | 2.72E-01 | no | | intergenic_variant | MODIFIER |
| 19 | 51550165 | AX-170343763 | 0 | T | C | 0.482 | 7.00E-02 | 6.44E-02 | 2.68E+00 | 2.77E+00 | 2.72E-01 | no | | intergenic_variant | MODIFIER |
| 19 | 51555910 | AX-170265622 | 0 | G | A | 0.482 | 7.00E-02 | 6.44E-02 | 2.68E+00 | 2.77E+00 | 2.72E-01 | no | | intergenic_variant | MODIFIER |
| 19 | 51566019 | AX-97836059 | 0 | G | A | 0.482 | 7.00E-02 | 6.44E-02 | 2.68E+00 | 2.77E+00 | 2.72E-01 | no | | intergenic_variant | MODIFIER |
| 19 | 51608778 | AX-97826652 | 0 | G | A | 0.482 | 7.00E-02 | 6.44E-02 | 2.68E+00 | 2.77E+00 | 2.72E-01 | no | | intergenic_variant | MODIFIER |
| 13 | 25458026 | AX-97830142 | 0 | C | T | 0.167 | -1.26E-01 | 1.12E-01 | 2.37E+00 | 2.43E+00 | 2.72E-01 | no | Tpi-rs8 | downstream_gene_variant | MODIFIER |
| 9 | 92530041 | AX-170208291 | 0 | A | G | 0.14 | -1.32E-01 | 1.22E-01 | 2.66E+00 | 2.74E+00 | 2.72E-01 | no | Gm25417 | upstream_gene_variant | MODIFIER |
| 9 | 92531765 | AX-170111579 | 0 | T | A | 0.14 | -1.32E-01 | 1.22E-01 | 2.66E+00 | 2.74E+00 | 2.72E-01 | no | Gm25417 | upstream_gene_variant | MODIFIER |
| 9 | 92548868 | AX-170270491 | 0 | A | G | 0.14 | -1.32E-01 | 1.22E-01 | 2.66E+00 | 2.74E+00 | 2.72E-01 | no | Plod2 | intron_variant | MODIFIER |
| 9 | 92550303 | AX-170083096 | 0 | A | G | 0.14 | -1.32E-01 | 1.22E-01 | 2.66E+00 | 2.74E+00 | 2.72E-01 | no | Plod2 | intron_variant | MODIFIER |
| 9 | 92566120 | AX-170118580 | 0 | G | T | 0.14 | -1.32E-01 | 1.22E-01 | 2.66E+00 | 2.74E+00 | 2.72E-01 | no | Plod2 | intron_variant | MODIFIER |
| 9 | 92571493 | AX-170315266 | 0 | A | G | 0.14 | -1.32E-01 | 1.22E-01 | 2.66E+00 | 2.74E+00 | 2.72E-01 | no | Plod2 | intron_variant | MODIFIER |
| 9 | 92571540 | AX-170185291 | 0 | A | G | 0.14 | -1.32E-01 | 1.22E-01 | 2.66E+00 | 2.74E+00 | 2.72E-01 | no | Plod2 | intron_variant | MODIFIER |
| 9 | 92588059 | AX-170502244 | 0 | A | G | 0.14 | -1.32E-01 | 1.22E-01 | 2.66E+00 | 2.74E+00 | 2.72E-01 | no | Plod2 | intron_variant | MODIFIER |
| 9 | 92588193 | AX-170354881 | 0 | T | C | 0.14 | -1.32E-01 | 1.22E-01 | 2.66E+00 | 2.74E+00 | 2.72E-01 | no | Plod2 | intron_variant | MODIFIER |
| 9 | 92591092 | AX-170267929 | 0 | C | T | 0.14 | -1.32E-01 | 1.22E-01 | 2.66E+00 | 2.74E+00 | 2.72E-01 | no | Plod2 | intron_variant | MODIFIER |
| 9 | 92593483 | AX-170201269 | 0 | G | A | 0.14 | -1.32E-01 | 1.22E-01 | 2.66E+00 | 2.74E+00 | 2.72E-01 | no | Plod2 | intron_variant | MODIFIER |
| 9 | 92594213 | AX-170250823 | 0 | G | T | 0.14 | -1.32E-01 | 1.22E-01 | 2.66E+00 | 2.74E+00 | 2.72E-01 | no | Plod2 | intron_variant | MODIFIER |
| 9 | 92598989 | AX-170382060 | 0 | C | A | 0.14 | -1.32E-01 | 1.22E-01 | 2.66E+00 | 2.74E+00 | 2.72E-01 | no | Plod2 | intron_variant | MODIFIER |
| 9 | 92604914 | AX-170510842 | 0 | A | C | 0.14 | -1.32E-01 | 1.22E-01 | 2.66E+00 | 2.74E+00 | 2.72E-01 | no | Plod2 | intron_variant | MODIFIER |
| 9 | 92605802 | AX-170509179 | 0 | T | C | 0.14 | -1.32E-01 | 1.22E-01 | 2.66E+00 | 2.74E+00 | 2.72E-01 | no | Plod2 | intron_variant | MODIFIER |
| 9 | 92606733 | AX-170408192 | 0 | G | A | 0.14 | -1.32E-01 | 1.22E-01 | 2.66E+00 | 2.74E+00 | 2.72E-01 | no | Plod2 | intron_variant | MODIFIER |
| 9 | 92609286 | AX-170368169 | 0 | A | G | 0.14 | -1.32E-01 | 1.22E-01 | 2.66E+00 | 2.74E+00 | 2.72E-01 | no | Plod2 | downstream_gene_variant | MODIFIER |
| 9 | 92609324 | AX-170207092 | 0 | T | G | 0.14 | -1.32E-01 | 1.22E-01 | 2.66E+00 | 2.74E+00 | 2.72E-01 | no | Plod2 | downstream_gene_variant | MODIFIER |
| 9 | 92610719 | AX-170135804 | 0 | G | C | 0.14 | -1.32E-01 | 1.22E-01 | 2.66E+00 | 2.74E+00 | 2.72E-01 | no | Plod2 | downstream_gene_variant | MODIFIER |
| 9 | 92613729 | AX-170083098 | 0 | G | C | 0.14 | -1.32E-01 | 1.22E-01 | 2.66E+00 | 2.74E+00 | 2.72E-01 | no | Gm2495 | upstream_gene_variant | MODIFIER |
| 9 | 92616627 | AX-170523275 | 0 | C | T | 0.14 | -1.32E-01 | 1.22E-01 | 2.66E+00 | 2.74E+00 | 2.72E-01 | no | Gm2495 | upstream_gene_variant | MODIFIER |

| | | | | | | | | | | | | | | | |
|---|---|---|---|---|---|---|---|---|---|---|---|---|---|---|---|
| 9 | 92640157 | AX-170445030 | 0 | A | G | 0.14 | -1.32E-01 | 1.22E-01 | 2.66E+00 | 2.74E+00 | 2.72E-01 | no | | intergenic_variant | MODIFIER |
| 9 | 92654694 | AX-170408517 | 0 | T | A | 0.14 | -1.32E-01 | 1.22E-01 | 2.66E+00 | 2.74E+00 | 2.72E-01 | no | | intergenic_variant | MODIFIER |
| 9 | 92654863 | AX-170256766 | 0 | A | G | 0.14 | -1.32E-01 | 1.22E-01 | 2.66E+00 | 2.74E+00 | 2.72E-01 | no | | intergenic_variant | MODIFIER |
| 9 | 92655317 | AX-170518024 | 0 | C | G | 0.14 | -1.32E-01 | 1.22E-01 | 2.66E+00 | 2.74E+00 | 2.72E-01 | no | | intergenic_variant | MODIFIER |
| 9 | 92655823 | AX-170171275 | 0 | C | A | 0.14 | -1.32E-01 | 1.22E-01 | 2.66E+00 | 2.74E+00 | 2.72E-01 | no | | intergenic_variant | MODIFIER |
| 9 | 92656101 | AX-170427033 | 0 | T | C | 0.14 | -1.32E-01 | 1.22E-01 | 2.66E+00 | 2.74E+00 | 2.72E-01 | no | | intergenic_variant | MODIFIER |
| 9 | 92757202 | AX-170083100 | 0 | T | A | 0.14 | -1.32E-01 | 1.22E-01 | 2.66E+00 | 2.74E+00 | 2.72E-01 | no | 4933400C23Rik | intron_variant,non_coding_transcript_variant | MODIFIER |
| 9 | 94261416 | AX-170433002 | 0 | G | C | 0.14 | -1.32E-01 | 1.22E-01 | 2.66E+00 | 2.74E+00 | 2.72E-01 | no | | intergenic_variant | MODIFIER |
| 9 | 94262273 | AX-170083132 | 0 | C | G | 0.14 | -1.32E-01 | 1.22E-01 | 2.66E+00 | 2.74E+00 | 2.72E-01 | no | | intergenic_variant | MODIFIER |
| 9 | 94262327 | AX-170083134 | 0 | G | T | 0.14 | -1.32E-01 | 1.22E-01 | 2.66E+00 | 2.74E+00 | 2.72E-01 | no | | intergenic_variant | MODIFIER |
| 9 | 94267571 | AX-97814770 | 0 | C | T | 0.14 | -1.32E-01 | 1.22E-01 | 2.66E+00 | 2.74E+00 | 2.72E-01 | no | | intergenic_variant | MODIFIER |
| 9 | 94284389 | AX-170357686 | 0 | C | T | 0.14 | -1.32E-01 | 1.22E-01 | 2.66E+00 | 2.74E+00 | 2.72E-01 | no | Gm24200 | upstream_gene_variant | MODIFIER |
| 9 | 94295404 | AX-170251911 | 0 | A | G | 0.14 | -1.32E-01 | 1.22E-01 | 2.66E+00 | 2.74E+00 | 2.72E-01 | no | | regulatory_region_variant | MODIFIER |
| 9 | 94301918 | AX-170187226 | 0 | A | T | 0.14 | -1.32E-01 | 1.22E-01 | 2.66E+00 | 2.74E+00 | 2.72E-01 | no | | intergenic_variant | MODIFIER |
| 9 | 94422231 | AX-97854279 | 0 | G | A | 0.14 | -1.32E-01 | 1.22E-01 | 2.66E+00 | 2.74E+00 | 2.72E-01 | no | | intergenic_variant | MODIFIER |
| 9 | 94426710 | AX-170117691 | 0 | G | A | 0.14 | -1.32E-01 | 1.22E-01 | 2.66E+00 | 2.74E+00 | 2.72E-01 | no | | intergenic_variant | MODIFIER |
| 9 | 94435847 | AX-170275392 | 0 | G | A | 0.14 | -1.32E-01 | 1.22E-01 | 2.66E+00 | 2.74E+00 | 2.72E-01 | no | | intergenic_variant | MODIFIER |
| 9 | 94447224 | AX-97782177 | 0 | C | A | 0.14 | -1.32E-01 | 1.22E-01 | 2.66E+00 | 2.74E+00 | 2.72E-01 | no | | intergenic_variant | MODIFIER |
| 9 | 94454518 | AX-170106965 | 0 | T | G | 0.14 | -1.32E-01 | 1.22E-01 | 2.66E+00 | 2.74E+00 | 2.72E-01 | no | | intergenic_variant | MODIFIER |
| 9 | 94461494 | AX-97854286 | 0 | A | G | 0.14 | -1.32E-01 | 1.22E-01 | 2.66E+00 | 2.74E+00 | 2.72E-01 | no | | intergenic_variant | MODIFIER |
| 9 | 94466878 | AX-170258503 | 0 | G | A | 0.14 | -1.32E-01 | 1.22E-01 | 2.66E+00 | 2.74E+00 | 2.72E-01 | no | | intergenic_variant | MODIFIER |
| 9 | 94466902 | AX-170410024 | 0 | C | T | 0.14 | -1.32E-01 | 1.22E-01 | 2.66E+00 | 2.74E+00 | 2.72E-01 | no | | intergenic_variant | MODIFIER |
| 9 | 94468212 | AX-170239927 | 0 | C | T | 0.14 | -1.32E-01 | 1.22E-01 | 2.66E+00 | 2.74E+00 | 2.72E-01 | no | | regulatory_region_variant | MODIFIER |
| 9 | 94469929 | AX-170528019 | 0 | A | G | 0.14 | -1.32E-01 | 1.22E-01 | 2.66E+00 | 2.74E+00 | 2.72E-01 | no | | intergenic_variant | MODIFIER |
| 9 | 94470329 | AX-170443000 | 0 | T | C | 0.14 | -1.32E-01 | 1.22E-01 | 2.66E+00 | 2.74E+00 | 2.72E-01 | no | | intergenic_variant | MODIFIER |
| 9 | 94476526 | AX-170473683 | 0 | A | G | 0.14 | -1.32E-01 | 1.22E-01 | 2.66E+00 | 2.74E+00 | 2.72E-01 | no | | intergenic_variant | MODIFIER |
| 9 | 94480022 | AX-170468127 | 0 | G | A | 0.14 | -1.32E-01 | 1.22E-01 | 2.66E+00 | 2.74E+00 | 2.72E-01 | no | | intergenic_variant | MODIFIER |
| 9 | 94498721 | AX-170186370 | 0 | C | T | 0.14 | -1.32E-01 | 1.22E-01 | 2.66E+00 | 2.74E+00 | 2.72E-01 | no | | intergenic_variant | MODIFIER |
| 9 | 94506982 | AX-170192618 | 0 | T | A | 0.14 | -1.32E-01 | 1.22E-01 | 2.66E+00 | 2.74E+00 | 2.72E-01 | no | | intergenic_variant | MODIFIER |
| 9 | 94530220 | AX-170090384 | 0 | A | G | 0.14 | -1.32E-01 | 1.22E-01 | 2.66E+00 | 2.74E+00 | 2.72E-01 | no | 1190002N15Rik | intron_variant | MODIFIER |
| 9 | 94541897 | AX-170033030 | 0 | G | C | 0.14 | -1.32E-01 | 1.22E-01 | 2.66E+00 | 2.74E+00 | 2.72E-01 | no | 1190002N15Rik | upstream_gene_variant | MODIFIER |
| 9 | 94552084 | AX-170339942 | 0 | T | C | 0.14 | -1.32E-01 | 1.22E-01 | 2.66E+00 | 2.74E+00 | 2.72E-01 | no | | intergenic_variant | MODIFIER |
| 9 | 94567931 | AX-97822662 | 0 | G | T | 0.14 | -1.32E-01 | 1.22E-01 | 2.66E+00 | 2.74E+00 | 2.72E-01 | no | | intergenic_variant | MODIFIER |
| 9 | 94633297 | AX-97799073 | 0 | T | A | 0.14 | -1.32E-01 | 1.22E-01 | 2.66E+00 | 2.74E+00 | 2.72E-01 | no | | intergenic_variant | MODIFIER |
| 9 | 94678395 | AX-97787192 | 0 | T | G | 0.14 | -1.32E-01 | 1.22E-01 | 2.66E+00 | 2.74E+00 | 2.72E-01 | no | Slc9a9 | intron_variant | MODIFIER |
| 9 | 94736238 | AX-97830369 | 0 | C | T | 0.14 | -1.32E-01 | 1.22E-01 | 2.66E+00 | 2.74E+00 | 2.72E-01 | no | Slc9a9 | intron_variant | MODIFIER |
| 16 | 56062583 | AX-170152854 | 1 | C | T | 0.473 | -9.11E-02 | 8.26E-02 | 2.52E+00 | 2.56E+00 | 2.72E-01 | no | Senp7 | intron_variant,non_coding_transcript_variant | MODIFIER |
| 7 | 39678149 | AX-170028866 | 0 | T | C | 0.36 | 9.84E-02 | 9.01E-02 | 2.60E+00 | 2.67E+00 | 2.72E-01 | no | | intergenic_variant | MODIFIER |
| 3 | 23851679 | AX-170371383 | 1 | A | G | 0.161 | -1.11E-01 | 1.02E-01 | 2.59E+00 | 2.67E+00 | 2.72E-01 | no | Naaladl2 | intron_variant,non_coding_transcript_variant | MODIFIER |
| 1 | 156100872 | AX-170002054 | 1 | C | T | 0.232 | 1.38E-01 | 1.27E-01 | 2.61E+00 | 2.68E+00 | 2.72E-01 | no | Fam163a | intron_variant | MODIFIER |
| 19 | 50695996 | AX-97806543 | 0 | A | G | 0.482 | 7.00E-02 | 6.44E-02 | 2.68E+00 | 2.78E+00 | 2.72E-01 | no | Gm26629 | intron_variant,non_coding_transcript_variant | MODIFIER |
| 6 | 119574570 | AX-170475854 | 0 | T | C | 0.07 | 2.21E-01 | 2.03E-01 | 2.83E+00 | 2.96E+00 | 2.73E-01 | Yes | Erc1 | 3_prime_UTR_variant | MODIFIER |
| 6 | 139121712 | AX-170208729 | 0 | A | G | 0.07 | 2.21E-01 | 2.03E-01 | 2.83E+00 | 2.96E+00 | 2.73E-01 | Yes | | intergenic_variant | MODIFIER |
| 4 | 80092177 | AX-170411623 | 1 | G | A | 0.446 | -9.35E-02 | 8.49E-02 | 2.52E+00 | 2.57E+00 | 2.73E-01 | no | | intergenic_variant | MODIFIER |
| 8 | 60630817 | AX-170269523 | 0 | A | G | 0.14 | -1.70E-01 | 1.53E-01 | 2.45E+00 | 2.50E+00 | 2.73E-01 | no | Mfap3l | upstream_gene_variant | MODIFIER |
| 8 | 60669181 | AX-170253788 | 0 | A | G | 0.14 | -1.70E-01 | 1.53E-01 | 2.45E+00 | 2.50E+00 | 2.73E-01 | no | Mfap3l | intron_variant | MODIFIER |
| 8 | 60694681 | AX-170192815 | 0 | C | A | 0.14 | -1.70E-01 | 1.53E-01 | 2.45E+00 | 2.50E+00 | 2.73E-01 | no | | intergenic_variant | MODIFIER |
| 8 | 60694899 | AX-170252994 | 0 | T | C | 0.14 | -1.70E-01 | 1.53E-01 | 2.45E+00 | 2.50E+00 | 2.73E-01 | no | | intergenic_variant | MODIFIER |
| 8 | 60712078 | AX-170210415 | 0 | G | C | 0.14 | -1.70E-01 | 1.53E-01 | 2.45E+00 | 2.50E+00 | 2.73E-01 | no | | intergenic_variant | MODIFIER |
| 8 | 60712861 | AX-170283216 | 0 | T | C | 0.14 | -1.70E-01 | 1.53E-01 | 2.45E+00 | 2.50E+00 | 2.73E-01 | no | | intergenic_variant | MODIFIER |
| 8 | 60736795 | AX-170412974 | 0 | G | T | 0.14 | -1.70E-01 | 1.53E-01 | 2.45E+00 | 2.50E+00 | 2.73E-01 | no | | intergenic_variant | MODIFIER |
| 8 | 60745468 | AX-170290070 | 0 | T | C | 0.14 | -1.70E-01 | 1.53E-01 | 2.45E+00 | 2.50E+00 | 2.73E-01 | no | | intergenic_variant | MODIFIER |
| 8 | 60747632 | AX-170167491 | 0 | T | C | 0.14 | -1.70E-01 | 1.53E-01 | 2.45E+00 | 2.50E+00 | 2.73E-01 | no | | intergenic_variant | MODIFIER |
| 8 | 60759962 | AX-170450610 | 0 | A | G | 0.14 | -1.70E-01 | 1.53E-01 | 2.45E+00 | 2.50E+00 | 2.73E-01 | no | Gm45651 | downstream_gene_variant | MODIFIER |

| | | | | | | | | | | | | | | | |
|---|---|---|---|---|---|---|---|---|---|---|---|---|---|---|---|
| 8 | 60760520 | AX-170329531 | 0 | C | T | 0.14 | -1.70E-01 | 1.53E-01 | 2.45E+00 | 2.50E+00 | 2.73E-01 | no | Gm45651 | downstream_gene_variant | MODIFIER |
| 8 | 60769055 | AX-170030848 | 0 | A | C | 0.14 | -1.70E-01 | 1.53E-01 | 2.45E+00 | 2.50E+00 | 2.73E-01 | no | | intergenic_variant | MODIFIER |
| 8 | 60770104 | AX-170362022 | 0 | A | G | 0.14 | -1.70E-01 | 1.53E-01 | 2.45E+00 | 2.50E+00 | 2.73E-01 | no | | intergenic_variant | MODIFIER |
| 8 | 60777519 | AX-170030849 | 0 | C | T | 0.14 | -1.70E-01 | 1.53E-01 | 2.45E+00 | 2.50E+00 | 2.73E-01 | no | | intergenic_variant | MODIFIER |
| 8 | 60789995 | AX-170030850 | 0 | G | A | 0.14 | -1.70E-01 | 1.53E-01 | 2.45E+00 | 2.50E+00 | 2.73E-01 | no | Stambp-ps1 | upstream_gene_variant | MODIFIER |
| 8 | 60795042 | AX-170030846 | 0 | T | C | 0.14 | -1.70E-01 | 1.53E-01 | 2.45E+00 | 2.50E+00 | 2.73E-01 | no | Stambp-ps1 | downstream_gene_variant | MODIFIER |
| 8 | 60811503 | AX-170266556 | 0 | A | C | 0.14 | -1.70E-01 | 1.53E-01 | 2.45E+00 | 2.50E+00 | 2.73E-01 | no | Gm23329 | upstream_gene_variant | MODIFIER |
| 8 | 60835466 | AX-170264914 | 0 | A | G | 0.14 | -1.70E-01 | 1.53E-01 | 2.45E+00 | 2.50E+00 | 2.73E-01 | no | | intergenic_variant | MODIFIER |
| 8 | 60845124 | AX-170490096 | 0 | A | G | 0.14 | -1.70E-01 | 1.53E-01 | 2.45E+00 | 2.50E+00 | 2.73E-01 | no | Gm45485 | upstream_gene_variant | MODIFIER |
| 8 | 60845204 | AX-170414456 | 0 | A | C | 0.14 | -1.70E-01 | 1.53E-01 | 2.45E+00 | 2.50E+00 | 2.73E-01 | no | Gm45485 | upstream_gene_variant | MODIFIER |
| 8 | 60846707 | AX-170411990 | 0 | T | C | 0.14 | -1.70E-01 | 1.53E-01 | 2.45E+00 | 2.50E+00 | 2.73E-01 | no | Gm45485 | downstream_gene_variant | MODIFIER |
| 8 | 60846941 | AX-170350916 | 0 | A | T | 0.14 | -1.70E-01 | 1.53E-01 | 2.45E+00 | 2.50E+00 | 2.73E-01 | no | Gm45485 | downstream_gene_variant | MODIFIER |
| 8 | 60862281 | AX-170096286 | 0 | A | G | 0.14 | -1.70E-01 | 1.53E-01 | 2.45E+00 | 2.50E+00 | 2.73E-01 | no | | intergenic_variant | MODIFIER |
| 8 | 61130651 | AX-168375220 | 0 | G | A | 0.14 | -1.70E-01 | 1.53E-01 | 2.45E+00 | 2.50E+00 | 2.73E-01 | no | Nek1 | 3_prime_UTR_variant | MODIFIER |
| 8 | 61160442 | AX-170503797 | 0 | C | T | 0.14 | -1.70E-01 | 1.53E-01 | 2.45E+00 | 2.50E+00 | 2.73E-01 | no | Gm16178 | intron_variant,non_coding_transcript_variant | MODIFIER |
| 8 | 61160995 | AX-170080218 | 0 | G | A | 0.14 | -1.70E-01 | 1.53E-01 | 2.45E+00 | 2.50E+00 | 2.73E-01 | no | Gm16178 | intron_variant,non_coding_transcript_variant | MODIFIER |
| 8 | 61177170 | AX-97822898 | 0 | C | T | 0.14 | -1.70E-01 | 1.53E-01 | 2.45E+00 | 2.50E+00 | 2.73E-01 | no | Gm16178 | intron_variant,non_coding_transcript_variant | MODIFIER |
| 8 | 61204704 | AX-170443362 | 0 | A | G | 0.14 | -1.70E-01 | 1.53E-01 | 2.45E+00 | 2.50E+00 | 2.73E-01 | no | | intergenic_variant | MODIFIER |
| 8 | 61205167 | AX-170460855 | 0 | T | G | 0.14 | -1.70E-01 | 1.53E-01 | 2.45E+00 | 2.50E+00 | 2.73E-01 | no | | intergenic_variant | MODIFIER |
| 8 | 61208871 | AX-170509882 | 0 | T | C | 0.14 | -1.70E-01 | 1.53E-01 | 2.45E+00 | 2.50E+00 | 2.73E-01 | no | | regulatory_region_variant | MODIFIER |
| 8 | 61281463 | AX-170030855 | 0 | T | G | 0.14 | -1.70E-01 | 1.53E-01 | 2.45E+00 | 2.50E+00 | 2.73E-01 | no | Sh3rf1 | intron_variant | MODIFIER |
| 8 | 61338191 | AX-170224167 | 0 | C | T | 0.14 | -1.70E-01 | 1.53E-01 | 2.45E+00 | 2.50E+00 | 2.73E-01 | no | Sh3rf1 | intron_variant | MODIFIER |
| 8 | 61348880 | AX-170030852 | 0 | C | T | 0.14 | -1.70E-01 | 1.53E-01 | 2.45E+00 | 2.50E+00 | 2.73E-01 | no | Sh3rf1 | intron_variant | MODIFIER |
| 8 | 61349143 | AX-170030851 | 0 | C | G | 0.14 | -1.70E-01 | 1.53E-01 | 2.45E+00 | 2.50E+00 | 2.73E-01 | no | Sh3rf1 | intron_variant | MODIFIER |
| 8 | 61354412 | AX-170080219 | 0 | A | C | 0.14 | -1.70E-01 | 1.53E-01 | 2.45E+00 | 2.50E+00 | 2.73E-01 | no | Sh3rf1 | intron_variant | MODIFIER |
| 8 | 61391865 | AX-170523531 | 0 | T | G | 0.14 | -1.70E-01 | 1.53E-01 | 2.45E+00 | 2.50E+00 | 2.73E-01 | no | Sh3rf1 | intron_variant | MODIFIER |
| 8 | 61410203 | AX-170468708 | 0 | A | G | 0.14 | -1.70E-01 | 1.53E-01 | 2.45E+00 | 2.50E+00 | 2.73E-01 | no | | intergenic_variant | MODIFIER |
| 8 | 61429960 | AX-170454430 | 0 | C | G | 0.14 | -1.70E-01 | 1.53E-01 | 2.45E+00 | 2.50E+00 | 2.73E-01 | no | | intergenic_variant | MODIFIER |
| 8 | 61435131 | AX-170385715 | 0 | C | T | 0.14 | -1.70E-01 | 1.53E-01 | 2.45E+00 | 2.50E+00 | 2.73E-01 | no | | intergenic_variant | MODIFIER |
| 8 | 61437941 | AX-97836070 | 0 | C | A | 0.14 | -1.70E-01 | 1.53E-01 | 2.45E+00 | 2.50E+00 | 2.73E-01 | no | | intergenic_variant | MODIFIER |
| 8 | 61466575 | AX-170336481 | 0 | C | T | 0.14 | -1.70E-01 | 1.53E-01 | 2.45E+00 | 2.50E+00 | 2.73E-01 | no | | intergenic_variant | MODIFIER |
| 8 | 61466785 | AX-170137755 | 0 | A | T | 0.14 | -1.70E-01 | 1.53E-01 | 2.45E+00 | 2.50E+00 | 2.73E-01 | no | | intergenic_variant | MODIFIER |
| 8 | 61472109 | AX-170080221 | 0 | T | G | 0.14 | -1.70E-01 | 1.53E-01 | 2.45E+00 | 2.50E+00 | 2.73E-01 | no | | intergenic_variant | MODIFIER |
| 8 | 61472597 | AX-97857185 | 0 | C | T | 0.14 | -1.70E-01 | 1.53E-01 | 2.45E+00 | 2.50E+00 | 2.73E-01 | no | | intergenic_variant | MODIFIER |
| 8 | 61500129 | AX-170522081 | 0 | C | G | 0.14 | -1.70E-01 | 1.53E-01 | 2.45E+00 | 2.50E+00 | 2.73E-01 | no | Cbr4 | intron_variant | MODIFIER |
| 8 | 61506773 | AX-170304217 | 0 | T | C | 0.14 | -1.70E-01 | 1.53E-01 | 2.45E+00 | 2.50E+00 | 2.73E-01 | no | Cbr4 | downstream_gene_variant | MODIFIER |
| 8 | 61506925 | AX-97822318 | 0 | G | A | 0.14 | -1.70E-01 | 1.53E-01 | 2.45E+00 | 2.50E+00 | 2.73E-01 | no | Cbr4 | downstream_gene_variant | MODIFIER |
| 8 | 61515604 | AX-170086037 | 0 | C | T | 0.14 | -1.70E-01 | 1.53E-01 | 2.45E+00 | 2.50E+00 | 2.73E-01 | no | Palld | intron_variant | MODIFIER |
| 8 | 61518024 | AX-170189044 | 0 | A | G | 0.14 | -1.70E-01 | 1.53E-01 | 2.45E+00 | 2.50E+00 | 2.73E-01 | no | Palld | intron_variant | MODIFIER |
| 8 | 61519051 | AX-97846124 | 0 | C | T | 0.14 | -1.70E-01 | 1.53E-01 | 2.45E+00 | 2.50E+00 | 2.73E-01 | no | Palld | intron_variant | MODIFIER |
| 8 | 61539721 | AX-97821904 | 0 | T | C | 0.14 | -1.70E-01 | 1.53E-01 | 2.45E+00 | 2.50E+00 | 2.73E-01 | no | Palld | intron_variant | MODIFIER |
| 8 | 61558300 | AX-170265826 | 0 | A | G | 0.14 | -1.70E-01 | 1.53E-01 | 2.45E+00 | 2.50E+00 | 2.73E-01 | no | Palld | intron_variant | MODIFIER |
| 8 | 61565208 | AX-170156264 | 0 | T | C | 0.14 | -1.70E-01 | 1.53E-01 | 2.45E+00 | 2.50E+00 | 2.73E-01 | no | Palld | intron_variant | MODIFIER |
| 8 | 61565738 | AX-170302080 | 0 | G | A | 0.14 | -1.70E-01 | 1.53E-01 | 2.45E+00 | 2.50E+00 | 2.73E-01 | no | Palld | intron_variant | MODIFIER |
| 8 | 61634454 | AX-170369496 | 0 | T | C | 0.14 | -1.70E-01 | 1.53E-01 | 2.45E+00 | 2.50E+00 | 2.73E-01 | no | Palld | intron_variant | MODIFIER |
| 8 | 61756194 | AX-170420766 | 0 | A | G | 0.14 | -1.70E-01 | 1.53E-01 | 2.45E+00 | 2.50E+00 | 2.73E-01 | no | Palld | intron_variant | MODIFIER |
| 8 | 61805481 | AX-170389218 | 0 | T | C | 0.14 | -1.70E-01 | 1.53E-01 | 2.45E+00 | 2.50E+00 | 2.73E-01 | no | Palld | intron_variant | MODIFIER |
| 8 | 61985227 | AX-170404938 | 0 | A | T | 0.14 | -1.70E-01 | 1.53E-01 | 2.45E+00 | 2.50E+00 | 2.73E-01 | no | Ddx60 | intron_variant | MODIFIER |
| 8 | 62010479 | AX-170150848 | 0 | C | T | 0.14 | -1.70E-01 | 1.53E-01 | 2.45E+00 | 2.50E+00 | 2.73E-01 | no | Ddx60 | intron_variant | MODIFIER |
| 8 | 62069191 | AX-170149797 | 0 | G | A | 0.14 | -1.70E-01 | 1.53E-01 | 2.45E+00 | 2.50E+00 | 2.73E-01 | no | Anxa10 | intron_variant | MODIFIER |
| 8 | 62469441 | AX-170030870 | 0 | A | G | 0.14 | -1.70E-01 | 1.53E-01 | 2.45E+00 | 2.50E+00 | 2.73E-01 | no | | intergenic_variant | MODIFIER |
| 8 | 62469707 | AX-170030871 | 0 | A | T | 0.14 | -1.70E-01 | 1.53E-01 | 2.45E+00 | 2.50E+00 | 2.73E-01 | no | | intergenic_variant | MODIFIER |
| 8 | 36060267 | AX-170388769 | 0 | G | A | 0.149 | -1.41E-01 | 1.30E-01 | 2.70E+00 | 2.80E+00 | 2.73E-01 | no | | intergenic_variant | MODIFIER |
| 16 | 81142362 | AX-170429193 | 0 | T | C | 0.474 | 8.13E-02 | 7.46E-02 | 2.62E+00 | 2.69E+00 | 2.73E-01 | no | | intergenic_variant | MODIFIER |

| | | | | | | | | | | | | | | | |
|---|---|---|---|---|---|---|---|---|---|---|---|---|---|---|---|
| 1 | 174847781 | AX-170432849 | 2 | G | T | 0.236 | 1.36E-01 | 1.24E-01 | 2.53E+00 | 2.59E+00 | 2.73E-01 | no | Grem2 | intron_variant | MODIFIER |
| 7 | 114563278 | AX-170492422 | 0 | C | T | 0.439 | 1.03E-01 | 9.52E-02 | 2.80E+00 | 2.86E+00 | 2.73E-01 | no | Cyp2r1 | upstream_gene_variant | MODIFIER |
| 13 | 11567562 | AX-170357587 | 1 | T | A | 0.205 | -1.16E-01 | 1.05E-01 | 2.42E+00 | 2.51E+00 | 2.73E-01 | no | Ryr2 | intron_variant | MODIFIER |
| 4 | 77297917 | AX-97846363 | 1 | A | G | 0.321 | 1.31E-01 | 1.21E-01 | 2.81E+00 | 2.87E+00 | 2.73E-01 | no | Ptprd | intron_variant,non_coding_transcript_variant | MODIFIER |
| 7 | 65701985 | AX-97834022 | 0 | C | T | 0.421 | -1.08E-01 | 9.91E-02 | 2.90E+00 | 2.99E+00 | 2.73E-01 | no | Tm2d3 | downstream_gene_variant | MODIFIER |
| 4 | 79156776 | AX-170172974 | 1 | T | A | 0.446 | -9.33E-02 | 8.48E-02 | 2.52E+00 | 2.57E+00 | 2.73E-01 | no | | intergenic_variant | MODIFIER |
| 16 | 37792420 | AX-170012844 | 0 | C | A | 0.482 | 8.90E-02 | 8.11E-02 | 2.54E+00 | 2.58E+00 | 2.73E-01 | no | Fstl1 | intron_variant | MODIFIER |
| 16 | 37821075 | AX-97793338 | 0 | T | C | 0.482 | 8.90E-02 | 8.11E-02 | 2.54E+00 | 2.58E+00 | 2.73E-01 | no | Fstl1 | intron_variant | MODIFIER |
| 16 | 37821615 | AX-170012848 | 0 | T | G | 0.482 | 8.90E-02 | 8.11E-02 | 2.54E+00 | 2.58E+00 | 2.73E-01 | no | Fstl1 | intron_variant | MODIFIER |
| 16 | 37978964 | AX-170275797 | 0 | A | G | 0.482 | 8.90E-02 | 8.11E-02 | 2.54E+00 | 2.58E+00 | 2.73E-01 | no | Gpr156 | intron_variant | MODIFIER |
| 16 | 38019377 | AX-170524663 | 0 | C | G | 0.482 | 8.90E-02 | 8.11E-02 | 2.54E+00 | 2.58E+00 | 2.73E-01 | no | | intergenic_variant | MODIFIER |
| 16 | 38237325 | AX-97817420 | 0 | G | T | 0.482 | 8.90E-02 | 8.11E-02 | 2.54E+00 | 2.58E+00 | 2.73E-01 | no | Gsk3b | intron_variant | MODIFIER |
| 16 | 38466013 | AX-170369540 | 0 | G | A | 0.482 | 8.90E-02 | 8.11E-02 | 2.54E+00 | 2.58E+00 | 2.73E-01 | no | Cd80 | intron_variant | MODIFIER |
| 16 | 38473861 | AX-97849596 | 0 | C | T | 0.482 | 8.90E-02 | 8.11E-02 | 2.54E+00 | 2.58E+00 | 2.73E-01 | no | Cd80 | splice_region_variant,intron_variant | LOW |
| 16 | 38475600 | AX-168390137 | 0 | T | C | 0.482 | 8.90E-02 | 8.11E-02 | 2.54E+00 | 2.58E+00 | 2.73E-01 | no | Cd80 | intron_variant | MODIFIER |
| 16 | 38476653 | AX-170057167 | 0 | G | T | 0.482 | 8.90E-02 | 8.11E-02 | 2.54E+00 | 2.58E+00 | 2.73E-01 | no | Cd80 | intron_variant | MODIFIER |
| 16 | 38477344 | AX-170057162 | 0 | C | T | 0.482 | 8.90E-02 | 8.11E-02 | 2.54E+00 | 2.58E+00 | 2.73E-01 | no | Cd80 | intron_variant | MODIFIER |
| 16 | 38482395 | AX-170057164 | 0 | T | C | 0.482 | 8.90E-02 | 8.11E-02 | 2.54E+00 | 2.58E+00 | 2.73E-01 | no | Cd80 | intron_variant | MODIFIER |
| 16 | 38482511 | AX-170057165 | 0 | A | G | 0.482 | 8.90E-02 | 8.11E-02 | 2.54E+00 | 2.58E+00 | 2.73E-01 | no | Cd80 | intron_variant | MODIFIER |
| 16 | 38518134 | AX-97789285 | 0 | A | G | 0.482 | 8.90E-02 | 8.11E-02 | 2.54E+00 | 2.58E+00 | 2.73E-01 | no | Timmdc1 | intron_variant | MODIFIER |
| 16 | 38531661 | AX-170433550 | 0 | C | A | 0.482 | 8.90E-02 | 8.11E-02 | 2.54E+00 | 2.58E+00 | 2.73E-01 | no | Poglut1 | intron_variant | MODIFIER |
| 16 | 38636624 | AX-97857380 | 0 | T | C | 0.482 | 8.90E-02 | 8.11E-02 | 2.54E+00 | 2.58E+00 | 2.73E-01 | no | Arhgap31 | intron_variant | MODIFIER |
| 16 | 38763411 | AX-97808834 | 0 | G | A | 0.482 | 8.90E-02 | 8.11E-02 | 2.54E+00 | 2.58E+00 | 2.73E-01 | no | B4galt4 | intron_variant | MODIFIER |
| 16 | 38816642 | AX-170012859 | 0 | A | G | 0.482 | 8.90E-02 | 8.11E-02 | 2.54E+00 | 2.58E+00 | 2.73E-01 | no | Gm15803 | downstream_gene_variant | MODIFIER |
| 16 | 38817198 | AX-170169755 | 0 | T | C | 0.482 | 8.90E-02 | 8.11E-02 | 2.54E+00 | 2.58E+00 | 2.73E-01 | no | Gm15803 | downstream_gene_variant | MODIFIER |
| 16 | 38948788 | AX-168396221 | 0 | T | C | 0.482 | 8.90E-02 | 8.11E-02 | 2.54E+00 | 2.58E+00 | 2.73E-01 | no | Igsf11 | intron_variant | MODIFIER |
| 16 | 38951069 | AX-170488661 | 0 | C | T | 0.482 | 8.90E-02 | 8.11E-02 | 2.54E+00 | 2.58E+00 | 2.73E-01 | no | Igsf11 | intron_variant | MODIFIER |
| 16 | 39003774 | AX-170354001 | 0 | T | C | 0.482 | 8.90E-02 | 8.11E-02 | 2.54E+00 | 2.58E+00 | 2.73E-01 | no | Igsf11 | intron_variant | MODIFIER |
| 16 | 39005887 | AX-170196483 | 0 | G | C | 0.482 | 8.90E-02 | 8.11E-02 | 2.54E+00 | 2.58E+00 | 2.73E-01 | no | Igsf11 | intron_variant | MODIFIER |
| 16 | 39096138 | AX-97828608 | 0 | A | C | 0.482 | 8.90E-02 | 8.11E-02 | 2.54E+00 | 2.58E+00 | 2.73E-01 | no | | intergenic_variant | MODIFIER |
| 16 | 39096837 | AX-170256120 | 0 | C | T | 0.482 | 8.90E-02 | 8.11E-02 | 2.54E+00 | 2.58E+00 | 2.73E-01 | no | | intergenic_variant | MODIFIER |
| 16 | 39097716 | AX-170302962 | 0 | T | C | 0.482 | 8.90E-02 | 8.11E-02 | 2.54E+00 | 2.58E+00 | 2.73E-01 | no | | intergenic_variant | MODIFIER |
| 16 | 39177276 | AX-170257644 | 0 | T | C | 0.482 | 8.90E-02 | 8.11E-02 | 2.54E+00 | 2.58E+00 | 2.73E-01 | no | | intergenic_variant | MODIFIER |
| 16 | 39177304 | AX-170420634 | 0 | T | C | 0.482 | 8.90E-02 | 8.11E-02 | 2.54E+00 | 2.58E+00 | 2.73E-01 | no | | intergenic_variant | MODIFIER |
| 16 | 39177382 | AX-170396786 | 0 | C | T | 0.482 | 8.90E-02 | 8.11E-02 | 2.54E+00 | 2.58E+00 | 2.73E-01 | no | | intergenic_variant | MODIFIER |
| 16 | 39200533 | AX-170327354 | 0 | C | T | 0.482 | 8.90E-02 | 8.11E-02 | 2.54E+00 | 2.58E+00 | 2.73E-01 | no | | intergenic_variant | MODIFIER |
| 16 | 39206742 | AX-170272567 | 0 | G | T | 0.482 | 8.90E-02 | 8.11E-02 | 2.54E+00 | 2.58E+00 | 2.73E-01 | no | | intergenic_variant | MODIFIER |
| 16 | 39223484 | AX-170314252 | 0 | G | A | 0.482 | 8.90E-02 | 8.11E-02 | 2.54E+00 | 2.58E+00 | 2.73E-01 | no | | intergenic_variant | MODIFIER |
| 16 | 39223575 | AX-97841036 | 0 | A | G | 0.482 | 8.90E-02 | 8.11E-02 | 2.54E+00 | 2.58E+00 | 2.73E-01 | no | | intergenic_variant | MODIFIER |
| 16 | 39242874 | AX-97837110 | 0 | T | G | 0.482 | 8.90E-02 | 8.11E-02 | 2.54E+00 | 2.58E+00 | 2.73E-01 | no | | intergenic_variant | MODIFIER |
| 16 | 39271733 | AX-170012864 | 0 | G | T | 0.482 | 8.90E-02 | 8.11E-02 | 2.54E+00 | 2.58E+00 | 2.73E-01 | no | | intergenic_variant | MODIFIER |
| 16 | 39272939 | AX-170012865 | 0 | T | C | 0.482 | 8.90E-02 | 8.11E-02 | 2.54E+00 | 2.58E+00 | 2.73E-01 | no | | intergenic_variant | MODIFIER |
| 16 | 39276928 | AX-170145538 | 0 | T | G | 0.482 | 8.90E-02 | 8.11E-02 | 2.54E+00 | 2.58E+00 | 2.73E-01 | no | | intergenic_variant | MODIFIER |
| 16 | 39302143 | AX-170012862 | 0 | T | C | 0.482 | 8.90E-02 | 8.11E-02 | 2.54E+00 | 2.58E+00 | 2.73E-01 | no | | regulatory_region_variant | MODIFIER |
| 16 | 39305465 | AX-170012866 | 0 | T | C | 0.482 | 8.90E-02 | 8.11E-02 | 2.54E+00 | 2.58E+00 | 2.73E-01 | no | | intergenic_variant | MODIFIER |
| 16 | 39306418 | AX-170355831 | 0 | T | C | 0.482 | 8.90E-02 | 8.11E-02 | 2.54E+00 | 2.58E+00 | 2.73E-01 | no | | intergenic_variant | MODIFIER |
| 16 | 39343264 | AX-97836845 | 0 | G | A | 0.482 | 8.90E-02 | 8.11E-02 | 2.54E+00 | 2.58E+00 | 2.73E-01 | no | | intergenic_variant | MODIFIER |
| 6 | 136258271 | AX-170028329 | 0 | T | C | 0.561 | 9.84E-02 | 8.80E-02 | 2.40E+00 | 2.45E+00 | 2.73E-01 | no | | intergenic_variant | MODIFIER |
| 4 | 80755560 | AX-170532153 | 1 | A | G | 0.446 | -9.32E-02 | 8.48E-02 | 2.52E+00 | 2.57E+00 | 2.73E-01 | no | | intergenic_variant | MODIFIER |
| 4 | 80073458 | AX-170023210 | 1 | A | G | 0.446 | -9.34E-02 | 8.50E-02 | 2.52E+00 | 2.57E+00 | 2.73E-01 | no | | intergenic_variant | MODIFIER |
| 5 | 50550716 | AX-170354219 | 1 | C | T | 0.509 | 1.15E-01 | 1.02E-01 | 2.35E+00 | 2.40E+00 | 2.73E-01 | no | Gm43016 | intron_variant,non_coding_transcript_variant | MODIFIER |
| 4 | 132622590 | AX-170023975 | 2 | G | A | 0.182 | 1.18E-01 | 1.08E-01 | 2.62E+00 | 2.67E+00 | 2.73E-01 | no | | regulatory_region_variant | MODIFIER |
| 4 | 128539874 | AX-170375312 | 0 | T | A | 0.167 | 1.12E-01 | 1.03E-01 | 2.63E+00 | 2.70E+00 | 2.73E-01 | no | Csmd2 | intron_variant | MODIFIER |
| 4 | 54664939 | AX-170426382 | 0 | T | C | 0.36 | -1.09E-01 | 9.99E-02 | 2.86E+00 | 2.96E+00 | 2.73E-01 | no | Gm12477 | downstream_gene_variant | MODIFIER |

| | | | | | | | | | | | | | | | |
|---|---|---|---|---|---|---|---|---|---|---|---|---|---|---|---|
| 4 | 54990334 | AX-170196757 | 0 | T | C | 0.36 | -1.09E-01 | 9.99E-02 | 2.86E+00 | 2.96E+00 | 2.73E-01 | no | Zfp462 | intron_variant | MODIFIER |
| 4 | 64048430 | AX-97798497 | 1 | A | G | 0.33 | 1.17E-01 | 1.08E-01 | 2.75E+00 | 2.83E+00 | 2.74E-01 | no | Tnc | upstream_gene_variant | MODIFIER |
| 11 | 7647735 | AX-170101390 | 1 | T | C | 0.036 | 1.45E-01 | 1.35E-01 | 2.66E+00 | 2.86E+00 | 2.74E-01 | Yes | | intergenic_variant | MODIFIER |
| 5 | 50321127 | AX-170488558 | 1 | A | G | 0.482 | 1.19E-01 | 1.06E-01 | 2.36E+00 | 2.40E+00 | 2.74E-01 | no | Gm7205 | intron_variant,non_coding_transcript_variant | MODIFIER |
| 18 | 53002236 | AX-170037815 | 0 | A | T | 0.237 | -1.14E-01 | 1.05E-01 | 2.63E+00 | 2.70E+00 | 2.74E-01 | no | | intergenic_variant | MODIFIER |
| 15 | 83135965 | AX-97816386 | 0 | T | C | 0.096 | -1.37E-01 | 1.03E-01 | 1.88E+00 | 1.96E+00 | 2.74E-01 | no | Poldip3 | intron_variant | MODIFIER |
| 15 | 83178526 | AX-97798527 | 0 | G | A | 0.096 | -1.37E-01 | 1.03E-01 | 1.88E+00 | 1.96E+00 | 2.74E-01 | no | 7530414M10Rik | upstream_gene_variant | MODIFIER |
| 15 | 83324873 | AX-97829189 | 0 | A | C | 0.096 | -1.37E-01 | 1.03E-01 | 1.88E+00 | 1.96E+00 | 2.74E-01 | no | Arfgap3 | intron_variant | MODIFIER |
| 15 | 83389653 | AX-170286044 | 0 | G | A | 0.096 | -1.37E-01 | 1.03E-01 | 1.88E+00 | 1.96E+00 | 2.74E-01 | no | Pacsin2 | intron_variant | MODIFIER |
| 15 | 83440523 | AX-97821840 | 0 | G | A | 0.096 | -1.37E-01 | 1.03E-01 | 1.88E+00 | 1.96E+00 | 2.74E-01 | no | Pacsin2 | intron_variant | MODIFIER |
| 15 | 83514154 | AX-170478171 | 0 | C | T | 0.096 | -1.37E-01 | 1.03E-01 | 1.88E+00 | 1.96E+00 | 2.74E-01 | no | Ttll1 | upstream_gene_variant | MODIFIER |
| 15 | 83983428 | AX-97782699 | 0 | C | T | 0.096 | -1.37E-01 | 1.03E-01 | 1.88E+00 | 1.96E+00 | 2.74E-01 | no | Efcab6 | intron_variant,non_coding_transcript_variant | MODIFIER |
| 15 | 84520402 | AX-170361023 | 0 | G | A | 0.096 | -1.37E-01 | 1.03E-01 | 1.88E+00 | 1.96E+00 | 2.74E-01 | no | | intergenic_variant | MODIFIER |
| 15 | 84523117 | AX-97826304 | 0 | A | G | 0.096 | -1.37E-01 | 1.03E-01 | 1.88E+00 | 1.96E+00 | 2.74E-01 | no | | intergenic_variant | MODIFIER |
| 15 | 84563633 | AX-170371482 | 0 | C | G | 0.096 | -1.37E-01 | 1.03E-01 | 1.88E+00 | 1.96E+00 | 2.74E-01 | no | | intergenic_variant | MODIFIER |
| 15 | 84563656 | AX-170362653 | 0 | T | C | 0.096 | -1.37E-01 | 1.03E-01 | 1.88E+00 | 1.96E+00 | 2.74E-01 | no | | intergenic_variant | MODIFIER |
| 15 | 84577442 | AX-170252468 | 0 | G | A | 0.096 | -1.37E-01 | 1.03E-01 | 1.88E+00 | 1.96E+00 | 2.74E-01 | no | Gm34095 | upstream_gene_variant | MODIFIER |
| 15 | 84578337 | AX-170112292 | 0 | T | C | 0.096 | -1.37E-01 | 1.03E-01 | 1.88E+00 | 1.96E+00 | 2.74E-01 | no | Gm34095 | upstream_gene_variant | MODIFIER |
| 15 | 84581829 | AX-170495894 | 0 | A | G | 0.096 | -1.37E-01 | 1.03E-01 | 1.88E+00 | 1.96E+00 | 2.74E-01 | no | Gm34095 | upstream_gene_variant | MODIFIER |
| 15 | 84582772 | AX-170178251 | 0 | G | C | 0.096 | -1.37E-01 | 1.03E-01 | 1.88E+00 | 1.96E+00 | 2.74E-01 | no | | intergenic_variant | MODIFIER |
| 15 | 84596395 | AX-170398749 | 0 | A | G | 0.096 | -1.37E-01 | 1.03E-01 | 1.88E+00 | 1.96E+00 | 2.74E-01 | no | | intergenic_variant | MODIFIER |
| 15 | 84597575 | AX-97837827 | 0 | A | G | 0.096 | -1.37E-01 | 1.03E-01 | 1.88E+00 | 1.96E+00 | 2.74E-01 | no | | intergenic_variant | MODIFIER |
| 15 | 84609771 | AX-170012151 | 0 | A | G | 0.096 | -1.37E-01 | 1.03E-01 | 1.88E+00 | 1.96E+00 | 2.74E-01 | no | | regulatory_region_variant | MODIFIER |
| 15 | 84609904 | AX-170231768 | 0 | C | A | 0.096 | -1.37E-01 | 1.03E-01 | 1.88E+00 | 1.96E+00 | 2.74E-01 | no | | regulatory_region_variant | MODIFIER |
| 15 | 84613587 | AX-170012155 | 0 | A | C | 0.096 | -1.37E-01 | 1.03E-01 | 1.88E+00 | 1.96E+00 | 2.74E-01 | no | | intergenic_variant | MODIFIER |
| 15 | 84613804 | AX-97829952 | 0 | T | C | 0.096 | -1.37E-01 | 1.03E-01 | 1.88E+00 | 1.96E+00 | 2.74E-01 | no | | intergenic_variant | MODIFIER |
| 15 | 84615326 | AX-170527107 | 0 | A | C | 0.096 | -1.37E-01 | 1.03E-01 | 1.88E+00 | 1.96E+00 | 2.74E-01 | no | | intergenic_variant | MODIFIER |
| 15 | 84654395 | AX-170012158 | 0 | T | C | 0.096 | -1.37E-01 | 1.03E-01 | 1.88E+00 | 1.96E+00 | 2.74E-01 | no | | regulatory_region_variant | MODIFIER |
| 15 | 84654676 | AX-170012156 | 0 | G | A | 0.096 | -1.37E-01 | 1.03E-01 | 1.88E+00 | 1.96E+00 | 2.74E-01 | no | | regulatory_region_variant | MODIFIER |
| 13 | 76013316 | AX-170505891 | 1 | T | C | 0.027 | -1.86E-01 | 1.68E-01 | 2.45E+00 | 2.57E+00 | 2.74E-01 | Yes | Rfesd | intron_variant | MODIFIER |
| 18 | 78332683 | AX-170158105 | 0 | C | T | 0.447 | 9.27E-02 | 8.43E-02 | 2.49E+00 | 2.57E+00 | 2.74E-01 | no | Slc14a2 | intron_variant | MODIFIER |
| 1 | 194801364 | AX-170253712 | 0 | T | C | 0.272 | 1.04E-01 | 9.24E-02 | 2.35E+00 | 2.43E+00 | 2.74E-01 | no | Plxna2 | intron_variant | MODIFIER |
| 7 | 145012842 | AX-170395776 | 1 | T | C | 0.205 | -7.92E-02 | 7.33E-02 | 2.68E+00 | 2.81E+00 | 2.75E-01 | no | | intergenic_variant | MODIFIER |
| 18 | 59166854 | AX-97834961 | 0 | T | C | 0.184 | -1.18E-01 | 1.09E-01 | 2.65E+00 | 2.71E+00 | 2.75E-01 | no | | intergenic_variant | MODIFIER |
| 10 | 101232400 | AX-97856639 | 2 | T | C | 0.291 | -9.27E-02 | 8.51E-02 | 2.58E+00 | 2.64E+00 | 2.75E-01 | no | | intergenic_variant | MODIFIER |
| 4 | 88092657 | AX-170423415 | 0 | A | G | 0.254 | 1.22E-01 | 1.11E-01 | 2.45E+00 | 2.50E+00 | 2.75E-01 | no | Focad | upstream_gene_variant | MODIFIER |
| 4 | 88098370 | AX-170132848 | 0 | C | T | 0.254 | 1.22E-01 | 1.11E-01 | 2.45E+00 | 2.50E+00 | 2.75E-01 | no | Focad | intron_variant | MODIFIER |
| 4 | 88098827 | AX-170499155 | 0 | A | G | 0.254 | 1.22E-01 | 1.11E-01 | 2.45E+00 | 2.50E+00 | 2.75E-01 | no | Focad | intron_variant | MODIFIER |
| 4 | 88128475 | AX-170069924 | 0 | T | G | 0.254 | 1.22E-01 | 1.11E-01 | 2.45E+00 | 2.50E+00 | 2.75E-01 | no | Focad | intron_variant | MODIFIER |
| 4 | 88147546 | AX-170471112 | 0 | T | C | 0.254 | 1.22E-01 | 1.11E-01 | 2.45E+00 | 2.50E+00 | 2.75E-01 | no | Focad | intron_variant | MODIFIER |
| 4 | 88914544 | AX-170102668 | 0 | G | A | 0.254 | 1.22E-01 | 1.11E-01 | 2.45E+00 | 2.50E+00 | 2.75E-01 | no | Mir31 | upstream_gene_variant | MODIFIER |
| 4 | 88914775 | AX-170069936 | 0 | T | C | 0.254 | 1.22E-01 | 1.11E-01 | 2.45E+00 | 2.50E+00 | 2.75E-01 | no | Mir31 | upstream_gene_variant | MODIFIER |
| 4 | 88914843 | AX-170495774 | 0 | C | A | 0.254 | 1.22E-01 | 1.11E-01 | 2.45E+00 | 2.50E+00 | 2.75E-01 | no | Mir31 | upstream_gene_variant | MODIFIER |
| 4 | 88916191 | AX-170324880 | 0 | C | G | 0.254 | 1.22E-01 | 1.11E-01 | 2.45E+00 | 2.50E+00 | 2.75E-01 | no | AL805899.1 | intron_variant,non_coding_transcript_variant | MODIFIER |
| 4 | 88925111 | AX-170257825 | 0 | A | T | 0.254 | 1.22E-01 | 1.11E-01 | 2.45E+00 | 2.50E+00 | 2.75E-01 | no | AL805899.1 | intron_variant,non_coding_transcript_variant | MODIFIER |
| 4 | 89113540 | AX-97856531 | 0 | A | G | 0.254 | 1.22E-01 | 1.11E-01 | 2.45E+00 | 2.50E+00 | 2.75E-01 | no | Gm12602 | intron_variant,non_coding_transcript_variant | MODIFIER |
| 4 | 90438609 | AX-170365983 | 0 | C | T | 0.254 | 1.22E-01 | 1.11E-01 | 2.45E+00 | 2.50E+00 | 2.75E-01 | no | | regulatory_region_variant | MODIFIER |
| 4 | 90447556 | AX-170095050 | 0 | A | G | 0.254 | 1.22E-01 | 1.11E-01 | 2.45E+00 | 2.50E+00 | 2.75E-01 | no | | intergenic_variant | MODIFIER |
| 4 | 90494928 | AX-170349613 | 0 | T | A | 0.254 | 1.22E-01 | 1.11E-01 | 2.45E+00 | 2.50E+00 | 2.75E-01 | no | | intergenic_variant | MODIFIER |
| 4 | 90642934 | AX-170234755 | 0 | C | T | 0.254 | 1.22E-01 | 1.11E-01 | 2.45E+00 | 2.50E+00 | 2.75E-01 | no | | intergenic_variant | MODIFIER |
| 1 | 20542727 | AX-170222962 | 0 | G | C | 0.404 | 9.23E-02 | 8.45E-02 | 2.55E+00 | 2.60E+00 | 2.75E-01 | no | Pkhd1 | intron_variant | MODIFIER |
| 5 | 51338915 | AX-170300752 | 1 | T | A | 0.5 | 1.16E-01 | 1.04E-01 | 2.41E+00 | 2.45E+00 | 2.75E-01 | no | | intergenic_variant | MODIFIER |
| 1 | 171236929 | AX-170177426 | 0 | C | T | 0.228 | 9.99E-02 | 9.27E-02 | 2.73E+00 | 2.88E+00 | 2.75E-01 | no | Ndufs2 | intron_variant | MODIFIER |
| 3 | 22826608 | AX-170303934 | 0 | T | C | 0.167 | -1.07E-01 | 9.85E-02 | 2.56E+00 | 2.64E+00 | 2.75E-01 | no | | intergenic_variant | MODIFIER |

| | | | | | | | | | | | | | | | |
|---|---|---|---|---|---|---|---|---|---|---|---|---|---|---|---|
| 8 | 21534493 | AX-170204750 | 2 | G | A | 0.355 | 1.12E-01 | 1.04E-01 | 2.75E+00 | 2.89E+00 | 2.75E-01 | no | Defa32 | downstream_gene_variant | MODIFIER |
| 4 | 55526631 | AX-170179262 | 1 | A | G | 0.357 | -1.07E-01 | 9.86E-02 | 2.89E+00 | 3.01E+00 | 2.75E-01 | no | Klf4 | downstream_gene_variant | MODIFIER |
| 4 | 130061408 | AX-170023945 | 1 | A | G | 0.161 | 1.28E-01 | 1.17E-01 | 2.56E+00 | 2.61E+00 | 2.75E-01 | no | Col16a1 | intron_variant | MODIFIER |
| 4 | 73822998 | AX-97804674 | 1 | A | G | 0.446 | -9.29E-02 | 8.48E-02 | 2.52E+00 | 2.57E+00 | 2.75E-01 | no | | intergenic_variant | MODIFIER |
| 5 | 50857290 | AX-170433346 | 0 | C | T | 0.5 | 1.15E-01 | 1.04E-01 | 2.43E+00 | 2.48E+00 | 2.75E-01 | no | | intergenic_variant | MODIFIER |
| 12 | 58471398 | AX-170191095 | 0 | C | T | 0.079 | -1.54E-01 | 1.44E-01 | 2.75E+00 | 2.93E+00 | 2.75E-01 | no | | intergenic_variant | MODIFIER |
| 4 | 49817660 | AX-97785931 | 0 | G | A | 0.36 | -1.00E-01 | 9.31E-02 | 2.75E+00 | 2.85E+00 | 2.75E-01 | no | Grin3a | intron_variant | MODIFIER |
| 4 | 117910431 | AX-97831484 | 1 | C | T | 0.241 | 1.14E-01 | 1.05E-01 | 2.56E+00 | 2.63E+00 | 2.75E-01 | no | Ipo13 | intron_variant | MODIFIER |
| 7 | 99630621 | AX-170029616 | 0 | C | A | 0.36 | -1.12E-01 | 1.03E-01 | 2.86E+00 | 2.95E+00 | 2.76E-01 | no | Tpbgl | upstream_gene_variant | MODIFIER |
| 7 | 99685485 | AX-170370016 | 0 | C | T | 0.36 | -1.12E-01 | 1.03E-01 | 2.86E+00 | 2.95E+00 | 2.76E-01 | no | Slco2b1 | missense_variant | MODERATE |
| 7 | 100175883 | AX-170439483 | 0 | T | C | 0.36 | -1.12E-01 | 1.03E-01 | 2.86E+00 | 2.95E+00 | 2.76E-01 | no | Kcne3 | upstream_gene_variant | MODIFIER |
| 7 | 100178085 | AX-170460283 | 0 | A | C | 0.36 | -1.12E-01 | 1.03E-01 | 2.86E+00 | 2.95E+00 | 2.76E-01 | no | Kcne3 | upstream_gene_variant | MODIFIER |
| 7 | 100178704 | AX-170100155 | 0 | C | T | 0.36 | -1.12E-01 | 1.03E-01 | 2.86E+00 | 2.95E+00 | 2.76E-01 | no | Kcne3 | 5_prime_UTR_variant | MODIFIER |
| 7 | 100294884 | AX-170212075 | 0 | C | T | 0.36 | -1.12E-01 | 1.03E-01 | 2.86E+00 | 2.95E+00 | 2.76E-01 | no | P4ha3 | intron_variant | MODIFIER |
| 7 | 100297978 | AX-170335349 | 0 | A | C | 0.36 | -1.12E-01 | 1.03E-01 | 2.86E+00 | 2.95E+00 | 2.76E-01 | no | P4ha3 | synonymous_variant | LOW |
| 7 | 100298377 | AX-97855011 | 0 | T | C | 0.36 | -1.12E-01 | 1.03E-01 | 2.86E+00 | 2.95E+00 | 2.76E-01 | no | P4ha3 | intron_variant | MODIFIER |
| 7 | 100303961 | AX-170237780 | 0 | T | A | 0.36 | -1.12E-01 | 1.03E-01 | 2.86E+00 | 2.95E+00 | 2.76E-01 | no | P4ha3 | intron_variant | MODIFIER |
| 7 | 100331589 | AX-170497922 | 0 | C | T | 0.36 | -1.12E-01 | 1.03E-01 | 2.86E+00 | 2.95E+00 | 2.76E-01 | no | Ppme1 | intron_variant | MODIFIER |
| 7 | 100345506 | AX-170324329 | 0 | T | A | 0.36 | -1.12E-01 | 1.03E-01 | 2.86E+00 | 2.95E+00 | 2.76E-01 | no | Ppme1 | intron_variant | MODIFIER |
| 7 | 100365194 | AX-97829513 | 0 | G | A | 0.36 | -1.12E-01 | 1.03E-01 | 2.86E+00 | 2.95E+00 | 2.76E-01 | no | Ppme1 | intron_variant | MODIFIER |
| 7 | 100426367 | AX-170491728 | 0 | C | A | 0.36 | -1.12E-01 | 1.03E-01 | 2.86E+00 | 2.95E+00 | 2.76E-01 | no | C2cd3 | intron_variant | MODIFIER |
| 7 | 100430801 | AX-168387855 | 0 | T | G | 0.36 | -1.12E-01 | 1.03E-01 | 2.86E+00 | 2.95E+00 | 2.76E-01 | no | C2cd3 | synonymous_variant | LOW |
| 7 | 100432682 | AX-170388025 | 0 | G | A | 0.36 | -1.12E-01 | 1.03E-01 | 2.86E+00 | 2.95E+00 | 2.76E-01 | no | C2cd3 | intron_variant | MODIFIER |
| 7 | 100437092 | AX-170333664 | 0 | G | A | 0.36 | -1.12E-01 | 1.03E-01 | 2.86E+00 | 2.95E+00 | 2.76E-01 | no | C2cd3 | intron_variant | MODIFIER |
| 7 | 100453246 | AX-170437408 | 0 | T | C | 0.36 | -1.12E-01 | 1.03E-01 | 2.86E+00 | 2.95E+00 | 2.76E-01 | no | C2cd3 | intron_variant | MODIFIER |
| 7 | 100463914 | AX-170442126 | 0 | T | G | 0.36 | -1.12E-01 | 1.03E-01 | 2.86E+00 | 2.95E+00 | 2.76E-01 | no | C2cd3 | intron_variant | MODIFIER |
| 7 | 100465284 | AX-170467889 | 0 | G | T | 0.36 | -1.12E-01 | 1.03E-01 | 2.86E+00 | 2.95E+00 | 2.76E-01 | no | C2cd3 | intron_variant | MODIFIER |
| 7 | 100518010 | AX-170339809 | 0 | A | G | 0.36 | -1.12E-01 | 1.03E-01 | 2.86E+00 | 2.95E+00 | 2.76E-01 | no | Dnajb13 | upstream_gene_variant | MODIFIER |
| 7 | 100535024 | AX-170131383 | 0 | T | C | 0.36 | -1.12E-01 | 1.03E-01 | 2.86E+00 | 2.95E+00 | 2.76E-01 | no | Coa4 | upstream_gene_variant | MODIFIER |
| 7 | 100535094 | AX-170493320 | 0 | T | C | 0.36 | -1.12E-01 | 1.03E-01 | 2.86E+00 | 2.95E+00 | 2.76E-01 | no | Coa4 | upstream_gene_variant | MODIFIER |
| 7 | 100536948 | AX-97803971 | 0 | A | C | 0.36 | -1.12E-01 | 1.03E-01 | 2.86E+00 | 2.95E+00 | 2.76E-01 | no | Coa4 | upstream_gene_variant | MODIFIER |
| 7 | 100620106 | AX-170419788 | 0 | A | G | 0.36 | -1.12E-01 | 1.03E-01 | 2.86E+00 | 2.95E+00 | 2.76E-01 | no | Rab6a | intron_variant | MODIFIER |
| 7 | 100633823 | AX-97803455 | 0 | G | A | 0.36 | -1.12E-01 | 1.03E-01 | 2.86E+00 | 2.95E+00 | 2.76E-01 | no | Rab6a | intron_variant | MODIFIER |
| 7 | 100637489 | AX-170210598 | 0 | C | T | 0.36 | -1.12E-01 | 1.03E-01 | 2.86E+00 | 2.95E+00 | 2.76E-01 | no | Rab6a | intron_variant | MODIFIER |
| 7 | 100645017 | AX-170029632 | 0 | T | G | 0.36 | -1.12E-01 | 1.03E-01 | 2.86E+00 | 2.95E+00 | 2.76E-01 | no | Rab6a | downstream_gene_variant | MODIFIER |
| 7 | 100664297 | AX-170162372 | 0 | C | G | 0.36 | -1.12E-01 | 1.03E-01 | 2.86E+00 | 2.95E+00 | 2.76E-01 | no | Plekhb1 | upstream_gene_variant | MODIFIER |
| 7 | 100673312 | AX-170029631 | 0 | T | C | 0.36 | -1.12E-01 | 1.03E-01 | 2.86E+00 | 2.95E+00 | 2.76E-01 | no | Gm35082 | upstream_gene_variant | MODIFIER |
| 7 | 100676692 | AX-170029630 | 0 | G | T | 0.36 | -1.12E-01 | 1.03E-01 | 2.86E+00 | 2.95E+00 | 2.76E-01 | no | Gm35082 | intron_variant,non_coding_transcript_variant | MODIFIER |
| 7 | 100688458 | AX-170355495 | 0 | T | C | 0.36 | -1.12E-01 | 1.03E-01 | 2.86E+00 | 2.95E+00 | 2.76E-01 | no | | intergenic_variant | MODIFIER |
| 7 | 100712791 | AX-170029633 | 0 | A | G | 0.36 | -1.12E-01 | 1.03E-01 | 2.86E+00 | 2.95E+00 | 2.76E-01 | no | Fam168a | intron_variant | MODIFIER |
| 7 | 100713704 | AX-170518817 | 0 | T | C | 0.36 | -1.12E-01 | 1.03E-01 | 2.86E+00 | 2.95E+00 | 2.76E-01 | no | Fam168a | intron_variant | MODIFIER |
| 7 | 100726843 | AX-170480861 | 0 | A | G | 0.36 | -1.12E-01 | 1.03E-01 | 2.86E+00 | 2.95E+00 | 2.76E-01 | no | Fam168a | intron_variant | MODIFIER |
| 7 | 100735852 | AX-97803453 | 0 | G | A | 0.36 | -1.12E-01 | 1.03E-01 | 2.86E+00 | 2.95E+00 | 2.76E-01 | no | Fam168a | intron_variant | MODIFIER |
| 7 | 100744426 | AX-170356186 | 0 | T | A | 0.36 | -1.12E-01 | 1.03E-01 | 2.86E+00 | 2.95E+00 | 2.76E-01 | no | Fam168a | intron_variant | MODIFIER |
| 7 | 100912996 | AX-170078243 | 0 | T | C | 0.36 | -1.12E-01 | 1.03E-01 | 2.86E+00 | 2.95E+00 | 2.76E-01 | no | Arhgef17 | intron_variant | MODIFIER |
| 7 | 100924136 | AX-170451337 | 0 | G | A | 0.36 | -1.12E-01 | 1.03E-01 | 2.86E+00 | 2.95E+00 | 2.76E-01 | no | Arhgef17 | intron_variant | MODIFIER |
| 7 | 100940710 | AX-170262506 | 0 | A | G | 0.36 | -1.12E-01 | 1.03E-01 | 2.86E+00 | 2.95E+00 | 2.76E-01 | no | P2ry6 | intron_variant | MODIFIER |
| 7 | 100940848 | AX-170369429 | 0 | A | G | 0.36 | -1.12E-01 | 1.03E-01 | 2.86E+00 | 2.95E+00 | 2.76E-01 | no | P2ry6 | intron_variant | MODIFIER |
| 7 | 100974347 | AX-170252275 | 0 | A | G | 0.36 | -1.12E-01 | 1.03E-01 | 2.86E+00 | 2.95E+00 | 2.76E-01 | no | P2ry6 | intron_variant,non_coding_transcript_variant | MODIFIER |
| 7 | 101114536 | AX-168419977 | 0 | T | C | 0.36 | -1.12E-01 | 1.03E-01 | 2.86E+00 | 2.95E+00 | 2.76E-01 | no | Fchsd2 | intron_variant | MODIFIER |
| 7 | 101644614 | AX-170112288 | 0 | T | C | 0.36 | -1.12E-01 | 1.03E-01 | 2.86E+00 | 2.95E+00 | 2.76E-01 | no | | intergenic_variant | MODIFIER |
| 18 | 59292201 | AX-170015794 | 0 | C | T | 0.184 | -1.30E-01 | 1.20E-01 | 2.69E+00 | 2.75E+00 | 2.76E-01 | no | Chsy3 | intron_variant | MODIFIER |
| 18 | 63974909 | AX-170219799 | 0 | C | T | 0.184 | -1.30E-01 | 1.20E-01 | 2.69E+00 | 2.75E+00 | 2.76E-01 | no | Wdr7 | intron_variant | MODIFIER |
| 18 | 65085116 | AX-170200273 | 0 | T | A | 0.184 | -1.30E-01 | 1.20E-01 | 2.69E+00 | 2.75E+00 | 2.76E-01 | no | Nedd4l | intron_variant | MODIFIER |

| | | | | | | | | | | | | | | | |
|---|---|---|---|---|---|---|---|---|---|---|---|---|---|---|---|
| 1 | 76147799 | AX-170450607 | 0 | G | A | 0.132 | 1.93E-01 | 1.79E-01 | 2.77E+00 | 2.89E+00 | 2.76E-01 | Yes | | intergenic_variant | MODIFIER |
| 16 | 45199848 | AX-170423668 | 0 | A | G | 0.5 | 8.29E-02 | 7.64E-02 | 2.58E+00 | 2.65E+00 | 2.76E-01 | no | | intergenic_variant | MODIFIER |
| 3 | 14849390 | AX-170349477 | 1 | G | A | 0.429 | -7.07E-02 | 6.56E-02 | 2.82E+00 | 2.97E+00 | 2.76E-01 | no | | intergenic_variant | MODIFIER |
| 4 | 48716499 | AX-170240419 | 0 | A | G | 0.36 | -1.01E-01 | 9.36E-02 | 2.80E+00 | 2.91E+00 | 2.76E-01 | no | | regulatory_region_variant | MODIFIER |
| 4 | 51210522 | AX-170128908 | 0 | C | T | 0.36 | -1.01E-01 | 9.36E-02 | 2.80E+00 | 2.91E+00 | 2.76E-01 | no | | intergenic_variant | MODIFIER |
| 1 | 172952599 | AX-170515311 | 0 | A | T | 0.237 | 1.29E-01 | 1.18E-01 | 2.54E+00 | 2.61E+00 | 2.76E-01 | no | Olfr16 | upstream_gene_variant | MODIFIER |
| 4 | 79492213 | AX-170023192 | 1 | A | C | 0.446 | -9.28E-02 | 8.49E-02 | 2.52E+00 | 2.57E+00 | 2.76E-01 | no | | intergenic_variant | MODIFIER |
| 3 | 24106545 | AX-170415899 | 1 | G | T | 0.152 | -1.10E-01 | 1.00E-01 | 2.46E+00 | 2.54E+00 | 2.76E-01 | no | Naaladl2 | intron_variant | MODIFIER |
| 9 | 6562976 | AX-170196634 | 0 | C | T | 0.482 | 9.14E-02 | 8.34E-02 | 2.48E+00 | 2.57E+00 | 2.76E-01 | no | | intergenic_variant | MODIFIER |
| 16 | 67381693 | AX-97837214 | 2 | A | C | 0.464 | 8.23E-02 | 7.58E-02 | 2.57E+00 | 2.65E+00 | 2.76E-01 | no | Cadm2 | intron_variant | MODIFIER |
| 18 | 67980512 | AX-170240415 | 0 | A | G | 0.175 | -1.34E-01 | 1.24E-01 | 2.67E+00 | 2.72E+00 | 2.77E-01 | no | Ldlrad4 | intron_variant | MODIFIER |
| 18 | 67986468 | AX-170088367 | 0 | G | A | 0.175 | -1.34E-01 | 1.24E-01 | 2.67E+00 | 2.72E+00 | 2.77E-01 | no | Ldlrad4 | intron_variant | MODIFIER |
| 18 | 68081403 | AX-170383796 | 0 | C | T | 0.175 | -1.34E-01 | 1.24E-01 | 2.67E+00 | 2.72E+00 | 2.77E-01 | no | Ldlrad4 | intron_variant | MODIFIER |
| 18 | 68087073 | AX-170088368 | 0 | G | A | 0.175 | -1.34E-01 | 1.24E-01 | 2.67E+00 | 2.72E+00 | 2.77E-01 | no | Ldlrad4 | intron_variant | MODIFIER |
| 18 | 68106767 | AX-170382543 | 0 | A | G | 0.175 | -1.34E-01 | 1.24E-01 | 2.67E+00 | 2.72E+00 | 2.77E-01 | no | Ldlrad4 | intron_variant | MODIFIER |
| 18 | 68106884 | AX-170297302 | 0 | A | G | 0.175 | -1.34E-01 | 1.24E-01 | 2.67E+00 | 2.72E+00 | 2.77E-01 | no | Ldlrad4 | intron_variant | MODIFIER |
| 18 | 68110526 | AX-97837339 | 0 | T | C | 0.175 | -1.34E-01 | 1.24E-01 | 2.67E+00 | 2.72E+00 | 2.77E-01 | no | Ldlrad4 | intron_variant | MODIFIER |
| 18 | 68115397 | AX-170277225 | 0 | G | T | 0.175 | -1.34E-01 | 1.24E-01 | 2.67E+00 | 2.72E+00 | 2.77E-01 | no | Ldlrad4 | intron_variant | MODIFIER |
| 18 | 68119488 | AX-170163788 | 0 | G | A | 0.175 | -1.34E-01 | 1.24E-01 | 2.67E+00 | 2.72E+00 | 2.77E-01 | no | Ldlrad4 | intron_variant | MODIFIER |
| 18 | 68133675 | AX-170271988 | 0 | T | C | 0.175 | -1.34E-01 | 1.24E-01 | 2.67E+00 | 2.72E+00 | 2.77E-01 | no | Ldlrad4 | intron_variant | MODIFIER |
| 18 | 68136517 | AX-170364002 | 0 | T | C | 0.175 | -1.34E-01 | 1.24E-01 | 2.67E+00 | 2.72E+00 | 2.77E-01 | no | Ldlrad4 | intron_variant | MODIFIER |
| 18 | 68137055 | AX-170136584 | 0 | T | C | 0.175 | -1.34E-01 | 1.24E-01 | 2.67E+00 | 2.72E+00 | 2.77E-01 | no | Ldlrad4 | intron_variant | MODIFIER |
| 18 | 68144435 | AX-170453780 | 0 | G | C | 0.175 | -1.34E-01 | 1.24E-01 | 2.67E+00 | 2.72E+00 | 2.77E-01 | no | Ldlrad4 | intron_variant | MODIFIER |
| 18 | 68147669 | AX-170228467 | 0 | G | C | 0.175 | -1.34E-01 | 1.24E-01 | 2.67E+00 | 2.72E+00 | 2.77E-01 | no | Ldlrad4 | intron_variant | MODIFIER |
| 18 | 68148911 | AX-170360770 | 0 | T | A | 0.175 | -1.34E-01 | 1.24E-01 | 2.67E+00 | 2.72E+00 | 2.77E-01 | no | Ldlrad4 | intron_variant | MODIFIER |
| 18 | 68154753 | AX-170450070 | 0 | A | G | 0.175 | -1.34E-01 | 1.24E-01 | 2.67E+00 | 2.72E+00 | 2.77E-01 | no | Ldlrad4 | intron_variant | MODIFIER |
| 18 | 68155446 | AX-170470914 | 0 | C | T | 0.175 | -1.34E-01 | 1.24E-01 | 2.67E+00 | 2.72E+00 | 2.77E-01 | no | Ldlrad4 | intron_variant | MODIFIER |
| 18 | 68155713 | AX-170213165 | 0 | A | G | 0.175 | -1.34E-01 | 1.24E-01 | 2.67E+00 | 2.72E+00 | 2.77E-01 | no | Ldlrad4 | intron_variant | MODIFIER |
| 18 | 68157021 | AX-170190418 | 0 | T | A | 0.175 | -1.34E-01 | 1.24E-01 | 2.67E+00 | 2.72E+00 | 2.77E-01 | no | Ldlrad4 | intron_variant | MODIFIER |
| 18 | 68157231 | AX-170061191 | 0 | T | C | 0.175 | -1.34E-01 | 1.24E-01 | 2.67E+00 | 2.72E+00 | 2.77E-01 | no | Ldlrad4 | intron_variant | MODIFIER |
| 18 | 68168778 | AX-170491014 | 0 | A | C | 0.175 | -1.34E-01 | 1.24E-01 | 2.67E+00 | 2.72E+00 | 2.77E-01 | no | Ldlrad4 | intron_variant | MODIFIER |
| 18 | 68175718 | AX-170189147 | 0 | A | G | 0.175 | -1.34E-01 | 1.24E-01 | 2.67E+00 | 2.72E+00 | 2.77E-01 | no | Ldlrad4 | intron_variant | MODIFIER |
| 18 | 68234368 | AX-170486628 | 0 | T | C | 0.175 | -1.34E-01 | 1.24E-01 | 2.67E+00 | 2.72E+00 | 2.77E-01 | no | Ldlrad4 | intron_variant | MODIFIER |
| 18 | 68571840 | AX-97791381 | 0 | A | G | 0.175 | -1.34E-01 | 1.24E-01 | 2.67E+00 | 2.72E+00 | 2.77E-01 | no | AC147020.2 | upstream_gene_variant | MODIFIER |
| 18 | 68586290 | AX-170147037 | 0 | G | A | 0.175 | -1.34E-01 | 1.24E-01 | 2.67E+00 | 2.72E+00 | 2.77E-01 | no | AC147020.3 | non_coding_transcript_exon_variant | MODIFIER |
| 18 | 68603409 | AX-170061198 | 0 | A | G | 0.175 | -1.34E-01 | 1.24E-01 | 2.67E+00 | 2.72E+00 | 2.77E-01 | no | | intergenic_variant | MODIFIER |
| 18 | 68626615 | AX-97851084 | 0 | T | G | 0.175 | -1.34E-01 | 1.24E-01 | 2.67E+00 | 2.72E+00 | 2.77E-01 | no | | intergenic_variant | MODIFIER |
| 18 | 68676255 | AX-97835046 | 0 | G | A | 0.175 | -1.34E-01 | 1.24E-01 | 2.67E+00 | 2.72E+00 | 2.77E-01 | no | | intergenic_variant | MODIFIER |
| 18 | 68698505 | AX-170528323 | 0 | T | C | 0.175 | -1.34E-01 | 1.24E-01 | 2.67E+00 | 2.72E+00 | 2.77E-01 | no | | intergenic_variant | MODIFIER |
| 18 | 68699800 | AX-170510074 | 0 | G | A | 0.175 | -1.34E-01 | 1.24E-01 | 2.67E+00 | 2.72E+00 | 2.77E-01 | no | | intergenic_variant | MODIFIER |
| 18 | 68700558 | AX-97840668 | 0 | C | A | 0.175 | -1.34E-01 | 1.24E-01 | 2.67E+00 | 2.72E+00 | 2.77E-01 | no | | intergenic_variant | MODIFIER |
| 18 | 68707658 | AX-170264102 | 0 | A | G | 0.175 | -1.34E-01 | 1.24E-01 | 2.67E+00 | 2.72E+00 | 2.77E-01 | no | | intergenic_variant | MODIFIER |
| 18 | 68744326 | AX-170265745 | 0 | T | C | 0.175 | -1.34E-01 | 1.24E-01 | 2.67E+00 | 2.72E+00 | 2.77E-01 | no | Mir6356 | downstream_gene_variant | MODIFIER |
| 18 | 68769099 | AX-170190110 | 0 | G | A | 0.175 | -1.34E-01 | 1.24E-01 | 2.67E+00 | 2.72E+00 | 2.77E-01 | no | | intergenic_variant | MODIFIER |
| 18 | 68769753 | AX-97835043 | 0 | G | A | 0.175 | -1.34E-01 | 1.24E-01 | 2.67E+00 | 2.72E+00 | 2.77E-01 | no | | intergenic_variant | MODIFIER |
| 18 | 68840850 | AX-170261173 | 0 | T | G | 0.175 | -1.34E-01 | 1.24E-01 | 2.67E+00 | 2.72E+00 | 2.77E-01 | no | | intergenic_variant | MODIFIER |
| 18 | 68841246 | AX-170242900 | 0 | C | T | 0.175 | -1.34E-01 | 1.24E-01 | 2.67E+00 | 2.72E+00 | 2.77E-01 | no | | regulatory_region_variant | MODIFIER |
| 18 | 68857094 | AX-170262905 | 0 | C | T | 0.175 | -1.34E-01 | 1.24E-01 | 2.67E+00 | 2.72E+00 | 2.77E-01 | no | | intergenic_variant | MODIFIER |
| 18 | 68858866 | AX-170407772 | 0 | A | C | 0.175 | -1.34E-01 | 1.24E-01 | 2.67E+00 | 2.72E+00 | 2.77E-01 | no | | intergenic_variant | MODIFIER |
| 18 | 68890046 | AX-170423675 | 0 | C | G | 0.175 | -1.34E-01 | 1.24E-01 | 2.67E+00 | 2.72E+00 | 2.77E-01 | no | 4930546C10Rik | intron_variant,non_coding_transcript_variant | MODIFIER |
| 18 | 68906654 | AX-97847650 | 0 | C | T | 0.175 | -1.34E-01 | 1.24E-01 | 2.67E+00 | 2.72E+00 | 2.77E-01 | no | 4930546C10Rik | intron_variant,non_coding_transcript_variant | MODIFIER |
| 18 | 68952236 | AX-168397499 | 0 | T | G | 0.175 | -1.34E-01 | 1.24E-01 | 2.67E+00 | 2.72E+00 | 2.77E-01 | no | 4930546C10Rik | upstream_gene_variant | MODIFIER |
| 18 | 68960340 | AX-170155106 | 0 | G | A | 0.175 | -1.34E-01 | 1.24E-01 | 2.67E+00 | 2.72E+00 | 2.77E-01 | no | | intergenic_variant | MODIFIER |
| 18 | 68960452 | AX-170404669 | 0 | C | T | 0.175 | -1.34E-01 | 1.24E-01 | 2.67E+00 | 2.72E+00 | 2.77E-01 | no | | intergenic_variant | MODIFIER |

| | | | | | | | | | | | | | | | |
|---|---|---|---|---|---|---|---|---|---|---|---|---|---|---|---|
| 18 | 68961456 | AX-170338258 | 0 | A | C | 0.175 | -1.34E-01 | 1.24E-01 | 2.67E+00 | 2.72E+00 | 2.77E-01 | no | | intergenic_variant | MODIFIER |
| 18 | 68961471 | AX-170523486 | 0 | A | G | 0.175 | -1.34E-01 | 1.24E-01 | 2.67E+00 | 2.72E+00 | 2.77E-01 | no | | intergenic_variant | MODIFIER |
| 18 | 68964326 | AX-170523421 | 0 | A | G | 0.175 | -1.34E-01 | 1.24E-01 | 2.67E+00 | 2.72E+00 | 2.77E-01 | no | | intergenic_variant | MODIFIER |
| 18 | 68973145 | AX-97782934 | 0 | A | G | 0.175 | -1.34E-01 | 1.24E-01 | 2.67E+00 | 2.72E+00 | 2.77E-01 | no | | intergenic_variant | MODIFIER |
| 4 | 71583622 | AX-170429306 | 0 | G | C | 0.447 | -9.27E-02 | 8.49E-02 | 2.52E+00 | 2.57E+00 | 2.77E-01 | no | | intergenic_variant | MODIFIER |
| 4 | 71585937 | AX-170379221 | 0 | A | G | 0.447 | -9.27E-02 | 8.49E-02 | 2.52E+00 | 2.57E+00 | 2.77E-01 | no | | intergenic_variant | MODIFIER |
| 4 | 71658628 | AX-170526796 | 0 | A | G | 0.447 | -9.27E-02 | 8.49E-02 | 2.52E+00 | 2.57E+00 | 2.77E-01 | no | | intergenic_variant | MODIFIER |
| 4 | 71696216 | AX-170502834 | 0 | A | G | 0.447 | -9.27E-02 | 8.49E-02 | 2.52E+00 | 2.57E+00 | 2.77E-01 | no | | intergenic_variant | MODIFIER |
| 4 | 71734620 | AX-170069650 | 0 | A | G | 0.447 | -9.27E-02 | 8.49E-02 | 2.52E+00 | 2.57E+00 | 2.77E-01 | no | | intergenic_variant | MODIFIER |
| 4 | 71936079 | AX-170159040 | 0 | G | C | 0.447 | -9.27E-02 | 8.49E-02 | 2.52E+00 | 2.57E+00 | 2.77E-01 | no | | intergenic_variant | MODIFIER |
| 4 | 71969908 | AX-170477522 | 0 | A | G | 0.447 | -9.27E-02 | 8.49E-02 | 2.52E+00 | 2.57E+00 | 2.77E-01 | no | | intergenic_variant | MODIFIER |
| 4 | 72014335 | AX-97800715 | 0 | G | A | 0.447 | -9.27E-02 | 8.49E-02 | 2.52E+00 | 2.57E+00 | 2.77E-01 | no | | regulatory_region_variant | MODIFIER |
| 4 | 72015022 | AX-170069653 | 0 | A | G | 0.447 | -9.27E-02 | 8.49E-02 | 2.52E+00 | 2.57E+00 | 2.77E-01 | no | | regulatory_region_variant | MODIFIER |
| 4 | 72129750 | AX-170254044 | 0 | C | T | 0.447 | -9.27E-02 | 8.49E-02 | 2.52E+00 | 2.57E+00 | 2.77E-01 | no | Tle1 | intron_variant | MODIFIER |
| 4 | 72138128 | AX-170069659 | 0 | C | T | 0.447 | -9.27E-02 | 8.49E-02 | 2.52E+00 | 2.57E+00 | 2.77E-01 | no | Tle1 | intron_variant | MODIFIER |
| 4 | 72166364 | AX-97836105 | 0 | G | A | 0.447 | -9.27E-02 | 8.49E-02 | 2.52E+00 | 2.57E+00 | 2.77E-01 | no | Tle1 | intron_variant | MODIFIER |
| 4 | 72177133 | AX-170366765 | 0 | T | C | 0.447 | -9.27E-02 | 8.49E-02 | 2.52E+00 | 2.57E+00 | 2.77E-01 | no | Tle1 | intron_variant | MODIFIER |
| 4 | 72189359 | AX-97806150 | 0 | A | G | 0.447 | -9.27E-02 | 8.49E-02 | 2.52E+00 | 2.57E+00 | 2.77E-01 | no | Tle1 | intron_variant | MODIFIER |
| 4 | 72232440 | AX-168402027 | 0 | T | C | 0.447 | -9.27E-02 | 8.49E-02 | 2.52E+00 | 2.57E+00 | 2.77E-01 | no | | intergenic_variant | MODIFIER |
| 4 | 72232669 | AX-170157749 | 0 | A | T | 0.447 | -9.27E-02 | 8.49E-02 | 2.52E+00 | 2.57E+00 | 2.77E-01 | no | | intergenic_variant | MODIFIER |
| 4 | 72232692 | AX-170239282 | 0 | C | T | 0.447 | -9.27E-02 | 8.49E-02 | 2.52E+00 | 2.57E+00 | 2.77E-01 | no | | intergenic_variant | MODIFIER |
| 4 | 72251444 | AX-170254745 | 0 | A | T | 0.447 | -9.27E-02 | 8.49E-02 | 2.52E+00 | 2.57E+00 | 2.77E-01 | no | | intergenic_variant | MODIFIER |
| 4 | 72557794 | AX-170346024 | 0 | G | A | 0.447 | -9.27E-02 | 8.49E-02 | 2.52E+00 | 2.57E+00 | 2.77E-01 | no | | intergenic_variant | MODIFIER |
| 4 | 72613878 | AX-170393316 | 0 | T | C | 0.447 | -9.27E-02 | 8.49E-02 | 2.52E+00 | 2.57E+00 | 2.77E-01 | no | | intergenic_variant | MODIFIER |
| 4 | 72626034 | AX-170212514 | 0 | A | G | 0.447 | -9.27E-02 | 8.49E-02 | 2.52E+00 | 2.57E+00 | 2.77E-01 | no | | intergenic_variant | MODIFIER |
| 4 | 72690959 | AX-170323701 | 0 | G | A | 0.447 | -9.27E-02 | 8.49E-02 | 2.52E+00 | 2.57E+00 | 2.77E-01 | no | | intergenic_variant | MODIFIER |
| 4 | 72788085 | AX-170069669 | 0 | T | G | 0.447 | -9.27E-02 | 8.49E-02 | 2.52E+00 | 2.57E+00 | 2.77E-01 | no | | intergenic_variant | MODIFIER |
| 4 | 72995863 | AX-170259155 | 0 | T | C | 0.447 | -9.27E-02 | 8.49E-02 | 2.52E+00 | 2.57E+00 | 2.77E-01 | no | | intergenic_variant | MODIFIER |
| 4 | 73173681 | AX-170394416 | 0 | G | A | 0.447 | -9.27E-02 | 8.49E-02 | 2.52E+00 | 2.57E+00 | 2.77E-01 | no | | intergenic_variant | MODIFIER |
| 4 | 73202099 | AX-97844442 | 0 | T | C | 0.447 | -9.27E-02 | 8.49E-02 | 2.52E+00 | 2.57E+00 | 2.77E-01 | no | | intergenic_variant | MODIFIER |
| 4 | 73241865 | AX-170222400 | 0 | T | C | 0.447 | -9.27E-02 | 8.49E-02 | 2.52E+00 | 2.57E+00 | 2.77E-01 | no | | intergenic_variant | MODIFIER |
| 4 | 73942950 | AX-97793088 | 0 | T | C | 0.447 | -9.27E-02 | 8.49E-02 | 2.52E+00 | 2.57E+00 | 2.77E-01 | no | 2310002L09Rik | missense_variant | MODERATE |
| 4 | 73943120 | AX-170282416 | 0 | A | G | 0.447 | -9.27E-02 | 8.49E-02 | 2.52E+00 | 2.57E+00 | 2.77E-01 | no | 2310002L09Rik | missense_variant | MODERATE |
| 4 | 74006678 | AX-97841922 | 0 | G | T | 0.447 | -9.27E-02 | 8.49E-02 | 2.52E+00 | 2.57E+00 | 2.77E-01 | no | | intergenic_variant | MODIFIER |
| 4 | 74053133 | AX-170226608 | 0 | A | C | 0.447 | -9.27E-02 | 8.49E-02 | 2.52E+00 | 2.57E+00 | 2.77E-01 | no | Frmd3 | intron_variant | MODIFIER |
| 4 | 74083946 | AX-97803218 | 0 | A | G | 0.447 | -9.27E-02 | 8.49E-02 | 2.52E+00 | 2.57E+00 | 2.77E-01 | no | Frmd3 | intron_variant | MODIFIER |
| 4 | 74119596 | AX-170023116 | 0 | A | G | 0.447 | -9.27E-02 | 8.49E-02 | 2.52E+00 | 2.57E+00 | 2.77E-01 | no | Frmd3 | intron_variant | MODIFIER |
| 4 | 74236623 | AX-170125209 | 0 | G | A | 0.447 | -9.27E-02 | 8.49E-02 | 2.52E+00 | 2.57E+00 | 2.77E-01 | no | | intergenic_variant | MODIFIER |
| 4 | 74480788 | AX-97854222 | 0 | G | T | 0.447 | -9.27E-02 | 8.49E-02 | 2.52E+00 | 2.57E+00 | 2.77E-01 | no | | intergenic_variant | MODIFIER |
| 4 | 74570991 | AX-170023135 | 0 | C | G | 0.447 | -9.27E-02 | 8.49E-02 | 2.52E+00 | 2.57E+00 | 2.77E-01 | no | | intergenic_variant | MODIFIER |
| 4 | 74587880 | AX-170023127 | 0 | T | C | 0.447 | -9.27E-02 | 8.49E-02 | 2.52E+00 | 2.57E+00 | 2.77E-01 | no | | intergenic_variant | MODIFIER |
| 4 | 74687998 | AX-97792697 | 0 | G | T | 0.447 | -9.27E-02 | 8.49E-02 | 2.52E+00 | 2.57E+00 | 2.77E-01 | no | Gm11258 | downstream_gene_variant | MODIFIER |
| 4 | 74784909 | AX-170230977 | 0 | T | A | 0.447 | -9.27E-02 | 8.49E-02 | 2.52E+00 | 2.57E+00 | 2.77E-01 | no | | intergenic_variant | MODIFIER |
| 4 | 74862955 | AX-97824491 | 0 | A | T | 0.447 | -9.27E-02 | 8.49E-02 | 2.52E+00 | 2.57E+00 | 2.77E-01 | no | | intergenic_variant | MODIFIER |
| 4 | 78928759 | AX-97813824 | 0 | G | T | 0.447 | -9.27E-02 | 8.49E-02 | 2.52E+00 | 2.57E+00 | 2.77E-01 | no | | intergenic_variant | MODIFIER |
| 4 | 79052161 | AX-170515877 | 0 | G | A | 0.447 | -9.27E-02 | 8.49E-02 | 2.52E+00 | 2.57E+00 | 2.77E-01 | no | | intergenic_variant | MODIFIER |
| 4 | 79119167 | AX-170263893 | 0 | T | C | 0.447 | -9.27E-02 | 8.49E-02 | 2.52E+00 | 2.57E+00 | 2.77E-01 | no | | intergenic_variant | MODIFIER |
| 4 | 79142592 | AX-97840077 | 0 | G | A | 0.447 | -9.27E-02 | 8.49E-02 | 2.52E+00 | 2.57E+00 | 2.77E-01 | no | | intergenic_variant | MODIFIER |
| 4 | 79190915 | AX-97813989 | 0 | A | G | 0.447 | -9.27E-02 | 8.49E-02 | 2.52E+00 | 2.57E+00 | 2.77E-01 | no | | intergenic_variant | MODIFIER |
| 4 | 79309648 | AX-170089286 | 0 | T | A | 0.447 | -9.27E-02 | 8.49E-02 | 2.52E+00 | 2.57E+00 | 2.77E-01 | no | | intergenic_variant | MODIFIER |
| 4 | 79360246 | AX-170481453 | 0 | A | C | 0.447 | -9.27E-02 | 8.49E-02 | 2.52E+00 | 2.57E+00 | 2.77E-01 | no | | intergenic_variant | MODIFIER |
| 4 | 79388000 | AX-97848171 | 0 | T | C | 0.447 | -9.27E-02 | 8.49E-02 | 2.52E+00 | 2.57E+00 | 2.77E-01 | no | | intergenic_variant | MODIFIER |
| 4 | 79404049 | AX-170407709 | 0 | G | C | 0.447 | -9.27E-02 | 8.49E-02 | 2.52E+00 | 2.57E+00 | 2.77E-01 | no | | intergenic_variant | MODIFIER |
| 4 | 79404264 | AX-170312796 | 0 | T | A | 0.447 | -9.27E-02 | 8.49E-02 | 2.52E+00 | 2.57E+00 | 2.77E-01 | no | | intergenic_variant | MODIFIER |

| | | | | | | | | | | | | | | | |
|---|---|---|---|---|---|---|---|---|---|---|---|---|---|---|---|
| 4 | 79458102 | AX-97808832 | 0 | C | T | 0.447 | -9.27E-02 | 8.49E-02 | 2.52E+00 | 2.57E+00 | 2.77E-01 | no | | intergenic_variant | MODIFIER |
| 4 | 79461867 | AX-170023189 | 0 | G | A | 0.447 | -9.27E-02 | 8.49E-02 | 2.52E+00 | 2.57E+00 | 2.77E-01 | no | | intergenic_variant | MODIFIER |
| 4 | 79495397 | AX-170415741 | 0 | T | C | 0.447 | -9.27E-02 | 8.49E-02 | 2.52E+00 | 2.57E+00 | 2.77E-01 | no | | intergenic_variant | MODIFIER |
| 4 | 79522706 | AX-170404962 | 0 | C | T | 0.447 | -9.27E-02 | 8.49E-02 | 2.52E+00 | 2.57E+00 | 2.77E-01 | no | | intergenic_variant | MODIFIER |
| 4 | 79692853 | AX-170350787 | 0 | C | A | 0.447 | -9.27E-02 | 8.49E-02 | 2.52E+00 | 2.57E+00 | 2.77E-01 | no | | intergenic_variant | MODIFIER |
| 4 | 79697967 | AX-97786234 | 0 | T | C | 0.447 | -9.27E-02 | 8.49E-02 | 2.52E+00 | 2.57E+00 | 2.77E-01 | no | | regulatory_region_variant | MODIFIER |
| 4 | 79699183 | AX-170308995 | 0 | A | T | 0.447 | -9.27E-02 | 8.49E-02 | 2.52E+00 | 2.57E+00 | 2.77E-01 | no | | intergenic_variant | MODIFIER |
| 4 | 79756841 | AX-97801569 | 0 | A | G | 0.447 | -9.27E-02 | 8.49E-02 | 2.52E+00 | 2.57E+00 | 2.77E-01 | no | | intergenic_variant | MODIFIER |
| 4 | 79811825 | AX-97805960 | 0 | G | A | 0.447 | -9.27E-02 | 8.49E-02 | 2.52E+00 | 2.57E+00 | 2.77E-01 | no | | intergenic_variant | MODIFIER |
| 4 | 79827383 | AX-170089287 | 0 | T | C | 0.447 | -9.27E-02 | 8.49E-02 | 2.52E+00 | 2.57E+00 | 2.77E-01 | no | | intergenic_variant | MODIFIER |
| 4 | 79873327 | AX-97834452 | 0 | G | T | 0.447 | -9.27E-02 | 8.49E-02 | 2.52E+00 | 2.57E+00 | 2.77E-01 | no | Gm11409 | upstream_gene_variant | MODIFIER |
| 4 | 79874537 | AX-170277491 | 0 | G | A | 0.447 | -9.27E-02 | 8.49E-02 | 2.52E+00 | 2.57E+00 | 2.77E-01 | no | Gm11409 | non_coding_transcript_exon_variant | MODIFIER |
| 4 | 79874623 | AX-170304337 | 0 | G | T | 0.447 | -9.27E-02 | 8.49E-02 | 2.52E+00 | 2.57E+00 | 2.77E-01 | no | Gm11409 | non_coding_transcript_exon_variant | MODIFIER |
| 4 | 79912551 | AX-170023198 | 0 | G | T | 0.447 | -9.27E-02 | 8.49E-02 | 2.52E+00 | 2.57E+00 | 2.77E-01 | no | | intergenic_variant | MODIFIER |
| 4 | 79943152 | AX-170212129 | 0 | T | C | 0.447 | -9.27E-02 | 8.49E-02 | 2.52E+00 | 2.57E+00 | 2.77E-01 | no | | intergenic_variant | MODIFIER |
| 4 | 79945741 | AX-97836622 | 0 | A | C | 0.447 | -9.27E-02 | 8.49E-02 | 2.52E+00 | 2.57E+00 | 2.77E-01 | no | | intergenic_variant | MODIFIER |
| 4 | 79949313 | AX-170023207 | 0 | C | A | 0.447 | -9.27E-02 | 8.49E-02 | 2.52E+00 | 2.57E+00 | 2.77E-01 | no | | intergenic_variant | MODIFIER |
| 4 | 79962556 | AX-97787136 | 0 | C | T | 0.447 | -9.27E-02 | 8.49E-02 | 2.52E+00 | 2.57E+00 | 2.77E-01 | no | | intergenic_variant | MODIFIER |
| 4 | 79978797 | AX-170023200 | 0 | C | A | 0.447 | -9.27E-02 | 8.49E-02 | 2.52E+00 | 2.57E+00 | 2.77E-01 | no | | intergenic_variant | MODIFIER |
| 4 | 79978852 | AX-170023208 | 0 | T | C | 0.447 | -9.27E-02 | 8.49E-02 | 2.52E+00 | 2.57E+00 | 2.77E-01 | no | | intergenic_variant | MODIFIER |
| 4 | 80017924 | AX-170380026 | 0 | C | G | 0.447 | -9.27E-02 | 8.49E-02 | 2.52E+00 | 2.57E+00 | 2.77E-01 | no | | intergenic_variant | MODIFIER |
| 4 | 80027148 | AX-170023199 | 0 | A | G | 0.447 | -9.27E-02 | 8.49E-02 | 2.52E+00 | 2.57E+00 | 2.77E-01 | no | | intergenic_variant | MODIFIER |
| 4 | 80073607 | AX-170023203 | 0 | T | C | 0.447 | -9.27E-02 | 8.49E-02 | 2.52E+00 | 2.57E+00 | 2.77E-01 | no | | intergenic_variant | MODIFIER |
| 4 | 80103051 | AX-97786307 | 0 | T | C | 0.447 | -9.27E-02 | 8.49E-02 | 2.52E+00 | 2.57E+00 | 2.77E-01 | no | | intergenic_variant | MODIFIER |
| 4 | 80152418 | AX-170023196 | 0 | G | A | 0.447 | -9.27E-02 | 8.49E-02 | 2.52E+00 | 2.57E+00 | 2.77E-01 | no | | intergenic_variant | MODIFIER |
| 4 | 80396637 | AX-170023220 | 0 | G | A | 0.447 | -9.27E-02 | 8.49E-02 | 2.52E+00 | 2.57E+00 | 2.77E-01 | no | | intergenic_variant | MODIFIER |
| 4 | 80684028 | AX-97797218 | 0 | G | A | 0.447 | -9.27E-02 | 8.49E-02 | 2.52E+00 | 2.57E+00 | 2.77E-01 | no | | intergenic_variant | MODIFIER |
| 4 | 80713991 | AX-97853943 | 0 | C | T | 0.447 | -9.27E-02 | 8.49E-02 | 2.52E+00 | 2.57E+00 | 2.77E-01 | no | | intergenic_variant | MODIFIER |
| 4 | 80739874 | AX-97837434 | 0 | C | T | 0.447 | -9.27E-02 | 8.49E-02 | 2.52E+00 | 2.57E+00 | 2.77E-01 | no | | intergenic_variant | MODIFIER |
| 4 | 80744933 | AX-170439076 | 0 | T | C | 0.447 | -9.27E-02 | 8.49E-02 | 2.52E+00 | 2.57E+00 | 2.77E-01 | no | | intergenic_variant | MODIFIER |
| 4 | 80745180 | AX-170258587 | 0 | T | G | 0.447 | -9.27E-02 | 8.49E-02 | 2.52E+00 | 2.57E+00 | 2.77E-01 | no | | intergenic_variant | MODIFIER |
| 4 | 80865216 | AX-170255920 | 0 | T | C | 0.447 | -9.27E-02 | 8.49E-02 | 2.52E+00 | 2.57E+00 | 2.77E-01 | no | | regulatory_region_variant | MODIFIER |
| 4 | 80866714 | AX-170226055 | 0 | G | A | 0.447 | -9.27E-02 | 8.49E-02 | 2.52E+00 | 2.57E+00 | 2.77E-01 | no | | intergenic_variant | MODIFIER |
| 4 | 80867797 | AX-97818198 | 0 | C | T | 0.447 | -9.27E-02 | 8.49E-02 | 2.52E+00 | 2.57E+00 | 2.77E-01 | no | | intergenic_variant | MODIFIER |
| 4 | 80874510 | AX-170402676 | 0 | A | G | 0.447 | -9.27E-02 | 8.49E-02 | 2.52E+00 | 2.57E+00 | 2.77E-01 | no | | regulatory_region_variant | MODIFIER |
| 4 | 80875000 | AX-170342944 | 0 | T | C | 0.447 | -9.27E-02 | 8.49E-02 | 2.52E+00 | 2.57E+00 | 2.77E-01 | no | | regulatory_region_variant | MODIFIER |
| 4 | 74243614 | AX-170414791 | 1 | T | C | 0.446 | -9.27E-02 | 8.49E-02 | 2.52E+00 | 2.57E+00 | 2.77E-01 | no | Kdm4c | intron_variant | MODIFIER |
| 16 | 80300838 | AX-170088074 | 1 | C | T | 0.464 | 7.99E-02 | 7.39E-02 | 2.61E+00 | 2.69E+00 | 2.77E-01 | no | | intergenic_variant | MODIFIER |
| 7 | 114562696 | AX-170029807 | 1 | C | T | 0.429 | 9.43E-02 | 8.74E-02 | 2.64E+00 | 2.72E+00 | 2.77E-01 | no | Cyp2r1 | intron_variant | MODIFIER |
| 4 | 48288922 | AX-97827549 | 0 | C | T | 0.36 | -1.03E-01 | 9.53E-02 | 2.80E+00 | 2.89E+00 | 2.77E-01 | no | Invs | intron_variant | MODIFIER |
| 4 | 100822940 | AX-170192609 | 2 | A | T | 0.427 | -8.10E-02 | 7.43E-02 | 2.52E+00 | 2.60E+00 | 2.77E-01 | no | Cachd1 | intron_variant | MODIFIER |
| 16 | 32700899 | AX-170012774 | 0 | G | A | 0.482 | 8.75E-02 | 7.95E-02 | 2.46E+00 | 2.51E+00 | 2.77E-01 | no | | intergenic_variant | MODIFIER |
| 4 | 134572131 | AX-170366736 | 1 | A | G | 0.17 | 1.15E-01 | 1.06E-01 | 2.55E+00 | 2.62E+00 | 2.77E-01 | no | Man1c1 | intron_variant | MODIFIER |
| 16 | 29995938 | AX-170161485 | 1 | C | T | 0.509 | 8.59E-02 | 7.93E-02 | 2.57E+00 | 2.63E+00 | 2.77E-01 | no | | regulatory_region_variant | MODIFIER |
| 16 | 31326257 | AX-170533646 | 1 | C | T | 0.509 | 8.59E-02 | 7.93E-02 | 2.57E+00 | 2.63E+00 | 2.77E-01 | no | AV205837 | intron_variant,non_coding_transcript_variant | MODIFIER |
| 5 | 43530003 | AX-97798202 | 0 | G | A | 0.184 | -1.19E-01 | 1.10E-01 | 2.63E+00 | 2.72E+00 | 2.77E-01 | no | C1qtnf7 | intron_variant | MODIFIER |
| 4 | 48063648 | AX-170408296 | 0 | C | T | 0.36 | -1.03E-01 | 9.56E-02 | 2.83E+00 | 2.93E+00 | 2.77E-01 | no | Nr4a3 | intron_variant | MODIFIER |
| 1 | 150714035 | AX-170216311 | 1 | T | C | 0.214 | 1.49E-01 | 1.38E-01 | 2.76E+00 | 2.81E+00 | 2.77E-01 | Yes | Hmcn1 | intron_variant | MODIFIER |
| 15 | 87672910 | AX-170366560 | 1 | G | A | 0.152 | 1.20E-01 | 1.12E-01 | 2.66E+00 | 2.79E+00 | 2.78E-01 | no | Fam19a5 | intron_variant | MODIFIER |
| 4 | 77361183 | AX-170466710 | 0 | T | A | 0.307 | 1.22E-01 | 1.13E-01 | 2.85E+00 | 2.92E+00 | 2.78E-01 | no | Ptprd | intron_variant,non_coding_transcript_variant | MODIFIER |
| 1 | 48706996 | AX-170000437 | 0 | T | A | 0.114 | 1.97E-01 | 1.83E-01 | 2.74E+00 | 2.84E+00 | 2.78E-01 | Yes | | intergenic_variant | MODIFIER |
| 18 | 86952591 | AX-170319929 | 0 | C | T | 0.167 | -1.16E-01 | 1.06E-01 | 2.51E+00 | 2.59E+00 | 2.78E-01 | no | | intergenic_variant | MODIFIER |
| 18 | 86964108 | AX-170315531 | 0 | A | G | 0.167 | -1.16E-01 | 1.06E-01 | 2.51E+00 | 2.59E+00 | 2.78E-01 | no | | intergenic_variant | MODIFIER |
| 18 | 87044226 | AX-97792984 | 0 | T | C | 0.167 | -1.16E-01 | 1.06E-01 | 2.51E+00 | 2.59E+00 | 2.78E-01 | no | | intergenic_variant | MODIFIER |

| | | | | | | | | | | | | | | | |
|---|---|---|---|---|---|---|---|---|---|---|---|---|---|---|---|
| 3 | 36963563 | AX-170286692 | 2 | C | T | 0.2 | 1.03E-01 | 9.50E-02 | 2.90E+00 | 3.00E+00 | 2.78E-01 | no | 4932438A13Rik | intron_variant | MODIFIER |
| 7 | 36917401 | AX-170452211 | 0 | G | A | 0.351 | 9.66E-02 | 8.92E-02 | 2.56E+00 | 2.65E+00 | 2.78E-01 | no | | intergenic_variant | MODIFIER |
| 6 | 136848563 | AX-170181276 | 0 | T | C | 0.561 | 9.66E-02 | 8.71E-02 | 2.40E+00 | 2.45E+00 | 2.78E-01 | no | Art4 | 3_prime_UTR_variant | MODIFIER |
| 8 | 95910756 | AX-170279329 | 0 | G | C | 0.228 | 1.20E-01 | 1.11E-01 | 2.85E+00 | 2.94E+00 | 2.78E-01 | no | Gm45895 | non_coding_transcript_exon_variant | MODIFIER |
| 8 | 98320431 | AX-170455109 | 0 | G | T | 0.228 | 1.20E-01 | 1.11E-01 | 2.85E+00 | 2.94E+00 | 2.78E-01 | no | | intergenic_variant | MODIFIER |
| 8 | 98619037 | AX-97837404 | 0 | G | T | 0.228 | 1.20E-01 | 1.11E-01 | 2.85E+00 | 2.94E+00 | 2.78E-01 | no | | intergenic_variant | MODIFIER |
| 14 | 111102329 | AX-170412232 | 1 | G | T | 0.304 | 8.03E-02 | 7.45E-02 | 2.60E+00 | 2.70E+00 | 2.78E-01 | no | | intergenic_variant | MODIFIER |
| 18 | 77934433 | AX-97835112 | 1 | G | A | 0.446 | 9.60E-02 | 8.70E-02 | 2.43E+00 | 2.49E+00 | 2.78E-01 | no | Epg5 | upstream_gene_variant | MODIFIER |
| 9 | 107223429 | AX-97810724 | 1 | C | T | 0.125 | -1.21E-01 | 1.13E-01 | 2.65E+00 | 2.74E+00 | 2.78E-01 | no | Dock3 | intron_variant | MODIFIER |
| 18 | 84666164 | AX-170354203 | 0 | T | C | 0.158 | -1.18E-01 | 1.08E-01 | 2.44E+00 | 2.52E+00 | 2.78E-01 | no | Cndp2 | downstream_gene_variant | MODIFIER |
| 7 | 100171172 | AX-170029627 | 1 | A | G | 0.348 | -1.09E-01 | 1.01E-01 | 2.81E+00 | 2.89E+00 | 2.78E-01 | no | Gm34821 | downstream_gene_variant | MODIFIER |
| 16 | 19767478 | AX-170383527 | 0 | C | T | 0.14 | 9.98E-02 | 9.31E-02 | 2.81E+00 | 2.95E+00 | 2.79E-01 | no | B3gnt5 | intron_variant | MODIFIER |
| 15 | 67757846 | AX-170302938 | 1 | A | G | 0.33 | -8.53E-02 | 7.58E-02 | 2.32E+00 | 2.41E+00 | 2.79E-01 | no | | intergenic_variant | MODIFIER |
| 1 | 166633734 | AX-170139360 | 0 | A | G | 0.246 | 1.16E-01 | 1.08E-01 | 2.66E+00 | 2.75E+00 | 2.79E-01 | no | Gm4847 | intron_variant | MODIFIER |
| 4 | 155974316 | AX-97780594 | 0 | G | A | 0.096 | 1.71E-01 | 1.60E-01 | 2.77E+00 | 2.90E+00 | 2.79E-01 | Yes | | intergenic_variant | MODIFIER |
| 5 | 49717091 | AX-170490811 | 1 | T | C | 0.482 | 1.18E-01 | 1.06E-01 | 2.37E+00 | 2.41E+00 | 2.79E-01 | no | | intergenic_variant | MODIFIER |
| 16 | 82065752 | AX-170349221 | 0 | G | A | 0.465 | 7.82E-02 | 7.23E-02 | 2.56E+00 | 2.63E+00 | 2.79E-01 | no | | intergenic_variant | MODIFIER |
| 1 | 190062935 | AX-170214200 | 1 | C | T | 0.259 | 1.08E-01 | 9.74E-02 | 2.38E+00 | 2.45E+00 | 2.79E-01 | no | | intergenic_variant | MODIFIER |
| 3 | 38625264 | AX-170470009 | 0 | G | C | 0.202 | 1.04E-01 | 9.65E-02 | 2.92E+00 | 3.02E+00 | 2.79E-01 | no | | intergenic_variant | MODIFIER |
| 4 | 145012389 | AX-97795580 | 1 | G | A | 0.116 | 1.84E-01 | 1.71E-01 | 2.77E+00 | 2.88E+00 | 2.79E-01 | Yes | Vps13d | intron_variant | MODIFIER |
| 2 | 169488187 | AX-170178530 | 0 | A | C | 0.175 | 1.66E-01 | 1.55E-01 | 2.59E+00 | 2.71E+00 | 2.79E-01 | Yes | Gm14249 | intron_variant,non_coding_transcript_variant | MODIFIER |
| 2 | 169874713 | AX-170260707 | 0 | A | G | 0.175 | 1.66E-01 | 1.55E-01 | 2.59E+00 | 2.71E+00 | 2.79E-01 | Yes | Tshz2 | intron_variant | MODIFIER |
| 18 | 35584812 | AX-170357150 | 1 | G | A | 0.098 | 1.73E-01 | 1.62E-01 | 2.77E+00 | 2.89E+00 | 2.79E-01 | Yes | Matr3 | synonymous_variant | LOW |
| 11 | 17638592 | AX-170314815 | 1 | G | A | 0.062 | 1.87E-01 | 1.74E-01 | 2.90E+00 | 3.07E+00 | 2.79E-01 | Yes | Gm12016 | non_coding_transcript_exon_variant | MODIFIER |
| 4 | 79681950 | AX-170212064 | 1 | A | G | 0.438 | -8.94E-02 | 8.26E-02 | 2.54E+00 | 2.62E+00 | 2.79E-01 | no | | intergenic_variant | MODIFIER |
| 18 | 88276489 | AX-170522022 | 1 | T | C | 0.161 | -1.15E-01 | 1.07E-01 | 2.53E+00 | 2.62E+00 | 2.79E-01 | no | | intergenic_variant | MODIFIER |
| 15 | 19316576 | AX-170470700 | 1 | C | T | 0.232 | -1.13E-01 | 1.05E-01 | 2.57E+00 | 2.63E+00 | 2.79E-01 | no | Gm49162 | intron_variant,non_coding_transcript_variant | MODIFIER |
| 18 | 88139228 | AX-170437227 | 1 | A | G | 0.17 | -1.10E-01 | 1.02E-01 | 2.57E+00 | 2.67E+00 | 2.80E-01 | no | | intergenic_variant | MODIFIER |
| 1 | 132471655 | AX-170166025 | 1 | G | A | 0.545 | 8.95E-02 | 8.32E-02 | 2.61E+00 | 2.68E+00 | 2.80E-01 | no | Dstyk | downstream_gene_variant | MODIFIER |
| 10 | 10554568 | AX-170514326 | 0 | A | G | 0.561 | 5.54E-02 | 5.17E-02 | 2.80E+00 | 2.90E+00 | 2.80E-01 | no | Rab32 | intron_variant | MODIFIER |
| 3 | 37789941 | AX-170328462 | 0 | T | C | 0.193 | 1.02E-01 | 9.51E-02 | 2.90E+00 | 3.00E+00 | 2.80E-01 | no | | regulatory_region_variant | MODIFIER |
| 5 | 38248613 | AX-170024718 | 2 | G | A | 0.327 | 1.18E-01 | 1.10E-01 | 2.85E+00 | 2.92E+00 | 2.80E-01 | no | | intergenic_variant | MODIFIER |
| 18 | 73124519 | AX-170186306 | 0 | C | T | 0.193 | -1.17E-01 | 1.10E-01 | 2.65E+00 | 2.71E+00 | 2.80E-01 | no | | intergenic_variant | MODIFIER |
| 7 | 36900537 | AX-170301506 | 1 | G | T | 0.357 | 9.68E-02 | 8.98E-02 | 2.55E+00 | 2.64E+00 | 2.80E-01 | no | | intergenic_variant | MODIFIER |
| 7 | 36736006 | AX-170077076 | 0 | T | C | 0.342 | 9.52E-02 | 8.83E-02 | 2.55E+00 | 2.64E+00 | 2.80E-01 | no | Tshz3 | intron_variant | MODIFIER |
| 16 | 61996676 | AX-97807866 | 0 | T | G | 0.474 | -9.25E-02 | 8.54E-02 | 2.52E+00 | 2.58E+00 | 2.80E-01 | no | | intergenic_variant | MODIFIER |
| 4 | 71826734 | AX-170496556 | 0 | A | T | 0.447 | -9.20E-02 | 8.49E-02 | 2.52E+00 | 2.57E+00 | 2.80E-01 | no | | intergenic_variant | MODIFIER |
| 5 | 52237360 | AX-170089476 | 0 | T | G | 0.509 | 1.12E-01 | 1.00E-01 | 2.34E+00 | 2.40E+00 | 2.80E-01 | no | Gm43180 | upstream_gene_variant | MODIFIER |
| 1 | 176921937 | AX-170219970 | 1 | T | C | 0.205 | 1.37E-01 | 1.28E-01 | 2.66E+00 | 2.74E+00 | 2.80E-01 | no | Sdccag8 | intron_variant | MODIFIER |
| 20 | 66797529 | AX-97800987 | 2 | C | T | 0.236 | 6.57E-02 | 6.16E-02 | 2.70E+00 | 2.81E+00 | 2.81E-01 | no | Mir463 | downstream_gene_variant | MODIFIER |
| 3 | 23851810 | AX-170339709 | 1 | T | C | 0.152 | -1.11E-01 | 1.03E-01 | 2.59E+00 | 2.66E+00 | 2.81E-01 | no | Naaladl2 | intron_variant,non_coding_transcript_variant | MODIFIER |
| 16 | 32350964 | AX-97812349 | 0 | C | A | 0.518 | 8.67E-02 | 8.05E-02 | 2.56E+00 | 2.61E+00 | 2.81E-01 | no | Ubxn7 | intron_variant | MODIFIER |
| 5 | 51552363 | AX-170024930 | 1 | G | T | 0.5 | 1.17E-01 | 1.06E-01 | 2.40E+00 | 2.44E+00 | 2.81E-01 | no | Ppargc1a | intron_variant,NMD_transcript_variant | MODIFIER |
| 2 | 169291315 | AX-170251541 | 0 | G | T | 0.175 | 1.67E-01 | 1.56E-01 | 2.58E+00 | 2.69E+00 | 2.81E-01 | Yes | | intergenic_variant | MODIFIER |
| 2 | 169547237 | AX-170495198 | 0 | G | A | 0.175 | 1.67E-01 | 1.56E-01 | 2.58E+00 | 2.69E+00 | 2.81E-01 | Yes | | intergenic_variant | MODIFIER |
| 3 | 20611335 | AX-97841045 | 0 | C | T | 0.149 | -1.07E-01 | 9.96E-02 | 2.62E+00 | 2.71E+00 | 2.82E-01 | no | | intergenic_variant | MODIFIER |
| 13 | 25247152 | AX-170535313 | 0 | C | G | 0.167 | -1.36E-01 | 1.22E-01 | 2.37E+00 | 2.42E+00 | 2.82E-01 | no | Nrsn1 | downstream_gene_variant | MODIFIER |
| 1 | 174675906 | AX-170422626 | 0 | A | G | 0.237 | 1.19E-01 | 1.11E-01 | 2.57E+00 | 2.64E+00 | 2.82E-01 | no | Fmn2 | intron_variant | MODIFIER |
| 9 | 102246421 | AX-97805501 | 0 | A | G | 0.132 | -1.24E-01 | 1.16E-01 | 2.62E+00 | 2.70E+00 | 2.82E-01 | no | Ephb1 | intron_variant | MODIFIER |
| 14 | 112657096 | AX-97785870 | 1 | C | T | 0.312 | 7.77E-02 | 7.28E-02 | 2.61E+00 | 2.71E+00 | 2.82E-01 | no | | intergenic_variant | MODIFIER |
| 13 | 10615253 | AX-97834847 | 0 | A | C | 0.193 | -1.28E-01 | 1.17E-01 | 2.43E+00 | 2.49E+00 | 2.82E-01 | no | Gm26861 | intron_variant,non_coding_transcript_variant | MODIFIER |
| 16 | 18647997 | AX-170172556 | 0 | T | C | 0.14 | 9.55E-02 | 9.00E-02 | 2.77E+00 | 2.90E+00 | 2.82E-01 | no | | intergenic_variant | MODIFIER |
| 5 | 49967198 | AX-170137769 | 0 | A | G | 0.5 | 1.12E-01 | 1.02E-01 | 2.43E+00 | 2.49E+00 | 2.83E-01 | no | Adgra3 | intron_variant | MODIFIER |
| 1 | 168267927 | AX-170261031 | 0 | G | A | 0.228 | 1.40E-01 | 1.31E-01 | 2.65E+00 | 2.71E+00 | 2.83E-01 | no | Pbx1 | intron_variant | MODIFIER |

| | | | | | | | | | | | | | | | |
|---|---|---|---|---|---|---|---|---|---|---|---|---|---|---|---|
| 13 | 4993968 | AX-170250707 | 0 | G | A | 0.219 | -1.23E-01 | 1.15E-01 | 2.71E+00 | 2.78E+00 | 2.83E-01 | no | | intergenic_variant | MODIFIER |
| 1 | 191252185 | AX-170464446 | 1 | T | C | 0.259 | 1.05E-01 | 9.45E-02 | 2.35E+00 | 2.43E+00 | 2.83E-01 | no | Gm37074 | downstream_gene_variant | MODIFIER |
| 1 | 76926559 | AX-170504057 | 1 | C | T | 0.143 | -1.78E-01 | 1.67E-01 | 2.85E+00 | 2.99E+00 | 2.83E-01 | Yes | Gm816 | intron_variant,non_coding_transcript_variant | MODIFIER |
| 7 | 70881133 | AX-170467788 | 2 | C | A | 0.382 | -1.09E-01 | 1.02E-01 | 2.92E+00 | 3.03E+00 | 2.83E-01 | no | Gm42397 | intron_variant,non_coding_transcript_variant | MODIFIER |
| 12 | 59656568 | AX-170006837 | 0 | T | C | 0.061 | -2.38E-01 | 2.23E-01 | 2.61E+00 | 2.69E+00 | 2.83E-01 | Yes | | intergenic_variant | MODIFIER |
| 4 | 77476479 | AX-170243659 | 0 | C | T | 0.325 | 1.16E-01 | 1.08E-01 | 2.91E+00 | 3.01E+00 | 2.83E-01 | no | Ptprd | intron_variant,non_coding_transcript_variant | MODIFIER |
| 4 | 128106764 | AX-170519275 | 0 | G | C | 0.14 | -1.26E-01 | 1.12E-01 | 3.28E+00 | 3.49E+00 | 2.83E-01 | no | Csmd2 | intron_variant,non_coding_transcript_variant | MODIFIER |
| 10 | 18226433 | AX-170232259 | 0 | G | A | 0.211 | 1.07E-01 | 9.88E-02 | 3.02E+00 | 3.14E+00 | 2.83E-01 | no | Ccdc28a | intron_variant | MODIFIER |
| 6 | 135056506 | AX-97856571 | 0 | C | T | 0.561 | 9.65E-02 | 8.80E-02 | 2.41E+00 | 2.45E+00 | 2.83E-01 | no | | intergenic_variant | MODIFIER |
| 6 | 135087667 | AX-170477302 | 0 | G | A | 0.561 | 9.65E-02 | 8.80E-02 | 2.41E+00 | 2.45E+00 | 2.83E-01 | no | Gprc5a | downstream_gene_variant | MODIFIER |
| 6 | 135141329 | AX-170443388 | 0 | C | T | 0.561 | 9.65E-02 | 8.80E-02 | 2.41E+00 | 2.45E+00 | 2.83E-01 | no | Hebp1 | intron_variant | MODIFIER |
| 6 | 135141342 | AX-170244429 | 0 | A | G | 0.561 | 9.65E-02 | 8.80E-02 | 2.41E+00 | 2.45E+00 | 2.83E-01 | no | Hebp1 | intron_variant | MODIFIER |
| 6 | 135153106 | AX-170343853 | 0 | G | A | 0.561 | 9.65E-02 | 8.80E-02 | 2.41E+00 | 2.45E+00 | 2.83E-01 | no | Hebp1 | intron_variant | MODIFIER |
| 6 | 135171639 | AX-97792351 | 0 | T | G | 0.561 | 9.65E-02 | 8.80E-02 | 2.41E+00 | 2.45E+00 | 2.83E-01 | no | Hebp1 | upstream_gene_variant | MODIFIER |
| 6 | 135225271 | AX-170268470 | 0 | T | C | 0.561 | 9.65E-02 | 8.80E-02 | 2.41E+00 | 2.45E+00 | 2.83E-01 | no | Fam234b | synonymous_variant | LOW |
| 6 | 135255453 | AX-97788646 | 0 | T | C | 0.561 | 9.65E-02 | 8.80E-02 | 2.41E+00 | 2.45E+00 | 2.83E-01 | no | Gsg1 | upstream_gene_variant | MODIFIER |
| 6 | 135275848 | AX-97805832 | 0 | A | G | 0.561 | 9.65E-02 | 8.80E-02 | 2.41E+00 | 2.45E+00 | 2.83E-01 | no | | intergenic_variant | MODIFIER |
| 6 | 135277248 | AX-170299386 | 0 | A | G | 0.561 | 9.65E-02 | 8.80E-02 | 2.41E+00 | 2.45E+00 | 2.83E-01 | no | | intergenic_variant | MODIFIER |
| 6 | 135280800 | AX-170497763 | 0 | T | C | 0.561 | 9.65E-02 | 8.80E-02 | 2.41E+00 | 2.45E+00 | 2.83E-01 | no | | regulatory_region_variant | MODIFIER |
| 6 | 135294873 | AX-170028314 | 0 | T | C | 0.561 | 9.65E-02 | 8.80E-02 | 2.41E+00 | 2.45E+00 | 2.83E-01 | no | Gm36640 | intron_variant,non_coding_transcript_variant | MODIFIER |
| 6 | 135296155 | AX-170320650 | 0 | C | T | 0.561 | 9.65E-02 | 8.80E-02 | 2.41E+00 | 2.45E+00 | 2.83E-01 | no | Gm36640 | intron_variant,non_coding_transcript_variant | MODIFIER |
| 6 | 135397736 | AX-170257453 | 0 | T | C | 0.561 | 9.65E-02 | 8.80E-02 | 2.41E+00 | 2.45E+00 | 2.83E-01 | no | | regulatory_region_variant | MODIFIER |
| 6 | 135400348 | AX-170503057 | 0 | T | C | 0.561 | 9.65E-02 | 8.80E-02 | 2.41E+00 | 2.45E+00 | 2.83E-01 | no | | regulatory_region_variant | MODIFIER |
| 6 | 135465133 | AX-170467091 | 0 | G | A | 0.561 | 9.65E-02 | 8.80E-02 | 2.41E+00 | 2.45E+00 | 2.83E-01 | no | | intergenic_variant | MODIFIER |
| 6 | 135465240 | AX-170365598 | 0 | T | C | 0.561 | 9.65E-02 | 8.80E-02 | 2.41E+00 | 2.45E+00 | 2.83E-01 | no | | intergenic_variant | MODIFIER |
| 6 | 135531836 | AX-170115953 | 0 | A | G | 0.561 | 9.65E-02 | 8.80E-02 | 2.41E+00 | 2.45E+00 | 2.83E-01 | no | | intergenic_variant | MODIFIER |
| 6 | 135571143 | AX-170219548 | 0 | G | C | 0.561 | 9.65E-02 | 8.80E-02 | 2.41E+00 | 2.45E+00 | 2.83E-01 | no | | intergenic_variant | MODIFIER |
| 6 | 135586352 | AX-170168561 | 0 | G | T | 0.561 | 9.65E-02 | 8.80E-02 | 2.41E+00 | 2.45E+00 | 2.83E-01 | no | Gm25136 | downstream_gene_variant | MODIFIER |
| 6 | 135613297 | AX-97809787 | 0 | A | G | 0.561 | 9.65E-02 | 8.80E-02 | 2.41E+00 | 2.45E+00 | 2.83E-01 | no | | intergenic_variant | MODIFIER |
| 6 | 135656173 | AX-170112158 | 0 | T | C | 0.561 | 9.65E-02 | 8.80E-02 | 2.41E+00 | 2.45E+00 | 2.83E-01 | no | 4930425L21Rik | upstream_gene_variant | MODIFIER |
| 6 | 135659424 | AX-97792884 | 0 | T | C | 0.561 | 9.65E-02 | 8.80E-02 | 2.41E+00 | 2.45E+00 | 2.83E-01 | no | 4930425L21Rik | intron_variant,non_coding_transcript_variant | MODIFIER |
| 6 | 135743739 | AX-170387685 | 0 | A | G | 0.561 | 9.65E-02 | 8.80E-02 | 2.41E+00 | 2.45E+00 | 2.83E-01 | no | Grin2b | intron_variant | MODIFIER |
| 6 | 135825710 | AX-170251658 | 0 | C | T | 0.561 | 9.65E-02 | 8.80E-02 | 2.41E+00 | 2.45E+00 | 2.83E-01 | no | Grin2b | intron_variant | MODIFIER |
| 6 | 136123869 | AX-170477775 | 0 | G | C | 0.561 | 9.65E-02 | 8.80E-02 | 2.41E+00 | 2.45E+00 | 2.83E-01 | no | Grin2b | intron_variant | MODIFIER |
| 6 | 136212096 | AX-170453445 | 0 | A | G | 0.561 | 9.65E-02 | 8.80E-02 | 2.41E+00 | 2.45E+00 | 2.83E-01 | no | | intergenic_variant | MODIFIER |
| 6 | 136218590 | AX-97793301 | 0 | A | G | 0.561 | 9.65E-02 | 8.80E-02 | 2.41E+00 | 2.45E+00 | 2.83E-01 | no | | intergenic_variant | MODIFIER |
| 6 | 136218616 | AX-170309477 | 0 | T | C | 0.561 | 9.65E-02 | 8.80E-02 | 2.41E+00 | 2.45E+00 | 2.83E-01 | no | | regulatory_region_variant | MODIFIER |
| 6 | 136259186 | AX-170028324 | 0 | A | G | 0.561 | 9.65E-02 | 8.80E-02 | 2.41E+00 | 2.45E+00 | 2.83E-01 | no | | intergenic_variant | MODIFIER |
| 6 | 136271722 | AX-170147989 | 0 | T | C | 0.561 | 9.65E-02 | 8.80E-02 | 2.41E+00 | 2.45E+00 | 2.83E-01 | no | | intergenic_variant | MODIFIER |
| 6 | 136276328 | AX-170181174 | 0 | T | C | 0.561 | 9.65E-02 | 8.80E-02 | 2.41E+00 | 2.45E+00 | 2.83E-01 | no | Gm26105 | upstream_gene_variant | MODIFIER |
| 6 | 136280761 | AX-170173830 | 0 | A | G | 0.561 | 9.65E-02 | 8.80E-02 | 2.41E+00 | 2.45E+00 | 2.83E-01 | no | Gm26105 | upstream_gene_variant | MODIFIER |
| 6 | 136282832 | AX-170028330 | 0 | T | C | 0.561 | 9.65E-02 | 8.80E-02 | 2.41E+00 | 2.45E+00 | 2.83E-01 | no | Gm26105 | downstream_gene_variant | MODIFIER |
| 6 | 136292038 | AX-170122039 | 0 | A | G | 0.561 | 9.65E-02 | 8.80E-02 | 2.41E+00 | 2.45E+00 | 2.83E-01 | no | | intergenic_variant | MODIFIER |
| 6 | 136416598 | AX-170380310 | 0 | C | G | 0.561 | 9.65E-02 | 8.80E-02 | 2.41E+00 | 2.45E+00 | 2.83E-01 | no | Gm25882 | upstream_gene_variant | MODIFIER |
| 6 | 136416707 | AX-170264719 | 0 | G | A | 0.561 | 9.65E-02 | 8.80E-02 | 2.41E+00 | 2.45E+00 | 2.83E-01 | no | Gm25882 | upstream_gene_variant | MODIFIER |
| 6 | 136417090 | AX-170095821 | 0 | A | G | 0.561 | 9.65E-02 | 8.80E-02 | 2.41E+00 | 2.45E+00 | 2.83E-01 | no | Gm25882 | upstream_gene_variant | MODIFIER |
| 6 | 136791003 | AX-170183135 | 0 | T | C | 0.561 | 9.65E-02 | 8.80E-02 | 2.41E+00 | 2.45E+00 | 2.83E-01 | no | | intergenic_variant | MODIFIER |
| 6 | 136801371 | AX-170473084 | 0 | A | G | 0.561 | 9.65E-02 | 8.80E-02 | 2.41E+00 | 2.45E+00 | 2.83E-01 | no | Hist4h4 | downstream_gene_variant | MODIFIER |
| 6 | 136847634 | AX-170466162 | 0 | A | G | 0.561 | 9.65E-02 | 8.80E-02 | 2.41E+00 | 2.45E+00 | 2.83E-01 | no | Art4 | downstream_gene_variant | MODIFIER |
| 6 | 136899505 | AX-170141737 | 0 | G | A | 0.561 | 9.65E-02 | 8.80E-02 | 2.41E+00 | 2.45E+00 | 2.83E-01 | no | | regulatory_region_variant | MODIFIER |
| 6 | 136899545 | AX-170361690 | 0 | C | T | 0.561 | 9.65E-02 | 8.80E-02 | 2.41E+00 | 2.45E+00 | 2.83E-01 | no | | regulatory_region_variant | MODIFIER |
| 6 | 136963430 | AX-170028334 | 0 | C | T | 0.561 | 9.65E-02 | 8.80E-02 | 2.41E+00 | 2.45E+00 | 2.83E-01 | no | Pde6h | non_coding_transcript_exon_variant | MODIFIER |
| 6 | 136966034 | AX-170334576 | 0 | A | G | 0.561 | 9.65E-02 | 8.80E-02 | 2.41E+00 | 2.45E+00 | 2.83E-01 | no | Pde6h | downstream_gene_variant | MODIFIER |
| 6 | 137027001 | AX-97802390 | 0 | A | G | 0.561 | 9.65E-02 | 8.80E-02 | 2.41E+00 | 2.45E+00 | 2.83E-01 | no | | intergenic_variant | MODIFIER |

| | | | | | | | | | | | | | | | |
|---|---|---|---|---|---|---|---|---|---|---|---|---|---|---|---|
| 6 | 137030480 | AX-170076359 | 0 | A | G | 0.561 | 9.65E-02 | 8.80E-02 | 2.41E+00 | 2.45E+00 | 2.83E-01 | no | | regulatory_region_variant | MODIFIER |
| 6 | 137666882 | AX-170386042 | 0 | G | A | 0.561 | 9.65E-02 | 8.80E-02 | 2.41E+00 | 2.45E+00 | 2.83E-01 | no | | intergenic_variant | MODIFIER |
| 6 | 137685958 | AX-97854242 | 0 | G | A | 0.561 | 9.65E-02 | 8.80E-02 | 2.41E+00 | 2.45E+00 | 2.83E-01 | no | | intergenic_variant | MODIFIER |
| 6 | 137687461 | AX-170076375 | 0 | A | G | 0.561 | 9.65E-02 | 8.80E-02 | 2.41E+00 | 2.45E+00 | 2.83E-01 | no | | intergenic_variant | MODIFIER |
| 6 | 137689058 | AX-168390206 | 0 | A | G | 0.561 | 9.65E-02 | 8.80E-02 | 2.41E+00 | 2.45E+00 | 2.83E-01 | no | | regulatory_region_variant | MODIFIER |
| 6 | 137695613 | AX-170327763 | 0 | A | C | 0.561 | 9.65E-02 | 8.80E-02 | 2.41E+00 | 2.45E+00 | 2.83E-01 | no | | intergenic_variant | MODIFIER |
| 6 | 137696425 | AX-170368104 | 0 | T | A | 0.561 | 9.65E-02 | 8.80E-02 | 2.41E+00 | 2.45E+00 | 2.83E-01 | no | | regulatory_region_variant | MODIFIER |
| 6 | 137700433 | AX-170124770 | 0 | T | G | 0.561 | 9.65E-02 | 8.80E-02 | 2.41E+00 | 2.45E+00 | 2.83E-01 | no | | intergenic_variant | MODIFIER |
| 6 | 137705714 | AX-97801523 | 0 | G | A | 0.561 | 9.65E-02 | 8.80E-02 | 2.41E+00 | 2.45E+00 | 2.83E-01 | no | | intergenic_variant | MODIFIER |
| 5 | 113882200 | AX-170119189 | 0 | G | A | 0.456 | 1.16E-01 | 1.09E-01 | 2.72E+00 | 2.78E+00 | 2.84E-01 | no | Coro1c | intron_variant | MODIFIER |
| 5 | 113900643 | AX-170505285 | 0 | G | A | 0.456 | 1.16E-01 | 1.09E-01 | 2.72E+00 | 2.78E+00 | 2.84E-01 | no | Coro1c | intron_variant | MODIFIER |
| 5 | 114227150 | AX-170376194 | 0 | G | A | 0.456 | 1.16E-01 | 1.09E-01 | 2.72E+00 | 2.78E+00 | 2.84E-01 | no | Acacb | intron_variant | MODIFIER |
| 12 | 101412653 | AX-170415192 | 0 | G | A | 0.088 | -1.67E-01 | 1.57E-01 | 2.84E+00 | 2.97E+00 | 2.84E-01 | no | Catsperb | intron_variant | MODIFIER |
| 18 | 57529642 | AX-170213223 | 1 | C | A | 0.42 | 1.11E-01 | 1.01E-01 | 2.39E+00 | 2.44E+00 | 2.84E-01 | no | Ccdc192 | upstream_gene_variant | MODIFIER |
| 9 | 112899131 | AX-170429454 | 0 | C | T | 0.123 | -9.93E-02 | 9.37E-02 | 2.71E+00 | 2.84E+00 | 2.84E-01 | no | | intergenic_variant | MODIFIER |
| 9 | 113368437 | AX-97797550 | 0 | T | G | 0.123 | -9.93E-02 | 9.37E-02 | 2.71E+00 | 2.84E+00 | 2.84E-01 | no | | intergenic_variant | MODIFIER |
| 15 | 5828542 | AX-170490328 | 1 | T | A | 0.241 | -1.13E-01 | 1.06E-01 | 2.65E+00 | 2.71E+00 | 2.84E-01 | no | | intergenic_variant | MODIFIER |
| 4 | 51198957 | AX-170486806 | 0 | C | T | 0.368 | -9.88E-02 | 9.28E-02 | 2.88E+00 | 2.99E+00 | 2.84E-01 | no | | intergenic_variant | MODIFIER |
| 6 | 139502385 | AX-170028384 | 0 | T | G | 0.053 | -2.29E-01 | 2.16E-01 | 2.76E+00 | 2.87E+00 | 2.84E-01 | Yes | Rergl | upstream_gene_variant | MODIFIER |
| 14 | 123405619 | AX-170010893 | 0 | G | A | 0.325 | 8.58E-02 | 8.08E-02 | 2.65E+00 | 2.77E+00 | 2.84E-01 | no | Nalcn | intron_variant | MODIFIER |
| 4 | 70098952 | AX-170415121 | 0 | C | T | 0.456 | -8.95E-02 | 8.32E-02 | 2.52E+00 | 2.57E+00 | 2.84E-01 | no | | intergenic_variant | MODIFIER |
| 4 | 70099153 | AX-97849592 | 0 | G | A | 0.456 | -8.95E-02 | 8.32E-02 | 2.52E+00 | 2.57E+00 | 2.84E-01 | no | | intergenic_variant | MODIFIER |
| 4 | 70104847 | AX-97799110 | 0 | T | C | 0.456 | -8.95E-02 | 8.32E-02 | 2.52E+00 | 2.57E+00 | 2.84E-01 | no | | intergenic_variant | MODIFIER |
| 4 | 70148180 | AX-170397077 | 0 | C | T | 0.456 | -8.95E-02 | 8.32E-02 | 2.52E+00 | 2.57E+00 | 2.84E-01 | no | | intergenic_variant | MODIFIER |
| 4 | 70153863 | AX-170145443 | 0 | A | G | 0.456 | -8.95E-02 | 8.32E-02 | 2.52E+00 | 2.57E+00 | 2.84E-01 | no | | intergenic_variant | MODIFIER |
| 4 | 70154055 | AX-170297293 | 0 | T | C | 0.456 | -8.95E-02 | 8.32E-02 | 2.52E+00 | 2.57E+00 | 2.84E-01 | no | | intergenic_variant | MODIFIER |
| 4 | 70162872 | AX-97797843 | 0 | G | T | 0.456 | -8.95E-02 | 8.32E-02 | 2.52E+00 | 2.57E+00 | 2.84E-01 | no | | intergenic_variant | MODIFIER |
| 4 | 70164296 | AX-170281242 | 0 | C | G | 0.456 | -8.95E-02 | 8.32E-02 | 2.52E+00 | 2.57E+00 | 2.84E-01 | no | | intergenic_variant | MODIFIER |
| 4 | 70180285 | AX-97845629 | 0 | T | C | 0.456 | -8.95E-02 | 8.32E-02 | 2.52E+00 | 2.57E+00 | 2.84E-01 | no | | intergenic_variant | MODIFIER |
| 4 | 70180627 | AX-170023053 | 0 | C | T | 0.456 | -8.95E-02 | 8.32E-02 | 2.52E+00 | 2.57E+00 | 2.84E-01 | no | | intergenic_variant | MODIFIER |
| 4 | 70182282 | AX-170023052 | 0 | A | G | 0.456 | -8.95E-02 | 8.32E-02 | 2.52E+00 | 2.57E+00 | 2.84E-01 | no | | regulatory_region_variant | MODIFIER |
| 4 | 70201797 | AX-170378329 | 0 | A | G | 0.456 | -8.95E-02 | 8.32E-02 | 2.52E+00 | 2.57E+00 | 2.84E-01 | no | | intergenic_variant | MODIFIER |
| 4 | 70201990 | AX-170226749 | 0 | A | G | 0.456 | -8.95E-02 | 8.32E-02 | 2.52E+00 | 2.57E+00 | 2.84E-01 | no | | intergenic_variant | MODIFIER |
| 4 | 70206636 | AX-170273752 | 0 | G | A | 0.456 | -8.95E-02 | 8.32E-02 | 2.52E+00 | 2.57E+00 | 2.84E-01 | no | | intergenic_variant | MODIFIER |
| 4 | 70234909 | AX-170286950 | 0 | C | T | 0.456 | -8.95E-02 | 8.32E-02 | 2.52E+00 | 2.57E+00 | 2.84E-01 | no | Cdk5rap2 | intron_variant,NMD_transcript_variant | MODIFIER |
| 4 | 70253122 | AX-170023058 | 0 | G | C | 0.456 | -8.95E-02 | 8.32E-02 | 2.52E+00 | 2.57E+00 | 2.84E-01 | no | Cdk5rap2 | intron_variant,NMD_transcript_variant | MODIFIER |
| 4 | 70266817 | AX-97798018 | 0 | T | C | 0.456 | -8.95E-02 | 8.32E-02 | 2.52E+00 | 2.57E+00 | 2.84E-01 | no | Cdk5rap2 | 3_prime_UTR_variant,NMD_transcript_variant | MODIFIER |
| 4 | 70268479 | AX-170023056 | 0 | T | C | 0.456 | -8.95E-02 | 8.32E-02 | 2.52E+00 | 2.57E+00 | 2.84E-01 | no | Cdk5rap2 | intron_variant,NMD_transcript_variant | MODIFIER |
| 4 | 70271383 | AX-170069625 | 0 | T | C | 0.456 | -8.95E-02 | 8.32E-02 | 2.52E+00 | 2.57E+00 | 2.84E-01 | no | Cdk5rap2 | intron_variant,NMD_transcript_variant | MODIFIER |
| 4 | 70279066 | AX-170069622 | 0 | C | T | 0.456 | -8.95E-02 | 8.32E-02 | 2.52E+00 | 2.57E+00 | 2.84E-01 | no | Cdk5rap2 | intron_variant,NMD_transcript_variant | MODIFIER |
| 4 | 70280703 | AX-170275300 | 0 | C | A | 0.456 | -8.95E-02 | 8.32E-02 | 2.52E+00 | 2.57E+00 | 2.84E-01 | no | Cdk5rap2 | intron_variant,NMD_transcript_variant | MODIFIER |
| 4 | 70281456 | AX-170355713 | 0 | G | A | 0.456 | -8.95E-02 | 8.32E-02 | 2.52E+00 | 2.57E+00 | 2.84E-01 | no | Cdk5rap2 | 3_prime_UTR_variant,NMD_transcript_variant | MODIFIER |
| 4 | 70282564 | AX-170023057 | 0 | C | T | 0.456 | -8.95E-02 | 8.32E-02 | 2.52E+00 | 2.57E+00 | 2.84E-01 | no | Cdk5rap2 | intron_variant,NMD_transcript_variant | MODIFIER |
| 4 | 70301042 | AX-170418270 | 0 | T | C | 0.456 | -8.95E-02 | 8.32E-02 | 2.52E+00 | 2.57E+00 | 2.84E-01 | no | Cdk5rap2 | intron_variant,NMD_transcript_variant | MODIFIER |
| 4 | 70303477 | AX-170095004 | 0 | T | C | 0.456 | -8.95E-02 | 8.32E-02 | 2.52E+00 | 2.57E+00 | 2.84E-01 | no | Cdk5rap2 | intron_variant,NMD_transcript_variant | MODIFIER |
| 4 | 70314740 | AX-170416365 | 0 | G | T | 0.456 | -8.95E-02 | 8.32E-02 | 2.52E+00 | 2.57E+00 | 2.84E-01 | no | Cdk5rap2 | intron_variant,NMD_transcript_variant | MODIFIER |
| 4 | 70330877 | AX-97803253 | 0 | C | A | 0.456 | -8.95E-02 | 8.32E-02 | 2.52E+00 | 2.57E+00 | 2.84E-01 | no | Cdk5rap2 | intron_variant,NMD_transcript_variant | MODIFIER |
| 4 | 70356810 | AX-170275731 | 0 | T | C | 0.456 | -8.95E-02 | 8.32E-02 | 2.52E+00 | 2.57E+00 | 2.84E-01 | no | Cdk5rap2 | intron_variant,NMD_transcript_variant | MODIFIER |
| 4 | 70375185 | AX-170330300 | 0 | C | G | 0.456 | -8.95E-02 | 8.32E-02 | 2.52E+00 | 2.57E+00 | 2.84E-01 | no | Cdk5rap2 | intron_variant,NMD_transcript_variant | MODIFIER |
| 4 | 70375246 | AX-170326553 | 0 | G | A | 0.456 | -8.95E-02 | 8.32E-02 | 2.52E+00 | 2.57E+00 | 2.84E-01 | no | Cdk5rap2 | intron_variant,NMD_transcript_variant | MODIFIER |
| 4 | 70376936 | AX-170175002 | 0 | A | G | 0.456 | -8.95E-02 | 8.32E-02 | 2.52E+00 | 2.57E+00 | 2.84E-01 | no | Cdk5rap2 | intron_variant,NMD_transcript_variant | MODIFIER |
| 4 | 70376965 | AX-170345366 | 0 | A | G | 0.456 | -8.95E-02 | 8.32E-02 | 2.52E+00 | 2.57E+00 | 2.84E-01 | no | Cdk5rap2 | intron_variant,NMD_transcript_variant | MODIFIER |
| 4 | 70377547 | AX-97781107 | 0 | A | G | 0.456 | -8.95E-02 | 8.32E-02 | 2.52E+00 | 2.57E+00 | 2.84E-01 | no | Cdk5rap2 | intron_variant,NMD_transcript_variant | MODIFIER |
| 4 | 70380873 | AX-170035685 | 0 | A | G | 0.456 | -8.95E-02 | 8.32E-02 | 2.52E+00 | 2.57E+00 | 2.84E-01 | no | Cdk5rap2 | intron_variant,NMD_transcript_variant | MODIFIER |

| | | | | | | | | | | | | | | | |
|---|---|---|---|---|---|---|---|---|---|---|---|---|---|---|---|
| 4 | 70385786 | AX-170305485 | 0 | A | G | 0.456 | -8.95E-02 | 8.32E-02 | 2.52E+00 | 2.57E+00 | 2.84E-01 | no | Cdk5rap2 | intron_variant,NMD_transcript_variant | MODIFIER |
| 4 | 71229693 | AX-170069640 | 0 | T | G | 0.456 | -8.95E-02 | 8.32E-02 | 2.52E+00 | 2.57E+00 | 2.84E-01 | no | | regulatory_region_variant | MODIFIER |
| 4 | 71583035 | AX-170211189 | 0 | C | T | 0.456 | -8.95E-02 | 8.32E-02 | 2.52E+00 | 2.57E+00 | 2.84E-01 | no | | intergenic_variant | MODIFIER |
| 4 | 71617177 | AX-170425071 | 0 | G | A | 0.456 | -8.95E-02 | 8.32E-02 | 2.52E+00 | 2.57E+00 | 2.84E-01 | no | | intergenic_variant | MODIFIER |
| 4 | 71646203 | AX-170396068 | 0 | G | A | 0.456 | -8.95E-02 | 8.32E-02 | 2.52E+00 | 2.57E+00 | 2.84E-01 | no | | intergenic_variant | MODIFIER |
| 4 | 71735177 | AX-97825464 | 0 | C | T | 0.456 | -8.95E-02 | 8.32E-02 | 2.52E+00 | 2.57E+00 | 2.84E-01 | no | | intergenic_variant | MODIFIER |
| 4 | 71756837 | AX-170371675 | 0 | A | C | 0.456 | -8.95E-02 | 8.32E-02 | 2.52E+00 | 2.57E+00 | 2.84E-01 | no | Gm11232 | missense_variant | MODERATE |
| 4 | 71759562 | AX-170137002 | 0 | C | T | 0.456 | -8.95E-02 | 8.32E-02 | 2.52E+00 | 2.57E+00 | 2.84E-01 | no | Gm11232 | upstream_gene_variant | MODIFIER |
| 4 | 71823147 | AX-170023083 | 0 | G | A | 0.456 | -8.95E-02 | 8.32E-02 | 2.52E+00 | 2.57E+00 | 2.84E-01 | no | | intergenic_variant | MODIFIER |
| 4 | 71846126 | AX-97806288 | 0 | A | G | 0.456 | -8.95E-02 | 8.32E-02 | 2.52E+00 | 2.57E+00 | 2.84E-01 | no | | intergenic_variant | MODIFIER |
| 4 | 71854468 | AX-170023084 | 0 | A | T | 0.456 | -8.95E-02 | 8.32E-02 | 2.52E+00 | 2.57E+00 | 2.84E-01 | no | | intergenic_variant | MODIFIER |
| 4 | 72136752 | AX-170069657 | 0 | C | T | 0.456 | -8.95E-02 | 8.32E-02 | 2.52E+00 | 2.57E+00 | 2.84E-01 | no | Tle1 | intron_variant | MODIFIER |
| 4 | 72170980 | AX-170398451 | 0 | G | A | 0.456 | -8.95E-02 | 8.32E-02 | 2.52E+00 | 2.57E+00 | 2.84E-01 | no | Tle1 | intron_variant | MODIFIER |
| 4 | 72232144 | AX-170469633 | 0 | G | A | 0.456 | -8.95E-02 | 8.32E-02 | 2.52E+00 | 2.57E+00 | 2.84E-01 | no | | intergenic_variant | MODIFIER |
| 4 | 72237352 | AX-170204214 | 0 | C | T | 0.456 | -8.95E-02 | 8.32E-02 | 2.52E+00 | 2.57E+00 | 2.84E-01 | no | | intergenic_variant | MODIFIER |
| 4 | 72353314 | AX-170258876 | 0 | C | T | 0.456 | -8.95E-02 | 8.32E-02 | 2.52E+00 | 2.57E+00 | 2.84E-01 | no | | intergenic_variant | MODIFIER |
| 4 | 72652029 | AX-170404989 | 0 | C | T | 0.456 | -8.95E-02 | 8.32E-02 | 2.52E+00 | 2.57E+00 | 2.84E-01 | no | | intergenic_variant | MODIFIER |
| 4 | 73109443 | AX-170143769 | 0 | C | T | 0.456 | -8.95E-02 | 8.32E-02 | 2.52E+00 | 2.57E+00 | 2.84E-01 | no | | intergenic_variant | MODIFIER |
| 4 | 74050840 | AX-170023121 | 0 | C | G | 0.456 | -8.95E-02 | 8.32E-02 | 2.52E+00 | 2.57E+00 | 2.84E-01 | no | Frmd3 | intron_variant | MODIFIER |
| 4 | 74252910 | AX-170502335 | 0 | G | A | 0.456 | -8.95E-02 | 8.32E-02 | 2.52E+00 | 2.57E+00 | 2.84E-01 | no | Kdm4c | intron_variant | MODIFIER |
| 4 | 74278661 | AX-170496862 | 0 | C | T | 0.456 | -8.95E-02 | 8.32E-02 | 2.52E+00 | 2.57E+00 | 2.84E-01 | no | Kdm4c | intron_variant | MODIFIER |
| 4 | 74740263 | AX-170023129 | 0 | C | T | 0.456 | -8.95E-02 | 8.32E-02 | 2.52E+00 | 2.57E+00 | 2.84E-01 | no | | intergenic_variant | MODIFIER |
| 4 | 74841162 | AX-170023133 | 0 | A | G | 0.456 | -8.95E-02 | 8.32E-02 | 2.52E+00 | 2.57E+00 | 2.84E-01 | no | | intergenic_variant | MODIFIER |
| 4 | 74954260 | AX-170103983 | 0 | C | G | 0.456 | -8.95E-02 | 8.32E-02 | 2.52E+00 | 2.57E+00 | 2.84E-01 | no | | intergenic_variant | MODIFIER |
| 13 | 6429914 | AX-170503108 | 1 | T | C | 0.205 | -1.29E-01 | 1.20E-01 | 2.53E+00 | 2.59E+00 | 2.84E-01 | no | | intergenic_variant | MODIFIER |
| 9 | 104539703 | AX-170259250 | 1 | A | G | 0.125 | -1.18E-01 | 1.11E-01 | 2.67E+00 | 2.76E+00 | 2.84E-01 | no | | intergenic_variant | MODIFIER |
| 5 | 117133851 | AX-170166269 | 0 | T | C | 0.456 | 1.21E-01 | 1.14E-01 | 2.63E+00 | 2.67E+00 | 2.84E-01 | no | Taok3 | intron_variant | MODIFIER |
| 1 | 120933699 | AX-170197061 | 0 | A | G | 0.57 | 8.61E-02 | 8.00E-02 | 2.50E+00 | 2.56E+00 | 2.84E-01 | no | | intergenic_variant | MODIFIER |
| 8 | 69011169 | AX-170381126 | 0 | A | G | 0.184 | -1.32E-01 | 1.24E-01 | 2.75E+00 | 2.85E+00 | 2.84E-01 | no | | intergenic_variant | MODIFIER |
| 9 | 15772382 | AX-170392380 | 1 | T | C | 0.464 | 8.91E-02 | 8.35E-02 | 2.57E+00 | 2.65E+00 | 2.85E-01 | no | | intergenic_variant | MODIFIER |
| 6 | 139403923 | AX-97852255 | 0 | G | A | 0.053 | -1.97E-01 | 1.83E-01 | 2.49E+00 | 2.61E+00 | 2.85E-01 | Yes | | intergenic_variant | MODIFIER |
| 10 | 92559511 | AX-170308872 | 0 | C | T | 0.298 | -9.02E-02 | 8.41E-02 | 2.53E+00 | 2.58E+00 | 2.85E-01 | no | | intergenic_variant | MODIFIER |
| 1 | 96842607 | AX-97845620 | 0 | G | A | 0.158 | 1.80E-01 | 1.70E-01 | 2.64E+00 | 2.73E+00 | 2.85E-01 | Yes | Slco4c1 | intron_variant | MODIFIER |
| 4 | 78674157 | AX-170378911 | 0 | T | C | 0.298 | 1.19E-01 | 1.12E-01 | 2.66E+00 | 2.74E+00 | 2.85E-01 | no | Gm22262 | downstream_gene_variant | MODIFIER |
| 5 | 4089976 | AX-170222084 | 1 | T | C | 0.411 | -7.06E-02 | 6.57E-02 | 2.50E+00 | 2.61E+00 | 2.85E-01 | no | Cyp51 | intron_variant | MODIFIER |
| 4 | 70143510 | AX-170207063 | 1 | C | T | 0.455 | -8.93E-02 | 8.32E-02 | 2.52E+00 | 2.57E+00 | 2.85E-01 | no | | intergenic_variant | MODIFIER |
| 8 | 65715809 | AX-170030928 | 0 | G | C | 0.132 | -1.42E-01 | 1.33E-01 | 2.51E+00 | 2.60E+00 | 2.85E-01 | no | MAR1 | intron_variant | MODIFIER |
| 8 | 34610185 | AX-170375688 | 0 | T | A | 0.14 | -1.39E-01 | 1.31E-01 | 2.66E+00 | 2.74E+00 | 2.85E-01 | no | Gm34096 | intron_variant,non_coding_transcript_variant | MODIFIER |
| 3 | 26325013 | AX-170521145 | 0 | G | A | 0.14 | -1.07E-01 | 1.01E-01 | 2.60E+00 | 2.70E+00 | 2.85E-01 | no | Nlgn1 | intron_variant,non_coding_transcript_variant | MODIFIER |
| 3 | 26983440 | AX-170255045 | 0 | G | A | 0.14 | -1.07E-01 | 1.01E-01 | 2.60E+00 | 2.70E+00 | 2.85E-01 | no | Spata16 | downstream_gene_variant | MODIFIER |
| 15 | 83805877 | AX-170126938 | 1 | T | A | 0.089 | -1.38E-01 | 1.05E-01 | 1.88E+00 | 1.96E+00 | 2.85E-01 | no | Mpped1 | intron_variant | MODIFIER |
| 1 | 139622173 | AX-170520900 | 1 | G | A | 0.214 | 1.26E-01 | 1.19E-01 | 2.61E+00 | 2.68E+00 | 2.85E-01 | no | Cfhr3 | non_coding_transcript_exon_variant | MODIFIER |
| 3 | 36824165 | AX-170351949 | 0 | T | C | 0.202 | 9.86E-02 | 9.27E-02 | 2.91E+00 | 3.02E+00 | 2.85E-01 | no | Gm12376 | downstream_gene_variant | MODIFIER |
| 10 | 66026504 | AX-170003368 | 0 | A | G | 0.412 | -8.41E-02 | 7.93E-02 | 2.63E+00 | 2.71E+00 | 2.85E-01 | no | | intergenic_variant | MODIFIER |
| 1 | 131612925 | AX-170001722 | 1 | G | C | 0.545 | 9.08E-02 | 8.57E-02 | 2.66E+00 | 2.72E+00 | 2.85E-01 | no | Avpr1b | downstream_gene_variant | MODIFIER |
| 18 | 66601686 | AX-170061155 | 0 | T | C | 0.281 | -1.01E-01 | 9.58E-02 | 2.80E+00 | 2.88E+00 | 2.85E-01 | no | AC100212.2 | downstream_gene_variant | MODIFIER |
| 4 | 88913416 | AX-170140913 | 0 | C | T | 0.263 | 1.17E-01 | 1.07E-01 | 2.40E+00 | 2.45E+00 | 2.85E-01 | no | Mir31 | upstream_gene_variant | MODIFIER |
| 5 | 49902205 | AX-170279639 | 1 | G | A | 0.473 | 1.12E-01 | 9.80E-02 | 2.24E+00 | 2.30E+00 | 2.86E-01 | no | Gm7988 | non_coding_transcript_exon_variant | MODIFIER |
| 1 | 72394390 | AX-170000784 | 0 | T | C | 0.114 | 1.94E-01 | 1.84E-01 | 2.78E+00 | 2.90E+00 | 2.86E-01 | Yes | Xrcc5 | intron_variant | MODIFIER |
| 4 | 51243733 | AX-170320043 | 1 | T | A | 0.366 | -1.01E-01 | 9.55E-02 | 2.82E+00 | 2.93E+00 | 2.86E-01 | no | | intergenic_variant | MODIFIER |
| 18 | 80743626 | AX-97835144 | 0 | T | C | 0.553 | 7.83E-02 | 7.19E-02 | 2.42E+00 | 2.50E+00 | 2.86E-01 | no | Atp9b | intron_variant | MODIFIER |
| 8 | 19256897 | AX-170498769 | 0 | C | A | 0.167 | -1.13E-01 | 1.06E-01 | 2.90E+00 | 3.04E+00 | 2.86E-01 | no | | intergenic_variant | MODIFIER |
| 8 | 21570179 | AX-170460437 | 0 | A | C | 0.167 | -1.13E-01 | 1.06E-01 | 2.90E+00 | 3.04E+00 | 2.86E-01 | no | Defa37 | downstream_gene_variant | MODIFIER |
| 8 | 21708384 | AX-170271789 | 0 | T | A | 0.167 | -1.13E-01 | 1.06E-01 | 2.90E+00 | 3.04E+00 | 2.86E-01 | no | AY761184 | upstream_gene_variant | MODIFIER |

| | | | | | | | | | | | | | | | |
|---|---|---|---|---|---|---|---|---|---|---|---|---|---|---|---|
| 9 | 92524801 | AX-170227613 | 0 | T | C | 0.132 | -1.34E-01 | 1.26E-01 | 2.67E+00 | 2.75E+00 | 2.86E-01 | no | Gm25417 | downstream_gene_variant | MODIFIER |
| 9 | 92565970 | AX-170280228 | 0 | G | T | 0.132 | -1.34E-01 | 1.26E-01 | 2.67E+00 | 2.75E+00 | 2.86E-01 | no | Plod2 | intron_variant | MODIFIER |
| 9 | 92579521 | AX-170330547 | 0 | T | C | 0.132 | -1.34E-01 | 1.26E-01 | 2.67E+00 | 2.75E+00 | 2.86E-01 | no | Plod2 | intron_variant | MODIFIER |
| 9 | 92592977 | AX-170258525 | 0 | C | T | 0.132 | -1.34E-01 | 1.26E-01 | 2.67E+00 | 2.75E+00 | 2.86E-01 | no | Plod2 | intron_variant | MODIFIER |
| 9 | 92612396 | AX-97828461 | 0 | A | C | 0.132 | -1.34E-01 | 1.26E-01 | 2.67E+00 | 2.75E+00 | 2.86E-01 | no | Plod2 | downstream_gene_variant | MODIFIER |
| 9 | 92686085 | AX-170229009 | 0 | T | C | 0.132 | -1.34E-01 | 1.26E-01 | 2.67E+00 | 2.75E+00 | 2.86E-01 | no | | intergenic_variant | MODIFIER |
| 9 | 92691597 | AX-170453978 | 0 | C | T | 0.132 | -1.34E-01 | 1.26E-01 | 2.67E+00 | 2.75E+00 | 2.86E-01 | no | Gm29396 | upstream_gene_variant | MODIFIER |
| 9 | 92704150 | AX-170096626 | 0 | T | C | 0.132 | -1.34E-01 | 1.26E-01 | 2.67E+00 | 2.75E+00 | 2.86E-01 | no | | intergenic_variant | MODIFIER |
| 9 | 94592460 | AX-97781313 | 0 | A | G | 0.132 | -1.34E-01 | 1.26E-01 | 2.67E+00 | 2.75E+00 | 2.86E-01 | no | Gm39404 | non_coding_transcript_exon_variant | MODIFIER |
| 9 | 94715658 | AX-170359452 | 0 | A | C | 0.132 | -1.34E-01 | 1.26E-01 | 2.67E+00 | 2.75E+00 | 2.86E-01 | no | Slc9a9 | intron_variant | MODIFIER |
| 9 | 94733507 | AX-170310927 | 0 | T | C | 0.132 | -1.34E-01 | 1.26E-01 | 2.67E+00 | 2.75E+00 | 2.86E-01 | no | Slc9a9 | intron_variant | MODIFIER |
| 4 | 84982018 | AX-170332914 | 1 | T | C | 0.33 | 1.20E-01 | 1.13E-01 | 2.80E+00 | 2.88E+00 | 2.86E-01 | no | Cntln | intron_variant | MODIFIER |
| 4 | 98512926 | AX-97811599 | 1 | C | T | 0.232 | 1.20E-01 | 1.13E-01 | 2.59E+00 | 2.66E+00 | 2.86E-01 | no | Patj | intron_variant | MODIFIER |
| 4 | 143468571 | AX-170353394 | 0 | G | A | 0.114 | 1.81E-01 | 1.72E-01 | 2.79E+00 | 2.90E+00 | 2.86E-01 | Yes | Gm13046 | upstream_gene_variant | MODIFIER |
| 18 | 55670523 | AX-97788082 | 1 | C | T | 0.455 | 1.04E-01 | 9.33E-02 | 2.31E+00 | 2.36E+00 | 2.86E-01 | no | | intergenic_variant | MODIFIER |
| 2 | 122452861 | AX-170358577 | 0 | A | C | 0.149 | 1.89E-01 | 1.79E-01 | 2.77E+00 | 2.86E+00 | 2.86E-01 | Yes | Slc28a2 | intron_variant | MODIFIER |
| 15 | 83103820 | AX-170012130 | 0 | C | T | 0.088 | -1.40E-01 | 1.06E-01 | 1.87E+00 | 1.96E+00 | 2.86E-01 | no | Serhl | intron_variant | MODIFIER |
| 15 | 83103992 | AX-170452072 | 0 | A | T | 0.088 | -1.40E-01 | 1.06E-01 | 1.87E+00 | 1.96E+00 | 2.86E-01 | no | Serhl | intron_variant | MODIFIER |
| 15 | 83359183 | AX-97832867 | 0 | A | G | 0.088 | -1.40E-01 | 1.06E-01 | 1.87E+00 | 1.96E+00 | 2.86E-01 | no | 1700001L05Rik | intron_variant,non_coding_transcript_variant | MODIFIER |
| 15 | 83382882 | AX-170225678 | 0 | T | C | 0.088 | -1.40E-01 | 1.06E-01 | 1.87E+00 | 1.96E+00 | 2.86E-01 | no | Pacsin2 | intron_variant | MODIFIER |
| 15 | 83808488 | AX-97855101 | 0 | T | C | 0.088 | -1.40E-01 | 1.06E-01 | 1.87E+00 | 1.96E+00 | 2.86E-01 | no | Mpped1 | intron_variant | MODIFIER |
| 15 | 83808756 | AX-170533207 | 0 | C | T | 0.088 | -1.40E-01 | 1.06E-01 | 1.87E+00 | 1.96E+00 | 2.86E-01 | no | Mpped1 | intron_variant | MODIFIER |
| 15 | 83899229 | AX-170165411 | 0 | C | T | 0.088 | -1.40E-01 | 1.06E-01 | 1.87E+00 | 1.96E+00 | 2.86E-01 | no | Efcab6 | intron_variant,non_coding_transcript_variant | MODIFIER |
| 15 | 83937937 | AX-97829929 | 0 | T | C | 0.088 | -1.40E-01 | 1.06E-01 | 1.87E+00 | 1.96E+00 | 2.86E-01 | no | Efcab6 | upstream_gene_variant | MODIFIER |
| 15 | 84003609 | AX-97829928 | 0 | C | T | 0.088 | -1.40E-01 | 1.06E-01 | 1.87E+00 | 1.96E+00 | 2.86E-01 | no | Efcab6 | intron_variant | MODIFIER |
| 15 | 84011097 | AX-170173556 | 0 | G | A | 0.088 | -1.40E-01 | 1.06E-01 | 1.87E+00 | 1.96E+00 | 2.86E-01 | no | Efcab6 | intron_variant | MODIFIER |
| 15 | 84011918 | AX-170210069 | 0 | T | A | 0.088 | -1.40E-01 | 1.06E-01 | 1.87E+00 | 1.96E+00 | 2.86E-01 | no | Efcab6 | intron_variant | MODIFIER |
| 15 | 84037487 | AX-97804787 | 0 | G | T | 0.088 | -1.40E-01 | 1.06E-01 | 1.87E+00 | 1.96E+00 | 2.86E-01 | no | Efcab6 | intron_variant,non_coding_transcript_variant | MODIFIER |
| 15 | 84162417 | AX-97789835 | 0 | A | G | 0.088 | -1.40E-01 | 1.06E-01 | 1.87E+00 | 1.96E+00 | 2.86E-01 | no | | intergenic_variant | MODIFIER |
| 15 | 84248407 | AX-170355763 | 0 | C | A | 0.088 | -1.40E-01 | 1.06E-01 | 1.87E+00 | 1.96E+00 | 2.86E-01 | no | Parvb | intron_variant | MODIFIER |
| 15 | 84303101 | AX-170151698 | 0 | G | A | 0.088 | -1.40E-01 | 1.06E-01 | 1.87E+00 | 1.96E+00 | 2.86E-01 | no | Parvb | intron_variant | MODIFIER |
| 15 | 84428171 | AX-170467649 | 0 | T | C | 0.088 | -1.40E-01 | 1.06E-01 | 1.87E+00 | 1.96E+00 | 2.86E-01 | no | Shisal1 | intron_variant | MODIFIER |
| 15 | 84619056 | AX-170012153 | 0 | G | C | 0.088 | -1.40E-01 | 1.06E-01 | 1.87E+00 | 1.96E+00 | 2.86E-01 | no | | intergenic_variant | MODIFIER |
| 15 | 84623297 | AX-170056165 | 0 | C | A | 0.088 | -1.40E-01 | 1.06E-01 | 1.87E+00 | 1.96E+00 | 2.86E-01 | no | | intergenic_variant | MODIFIER |
| 15 | 5189687 | AX-170037222 | 1 | A | C | 0.241 | -1.02E-01 | 9.65E-02 | 2.64E+00 | 2.71E+00 | 2.86E-01 | no | Ttc33 | missense_variant | MODERATE |
| 4 | 47369078 | AX-170220086 | 1 | G | A | 0.357 | -1.01E-01 | 9.56E-02 | 2.86E+00 | 2.96E+00 | 2.86E-01 | no | Tgfbr1 | intron_variant | MODIFIER |
| 5 | 50950329 | AX-170409524 | 0 | A | T | 0.491 | 1.04E-01 | 9.67E-02 | 2.47E+00 | 2.55E+00 | 2.86E-01 | no | | intergenic_variant | MODIFIER |
| 4 | 129733906 | AX-170426916 | 0 | G | A | 0.193 | 1.13E-01 | 1.06E-01 | 2.58E+00 | 2.64E+00 | 2.86E-01 | no | Khdrbs1 | intron_variant | MODIFIER |
| 1 | 21641028 | AX-170467601 | 0 | C | A | 0.395 | 8.33E-02 | 7.84E-02 | 2.58E+00 | 2.65E+00 | 2.86E-01 | no | Kcnq5 | intron_variant | MODIFIER |
| 15 | 93884200 | AX-170222317 | 1 | A | T | 0.152 | 1.64E-01 | 1.55E-01 | 2.68E+00 | 2.78E+00 | 2.86E-01 | Yes | | intergenic_variant | MODIFIER |
| 4 | 129525212 | AX-170023933 | 1 | C | T | 0.161 | 1.24E-01 | 1.17E-01 | 2.58E+00 | 2.62E+00 | 2.86E-01 | no | Hdac1 | intron_variant | MODIFIER |
| 15 | 6960902 | AX-170468754 | 1 | C | A | 0.241 | -1.12E-01 | 1.06E-01 | 2.67E+00 | 2.73E+00 | 2.87E-01 | no | | intergenic_variant | MODIFIER |
| 1 | 127692943 | AX-170086606 | 0 | T | A | 0.535 | 8.71E-02 | 8.24E-02 | 2.67E+00 | 2.73E+00 | 2.87E-01 | no | | intergenic_variant | MODIFIER |
| 11 | 9753728 | AX-170143513 | 0 | T | A | 0.07 | 1.81E-01 | 1.71E-01 | 2.85E+00 | 3.00E+00 | 2.87E-01 | Yes | | intergenic_variant | MODIFIER |
| 11 | 17640746 | AX-170107571 | 0 | T | C | 0.07 | 1.81E-01 | 1.71E-01 | 2.85E+00 | 3.00E+00 | 2.87E-01 | Yes | Gm12016 | upstream_gene_variant | MODIFIER |
| 18 | 69019406 | AX-170490815 | 1 | T | G | 0.161 | -1.29E-01 | 1.22E-01 | 2.59E+00 | 2.64E+00 | 2.87E-01 | no | Mir145b | downstream_gene_variant | MODIFIER |
| 1 | 50224000 | AX-170128912 | 0 | G | A | 0.114 | 1.89E-01 | 1.80E-01 | 2.81E+00 | 2.94E+00 | 2.87E-01 | Yes | Gm27330 | upstream_gene_variant | MODIFIER |
| 4 | 75439909 | AX-97802315 | 0 | G | A | 0.333 | 1.23E-01 | 1.17E-01 | 2.73E+00 | 2.79E+00 | 2.87E-01 | no | | intergenic_variant | MODIFIER |
| 3 | 37329947 | AX-97827327 | 2 | C | T | 0.2 | 9.89E-02 | 9.31E-02 | 2.92E+00 | 3.04E+00 | 2.87E-01 | no | Gm12540 | downstream_gene_variant | MODIFIER |
| 1 | 122830518 | AX-170357279 | 0 | C | T | 0.561 | 8.95E-02 | 8.41E-02 | 2.56E+00 | 2.61E+00 | 2.87E-01 | no | | intergenic_variant | MODIFIER |
| 4 | 63898025 | AX-97788071 | 1 | T | C | 0.348 | 1.21E-01 | 1.15E-01 | 2.77E+00 | 2.83E+00 | 2.87E-01 | no | Rpl17-ps4 | upstream_gene_variant | MODIFIER |
| 1 | 171696682 | AX-170288944 | 1 | T | C | 0.214 | 1.43E-01 | 1.35E-01 | 2.64E+00 | 2.69E+00 | 2.87E-01 | no | Cd48 | intron_variant | MODIFIER |
| 13 | 40482106 | AX-170401618 | 0 | T | C | 0.184 | -1.14E-01 | 1.05E-01 | 2.39E+00 | 2.48E+00 | 2.87E-01 | no | | intergenic_variant | MODIFIER |
| 8 | 93852242 | AX-170031274 | 0 | A | G | 0.211 | -1.19E-01 | 1.13E-01 | 2.80E+00 | 2.89E+00 | 2.87E-01 | no | Gnao1 | intron_variant | MODIFIER |

| | | | | | | | | | | | | | | | |
|---|---|---|---|---|---|---|---|---|---|---|---|---|---|---|---|
| 10 | 101219719 | AX-97848201 | 0 | G | T | 0.298 | -8.87E-02 | 8.36E-02 | 2.59E+00 | 2.66E+00 | 2.87E-01 | no | | intergenic_variant | MODIFIER |
| 4 | 69588674 | AX-170140481 | 0 | C | A | 0.333 | 1.26E-01 | 1.20E-01 | 2.75E+00 | 2.81E+00 | 2.87E-01 | no | | intergenic_variant | MODIFIER |
| 18 | 74634830 | AX-170061328 | 0 | T | G | 0.421 | 9.04E-02 | 8.48E-02 | 2.53E+00 | 2.60E+00 | 2.87E-01 | no | Myo5b | intron_variant | MODIFIER |
| 9 | 8350041 | AX-170031765 | 1 | G | A | 0.473 | 8.49E-02 | 7.86E-02 | 2.43E+00 | 2.52E+00 | 2.88E-01 | no | | intergenic_variant | MODIFIER |
| 16 | 29273989 | AX-170133280 | 0 | A | G | 0.5 | 8.30E-02 | 7.75E-02 | 2.50E+00 | 2.55E+00 | 2.88E-01 | no | Atp13a5 | intron_variant | MODIFIER |
| 16 | 80385836 | AX-170057920 | 1 | C | T | 0.473 | 7.97E-02 | 7.51E-02 | 2.57E+00 | 2.64E+00 | 2.88E-01 | no | | intergenic_variant | MODIFIER |
| 1 | 120271930 | AX-170001552 | 0 | A | G | 0.561 | 8.49E-02 | 7.93E-02 | 2.49E+00 | 2.57E+00 | 2.88E-01 | no | Steap3 | upstream_gene_variant | MODIFIER |
| 4 | 135208082 | AX-170258137 | 0 | C | T | 0.184 | 1.10E-01 | 1.04E-01 | 2.63E+00 | 2.69E+00 | 2.88E-01 | no | | intergenic_variant | MODIFIER |
| 18 | 66377879 | AX-170137781 | 1 | G | A | 0.277 | -9.24E-02 | 8.78E-02 | 2.67E+00 | 2.79E+00 | 2.88E-01 | no | | intergenic_variant | MODIFIER |
| 4 | 95947911 | AX-170023450 | 2 | A | T | 0.245 | 1.17E-01 | 1.10E-01 | 2.51E+00 | 2.58E+00 | 2.88E-01 | no | | intergenic_variant | MODIFIER |
| 5 | 84029807 | AX-170288106 | 0 | A | G | 0.447 | 1.07E-01 | 1.01E-01 | 2.79E+00 | 2.87E+00 | 2.88E-01 | no | | intergenic_variant | MODIFIER |
| 4 | 80152448 | AX-97827403 | 1 | A | G | 0.455 | -8.90E-02 | 8.36E-02 | 2.53E+00 | 2.60E+00 | 2.88E-01 | no | | intergenic_variant | MODIFIER |
| 9 | 89971481 | AX-170105454 | 0 | C | T | 0.132 | -1.16E-01 | 1.10E-01 | 2.60E+00 | 2.72E+00 | 2.88E-01 | no | Rasgrf1 | intron_variant | MODIFIER |
| 14 | 118999682 | AX-97828768 | 1 | A | G | 0.321 | 7.76E-02 | 7.35E-02 | 2.60E+00 | 2.71E+00 | 2.88E-01 | no | Uggt2 | intron_variant,non_coding_transcript_variant | MODIFIER |
| 18 | 71929245 | AX-170015956 | 0 | C | A | 0.184 | -1.18E-01 | 1.12E-01 | 2.62E+00 | 2.70E+00 | 2.88E-01 | no | Dcc | intron_variant | MODIFIER |
| 1 | 20803367 | AX-170323821 | 0 | G | A | 0.404 | 8.71E-02 | 8.22E-02 | 2.58E+00 | 2.63E+00 | 2.88E-01 | no | Mcm3 | 3_prime_UTR_variant | MODIFIER |
| 1 | 124562827 | AX-170042568 | 0 | C | T | 0.561 | 8.91E-02 | 8.39E-02 | 2.56E+00 | 2.61E+00 | 2.88E-01 | no | Gm28928 | intron_variant,non_coding_transcript_variant | MODIFIER |
| 15 | 5879916 | AX-170010949 | 1 | T | A | 0.125 | -7.44E-02 | 5.96E-02 | 1.98E+00 | 2.08E+00 | 2.89E-01 | no | | intergenic_variant | MODIFIER |
| 1 | 120971899 | AX-170433304 | 1 | A | G | 0.571 | 8.94E-02 | 8.38E-02 | 2.51E+00 | 2.57E+00 | 2.89E-01 | no | | intergenic_variant | MODIFIER |
| 15 | 66862876 | AX-170525251 | 1 | C | T | 0.339 | -8.42E-02 | 7.56E-02 | 2.29E+00 | 2.37E+00 | 2.89E-01 | no | | intergenic_variant | MODIFIER |
| 4 | 79978897 | AX-97789686 | 0 | G | T | 0.456 | -9.31E-02 | 8.80E-02 | 2.58E+00 | 2.63E+00 | 2.89E-01 | no | | intergenic_variant | MODIFIER |
| 14 | 111076630 | AX-97801339 | 0 | A | G | 0.316 | 7.71E-02 | 7.32E-02 | 2.60E+00 | 2.70E+00 | 2.89E-01 | no | | intergenic_variant | MODIFIER |
| 14 | 111076683 | AX-170534761 | 0 | A | G | 0.316 | 7.71E-02 | 7.32E-02 | 2.60E+00 | 2.70E+00 | 2.89E-01 | no | | intergenic_variant | MODIFIER |
| 14 | 111103641 | AX-170496479 | 0 | T | C | 0.316 | 7.71E-02 | 7.32E-02 | 2.60E+00 | 2.70E+00 | 2.89E-01 | no | | regulatory_region_variant | MODIFIER |
| 14 | 111112881 | AX-170220147 | 0 | G | C | 0.316 | 7.71E-02 | 7.32E-02 | 2.60E+00 | 2.70E+00 | 2.89E-01 | no | | intergenic_variant | MODIFIER |
| 14 | 111128759 | AX-170105478 | 0 | T | A | 0.316 | 7.71E-02 | 7.32E-02 | 2.60E+00 | 2.70E+00 | 2.89E-01 | no | | intergenic_variant | MODIFIER |
| 14 | 111136660 | AX-170203641 | 0 | C | T | 0.316 | 7.71E-02 | 7.32E-02 | 2.60E+00 | 2.70E+00 | 2.89E-01 | no | | intergenic_variant | MODIFIER |
| 14 | 111169330 | AX-97848316 | 0 | C | T | 0.316 | 7.71E-02 | 7.32E-02 | 2.60E+00 | 2.70E+00 | 2.89E-01 | no | | intergenic_variant | MODIFIER |
| 14 | 111175643 | AX-170523434 | 0 | G | C | 0.316 | 7.71E-02 | 7.32E-02 | 2.60E+00 | 2.70E+00 | 2.89E-01 | no | | intergenic_variant | MODIFIER |
| 14 | 111180492 | AX-170247346 | 0 | A | C | 0.316 | 7.71E-02 | 7.32E-02 | 2.60E+00 | 2.70E+00 | 2.89E-01 | no | | intergenic_variant | MODIFIER |
| 14 | 111228764 | AX-170479499 | 0 | G | T | 0.316 | 7.71E-02 | 7.32E-02 | 2.60E+00 | 2.70E+00 | 2.89E-01 | no | | intergenic_variant | MODIFIER |
| 14 | 111242893 | AX-170054460 | 0 | A | G | 0.316 | 7.71E-02 | 7.32E-02 | 2.60E+00 | 2.70E+00 | 2.89E-01 | no | | intergenic_variant | MODIFIER |
| 14 | 111247306 | AX-97837567 | 0 | T | G | 0.316 | 7.71E-02 | 7.32E-02 | 2.60E+00 | 2.70E+00 | 2.89E-01 | no | | intergenic_variant | MODIFIER |
| 14 | 111247337 | AX-170010692 | 0 | A | G | 0.316 | 7.71E-02 | 7.32E-02 | 2.60E+00 | 2.70E+00 | 2.89E-01 | no | | intergenic_variant | MODIFIER |
| 14 | 111392000 | AX-97827846 | 0 | G | A | 0.316 | 7.71E-02 | 7.32E-02 | 2.60E+00 | 2.70E+00 | 2.89E-01 | no | | intergenic_variant | MODIFIER |
| 14 | 111869859 | AX-97820703 | 0 | T | C | 0.316 | 7.71E-02 | 7.32E-02 | 2.60E+00 | 2.70E+00 | 2.89E-01 | no | | intergenic_variant | MODIFIER |
| 14 | 111870040 | AX-170149745 | 0 | G | A | 0.316 | 7.71E-02 | 7.32E-02 | 2.60E+00 | 2.70E+00 | 2.89E-01 | no | | intergenic_variant | MODIFIER |
| 14 | 111883834 | AX-97827585 | 0 | A | G | 0.316 | 7.71E-02 | 7.32E-02 | 2.60E+00 | 2.70E+00 | 2.89E-01 | no | | intergenic_variant | MODIFIER |
| 14 | 111944018 | AX-97846689 | 0 | T | G | 0.316 | 7.71E-02 | 7.32E-02 | 2.60E+00 | 2.70E+00 | 2.89E-01 | no | | intergenic_variant | MODIFIER |
| 14 | 111976016 | AX-170115344 | 0 | C | G | 0.316 | 7.71E-02 | 7.32E-02 | 2.60E+00 | 2.70E+00 | 2.89E-01 | no | | intergenic_variant | MODIFIER |
| 14 | 112003431 | AX-170129798 | 0 | A | T | 0.316 | 7.71E-02 | 7.32E-02 | 2.60E+00 | 2.70E+00 | 2.89E-01 | no | | intergenic_variant | MODIFIER |
| 14 | 112013399 | AX-170010705 | 0 | G | T | 0.316 | 7.71E-02 | 7.32E-02 | 2.60E+00 | 2.70E+00 | 2.89E-01 | no | | intergenic_variant | MODIFIER |
| 14 | 112106643 | AX-170413281 | 0 | A | T | 0.316 | 7.71E-02 | 7.32E-02 | 2.60E+00 | 2.70E+00 | 2.89E-01 | no | | intergenic_variant | MODIFIER |
| 14 | 112164168 | AX-170234339 | 0 | T | C | 0.316 | 7.71E-02 | 7.32E-02 | 2.60E+00 | 2.70E+00 | 2.89E-01 | no | | intergenic_variant | MODIFIER |
| 14 | 112165232 | AX-170223655 | 0 | G | C | 0.316 | 7.71E-02 | 7.32E-02 | 2.60E+00 | 2.70E+00 | 2.89E-01 | no | | intergenic_variant | MODIFIER |
| 14 | 112173472 | AX-170432684 | 0 | C | T | 0.316 | 7.71E-02 | 7.32E-02 | 2.60E+00 | 2.70E+00 | 2.89E-01 | no | | intergenic_variant | MODIFIER |
| 14 | 112225864 | AX-170273669 | 0 | T | C | 0.316 | 7.71E-02 | 7.32E-02 | 2.60E+00 | 2.70E+00 | 2.89E-01 | no | | intergenic_variant | MODIFIER |
| 14 | 112228376 | AX-170207103 | 0 | G | A | 0.316 | 7.71E-02 | 7.32E-02 | 2.60E+00 | 2.70E+00 | 2.89E-01 | no | | intergenic_variant | MODIFIER |
| 14 | 112229650 | AX-170310565 | 0 | C | T | 0.316 | 7.71E-02 | 7.32E-02 | 2.60E+00 | 2.70E+00 | 2.89E-01 | no | | intergenic_variant | MODIFIER |
| 14 | 112229822 | AX-170164295 | 0 | T | C | 0.316 | 7.71E-02 | 7.32E-02 | 2.60E+00 | 2.70E+00 | 2.89E-01 | no | | intergenic_variant | MODIFIER |
| 14 | 112233605 | AX-170278914 | 0 | G | A | 0.316 | 7.71E-02 | 7.32E-02 | 2.60E+00 | 2.70E+00 | 2.89E-01 | no | | intergenic_variant | MODIFIER |
| 14 | 112243035 | AX-170054476 | 0 | C | T | 0.316 | 7.71E-02 | 7.32E-02 | 2.60E+00 | 2.70E+00 | 2.89E-01 | no | | intergenic_variant | MODIFIER |
| 14 | 112243732 | AX-97795009 | 0 | T | A | 0.316 | 7.71E-02 | 7.32E-02 | 2.60E+00 | 2.70E+00 | 2.89E-01 | no | | intergenic_variant | MODIFIER |
| 14 | 112272629 | AX-170515527 | 0 | A | C | 0.316 | 7.71E-02 | 7.32E-02 | 2.60E+00 | 2.70E+00 | 2.89E-01 | no | | intergenic_variant | MODIFIER |

| | | | | | | | | | | | | | | | |
|---|---|---|---|---|---|---|---|---|---|---|---|---|---|---|---|
| 14 | 112293158 | AX-170054477 | 0 | C | A | 0.316 | 7.71E-02 | 7.32E-02 | 2.60E+00 | 2.70E+00 | 2.89E-01 | no | | intergenic_variant | MODIFIER |
| 14 | 112293219 | AX-170524273 | 0 | A | G | 0.316 | 7.71E-02 | 7.32E-02 | 2.60E+00 | 2.70E+00 | 2.89E-01 | no | | intergenic_variant | MODIFIER |
| 14 | 112293469 | AX-170349735 | 0 | G | A | 0.316 | 7.71E-02 | 7.32E-02 | 2.60E+00 | 2.70E+00 | 2.89E-01 | no | | intergenic_variant | MODIFIER |
| 14 | 112294270 | AX-170339861 | 0 | C | T | 0.316 | 7.71E-02 | 7.32E-02 | 2.60E+00 | 2.70E+00 | 2.89E-01 | no | | intergenic_variant | MODIFIER |
| 14 | 112295058 | AX-97854240 | 0 | T | C | 0.316 | 7.71E-02 | 7.32E-02 | 2.60E+00 | 2.70E+00 | 2.89E-01 | no | | intergenic_variant | MODIFIER |
| 14 | 112295167 | AX-170443850 | 0 | T | A | 0.316 | 7.71E-02 | 7.32E-02 | 2.60E+00 | 2.70E+00 | 2.89E-01 | no | | intergenic_variant | MODIFIER |
| 14 | 112386104 | AX-170511173 | 0 | A | G | 0.316 | 7.71E-02 | 7.32E-02 | 2.60E+00 | 2.70E+00 | 2.89E-01 | no | | intergenic_variant | MODIFIER |
| 14 | 112387172 | AX-170506118 | 0 | C | T | 0.316 | 7.71E-02 | 7.32E-02 | 2.60E+00 | 2.70E+00 | 2.89E-01 | no | | intergenic_variant | MODIFIER |
| 14 | 112387532 | AX-170159545 | 0 | C | T | 0.316 | 7.71E-02 | 7.32E-02 | 2.60E+00 | 2.70E+00 | 2.89E-01 | no | | intergenic_variant | MODIFIER |
| 14 | 112397264 | AX-170259110 | 0 | A | G | 0.316 | 7.71E-02 | 7.32E-02 | 2.60E+00 | 2.70E+00 | 2.89E-01 | no | | intergenic_variant | MODIFIER |
| 14 | 112397842 | AX-170523385 | 0 | T | C | 0.316 | 7.71E-02 | 7.32E-02 | 2.60E+00 | 2.70E+00 | 2.89E-01 | no | | intergenic_variant | MODIFIER |
| 14 | 112419990 | AX-97792064 | 0 | A | G | 0.316 | 7.71E-02 | 7.32E-02 | 2.60E+00 | 2.70E+00 | 2.89E-01 | no | | intergenic_variant | MODIFIER |
| 14 | 112421909 | AX-170100518 | 0 | A | G | 0.316 | 7.71E-02 | 7.32E-02 | 2.60E+00 | 2.70E+00 | 2.89E-01 | no | | intergenic_variant | MODIFIER |
| 14 | 112422116 | AX-170217618 | 0 | A | G | 0.316 | 7.71E-02 | 7.32E-02 | 2.60E+00 | 2.70E+00 | 2.89E-01 | no | | intergenic_variant | MODIFIER |
| 14 | 112447548 | AX-170010723 | 0 | T | G | 0.316 | 7.71E-02 | 7.32E-02 | 2.60E+00 | 2.70E+00 | 2.89E-01 | no | | intergenic_variant | MODIFIER |
| 14 | 112475110 | AX-170010720 | 0 | C | T | 0.316 | 7.71E-02 | 7.32E-02 | 2.60E+00 | 2.70E+00 | 2.89E-01 | no | | intergenic_variant | MODIFIER |
| 14 | 112580656 | AX-170010724 | 0 | A | C | 0.316 | 7.71E-02 | 7.32E-02 | 2.60E+00 | 2.70E+00 | 2.89E-01 | no | Gm19223 | upstream_gene_variant | MODIFIER |
| 14 | 112607403 | AX-170334668 | 0 | A | T | 0.316 | 7.71E-02 | 7.32E-02 | 2.60E+00 | 2.70E+00 | 2.89E-01 | no | | intergenic_variant | MODIFIER |
| 14 | 112612793 | AX-170010715 | 0 | G | A | 0.316 | 7.71E-02 | 7.32E-02 | 2.60E+00 | 2.70E+00 | 2.89E-01 | no | | intergenic_variant | MODIFIER |
| 14 | 112619334 | AX-170010719 | 0 | T | G | 0.316 | 7.71E-02 | 7.32E-02 | 2.60E+00 | 2.70E+00 | 2.89E-01 | no | | intergenic_variant | MODIFIER |
| 14 | 112635539 | AX-170010713 | 0 | C | T | 0.316 | 7.71E-02 | 7.32E-02 | 2.60E+00 | 2.70E+00 | 2.89E-01 | no | | intergenic_variant | MODIFIER |
| 14 | 112656658 | AX-170010716 | 0 | G | A | 0.316 | 7.71E-02 | 7.32E-02 | 2.60E+00 | 2.70E+00 | 2.89E-01 | no | | intergenic_variant | MODIFIER |
| 14 | 112753239 | AX-170171924 | 0 | T | C | 0.316 | 7.71E-02 | 7.32E-02 | 2.60E+00 | 2.70E+00 | 2.89E-01 | no | | intergenic_variant | MODIFIER |
| 14 | 113690226 | AX-170204896 | 0 | A | G | 0.316 | 7.71E-02 | 7.32E-02 | 2.60E+00 | 2.70E+00 | 2.89E-01 | no | | intergenic_variant | MODIFIER |
| 14 | 114049325 | AX-170291765 | 0 | C | G | 0.316 | 7.71E-02 | 7.32E-02 | 2.60E+00 | 2.70E+00 | 2.89E-01 | no | | intergenic_variant | MODIFIER |
| 14 | 115702035 | AX-97827992 | 0 | A | G | 0.316 | 7.71E-02 | 7.32E-02 | 2.60E+00 | 2.70E+00 | 2.89E-01 | no | Gpc5 | intron_variant | MODIFIER |
| 14 | 115722935 | AX-97848470 | 0 | C | A | 0.316 | 7.71E-02 | 7.32E-02 | 2.60E+00 | 2.70E+00 | 2.89E-01 | no | Gpc5 | intron_variant | MODIFIER |
| 14 | 116579512 | AX-170278503 | 0 | C | T | 0.316 | 7.71E-02 | 7.32E-02 | 2.60E+00 | 2.70E+00 | 2.89E-01 | no | | intergenic_variant | MODIFIER |
| 14 | 116620286 | AX-170260407 | 0 | A | T | 0.316 | 7.71E-02 | 7.32E-02 | 2.60E+00 | 2.70E+00 | 2.89E-01 | no | | intergenic_variant | MODIFIER |
| 14 | 116630202 | AX-97848752 | 0 | A | G | 0.316 | 7.71E-02 | 7.32E-02 | 2.60E+00 | 2.70E+00 | 2.89E-01 | no | | intergenic_variant | MODIFIER |
| 14 | 116654625 | AX-97844574 | 0 | T | C | 0.316 | 7.71E-02 | 7.32E-02 | 2.60E+00 | 2.70E+00 | 2.89E-01 | no | | intergenic_variant | MODIFIER |
| 14 | 116658557 | AX-168382048 | 0 | G | A | 0.316 | 7.71E-02 | 7.32E-02 | 2.60E+00 | 2.70E+00 | 2.89E-01 | no | | intergenic_variant | MODIFIER |
| 14 | 116692972 | AX-97854456 | 0 | T | G | 0.316 | 7.71E-02 | 7.32E-02 | 2.60E+00 | 2.70E+00 | 2.89E-01 | no | | intergenic_variant | MODIFIER |
| 14 | 116700482 | AX-170109764 | 0 | C | A | 0.316 | 7.71E-02 | 7.32E-02 | 2.60E+00 | 2.70E+00 | 2.89E-01 | no | | intergenic_variant | MODIFIER |
| 14 | 116711453 | AX-170191205 | 0 | T | C | 0.316 | 7.71E-02 | 7.32E-02 | 2.60E+00 | 2.70E+00 | 2.89E-01 | no | | intergenic_variant | MODIFIER |
| 14 | 116776694 | AX-170143518 | 0 | C | T | 0.316 | 7.71E-02 | 7.32E-02 | 2.60E+00 | 2.70E+00 | 2.89E-01 | no | | intergenic_variant | MODIFIER |
| 14 | 116846854 | AX-97785131 | 0 | C | T | 0.316 | 7.71E-02 | 7.32E-02 | 2.60E+00 | 2.70E+00 | 2.89E-01 | no | | intergenic_variant | MODIFIER |
| 14 | 116851052 | AX-170010783 | 0 | A | T | 0.316 | 7.71E-02 | 7.32E-02 | 2.60E+00 | 2.70E+00 | 2.89E-01 | no | | intergenic_variant | MODIFIER |
| 14 | 116899665 | AX-97796730 | 0 | T | C | 0.316 | 7.71E-02 | 7.32E-02 | 2.60E+00 | 2.70E+00 | 2.89E-01 | no | | intergenic_variant | MODIFIER |
| 14 | 116909890 | AX-170265164 | 0 | G | C | 0.316 | 7.71E-02 | 7.32E-02 | 2.60E+00 | 2.70E+00 | 2.89E-01 | no | | intergenic_variant | MODIFIER |
| 14 | 116933639 | AX-170407839 | 0 | A | G | 0.316 | 7.71E-02 | 7.32E-02 | 2.60E+00 | 2.70E+00 | 2.89E-01 | no | Gpc6 | intron_variant | MODIFIER |
| 14 | 116949626 | AX-97841634 | 0 | G | T | 0.316 | 7.71E-02 | 7.32E-02 | 2.60E+00 | 2.70E+00 | 2.89E-01 | no | Gpc6 | intron_variant | MODIFIER |
| 14 | 116951146 | AX-170263272 | 0 | C | G | 0.316 | 7.71E-02 | 7.32E-02 | 2.60E+00 | 2.70E+00 | 2.89E-01 | no | Gpc6 | intron_variant | MODIFIER |
| 14 | 116979303 | AX-170010776 | 0 | C | A | 0.316 | 7.71E-02 | 7.32E-02 | 2.60E+00 | 2.70E+00 | 2.89E-01 | no | Gpc6 | intron_variant | MODIFIER |
| 14 | 116982029 | AX-97835040 | 0 | G | C | 0.316 | 7.71E-02 | 7.32E-02 | 2.60E+00 | 2.70E+00 | 2.89E-01 | no | Gpc6 | intron_variant | MODIFIER |
| 14 | 117009591 | AX-170531184 | 0 | T | A | 0.316 | 7.71E-02 | 7.32E-02 | 2.60E+00 | 2.70E+00 | 2.89E-01 | no | Gpc6 | intron_variant | MODIFIER |
| 14 | 117097938 | AX-97837716 | 0 | C | T | 0.316 | 7.71E-02 | 7.32E-02 | 2.60E+00 | 2.70E+00 | 2.89E-01 | no | Gpc6 | intron_variant | MODIFIER |
| 14 | 117097953 | AX-170350570 | 0 | T | A | 0.316 | 7.71E-02 | 7.32E-02 | 2.60E+00 | 2.70E+00 | 2.89E-01 | no | Gpc6 | intron_variant | MODIFIER |
| 14 | 117137879 | AX-170398299 | 0 | T | A | 0.316 | 7.71E-02 | 7.32E-02 | 2.60E+00 | 2.70E+00 | 2.89E-01 | no | Gpc6 | intron_variant | MODIFIER |
| 14 | 117143301 | AX-97827738 | 0 | A | G | 0.316 | 7.71E-02 | 7.32E-02 | 2.60E+00 | 2.70E+00 | 2.89E-01 | no | Gpc6 | intron_variant | MODIFIER |
| 14 | 117190003 | AX-170183327 | 0 | G | A | 0.316 | 7.71E-02 | 7.32E-02 | 2.60E+00 | 2.70E+00 | 2.89E-01 | no | Gpc6 | intron_variant | MODIFIER |
| 14 | 117227556 | AX-170181039 | 0 | C | T | 0.316 | 7.71E-02 | 7.32E-02 | 2.60E+00 | 2.70E+00 | 2.89E-01 | no | Gpc6 | intron_variant | MODIFIER |
| 14 | 117848689 | AX-170010793 | 0 | T | A | 0.316 | 7.71E-02 | 7.32E-02 | 2.60E+00 | 2.70E+00 | 2.89E-01 | no | Gpc6 | intron_variant | MODIFIER |
| 14 | 118598853 | AX-170286751 | 0 | G | T | 0.316 | 7.71E-02 | 7.32E-02 | 2.60E+00 | 2.70E+00 | 2.89E-01 | no | Abcc4 | intron_variant | MODIFIER |

| | | | | | | | | | | | | | | | |
|---|---|---|---|---|---|---|---|---|---|---|---|---|---|---|---|
| 14 | 118607825 | AX-170412416 | 0 | T | G | 0.316 | 7.71E-02 | 7.32E-02 | 2.60E+00 | 2.70E+00 | 2.89E-01 | no | Abcc4 | intron_variant | MODIFIER |
| 14 | 118632104 | AX-170010811 | 0 | G | A | 0.316 | 7.71E-02 | 7.32E-02 | 2.60E+00 | 2.70E+00 | 2.89E-01 | no | Abcc4 | synonymous_variant | LOW |
| 14 | 118632116 | AX-170141144 | 0 | G | A | 0.316 | 7.71E-02 | 7.32E-02 | 2.60E+00 | 2.70E+00 | 2.89E-01 | no | Abcc4 | synonymous_variant | LOW |
| 14 | 119008882 | AX-170416578 | 0 | T | C | 0.316 | 7.71E-02 | 7.32E-02 | 2.60E+00 | 2.70E+00 | 2.89E-01 | no | Uggt2 | upstream_gene_variant | MODIFIER |
| 14 | 119107825 | AX-170453333 | 0 | G | A | 0.316 | 7.71E-02 | 7.32E-02 | 2.60E+00 | 2.70E+00 | 2.89E-01 | no | Gm16835 | intron_variant,non_coding_transcript_variant | MODIFIER |
| 14 | 119166560 | AX-97855483 | 0 | A | G | 0.316 | 7.71E-02 | 7.32E-02 | 2.60E+00 | 2.70E+00 | 2.89E-01 | no | Hs6st3 | intron_variant | MODIFIER |
| 14 | 119172320 | AX-170334906 | 0 | C | T | 0.316 | 7.71E-02 | 7.32E-02 | 2.60E+00 | 2.70E+00 | 2.89E-01 | no | Hs6st3 | intron_variant | MODIFIER |
| 14 | 119174003 | AX-170345101 | 0 | T | C | 0.316 | 7.71E-02 | 7.32E-02 | 2.60E+00 | 2.70E+00 | 2.89E-01 | no | Hs6st3 | intron_variant | MODIFIER |
| 14 | 119210365 | AX-170010821 | 0 | A | G | 0.316 | 7.71E-02 | 7.32E-02 | 2.60E+00 | 2.70E+00 | 2.89E-01 | no | Hs6st3 | intron_variant | MODIFIER |
| 14 | 119216192 | AX-97793677 | 0 | G | A | 0.316 | 7.71E-02 | 7.32E-02 | 2.60E+00 | 2.70E+00 | 2.89E-01 | no | Hs6st3 | intron_variant | MODIFIER |
| 4 | 69752917 | AX-170502360 | 0 | C | T | 0.325 | 1.18E-01 | 1.12E-01 | 2.80E+00 | 2.87E+00 | 2.89E-01 | no | | intergenic_variant | MODIFIER |
| 4 | 79404141 | AX-97812289 | 0 | T | C | 0.456 | -8.86E-02 | 8.34E-02 | 2.53E+00 | 2.60E+00 | 2.89E-01 | no | | intergenic_variant | MODIFIER |
| 13 | 5901762 | AX-170339575 | 1 | G | A | 0.205 | -1.31E-01 | 1.25E-01 | 2.63E+00 | 2.69E+00 | 2.89E-01 | no | Gm47507 | intron_variant,non_coding_transcript_variant | MODIFIER |
| 13 | 6455420 | AX-170195294 | 1 | C | T | 0.205 | -1.31E-01 | 1.25E-01 | 2.63E+00 | 2.69E+00 | 2.89E-01 | no | | intergenic_variant | MODIFIER |
| 4 | 120737593 | AX-97785771 | 0 | C | T | 0.228 | 1.16E-01 | 1.10E-01 | 2.59E+00 | 2.65E+00 | 2.89E-01 | no | Kcnq4 | intron_variant | MODIFIER |
| 6 | 135141445 | AX-170269302 | 1 | C | T | 0.562 | 9.54E-02 | 8.80E-02 | 2.41E+00 | 2.45E+00 | 2.89E-01 | no | Hebp1 | intron_variant | MODIFIER |
| 14 | 124107842 | AX-97790825 | 0 | G | A | 0.325 | 8.51E-02 | 8.11E-02 | 2.67E+00 | 2.78E+00 | 2.89E-01 | no | Fgf14 | intron_variant | MODIFIER |
| 9 | 6013561 | AX-97841108 | 1 | T | C | 0.464 | 8.32E-02 | 7.74E-02 | 2.44E+00 | 2.54E+00 | 2.89E-01 | no | | intergenic_variant | MODIFIER |
| 3 | 10688021 | AX-170019912 | 1 | T | C | 0.402 | -6.91E-02 | 6.61E-02 | 2.77E+00 | 2.92E+00 | 2.89E-01 | no | | intergenic_variant | MODIFIER |
| 4 | 89178448 | AX-170131884 | 1 | G | T | 0.25 | 1.19E-01 | 1.11E-01 | 2.44E+00 | 2.49E+00 | 2.89E-01 | no | Mtap | intron_variant | MODIFIER |
| 16 | 52383337 | AX-170339295 | 0 | G | T | 0.518 | 8.91E-02 | 8.34E-02 | 2.49E+00 | 2.53E+00 | 2.89E-01 | no | Alcam | intron_variant | MODIFIER |
| 8 | 49313211 | AX-170080023 | 0 | G | A | 0.149 | -1.27E-01 | 1.21E-01 | 2.57E+00 | 2.68E+00 | 2.89E-01 | no | | intergenic_variant | MODIFIER |
| 8 | 53370731 | AX-97848238 | 0 | T | G | 0.149 | -1.27E-01 | 1.21E-01 | 2.57E+00 | 2.68E+00 | 2.89E-01 | no | | intergenic_variant | MODIFIER |
| 8 | 54334922 | AX-97799815 | 0 | T | G | 0.149 | -1.27E-01 | 1.21E-01 | 2.57E+00 | 2.68E+00 | 2.89E-01 | no | | intergenic_variant | MODIFIER |
| 8 | 56291338 | AX-170214525 | 0 | A | G | 0.149 | -1.27E-01 | 1.21E-01 | 2.57E+00 | 2.68E+00 | 2.89E-01 | no | Hpgd | upstream_gene_variant | MODIFIER |
| 8 | 57699748 | AX-170330494 | 0 | T | C | 0.149 | -1.27E-01 | 1.21E-01 | 2.57E+00 | 2.68E+00 | 2.89E-01 | no | Gm45356 | upstream_gene_variant | MODIFIER |
| 10 | 18417250 | AX-170342519 | 1 | G | A | 0.196 | 1.08E-01 | 1.01E-01 | 3.06E+00 | 3.18E+00 | 2.89E-01 | no | Nhsl1 | intron_variant | MODIFIER |
| 4 | 75955815 | AX-170212326 | 0 | C | T | 0.325 | 1.23E-01 | 1.17E-01 | 2.79E+00 | 2.85E+00 | 2.89E-01 | no | Ptprd | intron_variant | MODIFIER |
| 17 | 79372592 | AX-170310959 | 1 | T | A | 0.062 | 1.42E-01 | 1.36E-01 | 2.79E+00 | 2.95E+00 | 2.89E-01 | no | | intergenic_variant | MODIFIER |
| 8 | 17582435 | AX-170420203 | 1 | C | T | 0.223 | -8.21E-02 | 7.76E-02 | 2.55E+00 | 2.64E+00 | 2.89E-01 | no | | intergenic_variant | MODIFIER |
| 8 | 19354686 | AX-170187423 | 0 | G | A | 0.167 | -1.11E-01 | 1.05E-01 | 2.91E+00 | 3.06E+00 | 2.90E-01 | no | | intergenic_variant | MODIFIER |
| 8 | 91130210 | AX-97792344 | 0 | T | C | 0.184 | -1.09E-01 | 1.04E-01 | 2.66E+00 | 2.78E+00 | 2.90E-01 | no | Aktip | intron_variant | MODIFIER |
| 18 | 80769329 | AX-170124586 | 0 | T | C | 0.553 | 7.77E-02 | 7.20E-02 | 2.42E+00 | 2.50E+00 | 2.90E-01 | no | Atp9b | intron_variant | MODIFIER |
| 18 | 80783063 | AX-170213900 | 0 | T | C | 0.553 | 7.77E-02 | 7.20E-02 | 2.42E+00 | 2.50E+00 | 2.90E-01 | no | Atp9b | intron_variant | MODIFIER |
| 18 | 80800827 | AX-97846639 | 0 | A | C | 0.553 | 7.77E-02 | 7.20E-02 | 2.42E+00 | 2.50E+00 | 2.90E-01 | no | Atp9b | intron_variant | MODIFIER |
| 18 | 80806310 | AX-170286671 | 0 | C | T | 0.553 | 7.77E-02 | 7.20E-02 | 2.42E+00 | 2.50E+00 | 2.90E-01 | no | Atp9b | intron_variant | MODIFIER |
| 18 | 80823231 | AX-170382847 | 0 | T | C | 0.553 | 7.77E-02 | 7.20E-02 | 2.42E+00 | 2.50E+00 | 2.90E-01 | no | Atp9b | intron_variant | MODIFIER |
| 18 | 80850894 | AX-97838454 | 0 | T | C | 0.553 | 7.77E-02 | 7.20E-02 | 2.42E+00 | 2.50E+00 | 2.90E-01 | no | Atp9b | intron_variant | MODIFIER |
| 18 | 80900458 | AX-170141580 | 0 | A | C | 0.553 | 7.77E-02 | 7.20E-02 | 2.42E+00 | 2.50E+00 | 2.90E-01 | no | Atp9b | intron_variant | MODIFIER |
| 18 | 80903162 | AX-170320387 | 0 | A | G | 0.553 | 7.77E-02 | 7.20E-02 | 2.42E+00 | 2.50E+00 | 2.90E-01 | no | Atp9b | intron_variant | MODIFIER |
| 18 | 80907436 | AX-170061466 | 0 | A | G | 0.553 | 7.77E-02 | 7.20E-02 | 2.42E+00 | 2.50E+00 | 2.90E-01 | no | Atp9b | intron_variant | MODIFIER |
| 18 | 80941220 | AX-170221104 | 0 | T | C | 0.553 | 7.77E-02 | 7.20E-02 | 2.42E+00 | 2.50E+00 | 2.90E-01 | no | | intergenic_variant | MODIFIER |
| 6 | 135401780 | AX-170316117 | 0 | A | G | 0.553 | 9.45E-02 | 8.78E-02 | 2.44E+00 | 2.49E+00 | 2.90E-01 | no | | intergenic_variant | MODIFIER |
| 16 | 39020936 | AX-170455675 | 0 | A | G | 0.5 | 8.63E-02 | 8.15E-02 | 2.55E+00 | 2.60E+00 | 2.90E-01 | no | Igsf11 | intron_variant | MODIFIER |
| 1 | 15702752 | AX-170391928 | 0 | G | A | 0.377 | 8.16E-02 | 7.71E-02 | 2.55E+00 | 2.61E+00 | 2.90E-01 | no | Kcnb2 | intron_variant | MODIFIER |
| 2 | 109907048 | AX-170088705 | 0 | C | T | 0.053 | 2.24E-01 | 2.14E-01 | 2.89E+00 | 3.04E+00 | 2.90E-01 | Yes | Platr8 | intron_variant,non_coding_transcript_variant | MODIFIER |
| 8 | 23714084 | AX-97837618 | 0 | A | C | 0.167 | -1.04E-01 | 1.00E-01 | 2.78E+00 | 2.92E+00 | 2.90E-01 | no | Zmat4 | intron_variant | MODIFIER |
| 3 | 30722869 | AX-170364733 | 0 | C | T | 0.175 | -1.00E-01 | 9.59E-02 | 2.82E+00 | 2.92E+00 | 2.90E-01 | no | | intergenic_variant | MODIFIER |
| 4 | 140826274 | AX-97821171 | 1 | T | A | 0.17 | 1.20E-01 | 1.14E-01 | 2.62E+00 | 2.67E+00 | 2.90E-01 | no | Padi1 | intron_variant | MODIFIER |
| 4 | 133783716 | AX-97854403 | 0 | C | T | 0.158 | 1.20E-01 | 1.12E-01 | 2.45E+00 | 2.50E+00 | 2.90E-01 | no | | intergenic_variant | MODIFIER |
| 18 | 86290475 | AX-170244725 | 0 | C | T | 0.158 | -1.11E-01 | 1.04E-01 | 2.51E+00 | 2.60E+00 | 2.91E-01 | no | | intergenic_variant | MODIFIER |
| 18 | 86574789 | AX-170152261 | 0 | G | A | 0.158 | -1.11E-01 | 1.04E-01 | 2.51E+00 | 2.60E+00 | 2.91E-01 | no | | intergenic_variant | MODIFIER |
| 18 | 86588974 | AX-170473432 | 0 | T | C | 0.158 | -1.11E-01 | 1.04E-01 | 2.51E+00 | 2.60E+00 | 2.91E-01 | no | | intergenic_variant | MODIFIER |
| 18 | 86590356 | AX-170473463 | 0 | A | G | 0.158 | -1.11E-01 | 1.04E-01 | 2.51E+00 | 2.60E+00 | 2.91E-01 | no | | intergenic_variant | MODIFIER |

| | | | | | | | | | | | | | | | |
|---|---|---|---|---|---|---|---|---|---|---|---|---|---|---|---|
| 18 | 59995520 | AX-170119046 | 1 | T | C | 0.411 | 1.01E-01 | 9.41E-02 | 2.45E+00 | 2.50E+00 | 2.91E-01 | no | | intergenic_variant | MODIFIER |
| 1 | 122482001 | AX-170212570 | 0 | C | T | 0.57 | 8.89E-02 | 8.42E-02 | 2.56E+00 | 2.61E+00 | 2.91E-01 | no | | intergenic_variant | MODIFIER |
| 4 | 54243059 | AX-97799439 | 0 | G | A | 0.368 | -1.08E-01 | 1.03E-01 | 2.84E+00 | 2.94E+00 | 2.91E-01 | no | | intergenic_variant | MODIFIER |
| 5 | 52170620 | AX-97794923 | 0 | G | A | 0.482 | 1.09E-01 | 1.02E-01 | 2.43E+00 | 2.48E+00 | 2.91E-01 | no | Dhx15 | intron_variant | MODIFIER |
| 1 | 131316540 | AX-97846251 | 0 | A | C | 0.544 | 8.98E-02 | 8.57E-02 | 2.66E+00 | 2.72E+00 | 2.91E-01 | no | Srgap2 | intron_variant | MODIFIER |
| 4 | 124064900 | AX-170531306 | 2 | C | T | 0.218 | 1.21E-01 | 1.14E-01 | 2.52E+00 | 2.59E+00 | 2.91E-01 | no | | intergenic_variant | MODIFIER |
| 13 | 3952584 | AX-170320818 | 0 | C | A | 0.202 | -1.30E-01 | 1.24E-01 | 2.62E+00 | 2.67E+00 | 2.91E-01 | no | Gm22460 | downstream_gene_variant | MODIFIER |
| 9 | 20472374 | AX-170524967 | 0 | C | T | 0.465 | 9.13E-02 | 8.66E-02 | 2.56E+00 | 2.63E+00 | 2.91E-01 | no | Zfp426 | intron_variant | MODIFIER |
| 9 | 21434941 | AX-170533966 | 0 | C | T | 0.465 | 9.13E-02 | 8.66E-02 | 2.56E+00 | 2.63E+00 | 2.91E-01 | no | Dnm2 | intron_variant | MODIFIER |
| 1 | 167332818 | AX-170115170 | 0 | G | A | 0.228 | 1.34E-01 | 1.28E-01 | 2.61E+00 | 2.68E+00 | 2.91E-01 | no | Tmco1 | 3_prime_UTR_variant,NMD_transcript_variant | MODIFIER |
| 16 | 29175847 | AX-170527728 | 1 | C | T | 0.509 | 8.33E-02 | 7.87E-02 | 2.53E+00 | 2.58E+00 | 2.92E-01 | no | | intergenic_variant | MODIFIER |
| 1 | 122431512 | AX-170497135 | 0 | G | A | 0.561 | 8.85E-02 | 8.40E-02 | 2.56E+00 | 2.61E+00 | 2.92E-01 | no | | intergenic_variant | MODIFIER |
| 1 | 122554923 | AX-170475796 | 0 | C | T | 0.561 | 8.85E-02 | 8.40E-02 | 2.56E+00 | 2.61E+00 | 2.92E-01 | no | Gm25771 | downstream_gene_variant | MODIFIER |
| 1 | 122667163 | AX-170371557 | 0 | G | A | 0.561 | 8.85E-02 | 8.40E-02 | 2.56E+00 | 2.61E+00 | 2.92E-01 | no | | intergenic_variant | MODIFIER |
| 1 | 122669834 | AX-170460500 | 0 | T | G | 0.561 | 8.85E-02 | 8.40E-02 | 2.56E+00 | 2.61E+00 | 2.92E-01 | no | | intergenic_variant | MODIFIER |
| 1 | 122698563 | AX-170479166 | 0 | G | A | 0.561 | 8.85E-02 | 8.40E-02 | 2.56E+00 | 2.61E+00 | 2.92E-01 | no | | intergenic_variant | MODIFIER |
| 1 | 122704101 | AX-97813183 | 0 | A | C | 0.561 | 8.85E-02 | 8.40E-02 | 2.56E+00 | 2.61E+00 | 2.92E-01 | no | | intergenic_variant | MODIFIER |
| 1 | 122743813 | AX-170244723 | 0 | A | C | 0.561 | 8.85E-02 | 8.40E-02 | 2.56E+00 | 2.61E+00 | 2.92E-01 | no | | intergenic_variant | MODIFIER |
| 1 | 122744556 | AX-170423285 | 0 | G | A | 0.561 | 8.85E-02 | 8.40E-02 | 2.56E+00 | 2.61E+00 | 2.92E-01 | no | | intergenic_variant | MODIFIER |
| 1 | 122763758 | AX-170042531 | 0 | A | G | 0.561 | 8.85E-02 | 8.40E-02 | 2.56E+00 | 2.61E+00 | 2.92E-01 | no | | intergenic_variant | MODIFIER |
| 1 | 122801117 | AX-170299024 | 0 | G | A | 0.561 | 8.85E-02 | 8.40E-02 | 2.56E+00 | 2.61E+00 | 2.92E-01 | no | | intergenic_variant | MODIFIER |
| 1 | 122835981 | AX-170380101 | 0 | A | G | 0.561 | 8.85E-02 | 8.40E-02 | 2.56E+00 | 2.61E+00 | 2.92E-01 | no | | intergenic_variant | MODIFIER |
| 1 | 122836318 | AX-170178099 | 0 | T | C | 0.561 | 8.85E-02 | 8.40E-02 | 2.56E+00 | 2.61E+00 | 2.92E-01 | no | | intergenic_variant | MODIFIER |
| 1 | 122850859 | AX-170223049 | 0 | A | G | 0.561 | 8.85E-02 | 8.40E-02 | 2.56E+00 | 2.61E+00 | 2.92E-01 | no | | intergenic_variant | MODIFIER |
| 1 | 122852627 | AX-97812354 | 0 | C | A | 0.561 | 8.85E-02 | 8.40E-02 | 2.56E+00 | 2.61E+00 | 2.92E-01 | no | | intergenic_variant | MODIFIER |
| 1 | 122853990 | AX-170367323 | 0 | G | C | 0.561 | 8.85E-02 | 8.40E-02 | 2.56E+00 | 2.61E+00 | 2.92E-01 | no | | intergenic_variant | MODIFIER |
| 1 | 122855070 | AX-170128254 | 0 | C | T | 0.561 | 8.85E-02 | 8.40E-02 | 2.56E+00 | 2.61E+00 | 2.92E-01 | no | | intergenic_variant | MODIFIER |
| 1 | 122860918 | AX-170469489 | 0 | T | C | 0.561 | 8.85E-02 | 8.40E-02 | 2.56E+00 | 2.61E+00 | 2.92E-01 | no | | intergenic_variant | MODIFIER |
| 1 | 122889449 | AX-170437687 | 0 | G | A | 0.561 | 8.85E-02 | 8.40E-02 | 2.56E+00 | 2.61E+00 | 2.92E-01 | no | | regulatory_region_variant | MODIFIER |
| 1 | 122890232 | AX-170086598 | 0 | C | A | 0.561 | 8.85E-02 | 8.40E-02 | 2.56E+00 | 2.61E+00 | 2.92E-01 | no | | intergenic_variant | MODIFIER |
| 1 | 122912105 | AX-170223957 | 0 | A | C | 0.561 | 8.85E-02 | 8.40E-02 | 2.56E+00 | 2.61E+00 | 2.92E-01 | no | | intergenic_variant | MODIFIER |
| 1 | 122912327 | AX-170340442 | 0 | A | G | 0.561 | 8.85E-02 | 8.40E-02 | 2.56E+00 | 2.61E+00 | 2.92E-01 | no | | intergenic_variant | MODIFIER |
| 1 | 122939512 | AX-170442529 | 0 | A | T | 0.561 | 8.85E-02 | 8.40E-02 | 2.56E+00 | 2.61E+00 | 2.92E-01 | no | | intergenic_variant | MODIFIER |
| 1 | 122963353 | AX-170310718 | 0 | T | G | 0.561 | 8.85E-02 | 8.40E-02 | 2.56E+00 | 2.61E+00 | 2.92E-01 | no | | intergenic_variant | MODIFIER |
| 1 | 122971657 | AX-170316414 | 0 | T | C | 0.561 | 8.85E-02 | 8.40E-02 | 2.56E+00 | 2.61E+00 | 2.92E-01 | no | | intergenic_variant | MODIFIER |
| 1 | 123042000 | AX-170464638 | 0 | G | A | 0.561 | 8.85E-02 | 8.40E-02 | 2.56E+00 | 2.61E+00 | 2.92E-01 | no | | intergenic_variant | MODIFIER |
| 1 | 123238397 | AX-170111686 | 0 | C | T | 0.561 | 8.85E-02 | 8.40E-02 | 2.56E+00 | 2.61E+00 | 2.92E-01 | no | | intergenic_variant | MODIFIER |
| 1 | 123385724 | AX-170217632 | 0 | T | C | 0.561 | 8.85E-02 | 8.40E-02 | 2.56E+00 | 2.61E+00 | 2.92E-01 | no | Gm24791 | downstream_gene_variant | MODIFIER |
| 1 | 123403487 | AX-170508512 | 0 | T | A | 0.561 | 8.85E-02 | 8.40E-02 | 2.56E+00 | 2.61E+00 | 2.92E-01 | no | Dpp10 | intron_variant | MODIFIER |
| 1 | 123410972 | AX-97786433 | 0 | C | T | 0.561 | 8.85E-02 | 8.40E-02 | 2.56E+00 | 2.61E+00 | 2.92E-01 | no | Dpp10 | intron_variant | MODIFIER |
| 1 | 123430177 | AX-170104300 | 0 | G | A | 0.561 | 8.85E-02 | 8.40E-02 | 2.56E+00 | 2.61E+00 | 2.92E-01 | no | Dpp10 | intron_variant | MODIFIER |
| 1 | 123432349 | AX-170098455 | 0 | C | A | 0.561 | 8.85E-02 | 8.40E-02 | 2.56E+00 | 2.61E+00 | 2.92E-01 | no | Dpp10 | intron_variant | MODIFIER |
| 1 | 123435154 | AX-170001594 | 0 | G | T | 0.561 | 8.85E-02 | 8.40E-02 | 2.56E+00 | 2.61E+00 | 2.92E-01 | no | Dpp10 | intron_variant | MODIFIER |
| 1 | 123436444 | AX-170527259 | 0 | T | G | 0.561 | 8.85E-02 | 8.40E-02 | 2.56E+00 | 2.61E+00 | 2.92E-01 | no | Dpp10 | intron_variant | MODIFIER |
| 1 | 123441940 | AX-170493230 | 0 | A | G | 0.561 | 8.85E-02 | 8.40E-02 | 2.56E+00 | 2.61E+00 | 2.92E-01 | no | Dpp10 | intron_variant | MODIFIER |
| 1 | 123487523 | AX-170261927 | 0 | A | T | 0.561 | 8.85E-02 | 8.40E-02 | 2.56E+00 | 2.61E+00 | 2.92E-01 | no | Dpp10 | intron_variant | MODIFIER |
| 1 | 123491814 | AX-170375559 | 0 | G | T | 0.561 | 8.85E-02 | 8.40E-02 | 2.56E+00 | 2.61E+00 | 2.92E-01 | no | Dpp10 | intron_variant | MODIFIER |
| 1 | 123492329 | AX-170370605 | 0 | A | T | 0.561 | 8.85E-02 | 8.40E-02 | 2.56E+00 | 2.61E+00 | 2.92E-01 | no | Dpp10 | intron_variant | MODIFIER |
| 1 | 123540167 | AX-170364686 | 0 | A | G | 0.561 | 8.85E-02 | 8.40E-02 | 2.56E+00 | 2.61E+00 | 2.92E-01 | no | Dpp10 | intron_variant | MODIFIER |
| 1 | 123540381 | AX-170414303 | 0 | G | C | 0.561 | 8.85E-02 | 8.40E-02 | 2.56E+00 | 2.61E+00 | 2.92E-01 | no | Dpp10 | intron_variant | MODIFIER |
| 1 | 123550442 | AX-170439265 | 0 | G | A | 0.561 | 8.85E-02 | 8.40E-02 | 2.56E+00 | 2.61E+00 | 2.92E-01 | no | Dpp10 | intron_variant | MODIFIER |
| 1 | 123557252 | AX-97790571 | 0 | T | C | 0.561 | 8.85E-02 | 8.40E-02 | 2.56E+00 | 2.61E+00 | 2.92E-01 | no | Dpp10 | intron_variant | MODIFIER |
| 1 | 123576033 | AX-170091673 | 0 | A | G | 0.561 | 8.85E-02 | 8.40E-02 | 2.56E+00 | 2.61E+00 | 2.92E-01 | no | Dpp10 | intron_variant | MODIFIER |
| 1 | 123578057 | AX-170119229 | 0 | A | C | 0.561 | 8.85E-02 | 8.40E-02 | 2.56E+00 | 2.61E+00 | 2.92E-01 | no | Dpp10 | intron_variant | MODIFIER |

| | | | | | | | | | | | | | | | |
|---|---|---|---|---|---|---|---|---|---|---|---|---|---|---|---|
| 1 | 123578369 | AX-170468871 | 0 | T | C | 0.561 | 8.85E-02 | 8.40E-02 | 2.56E+00 | 2.61E+00 | 2.92E-01 | no | Dpp10 | intron_variant | MODIFIER |
| 1 | 123613323 | AX-170309768 | 0 | A | C | 0.561 | 8.85E-02 | 8.40E-02 | 2.56E+00 | 2.61E+00 | 2.92E-01 | no | Dpp10 | intron_variant | MODIFIER |
| 1 | 123613427 | AX-170349167 | 0 | T | C | 0.561 | 8.85E-02 | 8.40E-02 | 2.56E+00 | 2.61E+00 | 2.92E-01 | no | Dpp10 | intron_variant | MODIFIER |
| 1 | 123613923 | AX-170136757 | 0 | C | T | 0.561 | 8.85E-02 | 8.40E-02 | 2.56E+00 | 2.61E+00 | 2.92E-01 | no | Dpp10 | intron_variant | MODIFIER |
| 1 | 123614455 | AX-170505172 | 0 | C | T | 0.561 | 8.85E-02 | 8.40E-02 | 2.56E+00 | 2.61E+00 | 2.92E-01 | no | Dpp10 | intron_variant | MODIFIER |
| 1 | 123703978 | AX-170199147 | 0 | A | T | 0.561 | 8.85E-02 | 8.40E-02 | 2.56E+00 | 2.61E+00 | 2.92E-01 | no | Dpp10 | intron_variant | MODIFIER |
| 1 | 123729503 | AX-170408132 | 0 | T | C | 0.561 | 8.85E-02 | 8.40E-02 | 2.56E+00 | 2.61E+00 | 2.92E-01 | no | Dpp10 | intron_variant | MODIFIER |
| 1 | 123734249 | AX-170158068 | 0 | C | A | 0.561 | 8.85E-02 | 8.40E-02 | 2.56E+00 | 2.61E+00 | 2.92E-01 | no | Dpp10 | intron_variant | MODIFIER |
| 1 | 123749934 | AX-170042554 | 0 | T | C | 0.561 | 8.85E-02 | 8.40E-02 | 2.56E+00 | 2.61E+00 | 2.92E-01 | no | Dpp10 | intron_variant | MODIFIER |
| 1 | 123751908 | AX-170263864 | 0 | C | G | 0.561 | 8.85E-02 | 8.40E-02 | 2.56E+00 | 2.61E+00 | 2.92E-01 | no | Dpp10 | intron_variant | MODIFIER |
| 1 | 123811384 | AX-170195935 | 0 | C | A | 0.561 | 8.85E-02 | 8.40E-02 | 2.56E+00 | 2.61E+00 | 2.92E-01 | no | Dpp10 | intron_variant | MODIFIER |
| 1 | 123816906 | AX-170220530 | 0 | A | G | 0.561 | 8.85E-02 | 8.40E-02 | 2.56E+00 | 2.61E+00 | 2.92E-01 | no | Dpp10 | intron_variant | MODIFIER |
| 1 | 123816928 | AX-170505755 | 0 | G | C | 0.561 | 8.85E-02 | 8.40E-02 | 2.56E+00 | 2.61E+00 | 2.92E-01 | no | Dpp10 | intron_variant | MODIFIER |
| 1 | 123836298 | AX-170462341 | 0 | T | C | 0.561 | 8.85E-02 | 8.40E-02 | 2.56E+00 | 2.61E+00 | 2.92E-01 | no | Dpp10 | intron_variant | MODIFIER |
| 1 | 123941083 | AX-97842652 | 0 | C | G | 0.561 | 8.85E-02 | 8.40E-02 | 2.56E+00 | 2.61E+00 | 2.92E-01 | no | Dpp10 | intron_variant | MODIFIER |
| 1 | 123982970 | AX-97849349 | 0 | G | T | 0.561 | 8.85E-02 | 8.40E-02 | 2.56E+00 | 2.61E+00 | 2.92E-01 | no | Dpp10 | intron_variant | MODIFIER |
| 1 | 124031083 | AX-170001601 | 0 | T | C | 0.561 | 8.85E-02 | 8.40E-02 | 2.56E+00 | 2.61E+00 | 2.92E-01 | no | Dpp10 | intron_variant | MODIFIER |
| 1 | 124148791 | AX-97824759 | 0 | C | T | 0.561 | 8.85E-02 | 8.40E-02 | 2.56E+00 | 2.61E+00 | 2.92E-01 | no | | intergenic_variant | MODIFIER |
| 1 | 124289705 | AX-170419156 | 0 | A | G | 0.561 | 8.85E-02 | 8.40E-02 | 2.56E+00 | 2.61E+00 | 2.92E-01 | no | | intergenic_variant | MODIFIER |
| 1 | 124290950 | AX-170503840 | 0 | A | G | 0.561 | 8.85E-02 | 8.40E-02 | 2.56E+00 | 2.61E+00 | 2.92E-01 | no | | intergenic_variant | MODIFIER |
| 1 | 124293149 | AX-170195195 | 0 | G | A | 0.561 | 8.85E-02 | 8.40E-02 | 2.56E+00 | 2.61E+00 | 2.92E-01 | no | | intergenic_variant | MODIFIER |
| 1 | 124313513 | AX-170101925 | 0 | C | T | 0.561 | 8.85E-02 | 8.40E-02 | 2.56E+00 | 2.61E+00 | 2.92E-01 | no | | intergenic_variant | MODIFIER |
| 1 | 124349673 | AX-170480130 | 0 | G | A | 0.561 | 8.85E-02 | 8.40E-02 | 2.56E+00 | 2.61E+00 | 2.92E-01 | no | | intergenic_variant | MODIFIER |
| 1 | 124357891 | AX-170086603 | 0 | A | G | 0.561 | 8.85E-02 | 8.40E-02 | 2.56E+00 | 2.61E+00 | 2.92E-01 | no | | intergenic_variant | MODIFIER |
| 1 | 124433247 | AX-170475587 | 0 | C | T | 0.561 | 8.85E-02 | 8.40E-02 | 2.56E+00 | 2.61E+00 | 2.92E-01 | no | | intergenic_variant | MODIFIER |
| 1 | 124485382 | AX-170291760 | 0 | A | T | 0.561 | 8.85E-02 | 8.40E-02 | 2.56E+00 | 2.61E+00 | 2.92E-01 | no | | intergenic_variant | MODIFIER |
| 1 | 124566254 | AX-170175829 | 0 | A | G | 0.561 | 8.85E-02 | 8.40E-02 | 2.56E+00 | 2.61E+00 | 2.92E-01 | no | Gm28928 | intron_variant,non_coding_transcript_variant | MODIFIER |
| 1 | 124634528 | AX-170245735 | 0 | G | A | 0.561 | 8.85E-02 | 8.40E-02 | 2.56E+00 | 2.61E+00 | 2.92E-01 | no | Gm28928 | intron_variant,non_coding_transcript_variant | MODIFIER |
| 1 | 124656590 | AX-170412316 | 0 | C | T | 0.561 | 8.85E-02 | 8.40E-02 | 2.56E+00 | 2.61E+00 | 2.92E-01 | no | Gm28928 | intron_variant,non_coding_transcript_variant | MODIFIER |
| 1 | 124678593 | AX-170229347 | 0 | G | A | 0.561 | 8.85E-02 | 8.40E-02 | 2.56E+00 | 2.61E+00 | 2.92E-01 | no | Gm28928 | intron_variant,non_coding_transcript_variant | MODIFIER |
| 1 | 124688454 | AX-97807974 | 0 | A | G | 0.561 | 8.85E-02 | 8.40E-02 | 2.56E+00 | 2.61E+00 | 2.92E-01 | no | Gm28928 | intron_variant,non_coding_transcript_variant | MODIFIER |
| 1 | 124691139 | AX-170489881 | 0 | A | G | 0.561 | 8.85E-02 | 8.40E-02 | 2.56E+00 | 2.61E+00 | 2.92E-01 | no | Gm28928 | intron_variant,non_coding_transcript_variant | MODIFIER |
| 1 | 124800758 | AX-170305962 | 0 | T | C | 0.561 | 8.85E-02 | 8.40E-02 | 2.56E+00 | 2.61E+00 | 2.92E-01 | no | Gm28928 | intron_variant,non_coding_transcript_variant | MODIFIER |
| 1 | 124800831 | AX-170516440 | 0 | A | C | 0.561 | 8.85E-02 | 8.40E-02 | 2.56E+00 | 2.61E+00 | 2.92E-01 | no | Gm28928 | intron_variant,non_coding_transcript_variant | MODIFIER |
| 1 | 124803067 | AX-170495326 | 0 | T | C | 0.561 | 8.85E-02 | 8.40E-02 | 2.56E+00 | 2.61E+00 | 2.92E-01 | no | Gm28928 | intron_variant,non_coding_transcript_variant | MODIFIER |
| 1 | 124822836 | AX-170199921 | 0 | T | A | 0.561 | 8.85E-02 | 8.40E-02 | 2.56E+00 | 2.61E+00 | 2.92E-01 | no | Gm28928 | intron_variant,non_coding_transcript_variant | MODIFIER |
| 1 | 124830611 | AX-170472612 | 0 | A | G | 0.561 | 8.85E-02 | 8.40E-02 | 2.56E+00 | 2.61E+00 | 2.92E-01 | no | Gm28928 | intron_variant,non_coding_transcript_variant | MODIFIER |
| 1 | 126619208 | AX-170042618 | 0 | G | T | 0.561 | 8.85E-02 | 8.40E-02 | 2.56E+00 | 2.61E+00 | 2.92E-01 | no | Nckap5 | intron_variant | MODIFIER |
| 1 | 126625736 | AX-170507254 | 0 | T | G | 0.561 | 8.85E-02 | 8.40E-02 | 2.56E+00 | 2.61E+00 | 2.92E-01 | no | Nckap5 | intron_variant | MODIFIER |
| 1 | 126646495 | AX-170260634 | 0 | A | G | 0.561 | 8.85E-02 | 8.40E-02 | 2.56E+00 | 2.61E+00 | 2.92E-01 | no | Nckap5 | intron_variant | MODIFIER |
| 1 | 126659671 | AX-170435235 | 0 | A | G | 0.561 | 8.85E-02 | 8.40E-02 | 2.56E+00 | 2.61E+00 | 2.92E-01 | no | Nckap5 | intron_variant | MODIFIER |
| 1 | 126659790 | AX-170526088 | 0 | T | C | 0.561 | 8.85E-02 | 8.40E-02 | 2.56E+00 | 2.61E+00 | 2.92E-01 | no | Nckap5 | intron_variant | MODIFIER |
| 18 | 59996779 | AX-170458338 | 1 | A | G | 0.402 | 1.01E-01 | 9.35E-02 | 2.42E+00 | 2.47E+00 | 2.92E-01 | no | | intergenic_variant | MODIFIER |
| 6 | 136537678 | AX-170318440 | 1 | A | C | 0.571 | 9.35E-02 | 8.69E-02 | 2.42E+00 | 2.47E+00 | 2.92E-01 | no | Atf7ip | intron_variant | MODIFIER |
| 8 | 90704001 | AX-170258048 | 1 | G | A | 0.196 | -1.18E-01 | 1.13E-01 | 2.66E+00 | 2.74E+00 | 2.92E-01 | no | | intergenic_variant | MODIFIER |
| 8 | 90502020 | AX-170031243 | 0 | G | A | 0.202 | -1.08E-01 | 1.04E-01 | 2.71E+00 | 2.83E+00 | 2.92E-01 | no | | intergenic_variant | MODIFIER |
| 15 | 6962752 | AX-170531055 | 1 | G | A | 0.241 | -1.11E-01 | 1.06E-01 | 2.63E+00 | 2.69E+00 | 2.92E-01 | no | | intergenic_variant | MODIFIER |
| 8 | 32152546 | AX-170090076 | 0 | A | G | 0.14 | -1.24E-01 | 1.19E-01 | 2.73E+00 | 2.84E+00 | 2.92E-01 | no | Nrg1 | intron_variant | MODIFIER |
| 18 | 60154620 | AX-170292332 | 0 | T | G | 0.404 | 9.77E-02 | 9.01E-02 | 2.38E+00 | 2.43E+00 | 2.92E-01 | no | | regulatory_region_variant | MODIFIER |
| 8 | 34422016 | AX-97788454 | 0 | G | A | 0.14 | -1.41E-01 | 1.35E-01 | 2.69E+00 | 2.75E+00 | 2.92E-01 | no | Gm45349 | intron_variant,non_coding_transcript_variant | MODIFIER |
| 8 | 34554504 | AX-170262342 | 0 | G | A | 0.14 | -1.41E-01 | 1.35E-01 | 2.69E+00 | 2.75E+00 | 2.92E-01 | no | | intergenic_variant | MODIFIER |
| 1 | 132473317 | AX-170303106 | 1 | A | T | 0.536 | 8.91E-02 | 8.53E-02 | 2.66E+00 | 2.72E+00 | 2.92E-01 | no | Rbbp5 | upstream_gene_variant | MODIFIER |
| 3 | 7421985 | AX-170105104 | 2 | C | G | 0.418 | -6.56E-02 | 6.32E-02 | 2.72E+00 | 2.87E+00 | 2.92E-01 | no | Pkia | intron_variant | MODIFIER |
| 14 | 119209126 | AX-170010816 | 0 | C | T | 0.342 | 7.70E-02 | 7.37E-02 | 2.62E+00 | 2.72E+00 | 2.92E-01 | no | Hs6st3 | intron_variant | MODIFIER |

| | | | | | | | | | | | | | | | |
|---|---|---|---|---|---|---|---|---|---|---|---|---|---|---|---|
| 11 | 71366922 | AX-170047389 | 1 | T | C | 0.241 | -9.98E-02 | 9.51E-02 | 2.56E+00 | 2.66E+00 | 2.93E-01 | no | | intergenic_variant | MODIFIER |
| 4 | 63273384 | AX-170466758 | 1 | T | C | 0.348 | 1.23E-01 | 1.18E-01 | 2.79E+00 | 2.84E+00 | 2.93E-01 | no | Col27a1 | intron_variant | MODIFIER |
| 4 | 68855000 | AX-170239333 | 1 | A | G | 0.348 | 1.23E-01 | 1.18E-01 | 2.79E+00 | 2.84E+00 | 2.93E-01 | no | Brinp1 | intron_variant | MODIFIER |
| 5 | 108602350 | AX-170263012 | 0 | T | A | 0.456 | 1.16E-01 | 1.11E-01 | 2.75E+00 | 2.81E+00 | 2.93E-01 | no | Gak | intron_variant | MODIFIER |
| 4 | 55435384 | AX-170524145 | 1 | C | T | 0.375 | -1.07E-01 | 1.03E-01 | 2.85E+00 | 2.95E+00 | 2.93E-01 | no | | intergenic_variant | MODIFIER |
| 9 | 17551328 | AX-97830391 | 1 | T | C | 0.491 | 8.87E-02 | 8.36E-02 | 2.49E+00 | 2.55E+00 | 2.93E-01 | no | | intergenic_variant | MODIFIER |
| 4 | 64529675 | AX-170224350 | 0 | C | T | 0.325 | 1.23E-01 | 1.18E-01 | 2.77E+00 | 2.83E+00 | 2.93E-01 | no | | intergenic_variant | MODIFIER |
| 10 | 114896222 | AX-97822635 | 1 | A | G | 0.348 | -7.73E-02 | 7.44E-02 | 2.78E+00 | 2.89E+00 | 2.93E-01 | no | | intergenic_variant | MODIFIER |
| 1 | 126561392 | AX-170001640 | 1 | T | C | 0.571 | 8.84E-02 | 8.41E-02 | 2.56E+00 | 2.62E+00 | 2.93E-01 | no | Nckap5 | intron_variant | MODIFIER |
| 4 | 48617092 | AX-170377181 | 1 | C | T | 0.366 | -1.01E-01 | 9.64E-02 | 2.87E+00 | 2.97E+00 | 2.93E-01 | no | Tmeff1 | intron_variant | MODIFIER |
| 4 | 48886560 | AX-170022736 | 1 | A | T | 0.366 | -1.01E-01 | 9.64E-02 | 2.87E+00 | 2.97E+00 | 2.93E-01 | no | | intergenic_variant | MODIFIER |
| 15 | 5819330 | AX-170010950 | 1 | G | C | 0.232 | -1.11E-01 | 1.06E-01 | 2.64E+00 | 2.69E+00 | 2.93E-01 | no | | intergenic_variant | MODIFIER |
| 5 | 54929983 | AX-170483951 | 0 | G | A | 0.482 | 1.02E-01 | 9.55E-02 | 2.45E+00 | 2.53E+00 | 2.93E-01 | no | | intergenic_variant | MODIFIER |
| 6 | 125268044 | AX-170481488 | 1 | A | G | 0.188 | -8.28E-02 | 7.98E-02 | 2.80E+00 | 2.96E+00 | 2.93E-01 | no | 4930417O13Rik | intron_variant,non_coding_transcript_variant | MODIFIER |
| 1 | 122415545 | AX-170001579 | 0 | T | G | 0.57 | 8.84E-02 | 8.42E-02 | 2.56E+00 | 2.62E+00 | 2.93E-01 | no | | intergenic_variant | MODIFIER |
| 1 | 122489277 | AX-97797534 | 0 | G | A | 0.57 | 8.84E-02 | 8.42E-02 | 2.56E+00 | 2.62E+00 | 2.93E-01 | no | | intergenic_variant | MODIFIER |
| 1 | 123389589 | AX-170244881 | 0 | T | C | 0.57 | 8.84E-02 | 8.42E-02 | 2.56E+00 | 2.62E+00 | 2.93E-01 | no | Gm24791 | downstream_gene_variant | MODIFIER |
| 1 | 124499602 | AX-170526425 | 0 | C | T | 0.57 | 8.84E-02 | 8.42E-02 | 2.56E+00 | 2.62E+00 | 2.93E-01 | no | | intergenic_variant | MODIFIER |
| 1 | 124508431 | AX-170380971 | 0 | C | T | 0.57 | 8.84E-02 | 8.42E-02 | 2.56E+00 | 2.62E+00 | 2.93E-01 | no | Gm28928 | intron_variant,non_coding_transcript_variant | MODIFIER |
| 1 | 124514997 | AX-170290960 | 0 | T | C | 0.57 | 8.84E-02 | 8.42E-02 | 2.56E+00 | 2.62E+00 | 2.93E-01 | no | Gm28928 | intron_variant,non_coding_transcript_variant | MODIFIER |
| 1 | 124518147 | AX-170319456 | 0 | T | C | 0.57 | 8.84E-02 | 8.42E-02 | 2.56E+00 | 2.62E+00 | 2.93E-01 | no | Gm28928 | intron_variant,non_coding_transcript_variant | MODIFIER |
| 1 | 124520934 | AX-170487470 | 0 | A | G | 0.57 | 8.84E-02 | 8.42E-02 | 2.56E+00 | 2.62E+00 | 2.93E-01 | no | Gm28928 | intron_variant,non_coding_transcript_variant | MODIFIER |
| 1 | 124521414 | AX-170152426 | 0 | C | T | 0.57 | 8.84E-02 | 8.42E-02 | 2.56E+00 | 2.62E+00 | 2.93E-01 | no | Gm28928 | intron_variant,non_coding_transcript_variant | MODIFIER |
| 1 | 124524344 | AX-170261560 | 0 | G | A | 0.57 | 8.84E-02 | 8.42E-02 | 2.56E+00 | 2.62E+00 | 2.93E-01 | no | Gm28928 | intron_variant,non_coding_transcript_variant | MODIFIER |
| 1 | 124529041 | AX-170379233 | 0 | T | C | 0.57 | 8.84E-02 | 8.42E-02 | 2.56E+00 | 2.62E+00 | 2.93E-01 | no | Gm28928 | intron_variant,non_coding_transcript_variant | MODIFIER |
| 1 | 124529439 | AX-170280264 | 0 | C | A | 0.57 | 8.84E-02 | 8.42E-02 | 2.56E+00 | 2.62E+00 | 2.93E-01 | no | Gm28928 | intron_variant,non_coding_transcript_variant | MODIFIER |
| 1 | 124541277 | AX-170465364 | 0 | A | T | 0.57 | 8.84E-02 | 8.42E-02 | 2.56E+00 | 2.62E+00 | 2.93E-01 | no | Gm28928 | intron_variant,non_coding_transcript_variant | MODIFIER |
| 1 | 124541752 | AX-170517249 | 0 | G | A | 0.57 | 8.84E-02 | 8.42E-02 | 2.56E+00 | 2.62E+00 | 2.93E-01 | no | Gm28928 | intron_variant,non_coding_transcript_variant | MODIFIER |
| 1 | 124543966 | AX-168412312 | 0 | T | C | 0.57 | 8.84E-02 | 8.42E-02 | 2.56E+00 | 2.62E+00 | 2.93E-01 | no | Gm28928 | intron_variant,non_coding_transcript_variant | MODIFIER |
| 1 | 124544719 | AX-170292872 | 0 | A | G | 0.57 | 8.84E-02 | 8.42E-02 | 2.56E+00 | 2.62E+00 | 2.93E-01 | no | Gm28928 | intron_variant,non_coding_transcript_variant | MODIFIER |
| 1 | 124555255 | AX-170001616 | 0 | C | A | 0.57 | 8.84E-02 | 8.42E-02 | 2.56E+00 | 2.62E+00 | 2.93E-01 | no | Gm28928 | intron_variant,non_coding_transcript_variant | MODIFIER |
| 1 | 124557802 | AX-170386913 | 0 | T | C | 0.57 | 8.84E-02 | 8.42E-02 | 2.56E+00 | 2.62E+00 | 2.93E-01 | no | Gm28928 | intron_variant,non_coding_transcript_variant | MODIFIER |
| 1 | 124566753 | AX-170042566 | 0 | A | G | 0.57 | 8.84E-02 | 8.42E-02 | 2.56E+00 | 2.62E+00 | 2.93E-01 | no | Gm28928 | intron_variant,non_coding_transcript_variant | MODIFIER |
| 1 | 124589421 | AX-170464537 | 0 | C | A | 0.57 | 8.84E-02 | 8.42E-02 | 2.56E+00 | 2.62E+00 | 2.93E-01 | no | Gm28928 | intron_variant,non_coding_transcript_variant | MODIFIER |
| 1 | 124591471 | AX-170218849 | 0 | G | C | 0.57 | 8.84E-02 | 8.42E-02 | 2.56E+00 | 2.62E+00 | 2.93E-01 | no | Gm28928 | intron_variant,non_coding_transcript_variant | MODIFIER |
| 1 | 124595808 | AX-170199071 | 0 | A | G | 0.57 | 8.84E-02 | 8.42E-02 | 2.56E+00 | 2.62E+00 | 2.93E-01 | no | Gm28928 | intron_variant,non_coding_transcript_variant | MODIFIER |
| 1 | 124596176 | AX-170151406 | 0 | G | T | 0.57 | 8.84E-02 | 8.42E-02 | 2.56E+00 | 2.62E+00 | 2.93E-01 | no | Gm28928 | intron_variant,non_coding_transcript_variant | MODIFIER |
| 1 | 124610303 | AX-170284403 | 0 | G | A | 0.57 | 8.84E-02 | 8.42E-02 | 2.56E+00 | 2.62E+00 | 2.93E-01 | no | Gm28928 | intron_variant,non_coding_transcript_variant | MODIFIER |
| 1 | 124612204 | AX-170492347 | 0 | C | T | 0.57 | 8.84E-02 | 8.42E-02 | 2.56E+00 | 2.62E+00 | 2.93E-01 | no | Gm28928 | intron_variant,non_coding_transcript_variant | MODIFIER |
| 1 | 124612776 | AX-97803471 | 0 | A | C | 0.57 | 8.84E-02 | 8.42E-02 | 2.56E+00 | 2.62E+00 | 2.93E-01 | no | Gm28928 | intron_variant,non_coding_transcript_variant | MODIFIER |
| 1 | 124618241 | AX-170515101 | 0 | T | G | 0.57 | 8.84E-02 | 8.42E-02 | 2.56E+00 | 2.62E+00 | 2.93E-01 | no | Gm28928 | intron_variant,non_coding_transcript_variant | MODIFIER |
| 1 | 124635295 | AX-170196972 | 0 | A | G | 0.57 | 8.84E-02 | 8.42E-02 | 2.56E+00 | 2.62E+00 | 2.93E-01 | no | Gm28928 | intron_variant,non_coding_transcript_variant | MODIFIER |
| 1 | 124645095 | AX-170232887 | 0 | A | C | 0.57 | 8.84E-02 | 8.42E-02 | 2.56E+00 | 2.62E+00 | 2.93E-01 | no | Gm28928 | intron_variant,non_coding_transcript_variant | MODIFIER |
| 1 | 124688490 | AX-170183368 | 0 | A | C | 0.57 | 8.84E-02 | 8.42E-02 | 2.56E+00 | 2.62E+00 | 2.93E-01 | no | Gm28928 | intron_variant,non_coding_transcript_variant | MODIFIER |
| 1 | 124695727 | AX-170501868 | 0 | A | G | 0.57 | 8.84E-02 | 8.42E-02 | 2.56E+00 | 2.62E+00 | 2.93E-01 | no | Gm28928 | intron_variant,non_coding_transcript_variant | MODIFIER |
| 1 | 124717449 | AX-97841681 | 0 | C | T | 0.57 | 8.84E-02 | 8.42E-02 | 2.56E+00 | 2.62E+00 | 2.93E-01 | no | Gm28928 | intron_variant,non_coding_transcript_variant | MODIFIER |
| 1 | 124739017 | AX-170232917 | 0 | A | G | 0.57 | 8.84E-02 | 8.42E-02 | 2.56E+00 | 2.62E+00 | 2.93E-01 | no | Gm28928 | intron_variant,non_coding_transcript_variant | MODIFIER |
| 1 | 124741652 | AX-170170864 | 0 | C | G | 0.57 | 8.84E-02 | 8.42E-02 | 2.56E+00 | 2.62E+00 | 2.93E-01 | no | Gm28928 | intron_variant,non_coding_transcript_variant | MODIFIER |
| 1 | 124741765 | AX-97788259 | 0 | G | A | 0.57 | 8.84E-02 | 8.42E-02 | 2.56E+00 | 2.62E+00 | 2.93E-01 | no | Gm28928 | intron_variant,non_coding_transcript_variant | MODIFIER |
| 1 | 124744823 | AX-170266075 | 0 | G | C | 0.57 | 8.84E-02 | 8.42E-02 | 2.56E+00 | 2.62E+00 | 2.93E-01 | no | Gm28928 | intron_variant,non_coding_transcript_variant | MODIFIER |
| 1 | 124748824 | AX-170472028 | 0 | T | C | 0.57 | 8.84E-02 | 8.42E-02 | 2.56E+00 | 2.62E+00 | 2.93E-01 | no | Gm28928 | intron_variant,non_coding_transcript_variant | MODIFIER |
| 1 | 124756533 | AX-170042571 | 0 | C | T | 0.57 | 8.84E-02 | 8.42E-02 | 2.56E+00 | 2.62E+00 | 2.93E-01 | no | Gm28928 | intron_variant,non_coding_transcript_variant | MODIFIER |
| 1 | 124794181 | AX-97829805 | 0 | C | T | 0.57 | 8.84E-02 | 8.42E-02 | 2.56E+00 | 2.62E+00 | 2.93E-01 | no | Gm28928 | intron_variant,non_coding_transcript_variant | MODIFIER |
| 1 | 124794791 | AX-170114879 | 0 | C | T | 0.57 | 8.84E-02 | 8.42E-02 | 2.56E+00 | 2.62E+00 | 2.93E-01 | no | Gm28928 | intron_variant,non_coding_transcript_variant | MODIFIER |

| | | | | | | | | | | | | | | | |
|---|---|---|---|---|---|---|---|---|---|---|---|---|---|---|---|
| 1 | 124796096 | AX-170206214 | 0 | C | T | 0.57 | 8.84E-02 | 8.42E-02 | 2.56E+00 | 2.62E+00 | 2.93E-01 | no | Gm28928 | intron_variant,non_coding_transcript_variant | MODIFIER |
| 1 | 124796359 | AX-170161606 | 0 | G | A | 0.57 | 8.84E-02 | 8.42E-02 | 2.56E+00 | 2.62E+00 | 2.93E-01 | no | Gm28928 | intron_variant,non_coding_transcript_variant | MODIFIER |
| 1 | 124852474 | AX-170468596 | 0 | G | A | 0.57 | 8.84E-02 | 8.42E-02 | 2.56E+00 | 2.62E+00 | 2.93E-01 | no | | intergenic_variant | MODIFIER |
| 1 | 124852781 | AX-170430430 | 0 | G | A | 0.57 | 8.84E-02 | 8.42E-02 | 2.56E+00 | 2.62E+00 | 2.93E-01 | no | | intergenic_variant | MODIFIER |
| 1 | 124866427 | AX-170156319 | 0 | A | C | 0.57 | 8.84E-02 | 8.42E-02 | 2.56E+00 | 2.62E+00 | 2.93E-01 | no | | intergenic_variant | MODIFIER |
| 1 | 124879073 | AX-170459292 | 0 | G | A | 0.57 | 8.84E-02 | 8.42E-02 | 2.56E+00 | 2.62E+00 | 2.93E-01 | no | | intergenic_variant | MODIFIER |
| 1 | 125938812 | AX-170364890 | 0 | T | C | 0.57 | 8.84E-02 | 8.42E-02 | 2.56E+00 | 2.62E+00 | 2.93E-01 | no | Nckap5 | intron_variant | MODIFIER |
| 1 | 125940733 | AX-170468113 | 0 | A | C | 0.57 | 8.84E-02 | 8.42E-02 | 2.56E+00 | 2.62E+00 | 2.93E-01 | no | Nckap5 | intron_variant | MODIFIER |
| 1 | 125942631 | AX-170187236 | 0 | C | T | 0.57 | 8.84E-02 | 8.42E-02 | 2.56E+00 | 2.62E+00 | 2.93E-01 | no | Nckap5 | intron_variant | MODIFIER |
| 1 | 125943028 | AX-170001633 | 0 | C | T | 0.57 | 8.84E-02 | 8.42E-02 | 2.56E+00 | 2.62E+00 | 2.93E-01 | no | Nckap5 | intron_variant | MODIFIER |
| 1 | 126357985 | AX-170185248 | 0 | C | A | 0.57 | 8.84E-02 | 8.42E-02 | 2.56E+00 | 2.62E+00 | 2.93E-01 | no | Nckap5 | intron_variant | MODIFIER |
| 1 | 126358073 | AX-170042611 | 0 | G | A | 0.57 | 8.84E-02 | 8.42E-02 | 2.56E+00 | 2.62E+00 | 2.93E-01 | no | Nckap5 | intron_variant | MODIFIER |
| 1 | 126360548 | AX-170091680 | 0 | T | C | 0.57 | 8.84E-02 | 8.42E-02 | 2.56E+00 | 2.62E+00 | 2.93E-01 | no | Nckap5 | intron_variant | MODIFIER |
| 1 | 126395977 | AX-97852459 | 0 | A | C | 0.57 | 8.84E-02 | 8.42E-02 | 2.56E+00 | 2.62E+00 | 2.93E-01 | no | Nckap5 | intron_variant | MODIFIER |
| 1 | 126396405 | AX-170250897 | 0 | T | C | 0.57 | 8.84E-02 | 8.42E-02 | 2.56E+00 | 2.62E+00 | 2.93E-01 | no | Nckap5 | intron_variant | MODIFIER |
| 1 | 126396691 | AX-170179105 | 0 | A | T | 0.57 | 8.84E-02 | 8.42E-02 | 2.56E+00 | 2.62E+00 | 2.93E-01 | no | Nckap5 | intron_variant | MODIFIER |
| 1 | 126396834 | AX-170432213 | 0 | C | T | 0.57 | 8.84E-02 | 8.42E-02 | 2.56E+00 | 2.62E+00 | 2.93E-01 | no | Nckap5 | intron_variant | MODIFIER |
| 1 | 126396864 | AX-170521327 | 0 | T | C | 0.57 | 8.84E-02 | 8.42E-02 | 2.56E+00 | 2.62E+00 | 2.93E-01 | no | Nckap5 | intron_variant | MODIFIER |
| 1 | 126397004 | AX-97783276 | 0 | G | A | 0.57 | 8.84E-02 | 8.42E-02 | 2.56E+00 | 2.62E+00 | 2.93E-01 | no | Nckap5 | intron_variant | MODIFIER |
| 1 | 126578596 | AX-170334332 | 0 | C | T | 0.57 | 8.84E-02 | 8.42E-02 | 2.56E+00 | 2.62E+00 | 2.93E-01 | no | Nckap5 | intron_variant | MODIFIER |
| 1 | 126579933 | AX-170167108 | 0 | T | A | 0.57 | 8.84E-02 | 8.42E-02 | 2.56E+00 | 2.62E+00 | 2.93E-01 | no | Nckap5 | intron_variant | MODIFIER |
| 1 | 126579953 | AX-170416978 | 0 | T | C | 0.57 | 8.84E-02 | 8.42E-02 | 2.56E+00 | 2.62E+00 | 2.93E-01 | no | Nckap5 | intron_variant | MODIFIER |
| 1 | 126582718 | AX-170324298 | 0 | T | G | 0.57 | 8.84E-02 | 8.42E-02 | 2.56E+00 | 2.62E+00 | 2.93E-01 | no | Nckap5 | intron_variant | MODIFIER |
| 1 | 126583869 | AX-170498053 | 0 | C | A | 0.57 | 8.84E-02 | 8.42E-02 | 2.56E+00 | 2.62E+00 | 2.93E-01 | no | Nckap5 | intron_variant | MODIFIER |
| 1 | 126616948 | AX-170223543 | 0 | G | A | 0.57 | 8.84E-02 | 8.42E-02 | 2.56E+00 | 2.62E+00 | 2.93E-01 | no | Nckap5 | intron_variant | MODIFIER |
| 1 | 126625683 | AX-97791524 | 0 | C | T | 0.57 | 8.84E-02 | 8.42E-02 | 2.56E+00 | 2.62E+00 | 2.93E-01 | no | Nckap5 | intron_variant | MODIFIER |
| 1 | 126651593 | AX-170240397 | 0 | A | C | 0.57 | 8.84E-02 | 8.42E-02 | 2.56E+00 | 2.62E+00 | 2.93E-01 | no | Nckap5 | intron_variant | MODIFIER |
| 2 | 164499043 | AX-170300777 | 1 | A | G | 0.161 | 2.02E-01 | 1.94E-01 | 2.82E+00 | 2.91E+00 | 2.93E-01 | Yes | Pigt | intron_variant | MODIFIER |
| 4 | 138078621 | AX-97835480 | 0 | C | T | 0.175 | 1.13E-01 | 1.08E-01 | 2.62E+00 | 2.69E+00 | 2.94E-01 | no | Eif4g3 | intron_variant | MODIFIER |
| 2 | 120767353 | AX-170520628 | 0 | A | T | 0.14 | 1.85E-01 | 1.78E-01 | 2.76E+00 | 2.86E+00 | 2.94E-01 | Yes | Ttbk2 | intron_variant | MODIFIER |
| 18 | 73099062 | AX-170015970 | 0 | G | T | 0.184 | -1.23E-01 | 1.17E-01 | 2.58E+00 | 2.63E+00 | 2.94E-01 | no | | intergenic_variant | MODIFIER |
| 4 | 144543601 | AX-170511837 | 0 | G | C | 0.114 | 1.82E-01 | 1.74E-01 | 2.84E+00 | 2.94E+00 | 2.94E-01 | Yes | Gm13126 | downstream_gene_variant | MODIFIER |
| 5 | 126448445 | AX-170354639 | 0 | T | C | 0.465 | 9.28E-02 | 8.93E-02 | 2.81E+00 | 2.93E+00 | 2.94E-01 | no | Gm37700 | upstream_gene_variant | MODIFIER |
| 1 | 129386361 | AX-170086610 | 1 | C | T | 0.545 | 8.91E-02 | 8.56E-02 | 2.66E+00 | 2.72E+00 | 2.94E-01 | no | Thsd7b | intron_variant | MODIFIER |
| 5 | 50140295 | AX-170103543 | 0 | C | T | 0.482 | 1.10E-01 | 1.03E-01 | 2.46E+00 | 2.52E+00 | 2.94E-01 | no | | intergenic_variant | MODIFIER |
| 5 | 50257827 | AX-97848291 | 0 | G | T | 0.482 | 1.10E-01 | 1.03E-01 | 2.46E+00 | 2.52E+00 | 2.94E-01 | no | Mir8117 | downstream_gene_variant | MODIFIER |
| 9 | 92678637 | AX-170140659 | 1 | G | A | 0.143 | -1.24E-01 | 1.19E-01 | 2.63E+00 | 2.71E+00 | 2.94E-01 | no | | intergenic_variant | MODIFIER |
| 15 | 5811689 | AX-170535092 | 1 | T | C | 0.232 | -1.10E-01 | 1.06E-01 | 2.63E+00 | 2.68E+00 | 2.94E-01 | no | | intergenic_variant | MODIFIER |
| 1 | 146909879 | AX-170267710 | 1 | G | A | 0.214 | 1.33E-01 | 1.28E-01 | 2.64E+00 | 2.71E+00 | 2.94E-01 | no | | intergenic_variant | MODIFIER |
| 2 | 163765903 | AX-170425322 | 0 | A | G | 0.158 | 2.02E-01 | 1.94E-01 | 2.82E+00 | 2.92E+00 | 2.94E-01 | Yes | | intergenic_variant | MODIFIER |
| 2 | 165045661 | AX-170191522 | 0 | C | T | 0.158 | 2.02E-01 | 1.94E-01 | 2.82E+00 | 2.92E+00 | 2.94E-01 | Yes | | intergenic_variant | MODIFIER |
| 1 | 130545586 | AX-170001706 | 0 | A | G | 0.535 | 8.88E-02 | 8.54E-02 | 2.65E+00 | 2.71E+00 | 2.94E-01 | no | Gm16083 | intron_variant,non_coding_transcript_variant | MODIFIER |
| 1 | 130547674 | AX-97813274 | 0 | A | C | 0.535 | 8.88E-02 | 8.54E-02 | 2.65E+00 | 2.71E+00 | 2.94E-01 | no | Gm16083 | intron_variant,non_coding_transcript_variant | MODIFIER |
| 1 | 130551830 | AX-170001705 | 0 | T | C | 0.535 | 8.88E-02 | 8.54E-02 | 2.65E+00 | 2.71E+00 | 2.94E-01 | no | Gm16083 | intron_variant,non_coding_transcript_variant | MODIFIER |
| 1 | 130592085 | AX-170001701 | 0 | G | A | 0.535 | 8.88E-02 | 8.54E-02 | 2.65E+00 | 2.71E+00 | 2.94E-01 | no | Zp3r | intron_variant | MODIFIER |
| 1 | 130592171 | AX-170001702 | 0 | A | C | 0.535 | 8.88E-02 | 8.54E-02 | 2.65E+00 | 2.71E+00 | 2.94E-01 | no | Zp3r | intron_variant | MODIFIER |
| 1 | 130598609 | AX-170318917 | 0 | T | C | 0.535 | 8.88E-02 | 8.54E-02 | 2.65E+00 | 2.71E+00 | 2.94E-01 | no | Zp3r | intron_variant | MODIFIER |
| 1 | 130608110 | AX-170205836 | 0 | G | A | 0.535 | 8.88E-02 | 8.54E-02 | 2.65E+00 | 2.71E+00 | 2.94E-01 | no | Zp3r | intron_variant | MODIFIER |
| 1 | 130608437 | AX-97808647 | 0 | A | G | 0.535 | 8.88E-02 | 8.54E-02 | 2.65E+00 | 2.71E+00 | 2.94E-01 | no | Zp3r | intron_variant | MODIFIER |
| 1 | 132068911 | AX-170042738 | 0 | A | G | 0.535 | 8.88E-02 | 8.54E-02 | 2.65E+00 | 2.71E+00 | 2.94E-01 | no | Mfsd4a | upstream_gene_variant | MODIFIER |
| 1 | 132100586 | AX-170370267 | 0 | T | G | 0.535 | 8.88E-02 | 8.54E-02 | 2.65E+00 | 2.71E+00 | 2.94E-01 | no | Gm26034 | upstream_gene_variant | MODIFIER |
| 1 | 132103818 | AX-170488039 | 0 | T | G | 0.535 | 8.88E-02 | 8.54E-02 | 2.65E+00 | 2.71E+00 | 2.94E-01 | no | Gm26034 | downstream_gene_variant | MODIFIER |
| 1 | 132126444 | AX-170150414 | 0 | T | C | 0.535 | 8.88E-02 | 8.54E-02 | 2.65E+00 | 2.71E+00 | 2.94E-01 | no | Cdk18 | intron_variant | MODIFIER |
| 1 | 132151269 | AX-170001730 | 0 | T | C | 0.535 | 8.88E-02 | 8.54E-02 | 2.65E+00 | 2.71E+00 | 2.94E-01 | no | | regulatory_region_variant | MODIFIER |

| | | | | | | | | | | | | | | | |
|---|---|---|---|---|---|---|---|---|---|---|---|---|---|---|---|
| 1 | 132156558 | AX-170367108 | 0 | C | T | 0.535 | 8.88E-02 | 8.54E-02 | 2.65E+00 | 2.71E+00 | 2.94E-01 | no | | intergenic_variant | MODIFIER |
| 1 | 132159349 | AX-170326134 | 0 | T | C | 0.535 | 8.88E-02 | 8.54E-02 | 2.65E+00 | 2.71E+00 | 2.94E-01 | no | | regulatory_region_variant | MODIFIER |
| 1 | 132162058 | AX-97812988 | 0 | T | C | 0.535 | 8.88E-02 | 8.54E-02 | 2.65E+00 | 2.71E+00 | 2.94E-01 | no | | intergenic_variant | MODIFIER |
| 1 | 132184201 | AX-97790817 | 0 | T | C | 0.535 | 8.88E-02 | 8.54E-02 | 2.65E+00 | 2.71E+00 | 2.94E-01 | no | | intergenic_variant | MODIFIER |
| 1 | 132197623 | AX-170001725 | 0 | A | G | 0.535 | 8.88E-02 | 8.54E-02 | 2.65E+00 | 2.71E+00 | 2.94E-01 | no | Mir135b | upstream_gene_variant | MODIFIER |
| 1 | 132199455 | AX-170042739 | 0 | C | A | 0.535 | 8.88E-02 | 8.54E-02 | 2.65E+00 | 2.71E+00 | 2.94E-01 | no | Mir135b | downstream_gene_variant | MODIFIER |
| 1 | 132242646 | AX-170440673 | 0 | C | G | 0.535 | 8.88E-02 | 8.54E-02 | 2.65E+00 | 2.71E+00 | 2.94E-01 | no | Lemd1 | intron_variant | MODIFIER |
| 1 | 132245442 | AX-170344404 | 0 | C | T | 0.535 | 8.88E-02 | 8.54E-02 | 2.65E+00 | 2.71E+00 | 2.94E-01 | no | Lemd1 | intron_variant | MODIFIER |
| 1 | 132256530 | AX-170171572 | 0 | G | A | 0.535 | 8.88E-02 | 8.54E-02 | 2.65E+00 | 2.71E+00 | 2.94E-01 | no | Lemd1 | intron_variant | MODIFIER |
| 1 | 132267892 | AX-170277408 | 0 | C | A | 0.535 | 8.88E-02 | 8.54E-02 | 2.65E+00 | 2.71E+00 | 2.94E-01 | no | | regulatory_region_variant | MODIFIER |
| 1 | 132271749 | AX-97824805 | 0 | G | T | 0.535 | 8.88E-02 | 8.54E-02 | 2.65E+00 | 2.71E+00 | 2.94E-01 | no | Gm29630 | downstream_gene_variant | MODIFIER |
| 1 | 132274761 | AX-170250861 | 0 | T | C | 0.535 | 8.88E-02 | 8.54E-02 | 2.65E+00 | 2.71E+00 | 2.94E-01 | no | Gm29630 | non_coding_transcript_exon_variant | MODIFIER |
| 1 | 132345061 | AX-170203872 | 0 | G | T | 0.535 | 8.88E-02 | 8.54E-02 | 2.65E+00 | 2.71E+00 | 2.94E-01 | no | F730311O21Rik | non_coding_transcript_exon_variant | MODIFIER |
| 1 | 132345586 | AX-170214447 | 0 | T | G | 0.535 | 8.88E-02 | 8.54E-02 | 2.65E+00 | 2.71E+00 | 2.94E-01 | no | F730311O21Rik | non_coding_transcript_exon_variant | MODIFIER |
| 1 | 132351839 | AX-170195865 | 0 | G | A | 0.535 | 8.88E-02 | 8.54E-02 | 2.65E+00 | 2.71E+00 | 2.94E-01 | no | Tmcc2 | downstream_gene_variant | MODIFIER |
| 1 | 132360178 | AX-170509816 | 0 | T | C | 0.535 | 8.88E-02 | 8.54E-02 | 2.65E+00 | 2.71E+00 | 2.94E-01 | no | Tmcc2 | intron_variant | MODIFIER |
| 1 | 132381909 | AX-168396035 | 0 | T | C | 0.535 | 8.88E-02 | 8.54E-02 | 2.65E+00 | 2.71E+00 | 2.94E-01 | no | Tmcc2 | intron_variant | MODIFIER |
| 1 | 132471705 | AX-170366455 | 0 | A | G | 0.535 | 8.88E-02 | 8.54E-02 | 2.65E+00 | 2.71E+00 | 2.94E-01 | no | Dstyk | downstream_gene_variant | MODIFIER |
| 1 | 132477668 | AX-170487764 | 0 | G | A | 0.535 | 8.88E-02 | 8.54E-02 | 2.65E+00 | 2.71E+00 | 2.94E-01 | no | Rbbp5 | intron_variant | MODIFIER |
| 1 | 132477802 | AX-170306609 | 0 | G | A | 0.535 | 8.88E-02 | 8.54E-02 | 2.65E+00 | 2.71E+00 | 2.94E-01 | no | Rbbp5 | intron_variant | MODIFIER |
| 1 | 132481090 | AX-97857159 | 0 | A | G | 0.535 | 8.88E-02 | 8.54E-02 | 2.65E+00 | 2.71E+00 | 2.94E-01 | no | Rbbp5 | intron_variant | MODIFIER |
| 1 | 132481276 | AX-170223323 | 0 | C | G | 0.535 | 8.88E-02 | 8.54E-02 | 2.65E+00 | 2.71E+00 | 2.94E-01 | no | Rbbp5 | intron_variant | MODIFIER |
| 1 | 132522484 | AX-170452681 | 0 | A | G | 0.535 | 8.88E-02 | 8.54E-02 | 2.65E+00 | 2.71E+00 | 2.94E-01 | no | Cntn2 | intron_variant | MODIFIER |
| 9 | 96362676 | AX-97809530 | 0 | G | A | 0.14 | -1.22E-01 | 1.17E-01 | 2.66E+00 | 2.74E+00 | 2.94E-01 | no | Atp1b3 | intron_variant | MODIFIER |
| 8 | 90435822 | AX-170254768 | 0 | G | A | 0.193 | -1.07E-01 | 1.03E-01 | 2.66E+00 | 2.78E+00 | 2.94E-01 | no | | intergenic_variant | MODIFIER |
| 1 | 171039894 | AX-170135035 | 0 | C | T | 0.211 | 1.43E-01 | 1.37E-01 | 2.62E+00 | 2.66E+00 | 2.94E-01 | no | | intergenic_variant | MODIFIER |
| 1 | 171057436 | AX-170116369 | 0 | T | G | 0.211 | 1.43E-01 | 1.37E-01 | 2.62E+00 | 2.66E+00 | 2.94E-01 | no | Fcgr3 | intron_variant | MODIFIER |
| 1 | 171057965 | AX-170141860 | 0 | A | G | 0.211 | 1.43E-01 | 1.37E-01 | 2.62E+00 | 2.66E+00 | 2.94E-01 | no | Fcgr3 | intron_variant | MODIFIER |
| 1 | 171061145 | AX-97799937 | 0 | A | G | 0.211 | 1.43E-01 | 1.37E-01 | 2.62E+00 | 2.66E+00 | 2.94E-01 | no | Fcgr3 | upstream_gene_variant | MODIFIER |
| 1 | 171061311 | AX-170121803 | 0 | A | G | 0.211 | 1.43E-01 | 1.37E-01 | 2.62E+00 | 2.66E+00 | 2.94E-01 | no | Fcgr3 | upstream_gene_variant | MODIFIER |
| 1 | 171090842 | AX-170517494 | 0 | G | A | 0.211 | 1.43E-01 | 1.37E-01 | 2.62E+00 | 2.66E+00 | 2.94E-01 | no | | intergenic_variant | MODIFIER |
| 1 | 171095442 | AX-170283689 | 0 | A | C | 0.211 | 1.43E-01 | 1.37E-01 | 2.62E+00 | 2.66E+00 | 2.94E-01 | no | | intergenic_variant | MODIFIER |
| 1 | 171146297 | AX-170002274 | 0 | T | C | 0.211 | 1.43E-01 | 1.37E-01 | 2.62E+00 | 2.66E+00 | 2.94E-01 | no | Mpz | upstream_gene_variant | MODIFIER |
| 1 | 171191690 | AX-170002278 | 0 | A | T | 0.211 | 1.43E-01 | 1.37E-01 | 2.62E+00 | 2.66E+00 | 2.94E-01 | no | Pcp4l1 | intron_variant | MODIFIER |
| 1 | 171195757 | AX-97845333 | 0 | G | A | 0.211 | 1.43E-01 | 1.37E-01 | 2.62E+00 | 2.66E+00 | 2.94E-01 | no | Pcp4l1 | intron_variant | MODIFIER |
| 1 | 171211317 | AX-97798100 | 0 | A | T | 0.211 | 1.43E-01 | 1.37E-01 | 2.62E+00 | 2.66E+00 | 2.94E-01 | no | Tomm40l | downstream_gene_variant | MODIFIER |
| 1 | 171222335 | AX-170002273 | 0 | C | A | 0.211 | 1.43E-01 | 1.37E-01 | 2.62E+00 | 2.66E+00 | 2.94E-01 | no | Tomm40l | intron_variant | MODIFIER |
| 1 | 171225381 | AX-170440145 | 0 | C | T | 0.211 | 1.43E-01 | 1.37E-01 | 2.62E+00 | 2.66E+00 | 2.94E-01 | no | Tomm40l | upstream_gene_variant | MODIFIER |
| 1 | 171238494 | AX-170266551 | 0 | T | C | 0.211 | 1.43E-01 | 1.37E-01 | 2.62E+00 | 2.66E+00 | 2.94E-01 | no | Ndufs2 | intron_variant | MODIFIER |
| 1 | 171244725 | AX-170216440 | 0 | G | A | 0.211 | 1.43E-01 | 1.37E-01 | 2.62E+00 | 2.66E+00 | 2.94E-01 | no | Ndufs2 | intron_variant | MODIFIER |
| 1 | 171246149 | AX-170485057 | 0 | C | G | 0.211 | 1.43E-01 | 1.37E-01 | 2.62E+00 | 2.66E+00 | 2.94E-01 | no | Ndufs2 | synonymous_variant | LOW |
| 1 | 171246239 | AX-170133455 | 0 | G | A | 0.211 | 1.43E-01 | 1.37E-01 | 2.62E+00 | 2.66E+00 | 2.94E-01 | no | Ndufs2 | intron_variant | MODIFIER |
| 1 | 171247008 | AX-170427451 | 0 | T | C | 0.211 | 1.43E-01 | 1.37E-01 | 2.62E+00 | 2.66E+00 | 2.94E-01 | no | Ndufs2 | synonymous_variant | LOW |
| 1 | 171248448 | AX-170141037 | 0 | A | G | 0.211 | 1.43E-01 | 1.37E-01 | 2.62E+00 | 2.66E+00 | 2.94E-01 | no | Ndufs2 | upstream_gene_variant | MODIFIER |
| 1 | 171249729 | AX-97810160 | 0 | T | C | 0.211 | 1.43E-01 | 1.37E-01 | 2.62E+00 | 2.66E+00 | 2.94E-01 | no | Ndufs2 | upstream_gene_variant | MODIFIER |
| 1 | 171258252 | AX-170002275 | 0 | T | C | 0.211 | 1.43E-01 | 1.37E-01 | 2.62E+00 | 2.66E+00 | 2.94E-01 | no | Adamts4 | intron_variant | MODIFIER |
| 1 | 171269985 | AX-170485262 | 0 | G | A | 0.211 | 1.43E-01 | 1.37E-01 | 2.62E+00 | 2.66E+00 | 2.94E-01 | no | B4galt3 | upstream_gene_variant | MODIFIER |
| 1 | 171276214 | AX-170345238 | 0 | C | T | 0.211 | 1.43E-01 | 1.37E-01 | 2.62E+00 | 2.66E+00 | 2.94E-01 | no | B4galt3 | synonymous_variant | LOW |
| 1 | 171288826 | AX-170387197 | 0 | A | G | 0.211 | 1.43E-01 | 1.37E-01 | 2.62E+00 | 2.66E+00 | 2.94E-01 | no | Usp21 | upstream_gene_variant | MODIFIER |
| 1 | 171289680 | AX-170043570 | 0 | T | C | 0.211 | 1.43E-01 | 1.37E-01 | 2.62E+00 | 2.66E+00 | 2.94E-01 | no | Usp21 | upstream_gene_variant | MODIFIER |
| 1 | 171290106 | AX-170091786 | 0 | G | A | 0.211 | 1.43E-01 | 1.37E-01 | 2.62E+00 | 2.66E+00 | 2.94E-01 | no | Usp21 | upstream_gene_variant | MODIFIER |
| 1 | 171292590 | AX-170232165 | 0 | G | A | 0.211 | 1.43E-01 | 1.37E-01 | 2.62E+00 | 2.66E+00 | 2.94E-01 | no | Usp21 | upstream_gene_variant | MODIFIER |
| 1 | 171296427 | AX-170476505 | 0 | T | C | 0.211 | 1.43E-01 | 1.37E-01 | 2.62E+00 | 2.66E+00 | 2.94E-01 | no | Ufc1 | upstream_gene_variant | MODIFIER |
| 1 | 171318820 | AX-170142960 | 0 | C | T | 0.211 | 1.43E-01 | 1.37E-01 | 2.62E+00 | 2.66E+00 | 2.94E-01 | no | Gm20045 | downstream_gene_variant | MODIFIER |

| | | | | | | | | | | | | | | | |
|---|---|---|---|---|---|---|---|---|---|---|---|---|---|---|---|
| 1 | 171334625 | AX-170373090 | 0 | G | C | 0.211 | 1.43E-01 | 1.37E-01 | 2.62E+00 | 2.66E+00 | 2.94E-01 | no | Dedd | intron_variant | MODIFIER |
| 1 | 171338636 | AX-170330852 | 0 | A | C | 0.211 | 1.43E-01 | 1.37E-01 | 2.62E+00 | 2.66E+00 | 2.94E-01 | no | Dedd | synonymous_variant | LOW |
| 1 | 171342027 | AX-170268254 | 0 | T | G | 0.211 | 1.43E-01 | 1.37E-01 | 2.62E+00 | 2.66E+00 | 2.94E-01 | no | Pfdn2 | upstream_gene_variant | MODIFIER |
| 1 | 171350073 | AX-170229066 | 0 | C | T | 0.211 | 1.43E-01 | 1.37E-01 | 2.62E+00 | 2.66E+00 | 2.94E-01 | no | Pfdn2 | intron_variant | MODIFIER |
| 1 | 171368654 | AX-170266696 | 0 | G | A | 0.211 | 1.43E-01 | 1.37E-01 | 2.62E+00 | 2.66E+00 | 2.94E-01 | no | Nectin4 | upstream_gene_variant | MODIFIER |
| 1 | 171369014 | AX-170176791 | 0 | A | G | 0.211 | 1.43E-01 | 1.37E-01 | 2.62E+00 | 2.66E+00 | 2.94E-01 | no | Nectin4 | upstream_gene_variant | MODIFIER |
| 1 | 171369609 | AX-170419945 | 0 | C | T | 0.211 | 1.43E-01 | 1.37E-01 | 2.62E+00 | 2.66E+00 | 2.94E-01 | no | Nectin4 | upstream_gene_variant | MODIFIER |
| 1 | 171373198 | AX-168406076 | 0 | C | T | 0.211 | 1.43E-01 | 1.37E-01 | 2.62E+00 | 2.66E+00 | 2.94E-01 | no | Nectin4 | intron_variant | MODIFIER |
| 1 | 171376218 | AX-97850049 | 0 | G | T | 0.211 | 1.43E-01 | 1.37E-01 | 2.62E+00 | 2.66E+00 | 2.94E-01 | no | Nectin4 | intron_variant | MODIFIER |
| 1 | 171384101 | AX-170160769 | 0 | G | C | 0.211 | 1.43E-01 | 1.37E-01 | 2.62E+00 | 2.66E+00 | 2.94E-01 | no | Nectin4 | intron_variant | MODIFIER |
| 1 | 171387257 | AX-170348963 | 0 | T | C | 0.211 | 1.43E-01 | 1.37E-01 | 2.62E+00 | 2.66E+00 | 2.94E-01 | no | Nectin4 | 3_prime_UTR_variant | MODIFIER |
| 1 | 171387752 | AX-170379379 | 0 | A | G | 0.211 | 1.43E-01 | 1.37E-01 | 2.62E+00 | 2.66E+00 | 2.94E-01 | no | Nectin4 | 3_prime_UTR_variant | MODIFIER |
| 1 | 171418823 | AX-170304495 | 0 | A | G | 0.211 | 1.43E-01 | 1.37E-01 | 2.62E+00 | 2.66E+00 | 2.94E-01 | no | Usf1 | downstream_gene_variant | MODIFIER |
| 1 | 171418926 | AX-170278189 | 0 | C | T | 0.211 | 1.43E-01 | 1.37E-01 | 2.62E+00 | 2.66E+00 | 2.94E-01 | no | Usf1 | downstream_gene_variant | MODIFIER |
| 1 | 171431737 | AX-97820006 | 0 | T | C | 0.211 | 1.43E-01 | 1.37E-01 | 2.62E+00 | 2.66E+00 | 2.94E-01 | no | Gm26641 | intron_variant,non_coding_transcript_variant | MODIFIER |
| 1 | 171435646 | AX-168391064 | 0 | C | T | 0.211 | 1.43E-01 | 1.37E-01 | 2.62E+00 | 2.66E+00 | 2.94E-01 | no | F11r | upstream_gene_variant | MODIFIER |
| 1 | 171438378 | AX-170170605 | 0 | A | G | 0.211 | 1.43E-01 | 1.37E-01 | 2.62E+00 | 2.66E+00 | 2.94E-01 | no | F11r | intron_variant | MODIFIER |
| 1 | 171460216 | AX-97830804 | 0 | T | C | 0.211 | 1.43E-01 | 1.37E-01 | 2.62E+00 | 2.66E+00 | 2.94E-01 | no | F11r | intron_variant | MODIFIER |
| 1 | 171461622 | AX-170339657 | 0 | A | G | 0.211 | 1.43E-01 | 1.37E-01 | 2.62E+00 | 2.66E+00 | 2.94E-01 | no | F11r | synonymous_variant | LOW |
| 1 | 171498421 | AX-170333686 | 0 | A | G | 0.211 | 1.43E-01 | 1.37E-01 | 2.62E+00 | 2.66E+00 | 2.94E-01 | no | Gm24871 | downstream_gene_variant | MODIFIER |
| 1 | 171503211 | AX-170428866 | 0 | C | A | 0.211 | 1.43E-01 | 1.37E-01 | 2.62E+00 | 2.66E+00 | 2.94E-01 | no | Alyref2 | upstream_gene_variant | MODIFIER |
| 1 | 171527620 | AX-170296119 | 0 | T | G | 0.211 | 1.43E-01 | 1.37E-01 | 2.62E+00 | 2.66E+00 | 2.94E-01 | no | Itln1 | intron_variant | MODIFIER |
| 1 | 171576905 | AX-170100051 | 0 | A | G | 0.211 | 1.43E-01 | 1.37E-01 | 2.62E+00 | 2.66E+00 | 2.94E-01 | no | Cd244a | intron_variant | MODIFIER |
| 1 | 171582944 | AX-170393050 | 0 | C | T | 0.211 | 1.43E-01 | 1.37E-01 | 2.62E+00 | 2.66E+00 | 2.94E-01 | no | Cd244a | 3_prime_UTR_variant | MODIFIER |
| 1 | 171589634 | AX-170043578 | 0 | G | A | 0.211 | 1.43E-01 | 1.37E-01 | 2.62E+00 | 2.66E+00 | 2.94E-01 | no | Ly9 | intron_variant | MODIFIER |
| 1 | 171593907 | AX-97818297 | 0 | C | T | 0.211 | 1.43E-01 | 1.37E-01 | 2.62E+00 | 2.66E+00 | 2.94E-01 | no | Ly9 | missense_variant | MODERATE |
| 1 | 171621022 | AX-97851133 | 0 | C | G | 0.211 | 1.43E-01 | 1.37E-01 | 2.62E+00 | 2.66E+00 | 2.94E-01 | no | | intergenic_variant | MODIFIER |
| 1 | 171664326 | AX-97782427 | 0 | A | G | 0.211 | 1.43E-01 | 1.37E-01 | 2.62E+00 | 2.66E+00 | 2.94E-01 | no | | intergenic_variant | MODIFIER |
| 1 | 171695858 | AX-170298886 | 0 | A | G | 0.211 | 1.43E-01 | 1.37E-01 | 2.62E+00 | 2.66E+00 | 2.94E-01 | no | Cd48 | missense_variant | MODERATE |
| 1 | 171696579 | AX-170002285 | 0 | G | A | 0.211 | 1.43E-01 | 1.37E-01 | 2.62E+00 | 2.66E+00 | 2.94E-01 | no | Cd48 | intron_variant | MODIFIER |
| 1 | 171701395 | AX-168374150 | 0 | T | C | 0.211 | 1.43E-01 | 1.37E-01 | 2.62E+00 | 2.66E+00 | 2.94E-01 | no | Cd48 | intron_variant | MODIFIER |
| 1 | 171729690 | AX-170152888 | 0 | A | G | 0.211 | 1.43E-01 | 1.37E-01 | 2.62E+00 | 2.66E+00 | 2.94E-01 | no | Gm22950 | downstream_gene_variant | MODIFIER |
| 1 | 171744758 | AX-170002284 | 0 | T | C | 0.211 | 1.43E-01 | 1.37E-01 | 2.62E+00 | 2.66E+00 | 2.94E-01 | no | | intergenic_variant | MODIFIER |
| 1 | 171746038 | AX-170002280 | 0 | T | C | 0.211 | 1.43E-01 | 1.37E-01 | 2.62E+00 | 2.66E+00 | 2.94E-01 | no | | intergenic_variant | MODIFIER |
| 1 | 171761892 | AX-97850331 | 0 | C | T | 0.211 | 1.43E-01 | 1.37E-01 | 2.62E+00 | 2.66E+00 | 2.94E-01 | no | | regulatory_region_variant | MODIFIER |
| 1 | 171778416 | AX-170002282 | 0 | T | G | 0.211 | 1.43E-01 | 1.37E-01 | 2.62E+00 | 2.66E+00 | 2.94E-01 | no | Slamf1 | intron_variant | MODIFIER |
| 1 | 171794289 | AX-97849582 | 0 | A | G | 0.211 | 1.43E-01 | 1.37E-01 | 2.62E+00 | 2.66E+00 | 2.94E-01 | no | Slamf1 | intron_variant | MODIFIER |
| 1 | 171799774 | AX-170446485 | 0 | G | A | 0.211 | 1.43E-01 | 1.37E-01 | 2.62E+00 | 2.66E+00 | 2.94E-01 | no | Slamf1 | 3_prime_UTR_variant | MODIFIER |
| 1 | 171845606 | AX-170150254 | 0 | T | C | 0.211 | 1.43E-01 | 1.37E-01 | 2.62E+00 | 2.66E+00 | 2.94E-01 | no | Cd84 | intron_variant | MODIFIER |
| 1 | 171845877 | AX-170126447 | 0 | G | T | 0.211 | 1.43E-01 | 1.37E-01 | 2.62E+00 | 2.66E+00 | 2.94E-01 | no | Cd84 | intron_variant | MODIFIER |
| 1 | 171848487 | AX-97800342 | 0 | G | A | 0.211 | 1.43E-01 | 1.37E-01 | 2.62E+00 | 2.66E+00 | 2.94E-01 | no | Cd84 | intron_variant | MODIFIER |
| 1 | 171849483 | AX-170424415 | 0 | T | G | 0.211 | 1.43E-01 | 1.37E-01 | 2.62E+00 | 2.66E+00 | 2.94E-01 | no | Cd84 | intron_variant | MODIFIER |
| 1 | 171849583 | AX-170143723 | 0 | G | A | 0.211 | 1.43E-01 | 1.37E-01 | 2.62E+00 | 2.66E+00 | 2.94E-01 | no | Cd84 | intron_variant | MODIFIER |
| 1 | 171946789 | AX-170209693 | 0 | C | T | 0.211 | 1.43E-01 | 1.37E-01 | 2.62E+00 | 2.66E+00 | 2.94E-01 | no | Slamf6 | downstream_gene_variant | MODIFIER |
| 1 | 171948902 | AX-170468315 | 0 | A | G | 0.211 | 1.43E-01 | 1.37E-01 | 2.62E+00 | 2.66E+00 | 2.94E-01 | no | Slamf6 | downstream_gene_variant | MODIFIER |
| 1 | 171949428 | AX-170426426 | 0 | T | C | 0.211 | 1.43E-01 | 1.37E-01 | 2.62E+00 | 2.66E+00 | 2.94E-01 | no | Slamf6 | downstream_gene_variant | MODIFIER |
| 1 | 171991896 | AX-97849655 | 0 | A | G | 0.211 | 1.43E-01 | 1.37E-01 | 2.62E+00 | 2.66E+00 | 2.94E-01 | no | | intergenic_variant | MODIFIER |
| 1 | 172004593 | AX-170271946 | 0 | C | T | 0.211 | 1.43E-01 | 1.37E-01 | 2.62E+00 | 2.66E+00 | 2.94E-01 | no | Vangl2 | 3_prime_UTR_variant | MODIFIER |
| 1 | 172012068 | AX-97844846 | 0 | G | T | 0.211 | 1.43E-01 | 1.37E-01 | 2.62E+00 | 2.66E+00 | 2.94E-01 | no | Vangl2 | intron_variant | MODIFIER |
| 1 | 172018615 | AX-170437939 | 0 | A | G | 0.211 | 1.43E-01 | 1.37E-01 | 2.62E+00 | 2.66E+00 | 2.94E-01 | no | Vangl2 | intron_variant | MODIFIER |
| 1 | 172032523 | AX-170493081 | 0 | A | G | 0.211 | 1.43E-01 | 1.37E-01 | 2.62E+00 | 2.66E+00 | 2.94E-01 | no | Vangl2 | upstream_gene_variant | MODIFIER |
| 1 | 172060216 | AX-170167161 | 0 | C | T | 0.211 | 1.43E-01 | 1.37E-01 | 2.62E+00 | 2.66E+00 | 2.94E-01 | no | Nhlh1 | upstream_gene_variant | MODIFIER |
| 1 | 172065168 | AX-97807414 | 0 | A | G | 0.211 | 1.43E-01 | 1.37E-01 | 2.62E+00 | 2.66E+00 | 2.94E-01 | no | Ncstn | downstream_gene_variant | MODIFIER |
| 1 | 172067996 | AX-170376011 | 0 | T | C | 0.211 | 1.43E-01 | 1.37E-01 | 2.62E+00 | 2.66E+00 | 2.94E-01 | no | Ncstn | intron_variant | MODIFIER |

| | | | | | | | | | | | | | | | |
|---|---|---|---|---|---|---|---|---|---|---|---|---|---|---|---|
| 1 | 172070053 | AX-170333532 | 0 | C | T | 0.211 | 1.43E-01 | 1.37E-01 | 2.62E+00 | 2.66E+00 | 2.94E-01 | no | Ncstn | synonymous_variant | LOW |
| 1 | 172070831 | AX-170420351 | 0 | G | A | 0.211 | 1.43E-01 | 1.37E-01 | 2.62E+00 | 2.66E+00 | 2.94E-01 | no | Ncstn | intron_variant | MODIFIER |
| 1 | 172073954 | AX-170408519 | 0 | C | T | 0.211 | 1.43E-01 | 1.37E-01 | 2.62E+00 | 2.66E+00 | 2.94E-01 | no | Ncstn | intron_variant | MODIFIER |
| 1 | 172091334 | AX-170176333 | 0 | T | C | 0.211 | 1.43E-01 | 1.37E-01 | 2.62E+00 | 2.66E+00 | 2.94E-01 | no | Copa | intron_variant | MODIFIER |
| 1 | 172098237 | AX-97853733 | 0 | A | G | 0.211 | 1.43E-01 | 1.37E-01 | 2.62E+00 | 2.66E+00 | 2.94E-01 | no | Copa | intron_variant | MODIFIER |
| 1 | 172098365 | AX-170104180 | 0 | A | G | 0.211 | 1.43E-01 | 1.37E-01 | 2.62E+00 | 2.66E+00 | 2.94E-01 | no | Copa | intron_variant | MODIFIER |
| 1 | 172098989 | AX-170531114 | 0 | C | T | 0.211 | 1.43E-01 | 1.37E-01 | 2.62E+00 | 2.66E+00 | 2.94E-01 | no | Copa | intron_variant | MODIFIER |
| 1 | 172113899 | AX-170216826 | 0 | C | A | 0.211 | 1.43E-01 | 1.37E-01 | 2.62E+00 | 2.66E+00 | 2.94E-01 | no | Copa | intron_variant | MODIFIER |
| 1 | 172138049 | AX-170355882 | 0 | T | A | 0.211 | 1.43E-01 | 1.37E-01 | 2.62E+00 | 2.66E+00 | 2.94E-01 | no | Pex19 | downstream_gene_variant | MODIFIER |
| 1 | 172138234 | AX-97814216 | 0 | G | A | 0.211 | 1.43E-01 | 1.37E-01 | 2.62E+00 | 2.66E+00 | 2.94E-01 | no | Pex19 | downstream_gene_variant | MODIFIER |
| 1 | 172160885 | AX-170043589 | 0 | G | C | 0.211 | 1.43E-01 | 1.37E-01 | 2.62E+00 | 2.66E+00 | 2.94E-01 | no | Dcaf8 | intron_variant | MODIFIER |
| 1 | 172196596 | AX-170043590 | 0 | T | A | 0.211 | 1.43E-01 | 1.37E-01 | 2.62E+00 | 2.66E+00 | 2.94E-01 | no | Pea15a | downstream_gene_variant | MODIFIER |
| 1 | 172203030 | AX-170470941 | 0 | G | A | 0.211 | 1.43E-01 | 1.37E-01 | 2.62E+00 | 2.66E+00 | 2.94E-01 | no | Pea15a | intron_variant | MODIFIER |
| 1 | 172203589 | AX-170508050 | 0 | C | T | 0.211 | 1.43E-01 | 1.37E-01 | 2.62E+00 | 2.66E+00 | 2.94E-01 | no | Pea15a | intron_variant | MODIFIER |
| 1 | 172204090 | AX-97845344 | 0 | A | G | 0.211 | 1.43E-01 | 1.37E-01 | 2.62E+00 | 2.66E+00 | 2.94E-01 | no | Pea15a | intron_variant | MODIFIER |
| 1 | 172219532 | AX-170002291 | 0 | G | A | 0.211 | 1.43E-01 | 1.37E-01 | 2.62E+00 | 2.66E+00 | 2.94E-01 | no | Casq1 | synonymous_variant | LOW |
| 1 | 172220439 | AX-170002290 | 0 | A | G | 0.211 | 1.43E-01 | 1.37E-01 | 2.62E+00 | 2.66E+00 | 2.94E-01 | no | Casq1 | upstream_gene_variant | MODIFIER |
| 1 | 172227674 | AX-170196550 | 0 | A | G | 0.211 | 1.43E-01 | 1.37E-01 | 2.62E+00 | 2.66E+00 | 2.94E-01 | no | Atp1a4 | intron_variant | MODIFIER |
| 1 | 172232788 | AX-170313007 | 0 | T | C | 0.211 | 1.43E-01 | 1.37E-01 | 2.62E+00 | 2.66E+00 | 2.94E-01 | no | Atp1a4 | intron_variant | MODIFIER |
| 1 | 172235364 | AX-170227947 | 0 | A | G | 0.211 | 1.43E-01 | 1.37E-01 | 2.62E+00 | 2.66E+00 | 2.94E-01 | no | Atp1a4 | intron_variant | MODIFIER |
| 1 | 172290161 | AX-170352547 | 0 | G | A | 0.211 | 1.43E-01 | 1.37E-01 | 2.62E+00 | 2.66E+00 | 2.94E-01 | no | Atp1a2 | intron_variant | MODIFIER |
| 1 | 172290559 | AX-170446809 | 0 | A | G | 0.211 | 1.43E-01 | 1.37E-01 | 2.62E+00 | 2.66E+00 | 2.94E-01 | no | Atp1a2 | intron_variant | MODIFIER |
| 1 | 172293063 | AX-170043594 | 0 | T | C | 0.211 | 1.43E-01 | 1.37E-01 | 2.62E+00 | 2.66E+00 | 2.94E-01 | no | Atp1a2 | intron_variant | MODIFIER |
| 1 | 172293943 | AX-170043596 | 0 | C | A | 0.211 | 1.43E-01 | 1.37E-01 | 2.62E+00 | 2.66E+00 | 2.94E-01 | no | Atp1a2 | intron_variant | MODIFIER |
| 1 | 172294541 | AX-170043595 | 0 | C | G | 0.211 | 1.43E-01 | 1.37E-01 | 2.62E+00 | 2.66E+00 | 2.94E-01 | no | Atp1a2 | intron_variant | MODIFIER |
| 1 | 172297040 | AX-170134836 | 0 | T | C | 0.211 | 1.43E-01 | 1.37E-01 | 2.62E+00 | 2.66E+00 | 2.94E-01 | no | Atp1a2 | intron_variant | MODIFIER |
| 1 | 172300552 | AX-170348535 | 0 | G | A | 0.211 | 1.43E-01 | 1.37E-01 | 2.62E+00 | 2.66E+00 | 2.94E-01 | no | Atp1a2 | upstream_gene_variant | MODIFIER |
| 1 | 172323525 | AX-170125483 | 0 | T | A | 0.211 | 1.43E-01 | 1.37E-01 | 2.62E+00 | 2.66E+00 | 2.94E-01 | no | Igsf8 | downstream_gene_variant | MODIFIER |
| 1 | 172325480 | AX-170488239 | 0 | C | T | 0.211 | 1.43E-01 | 1.37E-01 | 2.62E+00 | 2.66E+00 | 2.94E-01 | no | Kcnj9 | intron_variant | MODIFIER |
| 1 | 172325962 | AX-170207752 | 0 | C | T | 0.211 | 1.43E-01 | 1.37E-01 | 2.62E+00 | 2.66E+00 | 2.94E-01 | no | Kcnj9 | synonymous_variant | LOW |
| 1 | 172328115 | AX-170273535 | 0 | A | G | 0.211 | 1.43E-01 | 1.37E-01 | 2.62E+00 | 2.66E+00 | 2.94E-01 | no | Kcnj9 | intron_variant | MODIFIER |
| 1 | 172328453 | AX-170160807 | 0 | T | C | 0.211 | 1.43E-01 | 1.37E-01 | 2.62E+00 | 2.66E+00 | 2.94E-01 | no | Kcnj9 | intron_variant | MODIFIER |
| 1 | 172333385 | AX-170510730 | 0 | G | T | 0.211 | 1.43E-01 | 1.37E-01 | 2.62E+00 | 2.66E+00 | 2.94E-01 | no | Kcnj9 | upstream_gene_variant | MODIFIER |
| 1 | 172339456 | AX-170448693 | 0 | A | G | 0.211 | 1.43E-01 | 1.37E-01 | 2.62E+00 | 2.66E+00 | 2.94E-01 | no | Kcnj10 | upstream_gene_variant | MODIFIER |
| 10 | 18210806 | AX-170258922 | 1 | C | T | 0.205 | 1.07E-01 | 1.01E-01 | 3.05E+00 | 3.17E+00 | 2.95E-01 | no | Ccdc28a | downstream_gene_variant | MODIFIER |
| 2 | 72711958 | AX-170223364 | 0 | T | C | 0.044 | -2.00E-01 | 1.93E-01 | 2.67E+00 | 2.78E+00 | 2.95E-01 | Yes | 8430437L04Rik | intron_variant,non_coding_transcript_variant | MODIFIER |
| 4 | 54379540 | AX-170360704 | 1 | T | C | 0.357 | -1.01E-01 | 9.75E-02 | 2.85E+00 | 2.95E+00 | 2.95E-01 | no | | intergenic_variant | MODIFIER |
| 4 | 51243908 | AX-97845529 | 2 | T | C | 0.355 | -9.48E-02 | 9.15E-02 | 2.76E+00 | 2.87E+00 | 2.95E-01 | no | | intergenic_variant | MODIFIER |
| 4 | 83636833 | AX-170023271 | 0 | T | G | 0.333 | 1.18E-01 | 1.14E-01 | 2.70E+00 | 2.75E+00 | 2.95E-01 | no | Ccdc171 | intron_variant | MODIFIER |
| 1 | 152701710 | AX-170401268 | 0 | A | C | 0.202 | 1.45E-01 | 1.40E-01 | 2.71E+00 | 2.76E+00 | 2.95E-01 | Yes | Rgl1 | intron_variant | MODIFIER |
| 8 | 73513453 | AX-170220801 | 0 | T | C | 0.193 | -1.18E-01 | 1.14E-01 | 2.74E+00 | 2.86E+00 | 2.95E-01 | no | | intergenic_variant | MODIFIER |
| 7 | 17888061 | AX-170516344 | 0 | T | A | 0.36 | 8.35E-02 | 8.06E-02 | 2.64E+00 | 2.76E+00 | 2.95E-01 | no | Gm5155 | intron_variant | MODIFIER |
| 1 | 131292602 | AX-170001710 | 1 | T | C | 0.536 | 8.86E-02 | 8.52E-02 | 2.65E+00 | 2.71E+00 | 2.95E-01 | no | Srgap2 | synonymous_variant | LOW |
| 6 | 135909241 | AX-170181293 | 0 | T | C | 0.579 | 9.41E-02 | 8.83E-02 | 2.43E+00 | 2.48E+00 | 2.95E-01 | no | Grin2b | intron_variant | MODIFIER |
| 10 | 5315346 | AX-97823242 | 0 | A | T | 0.211 | -9.92E-02 | 9.59E-02 | 2.79E+00 | 2.92E+00 | 2.95E-01 | no | Syne1 | intron_variant | MODIFIER |
| 2 | 163828883 | AX-170349028 | 1 | C | T | 0.161 | 2.03E-01 | 1.96E-01 | 2.82E+00 | 2.91E+00 | 2.95E-01 | Yes | Ccn5 | synonymous_variant | LOW |
| 1 | 124586342 | AX-170523291 | 1 | T | C | 0.562 | 8.79E-02 | 8.40E-02 | 2.56E+00 | 2.61E+00 | 2.95E-01 | no | Gm28928 | intron_variant,non_coding_transcript_variant | MODIFIER |
| 4 | 46897719 | AX-170181800 | 0 | T | C | 0.36 | -1.00E-01 | 9.64E-02 | 2.85E+00 | 2.95E+00 | 2.95E-01 | no | Gabbr2 | intron_variant | MODIFIER |
| 4 | 46900792 | AX-170493356 | 0 | T | C | 0.36 | -1.00E-01 | 9.64E-02 | 2.85E+00 | 2.95E+00 | 2.95E-01 | no | Gabbr2 | intron_variant | MODIFIER |
| 4 | 46970599 | AX-170367994 | 0 | A | G | 0.36 | -1.00E-01 | 9.64E-02 | 2.85E+00 | 2.95E+00 | 2.95E-01 | no | Gabbr2 | intron_variant | MODIFIER |
| 4 | 46979041 | AX-170253948 | 0 | A | C | 0.36 | -1.00E-01 | 9.64E-02 | 2.85E+00 | 2.95E+00 | 2.95E-01 | no | Gabbr2 | intron_variant | MODIFIER |
| 4 | 47118700 | AX-97846898 | 0 | C | T | 0.36 | -1.00E-01 | 9.64E-02 | 2.85E+00 | 2.95E+00 | 2.95E-01 | no | Galnt12 | intron_variant | MODIFIER |
| 4 | 47144125 | AX-170191672 | 0 | G | A | 0.36 | -1.00E-01 | 9.64E-02 | 2.85E+00 | 2.95E+00 | 2.95E-01 | no | | intergenic_variant | MODIFIER |
| 4 | 47150633 | AX-170120646 | 0 | C | T | 0.36 | -1.00E-01 | 9.64E-02 | 2.85E+00 | 2.95E+00 | 2.95E-01 | no | | regulatory_region_variant | MODIFIER |

| | | | | | | | | | | | | | | | |
|---|---|---|---|---|---|---|---|---|---|---|---|---|---|---|---|
| 4 | 47158529 | AX-170022716 | 0 | G | A | 0.36 | -1.00E-01 | 9.64E-02 | 2.85E+00 | 2.95E+00 | 2.95E-01 | no | | intergenic_variant | MODIFIER |
| 4 | 47158811 | AX-170022715 | 0 | A | T | 0.36 | -1.00E-01 | 9.64E-02 | 2.85E+00 | 2.95E+00 | 2.95E-01 | no | | intergenic_variant | MODIFIER |
| 4 | 47162364 | AX-170022712 | 0 | C | T | 0.36 | -1.00E-01 | 9.64E-02 | 2.85E+00 | 2.95E+00 | 2.95E-01 | no | | regulatory_region_variant | MODIFIER |
| 4 | 47196907 | AX-170022713 | 0 | G | A | 0.36 | -1.00E-01 | 9.64E-02 | 2.85E+00 | 2.95E+00 | 2.95E-01 | no | Gm12426 | downstream_gene_variant | MODIFIER |
| 4 | 47292764 | AX-170278285 | 0 | C | G | 0.36 | -1.00E-01 | 9.64E-02 | 2.85E+00 | 2.95E+00 | 2.95E-01 | no | Col15a1 | intron_variant | MODIFIER |
| 4 | 47298043 | AX-170455437 | 0 | T | C | 0.36 | -1.00E-01 | 9.64E-02 | 2.85E+00 | 2.95E+00 | 2.95E-01 | no | Col15a1 | intron_variant | MODIFIER |
| 4 | 47298987 | AX-170351538 | 0 | A | G | 0.36 | -1.00E-01 | 9.64E-02 | 2.85E+00 | 2.95E+00 | 2.95E-01 | no | Col15a1 | intron_variant | MODIFIER |
| 4 | 47299615 | AX-170472620 | 0 | G | A | 0.36 | -1.00E-01 | 9.64E-02 | 2.85E+00 | 2.95E+00 | 2.95E-01 | no | Col15a1 | intron_variant | MODIFIER |
| 4 | 47305071 | AX-170265557 | 0 | C | T | 0.36 | -1.00E-01 | 9.64E-02 | 2.85E+00 | 2.95E+00 | 2.95E-01 | no | Col15a1 | intron_variant | MODIFIER |
| 4 | 47305222 | AX-170131254 | 0 | T | C | 0.36 | -1.00E-01 | 9.64E-02 | 2.85E+00 | 2.95E+00 | 2.95E-01 | no | Col15a1 | intron_variant | MODIFIER |
| 4 | 47332799 | AX-170130510 | 0 | C | T | 0.36 | -1.00E-01 | 9.64E-02 | 2.85E+00 | 2.95E+00 | 2.95E-01 | no | | intergenic_variant | MODIFIER |
| 4 | 47399570 | AX-170519789 | 0 | G | T | 0.36 | -1.00E-01 | 9.64E-02 | 2.85E+00 | 2.95E+00 | 2.95E-01 | no | Tgfbr1 | intron_variant | MODIFIER |
| 4 | 47430413 | AX-97795215 | 0 | A | G | 0.36 | -1.00E-01 | 9.64E-02 | 2.85E+00 | 2.95E+00 | 2.95E-01 | no | | intergenic_variant | MODIFIER |
| 4 | 47439973 | AX-170361516 | 0 | A | G | 0.36 | -1.00E-01 | 9.64E-02 | 2.85E+00 | 2.95E+00 | 2.95E-01 | no | | intergenic_variant | MODIFIER |
| 4 | 47443947 | AX-170176647 | 0 | G | T | 0.36 | -1.00E-01 | 9.64E-02 | 2.85E+00 | 2.95E+00 | 2.95E-01 | no | | intergenic_variant | MODIFIER |
| 4 | 47444369 | AX-170207045 | 0 | T | C | 0.36 | -1.00E-01 | 9.64E-02 | 2.85E+00 | 2.95E+00 | 2.95E-01 | no | | intergenic_variant | MODIFIER |
| 4 | 47447336 | AX-170482429 | 0 | C | T | 0.36 | -1.00E-01 | 9.64E-02 | 2.85E+00 | 2.95E+00 | 2.95E-01 | no | | intergenic_variant | MODIFIER |
| 4 | 47449824 | AX-170361179 | 0 | T | G | 0.36 | -1.00E-01 | 9.64E-02 | 2.85E+00 | 2.95E+00 | 2.95E-01 | no | Gm12430 | upstream_gene_variant | MODIFIER |
| 4 | 47465491 | AX-170380158 | 0 | T | C | 0.36 | -1.00E-01 | 9.64E-02 | 2.85E+00 | 2.95E+00 | 2.95E-01 | no | Alg2 | downstream_gene_variant | MODIFIER |
| 4 | 47471200 | AX-170323797 | 0 | T | C | 0.36 | -1.00E-01 | 9.64E-02 | 2.85E+00 | 2.95E+00 | 2.95E-01 | no | Alg2 | 3_prime_UTR_variant | MODIFIER |
| 4 | 47590398 | AX-97787700 | 0 | T | C | 0.36 | -1.00E-01 | 9.64E-02 | 2.85E+00 | 2.95E+00 | 2.95E-01 | no | Gm27601 | upstream_gene_variant | MODIFIER |
| 4 | 47699478 | AX-170035681 | 0 | G | A | 0.36 | -1.00E-01 | 9.64E-02 | 2.85E+00 | 2.95E+00 | 2.95E-01 | no | | regulatory_region_variant | MODIFIER |
| 4 | 47772603 | AX-97799440 | 0 | T | C | 0.36 | -1.00E-01 | 9.64E-02 | 2.85E+00 | 2.95E+00 | 2.95E-01 | no | | regulatory_region_variant | MODIFIER |
| 4 | 47772980 | AX-170022723 | 0 | C | T | 0.36 | -1.00E-01 | 9.64E-02 | 2.85E+00 | 2.95E+00 | 2.95E-01 | no | | regulatory_region_variant | MODIFIER |
| 4 | 47778482 | AX-170022720 | 0 | G | C | 0.36 | -1.00E-01 | 9.64E-02 | 2.85E+00 | 2.95E+00 | 2.95E-01 | no | | intergenic_variant | MODIFIER |
| 4 | 47963643 | AX-97844404 | 0 | G | A | 0.36 | -1.00E-01 | 9.64E-02 | 2.85E+00 | 2.95E+00 | 2.95E-01 | no | | intergenic_variant | MODIFIER |
| 4 | 47997613 | AX-170358966 | 0 | C | T | 0.36 | -1.00E-01 | 9.64E-02 | 2.85E+00 | 2.95E+00 | 2.95E-01 | no | Gm12434 | intron_variant,non_coding_transcript_variant | MODIFIER |
| 4 | 48051670 | AX-170267082 | 0 | G | T | 0.36 | -1.00E-01 | 9.64E-02 | 2.85E+00 | 2.95E+00 | 2.95E-01 | no | Nr4a3 | synonymous_variant | LOW |
| 4 | 48067429 | AX-170233592 | 0 | A | G | 0.36 | -1.00E-01 | 9.64E-02 | 2.85E+00 | 2.95E+00 | 2.95E-01 | no | Nr4a3 | intron_variant | MODIFIER |
| 4 | 48071457 | AX-170257834 | 0 | A | G | 0.36 | -1.00E-01 | 9.64E-02 | 2.85E+00 | 2.95E+00 | 2.95E-01 | no | Nr4a3 | intron_variant | MODIFIER |
| 4 | 48110540 | AX-170422558 | 0 | T | C | 0.36 | -1.00E-01 | 9.64E-02 | 2.85E+00 | 2.95E+00 | 2.95E-01 | no | | regulatory_region_variant | MODIFIER |
| 4 | 48153590 | AX-170270843 | 0 | G | A | 0.36 | -1.00E-01 | 9.64E-02 | 2.85E+00 | 2.95E+00 | 2.95E-01 | no | Stx17 | intron_variant | MODIFIER |
| 4 | 48164392 | AX-97806262 | 0 | T | G | 0.36 | -1.00E-01 | 9.64E-02 | 2.85E+00 | 2.95E+00 | 2.95E-01 | no | Stx17 | intron_variant | MODIFIER |
| 4 | 48180138 | AX-97785630 | 0 | C | T | 0.36 | -1.00E-01 | 9.64E-02 | 2.85E+00 | 2.95E+00 | 2.95E-01 | no | Stx17 | intron_variant | MODIFIER |
| 4 | 48187488 | AX-170069294 | 0 | A | T | 0.36 | -1.00E-01 | 9.64E-02 | 2.85E+00 | 2.95E+00 | 2.95E-01 | no | Stx17 | downstream_gene_variant | MODIFIER |
| 4 | 48236276 | AX-170022725 | 0 | C | A | 0.36 | -1.00E-01 | 9.64E-02 | 2.85E+00 | 2.95E+00 | 2.95E-01 | no | Erp44 | intron_variant | MODIFIER |
| 4 | 48264286 | AX-97797538 | 0 | A | G | 0.36 | -1.00E-01 | 9.64E-02 | 2.85E+00 | 2.95E+00 | 2.95E-01 | no | Erp44 | intron_variant | MODIFIER |
| 4 | 48267650 | AX-170069297 | 0 | T | C | 0.36 | -1.00E-01 | 9.64E-02 | 2.85E+00 | 2.95E+00 | 2.95E-01 | no | Erp44 | intron_variant | MODIFIER |
| 4 | 48380374 | AX-170153493 | 0 | T | C | 0.36 | -1.00E-01 | 9.64E-02 | 2.85E+00 | 2.95E+00 | 2.95E-01 | no | Invs | intron_variant | MODIFIER |
| 4 | 48380474 | AX-170022732 | 0 | C | A | 0.36 | -1.00E-01 | 9.64E-02 | 2.85E+00 | 2.95E+00 | 2.95E-01 | no | Invs | intron_variant | MODIFIER |
| 4 | 48388137 | AX-170022733 | 0 | C | A | 0.36 | -1.00E-01 | 9.64E-02 | 2.85E+00 | 2.95E+00 | 2.95E-01 | no | Invs | intron_variant | MODIFIER |
| 4 | 48394424 | AX-170382299 | 0 | A | G | 0.36 | -1.00E-01 | 9.64E-02 | 2.85E+00 | 2.95E+00 | 2.95E-01 | no | Invs | intron_variant | MODIFIER |
| 4 | 48397100 | AX-170369995 | 0 | C | T | 0.36 | -1.00E-01 | 9.64E-02 | 2.85E+00 | 2.95E+00 | 2.95E-01 | no | Invs | intron_variant | MODIFIER |
| 4 | 48405509 | AX-170270753 | 0 | A | G | 0.36 | -1.00E-01 | 9.64E-02 | 2.85E+00 | 2.95E+00 | 2.95E-01 | no | Invs | intron_variant | MODIFIER |
| 4 | 48430137 | AX-170235826 | 0 | G | A | 0.36 | -1.00E-01 | 9.64E-02 | 2.85E+00 | 2.95E+00 | 2.95E-01 | no | Invs | 3_prime_UTR_variant | MODIFIER |
| 4 | 48476625 | AX-170443078 | 0 | T | C | 0.36 | -1.00E-01 | 9.64E-02 | 2.85E+00 | 2.95E+00 | 2.95E-01 | no | Tex10 | upstream_gene_variant | MODIFIER |
| 4 | 48493032 | AX-168381158 | 0 | G | T | 0.36 | -1.00E-01 | 9.64E-02 | 2.85E+00 | 2.95E+00 | 2.95E-01 | no | | intergenic_variant | MODIFIER |
| 4 | 48545079 | AX-170127336 | 0 | G | A | 0.36 | -1.00E-01 | 9.64E-02 | 2.85E+00 | 2.95E+00 | 2.95E-01 | no | Msantd3 | intron_variant | MODIFIER |
| 4 | 48564208 | AX-170022726 | 0 | G | A | 0.36 | -1.00E-01 | 9.64E-02 | 2.85E+00 | 2.95E+00 | 2.95E-01 | no | Msantd3 | downstream_gene_variant | MODIFIER |
| 4 | 48589705 | AX-170210408 | 0 | T | C | 0.36 | -1.00E-01 | 9.64E-02 | 2.85E+00 | 2.95E+00 | 2.95E-01 | no | Tmeff1 | intron_variant | MODIFIER |
| 4 | 48589825 | AX-97838382 | 0 | G | A | 0.36 | -1.00E-01 | 9.64E-02 | 2.85E+00 | 2.95E+00 | 2.95E-01 | no | Tmeff1 | intron_variant | MODIFIER |
| 4 | 48596784 | AX-170488732 | 0 | T | C | 0.36 | -1.00E-01 | 9.64E-02 | 2.85E+00 | 2.95E+00 | 2.95E-01 | no | Tmeff1 | intron_variant | MODIFIER |
| 4 | 48608392 | AX-97805729 | 0 | A | G | 0.36 | -1.00E-01 | 9.64E-02 | 2.85E+00 | 2.95E+00 | 2.95E-01 | no | Tmeff1 | intron_variant | MODIFIER |
| 4 | 48617077 | AX-170275220 | 0 | A | C | 0.36 | -1.00E-01 | 9.64E-02 | 2.85E+00 | 2.95E+00 | 2.95E-01 | no | Tmeff1 | intron_variant | MODIFIER |

| | | | | | | | | | | | | | | | |
|---|---|---|---|---|---|---|---|---|---|---|---|---|---|---|---|
| 4 | 48618278 | AX-170245482 | 0 | T | C | 0.36 | -1.00E-01 | 9.64E-02 | 2.85E+00 | 2.95E+00 | 2.95E-01 | no | Tmeff1 | intron_variant | MODIFIER |
| 4 | 48645451 | AX-170217712 | 0 | A | G | 0.36 | -1.00E-01 | 9.64E-02 | 2.85E+00 | 2.95E+00 | 2.95E-01 | no | Tmeff1 | intron_variant | MODIFIER |
| 4 | 48645625 | AX-170328754 | 0 | C | T | 0.36 | -1.00E-01 | 9.64E-02 | 2.85E+00 | 2.95E+00 | 2.95E-01 | no | Tmeff1 | intron_variant | MODIFIER |
| 4 | 48650591 | AX-170345173 | 0 | T | A | 0.36 | -1.00E-01 | 9.64E-02 | 2.85E+00 | 2.95E+00 | 2.95E-01 | no | Tmeff1 | intron_variant | MODIFIER |
| 4 | 48651533 | AX-170219321 | 0 | A | G | 0.36 | -1.00E-01 | 9.64E-02 | 2.85E+00 | 2.95E+00 | 2.95E-01 | no | Tmeff1 | intron_variant | MODIFIER |
| 4 | 48658855 | AX-170069300 | 0 | T | C | 0.36 | -1.00E-01 | 9.64E-02 | 2.85E+00 | 2.95E+00 | 2.95E-01 | no | Tmeff1 | synonymous_variant | LOW |
| 4 | 48659006 | AX-170516970 | 0 | T | C | 0.36 | -1.00E-01 | 9.64E-02 | 2.85E+00 | 2.95E+00 | 2.95E-01 | no | Tmeff1 | intron_variant | MODIFIER |
| 4 | 48664040 | AX-170226473 | 0 | G | A | 0.36 | -1.00E-01 | 9.64E-02 | 2.85E+00 | 2.95E+00 | 2.95E-01 | no | Tmeff1 | downstream_gene_variant | MODIFIER |
| 4 | 48664313 | AX-170153935 | 0 | C | T | 0.36 | -1.00E-01 | 9.64E-02 | 2.85E+00 | 2.95E+00 | 2.95E-01 | no | Tmeff1 | downstream_gene_variant | MODIFIER |
| 4 | 48674022 | AX-170324629 | 0 | A | G | 0.36 | -1.00E-01 | 9.64E-02 | 2.85E+00 | 2.95E+00 | 2.95E-01 | no | Cavin4 | downstream_gene_variant | MODIFIER |
| 4 | 48718563 | AX-170216976 | 0 | C | A | 0.36 | -1.00E-01 | 9.64E-02 | 2.85E+00 | 2.95E+00 | 2.95E-01 | no | | intergenic_variant | MODIFIER |
| 4 | 48721230 | AX-170224941 | 0 | A | G | 0.36 | -1.00E-01 | 9.64E-02 | 2.85E+00 | 2.95E+00 | 2.95E-01 | no | | intergenic_variant | MODIFIER |
| 4 | 48722520 | AX-170346832 | 0 | G | T | 0.36 | -1.00E-01 | 9.64E-02 | 2.85E+00 | 2.95E+00 | 2.95E-01 | no | | intergenic_variant | MODIFIER |
| 4 | 48760435 | AX-170089224 | 0 | A | C | 0.36 | -1.00E-01 | 9.64E-02 | 2.85E+00 | 2.95E+00 | 2.95E-01 | no | Gm24573 | downstream_gene_variant | MODIFIER |
| 4 | 48772826 | AX-170155311 | 0 | C | T | 0.36 | -1.00E-01 | 9.64E-02 | 2.85E+00 | 2.95E+00 | 2.95E-01 | no | | intergenic_variant | MODIFIER |
| 4 | 48813874 | AX-170069303 | 0 | T | C | 0.36 | -1.00E-01 | 9.64E-02 | 2.85E+00 | 2.95E+00 | 2.95E-01 | no | | intergenic_variant | MODIFIER |
| 4 | 48817257 | AX-170022737 | 0 | C | T | 0.36 | -1.00E-01 | 9.64E-02 | 2.85E+00 | 2.95E+00 | 2.95E-01 | no | | intergenic_variant | MODIFIER |
| 4 | 48837825 | AX-170022735 | 0 | T | G | 0.36 | -1.00E-01 | 9.64E-02 | 2.85E+00 | 2.95E+00 | 2.95E-01 | no | | intergenic_variant | MODIFIER |
| 4 | 48837852 | AX-170069304 | 0 | T | A | 0.36 | -1.00E-01 | 9.64E-02 | 2.85E+00 | 2.95E+00 | 2.95E-01 | no | | intergenic_variant | MODIFIER |
| 4 | 48839035 | AX-170226285 | 0 | A | G | 0.36 | -1.00E-01 | 9.64E-02 | 2.85E+00 | 2.95E+00 | 2.95E-01 | no | | intergenic_variant | MODIFIER |
| 4 | 48840433 | AX-170022738 | 0 | C | T | 0.36 | -1.00E-01 | 9.64E-02 | 2.85E+00 | 2.95E+00 | 2.95E-01 | no | | regulatory_region_variant | MODIFIER |
| 4 | 48902503 | AX-97820855 | 0 | C | A | 0.36 | -1.00E-01 | 9.64E-02 | 2.85E+00 | 2.95E+00 | 2.95E-01 | no | Gm12436 | intron_variant,non_coding_transcript_variant | MODIFIER |
| 4 | 48915656 | AX-170145175 | 0 | C | T | 0.36 | -1.00E-01 | 9.64E-02 | 2.85E+00 | 2.95E+00 | 2.95E-01 | no | Gm12436 | intron_variant,non_coding_transcript_variant | MODIFIER |
| 4 | 48915702 | AX-170108942 | 0 | T | C | 0.36 | -1.00E-01 | 9.64E-02 | 2.85E+00 | 2.95E+00 | 2.95E-01 | no | Gm12436 | intron_variant,non_coding_transcript_variant | MODIFIER |
| 4 | 48924107 | AX-170405836 | 0 | C | G | 0.36 | -1.00E-01 | 9.64E-02 | 2.85E+00 | 2.95E+00 | 2.95E-01 | no | Gm12436 | intron_variant,non_coding_transcript_variant | MODIFIER |
| 4 | 48938442 | AX-170343044 | 0 | T | C | 0.36 | -1.00E-01 | 9.64E-02 | 2.85E+00 | 2.95E+00 | 2.95E-01 | no | Gm12436 | intron_variant,non_coding_transcript_variant | MODIFIER |
| 4 | 48938534 | AX-170203906 | 0 | G | T | 0.36 | -1.00E-01 | 9.64E-02 | 2.85E+00 | 2.95E+00 | 2.95E-01 | no | Gm12436 | intron_variant,non_coding_transcript_variant | MODIFIER |
| 4 | 48944165 | AX-170405291 | 0 | C | G | 0.36 | -1.00E-01 | 9.64E-02 | 2.85E+00 | 2.95E+00 | 2.95E-01 | no | Gm12436 | intron_variant,non_coding_transcript_variant | MODIFIER |
| 4 | 48945352 | AX-170380258 | 0 | T | G | 0.36 | -1.00E-01 | 9.64E-02 | 2.85E+00 | 2.95E+00 | 2.95E-01 | no | Gm12436 | intron_variant,non_coding_transcript_variant | MODIFIER |
| 4 | 48969275 | AX-170304847 | 0 | A | G | 0.36 | -1.00E-01 | 9.64E-02 | 2.85E+00 | 2.95E+00 | 2.95E-01 | no | Gm12442 | upstream_gene_variant | MODIFIER |
| 4 | 49097066 | AX-170122673 | 0 | A | G | 0.36 | -1.00E-01 | 9.64E-02 | 2.85E+00 | 2.95E+00 | 2.95E-01 | no | Plppr1 | intron_variant | MODIFIER |
| 4 | 49264649 | AX-170127443 | 0 | A | G | 0.36 | -1.00E-01 | 9.64E-02 | 2.85E+00 | 2.95E+00 | 2.95E-01 | no | Plppr1 | intron_variant | MODIFIER |
| 4 | 49276055 | AX-170469280 | 0 | C | G | 0.36 | -1.00E-01 | 9.64E-02 | 2.85E+00 | 2.95E+00 | 2.95E-01 | no | Plppr1 | intron_variant | MODIFIER |
| 4 | 49276345 | AX-170022742 | 0 | A | T | 0.36 | -1.00E-01 | 9.64E-02 | 2.85E+00 | 2.95E+00 | 2.95E-01 | no | Plppr1 | intron_variant | MODIFIER |
| 4 | 49306685 | AX-170022740 | 0 | G | A | 0.36 | -1.00E-01 | 9.64E-02 | 2.85E+00 | 2.95E+00 | 2.95E-01 | no | Plppr1 | intron_variant | MODIFIER |
| 4 | 49534945 | AX-170399192 | 0 | A | G | 0.36 | -1.00E-01 | 9.64E-02 | 2.85E+00 | 2.95E+00 | 2.95E-01 | no | Aldob | downstream_gene_variant | MODIFIER |
| 4 | 49534985 | AX-170150225 | 0 | G | A | 0.36 | -1.00E-01 | 9.64E-02 | 2.85E+00 | 2.95E+00 | 2.95E-01 | no | Aldob | downstream_gene_variant | MODIFIER |
| 4 | 49536874 | AX-170276086 | 0 | T | C | 0.36 | -1.00E-01 | 9.64E-02 | 2.85E+00 | 2.95E+00 | 2.95E-01 | no | Aldob | synonymous_variant | LOW |
| 4 | 49537035 | AX-170310251 | 0 | C | T | 0.36 | -1.00E-01 | 9.64E-02 | 2.85E+00 | 2.95E+00 | 2.95E-01 | no | Aldob | intron_variant | MODIFIER |
| 4 | 49537107 | AX-97852537 | 0 | C | T | 0.36 | -1.00E-01 | 9.64E-02 | 2.85E+00 | 2.95E+00 | 2.95E-01 | no | Aldob | intron_variant | MODIFIER |
| 4 | 49538835 | AX-170372359 | 0 | G | A | 0.36 | -1.00E-01 | 9.64E-02 | 2.85E+00 | 2.95E+00 | 2.95E-01 | no | Aldob | synonymous_variant | LOW |
| 4 | 49548930 | AX-170407157 | 0 | C | A | 0.36 | -1.00E-01 | 9.64E-02 | 2.85E+00 | 2.95E+00 | 2.95E-01 | no | Aldob | intron_variant | MODIFIER |
| 4 | 49550024 | AX-170327588 | 0 | C | T | 0.36 | -1.00E-01 | 9.64E-02 | 2.85E+00 | 2.95E+00 | 2.95E-01 | no | Aldob | upstream_gene_variant | MODIFIER |
| 4 | 49550480 | AX-170342429 | 0 | C | T | 0.36 | -1.00E-01 | 9.64E-02 | 2.85E+00 | 2.95E+00 | 2.95E-01 | no | Aldob | upstream_gene_variant | MODIFIER |
| 4 | 49563294 | AX-97785830 | 0 | T | C | 0.36 | -1.00E-01 | 9.64E-02 | 2.85E+00 | 2.95E+00 | 2.95E-01 | no | | intergenic_variant | MODIFIER |
| 4 | 49579852 | AX-170507835 | 0 | T | C | 0.36 | -1.00E-01 | 9.64E-02 | 2.85E+00 | 2.95E+00 | 2.95E-01 | no | Tmem246 | downstream_gene_variant | MODIFIER |
| 4 | 49582296 | AX-97788213 | 0 | G | A | 0.36 | -1.00E-01 | 9.64E-02 | 2.85E+00 | 2.95E+00 | 2.95E-01 | no | Tmem246 | downstream_gene_variant | MODIFIER |
| 4 | 49583192 | AX-170488610 | 0 | C | A | 0.36 | -1.00E-01 | 9.64E-02 | 2.85E+00 | 2.95E+00 | 2.95E-01 | no | Tmem246 | downstream_gene_variant | MODIFIER |
| 4 | 49583452 | AX-170159350 | 0 | C | T | 0.36 | -1.00E-01 | 9.64E-02 | 2.85E+00 | 2.95E+00 | 2.95E-01 | no | Tmem246 | downstream_gene_variant | MODIFIER |
| 4 | 49583480 | AX-170069320 | 0 | A | G | 0.36 | -1.00E-01 | 9.64E-02 | 2.85E+00 | 2.95E+00 | 2.95E-01 | no | Tmem246 | downstream_gene_variant | MODIFIER |
| 4 | 49590002 | AX-168400126 | 0 | C | A | 0.36 | -1.00E-01 | 9.64E-02 | 2.85E+00 | 2.95E+00 | 2.95E-01 | no | Tmem246 | intron_variant | MODIFIER |
| 4 | 49593629 | AX-170151965 | 0 | T | C | 0.36 | -1.00E-01 | 9.64E-02 | 2.85E+00 | 2.95E+00 | 2.95E-01 | no | Tmem246 | intron_variant | MODIFIER |
| 4 | 49660943 | AX-170520239 | 0 | G | A | 0.36 | -1.00E-01 | 9.64E-02 | 2.85E+00 | 2.95E+00 | 2.95E-01 | no | Rnf20 | downstream_gene_variant | MODIFIER |
| 4 | 49666686 | AX-170368720 | 0 | C | A | 0.36 | -1.00E-01 | 9.64E-02 | 2.85E+00 | 2.95E+00 | 2.95E-01 | no | Grin3a | intron_variant | MODIFIER |

| | | | | | | | | | | | | | | | |
|---|---|---|---|---|---|---|---|---|---|---|---|---|---|---|---|
| 4 | 49788610 | AX-170069322 | 0 | T | A | 0.36 | -1.00E-01 | 9.64E-02 | 2.85E+00 | 2.95E+00 | 2.95E-01 | no | Grin3a | intron_variant | MODIFIER |
| 4 | 49806137 | AX-170129279 | 0 | T | C | 0.36 | -1.00E-01 | 9.64E-02 | 2.85E+00 | 2.95E+00 | 2.95E-01 | no | Grin3a | intron_variant | MODIFIER |
| 4 | 49832554 | AX-170345728 | 0 | T | C | 0.36 | -1.00E-01 | 9.64E-02 | 2.85E+00 | 2.95E+00 | 2.95E-01 | no | Grin3a | intron_variant | MODIFIER |
| 4 | 49839951 | AX-170089227 | 0 | T | C | 0.36 | -1.00E-01 | 9.64E-02 | 2.85E+00 | 2.95E+00 | 2.95E-01 | no | Grin3a | intron_variant | MODIFIER |
| 4 | 50145585 | AX-97851579 | 0 | G | A | 0.36 | -1.00E-01 | 9.64E-02 | 2.85E+00 | 2.95E+00 | 2.95E-01 | no | | intergenic_variant | MODIFIER |
| 4 | 50145971 | AX-170125477 | 0 | T | C | 0.36 | -1.00E-01 | 9.64E-02 | 2.85E+00 | 2.95E+00 | 2.95E-01 | no | | intergenic_variant | MODIFIER |
| 4 | 50159132 | AX-170089228 | 0 | G | A | 0.36 | -1.00E-01 | 9.64E-02 | 2.85E+00 | 2.95E+00 | 2.95E-01 | no | | intergenic_variant | MODIFIER |
| 4 | 50175050 | AX-170214916 | 0 | G | A | 0.36 | -1.00E-01 | 9.64E-02 | 2.85E+00 | 2.95E+00 | 2.95E-01 | no | | intergenic_variant | MODIFIER |
| 4 | 50203071 | AX-170369020 | 0 | T | G | 0.36 | -1.00E-01 | 9.64E-02 | 2.85E+00 | 2.95E+00 | 2.95E-01 | no | | intergenic_variant | MODIFIER |
| 4 | 50203513 | AX-170501555 | 0 | C | T | 0.36 | -1.00E-01 | 9.64E-02 | 2.85E+00 | 2.95E+00 | 2.95E-01 | no | | intergenic_variant | MODIFIER |
| 4 | 50204044 | AX-170427118 | 0 | T | C | 0.36 | -1.00E-01 | 9.64E-02 | 2.85E+00 | 2.95E+00 | 2.95E-01 | no | | intergenic_variant | MODIFIER |
| 4 | 50204073 | AX-97855886 | 0 | T | C | 0.36 | -1.00E-01 | 9.64E-02 | 2.85E+00 | 2.95E+00 | 2.95E-01 | no | | intergenic_variant | MODIFIER |
| 4 | 50220344 | AX-170104066 | 0 | T | C | 0.36 | -1.00E-01 | 9.64E-02 | 2.85E+00 | 2.95E+00 | 2.95E-01 | no | | intergenic_variant | MODIFIER |
| 4 | 50291394 | AX-170220713 | 0 | G | A | 0.36 | -1.00E-01 | 9.64E-02 | 2.85E+00 | 2.95E+00 | 2.95E-01 | no | Gm25485 | downstream_gene_variant | MODIFIER |
| 4 | 50304571 | AX-170365733 | 0 | T | G | 0.36 | -1.00E-01 | 9.64E-02 | 2.85E+00 | 2.95E+00 | 2.95E-01 | no | | intergenic_variant | MODIFIER |
| 4 | 50324715 | AX-170408209 | 0 | C | T | 0.36 | -1.00E-01 | 9.64E-02 | 2.85E+00 | 2.95E+00 | 2.95E-01 | no | | intergenic_variant | MODIFIER |
| 4 | 50336063 | AX-97848593 | 0 | G | A | 0.36 | -1.00E-01 | 9.64E-02 | 2.85E+00 | 2.95E+00 | 2.95E-01 | no | | intergenic_variant | MODIFIER |
| 4 | 50350952 | AX-170022753 | 0 | A | G | 0.36 | -1.00E-01 | 9.64E-02 | 2.85E+00 | 2.95E+00 | 2.95E-01 | no | | intergenic_variant | MODIFIER |
| 4 | 50362250 | AX-170022756 | 0 | G | A | 0.36 | -1.00E-01 | 9.64E-02 | 2.85E+00 | 2.95E+00 | 2.95E-01 | no | | intergenic_variant | MODIFIER |
| 4 | 50369536 | AX-170195217 | 0 | T | C | 0.36 | -1.00E-01 | 9.64E-02 | 2.85E+00 | 2.95E+00 | 2.95E-01 | no | | intergenic_variant | MODIFIER |
| 4 | 50430582 | AX-97780586 | 0 | C | T | 0.36 | -1.00E-01 | 9.64E-02 | 2.85E+00 | 2.95E+00 | 2.95E-01 | no | | intergenic_variant | MODIFIER |
| 4 | 50437054 | AX-170022762 | 0 | G | A | 0.36 | -1.00E-01 | 9.64E-02 | 2.85E+00 | 2.95E+00 | 2.95E-01 | no | | intergenic_variant | MODIFIER |
| 4 | 50647592 | AX-168406803 | 0 | G | A | 0.36 | -1.00E-01 | 9.64E-02 | 2.85E+00 | 2.95E+00 | 2.95E-01 | no | | intergenic_variant | MODIFIER |
| 4 | 50884090 | AX-170022765 | 0 | G | C | 0.36 | -1.00E-01 | 9.64E-02 | 2.85E+00 | 2.95E+00 | 2.95E-01 | no | | intergenic_variant | MODIFIER |
| 4 | 51097974 | AX-170158382 | 0 | G | C | 0.36 | -1.00E-01 | 9.64E-02 | 2.85E+00 | 2.95E+00 | 2.95E-01 | no | | intergenic_variant | MODIFIER |
| 4 | 51105695 | AX-170312044 | 0 | A | G | 0.36 | -1.00E-01 | 9.64E-02 | 2.85E+00 | 2.95E+00 | 2.95E-01 | no | | intergenic_variant | MODIFIER |
| 4 | 51130000 | AX-170384650 | 0 | T | C | 0.36 | -1.00E-01 | 9.64E-02 | 2.85E+00 | 2.95E+00 | 2.95E-01 | no | | intergenic_variant | MODIFIER |
| 4 | 51147270 | AX-170022775 | 0 | C | T | 0.36 | -1.00E-01 | 9.64E-02 | 2.85E+00 | 2.95E+00 | 2.95E-01 | no | | intergenic_variant | MODIFIER |
| 4 | 51198843 | AX-97789437 | 0 | T | C | 0.36 | -1.00E-01 | 9.64E-02 | 2.85E+00 | 2.95E+00 | 2.95E-01 | no | | intergenic_variant | MODIFIER |
| 4 | 51290886 | AX-170225578 | 0 | T | C | 0.36 | -1.00E-01 | 9.64E-02 | 2.85E+00 | 2.95E+00 | 2.95E-01 | no | | intergenic_variant | MODIFIER |
| 4 | 51297590 | AX-97804809 | 0 | G | A | 0.36 | -1.00E-01 | 9.64E-02 | 2.85E+00 | 2.95E+00 | 2.95E-01 | no | | intergenic_variant | MODIFIER |
| 4 | 51311993 | AX-170287806 | 0 | T | G | 0.36 | -1.00E-01 | 9.64E-02 | 2.85E+00 | 2.95E+00 | 2.95E-01 | no | | intergenic_variant | MODIFIER |
| 4 | 51315614 | AX-97789258 | 0 | G | A | 0.36 | -1.00E-01 | 9.64E-02 | 2.85E+00 | 2.95E+00 | 2.95E-01 | no | | intergenic_variant | MODIFIER |
| 4 | 63746620 | AX-170435568 | 1 | G | C | 0.348 | 1.23E-01 | 1.18E-01 | 2.79E+00 | 2.85E+00 | 2.95E-01 | no | Tnfsf15 | upstream_gene_variant | MODIFIER |
| 8 | 36869959 | AX-170282473 | 0 | G | C | 0.158 | 1.45E-01 | 1.37E-01 | 3.04E+00 | 3.14E+00 | 2.95E-01 | Yes | Dlc1 | intron_variant,non_coding_transcript_variant | MODIFIER |
| 18 | 60181481 | AX-170210304 | 0 | C | A | 0.395 | 9.70E-02 | 9.15E-02 | 2.46E+00 | 2.51E+00 | 2.95E-01 | no | | intergenic_variant | MODIFIER |
| 15 | 4629998 | AX-170474025 | 0 | C | T | 0.254 | -1.04E-01 | 1.00E-01 | 2.60E+00 | 2.68E+00 | 2.96E-01 | no | | intergenic_variant | MODIFIER |
| 16 | 37618270 | AX-97782870 | 0 | C | T | 0.5 | 8.53E-02 | 8.15E-02 | 2.56E+00 | 2.60E+00 | 2.96E-01 | no | Hgd | intron_variant | MODIFIER |
| 16 | 37636700 | AX-97797732 | 0 | G | A | 0.5 | 8.53E-02 | 8.15E-02 | 2.56E+00 | 2.60E+00 | 2.96E-01 | no | Hgd | downstream_gene_variant | MODIFIER |
| 16 | 37646423 | AX-170012843 | 0 | A | G | 0.5 | 8.53E-02 | 8.15E-02 | 2.56E+00 | 2.60E+00 | 2.96E-01 | no | Ndufb4 | downstream_gene_variant | MODIFIER |
| 16 | 37847666 | AX-97857511 | 0 | G | A | 0.5 | 8.53E-02 | 8.15E-02 | 2.56E+00 | 2.60E+00 | 2.96E-01 | no | Gm36028 | downstream_gene_variant | MODIFIER |
| 16 | 37978502 | AX-170481658 | 0 | A | G | 0.5 | 8.53E-02 | 8.15E-02 | 2.56E+00 | 2.60E+00 | 2.96E-01 | no | Gpr156 | intron_variant | MODIFIER |
| 16 | 37978846 | AX-170409987 | 0 | C | T | 0.5 | 8.53E-02 | 8.15E-02 | 2.56E+00 | 2.60E+00 | 2.96E-01 | no | Gpr156 | intron_variant | MODIFIER |
| 16 | 37979192 | AX-170175643 | 0 | T | C | 0.5 | 8.53E-02 | 8.15E-02 | 2.56E+00 | 2.60E+00 | 2.96E-01 | no | Gpr156 | intron_variant | MODIFIER |
| 16 | 37982672 | AX-170345854 | 0 | A | G | 0.5 | 8.53E-02 | 8.15E-02 | 2.56E+00 | 2.60E+00 | 2.96E-01 | no | Gpr156 | intron_variant | MODIFIER |
| 16 | 38009627 | AX-170527671 | 0 | G | A | 0.5 | 8.53E-02 | 8.15E-02 | 2.56E+00 | 2.60E+00 | 2.96E-01 | no | Gpr156 | downstream_gene_variant | MODIFIER |
| 16 | 38014954 | AX-170087999 | 0 | A | G | 0.5 | 8.53E-02 | 8.15E-02 | 2.56E+00 | 2.60E+00 | 2.96E-01 | no | | intergenic_variant | MODIFIER |
| 16 | 38015216 | AX-170264691 | 0 | G | A | 0.5 | 8.53E-02 | 8.15E-02 | 2.56E+00 | 2.60E+00 | 2.96E-01 | no | | intergenic_variant | MODIFIER |
| 16 | 38051594 | AX-170121862 | 0 | A | G | 0.5 | 8.53E-02 | 8.15E-02 | 2.56E+00 | 2.60E+00 | 2.96E-01 | no | | intergenic_variant | MODIFIER |
| 16 | 38069124 | AX-170260021 | 0 | C | G | 0.5 | 8.53E-02 | 8.15E-02 | 2.56E+00 | 2.60E+00 | 2.96E-01 | no | | intergenic_variant | MODIFIER |
| 16 | 38257026 | AX-97831029 | 0 | A | G | 0.5 | 8.53E-02 | 8.15E-02 | 2.56E+00 | 2.60E+00 | 2.96E-01 | no | Nr1i2 | intron_variant | MODIFIER |
| 16 | 38416971 | AX-170309851 | 0 | A | G | 0.5 | 8.53E-02 | 8.15E-02 | 2.56E+00 | 2.60E+00 | 2.96E-01 | no | Pla1a | intron_variant | MODIFIER |
| 16 | 38418397 | AX-170335634 | 0 | T | C | 0.5 | 8.53E-02 | 8.15E-02 | 2.56E+00 | 2.60E+00 | 2.96E-01 | no | Pla1a | intron_variant | MODIFIER |
| 16 | 38421523 | AX-170387855 | 0 | A | G | 0.5 | 8.53E-02 | 8.15E-02 | 2.56E+00 | 2.60E+00 | 2.96E-01 | no | Pla1a | intron_variant | MODIFIER |

| | | | | | | | | | | | | | | |
|---|---|---|---|---|---|---|---|---|---|---|---|---|---|---|
| 16 | 38429826 | AX-170057163 | 0 | T | C | 0.5 | 8.53E-02 | 8.15E-02 | 2.56E+00 | 2.60E+00 | 2.96E-01 | no | Pla1a | intron_variant | MODIFIER |
| 16 | 38450538 | AX-170341191 | 0 | T | C | 0.5 | 8.53E-02 | 8.15E-02 | 2.56E+00 | 2.60E+00 | 2.96E-01 | no | Adprh | intron_variant | MODIFIER |
| 16 | 38873895 | AX-170131581 | 0 | G | A | 0.5 | 8.53E-02 | 8.15E-02 | 2.56E+00 | 2.60E+00 | 2.96E-01 | no | Gm15802 | intron_variant,non_coding_transcript_variant | MODIFIER |
| 16 | 38875710 | AX-170168814 | 0 | C | T | 0.5 | 8.53E-02 | 8.15E-02 | 2.56E+00 | 2.60E+00 | 2.96E-01 | no | Gm15802 | intron_variant,non_coding_transcript_variant | MODIFIER |
| 16 | 38888058 | AX-170473705 | 0 | G | T | 0.5 | 8.53E-02 | 8.15E-02 | 2.56E+00 | 2.60E+00 | 2.96E-01 | no | Gm15802 | intron_variant,non_coding_transcript_variant | MODIFIER |
| 16 | 38888773 | AX-170111915 | 0 | A | G | 0.5 | 8.53E-02 | 8.15E-02 | 2.56E+00 | 2.60E+00 | 2.96E-01 | no | Gm15802 | intron_variant,non_coding_transcript_variant | MODIFIER |
| 16 | 38908298 | AX-170463850 | 0 | C | T | 0.5 | 8.53E-02 | 8.15E-02 | 2.56E+00 | 2.60E+00 | 2.96E-01 | no | Igsf11 | intron_variant | MODIFIER |
| 16 | 38912726 | AX-97809785 | 0 | T | C | 0.5 | 8.53E-02 | 8.15E-02 | 2.56E+00 | 2.60E+00 | 2.96E-01 | no | Igsf11 | intron_variant | MODIFIER |
| 16 | 38914658 | AX-170217271 | 0 | T | C | 0.5 | 8.53E-02 | 8.15E-02 | 2.56E+00 | 2.60E+00 | 2.96E-01 | no | Igsf11 | intron_variant | MODIFIER |
| 16 | 38916619 | AX-170354663 | 0 | C | T | 0.5 | 8.53E-02 | 8.15E-02 | 2.56E+00 | 2.60E+00 | 2.96E-01 | no | Igsf11 | intron_variant | MODIFIER |
| 16 | 38945989 | AX-170338627 | 0 | C | T | 0.5 | 8.53E-02 | 8.15E-02 | 2.56E+00 | 2.60E+00 | 2.96E-01 | no | Igsf11 | intron_variant | MODIFIER |
| 16 | 38952547 | AX-170488733 | 0 | C | T | 0.5 | 8.53E-02 | 8.15E-02 | 2.56E+00 | 2.60E+00 | 2.96E-01 | no | Igsf11 | intron_variant | MODIFIER |
| 16 | 38953625 | AX-170182342 | 0 | A | G | 0.5 | 8.53E-02 | 8.15E-02 | 2.56E+00 | 2.60E+00 | 2.96E-01 | no | Igsf11 | intron_variant | MODIFIER |
| 16 | 38957477 | AX-170412873 | 0 | T | A | 0.5 | 8.53E-02 | 8.15E-02 | 2.56E+00 | 2.60E+00 | 2.96E-01 | no | Igsf11 | intron_variant | MODIFIER |
| 16 | 38959149 | AX-170329465 | 0 | C | T | 0.5 | 8.53E-02 | 8.15E-02 | 2.56E+00 | 2.60E+00 | 2.96E-01 | no | Igsf11 | intron_variant | MODIFIER |
| 16 | 38999310 | AX-170518824 | 0 | G | A | 0.5 | 8.53E-02 | 8.15E-02 | 2.56E+00 | 2.60E+00 | 2.96E-01 | no | Igsf11 | intron_variant | MODIFIER |
| 16 | 39004387 | AX-170527865 | 0 | G | A | 0.5 | 8.53E-02 | 8.15E-02 | 2.56E+00 | 2.60E+00 | 2.96E-01 | no | Igsf11 | intron_variant | MODIFIER |
| 16 | 39022940 | AX-170505386 | 0 | A | C | 0.5 | 8.53E-02 | 8.15E-02 | 2.56E+00 | 2.60E+00 | 2.96E-01 | no | Igsf11 | intron_variant | MODIFIER |
| 16 | 39039708 | AX-170278904 | 0 | T | C | 0.5 | 8.53E-02 | 8.15E-02 | 2.56E+00 | 2.60E+00 | 2.96E-01 | no | | intergenic_variant | MODIFIER |
| 16 | 39048843 | AX-170216432 | 0 | T | C | 0.5 | 8.53E-02 | 8.15E-02 | 2.56E+00 | 2.60E+00 | 2.96E-01 | no | | regulatory_region_variant | MODIFIER |
| 16 | 39049457 | AX-170288034 | 0 | A | G | 0.5 | 8.53E-02 | 8.15E-02 | 2.56E+00 | 2.60E+00 | 2.96E-01 | no | | intergenic_variant | MODIFIER |
| 16 | 39050334 | AX-170241124 | 0 | A | G | 0.5 | 8.53E-02 | 8.15E-02 | 2.56E+00 | 2.60E+00 | 2.96E-01 | no | | intergenic_variant | MODIFIER |
| 16 | 39056608 | AX-170334217 | 0 | G | C | 0.5 | 8.53E-02 | 8.15E-02 | 2.56E+00 | 2.60E+00 | 2.96E-01 | no | | regulatory_region_variant | MODIFIER |
| 16 | 39066090 | AX-170352580 | 0 | A | T | 0.5 | 8.53E-02 | 8.15E-02 | 2.56E+00 | 2.60E+00 | 2.96E-01 | no | | intergenic_variant | MODIFIER |
| 16 | 39097309 | AX-170450837 | 0 | G | T | 0.5 | 8.53E-02 | 8.15E-02 | 2.56E+00 | 2.60E+00 | 2.96E-01 | no | | intergenic_variant | MODIFIER |
| 16 | 39099427 | AX-170518885 | 0 | A | C | 0.5 | 8.53E-02 | 8.15E-02 | 2.56E+00 | 2.60E+00 | 2.96E-01 | no | | intergenic_variant | MODIFIER |
| 16 | 39127836 | AX-170057184 | 0 | T | A | 0.5 | 8.53E-02 | 8.15E-02 | 2.56E+00 | 2.60E+00 | 2.96E-01 | no | | intergenic_variant | MODIFIER |
| 16 | 39138028 | AX-170057182 | 0 | G | A | 0.5 | 8.53E-02 | 8.15E-02 | 2.56E+00 | 2.60E+00 | 2.96E-01 | no | | intergenic_variant | MODIFIER |
| 16 | 39139428 | AX-170221404 | 0 | G | T | 0.5 | 8.53E-02 | 8.15E-02 | 2.56E+00 | 2.60E+00 | 2.96E-01 | no | | intergenic_variant | MODIFIER |
| 16 | 39155680 | AX-170057181 | 0 | A | G | 0.5 | 8.53E-02 | 8.15E-02 | 2.56E+00 | 2.60E+00 | 2.96E-01 | no | | intergenic_variant | MODIFIER |
| 16 | 39164139 | AX-170161034 | 0 | A | G | 0.5 | 8.53E-02 | 8.15E-02 | 2.56E+00 | 2.60E+00 | 2.96E-01 | no | | intergenic_variant | MODIFIER |
| 16 | 39177144 | AX-170342787 | 0 | T | C | 0.5 | 8.53E-02 | 8.15E-02 | 2.56E+00 | 2.60E+00 | 2.96E-01 | no | | intergenic_variant | MODIFIER |
| 16 | 39196766 | AX-170190008 | 0 | C | T | 0.5 | 8.53E-02 | 8.15E-02 | 2.56E+00 | 2.60E+00 | 2.96E-01 | no | | intergenic_variant | MODIFIER |
| 16 | 39198653 | AX-170318338 | 0 | C | T | 0.5 | 8.53E-02 | 8.15E-02 | 2.56E+00 | 2.60E+00 | 2.96E-01 | no | | intergenic_variant | MODIFIER |
| 16 | 39202391 | AX-170521696 | 0 | T | C | 0.5 | 8.53E-02 | 8.15E-02 | 2.56E+00 | 2.60E+00 | 2.96E-01 | no | | intergenic_variant | MODIFIER |
| 16 | 39241946 | AX-170311121 | 0 | A | G | 0.5 | 8.53E-02 | 8.15E-02 | 2.56E+00 | 2.60E+00 | 2.96E-01 | no | | intergenic_variant | MODIFIER |
| 16 | 39283151 | AX-170515613 | 0 | G | A | 0.5 | 8.53E-02 | 8.15E-02 | 2.56E+00 | 2.60E+00 | 2.96E-01 | no | | intergenic_variant | MODIFIER |
| 16 | 39295428 | AX-170351639 | 0 | C | T | 0.5 | 8.53E-02 | 8.15E-02 | 2.56E+00 | 2.60E+00 | 2.96E-01 | no | | intergenic_variant | MODIFIER |
| 16 | 39298046 | AX-170326379 | 0 | G | A | 0.5 | 8.53E-02 | 8.15E-02 | 2.56E+00 | 2.60E+00 | 2.96E-01 | no | | intergenic_variant | MODIFIER |
| 16 | 39304390 | AX-170487675 | 0 | C | T | 0.5 | 8.53E-02 | 8.15E-02 | 2.56E+00 | 2.60E+00 | 2.96E-01 | no | | intergenic_variant | MODIFIER |
| 16 | 39320776 | AX-170213514 | 0 | G | A | 0.5 | 8.53E-02 | 8.15E-02 | 2.56E+00 | 2.60E+00 | 2.96E-01 | no | | intergenic_variant | MODIFIER |
| 16 | 39326077 | AX-170130911 | 0 | T | C | 0.5 | 8.53E-02 | 8.15E-02 | 2.56E+00 | 2.60E+00 | 2.96E-01 | no | | intergenic_variant | MODIFIER |
| 16 | 39326387 | AX-170123545 | 0 | C | T | 0.5 | 8.53E-02 | 8.15E-02 | 2.56E+00 | 2.60E+00 | 2.96E-01 | no | | intergenic_variant | MODIFIER |
| 16 | 39339852 | AX-170305608 | 0 | C | T | 0.5 | 8.53E-02 | 8.15E-02 | 2.56E+00 | 2.60E+00 | 2.96E-01 | no | | intergenic_variant | MODIFIER |
| 16 | 39345846 | AX-170266067 | 0 | C | T | 0.5 | 8.53E-02 | 8.15E-02 | 2.56E+00 | 2.60E+00 | 2.96E-01 | no | | intergenic_variant | MODIFIER |
| 16 | 39374320 | AX-97831554 | 0 | A | G | 0.5 | 8.53E-02 | 8.15E-02 | 2.56E+00 | 2.60E+00 | 2.96E-01 | no | | intergenic_variant | MODIFIER |
| 16 | 39396807 | AX-170396408 | 0 | C | T | 0.5 | 8.53E-02 | 8.15E-02 | 2.56E+00 | 2.60E+00 | 2.96E-01 | no | | intergenic_variant | MODIFIER |
| 16 | 39397606 | AX-97831553 | 0 | A | C | 0.5 | 8.53E-02 | 8.15E-02 | 2.56E+00 | 2.60E+00 | 2.96E-01 | no | | intergenic_variant | MODIFIER |
| 16 | 39399182 | AX-170469749 | 0 | G | T | 0.5 | 8.53E-02 | 8.15E-02 | 2.56E+00 | 2.60E+00 | 2.96E-01 | no | | intergenic_variant | MODIFIER |
| 16 | 39428299 | AX-170487739 | 0 | C | T | 0.5 | 8.53E-02 | 8.15E-02 | 2.56E+00 | 2.60E+00 | 2.96E-01 | no | | intergenic_variant | MODIFIER |
| 16 | 39442137 | AX-170414515 | 0 | T | C | 0.5 | 8.53E-02 | 8.15E-02 | 2.56E+00 | 2.60E+00 | 2.96E-01 | no | Gm36903 | upstream_gene_variant | MODIFIER |
| 16 | 39442187 | AX-170193251 | 0 | T | C | 0.5 | 8.53E-02 | 8.15E-02 | 2.56E+00 | 2.60E+00 | 2.96E-01 | no | Gm36903 | upstream_gene_variant | MODIFIER |
| 16 | 39452276 | AX-170277372 | 0 | C | A | 0.5 | 8.53E-02 | 8.15E-02 | 2.56E+00 | 2.60E+00 | 2.96E-01 | no | Gm24795 | upstream_gene_variant | MODIFIER |
| 16 | 39452364 | AX-170429462 | 0 | T | C | 0.5 | 8.53E-02 | 8.15E-02 | 2.56E+00 | 2.60E+00 | 2.96E-01 | no | Gm24795 | upstream_gene_variant | MODIFIER |

| | | | | | | | | | | | | | | |
|---|---|---|---|---|---|---|---|---|---|---|---|---|---|---|
| 16 | 39452579 | AX-170057189 | 0 | C | T | 0.5 | 8.53E-02 | 8.15E-02 | 2.56E+00 | 2.60E+00 | 2.96E-01 | no | Gm24795 | upstream_gene_variant | MODIFIER |
| 16 | 39493334 | AX-170212726 | 0 | G | A | 0.5 | 8.53E-02 | 8.15E-02 | 2.56E+00 | 2.60E+00 | 2.96E-01 | no | | intergenic_variant | MODIFIER |
| 16 | 39526874 | AX-170057188 | 0 | G | T | 0.5 | 8.53E-02 | 8.15E-02 | 2.56E+00 | 2.60E+00 | 2.96E-01 | no | | intergenic_variant | MODIFIER |
| 16 | 39532149 | AX-170057191 | 0 | A | G | 0.5 | 8.53E-02 | 8.15E-02 | 2.56E+00 | 2.60E+00 | 2.96E-01 | no | | intergenic_variant | MODIFIER |
| 16 | 39533055 | AX-170397701 | 0 | C | T | 0.5 | 8.53E-02 | 8.15E-02 | 2.56E+00 | 2.60E+00 | 2.96E-01 | no | | regulatory_region_variant | MODIFIER |
| 16 | 39580241 | AX-170263853 | 0 | G | T | 0.5 | 8.53E-02 | 8.15E-02 | 2.56E+00 | 2.60E+00 | 2.96E-01 | no | | intergenic_variant | MODIFIER |
| 16 | 39581757 | AX-170462459 | 0 | G | C | 0.5 | 8.53E-02 | 8.15E-02 | 2.56E+00 | 2.60E+00 | 2.96E-01 | no | | intergenic_variant | MODIFIER |
| 16 | 39585557 | AX-97781642 | 0 | A | G | 0.5 | 8.53E-02 | 8.15E-02 | 2.56E+00 | 2.60E+00 | 2.96E-01 | no | | intergenic_variant | MODIFIER |
| 16 | 39590039 | AX-170129275 | 0 | G | T | 0.5 | 8.53E-02 | 8.15E-02 | 2.56E+00 | 2.60E+00 | 2.96E-01 | no | | intergenic_variant | MODIFIER |
| 16 | 39600809 | AX-170423220 | 0 | T | C | 0.5 | 8.53E-02 | 8.15E-02 | 2.56E+00 | 2.60E+00 | 2.96E-01 | no | | intergenic_variant | MODIFIER |
| 16 | 39620311 | AX-170309466 | 0 | C | T | 0.5 | 8.53E-02 | 8.15E-02 | 2.56E+00 | 2.60E+00 | 2.96E-01 | no | | intergenic_variant | MODIFIER |
| 16 | 39727060 | AX-97844800 | 0 | T | C | 0.5 | 8.53E-02 | 8.15E-02 | 2.56E+00 | 2.60E+00 | 2.96E-01 | no | | intergenic_variant | MODIFIER |
| 16 | 39799432 | AX-97821783 | 0 | T | G | 0.5 | 8.53E-02 | 8.15E-02 | 2.56E+00 | 2.60E+00 | 2.96E-01 | no | | intergenic_variant | MODIFIER |
| 16 | 39820524 | AX-168399006 | 0 | T | C | 0.5 | 8.53E-02 | 8.15E-02 | 2.56E+00 | 2.60E+00 | 2.96E-01 | no | | intergenic_variant | MODIFIER |
| 16 | 39857171 | AX-170012868 | 0 | C | G | 0.5 | 8.53E-02 | 8.15E-02 | 2.56E+00 | 2.60E+00 | 2.96E-01 | no | | intergenic_variant | MODIFIER |
| 16 | 39971437 | AX-170012870 | 0 | G | A | 0.5 | 8.53E-02 | 8.15E-02 | 2.56E+00 | 2.60E+00 | 2.96E-01 | no | | intergenic_variant | MODIFIER |
| 16 | 39972168 | AX-170110206 | 0 | C | G | 0.5 | 8.53E-02 | 8.15E-02 | 2.56E+00 | 2.60E+00 | 2.96E-01 | no | | intergenic_variant | MODIFIER |
| 16 | 39972816 | AX-170093450 | 0 | T | C | 0.5 | 8.53E-02 | 8.15E-02 | 2.56E+00 | 2.60E+00 | 2.96E-01 | no | | intergenic_variant | MODIFIER |
| 16 | 40071287 | AX-97839338 | 0 | A | G | 0.5 | 8.53E-02 | 8.15E-02 | 2.56E+00 | 2.60E+00 | 2.96E-01 | no | Lsamp | intron_variant | MODIFIER |
| 9 | 16926088 | AX-97806791 | 0 | G | T | 0.482 | 9.29E-02 | 8.85E-02 | 2.53E+00 | 2.59E+00 | 2.96E-01 | no | | intergenic_variant | MODIFIER |
| 1 | 128543687 | AX-170495224 | 0 | G | A | 0.544 | 8.89E-02 | 8.56E-02 | 2.66E+00 | 2.72E+00 | 2.96E-01 | no | | intergenic_variant | MODIFIER |
| 1 | 128546304 | AX-170432597 | 0 | G | A | 0.544 | 8.89E-02 | 8.56E-02 | 2.66E+00 | 2.72E+00 | 2.96E-01 | no | | regulatory_region_variant | MODIFIER |
| 1 | 128571585 | AX-170128368 | 0 | G | C | 0.544 | 8.89E-02 | 8.56E-02 | 2.66E+00 | 2.72E+00 | 2.96E-01 | no | | intergenic_variant | MODIFIER |
| 1 | 128572434 | AX-170102713 | 0 | C | T | 0.544 | 8.89E-02 | 8.56E-02 | 2.66E+00 | 2.72E+00 | 2.96E-01 | no | | intergenic_variant | MODIFIER |
| 1 | 128586652 | AX-170108770 | 0 | G | C | 0.544 | 8.89E-02 | 8.56E-02 | 2.66E+00 | 2.72E+00 | 2.96E-01 | no | Cxcr4 | downstream_gene_variant | MODIFIER |
| 1 | 128606451 | AX-170486466 | 0 | G | C | 0.544 | 8.89E-02 | 8.56E-02 | 2.66E+00 | 2.72E+00 | 2.96E-01 | no | | regulatory_region_variant | MODIFIER |
| 1 | 128606474 | AX-170458582 | 0 | A | G | 0.544 | 8.89E-02 | 8.56E-02 | 2.66E+00 | 2.72E+00 | 2.96E-01 | no | | regulatory_region_variant | MODIFIER |
| 1 | 128608760 | AX-170168737 | 0 | T | C | 0.544 | 8.89E-02 | 8.56E-02 | 2.66E+00 | 2.72E+00 | 2.96E-01 | no | | intergenic_variant | MODIFIER |
| 1 | 128617917 | AX-170377081 | 0 | C | T | 0.544 | 8.89E-02 | 8.56E-02 | 2.66E+00 | 2.72E+00 | 2.96E-01 | no | | regulatory_region_variant | MODIFIER |
| 1 | 128618421 | AX-170091687 | 0 | G | A | 0.544 | 8.89E-02 | 8.56E-02 | 2.66E+00 | 2.72E+00 | 2.96E-01 | no | | intergenic_variant | MODIFIER |
| 1 | 128629623 | AX-97841787 | 0 | A | G | 0.544 | 8.89E-02 | 8.56E-02 | 2.66E+00 | 2.72E+00 | 2.96E-01 | no | | intergenic_variant | MODIFIER |
| 1 | 128638458 | AX-170001679 | 0 | C | T | 0.544 | 8.89E-02 | 8.56E-02 | 2.66E+00 | 2.72E+00 | 2.96E-01 | no | | intergenic_variant | MODIFIER |
| 1 | 128640652 | AX-170307827 | 0 | C | T | 0.544 | 8.89E-02 | 8.56E-02 | 2.66E+00 | 2.72E+00 | 2.96E-01 | no | | intergenic_variant | MODIFIER |
| 1 | 128662926 | AX-97823000 | 0 | C | T | 0.544 | 8.89E-02 | 8.56E-02 | 2.66E+00 | 2.72E+00 | 2.96E-01 | no | | intergenic_variant | MODIFIER |
| 1 | 128665903 | AX-170214168 | 0 | G | A | 0.544 | 8.89E-02 | 8.56E-02 | 2.66E+00 | 2.72E+00 | 2.96E-01 | no | | intergenic_variant | MODIFIER |
| 1 | 128666038 | AX-170042667 | 0 | T | C | 0.544 | 8.89E-02 | 8.56E-02 | 2.66E+00 | 2.72E+00 | 2.96E-01 | no | | intergenic_variant | MODIFIER |
| 1 | 128670917 | AX-170035629 | 0 | G | A | 0.544 | 8.89E-02 | 8.56E-02 | 2.66E+00 | 2.72E+00 | 2.96E-01 | no | | regulatory_region_variant | MODIFIER |
| 1 | 128674251 | AX-170001680 | 0 | C | T | 0.544 | 8.89E-02 | 8.56E-02 | 2.66E+00 | 2.72E+00 | 2.96E-01 | no | | intergenic_variant | MODIFIER |
| 1 | 128674616 | AX-170250309 | 0 | C | A | 0.544 | 8.89E-02 | 8.56E-02 | 2.66E+00 | 2.72E+00 | 2.96E-01 | no | | intergenic_variant | MODIFIER |
| 1 | 128682736 | AX-170042665 | 0 | C | T | 0.544 | 8.89E-02 | 8.56E-02 | 2.66E+00 | 2.72E+00 | 2.96E-01 | no | | regulatory_region_variant | MODIFIER |
| 1 | 128683331 | AX-97808610 | 0 | T | C | 0.544 | 8.89E-02 | 8.56E-02 | 2.66E+00 | 2.72E+00 | 2.96E-01 | no | | regulatory_region_variant | MODIFIER |
| 1 | 128705346 | AX-97826382 | 0 | C | T | 0.544 | 8.89E-02 | 8.56E-02 | 2.66E+00 | 2.72E+00 | 2.96E-01 | no | | regulatory_region_variant | MODIFIER |
| 1 | 128728831 | AX-170463247 | 0 | T | C | 0.544 | 8.89E-02 | 8.56E-02 | 2.66E+00 | 2.72E+00 | 2.96E-01 | no | | intergenic_variant | MODIFIER |
| 1 | 128738535 | AX-170329721 | 0 | C | T | 0.544 | 8.89E-02 | 8.56E-02 | 2.66E+00 | 2.72E+00 | 2.96E-01 | no | | intergenic_variant | MODIFIER |
| 1 | 128753782 | AX-170001676 | 0 | A | G | 0.544 | 8.89E-02 | 8.56E-02 | 2.66E+00 | 2.72E+00 | 2.96E-01 | no | Gm29599 | downstream_gene_variant | MODIFIER |
| 1 | 128754477 | AX-170001672 | 0 | C | T | 0.544 | 8.89E-02 | 8.56E-02 | 2.66E+00 | 2.72E+00 | 2.96E-01 | no | Gm29599 | downstream_gene_variant | MODIFIER |
| 1 | 128767193 | AX-170384816 | 0 | T | C | 0.544 | 8.89E-02 | 8.56E-02 | 2.66E+00 | 2.72E+00 | 2.96E-01 | no | Gm26686 | downstream_gene_variant | MODIFIER |
| 1 | 128782810 | AX-170253569 | 0 | G | A | 0.544 | 8.89E-02 | 8.56E-02 | 2.66E+00 | 2.72E+00 | 2.96E-01 | no | | intergenic_variant | MODIFIER |
| 1 | 128791270 | AX-170466856 | 0 | A | G | 0.544 | 8.89E-02 | 8.56E-02 | 2.66E+00 | 2.72E+00 | 2.96E-01 | no | | intergenic_variant | MODIFIER |
| 1 | 128793281 | AX-170001678 | 0 | T | C | 0.544 | 8.89E-02 | 8.56E-02 | 2.66E+00 | 2.72E+00 | 2.96E-01 | no | | intergenic_variant | MODIFIER |
| 1 | 130850953 | AX-170476894 | 0 | C | T | 0.544 | 8.89E-02 | 8.56E-02 | 2.66E+00 | 2.72E+00 | 2.96E-01 | no | Pigr | 3_prime_UTR_variant | MODIFIER |
| 1 | 130854208 | AX-97825614 | 0 | A | G | 0.544 | 8.89E-02 | 8.56E-02 | 2.66E+00 | 2.72E+00 | 2.96E-01 | no | Pigr | downstream_gene_variant | MODIFIER |
| 1 | 130863937 | AX-170198445 | 0 | C | T | 0.544 | 8.89E-02 | 8.56E-02 | 2.66E+00 | 2.72E+00 | 2.96E-01 | no | Fcmr | upstream_gene_variant | MODIFIER |
| 1 | 130864912 | AX-170116748 | 0 | G | A | 0.544 | 8.89E-02 | 8.56E-02 | 2.66E+00 | 2.72E+00 | 2.96E-01 | no | Fcmr | upstream_gene_variant | MODIFIER |

| | | | | | | | | | | | | | | | |
|---|---|---|---|---|---|---|---|---|---|---|---|---|---|---|---|
| 1 | 130917309 | AX-97781648 | 0 | A | G | 0.544 | 8.89E-02 | 8.56E-02 | 2.66E+00 | 2.72E+00 | 2.96E-01 | no | | intergenic_variant | MODIFIER |
| 1 | 130992372 | AX-170296701 | 0 | T | C | 0.544 | 8.89E-02 | 8.56E-02 | 2.66E+00 | 2.72E+00 | 2.96E-01 | no | | intergenic_variant | MODIFIER |
| 1 | 131007814 | AX-97801961 | 0 | A | G | 0.544 | 8.89E-02 | 8.56E-02 | 2.66E+00 | 2.72E+00 | 2.96E-01 | no | | intergenic_variant | MODIFIER |
| 1 | 131047652 | AX-97782609 | 0 | T | C | 0.544 | 8.89E-02 | 8.56E-02 | 2.66E+00 | 2.72E+00 | 2.96E-01 | no | | intergenic_variant | MODIFIER |
| 1 | 131067501 | AX-170001715 | 0 | A | G | 0.544 | 8.89E-02 | 8.56E-02 | 2.66E+00 | 2.72E+00 | 2.96E-01 | no | Mapkapk2 | intron_variant | MODIFIER |
| 1 | 131070466 | AX-170001716 | 0 | A | G | 0.544 | 8.89E-02 | 8.56E-02 | 2.66E+00 | 2.72E+00 | 2.96E-01 | no | Mapkapk2 | intron_variant | MODIFIER |
| 1 | 131073393 | AX-170001712 | 0 | G | A | 0.544 | 8.89E-02 | 8.56E-02 | 2.66E+00 | 2.72E+00 | 2.96E-01 | no | Mapkapk2 | intron_variant | MODIFIER |
| 1 | 131082815 | AX-97854965 | 0 | G | A | 0.544 | 8.89E-02 | 8.56E-02 | 2.66E+00 | 2.72E+00 | 2.96E-01 | no | Mapkapk2 | intron_variant | MODIFIER |
| 1 | 131130075 | AX-97823415 | 0 | T | C | 0.544 | 8.89E-02 | 8.56E-02 | 2.66E+00 | 2.72E+00 | 2.96E-01 | no | Dyrk3 | synonymous_variant | LOW |
| 1 | 131168383 | AX-170001717 | 0 | A | G | 0.544 | 8.89E-02 | 8.56E-02 | 2.66E+00 | 2.72E+00 | 2.96E-01 | no | Eif2d | intron_variant | MODIFIER |
| 1 | 131185993 | AX-97852377 | 0 | A | G | 0.544 | 8.89E-02 | 8.56E-02 | 2.66E+00 | 2.72E+00 | 2.96E-01 | no | Rassf5 | intron_variant | MODIFIER |
| 1 | 131189806 | AX-170001718 | 0 | G | C | 0.544 | 8.89E-02 | 8.56E-02 | 2.66E+00 | 2.72E+00 | 2.96E-01 | no | Rassf5 | intron_variant | MODIFIER |
| 1 | 131213416 | AX-97833646 | 0 | A | G | 0.544 | 8.89E-02 | 8.56E-02 | 2.66E+00 | 2.72E+00 | 2.96E-01 | no | Rassf5 | intron_variant | MODIFIER |
| 1 | 131230784 | AX-170001713 | 0 | G | A | 0.544 | 8.89E-02 | 8.56E-02 | 2.66E+00 | 2.72E+00 | 2.96E-01 | no | Rassf5 | intron_variant | MODIFIER |
| 1 | 131234205 | AX-97828574 | 0 | A | G | 0.544 | 8.89E-02 | 8.56E-02 | 2.66E+00 | 2.72E+00 | 2.96E-01 | no | Rassf5 | intron_variant | MODIFIER |
| 1 | 131247951 | AX-97841009 | 0 | G | A | 0.544 | 8.89E-02 | 8.56E-02 | 2.66E+00 | 2.72E+00 | 2.96E-01 | no | Rassf5 | upstream_gene_variant | MODIFIER |
| 1 | 131284742 | AX-97856466 | 0 | A | G | 0.544 | 8.89E-02 | 8.56E-02 | 2.66E+00 | 2.72E+00 | 2.96E-01 | no | Srgap2 | downstream_gene_variant | MODIFIER |
| 1 | 131298159 | AX-170103236 | 0 | A | G | 0.544 | 8.89E-02 | 8.56E-02 | 2.66E+00 | 2.72E+00 | 2.96E-01 | no | Srgap2 | intron_variant | MODIFIER |
| 1 | 131356554 | AX-97803684 | 0 | A | G | 0.544 | 8.89E-02 | 8.56E-02 | 2.66E+00 | 2.72E+00 | 2.96E-01 | no | Srgap2 | intron_variant | MODIFIER |
| 1 | 131361888 | AX-170110555 | 0 | A | G | 0.544 | 8.89E-02 | 8.56E-02 | 2.66E+00 | 2.72E+00 | 2.96E-01 | no | Srgap2 | intron_variant | MODIFIER |
| 1 | 131393810 | AX-170330909 | 0 | A | G | 0.544 | 8.89E-02 | 8.56E-02 | 2.66E+00 | 2.72E+00 | 2.96E-01 | no | Srgap2 | intron_variant | MODIFIER |
| 1 | 131394224 | AX-170361958 | 0 | C | T | 0.544 | 8.89E-02 | 8.56E-02 | 2.66E+00 | 2.72E+00 | 2.96E-01 | no | Srgap2 | intron_variant | MODIFIER |
| 1 | 131422959 | AX-170205629 | 0 | G | A | 0.544 | 8.89E-02 | 8.56E-02 | 2.66E+00 | 2.72E+00 | 2.96E-01 | no | Srgap2 | intron_variant | MODIFIER |
| 1 | 131435012 | AX-97813276 | 0 | C | A | 0.544 | 8.89E-02 | 8.56E-02 | 2.66E+00 | 2.72E+00 | 2.96E-01 | no | Srgap2 | intron_variant | MODIFIER |
| 1 | 131436250 | AX-170288321 | 0 | G | T | 0.544 | 8.89E-02 | 8.56E-02 | 2.66E+00 | 2.72E+00 | 2.96E-01 | no | Srgap2 | intron_variant | MODIFIER |
| 1 | 131448202 | AX-170140370 | 0 | G | C | 0.544 | 8.89E-02 | 8.56E-02 | 2.66E+00 | 2.72E+00 | 2.96E-01 | no | Srgap2 | intron_variant | MODIFIER |
| 1 | 131464885 | AX-170270870 | 0 | G | T | 0.544 | 8.89E-02 | 8.56E-02 | 2.66E+00 | 2.72E+00 | 2.96E-01 | no | Srgap2 | intron_variant | MODIFIER |
| 1 | 131468161 | AX-170456826 | 0 | G | A | 0.544 | 8.89E-02 | 8.56E-02 | 2.66E+00 | 2.72E+00 | 2.96E-01 | no | Srgap2 | intron_variant | MODIFIER |
| 1 | 131469582 | AX-97855593 | 0 | G | A | 0.544 | 8.89E-02 | 8.56E-02 | 2.66E+00 | 2.72E+00 | 2.96E-01 | no | Srgap2 | intron_variant | MODIFIER |
| 1 | 131504340 | AX-170159935 | 0 | T | C | 0.544 | 8.89E-02 | 8.56E-02 | 2.66E+00 | 2.72E+00 | 2.96E-01 | no | Srgap2 | intron_variant | MODIFIER |
| 1 | 131524089 | AX-170476210 | 0 | T | C | 0.544 | 8.89E-02 | 8.56E-02 | 2.66E+00 | 2.72E+00 | 2.96E-01 | no | Fam72a | upstream_gene_variant | MODIFIER |
| 1 | 131532284 | AX-170042720 | 0 | G | T | 0.544 | 8.89E-02 | 8.56E-02 | 2.66E+00 | 2.72E+00 | 2.96E-01 | no | Fam72a | intron_variant | MODIFIER |
| 1 | 131536584 | AX-170042722 | 0 | A | G | 0.544 | 8.89E-02 | 8.56E-02 | 2.66E+00 | 2.72E+00 | 2.96E-01 | no | Fam72a | intron_variant | MODIFIER |
| 1 | 131540443 | AX-170163206 | 0 | G | T | 0.544 | 8.89E-02 | 8.56E-02 | 2.66E+00 | 2.72E+00 | 2.96E-01 | no | Fam72a | downstream_gene_variant | MODIFIER |
| 1 | 131543963 | AX-97853817 | 0 | G | A | 0.544 | 8.89E-02 | 8.56E-02 | 2.66E+00 | 2.72E+00 | 2.96E-01 | no | Fam72a | downstream_gene_variant | MODIFIER |
| 1 | 131545206 | AX-170001723 | 0 | A | G | 0.544 | 8.89E-02 | 8.56E-02 | 2.66E+00 | 2.72E+00 | 2.96E-01 | no | | intergenic_variant | MODIFIER |
| 1 | 131552661 | AX-170302648 | 0 | A | G | 0.544 | 8.89E-02 | 8.56E-02 | 2.66E+00 | 2.72E+00 | 2.96E-01 | no | | intergenic_variant | MODIFIER |
| 1 | 131552855 | AX-170183810 | 0 | T | C | 0.544 | 8.89E-02 | 8.56E-02 | 2.66E+00 | 2.72E+00 | 2.96E-01 | no | | intergenic_variant | MODIFIER |
| 1 | 131553803 | AX-97827580 | 0 | G | A | 0.544 | 8.89E-02 | 8.56E-02 | 2.66E+00 | 2.72E+00 | 2.96E-01 | no | | intergenic_variant | MODIFIER |
| 1 | 131558624 | AX-170001724 | 0 | T | C | 0.544 | 8.89E-02 | 8.56E-02 | 2.66E+00 | 2.72E+00 | 2.96E-01 | no | | intergenic_variant | MODIFIER |
| 1 | 131561963 | AX-170166887 | 0 | T | G | 0.544 | 8.89E-02 | 8.56E-02 | 2.66E+00 | 2.72E+00 | 2.96E-01 | no | | intergenic_variant | MODIFIER |
| 1 | 131565566 | AX-170256082 | 0 | A | G | 0.544 | 8.89E-02 | 8.56E-02 | 2.66E+00 | 2.72E+00 | 2.96E-01 | no | | intergenic_variant | MODIFIER |
| 1 | 131598352 | AX-170345732 | 0 | T | C | 0.544 | 8.89E-02 | 8.56E-02 | 2.66E+00 | 2.72E+00 | 2.96E-01 | no | Avpr1b | upstream_gene_variant | MODIFIER |
| 1 | 131599282 | AX-170484404 | 0 | T | A | 0.544 | 8.89E-02 | 8.56E-02 | 2.66E+00 | 2.72E+00 | 2.96E-01 | no | Avpr1b | 5_prime_UTR_variant | MODIFIER |
| 1 | 131603798 | AX-170174284 | 0 | T | C | 0.544 | 8.89E-02 | 8.56E-02 | 2.66E+00 | 2.72E+00 | 2.96E-01 | no | Avpr1b | intron_variant | MODIFIER |
| 1 | 131614347 | AX-170042725 | 0 | T | G | 0.544 | 8.89E-02 | 8.56E-02 | 2.66E+00 | 2.72E+00 | 2.96E-01 | no | Avpr1b | downstream_gene_variant | MODIFIER |
| 1 | 131638481 | AX-170001719 | 0 | C | T | 0.544 | 8.89E-02 | 8.56E-02 | 2.66E+00 | 2.72E+00 | 2.96E-01 | no | Ctse | 5_prime_UTR_variant | MODIFIER |
| 1 | 131673977 | AX-170358963 | 0 | T | C | 0.544 | 8.89E-02 | 8.56E-02 | 2.66E+00 | 2.72E+00 | 2.96E-01 | no | Ctse | intron_variant | MODIFIER |
| 1 | 131688907 | AX-170243162 | 0 | A | G | 0.544 | 8.89E-02 | 8.56E-02 | 2.66E+00 | 2.72E+00 | 2.96E-01 | no | Rab7b | intron_variant | MODIFIER |
| 1 | 131700068 | AX-170329351 | 0 | C | A | 0.544 | 8.89E-02 | 8.56E-02 | 2.66E+00 | 2.72E+00 | 2.96E-01 | no | Rab7b | intron_variant | MODIFIER |
| 1 | 131702373 | AX-170399375 | 0 | G | A | 0.544 | 8.89E-02 | 8.56E-02 | 2.66E+00 | 2.72E+00 | 2.96E-01 | no | Rab7b | intron_variant | MODIFIER |
| 1 | 131704244 | AX-170524909 | 0 | A | G | 0.544 | 8.89E-02 | 8.56E-02 | 2.66E+00 | 2.72E+00 | 2.96E-01 | no | Rab7b | intron_variant | MODIFIER |
| 1 | 131709696 | AX-170187970 | 0 | T | C | 0.544 | 8.89E-02 | 8.56E-02 | 2.66E+00 | 2.72E+00 | 2.96E-01 | no | Rab7b | intron_variant | MODIFIER |
| 1 | 131710587 | AX-170368039 | 0 | C | T | 0.544 | 8.89E-02 | 8.56E-02 | 2.66E+00 | 2.72E+00 | 2.96E-01 | no | Rab7b | intron_variant | MODIFIER |

| | | | | | | | | | | | | | | | |
|---|---|---|---|---|---|---|---|---|---|---|---|---|---|---|---|
| 1 | 131713056 | AX-170396276 | 0 | C | T | 0.544 | 8.89E-02 | 8.56E-02 | 2.66E+00 | 2.72E+00 | 2.96E-01 | no | Rab7b | downstream_gene_variant | MODIFIER |
| 1 | 131740125 | AX-170248835 | 0 | G | C | 0.544 | 8.89E-02 | 8.56E-02 | 2.66E+00 | 2.72E+00 | 2.96E-01 | no | Slc26a9 | upstream_gene_variant | MODIFIER |
| 1 | 131752423 | AX-170446796 | 0 | A | G | 0.544 | 8.89E-02 | 8.56E-02 | 2.66E+00 | 2.72E+00 | 2.96E-01 | no | Slc26a9 | intron_variant | MODIFIER |
| 1 | 131752937 | AX-170294040 | 0 | C | T | 0.544 | 8.89E-02 | 8.56E-02 | 2.66E+00 | 2.72E+00 | 2.96E-01 | no | Slc26a9 | synonymous_variant | LOW |
| 1 | 131756584 | AX-170189899 | 0 | G | A | 0.544 | 8.89E-02 | 8.56E-02 | 2.66E+00 | 2.72E+00 | 2.96E-01 | no | Slc26a9 | intron_variant | MODIFIER |
| 1 | 131762231 | AX-170042728 | 0 | G | A | 0.544 | 8.89E-02 | 8.56E-02 | 2.66E+00 | 2.72E+00 | 2.96E-01 | no | Slc26a9 | intron_variant | MODIFIER |
| 1 | 131768657 | AX-170463293 | 0 | A | G | 0.544 | 8.89E-02 | 8.56E-02 | 2.66E+00 | 2.72E+00 | 2.96E-01 | no | Slc26a9 | intron_variant | MODIFIER |
| 1 | 131771281 | AX-97807948 | 0 | C | T | 0.544 | 8.89E-02 | 8.56E-02 | 2.66E+00 | 2.72E+00 | 2.96E-01 | no | Slc26a9 | 3_prime_UTR_variant | MODIFIER |
| 1 | 131819995 | AX-170104565 | 0 | G | A | 0.544 | 8.89E-02 | 8.56E-02 | 2.66E+00 | 2.72E+00 | 2.96E-01 | no | Pm20d1 | intron_variant | MODIFIER |
| 1 | 131872110 | AX-170393849 | 0 | G | A | 0.544 | 8.89E-02 | 8.56E-02 | 2.66E+00 | 2.72E+00 | 2.96E-01 | no | Rab29 | missense_variant | MODERATE |
| 1 | 132124172 | AX-97846905 | 0 | T | C | 0.544 | 8.89E-02 | 8.56E-02 | 2.66E+00 | 2.72E+00 | 2.96E-01 | no | Cdk18 | intron_variant | MODIFIER |
| 1 | 132245387 | AX-97839764 | 0 | C | T | 0.544 | 8.89E-02 | 8.56E-02 | 2.66E+00 | 2.72E+00 | 2.96E-01 | no | Lemd1 | intron_variant | MODIFIER |
| 1 | 132320426 | AX-170001732 | 0 | C | A | 0.544 | 8.89E-02 | 8.56E-02 | 2.66E+00 | 2.72E+00 | 2.96E-01 | no | Nuak2 | intron_variant | MODIFIER |
| 1 | 132345989 | AX-170487272 | 0 | T | C | 0.544 | 8.89E-02 | 8.56E-02 | 2.66E+00 | 2.72E+00 | 2.96E-01 | no | F730311O21Rik | non_coding_transcript_exon_variant | MODIFIER |
| 1 | 132385489 | AX-170450231 | 0 | G | A | 0.544 | 8.89E-02 | 8.56E-02 | 2.66E+00 | 2.72E+00 | 2.96E-01 | no | Tmcc2 | intron_variant | MODIFIER |
| 1 | 132413223 | AX-170098429 | 0 | A | G | 0.544 | 8.89E-02 | 8.56E-02 | 2.66E+00 | 2.72E+00 | 2.96E-01 | no | Dstyk | upstream_gene_variant | MODIFIER |
| 1 | 132413914 | AX-170510966 | 0 | C | T | 0.544 | 8.89E-02 | 8.56E-02 | 2.66E+00 | 2.72E+00 | 2.96E-01 | no | Dstyk | upstream_gene_variant | MODIFIER |
| 1 | 132438643 | AX-170228558 | 0 | C | T | 0.544 | 8.89E-02 | 8.56E-02 | 2.66E+00 | 2.72E+00 | 2.96E-01 | no | Dstyk | intron_variant | MODIFIER |
| 1 | 132441014 | AX-170452729 | 0 | A | G | 0.544 | 8.89E-02 | 8.56E-02 | 2.66E+00 | 2.72E+00 | 2.96E-01 | no | Dstyk | intron_variant | MODIFIER |
| 1 | 132453334 | AX-170147127 | 0 | A | G | 0.544 | 8.89E-02 | 8.56E-02 | 2.66E+00 | 2.72E+00 | 2.96E-01 | no | Dstyk | intron_variant | MODIFIER |
| 1 | 132505366 | AX-170205368 | 0 | C | T | 0.544 | 8.89E-02 | 8.56E-02 | 2.66E+00 | 2.72E+00 | 2.96E-01 | no | Rbbp5 | 3_prime_UTR_variant | MODIFIER |
| 1 | 132511515 | AX-170328433 | 0 | A | C | 0.544 | 8.89E-02 | 8.56E-02 | 2.66E+00 | 2.72E+00 | 2.96E-01 | no | Tmem81 | downstream_gene_variant | MODIFIER |
| 1 | 132516696 | AX-170219806 | 0 | G | T | 0.544 | 8.89E-02 | 8.56E-02 | 2.66E+00 | 2.72E+00 | 2.96E-01 | no | Cntn2 | intron_variant | MODIFIER |
| 7 | 12153357 | AX-97841167 | 0 | G | C | 0.351 | 8.31E-02 | 8.00E-02 | 2.59E+00 | 2.71E+00 | 2.96E-01 | no | Vmn1r78 | missense_variant | MODERATE |
| 7 | 12185127 | AX-170028595 | 0 | G | A | 0.351 | 8.31E-02 | 8.00E-02 | 2.59E+00 | 2.71E+00 | 2.96E-01 | no | Gm19201 | upstream_gene_variant | MODIFIER |
| 7 | 12195406 | AX-170028594 | 0 | T | C | 0.351 | 8.31E-02 | 8.00E-02 | 2.59E+00 | 2.71E+00 | 2.96E-01 | no | Vmn1r80 | downstream_gene_variant | MODIFIER |
| 7 | 12214849 | AX-97802532 | 0 | A | G | 0.351 | 8.31E-02 | 8.00E-02 | 2.59E+00 | 2.71E+00 | 2.96E-01 | no | Vmn1r-ps65 | non_coding_transcript_exon_variant | MODIFIER |
| 7 | 12469961 | AX-97824103 | 0 | G | A | 0.351 | 8.31E-02 | 8.00E-02 | 2.59E+00 | 2.71E+00 | 2.96E-01 | no | Vmn2r-ps53 | downstream_gene_variant | MODIFIER |
| 7 | 12824911 | AX-170367834 | 0 | T | C | 0.351 | 8.31E-02 | 8.00E-02 | 2.59E+00 | 2.71E+00 | 2.96E-01 | no | Gm45593 | upstream_gene_variant | MODIFIER |
| 7 | 12831359 | AX-170151174 | 0 | C | T | 0.351 | 8.31E-02 | 8.00E-02 | 2.59E+00 | 2.71E+00 | 2.96E-01 | no | Zfp110 | upstream_gene_variant | MODIFIER |
| 7 | 12950679 | AX-170341440 | 0 | G | T | 0.351 | 8.31E-02 | 8.00E-02 | 2.59E+00 | 2.71E+00 | 2.96E-01 | no | 1810019N24Rik | intron_variant,non_coding_transcript_variant | MODIFIER |
| 7 | 12950774 | AX-170076739 | 0 | T | C | 0.351 | 8.31E-02 | 8.00E-02 | 2.59E+00 | 2.71E+00 | 2.96E-01 | no | 1810019N24Rik | intron_variant,non_coding_transcript_variant | MODIFIER |
| 7 | 12958852 | AX-170377680 | 0 | T | C | 0.351 | 8.31E-02 | 8.00E-02 | 2.59E+00 | 2.71E+00 | 2.96E-01 | no | 1810019N24Rik | intron_variant,non_coding_transcript_variant | MODIFIER |
| 7 | 12959111 | AX-97832609 | 0 | T | C | 0.351 | 8.31E-02 | 8.00E-02 | 2.59E+00 | 2.71E+00 | 2.96E-01 | no | 1810019N24Rik | intron_variant,non_coding_transcript_variant | MODIFIER |
| 7 | 12983578 | AX-170442690 | 0 | A | G | 0.351 | 8.31E-02 | 8.00E-02 | 2.59E+00 | 2.71E+00 | 2.96E-01 | no | Slc27a5 | downstream_gene_variant | MODIFIER |
| 7 | 12990633 | AX-170235130 | 0 | C | T | 0.351 | 8.31E-02 | 8.00E-02 | 2.59E+00 | 2.71E+00 | 2.96E-01 | no | Slc27a5 | intron_variant | MODIFIER |
| 7 | 12996525 | AX-170187089 | 0 | A | G | 0.351 | 8.31E-02 | 8.00E-02 | 2.59E+00 | 2.71E+00 | 2.96E-01 | no | Slc27a5 | intron_variant | MODIFIER |
| 7 | 12998099 | AX-170350820 | 0 | A | G | 0.351 | 8.31E-02 | 8.00E-02 | 2.59E+00 | 2.71E+00 | 2.96E-01 | no | Slc27a5 | 5_prime_UTR_variant | MODIFIER |
| 1 | 131505059 | AX-97827485 | 1 | A | G | 0.545 | 8.87E-02 | 8.55E-02 | 2.66E+00 | 2.72E+00 | 2.96E-01 | no | Srgap2 | intron_variant | MODIFIER |
| 1 | 131548193 | AX-170320324 | 1 | C | T | 0.545 | 8.87E-02 | 8.55E-02 | 2.66E+00 | 2.72E+00 | 2.96E-01 | no | | intergenic_variant | MODIFIER |
| 16 | 25951035 | AX-97828037 | 1 | T | C | 0.25 | 8.52E-02 | 8.27E-02 | 2.70E+00 | 2.85E+00 | 2.96E-01 | no | | intergenic_variant | MODIFIER |
| 4 | 77367385 | AX-97854625 | 1 | T | G | 0.321 | 1.24E-01 | 1.19E-01 | 2.77E+00 | 2.82E+00 | 2.96E-01 | no | Ptprd | intron_variant,non_coding_transcript_variant | MODIFIER |
| 18 | 74656342 | AX-170359257 | 1 | T | A | 0.179 | -1.28E-01 | 1.24E-01 | 2.67E+00 | 2.72E+00 | 2.96E-01 | no | Myo5b | intron_variant | MODIFIER |
| 4 | 71695236 | AX-170288593 | 0 | G | T | 0.465 | -8.57E-02 | 8.18E-02 | 2.53E+00 | 2.60E+00 | 2.96E-01 | no | | intergenic_variant | MODIFIER |
| 7 | 90052835 | AX-170375202 | 2 | G | A | 0.118 | -1.83E-01 | 1.75E-01 | 2.93E+00 | 3.07E+00 | 2.96E-01 | Yes | E230029C05Rik | downstream_gene_variant | MODIFIER |
| 5 | 45688363 | AX-170024841 | 1 | A | G | 0.491 | 1.07E-01 | 9.95E-02 | 2.38E+00 | 2.44E+00 | 2.96E-01 | no | Ncapg | intron_variant | MODIFIER |
| 8 | 38226456 | AX-170291311 | 0 | T | C | 0.149 | 1.41E-01 | 1.35E-01 | 2.93E+00 | 3.02E+00 | 2.96E-01 | no | Sgcz | intron_variant | MODIFIER |
| 4 | 155187180 | AX-170486062 | 1 | C | A | 0.107 | 1.68E-01 | 1.63E-01 | 2.73E+00 | 2.84E+00 | 2.96E-01 | Yes | Ski | intron_variant | MODIFIER |
| 18 | 65183641 | AX-170157356 | 1 | A | C | 0.179 | -1.24E-01 | 1.20E-01 | 2.68E+00 | 2.73E+00 | 2.96E-01 | no | Nedd4l | intron_variant | MODIFIER |
| 17 | 7781949 | AX-170058331 | 0 | T | C | 0.044 | -1.84E-01 | 1.78E-01 | 2.69E+00 | 2.80E+00 | 2.96E-01 | Yes | Fndc1 | intron_variant | MODIFIER |
| 1 | 169489176 | AX-170307438 | 0 | A | G | 0.219 | 1.31E-01 | 1.26E-01 | 2.61E+00 | 2.68E+00 | 2.96E-01 | no | | intergenic_variant | MODIFIER |
| 10 | 93734348 | AX-170426353 | 1 | G | A | 0.295 | -8.68E-02 | 8.28E-02 | 2.51E+00 | 2.57E+00 | 2.97E-01 | no | Ntn4 | intron_variant | MODIFIER |
| 5 | 36285636 | AX-170166966 | 0 | A | G | 0.333 | 1.06E-01 | 1.02E-01 | 2.79E+00 | 2.87E+00 | 2.97E-01 | no | Sorcs2 | intron_variant | MODIFIER |
| 3 | 20602066 | AX-170135661 | 0 | A | G | 0.14 | -1.07E-01 | 1.03E-01 | 2.59E+00 | 2.67E+00 | 2.97E-01 | no | | intergenic_variant | MODIFIER |

| | | | | | | | | | | | | | | | |
|---|---|---|---|---|---|---|---|---|---|---|---|---|---|---|---|
| 3 | 20612442 | AX-170020082 | 0 | T | G | 0.14 | -1.07E-01 | 1.03E-01 | 2.59E+00 | 2.67E+00 | 2.97E-01 | no | | intergenic_variant | MODIFIER |
| 3 | 20646238 | AX-170211375 | 0 | G | A | 0.14 | -1.07E-01 | 1.03E-01 | 2.59E+00 | 2.67E+00 | 2.97E-01 | no | | intergenic_variant | MODIFIER |
| 3 | 20647091 | AX-170530523 | 0 | C | A | 0.14 | -1.07E-01 | 1.03E-01 | 2.59E+00 | 2.67E+00 | 2.97E-01 | no | | intergenic_variant | MODIFIER |
| 3 | 20663511 | AX-97801065 | 0 | T | C | 0.14 | -1.07E-01 | 1.03E-01 | 2.59E+00 | 2.67E+00 | 2.97E-01 | no | | intergenic_variant | MODIFIER |
| 3 | 20680930 | AX-170020081 | 0 | A | C | 0.14 | -1.07E-01 | 1.03E-01 | 2.59E+00 | 2.67E+00 | 2.97E-01 | no | | intergenic_variant | MODIFIER |
| 3 | 20684137 | AX-170201675 | 0 | T | C | 0.14 | -1.07E-01 | 1.03E-01 | 2.59E+00 | 2.67E+00 | 2.97E-01 | no | | intergenic_variant | MODIFIER |
| 3 | 20686875 | AX-97846877 | 0 | A | C | 0.14 | -1.07E-01 | 1.03E-01 | 2.59E+00 | 2.67E+00 | 2.97E-01 | no | | intergenic_variant | MODIFIER |
| 3 | 20695424 | AX-170217482 | 0 | G | A | 0.14 | -1.07E-01 | 1.03E-01 | 2.59E+00 | 2.67E+00 | 2.97E-01 | no | | intergenic_variant | MODIFIER |
| 3 | 20709149 | AX-170214989 | 0 | G | A | 0.14 | -1.07E-01 | 1.03E-01 | 2.59E+00 | 2.67E+00 | 2.97E-01 | no | | intergenic_variant | MODIFIER |
| 3 | 20739494 | AX-170094546 | 0 | T | C | 0.14 | -1.07E-01 | 1.03E-01 | 2.59E+00 | 2.67E+00 | 2.97E-01 | no | | intergenic_variant | MODIFIER |
| 3 | 20756861 | AX-170147987 | 0 | T | C | 0.14 | -1.07E-01 | 1.03E-01 | 2.59E+00 | 2.67E+00 | 2.97E-01 | no | | intergenic_variant | MODIFIER |
| 3 | 20757020 | AX-170066047 | 0 | T | G | 0.14 | -1.07E-01 | 1.03E-01 | 2.59E+00 | 2.67E+00 | 2.97E-01 | no | | intergenic_variant | MODIFIER |
| 3 | 20777695 | AX-97814179 | 0 | T | C | 0.14 | -1.07E-01 | 1.03E-01 | 2.59E+00 | 2.67E+00 | 2.97E-01 | no | Gm18491 | downstream_gene_variant | MODIFIER |
| 3 | 20820672 | AX-170403471 | 0 | C | T | 0.14 | -1.07E-01 | 1.03E-01 | 2.59E+00 | 2.67E+00 | 2.97E-01 | no | Gm37480 | intron_variant,non_coding_transcript_variant | MODIFIER |
| 3 | 20862312 | AX-170251335 | 0 | T | C | 0.14 | -1.07E-01 | 1.03E-01 | 2.59E+00 | 2.67E+00 | 2.97E-01 | no | | intergenic_variant | MODIFIER |
| 3 | 20893462 | AX-170479013 | 0 | T | C | 0.14 | -1.07E-01 | 1.03E-01 | 2.59E+00 | 2.67E+00 | 2.97E-01 | no | | intergenic_variant | MODIFIER |
| 3 | 20894529 | AX-170463079 | 0 | C | T | 0.14 | -1.07E-01 | 1.03E-01 | 2.59E+00 | 2.67E+00 | 2.97E-01 | no | | intergenic_variant | MODIFIER |
| 3 | 20894674 | AX-170427101 | 0 | G | A | 0.14 | -1.07E-01 | 1.03E-01 | 2.59E+00 | 2.67E+00 | 2.97E-01 | no | | intergenic_variant | MODIFIER |
| 3 | 20918541 | AX-170377046 | 0 | G | A | 0.14 | -1.07E-01 | 1.03E-01 | 2.59E+00 | 2.67E+00 | 2.97E-01 | no | | intergenic_variant | MODIFIER |
| 3 | 20926868 | AX-97857635 | 0 | T | G | 0.14 | -1.07E-01 | 1.03E-01 | 2.59E+00 | 2.67E+00 | 2.97E-01 | no | | intergenic_variant | MODIFIER |
| 3 | 20927025 | AX-170238520 | 0 | G | A | 0.14 | -1.07E-01 | 1.03E-01 | 2.59E+00 | 2.67E+00 | 2.97E-01 | no | | intergenic_variant | MODIFIER |
| 3 | 20940840 | AX-170309208 | 0 | G | A | 0.14 | -1.07E-01 | 1.03E-01 | 2.59E+00 | 2.67E+00 | 2.97E-01 | no | | intergenic_variant | MODIFIER |
| 3 | 20981821 | AX-97828384 | 0 | C | T | 0.14 | -1.07E-01 | 1.03E-01 | 2.59E+00 | 2.67E+00 | 2.97E-01 | no | | intergenic_variant | MODIFIER |
| 3 | 20986631 | AX-170066051 | 0 | G | A | 0.14 | -1.07E-01 | 1.03E-01 | 2.59E+00 | 2.67E+00 | 2.97E-01 | no | | regulatory_region_variant | MODIFIER |
| 3 | 20987754 | AX-97853551 | 0 | C | T | 0.14 | -1.07E-01 | 1.03E-01 | 2.59E+00 | 2.67E+00 | 2.97E-01 | no | | regulatory_region_variant | MODIFIER |
| 3 | 21107596 | AX-170404211 | 0 | A | G | 0.14 | -1.07E-01 | 1.03E-01 | 2.59E+00 | 2.67E+00 | 2.97E-01 | no | | intergenic_variant | MODIFIER |
| 3 | 21107691 | AX-170455488 | 0 | C | A | 0.14 | -1.07E-01 | 1.03E-01 | 2.59E+00 | 2.67E+00 | 2.97E-01 | no | | intergenic_variant | MODIFIER |
| 3 | 21108450 | AX-97821750 | 0 | T | C | 0.14 | -1.07E-01 | 1.03E-01 | 2.59E+00 | 2.67E+00 | 2.97E-01 | no | | intergenic_variant | MODIFIER |
| 3 | 21114896 | AX-170332748 | 0 | T | C | 0.14 | -1.07E-01 | 1.03E-01 | 2.59E+00 | 2.67E+00 | 2.97E-01 | no | Gm43160 | upstream_gene_variant | MODIFIER |
| 3 | 21119730 | AX-97812683 | 0 | A | G | 0.14 | -1.07E-01 | 1.03E-01 | 2.59E+00 | 2.67E+00 | 2.97E-01 | no | | intergenic_variant | MODIFIER |
| 3 | 21126248 | AX-170203460 | 0 | T | C | 0.14 | -1.07E-01 | 1.03E-01 | 2.59E+00 | 2.67E+00 | 2.97E-01 | no | | intergenic_variant | MODIFIER |
| 3 | 21126355 | AX-170375868 | 0 | C | G | 0.14 | -1.07E-01 | 1.03E-01 | 2.59E+00 | 2.67E+00 | 2.97E-01 | no | | intergenic_variant | MODIFIER |
| 3 | 21130598 | AX-170355446 | 0 | C | T | 0.14 | -1.07E-01 | 1.03E-01 | 2.59E+00 | 2.67E+00 | 2.97E-01 | no | | intergenic_variant | MODIFIER |
| 3 | 21149860 | AX-170088870 | 0 | C | T | 0.14 | -1.07E-01 | 1.03E-01 | 2.59E+00 | 2.67E+00 | 2.97E-01 | no | | intergenic_variant | MODIFIER |
| 3 | 21152537 | AX-170020094 | 0 | G | T | 0.14 | -1.07E-01 | 1.03E-01 | 2.59E+00 | 2.67E+00 | 2.97E-01 | no | | intergenic_variant | MODIFIER |
| 3 | 21161067 | AX-170020093 | 0 | G | C | 0.14 | -1.07E-01 | 1.03E-01 | 2.59E+00 | 2.67E+00 | 2.97E-01 | no | | intergenic_variant | MODIFIER |
| 3 | 21176799 | AX-170066052 | 0 | T | C | 0.14 | -1.07E-01 | 1.03E-01 | 2.59E+00 | 2.67E+00 | 2.97E-01 | no | | intergenic_variant | MODIFIER |
| 3 | 21179078 | AX-97842614 | 0 | A | G | 0.14 | -1.07E-01 | 1.03E-01 | 2.59E+00 | 2.67E+00 | 2.97E-01 | no | | intergenic_variant | MODIFIER |
| 3 | 21189291 | AX-170413079 | 0 | G | A | 0.14 | -1.07E-01 | 1.03E-01 | 2.59E+00 | 2.67E+00 | 2.97E-01 | no | | intergenic_variant | MODIFIER |
| 3 | 21221288 | AX-97846084 | 0 | A | G | 0.14 | -1.07E-01 | 1.03E-01 | 2.59E+00 | 2.67E+00 | 2.97E-01 | no | | intergenic_variant | MODIFIER |
| 3 | 21245341 | AX-170020090 | 0 | G | C | 0.14 | -1.07E-01 | 1.03E-01 | 2.59E+00 | 2.67E+00 | 2.97E-01 | no | | intergenic_variant | MODIFIER |
| 3 | 21938137 | AX-170362234 | 0 | A | G | 0.14 | -1.07E-01 | 1.03E-01 | 2.59E+00 | 2.67E+00 | 2.97E-01 | no | | regulatory_region_variant | MODIFIER |
| 1 | 130583339 | AX-170295137 | 1 | G | A | 0.536 | 8.84E-02 | 8.54E-02 | 2.66E+00 | 2.72E+00 | 2.97E-01 | no | Zp3r | intron_variant | MODIFIER |
| 15 | 25345083 | AX-97833236 | 0 | G | A | 0.246 | -1.07E-01 | 1.03E-01 | 2.61E+00 | 2.67E+00 | 2.97E-01 | no | | intergenic_variant | MODIFIER |
| 9 | 16931752 | AX-170212418 | 1 | C | T | 0.491 | 9.22E-02 | 8.78E-02 | 2.50E+00 | 2.57E+00 | 2.97E-01 | no | | intergenic_variant | MODIFIER |
| 4 | 98578825 | AX-97799094 | 0 | G | T | 0.237 | 1.16E-01 | 1.11E-01 | 2.50E+00 | 2.58E+00 | 2.97E-01 | no | Patj | intron_variant | MODIFIER |
| 5 | 66387479 | AX-170345822 | 1 | A | G | 0.473 | 1.00E-01 | 9.45E-02 | 2.43E+00 | 2.51E+00 | 2.97E-01 | no | Apbb2 | intron_variant | MODIFIER |
| 6 | 138482545 | AX-170414887 | 1 | A | G | 0.58 | 8.74E-02 | 8.26E-02 | 2.45E+00 | 2.52E+00 | 2.97E-01 | no | Lmo3 | intron_variant | MODIFIER |
| 5 | 9639095 | AX-170267947 | 0 | C | T | 0.325 | 9.75E-02 | 9.47E-02 | 2.73E+00 | 2.85E+00 | 2.97E-01 | no | Grm3 | intron_variant | MODIFIER |
| 15 | 5722353 | AX-170054702 | 1 | A | G | 0.232 | -1.11E-01 | 1.07E-01 | 2.66E+00 | 2.71E+00 | 2.97E-01 | no | | intergenic_variant | MODIFIER |
| 10 | 17899482 | AX-97802688 | 1 | G | A | 0.214 | 9.99E-02 | 9.60E-02 | 2.94E+00 | 3.06E+00 | 2.97E-01 | no | Heca | 3_prime_UTR_variant | MODIFIER |
| 18 | 78195143 | AX-170434316 | 0 | G | A | 0.439 | 9.13E-02 | 8.64E-02 | 2.46E+00 | 2.52E+00 | 2.97E-01 | no | Slc14a2 | intron_variant | MODIFIER |
| 6 | 138653632 | AX-170110505 | 0 | A | T | 0.561 | 8.42E-02 | 7.95E-02 | 2.43E+00 | 2.52E+00 | 2.98E-01 | no | Igbp1b | downstream_gene_variant | MODIFIER |
| 7 | 12597821 | AX-170440596 | 1 | T | A | 0.348 | 8.16E-02 | 7.88E-02 | 2.58E+00 | 2.70E+00 | 2.98E-01 | no | Vmn2r53 | intron_variant | MODIFIER |

| | | | | | | | | | | | | | | | |
|---|---|---|---|---|---|---|---|---|---|---|---|---|---|---|---|
| 4 | 129890008 | AX-97798112 | 2 | A | T | 0.164 | 1.22E-01 | 1.17E-01 | 2.55E+00 | 2.60E+00 | 2.98E-01 | no | | intergenic_variant | MODIFIER |
| 14 | 112159016 | AX-170344034 | 0 | T | A | 0.325 | 7.48E-02 | 7.21E-02 | 2.57E+00 | 2.67E+00 | 2.98E-01 | no | | intergenic_variant | MODIFIER |
| 4 | 134326710 | AX-170379545 | 1 | C | T | 0.161 | 1.21E-01 | 1.17E-01 | 2.57E+00 | 2.62E+00 | 2.98E-01 | no | Trim63 | intron_variant | MODIFIER |
| 16 | 45295583 | AX-170109553 | 1 | T | C | 0.473 | -8.32E-02 | 7.99E-02 | 2.55E+00 | 2.61E+00 | 2.98E-01 | no | | intergenic_variant | MODIFIER |
| 4 | 63810217 | AX-170022961 | 2 | C | G | 0.3 | 1.11E-01 | 1.08E-01 | 2.63E+00 | 2.73E+00 | 2.98E-01 | no | | intergenic_variant | MODIFIER |
| 9 | 19308927 | AX-170031904 | 1 | A | C | 0.464 | 8.67E-02 | 8.38E-02 | 2.60E+00 | 2.69E+00 | 2.98E-01 | no | | intergenic_variant | MODIFIER |
| 1 | 121211633 | AX-170380672 | 1 | G | A | 0.571 | 8.57E-02 | 8.19E-02 | 2.50E+00 | 2.57E+00 | 2.98E-01 | no | | intergenic_variant | MODIFIER |
| 4 | 75338487 | AX-97840409 | 2 | T | G | 0.318 | 1.24E-01 | 1.20E-01 | 2.78E+00 | 2.84E+00 | 2.98E-01 | no | | intergenic_variant | MODIFIER |
| 13 | 19373058 | AX-170477975 | 0 | G | A | 0.175 | -1.29E-01 | 1.23E-01 | 2.49E+00 | 2.55E+00 | 2.98E-01 | no | Stard3nl | intron_variant | MODIFIER |
| 8 | 19449171 | AX-170410728 | 0 | A | C | 0.175 | -1.07E-01 | 1.03E-01 | 2.92E+00 | 3.08E+00 | 2.98E-01 | no | Defb8 | upstream_gene_variant | MODIFIER |
| 13 | 3944024 | AX-170524235 | 0 | C | T | 0.211 | -1.21E-01 | 1.17E-01 | 2.62E+00 | 2.69E+00 | 2.98E-01 | no | Ucn3 | intron_variant | MODIFIER |
| 13 | 6261582 | AX-170337941 | 0 | G | A | 0.211 | -1.21E-01 | 1.17E-01 | 2.62E+00 | 2.69E+00 | 2.98E-01 | no | | intergenic_variant | MODIFIER |
| 6 | 137227323 | AX-170225456 | 0 | C | G | 0.579 | 9.36E-02 | 8.83E-02 | 2.43E+00 | 2.48E+00 | 2.98E-01 | no | | intergenic_variant | MODIFIER |
| 4 | 59122138 | AX-170233145 | 1 | T | C | 0.366 | 1.17E-01 | 1.13E-01 | 2.67E+00 | 2.73E+00 | 2.98E-01 | no | AI481877 | intron_variant | MODIFIER |
| 16 | 19285447 | AX-170037393 | 0 | A | T | 0.14 | 9.72E-02 | 9.47E-02 | 2.78E+00 | 2.92E+00 | 2.99E-01 | no | Olfr164 | downstream_gene_variant | MODIFIER |
| 16 | 32561240 | AX-170399584 | 0 | G | A | 0.482 | 8.37E-02 | 8.04E-02 | 2.55E+00 | 2.60E+00 | 2.99E-01 | no | Zdhhc19 | intron_variant,non_coding_transcript_variant | MODIFIER |
| 11 | 5738450 | AX-170285503 | 0 | A | G | 0.053 | 1.91E-01 | 1.86E-01 | 2.81E+00 | 2.96E+00 | 2.99E-01 | Yes | Urgcp | intron_variant | MODIFIER |
| 4 | 82349478 | AX-97845867 | 0 | C | T | 0.43 | -8.70E-02 | 8.23E-02 | 2.44E+00 | 2.50E+00 | 2.99E-01 | no | Nfib | intron_variant | MODIFIER |
| 10 | 92032195 | AX-97837212 | 0 | G | A | 0.289 | -8.54E-02 | 8.12E-02 | 2.47E+00 | 2.53E+00 | 2.99E-01 | no | | intergenic_variant | MODIFIER |
| 7 | 66179037 | AX-170199114 | 1 | G | A | 0.429 | -9.26E-02 | 9.01E-02 | 2.81E+00 | 2.92E+00 | 2.99E-01 | no | | regulatory_region_variant | MODIFIER |
| 8 | 41486348 | AX-170105463 | 0 | C | T | 0.158 | 1.45E-01 | 1.38E-01 | 3.03E+00 | 3.11E+00 | 2.99E-01 | Yes | | intergenic_variant | MODIFIER |
| 6 | 101501434 | AX-170493147 | 0 | A | T | 0.07 | -2.19E-01 | 2.12E-01 | 2.66E+00 | 2.74E+00 | 2.99E-01 | Yes | | intergenic_variant | MODIFIER |
| 7 | 71659936 | AX-170077685 | 1 | C | T | 0.375 | -1.08E-01 | 1.05E-01 | 2.88E+00 | 2.97E+00 | 2.99E-01 | no | | intergenic_variant | MODIFIER |
| 9 | 91981056 | AX-170209290 | 1 | T | A | 0.134 | -1.27E-01 | 1.24E-01 | 2.65E+00 | 2.73E+00 | 2.99E-01 | no | | intergenic_variant | MODIFIER |
| 16 | 54545876 | AX-97839158 | 0 | T | G | 0.518 | 8.63E-02 | 8.22E-02 | 2.48E+00 | 2.52E+00 | 2.99E-01 | no | | intergenic_variant | MODIFIER |
| 1 | 128500148 | AX-97847342 | 1 | G | T | 0.536 | 8.49E-02 | 8.24E-02 | 2.63E+00 | 2.70E+00 | 2.99E-01 | no | | intergenic_variant | MODIFIER |
| 1 | 68097777 | AX-170041531 | 1 | A | T | 0.116 | 1.74E-01 | 1.69E-01 | 2.73E+00 | 2.85E+00 | 2.99E-01 | Yes | Erbb4 | intron_variant | MODIFIER |
| 5 | 46842070 | AX-170071979 | 0 | C | T | 0.509 | 9.96E-02 | 9.42E-02 | 2.43E+00 | 2.50E+00 | 2.99E-01 | no | | intergenic_variant | MODIFIER |
| 16 | 51664778 | AX-97830773 | 2 | T | C | 0.509 | 8.44E-02 | 8.02E-02 | 2.46E+00 | 2.51E+00 | 2.99E-01 | no | | intergenic_variant | MODIFIER |
| 1 | 133988795 | AX-170522882 | 1 | G | A | 0.527 | 8.60E-02 | 8.32E-02 | 2.60E+00 | 2.66E+00 | 2.99E-01 | no | | regulatory_region_variant | MODIFIER |
| 9 | 4447130 | AX-170275650 | 1 | T | A | 0.464 | 7.75E-02 | 7.47E-02 | 2.53E+00 | 2.63E+00 | 3.00E-01 | no | Gria4 | intron_variant | MODIFIER |
| 13 | 4715064 | AX-170050416 | 1 | G | T | 0.205 | -1.14E-01 | 1.11E-01 | 2.65E+00 | 2.75E+00 | 3.00E-01 | no | Gm40658 | intron_variant,non_coding_transcript_variant | MODIFIER |
| 18 | 60499771 | AX-170257160 | 0 | T | C | 0.395 | 9.61E-02 | 9.14E-02 | 2.46E+00 | 2.51E+00 | 3.00E-01 | no | Smim3 | intron_variant | MODIFIER |
| 6 | 138612004 | AX-170342999 | 0 | G | T | 0.579 | 8.71E-02 | 8.27E-02 | 2.44E+00 | 2.52E+00 | 3.00E-01 | no | | intergenic_variant | MODIFIER |
| 18 | 85521392 | AX-170194915 | 0 | T | C | 0.149 | -1.13E-01 | 1.08E-01 | 2.52E+00 | 2.61E+00 | 3.00E-01 | no | | intergenic_variant | MODIFIER |
| 18 | 85529808 | AX-97846433 | 0 | G | A | 0.149 | -1.13E-01 | 1.08E-01 | 2.52E+00 | 2.61E+00 | 3.00E-01 | no | | intergenic_variant | MODIFIER |
| 18 | 85578657 | AX-170016127 | 0 | G | A | 0.149 | -1.13E-01 | 1.08E-01 | 2.52E+00 | 2.61E+00 | 3.00E-01 | no | | intergenic_variant | MODIFIER |
| 18 | 85765082 | AX-170240633 | 0 | T | C | 0.149 | -1.13E-01 | 1.08E-01 | 2.52E+00 | 2.61E+00 | 3.00E-01 | no | | intergenic_variant | MODIFIER |
| 18 | 85765424 | AX-170339459 | 0 | T | A | 0.149 | -1.13E-01 | 1.08E-01 | 2.52E+00 | 2.61E+00 | 3.00E-01 | no | | intergenic_variant | MODIFIER |
| 18 | 85771397 | AX-170369229 | 0 | T | G | 0.149 | -1.13E-01 | 1.08E-01 | 2.52E+00 | 2.61E+00 | 3.00E-01 | no | | intergenic_variant | MODIFIER |
| 18 | 85931422 | AX-97844332 | 0 | T | C | 0.149 | -1.13E-01 | 1.08E-01 | 2.52E+00 | 2.61E+00 | 3.00E-01 | no | | intergenic_variant | MODIFIER |
| 18 | 88009117 | AX-170401139 | 0 | C | T | 0.149 | -1.13E-01 | 1.08E-01 | 2.52E+00 | 2.61E+00 | 3.00E-01 | no | | intergenic_variant | MODIFIER |
| 14 | 111132732 | AX-170462744 | 1 | C | T | 0.321 | 7.17E-02 | 6.96E-02 | 2.60E+00 | 2.71E+00 | 3.00E-01 | no | | intergenic_variant | MODIFIER |
| 7 | 91688895 | AX-170405405 | 0 | G | A | 0.149 | 1.56E-01 | 1.52E-01 | 2.64E+00 | 2.77E+00 | 3.00E-01 | Yes | Dlg2 | intron_variant | MODIFIER |
| 6 | 126800729 | AX-170076157 | 1 | T | C | 0.527 | 8.53E-02 | 8.17E-02 | 2.49E+00 | 2.56E+00 | 3.00E-01 | no | | intergenic_variant | MODIFIER |
| 10 | 10748827 | AX-170308818 | 0 | C | G | 0.202 | -1.08E-01 | 1.05E-01 | 2.81E+00 | 2.91E+00 | 3.00E-01 | no | Grm1 | intron_variant | MODIFIER |
| 5 | 109674193 | AX-170441757 | 0 | C | A | 0.456 | 1.10E-01 | 1.07E-01 | 2.64E+00 | 2.70E+00 | 3.00E-01 | no | 5430403G16Rik | downstream_gene_variant | MODIFIER |
| 16 | 18872019 | AX-170056827 | 1 | C | T | 0.134 | 9.99E-02 | 9.76E-02 | 2.83E+00 | 2.97E+00 | 3.01E-01 | no | Mrpl40 | 3_prime_UTR_variant | MODIFIER |
| 4 | 125470349 | AX-170154219 | 0 | G | A | 0.237 | 1.12E-01 | 1.08E-01 | 2.60E+00 | 2.67E+00 | 3.01E-01 | no | | intergenic_variant | MODIFIER |
| 16 | 38962270 | AX-170355699 | 0 | C | A | 0.5 | 8.35E-02 | 8.02E-02 | 2.51E+00 | 2.56E+00 | 3.01E-01 | no | Igsf11 | intron_variant | MODIFIER |
| 13 | 24061144 | AX-170467011 | 2 | A | T | 0.173 | -1.14E-01 | 1.09E-01 | 2.50E+00 | 2.60E+00 | 3.01E-01 | no | Carmil1 | intron_variant | MODIFIER |
| 3 | 37249627 | AX-170088908 | 0 | A | G | 0.193 | 9.88E-02 | 9.59E-02 | 2.89E+00 | 2.99E+00 | 3.01E-01 | no | | intergenic_variant | MODIFIER |
| 1 | 124672894 | AX-170470756 | 1 | G | A | 0.562 | 8.70E-02 | 8.41E-02 | 2.56E+00 | 2.61E+00 | 3.01E-01 | no | Gm28928 | intron_variant,non_coding_transcript_variant | MODIFIER |
| 18 | 73188673 | AX-170404427 | 1 | G | A | 0.179 | -1.20E-01 | 1.16E-01 | 2.58E+00 | 2.64E+00 | 3.01E-01 | no | | intergenic_variant | MODIFIER |

| | | | | | | | | | | | | | | | |
|---|---|---|---|---|---|---|---|---|---|---|---|---|---|---|---|
| 1 | 131597284 | AX-170143666 | 1 | A | G | 0.545 | 8.79E-02 | 8.57E-02 | 2.65E+00 | 2.71E+00 | 3.01E-01 | no | Avpr1b | upstream_gene_variant | MODIFIER |
| 18 | 78204085 | AX-170446370 | 0 | A | G | 0.439 | 9.01E-02 | 8.61E-02 | 2.47E+00 | 2.53E+00 | 3.01E-01 | no | Slc14a2 | intron_variant | MODIFIER |
| 9 | 96755560 | AX-170033061 | 1 | C | T | 0.134 | -1.24E-01 | 1.21E-01 | 2.61E+00 | 2.69E+00 | 3.01E-01 | no | Zbtb38 | upstream_gene_variant | MODIFIER |
| 18 | 66927216 | AX-170015896 | 1 | C | T | 0.098 | 1.56E-01 | 1.53E-01 | 2.69E+00 | 2.82E+00 | 3.01E-01 | Yes | AC127234.1 | intron_variant,non_coding_transcript_variant | MODIFIER |
| 18 | 75427810 | AX-170107983 | 1 | A | G | 0.438 | 9.00E-02 | 8.62E-02 | 2.48E+00 | 2.54E+00 | 3.01E-01 | no | Ctif | downstream_gene_variant | MODIFIER |
| 16 | 19515797 | AX-170489187 | 0 | T | C | 0.14 | 9.82E-02 | 9.58E-02 | 2.87E+00 | 3.01E+00 | 3.01E-01 | no | Olfr167 | upstream_gene_variant | MODIFIER |
| 1 | 132268494 | AX-170261969 | 1 | C | T | 0.536 | 8.76E-02 | 8.54E-02 | 2.65E+00 | 2.71E+00 | 3.01E-01 | no | | intergenic_variant | MODIFIER |
| 5 | 80081078 | AX-170025384 | 0 | G | A | 0.439 | 1.07E-01 | 1.05E-01 | 2.73E+00 | 2.80E+00 | 3.01E-01 | no | | intergenic_variant | MODIFIER |
| 7 | 39770858 | AX-170174573 | 1 | G | T | 0.357 | 9.10E-02 | 8.89E-02 | 2.67E+00 | 2.76E+00 | 3.01E-01 | no | | intergenic_variant | MODIFIER |
| 10 | 17654928 | AX-97797590 | 2 | C | G | 0.218 | 9.98E-02 | 9.71E-02 | 2.91E+00 | 3.03E+00 | 3.02E-01 | no | Gm47771 | intron_variant,non_coding_transcript_variant | MODIFIER |
| 2 | 161769962 | AX-170345991 | 0 | A | G | 0.167 | 1.88E-01 | 1.84E-01 | 2.76E+00 | 2.87E+00 | 3.02E-01 | Yes | Ptprt | intron_variant | MODIFIER |
| 4 | 72209457 | AX-170164802 | 0 | T | C | 0.228 | -1.16E-01 | 1.06E-01 | 3.33E+00 | 3.51E+00 | 3.02E-01 | no | C630043F03Rik | intron_variant,non_coding_transcript_variant | MODIFIER |
| 4 | 149724554 | AX-170387670 | 0 | G | A | 0.096 | 1.69E-01 | 1.66E-01 | 2.77E+00 | 2.88E+00 | 3.02E-01 | Yes | Tmem201 | intron_variant | MODIFIER |
| 1 | 191197622 | AX-97855725 | 0 | C | G | 0.263 | 1.02E-01 | 9.52E-02 | 2.35E+00 | 2.43E+00 | 3.02E-01 | no | Gm24748 | downstream_gene_variant | MODIFIER |
| 1 | 191233484 | AX-97823923 | 0 | C | T | 0.263 | 1.02E-01 | 9.52E-02 | 2.35E+00 | 2.43E+00 | 3.02E-01 | no | | intergenic_variant | MODIFIER |
| 1 | 191234148 | AX-170002568 | 0 | T | G | 0.263 | 1.02E-01 | 9.52E-02 | 2.35E+00 | 2.43E+00 | 3.02E-01 | no | | intergenic_variant | MODIFIER |
| 1 | 191236114 | AX-170145672 | 0 | A | G | 0.263 | 1.02E-01 | 9.52E-02 | 2.35E+00 | 2.43E+00 | 3.02E-01 | no | | regulatory_region_variant | MODIFIER |
| 1 | 191254504 | AX-170533340 | 0 | C | T | 0.263 | 1.02E-01 | 9.52E-02 | 2.35E+00 | 2.43E+00 | 3.02E-01 | no | Gm37074 | downstream_gene_variant | MODIFIER |
| 1 | 191260725 | AX-97827325 | 0 | A | G | 0.263 | 1.02E-01 | 9.52E-02 | 2.35E+00 | 2.43E+00 | 3.02E-01 | no | | regulatory_region_variant | MODIFIER |
| 1 | 191263725 | AX-170148146 | 0 | T | C | 0.263 | 1.02E-01 | 9.52E-02 | 2.35E+00 | 2.43E+00 | 3.02E-01 | no | Gm37168 | upstream_gene_variant | MODIFIER |
| 1 | 191281590 | AX-170328038 | 0 | C | T | 0.263 | 1.02E-01 | 9.52E-02 | 2.35E+00 | 2.43E+00 | 3.02E-01 | no | Gm37168 | intron_variant,non_coding_transcript_variant | MODIFIER |
| 1 | 191283492 | AX-170206344 | 0 | A | G | 0.263 | 1.02E-01 | 9.52E-02 | 2.35E+00 | 2.43E+00 | 3.02E-01 | no | Gm37168 | intron_variant,non_coding_transcript_variant | MODIFIER |
| 1 | 191283542 | AX-97832439 | 0 | C | T | 0.263 | 1.02E-01 | 9.52E-02 | 2.35E+00 | 2.43E+00 | 3.02E-01 | no | Gm37168 | intron_variant,non_coding_transcript_variant | MODIFIER |
| 1 | 191365475 | AX-97791311 | 0 | A | G | 0.263 | 1.02E-01 | 9.52E-02 | 2.35E+00 | 2.43E+00 | 3.02E-01 | no | Ppp2r5a | intron_variant | MODIFIER |
| 1 | 191437971 | AX-170349784 | 0 | C | T | 0.263 | 1.02E-01 | 9.52E-02 | 2.35E+00 | 2.43E+00 | 3.02E-01 | no | Gm32200 | non_coding_transcript_exon_variant | MODIFIER |
| 1 | 191448099 | AX-170429280 | 0 | T | C | 0.263 | 1.02E-01 | 9.52E-02 | 2.35E+00 | 2.43E+00 | 3.02E-01 | no | Gm32200 | intron_variant,non_coding_transcript_variant | MODIFIER |
| 1 | 191449241 | AX-170148664 | 0 | C | A | 0.263 | 1.02E-01 | 9.52E-02 | 2.35E+00 | 2.43E+00 | 3.02E-01 | no | Gm32200 | intron_variant,non_coding_transcript_variant | MODIFIER |
| 1 | 191450222 | AX-170154554 | 0 | C | T | 0.263 | 1.02E-01 | 9.52E-02 | 2.35E+00 | 2.43E+00 | 3.02E-01 | no | Gm32200 | intron_variant,non_coding_transcript_variant | MODIFIER |
| 1 | 191454035 | AX-170182166 | 0 | A | G | 0.263 | 1.02E-01 | 9.52E-02 | 2.35E+00 | 2.43E+00 | 3.02E-01 | no | Gm32200 | intron_variant,non_coding_transcript_variant | MODIFIER |
| 1 | 191454735 | AX-170086092 | 0 | C | T | 0.263 | 1.02E-01 | 9.52E-02 | 2.35E+00 | 2.43E+00 | 3.02E-01 | no | Gm32200 | intron_variant,non_coding_transcript_variant | MODIFIER |
| 1 | 191459978 | AX-170357734 | 0 | A | G | 0.263 | 1.02E-01 | 9.52E-02 | 2.35E+00 | 2.43E+00 | 3.02E-01 | no | Gm32200 | intron_variant,non_coding_transcript_variant | MODIFIER |
| 1 | 191479703 | AX-170134390 | 0 | T | C | 0.263 | 1.02E-01 | 9.52E-02 | 2.35E+00 | 2.43E+00 | 3.02E-01 | no | Gm37688 | downstream_gene_variant | MODIFIER |
| 1 | 191490112 | AX-169999713 | 0 | C | T | 0.263 | 1.02E-01 | 9.52E-02 | 2.35E+00 | 2.43E+00 | 3.02E-01 | no | Gm37432 | downstream_gene_variant | MODIFIER |
| 1 | 191498216 | AX-170359120 | 0 | A | G | 0.263 | 1.02E-01 | 9.52E-02 | 2.35E+00 | 2.43E+00 | 3.02E-01 | no | Gm37432 | upstream_gene_variant | MODIFIER |
| 1 | 191521527 | AX-170268669 | 0 | A | G | 0.263 | 1.02E-01 | 9.52E-02 | 2.35E+00 | 2.43E+00 | 3.02E-01 | no | | regulatory_region_variant | MODIFIER |
| 1 | 191522126 | AX-170099859 | 0 | C | T | 0.263 | 1.02E-01 | 9.52E-02 | 2.35E+00 | 2.43E+00 | 3.02E-01 | no | | intergenic_variant | MODIFIER |
| 1 | 191532443 | AX-170043979 | 0 | G | C | 0.263 | 1.02E-01 | 9.52E-02 | 2.35E+00 | 2.43E+00 | 3.02E-01 | no | Dtl | downstream_gene_variant | MODIFIER |
| 1 | 191543820 | AX-97788106 | 0 | C | T | 0.263 | 1.02E-01 | 9.52E-02 | 2.35E+00 | 2.43E+00 | 3.02E-01 | no | Dtl | intron_variant | MODIFIER |
| 1 | 191560047 | AX-170335350 | 0 | C | T | 0.263 | 1.02E-01 | 9.52E-02 | 2.35E+00 | 2.43E+00 | 3.02E-01 | no | Dtl | intron_variant | MODIFIER |
| 1 | 191567045 | AX-170159579 | 0 | T | C | 0.263 | 1.02E-01 | 9.52E-02 | 2.35E+00 | 2.43E+00 | 3.02E-01 | no | Dtl | intron_variant | MODIFIER |
| 1 | 191575383 | AX-170272602 | 0 | T | C | 0.263 | 1.02E-01 | 9.52E-02 | 2.35E+00 | 2.43E+00 | 3.02E-01 | no | Dtl | synonymous_variant | LOW |
| 1 | 191576247 | AX-170467136 | 0 | A | G | 0.263 | 1.02E-01 | 9.52E-02 | 2.35E+00 | 2.43E+00 | 3.02E-01 | no | Dtl | upstream_gene_variant | MODIFIER |
| 1 | 191576262 | AX-170535536 | 0 | T | C | 0.263 | 1.02E-01 | 9.52E-02 | 2.35E+00 | 2.43E+00 | 3.02E-01 | no | Dtl | upstream_gene_variant | MODIFIER |
| 1 | 191579489 | AX-170503318 | 0 | G | A | 0.263 | 1.02E-01 | 9.52E-02 | 2.35E+00 | 2.43E+00 | 3.02E-01 | no | Dtl | upstream_gene_variant | MODIFIER |
| 1 | 191593452 | AX-170438951 | 0 | T | C | 0.263 | 1.02E-01 | 9.52E-02 | 2.35E+00 | 2.43E+00 | 3.02E-01 | no | Ints7 | intron_variant | MODIFIER |
| 1 | 191593827 | AX-170126970 | 0 | A | G | 0.263 | 1.02E-01 | 9.52E-02 | 2.35E+00 | 2.43E+00 | 3.02E-01 | no | Ints7 | intron_variant | MODIFIER |
| 1 | 191615215 | AX-168378553 | 0 | C | A | 0.263 | 1.02E-01 | 9.52E-02 | 2.35E+00 | 2.43E+00 | 3.02E-01 | no | Ints7 | intron_variant | MODIFIER |
| 1 | 191620358 | AX-170464083 | 0 | G | A | 0.263 | 1.02E-01 | 9.52E-02 | 2.35E+00 | 2.43E+00 | 3.02E-01 | no | Ints7 | intron_variant | MODIFIER |
| 1 | 191621003 | AX-170189063 | 0 | G | C | 0.263 | 1.02E-01 | 9.52E-02 | 2.35E+00 | 2.43E+00 | 3.02E-01 | no | Ints7 | intron_variant | MODIFIER |
| 1 | 191643505 | AX-170131947 | 0 | G | A | 0.263 | 1.02E-01 | 9.52E-02 | 2.35E+00 | 2.43E+00 | 3.02E-01 | no | | intergenic_variant | MODIFIER |
| 1 | 191644701 | AX-170002573 | 0 | T | C | 0.263 | 1.02E-01 | 9.52E-02 | 2.35E+00 | 2.43E+00 | 3.02E-01 | no | | intergenic_variant | MODIFIER |
| 1 | 191648529 | AX-170317271 | 0 | A | G | 0.263 | 1.02E-01 | 9.52E-02 | 2.35E+00 | 2.43E+00 | 3.02E-01 | no | | intergenic_variant | MODIFIER |
| 16 | 39155579 | AX-170228435 | 1 | G | A | 0.491 | 8.26E-02 | 8.00E-02 | 2.55E+00 | 2.60E+00 | 3.02E-01 | no | | intergenic_variant | MODIFIER |
| 1 | 171847215 | AX-170273620 | 0 | T | C | 0.211 | 1.39E-01 | 1.35E-01 | 2.63E+00 | 2.68E+00 | 3.02E-01 | no | Cd84 | intron_variant | MODIFIER |

| | | | | | | | | | | | | | | | |
|---|---|---|---|---|---|---|---|---|---|---|---|---|---|---|---|
| 5 | 46235342 | AX-170456070 | 0 | T | C | 0.5 | 1.08E-01 | 1.02E-01 | 2.40E+00 | 2.46E+00 | 3.02E-01 | no | | intergenic_variant | MODIFIER |
| 13 | 24505998 | AX-170413411 | 0 | C | G | 0.167 | -1.15E-01 | 1.05E-01 | 2.27E+00 | 2.34E+00 | 3.02E-01 | no | | intergenic_variant | MODIFIER |
| 6 | 136257959 | AX-97801236 | 1 | C | T | 0.589 | 9.43E-02 | 8.87E-02 | 2.38E+00 | 2.43E+00 | 3.02E-01 | no | | intergenic_variant | MODIFIER |
| 9 | 91336231 | AX-170032976 | 0 | G | A | 0.132 | -1.25E-01 | 1.21E-01 | 2.55E+00 | 2.64E+00 | 3.02E-01 | no | Gm28652 | upstream_gene_variant | MODIFIER |
| 16 | 39535369 | AX-170456945 | 2 | A | T | 0.5 | 8.39E-02 | 8.13E-02 | 2.56E+00 | 2.60E+00 | 3.02E-01 | no | | intergenic_variant | MODIFIER |
| 18 | 74524260 | AX-97844410 | 0 | T | C | 0.43 | 8.96E-02 | 8.63E-02 | 2.51E+00 | 2.56E+00 | 3.02E-01 | no | Myo5b | intron_variant | MODIFIER |
| 18 | 74591291 | AX-170459402 | 0 | G | A | 0.43 | 8.96E-02 | 8.63E-02 | 2.51E+00 | 2.56E+00 | 3.02E-01 | no | Myo5b | intron_variant | MODIFIER |
| 18 | 74591775 | AX-170335357 | 0 | G | A | 0.43 | 8.96E-02 | 8.63E-02 | 2.51E+00 | 2.56E+00 | 3.02E-01 | no | Myo5b | intron_variant | MODIFIER |
| 18 | 74594369 | AX-170245769 | 0 | T | C | 0.43 | 8.96E-02 | 8.63E-02 | 2.51E+00 | 2.56E+00 | 3.02E-01 | no | Myo5b | intron_variant | MODIFIER |
| 18 | 74609304 | AX-170397513 | 0 | G | A | 0.43 | 8.96E-02 | 8.63E-02 | 2.51E+00 | 2.56E+00 | 3.02E-01 | no | Myo5b | intron_variant | MODIFIER |
| 18 | 74615564 | AX-170509822 | 0 | C | G | 0.43 | 8.96E-02 | 8.63E-02 | 2.51E+00 | 2.56E+00 | 3.02E-01 | no | Myo5b | intron_variant | MODIFIER |
| 18 | 74615774 | AX-170264493 | 0 | C | T | 0.43 | 8.96E-02 | 8.63E-02 | 2.51E+00 | 2.56E+00 | 3.02E-01 | no | Myo5b | intron_variant | MODIFIER |
| 18 | 74619487 | AX-97808308 | 0 | C | T | 0.43 | 8.96E-02 | 8.63E-02 | 2.51E+00 | 2.56E+00 | 3.02E-01 | no | Myo5b | intron_variant | MODIFIER |
| 18 | 74637199 | AX-170121487 | 0 | T | C | 0.43 | 8.96E-02 | 8.63E-02 | 2.51E+00 | 2.56E+00 | 3.02E-01 | no | Myo5b | intron_variant | MODIFIER |
| 18 | 74650847 | AX-170291124 | 0 | G | A | 0.43 | 8.96E-02 | 8.63E-02 | 2.51E+00 | 2.56E+00 | 3.02E-01 | no | Myo5b | intron_variant | MODIFIER |
| 18 | 74652139 | AX-170234801 | 0 | A | G | 0.43 | 8.96E-02 | 8.63E-02 | 2.51E+00 | 2.56E+00 | 3.02E-01 | no | Myo5b | intron_variant | MODIFIER |
| 18 | 74654599 | AX-170113915 | 0 | G | T | 0.43 | 8.96E-02 | 8.63E-02 | 2.51E+00 | 2.56E+00 | 3.02E-01 | no | Myo5b | intron_variant | MODIFIER |
| 18 | 74681136 | AX-170322032 | 0 | C | T | 0.43 | 8.96E-02 | 8.63E-02 | 2.51E+00 | 2.56E+00 | 3.02E-01 | no | Myo5b | intron_variant | MODIFIER |
| 18 | 74685235 | AX-170015995 | 0 | G | A | 0.43 | 8.96E-02 | 8.63E-02 | 2.51E+00 | 2.56E+00 | 3.02E-01 | no | Myo5b | intron_variant | MODIFIER |
| 18 | 74714519 | AX-170507869 | 0 | G | A | 0.43 | 8.96E-02 | 8.63E-02 | 2.51E+00 | 2.56E+00 | 3.02E-01 | no | Myo5b | intron_variant | MODIFIER |
| 18 | 74729255 | AX-170061330 | 0 | G | A | 0.43 | 8.96E-02 | 8.63E-02 | 2.51E+00 | 2.56E+00 | 3.02E-01 | no | Myo5b | intron_variant | MODIFIER |
| 18 | 74818120 | AX-170088375 | 0 | G | A | 0.43 | 8.96E-02 | 8.63E-02 | 2.51E+00 | 2.56E+00 | 3.02E-01 | no | | intergenic_variant | MODIFIER |
| 18 | 75057880 | AX-97835868 | 0 | A | T | 0.43 | 8.96E-02 | 8.63E-02 | 2.51E+00 | 2.56E+00 | 3.02E-01 | no | Dym | intron_variant | MODIFIER |
| 18 | 75104327 | AX-97845772 | 0 | G | A | 0.43 | 8.96E-02 | 8.63E-02 | 2.51E+00 | 2.56E+00 | 3.02E-01 | no | Dym | intron_variant | MODIFIER |
| 18 | 75133495 | AX-170205848 | 0 | G | A | 0.43 | 8.96E-02 | 8.63E-02 | 2.51E+00 | 2.56E+00 | 3.02E-01 | no | Dym | intron_variant | MODIFIER |
| 18 | 75136383 | AX-170331359 | 0 | T | C | 0.43 | 8.96E-02 | 8.63E-02 | 2.51E+00 | 2.56E+00 | 3.02E-01 | no | Dym | intron_variant | MODIFIER |
| 18 | 75137418 | AX-170193935 | 0 | A | G | 0.43 | 8.96E-02 | 8.63E-02 | 2.51E+00 | 2.56E+00 | 3.02E-01 | no | Dym | intron_variant | MODIFIER |
| 18 | 75140924 | AX-170351976 | 0 | C | T | 0.43 | 8.96E-02 | 8.63E-02 | 2.51E+00 | 2.56E+00 | 3.02E-01 | no | Dym | intron_variant | MODIFIER |
| 18 | 75141354 | AX-97800065 | 0 | G | A | 0.43 | 8.96E-02 | 8.63E-02 | 2.51E+00 | 2.56E+00 | 3.02E-01 | no | Dym | intron_variant | MODIFIER |
| 18 | 75144320 | AX-170187380 | 0 | G | A | 0.43 | 8.96E-02 | 8.63E-02 | 2.51E+00 | 2.56E+00 | 3.02E-01 | no | Dym | intron_variant | MODIFIER |
| 18 | 75144587 | AX-170463224 | 0 | T | C | 0.43 | 8.96E-02 | 8.63E-02 | 2.51E+00 | 2.56E+00 | 3.02E-01 | no | Dym | intron_variant | MODIFIER |
| 18 | 75160211 | AX-170312476 | 0 | A | T | 0.43 | 8.96E-02 | 8.63E-02 | 2.51E+00 | 2.56E+00 | 3.02E-01 | no | Dym | intron_variant | MODIFIER |
| 18 | 75165254 | AX-170389225 | 0 | G | A | 0.43 | 8.96E-02 | 8.63E-02 | 2.51E+00 | 2.56E+00 | 3.02E-01 | no | Dym | intron_variant | MODIFIER |
| 18 | 75171731 | AX-170440069 | 0 | G | A | 0.43 | 8.96E-02 | 8.63E-02 | 2.51E+00 | 2.56E+00 | 3.02E-01 | no | Dym | intron_variant | MODIFIER |
| 18 | 75173076 | AX-170307885 | 0 | G | A | 0.43 | 8.96E-02 | 8.63E-02 | 2.51E+00 | 2.56E+00 | 3.02E-01 | no | Dym | intron_variant | MODIFIER |
| 18 | 75176550 | AX-97805813 | 0 | T | C | 0.43 | 8.96E-02 | 8.63E-02 | 2.51E+00 | 2.56E+00 | 3.02E-01 | no | Dym | intron_variant | MODIFIER |
| 18 | 75401036 | AX-170258546 | 0 | A | T | 0.43 | 8.96E-02 | 8.63E-02 | 2.51E+00 | 2.56E+00 | 3.02E-01 | no | | regulatory_region_variant | MODIFIER |
| 18 | 75401160 | AX-170347954 | 0 | C | A | 0.43 | 8.96E-02 | 8.63E-02 | 2.51E+00 | 2.56E+00 | 3.02E-01 | no | | regulatory_region_variant | MODIFIER |
| 18 | 75401755 | AX-170088377 | 0 | G | A | 0.43 | 8.96E-02 | 8.63E-02 | 2.51E+00 | 2.56E+00 | 3.02E-01 | no | | regulatory_region_variant | MODIFIER |
| 18 | 75416440 | AX-97835903 | 0 | T | C | 0.43 | 8.96E-02 | 8.63E-02 | 2.51E+00 | 2.56E+00 | 3.02E-01 | no | | intergenic_variant | MODIFIER |
| 18 | 75430589 | AX-170167247 | 0 | G | A | 0.43 | 8.96E-02 | 8.63E-02 | 2.51E+00 | 2.56E+00 | 3.02E-01 | no | Ctif | downstream_gene_variant | MODIFIER |
| 18 | 75436456 | AX-170398958 | 0 | A | G | 0.43 | 8.96E-02 | 8.63E-02 | 2.51E+00 | 2.56E+00 | 3.02E-01 | no | Ctif | intron_variant | MODIFIER |
| 18 | 75442396 | AX-170279253 | 0 | C | T | 0.43 | 8.96E-02 | 8.63E-02 | 2.51E+00 | 2.56E+00 | 3.02E-01 | no | Ctif | intron_variant | MODIFIER |
| 18 | 75445061 | AX-170178200 | 0 | A | G | 0.43 | 8.96E-02 | 8.63E-02 | 2.51E+00 | 2.56E+00 | 3.02E-01 | no | Ctif | intron_variant | MODIFIER |
| 18 | 75453182 | AX-170313455 | 0 | A | G | 0.43 | 8.96E-02 | 8.63E-02 | 2.51E+00 | 2.56E+00 | 3.02E-01 | no | Ctif | intron_variant | MODIFIER |
| 16 | 39445593 | AX-170057190 | 0 | A | G | 0.5 | 8.29E-02 | 7.99E-02 | 2.50E+00 | 2.56E+00 | 3.03E-01 | no | Gm36903 | upstream_gene_variant | MODIFIER |
| 4 | 137025927 | AX-170178928 | 1 | C | T | 0.179 | 1.14E-01 | 1.11E-01 | 2.57E+00 | 2.62E+00 | 3.03E-01 | no | Zbtb40 | intron_variant | MODIFIER |
| 5 | 113256236 | AX-170385379 | 1 | C | T | 0.464 | 1.21E-01 | 1.18E-01 | 2.74E+00 | 2.78E+00 | 3.03E-01 | no | Sgsm1 | intron_variant | MODIFIER |
| 18 | 73249967 | AX-170315309 | 0 | G | T | 0.193 | -1.23E-01 | 1.20E-01 | 2.61E+00 | 2.66E+00 | 3.03E-01 | no | | regulatory_region_variant | MODIFIER |
| 18 | 73250062 | AX-170406877 | 0 | A | T | 0.193 | -1.23E-01 | 1.20E-01 | 2.61E+00 | 2.66E+00 | 3.03E-01 | no | | regulatory_region_variant | MODIFIER |
| 18 | 73254037 | AX-97783403 | 0 | T | C | 0.193 | -1.23E-01 | 1.20E-01 | 2.61E+00 | 2.66E+00 | 3.03E-01 | no | | intergenic_variant | MODIFIER |
| 18 | 73438074 | AX-97802767 | 0 | T | C | 0.193 | -1.23E-01 | 1.20E-01 | 2.61E+00 | 2.66E+00 | 3.03E-01 | no | | intergenic_variant | MODIFIER |
| 18 | 73439154 | AX-170107854 | 0 | G | A | 0.193 | -1.23E-01 | 1.20E-01 | 2.61E+00 | 2.66E+00 | 3.03E-01 | no | | intergenic_variant | MODIFIER |
| 18 | 73456619 | AX-97835204 | 0 | C | T | 0.193 | -1.23E-01 | 1.20E-01 | 2.61E+00 | 2.66E+00 | 3.03E-01 | no | | intergenic_variant | MODIFIER |

| | | | | | | | | | | | | | | | |
|---|---|---|---|---|---|---|---|---|---|---|---|---|---|---|---|
| 18 | 73770662 | AX-170248071 | 0 | T | C | 0.193 | -1.23E-01 | 1.20E-01 | 2.61E+00 | 2.66E+00 | 3.03E-01 | no | Me2 | synonymous_variant | LOW |
| 18 | 73830964 | AX-97781520 | 0 | T | C | 0.193 | -1.23E-01 | 1.20E-01 | 2.61E+00 | 2.66E+00 | 3.03E-01 | no | | regulatory_region_variant | MODIFIER |
| 18 | 73856410 | AX-170271989 | 0 | C | T | 0.193 | -1.23E-01 | 1.20E-01 | 2.61E+00 | 2.66E+00 | 3.03E-01 | no | Mro | upstream_gene_variant | MODIFIER |
| 18 | 74406875 | AX-170405448 | 0 | A | G | 0.193 | -1.23E-01 | 1.20E-01 | 2.61E+00 | 2.66E+00 | 3.03E-01 | no | | intergenic_variant | MODIFIER |
| 18 | 74407021 | AX-170324444 | 0 | C | T | 0.193 | -1.23E-01 | 1.20E-01 | 2.61E+00 | 2.66E+00 | 3.03E-01 | no | | intergenic_variant | MODIFIER |
| 18 | 74431235 | AX-97805585 | 0 | C | T | 0.193 | -1.23E-01 | 1.20E-01 | 2.61E+00 | 2.66E+00 | 3.03E-01 | no | | regulatory_region_variant | MODIFIER |
| 18 | 74431425 | AX-170305990 | 0 | A | G | 0.193 | -1.23E-01 | 1.20E-01 | 2.61E+00 | 2.66E+00 | 3.03E-01 | no | | regulatory_region_variant | MODIFIER |
| 19 | 50744754 | AX-170016810 | 0 | A | T | 0.491 | 6.62E-02 | 6.48E-02 | 2.65E+00 | 2.74E+00 | 3.03E-01 | no | Gm26629 | intron_variant,non_coding_transcript_variant | MODIFIER |
| 4 | 134987673 | AX-170095158 | 0 | C | A | 0.158 | 1.19E-01 | 1.16E-01 | 2.52E+00 | 2.57E+00 | 3.03E-01 | no | | intergenic_variant | MODIFIER |
| 18 | 52198347 | AX-170190492 | 1 | C | T | 0.241 | -1.03E-01 | 1.00E-01 | 2.58E+00 | 2.65E+00 | 3.03E-01 | no | | intergenic_variant | MODIFIER |
| 13 | 31139799 | AX-97806296 | 0 | G | A | 0.193 | -1.08E-01 | 1.06E-01 | 2.65E+00 | 2.75E+00 | 3.03E-01 | no | | regulatory_region_variant | MODIFIER |
| 20 | 165004431 | AX-170085737 | 0 | G | A | 0.14 | 1.21E-01 | 1.19E-01 | 2.77E+00 | 2.90E+00 | 3.03E-01 | no | Gm17604 | intron_variant | MODIFIER |
| 6 | 137536744 | AX-170174266 | 2 | G | C | 0.555 | 8.98E-02 | 8.63E-02 | 2.48E+00 | 2.55E+00 | 3.03E-01 | no | Eps8 | intron_variant | MODIFIER |
| 1 | 132120419 | AX-170457511 | 1 | C | T | 0.545 | 8.76E-02 | 8.57E-02 | 2.65E+00 | 2.71E+00 | 3.03E-01 | no | Cdk18 | intron_variant | MODIFIER |
| 1 | 132481708 | AX-170386628 | 1 | C | T | 0.545 | 8.76E-02 | 8.57E-02 | 2.65E+00 | 2.71E+00 | 3.03E-01 | no | Rbbp5 | intron_variant | MODIFIER |
| 12 | 91944464 | AX-97811302 | 0 | G | A | 0.079 | 1.75E-01 | 1.72E-01 | 2.79E+00 | 2.93E+00 | 3.03E-01 | Yes | | intergenic_variant | MODIFIER |
| 5 | 106917178 | AX-168384187 | 0 | C | A | 0.439 | 1.12E-01 | 1.10E-01 | 2.69E+00 | 2.74E+00 | 3.03E-01 | no | Hfm1 | intron_variant | MODIFIER |
| 14 | 122953861 | AX-170261520 | 0 | T | C | 0.298 | 8.23E-02 | 8.06E-02 | 2.62E+00 | 2.72E+00 | 3.03E-01 | no | Tmtc4 | intron_variant | MODIFIER |
| 1 | 172326178 | AX-170171477 | 1 | C | G | 0.214 | 1.38E-01 | 1.34E-01 | 2.58E+00 | 2.62E+00 | 3.03E-01 | no | Kcnj9 | synonymous_variant | LOW |
| 1 | 172334273 | AX-170346236 | 1 | G | T | 0.214 | 1.38E-01 | 1.34E-01 | 2.58E+00 | 2.62E+00 | 3.03E-01 | no | Kcnj9 | upstream_gene_variant | MODIFIER |
| 7 | 12127199 | AX-97817786 | 1 | T | C | 0.339 | 8.00E-02 | 7.82E-02 | 2.58E+00 | 2.70E+00 | 3.03E-01 | no | Gm45498 | downstream_gene_variant | MODIFIER |
| 8 | 48689257 | AX-97792595 | 0 | G | T | 0.14 | -1.44E-01 | 1.40E-01 | 2.56E+00 | 2.63E+00 | 3.03E-01 | no | Tenm3 | intron_variant | MODIFIER |
| 15 | 4600121 | AX-170293594 | 0 | A | G | 0.246 | -1.03E-01 | 1.01E-01 | 2.66E+00 | 2.72E+00 | 3.04E-01 | no | | intergenic_variant | MODIFIER |
| 8 | 93550858 | AX-170142241 | 0 | A | G | 0.211 | -1.05E-01 | 1.03E-01 | 2.83E+00 | 2.96E+00 | 3.04E-01 | no | | intergenic_variant | MODIFIER |
| 1 | 120365894 | AX-170474289 | 1 | A | G | 0.562 | 8.61E-02 | 8.32E-02 | 2.51E+00 | 2.57E+00 | 3.04E-01 | no | | intergenic_variant | MODIFIER |
| 9 | 98487926 | AX-170218840 | 1 | G | A | 0.152 | -1.08E-01 | 1.06E-01 | 2.59E+00 | 2.69E+00 | 3.04E-01 | no | Rbp2 | upstream_gene_variant | MODIFIER |
| 1 | 132266540 | AX-170508531 | 1 | G | A | 0.536 | 8.71E-02 | 8.54E-02 | 2.65E+00 | 2.71E+00 | 3.04E-01 | no | | regulatory_region_variant | MODIFIER |
| 16 | 50063777 | AX-170181009 | 0 | A | G | 0.518 | 8.47E-02 | 8.12E-02 | 2.46E+00 | 2.50E+00 | 3.04E-01 | no | | intergenic_variant | MODIFIER |
| 1 | 171413911 | AX-170043573 | 1 | C | T | 0.214 | 1.37E-01 | 1.34E-01 | 2.58E+00 | 2.63E+00 | 3.04E-01 | no | Usf1 | intron_variant,NMD_transcript_variant | MODIFIER |
| 4 | 95791126 | AX-170489160 | 0 | C | T | 0.246 | 1.16E-01 | 1.12E-01 | 2.48E+00 | 2.53E+00 | 3.04E-01 | no | Fggy | intron_variant | MODIFIER |
| 4 | 95791541 | AX-170169810 | 0 | A | G | 0.246 | 1.16E-01 | 1.12E-01 | 2.48E+00 | 2.53E+00 | 3.04E-01 | no | Fggy | intron_variant | MODIFIER |
| 4 | 95798333 | AX-170395396 | 0 | C | T | 0.246 | 1.16E-01 | 1.12E-01 | 2.48E+00 | 2.53E+00 | 3.04E-01 | no | Fggy | intron_variant | MODIFIER |
| 4 | 95798422 | AX-170507610 | 0 | G | A | 0.246 | 1.16E-01 | 1.12E-01 | 2.48E+00 | 2.53E+00 | 3.04E-01 | no | Fggy | intron_variant | MODIFIER |
| 4 | 95810206 | AX-170267623 | 0 | A | G | 0.246 | 1.16E-01 | 1.12E-01 | 2.48E+00 | 2.53E+00 | 3.04E-01 | no | Fggy | intron_variant | MODIFIER |
| 4 | 95811222 | AX-170334903 | 0 | A | G | 0.246 | 1.16E-01 | 1.12E-01 | 2.48E+00 | 2.53E+00 | 3.04E-01 | no | Fggy | intron_variant | MODIFIER |
| 4 | 95814196 | AX-170371591 | 0 | T | A | 0.246 | 1.16E-01 | 1.12E-01 | 2.48E+00 | 2.53E+00 | 3.04E-01 | no | Fggy | intron_variant | MODIFIER |
| 4 | 95814276 | AX-170070064 | 0 | T | A | 0.246 | 1.16E-01 | 1.12E-01 | 2.48E+00 | 2.53E+00 | 3.04E-01 | no | Fggy | intron_variant | MODIFIER |
| 4 | 95847131 | AX-170147427 | 0 | A | G | 0.246 | 1.16E-01 | 1.12E-01 | 2.48E+00 | 2.53E+00 | 3.04E-01 | no | Fggy | intron_variant | MODIFIER |
| 4 | 95849499 | AX-170453673 | 0 | G | A | 0.246 | 1.16E-01 | 1.12E-01 | 2.48E+00 | 2.53E+00 | 3.04E-01 | no | Fggy | intron_variant | MODIFIER |
| 4 | 95859689 | AX-170166104 | 0 | T | C | 0.246 | 1.16E-01 | 1.12E-01 | 2.48E+00 | 2.53E+00 | 3.04E-01 | no | Fggy | intron_variant | MODIFIER |
| 4 | 95872927 | AX-170328521 | 0 | G | A | 0.246 | 1.16E-01 | 1.12E-01 | 2.48E+00 | 2.53E+00 | 3.04E-01 | no | Fggy | intron_variant | MODIFIER |
| 4 | 95885783 | AX-170487631 | 0 | A | G | 0.246 | 1.16E-01 | 1.12E-01 | 2.48E+00 | 2.53E+00 | 3.04E-01 | no | Fggy | intron_variant | MODIFIER |
| 4 | 95892200 | AX-170457072 | 0 | C | T | 0.246 | 1.16E-01 | 1.12E-01 | 2.48E+00 | 2.53E+00 | 3.04E-01 | no | Fggy | intron_variant | MODIFIER |
| 4 | 95899187 | AX-170305444 | 0 | A | G | 0.246 | 1.16E-01 | 1.12E-01 | 2.48E+00 | 2.53E+00 | 3.04E-01 | no | Fggy | intron_variant | MODIFIER |
| 4 | 95901751 | AX-170440278 | 0 | C | T | 0.246 | 1.16E-01 | 1.12E-01 | 2.48E+00 | 2.53E+00 | 3.04E-01 | no | Fggy | intron_variant | MODIFIER |
| 4 | 95931189 | AX-170023451 | 0 | C | A | 0.246 | 1.16E-01 | 1.12E-01 | 2.48E+00 | 2.53E+00 | 3.04E-01 | no | Fggy | downstream_gene_variant | MODIFIER |
| 4 | 95947288 | AX-170224530 | 0 | G | T | 0.246 | 1.16E-01 | 1.12E-01 | 2.48E+00 | 2.53E+00 | 3.04E-01 | no | | intergenic_variant | MODIFIER |
| 4 | 95947459 | AX-170170580 | 0 | G | T | 0.246 | 1.16E-01 | 1.12E-01 | 2.48E+00 | 2.53E+00 | 3.04E-01 | no | | intergenic_variant | MODIFIER |
| 4 | 95947933 | AX-170023456 | 0 | T | C | 0.246 | 1.16E-01 | 1.12E-01 | 2.48E+00 | 2.53E+00 | 3.04E-01 | no | | intergenic_variant | MODIFIER |
| 4 | 95948364 | AX-170214350 | 0 | G | A | 0.246 | 1.16E-01 | 1.12E-01 | 2.48E+00 | 2.53E+00 | 3.04E-01 | no | | intergenic_variant | MODIFIER |
| 4 | 95990791 | AX-170302887 | 0 | T | C | 0.246 | 1.16E-01 | 1.12E-01 | 2.48E+00 | 2.53E+00 | 3.04E-01 | no | Hook1 | intron_variant | MODIFIER |
| 4 | 96405169 | AX-97848277 | 0 | G | A | 0.246 | 1.16E-01 | 1.12E-01 | 2.48E+00 | 2.53E+00 | 3.04E-01 | no | | intergenic_variant | MODIFIER |
| 14 | 111242724 | AX-170054458 | 1 | T | C | 0.312 | 7.51E-02 | 7.36E-02 | 2.60E+00 | 2.70E+00 | 3.04E-01 | no | | intergenic_variant | MODIFIER |
| 9 | 9091491 | AX-170399921 | 1 | T | G | 0.464 | 8.41E-02 | 8.11E-02 | 2.47E+00 | 2.56E+00 | 3.04E-01 | no | Arhgap42 | intron_variant | MODIFIER |

| | | | | | | | | | | | | | | | |
|---|---|---|---|---|---|---|---|---|---|---|---|---|---|---|---|
| 3 | 36514604 | AX-170530055 | 0 | A | G | 0.193 | 9.65E-02 | 9.47E-02 | 2.86E+00 | 2.96E+00 | 3.05E-01 | no | Gm11549 | downstream_gene_variant | MODIFIER |
| 3 | 37519738 | AX-170101470 | 0 | A | G | 0.193 | 9.65E-02 | 9.47E-02 | 2.86E+00 | 2.96E+00 | 3.05E-01 | no | Spata5 | intron_variant | MODIFIER |
| 8 | 73918609 | AX-170116988 | 0 | G | A | 0.202 | 1.26E-01 | 1.23E-01 | 2.95E+00 | 3.06E+00 | 3.05E-01 | no | | intergenic_variant | MODIFIER |
| 4 | 128008726 | AX-170070634 | 1 | T | G | 0.143 | 1.24E-01 | 1.21E-01 | 2.60E+00 | 2.66E+00 | 3.05E-01 | no | Csmd2 | intron_variant,non_coding_transcript_variant | MODIFIER |
| 9 | 98465907 | AX-170425535 | 1 | G | T | 0.143 | -1.17E-01 | 1.14E-01 | 2.60E+00 | 2.67E+00 | 3.05E-01 | no | | intergenic_variant | MODIFIER |
| 9 | 4526789 | AX-97842822 | 0 | C | G | 0.465 | 8.31E-02 | 8.01E-02 | 2.46E+00 | 2.55E+00 | 3.05E-01 | no | Gria4 | intron_variant | MODIFIER |
| 18 | 59347113 | AX-170297283 | 1 | C | T | 0.179 | -1.24E-01 | 1.22E-01 | 2.72E+00 | 2.77E+00 | 3.06E-01 | no | Chsy3 | intron_variant | MODIFIER |
| 9 | 103998998 | AX-170436643 | 0 | A | C | 0.114 | -1.14E-01 | 1.11E-01 | 2.54E+00 | 2.64E+00 | 3.06E-01 | no | Nphp3 | upstream_gene_variant | MODIFIER |
| 16 | 33136849 | AX-170407252 | 0 | T | C | 0.491 | 7.97E-02 | 7.73E-02 | 2.50E+00 | 2.56E+00 | 3.06E-01 | no | Gm23916 | downstream_gene_variant | MODIFIER |
| 5 | 96364469 | AX-170525242 | 0 | A | T | 0.43 | 1.01E-01 | 9.88E-02 | 2.62E+00 | 2.70E+00 | 3.06E-01 | no | | regulatory_region_variant | MODIFIER |
| 1 | 22402809 | AX-168374878 | 1 | A | G | 0.411 | 8.36E-02 | 8.24E-02 | 2.66E+00 | 2.74E+00 | 3.06E-01 | no | Rims1 | intron_variant | MODIFIER |
| 18 | 85765116 | AX-170503249 | 0 | C | T | 0.158 | -1.09E-01 | 1.06E-01 | 2.52E+00 | 2.60E+00 | 3.06E-01 | no | | intergenic_variant | MODIFIER |
| 18 | 85781594 | AX-170061574 | 0 | G | A | 0.158 | -1.09E-01 | 1.06E-01 | 2.52E+00 | 2.60E+00 | 3.06E-01 | no | | intergenic_variant | MODIFIER |
| 18 | 85859950 | AX-170278644 | 0 | C | T | 0.158 | -1.09E-01 | 1.06E-01 | 2.52E+00 | 2.60E+00 | 3.06E-01 | no | | intergenic_variant | MODIFIER |
| 18 | 85896539 | AX-170416417 | 0 | C | T | 0.158 | -1.09E-01 | 1.06E-01 | 2.52E+00 | 2.60E+00 | 3.06E-01 | no | | intergenic_variant | MODIFIER |
| 18 | 86179662 | AX-170236709 | 0 | T | G | 0.158 | -1.09E-01 | 1.06E-01 | 2.52E+00 | 2.60E+00 | 3.06E-01 | no | | intergenic_variant | MODIFIER |
| 18 | 86185209 | AX-170409492 | 0 | T | C | 0.158 | -1.09E-01 | 1.06E-01 | 2.52E+00 | 2.60E+00 | 3.06E-01 | no | | intergenic_variant | MODIFIER |
| 18 | 86216951 | AX-170509328 | 0 | T | C | 0.158 | -1.09E-01 | 1.06E-01 | 2.52E+00 | 2.60E+00 | 3.06E-01 | no | | intergenic_variant | MODIFIER |
| 18 | 86220213 | AX-170420632 | 0 | G | A | 0.158 | -1.09E-01 | 1.06E-01 | 2.52E+00 | 2.60E+00 | 3.06E-01 | no | | intergenic_variant | MODIFIER |
| 18 | 86484171 | AX-170501369 | 0 | G | A | 0.158 | -1.09E-01 | 1.06E-01 | 2.52E+00 | 2.60E+00 | 3.06E-01 | no | Neto1 | intron_variant | MODIFIER |
| 18 | 86494610 | AX-170061589 | 0 | T | C | 0.158 | -1.09E-01 | 1.06E-01 | 2.52E+00 | 2.60E+00 | 3.06E-01 | no | Neto1 | intron_variant | MODIFIER |
| 18 | 86498345 | AX-170061590 | 0 | T | A | 0.158 | -1.09E-01 | 1.06E-01 | 2.52E+00 | 2.60E+00 | 3.06E-01 | no | Neto1 | intron_variant | MODIFIER |
| 18 | 86501408 | AX-170061591 | 0 | C | T | 0.158 | -1.09E-01 | 1.06E-01 | 2.52E+00 | 2.60E+00 | 3.06E-01 | no | Neto1 | 3_prime_UTR_variant | MODIFIER |
| 18 | 86525531 | AX-170349326 | 0 | A | G | 0.158 | -1.09E-01 | 1.06E-01 | 2.52E+00 | 2.60E+00 | 3.06E-01 | no | | intergenic_variant | MODIFIER |
| 18 | 86531739 | AX-170508429 | 0 | T | C | 0.158 | -1.09E-01 | 1.06E-01 | 2.52E+00 | 2.60E+00 | 3.06E-01 | no | | intergenic_variant | MODIFIER |
| 18 | 86538523 | AX-170464594 | 0 | T | C | 0.158 | -1.09E-01 | 1.06E-01 | 2.52E+00 | 2.60E+00 | 3.06E-01 | no | | intergenic_variant | MODIFIER |
| 18 | 86620376 | AX-97834866 | 0 | T | C | 0.158 | -1.09E-01 | 1.06E-01 | 2.52E+00 | 2.60E+00 | 3.06E-01 | no | AC113010.1 | downstream_gene_variant | MODIFIER |
| 18 | 86639799 | AX-97809999 | 0 | T | C | 0.158 | -1.09E-01 | 1.06E-01 | 2.52E+00 | 2.60E+00 | 3.06E-01 | no | | intergenic_variant | MODIFIER |
| 18 | 86640956 | AX-170160272 | 0 | T | C | 0.158 | -1.09E-01 | 1.06E-01 | 2.52E+00 | 2.60E+00 | 3.06E-01 | no | | intergenic_variant | MODIFIER |
| 18 | 87401586 | AX-170530957 | 0 | C | T | 0.158 | -1.09E-01 | 1.06E-01 | 2.52E+00 | 2.60E+00 | 3.06E-01 | no | | intergenic_variant | MODIFIER |
| 18 | 87410273 | AX-170155921 | 0 | T | C | 0.158 | -1.09E-01 | 1.06E-01 | 2.52E+00 | 2.60E+00 | 3.06E-01 | no | | intergenic_variant | MODIFIER |
| 18 | 87434094 | AX-170236571 | 0 | A | T | 0.158 | -1.09E-01 | 1.06E-01 | 2.52E+00 | 2.60E+00 | 3.06E-01 | no | | intergenic_variant | MODIFIER |
| 18 | 87486955 | AX-97835739 | 0 | A | G | 0.158 | -1.09E-01 | 1.06E-01 | 2.52E+00 | 2.60E+00 | 3.06E-01 | no | | intergenic_variant | MODIFIER |
| 18 | 87489567 | AX-170408965 | 0 | T | C | 0.158 | -1.09E-01 | 1.06E-01 | 2.52E+00 | 2.60E+00 | 3.06E-01 | no | | intergenic_variant | MODIFIER |
| 18 | 88040418 | AX-170207819 | 0 | T | C | 0.158 | -1.09E-01 | 1.06E-01 | 2.52E+00 | 2.60E+00 | 3.06E-01 | no | | regulatory_region_variant | MODIFIER |
| 18 | 88131237 | AX-170525615 | 0 | C | T | 0.158 | -1.09E-01 | 1.06E-01 | 2.52E+00 | 2.60E+00 | 3.06E-01 | no | | intergenic_variant | MODIFIER |
| 18 | 88146721 | AX-170149504 | 0 | A | C | 0.158 | -1.09E-01 | 1.06E-01 | 2.52E+00 | 2.60E+00 | 3.06E-01 | no | | intergenic_variant | MODIFIER |
| 18 | 88164838 | AX-170175995 | 0 | G | A | 0.158 | -1.09E-01 | 1.06E-01 | 2.52E+00 | 2.60E+00 | 3.06E-01 | no | | intergenic_variant | MODIFIER |
| 18 | 88209738 | AX-97848039 | 0 | G | A | 0.158 | -1.09E-01 | 1.06E-01 | 2.52E+00 | 2.60E+00 | 3.06E-01 | no | | intergenic_variant | MODIFIER |
| 18 | 88227819 | AX-97836175 | 0 | G | C | 0.158 | -1.09E-01 | 1.06E-01 | 2.52E+00 | 2.60E+00 | 3.06E-01 | no | | intergenic_variant | MODIFIER |
| 18 | 88259252 | AX-170434660 | 0 | C | T | 0.158 | -1.09E-01 | 1.06E-01 | 2.52E+00 | 2.60E+00 | 3.06E-01 | no | | intergenic_variant | MODIFIER |
| 18 | 88276319 | AX-170171002 | 0 | A | G | 0.158 | -1.09E-01 | 1.06E-01 | 2.52E+00 | 2.60E+00 | 3.06E-01 | no | | intergenic_variant | MODIFIER |
| 18 | 88285180 | AX-170099850 | 0 | A | G | 0.158 | -1.09E-01 | 1.06E-01 | 2.52E+00 | 2.60E+00 | 3.06E-01 | no | | intergenic_variant | MODIFIER |
| 18 | 88327418 | AX-170185794 | 0 | T | C | 0.158 | -1.09E-01 | 1.06E-01 | 2.52E+00 | 2.60E+00 | 3.06E-01 | no | | intergenic_variant | MODIFIER |
| 18 | 88446943 | AX-170236111 | 0 | T | C | 0.158 | -1.09E-01 | 1.06E-01 | 2.52E+00 | 2.60E+00 | 3.06E-01 | no | | intergenic_variant | MODIFIER |
| 18 | 88503729 | AX-170411778 | 0 | T | G | 0.158 | -1.09E-01 | 1.06E-01 | 2.52E+00 | 2.60E+00 | 3.06E-01 | no | | intergenic_variant | MODIFIER |
| 18 | 88516630 | AX-170158222 | 0 | G | A | 0.158 | -1.09E-01 | 1.06E-01 | 2.52E+00 | 2.60E+00 | 3.06E-01 | no | | intergenic_variant | MODIFIER |
| 18 | 88526996 | AX-170253591 | 0 | A | G | 0.158 | -1.09E-01 | 1.06E-01 | 2.52E+00 | 2.60E+00 | 3.06E-01 | no | | intergenic_variant | MODIFIER |
| 18 | 88536012 | AX-170301482 | 0 | G | A | 0.158 | -1.09E-01 | 1.06E-01 | 2.52E+00 | 2.60E+00 | 3.06E-01 | no | | intergenic_variant | MODIFIER |
| 18 | 88575153 | AX-170393589 | 0 | C | T | 0.158 | -1.09E-01 | 1.06E-01 | 2.52E+00 | 2.60E+00 | 3.06E-01 | no | | intergenic_variant | MODIFIER |
| 18 | 88586801 | AX-170308760 | 0 | C | T | 0.158 | -1.09E-01 | 1.06E-01 | 2.52E+00 | 2.60E+00 | 3.06E-01 | no | | intergenic_variant | MODIFIER |
| 18 | 88603206 | AX-170061632 | 0 | C | T | 0.158 | -1.09E-01 | 1.06E-01 | 2.52E+00 | 2.60E+00 | 3.06E-01 | no | | intergenic_variant | MODIFIER |
| 18 | 88640733 | AX-170219463 | 0 | G | A | 0.158 | -1.09E-01 | 1.06E-01 | 2.52E+00 | 2.60E+00 | 3.06E-01 | no | AC133941.2 | non_coding_transcript_exon_variant | MODIFIER |
| 18 | 88642161 | AX-170134721 | 0 | G | A | 0.158 | -1.09E-01 | 1.06E-01 | 2.52E+00 | 2.60E+00 | 3.06E-01 | no | AC133941.2 | intron_variant,non_coding_transcript_variant | MODIFIER |

| | | | | | | | | | | | | | | | |
|---|---|---|---|---|---|---|---|---|---|---|---|---|---|---|---|
| 18 | 88663678 | AX-170191086 | 0 | A | G | 0.158 | -1.09E-01 | 1.06E-01 | 2.52E+00 | 2.60E+00 | 3.06E-01 | no | Socs6 | downstream_gene_variant | MODIFIER |
| 18 | 88699505 | AX-170135814 | 0 | T | A | 0.158 | -1.09E-01 | 1.06E-01 | 2.52E+00 | 2.60E+00 | 3.06E-01 | no | Socs6 | intron_variant,non_coding_transcript_variant | MODIFIER |
| 18 | 88699788 | AX-170297749 | 0 | T | C | 0.158 | -1.09E-01 | 1.06E-01 | 2.52E+00 | 2.60E+00 | 3.06E-01 | no | Socs6 | intron_variant,non_coding_transcript_variant | MODIFIER |
| 18 | 88704069 | AX-97797011 | 0 | T | C | 0.158 | -1.09E-01 | 1.06E-01 | 2.52E+00 | 2.60E+00 | 3.06E-01 | no | Socs6 | intron_variant,non_coding_transcript_variant | MODIFIER |
| 18 | 88704586 | AX-170215510 | 0 | A | C | 0.158 | -1.09E-01 | 1.06E-01 | 2.52E+00 | 2.60E+00 | 3.06E-01 | no | Socs6 | intron_variant,non_coding_transcript_variant | MODIFIER |
| 18 | 88712798 | AX-170437740 | 0 | C | T | 0.158 | -1.09E-01 | 1.06E-01 | 2.52E+00 | 2.60E+00 | 3.06E-01 | no | Socs6 | intron_variant,non_coding_transcript_variant | MODIFIER |
| 18 | 88713207 | AX-170350967 | 0 | T | C | 0.158 | -1.09E-01 | 1.06E-01 | 2.52E+00 | 2.60E+00 | 3.06E-01 | no | Socs6 | intron_variant,non_coding_transcript_variant | MODIFIER |
| 18 | 88713468 | AX-170373898 | 0 | G | C | 0.158 | -1.09E-01 | 1.06E-01 | 2.52E+00 | 2.60E+00 | 3.06E-01 | no | Socs6 | intron_variant,non_coding_transcript_variant | MODIFIER |
| 18 | 88735809 | AX-170468979 | 0 | A | G | 0.158 | -1.09E-01 | 1.06E-01 | 2.52E+00 | 2.60E+00 | 3.06E-01 | no | Socs6 | intron_variant,non_coding_transcript_variant | MODIFIER |
| 18 | 88737489 | AX-170430920 | 0 | A | G | 0.158 | -1.09E-01 | 1.06E-01 | 2.52E+00 | 2.60E+00 | 3.06E-01 | no | Socs6 | intron_variant,non_coding_transcript_variant | MODIFIER |
| 18 | 88738275 | AX-170327685 | 0 | G | A | 0.158 | -1.09E-01 | 1.06E-01 | 2.52E+00 | 2.60E+00 | 3.06E-01 | no | Socs6 | intron_variant,non_coding_transcript_variant | MODIFIER |
| 18 | 88758106 | AX-170220926 | 0 | T | C | 0.158 | -1.09E-01 | 1.06E-01 | 2.52E+00 | 2.60E+00 | 3.06E-01 | no | Socs6 | intron_variant,non_coding_transcript_variant | MODIFIER |
| 18 | 88790228 | AX-170061637 | 0 | G | A | 0.158 | -1.09E-01 | 1.06E-01 | 2.52E+00 | 2.60E+00 | 3.06E-01 | no | Socs6 | intron_variant,non_coding_transcript_variant | MODIFIER |
| 18 | 88800304 | AX-170308688 | 0 | A | G | 0.158 | -1.09E-01 | 1.06E-01 | 2.52E+00 | 2.60E+00 | 3.06E-01 | no | Socs6 | intron_variant,non_coding_transcript_variant | MODIFIER |
| 18 | 88819933 | AX-170130966 | 0 | A | G | 0.158 | -1.09E-01 | 1.06E-01 | 2.52E+00 | 2.60E+00 | 3.06E-01 | no | Socs6 | intron_variant,non_coding_transcript_variant | MODIFIER |
| 1 | 135174138 | AX-170505275 | 0 | C | A | 0.544 | 8.41E-02 | 8.28E-02 | 2.66E+00 | 2.73E+00 | 3.06E-01 | no | | intergenic_variant | MODIFIER |
| 20 | 95903206 | AX-170272611 | 0 | A | T | 0.325 | 5.69E-02 | 5.63E-02 | 2.72E+00 | 2.84E+00 | 3.06E-01 | no | Zc3h12b | intron_variant | MODIFIER |
| 16 | 40798409 | AX-170312375 | 0 | A | C | 0.491 | 8.54E-02 | 8.34E-02 | 2.55E+00 | 2.59E+00 | 3.06E-01 | no | Lsamp | intron_variant | MODIFIER |
| 6 | 134421372 | AX-97827269 | 1 | C | T | 0.58 | 8.76E-02 | 8.39E-02 | 2.42E+00 | 2.49E+00 | 3.06E-01 | no | Bcl2l14 | intron_variant | MODIFIER |
| 2 | 168748725 | AX-170165124 | 1 | C | T | 0.17 | 1.84E-01 | 1.81E-01 | 2.67E+00 | 2.76E+00 | 3.06E-01 | Yes | Sall4 | 3_prime_UTR_variant | MODIFIER |
| 10 | 92192510 | AX-170281576 | 0 | G | T | 0.298 | -8.63E-02 | 8.39E-02 | 2.51E+00 | 2.57E+00 | 3.06E-01 | no | Gm20757 | intron_variant,non_coding_transcript_variant | MODIFIER |
| 1 | 155726264 | AX-170138221 | 0 | G | A | 0.228 | 1.36E-01 | 1.34E-01 | 2.70E+00 | 2.76E+00 | 3.07E-01 | no | Lhx4 | intron_variant | MODIFIER |
| 10 | 66152881 | AX-170309030 | 0 | A | T | 0.412 | -8.02E-02 | 7.91E-02 | 2.64E+00 | 2.72E+00 | 3.07E-01 | no | 1700023F02Rik | intron_variant,non_coding_transcript_variant | MODIFIER |
| 15 | 97470412 | AX-97837792 | 0 | C | T | 0.167 | 1.39E-01 | 1.36E-01 | 2.56E+00 | 2.67E+00 | 3.07E-01 | no | | regulatory_region_variant | MODIFIER |
| 5 | 36159107 | AX-97822716 | 0 | C | T | 0.342 | 1.08E-01 | 1.07E-01 | 2.80E+00 | 2.88E+00 | 3.07E-01 | no | Sorcs2 | intron_variant | MODIFIER |
| 7 | 93040857 | AX-170319798 | 0 | G | A | 0.14 | -1.68E-01 | 1.66E-01 | 2.85E+00 | 2.98E+00 | 3.07E-01 | no | Gm26862 | intron_variant,non_coding_transcript_variant | MODIFIER |
| 1 | 132274695 | AX-170266386 | 0 | A | G | 0.535 | 8.36E-02 | 8.20E-02 | 2.57E+00 | 2.64E+00 | 3.07E-01 | no | Gm29630 | downstream_gene_variant | MODIFIER |
| 1 | 124541291 | AX-170371504 | 0 | G | A | 0.561 | 8.45E-02 | 8.29E-02 | 2.58E+00 | 2.64E+00 | 3.07E-01 | no | Gm28928 | intron_variant,non_coding_transcript_variant | MODIFIER |
| 10 | 72810114 | AX-170355906 | 2 | T | C | 0.409 | -8.00E-02 | 7.88E-02 | 2.61E+00 | 2.70E+00 | 3.07E-01 | no | | regulatory_region_variant | MODIFIER |
| 4 | 139268471 | AX-170070852 | 1 | C | A | 0.179 | 1.10E-01 | 1.09E-01 | 2.64E+00 | 2.70E+00 | 3.07E-01 | no | Capzb | intron_variant | MODIFIER |
| 2 | 162814062 | AX-170390532 | 1 | G | A | 0.161 | 1.97E-01 | 1.95E-01 | 2.82E+00 | 2.91E+00 | 3.07E-01 | Yes | Gm24478 | downstream_gene_variant | MODIFIER |
| 7 | 144465118 | AX-170276832 | 0 | A | C | 0.193 | -7.83E-02 | 7.74E-02 | 2.62E+00 | 2.75E+00 | 3.07E-01 | no | Cttn | intron_variant | MODIFIER |
| 13 | 6304055 | AX-170382912 | 0 | A | G | 0.184 | -1.26E-01 | 1.24E-01 | 2.61E+00 | 2.67E+00 | 3.07E-01 | no | | intergenic_variant | MODIFIER |
| 7 | 37091363 | AX-170525064 | 0 | T | A | 0.342 | 9.35E-02 | 9.21E-02 | 2.59E+00 | 2.67E+00 | 3.08E-01 | no | | intergenic_variant | MODIFIER |
| 1 | 14669075 | AX-97795844 | 1 | T | C | 0.384 | 8.21E-02 | 8.09E-02 | 2.62E+00 | 2.67E+00 | 3.08E-01 | no | | intergenic_variant | MODIFIER |
| 16 | 32646574 | AX-97826951 | 0 | C | T | 0.482 | 8.22E-02 | 8.05E-02 | 2.55E+00 | 2.60E+00 | 3.08E-01 | no | Tnk2 | intron_variant | MODIFIER |
| 14 | 122285274 | AX-97828857 | 0 | C | T | 0.298 | 8.10E-02 | 7.99E-02 | 2.59E+00 | 2.70E+00 | 3.08E-01 | no | Clybl | intron_variant | MODIFIER |
| 4 | 50444564 | AX-170022760 | 1 | C | T | 0.366 | -9.88E-02 | 9.74E-02 | 2.89E+00 | 3.00E+00 | 3.08E-01 | no | | intergenic_variant | MODIFIER |
| 18 | 75950788 | AX-170061358 | 0 | G | A | 0.439 | 8.89E-02 | 8.64E-02 | 2.48E+00 | 2.54E+00 | 3.08E-01 | no | Zbtb7c | intron_variant | MODIFIER |
| 18 | 77365559 | AX-170309282 | 0 | C | T | 0.439 | 8.89E-02 | 8.64E-02 | 2.48E+00 | 2.54E+00 | 3.08E-01 | no | Loxhd1 | intron_variant,non_coding_transcript_variant | MODIFIER |
| 18 | 77548467 | AX-170325889 | 0 | C | A | 0.439 | 8.89E-02 | 8.64E-02 | 2.48E+00 | 2.54E+00 | 3.08E-01 | no | Rnf165 | intron_variant | MODIFIER |
| 18 | 77573334 | AX-170245262 | 0 | T | A | 0.439 | 8.89E-02 | 8.64E-02 | 2.48E+00 | 2.54E+00 | 3.08E-01 | no | | intergenic_variant | MODIFIER |
| 18 | 77781181 | AX-170088381 | 0 | G | A | 0.439 | 8.89E-02 | 8.64E-02 | 2.48E+00 | 2.54E+00 | 3.08E-01 | no | Atp5a1 | intron_variant | MODIFIER |
| 18 | 77781422 | AX-170016036 | 0 | T | C | 0.439 | 8.89E-02 | 8.64E-02 | 2.48E+00 | 2.54E+00 | 3.08E-01 | no | Atp5a1 | intron_variant | MODIFIER |
| 18 | 77784724 | AX-170160681 | 0 | C | T | 0.439 | 8.89E-02 | 8.64E-02 | 2.48E+00 | 2.54E+00 | 3.08E-01 | no | Atp5a1 | downstream_gene_variant | MODIFIER |
| 18 | 77785589 | AX-170061389 | 0 | C | T | 0.439 | 8.89E-02 | 8.64E-02 | 2.48E+00 | 2.54E+00 | 3.08E-01 | no | Atp5a1 | downstream_gene_variant | MODIFIER |
| 18 | 77786896 | AX-170061391 | 0 | C | G | 0.439 | 8.89E-02 | 8.64E-02 | 2.48E+00 | 2.54E+00 | 3.08E-01 | no | Atp5a1 | downstream_gene_variant | MODIFIER |
| 18 | 77800954 | AX-170105183 | 0 | T | C | 0.439 | 8.89E-02 | 8.64E-02 | 2.48E+00 | 2.54E+00 | 3.08E-01 | no | Pstpip2 | intron_variant | MODIFIER |
| 18 | 77804245 | AX-170456323 | 0 | T | C | 0.439 | 8.89E-02 | 8.64E-02 | 2.48E+00 | 2.54E+00 | 3.08E-01 | no | Pstpip2 | intron_variant | MODIFIER |
| 18 | 77814238 | AX-170360763 | 0 | C | A | 0.439 | 8.89E-02 | 8.64E-02 | 2.48E+00 | 2.54E+00 | 3.08E-01 | no | Pstpip2 | intron_variant | MODIFIER |
| 18 | 77815187 | AX-170334993 | 0 | T | G | 0.439 | 8.89E-02 | 8.64E-02 | 2.48E+00 | 2.54E+00 | 3.08E-01 | no | Pstpip2 | intron_variant | MODIFIER |
| 18 | 77816251 | AX-170316993 | 0 | A | C | 0.439 | 8.89E-02 | 8.64E-02 | 2.48E+00 | 2.54E+00 | 3.08E-01 | no | Pstpip2 | intron_variant | MODIFIER |
| 18 | 77816481 | AX-97799820 | 0 | C | T | 0.439 | 8.89E-02 | 8.64E-02 | 2.48E+00 | 2.54E+00 | 3.08E-01 | no | Pstpip2 | intron_variant | MODIFIER |
| 18 | 77847924 | AX-97803738 | 0 | T | C | 0.439 | 8.89E-02 | 8.64E-02 | 2.48E+00 | 2.54E+00 | 3.08E-01 | no | Pstpip2 | intron_variant | MODIFIER |

| | | | | | | | | | | | | | | | |
|---|---|---|---|---|---|---|---|---|---|---|---|---|---|---|---|
| 18 | 77871681 | AX-170386044 | 0 | G | A | 0.439 | 8.89E-02 | 8.64E-02 | 2.48E+00 | 2.54E+00 | 3.08E-01 | no | Pstpip2 | intron_variant | MODIFIER |
| 18 | 77872991 | AX-170535712 | 0 | T | C | 0.439 | 8.89E-02 | 8.64E-02 | 2.48E+00 | 2.54E+00 | 3.08E-01 | no | Pstpip2 | intron_variant | MODIFIER |
| 18 | 77934559 | AX-170452095 | 0 | A | G | 0.439 | 8.89E-02 | 8.64E-02 | 2.48E+00 | 2.54E+00 | 3.08E-01 | no | Epg5 | upstream_gene_variant | MODIFIER |
| 18 | 77939294 | AX-170061392 | 0 | T | C | 0.439 | 8.89E-02 | 8.64E-02 | 2.48E+00 | 2.54E+00 | 3.08E-01 | no | Epg5 | intron_variant | MODIFIER |
| 18 | 77948890 | AX-170103678 | 0 | C | T | 0.439 | 8.89E-02 | 8.64E-02 | 2.48E+00 | 2.54E+00 | 3.08E-01 | no | Epg5 | synonymous_variant | LOW |
| 18 | 77959807 | AX-170408589 | 0 | A | T | 0.439 | 8.89E-02 | 8.64E-02 | 2.48E+00 | 2.54E+00 | 3.08E-01 | no | Epg5 | intron_variant | MODIFIER |
| 18 | 77960007 | AX-97853169 | 0 | T | C | 0.439 | 8.89E-02 | 8.64E-02 | 2.48E+00 | 2.54E+00 | 3.08E-01 | no | Epg5 | intron_variant | MODIFIER |
| 18 | 77960089 | AX-170127640 | 0 | G | A | 0.439 | 8.89E-02 | 8.64E-02 | 2.48E+00 | 2.54E+00 | 3.08E-01 | no | Epg5 | intron_variant | MODIFIER |
| 18 | 77979154 | AX-170447324 | 0 | G | A | 0.439 | 8.89E-02 | 8.64E-02 | 2.48E+00 | 2.54E+00 | 3.08E-01 | no | Epg5 | synonymous_variant | LOW |
| 18 | 77981064 | AX-97806628 | 0 | G | A | 0.439 | 8.89E-02 | 8.64E-02 | 2.48E+00 | 2.54E+00 | 3.08E-01 | no | Epg5 | synonymous_variant | LOW |
| 18 | 77981532 | AX-170414399 | 0 | A | G | 0.439 | 8.89E-02 | 8.64E-02 | 2.48E+00 | 2.54E+00 | 3.08E-01 | no | Epg5 | intron_variant | MODIFIER |
| 18 | 77985611 | AX-170383153 | 0 | T | C | 0.439 | 8.89E-02 | 8.64E-02 | 2.48E+00 | 2.54E+00 | 3.08E-01 | no | Epg5 | intron_variant | MODIFIER |
| 18 | 78014182 | AX-112880240 | 0 | C | T | 0.439 | 8.89E-02 | 8.64E-02 | 2.48E+00 | 2.54E+00 | 3.08E-01 | no | Epg5 | synonymous_variant | LOW |
| 18 | 78092092 | AX-170401198 | 0 | C | T | 0.439 | 8.89E-02 | 8.64E-02 | 2.48E+00 | 2.54E+00 | 3.08E-01 | no | | intergenic_variant | MODIFIER |
| 18 | 78105760 | AX-170016047 | 0 | G | T | 0.439 | 8.89E-02 | 8.64E-02 | 2.48E+00 | 2.54E+00 | 3.08E-01 | no | Slc14a1 | intron_variant | MODIFIER |
| 18 | 78149075 | AX-97836757 | 0 | G | A | 0.439 | 8.89E-02 | 8.64E-02 | 2.48E+00 | 2.54E+00 | 3.08E-01 | no | Slc14a2 | intron_variant | MODIFIER |
| 18 | 78156407 | AX-170016048 | 0 | T | A | 0.439 | 8.89E-02 | 8.64E-02 | 2.48E+00 | 2.54E+00 | 3.08E-01 | no | Slc14a2 | intron_variant | MODIFIER |
| 18 | 78180890 | AX-170061401 | 0 | C | A | 0.439 | 8.89E-02 | 8.64E-02 | 2.48E+00 | 2.54E+00 | 3.08E-01 | no | Slc14a2 | intron_variant | MODIFIER |
| 18 | 78184054 | AX-170215774 | 0 | T | C | 0.439 | 8.89E-02 | 8.64E-02 | 2.48E+00 | 2.54E+00 | 3.08E-01 | no | Slc14a2 | intron_variant | MODIFIER |
| 18 | 78192899 | AX-170357257 | 0 | G | A | 0.439 | 8.89E-02 | 8.64E-02 | 2.48E+00 | 2.54E+00 | 3.08E-01 | no | Slc14a2 | intron_variant | MODIFIER |
| 18 | 78196307 | AX-170151409 | 0 | C | T | 0.439 | 8.89E-02 | 8.64E-02 | 2.48E+00 | 2.54E+00 | 3.08E-01 | no | Slc14a2 | intron_variant | MODIFIER |
| 18 | 78209177 | AX-97835336 | 0 | T | C | 0.439 | 8.89E-02 | 8.64E-02 | 2.48E+00 | 2.54E+00 | 3.08E-01 | no | Slc14a2 | upstream_gene_variant | MODIFIER |
| 18 | 78214348 | AX-170313351 | 0 | T | C | 0.439 | 8.89E-02 | 8.64E-02 | 2.48E+00 | 2.54E+00 | 3.08E-01 | no | Slc14a2 | intron_variant | MODIFIER |
| 18 | 78215007 | AX-170486673 | 0 | C | G | 0.439 | 8.89E-02 | 8.64E-02 | 2.48E+00 | 2.54E+00 | 3.08E-01 | no | Slc14a2 | intron_variant | MODIFIER |
| 18 | 78262785 | AX-170016045 | 0 | G | A | 0.439 | 8.89E-02 | 8.64E-02 | 2.48E+00 | 2.54E+00 | 3.08E-01 | no | Slc14a2 | intron_variant | MODIFIER |
| 18 | 78283047 | AX-170269402 | 0 | T | C | 0.439 | 8.89E-02 | 8.64E-02 | 2.48E+00 | 2.54E+00 | 3.08E-01 | no | Slc14a2 | intron_variant | MODIFIER |
| 18 | 78287667 | AX-170016046 | 0 | A | G | 0.439 | 8.89E-02 | 8.64E-02 | 2.48E+00 | 2.54E+00 | 3.08E-01 | no | Slc14a2 | intron_variant | MODIFIER |
| 18 | 78296341 | AX-170289441 | 0 | C | T | 0.439 | 8.89E-02 | 8.64E-02 | 2.48E+00 | 2.54E+00 | 3.08E-01 | no | Slc14a2 | intron_variant | MODIFIER |
| 18 | 78296906 | AX-97848267 | 0 | C | A | 0.439 | 8.89E-02 | 8.64E-02 | 2.48E+00 | 2.54E+00 | 3.08E-01 | no | Slc14a2 | intron_variant | MODIFIER |
| 2 | 123814144 | AX-170163943 | 1 | C | T | 0.152 | 1.76E-01 | 1.75E-01 | 2.75E+00 | 2.86E+00 | 3.08E-01 | Yes | | regulatory_region_variant | MODIFIER |
| 8 | 90849242 | AX-170304437 | 1 | C | A | 0.196 | -1.14E-01 | 1.13E-01 | 2.66E+00 | 2.75E+00 | 3.08E-01 | no | Chd9 | intron_variant | MODIFIER |
| 1 | 167866408 | AX-170175630 | 0 | T | C | 0.237 | 1.26E-01 | 1.25E-01 | 2.62E+00 | 2.69E+00 | 3.08E-01 | no | | intergenic_variant | MODIFIER |
| 7 | 113937890 | AX-97818240 | 0 | C | T | 0.43 | 9.38E-02 | 9.30E-02 | 2.74E+00 | 2.80E+00 | 3.08E-01 | no | Spon1 | intron_variant | MODIFIER |
| 7 | 37619938 | AX-170462349 | 0 | T | C | 0.342 | 9.40E-02 | 9.26E-02 | 2.58E+00 | 2.66E+00 | 3.08E-01 | no | Zfp536 | intron_variant | MODIFIER |
| 18 | 48957418 | AX-170462931 | 1 | C | A | 0.062 | 1.63E-01 | 1.62E-01 | 2.83E+00 | 3.00E+00 | 3.08E-01 | Yes | | intergenic_variant | MODIFIER |
| 1 | 148642583 | AX-170150228 | 1 | A | G | 0.205 | 1.25E-01 | 1.24E-01 | 2.71E+00 | 2.81E+00 | 3.09E-01 | no | | intergenic_variant | MODIFIER |
| 7 | 12982049 | AX-170242799 | 0 | G | T | 0.36 | 7.89E-02 | 7.76E-02 | 2.54E+00 | 2.66E+00 | 3.09E-01 | no | Zfp446 | intron_variant | MODIFIER |
| 2 | 89997458 | AX-170253335 | 0 | A | G | 0.061 | 2.13E-01 | 2.11E-01 | 2.83E+00 | 2.95E+00 | 3.09E-01 | Yes | Olfr1262 | upstream_gene_variant | MODIFIER |
| 5 | 24182164 | AX-170276114 | 0 | A | G | 0.061 | 2.13E-01 | 2.11E-01 | 2.83E+00 | 2.95E+00 | 3.09E-01 | Yes | Nupl2 | synonymous_variant | LOW |
| 13 | 39957677 | AX-170480533 | 0 | T | G | 0.167 | -1.12E-01 | 1.07E-01 | 2.41E+00 | 2.50E+00 | 3.09E-01 | no | | intergenic_variant | MODIFIER |
| 13 | 40691897 | AX-168410942 | 0 | C | T | 0.167 | -1.12E-01 | 1.07E-01 | 2.41E+00 | 2.50E+00 | 3.09E-01 | no | | intergenic_variant | MODIFIER |
| 18 | 88170284 | AX-170061624 | 1 | C | A | 0.161 | -1.09E-01 | 1.06E-01 | 2.53E+00 | 2.62E+00 | 3.09E-01 | no | | intergenic_variant | MODIFIER |
| 16 | 33113659 | AX-170514979 | 1 | T | A | 0.482 | 8.20E-02 | 8.05E-02 | 2.55E+00 | 2.60E+00 | 3.09E-01 | no | Lmln | intron_variant | MODIFIER |
| 3 | 27620836 | AX-97809031 | 0 | C | T | 0.132 | -1.04E-01 | 1.03E-01 | 2.62E+00 | 2.70E+00 | 3.09E-01 | no | Fndc3b | intron_variant | MODIFIER |
| 3 | 27621318 | AX-170113515 | 0 | A | G | 0.132 | -1.04E-01 | 1.03E-01 | 2.62E+00 | 2.70E+00 | 3.09E-01 | no | Fndc3b | intron_variant | MODIFIER |
| 9 | 72449804 | AX-170175483 | 0 | G | A | 0.193 | -1.29E-01 | 1.28E-01 | 2.79E+00 | 2.85E+00 | 3.09E-01 | no | Mns1 | intron_variant | MODIFIER |
| 7 | 145210883 | AX-170099156 | 1 | C | T | 0.205 | -7.56E-02 | 7.52E-02 | 2.65E+00 | 2.78E+00 | 3.09E-01 | no | 1810010D01Rik | upstream_gene_variant | MODIFIER |
| 9 | 64456672 | AX-170032546 | 0 | A | C | 0.193 | -1.16E-01 | 1.15E-01 | 2.82E+00 | 2.94E+00 | 3.10E-01 | no | Megf11 | intron_variant | MODIFIER |
| 9 | 77051520 | AX-170510541 | 0 | A | G | 0.193 | -1.16E-01 | 1.15E-01 | 2.82E+00 | 2.94E+00 | 3.10E-01 | no | Gm19541 | non_coding_transcript_exon_variant | MODIFIER |
| 9 | 77496363 | AX-170259037 | 0 | T | A | 0.193 | -1.16E-01 | 1.15E-01 | 2.82E+00 | 2.94E+00 | 3.10E-01 | no | Lrrc1 | intron_variant | MODIFIER |
| 15 | 15378847 | AX-97843165 | 2 | G | T | 0.227 | -1.06E-01 | 1.05E-01 | 2.63E+00 | 2.68E+00 | 3.10E-01 | no | | intergenic_variant | MODIFIER |
| 18 | 59564887 | AX-170442199 | 1 | T | G | 0.411 | 9.77E-02 | 9.38E-02 | 2.41E+00 | 2.45E+00 | 3.10E-01 | no | | intergenic_variant | MODIFIER |
| 18 | 67482743 | AX-170445666 | 0 | G | A | 0.281 | -9.99E-02 | 9.93E-02 | 2.78E+00 | 2.85E+00 | 3.10E-01 | no | Prelid3a | downstream_gene_variant | MODIFIER |
| 18 | 77379821 | AX-170325756 | 1 | C | T | 0.446 | 8.86E-02 | 8.58E-02 | 2.45E+00 | 2.51E+00 | 3.10E-01 | no | Loxhd1 | intron_variant,non_coding_transcript_variant | MODIFIER |

| | | | | | | | | | | | | | | | |
|---|---|---|---|---|---|---|---|---|---|---|---|---|---|---|---|
| 5 | 45841825 | AX-170281304 | 0 | T | C | 0.491 | 1.04E-01 | 9.97E-02 | 2.40E+00 | 2.45E+00 | 3.10E-01 | no | Lcorl | intron_variant | MODIFIER |
| 5 | 106577382 | AX-170120836 | 0 | A | G | 0.035 | -1.38E-01 | 1.38E-01 | 2.71E+00 | 2.90E+00 | 3.10E-01 | no | Gm28050 | upstream_gene_variant | MODIFIER |
| 5 | 46060814 | AX-170271177 | 0 | C | G | 0.482 | 1.07E-01 | 1.03E-01 | 2.39E+00 | 2.44E+00 | 3.10E-01 | no | | intergenic_variant | MODIFIER |
| 5 | 46158476 | AX-97804470 | 0 | C | T | 0.482 | 1.07E-01 | 1.03E-01 | 2.39E+00 | 2.44E+00 | 3.10E-01 | no | | intergenic_variant | MODIFIER |
| 5 | 46180392 | AX-97854401 | 0 | G | A | 0.482 | 1.07E-01 | 1.03E-01 | 2.39E+00 | 2.44E+00 | 3.10E-01 | no | | intergenic_variant | MODIFIER |
| 5 | 46206986 | AX-170476196 | 0 | G | A | 0.482 | 1.07E-01 | 1.03E-01 | 2.39E+00 | 2.44E+00 | 3.10E-01 | no | | intergenic_variant | MODIFIER |
| 5 | 46220435 | AX-170285667 | 0 | C | T | 0.482 | 1.07E-01 | 1.03E-01 | 2.39E+00 | 2.44E+00 | 3.10E-01 | no | | intergenic_variant | MODIFIER |
| 5 | 46220766 | AX-170159561 | 0 | A | T | 0.482 | 1.07E-01 | 1.03E-01 | 2.39E+00 | 2.44E+00 | 3.10E-01 | no | | intergenic_variant | MODIFIER |
| 5 | 46256444 | AX-170287758 | 0 | C | T | 0.482 | 1.07E-01 | 1.03E-01 | 2.39E+00 | 2.44E+00 | 3.10E-01 | no | | intergenic_variant | MODIFIER |
| 5 | 46293187 | AX-170532781 | 0 | A | T | 0.482 | 1.07E-01 | 1.03E-01 | 2.39E+00 | 2.44E+00 | 3.10E-01 | no | | intergenic_variant | MODIFIER |
| 5 | 46338230 | AX-170436878 | 0 | A | G | 0.482 | 1.07E-01 | 1.03E-01 | 2.39E+00 | 2.44E+00 | 3.10E-01 | no | | intergenic_variant | MODIFIER |
| 5 | 46351481 | AX-97790412 | 0 | A | G | 0.482 | 1.07E-01 | 1.03E-01 | 2.39E+00 | 2.44E+00 | 3.10E-01 | no | | intergenic_variant | MODIFIER |
| 5 | 46418442 | AX-97796179 | 0 | A | C | 0.482 | 1.07E-01 | 1.03E-01 | 2.39E+00 | 2.44E+00 | 3.10E-01 | no | Gm7931 | downstream_gene_variant | MODIFIER |
| 5 | 46422510 | AX-170089445 | 0 | A | G | 0.482 | 1.07E-01 | 1.03E-01 | 2.39E+00 | 2.44E+00 | 3.10E-01 | no | | intergenic_variant | MODIFIER |
| 5 | 46442742 | AX-170202360 | 0 | T | C | 0.482 | 1.07E-01 | 1.03E-01 | 2.39E+00 | 2.44E+00 | 3.10E-01 | no | | intergenic_variant | MODIFIER |
| 5 | 46449693 | AX-97815252 | 0 | A | G | 0.482 | 1.07E-01 | 1.03E-01 | 2.39E+00 | 2.44E+00 | 3.10E-01 | no | | intergenic_variant | MODIFIER |
| 5 | 46545073 | AX-170432939 | 0 | G | A | 0.482 | 1.07E-01 | 1.03E-01 | 2.39E+00 | 2.44E+00 | 3.10E-01 | no | | intergenic_variant | MODIFIER |
| 5 | 46575988 | AX-97783778 | 0 | G | A | 0.482 | 1.07E-01 | 1.03E-01 | 2.39E+00 | 2.44E+00 | 3.10E-01 | no | | intergenic_variant | MODIFIER |
| 5 | 46610595 | AX-170222617 | 0 | A | G | 0.482 | 1.07E-01 | 1.03E-01 | 2.39E+00 | 2.44E+00 | 3.10E-01 | no | | intergenic_variant | MODIFIER |
| 5 | 46642030 | AX-170095309 | 0 | A | G | 0.482 | 1.07E-01 | 1.03E-01 | 2.39E+00 | 2.44E+00 | 3.10E-01 | no | | intergenic_variant | MODIFIER |
| 5 | 46647996 | AX-170213825 | 0 | A | C | 0.482 | 1.07E-01 | 1.03E-01 | 2.39E+00 | 2.44E+00 | 3.10E-01 | no | | intergenic_variant | MODIFIER |
| 5 | 46660995 | AX-97843355 | 0 | C | T | 0.482 | 1.07E-01 | 1.03E-01 | 2.39E+00 | 2.44E+00 | 3.10E-01 | no | | intergenic_variant | MODIFIER |
| 5 | 46969179 | AX-170071983 | 0 | C | A | 0.482 | 1.07E-01 | 1.03E-01 | 2.39E+00 | 2.44E+00 | 3.10E-01 | no | | intergenic_variant | MODIFIER |
| 5 | 47023108 | AX-170223207 | 0 | A | T | 0.482 | 1.07E-01 | 1.03E-01 | 2.39E+00 | 2.44E+00 | 3.10E-01 | no | | intergenic_variant | MODIFIER |
| 5 | 47054726 | AX-170505237 | 0 | T | G | 0.482 | 1.07E-01 | 1.03E-01 | 2.39E+00 | 2.44E+00 | 3.10E-01 | no | | intergenic_variant | MODIFIER |
| 5 | 47111361 | AX-97842098 | 0 | C | A | 0.482 | 1.07E-01 | 1.03E-01 | 2.39E+00 | 2.44E+00 | 3.10E-01 | no | | intergenic_variant | MODIFIER |
| 5 | 47132871 | AX-170433362 | 0 | C | G | 0.482 | 1.07E-01 | 1.03E-01 | 2.39E+00 | 2.44E+00 | 3.10E-01 | no | | intergenic_variant | MODIFIER |
| 5 | 47133459 | AX-170071988 | 0 | A | G | 0.482 | 1.07E-01 | 1.03E-01 | 2.39E+00 | 2.44E+00 | 3.10E-01 | no | | intergenic_variant | MODIFIER |
| 5 | 47545554 | AX-170185756 | 0 | C | T | 0.482 | 1.07E-01 | 1.03E-01 | 2.39E+00 | 2.44E+00 | 3.10E-01 | no | | intergenic_variant | MODIFIER |
| 5 | 44799009 | AX-97846345 | 0 | C | T | 0.202 | -1.08E-01 | 1.07E-01 | 2.58E+00 | 2.66E+00 | 3.10E-01 | no | Ldb2 | intron_variant | MODIFIER |
| 1 | 124639534 | AX-170207554 | 1 | A | T | 0.571 | 7.92E-02 | 7.82E-02 | 2.57E+00 | 2.65E+00 | 3.10E-01 | no | Gm28928 | intron_variant,non_coding_transcript_variant | MODIFIER |
| 10 | 72810220 | AX-170395508 | 1 | G | C | 0.411 | -8.10E-02 | 8.05E-02 | 2.67E+00 | 2.74E+00 | 3.10E-01 | no | | regulatory_region_variant | MODIFIER |
| 6 | 135097298 | AX-170205406 | 0 | T | C | 0.579 | 9.14E-02 | 8.84E-02 | 2.44E+00 | 2.48E+00 | 3.10E-01 | no | | intergenic_variant | MODIFIER |
| 6 | 135099270 | AX-97831362 | 0 | G | A | 0.579 | 9.14E-02 | 8.84E-02 | 2.44E+00 | 2.48E+00 | 3.10E-01 | no | | intergenic_variant | MODIFIER |
| 6 | 135105083 | AX-170076317 | 0 | C | A | 0.579 | 9.14E-02 | 8.84E-02 | 2.44E+00 | 2.48E+00 | 3.10E-01 | no | Gprc5d | downstream_gene_variant | MODIFIER |
| 6 | 135117074 | AX-97839359 | 0 | T | C | 0.579 | 9.14E-02 | 8.84E-02 | 2.44E+00 | 2.48E+00 | 3.10E-01 | no | Gprc5d | intron_variant | MODIFIER |
| 6 | 135118752 | AX-170280573 | 0 | C | T | 0.579 | 9.14E-02 | 8.84E-02 | 2.44E+00 | 2.48E+00 | 3.10E-01 | no | Gprc5d | upstream_gene_variant | MODIFIER |
| 6 | 135135284 | AX-170309364 | 0 | A | G | 0.579 | 9.14E-02 | 8.84E-02 | 2.44E+00 | 2.48E+00 | 3.10E-01 | no | Hebp1 | downstream_gene_variant | MODIFIER |
| 6 | 135137961 | AX-170427559 | 0 | G | A | 0.579 | 9.14E-02 | 8.84E-02 | 2.44E+00 | 2.48E+00 | 3.10E-01 | no | Hebp1 | synonymous_variant | LOW |
| 6 | 135138751 | AX-170239153 | 0 | A | T | 0.579 | 9.14E-02 | 8.84E-02 | 2.44E+00 | 2.48E+00 | 3.10E-01 | no | Hebp1 | intron_variant | MODIFIER |
| 6 | 135139386 | AX-170279646 | 0 | T | C | 0.579 | 9.14E-02 | 8.84E-02 | 2.44E+00 | 2.48E+00 | 3.10E-01 | no | Hebp1 | intron_variant | MODIFIER |
| 6 | 135154659 | AX-170501075 | 0 | G | C | 0.579 | 9.14E-02 | 8.84E-02 | 2.44E+00 | 2.48E+00 | 3.10E-01 | no | Hebp1 | intron_variant | MODIFIER |
| 6 | 135161223 | AX-170533489 | 0 | T | C | 0.579 | 9.14E-02 | 8.84E-02 | 2.44E+00 | 2.48E+00 | 3.10E-01 | no | Hebp1 | intron_variant | MODIFIER |
| 6 | 135205309 | AX-170134563 | 0 | C | T | 0.579 | 9.14E-02 | 8.84E-02 | 2.44E+00 | 2.48E+00 | 3.10E-01 | no | Fam234b | intron_variant | MODIFIER |
| 6 | 135217254 | AX-97821944 | 0 | A | G | 0.579 | 9.14E-02 | 8.84E-02 | 2.44E+00 | 2.48E+00 | 3.10E-01 | no | Fam234b | intron_variant | MODIFIER |
| 6 | 135217423 | AX-170460776 | 0 | G | T | 0.579 | 9.14E-02 | 8.84E-02 | 2.44E+00 | 2.48E+00 | 3.10E-01 | no | Fam234b | intron_variant | MODIFIER |
| 6 | 135217647 | AX-170076318 | 0 | C | T | 0.579 | 9.14E-02 | 8.84E-02 | 2.44E+00 | 2.48E+00 | 3.10E-01 | no | Fam234b | intron_variant | MODIFIER |
| 6 | 135276891 | AX-170313312 | 0 | T | C | 0.579 | 9.14E-02 | 8.84E-02 | 2.44E+00 | 2.48E+00 | 3.10E-01 | no | | regulatory_region_variant | MODIFIER |
| 6 | 135294445 | AX-170309435 | 0 | A | G | 0.579 | 9.14E-02 | 8.84E-02 | 2.44E+00 | 2.48E+00 | 3.10E-01 | no | Gm36640 | intron_variant,non_coding_transcript_variant | MODIFIER |
| 6 | 135304543 | AX-170477058 | 0 | G | A | 0.579 | 9.14E-02 | 8.84E-02 | 2.44E+00 | 2.48E+00 | 3.10E-01 | no | Pbp2 | downstream_gene_variant | MODIFIER |
| 6 | 135306558 | AX-97852273 | 0 | G | A | 0.579 | 9.14E-02 | 8.84E-02 | 2.44E+00 | 2.48E+00 | 3.10E-01 | no | Pbp2 | downstream_gene_variant | MODIFIER |
| 6 | 135326258 | AX-170028316 | 0 | A | C | 0.579 | 9.14E-02 | 8.84E-02 | 2.44E+00 | 2.48E+00 | 3.10E-01 | no | Gm36640 | intron_variant,non_coding_transcript_variant | MODIFIER |
| 6 | 135336016 | AX-170518516 | 0 | C | T | 0.579 | 9.14E-02 | 8.84E-02 | 2.44E+00 | 2.48E+00 | 3.10E-01 | no | Gm36640 | intron_variant,non_coding_transcript_variant | MODIFIER |
| 6 | 135339201 | AX-170106543 | 0 | G | A | 0.579 | 9.14E-02 | 8.84E-02 | 2.44E+00 | 2.48E+00 | 3.10E-01 | no | Gm36640 | non_coding_transcript_exon_variant | MODIFIER |

| | | | | | | | | | | | | | | | |
|---|---|---|---|---|---|---|---|---|---|---|---|---|---|---|---|
| 6 | 135340348 | AX-170521297 | 0 | C | T | 0.579 | 9.14E-02 | 8.84E-02 | 2.44E+00 | 2.48E+00 | 3.10E-01 | no | Gm36640 | non_coding_transcript_exon_variant | MODIFIER |
| 6 | 135352845 | AX-170445647 | 0 | T | C | 0.579 | 9.14E-02 | 8.84E-02 | 2.44E+00 | 2.48E+00 | 3.10E-01 | no | Dynlt1-ps1 | upstream_gene_variant | MODIFIER |
| 6 | 135357898 | AX-170502290 | 0 | T | C | 0.579 | 9.14E-02 | 8.84E-02 | 2.44E+00 | 2.48E+00 | 3.10E-01 | no | Emp1 | upstream_gene_variant | MODIFIER |
| 6 | 135366673 | AX-170028309 | 0 | A | G | 0.579 | 9.14E-02 | 8.84E-02 | 2.44E+00 | 2.48E+00 | 3.10E-01 | no | Emp1 | intron_variant | MODIFIER |
| 6 | 135366819 | AX-170455481 | 0 | T | C | 0.579 | 9.14E-02 | 8.84E-02 | 2.44E+00 | 2.48E+00 | 3.10E-01 | no | Emp1 | intron_variant | MODIFIER |
| 6 | 135379748 | AX-170530729 | 0 | T | G | 0.579 | 9.14E-02 | 8.84E-02 | 2.44E+00 | 2.48E+00 | 3.10E-01 | no | Emp1 | intron_variant | MODIFIER |
| 6 | 135382175 | AX-170366668 | 0 | G | A | 0.579 | 9.14E-02 | 8.84E-02 | 2.44E+00 | 2.48E+00 | 3.10E-01 | no | Emp1 | 3_prime_UTR_variant | MODIFIER |
| 6 | 135382347 | AX-170124488 | 0 | A | G | 0.579 | 9.14E-02 | 8.84E-02 | 2.44E+00 | 2.48E+00 | 3.10E-01 | no | Emp1 | 3_prime_UTR_variant | MODIFIER |
| 6 | 135386007 | AX-170212269 | 0 | C | T | 0.579 | 9.14E-02 | 8.84E-02 | 2.44E+00 | 2.48E+00 | 3.10E-01 | no | Emp1 | downstream_gene_variant | MODIFIER |
| 6 | 135386541 | AX-170385924 | 0 | G | A | 0.579 | 9.14E-02 | 8.84E-02 | 2.44E+00 | 2.48E+00 | 3.10E-01 | no | Emp1 | downstream_gene_variant | MODIFIER |
| 6 | 135392528 | AX-170174472 | 0 | C | T | 0.579 | 9.14E-02 | 8.84E-02 | 2.44E+00 | 2.48E+00 | 3.10E-01 | no | | intergenic_variant | MODIFIER |
| 6 | 135402234 | AX-170028318 | 0 | A | G | 0.579 | 9.14E-02 | 8.84E-02 | 2.44E+00 | 2.48E+00 | 3.10E-01 | no | | intergenic_variant | MODIFIER |
| 6 | 135433462 | AX-170239729 | 0 | C | T | 0.579 | 9.14E-02 | 8.84E-02 | 2.44E+00 | 2.48E+00 | 3.10E-01 | no | | intergenic_variant | MODIFIER |
| 6 | 135435436 | AX-170497277 | 0 | C | T | 0.579 | 9.14E-02 | 8.84E-02 | 2.44E+00 | 2.48E+00 | 3.10E-01 | no | | intergenic_variant | MODIFIER |
| 6 | 135436426 | AX-170387258 | 0 | G | A | 0.579 | 9.14E-02 | 8.84E-02 | 2.44E+00 | 2.48E+00 | 3.10E-01 | no | | intergenic_variant | MODIFIER |
| 6 | 135444936 | AX-170368159 | 0 | T | C | 0.579 | 9.14E-02 | 8.84E-02 | 2.44E+00 | 2.48E+00 | 3.10E-01 | no | | regulatory_region_variant | MODIFIER |
| 6 | 135464866 | AX-170207370 | 0 | G | A | 0.579 | 9.14E-02 | 8.84E-02 | 2.44E+00 | 2.48E+00 | 3.10E-01 | no | | intergenic_variant | MODIFIER |
| 6 | 135478129 | AX-170076323 | 0 | C | A | 0.579 | 9.14E-02 | 8.84E-02 | 2.44E+00 | 2.48E+00 | 3.10E-01 | no | | intergenic_variant | MODIFIER |
| 6 | 135482415 | AX-170164478 | 0 | G | A | 0.579 | 9.14E-02 | 8.84E-02 | 2.44E+00 | 2.48E+00 | 3.10E-01 | no | | intergenic_variant | MODIFIER |
| 6 | 135501172 | AX-170283259 | 0 | G | C | 0.579 | 9.14E-02 | 8.84E-02 | 2.44E+00 | 2.48E+00 | 3.10E-01 | no | | intergenic_variant | MODIFIER |
| 6 | 135509882 | AX-97786161 | 0 | A | G | 0.579 | 9.14E-02 | 8.84E-02 | 2.44E+00 | 2.48E+00 | 3.10E-01 | no | | intergenic_variant | MODIFIER |
| 6 | 135520319 | AX-170332118 | 0 | T | C | 0.579 | 9.14E-02 | 8.84E-02 | 2.44E+00 | 2.48E+00 | 3.10E-01 | no | | regulatory_region_variant | MODIFIER |
| 6 | 135523080 | AX-170497492 | 0 | G | C | 0.579 | 9.14E-02 | 8.84E-02 | 2.44E+00 | 2.48E+00 | 3.10E-01 | no | | intergenic_variant | MODIFIER |
| 6 | 135533093 | AX-170217322 | 0 | T | C | 0.579 | 9.14E-02 | 8.84E-02 | 2.44E+00 | 2.48E+00 | 3.10E-01 | no | | intergenic_variant | MODIFIER |
| 6 | 135542053 | AX-97825067 | 0 | T | G | 0.579 | 9.14E-02 | 8.84E-02 | 2.44E+00 | 2.48E+00 | 3.10E-01 | no | | regulatory_region_variant | MODIFIER |
| 6 | 135546699 | AX-170114021 | 0 | A | C | 0.579 | 9.14E-02 | 8.84E-02 | 2.44E+00 | 2.48E+00 | 3.10E-01 | no | | intergenic_variant | MODIFIER |
| 6 | 135559032 | AX-170385153 | 0 | T | C | 0.579 | 9.14E-02 | 8.84E-02 | 2.44E+00 | 2.48E+00 | 3.10E-01 | no | | regulatory_region_variant | MODIFIER |
| 6 | 135567415 | AX-170490972 | 0 | A | G | 0.579 | 9.14E-02 | 8.84E-02 | 2.44E+00 | 2.48E+00 | 3.10E-01 | no | | intergenic_variant | MODIFIER |
| 6 | 135567465 | AX-170515868 | 0 | A | C | 0.579 | 9.14E-02 | 8.84E-02 | 2.44E+00 | 2.48E+00 | 3.10E-01 | no | | intergenic_variant | MODIFIER |
| 6 | 135577694 | AX-170243464 | 0 | C | T | 0.579 | 9.14E-02 | 8.84E-02 | 2.44E+00 | 2.48E+00 | 3.10E-01 | no | | intergenic_variant | MODIFIER |
| 6 | 135586158 | AX-170301484 | 0 | A | C | 0.579 | 9.14E-02 | 8.84E-02 | 2.44E+00 | 2.48E+00 | 3.10E-01 | no | Gm25136 | downstream_gene_variant | MODIFIER |
| 6 | 135604320 | AX-170328968 | 0 | T | A | 0.579 | 9.14E-02 | 8.84E-02 | 2.44E+00 | 2.48E+00 | 3.10E-01 | no | | intergenic_variant | MODIFIER |
| 6 | 135609111 | AX-170227253 | 0 | G | T | 0.579 | 9.14E-02 | 8.84E-02 | 2.44E+00 | 2.48E+00 | 3.10E-01 | no | | intergenic_variant | MODIFIER |
| 6 | 135627273 | AX-170076327 | 0 | T | C | 0.579 | 9.14E-02 | 8.84E-02 | 2.44E+00 | 2.48E+00 | 3.10E-01 | no | | intergenic_variant | MODIFIER |
| 6 | 135629891 | AX-170255089 | 0 | G | T | 0.579 | 9.14E-02 | 8.84E-02 | 2.44E+00 | 2.48E+00 | 3.10E-01 | no | | intergenic_variant | MODIFIER |
| 6 | 135657068 | AX-170119889 | 0 | T | C | 0.579 | 9.14E-02 | 8.84E-02 | 2.44E+00 | 2.48E+00 | 3.10E-01 | no | 4930425L21Rik | upstream_gene_variant | MODIFIER |
| 6 | 135657330 | AX-170156972 | 0 | T | C | 0.579 | 9.14E-02 | 8.84E-02 | 2.44E+00 | 2.48E+00 | 3.10E-01 | no | 4930425L21Rik | upstream_gene_variant | MODIFIER |
| 6 | 135659034 | AX-170107762 | 0 | C | T | 0.579 | 9.14E-02 | 8.84E-02 | 2.44E+00 | 2.48E+00 | 3.10E-01 | no | 4930425L21Rik | non_coding_transcript_exon_variant | MODIFIER |
| 6 | 135659453 | AX-170123184 | 0 | C | T | 0.579 | 9.14E-02 | 8.84E-02 | 2.44E+00 | 2.48E+00 | 3.10E-01 | no | 4930425L21Rik | intron_variant,non_coding_transcript_variant | MODIFIER |
| 6 | 135663038 | AX-170285527 | 0 | C | T | 0.579 | 9.14E-02 | 8.84E-02 | 2.44E+00 | 2.48E+00 | 3.10E-01 | no | 4930425L21Rik | intron_variant,non_coding_transcript_variant | MODIFIER |
| 6 | 135663550 | AX-170313932 | 0 | T | C | 0.579 | 9.14E-02 | 8.84E-02 | 2.44E+00 | 2.48E+00 | 3.10E-01 | no | 4930425L21Rik | intron_variant,non_coding_transcript_variant | MODIFIER |
| 6 | 135663683 | AX-170481292 | 0 | A | C | 0.579 | 9.14E-02 | 8.84E-02 | 2.44E+00 | 2.48E+00 | 3.10E-01 | no | 4930425L21Rik | intron_variant,non_coding_transcript_variant | MODIFIER |
| 6 | 135672788 | AX-170215686 | 0 | G | A | 0.579 | 9.14E-02 | 8.84E-02 | 2.44E+00 | 2.48E+00 | 3.10E-01 | no | 4930425L21Rik | intron_variant,non_coding_transcript_variant | MODIFIER |
| 6 | 135680160 | AX-170174243 | 0 | C | T | 0.579 | 9.14E-02 | 8.84E-02 | 2.44E+00 | 2.48E+00 | 3.10E-01 | no | 4930425L21Rik | non_coding_transcript_exon_variant | MODIFIER |
| 6 | 135690829 | AX-170248447 | 0 | T | C | 0.579 | 9.14E-02 | 8.84E-02 | 2.44E+00 | 2.48E+00 | 3.10E-01 | no | | intergenic_variant | MODIFIER |
| 6 | 135702532 | AX-170336962 | 0 | A | T | 0.579 | 9.14E-02 | 8.84E-02 | 2.44E+00 | 2.48E+00 | 3.10E-01 | no | | intergenic_variant | MODIFIER |
| 6 | 135714067 | AX-170180169 | 0 | G | A | 0.579 | 9.14E-02 | 8.84E-02 | 2.44E+00 | 2.48E+00 | 3.10E-01 | no | Grin2b | 3_prime_UTR_variant | MODIFIER |
| 6 | 135726527 | AX-170256029 | 0 | T | C | 0.579 | 9.14E-02 | 8.84E-02 | 2.44E+00 | 2.48E+00 | 3.10E-01 | no | Grin2b | 3_prime_UTR_variant | MODIFIER |
| 6 | 135738896 | AX-170492574 | 0 | C | G | 0.579 | 9.14E-02 | 8.84E-02 | 2.44E+00 | 2.48E+00 | 3.10E-01 | no | Grin2b | intron_variant | MODIFIER |
| 6 | 135749133 | AX-170432859 | 0 | A | G | 0.579 | 9.14E-02 | 8.84E-02 | 2.44E+00 | 2.48E+00 | 3.10E-01 | no | Grin2b | intron_variant | MODIFIER |
| 6 | 135751605 | AX-170028320 | 0 | C | G | 0.579 | 9.14E-02 | 8.84E-02 | 2.44E+00 | 2.48E+00 | 3.10E-01 | no | Grin2b | intron_variant | MODIFIER |
| 6 | 135756678 | AX-170214267 | 0 | A | G | 0.579 | 9.14E-02 | 8.84E-02 | 2.44E+00 | 2.48E+00 | 3.10E-01 | no | Grin2b | intron_variant | MODIFIER |
| 6 | 135771704 | AX-170028319 | 0 | T | C | 0.579 | 9.14E-02 | 8.84E-02 | 2.44E+00 | 2.48E+00 | 3.10E-01 | no | Grin2b | intron_variant | MODIFIER |
| 6 | 135771721 | AX-170028321 | 0 | T | A | 0.579 | 9.14E-02 | 8.84E-02 | 2.44E+00 | 2.48E+00 | 3.10E-01 | no | Grin2b | intron_variant | MODIFIER |

| | | | | | | | | | | | | | | | |
|---|---|---|---|---|---|---|---|---|---|---|---|---|---|---|---|
| 6 | 135780274 | AX-170209812 | 0 | T | A | 0.579 | 9.14E-02 | 8.84E-02 | 2.44E+00 | 2.48E+00 | 3.10E-01 | no | Grin2b | synonymous_variant | LOW |
| 6 | 135783515 | AX-97821551 | 0 | A | G | 0.579 | 9.14E-02 | 8.84E-02 | 2.44E+00 | 2.48E+00 | 3.10E-01 | no | Grin2b | intron_variant | MODIFIER |
| 6 | 135817843 | AX-170212950 | 0 | C | T | 0.579 | 9.14E-02 | 8.84E-02 | 2.44E+00 | 2.48E+00 | 3.10E-01 | no | Grin2b | intron_variant | MODIFIER |
| 6 | 135822391 | AX-170395282 | 0 | C | T | 0.579 | 9.14E-02 | 8.84E-02 | 2.44E+00 | 2.48E+00 | 3.10E-01 | no | Grin2b | intron_variant | MODIFIER |
| 6 | 135824437 | AX-170143000 | 0 | T | C | 0.579 | 9.14E-02 | 8.84E-02 | 2.44E+00 | 2.48E+00 | 3.10E-01 | no | Grin2b | intron_variant | MODIFIER |
| 6 | 135833623 | AX-170499342 | 0 | G | A | 0.579 | 9.14E-02 | 8.84E-02 | 2.44E+00 | 2.48E+00 | 3.10E-01 | no | Grin2b | intron_variant | MODIFIER |
| 6 | 135838129 | AX-170368505 | 0 | C | T | 0.579 | 9.14E-02 | 8.84E-02 | 2.44E+00 | 2.48E+00 | 3.10E-01 | no | Grin2b | intron_variant | MODIFIER |
| 6 | 135838765 | AX-170403126 | 0 | T | C | 0.579 | 9.14E-02 | 8.84E-02 | 2.44E+00 | 2.48E+00 | 3.10E-01 | no | Grin2b | intron_variant | MODIFIER |
| 6 | 135844143 | AX-170377254 | 0 | T | C | 0.579 | 9.14E-02 | 8.84E-02 | 2.44E+00 | 2.48E+00 | 3.10E-01 | no | Grin2b | intron_variant | MODIFIER |
| 6 | 135844424 | AX-170076334 | 0 | T | C | 0.579 | 9.14E-02 | 8.84E-02 | 2.44E+00 | 2.48E+00 | 3.10E-01 | no | Grin2b | intron_variant | MODIFIER |
| 6 | 135853395 | AX-170076331 | 0 | G | C | 0.579 | 9.14E-02 | 8.84E-02 | 2.44E+00 | 2.48E+00 | 3.10E-01 | no | Grin2b | intron_variant | MODIFIER |
| 6 | 135867103 | AX-170282436 | 0 | A | C | 0.579 | 9.14E-02 | 8.84E-02 | 2.44E+00 | 2.48E+00 | 3.10E-01 | no | Grin2b | intron_variant | MODIFIER |
| 6 | 135877608 | AX-170251684 | 0 | T | C | 0.579 | 9.14E-02 | 8.84E-02 | 2.44E+00 | 2.48E+00 | 3.10E-01 | no | Grin2b | intron_variant | MODIFIER |
| 6 | 135889585 | AX-170144528 | 0 | T | A | 0.579 | 9.14E-02 | 8.84E-02 | 2.44E+00 | 2.48E+00 | 3.10E-01 | no | Grin2b | intron_variant | MODIFIER |
| 6 | 135889708 | AX-170325012 | 0 | T | C | 0.579 | 9.14E-02 | 8.84E-02 | 2.44E+00 | 2.48E+00 | 3.10E-01 | no | Grin2b | intron_variant | MODIFIER |
| 6 | 135904994 | AX-170412510 | 0 | A | G | 0.579 | 9.14E-02 | 8.84E-02 | 2.44E+00 | 2.48E+00 | 3.10E-01 | no | Grin2b | intron_variant | MODIFIER |
| 6 | 135934182 | AX-97794777 | 0 | G | A | 0.579 | 9.14E-02 | 8.84E-02 | 2.44E+00 | 2.48E+00 | 3.10E-01 | no | Grin2b | intron_variant | MODIFIER |
| 6 | 135938086 | AX-170393078 | 0 | T | C | 0.579 | 9.14E-02 | 8.84E-02 | 2.44E+00 | 2.48E+00 | 3.10E-01 | no | Grin2b | intron_variant | MODIFIER |
| 6 | 135947674 | AX-170451541 | 0 | T | C | 0.579 | 9.14E-02 | 8.84E-02 | 2.44E+00 | 2.48E+00 | 3.10E-01 | no | Grin2b | intron_variant | MODIFIER |
| 6 | 135951859 | AX-170210476 | 0 | C | T | 0.579 | 9.14E-02 | 8.84E-02 | 2.44E+00 | 2.48E+00 | 3.10E-01 | no | Grin2b | intron_variant | MODIFIER |
| 6 | 135964679 | AX-170188844 | 0 | G | A | 0.579 | 9.14E-02 | 8.84E-02 | 2.44E+00 | 2.48E+00 | 3.10E-01 | no | Grin2b | intron_variant | MODIFIER |
| 6 | 135965077 | AX-170406150 | 0 | C | T | 0.579 | 9.14E-02 | 8.84E-02 | 2.44E+00 | 2.48E+00 | 3.10E-01 | no | Grin2b | intron_variant | MODIFIER |
| 6 | 136015058 | AX-97815062 | 0 | T | A | 0.579 | 9.14E-02 | 8.84E-02 | 2.44E+00 | 2.48E+00 | 3.10E-01 | no | Grin2b | intron_variant | MODIFIER |
| 6 | 136019491 | AX-170393829 | 0 | T | C | 0.579 | 9.14E-02 | 8.84E-02 | 2.44E+00 | 2.48E+00 | 3.10E-01 | no | Grin2b | intron_variant | MODIFIER |
| 6 | 136024469 | AX-170297139 | 0 | G | A | 0.579 | 9.14E-02 | 8.84E-02 | 2.44E+00 | 2.48E+00 | 3.10E-01 | no | Grin2b | intron_variant | MODIFIER |
| 6 | 136044399 | AX-170493654 | 0 | G | A | 0.579 | 9.14E-02 | 8.84E-02 | 2.44E+00 | 2.48E+00 | 3.10E-01 | no | Grin2b | intron_variant | MODIFIER |
| 6 | 136046506 | AX-170334771 | 0 | T | G | 0.579 | 9.14E-02 | 8.84E-02 | 2.44E+00 | 2.48E+00 | 3.10E-01 | no | Grin2b | intron_variant | MODIFIER |
| 6 | 136046695 | AX-170281208 | 0 | C | T | 0.579 | 9.14E-02 | 8.84E-02 | 2.44E+00 | 2.48E+00 | 3.10E-01 | no | Grin2b | intron_variant | MODIFIER |
| 6 | 136057926 | AX-97794678 | 0 | T | C | 0.579 | 9.14E-02 | 8.84E-02 | 2.44E+00 | 2.48E+00 | 3.10E-01 | no | Grin2b | intron_variant | MODIFIER |
| 6 | 136060670 | AX-170109182 | 0 | G | A | 0.579 | 9.14E-02 | 8.84E-02 | 2.44E+00 | 2.48E+00 | 3.10E-01 | no | Grin2b | intron_variant | MODIFIER |
| 6 | 136068700 | AX-170430523 | 0 | A | G | 0.579 | 9.14E-02 | 8.84E-02 | 2.44E+00 | 2.48E+00 | 3.10E-01 | no | Grin2b | intron_variant | MODIFIER |
| 6 | 136068852 | AX-170108108 | 0 | A | G | 0.579 | 9.14E-02 | 8.84E-02 | 2.44E+00 | 2.48E+00 | 3.10E-01 | no | Grin2b | intron_variant | MODIFIER |
| 6 | 136069059 | AX-170310512 | 0 | T | C | 0.579 | 9.14E-02 | 8.84E-02 | 2.44E+00 | 2.48E+00 | 3.10E-01 | no | Grin2b | intron_variant | MODIFIER |
| 6 | 136081377 | AX-170293688 | 0 | G | A | 0.579 | 9.14E-02 | 8.84E-02 | 2.44E+00 | 2.48E+00 | 3.10E-01 | no | Grin2b | intron_variant | MODIFIER |
| 6 | 136083382 | AX-170342260 | 0 | T | A | 0.579 | 9.14E-02 | 8.84E-02 | 2.44E+00 | 2.48E+00 | 3.10E-01 | no | Grin2b | intron_variant | MODIFIER |
| 6 | 136084707 | AX-170286746 | 0 | T | C | 0.579 | 9.14E-02 | 8.84E-02 | 2.44E+00 | 2.48E+00 | 3.10E-01 | no | Grin2b | intron_variant | MODIFIER |
| 6 | 136090676 | AX-170253308 | 0 | G | A | 0.579 | 9.14E-02 | 8.84E-02 | 2.44E+00 | 2.48E+00 | 3.10E-01 | no | Grin2b | intron_variant | MODIFIER |
| 6 | 136099750 | AX-170520633 | 0 | C | A | 0.579 | 9.14E-02 | 8.84E-02 | 2.44E+00 | 2.48E+00 | 3.10E-01 | no | Grin2b | intron_variant | MODIFIER |
| 6 | 136111282 | AX-170270560 | 0 | C | T | 0.579 | 9.14E-02 | 8.84E-02 | 2.44E+00 | 2.48E+00 | 3.10E-01 | no | Grin2b | intron_variant | MODIFIER |
| 6 | 136113283 | AX-170417607 | 0 | C | A | 0.579 | 9.14E-02 | 8.84E-02 | 2.44E+00 | 2.48E+00 | 3.10E-01 | no | Grin2b | intron_variant | MODIFIER |
| 6 | 136114360 | AX-170319871 | 0 | T | C | 0.579 | 9.14E-02 | 8.84E-02 | 2.44E+00 | 2.48E+00 | 3.10E-01 | no | Grin2b | intron_variant | MODIFIER |
| 6 | 136116842 | AX-170183037 | 0 | A | G | 0.579 | 9.14E-02 | 8.84E-02 | 2.44E+00 | 2.48E+00 | 3.10E-01 | no | Grin2b | intron_variant | MODIFIER |
| 6 | 136147931 | AX-170500741 | 0 | G | A | 0.579 | 9.14E-02 | 8.84E-02 | 2.44E+00 | 2.48E+00 | 3.10E-01 | no | Grin2b | intron_variant | MODIFIER |
| 6 | 136154501 | AX-170246448 | 0 | A | G | 0.579 | 9.14E-02 | 8.84E-02 | 2.44E+00 | 2.48E+00 | 3.10E-01 | no | Grin2b | intron_variant | MODIFIER |
| 6 | 136155982 | AX-170218230 | 0 | A | G | 0.579 | 9.14E-02 | 8.84E-02 | 2.44E+00 | 2.48E+00 | 3.10E-01 | no | Grin2b | intron_variant | MODIFIER |
| 6 | 136157978 | AX-170302161 | 0 | T | C | 0.579 | 9.14E-02 | 8.84E-02 | 2.44E+00 | 2.48E+00 | 3.10E-01 | no | Grin2b | intron_variant | MODIFIER |
| 6 | 136185109 | AX-97822791 | 0 | G | A | 0.579 | 9.14E-02 | 8.84E-02 | 2.44E+00 | 2.48E+00 | 3.10E-01 | no | | intergenic_variant | MODIFIER |
| 6 | 136212887 | AX-170344387 | 0 | C | T | 0.579 | 9.14E-02 | 8.84E-02 | 2.44E+00 | 2.48E+00 | 3.10E-01 | no | | intergenic_variant | MODIFIER |
| 6 | 136214628 | AX-170242250 | 0 | G | A | 0.579 | 9.14E-02 | 8.84E-02 | 2.44E+00 | 2.48E+00 | 3.10E-01 | no | | intergenic_variant | MODIFIER |
| 6 | 136247353 | AX-170226416 | 0 | C | A | 0.579 | 9.14E-02 | 8.84E-02 | 2.44E+00 | 2.48E+00 | 3.10E-01 | no | | intergenic_variant | MODIFIER |
| 6 | 136247555 | AX-170248168 | 0 | T | G | 0.579 | 9.14E-02 | 8.84E-02 | 2.44E+00 | 2.48E+00 | 3.10E-01 | no | | intergenic_variant | MODIFIER |
| 6 | 136261643 | AX-170521384 | 0 | C | T | 0.579 | 9.14E-02 | 8.84E-02 | 2.44E+00 | 2.48E+00 | 3.10E-01 | no | | intergenic_variant | MODIFIER |
| 6 | 136280848 | AX-170289439 | 0 | A | G | 0.579 | 9.14E-02 | 8.84E-02 | 2.44E+00 | 2.48E+00 | 3.10E-01 | no | Gm26105 | upstream_gene_variant | MODIFIER |
| 6 | 136282147 | AX-170378965 | 0 | T | C | 0.579 | 9.14E-02 | 8.84E-02 | 2.44E+00 | 2.48E+00 | 3.10E-01 | no | Gm26105 | downstream_gene_variant | MODIFIER |

| | | | | | | | | | | | | | | | |
|---|---|---|---|---|---|---|---|---|---|---|---|---|---|---|---|
| 6 | 136292568 | AX-170028328 | 0 | A | G | 0.579 | 9.14E-02 | 8.84E-02 | 2.44E+00 | 2.48E+00 | 3.10E-01 | no | | regulatory_region_variant | MODIFIER |
| 6 | 136302008 | AX-170440646 | 0 | T | C | 0.579 | 9.14E-02 | 8.84E-02 | 2.44E+00 | 2.48E+00 | 3.10E-01 | no | | intergenic_variant | MODIFIER |
| 6 | 136304372 | AX-170076344 | 0 | T | C | 0.579 | 9.14E-02 | 8.84E-02 | 2.44E+00 | 2.48E+00 | 3.10E-01 | no | | intergenic_variant | MODIFIER |
| 6 | 136305645 | AX-170345832 | 0 | A | C | 0.579 | 9.14E-02 | 8.84E-02 | 2.44E+00 | 2.48E+00 | 3.10E-01 | no | | intergenic_variant | MODIFIER |
| 6 | 136309123 | AX-170129482 | 0 | C | A | 0.579 | 9.14E-02 | 8.84E-02 | 2.44E+00 | 2.48E+00 | 3.10E-01 | no | | intergenic_variant | MODIFIER |
| 6 | 136320094 | AX-97846465 | 0 | C | T | 0.579 | 9.14E-02 | 8.84E-02 | 2.44E+00 | 2.48E+00 | 3.10E-01 | no | | intergenic_variant | MODIFIER |
| 6 | 136342039 | AX-170028325 | 0 | A | G | 0.579 | 9.14E-02 | 8.84E-02 | 2.44E+00 | 2.48E+00 | 3.10E-01 | no | | intergenic_variant | MODIFIER |
| 6 | 136345252 | AX-170483593 | 0 | G | A | 0.579 | 9.14E-02 | 8.84E-02 | 2.44E+00 | 2.48E+00 | 3.10E-01 | no | | intergenic_variant | MODIFIER |
| 6 | 136351210 | AX-97797248 | 0 | T | A | 0.579 | 9.14E-02 | 8.84E-02 | 2.44E+00 | 2.48E+00 | 3.10E-01 | no | | intergenic_variant | MODIFIER |
| 6 | 136356109 | AX-170325859 | 0 | T | C | 0.579 | 9.14E-02 | 8.84E-02 | 2.44E+00 | 2.48E+00 | 3.10E-01 | no | | intergenic_variant | MODIFIER |
| 6 | 136361185 | AX-170104342 | 0 | A | G | 0.579 | 9.14E-02 | 8.84E-02 | 2.44E+00 | 2.48E+00 | 3.10E-01 | no | | intergenic_variant | MODIFIER |
| 6 | 136361803 | AX-170352939 | 0 | T | C | 0.579 | 9.14E-02 | 8.84E-02 | 2.44E+00 | 2.48E+00 | 3.10E-01 | no | | intergenic_variant | MODIFIER |
| 6 | 136368539 | AX-97793887 | 0 | A | G | 0.579 | 9.14E-02 | 8.84E-02 | 2.44E+00 | 2.48E+00 | 3.10E-01 | no | | intergenic_variant | MODIFIER |
| 6 | 136390430 | AX-170028327 | 0 | C | T | 0.579 | 9.14E-02 | 8.84E-02 | 2.44E+00 | 2.48E+00 | 3.10E-01 | no | | intergenic_variant | MODIFIER |
| 6 | 136390540 | AX-170028326 | 0 | G | A | 0.579 | 9.14E-02 | 8.84E-02 | 2.44E+00 | 2.48E+00 | 3.10E-01 | no | | intergenic_variant | MODIFIER |
| 6 | 136452937 | AX-170142040 | 0 | T | C | 0.579 | 9.14E-02 | 8.84E-02 | 2.44E+00 | 2.48E+00 | 3.10E-01 | no | | regulatory_region_variant | MODIFIER |
| 6 | 136474702 | AX-170174352 | 0 | C | T | 0.579 | 9.14E-02 | 8.84E-02 | 2.44E+00 | 2.48E+00 | 3.10E-01 | no | | regulatory_region_variant | MODIFIER |
| 6 | 136500136 | AX-170312380 | 0 | A | G | 0.579 | 9.14E-02 | 8.84E-02 | 2.44E+00 | 2.48E+00 | 3.10E-01 | no | | regulatory_region_variant | MODIFIER |
| 6 | 136507869 | AX-97835379 | 0 | C | G | 0.579 | 9.14E-02 | 8.84E-02 | 2.44E+00 | 2.48E+00 | 3.10E-01 | no | Atf7ip | intron_variant | MODIFIER |
| 6 | 136516304 | AX-170245926 | 0 | T | C | 0.579 | 9.14E-02 | 8.84E-02 | 2.44E+00 | 2.48E+00 | 3.10E-01 | no | Atf7ip | upstream_gene_variant | MODIFIER |
| 6 | 136516595 | AX-170252739 | 0 | G | A | 0.579 | 9.14E-02 | 8.84E-02 | 2.44E+00 | 2.48E+00 | 3.10E-01 | no | Atf7ip | upstream_gene_variant | MODIFIER |
| 6 | 136537873 | AX-170115428 | 0 | A | G | 0.579 | 9.14E-02 | 8.84E-02 | 2.44E+00 | 2.48E+00 | 3.10E-01 | no | Atf7ip | intron_variant | MODIFIER |
| 6 | 136582001 | AX-170302629 | 0 | T | C | 0.579 | 9.14E-02 | 8.84E-02 | 2.44E+00 | 2.48E+00 | 3.10E-01 | no | Atf7ip | intron_variant | MODIFIER |
| 6 | 136582183 | AX-97797319 | 0 | T | G | 0.579 | 9.14E-02 | 8.84E-02 | 2.44E+00 | 2.48E+00 | 3.10E-01 | no | Atf7ip | intron_variant | MODIFIER |
| 6 | 136587815 | AX-170472750 | 0 | C | T | 0.579 | 9.14E-02 | 8.84E-02 | 2.44E+00 | 2.48E+00 | 3.10E-01 | no | Atf7ip | intron_variant | MODIFIER |
| 6 | 136591407 | AX-170324414 | 0 | A | G | 0.579 | 9.14E-02 | 8.84E-02 | 2.44E+00 | 2.48E+00 | 3.10E-01 | no | Atf7ip | intron_variant | MODIFIER |
| 6 | 136609682 | AX-170076352 | 0 | G | C | 0.579 | 9.14E-02 | 8.84E-02 | 2.44E+00 | 2.48E+00 | 3.10E-01 | no | Atf7ip | 3_prime_UTR_variant | MODIFIER |
| 6 | 136610606 | AX-170213542 | 0 | A | G | 0.579 | 9.14E-02 | 8.84E-02 | 2.44E+00 | 2.48E+00 | 3.10E-01 | no | Atf7ip | 3_prime_UTR_variant | MODIFIER |
| 6 | 136612247 | AX-170125270 | 0 | T | G | 0.579 | 9.14E-02 | 8.84E-02 | 2.44E+00 | 2.48E+00 | 3.10E-01 | no | Atf7ip | downstream_gene_variant | MODIFIER |
| 6 | 136616471 | AX-170352086 | 0 | C | T | 0.579 | 9.14E-02 | 8.84E-02 | 2.44E+00 | 2.48E+00 | 3.10E-01 | no | Plbd1 | intron_variant | MODIFIER |
| 6 | 136634081 | AX-170106830 | 0 | A | G | 0.579 | 9.14E-02 | 8.84E-02 | 2.44E+00 | 2.48E+00 | 3.10E-01 | no | Plbd1 | intron_variant | MODIFIER |
| 6 | 136644514 | AX-170531578 | 0 | T | G | 0.579 | 9.14E-02 | 8.84E-02 | 2.44E+00 | 2.48E+00 | 3.10E-01 | no | Plbd1 | intron_variant | MODIFIER |
| 6 | 136654971 | AX-97852671 | 0 | G | A | 0.579 | 9.14E-02 | 8.84E-02 | 2.44E+00 | 2.48E+00 | 3.10E-01 | no | Plbd1 | intron_variant | MODIFIER |
| 6 | 136679640 | AX-170274422 | 0 | G | T | 0.579 | 9.14E-02 | 8.84E-02 | 2.44E+00 | 2.48E+00 | 3.10E-01 | no | | intergenic_variant | MODIFIER |
| 6 | 136689300 | AX-170193787 | 0 | C | G | 0.579 | 9.14E-02 | 8.84E-02 | 2.44E+00 | 2.48E+00 | 3.10E-01 | no | | intergenic_variant | MODIFIER |
| 6 | 136691234 | AX-170356745 | 0 | C | A | 0.579 | 9.14E-02 | 8.84E-02 | 2.44E+00 | 2.48E+00 | 3.10E-01 | no | Gm4714 | downstream_gene_variant | MODIFIER |
| 6 | 136698834 | AX-170367654 | 0 | A | G | 0.579 | 9.14E-02 | 8.84E-02 | 2.44E+00 | 2.48E+00 | 3.10E-01 | no | Gucy2c | intron_variant | MODIFIER |
| 6 | 136712205 | AX-170120153 | 0 | G | A | 0.579 | 9.14E-02 | 8.84E-02 | 2.44E+00 | 2.48E+00 | 3.10E-01 | no | Gucy2c | intron_variant | MODIFIER |
| 6 | 136712838 | AX-170291196 | 0 | A | G | 0.579 | 9.14E-02 | 8.84E-02 | 2.44E+00 | 2.48E+00 | 3.10E-01 | no | Gucy2c | intron_variant | MODIFIER |
| 6 | 136760865 | AX-170249206 | 0 | A | C | 0.579 | 9.14E-02 | 8.84E-02 | 2.44E+00 | 2.48E+00 | 3.10E-01 | no | Gucy2c | intron_variant | MODIFIER |
| 6 | 136780434 | AX-170076356 | 0 | C | T | 0.579 | 9.14E-02 | 8.84E-02 | 2.44E+00 | 2.48E+00 | 3.10E-01 | no | Gucy2c | intron_variant | MODIFIER |
| 6 | 136824449 | AX-170410274 | 0 | G | C | 0.579 | 9.14E-02 | 8.84E-02 | 2.44E+00 | 2.48E+00 | 3.10E-01 | no | BC049715 | upstream_gene_variant | MODIFIER |
| 6 | 136839032 | AX-170372098 | 0 | T | C | 0.579 | 9.14E-02 | 8.84E-02 | 2.44E+00 | 2.48E+00 | 3.10E-01 | no | BC049715 | intron_variant | MODIFIER |
| 6 | 136839049 | AX-170521224 | 0 | A | G | 0.579 | 9.14E-02 | 8.84E-02 | 2.44E+00 | 2.48E+00 | 3.10E-01 | no | BC049715 | intron_variant | MODIFIER |
| 6 | 136854621 | AX-170156759 | 0 | T | C | 0.579 | 9.14E-02 | 8.84E-02 | 2.44E+00 | 2.48E+00 | 3.10E-01 | no | Art4 | missense_variant | MODERATE |
| 6 | 136855027 | AX-170511578 | 0 | T | C | 0.579 | 9.14E-02 | 8.84E-02 | 2.44E+00 | 2.48E+00 | 3.10E-01 | no | Art4 | missense_variant | MODERATE |
| 6 | 136856498 | AX-170201226 | 0 | A | G | 0.579 | 9.14E-02 | 8.84E-02 | 2.44E+00 | 2.48E+00 | 3.10E-01 | no | Art4 | intron_variant | MODIFIER |
| 6 | 136856704 | AX-170213561 | 0 | T | C | 0.579 | 9.14E-02 | 8.84E-02 | 2.44E+00 | 2.48E+00 | 3.10E-01 | no | Art4 | intron_variant | MODIFIER |
| 6 | 136857069 | AX-170237741 | 0 | T | A | 0.579 | 9.14E-02 | 8.84E-02 | 2.44E+00 | 2.48E+00 | 3.10E-01 | no | Art4 | intron_variant | MODIFIER |
| 6 | 136862416 | AX-170166729 | 0 | C | T | 0.579 | 9.14E-02 | 8.84E-02 | 2.44E+00 | 2.48E+00 | 3.10E-01 | no | Art4 | upstream_gene_variant | MODIFIER |
| 6 | 136865716 | AX-170303392 | 0 | G | A | 0.579 | 9.14E-02 | 8.84E-02 | 2.44E+00 | 2.48E+00 | 3.10E-01 | no | | regulatory_region_variant | MODIFIER |
| 6 | 136872487 | AX-170135693 | 0 | C | T | 0.579 | 9.14E-02 | 8.84E-02 | 2.44E+00 | 2.48E+00 | 3.10E-01 | no | Mgp | 3_prime_UTR_variant | MODIFIER |
| 6 | 136878008 | AX-170028336 | 0 | A | C | 0.579 | 9.14E-02 | 8.84E-02 | 2.44E+00 | 2.48E+00 | 3.10E-01 | no | Mgp | upstream_gene_variant | MODIFIER |
| 6 | 136894639 | AX-170227916 | 0 | A | G | 0.579 | 9.14E-02 | 8.84E-02 | 2.44E+00 | 2.48E+00 | 3.10E-01 | no | | regulatory_region_variant | MODIFIER |

| | | | | | | | | | | | | | | | | |
|---|---|---|---|---|---|---|---|---|---|---|---|---|---|---|---|---|
| 6 | 136896398 | AX-170282392 | 0 | G | T | 0.579 | 9.14E-02 | 8.84E-02 | 2.44E+00 | 2.48E+00 | 3.10E-01 | no | | | regulatory_region_variant | MODIFIER |
| 6 | 136897662 | AX-170432719 | 0 | C | T | 0.579 | 9.14E-02 | 8.84E-02 | 2.44E+00 | 2.48E+00 | 3.10E-01 | no | | | regulatory_region_variant | MODIFIER |
| 6 | 136898630 | AX-170445376 | 0 | G | A | 0.579 | 9.14E-02 | 8.84E-02 | 2.44E+00 | 2.48E+00 | 3.10E-01 | no | | | regulatory_region_variant | MODIFIER |
| 6 | 136898898 | AX-170269856 | 0 | C | T | 0.579 | 9.14E-02 | 8.84E-02 | 2.44E+00 | 2.48E+00 | 3.10E-01 | no | | | regulatory_region_variant | MODIFIER |
| 6 | 136899650 | AX-170398214 | 0 | A | G | 0.579 | 9.14E-02 | 8.84E-02 | 2.44E+00 | 2.48E+00 | 3.10E-01 | no | | | regulatory_region_variant | MODIFIER |
| 6 | 136900742 | AX-170028339 | 0 | C | T | 0.579 | 9.14E-02 | 8.84E-02 | 2.44E+00 | 2.48E+00 | 3.10E-01 | no | | | regulatory_region_variant | MODIFIER |
| 6 | 136925581 | AX-97807721 | 0 | T | G | 0.579 | 9.14E-02 | 8.84E-02 | 2.44E+00 | 2.48E+00 | 3.10E-01 | no | Erp27 | | upstream_gene_variant | MODIFIER |
| 6 | 136942575 | AX-170459699 | 0 | A | G | 0.579 | 9.14E-02 | 8.84E-02 | 2.44E+00 | 2.48E+00 | 3.10E-01 | no | Arhgdib | | upstream_gene_variant | MODIFIER |
| 6 | 136963925 | AX-170028337 | 0 | C | T | 0.579 | 9.14E-02 | 8.84E-02 | 2.44E+00 | 2.48E+00 | 3.10E-01 | no | Pde6h | | downstream_gene_variant | MODIFIER |
| 6 | 136981241 | AX-97794965 | 0 | C | T | 0.579 | 9.14E-02 | 8.84E-02 | 2.44E+00 | 2.48E+00 | 3.10E-01 | no | | | intergenic_variant | MODIFIER |
| 6 | 137013548 | AX-170028338 | 0 | C | T | 0.579 | 9.14E-02 | 8.84E-02 | 2.44E+00 | 2.48E+00 | 3.10E-01 | no | | | intergenic_variant | MODIFIER |
| 6 | 137063320 | AX-170127384 | 0 | A | G | 0.579 | 9.14E-02 | 8.84E-02 | 2.44E+00 | 2.48E+00 | 3.10E-01 | no | Rerg | | intron_variant | MODIFIER |
| 6 | 137067348 | AX-170234401 | 0 | G | A | 0.579 | 9.14E-02 | 8.84E-02 | 2.44E+00 | 2.48E+00 | 3.10E-01 | no | Rerg | | intron_variant | MODIFIER |
| 6 | 137070622 | AX-170450522 | 0 | C | T | 0.579 | 9.14E-02 | 8.84E-02 | 2.44E+00 | 2.48E+00 | 3.10E-01 | no | Rerg | | intron_variant | MODIFIER |
| 6 | 137076739 | AX-170183714 | 0 | T | C | 0.579 | 9.14E-02 | 8.84E-02 | 2.44E+00 | 2.48E+00 | 3.10E-01 | no | Rerg | | intron_variant | MODIFIER |
| 6 | 137098025 | AX-170395700 | 0 | A | G | 0.579 | 9.14E-02 | 8.84E-02 | 2.44E+00 | 2.48E+00 | 3.10E-01 | no | Rerg | | intron_variant | MODIFIER |
| 6 | 137113974 | AX-97816217 | 0 | A | G | 0.579 | 9.14E-02 | 8.84E-02 | 2.44E+00 | 2.48E+00 | 3.10E-01 | no | Rerg | | intron_variant | MODIFIER |
| 6 | 137115073 | AX-170244152 | 0 | T | C | 0.579 | 9.14E-02 | 8.84E-02 | 2.44E+00 | 2.48E+00 | 3.10E-01 | no | Rerg | | intron_variant | MODIFIER |
| 6 | 137146714 | AX-170410720 | 0 | A | C | 0.579 | 9.14E-02 | 8.84E-02 | 2.44E+00 | 2.48E+00 | 3.10E-01 | no | Rerg | | intron_variant | MODIFIER |
| 6 | 137149294 | AX-170161492 | 0 | T | C | 0.579 | 9.14E-02 | 8.84E-02 | 2.44E+00 | 2.48E+00 | 3.10E-01 | no | Rerg | | intron_variant | MODIFIER |
| 6 | 137151693 | AX-170506782 | 0 | T | C | 0.579 | 9.14E-02 | 8.84E-02 | 2.44E+00 | 2.48E+00 | 3.10E-01 | no | Rerg | | intron_variant | MODIFIER |
| 6 | 137190864 | AX-170280859 | 0 | G | C | 0.579 | 9.14E-02 | 8.84E-02 | 2.44E+00 | 2.48E+00 | 3.10E-01 | no | | | intergenic_variant | MODIFIER |
| 6 | 137237118 | AX-170333246 | 0 | T | C | 0.579 | 9.14E-02 | 8.84E-02 | 2.44E+00 | 2.48E+00 | 3.10E-01 | no | | | intergenic_variant | MODIFIER |
| 6 | 137286289 | AX-170327456 | 0 | T | C | 0.579 | 9.14E-02 | 8.84E-02 | 2.44E+00 | 2.48E+00 | 3.10E-01 | no | Ptpro | | intron_variant | MODIFIER |
| 6 | 137287176 | AX-170431883 | 0 | C | T | 0.579 | 9.14E-02 | 8.84E-02 | 2.44E+00 | 2.48E+00 | 3.10E-01 | no | Ptpro | | intron_variant | MODIFIER |
| 6 | 137288383 | AX-170356936 | 0 | G | C | 0.579 | 9.14E-02 | 8.84E-02 | 2.44E+00 | 2.48E+00 | 3.10E-01 | no | Ptpro | | intron_variant | MODIFIER |
| 6 | 137327554 | AX-170221926 | 0 | T | C | 0.579 | 9.14E-02 | 8.84E-02 | 2.44E+00 | 2.48E+00 | 3.10E-01 | no | Ptpro | | intron_variant | MODIFIER |
| 6 | 137329623 | AX-170257752 | 0 | G | T | 0.579 | 9.14E-02 | 8.84E-02 | 2.44E+00 | 2.48E+00 | 3.10E-01 | no | Ptpro | | intron_variant | MODIFIER |
| 6 | 137329730 | AX-170263638 | 0 | C | T | 0.579 | 9.14E-02 | 8.84E-02 | 2.44E+00 | 2.48E+00 | 3.10E-01 | no | Ptpro | | intron_variant | MODIFIER |
| 6 | 137334995 | AX-170182868 | 0 | G | A | 0.579 | 9.14E-02 | 8.84E-02 | 2.44E+00 | 2.48E+00 | 3.10E-01 | no | Ptpro | | intron_variant | MODIFIER |
| 6 | 137338515 | AX-170427710 | 0 | C | G | 0.579 | 9.14E-02 | 8.84E-02 | 2.44E+00 | 2.48E+00 | 3.10E-01 | no | Ptpro | | intron_variant | MODIFIER |
| 6 | 137385412 | AX-97794374 | 0 | C | G | 0.579 | 9.14E-02 | 8.84E-02 | 2.44E+00 | 2.48E+00 | 3.10E-01 | no | Ptpro | | intron_variant | MODIFIER |
| 6 | 137401073 | AX-170179070 | 0 | G | T | 0.579 | 9.14E-02 | 8.84E-02 | 2.44E+00 | 2.48E+00 | 3.10E-01 | no | Ptpro | | intron_variant | MODIFIER |
| 6 | 137401091 | AX-97791928 | 0 | C | T | 0.579 | 9.14E-02 | 8.84E-02 | 2.44E+00 | 2.48E+00 | 3.10E-01 | no | Ptpro | | intron_variant | MODIFIER |
| 6 | 137418189 | AX-170490699 | 0 | C | A | 0.579 | 9.14E-02 | 8.84E-02 | 2.44E+00 | 2.48E+00 | 3.10E-01 | no | Ptpro | | intron_variant | MODIFIER |
| 6 | 137423771 | AX-170452255 | 0 | T | C | 0.579 | 9.14E-02 | 8.84E-02 | 2.44E+00 | 2.48E+00 | 3.10E-01 | no | Ptpro | | intron_variant | MODIFIER |
| 6 | 137427577 | AX-170076366 | 0 | A | G | 0.579 | 9.14E-02 | 8.84E-02 | 2.44E+00 | 2.48E+00 | 3.10E-01 | no | Ptpro | | intron_variant | MODIFIER |
| 6 | 137432765 | AX-170028345 | 0 | A | G | 0.579 | 9.14E-02 | 8.84E-02 | 2.44E+00 | 2.48E+00 | 3.10E-01 | no | Ptpro | | intron_variant | MODIFIER |
| 6 | 137432805 | AX-97786107 | 0 | G | A | 0.579 | 9.14E-02 | 8.84E-02 | 2.44E+00 | 2.48E+00 | 3.10E-01 | no | Ptpro | | intron_variant | MODIFIER |
| 6 | 137448631 | AX-170266400 | 0 | T | G | 0.579 | 9.14E-02 | 8.84E-02 | 2.44E+00 | 2.48E+00 | 3.10E-01 | no | Ptpro | | intron_variant | MODIFIER |
| 6 | 137449249 | AX-170515296 | 0 | G | A | 0.579 | 9.14E-02 | 8.84E-02 | 2.44E+00 | 2.48E+00 | 3.10E-01 | no | Ptpro | | intron_variant | MODIFIER |
| 6 | 137449975 | AX-170115066 | 0 | G | A | 0.579 | 9.14E-02 | 8.84E-02 | 2.44E+00 | 2.48E+00 | 3.10E-01 | no | Ptpro | | synonymous_variant | LOW |
| 6 | 137455059 | AX-170442222 | 0 | A | G | 0.579 | 9.14E-02 | 8.84E-02 | 2.44E+00 | 2.48E+00 | 3.10E-01 | no | Ptpro | | intron_variant | MODIFIER |
| 6 | 137459295 | AX-170028346 | 0 | T | G | 0.579 | 9.14E-02 | 8.84E-02 | 2.44E+00 | 2.48E+00 | 3.10E-01 | no | Ptpro | | intron_variant | MODIFIER |
| 6 | 137460076 | AX-170533390 | 0 | A | G | 0.579 | 9.14E-02 | 8.84E-02 | 2.44E+00 | 2.48E+00 | 3.10E-01 | no | Ptpro | | intron_variant | MODIFIER |
| 6 | 137461866 | AX-170028341 | 0 | C | T | 0.579 | 9.14E-02 | 8.84E-02 | 2.44E+00 | 2.48E+00 | 3.10E-01 | no | Ptpro | | intron_variant | MODIFIER |
| 6 | 137462157 | AX-170450134 | 0 | C | T | 0.579 | 9.14E-02 | 8.84E-02 | 2.44E+00 | 2.48E+00 | 3.10E-01 | no | Ptpro | | 3_prime_UTR_variant | MODIFIER |
| 6 | 137463477 | AX-170176857 | 0 | A | C | 0.579 | 9.14E-02 | 8.84E-02 | 2.44E+00 | 2.48E+00 | 3.10E-01 | no | Ptpro | | downstream_gene_variant | MODIFIER |
| 6 | 137486557 | AX-170288055 | 0 | G | A | 0.579 | 9.14E-02 | 8.84E-02 | 2.44E+00 | 2.48E+00 | 3.10E-01 | no | Eps8 | | intron_variant | MODIFIER |
| 6 | 137530638 | AX-170115205 | 0 | A | G | 0.579 | 9.14E-02 | 8.84E-02 | 2.44E+00 | 2.48E+00 | 3.10E-01 | no | Eps8 | | intron_variant | MODIFIER |
| 6 | 137531158 | AX-170164126 | 0 | A | G | 0.579 | 9.14E-02 | 8.84E-02 | 2.44E+00 | 2.48E+00 | 3.10E-01 | no | Eps8 | | intron_variant | MODIFIER |
| 6 | 137532697 | AX-170322285 | 0 | C | T | 0.579 | 9.14E-02 | 8.84E-02 | 2.44E+00 | 2.48E+00 | 3.10E-01 | no | Eps8 | | intron_variant | MODIFIER |
| 6 | 137533514 | AX-170144993 | 0 | C | T | 0.579 | 9.14E-02 | 8.84E-02 | 2.44E+00 | 2.48E+00 | 3.10E-01 | no | Eps8 | | intron_variant | MODIFIER |
| 6 | 137533652 | AX-170035740 | 0 | G | A | 0.579 | 9.14E-02 | 8.84E-02 | 2.44E+00 | 2.48E+00 | 3.10E-01 | no | Eps8 | | intron_variant | MODIFIER |

| | | | | | | | | | | | | | | | |
|---|---|---|---|---|---|---|---|---|---|---|---|---|---|---|---|
| 6 | 137533920 | AX-170438386 | 0 | G | A | 0.579 | 9.14E-02 | 8.84E-02 | 2.44E+00 | 2.48E+00 | 3.10E-01 | no | Eps8 | intron_variant | MODIFIER |
| 6 | 137539621 | AX-170076367 | 0 | A | G | 0.579 | 9.14E-02 | 8.84E-02 | 2.44E+00 | 2.48E+00 | 3.10E-01 | no | Eps8 | intron_variant | MODIFIER |
| 6 | 137539661 | AX-97801640 | 0 | G | C | 0.579 | 9.14E-02 | 8.84E-02 | 2.44E+00 | 2.48E+00 | 3.10E-01 | no | Eps8 | intron_variant | MODIFIER |
| 6 | 137560252 | AX-170490739 | 0 | C | T | 0.579 | 9.14E-02 | 8.84E-02 | 2.44E+00 | 2.48E+00 | 3.10E-01 | no | Eps8 | intron_variant | MODIFIER |
| 6 | 137560625 | AX-170472736 | 0 | G | A | 0.579 | 9.14E-02 | 8.84E-02 | 2.44E+00 | 2.48E+00 | 3.10E-01 | no | Eps8 | intron_variant | MODIFIER |
| 6 | 137560842 | AX-170243044 | 0 | G | A | 0.579 | 9.14E-02 | 8.84E-02 | 2.44E+00 | 2.48E+00 | 3.10E-01 | no | Eps8 | intron_variant | MODIFIER |
| 6 | 137561809 | AX-170168153 | 0 | C | T | 0.579 | 9.14E-02 | 8.84E-02 | 2.44E+00 | 2.48E+00 | 3.10E-01 | no | Eps8 | intron_variant | MODIFIER |
| 6 | 137565058 | AX-170362741 | 0 | T | C | 0.579 | 9.14E-02 | 8.84E-02 | 2.44E+00 | 2.48E+00 | 3.10E-01 | no | Eps8 | intron_variant | MODIFIER |
| 6 | 137566234 | AX-170520423 | 0 | T | C | 0.579 | 9.14E-02 | 8.84E-02 | 2.44E+00 | 2.48E+00 | 3.10E-01 | no | Eps8 | intron_variant | MODIFIER |
| 6 | 137572587 | AX-170180301 | 0 | G | A | 0.579 | 9.14E-02 | 8.84E-02 | 2.44E+00 | 2.48E+00 | 3.10E-01 | no | Eps8 | intron_variant | MODIFIER |
| 6 | 137575603 | AX-170352408 | 0 | T | C | 0.579 | 9.14E-02 | 8.84E-02 | 2.44E+00 | 2.48E+00 | 3.10E-01 | no | Eps8 | intron_variant | MODIFIER |
| 6 | 137586997 | AX-170459439 | 0 | A | G | 0.579 | 9.14E-02 | 8.84E-02 | 2.44E+00 | 2.48E+00 | 3.10E-01 | no | Eps8 | intron_variant | MODIFIER |
| 6 | 137602357 | AX-170460563 | 0 | A | C | 0.579 | 9.14E-02 | 8.84E-02 | 2.44E+00 | 2.48E+00 | 3.10E-01 | no | Eps8 | intron_variant | MODIFIER |
| 6 | 137607373 | AX-170230365 | 0 | G | T | 0.579 | 9.14E-02 | 8.84E-02 | 2.44E+00 | 2.48E+00 | 3.10E-01 | no | Eps8 | intron_variant | MODIFIER |
| 6 | 137608841 | AX-170145561 | 0 | T | C | 0.579 | 9.14E-02 | 8.84E-02 | 2.44E+00 | 2.48E+00 | 3.10E-01 | no | Eps8 | intron_variant | MODIFIER |
| 6 | 137610577 | AX-170411309 | 0 | G | C | 0.579 | 9.14E-02 | 8.84E-02 | 2.44E+00 | 2.48E+00 | 3.10E-01 | no | Eps8 | intron_variant | MODIFIER |
| 6 | 137634232 | AX-170458604 | 0 | T | C | 0.579 | 9.14E-02 | 8.84E-02 | 2.44E+00 | 2.48E+00 | 3.10E-01 | no | Eps8 | intron_variant | MODIFIER |
| 6 | 137635023 | AX-170140652 | 0 | C | T | 0.579 | 9.14E-02 | 8.84E-02 | 2.44E+00 | 2.48E+00 | 3.10E-01 | no | Eps8 | intron_variant | MODIFIER |
| 1 | 151928196 | AX-170043149 | 1 | C | G | 0.205 | 1.38E-01 | 1.37E-01 | 2.72E+00 | 2.77E+00 | 3.10E-01 | no | 1700025G04Rik | intron_variant | MODIFIER |
| 5 | 89600148 | AX-170147530 | 2 | A | G | 0.427 | 1.07E-01 | 1.06E-01 | 2.93E+00 | 3.01E+00 | 3.10E-01 | no | | intergenic_variant | MODIFIER |
| 1 | 69725929 | AX-170000756 | 2 | C | T | 0.136 | -1.77E-01 | 1.75E-01 | 2.92E+00 | 3.06E+00 | 3.10E-01 | Yes | | intergenic_variant | MODIFIER |
| 1 | 170542428 | AX-97815083 | 1 | C | T | 0.214 | 1.38E-01 | 1.37E-01 | 2.70E+00 | 2.75E+00 | 3.10E-01 | no | Nos1ap | intron_variant,non_coding_transcript_variant | MODIFIER |
| 14 | 113108292 | AX-97838409 | 0 | A | G | 0.342 | 7.41E-02 | 7.36E-02 | 2.62E+00 | 2.72E+00 | 3.10E-01 | no | | intergenic_variant | MODIFIER |
| 14 | 113130125 | AX-170501989 | 0 | C | T | 0.342 | 7.41E-02 | 7.36E-02 | 2.62E+00 | 2.72E+00 | 3.10E-01 | no | Gm49005 | downstream_gene_variant | MODIFIER |
| 14 | 113153027 | AX-170122503 | 0 | T | A | 0.342 | 7.41E-02 | 7.36E-02 | 2.62E+00 | 2.72E+00 | 3.10E-01 | no | | intergenic_variant | MODIFIER |
| 14 | 113167150 | AX-170087731 | 0 | A | C | 0.342 | 7.41E-02 | 7.36E-02 | 2.62E+00 | 2.72E+00 | 3.10E-01 | no | | intergenic_variant | MODIFIER |
| 14 | 113183607 | AX-170355743 | 0 | T | G | 0.342 | 7.41E-02 | 7.36E-02 | 2.62E+00 | 2.72E+00 | 3.10E-01 | no | | intergenic_variant | MODIFIER |
| 14 | 113215817 | AX-170324545 | 0 | G | C | 0.342 | 7.41E-02 | 7.36E-02 | 2.62E+00 | 2.72E+00 | 3.10E-01 | no | | intergenic_variant | MODIFIER |
| 14 | 113222796 | AX-170489602 | 0 | A | C | 0.342 | 7.41E-02 | 7.36E-02 | 2.62E+00 | 2.72E+00 | 3.10E-01 | no | | intergenic_variant | MODIFIER |
| 14 | 113312880 | AX-170099214 | 0 | T | C | 0.342 | 7.41E-02 | 7.36E-02 | 2.62E+00 | 2.72E+00 | 3.10E-01 | no | Tpm3-rs7 | upstream_gene_variant | MODIFIER |
| 14 | 113320859 | AX-97788889 | 0 | A | G | 0.342 | 7.41E-02 | 7.36E-02 | 2.62E+00 | 2.72E+00 | 3.10E-01 | no | Tpm3-rs7 | downstream_gene_variant | MODIFIER |
| 14 | 113321160 | AX-170153481 | 0 | A | G | 0.342 | 7.41E-02 | 7.36E-02 | 2.62E+00 | 2.72E+00 | 3.10E-01 | no | Tpm3-rs7 | downstream_gene_variant | MODIFIER |
| 14 | 113484878 | AX-170010735 | 0 | C | T | 0.342 | 7.41E-02 | 7.36E-02 | 2.62E+00 | 2.72E+00 | 3.10E-01 | no | | intergenic_variant | MODIFIER |
| 14 | 117656471 | AX-170177447 | 0 | A | G | 0.342 | 7.41E-02 | 7.36E-02 | 2.62E+00 | 2.72E+00 | 3.10E-01 | no | Gpc6 | intron_variant | MODIFIER |
| 14 | 117657913 | AX-97806253 | 0 | G | A | 0.342 | 7.41E-02 | 7.36E-02 | 2.62E+00 | 2.72E+00 | 3.10E-01 | no | Gpc6 | intron_variant | MODIFIER |
| 14 | 117658073 | AX-170332800 | 0 | C | T | 0.342 | 7.41E-02 | 7.36E-02 | 2.62E+00 | 2.72E+00 | 3.10E-01 | no | Gpc6 | intron_variant | MODIFIER |
| 14 | 117658107 | AX-170146758 | 0 | C | T | 0.342 | 7.41E-02 | 7.36E-02 | 2.62E+00 | 2.72E+00 | 3.10E-01 | no | Gpc6 | intron_variant | MODIFIER |
| 14 | 117658507 | AX-170203105 | 0 | C | G | 0.342 | 7.41E-02 | 7.36E-02 | 2.62E+00 | 2.72E+00 | 3.10E-01 | no | Gpc6 | intron_variant | MODIFIER |
| 14 | 117658855 | AX-170054560 | 0 | T | C | 0.342 | 7.41E-02 | 7.36E-02 | 2.62E+00 | 2.72E+00 | 3.10E-01 | no | Gpc6 | intron_variant | MODIFIER |
| 14 | 117659939 | AX-170436398 | 0 | G | A | 0.342 | 7.41E-02 | 7.36E-02 | 2.62E+00 | 2.72E+00 | 3.10E-01 | no | Gpc6 | intron_variant | MODIFIER |
| 14 | 117707076 | AX-170131091 | 0 | G | C | 0.342 | 7.41E-02 | 7.36E-02 | 2.62E+00 | 2.72E+00 | 3.10E-01 | no | Gpc6 | intron_variant | MODIFIER |
| 14 | 117707194 | AX-170274227 | 0 | A | G | 0.342 | 7.41E-02 | 7.36E-02 | 2.62E+00 | 2.72E+00 | 3.10E-01 | no | Gpc6 | intron_variant | MODIFIER |
| 14 | 117731220 | AX-170506774 | 0 | C | T | 0.342 | 7.41E-02 | 7.36E-02 | 2.62E+00 | 2.72E+00 | 3.10E-01 | no | Gpc6 | intron_variant | MODIFIER |
| 14 | 117762284 | AX-170513233 | 0 | T | C | 0.342 | 7.41E-02 | 7.36E-02 | 2.62E+00 | 2.72E+00 | 3.10E-01 | no | Gpc6 | intron_variant | MODIFIER |
| 14 | 117762706 | AX-170479456 | 0 | A | G | 0.342 | 7.41E-02 | 7.36E-02 | 2.62E+00 | 2.72E+00 | 3.10E-01 | no | Gpc6 | intron_variant | MODIFIER |
| 14 | 117763371 | AX-170379932 | 0 | A | G | 0.342 | 7.41E-02 | 7.36E-02 | 2.62E+00 | 2.72E+00 | 3.10E-01 | no | Gpc6 | intron_variant | MODIFIER |
| 14 | 117765057 | AX-170473475 | 0 | A | G | 0.342 | 7.41E-02 | 7.36E-02 | 2.62E+00 | 2.72E+00 | 3.10E-01 | no | Gpc6 | intron_variant | MODIFIER |
| 14 | 117765329 | AX-170496351 | 0 | A | C | 0.342 | 7.41E-02 | 7.36E-02 | 2.62E+00 | 2.72E+00 | 3.10E-01 | no | Gpc6 | intron_variant | MODIFIER |
| 14 | 117767816 | AX-170231104 | 0 | G | T | 0.342 | 7.41E-02 | 7.36E-02 | 2.62E+00 | 2.72E+00 | 3.10E-01 | no | Gpc6 | intron_variant | MODIFIER |
| 14 | 117780798 | AX-170394140 | 0 | C | A | 0.342 | 7.41E-02 | 7.36E-02 | 2.62E+00 | 2.72E+00 | 3.10E-01 | no | Gpc6 | intron_variant | MODIFIER |
| 14 | 117789324 | AX-170529722 | 0 | G | A | 0.342 | 7.41E-02 | 7.36E-02 | 2.62E+00 | 2.72E+00 | 3.10E-01 | no | Gpc6 | intron_variant | MODIFIER |
| 14 | 118534181 | AX-170010810 | 0 | C | T | 0.342 | 7.41E-02 | 7.36E-02 | 2.62E+00 | 2.72E+00 | 3.10E-01 | no | Abcc4 | intron_variant | MODIFIER |
| 14 | 118535533 | AX-170010808 | 0 | A | C | 0.342 | 7.41E-02 | 7.36E-02 | 2.62E+00 | 2.72E+00 | 3.10E-01 | no | Abcc4 | intron_variant | MODIFIER |
| 14 | 118535862 | AX-97820784 | 0 | A | G | 0.342 | 7.41E-02 | 7.36E-02 | 2.62E+00 | 2.72E+00 | 3.10E-01 | no | Abcc4 | intron_variant | MODIFIER |

| | | | | | | | | | | | | | | | |
|---|---|---|---|---|---|---|---|---|---|---|---|---|---|---|---|
| 14 | 118550177 | AX-170010809 | 0 | A | G | 0.342 | 7.41E-02 | 7.36E-02 | 2.62E+00 | 2.72E+00 | 3.10E-01 | no | Abcc4 | intron_variant | MODIFIER |
| 14 | 118553478 | AX-170129486 | 0 | A | C | 0.342 | 7.41E-02 | 7.36E-02 | 2.62E+00 | 2.72E+00 | 3.10E-01 | no | Abcc4 | intron_variant | MODIFIER |
| 14 | 118571562 | AX-97848635 | 0 | G | T | 0.342 | 7.41E-02 | 7.36E-02 | 2.62E+00 | 2.72E+00 | 3.10E-01 | no | Abcc4 | intron_variant | MODIFIER |
| 14 | 118580382 | AX-170010812 | 0 | G | C | 0.342 | 7.41E-02 | 7.36E-02 | 2.62E+00 | 2.72E+00 | 3.10E-01 | no | Abcc4 | intron_variant | MODIFIER |
| 14 | 118594906 | AX-170457471 | 0 | G | A | 0.342 | 7.41E-02 | 7.36E-02 | 2.62E+00 | 2.72E+00 | 3.10E-01 | no | Abcc4 | splice_region_variant,synonymous_variant | LOW |
| 14 | 119000187 | AX-170132830 | 0 | G | A | 0.342 | 7.41E-02 | 7.36E-02 | 2.62E+00 | 2.72E+00 | 3.10E-01 | no | Uggt2 | intron_variant,non_coding_transcript_variant | MODIFIER |
| 14 | 119181567 | AX-170158883 | 0 | C | T | 0.342 | 7.41E-02 | 7.36E-02 | 2.62E+00 | 2.72E+00 | 3.10E-01 | no | Hs6st3 | intron_variant | MODIFIER |
| 14 | 119199865 | AX-170341914 | 0 | G | A | 0.342 | 7.41E-02 | 7.36E-02 | 2.62E+00 | 2.72E+00 | 3.10E-01 | no | Hs6st3 | intron_variant | MODIFIER |
| 14 | 119199895 | AX-170291404 | 0 | T | A | 0.342 | 7.41E-02 | 7.36E-02 | 2.62E+00 | 2.72E+00 | 3.10E-01 | no | Hs6st3 | intron_variant | MODIFIER |
| 15 | 83441528 | AX-170471632 | 0 | A | T | 0.105 | -1.20E-01 | 9.98E-02 | 1.98E+00 | 2.07E+00 | 3.11E-01 | no | Pacsin2 | intron_variant | MODIFIER |
| 18 | 51450465 | AX-97839622 | 0 | A | C | 0.237 | -1.06E-01 | 1.05E-01 | 2.58E+00 | 2.66E+00 | 3.11E-01 | no | | intergenic_variant | MODIFIER |
| 9 | 9425197 | AX-97845345 | 0 | C | T | 0.474 | 8.41E-02 | 8.22E-02 | 2.48E+00 | 2.57E+00 | 3.11E-01 | no | Gm46102 | intron_variant,non_coding_transcript_variant | MODIFIER |
| 14 | 119915529 | AX-170313117 | 0 | A | G | 0.342 | 7.33E-02 | 7.27E-02 | 2.59E+00 | 2.70E+00 | 3.11E-01 | no | | intergenic_variant | MODIFIER |
| 13 | 31259448 | AX-170105506 | 0 | T | C | 0.175 | -1.16E-01 | 1.15E-01 | 2.59E+00 | 2.68E+00 | 3.11E-01 | no | | intergenic_variant | MODIFIER |
| 13 | 32148754 | AX-170125780 | 0 | C | T | 0.175 | -1.16E-01 | 1.15E-01 | 2.59E+00 | 2.68E+00 | 3.11E-01 | no | Gmds | intron_variant | MODIFIER |
| 13 | 35040407 | AX-97783299 | 0 | A | G | 0.175 | -1.16E-01 | 1.15E-01 | 2.59E+00 | 2.68E+00 | 3.11E-01 | no | | intergenic_variant | MODIFIER |
| 6 | 136838939 | AX-170177185 | 1 | A | C | 0.58 | 9.12E-02 | 8.83E-02 | 2.44E+00 | 2.48E+00 | 3.11E-01 | no | BC049715 | intron_variant | MODIFIER |
| 9 | 92734552 | AX-170226879 | 1 | T | G | 0.152 | -1.08E-01 | 1.08E-01 | 2.72E+00 | 2.83E+00 | 3.11E-01 | no | 4933400C23Rik | intron_variant,non_coding_transcript_variant | MODIFIER |
| 2 | 162218346 | AX-170168704 | 0 | T | A | 0.149 | 1.86E-01 | 1.86E-01 | 2.81E+00 | 2.93E+00 | 3.11E-01 | Yes | Ptprt | intron_variant | MODIFIER |
| 1 | 88211750 | AX-170524064 | 0 | G | C | 0.237 | -1.07E-01 | 1.07E-01 | 2.63E+00 | 2.68E+00 | 3.11E-01 | no | Ugt1a6a | intron_variant | MODIFIER |
| 15 | 3134843 | AX-170518971 | 0 | T | C | 0.237 | -1.07E-01 | 1.07E-01 | 2.63E+00 | 2.68E+00 | 3.11E-01 | no | Gm49231 | intron_variant,non_coding_transcript_variant | MODIFIER |
| 15 | 3146128 | AX-170184285 | 0 | C | G | 0.237 | -1.07E-01 | 1.07E-01 | 2.63E+00 | 2.68E+00 | 3.11E-01 | no | Gm49231 | intron_variant,non_coding_transcript_variant | MODIFIER |
| 15 | 3175333 | AX-170293617 | 0 | A | G | 0.237 | -1.07E-01 | 1.07E-01 | 2.63E+00 | 2.68E+00 | 3.11E-01 | no | Gm49232 | upstream_gene_variant | MODIFIER |
| 15 | 3183557 | AX-170226837 | 0 | A | G | 0.237 | -1.07E-01 | 1.07E-01 | 2.63E+00 | 2.68E+00 | 3.11E-01 | no | Gm49232 | downstream_gene_variant | MODIFIER |
| 15 | 3203255 | AX-170233934 | 0 | T | C | 0.237 | -1.07E-01 | 1.07E-01 | 2.63E+00 | 2.68E+00 | 3.11E-01 | no | | intergenic_variant | MODIFIER |
| 15 | 3221625 | AX-170177824 | 0 | A | G | 0.237 | -1.07E-01 | 1.07E-01 | 2.63E+00 | 2.68E+00 | 3.11E-01 | no | Gm7962 | upstream_gene_variant | MODIFIER |
| 15 | 3256145 | AX-170054661 | 0 | A | G | 0.237 | -1.07E-01 | 1.07E-01 | 2.63E+00 | 2.68E+00 | 3.11E-01 | no | | intergenic_variant | MODIFIER |
| 15 | 3259168 | AX-97784139 | 0 | C | T | 0.237 | -1.07E-01 | 1.07E-01 | 2.63E+00 | 2.68E+00 | 3.11E-01 | no | | intergenic_variant | MODIFIER |
| 15 | 3281222 | AX-170196022 | 0 | T | C | 0.237 | -1.07E-01 | 1.07E-01 | 2.63E+00 | 2.68E+00 | 3.11E-01 | no | Selenop | downstream_gene_variant | MODIFIER |
| 15 | 3315456 | AX-170365761 | 0 | G | A | 0.237 | -1.07E-01 | 1.07E-01 | 2.63E+00 | 2.68E+00 | 3.11E-01 | no | Ghr | downstream_gene_variant | MODIFIER |
| 15 | 3331717 | AX-170054662 | 0 | A | G | 0.237 | -1.07E-01 | 1.07E-01 | 2.63E+00 | 2.68E+00 | 3.11E-01 | no | Ghr | intron_variant | MODIFIER |
| 15 | 3342748 | AX-170356043 | 0 | A | T | 0.237 | -1.07E-01 | 1.07E-01 | 2.63E+00 | 2.68E+00 | 3.11E-01 | no | Ghr | intron_variant | MODIFIER |
| 15 | 3349930 | AX-97840607 | 0 | G | A | 0.237 | -1.07E-01 | 1.07E-01 | 2.63E+00 | 2.68E+00 | 3.11E-01 | no | Ghr | intron_variant | MODIFIER |
| 15 | 3365013 | AX-170523459 | 0 | G | A | 0.237 | -1.07E-01 | 1.07E-01 | 2.63E+00 | 2.68E+00 | 3.11E-01 | no | Ghr | intron_variant | MODIFIER |
| 15 | 3374769 | AX-170054664 | 0 | T | C | 0.237 | -1.07E-01 | 1.07E-01 | 2.63E+00 | 2.68E+00 | 3.11E-01 | no | Ghr | intron_variant | MODIFIER |
| 15 | 3413867 | AX-170183281 | 0 | A | G | 0.237 | -1.07E-01 | 1.07E-01 | 2.63E+00 | 2.68E+00 | 3.11E-01 | no | Ghr | intron_variant | MODIFIER |
| 15 | 3425667 | AX-170380150 | 0 | G | A | 0.237 | -1.07E-01 | 1.07E-01 | 2.63E+00 | 2.68E+00 | 3.11E-01 | no | Ghr | intron_variant | MODIFIER |
| 15 | 3460611 | AX-170135497 | 0 | A | C | 0.237 | -1.07E-01 | 1.07E-01 | 2.63E+00 | 2.68E+00 | 3.11E-01 | no | Ghr | intron_variant | MODIFIER |
| 15 | 3468397 | AX-170359896 | 0 | C | T | 0.237 | -1.07E-01 | 1.07E-01 | 2.63E+00 | 2.68E+00 | 3.11E-01 | no | Ghr | intron_variant | MODIFIER |
| 15 | 3514502 | AX-170010916 | 0 | T | A | 0.237 | -1.07E-01 | 1.07E-01 | 2.63E+00 | 2.68E+00 | 3.11E-01 | no | Ghr | intron_variant | MODIFIER |
| 15 | 3536586 | AX-170413233 | 0 | T | A | 0.237 | -1.07E-01 | 1.07E-01 | 2.63E+00 | 2.68E+00 | 3.11E-01 | no | Ghr | intron_variant | MODIFIER |
| 15 | 3543386 | AX-170394695 | 0 | G | A | 0.237 | -1.07E-01 | 1.07E-01 | 2.63E+00 | 2.68E+00 | 3.11E-01 | no | Ghr | intron_variant | MODIFIER |
| 15 | 3568726 | AX-170161788 | 0 | G | T | 0.237 | -1.07E-01 | 1.07E-01 | 2.63E+00 | 2.68E+00 | 3.11E-01 | no | Ghr | intron_variant | MODIFIER |
| 15 | 3580446 | AX-170054667 | 0 | T | C | 0.237 | -1.07E-01 | 1.07E-01 | 2.63E+00 | 2.68E+00 | 3.11E-01 | no | Ghr | intron_variant | MODIFIER |
| 15 | 3607884 | AX-170416953 | 0 | A | G | 0.237 | -1.07E-01 | 1.07E-01 | 2.63E+00 | 2.68E+00 | 3.11E-01 | no | Gm22031 | upstream_gene_variant | MODIFIER |
| 15 | 3636931 | AX-170135776 | 0 | A | G | 0.237 | -1.07E-01 | 1.07E-01 | 2.63E+00 | 2.68E+00 | 3.11E-01 | no | | intergenic_variant | MODIFIER |
| 15 | 3640560 | AX-170359494 | 0 | T | G | 0.237 | -1.07E-01 | 1.07E-01 | 2.63E+00 | 2.68E+00 | 3.11E-01 | no | | regulatory_region_variant | MODIFIER |
| 15 | 3760898 | AX-170412752 | 0 | A | G | 0.237 | -1.07E-01 | 1.07E-01 | 2.63E+00 | 2.68E+00 | 3.11E-01 | no | | intergenic_variant | MODIFIER |
| 15 | 3787999 | AX-170413592 | 0 | A | G | 0.237 | -1.07E-01 | 1.07E-01 | 2.63E+00 | 2.68E+00 | 3.11E-01 | no | | intergenic_variant | MODIFIER |
| 15 | 3788342 | AX-170406501 | 0 | A | G | 0.237 | -1.07E-01 | 1.07E-01 | 2.63E+00 | 2.68E+00 | 3.11E-01 | no | | intergenic_variant | MODIFIER |
| 15 | 3801524 | AX-170144589 | 0 | G | A | 0.237 | -1.07E-01 | 1.07E-01 | 2.63E+00 | 2.68E+00 | 3.11E-01 | no | Gm23228 | downstream_gene_variant | MODIFIER |
| 15 | 3832985 | AX-170152047 | 0 | T | G | 0.237 | -1.07E-01 | 1.07E-01 | 2.63E+00 | 2.68E+00 | 3.11E-01 | no | | intergenic_variant | MODIFIER |
| 15 | 3855540 | AX-170204358 | 0 | T | C | 0.237 | -1.07E-01 | 1.07E-01 | 2.63E+00 | 2.68E+00 | 3.11E-01 | no | | intergenic_variant | MODIFIER |
| 15 | 3861175 | AX-170093140 | 0 | A | G | 0.237 | -1.07E-01 | 1.07E-01 | 2.63E+00 | 2.68E+00 | 3.11E-01 | no | | intergenic_variant | MODIFIER |

| | | | | | | | | | | | | | | | |
|---|---|---|---|---|---|---|---|---|---|---|---|---|---|---|---|
| 15 | 3878411 | AX-170362450 | 0 | C | A | 0.237 | -1.07E-01 | 1.07E-01 | 2.63E+00 | 2.68E+00 | 3.11E-01 | no | | intergenic_variant | MODIFIER |
| 15 | 3879227 | AX-170106704 | 0 | C | A | 0.237 | -1.07E-01 | 1.07E-01 | 2.63E+00 | 2.68E+00 | 3.11E-01 | no | | intergenic_variant | MODIFIER |
| 15 | 3879290 | AX-170436436 | 0 | A | C | 0.237 | -1.07E-01 | 1.07E-01 | 2.63E+00 | 2.68E+00 | 3.11E-01 | no | | intergenic_variant | MODIFIER |
| 15 | 3910442 | AX-97827922 | 0 | T | G | 0.237 | -1.07E-01 | 1.07E-01 | 2.63E+00 | 2.68E+00 | 3.11E-01 | no | | intergenic_variant | MODIFIER |
| 15 | 3969164 | AX-170168422 | 0 | A | T | 0.237 | -1.07E-01 | 1.07E-01 | 2.63E+00 | 2.68E+00 | 3.11E-01 | no | Fbxo4 | intron_variant | MODIFIER |
| 15 | 3983573 | AX-170418477 | 0 | C | T | 0.237 | -1.07E-01 | 1.07E-01 | 2.63E+00 | 2.68E+00 | 3.11E-01 | no | Fbxo4 | upstream_gene_variant | MODIFIER |
| 15 | 3984639 | AX-170509636 | 0 | C | T | 0.237 | -1.07E-01 | 1.07E-01 | 2.63E+00 | 2.68E+00 | 3.11E-01 | no | AW549877 | 3_prime_UTR_variant | MODIFIER |
| 15 | 4019432 | AX-170110364 | 0 | T | C | 0.237 | -1.07E-01 | 1.07E-01 | 2.63E+00 | 2.68E+00 | 3.11E-01 | no | BC037032 | downstream_gene_variant | MODIFIER |
| 15 | 4020334 | AX-170203635 | 0 | C | T | 0.237 | -1.07E-01 | 1.07E-01 | 2.63E+00 | 2.68E+00 | 3.11E-01 | no | BC037032 | downstream_gene_variant | MODIFIER |
| 15 | 4041128 | AX-170054672 | 0 | C | T | 0.237 | -1.07E-01 | 1.07E-01 | 2.63E+00 | 2.68E+00 | 3.11E-01 | no | Oxct1 | intron_variant | MODIFIER |
| 15 | 4044169 | AX-170518605 | 0 | A | G | 0.237 | -1.07E-01 | 1.07E-01 | 2.63E+00 | 2.68E+00 | 3.11E-01 | no | Oxct1 | intron_variant | MODIFIER |
| 15 | 4054587 | AX-170117401 | 0 | A | G | 0.237 | -1.07E-01 | 1.07E-01 | 2.63E+00 | 2.68E+00 | 3.11E-01 | no | Oxct1 | intron_variant | MODIFIER |
| 15 | 4054652 | AX-170387987 | 0 | A | C | 0.237 | -1.07E-01 | 1.07E-01 | 2.63E+00 | 2.68E+00 | 3.11E-01 | no | Oxct1 | intron_variant | MODIFIER |
| 15 | 4054681 | AX-170526851 | 0 | T | G | 0.237 | -1.07E-01 | 1.07E-01 | 2.63E+00 | 2.68E+00 | 3.11E-01 | no | Oxct1 | intron_variant | MODIFIER |
| 15 | 4056039 | AX-170172176 | 0 | A | G | 0.237 | -1.07E-01 | 1.07E-01 | 2.63E+00 | 2.68E+00 | 3.11E-01 | no | Oxct1 | intron_variant | MODIFIER |
| 15 | 4056502 | AX-170208511 | 0 | C | G | 0.237 | -1.07E-01 | 1.07E-01 | 2.63E+00 | 2.68E+00 | 3.11E-01 | no | Oxct1 | intron_variant | MODIFIER |
| 15 | 4057417 | AX-170229281 | 0 | T | A | 0.237 | -1.07E-01 | 1.07E-01 | 2.63E+00 | 2.68E+00 | 3.11E-01 | no | Oxct1 | intron_variant | MODIFIER |
| 15 | 4063033 | AX-170228737 | 0 | G | A | 0.237 | -1.07E-01 | 1.07E-01 | 2.63E+00 | 2.68E+00 | 3.11E-01 | no | Oxct1 | intron_variant | MODIFIER |
| 15 | 4068206 | AX-170010923 | 0 | C | G | 0.237 | -1.07E-01 | 1.07E-01 | 2.63E+00 | 2.68E+00 | 3.11E-01 | no | Oxct1 | intron_variant | MODIFIER |
| 15 | 4098747 | AX-170469885 | 0 | T | A | 0.237 | -1.07E-01 | 1.07E-01 | 2.63E+00 | 2.68E+00 | 3.11E-01 | no | Oxct1 | intron_variant | MODIFIER |
| 15 | 4126383 | AX-97846826 | 0 | G | A | 0.237 | -1.07E-01 | 1.07E-01 | 2.63E+00 | 2.68E+00 | 3.11E-01 | no | Oxct1 | intron_variant | MODIFIER |
| 15 | 4155333 | AX-97842037 | 0 | G | A | 0.237 | -1.07E-01 | 1.07E-01 | 2.63E+00 | 2.68E+00 | 3.11E-01 | no | Oxct1 | 3_prime_UTR_variant | MODIFIER |
| 15 | 4155354 | AX-170010922 | 0 | T | G | 0.237 | -1.07E-01 | 1.07E-01 | 2.63E+00 | 2.68E+00 | 3.11E-01 | no | Oxct1 | downstream_gene_variant | MODIFIER |
| 15 | 4162209 | AX-170527180 | 0 | G | A | 0.237 | -1.07E-01 | 1.07E-01 | 2.63E+00 | 2.68E+00 | 3.11E-01 | no | | intergenic_variant | MODIFIER |
| 15 | 4219766 | AX-170276422 | 0 | G | C | 0.237 | -1.07E-01 | 1.07E-01 | 2.63E+00 | 2.68E+00 | 3.11E-01 | no | | intergenic_variant | MODIFIER |
| 15 | 4224245 | AX-170054675 | 0 | G | T | 0.237 | -1.07E-01 | 1.07E-01 | 2.63E+00 | 2.68E+00 | 3.11E-01 | no | | intergenic_variant | MODIFIER |
| 15 | 4230681 | AX-170054674 | 0 | A | G | 0.237 | -1.07E-01 | 1.07E-01 | 2.63E+00 | 2.68E+00 | 3.11E-01 | no | | intergenic_variant | MODIFIER |
| 15 | 4231122 | AX-97853119 | 0 | G | A | 0.237 | -1.07E-01 | 1.07E-01 | 2.63E+00 | 2.68E+00 | 3.11E-01 | no | | intergenic_variant | MODIFIER |
| 15 | 4261498 | AX-170101802 | 0 | G | A | 0.237 | -1.07E-01 | 1.07E-01 | 2.63E+00 | 2.68E+00 | 3.11E-01 | no | | intergenic_variant | MODIFIER |
| 15 | 4265068 | AX-170503893 | 0 | T | G | 0.237 | -1.07E-01 | 1.07E-01 | 2.63E+00 | 2.68E+00 | 3.11E-01 | no | | intergenic_variant | MODIFIER |
| 15 | 4269570 | AX-97856862 | 0 | A | C | 0.237 | -1.07E-01 | 1.07E-01 | 2.63E+00 | 2.68E+00 | 3.11E-01 | no | | intergenic_variant | MODIFIER |
| 15 | 4275335 | AX-170478519 | 0 | C | T | 0.237 | -1.07E-01 | 1.07E-01 | 2.63E+00 | 2.68E+00 | 3.11E-01 | no | | intergenic_variant | MODIFIER |
| 15 | 4327673 | AX-170104197 | 0 | C | T | 0.237 | -1.07E-01 | 1.07E-01 | 2.63E+00 | 2.68E+00 | 3.11E-01 | no | | intergenic_variant | MODIFIER |
| 15 | 4347171 | AX-170109876 | 0 | A | C | 0.237 | -1.07E-01 | 1.07E-01 | 2.63E+00 | 2.68E+00 | 3.11E-01 | no | | intergenic_variant | MODIFIER |
| 15 | 4369428 | AX-170442110 | 0 | T | C | 0.237 | -1.07E-01 | 1.07E-01 | 2.63E+00 | 2.68E+00 | 3.11E-01 | no | | intergenic_variant | MODIFIER |
| 15 | 4389870 | AX-170279054 | 0 | T | C | 0.237 | -1.07E-01 | 1.07E-01 | 2.63E+00 | 2.68E+00 | 3.11E-01 | no | Plcxd3 | intron_variant | MODIFIER |
| 15 | 4390571 | AX-170374141 | 0 | T | C | 0.237 | -1.07E-01 | 1.07E-01 | 2.63E+00 | 2.68E+00 | 3.11E-01 | no | Plcxd3 | intron_variant | MODIFIER |
| 15 | 4402028 | AX-170370347 | 0 | G | T | 0.237 | -1.07E-01 | 1.07E-01 | 2.63E+00 | 2.68E+00 | 3.11E-01 | no | Plcxd3 | intron_variant | MODIFIER |
| 15 | 4429100 | AX-170366991 | 0 | G | A | 0.237 | -1.07E-01 | 1.07E-01 | 2.63E+00 | 2.68E+00 | 3.11E-01 | no | Plcxd3 | intron_variant | MODIFIER |
| 15 | 4443029 | AX-170317863 | 0 | A | G | 0.237 | -1.07E-01 | 1.07E-01 | 2.63E+00 | 2.68E+00 | 3.11E-01 | no | Plcxd3 | intron_variant | MODIFIER |
| 15 | 4452000 | AX-170157443 | 0 | G | A | 0.237 | -1.07E-01 | 1.07E-01 | 2.63E+00 | 2.68E+00 | 3.11E-01 | no | Plcxd3 | intron_variant | MODIFIER |
| 15 | 4462234 | AX-170157207 | 0 | A | T | 0.237 | -1.07E-01 | 1.07E-01 | 2.63E+00 | 2.68E+00 | 3.11E-01 | no | Plcxd3 | intron_variant | MODIFIER |
| 15 | 4462315 | AX-97817103 | 0 | T | C | 0.237 | -1.07E-01 | 1.07E-01 | 2.63E+00 | 2.68E+00 | 3.11E-01 | no | Plcxd3 | intron_variant | MODIFIER |
| 15 | 4507236 | AX-170206217 | 0 | T | C | 0.237 | -1.07E-01 | 1.07E-01 | 2.63E+00 | 2.68E+00 | 3.11E-01 | no | Plcxd3 | intron_variant | MODIFIER |
| 15 | 4507350 | AX-170532507 | 0 | A | G | 0.237 | -1.07E-01 | 1.07E-01 | 2.63E+00 | 2.68E+00 | 3.11E-01 | no | Plcxd3 | intron_variant | MODIFIER |
| 15 | 4523728 | AX-170157109 | 0 | G | A | 0.237 | -1.07E-01 | 1.07E-01 | 2.63E+00 | 2.68E+00 | 3.11E-01 | no | Plcxd3 | intron_variant | MODIFIER |
| 15 | 4523867 | AX-170054681 | 0 | A | G | 0.237 | -1.07E-01 | 1.07E-01 | 2.63E+00 | 2.68E+00 | 3.11E-01 | no | Plcxd3 | intron_variant | MODIFIER |
| 15 | 4537585 | AX-170054684 | 0 | A | G | 0.237 | -1.07E-01 | 1.07E-01 | 2.63E+00 | 2.68E+00 | 3.11E-01 | no | Plcxd3 | intron_variant | MODIFIER |
| 15 | 4537881 | AX-170323549 | 0 | G | A | 0.237 | -1.07E-01 | 1.07E-01 | 2.63E+00 | 2.68E+00 | 3.11E-01 | no | Plcxd3 | intron_variant | MODIFIER |
| 15 | 4540756 | AX-170204142 | 0 | T | C | 0.237 | -1.07E-01 | 1.07E-01 | 2.63E+00 | 2.68E+00 | 3.11E-01 | no | Plcxd3 | intron_variant | MODIFIER |
| 15 | 4559468 | AX-170144888 | 0 | A | C | 0.237 | -1.07E-01 | 1.07E-01 | 2.63E+00 | 2.68E+00 | 3.11E-01 | no | Plcxd3 | intron_variant | MODIFIER |
| 15 | 4579611 | AX-170010935 | 0 | T | C | 0.237 | -1.07E-01 | 1.07E-01 | 2.63E+00 | 2.68E+00 | 3.11E-01 | no | Plcxd3 | downstream_gene_variant | MODIFIER |
| 15 | 4591160 | AX-170010930 | 0 | G | A | 0.237 | -1.07E-01 | 1.07E-01 | 2.63E+00 | 2.68E+00 | 3.11E-01 | no | | intergenic_variant | MODIFIER |
| 15 | 4649783 | AX-170383677 | 0 | G | A | 0.237 | -1.07E-01 | 1.07E-01 | 2.63E+00 | 2.68E+00 | 3.11E-01 | no | | intergenic_variant | MODIFIER |

| | | | | | | | | | | | | | | | |
|---|---|---|---|---|---|---|---|---|---|---|---|---|---|---|---|
| 15 | 4663018 | AX-97803227 | 0 | T | C | 0.237 | -1.07E-01 | 1.07E-01 | 2.63E+00 | 2.68E+00 | 3.11E-01 | no | | intergenic_variant | MODIFIER |
| 15 | 4664143 | AX-170010934 | 0 | T | C | 0.237 | -1.07E-01 | 1.07E-01 | 2.63E+00 | 2.68E+00 | 3.11E-01 | no | | intergenic_variant | MODIFIER |
| 15 | 4701669 | AX-97839859 | 0 | T | A | 0.237 | -1.07E-01 | 1.07E-01 | 2.63E+00 | 2.68E+00 | 3.11E-01 | no | | intergenic_variant | MODIFIER |
| 15 | 4702277 | AX-170108055 | 0 | G | A | 0.237 | -1.07E-01 | 1.07E-01 | 2.63E+00 | 2.68E+00 | 3.11E-01 | no | | intergenic_variant | MODIFIER |
| 15 | 4752189 | AX-170126522 | 0 | G | A | 0.237 | -1.07E-01 | 1.07E-01 | 2.63E+00 | 2.68E+00 | 3.11E-01 | no | C6 | intron_variant | MODIFIER |
| 15 | 4755070 | AX-170054687 | 0 | T | C | 0.237 | -1.07E-01 | 1.07E-01 | 2.63E+00 | 2.68E+00 | 3.11E-01 | no | C6 | intron_variant | MODIFIER |
| 15 | 4755463 | AX-170054685 | 0 | G | A | 0.237 | -1.07E-01 | 1.07E-01 | 2.63E+00 | 2.68E+00 | 3.11E-01 | no | C6 | intron_variant | MODIFIER |
| 15 | 4761881 | AX-170384783 | 0 | A | G | 0.237 | -1.07E-01 | 1.07E-01 | 2.63E+00 | 2.68E+00 | 3.11E-01 | no | C6 | intron_variant | MODIFIER |
| 15 | 4762930 | AX-170125594 | 0 | G | T | 0.237 | -1.07E-01 | 1.07E-01 | 2.63E+00 | 2.68E+00 | 3.11E-01 | no | C6 | intron_variant | MODIFIER |
| 15 | 4766055 | AX-170229795 | 0 | C | T | 0.237 | -1.07E-01 | 1.07E-01 | 2.63E+00 | 2.68E+00 | 3.11E-01 | no | C6 | intron_variant | MODIFIER |
| 15 | 4782955 | AX-170370177 | 0 | T | C | 0.237 | -1.07E-01 | 1.07E-01 | 2.63E+00 | 2.68E+00 | 3.11E-01 | no | C6 | intron_variant | MODIFIER |
| 15 | 4790182 | AX-170349347 | 0 | C | T | 0.237 | -1.07E-01 | 1.07E-01 | 2.63E+00 | 2.68E+00 | 3.11E-01 | no | C6 | intron_variant | MODIFIER |
| 15 | 4790314 | AX-170382170 | 0 | T | C | 0.237 | -1.07E-01 | 1.07E-01 | 2.63E+00 | 2.68E+00 | 3.11E-01 | no | C6 | intron_variant | MODIFIER |
| 15 | 4815357 | AX-170249920 | 0 | C | G | 0.237 | -1.07E-01 | 1.07E-01 | 2.63E+00 | 2.68E+00 | 3.11E-01 | no | C6 | downstream_gene_variant | MODIFIER |
| 15 | 4839737 | AX-170303992 | 0 | C | T | 0.237 | -1.07E-01 | 1.07E-01 | 2.63E+00 | 2.68E+00 | 3.11E-01 | no | | intergenic_variant | MODIFIER |
| 15 | 4909078 | AX-170054689 | 0 | T | C | 0.237 | -1.07E-01 | 1.07E-01 | 2.63E+00 | 2.68E+00 | 3.11E-01 | no | Mroh2b | intron_variant | MODIFIER |
| 15 | 4916691 | AX-170295241 | 0 | G | A | 0.237 | -1.07E-01 | 1.07E-01 | 2.63E+00 | 2.68E+00 | 3.11E-01 | no | Mroh2b | intron_variant | MODIFIER |
| 15 | 4916913 | AX-170152894 | 0 | T | C | 0.237 | -1.07E-01 | 1.07E-01 | 2.63E+00 | 2.68E+00 | 3.11E-01 | no | Mroh2b | intron_variant | MODIFIER |
| 15 | 4950756 | AX-170113430 | 0 | T | A | 0.237 | -1.07E-01 | 1.07E-01 | 2.63E+00 | 2.68E+00 | 3.11E-01 | no | Mroh2b | intron_variant | MODIFIER |
| 15 | 4977892 | AX-170497958 | 0 | T | G | 0.237 | -1.07E-01 | 1.07E-01 | 2.63E+00 | 2.68E+00 | 3.11E-01 | no | | intergenic_variant | MODIFIER |
| 15 | 4993751 | AX-170130390 | 0 | G | A | 0.237 | -1.07E-01 | 1.07E-01 | 2.63E+00 | 2.68E+00 | 3.11E-01 | no | C7 | intron_variant | MODIFIER |
| 15 | 5009011 | AX-170318959 | 0 | C | T | 0.237 | -1.07E-01 | 1.07E-01 | 2.63E+00 | 2.68E+00 | 3.11E-01 | no | C7 | intron_variant | MODIFIER |
| 15 | 5014752 | AX-170098985 | 0 | C | T | 0.237 | -1.07E-01 | 1.07E-01 | 2.63E+00 | 2.68E+00 | 3.11E-01 | no | C7 | intron_variant | MODIFIER |
| 15 | 5022777 | AX-170214971 | 0 | T | C | 0.237 | -1.07E-01 | 1.07E-01 | 2.63E+00 | 2.68E+00 | 3.11E-01 | no | C7 | intron_variant | MODIFIER |
| 15 | 5023178 | AX-170366055 | 0 | T | G | 0.237 | -1.07E-01 | 1.07E-01 | 2.63E+00 | 2.68E+00 | 3.11E-01 | no | C7 | intron_variant | MODIFIER |
| 15 | 5046088 | AX-170180895 | 0 | A | C | 0.237 | -1.07E-01 | 1.07E-01 | 2.63E+00 | 2.68E+00 | 3.11E-01 | no | C7 | intron_variant | MODIFIER |
| 15 | 5047397 | AX-170093141 | 0 | G | T | 0.237 | -1.07E-01 | 1.07E-01 | 2.63E+00 | 2.68E+00 | 3.11E-01 | no | C7 | intron_variant | MODIFIER |
| 15 | 5059851 | AX-170368749 | 0 | T | C | 0.237 | -1.07E-01 | 1.07E-01 | 2.63E+00 | 2.68E+00 | 3.11E-01 | no | C7 | intron_variant | MODIFIER |
| 15 | 5083757 | AX-170037220 | 0 | A | G | 0.237 | -1.07E-01 | 1.07E-01 | 2.63E+00 | 2.68E+00 | 3.11E-01 | no | | intergenic_variant | MODIFIER |
| 15 | 5083780 | AX-170037223 | 0 | A | G | 0.237 | -1.07E-01 | 1.07E-01 | 2.63E+00 | 2.68E+00 | 3.11E-01 | no | | intergenic_variant | MODIFIER |
| 15 | 5114807 | AX-170010938 | 0 | C | A | 0.237 | -1.07E-01 | 1.07E-01 | 2.63E+00 | 2.68E+00 | 3.11E-01 | no | Rpl37 | upstream_gene_variant | MODIFIER |
| 15 | 5149694 | AX-170250347 | 0 | C | G | 0.237 | -1.07E-01 | 1.07E-01 | 2.63E+00 | 2.68E+00 | 3.11E-01 | no | Prkaa1 | intron_variant | MODIFIER |
| 15 | 5151112 | AX-170150725 | 0 | A | G | 0.237 | -1.07E-01 | 1.07E-01 | 2.63E+00 | 2.68E+00 | 3.11E-01 | no | Prkaa1 | intron_variant | MODIFIER |
| 15 | 5151159 | AX-170010936 | 0 | T | C | 0.237 | -1.07E-01 | 1.07E-01 | 2.63E+00 | 2.68E+00 | 3.11E-01 | no | Prkaa1 | intron_variant | MODIFIER |
| 15 | 5151205 | AX-170010940 | 0 | T | C | 0.237 | -1.07E-01 | 1.07E-01 | 2.63E+00 | 2.68E+00 | 3.11E-01 | no | Prkaa1 | intron_variant | MODIFIER |
| 15 | 5241128 | AX-170317175 | 0 | T | A | 0.237 | -1.07E-01 | 1.07E-01 | 2.63E+00 | 2.68E+00 | 3.11E-01 | no | Ptger4 | intron_variant | MODIFIER |
| 15 | 5343817 | AX-170010941 | 0 | C | T | 0.237 | -1.07E-01 | 1.07E-01 | 2.63E+00 | 2.68E+00 | 3.11E-01 | no | | regulatory_region_variant | MODIFIER |
| 15 | 5350278 | AX-170439216 | 0 | C | T | 0.237 | -1.07E-01 | 1.07E-01 | 2.63E+00 | 2.68E+00 | 3.11E-01 | no | | intergenic_variant | MODIFIER |
| 15 | 5355638 | AX-170248684 | 0 | T | C | 0.237 | -1.07E-01 | 1.07E-01 | 2.63E+00 | 2.68E+00 | 3.11E-01 | no | | intergenic_variant | MODIFIER |
| 15 | 5388922 | AX-170010942 | 0 | T | A | 0.237 | -1.07E-01 | 1.07E-01 | 2.63E+00 | 2.68E+00 | 3.11E-01 | no | | intergenic_variant | MODIFIER |
| 15 | 5389167 | AX-170010945 | 0 | C | T | 0.237 | -1.07E-01 | 1.07E-01 | 2.63E+00 | 2.68E+00 | 3.11E-01 | no | | intergenic_variant | MODIFIER |
| 15 | 5391040 | AX-97828021 | 0 | T | C | 0.237 | -1.07E-01 | 1.07E-01 | 2.63E+00 | 2.68E+00 | 3.11E-01 | no | | intergenic_variant | MODIFIER |
| 15 | 5403743 | AX-170054697 | 0 | C | T | 0.237 | -1.07E-01 | 1.07E-01 | 2.63E+00 | 2.68E+00 | 3.11E-01 | no | | intergenic_variant | MODIFIER |
| 15 | 5428131 | AX-170054696 | 0 | G | C | 0.237 | -1.07E-01 | 1.07E-01 | 2.63E+00 | 2.68E+00 | 3.11E-01 | no | | regulatory_region_variant | MODIFIER |
| 15 | 5473984 | AX-170195331 | 0 | T | C | 0.237 | -1.07E-01 | 1.07E-01 | 2.63E+00 | 2.68E+00 | 3.11E-01 | no | | intergenic_variant | MODIFIER |
| 15 | 5513560 | AX-97828018 | 0 | G | A | 0.237 | -1.07E-01 | 1.07E-01 | 2.63E+00 | 2.68E+00 | 3.11E-01 | no | 5430437J10Rik | intron_variant,non_coding_transcript_variant | MODIFIER |
| 15 | 5550156 | AX-170416068 | 0 | A | G | 0.237 | -1.07E-01 | 1.07E-01 | 2.63E+00 | 2.68E+00 | 3.11E-01 | no | 5430437J10Rik | intron_variant,non_coding_transcript_variant | MODIFIER |
| 15 | 5550212 | AX-170312345 | 0 | A | G | 0.237 | -1.07E-01 | 1.07E-01 | 2.63E+00 | 2.68E+00 | 3.11E-01 | no | 5430437J10Rik | intron_variant,non_coding_transcript_variant | MODIFIER |
| 15 | 5569165 | AX-170093142 | 0 | T | G | 0.237 | -1.07E-01 | 1.07E-01 | 2.63E+00 | 2.68E+00 | 3.11E-01 | no | 5430437J10Rik | intron_variant,non_coding_transcript_variant | MODIFIER |
| 15 | 5586574 | AX-170174782 | 0 | T | C | 0.237 | -1.07E-01 | 1.07E-01 | 2.63E+00 | 2.68E+00 | 3.11E-01 | no | 5430437J10Rik | intron_variant,non_coding_transcript_variant | MODIFIER |
| 15 | 5586592 | AX-170512362 | 0 | A | C | 0.237 | -1.07E-01 | 1.07E-01 | 2.63E+00 | 2.68E+00 | 3.11E-01 | no | 5430437J10Rik | intron_variant,non_coding_transcript_variant | MODIFIER |
| 15 | 5594756 | AX-170239627 | 0 | C | T | 0.237 | -1.07E-01 | 1.07E-01 | 2.63E+00 | 2.68E+00 | 3.11E-01 | no | 5430437J10Rik | non_coding_transcript_exon_variant | MODIFIER |
| 15 | 5619149 | AX-170511518 | 0 | C | T | 0.237 | -1.07E-01 | 1.07E-01 | 2.63E+00 | 2.68E+00 | 3.11E-01 | no | | intergenic_variant | MODIFIER |
| 15 | 5620420 | AX-170243924 | 0 | C | T | 0.237 | -1.07E-01 | 1.07E-01 | 2.63E+00 | 2.68E+00 | 3.11E-01 | no | | intergenic_variant | MODIFIER |

| | | | | | | | | | | | | | | | |
|---|---|---|---|---|---|---|---|---|---|---|---|---|---|---|---|
| 15 | 5620596 | AX-97789323 | 0 | C | T | 0.237 | -1.07E-01 | 1.07E-01 | 2.63E+00 | 2.68E+00 | 3.11E-01 | no | | intergenic_variant | MODIFIER |
| 15 | 5620988 | AX-170245626 | 0 | C | T | 0.237 | -1.07E-01 | 1.07E-01 | 2.63E+00 | 2.68E+00 | 3.11E-01 | no | | intergenic_variant | MODIFIER |
| 15 | 5675352 | AX-170531501 | 0 | G | C | 0.237 | -1.07E-01 | 1.07E-01 | 2.63E+00 | 2.68E+00 | 3.11E-01 | no | | intergenic_variant | MODIFIER |
| 15 | 5678347 | AX-170099957 | 0 | G | A | 0.237 | -1.07E-01 | 1.07E-01 | 2.63E+00 | 2.68E+00 | 3.11E-01 | no | | intergenic_variant | MODIFIER |
| 15 | 5691226 | AX-170128091 | 0 | T | A | 0.237 | -1.07E-01 | 1.07E-01 | 2.63E+00 | 2.68E+00 | 3.11E-01 | no | | intergenic_variant | MODIFIER |
| 15 | 5739647 | AX-170054703 | 0 | C | A | 0.237 | -1.07E-01 | 1.07E-01 | 2.63E+00 | 2.68E+00 | 3.11E-01 | no | | intergenic_variant | MODIFIER |
| 15 | 5740431 | AX-170242051 | 0 | C | T | 0.237 | -1.07E-01 | 1.07E-01 | 2.63E+00 | 2.68E+00 | 3.11E-01 | no | | intergenic_variant | MODIFIER |
| 15 | 5741479 | AX-170211978 | 0 | G | T | 0.237 | -1.07E-01 | 1.07E-01 | 2.63E+00 | 2.68E+00 | 3.11E-01 | no | | intergenic_variant | MODIFIER |
| 15 | 5778570 | AX-97837401 | 0 | C | G | 0.237 | -1.07E-01 | 1.07E-01 | 2.63E+00 | 2.68E+00 | 3.11E-01 | no | | intergenic_variant | MODIFIER |
| 15 | 5798666 | AX-97806412 | 0 | C | T | 0.237 | -1.07E-01 | 1.07E-01 | 2.63E+00 | 2.68E+00 | 3.11E-01 | no | | intergenic_variant | MODIFIER |
| 15 | 5805702 | AX-170286711 | 0 | G | A | 0.237 | -1.07E-01 | 1.07E-01 | 2.63E+00 | 2.68E+00 | 3.11E-01 | no | | intergenic_variant | MODIFIER |
| 15 | 5811345 | AX-170204374 | 0 | T | C | 0.237 | -1.07E-01 | 1.07E-01 | 2.63E+00 | 2.68E+00 | 3.11E-01 | no | | regulatory_region_variant | MODIFIER |
| 15 | 5811879 | AX-170421053 | 0 | A | G | 0.237 | -1.07E-01 | 1.07E-01 | 2.63E+00 | 2.68E+00 | 3.11E-01 | no | | intergenic_variant | MODIFIER |
| 15 | 5817965 | AX-170341024 | 0 | T | C | 0.237 | -1.07E-01 | 1.07E-01 | 2.63E+00 | 2.68E+00 | 3.11E-01 | no | | intergenic_variant | MODIFIER |
| 15 | 5819294 | AX-170010954 | 0 | C | G | 0.237 | -1.07E-01 | 1.07E-01 | 2.63E+00 | 2.68E+00 | 3.11E-01 | no | | intergenic_variant | MODIFIER |
| 15 | 5819353 | AX-170010948 | 0 | C | T | 0.237 | -1.07E-01 | 1.07E-01 | 2.63E+00 | 2.68E+00 | 3.11E-01 | no | | intergenic_variant | MODIFIER |
| 15 | 5892962 | AX-170195034 | 0 | T | C | 0.237 | -1.07E-01 | 1.07E-01 | 2.63E+00 | 2.68E+00 | 3.11E-01 | no | | intergenic_variant | MODIFIER |
| 15 | 5938770 | AX-170368980 | 0 | A | G | 0.237 | -1.07E-01 | 1.07E-01 | 2.63E+00 | 2.68E+00 | 3.11E-01 | no | | intergenic_variant | MODIFIER |
| 15 | 5974660 | AX-97809139 | 0 | C | T | 0.237 | -1.07E-01 | 1.07E-01 | 2.63E+00 | 2.68E+00 | 3.11E-01 | no | | intergenic_variant | MODIFIER |
| 15 | 6038503 | AX-170162894 | 0 | T | G | 0.237 | -1.07E-01 | 1.07E-01 | 2.63E+00 | 2.68E+00 | 3.11E-01 | no | Gm8047 | downstream_gene_variant | MODIFIER |
| 15 | 6054751 | AX-170204154 | 0 | A | G | 0.237 | -1.07E-01 | 1.07E-01 | 2.63E+00 | 2.68E+00 | 3.11E-01 | no | | intergenic_variant | MODIFIER |
| 15 | 6058961 | AX-170437855 | 0 | T | A | 0.237 | -1.07E-01 | 1.07E-01 | 2.63E+00 | 2.68E+00 | 3.11E-01 | no | | regulatory_region_variant | MODIFIER |
| 15 | 6060663 | AX-170301488 | 0 | A | C | 0.237 | -1.07E-01 | 1.07E-01 | 2.63E+00 | 2.68E+00 | 3.11E-01 | no | | regulatory_region_variant | MODIFIER |
| 15 | 6060932 | AX-170156741 | 0 | C | A | 0.237 | -1.07E-01 | 1.07E-01 | 2.63E+00 | 2.68E+00 | 3.11E-01 | no | | regulatory_region_variant | MODIFIER |
| 15 | 6067699 | AX-170152303 | 0 | G | A | 0.237 | -1.07E-01 | 1.07E-01 | 2.63E+00 | 2.68E+00 | 3.11E-01 | no | | intergenic_variant | MODIFIER |
| 15 | 6081941 | AX-170263620 | 0 | G | A | 0.237 | -1.07E-01 | 1.07E-01 | 2.63E+00 | 2.68E+00 | 3.11E-01 | no | | intergenic_variant | MODIFIER |
| 15 | 6102907 | AX-170532241 | 0 | A | G | 0.237 | -1.07E-01 | 1.07E-01 | 2.63E+00 | 2.68E+00 | 3.11E-01 | no | | intergenic_variant | MODIFIER |
| 15 | 6118794 | AX-170274498 | 0 | T | A | 0.237 | -1.07E-01 | 1.07E-01 | 2.63E+00 | 2.68E+00 | 3.11E-01 | no | | intergenic_variant | MODIFIER |
| 15 | 6123217 | AX-170337926 | 0 | T | C | 0.237 | -1.07E-01 | 1.07E-01 | 2.63E+00 | 2.68E+00 | 3.11E-01 | no | | intergenic_variant | MODIFIER |
| 15 | 6128226 | AX-170137093 | 0 | A | G | 0.237 | -1.07E-01 | 1.07E-01 | 2.63E+00 | 2.68E+00 | 3.11E-01 | no | | intergenic_variant | MODIFIER |
| 15 | 6181917 | AX-170054710 | 0 | A | G | 0.237 | -1.07E-01 | 1.07E-01 | 2.63E+00 | 2.68E+00 | 3.11E-01 | no | | intergenic_variant | MODIFIER |
| 15 | 6192339 | AX-170493182 | 0 | C | T | 0.237 | -1.07E-01 | 1.07E-01 | 2.63E+00 | 2.68E+00 | 3.11E-01 | no | | intergenic_variant | MODIFIER |
| 15 | 6219224 | AX-170054708 | 0 | A | G | 0.237 | -1.07E-01 | 1.07E-01 | 2.63E+00 | 2.68E+00 | 3.11E-01 | no | | intergenic_variant | MODIFIER |
| 15 | 6233863 | AX-170294700 | 0 | G | A | 0.237 | -1.07E-01 | 1.07E-01 | 2.63E+00 | 2.68E+00 | 3.11E-01 | no | | intergenic_variant | MODIFIER |
| 15 | 6245554 | AX-170297423 | 0 | A | T | 0.237 | -1.07E-01 | 1.07E-01 | 2.63E+00 | 2.68E+00 | 3.11E-01 | no | | intergenic_variant | MODIFIER |
| 15 | 6245623 | AX-170251561 | 0 | A | C | 0.237 | -1.07E-01 | 1.07E-01 | 2.63E+00 | 2.68E+00 | 3.11E-01 | no | | intergenic_variant | MODIFIER |
| 15 | 6249971 | AX-170474602 | 0 | T | C | 0.237 | -1.07E-01 | 1.07E-01 | 2.63E+00 | 2.68E+00 | 3.11E-01 | no | | intergenic_variant | MODIFIER |
| 15 | 6250069 | AX-170287590 | 0 | C | G | 0.237 | -1.07E-01 | 1.07E-01 | 2.63E+00 | 2.68E+00 | 3.11E-01 | no | | intergenic_variant | MODIFIER |
| 15 | 6250182 | AX-170190840 | 0 | G | A | 0.237 | -1.07E-01 | 1.07E-01 | 2.63E+00 | 2.68E+00 | 3.11E-01 | no | | intergenic_variant | MODIFIER |
| 15 | 6265501 | AX-170170053 | 0 | T | A | 0.237 | -1.07E-01 | 1.07E-01 | 2.63E+00 | 2.68E+00 | 3.11E-01 | no | | intergenic_variant | MODIFIER |
| 15 | 6279087 | AX-170283751 | 0 | T | C | 0.237 | -1.07E-01 | 1.07E-01 | 2.63E+00 | 2.68E+00 | 3.11E-01 | no | | regulatory_region_variant | MODIFIER |
| 15 | 6280480 | AX-170253713 | 0 | T | C | 0.237 | -1.07E-01 | 1.07E-01 | 2.63E+00 | 2.68E+00 | 3.11E-01 | no | | regulatory_region_variant | MODIFIER |
| 15 | 6310902 | AX-170105507 | 0 | A | C | 0.237 | -1.07E-01 | 1.07E-01 | 2.63E+00 | 2.68E+00 | 3.11E-01 | no | Dab2 | intron_variant | MODIFIER |
| 15 | 6335676 | AX-170109520 | 0 | T | C | 0.237 | -1.07E-01 | 1.07E-01 | 2.63E+00 | 2.68E+00 | 3.11E-01 | no | Dab2 | intron_variant | MODIFIER |
| 15 | 6336776 | AX-170336893 | 0 | G | C | 0.237 | -1.07E-01 | 1.07E-01 | 2.63E+00 | 2.68E+00 | 3.11E-01 | no | Dab2 | intron_variant | MODIFIER |
| 15 | 6359022 | AX-170195287 | 0 | T | C | 0.237 | -1.07E-01 | 1.07E-01 | 2.63E+00 | 2.68E+00 | 3.11E-01 | no | Dab2 | intron_variant | MODIFIER |
| 15 | 6414976 | AX-170054715 | 0 | A | G | 0.237 | -1.07E-01 | 1.07E-01 | 2.63E+00 | 2.68E+00 | 3.11E-01 | no | Dab2 | intron_variant | MODIFIER |
| 15 | 6440044 | AX-170105608 | 0 | T | C | 0.237 | -1.07E-01 | 1.07E-01 | 2.63E+00 | 2.68E+00 | 3.11E-01 | no | Dab2 | 3_prime_UTR_variant | MODIFIER |
| 15 | 6446260 | AX-170288045 | 0 | C | T | 0.237 | -1.07E-01 | 1.07E-01 | 2.63E+00 | 2.68E+00 | 3.11E-01 | no | C9 | intron_variant | MODIFIER |
| 15 | 6446283 | AX-170462507 | 0 | C | T | 0.237 | -1.07E-01 | 1.07E-01 | 2.63E+00 | 2.68E+00 | 3.11E-01 | no | C9 | intron_variant | MODIFIER |
| 15 | 6470316 | AX-170175476 | 0 | T | G | 0.237 | -1.07E-01 | 1.07E-01 | 2.63E+00 | 2.68E+00 | 3.11E-01 | no | C9 | intron_variant | MODIFIER |
| 15 | 6470374 | AX-170312519 | 0 | T | A | 0.237 | -1.07E-01 | 1.07E-01 | 2.63E+00 | 2.68E+00 | 3.11E-01 | no | C9 | intron_variant | MODIFIER |
| 15 | 6476566 | AX-170288660 | 0 | G | A | 0.237 | -1.07E-01 | 1.07E-01 | 2.63E+00 | 2.68E+00 | 3.11E-01 | no | C9 | intron_variant | MODIFIER |
| 15 | 6492469 | AX-170454638 | 0 | C | A | 0.237 | -1.07E-01 | 1.07E-01 | 2.63E+00 | 2.68E+00 | 3.11E-01 | no | C9 | intron_variant | MODIFIER |

| | | | | | | | | | | | | | | | |
|---|---|---|---|---|---|---|---|---|---|---|---|---|---|---|---|
| 15 | 6517774 | AX-170174259 | 0 | G | A | 0.237 | -1.07E-01 | 1.07E-01 | 2.63E+00 | 2.68E+00 | 3.11E-01 | no | | intergenic_variant | MODIFIER |
| 15 | 6518236 | AX-97820224 | 0 | T | G | 0.237 | -1.07E-01 | 1.07E-01 | 2.63E+00 | 2.68E+00 | 3.11E-01 | no | Fyb | upstream_gene_variant | MODIFIER |
| 15 | 6518500 | AX-170377284 | 0 | C | T | 0.237 | -1.07E-01 | 1.07E-01 | 2.63E+00 | 2.68E+00 | 3.11E-01 | no | Fyb | upstream_gene_variant | MODIFIER |
| 15 | 6520080 | AX-170184485 | 0 | C | T | 0.237 | -1.07E-01 | 1.07E-01 | 2.63E+00 | 2.68E+00 | 3.11E-01 | no | Fyb | upstream_gene_variant | MODIFIER |
| 15 | 6527096 | AX-170320967 | 0 | C | A | 0.237 | -1.07E-01 | 1.07E-01 | 2.63E+00 | 2.68E+00 | 3.11E-01 | no | Fyb | intron_variant | MODIFIER |
| 15 | 6534692 | AX-170438767 | 0 | C | G | 0.237 | -1.07E-01 | 1.07E-01 | 2.63E+00 | 2.68E+00 | 3.11E-01 | no | Fyb | intron_variant | MODIFIER |
| 15 | 6569602 | AX-170168453 | 0 | A | G | 0.237 | -1.07E-01 | 1.07E-01 | 2.63E+00 | 2.68E+00 | 3.11E-01 | no | Fyb | intron_variant | MODIFIER |
| 15 | 6571435 | AX-97814918 | 0 | G | A | 0.237 | -1.07E-01 | 1.07E-01 | 2.63E+00 | 2.68E+00 | 3.11E-01 | no | Fyb | intron_variant | MODIFIER |
| 15 | 6571464 | AX-170207680 | 0 | A | C | 0.237 | -1.07E-01 | 1.07E-01 | 2.63E+00 | 2.68E+00 | 3.11E-01 | no | Fyb | intron_variant | MODIFIER |
| 15 | 6585152 | AX-170441627 | 0 | G | A | 0.237 | -1.07E-01 | 1.07E-01 | 2.63E+00 | 2.68E+00 | 3.11E-01 | no | Fyb | intron_variant | MODIFIER |
| 15 | 6591690 | AX-170287608 | 0 | A | G | 0.237 | -1.07E-01 | 1.07E-01 | 2.63E+00 | 2.68E+00 | 3.11E-01 | no | Fyb | intron_variant | MODIFIER |
| 15 | 6591849 | AX-170145313 | 0 | C | T | 0.237 | -1.07E-01 | 1.07E-01 | 2.63E+00 | 2.68E+00 | 3.11E-01 | no | Fyb | intron_variant | MODIFIER |
| 15 | 6592083 | AX-170311308 | 0 | T | A | 0.237 | -1.07E-01 | 1.07E-01 | 2.63E+00 | 2.68E+00 | 3.11E-01 | no | Fyb | intron_variant | MODIFIER |
| 15 | 6592525 | AX-170340847 | 0 | A | G | 0.237 | -1.07E-01 | 1.07E-01 | 2.63E+00 | 2.68E+00 | 3.11E-01 | no | Fyb | intron_variant | MODIFIER |
| 15 | 6614362 | AX-170260669 | 0 | C | G | 0.237 | -1.07E-01 | 1.07E-01 | 2.63E+00 | 2.68E+00 | 3.11E-01 | no | Fyb | intron_variant | MODIFIER |
| 15 | 6625395 | AX-170054717 | 0 | C | G | 0.237 | -1.07E-01 | 1.07E-01 | 2.63E+00 | 2.68E+00 | 3.11E-01 | no | Fyb | intron_variant | MODIFIER |
| 15 | 6625479 | AX-97791042 | 0 | G | T | 0.237 | -1.07E-01 | 1.07E-01 | 2.63E+00 | 2.68E+00 | 3.11E-01 | no | Fyb | intron_variant | MODIFIER |
| 15 | 6626989 | AX-170054720 | 0 | A | G | 0.237 | -1.07E-01 | 1.07E-01 | 2.63E+00 | 2.68E+00 | 3.11E-01 | no | Fyb | intron_variant | MODIFIER |
| 15 | 6629443 | AX-170339812 | 0 | G | T | 0.237 | -1.07E-01 | 1.07E-01 | 2.63E+00 | 2.68E+00 | 3.11E-01 | no | Fyb | intron_variant | MODIFIER |
| 15 | 6664897 | AX-170394195 | 0 | G | A | 0.237 | -1.07E-01 | 1.07E-01 | 2.63E+00 | 2.68E+00 | 3.11E-01 | no | Fyb | downstream_gene_variant | MODIFIER |
| 15 | 6710888 | AX-170253495 | 0 | T | C | 0.237 | -1.07E-01 | 1.07E-01 | 2.63E+00 | 2.68E+00 | 3.11E-01 | no | Rictor | intron_variant | MODIFIER |
| 15 | 6713408 | AX-170371668 | 0 | C | A | 0.237 | -1.07E-01 | 1.07E-01 | 2.63E+00 | 2.68E+00 | 3.11E-01 | no | Rictor | intron_variant | MODIFIER |
| 15 | 6713873 | AX-170297809 | 0 | C | T | 0.237 | -1.07E-01 | 1.07E-01 | 2.63E+00 | 2.68E+00 | 3.11E-01 | no | Rictor | intron_variant | MODIFIER |
| 15 | 6714055 | AX-170222044 | 0 | A | G | 0.237 | -1.07E-01 | 1.07E-01 | 2.63E+00 | 2.68E+00 | 3.11E-01 | no | Rictor | intron_variant | MODIFIER |
| 15 | 6717084 | AX-170124781 | 0 | C | T | 0.237 | -1.07E-01 | 1.07E-01 | 2.63E+00 | 2.68E+00 | 3.11E-01 | no | Rictor | intron_variant | MODIFIER |
| 15 | 6717954 | AX-97795464 | 0 | C | T | 0.237 | -1.07E-01 | 1.07E-01 | 2.63E+00 | 2.68E+00 | 3.11E-01 | no | Rictor | intron_variant | MODIFIER |
| 15 | 6726053 | AX-170171334 | 0 | C | T | 0.237 | -1.07E-01 | 1.07E-01 | 2.63E+00 | 2.68E+00 | 3.11E-01 | no | Rictor | intron_variant | MODIFIER |
| 15 | 6753900 | AX-170514613 | 0 | C | A | 0.237 | -1.07E-01 | 1.07E-01 | 2.63E+00 | 2.68E+00 | 3.11E-01 | no | Rictor | intron_variant | MODIFIER |
| 15 | 6774970 | AX-170200444 | 0 | G | T | 0.237 | -1.07E-01 | 1.07E-01 | 2.63E+00 | 2.68E+00 | 3.11E-01 | no | Rictor | intron_variant | MODIFIER |
| 15 | 6775124 | AX-170350604 | 0 | T | C | 0.237 | -1.07E-01 | 1.07E-01 | 2.63E+00 | 2.68E+00 | 3.11E-01 | no | Rictor | intron_variant | MODIFIER |
| 15 | 6831116 | AX-170384099 | 0 | T | C | 0.237 | -1.07E-01 | 1.07E-01 | 2.63E+00 | 2.68E+00 | 3.11E-01 | no | Osmr | intron_variant | MODIFIER |
| 15 | 6840542 | AX-170258723 | 0 | G | A | 0.237 | -1.07E-01 | 1.07E-01 | 2.63E+00 | 2.68E+00 | 3.11E-01 | no | Osmr | intron_variant | MODIFIER |
| 15 | 6841845 | AX-170054724 | 0 | T | C | 0.237 | -1.07E-01 | 1.07E-01 | 2.63E+00 | 2.68E+00 | 3.11E-01 | no | Osmr | intron_variant | MODIFIER |
| 15 | 6888263 | AX-170375276 | 0 | C | T | 0.237 | -1.07E-01 | 1.07E-01 | 2.63E+00 | 2.68E+00 | 3.11E-01 | no | | intergenic_variant | MODIFIER |
| 15 | 6901954 | AX-170534163 | 0 | C | T | 0.237 | -1.07E-01 | 1.07E-01 | 2.63E+00 | 2.68E+00 | 3.11E-01 | no | | intergenic_variant | MODIFIER |
| 15 | 6922710 | AX-170486602 | 0 | G | A | 0.237 | -1.07E-01 | 1.07E-01 | 2.63E+00 | 2.68E+00 | 3.11E-01 | no | | intergenic_variant | MODIFIER |
| 15 | 6922942 | AX-170492012 | 0 | T | G | 0.237 | -1.07E-01 | 1.07E-01 | 2.63E+00 | 2.68E+00 | 3.11E-01 | no | | intergenic_variant | MODIFIER |
| 15 | 6923036 | AX-170322262 | 0 | T | C | 0.237 | -1.07E-01 | 1.07E-01 | 2.63E+00 | 2.68E+00 | 3.11E-01 | no | | intergenic_variant | MODIFIER |
| 15 | 6954485 | AX-170519303 | 0 | A | C | 0.237 | -1.07E-01 | 1.07E-01 | 2.63E+00 | 2.68E+00 | 3.11E-01 | no | | intergenic_variant | MODIFIER |
| 15 | 6957728 | AX-170208758 | 0 | C | G | 0.237 | -1.07E-01 | 1.07E-01 | 2.63E+00 | 2.68E+00 | 3.11E-01 | no | | intergenic_variant | MODIFIER |
| 15 | 6960918 | AX-170272516 | 0 | T | C | 0.237 | -1.07E-01 | 1.07E-01 | 2.63E+00 | 2.68E+00 | 3.11E-01 | no | | intergenic_variant | MODIFIER |
| 15 | 6992026 | AX-170534939 | 0 | A | G | 0.237 | -1.07E-01 | 1.07E-01 | 2.63E+00 | 2.68E+00 | 3.11E-01 | no | Gm38282 | upstream_gene_variant | MODIFIER |
| 15 | 7013666 | AX-170186047 | 0 | C | T | 0.237 | -1.07E-01 | 1.07E-01 | 2.63E+00 | 2.68E+00 | 3.11E-01 | no | | intergenic_variant | MODIFIER |
| 15 | 7036468 | AX-170514300 | 0 | A | C | 0.237 | -1.07E-01 | 1.07E-01 | 2.63E+00 | 2.68E+00 | 3.11E-01 | no | | intergenic_variant | MODIFIER |
| 15 | 7043011 | AX-170464463 | 0 | G | A | 0.237 | -1.07E-01 | 1.07E-01 | 2.63E+00 | 2.68E+00 | 3.11E-01 | no | | intergenic_variant | MODIFIER |
| 15 | 7043084 | AX-170120835 | 0 | A | G | 0.237 | -1.07E-01 | 1.07E-01 | 2.63E+00 | 2.68E+00 | 3.11E-01 | no | | intergenic_variant | MODIFIER |
| 15 | 7043493 | AX-97820767 | 0 | A | C | 0.237 | -1.07E-01 | 1.07E-01 | 2.63E+00 | 2.68E+00 | 3.11E-01 | no | | regulatory_region_variant | MODIFIER |
| 15 | 7047656 | AX-170422746 | 0 | G | C | 0.237 | -1.07E-01 | 1.07E-01 | 2.63E+00 | 2.68E+00 | 3.11E-01 | no | | intergenic_variant | MODIFIER |
| 15 | 7050652 | AX-170368172 | 0 | G | A | 0.237 | -1.07E-01 | 1.07E-01 | 2.63E+00 | 2.68E+00 | 3.11E-01 | no | | intergenic_variant | MODIFIER |
| 15 | 7050980 | AX-170126043 | 0 | A | C | 0.237 | -1.07E-01 | 1.07E-01 | 2.63E+00 | 2.68E+00 | 3.11E-01 | no | | intergenic_variant | MODIFIER |
| 15 | 7053193 | AX-170248324 | 0 | T | A | 0.237 | -1.07E-01 | 1.07E-01 | 2.63E+00 | 2.68E+00 | 3.11E-01 | no | | intergenic_variant | MODIFIER |
| 15 | 7060669 | AX-170312892 | 0 | A | G | 0.237 | -1.07E-01 | 1.07E-01 | 2.63E+00 | 2.68E+00 | 3.11E-01 | no | | regulatory_region_variant | MODIFIER |
| 15 | 7061089 | AX-170379242 | 0 | A | G | 0.237 | -1.07E-01 | 1.07E-01 | 2.63E+00 | 2.68E+00 | 3.11E-01 | no | | intergenic_variant | MODIFIER |
| 15 | 7064141 | AX-170488928 | 0 | C | T | 0.237 | -1.07E-01 | 1.07E-01 | 2.63E+00 | 2.68E+00 | 3.11E-01 | no | | intergenic_variant | MODIFIER |

| | | | | | | | | | | | | | | | |
|---|---|---|---|---|---|---|---|---|---|---|---|---|---|---|---|
| 15 | 7064793 | AX-170431845 | 0 | T | C | 0.237 | -1.07E-01 | 1.07E-01 | 2.63E+00 | 2.68E+00 | 3.11E-01 | no | | intergenic_variant | MODIFIER |
| 15 | 7065096 | AX-170375006 | 0 | T | C | 0.237 | -1.07E-01 | 1.07E-01 | 2.63E+00 | 2.68E+00 | 3.11E-01 | no | | intergenic_variant | MODIFIER |
| 15 | 7473654 | AX-97790059 | 0 | C | A | 0.237 | -1.07E-01 | 1.07E-01 | 2.63E+00 | 2.68E+00 | 3.11E-01 | no | | intergenic_variant | MODIFIER |
| 15 | 7497672 | AX-170010964 | 0 | T | G | 0.237 | -1.07E-01 | 1.07E-01 | 2.63E+00 | 2.68E+00 | 3.11E-01 | no | | intergenic_variant | MODIFIER |
| 15 | 7610880 | AX-170474057 | 0 | C | T | 0.237 | -1.07E-01 | 1.07E-01 | 2.63E+00 | 2.68E+00 | 3.11E-01 | no | | intergenic_variant | MODIFIER |
| 15 | 7618541 | AX-170292892 | 0 | T | C | 0.237 | -1.07E-01 | 1.07E-01 | 2.63E+00 | 2.68E+00 | 3.11E-01 | no | | intergenic_variant | MODIFIER |
| 15 | 7624373 | AX-170187424 | 0 | A | G | 0.237 | -1.07E-01 | 1.07E-01 | 2.63E+00 | 2.68E+00 | 3.11E-01 | no | | regulatory_region_variant | MODIFIER |
| 15 | 7626288 | AX-170260760 | 0 | T | C | 0.237 | -1.07E-01 | 1.07E-01 | 2.63E+00 | 2.68E+00 | 3.11E-01 | no | | intergenic_variant | MODIFIER |
| 15 | 7626888 | AX-170346600 | 0 | C | G | 0.237 | -1.07E-01 | 1.07E-01 | 2.63E+00 | 2.68E+00 | 3.11E-01 | no | | intergenic_variant | MODIFIER |
| 15 | 7629244 | AX-170514862 | 0 | C | T | 0.237 | -1.07E-01 | 1.07E-01 | 2.63E+00 | 2.68E+00 | 3.11E-01 | no | | intergenic_variant | MODIFIER |
| 15 | 7630505 | AX-170494600 | 0 | A | G | 0.237 | -1.07E-01 | 1.07E-01 | 2.63E+00 | 2.68E+00 | 3.11E-01 | no | | intergenic_variant | MODIFIER |
| 15 | 7634544 | AX-170138536 | 0 | A | C | 0.237 | -1.07E-01 | 1.07E-01 | 2.63E+00 | 2.68E+00 | 3.11E-01 | no | | intergenic_variant | MODIFIER |
| 15 | 7635278 | AX-170383353 | 0 | T | C | 0.237 | -1.07E-01 | 1.07E-01 | 2.63E+00 | 2.68E+00 | 3.11E-01 | no | | regulatory_region_variant | MODIFIER |
| 15 | 7637164 | AX-170374233 | 0 | G | C | 0.237 | -1.07E-01 | 1.07E-01 | 2.63E+00 | 2.68E+00 | 3.11E-01 | no | | intergenic_variant | MODIFIER |
| 15 | 7823941 | AX-170087756 | 0 | T | G | 0.237 | -1.07E-01 | 1.07E-01 | 2.63E+00 | 2.68E+00 | 3.11E-01 | no | Gdnf | intron_variant | MODIFIER |
| 15 | 7824824 | AX-170402501 | 0 | T | G | 0.237 | -1.07E-01 | 1.07E-01 | 2.63E+00 | 2.68E+00 | 3.11E-01 | no | Gdnf | intron_variant | MODIFIER |
| 15 | 7839004 | AX-97856883 | 0 | T | C | 0.237 | -1.07E-01 | 1.07E-01 | 2.63E+00 | 2.68E+00 | 3.11E-01 | no | Gdnf | downstream_gene_variant | MODIFIER |
| 15 | 7931407 | AX-97807371 | 0 | G | A | 0.237 | -1.07E-01 | 1.07E-01 | 2.63E+00 | 2.68E+00 | 3.11E-01 | no | Wdr70 | intron_variant | MODIFIER |
| 15 | 7955508 | AX-97828351 | 0 | C | T | 0.237 | -1.07E-01 | 1.07E-01 | 2.63E+00 | 2.68E+00 | 3.11E-01 | no | Wdr70 | intron_variant | MODIFIER |
| 15 | 7960120 | AX-170010973 | 0 | T | A | 0.237 | -1.07E-01 | 1.07E-01 | 2.63E+00 | 2.68E+00 | 3.11E-01 | no | Wdr70 | intron_variant | MODIFIER |
| 15 | 7978497 | AX-170087757 | 0 | C | T | 0.237 | -1.07E-01 | 1.07E-01 | 2.63E+00 | 2.68E+00 | 3.11E-01 | no | Wdr70 | intron_variant | MODIFIER |
| 15 | 8090036 | AX-170010969 | 0 | T | C | 0.237 | -1.07E-01 | 1.07E-01 | 2.63E+00 | 2.68E+00 | 3.11E-01 | no | Wdr70 | intron_variant | MODIFIER |
| 15 | 8090812 | AX-170269965 | 0 | A | T | 0.237 | -1.07E-01 | 1.07E-01 | 2.63E+00 | 2.68E+00 | 3.11E-01 | no | Wdr70 | intron_variant | MODIFIER |
| 15 | 8093533 | AX-97828354 | 0 | C | A | 0.237 | -1.07E-01 | 1.07E-01 | 2.63E+00 | 2.68E+00 | 3.11E-01 | no | Wdr70 | intron_variant | MODIFIER |
| 15 | 8098481 | AX-170098249 | 0 | C | A | 0.237 | -1.07E-01 | 1.07E-01 | 2.63E+00 | 2.68E+00 | 3.11E-01 | no | Wdr70 | intron_variant | MODIFIER |
| 15 | 8113200 | AX-170121619 | 0 | A | C | 0.237 | -1.07E-01 | 1.07E-01 | 2.63E+00 | 2.68E+00 | 3.11E-01 | no | Nup155 | intron_variant | MODIFIER |
| 15 | 8132660 | AX-170054755 | 0 | T | C | 0.237 | -1.07E-01 | 1.07E-01 | 2.63E+00 | 2.68E+00 | 3.11E-01 | no | Nup155 | intron_variant | MODIFIER |
| 15 | 8132871 | AX-170361340 | 0 | A | G | 0.237 | -1.07E-01 | 1.07E-01 | 2.63E+00 | 2.68E+00 | 3.11E-01 | no | Nup155 | intron_variant | MODIFIER |
| 15 | 8845050 | AX-97828377 | 0 | C | G | 0.237 | -1.07E-01 | 1.07E-01 | 2.63E+00 | 2.68E+00 | 3.11E-01 | no | | intergenic_variant | MODIFIER |
| 15 | 8978429 | AX-170336264 | 0 | C | T | 0.237 | -1.07E-01 | 1.07E-01 | 2.63E+00 | 2.68E+00 | 3.11E-01 | no | Ranbp3l | intron_variant | MODIFIER |
| 15 | 8979335 | AX-97828403 | 0 | G | A | 0.237 | -1.07E-01 | 1.07E-01 | 2.63E+00 | 2.68E+00 | 3.11E-01 | no | Ranbp3l | intron_variant | MODIFIER |
| 15 | 8991380 | AX-97828924 | 0 | G | A | 0.237 | -1.07E-01 | 1.07E-01 | 2.63E+00 | 2.68E+00 | 3.11E-01 | no | Ranbp3l | intron_variant | MODIFIER |
| 15 | 9196469 | AX-170273313 | 0 | C | T | 0.237 | -1.07E-01 | 1.07E-01 | 2.63E+00 | 2.68E+00 | 3.11E-01 | no | Lmbrd2 | intron_variant | MODIFIER |
| 15 | 9200530 | AX-170191709 | 0 | G | T | 0.237 | -1.07E-01 | 1.07E-01 | 2.63E+00 | 2.68E+00 | 3.11E-01 | no | Lmbrd2 | downstream_gene_variant | MODIFIER |
| 15 | 9203148 | AX-170316849 | 0 | C | T | 0.237 | -1.07E-01 | 1.07E-01 | 2.63E+00 | 2.68E+00 | 3.11E-01 | no | Lmbrd2 | downstream_gene_variant | MODIFIER |
| 15 | 9255336 | AX-170011000 | 0 | G | C | 0.237 | -1.07E-01 | 1.07E-01 | 2.63E+00 | 2.68E+00 | 3.11E-01 | no | Gm8139 | non_coding_transcript_exon_variant | MODIFIER |
| 15 | 9256494 | AX-170087762 | 0 | C | T | 0.237 | -1.07E-01 | 1.07E-01 | 2.63E+00 | 2.68E+00 | 3.11E-01 | no | Gm8139 | intron_variant,non_coding_transcript_variant | MODIFIER |
| 15 | 9336735 | AX-97831982 | 0 | T | C | 0.237 | -1.07E-01 | 1.07E-01 | 2.63E+00 | 2.68E+00 | 3.11E-01 | no | Ugt3a2 | intron_variant | MODIFIER |
| 15 | 9585361 | AX-168379283 | 0 | G | A | 0.237 | -1.07E-01 | 1.07E-01 | 2.63E+00 | 2.68E+00 | 3.11E-01 | no | Spef2 | intron_variant | MODIFIER |
| 15 | 9634190 | AX-170011003 | 0 | A | G | 0.237 | -1.07E-01 | 1.07E-01 | 2.63E+00 | 2.68E+00 | 3.11E-01 | no | Spef2 | intron_variant | MODIFIER |
| 15 | 9634294 | AX-170011007 | 0 | G | T | 0.237 | -1.07E-01 | 1.07E-01 | 2.63E+00 | 2.68E+00 | 3.11E-01 | no | Spef2 | intron_variant | MODIFIER |
| 15 | 9646634 | AX-97793977 | 0 | A | C | 0.237 | -1.07E-01 | 1.07E-01 | 2.63E+00 | 2.68E+00 | 3.11E-01 | no | Spef2 | intron_variant | MODIFIER |
| 15 | 9647669 | AX-170168934 | 0 | T | C | 0.237 | -1.07E-01 | 1.07E-01 | 2.63E+00 | 2.68E+00 | 3.11E-01 | no | Spef2 | intron_variant | MODIFIER |
| 15 | 9710532 | AX-170054772 | 0 | T | G | 0.237 | -1.07E-01 | 1.07E-01 | 2.63E+00 | 2.68E+00 | 3.11E-01 | no | Spef2 | intron_variant | MODIFIER |
| 15 | 9785056 | AX-170167745 | 0 | A | T | 0.237 | -1.07E-01 | 1.07E-01 | 2.63E+00 | 2.68E+00 | 3.11E-01 | no | | intergenic_variant | MODIFIER |
| 15 | 9836680 | AX-170248621 | 0 | T | C | 0.237 | -1.07E-01 | 1.07E-01 | 2.63E+00 | 2.68E+00 | 3.11E-01 | no | | intergenic_variant | MODIFIER |
| 15 | 9849738 | AX-170011004 | 0 | T | C | 0.237 | -1.07E-01 | 1.07E-01 | 2.63E+00 | 2.68E+00 | 3.11E-01 | no | | intergenic_variant | MODIFIER |
| 15 | 9890095 | AX-97828131 | 0 | G | A | 0.237 | -1.07E-01 | 1.07E-01 | 2.63E+00 | 2.68E+00 | 3.11E-01 | no | | intergenic_variant | MODIFIER |
| 15 | 10069685 | AX-97828432 | 0 | G | C | 0.237 | -1.07E-01 | 1.07E-01 | 2.63E+00 | 2.68E+00 | 3.11E-01 | no | | intergenic_variant | MODIFIER |
| 15 | 10120497 | AX-170424392 | 0 | A | G | 0.237 | -1.07E-01 | 1.07E-01 | 2.63E+00 | 2.68E+00 | 3.11E-01 | no | | intergenic_variant | MODIFIER |
| 15 | 10160923 | AX-170011021 | 0 | G | A | 0.237 | -1.07E-01 | 1.07E-01 | 2.63E+00 | 2.68E+00 | 3.11E-01 | no | | intergenic_variant | MODIFIER |
| 15 | 10161011 | AX-170011013 | 0 | G | A | 0.237 | -1.07E-01 | 1.07E-01 | 2.63E+00 | 2.68E+00 | 3.11E-01 | no | | intergenic_variant | MODIFIER |
| 15 | 10180647 | AX-97792081 | 0 | C | T | 0.237 | -1.07E-01 | 1.07E-01 | 2.63E+00 | 2.68E+00 | 3.11E-01 | no | Prlr | intron_variant,non_coding_transcript_variant | MODIFIER |
| 15 | 10194524 | AX-97827550 | 0 | C | T | 0.237 | -1.07E-01 | 1.07E-01 | 2.63E+00 | 2.68E+00 | 3.11E-01 | no | Prlr | intron_variant,non_coding_transcript_variant | MODIFIER |

| | | | | | | | | | | | | | | | |
|---|---|---|---|---|---|---|---|---|---|---|---|---|---|---|---|
| 15 | 10234967 | AX-170011020 | 0 | G | A | 0.237 | -1.07E-01 | 1.07E-01 | 2.63E+00 | 2.68E+00 | 3.11E-01 | no | Prlr | intron_variant,non_coding_transcript_variant | MODIFIER |
| 15 | 10267261 | AX-170093155 | 0 | G | A | 0.237 | -1.07E-01 | 1.07E-01 | 2.63E+00 | 2.68E+00 | 3.11E-01 | no | Prlr | intron_variant,non_coding_transcript_variant | MODIFIER |
| 15 | 10271069 | AX-170054783 | 0 | A | G | 0.237 | -1.07E-01 | 1.07E-01 | 2.63E+00 | 2.68E+00 | 3.11E-01 | no | Prlr | intron_variant,non_coding_transcript_variant | MODIFIER |
| 15 | 10274045 | AX-170011017 | 0 | A | G | 0.237 | -1.07E-01 | 1.07E-01 | 2.63E+00 | 2.68E+00 | 3.11E-01 | no | Prlr | intron_variant,non_coding_transcript_variant | MODIFIER |
| 15 | 10277694 | AX-170011015 | 0 | T | C | 0.237 | -1.07E-01 | 1.07E-01 | 2.63E+00 | 2.68E+00 | 3.11E-01 | no | Prlr | intron_variant,non_coding_transcript_variant | MODIFIER |
| 15 | 10280509 | AX-170054782 | 0 | G | T | 0.237 | -1.07E-01 | 1.07E-01 | 2.63E+00 | 2.68E+00 | 3.11E-01 | no | Prlr | intron_variant,non_coding_transcript_variant | MODIFIER |
| 15 | 10294297 | AX-170011019 | 0 | G | C | 0.237 | -1.07E-01 | 1.07E-01 | 2.63E+00 | 2.68E+00 | 3.11E-01 | no | Prlr | intron_variant,non_coding_transcript_variant | MODIFIER |
| 15 | 10297477 | AX-97802852 | 0 | A | G | 0.237 | -1.07E-01 | 1.07E-01 | 2.63E+00 | 2.68E+00 | 3.11E-01 | no | Prlr | intron_variant,non_coding_transcript_variant | MODIFIER |
| 15 | 10303721 | AX-170485292 | 0 | G | A | 0.237 | -1.07E-01 | 1.07E-01 | 2.63E+00 | 2.68E+00 | 3.11E-01 | no | Prlr | intron_variant,non_coding_transcript_variant | MODIFIER |
| 15 | 10328867 | AX-170162394 | 0 | G | A | 0.237 | -1.07E-01 | 1.07E-01 | 2.63E+00 | 2.68E+00 | 3.11E-01 | no | Prlr | non_coding_transcript_exon_variant | MODIFIER |
| 15 | 10426288 | AX-97814237 | 0 | C | T | 0.237 | -1.07E-01 | 1.07E-01 | 2.63E+00 | 2.68E+00 | 3.11E-01 | no | | intergenic_variant | MODIFIER |
| 15 | 10437817 | AX-97804092 | 0 | G | A | 0.237 | -1.07E-01 | 1.07E-01 | 2.63E+00 | 2.68E+00 | 3.11E-01 | no | | intergenic_variant | MODIFIER |
| 15 | 10506211 | AX-170289863 | 0 | A | G | 0.237 | -1.07E-01 | 1.07E-01 | 2.63E+00 | 2.68E+00 | 3.11E-01 | no | Ttc23l | intron_variant | MODIFIER |
| 15 | 10521491 | AX-97838882 | 0 | G | A | 0.237 | -1.07E-01 | 1.07E-01 | 2.63E+00 | 2.68E+00 | 3.11E-01 | no | Ttc23l | intron_variant | MODIFIER |
| 15 | 10521584 | AX-170292607 | 0 | A | G | 0.237 | -1.07E-01 | 1.07E-01 | 2.63E+00 | 2.68E+00 | 3.11E-01 | no | Ttc23l | intron_variant | MODIFIER |
| 15 | 10537828 | AX-170296016 | 0 | T | C | 0.237 | -1.07E-01 | 1.07E-01 | 2.63E+00 | 2.68E+00 | 3.11E-01 | no | Ttc23l | intron_variant | MODIFIER |
| 15 | 10545928 | AX-97838701 | 0 | A | G | 0.237 | -1.07E-01 | 1.07E-01 | 2.63E+00 | 2.68E+00 | 3.11E-01 | no | Ttc23l | intron_variant | MODIFIER |
| 15 | 10559418 | AX-168385523 | 0 | C | T | 0.237 | -1.07E-01 | 1.07E-01 | 2.63E+00 | 2.68E+00 | 3.11E-01 | no | Ttc23l | upstream_gene_variant | MODIFIER |
| 15 | 10563883 | AX-170105947 | 0 | C | T | 0.237 | -1.07E-01 | 1.07E-01 | 2.63E+00 | 2.68E+00 | 3.11E-01 | no | Gm8174 | upstream_gene_variant | MODIFIER |
| 15 | 10566880 | AX-170512762 | 0 | A | G | 0.237 | -1.07E-01 | 1.07E-01 | 2.63E+00 | 2.68E+00 | 3.11E-01 | no | Rai14 | downstream_gene_variant | MODIFIER |
| 15 | 10572664 | AX-97853649 | 0 | A | G | 0.237 | -1.07E-01 | 1.07E-01 | 2.63E+00 | 2.68E+00 | 3.11E-01 | no | Rai14 | intron_variant | MODIFIER |
| 15 | 10584809 | AX-170230836 | 0 | T | C | 0.237 | -1.07E-01 | 1.07E-01 | 2.63E+00 | 2.68E+00 | 3.11E-01 | no | Rai14 | intron_variant | MODIFIER |
| 15 | 10585359 | AX-97809023 | 0 | C | T | 0.237 | -1.07E-01 | 1.07E-01 | 2.63E+00 | 2.68E+00 | 3.11E-01 | no | Rai14 | intron_variant | MODIFIER |
| 15 | 10675971 | AX-170294821 | 0 | C | A | 0.237 | -1.07E-01 | 1.07E-01 | 2.63E+00 | 2.68E+00 | 3.11E-01 | no | Rai14 | intron_variant | MODIFIER |
| 15 | 10677081 | AX-170362992 | 0 | T | G | 0.237 | -1.07E-01 | 1.07E-01 | 2.63E+00 | 2.68E+00 | 3.11E-01 | no | Rai14 | intron_variant | MODIFIER |
| 15 | 10679416 | AX-170356462 | 0 | C | G | 0.237 | -1.07E-01 | 1.07E-01 | 2.63E+00 | 2.68E+00 | 3.11E-01 | no | Rai14 | intron_variant | MODIFIER |
| 15 | 10680888 | AX-170341184 | 0 | C | T | 0.237 | -1.07E-01 | 1.07E-01 | 2.63E+00 | 2.68E+00 | 3.11E-01 | no | Rai14 | intron_variant | MODIFIER |
| 15 | 10686241 | AX-170445172 | 0 | A | G | 0.237 | -1.07E-01 | 1.07E-01 | 2.63E+00 | 2.68E+00 | 3.11E-01 | no | Rai14 | intron_variant | MODIFIER |
| 15 | 10688111 | AX-170011023 | 0 | A | C | 0.237 | -1.07E-01 | 1.07E-01 | 2.63E+00 | 2.68E+00 | 3.11E-01 | no | Rai14 | intron_variant | MODIFIER |
| 15 | 10700578 | AX-170011025 | 0 | T | G | 0.237 | -1.07E-01 | 1.07E-01 | 2.63E+00 | 2.68E+00 | 3.11E-01 | no | Rai14 | intron_variant | MODIFIER |
| 15 | 10742102 | AX-97821371 | 0 | C | T | 0.237 | -1.07E-01 | 1.07E-01 | 2.63E+00 | 2.68E+00 | 3.11E-01 | no | 4930556M19Rik | intron_variant,non_coding_transcript_variant | MODIFIER |
| 15 | 10760350 | AX-170139202 | 0 | A | C | 0.237 | -1.07E-01 | 1.07E-01 | 2.63E+00 | 2.68E+00 | 3.11E-01 | no | 4930556M19Rik | intron_variant,non_coding_transcript_variant | MODIFIER |
| 15 | 10765679 | AX-170054794 | 0 | A | G | 0.237 | -1.07E-01 | 1.07E-01 | 2.63E+00 | 2.68E+00 | 3.11E-01 | no | 4930556M19Rik | intron_variant,non_coding_transcript_variant | MODIFIER |
| 15 | 10769171 | AX-170366071 | 0 | G | C | 0.237 | -1.07E-01 | 1.07E-01 | 2.63E+00 | 2.68E+00 | 3.11E-01 | no | 4930556M19Rik | non_coding_transcript_exon_variant | MODIFIER |
| 15 | 10769350 | AX-170397851 | 0 | G | A | 0.237 | -1.07E-01 | 1.07E-01 | 2.63E+00 | 2.68E+00 | 3.11E-01 | no | 4930556M19Rik | intron_variant,non_coding_transcript_variant | MODIFIER |
| 15 | 10769763 | AX-170011024 | 0 | G | C | 0.237 | -1.07E-01 | 1.07E-01 | 2.63E+00 | 2.68E+00 | 3.11E-01 | no | 4930556M19Rik | intron_variant,non_coding_transcript_variant | MODIFIER |
| 15 | 10770106 | AX-97813323 | 0 | C | T | 0.237 | -1.07E-01 | 1.07E-01 | 2.63E+00 | 2.68E+00 | 3.11E-01 | no | 4930556M19Rik | intron_variant,non_coding_transcript_variant | MODIFIER |
| 15 | 10780203 | AX-170011029 | 0 | C | T | 0.237 | -1.07E-01 | 1.07E-01 | 2.63E+00 | 2.68E+00 | 3.11E-01 | no | 4930556M19Rik | intron_variant,non_coding_transcript_variant | MODIFIER |
| 15 | 10813683 | AX-170011028 | 0 | T | A | 0.237 | -1.07E-01 | 1.07E-01 | 2.63E+00 | 2.68E+00 | 3.11E-01 | no | Gm19276 | upstream_gene_variant | MODIFIER |
| 15 | 10815076 | AX-170518760 | 0 | C | T | 0.237 | -1.07E-01 | 1.07E-01 | 2.63E+00 | 2.68E+00 | 3.11E-01 | no | Gm19276 | intron_variant,non_coding_transcript_variant | MODIFIER |
| 15 | 10815282 | AX-170175355 | 0 | C | A | 0.237 | -1.07E-01 | 1.07E-01 | 2.63E+00 | 2.68E+00 | 3.11E-01 | no | Gm19276 | intron_variant,non_coding_transcript_variant | MODIFIER |
| 15 | 10817773 | AX-170308217 | 0 | C | T | 0.237 | -1.07E-01 | 1.07E-01 | 2.63E+00 | 2.68E+00 | 3.11E-01 | no | Gm19276 | intron_variant,non_coding_transcript_variant | MODIFIER |
| 15 | 10818630 | AX-170424463 | 0 | C | T | 0.237 | -1.07E-01 | 1.07E-01 | 2.63E+00 | 2.68E+00 | 3.11E-01 | no | Gm19276 | intron_variant,non_coding_transcript_variant | MODIFIER |
| 15 | 10818784 | AX-170532387 | 0 | C | T | 0.237 | -1.07E-01 | 1.07E-01 | 2.63E+00 | 2.68E+00 | 3.11E-01 | no | Gm19276 | intron_variant,non_coding_transcript_variant | MODIFIER |
| 15 | 10819091 | AX-97803712 | 0 | T | C | 0.237 | -1.07E-01 | 1.07E-01 | 2.63E+00 | 2.68E+00 | 3.11E-01 | no | Gm19276 | intron_variant,non_coding_transcript_variant | MODIFIER |
| 15 | 10822570 | AX-170301605 | 0 | T | C | 0.237 | -1.07E-01 | 1.07E-01 | 2.63E+00 | 2.68E+00 | 3.11E-01 | no | Gm19276 | intron_variant,non_coding_transcript_variant | MODIFIER |
| 15 | 10830965 | AX-170285078 | 0 | T | C | 0.237 | -1.07E-01 | 1.07E-01 | 2.63E+00 | 2.68E+00 | 3.11E-01 | no | Gm19276 | intron_variant,non_coding_transcript_variant | MODIFIER |
| 15 | 10865610 | AX-170011022 | 0 | C | A | 0.237 | -1.07E-01 | 1.07E-01 | 2.63E+00 | 2.68E+00 | 3.11E-01 | no | | regulatory_region_variant | MODIFIER |
| 15 | 10869651 | AX-170150377 | 0 | A | G | 0.237 | -1.07E-01 | 1.07E-01 | 2.63E+00 | 2.68E+00 | 3.11E-01 | no | | regulatory_region_variant | MODIFIER |
| 15 | 10903982 | AX-170231085 | 0 | A | G | 0.237 | -1.07E-01 | 1.07E-01 | 2.63E+00 | 2.68E+00 | 3.11E-01 | no | | intergenic_variant | MODIFIER |
| 15 | 10911517 | AX-170356014 | 0 | A | T | 0.237 | -1.07E-01 | 1.07E-01 | 2.63E+00 | 2.68E+00 | 3.11E-01 | no | | intergenic_variant | MODIFIER |
| 15 | 10913444 | AX-170171868 | 0 | C | T | 0.237 | -1.07E-01 | 1.07E-01 | 2.63E+00 | 2.68E+00 | 3.11E-01 | no | Gm29742 | upstream_gene_variant | MODIFIER |
| 15 | 10918315 | AX-170489574 | 0 | T | G | 0.237 | -1.07E-01 | 1.07E-01 | 2.63E+00 | 2.68E+00 | 3.11E-01 | no | Gm29742 | downstream_gene_variant | MODIFIER |
| 15 | 10950081 | AX-170252416 | 0 | A | G | 0.237 | -1.07E-01 | 1.07E-01 | 2.63E+00 | 2.68E+00 | 3.11E-01 | no | C1qtnf3 | upstream_gene_variant | MODIFIER |

| | | | | | | | | | | | | | | | |
|---|---|---|---|---|---|---|---|---|---|---|---|---|---|---|---|
| 15 | 10950274 | AX-170271143 | 0 | G | A | 0.237 | -1.07E-01 | 1.07E-01 | 2.63E+00 | 2.68E+00 | 3.11E-01 | no | C1qtnf3 | upstream_gene_variant | MODIFIER |
| 15 | 10953405 | AX-170522833 | 0 | G | C | 0.237 | -1.07E-01 | 1.07E-01 | 2.63E+00 | 2.68E+00 | 3.11E-01 | no | C1qtnf3 | intron_variant | MODIFIER |
| 15 | 10955701 | AX-170161422 | 0 | G | A | 0.237 | -1.07E-01 | 1.07E-01 | 2.63E+00 | 2.68E+00 | 3.11E-01 | no | C1qtnf3 | intron_variant | MODIFIER |
| 15 | 10956266 | AX-170250493 | 0 | T | C | 0.237 | -1.07E-01 | 1.07E-01 | 2.63E+00 | 2.68E+00 | 3.11E-01 | no | C1qtnf3 | intron_variant | MODIFIER |
| 15 | 10958364 | AX-170427048 | 0 | C | T | 0.237 | -1.07E-01 | 1.07E-01 | 2.63E+00 | 2.68E+00 | 3.11E-01 | no | C1qtnf3 | intron_variant | MODIFIER |
| 15 | 10975856 | AX-170219829 | 0 | C | T | 0.237 | -1.07E-01 | 1.07E-01 | 2.63E+00 | 2.68E+00 | 3.11E-01 | no | C1qtnf3 | intron_variant | MODIFIER |
| 15 | 10979768 | AX-97854853 | 0 | A | G | 0.237 | -1.07E-01 | 1.07E-01 | 2.63E+00 | 2.68E+00 | 3.11E-01 | no | C1qtnf3 | 3_prime_UTR_variant | MODIFIER |
| 15 | 10979860 | AX-170273459 | 0 | G | A | 0.237 | -1.07E-01 | 1.07E-01 | 2.63E+00 | 2.68E+00 | 3.11E-01 | no | C1qtnf3 | 3_prime_UTR_variant | MODIFIER |
| 15 | 10988647 | AX-97844528 | 0 | A | G | 0.237 | -1.07E-01 | 1.07E-01 | 2.63E+00 | 2.68E+00 | 3.11E-01 | no | Amacr | intron_variant | MODIFIER |
| 15 | 10989162 | AX-170414248 | 0 | G | A | 0.237 | -1.07E-01 | 1.07E-01 | 2.63E+00 | 2.68E+00 | 3.11E-01 | no | Amacr | intron_variant | MODIFIER |
| 15 | 11030426 | AX-170347009 | 0 | T | C | 0.237 | -1.07E-01 | 1.07E-01 | 2.63E+00 | 2.68E+00 | 3.11E-01 | no | Rxfp3 | downstream_gene_variant | MODIFIER |
| 15 | 11100818 | AX-97827066 | 0 | A | G | 0.237 | -1.07E-01 | 1.07E-01 | 2.63E+00 | 2.68E+00 | 3.11E-01 | no | Adamts12 | intron_variant | MODIFIER |
| 15 | 11101184 | AX-170140870 | 0 | T | G | 0.237 | -1.07E-01 | 1.07E-01 | 2.63E+00 | 2.68E+00 | 3.11E-01 | no | Adamts12 | intron_variant | MODIFIER |
| 15 | 11107579 | AX-170159941 | 0 | C | T | 0.237 | -1.07E-01 | 1.07E-01 | 2.63E+00 | 2.68E+00 | 3.11E-01 | no | Adamts12 | intron_variant | MODIFIER |
| 15 | 11111376 | AX-170153331 | 0 | T | C | 0.237 | -1.07E-01 | 1.07E-01 | 2.63E+00 | 2.68E+00 | 3.11E-01 | no | Adamts12 | intron_variant | MODIFIER |
| 15 | 11117123 | AX-170178314 | 0 | C | T | 0.237 | -1.07E-01 | 1.07E-01 | 2.63E+00 | 2.68E+00 | 3.11E-01 | no | Adamts12 | intron_variant | MODIFIER |
| 15 | 11119834 | AX-97828444 | 0 | G | A | 0.237 | -1.07E-01 | 1.07E-01 | 2.63E+00 | 2.68E+00 | 3.11E-01 | no | Adamts12 | intron_variant | MODIFIER |
| 15 | 11178686 | AX-170333013 | 0 | G | A | 0.237 | -1.07E-01 | 1.07E-01 | 2.63E+00 | 2.68E+00 | 3.11E-01 | no | Adamts12 | intron_variant | MODIFIER |
| 15 | 11197403 | AX-170196389 | 0 | A | G | 0.237 | -1.07E-01 | 1.07E-01 | 2.63E+00 | 2.68E+00 | 3.11E-01 | no | Adamts12 | intron_variant | MODIFIER |
| 15 | 11198098 | AX-170423387 | 0 | T | A | 0.237 | -1.07E-01 | 1.07E-01 | 2.63E+00 | 2.68E+00 | 3.11E-01 | no | Adamts12 | intron_variant | MODIFIER |
| 15 | 11216436 | AX-170148272 | 0 | T | C | 0.237 | -1.07E-01 | 1.07E-01 | 2.63E+00 | 2.68E+00 | 3.11E-01 | no | Adamts12 | intron_variant | MODIFIER |
| 15 | 11217396 | AX-170532295 | 0 | G | C | 0.237 | -1.07E-01 | 1.07E-01 | 2.63E+00 | 2.68E+00 | 3.11E-01 | no | Adamts12 | intron_variant | MODIFIER |
| 15 | 11223906 | AX-170366367 | 0 | T | G | 0.237 | -1.07E-01 | 1.07E-01 | 2.63E+00 | 2.68E+00 | 3.11E-01 | no | Adamts12 | intron_variant | MODIFIER |
| 15 | 11233496 | AX-170498872 | 0 | G | C | 0.237 | -1.07E-01 | 1.07E-01 | 2.63E+00 | 2.68E+00 | 3.11E-01 | no | Adamts12 | intron_variant | MODIFIER |
| 15 | 11233668 | AX-170295773 | 0 | A | G | 0.237 | -1.07E-01 | 1.07E-01 | 2.63E+00 | 2.68E+00 | 3.11E-01 | no | Adamts12 | intron_variant | MODIFIER |
| 15 | 11288301 | AX-170087769 | 0 | C | A | 0.237 | -1.07E-01 | 1.07E-01 | 2.63E+00 | 2.68E+00 | 3.11E-01 | no | Adamts12 | intron_variant | MODIFIER |
| 15 | 11311464 | AX-97828445 | 0 | G | A | 0.237 | -1.07E-01 | 1.07E-01 | 2.63E+00 | 2.68E+00 | 3.11E-01 | no | Adamts12 | missense_variant | MODERATE |
| 15 | 11345159 | AX-170011032 | 0 | C | T | 0.237 | -1.07E-01 | 1.07E-01 | 2.63E+00 | 2.68E+00 | 3.11E-01 | no | Adamts12 | intron_variant | MODIFIER |
| 15 | 11348751 | AX-170054806 | 0 | G | A | 0.237 | -1.07E-01 | 1.07E-01 | 2.63E+00 | 2.68E+00 | 3.11E-01 | no | Adamts12 | 3_prime_UTR_variant | MODIFIER |
| 15 | 11349156 | AX-170011038 | 0 | G | A | 0.237 | -1.07E-01 | 1.07E-01 | 2.63E+00 | 2.68E+00 | 3.11E-01 | no | Adamts12 | 3_prime_UTR_variant | MODIFIER |
| 15 | 14777173 | AX-97831718 | 0 | C | A | 0.237 | -1.07E-01 | 1.07E-01 | 2.63E+00 | 2.68E+00 | 3.11E-01 | no | | intergenic_variant | MODIFIER |
| 15 | 14794649 | AX-97851716 | 0 | C | T | 0.237 | -1.07E-01 | 1.07E-01 | 2.63E+00 | 2.68E+00 | 3.11E-01 | no | | intergenic_variant | MODIFIER |
| 15 | 14828526 | AX-97835104 | 0 | T | C | 0.237 | -1.07E-01 | 1.07E-01 | 2.63E+00 | 2.68E+00 | 3.11E-01 | no | | intergenic_variant | MODIFIER |
| 15 | 14981649 | AX-170099211 | 0 | C | G | 0.237 | -1.07E-01 | 1.07E-01 | 2.63E+00 | 2.68E+00 | 3.11E-01 | no | | intergenic_variant | MODIFIER |
| 15 | 15159655 | AX-170087776 | 0 | G | A | 0.237 | -1.07E-01 | 1.07E-01 | 2.63E+00 | 2.68E+00 | 3.11E-01 | no | | intergenic_variant | MODIFIER |
| 15 | 15226055 | AX-170308928 | 0 | A | G | 0.237 | -1.07E-01 | 1.07E-01 | 2.63E+00 | 2.68E+00 | 3.11E-01 | no | | intergenic_variant | MODIFIER |
| 15 | 15251250 | AX-170011095 | 0 | T | C | 0.237 | -1.07E-01 | 1.07E-01 | 2.63E+00 | 2.68E+00 | 3.11E-01 | no | | intergenic_variant | MODIFIER |
| 15 | 15255205 | AX-170261012 | 0 | G | C | 0.237 | -1.07E-01 | 1.07E-01 | 2.63E+00 | 2.68E+00 | 3.11E-01 | no | | intergenic_variant | MODIFIER |
| 15 | 15276161 | AX-97791914 | 0 | G | A | 0.237 | -1.07E-01 | 1.07E-01 | 2.63E+00 | 2.68E+00 | 3.11E-01 | no | | intergenic_variant | MODIFIER |
| 15 | 15276446 | AX-170011103 | 0 | G | C | 0.237 | -1.07E-01 | 1.07E-01 | 2.63E+00 | 2.68E+00 | 3.11E-01 | no | | intergenic_variant | MODIFIER |
| 15 | 15276464 | AX-170011099 | 0 | T | C | 0.237 | -1.07E-01 | 1.07E-01 | 2.63E+00 | 2.68E+00 | 3.11E-01 | no | | intergenic_variant | MODIFIER |
| 15 | 15307834 | AX-170011107 | 0 | T | G | 0.237 | -1.07E-01 | 1.07E-01 | 2.63E+00 | 2.68E+00 | 3.11E-01 | no | | intergenic_variant | MODIFIER |
| 15 | 15310188 | AX-170087777 | 0 | T | C | 0.237 | -1.07E-01 | 1.07E-01 | 2.63E+00 | 2.68E+00 | 3.11E-01 | no | | intergenic_variant | MODIFIER |
| 15 | 15311319 | AX-170011104 | 0 | A | G | 0.237 | -1.07E-01 | 1.07E-01 | 2.63E+00 | 2.68E+00 | 3.11E-01 | no | | intergenic_variant | MODIFIER |
| 15 | 15348704 | AX-97814913 | 0 | A | G | 0.237 | -1.07E-01 | 1.07E-01 | 2.63E+00 | 2.68E+00 | 3.11E-01 | no | | intergenic_variant | MODIFIER |
| 15 | 15352496 | AX-170011100 | 0 | C | G | 0.237 | -1.07E-01 | 1.07E-01 | 2.63E+00 | 2.68E+00 | 3.11E-01 | no | | intergenic_variant | MODIFIER |
| 15 | 15373779 | AX-170393045 | 0 | C | G | 0.237 | -1.07E-01 | 1.07E-01 | 2.63E+00 | 2.68E+00 | 3.11E-01 | no | | intergenic_variant | MODIFIER |
| 15 | 15400457 | AX-170011106 | 0 | C | G | 0.237 | -1.07E-01 | 1.07E-01 | 2.63E+00 | 2.68E+00 | 3.11E-01 | no | | intergenic_variant | MODIFIER |
| 15 | 15412059 | AX-170515437 | 0 | A | G | 0.237 | -1.07E-01 | 1.07E-01 | 2.63E+00 | 2.68E+00 | 3.11E-01 | no | | intergenic_variant | MODIFIER |
| 15 | 15426631 | AX-170011096 | 0 | C | T | 0.237 | -1.07E-01 | 1.07E-01 | 2.63E+00 | 2.68E+00 | 3.11E-01 | no | | intergenic_variant | MODIFIER |
| 15 | 15456535 | AX-170483971 | 0 | G | T | 0.237 | -1.07E-01 | 1.07E-01 | 2.63E+00 | 2.68E+00 | 3.11E-01 | no | | intergenic_variant | MODIFIER |
| 15 | 19028353 | AX-170246556 | 0 | G | A | 0.237 | -1.07E-01 | 1.07E-01 | 2.63E+00 | 2.68E+00 | 3.11E-01 | no | | intergenic_variant | MODIFIER |
| 15 | 19044037 | AX-170529519 | 0 | C | A | 0.237 | -1.07E-01 | 1.07E-01 | 2.63E+00 | 2.68E+00 | 3.11E-01 | no | | intergenic_variant | MODIFIER |
| 15 | 19047685 | AX-97790419 | 0 | G | A | 0.237 | -1.07E-01 | 1.07E-01 | 2.63E+00 | 2.68E+00 | 3.11E-01 | no | | intergenic_variant | MODIFIER |

| | | | | | | | | | | | | | | | |
|---|---|---|---|---|---|---|---|---|---|---|---|---|---|---|---|
| 15 | 19066402 | AX-170054942 | 0 | T | C | 0.237 | -1.07E-01 | 1.07E-01 | 2.63E+00 | 2.68E+00 | 3.11E-01 | no | Gm8328 | non_coding_transcript_exon_variant | MODIFIER |
| 15 | 19067603 | AX-170492433 | 0 | G | A | 0.237 | -1.07E-01 | 1.07E-01 | 2.63E+00 | 2.68E+00 | 3.11E-01 | no | Gm8328 | upstream_gene_variant | MODIFIER |
| 15 | 19069512 | AX-170337717 | 0 | T | C | 0.237 | -1.07E-01 | 1.07E-01 | 2.63E+00 | 2.68E+00 | 3.11E-01 | no | Gm8328 | upstream_gene_variant | MODIFIER |
| 15 | 19070318 | AX-170497330 | 0 | T | C | 0.237 | -1.07E-01 | 1.07E-01 | 2.63E+00 | 2.68E+00 | 3.11E-01 | no | Gm8328 | upstream_gene_variant | MODIFIER |
| 15 | 19070951 | AX-170128337 | 0 | G | A | 0.237 | -1.07E-01 | 1.07E-01 | 2.63E+00 | 2.68E+00 | 3.11E-01 | no | Gm8328 | upstream_gene_variant | MODIFIER |
| 15 | 19071247 | AX-97853929 | 0 | A | C | 0.237 | -1.07E-01 | 1.07E-01 | 2.63E+00 | 2.68E+00 | 3.11E-01 | no | Gm8328 | upstream_gene_variant | MODIFIER |
| 15 | 19105543 | AX-97828430 | 0 | T | C | 0.237 | -1.07E-01 | 1.07E-01 | 2.63E+00 | 2.68E+00 | 3.11E-01 | no | | intergenic_variant | MODIFIER |
| 15 | 19114801 | AX-170011167 | 0 | A | G | 0.237 | -1.07E-01 | 1.07E-01 | 2.63E+00 | 2.68E+00 | 3.11E-01 | no | | intergenic_variant | MODIFIER |
| 15 | 19118800 | AX-97831724 | 0 | A | G | 0.237 | -1.07E-01 | 1.07E-01 | 2.63E+00 | 2.68E+00 | 3.11E-01 | no | | intergenic_variant | MODIFIER |
| 15 | 19160564 | AX-170011172 | 0 | C | T | 0.237 | -1.07E-01 | 1.07E-01 | 2.63E+00 | 2.68E+00 | 3.11E-01 | no | | intergenic_variant | MODIFIER |
| 15 | 19173397 | AX-97798426 | 0 | G | A | 0.237 | -1.07E-01 | 1.07E-01 | 2.63E+00 | 2.68E+00 | 3.11E-01 | no | | intergenic_variant | MODIFIER |
| 15 | 19204979 | AX-97822650 | 0 | C | A | 0.237 | -1.07E-01 | 1.07E-01 | 2.63E+00 | 2.68E+00 | 3.11E-01 | no | Gm35496 | downstream_gene_variant | MODIFIER |
| 15 | 19210013 | AX-97796981 | 0 | G | C | 0.237 | -1.07E-01 | 1.07E-01 | 2.63E+00 | 2.68E+00 | 3.11E-01 | no | Gm18533 | downstream_gene_variant | MODIFIER |
| 15 | 19252074 | AX-97838332 | 0 | T | G | 0.237 | -1.07E-01 | 1.07E-01 | 2.63E+00 | 2.68E+00 | 3.11E-01 | no | | intergenic_variant | MODIFIER |
| 15 | 19269884 | AX-170451647 | 0 | T | A | 0.237 | -1.07E-01 | 1.07E-01 | 2.63E+00 | 2.68E+00 | 3.11E-01 | no | | intergenic_variant | MODIFIER |
| 15 | 19308215 | AX-97799662 | 0 | T | C | 0.237 | -1.07E-01 | 1.07E-01 | 2.63E+00 | 2.68E+00 | 3.11E-01 | no | Gm49161 | downstream_gene_variant | MODIFIER |
| 15 | 19308935 | AX-170158048 | 0 | T | C | 0.237 | -1.07E-01 | 1.07E-01 | 2.63E+00 | 2.68E+00 | 3.11E-01 | no | Gm49161 | downstream_gene_variant | MODIFIER |
| 15 | 19342279 | AX-170434945 | 0 | A | G | 0.237 | -1.07E-01 | 1.07E-01 | 2.63E+00 | 2.68E+00 | 3.11E-01 | no | Gm49162 | intron_variant,non_coding_transcript_variant | MODIFIER |
| 15 | 19357374 | AX-170215844 | 0 | A | G | 0.237 | -1.07E-01 | 1.07E-01 | 2.63E+00 | 2.68E+00 | 3.11E-01 | no | Gm49162 | intron_variant,non_coding_transcript_variant | MODIFIER |
| 15 | 19373427 | AX-170087787 | 0 | T | C | 0.237 | -1.07E-01 | 1.07E-01 | 2.63E+00 | 2.68E+00 | 3.11E-01 | no | Gm49162 | intron_variant,non_coding_transcript_variant | MODIFIER |
| 15 | 19409671 | AX-170378938 | 0 | A | C | 0.237 | -1.07E-01 | 1.07E-01 | 2.63E+00 | 2.68E+00 | 3.11E-01 | no | | intergenic_variant | MODIFIER |
| 15 | 19418398 | AX-170454199 | 0 | A | G | 0.237 | -1.07E-01 | 1.07E-01 | 2.63E+00 | 2.68E+00 | 3.11E-01 | no | | intergenic_variant | MODIFIER |
| 15 | 19422161 | AX-170054951 | 0 | A | G | 0.237 | -1.07E-01 | 1.07E-01 | 2.63E+00 | 2.68E+00 | 3.11E-01 | no | | intergenic_variant | MODIFIER |
| 15 | 19422380 | AX-170461702 | 0 | T | C | 0.237 | -1.07E-01 | 1.07E-01 | 2.63E+00 | 2.68E+00 | 3.11E-01 | no | | intergenic_variant | MODIFIER |
| 15 | 19437520 | AX-170428398 | 0 | A | G | 0.237 | -1.07E-01 | 1.07E-01 | 2.63E+00 | 2.68E+00 | 3.11E-01 | no | | intergenic_variant | MODIFIER |
| 15 | 19437551 | AX-170300213 | 0 | T | G | 0.237 | -1.07E-01 | 1.07E-01 | 2.63E+00 | 2.68E+00 | 3.11E-01 | no | | intergenic_variant | MODIFIER |
| 15 | 19438632 | AX-97824917 | 0 | A | G | 0.237 | -1.07E-01 | 1.07E-01 | 2.63E+00 | 2.68E+00 | 3.11E-01 | no | | intergenic_variant | MODIFIER |
| 15 | 19443822 | AX-170320170 | 0 | G | A | 0.237 | -1.07E-01 | 1.07E-01 | 2.63E+00 | 2.68E+00 | 3.11E-01 | no | | intergenic_variant | MODIFIER |
| 15 | 19686836 | AX-170011180 | 0 | G | A | 0.237 | -1.07E-01 | 1.07E-01 | 2.63E+00 | 2.68E+00 | 3.11E-01 | no | Gm49166 | intron_variant,non_coding_transcript_variant | MODIFIER |
| 15 | 19752153 | AX-170306523 | 0 | C | T | 0.237 | -1.07E-01 | 1.07E-01 | 2.63E+00 | 2.68E+00 | 3.11E-01 | no | Gm46498 | upstream_gene_variant | MODIFIER |
| 15 | 19779948 | AX-97818884 | 0 | G | A | 0.237 | -1.07E-01 | 1.07E-01 | 2.63E+00 | 2.68E+00 | 3.11E-01 | no | | intergenic_variant | MODIFIER |
| 15 | 19810373 | AX-170257434 | 0 | C | G | 0.237 | -1.07E-01 | 1.07E-01 | 2.63E+00 | 2.68E+00 | 3.11E-01 | no | | intergenic_variant | MODIFIER |
| 15 | 19810551 | AX-170531484 | 0 | G | T | 0.237 | -1.07E-01 | 1.07E-01 | 2.63E+00 | 2.68E+00 | 3.11E-01 | no | | intergenic_variant | MODIFIER |
| 15 | 19814428 | AX-97844724 | 0 | A | G | 0.237 | -1.07E-01 | 1.07E-01 | 2.63E+00 | 2.68E+00 | 3.11E-01 | no | | intergenic_variant | MODIFIER |
| 15 | 19854229 | AX-97795807 | 0 | T | C | 0.237 | -1.07E-01 | 1.07E-01 | 2.63E+00 | 2.68E+00 | 3.11E-01 | no | | intergenic_variant | MODIFIER |
| 15 | 20091588 | AX-170113148 | 0 | A | G | 0.237 | -1.07E-01 | 1.07E-01 | 2.63E+00 | 2.68E+00 | 3.11E-01 | no | | intergenic_variant | MODIFIER |
| 15 | 20139226 | AX-168410991 | 0 | T | C | 0.237 | -1.07E-01 | 1.07E-01 | 2.63E+00 | 2.68E+00 | 3.11E-01 | no | | intergenic_variant | MODIFIER |
| 15 | 20140068 | AX-97796216 | 0 | T | C | 0.237 | -1.07E-01 | 1.07E-01 | 2.63E+00 | 2.68E+00 | 3.11E-01 | no | | intergenic_variant | MODIFIER |
| 15 | 20140274 | AX-170289357 | 0 | C | T | 0.237 | -1.07E-01 | 1.07E-01 | 2.63E+00 | 2.68E+00 | 3.11E-01 | no | | intergenic_variant | MODIFIER |
| 15 | 20208202 | AX-97821035 | 0 | A | G | 0.237 | -1.07E-01 | 1.07E-01 | 2.63E+00 | 2.68E+00 | 3.11E-01 | no | | intergenic_variant | MODIFIER |
| 15 | 20255372 | AX-170421849 | 0 | T | C | 0.237 | -1.07E-01 | 1.07E-01 | 2.63E+00 | 2.68E+00 | 3.11E-01 | no | | intergenic_variant | MODIFIER |
| 15 | 20408453 | AX-170415944 | 0 | G | T | 0.237 | -1.07E-01 | 1.07E-01 | 2.63E+00 | 2.68E+00 | 3.11E-01 | no | | intergenic_variant | MODIFIER |
| 15 | 20413899 | AX-170403072 | 0 | C | T | 0.237 | -1.07E-01 | 1.07E-01 | 2.63E+00 | 2.68E+00 | 3.11E-01 | no | | intergenic_variant | MODIFIER |
| 15 | 20469743 | AX-97842537 | 0 | T | C | 0.237 | -1.07E-01 | 1.07E-01 | 2.63E+00 | 2.68E+00 | 3.11E-01 | no | Cdh12 | intron_variant,non_coding_transcript_variant | MODIFIER |
| 15 | 20904375 | AX-97812832 | 0 | T | C | 0.237 | -1.07E-01 | 1.07E-01 | 2.63E+00 | 2.68E+00 | 3.11E-01 | no | Gm25711 | upstream_gene_variant | MODIFIER |
| 15 | 20911557 | AX-170455013 | 0 | G | A | 0.237 | -1.07E-01 | 1.07E-01 | 2.63E+00 | 2.68E+00 | 3.11E-01 | no | Gm25711 | downstream_gene_variant | MODIFIER |
| 15 | 20958642 | AX-170208480 | 0 | T | C | 0.237 | -1.07E-01 | 1.07E-01 | 2.63E+00 | 2.68E+00 | 3.11E-01 | no | Cdh12 | intron_variant,non_coding_transcript_variant | MODIFIER |
| 15 | 20982336 | AX-170333834 | 0 | G | T | 0.237 | -1.07E-01 | 1.07E-01 | 2.63E+00 | 2.68E+00 | 3.11E-01 | no | Cdh12 | intron_variant,non_coding_transcript_variant | MODIFIER |
| 15 | 20986072 | AX-170298377 | 0 | T | C | 0.237 | -1.07E-01 | 1.07E-01 | 2.63E+00 | 2.68E+00 | 3.11E-01 | no | Cdh12 | intron_variant,non_coding_transcript_variant | MODIFIER |
| 15 | 21125603 | AX-170458074 | 0 | C | G | 0.237 | -1.07E-01 | 1.07E-01 | 2.63E+00 | 2.68E+00 | 3.11E-01 | no | Cdh12 | intron_variant | MODIFIER |
| 15 | 21134789 | AX-170117792 | 0 | A | G | 0.237 | -1.07E-01 | 1.07E-01 | 2.63E+00 | 2.68E+00 | 3.11E-01 | no | Cdh12 | intron_variant | MODIFIER |
| 15 | 21278154 | AX-170179914 | 0 | T | C | 0.237 | -1.07E-01 | 1.07E-01 | 2.63E+00 | 2.68E+00 | 3.11E-01 | no | Cdh12 | intron_variant | MODIFIER |
| 15 | 21557563 | AX-170405107 | 0 | G | A | 0.237 | -1.07E-01 | 1.07E-01 | 2.63E+00 | 2.68E+00 | 3.11E-01 | no | Cdh12 | intron_variant | MODIFIER |
| 15 | 21557893 | AX-170102065 | 0 | G | A | 0.237 | -1.07E-01 | 1.07E-01 | 2.63E+00 | 2.68E+00 | 3.11E-01 | no | Cdh12 | intron_variant | MODIFIER |

| | | | | | | | | | | | | | | | |
|---|---|---|---|---|---|---|---|---|---|---|---|---|---|---|---|
| 15 | 21609072 | AX-170340170 | 0 | T | C | 0.237 | -1.07E-01 | 1.07E-01 | 2.63E+00 | 2.68E+00 | 3.11E-01 | no | | intergenic_variant | MODIFIER |
| 15 | 21611565 | AX-170483526 | 0 | T | C | 0.237 | -1.07E-01 | 1.07E-01 | 2.63E+00 | 2.68E+00 | 3.11E-01 | no | | intergenic_variant | MODIFIER |
| 15 | 21633374 | AX-170285272 | 0 | T | C | 0.237 | -1.07E-01 | 1.07E-01 | 2.63E+00 | 2.68E+00 | 3.11E-01 | no | | intergenic_variant | MODIFIER |
| 15 | 21633899 | AX-170403807 | 0 | C | G | 0.237 | -1.07E-01 | 1.07E-01 | 2.63E+00 | 2.68E+00 | 3.11E-01 | no | | intergenic_variant | MODIFIER |
| 15 | 21714033 | AX-170239352 | 0 | C | A | 0.237 | -1.07E-01 | 1.07E-01 | 2.63E+00 | 2.68E+00 | 3.11E-01 | no | | intergenic_variant | MODIFIER |
| 15 | 22198582 | AX-97799985 | 0 | T | C | 0.237 | -1.07E-01 | 1.07E-01 | 2.63E+00 | 2.68E+00 | 3.11E-01 | no | | intergenic_variant | MODIFIER |
| 15 | 22237878 | AX-170395622 | 0 | G | T | 0.237 | -1.07E-01 | 1.07E-01 | 2.63E+00 | 2.68E+00 | 3.11E-01 | no | | intergenic_variant | MODIFIER |
| 15 | 22594293 | AX-170055012 | 0 | C | T | 0.237 | -1.07E-01 | 1.07E-01 | 2.63E+00 | 2.68E+00 | 3.11E-01 | no | Cdh18 | intron_variant | MODIFIER |
| 15 | 22679199 | AX-97828973 | 0 | A | C | 0.237 | -1.07E-01 | 1.07E-01 | 2.63E+00 | 2.68E+00 | 3.11E-01 | no | Cdh18 | intron_variant | MODIFIER |
| 15 | 22822361 | AX-170222491 | 0 | A | G | 0.237 | -1.07E-01 | 1.07E-01 | 2.63E+00 | 2.68E+00 | 3.11E-01 | no | Cdh18 | intron_variant | MODIFIER |
| 15 | 22825761 | AX-170429054 | 0 | A | G | 0.237 | -1.07E-01 | 1.07E-01 | 2.63E+00 | 2.68E+00 | 3.11E-01 | no | Cdh18 | intron_variant | MODIFIER |
| 15 | 22937154 | AX-170011221 | 0 | T | C | 0.237 | -1.07E-01 | 1.07E-01 | 2.63E+00 | 2.68E+00 | 3.11E-01 | no | Cdh18 | intron_variant | MODIFIER |
| 15 | 23111856 | AX-170185991 | 0 | G | T | 0.237 | -1.07E-01 | 1.07E-01 | 2.63E+00 | 2.68E+00 | 3.11E-01 | no | Cdh18 | intron_variant,non_coding_transcript_variant | MODIFIER |
| 15 | 23112034 | AX-170148409 | 0 | C | T | 0.237 | -1.07E-01 | 1.07E-01 | 2.63E+00 | 2.68E+00 | 3.11E-01 | no | Cdh18 | intron_variant,non_coding_transcript_variant | MODIFIER |
| 15 | 23112446 | AX-170506450 | 0 | A | G | 0.237 | -1.07E-01 | 1.07E-01 | 2.63E+00 | 2.68E+00 | 3.11E-01 | no | Cdh18 | intron_variant,non_coding_transcript_variant | MODIFIER |
| 15 | 23121963 | AX-170087796 | 0 | A | G | 0.237 | -1.07E-01 | 1.07E-01 | 2.63E+00 | 2.68E+00 | 3.11E-01 | no | Cdh18 | intron_variant,non_coding_transcript_variant | MODIFIER |
| 15 | 23122022 | AX-170350454 | 0 | T | A | 0.237 | -1.07E-01 | 1.07E-01 | 2.63E+00 | 2.68E+00 | 3.11E-01 | no | Cdh18 | intron_variant,non_coding_transcript_variant | MODIFIER |
| 8 | 38541244 | AX-170265977 | 0 | C | A | 0.149 | 1.53E-01 | 1.48E-01 | 3.09E+00 | 3.16E+00 | 3.11E-01 | Yes | Sgcz | intron_variant | MODIFIER |
| 8 | 39459515 | AX-170208232 | 0 | A | C | 0.149 | 1.53E-01 | 1.48E-01 | 3.09E+00 | 3.16E+00 | 3.11E-01 | Yes | Gm5609 | downstream_gene_variant | MODIFIER |
| 8 | 41377872 | AX-170354049 | 0 | G | T | 0.149 | 1.53E-01 | 1.48E-01 | 3.09E+00 | 3.16E+00 | 3.11E-01 | Yes | Asah1 | upstream_gene_variant | MODIFIER |
| 4 | 74772721 | AX-170023131 | 1 | T | G | 0.438 | -8.54E-02 | 8.32E-02 | 2.46E+00 | 2.53E+00 | 3.11E-01 | no | | intergenic_variant | MODIFIER |
| 18 | 78545790 | AX-170414474 | 0 | T | C | 0.237 | -6.34E-02 | 6.34E-02 | 2.72E+00 | 2.85E+00 | 3.11E-01 | no | Slc14a2 | intron_variant | MODIFIER |
| 7 | 145311961 | AX-170090795 | 1 | A | G | 0.196 | -7.71E-02 | 7.69E-02 | 2.64E+00 | 2.76E+00 | 3.11E-01 | no | Mrgprf | downstream_gene_variant | MODIFIER |
| 15 | 6011761 | AX-170525159 | 1 | G | A | 0.241 | -1.07E-01 | 1.07E-01 | 2.65E+00 | 2.71E+00 | 3.11E-01 | no | | intergenic_variant | MODIFIER |
| 18 | 59256820 | AX-170108554 | 0 | A | G | 0.404 | 9.71E-02 | 9.41E-02 | 2.44E+00 | 2.48E+00 | 3.11E-01 | no | Chsy3 | intron_variant | MODIFIER |
| 18 | 59286771 | AX-170140474 | 0 | C | T | 0.404 | 9.71E-02 | 9.41E-02 | 2.44E+00 | 2.48E+00 | 3.11E-01 | no | Chsy3 | intron_variant | MODIFIER |
| 18 | 59320582 | AX-170129667 | 0 | C | A | 0.404 | 9.71E-02 | 9.41E-02 | 2.44E+00 | 2.48E+00 | 3.11E-01 | no | Chsy3 | intron_variant | MODIFIER |
| 18 | 59328545 | AX-97822255 | 0 | T | C | 0.404 | 9.71E-02 | 9.41E-02 | 2.44E+00 | 2.48E+00 | 3.11E-01 | no | Chsy3 | intron_variant | MODIFIER |
| 18 | 59330124 | AX-170335565 | 0 | G | T | 0.404 | 9.71E-02 | 9.41E-02 | 2.44E+00 | 2.48E+00 | 3.11E-01 | no | Chsy3 | intron_variant | MODIFIER |
| 18 | 59330797 | AX-170229245 | 0 | A | G | 0.404 | 9.71E-02 | 9.41E-02 | 2.44E+00 | 2.48E+00 | 3.11E-01 | no | Chsy3 | intron_variant | MODIFIER |
| 18 | 59341720 | AX-170232083 | 0 | A | C | 0.404 | 9.71E-02 | 9.41E-02 | 2.44E+00 | 2.48E+00 | 3.11E-01 | no | Chsy3 | intron_variant | MODIFIER |
| 18 | 59351472 | AX-170470298 | 0 | C | T | 0.404 | 9.71E-02 | 9.41E-02 | 2.44E+00 | 2.48E+00 | 3.11E-01 | no | Chsy3 | intron_variant | MODIFIER |
| 18 | 59352104 | AX-170159785 | 0 | T | C | 0.404 | 9.71E-02 | 9.41E-02 | 2.44E+00 | 2.48E+00 | 3.11E-01 | no | Chsy3 | intron_variant | MODIFIER |
| 18 | 59352609 | AX-170319247 | 0 | T | A | 0.404 | 9.71E-02 | 9.41E-02 | 2.44E+00 | 2.48E+00 | 3.11E-01 | no | Chsy3 | intron_variant | MODIFIER |
| 18 | 59382683 | AX-170060992 | 0 | T | G | 0.404 | 9.71E-02 | 9.41E-02 | 2.44E+00 | 2.48E+00 | 3.11E-01 | no | Chsy3 | intron_variant | MODIFIER |
| 18 | 59385276 | AX-170480873 | 0 | G | C | 0.404 | 9.71E-02 | 9.41E-02 | 2.44E+00 | 2.48E+00 | 3.11E-01 | no | Chsy3 | intron_variant | MODIFIER |
| 18 | 59445335 | AX-170481897 | 0 | G | C | 0.404 | 9.71E-02 | 9.41E-02 | 2.44E+00 | 2.48E+00 | 3.11E-01 | no | Chsy3 | intron_variant,NMD_transcript_variant | MODIFIER |
| 18 | 59445475 | AX-97846994 | 0 | C | T | 0.404 | 9.71E-02 | 9.41E-02 | 2.44E+00 | 2.48E+00 | 3.11E-01 | no | Chsy3 | intron_variant,NMD_transcript_variant | MODIFIER |
| 18 | 59479388 | AX-170108671 | 0 | T | A | 0.404 | 9.71E-02 | 9.41E-02 | 2.44E+00 | 2.48E+00 | 3.11E-01 | no | Chsy3 | intron_variant,NMD_transcript_variant | MODIFIER |
| 18 | 59488328 | AX-170215405 | 0 | T | C | 0.404 | 9.71E-02 | 9.41E-02 | 2.44E+00 | 2.48E+00 | 3.11E-01 | no | Chsy3 | intron_variant,NMD_transcript_variant | MODIFIER |
| 18 | 59530699 | AX-170155204 | 0 | T | G | 0.404 | 9.71E-02 | 9.41E-02 | 2.44E+00 | 2.48E+00 | 3.11E-01 | no | Chsy3 | intron_variant,NMD_transcript_variant | MODIFIER |
| 18 | 59557539 | AX-170331561 | 0 | G | A | 0.404 | 9.71E-02 | 9.41E-02 | 2.44E+00 | 2.48E+00 | 3.11E-01 | no | | intergenic_variant | MODIFIER |
| 18 | 59568282 | AX-170325922 | 0 | T | C | 0.404 | 9.71E-02 | 9.41E-02 | 2.44E+00 | 2.48E+00 | 3.11E-01 | no | | intergenic_variant | MODIFIER |
| 18 | 59591399 | AX-170411414 | 0 | G | A | 0.404 | 9.71E-02 | 9.41E-02 | 2.44E+00 | 2.48E+00 | 3.11E-01 | no | | intergenic_variant | MODIFIER |
| 18 | 59599225 | AX-170425271 | 0 | T | C | 0.404 | 9.71E-02 | 9.41E-02 | 2.44E+00 | 2.48E+00 | 3.11E-01 | no | | intergenic_variant | MODIFIER |
| 18 | 59642770 | AX-170400076 | 0 | T | C | 0.404 | 9.71E-02 | 9.41E-02 | 2.44E+00 | 2.48E+00 | 3.11E-01 | no | | intergenic_variant | MODIFIER |
| 18 | 59696722 | AX-170098550 | 0 | A | T | 0.404 | 9.71E-02 | 9.41E-02 | 2.44E+00 | 2.48E+00 | 3.11E-01 | no | | intergenic_variant | MODIFIER |
| 18 | 59697366 | AX-170252632 | 0 | G | T | 0.404 | 9.71E-02 | 9.41E-02 | 2.44E+00 | 2.48E+00 | 3.11E-01 | no | | intergenic_variant | MODIFIER |
| 18 | 59697945 | AX-170265848 | 0 | A | G | 0.404 | 9.71E-02 | 9.41E-02 | 2.44E+00 | 2.48E+00 | 3.11E-01 | no | | intergenic_variant | MODIFIER |
| 18 | 59724216 | AX-170220029 | 0 | C | T | 0.404 | 9.71E-02 | 9.41E-02 | 2.44E+00 | 2.48E+00 | 3.11E-01 | no | | intergenic_variant | MODIFIER |
| 18 | 59751443 | AX-170505775 | 0 | C | G | 0.404 | 9.71E-02 | 9.41E-02 | 2.44E+00 | 2.48E+00 | 3.11E-01 | no | | intergenic_variant | MODIFIER |
| 18 | 59751918 | AX-170510227 | 0 | C | T | 0.404 | 9.71E-02 | 9.41E-02 | 2.44E+00 | 2.48E+00 | 3.11E-01 | no | | intergenic_variant | MODIFIER |
| 18 | 59758169 | AX-170459195 | 0 | G | A | 0.404 | 9.71E-02 | 9.41E-02 | 2.44E+00 | 2.48E+00 | 3.11E-01 | no | | intergenic_variant | MODIFIER |
| 18 | 59762389 | AX-170456062 | 0 | C | G | 0.404 | 9.71E-02 | 9.41E-02 | 2.44E+00 | 2.48E+00 | 3.11E-01 | no | | intergenic_variant | MODIFIER |

| | | | | | | | | | | | | | | | |
|---|---|---|---|---|---|---|---|---|---|---|---|---|---|---|---|
| 18 | 59771154 | AX-97825076 | 0 | T | C | 0.404 | 9.71E-02 | 9.41E-02 | 2.44E+00 | 2.48E+00 | 3.11E-01 | no | AC138134.2 | downstream_gene_variant | MODIFIER |
| 18 | 59785212 | AX-170486225 | 0 | C | T | 0.404 | 9.71E-02 | 9.41E-02 | 2.44E+00 | 2.48E+00 | 3.11E-01 | no | | intergenic_variant | MODIFIER |
| 18 | 59791873 | AX-170129895 | 0 | C | A | 0.404 | 9.71E-02 | 9.41E-02 | 2.44E+00 | 2.48E+00 | 3.11E-01 | no | | intergenic_variant | MODIFIER |
| 18 | 59794994 | AX-170261040 | 0 | T | C | 0.404 | 9.71E-02 | 9.41E-02 | 2.44E+00 | 2.48E+00 | 3.11E-01 | no | | intergenic_variant | MODIFIER |
| 18 | 59801198 | AX-170242262 | 0 | A | T | 0.404 | 9.71E-02 | 9.41E-02 | 2.44E+00 | 2.48E+00 | 3.11E-01 | no | | intergenic_variant | MODIFIER |
| 18 | 59801442 | AX-170479557 | 0 | C | G | 0.404 | 9.71E-02 | 9.41E-02 | 2.44E+00 | 2.48E+00 | 3.11E-01 | no | | intergenic_variant | MODIFIER |
| 18 | 59835576 | AX-170182056 | 0 | A | G | 0.404 | 9.71E-02 | 9.41E-02 | 2.44E+00 | 2.48E+00 | 3.11E-01 | no | AC138134.1 | intron_variant,non_coding_transcript_variant | MODIFIER |
| 18 | 59861419 | AX-97821090 | 0 | C | T | 0.404 | 9.71E-02 | 9.41E-02 | 2.44E+00 | 2.48E+00 | 3.11E-01 | no | | intergenic_variant | MODIFIER |
| 18 | 59869439 | AX-170060995 | 0 | A | G | 0.404 | 9.71E-02 | 9.41E-02 | 2.44E+00 | 2.48E+00 | 3.11E-01 | no | | intergenic_variant | MODIFIER |
| 18 | 59873083 | AX-170060997 | 0 | C | T | 0.404 | 9.71E-02 | 9.41E-02 | 2.44E+00 | 2.48E+00 | 3.11E-01 | no | | intergenic_variant | MODIFIER |
| 18 | 59886794 | AX-170300037 | 0 | A | G | 0.404 | 9.71E-02 | 9.41E-02 | 2.44E+00 | 2.48E+00 | 3.11E-01 | no | | intergenic_variant | MODIFIER |
| 18 | 59886862 | AX-170112612 | 0 | A | G | 0.404 | 9.71E-02 | 9.41E-02 | 2.44E+00 | 2.48E+00 | 3.11E-01 | no | | intergenic_variant | MODIFIER |
| 18 | 59897940 | AX-170015806 | 0 | G | T | 0.404 | 9.71E-02 | 9.41E-02 | 2.44E+00 | 2.48E+00 | 3.11E-01 | no | AC102091.2 | non_coding_transcript_exon_variant | MODIFIER |
| 18 | 59898353 | AX-97793615 | 0 | C | T | 0.404 | 9.71E-02 | 9.41E-02 | 2.44E+00 | 2.48E+00 | 3.11E-01 | no | AC102091.2 | downstream_gene_variant | MODIFIER |
| 18 | 59927665 | AX-170015805 | 0 | A | C | 0.404 | 9.71E-02 | 9.41E-02 | 2.44E+00 | 2.48E+00 | 3.11E-01 | no | | intergenic_variant | MODIFIER |
| 18 | 59956505 | AX-170120339 | 0 | C | T | 0.404 | 9.71E-02 | 9.41E-02 | 2.44E+00 | 2.48E+00 | 3.11E-01 | no | | intergenic_variant | MODIFIER |
| 18 | 59970774 | AX-170226474 | 0 | C | T | 0.404 | 9.71E-02 | 9.41E-02 | 2.44E+00 | 2.48E+00 | 3.11E-01 | no | | intergenic_variant | MODIFIER |
| 18 | 59973905 | AX-170318710 | 0 | T | C | 0.404 | 9.71E-02 | 9.41E-02 | 2.44E+00 | 2.48E+00 | 3.11E-01 | no | | intergenic_variant | MODIFIER |
| 18 | 59977531 | AX-170493504 | 0 | A | G | 0.404 | 9.71E-02 | 9.41E-02 | 2.44E+00 | 2.48E+00 | 3.11E-01 | no | | intergenic_variant | MODIFIER |
| 18 | 59995605 | AX-170206715 | 0 | C | G | 0.404 | 9.71E-02 | 9.41E-02 | 2.44E+00 | 2.48E+00 | 3.11E-01 | no | | intergenic_variant | MODIFIER |
| 18 | 59995622 | AX-170171671 | 0 | A | G | 0.404 | 9.71E-02 | 9.41E-02 | 2.44E+00 | 2.48E+00 | 3.11E-01 | no | | intergenic_variant | MODIFIER |
| 18 | 59995892 | AX-170140543 | 0 | T | C | 0.404 | 9.71E-02 | 9.41E-02 | 2.44E+00 | 2.48E+00 | 3.11E-01 | no | | intergenic_variant | MODIFIER |
| 18 | 60000955 | AX-170268368 | 0 | A | C | 0.404 | 9.71E-02 | 9.41E-02 | 2.44E+00 | 2.48E+00 | 3.11E-01 | no | | intergenic_variant | MODIFIER |
| 18 | 60001008 | AX-170191447 | 0 | A | G | 0.404 | 9.71E-02 | 9.41E-02 | 2.44E+00 | 2.48E+00 | 3.11E-01 | no | | intergenic_variant | MODIFIER |
| 18 | 60013160 | AX-170259262 | 0 | G | A | 0.404 | 9.71E-02 | 9.41E-02 | 2.44E+00 | 2.48E+00 | 3.11E-01 | no | | intergenic_variant | MODIFIER |
| 18 | 60034057 | AX-170416542 | 0 | C | G | 0.404 | 9.71E-02 | 9.41E-02 | 2.44E+00 | 2.48E+00 | 3.11E-01 | no | | intergenic_variant | MODIFIER |
| 18 | 60037899 | AX-170061000 | 0 | C | T | 0.404 | 9.71E-02 | 9.41E-02 | 2.44E+00 | 2.48E+00 | 3.11E-01 | no | | intergenic_variant | MODIFIER |
| 18 | 60046491 | AX-170341170 | 0 | T | G | 0.404 | 9.71E-02 | 9.41E-02 | 2.44E+00 | 2.48E+00 | 3.11E-01 | no | | intergenic_variant | MODIFIER |
| 18 | 60048127 | AX-170522058 | 0 | C | G | 0.404 | 9.71E-02 | 9.41E-02 | 2.44E+00 | 2.48E+00 | 3.11E-01 | no | | intergenic_variant | MODIFIER |
| 18 | 60083156 | AX-97835601 | 0 | C | T | 0.404 | 9.71E-02 | 9.41E-02 | 2.44E+00 | 2.48E+00 | 3.11E-01 | no | | intergenic_variant | MODIFIER |
| 18 | 60124955 | AX-170203936 | 0 | G | A | 0.404 | 9.71E-02 | 9.41E-02 | 2.44E+00 | 2.48E+00 | 3.11E-01 | no | | intergenic_variant | MODIFIER |
| 18 | 60142798 | AX-170379595 | 0 | T | C | 0.404 | 9.71E-02 | 9.41E-02 | 2.44E+00 | 2.48E+00 | 3.11E-01 | no | | intergenic_variant | MODIFIER |
| 18 | 60154642 | AX-170447450 | 0 | C | T | 0.404 | 9.71E-02 | 9.41E-02 | 2.44E+00 | 2.48E+00 | 3.11E-01 | no | | regulatory_region_variant | MODIFIER |
| 18 | 60155438 | AX-170439518 | 0 | C | T | 0.404 | 9.71E-02 | 9.41E-02 | 2.44E+00 | 2.48E+00 | 3.11E-01 | no | | regulatory_region_variant | MODIFIER |
| 18 | 60157486 | AX-170061007 | 0 | T | C | 0.404 | 9.71E-02 | 9.41E-02 | 2.44E+00 | 2.48E+00 | 3.11E-01 | no | | intergenic_variant | MODIFIER |
| 18 | 60157547 | AX-170061005 | 0 | T | C | 0.404 | 9.71E-02 | 9.41E-02 | 2.44E+00 | 2.48E+00 | 3.11E-01 | no | | intergenic_variant | MODIFIER |
| 18 | 60184525 | AX-170236484 | 0 | A | G | 0.404 | 9.71E-02 | 9.41E-02 | 2.44E+00 | 2.48E+00 | 3.11E-01 | no | | intergenic_variant | MODIFIER |
| 18 | 60185028 | AX-170170956 | 0 | G | A | 0.404 | 9.71E-02 | 9.41E-02 | 2.44E+00 | 2.48E+00 | 3.11E-01 | no | | intergenic_variant | MODIFIER |
| 18 | 60196105 | AX-97823649 | 0 | A | G | 0.404 | 9.71E-02 | 9.41E-02 | 2.44E+00 | 2.48E+00 | 3.11E-01 | no | | regulatory_region_variant | MODIFIER |
| 18 | 60220839 | AX-97853299 | 0 | G | A | 0.404 | 9.71E-02 | 9.41E-02 | 2.44E+00 | 2.48E+00 | 3.11E-01 | no | Gm4951 | intron_variant | MODIFIER |
| 18 | 60298645 | AX-170099537 | 0 | A | G | 0.404 | 9.71E-02 | 9.41E-02 | 2.44E+00 | 2.48E+00 | 3.11E-01 | no | F830016B08Rik | intron_variant | MODIFIER |
| 18 | 60356047 | AX-170341991 | 0 | G | T | 0.404 | 9.71E-02 | 9.41E-02 | 2.44E+00 | 2.48E+00 | 3.11E-01 | no | AC132320.1 | non_coding_transcript_exon_variant | MODIFIER |
| 18 | 60359769 | AX-170450841 | 0 | C | T | 0.404 | 9.71E-02 | 9.41E-02 | 2.44E+00 | 2.48E+00 | 3.11E-01 | no | AC151990.1 | upstream_gene_variant | MODIFIER |
| 18 | 60362099 | AX-170120845 | 0 | T | C | 0.404 | 9.71E-02 | 9.41E-02 | 2.44E+00 | 2.48E+00 | 3.11E-01 | no | AC151990.1 | non_coding_transcript_exon_variant | MODIFIER |
| 18 | 60362231 | AX-170061009 | 0 | C | T | 0.404 | 9.71E-02 | 9.41E-02 | 2.44E+00 | 2.48E+00 | 3.11E-01 | no | AC151990.1 | non_coding_transcript_exon_variant | MODIFIER |
| 18 | 60362349 | AX-170061010 | 0 | A | G | 0.404 | 9.71E-02 | 9.41E-02 | 2.44E+00 | 2.48E+00 | 3.11E-01 | no | AC151990.1 | non_coding_transcript_exon_variant | MODIFIER |
| 18 | 60362990 | AX-170061008 | 0 | C | A | 0.404 | 9.71E-02 | 9.41E-02 | 2.44E+00 | 2.48E+00 | 3.11E-01 | no | AC151990.1 | downstream_gene_variant | MODIFIER |
| 18 | 60463266 | AX-170365887 | 0 | G | A | 0.404 | 9.71E-02 | 9.41E-02 | 2.44E+00 | 2.48E+00 | 3.11E-01 | no | | intergenic_variant | MODIFIER |
| 18 | 60483450 | AX-170275482 | 0 | T | C | 0.404 | 9.71E-02 | 9.41E-02 | 2.44E+00 | 2.48E+00 | 3.11E-01 | no | Smim3 | intron_variant | MODIFIER |
| 18 | 60522165 | AX-170357915 | 0 | C | T | 0.404 | 9.71E-02 | 9.41E-02 | 2.44E+00 | 2.48E+00 | 3.11E-01 | no | Dctn4 | upstream_gene_variant | MODIFIER |
| 18 | 60522467 | AX-170184075 | 0 | G | A | 0.404 | 9.71E-02 | 9.41E-02 | 2.44E+00 | 2.48E+00 | 3.11E-01 | no | Dctn4 | upstream_gene_variant | MODIFIER |
| 18 | 60522613 | AX-170192926 | 0 | T | G | 0.404 | 9.71E-02 | 9.41E-02 | 2.44E+00 | 2.48E+00 | 3.11E-01 | no | Dctn4 | upstream_gene_variant | MODIFIER |
| 18 | 60523210 | AX-170061012 | 0 | A | G | 0.404 | 9.71E-02 | 9.41E-02 | 2.44E+00 | 2.48E+00 | 3.11E-01 | no | Dctn4 | upstream_gene_variant | MODIFIER |
| 18 | 60526121 | AX-170346825 | 0 | C | G | 0.404 | 9.71E-02 | 9.41E-02 | 2.44E+00 | 2.48E+00 | 3.11E-01 | no | Dctn4 | upstream_gene_variant | MODIFIER |

| | | | | | | | | | | | | | | | |
|---|---|---|---|---|---|---|---|---|---|---|---|---|---|---|---|
| 18 | 60535182 | AX-170297648 | 0 | C | T | 0.404 | 9.71E-02 | 9.41E-02 | 2.44E+00 | 2.48E+00 | 3.11E-01 | no | Dctn4 | intron_variant | MODIFIER |
| 18 | 60549571 | AX-170153946 | 0 | C | A | 0.404 | 9.71E-02 | 9.41E-02 | 2.44E+00 | 2.48E+00 | 3.11E-01 | no | Dctn4 | intron_variant | MODIFIER |
| 18 | 60551562 | AX-170151522 | 0 | A | G | 0.404 | 9.71E-02 | 9.41E-02 | 2.44E+00 | 2.48E+00 | 3.11E-01 | no | Dctn4 | intron_variant | MODIFIER |
| 18 | 60557667 | AX-170496506 | 0 | G | A | 0.404 | 9.71E-02 | 9.41E-02 | 2.44E+00 | 2.48E+00 | 3.11E-01 | no | Dctn4 | 3_prime_UTR_variant | MODIFIER |
| 18 | 60574955 | AX-170015811 | 0 | A | G | 0.404 | 9.71E-02 | 9.41E-02 | 2.44E+00 | 2.48E+00 | 3.11E-01 | no | Rbm22 | downstream_gene_variant | MODIFIER |
| 18 | 60574998 | AX-170015814 | 0 | T | C | 0.404 | 9.71E-02 | 9.41E-02 | 2.44E+00 | 2.48E+00 | 3.11E-01 | no | Rbm22 | downstream_gene_variant | MODIFIER |
| 18 | 60575098 | AX-97843821 | 0 | T | C | 0.404 | 9.71E-02 | 9.41E-02 | 2.44E+00 | 2.48E+00 | 3.11E-01 | no | Rbm22 | downstream_gene_variant | MODIFIER |
| 18 | 60589841 | AX-170525096 | 0 | G | T | 0.404 | 9.71E-02 | 9.41E-02 | 2.44E+00 | 2.48E+00 | 3.11E-01 | no | Myoz3 | intron_variant | MODIFIER |
| 18 | 61093056 | AX-170282806 | 0 | C | T | 0.404 | 9.71E-02 | 9.41E-02 | 2.44E+00 | 2.48E+00 | 3.11E-01 | no | | regulatory_region_variant | MODIFIER |
| 18 | 61270800 | AX-97832532 | 0 | C | A | 0.404 | 9.71E-02 | 9.41E-02 | 2.44E+00 | 2.48E+00 | 3.11E-01 | no | Pde6a | intron_variant | MODIFIER |
| 18 | 61285414 | AX-170182267 | 0 | C | G | 0.404 | 9.71E-02 | 9.41E-02 | 2.44E+00 | 2.48E+00 | 3.11E-01 | no | Pde6a | intron_variant | MODIFIER |
| 18 | 61349157 | AX-97852137 | 0 | C | T | 0.404 | 9.71E-02 | 9.41E-02 | 2.44E+00 | 2.48E+00 | 3.11E-01 | no | Ppargc1b | intron_variant | MODIFIER |
| 18 | 61357915 | AX-170262504 | 0 | A | G | 0.404 | 9.71E-02 | 9.41E-02 | 2.44E+00 | 2.48E+00 | 3.11E-01 | no | Ppargc1b | intron_variant | MODIFIER |
| 18 | 61381857 | AX-97834960 | 0 | G | A | 0.404 | 9.71E-02 | 9.41E-02 | 2.44E+00 | 2.48E+00 | 3.11E-01 | no | Ppargc1b | intron_variant | MODIFIER |
| 18 | 61391650 | AX-97835604 | 0 | T | C | 0.404 | 9.71E-02 | 9.41E-02 | 2.44E+00 | 2.48E+00 | 3.11E-01 | no | Ppargc1b | intron_variant | MODIFIER |
| 18 | 61437443 | AX-170140947 | 0 | C | T | 0.404 | 9.71E-02 | 9.41E-02 | 2.44E+00 | 2.48E+00 | 3.11E-01 | no | AC116595.1 | upstream_gene_variant | MODIFIER |
| 18 | 61677051 | AX-170395277 | 0 | G | C | 0.404 | 9.71E-02 | 9.41E-02 | 2.44E+00 | 2.48E+00 | 3.11E-01 | no | | regulatory_region_variant | MODIFIER |
| 18 | 61684675 | AX-97828035 | 0 | C | T | 0.404 | 9.71E-02 | 9.41E-02 | 2.44E+00 | 2.48E+00 | 3.11E-01 | no | Il17b | upstream_gene_variant | MODIFIER |
| 18 | 61684924 | AX-170271289 | 0 | T | C | 0.404 | 9.71E-02 | 9.41E-02 | 2.44E+00 | 2.48E+00 | 3.11E-01 | no | Il17b | upstream_gene_variant | MODIFIER |
| 18 | 61731078 | AX-170344633 | 0 | T | C | 0.404 | 9.71E-02 | 9.41E-02 | 2.44E+00 | 2.48E+00 | 3.11E-01 | no | Grpel2 | upstream_gene_variant | MODIFIER |
| 18 | 61737739 | AX-170238929 | 0 | C | T | 0.404 | 9.71E-02 | 9.41E-02 | 2.44E+00 | 2.48E+00 | 3.11E-01 | no | Afap1l1 | intron_variant | MODIFIER |
| 18 | 61738327 | AX-170412095 | 0 | C | T | 0.404 | 9.71E-02 | 9.41E-02 | 2.44E+00 | 2.48E+00 | 3.11E-01 | no | Afap1l1 | intron_variant | MODIFIER |
| 18 | 61781960 | AX-97834954 | 0 | A | C | 0.404 | 9.71E-02 | 9.41E-02 | 2.44E+00 | 2.48E+00 | 3.11E-01 | no | Afap1l1 | intron_variant | MODIFIER |
| 18 | 61784500 | AX-170371389 | 0 | T | C | 0.404 | 9.71E-02 | 9.41E-02 | 2.44E+00 | 2.48E+00 | 3.11E-01 | no | Afap1l1 | intron_variant | MODIFIER |
| 18 | 61793610 | AX-170282768 | 0 | T | C | 0.404 | 9.71E-02 | 9.41E-02 | 2.44E+00 | 2.48E+00 | 3.11E-01 | no | | regulatory_region_variant | MODIFIER |
| 18 | 61804350 | AX-170171555 | 0 | T | A | 0.404 | 9.71E-02 | 9.41E-02 | 2.44E+00 | 2.48E+00 | 3.11E-01 | no | Ablim3 | intron_variant | MODIFIER |
| 18 | 61804777 | AX-97826897 | 0 | T | C | 0.404 | 9.71E-02 | 9.41E-02 | 2.44E+00 | 2.48E+00 | 3.11E-01 | no | Ablim3 | intron_variant | MODIFIER |
| 18 | 61804824 | AX-170380353 | 0 | T | C | 0.404 | 9.71E-02 | 9.41E-02 | 2.44E+00 | 2.48E+00 | 3.11E-01 | no | Ablim3 | intron_variant | MODIFIER |
| 18 | 61807346 | AX-170181618 | 0 | A | C | 0.404 | 9.71E-02 | 9.41E-02 | 2.44E+00 | 2.48E+00 | 3.11E-01 | no | Ablim3 | intron_variant | MODIFIER |
| 18 | 61809071 | AX-170113177 | 0 | A | G | 0.404 | 9.71E-02 | 9.41E-02 | 2.44E+00 | 2.48E+00 | 3.11E-01 | no | Ablim3 | intron_variant | MODIFIER |
| 18 | 61905716 | AX-170351156 | 0 | C | T | 0.404 | 9.71E-02 | 9.41E-02 | 2.44E+00 | 2.48E+00 | 3.11E-01 | no | Ablim3 | intron_variant | MODIFIER |
| 18 | 61941167 | AX-170474957 | 0 | G | A | 0.404 | 9.71E-02 | 9.41E-02 | 2.44E+00 | 2.48E+00 | 3.11E-01 | no | | intergenic_variant | MODIFIER |
| 18 | 61944953 | AX-170446803 | 0 | G | T | 0.404 | 9.71E-02 | 9.41E-02 | 2.44E+00 | 2.48E+00 | 3.11E-01 | no | | regulatory_region_variant | MODIFIER |
| 18 | 61975964 | AX-170257702 | 0 | T | C | 0.404 | 9.71E-02 | 9.41E-02 | 2.44E+00 | 2.48E+00 | 3.11E-01 | no | Sh3tc2 | intron_variant | MODIFIER |
| 18 | 61976851 | AX-170405619 | 0 | C | A | 0.404 | 9.71E-02 | 9.41E-02 | 2.44E+00 | 2.48E+00 | 3.11E-01 | no | Sh3tc2 | intron_variant | MODIFIER |
| 18 | 61981552 | AX-170447453 | 0 | T | C | 0.404 | 9.71E-02 | 9.41E-02 | 2.44E+00 | 2.48E+00 | 3.11E-01 | no | Sh3tc2 | intron_variant | MODIFIER |
| 18 | 61981568 | AX-170516722 | 0 | C | T | 0.404 | 9.71E-02 | 9.41E-02 | 2.44E+00 | 2.48E+00 | 3.11E-01 | no | Sh3tc2 | intron_variant | MODIFIER |
| 18 | 61986244 | AX-170284143 | 0 | G | T | 0.404 | 9.71E-02 | 9.41E-02 | 2.44E+00 | 2.48E+00 | 3.11E-01 | no | Sh3tc2 | intron_variant | MODIFIER |
| 18 | 61995817 | AX-170176189 | 0 | C | G | 0.404 | 9.71E-02 | 9.41E-02 | 2.44E+00 | 2.48E+00 | 3.11E-01 | no | Sh3tc2 | intron_variant | MODIFIER |
| 18 | 61996937 | AX-170223671 | 0 | C | T | 0.404 | 9.71E-02 | 9.41E-02 | 2.44E+00 | 2.48E+00 | 3.11E-01 | no | Sh3tc2 | intron_variant | MODIFIER |
| 18 | 62002218 | AX-170415670 | 0 | A | G | 0.404 | 9.71E-02 | 9.41E-02 | 2.44E+00 | 2.48E+00 | 3.11E-01 | no | Sh3tc2 | intron_variant | MODIFIER |
| 18 | 62126154 | AX-170198376 | 0 | A | G | 0.404 | 9.71E-02 | 9.41E-02 | 2.44E+00 | 2.48E+00 | 3.11E-01 | no | | intergenic_variant | MODIFIER |
| 18 | 62204063 | AX-170525650 | 0 | G | A | 0.404 | 9.71E-02 | 9.41E-02 | 2.44E+00 | 2.48E+00 | 3.11E-01 | no | | regulatory_region_variant | MODIFIER |
| 18 | 62998403 | AX-170362402 | 0 | G | A | 0.404 | 9.71E-02 | 9.41E-02 | 2.44E+00 | 2.48E+00 | 3.11E-01 | no | Napg | 3_prime_UTR_variant | MODIFIER |
| 18 | 62998547 | AX-170422832 | 0 | C | T | 0.404 | 9.71E-02 | 9.41E-02 | 2.44E+00 | 2.48E+00 | 3.11E-01 | no | Napg | 3_prime_UTR_variant | MODIFIER |
| 18 | 62998811 | AX-170434531 | 0 | C | T | 0.404 | 9.71E-02 | 9.41E-02 | 2.44E+00 | 2.48E+00 | 3.11E-01 | no | Napg | 3_prime_UTR_variant | MODIFIER |
| 18 | 63126930 | AX-170340957 | 0 | A | C | 0.404 | 9.71E-02 | 9.41E-02 | 2.44E+00 | 2.48E+00 | 3.11E-01 | no | Piezo2 | intron_variant | MODIFIER |
| 18 | 63162234 | AX-97798635 | 0 | C | T | 0.404 | 9.71E-02 | 9.41E-02 | 2.44E+00 | 2.48E+00 | 3.11E-01 | no | Piezo2 | intron_variant | MODIFIER |
| 18 | 63245586 | AX-170175751 | 0 | C | T | 0.404 | 9.71E-02 | 9.41E-02 | 2.44E+00 | 2.48E+00 | 3.11E-01 | no | Piezo2 | missense_variant | MODERATE |
| 18 | 63246577 | AX-170209219 | 0 | A | G | 0.404 | 9.71E-02 | 9.41E-02 | 2.44E+00 | 2.48E+00 | 3.11E-01 | no | Piezo2 | intron_variant | MODIFIER |
| 7 | 71634052 | AX-170212756 | 0 | A | G | 0.368 | -1.03E-01 | 1.02E-01 | 2.83E+00 | 2.92E+00 | 3.11E-01 | no | | intergenic_variant | MODIFIER |
| 7 | 71634908 | AX-170145528 | 0 | A | G | 0.368 | -1.03E-01 | 1.02E-01 | 2.83E+00 | 2.92E+00 | 3.11E-01 | no | | intergenic_variant | MODIFIER |
| 7 | 71636953 | AX-97843883 | 0 | A | C | 0.368 | -1.03E-01 | 1.02E-01 | 2.83E+00 | 2.92E+00 | 3.11E-01 | no | | intergenic_variant | MODIFIER |
| 7 | 71657854 | AX-170386497 | 0 | C | T | 0.368 | -1.03E-01 | 1.02E-01 | 2.83E+00 | 2.92E+00 | 3.11E-01 | no | | intergenic_variant | MODIFIER |

| | | | | | | | | | | | | | | | |
|---|---|---|---|---|---|---|---|---|---|---|---|---|---|---|---|
| 7 | 71670245 | AX-170292791 | 0 | C | A | 0.368 | -1.03E-01 | 1.02E-01 | 2.83E+00 | 2.92E+00 | 3.11E-01 | no | | intergenic_variant | MODIFIER |
| 7 | 71696852 | AX-170371056 | 0 | T | C | 0.368 | -1.03E-01 | 1.02E-01 | 2.83E+00 | 2.92E+00 | 3.11E-01 | no | Gm7675 | non_coding_transcript_exon_variant | MODIFIER |
| 7 | 71776507 | AX-97852297 | 0 | G | A | 0.368 | -1.03E-01 | 1.02E-01 | 2.83E+00 | 2.92E+00 | 3.11E-01 | no | | intergenic_variant | MODIFIER |
| 7 | 71816909 | AX-170128931 | 0 | G | C | 0.368 | -1.03E-01 | 1.02E-01 | 2.83E+00 | 2.92E+00 | 3.11E-01 | no | | intergenic_variant | MODIFIER |
| 7 | 72067583 | AX-170322453 | 0 | G | A | 0.368 | -1.03E-01 | 1.02E-01 | 2.83E+00 | 2.92E+00 | 3.11E-01 | no | | intergenic_variant | MODIFIER |
| 7 | 72103936 | AX-97810043 | 0 | A | G | 0.368 | -1.03E-01 | 1.02E-01 | 2.83E+00 | 2.92E+00 | 3.11E-01 | no | Mctp2 | intron_variant | MODIFIER |
| 7 | 72110157 | AX-170520837 | 0 | G | T | 0.368 | -1.03E-01 | 1.02E-01 | 2.83E+00 | 2.92E+00 | 3.11E-01 | no | Mctp2 | intron_variant | MODIFIER |
| 7 | 72113385 | AX-170278214 | 0 | T | G | 0.368 | -1.03E-01 | 1.02E-01 | 2.83E+00 | 2.92E+00 | 3.11E-01 | no | Mctp2 | intron_variant | MODIFIER |
| 7 | 72115604 | AX-170029269 | 0 | A | T | 0.368 | -1.03E-01 | 1.02E-01 | 2.83E+00 | 2.92E+00 | 3.11E-01 | no | Mctp2 | intron_variant | MODIFIER |
| 7 | 72121626 | AX-170504444 | 0 | G | A | 0.368 | -1.03E-01 | 1.02E-01 | 2.83E+00 | 2.92E+00 | 3.11E-01 | no | Mctp2 | intron_variant | MODIFIER |
| 7 | 72147183 | AX-97790370 | 0 | C | A | 0.368 | -1.03E-01 | 1.02E-01 | 2.83E+00 | 2.92E+00 | 3.11E-01 | no | Mctp2 | intron_variant | MODIFIER |
| 7 | 72149624 | AX-97813282 | 0 | G | A | 0.368 | -1.03E-01 | 1.02E-01 | 2.83E+00 | 2.92E+00 | 3.11E-01 | no | Mctp2 | intron_variant | MODIFIER |
| 7 | 72350401 | AX-170029267 | 0 | G | A | 0.368 | -1.03E-01 | 1.02E-01 | 2.83E+00 | 2.92E+00 | 3.11E-01 | no | | regulatory_region_variant | MODIFIER |
| 7 | 72359934 | AX-170193187 | 0 | T | C | 0.368 | -1.03E-01 | 1.02E-01 | 2.83E+00 | 2.92E+00 | 3.11E-01 | no | | intergenic_variant | MODIFIER |
| 7 | 72543649 | AX-170266754 | 0 | A | G | 0.368 | -1.03E-01 | 1.02E-01 | 2.83E+00 | 2.92E+00 | 3.11E-01 | no | | regulatory_region_variant | MODIFIER |
| 7 | 72577532 | AX-170248187 | 0 | G | A | 0.368 | -1.03E-01 | 1.02E-01 | 2.83E+00 | 2.92E+00 | 3.11E-01 | no | | intergenic_variant | MODIFIER |
| 7 | 72579250 | AX-170320833 | 0 | C | T | 0.368 | -1.03E-01 | 1.02E-01 | 2.83E+00 | 2.92E+00 | 3.11E-01 | no | | intergenic_variant | MODIFIER |
| 7 | 72579285 | AX-170431037 | 0 | G | A | 0.368 | -1.03E-01 | 1.02E-01 | 2.83E+00 | 2.92E+00 | 3.11E-01 | no | | intergenic_variant | MODIFIER |
| 7 | 72580556 | AX-170340619 | 0 | C | T | 0.368 | -1.03E-01 | 1.02E-01 | 2.83E+00 | 2.92E+00 | 3.11E-01 | no | | intergenic_variant | MODIFIER |
| 7 | 72598621 | AX-170247700 | 0 | G | A | 0.368 | -1.03E-01 | 1.02E-01 | 2.83E+00 | 2.92E+00 | 3.11E-01 | no | Gm37620 | upstream_gene_variant | MODIFIER |
| 7 | 72627640 | AX-170485266 | 0 | C | A | 0.368 | -1.03E-01 | 1.02E-01 | 2.83E+00 | 2.92E+00 | 3.11E-01 | no | | regulatory_region_variant | MODIFIER |
| 7 | 72665038 | AX-170281596 | 0 | G | A | 0.368 | -1.03E-01 | 1.02E-01 | 2.83E+00 | 2.92E+00 | 3.11E-01 | no | Gm28744 | downstream_gene_variant | MODIFIER |
| 7 | 72686000 | AX-170029281 | 0 | T | C | 0.368 | -1.03E-01 | 1.02E-01 | 2.83E+00 | 2.92E+00 | 3.11E-01 | no | | intergenic_variant | MODIFIER |
| 7 | 72686265 | AX-170029278 | 0 | T | C | 0.368 | -1.03E-01 | 1.02E-01 | 2.83E+00 | 2.92E+00 | 3.11E-01 | no | | intergenic_variant | MODIFIER |
| 7 | 72686404 | AX-170029282 | 0 | A | G | 0.368 | -1.03E-01 | 1.02E-01 | 2.83E+00 | 2.92E+00 | 3.11E-01 | no | | intergenic_variant | MODIFIER |
| 7 | 72686667 | AX-170290269 | 0 | G | A | 0.368 | -1.03E-01 | 1.02E-01 | 2.83E+00 | 2.92E+00 | 3.11E-01 | no | | intergenic_variant | MODIFIER |
| 7 | 72696850 | AX-170029279 | 0 | G | A | 0.368 | -1.03E-01 | 1.02E-01 | 2.83E+00 | 2.92E+00 | 3.11E-01 | no | | intergenic_variant | MODIFIER |
| 7 | 72702182 | AX-170029277 | 0 | T | C | 0.368 | -1.03E-01 | 1.02E-01 | 2.83E+00 | 2.92E+00 | 3.11E-01 | no | | intergenic_variant | MODIFIER |
| 7 | 72702907 | AX-170298593 | 0 | G | A | 0.368 | -1.03E-01 | 1.02E-01 | 2.83E+00 | 2.92E+00 | 3.11E-01 | no | | intergenic_variant | MODIFIER |
| 7 | 72725941 | AX-170532291 | 0 | A | C | 0.368 | -1.03E-01 | 1.02E-01 | 2.83E+00 | 2.92E+00 | 3.11E-01 | no | | intergenic_variant | MODIFIER |
| 7 | 72808034 | AX-170310453 | 0 | T | G | 0.368 | -1.03E-01 | 1.02E-01 | 2.83E+00 | 2.92E+00 | 3.11E-01 | no | | intergenic_variant | MODIFIER |
| 7 | 72810895 | AX-170155760 | 0 | T | C | 0.368 | -1.03E-01 | 1.02E-01 | 2.83E+00 | 2.92E+00 | 3.11E-01 | no | | intergenic_variant | MODIFIER |
| 7 | 72826512 | AX-170077704 | 0 | T | G | 0.368 | -1.03E-01 | 1.02E-01 | 2.83E+00 | 2.92E+00 | 3.11E-01 | no | | intergenic_variant | MODIFIER |
| 7 | 72945387 | AX-170513736 | 0 | G | A | 0.368 | -1.03E-01 | 1.02E-01 | 2.83E+00 | 2.92E+00 | 3.11E-01 | no | | intergenic_variant | MODIFIER |
| 7 | 72956594 | AX-97837599 | 0 | C | G | 0.368 | -1.03E-01 | 1.02E-01 | 2.83E+00 | 2.92E+00 | 3.11E-01 | no | | intergenic_variant | MODIFIER |
| 5 | 89301465 | AX-97798094 | 0 | A | G | 0.43 | 1.07E-01 | 1.06E-01 | 2.91E+00 | 2.99E+00 | 3.11E-01 | no | | intergenic_variant | MODIFIER |
| 18 | 77938436 | AX-170380046 | 1 | C | G | 0.438 | 8.79E-02 | 8.60E-02 | 2.49E+00 | 2.55E+00 | 3.11E-01 | no | Epg5 | upstream_gene_variant | MODIFIER |
| 15 | 53425071 | AX-170218634 | 0 | A | T | 0.061 | -1.19E-01 | 9.93E-02 | 1.99E+00 | 2.11E+00 | 3.11E-01 | no | | intergenic_variant | MODIFIER |
| 4 | 155860646 | AX-170024311 | 0 | G | A | 0.123 | 1.34E-01 | 1.34E-01 | 2.64E+00 | 2.78E+00 | 3.11E-01 | no | Dvl1 | downstream_gene_variant | MODIFIER |
| 1 | 166167682 | AX-170341764 | 1 | C | T | 0.223 | 1.36E-01 | 1.36E-01 | 2.76E+00 | 2.82E+00 | 3.12E-01 | no | Gpa33 | downstream_gene_variant | MODIFIER |
| 1 | 138156067 | AX-170344762 | 0 | G | C | 0.211 | 1.27E-01 | 1.27E-01 | 2.74E+00 | 2.81E+00 | 3.12E-01 | no | Ptprc | intron_variant,non_coding_transcript_variant | MODIFIER |
| 14 | 111349317 | AX-170192064 | 0 | G | T | 0.026 | 1.42E-01 | 1.41E-01 | 2.55E+00 | 2.73E+00 | 3.12E-01 | no | Gm48988 | downstream_gene_variant | MODIFIER |
| 16 | 24562399 | AX-97800090 | 0 | T | G | 0.246 | 8.65E-02 | 8.67E-02 | 2.72E+00 | 2.85E+00 | 3.12E-01 | no | Lpp | intron_variant | MODIFIER |
| 15 | 4429072 | AX-170191586 | 1 | G | A | 0.241 | -1.07E-01 | 1.07E-01 | 2.65E+00 | 2.70E+00 | 3.12E-01 | no | Plcxd3 | intron_variant | MODIFIER |
| 15 | 22908630 | AX-170011228 | 1 | G | A | 0.241 | -1.07E-01 | 1.07E-01 | 2.65E+00 | 2.70E+00 | 3.12E-01 | no | Cdh18 | intron_variant | MODIFIER |
| 9 | 72122573 | AX-170409589 | 0 | G | A | 0.184 | -1.08E-01 | 1.08E-01 | 2.73E+00 | 2.85E+00 | 3.12E-01 | no | Gm25045 | upstream_gene_variant | MODIFIER |
| 5 | 42939301 | AX-170173635 | 0 | A | T | 0.202 | -1.07E-01 | 1.06E-01 | 2.56E+00 | 2.65E+00 | 3.12E-01 | no | | intergenic_variant | MODIFIER |
| 7 | 72350281 | AX-170029271 | 0 | A | G | 0.377 | -1.02E-01 | 1.02E-01 | 2.86E+00 | 2.96E+00 | 3.12E-01 | no | | regulatory_region_variant | MODIFIER |
| 16 | 40744291 | AX-170352502 | 1 | A | G | 0.491 | 8.44E-02 | 8.35E-02 | 2.56E+00 | 2.59E+00 | 3.12E-01 | no | Lsamp | intron_variant | MODIFIER |
| 12 | 91537729 | AX-170292843 | 0 | A | G | 0.088 | 1.57E-01 | 1.58E-01 | 2.80E+00 | 2.92E+00 | 3.12E-01 | Yes | Tshr | synonymous_variant | LOW |
| 18 | 60002064 | AX-170093904 | 1 | T | C | 0.402 | 9.67E-02 | 9.42E-02 | 2.45E+00 | 2.49E+00 | 3.12E-01 | no | | intergenic_variant | MODIFIER |
| 18 | 62307328 | AX-170292497 | 1 | A | G | 0.402 | 9.67E-02 | 9.42E-02 | 2.45E+00 | 2.49E+00 | 3.12E-01 | no | | intergenic_variant | MODIFIER |
| 8 | 62977751 | AX-97830794 | 0 | A | T | 0.14 | -1.56E-01 | 1.54E-01 | 2.53E+00 | 2.58E+00 | 3.13E-01 | no | Spock3 | intron_variant | MODIFIER |
| 8 | 65719396 | AX-97852084 | 0 | C | T | 0.14 | -1.56E-01 | 1.54E-01 | 2.53E+00 | 2.58E+00 | 3.13E-01 | no | MAR1 | intron_variant | MODIFIER |

| | | | | | | | | | | | | | | | |
|---|---|---|---|---|---|---|---|---|---|---|---|---|---|---|---|
| 2 | 159844005 | AX-170473387 | 0 | G | T | 0.158 | -1.68E-01 | 1.67E-01 | 2.92E+00 | 3.05E+00 | 3.13E-01 | no | | intergenic_variant | MODIFIER |
| 15 | 4246392 | AX-170132548 | 1 | C | T | 0.241 | -1.07E-01 | 1.07E-01 | 2.65E+00 | 2.70E+00 | 3.13E-01 | no | Gm49301 | upstream_gene_variant | MODIFIER |
| 15 | 5805302 | AX-170363369 | 1 | A | G | 0.241 | -1.07E-01 | 1.07E-01 | 2.65E+00 | 2.70E+00 | 3.13E-01 | no | | intergenic_variant | MODIFIER |
| 15 | 10951451 | AX-170300240 | 1 | G | A | 0.241 | -1.07E-01 | 1.07E-01 | 2.65E+00 | 2.70E+00 | 3.13E-01 | no | C1qtnf3 | upstream_gene_variant | MODIFIER |
| 15 | 21726325 | AX-170241157 | 1 | C | T | 0.241 | -1.07E-01 | 1.07E-01 | 2.65E+00 | 2.70E+00 | 3.13E-01 | no | | intergenic_variant | MODIFIER |
| 5 | 46969427 | AX-170071986 | 1 | G | T | 0.491 | 1.06E-01 | 1.01E-01 | 2.35E+00 | 2.40E+00 | 3.13E-01 | no | | intergenic_variant | MODIFIER |
| 1 | 136326087 | AX-170463149 | 0 | C | T | 0.219 | 1.15E-01 | 1.16E-01 | 2.67E+00 | 2.76E+00 | 3.13E-01 | no | Camsap2 | intron_variant | MODIFIER |
| 1 | 137539440 | AX-170476298 | 0 | A | T | 0.219 | 1.15E-01 | 1.16E-01 | 2.67E+00 | 2.76E+00 | 3.13E-01 | no | | regulatory_region_variant | MODIFIER |
| 15 | 6640743 | AX-97840073 | 1 | C | G | 0.089 | -1.20E-01 | 9.93E-02 | 1.97E+00 | 2.07E+00 | 3.13E-01 | no | Fyb | intron_variant | MODIFIER |
| 16 | 18418039 | AX-170493350 | 1 | G | A | 0.134 | 9.76E-02 | 9.78E-02 | 2.83E+00 | 2.97E+00 | 3.13E-01 | no | Comt | intron_variant | MODIFIER |
| 16 | 18645842 | AX-97806484 | 1 | A | G | 0.134 | 9.76E-02 | 9.78E-02 | 2.83E+00 | 2.97E+00 | 3.13E-01 | no | | intergenic_variant | MODIFIER |
| 16 | 18959917 | AX-170279699 | 1 | A | G | 0.134 | 9.76E-02 | 9.78E-02 | 2.83E+00 | 2.97E+00 | 3.13E-01 | no | Hira | intron_variant | MODIFIER |
| 3 | 32183964 | AX-170496596 | 0 | G | C | 0.158 | -9.70E-02 | 9.70E-02 | 2.82E+00 | 2.92E+00 | 3.13E-01 | no | Kcnmb2 | intron_variant | MODIFIER |
| 14 | 113217915 | AX-97827947 | 0 | A | G | 0.342 | 7.27E-02 | 7.24E-02 | 2.58E+00 | 2.68E+00 | 3.13E-01 | no | | intergenic_variant | MODIFIER |
| 1 | 191715097 | AX-170002576 | 0 | G | C | 0.272 | 9.73E-02 | 9.27E-02 | 2.34E+00 | 2.42E+00 | 3.13E-01 | no | Lpgat1 | upstream_gene_variant | MODIFIER |
| 2 | 168881302 | AX-97809933 | 1 | G | T | 0.17 | 1.79E-01 | 1.79E-01 | 2.66E+00 | 2.75E+00 | 3.14E-01 | Yes | Gm25214 | downstream_gene_variant | MODIFIER |
| 4 | 138204020 | AX-97795767 | 1 | G | A | 0.17 | 1.15E-01 | 1.15E-01 | 2.62E+00 | 2.67E+00 | 3.14E-01 | no | Eif4g3 | intron_variant | MODIFIER |
| 4 | 95794977 | AX-170252773 | 0 | T | C | 0.254 | 1.12E-01 | 1.10E-01 | 2.49E+00 | 2.55E+00 | 3.14E-01 | no | Fggy | intron_variant | MODIFIER |
| 4 | 95863428 | AX-170224663 | 0 | G | A | 0.254 | 1.12E-01 | 1.10E-01 | 2.49E+00 | 2.55E+00 | 3.14E-01 | no | Fggy | intron_variant | MODIFIER |
| 4 | 95866344 | AX-170463902 | 0 | G | A | 0.254 | 1.12E-01 | 1.10E-01 | 2.49E+00 | 2.55E+00 | 3.14E-01 | no | Fggy | intron_variant | MODIFIER |
| 4 | 95868488 | AX-170272918 | 0 | G | A | 0.254 | 1.12E-01 | 1.10E-01 | 2.49E+00 | 2.55E+00 | 3.14E-01 | no | Fggy | intron_variant | MODIFIER |
| 4 | 95885991 | AX-170023452 | 0 | A | C | 0.254 | 1.12E-01 | 1.10E-01 | 2.49E+00 | 2.55E+00 | 3.14E-01 | no | Fggy | intron_variant | MODIFIER |
| 4 | 95886242 | AX-170023453 | 0 | A | G | 0.254 | 1.12E-01 | 1.10E-01 | 2.49E+00 | 2.55E+00 | 3.14E-01 | no | Fggy | intron_variant | MODIFIER |
| 4 | 95900450 | AX-170070067 | 0 | T | C | 0.254 | 1.12E-01 | 1.10E-01 | 2.49E+00 | 2.55E+00 | 3.14E-01 | no | Fggy | intron_variant | MODIFIER |
| 4 | 95900723 | AX-170070066 | 0 | G | T | 0.254 | 1.12E-01 | 1.10E-01 | 2.49E+00 | 2.55E+00 | 3.14E-01 | no | Fggy | intron_variant | MODIFIER |
| 4 | 95901576 | AX-170228363 | 0 | G | A | 0.254 | 1.12E-01 | 1.10E-01 | 2.49E+00 | 2.55E+00 | 3.14E-01 | no | Fggy | intron_variant | MODIFIER |
| 4 | 95907331 | AX-170241875 | 0 | C | A | 0.254 | 1.12E-01 | 1.10E-01 | 2.49E+00 | 2.55E+00 | 3.14E-01 | no | Fggy | intron_variant | MODIFIER |
| 4 | 95917655 | AX-170124129 | 0 | T | G | 0.254 | 1.12E-01 | 1.10E-01 | 2.49E+00 | 2.55E+00 | 3.14E-01 | no | Fggy | intron_variant | MODIFIER |
| 4 | 95919565 | AX-170447173 | 0 | A | G | 0.254 | 1.12E-01 | 1.10E-01 | 2.49E+00 | 2.55E+00 | 3.14E-01 | no | Fggy | intron_variant | MODIFIER |
| 4 | 96022491 | AX-97821599 | 0 | G | C | 0.254 | 1.12E-01 | 1.10E-01 | 2.49E+00 | 2.55E+00 | 3.14E-01 | no | Hook1 | 3_prime_UTR_variant | MODIFIER |
| 4 | 96069209 | AX-97802602 | 0 | A | T | 0.254 | 1.12E-01 | 1.10E-01 | 2.49E+00 | 2.55E+00 | 3.14E-01 | no | Cyp2j13 | intron_variant | MODIFIER |
| 4 | 96174408 | AX-170301201 | 0 | C | T | 0.254 | 1.12E-01 | 1.10E-01 | 2.49E+00 | 2.55E+00 | 3.14E-01 | no | Cyp2j7-ps2 | upstream_gene_variant | MODIFIER |
| 4 | 96200139 | AX-97843476 | 0 | T | C | 0.254 | 1.12E-01 | 1.10E-01 | 2.49E+00 | 2.55E+00 | 3.14E-01 | no | Cyp2j7 | intron_variant | MODIFIER |
| 4 | 96200781 | AX-170439116 | 0 | C | T | 0.254 | 1.12E-01 | 1.10E-01 | 2.49E+00 | 2.55E+00 | 3.14E-01 | no | Cyp2j7 | intron_variant | MODIFIER |
| 4 | 96203525 | AX-170220849 | 0 | A | G | 0.254 | 1.12E-01 | 1.10E-01 | 2.49E+00 | 2.55E+00 | 3.14E-01 | no | Cyp2j7 | intron_variant | MODIFIER |
| 9 | 16694504 | AX-170096481 | 1 | T | C | 0.473 | 8.98E-02 | 8.91E-02 | 2.56E+00 | 2.63E+00 | 3.14E-01 | no | | intergenic_variant | MODIFIER |
| 9 | 17586852 | AX-170314774 | 1 | T | G | 0.473 | 8.98E-02 | 8.91E-02 | 2.56E+00 | 2.63E+00 | 3.14E-01 | no | | intergenic_variant | MODIFIER |
| 18 | 74030024 | AX-170470596 | 1 | G | A | 0.179 | -1.22E-01 | 1.22E-01 | 2.64E+00 | 2.69E+00 | 3.14E-01 | no | Mapk4 | intron_variant | MODIFIER |
| 16 | 39493297 | AX-170250930 | 0 | G | A | 0.509 | 7.90E-02 | 7.80E-02 | 2.51E+00 | 2.57E+00 | 3.14E-01 | no | | intergenic_variant | MODIFIER |
| 4 | 155567444 | AX-170202104 | 1 | C | T | 0.125 | 1.61E-01 | 1.61E-01 | 2.78E+00 | 2.89E+00 | 3.14E-01 | Yes | Nadk | intron_variant | MODIFIER |
| 6 | 135607827 | AX-170125476 | 1 | A | C | 0.58 | 9.06E-02 | 8.82E-02 | 2.44E+00 | 2.48E+00 | 3.14E-01 | no | | intergenic_variant | MODIFIER |
| 9 | 7617732 | AX-170301738 | 2 | T | C | 0.445 | 7.98E-02 | 7.84E-02 | 2.47E+00 | 2.57E+00 | 3.14E-01 | no | | intergenic_variant | MODIFIER |
| 4 | 73168332 | AX-97789765 | 2 | T | C | 0.436 | -8.50E-02 | 8.32E-02 | 2.46E+00 | 2.53E+00 | 3.14E-01 | no | | intergenic_variant | MODIFIER |
| 18 | 68155823 | AX-170521010 | 0 | C | A | 0.184 | -1.15E-01 | 1.16E-01 | 2.67E+00 | 2.74E+00 | 3.14E-01 | no | Ldlrad4 | intron_variant | MODIFIER |
| 16 | 37212919 | AX-97836827 | 0 | C | T | 0.482 | 8.11E-02 | 8.05E-02 | 2.55E+00 | 2.60E+00 | 3.14E-01 | no | Stxbp5l | intron_variant,non_coding_transcript_variant | MODIFIER |
| 4 | 81956690 | AX-97847303 | 0 | C | A | 0.342 | 1.18E-01 | 1.19E-01 | 2.73E+00 | 2.79E+00 | 3.14E-01 | no | | regulatory_region_variant | MODIFIER |
| 3 | 38140776 | AX-170482090 | 0 | T | C | 0.202 | 8.75E-02 | 8.75E-02 | 2.88E+00 | 2.99E+00 | 3.14E-01 | no | | intergenic_variant | MODIFIER |
| 10 | 19018163 | AX-170305715 | 1 | C | T | 0.205 | 1.04E-01 | 1.02E-01 | 3.05E+00 | 3.16E+00 | 3.14E-01 | no | Tnfaip3 | upstream_gene_variant | MODIFIER |
| 11 | 53379428 | AX-170532172 | 0 | T | C | 0.035 | -1.66E-01 | 1.65E-01 | 2.54E+00 | 2.67E+00 | 3.14E-01 | no | Aff4 | intron_variant | MODIFIER |
| 13 | 8194967 | AX-170436930 | 1 | G | A | 0.188 | -1.18E-01 | 1.16E-01 | 2.52E+00 | 2.59E+00 | 3.14E-01 | no | Gm48257 | downstream_gene_variant | MODIFIER |
| 8 | 78678480 | AX-170428480 | 0 | T | G | 0.202 | 1.21E-01 | 1.21E-01 | 2.89E+00 | 2.99E+00 | 3.14E-01 | no | Slc10a7 | intron_variant | MODIFIER |
| 1 | 126650096 | AX-170036020 | 2 | A | G | 0.555 | 8.42E-02 | 8.33E-02 | 2.52E+00 | 2.57E+00 | 3.14E-01 | no | Nckap5 | intron_variant | MODIFIER |
| 14 | 81017420 | AX-97855020 | 0 | C | T | 0.044 | -1.70E-01 | 1.71E-01 | 2.76E+00 | 2.89E+00 | 3.14E-01 | no | | intergenic_variant | MODIFIER |
| 16 | 32509147 | AX-170358726 | 1 | T | C | 0.491 | 8.04E-02 | 8.01E-02 | 2.58E+00 | 2.64E+00 | 3.14E-01 | no | Zdhhc19 | downstream_gene_variant | MODIFIER |

| | | | | | | | | | | | | | | | |
|---|---|---|---|---|---|---|---|---|---|---|---|---|---|---|---|
| 15 | 21730051 | AX-170055001 | 1 | T | G | 0.232 | -1.06E-01 | 1.06E-01 | 2.64E+00 | 2.68E+00 | 3.14E-01 | no | | intergenic_variant | MODIFIER |
| 6 | 137093801 | AX-170414661 | 1 | C | T | 0.58 | 9.06E-02 | 8.84E-02 | 2.44E+00 | 2.49E+00 | 3.15E-01 | no | Rerg | intron_variant | MODIFIER |
| 6 | 137417691 | AX-170353084 | 0 | T | C | 0.57 | 9.02E-02 | 8.85E-02 | 2.47E+00 | 2.52E+00 | 3.15E-01 | no | Ptpro | intron_variant | MODIFIER |
| 4 | 78348435 | AX-170023174 | 1 | A | G | 0.312 | 1.12E-01 | 1.13E-01 | 2.77E+00 | 2.85E+00 | 3.15E-01 | no | | intergenic_variant | MODIFIER |
| 5 | 48506063 | AX-170214509 | 1 | C | G | 0.5 | 1.09E-01 | 1.05E-01 | 2.41E+00 | 2.45E+00 | 3.15E-01 | no | Kcnip4 | intron_variant | MODIFIER |
| 10 | 77161272 | AX-170154515 | 0 | T | C | 0.421 | -7.83E-02 | 7.86E-02 | 2.67E+00 | 2.75E+00 | 3.15E-01 | no | Col18a1 | intron_variant | MODIFIER |
| 1 | 121834316 | AX-170515837 | 0 | G | A | 0.57 | 8.35E-02 | 8.28E-02 | 2.53E+00 | 2.59E+00 | 3.15E-01 | no | | intergenic_variant | MODIFIER |
| 13 | 39343336 | AX-170344545 | 1 | A | T | 0.179 | -9.86E-02 | 9.71E-02 | 2.47E+00 | 2.59E+00 | 3.15E-01 | no | | intergenic_variant | MODIFIER |
| 7 | 114423864 | AX-170260095 | 0 | A | G | 0.447 | 9.29E-02 | 9.33E-02 | 2.74E+00 | 2.80E+00 | 3.15E-01 | no | Pde3b | intron_variant | MODIFIER |
| 13 | 5719239 | AX-170513945 | 1 | C | A | 0.205 | -1.26E-01 | 1.26E-01 | 2.61E+00 | 2.66E+00 | 3.15E-01 | no | Gm35330 | intron_variant,non_coding_transcript_variant | MODIFIER |
| 5 | 51617691 | AX-170024942 | 0 | T | C | 0.5 | 1.06E-01 | 1.04E-01 | 2.47E+00 | 2.53E+00 | 3.15E-01 | no | | intergenic_variant | MODIFIER |
| 7 | 37895492 | AX-170163733 | 1 | G | A | 0.357 | 9.58E-02 | 9.52E-02 | 2.55E+00 | 2.62E+00 | 3.15E-01 | no | | intergenic_variant | MODIFIER |
| 1 | 151633898 | AX-170260041 | 1 | C | T | 0.223 | 1.26E-01 | 1.26E-01 | 2.69E+00 | 2.77E+00 | 3.15E-01 | no | Fam129a | intron_variant,NMD_transcript_variant | MODIFIER |
| 6 | 135117058 | AX-170321977 | 1 | A | G | 0.571 | 8.91E-02 | 8.69E-02 | 2.43E+00 | 2.48E+00 | 3.15E-01 | no | Gprc5d | intron_variant | MODIFIER |
| 6 | 134786583 | AX-170445174 | 1 | G | A | 0.58 | 9.06E-02 | 8.85E-02 | 2.44E+00 | 2.49E+00 | 3.15E-01 | no | Dusp16 | intron_variant | MODIFIER |
| 16 | 40121268 | AX-97789445 | 0 | C | A | 0.491 | 8.39E-02 | 8.34E-02 | 2.56E+00 | 2.59E+00 | 3.15E-01 | no | Lsamp | intron_variant | MODIFIER |
| 16 | 40140268 | AX-170307270 | 0 | A | T | 0.491 | 8.39E-02 | 8.34E-02 | 2.56E+00 | 2.59E+00 | 3.15E-01 | no | Lsamp | intron_variant | MODIFIER |
| 16 | 40142350 | AX-97827701 | 0 | T | C | 0.491 | 8.39E-02 | 8.34E-02 | 2.56E+00 | 2.59E+00 | 3.15E-01 | no | Lsamp | intron_variant | MODIFIER |
| 16 | 40144122 | AX-170057205 | 0 | A | T | 0.491 | 8.39E-02 | 8.34E-02 | 2.56E+00 | 2.59E+00 | 3.15E-01 | no | Lsamp | intron_variant | MODIFIER |
| 16 | 40144614 | AX-170494856 | 0 | A | T | 0.491 | 8.39E-02 | 8.34E-02 | 2.56E+00 | 2.59E+00 | 3.15E-01 | no | Lsamp | intron_variant | MODIFIER |
| 16 | 40164307 | AX-97849489 | 0 | A | C | 0.491 | 8.39E-02 | 8.34E-02 | 2.56E+00 | 2.59E+00 | 3.15E-01 | no | Lsamp | intron_variant | MODIFIER |
| 16 | 40217638 | AX-97831555 | 0 | C | T | 0.491 | 8.39E-02 | 8.34E-02 | 2.56E+00 | 2.59E+00 | 3.15E-01 | no | Lsamp | intron_variant | MODIFIER |
| 16 | 40234938 | AX-170294642 | 0 | T | G | 0.491 | 8.39E-02 | 8.34E-02 | 2.56E+00 | 2.59E+00 | 3.15E-01 | no | Lsamp | intron_variant | MODIFIER |
| 16 | 40238220 | AX-170444154 | 0 | C | G | 0.491 | 8.39E-02 | 8.34E-02 | 2.56E+00 | 2.59E+00 | 3.15E-01 | no | Lsamp | intron_variant | MODIFIER |
| 16 | 40239795 | AX-170515539 | 0 | A | G | 0.491 | 8.39E-02 | 8.34E-02 | 2.56E+00 | 2.59E+00 | 3.15E-01 | no | Lsamp | intron_variant | MODIFIER |
| 16 | 40240795 | AX-170130892 | 0 | T | G | 0.491 | 8.39E-02 | 8.34E-02 | 2.56E+00 | 2.59E+00 | 3.15E-01 | no | Lsamp | intron_variant | MODIFIER |
| 16 | 40321333 | AX-170093452 | 0 | T | C | 0.491 | 8.39E-02 | 8.34E-02 | 2.56E+00 | 2.59E+00 | 3.15E-01 | no | Lsamp | intron_variant | MODIFIER |
| 16 | 40449922 | AX-97821841 | 0 | C | T | 0.491 | 8.39E-02 | 8.34E-02 | 2.56E+00 | 2.59E+00 | 3.15E-01 | no | Lsamp | intron_variant | MODIFIER |
| 16 | 40450551 | AX-170227668 | 0 | G | C | 0.491 | 8.39E-02 | 8.34E-02 | 2.56E+00 | 2.59E+00 | 3.15E-01 | no | Lsamp | intron_variant | MODIFIER |
| 16 | 40451202 | AX-170012882 | 0 | G | A | 0.491 | 8.39E-02 | 8.34E-02 | 2.56E+00 | 2.59E+00 | 3.15E-01 | no | Lsamp | intron_variant | MODIFIER |
| 16 | 40456499 | AX-170482035 | 0 | A | T | 0.491 | 8.39E-02 | 8.34E-02 | 2.56E+00 | 2.59E+00 | 3.15E-01 | no | Lsamp | intron_variant | MODIFIER |
| 16 | 40463607 | AX-170186772 | 0 | G | C | 0.491 | 8.39E-02 | 8.34E-02 | 2.56E+00 | 2.59E+00 | 3.15E-01 | no | Lsamp | intron_variant | MODIFIER |
| 16 | 40480046 | AX-97828160 | 0 | T | G | 0.491 | 8.39E-02 | 8.34E-02 | 2.56E+00 | 2.59E+00 | 3.15E-01 | no | Lsamp | intron_variant | MODIFIER |
| 16 | 40590385 | AX-97853236 | 0 | A | G | 0.491 | 8.39E-02 | 8.34E-02 | 2.56E+00 | 2.59E+00 | 3.15E-01 | no | Lsamp | intron_variant | MODIFIER |
| 16 | 40615096 | AX-170196369 | 0 | T | C | 0.491 | 8.39E-02 | 8.34E-02 | 2.56E+00 | 2.59E+00 | 3.15E-01 | no | Lsamp | intron_variant | MODIFIER |
| 16 | 40616094 | AX-170128379 | 0 | G | C | 0.491 | 8.39E-02 | 8.34E-02 | 2.56E+00 | 2.59E+00 | 3.15E-01 | no | Lsamp | intron_variant | MODIFIER |
| 16 | 40616268 | AX-170180610 | 0 | G | T | 0.491 | 8.39E-02 | 8.34E-02 | 2.56E+00 | 2.59E+00 | 3.15E-01 | no | Lsamp | intron_variant | MODIFIER |
| 16 | 40625432 | AX-170422656 | 0 | A | G | 0.491 | 8.39E-02 | 8.34E-02 | 2.56E+00 | 2.59E+00 | 3.15E-01 | no | Lsamp | intron_variant | MODIFIER |
| 16 | 40634765 | AX-170143270 | 0 | A | G | 0.491 | 8.39E-02 | 8.34E-02 | 2.56E+00 | 2.59E+00 | 3.15E-01 | no | Lsamp | intron_variant | MODIFIER |
| 16 | 40645747 | AX-170511188 | 0 | C | T | 0.491 | 8.39E-02 | 8.34E-02 | 2.56E+00 | 2.59E+00 | 3.15E-01 | no | Lsamp | intron_variant | MODIFIER |
| 16 | 40647182 | AX-170057214 | 0 | C | A | 0.491 | 8.39E-02 | 8.34E-02 | 2.56E+00 | 2.59E+00 | 3.15E-01 | no | Lsamp | intron_variant | MODIFIER |
| 16 | 40647228 | AX-170057213 | 0 | A | T | 0.491 | 8.39E-02 | 8.34E-02 | 2.56E+00 | 2.59E+00 | 3.15E-01 | no | Lsamp | intron_variant | MODIFIER |
| 16 | 40707558 | AX-170185482 | 0 | G | C | 0.491 | 8.39E-02 | 8.34E-02 | 2.56E+00 | 2.59E+00 | 3.15E-01 | no | Lsamp | intron_variant | MODIFIER |
| 16 | 40708687 | AX-97830683 | 0 | C | T | 0.491 | 8.39E-02 | 8.34E-02 | 2.56E+00 | 2.59E+00 | 3.15E-01 | no | Lsamp | intron_variant | MODIFIER |
| 16 | 40744335 | AX-170115582 | 0 | C | T | 0.491 | 8.39E-02 | 8.34E-02 | 2.56E+00 | 2.59E+00 | 3.15E-01 | no | Lsamp | intron_variant | MODIFIER |
| 16 | 40753746 | AX-170219162 | 0 | G | A | 0.491 | 8.39E-02 | 8.34E-02 | 2.56E+00 | 2.59E+00 | 3.15E-01 | no | Lsamp | intron_variant | MODIFIER |
| 16 | 40755031 | AX-170247734 | 0 | C | T | 0.491 | 8.39E-02 | 8.34E-02 | 2.56E+00 | 2.59E+00 | 3.15E-01 | no | Lsamp | intron_variant | MODIFIER |
| 16 | 40755769 | AX-170293559 | 0 | G | A | 0.491 | 8.39E-02 | 8.34E-02 | 2.56E+00 | 2.59E+00 | 3.15E-01 | no | Lsamp | intron_variant | MODIFIER |
| 16 | 40788399 | AX-170355554 | 0 | C | G | 0.491 | 8.39E-02 | 8.34E-02 | 2.56E+00 | 2.59E+00 | 3.15E-01 | no | Lsamp | intron_variant | MODIFIER |
| 16 | 40790314 | AX-170112295 | 0 | C | T | 0.491 | 8.39E-02 | 8.34E-02 | 2.56E+00 | 2.59E+00 | 3.15E-01 | no | Lsamp | intron_variant | MODIFIER |
| 16 | 40806331 | AX-170012891 | 0 | A | C | 0.491 | 8.39E-02 | 8.34E-02 | 2.56E+00 | 2.59E+00 | 3.15E-01 | no | Lsamp | intron_variant | MODIFIER |
| 16 | 40914655 | AX-170057219 | 0 | T | C | 0.491 | 8.39E-02 | 8.34E-02 | 2.56E+00 | 2.59E+00 | 3.15E-01 | no | Lsamp | intron_variant | MODIFIER |
| 16 | 40979980 | AX-170456685 | 0 | C | G | 0.491 | 8.39E-02 | 8.34E-02 | 2.56E+00 | 2.59E+00 | 3.15E-01 | no | Lsamp | intron_variant | MODIFIER |
| 16 | 41013158 | AX-170212159 | 0 | G | A | 0.491 | 8.39E-02 | 8.34E-02 | 2.56E+00 | 2.59E+00 | 3.15E-01 | no | Lsamp | intron_variant | MODIFIER |

| | | | | | | | | | | | | | | | |
|---|---|---|---|---|---|---|---|---|---|---|---|---|---|---|---|
| 4 | 85396568 | AX-170152382 | 1 | T | G | 0.339 | 1.05E-01 | 1.06E-01 | 2.75E+00 | 2.84E+00 | 3.15E-01 | no | Sh3gl2 | intron_variant | MODIFIER |
| 2 | 169773139 | AX-170496235 | 1 | G | C | 0.17 | 1.79E-01 | 1.80E-01 | 2.66E+00 | 2.75E+00 | 3.15E-01 | Yes | Tshz2 | intron_variant | MODIFIER |
| 14 | 119919161 | AX-170528390 | 1 | G | A | 0.33 | 7.43E-02 | 7.46E-02 | 2.62E+00 | 2.73E+00 | 3.15E-01 | no | | intergenic_variant | MODIFIER |
| 1 | 10029501 | AX-170101733 | 0 | C | T | 0.096 | -1.52E-01 | 1.53E-01 | 2.81E+00 | 2.94E+00 | 3.15E-01 | no | Cops5 | intron_variant | MODIFIER |
| 4 | 85953308 | AX-170136668 | 1 | C | T | 0.357 | 1.13E-01 | 1.13E-01 | 2.69E+00 | 2.77E+00 | 3.15E-01 | no | Adamtsl1 | intron_variant | MODIFIER |
| 2 | 161915859 | AX-170460105 | 1 | T | C | 0.152 | 1.91E-01 | 1.93E-01 | 2.81E+00 | 2.92E+00 | 3.15E-01 | Yes | Ptprt | intron_variant | MODIFIER |
| 2 | 164925017 | AX-97810378 | 1 | C | T | 0.152 | 1.91E-01 | 1.93E-01 | 2.81E+00 | 2.92E+00 | 3.15E-01 | Yes | | regulatory_region_variant | MODIFIER |
| 2 | 165686302 | AX-170019598 | 1 | A | T | 0.152 | 1.91E-01 | 1.93E-01 | 2.81E+00 | 2.92E+00 | 3.15E-01 | Yes | Eya2 | intron_variant | MODIFIER |
| 4 | 46914993 | AX-170089219 | 1 | A | G | 0.357 | -9.67E-02 | 9.71E-02 | 2.84E+00 | 2.93E+00 | 3.16E-01 | no | Gabbr2 | intron_variant | MODIFIER |
| 6 | 135336041 | AX-170176663 | 1 | G | C | 0.58 | 9.03E-02 | 8.83E-02 | 2.44E+00 | 2.48E+00 | 3.16E-01 | no | Gm36640 | intron_variant,non_coding_transcript_variant | MODIFIER |
| 8 | 35996618 | AX-170030505 | 0 | A | G | 0.149 | -1.31E-01 | 1.31E-01 | 2.65E+00 | 2.73E+00 | 3.16E-01 | no | | regulatory_region_variant | MODIFIER |
| 5 | 38079848 | AX-170502684 | 1 | G | A | 0.348 | 1.09E-01 | 1.09E-01 | 2.90E+00 | 2.99E+00 | 3.16E-01 | no | Stx18 | intron_variant | MODIFIER |
| 4 | 72241642 | AX-170362379 | 1 | A | G | 0.232 | 1.00E-01 | 9.93E-02 | 2.52E+00 | 2.60E+00 | 3.16E-01 | no | | intergenic_variant | MODIFIER |
| 5 | 33825612 | AX-170024653 | 0 | G | A | 0.333 | 1.04E-01 | 1.05E-01 | 2.73E+00 | 2.79E+00 | 3.16E-01 | no | Nsd2 | intron_variant | MODIFIER |
| 16 | 32578099 | AX-170322136 | 0 | T | C | 0.482 | 8.09E-02 | 8.05E-02 | 2.55E+00 | 2.60E+00 | 3.16E-01 | no | | intergenic_variant | MODIFIER |
| 16 | 32699967 | AX-97850555 | 0 | T | C | 0.482 | 8.09E-02 | 8.05E-02 | 2.55E+00 | 2.60E+00 | 3.16E-01 | no | | regulatory_region_variant | MODIFIER |
| 16 | 32769607 | AX-170012777 | 0 | A | T | 0.482 | 8.09E-02 | 8.05E-02 | 2.55E+00 | 2.60E+00 | 3.16E-01 | no | Muc4 | intron_variant | MODIFIER |
| 16 | 32770006 | AX-170448972 | 0 | T | C | 0.482 | 8.09E-02 | 8.05E-02 | 2.55E+00 | 2.60E+00 | 3.16E-01 | no | Muc4 | intron_variant | MODIFIER |
| 16 | 32770526 | AX-170122700 | 0 | T | C | 0.482 | 8.09E-02 | 8.05E-02 | 2.55E+00 | 2.60E+00 | 3.16E-01 | no | Muc4 | intron_variant | MODIFIER |
| 16 | 32771082 | AX-170195750 | 0 | T | C | 0.482 | 8.09E-02 | 8.05E-02 | 2.55E+00 | 2.60E+00 | 3.16E-01 | no | Muc4 | intron_variant | MODIFIER |
| 16 | 32771703 | AX-170259761 | 0 | C | A | 0.482 | 8.09E-02 | 8.05E-02 | 2.55E+00 | 2.60E+00 | 3.16E-01 | no | Muc4 | missense_variant | MODERATE |
| 16 | 33111835 | AX-97851724 | 0 | A | G | 0.482 | 8.09E-02 | 8.05E-02 | 2.55E+00 | 2.60E+00 | 3.16E-01 | no | Lmln | intron_variant | MODIFIER |
| 16 | 33123249 | AX-170299085 | 0 | G | A | 0.482 | 8.09E-02 | 8.05E-02 | 2.55E+00 | 2.60E+00 | 3.16E-01 | no | Lmln | intron_variant | MODIFIER |
| 16 | 33124626 | AX-170355182 | 0 | A | G | 0.482 | 8.09E-02 | 8.05E-02 | 2.55E+00 | 2.60E+00 | 3.16E-01 | no | Lmln | 3_prime_UTR_variant | MODIFIER |
| 16 | 33136766 | AX-170289847 | 0 | T | C | 0.482 | 8.09E-02 | 8.05E-02 | 2.55E+00 | 2.60E+00 | 3.16E-01 | no | Gm23916 | downstream_gene_variant | MODIFIER |
| 16 | 33140496 | AX-97825205 | 0 | T | G | 0.482 | 8.09E-02 | 8.05E-02 | 2.55E+00 | 2.60E+00 | 3.16E-01 | no | Gm23916 | upstream_gene_variant | MODIFIER |
| 16 | 33165245 | AX-97830890 | 0 | T | C | 0.482 | 8.09E-02 | 8.05E-02 | 2.55E+00 | 2.60E+00 | 3.16E-01 | no | | intergenic_variant | MODIFIER |
| 16 | 33168384 | AX-170418277 | 0 | A | G | 0.482 | 8.09E-02 | 8.05E-02 | 2.55E+00 | 2.60E+00 | 3.16E-01 | no | | intergenic_variant | MODIFIER |
| 16 | 33242133 | AX-97831426 | 0 | A | C | 0.482 | 8.09E-02 | 8.05E-02 | 2.55E+00 | 2.60E+00 | 3.16E-01 | no | Osbpl11 | 3_prime_UTR_variant | MODIFIER |
| 16 | 75508011 | AX-170013401 | 0 | T | A | 0.491 | 7.49E-02 | 7.40E-02 | 2.48E+00 | 2.57E+00 | 3.16E-01 | no | | intergenic_variant | MODIFIER |
| 4 | 46897160 | AX-170326788 | 1 | G | A | 0.357 | -9.58E-02 | 9.64E-02 | 2.83E+00 | 2.92E+00 | 3.16E-01 | no | Gabbr2 | intron_variant | MODIFIER |
| 2 | 169555838 | AX-170404515 | 1 | G | A | 0.17 | 1.79E-01 | 1.80E-01 | 2.64E+00 | 2.72E+00 | 3.16E-01 | Yes | | intergenic_variant | MODIFIER |
| 7 | 144579810 | AX-170217111 | 2 | C | T | 0.2 | -7.52E-02 | 7.58E-02 | 2.62E+00 | 2.75E+00 | 3.16E-01 | no | Fadd | 3_prime_UTR_variant | MODIFIER |
| 4 | 98092404 | AX-170393146 | 1 | C | T | 0.241 | 1.15E-01 | 1.15E-01 | 2.59E+00 | 2.65E+00 | 3.16E-01 | no | Nfia | intron_variant | MODIFIER |
| 1 | 128730859 | AX-170168025 | 1 | C | T | 0.554 | 8.28E-02 | 8.34E-02 | 2.67E+00 | 2.75E+00 | 3.16E-01 | no | | intergenic_variant | MODIFIER |
| 4 | 49829869 | AX-170465126 | 0 | C | T | 0.368 | -9.93E-02 | 1.00E-01 | 2.85E+00 | 2.96E+00 | 3.16E-01 | no | Grin3a | intron_variant | MODIFIER |
| 10 | 94015282 | AX-170402386 | 0 | T | C | 0.289 | -8.77E-02 | 8.71E-02 | 2.52E+00 | 2.57E+00 | 3.16E-01 | no | Vezt | intron_variant | MODIFIER |
| 4 | 98513155 | AX-170070123 | 1 | C | T | 0.232 | 1.18E-01 | 1.18E-01 | 2.57E+00 | 2.63E+00 | 3.17E-01 | no | Patj | intron_variant | MODIFIER |
| 12 | 117358762 | AX-170281010 | 0 | G | A | 0.14 | -1.59E-01 | 1.61E-01 | 2.69E+00 | 2.80E+00 | 3.17E-01 | no | | intergenic_variant | MODIFIER |
| 1 | 177310900 | AX-170533541 | 1 | A | G | 0.223 | 1.13E-01 | 1.13E-01 | 2.62E+00 | 2.70E+00 | 3.17E-01 | no | | intergenic_variant | MODIFIER |
| 18 | 63482778 | AX-170459354 | 0 | C | A | 0.184 | -1.21E-01 | 1.21E-01 | 2.66E+00 | 2.70E+00 | 3.17E-01 | no | | intergenic_variant | MODIFIER |
| 10 | 4964543 | AX-170146434 | 0 | T | C | 0.193 | -9.61E-02 | 9.72E-02 | 2.79E+00 | 2.93E+00 | 3.17E-01 | no | Esr1 | intron_variant | MODIFIER |
| 13 | 20918266 | AX-170176782 | 0 | C | A | 0.175 | -1.21E-01 | 1.21E-01 | 2.54E+00 | 2.61E+00 | 3.17E-01 | no | Aoah | intron_variant | MODIFIER |
| 13 | 20977434 | AX-170398487 | 0 | C | A | 0.175 | -1.21E-01 | 1.21E-01 | 2.54E+00 | 2.61E+00 | 3.17E-01 | no | Aoah | intron_variant | MODIFIER |
| 1 | 132161077 | AX-170387326 | 0 | C | T | 0.535 | 8.40E-02 | 8.47E-02 | 2.68E+00 | 2.75E+00 | 3.17E-01 | no | | intergenic_variant | MODIFIER |
| 18 | 79680692 | AX-170243325 | 1 | T | C | 0.455 | 8.38E-02 | 8.14E-02 | 2.39E+00 | 2.47E+00 | 3.17E-01 | no | | intergenic_variant | MODIFIER |
| 18 | 59140958 | AX-170350807 | 0 | G | A | 0.404 | -6.04E-02 | 5.86E-02 | 2.38E+00 | 2.47E+00 | 3.17E-01 | no | | intergenic_variant | MODIFIER |
| 4 | 137172311 | AX-170070821 | 1 | G | A | 0.161 | 1.16E-01 | 1.16E-01 | 2.62E+00 | 2.67E+00 | 3.17E-01 | no | | intergenic_variant | MODIFIER |
| 16 | 81136655 | AX-170499967 | 0 | G | A | 0.474 | 7.18E-02 | 7.24E-02 | 2.63E+00 | 2.72E+00 | 3.17E-01 | no | | intergenic_variant | MODIFIER |
| 10 | 71168761 | AX-97815834 | 1 | G | T | 0.402 | -8.14E-02 | 8.20E-02 | 2.63E+00 | 2.70E+00 | 3.17E-01 | no | C730027H18Rik | non_coding_transcript_exon_variant | MODIFIER |
| 18 | 72932510 | AX-170478899 | 0 | T | C | 0.184 | -1.17E-01 | 1.17E-01 | 2.61E+00 | 2.66E+00 | 3.17E-01 | no | | intergenic_variant | MODIFIER |
| 18 | 73099285 | AX-170015971 | 0 | C | G | 0.184 | -1.17E-01 | 1.17E-01 | 2.61E+00 | 2.66E+00 | 3.17E-01 | no | | intergenic_variant | MODIFIER |
| 18 | 73117436 | AX-170199618 | 0 | A | G | 0.184 | -1.17E-01 | 1.17E-01 | 2.61E+00 | 2.66E+00 | 3.17E-01 | no | | intergenic_variant | MODIFIER |
| 18 | 73118508 | AX-170517032 | 0 | C | G | 0.184 | -1.17E-01 | 1.17E-01 | 2.61E+00 | 2.66E+00 | 3.17E-01 | no | | intergenic_variant | MODIFIER |

| | | | | | | | | | | | | | | | |
|---|---|---|---|---|---|---|---|---|---|---|---|---|---|---|---|
| 18 | 73126999 | AX-170526212 | 0 | A | C | 0.184 | -1.17E-01 | 1.17E-01 | 2.61E+00 | 2.66E+00 | 3.17E-01 | no | | intergenic_variant | MODIFIER |
| 18 | 73131958 | AX-170514433 | 0 | T | A | 0.184 | -1.17E-01 | 1.17E-01 | 2.61E+00 | 2.66E+00 | 3.17E-01 | no | | regulatory_region_variant | MODIFIER |
| 18 | 73133118 | AX-170015975 | 0 | T | C | 0.184 | -1.17E-01 | 1.17E-01 | 2.61E+00 | 2.66E+00 | 3.17E-01 | no | | regulatory_region_variant | MODIFIER |
| 18 | 73137208 | AX-170061299 | 0 | T | C | 0.184 | -1.17E-01 | 1.17E-01 | 2.61E+00 | 2.66E+00 | 3.17E-01 | no | | intergenic_variant | MODIFIER |
| 18 | 73145796 | AX-170504581 | 0 | C | T | 0.184 | -1.17E-01 | 1.17E-01 | 2.61E+00 | 2.66E+00 | 3.17E-01 | no | | intergenic_variant | MODIFIER |
| 18 | 73167566 | AX-170451014 | 0 | T | C | 0.184 | -1.17E-01 | 1.17E-01 | 2.61E+00 | 2.66E+00 | 3.17E-01 | no | | intergenic_variant | MODIFIER |
| 18 | 73171143 | AX-170510028 | 0 | A | G | 0.184 | -1.17E-01 | 1.17E-01 | 2.61E+00 | 2.66E+00 | 3.17E-01 | no | | intergenic_variant | MODIFIER |
| 18 | 73175617 | AX-97810369 | 0 | C | T | 0.184 | -1.17E-01 | 1.17E-01 | 2.61E+00 | 2.66E+00 | 3.17E-01 | no | | intergenic_variant | MODIFIER |
| 18 | 73182891 | AX-170380619 | 0 | A | C | 0.184 | -1.17E-01 | 1.17E-01 | 2.61E+00 | 2.66E+00 | 3.17E-01 | no | | regulatory_region_variant | MODIFIER |
| 18 | 73183142 | AX-170373362 | 0 | A | G | 0.184 | -1.17E-01 | 1.17E-01 | 2.61E+00 | 2.66E+00 | 3.17E-01 | no | | regulatory_region_variant | MODIFIER |
| 18 | 73189268 | AX-170191712 | 0 | T | C | 0.184 | -1.17E-01 | 1.17E-01 | 2.61E+00 | 2.66E+00 | 3.17E-01 | no | | regulatory_region_variant | MODIFIER |
| 18 | 73189684 | AX-170419652 | 0 | A | G | 0.184 | -1.17E-01 | 1.17E-01 | 2.61E+00 | 2.66E+00 | 3.17E-01 | no | | regulatory_region_variant | MODIFIER |
| 16 | 78900399 | AX-170186994 | 0 | A | C | 0.447 | 7.41E-02 | 7.42E-02 | 2.55E+00 | 2.63E+00 | 3.18E-01 | no | | intergenic_variant | MODIFIER |
| 3 | 36611943 | AX-170308462 | 1 | G | A | 0.196 | 9.39E-02 | 9.47E-02 | 2.87E+00 | 2.96E+00 | 3.18E-01 | no | Bbs7 | intron_variant | MODIFIER |
| 4 | 136179753 | AX-170070796 | 1 | C | T | 0.17 | 1.16E-01 | 1.16E-01 | 2.55E+00 | 2.59E+00 | 3.18E-01 | no | E2f2 | intron_variant | MODIFIER |
| 9 | 112568599 | AX-170475904 | 0 | T | G | 0.132 | -8.64E-02 | 8.77E-02 | 2.75E+00 | 2.88E+00 | 3.18E-01 | no | | intergenic_variant | MODIFIER |
| 1 | 82240027 | AX-97784645 | 1 | A | C | 0.161 | 1.64E-01 | 1.66E-01 | 2.64E+00 | 2.76E+00 | 3.18E-01 | Yes | Gm9747 | downstream_gene_variant | MODIFIER |
| 7 | 71731908 | AX-170129768 | 0 | A | G | 0.36 | -9.28E-02 | 9.37E-02 | 2.85E+00 | 2.94E+00 | 3.18E-01 | no | | intergenic_variant | MODIFIER |
| 4 | 138852267 | AX-97822679 | 0 | C | T | 0.167 | 1.10E-01 | 1.10E-01 | 2.54E+00 | 2.59E+00 | 3.18E-01 | no | Pla2g5 | intron_variant | MODIFIER |
| 5 | 5159272 | AX-170071197 | 0 | A | T | 0.439 | -6.37E-02 | 6.35E-02 | 2.50E+00 | 2.61E+00 | 3.18E-01 | no | Cdk14 | intron_variant | MODIFIER |
| 5 | 37332160 | AX-170233834 | 1 | G | A | 0.33 | 1.11E-01 | 1.12E-01 | 2.80E+00 | 2.86E+00 | 3.18E-01 | no | Evc | intron_variant | MODIFIER |
| 13 | 32312309 | AX-170449291 | 1 | A | T | 0.188 | -1.11E-01 | 1.12E-01 | 2.62E+00 | 2.71E+00 | 3.18E-01 | no | Gmds | intron_variant | MODIFIER |
| 18 | 59385896 | AX-170278473 | 1 | T | C | 0.402 | 9.56E-02 | 9.38E-02 | 2.43E+00 | 2.47E+00 | 3.19E-01 | no | Chsy3 | intron_variant | MODIFIER |
| 2 | 107123062 | AX-170125768 | 0 | C | T | 0.053 | -1.86E-01 | 1.86E-01 | 2.54E+00 | 2.64E+00 | 3.19E-01 | Yes | Gm13903 | downstream_gene_variant | MODIFIER |
| 1 | 170128098 | AX-170002265 | 0 | A | G | 0.219 | 1.28E-01 | 1.30E-01 | 2.68E+00 | 2.75E+00 | 3.19E-01 | no | | intergenic_variant | MODIFIER |
| 1 | 170671193 | AX-170043556 | 0 | G | A | 0.219 | 1.28E-01 | 1.30E-01 | 2.68E+00 | 2.75E+00 | 3.19E-01 | no | Olfml2b | intron_variant | MODIFIER |
| 4 | 127705442 | AX-170137225 | 0 | G | A | 0.14 | 1.20E-01 | 1.20E-01 | 2.54E+00 | 2.60E+00 | 3.19E-01 | no | | intergenic_variant | MODIFIER |
| 9 | 60171802 | AX-170145424 | 0 | T | C | 0.202 | -1.11E-01 | 1.13E-01 | 2.80E+00 | 2.92E+00 | 3.19E-01 | no | Thsd4 | intron_variant | MODIFIER |
| 10 | 17791438 | AX-170002840 | 1 | G | A | 0.205 | 1.01E-01 | 1.00E-01 | 3.04E+00 | 3.16E+00 | 3.19E-01 | no | Txlnb | upstream_gene_variant | MODIFIER |
| 4 | 156004536 | AX-170486324 | 1 | C | T | 0.107 | 1.60E-01 | 1.62E-01 | 2.71E+00 | 2.83E+00 | 3.19E-01 | Yes | Sdf4 | intron_variant | MODIFIER |
| 1 | 126648580 | AX-170291081 | 0 | T | C | 0.561 | 7.99E-02 | 8.05E-02 | 2.59E+00 | 2.65E+00 | 3.19E-01 | no | Nckap5 | intron_variant | MODIFIER |
| 1 | 151289774 | AX-170244746 | 1 | A | G | 0.214 | 1.33E-01 | 1.35E-01 | 2.73E+00 | 2.78E+00 | 3.19E-01 | no | | regulatory_region_variant | MODIFIER |
| 4 | 48202319 | AX-170022730 | 0 | C | A | 0.368 | -9.35E-02 | 9.50E-02 | 2.74E+00 | 2.84E+00 | 3.19E-01 | no | Erp44 | intron_variant | MODIFIER |
| 19 | 50837325 | AX-170472515 | 2 | G | C | 0.464 | 6.29E-02 | 6.40E-02 | 2.71E+00 | 2.81E+00 | 3.19E-01 | no | | intergenic_variant | MODIFIER |
| 18 | 68263932 | AX-170181502 | 0 | A | G | 0.272 | -9.28E-02 | 9.41E-02 | 2.70E+00 | 2.78E+00 | 3.19E-01 | no | Fam210a | 3_prime_UTR_variant | MODIFIER |
| 16 | 65499882 | AX-170013264 | 1 | C | A | 0.509 | 8.27E-02 | 8.26E-02 | 2.51E+00 | 2.57E+00 | 3.19E-01 | no | | intergenic_variant | MODIFIER |
| 6 | 134460079 | AX-170448167 | 0 | T | C | 0.561 | 8.55E-02 | 8.55E-02 | 2.52E+00 | 2.58E+00 | 3.20E-01 | no | Lrp6 | intron_variant | MODIFIER |
| 4 | 54520077 | AX-170228425 | 1 | G | A | 0.375 | -1.01E-01 | 1.03E-01 | 2.79E+00 | 2.89E+00 | 3.20E-01 | no | | intergenic_variant | MODIFIER |
| 4 | 48075143 | AX-170464013 | 0 | A | G | 0.351 | -9.33E-02 | 9.47E-02 | 2.82E+00 | 2.93E+00 | 3.20E-01 | no | Nr4a3 | intron_variant | MODIFIER |
| 8 | 34926985 | AX-170333492 | 1 | A | G | 0.143 | -1.14E-01 | 1.15E-01 | 2.60E+00 | 2.71E+00 | 3.20E-01 | no | Tnks | intron_variant | MODIFIER |
| 6 | 137088395 | AX-170503772 | 1 | A | G | 0.58 | 8.98E-02 | 8.85E-02 | 2.44E+00 | 2.48E+00 | 3.20E-01 | no | Rerg | intron_variant | MODIFIER |
| 10 | 27777283 | AX-170188672 | 0 | C | T | 0.351 | -4.68E-02 | 4.77E-02 | 2.70E+00 | 2.83E+00 | 3.20E-01 | no | Gm18041 | upstream_gene_variant | MODIFIER |
| 4 | 89998562 | AX-170108499 | 1 | T | C | 0.33 | 8.49E-02 | 8.60E-02 | 2.59E+00 | 2.72E+00 | 3.20E-01 | no | | intergenic_variant | MODIFIER |
| 18 | 77813083 | AX-170412486 | 0 | G | A | 0.439 | 8.51E-02 | 8.48E-02 | 2.50E+00 | 2.57E+00 | 3.20E-01 | no | Pstpip2 | intron_variant | MODIFIER |
| 9 | 105436314 | AX-170033197 | 0 | T | C | 0.123 | -1.00E-01 | 1.01E-01 | 2.60E+00 | 2.72E+00 | 3.20E-01 | no | Atp2c1 | intron_variant | MODIFIER |
| 9 | 106684628 | AX-170211425 | 0 | A | G | 0.123 | -1.00E-01 | 1.01E-01 | 2.60E+00 | 2.72E+00 | 3.20E-01 | no | Rad54l2 | downstream_gene_variant | MODIFIER |
| 7 | 36501449 | AX-170211591 | 0 | G | A | 0.351 | 8.58E-02 | 8.69E-02 | 2.62E+00 | 2.72E+00 | 3.20E-01 | no | | intergenic_variant | MODIFIER |
| 15 | 3466683 | AX-170523427 | 1 | G | T | 0.241 | -1.03E-01 | 1.04E-01 | 2.60E+00 | 2.66E+00 | 3.20E-01 | no | Ghr | intron_variant | MODIFIER |
| 5 | 36103059 | AX-97839129 | 0 | C | T | 0.325 | 1.03E-01 | 1.05E-01 | 2.73E+00 | 2.80E+00 | 3.20E-01 | no | Sorcs2 | intron_variant | MODIFIER |
| 15 | 81422249 | AX-97813545 | 0 | C | T | 0.561 | 7.66E-02 | 7.71E-02 | 2.56E+00 | 2.64E+00 | 3.20E-01 | no | Xpnpep3 | intron_variant | MODIFIER |
| 18 | 59339100 | AX-170135240 | 1 | C | A | 0.339 | -6.43E-02 | 6.52E-02 | 2.86E+00 | 2.96E+00 | 3.20E-01 | no | Chsy3 | intron_variant | MODIFIER |
| 3 | 27621548 | AX-170086109 | 0 | A | C | 0.149 | -8.31E-02 | 8.46E-02 | 2.66E+00 | 2.77E+00 | 3.20E-01 | no | Fndc3b | intron_variant | MODIFIER |
| 4 | 95576310 | AX-170382033 | 0 | C | T | 0.342 | 1.03E-01 | 1.05E-01 | 2.71E+00 | 2.80E+00 | 3.20E-01 | no | Fggy | upstream_gene_variant | MODIFIER |
| 6 | 130735853 | AX-170143736 | 1 | C | T | 0.545 | 8.19E-02 | 8.18E-02 | 2.51E+00 | 2.58E+00 | 3.20E-01 | no | Gm44527 | intron_variant,non_coding_transcript_variant | MODIFIER |

| | | | | | | | | | | | | | | | |
|---|---|---|---|---|---|---|---|---|---|---|---|---|---|---|---|
| 5 | 107107163 | AX-170318916 | 0 | A | T | 0.465 | 1.08E-01 | 1.10E-01 | 2.66E+00 | 2.71E+00 | 3.20E-01 | no | Tgfbr3 | 3_prime_UTR_variant | MODIFIER |
| 18 | 75172075 | AX-170498641 | 1 | A | G | 0.42 | 8.35E-02 | 8.39E-02 | 2.54E+00 | 2.62E+00 | 3.21E-01 | no | Dym | intron_variant | MODIFIER |
| 4 | 140195687 | AX-170360828 | 1 | C | T | 0.161 | 1.14E-01 | 1.15E-01 | 2.58E+00 | 2.63E+00 | 3.21E-01 | no | Igsf21 | intron_variant | MODIFIER |
| 5 | 56520662 | AX-170380350 | 2 | C | T | 0.436 | 9.64E-02 | 9.65E-02 | 2.51E+00 | 2.60E+00 | 3.21E-01 | no | | intergenic_variant | MODIFIER |
| 18 | 67992722 | AX-170143900 | 1 | C | T | 0.179 | -1.21E-01 | 1.23E-01 | 2.65E+00 | 2.70E+00 | 3.21E-01 | no | Ldlrad4 | intron_variant | MODIFIER |
| 3 | 10717095 | AX-97803098 | 1 | C | T | 0.402 | -6.33E-02 | 6.47E-02 | 2.74E+00 | 2.89E+00 | 3.21E-01 | no | | intergenic_variant | MODIFIER |
| 14 | 119750306 | AX-97801882 | 0 | T | G | 0.333 | 7.41E-02 | 7.53E-02 | 2.62E+00 | 2.72E+00 | 3.21E-01 | no | Hs6st3 | intron_variant | MODIFIER |
| 14 | 119806246 | AX-97796609 | 0 | T | A | 0.333 | 7.41E-02 | 7.53E-02 | 2.62E+00 | 2.72E+00 | 3.21E-01 | no | Hs6st3 | intron_variant | MODIFIER |
| 14 | 119911596 | AX-170156885 | 0 | G | A | 0.333 | 7.41E-02 | 7.53E-02 | 2.62E+00 | 2.72E+00 | 3.21E-01 | no | | intergenic_variant | MODIFIER |
| 14 | 119915078 | AX-170430213 | 0 | A | G | 0.333 | 7.41E-02 | 7.53E-02 | 2.62E+00 | 2.72E+00 | 3.21E-01 | no | | intergenic_variant | MODIFIER |
| 14 | 119915467 | AX-170322153 | 0 | T | G | 0.333 | 7.41E-02 | 7.53E-02 | 2.62E+00 | 2.72E+00 | 3.21E-01 | no | | intergenic_variant | MODIFIER |
| 14 | 119923521 | AX-97810094 | 0 | A | C | 0.333 | 7.41E-02 | 7.53E-02 | 2.62E+00 | 2.72E+00 | 3.21E-01 | no | | intergenic_variant | MODIFIER |
| 14 | 119955562 | AX-170526746 | 0 | A | C | 0.333 | 7.41E-02 | 7.53E-02 | 2.62E+00 | 2.72E+00 | 3.21E-01 | no | | intergenic_variant | MODIFIER |
| 14 | 119955894 | AX-170113300 | 0 | A | G | 0.333 | 7.41E-02 | 7.53E-02 | 2.62E+00 | 2.72E+00 | 3.21E-01 | no | | intergenic_variant | MODIFIER |
| 14 | 119963711 | AX-170366558 | 0 | C | T | 0.333 | 7.41E-02 | 7.53E-02 | 2.62E+00 | 2.72E+00 | 3.21E-01 | no | | intergenic_variant | MODIFIER |
| 14 | 119978986 | AX-170010831 | 0 | G | A | 0.333 | 7.41E-02 | 7.53E-02 | 2.62E+00 | 2.72E+00 | 3.21E-01 | no | | intergenic_variant | MODIFIER |
| 14 | 120051254 | AX-97854260 | 0 | C | T | 0.333 | 7.41E-02 | 7.53E-02 | 2.62E+00 | 2.72E+00 | 3.21E-01 | no | | intergenic_variant | MODIFIER |
| 14 | 120072642 | AX-97855461 | 0 | G | A | 0.333 | 7.41E-02 | 7.53E-02 | 2.62E+00 | 2.72E+00 | 3.21E-01 | no | Gm20167 | upstream_gene_variant | MODIFIER |
| 14 | 120116933 | AX-97828137 | 0 | G | T | 0.333 | 7.41E-02 | 7.53E-02 | 2.62E+00 | 2.72E+00 | 3.21E-01 | no | Gm4681 | intron_variant,non_coding_transcript_variant | MODIFIER |
| 14 | 120145420 | AX-170503666 | 0 | A | G | 0.333 | 7.41E-02 | 7.53E-02 | 2.62E+00 | 2.72E+00 | 3.21E-01 | no | Gm4681 | intron_variant,non_coding_transcript_variant | MODIFIER |
| 14 | 120281294 | AX-170010837 | 0 | G | T | 0.333 | 7.41E-02 | 7.53E-02 | 2.62E+00 | 2.72E+00 | 3.21E-01 | no | Mbnl2 | intron_variant | MODIFIER |
| 14 | 120281761 | AX-170010838 | 0 | A | G | 0.333 | 7.41E-02 | 7.53E-02 | 2.62E+00 | 2.72E+00 | 3.21E-01 | no | Mbnl2 | intron_variant | MODIFIER |
| 14 | 120281795 | AX-170010835 | 0 | A | C | 0.333 | 7.41E-02 | 7.53E-02 | 2.62E+00 | 2.72E+00 | 3.21E-01 | no | Mbnl2 | intron_variant | MODIFIER |
| 16 | 32769839 | AX-170255848 | 1 | T | A | 0.482 | 8.00E-02 | 8.05E-02 | 2.55E+00 | 2.60E+00 | 3.21E-01 | no | Muc4 | missense_variant | MODERATE |
| 1 | 120191934 | AX-170419605 | 0 | C | T | 0.57 | 8.10E-02 | 8.12E-02 | 2.52E+00 | 2.58E+00 | 3.21E-01 | no | 3110009E18Rik | downstream_gene_variant | MODIFIER |
| 1 | 138840154 | AX-170406970 | 0 | C | T | 0.219 | 1.14E-01 | 1.16E-01 | 2.68E+00 | 2.75E+00 | 3.21E-01 | no | Lhx9 | intron_variant | MODIFIER |
| 8 | 32346403 | AX-170175795 | 0 | A | G | 0.14 | -1.15E-01 | 1.17E-01 | 2.72E+00 | 2.82E+00 | 3.21E-01 | no | Nrg1 | intron_variant | MODIFIER |
| 18 | 67781242 | AX-170294772 | 2 | C | T | 0.191 | -1.19E-01 | 1.21E-01 | 2.75E+00 | 2.81E+00 | 3.21E-01 | no | Seh1l | intron_variant | MODIFIER |
| 2 | 90544645 | AX-97797094 | 0 | A | G | 0.053 | -2.12E-01 | 2.16E-01 | 2.72E+00 | 2.82E+00 | 3.21E-01 | Yes | Ptprj | intron_variant | MODIFIER |
| 2 | 99934807 | AX-170018470 | 0 | C | T | 0.053 | -2.12E-01 | 2.16E-01 | 2.72E+00 | 2.82E+00 | 3.21E-01 | Yes | | intergenic_variant | MODIFIER |
| 5 | 27482724 | AX-170303279 | 0 | A | G | 0.053 | -2.12E-01 | 2.16E-01 | 2.72E+00 | 2.82E+00 | 3.21E-01 | Yes | Dpp6 | intron_variant | MODIFIER |
| 5 | 117530154 | AX-170444684 | 0 | C | T | 0.456 | 1.07E-01 | 1.08E-01 | 2.80E+00 | 2.86E+00 | 3.21E-01 | no | Ksr2 | intron_variant,non_coding_transcript_variant | MODIFIER |
| 4 | 117242301 | AX-170023772 | 0 | G | C | 0.211 | 1.16E-01 | 1.18E-01 | 2.59E+00 | 2.66E+00 | 3.21E-01 | no | Armh1 | intron_variant,non_coding_transcript_variant | MODIFIER |
| 7 | 145112342 | AX-170472172 | 1 | G | C | 0.205 | -7.39E-02 | 7.53E-02 | 2.65E+00 | 2.78E+00 | 3.21E-01 | no | | regulatory_region_variant | MODIFIER |
| 5 | 47054857 | AX-97835764 | 2 | A | G | 0.482 | 1.04E-01 | 1.03E-01 | 2.41E+00 | 2.46E+00 | 3.21E-01 | no | | intergenic_variant | MODIFIER |
| 4 | 129790757 | AX-170286554 | 1 | A | G | 0.161 | 1.18E-01 | 1.19E-01 | 2.57E+00 | 2.61E+00 | 3.21E-01 | no | | regulatory_region_variant | MODIFIER |
| 4 | 132615085 | AX-170189277 | 1 | A | T | 0.161 | 1.18E-01 | 1.19E-01 | 2.57E+00 | 2.61E+00 | 3.21E-01 | no | | intergenic_variant | MODIFIER |
| 4 | 133303156 | AX-170143103 | 1 | T | A | 0.161 | 1.18E-01 | 1.19E-01 | 2.57E+00 | 2.61E+00 | 3.21E-01 | no | Wdtc1 | intron_variant | MODIFIER |
| 4 | 134073801 | AX-170262601 | 1 | T | C | 0.161 | 1.18E-01 | 1.19E-01 | 2.57E+00 | 2.61E+00 | 3.21E-01 | no | Crybg2 | synonymous_variant | LOW |
| 4 | 72939389 | AX-97798050 | 1 | C | A | 0.455 | -8.45E-02 | 8.43E-02 | 2.48E+00 | 2.54E+00 | 3.21E-01 | no | | intergenic_variant | MODIFIER |
| 4 | 49662330 | AX-97857120 | 0 | T | C | 0.368 | -8.96E-02 | 9.12E-02 | 2.85E+00 | 2.96E+00 | 3.22E-01 | no | Grin3a | 3_prime_UTR_variant | MODIFIER |
| 15 | 66902078 | AX-170233137 | 0 | A | G | 0.333 | -7.78E-02 | 7.61E-02 | 2.38E+00 | 2.46E+00 | 3.22E-01 | no | Ccn4 | intron_variant | MODIFIER |
| 7 | 144693763 | AX-97800737 | 1 | G | A | 0.205 | -7.38E-02 | 7.54E-02 | 2.66E+00 | 2.79E+00 | 3.22E-01 | no | Ano1 | intron_variant | MODIFIER |
| 7 | 144908318 | AX-170079132 | 1 | G | T | 0.205 | -7.38E-02 | 7.54E-02 | 2.66E+00 | 2.79E+00 | 3.22E-01 | no | Gm39119 | intron_variant,non_coding_transcript_variant | MODIFIER |
| 15 | 96159770 | AX-97830100 | 1 | G | A | 0.161 | 1.49E-01 | 1.52E-01 | 2.66E+00 | 2.75E+00 | 3.22E-01 | Yes | 2610037D02Rik | intron_variant,non_coding_transcript_variant | MODIFIER |
| 7 | 143845618 | AX-97827481 | 0 | A | G | 0.211 | -7.35E-02 | 7.50E-02 | 2.64E+00 | 2.77E+00 | 3.22E-01 | no | Dhcr7 | intron_variant | MODIFIER |
| 7 | 144017443 | AX-170332573 | 0 | G | T | 0.211 | -7.35E-02 | 7.50E-02 | 2.64E+00 | 2.77E+00 | 3.22E-01 | no | Shank2 | intron_variant | MODIFIER |
| 7 | 144020043 | AX-170242124 | 0 | C | T | 0.211 | -7.35E-02 | 7.50E-02 | 2.64E+00 | 2.77E+00 | 3.22E-01 | no | Shank2 | intron_variant | MODIFIER |
| 7 | 145212467 | AX-170272641 | 0 | G | A | 0.211 | -7.35E-02 | 7.50E-02 | 2.64E+00 | 2.77E+00 | 3.22E-01 | no | 1810010D01Rik | upstream_gene_variant | MODIFIER |
| 2 | 160104755 | AX-170088815 | 2 | G | A | 0.145 | 1.57E-01 | 1.61E-01 | 2.79E+00 | 2.90E+00 | 3.22E-01 | Yes | | intergenic_variant | MODIFIER |
| 4 | 129003123 | AX-170373005 | 0 | T | C | 0.158 | 1.17E-01 | 1.18E-01 | 2.57E+00 | 2.61E+00 | 3.22E-01 | no | Ak2 | intron_variant | MODIFIER |
| 4 | 129524852 | AX-97811288 | 0 | A | G | 0.158 | 1.17E-01 | 1.18E-01 | 2.57E+00 | 2.61E+00 | 3.22E-01 | no | Hdac1 | intron_variant | MODIFIER |
| 4 | 129525291 | AX-170023937 | 0 | G | A | 0.158 | 1.17E-01 | 1.18E-01 | 2.57E+00 | 2.61E+00 | 3.22E-01 | no | Hdac1 | intron_variant | MODIFIER |
| 4 | 129533391 | AX-97786395 | 0 | G | C | 0.158 | 1.17E-01 | 1.18E-01 | 2.57E+00 | 2.61E+00 | 3.22E-01 | no | Hdac1 | intron_variant | MODIFIER |

| | | | | | | | | | | | | | | | |
|---|---|---|---|---|---|---|---|---|---|---|---|---|---|---|---|
| 4 | 129561319 | AX-170023934 | 0 | T | G | 0.158 | 1.17E-01 | 1.18E-01 | 2.57E+00 | 2.61E+00 | 3.22E-01 | no | Lck | upstream_gene_variant | MODIFIER |
| 4 | 129577499 | AX-170121217 | 0 | G | T | 0.158 | 1.17E-01 | 1.18E-01 | 2.57E+00 | 2.61E+00 | 3.22E-01 | no | Fam167b | intron_variant | MODIFIER |
| 4 | 129589140 | AX-170023936 | 0 | A | T | 0.158 | 1.17E-01 | 1.18E-01 | 2.57E+00 | 2.61E+00 | 3.22E-01 | no | Eif3i | downstream_gene_variant | MODIFIER |
| 4 | 129594069 | AX-170089369 | 0 | G | A | 0.158 | 1.17E-01 | 1.18E-01 | 2.57E+00 | 2.61E+00 | 3.22E-01 | no | Eif3i | intron_variant | MODIFIER |
| 4 | 129596789 | AX-170514182 | 0 | C | T | 0.158 | 1.17E-01 | 1.18E-01 | 2.57E+00 | 2.61E+00 | 3.22E-01 | no | Tmem234 | upstream_gene_variant | MODIFIER |
| 4 | 129616164 | AX-170179665 | 0 | G | C | 0.158 | 1.17E-01 | 1.18E-01 | 2.57E+00 | 2.61E+00 | 3.22E-01 | no | Ccdc28b | downstream_gene_variant | MODIFIER |
| 4 | 129623415 | AX-170023927 | 0 | T | C | 0.158 | 1.17E-01 | 1.18E-01 | 2.57E+00 | 2.61E+00 | 3.22E-01 | no | Ccdc28b | intron_variant | MODIFIER |
| 4 | 129641173 | AX-97819205 | 0 | G | A | 0.158 | 1.17E-01 | 1.18E-01 | 2.57E+00 | 2.61E+00 | 3.22E-01 | no | Txlna | upstream_gene_variant | MODIFIER |
| 4 | 129654873 | AX-170023931 | 0 | G | A | 0.158 | 1.17E-01 | 1.18E-01 | 2.57E+00 | 2.61E+00 | 3.22E-01 | no | Kpna6 | synonymous_variant | LOW |
| 4 | 129684417 | AX-170023930 | 0 | T | C | 0.158 | 1.17E-01 | 1.18E-01 | 2.57E+00 | 2.61E+00 | 3.22E-01 | no | Tmem39b | missense_variant | MODERATE |
| 4 | 129688198 | AX-97850280 | 0 | G | T | 0.158 | 1.17E-01 | 1.18E-01 | 2.57E+00 | 2.61E+00 | 3.22E-01 | no | Tmem39b | intron_variant | MODIFIER |
| 4 | 129693877 | AX-170023929 | 0 | G | T | 0.158 | 1.17E-01 | 1.18E-01 | 2.57E+00 | 2.61E+00 | 3.22E-01 | no | Tmem39b | intron_variant | MODIFIER |
| 4 | 129696010 | AX-170023938 | 0 | G | T | 0.158 | 1.17E-01 | 1.18E-01 | 2.57E+00 | 2.61E+00 | 3.22E-01 | no | Tmem39b | intron_variant | MODIFIER |
| 4 | 129713263 | AX-170361151 | 0 | A | C | 0.158 | 1.17E-01 | 1.18E-01 | 2.57E+00 | 2.61E+00 | 3.22E-01 | no | Khdrbs1 | downstream_gene_variant | MODIFIER |
| 4 | 129713375 | AX-97845396 | 0 | A | C | 0.158 | 1.17E-01 | 1.18E-01 | 2.57E+00 | 2.61E+00 | 3.22E-01 | no | Khdrbs1 | downstream_gene_variant | MODIFIER |
| 4 | 129717574 | AX-170309750 | 0 | C | A | 0.158 | 1.17E-01 | 1.18E-01 | 2.57E+00 | 2.61E+00 | 3.22E-01 | no | Khdrbs1 | intron_variant | MODIFIER |
| 4 | 129720019 | AX-170508674 | 0 | T | C | 0.158 | 1.17E-01 | 1.18E-01 | 2.57E+00 | 2.61E+00 | 3.22E-01 | no | Khdrbs1 | intron_variant | MODIFIER |
| 4 | 129728090 | AX-170023932 | 0 | G | T | 0.158 | 1.17E-01 | 1.18E-01 | 2.57E+00 | 2.61E+00 | 3.22E-01 | no | Khdrbs1 | intron_variant | MODIFIER |
| 4 | 129731515 | AX-97780408 | 0 | C | T | 0.158 | 1.17E-01 | 1.18E-01 | 2.57E+00 | 2.61E+00 | 3.22E-01 | no | Khdrbs1 | intron_variant | MODIFIER |
| 4 | 129759808 | AX-97814204 | 0 | T | C | 0.158 | 1.17E-01 | 1.18E-01 | 2.57E+00 | 2.61E+00 | 3.22E-01 | no | Gm12967 | downstream_gene_variant | MODIFIER |
| 4 | 129783426 | AX-170165849 | 0 | T | A | 0.158 | 1.17E-01 | 1.18E-01 | 2.57E+00 | 2.61E+00 | 3.22E-01 | no | | regulatory_region_variant | MODIFIER |
| 4 | 129798509 | AX-170461562 | 0 | G | T | 0.158 | 1.17E-01 | 1.18E-01 | 2.57E+00 | 2.61E+00 | 3.22E-01 | no | | regulatory_region_variant | MODIFIER |
| 4 | 129808034 | AX-170222913 | 0 | G | C | 0.158 | 1.17E-01 | 1.18E-01 | 2.57E+00 | 2.61E+00 | 3.22E-01 | no | Ptp4a2 | upstream_gene_variant | MODIFIER |
| 4 | 129826461 | AX-97796332 | 0 | C | T | 0.158 | 1.17E-01 | 1.18E-01 | 2.57E+00 | 2.61E+00 | 3.22E-01 | no | Ptp4a2 | intron_variant | MODIFIER |
| 4 | 129855394 | AX-170335412 | 0 | C | T | 0.158 | 1.17E-01 | 1.18E-01 | 2.57E+00 | 2.61E+00 | 3.22E-01 | no | Gm10571 | non_coding_transcript_exon_variant | MODIFIER |
| 4 | 129864982 | AX-170191489 | 0 | G | A | 0.158 | 1.17E-01 | 1.18E-01 | 2.57E+00 | 2.61E+00 | 3.22E-01 | no | | intergenic_variant | MODIFIER |
| 4 | 129873514 | AX-170070669 | 0 | A | G | 0.158 | 1.17E-01 | 1.18E-01 | 2.57E+00 | 2.61E+00 | 3.22E-01 | no | | regulatory_region_variant | MODIFIER |
| 4 | 129912883 | AX-170274477 | 0 | T | A | 0.158 | 1.17E-01 | 1.18E-01 | 2.57E+00 | 2.61E+00 | 3.22E-01 | no | E330017L17Rik | downstream_gene_variant | MODIFIER |
| 4 | 129912930 | AX-170233301 | 0 | C | T | 0.158 | 1.17E-01 | 1.18E-01 | 2.57E+00 | 2.61E+00 | 3.22E-01 | no | E330017L17Rik | downstream_gene_variant | MODIFIER |
| 4 | 129924391 | AX-97793012 | 0 | C | A | 0.158 | 1.17E-01 | 1.18E-01 | 2.57E+00 | 2.61E+00 | 3.22E-01 | no | Spocd1 | upstream_gene_variant | MODIFIER |
| 4 | 129925230 | AX-170258727 | 0 | T | C | 0.158 | 1.17E-01 | 1.18E-01 | 2.57E+00 | 2.61E+00 | 3.22E-01 | no | Spocd1 | upstream_gene_variant | MODIFIER |
| 4 | 129930242 | AX-170191631 | 0 | T | C | 0.158 | 1.17E-01 | 1.18E-01 | 2.57E+00 | 2.61E+00 | 3.22E-01 | no | Spocd1 | intron_variant | MODIFIER |
| 4 | 129936135 | AX-97783989 | 0 | T | A | 0.158 | 1.17E-01 | 1.18E-01 | 2.57E+00 | 2.61E+00 | 3.22E-01 | no | Spocd1 | intron_variant | MODIFIER |
| 4 | 129955957 | AX-170426467 | 0 | A | G | 0.158 | 1.17E-01 | 1.18E-01 | 2.57E+00 | 2.61E+00 | 3.22E-01 | no | Spocd1 | intron_variant | MODIFIER |
| 4 | 129968813 | AX-170325009 | 0 | G | A | 0.158 | 1.17E-01 | 1.18E-01 | 2.57E+00 | 2.61E+00 | 3.22E-01 | no | | regulatory_region_variant | MODIFIER |
| 4 | 129973306 | AX-170461956 | 0 | A | G | 0.158 | 1.17E-01 | 1.18E-01 | 2.57E+00 | 2.61E+00 | 3.22E-01 | no | | intergenic_variant | MODIFIER |
| 4 | 129976197 | AX-170339479 | 0 | C | A | 0.158 | 1.17E-01 | 1.18E-01 | 2.57E+00 | 2.61E+00 | 3.22E-01 | no | | intergenic_variant | MODIFIER |
| 4 | 129990469 | AX-97790607 | 0 | C | T | 0.158 | 1.17E-01 | 1.18E-01 | 2.57E+00 | 2.61E+00 | 3.22E-01 | no | Adgrb2 | upstream_gene_variant | MODIFIER |
| 4 | 130014604 | AX-170216539 | 0 | C | T | 0.158 | 1.17E-01 | 1.18E-01 | 2.57E+00 | 2.61E+00 | 3.22E-01 | no | Adgrb2 | synonymous_variant | LOW |
| 4 | 130029938 | AX-170360040 | 0 | T | G | 0.158 | 1.17E-01 | 1.18E-01 | 2.57E+00 | 2.61E+00 | 3.22E-01 | no | Gm12963 | intron_variant,non_coding_transcript_variant | MODIFIER |
| 4 | 130041394 | AX-170089371 | 0 | G | A | 0.158 | 1.17E-01 | 1.18E-01 | 2.57E+00 | 2.61E+00 | 3.22E-01 | no | Gm12963 | intron_variant,non_coding_transcript_variant | MODIFIER |
| 4 | 130058731 | AX-97809462 | 0 | C | T | 0.158 | 1.17E-01 | 1.18E-01 | 2.57E+00 | 2.61E+00 | 3.22E-01 | no | Col16a1 | intron_variant | MODIFIER |
| 4 | 130061678 | AX-170402056 | 0 | C | T | 0.158 | 1.17E-01 | 1.18E-01 | 2.57E+00 | 2.61E+00 | 3.22E-01 | no | Col16a1 | intron_variant | MODIFIER |
| 4 | 130076718 | AX-170262740 | 0 | G | T | 0.158 | 1.17E-01 | 1.18E-01 | 2.57E+00 | 2.61E+00 | 3.22E-01 | no | Col16a1 | intron_variant | MODIFIER |
| 4 | 130081799 | AX-170298098 | 0 | A | G | 0.158 | 1.17E-01 | 1.18E-01 | 2.57E+00 | 2.61E+00 | 3.22E-01 | no | Col16a1 | intron_variant | MODIFIER |
| 4 | 130082283 | AX-97782274 | 0 | C | T | 0.158 | 1.17E-01 | 1.18E-01 | 2.57E+00 | 2.61E+00 | 3.22E-01 | no | Col16a1 | intron_variant | MODIFIER |
| 4 | 131598175 | AX-170089372 | 0 | G | A | 0.158 | 1.17E-01 | 1.18E-01 | 2.57E+00 | 2.61E+00 | 3.22E-01 | no | | intergenic_variant | MODIFIER |
| 4 | 131598359 | AX-170282759 | 0 | T | C | 0.158 | 1.17E-01 | 1.18E-01 | 2.57E+00 | 2.61E+00 | 3.22E-01 | no | | intergenic_variant | MODIFIER |
| 4 | 131598457 | AX-170287605 | 0 | C | T | 0.158 | 1.17E-01 | 1.18E-01 | 2.57E+00 | 2.61E+00 | 3.22E-01 | no | | intergenic_variant | MODIFIER |
| 4 | 131623219 | AX-97804706 | 0 | G | A | 0.158 | 1.17E-01 | 1.18E-01 | 2.57E+00 | 2.61E+00 | 3.22E-01 | no | | intergenic_variant | MODIFIER |
| 4 | 131625769 | AX-170023961 | 0 | G | A | 0.158 | 1.17E-01 | 1.18E-01 | 2.57E+00 | 2.61E+00 | 3.22E-01 | no | | intergenic_variant | MODIFIER |
| 4 | 131999242 | AX-170257502 | 0 | A | G | 0.158 | 1.17E-01 | 1.18E-01 | 2.57E+00 | 2.61E+00 | 3.22E-01 | no | Epb41 | intron_variant | MODIFIER |
| 4 | 132166285 | AX-97790723 | 0 | T | C | 0.158 | 1.17E-01 | 1.18E-01 | 2.57E+00 | 2.61E+00 | 3.22E-01 | no | | regulatory_region_variant | MODIFIER |
| 4 | 132180308 | AX-170169502 | 0 | A | G | 0.158 | 1.17E-01 | 1.18E-01 | 2.57E+00 | 2.61E+00 | 3.22E-01 | no | Ythdf2 | downstream_gene_variant | MODIFIER |

| | | | | | | | | | | | | | | | |
|---|---|---|---|---|---|---|---|---|---|---|---|---|---|---|---|
| 4 | 132181255 | AX-170507790 | 0 | T | C | 0.158 | 1.17E-01 | 1.18E-01 | 2.57E+00 | 2.61E+00 | 3.22E-01 | no | Ythdf2 | downstream_gene_variant | MODIFIER |
| 4 | 132247951 | AX-170378095 | 0 | T | C | 0.158 | 1.17E-01 | 1.18E-01 | 2.57E+00 | 2.61E+00 | 3.22E-01 | no | Gmeb1 | intron_variant | MODIFIER |
| 4 | 132379057 | AX-170155731 | 0 | C | A | 0.158 | 1.17E-01 | 1.18E-01 | 2.57E+00 | 2.61E+00 | 3.22E-01 | no | Phactr4 | intron_variant | MODIFIER |
| 4 | 132415160 | AX-97849460 | 0 | G | C | 0.158 | 1.17E-01 | 1.18E-01 | 2.57E+00 | 2.61E+00 | 3.22E-01 | no | Phactr4 | intron_variant | MODIFIER |
| 4 | 132428395 | AX-170327194 | 0 | G | A | 0.158 | 1.17E-01 | 1.18E-01 | 2.57E+00 | 2.61E+00 | 3.22E-01 | no | | regulatory_region_variant | MODIFIER |
| 4 | 132433241 | AX-170245180 | 0 | G | T | 0.158 | 1.17E-01 | 1.18E-01 | 2.57E+00 | 2.61E+00 | 3.22E-01 | no | | regulatory_region_variant | MODIFIER |
| 4 | 132702492 | AX-170130842 | 0 | T | C | 0.158 | 1.17E-01 | 1.18E-01 | 2.57E+00 | 2.61E+00 | 3.22E-01 | no | Eya3 | intron_variant | MODIFIER |
| 4 | 132703024 | AX-170142216 | 0 | C | G | 0.158 | 1.17E-01 | 1.18E-01 | 2.57E+00 | 2.61E+00 | 3.22E-01 | no | Eya3 | intron_variant | MODIFIER |
| 4 | 132708992 | AX-97817444 | 0 | C | T | 0.158 | 1.17E-01 | 1.18E-01 | 2.57E+00 | 2.61E+00 | 3.22E-01 | no | Eya3 | intron_variant | MODIFIER |
| 4 | 132711110 | AX-170410645 | 0 | C | A | 0.158 | 1.17E-01 | 1.18E-01 | 2.57E+00 | 2.61E+00 | 3.22E-01 | no | Eya3 | intron_variant | MODIFIER |
| 4 | 132718299 | AX-170249891 | 0 | C | G | 0.158 | 1.17E-01 | 1.18E-01 | 2.57E+00 | 2.61E+00 | 3.22E-01 | no | Eya3 | intron_variant | MODIFIER |
| 4 | 132729142 | AX-168407119 | 0 | G | A | 0.158 | 1.17E-01 | 1.18E-01 | 2.57E+00 | 2.61E+00 | 3.22E-01 | no | Eya3 | downstream_gene_variant | MODIFIER |
| 4 | 132731658 | AX-170448452 | 0 | C | T | 0.158 | 1.17E-01 | 1.18E-01 | 2.57E+00 | 2.61E+00 | 3.22E-01 | no | Smpdl3b | downstream_gene_variant | MODIFIER |
| 4 | 132783760 | AX-97811815 | 0 | A | C | 0.158 | 1.17E-01 | 1.18E-01 | 2.57E+00 | 2.61E+00 | 3.22E-01 | no | Themis2 | intron_variant | MODIFIER |
| 4 | 132792878 | AX-170451330 | 0 | A | G | 0.158 | 1.17E-01 | 1.18E-01 | 2.57E+00 | 2.61E+00 | 3.22E-01 | no | Themis2 | intron_variant | MODIFIER |
| 4 | 132936651 | AX-170529727 | 0 | G | T | 0.158 | 1.17E-01 | 1.18E-01 | 2.57E+00 | 2.61E+00 | 3.22E-01 | no | | intergenic_variant | MODIFIER |
| 4 | 132937009 | AX-170485025 | 0 | G | A | 0.158 | 1.17E-01 | 1.18E-01 | 2.57E+00 | 2.61E+00 | 3.22E-01 | no | | intergenic_variant | MODIFIER |
| 4 | 132950161 | AX-170322896 | 0 | A | G | 0.158 | 1.17E-01 | 1.18E-01 | 2.57E+00 | 2.61E+00 | 3.22E-01 | no | | regulatory_region_variant | MODIFIER |
| 4 | 132950916 | AX-170198981 | 0 | A | C | 0.158 | 1.17E-01 | 1.18E-01 | 2.57E+00 | 2.61E+00 | 3.22E-01 | no | | regulatory_region_variant | MODIFIER |
| 4 | 132982560 | AX-170497837 | 0 | C | T | 0.158 | 1.17E-01 | 1.18E-01 | 2.57E+00 | 2.61E+00 | 3.22E-01 | no | Fgr | intron_variant | MODIFIER |
| 4 | 132987662 | AX-170194212 | 0 | G | T | 0.158 | 1.17E-01 | 1.18E-01 | 2.57E+00 | 2.61E+00 | 3.22E-01 | no | Fgr | intron_variant | MODIFIER |
| 4 | 132987720 | AX-170414861 | 0 | G | A | 0.158 | 1.17E-01 | 1.18E-01 | 2.57E+00 | 2.61E+00 | 3.22E-01 | no | Fgr | intron_variant | MODIFIER |
| 4 | 132991683 | AX-170337375 | 0 | A | G | 0.158 | 1.17E-01 | 1.18E-01 | 2.57E+00 | 2.61E+00 | 3.22E-01 | no | Fgr | intron_variant | MODIFIER |
| 4 | 132991759 | AX-170381380 | 0 | G | A | 0.158 | 1.17E-01 | 1.18E-01 | 2.57E+00 | 2.61E+00 | 3.22E-01 | no | Fgr | intron_variant | MODIFIER |
| 4 | 132995922 | AX-170491115 | 0 | G | A | 0.158 | 1.17E-01 | 1.18E-01 | 2.57E+00 | 2.61E+00 | 3.22E-01 | no | Fgr | intron_variant | MODIFIER |
| 4 | 133001361 | AX-170309164 | 0 | A | C | 0.158 | 1.17E-01 | 1.18E-01 | 2.57E+00 | 2.61E+00 | 3.22E-01 | no | Fgr | 3_prime_UTR_variant | MODIFIER |
| 4 | 133002842 | AX-170446381 | 0 | C | T | 0.158 | 1.17E-01 | 1.18E-01 | 2.57E+00 | 2.61E+00 | 3.22E-01 | no | Fgr | downstream_gene_variant | MODIFIER |
| 4 | 133004638 | AX-170394185 | 0 | C | T | 0.158 | 1.17E-01 | 1.18E-01 | 2.57E+00 | 2.61E+00 | 3.22E-01 | no | Fgr | downstream_gene_variant | MODIFIER |
| 4 | 133053672 | AX-170455191 | 0 | C | T | 0.158 | 1.17E-01 | 1.18E-01 | 2.57E+00 | 2.61E+00 | 3.22E-01 | no | Ahdc1 | intron_variant | MODIFIER |
| 4 | 133060312 | AX-97833049 | 0 | T | C | 0.158 | 1.17E-01 | 1.18E-01 | 2.57E+00 | 2.61E+00 | 3.22E-01 | no | Ahdc1 | intron_variant | MODIFIER |
| 4 | 133065121 | AX-170070731 | 0 | C | T | 0.158 | 1.17E-01 | 1.18E-01 | 2.57E+00 | 2.61E+00 | 3.22E-01 | no | Ahdc1 | synonymous_variant | LOW |
| 4 | 133069194 | AX-170070732 | 0 | A | G | 0.158 | 1.17E-01 | 1.18E-01 | 2.57E+00 | 2.61E+00 | 3.22E-01 | no | Ahdc1 | intron_variant | MODIFIER |
| 4 | 133069349 | AX-170171929 | 0 | C | T | 0.158 | 1.17E-01 | 1.18E-01 | 2.57E+00 | 2.61E+00 | 3.22E-01 | no | Ahdc1 | intron_variant | MODIFIER |
| 4 | 133079350 | AX-170252980 | 0 | C | T | 0.158 | 1.17E-01 | 1.18E-01 | 2.57E+00 | 2.61E+00 | 3.22E-01 | no | Ahdc1 | downstream_gene_variant | MODIFIER |
| 4 | 133079646 | AX-170354190 | 0 | A | G | 0.158 | 1.17E-01 | 1.18E-01 | 2.57E+00 | 2.61E+00 | 3.22E-01 | no | Ahdc1 | downstream_gene_variant | MODIFIER |
| 4 | 133086469 | AX-170376639 | 0 | A | G | 0.158 | 1.17E-01 | 1.18E-01 | 2.57E+00 | 2.61E+00 | 3.22E-01 | no | Gm26615 | non_coding_transcript_exon_variant | MODIFIER |
| 4 | 133093032 | AX-97836364 | 0 | T | C | 0.158 | 1.17E-01 | 1.18E-01 | 2.57E+00 | 2.61E+00 | 3.22E-01 | no | Gm26615 | downstream_gene_variant | MODIFIER |
| 4 | 133102026 | AX-170458131 | 0 | C | T | 0.158 | 1.17E-01 | 1.18E-01 | 2.57E+00 | 2.61E+00 | 3.22E-01 | no | | intergenic_variant | MODIFIER |
| 4 | 133105951 | AX-97791457 | 0 | T | G | 0.158 | 1.17E-01 | 1.18E-01 | 2.57E+00 | 2.61E+00 | 3.22E-01 | no | | regulatory_region_variant | MODIFIER |
| 4 | 133108124 | AX-170327580 | 0 | T | C | 0.158 | 1.17E-01 | 1.18E-01 | 2.57E+00 | 2.61E+00 | 3.22E-01 | no | | regulatory_region_variant | MODIFIER |
| 4 | 133161147 | AX-170023980 | 0 | T | C | 0.158 | 1.17E-01 | 1.18E-01 | 2.57E+00 | 2.61E+00 | 3.22E-01 | no | Wasf2 | intron_variant | MODIFIER |
| 4 | 133176623 | AX-170392964 | 0 | G | C | 0.158 | 1.17E-01 | 1.18E-01 | 2.57E+00 | 2.61E+00 | 3.22E-01 | no | Wasf2 | intron_variant | MODIFIER |
| 4 | 133205928 | AX-170353600 | 0 | A | G | 0.158 | 1.17E-01 | 1.18E-01 | 2.57E+00 | 2.61E+00 | 3.22E-01 | no | Gpr3 | downstream_gene_variant | MODIFIER |
| 4 | 133209321 | AX-170266739 | 0 | G | A | 0.158 | 1.17E-01 | 1.18E-01 | 2.57E+00 | 2.61E+00 | 3.22E-01 | no | Gpr3 | downstream_gene_variant | MODIFIER |
| 4 | 133218427 | AX-170327300 | 0 | T | C | 0.158 | 1.17E-01 | 1.18E-01 | 2.57E+00 | 2.61E+00 | 3.22E-01 | no | Cd164l2 | upstream_gene_variant | MODIFIER |
| 4 | 133226063 | AX-170023985 | 0 | A | G | 0.158 | 1.17E-01 | 1.18E-01 | 2.57E+00 | 2.61E+00 | 3.22E-01 | no | Cd164l2 | downstream_gene_variant | MODIFIER |
| 4 | 133226213 | AX-170023984 | 0 | G | T | 0.158 | 1.17E-01 | 1.18E-01 | 2.57E+00 | 2.61E+00 | 3.22E-01 | no | Cd164l2 | downstream_gene_variant | MODIFIER |
| 4 | 133251429 | AX-170070738 | 0 | A | C | 0.158 | 1.17E-01 | 1.18E-01 | 2.57E+00 | 2.61E+00 | 3.22E-01 | no | Sytl1 | downstream_gene_variant | MODIFIER |
| 4 | 133252016 | AX-170070735 | 0 | A | G | 0.158 | 1.17E-01 | 1.18E-01 | 2.57E+00 | 2.61E+00 | 3.22E-01 | no | Sytl1 | downstream_gene_variant | MODIFIER |
| 4 | 133270585 | AX-170449425 | 0 | G | C | 0.158 | 1.17E-01 | 1.18E-01 | 2.57E+00 | 2.61E+00 | 3.22E-01 | no | Tmem222 | intron_variant | MODIFIER |
| 4 | 133280316 | AX-170273403 | 0 | C | T | 0.158 | 1.17E-01 | 1.18E-01 | 2.57E+00 | 2.61E+00 | 3.22E-01 | no | Tmem222 | upstream_gene_variant | MODIFIER |
| 4 | 133289417 | AX-170122070 | 0 | T | C | 0.158 | 1.17E-01 | 1.18E-01 | 2.57E+00 | 2.61E+00 | 3.22E-01 | no | Wdtc1 | downstream_gene_variant | MODIFIER |
| 4 | 133297690 | AX-170276289 | 0 | A | G | 0.158 | 1.17E-01 | 1.18E-01 | 2.57E+00 | 2.61E+00 | 3.22E-01 | no | Wdtc1 | intron_variant | MODIFIER |
| 4 | 133308688 | AX-170511138 | 0 | A | G | 0.158 | 1.17E-01 | 1.18E-01 | 2.57E+00 | 2.61E+00 | 3.22E-01 | no | Wdtc1 | intron_variant | MODIFIER |

| | | | | | | | | | | | | | | | |
|---|---|---|---|---|---|---|---|---|---|---|---|---|---|---|---|
| 4 | 133333908 | AX-170408407 | 0 | G | A | 0.158 | 1.17E-01 | 1.18E-01 | 2.57E+00 | 2.61E+00 | 3.22E-01 | no | Wdtc1 | intron_variant | MODIFIER |
| 4 | 133343051 | AX-170153290 | 0 | T | C | 0.158 | 1.17E-01 | 1.18E-01 | 2.57E+00 | 2.61E+00 | 3.22E-01 | no | Wdtc1 | upstream_gene_variant | MODIFIER |
| 4 | 133344756 | AX-170125521 | 0 | G | A | 0.158 | 1.17E-01 | 1.18E-01 | 2.57E+00 | 2.61E+00 | 3.22E-01 | no | Wdtc1 | intron_variant,non_coding_transcript_variant | MODIFIER |
| 4 | 133367920 | AX-170515237 | 0 | T | C | 0.158 | 1.17E-01 | 1.18E-01 | 2.57E+00 | 2.61E+00 | 3.22E-01 | no | Slc9a1 | upstream_gene_variant | MODIFIER |
| 4 | 133398368 | AX-170338086 | 0 | T | C | 0.158 | 1.17E-01 | 1.18E-01 | 2.57E+00 | 2.61E+00 | 3.22E-01 | no | Slc9a1 | intron_variant | MODIFIER |
| 4 | 133414727 | AX-170377812 | 0 | T | C | 0.158 | 1.17E-01 | 1.18E-01 | 2.57E+00 | 2.61E+00 | 3.22E-01 | no | Slc9a1 | intron_variant | MODIFIER |
| 4 | 133443185 | AX-170070739 | 0 | G | A | 0.158 | 1.17E-01 | 1.18E-01 | 2.57E+00 | 2.61E+00 | 3.22E-01 | no | | regulatory_region_variant | MODIFIER |
| 4 | 133447742 | AX-170246486 | 0 | A | G | 0.158 | 1.17E-01 | 1.18E-01 | 2.57E+00 | 2.61E+00 | 3.22E-01 | no | | intergenic_variant | MODIFIER |
| 4 | 133462982 | AX-170297895 | 0 | T | C | 0.158 | 1.17E-01 | 1.18E-01 | 2.57E+00 | 2.61E+00 | 3.22E-01 | no | | regulatory_region_variant | MODIFIER |
| 4 | 133487014 | AX-170270942 | 0 | G | A | 0.158 | 1.17E-01 | 1.18E-01 | 2.57E+00 | 2.61E+00 | 3.22E-01 | no | Tent5b | missense_variant | MODERATE |
| 4 | 133499338 | AX-170421818 | 0 | C | T | 0.158 | 1.17E-01 | 1.18E-01 | 2.57E+00 | 2.61E+00 | 3.22E-01 | no | Trnp1 | upstream_gene_variant | MODIFIER |
| 4 | 133503397 | AX-170341379 | 0 | A | G | 0.158 | 1.17E-01 | 1.18E-01 | 2.57E+00 | 2.61E+00 | 3.22E-01 | no | Trnp1 | upstream_gene_variant | MODIFIER |
| 4 | 133503795 | AX-170502421 | 0 | T | C | 0.158 | 1.17E-01 | 1.18E-01 | 2.57E+00 | 2.61E+00 | 3.22E-01 | no | | intergenic_variant | MODIFIER |
| 4 | 133554441 | AX-170438426 | 0 | A | G | 0.158 | 1.17E-01 | 1.18E-01 | 2.57E+00 | 2.61E+00 | 3.22E-01 | no | Nr0b2 | intron_variant | MODIFIER |
| 4 | 133559384 | AX-170210820 | 0 | T | C | 0.158 | 1.17E-01 | 1.18E-01 | 2.57E+00 | 2.61E+00 | 3.22E-01 | no | Nr0b2 | downstream_gene_variant | MODIFIER |
| 4 | 133563364 | AX-97811701 | 0 | A | G | 0.158 | 1.17E-01 | 1.18E-01 | 2.57E+00 | 2.61E+00 | 3.22E-01 | no | Gm23158 | upstream_gene_variant | MODIFIER |
| 4 | 133563483 | AX-170369011 | 0 | A | G | 0.158 | 1.17E-01 | 1.18E-01 | 2.57E+00 | 2.61E+00 | 3.22E-01 | no | Gm23158 | upstream_gene_variant | MODIFIER |
| 4 | 133583263 | AX-170095156 | 0 | T | C | 0.158 | 1.17E-01 | 1.18E-01 | 2.57E+00 | 2.61E+00 | 3.22E-01 | no | Gpn2 | upstream_gene_variant | MODIFIER |
| 4 | 133602600 | AX-170358657 | 0 | T | C | 0.158 | 1.17E-01 | 1.18E-01 | 2.57E+00 | 2.61E+00 | 3.22E-01 | no | Sfn | upstream_gene_variant | MODIFIER |
| 4 | 133605582 | AX-170475374 | 0 | A | G | 0.158 | 1.17E-01 | 1.18E-01 | 2.57E+00 | 2.61E+00 | 3.22E-01 | no | Sfn | upstream_gene_variant | MODIFIER |
| 4 | 133613402 | AX-170211000 | 0 | A | G | 0.158 | 1.17E-01 | 1.18E-01 | 2.57E+00 | 2.61E+00 | 3.22E-01 | no | Zdhhc18 | intron_variant | MODIFIER |
| 4 | 133615047 | AX-170214515 | 0 | G | C | 0.158 | 1.17E-01 | 1.18E-01 | 2.57E+00 | 2.61E+00 | 3.22E-01 | no | Zdhhc18 | intron_variant | MODIFIER |
| 4 | 133626682 | AX-170023996 | 0 | C | T | 0.158 | 1.17E-01 | 1.18E-01 | 2.57E+00 | 2.61E+00 | 3.22E-01 | no | Zdhhc18 | intron_variant | MODIFIER |
| 4 | 133639330 | AX-170023991 | 0 | T | C | 0.158 | 1.17E-01 | 1.18E-01 | 2.57E+00 | 2.61E+00 | 3.22E-01 | no | Gm13213 | upstream_gene_variant | MODIFIER |
| 4 | 133644737 | AX-170441478 | 0 | G | C | 0.158 | 1.17E-01 | 1.18E-01 | 2.57E+00 | 2.61E+00 | 3.22E-01 | no | Zdhhc18 | intron_variant,non_coding_transcript_variant | MODIFIER |
| 4 | 133651651 | AX-170393937 | 0 | A | G | 0.158 | 1.17E-01 | 1.18E-01 | 2.57E+00 | 2.61E+00 | 3.22E-01 | no | Zdhhc18 | upstream_gene_variant | MODIFIER |
| 4 | 133663018 | AX-170023990 | 0 | A | T | 0.158 | 1.17E-01 | 1.18E-01 | 2.57E+00 | 2.61E+00 | 3.22E-01 | no | Pigv | intron_variant | MODIFIER |
| 4 | 133663089 | AX-170070746 | 0 | G | C | 0.158 | 1.17E-01 | 1.18E-01 | 2.57E+00 | 2.61E+00 | 3.22E-01 | no | Pigv | intron_variant | MODIFIER |
| 4 | 133663325 | AX-97828830 | 0 | G | A | 0.158 | 1.17E-01 | 1.18E-01 | 2.57E+00 | 2.61E+00 | 3.22E-01 | no | Pigv | intron_variant | MODIFIER |
| 4 | 133694392 | AX-170463885 | 0 | G | C | 0.158 | 1.17E-01 | 1.18E-01 | 2.57E+00 | 2.61E+00 | 3.22E-01 | no | Arid1a | intron_variant | MODIFIER |
| 4 | 133694943 | AX-97852453 | 0 | T | C | 0.158 | 1.17E-01 | 1.18E-01 | 2.57E+00 | 2.61E+00 | 3.22E-01 | no | Arid1a | intron_variant | MODIFIER |
| 4 | 133706746 | AX-170023994 | 0 | T | C | 0.158 | 1.17E-01 | 1.18E-01 | 2.57E+00 | 2.61E+00 | 3.22E-01 | no | Arid1a | intron_variant | MODIFIER |
| 4 | 133707961 | AX-170376398 | 0 | G | A | 0.158 | 1.17E-01 | 1.18E-01 | 2.57E+00 | 2.61E+00 | 3.22E-01 | no | Arid1a | intron_variant | MODIFIER |
| 4 | 133716120 | AX-97798714 | 0 | G | A | 0.158 | 1.17E-01 | 1.18E-01 | 2.57E+00 | 2.61E+00 | 3.22E-01 | no | Arid1a | intron_variant | MODIFIER |
| 4 | 133726298 | AX-170300606 | 0 | A | G | 0.158 | 1.17E-01 | 1.18E-01 | 2.57E+00 | 2.61E+00 | 3.22E-01 | no | Arid1a | intron_variant | MODIFIER |
| 4 | 133734892 | AX-170379819 | 0 | T | C | 0.158 | 1.17E-01 | 1.18E-01 | 2.57E+00 | 2.61E+00 | 3.22E-01 | no | Arid1a | intron_variant | MODIFIER |
| 4 | 133759746 | AX-170269827 | 0 | A | G | 0.158 | 1.17E-01 | 1.18E-01 | 2.57E+00 | 2.61E+00 | 3.22E-01 | no | Arid1a | upstream_gene_variant | MODIFIER |
| 4 | 133762330 | AX-97828846 | 0 | A | G | 0.158 | 1.17E-01 | 1.18E-01 | 2.57E+00 | 2.61E+00 | 3.22E-01 | no | | intergenic_variant | MODIFIER |
| 4 | 133816755 | AX-170023998 | 0 | T | A | 0.158 | 1.17E-01 | 1.18E-01 | 2.57E+00 | 2.61E+00 | 3.22E-01 | no | Gm12974 | intron_variant,non_coding_transcript_variant | MODIFIER |
| 4 | 133816991 | AX-97782495 | 0 | G | A | 0.158 | 1.17E-01 | 1.18E-01 | 2.57E+00 | 2.61E+00 | 3.22E-01 | no | Gm12974 | intron_variant,non_coding_transcript_variant | MODIFIER |
| 4 | 133820128 | AX-170236173 | 0 | C | T | 0.158 | 1.17E-01 | 1.18E-01 | 2.57E+00 | 2.61E+00 | 3.22E-01 | no | Gm12974 | downstream_gene_variant | MODIFIER |
| 4 | 133846877 | AX-170102176 | 0 | C | T | 0.158 | 1.17E-01 | 1.18E-01 | 2.57E+00 | 2.61E+00 | 3.22E-01 | no | Rps6ka1 | downstream_gene_variant | MODIFIER |
| 4 | 133872514 | AX-170459272 | 0 | A | C | 0.158 | 1.17E-01 | 1.18E-01 | 2.57E+00 | 2.61E+00 | 3.22E-01 | no | Rps6ka1 | intron_variant | MODIFIER |
| 4 | 133876216 | AX-170175059 | 0 | T | G | 0.158 | 1.17E-01 | 1.18E-01 | 2.57E+00 | 2.61E+00 | 3.22E-01 | no | Rps6ka1 | intron_variant | MODIFIER |
| 4 | 133879171 | AX-170505519 | 0 | C | T | 0.158 | 1.17E-01 | 1.18E-01 | 2.57E+00 | 2.61E+00 | 3.22E-01 | no | Rps6ka1 | intron_variant | MODIFIER |
| 4 | 133879347 | AX-170103443 | 0 | A | C | 0.158 | 1.17E-01 | 1.18E-01 | 2.57E+00 | 2.61E+00 | 3.22E-01 | no | Rps6ka1 | intron_variant | MODIFIER |
| 4 | 133908158 | AX-97817756 | 0 | A | C | 0.158 | 1.17E-01 | 1.18E-01 | 2.57E+00 | 2.61E+00 | 3.22E-01 | no | | regulatory_region_variant | MODIFIER |
| 4 | 133908567 | AX-170382692 | 0 | T | A | 0.158 | 1.17E-01 | 1.18E-01 | 2.57E+00 | 2.61E+00 | 3.22E-01 | no | | regulatory_region_variant | MODIFIER |
| 4 | 133941432 | AX-170070750 | 0 | C | G | 0.158 | 1.17E-01 | 1.18E-01 | 2.57E+00 | 2.61E+00 | 3.22E-01 | no | | regulatory_region_variant | MODIFIER |
| 4 | 133977667 | AX-97799521 | 0 | A | G | 0.158 | 1.17E-01 | 1.18E-01 | 2.57E+00 | 2.61E+00 | 3.22E-01 | no | Dhdds | intron_variant | MODIFIER |
| 4 | 133977766 | AX-170311317 | 0 | T | C | 0.158 | 1.17E-01 | 1.18E-01 | 2.57E+00 | 2.61E+00 | 3.22E-01 | no | Dhdds | intron_variant | MODIFIER |
| 4 | 133985768 | AX-170211604 | 0 | T | C | 0.158 | 1.17E-01 | 1.18E-01 | 2.57E+00 | 2.61E+00 | 3.22E-01 | no | Dhdds | intron_variant | MODIFIER |
| 4 | 133985942 | AX-170188901 | 0 | A | G | 0.158 | 1.17E-01 | 1.18E-01 | 2.57E+00 | 2.61E+00 | 3.22E-01 | no | Dhdds | intron_variant | MODIFIER |
| 4 | 133991692 | AX-170427184 | 0 | A | G | 0.158 | 1.17E-01 | 1.18E-01 | 2.57E+00 | 2.61E+00 | 3.22E-01 | no | Dhdds | intron_variant | MODIFIER |

| | | | | | | | | | | | | | | | |
|---|---|---|---|---|---|---|---|---|---|---|---|---|---|---|---|
| 4 | 134007869 | AX-170229979 | 0 | G | A | 0.158 | 1.17E-01 | 1.18E-01 | 2.57E+00 | 2.61E+00 | 3.22E-01 | no | Lin28a | intron_variant | MODIFIER |
| 4 | 134007967 | AX-170513543 | 0 | C | T | 0.158 | 1.17E-01 | 1.18E-01 | 2.57E+00 | 2.61E+00 | 3.22E-01 | no | Lin28a | synonymous_variant | LOW |
| 4 | 134010713 | AX-170226413 | 0 | A | C | 0.158 | 1.17E-01 | 1.18E-01 | 2.57E+00 | 2.61E+00 | 3.22E-01 | no | Lin28a | intron_variant | MODIFIER |
| 4 | 134011445 | AX-170526321 | 0 | T | C | 0.158 | 1.17E-01 | 1.18E-01 | 2.57E+00 | 2.61E+00 | 3.22E-01 | no | Lin28a | intron_variant | MODIFIER |
| 4 | 134065057 | AX-97809532 | 0 | T | C | 0.158 | 1.17E-01 | 1.18E-01 | 2.57E+00 | 2.61E+00 | 3.22E-01 | no | Crybg2 | upstream_gene_variant | MODIFIER |
| 4 | 134072257 | AX-170387686 | 0 | G | A | 0.158 | 1.17E-01 | 1.18E-01 | 2.57E+00 | 2.61E+00 | 3.22E-01 | no | Crybg2 | intron_variant | MODIFIER |
| 4 | 134081140 | AX-170235335 | 0 | A | G | 0.158 | 1.17E-01 | 1.18E-01 | 2.57E+00 | 2.61E+00 | 3.22E-01 | no | Crybg2 | missense_variant | MODERATE |
| 4 | 134091341 | AX-170169419 | 0 | C | T | 0.158 | 1.17E-01 | 1.18E-01 | 2.57E+00 | 2.61E+00 | 3.22E-01 | no | Cd52 | downstream_gene_variant | MODIFIER |
| 4 | 134116273 | AX-170227241 | 0 | C | T | 0.158 | 1.17E-01 | 1.18E-01 | 2.57E+00 | 2.61E+00 | 3.22E-01 | no | Ubxn11 | intron_variant | MODIFIER |
| 4 | 134121005 | AX-170334224 | 0 | T | A | 0.158 | 1.17E-01 | 1.18E-01 | 2.57E+00 | 2.61E+00 | 3.22E-01 | no | Ubxn11 | intron_variant | MODIFIER |
| 4 | 134137775 | AX-170423109 | 0 | A | G | 0.158 | 1.17E-01 | 1.18E-01 | 2.57E+00 | 2.61E+00 | 3.22E-01 | no | Cep85 | intron_variant | MODIFIER |
| 4 | 134146703 | AX-170203859 | 0 | C | T | 0.158 | 1.17E-01 | 1.18E-01 | 2.57E+00 | 2.61E+00 | 3.22E-01 | no | Cep85 | intron_variant | MODIFIER |
| 4 | 134147389 | AX-170388787 | 0 | G | C | 0.158 | 1.17E-01 | 1.18E-01 | 2.57E+00 | 2.61E+00 | 3.22E-01 | no | Cep85 | intron_variant | MODIFIER |
| 4 | 134147980 | AX-97803264 | 0 | C | T | 0.158 | 1.17E-01 | 1.18E-01 | 2.57E+00 | 2.61E+00 | 3.22E-01 | no | Cep85 | intron_variant | MODIFIER |
| 4 | 134152757 | AX-170345661 | 0 | C | T | 0.158 | 1.17E-01 | 1.18E-01 | 2.57E+00 | 2.61E+00 | 3.22E-01 | no | Cep85 | intron_variant | MODIFIER |
| 4 | 134153658 | AX-170146269 | 0 | T | G | 0.158 | 1.17E-01 | 1.18E-01 | 2.57E+00 | 2.61E+00 | 3.22E-01 | no | Cep85 | intron_variant | MODIFIER |
| 4 | 134169169 | AX-170271041 | 0 | C | T | 0.158 | 1.17E-01 | 1.18E-01 | 2.57E+00 | 2.61E+00 | 3.22E-01 | no | Cep85 | intron_variant | MODIFIER |
| 4 | 134176845 | AX-170166122 | 0 | A | G | 0.158 | 1.17E-01 | 1.18E-01 | 2.57E+00 | 2.61E+00 | 3.22E-01 | no | Cep85 | intron_variant | MODIFIER |
| 4 | 134187313 | AX-170394542 | 0 | G | A | 0.158 | 1.17E-01 | 1.18E-01 | 2.57E+00 | 2.61E+00 | 3.22E-01 | no | Cep85 | upstream_gene_variant | MODIFIER |
| 4 | 134187989 | AX-170128626 | 0 | C | G | 0.158 | 1.17E-01 | 1.18E-01 | 2.57E+00 | 2.61E+00 | 3.22E-01 | no | Cep85 | upstream_gene_variant | MODIFIER |
| 4 | 134188222 | AX-170024001 | 0 | G | C | 0.158 | 1.17E-01 | 1.18E-01 | 2.57E+00 | 2.61E+00 | 3.22E-01 | no | Cep85 | upstream_gene_variant | MODIFIER |
| 4 | 134200924 | AX-170277354 | 0 | C | G | 0.158 | 1.17E-01 | 1.18E-01 | 2.57E+00 | 2.61E+00 | 3.22E-01 | no | Gm7534 | intron_variant | MODIFIER |
| 4 | 134218795 | AX-170024002 | 0 | A | G | 0.158 | 1.17E-01 | 1.18E-01 | 2.57E+00 | 2.61E+00 | 3.22E-01 | no | Catsper4 | intron_variant | MODIFIER |
| 4 | 134223863 | AX-170070755 | 0 | A | G | 0.158 | 1.17E-01 | 1.18E-01 | 2.57E+00 | 2.61E+00 | 3.22E-01 | no | Cnksr1 | downstream_gene_variant | MODIFIER |
| 4 | 134237520 | AX-170070756 | 0 | C | T | 0.158 | 1.17E-01 | 1.18E-01 | 2.57E+00 | 2.61E+00 | 3.22E-01 | no | Cnksr1 | intron_variant | MODIFIER |
| 4 | 134267046 | AX-170477209 | 0 | T | A | 0.158 | 1.17E-01 | 1.18E-01 | 2.57E+00 | 2.61E+00 | 3.22E-01 | no | | intergenic_variant | MODIFIER |
| 4 | 134271476 | AX-170286135 | 0 | A | T | 0.158 | 1.17E-01 | 1.18E-01 | 2.57E+00 | 2.61E+00 | 3.22E-01 | no | Pdik1l | downstream_gene_variant | MODIFIER |
| 4 | 134294205 | AX-170385513 | 0 | T | C | 0.158 | 1.17E-01 | 1.18E-01 | 2.57E+00 | 2.61E+00 | 3.22E-01 | no | | regulatory_region_variant | MODIFIER |
| 4 | 134309493 | AX-170320227 | 0 | G | T | 0.158 | 1.17E-01 | 1.18E-01 | 2.57E+00 | 2.61E+00 | 3.22E-01 | no | | intergenic_variant | MODIFIER |
| 4 | 134316322 | AX-97813689 | 0 | A | G | 0.158 | 1.17E-01 | 1.18E-01 | 2.57E+00 | 2.61E+00 | 3.22E-01 | no | Trim63 | intron_variant | MODIFIER |
| 4 | 134328918 | AX-170402226 | 0 | A | G | 0.158 | 1.17E-01 | 1.18E-01 | 2.57E+00 | 2.61E+00 | 3.22E-01 | no | Trim63 | 3_prime_UTR_variant | MODIFIER |
| 4 | 134347705 | AX-170406245 | 0 | T | C | 0.158 | 1.17E-01 | 1.18E-01 | 2.57E+00 | 2.61E+00 | 3.22E-01 | no | Slc30a2 | intron_variant | MODIFIER |
| 4 | 134347740 | AX-170461924 | 0 | T | C | 0.158 | 1.17E-01 | 1.18E-01 | 2.57E+00 | 2.61E+00 | 3.22E-01 | no | Slc30a2 | intron_variant | MODIFIER |
| 4 | 134348359 | AX-170352249 | 0 | C | G | 0.158 | 1.17E-01 | 1.18E-01 | 2.57E+00 | 2.61E+00 | 3.22E-01 | no | Slc30a2 | intron_variant | MODIFIER |
| 4 | 134371151 | AX-170231282 | 0 | A | T | 0.158 | 1.17E-01 | 1.18E-01 | 2.57E+00 | 2.61E+00 | 3.22E-01 | no | Extl1 | missense_variant | MODERATE |
| 4 | 134372548 | AX-170174201 | 0 | A | T | 0.158 | 1.17E-01 | 1.18E-01 | 2.57E+00 | 2.61E+00 | 3.22E-01 | no | Extl1 | 5_prime_UTR_variant | MODIFIER |
| 4 | 134373468 | AX-170324976 | 0 | C | A | 0.158 | 1.17E-01 | 1.18E-01 | 2.57E+00 | 2.61E+00 | 3.22E-01 | no | Extl1 | upstream_gene_variant | MODIFIER |
| 4 | 134404469 | AX-170520051 | 0 | G | T | 0.158 | 1.17E-01 | 1.18E-01 | 2.57E+00 | 2.61E+00 | 3.22E-01 | no | Pafah2 | intron_variant | MODIFIER |
| 4 | 134404704 | AX-170155607 | 0 | A | C | 0.158 | 1.17E-01 | 1.18E-01 | 2.57E+00 | 2.61E+00 | 3.22E-01 | no | Pafah2 | intron_variant | MODIFIER |
| 4 | 134432536 | AX-170141058 | 0 | T | C | 0.158 | 1.17E-01 | 1.18E-01 | 2.57E+00 | 2.61E+00 | 3.22E-01 | no | | regulatory_region_variant | MODIFIER |
| 4 | 134440589 | AX-170354678 | 0 | G | C | 0.158 | 1.17E-01 | 1.18E-01 | 2.57E+00 | 2.61E+00 | 3.22E-01 | no | | intergenic_variant | MODIFIER |
| 4 | 134458997 | AX-170151596 | 0 | C | A | 0.158 | 1.17E-01 | 1.18E-01 | 2.57E+00 | 2.61E+00 | 3.22E-01 | no | | intergenic_variant | MODIFIER |
| 4 | 134459736 | AX-170342195 | 0 | A | G | 0.158 | 1.17E-01 | 1.18E-01 | 2.57E+00 | 2.61E+00 | 3.22E-01 | no | | intergenic_variant | MODIFIER |
| 4 | 134465931 | AX-170205111 | 0 | G | A | 0.158 | 1.17E-01 | 1.18E-01 | 2.57E+00 | 2.61E+00 | 3.22E-01 | no | Stmn1 | upstream_gene_variant | MODIFIER |
| 4 | 134466540 | AX-170097939 | 0 | T | C | 0.158 | 1.17E-01 | 1.18E-01 | 2.57E+00 | 2.61E+00 | 3.22E-01 | no | Stmn1 | upstream_gene_variant | MODIFIER |
| 4 | 134474925 | AX-170385043 | 0 | C | T | 0.158 | 1.17E-01 | 1.18E-01 | 2.57E+00 | 2.61E+00 | 3.22E-01 | no | Stmn1 | downstream_gene_variant | MODIFIER |
| 4 | 134490309 | AX-170070762 | 0 | A | G | 0.158 | 1.17E-01 | 1.18E-01 | 2.57E+00 | 2.61E+00 | 3.22E-01 | no | Gm13250 | upstream_gene_variant | MODIFIER |
| 4 | 134491308 | AX-170070761 | 0 | T | C | 0.158 | 1.17E-01 | 1.18E-01 | 2.57E+00 | 2.61E+00 | 3.22E-01 | no | Gm13250 | upstream_gene_variant | MODIFIER |
| 4 | 134491943 | AX-97803531 | 0 | G | A | 0.158 | 1.17E-01 | 1.18E-01 | 2.57E+00 | 2.61E+00 | 3.22E-01 | no | Paqr7 | upstream_gene_variant | MODIFIER |
| 4 | 134501332 | AX-170527618 | 0 | A | T | 0.158 | 1.17E-01 | 1.18E-01 | 2.57E+00 | 2.61E+00 | 3.22E-01 | no | Paqr7 | intron_variant | MODIFIER |
| 4 | 134503596 | AX-170268818 | 0 | A | C | 0.158 | 1.17E-01 | 1.18E-01 | 2.57E+00 | 2.61E+00 | 3.22E-01 | no | Paqr7 | intron_variant | MODIFIER |
| 4 | 134634897 | AX-170070768 | 0 | T | C | 0.158 | 1.17E-01 | 1.18E-01 | 2.57E+00 | 2.61E+00 | 3.22E-01 | no | Man1c1 | intron_variant | MODIFIER |
| 4 | 134643205 | AX-170070763 | 0 | C | T | 0.158 | 1.17E-01 | 1.18E-01 | 2.57E+00 | 2.61E+00 | 3.22E-01 | no | Man1c1 | intron_variant | MODIFIER |
| 4 | 134643699 | AX-170328543 | 0 | A | G | 0.158 | 1.17E-01 | 1.18E-01 | 2.57E+00 | 2.61E+00 | 3.22E-01 | no | Man1c1 | intron_variant | MODIFIER |

| | | | | | | | | | | | | | | | |
|---|---|---|---|---|---|---|---|---|---|---|---|---|---|---|---|
| 4 | 134665320 | AX-97840447 | 0 | C | A | 0.158 | 1.17E-01 | 1.18E-01 | 2.57E+00 | 2.61E+00 | 3.22E-01 | no | Man1c1 | intron_variant | MODIFIER |
| 4 | 134666169 | AX-170210052 | 0 | C | G | 0.158 | 1.17E-01 | 1.18E-01 | 2.57E+00 | 2.61E+00 | 3.22E-01 | no | Man1c1 | intron_variant | MODIFIER |
| 4 | 134677102 | AX-170451678 | 0 | A | G | 0.158 | 1.17E-01 | 1.18E-01 | 2.57E+00 | 2.61E+00 | 3.22E-01 | no | Man1c1 | intron_variant | MODIFIER |
| 4 | 134677217 | AX-170247268 | 0 | T | G | 0.158 | 1.17E-01 | 1.18E-01 | 2.57E+00 | 2.61E+00 | 3.22E-01 | no | Man1c1 | intron_variant | MODIFIER |
| 4 | 134684065 | AX-170454962 | 0 | A | G | 0.158 | 1.17E-01 | 1.18E-01 | 2.57E+00 | 2.61E+00 | 3.22E-01 | no | Man1c1 | intron_variant | MODIFIER |
| 4 | 134684453 | AX-170258479 | 0 | T | C | 0.158 | 1.17E-01 | 1.18E-01 | 2.57E+00 | 2.61E+00 | 3.22E-01 | no | Man1c1 | intron_variant | MODIFIER |
| 4 | 134685919 | AX-170177702 | 0 | C | T | 0.158 | 1.17E-01 | 1.18E-01 | 2.57E+00 | 2.61E+00 | 3.22E-01 | no | Man1c1 | intron_variant | MODIFIER |
| 4 | 134905506 | AX-170474875 | 0 | A | C | 0.158 | 1.17E-01 | 1.18E-01 | 2.57E+00 | 2.61E+00 | 3.22E-01 | no | Tmem50a | intron_variant | MODIFIER |
| 4 | 134989185 | AX-170070775 | 0 | G | T | 0.158 | 1.17E-01 | 1.18E-01 | 2.57E+00 | 2.61E+00 | 3.22E-01 | no | | intergenic_variant | MODIFIER |
| 4 | 135124118 | AX-170240558 | 0 | A | G | 0.158 | 1.17E-01 | 1.18E-01 | 2.57E+00 | 2.61E+00 | 3.22E-01 | no | Runx3 | intron_variant | MODIFIER |
| 4 | 135168655 | AX-170379589 | 0 | G | A | 0.158 | 1.17E-01 | 1.18E-01 | 2.57E+00 | 2.61E+00 | 3.22E-01 | no | Runx3 | intron_variant | MODIFIER |
| 4 | 135267818 | AX-170526584 | 0 | C | A | 0.158 | 1.17E-01 | 1.18E-01 | 2.57E+00 | 2.61E+00 | 3.22E-01 | no | Clic4 | intron_variant | MODIFIER |
| 4 | 135371261 | AX-170134093 | 0 | T | A | 0.158 | 1.17E-01 | 1.18E-01 | 2.57E+00 | 2.61E+00 | 3.22E-01 | no | Ncmap | intron_variant | MODIFIER |
| 4 | 135375329 | AX-170095160 | 0 | A | G | 0.158 | 1.17E-01 | 1.18E-01 | 2.57E+00 | 2.61E+00 | 3.22E-01 | no | Ncmap | intron_variant | MODIFIER |
| 4 | 135403238 | AX-170250441 | 0 | G | A | 0.158 | 1.17E-01 | 1.18E-01 | 2.57E+00 | 2.61E+00 | 3.22E-01 | no | Gm12990 | non_coding_transcript_exon_variant | MODIFIER |
| 4 | 135404358 | AX-170024018 | 0 | A | G | 0.158 | 1.17E-01 | 1.18E-01 | 2.57E+00 | 2.61E+00 | 3.22E-01 | no | Gm12990 | upstream_gene_variant | MODIFIER |
| 4 | 135414168 | AX-170234307 | 0 | C | T | 0.158 | 1.17E-01 | 1.18E-01 | 2.57E+00 | 2.61E+00 | 3.22E-01 | no | Rcan3 | 3_prime_UTR_variant | MODIFIER |
| 4 | 135414980 | AX-170212659 | 0 | T | C | 0.158 | 1.17E-01 | 1.18E-01 | 2.57E+00 | 2.61E+00 | 3.22E-01 | no | Rcan3 | 3_prime_UTR_variant | MODIFIER |
| 4 | 135418900 | AX-97816294 | 0 | G | A | 0.158 | 1.17E-01 | 1.18E-01 | 2.57E+00 | 2.61E+00 | 3.22E-01 | no | Rcan3 | intron_variant | MODIFIER |
| 4 | 135427104 | AX-170024013 | 0 | C | T | 0.158 | 1.17E-01 | 1.18E-01 | 2.57E+00 | 2.61E+00 | 3.22E-01 | no | Rcan3 | intron_variant | MODIFIER |
| 4 | 135427211 | AX-97810547 | 0 | A | G | 0.158 | 1.17E-01 | 1.18E-01 | 2.57E+00 | 2.61E+00 | 3.22E-01 | no | Rcan3 | intron_variant | MODIFIER |
| 4 | 136442526 | AX-170279231 | 0 | T | C | 0.158 | 1.17E-01 | 1.18E-01 | 2.57E+00 | 2.61E+00 | 3.22E-01 | no | Htr1d | missense_variant | MODERATE |
| 4 | 136468696 | AX-170145263 | 0 | A | T | 0.158 | 1.17E-01 | 1.18E-01 | 2.57E+00 | 2.61E+00 | 3.22E-01 | no | 6030445D17Rik | downstream_gene_variant | MODIFIER |
| 4 | 136483074 | AX-170095163 | 0 | C | T | 0.158 | 1.17E-01 | 1.18E-01 | 2.57E+00 | 2.61E+00 | 3.22E-01 | no | Luzp1 | intron_variant | MODIFIER |
| 4 | 136484700 | AX-170213540 | 0 | A | G | 0.158 | 1.17E-01 | 1.18E-01 | 2.57E+00 | 2.61E+00 | 3.22E-01 | no | Luzp1 | intron_variant | MODIFIER |
| 4 | 136502427 | AX-170024030 | 0 | A | C | 0.158 | 1.17E-01 | 1.18E-01 | 2.57E+00 | 2.61E+00 | 3.22E-01 | no | Luzp1 | intron_variant | MODIFIER |
| 4 | 136502749 | AX-170426953 | 0 | T | G | 0.158 | 1.17E-01 | 1.18E-01 | 2.57E+00 | 2.61E+00 | 3.22E-01 | no | Luzp1 | intron_variant | MODIFIER |
| 4 | 136521759 | AX-170024037 | 0 | C | T | 0.158 | 1.17E-01 | 1.18E-01 | 2.57E+00 | 2.61E+00 | 3.22E-01 | no | Luzp1 | intron_variant | MODIFIER |
| 4 | 136544478 | AX-97798743 | 0 | G | A | 0.158 | 1.17E-01 | 1.18E-01 | 2.57E+00 | 2.61E+00 | 3.22E-01 | no | Luzp1 | intron_variant | MODIFIER |
| 4 | 136548079 | AX-170024033 | 0 | T | C | 0.158 | 1.17E-01 | 1.18E-01 | 2.57E+00 | 2.61E+00 | 3.22E-01 | no | Luzp1 | 3_prime_UTR_variant | MODIFIER |
| 4 | 136553613 | AX-170320649 | 0 | G | T | 0.158 | 1.17E-01 | 1.18E-01 | 2.57E+00 | 2.61E+00 | 3.22E-01 | no | Luzp1 | downstream_gene_variant | MODIFIER |
| 4 | 136558304 | AX-170178703 | 0 | A | G | 0.158 | 1.17E-01 | 1.18E-01 | 2.57E+00 | 2.61E+00 | 3.22E-01 | no | Kdm1a | intron_variant | MODIFIER |
| 4 | 136563596 | AX-170467431 | 0 | C | A | 0.158 | 1.17E-01 | 1.18E-01 | 2.57E+00 | 2.61E+00 | 3.22E-01 | no | Kdm1a | intron_variant | MODIFIER |
| 4 | 136568104 | AX-170024032 | 0 | C | T | 0.158 | 1.17E-01 | 1.18E-01 | 2.57E+00 | 2.61E+00 | 3.22E-01 | no | Kdm1a | intron_variant | MODIFIER |
| 4 | 136583324 | AX-170385695 | 0 | C | T | 0.158 | 1.17E-01 | 1.18E-01 | 2.57E+00 | 2.61E+00 | 3.22E-01 | no | Kdm1a | intron_variant | MODIFIER |
| 4 | 136626250 | AX-170070806 | 0 | A | G | 0.158 | 1.17E-01 | 1.18E-01 | 2.57E+00 | 2.61E+00 | 3.22E-01 | no | Lactbl1 | intron_variant | MODIFIER |
| 4 | 136648400 | AX-97824045 | 0 | T | C | 0.158 | 1.17E-01 | 1.18E-01 | 2.57E+00 | 2.61E+00 | 3.22E-01 | no | Ephb2 | 3_prime_UTR_variant | MODIFIER |
| 4 | 136660088 | AX-170374746 | 0 | T | C | 0.158 | 1.17E-01 | 1.18E-01 | 2.57E+00 | 2.61E+00 | 3.22E-01 | no | Ephb2 | intron_variant | MODIFIER |
| 4 | 136661401 | AX-170211953 | 0 | G | A | 0.158 | 1.17E-01 | 1.18E-01 | 2.57E+00 | 2.61E+00 | 3.22E-01 | no | Ephb2 | synonymous_variant | LOW |
| 4 | 136661859 | AX-170514810 | 0 | G | T | 0.158 | 1.17E-01 | 1.18E-01 | 2.57E+00 | 2.61E+00 | 3.22E-01 | no | Ephb2 | intron_variant | MODIFIER |
| 4 | 136666318 | AX-97804176 | 0 | T | G | 0.158 | 1.17E-01 | 1.18E-01 | 2.57E+00 | 2.61E+00 | 3.22E-01 | no | Ephb2 | intron_variant | MODIFIER |
| 4 | 136685347 | AX-170173157 | 0 | T | C | 0.158 | 1.17E-01 | 1.18E-01 | 2.57E+00 | 2.61E+00 | 3.22E-01 | no | Ephb2 | intron_variant | MODIFIER |
| 4 | 136687738 | AX-170155035 | 0 | C | T | 0.158 | 1.17E-01 | 1.18E-01 | 2.57E+00 | 2.61E+00 | 3.22E-01 | no | Ephb2 | intron_variant | MODIFIER |
| 4 | 136688588 | AX-97798746 | 0 | T | G | 0.158 | 1.17E-01 | 1.18E-01 | 2.57E+00 | 2.61E+00 | 3.22E-01 | no | Ephb2 | intron_variant | MODIFIER |
| 4 | 136706464 | AX-97809941 | 0 | A | G | 0.158 | 1.17E-01 | 1.18E-01 | 2.57E+00 | 2.61E+00 | 3.22E-01 | no | Ephb2 | intron_variant | MODIFIER |
| 4 | 136724982 | AX-170356708 | 0 | C | T | 0.158 | 1.17E-01 | 1.18E-01 | 2.57E+00 | 2.61E+00 | 3.22E-01 | no | Ephb2 | intron_variant | MODIFIER |
| 4 | 136744356 | AX-97833417 | 0 | T | C | 0.158 | 1.17E-01 | 1.18E-01 | 2.57E+00 | 2.61E+00 | 3.22E-01 | no | Ephb2 | intron_variant | MODIFIER |
| 4 | 136752145 | AX-170355510 | 0 | C | T | 0.158 | 1.17E-01 | 1.18E-01 | 2.57E+00 | 2.61E+00 | 3.22E-01 | no | Ephb2 | intron_variant | MODIFIER |
| 4 | 136754142 | AX-170310472 | 0 | G | A | 0.158 | 1.17E-01 | 1.18E-01 | 2.57E+00 | 2.61E+00 | 3.22E-01 | no | Ephb2 | intron_variant | MODIFIER |
| 4 | 136759236 | AX-170372208 | 0 | T | G | 0.158 | 1.17E-01 | 1.18E-01 | 2.57E+00 | 2.61E+00 | 3.22E-01 | no | Ephb2 | intron_variant | MODIFIER |
| 4 | 136759278 | AX-170281245 | 0 | C | A | 0.158 | 1.17E-01 | 1.18E-01 | 2.57E+00 | 2.61E+00 | 3.22E-01 | no | Ephb2 | intron_variant | MODIFIER |
| 4 | 136775606 | AX-97811127 | 0 | T | C | 0.158 | 1.17E-01 | 1.18E-01 | 2.57E+00 | 2.61E+00 | 3.22E-01 | no | Ephb2 | intron_variant | MODIFIER |
| 4 | 136775635 | AX-170374504 | 0 | A | G | 0.158 | 1.17E-01 | 1.18E-01 | 2.57E+00 | 2.61E+00 | 3.22E-01 | no | Ephb2 | intron_variant | MODIFIER |
| 4 | 136777252 | AX-170374464 | 0 | T | C | 0.158 | 1.17E-01 | 1.18E-01 | 2.57E+00 | 2.61E+00 | 3.22E-01 | no | Ephb2 | intron_variant | MODIFIER |

| | | | | | | | | | | | | | | | |
|---|---|---|---|---|---|---|---|---|---|---|---|---|---|---|---|
| 4 | 136781708 | AX-170095165 | 0 | A | G | 0.158 | 1.17E-01 | 1.18E-01 | 2.57E+00 | 2.61E+00 | 3.22E-01 | no | Ephb2 | intron_variant | MODIFIER |
| 4 | 136806790 | AX-170157061 | 0 | C | T | 0.158 | 1.17E-01 | 1.18E-01 | 2.57E+00 | 2.61E+00 | 3.22E-01 | no | Ephb2 | intron_variant | MODIFIER |
| 4 | 136817350 | AX-170173553 | 0 | G | A | 0.158 | 1.17E-01 | 1.18E-01 | 2.57E+00 | 2.61E+00 | 3.22E-01 | no | Ephb2 | intron_variant | MODIFIER |
| 4 | 136821276 | AX-170070808 | 0 | A | G | 0.158 | 1.17E-01 | 1.18E-01 | 2.57E+00 | 2.61E+00 | 3.22E-01 | no | Ephb2 | intron_variant | MODIFIER |
| 4 | 136823293 | AX-170070807 | 0 | G | A | 0.158 | 1.17E-01 | 1.18E-01 | 2.57E+00 | 2.61E+00 | 3.22E-01 | no | Ephb2 | intron_variant | MODIFIER |
| 4 | 136824217 | AX-170281943 | 0 | G | A | 0.158 | 1.17E-01 | 1.18E-01 | 2.57E+00 | 2.61E+00 | 3.22E-01 | no | Ephb2 | intron_variant | MODIFIER |
| 4 | 136861564 | AX-170443701 | 0 | A | G | 0.158 | 1.17E-01 | 1.18E-01 | 2.57E+00 | 2.61E+00 | 3.22E-01 | no | | regulatory_region_variant | MODIFIER |
| 4 | 136869390 | AX-170242391 | 0 | C | T | 0.158 | 1.17E-01 | 1.18E-01 | 2.57E+00 | 2.61E+00 | 3.22E-01 | no | | intergenic_variant | MODIFIER |
| 4 | 136886057 | AX-170449897 | 0 | C | T | 0.158 | 1.17E-01 | 1.18E-01 | 2.57E+00 | 2.61E+00 | 3.22E-01 | no | C1qc | downstream_gene_variant | MODIFIER |
| 4 | 136930604 | AX-170340501 | 0 | C | T | 0.158 | 1.17E-01 | 1.18E-01 | 2.57E+00 | 2.61E+00 | 3.22E-01 | no | Epha8 | 3_prime_UTR_variant | MODIFIER |
| 4 | 136937660 | AX-97856502 | 0 | A | G | 0.158 | 1.17E-01 | 1.18E-01 | 2.57E+00 | 2.61E+00 | 3.22E-01 | no | Epha8 | intron_variant | MODIFIER |
| 4 | 136960167 | AX-170536073 | 0 | C | T | 0.158 | 1.17E-01 | 1.18E-01 | 2.57E+00 | 2.61E+00 | 3.22E-01 | no | Epha8 | upstream_gene_variant | MODIFIER |
| 4 | 136967020 | AX-170397822 | 0 | C | T | 0.158 | 1.17E-01 | 1.18E-01 | 2.57E+00 | 2.61E+00 | 3.22E-01 | no | | regulatory_region_variant | MODIFIER |
| 4 | 136977125 | AX-170137098 | 0 | T | C | 0.158 | 1.17E-01 | 1.18E-01 | 2.57E+00 | 2.61E+00 | 3.22E-01 | no | Zbtb40 | downstream_gene_variant | MODIFIER |
| 4 | 136979516 | AX-170320404 | 0 | G | C | 0.158 | 1.17E-01 | 1.18E-01 | 2.57E+00 | 2.61E+00 | 3.22E-01 | no | Zbtb40 | downstream_gene_variant | MODIFIER |
| 4 | 136992894 | AX-170298424 | 0 | T | G | 0.158 | 1.17E-01 | 1.18E-01 | 2.57E+00 | 2.61E+00 | 3.22E-01 | no | Zbtb40 | intron_variant | MODIFIER |
| 4 | 136995258 | AX-170286564 | 0 | A | G | 0.158 | 1.17E-01 | 1.18E-01 | 2.57E+00 | 2.61E+00 | 3.22E-01 | no | Zbtb40 | synonymous_variant | LOW |
| 4 | 137002062 | AX-170475881 | 0 | A | G | 0.158 | 1.17E-01 | 1.18E-01 | 2.57E+00 | 2.61E+00 | 3.22E-01 | no | Zbtb40 | intron_variant | MODIFIER |
| 4 | 137004170 | AX-170513408 | 0 | T | C | 0.158 | 1.17E-01 | 1.18E-01 | 2.57E+00 | 2.61E+00 | 3.22E-01 | no | Zbtb40 | intron_variant | MODIFIER |
| 4 | 137022149 | AX-170224908 | 0 | C | G | 0.158 | 1.17E-01 | 1.18E-01 | 2.57E+00 | 2.61E+00 | 3.22E-01 | no | Zbtb40 | intron_variant | MODIFIER |
| 4 | 137031711 | AX-170347346 | 0 | A | G | 0.158 | 1.17E-01 | 1.18E-01 | 2.57E+00 | 2.61E+00 | 3.22E-01 | no | Zbtb40 | intron_variant | MODIFIER |
| 4 | 137038700 | AX-170024039 | 0 | C | A | 0.158 | 1.17E-01 | 1.18E-01 | 2.57E+00 | 2.61E+00 | 3.22E-01 | no | Zbtb40 | intron_variant | MODIFIER |
| 4 | 137040406 | AX-170099970 | 0 | T | C | 0.158 | 1.17E-01 | 1.18E-01 | 2.57E+00 | 2.61E+00 | 3.22E-01 | no | Zbtb40 | intron_variant | MODIFIER |
| 4 | 137049413 | AX-170247395 | 0 | T | C | 0.158 | 1.17E-01 | 1.18E-01 | 2.57E+00 | 2.61E+00 | 3.22E-01 | no | Zbtb40 | upstream_gene_variant | MODIFIER |
| 4 | 137052296 | AX-170230937 | 0 | A | G | 0.158 | 1.17E-01 | 1.18E-01 | 2.57E+00 | 2.61E+00 | 3.22E-01 | no | Zbtb40 | upstream_gene_variant | MODIFIER |
| 4 | 137059139 | AX-170234961 | 0 | G | C | 0.158 | 1.17E-01 | 1.18E-01 | 2.57E+00 | 2.61E+00 | 3.22E-01 | no | | intergenic_variant | MODIFIER |
| 4 | 137065181 | AX-170089383 | 0 | G | A | 0.158 | 1.17E-01 | 1.18E-01 | 2.57E+00 | 2.61E+00 | 3.22E-01 | no | | intergenic_variant | MODIFIER |
| 4 | 137072883 | AX-170024040 | 0 | A | G | 0.158 | 1.17E-01 | 1.18E-01 | 2.57E+00 | 2.61E+00 | 3.22E-01 | no | | regulatory_region_variant | MODIFIER |
| 15 | 67685357 | AX-170362739 | 1 | A | T | 0.33 | -7.92E-02 | 7.68E-02 | 2.34E+00 | 2.42E+00 | 3.22E-01 | no | | intergenic_variant | MODIFIER |
| 1 | 14797138 | AX-170445353 | 0 | G | A | 0.395 | 7.53E-02 | 7.63E-02 | 2.59E+00 | 2.65E+00 | 3.22E-01 | no | Gm37412 | downstream_gene_variant | MODIFIER |
| 13 | 5741506 | AX-170385318 | 2 | C | T | 0.209 | -1.17E-01 | 1.19E-01 | 2.59E+00 | 2.66E+00 | 3.23E-01 | no | | intergenic_variant | MODIFIER |
| 9 | 96402442 | AX-97830725 | 0 | T | C | 0.14 | -1.13E-01 | 1.14E-01 | 2.58E+00 | 2.66E+00 | 3.23E-01 | no | | intergenic_variant | MODIFIER |
| 9 | 96416036 | AX-170398911 | 0 | C | T | 0.14 | -1.13E-01 | 1.14E-01 | 2.58E+00 | 2.66E+00 | 3.23E-01 | no | | regulatory_region_variant | MODIFIER |
| 9 | 96423264 | AX-170378458 | 0 | A | T | 0.14 | -1.13E-01 | 1.14E-01 | 2.58E+00 | 2.66E+00 | 3.23E-01 | no | | intergenic_variant | MODIFIER |
| 9 | 96425051 | AX-170329644 | 0 | A | G | 0.14 | -1.13E-01 | 1.14E-01 | 2.58E+00 | 2.66E+00 | 3.23E-01 | no | | intergenic_variant | MODIFIER |
| 9 | 96425959 | AX-170411222 | 0 | C | T | 0.14 | -1.13E-01 | 1.14E-01 | 2.58E+00 | 2.66E+00 | 3.23E-01 | no | | intergenic_variant | MODIFIER |
| 9 | 96431238 | AX-170185420 | 0 | T | C | 0.14 | -1.13E-01 | 1.14E-01 | 2.58E+00 | 2.66E+00 | 3.23E-01 | no | BC043934 | downstream_gene_variant | MODIFIER |
| 9 | 96431389 | AX-170382668 | 0 | T | G | 0.14 | -1.13E-01 | 1.14E-01 | 2.58E+00 | 2.66E+00 | 3.23E-01 | no | BC043934 | downstream_gene_variant | MODIFIER |
| 9 | 96652344 | AX-97819637 | 0 | T | G | 0.14 | -1.13E-01 | 1.14E-01 | 2.58E+00 | 2.66E+00 | 3.23E-01 | no | | intergenic_variant | MODIFIER |
| 9 | 96664006 | AX-97806034 | 0 | A | G | 0.14 | -1.13E-01 | 1.14E-01 | 2.58E+00 | 2.66E+00 | 3.23E-01 | no | | intergenic_variant | MODIFIER |
| 9 | 96680240 | AX-170033065 | 0 | T | C | 0.14 | -1.13E-01 | 1.14E-01 | 2.58E+00 | 2.66E+00 | 3.23E-01 | no | Zbtb38 | downstream_gene_variant | MODIFIER |
| 9 | 96684293 | AX-170485017 | 0 | G | T | 0.14 | -1.13E-01 | 1.14E-01 | 2.58E+00 | 2.66E+00 | 3.23E-01 | no | Zbtb38 | downstream_gene_variant | MODIFIER |
| 9 | 96687416 | AX-170033066 | 0 | G | A | 0.14 | -1.13E-01 | 1.14E-01 | 2.58E+00 | 2.66E+00 | 3.23E-01 | no | Zbtb38 | synonymous_variant | LOW |
| 9 | 96883578 | AX-97816781 | 0 | G | A | 0.14 | -1.13E-01 | 1.14E-01 | 2.58E+00 | 2.66E+00 | 3.23E-01 | no | Pxylp1 | intron_variant | MODIFIER |
| 9 | 96892408 | AX-170033069 | 0 | G | C | 0.14 | -1.13E-01 | 1.14E-01 | 2.58E+00 | 2.66E+00 | 3.23E-01 | no | Gm10123 | upstream_gene_variant | MODIFIER |
| 9 | 96897577 | AX-97832357 | 0 | G | A | 0.14 | -1.13E-01 | 1.14E-01 | 2.58E+00 | 2.66E+00 | 3.23E-01 | no | Gm10123 | downstream_gene_variant | MODIFIER |
| 9 | 96898272 | AX-170149284 | 0 | C | T | 0.14 | -1.13E-01 | 1.14E-01 | 2.58E+00 | 2.66E+00 | 3.23E-01 | no | Gm10123 | downstream_gene_variant | MODIFIER |
| 9 | 96933259 | AX-97847880 | 0 | C | T | 0.14 | -1.13E-01 | 1.14E-01 | 2.58E+00 | 2.66E+00 | 3.23E-01 | no | | intergenic_variant | MODIFIER |
| 9 | 96949513 | AX-170090387 | 0 | C | T | 0.14 | -1.13E-01 | 1.14E-01 | 2.58E+00 | 2.66E+00 | 3.23E-01 | no | Spsb4 | intron_variant | MODIFIER |
| 9 | 97203515 | AX-97793935 | 0 | G | A | 0.14 | -1.13E-01 | 1.14E-01 | 2.58E+00 | 2.66E+00 | 3.23E-01 | no | | intergenic_variant | MODIFIER |
| 9 | 97289338 | AX-97793019 | 0 | C | G | 0.14 | -1.13E-01 | 1.14E-01 | 2.58E+00 | 2.66E+00 | 3.23E-01 | no | A030012G06Rik | non_coding_transcript_exon_variant | MODIFIER |
| 9 | 97289644 | AX-170033077 | 0 | G | C | 0.14 | -1.13E-01 | 1.14E-01 | 2.58E+00 | 2.66E+00 | 3.23E-01 | no | A030012G06Rik | non_coding_transcript_exon_variant | MODIFIER |
| 9 | 97307008 | AX-170175851 | 0 | C | G | 0.14 | -1.13E-01 | 1.14E-01 | 2.58E+00 | 2.66E+00 | 3.23E-01 | no | | intergenic_variant | MODIFIER |
| 9 | 97320583 | AX-170313409 | 0 | A | C | 0.14 | -1.13E-01 | 1.14E-01 | 2.58E+00 | 2.66E+00 | 3.23E-01 | no | | intergenic_variant | MODIFIER |

| | | | | | | | | | | | | | | | |
|---|---|---|---|---|---|---|---|---|---|---|---|---|---|---|---|
| 9 | 97321184 | AX-170033074 | 0 | T | C | 0.14 | -1.13E-01 | 1.14E-01 | 2.58E+00 | 2.66E+00 | 3.23E-01 | no | | intergenic_variant | MODIFIER |
| 9 | 97354208 | AX-170230799 | 0 | C | T | 0.14 | -1.13E-01 | 1.14E-01 | 2.58E+00 | 2.66E+00 | 3.23E-01 | no | Trim42 | intron_variant | MODIFIER |
| 9 | 97355769 | AX-170280265 | 0 | A | G | 0.14 | -1.13E-01 | 1.14E-01 | 2.58E+00 | 2.66E+00 | 3.23E-01 | no | Trim42 | intron_variant | MODIFIER |
| 9 | 97367844 | AX-97823363 | 0 | C | T | 0.14 | -1.13E-01 | 1.14E-01 | 2.58E+00 | 2.66E+00 | 3.23E-01 | no | Trim42 | intron_variant | MODIFIER |
| 9 | 97415839 | AX-97816035 | 0 | G | A | 0.14 | -1.13E-01 | 1.14E-01 | 2.58E+00 | 2.66E+00 | 3.23E-01 | no | | intergenic_variant | MODIFIER |
| 9 | 97422262 | AX-170033075 | 0 | T | C | 0.14 | -1.13E-01 | 1.14E-01 | 2.58E+00 | 2.66E+00 | 3.23E-01 | no | | intergenic_variant | MODIFIER |
| 9 | 97423055 | AX-170452535 | 0 | A | G | 0.14 | -1.13E-01 | 1.14E-01 | 2.58E+00 | 2.66E+00 | 3.23E-01 | no | | intergenic_variant | MODIFIER |
| 9 | 97446045 | AX-170096639 | 0 | A | G | 0.14 | -1.13E-01 | 1.14E-01 | 2.58E+00 | 2.66E+00 | 3.23E-01 | no | Clstn2 | intron_variant | MODIFIER |
| 9 | 97460124 | AX-170473964 | 0 | A | C | 0.14 | -1.13E-01 | 1.14E-01 | 2.58E+00 | 2.66E+00 | 3.23E-01 | no | Clstn2 | intron_variant | MODIFIER |
| 9 | 97463410 | AX-170178190 | 0 | C | A | 0.14 | -1.13E-01 | 1.14E-01 | 2.58E+00 | 2.66E+00 | 3.23E-01 | no | Clstn2 | intron_variant | MODIFIER |
| 9 | 97464546 | AX-170370324 | 0 | C | A | 0.14 | -1.13E-01 | 1.14E-01 | 2.58E+00 | 2.66E+00 | 3.23E-01 | no | Clstn2 | intron_variant | MODIFIER |
| 9 | 97484541 | AX-170385320 | 0 | C | G | 0.14 | -1.13E-01 | 1.14E-01 | 2.58E+00 | 2.66E+00 | 3.23E-01 | no | Clstn2 | intron_variant | MODIFIER |
| 9 | 97484751 | AX-170312308 | 0 | T | C | 0.14 | -1.13E-01 | 1.14E-01 | 2.58E+00 | 2.66E+00 | 3.23E-01 | no | Clstn2 | intron_variant | MODIFIER |
| 9 | 97487161 | AX-170083185 | 0 | T | C | 0.14 | -1.13E-01 | 1.14E-01 | 2.58E+00 | 2.66E+00 | 3.23E-01 | no | Clstn2 | intron_variant | MODIFIER |
| 9 | 97487567 | AX-170083188 | 0 | G | T | 0.14 | -1.13E-01 | 1.14E-01 | 2.58E+00 | 2.66E+00 | 3.23E-01 | no | Clstn2 | intron_variant | MODIFIER |
| 9 | 97488444 | AX-170083186 | 0 | A | T | 0.14 | -1.13E-01 | 1.14E-01 | 2.58E+00 | 2.66E+00 | 3.23E-01 | no | Clstn2 | intron_variant | MODIFIER |
| 9 | 97488609 | AX-97844381 | 0 | A | G | 0.14 | -1.13E-01 | 1.14E-01 | 2.58E+00 | 2.66E+00 | 3.23E-01 | no | Clstn2 | intron_variant | MODIFIER |
| 9 | 97488755 | AX-170450877 | 0 | G | A | 0.14 | -1.13E-01 | 1.14E-01 | 2.58E+00 | 2.66E+00 | 3.23E-01 | no | Clstn2 | intron_variant | MODIFIER |
| 9 | 97490179 | AX-170322158 | 0 | T | C | 0.14 | -1.13E-01 | 1.14E-01 | 2.58E+00 | 2.66E+00 | 3.23E-01 | no | Clstn2 | intron_variant | MODIFIER |
| 9 | 97490202 | AX-170083187 | 0 | A | G | 0.14 | -1.13E-01 | 1.14E-01 | 2.58E+00 | 2.66E+00 | 3.23E-01 | no | Clstn2 | intron_variant | MODIFIER |
| 9 | 97510810 | AX-170362244 | 0 | G | A | 0.14 | -1.13E-01 | 1.14E-01 | 2.58E+00 | 2.66E+00 | 3.23E-01 | no | Clstn2 | intron_variant | MODIFIER |
| 9 | 97511330 | AX-170412743 | 0 | C | T | 0.14 | -1.13E-01 | 1.14E-01 | 2.58E+00 | 2.66E+00 | 3.23E-01 | no | Clstn2 | intron_variant | MODIFIER |
| 9 | 97616951 | AX-170233493 | 0 | T | A | 0.14 | -1.13E-01 | 1.14E-01 | 2.58E+00 | 2.66E+00 | 3.23E-01 | no | Clstn2 | intron_variant | MODIFIER |
| 9 | 97655852 | AX-170110385 | 0 | A | G | 0.14 | -1.13E-01 | 1.14E-01 | 2.58E+00 | 2.66E+00 | 3.23E-01 | no | Clstn2 | intron_variant | MODIFIER |
| 9 | 97669604 | AX-170361257 | 0 | A | G | 0.14 | -1.13E-01 | 1.14E-01 | 2.58E+00 | 2.66E+00 | 3.23E-01 | no | Clstn2 | intron_variant | MODIFIER |
| 9 | 97730183 | AX-170287837 | 0 | A | G | 0.14 | -1.13E-01 | 1.14E-01 | 2.58E+00 | 2.66E+00 | 3.23E-01 | no | Clstn2 | intron_variant | MODIFIER |
| 9 | 97826419 | AX-170165864 | 0 | T | C | 0.14 | -1.13E-01 | 1.14E-01 | 2.58E+00 | 2.66E+00 | 3.23E-01 | no | Clstn2 | intron_variant | MODIFIER |
| 9 | 97826659 | AX-170443797 | 0 | C | T | 0.14 | -1.13E-01 | 1.14E-01 | 2.58E+00 | 2.66E+00 | 3.23E-01 | no | Clstn2 | intron_variant | MODIFIER |
| 9 | 97879890 | AX-170083197 | 0 | A | T | 0.14 | -1.13E-01 | 1.14E-01 | 2.58E+00 | 2.66E+00 | 3.23E-01 | no | Clstn2 | intron_variant | MODIFIER |
| 9 | 97883136 | AX-170083192 | 0 | A | G | 0.14 | -1.13E-01 | 1.14E-01 | 2.58E+00 | 2.66E+00 | 3.23E-01 | no | Clstn2 | intron_variant | MODIFIER |
| 9 | 97909643 | AX-170473569 | 0 | G | A | 0.14 | -1.13E-01 | 1.14E-01 | 2.58E+00 | 2.66E+00 | 3.23E-01 | no | Clstn2 | intron_variant | MODIFIER |
| 9 | 97920328 | AX-170295769 | 0 | C | T | 0.14 | -1.13E-01 | 1.14E-01 | 2.58E+00 | 2.66E+00 | 3.23E-01 | no | Clstn2 | intron_variant | MODIFIER |
| 9 | 97976997 | AX-170306194 | 0 | C | T | 0.14 | -1.13E-01 | 1.14E-01 | 2.58E+00 | 2.66E+00 | 3.23E-01 | no | Clstn2 | intron_variant | MODIFIER |
| 9 | 98037197 | AX-170115241 | 0 | G | C | 0.14 | -1.13E-01 | 1.14E-01 | 2.58E+00 | 2.66E+00 | 3.23E-01 | no | Clstn2 | upstream_gene_variant | MODIFIER |
| 9 | 98047455 | AX-97798312 | 0 | T | C | 0.14 | -1.13E-01 | 1.14E-01 | 2.58E+00 | 2.66E+00 | 3.23E-01 | no | | intergenic_variant | MODIFIER |
| 9 | 98053342 | AX-170098513 | 0 | G | A | 0.14 | -1.13E-01 | 1.14E-01 | 2.58E+00 | 2.66E+00 | 3.23E-01 | no | | intergenic_variant | MODIFIER |
| 9 | 98077963 | AX-170291080 | 0 | C | T | 0.14 | -1.13E-01 | 1.14E-01 | 2.58E+00 | 2.66E+00 | 3.23E-01 | no | | intergenic_variant | MODIFIER |
| 9 | 98108465 | AX-170440159 | 0 | G | A | 0.14 | -1.13E-01 | 1.14E-01 | 2.58E+00 | 2.66E+00 | 3.23E-01 | no | | intergenic_variant | MODIFIER |
| 9 | 98116595 | AX-170083201 | 0 | G | C | 0.14 | -1.13E-01 | 1.14E-01 | 2.58E+00 | 2.66E+00 | 3.23E-01 | no | | intergenic_variant | MODIFIER |
| 9 | 98117104 | AX-170083199 | 0 | T | C | 0.14 | -1.13E-01 | 1.14E-01 | 2.58E+00 | 2.66E+00 | 3.23E-01 | no | | intergenic_variant | MODIFIER |
| 9 | 98128974 | AX-170393034 | 0 | A | G | 0.14 | -1.13E-01 | 1.14E-01 | 2.58E+00 | 2.66E+00 | 3.23E-01 | no | | intergenic_variant | MODIFIER |
| 9 | 98152870 | AX-170083200 | 0 | A | G | 0.14 | -1.13E-01 | 1.14E-01 | 2.58E+00 | 2.66E+00 | 3.23E-01 | no | | intergenic_variant | MODIFIER |
| 9 | 98153959 | AX-170083202 | 0 | A | G | 0.14 | -1.13E-01 | 1.14E-01 | 2.58E+00 | 2.66E+00 | 3.23E-01 | no | | intergenic_variant | MODIFIER |
| 9 | 98161167 | AX-170106921 | 0 | C | T | 0.14 | -1.13E-01 | 1.14E-01 | 2.58E+00 | 2.66E+00 | 3.23E-01 | no | | intergenic_variant | MODIFIER |
| 9 | 98172981 | AX-170275864 | 0 | C | T | 0.14 | -1.13E-01 | 1.14E-01 | 2.58E+00 | 2.66E+00 | 3.23E-01 | no | | intergenic_variant | MODIFIER |
| 9 | 98192749 | AX-170289286 | 0 | T | C | 0.14 | -1.13E-01 | 1.14E-01 | 2.58E+00 | 2.66E+00 | 3.23E-01 | no | | intergenic_variant | MODIFIER |
| 9 | 98214880 | AX-97810970 | 0 | C | T | 0.14 | -1.13E-01 | 1.14E-01 | 2.58E+00 | 2.66E+00 | 3.23E-01 | no | | intergenic_variant | MODIFIER |
| 9 | 98228170 | AX-170239142 | 0 | C | G | 0.14 | -1.13E-01 | 1.14E-01 | 2.58E+00 | 2.66E+00 | 3.23E-01 | no | | intergenic_variant | MODIFIER |
| 9 | 98247310 | AX-170452988 | 0 | A | G | 0.14 | -1.13E-01 | 1.14E-01 | 2.58E+00 | 2.66E+00 | 3.23E-01 | no | | intergenic_variant | MODIFIER |
| 9 | 98247933 | AX-170033090 | 0 | A | C | 0.14 | -1.13E-01 | 1.14E-01 | 2.58E+00 | 2.66E+00 | 3.23E-01 | no | | intergenic_variant | MODIFIER |
| 9 | 98247974 | AX-97783602 | 0 | G | T | 0.14 | -1.13E-01 | 1.14E-01 | 2.58E+00 | 2.66E+00 | 3.23E-01 | no | | intergenic_variant | MODIFIER |
| 9 | 98248726 | AX-170033091 | 0 | C | T | 0.14 | -1.13E-01 | 1.14E-01 | 2.58E+00 | 2.66E+00 | 3.23E-01 | no | | intergenic_variant | MODIFIER |
| 9 | 98270694 | AX-170125291 | 0 | A | T | 0.14 | -1.13E-01 | 1.14E-01 | 2.58E+00 | 2.66E+00 | 3.23E-01 | no | | intergenic_variant | MODIFIER |
| 9 | 98282167 | AX-97781760 | 0 | C | T | 0.14 | -1.13E-01 | 1.14E-01 | 2.58E+00 | 2.66E+00 | 3.23E-01 | no | | intergenic_variant | MODIFIER |

| | | | | | | | | | | | | | | | |
|---|---|---|---|---|---|---|---|---|---|---|---|---|---|---|---|
| 9 | 98291382 | AX-170033097 | 0 | C | A | 0.14 | -1.13E-01 | 1.14E-01 | 2.58E+00 | 2.66E+00 | 3.23E-01 | no | Nmnat3 | intron_variant,non_coding_transcript_variant | MODIFIER |
| 9 | 98292168 | AX-170033096 | 0 | A | G | 0.14 | -1.13E-01 | 1.14E-01 | 2.58E+00 | 2.66E+00 | 3.23E-01 | no | Nmnat3 | upstream_gene_variant | MODIFIER |
| 9 | 98306199 | AX-97812831 | 0 | G | A | 0.14 | -1.13E-01 | 1.14E-01 | 2.58E+00 | 2.66E+00 | 3.23E-01 | no | Nmnat3 | intron_variant | MODIFIER |
| 9 | 98308005 | AX-170455195 | 0 | A | G | 0.14 | -1.13E-01 | 1.14E-01 | 2.58E+00 | 2.66E+00 | 3.23E-01 | no | Nmnat3 | intron_variant | MODIFIER |
| 9 | 98336462 | AX-170212336 | 0 | A | T | 0.14 | -1.13E-01 | 1.14E-01 | 2.58E+00 | 2.66E+00 | 3.23E-01 | no | Nmnat3 | intron_variant | MODIFIER |
| 9 | 98347468 | AX-170211789 | 0 | T | C | 0.14 | -1.13E-01 | 1.14E-01 | 2.58E+00 | 2.66E+00 | 3.23E-01 | no | Nmnat3 | intron_variant | MODIFIER |
| 9 | 98350342 | AX-170525832 | 0 | G | A | 0.14 | -1.13E-01 | 1.14E-01 | 2.58E+00 | 2.66E+00 | 3.23E-01 | no | Nmnat3 | intron_variant | MODIFIER |
| 9 | 98398516 | AX-170225481 | 0 | A | G | 0.14 | -1.13E-01 | 1.14E-01 | 2.58E+00 | 2.66E+00 | 3.23E-01 | no | Nmnat3 | intron_variant | MODIFIER |
| 9 | 98434952 | AX-170083204 | 0 | C | T | 0.14 | -1.13E-01 | 1.14E-01 | 2.58E+00 | 2.66E+00 | 3.23E-01 | no | Rbp1 | intron_variant | MODIFIER |
| 9 | 98438774 | AX-170175807 | 0 | T | G | 0.14 | -1.13E-01 | 1.14E-01 | 2.58E+00 | 2.66E+00 | 3.23E-01 | no | Rbp1 | intron_variant | MODIFIER |
| 9 | 98446334 | AX-97798386 | 0 | T | C | 0.14 | -1.13E-01 | 1.14E-01 | 2.58E+00 | 2.66E+00 | 3.23E-01 | no | Rbp1 | synonymous_variant | LOW |
| 9 | 98465702 | AX-97784182 | 0 | C | T | 0.14 | -1.13E-01 | 1.14E-01 | 2.58E+00 | 2.66E+00 | 3.23E-01 | no | | intergenic_variant | MODIFIER |
| 9 | 98491452 | AX-170503146 | 0 | G | A | 0.14 | -1.13E-01 | 1.14E-01 | 2.58E+00 | 2.66E+00 | 3.23E-01 | no | Rbp2 | intron_variant | MODIFIER |
| 9 | 98496367 | AX-170484069 | 0 | A | G | 0.14 | -1.13E-01 | 1.14E-01 | 2.58E+00 | 2.66E+00 | 3.23E-01 | no | Rbp2 | intron_variant | MODIFIER |
| 18 | 58442841 | AX-97789126 | 2 | C | T | 0.418 | 9.70E-02 | 9.52E-02 | 2.39E+00 | 2.44E+00 | 3.23E-01 | no | | regulatory_region_variant | MODIFIER |
| 3 | 37344032 | AX-170121192 | 0 | T | C | 0.202 | 8.99E-02 | 9.16E-02 | 2.86E+00 | 2.96E+00 | 3.23E-01 | no | Fgf2 | upstream_gene_variant | MODIFIER |
| 7 | 70829237 | AX-97847546 | 0 | A | C | 0.36 | -9.80E-02 | 9.97E-02 | 2.88E+00 | 2.98E+00 | 3.23E-01 | no | | intergenic_variant | MODIFIER |
| 6 | 126797674 | AX-170166567 | 1 | G | A | 0.536 | 8.19E-02 | 8.19E-02 | 2.48E+00 | 2.55E+00 | 3.23E-01 | no | | intergenic_variant | MODIFIER |
| 5 | 30640643 | AX-170136005 | 0 | C | A | 0.36 | -1.01E-01 | 1.02E-01 | 2.95E+00 | 3.05E+00 | 3.23E-01 | no | | intergenic_variant | MODIFIER |
| 7 | 140247394 | AX-97836555 | 0 | T | C | 0.202 | -7.33E-02 | 7.51E-02 | 2.65E+00 | 2.78E+00 | 3.23E-01 | no | 5830411N06Rik | non_coding_transcript_exon_variant | MODIFIER |
| 7 | 140298293 | AX-97815733 | 0 | G | A | 0.202 | -7.33E-02 | 7.51E-02 | 2.65E+00 | 2.78E+00 | 3.23E-01 | no | 5830411N06Rik | downstream_gene_variant | MODIFIER |
| 7 | 140299658 | AX-97837064 | 0 | G | A | 0.202 | -7.33E-02 | 7.51E-02 | 2.65E+00 | 2.78E+00 | 3.23E-01 | no | 5830411N06Rik | downstream_gene_variant | MODIFIER |
| 7 | 144068370 | AX-170350135 | 0 | A | G | 0.202 | -7.33E-02 | 7.51E-02 | 2.65E+00 | 2.78E+00 | 3.23E-01 | no | Shank2 | intron_variant | MODIFIER |
| 7 | 144068418 | AX-170230299 | 0 | T | C | 0.202 | -7.33E-02 | 7.51E-02 | 2.65E+00 | 2.78E+00 | 3.23E-01 | no | Shank2 | intron_variant | MODIFIER |
| 7 | 144068497 | AX-170364771 | 0 | T | C | 0.202 | -7.33E-02 | 7.51E-02 | 2.65E+00 | 2.78E+00 | 3.23E-01 | no | Shank2 | intron_variant | MODIFIER |
| 7 | 144074859 | AX-170177246 | 0 | G | A | 0.202 | -7.33E-02 | 7.51E-02 | 2.65E+00 | 2.78E+00 | 3.23E-01 | no | Shank2 | intron_variant | MODIFIER |
| 7 | 144090838 | AX-170205484 | 0 | A | G | 0.202 | -7.33E-02 | 7.51E-02 | 2.65E+00 | 2.78E+00 | 3.23E-01 | no | Shank2 | intron_variant | MODIFIER |
| 7 | 144137998 | AX-170518018 | 0 | T | C | 0.202 | -7.33E-02 | 7.51E-02 | 2.65E+00 | 2.78E+00 | 3.23E-01 | no | Shank2 | intron_variant | MODIFIER |
| 7 | 144138546 | AX-170484946 | 0 | T | C | 0.202 | -7.33E-02 | 7.51E-02 | 2.65E+00 | 2.78E+00 | 3.23E-01 | no | Shank2 | intron_variant | MODIFIER |
| 7 | 144139231 | AX-170244965 | 0 | A | G | 0.202 | -7.33E-02 | 7.51E-02 | 2.65E+00 | 2.78E+00 | 3.23E-01 | no | Shank2 | intron_variant | MODIFIER |
| 7 | 144139286 | AX-170416533 | 0 | C | T | 0.202 | -7.33E-02 | 7.51E-02 | 2.65E+00 | 2.78E+00 | 3.23E-01 | no | Shank2 | intron_variant | MODIFIER |
| 7 | 144139830 | AX-170376911 | 0 | T | C | 0.202 | -7.33E-02 | 7.51E-02 | 2.65E+00 | 2.78E+00 | 3.23E-01 | no | Shank2 | intron_variant | MODIFIER |
| 7 | 144140494 | AX-170374250 | 0 | C | T | 0.202 | -7.33E-02 | 7.51E-02 | 2.65E+00 | 2.78E+00 | 3.23E-01 | no | Shank2 | intron_variant | MODIFIER |
| 7 | 144141103 | AX-170330444 | 0 | C | T | 0.202 | -7.33E-02 | 7.51E-02 | 2.65E+00 | 2.78E+00 | 3.23E-01 | no | Shank2 | intron_variant | MODIFIER |
| 7 | 144141163 | AX-170372629 | 0 | C | G | 0.202 | -7.33E-02 | 7.51E-02 | 2.65E+00 | 2.78E+00 | 3.23E-01 | no | Shank2 | intron_variant | MODIFIER |
| 7 | 144141243 | AX-170348817 | 0 | A | G | 0.202 | -7.33E-02 | 7.51E-02 | 2.65E+00 | 2.78E+00 | 3.23E-01 | no | Shank2 | intron_variant | MODIFIER |
| 7 | 144142556 | AX-170408948 | 0 | T | C | 0.202 | -7.33E-02 | 7.51E-02 | 2.65E+00 | 2.78E+00 | 3.23E-01 | no | Shank2 | intron_variant | MODIFIER |
| 7 | 144147699 | AX-170378362 | 0 | T | C | 0.202 | -7.33E-02 | 7.51E-02 | 2.65E+00 | 2.78E+00 | 3.23E-01 | no | Shank2 | intron_variant | MODIFIER |
| 7 | 144153078 | AX-170494348 | 0 | G | A | 0.202 | -7.33E-02 | 7.51E-02 | 2.65E+00 | 2.78E+00 | 3.23E-01 | no | Shank2 | intron_variant | MODIFIER |
| 7 | 144158430 | AX-170079112 | 0 | T | C | 0.202 | -7.33E-02 | 7.51E-02 | 2.65E+00 | 2.78E+00 | 3.23E-01 | no | Shank2 | intron_variant | MODIFIER |
| 7 | 144161535 | AX-170533627 | 0 | C | T | 0.202 | -7.33E-02 | 7.51E-02 | 2.65E+00 | 2.78E+00 | 3.23E-01 | no | Shank2 | intron_variant | MODIFIER |
| 7 | 144168083 | AX-97808175 | 0 | G | A | 0.202 | -7.33E-02 | 7.51E-02 | 2.65E+00 | 2.78E+00 | 3.23E-01 | no | Shank2 | intron_variant | MODIFIER |
| 7 | 144168559 | AX-170240408 | 0 | G | A | 0.202 | -7.33E-02 | 7.51E-02 | 2.65E+00 | 2.78E+00 | 3.23E-01 | no | Shank2 | intron_variant | MODIFIER |
| 7 | 144170433 | AX-170112051 | 0 | A | G | 0.202 | -7.33E-02 | 7.51E-02 | 2.65E+00 | 2.78E+00 | 3.23E-01 | no | Shank2 | intron_variant | MODIFIER |
| 7 | 144171844 | AX-170286850 | 0 | T | C | 0.202 | -7.33E-02 | 7.51E-02 | 2.65E+00 | 2.78E+00 | 3.23E-01 | no | Shank2 | upstream_gene_variant | MODIFIER |
| 7 | 144171939 | AX-170283331 | 0 | G | A | 0.202 | -7.33E-02 | 7.51E-02 | 2.65E+00 | 2.78E+00 | 3.23E-01 | no | Shank2 | upstream_gene_variant | MODIFIER |
| 7 | 144408898 | AX-170242701 | 0 | T | C | 0.202 | -7.33E-02 | 7.51E-02 | 2.65E+00 | 2.78E+00 | 3.23E-01 | no | Shank2 | intron_variant | MODIFIER |
| 7 | 144412533 | AX-170030173 | 0 | C | T | 0.202 | -7.33E-02 | 7.51E-02 | 2.65E+00 | 2.78E+00 | 3.23E-01 | no | Shank2 | intron_variant | MODIFIER |
| 7 | 144412671 | AX-97828062 | 0 | G | A | 0.202 | -7.33E-02 | 7.51E-02 | 2.65E+00 | 2.78E+00 | 3.23E-01 | no | Shank2 | intron_variant | MODIFIER |
| 7 | 144445141 | AX-170030166 | 0 | A | G | 0.202 | -7.33E-02 | 7.51E-02 | 2.65E+00 | 2.78E+00 | 3.23E-01 | no | Cttn | intron_variant | MODIFIER |
| 7 | 144458041 | AX-97845603 | 0 | A | C | 0.202 | -7.33E-02 | 7.51E-02 | 2.65E+00 | 2.78E+00 | 3.23E-01 | no | Cttn | intron_variant | MODIFIER |
| 7 | 144482632 | AX-170030168 | 0 | G | C | 0.202 | -7.33E-02 | 7.51E-02 | 2.65E+00 | 2.78E+00 | 3.23E-01 | no | Ppfia1 | intron_variant | MODIFIER |
| 7 | 144490135 | AX-97836222 | 0 | A | G | 0.202 | -7.33E-02 | 7.51E-02 | 2.65E+00 | 2.78E+00 | 3.23E-01 | no | Ppfia1 | intron_variant | MODIFIER |
| 7 | 144502815 | AX-170304120 | 0 | A | T | 0.202 | -7.33E-02 | 7.51E-02 | 2.65E+00 | 2.78E+00 | 3.23E-01 | no | Ppfia1 | intron_variant | MODIFIER |

| | | | | | | | | | | | | | | | |
|---|---|---|---|---|---|---|---|---|---|---|---|---|---|---|---|
| 7 | 144503825 | AX-97789771 | 0 | A | C | 0.202 | -7.33E-02 | 7.51E-02 | 2.65E+00 | 2.78E+00 | 3.23E-01 | no | Ppfia1 | intron_variant | MODIFIER |
| 7 | 144529901 | AX-97836550 | 0 | C | T | 0.202 | -7.33E-02 | 7.51E-02 | 2.65E+00 | 2.78E+00 | 3.23E-01 | no | Ppfia1 | intron_variant | MODIFIER |
| 7 | 144547830 | AX-170030172 | 0 | A | G | 0.202 | -7.33E-02 | 7.51E-02 | 2.65E+00 | 2.78E+00 | 3.23E-01 | no | Ppfia1 | intron_variant | MODIFIER |
| 7 | 144579845 | AX-170247195 | 0 | A | G | 0.202 | -7.33E-02 | 7.51E-02 | 2.65E+00 | 2.78E+00 | 3.23E-01 | no | Fadd | 3_prime_UTR_variant | MODIFIER |
| 7 | 144595703 | AX-170377864 | 0 | A | G | 0.202 | -7.33E-02 | 7.51E-02 | 2.65E+00 | 2.78E+00 | 3.23E-01 | no | Ano1 | intron_variant | MODIFIER |
| 7 | 144645153 | AX-170096148 | 0 | G | T | 0.202 | -7.33E-02 | 7.51E-02 | 2.65E+00 | 2.78E+00 | 3.23E-01 | no | Ano1 | intron_variant | MODIFIER |
| 7 | 144646119 | AX-170519773 | 0 | T | C | 0.202 | -7.33E-02 | 7.51E-02 | 2.65E+00 | 2.78E+00 | 3.23E-01 | no | Ano1 | intron_variant | MODIFIER |
| 7 | 144656386 | AX-97807499 | 0 | G | A | 0.202 | -7.33E-02 | 7.51E-02 | 2.65E+00 | 2.78E+00 | 3.23E-01 | no | Ano1 | intron_variant | MODIFIER |
| 7 | 144691535 | AX-170486326 | 0 | C | A | 0.202 | -7.33E-02 | 7.51E-02 | 2.65E+00 | 2.78E+00 | 3.23E-01 | no | Ano1 | intron_variant | MODIFIER |
| 7 | 144704904 | AX-170353151 | 0 | G | A | 0.202 | -7.33E-02 | 7.51E-02 | 2.65E+00 | 2.78E+00 | 3.23E-01 | no | Ano1 | intron_variant | MODIFIER |
| 7 | 144705001 | AX-170330068 | 0 | G | A | 0.202 | -7.33E-02 | 7.51E-02 | 2.65E+00 | 2.78E+00 | 3.23E-01 | no | Ano1 | intron_variant | MODIFIER |
| 7 | 144707824 | AX-170079127 | 0 | C | G | 0.202 | -7.33E-02 | 7.51E-02 | 2.65E+00 | 2.78E+00 | 3.23E-01 | no | Ano1 | intron_variant | MODIFIER |
| 7 | 144708150 | AX-170079126 | 0 | T | C | 0.202 | -7.33E-02 | 7.51E-02 | 2.65E+00 | 2.78E+00 | 3.23E-01 | no | Ano1 | intron_variant | MODIFIER |
| 7 | 144709143 | AX-170153351 | 0 | T | C | 0.202 | -7.33E-02 | 7.51E-02 | 2.65E+00 | 2.78E+00 | 3.23E-01 | no | Ano1 | intron_variant | MODIFIER |
| 7 | 144711789 | AX-170187810 | 0 | G | A | 0.202 | -7.33E-02 | 7.51E-02 | 2.65E+00 | 2.78E+00 | 3.23E-01 | no | Ano1 | intron_variant | MODIFIER |
| 7 | 144751656 | AX-170175856 | 0 | G | A | 0.202 | -7.33E-02 | 7.51E-02 | 2.65E+00 | 2.78E+00 | 3.23E-01 | no | Ano1 | intron_variant | MODIFIER |
| 7 | 144794732 | AX-170090040 | 0 | T | G | 0.202 | -7.33E-02 | 7.51E-02 | 2.65E+00 | 2.78E+00 | 3.23E-01 | no | Gm34964 | upstream_gene_variant | MODIFIER |
| 7 | 144796040 | AX-170148860 | 0 | A | G | 0.202 | -7.33E-02 | 7.51E-02 | 2.65E+00 | 2.78E+00 | 3.23E-01 | no | Gm34964 | upstream_gene_variant | MODIFIER |
| 7 | 144796070 | AX-170398059 | 0 | T | C | 0.202 | -7.33E-02 | 7.51E-02 | 2.65E+00 | 2.78E+00 | 3.23E-01 | no | Gm34964 | upstream_gene_variant | MODIFIER |
| 7 | 144796966 | AX-170294342 | 0 | C | T | 0.202 | -7.33E-02 | 7.51E-02 | 2.65E+00 | 2.78E+00 | 3.23E-01 | no | Gm9711 | downstream_gene_variant | MODIFIER |
| 7 | 144828115 | AX-97848693 | 0 | G | A | 0.202 | -7.33E-02 | 7.51E-02 | 2.65E+00 | 2.78E+00 | 3.23E-01 | no | | intergenic_variant | MODIFIER |
| 7 | 144878363 | AX-170260323 | 0 | A | G | 0.202 | -7.33E-02 | 7.51E-02 | 2.65E+00 | 2.78E+00 | 3.23E-01 | no | | intergenic_variant | MODIFIER |
| 7 | 144906992 | AX-170079133 | 0 | A | C | 0.202 | -7.33E-02 | 7.51E-02 | 2.65E+00 | 2.78E+00 | 3.23E-01 | no | Gm39119 | intron_variant,non_coding_transcript_variant | MODIFIER |
| 7 | 144907056 | AX-170079128 | 0 | C | G | 0.202 | -7.33E-02 | 7.51E-02 | 2.65E+00 | 2.78E+00 | 3.23E-01 | no | Gm39119 | intron_variant,non_coding_transcript_variant | MODIFIER |
| 7 | 144914004 | AX-170079129 | 0 | T | C | 0.202 | -7.33E-02 | 7.51E-02 | 2.65E+00 | 2.78E+00 | 3.23E-01 | no | LTO1 | upstream_gene_variant | MODIFIER |
| 7 | 144919118 | AX-170428696 | 0 | C | T | 0.202 | -7.33E-02 | 7.51E-02 | 2.65E+00 | 2.78E+00 | 3.23E-01 | no | LTO1 | intron_variant | MODIFIER |
| 7 | 144934224 | AX-170354551 | 0 | C | T | 0.202 | -7.33E-02 | 7.51E-02 | 2.65E+00 | 2.78E+00 | 3.23E-01 | no | Ccnd1 | intron_variant | MODIFIER |
| 7 | 144945983 | AX-170030174 | 0 | G | A | 0.202 | -7.33E-02 | 7.51E-02 | 2.65E+00 | 2.78E+00 | 3.23E-01 | no | | intergenic_variant | MODIFIER |
| 7 | 144947541 | AX-170530056 | 0 | A | G | 0.202 | -7.33E-02 | 7.51E-02 | 2.65E+00 | 2.78E+00 | 3.23E-01 | no | | intergenic_variant | MODIFIER |
| 7 | 144953376 | AX-97794722 | 0 | C | T | 0.202 | -7.33E-02 | 7.51E-02 | 2.65E+00 | 2.78E+00 | 3.23E-01 | no | | intergenic_variant | MODIFIER |
| 7 | 144976540 | AX-170030177 | 0 | C | G | 0.202 | -7.33E-02 | 7.51E-02 | 2.65E+00 | 2.78E+00 | 3.23E-01 | no | | intergenic_variant | MODIFIER |
| 7 | 144979597 | AX-170470113 | 0 | A | G | 0.202 | -7.33E-02 | 7.51E-02 | 2.65E+00 | 2.78E+00 | 3.23E-01 | no | | intergenic_variant | MODIFIER |
| 7 | 144991295 | AX-97807211 | 0 | T | C | 0.202 | -7.33E-02 | 7.51E-02 | 2.65E+00 | 2.78E+00 | 3.23E-01 | no | | intergenic_variant | MODIFIER |
| 7 | 145008163 | AX-170030180 | 0 | G | A | 0.202 | -7.33E-02 | 7.51E-02 | 2.65E+00 | 2.78E+00 | 3.23E-01 | no | | intergenic_variant | MODIFIER |
| 7 | 145025153 | AX-170030178 | 0 | T | C | 0.202 | -7.33E-02 | 7.51E-02 | 2.65E+00 | 2.78E+00 | 3.23E-01 | no | | regulatory_region_variant | MODIFIER |
| 7 | 145043373 | AX-97812853 | 0 | C | A | 0.202 | -7.33E-02 | 7.51E-02 | 2.65E+00 | 2.78E+00 | 3.23E-01 | no | | regulatory_region_variant | MODIFIER |
| 7 | 145050651 | AX-170030175 | 0 | G | T | 0.202 | -7.33E-02 | 7.51E-02 | 2.65E+00 | 2.78E+00 | 3.23E-01 | no | | intergenic_variant | MODIFIER |
| 7 | 145066670 | AX-170351247 | 0 | G | A | 0.202 | -7.33E-02 | 7.51E-02 | 2.65E+00 | 2.78E+00 | 3.23E-01 | no | | intergenic_variant | MODIFIER |
| 7 | 145067526 | AX-170229802 | 0 | A | G | 0.202 | -7.33E-02 | 7.51E-02 | 2.65E+00 | 2.78E+00 | 3.23E-01 | no | | intergenic_variant | MODIFIER |
| 7 | 145069272 | AX-170137526 | 0 | T | C | 0.202 | -7.33E-02 | 7.51E-02 | 2.65E+00 | 2.78E+00 | 3.23E-01 | no | | intergenic_variant | MODIFIER |
| 7 | 145092684 | AX-170125307 | 0 | C | T | 0.202 | -7.33E-02 | 7.51E-02 | 2.65E+00 | 2.78E+00 | 3.23E-01 | no | | regulatory_region_variant | MODIFIER |
| 7 | 145097887 | AX-170368987 | 0 | A | T | 0.202 | -7.33E-02 | 7.51E-02 | 2.65E+00 | 2.78E+00 | 3.23E-01 | no | | intergenic_variant | MODIFIER |
| 7 | 145109128 | AX-170388244 | 0 | A | T | 0.202 | -7.33E-02 | 7.51E-02 | 2.65E+00 | 2.78E+00 | 3.23E-01 | no | | intergenic_variant | MODIFIER |
| 7 | 145137386 | AX-170234549 | 0 | T | C | 0.202 | -7.33E-02 | 7.51E-02 | 2.65E+00 | 2.78E+00 | 3.23E-01 | no | | intergenic_variant | MODIFIER |
| 7 | 145155303 | AX-170270259 | 0 | C | T | 0.202 | -7.33E-02 | 7.51E-02 | 2.65E+00 | 2.78E+00 | 3.23E-01 | no | | intergenic_variant | MODIFIER |
| 7 | 145159101 | AX-170518799 | 0 | A | G | 0.202 | -7.33E-02 | 7.51E-02 | 2.65E+00 | 2.78E+00 | 3.23E-01 | no | Gm45181 | upstream_gene_variant | MODIFIER |
| 7 | 145161526 | AX-97842406 | 0 | G | A | 0.202 | -7.33E-02 | 7.51E-02 | 2.65E+00 | 2.78E+00 | 3.23E-01 | no | Gm45181 | upstream_gene_variant | MODIFIER |
| 7 | 145175519 | AX-170347038 | 0 | C | T | 0.202 | -7.33E-02 | 7.51E-02 | 2.65E+00 | 2.78E+00 | 3.23E-01 | no | | intergenic_variant | MODIFIER |
| 7 | 145184705 | AX-170268673 | 0 | G | C | 0.202 | -7.33E-02 | 7.51E-02 | 2.65E+00 | 2.78E+00 | 3.23E-01 | no | Tpcn2 | downstream_gene_variant | MODIFIER |
| 7 | 145186653 | AX-97833947 | 0 | C | T | 0.202 | -7.33E-02 | 7.51E-02 | 2.65E+00 | 2.78E+00 | 3.23E-01 | no | Tpcn2 | 3_prime_UTR_variant,NMD_transcript_variant | MODIFIER |
| 7 | 145201837 | AX-170528519 | 0 | T | C | 0.202 | -7.33E-02 | 7.51E-02 | 2.65E+00 | 2.78E+00 | 3.23E-01 | no | 1810010D01Rik | downstream_gene_variant | MODIFIER |
| 7 | 145216089 | AX-170259732 | 0 | A | G | 0.202 | -7.33E-02 | 7.51E-02 | 2.65E+00 | 2.78E+00 | 3.23E-01 | no | Tpcn2 | intron_variant,NMD_transcript_variant | MODIFIER |
| 7 | 145228052 | AX-170348656 | 0 | G | C | 0.202 | -7.33E-02 | 7.51E-02 | 2.65E+00 | 2.78E+00 | 3.23E-01 | no | Tpcn2 | intron_variant,NMD_transcript_variant | MODIFIER |
| 7 | 145248634 | AX-170098973 | 0 | A | G | 0.202 | -7.33E-02 | 7.51E-02 | 2.65E+00 | 2.78E+00 | 3.23E-01 | no | Tpcn2 | intron_variant,NMD_transcript_variant | MODIFIER |

| | | | | | | | | | | | | | | | |
|---|---|---|---|---|---|---|---|---|---|---|---|---|---|---|---|
| 7 | 145254764 | AX-170441690 | 0 | G | A | 0.202 | -7.33E-02 | 7.51E-02 | 2.65E+00 | 2.78E+00 | 3.23E-01 | no | Tpcn2 | intron_variant | MODIFIER |
| 7 | 145263752 | AX-170240760 | 0 | T | C | 0.202 | -7.33E-02 | 7.51E-02 | 2.65E+00 | 2.78E+00 | 3.23E-01 | no | Tpcn2 | intron_variant | MODIFIER |
| 7 | 145264731 | AX-170238368 | 0 | A | G | 0.202 | -7.33E-02 | 7.51E-02 | 2.65E+00 | 2.78E+00 | 3.23E-01 | no | Tpcn2 | intron_variant | MODIFIER |
| 7 | 145314521 | AX-170079139 | 0 | G | A | 0.202 | -7.33E-02 | 7.51E-02 | 2.65E+00 | 2.78E+00 | 3.23E-01 | no | Mrgprf | downstream_gene_variant | MODIFIER |
| 7 | 145315401 | AX-170463074 | 0 | T | C | 0.202 | -7.33E-02 | 7.51E-02 | 2.65E+00 | 2.78E+00 | 3.23E-01 | no | Mrgprd | intron_variant | MODIFIER |
| 7 | 145321555 | AX-170370977 | 0 | T | C | 0.202 | -7.33E-02 | 7.51E-02 | 2.65E+00 | 2.78E+00 | 3.23E-01 | no | Mrgprd | synonymous_variant | LOW |
| 7 | 145322194 | AX-170181179 | 0 | G | A | 0.202 | -7.33E-02 | 7.51E-02 | 2.65E+00 | 2.78E+00 | 3.23E-01 | no | Mrgprd | synonymous_variant | LOW |
| 7 | 145323643 | AX-170458605 | 0 | C | T | 0.202 | -7.33E-02 | 7.51E-02 | 2.65E+00 | 2.78E+00 | 3.23E-01 | no | Mrgprd | 3_prime_UTR_variant | MODIFIER |
| 7 | 145324205 | AX-97840630 | 0 | G | A | 0.202 | -7.33E-02 | 7.51E-02 | 2.65E+00 | 2.78E+00 | 3.23E-01 | no | Mrgprd | downstream_gene_variant | MODIFIER |
| 13 | 5957623 | AX-170238694 | 0 | C | T | 0.211 | -1.19E-01 | 1.21E-01 | 2.65E+00 | 2.72E+00 | 3.24E-01 | no | Gm46401 | intron_variant,non_coding_transcript_variant | MODIFIER |
| 5 | 81724054 | AX-170025413 | 0 | A | G | 0.439 | 1.05E-01 | 1.07E-01 | 2.74E+00 | 2.80E+00 | 3.24E-01 | no | Adgrl3 | splice_region_variant,intron_variant | LOW |
| 4 | 134078397 | AX-170450084 | 1 | T | C | 0.152 | 1.16E-01 | 1.18E-01 | 2.57E+00 | 2.62E+00 | 3.24E-01 | no | Crybg2 | intron_variant | MODIFIER |
| 4 | 97224594 | AX-170433343 | 1 | A | G | 0.223 | 1.15E-01 | 1.17E-01 | 2.57E+00 | 2.63E+00 | 3.24E-01 | no | | intergenic_variant | MODIFIER |
| 9 | 98174413 | AX-170116436 | 1 | A | G | 0.143 | -1.12E-01 | 1.14E-01 | 2.59E+00 | 2.67E+00 | 3.24E-01 | no | | intergenic_variant | MODIFIER |
| 4 | 84786005 | AX-97842212 | 1 | G | A | 0.348 | 1.10E-01 | 1.12E-01 | 2.72E+00 | 2.80E+00 | 3.24E-01 | no | | intergenic_variant | MODIFIER |
| 10 | 17592787 | AX-170164782 | 0 | C | T | 0.211 | 1.02E-01 | 1.02E-01 | 3.04E+00 | 3.15E+00 | 3.24E-01 | no | Gm47771 | intron_variant,non_coding_transcript_variant | MODIFIER |
| 10 | 17643207 | AX-97825107 | 0 | T | C | 0.211 | 1.02E-01 | 1.02E-01 | 3.04E+00 | 3.15E+00 | 3.24E-01 | no | Gm47771 | intron_variant,non_coding_transcript_variant | MODIFIER |
| 10 | 17739339 | AX-97857119 | 0 | C | T | 0.211 | 1.02E-01 | 1.02E-01 | 3.04E+00 | 3.15E+00 | 3.24E-01 | no | | intergenic_variant | MODIFIER |
| 10 | 17758312 | AX-170002839 | 0 | G | A | 0.211 | 1.02E-01 | 1.02E-01 | 3.04E+00 | 3.15E+00 | 3.24E-01 | no | | intergenic_variant | MODIFIER |
| 10 | 17769305 | AX-97841740 | 0 | A | G | 0.211 | 1.02E-01 | 1.02E-01 | 3.04E+00 | 3.15E+00 | 3.24E-01 | no | | intergenic_variant | MODIFIER |
| 10 | 100870952 | AX-97789444 | 0 | C | A | 0.289 | -8.37E-02 | 8.50E-02 | 2.56E+00 | 2.62E+00 | 3.24E-01 | no | | intergenic_variant | MODIFIER |
| 9 | 94395286 | AX-97838712 | 1 | C | T | 0.143 | -1.20E-01 | 1.23E-01 | 2.68E+00 | 2.76E+00 | 3.24E-01 | no | | regulatory_region_variant | MODIFIER |
| 15 | 20100481 | AX-170415522 | 1 | G | A | 0.241 | -1.00E-01 | 1.02E-01 | 2.61E+00 | 2.67E+00 | 3.24E-01 | no | | intergenic_variant | MODIFIER |
| 5 | 104862575 | AX-97780555 | 0 | C | T | 0.456 | 1.09E-01 | 1.12E-01 | 2.71E+00 | 2.76E+00 | 3.24E-01 | no | Zfp951 | upstream_gene_variant | MODIFIER |
| 5 | 114252982 | AX-170385522 | 0 | G | A | 0.456 | 1.09E-01 | 1.12E-01 | 2.71E+00 | 2.76E+00 | 3.24E-01 | no | Acacb | downstream_gene_variant | MODIFIER |
| 5 | 114722598 | AX-170174212 | 0 | C | T | 0.456 | 1.09E-01 | 1.12E-01 | 2.71E+00 | 2.76E+00 | 3.24E-01 | no | Git2 | downstream_gene_variant | MODIFIER |
| 15 | 6171181 | AX-170054711 | 0 | C | T | 0.246 | -1.02E-01 | 1.04E-01 | 2.66E+00 | 2.73E+00 | 3.24E-01 | no | | intergenic_variant | MODIFIER |
| 3 | 151609164 | AX-170488017 | 1 | G | A | 0.339 | 7.20E-02 | 7.38E-02 | 2.81E+00 | 2.90E+00 | 3.24E-01 | no | Gm24458 | downstream_gene_variant | MODIFIER |
| 4 | 49532394 | AX-170162367 | 0 | A | G | 0.351 | -9.36E-02 | 9.58E-02 | 2.85E+00 | 2.95E+00 | 3.24E-01 | no | Aldob | downstream_gene_variant | MODIFIER |
| 15 | 10919775 | AX-170501814 | 0 | C | T | 0.246 | -1.02E-01 | 1.04E-01 | 2.65E+00 | 2.71E+00 | 3.25E-01 | no | Gm29742 | downstream_gene_variant | MODIFIER |
| 16 | 72218831 | AX-170166265 | 1 | A | T | 0.473 | 7.62E-02 | 7.79E-02 | 2.61E+00 | 2.69E+00 | 3.25E-01 | no | | intergenic_variant | MODIFIER |
| 18 | 72866489 | AX-170261596 | 0 | G | A | 0.175 | -1.21E-01 | 1.24E-01 | 2.63E+00 | 2.67E+00 | 3.25E-01 | no | | intergenic_variant | MODIFIER |
| 18 | 72891475 | AX-97835075 | 0 | T | G | 0.175 | -1.21E-01 | 1.24E-01 | 2.63E+00 | 2.67E+00 | 3.25E-01 | no | | intergenic_variant | MODIFIER |
| 18 | 72961843 | AX-170187815 | 0 | A | G | 0.175 | -1.21E-01 | 1.24E-01 | 2.63E+00 | 2.67E+00 | 3.25E-01 | no | | intergenic_variant | MODIFIER |
| 18 | 72966170 | AX-170118565 | 0 | G | A | 0.175 | -1.21E-01 | 1.24E-01 | 2.63E+00 | 2.67E+00 | 3.25E-01 | no | | intergenic_variant | MODIFIER |
| 18 | 72966796 | AX-97857233 | 0 | C | T | 0.175 | -1.21E-01 | 1.24E-01 | 2.63E+00 | 2.67E+00 | 3.25E-01 | no | | intergenic_variant | MODIFIER |
| 18 | 72985373 | AX-170329458 | 0 | G | A | 0.175 | -1.21E-01 | 1.24E-01 | 2.63E+00 | 2.67E+00 | 3.25E-01 | no | | intergenic_variant | MODIFIER |
| 18 | 72985425 | AX-170272467 | 0 | G | A | 0.175 | -1.21E-01 | 1.24E-01 | 2.63E+00 | 2.67E+00 | 3.25E-01 | no | | intergenic_variant | MODIFIER |
| 18 | 72985521 | AX-170341175 | 0 | A | G | 0.175 | -1.21E-01 | 1.24E-01 | 2.63E+00 | 2.67E+00 | 3.25E-01 | no | | intergenic_variant | MODIFIER |
| 18 | 72985750 | AX-170408703 | 0 | C | T | 0.175 | -1.21E-01 | 1.24E-01 | 2.63E+00 | 2.67E+00 | 3.25E-01 | no | | intergenic_variant | MODIFIER |
| 18 | 72986154 | AX-170461413 | 0 | A | G | 0.175 | -1.21E-01 | 1.24E-01 | 2.63E+00 | 2.67E+00 | 3.25E-01 | no | | intergenic_variant | MODIFIER |
| 18 | 72986344 | AX-170124026 | 0 | A | G | 0.175 | -1.21E-01 | 1.24E-01 | 2.63E+00 | 2.67E+00 | 3.25E-01 | no | | intergenic_variant | MODIFIER |
| 18 | 73034844 | AX-170015969 | 0 | T | G | 0.175 | -1.21E-01 | 1.24E-01 | 2.63E+00 | 2.67E+00 | 3.25E-01 | no | | intergenic_variant | MODIFIER |
| 18 | 73061527 | AX-97835074 | 0 | C | T | 0.175 | -1.21E-01 | 1.24E-01 | 2.63E+00 | 2.67E+00 | 3.25E-01 | no | | intergenic_variant | MODIFIER |
| 18 | 73656308 | AX-170015985 | 0 | C | T | 0.175 | -1.21E-01 | 1.24E-01 | 2.63E+00 | 2.67E+00 | 3.25E-01 | no | Smad4 | intron_variant | MODIFIER |
| 18 | 73662923 | AX-170015976 | 0 | G | A | 0.175 | -1.21E-01 | 1.24E-01 | 2.63E+00 | 2.67E+00 | 3.25E-01 | no | Smad4 | intron_variant | MODIFIER |
| 18 | 73667090 | AX-97831779 | 0 | T | C | 0.175 | -1.21E-01 | 1.24E-01 | 2.63E+00 | 2.67E+00 | 3.25E-01 | no | Smad4 | intron_variant | MODIFIER |
| 18 | 73752885 | AX-97835844 | 0 | G | A | 0.175 | -1.21E-01 | 1.24E-01 | 2.63E+00 | 2.67E+00 | 3.25E-01 | no | Elac1 | intron_variant | MODIFIER |
| 18 | 73795066 | AX-170316877 | 0 | A | G | 0.175 | -1.21E-01 | 1.24E-01 | 2.63E+00 | 2.67E+00 | 3.25E-01 | no | Me2 | intron_variant | MODIFIER |
| 18 | 73824531 | AX-170453805 | 0 | A | G | 0.175 | -1.21E-01 | 1.24E-01 | 2.63E+00 | 2.67E+00 | 3.25E-01 | no | | intergenic_variant | MODIFIER |
| 18 | 73851007 | AX-170191954 | 0 | A | C | 0.175 | -1.21E-01 | 1.24E-01 | 2.63E+00 | 2.67E+00 | 3.25E-01 | no | | intergenic_variant | MODIFIER |
| 18 | 73862594 | AX-170061311 | 0 | C | T | 0.175 | -1.21E-01 | 1.24E-01 | 2.63E+00 | 2.67E+00 | 3.25E-01 | no | Mro | intron_variant | MODIFIER |
| 18 | 73898397 | AX-97840338 | 0 | T | C | 0.175 | -1.21E-01 | 1.24E-01 | 2.63E+00 | 2.67E+00 | 3.25E-01 | no | | regulatory_region_variant | MODIFIER |
| 18 | 73909755 | AX-170183713 | 0 | A | G | 0.175 | -1.21E-01 | 1.24E-01 | 2.63E+00 | 2.67E+00 | 3.25E-01 | no | | intergenic_variant | MODIFIER |

| | | | | | | | | | | | | | | | |
|---|---|---|---|---|---|---|---|---|---|---|---|---|---|---|---|
| 18 | 73910929 | AX-170233065 | 0 | G | A | 0.175 | -1.21E-01 | 1.24E-01 | 2.63E+00 | 2.67E+00 | 3.25E-01 | no | | intergenic_variant | MODIFIER |
| 18 | 73911525 | AX-170262927 | 0 | C | A | 0.175 | -1.21E-01 | 1.24E-01 | 2.63E+00 | 2.67E+00 | 3.25E-01 | no | | intergenic_variant | MODIFIER |
| 18 | 73911809 | AX-170398399 | 0 | T | C | 0.175 | -1.21E-01 | 1.24E-01 | 2.63E+00 | 2.67E+00 | 3.25E-01 | no | | intergenic_variant | MODIFIER |
| 18 | 73912802 | AX-170518505 | 0 | T | C | 0.175 | -1.21E-01 | 1.24E-01 | 2.63E+00 | 2.67E+00 | 3.25E-01 | no | | intergenic_variant | MODIFIER |
| 18 | 73912878 | AX-170208555 | 0 | G | C | 0.175 | -1.21E-01 | 1.24E-01 | 2.63E+00 | 2.67E+00 | 3.25E-01 | no | | intergenic_variant | MODIFIER |
| 18 | 73913515 | AX-170153291 | 0 | A | C | 0.175 | -1.21E-01 | 1.24E-01 | 2.63E+00 | 2.67E+00 | 3.25E-01 | no | | intergenic_variant | MODIFIER |
| 18 | 73922723 | AX-170328958 | 0 | T | C | 0.175 | -1.21E-01 | 1.24E-01 | 2.63E+00 | 2.67E+00 | 3.25E-01 | no | | intergenic_variant | MODIFIER |
| 18 | 73922774 | AX-170093949 | 0 | T | C | 0.175 | -1.21E-01 | 1.24E-01 | 2.63E+00 | 2.67E+00 | 3.25E-01 | no | | intergenic_variant | MODIFIER |
| 18 | 74010820 | AX-170294575 | 0 | G | C | 0.175 | -1.21E-01 | 1.24E-01 | 2.63E+00 | 2.67E+00 | 3.25E-01 | no | Mapk4 | intron_variant | MODIFIER |
| 18 | 74010876 | AX-97831783 | 0 | G | A | 0.175 | -1.21E-01 | 1.24E-01 | 2.63E+00 | 2.67E+00 | 3.25E-01 | no | Mapk4 | intron_variant | MODIFIER |
| 18 | 74021079 | AX-170298223 | 0 | G | A | 0.175 | -1.21E-01 | 1.24E-01 | 2.63E+00 | 2.67E+00 | 3.25E-01 | no | Mapk4 | intron_variant | MODIFIER |
| 18 | 74023623 | AX-170136481 | 0 | G | A | 0.175 | -1.21E-01 | 1.24E-01 | 2.63E+00 | 2.67E+00 | 3.25E-01 | no | Mapk4 | intron_variant | MODIFIER |
| 18 | 74030063 | AX-170266888 | 0 | C | T | 0.175 | -1.21E-01 | 1.24E-01 | 2.63E+00 | 2.67E+00 | 3.25E-01 | no | Mapk4 | intron_variant | MODIFIER |
| 18 | 74135648 | AX-97831778 | 0 | A | G | 0.175 | -1.21E-01 | 1.24E-01 | 2.63E+00 | 2.67E+00 | 3.25E-01 | no | | intergenic_variant | MODIFIER |
| 18 | 74147901 | AX-170015989 | 0 | G | C | 0.175 | -1.21E-01 | 1.24E-01 | 2.63E+00 | 2.67E+00 | 3.25E-01 | no | | intergenic_variant | MODIFIER |
| 18 | 74168284 | AX-97824423 | 0 | C | T | 0.175 | -1.21E-01 | 1.24E-01 | 2.63E+00 | 2.67E+00 | 3.25E-01 | no | | intergenic_variant | MODIFIER |
| 18 | 74201892 | AX-170191219 | 0 | T | C | 0.175 | -1.21E-01 | 1.24E-01 | 2.63E+00 | 2.67E+00 | 3.25E-01 | no | Ska1 | intron_variant | MODIFIER |
| 18 | 74203465 | AX-170282604 | 0 | G | C | 0.175 | -1.21E-01 | 1.24E-01 | 2.63E+00 | 2.67E+00 | 3.25E-01 | no | Ska1 | intron_variant | MODIFIER |
| 18 | 74208368 | AX-170125853 | 0 | T | C | 0.175 | -1.21E-01 | 1.24E-01 | 2.63E+00 | 2.67E+00 | 3.25E-01 | no | Ska1 | upstream_gene_variant | MODIFIER |
| 18 | 74209054 | AX-170485838 | 0 | A | C | 0.175 | -1.21E-01 | 1.24E-01 | 2.63E+00 | 2.67E+00 | 3.25E-01 | no | Ska1 | upstream_gene_variant | MODIFIER |
| 18 | 74214257 | AX-170015992 | 0 | G | C | 0.175 | -1.21E-01 | 1.24E-01 | 2.63E+00 | 2.67E+00 | 3.25E-01 | no | Cxxc1 | upstream_gene_variant | MODIFIER |
| 18 | 74215068 | AX-97804294 | 0 | T | C | 0.175 | -1.21E-01 | 1.24E-01 | 2.63E+00 | 2.67E+00 | 3.25E-01 | no | Cxxc1 | upstream_gene_variant | MODIFIER |
| 18 | 74246207 | AX-97835871 | 0 | C | T | 0.175 | -1.21E-01 | 1.24E-01 | 2.63E+00 | 2.67E+00 | 3.25E-01 | no | AC133186.1 | downstream_gene_variant | MODIFIER |
| 18 | 74302796 | AX-97842567 | 0 | G | A | 0.175 | -1.21E-01 | 1.24E-01 | 2.63E+00 | 2.67E+00 | 3.25E-01 | no | Cfap53 | intron_variant | MODIFIER |
| 18 | 74333038 | AX-170088374 | 0 | C | T | 0.175 | -1.21E-01 | 1.24E-01 | 2.63E+00 | 2.67E+00 | 3.25E-01 | no | Cfap53 | intron_variant | MODIFIER |
| 18 | 74347536 | AX-97835091 | 0 | T | C | 0.175 | -1.21E-01 | 1.24E-01 | 2.63E+00 | 2.67E+00 | 3.25E-01 | no | Cfap53 | intron_variant | MODIFIER |
| 18 | 74374964 | AX-170427261 | 0 | A | G | 0.175 | -1.21E-01 | 1.24E-01 | 2.63E+00 | 2.67E+00 | 3.25E-01 | no | | intergenic_variant | MODIFIER |
| 18 | 74387100 | AX-170132450 | 0 | G | A | 0.175 | -1.21E-01 | 1.24E-01 | 2.63E+00 | 2.67E+00 | 3.25E-01 | no | | intergenic_variant | MODIFIER |
| 18 | 74423545 | AX-170226781 | 0 | A | G | 0.175 | -1.21E-01 | 1.24E-01 | 2.63E+00 | 2.67E+00 | 3.25E-01 | no | AC147992.2 | downstream_gene_variant | MODIFIER |
| 18 | 74487158 | AX-170061324 | 0 | C | T | 0.175 | -1.21E-01 | 1.24E-01 | 2.63E+00 | 2.67E+00 | 3.25E-01 | no | Myo5b | intron_variant | MODIFIER |
| 18 | 74487259 | AX-170355717 | 0 | C | T | 0.175 | -1.21E-01 | 1.24E-01 | 2.63E+00 | 2.67E+00 | 3.25E-01 | no | Myo5b | intron_variant | MODIFIER |
| 18 | 74487338 | AX-170250101 | 0 | T | C | 0.175 | -1.21E-01 | 1.24E-01 | 2.63E+00 | 2.67E+00 | 3.25E-01 | no | Myo5b | intron_variant | MODIFIER |
| 18 | 74487400 | AX-170483725 | 0 | A | G | 0.175 | -1.21E-01 | 1.24E-01 | 2.63E+00 | 2.67E+00 | 3.25E-01 | no | Myo5b | intron_variant | MODIFIER |
| 18 | 74492266 | AX-170197141 | 0 | A | G | 0.175 | -1.21E-01 | 1.24E-01 | 2.63E+00 | 2.67E+00 | 3.25E-01 | no | Myo5b | intron_variant | MODIFIER |
| 18 | 74499542 | AX-170417917 | 0 | G | C | 0.175 | -1.21E-01 | 1.24E-01 | 2.63E+00 | 2.67E+00 | 3.25E-01 | no | Myo5b | intron_variant | MODIFIER |
| 18 | 74526079 | AX-170423412 | 0 | A | G | 0.175 | -1.21E-01 | 1.24E-01 | 2.63E+00 | 2.67E+00 | 3.25E-01 | no | Myo5b | intron_variant | MODIFIER |
| 18 | 74536408 | AX-170387121 | 0 | A | G | 0.175 | -1.21E-01 | 1.24E-01 | 2.63E+00 | 2.67E+00 | 3.25E-01 | no | Myo5b | intron_variant | MODIFIER |
| 18 | 74560330 | AX-170167543 | 0 | T | A | 0.175 | -1.21E-01 | 1.24E-01 | 2.63E+00 | 2.67E+00 | 3.25E-01 | no | Myo5b | intron_variant | MODIFIER |
| 18 | 74591143 | AX-170441372 | 0 | T | C | 0.175 | -1.21E-01 | 1.24E-01 | 2.63E+00 | 2.67E+00 | 3.25E-01 | no | Myo5b | intron_variant | MODIFIER |
| 18 | 74655910 | AX-170113855 | 0 | A | C | 0.175 | -1.21E-01 | 1.24E-01 | 2.63E+00 | 2.67E+00 | 3.25E-01 | no | Myo5b | intron_variant | MODIFIER |
| 18 | 74655955 | AX-170233728 | 0 | A | G | 0.175 | -1.21E-01 | 1.24E-01 | 2.63E+00 | 2.67E+00 | 3.25E-01 | no | Myo5b | intron_variant | MODIFIER |
| 18 | 74661834 | AX-170373871 | 0 | C | T | 0.175 | -1.21E-01 | 1.24E-01 | 2.63E+00 | 2.67E+00 | 3.25E-01 | no | Myo5b | intron_variant | MODIFIER |
| 18 | 74663231 | AX-170015998 | 0 | T | C | 0.175 | -1.21E-01 | 1.24E-01 | 2.63E+00 | 2.67E+00 | 3.25E-01 | no | Myo5b | intron_variant | MODIFIER |
| 18 | 74664463 | AX-170371763 | 0 | C | G | 0.175 | -1.21E-01 | 1.24E-01 | 2.63E+00 | 2.67E+00 | 3.25E-01 | no | Myo5b | intron_variant | MODIFIER |
| 18 | 74820097 | AX-170340664 | 0 | G | C | 0.175 | -1.21E-01 | 1.24E-01 | 2.63E+00 | 2.67E+00 | 3.25E-01 | no | | intergenic_variant | MODIFIER |
| 18 | 74823039 | AX-170243546 | 0 | C | T | 0.175 | -1.21E-01 | 1.24E-01 | 2.63E+00 | 2.67E+00 | 3.25E-01 | no | | regulatory_region_variant | MODIFIER |
| 18 | 74850773 | AX-170369473 | 0 | T | C | 0.175 | -1.21E-01 | 1.24E-01 | 2.63E+00 | 2.67E+00 | 3.25E-01 | no | | regulatory_region_variant | MODIFIER |
| 18 | 74861095 | AX-170417719 | 0 | C | T | 0.175 | -1.21E-01 | 1.24E-01 | 2.63E+00 | 2.67E+00 | 3.25E-01 | no | | intergenic_variant | MODIFIER |
| 18 | 74866034 | AX-170462995 | 0 | G | A | 0.175 | -1.21E-01 | 1.24E-01 | 2.63E+00 | 2.67E+00 | 3.25E-01 | no | | intergenic_variant | MODIFIER |
| 18 | 74875248 | AX-170512588 | 0 | C | G | 0.175 | -1.21E-01 | 1.24E-01 | 2.63E+00 | 2.67E+00 | 3.25E-01 | no | | intergenic_variant | MODIFIER |
| 18 | 74878171 | AX-170340258 | 0 | T | C | 0.175 | -1.21E-01 | 1.24E-01 | 2.63E+00 | 2.67E+00 | 3.25E-01 | no | | intergenic_variant | MODIFIER |
| 18 | 74971328 | AX-97842891 | 0 | C | T | 0.175 | -1.21E-01 | 1.24E-01 | 2.63E+00 | 2.67E+00 | 3.25E-01 | no | 9030625G05Rik | downstream_gene_variant | MODIFIER |
| 18 | 74982974 | AX-170061336 | 0 | T | C | 0.175 | -1.21E-01 | 1.24E-01 | 2.63E+00 | 2.67E+00 | 3.25E-01 | no | 9030625G05Rik | upstream_gene_variant | MODIFIER |
| 18 | 75001858 | AX-170496776 | 0 | T | C | 0.175 | -1.21E-01 | 1.24E-01 | 2.63E+00 | 2.67E+00 | 3.25E-01 | no | BC031181 | upstream_gene_variant | MODIFIER |

| | | | | | | | | | | | | | | | |
|---|---|---|---|---|---|---|---|---|---|---|---|---|---|---|---|
| 18 | 75001932 | AX-170503292 | 0 | A | T | 0.175 | -1.21E-01 | 1.24E-01 | 2.63E+00 | 2.67E+00 | 3.25E-01 | no | BC031181 | upstream_gene_variant | MODIFIER |
| 18 | 75007619 | AX-97852433 | 0 | A | G | 0.175 | -1.21E-01 | 1.24E-01 | 2.63E+00 | 2.67E+00 | 3.25E-01 | no | BC031181 | intron_variant | MODIFIER |
| 18 | 75015638 | AX-170136093 | 0 | C | T | 0.175 | -1.21E-01 | 1.24E-01 | 2.63E+00 | 2.67E+00 | 3.25E-01 | no | Dym | upstream_gene_variant | MODIFIER |
| 18 | 75042879 | AX-170247870 | 0 | C | T | 0.175 | -1.21E-01 | 1.24E-01 | 2.63E+00 | 2.67E+00 | 3.25E-01 | no | Dym | intron_variant | MODIFIER |
| 18 | 75053778 | AX-170447980 | 0 | C | G | 0.175 | -1.21E-01 | 1.24E-01 | 2.63E+00 | 2.67E+00 | 3.25E-01 | no | Dym | intron_variant | MODIFIER |
| 18 | 75053980 | AX-170192794 | 0 | T | G | 0.175 | -1.21E-01 | 1.24E-01 | 2.63E+00 | 2.67E+00 | 3.25E-01 | no | Dym | intron_variant | MODIFIER |
| 18 | 75054041 | AX-170292069 | 0 | T | C | 0.175 | -1.21E-01 | 1.24E-01 | 2.63E+00 | 2.67E+00 | 3.25E-01 | no | Dym | intron_variant | MODIFIER |
| 18 | 75127530 | AX-170355790 | 0 | G | A | 0.175 | -1.21E-01 | 1.24E-01 | 2.63E+00 | 2.67E+00 | 3.25E-01 | no | Dym | intron_variant | MODIFIER |
| 18 | 75189211 | AX-170513550 | 0 | A | G | 0.175 | -1.21E-01 | 1.24E-01 | 2.63E+00 | 2.67E+00 | 3.25E-01 | no | Dym | intron_variant | MODIFIER |
| 18 | 75196077 | AX-170016004 | 0 | C | A | 0.175 | -1.21E-01 | 1.24E-01 | 2.63E+00 | 2.67E+00 | 3.25E-01 | no | Dym | intron_variant | MODIFIER |
| 18 | 75208062 | AX-170093954 | 0 | C | T | 0.175 | -1.21E-01 | 1.24E-01 | 2.63E+00 | 2.67E+00 | 3.25E-01 | no | Dym | intron_variant | MODIFIER |
| 18 | 75231133 | AX-170016005 | 0 | G | A | 0.175 | -1.21E-01 | 1.24E-01 | 2.63E+00 | 2.67E+00 | 3.25E-01 | no | Dym | intron_variant | MODIFIER |
| 18 | 75259621 | AX-170016003 | 0 | C | T | 0.175 | -1.21E-01 | 1.24E-01 | 2.63E+00 | 2.67E+00 | 3.25E-01 | no | Dym | intron_variant | MODIFIER |
| 18 | 75274453 | AX-170016008 | 0 | G | C | 0.175 | -1.21E-01 | 1.24E-01 | 2.63E+00 | 2.67E+00 | 3.25E-01 | no | Dym | intron_variant | MODIFIER |
| 18 | 75276253 | AX-170492644 | 0 | G | A | 0.175 | -1.21E-01 | 1.24E-01 | 2.63E+00 | 2.67E+00 | 3.25E-01 | no | Dym | intron_variant | MODIFIER |
| 18 | 75280384 | AX-170325857 | 0 | C | T | 0.175 | -1.21E-01 | 1.24E-01 | 2.63E+00 | 2.67E+00 | 3.25E-01 | no | Dym | intron_variant | MODIFIER |
| 18 | 75281334 | AX-97827749 | 0 | A | G | 0.175 | -1.21E-01 | 1.24E-01 | 2.63E+00 | 2.67E+00 | 3.25E-01 | no | Dym | intron_variant | MODIFIER |
| 8 | 47174982 | AX-170079989 | 0 | T | C | 0.149 | -1.16E-01 | 1.19E-01 | 2.68E+00 | 2.79E+00 | 3.25E-01 | no | | intergenic_variant | MODIFIER |
| 7 | 70848905 | AX-170159539 | 0 | T | G | 0.377 | -9.51E-02 | 9.78E-02 | 2.76E+00 | 2.86E+00 | 3.25E-01 | no | | intergenic_variant | MODIFIER |
| 8 | 61408442 | AX-170109808 | 0 | T | A | 0.158 | -1.15E-01 | 1.16E-01 | 2.55E+00 | 2.64E+00 | 3.25E-01 | no | | intergenic_variant | MODIFIER |
| 16 | 80335128 | AX-170114489 | 0 | T | C | 0.447 | 7.27E-02 | 7.42E-02 | 2.59E+00 | 2.67E+00 | 3.25E-01 | no | | intergenic_variant | MODIFIER |
| 16 | 29259646 | AX-170500035 | 0 | A | G | 0.5 | 7.86E-02 | 7.93E-02 | 2.51E+00 | 2.56E+00 | 3.25E-01 | no | Atp13a5 | intron_variant | MODIFIER |
| 5 | 115941282 | AX-170187999 | 1 | A | T | 0.455 | 1.02E-01 | 1.05E-01 | 2.74E+00 | 2.82E+00 | 3.25E-01 | no | Cit | intron_variant | MODIFIER |
| 1 | 143275595 | AX-170001881 | 1 | G | A | 0.214 | 1.34E-01 | 1.37E-01 | 2.74E+00 | 2.79E+00 | 3.25E-01 | no | | intergenic_variant | MODIFIER |
| 10 | 17600274 | AX-97814878 | 0 | A | G | 0.202 | 1.01E-01 | 1.02E-01 | 3.02E+00 | 3.13E+00 | 3.25E-01 | no | Gm47771 | intron_variant,non_coding_transcript_variant | MODIFIER |
| 10 | 17623867 | AX-170143508 | 0 | T | C | 0.202 | 1.01E-01 | 1.02E-01 | 3.02E+00 | 3.13E+00 | 3.25E-01 | no | Gm47771 | intron_variant,non_coding_transcript_variant | MODIFIER |
| 10 | 17625114 | AX-97814000 | 0 | T | C | 0.202 | 1.01E-01 | 1.02E-01 | 3.02E+00 | 3.13E+00 | 3.25E-01 | no | Gm47771 | intron_variant,non_coding_transcript_variant | MODIFIER |
| 10 | 17628488 | AX-170436241 | 0 | T | G | 0.202 | 1.01E-01 | 1.02E-01 | 3.02E+00 | 3.13E+00 | 3.25E-01 | no | Gm47771 | intron_variant,non_coding_transcript_variant | MODIFIER |
| 10 | 17628603 | AX-170478452 | 0 | G | C | 0.202 | 1.01E-01 | 1.02E-01 | 3.02E+00 | 3.13E+00 | 3.25E-01 | no | Gm47771 | intron_variant,non_coding_transcript_variant | MODIFIER |
| 10 | 17732585 | AX-170002842 | 0 | G | A | 0.202 | 1.01E-01 | 1.02E-01 | 3.02E+00 | 3.13E+00 | 3.25E-01 | no | | regulatory_region_variant | MODIFIER |
| 10 | 17752489 | AX-170158707 | 0 | T | C | 0.202 | 1.01E-01 | 1.02E-01 | 3.02E+00 | 3.13E+00 | 3.25E-01 | no | | intergenic_variant | MODIFIER |
| 10 | 17785764 | AX-170463379 | 0 | T | C | 0.202 | 1.01E-01 | 1.02E-01 | 3.02E+00 | 3.13E+00 | 3.25E-01 | no | Gm20655 | downstream_gene_variant | MODIFIER |
| 10 | 17786786 | AX-170002844 | 0 | G | C | 0.202 | 1.01E-01 | 1.02E-01 | 3.02E+00 | 3.13E+00 | 3.25E-01 | no | Gm20655 | downstream_gene_variant | MODIFIER |
| 10 | 17790979 | AX-170002843 | 0 | C | T | 0.202 | 1.01E-01 | 1.02E-01 | 3.02E+00 | 3.13E+00 | 3.25E-01 | no | Gm20655 | downstream_gene_variant | MODIFIER |
| 10 | 17852476 | AX-97815016 | 0 | T | G | 0.202 | 1.01E-01 | 1.02E-01 | 3.02E+00 | 3.13E+00 | 3.25E-01 | no | | intergenic_variant | MODIFIER |
| 10 | 17903519 | AX-97795395 | 0 | A | G | 0.202 | 1.01E-01 | 1.02E-01 | 3.02E+00 | 3.13E+00 | 3.25E-01 | no | Heca | intron_variant | MODIFIER |
| 10 | 17919675 | AX-170310010 | 0 | T | A | 0.202 | 1.01E-01 | 1.02E-01 | 3.02E+00 | 3.13E+00 | 3.25E-01 | no | Heca | intron_variant | MODIFIER |
| 10 | 17920709 | AX-170457880 | 0 | G | C | 0.202 | 1.01E-01 | 1.02E-01 | 3.02E+00 | 3.13E+00 | 3.25E-01 | no | Heca | intron_variant | MODIFIER |
| 10 | 17930520 | AX-97793658 | 0 | A | G | 0.202 | 1.01E-01 | 1.02E-01 | 3.02E+00 | 3.13E+00 | 3.25E-01 | no | Heca | intron_variant | MODIFIER |
| 4 | 120800108 | AX-170427272 | 1 | A | T | 0.232 | 1.03E-01 | 1.05E-01 | 2.60E+00 | 2.67E+00 | 3.26E-01 | no | Nfyc | intron_variant | MODIFIER |
| 5 | 43551797 | AX-170404507 | 0 | A | C | 0.184 | -9.81E-02 | 1.00E-01 | 2.60E+00 | 2.69E+00 | 3.26E-01 | no | C1qtnf7 | intron_variant | MODIFIER |
| 16 | 85356203 | AX-170290779 | 0 | A | G | 0.447 | 7.64E-02 | 7.85E-02 | 2.70E+00 | 2.78E+00 | 3.26E-01 | no | | intergenic_variant | MODIFIER |
| 16 | 85372608 | AX-170378927 | 0 | G | C | 0.447 | 7.64E-02 | 7.85E-02 | 2.70E+00 | 2.78E+00 | 3.26E-01 | no | | intergenic_variant | MODIFIER |
| 1 | 106600325 | AX-170110880 | 0 | A | G | 0.167 | 1.54E-01 | 1.56E-01 | 2.53E+00 | 2.64E+00 | 3.26E-01 | Yes | Bcl2 | intron_variant | MODIFIER |
| 1 | 113980617 | AX-170001429 | 0 | C | A | 0.167 | 1.54E-01 | 1.56E-01 | 2.53E+00 | 2.64E+00 | 3.26E-01 | Yes | | intergenic_variant | MODIFIER |
| 9 | 61196230 | AX-97847731 | 1 | C | T | 0.205 | 1.14E-01 | 1.17E-01 | 2.85E+00 | 2.94E+00 | 3.26E-01 | no | | intergenic_variant | MODIFIER |
| 9 | 92679979 | AX-97823049 | 1 | T | C | 0.143 | -1.17E-01 | 1.20E-01 | 2.64E+00 | 2.72E+00 | 3.26E-01 | no | | intergenic_variant | MODIFIER |
| 6 | 135137729 | AX-170528592 | 0 | C | T | 0.579 | 8.61E-02 | 8.63E-02 | 2.46E+00 | 2.52E+00 | 3.26E-01 | no | Hebp1 | 3_prime_UTR_variant | MODIFIER |
| 10 | 18249627 | AX-170035477 | 0 | T | C | 0.184 | 1.03E-01 | 1.04E-01 | 3.00E+00 | 3.11E+00 | 3.26E-01 | no | Gm10827 | downstream_gene_variant | MODIFIER |
| 10 | 26430722 | AX-170268690 | 0 | T | C | 0.184 | 1.03E-01 | 1.04E-01 | 3.00E+00 | 3.11E+00 | 3.26E-01 | no | | intergenic_variant | MODIFIER |
| 10 | 26445712 | AX-170002942 | 0 | A | G | 0.184 | 1.03E-01 | 1.04E-01 | 3.00E+00 | 3.11E+00 | 3.26E-01 | no | | intergenic_variant | MODIFIER |
| 10 | 26458139 | AX-97845010 | 0 | C | T | 0.184 | 1.03E-01 | 1.04E-01 | 3.00E+00 | 3.11E+00 | 3.26E-01 | no | | intergenic_variant | MODIFIER |
| 10 | 26477163 | AX-170254912 | 0 | C | T | 0.184 | 1.03E-01 | 1.04E-01 | 3.00E+00 | 3.11E+00 | 3.26E-01 | no | | intergenic_variant | MODIFIER |
| 10 | 26487161 | AX-170152122 | 0 | T | C | 0.184 | 1.03E-01 | 1.04E-01 | 3.00E+00 | 3.11E+00 | 3.26E-01 | no | | intergenic_variant | MODIFIER |

| | | | | | | | | | | | | | | | |
|---|---|---|---|---|---|---|---|---|---|---|---|---|---|---|---|
| 10 | 26497517 | AX-170116847 | 0 | C | G | 0.184 | 1.03E-01 | 1.04E-01 | 3.00E+00 | 3.11E+00 | 3.26E-01 | no | Gm30034 | intron_variant,non_coding_transcript_variant | MODIFIER |
| 10 | 26497648 | AX-97839771 | 0 | A | G | 0.184 | 1.03E-01 | 1.04E-01 | 3.00E+00 | 3.11E+00 | 3.26E-01 | no | Gm30034 | intron_variant,non_coding_transcript_variant | MODIFIER |
| 10 | 26499980 | AX-170141055 | 0 | A | G | 0.184 | 1.03E-01 | 1.04E-01 | 3.00E+00 | 3.11E+00 | 3.26E-01 | no | Gm30034 | downstream_gene_variant | MODIFIER |
| 10 | 26510603 | AX-170496179 | 0 | C | A | 0.184 | 1.03E-01 | 1.04E-01 | 3.00E+00 | 3.11E+00 | 3.26E-01 | no | | intergenic_variant | MODIFIER |
| 10 | 26539661 | AX-97850672 | 0 | G | T | 0.184 | 1.03E-01 | 1.04E-01 | 3.00E+00 | 3.11E+00 | 3.26E-01 | no | | intergenic_variant | MODIFIER |
| 10 | 26558060 | AX-170318628 | 0 | T | G | 0.184 | 1.03E-01 | 1.04E-01 | 3.00E+00 | 3.11E+00 | 3.26E-01 | no | | intergenic_variant | MODIFIER |
| 10 | 26569968 | AX-170320289 | 0 | C | T | 0.184 | 1.03E-01 | 1.04E-01 | 3.00E+00 | 3.11E+00 | 3.26E-01 | no | | intergenic_variant | MODIFIER |
| 10 | 26570730 | AX-170356238 | 0 | A | C | 0.184 | 1.03E-01 | 1.04E-01 | 3.00E+00 | 3.11E+00 | 3.26E-01 | no | | regulatory_region_variant | MODIFIER |
| 10 | 26574487 | AX-170044412 | 0 | A | G | 0.184 | 1.03E-01 | 1.04E-01 | 3.00E+00 | 3.11E+00 | 3.26E-01 | no | | intergenic_variant | MODIFIER |
| 10 | 26587350 | AX-170353830 | 0 | C | A | 0.184 | 1.03E-01 | 1.04E-01 | 3.00E+00 | 3.11E+00 | 3.26E-01 | no | | intergenic_variant | MODIFIER |
| 10 | 26589956 | AX-170327575 | 0 | T | G | 0.184 | 1.03E-01 | 1.04E-01 | 3.00E+00 | 3.11E+00 | 3.26E-01 | no | | intergenic_variant | MODIFIER |
| 10 | 26609977 | AX-170440284 | 0 | A | C | 0.184 | 1.03E-01 | 1.04E-01 | 3.00E+00 | 3.11E+00 | 3.26E-01 | no | | intergenic_variant | MODIFIER |
| 10 | 26610628 | AX-170482438 | 0 | C | T | 0.184 | 1.03E-01 | 1.04E-01 | 3.00E+00 | 3.11E+00 | 3.26E-01 | no | | intergenic_variant | MODIFIER |
| 10 | 26632222 | AX-97795877 | 0 | A | G | 0.184 | 1.03E-01 | 1.04E-01 | 3.00E+00 | 3.11E+00 | 3.26E-01 | no | Gm48864 | downstream_gene_variant | MODIFIER |
| 10 | 26648618 | AX-170349509 | 0 | T | C | 0.184 | 1.03E-01 | 1.04E-01 | 3.00E+00 | 3.11E+00 | 3.26E-01 | no | | intergenic_variant | MODIFIER |
| 10 | 26675240 | AX-170322339 | 0 | T | C | 0.184 | 1.03E-01 | 1.04E-01 | 3.00E+00 | 3.11E+00 | 3.26E-01 | no | Gm8709 | upstream_gene_variant | MODIFIER |
| 10 | 26679465 | AX-170086769 | 0 | C | T | 0.184 | 1.03E-01 | 1.04E-01 | 3.00E+00 | 3.11E+00 | 3.26E-01 | no | Gm8709 | upstream_gene_variant | MODIFIER |
| 10 | 26680149 | AX-170434347 | 0 | C | A | 0.184 | 1.03E-01 | 1.04E-01 | 3.00E+00 | 3.11E+00 | 3.26E-01 | no | | regulatory_region_variant | MODIFIER |
| 10 | 26686258 | AX-170259473 | 0 | C | T | 0.184 | 1.03E-01 | 1.04E-01 | 3.00E+00 | 3.11E+00 | 3.26E-01 | no | | intergenic_variant | MODIFIER |
| 10 | 26686427 | AX-170499706 | 0 | T | C | 0.184 | 1.03E-01 | 1.04E-01 | 3.00E+00 | 3.11E+00 | 3.26E-01 | no | | intergenic_variant | MODIFIER |
| 10 | 26937675 | AX-170322172 | 0 | T | C | 0.184 | 1.03E-01 | 1.04E-01 | 3.00E+00 | 3.11E+00 | 3.26E-01 | no | | intergenic_variant | MODIFIER |
| 10 | 26950277 | AX-97791338 | 0 | A | G | 0.184 | 1.03E-01 | 1.04E-01 | 3.00E+00 | 3.11E+00 | 3.26E-01 | no | Gm48084 | intron_variant,non_coding_transcript_variant | MODIFIER |
| 10 | 26985086 | AX-170241635 | 0 | T | C | 0.184 | 1.03E-01 | 1.04E-01 | 3.00E+00 | 3.11E+00 | 3.26E-01 | no | Lama2 | intron_variant | MODIFIER |
| 10 | 26986761 | AX-170274020 | 0 | A | C | 0.184 | 1.03E-01 | 1.04E-01 | 3.00E+00 | 3.11E+00 | 3.26E-01 | no | Lama2 | intron_variant | MODIFIER |
| 10 | 26988093 | AX-170372190 | 0 | G | A | 0.184 | 1.03E-01 | 1.04E-01 | 3.00E+00 | 3.11E+00 | 3.26E-01 | no | Lama2 | intron_variant | MODIFIER |
| 10 | 26992397 | AX-97793646 | 0 | C | T | 0.184 | 1.03E-01 | 1.04E-01 | 3.00E+00 | 3.11E+00 | 3.26E-01 | no | Lama2 | intron_variant | MODIFIER |
| 10 | 26993405 | AX-170175953 | 0 | A | G | 0.184 | 1.03E-01 | 1.04E-01 | 3.00E+00 | 3.11E+00 | 3.26E-01 | no | Lama2 | splice_region_variant,synonymous_variant | LOW |
| 10 | 26994453 | AX-170176979 | 0 | A | G | 0.184 | 1.03E-01 | 1.04E-01 | 3.00E+00 | 3.11E+00 | 3.26E-01 | no | Lama2 | intron_variant | MODIFIER |
| 10 | 27002201 | AX-170044424 | 0 | T | C | 0.184 | 1.03E-01 | 1.04E-01 | 3.00E+00 | 3.11E+00 | 3.26E-01 | no | Lama2 | intron_variant | MODIFIER |
| 10 | 27004950 | AX-170044422 | 0 | G | A | 0.184 | 1.03E-01 | 1.04E-01 | 3.00E+00 | 3.11E+00 | 3.26E-01 | no | Lama2 | intron_variant | MODIFIER |
| 10 | 27005498 | AX-170361507 | 0 | C | T | 0.184 | 1.03E-01 | 1.04E-01 | 3.00E+00 | 3.11E+00 | 3.26E-01 | no | Lama2 | intron_variant | MODIFIER |
| 10 | 27014522 | AX-170147780 | 0 | C | T | 0.184 | 1.03E-01 | 1.04E-01 | 3.00E+00 | 3.11E+00 | 3.26E-01 | no | Lama2 | intron_variant | MODIFIER |
| 14 | 119967163 | AX-170533650 | 0 | G | A | 0.325 | 7.08E-02 | 7.26E-02 | 2.60E+00 | 2.71E+00 | 3.26E-01 | no | | intergenic_variant | MODIFIER |
| 7 | 77241101 | AX-170344365 | 2 | T | C | 0.355 | -8.55E-02 | 8.81E-02 | 2.80E+00 | 2.92E+00 | 3.26E-01 | no | | intergenic_variant | MODIFIER |
| 1 | 121570158 | AX-170323699 | 0 | G | A | 0.579 | 8.02E-02 | 8.12E-02 | 2.52E+00 | 2.59E+00 | 3.26E-01 | no | Ddx18 | upstream_gene_variant | MODIFIER |
| 1 | 128461016 | AX-170189263 | 0 | A | T | 0.544 | 8.18E-02 | 8.30E-02 | 2.53E+00 | 2.59E+00 | 3.26E-01 | no | | intergenic_variant | MODIFIER |
| 3 | 28339864 | AX-170440548 | 0 | T | C | 0.175 | -9.34E-02 | 9.62E-02 | 2.76E+00 | 2.85E+00 | 3.26E-01 | no | Tnik | intron_variant | MODIFIER |
| 3 | 29419883 | AX-170066214 | 0 | T | C | 0.175 | -9.34E-02 | 9.62E-02 | 2.76E+00 | 2.85E+00 | 3.26E-01 | no | Mir551b | downstream_gene_variant | MODIFIER |
| 8 | 65375902 | AX-170275835 | 1 | C | T | 0.134 | -1.38E-01 | 1.40E-01 | 2.53E+00 | 2.61E+00 | 3.26E-01 | no | | intergenic_variant | MODIFIER |
| 7 | 19704519 | AX-170508806 | 0 | T | C | 0.412 | 8.82E-02 | 9.00E-02 | 2.55E+00 | 2.65E+00 | 3.26E-01 | no | Tomm40 | intron_variant | MODIFIER |
| 7 | 19756017 | AX-97805099 | 0 | C | T | 0.412 | 8.82E-02 | 9.00E-02 | 2.55E+00 | 2.65E+00 | 3.26E-01 | no | Bcam | downstream_gene_variant | MODIFIER |
| 7 | 19757059 | AX-170469302 | 0 | G | A | 0.412 | 8.82E-02 | 9.00E-02 | 2.55E+00 | 2.65E+00 | 3.26E-01 | no | Bcam | intron_variant | MODIFIER |
| 7 | 19768347 | AX-170180390 | 0 | A | G | 0.412 | 8.82E-02 | 9.00E-02 | 2.55E+00 | 2.65E+00 | 3.26E-01 | no | Bcam | intron_variant | MODIFIER |
| 7 | 19801074 | AX-170154447 | 0 | C | T | 0.412 | 8.82E-02 | 9.00E-02 | 2.55E+00 | 2.65E+00 | 3.26E-01 | no | Cblc | upstream_gene_variant | MODIFIER |
| 7 | 19803956 | AX-97792569 | 0 | C | A | 0.412 | 8.82E-02 | 9.00E-02 | 2.55E+00 | 2.65E+00 | 3.26E-01 | no | Bcl3 | downstream_gene_variant | MODIFIER |
| 7 | 19829746 | AX-170408654 | 0 | C | T | 0.412 | 8.82E-02 | 9.00E-02 | 2.55E+00 | 2.65E+00 | 3.26E-01 | no | | regulatory_region_variant | MODIFIER |
| 7 | 19829851 | AX-97821511 | 0 | T | C | 0.412 | 8.82E-02 | 9.00E-02 | 2.55E+00 | 2.65E+00 | 3.26E-01 | no | | regulatory_region_variant | MODIFIER |
| 7 | 19932164 | AX-170076832 | 0 | G | A | 0.412 | 8.82E-02 | 9.00E-02 | 2.55E+00 | 2.65E+00 | 3.26E-01 | no | Gm16251 | downstream_gene_variant | MODIFIER |
| 7 | 19932326 | AX-170274972 | 0 | G | A | 0.412 | 8.82E-02 | 9.00E-02 | 2.55E+00 | 2.65E+00 | 3.26E-01 | no | Igsf23 | downstream_gene_variant | MODIFIER |
| 7 | 19938326 | AX-170368778 | 0 | A | G | 0.412 | 8.82E-02 | 9.00E-02 | 2.55E+00 | 2.65E+00 | 3.26E-01 | no | Igsf23 | intron_variant | MODIFIER |
| 7 | 19939082 | AX-170395925 | 0 | G | T | 0.412 | 8.82E-02 | 9.00E-02 | 2.55E+00 | 2.65E+00 | 3.26E-01 | no | Igsf23 | intron_variant | MODIFIER |
| 7 | 20024672 | AX-170319754 | 0 | C | T | 0.412 | 8.82E-02 | 9.00E-02 | 2.55E+00 | 2.65E+00 | 3.26E-01 | no | Nlrp9b | synonymous_variant | LOW |
| 7 | 23367851 | AX-97851435 | 0 | G | A | 0.412 | 8.82E-02 | 9.00E-02 | 2.55E+00 | 2.65E+00 | 3.26E-01 | no | | intergenic_variant | MODIFIER |
| 7 | 23386383 | AX-97842691 | 0 | G | T | 0.412 | 8.82E-02 | 9.00E-02 | 2.55E+00 | 2.65E+00 | 3.26E-01 | no | Nlrp5 | intron_variant | MODIFIER |

| | | | | | | | | | | | | | | | |
|---|---|---|---|---|---|---|---|---|---|---|---|---|---|---|---|
| 7 | 23399149 | AX-170112113 | 0 | C | T | 0.412 | 8.82E-02 | 9.00E-02 | 2.55E+00 | 2.65E+00 | 3.26E-01 | no | Nlrp5 | intron_variant | MODIFIER |
| 7 | 23400363 | AX-170309379 | 0 | A | C | 0.412 | 8.82E-02 | 9.00E-02 | 2.55E+00 | 2.65E+00 | 3.26E-01 | no | Nlrp5 | intron_variant | MODIFIER |
| 7 | 23400744 | AX-170399165 | 0 | G | T | 0.412 | 8.82E-02 | 9.00E-02 | 2.55E+00 | 2.65E+00 | 3.26E-01 | no | Nlrp5 | intron_variant | MODIFIER |
| 7 | 24293421 | AX-170095873 | 0 | A | G | 0.412 | 8.82E-02 | 9.00E-02 | 2.55E+00 | 2.65E+00 | 3.26E-01 | no | Zfp61 | synonymous_variant | LOW |
| 7 | 24315657 | AX-97801371 | 0 | C | T | 0.412 | 8.82E-02 | 9.00E-02 | 2.55E+00 | 2.65E+00 | 3.26E-01 | no | Zfp94 | intron_variant | MODIFIER |
| 7 | 24315717 | AX-170076848 | 0 | A | G | 0.412 | 8.82E-02 | 9.00E-02 | 2.55E+00 | 2.65E+00 | 3.26E-01 | no | Zfp94 | intron_variant | MODIFIER |
| 7 | 24327917 | AX-97809505 | 0 | G | A | 0.412 | 8.82E-02 | 9.00E-02 | 2.55E+00 | 2.65E+00 | 3.26E-01 | no | | intergenic_variant | MODIFIER |
| 7 | 24355259 | AX-170300199 | 0 | T | C | 0.412 | 8.82E-02 | 9.00E-02 | 2.55E+00 | 2.65E+00 | 3.26E-01 | no | Lypd5 | downstream_gene_variant | MODIFIER |
| 7 | 24355448 | AX-170098230 | 0 | C | G | 0.412 | 8.82E-02 | 9.00E-02 | 2.55E+00 | 2.65E+00 | 3.26E-01 | no | Lypd5 | downstream_gene_variant | MODIFIER |
| 7 | 24356539 | AX-170140690 | 0 | G | A | 0.412 | 8.82E-02 | 9.00E-02 | 2.55E+00 | 2.65E+00 | 3.26E-01 | no | Lypd5 | downstream_gene_variant | MODIFIER |
| 7 | 24359357 | AX-97850503 | 0 | A | G | 0.412 | 8.82E-02 | 9.00E-02 | 2.55E+00 | 2.65E+00 | 3.26E-01 | no | Lypd5 | downstream_gene_variant | MODIFIER |
| 7 | 24387577 | AX-97818854 | 0 | C | T | 0.412 | 8.82E-02 | 9.00E-02 | 2.55E+00 | 2.65E+00 | 3.26E-01 | no | Kcnn4 | downstream_gene_variant | MODIFIER |
| 7 | 24407400 | AX-97832662 | 0 | T | C | 0.412 | 8.82E-02 | 9.00E-02 | 2.55E+00 | 2.65E+00 | 3.26E-01 | no | Smg9 | intron_variant | MODIFIER |
| 7 | 24473721 | AX-97848155 | 0 | C | G | 0.412 | 8.82E-02 | 9.00E-02 | 2.55E+00 | 2.65E+00 | 3.26E-01 | no | Plaur | intron_variant | MODIFIER |
| 7 | 24513079 | AX-170028693 | 0 | A | G | 0.412 | 8.82E-02 | 9.00E-02 | 2.55E+00 | 2.65E+00 | 3.26E-01 | no | Zfp428 | intron_variant | MODIFIER |
| 18 | 60298683 | AX-97815125 | 1 | A | G | 0.411 | 9.35E-02 | 9.33E-02 | 2.43E+00 | 2.48E+00 | 3.26E-01 | no | F830016B08Rik | intron_variant | MODIFIER |
| 1 | 15718815 | AX-170493563 | 0 | A | G | 0.377 | 7.79E-02 | 7.97E-02 | 2.60E+00 | 2.65E+00 | 3.27E-01 | no | Kcnb2 | 3_prime_UTR_variant | MODIFIER |
| 16 | 27937166 | AX-97787163 | 0 | G | A | 0.254 | 8.31E-02 | 8.55E-02 | 2.61E+00 | 2.74E+00 | 3.27E-01 | no | | intergenic_variant | MODIFIER |
| 13 | 23340680 | AX-170219664 | 0 | C | A | 0.167 | -1.24E-01 | 1.24E-01 | 2.43E+00 | 2.47E+00 | 3.27E-01 | no | 4930586N03Rik | intron_variant,non_coding_transcript_variant | MODIFIER |
| 10 | 66112636 | AX-170331113 | 0 | C | T | 0.412 | -7.71E-02 | 7.92E-02 | 2.65E+00 | 2.72E+00 | 3.27E-01 | no | | intergenic_variant | MODIFIER |
| 4 | 135738433 | AX-97813887 | 1 | T | C | 0.179 | 1.06E-01 | 1.09E-01 | 2.60E+00 | 2.65E+00 | 3.27E-01 | no | Il22ra1 | intron_variant | MODIFIER |
| 1 | 138163449 | AX-170509374 | 0 | G | A | 0.211 | 1.21E-01 | 1.25E-01 | 2.70E+00 | 2.78E+00 | 3.27E-01 | no | Ptprc | intron_variant,non_coding_transcript_variant | MODIFIER |
| 18 | 88699553 | AX-170114558 | 1 | G | A | 0.161 | -1.03E-01 | 1.05E-01 | 2.51E+00 | 2.61E+00 | 3.27E-01 | no | Socs6 | intron_variant,non_coding_transcript_variant | MODIFIER |
| 14 | 113383270 | AX-170509513 | 1 | C | T | 0.348 | 6.98E-02 | 7.18E-02 | 2.61E+00 | 2.71E+00 | 3.27E-01 | no | | intergenic_variant | MODIFIER |
| 18 | 80172951 | AX-170061445 | 2 | T | G | 0.373 | -7.97E-02 | 8.22E-02 | 2.66E+00 | 2.76E+00 | 3.27E-01 | no | | regulatory_region_variant | MODIFIER |
| 8 | 24628890 | AX-170449183 | 0 | A | G | 0.342 | -6.21E-02 | 6.34E-02 | 2.54E+00 | 2.64E+00 | 3.27E-01 | no | Adam18 | intron_variant | MODIFIER |
| 1 | 187423517 | AX-170117262 | 1 | G | T | 0.223 | 1.10E-01 | 1.10E-01 | 2.44E+00 | 2.53E+00 | 3.27E-01 | no | | intergenic_variant | MODIFIER |
| 10 | 27005048 | AX-170132381 | 1 | C | T | 0.188 | 1.03E-01 | 1.04E-01 | 2.98E+00 | 3.08E+00 | 3.27E-01 | no | Lama2 | intron_variant | MODIFIER |
| 16 | 29988106 | AX-97782650 | 0 | C | T | 0.5 | 7.89E-02 | 8.06E-02 | 2.56E+00 | 2.61E+00 | 3.28E-01 | no | | intergenic_variant | MODIFIER |
| 16 | 19154417 | AX-170493374 | 2 | T | C | 0.136 | 9.33E-02 | 9.65E-02 | 2.81E+00 | 2.94E+00 | 3.28E-01 | no | | intergenic_variant | MODIFIER |
| 1 | 123609177 | AX-170042553 | 1 | C | T | 0.554 | 8.04E-02 | 8.21E-02 | 2.56E+00 | 2.62E+00 | 3.28E-01 | no | Dpp10 | intron_variant | MODIFIER |
| 3 | 7569827 | AX-170170192 | 0 | G | A | 0.412 | -6.12E-02 | 6.36E-02 | 2.76E+00 | 2.91E+00 | 3.28E-01 | no | Il7 | downstream_gene_variant | MODIFIER |
| 4 | 84284694 | AX-170388923 | 0 | C | T | 0.333 | 1.06E-01 | 1.09E-01 | 2.71E+00 | 2.79E+00 | 3.28E-01 | no | Bnc2 | intron_variant | MODIFIER |
| 16 | 79137272 | AX-170320577 | 0 | A | C | 0.465 | 7.38E-02 | 7.57E-02 | 2.60E+00 | 2.66E+00 | 3.28E-01 | no | | intergenic_variant | MODIFIER |
| 5 | 97814198 | AX-170254474 | 0 | G | A | 0.447 | 1.09E-01 | 1.13E-01 | 2.82E+00 | 2.88E+00 | 3.28E-01 | no | | intergenic_variant | MODIFIER |
| 4 | 155008206 | AX-170095204 | 0 | G | A | 0.105 | 1.57E-01 | 1.62E-01 | 2.71E+00 | 2.82E+00 | 3.28E-01 | Yes | Plch2 | intron_variant | MODIFIER |
| 4 | 155155860 | AX-170261694 | 0 | T | C | 0.105 | 1.57E-01 | 1.62E-01 | 2.71E+00 | 2.82E+00 | 3.28E-01 | Yes | Ski | 3_prime_UTR_variant | MODIFIER |
| 4 | 155184627 | AX-97809121 | 0 | C | T | 0.105 | 1.57E-01 | 1.62E-01 | 2.71E+00 | 2.82E+00 | 3.28E-01 | Yes | Ski | intron_variant | MODIFIER |
| 4 | 155184760 | AX-170071134 | 0 | T | C | 0.105 | 1.57E-01 | 1.62E-01 | 2.71E+00 | 2.82E+00 | 3.28E-01 | Yes | Ski | intron_variant | MODIFIER |
| 4 | 155225673 | AX-168394690 | 0 | A | G | 0.105 | 1.57E-01 | 1.62E-01 | 2.71E+00 | 2.82E+00 | 3.28E-01 | Yes | Ski | upstream_gene_variant | MODIFIER |
| 4 | 155627729 | AX-170125463 | 0 | G | A | 0.105 | 1.57E-01 | 1.62E-01 | 2.71E+00 | 2.82E+00 | 3.28E-01 | Yes | Cdk11b | intron_variant | MODIFIER |
| 4 | 155771368 | AX-170024307 | 0 | C | T | 0.105 | 1.57E-01 | 1.62E-01 | 2.71E+00 | 2.82E+00 | 3.28E-01 | Yes | Vwa1 | intron_variant | MODIFIER |
| 4 | 155771496 | AX-170024313 | 0 | A | C | 0.105 | 1.57E-01 | 1.62E-01 | 2.71E+00 | 2.82E+00 | 3.28E-01 | Yes | Vwa1 | intron_variant | MODIFIER |
| 4 | 155772058 | AX-170024310 | 0 | T | C | 0.105 | 1.57E-01 | 1.62E-01 | 2.71E+00 | 2.82E+00 | 3.28E-01 | Yes | Vwa1 | intron_variant | MODIFIER |
| 4 | 155791054 | AX-97832218 | 0 | T | C | 0.105 | 1.57E-01 | 1.62E-01 | 2.71E+00 | 2.82E+00 | 3.28E-01 | Yes | Ankrd65 | upstream_gene_variant | MODIFIER |
| 4 | 155824952 | AX-97795992 | 0 | A | G | 0.105 | 1.57E-01 | 1.62E-01 | 2.71E+00 | 2.82E+00 | 3.28E-01 | Yes | Ccnl2 | downstream_gene_variant | MODIFIER |
| 4 | 155850695 | AX-170024308 | 0 | G | T | 0.105 | 1.57E-01 | 1.62E-01 | 2.71E+00 | 2.82E+00 | 3.28E-01 | Yes | Dvl1 | intron_variant | MODIFIER |
| 4 | 155874988 | AX-97794282 | 0 | G | A | 0.105 | 1.57E-01 | 1.62E-01 | 2.71E+00 | 2.82E+00 | 3.28E-01 | Yes | Ints11 | intron_variant | MODIFIER |
| 4 | 155878647 | AX-170209476 | 0 | G | A | 0.105 | 1.57E-01 | 1.62E-01 | 2.71E+00 | 2.82E+00 | 3.28E-01 | Yes | Ints11 | intron_variant | MODIFIER |
| 4 | 155886688 | AX-170183269 | 0 | A | G | 0.105 | 1.57E-01 | 1.62E-01 | 2.71E+00 | 2.82E+00 | 3.28E-01 | Yes | Ints11 | intron_variant | MODIFIER |
| 4 | 155886963 | AX-170529662 | 0 | G | A | 0.105 | 1.57E-01 | 1.62E-01 | 2.71E+00 | 2.82E+00 | 3.28E-01 | Yes | Ints11 | synonymous_variant | LOW |
| 4 | 155887172 | AX-170210005 | 0 | G | A | 0.105 | 1.57E-01 | 1.62E-01 | 2.71E+00 | 2.82E+00 | 3.28E-01 | Yes | Ints11 | synonymous_variant | LOW |
| 4 | 155888094 | AX-170322694 | 0 | C | A | 0.105 | 1.57E-01 | 1.62E-01 | 2.71E+00 | 2.82E+00 | 3.28E-01 | Yes | Ints11 | synonymous_variant | LOW |
| 4 | 155888658 | AX-97820412 | 0 | G | A | 0.105 | 1.57E-01 | 1.62E-01 | 2.71E+00 | 2.82E+00 | 3.28E-01 | Yes | Ints11 | synonymous_variant | LOW |

| | | | | | | | | | | | | | | | |
|---|---|---|---|---|---|---|---|---|---|---|---|---|---|---|---|
| 4 | 155889902 | AX-170405602 | 0 | T | C | 0.105 | 1.57E-01 | 1.62E-01 | 2.71E+00 | 2.82E+00 | 3.28E-01 | Yes | Ints11 | downstream_gene_variant | MODIFIER |
| 4 | 155894026 | AX-170024312 | 0 | G | A | 0.105 | 1.57E-01 | 1.62E-01 | 2.71E+00 | 2.82E+00 | 3.28E-01 | Yes | Ints11 | downstream_gene_variant | MODIFIER |
| 4 | 155903497 | AX-170024309 | 0 | C | T | 0.105 | 1.57E-01 | 1.62E-01 | 2.71E+00 | 2.82E+00 | 3.28E-01 | Yes | Acap3 | synonymous_variant | LOW |
| 4 | 155907594 | AX-170478464 | 0 | T | C | 0.105 | 1.57E-01 | 1.62E-01 | 2.71E+00 | 2.82E+00 | 3.28E-01 | Yes | Acap3 | downstream_gene_variant | MODIFIER |
| 4 | 155910781 | AX-97834077 | 0 | T | G | 0.105 | 1.57E-01 | 1.62E-01 | 2.71E+00 | 2.82E+00 | 3.28E-01 | Yes | Acap3 | downstream_gene_variant | MODIFIER |
| 4 | 155911120 | AX-170071153 | 0 | C | G | 0.105 | 1.57E-01 | 1.62E-01 | 2.71E+00 | 2.82E+00 | 3.28E-01 | Yes | Acap3 | downstream_gene_variant | MODIFIER |
| 4 | 155941221 | AX-97850600 | 0 | T | C | 0.105 | 1.57E-01 | 1.62E-01 | 2.71E+00 | 2.82E+00 | 3.28E-01 | Yes | Ube2j2 | upstream_gene_variant | MODIFIER |
| 4 | 155941330 | AX-170024314 | 0 | C | T | 0.105 | 1.57E-01 | 1.62E-01 | 2.71E+00 | 2.82E+00 | 3.28E-01 | Yes | Ube2j2 | upstream_gene_variant | MODIFIER |
| 4 | 155941789 | AX-170353669 | 0 | T | A | 0.105 | 1.57E-01 | 1.62E-01 | 2.71E+00 | 2.82E+00 | 3.28E-01 | Yes | Ube2j2 | upstream_gene_variant | MODIFIER |
| 4 | 155970709 | AX-170174033 | 0 | A | G | 0.105 | 1.57E-01 | 1.62E-01 | 2.71E+00 | 2.82E+00 | 3.28E-01 | Yes | C1qtnf12 | downstream_gene_variant | MODIFIER |
| 4 | 155970754 | AX-170389282 | 0 | T | C | 0.105 | 1.57E-01 | 1.62E-01 | 2.71E+00 | 2.82E+00 | 3.28E-01 | Yes | C1qtnf12 | downstream_gene_variant | MODIFIER |
| 4 | 156014280 | AX-170282425 | 0 | A | G | 0.105 | 1.57E-01 | 1.62E-01 | 2.71E+00 | 2.82E+00 | 3.28E-01 | Yes | Tnfrsf4 | synonymous_variant | LOW |
| 4 | 156045338 | AX-97827806 | 0 | A | G | 0.105 | 1.57E-01 | 1.62E-01 | 2.71E+00 | 2.82E+00 | 3.28E-01 | Yes | Ttll10 | downstream_gene_variant | MODIFIER |
| 4 | 156053729 | AX-170180946 | 0 | T | C | 0.105 | 1.57E-01 | 1.62E-01 | 2.71E+00 | 2.82E+00 | 3.28E-01 | Yes | Ttll10 | upstream_gene_variant | MODIFIER |
| 8 | 65605681 | AX-170527959 | 1 | T | G | 0.143 | -1.40E-01 | 1.42E-01 | 2.49E+00 | 2.56E+00 | 3.28E-01 | no | | intergenic_variant | MODIFIER |
| 13 | 21008754 | AX-170232634 | 0 | G | T | 0.167 | -1.27E-01 | 1.29E-01 | 2.49E+00 | 2.54E+00 | 3.28E-01 | no | Aoah | intron_variant | MODIFIER |
| 4 | 79455404 | AX-170023187 | 1 | C | T | 0.455 | -8.36E-02 | 8.49E-02 | 2.51E+00 | 2.56E+00 | 3.28E-01 | no | | intergenic_variant | MODIFIER |
| 4 | 80010461 | AX-97795702 | 1 | C | T | 0.455 | -8.36E-02 | 8.49E-02 | 2.51E+00 | 2.56E+00 | 3.28E-01 | no | | intergenic_variant | MODIFIER |
| 9 | 97209562 | AX-170033076 | 0 | A | G | 0.14 | -1.10E-01 | 1.13E-01 | 2.60E+00 | 2.68E+00 | 3.28E-01 | no | Gm5371 | non_coding_transcript_exon_variant | MODIFIER |
| 1 | 191600903 | AX-170289143 | 1 | A | G | 0.259 | 9.42E-02 | 9.25E-02 | 2.34E+00 | 2.43E+00 | 3.28E-01 | no | Ints7 | intron_variant | MODIFIER |
| 1 | 126901567 | AX-170042629 | 1 | T | C | 0.536 | 8.08E-02 | 8.31E-02 | 2.60E+00 | 2.67E+00 | 3.29E-01 | no | | intergenic_variant | MODIFIER |
| 1 | 128164264 | AX-170363459 | 1 | G | A | 0.536 | 8.08E-02 | 8.31E-02 | 2.60E+00 | 2.67E+00 | 3.29E-01 | no | R3hdm1 | intron_variant | MODIFIER |
| 1 | 128279003 | AX-97796560 | 1 | T | C | 0.536 | 8.08E-02 | 8.31E-02 | 2.60E+00 | 2.67E+00 | 3.29E-01 | no | Ubxn4 | 3_prime_UTR_variant | MODIFIER |
| 2 | 169916883 | AX-170181968 | 1 | C | T | 0.17 | 1.75E-01 | 1.81E-01 | 2.64E+00 | 2.73E+00 | 3.29E-01 | Yes | Tshz2 | downstream_gene_variant | MODIFIER |
| 1 | 131712664 | AX-170324059 | 1 | T | G | 0.536 | 8.17E-02 | 8.43E-02 | 2.65E+00 | 2.72E+00 | 3.29E-01 | no | Rab7b | 3_prime_UTR_variant | MODIFIER |
| 6 | 137574105 | AX-170274772 | 2 | C | T | 0.564 | 8.40E-02 | 8.44E-02 | 2.44E+00 | 2.50E+00 | 3.29E-01 | no | Eps8 | intron_variant | MODIFIER |
| 18 | 57289386 | AX-170309076 | 0 | C | T | 0.404 | 9.19E-02 | 9.21E-02 | 2.43E+00 | 2.49E+00 | 3.29E-01 | no | Megf10 | synonymous_variant | LOW |
| 18 | 57495254 | AX-170093896 | 0 | C | T | 0.404 | 9.19E-02 | 9.21E-02 | 2.43E+00 | 2.49E+00 | 3.29E-01 | no | 4930511M06Rik | intron_variant,non_coding_transcript_variant | MODIFIER |
| 15 | 3462060 | AX-170497761 | 0 | G | A | 0.246 | -1.00E-01 | 1.03E-01 | 2.65E+00 | 2.71E+00 | 3.29E-01 | no | Ghr | intron_variant | MODIFIER |
| 15 | 3537029 | AX-170480495 | 0 | C | T | 0.246 | -1.00E-01 | 1.03E-01 | 2.65E+00 | 2.71E+00 | 3.29E-01 | no | Ghr | intron_variant | MODIFIER |
| 15 | 4729579 | AX-97843870 | 0 | G | A | 0.246 | -1.00E-01 | 1.03E-01 | 2.65E+00 | 2.71E+00 | 3.29E-01 | no | C6 | intron_variant | MODIFIER |
| 15 | 4790272 | AX-170436130 | 0 | G | C | 0.246 | -1.00E-01 | 1.03E-01 | 2.65E+00 | 2.71E+00 | 3.29E-01 | no | C6 | intron_variant | MODIFIER |
| 15 | 6962646 | AX-170529587 | 0 | G | A | 0.246 | -1.00E-01 | 1.03E-01 | 2.65E+00 | 2.71E+00 | 3.29E-01 | no | | intergenic_variant | MODIFIER |
| 15 | 7052751 | AX-170457926 | 0 | A | G | 0.246 | -1.00E-01 | 1.03E-01 | 2.65E+00 | 2.71E+00 | 3.29E-01 | no | | intergenic_variant | MODIFIER |
| 15 | 51258840 | AX-170118683 | 1 | C | T | 0.018 | -1.30E-01 | 1.35E-01 | 2.60E+00 | 2.77E+00 | 3.29E-01 | no | Gm19303 | intron_variant,non_coding_transcript_variant | MODIFIER |
| 10 | 69242755 | AX-170245056 | 0 | G | A | 0.412 | -7.75E-02 | 8.02E-02 | 2.67E+00 | 2.74E+00 | 3.29E-01 | no | Rhobtb1 | intron_variant | MODIFIER |
| 10 | 69333499 | AX-170524336 | 0 | A | G | 0.412 | -7.75E-02 | 8.02E-02 | 2.67E+00 | 2.74E+00 | 3.29E-01 | no | Cdk1 | downstream_gene_variant | MODIFIER |
| 10 | 69334469 | AX-170473752 | 0 | G | A | 0.412 | -7.75E-02 | 8.02E-02 | 2.67E+00 | 2.74E+00 | 3.29E-01 | no | Cdk1 | downstream_gene_variant | MODIFIER |
| 10 | 69335575 | AX-97793438 | 0 | G | A | 0.412 | -7.75E-02 | 8.02E-02 | 2.67E+00 | 2.74E+00 | 3.29E-01 | no | Cdk1 | 3_prime_UTR_variant | MODIFIER |
| 10 | 69346633 | AX-170501043 | 0 | A | C | 0.412 | -7.75E-02 | 8.02E-02 | 2.67E+00 | 2.74E+00 | 3.29E-01 | no | Cdk1 | intron_variant | MODIFIER |
| 10 | 69444667 | AX-170163408 | 0 | G | T | 0.412 | -7.75E-02 | 8.02E-02 | 2.67E+00 | 2.74E+00 | 3.29E-01 | no | Ank3 | intron_variant,non_coding_transcript_variant | MODIFIER |
| 10 | 69453330 | AX-170261541 | 0 | T | C | 0.412 | -7.75E-02 | 8.02E-02 | 2.67E+00 | 2.74E+00 | 3.29E-01 | no | Ank3 | intron_variant,non_coding_transcript_variant | MODIFIER |
| 10 | 69476670 | AX-170296839 | 0 | A | G | 0.412 | -7.75E-02 | 8.02E-02 | 2.67E+00 | 2.74E+00 | 3.29E-01 | no | Ank3 | intron_variant,non_coding_transcript_variant | MODIFIER |
| 10 | 69488407 | AX-170045147 | 0 | A | C | 0.412 | -7.75E-02 | 8.02E-02 | 2.67E+00 | 2.74E+00 | 3.29E-01 | no | Ank3 | downstream_gene_variant | MODIFIER |
| 10 | 69488867 | AX-97838331 | 0 | A | G | 0.412 | -7.75E-02 | 8.02E-02 | 2.67E+00 | 2.74E+00 | 3.29E-01 | no | Ank3 | downstream_gene_variant | MODIFIER |
| 10 | 69499253 | AX-170513853 | 0 | A | C | 0.412 | -7.75E-02 | 8.02E-02 | 2.67E+00 | 2.74E+00 | 3.29E-01 | no | Ank3 | intron_variant,non_coding_transcript_variant | MODIFIER |
| 10 | 69502623 | AX-170132929 | 0 | G | A | 0.412 | -7.75E-02 | 8.02E-02 | 2.67E+00 | 2.74E+00 | 3.29E-01 | no | Ank3 | intron_variant,non_coding_transcript_variant | MODIFIER |
| 10 | 69505928 | AX-170442557 | 0 | A | T | 0.412 | -7.75E-02 | 8.02E-02 | 2.67E+00 | 2.74E+00 | 3.29E-01 | no | Ank3 | intron_variant,non_coding_transcript_variant | MODIFIER |
| 10 | 69507742 | AX-170311963 | 0 | C | T | 0.412 | -7.75E-02 | 8.02E-02 | 2.67E+00 | 2.74E+00 | 3.29E-01 | no | Ank3 | intron_variant,non_coding_transcript_variant | MODIFIER |
| 10 | 69516718 | AX-170237625 | 0 | T | C | 0.412 | -7.75E-02 | 8.02E-02 | 2.67E+00 | 2.74E+00 | 3.29E-01 | no | Ank3 | intron_variant,non_coding_transcript_variant | MODIFIER |
| 10 | 69546160 | AX-170416538 | 0 | C | G | 0.412 | -7.75E-02 | 8.02E-02 | 2.67E+00 | 2.74E+00 | 3.29E-01 | no | Ank3 | intron_variant | MODIFIER |
| 10 | 69547019 | AX-170298434 | 0 | T | C | 0.412 | -7.75E-02 | 8.02E-02 | 2.67E+00 | 2.74E+00 | 3.29E-01 | no | Ank3 | intron_variant | MODIFIER |
| 10 | 69559371 | AX-170466707 | 0 | A | T | 0.412 | -7.75E-02 | 8.02E-02 | 2.67E+00 | 2.74E+00 | 3.29E-01 | no | Ank3 | intron_variant | MODIFIER |
| 10 | 69559920 | AX-170356603 | 0 | A | G | 0.412 | -7.75E-02 | 8.02E-02 | 2.67E+00 | 2.74E+00 | 3.29E-01 | no | Ank3 | intron_variant | MODIFIER |

| | | | | | | | | | | | | | | | |
|---|---|---|---|---|---|---|---|---|---|---|---|---|---|---|---|
| 10 | 69560586 | AX-170269839 | 0 | T | C | 0.412 | -7.75E-02 | 8.02E-02 | 2.67E+00 | 2.74E+00 | 3.29E-01 | no | Ank3 | intron_variant | MODIFIER |
| 10 | 69570000 | AX-170331964 | 0 | G | A | 0.412 | -7.75E-02 | 8.02E-02 | 2.67E+00 | 2.74E+00 | 3.29E-01 | no | Ank3 | intron_variant | MODIFIER |
| 10 | 69578222 | AX-170521345 | 0 | C | T | 0.412 | -7.75E-02 | 8.02E-02 | 2.67E+00 | 2.74E+00 | 3.29E-01 | no | Ank3 | intron_variant | MODIFIER |
| 10 | 69578403 | AX-170406197 | 0 | T | C | 0.412 | -7.75E-02 | 8.02E-02 | 2.67E+00 | 2.74E+00 | 3.29E-01 | no | Ank3 | intron_variant | MODIFIER |
| 10 | 69580046 | AX-97801942 | 0 | A | G | 0.412 | -7.75E-02 | 8.02E-02 | 2.67E+00 | 2.74E+00 | 3.29E-01 | no | Ank3 | intron_variant | MODIFIER |
| 10 | 69604242 | AX-97790567 | 0 | C | T | 0.412 | -7.75E-02 | 8.02E-02 | 2.67E+00 | 2.74E+00 | 3.29E-01 | no | Ank3 | intron_variant | MODIFIER |
| 10 | 69636980 | AX-170045151 | 0 | T | C | 0.412 | -7.75E-02 | 8.02E-02 | 2.67E+00 | 2.74E+00 | 3.29E-01 | no | Ank3 | intron_variant | MODIFIER |
| 10 | 69744902 | AX-97808962 | 0 | C | T | 0.412 | -7.75E-02 | 8.02E-02 | 2.67E+00 | 2.74E+00 | 3.29E-01 | no | Ank3 | intron_variant | MODIFIER |
| 10 | 69745163 | AX-170255031 | 0 | C | G | 0.412 | -7.75E-02 | 8.02E-02 | 2.67E+00 | 2.74E+00 | 3.29E-01 | no | Ank3 | intron_variant | MODIFIER |
| 10 | 69746326 | AX-170104036 | 0 | G | A | 0.412 | -7.75E-02 | 8.02E-02 | 2.67E+00 | 2.74E+00 | 3.29E-01 | no | Ank3 | intron_variant | MODIFIER |
| 10 | 69747976 | AX-170045155 | 0 | T | C | 0.412 | -7.75E-02 | 8.02E-02 | 2.67E+00 | 2.74E+00 | 3.29E-01 | no | Ank3 | intron_variant | MODIFIER |
| 10 | 69750539 | AX-170170474 | 0 | G | C | 0.412 | -7.75E-02 | 8.02E-02 | 2.67E+00 | 2.74E+00 | 3.29E-01 | no | Ank3 | intron_variant | MODIFIER |
| 10 | 69752684 | AX-170419973 | 0 | T | C | 0.412 | -7.75E-02 | 8.02E-02 | 2.67E+00 | 2.74E+00 | 3.29E-01 | no | Ank3 | intron_variant | MODIFIER |
| 10 | 69753028 | AX-170471525 | 0 | G | A | 0.412 | -7.75E-02 | 8.02E-02 | 2.67E+00 | 2.74E+00 | 3.29E-01 | no | Ank3 | intron_variant | MODIFIER |
| 10 | 69769061 | AX-97785868 | 0 | C | T | 0.412 | -7.75E-02 | 8.02E-02 | 2.67E+00 | 2.74E+00 | 3.29E-01 | no | Ank3 | intron_variant | MODIFIER |
| 10 | 69771359 | AX-170279690 | 0 | G | A | 0.412 | -7.75E-02 | 8.02E-02 | 2.67E+00 | 2.74E+00 | 3.29E-01 | no | Ank3 | intron_variant | MODIFIER |
| 10 | 69771801 | AX-170468375 | 0 | T | C | 0.412 | -7.75E-02 | 8.02E-02 | 2.67E+00 | 2.74E+00 | 3.29E-01 | no | Ank3 | intron_variant | MODIFIER |
| 10 | 69771854 | AX-170175575 | 0 | G | A | 0.412 | -7.75E-02 | 8.02E-02 | 2.67E+00 | 2.74E+00 | 3.29E-01 | no | Ank3 | intron_variant | MODIFIER |
| 10 | 69781853 | AX-170171515 | 0 | A | C | 0.412 | -7.75E-02 | 8.02E-02 | 2.67E+00 | 2.74E+00 | 3.29E-01 | no | Ank3 | intron_variant | MODIFIER |
| 10 | 69782055 | AX-170493685 | 0 | T | C | 0.412 | -7.75E-02 | 8.02E-02 | 2.67E+00 | 2.74E+00 | 3.29E-01 | no | Ank3 | intron_variant | MODIFIER |
| 10 | 69786211 | AX-170446497 | 0 | A | T | 0.412 | -7.75E-02 | 8.02E-02 | 2.67E+00 | 2.74E+00 | 3.29E-01 | no | Ank3 | intron_variant | MODIFIER |
| 10 | 69786353 | AX-97790396 | 0 | T | C | 0.412 | -7.75E-02 | 8.02E-02 | 2.67E+00 | 2.74E+00 | 3.29E-01 | no | Ank3 | intron_variant | MODIFIER |
| 10 | 69799651 | AX-170003424 | 0 | A | G | 0.412 | -7.75E-02 | 8.02E-02 | 2.67E+00 | 2.74E+00 | 3.29E-01 | no | Ank3 | intron_variant | MODIFIER |
| 10 | 69813377 | AX-97848134 | 0 | C | T | 0.412 | -7.75E-02 | 8.02E-02 | 2.67E+00 | 2.74E+00 | 3.29E-01 | no | Ank3 | intron_variant | MODIFIER |
| 10 | 69829723 | AX-170003420 | 0 | A | G | 0.412 | -7.75E-02 | 8.02E-02 | 2.67E+00 | 2.74E+00 | 3.29E-01 | no | Ank3 | intron_variant | MODIFIER |
| 10 | 69906707 | AX-97815783 | 0 | G | A | 0.412 | -7.75E-02 | 8.02E-02 | 2.67E+00 | 2.74E+00 | 3.29E-01 | no | Ank3 | intron_variant | MODIFIER |
| 10 | 69919458 | AX-170003421 | 0 | C | T | 0.412 | -7.75E-02 | 8.02E-02 | 2.67E+00 | 2.74E+00 | 3.29E-01 | no | Ank3 | intron_variant | MODIFIER |
| 10 | 69921387 | AX-170230101 | 0 | G | A | 0.412 | -7.75E-02 | 8.02E-02 | 2.67E+00 | 2.74E+00 | 3.29E-01 | no | Ank3 | intron_variant | MODIFIER |
| 10 | 69929903 | AX-170236797 | 0 | G | A | 0.412 | -7.75E-02 | 8.02E-02 | 2.67E+00 | 2.74E+00 | 3.29E-01 | no | Ank3 | intron_variant | MODIFIER |
| 10 | 69930659 | AX-170373350 | 0 | A | G | 0.412 | -7.75E-02 | 8.02E-02 | 2.67E+00 | 2.74E+00 | 3.29E-01 | no | Ank3 | intron_variant | MODIFIER |
| 10 | 69930798 | AX-170165016 | 0 | A | G | 0.412 | -7.75E-02 | 8.02E-02 | 2.67E+00 | 2.74E+00 | 3.29E-01 | no | Ank3 | intron_variant | MODIFIER |
| 10 | 69936512 | AX-170449953 | 0 | C | A | 0.412 | -7.75E-02 | 8.02E-02 | 2.67E+00 | 2.74E+00 | 3.29E-01 | no | Ank3 | intron_variant | MODIFIER |
| 10 | 69937947 | AX-170391411 | 0 | C | T | 0.412 | -7.75E-02 | 8.02E-02 | 2.67E+00 | 2.74E+00 | 3.29E-01 | no | Ank3 | intron_variant | MODIFIER |
| 10 | 69939143 | AX-97822972 | 0 | A | G | 0.412 | -7.75E-02 | 8.02E-02 | 2.67E+00 | 2.74E+00 | 3.29E-01 | no | Ank3 | intron_variant | MODIFIER |
| 10 | 69939492 | AX-170427090 | 0 | A | C | 0.412 | -7.75E-02 | 8.02E-02 | 2.67E+00 | 2.74E+00 | 3.29E-01 | no | Ank3 | intron_variant | MODIFIER |
| 10 | 69982862 | AX-170273652 | 0 | G | A | 0.412 | -7.75E-02 | 8.02E-02 | 2.67E+00 | 2.74E+00 | 3.29E-01 | no | Ank3 | intron_variant | MODIFIER |
| 10 | 69983408 | AX-97815785 | 0 | T | C | 0.412 | -7.75E-02 | 8.02E-02 | 2.67E+00 | 2.74E+00 | 3.29E-01 | no | Ank3 | intron_variant | MODIFIER |
| 10 | 70028591 | AX-170498768 | 0 | A | G | 0.412 | -7.75E-02 | 8.02E-02 | 2.67E+00 | 2.74E+00 | 3.29E-01 | no | Ank3 | downstream_gene_variant | MODIFIER |
| 10 | 70040176 | AX-170306359 | 0 | A | G | 0.412 | -7.75E-02 | 8.02E-02 | 2.67E+00 | 2.74E+00 | 3.29E-01 | no | | regulatory_region_variant | MODIFIER |
| 10 | 70047048 | AX-170137165 | 0 | G | C | 0.412 | -7.75E-02 | 8.02E-02 | 2.67E+00 | 2.74E+00 | 3.29E-01 | no | | intergenic_variant | MODIFIER |
| 10 | 70055064 | AX-97801503 | 0 | T | C | 0.412 | -7.75E-02 | 8.02E-02 | 2.67E+00 | 2.74E+00 | 3.29E-01 | no | | intergenic_variant | MODIFIER |
| 10 | 70055334 | AX-170204397 | 0 | C | T | 0.412 | -7.75E-02 | 8.02E-02 | 2.67E+00 | 2.74E+00 | 3.29E-01 | no | | intergenic_variant | MODIFIER |
| 10 | 70121570 | AX-170045163 | 0 | C | G | 0.412 | -7.75E-02 | 8.02E-02 | 2.67E+00 | 2.74E+00 | 3.29E-01 | no | Ccdc6 | intron_variant | MODIFIER |
| 10 | 70129817 | AX-170045162 | 0 | C | T | 0.412 | -7.75E-02 | 8.02E-02 | 2.67E+00 | 2.74E+00 | 3.29E-01 | no | Ccdc6 | intron_variant | MODIFIER |
| 10 | 70147310 | AX-170139968 | 0 | C | T | 0.412 | -7.75E-02 | 8.02E-02 | 2.67E+00 | 2.74E+00 | 3.29E-01 | no | Ccdc6 | intron_variant | MODIFIER |
| 10 | 70147324 | AX-170180385 | 0 | C | G | 0.412 | -7.75E-02 | 8.02E-02 | 2.67E+00 | 2.74E+00 | 3.29E-01 | no | Ccdc6 | intron_variant | MODIFIER |
| 10 | 70149411 | AX-170367458 | 0 | C | T | 0.412 | -7.75E-02 | 8.02E-02 | 2.67E+00 | 2.74E+00 | 3.29E-01 | no | Ccdc6 | intron_variant | MODIFIER |
| 10 | 70159016 | AX-170282590 | 0 | G | A | 0.412 | -7.75E-02 | 8.02E-02 | 2.67E+00 | 2.74E+00 | 3.29E-01 | no | Ccdc6 | intron_variant | MODIFIER |
| 10 | 70230789 | AX-170206438 | 0 | A | G | 0.412 | -7.75E-02 | 8.02E-02 | 2.67E+00 | 2.74E+00 | 3.29E-01 | no | | intergenic_variant | MODIFIER |
| 10 | 70233646 | AX-170439789 | 0 | C | T | 0.412 | -7.75E-02 | 8.02E-02 | 2.67E+00 | 2.74E+00 | 3.29E-01 | no | | intergenic_variant | MODIFIER |
| 10 | 70261451 | AX-170430832 | 0 | T | C | 0.412 | -7.75E-02 | 8.02E-02 | 2.67E+00 | 2.74E+00 | 3.29E-01 | no | Slc16a9 | intron_variant | MODIFIER |
| 10 | 70263256 | AX-170448293 | 0 | A | G | 0.412 | -7.75E-02 | 8.02E-02 | 2.67E+00 | 2.74E+00 | 3.29E-01 | no | Slc16a9 | intron_variant | MODIFIER |
| 10 | 70265714 | AX-170045166 | 0 | C | G | 0.412 | -7.75E-02 | 8.02E-02 | 2.67E+00 | 2.74E+00 | 3.29E-01 | no | Slc16a9 | intron_variant | MODIFIER |
| 10 | 70276031 | AX-97836899 | 0 | T | C | 0.412 | -7.75E-02 | 8.02E-02 | 2.67E+00 | 2.74E+00 | 3.29E-01 | no | Slc16a9 | intron_variant | MODIFIER |

| | | | | | | | | | | | | | | | |
|---|---|---|---|---|---|---|---|---|---|---|---|---|---|---|---|
| 10 | 70276796 | AX-170045165 | 0 | A | G | 0.412 | -7.75E-02 | 8.02E-02 | 2.67E+00 | 2.74E+00 | 3.29E-01 | no | Slc16a9 | intron_variant | MODIFIER |
| 10 | 70277520 | AX-170521993 | 0 | C | T | 0.412 | -7.75E-02 | 8.02E-02 | 2.67E+00 | 2.74E+00 | 3.29E-01 | no | Slc16a9 | intron_variant | MODIFIER |
| 10 | 70283327 | AX-97835907 | 0 | A | G | 0.412 | -7.75E-02 | 8.02E-02 | 2.67E+00 | 2.74E+00 | 3.29E-01 | no | Slc16a9 | intron_variant | MODIFIER |
| 10 | 70313551 | AX-170003433 | 0 | C | T | 0.412 | -7.75E-02 | 8.02E-02 | 2.67E+00 | 2.74E+00 | 3.29E-01 | no | | intergenic_variant | MODIFIER |
| 10 | 70323288 | AX-170301158 | 0 | A | G | 0.412 | -7.75E-02 | 8.02E-02 | 2.67E+00 | 2.74E+00 | 3.29E-01 | no | | intergenic_variant | MODIFIER |
| 10 | 70337397 | AX-170277711 | 0 | A | G | 0.412 | -7.75E-02 | 8.02E-02 | 2.67E+00 | 2.74E+00 | 3.29E-01 | no | Gm20108 | upstream_gene_variant | MODIFIER |
| 10 | 70341599 | AX-170003431 | 0 | C | T | 0.412 | -7.75E-02 | 8.02E-02 | 2.67E+00 | 2.74E+00 | 3.29E-01 | no | Gm20108 | downstream_gene_variant | MODIFIER |
| 10 | 70343634 | AX-170003432 | 0 | G | A | 0.412 | -7.75E-02 | 8.02E-02 | 2.67E+00 | 2.74E+00 | 3.29E-01 | no | | intergenic_variant | MODIFIER |
| 10 | 70355706 | AX-170426171 | 0 | T | G | 0.412 | -7.75E-02 | 8.02E-02 | 2.67E+00 | 2.74E+00 | 3.29E-01 | no | | intergenic_variant | MODIFIER |
| 10 | 70356983 | AX-170248072 | 0 | A | G | 0.412 | -7.75E-02 | 8.02E-02 | 2.67E+00 | 2.74E+00 | 3.29E-01 | no | | intergenic_variant | MODIFIER |
| 10 | 70362484 | AX-170231495 | 0 | T | C | 0.412 | -7.75E-02 | 8.02E-02 | 2.67E+00 | 2.74E+00 | 3.29E-01 | no | | intergenic_variant | MODIFIER |
| 10 | 70366150 | AX-97793633 | 0 | C | T | 0.412 | -7.75E-02 | 8.02E-02 | 2.67E+00 | 2.74E+00 | 3.29E-01 | no | | intergenic_variant | MODIFIER |
| 10 | 70393393 | AX-170003430 | 0 | C | T | 0.412 | -7.75E-02 | 8.02E-02 | 2.67E+00 | 2.74E+00 | 3.29E-01 | no | | intergenic_variant | MODIFIER |
| 10 | 70403017 | AX-170003428 | 0 | C | T | 0.412 | -7.75E-02 | 8.02E-02 | 2.67E+00 | 2.74E+00 | 3.29E-01 | no | | intergenic_variant | MODIFIER |
| 10 | 70407805 | AX-170003427 | 0 | A | G | 0.412 | -7.75E-02 | 8.02E-02 | 2.67E+00 | 2.74E+00 | 3.29E-01 | no | Gm47946 | downstream_gene_variant | MODIFIER |
| 10 | 70446894 | AX-170447693 | 0 | T | C | 0.412 | -7.75E-02 | 8.02E-02 | 2.67E+00 | 2.74E+00 | 3.29E-01 | no | Fam13c | intron_variant | MODIFIER |
| 10 | 70463581 | AX-170086837 | 0 | C | T | 0.412 | -7.75E-02 | 8.02E-02 | 2.67E+00 | 2.74E+00 | 3.29E-01 | no | Fam13c | intron_variant | MODIFIER |
| 10 | 70465015 | AX-170003434 | 0 | C | T | 0.412 | -7.75E-02 | 8.02E-02 | 2.67E+00 | 2.74E+00 | 3.29E-01 | no | Fam13c | intron_variant | MODIFIER |
| 10 | 70465613 | AX-170370759 | 0 | C | T | 0.412 | -7.75E-02 | 8.02E-02 | 2.67E+00 | 2.74E+00 | 3.29E-01 | no | Fam13c | intron_variant | MODIFIER |
| 10 | 70562348 | AX-170385737 | 0 | G | A | 0.412 | -7.75E-02 | 8.02E-02 | 2.67E+00 | 2.74E+00 | 3.29E-01 | no | Phyhipl | intron_variant | MODIFIER |
| 10 | 70753796 | AX-170506457 | 0 | A | G | 0.412 | -7.75E-02 | 8.02E-02 | 2.67E+00 | 2.74E+00 | 3.29E-01 | no | | intergenic_variant | MODIFIER |
| 10 | 70941317 | AX-170186752 | 0 | C | T | 0.412 | -7.75E-02 | 8.02E-02 | 2.67E+00 | 2.74E+00 | 3.29E-01 | no | Bicc1 | intron_variant | MODIFIER |
| 10 | 70941339 | AX-170003444 | 0 | A | C | 0.412 | -7.75E-02 | 8.02E-02 | 2.67E+00 | 2.74E+00 | 3.29E-01 | no | Bicc1 | intron_variant | MODIFIER |
| 10 | 70943629 | AX-170003443 | 0 | T | A | 0.412 | -7.75E-02 | 8.02E-02 | 2.67E+00 | 2.74E+00 | 3.29E-01 | no | Bicc1 | intron_variant | MODIFIER |
| 10 | 70957632 | AX-170045180 | 0 | G | A | 0.412 | -7.75E-02 | 8.02E-02 | 2.67E+00 | 2.74E+00 | 3.29E-01 | no | Bicc1 | intron_variant | MODIFIER |
| 10 | 70957866 | AX-170442114 | 0 | G | T | 0.412 | -7.75E-02 | 8.02E-02 | 2.67E+00 | 2.74E+00 | 3.29E-01 | no | Bicc1 | intron_variant | MODIFIER |
| 10 | 70958068 | AX-170431620 | 0 | T | C | 0.412 | -7.75E-02 | 8.02E-02 | 2.67E+00 | 2.74E+00 | 3.29E-01 | no | Bicc1 | intron_variant | MODIFIER |
| 10 | 70958917 | AX-97839862 | 0 | G | A | 0.412 | -7.75E-02 | 8.02E-02 | 2.67E+00 | 2.74E+00 | 3.29E-01 | no | Bicc1 | synonymous_variant | LOW |
| 10 | 70962855 | AX-170222401 | 0 | C | A | 0.412 | -7.75E-02 | 8.02E-02 | 2.67E+00 | 2.74E+00 | 3.29E-01 | no | Bicc1 | intron_variant | MODIFIER |
| 10 | 70963916 | AX-170483573 | 0 | T | G | 0.412 | -7.75E-02 | 8.02E-02 | 2.67E+00 | 2.74E+00 | 3.29E-01 | no | Bicc1 | intron_variant | MODIFIER |
| 10 | 70968432 | AX-170258239 | 0 | A | G | 0.412 | -7.75E-02 | 8.02E-02 | 2.67E+00 | 2.74E+00 | 3.29E-01 | no | Bicc1 | intron_variant | MODIFIER |
| 10 | 70969038 | AX-170172076 | 0 | A | C | 0.412 | -7.75E-02 | 8.02E-02 | 2.67E+00 | 2.74E+00 | 3.29E-01 | no | Bicc1 | intron_variant | MODIFIER |
| 10 | 71031613 | AX-170240576 | 0 | A | G | 0.412 | -7.75E-02 | 8.02E-02 | 2.67E+00 | 2.74E+00 | 3.29E-01 | no | Bicc1 | intron_variant | MODIFIER |
| 10 | 71031757 | AX-97823358 | 0 | T | G | 0.412 | -7.75E-02 | 8.02E-02 | 2.67E+00 | 2.74E+00 | 3.29E-01 | no | Bicc1 | intron_variant | MODIFIER |
| 10 | 71031800 | AX-170474532 | 0 | A | G | 0.412 | -7.75E-02 | 8.02E-02 | 2.67E+00 | 2.74E+00 | 3.29E-01 | no | Bicc1 | intron_variant | MODIFIER |
| 10 | 71035579 | AX-170480892 | 0 | T | C | 0.412 | -7.75E-02 | 8.02E-02 | 2.67E+00 | 2.74E+00 | 3.29E-01 | no | Bicc1 | intron_variant | MODIFIER |
| 10 | 71036828 | AX-170260542 | 0 | T | C | 0.412 | -7.75E-02 | 8.02E-02 | 2.67E+00 | 2.74E+00 | 3.29E-01 | no | Bicc1 | intron_variant | MODIFIER |
| 10 | 71037113 | AX-170234200 | 0 | C | A | 0.412 | -7.75E-02 | 8.02E-02 | 2.67E+00 | 2.74E+00 | 3.29E-01 | no | Bicc1 | intron_variant | MODIFIER |
| 10 | 71107626 | AX-97848713 | 0 | A | G | 0.412 | -7.75E-02 | 8.02E-02 | 2.67E+00 | 2.74E+00 | 3.29E-01 | no | Bicc1 | intron_variant | MODIFIER |
| 10 | 71109880 | AX-170218411 | 0 | C | G | 0.412 | -7.75E-02 | 8.02E-02 | 2.67E+00 | 2.74E+00 | 3.29E-01 | no | Bicc1 | intron_variant | MODIFIER |
| 10 | 71126373 | AX-170252136 | 0 | A | T | 0.412 | -7.75E-02 | 8.02E-02 | 2.67E+00 | 2.74E+00 | 3.29E-01 | no | Bicc1 | intron_variant | MODIFIER |
| 10 | 71132675 | AX-170106183 | 0 | G | A | 0.412 | -7.75E-02 | 8.02E-02 | 2.67E+00 | 2.74E+00 | 3.29E-01 | no | Bicc1 | intron_variant | MODIFIER |
| 10 | 71136618 | AX-170272315 | 0 | G | A | 0.412 | -7.75E-02 | 8.02E-02 | 2.67E+00 | 2.74E+00 | 3.29E-01 | no | Bicc1 | intron_variant | MODIFIER |
| 10 | 71136884 | AX-170150712 | 0 | A | G | 0.412 | -7.75E-02 | 8.02E-02 | 2.67E+00 | 2.74E+00 | 3.29E-01 | no | Bicc1 | intron_variant | MODIFIER |
| 10 | 71149241 | AX-97815888 | 0 | A | G | 0.412 | -7.75E-02 | 8.02E-02 | 2.67E+00 | 2.74E+00 | 3.29E-01 | no | Bicc1 | intron_variant | MODIFIER |
| 10 | 71149296 | AX-170270883 | 0 | C | T | 0.412 | -7.75E-02 | 8.02E-02 | 2.67E+00 | 2.74E+00 | 3.29E-01 | no | Bicc1 | intron_variant | MODIFIER |
| 10 | 71149633 | AX-170448757 | 0 | G | A | 0.412 | -7.75E-02 | 8.02E-02 | 2.67E+00 | 2.74E+00 | 3.29E-01 | no | Bicc1 | intron_variant | MODIFIER |
| 10 | 71149733 | AX-170455060 | 0 | C | A | 0.412 | -7.75E-02 | 8.02E-02 | 2.67E+00 | 2.74E+00 | 3.29E-01 | no | Bicc1 | intron_variant | MODIFIER |
| 10 | 71157674 | AX-170390419 | 0 | G | A | 0.412 | -7.75E-02 | 8.02E-02 | 2.67E+00 | 2.74E+00 | 3.29E-01 | no | Bicc1 | intron_variant | MODIFIER |
| 10 | 71168282 | AX-170513631 | 0 | T | C | 0.412 | -7.75E-02 | 8.02E-02 | 2.67E+00 | 2.74E+00 | 3.29E-01 | no | Gm6331 | downstream_gene_variant | MODIFIER |
| 10 | 71169306 | AX-170240168 | 0 | T | C | 0.412 | -7.75E-02 | 8.02E-02 | 2.67E+00 | 2.74E+00 | 3.29E-01 | no | C730027H18Rik | non_coding_transcript_exon_variant | MODIFIER |
| 10 | 71169718 | AX-170374918 | 0 | G | A | 0.412 | -7.75E-02 | 8.02E-02 | 2.67E+00 | 2.74E+00 | 3.29E-01 | no | C730027H18Rik | non_coding_transcript_exon_variant | MODIFIER |
| 10 | 71216245 | AX-170035482 | 0 | A | C | 0.412 | -7.75E-02 | 8.02E-02 | 2.67E+00 | 2.74E+00 | 3.29E-01 | no | | intergenic_variant | MODIFIER |
| 10 | 71328934 | AX-97815863 | 0 | G | A | 0.412 | -7.75E-02 | 8.02E-02 | 2.67E+00 | 2.74E+00 | 3.29E-01 | no | Cisd1 | downstream_gene_variant | MODIFIER |

| | | | | | | | | | | | | | | | |
|---|---|---|---|---|---|---|---|---|---|---|---|---|---|---|---|
| 10 | 71387539 | AX-97823436 | 0 | C | T | 0.412 | -7.75E-02 | 8.02E-02 | 2.67E+00 | 2.74E+00 | 3.29E-01 | no | Ipmk | downstream_gene_variant | MODIFIER |
| 10 | 71467755 | AX-170178117 | 0 | T | A | 0.412 | -7.75E-02 | 8.02E-02 | 2.67E+00 | 2.74E+00 | 3.29E-01 | no | | intergenic_variant | MODIFIER |
| 10 | 71480032 | AX-97816993 | 0 | C | A | 0.412 | -7.75E-02 | 8.02E-02 | 2.67E+00 | 2.74E+00 | 3.29E-01 | no | Gm46233 | upstream_gene_variant | MODIFIER |
| 10 | 71498018 | AX-170255010 | 0 | T | C | 0.412 | -7.75E-02 | 8.02E-02 | 2.67E+00 | 2.74E+00 | 3.29E-01 | no | | intergenic_variant | MODIFIER |
| 10 | 71507361 | AX-97815937 | 0 | C | T | 0.412 | -7.75E-02 | 8.02E-02 | 2.67E+00 | 2.74E+00 | 3.29E-01 | no | | intergenic_variant | MODIFIER |
| 10 | 71546127 | AX-97835685 | 0 | G | T | 0.412 | -7.75E-02 | 8.02E-02 | 2.67E+00 | 2.74E+00 | 3.29E-01 | no | Gm34544 | intron_variant,non_coding_transcript_variant | MODIFIER |
| 10 | 71566394 | AX-170383577 | 0 | G | A | 0.412 | -7.75E-02 | 8.02E-02 | 2.67E+00 | 2.74E+00 | 3.29E-01 | no | Gm34544 | intron_variant,non_coding_transcript_variant | MODIFIER |
| 10 | 71566838 | AX-97819879 | 0 | C | T | 0.412 | -7.75E-02 | 8.02E-02 | 2.67E+00 | 2.74E+00 | 3.29E-01 | no | Gm34544 | intron_variant,non_coding_transcript_variant | MODIFIER |
| 10 | 71985862 | AX-170003456 | 0 | G | T | 0.412 | -7.75E-02 | 8.02E-02 | 2.67E+00 | 2.74E+00 | 3.29E-01 | no | | intergenic_variant | MODIFIER |
| 10 | 71990414 | AX-97856329 | 0 | G | T | 0.412 | -7.75E-02 | 8.02E-02 | 2.67E+00 | 2.74E+00 | 3.29E-01 | no | | intergenic_variant | MODIFIER |
| 10 | 72040227 | AX-170504686 | 0 | A | C | 0.412 | -7.75E-02 | 8.02E-02 | 2.67E+00 | 2.74E+00 | 3.29E-01 | no | Gm34609 | intron_variant,non_coding_transcript_variant | MODIFIER |
| 10 | 72049387 | AX-97823446 | 0 | C | T | 0.412 | -7.75E-02 | 8.02E-02 | 2.67E+00 | 2.74E+00 | 3.29E-01 | no | Gm34609 | intron_variant,non_coding_transcript_variant | MODIFIER |
| 10 | 72124603 | AX-170045207 | 0 | T | G | 0.412 | -7.75E-02 | 8.02E-02 | 2.67E+00 | 2.74E+00 | 3.29E-01 | no | | intergenic_variant | MODIFIER |
| 10 | 72143023 | AX-170133849 | 0 | T | C | 0.412 | -7.75E-02 | 8.02E-02 | 2.67E+00 | 2.74E+00 | 3.29E-01 | no | | intergenic_variant | MODIFIER |
| 10 | 72144584 | AX-170136137 | 0 | G | C | 0.412 | -7.75E-02 | 8.02E-02 | 2.67E+00 | 2.74E+00 | 3.29E-01 | no | | intergenic_variant | MODIFIER |
| 10 | 72157319 | AX-170045202 | 0 | C | T | 0.412 | -7.75E-02 | 8.02E-02 | 2.67E+00 | 2.74E+00 | 3.29E-01 | no | | intergenic_variant | MODIFIER |
| 10 | 72159695 | AX-170117315 | 0 | G | A | 0.412 | -7.75E-02 | 8.02E-02 | 2.67E+00 | 2.74E+00 | 3.29E-01 | no | | intergenic_variant | MODIFIER |
| 10 | 72160874 | AX-170045208 | 0 | C | A | 0.412 | -7.75E-02 | 8.02E-02 | 2.67E+00 | 2.74E+00 | 3.29E-01 | no | | intergenic_variant | MODIFIER |
| 10 | 72161277 | AX-170045201 | 0 | C | A | 0.412 | -7.75E-02 | 8.02E-02 | 2.67E+00 | 2.74E+00 | 3.29E-01 | no | | intergenic_variant | MODIFIER |
| 10 | 72186736 | AX-170280449 | 0 | A | C | 0.412 | -7.75E-02 | 8.02E-02 | 2.67E+00 | 2.74E+00 | 3.29E-01 | no | | intergenic_variant | MODIFIER |
| 10 | 72187258 | AX-170267611 | 0 | A | G | 0.412 | -7.75E-02 | 8.02E-02 | 2.67E+00 | 2.74E+00 | 3.29E-01 | no | | intergenic_variant | MODIFIER |
| 10 | 72214153 | AX-170253445 | 0 | T | C | 0.412 | -7.75E-02 | 8.02E-02 | 2.67E+00 | 2.74E+00 | 3.29E-01 | no | | intergenic_variant | MODIFIER |
| 10 | 72222004 | AX-170408224 | 0 | A | T | 0.412 | -7.75E-02 | 8.02E-02 | 2.67E+00 | 2.74E+00 | 3.29E-01 | no | | intergenic_variant | MODIFIER |
| 10 | 72256565 | AX-170092025 | 0 | C | G | 0.412 | -7.75E-02 | 8.02E-02 | 2.67E+00 | 2.74E+00 | 3.29E-01 | no | | intergenic_variant | MODIFIER |
| 10 | 72281295 | AX-170337906 | 0 | A | C | 0.412 | -7.75E-02 | 8.02E-02 | 2.67E+00 | 2.74E+00 | 3.29E-01 | no | | intergenic_variant | MODIFIER |
| 10 | 72291719 | AX-170459218 | 0 | T | G | 0.412 | -7.75E-02 | 8.02E-02 | 2.67E+00 | 2.74E+00 | 3.29E-01 | no | | intergenic_variant | MODIFIER |
| 10 | 72295954 | AX-170045210 | 0 | A | C | 0.412 | -7.75E-02 | 8.02E-02 | 2.67E+00 | 2.74E+00 | 3.29E-01 | no | | intergenic_variant | MODIFIER |
| 10 | 72330012 | AX-170132440 | 0 | T | G | 0.412 | -7.75E-02 | 8.02E-02 | 2.67E+00 | 2.74E+00 | 3.29E-01 | no | | intergenic_variant | MODIFIER |
| 10 | 72333479 | AX-170477065 | 0 | T | C | 0.412 | -7.75E-02 | 8.02E-02 | 2.67E+00 | 2.74E+00 | 3.29E-01 | no | | intergenic_variant | MODIFIER |
| 10 | 72363286 | AX-170306998 | 0 | T | A | 0.412 | -7.75E-02 | 8.02E-02 | 2.67E+00 | 2.74E+00 | 3.29E-01 | no | | intergenic_variant | MODIFIER |
| 10 | 72409846 | AX-170461953 | 0 | A | T | 0.412 | -7.75E-02 | 8.02E-02 | 2.67E+00 | 2.74E+00 | 3.29E-01 | no | | intergenic_variant | MODIFIER |
| 10 | 72424824 | AX-170416614 | 0 | G | A | 0.412 | -7.75E-02 | 8.02E-02 | 2.67E+00 | 2.74E+00 | 3.29E-01 | no | | intergenic_variant | MODIFIER |
| 10 | 72486142 | AX-170415340 | 0 | G | A | 0.412 | -7.75E-02 | 8.02E-02 | 2.67E+00 | 2.74E+00 | 3.29E-01 | no | | intergenic_variant | MODIFIER |
| 10 | 72487170 | AX-170481903 | 0 | C | T | 0.412 | -7.75E-02 | 8.02E-02 | 2.67E+00 | 2.74E+00 | 3.29E-01 | no | | intergenic_variant | MODIFIER |
| 10 | 72489657 | AX-97811454 | 0 | G | A | 0.412 | -7.75E-02 | 8.02E-02 | 2.67E+00 | 2.74E+00 | 3.29E-01 | no | | intergenic_variant | MODIFIER |
| 10 | 72489712 | AX-170102458 | 0 | G | A | 0.412 | -7.75E-02 | 8.02E-02 | 2.67E+00 | 2.74E+00 | 3.29E-01 | no | | intergenic_variant | MODIFIER |
| 10 | 72623817 | AX-170221018 | 0 | G | A | 0.412 | -7.75E-02 | 8.02E-02 | 2.67E+00 | 2.74E+00 | 3.29E-01 | no | | regulatory_region_variant | MODIFIER |
| 10 | 72682616 | AX-170426758 | 0 | G | C | 0.412 | -7.75E-02 | 8.02E-02 | 2.67E+00 | 2.74E+00 | 3.29E-01 | no | | intergenic_variant | MODIFIER |
| 10 | 72780332 | AX-170138347 | 0 | G | A | 0.412 | -7.75E-02 | 8.02E-02 | 2.67E+00 | 2.74E+00 | 3.29E-01 | no | | intergenic_variant | MODIFIER |
| 10 | 72795663 | AX-97803487 | 0 | G | A | 0.412 | -7.75E-02 | 8.02E-02 | 2.67E+00 | 2.74E+00 | 3.29E-01 | no | | intergenic_variant | MODIFIER |
| 10 | 72796349 | AX-170226031 | 0 | A | G | 0.412 | -7.75E-02 | 8.02E-02 | 2.67E+00 | 2.74E+00 | 3.29E-01 | no | | intergenic_variant | MODIFIER |
| 10 | 72806017 | AX-170264488 | 0 | A | G | 0.412 | -7.75E-02 | 8.02E-02 | 2.67E+00 | 2.74E+00 | 3.29E-01 | no | | intergenic_variant | MODIFIER |
| 10 | 72810064 | AX-170531571 | 0 | T | G | 0.412 | -7.75E-02 | 8.02E-02 | 2.67E+00 | 2.74E+00 | 3.29E-01 | no | | regulatory_region_variant | MODIFIER |
| 10 | 72981462 | AX-170045223 | 0 | C | T | 0.412 | -7.75E-02 | 8.02E-02 | 2.67E+00 | 2.74E+00 | 3.29E-01 | no | A330049N07Rik | intron_variant,non_coding_transcript_variant | MODIFIER |
| 10 | 73022107 | AX-170450775 | 0 | C | T | 0.412 | -7.75E-02 | 8.02E-02 | 2.67E+00 | 2.74E+00 | 3.29E-01 | no | A330049N07Rik | intron_variant,non_coding_transcript_variant | MODIFIER |
| 10 | 73026245 | AX-170086840 | 0 | G | A | 0.412 | -7.75E-02 | 8.02E-02 | 2.67E+00 | 2.74E+00 | 3.29E-01 | no | A330049N07Rik | intron_variant,non_coding_transcript_variant | MODIFIER |
| 10 | 73035336 | AX-170394482 | 0 | G | A | 0.412 | -7.75E-02 | 8.02E-02 | 2.67E+00 | 2.74E+00 | 3.29E-01 | no | A330049N07Rik | intron_variant,non_coding_transcript_variant | MODIFIER |
| 10 | 73063527 | AX-170218662 | 0 | A | G | 0.412 | -7.75E-02 | 8.02E-02 | 2.67E+00 | 2.74E+00 | 3.29E-01 | no | A330049N07Rik | intron_variant,non_coding_transcript_variant | MODIFIER |
| 10 | 73126463 | AX-170045226 | 0 | A | G | 0.412 | -7.75E-02 | 8.02E-02 | 2.67E+00 | 2.74E+00 | 3.29E-01 | no | Pcdh15 | intron_variant,non_coding_transcript_variant | MODIFIER |
| 4 | 134025871 | AX-170354134 | 0 | G | A | 0.167 | 1.15E-01 | 1.18E-01 | 2.58E+00 | 2.62E+00 | 3.29E-01 | no | | intergenic_variant | MODIFIER |
| 4 | 134053095 | AX-170150530 | 0 | C | T | 0.167 | 1.15E-01 | 1.18E-01 | 2.58E+00 | 2.62E+00 | 3.29E-01 | no | Zfp683 | upstream_gene_variant | MODIFIER |
| 4 | 134071346 | AX-170379616 | 0 | G | A | 0.167 | 1.15E-01 | 1.18E-01 | 2.58E+00 | 2.62E+00 | 3.29E-01 | no | Crybg2 | intron_variant | MODIFIER |
| 4 | 134075927 | AX-170356247 | 0 | C | T | 0.167 | 1.15E-01 | 1.18E-01 | 2.58E+00 | 2.62E+00 | 3.29E-01 | no | Crybg2 | intron_variant | MODIFIER |
| 4 | 134076670 | AX-170070753 | 0 | A | C | 0.167 | 1.15E-01 | 1.18E-01 | 2.58E+00 | 2.62E+00 | 3.29E-01 | no | Crybg2 | intron_variant | MODIFIER |

| | | | | | | | | | | | | | | | |
|---|---|---|---|---|---|---|---|---|---|---|---|---|---|---|---|
| 4 | 134238248 | AX-97850370 | 0 | T | G | 0.167 | 1.15E-01 | 1.18E-01 | 2.58E+00 | 2.62E+00 | 3.29E-01 | no | Cnksr1 | intron_variant | MODIFIER |
| 4 | 134471307 | AX-170410486 | 0 | C | T | 0.167 | 1.15E-01 | 1.18E-01 | 2.58E+00 | 2.62E+00 | 3.29E-01 | no | Stmn1 | intron_variant | MODIFIER |
| 4 | 134544212 | AX-170267217 | 0 | T | C | 0.167 | 1.15E-01 | 1.18E-01 | 2.58E+00 | 2.62E+00 | 3.29E-01 | no | Selenon | intron_variant | MODIFIER |
| 4 | 134571173 | AX-97820552 | 0 | T | G | 0.167 | 1.15E-01 | 1.18E-01 | 2.58E+00 | 2.62E+00 | 3.29E-01 | no | Man1c1 | intron_variant | MODIFIER |
| 4 | 134571881 | AX-170469475 | 0 | G | A | 0.167 | 1.15E-01 | 1.18E-01 | 2.58E+00 | 2.62E+00 | 3.29E-01 | no | Man1c1 | intron_variant | MODIFIER |
| 4 | 135328906 | AX-97801572 | 0 | A | C | 0.167 | 1.15E-01 | 1.18E-01 | 2.58E+00 | 2.62E+00 | 3.29E-01 | no | Srrm1 | intron_variant | MODIFIER |
| 4 | 135336184 | AX-170024015 | 0 | T | C | 0.167 | 1.15E-01 | 1.18E-01 | 2.58E+00 | 2.62E+00 | 3.29E-01 | no | Srrm1 | intron_variant | MODIFIER |
| 4 | 135336456 | AX-170152742 | 0 | A | G | 0.167 | 1.15E-01 | 1.18E-01 | 2.58E+00 | 2.62E+00 | 3.29E-01 | no | Srrm1 | intron_variant | MODIFIER |
| 4 | 135452136 | AX-97815324 | 0 | T | C | 0.167 | 1.15E-01 | 1.18E-01 | 2.58E+00 | 2.62E+00 | 3.29E-01 | no | Nipal3 | 3_prime_UTR_variant | MODIFIER |
| 4 | 135452790 | AX-170380890 | 0 | G | A | 0.167 | 1.15E-01 | 1.18E-01 | 2.58E+00 | 2.62E+00 | 3.29E-01 | no | Nipal3 | intron_variant | MODIFIER |
| 4 | 135452807 | AX-170327043 | 0 | G | T | 0.167 | 1.15E-01 | 1.18E-01 | 2.58E+00 | 2.62E+00 | 3.29E-01 | no | Nipal3 | intron_variant | MODIFIER |
| 4 | 135466700 | AX-170281321 | 0 | T | C | 0.167 | 1.15E-01 | 1.18E-01 | 2.58E+00 | 2.62E+00 | 3.29E-01 | no | Nipal3 | intron_variant | MODIFIER |
| 4 | 135466777 | AX-170150838 | 0 | G | A | 0.167 | 1.15E-01 | 1.18E-01 | 2.58E+00 | 2.62E+00 | 3.29E-01 | no | Nipal3 | intron_variant | MODIFIER |
| 4 | 135469906 | AX-170024012 | 0 | T | G | 0.167 | 1.15E-01 | 1.18E-01 | 2.58E+00 | 2.62E+00 | 3.29E-01 | no | Nipal3 | intron_variant | MODIFIER |
| 4 | 135478511 | AX-170221689 | 0 | A | G | 0.167 | 1.15E-01 | 1.18E-01 | 2.58E+00 | 2.62E+00 | 3.29E-01 | no | Nipal3 | intron_variant | MODIFIER |
| 4 | 135547923 | AX-97781343 | 0 | G | A | 0.167 | 1.15E-01 | 1.18E-01 | 2.58E+00 | 2.62E+00 | 3.29E-01 | no | Grhl3 | intron_variant | MODIFIER |
| 4 | 135704092 | AX-170424097 | 0 | T | C | 0.167 | 1.15E-01 | 1.18E-01 | 2.58E+00 | 2.62E+00 | 3.29E-01 | no | Ifnlr1 | intron_variant | MODIFIER |
| 4 | 135714651 | AX-97833991 | 0 | G | A | 0.167 | 1.15E-01 | 1.18E-01 | 2.58E+00 | 2.62E+00 | 3.29E-01 | no | Gm12989 | downstream_gene_variant | MODIFIER |
| 4 | 135715002 | AX-170273849 | 0 | G | A | 0.167 | 1.15E-01 | 1.18E-01 | 2.58E+00 | 2.62E+00 | 3.29E-01 | no | Gm12989 | downstream_gene_variant | MODIFIER |
| 4 | 135725234 | AX-170407044 | 0 | A | G | 0.167 | 1.15E-01 | 1.18E-01 | 2.58E+00 | 2.62E+00 | 3.29E-01 | no | Il22ra1 | upstream_gene_variant | MODIFIER |
| 4 | 136034669 | AX-170024021 | 0 | G | C | 0.167 | 1.15E-01 | 1.18E-01 | 2.58E+00 | 2.62E+00 | 3.29E-01 | no | Rpl11 | intron_variant,non_coding_transcript_variant | MODIFIER |
| 4 | 136067146 | AX-97785010 | 0 | C | T | 0.167 | 1.15E-01 | 1.18E-01 | 2.58E+00 | 2.62E+00 | 3.29E-01 | no | | intergenic_variant | MODIFIER |
| 4 | 136118806 | AX-170315606 | 0 | A | C | 0.167 | 1.15E-01 | 1.18E-01 | 2.58E+00 | 2.62E+00 | 3.29E-01 | no | | regulatory_region_variant | MODIFIER |
| 4 | 136121759 | AX-170095162 | 0 | C | T | 0.167 | 1.15E-01 | 1.18E-01 | 2.58E+00 | 2.62E+00 | 3.29E-01 | no | | regulatory_region_variant | MODIFIER |
| 4 | 136176383 | AX-97809086 | 0 | T | C | 0.167 | 1.15E-01 | 1.18E-01 | 2.58E+00 | 2.62E+00 | 3.29E-01 | no | E2f2 | intron_variant | MODIFIER |
| 4 | 136179444 | AX-170070797 | 0 | G | T | 0.167 | 1.15E-01 | 1.18E-01 | 2.58E+00 | 2.62E+00 | 3.29E-01 | no | E2f2 | intron_variant | MODIFIER |
| 4 | 136188467 | AX-97852554 | 0 | G | A | 0.167 | 1.15E-01 | 1.18E-01 | 2.58E+00 | 2.62E+00 | 3.29E-01 | no | E2f2 | intron_variant | MODIFIER |
| 4 | 136223324 | AX-170374847 | 0 | T | C | 0.167 | 1.15E-01 | 1.18E-01 | 2.58E+00 | 2.62E+00 | 3.29E-01 | no | Asap3 | intron_variant | MODIFIER |
| 4 | 136233426 | AX-170511164 | 0 | A | G | 0.167 | 1.15E-01 | 1.18E-01 | 2.58E+00 | 2.62E+00 | 3.29E-01 | no | Asap3 | intron_variant | MODIFIER |
| 4 | 136275949 | AX-170242502 | 0 | A | G | 0.167 | 1.15E-01 | 1.18E-01 | 2.58E+00 | 2.62E+00 | 3.29E-01 | no | Tcea3 | downstream_gene_variant | MODIFIER |
| 4 | 136325052 | AX-97809743 | 0 | C | T | 0.167 | 1.15E-01 | 1.18E-01 | 2.58E+00 | 2.62E+00 | 3.29E-01 | no | Hnrnpr | intron_variant | MODIFIER |
| 4 | 136359234 | AX-170133120 | 0 | C | T | 0.167 | 1.15E-01 | 1.18E-01 | 2.58E+00 | 2.62E+00 | 3.29E-01 | no | Hnrnpr | 3_prime_UTR_variant | MODIFIER |
| 4 | 136442401 | AX-170089382 | 0 | G | A | 0.167 | 1.15E-01 | 1.18E-01 | 2.58E+00 | 2.62E+00 | 3.29E-01 | no | Htr1d | 5_prime_UTR_variant | MODIFIER |
| 4 | 136442626 | AX-170392928 | 0 | C | T | 0.167 | 1.15E-01 | 1.18E-01 | 2.58E+00 | 2.62E+00 | 3.29E-01 | no | Htr1d | synonymous_variant | LOW |
| 4 | 136566466 | AX-170416743 | 0 | C | T | 0.167 | 1.15E-01 | 1.18E-01 | 2.58E+00 | 2.62E+00 | 3.29E-01 | no | Kdm1a | intron_variant | MODIFIER |
| 4 | 136573333 | AX-170101844 | 0 | T | A | 0.167 | 1.15E-01 | 1.18E-01 | 2.58E+00 | 2.62E+00 | 3.29E-01 | no | Kdm1a | intron_variant | MODIFIER |
| 4 | 136583368 | AX-170095164 | 0 | G | A | 0.167 | 1.15E-01 | 1.18E-01 | 2.58E+00 | 2.62E+00 | 3.29E-01 | no | Kdm1a | intron_variant | MODIFIER |
| 4 | 136621551 | AX-97855154 | 0 | A | G | 0.167 | 1.15E-01 | 1.18E-01 | 2.58E+00 | 2.62E+00 | 3.29E-01 | no | Lactbl1 | upstream_gene_variant | MODIFIER |
| 4 | 136621713 | AX-170070804 | 0 | T | C | 0.167 | 1.15E-01 | 1.18E-01 | 2.58E+00 | 2.62E+00 | 3.29E-01 | no | Lactbl1 | upstream_gene_variant | MODIFIER |
| 4 | 136824585 | AX-170318632 | 0 | A | T | 0.167 | 1.15E-01 | 1.18E-01 | 2.58E+00 | 2.62E+00 | 3.29E-01 | no | Ephb2 | intron_variant | MODIFIER |
| 4 | 136868638 | AX-97849796 | 0 | G | A | 0.167 | 1.15E-01 | 1.18E-01 | 2.58E+00 | 2.62E+00 | 3.29E-01 | no | | regulatory_region_variant | MODIFIER |
| 4 | 136923804 | AX-97834492 | 0 | A | G | 0.167 | 1.15E-01 | 1.18E-01 | 2.58E+00 | 2.62E+00 | 3.29E-01 | no | | intergenic_variant | MODIFIER |
| 4 | 136925971 | AX-170521538 | 0 | C | T | 0.167 | 1.15E-01 | 1.18E-01 | 2.58E+00 | 2.62E+00 | 3.29E-01 | no | Epha8 | downstream_gene_variant | MODIFIER |
| 4 | 136958690 | AX-170378577 | 0 | T | C | 0.167 | 1.15E-01 | 1.18E-01 | 2.58E+00 | 2.62E+00 | 3.29E-01 | no | Epha8 | upstream_gene_variant | MODIFIER |
| 4 | 136958785 | AX-170484498 | 0 | C | T | 0.167 | 1.15E-01 | 1.18E-01 | 2.58E+00 | 2.62E+00 | 3.29E-01 | no | Epha8 | upstream_gene_variant | MODIFIER |
| 4 | 136963250 | AX-170070812 | 0 | A | G | 0.167 | 1.15E-01 | 1.18E-01 | 2.58E+00 | 2.62E+00 | 3.29E-01 | no | | intergenic_variant | MODIFIER |
| 4 | 136974903 | AX-97843716 | 0 | C | T | 0.167 | 1.15E-01 | 1.18E-01 | 2.58E+00 | 2.62E+00 | 3.29E-01 | no | Zbtb40 | downstream_gene_variant | MODIFIER |
| 4 | 136987928 | AX-170149374 | 0 | A | G | 0.167 | 1.15E-01 | 1.18E-01 | 2.58E+00 | 2.62E+00 | 3.29E-01 | no | Zbtb40 | intron_variant | MODIFIER |
| 4 | 136988860 | AX-170236527 | 0 | C | T | 0.167 | 1.15E-01 | 1.18E-01 | 2.58E+00 | 2.62E+00 | 3.29E-01 | no | Zbtb40 | intron_variant | MODIFIER |
| 4 | 136995155 | AX-170514599 | 0 | G | A | 0.167 | 1.15E-01 | 1.18E-01 | 2.58E+00 | 2.62E+00 | 3.29E-01 | no | Zbtb40 | intron_variant | MODIFIER |
| 4 | 137009590 | AX-97822527 | 0 | A | C | 0.167 | 1.15E-01 | 1.18E-01 | 2.58E+00 | 2.62E+00 | 3.29E-01 | no | Zbtb40 | intron_variant | MODIFIER |
| 4 | 137103983 | AX-170070817 | 0 | G | A | 0.167 | 1.15E-01 | 1.18E-01 | 2.58E+00 | 2.62E+00 | 3.29E-01 | no | | regulatory_region_variant | MODIFIER |
| 4 | 137105319 | AX-170239817 | 0 | T | C | 0.167 | 1.15E-01 | 1.18E-01 | 2.58E+00 | 2.62E+00 | 3.29E-01 | no | | regulatory_region_variant | MODIFIER |
| 4 | 137108141 | AX-170252306 | 0 | C | T | 0.167 | 1.15E-01 | 1.18E-01 | 2.58E+00 | 2.62E+00 | 3.29E-01 | no | | intergenic_variant | MODIFIER |

| | | | | | | | | | | | | | | | |
|---|---|---|---|---|---|---|---|---|---|---|---|---|---|---|---|
| 4 | 137119830 | AX-170024044 | 0 | C | T | 0.167 | 1.15E-01 | 1.18E-01 | 2.58E+00 | 2.62E+00 | 3.29E-01 | no | 1700037C06Rik | upstream_gene_variant | MODIFIER |
| 4 | 137128986 | AX-170111355 | 0 | T | C | 0.167 | 1.15E-01 | 1.18E-01 | 2.58E+00 | 2.62E+00 | 3.29E-01 | no | | intergenic_variant | MODIFIER |
| 4 | 137136727 | AX-170327938 | 0 | A | G | 0.167 | 1.15E-01 | 1.18E-01 | 2.58E+00 | 2.62E+00 | 3.29E-01 | no | | intergenic_variant | MODIFIER |
| 4 | 137138689 | AX-97788980 | 0 | A | C | 0.167 | 1.15E-01 | 1.18E-01 | 2.58E+00 | 2.62E+00 | 3.29E-01 | no | | intergenic_variant | MODIFIER |
| 4 | 137142041 | AX-170367130 | 0 | T | C | 0.167 | 1.15E-01 | 1.18E-01 | 2.58E+00 | 2.62E+00 | 3.29E-01 | no | | regulatory_region_variant | MODIFIER |
| 4 | 137142870 | AX-170291701 | 0 | A | G | 0.167 | 1.15E-01 | 1.18E-01 | 2.58E+00 | 2.62E+00 | 3.29E-01 | no | | regulatory_region_variant | MODIFIER |
| 4 | 137149854 | AX-170236650 | 0 | T | C | 0.167 | 1.15E-01 | 1.18E-01 | 2.58E+00 | 2.62E+00 | 3.29E-01 | no | | intergenic_variant | MODIFIER |
| 4 | 137153594 | AX-170024041 | 0 | A | C | 0.167 | 1.15E-01 | 1.18E-01 | 2.58E+00 | 2.62E+00 | 3.29E-01 | no | | intergenic_variant | MODIFIER |
| 4 | 137154149 | AX-170510507 | 0 | G | A | 0.167 | 1.15E-01 | 1.18E-01 | 2.58E+00 | 2.62E+00 | 3.29E-01 | no | Gm13003 | downstream_gene_variant | MODIFIER |
| 4 | 137156515 | AX-170024043 | 0 | T | C | 0.167 | 1.15E-01 | 1.18E-01 | 2.58E+00 | 2.62E+00 | 3.29E-01 | no | Gm13003 | downstream_gene_variant | MODIFIER |
| 4 | 137163976 | AX-170101094 | 0 | A | T | 0.167 | 1.15E-01 | 1.18E-01 | 2.58E+00 | 2.62E+00 | 3.29E-01 | no | Gm13003 | non_coding_transcript_exon_variant | MODIFIER |
| 4 | 137172297 | AX-170070818 | 0 | T | G | 0.167 | 1.15E-01 | 1.18E-01 | 2.58E+00 | 2.62E+00 | 3.29E-01 | no | | intergenic_variant | MODIFIER |
| 4 | 98936759 | AX-170023481 | 0 | A | C | 0.237 | 1.10E-01 | 1.11E-01 | 2.48E+00 | 2.56E+00 | 3.29E-01 | no | Dock7 | 3_prime_UTR_variant | MODIFIER |
| 1 | 75276632 | AX-170346453 | 1 | C | T | 0.098 | -1.51E-01 | 1.55E-01 | 2.92E+00 | 3.06E+00 | 3.30E-01 | no | Resp18 | intron_variant | MODIFIER |
| 15 | 19581652 | AX-170087788 | 1 | A | T | 0.25 | -9.93E-02 | 1.03E-01 | 2.69E+00 | 2.76E+00 | 3.30E-01 | no | | intergenic_variant | MODIFIER |
| 18 | 88759700 | AX-170061633 | 1 | C | G | 0.161 | -1.02E-01 | 1.05E-01 | 2.55E+00 | 2.64E+00 | 3.30E-01 | no | Socs6 | intron_variant,non_coding_transcript_variant | MODIFIER |
| 3 | 20965713 | AX-170118415 | 1 | G | A | 0.143 | -9.93E-02 | 1.02E-01 | 2.58E+00 | 2.66E+00 | 3.30E-01 | no | | intergenic_variant | MODIFIER |
| 1 | 136673243 | AX-170001788 | 0 | G | A | 0.211 | 1.20E-01 | 1.24E-01 | 2.70E+00 | 2.77E+00 | 3.30E-01 | no | Gm19705 | intron_variant,non_coding_transcript_variant | MODIFIER |
| 5 | 37423601 | AX-97805460 | 0 | T | G | 0.316 | 1.08E-01 | 1.12E-01 | 2.84E+00 | 2.91E+00 | 3.30E-01 | no | Evc2 | intron_variant | MODIFIER |
| 6 | 135947829 | AX-170294097 | 1 | T | C | 0.571 | 8.86E-02 | 8.82E-02 | 2.39E+00 | 2.44E+00 | 3.30E-01 | no | Grin2b | intron_variant | MODIFIER |
| 5 | 85469991 | AX-170025487 | 0 | G | A | 0.43 | 1.05E-01 | 1.08E-01 | 2.94E+00 | 3.01E+00 | 3.30E-01 | no | | intergenic_variant | MODIFIER |
| 15 | 7629139 | AX-170263154 | 1 | C | T | 0.232 | -1.03E-01 | 1.06E-01 | 2.62E+00 | 2.66E+00 | 3.30E-01 | no | | intergenic_variant | MODIFIER |
| 15 | 22906729 | AX-170093181 | 1 | A | G | 0.232 | -1.03E-01 | 1.06E-01 | 2.62E+00 | 2.66E+00 | 3.30E-01 | no | Cdh18 | intron_variant | MODIFIER |
| 5 | 64518057 | AX-170295792 | 0 | G | T | 0.482 | 9.77E-02 | 9.88E-02 | 2.46E+00 | 2.53E+00 | 3.30E-01 | no | | regulatory_region_variant | MODIFIER |
| 5 | 92861086 | AX-170304660 | 0 | C | T | 0.421 | 9.82E-02 | 1.02E-01 | 2.77E+00 | 2.85E+00 | 3.30E-01 | no | Shroom3 | intron_variant | MODIFIER |
| 8 | 14824940 | AX-170030294 | 0 | C | A | 0.237 | -7.42E-02 | 7.57E-02 | 2.50E+00 | 2.59E+00 | 3.30E-01 | no | Dlgap2 | intron_variant | MODIFIER |
| 1 | 176981903 | AX-170407318 | 0 | G | A | 0.228 | 1.12E-01 | 1.16E-01 | 2.61E+00 | 2.68E+00 | 3.31E-01 | no | Sdccag8 | intron_variant | MODIFIER |
| 1 | 176986726 | AX-170299608 | 0 | C | G | 0.228 | 1.12E-01 | 1.16E-01 | 2.61E+00 | 2.68E+00 | 3.31E-01 | no | Sdccag8 | intron_variant | MODIFIER |
| 1 | 176987729 | AX-170480008 | 0 | G | C | 0.228 | 1.12E-01 | 1.16E-01 | 2.61E+00 | 2.68E+00 | 3.31E-01 | no | Sdccag8 | intron_variant | MODIFIER |
| 1 | 176988034 | AX-170264076 | 0 | G | A | 0.228 | 1.12E-01 | 1.16E-01 | 2.61E+00 | 2.68E+00 | 3.31E-01 | no | Sdccag8 | intron_variant | MODIFIER |
| 1 | 176988141 | AX-170426777 | 0 | G | C | 0.228 | 1.12E-01 | 1.16E-01 | 2.61E+00 | 2.68E+00 | 3.31E-01 | no | Sdccag8 | intron_variant | MODIFIER |
| 1 | 177025106 | AX-170444776 | 0 | G | A | 0.228 | 1.12E-01 | 1.16E-01 | 2.61E+00 | 2.68E+00 | 3.31E-01 | no | Akt3 | intron_variant | MODIFIER |
| 1 | 177026139 | AX-170504067 | 0 | T | G | 0.228 | 1.12E-01 | 1.16E-01 | 2.61E+00 | 2.68E+00 | 3.31E-01 | no | Akt3 | intron_variant | MODIFIER |
| 1 | 177027375 | AX-170221338 | 0 | A | C | 0.228 | 1.12E-01 | 1.16E-01 | 2.61E+00 | 2.68E+00 | 3.31E-01 | no | Akt3 | intron_variant | MODIFIER |
| 1 | 177027584 | AX-170086697 | 0 | G | A | 0.228 | 1.12E-01 | 1.16E-01 | 2.61E+00 | 2.68E+00 | 3.31E-01 | no | Akt3 | intron_variant | MODIFIER |
| 1 | 177082062 | AX-97824881 | 0 | T | C | 0.228 | 1.12E-01 | 1.16E-01 | 2.61E+00 | 2.68E+00 | 3.31E-01 | no | Akt3 | intron_variant | MODIFIER |
| 1 | 177296065 | AX-170352526 | 0 | A | G | 0.228 | 1.12E-01 | 1.16E-01 | 2.61E+00 | 2.68E+00 | 3.31E-01 | no | | intergenic_variant | MODIFIER |
| 1 | 177314697 | AX-170414695 | 0 | C | T | 0.228 | 1.12E-01 | 1.16E-01 | 2.61E+00 | 2.68E+00 | 3.31E-01 | no | | intergenic_variant | MODIFIER |
| 1 | 177314948 | AX-170533144 | 0 | A | C | 0.228 | 1.12E-01 | 1.16E-01 | 2.61E+00 | 2.68E+00 | 3.31E-01 | no | | intergenic_variant | MODIFIER |
| 1 | 177317825 | AX-170131902 | 0 | C | T | 0.228 | 1.12E-01 | 1.16E-01 | 2.61E+00 | 2.68E+00 | 3.31E-01 | no | | intergenic_variant | MODIFIER |
| 1 | 177320056 | AX-170312689 | 0 | G | A | 0.228 | 1.12E-01 | 1.16E-01 | 2.61E+00 | 2.68E+00 | 3.31E-01 | no | | intergenic_variant | MODIFIER |
| 1 | 177320664 | AX-170387551 | 0 | T | C | 0.228 | 1.12E-01 | 1.16E-01 | 2.61E+00 | 2.68E+00 | 3.31E-01 | no | | intergenic_variant | MODIFIER |
| 1 | 177331101 | AX-170280588 | 0 | A | G | 0.228 | 1.12E-01 | 1.16E-01 | 2.61E+00 | 2.68E+00 | 3.31E-01 | no | | intergenic_variant | MODIFIER |
| 1 | 177333299 | AX-170424672 | 0 | G | C | 0.228 | 1.12E-01 | 1.16E-01 | 2.61E+00 | 2.68E+00 | 3.31E-01 | no | | intergenic_variant | MODIFIER |
| 1 | 177335446 | AX-170380219 | 0 | C | T | 0.228 | 1.12E-01 | 1.16E-01 | 2.61E+00 | 2.68E+00 | 3.31E-01 | no | | intergenic_variant | MODIFIER |
| 1 | 134831643 | AX-170129007 | 0 | C | T | 0.544 | 7.63E-02 | 7.93E-02 | 2.69E+00 | 2.77E+00 | 3.31E-01 | no | Ppp1r12b | intron_variant | MODIFIER |
| 2 | 164926802 | AX-170205088 | 1 | G | A | 0.161 | 1.88E-01 | 1.96E-01 | 2.79E+00 | 2.87E+00 | 3.31E-01 | Yes | | intergenic_variant | MODIFIER |
| 2 | 167166185 | AX-97831577 | 1 | C | T | 0.161 | 1.88E-01 | 1.96E-01 | 2.79E+00 | 2.87E+00 | 3.31E-01 | Yes | Kcnb1 | intron_variant | MODIFIER |
| 2 | 169947937 | AX-170512091 | 1 | T | C | 0.17 | 1.73E-01 | 1.79E-01 | 2.64E+00 | 2.73E+00 | 3.31E-01 | Yes | Tshz2 | intron_variant | MODIFIER |
| 15 | 20094248 | AX-168381968 | 1 | T | C | 0.241 | -1.02E-01 | 1.06E-01 | 2.63E+00 | 2.68E+00 | 3.31E-01 | no | | intergenic_variant | MODIFIER |
| 5 | 78544367 | AX-170025356 | 0 | T | G | 0.43 | 1.01E-01 | 1.04E-01 | 2.77E+00 | 2.83E+00 | 3.31E-01 | no | Gm43232 | downstream_gene_variant | MODIFIER |
| 3 | 149493096 | AX-170455767 | 0 | G | A | 0.36 | 6.88E-02 | 7.14E-02 | 2.82E+00 | 2.91E+00 | 3.31E-01 | no | | intergenic_variant | MODIFIER |
| 16 | 52181435 | AX-170113126 | 1 | A | G | 0.518 | 8.03E-02 | 8.22E-02 | 2.53E+00 | 2.58E+00 | 3.31E-01 | no | Cblb | intron_variant | MODIFIER |
| 16 | 52652338 | AX-170013051 | 1 | G | C | 0.518 | 8.03E-02 | 8.22E-02 | 2.53E+00 | 2.58E+00 | 3.31E-01 | no | | intergenic_variant | MODIFIER |

| | | | | | | | | | | | | | | |
|---|---|---|---|---|---|---|---|---|---|---|---|---|---|---|
| 5 | 5305596 | AX-170394304 | 1 | T | C | 0.429 | -6.38E-02 | 6.51E-02 | 2.49E+00 | 2.60E+00 | 3.31E-01 | no | Cdk14 | intron_variant | MODIFIER |
| 3 | 23979574 | AX-97796381 | 0 | C | T | 0.167 | -9.77E-02 | 9.99E-02 | 2.52E+00 | 2.59E+00 | 3.31E-01 | no | Naaladl2 | intron_variant | MODIFIER |
| 8 | 97417401 | AX-170472901 | 0 | C | G | 0.228 | 1.07E-01 | 1.11E-01 | 2.81E+00 | 2.89E+00 | 3.31E-01 | no | | regulatory_region_variant | MODIFIER |
| 1 | 82246553 | AX-170524076 | 0 | C | T | 0.167 | 1.65E-01 | 1.72E-01 | 2.66E+00 | 2.77E+00 | 3.31E-01 | Yes | Irs1 | intron_variant | MODIFIER |
| 7 | 24606264 | AX-170372609 | 0 | T | C | 0.404 | 8.75E-02 | 9.03E-02 | 2.57E+00 | 2.68E+00 | 3.31E-01 | no | Ethe1 | synonymous_variant | LOW |
| 7 | 24622528 | AX-97831539 | 0 | G | A | 0.404 | 8.75E-02 | 9.03E-02 | 2.57E+00 | 2.68E+00 | 3.31E-01 | no | Phldb3 | intron_variant | MODIFIER |
| 7 | 24643061 | AX-97843244 | 0 | A | G | 0.404 | 8.75E-02 | 9.03E-02 | 2.57E+00 | 2.68E+00 | 3.31E-01 | no | Lypd3 | downstream_gene_variant | MODIFIER |
| 7 | 24643890 | AX-170076857 | 0 | C | T | 0.404 | 8.75E-02 | 9.03E-02 | 2.57E+00 | 2.68E+00 | 3.31E-01 | no | Lypd3 | downstream_gene_variant | MODIFIER |
| 7 | 24645428 | AX-170076856 | 0 | G | A | 0.404 | 8.75E-02 | 9.03E-02 | 2.57E+00 | 2.68E+00 | 3.31E-01 | no | Lypd3 | downstream_gene_variant | MODIFIER |
| 7 | 24650612 | AX-170076861 | 0 | T | C | 0.404 | 8.75E-02 | 9.03E-02 | 2.57E+00 | 2.68E+00 | 3.31E-01 | no | | intergenic_variant | MODIFIER |
| 7 | 24652437 | AX-97785371 | 0 | G | A | 0.404 | 8.75E-02 | 9.03E-02 | 2.57E+00 | 2.68E+00 | 3.31E-01 | no | | intergenic_variant | MODIFIER |
| 7 | 24668391 | AX-170340656 | 0 | A | C | 0.404 | 8.75E-02 | 9.03E-02 | 2.57E+00 | 2.68E+00 | 3.31E-01 | no | Tex101 | missense_variant | MODERATE |
| 7 | 24693163 | AX-97791982 | 0 | G | A | 0.404 | 8.75E-02 | 9.03E-02 | 2.57E+00 | 2.68E+00 | 3.31E-01 | no | | intergenic_variant | MODIFIER |
| 9 | 97702708 | AX-170364368 | 1 | C | A | 0.143 | -1.09E-01 | 1.12E-01 | 2.60E+00 | 2.67E+00 | 3.31E-01 | no | Clstn2 | intron_variant | MODIFIER |
| 1 | 172194866 | AX-170043588 | 1 | A | C | 0.214 | 1.30E-01 | 1.35E-01 | 2.64E+00 | 2.69E+00 | 3.31E-01 | no | Pea15a | downstream_gene_variant | MODIFIER |
| 4 | 137668398 | AX-170024048 | 1 | C | T | 0.17 | 1.11E-01 | 1.15E-01 | 2.61E+00 | 2.66E+00 | 3.31E-01 | no | Rap1gap | intron_variant,non_coding_transcript_variant | MODIFIER |
| 1 | 16049333 | AX-170386588 | 0 | G | A | 0.386 | 7.65E-02 | 7.84E-02 | 2.53E+00 | 2.59E+00 | 3.31E-01 | no | Gm7634 | upstream_gene_variant | MODIFIER |
| 4 | 154233216 | AX-97795321 | 0 | C | T | 0.123 | 1.52E-01 | 1.59E-01 | 2.76E+00 | 2.88E+00 | 3.32E-01 | Yes | Megf6 | intron_variant | MODIFIER |
| 11 | 18344218 | AX-170381323 | 0 | C | T | 0.07 | 1.68E-01 | 1.75E-01 | 2.82E+00 | 2.95E+00 | 3.32E-01 | Yes | | intergenic_variant | MODIFIER |
| 16 | 29905094 | AX-170305263 | 0 | T | C | 0.5 | 7.82E-02 | 8.06E-02 | 2.57E+00 | 2.62E+00 | 3.32E-01 | no | Gm26569 | downstream_gene_variant | MODIFIER |
| 9 | 92490843 | AX-170179613 | 0 | T | A | 0.123 | -1.26E-01 | 1.31E-01 | 2.61E+00 | 2.68E+00 | 3.32E-01 | no | Plscr4 | 3_prime_UTR_variant | MODIFIER |
| 6 | 137479037 | AX-170365591 | 0 | A | G | 0.579 | 8.41E-02 | 8.33E-02 | 2.36E+00 | 2.41E+00 | 3.32E-01 | no | Eps8 | synonymous_variant | LOW |
| 5 | 106790156 | AX-97803825 | 1 | C | T | 0.446 | 1.07E-01 | 1.11E-01 | 2.64E+00 | 2.69E+00 | 3.32E-01 | no | | intergenic_variant | MODIFIER |
| 18 | 88552272 | AX-170351158 | 1 | C | A | 0.161 | -1.02E-01 | 1.05E-01 | 2.52E+00 | 2.61E+00 | 3.32E-01 | no | | intergenic_variant | MODIFIER |
| 14 | 117699727 | AX-170390686 | 0 | G | A | 0.325 | 7.03E-02 | 7.28E-02 | 2.58E+00 | 2.67E+00 | 3.32E-01 | no | Gpc6 | intron_variant | MODIFIER |
| 9 | 118151490 | AX-170261868 | 0 | G | A | 0.123 | -9.31E-02 | 9.71E-02 | 2.65E+00 | 2.77E+00 | 3.32E-01 | no | Cmc1 | upstream_gene_variant | MODIFIER |
| 4 | 139299581 | AX-170244717 | 1 | G | A | 0.161 | 1.13E-01 | 1.17E-01 | 2.61E+00 | 2.65E+00 | 3.32E-01 | no | Pqlc2 | intron_variant | MODIFIER |
| 4 | 80756624 | AX-97856155 | 1 | G | T | 0.455 | -8.29E-02 | 8.48E-02 | 2.51E+00 | 2.56E+00 | 3.32E-01 | no | | intergenic_variant | MODIFIER |
| 10 | 72410184 | AX-170176685 | 1 | A | T | 0.411 | -7.67E-02 | 8.00E-02 | 2.67E+00 | 2.75E+00 | 3.33E-01 | no | | regulatory_region_variant | MODIFIER |
| 10 | 17625918 | AX-170207119 | 1 | G | A | 0.205 | 9.76E-02 | 1.00E-01 | 2.98E+00 | 3.09E+00 | 3.33E-01 | no | Gm47771 | intron_variant,non_coding_transcript_variant | MODIFIER |
| 5 | 106470036 | AX-170299031 | 0 | T | A | 0.456 | 1.02E-01 | 1.06E-01 | 2.71E+00 | 2.78E+00 | 3.33E-01 | no | Gm26872 | intron_variant,non_coding_transcript_variant | MODIFIER |
| 4 | 75598356 | AX-97825335 | 1 | C | T | 0.312 | 1.09E-01 | 1.14E-01 | 2.71E+00 | 2.77E+00 | 3.33E-01 | no | | intergenic_variant | MODIFIER |
| 4 | 132778855 | AX-170283398 | 0 | A | G | 0.158 | 1.12E-01 | 1.17E-01 | 2.60E+00 | 2.65E+00 | 3.33E-01 | no | Themis2 | downstream_gene_variant | MODIFIER |
| 4 | 138590263 | AX-97786049 | 0 | T | C | 0.158 | 1.12E-01 | 1.17E-01 | 2.60E+00 | 2.65E+00 | 3.33E-01 | no | Vwa5b1 | intron_variant | MODIFIER |
| 4 | 138602422 | AX-170421439 | 0 | C | G | 0.158 | 1.12E-01 | 1.17E-01 | 2.60E+00 | 2.65E+00 | 3.33E-01 | no | Vwa5b1 | intron_variant | MODIFIER |
| 4 | 138604214 | AX-170282029 | 0 | G | A | 0.158 | 1.12E-01 | 1.17E-01 | 2.60E+00 | 2.65E+00 | 3.33E-01 | no | Vwa5b1 | intron_variant | MODIFIER |
| 4 | 138605784 | AX-170487266 | 0 | A | G | 0.158 | 1.12E-01 | 1.17E-01 | 2.60E+00 | 2.65E+00 | 3.33E-01 | no | Vwa5b1 | intron_variant | MODIFIER |
| 4 | 138605803 | AX-170315264 | 0 | G | A | 0.158 | 1.12E-01 | 1.17E-01 | 2.60E+00 | 2.65E+00 | 3.33E-01 | no | Vwa5b1 | intron_variant | MODIFIER |
| 4 | 138623656 | AX-170089388 | 0 | T | C | 0.158 | 1.12E-01 | 1.17E-01 | 2.60E+00 | 2.65E+00 | 3.33E-01 | no | Vwa5b1 | intron_variant | MODIFIER |
| 4 | 138629996 | AX-170024082 | 0 | T | C | 0.158 | 1.12E-01 | 1.17E-01 | 2.60E+00 | 2.65E+00 | 3.33E-01 | no | Vwa5b1 | intron_variant,NMD_transcript_variant | MODIFIER |
| 4 | 138630086 | AX-97809819 | 0 | A | G | 0.158 | 1.12E-01 | 1.17E-01 | 2.60E+00 | 2.65E+00 | 3.33E-01 | no | Vwa5b1 | intron_variant,NMD_transcript_variant | MODIFIER |
| 4 | 138655777 | AX-170095170 | 0 | T | G | 0.158 | 1.12E-01 | 1.17E-01 | 2.60E+00 | 2.65E+00 | 3.33E-01 | no | | intergenic_variant | MODIFIER |
| 4 | 138669150 | AX-170024078 | 0 | T | G | 0.158 | 1.12E-01 | 1.17E-01 | 2.60E+00 | 2.65E+00 | 3.33E-01 | no | | intergenic_variant | MODIFIER |
| 4 | 138669273 | AX-170024073 | 0 | C | T | 0.158 | 1.12E-01 | 1.17E-01 | 2.60E+00 | 2.65E+00 | 3.33E-01 | no | | intergenic_variant | MODIFIER |
| 4 | 138669485 | AX-170024079 | 0 | C | T | 0.158 | 1.12E-01 | 1.17E-01 | 2.60E+00 | 2.65E+00 | 3.33E-01 | no | | intergenic_variant | MODIFIER |
| 4 | 138673766 | AX-170131463 | 0 | A | G | 0.158 | 1.12E-01 | 1.17E-01 | 2.60E+00 | 2.65E+00 | 3.33E-01 | no | | intergenic_variant | MODIFIER |
| 4 | 138705855 | AX-97803460 | 0 | C | A | 0.158 | 1.12E-01 | 1.17E-01 | 2.60E+00 | 2.65E+00 | 3.33E-01 | no | Ubxn10 | downstream_gene_variant | MODIFIER |
| 4 | 138726830 | AX-170482826 | 0 | A | G | 0.158 | 1.12E-01 | 1.17E-01 | 2.60E+00 | 2.65E+00 | 3.33E-01 | no | Pla2g2c | intron_variant | MODIFIER |
| 4 | 138727362 | AX-170024074 | 0 | C | G | 0.158 | 1.12E-01 | 1.17E-01 | 2.60E+00 | 2.65E+00 | 3.33E-01 | no | Pla2g2c | intron_variant | MODIFIER |
| 4 | 138735132 | AX-170024080 | 0 | G | A | 0.158 | 1.12E-01 | 1.17E-01 | 2.60E+00 | 2.65E+00 | 3.33E-01 | no | Pla2g2c | intron_variant | MODIFIER |
| 4 | 138749865 | AX-97821114 | 0 | A | G | 0.158 | 1.12E-01 | 1.17E-01 | 2.60E+00 | 2.65E+00 | 3.33E-01 | no | Pla2g2f | downstream_gene_variant | MODIFIER |
| 4 | 138753420 | AX-170024075 | 0 | C | T | 0.158 | 1.12E-01 | 1.17E-01 | 2.60E+00 | 2.65E+00 | 3.33E-01 | no | Pla2g2f | intron_variant | MODIFIER |
| 4 | 138755871 | AX-170070843 | 0 | T | C | 0.158 | 1.12E-01 | 1.17E-01 | 2.60E+00 | 2.65E+00 | 3.33E-01 | no | Pla2g2f | intron_variant | MODIFIER |
| 4 | 138757625 | AX-170070844 | 0 | A | G | 0.158 | 1.12E-01 | 1.17E-01 | 2.60E+00 | 2.65E+00 | 3.33E-01 | no | Pla2g2f | 5_prime_UTR_variant | MODIFIER |

| | | | | | | | | | | | | | | | |
|---|---|---|---|---|---|---|---|---|---|---|---|---|---|---|---|
| 4 | 138766944 | AX-170070845 | 0 | T | C | 0.158 | 1.12E-01 | 1.17E-01 | 2.60E+00 | 2.65E+00 | 3.33E-01 | no | | intergenic_variant | MODIFIER |
| 4 | 138767968 | AX-97824890 | 0 | T | C | 0.158 | 1.12E-01 | 1.17E-01 | 2.60E+00 | 2.65E+00 | 3.33E-01 | no | | intergenic_variant | MODIFIER |
| 4 | 138771466 | AX-170024077 | 0 | C | T | 0.158 | 1.12E-01 | 1.17E-01 | 2.60E+00 | 2.65E+00 | 3.33E-01 | no | Pla2g2d | upstream_gene_variant | MODIFIER |
| 4 | 138785418 | AX-170024083 | 0 | A | G | 0.158 | 1.12E-01 | 1.17E-01 | 2.60E+00 | 2.65E+00 | 3.33E-01 | no | Pla2g2d | downstream_gene_variant | MODIFIER |
| 4 | 138785785 | AX-97798600 | 0 | A | G | 0.158 | 1.12E-01 | 1.17E-01 | 2.60E+00 | 2.65E+00 | 3.33E-01 | no | Pla2g2d | downstream_gene_variant | MODIFIER |
| 4 | 138785835 | AX-170529445 | 0 | C | T | 0.158 | 1.12E-01 | 1.17E-01 | 2.60E+00 | 2.65E+00 | 3.33E-01 | no | Pla2g2d | downstream_gene_variant | MODIFIER |
| 4 | 138788802 | AX-170024076 | 0 | A | G | 0.158 | 1.12E-01 | 1.17E-01 | 2.60E+00 | 2.65E+00 | 3.33E-01 | no | | intergenic_variant | MODIFIER |
| 4 | 138803673 | AX-170298299 | 0 | A | G | 0.158 | 1.12E-01 | 1.17E-01 | 2.60E+00 | 2.65E+00 | 3.33E-01 | no | Pla2g5 | intron_variant | MODIFIER |
| 4 | 138831905 | AX-170312155 | 0 | A | C | 0.158 | 1.12E-01 | 1.17E-01 | 2.60E+00 | 2.65E+00 | 3.33E-01 | no | Pla2g5 | intron_variant | MODIFIER |
| 4 | 138832336 | AX-97839890 | 0 | A | G | 0.158 | 1.12E-01 | 1.17E-01 | 2.60E+00 | 2.65E+00 | 3.33E-01 | no | Pla2g5 | intron_variant | MODIFIER |
| 4 | 138848941 | AX-170417230 | 0 | G | A | 0.158 | 1.12E-01 | 1.17E-01 | 2.60E+00 | 2.65E+00 | 3.33E-01 | no | Pla2g5 | intron_variant | MODIFIER |
| 4 | 138898245 | AX-170271776 | 0 | G | A | 0.158 | 1.12E-01 | 1.17E-01 | 2.60E+00 | 2.65E+00 | 3.33E-01 | no | Otud3 | intron_variant | MODIFIER |
| 4 | 138898436 | AX-97803631 | 0 | T | G | 0.158 | 1.12E-01 | 1.17E-01 | 2.60E+00 | 2.65E+00 | 3.33E-01 | no | Otud3 | intron_variant | MODIFIER |
| 4 | 138923315 | AX-170186640 | 0 | A | C | 0.158 | 1.12E-01 | 1.17E-01 | 2.60E+00 | 2.65E+00 | 3.33E-01 | no | | intergenic_variant | MODIFIER |
| 4 | 138953002 | AX-170492789 | 0 | A | G | 0.158 | 1.12E-01 | 1.17E-01 | 2.60E+00 | 2.65E+00 | 3.33E-01 | no | | intergenic_variant | MODIFIER |
| 4 | 138953176 | AX-170488790 | 0 | C | T | 0.158 | 1.12E-01 | 1.17E-01 | 2.60E+00 | 2.65E+00 | 3.33E-01 | no | | intergenic_variant | MODIFIER |
| 4 | 138953267 | AX-170500001 | 0 | C | G | 0.158 | 1.12E-01 | 1.17E-01 | 2.60E+00 | 2.65E+00 | 3.33E-01 | no | | intergenic_variant | MODIFIER |
| 4 | 138953551 | AX-97817064 | 0 | A | G | 0.158 | 1.12E-01 | 1.17E-01 | 2.60E+00 | 2.65E+00 | 3.33E-01 | no | | intergenic_variant | MODIFIER |
| 4 | 138989441 | AX-97840124 | 0 | T | C | 0.158 | 1.12E-01 | 1.17E-01 | 2.60E+00 | 2.65E+00 | 3.33E-01 | no | Tmco4 | intron_variant | MODIFIER |
| 4 | 139004345 | AX-170070851 | 0 | T | C | 0.158 | 1.12E-01 | 1.17E-01 | 2.60E+00 | 2.65E+00 | 3.33E-01 | no | Tmco4 | intron_variant | MODIFIER |
| 4 | 139018345 | AX-170070850 | 0 | T | C | 0.158 | 1.12E-01 | 1.17E-01 | 2.60E+00 | 2.65E+00 | 3.33E-01 | no | Tmco4 | intron_variant | MODIFIER |
| 4 | 139029633 | AX-97813737 | 0 | C | T | 0.158 | 1.12E-01 | 1.17E-01 | 2.60E+00 | 2.65E+00 | 3.33E-01 | no | Tmco4 | intron_variant | MODIFIER |
| 4 | 139064097 | AX-170516581 | 0 | C | T | 0.158 | 1.12E-01 | 1.17E-01 | 2.60E+00 | 2.65E+00 | 3.33E-01 | no | Tmco4 | downstream_gene_variant | MODIFIER |
| 4 | 139086190 | AX-97808008 | 0 | T | C | 0.158 | 1.12E-01 | 1.17E-01 | 2.60E+00 | 2.65E+00 | 3.33E-01 | no | Nbl1 | intron_variant | MODIFIER |
| 4 | 139115017 | AX-97813743 | 0 | T | C | 0.158 | 1.12E-01 | 1.17E-01 | 2.60E+00 | 2.65E+00 | 3.33E-01 | no | Minos1 | upstream_gene_variant | MODIFIER |
| 4 | 139127565 | AX-170455748 | 0 | T | C | 0.158 | 1.12E-01 | 1.17E-01 | 2.60E+00 | 2.65E+00 | 3.33E-01 | no | Minos1 | intron_variant | MODIFIER |
| 4 | 139155261 | AX-97822406 | 0 | G | A | 0.158 | 1.12E-01 | 1.17E-01 | 2.60E+00 | 2.65E+00 | 3.33E-01 | no | Hspe1-ps4 | downstream_gene_variant | MODIFIER |
| 4 | 139177951 | AX-170528932 | 0 | A | G | 0.158 | 1.12E-01 | 1.17E-01 | 2.60E+00 | 2.65E+00 | 3.33E-01 | no | Gm16287 | intron_variant,non_coding_transcript_variant | MODIFIER |
| 4 | 139183456 | AX-97811818 | 0 | T | C | 0.158 | 1.12E-01 | 1.17E-01 | 2.60E+00 | 2.65E+00 | 3.33E-01 | no | Gm16287 | upstream_gene_variant | MODIFIER |
| 4 | 139268767 | AX-97794983 | 0 | A | G | 0.158 | 1.12E-01 | 1.17E-01 | 2.60E+00 | 2.65E+00 | 3.33E-01 | no | Capzb | intron_variant | MODIFIER |
| 4 | 139314783 | AX-170348821 | 0 | A | G | 0.158 | 1.12E-01 | 1.17E-01 | 2.60E+00 | 2.65E+00 | 3.33E-01 | no | Pqlc2 | upstream_gene_variant | MODIFIER |
| 4 | 139323867 | AX-170507766 | 0 | G | C | 0.158 | 1.12E-01 | 1.17E-01 | 2.60E+00 | 2.65E+00 | 3.33E-01 | no | | intergenic_variant | MODIFIER |
| 4 | 139351378 | AX-97856192 | 0 | T | C | 0.158 | 1.12E-01 | 1.17E-01 | 2.60E+00 | 2.65E+00 | 3.33E-01 | no | Mrto4 | intron_variant | MODIFIER |
| 4 | 139354411 | AX-170288804 | 0 | C | A | 0.158 | 1.12E-01 | 1.17E-01 | 2.60E+00 | 2.65E+00 | 3.33E-01 | no | Mrto4 | upstream_gene_variant | MODIFIER |
| 4 | 139354682 | AX-97828623 | 0 | C | A | 0.158 | 1.12E-01 | 1.17E-01 | 2.60E+00 | 2.65E+00 | 3.33E-01 | no | Mrto4 | upstream_gene_variant | MODIFIER |
| 4 | 139357190 | AX-170024085 | 0 | A | G | 0.158 | 1.12E-01 | 1.17E-01 | 2.60E+00 | 2.65E+00 | 3.33E-01 | no | Mrto4 | upstream_gene_variant | MODIFIER |
| 4 | 139360570 | AX-170353214 | 0 | C | T | 0.158 | 1.12E-01 | 1.17E-01 | 2.60E+00 | 2.65E+00 | 3.33E-01 | no | Emc1 | synonymous_variant | LOW |
| 4 | 139398742 | AX-170024092 | 0 | A | G | 0.158 | 1.12E-01 | 1.17E-01 | 2.60E+00 | 2.65E+00 | 3.33E-01 | no | Ubr4 | intron_variant | MODIFIER |
| 4 | 139400738 | AX-97856788 | 0 | C | T | 0.158 | 1.12E-01 | 1.17E-01 | 2.60E+00 | 2.65E+00 | 3.33E-01 | no | Ubr4 | intron_variant | MODIFIER |
| 4 | 139403892 | AX-97813741 | 0 | A | G | 0.158 | 1.12E-01 | 1.17E-01 | 2.60E+00 | 2.65E+00 | 3.33E-01 | no | Ubr4 | intron_variant | MODIFIER |
| 4 | 139426376 | AX-170024095 | 0 | A | G | 0.158 | 1.12E-01 | 1.17E-01 | 2.60E+00 | 2.65E+00 | 3.33E-01 | no | Ubr4 | intron_variant | MODIFIER |
| 4 | 139448795 | AX-170070855 | 0 | G | C | 0.158 | 1.12E-01 | 1.17E-01 | 2.60E+00 | 2.65E+00 | 3.33E-01 | no | Ubr4 | intron_variant | MODIFIER |
| 4 | 139467848 | AX-97812000 | 0 | G | A | 0.158 | 1.12E-01 | 1.17E-01 | 2.60E+00 | 2.65E+00 | 3.33E-01 | no | Ubr4 | intron_variant | MODIFIER |
| 4 | 139476420 | AX-170024088 | 0 | A | G | 0.158 | 1.12E-01 | 1.17E-01 | 2.60E+00 | 2.65E+00 | 3.33E-01 | no | Ubr4 | intron_variant | MODIFIER |
| 4 | 139506356 | AX-170099797 | 0 | T | C | 0.158 | 1.12E-01 | 1.17E-01 | 2.60E+00 | 2.65E+00 | 3.33E-01 | no | | regulatory_region_variant | MODIFIER |
| 4 | 139506749 | AX-97849439 | 0 | A | G | 0.158 | 1.12E-01 | 1.17E-01 | 2.60E+00 | 2.65E+00 | 3.33E-01 | no | | regulatory_region_variant | MODIFIER |
| 4 | 139556491 | AX-170089389 | 0 | G | A | 0.158 | 1.12E-01 | 1.17E-01 | 2.60E+00 | 2.65E+00 | 3.33E-01 | no | Iffo2 | intron_variant | MODIFIER |
| 4 | 139562406 | AX-170535932 | 0 | G | A | 0.158 | 1.12E-01 | 1.17E-01 | 2.60E+00 | 2.65E+00 | 3.33E-01 | no | Iffo2 | intron_variant | MODIFIER |
| 4 | 139569906 | AX-170141099 | 0 | T | C | 0.158 | 1.12E-01 | 1.17E-01 | 2.60E+00 | 2.65E+00 | 3.33E-01 | no | Iffo2 | intron_variant | MODIFIER |
| 4 | 139570172 | AX-170211210 | 0 | T | C | 0.158 | 1.12E-01 | 1.17E-01 | 2.60E+00 | 2.65E+00 | 3.33E-01 | no | Iffo2 | intron_variant | MODIFIER |
| 4 | 139581391 | AX-170209069 | 0 | T | C | 0.158 | 1.12E-01 | 1.17E-01 | 2.60E+00 | 2.65E+00 | 3.33E-01 | no | Iffo2 | intron_variant | MODIFIER |
| 4 | 139583460 | AX-170070857 | 0 | G | A | 0.158 | 1.12E-01 | 1.17E-01 | 2.60E+00 | 2.65E+00 | 3.33E-01 | no | Iffo2 | intron_variant | MODIFIER |
| 4 | 139585068 | AX-170070859 | 0 | C | T | 0.158 | 1.12E-01 | 1.17E-01 | 2.60E+00 | 2.65E+00 | 3.33E-01 | no | Iffo2 | intron_variant | MODIFIER |
| 4 | 139585933 | AX-170274785 | 0 | A | G | 0.158 | 1.12E-01 | 1.17E-01 | 2.60E+00 | 2.65E+00 | 3.33E-01 | no | Iffo2 | intron_variant | MODIFIER |

| | | | | | | | | | | | | | | | |
|---|---|---|---|---|---|---|---|---|---|---|---|---|---|---|---|
| 4 | 139591236 | AX-97813785 | 0 | A | G | 0.158 | 1.12E-01 | 1.17E-01 | 2.60E+00 | 2.65E+00 | 3.33E-01 | no | Iffo2 | intron_variant | MODIFIER |
| 4 | 139591308 | AX-170421244 | 0 | G | A | 0.158 | 1.12E-01 | 1.17E-01 | 2.60E+00 | 2.65E+00 | 3.33E-01 | no | Iffo2 | intron_variant | MODIFIER |
| 4 | 139667042 | AX-97848192 | 0 | A | G | 0.158 | 1.12E-01 | 1.17E-01 | 2.60E+00 | 2.65E+00 | 3.33E-01 | no | Tas1r2 | intron_variant | MODIFIER |
| 4 | 139680934 | AX-97854987 | 0 | C | T | 0.158 | 1.12E-01 | 1.17E-01 | 2.60E+00 | 2.65E+00 | 3.33E-01 | no | | intergenic_variant | MODIFIER |
| 4 | 139710884 | AX-170383291 | 0 | T | C | 0.158 | 1.12E-01 | 1.17E-01 | 2.60E+00 | 2.65E+00 | 3.33E-01 | no | | regulatory_region_variant | MODIFIER |
| 4 | 139713478 | AX-170395045 | 0 | A | T | 0.158 | 1.12E-01 | 1.17E-01 | 2.60E+00 | 2.65E+00 | 3.33E-01 | no | | intergenic_variant | MODIFIER |
| 4 | 139713528 | AX-97838765 | 0 | G | A | 0.158 | 1.12E-01 | 1.17E-01 | 2.60E+00 | 2.65E+00 | 3.33E-01 | no | | intergenic_variant | MODIFIER |
| 4 | 139732154 | AX-170383214 | 0 | G | A | 0.158 | 1.12E-01 | 1.17E-01 | 2.60E+00 | 2.65E+00 | 3.33E-01 | no | Pax7 | downstream_gene_variant | MODIFIER |
| 4 | 139742048 | AX-97847818 | 0 | T | G | 0.158 | 1.12E-01 | 1.17E-01 | 2.60E+00 | 2.65E+00 | 3.33E-01 | no | Pax7 | intron_variant | MODIFIER |
| 4 | 139745376 | AX-170024103 | 0 | T | C | 0.158 | 1.12E-01 | 1.17E-01 | 2.60E+00 | 2.65E+00 | 3.33E-01 | no | Pax7 | intron_variant | MODIFIER |
| 4 | 139751636 | AX-170024100 | 0 | A | G | 0.158 | 1.12E-01 | 1.17E-01 | 2.60E+00 | 2.65E+00 | 3.33E-01 | no | Pax7 | intron_variant | MODIFIER |
| 4 | 139752706 | AX-170302889 | 0 | G | A | 0.158 | 1.12E-01 | 1.17E-01 | 2.60E+00 | 2.65E+00 | 3.33E-01 | no | Pax7 | intron_variant | MODIFIER |
| 4 | 139756037 | AX-170118809 | 0 | C | T | 0.158 | 1.12E-01 | 1.17E-01 | 2.60E+00 | 2.65E+00 | 3.33E-01 | no | Pax7 | intron_variant | MODIFIER |
| 4 | 139756113 | AX-97835479 | 0 | C | A | 0.158 | 1.12E-01 | 1.17E-01 | 2.60E+00 | 2.65E+00 | 3.33E-01 | no | Pax7 | intron_variant | MODIFIER |
| 4 | 139784678 | AX-97835570 | 0 | G | A | 0.158 | 1.12E-01 | 1.17E-01 | 2.60E+00 | 2.65E+00 | 3.33E-01 | no | Pax7 | synonymous_variant | LOW |
| 4 | 139785368 | AX-170485830 | 0 | A | G | 0.158 | 1.12E-01 | 1.17E-01 | 2.60E+00 | 2.65E+00 | 3.33E-01 | no | Pax7 | intron_variant | MODIFIER |
| 4 | 139842551 | AX-170024097 | 0 | T | C | 0.158 | 1.12E-01 | 1.17E-01 | 2.60E+00 | 2.65E+00 | 3.33E-01 | no | | intergenic_variant | MODIFIER |
| 4 | 139850542 | AX-97801134 | 0 | G | A | 0.158 | 1.12E-01 | 1.17E-01 | 2.60E+00 | 2.65E+00 | 3.33E-01 | no | | intergenic_variant | MODIFIER |
| 4 | 139867316 | AX-97831285 | 0 | A | G | 0.158 | 1.12E-01 | 1.17E-01 | 2.60E+00 | 2.65E+00 | 3.33E-01 | no | | intergenic_variant | MODIFIER |
| 4 | 139867433 | AX-170024098 | 0 | T | C | 0.158 | 1.12E-01 | 1.17E-01 | 2.60E+00 | 2.65E+00 | 3.33E-01 | no | | intergenic_variant | MODIFIER |
| 4 | 139890847 | AX-97813783 | 0 | A | G | 0.158 | 1.12E-01 | 1.17E-01 | 2.60E+00 | 2.65E+00 | 3.33E-01 | no | | intergenic_variant | MODIFIER |
| 4 | 139896828 | AX-170323907 | 0 | T | C | 0.158 | 1.12E-01 | 1.17E-01 | 2.60E+00 | 2.65E+00 | 3.33E-01 | no | | intergenic_variant | MODIFIER |
| 4 | 139907767 | AX-97829609 | 0 | A | C | 0.158 | 1.12E-01 | 1.17E-01 | 2.60E+00 | 2.65E+00 | 3.33E-01 | no | Gm1667 | upstream_gene_variant | MODIFIER |
| 4 | 139951049 | AX-170095172 | 0 | C | T | 0.158 | 1.12E-01 | 1.17E-01 | 2.60E+00 | 2.65E+00 | 3.33E-01 | no | | regulatory_region_variant | MODIFIER |
| 4 | 139952747 | AX-170277363 | 0 | T | C | 0.158 | 1.12E-01 | 1.17E-01 | 2.60E+00 | 2.65E+00 | 3.33E-01 | no | | intergenic_variant | MODIFIER |
| 4 | 139958965 | AX-170425751 | 0 | A | G | 0.158 | 1.12E-01 | 1.17E-01 | 2.60E+00 | 2.65E+00 | 3.33E-01 | no | Klhdc7a | downstream_gene_variant | MODIFIER |
| 4 | 139959354 | AX-170120443 | 0 | G | A | 0.158 | 1.12E-01 | 1.17E-01 | 2.60E+00 | 2.65E+00 | 3.33E-01 | no | Klhdc7a | downstream_gene_variant | MODIFIER |
| 4 | 139959632 | AX-170469228 | 0 | T | G | 0.158 | 1.12E-01 | 1.17E-01 | 2.60E+00 | 2.65E+00 | 3.33E-01 | no | Klhdc7a | downstream_gene_variant | MODIFIER |
| 4 | 139959659 | AX-170024101 | 0 | A | G | 0.158 | 1.12E-01 | 1.17E-01 | 2.60E+00 | 2.65E+00 | 3.33E-01 | no | Klhdc7a | downstream_gene_variant | MODIFIER |
| 4 | 139960081 | AX-170024106 | 0 | A | G | 0.158 | 1.12E-01 | 1.17E-01 | 2.60E+00 | 2.65E+00 | 3.33E-01 | no | Klhdc7a | downstream_gene_variant | MODIFIER |
| 4 | 139965973 | AX-170024108 | 0 | G | A | 0.158 | 1.12E-01 | 1.17E-01 | 2.60E+00 | 2.65E+00 | 3.33E-01 | no | Klhdc7a | synonymous_variant | LOW |
| 4 | 139968808 | AX-170095173 | 0 | G | A | 0.158 | 1.12E-01 | 1.17E-01 | 2.60E+00 | 2.65E+00 | 3.33E-01 | no | Klhdc7a | upstream_gene_variant | MODIFIER |
| 4 | 139971984 | AX-168408300 | 0 | A | G | 0.158 | 1.12E-01 | 1.17E-01 | 2.60E+00 | 2.65E+00 | 3.33E-01 | no | Klhdc7a | upstream_gene_variant | MODIFIER |
| 4 | 139978369 | AX-97813786 | 0 | T | C | 0.158 | 1.12E-01 | 1.17E-01 | 2.60E+00 | 2.65E+00 | 3.33E-01 | no | | regulatory_region_variant | MODIFIER |
| 4 | 140003130 | AX-170024099 | 0 | A | G | 0.158 | 1.12E-01 | 1.17E-01 | 2.60E+00 | 2.65E+00 | 3.33E-01 | no | | intergenic_variant | MODIFIER |
| 4 | 140030005 | AX-170287702 | 0 | G | A | 0.158 | 1.12E-01 | 1.17E-01 | 2.60E+00 | 2.65E+00 | 3.33E-01 | no | Igsf21 | intron_variant | MODIFIER |
| 4 | 140052952 | AX-170070864 | 0 | A | G | 0.158 | 1.12E-01 | 1.17E-01 | 2.60E+00 | 2.65E+00 | 3.33E-01 | no | Igsf21 | intron_variant | MODIFIER |
| 4 | 140061822 | AX-170070865 | 0 | C | T | 0.158 | 1.12E-01 | 1.17E-01 | 2.60E+00 | 2.65E+00 | 3.33E-01 | no | Igsf21 | intron_variant | MODIFIER |
| 4 | 140064930 | AX-170070866 | 0 | C | T | 0.158 | 1.12E-01 | 1.17E-01 | 2.60E+00 | 2.65E+00 | 3.33E-01 | no | Igsf21 | intron_variant | MODIFIER |
| 4 | 140065223 | AX-97795594 | 0 | A | C | 0.158 | 1.12E-01 | 1.17E-01 | 2.60E+00 | 2.65E+00 | 3.33E-01 | no | Igsf21 | intron_variant | MODIFIER |
| 4 | 140085139 | AX-97839495 | 0 | T | C | 0.158 | 1.12E-01 | 1.17E-01 | 2.60E+00 | 2.65E+00 | 3.33E-01 | no | Igsf21 | intron_variant | MODIFIER |
| 4 | 140109999 | AX-97835487 | 0 | A | G | 0.158 | 1.12E-01 | 1.17E-01 | 2.60E+00 | 2.65E+00 | 3.33E-01 | no | Igsf21 | intron_variant | MODIFIER |
| 4 | 140112682 | AX-170182884 | 0 | G | T | 0.158 | 1.12E-01 | 1.17E-01 | 2.60E+00 | 2.65E+00 | 3.33E-01 | no | Igsf21 | intron_variant | MODIFIER |
| 4 | 140112748 | AX-170294437 | 0 | T | C | 0.158 | 1.12E-01 | 1.17E-01 | 2.60E+00 | 2.65E+00 | 3.33E-01 | no | Igsf21 | intron_variant | MODIFIER |
| 4 | 140117640 | AX-170139997 | 0 | C | T | 0.158 | 1.12E-01 | 1.17E-01 | 2.60E+00 | 2.65E+00 | 3.33E-01 | no | Igsf21 | intron_variant | MODIFIER |
| 4 | 140121588 | AX-170340395 | 0 | A | G | 0.158 | 1.12E-01 | 1.17E-01 | 2.60E+00 | 2.65E+00 | 3.33E-01 | no | Igsf21 | intron_variant | MODIFIER |
| 4 | 140131695 | AX-170523227 | 0 | C | T | 0.158 | 1.12E-01 | 1.17E-01 | 2.60E+00 | 2.65E+00 | 3.33E-01 | no | Igsf21 | intron_variant | MODIFIER |
| 4 | 140137387 | AX-97810259 | 0 | G | A | 0.158 | 1.12E-01 | 1.17E-01 | 2.60E+00 | 2.65E+00 | 3.33E-01 | no | Igsf21 | intron_variant | MODIFIER |
| 4 | 140151116 | AX-170514446 | 0 | T | A | 0.158 | 1.12E-01 | 1.17E-01 | 2.60E+00 | 2.65E+00 | 3.33E-01 | no | Igsf21 | intron_variant | MODIFIER |
| 4 | 140163325 | AX-97793816 | 0 | A | G | 0.158 | 1.12E-01 | 1.17E-01 | 2.60E+00 | 2.65E+00 | 3.33E-01 | no | Igsf21 | intron_variant | MODIFIER |
| 4 | 140186173 | AX-170348225 | 0 | T | C | 0.158 | 1.12E-01 | 1.17E-01 | 2.60E+00 | 2.65E+00 | 3.33E-01 | no | Igsf21 | intron_variant | MODIFIER |
| 4 | 140186354 | AX-97849997 | 0 | A | C | 0.158 | 1.12E-01 | 1.17E-01 | 2.60E+00 | 2.65E+00 | 3.33E-01 | no | Igsf21 | intron_variant | MODIFIER |
| 4 | 140208974 | AX-97820563 | 0 | T | C | 0.158 | 1.12E-01 | 1.17E-01 | 2.60E+00 | 2.65E+00 | 3.33E-01 | no | Igsf21 | intron_variant | MODIFIER |
| 4 | 140239006 | AX-170340762 | 0 | G | C | 0.158 | 1.12E-01 | 1.17E-01 | 2.60E+00 | 2.65E+00 | 3.33E-01 | no | Igsf21 | intron_variant | MODIFIER |

| | | | | | | | | | | | | | | | |
|---|---|---|---|---|---|---|---|---|---|---|---|---|---|---|---|
| 4 | 140249845 | AX-170380836 | 0 | G | A | 0.158 | 1.12E-01 | 1.17E-01 | 2.60E+00 | 2.65E+00 | 3.33E-01 | no | Igsf21 | upstream_gene_variant | MODIFIER |
| 4 | 140284947 | AX-170024116 | 0 | G | A | 0.158 | 1.12E-01 | 1.17E-01 | 2.60E+00 | 2.65E+00 | 3.33E-01 | no | | intergenic_variant | MODIFIER |
| 4 | 140285019 | AX-170024112 | 0 | T | G | 0.158 | 1.12E-01 | 1.17E-01 | 2.60E+00 | 2.65E+00 | 3.33E-01 | no | | regulatory_region_variant | MODIFIER |
| 4 | 140286763 | AX-97824026 | 0 | G | A | 0.158 | 1.12E-01 | 1.17E-01 | 2.60E+00 | 2.65E+00 | 3.33E-01 | no | | intergenic_variant | MODIFIER |
| 4 | 140292351 | AX-170423983 | 0 | C | G | 0.158 | 1.12E-01 | 1.17E-01 | 2.60E+00 | 2.65E+00 | 3.33E-01 | no | | intergenic_variant | MODIFIER |
| 4 | 140308704 | AX-97851414 | 0 | G | A | 0.158 | 1.12E-01 | 1.17E-01 | 2.60E+00 | 2.65E+00 | 3.33E-01 | no | | intergenic_variant | MODIFIER |
| 4 | 140313063 | AX-170070870 | 0 | G | A | 0.158 | 1.12E-01 | 1.17E-01 | 2.60E+00 | 2.65E+00 | 3.33E-01 | no | | regulatory_region_variant | MODIFIER |
| 4 | 140313282 | AX-170024118 | 0 | A | G | 0.158 | 1.12E-01 | 1.17E-01 | 2.60E+00 | 2.65E+00 | 3.33E-01 | no | | regulatory_region_variant | MODIFIER |
| 4 | 140321297 | AX-170070868 | 0 | T | A | 0.158 | 1.12E-01 | 1.17E-01 | 2.60E+00 | 2.65E+00 | 3.33E-01 | no | Gm9867 | non_coding_transcript_exon_variant | MODIFIER |
| 4 | 140331038 | AX-170024115 | 0 | T | C | 0.158 | 1.12E-01 | 1.17E-01 | 2.60E+00 | 2.65E+00 | 3.33E-01 | no | | intergenic_variant | MODIFIER |
| 4 | 140346282 | AX-170222502 | 0 | G | A | 0.158 | 1.12E-01 | 1.17E-01 | 2.60E+00 | 2.65E+00 | 3.33E-01 | no | | regulatory_region_variant | MODIFIER |
| 4 | 140360793 | AX-170024117 | 0 | T | C | 0.158 | 1.12E-01 | 1.17E-01 | 2.60E+00 | 2.65E+00 | 3.33E-01 | no | | intergenic_variant | MODIFIER |
| 4 | 140401947 | AX-170024114 | 0 | C | T | 0.158 | 1.12E-01 | 1.17E-01 | 2.60E+00 | 2.65E+00 | 3.33E-01 | no | | intergenic_variant | MODIFIER |
| 4 | 140409944 | AX-170024110 | 0 | C | T | 0.158 | 1.12E-01 | 1.17E-01 | 2.60E+00 | 2.65E+00 | 3.33E-01 | no | | intergenic_variant | MODIFIER |
| 4 | 140423385 | AX-170216954 | 0 | A | G | 0.158 | 1.12E-01 | 1.17E-01 | 2.60E+00 | 2.65E+00 | 3.33E-01 | no | Gm13021 | intron_variant,non_coding_transcript_variant | MODIFIER |
| 4 | 140433829 | AX-170366965 | 0 | C | T | 0.158 | 1.12E-01 | 1.17E-01 | 2.60E+00 | 2.65E+00 | 3.33E-01 | no | Gm13021 | downstream_gene_variant | MODIFIER |
| 4 | 140445495 | AX-170341938 | 0 | A | G | 0.158 | 1.12E-01 | 1.17E-01 | 2.60E+00 | 2.65E+00 | 3.33E-01 | no | Gm13026 | intron_variant,non_coding_transcript_variant | MODIFIER |
| 4 | 140446072 | AX-170335526 | 0 | C | T | 0.158 | 1.12E-01 | 1.17E-01 | 2.60E+00 | 2.65E+00 | 3.33E-01 | no | Gm13026 | intron_variant,non_coding_transcript_variant | MODIFIER |
| 4 | 140466293 | AX-97804998 | 0 | C | T | 0.158 | 1.12E-01 | 1.17E-01 | 2.60E+00 | 2.65E+00 | 3.33E-01 | no | | intergenic_variant | MODIFIER |
| 4 | 140471848 | AX-170462194 | 0 | C | T | 0.158 | 1.12E-01 | 1.17E-01 | 2.60E+00 | 2.65E+00 | 3.33E-01 | no | | intergenic_variant | MODIFIER |
| 4 | 140506462 | AX-170245321 | 0 | G | A | 0.158 | 1.12E-01 | 1.17E-01 | 2.60E+00 | 2.65E+00 | 3.33E-01 | no | | regulatory_region_variant | MODIFIER |
| 4 | 140506829 | AX-170369426 | 0 | G | C | 0.158 | 1.12E-01 | 1.17E-01 | 2.60E+00 | 2.65E+00 | 3.33E-01 | no | | intergenic_variant | MODIFIER |
| 4 | 140556040 | AX-170421664 | 0 | T | G | 0.158 | 1.12E-01 | 1.17E-01 | 2.60E+00 | 2.65E+00 | 3.33E-01 | no | Arhgef10l | intron_variant | MODIFIER |
| 4 | 140557307 | AX-170293443 | 0 | G | T | 0.158 | 1.12E-01 | 1.17E-01 | 2.60E+00 | 2.65E+00 | 3.33E-01 | no | Arhgef10l | intron_variant | MODIFIER |
| 4 | 140565445 | AX-97794196 | 0 | A | G | 0.158 | 1.12E-01 | 1.17E-01 | 2.60E+00 | 2.65E+00 | 3.33E-01 | no | Arhgef10l | intron_variant | MODIFIER |
| 4 | 140569204 | AX-170481788 | 0 | G | A | 0.158 | 1.12E-01 | 1.17E-01 | 2.60E+00 | 2.65E+00 | 3.33E-01 | no | Arhgef10l | intron_variant | MODIFIER |
| 4 | 140582041 | AX-97798791 | 0 | T | C | 0.158 | 1.12E-01 | 1.17E-01 | 2.60E+00 | 2.65E+00 | 3.33E-01 | no | Arhgef10l | intron_variant | MODIFIER |
| 4 | 140605784 | AX-170435304 | 0 | T | C | 0.158 | 1.12E-01 | 1.17E-01 | 2.60E+00 | 2.65E+00 | 3.33E-01 | no | Arhgef10l | intron_variant | MODIFIER |
| 4 | 140608130 | AX-170274285 | 0 | A | G | 0.158 | 1.12E-01 | 1.17E-01 | 2.60E+00 | 2.65E+00 | 3.33E-01 | no | Arhgef10l | intron_variant | MODIFIER |
| 4 | 140609530 | AX-170070876 | 0 | G | T | 0.158 | 1.12E-01 | 1.17E-01 | 2.60E+00 | 2.65E+00 | 3.33E-01 | no | Arhgef10l | intron_variant | MODIFIER |
| 4 | 140610380 | AX-97830980 | 0 | A | G | 0.158 | 1.12E-01 | 1.17E-01 | 2.60E+00 | 2.65E+00 | 3.33E-01 | no | Arhgef10l | intron_variant | MODIFIER |
| 4 | 140635072 | AX-170394163 | 0 | A | G | 0.158 | 1.12E-01 | 1.17E-01 | 2.60E+00 | 2.65E+00 | 3.33E-01 | no | Arhgef10l | intron_variant | MODIFIER |
| 4 | 140640361 | AX-170118738 | 0 | G | C | 0.158 | 1.12E-01 | 1.17E-01 | 2.60E+00 | 2.65E+00 | 3.33E-01 | no | Arhgef10l | intron_variant | MODIFIER |
| 4 | 140666594 | AX-170361425 | 0 | T | C | 0.158 | 1.12E-01 | 1.17E-01 | 2.60E+00 | 2.65E+00 | 3.33E-01 | no | Arhgef10l | upstream_gene_variant | MODIFIER |
| 4 | 140687056 | AX-170292808 | 0 | A | G | 0.158 | 1.12E-01 | 1.17E-01 | 2.60E+00 | 2.65E+00 | 3.33E-01 | no | Gm13025 | downstream_gene_variant | MODIFIER |
| 4 | 140704647 | AX-170401070 | 0 | T | C | 0.158 | 1.12E-01 | 1.17E-01 | 2.60E+00 | 2.65E+00 | 3.33E-01 | no | Rcc2 | intron_variant | MODIFIER |
| 4 | 140709219 | AX-170367825 | 0 | G | T | 0.158 | 1.12E-01 | 1.17E-01 | 2.60E+00 | 2.65E+00 | 3.33E-01 | no | Rcc2 | intron_variant | MODIFIER |
| 4 | 140748806 | AX-170354702 | 0 | C | A | 0.158 | 1.12E-01 | 1.17E-01 | 2.60E+00 | 2.65E+00 | 3.33E-01 | no | Padi4 | intron_variant | MODIFIER |
| 18 | 69255987 | AX-97825994 | 0 | T | C | 0.175 | -1.11E-01 | 1.16E-01 | 2.59E+00 | 2.66E+00 | 3.33E-01 | no | Gm4842 | upstream_gene_variant | MODIFIER |
| 18 | 59864406 | AX-97798148 | 1 | G | A | 0.402 | 9.25E-02 | 9.36E-02 | 2.44E+00 | 2.48E+00 | 3.33E-01 | no | | intergenic_variant | MODIFIER |
| 15 | 3750616 | AX-170247009 | 0 | G | T | 0.237 | -1.01E-01 | 1.05E-01 | 2.59E+00 | 2.66E+00 | 3.33E-01 | no | Gm4823 | upstream_gene_variant | MODIFIER |
| 2 | 114149976 | AX-170280323 | 1 | T | C | 0.152 | -1.26E-01 | 1.32E-01 | 2.84E+00 | 2.98E+00 | 3.33E-01 | no | Aqr | intron_variant | MODIFIER |
| 4 | 133964175 | AX-97812720 | 1 | G | A | 0.161 | 1.15E-01 | 1.19E-01 | 2.55E+00 | 2.59E+00 | 3.33E-01 | no | Dhdds | downstream_gene_variant | MODIFIER |
| 6 | 130032071 | AX-170339202 | 1 | C | A | 0.545 | 7.91E-02 | 8.13E-02 | 2.52E+00 | 2.59E+00 | 3.33E-01 | no | | intergenic_variant | MODIFIER |
| 13 | 6893260 | AX-97785642 | 1 | C | T | 0.205 | -1.20E-01 | 1.24E-01 | 2.59E+00 | 2.65E+00 | 3.33E-01 | no | | intergenic_variant | MODIFIER |
| 12 | 12554828 | AX-170251109 | 1 | C | T | 0.036 | -1.65E-01 | 1.70E-01 | 2.51E+00 | 2.61E+00 | 3.34E-01 | no | | intergenic_variant | MODIFIER |
| 6 | 124518033 | AX-97783286 | 1 | C | A | 0.179 | -7.89E-02 | 8.28E-02 | 2.76E+00 | 2.90E+00 | 3.34E-01 | no | C1ra | intron_variant | MODIFIER |
| 6 | 129687757 | AX-170028245 | 0 | A | G | 0.553 | 7.71E-02 | 7.99E-02 | 2.57E+00 | 2.65E+00 | 3.34E-01 | no | Mir680-1 | upstream_gene_variant | MODIFIER |
| 4 | 69169833 | AX-170023043 | 1 | C | T | 0.33 | 1.11E-01 | 1.16E-01 | 2.71E+00 | 2.77E+00 | 3.34E-01 | no | | intergenic_variant | MODIFIER |
| 6 | 127970348 | AX-170184708 | 0 | A | G | 0.535 | 8.05E-02 | 8.23E-02 | 2.48E+00 | 2.54E+00 | 3.34E-01 | no | Tspan9 | intron_variant | MODIFIER |
| 7 | 23386726 | AX-170512288 | 0 | G | A | 0.421 | 8.55E-02 | 8.80E-02 | 2.50E+00 | 2.61E+00 | 3.34E-01 | no | Nlrp5 | intron_variant | MODIFIER |
| 9 | 97883050 | AX-170033088 | 1 | A | C | 0.143 | -1.08E-01 | 1.12E-01 | 2.59E+00 | 2.67E+00 | 3.34E-01 | no | Clstn2 | intron_variant | MODIFIER |
| 16 | 29477308 | AX-170231665 | 1 | A | T | 0.482 | 7.51E-02 | 7.78E-02 | 2.56E+00 | 2.62E+00 | 3.34E-01 | no | Atp13a4 | intron_variant | MODIFIER |
| 1 | 171214250 | AX-170518954 | 0 | G | A | 0.211 | 1.22E-01 | 1.27E-01 | 2.62E+00 | 2.69E+00 | 3.34E-01 | no | Tomm40l | downstream_gene_variant | MODIFIER |

| | | | | | | | | | | | | | | | |
|---|---|---|---|---|---|---|---|---|---|---|---|---|---|---|---|
| 13 | 6375171 | AX-170163546 | 1 | C | T | 0.205 | -1.19E-01 | 1.23E-01 | 2.60E+00 | 2.65E+00 | 3.35E-01 | no | | intergenic_variant | MODIFIER |
| 18 | 69775509 | AX-97788434 | 1 | T | C | 0.25 | -9.53E-02 | 9.98E-02 | 2.68E+00 | 2.75E+00 | 3.35E-01 | no | Gm22508 | downstream_gene_variant | MODIFIER |
| 16 | 18690344 | AX-170117343 | 0 | G | A | 0.14 | 8.91E-02 | 9.39E-02 | 2.75E+00 | 2.89E+00 | 3.35E-01 | no | | intergenic_variant | MODIFIER |
| 5 | 4246250 | AX-170232253 | 1 | A | G | 0.411 | -6.47E-02 | 6.58E-02 | 2.43E+00 | 2.54E+00 | 3.35E-01 | no | | intergenic_variant | MODIFIER |
| 12 | 50342499 | AX-97821140 | 1 | G | A | 0.045 | -1.57E-01 | 1.65E-01 | 2.69E+00 | 2.82E+00 | 3.35E-01 | no | Prkd1 | intron_variant | MODIFIER |
| 18 | 73796953 | AX-170402600 | 0 | C | T | 0.193 | -1.14E-01 | 1.19E-01 | 2.66E+00 | 2.71E+00 | 3.35E-01 | no | Me2 | intron_variant | MODIFIER |
| 11 | 18876275 | AX-170268108 | 0 | C | T | 0.061 | 1.67E-01 | 1.76E-01 | 2.81E+00 | 2.95E+00 | 3.35E-01 | Yes | Meis1 | downstream_gene_variant | MODIFIER |
| 10 | 71468458 | AX-170292035 | 1 | C | G | 0.42 | -7.58E-02 | 7.95E-02 | 2.68E+00 | 2.76E+00 | 3.35E-01 | no | | intergenic_variant | MODIFIER |
| 10 | 72768870 | AX-97845911 | 1 | G | T | 0.42 | -7.58E-02 | 7.95E-02 | 2.68E+00 | 2.76E+00 | 3.35E-01 | no | | intergenic_variant | MODIFIER |
| 4 | 140748882 | AX-97856340 | 1 | A | G | 0.152 | 1.12E-01 | 1.17E-01 | 2.62E+00 | 2.67E+00 | 3.36E-01 | no | Padi4 | intron_variant | MODIFIER |
| 1 | 128669564 | AX-170042666 | 0 | C | T | 0.544 | 8.10E-02 | 8.51E-02 | 2.71E+00 | 2.78E+00 | 3.36E-01 | no | | regulatory_region_variant | MODIFIER |
| 16 | 79549389 | AX-170437438 | 0 | C | G | 0.465 | 7.23E-02 | 7.54E-02 | 2.59E+00 | 2.66E+00 | 3.36E-01 | no | | intergenic_variant | MODIFIER |
| 6 | 129034330 | AX-170076207 | 0 | T | C | 0.544 | 7.91E-02 | 8.17E-02 | 2.52E+00 | 2.59E+00 | 3.36E-01 | no | Gm16186 | upstream_gene_variant | MODIFIER |
| 15 | 15476449 | AX-97828301 | 1 | G | T | 0.241 | -1.03E-01 | 1.08E-01 | 2.60E+00 | 2.65E+00 | 3.36E-01 | no | | intergenic_variant | MODIFIER |
| 10 | 126866036 | AX-170137360 | 1 | A | G | 0.312 | -6.97E-02 | 7.33E-02 | 2.66E+00 | 2.77E+00 | 3.36E-01 | no | Atp23 | downstream_gene_variant | MODIFIER |
| 10 | 127497912 | AX-170004126 | 1 | A | G | 0.312 | -6.97E-02 | 7.33E-02 | 2.66E+00 | 2.77E+00 | 3.36E-01 | no | Stac3 | upstream_gene_variant | MODIFIER |
| 1 | 126688535 | AX-170120587 | 1 | A | G | 0.536 | 7.72E-02 | 8.06E-02 | 2.60E+00 | 2.68E+00 | 3.36E-01 | no | Nckap5 | intron_variant | MODIFIER |
| 8 | 16274090 | AX-170472422 | 1 | C | A | 0.232 | -7.37E-02 | 7.68E-02 | 2.56E+00 | 2.66E+00 | 3.36E-01 | no | Csmd1 | intron_variant | MODIFIER |
| 10 | 69822189 | AX-170003422 | 1 | C | T | 0.411 | -7.62E-02 | 8.00E-02 | 2.66E+00 | 2.73E+00 | 3.36E-01 | no | Ank3 | intron_variant | MODIFIER |
| 7 | 70132661 | AX-97799186 | 0 | T | C | 0.36 | -9.92E-02 | 1.03E-01 | 2.94E+00 | 3.03E+00 | 3.36E-01 | no | | intergenic_variant | MODIFIER |
| 16 | 73052945 | AX-170103560 | 1 | A | G | 0.491 | 7.02E-02 | 7.29E-02 | 2.54E+00 | 2.62E+00 | 3.36E-01 | no | | intergenic_variant | MODIFIER |
| 3 | 150891893 | AX-170189164 | 0 | A | T | 0.351 | 6.68E-02 | 7.03E-02 | 2.78E+00 | 2.88E+00 | 3.36E-01 | no | | intergenic_variant | MODIFIER |
| 13 | 8962074 | AX-170414892 | 0 | C | T | 0.193 | -1.18E-01 | 1.20E-01 | 2.44E+00 | 2.50E+00 | 3.36E-01 | no | Idi2 | downstream_gene_variant | MODIFIER |
| 13 | 11117023 | AX-170466999 | 0 | G | A | 0.193 | -1.18E-01 | 1.20E-01 | 2.44E+00 | 2.50E+00 | 3.36E-01 | no | | intergenic_variant | MODIFIER |
| 5 | 4349483 | AX-170159993 | 1 | C | T | 0.438 | -6.34E-02 | 6.53E-02 | 2.48E+00 | 2.59E+00 | 3.36E-01 | no | | intergenic_variant | MODIFIER |
| 18 | 86640842 | AX-170375464 | 0 | G | T | 0.167 | -9.69E-02 | 1.01E-01 | 2.54E+00 | 2.65E+00 | 3.37E-01 | no | | intergenic_variant | MODIFIER |
| 7 | 75443720 | AX-170454324 | 0 | T | C | 0.368 | -8.98E-02 | 9.46E-02 | 2.68E+00 | 2.78E+00 | 3.37E-01 | no | | intergenic_variant | MODIFIER |
| 16 | 50300886 | AX-97852128 | 0 | G | A | 0.518 | 8.04E-02 | 8.34E-02 | 2.55E+00 | 2.59E+00 | 3.37E-01 | no | Bbx | intron_variant | MODIFIER |
| 10 | 71107161 | AX-170003440 | 0 | C | T | 0.412 | -7.64E-02 | 8.03E-02 | 2.68E+00 | 2.75E+00 | 3.37E-01 | no | Bicc1 | intron_variant | MODIFIER |
| 10 | 75681514 | AX-170529943 | 0 | G | A | 0.43 | -7.58E-02 | 7.97E-02 | 2.70E+00 | 2.77E+00 | 3.37E-01 | no | Cabin1 | intron_variant | MODIFIER |
| 10 | 75684292 | AX-170171985 | 0 | A | G | 0.43 | -7.58E-02 | 7.97E-02 | 2.70E+00 | 2.77E+00 | 3.37E-01 | no | Cabin1 | splice_region_variant,synonymous_variant | LOW |
| 10 | 75690840 | AX-97790936 | 0 | A | G | 0.43 | -7.58E-02 | 7.97E-02 | 2.70E+00 | 2.77E+00 | 3.37E-01 | no | Cabin1 | intron_variant | MODIFIER |
| 10 | 75832585 | AX-97816016 | 0 | C | A | 0.43 | -7.58E-02 | 7.97E-02 | 2.70E+00 | 2.77E+00 | 3.37E-01 | no | Gstt2 | intron_variant | MODIFIER |
| 9 | 15613163 | AX-97836623 | 0 | C | T | 0.456 | 8.24E-02 | 8.53E-02 | 2.52E+00 | 2.59E+00 | 3.37E-01 | no | Deup1 | intron_variant | MODIFIER |
| 9 | 15637107 | AX-97801536 | 0 | C | T | 0.456 | 8.24E-02 | 8.53E-02 | 2.52E+00 | 2.59E+00 | 3.37E-01 | no | 4930540M03Rik | intron_variant,non_coding_transcript_variant | MODIFIER |
| 9 | 15686867 | AX-97816838 | 0 | T | G | 0.456 | 8.24E-02 | 8.53E-02 | 2.52E+00 | 2.59E+00 | 3.37E-01 | no | Gm5119 | downstream_gene_variant | MODIFIER |
| 12 | 109985644 | AX-170353501 | 0 | C | T | 0.132 | -1.53E-01 | 1.61E-01 | 2.75E+00 | 2.84E+00 | 3.37E-01 | no | | intergenic_variant | MODIFIER |
| 15 | 98450354 | AX-97835937 | 0 | G | A | 0.132 | -1.07E-01 | 1.13E-01 | 2.72E+00 | 2.84E+00 | 3.37E-01 | no | Olfr281 | upstream_gene_variant | MODIFIER |
| 13 | 20771638 | AX-170137901 | 1 | A | C | 0.179 | -1.18E-01 | 1.20E-01 | 2.45E+00 | 2.51E+00 | 3.37E-01 | no | | intergenic_variant | MODIFIER |
| 15 | 4728626 | AX-170473038 | 0 | T | A | 0.237 | -9.54E-02 | 9.99E-02 | 2.61E+00 | 2.68E+00 | 3.37E-01 | no | C6 | intron_variant | MODIFIER |
| 1 | 141064336 | AX-170249989 | 0 | G | A | 0.219 | 1.10E-01 | 1.16E-01 | 2.79E+00 | 2.91E+00 | 3.37E-01 | no | | intergenic_variant | MODIFIER |
| 7 | 82372222 | AX-170195235 | 0 | C | T | 0.123 | 1.64E-01 | 1.73E-01 | 2.72E+00 | 2.84E+00 | 3.37E-01 | Yes | Adamtsl3 | intron_variant | MODIFIER |
| 7 | 82842508 | AX-170475585 | 0 | T | A | 0.123 | 1.64E-01 | 1.73E-01 | 2.72E+00 | 2.84E+00 | 3.37E-01 | Yes | | regulatory_region_variant | MODIFIER |
| 8 | 15532176 | AX-170413573 | 0 | T | G | 0.228 | -7.33E-02 | 7.64E-02 | 2.55E+00 | 2.64E+00 | 3.37E-01 | no | | intergenic_variant | MODIFIER |
| 8 | 15549472 | AX-170454418 | 0 | C | T | 0.228 | -7.33E-02 | 7.64E-02 | 2.55E+00 | 2.64E+00 | 3.37E-01 | no | | intergenic_variant | MODIFIER |
| 8 | 15587645 | AX-170103547 | 0 | G | A | 0.228 | -7.33E-02 | 7.64E-02 | 2.55E+00 | 2.64E+00 | 3.37E-01 | no | Gm35453 | upstream_gene_variant | MODIFIER |
| 8 | 15587838 | AX-170256292 | 0 | A | G | 0.228 | -7.33E-02 | 7.64E-02 | 2.55E+00 | 2.64E+00 | 3.37E-01 | no | Gm35453 | upstream_gene_variant | MODIFIER |
| 8 | 15588557 | AX-170274426 | 0 | A | G | 0.228 | -7.33E-02 | 7.64E-02 | 2.55E+00 | 2.64E+00 | 3.37E-01 | no | Gm35453 | upstream_gene_variant | MODIFIER |
| 8 | 15604763 | AX-170249885 | 0 | T | C | 0.228 | -7.33E-02 | 7.64E-02 | 2.55E+00 | 2.64E+00 | 3.37E-01 | no | Gm45408 | intron_variant,non_coding_transcript_variant | MODIFIER |
| 8 | 15625995 | AX-170480690 | 0 | G | A | 0.228 | -7.33E-02 | 7.64E-02 | 2.55E+00 | 2.64E+00 | 3.37E-01 | no | Gm45408 | intron_variant,non_coding_transcript_variant | MODIFIER |
| 8 | 15628963 | AX-170309162 | 0 | A | G | 0.228 | -7.33E-02 | 7.64E-02 | 2.55E+00 | 2.64E+00 | 3.37E-01 | no | Gm45408 | intron_variant,non_coding_transcript_variant | MODIFIER |
| 8 | 15635407 | AX-170309814 | 0 | G | T | 0.228 | -7.33E-02 | 7.64E-02 | 2.55E+00 | 2.64E+00 | 3.37E-01 | no | Gm45408 | intron_variant,non_coding_transcript_variant | MODIFIER |
| 8 | 15659807 | AX-170173809 | 0 | C | T | 0.228 | -7.33E-02 | 7.64E-02 | 2.55E+00 | 2.64E+00 | 3.37E-01 | no | Gm45408 | intron_variant,non_coding_transcript_variant | MODIFIER |
| 8 | 15667095 | AX-170449679 | 0 | T | C | 0.228 | -7.33E-02 | 7.64E-02 | 2.55E+00 | 2.64E+00 | 3.37E-01 | no | Gm45408 | intron_variant,non_coding_transcript_variant | MODIFIER |

| | | | | | | | | | | | | | | | |
|---|---|---|---|---|---|---|---|---|---|---|---|---|---|---|---|
| 8 | 15670589 | AX-170351634 | 0 | A | G | 0.228 | -7.33E-02 | 7.64E-02 | 2.55E+00 | 2.64E+00 | 3.37E-01 | no | Gm45408 | downstream_gene_variant | MODIFIER |
| 8 | 15863591 | AX-170331979 | 0 | T | A | 0.228 | -7.33E-02 | 7.64E-02 | 2.55E+00 | 2.64E+00 | 3.37E-01 | no | | intergenic_variant | MODIFIER |
| 8 | 15863820 | AX-170258581 | 0 | T | C | 0.228 | -7.33E-02 | 7.64E-02 | 2.55E+00 | 2.64E+00 | 3.37E-01 | no | | intergenic_variant | MODIFIER |
| 8 | 15868906 | AX-170138718 | 0 | C | T | 0.228 | -7.33E-02 | 7.64E-02 | 2.55E+00 | 2.64E+00 | 3.37E-01 | no | | intergenic_variant | MODIFIER |
| 8 | 15870524 | AX-97824369 | 0 | C | T | 0.228 | -7.33E-02 | 7.64E-02 | 2.55E+00 | 2.64E+00 | 3.37E-01 | no | | intergenic_variant | MODIFIER |
| 8 | 15888705 | AX-170079397 | 0 | G | A | 0.228 | -7.33E-02 | 7.64E-02 | 2.55E+00 | 2.64E+00 | 3.37E-01 | no | Csmd1 | downstream_gene_variant | MODIFIER |
| 8 | 15911814 | AX-170176879 | 0 | G | A | 0.228 | -7.33E-02 | 7.64E-02 | 2.55E+00 | 2.64E+00 | 3.37E-01 | no | Csmd1 | intron_variant | MODIFIER |
| 8 | 15912055 | AX-170467820 | 0 | C | G | 0.228 | -7.33E-02 | 7.64E-02 | 2.55E+00 | 2.64E+00 | 3.37E-01 | no | Csmd1 | intron_variant | MODIFIER |
| 8 | 15914320 | AX-97825381 | 0 | C | T | 0.228 | -7.33E-02 | 7.64E-02 | 2.55E+00 | 2.64E+00 | 3.37E-01 | no | Csmd1 | intron_variant | MODIFIER |
| 8 | 15914448 | AX-170431975 | 0 | C | A | 0.228 | -7.33E-02 | 7.64E-02 | 2.55E+00 | 2.64E+00 | 3.37E-01 | no | Csmd1 | intron_variant | MODIFIER |
| 8 | 15919471 | AX-170481376 | 0 | C | T | 0.228 | -7.33E-02 | 7.64E-02 | 2.55E+00 | 2.64E+00 | 3.37E-01 | no | Csmd1 | intron_variant | MODIFIER |
| 8 | 15924782 | AX-170346219 | 0 | T | C | 0.228 | -7.33E-02 | 7.64E-02 | 2.55E+00 | 2.64E+00 | 3.37E-01 | no | Csmd1 | intron_variant | MODIFIER |
| 8 | 15931391 | AX-170519085 | 0 | T | C | 0.228 | -7.33E-02 | 7.64E-02 | 2.55E+00 | 2.64E+00 | 3.37E-01 | no | Csmd1 | intron_variant | MODIFIER |
| 8 | 15942257 | AX-170376689 | 0 | G | A | 0.228 | -7.33E-02 | 7.64E-02 | 2.55E+00 | 2.64E+00 | 3.37E-01 | no | Csmd1 | intron_variant | MODIFIER |
| 8 | 15942782 | AX-170495848 | 0 | C | T | 0.228 | -7.33E-02 | 7.64E-02 | 2.55E+00 | 2.64E+00 | 3.37E-01 | no | Csmd1 | intron_variant | MODIFIER |
| 8 | 15945622 | AX-170352923 | 0 | C | T | 0.228 | -7.33E-02 | 7.64E-02 | 2.55E+00 | 2.64E+00 | 3.37E-01 | no | Csmd1 | intron_variant | MODIFIER |
| 8 | 15966024 | AX-170119515 | 0 | G | A | 0.228 | -7.33E-02 | 7.64E-02 | 2.55E+00 | 2.64E+00 | 3.37E-01 | no | Csmd1 | intron_variant | MODIFIER |
| 8 | 15996674 | AX-170184697 | 0 | T | C | 0.228 | -7.33E-02 | 7.64E-02 | 2.55E+00 | 2.64E+00 | 3.37E-01 | no | Csmd1 | intron_variant | MODIFIER |
| 8 | 15997291 | AX-170196764 | 0 | C | T | 0.228 | -7.33E-02 | 7.64E-02 | 2.55E+00 | 2.64E+00 | 3.37E-01 | no | Csmd1 | intron_variant | MODIFIER |
| 8 | 15999502 | AX-170145807 | 0 | T | G | 0.228 | -7.33E-02 | 7.64E-02 | 2.55E+00 | 2.64E+00 | 3.37E-01 | no | Csmd1 | intron_variant | MODIFIER |
| 8 | 16006559 | AX-170468611 | 0 | A | C | 0.228 | -7.33E-02 | 7.64E-02 | 2.55E+00 | 2.64E+00 | 3.37E-01 | no | Csmd1 | intron_variant | MODIFIER |
| 8 | 16006601 | AX-170425382 | 0 | T | C | 0.228 | -7.33E-02 | 7.64E-02 | 2.55E+00 | 2.64E+00 | 3.37E-01 | no | Csmd1 | intron_variant | MODIFIER |
| 8 | 16011153 | AX-170484453 | 0 | G | A | 0.228 | -7.33E-02 | 7.64E-02 | 2.55E+00 | 2.64E+00 | 3.37E-01 | no | Csmd1 | intron_variant | MODIFIER |
| 8 | 16037174 | AX-170235376 | 0 | C | T | 0.228 | -7.33E-02 | 7.64E-02 | 2.55E+00 | 2.64E+00 | 3.37E-01 | no | Csmd1 | intron_variant | MODIFIER |
| 8 | 16051456 | AX-170297673 | 0 | C | G | 0.228 | -7.33E-02 | 7.64E-02 | 2.55E+00 | 2.64E+00 | 3.37E-01 | no | Csmd1 | intron_variant | MODIFIER |
| 8 | 16073160 | AX-97802335 | 0 | G | T | 0.228 | -7.33E-02 | 7.64E-02 | 2.55E+00 | 2.64E+00 | 3.37E-01 | no | Csmd1 | intron_variant | MODIFIER |
| 8 | 16074102 | AX-170259297 | 0 | G | A | 0.228 | -7.33E-02 | 7.64E-02 | 2.55E+00 | 2.64E+00 | 3.37E-01 | no | Csmd1 | intron_variant | MODIFIER |
| 8 | 16075765 | AX-170400019 | 0 | A | G | 0.228 | -7.33E-02 | 7.64E-02 | 2.55E+00 | 2.64E+00 | 3.37E-01 | no | Csmd1 | intron_variant | MODIFIER |
| 8 | 16085057 | AX-170157559 | 0 | A | G | 0.228 | -7.33E-02 | 7.64E-02 | 2.55E+00 | 2.64E+00 | 3.37E-01 | no | Csmd1 | intron_variant | MODIFIER |
| 8 | 16088440 | AX-170249959 | 0 | T | C | 0.228 | -7.33E-02 | 7.64E-02 | 2.55E+00 | 2.64E+00 | 3.37E-01 | no | Csmd1 | intron_variant | MODIFIER |
| 8 | 16098741 | AX-170420064 | 0 | T | C | 0.228 | -7.33E-02 | 7.64E-02 | 2.55E+00 | 2.64E+00 | 3.37E-01 | no | Csmd1 | intron_variant | MODIFIER |
| 8 | 16211447 | AX-170433492 | 0 | A | T | 0.228 | -7.33E-02 | 7.64E-02 | 2.55E+00 | 2.64E+00 | 3.37E-01 | no | Csmd1 | intron_variant | MODIFIER |
| 8 | 16248655 | AX-170488356 | 0 | A | G | 0.228 | -7.33E-02 | 7.64E-02 | 2.55E+00 | 2.64E+00 | 3.37E-01 | no | Csmd1 | intron_variant | MODIFIER |
| 8 | 16257621 | AX-170232269 | 0 | T | C | 0.228 | -7.33E-02 | 7.64E-02 | 2.55E+00 | 2.64E+00 | 3.37E-01 | no | Csmd1 | intron_variant | MODIFIER |
| 8 | 16272987 | AX-170496632 | 0 | T | A | 0.228 | -7.33E-02 | 7.64E-02 | 2.55E+00 | 2.64E+00 | 3.37E-01 | no | Csmd1 | intron_variant | MODIFIER |
| 8 | 16280066 | AX-170101206 | 0 | T | C | 0.228 | -7.33E-02 | 7.64E-02 | 2.55E+00 | 2.64E+00 | 3.37E-01 | no | Csmd1 | intron_variant | MODIFIER |
| 8 | 16282080 | AX-170387922 | 0 | A | T | 0.228 | -7.33E-02 | 7.64E-02 | 2.55E+00 | 2.64E+00 | 3.37E-01 | no | Csmd1 | intron_variant | MODIFIER |
| 8 | 16282227 | AX-170344275 | 0 | G | A | 0.228 | -7.33E-02 | 7.64E-02 | 2.55E+00 | 2.64E+00 | 3.37E-01 | no | Csmd1 | intron_variant | MODIFIER |
| 8 | 16424436 | AX-170030315 | 0 | T | C | 0.228 | -7.33E-02 | 7.64E-02 | 2.55E+00 | 2.64E+00 | 3.37E-01 | no | Csmd1 | intron_variant | MODIFIER |
| 8 | 16424991 | AX-170030312 | 0 | A | G | 0.228 | -7.33E-02 | 7.64E-02 | 2.55E+00 | 2.64E+00 | 3.37E-01 | no | Csmd1 | intron_variant | MODIFIER |
| 8 | 16452177 | AX-170167184 | 0 | A | G | 0.228 | -7.33E-02 | 7.64E-02 | 2.55E+00 | 2.64E+00 | 3.37E-01 | no | Csmd1 | intron_variant | MODIFIER |
| 8 | 16452269 | AX-97856038 | 0 | G | T | 0.228 | -7.33E-02 | 7.64E-02 | 2.55E+00 | 2.64E+00 | 3.37E-01 | no | Csmd1 | intron_variant | MODIFIER |
| 8 | 16457748 | AX-170534645 | 0 | A | G | 0.228 | -7.33E-02 | 7.64E-02 | 2.55E+00 | 2.64E+00 | 3.37E-01 | no | Csmd1 | intron_variant | MODIFIER |
| 8 | 16461499 | AX-170416163 | 0 | A | G | 0.228 | -7.33E-02 | 7.64E-02 | 2.55E+00 | 2.64E+00 | 3.37E-01 | no | Csmd1 | intron_variant | MODIFIER |
| 8 | 16461519 | AX-170119544 | 0 | T | C | 0.228 | -7.33E-02 | 7.64E-02 | 2.55E+00 | 2.64E+00 | 3.37E-01 | no | Csmd1 | intron_variant | MODIFIER |
| 8 | 16471482 | AX-170237781 | 0 | T | C | 0.228 | -7.33E-02 | 7.64E-02 | 2.55E+00 | 2.64E+00 | 3.37E-01 | no | Csmd1 | intron_variant | MODIFIER |
| 8 | 16481803 | AX-170079412 | 0 | A | G | 0.228 | -7.33E-02 | 7.64E-02 | 2.55E+00 | 2.64E+00 | 3.37E-01 | no | Csmd1 | intron_variant | MODIFIER |
| 8 | 16485309 | AX-170147464 | 0 | G | A | 0.228 | -7.33E-02 | 7.64E-02 | 2.55E+00 | 2.64E+00 | 3.37E-01 | no | Csmd1 | intron_variant | MODIFIER |
| 8 | 16485336 | AX-170413553 | 0 | A | G | 0.228 | -7.33E-02 | 7.64E-02 | 2.55E+00 | 2.64E+00 | 3.37E-01 | no | Csmd1 | intron_variant | MODIFIER |
| 8 | 16485873 | AX-170144454 | 0 | T | C | 0.228 | -7.33E-02 | 7.64E-02 | 2.55E+00 | 2.64E+00 | 3.37E-01 | no | Csmd1 | intron_variant | MODIFIER |
| 8 | 16487663 | AX-170460541 | 0 | C | A | 0.228 | -7.33E-02 | 7.64E-02 | 2.55E+00 | 2.64E+00 | 3.37E-01 | no | Csmd1 | intron_variant | MODIFIER |
| 8 | 16499694 | AX-97827425 | 0 | T | C | 0.228 | -7.33E-02 | 7.64E-02 | 2.55E+00 | 2.64E+00 | 3.37E-01 | no | Csmd1 | intron_variant | MODIFIER |
| 8 | 16505012 | AX-170270676 | 0 | A | G | 0.228 | -7.33E-02 | 7.64E-02 | 2.55E+00 | 2.64E+00 | 3.37E-01 | no | Csmd1 | intron_variant | MODIFIER |
| 8 | 16505430 | AX-170487381 | 0 | T | C | 0.228 | -7.33E-02 | 7.64E-02 | 2.55E+00 | 2.64E+00 | 3.37E-01 | no | Csmd1 | intron_variant | MODIFIER |

| | | | | | | | | | | | | | | | |
|---|---|---|---|---|---|---|---|---|---|---|---|---|---|---|---|
| 8 | 16521385 | AX-97856218 | 0 | G | T | 0.228 | -7.33E-02 | 7.64E-02 | 2.55E+00 | 2.64E+00 | 3.37E-01 | no | Csmd1 | intron_variant | MODIFIER |
| 8 | 16536623 | AX-170086231 | 0 | A | C | 0.228 | -7.33E-02 | 7.64E-02 | 2.55E+00 | 2.64E+00 | 3.37E-01 | no | Csmd1 | intron_variant | MODIFIER |
| 8 | 16537958 | AX-170270704 | 0 | A | T | 0.228 | -7.33E-02 | 7.64E-02 | 2.55E+00 | 2.64E+00 | 3.37E-01 | no | Csmd1 | intron_variant | MODIFIER |
| 8 | 16549710 | AX-170316955 | 0 | C | G | 0.228 | -7.33E-02 | 7.64E-02 | 2.55E+00 | 2.64E+00 | 3.37E-01 | no | Csmd1 | intron_variant | MODIFIER |
| 8 | 16566800 | AX-170317159 | 0 | T | C | 0.228 | -7.33E-02 | 7.64E-02 | 2.55E+00 | 2.64E+00 | 3.37E-01 | no | Csmd1 | intron_variant | MODIFIER |
| 8 | 16572203 | AX-170210914 | 0 | G | C | 0.228 | -7.33E-02 | 7.64E-02 | 2.55E+00 | 2.64E+00 | 3.37E-01 | no | Csmd1 | intron_variant | MODIFIER |
| 8 | 16580115 | AX-170481348 | 0 | G | A | 0.228 | -7.33E-02 | 7.64E-02 | 2.55E+00 | 2.64E+00 | 3.37E-01 | no | Csmd1 | intron_variant | MODIFIER |
| 8 | 16589949 | AX-170079416 | 0 | A | G | 0.228 | -7.33E-02 | 7.64E-02 | 2.55E+00 | 2.64E+00 | 3.37E-01 | no | Csmd1 | intron_variant | MODIFIER |
| 8 | 16590104 | AX-97837325 | 0 | T | C | 0.228 | -7.33E-02 | 7.64E-02 | 2.55E+00 | 2.64E+00 | 3.37E-01 | no | Csmd1 | intron_variant | MODIFIER |
| 8 | 16610566 | AX-170358221 | 0 | C | T | 0.228 | -7.33E-02 | 7.64E-02 | 2.55E+00 | 2.64E+00 | 3.37E-01 | no | Csmd1 | intron_variant | MODIFIER |
| 8 | 16619318 | AX-170514311 | 0 | G | A | 0.228 | -7.33E-02 | 7.64E-02 | 2.55E+00 | 2.64E+00 | 3.37E-01 | no | Csmd1 | intron_variant | MODIFIER |
| 8 | 16619522 | AX-97781745 | 0 | G | A | 0.228 | -7.33E-02 | 7.64E-02 | 2.55E+00 | 2.64E+00 | 3.37E-01 | no | Csmd1 | intron_variant | MODIFIER |
| 8 | 16623937 | AX-170382880 | 0 | T | C | 0.228 | -7.33E-02 | 7.64E-02 | 2.55E+00 | 2.64E+00 | 3.37E-01 | no | Csmd1 | intron_variant | MODIFIER |
| 8 | 16673836 | AX-170333851 | 0 | T | C | 0.228 | -7.33E-02 | 7.64E-02 | 2.55E+00 | 2.64E+00 | 3.37E-01 | no | Csmd1 | intron_variant | MODIFIER |
| 8 | 16694359 | AX-97817594 | 0 | C | T | 0.228 | -7.33E-02 | 7.64E-02 | 2.55E+00 | 2.64E+00 | 3.37E-01 | no | Csmd1 | intron_variant | MODIFIER |
| 8 | 16741566 | AX-170281553 | 0 | T | C | 0.228 | -7.33E-02 | 7.64E-02 | 2.55E+00 | 2.64E+00 | 3.37E-01 | no | Csmd1 | intron_variant | MODIFIER |
| 8 | 16952317 | AX-170030316 | 0 | C | T | 0.228 | -7.33E-02 | 7.64E-02 | 2.55E+00 | 2.64E+00 | 3.37E-01 | no | Csmd1 | intron_variant | MODIFIER |
| 8 | 16953253 | AX-170302429 | 0 | G | A | 0.228 | -7.33E-02 | 7.64E-02 | 2.55E+00 | 2.64E+00 | 3.37E-01 | no | Csmd1 | intron_variant | MODIFIER |
| 8 | 16956175 | AX-170030317 | 0 | A | G | 0.228 | -7.33E-02 | 7.64E-02 | 2.55E+00 | 2.64E+00 | 3.37E-01 | no | Csmd1 | intron_variant | MODIFIER |
| 8 | 16956241 | AX-170030318 | 0 | C | T | 0.228 | -7.33E-02 | 7.64E-02 | 2.55E+00 | 2.64E+00 | 3.37E-01 | no | Csmd1 | intron_variant | MODIFIER |
| 8 | 16960141 | AX-170233494 | 0 | G | A | 0.228 | -7.33E-02 | 7.64E-02 | 2.55E+00 | 2.64E+00 | 3.37E-01 | no | Csmd1 | intron_variant | MODIFIER |
| 8 | 16965112 | AX-170030320 | 0 | A | T | 0.228 | -7.33E-02 | 7.64E-02 | 2.55E+00 | 2.64E+00 | 3.37E-01 | no | Csmd1 | intron_variant | MODIFIER |
| 8 | 16965247 | AX-170461232 | 0 | G | T | 0.228 | -7.33E-02 | 7.64E-02 | 2.55E+00 | 2.64E+00 | 3.37E-01 | no | Csmd1 | intron_variant | MODIFIER |
| 8 | 16974056 | AX-170114673 | 0 | T | C | 0.228 | -7.33E-02 | 7.64E-02 | 2.55E+00 | 2.64E+00 | 3.37E-01 | no | Csmd1 | intron_variant | MODIFIER |
| 8 | 17031930 | AX-97826918 | 0 | C | T | 0.228 | -7.33E-02 | 7.64E-02 | 2.55E+00 | 2.64E+00 | 3.37E-01 | no | Csmd1 | intron_variant | MODIFIER |
| 8 | 17033700 | AX-170257886 | 0 | C | T | 0.228 | -7.33E-02 | 7.64E-02 | 2.55E+00 | 2.64E+00 | 3.37E-01 | no | Csmd1 | intron_variant | MODIFIER |
| 8 | 17037940 | AX-97851888 | 0 | C | T | 0.228 | -7.33E-02 | 7.64E-02 | 2.55E+00 | 2.64E+00 | 3.37E-01 | no | Csmd1 | intron_variant | MODIFIER |
| 8 | 17044636 | AX-170385148 | 0 | T | C | 0.228 | -7.33E-02 | 7.64E-02 | 2.55E+00 | 2.64E+00 | 3.37E-01 | no | Csmd1 | intron_variant | MODIFIER |
| 8 | 17053478 | AX-170266674 | 0 | C | T | 0.228 | -7.33E-02 | 7.64E-02 | 2.55E+00 | 2.64E+00 | 3.37E-01 | no | Csmd1 | intron_variant | MODIFIER |
| 8 | 17154523 | AX-170079430 | 0 | G | T | 0.228 | -7.33E-02 | 7.64E-02 | 2.55E+00 | 2.64E+00 | 3.37E-01 | no | Csmd1 | intron_variant | MODIFIER |
| 8 | 17159295 | AX-170272663 | 0 | G | A | 0.228 | -7.33E-02 | 7.64E-02 | 2.55E+00 | 2.64E+00 | 3.37E-01 | no | Csmd1 | intron_variant | MODIFIER |
| 8 | 17163218 | AX-170235033 | 0 | C | G | 0.228 | -7.33E-02 | 7.64E-02 | 2.55E+00 | 2.64E+00 | 3.37E-01 | no | Csmd1 | intron_variant | MODIFIER |
| 8 | 17164917 | AX-170308146 | 0 | C | G | 0.228 | -7.33E-02 | 7.64E-02 | 2.55E+00 | 2.64E+00 | 3.37E-01 | no | Csmd1 | intron_variant | MODIFIER |
| 8 | 17311541 | AX-170432153 | 0 | A | G | 0.228 | -7.33E-02 | 7.64E-02 | 2.55E+00 | 2.64E+00 | 3.37E-01 | no | Csmd1 | intron_variant | MODIFIER |
| 8 | 17312409 | AX-170517462 | 0 | T | A | 0.228 | -7.33E-02 | 7.64E-02 | 2.55E+00 | 2.64E+00 | 3.37E-01 | no | Csmd1 | intron_variant | MODIFIER |
| 8 | 17313890 | AX-170180254 | 0 | T | C | 0.228 | -7.33E-02 | 7.64E-02 | 2.55E+00 | 2.64E+00 | 3.37E-01 | no | Csmd1 | intron_variant | MODIFIER |
| 8 | 17313976 | AX-170416724 | 0 | C | T | 0.228 | -7.33E-02 | 7.64E-02 | 2.55E+00 | 2.64E+00 | 3.37E-01 | no | Csmd1 | intron_variant | MODIFIER |
| 8 | 17320545 | AX-170523130 | 0 | C | T | 0.228 | -7.33E-02 | 7.64E-02 | 2.55E+00 | 2.64E+00 | 3.37E-01 | no | Csmd1 | intron_variant | MODIFIER |
| 8 | 17328897 | AX-170313176 | 0 | C | T | 0.228 | -7.33E-02 | 7.64E-02 | 2.55E+00 | 2.64E+00 | 3.37E-01 | no | Csmd1 | intron_variant | MODIFIER |
| 8 | 17329577 | AX-170255107 | 0 | A | G | 0.228 | -7.33E-02 | 7.64E-02 | 2.55E+00 | 2.64E+00 | 3.37E-01 | no | Csmd1 | intron_variant | MODIFIER |
| 8 | 17347125 | AX-170305867 | 0 | G | A | 0.228 | -7.33E-02 | 7.64E-02 | 2.55E+00 | 2.64E+00 | 3.37E-01 | no | Csmd1 | intron_variant | MODIFIER |
| 8 | 17354027 | AX-170357429 | 0 | T | C | 0.228 | -7.33E-02 | 7.64E-02 | 2.55E+00 | 2.64E+00 | 3.37E-01 | no | Csmd1 | intron_variant | MODIFIER |
| 8 | 17359084 | AX-170111873 | 0 | G | A | 0.228 | -7.33E-02 | 7.64E-02 | 2.55E+00 | 2.64E+00 | 3.37E-01 | no | Csmd1 | intron_variant | MODIFIER |
| 8 | 17364402 | AX-97851355 | 0 | A | G | 0.228 | -7.33E-02 | 7.64E-02 | 2.55E+00 | 2.64E+00 | 3.37E-01 | no | Csmd1 | intron_variant | MODIFIER |
| 8 | 17378623 | AX-170079435 | 0 | A | G | 0.228 | -7.33E-02 | 7.64E-02 | 2.55E+00 | 2.64E+00 | 3.37E-01 | no | Csmd1 | intron_variant | MODIFIER |
| 8 | 17395755 | AX-170108204 | 0 | C | T | 0.228 | -7.33E-02 | 7.64E-02 | 2.55E+00 | 2.64E+00 | 3.37E-01 | no | Csmd1 | intron_variant | MODIFIER |
| 8 | 17458800 | AX-170134161 | 0 | A | T | 0.228 | -7.33E-02 | 7.64E-02 | 2.55E+00 | 2.64E+00 | 3.37E-01 | no | Csmd1 | intron_variant | MODIFIER |
| 8 | 17475484 | AX-170030323 | 0 | A | G | 0.228 | -7.33E-02 | 7.64E-02 | 2.55E+00 | 2.64E+00 | 3.37E-01 | no | Csmd1 | intron_variant | MODIFIER |
| 8 | 17479625 | AX-170030322 | 0 | A | G | 0.228 | -7.33E-02 | 7.64E-02 | 2.55E+00 | 2.64E+00 | 3.37E-01 | no | Csmd1 | intron_variant | MODIFIER |
| 8 | 17481914 | AX-170030321 | 0 | T | C | 0.228 | -7.33E-02 | 7.64E-02 | 2.55E+00 | 2.64E+00 | 3.37E-01 | no | Csmd1 | intron_variant | MODIFIER |
| 8 | 17528527 | AX-170361542 | 0 | A | G | 0.228 | -7.33E-02 | 7.64E-02 | 2.55E+00 | 2.64E+00 | 3.37E-01 | no | Csmd1 | intron_variant | MODIFIER |
| 8 | 17572525 | AX-97851645 | 0 | C | T | 0.228 | -7.33E-02 | 7.64E-02 | 2.55E+00 | 2.64E+00 | 3.37E-01 | no | | intergenic_variant | MODIFIER |
| 8 | 17573647 | AX-170463686 | 0 | C | T | 0.228 | -7.33E-02 | 7.64E-02 | 2.55E+00 | 2.64E+00 | 3.37E-01 | no | | intergenic_variant | MODIFIER |
| 8 | 17574092 | AX-170158575 | 0 | T | G | 0.228 | -7.33E-02 | 7.64E-02 | 2.55E+00 | 2.64E+00 | 3.37E-01 | no | | regulatory_region_variant | MODIFIER |

| | | | | | | | | | | | | | | | |
|---|---|---|---|---|---|---|---|---|---|---|---|---|---|---|---|
| 8 | 17579307 | AX-170204588 | 0 | C | T | 0.228 | -7.33E-02 | 7.64E-02 | 2.55E+00 | 2.64E+00 | 3.37E-01 | no | | intergenic_variant | MODIFIER |
| 8 | 17580228 | AX-170387156 | 0 | G | A | 0.228 | -7.33E-02 | 7.64E-02 | 2.55E+00 | 2.64E+00 | 3.37E-01 | no | | intergenic_variant | MODIFIER |
| 8 | 17595513 | AX-170473892 | 0 | G | C | 0.228 | -7.33E-02 | 7.64E-02 | 2.55E+00 | 2.64E+00 | 3.37E-01 | no | | intergenic_variant | MODIFIER |
| 8 | 17681947 | AX-170370354 | 0 | A | G | 0.228 | -7.33E-02 | 7.64E-02 | 2.55E+00 | 2.64E+00 | 3.37E-01 | no | | intergenic_variant | MODIFIER |
| 8 | 17940159 | AX-97786608 | 0 | G | A | 0.228 | -7.33E-02 | 7.64E-02 | 2.55E+00 | 2.64E+00 | 3.37E-01 | no | | intergenic_variant | MODIFIER |
| 8 | 17944548 | AX-170424606 | 0 | G | A | 0.228 | -7.33E-02 | 7.64E-02 | 2.55E+00 | 2.64E+00 | 3.37E-01 | no | | intergenic_variant | MODIFIER |
| 8 | 17948087 | AX-170237235 | 0 | G | A | 0.228 | -7.33E-02 | 7.64E-02 | 2.55E+00 | 2.64E+00 | 3.37E-01 | no | | regulatory_region_variant | MODIFIER |
| 8 | 17951389 | AX-170203285 | 0 | T | G | 0.228 | -7.33E-02 | 7.64E-02 | 2.55E+00 | 2.64E+00 | 3.37E-01 | no | | intergenic_variant | MODIFIER |
| 8 | 17951459 | AX-170442694 | 0 | G | A | 0.228 | -7.33E-02 | 7.64E-02 | 2.55E+00 | 2.64E+00 | 3.37E-01 | no | | intergenic_variant | MODIFIER |
| 10 | 7005489 | AX-170284438 | 0 | A | G | 0.386 | -4.78E-02 | 5.04E-02 | 2.75E+00 | 2.87E+00 | 3.37E-01 | no | Oprm1 | intron_variant | MODIFIER |
| 7 | 145143351 | AX-97817947 | 1 | G | T | 0.205 | -7.27E-02 | 7.66E-02 | 2.63E+00 | 2.76E+00 | 3.37E-01 | no | | intergenic_variant | MODIFIER |
| 6 | 94603924 | AX-170315492 | 0 | G | A | 0.43 | -5.66E-02 | 5.89E-02 | 2.53E+00 | 2.65E+00 | 3.37E-01 | no | Lrig1 | downstream_gene_variant | MODIFIER |
| 14 | 48026162 | AX-170053310 | 1 | T | C | 0.054 | 1.52E-01 | 1.61E-01 | 2.76E+00 | 2.92E+00 | 3.37E-01 | Yes | | intergenic_variant | MODIFIER |
| 10 | 69545520 | AX-97814008 | 0 | A | C | 0.412 | -7.61E-02 | 8.00E-02 | 2.67E+00 | 2.74E+00 | 3.37E-01 | no | Ank3 | intron_variant | MODIFIER |
| 1 | 150910472 | AX-170304958 | 1 | G | C | 0.214 | -1.25E-01 | 1.27E-01 | 3.13E+00 | 3.26E+00 | 3.37E-01 | no | Hmcn1 | intron_variant | MODIFIER |
| 9 | 96484275 | AX-170090386 | 0 | A | G | 0.132 | -1.13E-01 | 1.19E-01 | 2.59E+00 | 2.67E+00 | 3.37E-01 | no | Gm28729 | downstream_gene_variant | MODIFIER |
| 9 | 96548626 | AX-97786986 | 0 | C | G | 0.132 | -1.13E-01 | 1.19E-01 | 2.59E+00 | 2.67E+00 | 3.37E-01 | no | Rasa2 | intron_variant | MODIFIER |
| 9 | 96737712 | AX-170033059 | 0 | G | A | 0.132 | -1.13E-01 | 1.19E-01 | 2.59E+00 | 2.67E+00 | 3.37E-01 | no | Zbtb38 | intron_variant,non_coding_transcript_variant | MODIFIER |
| 9 | 96744525 | AX-170033067 | 0 | G | A | 0.132 | -1.13E-01 | 1.19E-01 | 2.59E+00 | 2.67E+00 | 3.37E-01 | no | Zbtb38 | intron_variant,non_coding_transcript_variant | MODIFIER |
| 9 | 96744816 | AX-97834174 | 0 | T | C | 0.132 | -1.13E-01 | 1.19E-01 | 2.59E+00 | 2.67E+00 | 3.37E-01 | no | Zbtb38 | intron_variant,non_coding_transcript_variant | MODIFIER |
| 9 | 96744858 | AX-170033068 | 0 | A | C | 0.132 | -1.13E-01 | 1.19E-01 | 2.59E+00 | 2.67E+00 | 3.37E-01 | no | Zbtb38 | intron_variant,non_coding_transcript_variant | MODIFIER |
| 9 | 96771115 | AX-97806067 | 0 | C | T | 0.132 | -1.13E-01 | 1.19E-01 | 2.59E+00 | 2.67E+00 | 3.37E-01 | no | C430002N11Rik | intron_variant,non_coding_transcript_variant | MODIFIER |
| 9 | 96771575 | AX-170033062 | 0 | T | C | 0.132 | -1.13E-01 | 1.19E-01 | 2.59E+00 | 2.67E+00 | 3.37E-01 | no | C430002N11Rik | non_coding_transcript_exon_variant | MODIFIER |
| 9 | 96888558 | AX-170033063 | 0 | A | T | 0.132 | -1.13E-01 | 1.19E-01 | 2.59E+00 | 2.67E+00 | 3.37E-01 | no | Pxylp1 | intron_variant | MODIFIER |
| 9 | 96908299 | AX-170203027 | 0 | A | G | 0.132 | -1.13E-01 | 1.19E-01 | 2.59E+00 | 2.67E+00 | 3.37E-01 | no | Gm8524 | downstream_gene_variant | MODIFIER |
| 9 | 96925609 | AX-170209886 | 0 | A | G | 0.132 | -1.13E-01 | 1.19E-01 | 2.59E+00 | 2.67E+00 | 3.37E-01 | no | | intergenic_variant | MODIFIER |
| 9 | 96929095 | AX-170371002 | 0 | A | G | 0.132 | -1.13E-01 | 1.19E-01 | 2.59E+00 | 2.67E+00 | 3.37E-01 | no | | intergenic_variant | MODIFIER |
| 9 | 97616874 | AX-170137248 | 0 | A | G | 0.132 | -1.13E-01 | 1.19E-01 | 2.59E+00 | 2.67E+00 | 3.37E-01 | no | Clstn2 | intron_variant | MODIFIER |
| 9 | 97635836 | AX-170219068 | 0 | A | G | 0.132 | -1.13E-01 | 1.19E-01 | 2.59E+00 | 2.67E+00 | 3.37E-01 | no | Clstn2 | intron_variant | MODIFIER |
| 9 | 97651191 | AX-97846613 | 0 | A | T | 0.132 | -1.13E-01 | 1.19E-01 | 2.59E+00 | 2.67E+00 | 3.37E-01 | no | Clstn2 | intron_variant | MODIFIER |
| 9 | 97669910 | AX-170129145 | 0 | T | C | 0.132 | -1.13E-01 | 1.19E-01 | 2.59E+00 | 2.67E+00 | 3.37E-01 | no | Clstn2 | intron_variant | MODIFIER |
| 9 | 97677625 | AX-170261271 | 0 | T | G | 0.132 | -1.13E-01 | 1.19E-01 | 2.59E+00 | 2.67E+00 | 3.37E-01 | no | Clstn2 | intron_variant | MODIFIER |
| 9 | 97679027 | AX-170451955 | 0 | C | G | 0.132 | -1.13E-01 | 1.19E-01 | 2.59E+00 | 2.67E+00 | 3.37E-01 | no | Clstn2 | intron_variant | MODIFIER |
| 9 | 97691551 | AX-97811732 | 0 | A | G | 0.132 | -1.13E-01 | 1.19E-01 | 2.59E+00 | 2.67E+00 | 3.37E-01 | no | Clstn2 | intron_variant | MODIFIER |
| 9 | 97703808 | AX-170395675 | 0 | A | G | 0.132 | -1.13E-01 | 1.19E-01 | 2.59E+00 | 2.67E+00 | 3.37E-01 | no | Clstn2 | intron_variant | MODIFIER |
| 9 | 97715303 | AX-170231189 | 0 | C | T | 0.132 | -1.13E-01 | 1.19E-01 | 2.59E+00 | 2.67E+00 | 3.37E-01 | no | Clstn2 | intron_variant | MODIFIER |
| 9 | 97731269 | AX-170090389 | 0 | T | G | 0.132 | -1.13E-01 | 1.19E-01 | 2.59E+00 | 2.67E+00 | 3.37E-01 | no | Clstn2 | intron_variant | MODIFIER |
| 9 | 97740610 | AX-170423715 | 0 | G | A | 0.132 | -1.13E-01 | 1.19E-01 | 2.59E+00 | 2.67E+00 | 3.37E-01 | no | Clstn2 | intron_variant | MODIFIER |
| 9 | 97791640 | AX-170531259 | 0 | A | C | 0.132 | -1.13E-01 | 1.19E-01 | 2.59E+00 | 2.67E+00 | 3.37E-01 | no | Clstn2 | intron_variant | MODIFIER |
| 9 | 97793080 | AX-97831445 | 0 | T | C | 0.132 | -1.13E-01 | 1.19E-01 | 2.59E+00 | 2.67E+00 | 3.37E-01 | no | Clstn2 | intron_variant | MODIFIER |
| 9 | 97795147 | AX-170329811 | 0 | T | C | 0.132 | -1.13E-01 | 1.19E-01 | 2.59E+00 | 2.67E+00 | 3.37E-01 | no | Clstn2 | intron_variant | MODIFIER |
| 9 | 97823082 | AX-97812157 | 0 | A | G | 0.132 | -1.13E-01 | 1.19E-01 | 2.59E+00 | 2.67E+00 | 3.37E-01 | no | Clstn2 | intron_variant | MODIFIER |
| 9 | 97858780 | AX-170083198 | 0 | T | C | 0.132 | -1.13E-01 | 1.19E-01 | 2.59E+00 | 2.67E+00 | 3.37E-01 | no | Clstn2 | intron_variant | MODIFIER |
| 9 | 97862298 | AX-168394201 | 0 | A | G | 0.132 | -1.13E-01 | 1.19E-01 | 2.59E+00 | 2.67E+00 | 3.37E-01 | no | Clstn2 | intron_variant | MODIFIER |
| 9 | 97862984 | AX-170083193 | 0 | T | C | 0.132 | -1.13E-01 | 1.19E-01 | 2.59E+00 | 2.67E+00 | 3.37E-01 | no | Clstn2 | intron_variant | MODIFIER |
| 9 | 97864681 | AX-170083195 | 0 | T | A | 0.132 | -1.13E-01 | 1.19E-01 | 2.59E+00 | 2.67E+00 | 3.37E-01 | no | Clstn2 | intron_variant | MODIFIER |
| 9 | 97919272 | AX-170271795 | 0 | T | C | 0.132 | -1.13E-01 | 1.19E-01 | 2.59E+00 | 2.67E+00 | 3.37E-01 | no | Clstn2 | intron_variant | MODIFIER |
| 9 | 97920717 | AX-170185661 | 0 | G | A | 0.132 | -1.13E-01 | 1.19E-01 | 2.59E+00 | 2.67E+00 | 3.37E-01 | no | Clstn2 | intron_variant | MODIFIER |
| 9 | 97921418 | AX-170459745 | 0 | G | A | 0.132 | -1.13E-01 | 1.19E-01 | 2.59E+00 | 2.67E+00 | 3.37E-01 | no | Clstn2 | intron_variant | MODIFIER |
| 9 | 97933702 | AX-170495430 | 0 | T | A | 0.132 | -1.13E-01 | 1.19E-01 | 2.59E+00 | 2.67E+00 | 3.37E-01 | no | Clstn2 | intron_variant | MODIFIER |
| 9 | 97966906 | AX-170433679 | 0 | C | G | 0.132 | -1.13E-01 | 1.19E-01 | 2.59E+00 | 2.67E+00 | 3.37E-01 | no | Clstn2 | intron_variant | MODIFIER |
| 9 | 97977477 | AX-170232500 | 0 | A | G | 0.132 | -1.13E-01 | 1.19E-01 | 2.59E+00 | 2.67E+00 | 3.37E-01 | no | Clstn2 | intron_variant | MODIFIER |
| 9 | 98180487 | AX-97818491 | 0 | A | C | 0.132 | -1.13E-01 | 1.19E-01 | 2.59E+00 | 2.67E+00 | 3.37E-01 | no | | intergenic_variant | MODIFIER |
| 9 | 98314464 | AX-170265399 | 0 | T | A | 0.132 | -1.13E-01 | 1.19E-01 | 2.59E+00 | 2.67E+00 | 3.37E-01 | no | Nmnat3 | intron_variant | MODIFIER |

| | | | | | | | | | | | | | | | |
|---|---|---|---|---|---|---|---|---|---|---|---|---|---|---|---|
| 9 | 98333273 | AX-170452881 | 0 | A | G | 0.132 | -1.13E-01 | 1.19E-01 | 2.59E+00 | 2.67E+00 | 3.37E-01 | no | Nmnat3 | intron_variant | MODIFIER |
| 9 | 98373402 | AX-170033094 | 0 | A | G | 0.132 | -1.13E-01 | 1.19E-01 | 2.59E+00 | 2.67E+00 | 3.37E-01 | no | Nmnat3 | intron_variant | MODIFIER |
| 9 | 98429637 | AX-170267498 | 0 | C | A | 0.132 | -1.13E-01 | 1.19E-01 | 2.59E+00 | 2.67E+00 | 3.37E-01 | no | Rbp1 | intron_variant | MODIFIER |
| 9 | 98431198 | AX-170356820 | 0 | A | C | 0.132 | -1.13E-01 | 1.19E-01 | 2.59E+00 | 2.67E+00 | 3.37E-01 | no | Rbp1 | intron_variant | MODIFIER |
| 9 | 98450403 | AX-170306737 | 0 | T | C | 0.132 | -1.13E-01 | 1.19E-01 | 2.59E+00 | 2.67E+00 | 3.37E-01 | no | Rbp1 | downstream_gene_variant | MODIFIER |
| 9 | 98487779 | AX-170172701 | 0 | C | T | 0.132 | -1.13E-01 | 1.19E-01 | 2.59E+00 | 2.67E+00 | 3.37E-01 | no | Rbp2 | upstream_gene_variant | MODIFIER |
| 9 | 98488094 | AX-170527778 | 0 | T | G | 0.132 | -1.13E-01 | 1.19E-01 | 2.59E+00 | 2.67E+00 | 3.37E-01 | no | Rbp2 | upstream_gene_variant | MODIFIER |
| 9 | 98491651 | AX-170411852 | 0 | C | T | 0.132 | -1.13E-01 | 1.19E-01 | 2.59E+00 | 2.67E+00 | 3.37E-01 | no | Rbp2 | intron_variant | MODIFIER |
| 4 | 155844089 | AX-170365611 | 1 | C | T | 0.107 | 1.53E-01 | 1.62E-01 | 2.69E+00 | 2.80E+00 | 3.38E-01 | Yes | Mxra8 | downstream_gene_variant | MODIFIER |
| 1 | 149557012 | AX-170320801 | 0 | T | C | 0.211 | 1.20E-01 | 1.26E-01 | 2.63E+00 | 2.69E+00 | 3.38E-01 | no | | regulatory_region_variant | MODIFIER |
| 3 | 22395981 | AX-170066069 | 0 | C | T | 0.044 | -1.09E-01 | 1.16E-01 | 2.68E+00 | 2.86E+00 | 3.38E-01 | no | | intergenic_variant | MODIFIER |
| 10 | 69546047 | AX-170180434 | 1 | A | C | 0.411 | -7.64E-02 | 8.04E-02 | 2.66E+00 | 2.73E+00 | 3.38E-01 | no | Ank3 | intron_variant | MODIFIER |
| 7 | 70832514 | AX-170035755 | 0 | C | T | 0.377 | -9.89E-02 | 1.04E-01 | 2.88E+00 | 2.97E+00 | 3.38E-01 | no | | intergenic_variant | MODIFIER |
| 7 | 70841706 | AX-170527549 | 0 | C | T | 0.377 | -9.89E-02 | 1.04E-01 | 2.88E+00 | 2.97E+00 | 3.38E-01 | no | | intergenic_variant | MODIFIER |
| 7 | 70848220 | AX-170527121 | 0 | G | A | 0.377 | -9.89E-02 | 1.04E-01 | 2.88E+00 | 2.97E+00 | 3.38E-01 | no | | regulatory_region_variant | MODIFIER |
| 7 | 70850199 | AX-170389032 | 0 | A | G | 0.377 | -9.89E-02 | 1.04E-01 | 2.88E+00 | 2.97E+00 | 3.38E-01 | no | | intergenic_variant | MODIFIER |
| 7 | 70852124 | AX-170187751 | 0 | T | C | 0.377 | -9.89E-02 | 1.04E-01 | 2.88E+00 | 2.97E+00 | 3.38E-01 | no | | intergenic_variant | MODIFIER |
| 7 | 70917184 | AX-170200818 | 0 | C | T | 0.377 | -9.89E-02 | 1.04E-01 | 2.88E+00 | 2.97E+00 | 3.38E-01 | no | Gm42397 | downstream_gene_variant | MODIFIER |
| 7 | 70919619 | AX-170285976 | 0 | A | G | 0.377 | -9.89E-02 | 1.04E-01 | 2.88E+00 | 2.97E+00 | 3.38E-01 | no | Gm42397 | downstream_gene_variant | MODIFIER |
| 7 | 70919698 | AX-170224585 | 0 | A | G | 0.377 | -9.89E-02 | 1.04E-01 | 2.88E+00 | 2.97E+00 | 3.38E-01 | no | Gm42397 | downstream_gene_variant | MODIFIER |
| 7 | 71389202 | AX-170203509 | 0 | G | T | 0.377 | -9.89E-02 | 1.04E-01 | 2.88E+00 | 2.97E+00 | 3.38E-01 | no | Gm29295 | intron_variant,non_coding_transcript_variant | MODIFIER |
| 7 | 71389299 | AX-97784455 | 0 | C | A | 0.377 | -9.89E-02 | 1.04E-01 | 2.88E+00 | 2.97E+00 | 3.38E-01 | no | Gm29295 | intron_variant,non_coding_transcript_variant | MODIFIER |
| 7 | 71389751 | AX-170292882 | 0 | A | G | 0.377 | -9.89E-02 | 1.04E-01 | 2.88E+00 | 2.97E+00 | 3.38E-01 | no | Gm29295 | intron_variant,non_coding_transcript_variant | MODIFIER |
| 7 | 71389942 | AX-170137334 | 0 | T | A | 0.377 | -9.89E-02 | 1.04E-01 | 2.88E+00 | 2.97E+00 | 3.38E-01 | no | Gm29295 | intron_variant,non_coding_transcript_variant | MODIFIER |
| 7 | 71394212 | AX-170120068 | 0 | G | A | 0.377 | -9.89E-02 | 1.04E-01 | 2.88E+00 | 2.97E+00 | 3.38E-01 | no | Gm29295 | downstream_gene_variant | MODIFIER |
| 7 | 71395168 | AX-170287130 | 0 | C | T | 0.377 | -9.89E-02 | 1.04E-01 | 2.88E+00 | 2.97E+00 | 3.38E-01 | no | Gm29295 | downstream_gene_variant | MODIFIER |
| 7 | 71401886 | AX-170261064 | 0 | A | G | 0.377 | -9.89E-02 | 1.04E-01 | 2.88E+00 | 2.97E+00 | 3.38E-01 | no | | intergenic_variant | MODIFIER |
| 7 | 71441564 | AX-170531290 | 0 | G | A | 0.377 | -9.89E-02 | 1.04E-01 | 2.88E+00 | 2.97E+00 | 3.38E-01 | no | | intergenic_variant | MODIFIER |
| 7 | 71460796 | AX-170228311 | 0 | C | A | 0.377 | -9.89E-02 | 1.04E-01 | 2.88E+00 | 2.97E+00 | 3.38E-01 | no | | intergenic_variant | MODIFIER |
| 7 | 71467612 | AX-97847509 | 0 | C | T | 0.377 | -9.89E-02 | 1.04E-01 | 2.88E+00 | 2.97E+00 | 3.38E-01 | no | | regulatory_region_variant | MODIFIER |
| 7 | 71467817 | AX-170138586 | 0 | G | A | 0.377 | -9.89E-02 | 1.04E-01 | 2.88E+00 | 2.97E+00 | 3.38E-01 | no | | regulatory_region_variant | MODIFIER |
| 7 | 71516345 | AX-170383427 | 0 | T | C | 0.377 | -9.89E-02 | 1.04E-01 | 2.88E+00 | 2.97E+00 | 3.38E-01 | no | | intergenic_variant | MODIFIER |
| 3 | 32831651 | AX-170190605 | 0 | T | C | 0.175 | 9.06E-02 | 9.54E-02 | 2.83E+00 | 2.93E+00 | 3.38E-01 | no | Usp13 | intron_variant | MODIFIER |
| 3 | 32871876 | AX-170232221 | 0 | C | T | 0.175 | 9.06E-02 | 9.54E-02 | 2.83E+00 | 2.93E+00 | 3.38E-01 | no | Usp13 | intron_variant | MODIFIER |
| 5 | 37023512 | AX-97786376 | 0 | A | G | 0.325 | -1.07E-01 | 1.11E-01 | 3.01E+00 | 3.12E+00 | 3.38E-01 | no | Jakmip1 | upstream_gene_variant | MODIFIER |
| 7 | 145262015 | AX-170346606 | 1 | G | A | 0.196 | -7.12E-02 | 7.51E-02 | 2.63E+00 | 2.76E+00 | 3.38E-01 | no | Tpcn2 | intron_variant | MODIFIER |
| 15 | 93654977 | AX-170012296 | 0 | G | A | 0.149 | 1.48E-01 | 1.56E-01 | 2.67E+00 | 2.77E+00 | 3.38E-01 | Yes | | intergenic_variant | MODIFIER |
| 16 | 41004695 | AX-170106040 | 1 | C | A | 0.482 | 7.62E-02 | 7.95E-02 | 2.55E+00 | 2.61E+00 | 3.38E-01 | no | Lsamp | intron_variant | MODIFIER |
| 4 | 115941253 | AX-170223116 | 1 | A | G | 0.205 | 1.07E-01 | 1.12E-01 | 2.60E+00 | 2.68E+00 | 3.38E-01 | no | Dmbx1 | upstream_gene_variant | MODIFIER |
| 9 | 101504101 | AX-170033144 | 0 | C | T | 0.132 | -1.12E-01 | 1.18E-01 | 2.61E+00 | 2.68E+00 | 3.38E-01 | no | | intergenic_variant | MODIFIER |
| 16 | 78941198 | AX-170037518 | 0 | G | C | 0.465 | 7.22E-02 | 7.57E-02 | 2.60E+00 | 2.66E+00 | 3.38E-01 | no | Chodl | intron_variant | MODIFIER |
| 16 | 78950133 | AX-170253484 | 0 | G | A | 0.465 | 7.22E-02 | 7.57E-02 | 2.60E+00 | 2.66E+00 | 3.38E-01 | no | Tmprss15 | downstream_gene_variant | MODIFIER |
| 16 | 78964000 | AX-170057896 | 0 | A | G | 0.465 | 7.22E-02 | 7.57E-02 | 2.60E+00 | 2.66E+00 | 3.38E-01 | no | Tmprss15 | intron_variant | MODIFIER |
| 16 | 78968107 | AX-170057897 | 0 | T | C | 0.465 | 7.22E-02 | 7.57E-02 | 2.60E+00 | 2.66E+00 | 3.38E-01 | no | Tmprss15 | missense_variant | MODERATE |
| 16 | 78969244 | AX-170136374 | 0 | A | C | 0.465 | 7.22E-02 | 7.57E-02 | 2.60E+00 | 2.66E+00 | 3.38E-01 | no | Tmprss15 | intron_variant | MODIFIER |
| 16 | 78988059 | AX-97792087 | 0 | C | A | 0.465 | 7.22E-02 | 7.57E-02 | 2.60E+00 | 2.66E+00 | 3.38E-01 | no | Tmprss15 | intron_variant | MODIFIER |
| 16 | 79015650 | AX-97799551 | 0 | G | A | 0.465 | 7.22E-02 | 7.57E-02 | 2.60E+00 | 2.66E+00 | 3.38E-01 | no | Tmprss15 | intron_variant | MODIFIER |
| 16 | 79017211 | AX-170532450 | 0 | A | G | 0.465 | 7.22E-02 | 7.57E-02 | 2.60E+00 | 2.66E+00 | 3.38E-01 | no | Tmprss15 | intron_variant | MODIFIER |
| 16 | 79021110 | AX-170463999 | 0 | G | A | 0.465 | 7.22E-02 | 7.57E-02 | 2.60E+00 | 2.66E+00 | 3.38E-01 | no | Tmprss15 | intron_variant | MODIFIER |
| 16 | 79055499 | AX-170187000 | 0 | T | C | 0.465 | 7.22E-02 | 7.57E-02 | 2.60E+00 | 2.66E+00 | 3.38E-01 | no | Tmprss15 | intron_variant | MODIFIER |
| 16 | 79085725 | AX-170462320 | 0 | T | C | 0.465 | 7.22E-02 | 7.57E-02 | 2.60E+00 | 2.66E+00 | 3.38E-01 | no | Tmprss15 | intron_variant | MODIFIER |
| 16 | 79086452 | AX-97845389 | 0 | A | C | 0.465 | 7.22E-02 | 7.57E-02 | 2.60E+00 | 2.66E+00 | 3.38E-01 | no | Tmprss15 | intron_variant | MODIFIER |
| 16 | 79133675 | AX-170354449 | 0 | C | T | 0.465 | 7.22E-02 | 7.57E-02 | 2.60E+00 | 2.66E+00 | 3.38E-01 | no | | intergenic_variant | MODIFIER |
| 16 | 79133899 | AX-170157133 | 0 | A | C | 0.465 | 7.22E-02 | 7.57E-02 | 2.60E+00 | 2.66E+00 | 3.38E-01 | no | | intergenic_variant | MODIFIER |

| | | | | | | | | | | | | | | | |
|---|---|---|---|---|---|---|---|---|---|---|---|---|---|---|---|
| 16 | 79185000 | AX-170088072 | 0 | T | C | 0.465 | 7.22E-02 | 7.57E-02 | 2.60E+00 | 2.66E+00 | 3.38E-01 | no | | intergenic_variant | MODIFIER |
| 16 | 79228912 | AX-170125953 | 0 | C | A | 0.465 | 7.22E-02 | 7.57E-02 | 2.60E+00 | 2.66E+00 | 3.38E-01 | no | | intergenic_variant | MODIFIER |
| 16 | 79228985 | AX-170128371 | 0 | C | T | 0.465 | 7.22E-02 | 7.57E-02 | 2.60E+00 | 2.66E+00 | 3.38E-01 | no | | intergenic_variant | MODIFIER |
| 16 | 79255758 | AX-170013451 | 0 | C | T | 0.465 | 7.22E-02 | 7.57E-02 | 2.60E+00 | 2.66E+00 | 3.38E-01 | no | | intergenic_variant | MODIFIER |
| 16 | 79256372 | AX-170013457 | 0 | T | C | 0.465 | 7.22E-02 | 7.57E-02 | 2.60E+00 | 2.66E+00 | 3.38E-01 | no | | intergenic_variant | MODIFIER |
| 16 | 79272539 | AX-170351405 | 0 | G | A | 0.465 | 7.22E-02 | 7.57E-02 | 2.60E+00 | 2.66E+00 | 3.38E-01 | no | | intergenic_variant | MODIFIER |
| 16 | 79273876 | AX-97831171 | 0 | C | T | 0.465 | 7.22E-02 | 7.57E-02 | 2.60E+00 | 2.66E+00 | 3.38E-01 | no | | intergenic_variant | MODIFIER |
| 16 | 79316309 | AX-97851532 | 0 | A | G | 0.465 | 7.22E-02 | 7.57E-02 | 2.60E+00 | 2.66E+00 | 3.38E-01 | no | | intergenic_variant | MODIFIER |
| 16 | 79327673 | AX-170529656 | 0 | C | T | 0.465 | 7.22E-02 | 7.57E-02 | 2.60E+00 | 2.66E+00 | 3.38E-01 | no | | intergenic_variant | MODIFIER |
| 16 | 79380334 | AX-97802948 | 0 | C | A | 0.465 | 7.22E-02 | 7.57E-02 | 2.60E+00 | 2.66E+00 | 3.38E-01 | no | | intergenic_variant | MODIFIER |
| 16 | 79408054 | AX-170013454 | 0 | A | G | 0.465 | 7.22E-02 | 7.57E-02 | 2.60E+00 | 2.66E+00 | 3.38E-01 | no | | intergenic_variant | MODIFIER |
| 16 | 79445575 | AX-170013453 | 0 | T | C | 0.465 | 7.22E-02 | 7.57E-02 | 2.60E+00 | 2.66E+00 | 3.38E-01 | no | | intergenic_variant | MODIFIER |
| 16 | 79518061 | AX-97791322 | 0 | T | C | 0.465 | 7.22E-02 | 7.57E-02 | 2.60E+00 | 2.66E+00 | 3.38E-01 | no | | intergenic_variant | MODIFIER |
| 16 | 79549582 | AX-170232605 | 0 | A | G | 0.465 | 7.22E-02 | 7.57E-02 | 2.60E+00 | 2.66E+00 | 3.38E-01 | no | | intergenic_variant | MODIFIER |
| 16 | 79570256 | AX-170310448 | 0 | A | C | 0.465 | 7.22E-02 | 7.57E-02 | 2.60E+00 | 2.66E+00 | 3.38E-01 | no | | intergenic_variant | MODIFIER |
| 16 | 79583861 | AX-170238102 | 0 | C | T | 0.465 | 7.22E-02 | 7.57E-02 | 2.60E+00 | 2.66E+00 | 3.38E-01 | no | | regulatory_region_variant | MODIFIER |
| 16 | 79644924 | AX-97832472 | 0 | T | G | 0.465 | 7.22E-02 | 7.57E-02 | 2.60E+00 | 2.66E+00 | 3.38E-01 | no | | intergenic_variant | MODIFIER |
| 16 | 79658273 | AX-170057908 | 0 | C | T | 0.465 | 7.22E-02 | 7.57E-02 | 2.60E+00 | 2.66E+00 | 3.38E-01 | no | | intergenic_variant | MODIFIER |
| 16 | 79738238 | AX-170310999 | 0 | C | T | 0.465 | 7.22E-02 | 7.57E-02 | 2.60E+00 | 2.66E+00 | 3.38E-01 | no | | intergenic_variant | MODIFIER |
| 16 | 79748702 | AX-170479144 | 0 | G | A | 0.465 | 7.22E-02 | 7.57E-02 | 2.60E+00 | 2.66E+00 | 3.38E-01 | no | | intergenic_variant | MODIFIER |
| 16 | 79851928 | AX-170336801 | 0 | G | A | 0.465 | 7.22E-02 | 7.57E-02 | 2.60E+00 | 2.66E+00 | 3.38E-01 | no | | intergenic_variant | MODIFIER |
| 16 | 79909144 | AX-170114510 | 0 | T | C | 0.465 | 7.22E-02 | 7.57E-02 | 2.60E+00 | 2.66E+00 | 3.38E-01 | no | | intergenic_variant | MODIFIER |
| 16 | 79921347 | AX-170162471 | 0 | C | T | 0.465 | 7.22E-02 | 7.57E-02 | 2.60E+00 | 2.66E+00 | 3.38E-01 | no | | intergenic_variant | MODIFIER |
| 16 | 80073922 | AX-170328028 | 0 | T | C | 0.465 | 7.22E-02 | 7.57E-02 | 2.60E+00 | 2.66E+00 | 3.38E-01 | no | | regulatory_region_variant | MODIFIER |
| 16 | 80137832 | AX-170057919 | 0 | A | C | 0.465 | 7.22E-02 | 7.57E-02 | 2.60E+00 | 2.66E+00 | 3.38E-01 | no | Gm23083 | downstream_gene_variant | MODIFIER |
| 16 | 80187412 | AX-170184890 | 0 | T | C | 0.465 | 7.22E-02 | 7.57E-02 | 2.60E+00 | 2.66E+00 | 3.38E-01 | no | | intergenic_variant | MODIFIER |
| 16 | 80189758 | AX-97793435 | 0 | A | G | 0.465 | 7.22E-02 | 7.57E-02 | 2.60E+00 | 2.66E+00 | 3.38E-01 | no | | intergenic_variant | MODIFIER |
| 16 | 80218385 | AX-170205534 | 0 | A | G | 0.465 | 7.22E-02 | 7.57E-02 | 2.60E+00 | 2.66E+00 | 3.38E-01 | no | | intergenic_variant | MODIFIER |
| 16 | 80219189 | AX-170156181 | 0 | G | A | 0.465 | 7.22E-02 | 7.57E-02 | 2.60E+00 | 2.66E+00 | 3.38E-01 | no | | intergenic_variant | MODIFIER |
| 16 | 80220827 | AX-97790148 | 0 | T | C | 0.465 | 7.22E-02 | 7.57E-02 | 2.60E+00 | 2.66E+00 | 3.38E-01 | no | | intergenic_variant | MODIFIER |
| 16 | 80284459 | AX-170214225 | 0 | T | C | 0.465 | 7.22E-02 | 7.57E-02 | 2.60E+00 | 2.66E+00 | 3.38E-01 | no | | intergenic_variant | MODIFIER |
| 16 | 80366405 | AX-170319113 | 0 | G | A | 0.465 | 7.22E-02 | 7.57E-02 | 2.60E+00 | 2.66E+00 | 3.38E-01 | no | | intergenic_variant | MODIFIER |
| 16 | 80370513 | AX-170102761 | 0 | T | C | 0.465 | 7.22E-02 | 7.57E-02 | 2.60E+00 | 2.66E+00 | 3.38E-01 | no | | intergenic_variant | MODIFIER |
| 16 | 80371085 | AX-170362789 | 0 | C | T | 0.465 | 7.22E-02 | 7.57E-02 | 2.60E+00 | 2.66E+00 | 3.38E-01 | no | | intergenic_variant | MODIFIER |
| 16 | 80388582 | AX-97822271 | 0 | G | A | 0.465 | 7.22E-02 | 7.57E-02 | 2.60E+00 | 2.66E+00 | 3.38E-01 | no | | intergenic_variant | MODIFIER |
| 16 | 80398308 | AX-170364475 | 0 | G | A | 0.465 | 7.22E-02 | 7.57E-02 | 2.60E+00 | 2.66E+00 | 3.38E-01 | no | | intergenic_variant | MODIFIER |
| 16 | 80405668 | AX-97844798 | 0 | C | A | 0.465 | 7.22E-02 | 7.57E-02 | 2.60E+00 | 2.66E+00 | 3.38E-01 | no | | intergenic_variant | MODIFIER |
| 16 | 80412125 | AX-170013479 | 0 | C | T | 0.465 | 7.22E-02 | 7.57E-02 | 2.60E+00 | 2.66E+00 | 3.38E-01 | no | | intergenic_variant | MODIFIER |
| 16 | 80468011 | AX-170013474 | 0 | C | A | 0.465 | 7.22E-02 | 7.57E-02 | 2.60E+00 | 2.66E+00 | 3.38E-01 | no | | intergenic_variant | MODIFIER |
| 16 | 80477686 | AX-170013470 | 0 | C | A | 0.465 | 7.22E-02 | 7.57E-02 | 2.60E+00 | 2.66E+00 | 3.38E-01 | no | | intergenic_variant | MODIFIER |
| 16 | 80490164 | AX-170365294 | 0 | T | G | 0.465 | 7.22E-02 | 7.57E-02 | 2.60E+00 | 2.66E+00 | 3.38E-01 | no | | intergenic_variant | MODIFIER |
| 16 | 80565798 | AX-97782305 | 0 | T | C | 0.465 | 7.22E-02 | 7.57E-02 | 2.60E+00 | 2.66E+00 | 3.38E-01 | no | | intergenic_variant | MODIFIER |
| 16 | 80586134 | AX-170013472 | 0 | T | G | 0.465 | 7.22E-02 | 7.57E-02 | 2.60E+00 | 2.66E+00 | 3.38E-01 | no | | intergenic_variant | MODIFIER |
| 16 | 80586250 | AX-97832496 | 0 | T | C | 0.465 | 7.22E-02 | 7.57E-02 | 2.60E+00 | 2.66E+00 | 3.38E-01 | no | | intergenic_variant | MODIFIER |
| 16 | 80589484 | AX-170375221 | 0 | A | G | 0.465 | 7.22E-02 | 7.57E-02 | 2.60E+00 | 2.66E+00 | 3.38E-01 | no | | intergenic_variant | MODIFIER |
| 16 | 80605428 | AX-170148175 | 0 | G | A | 0.465 | 7.22E-02 | 7.57E-02 | 2.60E+00 | 2.66E+00 | 3.38E-01 | no | | intergenic_variant | MODIFIER |
| 16 | 80614656 | AX-170439695 | 0 | T | G | 0.465 | 7.22E-02 | 7.57E-02 | 2.60E+00 | 2.66E+00 | 3.38E-01 | no | | intergenic_variant | MODIFIER |
| 16 | 80666317 | AX-170266643 | 0 | T | C | 0.465 | 7.22E-02 | 7.57E-02 | 2.60E+00 | 2.66E+00 | 3.38E-01 | no | | intergenic_variant | MODIFIER |
| 16 | 80666703 | AX-170296603 | 0 | G | A | 0.465 | 7.22E-02 | 7.57E-02 | 2.60E+00 | 2.66E+00 | 3.38E-01 | no | | intergenic_variant | MODIFIER |
| 16 | 80697547 | AX-170215552 | 0 | C | T | 0.465 | 7.22E-02 | 7.57E-02 | 2.60E+00 | 2.66E+00 | 3.38E-01 | no | | intergenic_variant | MODIFIER |
| 16 | 80705439 | AX-170433062 | 0 | A | G | 0.465 | 7.22E-02 | 7.57E-02 | 2.60E+00 | 2.66E+00 | 3.38E-01 | no | | intergenic_variant | MODIFIER |
| 16 | 80726623 | AX-170456033 | 0 | T | G | 0.465 | 7.22E-02 | 7.57E-02 | 2.60E+00 | 2.66E+00 | 3.38E-01 | no | Gm19867 | upstream_gene_variant | MODIFIER |
| 16 | 80731851 | AX-170093540 | 0 | C | T | 0.465 | 7.22E-02 | 7.57E-02 | 2.60E+00 | 2.66E+00 | 3.38E-01 | no | Gm19867 | downstream_gene_variant | MODIFIER |
| 16 | 80734811 | AX-170199789 | 0 | A | G | 0.465 | 7.22E-02 | 7.57E-02 | 2.60E+00 | 2.66E+00 | 3.38E-01 | no | | intergenic_variant | MODIFIER |

| | | | | | | | | | | | | | | | |
|---|---|---|---|---|---|---|---|---|---|---|---|---|---|---|---|
| 16 | 80735020 | AX-170247628 | 0 | T | C | 0.465 | 7.22E-02 | 7.57E-02 | 2.60E+00 | 2.66E+00 | 3.38E-01 | no | | intergenic_variant | MODIFIER |
| 16 | 80737815 | AX-170512644 | 0 | T | C | 0.465 | 7.22E-02 | 7.57E-02 | 2.60E+00 | 2.66E+00 | 3.38E-01 | no | | intergenic_variant | MODIFIER |
| 16 | 80772571 | AX-170127671 | 0 | C | G | 0.465 | 7.22E-02 | 7.57E-02 | 2.60E+00 | 2.66E+00 | 3.38E-01 | no | | intergenic_variant | MODIFIER |
| 16 | 80791510 | AX-97791351 | 0 | G | A | 0.465 | 7.22E-02 | 7.57E-02 | 2.60E+00 | 2.66E+00 | 3.38E-01 | no | | intergenic_variant | MODIFIER |
| 16 | 80813221 | AX-97809582 | 0 | T | C | 0.465 | 7.22E-02 | 7.57E-02 | 2.60E+00 | 2.66E+00 | 3.38E-01 | no | | intergenic_variant | MODIFIER |
| 16 | 80821587 | AX-170242756 | 0 | A | T | 0.465 | 7.22E-02 | 7.57E-02 | 2.60E+00 | 2.66E+00 | 3.38E-01 | no | | intergenic_variant | MODIFIER |
| 16 | 80830072 | AX-97796287 | 0 | C | A | 0.465 | 7.22E-02 | 7.57E-02 | 2.60E+00 | 2.66E+00 | 3.38E-01 | no | | intergenic_variant | MODIFIER |
| 16 | 80836772 | AX-170351110 | 0 | A | C | 0.465 | 7.22E-02 | 7.57E-02 | 2.60E+00 | 2.66E+00 | 3.38E-01 | no | | intergenic_variant | MODIFIER |
| 16 | 80837909 | AX-170470665 | 0 | G | A | 0.465 | 7.22E-02 | 7.57E-02 | 2.60E+00 | 2.66E+00 | 3.38E-01 | no | | intergenic_variant | MODIFIER |
| 16 | 80861898 | AX-170333543 | 0 | T | A | 0.465 | 7.22E-02 | 7.57E-02 | 2.60E+00 | 2.66E+00 | 3.38E-01 | no | | intergenic_variant | MODIFIER |
| 16 | 80873163 | AX-170360845 | 0 | T | C | 0.465 | 7.22E-02 | 7.57E-02 | 2.60E+00 | 2.66E+00 | 3.38E-01 | no | | intergenic_variant | MODIFIER |
| 16 | 80898997 | AX-97804546 | 0 | G | A | 0.465 | 7.22E-02 | 7.57E-02 | 2.60E+00 | 2.66E+00 | 3.38E-01 | no | | intergenic_variant | MODIFIER |
| 16 | 80933342 | AX-170093542 | 0 | A | G | 0.465 | 7.22E-02 | 7.57E-02 | 2.60E+00 | 2.66E+00 | 3.38E-01 | no | | intergenic_variant | MODIFIER |
| 16 | 80973468 | AX-170091005 | 0 | A | G | 0.465 | 7.22E-02 | 7.57E-02 | 2.60E+00 | 2.66E+00 | 3.38E-01 | no | | intergenic_variant | MODIFIER |
| 16 | 80978081 | AX-170148447 | 0 | C | T | 0.465 | 7.22E-02 | 7.57E-02 | 2.60E+00 | 2.66E+00 | 3.38E-01 | no | | intergenic_variant | MODIFIER |
| 16 | 80983160 | AX-170375710 | 0 | C | A | 0.465 | 7.22E-02 | 7.57E-02 | 2.60E+00 | 2.66E+00 | 3.38E-01 | no | | intergenic_variant | MODIFIER |
| 16 | 81024620 | AX-97817162 | 0 | C | T | 0.465 | 7.22E-02 | 7.57E-02 | 2.60E+00 | 2.66E+00 | 3.38E-01 | no | | intergenic_variant | MODIFIER |
| 16 | 81037615 | AX-170425286 | 0 | C | T | 0.465 | 7.22E-02 | 7.57E-02 | 2.60E+00 | 2.66E+00 | 3.38E-01 | no | | intergenic_variant | MODIFIER |
| 16 | 81041861 | AX-170396375 | 0 | A | G | 0.465 | 7.22E-02 | 7.57E-02 | 2.60E+00 | 2.66E+00 | 3.38E-01 | no | | intergenic_variant | MODIFIER |
| 16 | 81142526 | AX-170435338 | 0 | A | G | 0.465 | 7.22E-02 | 7.57E-02 | 2.60E+00 | 2.66E+00 | 3.38E-01 | no | | intergenic_variant | MODIFIER |
| 16 | 81879955 | AX-170422260 | 0 | C | G | 0.465 | 7.22E-02 | 7.57E-02 | 2.60E+00 | 2.66E+00 | 3.38E-01 | no | | intergenic_variant | MODIFIER |
| 16 | 81905582 | AX-97781442 | 0 | C | T | 0.465 | 7.22E-02 | 7.57E-02 | 2.60E+00 | 2.66E+00 | 3.38E-01 | no | Gm49560 | intron_variant,non_coding_transcript_variant | MODIFIER |
| 16 | 81910780 | AX-170215883 | 0 | T | C | 0.465 | 7.22E-02 | 7.57E-02 | 2.60E+00 | 2.66E+00 | 3.38E-01 | no | Gm49560 | intron_variant,non_coding_transcript_variant | MODIFIER |
| 16 | 81933892 | AX-170339513 | 0 | T | A | 0.465 | 7.22E-02 | 7.57E-02 | 2.60E+00 | 2.66E+00 | 3.38E-01 | no | Gm49560 | upstream_gene_variant | MODIFIER |
| 16 | 81970966 | AX-170057944 | 0 | A | G | 0.465 | 7.22E-02 | 7.57E-02 | 2.60E+00 | 2.66E+00 | 3.38E-01 | no | | intergenic_variant | MODIFIER |
| 16 | 81974296 | AX-170186678 | 0 | G | T | 0.465 | 7.22E-02 | 7.57E-02 | 2.60E+00 | 2.66E+00 | 3.38E-01 | no | | intergenic_variant | MODIFIER |
| 4 | 116804401 | AX-97852331 | 0 | T | C | 0.219 | 1.06E-01 | 1.11E-01 | 2.61E+00 | 2.68E+00 | 3.38E-01 | no | Toe1 | 3_prime_UTR_variant | MODIFIER |
| 16 | 32390552 | AX-170331829 | 1 | G | C | 0.509 | 7.69E-02 | 8.01E-02 | 2.54E+00 | 2.59E+00 | 3.38E-01 | no | Ubxn7 | 3_prime_UTR_variant | MODIFIER |
| 4 | 58891621 | AX-97798443 | 1 | C | A | 0.375 | 1.01E-01 | 1.07E-01 | 2.67E+00 | 2.76E+00 | 3.39E-01 | no | Ecpas | intron_variant | MODIFIER |
| 4 | 70130803 | AX-170246497 | 0 | C | A | 0.465 | -8.01E-02 | 8.30E-02 | 2.51E+00 | 2.57E+00 | 3.39E-01 | no | | intergenic_variant | MODIFIER |
| 4 | 71872049 | AX-168419997 | 0 | C | T | 0.465 | -8.01E-02 | 8.30E-02 | 2.51E+00 | 2.57E+00 | 3.39E-01 | no | Gm11233 | downstream_gene_variant | MODIFIER |
| 3 | 26341091 | AX-170190100 | 1 | C | T | 0.134 | -9.89E-02 | 1.04E-01 | 2.58E+00 | 2.66E+00 | 3.39E-01 | no | | intergenic_variant | MODIFIER |
| 7 | 23339502 | AX-170076834 | 0 | C | T | 0.36 | 8.47E-02 | 8.88E-02 | 2.55E+00 | 2.68E+00 | 3.39E-01 | no | Nlrp4e | intron_variant | MODIFIER |
| 7 | 94324056 | AX-170320776 | 2 | C | T | 0.127 | 1.29E-01 | 1.36E-01 | 2.59E+00 | 2.74E+00 | 3.39E-01 | no | Gm32647 | intron_variant,non_coding_transcript_variant | MODIFIER |
| 4 | 133628315 | AX-170245131 | 1 | G | A | 0.161 | 1.11E-01 | 1.16E-01 | 2.57E+00 | 2.61E+00 | 3.39E-01 | no | Zdhhc18 | intron_variant | MODIFIER |
| 7 | 16259512 | AX-170451988 | 0 | T | C | 0.368 | 7.67E-02 | 8.09E-02 | 2.61E+00 | 2.73E+00 | 3.39E-01 | no | C5ar1 | splice_donor_variant | HIGH |
| 14 | 117848342 | AX-170010797 | 1 | C | A | 0.339 | 7.02E-02 | 7.40E-02 | 2.62E+00 | 2.72E+00 | 3.39E-01 | no | Gpc6 | intron_variant | MODIFIER |
| 1 | 14717847 | AX-97794053 | 1 | C | T | 0.393 | 7.41E-02 | 7.70E-02 | 2.52E+00 | 2.58E+00 | 3.39E-01 | no | | intergenic_variant | MODIFIER |
| 20 | 57778783 | AX-170033930 | 0 | T | C | 0.246 | 5.81E-02 | 6.16E-02 | 2.71E+00 | 2.83E+00 | 3.39E-01 | no | | intergenic_variant | MODIFIER |
| 1 | 172383428 | AX-170234211 | 0 | G | A | 0.219 | 1.06E-01 | 1.11E-01 | 2.57E+00 | 2.66E+00 | 3.39E-01 | no | Pigm | 3_prime_UTR_variant | MODIFIER |
| 18 | 67787746 | AX-170484815 | 2 | T | A | 0.173 | -1.13E-01 | 1.19E-01 | 2.65E+00 | 2.71E+00 | 3.39E-01 | no | Seh1l | intron_variant | MODIFIER |
| 1 | 121799073 | AX-170224121 | 1 | G | A | 0.571 | 7.98E-02 | 8.27E-02 | 2.50E+00 | 2.56E+00 | 3.39E-01 | no | | intergenic_variant | MODIFIER |
| 1 | 82205953 | AX-170175488 | 0 | A | T | 0.167 | 1.83E-01 | 1.93E-01 | 2.70E+00 | 2.77E+00 | 3.39E-01 | Yes | | intergenic_variant | MODIFIER |
| 10 | 26434515 | AX-97839968 | 0 | C | T | 0.175 | 9.64E-02 | 1.01E-01 | 2.97E+00 | 3.10E+00 | 3.39E-01 | no | | intergenic_variant | MODIFIER |
| 5 | 47023893 | AX-170159804 | 0 | C | A | 0.491 | 1.02E-01 | 1.01E-01 | 2.31E+00 | 2.37E+00 | 3.39E-01 | no | | intergenic_variant | MODIFIER |
| 6 | 105830510 | AX-170027884 | 0 | G | C | 0.07 | -1.91E-01 | 2.03E-01 | 2.72E+00 | 2.83E+00 | 3.39E-01 | Yes | Cntn4 | intron_variant | MODIFIER |
| 8 | 66167794 | AX-170137308 | 0 | G | A | 0.149 | -1.26E-01 | 1.32E-01 | 2.60E+00 | 2.69E+00 | 3.40E-01 | no | MAR1 | intron_variant | MODIFIER |
| 1 | 63600176 | AX-97811760 | 1 | G | T | 0.143 | 1.57E-01 | 1.66E-01 | 2.66E+00 | 2.78E+00 | 3.40E-01 | Yes | Adam23 | downstream_gene_variant | MODIFIER |
| 13 | 20035548 | AX-170167286 | 0 | G | A | 0.175 | -1.14E-01 | 1.18E-01 | 2.49E+00 | 2.56E+00 | 3.40E-01 | no | | intergenic_variant | MODIFIER |
| 5 | 42308236 | AX-97839757 | 0 | A | G | 0.202 | -9.61E-02 | 1.01E-01 | 2.56E+00 | 2.64E+00 | 3.40E-01 | no | | intergenic_variant | MODIFIER |
| 2 | 92065028 | AX-170223005 | 0 | A | C | 0.053 | -1.96E-01 | 2.08E-01 | 2.70E+00 | 2.82E+00 | 3.40E-01 | Yes | Gm13780 | intron_variant,non_coding_transcript_variant | MODIFIER |
| 10 | 68199629 | AX-170437497 | 0 | C | G | 0.404 | -7.68E-02 | 8.09E-02 | 2.60E+00 | 2.67E+00 | 3.40E-01 | no | Arid5b | intron_variant,NMD_transcript_variant | MODIFIER |
| 5 | 36683534 | AX-170089449 | 1 | A | T | 0.33 | 9.75E-02 | 1.03E-01 | 2.81E+00 | 2.89E+00 | 3.40E-01 | no | D5Ertd579e | intron_variant | MODIFIER |
| 16 | 79851968 | AX-170339544 | 1 | T | A | 0.473 | 7.17E-02 | 7.52E-02 | 2.57E+00 | 2.64E+00 | 3.40E-01 | no | | intergenic_variant | MODIFIER |

| | | | | | | | | | | | | | | | |
|---|---|---|---|---|---|---|---|---|---|---|---|---|---|---|---|
| 7 | 90142045 | AX-170078026 | 0 | A | T | 0.14 | 1.61E-01 | 1.71E-01 | 2.77E+00 | 2.90E+00 | 3.40E-01 | Yes | Picalm | intron_variant | MODIFIER |
| 4 | 155835050 | AX-170024316 | 0 | G | T | 0.105 | 1.52E-01 | 1.61E-01 | 2.72E+00 | 2.83E+00 | 3.40E-01 | Yes | Mxra8 | upstream_gene_variant | MODIFIER |
| 7 | 19850776 | AX-170356308 | 1 | T | C | 0.402 | 8.46E-02 | 8.91E-02 | 2.57E+00 | 2.69E+00 | 3.40E-01 | no | Ceacam16 | downstream_gene_variant | MODIFIER |
| 14 | 113484668 | AX-170527232 | 1 | A | T | 0.348 | 6.80E-02 | 7.18E-02 | 2.61E+00 | 2.73E+00 | 3.40E-01 | no | | intergenic_variant | MODIFIER |
| 4 | 139300934 | AX-170148404 | 1 | G | A | 0.179 | 1.02E-01 | 1.08E-01 | 2.61E+00 | 2.66E+00 | 3.40E-01 | no | Pqlc2 | intron_variant | MODIFIER |
| 16 | 40321521 | AX-170439033 | 0 | C | T | 0.509 | 7.57E-02 | 7.92E-02 | 2.55E+00 | 2.61E+00 | 3.40E-01 | no | Lsamp | intron_variant | MODIFIER |
| 4 | 137231749 | AX-170449506 | 0 | C | T | 0.167 | 1.10E-01 | 1.16E-01 | 2.61E+00 | 2.66E+00 | 3.40E-01 | no | | regulatory_region_variant | MODIFIER |
| 4 | 137252490 | AX-170289852 | 0 | T | C | 0.167 | 1.10E-01 | 1.16E-01 | 2.61E+00 | 2.66E+00 | 3.40E-01 | no | | intergenic_variant | MODIFIER |
| 4 | 137266509 | AX-170446060 | 0 | G | C | 0.167 | 1.10E-01 | 1.16E-01 | 2.61E+00 | 2.66E+00 | 3.40E-01 | no | | intergenic_variant | MODIFIER |
| 4 | 137293684 | AX-170285846 | 0 | T | C | 0.167 | 1.10E-01 | 1.16E-01 | 2.61E+00 | 2.66E+00 | 3.40E-01 | no | Wnt4 | intron_variant | MODIFIER |
| 4 | 137297424 | AX-170198882 | 0 | T | C | 0.167 | 1.10E-01 | 1.16E-01 | 2.61E+00 | 2.66E+00 | 3.40E-01 | no | Wnt4 | 3_prime_UTR_variant | MODIFIER |
| 4 | 137326545 | AX-170267319 | 0 | A | G | 0.167 | 1.10E-01 | 1.16E-01 | 2.61E+00 | 2.66E+00 | 3.40E-01 | no | Cdc42 | intron_variant | MODIFIER |
| 4 | 137328671 | AX-97795557 | 0 | A | G | 0.167 | 1.10E-01 | 1.16E-01 | 2.61E+00 | 2.66E+00 | 3.40E-01 | no | Cdc42 | intron_variant | MODIFIER |
| 4 | 137346219 | AX-170439521 | 0 | T | A | 0.167 | 1.10E-01 | 1.16E-01 | 2.61E+00 | 2.66E+00 | 3.40E-01 | no | Cdc42 | intron_variant | MODIFIER |
| 4 | 137360734 | AX-170150570 | 0 | T | C | 0.167 | 1.10E-01 | 1.16E-01 | 2.61E+00 | 2.66E+00 | 3.40E-01 | no | Cdc42 | upstream_gene_variant | MODIFIER |
| 4 | 137360805 | AX-170499574 | 0 | T | C | 0.167 | 1.10E-01 | 1.16E-01 | 2.61E+00 | 2.66E+00 | 3.40E-01 | no | Cdc42 | upstream_gene_variant | MODIFIER |
| 4 | 137389073 | AX-170526473 | 0 | T | G | 0.167 | 1.10E-01 | 1.16E-01 | 2.61E+00 | 2.66E+00 | 3.40E-01 | no | 2810405F17Rik | upstream_gene_variant | MODIFIER |
| 4 | 137416016 | AX-170186409 | 0 | C | T | 0.167 | 1.10E-01 | 1.16E-01 | 2.61E+00 | 2.66E+00 | 3.40E-01 | no | Cela3b | downstream_gene_variant | MODIFIER |
| 4 | 137416195 | AX-170432401 | 0 | T | C | 0.167 | 1.10E-01 | 1.16E-01 | 2.61E+00 | 2.66E+00 | 3.40E-01 | no | Cela3b | downstream_gene_variant | MODIFIER |
| 4 | 137439892 | AX-170070825 | 0 | A | T | 0.167 | 1.10E-01 | 1.16E-01 | 2.61E+00 | 2.66E+00 | 3.40E-01 | no | Gm13010 | upstream_gene_variant | MODIFIER |
| 4 | 137443787 | AX-170484630 | 0 | C | T | 0.167 | 1.10E-01 | 1.16E-01 | 2.61E+00 | 2.66E+00 | 3.40E-01 | no | | regulatory_region_variant | MODIFIER |
| 4 | 137456998 | AX-170292205 | 0 | G | A | 0.167 | 1.10E-01 | 1.16E-01 | 2.61E+00 | 2.66E+00 | 3.40E-01 | no | 1700013G24Rik | downstream_gene_variant | MODIFIER |
| 4 | 137464225 | AX-170457212 | 0 | A | G | 0.167 | 1.10E-01 | 1.16E-01 | 2.61E+00 | 2.66E+00 | 3.40E-01 | no | Hspg2 | upstream_gene_variant | MODIFIER |
| 4 | 137464898 | AX-170530478 | 0 | A | C | 0.167 | 1.10E-01 | 1.16E-01 | 2.61E+00 | 2.66E+00 | 3.40E-01 | no | Hspg2 | upstream_gene_variant | MODIFIER |
| 4 | 137465152 | AX-97798744 | 0 | T | C | 0.167 | 1.10E-01 | 1.16E-01 | 2.61E+00 | 2.66E+00 | 3.40E-01 | no | Hspg2 | upstream_gene_variant | MODIFIER |
| 4 | 137492548 | AX-97813687 | 0 | C | A | 0.167 | 1.10E-01 | 1.16E-01 | 2.61E+00 | 2.66E+00 | 3.40E-01 | no | Hspg2 | intron_variant | MODIFIER |
| 4 | 137506112 | AX-170089385 | 0 | A | G | 0.167 | 1.10E-01 | 1.16E-01 | 2.61E+00 | 2.66E+00 | 3.40E-01 | no | Hspg2 | intron_variant | MODIFIER |
| 4 | 137508498 | AX-170184506 | 0 | T | C | 0.167 | 1.10E-01 | 1.16E-01 | 2.61E+00 | 2.66E+00 | 3.40E-01 | no | Hspg2 | intron_variant | MODIFIER |
| 4 | 137543897 | AX-170024050 | 0 | G | A | 0.167 | 1.10E-01 | 1.16E-01 | 2.61E+00 | 2.66E+00 | 3.40E-01 | no | Hspg2 | synonymous_variant | LOW |
| 4 | 137551036 | AX-170368202 | 0 | C | T | 0.167 | 1.10E-01 | 1.16E-01 | 2.61E+00 | 2.66E+00 | 3.40E-01 | no | Hspg2 | intron_variant | MODIFIER |
| 4 | 137582737 | AX-170024059 | 0 | C | T | 0.167 | 1.10E-01 | 1.16E-01 | 2.61E+00 | 2.66E+00 | 3.40E-01 | no | | intergenic_variant | MODIFIER |
| 4 | 137618509 | AX-170024052 | 0 | C | G | 0.167 | 1.10E-01 | 1.16E-01 | 2.61E+00 | 2.66E+00 | 3.40E-01 | no | Usp48 | intron_variant | MODIFIER |
| 4 | 137622835 | AX-97816727 | 0 | T | C | 0.167 | 1.10E-01 | 1.16E-01 | 2.61E+00 | 2.66E+00 | 3.40E-01 | no | Usp48 | intron_variant | MODIFIER |
| 4 | 137648937 | AX-97831665 | 0 | T | C | 0.167 | 1.10E-01 | 1.16E-01 | 2.61E+00 | 2.66E+00 | 3.40E-01 | no | Usp48 | intron_variant | MODIFIER |
| 4 | 137668206 | AX-97833638 | 0 | T | C | 0.167 | 1.10E-01 | 1.16E-01 | 2.61E+00 | 2.66E+00 | 3.40E-01 | no | Rap1gap | intron_variant,non_coding_transcript_variant | MODIFIER |
| 4 | 137690148 | AX-170504108 | 0 | T | G | 0.167 | 1.10E-01 | 1.16E-01 | 2.61E+00 | 2.66E+00 | 3.40E-01 | no | Rap1gap | intron_variant | MODIFIER |
| 4 | 137699097 | AX-97845712 | 0 | T | C | 0.167 | 1.10E-01 | 1.16E-01 | 2.61E+00 | 2.66E+00 | 3.40E-01 | no | Rap1gap | intron_variant | MODIFIER |
| 4 | 137704357 | AX-97828187 | 0 | T | C | 0.167 | 1.10E-01 | 1.16E-01 | 2.61E+00 | 2.66E+00 | 3.40E-01 | no | Rap1gap | intron_variant | MODIFIER |
| 4 | 137757285 | AX-97823623 | 0 | A | G | 0.167 | 1.10E-01 | 1.16E-01 | 2.61E+00 | 2.66E+00 | 3.40E-01 | no | Alpl | intron_variant | MODIFIER |
| 4 | 137772838 | AX-170457262 | 0 | A | G | 0.167 | 1.10E-01 | 1.16E-01 | 2.61E+00 | 2.66E+00 | 3.40E-01 | no | Alpl | intron_variant | MODIFIER |
| 4 | 137774258 | AX-170267687 | 0 | T | C | 0.167 | 1.10E-01 | 1.16E-01 | 2.61E+00 | 2.66E+00 | 3.40E-01 | no | Alpl | intron_variant | MODIFIER |
| 4 | 137777332 | AX-97811315 | 0 | T | G | 0.167 | 1.10E-01 | 1.16E-01 | 2.61E+00 | 2.66E+00 | 3.40E-01 | no | Alpl | intron_variant | MODIFIER |
| 4 | 137779860 | AX-170338831 | 0 | A | G | 0.167 | 1.10E-01 | 1.16E-01 | 2.61E+00 | 2.66E+00 | 3.40E-01 | no | Alpl | intron_variant | MODIFIER |
| 4 | 137785533 | AX-170024049 | 0 | G | A | 0.167 | 1.10E-01 | 1.16E-01 | 2.61E+00 | 2.66E+00 | 3.40E-01 | no | Alpl | intron_variant | MODIFIER |
| 4 | 137788768 | AX-170377708 | 0 | G | A | 0.167 | 1.10E-01 | 1.16E-01 | 2.61E+00 | 2.66E+00 | 3.40E-01 | no | Alpl | intron_variant | MODIFIER |
| 4 | 137853363 | AX-170223946 | 0 | C | G | 0.167 | 1.10E-01 | 1.16E-01 | 2.61E+00 | 2.66E+00 | 3.40E-01 | no | | intergenic_variant | MODIFIER |
| 4 | 137867051 | AX-170024055 | 0 | T | C | 0.167 | 1.10E-01 | 1.16E-01 | 2.61E+00 | 2.66E+00 | 3.40E-01 | no | Ece1 | intron_variant | MODIFIER |
| 4 | 137867164 | AX-97784375 | 0 | A | G | 0.167 | 1.10E-01 | 1.16E-01 | 2.61E+00 | 2.66E+00 | 3.40E-01 | no | Ece1 | intron_variant | MODIFIER |
| 4 | 137913392 | AX-97854124 | 0 | A | G | 0.167 | 1.10E-01 | 1.16E-01 | 2.61E+00 | 2.66E+00 | 3.40E-01 | no | Ece1 | intron_variant | MODIFIER |
| 4 | 137922847 | AX-170334681 | 0 | T | C | 0.167 | 1.10E-01 | 1.16E-01 | 2.61E+00 | 2.66E+00 | 3.40E-01 | no | Ece1 | intron_variant | MODIFIER |
| 4 | 137923419 | AX-170281384 | 0 | G | A | 0.167 | 1.10E-01 | 1.16E-01 | 2.61E+00 | 2.66E+00 | 3.40E-01 | no | Ece1 | intron_variant | MODIFIER |
| 4 | 137924452 | AX-170382581 | 0 | A | G | 0.167 | 1.10E-01 | 1.16E-01 | 2.61E+00 | 2.66E+00 | 3.40E-01 | no | Ece1 | intron_variant | MODIFIER |
| 4 | 137926133 | AX-170295336 | 0 | T | C | 0.167 | 1.10E-01 | 1.16E-01 | 2.61E+00 | 2.66E+00 | 3.40E-01 | no | Ece1 | intron_variant | MODIFIER |
| 4 | 137942396 | AX-170095168 | 0 | C | T | 0.167 | 1.10E-01 | 1.16E-01 | 2.61E+00 | 2.66E+00 | 3.40E-01 | no | Ece1 | intron_variant | MODIFIER |

| | | | | | | | | | | | | | | | |
|---|---|---|---|---|---|---|---|---|---|---|---|---|---|---|---|
| 4 | 137945455 | AX-170099428 | 0 | T | C | 0.167 | 1.10E-01 | 1.16E-01 | 2.61E+00 | 2.66E+00 | 3.40E-01 | no | Ece1 | intron_variant | MODIFIER |
| 4 | 137949714 | AX-170484294 | 0 | A | G | 0.167 | 1.10E-01 | 1.16E-01 | 2.61E+00 | 2.66E+00 | 3.40E-01 | no | Ece1 | intron_variant | MODIFIER |
| 4 | 137949848 | AX-170272212 | 0 | G | A | 0.167 | 1.10E-01 | 1.16E-01 | 2.61E+00 | 2.66E+00 | 3.40E-01 | no | Ece1 | intron_variant | MODIFIER |
| 4 | 137996341 | AX-97846825 | 0 | G | T | 0.167 | 1.10E-01 | 1.16E-01 | 2.61E+00 | 2.66E+00 | 3.40E-01 | no | Eif4g3 | intron_variant | MODIFIER |
| 4 | 137996971 | AX-170282391 | 0 | A | G | 0.167 | 1.10E-01 | 1.16E-01 | 2.61E+00 | 2.66E+00 | 3.40E-01 | no | Eif4g3 | intron_variant | MODIFIER |
| 4 | 138016596 | AX-97852402 | 0 | A | G | 0.167 | 1.10E-01 | 1.16E-01 | 2.61E+00 | 2.66E+00 | 3.40E-01 | no | Eif4g3 | intron_variant | MODIFIER |
| 4 | 138016618 | AX-170505407 | 0 | C | G | 0.167 | 1.10E-01 | 1.16E-01 | 2.61E+00 | 2.66E+00 | 3.40E-01 | no | Eif4g3 | intron_variant | MODIFIER |
| 4 | 138016806 | AX-170465662 | 0 | A | G | 0.167 | 1.10E-01 | 1.16E-01 | 2.61E+00 | 2.66E+00 | 3.40E-01 | no | Eif4g3 | intron_variant | MODIFIER |
| 4 | 138026032 | AX-170334992 | 0 | C | T | 0.167 | 1.10E-01 | 1.16E-01 | 2.61E+00 | 2.66E+00 | 3.40E-01 | no | Eif4g3 | intron_variant | MODIFIER |
| 4 | 138032779 | AX-170269032 | 0 | G | A | 0.167 | 1.10E-01 | 1.16E-01 | 2.61E+00 | 2.66E+00 | 3.40E-01 | no | Eif4g3 | intron_variant | MODIFIER |
| 4 | 138035130 | AX-170263243 | 0 | A | G | 0.167 | 1.10E-01 | 1.16E-01 | 2.61E+00 | 2.66E+00 | 3.40E-01 | no | Eif4g3 | intron_variant | MODIFIER |
| 4 | 138035227 | AX-97796988 | 0 | T | C | 0.167 | 1.10E-01 | 1.16E-01 | 2.61E+00 | 2.66E+00 | 3.40E-01 | no | Eif4g3 | intron_variant | MODIFIER |
| 4 | 138066502 | AX-170199160 | 0 | G | T | 0.167 | 1.10E-01 | 1.16E-01 | 2.61E+00 | 2.66E+00 | 3.40E-01 | no | Eif4g3 | intron_variant | MODIFIER |
| 4 | 138098696 | AX-170024064 | 0 | A | G | 0.167 | 1.10E-01 | 1.16E-01 | 2.61E+00 | 2.66E+00 | 3.40E-01 | no | Eif4g3 | intron_variant | MODIFIER |
| 4 | 138105958 | AX-170024071 | 0 | G | T | 0.167 | 1.10E-01 | 1.16E-01 | 2.61E+00 | 2.66E+00 | 3.40E-01 | no | Eif4g3 | intron_variant | MODIFIER |
| 4 | 138107180 | AX-170343662 | 0 | G | A | 0.167 | 1.10E-01 | 1.16E-01 | 2.61E+00 | 2.66E+00 | 3.40E-01 | no | Eif4g3 | intron_variant | MODIFIER |
| 4 | 138107841 | AX-170095169 | 0 | G | A | 0.167 | 1.10E-01 | 1.16E-01 | 2.61E+00 | 2.66E+00 | 3.40E-01 | no | Eif4g3 | intron_variant | MODIFIER |
| 4 | 138123656 | AX-170520780 | 0 | A | C | 0.167 | 1.10E-01 | 1.16E-01 | 2.61E+00 | 2.66E+00 | 3.40E-01 | no | Eif4g3 | intron_variant | MODIFIER |
| 4 | 138150383 | AX-170024072 | 0 | A | G | 0.167 | 1.10E-01 | 1.16E-01 | 2.61E+00 | 2.66E+00 | 3.40E-01 | no | Eif4g3 | intron_variant | MODIFIER |
| 4 | 138158616 | AX-170179068 | 0 | G | A | 0.167 | 1.10E-01 | 1.16E-01 | 2.61E+00 | 2.66E+00 | 3.40E-01 | no | Eif4g3 | intron_variant | MODIFIER |
| 4 | 138166026 | AX-97835483 | 0 | G | A | 0.167 | 1.10E-01 | 1.16E-01 | 2.61E+00 | 2.66E+00 | 3.40E-01 | no | Eif4g3 | intron_variant | MODIFIER |
| 4 | 138172638 | AX-170331169 | 0 | A | G | 0.167 | 1.10E-01 | 1.16E-01 | 2.61E+00 | 2.66E+00 | 3.40E-01 | no | Eif4g3 | synonymous_variant | LOW |
| 4 | 138207868 | AX-170296696 | 0 | A | G | 0.167 | 1.10E-01 | 1.16E-01 | 2.61E+00 | 2.66E+00 | 3.40E-01 | no | Eif4g3 | downstream_gene_variant | MODIFIER |
| 4 | 138228924 | AX-170024061 | 0 | C | T | 0.167 | 1.10E-01 | 1.16E-01 | 2.61E+00 | 2.66E+00 | 3.40E-01 | no | Hp1bp3 | intron_variant | MODIFIER |
| 4 | 138242553 | AX-97810980 | 0 | T | G | 0.167 | 1.10E-01 | 1.16E-01 | 2.61E+00 | 2.66E+00 | 3.40E-01 | no | Hp1bp3 | 3_prime_UTR_variant | MODIFIER |
| 4 | 138247077 | AX-170254679 | 0 | C | T | 0.167 | 1.10E-01 | 1.16E-01 | 2.61E+00 | 2.66E+00 | 3.40E-01 | no | Hp1bp3 | downstream_gene_variant | MODIFIER |
| 4 | 138261366 | AX-97785004 | 0 | C | T | 0.167 | 1.10E-01 | 1.16E-01 | 2.61E+00 | 2.66E+00 | 3.40E-01 | no | Kif17 | upstream_gene_variant | MODIFIER |
| 4 | 138277821 | AX-170306285 | 0 | T | C | 0.167 | 1.10E-01 | 1.16E-01 | 2.61E+00 | 2.66E+00 | 3.40E-01 | no | Kif17 | synonymous_variant | LOW |
| 4 | 138278614 | AX-170173781 | 0 | G | A | 0.167 | 1.10E-01 | 1.16E-01 | 2.61E+00 | 2.66E+00 | 3.40E-01 | no | Kif17 | intron_variant | MODIFIER |
| 4 | 138283661 | AX-170351589 | 0 | A | G | 0.167 | 1.10E-01 | 1.16E-01 | 2.61E+00 | 2.66E+00 | 3.40E-01 | no | Kif17 | intron_variant | MODIFIER |
| 4 | 138290783 | AX-97805199 | 0 | T | C | 0.167 | 1.10E-01 | 1.16E-01 | 2.61E+00 | 2.66E+00 | 3.40E-01 | no | Kif17 | intron_variant | MODIFIER |
| 4 | 138307356 | AX-170070838 | 0 | T | C | 0.167 | 1.10E-01 | 1.16E-01 | 2.61E+00 | 2.66E+00 | 3.40E-01 | no | Ddost | intron_variant | MODIFIER |
| 4 | 138307845 | AX-170024068 | 0 | T | A | 0.167 | 1.10E-01 | 1.16E-01 | 2.61E+00 | 2.66E+00 | 3.40E-01 | no | Ddost | intron_variant | MODIFIER |
| 4 | 138307932 | AX-170380894 | 0 | C | T | 0.167 | 1.10E-01 | 1.16E-01 | 2.61E+00 | 2.66E+00 | 3.40E-01 | no | Ddost | intron_variant | MODIFIER |
| 4 | 138308043 | AX-97818164 | 0 | C | A | 0.167 | 1.10E-01 | 1.16E-01 | 2.61E+00 | 2.66E+00 | 3.40E-01 | no | Ddost | intron_variant | MODIFIER |
| 4 | 138315709 | AX-170263179 | 0 | T | C | 0.167 | 1.10E-01 | 1.16E-01 | 2.61E+00 | 2.66E+00 | 3.40E-01 | no | Pink1 | missense_variant | MODERATE |
| 4 | 138339179 | AX-170024063 | 0 | T | C | 0.167 | 1.10E-01 | 1.16E-01 | 2.61E+00 | 2.66E+00 | 3.40E-01 | no | Cda | intron_variant | MODIFIER |
| 4 | 138346433 | AX-170024067 | 0 | T | C | 0.167 | 1.10E-01 | 1.16E-01 | 2.61E+00 | 2.66E+00 | 3.40E-01 | no | Cda | intron_variant | MODIFIER |
| 4 | 138348502 | AX-97844594 | 0 | T | G | 0.167 | 1.10E-01 | 1.16E-01 | 2.61E+00 | 2.66E+00 | 3.40E-01 | no | Cda | intron_variant | MODIFIER |
| 4 | 138369515 | AX-170248227 | 0 | G | A | 0.167 | 1.10E-01 | 1.16E-01 | 2.61E+00 | 2.66E+00 | 3.40E-01 | no | Cda | upstream_gene_variant | MODIFIER |
| 4 | 138404710 | AX-97788026 | 0 | C | T | 0.167 | 1.10E-01 | 1.16E-01 | 2.61E+00 | 2.66E+00 | 3.40E-01 | no | | regulatory_region_variant | MODIFIER |
| 4 | 138404770 | AX-170342124 | 0 | A | G | 0.167 | 1.10E-01 | 1.16E-01 | 2.61E+00 | 2.66E+00 | 3.40E-01 | no | | regulatory_region_variant | MODIFIER |
| 4 | 138404850 | AX-170421538 | 0 | T | C | 0.167 | 1.10E-01 | 1.16E-01 | 2.61E+00 | 2.66E+00 | 3.40E-01 | no | | regulatory_region_variant | MODIFIER |
| 4 | 138427554 | AX-170241744 | 0 | T | C | 0.167 | 1.10E-01 | 1.16E-01 | 2.61E+00 | 2.66E+00 | 3.40E-01 | no | Rpl38-ps1 | downstream_gene_variant | MODIFIER |
| 4 | 138437400 | AX-170168498 | 0 | T | G | 0.167 | 1.10E-01 | 1.16E-01 | 2.61E+00 | 2.66E+00 | 3.40E-01 | no | Mul1 | intron_variant | MODIFIER |
| 4 | 138437702 | AX-170218748 | 0 | T | C | 0.167 | 1.10E-01 | 1.16E-01 | 2.61E+00 | 2.66E+00 | 3.40E-01 | no | Mul1 | synonymous_variant | LOW |
| 4 | 138438128 | AX-97811999 | 0 | T | C | 0.167 | 1.10E-01 | 1.16E-01 | 2.61E+00 | 2.66E+00 | 3.40E-01 | no | Mul1 | intron_variant | MODIFIER |
| 4 | 138451393 | AX-170388291 | 0 | A | G | 0.167 | 1.10E-01 | 1.16E-01 | 2.61E+00 | 2.66E+00 | 3.40E-01 | no | Camk2n1 | upstream_gene_variant | MODIFIER |
| 4 | 138471784 | AX-97835207 | 0 | C | T | 0.167 | 1.10E-01 | 1.16E-01 | 2.61E+00 | 2.66E+00 | 3.40E-01 | no | | intergenic_variant | MODIFIER |
| 4 | 138506363 | AX-97838159 | 0 | G | A | 0.167 | 1.10E-01 | 1.16E-01 | 2.61E+00 | 2.66E+00 | 3.40E-01 | no | | intergenic_variant | MODIFIER |
| 4 | 138536925 | AX-170200610 | 0 | A | T | 0.167 | 1.10E-01 | 1.16E-01 | 2.61E+00 | 2.66E+00 | 3.40E-01 | no | | regulatory_region_variant | MODIFIER |
| 4 | 138537722 | AX-170089387 | 0 | T | C | 0.167 | 1.10E-01 | 1.16E-01 | 2.61E+00 | 2.66E+00 | 3.40E-01 | no | | regulatory_region_variant | MODIFIER |
| 4 | 138544699 | AX-170377599 | 0 | G | C | 0.167 | 1.10E-01 | 1.16E-01 | 2.61E+00 | 2.66E+00 | 3.40E-01 | no | | intergenic_variant | MODIFIER |
| 4 | 138555341 | AX-97845324 | 0 | A | G | 0.167 | 1.10E-01 | 1.16E-01 | 2.61E+00 | 2.66E+00 | 3.40E-01 | no | | regulatory_region_variant | MODIFIER |

| | | | | | | | | | | | | | | | |
|---|---|---|---|---|---|---|---|---|---|---|---|---|---|---|---|
| 4 | 140666565 | AX-97830802 | 0 | G | A | 0.167 | 1.10E-01 | 1.16E-01 | 2.61E+00 | 2.66E+00 | 3.40E-01 | no | Arhgef10l | upstream_gene_variant | MODIFIER |
| 4 | 140666634 | AX-170312932 | 0 | G | A | 0.167 | 1.10E-01 | 1.16E-01 | 2.61E+00 | 2.66E+00 | 3.40E-01 | no | Arhgef10l | upstream_gene_variant | MODIFIER |
| 4 | 140718527 | AX-170283326 | 0 | T | C | 0.167 | 1.10E-01 | 1.16E-01 | 2.61E+00 | 2.66E+00 | 3.40E-01 | no | Rcc2 | intron_variant | MODIFIER |
| 4 | 140724110 | AX-170263107 | 0 | A | G | 0.167 | 1.10E-01 | 1.16E-01 | 2.61E+00 | 2.66E+00 | 3.40E-01 | no | Padi6 | downstream_gene_variant | MODIFIER |
| 4 | 140752018 | AX-170257187 | 0 | G | C | 0.167 | 1.10E-01 | 1.16E-01 | 2.61E+00 | 2.66E+00 | 3.40E-01 | no | Padi4 | intron_variant | MODIFIER |
| 4 | 140762806 | AX-170225835 | 0 | G | A | 0.167 | 1.10E-01 | 1.16E-01 | 2.61E+00 | 2.66E+00 | 3.40E-01 | no | Padi4 | intron_variant | MODIFIER |
| 4 | 140799116 | AX-170422248 | 0 | G | A | 0.167 | 1.10E-01 | 1.16E-01 | 2.61E+00 | 2.66E+00 | 3.40E-01 | no | Padi3 | intron_variant | MODIFIER |
| 4 | 140803087 | AX-170356509 | 0 | A | G | 0.167 | 1.10E-01 | 1.16E-01 | 2.61E+00 | 2.66E+00 | 3.40E-01 | no | Padi3 | intron_variant | MODIFIER |
| 4 | 140814094 | AX-170369293 | 0 | G | A | 0.167 | 1.10E-01 | 1.16E-01 | 2.61E+00 | 2.66E+00 | 3.40E-01 | no | Padi3 | upstream_gene_variant | MODIFIER |
| 4 | 140826391 | AX-170480825 | 0 | T | C | 0.167 | 1.10E-01 | 1.16E-01 | 2.61E+00 | 2.66E+00 | 3.40E-01 | no | Padi1 | intron_variant | MODIFIER |
| 4 | 140831667 | AX-170095175 | 0 | G | A | 0.167 | 1.10E-01 | 1.16E-01 | 2.61E+00 | 2.66E+00 | 3.40E-01 | no | Padi1 | intron_variant | MODIFIER |
| 4 | 140845088 | AX-170024127 | 0 | G | A | 0.167 | 1.10E-01 | 1.16E-01 | 2.61E+00 | 2.66E+00 | 3.40E-01 | no | Padi1 | intron_variant | MODIFIER |
| 4 | 140859791 | AX-170024120 | 0 | T | C | 0.167 | 1.10E-01 | 1.16E-01 | 2.61E+00 | 2.66E+00 | 3.40E-01 | no | | regulatory_region_variant | MODIFIER |
| 4 | 140872123 | AX-170328792 | 0 | G | A | 0.167 | 1.10E-01 | 1.16E-01 | 2.61E+00 | 2.66E+00 | 3.40E-01 | no | 4930515B02Rik | upstream_gene_variant | MODIFIER |
| 4 | 140879221 | AX-170515672 | 0 | T | C | 0.167 | 1.10E-01 | 1.16E-01 | 2.61E+00 | 2.66E+00 | 3.40E-01 | no | 4930515B02Rik | downstream_gene_variant | MODIFIER |
| 4 | 140892350 | AX-170405716 | 0 | C | T | 0.167 | 1.10E-01 | 1.16E-01 | 2.61E+00 | 2.66E+00 | 3.40E-01 | no | | intergenic_variant | MODIFIER |
| 10 | 12230151 | AX-97783141 | 1 | G | A | 0.205 | -9.83E-02 | 1.04E-01 | 2.77E+00 | 2.88E+00 | 3.40E-01 | no | | intergenic_variant | MODIFIER |
| 1 | 169897036 | AX-170343241 | 0 | A | G | 0.211 | 1.22E-01 | 1.29E-01 | 2.64E+00 | 2.71E+00 | 3.40E-01 | no | | intergenic_variant | MODIFIER |
| 1 | 170898460 | AX-170346683 | 0 | A | C | 0.211 | 1.22E-01 | 1.29E-01 | 2.64E+00 | 2.71E+00 | 3.40E-01 | no | | regulatory_region_variant | MODIFIER |
| 10 | 72682586 | AX-97794787 | 0 | G | A | 0.421 | -7.42E-02 | 7.87E-02 | 2.68E+00 | 2.76E+00 | 3.40E-01 | no | | intergenic_variant | MODIFIER |
| 16 | 78950249 | AX-170088071 | 1 | A | G | 0.455 | 7.11E-02 | 7.49E-02 | 2.59E+00 | 2.66E+00 | 3.40E-01 | no | Tmprss15 | downstream_gene_variant | MODIFIER |
| 4 | 47201540 | AX-170473726 | 1 | C | G | 0.348 | -8.65E-02 | 9.19E-02 | 2.80E+00 | 2.91E+00 | 3.41E-01 | no | Gm12426 | intron_variant,non_coding_transcript_variant | MODIFIER |
| 18 | 84630558 | AX-170294568 | 0 | C | T | 0.158 | -1.09E-01 | 1.12E-01 | 2.44E+00 | 2.51E+00 | 3.41E-01 | no | Cndp1 | intron_variant | MODIFIER |
| 15 | 3544411 | AX-170181013 | 1 | C | A | 0.241 | -9.89E-02 | 1.05E-01 | 2.65E+00 | 2.71E+00 | 3.41E-01 | no | Ghr | intron_variant | MODIFIER |
| 15 | 5722648 | AX-170054701 | 1 | A | G | 0.241 | -9.89E-02 | 1.05E-01 | 2.65E+00 | 2.71E+00 | 3.41E-01 | no | | intergenic_variant | MODIFIER |
| 3 | 26868519 | AX-170395171 | 1 | A | G | 0.143 | -9.60E-02 | 1.02E-01 | 2.63E+00 | 2.71E+00 | 3.41E-01 | no | Spata16 | intron_variant | MODIFIER |
| 8 | 16725153 | AX-97805576 | 0 | T | G | 0.237 | -6.85E-02 | 7.22E-02 | 2.57E+00 | 2.67E+00 | 3.41E-01 | no | Csmd1 | intron_variant | MODIFIER |
| 16 | 29069094 | AX-170468785 | 0 | T | A | 0.5 | 7.68E-02 | 8.07E-02 | 2.57E+00 | 2.62E+00 | 3.41E-01 | no | | intergenic_variant | MODIFIER |
| 16 | 29177683 | AX-170176812 | 0 | C | T | 0.5 | 7.68E-02 | 8.07E-02 | 2.57E+00 | 2.62E+00 | 3.41E-01 | no | | intergenic_variant | MODIFIER |
| 16 | 29180007 | AX-170494860 | 0 | A | G | 0.5 | 7.68E-02 | 8.07E-02 | 2.57E+00 | 2.62E+00 | 3.41E-01 | no | | intergenic_variant | MODIFIER |
| 16 | 29185801 | AX-170343924 | 0 | C | T | 0.5 | 7.68E-02 | 8.07E-02 | 2.57E+00 | 2.62E+00 | 3.41E-01 | no | | intergenic_variant | MODIFIER |
| 16 | 29186505 | AX-170489998 | 0 | G | T | 0.5 | 7.68E-02 | 8.07E-02 | 2.57E+00 | 2.62E+00 | 3.41E-01 | no | | intergenic_variant | MODIFIER |
| 16 | 29196780 | AX-170348432 | 0 | A | G | 0.5 | 7.68E-02 | 8.07E-02 | 2.57E+00 | 2.62E+00 | 3.41E-01 | no | | intergenic_variant | MODIFIER |
| 16 | 29196795 | AX-170057019 | 0 | A | G | 0.5 | 7.68E-02 | 8.07E-02 | 2.57E+00 | 2.62E+00 | 3.41E-01 | no | | intergenic_variant | MODIFIER |
| 16 | 29199428 | AX-170057018 | 0 | C | T | 0.5 | 7.68E-02 | 8.07E-02 | 2.57E+00 | 2.62E+00 | 3.41E-01 | no | | intergenic_variant | MODIFIER |
| 16 | 29217983 | AX-170468721 | 0 | A | G | 0.5 | 7.68E-02 | 8.07E-02 | 2.57E+00 | 2.62E+00 | 3.41E-01 | no | Hrasls | intron_variant | MODIFIER |
| 16 | 29218664 | AX-170087983 | 0 | T | G | 0.5 | 7.68E-02 | 8.07E-02 | 2.57E+00 | 2.62E+00 | 3.41E-01 | no | Hrasls | intron_variant | MODIFIER |
| 16 | 29218760 | AX-170211352 | 0 | A | T | 0.5 | 7.68E-02 | 8.07E-02 | 2.57E+00 | 2.62E+00 | 3.41E-01 | no | Hrasls | intron_variant | MODIFIER |
| 16 | 29223655 | AX-170103488 | 0 | A | G | 0.5 | 7.68E-02 | 8.07E-02 | 2.57E+00 | 2.62E+00 | 3.41E-01 | no | Hrasls | intron_variant | MODIFIER |
| 16 | 29223742 | AX-170445485 | 0 | G | C | 0.5 | 7.68E-02 | 8.07E-02 | 2.57E+00 | 2.62E+00 | 3.41E-01 | no | Hrasls | intron_variant | MODIFIER |
| 16 | 29261320 | AX-170227124 | 0 | G | A | 0.5 | 7.68E-02 | 8.07E-02 | 2.57E+00 | 2.62E+00 | 3.41E-01 | no | Atp13a5 | intron_variant | MODIFIER |
| 16 | 29274167 | AX-170105482 | 0 | T | G | 0.5 | 7.68E-02 | 8.07E-02 | 2.57E+00 | 2.62E+00 | 3.41E-01 | no | Atp13a5 | intron_variant | MODIFIER |
| 16 | 29421099 | AX-97831797 | 0 | T | C | 0.5 | 7.68E-02 | 8.07E-02 | 2.57E+00 | 2.62E+00 | 3.41E-01 | no | Atp13a4 | intron_variant | MODIFIER |
| 16 | 29421175 | AX-170012704 | 0 | A | G | 0.5 | 7.68E-02 | 8.07E-02 | 2.57E+00 | 2.62E+00 | 3.41E-01 | no | Atp13a4 | intron_variant | MODIFIER |
| 16 | 29421429 | AX-170108924 | 0 | A | G | 0.5 | 7.68E-02 | 8.07E-02 | 2.57E+00 | 2.62E+00 | 3.41E-01 | no | Atp13a4 | intron_variant | MODIFIER |
| 16 | 29421801 | AX-170214610 | 0 | G | C | 0.5 | 7.68E-02 | 8.07E-02 | 2.57E+00 | 2.62E+00 | 3.41E-01 | no | Atp13a4 | intron_variant | MODIFIER |
| 16 | 29421836 | AX-170307512 | 0 | G | A | 0.5 | 7.68E-02 | 8.07E-02 | 2.57E+00 | 2.62E+00 | 3.41E-01 | no | Atp13a4 | intron_variant | MODIFIER |
| 16 | 29476440 | AX-170230981 | 0 | A | G | 0.5 | 7.68E-02 | 8.07E-02 | 2.57E+00 | 2.62E+00 | 3.41E-01 | no | Atp13a4 | intron_variant | MODIFIER |
| 16 | 29476590 | AX-170349023 | 0 | T | C | 0.5 | 7.68E-02 | 8.07E-02 | 2.57E+00 | 2.62E+00 | 3.41E-01 | no | Atp13a4 | intron_variant | MODIFIER |
| 16 | 29477508 | AX-170176214 | 0 | G | T | 0.5 | 7.68E-02 | 8.07E-02 | 2.57E+00 | 2.62E+00 | 3.41E-01 | no | Atp13a4 | intron_variant | MODIFIER |
| 16 | 29499227 | AX-170012703 | 0 | T | G | 0.5 | 7.68E-02 | 8.07E-02 | 2.57E+00 | 2.62E+00 | 3.41E-01 | no | Atp13a4 | intron_variant | MODIFIER |
| 16 | 29508048 | AX-168378928 | 0 | C | T | 0.5 | 7.68E-02 | 8.07E-02 | 2.57E+00 | 2.62E+00 | 3.41E-01 | no | Atp13a4 | intron_variant | MODIFIER |
| 16 | 29523957 | AX-97788151 | 0 | G | A | 0.5 | 7.68E-02 | 8.07E-02 | 2.57E+00 | 2.62E+00 | 3.41E-01 | no | Atp13a4 | intron_variant | MODIFIER |
| 16 | 29524012 | AX-170208303 | 0 | G | A | 0.5 | 7.68E-02 | 8.07E-02 | 2.57E+00 | 2.62E+00 | 3.41E-01 | no | Atp13a4 | intron_variant | MODIFIER |

| | | | | | | | | | | | | | | | |
|---|---|---|---|---|---|---|---|---|---|---|---|---|---|---|---|
| 16 | 29525415 | AX-170427711 | 0 | T | C | 0.5 | 7.68E-02 | 8.07E-02 | 2.57E+00 | 2.62E+00 | 3.41E-01 | no | Atp13a4 | intron_variant | MODIFIER |
| 16 | 29530223 | AX-170423023 | 0 | G | A | 0.5 | 7.68E-02 | 8.07E-02 | 2.57E+00 | 2.62E+00 | 3.41E-01 | no | Atp13a4 | intron_variant | MODIFIER |
| 16 | 29530310 | AX-170179010 | 0 | T | A | 0.5 | 7.68E-02 | 8.07E-02 | 2.57E+00 | 2.62E+00 | 3.41E-01 | no | Atp13a4 | intron_variant | MODIFIER |
| 16 | 29533389 | AX-170302868 | 0 | G | A | 0.5 | 7.68E-02 | 8.07E-02 | 2.57E+00 | 2.62E+00 | 3.41E-01 | no | Atp13a4 | intron_variant | MODIFIER |
| 16 | 29535499 | AX-168413729 | 0 | G | A | 0.5 | 7.68E-02 | 8.07E-02 | 2.57E+00 | 2.62E+00 | 3.41E-01 | no | Atp13a4 | intron_variant | MODIFIER |
| 16 | 29545204 | AX-170505844 | 0 | G | A | 0.5 | 7.68E-02 | 8.07E-02 | 2.57E+00 | 2.62E+00 | 3.41E-01 | no | Atp13a4 | upstream_gene_variant | MODIFIER |
| 16 | 29550783 | AX-170239726 | 0 | C | A | 0.5 | 7.68E-02 | 8.07E-02 | 2.57E+00 | 2.62E+00 | 3.41E-01 | no | | intergenic_variant | MODIFIER |
| 16 | 29636533 | AX-168394691 | 0 | T | C | 0.5 | 7.68E-02 | 8.07E-02 | 2.57E+00 | 2.62E+00 | 3.41E-01 | no | Opa1 | intron_variant | MODIFIER |
| 16 | 29750421 | AX-170148127 | 0 | A | G | 0.5 | 7.68E-02 | 8.07E-02 | 2.57E+00 | 2.62E+00 | 3.41E-01 | no | Gm49636 | downstream_gene_variant | MODIFIER |
| 16 | 29757940 | AX-170457835 | 0 | C | T | 0.5 | 7.68E-02 | 8.07E-02 | 2.57E+00 | 2.62E+00 | 3.41E-01 | no | Gm49636 | upstream_gene_variant | MODIFIER |
| 16 | 29760935 | AX-97815040 | 0 | A | G | 0.5 | 7.68E-02 | 8.07E-02 | 2.57E+00 | 2.62E+00 | 3.41E-01 | no | | regulatory_region_variant | MODIFIER |
| 16 | 29769107 | AX-170194354 | 0 | T | A | 0.5 | 7.68E-02 | 8.07E-02 | 2.57E+00 | 2.62E+00 | 3.41E-01 | no | | intergenic_variant | MODIFIER |
| 16 | 29785761 | AX-170402617 | 0 | C | T | 0.5 | 7.68E-02 | 8.07E-02 | 2.57E+00 | 2.62E+00 | 3.41E-01 | no | | intergenic_variant | MODIFIER |
| 16 | 29785796 | AX-170457821 | 0 | T | G | 0.5 | 7.68E-02 | 8.07E-02 | 2.57E+00 | 2.62E+00 | 3.41E-01 | no | | intergenic_variant | MODIFIER |
| 16 | 29788274 | AX-97809077 | 0 | C | T | 0.5 | 7.68E-02 | 8.07E-02 | 2.57E+00 | 2.62E+00 | 3.41E-01 | no | | regulatory_region_variant | MODIFIER |
| 16 | 29833700 | AX-170359078 | 0 | G | C | 0.5 | 7.68E-02 | 8.07E-02 | 2.57E+00 | 2.62E+00 | 3.41E-01 | no | Gm26834 | intron_variant,non_coding_transcript_variant | MODIFIER |
| 16 | 29845147 | AX-170140190 | 0 | G | C | 0.5 | 7.68E-02 | 8.07E-02 | 2.57E+00 | 2.62E+00 | 3.41E-01 | no | | regulatory_region_variant | MODIFIER |
| 16 | 29845528 | AX-170457887 | 0 | A | G | 0.5 | 7.68E-02 | 8.07E-02 | 2.57E+00 | 2.62E+00 | 3.41E-01 | no | | regulatory_region_variant | MODIFIER |
| 16 | 29845755 | AX-170332135 | 0 | T | C | 0.5 | 7.68E-02 | 8.07E-02 | 2.57E+00 | 2.62E+00 | 3.41E-01 | no | | regulatory_region_variant | MODIFIER |
| 16 | 29848023 | AX-170057034 | 0 | G | A | 0.5 | 7.68E-02 | 8.07E-02 | 2.57E+00 | 2.62E+00 | 3.41E-01 | no | | regulatory_region_variant | MODIFIER |
| 16 | 29849073 | AX-97851416 | 0 | T | C | 0.5 | 7.68E-02 | 8.07E-02 | 2.57E+00 | 2.62E+00 | 3.41E-01 | no | | regulatory_region_variant | MODIFIER |
| 16 | 29858502 | AX-170087980 | 0 | C | T | 0.5 | 7.68E-02 | 8.07E-02 | 2.57E+00 | 2.62E+00 | 3.41E-01 | no | | intergenic_variant | MODIFIER |
| 16 | 29859372 | AX-170012719 | 0 | A | T | 0.5 | 7.68E-02 | 8.07E-02 | 2.57E+00 | 2.62E+00 | 3.41E-01 | no | | intergenic_variant | MODIFIER |
| 16 | 29864968 | AX-170212085 | 0 | C | T | 0.5 | 7.68E-02 | 8.07E-02 | 2.57E+00 | 2.62E+00 | 3.41E-01 | no | | regulatory_region_variant | MODIFIER |
| 16 | 29881200 | AX-170012714 | 0 | C | T | 0.5 | 7.68E-02 | 8.07E-02 | 2.57E+00 | 2.62E+00 | 3.41E-01 | no | | regulatory_region_variant | MODIFIER |
| 16 | 29898139 | AX-97830763 | 0 | C | T | 0.5 | 7.68E-02 | 8.07E-02 | 2.57E+00 | 2.62E+00 | 3.41E-01 | no | | intergenic_variant | MODIFIER |
| 16 | 29905153 | AX-170454175 | 0 | A | G | 0.5 | 7.68E-02 | 8.07E-02 | 2.57E+00 | 2.62E+00 | 3.41E-01 | no | Gm26569 | downstream_gene_variant | MODIFIER |
| 16 | 29936950 | AX-170012716 | 0 | G | A | 0.5 | 7.68E-02 | 8.07E-02 | 2.57E+00 | 2.62E+00 | 3.41E-01 | no | Gm26569 | intron_variant,non_coding_transcript_variant | MODIFIER |
| 16 | 30058454 | AX-97829213 | 0 | C | T | 0.5 | 7.68E-02 | 8.07E-02 | 2.57E+00 | 2.62E+00 | 3.41E-01 | no | | intergenic_variant | MODIFIER |
| 16 | 30085429 | AX-170012712 | 0 | T | C | 0.5 | 7.68E-02 | 8.07E-02 | 2.57E+00 | 2.62E+00 | 3.41E-01 | no | | intergenic_variant | MODIFIER |
| 16 | 30089189 | AX-170012717 | 0 | C | T | 0.5 | 7.68E-02 | 8.07E-02 | 2.57E+00 | 2.62E+00 | 3.41E-01 | no | Gm20040 | downstream_gene_variant | MODIFIER |
| 16 | 30130221 | AX-170012711 | 0 | G | T | 0.5 | 7.68E-02 | 8.07E-02 | 2.57E+00 | 2.62E+00 | 3.41E-01 | no | | regulatory_region_variant | MODIFIER |
| 16 | 30157847 | AX-170012715 | 0 | T | A | 0.5 | 7.68E-02 | 8.07E-02 | 2.57E+00 | 2.62E+00 | 3.41E-01 | no | | intergenic_variant | MODIFIER |
| 16 | 30270652 | AX-168414937 | 0 | G | C | 0.5 | 7.68E-02 | 8.07E-02 | 2.57E+00 | 2.62E+00 | 3.41E-01 | no | Lrrc15 | 3_prime_UTR_variant | MODIFIER |
| 16 | 31207237 | AX-170012743 | 0 | A | C | 0.5 | 7.68E-02 | 8.07E-02 | 2.57E+00 | 2.62E+00 | 3.41E-01 | no | | intergenic_variant | MODIFIER |
| 16 | 31212627 | AX-97804064 | 0 | G | T | 0.5 | 7.68E-02 | 8.07E-02 | 2.57E+00 | 2.62E+00 | 3.41E-01 | no | | regulatory_region_variant | MODIFIER |
| 16 | 31255256 | AX-97790754 | 0 | T | C | 0.5 | 7.68E-02 | 8.07E-02 | 2.57E+00 | 2.62E+00 | 3.41E-01 | no | Ppp1r2 | intron_variant | MODIFIER |
| 16 | 31256470 | AX-170012739 | 0 | C | T | 0.5 | 7.68E-02 | 8.07E-02 | 2.57E+00 | 2.62E+00 | 3.41E-01 | no | Ppp1r2 | intron_variant | MODIFIER |
| 16 | 31298504 | AX-97830819 | 0 | G | A | 0.5 | 7.68E-02 | 8.07E-02 | 2.57E+00 | 2.62E+00 | 3.41E-01 | no | Apod | intron_variant,non_coding_transcript_variant | MODIFIER |
| 16 | 31299240 | AX-170012742 | 0 | A | G | 0.5 | 7.68E-02 | 8.07E-02 | 2.57E+00 | 2.62E+00 | 3.41E-01 | no | Apod | intron_variant,non_coding_transcript_variant | MODIFIER |
| 16 | 31302218 | AX-170338065 | 0 | T | C | 0.5 | 7.68E-02 | 8.07E-02 | 2.57E+00 | 2.62E+00 | 3.41E-01 | no | Apod | intron_variant,non_coding_transcript_variant | MODIFIER |
| 16 | 31326083 | AX-97823326 | 0 | C | T | 0.5 | 7.68E-02 | 8.07E-02 | 2.57E+00 | 2.62E+00 | 3.41E-01 | no | AV205837 | intron_variant,non_coding_transcript_variant | MODIFIER |
| 16 | 31338593 | AX-170426469 | 0 | C | T | 0.5 | 7.68E-02 | 8.07E-02 | 2.57E+00 | 2.62E+00 | 3.41E-01 | no | | intergenic_variant | MODIFIER |
| 16 | 31342379 | AX-97782002 | 0 | A | G | 0.5 | 7.68E-02 | 8.07E-02 | 2.57E+00 | 2.62E+00 | 3.41E-01 | no | Gm49404 | upstream_gene_variant | MODIFIER |
| 16 | 31403109 | AX-97832262 | 0 | A | C | 0.5 | 7.68E-02 | 8.07E-02 | 2.57E+00 | 2.62E+00 | 3.41E-01 | no | | regulatory_region_variant | MODIFIER |
| 16 | 31417774 | AX-97793978 | 0 | T | C | 0.5 | 7.68E-02 | 8.07E-02 | 2.57E+00 | 2.62E+00 | 3.41E-01 | no | Bdh1 | upstream_gene_variant | MODIFIER |
| 16 | 31418006 | AX-170210511 | 0 | G | A | 0.5 | 7.68E-02 | 8.07E-02 | 2.57E+00 | 2.62E+00 | 3.41E-01 | no | Bdh1 | upstream_gene_variant | MODIFIER |
| 16 | 31423453 | AX-170362920 | 0 | G | C | 0.5 | 7.68E-02 | 8.07E-02 | 2.57E+00 | 2.62E+00 | 3.41E-01 | no | Bdh1 | intron_variant | MODIFIER |
| 16 | 31426351 | AX-170411025 | 0 | G | A | 0.5 | 7.68E-02 | 8.07E-02 | 2.57E+00 | 2.62E+00 | 3.41E-01 | no | Bdh1 | intron_variant | MODIFIER |
| 16 | 31448600 | AX-97830824 | 0 | A | G | 0.5 | 7.68E-02 | 8.07E-02 | 2.57E+00 | 2.62E+00 | 3.41E-01 | no | Bdh1 | intron_variant | MODIFIER |
| 16 | 31448707 | AX-170331771 | 0 | C | T | 0.5 | 7.68E-02 | 8.07E-02 | 2.57E+00 | 2.62E+00 | 3.41E-01 | no | Bdh1 | intron_variant | MODIFIER |
| 16 | 31494773 | AX-97824696 | 0 | G | A | 0.5 | 7.68E-02 | 8.07E-02 | 2.57E+00 | 2.62E+00 | 3.41E-01 | no | | regulatory_region_variant | MODIFIER |
| 16 | 31499244 | AX-170273227 | 0 | G | A | 0.5 | 7.68E-02 | 8.07E-02 | 2.57E+00 | 2.62E+00 | 3.41E-01 | no | | regulatory_region_variant | MODIFIER |
| 16 | 31511112 | AX-97831402 | 0 | G | A | 0.5 | 7.68E-02 | 8.07E-02 | 2.57E+00 | 2.62E+00 | 3.41E-01 | no | Gm46560 | intron_variant,non_coding_transcript_variant | MODIFIER |

| | | | | | | | | | | | | | | | |
|---|---|---|---|---|---|---|---|---|---|---|---|---|---|---|---|
| 16 | 31529652 | AX-170178225 | 0 | C | T | 0.5 | 7.68E-02 | 8.07E-02 | 2.57E+00 | 2.62E+00 | 3.41E-01 | no | | regulatory_region_variant | MODIFIER |
| 16 | 31531989 | AX-170518231 | 0 | A | G | 0.5 | 7.68E-02 | 8.07E-02 | 2.57E+00 | 2.62E+00 | 3.41E-01 | no | | intergenic_variant | MODIFIER |
| 16 | 31533498 | AX-97834896 | 0 | A | G | 0.5 | 7.68E-02 | 8.07E-02 | 2.57E+00 | 2.62E+00 | 3.41E-01 | no | | regulatory_region_variant | MODIFIER |
| 16 | 31534208 | AX-170479045 | 0 | C | G | 0.5 | 7.68E-02 | 8.07E-02 | 2.57E+00 | 2.62E+00 | 3.41E-01 | no | | regulatory_region_variant | MODIFIER |
| 16 | 31534349 | AX-170480688 | 0 | G | A | 0.5 | 7.68E-02 | 8.07E-02 | 2.57E+00 | 2.62E+00 | 3.41E-01 | no | | regulatory_region_variant | MODIFIER |
| 16 | 31573611 | AX-97857197 | 0 | G | A | 0.5 | 7.68E-02 | 8.07E-02 | 2.57E+00 | 2.62E+00 | 3.41E-01 | no | | intergenic_variant | MODIFIER |
| 16 | 31577171 | AX-170012751 | 0 | A | G | 0.5 | 7.68E-02 | 8.07E-02 | 2.57E+00 | 2.62E+00 | 3.41E-01 | no | | intergenic_variant | MODIFIER |
| 16 | 31583683 | AX-170012755 | 0 | A | G | 0.5 | 7.68E-02 | 8.07E-02 | 2.57E+00 | 2.62E+00 | 3.41E-01 | no | | intergenic_variant | MODIFIER |
| 16 | 31591135 | AX-97822078 | 0 | T | C | 0.5 | 7.68E-02 | 8.07E-02 | 2.57E+00 | 2.62E+00 | 3.41E-01 | no | Gm34256 | non_coding_transcript_exon_variant | MODIFIER |
| 16 | 31689734 | AX-170215014 | 0 | C | T | 0.5 | 7.68E-02 | 8.07E-02 | 2.57E+00 | 2.62E+00 | 3.41E-01 | no | Dlg1 | intron_variant | MODIFIER |
| 16 | 31862839 | AX-97793631 | 0 | T | G | 0.5 | 7.68E-02 | 8.07E-02 | 2.57E+00 | 2.62E+00 | 3.41E-01 | no | Dlg1 | intron_variant | MODIFIER |
| 16 | 32142494 | AX-170057069 | 0 | G | A | 0.5 | 7.68E-02 | 8.07E-02 | 2.57E+00 | 2.62E+00 | 3.41E-01 | no | Nrros | downstream_gene_variant | MODIFIER |
| 16 | 32179133 | AX-170012766 | 0 | A | T | 0.5 | 7.68E-02 | 8.07E-02 | 2.57E+00 | 2.62E+00 | 3.41E-01 | no | Bex6 | upstream_gene_variant | MODIFIER |
| 16 | 32214659 | AX-170516640 | 0 | A | G | 0.5 | 7.68E-02 | 8.07E-02 | 2.57E+00 | 2.62E+00 | 3.41E-01 | no | | regulatory_region_variant | MODIFIER |
| 16 | 32216578 | AX-170012764 | 0 | T | C | 0.5 | 7.68E-02 | 8.07E-02 | 2.57E+00 | 2.62E+00 | 3.41E-01 | no | | intergenic_variant | MODIFIER |
| 16 | 32217110 | AX-97853489 | 0 | C | T | 0.5 | 7.68E-02 | 8.07E-02 | 2.57E+00 | 2.62E+00 | 3.41E-01 | no | | intergenic_variant | MODIFIER |
| 16 | 32282957 | AX-170012769 | 0 | C | A | 0.5 | 7.68E-02 | 8.07E-02 | 2.57E+00 | 2.62E+00 | 3.41E-01 | no | Rnf168 | intron_variant | MODIFIER |
| 16 | 32286081 | AX-170220134 | 0 | A | G | 0.5 | 7.68E-02 | 8.07E-02 | 2.57E+00 | 2.62E+00 | 3.41E-01 | no | Rnf168 | intron_variant | MODIFIER |
| 16 | 32314224 | AX-97822771 | 0 | G | T | 0.5 | 7.68E-02 | 8.07E-02 | 2.57E+00 | 2.62E+00 | 3.41E-01 | no | | regulatory_region_variant | MODIFIER |
| 16 | 32385603 | AX-97831425 | 0 | T | C | 0.5 | 7.68E-02 | 8.07E-02 | 2.57E+00 | 2.62E+00 | 3.41E-01 | no | Ubxn7 | 3_prime_UTR_variant | MODIFIER |
| 16 | 32407365 | AX-170466017 | 0 | G | A | 0.5 | 7.68E-02 | 8.07E-02 | 2.57E+00 | 2.62E+00 | 3.41E-01 | no | Tm4sf19 | intron_variant | MODIFIER |
| 16 | 32408178 | AX-170057071 | 0 | G | A | 0.5 | 7.68E-02 | 8.07E-02 | 2.57E+00 | 2.62E+00 | 3.41E-01 | no | Tm4sf19 | 3_prime_UTR_variant | MODIFIER |
| 16 | 32422403 | AX-170106651 | 0 | C | T | 0.5 | 7.68E-02 | 8.07E-02 | 2.57E+00 | 2.62E+00 | 3.41E-01 | no | Tctex1d2 | intron_variant | MODIFIER |
| 16 | 32455776 | AX-170250248 | 0 | G | A | 0.5 | 7.68E-02 | 8.07E-02 | 2.57E+00 | 2.62E+00 | 3.41E-01 | no | Pcyt1a | intron_variant | MODIFIER |
| 16 | 32460749 | AX-170360399 | 0 | C | T | 0.5 | 7.68E-02 | 8.07E-02 | 2.57E+00 | 2.62E+00 | 3.41E-01 | no | Pcyt1a | intron_variant | MODIFIER |
| 16 | 32461341 | AX-170087988 | 0 | A | G | 0.5 | 7.68E-02 | 8.07E-02 | 2.57E+00 | 2.62E+00 | 3.41E-01 | no | Pcyt1a | intron_variant | MODIFIER |
| 16 | 32461426 | AX-170287227 | 0 | C | A | 0.5 | 7.68E-02 | 8.07E-02 | 2.57E+00 | 2.62E+00 | 3.41E-01 | no | Pcyt1a | intron_variant | MODIFIER |
| 16 | 32498712 | AX-170348674 | 0 | G | T | 0.5 | 7.68E-02 | 8.07E-02 | 2.57E+00 | 2.62E+00 | 3.41E-01 | no | Zdhhc19 | intron_variant | MODIFIER |
| 16 | 32500441 | AX-170169744 | 0 | A | G | 0.5 | 7.68E-02 | 8.07E-02 | 2.57E+00 | 2.62E+00 | 3.41E-01 | no | Zdhhc19 | intron_variant | MODIFIER |
| 16 | 32504399 | AX-170353587 | 0 | G | T | 0.5 | 7.68E-02 | 8.07E-02 | 2.57E+00 | 2.62E+00 | 3.41E-01 | no | Zdhhc19 | intron_variant | MODIFIER |
| 16 | 32507949 | AX-170283458 | 0 | G | A | 0.5 | 7.68E-02 | 8.07E-02 | 2.57E+00 | 2.62E+00 | 3.41E-01 | no | Zdhhc19 | downstream_gene_variant | MODIFIER |
| 16 | 32768729 | AX-170248131 | 0 | T | C | 0.5 | 7.68E-02 | 8.07E-02 | 2.57E+00 | 2.62E+00 | 3.41E-01 | no | Muc4 | synonymous_variant | LOW |
| 16 | 32793353 | AX-97829633 | 0 | T | C | 0.5 | 7.68E-02 | 8.07E-02 | 2.57E+00 | 2.62E+00 | 3.41E-01 | no | Muc20 | synonymous_variant | LOW |
| 16 | 35379627 | AX-170012815 | 0 | G | A | 0.5 | 7.68E-02 | 8.07E-02 | 2.57E+00 | 2.62E+00 | 3.41E-01 | no | Gm25079 | upstream_gene_variant | MODIFIER |
| 16 | 35431333 | AX-97831503 | 0 | A | G | 0.5 | 7.68E-02 | 8.07E-02 | 2.57E+00 | 2.62E+00 | 3.41E-01 | no | Pdia5 | intron_variant | MODIFIER |
| 16 | 35439126 | AX-170247150 | 0 | C | T | 0.5 | 7.68E-02 | 8.07E-02 | 2.57E+00 | 2.62E+00 | 3.41E-01 | no | Pdia5 | intron_variant | MODIFIER |
| 16 | 39429057 | AX-170226403 | 0 | T | G | 0.509 | 7.78E-02 | 8.14E-02 | 2.53E+00 | 2.57E+00 | 3.41E-01 | no | | intergenic_variant | MODIFIER |
| 1 | 190910312 | AX-170443687 | 0 | G | C | 0.254 | 9.51E-02 | 9.65E-02 | 2.38E+00 | 2.45E+00 | 3.41E-01 | no | Rps6kc1 | intron_variant | MODIFIER |
| 2 | 102210411 | AX-97808605 | 1 | G | A | 0.062 | 2.07E-01 | 2.20E-01 | 2.82E+00 | 2.92E+00 | 3.41E-01 | Yes | Gm13866 | upstream_gene_variant | MODIFIER |
| 1 | 189434259 | AX-170105882 | 0 | A | C | 0.254 | 9.72E-02 | 9.96E-02 | 2.41E+00 | 2.50E+00 | 3.41E-01 | no | | intergenic_variant | MODIFIER |
| 1 | 189437329 | AX-170141875 | 0 | G | A | 0.254 | 9.72E-02 | 9.96E-02 | 2.41E+00 | 2.50E+00 | 3.41E-01 | no | | intergenic_variant | MODIFIER |
| 1 | 189438728 | AX-170438501 | 0 | A | G | 0.254 | 9.72E-02 | 9.96E-02 | 2.41E+00 | 2.50E+00 | 3.41E-01 | no | | intergenic_variant | MODIFIER |
| 1 | 189484607 | AX-170439081 | 0 | T | C | 0.254 | 9.72E-02 | 9.96E-02 | 2.41E+00 | 2.50E+00 | 3.41E-01 | no | | intergenic_variant | MODIFIER |
| 1 | 189524226 | AX-170259469 | 0 | A | G | 0.254 | 9.72E-02 | 9.96E-02 | 2.41E+00 | 2.50E+00 | 3.41E-01 | no | | intergenic_variant | MODIFIER |
| 1 | 189553019 | AX-170176966 | 0 | G | A | 0.254 | 9.72E-02 | 9.96E-02 | 2.41E+00 | 2.50E+00 | 3.41E-01 | no | | intergenic_variant | MODIFIER |
| 1 | 189553244 | AX-170139311 | 0 | G | A | 0.254 | 9.72E-02 | 9.96E-02 | 2.41E+00 | 2.50E+00 | 3.41E-01 | no | | intergenic_variant | MODIFIER |
| 1 | 189580328 | AX-170195419 | 0 | G | A | 0.254 | 9.72E-02 | 9.96E-02 | 2.41E+00 | 2.50E+00 | 3.41E-01 | no | | intergenic_variant | MODIFIER |
| 1 | 189587365 | AX-170043943 | 0 | A | G | 0.254 | 9.72E-02 | 9.96E-02 | 2.41E+00 | 2.50E+00 | 3.41E-01 | no | | intergenic_variant | MODIFIER |
| 1 | 189591167 | AX-170002542 | 0 | T | C | 0.254 | 9.72E-02 | 9.96E-02 | 2.41E+00 | 2.50E+00 | 3.41E-01 | no | | intergenic_variant | MODIFIER |
| 1 | 189591509 | AX-170002541 | 0 | G | T | 0.254 | 9.72E-02 | 9.96E-02 | 2.41E+00 | 2.50E+00 | 3.41E-01 | no | | intergenic_variant | MODIFIER |
| 1 | 189600067 | AX-170002538 | 0 | A | G | 0.254 | 9.72E-02 | 9.96E-02 | 2.41E+00 | 2.50E+00 | 3.41E-01 | no | | regulatory_region_variant | MODIFIER |
| 1 | 189603174 | AX-97789353 | 0 | A | G | 0.254 | 9.72E-02 | 9.96E-02 | 2.41E+00 | 2.50E+00 | 3.41E-01 | no | | intergenic_variant | MODIFIER |
| 1 | 189604680 | AX-170243734 | 0 | C | G | 0.254 | 9.72E-02 | 9.96E-02 | 2.41E+00 | 2.50E+00 | 3.41E-01 | no | | intergenic_variant | MODIFIER |
| 1 | 189616486 | AX-170227510 | 0 | A | C | 0.254 | 9.72E-02 | 9.96E-02 | 2.41E+00 | 2.50E+00 | 3.41E-01 | no | | intergenic_variant | MODIFIER |

| | | | | | | | | | | | | | | | |
|---|---|---|---|---|---|---|---|---|---|---|---|---|---|---|---|
| 1 | 189632680 | AX-170352275 | 0 | A | C | 0.254 | 9.72E-02 | 9.96E-02 | 2.41E+00 | 2.50E+00 | 3.41E-01 | no | | intergenic_variant | MODIFIER |
| 1 | 189686196 | AX-170372234 | 0 | T | C | 0.254 | 9.72E-02 | 9.96E-02 | 2.41E+00 | 2.50E+00 | 3.41E-01 | no | Cenpf | intron_variant,NMD_transcript_variant | MODIFIER |
| 1 | 189708632 | AX-170409160 | 0 | G | A | 0.254 | 9.72E-02 | 9.96E-02 | 2.41E+00 | 2.50E+00 | 3.41E-01 | no | | regulatory_region_variant | MODIFIER |
| 1 | 189714737 | AX-170002534 | 0 | A | C | 0.254 | 9.72E-02 | 9.96E-02 | 2.41E+00 | 2.50E+00 | 3.41E-01 | no | | intergenic_variant | MODIFIER |
| 1 | 189714772 | AX-170130271 | 0 | G | A | 0.254 | 9.72E-02 | 9.96E-02 | 2.41E+00 | 2.50E+00 | 3.41E-01 | no | | intergenic_variant | MODIFIER |
| 1 | 189755823 | AX-170471129 | 0 | A | G | 0.254 | 9.72E-02 | 9.96E-02 | 2.41E+00 | 2.50E+00 | 3.41E-01 | no | Ptpn14 | intron_variant | MODIFIER |
| 1 | 189836099 | AX-97814571 | 0 | A | C | 0.254 | 9.72E-02 | 9.96E-02 | 2.41E+00 | 2.50E+00 | 3.41E-01 | no | Ptpn14 | intron_variant | MODIFIER |
| 1 | 189849718 | AX-170411674 | 0 | T | C | 0.254 | 9.72E-02 | 9.96E-02 | 2.41E+00 | 2.50E+00 | 3.41E-01 | no | Ptpn14 | intron_variant | MODIFIER |
| 1 | 189856693 | AX-170322426 | 0 | T | C | 0.254 | 9.72E-02 | 9.96E-02 | 2.41E+00 | 2.50E+00 | 3.41E-01 | no | Ptpn14 | synonymous_variant | LOW |
| 1 | 189856744 | AX-170115115 | 0 | T | C | 0.254 | 9.72E-02 | 9.96E-02 | 2.41E+00 | 2.50E+00 | 3.41E-01 | no | Ptpn14 | synonymous_variant | LOW |
| 1 | 189880901 | AX-170273554 | 0 | G | C | 0.254 | 9.72E-02 | 9.96E-02 | 2.41E+00 | 2.50E+00 | 3.41E-01 | no | Smyd2 | intron_variant | MODIFIER |
| 1 | 189903784 | AX-170099791 | 0 | A | G | 0.254 | 9.72E-02 | 9.96E-02 | 2.41E+00 | 2.50E+00 | 3.41E-01 | no | Smyd2 | intron_variant | MODIFIER |
| 1 | 189927912 | AX-170244829 | 0 | T | C | 0.254 | 9.72E-02 | 9.96E-02 | 2.41E+00 | 2.50E+00 | 3.41E-01 | no | | regulatory_region_variant | MODIFIER |
| 1 | 189948817 | AX-170313880 | 0 | T | A | 0.254 | 9.72E-02 | 9.96E-02 | 2.41E+00 | 2.50E+00 | 3.41E-01 | no | | intergenic_variant | MODIFIER |
| 1 | 189957359 | AX-170157986 | 0 | C | T | 0.254 | 9.72E-02 | 9.96E-02 | 2.41E+00 | 2.50E+00 | 3.41E-01 | no | | intergenic_variant | MODIFIER |
| 1 | 189974699 | AX-170114190 | 0 | T | C | 0.254 | 9.72E-02 | 9.96E-02 | 2.41E+00 | 2.50E+00 | 3.41E-01 | no | | intergenic_variant | MODIFIER |
| 1 | 189992453 | AX-170167696 | 0 | G | A | 0.254 | 9.72E-02 | 9.96E-02 | 2.41E+00 | 2.50E+00 | 3.41E-01 | no | | intergenic_variant | MODIFIER |
| 1 | 189993099 | AX-170486288 | 0 | A | G | 0.254 | 9.72E-02 | 9.96E-02 | 2.41E+00 | 2.50E+00 | 3.41E-01 | no | | intergenic_variant | MODIFIER |
| 7 | 73878952 | AX-170106711 | 0 | C | T | 0.36 | -8.79E-02 | 9.35E-02 | 2.80E+00 | 2.90E+00 | 3.42E-01 | no | | intergenic_variant | MODIFIER |
| 8 | 16489614 | AX-170406465 | 0 | C | T | 0.219 | -7.24E-02 | 7.61E-02 | 2.55E+00 | 2.64E+00 | 3.42E-01 | no | Csmd1 | intron_variant | MODIFIER |
| 1 | 190063238 | AX-170416299 | 1 | G | A | 0.259 | 9.61E-02 | 9.72E-02 | 2.36E+00 | 2.44E+00 | 3.42E-01 | no | | intergenic_variant | MODIFIER |
| 4 | 137954683 | AX-97792382 | 1 | G | T | 0.17 | 1.09E-01 | 1.15E-01 | 2.60E+00 | 2.65E+00 | 3.42E-01 | no | Ece1 | intron_variant | MODIFIER |
| 7 | 78103284 | AX-170029364 | 1 | T | C | 0.375 | -9.38E-02 | 9.97E-02 | 2.81E+00 | 2.91E+00 | 3.42E-01 | no | | intergenic_variant | MODIFIER |
| 4 | 55278284 | AX-170103332 | 1 | A | G | 0.375 | -9.85E-02 | 1.05E-01 | 2.81E+00 | 2.91E+00 | 3.42E-01 | no | Gm12512 | downstream_gene_variant | MODIFIER |
| 4 | 140552360 | AX-170253624 | 0 | C | A | 0.158 | 1.11E-01 | 1.17E-01 | 2.62E+00 | 2.67E+00 | 3.42E-01 | no | Arhgef10l | intron_variant | MODIFIER |
| 1 | 179994547 | AX-170198579 | 1 | A | T | 0.232 | 1.08E-01 | 1.13E-01 | 2.55E+00 | 2.62E+00 | 3.42E-01 | no | Cdc42bpa | intron_variant | MODIFIER |
| 8 | 41290633 | AX-170030561 | 0 | C | T | 0.158 | 1.20E-01 | 1.27E-01 | 2.91E+00 | 3.02E+00 | 3.42E-01 | no | Pcm1 | intron_variant | MODIFIER |
| 9 | 90488216 | AX-170083077 | 0 | A | C | 0.132 | -1.17E-01 | 1.24E-01 | 2.62E+00 | 2.69E+00 | 3.42E-01 | no | | intergenic_variant | MODIFIER |
| 9 | 90507142 | AX-97822146 | 0 | A | G | 0.132 | -1.17E-01 | 1.24E-01 | 2.62E+00 | 2.69E+00 | 3.42E-01 | no | | intergenic_variant | MODIFIER |
| 9 | 90507207 | AX-170209010 | 0 | A | G | 0.132 | -1.17E-01 | 1.24E-01 | 2.62E+00 | 2.69E+00 | 3.42E-01 | no | | intergenic_variant | MODIFIER |
| 9 | 90523854 | AX-170249641 | 0 | T | C | 0.132 | -1.17E-01 | 1.24E-01 | 2.62E+00 | 2.69E+00 | 3.42E-01 | no | | intergenic_variant | MODIFIER |
| 9 | 90524090 | AX-97829886 | 0 | T | G | 0.132 | -1.17E-01 | 1.24E-01 | 2.62E+00 | 2.69E+00 | 3.42E-01 | no | | intergenic_variant | MODIFIER |
| 9 | 90641003 | AX-170032967 | 0 | A | C | 0.132 | -1.17E-01 | 1.24E-01 | 2.62E+00 | 2.69E+00 | 3.42E-01 | no | | intergenic_variant | MODIFIER |
| 9 | 90658950 | AX-170032960 | 0 | T | C | 0.132 | -1.17E-01 | 1.24E-01 | 2.62E+00 | 2.69E+00 | 3.42E-01 | no | | intergenic_variant | MODIFIER |
| 9 | 90673429 | AX-170454076 | 0 | C | G | 0.132 | -1.17E-01 | 1.24E-01 | 2.62E+00 | 2.69E+00 | 3.42E-01 | no | | intergenic_variant | MODIFIER |
| 9 | 90677892 | AX-170032964 | 0 | G | A | 0.132 | -1.17E-01 | 1.24E-01 | 2.62E+00 | 2.69E+00 | 3.42E-01 | no | | intergenic_variant | MODIFIER |
| 9 | 90707340 | AX-170090369 | 0 | G | A | 0.132 | -1.17E-01 | 1.24E-01 | 2.62E+00 | 2.69E+00 | 3.42E-01 | no | | intergenic_variant | MODIFIER |
| 9 | 90715902 | AX-170215555 | 0 | C | T | 0.132 | -1.17E-01 | 1.24E-01 | 2.62E+00 | 2.69E+00 | 3.42E-01 | no | | intergenic_variant | MODIFIER |
| 9 | 90726419 | AX-170408105 | 0 | A | G | 0.132 | -1.17E-01 | 1.24E-01 | 2.62E+00 | 2.69E+00 | 3.42E-01 | no | | intergenic_variant | MODIFIER |
| 9 | 91093845 | AX-97801283 | 0 | T | G | 0.132 | -1.17E-01 | 1.24E-01 | 2.62E+00 | 2.69E+00 | 3.42E-01 | no | Gm5620 | upstream_gene_variant | MODIFIER |
| 9 | 91138599 | AX-170032975 | 0 | C | A | 0.132 | -1.17E-01 | 1.24E-01 | 2.62E+00 | 2.69E+00 | 3.42E-01 | no | | intergenic_variant | MODIFIER |
| 9 | 91138762 | AX-170032974 | 0 | A | G | 0.132 | -1.17E-01 | 1.24E-01 | 2.62E+00 | 2.69E+00 | 3.42E-01 | no | | intergenic_variant | MODIFIER |
| 9 | 91152180 | AX-170174017 | 0 | C | T | 0.132 | -1.17E-01 | 1.24E-01 | 2.62E+00 | 2.69E+00 | 3.42E-01 | no | | regulatory_region_variant | MODIFIER |
| 9 | 91171392 | AX-97790053 | 0 | G | A | 0.132 | -1.17E-01 | 1.24E-01 | 2.62E+00 | 2.69E+00 | 3.42E-01 | no | | intergenic_variant | MODIFIER |
| 9 | 91199876 | AX-170162845 | 0 | T | C | 0.132 | -1.17E-01 | 1.24E-01 | 2.62E+00 | 2.69E+00 | 3.42E-01 | no | | intergenic_variant | MODIFIER |
| 9 | 91335891 | AX-170032972 | 0 | A | G | 0.132 | -1.17E-01 | 1.24E-01 | 2.62E+00 | 2.69E+00 | 3.42E-01 | no | Gm28652 | upstream_gene_variant | MODIFIER |
| 9 | 91520226 | AX-170083092 | 0 | T | A | 0.132 | -1.17E-01 | 1.24E-01 | 2.62E+00 | 2.69E+00 | 3.42E-01 | no | Gm29478 | intron_variant,non_coding_transcript_variant | MODIFIER |
| 9 | 91672901 | AX-97834907 | 0 | C | A | 0.132 | -1.17E-01 | 1.24E-01 | 2.62E+00 | 2.69E+00 | 3.42E-01 | no | | intergenic_variant | MODIFIER |
| 9 | 91681258 | AX-170032981 | 0 | G | C | 0.132 | -1.17E-01 | 1.24E-01 | 2.62E+00 | 2.69E+00 | 3.42E-01 | no | | regulatory_region_variant | MODIFIER |
| 9 | 91730651 | AX-170032986 | 0 | A | G | 0.132 | -1.17E-01 | 1.24E-01 | 2.62E+00 | 2.69E+00 | 3.42E-01 | no | | intergenic_variant | MODIFIER |
| 9 | 91753684 | AX-97808713 | 0 | T | A | 0.132 | -1.17E-01 | 1.24E-01 | 2.62E+00 | 2.69E+00 | 3.42E-01 | no | Gm37987 | downstream_gene_variant | MODIFIER |
| 9 | 91768496 | AX-170032982 | 0 | T | C | 0.132 | -1.17E-01 | 1.24E-01 | 2.62E+00 | 2.69E+00 | 3.42E-01 | no | | intergenic_variant | MODIFIER |
| 9 | 91774308 | AX-170032987 | 0 | A | C | 0.132 | -1.17E-01 | 1.24E-01 | 2.62E+00 | 2.69E+00 | 3.42E-01 | no | | intergenic_variant | MODIFIER |
| 9 | 91804379 | AX-170032988 | 0 | T | C | 0.132 | -1.17E-01 | 1.24E-01 | 2.62E+00 | 2.69E+00 | 3.42E-01 | no | | intergenic_variant | MODIFIER |

| | | | | | | | | | | | | | | | |
|---|---|---|---|---|---|---|---|---|---|---|---|---|---|---|---|
| 9 | 91909732 | AX-170445578 | 0 | A | G | 0.132 | -1.17E-01 | 1.24E-01 | 2.62E+00 | 2.69E+00 | 3.42E-01 | no | | intergenic_variant | MODIFIER |
| 9 | 91937357 | AX-170139330 | 0 | G | T | 0.132 | -1.17E-01 | 1.24E-01 | 2.62E+00 | 2.69E+00 | 3.42E-01 | no | Gm31409 | intron_variant,non_coding_transcript_variant | MODIFIER |
| 9 | 91957316 | AX-170083093 | 0 | T | C | 0.132 | -1.17E-01 | 1.24E-01 | 2.62E+00 | 2.69E+00 | 3.42E-01 | no | Gm31409 | intron_variant,non_coding_transcript_variant | MODIFIER |
| 9 | 91981211 | AX-97819137 | 0 | A | G | 0.132 | -1.17E-01 | 1.24E-01 | 2.62E+00 | 2.69E+00 | 3.42E-01 | no | | intergenic_variant | MODIFIER |
| 9 | 92079417 | AX-170238779 | 0 | A | G | 0.132 | -1.17E-01 | 1.24E-01 | 2.62E+00 | 2.69E+00 | 3.42E-01 | no | | intergenic_variant | MODIFIER |
| 9 | 92095720 | AX-170256914 | 0 | A | G | 0.132 | -1.17E-01 | 1.24E-01 | 2.62E+00 | 2.69E+00 | 3.42E-01 | no | | intergenic_variant | MODIFIER |
| 9 | 92128776 | AX-170205390 | 0 | A | G | 0.132 | -1.17E-01 | 1.24E-01 | 2.62E+00 | 2.69E+00 | 3.42E-01 | no | | intergenic_variant | MODIFIER |
| 9 | 92165899 | AX-170504512 | 0 | T | C | 0.132 | -1.17E-01 | 1.24E-01 | 2.62E+00 | 2.69E+00 | 3.42E-01 | no | | intergenic_variant | MODIFIER |
| 9 | 92169617 | AX-170162237 | 0 | G | T | 0.132 | -1.17E-01 | 1.24E-01 | 2.62E+00 | 2.69E+00 | 3.42E-01 | no | | intergenic_variant | MODIFIER |
| 9 | 92182509 | AX-170032993 | 0 | A | G | 0.132 | -1.17E-01 | 1.24E-01 | 2.62E+00 | 2.69E+00 | 3.42E-01 | no | | intergenic_variant | MODIFIER |
| 9 | 92186699 | AX-170395602 | 0 | T | C | 0.132 | -1.17E-01 | 1.24E-01 | 2.62E+00 | 2.69E+00 | 3.42E-01 | no | | intergenic_variant | MODIFIER |
| 9 | 92237091 | AX-170472887 | 0 | G | A | 0.132 | -1.17E-01 | 1.24E-01 | 2.62E+00 | 2.69E+00 | 3.42E-01 | no | | intergenic_variant | MODIFIER |
| 9 | 92243335 | AX-170177783 | 0 | G | A | 0.132 | -1.17E-01 | 1.24E-01 | 2.62E+00 | 2.69E+00 | 3.42E-01 | no | | intergenic_variant | MODIFIER |
| 9 | 92260914 | AX-170240383 | 0 | G | T | 0.132 | -1.17E-01 | 1.24E-01 | 2.62E+00 | 2.69E+00 | 3.42E-01 | no | Plscr1 | intron_variant | MODIFIER |
| 9 | 92260985 | AX-170338463 | 0 | T | C | 0.132 | -1.17E-01 | 1.24E-01 | 2.62E+00 | 2.69E+00 | 3.42E-01 | no | Plscr1 | intron_variant | MODIFIER |
| 9 | 92261427 | AX-170032994 | 0 | G | C | 0.132 | -1.17E-01 | 1.24E-01 | 2.62E+00 | 2.69E+00 | 3.42E-01 | no | Plscr1 | intron_variant | MODIFIER |
| 9 | 92263018 | AX-170132354 | 0 | T | C | 0.132 | -1.17E-01 | 1.24E-01 | 2.62E+00 | 2.69E+00 | 3.42E-01 | no | Plscr1 | intron_variant | MODIFIER |
| 9 | 92281908 | AX-170083095 | 0 | A | C | 0.132 | -1.17E-01 | 1.24E-01 | 2.62E+00 | 2.69E+00 | 3.42E-01 | no | Plscr2 | intron_variant | MODIFIER |
| 9 | 92338763 | AX-170390613 | 0 | A | G | 0.132 | -1.17E-01 | 1.24E-01 | 2.62E+00 | 2.69E+00 | 3.42E-01 | no | 1700057G04Rik | intron_variant | MODIFIER |
| 9 | 92382324 | AX-170269039 | 0 | T | C | 0.132 | -1.17E-01 | 1.24E-01 | 2.62E+00 | 2.69E+00 | 3.42E-01 | no | | intergenic_variant | MODIFIER |
| 9 | 92384080 | AX-170139505 | 0 | A | C | 0.132 | -1.17E-01 | 1.24E-01 | 2.62E+00 | 2.69E+00 | 3.42E-01 | no | | regulatory_region_variant | MODIFIER |
| 9 | 92413234 | AX-170112304 | 0 | G | A | 0.132 | -1.17E-01 | 1.24E-01 | 2.62E+00 | 2.69E+00 | 3.42E-01 | no | | intergenic_variant | MODIFIER |
| 9 | 92444426 | AX-170473424 | 0 | G | A | 0.132 | -1.17E-01 | 1.24E-01 | 2.62E+00 | 2.69E+00 | 3.42E-01 | no | | intergenic_variant | MODIFIER |
| 9 | 92465318 | AX-170381178 | 0 | C | T | 0.132 | -1.17E-01 | 1.24E-01 | 2.62E+00 | 2.69E+00 | 3.42E-01 | no | Plscr4 | intron_variant | MODIFIER |
| 9 | 92465368 | AX-170106985 | 0 | G | A | 0.132 | -1.17E-01 | 1.24E-01 | 2.62E+00 | 2.69E+00 | 3.42E-01 | no | Plscr4 | intron_variant | MODIFIER |
| 9 | 92476650 | AX-170450937 | 0 | C | T | 0.132 | -1.17E-01 | 1.24E-01 | 2.62E+00 | 2.69E+00 | 3.42E-01 | no | Plscr4 | intron_variant | MODIFIER |
| 9 | 92490603 | AX-170129095 | 0 | A | G | 0.132 | -1.17E-01 | 1.24E-01 | 2.62E+00 | 2.69E+00 | 3.42E-01 | no | Plscr4 | intron_variant | MODIFIER |
| 9 | 92496399 | AX-170307689 | 0 | A | C | 0.132 | -1.17E-01 | 1.24E-01 | 2.62E+00 | 2.69E+00 | 3.42E-01 | no | Plscr4 | downstream_gene_variant | MODIFIER |
| 9 | 92511972 | AX-97793753 | 0 | G | A | 0.132 | -1.17E-01 | 1.24E-01 | 2.62E+00 | 2.69E+00 | 3.42E-01 | no | | intergenic_variant | MODIFIER |
| 10 | 92072038 | AX-170322613 | 0 | C | T | 0.298 | -8.10E-02 | 8.45E-02 | 2.51E+00 | 2.56E+00 | 3.42E-01 | no | Mir135a-2 | downstream_gene_variant | MODIFIER |
| 16 | 80727025 | AX-170383714 | 1 | A | G | 0.464 | 7.15E-02 | 7.56E-02 | 2.60E+00 | 2.67E+00 | 3.42E-01 | no | Gm19867 | non_coding_transcript_exon_variant | MODIFIER |
| 8 | 25161836 | AX-170090064 | 0 | A | T | 0.351 | -6.19E-02 | 6.55E-02 | 2.59E+00 | 2.68E+00 | 3.42E-01 | no | Tacc1 | intron_variant | MODIFIER |
| 1 | 190913438 | AX-170372448 | 1 | T | C | 0.25 | 9.38E-02 | 9.62E-02 | 2.41E+00 | 2.49E+00 | 3.43E-01 | no | Rps6kc1 | upstream_gene_variant | MODIFIER |
| 13 | 25551343 | AX-170227719 | 1 | T | C | 0.161 | -1.19E-01 | 1.22E-01 | 2.40E+00 | 2.45E+00 | 3.43E-01 | no | | intergenic_variant | MODIFIER |
| 13 | 25551494 | AX-170153834 | 1 | C | T | 0.161 | -1.19E-01 | 1.22E-01 | 2.40E+00 | 2.45E+00 | 3.43E-01 | no | | intergenic_variant | MODIFIER |
| 3 | 24630028 | AX-170518742 | 1 | A | T | 0.152 | -9.79E-02 | 1.04E-01 | 2.60E+00 | 2.67E+00 | 3.43E-01 | no | Naaladl2 | intron_variant | MODIFIER |
| 7 | 70600786 | AX-170337183 | 0 | A | G | 0.377 | -9.07E-02 | 9.62E-02 | 2.90E+00 | 3.04E+00 | 3.43E-01 | no | | intergenic_variant | MODIFIER |
| 2 | 120204893 | AX-170178249 | 0 | A | G | 0.14 | -1.75E-01 | 1.85E-01 | 2.92E+00 | 3.02E+00 | 3.43E-01 | Yes | Pla2g4e | intron_variant | MODIFIER |
| 13 | 37393428 | AX-170050986 | 0 | A | G | 0.175 | -9.80E-02 | 1.04E-01 | 2.59E+00 | 2.69E+00 | 3.43E-01 | no | Ly86 | intron_variant | MODIFIER |
| 13 | 38118993 | AX-170051007 | 0 | C | T | 0.175 | -9.80E-02 | 1.04E-01 | 2.59E+00 | 2.69E+00 | 3.43E-01 | no | | intergenic_variant | MODIFIER |
| 1 | 187635404 | AX-170111821 | 1 | T | A | 0.25 | 1.02E-01 | 1.04E-01 | 2.37E+00 | 2.45E+00 | 3.43E-01 | no | Esrrg | intron_variant | MODIFIER |
| 16 | 71769191 | AX-97790931 | 1 | G | A | 0.491 | 7.31E-02 | 7.70E-02 | 2.55E+00 | 2.62E+00 | 3.43E-01 | no | | intergenic_variant | MODIFIER |
| 15 | 84582523 | AX-170139334 | 1 | C | T | 0.098 | -1.21E-01 | 1.03E-01 | 1.94E+00 | 2.03E+00 | 3.43E-01 | no | | intergenic_variant | MODIFIER |
| 13 | 19813353 | AX-97783802 | 0 | G | T | 0.184 | -1.11E-01 | 1.15E-01 | 2.47E+00 | 2.55E+00 | 3.43E-01 | no | Gpr141 | intron_variant | MODIFIER |
| 2 | 146726455 | AX-170065191 | 0 | A | G | 0.14 | -1.77E-01 | 1.88E-01 | 2.86E+00 | 2.94E+00 | 3.43E-01 | Yes | | intergenic_variant | MODIFIER |
| 3 | 36770559 | AX-170193533 | 1 | G | A | 0.196 | 8.87E-02 | 9.42E-02 | 2.86E+00 | 2.95E+00 | 3.43E-01 | no | | intergenic_variant | MODIFIER |
| 6 | 135940983 | AX-170388693 | 0 | A | T | 0.588 | 8.57E-02 | 8.83E-02 | 2.43E+00 | 2.48E+00 | 3.43E-01 | no | Grin2b | intron_variant | MODIFIER |
| 4 | 132713789 | AX-170023977 | 0 | A | T | 0.167 | 1.07E-01 | 1.13E-01 | 2.52E+00 | 2.57E+00 | 3.43E-01 | no | Eya3 | intron_variant | MODIFIER |
| 4 | 133649850 | AX-170023989 | 0 | G | A | 0.167 | 1.07E-01 | 1.13E-01 | 2.52E+00 | 2.57E+00 | 3.43E-01 | no | Zdhhc18 | intron_variant,non_coding_transcript_variant | MODIFIER |
| 2 | 161895605 | AX-170337479 | 1 | C | T | 0.161 | 1.83E-01 | 1.95E-01 | 2.79E+00 | 2.88E+00 | 3.43E-01 | Yes | Ptprt | intron_variant | MODIFIER |
| 10 | 69458589 | AX-170500796 | 1 | C | A | 0.402 | -7.36E-02 | 7.84E-02 | 2.66E+00 | 2.74E+00 | 3.43E-01 | no | Ank3 | intron_variant,non_coding_transcript_variant | MODIFIER |
| 1 | 187279610 | AX-170364702 | 0 | A | C | 0.237 | 9.50E-02 | 9.84E-02 | 2.45E+00 | 2.53E+00 | 3.44E-01 | no | Gpatch2 | intron_variant | MODIFIER |
| 1 | 121552559 | AX-170202020 | 1 | T | C | 0.58 | 7.93E-02 | 8.30E-02 | 2.50E+00 | 2.56E+00 | 3.44E-01 | no | Ddx18 | downstream_gene_variant | MODIFIER |
| 14 | 93451834 | AX-170214401 | 0 | C | T | 0.044 | 1.61E-01 | 1.72E-01 | 2.76E+00 | 2.91E+00 | 3.44E-01 | Yes | Pcdh9 | intron_variant | MODIFIER |

| | | | | | | | | | | | | | | | |
|---|---|---|---|---|---|---|---|---|---|---|---|---|---|---|---|
| 8 | 15628653 | AX-170503265 | 1 | G | T | 0.232 | -7.10E-02 | 7.51E-02 | 2.56E+00 | 2.66E+00 | 3.44E-01 | no | Gm45408 | intron_variant,non_coding_transcript_variant | MODIFIER |
| 1 | 169053013 | AX-170112220 | 2 | G | A | 0.218 | 1.31E-01 | 1.39E-01 | 2.64E+00 | 2.69E+00 | 3.44E-01 | no | | intergenic_variant | MODIFIER |
| 15 | 67404554 | AX-170386774 | 1 | G | A | 0.339 | -7.69E-02 | 7.70E-02 | 2.32E+00 | 2.39E+00 | 3.44E-01 | no | | intergenic_variant | MODIFIER |
| 6 | 126797056 | AX-170246501 | 1 | G | A | 0.536 | 7.86E-02 | 8.21E-02 | 2.49E+00 | 2.55E+00 | 3.44E-01 | no | | intergenic_variant | MODIFIER |
| 13 | 32955831 | AX-170008058 | 1 | T | G | 0.17 | -1.09E-01 | 1.16E-01 | 2.57E+00 | 2.65E+00 | 3.44E-01 | no | | intergenic_variant | MODIFIER |
| 5 | 51088177 | AX-170490831 | 1 | C | T | 0.509 | 9.95E-02 | 1.02E-01 | 2.41E+00 | 2.47E+00 | 3.44E-01 | no | | intergenic_variant | MODIFIER |
| 16 | 30056931 | AX-170012721 | 0 | T | A | 0.509 | 7.30E-02 | 7.73E-02 | 2.58E+00 | 2.64E+00 | 3.44E-01 | no | | intergenic_variant | MODIFIER |
| 9 | 17423883 | AX-170234319 | 0 | G | A | 0.474 | 8.10E-02 | 8.52E-02 | 2.52E+00 | 2.59E+00 | 3.44E-01 | no | | intergenic_variant | MODIFIER |
| 2 | 36128488 | AX-97837764 | 2 | T | C | 0.045 | 1.85E-01 | 1.97E-01 | 2.87E+00 | 3.01E+00 | 3.44E-01 | Yes | Rbm18 | intron_variant | MODIFIER |
| 18 | 54058291 | AX-170519450 | 1 | G | A | 0.455 | 9.25E-02 | 9.32E-02 | 2.34E+00 | 2.40E+00 | 3.44E-01 | no | | regulatory_region_variant | MODIFIER |
| 18 | 55162051 | AX-170088345 | 1 | G | C | 0.455 | 9.25E-02 | 9.32E-02 | 2.34E+00 | 2.40E+00 | 3.44E-01 | no | | intergenic_variant | MODIFIER |
| 18 | 55546792 | AX-170105820 | 1 | G | A | 0.455 | 9.25E-02 | 9.32E-02 | 2.34E+00 | 2.40E+00 | 3.44E-01 | no | | regulatory_region_variant | MODIFIER |
| 4 | 129791891 | AX-97833803 | 0 | G | T | 0.175 | 1.03E-01 | 1.08E-01 | 2.56E+00 | 2.61E+00 | 3.44E-01 | no | | regulatory_region_variant | MODIFIER |
| 4 | 132623795 | AX-170070720 | 0 | C | T | 0.175 | 1.03E-01 | 1.08E-01 | 2.56E+00 | 2.61E+00 | 3.44E-01 | no | | regulatory_region_variant | MODIFIER |
| 18 | 69013194 | AX-170355452 | 0 | C | A | 0.167 | -1.17E-01 | 1.24E-01 | 2.61E+00 | 2.66E+00 | 3.44E-01 | no | | intergenic_variant | MODIFIER |
| 18 | 69016727 | AX-170512918 | 0 | G | A | 0.167 | -1.17E-01 | 1.24E-01 | 2.61E+00 | 2.66E+00 | 3.44E-01 | no | | regulatory_region_variant | MODIFIER |
| 18 | 69019636 | AX-170236587 | 0 | C | G | 0.167 | -1.17E-01 | 1.24E-01 | 2.61E+00 | 2.66E+00 | 3.44E-01 | no | Mir145b | downstream_gene_variant | MODIFIER |
| 18 | 69019688 | AX-170445638 | 0 | G | A | 0.167 | -1.17E-01 | 1.24E-01 | 2.61E+00 | 2.66E+00 | 3.44E-01 | no | Mir145b | downstream_gene_variant | MODIFIER |
| 18 | 69030662 | AX-170215195 | 0 | A | T | 0.167 | -1.17E-01 | 1.24E-01 | 2.61E+00 | 2.66E+00 | 3.44E-01 | no | | intergenic_variant | MODIFIER |
| 18 | 69071666 | AX-170331904 | 0 | A | G | 0.167 | -1.17E-01 | 1.24E-01 | 2.61E+00 | 2.66E+00 | 3.44E-01 | no | | intergenic_variant | MODIFIER |
| 18 | 69071943 | AX-170213458 | 0 | A | G | 0.167 | -1.17E-01 | 1.24E-01 | 2.61E+00 | 2.66E+00 | 3.44E-01 | no | | regulatory_region_variant | MODIFIER |
| 18 | 69078530 | AX-170514887 | 0 | T | C | 0.167 | -1.17E-01 | 1.24E-01 | 2.61E+00 | 2.66E+00 | 3.44E-01 | no | | intergenic_variant | MODIFIER |
| 18 | 69086952 | AX-170153317 | 0 | T | C | 0.167 | -1.17E-01 | 1.24E-01 | 2.61E+00 | 2.66E+00 | 3.44E-01 | no | | regulatory_region_variant | MODIFIER |
| 18 | 69168481 | AX-170061216 | 0 | T | C | 0.167 | -1.17E-01 | 1.24E-01 | 2.61E+00 | 2.66E+00 | 3.44E-01 | no | | regulatory_region_variant | MODIFIER |
| 18 | 69190508 | AX-170445062 | 0 | A | C | 0.167 | -1.17E-01 | 1.24E-01 | 2.61E+00 | 2.66E+00 | 3.44E-01 | no | Gm31294 | downstream_gene_variant | MODIFIER |
| 18 | 69220613 | AX-170154411 | 0 | T | C | 0.167 | -1.17E-01 | 1.24E-01 | 2.61E+00 | 2.66E+00 | 3.44E-01 | no | AC121263.2 | non_coding_transcript_exon_variant | MODIFIER |
| 18 | 69245597 | AX-170215935 | 0 | G | A | 0.167 | -1.17E-01 | 1.24E-01 | 2.61E+00 | 2.66E+00 | 3.44E-01 | no | | intergenic_variant | MODIFIER |
| 18 | 69246445 | AX-170500990 | 0 | G | A | 0.167 | -1.17E-01 | 1.24E-01 | 2.61E+00 | 2.66E+00 | 3.44E-01 | no | | intergenic_variant | MODIFIER |
| 18 | 69306578 | AX-170131449 | 0 | A | T | 0.167 | -1.17E-01 | 1.24E-01 | 2.61E+00 | 2.66E+00 | 3.44E-01 | no | | intergenic_variant | MODIFIER |
| 18 | 69395664 | AX-170088370 | 0 | T | C | 0.167 | -1.17E-01 | 1.24E-01 | 2.61E+00 | 2.66E+00 | 3.44E-01 | no | Tcf4 | intron_variant | MODIFIER |
| 18 | 69403772 | AX-170369344 | 0 | G | A | 0.167 | -1.17E-01 | 1.24E-01 | 2.61E+00 | 2.66E+00 | 3.44E-01 | no | Tcf4 | intron_variant | MODIFIER |
| 18 | 69403803 | AX-170485371 | 0 | G | C | 0.167 | -1.17E-01 | 1.24E-01 | 2.61E+00 | 2.66E+00 | 3.44E-01 | no | Tcf4 | intron_variant | MODIFIER |
| 18 | 69409335 | AX-170389464 | 0 | G | C | 0.167 | -1.17E-01 | 1.24E-01 | 2.61E+00 | 2.66E+00 | 3.44E-01 | no | Tcf4 | intron_variant | MODIFIER |
| 18 | 69413407 | AX-170468383 | 0 | G | T | 0.167 | -1.17E-01 | 1.24E-01 | 2.61E+00 | 2.66E+00 | 3.44E-01 | no | Tcf4 | intron_variant | MODIFIER |
| 18 | 69439162 | AX-97797107 | 0 | C | T | 0.167 | -1.17E-01 | 1.24E-01 | 2.61E+00 | 2.66E+00 | 3.44E-01 | no | Tcf4 | intron_variant | MODIFIER |
| 18 | 69470125 | AX-170061223 | 0 | A | G | 0.167 | -1.17E-01 | 1.24E-01 | 2.61E+00 | 2.66E+00 | 3.44E-01 | no | Tcf4 | intron_variant | MODIFIER |
| 18 | 69475486 | AX-97849478 | 0 | A | G | 0.167 | -1.17E-01 | 1.24E-01 | 2.61E+00 | 2.66E+00 | 3.44E-01 | no | Tcf4 | intron_variant | MODIFIER |
| 18 | 69476336 | AX-170167936 | 0 | A | G | 0.167 | -1.17E-01 | 1.24E-01 | 2.61E+00 | 2.66E+00 | 3.44E-01 | no | Tcf4 | intron_variant | MODIFIER |
| 18 | 69477627 | AX-170221509 | 0 | C | T | 0.167 | -1.17E-01 | 1.24E-01 | 2.61E+00 | 2.66E+00 | 3.44E-01 | no | Tcf4 | intron_variant | MODIFIER |
| 18 | 69478797 | AX-170400620 | 0 | A | G | 0.167 | -1.17E-01 | 1.24E-01 | 2.61E+00 | 2.66E+00 | 3.44E-01 | no | Tcf4 | intron_variant | MODIFIER |
| 18 | 69774093 | AX-170370733 | 0 | T | C | 0.167 | -1.17E-01 | 1.24E-01 | 2.61E+00 | 2.66E+00 | 3.44E-01 | no | Gm22508 | downstream_gene_variant | MODIFIER |
| 18 | 69775189 | AX-170514885 | 0 | A | G | 0.167 | -1.17E-01 | 1.24E-01 | 2.61E+00 | 2.66E+00 | 3.44E-01 | no | Gm22508 | downstream_gene_variant | MODIFIER |
| 18 | 69812557 | AX-170508364 | 0 | T | C | 0.167 | -1.17E-01 | 1.24E-01 | 2.61E+00 | 2.66E+00 | 3.44E-01 | no | | intergenic_variant | MODIFIER |
| 18 | 69831402 | AX-97846241 | 0 | T | C | 0.167 | -1.17E-01 | 1.24E-01 | 2.61E+00 | 2.66E+00 | 3.44E-01 | no | | intergenic_variant | MODIFIER |
| 18 | 69856151 | AX-97823079 | 0 | A | C | 0.167 | -1.17E-01 | 1.24E-01 | 2.61E+00 | 2.66E+00 | 3.44E-01 | no | | intergenic_variant | MODIFIER |
| 5 | 49762317 | AX-170408414 | 0 | T | G | 0.474 | 1.00E-01 | 1.02E-01 | 2.36E+00 | 2.42E+00 | 3.45E-01 | no | | intergenic_variant | MODIFIER |
| 5 | 50678001 | AX-170072060 | 0 | A | G | 0.474 | 1.00E-01 | 1.02E-01 | 2.36E+00 | 2.42E+00 | 3.45E-01 | no | Gm43016 | intron_variant,non_coding_transcript_variant | MODIFIER |
| 13 | 24538099 | AX-170087387 | 0 | C | T | 0.167 | -1.04E-01 | 1.08E-01 | 2.42E+00 | 2.51E+00 | 3.45E-01 | no | | regulatory_region_variant | MODIFIER |
| 5 | 5316772 | AX-170208867 | 1 | C | T | 0.438 | -6.23E-02 | 6.53E-02 | 2.48E+00 | 2.59E+00 | 3.45E-01 | no | Cdk14 | intron_variant | MODIFIER |
| 16 | 32540701 | AX-170301246 | 1 | C | T | 0.5 | 7.63E-02 | 8.08E-02 | 2.57E+00 | 2.62E+00 | 3.45E-01 | no | Zdhhc19 | intron_variant,non_coding_transcript_variant | MODIFIER |
| 16 | 56057455 | AX-170013109 | 0 | G | T | 0.474 | -7.93E-02 | 8.33E-02 | 2.52E+00 | 2.57E+00 | 3.45E-01 | no | Senp7 | intron_variant,non_coding_transcript_variant | MODIFIER |
| 5 | 3941195 | AX-170409280 | 0 | C | T | 0.439 | -6.23E-02 | 6.53E-02 | 2.48E+00 | 2.59E+00 | 3.45E-01 | no | Akap9 | intron_variant | MODIFIER |
| 5 | 3950228 | AX-170071168 | 0 | C | A | 0.439 | -6.23E-02 | 6.53E-02 | 2.48E+00 | 2.59E+00 | 3.45E-01 | no | Akap9 | intron_variant | MODIFIER |
| 5 | 3989136 | AX-170336123 | 0 | C | T | 0.439 | -6.23E-02 | 6.53E-02 | 2.48E+00 | 2.59E+00 | 3.45E-01 | no | Akap9 | intron_variant | MODIFIER |

| | | | | | | | | | | | | | | | |
|---|---|---|---|---|---|---|---|---|---|---|---|---|---|---|---|
| 5 | 4003922 | AX-170487653 | 0 | G | A | 0.439 | -6.23E-02 | 6.53E-02 | 2.48E+00 | 2.59E+00 | 3.45E-01 | no | Akap9 | intron_variant | MODIFIER |
| 5 | 4011623 | AX-170301262 | 0 | G | C | 0.439 | -6.23E-02 | 6.53E-02 | 2.48E+00 | 2.59E+00 | 3.45E-01 | no | Akap9 | intron_variant | MODIFIER |
| 5 | 4013806 | AX-170409014 | 0 | A | C | 0.439 | -6.23E-02 | 6.53E-02 | 2.48E+00 | 2.59E+00 | 3.45E-01 | no | Akap9 | intron_variant | MODIFIER |
| 5 | 4031192 | AX-170241420 | 0 | G | A | 0.439 | -6.23E-02 | 6.53E-02 | 2.48E+00 | 2.59E+00 | 3.45E-01 | no | Akap9 | intron_variant | MODIFIER |
| 5 | 4034294 | AX-170356232 | 0 | A | G | 0.439 | -6.23E-02 | 6.53E-02 | 2.48E+00 | 2.59E+00 | 3.45E-01 | no | Akap9 | intron_variant | MODIFIER |
| 5 | 4069159 | AX-170106996 | 0 | C | T | 0.439 | -6.23E-02 | 6.53E-02 | 2.48E+00 | 2.59E+00 | 3.45E-01 | no | Akap9 | synonymous_variant | LOW |
| 5 | 4426592 | AX-170035705 | 0 | A | T | 0.439 | -6.23E-02 | 6.53E-02 | 2.48E+00 | 2.59E+00 | 3.45E-01 | no | | intergenic_variant | MODIFIER |
| 5 | 4944497 | AX-170192310 | 0 | G | A | 0.439 | -6.23E-02 | 6.53E-02 | 2.48E+00 | 2.59E+00 | 3.45E-01 | no | Cdk14 | intron_variant | MODIFIER |
| 5 | 5034059 | AX-170127477 | 0 | T | G | 0.439 | -6.23E-02 | 6.53E-02 | 2.48E+00 | 2.59E+00 | 3.45E-01 | no | Cdk14 | intron_variant | MODIFIER |
| 5 | 5056511 | AX-170314038 | 0 | C | T | 0.439 | -6.23E-02 | 6.53E-02 | 2.48E+00 | 2.59E+00 | 3.45E-01 | no | Cdk14 | intron_variant | MODIFIER |
| 5 | 5097369 | AX-170134609 | 0 | A | T | 0.439 | -6.23E-02 | 6.53E-02 | 2.48E+00 | 2.59E+00 | 3.45E-01 | no | Cdk14 | intron_variant | MODIFIER |
| 5 | 5152040 | AX-170170379 | 0 | G | A | 0.439 | -6.23E-02 | 6.53E-02 | 2.48E+00 | 2.59E+00 | 3.45E-01 | no | Cdk14 | intron_variant | MODIFIER |
| 5 | 5153983 | AX-170499539 | 0 | G | C | 0.439 | -6.23E-02 | 6.53E-02 | 2.48E+00 | 2.59E+00 | 3.45E-01 | no | Cdk14 | intron_variant | MODIFIER |
| 5 | 5158410 | AX-170287500 | 0 | T | C | 0.439 | -6.23E-02 | 6.53E-02 | 2.48E+00 | 2.59E+00 | 3.45E-01 | no | Cdk14 | intron_variant | MODIFIER |
| 5 | 5158630 | AX-170442916 | 0 | A | G | 0.439 | -6.23E-02 | 6.53E-02 | 2.48E+00 | 2.59E+00 | 3.45E-01 | no | Cdk14 | intron_variant | MODIFIER |
| 5 | 5196004 | AX-170185678 | 0 | G | C | 0.439 | -6.23E-02 | 6.53E-02 | 2.48E+00 | 2.59E+00 | 3.45E-01 | no | Cdk14 | intron_variant | MODIFIER |
| 5 | 5197943 | AX-170414102 | 0 | T | G | 0.439 | -6.23E-02 | 6.53E-02 | 2.48E+00 | 2.59E+00 | 3.45E-01 | no | Cdk14 | intron_variant | MODIFIER |
| 5 | 5255830 | AX-170534185 | 0 | C | T | 0.439 | -6.23E-02 | 6.53E-02 | 2.48E+00 | 2.59E+00 | 3.45E-01 | no | Cdk14 | intron_variant | MODIFIER |
| 5 | 5259539 | AX-170121164 | 0 | G | C | 0.439 | -6.23E-02 | 6.53E-02 | 2.48E+00 | 2.59E+00 | 3.45E-01 | no | Cdk14 | intron_variant | MODIFIER |
| 5 | 5266128 | AX-170105348 | 0 | G | A | 0.439 | -6.23E-02 | 6.53E-02 | 2.48E+00 | 2.59E+00 | 3.45E-01 | no | Cdk14 | intron_variant | MODIFIER |
| 5 | 5269035 | AX-170248953 | 0 | C | G | 0.439 | -6.23E-02 | 6.53E-02 | 2.48E+00 | 2.59E+00 | 3.45E-01 | no | Cdk14 | intron_variant | MODIFIER |
| 5 | 5307851 | AX-97840335 | 0 | G | A | 0.439 | -6.23E-02 | 6.53E-02 | 2.48E+00 | 2.59E+00 | 3.45E-01 | no | Cdk14 | intron_variant | MODIFIER |
| 5 | 5362888 | AX-170136712 | 0 | G | C | 0.439 | -6.23E-02 | 6.53E-02 | 2.48E+00 | 2.59E+00 | 3.45E-01 | no | Cdk14 | intron_variant | MODIFIER |
| 5 | 5363702 | AX-170450333 | 0 | A | G | 0.439 | -6.23E-02 | 6.53E-02 | 2.48E+00 | 2.59E+00 | 3.45E-01 | no | Cdk14 | intron_variant | MODIFIER |
| 5 | 5364241 | AX-170129514 | 0 | A | G | 0.439 | -6.23E-02 | 6.53E-02 | 2.48E+00 | 2.59E+00 | 3.45E-01 | no | Cdk14 | intron_variant | MODIFIER |
| 5 | 5365277 | AX-170145858 | 0 | G | A | 0.439 | -6.23E-02 | 6.53E-02 | 2.48E+00 | 2.59E+00 | 3.45E-01 | no | Cdk14 | intron_variant | MODIFIER |
| 5 | 89288461 | AX-170095393 | 1 | T | C | 0.446 | 9.25E-02 | 9.88E-02 | 2.85E+00 | 2.94E+00 | 3.45E-01 | no | | intergenic_variant | MODIFIER |
| 10 | 99678204 | AX-170045721 | 0 | C | T | 0.272 | -8.05E-02 | 8.53E-02 | 2.57E+00 | 2.64E+00 | 3.45E-01 | no | Gm35101 | downstream_gene_variant | MODIFIER |
| 5 | 43846339 | AX-170405372 | 1 | A | G | 0.179 | -1.03E-01 | 1.10E-01 | 2.61E+00 | 2.70E+00 | 3.45E-01 | no | Bst1 | downstream_gene_variant | MODIFIER |
| 7 | 5913690 | AX-97809325 | 2 | C | A | 0.182 | 7.42E-02 | 7.96E-02 | 2.81E+00 | 2.95E+00 | 3.45E-01 | no | | intergenic_variant | MODIFIER |
| 16 | 97175760 | AX-170377460 | 0 | A | G | 0.465 | 7.10E-02 | 7.54E-02 | 2.93E+00 | 3.07E+00 | 3.45E-01 | no | | intergenic_variant | MODIFIER |
| 13 | 7149550 | AX-97802527 | 1 | T | C | 0.205 | -1.14E-01 | 1.21E-01 | 2.57E+00 | 2.63E+00 | 3.45E-01 | no | Gm40663 | downstream_gene_variant | MODIFIER |
| 14 | 123276988 | AX-170328760 | 0 | A | G | 0.298 | 7.40E-02 | 7.87E-02 | 2.58E+00 | 2.69E+00 | 3.45E-01 | no | Nalcn | 3_prime_UTR_variant | MODIFIER |
| 14 | 92416202 | AX-97827591 | 1 | A | G | 0.054 | -1.31E-01 | 1.38E-01 | 2.52E+00 | 2.63E+00 | 3.46E-01 | no | | intergenic_variant | MODIFIER |
| 18 | 60360563 | AX-97783064 | 1 | A | G | 0.402 | 9.02E-02 | 9.39E-02 | 2.46E+00 | 2.50E+00 | 3.46E-01 | no | AC151990.1 | upstream_gene_variant | MODIFIER |
| 4 | 133390344 | AX-170469218 | 0 | G | C | 0.167 | 1.05E-01 | 1.11E-01 | 2.54E+00 | 2.59E+00 | 3.46E-01 | no | Slc9a1 | intron_variant | MODIFIER |
| 10 | 76712846 | AX-170470880 | 0 | G | T | 0.439 | -7.32E-02 | 7.86E-02 | 2.71E+00 | 2.80E+00 | 3.46E-01 | no | Col6a1 | intron_variant | MODIFIER |
| 6 | 127729644 | AX-170349930 | 0 | G | A | 0.535 | 7.88E-02 | 8.25E-02 | 2.48E+00 | 2.54E+00 | 3.46E-01 | no | Prmt8 | intron_variant | MODIFIER |
| 12 | 79151974 | AX-170150318 | 1 | T | C | 0.054 | 1.89E-01 | 2.02E-01 | 2.88E+00 | 3.00E+00 | 3.46E-01 | Yes | Arg2 | synonymous_variant | LOW |
| 5 | 78633251 | AX-97822529 | 0 | C | A | 0.43 | 9.44E-02 | 1.01E-01 | 2.72E+00 | 2.78E+00 | 3.46E-01 | no | | intergenic_variant | MODIFIER |
| 1 | 128318803 | AX-170449559 | 1 | T | A | 0.545 | 7.89E-02 | 8.43E-02 | 2.62E+00 | 2.68E+00 | 3.46E-01 | no | Lct | intron_variant | MODIFIER |
| 7 | 24490566 | AX-170414917 | 0 | T | C | 0.421 | 8.39E-02 | 8.92E-02 | 2.55E+00 | 2.66E+00 | 3.46E-01 | no | Cadm4 | intron_variant | MODIFIER |
| 1 | 21556327 | AX-170501133 | 1 | T | A | 0.411 | 7.60E-02 | 8.02E-02 | 2.52E+00 | 2.59E+00 | 3.46E-01 | no | Kcnq5 | intron_variant | MODIFIER |
| 5 | 82289197 | AX-170025423 | 0 | A | T | 0.439 | 1.02E-01 | 1.09E-01 | 2.79E+00 | 2.84E+00 | 3.46E-01 | no | | intergenic_variant | MODIFIER |
| 7 | 57668031 | AX-170176076 | 0 | T | G | 0.43 | 8.91E-02 | 9.55E-02 | 2.65E+00 | 2.74E+00 | 3.46E-01 | no | Gabrb3 | intron_variant | MODIFIER |
| 8 | 109442552 | AX-170430655 | 0 | G | T | 0.246 | 8.69E-02 | 9.36E-02 | 2.79E+00 | 2.92E+00 | 3.47E-01 | no | | intergenic_variant | MODIFIER |
| 16 | 39812293 | AX-170100577 | 0 | A | G | 0.491 | 7.30E-02 | 7.79E-02 | 2.60E+00 | 2.66E+00 | 3.47E-01 | no | | intergenic_variant | MODIFIER |
| 9 | 104207633 | AX-170033177 | 1 | C | A | 0.125 | -1.10E-01 | 1.18E-01 | 2.62E+00 | 2.70E+00 | 3.47E-01 | no | Dnajc13 | intron_variant | MODIFIER |
| 5 | 71111887 | AX-170072427 | 0 | T | G | 0.447 | 8.74E-02 | 9.42E-02 | 2.70E+00 | 2.81E+00 | 3.47E-01 | no | | intergenic_variant | MODIFIER |
| 5 | 89804366 | AX-170442207 | 1 | A | T | 0.438 | 1.08E-01 | 1.15E-01 | 2.91E+00 | 2.97E+00 | 3.48E-01 | no | Adamts3 | intron_variant | MODIFIER |
| 13 | 19370215 | AX-97792550 | 0 | G | A | 0.175 | -1.12E-01 | 1.17E-01 | 2.46E+00 | 2.53E+00 | 3.48E-01 | no | Stard3nl | intron_variant | MODIFIER |
| 4 | 129611885 | AX-170095148 | 1 | A | G | 0.152 | 1.10E-01 | 1.18E-01 | 2.58E+00 | 2.63E+00 | 3.48E-01 | no | Iqcc | downstream_gene_variant | MODIFIER |
| 15 | 9205948 | AX-170010996 | 1 | T | C | 0.241 | -1.00E-01 | 1.07E-01 | 2.64E+00 | 2.69E+00 | 3.48E-01 | no | Lmbrd2 | downstream_gene_variant | MODIFIER |
| 1 | 186207177 | AX-170285880 | 1 | A | G | 0.223 | 1.06E-01 | 1.09E-01 | 2.41E+00 | 2.48E+00 | 3.48E-01 | no | Gm37134 | intron_variant,non_coding_transcript_variant | MODIFIER |

| | | | | | | | | | | | | | | | |
|---|---|---|---|---|---|---|---|---|---|---|---|---|---|---|---|
| 4 | 127326192 | AX-170356307 | 0 | C | A | 0.158 | 1.08E-01 | 1.15E-01 | 2.56E+00 | 2.63E+00 | 3.48E-01 | no | Gjb3 | synonymous_variant | LOW |
| 8 | 63397327 | AX-97810230 | 1 | A | G | 0.143 | -1.33E-01 | 1.42E-01 | 2.59E+00 | 2.67E+00 | 3.48E-01 | no | | intergenic_variant | MODIFIER |
| 8 | 65869305 | AX-170457347 | 1 | A | G | 0.143 | -1.33E-01 | 1.42E-01 | 2.59E+00 | 2.67E+00 | 3.48E-01 | no | MAR1 | intron_variant | MODIFIER |
| 8 | 66311402 | AX-170330374 | 1 | A | G | 0.143 | -1.33E-01 | 1.42E-01 | 2.59E+00 | 2.67E+00 | 3.48E-01 | no | MAR1 | intron_variant | MODIFIER |
| 6 | 125324530 | AX-170206928 | 0 | T | C | 0.184 | -7.36E-02 | 7.97E-02 | 2.75E+00 | 2.90E+00 | 3.48E-01 | no | Scnn1a | intron_variant | MODIFIER |
| 6 | 125994998 | AX-97801830 | 0 | T | A | 0.184 | -7.36E-02 | 7.97E-02 | 2.75E+00 | 2.90E+00 | 3.48E-01 | no | Ano2 | intron_variant,NMD_transcript_variant | MODIFIER |
| 13 | 24398450 | AX-170300264 | 1 | T | A | 0.17 | -1.16E-01 | 1.19E-01 | 2.36E+00 | 2.41E+00 | 3.48E-01 | no | Cmah | intron_variant | MODIFIER |
| 8 | 16485388 | AX-170229513 | 1 | G | T | 0.232 | -7.20E-02 | 7.70E-02 | 2.57E+00 | 2.66E+00 | 3.48E-01 | no | Csmd1 | intron_variant | MODIFIER |
| 1 | 126684688 | AX-170463308 | 0 | G | A | 0.544 | 7.87E-02 | 8.43E-02 | 2.62E+00 | 2.68E+00 | 3.48E-01 | no | Nckap5 | intron_variant | MODIFIER |
| 1 | 126687507 | AX-170148766 | 0 | C | A | 0.544 | 7.87E-02 | 8.43E-02 | 2.62E+00 | 2.68E+00 | 3.48E-01 | no | Nckap5 | intron_variant | MODIFIER |
| 1 | 126687806 | AX-170310092 | 0 | A | T | 0.544 | 7.87E-02 | 8.43E-02 | 2.62E+00 | 2.68E+00 | 3.48E-01 | no | Nckap5 | intron_variant | MODIFIER |
| 1 | 126706226 | AX-170042621 | 0 | A | G | 0.544 | 7.87E-02 | 8.43E-02 | 2.62E+00 | 2.68E+00 | 3.48E-01 | no | Nckap5 | intron_variant | MODIFIER |
| 1 | 126712775 | AX-170453272 | 0 | G | A | 0.544 | 7.87E-02 | 8.43E-02 | 2.62E+00 | 2.68E+00 | 3.48E-01 | no | Nckap5 | intron_variant | MODIFIER |
| 1 | 126740423 | AX-170469738 | 0 | G | A | 0.544 | 7.87E-02 | 8.43E-02 | 2.62E+00 | 2.68E+00 | 3.48E-01 | no | Nckap5 | intron_variant | MODIFIER |
| 1 | 126743280 | AX-170421379 | 0 | T | C | 0.544 | 7.87E-02 | 8.43E-02 | 2.62E+00 | 2.68E+00 | 3.48E-01 | no | Nckap5 | intron_variant | MODIFIER |
| 1 | 126743442 | AX-170290386 | 0 | G | A | 0.544 | 7.87E-02 | 8.43E-02 | 2.62E+00 | 2.68E+00 | 3.48E-01 | no | Nckap5 | intron_variant | MODIFIER |
| 1 | 126825792 | AX-170210721 | 0 | A | G | 0.544 | 7.87E-02 | 8.43E-02 | 2.62E+00 | 2.68E+00 | 3.48E-01 | no | Nckap5 | intron_variant | MODIFIER |
| 1 | 126841509 | AX-170343309 | 0 | C | T | 0.544 | 7.87E-02 | 8.43E-02 | 2.62E+00 | 2.68E+00 | 3.48E-01 | no | | intergenic_variant | MODIFIER |
| 1 | 126842546 | AX-170223639 | 0 | A | T | 0.544 | 7.87E-02 | 8.43E-02 | 2.62E+00 | 2.68E+00 | 3.48E-01 | no | | intergenic_variant | MODIFIER |
| 1 | 126842587 | AX-170102125 | 0 | A | G | 0.544 | 7.87E-02 | 8.43E-02 | 2.62E+00 | 2.68E+00 | 3.48E-01 | no | | intergenic_variant | MODIFIER |
| 1 | 126844261 | AX-170237347 | 0 | G | C | 0.544 | 7.87E-02 | 8.43E-02 | 2.62E+00 | 2.68E+00 | 3.48E-01 | no | | intergenic_variant | MODIFIER |
| 1 | 126874126 | AX-170468047 | 0 | T | G | 0.544 | 7.87E-02 | 8.43E-02 | 2.62E+00 | 2.68E+00 | 3.48E-01 | no | 4930599A14Rik | intron_variant,non_coding_transcript_variant | MODIFIER |
| 1 | 126877317 | AX-170375067 | 0 | T | C | 0.544 | 7.87E-02 | 8.43E-02 | 2.62E+00 | 2.68E+00 | 3.48E-01 | no | 4930599A14Rik | upstream_gene_variant | MODIFIER |
| 1 | 126877907 | AX-97848727 | 0 | T | C | 0.544 | 7.87E-02 | 8.43E-02 | 2.62E+00 | 2.68E+00 | 3.48E-01 | no | 4930599A14Rik | upstream_gene_variant | MODIFIER |
| 1 | 126897766 | AX-170258800 | 0 | G | A | 0.544 | 7.87E-02 | 8.43E-02 | 2.62E+00 | 2.68E+00 | 3.48E-01 | no | | intergenic_variant | MODIFIER |
| 1 | 126901519 | AX-170042627 | 0 | G | A | 0.544 | 7.87E-02 | 8.43E-02 | 2.62E+00 | 2.68E+00 | 3.48E-01 | no | | intergenic_variant | MODIFIER |
| 1 | 126909509 | AX-97830865 | 0 | T | C | 0.544 | 7.87E-02 | 8.43E-02 | 2.62E+00 | 2.68E+00 | 3.48E-01 | no | | intergenic_variant | MODIFIER |
| 1 | 126945129 | AX-97838676 | 0 | C | T | 0.544 | 7.87E-02 | 8.43E-02 | 2.62E+00 | 2.68E+00 | 3.48E-01 | no | | intergenic_variant | MODIFIER |
| 1 | 126952532 | AX-97818996 | 0 | C | T | 0.544 | 7.87E-02 | 8.43E-02 | 2.62E+00 | 2.68E+00 | 3.48E-01 | no | | intergenic_variant | MODIFIER |
| 1 | 127064708 | AX-170322048 | 0 | T | C | 0.544 | 7.87E-02 | 8.43E-02 | 2.62E+00 | 2.68E+00 | 3.48E-01 | no | | intergenic_variant | MODIFIER |
| 1 | 127079881 | AX-170215782 | 0 | G | A | 0.544 | 7.87E-02 | 8.43E-02 | 2.62E+00 | 2.68E+00 | 3.48E-01 | no | | intergenic_variant | MODIFIER |
| 1 | 127098781 | AX-170001645 | 0 | A | G | 0.544 | 7.87E-02 | 8.43E-02 | 2.62E+00 | 2.68E+00 | 3.48E-01 | no | | intergenic_variant | MODIFIER |
| 1 | 127104825 | AX-170001650 | 0 | G | A | 0.544 | 7.87E-02 | 8.43E-02 | 2.62E+00 | 2.68E+00 | 3.48E-01 | no | | intergenic_variant | MODIFIER |
| 1 | 127188500 | AX-170086171 | 0 | G | A | 0.544 | 7.87E-02 | 8.43E-02 | 2.62E+00 | 2.68E+00 | 3.48E-01 | no | | intergenic_variant | MODIFIER |
| 1 | 127188907 | AX-170332198 | 0 | T | A | 0.544 | 7.87E-02 | 8.43E-02 | 2.62E+00 | 2.68E+00 | 3.48E-01 | no | | intergenic_variant | MODIFIER |
| 1 | 127216866 | AX-170429638 | 0 | T | A | 0.544 | 7.87E-02 | 8.43E-02 | 2.62E+00 | 2.68E+00 | 3.48E-01 | no | Mgat5 | intron_variant | MODIFIER |
| 1 | 127218638 | AX-170527635 | 0 | A | G | 0.544 | 7.87E-02 | 8.43E-02 | 2.62E+00 | 2.68E+00 | 3.48E-01 | no | Mgat5 | intron_variant | MODIFIER |
| 1 | 127229093 | AX-170149641 | 0 | A | G | 0.544 | 7.87E-02 | 8.43E-02 | 2.62E+00 | 2.68E+00 | 3.48E-01 | no | Mgat5 | intron_variant | MODIFIER |
| 1 | 127391744 | AX-170530589 | 0 | G | A | 0.544 | 7.87E-02 | 8.43E-02 | 2.62E+00 | 2.68E+00 | 3.48E-01 | no | Mgat5 | intron_variant | MODIFIER |
| 1 | 127396598 | AX-170422682 | 0 | T | A | 0.544 | 7.87E-02 | 8.43E-02 | 2.62E+00 | 2.68E+00 | 3.48E-01 | no | Mgat5 | intron_variant | MODIFIER |
| 1 | 127419037 | AX-97813188 | 0 | A | G | 0.544 | 7.87E-02 | 8.43E-02 | 2.62E+00 | 2.68E+00 | 3.48E-01 | no | Mgat5 | intron_variant | MODIFIER |
| 1 | 127489056 | AX-170220719 | 0 | C | T | 0.544 | 7.87E-02 | 8.43E-02 | 2.62E+00 | 2.68E+00 | 3.48E-01 | no | Tmem163 | intron_variant,NMD_transcript_variant | MODIFIER |
| 1 | 127574769 | AX-170001658 | 0 | C | T | 0.544 | 7.87E-02 | 8.43E-02 | 2.62E+00 | 2.68E+00 | 3.48E-01 | no | Tmem163 | intron_variant,NMD_transcript_variant | MODIFIER |
| 1 | 127578276 | AX-170001652 | 0 | A | G | 0.544 | 7.87E-02 | 8.43E-02 | 2.62E+00 | 2.68E+00 | 3.48E-01 | no | Tmem163 | intron_variant,NMD_transcript_variant | MODIFIER |
| 1 | 127628338 | AX-97809782 | 0 | A | G | 0.544 | 7.87E-02 | 8.43E-02 | 2.62E+00 | 2.68E+00 | 3.48E-01 | no | Tmem163 | intron_variant,NMD_transcript_variant | MODIFIER |
| 1 | 127630768 | AX-170001653 | 0 | A | G | 0.544 | 7.87E-02 | 8.43E-02 | 2.62E+00 | 2.68E+00 | 3.48E-01 | no | Tmem163 | intron_variant,NMD_transcript_variant | MODIFIER |
| 1 | 127634301 | AX-170042647 | 0 | T | G | 0.544 | 7.87E-02 | 8.43E-02 | 2.62E+00 | 2.68E+00 | 3.48E-01 | no | Tmem163 | intron_variant,NMD_transcript_variant | MODIFIER |
| 1 | 127635099 | AX-170001655 | 0 | A | G | 0.544 | 7.87E-02 | 8.43E-02 | 2.62E+00 | 2.68E+00 | 3.48E-01 | no | Tmem163 | intron_variant,NMD_transcript_variant | MODIFIER |
| 1 | 127646636 | AX-170469635 | 0 | A | G | 0.544 | 7.87E-02 | 8.43E-02 | 2.62E+00 | 2.68E+00 | 3.48E-01 | no | Tmem163 | intron_variant,NMD_transcript_variant | MODIFIER |
| 1 | 127646846 | AX-170001657 | 0 | A | C | 0.544 | 7.87E-02 | 8.43E-02 | 2.62E+00 | 2.68E+00 | 3.48E-01 | no | Tmem163 | intron_variant,NMD_transcript_variant | MODIFIER |
| 1 | 127647255 | AX-170035630 | 0 | C | T | 0.544 | 7.87E-02 | 8.43E-02 | 2.62E+00 | 2.68E+00 | 3.48E-01 | no | Tmem163 | intron_variant,NMD_transcript_variant | MODIFIER |
| 1 | 127649973 | AX-170371473 | 0 | T | A | 0.544 | 7.87E-02 | 8.43E-02 | 2.62E+00 | 2.68E+00 | 3.48E-01 | no | Tmem163 | intron_variant,NMD_transcript_variant | MODIFIER |
| 1 | 127666610 | AX-170001656 | 0 | T | C | 0.544 | 7.87E-02 | 8.43E-02 | 2.62E+00 | 2.68E+00 | 3.48E-01 | no | Tmem163 | intron_variant,NMD_transcript_variant | MODIFIER |
| 1 | 127863321 | AX-170042652 | 0 | A | T | 0.544 | 7.87E-02 | 8.43E-02 | 2.62E+00 | 2.68E+00 | 3.48E-01 | no | | intergenic_variant | MODIFIER |

| | | | | | | | | | | | | | | | |
|---|---|---|---|---|---|---|---|---|---|---|---|---|---|---|---|
| 1 | 127871403 | AX-170120655 | 0 | G | A | 0.544 | 7.87E-02 | 8.43E-02 | 2.62E+00 | 2.68E+00 | 3.48E-01 | no | Rab3gap1 | intron_variant | MODIFIER |
| 1 | 127871436 | AX-170194196 | 0 | G | A | 0.544 | 7.87E-02 | 8.43E-02 | 2.62E+00 | 2.68E+00 | 3.48E-01 | no | Rab3gap1 | intron_variant | MODIFIER |
| 1 | 127872245 | AX-170334179 | 0 | C | T | 0.544 | 7.87E-02 | 8.43E-02 | 2.62E+00 | 2.68E+00 | 3.48E-01 | no | Rab3gap1 | intron_variant | MODIFIER |
| 1 | 127885582 | AX-97802795 | 0 | G | A | 0.544 | 7.87E-02 | 8.43E-02 | 2.62E+00 | 2.68E+00 | 3.48E-01 | no | Rab3gap1 | intron_variant | MODIFIER |
| 1 | 127918808 | AX-170464959 | 0 | C | T | 0.544 | 7.87E-02 | 8.43E-02 | 2.62E+00 | 2.68E+00 | 3.48E-01 | no | Rab3gap1 | intron_variant | MODIFIER |
| 1 | 127950134 | AX-170174057 | 0 | G | A | 0.544 | 7.87E-02 | 8.43E-02 | 2.62E+00 | 2.68E+00 | 3.48E-01 | no | Zranb3 | downstream_gene_variant | MODIFIER |
| 1 | 127950217 | AX-170373666 | 0 | A | G | 0.544 | 7.87E-02 | 8.43E-02 | 2.62E+00 | 2.68E+00 | 3.48E-01 | no | Zranb3 | downstream_gene_variant | MODIFIER |
| 1 | 127950355 | AX-170236963 | 0 | T | A | 0.544 | 7.87E-02 | 8.43E-02 | 2.62E+00 | 2.68E+00 | 3.48E-01 | no | Zranb3 | downstream_gene_variant | MODIFIER |
| 1 | 127952750 | AX-170379551 | 0 | G | A | 0.544 | 7.87E-02 | 8.43E-02 | 2.62E+00 | 2.68E+00 | 3.48E-01 | no | Zranb3 | downstream_gene_variant | MODIFIER |
| 1 | 127953181 | AX-170401590 | 0 | T | C | 0.544 | 7.87E-02 | 8.43E-02 | 2.62E+00 | 2.68E+00 | 3.48E-01 | no | Zranb3 | downstream_gene_variant | MODIFIER |
| 1 | 127956469 | AX-170147855 | 0 | C | T | 0.544 | 7.87E-02 | 8.43E-02 | 2.62E+00 | 2.68E+00 | 3.48E-01 | no | Zranb3 | intron_variant | MODIFIER |
| 1 | 127959819 | AX-170450289 | 0 | C | T | 0.544 | 7.87E-02 | 8.43E-02 | 2.62E+00 | 2.68E+00 | 3.48E-01 | no | Zranb3 | missense_variant | MODERATE |
| 1 | 128003008 | AX-97803062 | 0 | A | G | 0.544 | 7.87E-02 | 8.43E-02 | 2.62E+00 | 2.68E+00 | 3.48E-01 | no | Zranb3 | intron_variant | MODIFIER |
| 1 | 128008813 | AX-170452135 | 0 | A | T | 0.544 | 7.87E-02 | 8.43E-02 | 2.62E+00 | 2.68E+00 | 3.48E-01 | no | Zranb3 | intron_variant | MODIFIER |
| 1 | 128015918 | AX-170406417 | 0 | G | C | 0.544 | 7.87E-02 | 8.43E-02 | 2.62E+00 | 2.68E+00 | 3.48E-01 | no | Zranb3 | intron_variant | MODIFIER |
| 1 | 128062364 | AX-97843388 | 0 | T | C | 0.544 | 7.87E-02 | 8.43E-02 | 2.62E+00 | 2.68E+00 | 3.48E-01 | no | Zranb3 | intron_variant | MODIFIER |
| 1 | 128084987 | AX-170001663 | 0 | T | A | 0.544 | 7.87E-02 | 8.43E-02 | 2.62E+00 | 2.68E+00 | 3.48E-01 | no | Zranb3 | intron_variant | MODIFIER |
| 1 | 128090858 | AX-170001668 | 0 | G | A | 0.544 | 7.87E-02 | 8.43E-02 | 2.62E+00 | 2.68E+00 | 3.48E-01 | no | Zranb3 | intron_variant | MODIFIER |
| 1 | 128118229 | AX-97813227 | 0 | G | T | 0.544 | 7.87E-02 | 8.43E-02 | 2.62E+00 | 2.68E+00 | 3.48E-01 | no | R3hdm1 | intron_variant | MODIFIER |
| 1 | 128148331 | AX-170001667 | 0 | G | C | 0.544 | 7.87E-02 | 8.43E-02 | 2.62E+00 | 2.68E+00 | 3.48E-01 | no | R3hdm1 | intron_variant | MODIFIER |
| 1 | 128184968 | AX-170042656 | 0 | A | G | 0.544 | 7.87E-02 | 8.43E-02 | 2.62E+00 | 2.68E+00 | 3.48E-01 | no | R3hdm1 | intron_variant | MODIFIER |
| 1 | 128190198 | AX-170461732 | 0 | C | G | 0.544 | 7.87E-02 | 8.43E-02 | 2.62E+00 | 2.68E+00 | 3.48E-01 | no | R3hdm1 | intron_variant | MODIFIER |
| 1 | 128239956 | AX-170001670 | 0 | A | G | 0.544 | 7.87E-02 | 8.43E-02 | 2.62E+00 | 2.68E+00 | 3.48E-01 | no | Ubxn4 | upstream_gene_variant | MODIFIER |
| 1 | 128313152 | AX-97822798 | 0 | A | G | 0.544 | 7.87E-02 | 8.43E-02 | 2.62E+00 | 2.68E+00 | 3.48E-01 | no | Lct | intron_variant | MODIFIER |
| 1 | 128323954 | AX-170534246 | 0 | T | C | 0.544 | 7.87E-02 | 8.43E-02 | 2.62E+00 | 2.68E+00 | 3.48E-01 | no | Lct | intron_variant | MODIFIER |
| 1 | 128327768 | AX-97818577 | 0 | C | T | 0.544 | 7.87E-02 | 8.43E-02 | 2.62E+00 | 2.68E+00 | 3.48E-01 | no | Mcm6 | downstream_gene_variant | MODIFIER |
| 1 | 128351928 | AX-97821713 | 0 | G | A | 0.544 | 7.87E-02 | 8.43E-02 | 2.62E+00 | 2.68E+00 | 3.48E-01 | no | Mcm6 | intron_variant | MODIFIER |
| 1 | 128388586 | AX-170105766 | 0 | A | G | 0.544 | 7.87E-02 | 8.43E-02 | 2.62E+00 | 2.68E+00 | 3.48E-01 | no | Dars | intron_variant | MODIFIER |
| 1 | 128389061 | AX-170001661 | 0 | A | G | 0.544 | 7.87E-02 | 8.43E-02 | 2.62E+00 | 2.68E+00 | 3.48E-01 | no | Dars | intron_variant | MODIFIER |
| 1 | 128404893 | AX-170086609 | 0 | C | T | 0.544 | 7.87E-02 | 8.43E-02 | 2.62E+00 | 2.68E+00 | 3.48E-01 | no | Dars | intron_variant | MODIFIER |
| 1 | 128424016 | AX-170530824 | 0 | C | G | 0.544 | 7.87E-02 | 8.43E-02 | 2.62E+00 | 2.68E+00 | 3.48E-01 | no | | intergenic_variant | MODIFIER |
| 1 | 128456191 | AX-170121281 | 0 | G | A | 0.544 | 7.87E-02 | 8.43E-02 | 2.62E+00 | 2.68E+00 | 3.48E-01 | no | | intergenic_variant | MODIFIER |
| 1 | 128459049 | AX-170396011 | 0 | T | A | 0.544 | 7.87E-02 | 8.43E-02 | 2.62E+00 | 2.68E+00 | 3.48E-01 | no | | intergenic_variant | MODIFIER |
| 1 | 128459582 | AX-170422771 | 0 | C | G | 0.544 | 7.87E-02 | 8.43E-02 | 2.62E+00 | 2.68E+00 | 3.48E-01 | no | | intergenic_variant | MODIFIER |
| 1 | 128472297 | AX-170356541 | 0 | A | G | 0.544 | 7.87E-02 | 8.43E-02 | 2.62E+00 | 2.68E+00 | 3.48E-01 | no | | regulatory_region_variant | MODIFIER |
| 1 | 128477356 | AX-97824102 | 0 | G | A | 0.544 | 7.87E-02 | 8.43E-02 | 2.62E+00 | 2.68E+00 | 3.48E-01 | no | | intergenic_variant | MODIFIER |
| 1 | 128477889 | AX-170378533 | 0 | A | G | 0.544 | 7.87E-02 | 8.43E-02 | 2.62E+00 | 2.68E+00 | 3.48E-01 | no | | intergenic_variant | MODIFIER |
| 1 | 128483084 | AX-170180599 | 0 | G | A | 0.544 | 7.87E-02 | 8.43E-02 | 2.62E+00 | 2.68E+00 | 3.48E-01 | no | | regulatory_region_variant | MODIFIER |
| 18 | 80085318 | AX-170438691 | 1 | G | A | 0.152 | -1.10E-01 | 1.17E-01 | 2.52E+00 | 2.59E+00 | 3.48E-01 | no | Pard6g | intron_variant | MODIFIER |
| 1 | 150903228 | AX-170107558 | 0 | A | G | 0.202 | 1.29E-01 | 1.38E-01 | 2.68E+00 | 2.73E+00 | 3.48E-01 | no | Hmcn1 | intron_variant | MODIFIER |
| 3 | 151615405 | AX-97782182 | 1 | C | A | 0.321 | 6.74E-02 | 7.28E-02 | 2.78E+00 | 2.88E+00 | 3.49E-01 | no | | intergenic_variant | MODIFIER |
| 3 | 151738792 | AX-170151167 | 1 | T | C | 0.339 | 6.89E-02 | 7.43E-02 | 2.79E+00 | 2.88E+00 | 3.49E-01 | no | Ifi44 | intron_variant | MODIFIER |
| 18 | 84999855 | AX-170016124 | 1 | G | A | 0.161 | -1.05E-01 | 1.11E-01 | 2.49E+00 | 2.56E+00 | 3.49E-01 | no | | regulatory_region_variant | MODIFIER |
| 18 | 85299256 | AX-170386915 | 1 | T | C | 0.161 | -1.05E-01 | 1.11E-01 | 2.49E+00 | 2.56E+00 | 3.49E-01 | no | | intergenic_variant | MODIFIER |
| 5 | 3938277 | AX-168376096 | 0 | T | C | 0.43 | -6.16E-02 | 6.52E-02 | 2.49E+00 | 2.60E+00 | 3.49E-01 | no | Akap9 | intron_variant | MODIFIER |
| 5 | 3939231 | AX-170344700 | 0 | C | G | 0.43 | -6.16E-02 | 6.52E-02 | 2.49E+00 | 2.60E+00 | 3.49E-01 | no | Akap9 | intron_variant | MODIFIER |
| 5 | 3950304 | AX-170071167 | 0 | G | A | 0.43 | -6.16E-02 | 6.52E-02 | 2.49E+00 | 2.60E+00 | 3.49E-01 | no | Akap9 | intron_variant | MODIFIER |
| 5 | 3950367 | AX-97809251 | 0 | G | A | 0.43 | -6.16E-02 | 6.52E-02 | 2.49E+00 | 2.60E+00 | 3.49E-01 | no | Akap9 | intron_variant | MODIFIER |
| 5 | 3990513 | AX-170322955 | 0 | A | G | 0.43 | -6.16E-02 | 6.52E-02 | 2.49E+00 | 2.60E+00 | 3.49E-01 | no | Akap9 | intron_variant | MODIFIER |
| 5 | 4005323 | AX-97812658 | 0 | G | A | 0.43 | -6.16E-02 | 6.52E-02 | 2.49E+00 | 2.60E+00 | 3.49E-01 | no | Akap9 | intron_variant | MODIFIER |
| 5 | 4034771 | AX-170215179 | 0 | T | C | 0.43 | -6.16E-02 | 6.52E-02 | 2.49E+00 | 2.60E+00 | 3.49E-01 | no | Akap9 | synonymous_variant | LOW |
| 5 | 4039309 | AX-97791195 | 0 | C | T | 0.43 | -6.16E-02 | 6.52E-02 | 2.49E+00 | 2.60E+00 | 3.49E-01 | no | Akap9 | missense_variant | MODERATE |
| 5 | 4049610 | AX-170534317 | 0 | A | G | 0.43 | -6.16E-02 | 6.52E-02 | 2.49E+00 | 2.60E+00 | 3.49E-01 | no | Akap9 | intron_variant | MODIFIER |
| 5 | 4071106 | AX-170437455 | 0 | G | T | 0.43 | -6.16E-02 | 6.52E-02 | 2.49E+00 | 2.60E+00 | 3.49E-01 | no | Akap9 | intron_variant | MODIFIER |

| | | | | | | | | | | | | | | | |
|---|---|---|---|---|---|---|---|---|---|---|---|---|---|---|---|
| 5 | 4427812 | AX-170024329 | 0 | T | C | 0.43 | -6.16E-02 | 6.52E-02 | 2.49E+00 | 2.60E+00 | 3.49E-01 | no | | intergenic_variant | MODIFIER |
| 5 | 4674287 | AX-170522916 | 0 | C | A | 0.43 | -6.16E-02 | 6.52E-02 | 2.49E+00 | 2.60E+00 | 3.49E-01 | no | | intergenic_variant | MODIFIER |
| 5 | 5019104 | AX-97843788 | 0 | T | C | 0.43 | -6.16E-02 | 6.52E-02 | 2.49E+00 | 2.60E+00 | 3.49E-01 | no | Cdk14 | intron_variant | MODIFIER |
| 5 | 5044004 | AX-170404786 | 0 | A | C | 0.43 | -6.16E-02 | 6.52E-02 | 2.49E+00 | 2.60E+00 | 3.49E-01 | no | Cdk14 | intron_variant | MODIFIER |
| 5 | 5049259 | AX-97802379 | 0 | T | G | 0.43 | -6.16E-02 | 6.52E-02 | 2.49E+00 | 2.60E+00 | 3.49E-01 | no | Cdk14 | intron_variant | MODIFIER |
| 5 | 5068694 | AX-170195417 | 0 | C | T | 0.43 | -6.16E-02 | 6.52E-02 | 2.49E+00 | 2.60E+00 | 3.49E-01 | no | Cdk14 | intron_variant | MODIFIER |
| 5 | 5130510 | AX-168395114 | 0 | T | C | 0.43 | -6.16E-02 | 6.52E-02 | 2.49E+00 | 2.60E+00 | 3.49E-01 | no | Cdk14 | intron_variant | MODIFIER |
| 5 | 5154734 | AX-170138918 | 0 | G | C | 0.43 | -6.16E-02 | 6.52E-02 | 2.49E+00 | 2.60E+00 | 3.49E-01 | no | Cdk14 | intron_variant | MODIFIER |
| 5 | 5184452 | AX-170475615 | 0 | A | G | 0.43 | -6.16E-02 | 6.52E-02 | 2.49E+00 | 2.60E+00 | 3.49E-01 | no | Cdk14 | intron_variant | MODIFIER |
| 5 | 5211438 | AX-170416655 | 0 | T | C | 0.43 | -6.16E-02 | 6.52E-02 | 2.49E+00 | 2.60E+00 | 3.49E-01 | no | Cdk14 | intron_variant | MODIFIER |
| 5 | 5244884 | AX-170321577 | 0 | A | G | 0.43 | -6.16E-02 | 6.52E-02 | 2.49E+00 | 2.60E+00 | 3.49E-01 | no | Cdk14 | intron_variant | MODIFIER |
| 5 | 5261926 | AX-170260490 | 0 | A | G | 0.43 | -6.16E-02 | 6.52E-02 | 2.49E+00 | 2.60E+00 | 3.49E-01 | no | Cdk14 | intron_variant | MODIFIER |
| 5 | 5262094 | AX-170272055 | 0 | C | T | 0.43 | -6.16E-02 | 6.52E-02 | 2.49E+00 | 2.60E+00 | 3.49E-01 | no | Cdk14 | intron_variant | MODIFIER |
| 5 | 5262226 | AX-170130742 | 0 | C | G | 0.43 | -6.16E-02 | 6.52E-02 | 2.49E+00 | 2.60E+00 | 3.49E-01 | no | Cdk14 | intron_variant | MODIFIER |
| 5 | 5263710 | AX-170182447 | 0 | G | T | 0.43 | -6.16E-02 | 6.52E-02 | 2.49E+00 | 2.60E+00 | 3.49E-01 | no | Cdk14 | intron_variant | MODIFIER |
| 5 | 5285050 | AX-170487562 | 0 | A | G | 0.43 | -6.16E-02 | 6.52E-02 | 2.49E+00 | 2.60E+00 | 3.49E-01 | no | Cdk14 | intron_variant | MODIFIER |
| 5 | 5296507 | AX-170244869 | 0 | T | C | 0.43 | -6.16E-02 | 6.52E-02 | 2.49E+00 | 2.60E+00 | 3.49E-01 | no | Cdk14 | intron_variant | MODIFIER |
| 7 | 74434168 | AX-97817106 | 0 | G | A | 0.377 | -9.06E-02 | 9.79E-02 | 2.78E+00 | 2.89E+00 | 3.49E-01 | no | Slco3a1 | intron_variant | MODIFIER |
| 6 | 127013233 | AX-170028200 | 1 | C | A | 0.536 | 7.80E-02 | 8.23E-02 | 2.49E+00 | 2.55E+00 | 3.49E-01 | no | Fgf6 | upstream_gene_variant | MODIFIER |
| 16 | 80135369 | AX-97788396 | 0 | A | C | 0.456 | 7.28E-02 | 7.84E-02 | 2.65E+00 | 2.72E+00 | 3.49E-01 | no | Gm23083 | upstream_gene_variant | MODIFIER |
| 16 | 80540019 | AX-170013478 | 0 | A | G | 0.456 | 7.28E-02 | 7.84E-02 | 2.65E+00 | 2.72E+00 | 3.49E-01 | no | | intergenic_variant | MODIFIER |
| 16 | 85316584 | AX-97824035 | 0 | T | C | 0.456 | 7.28E-02 | 7.84E-02 | 2.65E+00 | 2.72E+00 | 3.49E-01 | no | | intergenic_variant | MODIFIER |
| 16 | 85317027 | AX-170429714 | 0 | G | A | 0.456 | 7.28E-02 | 7.84E-02 | 2.65E+00 | 2.72E+00 | 3.49E-01 | no | | intergenic_variant | MODIFIER |
| 16 | 85317891 | AX-170463073 | 0 | C | T | 0.456 | 7.28E-02 | 7.84E-02 | 2.65E+00 | 2.72E+00 | 3.49E-01 | no | | intergenic_variant | MODIFIER |
| 16 | 85324054 | AX-170369463 | 0 | C | T | 0.456 | 7.28E-02 | 7.84E-02 | 2.65E+00 | 2.72E+00 | 3.49E-01 | no | | intergenic_variant | MODIFIER |
| 16 | 85335439 | AX-170353101 | 0 | A | G | 0.456 | 7.28E-02 | 7.84E-02 | 2.65E+00 | 2.72E+00 | 3.49E-01 | no | | intergenic_variant | MODIFIER |
| 18 | 65870379 | AX-170194343 | 2 | A | C | 0.182 | -1.12E-01 | 1.21E-01 | 2.70E+00 | 2.75E+00 | 3.49E-01 | no | Grp | upstream_gene_variant | MODIFIER |
| 15 | 82132381 | AX-170056125 | 0 | C | T | 0.553 | 7.43E-02 | 7.91E-02 | 2.54E+00 | 2.62E+00 | 3.49E-01 | no | Ccdc134 | intron_variant | MODIFIER |
| 15 | 82777081 | AX-170151488 | 0 | G | A | 0.553 | 7.43E-02 | 7.91E-02 | 2.54E+00 | 2.62E+00 | 3.49E-01 | no | Cyp2d41-ps | downstream_gene_variant | MODIFIER |
| 7 | 99256382 | AX-170138180 | 0 | A | T | 0.377 | -9.48E-02 | 1.02E-01 | 2.81E+00 | 2.91E+00 | 3.49E-01 | no | Gm26705 | downstream_gene_variant | MODIFIER |
| 1 | 168443306 | AX-170043486 | 1 | T | C | 0.214 | 1.30E-01 | 1.40E-01 | 2.68E+00 | 2.73E+00 | 3.49E-01 | no | | intergenic_variant | MODIFIER |
| 15 | 64049138 | AX-170190234 | 0 | C | T | 0.333 | -7.93E-02 | 7.59E-02 | 2.17E+00 | 2.25E+00 | 3.49E-01 | no | Fam49b | intron_variant | MODIFIER |
| 16 | 72697178 | AX-170166381 | 0 | C | T | 0.474 | 6.90E-02 | 7.43E-02 | 2.64E+00 | 2.74E+00 | 3.49E-01 | no | Robo1 | intron_variant | MODIFIER |
| 7 | 37240665 | AX-170143166 | 0 | T | C | 0.342 | 8.60E-02 | 9.18E-02 | 2.56E+00 | 2.64E+00 | 3.49E-01 | no | | intergenic_variant | MODIFIER |
| 18 | 73118560 | AX-170523769 | 0 | G | A | 0.193 | -1.04E-01 | 1.12E-01 | 2.62E+00 | 2.69E+00 | 3.49E-01 | no | | intergenic_variant | MODIFIER |
| 1 | 150813160 | AX-170192239 | 0 | A | G | 0.211 | 1.22E-01 | 1.31E-01 | 2.59E+00 | 2.65E+00 | 3.49E-01 | no | Hmcn1 | intron_variant | MODIFIER |
| 16 | 70525931 | AX-170409925 | 0 | C | T | 0.491 | 7.05E-02 | 7.52E-02 | 2.55E+00 | 2.63E+00 | 3.49E-01 | no | Gbe1 | intron_variant | MODIFIER |
| 1 | 128389220 | AX-170001660 | 1 | A | C | 0.545 | 7.83E-02 | 8.42E-02 | 2.62E+00 | 2.68E+00 | 3.49E-01 | no | Dars | intron_variant | MODIFIER |
| 3 | 149771177 | AX-170021906 | 0 | C | T | 0.342 | 6.91E-02 | 7.46E-02 | 2.79E+00 | 2.89E+00 | 3.49E-01 | no | | intergenic_variant | MODIFIER |
| 16 | 78226749 | AX-97814932 | 0 | G | A | 0.456 | 7.07E-02 | 7.58E-02 | 2.59E+00 | 2.66E+00 | 3.49E-01 | no | | intergenic_variant | MODIFIER |
| 1 | 126839747 | AX-168420033 | 0 | G | A | 0.553 | 7.88E-02 | 8.47E-02 | 2.62E+00 | 2.68E+00 | 3.49E-01 | no | | intergenic_variant | MODIFIER |
| 1 | 126844645 | AX-170198008 | 0 | G | C | 0.553 | 7.88E-02 | 8.47E-02 | 2.62E+00 | 2.68E+00 | 3.49E-01 | no | | intergenic_variant | MODIFIER |
| 1 | 126918418 | AX-168400119 | 0 | T | C | 0.553 | 7.88E-02 | 8.47E-02 | 2.62E+00 | 2.68E+00 | 3.49E-01 | no | | regulatory_region_variant | MODIFIER |
| 1 | 126947911 | AX-170167658 | 0 | A | G | 0.553 | 7.88E-02 | 8.47E-02 | 2.62E+00 | 2.68E+00 | 3.49E-01 | no | | regulatory_region_variant | MODIFIER |
| 1 | 126952466 | AX-170350345 | 0 | A | G | 0.553 | 7.88E-02 | 8.47E-02 | 2.62E+00 | 2.68E+00 | 3.49E-01 | no | | intergenic_variant | MODIFIER |
| 1 | 127049404 | AX-170001642 | 0 | C | T | 0.553 | 7.88E-02 | 8.47E-02 | 2.62E+00 | 2.68E+00 | 3.49E-01 | no | Gm5261 | downstream_gene_variant | MODIFIER |
| 1 | 127060245 | AX-97820346 | 0 | T | G | 0.553 | 7.88E-02 | 8.47E-02 | 2.62E+00 | 2.68E+00 | 3.49E-01 | no | | intergenic_variant | MODIFIER |
| 1 | 127179958 | AX-170001643 | 0 | A | C | 0.553 | 7.88E-02 | 8.47E-02 | 2.62E+00 | 2.68E+00 | 3.49E-01 | no | | intergenic_variant | MODIFIER |
| 1 | 127184379 | AX-97827245 | 0 | C | T | 0.553 | 7.88E-02 | 8.47E-02 | 2.62E+00 | 2.68E+00 | 3.49E-01 | no | | intergenic_variant | MODIFIER |
| 1 | 127215814 | AX-97825923 | 0 | C | T | 0.553 | 7.88E-02 | 8.47E-02 | 2.62E+00 | 2.68E+00 | 3.49E-01 | no | Mgat5 | intron_variant | MODIFIER |
| 1 | 127226840 | AX-170403085 | 0 | C | T | 0.553 | 7.88E-02 | 8.47E-02 | 2.62E+00 | 2.68E+00 | 3.49E-01 | no | Mgat5 | intron_variant | MODIFIER |
| 1 | 127228495 | AX-170424998 | 0 | T | G | 0.553 | 7.88E-02 | 8.47E-02 | 2.62E+00 | 2.68E+00 | 3.49E-01 | no | Mgat5 | intron_variant | MODIFIER |
| 1 | 127352365 | AX-170141866 | 0 | A | G | 0.553 | 7.88E-02 | 8.47E-02 | 2.62E+00 | 2.68E+00 | 3.49E-01 | no | Mgat5 | intron_variant | MODIFIER |
| 1 | 127369352 | AX-168377959 | 0 | T | C | 0.553 | 7.88E-02 | 8.47E-02 | 2.62E+00 | 2.68E+00 | 3.49E-01 | no | Mgat5 | intron_variant | MODIFIER |

| | | | | | | | | | | | | | | | |
|---|---|---|---|---|---|---|---|---|---|---|---|---|---|---|---|
| 1 | 127371548 | AX-170482923 | 0 | C | T | 0.553 | 7.88E-02 | 8.47E-02 | 2.62E+00 | 2.68E+00 | 3.49E-01 | no | Mgat5 | intron_variant | MODIFIER |
| 1 | 127459710 | AX-170513533 | 0 | T | C | 0.553 | 7.88E-02 | 8.47E-02 | 2.62E+00 | 2.68E+00 | 3.49E-01 | no | Mgat5 | intron_variant | MODIFIER |
| 1 | 127482421 | AX-97801017 | 0 | T | G | 0.553 | 7.88E-02 | 8.47E-02 | 2.62E+00 | 2.68E+00 | 3.49E-01 | no | Tmem163 | downstream_gene_variant | MODIFIER |
| 1 | 127515029 | AX-97820253 | 0 | G | A | 0.553 | 7.88E-02 | 8.47E-02 | 2.62E+00 | 2.68E+00 | 3.49E-01 | no | Tmem163 | intron_variant,NMD_transcript_variant | MODIFIER |
| 1 | 127555388 | AX-170184266 | 0 | T | G | 0.553 | 7.88E-02 | 8.47E-02 | 2.62E+00 | 2.68E+00 | 3.49E-01 | no | Tmem163 | intron_variant,NMD_transcript_variant | MODIFIER |
| 1 | 127555606 | AX-170515090 | 0 | A | C | 0.553 | 7.88E-02 | 8.47E-02 | 2.62E+00 | 2.68E+00 | 3.49E-01 | no | Tmem163 | intron_variant,NMD_transcript_variant | MODIFIER |
| 1 | 127556255 | AX-170001651 | 0 | T | C | 0.553 | 7.88E-02 | 8.47E-02 | 2.62E+00 | 2.68E+00 | 3.49E-01 | no | Tmem163 | intron_variant,NMD_transcript_variant | MODIFIER |
| 1 | 127623408 | AX-170042645 | 0 | G | A | 0.553 | 7.88E-02 | 8.47E-02 | 2.62E+00 | 2.68E+00 | 3.49E-01 | no | Tmem163 | intron_variant,NMD_transcript_variant | MODIFIER |
| 1 | 127666562 | AX-97790823 | 0 | T | C | 0.553 | 7.88E-02 | 8.47E-02 | 2.62E+00 | 2.68E+00 | 3.49E-01 | no | Tmem163 | intron_variant,NMD_transcript_variant | MODIFIER |
| 1 | 127747026 | AX-170270780 | 0 | C | T | 0.553 | 7.88E-02 | 8.47E-02 | 2.62E+00 | 2.68E+00 | 3.49E-01 | no | Acmsd | intron_variant | MODIFIER |
| 1 | 127832206 | AX-170128496 | 0 | G | A | 0.553 | 7.88E-02 | 8.47E-02 | 2.62E+00 | 2.68E+00 | 3.49E-01 | no | Map3k19 | intron_variant | MODIFIER |
| 1 | 127943084 | AX-170288371 | 0 | T | C | 0.553 | 7.88E-02 | 8.47E-02 | 2.62E+00 | 2.68E+00 | 3.49E-01 | no | Rab3gap1 | 3_prime_UTR_variant | MODIFIER |
| 1 | 128063201 | AX-170285768 | 0 | T | C | 0.553 | 7.88E-02 | 8.47E-02 | 2.62E+00 | 2.68E+00 | 3.49E-01 | no | Zranb3 | intron_variant | MODIFIER |
| 1 | 128271901 | AX-170001671 | 0 | A | T | 0.553 | 7.88E-02 | 8.47E-02 | 2.62E+00 | 2.68E+00 | 3.49E-01 | no | Ubxn4 | intron_variant | MODIFIER |
| 1 | 128529692 | AX-170298503 | 0 | A | G | 0.553 | 7.88E-02 | 8.47E-02 | 2.62E+00 | 2.68E+00 | 3.49E-01 | no | | regulatory_region_variant | MODIFIER |
| 1 | 128529718 | AX-97851897 | 0 | C | T | 0.553 | 7.88E-02 | 8.47E-02 | 2.62E+00 | 2.68E+00 | 3.49E-01 | no | | regulatory_region_variant | MODIFIER |
| 8 | 17613940 | AX-170292576 | 1 | T | C | 0.214 | -7.03E-02 | 7.52E-02 | 2.55E+00 | 2.65E+00 | 3.50E-01 | no | | intergenic_variant | MODIFIER |
| 5 | 45338084 | AX-170071951 | 0 | C | T | 0.5 | 9.79E-02 | 1.02E-01 | 2.42E+00 | 2.47E+00 | 3.50E-01 | no | D5Ertd615e | intron_variant,non_coding_transcript_variant | MODIFIER |
| 5 | 45338287 | AX-170219266 | 0 | T | C | 0.5 | 9.79E-02 | 1.02E-01 | 2.42E+00 | 2.47E+00 | 3.50E-01 | no | D5Ertd615e | intron_variant,non_coding_transcript_variant | MODIFIER |
| 5 | 45341811 | AX-170350645 | 0 | G | A | 0.5 | 9.79E-02 | 1.02E-01 | 2.42E+00 | 2.47E+00 | 3.50E-01 | no | D5Ertd615e | intron_variant,non_coding_transcript_variant | MODIFIER |
| 5 | 45342937 | AX-170406902 | 0 | G | A | 0.5 | 9.79E-02 | 1.02E-01 | 2.42E+00 | 2.47E+00 | 3.50E-01 | no | D5Ertd615e | intron_variant,non_coding_transcript_variant | MODIFIER |
| 5 | 45343020 | AX-170402525 | 0 | G | A | 0.5 | 9.79E-02 | 1.02E-01 | 2.42E+00 | 2.47E+00 | 3.50E-01 | no | D5Ertd615e | intron_variant,non_coding_transcript_variant | MODIFIER |
| 5 | 45345150 | AX-170099007 | 0 | A | G | 0.5 | 9.79E-02 | 1.02E-01 | 2.42E+00 | 2.47E+00 | 3.50E-01 | no | D5Ertd615e | intron_variant,non_coding_transcript_variant | MODIFIER |
| 5 | 45391798 | AX-97832291 | 0 | C | T | 0.5 | 9.79E-02 | 1.02E-01 | 2.42E+00 | 2.47E+00 | 3.50E-01 | no | D5Ertd615e | intron_variant,non_coding_transcript_variant | MODIFIER |
| 5 | 45402534 | AX-170152518 | 0 | T | C | 0.5 | 9.79E-02 | 1.02E-01 | 2.42E+00 | 2.47E+00 | 3.50E-01 | no | D5Ertd615e | intron_variant,non_coding_transcript_variant | MODIFIER |
| 5 | 45404591 | AX-170326758 | 0 | G | T | 0.5 | 9.79E-02 | 1.02E-01 | 2.42E+00 | 2.47E+00 | 3.50E-01 | no | D5Ertd615e | intron_variant,non_coding_transcript_variant | MODIFIER |
| 5 | 45443252 | AX-170244502 | 0 | G | A | 0.5 | 9.79E-02 | 1.02E-01 | 2.42E+00 | 2.47E+00 | 3.50E-01 | no | Qdpr | intron_variant | MODIFIER |
| 5 | 45499248 | AX-170431573 | 0 | T | C | 0.5 | 9.79E-02 | 1.02E-01 | 2.42E+00 | 2.47E+00 | 3.50E-01 | no | Lap3 | intron_variant | MODIFIER |
| 5 | 45529975 | AX-170071955 | 0 | G | A | 0.5 | 9.79E-02 | 1.02E-01 | 2.42E+00 | 2.47E+00 | 3.50E-01 | no | Med28 | downstream_gene_variant | MODIFIER |
| 5 | 45533334 | AX-170450586 | 0 | A | G | 0.5 | 9.79E-02 | 1.02E-01 | 2.42E+00 | 2.47E+00 | 3.50E-01 | no | Fam184b | intron_variant | MODIFIER |
| 5 | 45563777 | AX-170381933 | 0 | T | G | 0.5 | 9.79E-02 | 1.02E-01 | 2.42E+00 | 2.47E+00 | 3.50E-01 | no | Fam184b | intron_variant | MODIFIER |
| 5 | 45565561 | AX-170392116 | 0 | A | G | 0.5 | 9.79E-02 | 1.02E-01 | 2.42E+00 | 2.47E+00 | 3.50E-01 | no | Fam184b | intron_variant | MODIFIER |
| 5 | 45568716 | AX-170501215 | 0 | A | C | 0.5 | 9.79E-02 | 1.02E-01 | 2.42E+00 | 2.47E+00 | 3.50E-01 | no | Fam184b | intron_variant | MODIFIER |
| 5 | 45670318 | AX-170024840 | 0 | G | A | 0.5 | 9.79E-02 | 1.02E-01 | 2.42E+00 | 2.47E+00 | 3.50E-01 | no | Ncapg | intron_variant | MODIFIER |
| 5 | 45695402 | AX-170024844 | 0 | G | A | 0.5 | 9.79E-02 | 1.02E-01 | 2.42E+00 | 2.47E+00 | 3.50E-01 | no | Lcorl | downstream_gene_variant | MODIFIER |
| 5 | 45722344 | AX-97837130 | 0 | T | C | 0.5 | 9.79E-02 | 1.02E-01 | 2.42E+00 | 2.47E+00 | 3.50E-01 | no | Lcorl | intron_variant | MODIFIER |
| 5 | 45769957 | AX-170403283 | 0 | T | C | 0.5 | 9.79E-02 | 1.02E-01 | 2.42E+00 | 2.47E+00 | 3.50E-01 | no | Lcorl | intron_variant | MODIFIER |
| 5 | 45783594 | AX-170112365 | 0 | T | C | 0.5 | 9.79E-02 | 1.02E-01 | 2.42E+00 | 2.47E+00 | 3.50E-01 | no | Lcorl | intron_variant | MODIFIER |
| 5 | 45797975 | AX-170521199 | 0 | A | G | 0.5 | 9.79E-02 | 1.02E-01 | 2.42E+00 | 2.47E+00 | 3.50E-01 | no | Lcorl | intron_variant | MODIFIER |
| 5 | 45807797 | AX-170492308 | 0 | G | A | 0.5 | 9.79E-02 | 1.02E-01 | 2.42E+00 | 2.47E+00 | 3.50E-01 | no | Lcorl | intron_variant | MODIFIER |
| 5 | 45834601 | AX-170224328 | 0 | C | A | 0.5 | 9.79E-02 | 1.02E-01 | 2.42E+00 | 2.47E+00 | 3.50E-01 | no | Lcorl | intron_variant | MODIFIER |
| 5 | 45835074 | AX-170430771 | 0 | T | G | 0.5 | 9.79E-02 | 1.02E-01 | 2.42E+00 | 2.47E+00 | 3.50E-01 | no | Lcorl | intron_variant | MODIFIER |
| 5 | 45840908 | AX-170413941 | 0 | C | G | 0.5 | 9.79E-02 | 1.02E-01 | 2.42E+00 | 2.47E+00 | 3.50E-01 | no | Lcorl | intron_variant | MODIFIER |
| 5 | 45841694 | AX-97807197 | 0 | C | T | 0.5 | 9.79E-02 | 1.02E-01 | 2.42E+00 | 2.47E+00 | 3.50E-01 | no | Lcorl | intron_variant | MODIFIER |
| 5 | 45871885 | AX-170527238 | 0 | C | A | 0.5 | 9.79E-02 | 1.02E-01 | 2.42E+00 | 2.47E+00 | 3.50E-01 | no | | intergenic_variant | MODIFIER |
| 5 | 45872017 | AX-170136770 | 0 | T | C | 0.5 | 9.79E-02 | 1.02E-01 | 2.42E+00 | 2.47E+00 | 3.50E-01 | no | | intergenic_variant | MODIFIER |
| 5 | 45882315 | AX-97831478 | 0 | T | C | 0.5 | 9.79E-02 | 1.02E-01 | 2.42E+00 | 2.47E+00 | 3.50E-01 | no | | intergenic_variant | MODIFIER |
| 5 | 45882782 | AX-170199765 | 0 | C | T | 0.5 | 9.79E-02 | 1.02E-01 | 2.42E+00 | 2.47E+00 | 3.50E-01 | no | | intergenic_variant | MODIFIER |
| 5 | 45894716 | AX-170226427 | 0 | T | C | 0.5 | 9.79E-02 | 1.02E-01 | 2.42E+00 | 2.47E+00 | 3.50E-01 | no | | intergenic_variant | MODIFIER |
| 5 | 45895347 | AX-170474714 | 0 | A | T | 0.5 | 9.79E-02 | 1.02E-01 | 2.42E+00 | 2.47E+00 | 3.50E-01 | no | | intergenic_variant | MODIFIER |
| 5 | 45916808 | AX-170364235 | 0 | A | G | 0.5 | 9.79E-02 | 1.02E-01 | 2.42E+00 | 2.47E+00 | 3.50E-01 | no | | intergenic_variant | MODIFIER |
| 5 | 45920105 | AX-170396031 | 0 | C | T | 0.5 | 9.79E-02 | 1.02E-01 | 2.42E+00 | 2.47E+00 | 3.50E-01 | no | | intergenic_variant | MODIFIER |
| 5 | 45920220 | AX-170357132 | 0 | A | T | 0.5 | 9.79E-02 | 1.02E-01 | 2.42E+00 | 2.47E+00 | 3.50E-01 | no | | intergenic_variant | MODIFIER |
| 5 | 45975630 | AX-170416957 | 0 | A | G | 0.5 | 9.79E-02 | 1.02E-01 | 2.42E+00 | 2.47E+00 | 3.50E-01 | no | | intergenic_variant | MODIFIER |

| | | | | | | | | | | | | | | | |
|---|---|---|---|---|---|---|---|---|---|---|---|---|---|---|---|
| 5 | 45988110 | AX-170109291 | 0 | C | T | 0.5 | 9.79E-02 | 1.02E-01 | 2.42E+00 | 2.47E+00 | 3.50E-01 | no | | intergenic_variant | MODIFIER |
| 5 | 45996507 | AX-170341006 | 0 | C | T | 0.5 | 9.79E-02 | 1.02E-01 | 2.42E+00 | 2.47E+00 | 3.50E-01 | no | | intergenic_variant | MODIFIER |
| 5 | 46010511 | AX-170471047 | 0 | C | T | 0.5 | 9.79E-02 | 1.02E-01 | 2.42E+00 | 2.47E+00 | 3.50E-01 | no | | intergenic_variant | MODIFIER |
| 5 | 46024032 | AX-170496096 | 0 | A | G | 0.5 | 9.79E-02 | 1.02E-01 | 2.42E+00 | 2.47E+00 | 3.50E-01 | no | | intergenic_variant | MODIFIER |
| 5 | 46024154 | AX-170223393 | 0 | C | T | 0.5 | 9.79E-02 | 1.02E-01 | 2.42E+00 | 2.47E+00 | 3.50E-01 | no | | intergenic_variant | MODIFIER |
| 5 | 46095969 | AX-170455062 | 0 | G | A | 0.5 | 9.79E-02 | 1.02E-01 | 2.42E+00 | 2.47E+00 | 3.50E-01 | no | | intergenic_variant | MODIFIER |
| 5 | 46108845 | AX-170118959 | 0 | G | A | 0.5 | 9.79E-02 | 1.02E-01 | 2.42E+00 | 2.47E+00 | 3.50E-01 | no | | intergenic_variant | MODIFIER |
| 5 | 46220451 | AX-170422548 | 0 | G | A | 0.5 | 9.79E-02 | 1.02E-01 | 2.42E+00 | 2.47E+00 | 3.50E-01 | no | | intergenic_variant | MODIFIER |
| 5 | 46230447 | AX-170483784 | 0 | C | T | 0.5 | 9.79E-02 | 1.02E-01 | 2.42E+00 | 2.47E+00 | 3.50E-01 | no | | intergenic_variant | MODIFIER |
| 5 | 46233095 | AX-97804797 | 0 | A | G | 0.5 | 9.79E-02 | 1.02E-01 | 2.42E+00 | 2.47E+00 | 3.50E-01 | no | | intergenic_variant | MODIFIER |
| 5 | 46253131 | AX-170497214 | 0 | G | A | 0.5 | 9.79E-02 | 1.02E-01 | 2.42E+00 | 2.47E+00 | 3.50E-01 | no | | intergenic_variant | MODIFIER |
| 5 | 46292509 | AX-170403701 | 0 | C | T | 0.5 | 9.79E-02 | 1.02E-01 | 2.42E+00 | 2.47E+00 | 3.50E-01 | no | | intergenic_variant | MODIFIER |
| 5 | 46363511 | AX-170246789 | 0 | C | A | 0.5 | 9.79E-02 | 1.02E-01 | 2.42E+00 | 2.47E+00 | 3.50E-01 | no | | intergenic_variant | MODIFIER |
| 5 | 46366934 | AX-170108194 | 0 | A | G | 0.5 | 9.79E-02 | 1.02E-01 | 2.42E+00 | 2.47E+00 | 3.50E-01 | no | | intergenic_variant | MODIFIER |
| 5 | 46367950 | AX-170363114 | 0 | C | T | 0.5 | 9.79E-02 | 1.02E-01 | 2.42E+00 | 2.47E+00 | 3.50E-01 | no | | intergenic_variant | MODIFIER |
| 5 | 46592249 | AX-170380696 | 0 | G | A | 0.5 | 9.79E-02 | 1.02E-01 | 2.42E+00 | 2.47E+00 | 3.50E-01 | no | | intergenic_variant | MODIFIER |
| 5 | 46600234 | AX-170478257 | 0 | G | A | 0.5 | 9.79E-02 | 1.02E-01 | 2.42E+00 | 2.47E+00 | 3.50E-01 | no | | intergenic_variant | MODIFIER |
| 5 | 46600309 | AX-170260144 | 0 | C | T | 0.5 | 9.79E-02 | 1.02E-01 | 2.42E+00 | 2.47E+00 | 3.50E-01 | no | | intergenic_variant | MODIFIER |
| 5 | 46608821 | AX-170473393 | 0 | G | A | 0.5 | 9.79E-02 | 1.02E-01 | 2.42E+00 | 2.47E+00 | 3.50E-01 | no | | intergenic_variant | MODIFIER |
| 5 | 46642131 | AX-170152847 | 0 | G | A | 0.5 | 9.79E-02 | 1.02E-01 | 2.42E+00 | 2.47E+00 | 3.50E-01 | no | | intergenic_variant | MODIFIER |
| 5 | 46642711 | AX-170206180 | 0 | C | T | 0.5 | 9.79E-02 | 1.02E-01 | 2.42E+00 | 2.47E+00 | 3.50E-01 | no | | intergenic_variant | MODIFIER |
| 5 | 46669884 | AX-170431266 | 0 | T | A | 0.5 | 9.79E-02 | 1.02E-01 | 2.42E+00 | 2.47E+00 | 3.50E-01 | no | | intergenic_variant | MODIFIER |
| 5 | 46670321 | AX-170180819 | 0 | T | G | 0.5 | 9.79E-02 | 1.02E-01 | 2.42E+00 | 2.47E+00 | 3.50E-01 | no | | intergenic_variant | MODIFIER |
| 5 | 46671193 | AX-170499408 | 0 | T | G | 0.5 | 9.79E-02 | 1.02E-01 | 2.42E+00 | 2.47E+00 | 3.50E-01 | no | | intergenic_variant | MODIFIER |
| 5 | 46726856 | AX-170024854 | 0 | C | A | 0.5 | 9.79E-02 | 1.02E-01 | 2.42E+00 | 2.47E+00 | 3.50E-01 | no | | intergenic_variant | MODIFIER |
| 5 | 46728249 | AX-170186429 | 0 | A | T | 0.5 | 9.79E-02 | 1.02E-01 | 2.42E+00 | 2.47E+00 | 3.50E-01 | no | | intergenic_variant | MODIFIER |
| 5 | 46731967 | AX-97799183 | 0 | T | C | 0.5 | 9.79E-02 | 1.02E-01 | 2.42E+00 | 2.47E+00 | 3.50E-01 | no | | intergenic_variant | MODIFIER |
| 5 | 46737339 | AX-170423224 | 0 | G | A | 0.5 | 9.79E-02 | 1.02E-01 | 2.42E+00 | 2.47E+00 | 3.50E-01 | no | | intergenic_variant | MODIFIER |
| 5 | 46773925 | AX-170024856 | 0 | A | T | 0.5 | 9.79E-02 | 1.02E-01 | 2.42E+00 | 2.47E+00 | 3.50E-01 | no | | intergenic_variant | MODIFIER |
| 5 | 46774766 | AX-170110089 | 0 | G | T | 0.5 | 9.79E-02 | 1.02E-01 | 2.42E+00 | 2.47E+00 | 3.50E-01 | no | | intergenic_variant | MODIFIER |
| 5 | 46782814 | AX-170024861 | 0 | A | G | 0.5 | 9.79E-02 | 1.02E-01 | 2.42E+00 | 2.47E+00 | 3.50E-01 | no | | intergenic_variant | MODIFIER |
| 5 | 46797501 | AX-170024857 | 0 | T | C | 0.5 | 9.79E-02 | 1.02E-01 | 2.42E+00 | 2.47E+00 | 3.50E-01 | no | Gm43092 | downstream_gene_variant | MODIFIER |
| 5 | 46842044 | AX-170071980 | 0 | T | G | 0.5 | 9.79E-02 | 1.02E-01 | 2.42E+00 | 2.47E+00 | 3.50E-01 | no | | intergenic_variant | MODIFIER |
| 5 | 46851892 | AX-170134018 | 0 | A | G | 0.5 | 9.79E-02 | 1.02E-01 | 2.42E+00 | 2.47E+00 | 3.50E-01 | no | | intergenic_variant | MODIFIER |
| 5 | 46857493 | AX-170024858 | 0 | A | G | 0.5 | 9.79E-02 | 1.02E-01 | 2.42E+00 | 2.47E+00 | 3.50E-01 | no | | intergenic_variant | MODIFIER |
| 5 | 46864947 | AX-170462986 | 0 | G | A | 0.5 | 9.79E-02 | 1.02E-01 | 2.42E+00 | 2.47E+00 | 3.50E-01 | no | | intergenic_variant | MODIFIER |
| 5 | 46869660 | AX-97848112 | 0 | G | A | 0.5 | 9.79E-02 | 1.02E-01 | 2.42E+00 | 2.47E+00 | 3.50E-01 | no | | intergenic_variant | MODIFIER |
| 5 | 46900452 | AX-170024860 | 0 | G | A | 0.5 | 9.79E-02 | 1.02E-01 | 2.42E+00 | 2.47E+00 | 3.50E-01 | no | | intergenic_variant | MODIFIER |
| 5 | 46927993 | AX-97846680 | 0 | G | A | 0.5 | 9.79E-02 | 1.02E-01 | 2.42E+00 | 2.47E+00 | 3.50E-01 | no | | intergenic_variant | MODIFIER |
| 5 | 46930956 | AX-170024853 | 0 | G | A | 0.5 | 9.79E-02 | 1.02E-01 | 2.42E+00 | 2.47E+00 | 3.50E-01 | no | | intergenic_variant | MODIFIER |
| 5 | 46940309 | AX-170172333 | 0 | C | T | 0.5 | 9.79E-02 | 1.02E-01 | 2.42E+00 | 2.47E+00 | 3.50E-01 | no | | intergenic_variant | MODIFIER |
| 5 | 46947864 | AX-170292747 | 0 | G | A | 0.5 | 9.79E-02 | 1.02E-01 | 2.42E+00 | 2.47E+00 | 3.50E-01 | no | | intergenic_variant | MODIFIER |
| 5 | 46948144 | AX-170491604 | 0 | A | T | 0.5 | 9.79E-02 | 1.02E-01 | 2.42E+00 | 2.47E+00 | 3.50E-01 | no | | intergenic_variant | MODIFIER |
| 5 | 46963651 | AX-170451804 | 0 | T | G | 0.5 | 9.79E-02 | 1.02E-01 | 2.42E+00 | 2.47E+00 | 3.50E-01 | no | | intergenic_variant | MODIFIER |
| 5 | 47005075 | AX-170511681 | 0 | A | G | 0.5 | 9.79E-02 | 1.02E-01 | 2.42E+00 | 2.47E+00 | 3.50E-01 | no | | intergenic_variant | MODIFIER |
| 5 | 47034355 | AX-170521812 | 0 | G | T | 0.5 | 9.79E-02 | 1.02E-01 | 2.42E+00 | 2.47E+00 | 3.50E-01 | no | | regulatory_region_variant | MODIFIER |
| 5 | 47133065 | AX-170196202 | 0 | A | C | 0.5 | 9.79E-02 | 1.02E-01 | 2.42E+00 | 2.47E+00 | 3.50E-01 | no | | intergenic_variant | MODIFIER |
| 5 | 47191935 | AX-170149601 | 0 | A | T | 0.5 | 9.79E-02 | 1.02E-01 | 2.42E+00 | 2.47E+00 | 3.50E-01 | no | | intergenic_variant | MODIFIER |
| 5 | 47557133 | AX-170343611 | 0 | T | A | 0.5 | 9.79E-02 | 1.02E-01 | 2.42E+00 | 2.47E+00 | 3.50E-01 | no | | regulatory_region_variant | MODIFIER |
| 5 | 47676618 | AX-170407012 | 0 | C | T | 0.5 | 9.79E-02 | 1.02E-01 | 2.42E+00 | 2.47E+00 | 3.50E-01 | no | | intergenic_variant | MODIFIER |
| 5 | 47699845 | AX-170250538 | 0 | C | A | 0.5 | 9.79E-02 | 1.02E-01 | 2.42E+00 | 2.47E+00 | 3.50E-01 | no | Gm3010 | non_coding_transcript_exon_variant | MODIFIER |
| 5 | 47700186 | AX-170188680 | 0 | G | A | 0.5 | 9.79E-02 | 1.02E-01 | 2.42E+00 | 2.47E+00 | 3.50E-01 | no | Gm3010 | non_coding_transcript_exon_variant | MODIFIER |
| 5 | 47731817 | AX-97797501 | 0 | A | G | 0.5 | 9.79E-02 | 1.02E-01 | 2.42E+00 | 2.47E+00 | 3.50E-01 | no | | intergenic_variant | MODIFIER |
| 4 | 139549836 | AX-97811707 | 1 | C | T | 0.161 | 1.07E-01 | 1.15E-01 | 2.60E+00 | 2.65E+00 | 3.50E-01 | no | Iffo2 | intron_variant | MODIFIER |

| | | | | | | | | | | | | | | | |
|---|---|---|---|---|---|---|---|---|---|---|---|---|---|---|---|
| 4 | 140089410 | AX-170159507 | 1 | G | C | 0.161 | 1.07E-01 | 1.15E-01 | 2.60E+00 | 2.65E+00 | 3.50E-01 | no | Igsf21 | intron_variant | MODIFIER |
| 1 | 19705448 | AX-170149511 | 2 | T | A | 0.418 | 7.93E-02 | 8.51E-02 | 2.61E+00 | 2.66E+00 | 3.50E-01 | no | | intergenic_variant | MODIFIER |
| 10 | 18904885 | AX-170002860 | 0 | G | A | 0.211 | 9.30E-02 | 9.96E-02 | 2.93E+00 | 3.02E+00 | 3.50E-01 | no | | intergenic_variant | MODIFIER |
| 3 | 10761065 | AX-170019907 | 1 | A | G | 0.429 | -5.97E-02 | 6.49E-02 | 2.73E+00 | 2.89E+00 | 3.50E-01 | no | | intergenic_variant | MODIFIER |
| 16 | 32232384 | AX-97830841 | 0 | G | C | 0.509 | 7.42E-02 | 7.88E-02 | 2.52E+00 | 2.58E+00 | 3.50E-01 | no | Fbxo45 | 3_prime_UTR_variant | MODIFIER |
| 9 | 57792917 | AX-170164044 | 0 | T | G | 0.219 | -9.73E-02 | 1.05E-01 | 2.81E+00 | 2.93E+00 | 3.50E-01 | no | Arid3b | intron_variant | MODIFIER |
| 3 | 14807268 | AX-170019985 | 0 | T | C | 0.412 | -5.76E-02 | 6.26E-02 | 2.74E+00 | 2.90E+00 | 3.50E-01 | no | Car1 | intron_variant | MODIFIER |
| 6 | 127092933 | AX-170510152 | 1 | A | G | 0.536 | 7.75E-02 | 8.19E-02 | 2.48E+00 | 2.55E+00 | 3.50E-01 | no | Tigar | intron_variant | MODIFIER |
| 1 | 127169741 | AX-170001649 | 1 | T | C | 0.545 | 7.83E-02 | 8.44E-02 | 2.62E+00 | 2.68E+00 | 3.50E-01 | no | | intergenic_variant | MODIFIER |
| 2 | 161418112 | AX-170356664 | 1 | C | T | 0.161 | 1.84E-01 | 1.99E-01 | 2.79E+00 | 2.87E+00 | 3.50E-01 | Yes | | intergenic_variant | MODIFIER |
| 10 | 92232532 | AX-170505957 | 1 | T | A | 0.295 | -7.88E-02 | 8.39E-02 | 2.52E+00 | 2.58E+00 | 3.50E-01 | no | Gm20757 | intron_variant,non_coding_transcript_variant | MODIFIER |
| 1 | 173205930 | AX-170385389 | 1 | A | T | 0.241 | 9.74E-02 | 1.04E-01 | 2.56E+00 | 2.65E+00 | 3.51E-01 | no | Olfr218 | downstream_gene_variant | MODIFIER |
| 9 | 5091498 | AX-170309154 | 0 | C | A | 0.474 | 7.35E-02 | 7.84E-02 | 2.52E+00 | 2.62E+00 | 3.51E-01 | no | | intergenic_variant | MODIFIER |
| 8 | 37832642 | AX-170086036 | 0 | A | G | 0.132 | -1.17E-01 | 1.27E-01 | 2.66E+00 | 2.76E+00 | 3.51E-01 | no | Sgcz | intron_variant | MODIFIER |
| 5 | 36362227 | AX-170406643 | 0 | A | G | 0.333 | 9.64E-02 | 1.04E-01 | 2.74E+00 | 2.81E+00 | 3.51E-01 | no | Sorcs2 | intron_variant | MODIFIER |
| 7 | 77212544 | AX-170416940 | 0 | A | G | 0.368 | -9.04E-02 | 9.81E-02 | 2.77E+00 | 2.87E+00 | 3.51E-01 | no | | intergenic_variant | MODIFIER |
| 14 | 119914870 | AX-170116414 | 0 | A | C | 0.342 | 6.94E-02 | 7.48E-02 | 2.60E+00 | 2.70E+00 | 3.51E-01 | no | | regulatory_region_variant | MODIFIER |
| 14 | 96010434 | AX-170513494 | 2 | C | T | 0.036 | 1.24E-01 | 1.35E-01 | 2.68E+00 | 2.86E+00 | 3.51E-01 | no | | intergenic_variant | MODIFIER |
| 4 | 124051418 | AX-170174432 | 0 | T | C | 0.219 | 1.02E-01 | 1.09E-01 | 2.55E+00 | 2.62E+00 | 3.51E-01 | no | | intergenic_variant | MODIFIER |
| 16 | 45300418 | AX-170122859 | 1 | G | C | 0.473 | -7.53E-02 | 8.02E-02 | 2.52E+00 | 2.57E+00 | 3.51E-01 | no | | intergenic_variant | MODIFIER |
| 18 | 88070830 | AX-170342977 | 0 | A | C | 0.158 | -9.75E-02 | 1.04E-01 | 2.53E+00 | 2.62E+00 | 3.51E-01 | no | | intergenic_variant | MODIFIER |
| 16 | 80079488 | AX-170529110 | 0 | C | T | 0.474 | 6.83E-02 | 7.33E-02 | 2.56E+00 | 2.63E+00 | 3.51E-01 | no | | intergenic_variant | MODIFIER |
| 3 | 30394714 | AX-170358011 | 0 | T | G | 0.158 | -8.97E-02 | 9.73E-02 | 2.76E+00 | 2.86E+00 | 3.51E-01 | no | Mecom | intron_variant | MODIFIER |
| 14 | 117649981 | AX-170189788 | 0 | G | A | 0.342 | 6.69E-02 | 7.26E-02 | 2.66E+00 | 2.78E+00 | 3.51E-01 | no | Gpc6 | intron_variant | MODIFIER |
| 5 | 80890831 | AX-170142002 | 0 | T | C | 0.439 | 9.69E-02 | 1.05E-01 | 2.64E+00 | 2.71E+00 | 3.51E-01 | no | Gm22974 | upstream_gene_variant | MODIFIER |
| 8 | 32591064 | AX-170030458 | 0 | A | G | 0.14 | -1.03E-01 | 1.11E-01 | 2.64E+00 | 2.73E+00 | 3.51E-01 | no | Nrg1 | intron_variant | MODIFIER |
| 5 | 114878955 | AX-170073195 | 1 | A | G | 0.455 | 1.03E-01 | 1.11E-01 | 2.65E+00 | 2.70E+00 | 3.51E-01 | no | 4930519G04Rik | intron_variant | MODIFIER |
| 8 | 121211859 | AX-170272184 | 0 | T | C | 0.044 | -1.56E-01 | 1.70E-01 | 2.70E+00 | 2.82E+00 | 3.51E-01 | no | Gm20388 | intron_variant | MODIFIER |
| 16 | 80512686 | AX-97822379 | 1 | T | G | 0.464 | 7.00E-02 | 7.53E-02 | 2.59E+00 | 2.66E+00 | 3.52E-01 | no | | regulatory_region_variant | MODIFIER |
| 4 | 136215120 | AX-97819654 | 1 | A | G | 0.17 | 1.01E-01 | 1.09E-01 | 2.63E+00 | 2.68E+00 | 3.52E-01 | no | Asap3 | intron_variant | MODIFIER |
| 9 | 16228057 | AX-97844941 | 0 | C | A | 0.482 | 8.27E-02 | 8.84E-02 | 2.53E+00 | 2.60E+00 | 3.52E-01 | no | Fat3 | intron_variant | MODIFIER |
| 9 | 16228430 | AX-170120808 | 0 | A | C | 0.482 | 8.27E-02 | 8.84E-02 | 2.53E+00 | 2.60E+00 | 3.52E-01 | no | Fat3 | intron_variant | MODIFIER |
| 9 | 16246492 | AX-97850004 | 0 | T | G | 0.482 | 8.27E-02 | 8.84E-02 | 2.53E+00 | 2.60E+00 | 3.52E-01 | no | Fat3 | intron_variant | MODIFIER |
| 9 | 16250942 | AX-170031845 | 0 | A | G | 0.482 | 8.27E-02 | 8.84E-02 | 2.53E+00 | 2.60E+00 | 3.52E-01 | no | Fat3 | intron_variant | MODIFIER |
| 9 | 16257946 | AX-170311134 | 0 | T | C | 0.482 | 8.27E-02 | 8.84E-02 | 2.53E+00 | 2.60E+00 | 3.52E-01 | no | Fat3 | intron_variant | MODIFIER |
| 9 | 16279947 | AX-97844121 | 0 | G | A | 0.482 | 8.27E-02 | 8.84E-02 | 2.53E+00 | 2.60E+00 | 3.52E-01 | no | Fat3 | intron_variant | MODIFIER |
| 9 | 16287998 | AX-170031846 | 0 | T | C | 0.482 | 8.27E-02 | 8.84E-02 | 2.53E+00 | 2.60E+00 | 3.52E-01 | no | Fat3 | intron_variant | MODIFIER |
| 9 | 16291085 | AX-170366396 | 0 | T | A | 0.482 | 8.27E-02 | 8.84E-02 | 2.53E+00 | 2.60E+00 | 3.52E-01 | no | Fat3 | intron_variant | MODIFIER |
| 9 | 16298797 | AX-170096479 | 0 | G | A | 0.482 | 8.27E-02 | 8.84E-02 | 2.53E+00 | 2.60E+00 | 3.52E-01 | no | Fat3 | intron_variant | MODIFIER |
| 9 | 16298850 | AX-170107433 | 0 | C | T | 0.482 | 8.27E-02 | 8.84E-02 | 2.53E+00 | 2.60E+00 | 3.52E-01 | no | Fat3 | intron_variant | MODIFIER |
| 9 | 16299119 | AX-170453402 | 0 | C | T | 0.482 | 8.27E-02 | 8.84E-02 | 2.53E+00 | 2.60E+00 | 3.52E-01 | no | Fat3 | intron_variant | MODIFIER |
| 9 | 16303665 | AX-170031847 | 0 | T | G | 0.482 | 8.27E-02 | 8.84E-02 | 2.53E+00 | 2.60E+00 | 3.52E-01 | no | Fat3 | intron_variant | MODIFIER |
| 9 | 16306080 | AX-170031844 | 0 | A | G | 0.482 | 8.27E-02 | 8.84E-02 | 2.53E+00 | 2.60E+00 | 3.52E-01 | no | Fat3 | intron_variant | MODIFIER |
| 9 | 16332169 | AX-97848012 | 0 | C | T | 0.482 | 8.27E-02 | 8.84E-02 | 2.53E+00 | 2.60E+00 | 3.52E-01 | no | Fat3 | intron_variant | MODIFIER |
| 9 | 16340441 | AX-170446654 | 0 | T | G | 0.482 | 8.27E-02 | 8.84E-02 | 2.53E+00 | 2.60E+00 | 3.52E-01 | no | Fat3 | intron_variant | MODIFIER |
| 9 | 16341502 | AX-170137689 | 0 | C | T | 0.482 | 8.27E-02 | 8.84E-02 | 2.53E+00 | 2.60E+00 | 3.52E-01 | no | Fat3 | intron_variant | MODIFIER |
| 9 | 16344551 | AX-170197340 | 0 | T | G | 0.482 | 8.27E-02 | 8.84E-02 | 2.53E+00 | 2.60E+00 | 3.52E-01 | no | Fat3 | intron_variant | MODIFIER |
| 9 | 16344630 | AX-170144309 | 0 | A | C | 0.482 | 8.27E-02 | 8.84E-02 | 2.53E+00 | 2.60E+00 | 3.52E-01 | no | Fat3 | intron_variant | MODIFIER |
| 9 | 16346249 | AX-97793520 | 0 | T | G | 0.482 | 8.27E-02 | 8.84E-02 | 2.53E+00 | 2.60E+00 | 3.52E-01 | no | Fat3 | intron_variant | MODIFIER |
| 9 | 16357398 | AX-170081772 | 0 | C | T | 0.482 | 8.27E-02 | 8.84E-02 | 2.53E+00 | 2.60E+00 | 3.52E-01 | no | Fat3 | intron_variant | MODIFIER |
| 9 | 16376022 | AX-97806326 | 0 | G | A | 0.482 | 8.27E-02 | 8.84E-02 | 2.53E+00 | 2.60E+00 | 3.52E-01 | no | Fat3 | missense_variant | MODERATE |
| 9 | 16380565 | AX-170476295 | 0 | G | A | 0.482 | 8.27E-02 | 8.84E-02 | 2.53E+00 | 2.60E+00 | 3.52E-01 | no | Fat3 | upstream_gene_variant | MODIFIER |
| 9 | 16402379 | AX-170096480 | 0 | A | G | 0.482 | 8.27E-02 | 8.84E-02 | 2.53E+00 | 2.60E+00 | 3.52E-01 | no | Fat3 | intron_variant | MODIFIER |
| 9 | 16574474 | AX-97816515 | 0 | G | A | 0.482 | 8.27E-02 | 8.84E-02 | 2.53E+00 | 2.60E+00 | 3.52E-01 | no | | intergenic_variant | MODIFIER |

| | | | | | | | | | | | | | | | |
|---|---|---|---|---|---|---|---|---|---|---|---|---|---|---|---|
| 9 | 16579748 | AX-170081778 | 0 | C | T | 0.482 | 8.27E-02 | 8.84E-02 | 2.53E+00 | 2.60E+00 | 3.52E-01 | no | | intergenic_variant | MODIFIER |
| 9 | 16580538 | AX-170291524 | 0 | G | C | 0.482 | 8.27E-02 | 8.84E-02 | 2.53E+00 | 2.60E+00 | 3.52E-01 | no | | intergenic_variant | MODIFIER |
| 9 | 16708187 | AX-170499980 | 0 | A | C | 0.482 | 8.27E-02 | 8.84E-02 | 2.53E+00 | 2.60E+00 | 3.52E-01 | no | | intergenic_variant | MODIFIER |
| 9 | 16708401 | AX-170401255 | 0 | A | C | 0.482 | 8.27E-02 | 8.84E-02 | 2.53E+00 | 2.60E+00 | 3.52E-01 | no | | intergenic_variant | MODIFIER |
| 9 | 16708855 | AX-170125726 | 0 | T | C | 0.482 | 8.27E-02 | 8.84E-02 | 2.53E+00 | 2.60E+00 | 3.52E-01 | no | | intergenic_variant | MODIFIER |
| 9 | 16728045 | AX-170127753 | 0 | C | G | 0.482 | 8.27E-02 | 8.84E-02 | 2.53E+00 | 2.60E+00 | 3.52E-01 | no | | intergenic_variant | MODIFIER |
| 9 | 16729863 | AX-170256177 | 0 | C | T | 0.482 | 8.27E-02 | 8.84E-02 | 2.53E+00 | 2.60E+00 | 3.52E-01 | no | | intergenic_variant | MODIFIER |
| 9 | 16729968 | AX-170090249 | 0 | C | A | 0.482 | 8.27E-02 | 8.84E-02 | 2.53E+00 | 2.60E+00 | 3.52E-01 | no | | intergenic_variant | MODIFIER |
| 9 | 16730820 | AX-170490172 | 0 | T | C | 0.482 | 8.27E-02 | 8.84E-02 | 2.53E+00 | 2.60E+00 | 3.52E-01 | no | | intergenic_variant | MODIFIER |
| 9 | 16814704 | AX-97782214 | 0 | A | T | 0.482 | 8.27E-02 | 8.84E-02 | 2.53E+00 | 2.60E+00 | 3.52E-01 | no | | intergenic_variant | MODIFIER |
| 9 | 16926358 | AX-170031852 | 0 | C | T | 0.482 | 8.27E-02 | 8.84E-02 | 2.53E+00 | 2.60E+00 | 3.52E-01 | no | | intergenic_variant | MODIFIER |
| 9 | 16933544 | AX-170292512 | 0 | T | C | 0.482 | 8.27E-02 | 8.84E-02 | 2.53E+00 | 2.60E+00 | 3.52E-01 | no | | intergenic_variant | MODIFIER |
| 9 | 16957514 | AX-97809062 | 0 | T | C | 0.482 | 8.27E-02 | 8.84E-02 | 2.53E+00 | 2.60E+00 | 3.52E-01 | no | | intergenic_variant | MODIFIER |
| 9 | 17028511 | AX-97848121 | 0 | A | G | 0.482 | 8.27E-02 | 8.84E-02 | 2.53E+00 | 2.60E+00 | 3.52E-01 | no | Gm5611 | upstream_gene_variant | MODIFIER |
| 9 | 17029297 | AX-170031851 | 0 | G | C | 0.482 | 8.27E-02 | 8.84E-02 | 2.53E+00 | 2.60E+00 | 3.52E-01 | no | Gm5611 | upstream_gene_variant | MODIFIER |
| 9 | 17029373 | AX-170325111 | 0 | C | T | 0.482 | 8.27E-02 | 8.84E-02 | 2.53E+00 | 2.60E+00 | 3.52E-01 | no | Gm5611 | upstream_gene_variant | MODIFIER |
| 9 | 17058588 | AX-170031861 | 0 | T | C | 0.482 | 8.27E-02 | 8.84E-02 | 2.53E+00 | 2.60E+00 | 3.52E-01 | no | | intergenic_variant | MODIFIER |
| 9 | 17067095 | AX-97851852 | 0 | C | T | 0.482 | 8.27E-02 | 8.84E-02 | 2.53E+00 | 2.60E+00 | 3.52E-01 | no | | intergenic_variant | MODIFIER |
| 9 | 17117851 | AX-170031855 | 0 | A | T | 0.482 | 8.27E-02 | 8.84E-02 | 2.53E+00 | 2.60E+00 | 3.52E-01 | no | | intergenic_variant | MODIFIER |
| 9 | 17122900 | AX-170230052 | 0 | C | G | 0.482 | 8.27E-02 | 8.84E-02 | 2.53E+00 | 2.60E+00 | 3.52E-01 | no | | intergenic_variant | MODIFIER |
| 9 | 17161530 | AX-170199893 | 0 | A | C | 0.482 | 8.27E-02 | 8.84E-02 | 2.53E+00 | 2.60E+00 | 3.52E-01 | no | | intergenic_variant | MODIFIER |
| 9 | 17172798 | AX-170196652 | 0 | G | C | 0.482 | 8.27E-02 | 8.84E-02 | 2.53E+00 | 2.60E+00 | 3.52E-01 | no | | intergenic_variant | MODIFIER |
| 9 | 17215098 | AX-170136917 | 0 | A | G | 0.482 | 8.27E-02 | 8.84E-02 | 2.53E+00 | 2.60E+00 | 3.52E-01 | no | | intergenic_variant | MODIFIER |
| 9 | 17231747 | AX-170147475 | 0 | A | G | 0.482 | 8.27E-02 | 8.84E-02 | 2.53E+00 | 2.60E+00 | 3.52E-01 | no | | intergenic_variant | MODIFIER |
| 9 | 17245829 | AX-97797021 | 0 | T | G | 0.482 | 8.27E-02 | 8.84E-02 | 2.53E+00 | 2.60E+00 | 3.52E-01 | no | | intergenic_variant | MODIFIER |
| 9 | 17250280 | AX-170228643 | 0 | T | C | 0.482 | 8.27E-02 | 8.84E-02 | 2.53E+00 | 2.60E+00 | 3.52E-01 | no | | intergenic_variant | MODIFIER |
| 9 | 17257007 | AX-170192469 | 0 | T | A | 0.482 | 8.27E-02 | 8.84E-02 | 2.53E+00 | 2.60E+00 | 3.52E-01 | no | | intergenic_variant | MODIFIER |
| 9 | 17260408 | AX-170322189 | 0 | T | G | 0.482 | 8.27E-02 | 8.84E-02 | 2.53E+00 | 2.60E+00 | 3.52E-01 | no | | intergenic_variant | MODIFIER |
| 9 | 17276109 | AX-170369329 | 0 | G | A | 0.482 | 8.27E-02 | 8.84E-02 | 2.53E+00 | 2.60E+00 | 3.52E-01 | no | | intergenic_variant | MODIFIER |
| 9 | 17277846 | AX-170447134 | 0 | G | C | 0.482 | 8.27E-02 | 8.84E-02 | 2.53E+00 | 2.60E+00 | 3.52E-01 | no | | intergenic_variant | MODIFIER |
| 9 | 17278021 | AX-97788807 | 0 | G | A | 0.482 | 8.27E-02 | 8.84E-02 | 2.53E+00 | 2.60E+00 | 3.52E-01 | no | | intergenic_variant | MODIFIER |
| 9 | 17288288 | AX-97786461 | 0 | C | T | 0.482 | 8.27E-02 | 8.84E-02 | 2.53E+00 | 2.60E+00 | 3.52E-01 | no | | intergenic_variant | MODIFIER |
| 9 | 17307606 | AX-170388047 | 0 | G | A | 0.482 | 8.27E-02 | 8.84E-02 | 2.53E+00 | 2.60E+00 | 3.52E-01 | no | | intergenic_variant | MODIFIER |
| 9 | 17318645 | AX-170343292 | 0 | A | G | 0.482 | 8.27E-02 | 8.84E-02 | 2.53E+00 | 2.60E+00 | 3.52E-01 | no | | intergenic_variant | MODIFIER |
| 9 | 17321314 | AX-170447723 | 0 | G | A | 0.482 | 8.27E-02 | 8.84E-02 | 2.53E+00 | 2.60E+00 | 3.52E-01 | no | | intergenic_variant | MODIFIER |
| 9 | 17321749 | AX-97785102 | 0 | T | G | 0.482 | 8.27E-02 | 8.84E-02 | 2.53E+00 | 2.60E+00 | 3.52E-01 | no | | intergenic_variant | MODIFIER |
| 9 | 17327723 | AX-170257442 | 0 | G | T | 0.482 | 8.27E-02 | 8.84E-02 | 2.53E+00 | 2.60E+00 | 3.52E-01 | no | | intergenic_variant | MODIFIER |
| 9 | 17351506 | AX-170327135 | 0 | T | C | 0.482 | 8.27E-02 | 8.84E-02 | 2.53E+00 | 2.60E+00 | 3.52E-01 | no | | intergenic_variant | MODIFIER |
| 9 | 17359985 | AX-170081787 | 0 | A | C | 0.482 | 8.27E-02 | 8.84E-02 | 2.53E+00 | 2.60E+00 | 3.52E-01 | no | | intergenic_variant | MODIFIER |
| 9 | 17416067 | AX-170326030 | 0 | A | C | 0.482 | 8.27E-02 | 8.84E-02 | 2.53E+00 | 2.60E+00 | 3.52E-01 | no | | intergenic_variant | MODIFIER |
| 9 | 17417022 | AX-170247113 | 0 | A | C | 0.482 | 8.27E-02 | 8.84E-02 | 2.53E+00 | 2.60E+00 | 3.52E-01 | no | | intergenic_variant | MODIFIER |
| 9 | 17442611 | AX-170270539 | 0 | G | A | 0.482 | 8.27E-02 | 8.84E-02 | 2.53E+00 | 2.60E+00 | 3.52E-01 | no | | intergenic_variant | MODIFIER |
| 9 | 17444813 | AX-97844914 | 0 | T | G | 0.482 | 8.27E-02 | 8.84E-02 | 2.53E+00 | 2.60E+00 | 3.52E-01 | no | | intergenic_variant | MODIFIER |
| 9 | 17499453 | AX-97796729 | 0 | C | A | 0.482 | 8.27E-02 | 8.84E-02 | 2.53E+00 | 2.60E+00 | 3.52E-01 | no | Gm2594 | downstream_gene_variant | MODIFIER |
| 9 | 17512078 | AX-170277837 | 0 | C | T | 0.482 | 8.27E-02 | 8.84E-02 | 2.53E+00 | 2.60E+00 | 3.52E-01 | no | | intergenic_variant | MODIFIER |
| 9 | 17560320 | AX-170081791 | 0 | G | A | 0.482 | 8.27E-02 | 8.84E-02 | 2.53E+00 | 2.60E+00 | 3.52E-01 | no | | intergenic_variant | MODIFIER |
| 9 | 17565825 | AX-170129994 | 0 | C | A | 0.482 | 8.27E-02 | 8.84E-02 | 2.53E+00 | 2.60E+00 | 3.52E-01 | no | | intergenic_variant | MODIFIER |
| 9 | 17574519 | AX-97813623 | 0 | C | A | 0.482 | 8.27E-02 | 8.84E-02 | 2.53E+00 | 2.60E+00 | 3.52E-01 | no | | intergenic_variant | MODIFIER |
| 9 | 17577907 | AX-170031866 | 0 | G | A | 0.482 | 8.27E-02 | 8.84E-02 | 2.53E+00 | 2.60E+00 | 3.52E-01 | no | | intergenic_variant | MODIFIER |
| 9 | 17578113 | AX-170240668 | 0 | G | A | 0.482 | 8.27E-02 | 8.84E-02 | 2.53E+00 | 2.60E+00 | 3.52E-01 | no | | intergenic_variant | MODIFIER |
| 9 | 17581008 | AX-170215164 | 0 | G | T | 0.482 | 8.27E-02 | 8.84E-02 | 2.53E+00 | 2.60E+00 | 3.52E-01 | no | | intergenic_variant | MODIFIER |
| 9 | 17585764 | AX-170039877 | 0 | G | T | 0.482 | 8.27E-02 | 8.84E-02 | 2.53E+00 | 2.60E+00 | 3.52E-01 | no | | intergenic_variant | MODIFIER |
| 9 | 17586328 | AX-170090253 | 0 | C | T | 0.482 | 8.27E-02 | 8.84E-02 | 2.53E+00 | 2.60E+00 | 3.52E-01 | no | | intergenic_variant | MODIFIER |
| 9 | 17622486 | AX-97792224 | 0 | A | G | 0.482 | 8.27E-02 | 8.84E-02 | 2.53E+00 | 2.60E+00 | 3.52E-01 | no | | intergenic_variant | MODIFIER |

| | | | | | | | | | | | | | | | |
|---|---|---|---|---|---|---|---|---|---|---|---|---|---|---|---|
| 9 | 17623646 | AX-170109807 | 0 | C | T | 0.482 | 8.27E-02 | 8.84E-02 | 2.53E+00 | 2.60E+00 | 3.52E-01 | no | | intergenic_variant | MODIFIER |
| 9 | 17624505 | AX-170396333 | 0 | G | A | 0.482 | 8.27E-02 | 8.84E-02 | 2.53E+00 | 2.60E+00 | 3.52E-01 | no | | intergenic_variant | MODIFIER |
| 9 | 17642741 | AX-170282717 | 0 | T | G | 0.482 | 8.27E-02 | 8.84E-02 | 2.53E+00 | 2.60E+00 | 3.52E-01 | no | | intergenic_variant | MODIFIER |
| 9 | 17661354 | AX-170081793 | 0 | T | C | 0.482 | 8.27E-02 | 8.84E-02 | 2.53E+00 | 2.60E+00 | 3.52E-01 | no | | intergenic_variant | MODIFIER |
| 9 | 17666990 | AX-170511947 | 0 | C | T | 0.482 | 8.27E-02 | 8.84E-02 | 2.53E+00 | 2.60E+00 | 3.52E-01 | no | | intergenic_variant | MODIFIER |
| 9 | 17669424 | AX-170467053 | 0 | T | C | 0.482 | 8.27E-02 | 8.84E-02 | 2.53E+00 | 2.60E+00 | 3.52E-01 | no | | intergenic_variant | MODIFIER |
| 9 | 17670262 | AX-170303448 | 0 | T | C | 0.482 | 8.27E-02 | 8.84E-02 | 2.53E+00 | 2.60E+00 | 3.52E-01 | no | | intergenic_variant | MODIFIER |
| 6 | 126464383 | AX-97829761 | 0 | T | C | 0.535 | 7.76E-02 | 8.23E-02 | 2.48E+00 | 2.55E+00 | 3.52E-01 | no | | intergenic_variant | MODIFIER |
| 6 | 126614137 | AX-170182798 | 0 | A | G | 0.535 | 7.76E-02 | 8.23E-02 | 2.48E+00 | 2.55E+00 | 3.52E-01 | no | | intergenic_variant | MODIFIER |
| 6 | 126790284 | AX-97848584 | 0 | A | G | 0.535 | 7.76E-02 | 8.23E-02 | 2.48E+00 | 2.55E+00 | 3.52E-01 | no | | intergenic_variant | MODIFIER |
| 6 | 126796931 | AX-170362677 | 0 | T | C | 0.535 | 7.76E-02 | 8.23E-02 | 2.48E+00 | 2.55E+00 | 3.52E-01 | no | | intergenic_variant | MODIFIER |
| 6 | 126861989 | AX-170322065 | 0 | T | C | 0.535 | 7.76E-02 | 8.23E-02 | 2.48E+00 | 2.55E+00 | 3.52E-01 | no | Akap3 | intron_variant | MODIFIER |
| 6 | 126865370 | AX-97815188 | 0 | A | G | 0.535 | 7.76E-02 | 8.23E-02 | 2.48E+00 | 2.55E+00 | 3.52E-01 | no | Akap3 | synonymous_variant | LOW |
| 6 | 126875999 | AX-170305399 | 0 | A | G | 0.535 | 7.76E-02 | 8.23E-02 | 2.48E+00 | 2.55E+00 | 3.52E-01 | no | Dyrk4 | downstream_gene_variant | MODIFIER |
| 6 | 126878039 | AX-170028196 | 0 | A | C | 0.535 | 7.76E-02 | 8.23E-02 | 2.48E+00 | 2.55E+00 | 3.52E-01 | no | Dyrk4 | intron_variant,non_coding_transcript_variant | MODIFIER |
| 6 | 126878594 | AX-170028203 | 0 | G | A | 0.535 | 7.76E-02 | 8.23E-02 | 2.48E+00 | 2.55E+00 | 3.52E-01 | no | Dyrk4 | intron_variant,non_coding_transcript_variant | MODIFIER |
| 6 | 126890802 | AX-170382425 | 0 | T | C | 0.535 | 7.76E-02 | 8.23E-02 | 2.48E+00 | 2.55E+00 | 3.52E-01 | no | Dyrk4 | intron_variant,non_coding_transcript_variant | MODIFIER |
| 6 | 126912755 | AX-170102160 | 0 | T | A | 0.535 | 7.76E-02 | 8.23E-02 | 2.48E+00 | 2.55E+00 | 3.52E-01 | no | Dyrk4 | intron_variant | MODIFIER |
| 6 | 126939360 | AX-170028201 | 0 | G | A | 0.535 | 7.76E-02 | 8.23E-02 | 2.48E+00 | 2.55E+00 | 3.52E-01 | no | D6Wsu163e | upstream_gene_variant | MODIFIER |
| 6 | 126940734 | AX-170028199 | 0 | T | C | 0.535 | 7.76E-02 | 8.23E-02 | 2.48E+00 | 2.55E+00 | 3.52E-01 | no | D6Wsu163e | intron_variant | MODIFIER |
| 6 | 126957271 | AX-97803405 | 0 | C | T | 0.535 | 7.76E-02 | 8.23E-02 | 2.48E+00 | 2.55E+00 | 3.52E-01 | no | D6Wsu163e | intron_variant | MODIFIER |
| 6 | 126958131 | AX-170338695 | 0 | G | A | 0.535 | 7.76E-02 | 8.23E-02 | 2.48E+00 | 2.55E+00 | 3.52E-01 | no | D6Wsu163e | intron_variant | MODIFIER |
| 6 | 127007049 | AX-170406119 | 0 | A | G | 0.535 | 7.76E-02 | 8.23E-02 | 2.48E+00 | 2.55E+00 | 3.52E-01 | no | | intergenic_variant | MODIFIER |
| 6 | 127044467 | AX-170113963 | 0 | T | C | 0.535 | 7.76E-02 | 8.23E-02 | 2.48E+00 | 2.55E+00 | 3.52E-01 | no | | intergenic_variant | MODIFIER |
| 6 | 127046118 | AX-170028202 | 0 | T | A | 0.535 | 7.76E-02 | 8.23E-02 | 2.48E+00 | 2.55E+00 | 3.52E-01 | no | | intergenic_variant | MODIFIER |
| 6 | 127051987 | AX-170028205 | 0 | C | T | 0.535 | 7.76E-02 | 8.23E-02 | 2.48E+00 | 2.55E+00 | 3.52E-01 | no | | regulatory_region_variant | MODIFIER |
| 6 | 127052014 | AX-97855842 | 0 | A | C | 0.535 | 7.76E-02 | 8.23E-02 | 2.48E+00 | 2.55E+00 | 3.52E-01 | no | | regulatory_region_variant | MODIFIER |
| 6 | 127060126 | AX-170152297 | 0 | G | A | 0.535 | 7.76E-02 | 8.23E-02 | 2.48E+00 | 2.55E+00 | 3.52E-01 | no | | intergenic_variant | MODIFIER |
| 6 | 127063520 | AX-170405131 | 0 | T | C | 0.535 | 7.76E-02 | 8.23E-02 | 2.48E+00 | 2.55E+00 | 3.52E-01 | no | | intergenic_variant | MODIFIER |
| 6 | 127089942 | AX-170209876 | 0 | A | G | 0.535 | 7.76E-02 | 8.23E-02 | 2.48E+00 | 2.55E+00 | 3.52E-01 | no | Tigar | intron_variant | MODIFIER |
| 6 | 127092581 | AX-170423896 | 0 | G | A | 0.535 | 7.76E-02 | 8.23E-02 | 2.48E+00 | 2.55E+00 | 3.52E-01 | no | Tigar | intron_variant | MODIFIER |
| 6 | 127105652 | AX-97801962 | 0 | G | T | 0.535 | 7.76E-02 | 8.23E-02 | 2.48E+00 | 2.55E+00 | 3.52E-01 | no | Tigar | intron_variant | MODIFIER |
| 6 | 127134432 | AX-170286994 | 0 | A | G | 0.535 | 7.76E-02 | 8.23E-02 | 2.48E+00 | 2.55E+00 | 3.52E-01 | no | Ccnd2 | intron_variant | MODIFIER |
| 6 | 127145920 | AX-170421698 | 0 | T | C | 0.535 | 7.76E-02 | 8.23E-02 | 2.48E+00 | 2.55E+00 | 3.52E-01 | no | Ccnd2 | intron_variant | MODIFIER |
| 6 | 127164799 | AX-97803623 | 0 | A | G | 0.535 | 7.76E-02 | 8.23E-02 | 2.48E+00 | 2.55E+00 | 3.52E-01 | no | 9330179D12Rik | intron_variant,non_coding_transcript_variant | MODIFIER |
| 6 | 127200609 | AX-170270511 | 0 | T | C | 0.535 | 7.76E-02 | 8.23E-02 | 2.48E+00 | 2.55E+00 | 3.52E-01 | no | 9330179D12Rik | intron_variant,non_coding_transcript_variant | MODIFIER |
| 6 | 127274814 | AX-170251909 | 0 | A | G | 0.535 | 7.76E-02 | 8.23E-02 | 2.48E+00 | 2.55E+00 | 3.52E-01 | no | | regulatory_region_variant | MODIFIER |
| 6 | 127277512 | AX-170076167 | 0 | C | T | 0.535 | 7.76E-02 | 8.23E-02 | 2.48E+00 | 2.55E+00 | 3.52E-01 | no | Gm7308 | downstream_gene_variant | MODIFIER |
| 6 | 127317092 | AX-170189465 | 0 | T | C | 0.535 | 7.76E-02 | 8.23E-02 | 2.48E+00 | 2.55E+00 | 3.52E-01 | no | Gm43636 | upstream_gene_variant | MODIFIER |
| 6 | 127318272 | AX-170181058 | 0 | C | T | 0.535 | 7.76E-02 | 8.23E-02 | 2.48E+00 | 2.55E+00 | 3.52E-01 | no | Gm43636 | upstream_gene_variant | MODIFIER |
| 6 | 127335376 | AX-170206583 | 0 | G | C | 0.535 | 7.76E-02 | 8.23E-02 | 2.48E+00 | 2.55E+00 | 3.52E-01 | no | | regulatory_region_variant | MODIFIER |
| 6 | 127341022 | AX-97832684 | 0 | T | C | 0.535 | 7.76E-02 | 8.23E-02 | 2.48E+00 | 2.55E+00 | 3.52E-01 | no | | regulatory_region_variant | MODIFIER |
| 6 | 127345345 | AX-170124944 | 0 | G | C | 0.535 | 7.76E-02 | 8.23E-02 | 2.48E+00 | 2.55E+00 | 3.52E-01 | no | | regulatory_region_variant | MODIFIER |
| 6 | 127345675 | AX-170191536 | 0 | G | A | 0.535 | 7.76E-02 | 8.23E-02 | 2.48E+00 | 2.55E+00 | 3.52E-01 | no | | regulatory_region_variant | MODIFIER |
| 6 | 127354351 | AX-170089835 | 0 | C | T | 0.535 | 7.76E-02 | 8.23E-02 | 2.48E+00 | 2.55E+00 | 3.52E-01 | no | | intergenic_variant | MODIFIER |
| 6 | 127357570 | AX-170233176 | 0 | G | C | 0.535 | 7.76E-02 | 8.23E-02 | 2.48E+00 | 2.55E+00 | 3.52E-01 | no | | intergenic_variant | MODIFIER |
| 6 | 127359507 | AX-170156351 | 0 | C | T | 0.535 | 7.76E-02 | 8.23E-02 | 2.48E+00 | 2.55E+00 | 3.52E-01 | no | | regulatory_region_variant | MODIFIER |
| 6 | 127378432 | AX-170365224 | 0 | A | G | 0.535 | 7.76E-02 | 8.23E-02 | 2.48E+00 | 2.55E+00 | 3.52E-01 | no | | intergenic_variant | MODIFIER |
| 6 | 127384084 | AX-170407043 | 0 | C | T | 0.535 | 7.76E-02 | 8.23E-02 | 2.48E+00 | 2.55E+00 | 3.52E-01 | no | | intergenic_variant | MODIFIER |
| 6 | 127387494 | AX-170292207 | 0 | T | G | 0.535 | 7.76E-02 | 8.23E-02 | 2.48E+00 | 2.55E+00 | 3.52E-01 | no | | intergenic_variant | MODIFIER |
| 6 | 127390698 | AX-170209215 | 0 | T | C | 0.535 | 7.76E-02 | 8.23E-02 | 2.48E+00 | 2.55E+00 | 3.52E-01 | no | | intergenic_variant | MODIFIER |
| 6 | 127392226 | AX-170123001 | 0 | C | T | 0.535 | 7.76E-02 | 8.23E-02 | 2.48E+00 | 2.55E+00 | 3.52E-01 | no | | regulatory_region_variant | MODIFIER |
| 6 | 127392464 | AX-170120997 | 0 | T | C | 0.535 | 7.76E-02 | 8.23E-02 | 2.48E+00 | 2.55E+00 | 3.52E-01 | no | | regulatory_region_variant | MODIFIER |
| 6 | 127399126 | AX-170358598 | 0 | T | G | 0.535 | 7.76E-02 | 8.23E-02 | 2.48E+00 | 2.55E+00 | 3.52E-01 | no | | intergenic_variant | MODIFIER |

| | | | | | | | | | | | | | | | |
|---|---|---|---|---|---|---|---|---|---|---|---|---|---|---|---|
| 6 | 127413247 | AX-170197293 | 0 | T | C | 0.535 | 7.76E-02 | 8.23E-02 | 2.48E+00 | 2.55E+00 | 3.52E-01 | no | Gm34091 | downstream_gene_variant | MODIFIER |
| 6 | 127414210 | AX-170522050 | 0 | C | T | 0.535 | 7.76E-02 | 8.23E-02 | 2.48E+00 | 2.55E+00 | 3.52E-01 | no | Gm34091 | downstream_gene_variant | MODIFIER |
| 6 | 127441723 | AX-170328498 | 0 | G | A | 0.535 | 7.76E-02 | 8.23E-02 | 2.48E+00 | 2.55E+00 | 3.52E-01 | no | | regulatory_region_variant | MODIFIER |
| 6 | 127443889 | AX-170136132 | 0 | A | G | 0.535 | 7.76E-02 | 8.23E-02 | 2.48E+00 | 2.55E+00 | 3.52E-01 | no | Parp11 | upstream_gene_variant | MODIFIER |
| 6 | 127460860 | AX-97838529 | 0 | A | G | 0.535 | 7.76E-02 | 8.23E-02 | 2.48E+00 | 2.55E+00 | 3.52E-01 | no | Parp11 | intron_variant | MODIFIER |
| 6 | 127464431 | AX-170028215 | 0 | C | T | 0.535 | 7.76E-02 | 8.23E-02 | 2.48E+00 | 2.55E+00 | 3.52E-01 | no | Parp11 | intron_variant | MODIFIER |
| 6 | 127488500 | AX-97788475 | 0 | T | C | 0.535 | 7.76E-02 | 8.23E-02 | 2.48E+00 | 2.55E+00 | 3.52E-01 | no | Parp11 | intron_variant | MODIFIER |
| 6 | 127489475 | AX-170076172 | 0 | T | C | 0.535 | 7.76E-02 | 8.23E-02 | 2.48E+00 | 2.55E+00 | 3.52E-01 | no | Parp11 | intron_variant | MODIFIER |
| 6 | 127497015 | AX-170028206 | 0 | G | T | 0.535 | 7.76E-02 | 8.23E-02 | 2.48E+00 | 2.55E+00 | 3.52E-01 | no | Parp11 | downstream_gene_variant | MODIFIER |
| 6 | 127497140 | AX-97816769 | 0 | G | T | 0.535 | 7.76E-02 | 8.23E-02 | 2.48E+00 | 2.55E+00 | 3.52E-01 | no | Parp11 | downstream_gene_variant | MODIFIER |
| 6 | 127499504 | AX-170076169 | 0 | A | G | 0.535 | 7.76E-02 | 8.23E-02 | 2.48E+00 | 2.55E+00 | 3.52E-01 | no | | intergenic_variant | MODIFIER |
| 6 | 127533606 | AX-170028209 | 0 | A | G | 0.535 | 7.76E-02 | 8.23E-02 | 2.48E+00 | 2.55E+00 | 3.52E-01 | no | | intergenic_variant | MODIFIER |
| 6 | 127533633 | AX-170028207 | 0 | A | G | 0.535 | 7.76E-02 | 8.23E-02 | 2.48E+00 | 2.55E+00 | 3.52E-01 | no | | intergenic_variant | MODIFIER |
| 6 | 127533660 | AX-170028208 | 0 | T | C | 0.535 | 7.76E-02 | 8.23E-02 | 2.48E+00 | 2.55E+00 | 3.52E-01 | no | | intergenic_variant | MODIFIER |
| 6 | 127542458 | AX-170200945 | 0 | C | G | 0.535 | 7.76E-02 | 8.23E-02 | 2.48E+00 | 2.55E+00 | 3.52E-01 | no | | regulatory_region_variant | MODIFIER |
| 6 | 127544932 | AX-170461599 | 0 | C | A | 0.535 | 7.76E-02 | 8.23E-02 | 2.48E+00 | 2.55E+00 | 3.52E-01 | no | | intergenic_variant | MODIFIER |
| 6 | 127550135 | AX-97800923 | 0 | C | T | 0.535 | 7.76E-02 | 8.23E-02 | 2.48E+00 | 2.55E+00 | 3.52E-01 | no | | intergenic_variant | MODIFIER |
| 6 | 127568278 | AX-170448559 | 0 | C | T | 0.535 | 7.76E-02 | 8.23E-02 | 2.48E+00 | 2.55E+00 | 3.52E-01 | no | Cracr2a | intron_variant | MODIFIER |
| 6 | 127649547 | AX-97811414 | 0 | C | T | 0.535 | 7.76E-02 | 8.23E-02 | 2.48E+00 | 2.55E+00 | 3.52E-01 | no | Gm43634 | intron_variant,non_coding_transcript_variant | MODIFIER |
| 6 | 127652950 | AX-170262267 | 0 | A | G | 0.535 | 7.76E-02 | 8.23E-02 | 2.48E+00 | 2.55E+00 | 3.52E-01 | no | Gm43634 | intron_variant,non_coding_transcript_variant | MODIFIER |
| 6 | 127655309 | AX-170028210 | 0 | A | G | 0.535 | 7.76E-02 | 8.23E-02 | 2.48E+00 | 2.55E+00 | 3.52E-01 | no | Gm43634 | intron_variant,non_coding_transcript_variant | MODIFIER |
| 6 | 127657362 | AX-170028211 | 0 | A | G | 0.535 | 7.76E-02 | 8.23E-02 | 2.48E+00 | 2.55E+00 | 3.52E-01 | no | Gm43634 | intron_variant,non_coding_transcript_variant | MODIFIER |
| 6 | 127658801 | AX-170148350 | 0 | C | T | 0.535 | 7.76E-02 | 8.23E-02 | 2.48E+00 | 2.55E+00 | 3.52E-01 | no | Gm43634 | downstream_gene_variant | MODIFIER |
| 6 | 127669076 | AX-170184957 | 0 | G | C | 0.535 | 7.76E-02 | 8.23E-02 | 2.48E+00 | 2.55E+00 | 3.52E-01 | no | Cracr2a | missense_variant | MODERATE |
| 6 | 127671713 | AX-170076174 | 0 | C | G | 0.535 | 7.76E-02 | 8.23E-02 | 2.48E+00 | 2.55E+00 | 3.52E-01 | no | Cracr2a | 3_prime_UTR_variant | MODIFIER |
| 6 | 127683398 | AX-170302912 | 0 | G | A | 0.535 | 7.76E-02 | 8.23E-02 | 2.48E+00 | 2.55E+00 | 3.52E-01 | no | | regulatory_region_variant | MODIFIER |
| 6 | 127689938 | AX-170395247 | 0 | C | T | 0.535 | 7.76E-02 | 8.23E-02 | 2.48E+00 | 2.55E+00 | 3.52E-01 | no | Prmt8 | intron_variant | MODIFIER |
| 6 | 127712399 | AX-170095801 | 0 | G | A | 0.535 | 7.76E-02 | 8.23E-02 | 2.48E+00 | 2.55E+00 | 3.52E-01 | no | Prmt8 | intron_variant | MODIFIER |
| 6 | 127714249 | AX-170488447 | 0 | G | A | 0.535 | 7.76E-02 | 8.23E-02 | 2.48E+00 | 2.55E+00 | 3.52E-01 | no | Prmt8 | intron_variant | MODIFIER |
| 6 | 127716973 | AX-170483612 | 0 | G | A | 0.535 | 7.76E-02 | 8.23E-02 | 2.48E+00 | 2.55E+00 | 3.52E-01 | no | Prmt8 | intron_variant | MODIFIER |
| 6 | 127727523 | AX-170442227 | 0 | G | T | 0.535 | 7.76E-02 | 8.23E-02 | 2.48E+00 | 2.55E+00 | 3.52E-01 | no | Prmt8 | intron_variant | MODIFIER |
| 6 | 127728441 | AX-170271584 | 0 | A | C | 0.535 | 7.76E-02 | 8.23E-02 | 2.48E+00 | 2.55E+00 | 3.52E-01 | no | Prmt8 | intron_variant | MODIFIER |
| 6 | 127729680 | AX-170444431 | 0 | A | G | 0.535 | 7.76E-02 | 8.23E-02 | 2.48E+00 | 2.55E+00 | 3.52E-01 | no | Prmt8 | intron_variant | MODIFIER |
| 6 | 127735762 | AX-170312120 | 0 | T | C | 0.535 | 7.76E-02 | 8.23E-02 | 2.48E+00 | 2.55E+00 | 3.52E-01 | no | Prmt8 | intron_variant | MODIFIER |
| 6 | 127754108 | AX-170253436 | 0 | T | C | 0.535 | 7.76E-02 | 8.23E-02 | 2.48E+00 | 2.55E+00 | 3.52E-01 | no | Prmt8 | intron_variant | MODIFIER |
| 6 | 127762663 | AX-170532238 | 0 | A | G | 0.535 | 7.76E-02 | 8.23E-02 | 2.48E+00 | 2.55E+00 | 3.52E-01 | no | Prmt8 | intron_variant | MODIFIER |
| 6 | 127777800 | AX-170124313 | 0 | C | T | 0.535 | 7.76E-02 | 8.23E-02 | 2.48E+00 | 2.55E+00 | 3.52E-01 | no | Gm42741 | upstream_gene_variant | MODIFIER |
| 6 | 127783438 | AX-170255721 | 0 | C | T | 0.535 | 7.76E-02 | 8.23E-02 | 2.48E+00 | 2.55E+00 | 3.52E-01 | no | Mir7233 | downstream_gene_variant | MODIFIER |
| 6 | 127788846 | AX-170368978 | 0 | C | A | 0.535 | 7.76E-02 | 8.23E-02 | 2.48E+00 | 2.55E+00 | 3.52E-01 | no | Mir7233 | upstream_gene_variant | MODIFIER |
| 6 | 127824793 | AX-97834786 | 0 | T | G | 0.535 | 7.76E-02 | 8.23E-02 | 2.48E+00 | 2.55E+00 | 3.52E-01 | no | | intergenic_variant | MODIFIER |
| 6 | 127838014 | AX-170327273 | 0 | C | T | 0.535 | 7.76E-02 | 8.23E-02 | 2.48E+00 | 2.55E+00 | 3.52E-01 | no | | intergenic_variant | MODIFIER |
| 6 | 127850668 | AX-170302433 | 0 | T | C | 0.535 | 7.76E-02 | 8.23E-02 | 2.48E+00 | 2.55E+00 | 3.52E-01 | no | Gm23098 | downstream_gene_variant | MODIFIER |
| 6 | 127854150 | AX-170465971 | 0 | A | G | 0.535 | 7.76E-02 | 8.23E-02 | 2.48E+00 | 2.55E+00 | 3.52E-01 | no | Gm23098 | downstream_gene_variant | MODIFIER |
| 6 | 127858388 | AX-97839926 | 0 | T | G | 0.535 | 7.76E-02 | 8.23E-02 | 2.48E+00 | 2.55E+00 | 3.52E-01 | no | | intergenic_variant | MODIFIER |
| 6 | 127877187 | AX-170351764 | 0 | G | A | 0.535 | 7.76E-02 | 8.23E-02 | 2.48E+00 | 2.55E+00 | 3.52E-01 | no | Gm38901 | non_coding_transcript_exon_variant | MODIFIER |
| 6 | 127953022 | AX-97830930 | 0 | T | G | 0.535 | 7.76E-02 | 8.23E-02 | 2.48E+00 | 2.55E+00 | 3.52E-01 | no | Tspan11 | 3_prime_UTR_variant | MODIFIER |
| 6 | 127972812 | AX-170443061 | 0 | T | C | 0.535 | 7.76E-02 | 8.23E-02 | 2.48E+00 | 2.55E+00 | 3.52E-01 | no | Tspan9 | intron_variant | MODIFIER |
| 6 | 127976223 | AX-170436321 | 0 | T | C | 0.535 | 7.76E-02 | 8.23E-02 | 2.48E+00 | 2.55E+00 | 3.52E-01 | no | Tspan9 | intron_variant | MODIFIER |
| 6 | 127987471 | AX-170101982 | 0 | A | G | 0.535 | 7.76E-02 | 8.23E-02 | 2.48E+00 | 2.55E+00 | 3.52E-01 | no | Tspan9 | intron_variant | MODIFIER |
| 6 | 127987729 | AX-170076184 | 0 | C | T | 0.535 | 7.76E-02 | 8.23E-02 | 2.48E+00 | 2.55E+00 | 3.52E-01 | no | Tspan9 | intron_variant | MODIFIER |
| 6 | 127998280 | AX-170028217 | 0 | G | A | 0.535 | 7.76E-02 | 8.23E-02 | 2.48E+00 | 2.55E+00 | 3.52E-01 | no | Tspan9 | intron_variant | MODIFIER |
| 6 | 129713720 | AX-97844384 | 0 | A | C | 0.535 | 7.76E-02 | 8.23E-02 | 2.48E+00 | 2.55E+00 | 3.52E-01 | no | Klri1 | intron_variant | MODIFIER |
| 6 | 130060336 | AX-170422230 | 0 | G | A | 0.535 | 7.76E-02 | 8.23E-02 | 2.48E+00 | 2.55E+00 | 3.52E-01 | no | Klra4 | intron_variant | MODIFIER |
| 6 | 130222754 | AX-97818492 | 0 | T | G | 0.535 | 7.76E-02 | 8.23E-02 | 2.48E+00 | 2.55E+00 | 3.52E-01 | no | Klra7 | intron_variant | MODIFIER |

| | | | | | | | | | | | | | | | |
|---|---|---|---|---|---|---|---|---|---|---|---|---|---|---|---|
| 6 | 130477147 | AX-170253690 | 0 | G | A | 0.535 | 7.76E-02 | 8.23E-02 | 2.48E+00 | 2.55E+00 | 3.52E-01 | no | Gm6590 | upstream_gene_variant | MODIFIER |
| 6 | 130825300 | AX-170290636 | 0 | C | T | 0.535 | 7.76E-02 | 8.23E-02 | 2.48E+00 | 2.55E+00 | 3.52E-01 | no | Gm44118 | upstream_gene_variant | MODIFIER |
| 9 | 15703854 | AX-170292329 | 0 | C | T | 0.474 | 8.21E-02 | 8.78E-02 | 2.53E+00 | 2.60E+00 | 3.52E-01 | no | | intergenic_variant | MODIFIER |
| 9 | 15721672 | AX-97791155 | 0 | G | A | 0.474 | 8.21E-02 | 8.78E-02 | 2.53E+00 | 2.60E+00 | 3.52E-01 | no | Slc36a4 | intron_variant | MODIFIER |
| 9 | 15730009 | AX-170149997 | 0 | A | G | 0.474 | 8.21E-02 | 8.78E-02 | 2.53E+00 | 2.60E+00 | 3.52E-01 | no | Slc36a4 | intron_variant | MODIFIER |
| 9 | 15735197 | AX-170465129 | 0 | C | T | 0.474 | 8.21E-02 | 8.78E-02 | 2.53E+00 | 2.60E+00 | 3.52E-01 | no | Slc36a4 | intron_variant | MODIFIER |
| 9 | 15755226 | AX-170424670 | 0 | T | C | 0.474 | 8.21E-02 | 8.78E-02 | 2.53E+00 | 2.60E+00 | 3.52E-01 | no | | intergenic_variant | MODIFIER |
| 9 | 15778985 | AX-170260636 | 0 | G | A | 0.474 | 8.21E-02 | 8.78E-02 | 2.53E+00 | 2.60E+00 | 3.52E-01 | no | | intergenic_variant | MODIFIER |
| 9 | 15789126 | AX-170031840 | 0 | T | C | 0.474 | 8.21E-02 | 8.78E-02 | 2.53E+00 | 2.60E+00 | 3.52E-01 | no | | intergenic_variant | MODIFIER |
| 9 | 15789228 | AX-170278386 | 0 | T | C | 0.474 | 8.21E-02 | 8.78E-02 | 2.53E+00 | 2.60E+00 | 3.52E-01 | no | | intergenic_variant | MODIFIER |
| 9 | 15792727 | AX-170338043 | 0 | A | G | 0.474 | 8.21E-02 | 8.78E-02 | 2.53E+00 | 2.60E+00 | 3.52E-01 | no | | intergenic_variant | MODIFIER |
| 9 | 15792890 | AX-170365346 | 0 | G | T | 0.474 | 8.21E-02 | 8.78E-02 | 2.53E+00 | 2.60E+00 | 3.52E-01 | no | | intergenic_variant | MODIFIER |
| 9 | 15793000 | AX-97850696 | 0 | T | G | 0.474 | 8.21E-02 | 8.78E-02 | 2.53E+00 | 2.60E+00 | 3.52E-01 | no | | intergenic_variant | MODIFIER |
| 9 | 15806281 | AX-170081759 | 0 | T | C | 0.474 | 8.21E-02 | 8.78E-02 | 2.53E+00 | 2.60E+00 | 3.52E-01 | no | | intergenic_variant | MODIFIER |
| 9 | 15824837 | AX-170294230 | 0 | G | C | 0.474 | 8.21E-02 | 8.78E-02 | 2.53E+00 | 2.60E+00 | 3.52E-01 | no | Mtnr1b | 3_prime_UTR_variant,NMD_transcript_variant | MODIFIER |
| 9 | 15825910 | AX-170188389 | 0 | C | T | 0.474 | 8.21E-02 | 8.78E-02 | 2.53E+00 | 2.60E+00 | 3.52E-01 | no | Mtnr1b | intron_variant,NMD_transcript_variant | MODIFIER |
| 1 | 150508166 | AX-97828911 | 2 | G | A | 0.2 | 1.22E-01 | 1.32E-01 | 2.68E+00 | 2.75E+00 | 3.52E-01 | no | | intergenic_variant | MODIFIER |
| 16 | 35926716 | AX-97801195 | 0 | A | G | 0.491 | 7.37E-02 | 7.85E-02 | 2.51E+00 | 2.56E+00 | 3.52E-01 | no | Dtx3l | 3_prime_UTR_variant | MODIFIER |
| 16 | 85330121 | AX-170107985 | 1 | C | A | 0.446 | 7.15E-02 | 7.74E-02 | 2.64E+00 | 2.71E+00 | 3.52E-01 | no | | intergenic_variant | MODIFIER |
| 5 | 86344118 | AX-170414239 | 0 | C | T | 0.43 | 9.52E-02 | 1.02E-01 | 2.95E+00 | 3.04E+00 | 3.52E-01 | no | Tmprss11d | intron_variant | MODIFIER |
| 18 | 65591560 | AX-170061133 | 0 | G | C | 0.412 | 8.19E-02 | 8.60E-02 | 2.43E+00 | 2.48E+00 | 3.53E-01 | no | Zfp532 | intron_variant | MODIFIER |
| 7 | 24598865 | AX-170229071 | 1 | G | A | 0.402 | 8.38E-02 | 9.05E-02 | 2.57E+00 | 2.69E+00 | 3.53E-01 | no | Ethe1 | intron_variant | MODIFIER |
| 4 | 135005260 | AX-97833392 | 1 | G | A | 0.161 | 1.10E-01 | 1.18E-01 | 2.56E+00 | 2.60E+00 | 3.53E-01 | no | | intergenic_variant | MODIFIER |
| 5 | 104563518 | AX-97852375 | 1 | G | A | 0.464 | 9.18E-02 | 9.99E-02 | 2.71E+00 | 2.79E+00 | 3.53E-01 | no | | intergenic_variant | MODIFIER |
| 4 | 77490112 | AX-97806071 | 0 | C | T | 0.316 | 1.03E-01 | 1.12E-01 | 2.76E+00 | 2.83E+00 | 3.53E-01 | no | Ptprd | intron_variant,non_coding_transcript_variant | MODIFIER |
| 18 | 79121733 | AX-170408544 | 0 | G | A | 0.456 | 8.20E-02 | 8.67E-02 | 2.46E+00 | 2.53E+00 | 3.53E-01 | no | | intergenic_variant | MODIFIER |
| 18 | 79298545 | AX-170397922 | 0 | A | G | 0.456 | 8.20E-02 | 8.67E-02 | 2.46E+00 | 2.53E+00 | 3.53E-01 | no | | intergenic_variant | MODIFIER |
| 10 | 107818505 | AX-170302120 | 0 | C | T | 0.316 | -7.58E-02 | 8.25E-02 | 2.68E+00 | 2.76E+00 | 3.53E-01 | no | Otogl | intron_variant | MODIFIER |
| 8 | 109291756 | AX-170402891 | 2 | G | A | 0.545 | 4.88E-02 | 5.30E-02 | 2.85E+00 | 2.96E+00 | 3.53E-01 | no | | intergenic_variant | MODIFIER |
| 8 | 16485834 | AX-170456993 | 0 | G | A | 0.228 | -7.13E-02 | 7.71E-02 | 2.58E+00 | 2.68E+00 | 3.53E-01 | no | Csmd1 | intron_variant | MODIFIER |
| 5 | 5316751 | AX-170268376 | 1 | C | T | 0.429 | -6.11E-02 | 6.52E-02 | 2.48E+00 | 2.59E+00 | 3.53E-01 | no | Cdk14 | intron_variant | MODIFIER |
| 1 | 91153924 | AX-170507606 | 0 | G | A | 0.167 | 1.56E-01 | 1.70E-01 | 2.67E+00 | 2.78E+00 | 3.53E-01 | Yes | Rbm44 | missense_variant | MODERATE |
| 9 | 16668728 | AX-170250295 | 1 | T | C | 0.473 | 8.24E-02 | 8.87E-02 | 2.56E+00 | 2.64E+00 | 3.53E-01 | no | | intergenic_variant | MODIFIER |
| 7 | 75300801 | AX-97792099 | 1 | C | T | 0.366 | -8.89E-02 | 9.66E-02 | 2.85E+00 | 2.96E+00 | 3.53E-01 | no | Sv2b | intron_variant | MODIFIER |
| 16 | 18333787 | AX-170482741 | 0 | T | C | 0.132 | 8.87E-02 | 9.69E-02 | 2.80E+00 | 2.93E+00 | 3.53E-01 | no | Tango2 | intron_variant | MODIFIER |
| 16 | 18339412 | AX-170327619 | 0 | C | T | 0.132 | 8.87E-02 | 9.69E-02 | 2.80E+00 | 2.93E+00 | 3.53E-01 | no | Tango2 | intron_variant | MODIFIER |
| 16 | 18353929 | AX-170400638 | 0 | C | G | 0.132 | 8.87E-02 | 9.69E-02 | 2.80E+00 | 2.93E+00 | 3.53E-01 | no | Arvcf | intron_variant | MODIFIER |
| 16 | 18383481 | AX-170279592 | 0 | G | A | 0.132 | 8.87E-02 | 9.69E-02 | 2.80E+00 | 2.93E+00 | 3.53E-01 | no | Arvcf | intron_variant | MODIFIER |
| 16 | 18406050 | AX-170012594 | 0 | G | A | 0.132 | 8.87E-02 | 9.69E-02 | 2.80E+00 | 2.93E+00 | 3.53E-01 | no | Comt | downstream_gene_variant | MODIFIER |
| 16 | 18425980 | AX-97851980 | 0 | T | C | 0.132 | 8.87E-02 | 9.69E-02 | 2.80E+00 | 2.93E+00 | 3.53E-01 | no | Comt | intron_variant | MODIFIER |
| 16 | 18427548 | AX-170012597 | 0 | A | G | 0.132 | 8.87E-02 | 9.69E-02 | 2.80E+00 | 2.93E+00 | 3.53E-01 | no | Comt | upstream_gene_variant | MODIFIER |
| 16 | 18450552 | AX-170458828 | 0 | A | G | 0.132 | 8.87E-02 | 9.69E-02 | 2.80E+00 | 2.93E+00 | 3.53E-01 | no | Txnrd2 | intron_variant | MODIFIER |
| 16 | 18471506 | AX-170479064 | 0 | G | C | 0.132 | 8.87E-02 | 9.69E-02 | 2.80E+00 | 2.93E+00 | 3.53E-01 | no | Txnrd2 | intron_variant,NMD_transcript_variant | MODIFIER |
| 16 | 18492046 | AX-170515881 | 0 | C | T | 0.132 | 8.87E-02 | 9.69E-02 | 2.80E+00 | 2.93E+00 | 3.53E-01 | no | | intergenic_variant | MODIFIER |
| 16 | 18507627 | AX-170214658 | 0 | C | T | 0.132 | 8.87E-02 | 9.69E-02 | 2.80E+00 | 2.93E+00 | 3.53E-01 | no | Gnb1l | intron_variant,NMD_transcript_variant | MODIFIER |
| 16 | 18521283 | AX-170238026 | 0 | G | C | 0.132 | 8.87E-02 | 9.69E-02 | 2.80E+00 | 2.93E+00 | 3.53E-01 | no | Gnb1l | intron_variant,NMD_transcript_variant | MODIFIER |
| 16 | 18540955 | AX-170093404 | 0 | C | T | 0.132 | 8.87E-02 | 9.69E-02 | 2.80E+00 | 2.93E+00 | 3.53E-01 | no | Gnb1l | intron_variant,NMD_transcript_variant | MODIFIER |
| 16 | 18544827 | AX-170247594 | 0 | T | A | 0.132 | 8.87E-02 | 9.69E-02 | 2.80E+00 | 2.93E+00 | 3.53E-01 | no | Gnb1l | intron_variant,NMD_transcript_variant | MODIFIER |
| 16 | 18546889 | AX-170172087 | 0 | T | C | 0.132 | 8.87E-02 | 9.69E-02 | 2.80E+00 | 2.93E+00 | 3.53E-01 | no | Gnb1l | intron_variant,NMD_transcript_variant | MODIFIER |
| 16 | 18552019 | AX-170056821 | 0 | G | A | 0.132 | 8.87E-02 | 9.69E-02 | 2.80E+00 | 2.93E+00 | 3.53E-01 | no | Gnb1l | intron_variant,NMD_transcript_variant | MODIFIER |
| 16 | 18557362 | AX-170056823 | 0 | A | T | 0.132 | 8.87E-02 | 9.69E-02 | 2.80E+00 | 2.93E+00 | 3.53E-01 | no | Gnb1l | intron_variant,NMD_transcript_variant | MODIFIER |
| 16 | 18567013 | AX-97832691 | 0 | G | A | 0.132 | 8.87E-02 | 9.69E-02 | 2.80E+00 | 2.93E+00 | 3.53E-01 | no | Gnb1l | downstream_gene_variant | MODIFIER |
| 16 | 18574458 | AX-170354011 | 0 | T | C | 0.132 | 8.87E-02 | 9.69E-02 | 2.80E+00 | 2.93E+00 | 3.53E-01 | no | | regulatory_region_variant | MODIFIER |
| 16 | 18579292 | AX-170156789 | 0 | T | C | 0.132 | 8.87E-02 | 9.69E-02 | 2.80E+00 | 2.93E+00 | 3.53E-01 | no | Tbx1 | downstream_gene_variant | MODIFIER |

| | | | | | | | | | | | | | | | |
|---|---|---|---|---|---|---|---|---|---|---|---|---|---|---|---|
| 16 | 18606427 | AX-170150539 | 0 | T | C | 0.132 | 8.87E-02 | 9.69E-02 | 2.80E+00 | 2.93E+00 | 3.53E-01 | no | 4930588K23Rik | intron_variant,non_coding_transcript_variant | MODIFIER |
| 16 | 18608818 | AX-170422321 | 0 | C | T | 0.132 | 8.87E-02 | 9.69E-02 | 2.80E+00 | 2.93E+00 | 3.53E-01 | no | 4930588K23Rik | non_coding_transcript_exon_variant | MODIFIER |
| 16 | 18609779 | AX-170498531 | 0 | C | T | 0.132 | 8.87E-02 | 9.69E-02 | 2.80E+00 | 2.93E+00 | 3.53E-01 | no | 4930588K23Rik | downstream_gene_variant | MODIFIER |
| 16 | 18635570 | AX-170421509 | 0 | T | A | 0.132 | 8.87E-02 | 9.69E-02 | 2.80E+00 | 2.93E+00 | 3.53E-01 | no | | intergenic_variant | MODIFIER |
| 16 | 18648800 | AX-170162207 | 0 | A | G | 0.132 | 8.87E-02 | 9.69E-02 | 2.80E+00 | 2.93E+00 | 3.53E-01 | no | | intergenic_variant | MODIFIER |
| 16 | 18649348 | AX-170482597 | 0 | C | G | 0.132 | 8.87E-02 | 9.69E-02 | 2.80E+00 | 2.93E+00 | 3.53E-01 | no | | regulatory_region_variant | MODIFIER |
| 16 | 18691716 | AX-97816413 | 0 | C | T | 0.132 | 8.87E-02 | 9.69E-02 | 2.80E+00 | 2.93E+00 | 3.53E-01 | no | | intergenic_variant | MODIFIER |
| 16 | 18704832 | AX-170192190 | 0 | T | A | 0.132 | 8.87E-02 | 9.69E-02 | 2.80E+00 | 2.93E+00 | 3.53E-01 | no | | regulatory_region_variant | MODIFIER |
| 16 | 18706251 | AX-170407753 | 0 | T | G | 0.132 | 8.87E-02 | 9.69E-02 | 2.80E+00 | 2.93E+00 | 3.53E-01 | no | | regulatory_region_variant | MODIFIER |
| 16 | 18707606 | AX-170420747 | 0 | A | T | 0.132 | 8.87E-02 | 9.69E-02 | 2.80E+00 | 2.93E+00 | 3.53E-01 | no | | intergenic_variant | MODIFIER |
| 16 | 18714355 | AX-170322749 | 0 | T | G | 0.132 | 8.87E-02 | 9.69E-02 | 2.80E+00 | 2.93E+00 | 3.53E-01 | no | | intergenic_variant | MODIFIER |
| 16 | 18715361 | AX-97853993 | 0 | A | G | 0.132 | 8.87E-02 | 9.69E-02 | 2.80E+00 | 2.93E+00 | 3.53E-01 | no | | intergenic_variant | MODIFIER |
| 16 | 18740906 | AX-170250726 | 0 | G | A | 0.132 | 8.87E-02 | 9.69E-02 | 2.80E+00 | 2.93E+00 | 3.53E-01 | no | | intergenic_variant | MODIFIER |
| 16 | 18745661 | AX-170327004 | 0 | A | G | 0.132 | 8.87E-02 | 9.69E-02 | 2.80E+00 | 2.93E+00 | 3.53E-01 | no | | intergenic_variant | MODIFIER |
| 16 | 18745756 | AX-170426106 | 0 | A | T | 0.132 | 8.87E-02 | 9.69E-02 | 2.80E+00 | 2.93E+00 | 3.53E-01 | no | | intergenic_variant | MODIFIER |
| 16 | 18754127 | AX-170487113 | 0 | G | A | 0.132 | 8.87E-02 | 9.69E-02 | 2.80E+00 | 2.93E+00 | 3.53E-01 | no | | intergenic_variant | MODIFIER |
| 16 | 18767257 | AX-170090984 | 0 | C | T | 0.132 | 8.87E-02 | 9.69E-02 | 2.80E+00 | 2.93E+00 | 3.53E-01 | no | | intergenic_variant | MODIFIER |
| 16 | 18767468 | AX-170100606 | 0 | A | G | 0.132 | 8.87E-02 | 9.69E-02 | 2.80E+00 | 2.93E+00 | 3.53E-01 | no | | intergenic_variant | MODIFIER |
| 16 | 18784522 | AX-170487254 | 0 | T | G | 0.132 | 8.87E-02 | 9.69E-02 | 2.80E+00 | 2.93E+00 | 3.53E-01 | no | Cdc45 | intron_variant | MODIFIER |
| 16 | 18787867 | AX-170176353 | 0 | C | T | 0.132 | 8.87E-02 | 9.69E-02 | 2.80E+00 | 2.93E+00 | 3.53E-01 | no | Cdc45 | intron_variant | MODIFIER |
| 16 | 18793994 | AX-170090985 | 0 | G | A | 0.132 | 8.87E-02 | 9.69E-02 | 2.80E+00 | 2.93E+00 | 3.53E-01 | no | Cdc45 | intron_variant | MODIFIER |
| 16 | 18800813 | AX-170118448 | 0 | A | G | 0.132 | 8.87E-02 | 9.69E-02 | 2.80E+00 | 2.93E+00 | 3.53E-01 | no | Cdc45 | intron_variant | MODIFIER |
| 16 | 18828250 | AX-170426684 | 0 | G | A | 0.132 | 8.87E-02 | 9.69E-02 | 2.80E+00 | 2.93E+00 | 3.53E-01 | no | Ufd1 | intron_variant | MODIFIER |
| 16 | 18864991 | AX-170012601 | 0 | C | T | 0.132 | 8.87E-02 | 9.69E-02 | 2.80E+00 | 2.93E+00 | 3.53E-01 | no | Gm28539 | intron_variant,NMD_transcript_variant | MODIFIER |
| 16 | 18905077 | AX-97830663 | 0 | T | G | 0.132 | 8.87E-02 | 9.69E-02 | 2.80E+00 | 2.93E+00 | 3.53E-01 | no | Hira | intron_variant | MODIFIER |
| 16 | 18939900 | AX-170254871 | 0 | A | G | 0.132 | 8.87E-02 | 9.69E-02 | 2.80E+00 | 2.93E+00 | 3.53E-01 | no | Hira | intron_variant | MODIFIER |
| 16 | 18960334 | AX-170037392 | 0 | C | T | 0.132 | 8.87E-02 | 9.69E-02 | 2.80E+00 | 2.93E+00 | 3.53E-01 | no | Hira | intron_variant | MODIFIER |
| 16 | 18968816 | AX-170383099 | 0 | A | C | 0.132 | 8.87E-02 | 9.69E-02 | 2.80E+00 | 2.93E+00 | 3.53E-01 | no | Hira | intron_variant | MODIFIER |
| 16 | 18974696 | AX-170012599 | 0 | A | G | 0.132 | 8.87E-02 | 9.69E-02 | 2.80E+00 | 2.93E+00 | 3.53E-01 | no | Hira | downstream_gene_variant | MODIFIER |
| 16 | 18974758 | AX-97831131 | 0 | G | A | 0.132 | 8.87E-02 | 9.69E-02 | 2.80E+00 | 2.93E+00 | 3.53E-01 | no | Hira | downstream_gene_variant | MODIFIER |
| 16 | 19006082 | AX-170012604 | 0 | C | T | 0.132 | 8.87E-02 | 9.69E-02 | 2.80E+00 | 2.93E+00 | 3.53E-01 | no | | intergenic_variant | MODIFIER |
| 16 | 19030846 | AX-170399773 | 0 | C | T | 0.132 | 8.87E-02 | 9.69E-02 | 2.80E+00 | 2.93E+00 | 3.53E-01 | no | Gm10088 | downstream_gene_variant | MODIFIER |
| 16 | 19052728 | AX-170525604 | 0 | C | T | 0.132 | 8.87E-02 | 9.69E-02 | 2.80E+00 | 2.93E+00 | 3.53E-01 | no | 2010309G21Rik | intron_variant,non_coding_transcript_variant | MODIFIER |
| 16 | 19085742 | AX-170153553 | 0 | A | G | 0.132 | 8.87E-02 | 9.69E-02 | 2.80E+00 | 2.93E+00 | 3.53E-01 | no | Iglv1 | upstream_gene_variant | MODIFIER |
| 16 | 19153231 | AX-170457374 | 0 | A | G | 0.132 | 8.87E-02 | 9.69E-02 | 2.80E+00 | 2.93E+00 | 3.53E-01 | no | | regulatory_region_variant | MODIFIER |
| 16 | 19161712 | AX-97847459 | 0 | T | C | 0.132 | 8.87E-02 | 9.69E-02 | 2.80E+00 | 2.93E+00 | 3.53E-01 | no | Gm18748 | non_coding_transcript_exon_variant | MODIFIER |
| 16 | 19200575 | AX-170217809 | 0 | T | C | 0.132 | 8.87E-02 | 9.69E-02 | 2.80E+00 | 2.93E+00 | 3.53E-01 | no | Iglc2 | upstream_gene_variant | MODIFIER |
| 16 | 19232940 | AX-170056833 | 0 | G | A | 0.132 | 8.87E-02 | 9.69E-02 | 2.80E+00 | 2.93E+00 | 3.53E-01 | no | Gm43388 | intron_variant,non_coding_transcript_variant | MODIFIER |
| 16 | 19262988 | AX-170338708 | 0 | T | C | 0.132 | 8.87E-02 | 9.69E-02 | 2.80E+00 | 2.93E+00 | 3.53E-01 | no | Iglv2 | upstream_gene_variant | MODIFIER |
| 16 | 19285289 | AX-97851680 | 0 | T | C | 0.132 | 8.87E-02 | 9.69E-02 | 2.80E+00 | 2.93E+00 | 3.53E-01 | no | Olfr164 | downstream_gene_variant | MODIFIER |
| 16 | 19308028 | AX-170464565 | 0 | A | T | 0.132 | 8.87E-02 | 9.69E-02 | 2.80E+00 | 2.93E+00 | 3.53E-01 | no | Gm22158 | downstream_gene_variant | MODIFIER |
| 16 | 19351555 | AX-170522232 | 0 | T | C | 0.132 | 8.87E-02 | 9.69E-02 | 2.80E+00 | 2.93E+00 | 3.53E-01 | no | | intergenic_variant | MODIFIER |
| 16 | 19351644 | AX-170338628 | 0 | G | A | 0.132 | 8.87E-02 | 9.69E-02 | 2.80E+00 | 2.93E+00 | 3.53E-01 | no | | intergenic_variant | MODIFIER |
| 16 | 19351753 | AX-170323974 | 0 | T | C | 0.132 | 8.87E-02 | 9.69E-02 | 2.80E+00 | 2.93E+00 | 3.53E-01 | no | | intergenic_variant | MODIFIER |
| 16 | 19403943 | AX-170292732 | 0 | C | A | 0.132 | 8.87E-02 | 9.69E-02 | 2.80E+00 | 2.93E+00 | 3.53E-01 | no | Olfr165 | downstream_gene_variant | MODIFIER |
| 16 | 19405670 | AX-170206636 | 0 | A | T | 0.132 | 8.87E-02 | 9.69E-02 | 2.80E+00 | 2.93E+00 | 3.53E-01 | no | Olfr165 | downstream_gene_variant | MODIFIER |
| 16 | 19437738 | AX-97793474 | 0 | C | A | 0.132 | 8.87E-02 | 9.69E-02 | 2.80E+00 | 2.93E+00 | 3.53E-01 | no | | intergenic_variant | MODIFIER |
| 16 | 19465453 | AX-170012606 | 0 | T | G | 0.132 | 8.87E-02 | 9.69E-02 | 2.80E+00 | 2.93E+00 | 3.53E-01 | no | | intergenic_variant | MODIFIER |
| 16 | 19501512 | AX-170037387 | 0 | A | G | 0.132 | 8.87E-02 | 9.69E-02 | 2.80E+00 | 2.93E+00 | 3.53E-01 | no | | intergenic_variant | MODIFIER |
| 16 | 19508382 | AX-170012610 | 0 | G | C | 0.132 | 8.87E-02 | 9.69E-02 | 2.80E+00 | 2.93E+00 | 3.53E-01 | no | Olfr167 | downstream_gene_variant | MODIFIER |
| 16 | 19508761 | AX-170012611 | 0 | T | G | 0.132 | 8.87E-02 | 9.69E-02 | 2.80E+00 | 2.93E+00 | 3.53E-01 | no | Olfr167 | downstream_gene_variant | MODIFIER |
| 16 | 19543214 | AX-170056835 | 0 | T | C | 0.132 | 8.87E-02 | 9.69E-02 | 2.80E+00 | 2.93E+00 | 3.53E-01 | no | | intergenic_variant | MODIFIER |
| 16 | 19569138 | AX-170135745 | 0 | A | G | 0.132 | 8.87E-02 | 9.69E-02 | 2.80E+00 | 2.93E+00 | 3.53E-01 | no | Olfr169 | upstream_gene_variant | MODIFIER |
| 16 | 19570572 | AX-170316321 | 0 | G | A | 0.132 | 8.87E-02 | 9.69E-02 | 2.80E+00 | 2.93E+00 | 3.53E-01 | no | Olfr169 | upstream_gene_variant | MODIFIER |

| | | | | | | | | | | | | | | | |
|---|---|---|---|---|---|---|---|---|---|---|---|---|---|---|---|
| 16 | 19628163 | AX-170506460 | 0 | T | C | 0.132 | 8.87E-02 | 9.69E-02 | 2.80E+00 | 2.93E+00 | 3.53E-01 | no | Olfr171 | upstream_gene_variant | MODIFIER |
| 16 | 19666940 | AX-170226861 | 0 | A | G | 0.132 | 8.87E-02 | 9.69E-02 | 2.80E+00 | 2.93E+00 | 3.53E-01 | no | Lamp3 | intron_variant | MODIFIER |
| 16 | 19676389 | AX-97821621 | 0 | A | C | 0.132 | 8.87E-02 | 9.69E-02 | 2.80E+00 | 2.93E+00 | 3.53E-01 | no | Lamp3 | intron_variant | MODIFIER |
| 16 | 19721434 | AX-170336341 | 0 | T | G | 0.132 | 8.87E-02 | 9.69E-02 | 2.80E+00 | 2.93E+00 | 3.53E-01 | no | Gm17799 | upstream_gene_variant | MODIFIER |
| 16 | 19727322 | AX-170486291 | 0 | A | G | 0.132 | 8.87E-02 | 9.69E-02 | 2.80E+00 | 2.93E+00 | 3.53E-01 | no | Gm17799 | downstream_gene_variant | MODIFIER |
| 16 | 19769088 | AX-170119659 | 0 | T | C | 0.132 | 8.87E-02 | 9.69E-02 | 2.80E+00 | 2.93E+00 | 3.53E-01 | no | B3gnt5 | missense_variant | MODERATE |
| 16 | 19773435 | AX-170180778 | 0 | G | C | 0.132 | 8.87E-02 | 9.69E-02 | 2.80E+00 | 2.93E+00 | 3.53E-01 | no | B3gnt5 | downstream_gene_variant | MODIFIER |
| 16 | 19786553 | AX-170137198 | 0 | A | G | 0.132 | 8.87E-02 | 9.69E-02 | 2.80E+00 | 2.93E+00 | 3.53E-01 | no | | regulatory_region_variant | MODIFIER |
| 16 | 19800471 | AX-170475142 | 0 | T | C | 0.132 | 8.87E-02 | 9.69E-02 | 2.80E+00 | 2.93E+00 | 3.53E-01 | no | | intergenic_variant | MODIFIER |
| 16 | 19834489 | AX-170056843 | 0 | G | C | 0.132 | 8.87E-02 | 9.69E-02 | 2.80E+00 | 2.93E+00 | 3.53E-01 | no | | intergenic_variant | MODIFIER |
| 16 | 19835655 | AX-170056839 | 0 | A | G | 0.132 | 8.87E-02 | 9.69E-02 | 2.80E+00 | 2.93E+00 | 3.53E-01 | no | | intergenic_variant | MODIFIER |
| 16 | 19895707 | AX-170190194 | 0 | C | T | 0.132 | 8.87E-02 | 9.69E-02 | 2.80E+00 | 2.93E+00 | 3.53E-01 | no | | intergenic_variant | MODIFIER |
| 16 | 21339004 | AX-170087972 | 0 | A | G | 0.132 | 8.87E-02 | 9.69E-02 | 2.80E+00 | 2.93E+00 | 3.53E-01 | no | | regulatory_region_variant | MODIFIER |
| 16 | 21678400 | AX-170349216 | 0 | C | G | 0.132 | 8.87E-02 | 9.69E-02 | 2.80E+00 | 2.93E+00 | 3.53E-01 | no | 2510009E07Rik | intron_variant | MODIFIER |
| 8 | 122467223 | AX-170166637 | 0 | A | G | 0.044 | 1.80E-01 | 1.95E-01 | 2.87E+00 | 3.01E+00 | 3.53E-01 | Yes | Rnf166 | synonymous_variant | LOW |
| 6 | 129532037 | AX-170129314 | 0 | G | A | 0.553 | 7.77E-02 | 8.29E-02 | 2.50E+00 | 2.56E+00 | 3.53E-01 | no | Gabarapl1 | upstream_gene_variant | MODIFIER |
| 5 | 97149556 | AX-170072825 | 0 | C | T | 0.439 | 9.54E-02 | 1.04E-01 | 2.78E+00 | 2.85E+00 | 3.53E-01 | no | Gm33609 | non_coding_transcript_exon_variant | MODIFIER |
| 14 | 116569213 | AX-97828019 | 1 | C | T | 0.312 | 6.73E-02 | 7.30E-02 | 2.60E+00 | 2.70E+00 | 3.53E-01 | no | | intergenic_variant | MODIFIER |
| 16 | 18448807 | AX-97844656 | 0 | C | A | 0.132 | 8.86E-02 | 9.68E-02 | 2.81E+00 | 2.94E+00 | 3.53E-01 | no | Txnrd2 | intron_variant | MODIFIER |
| 7 | 37045912 | AX-170485561 | 1 | T | A | 0.33 | 8.31E-02 | 9.02E-02 | 2.61E+00 | 2.70E+00 | 3.53E-01 | no | | intergenic_variant | MODIFIER |
| 7 | 37196407 | AX-97803923 | 1 | T | C | 0.33 | 8.31E-02 | 9.02E-02 | 2.61E+00 | 2.70E+00 | 3.53E-01 | no | | intergenic_variant | MODIFIER |
| 15 | 3453497 | AX-97827784 | 1 | G | A | 0.232 | -9.36E-02 | 1.01E-01 | 2.59E+00 | 2.65E+00 | 3.54E-01 | no | Ghr | intron_variant | MODIFIER |
| 7 | 100024416 | AX-170078220 | 2 | T | C | 0.345 | -8.77E-02 | 9.57E-02 | 2.70E+00 | 2.80E+00 | 3.54E-01 | no | Chrdl2 | intron_variant | MODIFIER |
| 18 | 77867703 | AX-97826252 | 0 | G | A | 0.43 | 8.36E-02 | 8.87E-02 | 2.47E+00 | 2.53E+00 | 3.54E-01 | no | Pstpip2 | intron_variant | MODIFIER |
| 4 | 133660829 | AX-170223576 | 0 | C | T | 0.149 | 1.13E-01 | 1.22E-01 | 2.58E+00 | 2.63E+00 | 3.54E-01 | no | Pigv | 3_prime_UTR_variant | MODIFIER |
| 20 | 48547068 | AX-170445909 | 1 | C | T | 0.241 | 5.33E-02 | 5.79E-02 | 2.58E+00 | 2.72E+00 | 3.54E-01 | no | Zfp280c | intron_variant | MODIFIER |
| 8 | 65585655 | AX-170278521 | 1 | T | C | 0.134 | -1.31E-01 | 1.42E-01 | 2.58E+00 | 2.66E+00 | 3.54E-01 | no | | intergenic_variant | MODIFIER |
| 1 | 128717994 | AX-170225687 | 1 | G | A | 0.536 | 7.67E-02 | 8.36E-02 | 2.68E+00 | 2.75E+00 | 3.54E-01 | no | | intergenic_variant | MODIFIER |
| 4 | 137405600 | AX-97807711 | 0 | A | G | 0.167 | 1.07E-01 | 1.16E-01 | 2.65E+00 | 2.70E+00 | 3.54E-01 | no | Cela3a | missense_variant | MODERATE |
| 18 | 54842826 | AX-170301655 | 1 | C | A | 0.455 | 9.15E-02 | 9.42E-02 | 2.35E+00 | 2.40E+00 | 3.54E-01 | no | AC129179.1 | downstream_gene_variant | MODIFIER |
| 1 | 188734170 | AX-97802542 | 0 | G | T | 0.254 | 9.62E-02 | 1.01E-01 | 2.40E+00 | 2.49E+00 | 3.54E-01 | no | Ush2a | intron_variant | MODIFIER |
| 1 | 141067649 | AX-170168061 | 0 | A | C | 0.211 | 1.14E-01 | 1.24E-01 | 2.69E+00 | 2.78E+00 | 3.54E-01 | no | | regulatory_region_variant | MODIFIER |
| 15 | 75135935 | AX-170301584 | 0 | T | A | 0.553 | 7.36E-02 | 7.98E-02 | 2.60E+00 | 2.69E+00 | 3.54E-01 | no | I830127L07Rik | upstream_gene_variant | MODIFIER |
| 6 | 103386900 | AX-170117703 | 0 | G | A | 0.061 | 2.02E-01 | 2.19E-01 | 2.56E+00 | 2.68E+00 | 3.54E-01 | Yes | | regulatory_region_variant | MODIFIER |
| 18 | 65686640 | AX-170279240 | 2 | G | A | 0.418 | 8.08E-02 | 8.42E-02 | 2.39E+00 | 2.45E+00 | 3.54E-01 | no | Zfp532 | intron_variant | MODIFIER |
| 18 | 69776365 | AX-170061232 | 0 | G | A | 0.263 | -8.87E-02 | 9.66E-02 | 2.65E+00 | 2.73E+00 | 3.54E-01 | no | Gm22508 | downstream_gene_variant | MODIFIER |
| 1 | 127357527 | AX-170109571 | 1 | G | A | 0.545 | 7.76E-02 | 8.43E-02 | 2.62E+00 | 2.67E+00 | 3.55E-01 | no | Mgat5 | intron_variant | MODIFIER |
| 16 | 78946166 | AX-170037517 | 1 | C | G | 0.473 | 6.91E-02 | 7.49E-02 | 2.60E+00 | 2.67E+00 | 3.55E-01 | no | Chodl | intron_variant | MODIFIER |
| 15 | 71408553 | AX-170011960 | 0 | T | C | 0.561 | 7.13E-02 | 7.73E-02 | 2.59E+00 | 2.67E+00 | 3.55E-01 | no | | intergenic_variant | MODIFIER |
| 10 | 93556696 | AX-170003674 | 0 | T | A | 0.289 | -7.76E-02 | 8.29E-02 | 2.49E+00 | 2.55E+00 | 3.55E-01 | no | Ccdc38 | intron_variant | MODIFIER |
| 16 | 58538223 | AX-170257544 | 0 | A | G | 0.456 | -8.14E-02 | 8.62E-02 | 2.45E+00 | 2.49E+00 | 3.55E-01 | no | | intergenic_variant | MODIFIER |
| 10 | 18992572 | AX-170417267 | 1 | T | C | 0.196 | 9.38E-02 | 1.01E-01 | 2.95E+00 | 3.05E+00 | 3.55E-01 | no | | intergenic_variant | MODIFIER |
| 4 | 135059737 | AX-170494822 | 1 | C | T | 0.179 | 1.01E-01 | 1.10E-01 | 2.59E+00 | 2.64E+00 | 3.55E-01 | no | | intergenic_variant | MODIFIER |
| 1 | 153013524 | AX-170239936 | 0 | G | A | 0.219 | 1.10E-01 | 1.19E-01 | 2.62E+00 | 2.71E+00 | 3.55E-01 | no | Nmnat2 | intron_variant | MODIFIER |
| 1 | 187567509 | AX-170002508 | 1 | C | A | 0.25 | 1.02E-01 | 1.05E-01 | 2.35E+00 | 2.42E+00 | 3.55E-01 | no | | intergenic_variant | MODIFIER |
| 1 | 173924840 | AX-170043639 | 1 | G | A | 0.223 | -1.15E-01 | 1.21E-01 | 3.13E+00 | 3.30E+00 | 3.55E-01 | no | Ifi203 | intron_variant | MODIFIER |
| 15 | 20942406 | AX-170117568 | 1 | C | T | 0.241 | -9.74E-02 | 1.06E-01 | 2.61E+00 | 2.66E+00 | 3.55E-01 | no | Cdh12 | intron_variant,non_coding_transcript_variant | MODIFIER |
| 4 | 49264662 | AX-170216396 | 1 | G | A | 0.366 | -8.82E-02 | 9.65E-02 | 2.82E+00 | 2.92E+00 | 3.55E-01 | no | Plppr1 | intron_variant | MODIFIER |
| 4 | 155005036 | AX-170498028 | 1 | T | C | 0.125 | 1.47E-01 | 1.61E-01 | 2.75E+00 | 2.85E+00 | 3.56E-01 | Yes | Plch2 | intron_variant | MODIFIER |
| 1 | 14763252 | AX-170532721 | 2 | A | T | 0.382 | 7.19E-02 | 7.85E-02 | 2.68E+00 | 2.74E+00 | 3.56E-01 | no | Gm9947 | downstream_gene_variant | MODIFIER |
| 7 | 93773378 | AX-170029540 | 0 | A | G | 0.14 | 1.55E-01 | 1.70E-01 | 2.78E+00 | 2.91E+00 | 3.56E-01 | Yes | | intergenic_variant | MODIFIER |
| 18 | 69013699 | AX-170100074 | 1 | C | T | 0.161 | -1.11E-01 | 1.21E-01 | 2.61E+00 | 2.67E+00 | 3.56E-01 | no | | intergenic_variant | MODIFIER |
| 2 | 166399938 | AX-170233643 | 1 | G | A | 0.161 | 1.79E-01 | 1.96E-01 | 2.75E+00 | 2.83E+00 | 3.56E-01 | Yes | Gm14268 | upstream_gene_variant | MODIFIER |
| 8 | 61221903 | AX-97803794 | 1 | G | T | 0.152 | -1.35E-01 | 1.46E-01 | 2.54E+00 | 2.62E+00 | 3.56E-01 | no | Sh3rf1 | upstream_gene_variant | MODIFIER |

| | | | | | | | | | | | | | | | |
|---|---|---|---|---|---|---|---|---|---|---|---|---|---|---|---|
| 18 | 60183841 | AX-170221639 | 1 | T | C | 0.402 | 8.88E-02 | 9.39E-02 | 2.44E+00 | 2.48E+00 | 3.56E-01 | no | | intergenic_variant | MODIFIER |
| 1 | 25940392 | AX-170105339 | 0 | T | C | 0.404 | 7.60E-02 | 8.33E-02 | 2.73E+00 | 2.81E+00 | 3.56E-01 | no | | intergenic_variant | MODIFIER |
| 7 | 71543469 | AX-97843058 | 0 | C | T | 0.351 | -9.12E-02 | 9.97E-02 | 2.82E+00 | 2.91E+00 | 3.56E-01 | no | | intergenic_variant | MODIFIER |
| 7 | 71547054 | AX-170365922 | 0 | T | C | 0.351 | -9.12E-02 | 9.97E-02 | 2.82E+00 | 2.91E+00 | 3.56E-01 | no | | intergenic_variant | MODIFIER |
| 7 | 71586027 | AX-97791003 | 0 | A | G | 0.351 | -9.12E-02 | 9.97E-02 | 2.82E+00 | 2.91E+00 | 3.56E-01 | no | | intergenic_variant | MODIFIER |
| 7 | 72083506 | AX-170382752 | 0 | A | G | 0.351 | -9.12E-02 | 9.97E-02 | 2.82E+00 | 2.91E+00 | 3.56E-01 | no | Mctp2 | intron_variant | MODIFIER |
| 7 | 72551345 | AX-97846233 | 0 | A | G | 0.351 | -9.12E-02 | 9.97E-02 | 2.82E+00 | 2.91E+00 | 3.56E-01 | no | | intergenic_variant | MODIFIER |
| 7 | 72567359 | AX-170321392 | 0 | G | C | 0.351 | -9.12E-02 | 9.97E-02 | 2.82E+00 | 2.91E+00 | 3.56E-01 | no | | intergenic_variant | MODIFIER |
| 7 | 72572344 | AX-170324534 | 0 | T | A | 0.351 | -9.12E-02 | 9.97E-02 | 2.82E+00 | 2.91E+00 | 3.56E-01 | no | | intergenic_variant | MODIFIER |
| 7 | 72585546 | AX-97842506 | 0 | G | T | 0.351 | -9.12E-02 | 9.97E-02 | 2.82E+00 | 2.91E+00 | 3.56E-01 | no | | intergenic_variant | MODIFIER |
| 7 | 72770960 | AX-170269255 | 0 | T | C | 0.351 | -9.12E-02 | 9.97E-02 | 2.82E+00 | 2.91E+00 | 3.56E-01 | no | | intergenic_variant | MODIFIER |
| 7 | 72792763 | AX-170143829 | 0 | A | G | 0.351 | -9.12E-02 | 9.97E-02 | 2.82E+00 | 2.91E+00 | 3.56E-01 | no | | intergenic_variant | MODIFIER |
| 7 | 72823202 | AX-170196717 | 0 | T | C | 0.351 | -9.12E-02 | 9.97E-02 | 2.82E+00 | 2.91E+00 | 3.56E-01 | no | | intergenic_variant | MODIFIER |
| 7 | 72852678 | AX-170116228 | 0 | C | T | 0.351 | -9.12E-02 | 9.97E-02 | 2.82E+00 | 2.91E+00 | 3.56E-01 | no | | intergenic_variant | MODIFIER |
| 7 | 72871343 | AX-97845053 | 0 | A | G | 0.351 | -9.12E-02 | 9.97E-02 | 2.82E+00 | 2.91E+00 | 3.56E-01 | no | | intergenic_variant | MODIFIER |
| 7 | 72927003 | AX-170194771 | 0 | A | G | 0.351 | -9.12E-02 | 9.97E-02 | 2.82E+00 | 2.91E+00 | 3.56E-01 | no | | regulatory_region_variant | MODIFIER |
| 7 | 72935652 | AX-97787710 | 0 | A | G | 0.351 | -9.12E-02 | 9.97E-02 | 2.82E+00 | 2.91E+00 | 3.56E-01 | no | | intergenic_variant | MODIFIER |
| 7 | 73086564 | AX-97810397 | 0 | C | T | 0.351 | -9.12E-02 | 9.97E-02 | 2.82E+00 | 2.91E+00 | 3.56E-01 | no | Gm44725 | upstream_gene_variant | MODIFIER |
| 15 | 4019292 | AX-170208420 | 0 | T | C | 0.228 | -9.26E-02 | 1.01E-01 | 2.60E+00 | 2.67E+00 | 3.56E-01 | no | BC037032 | downstream_gene_variant | MODIFIER |
| 9 | 97669802 | AX-170436089 | 1 | G | A | 0.134 | -1.09E-01 | 1.19E-01 | 2.62E+00 | 2.70E+00 | 3.56E-01 | no | Clstn2 | intron_variant | MODIFIER |
| 7 | 38173649 | AX-170312379 | 1 | C | G | 0.357 | 8.96E-02 | 9.69E-02 | 2.55E+00 | 2.62E+00 | 3.56E-01 | no | Gm22203 | downstream_gene_variant | MODIFIER |
| 10 | 18156096 | AX-170485440 | 0 | A | G | 0.202 | 9.29E-02 | 1.01E-01 | 2.93E+00 | 3.03E+00 | 3.56E-01 | no | Ect2l | intron_variant | MODIFIER |
| 4 | 118213556 | AX-170293277 | 0 | C | T | 0.211 | 1.09E-01 | 1.18E-01 | 2.57E+00 | 2.64E+00 | 3.56E-01 | no | Ptprf | synonymous_variant | LOW |
| 18 | 84464809 | AX-170126358 | 1 | A | G | 0.152 | -1.03E-01 | 1.09E-01 | 2.44E+00 | 2.52E+00 | 3.56E-01 | no | Zfp407 | intron_variant | MODIFIER |
| 15 | 23438157 | AX-170170296 | 0 | G | C | 0.254 | -9.43E-02 | 1.03E-01 | 2.64E+00 | 2.72E+00 | 3.56E-01 | no | Cdh18 | intron_variant,non_coding_transcript_variant | MODIFIER |
| 1 | 91527926 | AX-97805121 | 1 | C | A | 0.125 | 1.56E-01 | 1.71E-01 | 2.65E+00 | 2.75E+00 | 3.56E-01 | Yes | Traf3ip1 | 3_prime_UTR_variant | MODIFIER |
| 16 | 31501257 | AX-170246930 | 0 | C | T | 0.509 | 7.19E-02 | 7.76E-02 | 2.53E+00 | 2.59E+00 | 3.57E-01 | no | | regulatory_region_variant | MODIFIER |
| 16 | 19705723 | AX-170398491 | 0 | C | T | 0.149 | 8.19E-02 | 9.03E-02 | 2.72E+00 | 2.85E+00 | 3.57E-01 | no | Lamp3 | intron_variant | MODIFIER |
| 4 | 128417120 | AX-170203550 | 2 | C | T | 0.145 | 1.12E-01 | 1.22E-01 | 2.60E+00 | 2.66E+00 | 3.57E-01 | no | Csmd2 | intron_variant,non_coding_transcript_variant | MODIFIER |
| 5 | 4281118 | AX-170490115 | 1 | A | G | 0.42 | -6.14E-02 | 6.60E-02 | 2.49E+00 | 2.60E+00 | 3.57E-01 | no | | intergenic_variant | MODIFIER |
| 16 | 18922684 | AX-170056824 | 1 | A | T | 0.116 | 8.38E-02 | 9.24E-02 | 2.77E+00 | 2.92E+00 | 3.57E-01 | no | Hira | intron_variant | MODIFIER |
| 2 | 88679280 | AX-170403836 | 1 | A | T | 0.054 | -1.99E-01 | 2.19E-01 | 2.74E+00 | 2.84E+00 | 3.57E-01 | Yes | Olfr1193 | downstream_gene_variant | MODIFIER |
| 4 | 47783882 | AX-170022717 | 0 | A | G | 0.36 | -8.80E-02 | 9.65E-02 | 2.81E+00 | 2.91E+00 | 3.57E-01 | no | | intergenic_variant | MODIFIER |
| 4 | 48547577 | AX-97845014 | 0 | T | C | 0.36 | -8.80E-02 | 9.65E-02 | 2.81E+00 | 2.91E+00 | 3.57E-01 | no | Msantd3 | intron_variant | MODIFIER |
| 2 | 161926892 | AX-97795451 | 1 | G | A | 0.152 | 1.48E-01 | 1.63E-01 | 2.70E+00 | 2.82E+00 | 3.57E-01 | Yes | Ptprt | intron_variant | MODIFIER |
| 5 | 109527524 | AX-97806679 | 1 | G | T | 0.464 | 1.03E-01 | 1.12E-01 | 2.66E+00 | 2.70E+00 | 3.57E-01 | no | Gm8488 | non_coding_transcript_exon_variant | MODIFIER |
| 5 | 83972900 | AX-170095382 | 0 | A | C | 0.447 | 9.58E-02 | 1.05E-01 | 2.87E+00 | 2.96E+00 | 3.57E-01 | no | | intergenic_variant | MODIFIER |
| 20 | 69103195 | AX-170034089 | 0 | A | T | 0.254 | 5.70E-02 | 6.27E-02 | 2.73E+00 | 2.83E+00 | 3.57E-01 | no | | intergenic_variant | MODIFIER |
| 3 | 26927355 | AX-170020169 | 0 | G | A | 0.123 | -9.58E-02 | 1.03E-01 | 2.50E+00 | 2.59E+00 | 3.57E-01 | no | Spata16 | missense_variant | MODERATE |
| 2 | 169301271 | AX-170270848 | 1 | G | A | 0.17 | 1.64E-01 | 1.80E-01 | 2.62E+00 | 2.70E+00 | 3.57E-01 | Yes | | regulatory_region_variant | MODIFIER |
| 9 | 96693401 | AX-97799621 | 0 | G | A | 0.14 | -1.03E-01 | 1.12E-01 | 2.55E+00 | 2.63E+00 | 3.58E-01 | no | Zbtb38 | intron_variant | MODIFIER |
| 1 | 149475124 | AX-170506342 | 1 | T | C | 0.196 | 1.23E-01 | 1.36E-01 | 2.70E+00 | 2.76E+00 | 3.58E-01 | no | | intergenic_variant | MODIFIER |
| 1 | 152260073 | AX-170352326 | 1 | T | C | 0.196 | 1.23E-01 | 1.36E-01 | 2.70E+00 | 2.76E+00 | 3.58E-01 | no | | intergenic_variant | MODIFIER |
| 13 | 39505598 | AX-170183933 | 0 | C | T | 0.167 | -1.04E-01 | 1.10E-01 | 2.40E+00 | 2.48E+00 | 3.58E-01 | no | | intergenic_variant | MODIFIER |
| 1 | 119326600 | AX-170116448 | 1 | G | A | 0.571 | 7.40E-02 | 8.01E-02 | 2.52E+00 | 2.60E+00 | 3.58E-01 | no | Gm29455 | upstream_gene_variant | MODIFIER |
| 4 | 95868149 | AX-170333691 | 0 | G | C | 0.263 | 9.85E-02 | 1.05E-01 | 2.44E+00 | 2.51E+00 | 3.58E-01 | no | Fggy | intron_variant | MODIFIER |
| 5 | 3361605 | AX-170399222 | 1 | A | T | 0.42 | -6.13E-02 | 6.61E-02 | 2.49E+00 | 2.60E+00 | 3.58E-01 | no | Cdk6 | intron_variant | MODIFIER |
| 13 | 4820812 | AX-170460787 | 1 | G | A | 0.205 | -1.14E-01 | 1.24E-01 | 2.56E+00 | 2.62E+00 | 3.58E-01 | no | Gm5444 | intron_variant,non_coding_transcript_variant | MODIFIER |
| 16 | 79131231 | AX-170401762 | 0 | G | A | 0.474 | 6.83E-02 | 7.46E-02 | 2.60E+00 | 2.67E+00 | 3.58E-01 | no | | intergenic_variant | MODIFIER |
| 1 | 168164964 | AX-170411432 | 0 | A | G | 0.219 | 1.30E-01 | 1.43E-01 | 2.70E+00 | 2.75E+00 | 3.58E-01 | no | Pbx1 | intron_variant | MODIFIER |
| 1 | 174878069 | AX-170215519 | 0 | T | A | 0.228 | 1.13E-01 | 1.22E-01 | 2.48E+00 | 2.54E+00 | 3.58E-01 | no | Grem2 | intron_variant | MODIFIER |
| 4 | 140642724 | AX-97809203 | 1 | G | A | 0.161 | 1.07E-01 | 1.17E-01 | 2.59E+00 | 2.64E+00 | 3.58E-01 | no | Arhgef10l | intron_variant | MODIFIER |
| 7 | 20005695 | AX-170403717 | 1 | A | T | 0.411 | 8.25E-02 | 8.98E-02 | 2.54E+00 | 2.65E+00 | 3.58E-01 | no | Nlrp9b | upstream_gene_variant | MODIFIER |
| 9 | 50396817 | AX-168419945 | 2 | A | G | 0.191 | -9.80E-02 | 1.08E-01 | 2.72E+00 | 2.82E+00 | 3.58E-01 | no | | intergenic_variant | MODIFIER |

| | | | | | | | | | | | | | | | |
|---|---|---|---|---|---|---|---|---|---|---|---|---|---|---|---|
| 4 | 94742244 | AX-170270596 | 0 | A | T | 0.342 | 9.46E-02 | 1.04E-01 | 2.69E+00 | 2.77E+00 | 3.58E-01 | no | Tek | intron_variant | MODIFIER |
| 16 | 50362041 | AX-168397560 | 0 | T | G | 0.518 | 7.73E-02 | 8.30E-02 | 2.49E+00 | 2.52E+00 | 3.58E-01 | no | Bbx | intron_variant | MODIFIER |
| 16 | 50387228 | AX-170013017 | 0 | A | C | 0.518 | 7.73E-02 | 8.30E-02 | 2.49E+00 | 2.52E+00 | 3.58E-01 | no | Bbx | intron_variant | MODIFIER |
| 16 | 50391800 | AX-170258397 | 0 | G | A | 0.518 | 7.73E-02 | 8.30E-02 | 2.49E+00 | 2.52E+00 | 3.58E-01 | no | Bbx | intron_variant | MODIFIER |
| 16 | 50403757 | AX-170288588 | 0 | A | G | 0.518 | 7.73E-02 | 8.30E-02 | 2.49E+00 | 2.52E+00 | 3.58E-01 | no | Bbx | intron_variant | MODIFIER |
| 16 | 50410914 | AX-170160092 | 0 | T | G | 0.518 | 7.73E-02 | 8.30E-02 | 2.49E+00 | 2.52E+00 | 3.58E-01 | no | Bbx | intron_variant | MODIFIER |
| 16 | 50449689 | AX-170013021 | 0 | G | A | 0.518 | 7.73E-02 | 8.30E-02 | 2.49E+00 | 2.52E+00 | 3.58E-01 | no | | intergenic_variant | MODIFIER |
| 16 | 50469419 | AX-170334184 | 0 | G | A | 0.518 | 7.73E-02 | 8.30E-02 | 2.49E+00 | 2.52E+00 | 3.58E-01 | no | | intergenic_variant | MODIFIER |
| 16 | 50472602 | AX-97802048 | 0 | C | T | 0.518 | 7.73E-02 | 8.30E-02 | 2.49E+00 | 2.52E+00 | 3.58E-01 | no | | intergenic_variant | MODIFIER |
| 16 | 50485219 | AX-170329304 | 0 | T | C | 0.518 | 7.73E-02 | 8.30E-02 | 2.49E+00 | 2.52E+00 | 3.58E-01 | no | | intergenic_variant | MODIFIER |
| 16 | 50485236 | AX-170279225 | 0 | T | A | 0.518 | 7.73E-02 | 8.30E-02 | 2.49E+00 | 2.52E+00 | 3.58E-01 | no | | intergenic_variant | MODIFIER |
| 16 | 50486326 | AX-97833216 | 0 | G | A | 0.482 | -7.73E-02 | 8.30E-02 | 2.49E+00 | 2.52E+00 | 3.58E-01 | no | | intergenic_variant | MODIFIER |
| 16 | 50498904 | AX-170013022 | 0 | G | A | 0.518 | 7.73E-02 | 8.30E-02 | 2.49E+00 | 2.52E+00 | 3.58E-01 | no | | intergenic_variant | MODIFIER |
| 16 | 50506661 | AX-97850130 | 0 | T | A | 0.518 | 7.73E-02 | 8.30E-02 | 2.49E+00 | 2.52E+00 | 3.58E-01 | no | | intergenic_variant | MODIFIER |
| 16 | 50637034 | AX-170455480 | 0 | C | T | 0.518 | 7.73E-02 | 8.30E-02 | 2.49E+00 | 2.52E+00 | 3.58E-01 | no | 4930542D17Rik | intron_variant,non_coding_transcript_variant | MODIFIER |
| 16 | 50663047 | AX-170127772 | 0 | T | A | 0.518 | 7.73E-02 | 8.30E-02 | 2.49E+00 | 2.52E+00 | 3.58E-01 | no | Gm9575 | downstream_gene_variant | MODIFIER |
| 16 | 50687937 | AX-170454652 | 0 | A | G | 0.518 | 7.73E-02 | 8.30E-02 | 2.49E+00 | 2.52E+00 | 3.58E-01 | no | | intergenic_variant | MODIFIER |
| 16 | 50704600 | AX-170431745 | 0 | C | A | 0.518 | 7.73E-02 | 8.30E-02 | 2.49E+00 | 2.52E+00 | 3.58E-01 | no | | intergenic_variant | MODIFIER |
| 16 | 50706026 | AX-170480331 | 0 | C | T | 0.518 | 7.73E-02 | 8.30E-02 | 2.49E+00 | 2.52E+00 | 3.58E-01 | no | | intergenic_variant | MODIFIER |
| 16 | 50706243 | AX-170315622 | 0 | C | G | 0.518 | 7.73E-02 | 8.30E-02 | 2.49E+00 | 2.52E+00 | 3.58E-01 | no | | intergenic_variant | MODIFIER |
| 16 | 50713574 | AX-170057409 | 0 | G | T | 0.518 | 7.73E-02 | 8.30E-02 | 2.49E+00 | 2.52E+00 | 3.58E-01 | no | | intergenic_variant | MODIFIER |
| 16 | 50725348 | AX-170354970 | 0 | A | G | 0.518 | 7.73E-02 | 8.30E-02 | 2.49E+00 | 2.52E+00 | 3.58E-01 | no | Dubr | downstream_gene_variant | MODIFIER |
| 16 | 50738818 | AX-170397956 | 0 | A | C | 0.518 | 7.73E-02 | 8.30E-02 | 2.49E+00 | 2.52E+00 | 3.58E-01 | no | | intergenic_variant | MODIFIER |
| 16 | 50743763 | AX-170375704 | 0 | A | G | 0.518 | 7.73E-02 | 8.30E-02 | 2.49E+00 | 2.52E+00 | 3.58E-01 | no | | intergenic_variant | MODIFIER |
| 16 | 50745515 | AX-168416228 | 0 | C | T | 0.518 | 7.73E-02 | 8.30E-02 | 2.49E+00 | 2.52E+00 | 3.58E-01 | no | | intergenic_variant | MODIFIER |
| 16 | 50748416 | AX-170140014 | 0 | G | A | 0.518 | 7.73E-02 | 8.30E-02 | 2.49E+00 | 2.52E+00 | 3.58E-01 | no | | intergenic_variant | MODIFIER |
| 16 | 50749792 | AX-170102002 | 0 | A | G | 0.518 | 7.73E-02 | 8.30E-02 | 2.49E+00 | 2.52E+00 | 3.58E-01 | no | | intergenic_variant | MODIFIER |
| 16 | 50749855 | AX-170128184 | 0 | A | G | 0.518 | 7.73E-02 | 8.30E-02 | 2.49E+00 | 2.52E+00 | 3.58E-01 | no | | intergenic_variant | MODIFIER |
| 16 | 50800804 | AX-170057410 | 0 | C | T | 0.518 | 7.73E-02 | 8.30E-02 | 2.49E+00 | 2.52E+00 | 3.58E-01 | no | | intergenic_variant | MODIFIER |
| 16 | 50811875 | AX-170255745 | 0 | C | A | 0.518 | 7.73E-02 | 8.30E-02 | 2.49E+00 | 2.52E+00 | 3.58E-01 | no | | intergenic_variant | MODIFIER |
| 16 | 50812008 | AX-170057411 | 0 | T | C | 0.518 | 7.73E-02 | 8.30E-02 | 2.49E+00 | 2.52E+00 | 3.58E-01 | no | | intergenic_variant | MODIFIER |
| 16 | 50826348 | AX-170412450 | 0 | G | A | 0.518 | 7.73E-02 | 8.30E-02 | 2.49E+00 | 2.52E+00 | 3.58E-01 | no | | intergenic_variant | MODIFIER |
| 16 | 50826508 | AX-170230791 | 0 | G | T | 0.518 | 7.73E-02 | 8.30E-02 | 2.49E+00 | 2.52E+00 | 3.58E-01 | no | | intergenic_variant | MODIFIER |
| 16 | 50826692 | AX-170311814 | 0 | A | G | 0.518 | 7.73E-02 | 8.30E-02 | 2.49E+00 | 2.52E+00 | 3.58E-01 | no | | intergenic_variant | MODIFIER |
| 16 | 50830601 | AX-170465976 | 0 | T | C | 0.518 | 7.73E-02 | 8.30E-02 | 2.49E+00 | 2.52E+00 | 3.58E-01 | no | | intergenic_variant | MODIFIER |
| 16 | 50830672 | AX-170441955 | 0 | G | T | 0.518 | 7.73E-02 | 8.30E-02 | 2.49E+00 | 2.52E+00 | 3.58E-01 | no | | intergenic_variant | MODIFIER |
| 16 | 50831074 | AX-170131204 | 0 | C | T | 0.518 | 7.73E-02 | 8.30E-02 | 2.49E+00 | 2.52E+00 | 3.58E-01 | no | | intergenic_variant | MODIFIER |
| 16 | 50977118 | AX-170500540 | 0 | C | A | 0.518 | 7.73E-02 | 8.30E-02 | 2.49E+00 | 2.52E+00 | 3.58E-01 | no | | intergenic_variant | MODIFIER |
| 16 | 51008384 | AX-97815565 | 0 | C | G | 0.518 | 7.73E-02 | 8.30E-02 | 2.49E+00 | 2.52E+00 | 3.58E-01 | no | | intergenic_variant | MODIFIER |
| 16 | 51018559 | AX-170013028 | 0 | C | T | 0.518 | 7.73E-02 | 8.30E-02 | 2.49E+00 | 2.52E+00 | 3.58E-01 | no | | intergenic_variant | MODIFIER |
| 16 | 51069947 | AX-170013029 | 0 | T | C | 0.518 | 7.73E-02 | 8.30E-02 | 2.49E+00 | 2.52E+00 | 3.58E-01 | no | | intergenic_variant | MODIFIER |
| 16 | 51077931 | AX-170504216 | 0 | T | C | 0.518 | 7.73E-02 | 8.30E-02 | 2.49E+00 | 2.52E+00 | 3.58E-01 | no | | intergenic_variant | MODIFIER |
| 16 | 51086919 | AX-170013031 | 0 | T | C | 0.518 | 7.73E-02 | 8.30E-02 | 2.49E+00 | 2.52E+00 | 3.58E-01 | no | | intergenic_variant | MODIFIER |
| 16 | 51105876 | AX-170298391 | 0 | T | G | 0.518 | 7.73E-02 | 8.30E-02 | 2.49E+00 | 2.52E+00 | 3.58E-01 | no | | intergenic_variant | MODIFIER |
| 16 | 51112600 | AX-170289121 | 0 | T | C | 0.518 | 7.73E-02 | 8.30E-02 | 2.49E+00 | 2.52E+00 | 3.58E-01 | no | | intergenic_variant | MODIFIER |
| 16 | 51117498 | AX-170323730 | 0 | A | C | 0.518 | 7.73E-02 | 8.30E-02 | 2.49E+00 | 2.52E+00 | 3.58E-01 | no | | intergenic_variant | MODIFIER |
| 16 | 51171956 | AX-170395047 | 0 | T | C | 0.518 | 7.73E-02 | 8.30E-02 | 2.49E+00 | 2.52E+00 | 3.58E-01 | no | | intergenic_variant | MODIFIER |
| 16 | 51216678 | AX-170395495 | 0 | T | A | 0.518 | 7.73E-02 | 8.30E-02 | 2.49E+00 | 2.52E+00 | 3.58E-01 | no | | intergenic_variant | MODIFIER |
| 16 | 51216720 | AX-170118726 | 0 | T | C | 0.518 | 7.73E-02 | 8.30E-02 | 2.49E+00 | 2.52E+00 | 3.58E-01 | no | | intergenic_variant | MODIFIER |
| 16 | 51219034 | AX-170438892 | 0 | T | C | 0.518 | 7.73E-02 | 8.30E-02 | 2.49E+00 | 2.52E+00 | 3.58E-01 | no | | intergenic_variant | MODIFIER |
| 16 | 51221381 | AX-170508529 | 0 | G | A | 0.518 | 7.73E-02 | 8.30E-02 | 2.49E+00 | 2.52E+00 | 3.58E-01 | no | | intergenic_variant | MODIFIER |
| 16 | 51223314 | AX-170251047 | 0 | A | G | 0.518 | 7.73E-02 | 8.30E-02 | 2.49E+00 | 2.52E+00 | 3.58E-01 | no | | intergenic_variant | MODIFIER |
| 16 | 51223662 | AX-170351064 | 0 | A | G | 0.518 | 7.73E-02 | 8.30E-02 | 2.49E+00 | 2.52E+00 | 3.58E-01 | no | | intergenic_variant | MODIFIER |
| 16 | 51314552 | AX-170118261 | 0 | A | G | 0.518 | 7.73E-02 | 8.30E-02 | 2.49E+00 | 2.52E+00 | 3.58E-01 | no | | intergenic_variant | MODIFIER |

| | | | | | | | | | | | | | | | |
|---|---|---|---|---|---|---|---|---|---|---|---|---|---|---|---|
| 16 | 51345167 | AX-170297552 | 0 | T | C | 0.518 | 7.73E-02 | 8.30E-02 | 2.49E+00 | 2.52E+00 | 3.58E-01 | no | | intergenic_variant | MODIFIER |
| 16 | 51394037 | AX-170228297 | 0 | T | C | 0.518 | 7.73E-02 | 8.30E-02 | 2.49E+00 | 2.52E+00 | 3.58E-01 | no | | intergenic_variant | MODIFIER |
| 16 | 51401689 | AX-170394485 | 0 | A | G | 0.518 | 7.73E-02 | 8.30E-02 | 2.49E+00 | 2.52E+00 | 3.58E-01 | no | | intergenic_variant | MODIFIER |
| 16 | 51409412 | AX-170185955 | 0 | C | T | 0.518 | 7.73E-02 | 8.30E-02 | 2.49E+00 | 2.52E+00 | 3.58E-01 | no | | intergenic_variant | MODIFIER |
| 16 | 51442133 | AX-97831781 | 0 | C | G | 0.518 | 7.73E-02 | 8.30E-02 | 2.49E+00 | 2.52E+00 | 3.58E-01 | no | | intergenic_variant | MODIFIER |
| 16 | 51476083 | AX-170424215 | 0 | T | C | 0.518 | 7.73E-02 | 8.30E-02 | 2.49E+00 | 2.52E+00 | 3.58E-01 | no | | intergenic_variant | MODIFIER |
| 16 | 51476151 | AX-170432816 | 0 | G | C | 0.518 | 7.73E-02 | 8.30E-02 | 2.49E+00 | 2.52E+00 | 3.58E-01 | no | | intergenic_variant | MODIFIER |
| 16 | 51501878 | AX-170098615 | 0 | A | G | 0.518 | 7.73E-02 | 8.30E-02 | 2.49E+00 | 2.52E+00 | 3.58E-01 | no | | intergenic_variant | MODIFIER |
| 16 | 51502327 | AX-170013041 | 0 | C | T | 0.518 | 7.73E-02 | 8.30E-02 | 2.49E+00 | 2.52E+00 | 3.58E-01 | no | | intergenic_variant | MODIFIER |
| 16 | 51545111 | AX-97831179 | 0 | C | T | 0.518 | 7.73E-02 | 8.30E-02 | 2.49E+00 | 2.52E+00 | 3.58E-01 | no | | intergenic_variant | MODIFIER |
| 16 | 51563185 | AX-97845446 | 0 | C | T | 0.518 | 7.73E-02 | 8.30E-02 | 2.49E+00 | 2.52E+00 | 3.58E-01 | no | | intergenic_variant | MODIFIER |
| 16 | 51599543 | AX-170117023 | 0 | A | G | 0.518 | 7.73E-02 | 8.30E-02 | 2.49E+00 | 2.52E+00 | 3.58E-01 | no | | intergenic_variant | MODIFIER |
| 16 | 51623605 | AX-170164047 | 0 | A | G | 0.518 | 7.73E-02 | 8.30E-02 | 2.49E+00 | 2.52E+00 | 3.58E-01 | no | | intergenic_variant | MODIFIER |
| 16 | 51624126 | AX-170013040 | 0 | C | T | 0.518 | 7.73E-02 | 8.30E-02 | 2.49E+00 | 2.52E+00 | 3.58E-01 | no | | intergenic_variant | MODIFIER |
| 16 | 51630029 | AX-170013039 | 0 | C | T | 0.518 | 7.73E-02 | 8.30E-02 | 2.49E+00 | 2.52E+00 | 3.58E-01 | no | | intergenic_variant | MODIFIER |
| 16 | 51648060 | AX-170431009 | 0 | A | G | 0.518 | 7.73E-02 | 8.30E-02 | 2.49E+00 | 2.52E+00 | 3.58E-01 | no | | intergenic_variant | MODIFIER |
| 9 | 7207171 | AX-97781578 | 0 | G | A | 0.474 | 7.39E-02 | 7.88E-02 | 2.44E+00 | 2.54E+00 | 3.59E-01 | no | Dcun1d5 | downstream_gene_variant | MODIFIER |
| 1 | 170663219 | AX-170445878 | 0 | G | A | 0.219 | 1.20E-01 | 1.32E-01 | 2.68E+00 | 2.73E+00 | 3.59E-01 | no | Olfml2b | intron_variant | MODIFIER |
| 18 | 83892300 | AX-170347962 | 0 | C | A | 0.158 | -1.01E-01 | 1.09E-01 | 2.46E+00 | 2.54E+00 | 3.59E-01 | no | | intergenic_variant | MODIFIER |
| 18 | 83897333 | AX-170506796 | 0 | T | C | 0.158 | -1.01E-01 | 1.09E-01 | 2.46E+00 | 2.54E+00 | 3.59E-01 | no | | intergenic_variant | MODIFIER |
| 18 | 83898787 | AX-97828047 | 0 | A | G | 0.158 | -1.01E-01 | 1.09E-01 | 2.46E+00 | 2.54E+00 | 3.59E-01 | no | | intergenic_variant | MODIFIER |
| 18 | 83945713 | AX-170338910 | 0 | T | C | 0.158 | -1.01E-01 | 1.09E-01 | 2.46E+00 | 2.54E+00 | 3.59E-01 | no | | intergenic_variant | MODIFIER |
| 18 | 83945826 | AX-170431136 | 0 | G | C | 0.158 | -1.01E-01 | 1.09E-01 | 2.46E+00 | 2.54E+00 | 3.59E-01 | no | | intergenic_variant | MODIFIER |
| 18 | 83950382 | AX-170161625 | 0 | T | G | 0.158 | -1.01E-01 | 1.09E-01 | 2.46E+00 | 2.54E+00 | 3.59E-01 | no | | intergenic_variant | MODIFIER |
| 18 | 83950512 | AX-170370045 | 0 | G | A | 0.158 | -1.01E-01 | 1.09E-01 | 2.46E+00 | 2.54E+00 | 3.59E-01 | no | | intergenic_variant | MODIFIER |
| 18 | 83952264 | AX-170335594 | 0 | C | T | 0.158 | -1.01E-01 | 1.09E-01 | 2.46E+00 | 2.54E+00 | 3.59E-01 | no | | intergenic_variant | MODIFIER |
| 18 | 83954405 | AX-170508884 | 0 | A | G | 0.158 | -1.01E-01 | 1.09E-01 | 2.46E+00 | 2.54E+00 | 3.59E-01 | no | | regulatory_region_variant | MODIFIER |
| 18 | 83954466 | AX-97840231 | 0 | T | C | 0.158 | -1.01E-01 | 1.09E-01 | 2.46E+00 | 2.54E+00 | 3.59E-01 | no | | regulatory_region_variant | MODIFIER |
| 18 | 83954755 | AX-170207088 | 0 | T | C | 0.158 | -1.01E-01 | 1.09E-01 | 2.46E+00 | 2.54E+00 | 3.59E-01 | no | | regulatory_region_variant | MODIFIER |
| 18 | 84033542 | AX-170269710 | 0 | T | A | 0.158 | -1.01E-01 | 1.09E-01 | 2.46E+00 | 2.54E+00 | 3.59E-01 | no | Tshz1 | intron_variant | MODIFIER |
| 18 | 84040928 | AX-170510201 | 0 | T | G | 0.158 | -1.01E-01 | 1.09E-01 | 2.46E+00 | 2.54E+00 | 3.59E-01 | no | Tshz1 | intron_variant | MODIFIER |
| 18 | 84055350 | AX-97788325 | 0 | G | A | 0.158 | -1.01E-01 | 1.09E-01 | 2.46E+00 | 2.54E+00 | 3.59E-01 | no | Tshz1 | intron_variant | MODIFIER |
| 18 | 84064027 | AX-170335672 | 0 | G | A | 0.158 | -1.01E-01 | 1.09E-01 | 2.46E+00 | 2.54E+00 | 3.59E-01 | no | Tshz1 | intron_variant | MODIFIER |
| 18 | 84096424 | AX-170355201 | 0 | T | C | 0.158 | -1.01E-01 | 1.09E-01 | 2.46E+00 | 2.54E+00 | 3.59E-01 | no | Zadh2 | 3_prime_UTR_variant | MODIFIER |
| 18 | 84098181 | AX-170522399 | 0 | T | G | 0.158 | -1.01E-01 | 1.09E-01 | 2.46E+00 | 2.54E+00 | 3.59E-01 | no | Zadh2 | downstream_gene_variant | MODIFIER |
| 18 | 84102467 | AX-170346869 | 0 | A | G | 0.158 | -1.01E-01 | 1.09E-01 | 2.46E+00 | 2.54E+00 | 3.59E-01 | no | Zadh2 | downstream_gene_variant | MODIFIER |
| 18 | 84102776 | AX-170407331 | 0 | A | G | 0.158 | -1.01E-01 | 1.09E-01 | 2.46E+00 | 2.54E+00 | 3.59E-01 | no | AC124012.2 | upstream_gene_variant | MODIFIER |
| 18 | 84115757 | AX-97790956 | 0 | C | T | 0.158 | -1.01E-01 | 1.09E-01 | 2.46E+00 | 2.54E+00 | 3.59E-01 | no | AC124012.1 | upstream_gene_variant | MODIFIER |
| 18 | 84117974 | AX-170507121 | 0 | C | T | 0.158 | -1.01E-01 | 1.09E-01 | 2.46E+00 | 2.54E+00 | 3.59E-01 | no | | intergenic_variant | MODIFIER |
| 18 | 84121372 | AX-170358445 | 0 | T | G | 0.158 | -1.01E-01 | 1.09E-01 | 2.46E+00 | 2.54E+00 | 3.59E-01 | no | | regulatory_region_variant | MODIFIER |
| 18 | 84122149 | AX-170176088 | 0 | A | G | 0.158 | -1.01E-01 | 1.09E-01 | 2.46E+00 | 2.54E+00 | 3.59E-01 | no | | regulatory_region_variant | MODIFIER |
| 18 | 84150880 | AX-170367317 | 0 | A | G | 0.158 | -1.01E-01 | 1.09E-01 | 2.46E+00 | 2.54E+00 | 3.59E-01 | no | Zfp407 | intron_variant | MODIFIER |
| 18 | 84151359 | AX-170533730 | 0 | A | G | 0.158 | -1.01E-01 | 1.09E-01 | 2.46E+00 | 2.54E+00 | 3.59E-01 | no | Zfp407 | intron_variant | MODIFIER |
| 18 | 84154926 | AX-170105438 | 0 | T | C | 0.158 | -1.01E-01 | 1.09E-01 | 2.46E+00 | 2.54E+00 | 3.59E-01 | no | Zfp407 | intron_variant | MODIFIER |
| 18 | 84155562 | AX-170275514 | 0 | T | C | 0.158 | -1.01E-01 | 1.09E-01 | 2.46E+00 | 2.54E+00 | 3.59E-01 | no | Zfp407 | intron_variant | MODIFIER |
| 18 | 84155866 | AX-170153246 | 0 | A | C | 0.158 | -1.01E-01 | 1.09E-01 | 2.46E+00 | 2.54E+00 | 3.59E-01 | no | Zfp407 | intron_variant | MODIFIER |
| 18 | 84157249 | AX-170424341 | 0 | A | G | 0.158 | -1.01E-01 | 1.09E-01 | 2.46E+00 | 2.54E+00 | 3.59E-01 | no | Zfp407 | intron_variant | MODIFIER |
| 18 | 84165633 | AX-97835170 | 0 | G | A | 0.158 | -1.01E-01 | 1.09E-01 | 2.46E+00 | 2.54E+00 | 3.59E-01 | no | Zfp407 | intron_variant | MODIFIER |
| 18 | 84188863 | AX-170277948 | 0 | A | G | 0.158 | -1.01E-01 | 1.09E-01 | 2.46E+00 | 2.54E+00 | 3.59E-01 | no | Zfp407 | intron_variant | MODIFIER |
| 18 | 84192912 | AX-170145119 | 0 | A | G | 0.158 | -1.01E-01 | 1.09E-01 | 2.46E+00 | 2.54E+00 | 3.59E-01 | no | Zfp407 | intron_variant | MODIFIER |
| 18 | 84195707 | AX-97789350 | 0 | T | C | 0.158 | -1.01E-01 | 1.09E-01 | 2.46E+00 | 2.54E+00 | 3.59E-01 | no | Zfp407 | intron_variant | MODIFIER |
| 18 | 84198656 | AX-170303496 | 0 | A | G | 0.158 | -1.01E-01 | 1.09E-01 | 2.46E+00 | 2.54E+00 | 3.59E-01 | no | Zfp407 | intron_variant | MODIFIER |
| 18 | 84198940 | AX-170363193 | 0 | G | A | 0.158 | -1.01E-01 | 1.09E-01 | 2.46E+00 | 2.54E+00 | 3.59E-01 | no | Zfp407 | intron_variant | MODIFIER |
| 18 | 84203737 | AX-170520555 | 0 | T | C | 0.158 | -1.01E-01 | 1.09E-01 | 2.46E+00 | 2.54E+00 | 3.59E-01 | no | Zfp407 | intron_variant | MODIFIER |

| | | | | | | | | | | | | | | | |
|---|---|---|---|---|---|---|---|---|---|---|---|---|---|---|---|
| 18 | 84204046 | AX-170440825 | 0 | A | G | 0.158 | -1.01E-01 | 1.09E-01 | 2.46E+00 | 2.54E+00 | 3.59E-01 | no | Zfp407 | intron_variant | MODIFIER |
| 18 | 84209746 | AX-170302199 | 0 | C | T | 0.158 | -1.01E-01 | 1.09E-01 | 2.46E+00 | 2.54E+00 | 3.59E-01 | no | Zfp407 | intron_variant | MODIFIER |
| 18 | 84213879 | AX-170180087 | 0 | C | G | 0.158 | -1.01E-01 | 1.09E-01 | 2.46E+00 | 2.54E+00 | 3.59E-01 | no | Zfp407 | intron_variant | MODIFIER |
| 18 | 84227977 | AX-170254457 | 0 | C | T | 0.158 | -1.01E-01 | 1.09E-01 | 2.46E+00 | 2.54E+00 | 3.59E-01 | no | Zfp407 | intron_variant | MODIFIER |
| 18 | 84236916 | AX-170356929 | 0 | T | G | 0.158 | -1.01E-01 | 1.09E-01 | 2.46E+00 | 2.54E+00 | 3.59E-01 | no | Zfp407 | intron_variant | MODIFIER |
| 18 | 84240479 | AX-170117813 | 0 | G | A | 0.158 | -1.01E-01 | 1.09E-01 | 2.46E+00 | 2.54E+00 | 3.59E-01 | no | Zfp407 | intron_variant | MODIFIER |
| 18 | 84241167 | AX-170204464 | 0 | G | T | 0.158 | -1.01E-01 | 1.09E-01 | 2.46E+00 | 2.54E+00 | 3.59E-01 | no | Zfp407 | intron_variant | MODIFIER |
| 18 | 84241405 | AX-170116257 | 0 | T | C | 0.158 | -1.01E-01 | 1.09E-01 | 2.46E+00 | 2.54E+00 | 3.59E-01 | no | Zfp407 | intron_variant | MODIFIER |
| 18 | 84261165 | AX-170309106 | 0 | T | C | 0.158 | -1.01E-01 | 1.09E-01 | 2.46E+00 | 2.54E+00 | 3.59E-01 | no | Zfp407 | intron_variant | MODIFIER |
| 18 | 84264884 | AX-170135498 | 0 | T | C | 0.158 | -1.01E-01 | 1.09E-01 | 2.46E+00 | 2.54E+00 | 3.59E-01 | no | Zfp407 | intron_variant | MODIFIER |
| 18 | 84266857 | AX-170326729 | 0 | T | C | 0.158 | -1.01E-01 | 1.09E-01 | 2.46E+00 | 2.54E+00 | 3.59E-01 | no | Zfp407 | intron_variant | MODIFIER |
| 18 | 84287382 | AX-170205550 | 0 | T | G | 0.158 | -1.01E-01 | 1.09E-01 | 2.46E+00 | 2.54E+00 | 3.59E-01 | no | Zfp407 | intron_variant | MODIFIER |
| 18 | 84316559 | AX-170177318 | 0 | C | T | 0.158 | -1.01E-01 | 1.09E-01 | 2.46E+00 | 2.54E+00 | 3.59E-01 | no | Zfp407 | intron_variant | MODIFIER |
| 18 | 84330057 | AX-170250063 | 0 | A | G | 0.158 | -1.01E-01 | 1.09E-01 | 2.46E+00 | 2.54E+00 | 3.59E-01 | no | Zfp407 | intron_variant | MODIFIER |
| 18 | 84331245 | AX-170523323 | 0 | A | C | 0.158 | -1.01E-01 | 1.09E-01 | 2.46E+00 | 2.54E+00 | 3.59E-01 | no | Zfp407 | intron_variant | MODIFIER |
| 18 | 84332214 | AX-170471578 | 0 | T | C | 0.158 | -1.01E-01 | 1.09E-01 | 2.46E+00 | 2.54E+00 | 3.59E-01 | no | Zfp407 | intron_variant | MODIFIER |
| 18 | 84344217 | AX-170233781 | 0 | G | A | 0.158 | -1.01E-01 | 1.09E-01 | 2.46E+00 | 2.54E+00 | 3.59E-01 | no | Zfp407 | intron_variant | MODIFIER |
| 18 | 84405252 | AX-170016117 | 0 | T | C | 0.158 | -1.01E-01 | 1.09E-01 | 2.46E+00 | 2.54E+00 | 3.59E-01 | no | Zfp407 | intron_variant | MODIFIER |
| 18 | 84405591 | AX-97807916 | 0 | C | T | 0.158 | -1.01E-01 | 1.09E-01 | 2.46E+00 | 2.54E+00 | 3.59E-01 | no | Zfp407 | intron_variant | MODIFIER |
| 18 | 84405814 | AX-170260171 | 0 | A | T | 0.158 | -1.01E-01 | 1.09E-01 | 2.46E+00 | 2.54E+00 | 3.59E-01 | no | Zfp407 | intron_variant | MODIFIER |
| 18 | 84436470 | AX-170016113 | 0 | A | T | 0.158 | -1.01E-01 | 1.09E-01 | 2.46E+00 | 2.54E+00 | 3.59E-01 | no | Zfp407 | intron_variant | MODIFIER |
| 18 | 84464450 | AX-97824662 | 0 | T | C | 0.158 | -1.01E-01 | 1.09E-01 | 2.46E+00 | 2.54E+00 | 3.59E-01 | no | Zfp407 | intron_variant | MODIFIER |
| 18 | 84529309 | AX-170208589 | 0 | G | A | 0.158 | -1.01E-01 | 1.09E-01 | 2.46E+00 | 2.54E+00 | 3.59E-01 | no | Zfp407 | intron_variant | MODIFIER |
| 18 | 84540848 | AX-170061549 | 0 | G | A | 0.158 | -1.01E-01 | 1.09E-01 | 2.46E+00 | 2.54E+00 | 3.59E-01 | no | Zfp407 | intron_variant | MODIFIER |
| 18 | 84548940 | AX-170208922 | 0 | G | T | 0.158 | -1.01E-01 | 1.09E-01 | 2.46E+00 | 2.54E+00 | 3.59E-01 | no | Zfp407 | intron_variant | MODIFIER |
| 18 | 84555159 | AX-170102204 | 0 | G | T | 0.158 | -1.01E-01 | 1.09E-01 | 2.46E+00 | 2.54E+00 | 3.59E-01 | no | Zfp407 | intron_variant | MODIFIER |
| 18 | 84561006 | AX-170282209 | 0 | G | T | 0.158 | -1.01E-01 | 1.09E-01 | 2.46E+00 | 2.54E+00 | 3.59E-01 | no | Zfp407 | upstream_gene_variant | MODIFIER |
| 18 | 84644638 | AX-170383237 | 0 | C | T | 0.158 | -1.01E-01 | 1.09E-01 | 2.46E+00 | 2.54E+00 | 3.59E-01 | no | Cndp1 | intron_variant | MODIFIER |
| 18 | 84644670 | AX-170310880 | 0 | G | A | 0.158 | -1.01E-01 | 1.09E-01 | 2.46E+00 | 2.54E+00 | 3.59E-01 | no | Cndp1 | intron_variant | MODIFIER |
| 18 | 84646518 | AX-97792691 | 0 | T | C | 0.158 | -1.01E-01 | 1.09E-01 | 2.46E+00 | 2.54E+00 | 3.59E-01 | no | Cndp1 | intron_variant | MODIFIER |
| 18 | 84681140 | AX-170358806 | 0 | C | A | 0.158 | -1.01E-01 | 1.09E-01 | 2.46E+00 | 2.54E+00 | 3.59E-01 | no | Cndp2 | intron_variant | MODIFIER |
| 18 | 84694488 | AX-170359595 | 0 | T | C | 0.158 | -1.01E-01 | 1.09E-01 | 2.46E+00 | 2.54E+00 | 3.59E-01 | no | | intergenic_variant | MODIFIER |
| 18 | 84723743 | AX-170243401 | 0 | C | G | 0.158 | -1.01E-01 | 1.09E-01 | 2.46E+00 | 2.54E+00 | 3.59E-01 | no | Fam69c | intron_variant | MODIFIER |
| 18 | 84741509 | AX-170318212 | 0 | A | T | 0.158 | -1.01E-01 | 1.09E-01 | 2.46E+00 | 2.54E+00 | 3.59E-01 | no | Fam69c | downstream_gene_variant | MODIFIER |
| 18 | 84757335 | AX-97824666 | 0 | A | G | 0.158 | -1.01E-01 | 1.09E-01 | 2.46E+00 | 2.54E+00 | 3.59E-01 | no | AC142448.2 | downstream_gene_variant | MODIFIER |
| 18 | 84758877 | AX-170467462 | 0 | T | G | 0.158 | -1.01E-01 | 1.09E-01 | 2.46E+00 | 2.54E+00 | 3.59E-01 | no | AC142448.2 | downstream_gene_variant | MODIFIER |
| 18 | 84765795 | AX-170396402 | 0 | T | C | 0.158 | -1.01E-01 | 1.09E-01 | 2.46E+00 | 2.54E+00 | 3.59E-01 | no | | regulatory_region_variant | MODIFIER |
| 18 | 84766275 | AX-170167515 | 0 | T | C | 0.158 | -1.01E-01 | 1.09E-01 | 2.46E+00 | 2.54E+00 | 3.59E-01 | no | | intergenic_variant | MODIFIER |
| 18 | 84766300 | AX-170238237 | 0 | T | C | 0.158 | -1.01E-01 | 1.09E-01 | 2.46E+00 | 2.54E+00 | 3.59E-01 | no | | intergenic_variant | MODIFIER |
| 18 | 84766524 | AX-170247971 | 0 | T | C | 0.158 | -1.01E-01 | 1.09E-01 | 2.46E+00 | 2.54E+00 | 3.59E-01 | no | | intergenic_variant | MODIFIER |
| 18 | 84767010 | AX-170178335 | 0 | A | G | 0.158 | -1.01E-01 | 1.09E-01 | 2.46E+00 | 2.54E+00 | 3.59E-01 | no | | intergenic_variant | MODIFIER |
| 18 | 84768190 | AX-97824659 | 0 | G | A | 0.158 | -1.01E-01 | 1.09E-01 | 2.46E+00 | 2.54E+00 | 3.59E-01 | no | | intergenic_variant | MODIFIER |
| 18 | 84768734 | AX-170061554 | 0 | C | A | 0.158 | -1.01E-01 | 1.09E-01 | 2.46E+00 | 2.54E+00 | 3.59E-01 | no | | intergenic_variant | MODIFIER |
| 18 | 84803228 | AX-97835174 | 0 | A | C | 0.158 | -1.01E-01 | 1.09E-01 | 2.46E+00 | 2.54E+00 | 3.59E-01 | no | C18orf63 | intron_variant | MODIFIER |
| 18 | 84817838 | AX-97824379 | 0 | A | T | 0.158 | -1.01E-01 | 1.09E-01 | 2.46E+00 | 2.54E+00 | 3.59E-01 | no | Gm22819 | upstream_gene_variant | MODIFIER |
| 18 | 84848797 | AX-97844354 | 0 | A | G | 0.158 | -1.01E-01 | 1.09E-01 | 2.46E+00 | 2.54E+00 | 3.59E-01 | no | Cyb5a | upstream_gene_variant | MODIFIER |
| 18 | 84871118 | AX-170255274 | 0 | C | T | 0.158 | -1.01E-01 | 1.09E-01 | 2.46E+00 | 2.54E+00 | 3.59E-01 | no | Cyb5a | intron_variant,NMD_transcript_variant | MODIFIER |
| 18 | 84872325 | AX-97795793 | 0 | C | T | 0.158 | -1.01E-01 | 1.09E-01 | 2.46E+00 | 2.54E+00 | 3.59E-01 | no | Cyb5a | intron_variant,NMD_transcript_variant | MODIFIER |
| 18 | 84872524 | AX-170368533 | 0 | G | A | 0.158 | -1.01E-01 | 1.09E-01 | 2.46E+00 | 2.54E+00 | 3.59E-01 | no | Cyb5a | intron_variant,NMD_transcript_variant | MODIFIER |
| 18 | 84877793 | AX-170142318 | 0 | G | T | 0.158 | -1.01E-01 | 1.09E-01 | 2.46E+00 | 2.54E+00 | 3.59E-01 | no | Cyb5a | intron_variant,NMD_transcript_variant | MODIFIER |
| 18 | 84877847 | AX-170464422 | 0 | G | C | 0.158 | -1.01E-01 | 1.09E-01 | 2.46E+00 | 2.54E+00 | 3.59E-01 | no | Cyb5a | 3_prime_UTR_variant,NMD_transcript_variant | MODIFIER |
| 18 | 84879583 | AX-170175862 | 0 | G | A | 0.158 | -1.01E-01 | 1.09E-01 | 2.46E+00 | 2.54E+00 | 3.59E-01 | no | Cyb5a | 3_prime_UTR_variant,NMD_transcript_variant | MODIFIER |
| 18 | 84880596 | AX-170393196 | 0 | G | A | 0.158 | -1.01E-01 | 1.09E-01 | 2.46E+00 | 2.54E+00 | 3.59E-01 | no | Cyb5a | downstream_gene_variant | MODIFIER |
| 18 | 84886425 | AX-170229929 | 0 | A | G | 0.158 | -1.01E-01 | 1.09E-01 | 2.46E+00 | 2.54E+00 | 3.59E-01 | no | | regulatory_region_variant | MODIFIER |

| | | | | | | | | | | | | | | | |
|---|---|---|---|---|---|---|---|---|---|---|---|---|---|---|---|
| 18 | 84890867 | AX-97783552 | 0 | T | C | 0.158 | -1.01E-01 | 1.09E-01 | 2.46E+00 | 2.54E+00 | 3.59E-01 | no | | regulatory_region_variant | MODIFIER |
| 18 | 84916477 | AX-97801335 | 0 | G | A | 0.158 | -1.01E-01 | 1.09E-01 | 2.46E+00 | 2.54E+00 | 3.59E-01 | no | | intergenic_variant | MODIFIER |
| 18 | 84934465 | AX-170149324 | 0 | C | T | 0.158 | -1.01E-01 | 1.09E-01 | 2.46E+00 | 2.54E+00 | 3.59E-01 | no | Fbxo15 | upstream_gene_variant | MODIFIER |
| 18 | 84938247 | AX-170440881 | 0 | C | G | 0.158 | -1.01E-01 | 1.09E-01 | 2.46E+00 | 2.54E+00 | 3.59E-01 | no | Fbxo15 | intron_variant | MODIFIER |
| 18 | 84968400 | AX-170016122 | 0 | C | A | 0.158 | -1.01E-01 | 1.09E-01 | 2.46E+00 | 2.54E+00 | 3.59E-01 | no | Fbxo15 | intron_variant | MODIFIER |
| 18 | 84978284 | AX-170177611 | 0 | T | G | 0.158 | -1.01E-01 | 1.09E-01 | 2.46E+00 | 2.54E+00 | 3.59E-01 | no | Fbxo15 | intron_variant | MODIFIER |
| 18 | 84994635 | AX-170416064 | 0 | C | T | 0.158 | -1.01E-01 | 1.09E-01 | 2.46E+00 | 2.54E+00 | 3.59E-01 | no | | intergenic_variant | MODIFIER |
| 18 | 84995224 | AX-97824656 | 0 | C | T | 0.158 | -1.01E-01 | 1.09E-01 | 2.46E+00 | 2.54E+00 | 3.59E-01 | no | | intergenic_variant | MODIFIER |
| 18 | 85016340 | AX-170279794 | 0 | A | C | 0.158 | -1.01E-01 | 1.09E-01 | 2.46E+00 | 2.54E+00 | 3.59E-01 | no | AC128672.2 | upstream_gene_variant | MODIFIER |
| 18 | 85022535 | AX-170230876 | 0 | C | T | 0.158 | -1.01E-01 | 1.09E-01 | 2.46E+00 | 2.54E+00 | 3.59E-01 | no | | intergenic_variant | MODIFIER |
| 18 | 85063531 | AX-170434164 | 0 | T | C | 0.158 | -1.01E-01 | 1.09E-01 | 2.46E+00 | 2.54E+00 | 3.59E-01 | no | | intergenic_variant | MODIFIER |
| 18 | 85063706 | AX-170140862 | 0 | C | A | 0.158 | -1.01E-01 | 1.09E-01 | 2.46E+00 | 2.54E+00 | 3.59E-01 | no | | intergenic_variant | MODIFIER |
| 18 | 85068714 | AX-170397662 | 0 | A | T | 0.158 | -1.01E-01 | 1.09E-01 | 2.46E+00 | 2.54E+00 | 3.59E-01 | no | AC128672.1 | upstream_gene_variant | MODIFIER |
| 18 | 85088127 | AX-97830655 | 0 | T | C | 0.158 | -1.01E-01 | 1.09E-01 | 2.46E+00 | 2.54E+00 | 3.59E-01 | no | AC128672.1 | downstream_gene_variant | MODIFIER |
| 18 | 85152411 | AX-170061562 | 0 | C | A | 0.158 | -1.01E-01 | 1.09E-01 | 2.46E+00 | 2.54E+00 | 3.59E-01 | no | | intergenic_variant | MODIFIER |
| 18 | 85253624 | AX-97846938 | 0 | T | C | 0.158 | -1.01E-01 | 1.09E-01 | 2.46E+00 | 2.54E+00 | 3.59E-01 | no | | intergenic_variant | MODIFIER |
| 18 | 85272781 | AX-170188180 | 0 | G | T | 0.158 | -1.01E-01 | 1.09E-01 | 2.46E+00 | 2.54E+00 | 3.59E-01 | no | | intergenic_variant | MODIFIER |
| 18 | 85272815 | AX-170464980 | 0 | G | A | 0.158 | -1.01E-01 | 1.09E-01 | 2.46E+00 | 2.54E+00 | 3.59E-01 | no | | intergenic_variant | MODIFIER |
| 18 | 85277651 | AX-170170629 | 0 | C | G | 0.158 | -1.01E-01 | 1.09E-01 | 2.46E+00 | 2.54E+00 | 3.59E-01 | no | | intergenic_variant | MODIFIER |
| 18 | 85277755 | AX-170225692 | 0 | G | A | 0.158 | -1.01E-01 | 1.09E-01 | 2.46E+00 | 2.54E+00 | 3.59E-01 | no | | intergenic_variant | MODIFIER |
| 18 | 85288246 | AX-170495260 | 0 | C | T | 0.158 | -1.01E-01 | 1.09E-01 | 2.46E+00 | 2.54E+00 | 3.59E-01 | no | | intergenic_variant | MODIFIER |
| 18 | 85301770 | AX-170181660 | 0 | A | C | 0.158 | -1.01E-01 | 1.09E-01 | 2.46E+00 | 2.54E+00 | 3.59E-01 | no | | intergenic_variant | MODIFIER |
| 18 | 85302462 | AX-97804351 | 0 | G | A | 0.158 | -1.01E-01 | 1.09E-01 | 2.46E+00 | 2.54E+00 | 3.59E-01 | no | | intergenic_variant | MODIFIER |
| 18 | 85306598 | AX-170061566 | 0 | A | G | 0.158 | -1.01E-01 | 1.09E-01 | 2.46E+00 | 2.54E+00 | 3.59E-01 | no | | intergenic_variant | MODIFIER |
| 18 | 85306811 | AX-170061567 | 0 | T | C | 0.158 | -1.01E-01 | 1.09E-01 | 2.46E+00 | 2.54E+00 | 3.59E-01 | no | | intergenic_variant | MODIFIER |
| 18 | 85307241 | AX-170296059 | 0 | C | T | 0.158 | -1.01E-01 | 1.09E-01 | 2.46E+00 | 2.54E+00 | 3.59E-01 | no | | intergenic_variant | MODIFIER |
| 18 | 85339792 | AX-170505496 | 0 | T | C | 0.158 | -1.01E-01 | 1.09E-01 | 2.46E+00 | 2.54E+00 | 3.59E-01 | no | | intergenic_variant | MODIFIER |
| 18 | 85346962 | AX-170173741 | 0 | A | C | 0.158 | -1.01E-01 | 1.09E-01 | 2.46E+00 | 2.54E+00 | 3.59E-01 | no | | intergenic_variant | MODIFIER |
| 18 | 85363944 | AX-97847221 | 0 | A | G | 0.158 | -1.01E-01 | 1.09E-01 | 2.46E+00 | 2.54E+00 | 3.59E-01 | no | | intergenic_variant | MODIFIER |
| 18 | 85391106 | AX-170308718 | 0 | C | T | 0.158 | -1.01E-01 | 1.09E-01 | 2.46E+00 | 2.54E+00 | 3.59E-01 | no | | intergenic_variant | MODIFIER |
| 18 | 85391382 | AX-170147436 | 0 | T | G | 0.158 | -1.01E-01 | 1.09E-01 | 2.46E+00 | 2.54E+00 | 3.59E-01 | no | | intergenic_variant | MODIFIER |
| 18 | 85404319 | AX-97836092 | 0 | A | G | 0.158 | -1.01E-01 | 1.09E-01 | 2.46E+00 | 2.54E+00 | 3.59E-01 | no | | intergenic_variant | MODIFIER |
| 18 | 85405193 | AX-170126079 | 0 | A | G | 0.158 | -1.01E-01 | 1.09E-01 | 2.46E+00 | 2.54E+00 | 3.59E-01 | no | | intergenic_variant | MODIFIER |
| 8 | 98858354 | AX-170264195 | 2 | C | G | 0.227 | 1.02E-01 | 1.13E-01 | 2.82E+00 | 2.90E+00 | 3.59E-01 | no | | intergenic_variant | MODIFIER |
| 9 | 17461439 | AX-170090252 | 1 | C | A | 0.473 | 7.89E-02 | 8.64E-02 | 2.61E+00 | 2.69E+00 | 3.59E-01 | no | | intergenic_variant | MODIFIER |
| 9 | 90762286 | AX-97815280 | 0 | T | G | 0.123 | -1.18E-01 | 1.29E-01 | 2.63E+00 | 2.70E+00 | 3.59E-01 | no | | intergenic_variant | MODIFIER |
| 9 | 90781767 | AX-170209426 | 0 | T | A | 0.123 | -1.18E-01 | 1.29E-01 | 2.63E+00 | 2.70E+00 | 3.59E-01 | no | | intergenic_variant | MODIFIER |
| 9 | 90793022 | AX-97787376 | 0 | C | T | 0.123 | -1.18E-01 | 1.29E-01 | 2.63E+00 | 2.70E+00 | 3.59E-01 | no | | intergenic_variant | MODIFIER |
| 9 | 90799778 | AX-170083082 | 0 | G | C | 0.123 | -1.18E-01 | 1.29E-01 | 2.63E+00 | 2.70E+00 | 3.59E-01 | no | | intergenic_variant | MODIFIER |
| 9 | 90822969 | AX-170343081 | 0 | A | G | 0.123 | -1.18E-01 | 1.29E-01 | 2.63E+00 | 2.70E+00 | 3.59E-01 | no | Gm28703 | downstream_gene_variant | MODIFIER |
| 9 | 90924410 | AX-170198769 | 0 | A | C | 0.123 | -1.18E-01 | 1.29E-01 | 2.63E+00 | 2.70E+00 | 3.59E-01 | no | | intergenic_variant | MODIFIER |
| 9 | 90971535 | AX-170340080 | 0 | T | A | 0.123 | -1.18E-01 | 1.29E-01 | 2.63E+00 | 2.70E+00 | 3.59E-01 | no | | intergenic_variant | MODIFIER |
| 9 | 91021913 | AX-168400241 | 0 | T | C | 0.123 | -1.18E-01 | 1.29E-01 | 2.63E+00 | 2.70E+00 | 3.59E-01 | no | | intergenic_variant | MODIFIER |
| 9 | 91217127 | AX-97821343 | 0 | C | T | 0.123 | -1.18E-01 | 1.29E-01 | 2.63E+00 | 2.70E+00 | 3.59E-01 | no | Gm29602 | downstream_gene_variant | MODIFIER |
| 9 | 91223998 | AX-170032971 | 0 | G | T | 0.123 | -1.18E-01 | 1.29E-01 | 2.63E+00 | 2.70E+00 | 3.59E-01 | no | | intergenic_variant | MODIFIER |
| 9 | 91298261 | AX-170462631 | 0 | C | A | 0.123 | -1.18E-01 | 1.29E-01 | 2.63E+00 | 2.70E+00 | 3.59E-01 | no | | intergenic_variant | MODIFIER |
| 9 | 91302303 | AX-170165003 | 0 | T | C | 0.123 | -1.18E-01 | 1.29E-01 | 2.63E+00 | 2.70E+00 | 3.59E-01 | no | | intergenic_variant | MODIFIER |
| 9 | 91302383 | AX-170241512 | 0 | C | G | 0.123 | -1.18E-01 | 1.29E-01 | 2.63E+00 | 2.70E+00 | 3.59E-01 | no | | intergenic_variant | MODIFIER |
| 9 | 91302446 | AX-170032968 | 0 | G | A | 0.123 | -1.18E-01 | 1.29E-01 | 2.63E+00 | 2.70E+00 | 3.59E-01 | no | | intergenic_variant | MODIFIER |
| 9 | 91317257 | AX-170468869 | 0 | G | T | 0.123 | -1.18E-01 | 1.29E-01 | 2.63E+00 | 2.70E+00 | 3.59E-01 | no | | regulatory_region_variant | MODIFIER |
| 9 | 91537865 | AX-97785348 | 0 | C | T | 0.123 | -1.18E-01 | 1.29E-01 | 2.63E+00 | 2.70E+00 | 3.59E-01 | no | Gm29478 | intron_variant,non_coding_transcript_variant | MODIFIER |
| 9 | 91544749 | AX-170366739 | 0 | C | T | 0.123 | -1.18E-01 | 1.29E-01 | 2.63E+00 | 2.70E+00 | 3.59E-01 | no | Gm29478 | intron_variant,non_coding_transcript_variant | MODIFIER |
| 9 | 91830452 | AX-97812460 | 0 | A | G | 0.123 | -1.18E-01 | 1.29E-01 | 2.63E+00 | 2.70E+00 | 3.59E-01 | no | | intergenic_variant | MODIFIER |
| 9 | 92291332 | AX-170032995 | 0 | T | G | 0.123 | -1.18E-01 | 1.29E-01 | 2.63E+00 | 2.70E+00 | 3.59E-01 | no | Plscr2 | intron_variant | MODIFIER |

| | | | | | | | | | | | | | | | |
|---|---|---|---|---|---|---|---|---|---|---|---|---|---|---|---|
| 9 | 92323681 | AX-170083094 | 0 | C | A | 0.123 | -1.18E-01 | 1.29E-01 | 2.63E+00 | 2.70E+00 | 3.59E-01 | no | 1700057G04Rik | intron_variant | MODIFIER |
| 9 | 92355736 | AX-97782730 | 0 | T | C | 0.123 | -1.18E-01 | 1.29E-01 | 2.63E+00 | 2.70E+00 | 3.59E-01 | no | 1700057G04Rik | intron_variant | MODIFIER |
| 9 | 92373337 | AX-97800790 | 0 | A | G | 0.123 | -1.18E-01 | 1.29E-01 | 2.63E+00 | 2.70E+00 | 3.59E-01 | no | | intergenic_variant | MODIFIER |
| 9 | 92378237 | AX-170123480 | 0 | T | C | 0.123 | -1.18E-01 | 1.29E-01 | 2.63E+00 | 2.70E+00 | 3.59E-01 | no | | intergenic_variant | MODIFIER |
| 9 | 92378451 | AX-170032991 | 0 | C | G | 0.123 | -1.18E-01 | 1.29E-01 | 2.63E+00 | 2.70E+00 | 3.59E-01 | no | | intergenic_variant | MODIFIER |
| 9 | 92386066 | AX-170251753 | 0 | T | C | 0.123 | -1.18E-01 | 1.29E-01 | 2.63E+00 | 2.70E+00 | 3.59E-01 | no | | intergenic_variant | MODIFIER |
| 9 | 92445387 | AX-170032990 | 0 | A | T | 0.123 | -1.18E-01 | 1.29E-01 | 2.63E+00 | 2.70E+00 | 3.59E-01 | no | | intergenic_variant | MODIFIER |
| 9 | 92491443 | AX-170262996 | 0 | T | G | 0.123 | -1.18E-01 | 1.29E-01 | 2.63E+00 | 2.70E+00 | 3.59E-01 | no | Plscr4 | 3_prime_UTR_variant | MODIFIER |
| 2 | 169533160 | AX-170200707 | 1 | C | G | 0.17 | 1.63E-01 | 1.79E-01 | 2.63E+00 | 2.72E+00 | 3.59E-01 | Yes | | intergenic_variant | MODIFIER |
| 7 | 37958373 | AX-170422930 | 1 | A | T | 0.339 | 8.42E-02 | 9.14E-02 | 2.53E+00 | 2.61E+00 | 3.59E-01 | no | Uri1 | downstream_gene_variant | MODIFIER |
| 18 | 80055405 | AX-170486715 | 0 | G | T | 0.149 | -1.09E-01 | 1.17E-01 | 2.49E+00 | 2.55E+00 | 3.59E-01 | no | Pard6g | intron_variant | MODIFIER |
| 18 | 80055885 | AX-170505880 | 0 | T | C | 0.149 | -1.09E-01 | 1.17E-01 | 2.49E+00 | 2.55E+00 | 3.59E-01 | no | Pard6g | intron_variant | MODIFIER |
| 18 | 80060663 | AX-170461222 | 0 | T | C | 0.149 | -1.09E-01 | 1.17E-01 | 2.49E+00 | 2.55E+00 | 3.59E-01 | no | Pard6g | intron_variant | MODIFIER |
| 18 | 80079793 | AX-170333822 | 0 | C | T | 0.149 | -1.09E-01 | 1.17E-01 | 2.49E+00 | 2.55E+00 | 3.59E-01 | no | Pard6g | intron_variant | MODIFIER |
| 18 | 80083438 | AX-170533919 | 0 | A | G | 0.149 | -1.09E-01 | 1.17E-01 | 2.49E+00 | 2.55E+00 | 3.59E-01 | no | Pard6g | intron_variant | MODIFIER |
| 18 | 80265586 | AX-170397440 | 0 | C | T | 0.149 | -1.09E-01 | 1.17E-01 | 2.49E+00 | 2.55E+00 | 3.59E-01 | no | Pqlc1 | intron_variant | MODIFIER |
| 18 | 80265799 | AX-170380450 | 0 | G | A | 0.149 | -1.09E-01 | 1.17E-01 | 2.49E+00 | 2.55E+00 | 3.59E-01 | no | Pqlc1 | intron_variant | MODIFIER |
| 18 | 80266011 | AX-170472686 | 0 | C | T | 0.149 | -1.09E-01 | 1.17E-01 | 2.49E+00 | 2.55E+00 | 3.59E-01 | no | Pqlc1 | intron_variant | MODIFIER |
| 18 | 80266362 | AX-170463633 | 0 | T | A | 0.149 | -1.09E-01 | 1.17E-01 | 2.49E+00 | 2.55E+00 | 3.59E-01 | no | Pqlc1 | intron_variant | MODIFIER |
| 18 | 80328492 | AX-97853981 | 0 | C | T | 0.149 | -1.09E-01 | 1.17E-01 | 2.49E+00 | 2.55E+00 | 3.59E-01 | no | Kcng2 | intron_variant | MODIFIER |
| 18 | 80339062 | AX-170244472 | 0 | G | C | 0.149 | -1.09E-01 | 1.17E-01 | 2.49E+00 | 2.55E+00 | 3.59E-01 | no | Kcng2 | intron_variant | MODIFIER |
| 18 | 80339215 | AX-170210473 | 0 | G | A | 0.149 | -1.09E-01 | 1.17E-01 | 2.49E+00 | 2.55E+00 | 3.59E-01 | no | Kcng2 | intron_variant | MODIFIER |
| 18 | 80346202 | AX-97786731 | 0 | G | A | 0.149 | -1.09E-01 | 1.17E-01 | 2.49E+00 | 2.55E+00 | 3.59E-01 | no | Kcng2 | intron_variant | MODIFIER |
| 18 | 80383370 | AX-97837200 | 0 | T | C | 0.149 | -1.09E-01 | 1.17E-01 | 2.49E+00 | 2.55E+00 | 3.59E-01 | no | | regulatory_region_variant | MODIFIER |
| 18 | 80403819 | AX-170424501 | 0 | T | A | 0.149 | -1.09E-01 | 1.17E-01 | 2.49E+00 | 2.55E+00 | 3.59E-01 | no | Ctdp1 | downstream_gene_variant | MODIFIER |
| 18 | 80429337 | AX-170336247 | 0 | T | C | 0.149 | -1.09E-01 | 1.17E-01 | 2.49E+00 | 2.55E+00 | 3.59E-01 | no | Ctdp1 | intron_variant | MODIFIER |
| 18 | 80429868 | AX-170315366 | 0 | T | C | 0.149 | -1.09E-01 | 1.17E-01 | 2.49E+00 | 2.55E+00 | 3.59E-01 | no | Ctdp1 | intron_variant | MODIFIER |
| 18 | 80433737 | AX-170145393 | 0 | C | A | 0.149 | -1.09E-01 | 1.17E-01 | 2.49E+00 | 2.55E+00 | 3.59E-01 | no | Ctdp1 | intron_variant | MODIFIER |
| 18 | 80446204 | AX-170372134 | 0 | G | A | 0.149 | -1.09E-01 | 1.17E-01 | 2.49E+00 | 2.55E+00 | 3.59E-01 | no | Ctdp1 | intron_variant | MODIFIER |
| 18 | 80446299 | AX-170240494 | 0 | T | G | 0.149 | -1.09E-01 | 1.17E-01 | 2.49E+00 | 2.55E+00 | 3.59E-01 | no | Ctdp1 | intron_variant | MODIFIER |
| 18 | 80449491 | AX-97850678 | 0 | T | C | 0.149 | -1.09E-01 | 1.17E-01 | 2.49E+00 | 2.55E+00 | 3.59E-01 | no | Ctdp1 | synonymous_variant | LOW |
| 18 | 80543668 | AX-170282654 | 0 | T | C | 0.149 | -1.09E-01 | 1.17E-01 | 2.49E+00 | 2.55E+00 | 3.59E-01 | no | AC127256.2 | non_coding_transcript_exon_variant | MODIFIER |
| 18 | 80543785 | AX-170264375 | 0 | C | T | 0.149 | -1.09E-01 | 1.17E-01 | 2.49E+00 | 2.55E+00 | 3.59E-01 | no | AC127256.2 | non_coding_transcript_exon_variant | MODIFIER |
| 18 | 80597871 | AX-170455113 | 0 | A | G | 0.149 | -1.09E-01 | 1.17E-01 | 2.49E+00 | 2.55E+00 | 3.59E-01 | no | AC149054.1 | intron_variant,non_coding_transcript_variant | MODIFIER |
| 18 | 80607077 | AX-170472318 | 0 | T | G | 0.149 | -1.09E-01 | 1.17E-01 | 2.49E+00 | 2.55E+00 | 3.59E-01 | no | Nfatc1 | 3_prime_UTR_variant | MODIFIER |
| 18 | 80611637 | AX-170151556 | 0 | A | G | 0.149 | -1.09E-01 | 1.17E-01 | 2.49E+00 | 2.55E+00 | 3.59E-01 | no | Nfatc1 | intron_variant | MODIFIER |
| 18 | 80616226 | AX-170175086 | 0 | A | G | 0.149 | -1.09E-01 | 1.17E-01 | 2.49E+00 | 2.55E+00 | 3.59E-01 | no | Nfatc1 | intron_variant | MODIFIER |
| 18 | 80768222 | AX-170061463 | 0 | A | G | 0.149 | -1.09E-01 | 1.17E-01 | 2.49E+00 | 2.55E+00 | 3.59E-01 | no | Atp9b | intron_variant | MODIFIER |
| 18 | 80768463 | AX-170289542 | 0 | C | T | 0.149 | -1.09E-01 | 1.17E-01 | 2.49E+00 | 2.55E+00 | 3.59E-01 | no | Atp9b | intron_variant | MODIFIER |
| 18 | 80800259 | AX-170243601 | 0 | A | C | 0.149 | -1.09E-01 | 1.17E-01 | 2.49E+00 | 2.55E+00 | 3.59E-01 | no | Atp9b | intron_variant | MODIFIER |
| 18 | 80802800 | AX-170233864 | 0 | A | G | 0.149 | -1.09E-01 | 1.17E-01 | 2.49E+00 | 2.55E+00 | 3.59E-01 | no | Atp9b | intron_variant | MODIFIER |
| 18 | 80805404 | AX-170130250 | 0 | A | G | 0.149 | -1.09E-01 | 1.17E-01 | 2.49E+00 | 2.55E+00 | 3.59E-01 | no | Atp9b | intron_variant | MODIFIER |
| 18 | 80831870 | AX-97820454 | 0 | T | C | 0.149 | -1.09E-01 | 1.17E-01 | 2.49E+00 | 2.55E+00 | 3.59E-01 | no | Atp9b | intron_variant | MODIFIER |
| 18 | 80841091 | AX-170357356 | 0 | G | A | 0.149 | -1.09E-01 | 1.17E-01 | 2.49E+00 | 2.55E+00 | 3.59E-01 | no | Atp9b | intron_variant | MODIFIER |
| 18 | 80888928 | AX-170262270 | 0 | A | G | 0.149 | -1.09E-01 | 1.17E-01 | 2.49E+00 | 2.55E+00 | 3.59E-01 | no | Atp9b | intron_variant | MODIFIER |
| 18 | 80940106 | AX-170126594 | 0 | C | T | 0.149 | -1.09E-01 | 1.17E-01 | 2.49E+00 | 2.55E+00 | 3.59E-01 | no | | intergenic_variant | MODIFIER |
| 18 | 80943936 | AX-97826404 | 0 | A | C | 0.149 | -1.09E-01 | 1.17E-01 | 2.49E+00 | 2.55E+00 | 3.59E-01 | no | | intergenic_variant | MODIFIER |
| 18 | 80990693 | AX-170280579 | 0 | C | A | 0.149 | -1.09E-01 | 1.17E-01 | 2.49E+00 | 2.55E+00 | 3.59E-01 | no | Sall3 | upstream_gene_variant | MODIFIER |
| 10 | 126887563 | AX-170450018 | 1 | C | T | 0.33 | -6.63E-02 | 7.31E-02 | 2.66E+00 | 2.77E+00 | 3.59E-01 | no | Atp23 | intron_variant | MODIFIER |
| 6 | 130736615 | AX-170410541 | 1 | A | C | 0.545 | 7.72E-02 | 8.31E-02 | 2.49E+00 | 2.55E+00 | 3.59E-01 | no | Gm44527 | intron_variant,non_coding_transcript_variant | MODIFIER |
| 1 | 90213506 | AX-97848975 | 0 | G | A | 0.167 | 1.57E-01 | 1.73E-01 | 2.59E+00 | 2.69E+00 | 3.59E-01 | Yes | Ackr3 | intron_variant | MODIFIER |
| 10 | 92570580 | AX-170364541 | 1 | C | T | 0.304 | -7.69E-02 | 8.36E-02 | 2.54E+00 | 2.60E+00 | 3.59E-01 | no | | intergenic_variant | MODIFIER |
| 18 | 88851296 | AX-170291540 | 0 | C | T | 0.333 | 5.47E-02 | 6.05E-02 | 2.68E+00 | 2.82E+00 | 3.59E-01 | no | Socs6 | intron_variant,non_coding_transcript_variant | MODIFIER |
| 7 | 72736983 | AX-170411935 | 0 | T | C | 0.36 | -8.91E-02 | 9.83E-02 | 2.81E+00 | 2.90E+00 | 3.60E-01 | no | | intergenic_variant | MODIFIER |

| | | | | | | | | | | | | | | | |
|---|---|---|---|---|---|---|---|---|---|---|---|---|---|---|---|
| 16 | 45500006 | AX-170286664 | 0 | A | T | 0.518 | 7.74E-02 | 8.39E-02 | 2.53E+00 | 2.56E+00 | 3.60E-01 | no | | intergenic_variant | MODIFIER |
| 16 | 46114724 | AX-170057318 | 0 | A | G | 0.518 | 7.74E-02 | 8.39E-02 | 2.53E+00 | 2.56E+00 | 3.60E-01 | no | Cd96 | intron_variant | MODIFIER |
| 5 | 28524190 | AX-170199436 | 1 | T | C | 0.357 | 1.00E-01 | 1.10E-01 | 2.83E+00 | 2.90E+00 | 3.60E-01 | no | 9530036O11Rik | intron_variant,non_coding_transcript_variant | MODIFIER |
| 2 | 91209411 | AX-170018310 | 0 | C | T | 0.061 | 1.99E-01 | 2.20E-01 | 2.81E+00 | 2.91E+00 | 3.60E-01 | Yes | Acp2 | intron_variant | MODIFIER |
| 2 | 94355051 | AX-97851823 | 0 | T | C | 0.061 | 1.99E-01 | 2.20E-01 | 2.81E+00 | 2.91E+00 | 3.60E-01 | Yes | Ttc17 | intron_variant | MODIFIER |
| 2 | 110471043 | AX-170109822 | 0 | T | A | 0.061 | 1.99E-01 | 2.20E-01 | 2.81E+00 | 2.91E+00 | 3.60E-01 | Yes | | intergenic_variant | MODIFIER |
| 1 | 169496755 | AX-170126405 | 1 | C | T | 0.214 | 1.26E-01 | 1.39E-01 | 2.67E+00 | 2.71E+00 | 3.60E-01 | no | Nuf2 | downstream_gene_variant | MODIFIER |
| 1 | 169903870 | AX-170439889 | 1 | G | A | 0.214 | 1.26E-01 | 1.39E-01 | 2.67E+00 | 2.71E+00 | 3.60E-01 | no | | intergenic_variant | MODIFIER |
| 3 | 37471067 | AX-170066379 | 1 | G | A | 0.196 | 8.53E-02 | 9.41E-02 | 2.82E+00 | 2.91E+00 | 3.60E-01 | no | Spata5 | intron_variant | MODIFIER |
| 4 | 96588037 | AX-97855416 | 0 | A | G | 0.228 | 1.08E-01 | 1.18E-01 | 2.57E+00 | 2.62E+00 | 3.60E-01 | no | Cyp2j9 | intron_variant | MODIFIER |
| 4 | 96592162 | AX-170503503 | 0 | C | T | 0.228 | 1.08E-01 | 1.18E-01 | 2.57E+00 | 2.62E+00 | 3.60E-01 | no | Cyp2j9 | upstream_gene_variant | MODIFIER |
| 4 | 97093336 | AX-97830229 | 0 | A | G | 0.228 | 1.08E-01 | 1.18E-01 | 2.57E+00 | 2.62E+00 | 3.60E-01 | no | | intergenic_variant | MODIFIER |
| 4 | 97204464 | AX-97808942 | 0 | T | C | 0.228 | 1.08E-01 | 1.18E-01 | 2.57E+00 | 2.62E+00 | 3.60E-01 | no | | intergenic_variant | MODIFIER |
| 4 | 97255579 | AX-170253060 | 0 | A | G | 0.228 | 1.08E-01 | 1.18E-01 | 2.57E+00 | 2.62E+00 | 3.60E-01 | no | | intergenic_variant | MODIFIER |
| 4 | 97255871 | AX-170143471 | 0 | A | T | 0.228 | 1.08E-01 | 1.18E-01 | 2.57E+00 | 2.62E+00 | 3.60E-01 | no | | intergenic_variant | MODIFIER |
| 4 | 97264475 | AX-97850820 | 0 | G | A | 0.228 | 1.08E-01 | 1.18E-01 | 2.57E+00 | 2.62E+00 | 3.60E-01 | no | | intergenic_variant | MODIFIER |
| 4 | 97269665 | AX-170330639 | 0 | T | C | 0.228 | 1.08E-01 | 1.18E-01 | 2.57E+00 | 2.62E+00 | 3.60E-01 | no | | intergenic_variant | MODIFIER |
| 4 | 97761313 | AX-170455637 | 0 | T | C | 0.228 | 1.08E-01 | 1.18E-01 | 2.57E+00 | 2.62E+00 | 3.60E-01 | no | E130114P18Rik | intron_variant | MODIFIER |
| 4 | 97768679 | AX-170115358 | 0 | T | C | 0.228 | 1.08E-01 | 1.18E-01 | 2.57E+00 | 2.62E+00 | 3.60E-01 | no | E130114P18Rik | intron_variant | MODIFIER |
| 4 | 97796532 | AX-170130818 | 0 | C | T | 0.228 | 1.08E-01 | 1.18E-01 | 2.57E+00 | 2.62E+00 | 3.60E-01 | no | Nfia | intron_variant | MODIFIER |
| 4 | 97799956 | AX-170414534 | 0 | C | T | 0.228 | 1.08E-01 | 1.18E-01 | 2.57E+00 | 2.62E+00 | 3.60E-01 | no | Nfia | intron_variant | MODIFIER |
| 4 | 97807667 | AX-170415390 | 0 | G | A | 0.228 | 1.08E-01 | 1.18E-01 | 2.57E+00 | 2.62E+00 | 3.60E-01 | no | Nfia | intron_variant | MODIFIER |
| 4 | 97841169 | AX-170492865 | 0 | G | C | 0.228 | 1.08E-01 | 1.18E-01 | 2.57E+00 | 2.62E+00 | 3.60E-01 | no | Nfia | intron_variant | MODIFIER |
| 4 | 97842644 | AX-170460406 | 0 | A | C | 0.228 | 1.08E-01 | 1.18E-01 | 2.57E+00 | 2.62E+00 | 3.60E-01 | no | Nfia | intron_variant | MODIFIER |
| 4 | 97864095 | AX-97820429 | 0 | T | G | 0.228 | 1.08E-01 | 1.18E-01 | 2.57E+00 | 2.62E+00 | 3.60E-01 | no | Nfia | intron_variant | MODIFIER |
| 4 | 97870862 | AX-170355424 | 0 | A | G | 0.228 | 1.08E-01 | 1.18E-01 | 2.57E+00 | 2.62E+00 | 3.60E-01 | no | Nfia | intron_variant | MODIFIER |
| 4 | 97872138 | AX-170213438 | 0 | T | C | 0.228 | 1.08E-01 | 1.18E-01 | 2.57E+00 | 2.62E+00 | 3.60E-01 | no | Nfia | intron_variant | MODIFIER |
| 4 | 97877913 | AX-170345988 | 0 | A | G | 0.228 | 1.08E-01 | 1.18E-01 | 2.57E+00 | 2.62E+00 | 3.60E-01 | no | Nfia | intron_variant | MODIFIER |
| 4 | 97897486 | AX-170395640 | 0 | A | G | 0.228 | 1.08E-01 | 1.18E-01 | 2.57E+00 | 2.62E+00 | 3.60E-01 | no | Nfia | intron_variant | MODIFIER |
| 4 | 97914387 | AX-170290171 | 0 | T | C | 0.228 | 1.08E-01 | 1.18E-01 | 2.57E+00 | 2.62E+00 | 3.60E-01 | no | Nfia | intron_variant | MODIFIER |
| 4 | 97946940 | AX-170365410 | 0 | T | C | 0.228 | 1.08E-01 | 1.18E-01 | 2.57E+00 | 2.62E+00 | 3.60E-01 | no | Nfia | intron_variant | MODIFIER |
| 4 | 97950876 | AX-170333998 | 0 | T | C | 0.228 | 1.08E-01 | 1.18E-01 | 2.57E+00 | 2.62E+00 | 3.60E-01 | no | Nfia | intron_variant | MODIFIER |
| 4 | 97963542 | AX-97822077 | 0 | T | C | 0.228 | 1.08E-01 | 1.18E-01 | 2.57E+00 | 2.62E+00 | 3.60E-01 | no | Nfia | intron_variant | MODIFIER |
| 4 | 97977064 | AX-97805272 | 0 | C | T | 0.228 | 1.08E-01 | 1.18E-01 | 2.57E+00 | 2.62E+00 | 3.60E-01 | no | Nfia | intron_variant | MODIFIER |
| 4 | 97981687 | AX-170122289 | 0 | A | T | 0.228 | 1.08E-01 | 1.18E-01 | 2.57E+00 | 2.62E+00 | 3.60E-01 | no | Nfia | intron_variant | MODIFIER |
| 4 | 98007742 | AX-170220221 | 0 | A | G | 0.228 | 1.08E-01 | 1.18E-01 | 2.57E+00 | 2.62E+00 | 3.60E-01 | no | Nfia | intron_variant | MODIFIER |
| 4 | 98015234 | AX-170089322 | 0 | A | G | 0.228 | 1.08E-01 | 1.18E-01 | 2.57E+00 | 2.62E+00 | 3.60E-01 | no | Nfia | intron_variant | MODIFIER |
| 4 | 98021053 | AX-170023480 | 0 | T | C | 0.228 | 1.08E-01 | 1.18E-01 | 2.57E+00 | 2.62E+00 | 3.60E-01 | no | Nfia | intron_variant | MODIFIER |
| 4 | 98028056 | AX-170280397 | 0 | A | G | 0.228 | 1.08E-01 | 1.18E-01 | 2.57E+00 | 2.62E+00 | 3.60E-01 | no | Nfia | intron_variant | MODIFIER |
| 4 | 98072350 | AX-97792232 | 0 | C | T | 0.228 | 1.08E-01 | 1.18E-01 | 2.57E+00 | 2.62E+00 | 3.60E-01 | no | Nfia | intron_variant | MODIFIER |
| 4 | 98255368 | AX-170509790 | 0 | A | C | 0.228 | 1.08E-01 | 1.18E-01 | 2.57E+00 | 2.62E+00 | 3.60E-01 | no | | intergenic_variant | MODIFIER |
| 4 | 98261146 | AX-97786787 | 0 | T | C | 0.228 | 1.08E-01 | 1.18E-01 | 2.57E+00 | 2.62E+00 | 3.60E-01 | no | | intergenic_variant | MODIFIER |
| 4 | 98266528 | AX-170110504 | 0 | T | G | 0.228 | 1.08E-01 | 1.18E-01 | 2.57E+00 | 2.62E+00 | 3.60E-01 | no | Gm12787 | downstream_gene_variant | MODIFIER |
| 4 | 98271595 | AX-170177915 | 0 | A | G | 0.228 | 1.08E-01 | 1.18E-01 | 2.57E+00 | 2.62E+00 | 3.60E-01 | no | Gm24203 | upstream_gene_variant | MODIFIER |
| 4 | 98297639 | AX-170501051 | 0 | A | G | 0.228 | 1.08E-01 | 1.18E-01 | 2.57E+00 | 2.62E+00 | 3.60E-01 | no | | intergenic_variant | MODIFIER |
| 4 | 98317577 | AX-97790258 | 0 | G | A | 0.228 | 1.08E-01 | 1.18E-01 | 2.57E+00 | 2.62E+00 | 3.60E-01 | no | 0610025J13Rik | upstream_gene_variant | MODIFIER |
| 4 | 98321093 | AX-170070118 | 0 | A | G | 0.228 | 1.08E-01 | 1.18E-01 | 2.57E+00 | 2.62E+00 | 3.60E-01 | no | 0610025J13Rik | upstream_gene_variant | MODIFIER |
| 4 | 98513464 | AX-170070121 | 0 | A | G | 0.228 | 1.08E-01 | 1.18E-01 | 2.57E+00 | 2.62E+00 | 3.60E-01 | no | Patj | intron_variant | MODIFIER |
| 4 | 98514547 | AX-170228741 | 0 | A | G | 0.228 | 1.08E-01 | 1.18E-01 | 2.57E+00 | 2.62E+00 | 3.60E-01 | no | Patj | intron_variant | MODIFIER |
| 4 | 98518751 | AX-170337806 | 0 | C | A | 0.228 | 1.08E-01 | 1.18E-01 | 2.57E+00 | 2.62E+00 | 3.60E-01 | no | Patj | intron_variant | MODIFIER |
| 4 | 98541775 | AX-170407106 | 0 | T | C | 0.228 | 1.08E-01 | 1.18E-01 | 2.57E+00 | 2.62E+00 | 3.60E-01 | no | Patj | intron_variant | MODIFIER |
| 4 | 98554884 | AX-170365635 | 0 | A | C | 0.228 | 1.08E-01 | 1.18E-01 | 2.57E+00 | 2.62E+00 | 3.60E-01 | no | Patj | intron_variant | MODIFIER |
| 4 | 98555252 | AX-170103168 | 0 | T | C | 0.228 | 1.08E-01 | 1.18E-01 | 2.57E+00 | 2.62E+00 | 3.60E-01 | no | Patj | intron_variant | MODIFIER |
| 4 | 98569264 | AX-97799449 | 0 | G | A | 0.228 | 1.08E-01 | 1.18E-01 | 2.57E+00 | 2.62E+00 | 3.60E-01 | no | Patj | intron_variant | MODIFIER |

| | | | | | | | | | | | | | | | |
|---|---|---|---|---|---|---|---|---|---|---|---|---|---|---|---|
| 4 | 133884986 | AX-170070752 | 1 | T | C | 0.17 | 1.03E-01 | 1.12E-01 | 2.54E+00 | 2.59E+00 | 3.60E-01 | no | Rps6ka1 | intron_variant | MODIFIER |
| 5 | 3333763 | AX-97799492 | 1 | G | A | 0.429 | -6.02E-02 | 6.53E-02 | 2.49E+00 | 2.60E+00 | 3.60E-01 | no | | intergenic_variant | MODIFIER |
| 3 | 150533486 | AX-170395909 | 0 | G | A | 0.36 | 6.62E-02 | 7.31E-02 | 2.78E+00 | 2.87E+00 | 3.60E-01 | no | | intergenic_variant | MODIFIER |
| 3 | 150834117 | AX-170068441 | 0 | C | T | 0.36 | 6.62E-02 | 7.31E-02 | 2.78E+00 | 2.87E+00 | 3.60E-01 | no | | intergenic_variant | MODIFIER |
| 8 | 33876980 | AX-170232955 | 0 | G | C | 0.14 | -1.22E-01 | 1.35E-01 | 2.65E+00 | 2.71E+00 | 3.60E-01 | no | Rbpms | intron_variant | MODIFIER |
| 10 | 93878100 | AX-170105453 | 0 | A | C | 0.298 | -7.87E-02 | 8.60E-02 | 2.56E+00 | 2.61E+00 | 3.61E-01 | no | Metap2 | intron_variant | MODIFIER |
| 7 | 39413286 | AX-170392508 | 0 | C | A | 0.342 | 8.58E-02 | 9.36E-02 | 2.54E+00 | 2.61E+00 | 3.61E-01 | no | Gm5114 | upstream_gene_variant | MODIFIER |
| 6 | 127872873 | AX-170383669 | 1 | G | T | 0.536 | 7.60E-02 | 8.21E-02 | 2.48E+00 | 2.55E+00 | 3.61E-01 | no | Gm38901 | downstream_gene_variant | MODIFIER |
| 7 | 114196237 | AX-170136792 | 0 | G | A | 0.43 | 8.31E-02 | 9.19E-02 | 2.71E+00 | 2.78E+00 | 3.61E-01 | no | | intergenic_variant | MODIFIER |
| 8 | 21159531 | AX-170239872 | 2 | A | G | 0.173 | -9.53E-02 | 1.06E-01 | 2.83E+00 | 2.97E+00 | 3.61E-01 | no | Defa22 | upstream_gene_variant | MODIFIER |
| 5 | 49987983 | AX-170238302 | 2 | T | C | 0.5 | 9.35E-02 | 9.70E-02 | 2.33E+00 | 2.40E+00 | 3.61E-01 | no | Adgra3 | intron_variant | MODIFIER |
| 16 | 78095242 | AX-170131918 | 0 | A | G | 0.456 | 6.89E-02 | 7.57E-02 | 2.59E+00 | 2.66E+00 | 3.61E-01 | no | | intergenic_variant | MODIFIER |
| 16 | 78102128 | AX-170093532 | 0 | T | C | 0.456 | 6.89E-02 | 7.57E-02 | 2.59E+00 | 2.66E+00 | 3.61E-01 | no | | regulatory_region_variant | MODIFIER |
| 16 | 78145467 | AX-170196434 | 0 | T | C | 0.456 | 6.89E-02 | 7.57E-02 | 2.59E+00 | 2.66E+00 | 3.61E-01 | no | | regulatory_region_variant | MODIFIER |
| 16 | 78147742 | AX-170193041 | 0 | T | C | 0.456 | 6.89E-02 | 7.57E-02 | 2.59E+00 | 2.66E+00 | 3.61E-01 | no | | intergenic_variant | MODIFIER |
| 16 | 78149398 | AX-170403335 | 0 | T | C | 0.456 | 6.89E-02 | 7.57E-02 | 2.59E+00 | 2.66E+00 | 3.61E-01 | no | | intergenic_variant | MODIFIER |
| 16 | 78153297 | AX-170401479 | 0 | G | A | 0.456 | 6.89E-02 | 7.57E-02 | 2.59E+00 | 2.66E+00 | 3.61E-01 | no | | intergenic_variant | MODIFIER |
| 16 | 78182735 | AX-97789857 | 0 | G | A | 0.456 | 6.89E-02 | 7.57E-02 | 2.59E+00 | 2.66E+00 | 3.61E-01 | no | | intergenic_variant | MODIFIER |
| 16 | 78196553 | AX-170099308 | 0 | G | A | 0.456 | 6.89E-02 | 7.57E-02 | 2.59E+00 | 2.66E+00 | 3.61E-01 | no | | intergenic_variant | MODIFIER |
| 16 | 78202814 | AX-170383937 | 0 | C | T | 0.456 | 6.89E-02 | 7.57E-02 | 2.59E+00 | 2.66E+00 | 3.61E-01 | no | | intergenic_variant | MODIFIER |
| 16 | 78203377 | AX-170141617 | 0 | A | G | 0.456 | 6.89E-02 | 7.57E-02 | 2.59E+00 | 2.66E+00 | 3.61E-01 | no | | intergenic_variant | MODIFIER |
| 16 | 78219820 | AX-170317726 | 0 | C | T | 0.456 | 6.89E-02 | 7.57E-02 | 2.59E+00 | 2.66E+00 | 3.61E-01 | no | | intergenic_variant | MODIFIER |
| 16 | 78219940 | AX-170378211 | 0 | T | C | 0.456 | 6.89E-02 | 7.57E-02 | 2.59E+00 | 2.66E+00 | 3.61E-01 | no | | intergenic_variant | MODIFIER |
| 16 | 78225876 | AX-170057885 | 0 | A | G | 0.456 | 6.89E-02 | 7.57E-02 | 2.59E+00 | 2.66E+00 | 3.61E-01 | no | | intergenic_variant | MODIFIER |
| 16 | 78248683 | AX-170488918 | 0 | C | A | 0.456 | 6.89E-02 | 7.57E-02 | 2.59E+00 | 2.66E+00 | 3.61E-01 | no | E330011O21Rik | downstream_gene_variant | MODIFIER |
| 16 | 78259842 | AX-170269943 | 0 | C | G | 0.456 | 6.89E-02 | 7.57E-02 | 2.59E+00 | 2.66E+00 | 3.61E-01 | no | E330011O21Rik | intron_variant,non_coding_transcript_variant | MODIFIER |
| 16 | 78261101 | AX-170180382 | 0 | A | C | 0.456 | 6.89E-02 | 7.57E-02 | 2.59E+00 | 2.66E+00 | 3.61E-01 | no | E330011O21Rik | intron_variant,non_coding_transcript_variant | MODIFIER |
| 16 | 78261127 | AX-170453802 | 0 | A | G | 0.456 | 6.89E-02 | 7.57E-02 | 2.59E+00 | 2.66E+00 | 3.61E-01 | no | E330011O21Rik | intron_variant,non_coding_transcript_variant | MODIFIER |
| 16 | 78263621 | AX-170392309 | 0 | C | T | 0.456 | 6.89E-02 | 7.57E-02 | 2.59E+00 | 2.66E+00 | 3.61E-01 | no | E330011O21Rik | intron_variant,non_coding_transcript_variant | MODIFIER |
| 16 | 78264276 | AX-170143172 | 0 | A | G | 0.456 | 6.89E-02 | 7.57E-02 | 2.59E+00 | 2.66E+00 | 3.61E-01 | no | E330011O21Rik | intron_variant,non_coding_transcript_variant | MODIFIER |
| 16 | 78266948 | AX-170426176 | 0 | C | A | 0.456 | 6.89E-02 | 7.57E-02 | 2.59E+00 | 2.66E+00 | 3.61E-01 | no | E330011O21Rik | intron_variant,non_coding_transcript_variant | MODIFIER |
| 16 | 78281183 | AX-170400215 | 0 | G | T | 0.456 | 6.89E-02 | 7.57E-02 | 2.59E+00 | 2.66E+00 | 3.61E-01 | no | E330011O21Rik | intron_variant,non_coding_transcript_variant | MODIFIER |
| 16 | 78282804 | AX-97832445 | 0 | G | A | 0.456 | 6.89E-02 | 7.57E-02 | 2.59E+00 | 2.66E+00 | 3.61E-01 | no | E330011O21Rik | intron_variant,non_coding_transcript_variant | MODIFIER |
| 16 | 78313358 | AX-170330536 | 0 | A | G | 0.456 | 6.89E-02 | 7.57E-02 | 2.59E+00 | 2.66E+00 | 3.61E-01 | no | Cxadr | intron_variant | MODIFIER |
| 16 | 78325948 | AX-97808399 | 0 | G | A | 0.456 | 6.89E-02 | 7.57E-02 | 2.59E+00 | 2.66E+00 | 3.61E-01 | no | Cxadr | intron_variant | MODIFIER |
| 16 | 78339194 | AX-97817659 | 0 | T | C | 0.456 | 6.89E-02 | 7.57E-02 | 2.59E+00 | 2.66E+00 | 3.61E-01 | no | Cxadr | 3_prime_UTR_variant | MODIFIER |
| 16 | 78342544 | AX-170516614 | 0 | G | A | 0.456 | 6.89E-02 | 7.57E-02 | 2.59E+00 | 2.66E+00 | 3.61E-01 | no | Cxadr | downstream_gene_variant | MODIFIER |
| 16 | 78379323 | AX-170440285 | 0 | T | C | 0.456 | 6.89E-02 | 7.57E-02 | 2.59E+00 | 2.66E+00 | 3.61E-01 | no | Btg3 | upstream_gene_variant | MODIFIER |
| 16 | 78379435 | AX-170414785 | 0 | A | T | 0.456 | 6.89E-02 | 7.57E-02 | 2.59E+00 | 2.66E+00 | 3.61E-01 | no | Btg3 | upstream_gene_variant | MODIFIER |
| 16 | 78393688 | AX-170460122 | 0 | T | C | 0.456 | 6.89E-02 | 7.57E-02 | 2.59E+00 | 2.66E+00 | 3.61E-01 | no | | intergenic_variant | MODIFIER |
| 16 | 78413900 | AX-170175402 | 0 | C | T | 0.456 | 6.89E-02 | 7.57E-02 | 2.59E+00 | 2.66E+00 | 3.61E-01 | no | | intergenic_variant | MODIFIER |
| 16 | 78414049 | AX-170455941 | 0 | G | A | 0.456 | 6.89E-02 | 7.57E-02 | 2.59E+00 | 2.66E+00 | 3.61E-01 | no | | intergenic_variant | MODIFIER |
| 16 | 78414217 | AX-170209882 | 0 | G | A | 0.456 | 6.89E-02 | 7.57E-02 | 2.59E+00 | 2.66E+00 | 3.61E-01 | no | | intergenic_variant | MODIFIER |
| 16 | 78415078 | AX-97832443 | 0 | G | C | 0.456 | 6.89E-02 | 7.57E-02 | 2.59E+00 | 2.66E+00 | 3.61E-01 | no | | intergenic_variant | MODIFIER |
| 16 | 78418931 | AX-170057888 | 0 | T | C | 0.456 | 6.89E-02 | 7.57E-02 | 2.59E+00 | 2.66E+00 | 3.61E-01 | no | Gm25038 | downstream_gene_variant | MODIFIER |
| 16 | 78438669 | AX-170140337 | 0 | A | C | 0.456 | 6.89E-02 | 7.57E-02 | 2.59E+00 | 2.66E+00 | 3.61E-01 | no | | intergenic_variant | MODIFIER |
| 16 | 78442597 | AX-170410512 | 0 | A | C | 0.456 | 6.89E-02 | 7.57E-02 | 2.59E+00 | 2.66E+00 | 3.61E-01 | no | | intergenic_variant | MODIFIER |
| 16 | 78443568 | AX-170171826 | 0 | G | C | 0.456 | 6.89E-02 | 7.57E-02 | 2.59E+00 | 2.66E+00 | 3.61E-01 | no | | intergenic_variant | MODIFIER |
| 16 | 78450278 | AX-170407555 | 0 | T | C | 0.456 | 6.89E-02 | 7.57E-02 | 2.59E+00 | 2.66E+00 | 3.61E-01 | no | | intergenic_variant | MODIFIER |
| 16 | 78463177 | AX-170203884 | 0 | T | A | 0.456 | 6.89E-02 | 7.57E-02 | 2.59E+00 | 2.66E+00 | 3.61E-01 | no | | intergenic_variant | MODIFIER |
| 16 | 78480609 | AX-170468232 | 0 | T | C | 0.456 | 6.89E-02 | 7.57E-02 | 2.59E+00 | 2.66E+00 | 3.61E-01 | no | Gm49603 | upstream_gene_variant | MODIFIER |
| 16 | 78483094 | AX-170355352 | 0 | T | A | 0.456 | 6.89E-02 | 7.57E-02 | 2.59E+00 | 2.66E+00 | 3.61E-01 | no | Gm49603 | upstream_gene_variant | MODIFIER |
| 16 | 78490676 | AX-170492907 | 0 | G | A | 0.456 | 6.89E-02 | 7.57E-02 | 2.59E+00 | 2.66E+00 | 3.61E-01 | no | Gm49604 | non_coding_transcript_exon_variant | MODIFIER |
| 16 | 78491010 | AX-170295591 | 0 | A | G | 0.456 | 6.89E-02 | 7.57E-02 | 2.59E+00 | 2.66E+00 | 3.61E-01 | no | Gm49604 | non_coding_transcript_exon_variant | MODIFIER |

| | | | | | | | | | | | | | | | |
|---|---|---|---|---|---|---|---|---|---|---|---|---|---|---|---|
| 16 | 78501212 | AX-170198985 | 0 | C | T | 0.456 | 6.89E-02 | 7.57E-02 | 2.59E+00 | 2.66E+00 | 3.61E-01 | no | | intergenic_variant | MODIFIER |
| 16 | 78516113 | AX-170438871 | 0 | C | T | 0.456 | 6.89E-02 | 7.57E-02 | 2.59E+00 | 2.66E+00 | 3.61E-01 | no | | intergenic_variant | MODIFIER |
| 16 | 78519431 | AX-170125389 | 0 | T | C | 0.456 | 6.89E-02 | 7.57E-02 | 2.59E+00 | 2.66E+00 | 3.61E-01 | no | | intergenic_variant | MODIFIER |
| 16 | 78526114 | AX-170391494 | 0 | A | G | 0.456 | 6.89E-02 | 7.57E-02 | 2.59E+00 | 2.66E+00 | 3.61E-01 | no | | intergenic_variant | MODIFIER |
| 16 | 78527242 | AX-97822903 | 0 | C | T | 0.456 | 6.89E-02 | 7.57E-02 | 2.59E+00 | 2.66E+00 | 3.61E-01 | no | | intergenic_variant | MODIFIER |
| 16 | 78541576 | AX-170436942 | 0 | A | G | 0.456 | 6.89E-02 | 7.57E-02 | 2.59E+00 | 2.66E+00 | 3.61E-01 | no | D16Ertd472e | downstream_gene_variant | MODIFIER |
| 16 | 78547708 | AX-170259629 | 0 | C | G | 0.456 | 6.89E-02 | 7.57E-02 | 2.59E+00 | 2.66E+00 | 3.61E-01 | no | D16Ertd472e | missense_variant | MODERATE |
| 16 | 78565601 | AX-97805396 | 0 | T | G | 0.456 | 6.89E-02 | 7.57E-02 | 2.59E+00 | 2.66E+00 | 3.61E-01 | no | D16Ertd472e | intron_variant | MODIFIER |
| 16 | 78622720 | AX-97818232 | 0 | T | C | 0.456 | 6.89E-02 | 7.57E-02 | 2.59E+00 | 2.66E+00 | 3.61E-01 | no | Gm24861 | upstream_gene_variant | MODIFIER |
| 16 | 78624382 | AX-170057893 | 0 | T | C | 0.456 | 6.89E-02 | 7.57E-02 | 2.59E+00 | 2.66E+00 | 3.61E-01 | no | Gm24861 | upstream_gene_variant | MODIFIER |
| 16 | 78632226 | AX-170283320 | 0 | C | T | 0.456 | 6.89E-02 | 7.57E-02 | 2.59E+00 | 2.66E+00 | 3.61E-01 | no | | regulatory_region_variant | MODIFIER |
| 16 | 78641125 | AX-97856838 | 0 | C | T | 0.456 | 6.89E-02 | 7.57E-02 | 2.59E+00 | 2.66E+00 | 3.61E-01 | no | | intergenic_variant | MODIFIER |
| 16 | 78644529 | AX-170057891 | 0 | A | C | 0.456 | 6.89E-02 | 7.57E-02 | 2.59E+00 | 2.66E+00 | 3.61E-01 | no | | intergenic_variant | MODIFIER |
| 16 | 78672728 | AX-170425497 | 0 | T | A | 0.456 | 6.89E-02 | 7.57E-02 | 2.59E+00 | 2.66E+00 | 3.61E-01 | no | | intergenic_variant | MODIFIER |
| 16 | 78693836 | AX-170013448 | 0 | C | T | 0.456 | 6.89E-02 | 7.57E-02 | 2.59E+00 | 2.66E+00 | 3.61E-01 | no | 4930570E03Rik | upstream_gene_variant | MODIFIER |
| 16 | 78697998 | AX-170013442 | 0 | C | T | 0.456 | 6.89E-02 | 7.57E-02 | 2.59E+00 | 2.66E+00 | 3.61E-01 | no | 4930570E03Rik | intron_variant,non_coding_transcript_variant | MODIFIER |
| 16 | 78706771 | AX-97792391 | 0 | C | A | 0.456 | 6.89E-02 | 7.57E-02 | 2.59E+00 | 2.66E+00 | 3.61E-01 | no | 4930570E03Rik | intron_variant,non_coding_transcript_variant | MODIFIER |
| 16 | 78717633 | AX-170194544 | 0 | A | G | 0.456 | 6.89E-02 | 7.57E-02 | 2.59E+00 | 2.66E+00 | 3.61E-01 | no | | intergenic_variant | MODIFIER |
| 16 | 78730580 | AX-170013449 | 0 | G | T | 0.456 | 6.89E-02 | 7.57E-02 | 2.59E+00 | 2.66E+00 | 3.61E-01 | no | | intergenic_variant | MODIFIER |
| 16 | 78736854 | AX-170013446 | 0 | C | T | 0.456 | 6.89E-02 | 7.57E-02 | 2.59E+00 | 2.66E+00 | 3.61E-01 | no | | intergenic_variant | MODIFIER |
| 16 | 78741547 | AX-97782151 | 0 | G | A | 0.456 | 6.89E-02 | 7.57E-02 | 2.59E+00 | 2.66E+00 | 3.61E-01 | no | | intergenic_variant | MODIFIER |
| 16 | 78749020 | AX-170455454 | 0 | C | G | 0.456 | 6.89E-02 | 7.57E-02 | 2.59E+00 | 2.66E+00 | 3.61E-01 | no | | intergenic_variant | MODIFIER |
| 16 | 78749093 | AX-170472639 | 0 | C | T | 0.456 | 6.89E-02 | 7.57E-02 | 2.59E+00 | 2.66E+00 | 3.61E-01 | no | | intergenic_variant | MODIFIER |
| 16 | 78781929 | AX-170013445 | 0 | A | C | 0.456 | 6.89E-02 | 7.57E-02 | 2.59E+00 | 2.66E+00 | 3.61E-01 | no | | intergenic_variant | MODIFIER |
| 16 | 78801732 | AX-170198035 | 0 | G | C | 0.456 | 6.89E-02 | 7.57E-02 | 2.59E+00 | 2.66E+00 | 3.61E-01 | no | | intergenic_variant | MODIFIER |
| 16 | 78802066 | AX-170353203 | 0 | T | C | 0.456 | 6.89E-02 | 7.57E-02 | 2.59E+00 | 2.66E+00 | 3.61E-01 | no | | intergenic_variant | MODIFIER |
| 16 | 78835982 | AX-170216044 | 0 | C | T | 0.456 | 6.89E-02 | 7.57E-02 | 2.59E+00 | 2.66E+00 | 3.61E-01 | no | | intergenic_variant | MODIFIER |
| 16 | 78840752 | AX-170210484 | 0 | C | T | 0.456 | 6.89E-02 | 7.57E-02 | 2.59E+00 | 2.66E+00 | 3.61E-01 | no | | intergenic_variant | MODIFIER |
| 16 | 78894254 | AX-170257055 | 0 | T | G | 0.456 | 6.89E-02 | 7.57E-02 | 2.59E+00 | 2.66E+00 | 3.61E-01 | no | | intergenic_variant | MODIFIER |
| 5 | 73585133 | AX-170072470 | 1 | T | A | 0.429 | 9.01E-02 | 9.80E-02 | 2.52E+00 | 2.60E+00 | 3.61E-01 | no | | regulatory_region_variant | MODIFIER |
| 18 | 62811787 | AX-170485881 | 1 | G | A | 0.188 | -1.09E-01 | 1.20E-01 | 2.68E+00 | 2.73E+00 | 3.61E-01 | no | | regulatory_region_variant | MODIFIER |
| 1 | 152362663 | AX-170255399 | 2 | A | T | 0.2 | 1.25E-01 | 1.38E-01 | 2.66E+00 | 2.71E+00 | 3.61E-01 | no | | intergenic_variant | MODIFIER |
| 4 | 134244661 | AX-170023999 | 0 | C | T | 0.175 | 1.01E-01 | 1.09E-01 | 2.51E+00 | 2.57E+00 | 3.61E-01 | no | Zfp593 | 3_prime_UTR_variant | MODIFIER |
| 13 | 20599875 | AX-170501607 | 0 | C | T | 0.193 | -1.04E-01 | 1.13E-01 | 2.51E+00 | 2.59E+00 | 3.61E-01 | no | Elmo1 | intron_variant | MODIFIER |
| 1 | 145567531 | AX-170137861 | 0 | C | A | 0.211 | 1.18E-01 | 1.31E-01 | 2.69E+00 | 2.75E+00 | 3.61E-01 | no | | intergenic_variant | MODIFIER |
| 5 | 3575851 | AX-170155704 | 0 | A | G | 0.421 | -6.09E-02 | 6.62E-02 | 2.49E+00 | 2.60E+00 | 3.61E-01 | no | 1700109H08Rik | intron_variant | MODIFIER |
| 5 | 3930563 | AX-97856868 | 0 | G | A | 0.421 | -6.09E-02 | 6.62E-02 | 2.49E+00 | 2.60E+00 | 3.61E-01 | no | Akap9 | intron_variant | MODIFIER |
| 5 | 4127019 | AX-170505010 | 0 | G | T | 0.421 | -6.09E-02 | 6.62E-02 | 2.49E+00 | 2.60E+00 | 3.61E-01 | no | | intergenic_variant | MODIFIER |
| 5 | 4188176 | AX-170102619 | 0 | C | A | 0.421 | -6.09E-02 | 6.62E-02 | 2.49E+00 | 2.60E+00 | 3.61E-01 | no | Mterf1b | upstream_gene_variant | MODIFIER |
| 5 | 4194084 | AX-170461362 | 0 | T | C | 0.421 | -6.09E-02 | 6.62E-02 | 2.49E+00 | 2.60E+00 | 3.61E-01 | no | Mterf1b | intron_variant | MODIFIER |
| 5 | 4217842 | AX-170203580 | 0 | A | G | 0.421 | -6.09E-02 | 6.62E-02 | 2.49E+00 | 2.60E+00 | 3.61E-01 | no | | intergenic_variant | MODIFIER |
| 5 | 4245908 | AX-170071177 | 0 | T | C | 0.421 | -6.09E-02 | 6.62E-02 | 2.49E+00 | 2.60E+00 | 3.61E-01 | no | | intergenic_variant | MODIFIER |
| 5 | 4246169 | AX-170188457 | 0 | A | G | 0.421 | -6.09E-02 | 6.62E-02 | 2.49E+00 | 2.60E+00 | 3.61E-01 | no | | intergenic_variant | MODIFIER |
| 5 | 4254439 | AX-170171862 | 0 | A | G | 0.421 | -6.09E-02 | 6.62E-02 | 2.49E+00 | 2.60E+00 | 3.61E-01 | no | | intergenic_variant | MODIFIER |
| 5 | 4273409 | AX-170310517 | 0 | C | G | 0.421 | -6.09E-02 | 6.62E-02 | 2.49E+00 | 2.60E+00 | 3.61E-01 | no | | regulatory_region_variant | MODIFIER |
| 5 | 4293198 | AX-170162422 | 0 | T | C | 0.421 | -6.09E-02 | 6.62E-02 | 2.49E+00 | 2.60E+00 | 3.61E-01 | no | | intergenic_variant | MODIFIER |
| 5 | 4364099 | AX-170218716 | 0 | T | C | 0.421 | -6.09E-02 | 6.62E-02 | 2.49E+00 | 2.60E+00 | 3.61E-01 | no | | intergenic_variant | MODIFIER |
| 5 | 4488907 | AX-170313194 | 0 | A | T | 0.421 | -6.09E-02 | 6.62E-02 | 2.49E+00 | 2.60E+00 | 3.61E-01 | no | | regulatory_region_variant | MODIFIER |
| 5 | 4650534 | AX-170487721 | 0 | T | G | 0.421 | -6.09E-02 | 6.62E-02 | 2.49E+00 | 2.60E+00 | 3.61E-01 | no | | regulatory_region_variant | MODIFIER |
| 5 | 4844004 | AX-170448773 | 0 | C | T | 0.421 | -6.09E-02 | 6.62E-02 | 2.49E+00 | 2.60E+00 | 3.61E-01 | no | Cdk14 | intron_variant | MODIFIER |
| 5 | 4844068 | AX-170268502 | 0 | A | C | 0.421 | -6.09E-02 | 6.62E-02 | 2.49E+00 | 2.60E+00 | 3.61E-01 | no | Cdk14 | intron_variant | MODIFIER |
| 5 | 4887790 | AX-170071190 | 0 | T | G | 0.421 | -6.09E-02 | 6.62E-02 | 2.49E+00 | 2.60E+00 | 3.61E-01 | no | Cdk14 | intron_variant | MODIFIER |
| 3 | 150277436 | AX-170316475 | 1 | C | T | 0.357 | 6.61E-02 | 7.32E-02 | 2.80E+00 | 2.89E+00 | 3.61E-01 | no | | intergenic_variant | MODIFIER |
| 15 | 100309471 | AX-170375956 | 2 | C | A | 0.082 | -1.19E-01 | 1.32E-01 | 2.71E+00 | 2.83E+00 | 3.61E-01 | no | Mettl7a1 | intron_variant | MODIFIER |

| | | | | | | | | | | | | | | | |
|---|---|---|---|---|---|---|---|---|---|---|---|---|---|---|---|
| 11 | 32072583 | AX-170004648 | 1 | C | T | 0.098 | 1.37E-01 | 1.51E-01 | 2.59E+00 | 2.72E+00 | 3.61E-01 | no | | intergenic_variant | MODIFIER |
| 4 | 90646869 | AX-170023386 | 0 | C | G | 0.421 | -7.61E-02 | 8.22E-02 | 2.48E+00 | 2.54E+00 | 3.62E-01 | no | | intergenic_variant | MODIFIER |
| 8 | 25190609 | AX-170237909 | 1 | G | A | 0.357 | -5.98E-02 | 6.56E-02 | 2.56E+00 | 2.65E+00 | 3.62E-01 | no | Tacc1 | intron_variant | MODIFIER |
| 6 | 101428225 | AX-170027837 | 0 | A | C | 0.079 | 1.81E-01 | 2.01E-01 | 2.71E+00 | 2.82E+00 | 3.62E-01 | Yes | | intergenic_variant | MODIFIER |
| 9 | 77147757 | AX-170397032 | 0 | G | A | 0.193 | -1.17E-01 | 1.30E-01 | 2.78E+00 | 2.84E+00 | 3.62E-01 | no | Mlip | intron_variant | MODIFIER |
| 4 | 128020010 | AX-170122204 | 1 | C | T | 0.143 | 1.11E-01 | 1.22E-01 | 2.59E+00 | 2.65E+00 | 3.62E-01 | no | Csmd2 | intron_variant,non_coding_transcript_variant | MODIFIER |
| 10 | 72026572 | AX-170485048 | 0 | A | C | 0.412 | -7.02E-02 | 7.76E-02 | 2.63E+00 | 2.71E+00 | 3.62E-01 | no | Gm34609 | upstream_gene_variant | MODIFIER |
| 6 | 36137755 | AX-170467721 | 0 | A | G | 0.035 | -1.76E-01 | 1.94E-01 | 2.61E+00 | 2.72E+00 | 3.62E-01 | Yes | | intergenic_variant | MODIFIER |
| 6 | 127716243 | AX-170132577 | 1 | T | C | 0.536 | 7.60E-02 | 8.23E-02 | 2.48E+00 | 2.55E+00 | 3.62E-01 | no | Prmt8 | intron_variant | MODIFIER |
| 7 | 74518778 | AX-170474142 | 0 | A | G | 0.377 | -8.54E-02 | 9.49E-02 | 2.74E+00 | 2.85E+00 | 3.62E-01 | no | Slco3a1 | intron_variant | MODIFIER |
| 6 | 129528567 | AX-97837454 | 1 | G | A | 0.554 | 7.63E-02 | 8.30E-02 | 2.50E+00 | 2.57E+00 | 3.62E-01 | no | Gabarapl1 | upstream_gene_variant | MODIFIER |
| 9 | 61715828 | AX-97843370 | 1 | T | C | 0.214 | 1.02E-01 | 1.13E-01 | 2.84E+00 | 2.93E+00 | 3.63E-01 | no | | intergenic_variant | MODIFIER |
| 1 | 127215997 | AX-170470380 | 1 | T | C | 0.545 | 7.64E-02 | 8.43E-02 | 2.61E+00 | 2.67E+00 | 3.63E-01 | no | Mgat5 | intron_variant | MODIFIER |
| 4 | 118041183 | AX-170376561 | 0 | A | C | 0.219 | 1.03E-01 | 1.13E-01 | 2.60E+00 | 2.67E+00 | 3.63E-01 | no | St3gal3 | intron_variant | MODIFIER |
| 8 | 26071138 | AX-170030375 | 0 | A | G | 0.351 | -5.93E-02 | 6.54E-02 | 2.59E+00 | 2.68E+00 | 3.63E-01 | no | Hook3 | synonymous_variant | LOW |
| 8 | 74210739 | AX-170031011 | 0 | A | T | 0.193 | -1.11E-01 | 1.24E-01 | 2.77E+00 | 2.88E+00 | 3.63E-01 | no | | intergenic_variant | MODIFIER |
| 6 | 118374495 | AX-170432334 | 1 | C | T | 0.08 | 1.87E-01 | 2.08E-01 | 2.79E+00 | 2.90E+00 | 3.63E-01 | Yes | | intergenic_variant | MODIFIER |
| 8 | 41879486 | AX-97786971 | 0 | A | G | 0.158 | 1.20E-01 | 1.32E-01 | 2.89E+00 | 2.97E+00 | 3.63E-01 | no | 2810404M03Rik | intron_variant,non_coding_transcript_variant | MODIFIER |
| 3 | 23933045 | AX-170159245 | 1 | C | A | 0.161 | -9.29E-02 | 1.02E-01 | 2.56E+00 | 2.63E+00 | 3.63E-01 | no | Naaladl2 | intron_variant,non_coding_transcript_variant | MODIFIER |
| 18 | 84797193 | AX-170482256 | 1 | T | C | 0.152 | -1.00E-01 | 1.08E-01 | 2.47E+00 | 2.55E+00 | 3.63E-01 | no | C18orf63 | downstream_gene_variant | MODIFIER |
| 8 | 93410787 | AX-170463091 | 0 | T | G | 0.211 | -9.33E-02 | 1.04E-01 | 2.75E+00 | 2.86E+00 | 3.63E-01 | no | Gm45765 | intron_variant,non_coding_transcript_variant | MODIFIER |
| 1 | 144651531 | AX-170252732 | 0 | G | A | 0.219 | 1.16E-01 | 1.29E-01 | 2.60E+00 | 2.66E+00 | 3.63E-01 | no | | intergenic_variant | MODIFIER |
| 10 | 92871904 | AX-170294910 | 0 | G | A | 0.307 | -7.39E-02 | 8.08E-02 | 2.52E+00 | 2.60E+00 | 3.63E-01 | no | Cfap54 | intron_variant | MODIFIER |
| 7 | 39646172 | AX-97808917 | 1 | G | A | 0.357 | 8.62E-02 | 9.48E-02 | 2.56E+00 | 2.63E+00 | 3.63E-01 | no | | intergenic_variant | MODIFIER |
| 8 | 75490348 | AX-97834834 | 0 | T | C | 0.193 | -1.02E-01 | 1.14E-01 | 2.72E+00 | 2.83E+00 | 3.63E-01 | no | Iqcm | intron_variant | MODIFIER |
| 11 | 107867220 | AX-170281014 | 0 | A | G | 0.044 | -1.78E-01 | 1.97E-01 | 2.61E+00 | 2.71E+00 | 3.64E-01 | Yes | | regulatory_region_variant | MODIFIER |
| 5 | 3935452 | AX-170024327 | 1 | G | A | 0.446 | -5.98E-02 | 6.48E-02 | 2.46E+00 | 2.57E+00 | 3.64E-01 | no | Akap9 | intron_variant | MODIFIER |
| 5 | 115938522 | AX-170206908 | 1 | C | T | 0.473 | 8.98E-02 | 9.96E-02 | 2.63E+00 | 2.71E+00 | 3.64E-01 | no | Cit | intron_variant | MODIFIER |
| 9 | 94907634 | AX-170260126 | 0 | C | G | 0.14 | -1.04E-01 | 1.16E-01 | 2.66E+00 | 2.74E+00 | 3.64E-01 | no | Slc9a9 | intron_variant | MODIFIER |
| 9 | 94910262 | AX-170184829 | 0 | T | C | 0.14 | -1.04E-01 | 1.16E-01 | 2.66E+00 | 2.74E+00 | 3.64E-01 | no | Slc9a9 | intron_variant | MODIFIER |
| 9 | 94910646 | AX-170533849 | 0 | C | G | 0.14 | -1.04E-01 | 1.16E-01 | 2.66E+00 | 2.74E+00 | 3.64E-01 | no | Slc9a9 | intron_variant | MODIFIER |
| 9 | 94910735 | AX-170227272 | 0 | A | T | 0.14 | -1.04E-01 | 1.16E-01 | 2.66E+00 | 2.74E+00 | 3.64E-01 | no | Slc9a9 | intron_variant | MODIFIER |
| 9 | 94913935 | AX-97782411 | 0 | A | G | 0.14 | -1.04E-01 | 1.16E-01 | 2.66E+00 | 2.74E+00 | 3.64E-01 | no | Slc9a9 | intron_variant | MODIFIER |
| 9 | 94947572 | AX-170107908 | 0 | A | G | 0.14 | -1.04E-01 | 1.16E-01 | 2.66E+00 | 2.74E+00 | 3.64E-01 | no | Slc9a9 | intron_variant | MODIFIER |
| 9 | 94967982 | AX-170090385 | 0 | C | A | 0.14 | -1.04E-01 | 1.16E-01 | 2.66E+00 | 2.74E+00 | 3.64E-01 | no | Slc9a9 | intron_variant | MODIFIER |
| 9 | 94985094 | AX-170098134 | 0 | A | T | 0.14 | -1.04E-01 | 1.16E-01 | 2.66E+00 | 2.74E+00 | 3.64E-01 | no | Slc9a9 | intron_variant | MODIFIER |
| 9 | 95132242 | AX-97796129 | 0 | A | C | 0.14 | -1.04E-01 | 1.16E-01 | 2.66E+00 | 2.74E+00 | 3.64E-01 | no | Slc9a9 | intron_variant | MODIFIER |
| 9 | 95139277 | AX-97785744 | 0 | A | G | 0.14 | -1.04E-01 | 1.16E-01 | 2.66E+00 | 2.74E+00 | 3.64E-01 | no | Slc9a9 | intron_variant | MODIFIER |
| 9 | 95154474 | AX-170501909 | 0 | T | C | 0.14 | -1.04E-01 | 1.16E-01 | 2.66E+00 | 2.74E+00 | 3.64E-01 | no | Slc9a9 | intron_variant | MODIFIER |
| 9 | 95154543 | AX-170534882 | 0 | C | T | 0.14 | -1.04E-01 | 1.16E-01 | 2.66E+00 | 2.74E+00 | 3.64E-01 | no | Slc9a9 | intron_variant | MODIFIER |
| 9 | 95155834 | AX-170381897 | 0 | C | T | 0.14 | -1.04E-01 | 1.16E-01 | 2.66E+00 | 2.74E+00 | 3.64E-01 | no | Slc9a9 | intron_variant | MODIFIER |
| 9 | 95162472 | AX-170033035 | 0 | C | T | 0.14 | -1.04E-01 | 1.16E-01 | 2.66E+00 | 2.74E+00 | 3.64E-01 | no | Slc9a9 | intron_variant | MODIFIER |
| 9 | 95168703 | AX-170033037 | 0 | C | T | 0.14 | -1.04E-01 | 1.16E-01 | 2.66E+00 | 2.74E+00 | 3.64E-01 | no | Slc9a9 | intron_variant | MODIFIER |
| 9 | 95169310 | AX-97794512 | 0 | T | G | 0.14 | -1.04E-01 | 1.16E-01 | 2.66E+00 | 2.74E+00 | 3.64E-01 | no | Slc9a9 | intron_variant | MODIFIER |
| 9 | 95169574 | AX-170429076 | 0 | C | T | 0.14 | -1.04E-01 | 1.16E-01 | 2.66E+00 | 2.74E+00 | 3.64E-01 | no | Slc9a9 | intron_variant | MODIFIER |
| 9 | 95182866 | AX-170518747 | 0 | T | C | 0.14 | -1.04E-01 | 1.16E-01 | 2.66E+00 | 2.74E+00 | 3.64E-01 | no | Slc9a9 | intron_variant | MODIFIER |
| 9 | 95334029 | AX-97789301 | 0 | T | A | 0.14 | -1.04E-01 | 1.16E-01 | 2.66E+00 | 2.74E+00 | 3.64E-01 | no | | intergenic_variant | MODIFIER |
| 9 | 95337921 | AX-97843717 | 0 | T | C | 0.14 | -1.04E-01 | 1.16E-01 | 2.66E+00 | 2.74E+00 | 3.64E-01 | no | | intergenic_variant | MODIFIER |
| 9 | 95359562 | AX-170474702 | 0 | C | T | 0.14 | -1.04E-01 | 1.16E-01 | 2.66E+00 | 2.74E+00 | 3.64E-01 | no | | intergenic_variant | MODIFIER |
| 9 | 95366011 | AX-170471878 | 0 | A | G | 0.14 | -1.04E-01 | 1.16E-01 | 2.66E+00 | 2.74E+00 | 3.64E-01 | no | | intergenic_variant | MODIFIER |
| 9 | 95366175 | AX-97787994 | 0 | C | A | 0.14 | -1.04E-01 | 1.16E-01 | 2.66E+00 | 2.74E+00 | 3.64E-01 | no | | intergenic_variant | MODIFIER |
| 9 | 95381918 | AX-170146711 | 0 | G | A | 0.14 | -1.04E-01 | 1.16E-01 | 2.66E+00 | 2.74E+00 | 3.64E-01 | no | | intergenic_variant | MODIFIER |
| 9 | 95390714 | AX-97798749 | 0 | C | T | 0.14 | -1.04E-01 | 1.16E-01 | 2.66E+00 | 2.74E+00 | 3.64E-01 | no | | intergenic_variant | MODIFIER |
| 9 | 95402986 | AX-170535813 | 0 | T | C | 0.14 | -1.04E-01 | 1.16E-01 | 2.66E+00 | 2.74E+00 | 3.64E-01 | no | Chst2 | 3_prime_UTR_variant | MODIFIER |

| | | | | | | | | | | | | | | | |
|---|---|---|---|---|---|---|---|---|---|---|---|---|---|---|---|
| 9 | 95418755 | AX-170353486 | 0 | C | A | 0.14 | -1.04E-01 | 1.16E-01 | 2.66E+00 | 2.74E+00 | 3.64E-01 | no | | intergenic_variant | MODIFIER |
| 9 | 95427286 | AX-97822613 | 0 | C | T | 0.14 | -1.04E-01 | 1.16E-01 | 2.66E+00 | 2.74E+00 | 3.64E-01 | no | Gm19193 | downstream_gene_variant | MODIFIER |
| 9 | 95428304 | AX-170083152 | 0 | T | C | 0.14 | -1.04E-01 | 1.16E-01 | 2.66E+00 | 2.74E+00 | 3.64E-01 | no | Gm19193 | downstream_gene_variant | MODIFIER |
| 9 | 95446907 | AX-97809110 | 0 | A | G | 0.14 | -1.04E-01 | 1.16E-01 | 2.66E+00 | 2.74E+00 | 3.64E-01 | no | | intergenic_variant | MODIFIER |
| 9 | 95450390 | AX-170136534 | 0 | A | G | 0.14 | -1.04E-01 | 1.16E-01 | 2.66E+00 | 2.74E+00 | 3.64E-01 | no | | intergenic_variant | MODIFIER |
| 9 | 95450691 | AX-170229168 | 0 | C | T | 0.14 | -1.04E-01 | 1.16E-01 | 2.66E+00 | 2.74E+00 | 3.64E-01 | no | | regulatory_region_variant | MODIFIER |
| 9 | 95456796 | AX-170209832 | 0 | T | C | 0.14 | -1.04E-01 | 1.16E-01 | 2.66E+00 | 2.74E+00 | 3.64E-01 | no | U2surp | downstream_gene_variant | MODIFIER |
| 9 | 95459525 | AX-97795467 | 0 | G | A | 0.14 | -1.04E-01 | 1.16E-01 | 2.66E+00 | 2.74E+00 | 3.64E-01 | no | U2surp | downstream_gene_variant | MODIFIER |
| 9 | 95488315 | AX-170461536 | 0 | G | A | 0.14 | -1.04E-01 | 1.16E-01 | 2.66E+00 | 2.74E+00 | 3.64E-01 | no | U2surp | intron_variant | MODIFIER |
| 9 | 95490322 | AX-170135957 | 0 | A | G | 0.14 | -1.04E-01 | 1.16E-01 | 2.66E+00 | 2.74E+00 | 3.64E-01 | no | U2surp | intron_variant | MODIFIER |
| 9 | 95497009 | AX-97790056 | 0 | T | C | 0.14 | -1.04E-01 | 1.16E-01 | 2.66E+00 | 2.74E+00 | 3.64E-01 | no | U2surp | intron_variant | MODIFIER |
| 9 | 95502880 | AX-170349481 | 0 | G | A | 0.14 | -1.04E-01 | 1.16E-01 | 2.66E+00 | 2.74E+00 | 3.64E-01 | no | U2surp | intron_variant | MODIFIER |
| 9 | 95512282 | AX-170397660 | 0 | T | A | 0.14 | -1.04E-01 | 1.16E-01 | 2.66E+00 | 2.74E+00 | 3.64E-01 | no | U2surp | upstream_gene_variant | MODIFIER |
| 9 | 95757081 | AX-97834536 | 0 | C | T | 0.14 | -1.04E-01 | 1.16E-01 | 2.66E+00 | 2.74E+00 | 3.64E-01 | no | Pls1 | intron_variant | MODIFIER |
| 9 | 95760816 | AX-170421825 | 0 | A | G | 0.14 | -1.04E-01 | 1.16E-01 | 2.66E+00 | 2.74E+00 | 3.64E-01 | no | Pls1 | intron_variant | MODIFIER |
| 9 | 95767970 | AX-97826156 | 0 | C | T | 0.14 | -1.04E-01 | 1.16E-01 | 2.66E+00 | 2.74E+00 | 3.64E-01 | no | Pls1 | intron_variant | MODIFIER |
| 9 | 95773352 | AX-170481008 | 0 | C | T | 0.14 | -1.04E-01 | 1.16E-01 | 2.66E+00 | 2.74E+00 | 3.64E-01 | no | Pls1 | intron_variant | MODIFIER |
| 9 | 95774556 | AX-170185534 | 0 | T | A | 0.14 | -1.04E-01 | 1.16E-01 | 2.66E+00 | 2.74E+00 | 3.64E-01 | no | Pls1 | intron_variant | MODIFIER |
| 9 | 95777277 | AX-170353972 | 0 | C | T | 0.14 | -1.04E-01 | 1.16E-01 | 2.66E+00 | 2.74E+00 | 3.64E-01 | no | Pls1 | intron_variant | MODIFIER |
| 9 | 95869194 | AX-170465173 | 0 | C | T | 0.14 | -1.04E-01 | 1.16E-01 | 2.66E+00 | 2.74E+00 | 3.64E-01 | no | Atr | intron_variant | MODIFIER |
| 9 | 96252922 | AX-170033051 | 0 | A | G | 0.14 | -1.04E-01 | 1.16E-01 | 2.66E+00 | 2.74E+00 | 3.64E-01 | no | Tfdp2 | intron_variant | MODIFIER |
| 9 | 96260091 | AX-170369469 | 0 | C | T | 0.14 | -1.04E-01 | 1.16E-01 | 2.66E+00 | 2.74E+00 | 3.64E-01 | no | Tfdp2 | intron_variant | MODIFIER |
| 9 | 96303001 | AX-170379352 | 0 | A | G | 0.14 | -1.04E-01 | 1.16E-01 | 2.66E+00 | 2.74E+00 | 3.64E-01 | no | Tfdp2 | intron_variant | MODIFIER |
| 9 | 96304029 | AX-170497960 | 0 | T | C | 0.14 | -1.04E-01 | 1.16E-01 | 2.66E+00 | 2.74E+00 | 3.64E-01 | no | Tfdp2 | intron_variant | MODIFIER |
| 9 | 96304180 | AX-170482410 | 0 | G | A | 0.14 | -1.04E-01 | 1.16E-01 | 2.66E+00 | 2.74E+00 | 3.64E-01 | no | Tfdp2 | intron_variant | MODIFIER |
| 9 | 96306151 | AX-170354454 | 0 | G | A | 0.14 | -1.04E-01 | 1.16E-01 | 2.66E+00 | 2.74E+00 | 3.64E-01 | no | Tfdp2 | intron_variant | MODIFIER |
| 9 | 96317779 | AX-97850359 | 0 | G | A | 0.14 | -1.04E-01 | 1.16E-01 | 2.66E+00 | 2.74E+00 | 3.64E-01 | no | Tfdp2 | intron_variant | MODIFIER |
| 9 | 96321339 | AX-170518494 | 0 | C | T | 0.14 | -1.04E-01 | 1.16E-01 | 2.66E+00 | 2.74E+00 | 3.64E-01 | no | Tfdp2 | 3_prime_UTR_variant | MODIFIER |
| 9 | 96332674 | AX-170251232 | 0 | T | G | 0.14 | -1.04E-01 | 1.16E-01 | 2.66E+00 | 2.74E+00 | 3.64E-01 | no | Atp1b3 | 3_prime_UTR_variant | MODIFIER |
| 9 | 96349506 | AX-97836704 | 0 | C | T | 0.14 | -1.04E-01 | 1.16E-01 | 2.66E+00 | 2.74E+00 | 3.64E-01 | no | Atp1b3 | intron_variant | MODIFIER |
| 9 | 96356392 | AX-170328986 | 0 | T | C | 0.14 | -1.04E-01 | 1.16E-01 | 2.66E+00 | 2.74E+00 | 3.64E-01 | no | Atp1b3 | intron_variant | MODIFIER |
| 9 | 91559358 | AX-170528054 | 1 | T | C | 0.134 | -1.11E-01 | 1.23E-01 | 2.62E+00 | 2.71E+00 | 3.64E-01 | no | Gm29478 | intron_variant,non_coding_transcript_variant | MODIFIER |
| 2 | 154809368 | AX-170448491 | 1 | A | G | 0.134 | 1.50E-01 | 1.67E-01 | 2.72E+00 | 2.83E+00 | 3.64E-01 | Yes | Raly | intron_variant | MODIFIER |
| 6 | 127322622 | AX-170318217 | 1 | T | C | 0.536 | 7.55E-02 | 8.20E-02 | 2.49E+00 | 2.55E+00 | 3.64E-01 | no | Gm43636 | downstream_gene_variant | MODIFIER |
| 18 | 58270498 | AX-170369804 | 0 | T | C | 0.289 | -5.92E-02 | 6.58E-02 | 2.85E+00 | 2.97E+00 | 3.64E-01 | no | | intergenic_variant | MODIFIER |
| 4 | 88065369 | AX-97846162 | 2 | A | T | 0.255 | 1.00E-01 | 1.08E-01 | 2.45E+00 | 2.51E+00 | 3.64E-01 | no | | intergenic_variant | MODIFIER |
| 2 | 162363170 | AX-170261763 | 1 | G | A | 0.161 | 1.79E-01 | 2.00E-01 | 2.77E+00 | 2.86E+00 | 3.64E-01 | Yes | Ptprt | intron_variant | MODIFIER |
| 8 | 65932037 | AX-170316081 | 0 | C | T | 0.149 | -1.22E-01 | 1.35E-01 | 2.62E+00 | 2.71E+00 | 3.64E-01 | no | MAR1 | intron_variant | MODIFIER |
| 5 | 3355184 | AX-170194134 | 0 | A | G | 0.43 | -5.98E-02 | 6.54E-02 | 2.49E+00 | 2.60E+00 | 3.64E-01 | no | Cdk6 | intron_variant | MODIFIER |
| 5 | 3363054 | AX-97800265 | 0 | T | C | 0.43 | -5.98E-02 | 6.54E-02 | 2.49E+00 | 2.60E+00 | 3.64E-01 | no | Cdk6 | intron_variant | MODIFIER |
| 5 | 3501668 | AX-97829096 | 0 | C | T | 0.43 | -5.98E-02 | 6.54E-02 | 2.49E+00 | 2.60E+00 | 3.64E-01 | no | Cdk6 | intron_variant | MODIFIER |
| 5 | 3560625 | AX-170401145 | 0 | C | T | 0.43 | -5.98E-02 | 6.54E-02 | 2.49E+00 | 2.60E+00 | 3.64E-01 | no | n-R5s169 | upstream_gene_variant | MODIFIER |
| 5 | 3560712 | AX-170122939 | 0 | T | C | 0.43 | -5.98E-02 | 6.54E-02 | 2.49E+00 | 2.60E+00 | 3.64E-01 | no | n-R5s169 | upstream_gene_variant | MODIFIER |
| 5 | 3922341 | AX-170516090 | 0 | G | A | 0.43 | -5.98E-02 | 6.54E-02 | 2.49E+00 | 2.60E+00 | 3.64E-01 | no | | intergenic_variant | MODIFIER |
| 5 | 4161411 | AX-170501785 | 0 | T | G | 0.43 | -5.98E-02 | 6.54E-02 | 2.49E+00 | 2.60E+00 | 3.64E-01 | no | | intergenic_variant | MODIFIER |
| 5 | 4191342 | AX-170119482 | 0 | G | A | 0.43 | -5.98E-02 | 6.54E-02 | 2.49E+00 | 2.60E+00 | 3.64E-01 | no | Mterf1b | upstream_gene_variant | MODIFIER |
| 5 | 4200587 | AX-97820557 | 0 | A | C | 0.43 | -5.98E-02 | 6.54E-02 | 2.49E+00 | 2.60E+00 | 3.64E-01 | no | Mterf1b | downstream_gene_variant | MODIFIER |
| 5 | 4200809 | AX-170528810 | 0 | C | T | 0.43 | -5.98E-02 | 6.54E-02 | 2.49E+00 | 2.60E+00 | 3.64E-01 | no | Mterf1b | downstream_gene_variant | MODIFIER |
| 5 | 4237242 | AX-170381728 | 0 | C | T | 0.43 | -5.98E-02 | 6.54E-02 | 2.49E+00 | 2.60E+00 | 3.64E-01 | no | | intergenic_variant | MODIFIER |
| 5 | 4245937 | AX-170071175 | 0 | G | A | 0.43 | -5.98E-02 | 6.54E-02 | 2.49E+00 | 2.60E+00 | 3.64E-01 | no | | intergenic_variant | MODIFIER |
| 5 | 4254356 | AX-170496421 | 0 | A | C | 0.43 | -5.98E-02 | 6.54E-02 | 2.49E+00 | 2.60E+00 | 3.64E-01 | no | | intergenic_variant | MODIFIER |
| 5 | 4265142 | AX-170095210 | 0 | C | T | 0.43 | -5.98E-02 | 6.54E-02 | 2.49E+00 | 2.60E+00 | 3.64E-01 | no | | intergenic_variant | MODIFIER |
| 5 | 4271701 | AX-97847150 | 0 | T | A | 0.43 | -5.98E-02 | 6.54E-02 | 2.49E+00 | 2.60E+00 | 3.64E-01 | no | | intergenic_variant | MODIFIER |
| 5 | 4273122 | AX-170440059 | 0 | C | T | 0.43 | -5.98E-02 | 6.54E-02 | 2.49E+00 | 2.60E+00 | 3.64E-01 | no | | regulatory_region_variant | MODIFIER |

| | | | | | | | | | | | | | | | |
|---|---|---|---|---|---|---|---|---|---|---|---|---|---|---|---|
| 5 | 4275382 | AX-170533757 | 0 | T | C | 0.43 | -5.98E-02 | 6.54E-02 | 2.49E+00 | 2.60E+00 | 3.64E-01 | no | | intergenic_variant | MODIFIER |
| 5 | 4476402 | AX-170024333 | 0 | G | T | 0.43 | -5.98E-02 | 6.54E-02 | 2.49E+00 | 2.60E+00 | 3.64E-01 | no | | intergenic_variant | MODIFIER |
| 5 | 4888681 | AX-170311203 | 0 | A | T | 0.43 | -5.98E-02 | 6.54E-02 | 2.49E+00 | 2.60E+00 | 3.64E-01 | no | Cdk14 | intron_variant | MODIFIER |
| 5 | 4888800 | AX-170320145 | 0 | C | T | 0.43 | -5.98E-02 | 6.54E-02 | 2.49E+00 | 2.60E+00 | 3.64E-01 | no | Cdk14 | intron_variant | MODIFIER |
| 5 | 4890956 | AX-170115387 | 0 | A | C | 0.43 | -5.98E-02 | 6.54E-02 | 2.49E+00 | 2.60E+00 | 3.64E-01 | no | Cdk14 | intron_variant | MODIFIER |
| 5 | 4890990 | AX-170272520 | 0 | C | G | 0.43 | -5.98E-02 | 6.54E-02 | 2.49E+00 | 2.60E+00 | 3.64E-01 | no | Cdk14 | intron_variant | MODIFIER |
| 5 | 4967351 | AX-170458288 | 0 | C | A | 0.43 | -5.98E-02 | 6.54E-02 | 2.49E+00 | 2.60E+00 | 3.64E-01 | no | Cdk14 | intron_variant | MODIFIER |
| 5 | 4974874 | AX-170024335 | 0 | T | C | 0.43 | -5.98E-02 | 6.54E-02 | 2.49E+00 | 2.60E+00 | 3.64E-01 | no | Cdk14 | intron_variant | MODIFIER |
| 5 | 4988697 | AX-170024338 | 0 | C | T | 0.43 | -5.98E-02 | 6.54E-02 | 2.49E+00 | 2.60E+00 | 3.64E-01 | no | Cdk14 | intron_variant | MODIFIER |
| 5 | 5007703 | AX-170024336 | 0 | C | T | 0.43 | -5.98E-02 | 6.54E-02 | 2.49E+00 | 2.60E+00 | 3.64E-01 | no | Cdk14 | intron_variant | MODIFIER |
| 12 | 68702187 | AX-170471934 | 1 | C | T | 0.071 | 1.93E-01 | 2.14E-01 | 2.90E+00 | 3.02E+00 | 3.64E-01 | Yes | | intergenic_variant | MODIFIER |
| 1 | 20898081 | AX-170040571 | 1 | G | A | 0.393 | 7.27E-02 | 8.04E-02 | 2.59E+00 | 2.66E+00 | 3.64E-01 | no | Paqr8 | intron_variant | MODIFIER |
| 5 | 89863601 | AX-170442525 | 0 | C | T | 0.43 | 9.79E-02 | 1.08E-01 | 2.88E+00 | 2.95E+00 | 3.65E-01 | no | Adamts3 | intron_variant | MODIFIER |
| 16 | 78835890 | AX-170515144 | 1 | T | C | 0.464 | 6.79E-02 | 7.49E-02 | 2.56E+00 | 2.64E+00 | 3.65E-01 | no | | intergenic_variant | MODIFIER |
| 10 | 26740130 | AX-170104932 | 1 | A | G | 0.188 | 9.47E-02 | 1.04E-01 | 2.95E+00 | 3.06E+00 | 3.65E-01 | no | Gm48893 | intron_variant,non_coding_transcript_variant | MODIFIER |
| 2 | 164429337 | AX-170065514 | 2 | G | A | 0.164 | 1.73E-01 | 1.93E-01 | 2.74E+00 | 2.82E+00 | 3.65E-01 | Yes | Sdc4 | intron_variant | MODIFIER |
| 1 | 101566265 | AX-170120188 | 1 | T | C | 0.152 | 1.41E-01 | 1.58E-01 | 2.66E+00 | 2.77E+00 | 3.65E-01 | no | | intergenic_variant | MODIFIER |
| 1 | 25673482 | AX-170511817 | 1 | G | A | 0.411 | 7.27E-02 | 8.07E-02 | 2.60E+00 | 2.69E+00 | 3.65E-01 | no | Adgrb3 | intron_variant | MODIFIER |
| 15 | 19905373 | AX-170199848 | 1 | G | A | 0.241 | -9.45E-02 | 1.05E-01 | 2.63E+00 | 2.69E+00 | 3.65E-01 | no | | intergenic_variant | MODIFIER |
| 5 | 67331841 | AX-97809662 | 0 | G | A | 0.482 | 9.12E-02 | 1.00E-01 | 2.53E+00 | 2.60E+00 | 3.65E-01 | no | Slc30a9 | intron_variant | MODIFIER |
| 14 | 112480120 | AX-170087728 | 0 | A | T | 0.307 | 6.43E-02 | 7.13E-02 | 2.57E+00 | 2.68E+00 | 3.65E-01 | no | Gm4822 | upstream_gene_variant | MODIFIER |
| 4 | 144519321 | AX-170310424 | 0 | G | A | 0.114 | 1.52E-01 | 1.70E-01 | 2.69E+00 | 2.79E+00 | 3.65E-01 | Yes | 9430007A20Rik | upstream_gene_variant | MODIFIER |
| 4 | 129720504 | AX-170309644 | 0 | C | T | 0.175 | 9.97E-02 | 1.10E-01 | 2.59E+00 | 2.64E+00 | 3.66E-01 | no | Khdrbs1 | intron_variant | MODIFIER |
| 4 | 129953532 | AX-97855686 | 0 | G | A | 0.175 | 9.97E-02 | 1.10E-01 | 2.59E+00 | 2.64E+00 | 3.66E-01 | no | Spocd1 | missense_variant,splice_region_variant | MODERATE |
| 4 | 132227760 | AX-170097954 | 0 | G | A | 0.175 | 9.97E-02 | 1.10E-01 | 2.59E+00 | 2.64E+00 | 3.66E-01 | no | Gmeb1 | intron_variant | MODIFIER |
| 4 | 132302298 | AX-170227113 | 0 | A | G | 0.175 | 9.97E-02 | 1.10E-01 | 2.59E+00 | 2.64E+00 | 3.66E-01 | no | Rab42 | synonymous_variant | LOW |
| 4 | 132339357 | AX-170140256 | 0 | A | G | 0.175 | 9.97E-02 | 1.10E-01 | 2.59E+00 | 2.64E+00 | 3.66E-01 | no | Rcc1 | intron_variant | MODIFIER |
| 4 | 132365126 | AX-170070716 | 0 | C | G | 0.175 | 9.97E-02 | 1.10E-01 | 2.59E+00 | 2.64E+00 | 3.66E-01 | no | Phactr4 | intron_variant | MODIFIER |
| 4 | 132365279 | AX-170458999 | 0 | G | C | 0.175 | 9.97E-02 | 1.10E-01 | 2.59E+00 | 2.64E+00 | 3.66E-01 | no | Phactr4 | intron_variant | MODIFIER |
| 4 | 132556485 | AX-170241941 | 0 | C | T | 0.175 | 9.97E-02 | 1.10E-01 | 2.59E+00 | 2.64E+00 | 3.66E-01 | no | Dnajc8 | downstream_gene_variant | MODIFIER |
| 4 | 132562904 | AX-170294610 | 0 | T | C | 0.175 | 9.97E-02 | 1.10E-01 | 2.59E+00 | 2.64E+00 | 3.66E-01 | no | Ptafr | upstream_gene_variant | MODIFIER |
| 4 | 132565467 | AX-170280926 | 0 | A | G | 0.175 | 9.97E-02 | 1.10E-01 | 2.59E+00 | 2.64E+00 | 3.66E-01 | no | Ptafr | intron_variant | MODIFIER |
| 4 | 132577756 | AX-170179592 | 0 | C | T | 0.175 | 9.97E-02 | 1.10E-01 | 2.59E+00 | 2.64E+00 | 3.66E-01 | no | Ptafr | intron_variant | MODIFIER |
| 4 | 132615048 | AX-170319781 | 0 | G | C | 0.175 | 9.97E-02 | 1.10E-01 | 2.59E+00 | 2.64E+00 | 3.66E-01 | no | | intergenic_variant | MODIFIER |
| 4 | 132617161 | AX-170187731 | 0 | C | T | 0.175 | 9.97E-02 | 1.10E-01 | 2.59E+00 | 2.64E+00 | 3.66E-01 | no | | regulatory_region_variant | MODIFIER |
| 4 | 132617330 | AX-170210216 | 0 | C | T | 0.175 | 9.97E-02 | 1.10E-01 | 2.59E+00 | 2.64E+00 | 3.66E-01 | no | | regulatory_region_variant | MODIFIER |
| 4 | 132618074 | AX-97786656 | 0 | A | G | 0.175 | 9.97E-02 | 1.10E-01 | 2.59E+00 | 2.64E+00 | 3.66E-01 | no | | regulatory_region_variant | MODIFIER |
| 4 | 132628205 | AX-170070724 | 0 | C | T | 0.175 | 9.97E-02 | 1.10E-01 | 2.59E+00 | 2.64E+00 | 3.66E-01 | no | | intergenic_variant | MODIFIER |
| 4 | 132628563 | AX-170023972 | 0 | C | A | 0.175 | 9.97E-02 | 1.10E-01 | 2.59E+00 | 2.64E+00 | 3.66E-01 | no | | intergenic_variant | MODIFIER |
| 4 | 132673187 | AX-97799368 | 0 | C | T | 0.175 | 9.97E-02 | 1.10E-01 | 2.59E+00 | 2.64E+00 | 3.66E-01 | no | Eya3 | intron_variant | MODIFIER |
| 4 | 132681532 | AX-170089376 | 0 | G | A | 0.175 | 9.97E-02 | 1.10E-01 | 2.59E+00 | 2.64E+00 | 3.66E-01 | no | Eya3 | intron_variant | MODIFIER |
| 4 | 133177300 | AX-170430882 | 0 | T | G | 0.175 | 9.97E-02 | 1.10E-01 | 2.59E+00 | 2.64E+00 | 3.66E-01 | no | Wasf2 | intron_variant | MODIFIER |
| 4 | 133180368 | AX-170473981 | 0 | C | G | 0.175 | 9.97E-02 | 1.10E-01 | 2.59E+00 | 2.64E+00 | 3.66E-01 | no | Wasf2 | intron_variant | MODIFIER |
| 4 | 133188967 | AX-97821067 | 0 | C | T | 0.175 | 9.97E-02 | 1.10E-01 | 2.59E+00 | 2.64E+00 | 3.66E-01 | no | Wasf2 | intron_variant | MODIFIER |
| 4 | 133205292 | AX-170314509 | 0 | G | A | 0.175 | 9.97E-02 | 1.10E-01 | 2.59E+00 | 2.64E+00 | 3.66E-01 | no | Gpr3 | downstream_gene_variant | MODIFIER |
| 4 | 133257376 | AX-170070734 | 0 | G | A | 0.175 | 9.97E-02 | 1.10E-01 | 2.59E+00 | 2.64E+00 | 3.66E-01 | no | Sytl1 | intron_variant | MODIFIER |
| 4 | 133292663 | AX-170377330 | 0 | G | T | 0.175 | 9.97E-02 | 1.10E-01 | 2.59E+00 | 2.64E+00 | 3.66E-01 | no | Wdtc1 | 3_prime_UTR_variant | MODIFIER |
| 4 | 133366803 | AX-170188853 | 0 | T | C | 0.175 | 9.97E-02 | 1.10E-01 | 2.59E+00 | 2.64E+00 | 3.66E-01 | no | Slc9a1 | upstream_gene_variant | MODIFIER |
| 4 | 133389171 | AX-97841769 | 0 | A | G | 0.175 | 9.97E-02 | 1.10E-01 | 2.59E+00 | 2.64E+00 | 3.66E-01 | no | Slc9a1 | intron_variant | MODIFIER |
| 4 | 133405917 | AX-170288684 | 0 | C | G | 0.175 | 9.97E-02 | 1.10E-01 | 2.59E+00 | 2.64E+00 | 3.66E-01 | no | Slc9a1 | intron_variant | MODIFIER |
| 4 | 133431761 | AX-97805045 | 0 | A | G | 0.175 | 9.97E-02 | 1.10E-01 | 2.59E+00 | 2.64E+00 | 3.66E-01 | no | | intergenic_variant | MODIFIER |
| 4 | 133755539 | AX-170415713 | 0 | G | A | 0.175 | 9.97E-02 | 1.10E-01 | 2.59E+00 | 2.64E+00 | 3.66E-01 | no | Arid1a | intron_variant | MODIFIER |
| 4 | 133846486 | AX-97849615 | 0 | G | A | 0.175 | 9.97E-02 | 1.10E-01 | 2.59E+00 | 2.64E+00 | 3.66E-01 | no | Rps6ka1 | downstream_gene_variant | MODIFIER |
| 4 | 133864746 | AX-170288469 | 0 | C | T | 0.175 | 9.97E-02 | 1.10E-01 | 2.59E+00 | 2.64E+00 | 3.66E-01 | no | Rps6ka1 | intron_variant | MODIFIER |

| | | | | | | | | | | | | | | | |
|---|---|---|---|---|---|---|---|---|---|---|---|---|---|---|---|
| 4 | 133882278 | AX-170348435 | 0 | C | T | 0.175 | 9.97E-02 | 1.10E-01 | 2.59E+00 | 2.64E+00 | 3.66E-01 | no | Rps6ka1 | intron_variant | MODIFIER |
| 4 | 134333879 | AX-170213279 | 0 | G | A | 0.175 | 9.97E-02 | 1.10E-01 | 2.59E+00 | 2.64E+00 | 3.66E-01 | no | Trim63 | downstream_gene_variant | MODIFIER |
| 4 | 134336464 | AX-170369415 | 0 | C | T | 0.175 | 9.97E-02 | 1.10E-01 | 2.59E+00 | 2.64E+00 | 3.66E-01 | no | Gm13195 | upstream_gene_variant | MODIFIER |
| 4 | 134470833 | AX-170319718 | 0 | A | G | 0.175 | 9.97E-02 | 1.10E-01 | 2.59E+00 | 2.64E+00 | 3.66E-01 | no | Stmn1 | synonymous_variant | LOW |
| 4 | 134474755 | AX-170469723 | 0 | G | A | 0.175 | 9.97E-02 | 1.10E-01 | 2.59E+00 | 2.64E+00 | 3.66E-01 | no | Stmn1 | downstream_gene_variant | MODIFIER |
| 4 | 134475162 | AX-170176277 | 0 | T | C | 0.175 | 9.97E-02 | 1.10E-01 | 2.59E+00 | 2.64E+00 | 3.66E-01 | no | Stmn1 | downstream_gene_variant | MODIFIER |
| 4 | 134589370 | AX-170247932 | 0 | G | C | 0.175 | 9.97E-02 | 1.10E-01 | 2.59E+00 | 2.64E+00 | 3.66E-01 | no | Man1c1 | intron_variant | MODIFIER |
| 4 | 134609087 | AX-170144155 | 0 | T | C | 0.175 | 9.97E-02 | 1.10E-01 | 2.59E+00 | 2.64E+00 | 3.66E-01 | no | Man1c1 | intron_variant | MODIFIER |
| 4 | 134611465 | AX-170243539 | 0 | A | G | 0.175 | 9.97E-02 | 1.10E-01 | 2.59E+00 | 2.64E+00 | 3.66E-01 | no | Man1c1 | intron_variant | MODIFIER |
| 4 | 134621911 | AX-170204184 | 0 | G | A | 0.175 | 9.97E-02 | 1.10E-01 | 2.59E+00 | 2.64E+00 | 3.66E-01 | no | Man1c1 | intron_variant | MODIFIER |
| 4 | 134627377 | AX-170354846 | 0 | C | T | 0.175 | 9.97E-02 | 1.10E-01 | 2.59E+00 | 2.64E+00 | 3.66E-01 | no | Man1c1 | intron_variant | MODIFIER |
| 4 | 134747894 | AX-170373413 | 0 | G | A | 0.175 | 9.97E-02 | 1.10E-01 | 2.59E+00 | 2.64E+00 | 3.66E-01 | no | Ldlrap1 | intron_variant | MODIFIER |
| 4 | 134760306 | AX-97827836 | 0 | C | T | 0.175 | 9.97E-02 | 1.10E-01 | 2.59E+00 | 2.64E+00 | 3.66E-01 | no | Ldlrap1 | intron_variant | MODIFIER |
| 4 | 134782422 | AX-170024008 | 0 | A | G | 0.175 | 9.97E-02 | 1.10E-01 | 2.59E+00 | 2.64E+00 | 3.66E-01 | no | | regulatory_region_variant | MODIFIER |
| 4 | 134782448 | AX-170469946 | 0 | T | C | 0.175 | 9.97E-02 | 1.10E-01 | 2.59E+00 | 2.64E+00 | 3.66E-01 | no | | regulatory_region_variant | MODIFIER |
| 4 | 134790766 | AX-97835574 | 0 | A | G | 0.175 | 9.97E-02 | 1.10E-01 | 2.59E+00 | 2.64E+00 | 3.66E-01 | no | | intergenic_variant | MODIFIER |
| 4 | 134919982 | AX-97833920 | 0 | G | A | 0.175 | 9.97E-02 | 1.10E-01 | 2.59E+00 | 2.64E+00 | 3.66E-01 | no | Tmem50a | upstream_gene_variant | MODIFIER |
| 4 | 135039978 | AX-97807191 | 0 | C | T | 0.175 | 9.97E-02 | 1.10E-01 | 2.59E+00 | 2.64E+00 | 3.66E-01 | no | | intergenic_variant | MODIFIER |
| 4 | 135047966 | AX-170322632 | 0 | G | A | 0.175 | 9.97E-02 | 1.10E-01 | 2.59E+00 | 2.64E+00 | 3.66E-01 | no | | regulatory_region_variant | MODIFIER |
| 4 | 135053084 | AX-97809884 | 0 | C | T | 0.175 | 9.97E-02 | 1.10E-01 | 2.59E+00 | 2.64E+00 | 3.66E-01 | no | | intergenic_variant | MODIFIER |
| 4 | 135054316 | AX-170367163 | 0 | G | A | 0.175 | 9.97E-02 | 1.10E-01 | 2.59E+00 | 2.64E+00 | 3.66E-01 | no | | regulatory_region_variant | MODIFIER |
| 4 | 135058976 | AX-170405482 | 0 | C | A | 0.175 | 9.97E-02 | 1.10E-01 | 2.59E+00 | 2.64E+00 | 3.66E-01 | no | | intergenic_variant | MODIFIER |
| 4 | 135059487 | AX-170519841 | 0 | C | T | 0.175 | 9.97E-02 | 1.10E-01 | 2.59E+00 | 2.64E+00 | 3.66E-01 | no | | intergenic_variant | MODIFIER |
| 4 | 135091854 | AX-97830983 | 0 | T | C | 0.175 | 9.97E-02 | 1.10E-01 | 2.59E+00 | 2.64E+00 | 3.66E-01 | no | | intergenic_variant | MODIFIER |
| 4 | 135095754 | AX-170110695 | 0 | T | C | 0.175 | 9.97E-02 | 1.10E-01 | 2.59E+00 | 2.64E+00 | 3.66E-01 | no | | intergenic_variant | MODIFIER |
| 4 | 135118219 | AX-170376944 | 0 | G | A | 0.175 | 9.97E-02 | 1.10E-01 | 2.59E+00 | 2.64E+00 | 3.66E-01 | no | Runx3 | upstream_gene_variant | MODIFIER |
| 4 | 135136466 | AX-170266992 | 0 | G | A | 0.175 | 9.97E-02 | 1.10E-01 | 2.59E+00 | 2.64E+00 | 3.66E-01 | no | Runx3 | intron_variant | MODIFIER |
| 4 | 135136736 | AX-170180424 | 0 | T | A | 0.175 | 9.97E-02 | 1.10E-01 | 2.59E+00 | 2.64E+00 | 3.66E-01 | no | Runx3 | intron_variant | MODIFIER |
| 4 | 135138671 | AX-170095159 | 0 | A | G | 0.175 | 9.97E-02 | 1.10E-01 | 2.59E+00 | 2.64E+00 | 3.66E-01 | no | Runx3 | intron_variant | MODIFIER |
| 4 | 135145501 | AX-170178027 | 0 | G | C | 0.175 | 9.97E-02 | 1.10E-01 | 2.59E+00 | 2.64E+00 | 3.66E-01 | no | Runx3 | intron_variant | MODIFIER |
| 4 | 135192707 | AX-97782592 | 0 | C | T | 0.175 | 9.97E-02 | 1.10E-01 | 2.59E+00 | 2.64E+00 | 3.66E-01 | no | | intergenic_variant | MODIFIER |
| 4 | 135211573 | AX-97842417 | 0 | A | G | 0.175 | 9.97E-02 | 1.10E-01 | 2.59E+00 | 2.64E+00 | 3.66E-01 | no | Clic4 | downstream_gene_variant | MODIFIER |
| 4 | 135741090 | AX-170480250 | 0 | C | A | 0.175 | 9.97E-02 | 1.10E-01 | 2.59E+00 | 2.64E+00 | 3.66E-01 | no | Il22ra1 | intron_variant | MODIFIER |
| 4 | 135742891 | AX-170306164 | 0 | T | C | 0.175 | 9.97E-02 | 1.10E-01 | 2.59E+00 | 2.64E+00 | 3.66E-01 | no | Il22ra1 | intron_variant | MODIFIER |
| 4 | 135848354 | AX-170215122 | 0 | C | A | 0.175 | 9.97E-02 | 1.10E-01 | 2.59E+00 | 2.64E+00 | 3.66E-01 | no | | intergenic_variant | MODIFIER |
| 4 | 135849002 | AX-97857561 | 0 | A | G | 0.175 | 9.97E-02 | 1.10E-01 | 2.59E+00 | 2.64E+00 | 3.66E-01 | no | Gm13000 | upstream_gene_variant | MODIFIER |
| 4 | 135955595 | AX-170285011 | 0 | G | A | 0.175 | 9.97E-02 | 1.10E-01 | 2.59E+00 | 2.64E+00 | 3.66E-01 | no | Hmgcl | synonymous_variant | LOW |
| 4 | 135958824 | AX-170402517 | 0 | C | T | 0.175 | 9.97E-02 | 1.10E-01 | 2.59E+00 | 2.64E+00 | 3.66E-01 | no | Hmgcl | intron_variant | MODIFIER |
| 4 | 136014429 | AX-97852767 | 0 | T | C | 0.175 | 9.97E-02 | 1.10E-01 | 2.59E+00 | 2.64E+00 | 3.66E-01 | no | Eloa | missense_variant | MODERATE |
| 4 | 136134142 | AX-97843739 | 0 | C | T | 0.175 | 9.97E-02 | 1.10E-01 | 2.59E+00 | 2.64E+00 | 3.66E-01 | no | Gm26001 | upstream_gene_variant | MODIFIER |
| 4 | 136226801 | AX-170279968 | 0 | T | C | 0.175 | 9.97E-02 | 1.10E-01 | 2.59E+00 | 2.64E+00 | 3.66E-01 | no | Asap3 | intron_variant | MODIFIER |
| 4 | 136979157 | AX-170311737 | 0 | A | G | 0.175 | 9.97E-02 | 1.10E-01 | 2.59E+00 | 2.64E+00 | 3.66E-01 | no | Zbtb40 | downstream_gene_variant | MODIFIER |
| 4 | 136997715 | AX-170319038 | 0 | G | A | 0.175 | 9.97E-02 | 1.10E-01 | 2.59E+00 | 2.64E+00 | 3.66E-01 | no | Zbtb40 | intron_variant | MODIFIER |
| 4 | 136998031 | AX-170220422 | 0 | G | A | 0.175 | 9.97E-02 | 1.10E-01 | 2.59E+00 | 2.64E+00 | 3.66E-01 | no | Zbtb40 | intron_variant | MODIFIER |
| 4 | 136998113 | AX-170325339 | 0 | T | C | 0.175 | 9.97E-02 | 1.10E-01 | 2.59E+00 | 2.64E+00 | 3.66E-01 | no | Zbtb40 | intron_variant | MODIFIER |
| 4 | 136998489 | AX-170518124 | 0 | C | T | 0.175 | 9.97E-02 | 1.10E-01 | 2.59E+00 | 2.64E+00 | 3.66E-01 | no | Zbtb40 | splice_region_variant,intron_variant | LOW |
| 4 | 137022919 | AX-170306511 | 0 | G | A | 0.175 | 9.97E-02 | 1.10E-01 | 2.59E+00 | 2.64E+00 | 3.66E-01 | no | Zbtb40 | intron_variant | MODIFIER |
| 4 | 137029389 | AX-170314563 | 0 | G | A | 0.175 | 9.97E-02 | 1.10E-01 | 2.59E+00 | 2.64E+00 | 3.66E-01 | no | Zbtb40 | intron_variant | MODIFIER |
| 4 | 137029805 | AX-170095166 | 0 | G | T | 0.175 | 9.97E-02 | 1.10E-01 | 2.59E+00 | 2.64E+00 | 3.66E-01 | no | Zbtb40 | intron_variant | MODIFIER |
| 4 | 137029878 | AX-170355637 | 0 | A | G | 0.175 | 9.97E-02 | 1.10E-01 | 2.59E+00 | 2.64E+00 | 3.66E-01 | no | Zbtb40 | intron_variant | MODIFIER |
| 4 | 137091175 | AX-170302561 | 0 | A | T | 0.175 | 9.97E-02 | 1.10E-01 | 2.59E+00 | 2.64E+00 | 3.66E-01 | no | | intergenic_variant | MODIFIER |
| 4 | 137094894 | AX-170132346 | 0 | T | G | 0.175 | 9.97E-02 | 1.10E-01 | 2.59E+00 | 2.64E+00 | 3.66E-01 | no | | intergenic_variant | MODIFIER |
| 4 | 137106830 | AX-170024046 | 0 | A | C | 0.175 | 9.97E-02 | 1.10E-01 | 2.59E+00 | 2.64E+00 | 3.66E-01 | no | | intergenic_variant | MODIFIER |
| 4 | 137110519 | AX-170024045 | 0 | C | T | 0.175 | 9.97E-02 | 1.10E-01 | 2.59E+00 | 2.64E+00 | 3.66E-01 | no | | intergenic_variant | MODIFIER |

| | | | | | | | | | | | | | | | |
|---|---|---|---|---|---|---|---|---|---|---|---|---|---|---|---|
| 4 | 137164094 | AX-170167078 | 0 | T | C | 0.175 | 9.97E-02 | 1.10E-01 | 2.59E+00 | 2.64E+00 | 3.66E-01 | no | Gm13003 | non_coding_transcript_exon_variant | MODIFIER |
| 4 | 137189273 | AX-170150430 | 0 | A | G | 0.175 | 9.97E-02 | 1.10E-01 | 2.59E+00 | 2.64E+00 | 3.66E-01 | no | | regulatory_region_variant | MODIFIER |
| 7 | 17887062 | AX-170466169 | 0 | G | A | 0.36 | 7.43E-02 | 8.28E-02 | 2.62E+00 | 2.74E+00 | 3.66E-01 | no | Gm5155 | intron_variant | MODIFIER |
| 7 | 19265253 | AX-170286565 | 0 | G | A | 0.36 | 7.43E-02 | 8.28E-02 | 2.62E+00 | 2.74E+00 | 3.66E-01 | no | Vasp | intron_variant | MODIFIER |
| 16 | 18343079 | AX-97830635 | 1 | G | A | 0.134 | 8.63E-02 | 9.68E-02 | 2.77E+00 | 2.90E+00 | 3.66E-01 | no | Tango2 | intron_variant | MODIFIER |
| 1 | 59580906 | AX-170352845 | 0 | T | C | 0.105 | -1.66E-01 | 1.86E-01 | 2.75E+00 | 2.85E+00 | 3.66E-01 | no | Gm973 | intron_variant | MODIFIER |
| 1 | 148643720 | AX-170410778 | 0 | C | A | 0.211 | 1.16E-01 | 1.29E-01 | 2.64E+00 | 2.71E+00 | 3.66E-01 | no | | intergenic_variant | MODIFIER |
| 3 | 26832545 | AX-170020174 | 0 | A | G | 0.132 | -9.42E-02 | 1.04E-01 | 2.57E+00 | 2.65E+00 | 3.66E-01 | no | Spata16 | intron_variant | MODIFIER |
| 4 | 135959081 | AX-97817273 | 1 | C | A | 0.17 | 9.92E-02 | 1.10E-01 | 2.60E+00 | 2.65E+00 | 3.66E-01 | no | Hmgcl | intron_variant | MODIFIER |
| 14 | 111122157 | AX-170087724 | 1 | T | C | 0.304 | 6.57E-02 | 7.30E-02 | 2.59E+00 | 2.69E+00 | 3.66E-01 | no | | intergenic_variant | MODIFIER |
| 3 | 55575179 | AX-170229157 | 0 | C | T | 0.044 | -1.56E-01 | 1.73E-01 | 2.60E+00 | 2.72E+00 | 3.66E-01 | no | Gm43548 | non_coding_transcript_exon_variant | MODIFIER |
| 8 | 19610657 | AX-170129772 | 0 | A | T | 0.184 | 8.84E-02 | 9.92E-02 | 2.74E+00 | 2.85E+00 | 3.66E-01 | no | Gm35998 | intron_variant,non_coding_transcript_variant | MODIFIER |
| 13 | 24697258 | AX-170370960 | 0 | G | A | 0.167 | -1.08E-01 | 1.11E-01 | 2.25E+00 | 2.31E+00 | 3.66E-01 | no | Ripor2 | intron_variant | MODIFIER |
| 1 | 20557001 | AX-170367818 | 1 | A | T | 0.402 | 7.24E-02 | 8.06E-02 | 2.62E+00 | 2.68E+00 | 3.66E-01 | no | Pkhd1 | intron_variant | MODIFIER |
| 2 | 120176676 | AX-170264021 | 0 | G | A | 0.167 | 1.25E-01 | 1.40E-01 | 2.60E+00 | 2.74E+00 | 3.66E-01 | no | Pla2g4e | intron_variant | MODIFIER |
| 18 | 84214999 | AX-170304547 | 1 | G | A | 0.17 | -9.69E-02 | 1.06E-01 | 2.49E+00 | 2.57E+00 | 3.66E-01 | no | Zfp407 | intron_variant | MODIFIER |
| 8 | 17376060 | AX-170335427 | 0 | C | A | 0.237 | -6.88E-02 | 7.65E-02 | 2.59E+00 | 2.69E+00 | 3.66E-01 | no | Csmd1 | intron_variant | MODIFIER |
| 6 | 137334963 | AX-170323155 | 0 | C | G | 0.579 | 7.93E-02 | 8.70E-02 | 2.51E+00 | 2.57E+00 | 3.66E-01 | no | Ptpro | intron_variant | MODIFIER |
| 6 | 137359036 | AX-170089851 | 0 | C | A | 0.579 | 7.93E-02 | 8.70E-02 | 2.51E+00 | 2.57E+00 | 3.66E-01 | no | Ptpro | intron_variant | MODIFIER |
| 4 | 138341727 | AX-170230035 | 1 | G | A | 0.17 | 1.05E-01 | 1.16E-01 | 2.60E+00 | 2.64E+00 | 3.67E-01 | no | Cda | intron_variant | MODIFIER |
| 1 | 190047789 | AX-170363076 | 0 | C | T | 0.254 | 9.04E-02 | 9.69E-02 | 2.40E+00 | 2.48E+00 | 3.67E-01 | no | | intergenic_variant | MODIFIER |
| 1 | 190048180 | AX-170319884 | 0 | C | T | 0.254 | 9.04E-02 | 9.69E-02 | 2.40E+00 | 2.48E+00 | 3.67E-01 | no | | intergenic_variant | MODIFIER |
| 1 | 190054649 | AX-170332868 | 0 | G | A | 0.254 | 9.04E-02 | 9.69E-02 | 2.40E+00 | 2.48E+00 | 3.67E-01 | no | | intergenic_variant | MODIFIER |
| 1 | 190062247 | AX-170269282 | 0 | A | G | 0.254 | 9.04E-02 | 9.69E-02 | 2.40E+00 | 2.48E+00 | 3.67E-01 | no | | regulatory_region_variant | MODIFIER |
| 1 | 190068254 | AX-170118524 | 0 | C | T | 0.254 | 9.04E-02 | 9.69E-02 | 2.40E+00 | 2.48E+00 | 3.67E-01 | no | | intergenic_variant | MODIFIER |
| 1 | 190069719 | AX-170002549 | 0 | T | C | 0.254 | 9.04E-02 | 9.69E-02 | 2.40E+00 | 2.48E+00 | 3.67E-01 | no | | intergenic_variant | MODIFIER |
| 1 | 190069956 | AX-170279661 | 0 | G | T | 0.254 | 9.04E-02 | 9.69E-02 | 2.40E+00 | 2.48E+00 | 3.67E-01 | no | | intergenic_variant | MODIFIER |
| 1 | 190073300 | AX-170383826 | 0 | C | A | 0.254 | 9.04E-02 | 9.69E-02 | 2.40E+00 | 2.48E+00 | 3.67E-01 | no | | regulatory_region_variant | MODIFIER |
| 1 | 190078444 | AX-170520183 | 0 | C | T | 0.254 | 9.04E-02 | 9.69E-02 | 2.40E+00 | 2.48E+00 | 3.67E-01 | no | | intergenic_variant | MODIFIER |
| 1 | 190081121 | AX-170221472 | 0 | T | C | 0.254 | 9.04E-02 | 9.69E-02 | 2.40E+00 | 2.48E+00 | 3.67E-01 | no | | regulatory_region_variant | MODIFIER |
| 1 | 190081651 | AX-170002550 | 0 | C | T | 0.254 | 9.04E-02 | 9.69E-02 | 2.40E+00 | 2.48E+00 | 3.67E-01 | no | | intergenic_variant | MODIFIER |
| 1 | 190083306 | AX-170155295 | 0 | G | T | 0.254 | 9.04E-02 | 9.69E-02 | 2.40E+00 | 2.48E+00 | 3.67E-01 | no | | intergenic_variant | MODIFIER |
| 1 | 190085535 | AX-170444069 | 0 | C | T | 0.254 | 9.04E-02 | 9.69E-02 | 2.40E+00 | 2.48E+00 | 3.67E-01 | no | | regulatory_region_variant | MODIFIER |
| 1 | 190086431 | AX-170002552 | 0 | C | T | 0.254 | 9.04E-02 | 9.69E-02 | 2.40E+00 | 2.48E+00 | 3.67E-01 | no | | regulatory_region_variant | MODIFIER |
| 1 | 190097168 | AX-170043948 | 0 | G | A | 0.254 | 9.04E-02 | 9.69E-02 | 2.40E+00 | 2.48E+00 | 3.67E-01 | no | | intergenic_variant | MODIFIER |
| 1 | 190141938 | AX-170236141 | 0 | C | G | 0.254 | 9.04E-02 | 9.69E-02 | 2.40E+00 | 2.48E+00 | 3.67E-01 | no | Prox1 | intron_variant | MODIFIER |
| 1 | 190142015 | AX-170421373 | 0 | T | C | 0.254 | 9.04E-02 | 9.69E-02 | 2.40E+00 | 2.48E+00 | 3.67E-01 | no | Prox1 | intron_variant | MODIFIER |
| 1 | 190145951 | AX-97836807 | 0 | A | G | 0.254 | 9.04E-02 | 9.69E-02 | 2.40E+00 | 2.48E+00 | 3.67E-01 | no | Prox1 | intron_variant | MODIFIER |
| 1 | 190182341 | AX-97809115 | 0 | C | T | 0.254 | 9.04E-02 | 9.69E-02 | 2.40E+00 | 2.48E+00 | 3.67E-01 | no | Prox1os | intron_variant,non_coding_transcript_variant | MODIFIER |
| 1 | 190194673 | AX-170002554 | 0 | C | T | 0.254 | 9.04E-02 | 9.69E-02 | 2.40E+00 | 2.48E+00 | 3.67E-01 | no | Prox1os | intron_variant,non_coding_transcript_variant | MODIFIER |
| 1 | 190194841 | AX-170236533 | 0 | G | A | 0.254 | 9.04E-02 | 9.69E-02 | 2.40E+00 | 2.48E+00 | 3.67E-01 | no | Prox1os | intron_variant,non_coding_transcript_variant | MODIFIER |
| 1 | 190199717 | AX-170489339 | 0 | G | C | 0.254 | 9.04E-02 | 9.69E-02 | 2.40E+00 | 2.48E+00 | 3.67E-01 | no | Prox1os | intron_variant,non_coding_transcript_variant | MODIFIER |
| 1 | 190200406 | AX-170529700 | 0 | C | T | 0.254 | 9.04E-02 | 9.69E-02 | 2.40E+00 | 2.48E+00 | 3.67E-01 | no | Prox1os | intron_variant,non_coding_transcript_variant | MODIFIER |
| 1 | 190297516 | AX-170311463 | 0 | T | C | 0.254 | 9.04E-02 | 9.69E-02 | 2.40E+00 | 2.48E+00 | 3.67E-01 | no | | intergenic_variant | MODIFIER |
| 1 | 190324569 | AX-170191376 | 0 | T | C | 0.254 | 9.04E-02 | 9.69E-02 | 2.40E+00 | 2.48E+00 | 3.67E-01 | no | | intergenic_variant | MODIFIER |
| 1 | 190324641 | AX-170259211 | 0 | C | G | 0.254 | 9.04E-02 | 9.69E-02 | 2.40E+00 | 2.48E+00 | 3.67E-01 | no | | regulatory_region_variant | MODIFIER |
| 1 | 190324701 | AX-170274694 | 0 | T | C | 0.254 | 9.04E-02 | 9.69E-02 | 2.40E+00 | 2.48E+00 | 3.67E-01 | no | | regulatory_region_variant | MODIFIER |
| 1 | 190329334 | AX-97835797 | 0 | A | G | 0.254 | 9.04E-02 | 9.69E-02 | 2.40E+00 | 2.48E+00 | 3.67E-01 | no | | intergenic_variant | MODIFIER |
| 1 | 190340095 | AX-170446541 | 0 | G | A | 0.254 | 9.04E-02 | 9.69E-02 | 2.40E+00 | 2.48E+00 | 3.67E-01 | no | | intergenic_variant | MODIFIER |
| 1 | 190343081 | AX-170267911 | 0 | C | A | 0.254 | 9.04E-02 | 9.69E-02 | 2.40E+00 | 2.48E+00 | 3.67E-01 | no | | intergenic_variant | MODIFIER |
| 1 | 190468772 | AX-170470994 | 0 | A | C | 0.254 | 9.04E-02 | 9.69E-02 | 2.40E+00 | 2.48E+00 | 3.67E-01 | no | | intergenic_variant | MODIFIER |
| 1 | 190469788 | AX-97843752 | 0 | T | C | 0.254 | 9.04E-02 | 9.69E-02 | 2.40E+00 | 2.48E+00 | 3.67E-01 | no | | intergenic_variant | MODIFIER |
| 1 | 190474850 | AX-170315966 | 0 | T | G | 0.254 | 9.04E-02 | 9.69E-02 | 2.40E+00 | 2.48E+00 | 3.67E-01 | no | | intergenic_variant | MODIFIER |
| 1 | 190475126 | AX-170367507 | 0 | A | G | 0.254 | 9.04E-02 | 9.69E-02 | 2.40E+00 | 2.48E+00 | 3.67E-01 | no | | intergenic_variant | MODIFIER |

| | | | | | | | | | | | | | | | |
|---|---|---|---|---|---|---|---|---|---|---|---|---|---|---|---|
| 1 | 190487419 | AX-170086722 | 0 | G | C | 0.254 | 9.04E-02 | 9.69E-02 | 2.40E+00 | 2.48E+00 | 3.67E-01 | no | Gm23153 | upstream_gene_variant | MODIFIER |
| 1 | 190537302 | AX-170507327 | 0 | C | A | 0.254 | 9.04E-02 | 9.69E-02 | 2.40E+00 | 2.48E+00 | 3.67E-01 | no | | intergenic_variant | MODIFIER |
| 1 | 190557548 | AX-170002561 | 0 | T | C | 0.254 | 9.04E-02 | 9.69E-02 | 2.40E+00 | 2.48E+00 | 3.67E-01 | no | | intergenic_variant | MODIFIER |
| 1 | 190597307 | AX-97842608 | 0 | G | A | 0.254 | 9.04E-02 | 9.69E-02 | 2.40E+00 | 2.48E+00 | 3.67E-01 | no | Gm30932 | intron_variant,non_coding_transcript_variant | MODIFIER |
| 1 | 190609897 | AX-170043958 | 0 | T | C | 0.254 | 9.04E-02 | 9.69E-02 | 2.40E+00 | 2.48E+00 | 3.67E-01 | no | | intergenic_variant | MODIFIER |
| 1 | 190609938 | AX-170043955 | 0 | G | C | 0.254 | 9.04E-02 | 9.69E-02 | 2.40E+00 | 2.48E+00 | 3.67E-01 | no | | intergenic_variant | MODIFIER |
| 1 | 190625563 | AX-170043960 | 0 | A | G | 0.254 | 9.04E-02 | 9.69E-02 | 2.40E+00 | 2.48E+00 | 3.67E-01 | no | | intergenic_variant | MODIFIER |
| 1 | 190630138 | AX-170043956 | 0 | G | A | 0.254 | 9.04E-02 | 9.69E-02 | 2.40E+00 | 2.48E+00 | 3.67E-01 | no | | intergenic_variant | MODIFIER |
| 1 | 190630420 | AX-170043957 | 0 | A | C | 0.254 | 9.04E-02 | 9.69E-02 | 2.40E+00 | 2.48E+00 | 3.67E-01 | no | | intergenic_variant | MODIFIER |
| 1 | 190640000 | AX-170002560 | 0 | G | A | 0.254 | 9.04E-02 | 9.69E-02 | 2.40E+00 | 2.48E+00 | 3.67E-01 | no | | intergenic_variant | MODIFIER |
| 1 | 190643563 | AX-97834300 | 0 | G | A | 0.254 | 9.04E-02 | 9.69E-02 | 2.40E+00 | 2.48E+00 | 3.67E-01 | no | | intergenic_variant | MODIFIER |
| 1 | 190655252 | AX-170352584 | 0 | G | A | 0.254 | 9.04E-02 | 9.69E-02 | 2.40E+00 | 2.48E+00 | 3.67E-01 | no | | regulatory_region_variant | MODIFIER |
| 1 | 190685102 | AX-170530935 | 0 | G | A | 0.254 | 9.04E-02 | 9.69E-02 | 2.40E+00 | 2.48E+00 | 3.67E-01 | no | | intergenic_variant | MODIFIER |
| 1 | 190748177 | AX-170487924 | 0 | A | G | 0.254 | 9.04E-02 | 9.69E-02 | 2.40E+00 | 2.48E+00 | 3.67E-01 | no | Rps6kc1 | intron_variant,NMD_transcript_variant | MODIFIER |
| 1 | 190786820 | AX-97812077 | 0 | T | C | 0.254 | 9.04E-02 | 9.69E-02 | 2.40E+00 | 2.48E+00 | 3.67E-01 | no | Rps6kc1 | intron_variant | MODIFIER |
| 1 | 190809861 | AX-97788717 | 0 | T | C | 0.254 | 9.04E-02 | 9.69E-02 | 2.40E+00 | 2.48E+00 | 3.67E-01 | no | Rps6kc1 | intron_variant | MODIFIER |
| 1 | 190824909 | AX-170043961 | 0 | T | C | 0.254 | 9.04E-02 | 9.69E-02 | 2.40E+00 | 2.48E+00 | 3.67E-01 | no | Rps6kc1 | intron_variant | MODIFIER |
| 1 | 190825285 | AX-170043963 | 0 | A | G | 0.254 | 9.04E-02 | 9.69E-02 | 2.40E+00 | 2.48E+00 | 3.67E-01 | no | Rps6kc1 | intron_variant | MODIFIER |
| 1 | 190848827 | AX-97809418 | 0 | A | G | 0.254 | 9.04E-02 | 9.69E-02 | 2.40E+00 | 2.48E+00 | 3.67E-01 | no | Rps6kc1 | intron_variant | MODIFIER |
| 1 | 190903962 | AX-170320270 | 0 | A | G | 0.254 | 9.04E-02 | 9.69E-02 | 2.40E+00 | 2.48E+00 | 3.67E-01 | no | Rps6kc1 | intron_variant | MODIFIER |
| 1 | 190909101 | AX-170497511 | 0 | A | G | 0.254 | 9.04E-02 | 9.69E-02 | 2.40E+00 | 2.48E+00 | 3.67E-01 | no | Rps6kc1 | intron_variant | MODIFIER |
| 1 | 190910525 | AX-170104053 | 0 | A | G | 0.254 | 9.04E-02 | 9.69E-02 | 2.40E+00 | 2.48E+00 | 3.67E-01 | no | Rps6kc1 | intron_variant | MODIFIER |
| 1 | 190910625 | AX-170406137 | 0 | T | C | 0.254 | 9.04E-02 | 9.69E-02 | 2.40E+00 | 2.48E+00 | 3.67E-01 | no | Rps6kc1 | intron_variant | MODIFIER |
| 1 | 190919717 | AX-170424822 | 0 | C | T | 0.254 | 9.04E-02 | 9.69E-02 | 2.40E+00 | 2.48E+00 | 3.67E-01 | no | | intergenic_variant | MODIFIER |
| 1 | 190934638 | AX-170120515 | 0 | A | G | 0.254 | 9.04E-02 | 9.69E-02 | 2.40E+00 | 2.48E+00 | 3.67E-01 | no | Angel2 | intron_variant | MODIFIER |
| 1 | 190935335 | AX-170440482 | 0 | T | C | 0.254 | 9.04E-02 | 9.69E-02 | 2.40E+00 | 2.48E+00 | 3.67E-01 | no | Angel2 | intron_variant | MODIFIER |
| 1 | 190940240 | AX-170262178 | 0 | C | T | 0.254 | 9.04E-02 | 9.69E-02 | 2.40E+00 | 2.48E+00 | 3.67E-01 | no | Angel2 | intron_variant | MODIFIER |
| 1 | 190958241 | AX-170086723 | 0 | G | A | 0.254 | 9.04E-02 | 9.69E-02 | 2.40E+00 | 2.48E+00 | 3.67E-01 | no | Vash2 | intron_variant | MODIFIER |
| 1 | 191008195 | AX-170430435 | 0 | G | T | 0.254 | 9.04E-02 | 9.69E-02 | 2.40E+00 | 2.48E+00 | 3.67E-01 | no | Flvcr1 | synonymous_variant | LOW |
| 1 | 191017995 | AX-170340893 | 0 | A | G | 0.254 | 9.04E-02 | 9.69E-02 | 2.40E+00 | 2.48E+00 | 3.67E-01 | no | Flvcr1 | intron_variant | MODIFIER |
| 1 | 191027776 | AX-170413967 | 0 | T | C | 0.254 | 9.04E-02 | 9.69E-02 | 2.40E+00 | 2.48E+00 | 3.67E-01 | no | Flvcr1 | upstream_gene_variant | MODIFIER |
| 1 | 191081757 | AX-170043966 | 0 | T | A | 0.254 | 9.04E-02 | 9.69E-02 | 2.40E+00 | 2.48E+00 | 3.67E-01 | no | Nsl1 | intron_variant | MODIFIER |
| 1 | 191091564 | AX-170002564 | 0 | T | C | 0.254 | 9.04E-02 | 9.69E-02 | 2.40E+00 | 2.48E+00 | 3.67E-01 | no | | intergenic_variant | MODIFIER |
| 4 | 76530392 | AX-170310375 | 0 | C | A | 0.325 | 9.93E-02 | 1.11E-01 | 2.75E+00 | 2.82E+00 | 3.67E-01 | no | Ptprd | intron_variant,non_coding_transcript_variant | MODIFIER |
| 16 | 78228186 | AX-170298025 | 0 | G | A | 0.465 | 6.74E-02 | 7.46E-02 | 2.56E+00 | 2.63E+00 | 3.67E-01 | no | | intergenic_variant | MODIFIER |
| 16 | 78664402 | AX-170401440 | 0 | G | C | 0.465 | 6.74E-02 | 7.46E-02 | 2.56E+00 | 2.63E+00 | 3.67E-01 | no | | intergenic_variant | MODIFIER |
| 8 | 99029209 | AX-170534390 | 0 | A | C | 0.228 | 1.02E-01 | 1.14E-01 | 2.79E+00 | 2.87E+00 | 3.67E-01 | no | Cdh8 | 3_prime_UTR_variant | MODIFIER |
| 1 | 135748325 | AX-170316851 | 0 | C | T | 0.544 | 7.42E-02 | 8.25E-02 | 2.59E+00 | 2.66E+00 | 3.67E-01 | no | Csrp1 | intron_variant | MODIFIER |
| 18 | 83873010 | AX-170016111 | 1 | G | A | 0.161 | -9.83E-02 | 1.07E-01 | 2.48E+00 | 2.56E+00 | 3.67E-01 | no | | intergenic_variant | MODIFIER |
| 18 | 85335227 | AX-170463328 | 1 | C | A | 0.161 | -9.83E-02 | 1.07E-01 | 2.48E+00 | 2.56E+00 | 3.67E-01 | no | | intergenic_variant | MODIFIER |
| 2 | 162275840 | AX-170088825 | 0 | G | T | 0.167 | 1.63E-01 | 1.82E-01 | 2.65E+00 | 2.73E+00 | 3.67E-01 | Yes | Ptprt | intron_variant | MODIFIER |
| 2 | 168424389 | AX-170319079 | 0 | C | T | 0.167 | 1.63E-01 | 1.82E-01 | 2.65E+00 | 2.73E+00 | 3.67E-01 | Yes | | regulatory_region_variant | MODIFIER |
| 2 | 168436299 | AX-170094481 | 0 | C | T | 0.167 | 1.63E-01 | 1.82E-01 | 2.65E+00 | 2.73E+00 | 3.67E-01 | Yes | | intergenic_variant | MODIFIER |
| 2 | 168442355 | AX-170019639 | 0 | A | G | 0.167 | 1.63E-01 | 1.82E-01 | 2.65E+00 | 2.73E+00 | 3.67E-01 | Yes | | intergenic_variant | MODIFIER |
| 2 | 168445946 | AX-97833132 | 0 | A | G | 0.167 | 1.63E-01 | 1.82E-01 | 2.65E+00 | 2.73E+00 | 3.67E-01 | Yes | | regulatory_region_variant | MODIFIER |
| 2 | 168455439 | AX-170019643 | 0 | C | G | 0.167 | 1.63E-01 | 1.82E-01 | 2.65E+00 | 2.73E+00 | 3.67E-01 | Yes | | intergenic_variant | MODIFIER |
| 2 | 168471521 | AX-170348840 | 0 | T | G | 0.167 | 1.63E-01 | 1.82E-01 | 2.65E+00 | 2.73E+00 | 3.67E-01 | Yes | Nfatc2 | downstream_gene_variant | MODIFIER |
| 2 | 168478014 | AX-170268522 | 0 | C | G | 0.167 | 1.63E-01 | 1.82E-01 | 2.65E+00 | 2.73E+00 | 3.67E-01 | Yes | Nfatc2 | 3_prime_UTR_variant | MODIFIER |
| 2 | 168478096 | AX-170235970 | 0 | C | T | 0.167 | 1.63E-01 | 1.82E-01 | 2.65E+00 | 2.73E+00 | 3.67E-01 | Yes | Nfatc2 | 3_prime_UTR_variant | MODIFIER |
| 2 | 168482135 | AX-97835344 | 0 | A | G | 0.167 | 1.63E-01 | 1.82E-01 | 2.65E+00 | 2.73E+00 | 3.67E-01 | Yes | Nfatc2 | intron_variant | MODIFIER |
| 2 | 168485925 | AX-170019641 | 0 | C | T | 0.167 | 1.63E-01 | 1.82E-01 | 2.65E+00 | 2.73E+00 | 3.67E-01 | Yes | Nfatc2 | intron_variant | MODIFIER |
| 2 | 168486902 | AX-170019644 | 0 | T | C | 0.167 | 1.63E-01 | 1.82E-01 | 2.65E+00 | 2.73E+00 | 3.67E-01 | Yes | Nfatc2 | intron_variant | MODIFIER |
| 2 | 168487628 | AX-170019647 | 0 | T | C | 0.167 | 1.63E-01 | 1.82E-01 | 2.65E+00 | 2.73E+00 | 3.67E-01 | Yes | Nfatc2 | intron_variant | MODIFIER |
| 2 | 168496711 | AX-170239264 | 0 | T | C | 0.167 | 1.63E-01 | 1.82E-01 | 2.65E+00 | 2.73E+00 | 3.67E-01 | Yes | Nfatc2 | intron_variant | MODIFIER |

| | | | | | | | | | | | | | | | |
|---|---|---|---|---|---|---|---|---|---|---|---|---|---|---|---|
| 2 | 168519571 | AX-170142652 | 0 | A | G | 0.167 | 1.63E-01 | 1.82E-01 | 2.65E+00 | 2.73E+00 | 3.67E-01 | Yes | Nfatc2 | intron_variant | MODIFIER |
| 2 | 168521342 | AX-170489340 | 0 | C | T | 0.167 | 1.63E-01 | 1.82E-01 | 2.65E+00 | 2.73E+00 | 3.67E-01 | Yes | Nfatc2 | intron_variant | MODIFIER |
| 2 | 168573811 | AX-170019638 | 0 | A | G | 0.167 | 1.63E-01 | 1.82E-01 | 2.65E+00 | 2.73E+00 | 3.67E-01 | Yes | Nfatc2 | intron_variant | MODIFIER |
| 2 | 168575477 | AX-97839853 | 0 | C | T | 0.167 | 1.63E-01 | 1.82E-01 | 2.65E+00 | 2.73E+00 | 3.67E-01 | Yes | Nfatc2 | intron_variant | MODIFIER |
| 2 | 168579902 | AX-170019642 | 0 | G | T | 0.167 | 1.63E-01 | 1.82E-01 | 2.65E+00 | 2.73E+00 | 3.67E-01 | Yes | Nfatc2 | intron_variant | MODIFIER |
| 2 | 168593280 | AX-170019640 | 0 | A | G | 0.167 | 1.63E-01 | 1.82E-01 | 2.65E+00 | 2.73E+00 | 3.67E-01 | Yes | Nfatc2 | upstream_gene_variant | MODIFIER |
| 2 | 168604903 | AX-170355285 | 0 | A | G | 0.167 | 1.63E-01 | 1.82E-01 | 2.65E+00 | 2.73E+00 | 3.67E-01 | Yes | Nfatc2 | upstream_gene_variant | MODIFIER |
| 2 | 168613682 | AX-170088830 | 0 | G | A | 0.167 | 1.63E-01 | 1.82E-01 | 2.65E+00 | 2.73E+00 | 3.67E-01 | Yes | | regulatory_region_variant | MODIFIER |
| 2 | 168621441 | AX-170190425 | 0 | A | G | 0.167 | 1.63E-01 | 1.82E-01 | 2.65E+00 | 2.73E+00 | 3.67E-01 | Yes | | intergenic_variant | MODIFIER |
| 2 | 168631682 | AX-97818585 | 0 | G | A | 0.167 | 1.63E-01 | 1.82E-01 | 2.65E+00 | 2.73E+00 | 3.67E-01 | Yes | Atp9a | downstream_gene_variant | MODIFIER |
| 2 | 168634405 | AX-170500008 | 0 | G | A | 0.167 | 1.63E-01 | 1.82E-01 | 2.65E+00 | 2.73E+00 | 3.67E-01 | Yes | Atp9a | downstream_gene_variant | MODIFIER |
| 2 | 168634918 | AX-170467544 | 0 | A | T | 0.167 | 1.63E-01 | 1.82E-01 | 2.65E+00 | 2.73E+00 | 3.67E-01 | Yes | Atp9a | intron_variant | MODIFIER |
| 2 | 168635263 | AX-170019646 | 0 | A | G | 0.167 | 1.63E-01 | 1.82E-01 | 2.65E+00 | 2.73E+00 | 3.67E-01 | Yes | Atp9a | intron_variant | MODIFIER |
| 2 | 168643559 | AX-170446248 | 0 | T | C | 0.167 | 1.63E-01 | 1.82E-01 | 2.65E+00 | 2.73E+00 | 3.67E-01 | Yes | Atp9a | intron_variant | MODIFIER |
| 2 | 168649131 | AX-170352667 | 0 | C | T | 0.167 | 1.63E-01 | 1.82E-01 | 2.65E+00 | 2.73E+00 | 3.67E-01 | Yes | Atp9a | intron_variant | MODIFIER |
| 2 | 168656997 | AX-170366247 | 0 | A | G | 0.167 | 1.63E-01 | 1.82E-01 | 2.65E+00 | 2.73E+00 | 3.67E-01 | Yes | Atp9a | intron_variant | MODIFIER |
| 2 | 168662266 | AX-170473903 | 0 | C | T | 0.167 | 1.63E-01 | 1.82E-01 | 2.65E+00 | 2.73E+00 | 3.67E-01 | Yes | Atp9a | intron_variant | MODIFIER |
| 2 | 168700141 | AX-170144362 | 0 | T | C | 0.167 | 1.63E-01 | 1.82E-01 | 2.65E+00 | 2.73E+00 | 3.67E-01 | Yes | Atp9a | intron_variant | MODIFIER |
| 2 | 168751638 | AX-170142345 | 0 | T | C | 0.167 | 1.63E-01 | 1.82E-01 | 2.65E+00 | 2.73E+00 | 3.67E-01 | Yes | Sall4 | intron_variant | MODIFIER |
| 2 | 168755443 | AX-97850448 | 0 | C | T | 0.167 | 1.63E-01 | 1.82E-01 | 2.65E+00 | 2.73E+00 | 3.67E-01 | Yes | Sall4 | synonymous_variant | LOW |
| 2 | 168759382 | AX-170065585 | 0 | G | A | 0.167 | 1.63E-01 | 1.82E-01 | 2.65E+00 | 2.73E+00 | 3.67E-01 | Yes | Sall4 | intron_variant | MODIFIER |
| 2 | 168777343 | AX-97781502 | 0 | C | T | 0.167 | 1.63E-01 | 1.82E-01 | 2.65E+00 | 2.73E+00 | 3.67E-01 | Yes | | regulatory_region_variant | MODIFIER |
| 2 | 168794745 | AX-170429389 | 0 | C | T | 0.167 | 1.63E-01 | 1.82E-01 | 2.65E+00 | 2.73E+00 | 3.67E-01 | Yes | | intergenic_variant | MODIFIER |
| 2 | 168819094 | AX-170287691 | 0 | T | C | 0.167 | 1.63E-01 | 1.82E-01 | 2.65E+00 | 2.73E+00 | 3.67E-01 | Yes | | intergenic_variant | MODIFIER |
| 2 | 168861222 | AX-170260790 | 0 | A | G | 0.167 | 1.63E-01 | 1.82E-01 | 2.65E+00 | 2.73E+00 | 3.67E-01 | Yes | | regulatory_region_variant | MODIFIER |
| 2 | 168879257 | AX-170381176 | 0 | A | C | 0.167 | 1.63E-01 | 1.82E-01 | 2.65E+00 | 2.73E+00 | 3.67E-01 | Yes | Gm25214 | downstream_gene_variant | MODIFIER |
| 2 | 168920385 | AX-170409131 | 0 | A | C | 0.167 | 1.63E-01 | 1.82E-01 | 2.65E+00 | 2.73E+00 | 3.67E-01 | Yes | Zfp64 | downstream_gene_variant | MODIFIER |
| 2 | 168928434 | AX-170473848 | 0 | C | T | 0.167 | 1.63E-01 | 1.82E-01 | 2.65E+00 | 2.73E+00 | 3.67E-01 | Yes | Zfp64 | intron_variant | MODIFIER |
| 2 | 168933354 | AX-97798382 | 0 | T | C | 0.167 | 1.63E-01 | 1.82E-01 | 2.65E+00 | 2.73E+00 | 3.67E-01 | Yes | Zfp64 | intron_variant | MODIFIER |
| 2 | 168947735 | AX-97837961 | 0 | A | G | 0.167 | 1.63E-01 | 1.82E-01 | 2.65E+00 | 2.73E+00 | 3.67E-01 | Yes | Zfp64 | intron_variant | MODIFIER |
| 2 | 168949359 | AX-170293370 | 0 | T | C | 0.167 | 1.63E-01 | 1.82E-01 | 2.65E+00 | 2.73E+00 | 3.67E-01 | Yes | Zfp64 | intron_variant | MODIFIER |
| 2 | 168978868 | AX-97833154 | 0 | G | T | 0.167 | 1.63E-01 | 1.82E-01 | 2.65E+00 | 2.73E+00 | 3.67E-01 | Yes | 1700017J07Rik | non_coding_transcript_exon_variant | MODIFIER |
| 2 | 169026789 | AX-170108745 | 0 | C | T | 0.167 | 1.63E-01 | 1.82E-01 | 2.65E+00 | 2.73E+00 | 3.67E-01 | Yes | | intergenic_variant | MODIFIER |
| 2 | 169029953 | AX-170019653 | 0 | T | A | 0.167 | 1.63E-01 | 1.82E-01 | 2.65E+00 | 2.73E+00 | 3.67E-01 | Yes | | intergenic_variant | MODIFIER |
| 2 | 169054028 | AX-170019652 | 0 | T | G | 0.167 | 1.63E-01 | 1.82E-01 | 2.65E+00 | 2.73E+00 | 3.67E-01 | Yes | | intergenic_variant | MODIFIER |
| 2 | 169054587 | AX-170284200 | 0 | A | T | 0.167 | 1.63E-01 | 1.82E-01 | 2.65E+00 | 2.73E+00 | 3.67E-01 | Yes | | intergenic_variant | MODIFIER |
| 2 | 169059122 | AX-97829848 | 0 | A | G | 0.167 | 1.63E-01 | 1.82E-01 | 2.65E+00 | 2.73E+00 | 3.67E-01 | Yes | Gm26469 | upstream_gene_variant | MODIFIER |
| 2 | 169062462 | AX-170440142 | 0 | C | T | 0.167 | 1.63E-01 | 1.82E-01 | 2.65E+00 | 2.73E+00 | 3.67E-01 | Yes | Gm26469 | downstream_gene_variant | MODIFIER |
| 2 | 169062593 | AX-170301148 | 0 | G | A | 0.167 | 1.63E-01 | 1.82E-01 | 2.65E+00 | 2.73E+00 | 3.67E-01 | Yes | Gm26469 | downstream_gene_variant | MODIFIER |
| 2 | 169067782 | AX-170258178 | 0 | T | C | 0.167 | 1.63E-01 | 1.82E-01 | 2.65E+00 | 2.73E+00 | 3.67E-01 | Yes | | intergenic_variant | MODIFIER |
| 2 | 169068475 | AX-170467499 | 0 | T | C | 0.167 | 1.63E-01 | 1.82E-01 | 2.65E+00 | 2.73E+00 | 3.67E-01 | Yes | | intergenic_variant | MODIFIER |
| 2 | 169070517 | AX-170019648 | 0 | G | A | 0.167 | 1.63E-01 | 1.82E-01 | 2.65E+00 | 2.73E+00 | 3.67E-01 | Yes | | intergenic_variant | MODIFIER |
| 2 | 169087753 | AX-170019651 | 0 | A | T | 0.167 | 1.63E-01 | 1.82E-01 | 2.65E+00 | 2.73E+00 | 3.67E-01 | Yes | | intergenic_variant | MODIFIER |
| 2 | 169088185 | AX-97785980 | 0 | C | T | 0.167 | 1.63E-01 | 1.82E-01 | 2.65E+00 | 2.73E+00 | 3.67E-01 | Yes | | intergenic_variant | MODIFIER |
| 2 | 169088415 | AX-170149241 | 0 | A | G | 0.167 | 1.63E-01 | 1.82E-01 | 2.65E+00 | 2.73E+00 | 3.67E-01 | Yes | | regulatory_region_variant | MODIFIER |
| 2 | 169113614 | AX-97806782 | 0 | A | G | 0.167 | 1.63E-01 | 1.82E-01 | 2.65E+00 | 2.73E+00 | 3.67E-01 | Yes | Gm14258 | downstream_gene_variant | MODIFIER |
| 2 | 169126766 | AX-170359867 | 0 | T | G | 0.167 | 1.63E-01 | 1.82E-01 | 2.65E+00 | 2.73E+00 | 3.67E-01 | Yes | | intergenic_variant | MODIFIER |
| 2 | 169133884 | AX-97851124 | 0 | A | G | 0.167 | 1.63E-01 | 1.82E-01 | 2.65E+00 | 2.73E+00 | 3.67E-01 | Yes | | regulatory_region_variant | MODIFIER |
| 2 | 169203174 | AX-97833731 | 0 | T | C | 0.167 | 1.63E-01 | 1.82E-01 | 2.65E+00 | 2.73E+00 | 3.67E-01 | Yes | | intergenic_variant | MODIFIER |
| 2 | 169206507 | AX-170522569 | 0 | A | G | 0.167 | 1.63E-01 | 1.82E-01 | 2.65E+00 | 2.73E+00 | 3.67E-01 | Yes | | intergenic_variant | MODIFIER |
| 2 | 169221274 | AX-170136337 | 0 | A | C | 0.167 | 1.63E-01 | 1.82E-01 | 2.65E+00 | 2.73E+00 | 3.67E-01 | Yes | | intergenic_variant | MODIFIER |
| 2 | 169223535 | AX-170204390 | 0 | C | A | 0.167 | 1.63E-01 | 1.82E-01 | 2.65E+00 | 2.73E+00 | 3.67E-01 | Yes | | intergenic_variant | MODIFIER |
| 2 | 169228524 | AX-170252786 | 0 | G | A | 0.167 | 1.63E-01 | 1.82E-01 | 2.65E+00 | 2.73E+00 | 3.67E-01 | Yes | | intergenic_variant | MODIFIER |
| 2 | 169232019 | AX-97846641 | 0 | G | T | 0.167 | 1.63E-01 | 1.82E-01 | 2.65E+00 | 2.73E+00 | 3.67E-01 | Yes | 9430093N23Rik | upstream_gene_variant | MODIFIER |

| | | | | | | | | | | | | | | | |
|---|---|---|---|---|---|---|---|---|---|---|---|---|---|---|---|
| 2 | 169237429 | AX-170236423 | 0 | C | T | 0.167 | 1.63E-01 | 1.82E-01 | 2.65E+00 | 2.73E+00 | 3.67E-01 | Yes | 9430093N23Rik | intron_variant,non_coding_transcript_variant | MODIFIER |
| 2 | 169237645 | AX-170281201 | 0 | A | T | 0.167 | 1.63E-01 | 1.82E-01 | 2.65E+00 | 2.73E+00 | 3.67E-01 | Yes | 9430093N23Rik | non_coding_transcript_exon_variant | MODIFIER |
| 2 | 169244850 | AX-170122552 | 0 | C | T | 0.167 | 1.63E-01 | 1.82E-01 | 2.65E+00 | 2.73E+00 | 3.67E-01 | Yes | | intergenic_variant | MODIFIER |
| 2 | 169265215 | AX-170094483 | 0 | T | C | 0.167 | 1.63E-01 | 1.82E-01 | 2.65E+00 | 2.73E+00 | 3.67E-01 | Yes | | intergenic_variant | MODIFIER |
| 2 | 169267169 | AX-170178134 | 0 | G | C | 0.167 | 1.63E-01 | 1.82E-01 | 2.65E+00 | 2.73E+00 | 3.67E-01 | Yes | | intergenic_variant | MODIFIER |
| 2 | 169276219 | AX-97804178 | 0 | A | G | 0.167 | 1.63E-01 | 1.82E-01 | 2.65E+00 | 2.73E+00 | 3.67E-01 | Yes | | intergenic_variant | MODIFIER |
| 2 | 169366399 | AX-97820658 | 0 | C | T | 0.167 | 1.63E-01 | 1.82E-01 | 2.65E+00 | 2.73E+00 | 3.67E-01 | Yes | | intergenic_variant | MODIFIER |
| 2 | 169368690 | AX-170134248 | 0 | A | T | 0.167 | 1.63E-01 | 1.82E-01 | 2.65E+00 | 2.73E+00 | 3.67E-01 | Yes | | intergenic_variant | MODIFIER |
| 2 | 169369106 | AX-170509933 | 0 | C | T | 0.167 | 1.63E-01 | 1.82E-01 | 2.65E+00 | 2.73E+00 | 3.67E-01 | Yes | | intergenic_variant | MODIFIER |
| 2 | 169395015 | AX-170100912 | 0 | G | C | 0.167 | 1.63E-01 | 1.82E-01 | 2.65E+00 | 2.73E+00 | 3.67E-01 | Yes | 4930529I22Rik | downstream_gene_variant | MODIFIER |
| 2 | 169400879 | AX-170232441 | 0 | A | G | 0.167 | 1.63E-01 | 1.82E-01 | 2.65E+00 | 2.73E+00 | 3.67E-01 | Yes | 4930529I22Rik | intron_variant,non_coding_transcript_variant | MODIFIER |
| 2 | 169402511 | AX-170452171 | 0 | G | C | 0.167 | 1.63E-01 | 1.82E-01 | 2.65E+00 | 2.73E+00 | 3.67E-01 | Yes | 4930529I22Rik | intron_variant,non_coding_transcript_variant | MODIFIER |
| 2 | 169404077 | AX-170275043 | 0 | C | A | 0.167 | 1.63E-01 | 1.82E-01 | 2.65E+00 | 2.73E+00 | 3.67E-01 | Yes | 4930529I22Rik | intron_variant,non_coding_transcript_variant | MODIFIER |
| 2 | 169404950 | AX-97830904 | 0 | A | G | 0.167 | 1.63E-01 | 1.82E-01 | 2.65E+00 | 2.73E+00 | 3.67E-01 | Yes | 4930529I22Rik | intron_variant,non_coding_transcript_variant | MODIFIER |
| 2 | 169430186 | AX-97805833 | 0 | C | A | 0.167 | 1.63E-01 | 1.82E-01 | 2.65E+00 | 2.73E+00 | 3.67E-01 | Yes | | intergenic_variant | MODIFIER |
| 2 | 169442569 | AX-170353971 | 0 | T | C | 0.167 | 1.63E-01 | 1.82E-01 | 2.65E+00 | 2.73E+00 | 3.67E-01 | Yes | | intergenic_variant | MODIFIER |
| 2 | 169443351 | AX-170293109 | 0 | A | G | 0.167 | 1.63E-01 | 1.82E-01 | 2.65E+00 | 2.73E+00 | 3.67E-01 | Yes | | intergenic_variant | MODIFIER |
| 2 | 169446630 | AX-170380854 | 0 | C | T | 0.167 | 1.63E-01 | 1.82E-01 | 2.65E+00 | 2.73E+00 | 3.67E-01 | Yes | | intergenic_variant | MODIFIER |
| 2 | 169446665 | AX-170508101 | 0 | T | C | 0.167 | 1.63E-01 | 1.82E-01 | 2.65E+00 | 2.73E+00 | 3.67E-01 | Yes | | intergenic_variant | MODIFIER |
| 2 | 169449322 | AX-170285630 | 0 | C | T | 0.167 | 1.63E-01 | 1.82E-01 | 2.65E+00 | 2.73E+00 | 3.67E-01 | Yes | | intergenic_variant | MODIFIER |
| 2 | 169452952 | AX-170171668 | 0 | A | C | 0.167 | 1.63E-01 | 1.82E-01 | 2.65E+00 | 2.73E+00 | 3.67E-01 | Yes | | intergenic_variant | MODIFIER |
| 2 | 169455428 | AX-97856662 | 0 | G | A | 0.167 | 1.63E-01 | 1.82E-01 | 2.65E+00 | 2.73E+00 | 3.67E-01 | Yes | | intergenic_variant | MODIFIER |
| 2 | 169458092 | AX-170240904 | 0 | G | T | 0.167 | 1.63E-01 | 1.82E-01 | 2.65E+00 | 2.73E+00 | 3.67E-01 | Yes | | intergenic_variant | MODIFIER |
| 2 | 169482555 | AX-97833990 | 0 | A | G | 0.167 | 1.63E-01 | 1.82E-01 | 2.65E+00 | 2.73E+00 | 3.67E-01 | Yes | Gm14249 | intron_variant,non_coding_transcript_variant | MODIFIER |
| 2 | 169487900 | AX-170304768 | 0 | T | C | 0.167 | 1.63E-01 | 1.82E-01 | 2.65E+00 | 2.73E+00 | 3.67E-01 | Yes | Gm14249 | intron_variant,non_coding_transcript_variant | MODIFIER |
| 2 | 169488272 | AX-170168330 | 0 | G | C | 0.167 | 1.63E-01 | 1.82E-01 | 2.65E+00 | 2.73E+00 | 3.67E-01 | Yes | Gm14249 | intron_variant,non_coding_transcript_variant | MODIFIER |
| 2 | 169524103 | AX-170065598 | 0 | A | G | 0.167 | 1.63E-01 | 1.82E-01 | 2.65E+00 | 2.73E+00 | 3.67E-01 | Yes | | intergenic_variant | MODIFIER |
| 2 | 169527233 | AX-170490367 | 0 | C | T | 0.167 | 1.63E-01 | 1.82E-01 | 2.65E+00 | 2.73E+00 | 3.67E-01 | Yes | | intergenic_variant | MODIFIER |
| 2 | 169532704 | AX-97788416 | 0 | C | T | 0.167 | 1.63E-01 | 1.82E-01 | 2.65E+00 | 2.73E+00 | 3.67E-01 | Yes | | intergenic_variant | MODIFIER |
| 2 | 169533080 | AX-170384472 | 0 | C | T | 0.167 | 1.63E-01 | 1.82E-01 | 2.65E+00 | 2.73E+00 | 3.67E-01 | Yes | | intergenic_variant | MODIFIER |
| 2 | 169534213 | AX-170284444 | 0 | G | C | 0.167 | 1.63E-01 | 1.82E-01 | 2.65E+00 | 2.73E+00 | 3.67E-01 | Yes | | intergenic_variant | MODIFIER |
| 2 | 169535353 | AX-170372815 | 0 | T | C | 0.167 | 1.63E-01 | 1.82E-01 | 2.65E+00 | 2.73E+00 | 3.67E-01 | Yes | | intergenic_variant | MODIFIER |
| 2 | 169554914 | AX-170269863 | 0 | C | T | 0.167 | 1.63E-01 | 1.82E-01 | 2.65E+00 | 2.73E+00 | 3.67E-01 | Yes | | intergenic_variant | MODIFIER |
| 2 | 169555319 | AX-170186020 | 0 | T | C | 0.167 | 1.63E-01 | 1.82E-01 | 2.65E+00 | 2.73E+00 | 3.67E-01 | Yes | | intergenic_variant | MODIFIER |
| 2 | 169561562 | AX-170373534 | 0 | C | T | 0.167 | 1.63E-01 | 1.82E-01 | 2.65E+00 | 2.73E+00 | 3.67E-01 | Yes | | intergenic_variant | MODIFIER |
| 2 | 169562913 | AX-170135513 | 0 | C | T | 0.167 | 1.63E-01 | 1.82E-01 | 2.65E+00 | 2.73E+00 | 3.67E-01 | Yes | | intergenic_variant | MODIFIER |
| 2 | 169570760 | AX-170091113 | 0 | T | C | 0.167 | 1.63E-01 | 1.82E-01 | 2.65E+00 | 2.73E+00 | 3.67E-01 | Yes | | intergenic_variant | MODIFIER |
| 2 | 169596381 | AX-170259980 | 0 | C | T | 0.167 | 1.63E-01 | 1.82E-01 | 2.65E+00 | 2.73E+00 | 3.67E-01 | Yes | | intergenic_variant | MODIFIER |
| 2 | 169597084 | AX-170210923 | 0 | C | T | 0.167 | 1.63E-01 | 1.82E-01 | 2.65E+00 | 2.73E+00 | 3.67E-01 | Yes | | intergenic_variant | MODIFIER |
| 2 | 169602171 | AX-170178748 | 0 | G | A | 0.167 | 1.63E-01 | 1.82E-01 | 2.65E+00 | 2.73E+00 | 3.67E-01 | Yes | | intergenic_variant | MODIFIER |
| 2 | 169616951 | AX-97826428 | 0 | C | T | 0.167 | 1.63E-01 | 1.82E-01 | 2.65E+00 | 2.73E+00 | 3.67E-01 | Yes | | intergenic_variant | MODIFIER |
| 2 | 169616996 | AX-170450381 | 0 | A | G | 0.167 | 1.63E-01 | 1.82E-01 | 2.65E+00 | 2.73E+00 | 3.67E-01 | Yes | | intergenic_variant | MODIFIER |
| 2 | 169622577 | AX-170385600 | 0 | C | T | 0.167 | 1.63E-01 | 1.82E-01 | 2.65E+00 | 2.73E+00 | 3.67E-01 | Yes | Gm34294 | downstream_gene_variant | MODIFIER |
| 2 | 169622659 | AX-97833283 | 0 | A | G | 0.167 | 1.63E-01 | 1.82E-01 | 2.65E+00 | 2.73E+00 | 3.67E-01 | Yes | Gm34294 | downstream_gene_variant | MODIFIER |
| 2 | 169660186 | AX-97850338 | 0 | G | A | 0.167 | 1.63E-01 | 1.82E-01 | 2.65E+00 | 2.73E+00 | 3.67E-01 | Yes | Tshz2 | intron_variant | MODIFIER |
| 2 | 169660842 | AX-170512788 | 0 | A | G | 0.167 | 1.63E-01 | 1.82E-01 | 2.65E+00 | 2.73E+00 | 3.67E-01 | Yes | Tshz2 | intron_variant | MODIFIER |
| 2 | 169661045 | AX-170384054 | 0 | A | C | 0.167 | 1.63E-01 | 1.82E-01 | 2.65E+00 | 2.73E+00 | 3.67E-01 | Yes | Tshz2 | intron_variant | MODIFIER |
| 2 | 169675019 | AX-170399975 | 0 | A | G | 0.167 | 1.63E-01 | 1.82E-01 | 2.65E+00 | 2.73E+00 | 3.67E-01 | Yes | Tshz2 | intron_variant | MODIFIER |
| 2 | 169696560 | AX-170181740 | 0 | G | T | 0.167 | 1.63E-01 | 1.82E-01 | 2.65E+00 | 2.73E+00 | 3.67E-01 | Yes | Tshz2 | intron_variant | MODIFIER |
| 2 | 169715422 | AX-170151701 | 0 | T | A | 0.167 | 1.63E-01 | 1.82E-01 | 2.65E+00 | 2.73E+00 | 3.67E-01 | Yes | Tshz2 | intron_variant | MODIFIER |
| 2 | 169719117 | AX-170328860 | 0 | G | A | 0.167 | 1.63E-01 | 1.82E-01 | 2.65E+00 | 2.73E+00 | 3.67E-01 | Yes | Tshz2 | intron_variant | MODIFIER |
| 2 | 169728485 | AX-170187023 | 0 | A | G | 0.167 | 1.63E-01 | 1.82E-01 | 2.65E+00 | 2.73E+00 | 3.67E-01 | Yes | Tshz2 | intron_variant | MODIFIER |
| 2 | 169729468 | AX-170117839 | 0 | T | C | 0.167 | 1.63E-01 | 1.82E-01 | 2.65E+00 | 2.73E+00 | 3.67E-01 | Yes | Tshz2 | intron_variant | MODIFIER |
| 2 | 169734102 | AX-170165278 | 0 | C | T | 0.167 | 1.63E-01 | 1.82E-01 | 2.65E+00 | 2.73E+00 | 3.67E-01 | Yes | Tshz2 | intron_variant | MODIFIER |

| | | | | | | | | | | | | | | | |
|---|---|---|---|---|---|---|---|---|---|---|---|---|---|---|---|
| 2 | 169738134 | AX-170506312 | 0 | G | A | 0.167 | 1.63E-01 | 1.82E-01 | 2.65E+00 | 2.73E+00 | 3.67E-01 | Yes | Tshz2 | intron_variant | MODIFIER |
| 2 | 169744399 | AX-97848463 | 0 | C | T | 0.167 | 1.63E-01 | 1.82E-01 | 2.65E+00 | 2.73E+00 | 3.67E-01 | Yes | Tshz2 | intron_variant | MODIFIER |
| 2 | 169749423 | AX-170535137 | 0 | C | T | 0.167 | 1.63E-01 | 1.82E-01 | 2.65E+00 | 2.73E+00 | 3.67E-01 | Yes | Tshz2 | intron_variant | MODIFIER |
| 2 | 169753150 | AX-97828096 | 0 | G | A | 0.167 | 1.63E-01 | 1.82E-01 | 2.65E+00 | 2.73E+00 | 3.67E-01 | Yes | Tshz2 | intron_variant | MODIFIER |
| 2 | 169764005 | AX-170353249 | 0 | G | A | 0.167 | 1.63E-01 | 1.82E-01 | 2.65E+00 | 2.73E+00 | 3.67E-01 | Yes | Tshz2 | intron_variant | MODIFIER |
| 2 | 169767848 | AX-170401090 | 0 | T | C | 0.167 | 1.63E-01 | 1.82E-01 | 2.65E+00 | 2.73E+00 | 3.67E-01 | Yes | Tshz2 | intron_variant | MODIFIER |
| 2 | 169768032 | AX-170497683 | 0 | A | G | 0.167 | 1.63E-01 | 1.82E-01 | 2.65E+00 | 2.73E+00 | 3.67E-01 | Yes | Tshz2 | intron_variant | MODIFIER |
| 2 | 169773955 | AX-97840426 | 0 | C | A | 0.167 | 1.63E-01 | 1.82E-01 | 2.65E+00 | 2.73E+00 | 3.67E-01 | Yes | Tshz2 | intron_variant | MODIFIER |
| 2 | 169779862 | AX-170152046 | 0 | T | A | 0.167 | 1.63E-01 | 1.82E-01 | 2.65E+00 | 2.73E+00 | 3.67E-01 | Yes | Tshz2 | intron_variant | MODIFIER |
| 2 | 169783476 | AX-170362394 | 0 | G | A | 0.167 | 1.63E-01 | 1.82E-01 | 2.65E+00 | 2.73E+00 | 3.67E-01 | Yes | Tshz2 | intron_variant | MODIFIER |
| 2 | 169836130 | AX-97817674 | 0 | G | A | 0.167 | 1.63E-01 | 1.82E-01 | 2.65E+00 | 2.73E+00 | 3.67E-01 | Yes | Tshz2 | intron_variant | MODIFIER |
| 2 | 169836242 | AX-170476203 | 0 | C | G | 0.167 | 1.63E-01 | 1.82E-01 | 2.65E+00 | 2.73E+00 | 3.67E-01 | Yes | Tshz2 | intron_variant | MODIFIER |
| 2 | 169841223 | AX-170481322 | 0 | C | T | 0.167 | 1.63E-01 | 1.82E-01 | 2.65E+00 | 2.73E+00 | 3.67E-01 | Yes | Tshz2 | intron_variant | MODIFIER |
| 2 | 169847756 | AX-170325919 | 0 | C | G | 0.167 | 1.63E-01 | 1.82E-01 | 2.65E+00 | 2.73E+00 | 3.67E-01 | Yes | Tshz2 | intron_variant | MODIFIER |
| 2 | 169858154 | AX-170088831 | 0 | A | C | 0.167 | 1.63E-01 | 1.82E-01 | 2.65E+00 | 2.73E+00 | 3.67E-01 | Yes | Tshz2 | intron_variant | MODIFIER |
| 2 | 169870723 | AX-170176026 | 0 | A | C | 0.167 | 1.63E-01 | 1.82E-01 | 2.65E+00 | 2.73E+00 | 3.67E-01 | Yes | Tshz2 | intron_variant | MODIFIER |
| 2 | 169871722 | AX-170388822 | 0 | G | A | 0.167 | 1.63E-01 | 1.82E-01 | 2.65E+00 | 2.73E+00 | 3.67E-01 | Yes | Tshz2 | intron_variant | MODIFIER |
| 2 | 169875757 | AX-170471714 | 0 | T | C | 0.167 | 1.63E-01 | 1.82E-01 | 2.65E+00 | 2.73E+00 | 3.67E-01 | Yes | Tshz2 | intron_variant | MODIFIER |
| 2 | 169875800 | AX-170094485 | 0 | C | T | 0.167 | 1.63E-01 | 1.82E-01 | 2.65E+00 | 2.73E+00 | 3.67E-01 | Yes | Tshz2 | intron_variant | MODIFIER |
| 2 | 169897347 | AX-170242956 | 0 | G | C | 0.167 | 1.63E-01 | 1.82E-01 | 2.65E+00 | 2.73E+00 | 3.67E-01 | Yes | Tshz2 | intron_variant | MODIFIER |
| 2 | 169898504 | AX-170180306 | 0 | A | G | 0.167 | 1.63E-01 | 1.82E-01 | 2.65E+00 | 2.73E+00 | 3.67E-01 | Yes | Tshz2 | intron_variant | MODIFIER |
| 2 | 169899430 | AX-170304689 | 0 | A | G | 0.167 | 1.63E-01 | 1.82E-01 | 2.65E+00 | 2.73E+00 | 3.67E-01 | Yes | Tshz2 | intron_variant | MODIFIER |
| 2 | 169903673 | AX-170396001 | 0 | C | T | 0.167 | 1.63E-01 | 1.82E-01 | 2.65E+00 | 2.73E+00 | 3.67E-01 | Yes | Tshz2 | intron_variant | MODIFIER |
| 2 | 169916707 | AX-97856356 | 0 | C | T | 0.167 | 1.63E-01 | 1.82E-01 | 2.65E+00 | 2.73E+00 | 3.67E-01 | Yes | Tshz2 | downstream_gene_variant | MODIFIER |
| 2 | 169927019 | AX-97829630 | 0 | G | A | 0.167 | 1.63E-01 | 1.82E-01 | 2.65E+00 | 2.73E+00 | 3.67E-01 | Yes | Tshz2 | intron_variant | MODIFIER |
| 2 | 169948325 | AX-170331852 | 0 | T | C | 0.167 | 1.63E-01 | 1.82E-01 | 2.65E+00 | 2.73E+00 | 3.67E-01 | Yes | Tshz2 | intron_variant | MODIFIER |
| 2 | 169955213 | AX-170323051 | 0 | G | A | 0.167 | 1.63E-01 | 1.82E-01 | 2.65E+00 | 2.73E+00 | 3.67E-01 | Yes | Tshz2 | intron_variant | MODIFIER |
| 2 | 169960291 | AX-170235134 | 0 | A | G | 0.167 | 1.63E-01 | 1.82E-01 | 2.65E+00 | 2.73E+00 | 3.67E-01 | Yes | Tshz2 | intron_variant | MODIFIER |
| 2 | 170101311 | AX-97790942 | 0 | C | T | 0.167 | 1.63E-01 | 1.82E-01 | 2.65E+00 | 2.73E+00 | 3.67E-01 | Yes | | regulatory_region_variant | MODIFIER |
| 2 | 170277878 | AX-97839902 | 0 | T | C | 0.167 | 1.63E-01 | 1.82E-01 | 2.65E+00 | 2.73E+00 | 3.67E-01 | Yes | | regulatory_region_variant | MODIFIER |
| 2 | 170390902 | AX-97786786 | 0 | A | T | 0.167 | 1.63E-01 | 1.82E-01 | 2.65E+00 | 2.73E+00 | 3.67E-01 | Yes | Bcas1 | intron_variant | MODIFIER |
| 2 | 170406608 | AX-170399018 | 0 | A | G | 0.167 | 1.63E-01 | 1.82E-01 | 2.65E+00 | 2.73E+00 | 3.67E-01 | Yes | Bcas1 | missense_variant | MODERATE |
| 2 | 170406637 | AX-97849093 | 0 | C | T | 0.167 | 1.63E-01 | 1.82E-01 | 2.65E+00 | 2.73E+00 | 3.67E-01 | Yes | Bcas1 | synonymous_variant | LOW |
| 2 | 170407914 | AX-170137992 | 0 | G | A | 0.167 | 1.63E-01 | 1.82E-01 | 2.65E+00 | 2.73E+00 | 3.67E-01 | Yes | Bcas1 | intron_variant | MODIFIER |
| 16 | 98017278 | AX-170093581 | 0 | A | T | 0.465 | 6.77E-02 | 7.56E-02 | 2.88E+00 | 3.01E+00 | 3.67E-01 | no | B230307C23Rik | downstream_gene_variant | MODIFIER |
| 16 | 35438075 | AX-170395647 | 1 | G | C | 0.509 | 7.10E-02 | 7.87E-02 | 2.56E+00 | 2.62E+00 | 3.67E-01 | no | Pdia5 | intron_variant | MODIFIER |
| 7 | 144648821 | AX-170269672 | 0 | T | G | 0.211 | -6.14E-02 | 6.90E-02 | 2.65E+00 | 2.79E+00 | 3.67E-01 | no | Ano1 | intron_variant | MODIFIER |
| 1 | 150681134 | AX-170128616 | 1 | G | A | 0.205 | 1.22E-01 | 1.37E-01 | 2.68E+00 | 2.73E+00 | 3.67E-01 | no | Hmcn1 | intron_variant | MODIFIER |
| 3 | 20946633 | AX-170519458 | 1 | C | T | 0.143 | -9.19E-02 | 1.02E-01 | 2.56E+00 | 2.65E+00 | 3.67E-01 | no | | intergenic_variant | MODIFIER |
| 10 | 18159754 | AX-97810758 | 0 | C | T | 0.202 | 9.33E-02 | 1.03E-01 | 2.99E+00 | 3.09E+00 | 3.67E-01 | no | Ect2l | missense_variant | MODERATE |
| 10 | 18160721 | AX-170348221 | 0 | A | G | 0.202 | 9.33E-02 | 1.03E-01 | 2.99E+00 | 3.09E+00 | 3.67E-01 | no | Ect2l | intron_variant | MODIFIER |
| 10 | 18205035 | AX-170392252 | 0 | C | T | 0.202 | 9.33E-02 | 1.03E-01 | 2.99E+00 | 3.09E+00 | 3.67E-01 | no | Ect2l | intron_variant | MODIFIER |
| 10 | 18220847 | AX-170144079 | 0 | T | G | 0.202 | 9.33E-02 | 1.03E-01 | 2.99E+00 | 3.09E+00 | 3.67E-01 | no | Ccdc28a | intron_variant | MODIFIER |
| 10 | 18223621 | AX-170134086 | 0 | G | A | 0.202 | 9.33E-02 | 1.03E-01 | 2.99E+00 | 3.09E+00 | 3.67E-01 | no | Ccdc28a | intron_variant | MODIFIER |
| 10 | 18237710 | AX-170190927 | 0 | A | G | 0.202 | 9.33E-02 | 1.03E-01 | 2.99E+00 | 3.09E+00 | 3.67E-01 | no | Ccdc28a | upstream_gene_variant | MODIFIER |
| 10 | 18299595 | AX-170522067 | 0 | T | C | 0.202 | 9.33E-02 | 1.03E-01 | 2.99E+00 | 3.09E+00 | 3.67E-01 | no | Gm48229 | downstream_gene_variant | MODIFIER |
| 10 | 18299788 | AX-170311061 | 0 | G | A | 0.202 | 9.33E-02 | 1.03E-01 | 2.99E+00 | 3.09E+00 | 3.67E-01 | no | Gm48229 | downstream_gene_variant | MODIFIER |
| 10 | 18352983 | AX-170002848 | 0 | C | T | 0.202 | 9.33E-02 | 1.03E-01 | 2.99E+00 | 3.09E+00 | 3.67E-01 | no | Nhsl1 | intron_variant | MODIFIER |
| 10 | 18400820 | AX-170321009 | 0 | C | T | 0.202 | 9.33E-02 | 1.03E-01 | 2.99E+00 | 3.09E+00 | 3.67E-01 | no | Nhsl1 | intron_variant,non_coding_transcript_variant | MODIFIER |
| 10 | 18409771 | AX-170002847 | 0 | G | A | 0.202 | 9.33E-02 | 1.03E-01 | 2.99E+00 | 3.09E+00 | 3.67E-01 | no | Nhsl1 | intron_variant | MODIFIER |
| 10 | 18416923 | AX-170320085 | 0 | C | T | 0.202 | 9.33E-02 | 1.03E-01 | 2.99E+00 | 3.09E+00 | 3.67E-01 | no | Nhsl1 | intron_variant | MODIFIER |
| 10 | 18420472 | AX-170396828 | 0 | T | G | 0.202 | 9.33E-02 | 1.03E-01 | 2.99E+00 | 3.09E+00 | 3.67E-01 | no | Nhsl1 | intron_variant | MODIFIER |
| 10 | 18438898 | AX-97814930 | 0 | G | A | 0.202 | 9.33E-02 | 1.03E-01 | 2.99E+00 | 3.09E+00 | 3.67E-01 | no | Nhsl1 | intron_variant | MODIFIER |
| 10 | 18471890 | AX-97834570 | 0 | A | C | 0.202 | 9.33E-02 | 1.03E-01 | 2.99E+00 | 3.09E+00 | 3.67E-01 | no | Nhsl1 | intron_variant | MODIFIER |

| | | | | | | | | | | | | | | | |
|---|---|---|---|---|---|---|---|---|---|---|---|---|---|---|---|
| 10 | 18539355 | AX-170247972 | 0 | A | T | 0.202 | 9.33E-02 | 1.03E-01 | 2.99E+00 | 3.09E+00 | 3.67E-01 | no | Hebp2 | downstream_gene_variant | MODIFIER |
| 10 | 18541336 | AX-97853355 | 0 | C | T | 0.202 | 9.33E-02 | 1.03E-01 | 2.99E+00 | 3.09E+00 | 3.67E-01 | no | Hebp2 | splice_region_variant,intron_variant | LOW |
| 10 | 18541558 | AX-170166742 | 0 | C | G | 0.202 | 9.33E-02 | 1.03E-01 | 2.99E+00 | 3.09E+00 | 3.67E-01 | no | Hebp2 | intron_variant | MODIFIER |
| 10 | 18542178 | AX-170426009 | 0 | T | A | 0.202 | 9.33E-02 | 1.03E-01 | 2.99E+00 | 3.09E+00 | 3.67E-01 | no | Hebp2 | intron_variant | MODIFIER |
| 10 | 18590003 | AX-170166406 | 0 | C | T | 0.202 | 9.33E-02 | 1.03E-01 | 2.99E+00 | 3.09E+00 | 3.67E-01 | no | Arfgef3 | intron_variant | MODIFIER |
| 10 | 18590358 | AX-170086758 | 0 | T | C | 0.202 | 9.33E-02 | 1.03E-01 | 2.99E+00 | 3.09E+00 | 3.67E-01 | no | Arfgef3 | intron_variant | MODIFIER |
| 10 | 18591154 | AX-170244405 | 0 | T | C | 0.202 | 9.33E-02 | 1.03E-01 | 2.99E+00 | 3.09E+00 | 3.67E-01 | no | Arfgef3 | intron_variant | MODIFIER |
| 10 | 18597496 | AX-170427474 | 0 | T | C | 0.202 | 9.33E-02 | 1.03E-01 | 2.99E+00 | 3.09E+00 | 3.67E-01 | no | Arfgef3 | intron_variant | MODIFIER |
| 10 | 18607255 | AX-97799979 | 0 | A | C | 0.202 | 9.33E-02 | 1.03E-01 | 2.99E+00 | 3.09E+00 | 3.67E-01 | no | Arfgef3 | intron_variant | MODIFIER |
| 10 | 18632942 | AX-170302356 | 0 | A | G | 0.202 | 9.33E-02 | 1.03E-01 | 2.99E+00 | 3.09E+00 | 3.67E-01 | no | Arfgef3 | intron_variant | MODIFIER |
| 10 | 18644861 | AX-170374253 | 0 | C | T | 0.202 | 9.33E-02 | 1.03E-01 | 2.99E+00 | 3.09E+00 | 3.67E-01 | no | Arfgef3 | intron_variant | MODIFIER |
| 10 | 18720326 | AX-97814956 | 0 | C | T | 0.202 | 9.33E-02 | 1.03E-01 | 2.99E+00 | 3.09E+00 | 3.67E-01 | no | Arfgef3 | intron_variant | MODIFIER |
| 10 | 18720711 | AX-170172233 | 0 | A | C | 0.202 | 9.33E-02 | 1.03E-01 | 2.99E+00 | 3.09E+00 | 3.67E-01 | no | Arfgef3 | intron_variant | MODIFIER |
| 10 | 18789652 | AX-170272765 | 0 | A | C | 0.202 | 9.33E-02 | 1.03E-01 | 2.99E+00 | 3.09E+00 | 3.67E-01 | no | Gm4922 | upstream_gene_variant | MODIFIER |
| 10 | 18796850 | AX-170185561 | 0 | A | G | 0.202 | 9.33E-02 | 1.03E-01 | 2.99E+00 | 3.09E+00 | 3.67E-01 | no | Gm32926 | intron_variant,non_coding_transcript_variant | MODIFIER |
| 10 | 18799201 | AX-170168116 | 0 | C | T | 0.202 | 9.33E-02 | 1.03E-01 | 2.99E+00 | 3.09E+00 | 3.67E-01 | no | Gm32926 | intron_variant,non_coding_transcript_variant | MODIFIER |
| 10 | 18800800 | AX-170218323 | 0 | T | C | 0.202 | 9.33E-02 | 1.03E-01 | 2.99E+00 | 3.09E+00 | 3.67E-01 | no | Gm32926 | intron_variant,non_coding_transcript_variant | MODIFIER |
| 10 | 18803668 | AX-97820235 | 0 | C | A | 0.202 | 9.33E-02 | 1.03E-01 | 2.99E+00 | 3.09E+00 | 3.67E-01 | no | Gm32926 | intron_variant,non_coding_transcript_variant | MODIFIER |
| 10 | 18855988 | AX-97814958 | 0 | T | C | 0.202 | 9.33E-02 | 1.03E-01 | 2.99E+00 | 3.09E+00 | 3.67E-01 | no | Perp | 3_prime_UTR_variant | MODIFIER |
| 10 | 18862528 | AX-170404413 | 0 | T | C | 0.202 | 9.33E-02 | 1.03E-01 | 2.99E+00 | 3.09E+00 | 3.67E-01 | no | | intergenic_variant | MODIFIER |
| 10 | 18869683 | AX-170002859 | 0 | T | C | 0.202 | 9.33E-02 | 1.03E-01 | 2.99E+00 | 3.09E+00 | 3.67E-01 | no | | intergenic_variant | MODIFIER |
| 10 | 18877313 | AX-170254707 | 0 | G | A | 0.202 | 9.33E-02 | 1.03E-01 | 2.99E+00 | 3.09E+00 | 3.67E-01 | no | Gm48483 | downstream_gene_variant | MODIFIER |
| 10 | 18877535 | AX-170264475 | 0 | C | A | 0.202 | 9.33E-02 | 1.03E-01 | 2.99E+00 | 3.09E+00 | 3.67E-01 | no | Gm48483 | downstream_gene_variant | MODIFIER |
| 10 | 18879255 | AX-170327192 | 0 | G | A | 0.202 | 9.33E-02 | 1.03E-01 | 2.99E+00 | 3.09E+00 | 3.67E-01 | no | Gm48483 | downstream_gene_variant | MODIFIER |
| 10 | 18890676 | AX-170002858 | 0 | G | A | 0.202 | 9.33E-02 | 1.03E-01 | 2.99E+00 | 3.09E+00 | 3.67E-01 | no | Gm48483 | upstream_gene_variant | MODIFIER |
| 10 | 18890826 | AX-170219961 | 0 | C | T | 0.202 | 9.33E-02 | 1.03E-01 | 2.99E+00 | 3.09E+00 | 3.67E-01 | no | Gm48483 | upstream_gene_variant | MODIFIER |
| 10 | 18896257 | AX-97789106 | 0 | G | A | 0.202 | 9.33E-02 | 1.03E-01 | 2.99E+00 | 3.09E+00 | 3.67E-01 | no | | intergenic_variant | MODIFIER |
| 10 | 18924817 | AX-170198309 | 0 | T | C | 0.202 | 9.33E-02 | 1.03E-01 | 2.99E+00 | 3.09E+00 | 3.67E-01 | no | | intergenic_variant | MODIFIER |
| 10 | 18927875 | AX-170044253 | 0 | A | G | 0.202 | 9.33E-02 | 1.03E-01 | 2.99E+00 | 3.09E+00 | 3.67E-01 | no | | intergenic_variant | MODIFIER |
| 10 | 18939710 | AX-170467496 | 0 | A | T | 0.202 | 9.33E-02 | 1.03E-01 | 2.99E+00 | 3.09E+00 | 3.67E-01 | no | | intergenic_variant | MODIFIER |
| 10 | 18940727 | AX-97847538 | 0 | T | C | 0.202 | 9.33E-02 | 1.03E-01 | 2.99E+00 | 3.09E+00 | 3.67E-01 | no | | intergenic_variant | MODIFIER |
| 10 | 18943616 | AX-170161056 | 0 | A | C | 0.202 | 9.33E-02 | 1.03E-01 | 2.99E+00 | 3.09E+00 | 3.67E-01 | no | | intergenic_variant | MODIFIER |
| 10 | 18991691 | AX-170181552 | 0 | T | C | 0.202 | 9.33E-02 | 1.03E-01 | 2.99E+00 | 3.09E+00 | 3.67E-01 | no | | intergenic_variant | MODIFIER |
| 10 | 18994514 | AX-97812223 | 0 | G | T | 0.202 | 9.33E-02 | 1.03E-01 | 2.99E+00 | 3.09E+00 | 3.67E-01 | no | | intergenic_variant | MODIFIER |
| 10 | 19008783 | AX-170238346 | 0 | A | G | 0.202 | 9.33E-02 | 1.03E-01 | 2.99E+00 | 3.09E+00 | 3.67E-01 | no | Tnfaip3 | intron_variant | MODIFIER |
| 10 | 19011361 | AX-170230350 | 0 | T | C | 0.202 | 9.33E-02 | 1.03E-01 | 2.99E+00 | 3.09E+00 | 3.67E-01 | no | Tnfaip3 | intron_variant | MODIFIER |
| 10 | 19012967 | AX-97845615 | 0 | G | A | 0.202 | 9.33E-02 | 1.03E-01 | 2.99E+00 | 3.09E+00 | 3.67E-01 | no | Tnfaip3 | intron_variant | MODIFIER |
| 10 | 19022892 | AX-170528726 | 0 | T | A | 0.202 | 9.33E-02 | 1.03E-01 | 2.99E+00 | 3.09E+00 | 3.67E-01 | no | | intergenic_variant | MODIFIER |
| 10 | 19023012 | AX-170103153 | 0 | C | T | 0.202 | 9.33E-02 | 1.03E-01 | 2.99E+00 | 3.09E+00 | 3.67E-01 | no | | intergenic_variant | MODIFIER |
| 10 | 19023089 | AX-170128897 | 0 | G | A | 0.202 | 9.33E-02 | 1.03E-01 | 2.99E+00 | 3.09E+00 | 3.67E-01 | no | | intergenic_variant | MODIFIER |
| 10 | 19023673 | AX-170512171 | 0 | T | C | 0.202 | 9.33E-02 | 1.03E-01 | 2.99E+00 | 3.09E+00 | 3.67E-01 | no | | regulatory_region_variant | MODIFIER |
| 10 | 19027093 | AX-170202661 | 0 | C | T | 0.202 | 9.33E-02 | 1.03E-01 | 2.99E+00 | 3.09E+00 | 3.67E-01 | no | | intergenic_variant | MODIFIER |
| 10 | 19033597 | AX-170455741 | 0 | T | C | 0.202 | 9.33E-02 | 1.03E-01 | 2.99E+00 | 3.09E+00 | 3.67E-01 | no | | intergenic_variant | MODIFIER |
| 10 | 19034526 | AX-170044254 | 0 | C | T | 0.202 | 9.33E-02 | 1.03E-01 | 2.99E+00 | 3.09E+00 | 3.67E-01 | no | | regulatory_region_variant | MODIFIER |
| 10 | 19039300 | AX-97856349 | 0 | A | G | 0.202 | 9.33E-02 | 1.03E-01 | 2.99E+00 | 3.09E+00 | 3.67E-01 | no | | intergenic_variant | MODIFIER |
| 10 | 19109935 | AX-170173118 | 0 | A | G | 0.202 | 9.33E-02 | 1.03E-01 | 2.99E+00 | 3.09E+00 | 3.67E-01 | no | | intergenic_variant | MODIFIER |
| 10 | 19130964 | AX-170121839 | 0 | T | A | 0.202 | 9.33E-02 | 1.03E-01 | 2.99E+00 | 3.09E+00 | 3.67E-01 | no | | intergenic_variant | MODIFIER |
| 3 | 154801343 | AX-97805471 | 0 | T | C | 0.36 | 6.33E-02 | 7.09E-02 | 2.82E+00 | 2.92E+00 | 3.67E-01 | no | Tnni3k | intron_variant | MODIFIER |
| 10 | 54877090 | AX-97832938 | 1 | G | T | 0.304 | 6.99E-02 | 7.65E-02 | 3.03E+00 | 3.18E+00 | 3.67E-01 | no | | intergenic_variant | MODIFIER |
| 15 | 4827984 | AX-170485210 | 1 | G | A | 0.241 | -9.53E-02 | 1.06E-01 | 2.63E+00 | 2.68E+00 | 3.68E-01 | no | Gm49073 | upstream_gene_variant | MODIFIER |
| 15 | 6472757 | AX-170476471 | 1 | G | T | 0.241 | -9.53E-02 | 1.06E-01 | 2.63E+00 | 2.68E+00 | 3.68E-01 | no | C9 | intron_variant | MODIFIER |
| 15 | 6963369 | AX-170159221 | 1 | C | T | 0.241 | -9.53E-02 | 1.06E-01 | 2.63E+00 | 2.68E+00 | 3.68E-01 | no | | intergenic_variant | MODIFIER |
| 15 | 19080441 | AX-170011173 | 1 | G | A | 0.241 | -9.53E-02 | 1.06E-01 | 2.63E+00 | 2.68E+00 | 3.68E-01 | no | | intergenic_variant | MODIFIER |
| 15 | 20407107 | AX-170323377 | 1 | C | T | 0.241 | -9.53E-02 | 1.06E-01 | 2.63E+00 | 2.68E+00 | 3.68E-01 | no | | intergenic_variant | MODIFIER |

| | | | | | | | | | | | | | | | |
|---|---|---|---|---|---|---|---|---|---|---|---|---|---|---|---|
| 9 | 56586872 | AX-170082505 | 1 | G | T | 0.205 | -1.00E-01 | 1.12E-01 | 2.81E+00 | 2.90E+00 | 3.68E-01 | no | | intergenic_variant | MODIFIER |
| 1 | 128230646 | AX-97805723 | 0 | A | G | 0.553 | 7.49E-02 | 8.35E-02 | 2.60E+00 | 2.67E+00 | 3.68E-01 | no | R3hdm1 | intron_variant | MODIFIER |
| 7 | 70914351 | AX-97839044 | 0 | C | T | 0.36 | -8.86E-02 | 9.93E-02 | 2.84E+00 | 2.94E+00 | 3.68E-01 | no | Gm42397 | non_coding_transcript_exon_variant | MODIFIER |
| 5 | 50936576 | AX-170229665 | 0 | G | A | 0.482 | 9.79E-02 | 1.05E-01 | 2.40E+00 | 2.45E+00 | 3.68E-01 | no | | intergenic_variant | MODIFIER |
| 4 | 79134994 | AX-170294928 | 0 | T | C | 0.447 | -7.52E-02 | 8.35E-02 | 2.57E+00 | 2.64E+00 | 3.68E-01 | no | | intergenic_variant | MODIFIER |
| 4 | 108572681 | AX-170023636 | 1 | C | T | 0.045 | 1.45E-01 | 1.64E-01 | 2.64E+00 | 2.79E+00 | 3.68E-01 | Yes | Prpf38a | intron_variant | MODIFIER |
| 2 | 168705972 | AX-97833126 | 2 | T | C | 0.145 | 1.34E-01 | 1.50E-01 | 2.65E+00 | 2.76E+00 | 3.68E-01 | no | Atp9a | intron_variant | MODIFIER |
| 2 | 169692343 | AX-170227917 | 2 | G | A | 0.145 | 1.34E-01 | 1.50E-01 | 2.65E+00 | 2.76E+00 | 3.68E-01 | no | Tshz2 | intron_variant | MODIFIER |
| 9 | 94856708 | AX-170431637 | 0 | A | G | 0.123 | -1.11E-01 | 1.24E-01 | 2.61E+00 | 2.69E+00 | 3.68E-01 | no | Slc9a9 | intron_variant | MODIFIER |
| 5 | 45056950 | AX-170272927 | 0 | G | A | 0.202 | -9.71E-02 | 1.08E-01 | 2.58E+00 | 2.66E+00 | 3.68E-01 | no | D5Ertd615e | intron_variant,non_coding_transcript_variant | MODIFIER |
| 1 | 169301062 | AX-97805780 | 2 | C | A | 0.209 | 1.19E-01 | 1.33E-01 | 2.65E+00 | 2.71E+00 | 3.68E-01 | no | | intergenic_variant | MODIFIER |
| 9 | 96430940 | AX-170117365 | 2 | G | A | 0.145 | -1.00E-01 | 1.11E-01 | 2.57E+00 | 2.65E+00 | 3.68E-01 | no | BC043934 | downstream_gene_variant | MODIFIER |
| 2 | 169423691 | AX-170118383 | 1 | A | T | 0.17 | 1.62E-01 | 1.81E-01 | 2.63E+00 | 2.71E+00 | 3.68E-01 | Yes | | intergenic_variant | MODIFIER |
| 4 | 98518727 | AX-170152314 | 1 | T | C | 0.223 | 1.04E-01 | 1.16E-01 | 2.57E+00 | 2.64E+00 | 3.68E-01 | no | Patj | intron_variant | MODIFIER |
| 15 | 79570673 | AX-170087919 | 0 | A | T | 0.561 | 7.17E-02 | 7.95E-02 | 2.55E+00 | 2.63E+00 | 3.68E-01 | no | Dmc1 | intron_variant | MODIFIER |
| 15 | 80251623 | AX-170012081 | 0 | G | A | 0.561 | 7.17E-02 | 7.95E-02 | 2.55E+00 | 2.63E+00 | 3.68E-01 | no | Mief1 | 3_prime_UTR_variant | MODIFIER |
| 5 | 51026694 | AX-97799645 | 0 | C | T | 0.491 | 9.35E-02 | 1.02E-01 | 2.45E+00 | 2.50E+00 | 3.68E-01 | no | | intergenic_variant | MODIFIER |
| 5 | 128250723 | AX-170137046 | 0 | C | T | 0.149 | -8.68E-02 | 8.93E-02 | 2.27E+00 | 2.35E+00 | 3.68E-01 | no | Tmem132d | intron_variant | MODIFIER |
| 5 | 128250911 | AX-170234180 | 0 | C | A | 0.149 | -8.68E-02 | 8.93E-02 | 2.27E+00 | 2.35E+00 | 3.68E-01 | no | Tmem132d | intron_variant | MODIFIER |
| 5 | 128251310 | AX-170204673 | 0 | C | T | 0.149 | -8.68E-02 | 8.93E-02 | 2.27E+00 | 2.35E+00 | 3.68E-01 | no | Tmem132d | intron_variant | MODIFIER |
| 5 | 128252470 | AX-170153886 | 0 | C | G | 0.149 | -8.68E-02 | 8.93E-02 | 2.27E+00 | 2.35E+00 | 3.68E-01 | no | Tmem132d | intron_variant | MODIFIER |
| 5 | 128252590 | AX-97848654 | 0 | A | G | 0.149 | -8.68E-02 | 8.93E-02 | 2.27E+00 | 2.35E+00 | 3.68E-01 | no | Tmem132d | intron_variant | MODIFIER |
| 5 | 128260035 | AX-170026137 | 0 | C | T | 0.149 | -8.68E-02 | 8.93E-02 | 2.27E+00 | 2.35E+00 | 3.68E-01 | no | Tmem132d | intron_variant | MODIFIER |
| 5 | 128261496 | AX-170026134 | 0 | C | T | 0.149 | -8.68E-02 | 8.93E-02 | 2.27E+00 | 2.35E+00 | 3.68E-01 | no | Tmem132d | intron_variant | MODIFIER |
| 5 | 128270527 | AX-170140629 | 0 | G | A | 0.149 | -8.68E-02 | 8.93E-02 | 2.27E+00 | 2.35E+00 | 3.68E-01 | no | Tmem132d | intron_variant | MODIFIER |
| 5 | 128275402 | AX-170073468 | 0 | G | A | 0.149 | -8.68E-02 | 8.93E-02 | 2.27E+00 | 2.35E+00 | 3.68E-01 | no | Tmem132d | intron_variant | MODIFIER |
| 5 | 128278387 | AX-97788550 | 0 | T | C | 0.149 | -8.68E-02 | 8.93E-02 | 2.27E+00 | 2.35E+00 | 3.68E-01 | no | Tmem132d | intron_variant | MODIFIER |
| 5 | 128287816 | AX-170026135 | 0 | T | C | 0.149 | -8.68E-02 | 8.93E-02 | 2.27E+00 | 2.35E+00 | 3.68E-01 | no | Tmem132d | intron_variant | MODIFIER |
| 5 | 128291411 | AX-170026139 | 0 | A | G | 0.149 | -8.68E-02 | 8.93E-02 | 2.27E+00 | 2.35E+00 | 3.68E-01 | no | Tmem132d | intron_variant | MODIFIER |
| 5 | 128349299 | AX-170129035 | 0 | T | C | 0.149 | -8.68E-02 | 8.93E-02 | 2.27E+00 | 2.35E+00 | 3.68E-01 | no | Tmem132d | intron_variant | MODIFIER |
| 5 | 128407970 | AX-97811989 | 0 | A | T | 0.149 | -8.68E-02 | 8.93E-02 | 2.27E+00 | 2.35E+00 | 3.68E-01 | no | Tmem132d | intron_variant | MODIFIER |
| 5 | 128450420 | AX-170182027 | 0 | A | G | 0.149 | -8.68E-02 | 8.93E-02 | 2.27E+00 | 2.35E+00 | 3.68E-01 | no | Gm42500 | downstream_gene_variant | MODIFIER |
| 5 | 128450467 | AX-170411443 | 0 | G | A | 0.149 | -8.68E-02 | 8.93E-02 | 2.27E+00 | 2.35E+00 | 3.68E-01 | no | Gm42500 | downstream_gene_variant | MODIFIER |
| 5 | 128450531 | AX-170360061 | 0 | A | G | 0.149 | -8.68E-02 | 8.93E-02 | 2.27E+00 | 2.35E+00 | 3.68E-01 | no | Gm42500 | downstream_gene_variant | MODIFIER |
| 5 | 128455138 | AX-170448539 | 0 | A | G | 0.149 | -8.68E-02 | 8.93E-02 | 2.27E+00 | 2.35E+00 | 3.68E-01 | no | | intergenic_variant | MODIFIER |
| 5 | 128455627 | AX-170285860 | 0 | A | G | 0.149 | -8.68E-02 | 8.93E-02 | 2.27E+00 | 2.35E+00 | 3.68E-01 | no | | intergenic_variant | MODIFIER |
| 5 | 128455885 | AX-170198420 | 0 | T | A | 0.149 | -8.68E-02 | 8.93E-02 | 2.27E+00 | 2.35E+00 | 3.68E-01 | no | | intergenic_variant | MODIFIER |
| 5 | 128456556 | AX-170418833 | 0 | A | T | 0.149 | -8.68E-02 | 8.93E-02 | 2.27E+00 | 2.35E+00 | 3.68E-01 | no | | intergenic_variant | MODIFIER |
| 5 | 128472892 | AX-170201366 | 0 | A | G | 0.149 | -8.68E-02 | 8.93E-02 | 2.27E+00 | 2.35E+00 | 3.68E-01 | no | | intergenic_variant | MODIFIER |
| 5 | 128482387 | AX-170219537 | 0 | G | A | 0.149 | -8.68E-02 | 8.93E-02 | 2.27E+00 | 2.35E+00 | 3.68E-01 | no | | intergenic_variant | MODIFIER |
| 5 | 128482424 | AX-170112534 | 0 | G | A | 0.149 | -8.68E-02 | 8.93E-02 | 2.27E+00 | 2.35E+00 | 3.68E-01 | no | | regulatory_region_variant | MODIFIER |
| 5 | 128517527 | AX-170265629 | 0 | G | A | 0.149 | -8.68E-02 | 8.93E-02 | 2.27E+00 | 2.35E+00 | 3.68E-01 | no | Gm42499 | upstream_gene_variant | MODIFIER |
| 5 | 128533981 | AX-97850859 | 0 | G | A | 0.149 | -8.68E-02 | 8.93E-02 | 2.27E+00 | 2.35E+00 | 3.68E-01 | no | | intergenic_variant | MODIFIER |
| 5 | 128553228 | AX-97800232 | 0 | C | T | 0.149 | -8.68E-02 | 8.93E-02 | 2.27E+00 | 2.35E+00 | 3.68E-01 | no | | regulatory_region_variant | MODIFIER |
| 5 | 128553368 | AX-170452705 | 0 | C | G | 0.149 | -8.68E-02 | 8.93E-02 | 2.27E+00 | 2.35E+00 | 3.68E-01 | no | | regulatory_region_variant | MODIFIER |
| 5 | 128560459 | AX-170403405 | 0 | A | G | 0.149 | -8.68E-02 | 8.93E-02 | 2.27E+00 | 2.35E+00 | 3.68E-01 | no | | intergenic_variant | MODIFIER |
| 5 | 128566339 | AX-170367816 | 0 | A | C | 0.149 | -8.68E-02 | 8.93E-02 | 2.27E+00 | 2.35E+00 | 3.68E-01 | no | | intergenic_variant | MODIFIER |
| 5 | 128572496 | AX-170285677 | 0 | G | A | 0.149 | -8.68E-02 | 8.93E-02 | 2.27E+00 | 2.35E+00 | 3.68E-01 | no | | regulatory_region_variant | MODIFIER |
| 5 | 128573768 | AX-170145614 | 0 | A | G | 0.149 | -8.68E-02 | 8.93E-02 | 2.27E+00 | 2.35E+00 | 3.68E-01 | no | | intergenic_variant | MODIFIER |
| 5 | 128593981 | AX-170353037 | 0 | G | A | 0.149 | -8.68E-02 | 8.93E-02 | 2.27E+00 | 2.35E+00 | 3.68E-01 | no | 5930412G12Rik | intron_variant,non_coding_transcript_variant | MODIFIER |
| 5 | 128595069 | AX-170073478 | 0 | C | T | 0.149 | -8.68E-02 | 8.93E-02 | 2.27E+00 | 2.35E+00 | 3.68E-01 | no | 5930412G12Rik | non_coding_transcript_exon_variant | MODIFIER |
| 5 | 128595348 | AX-170405874 | 0 | G | A | 0.149 | -8.68E-02 | 8.93E-02 | 2.27E+00 | 2.35E+00 | 3.68E-01 | no | 5930412G12Rik | non_coding_transcript_exon_variant | MODIFIER |
| 5 | 128595804 | AX-97791744 | 0 | G | A | 0.149 | -8.68E-02 | 8.93E-02 | 2.27E+00 | 2.35E+00 | 3.68E-01 | no | 5930412G12Rik | intron_variant,non_coding_transcript_variant | MODIFIER |
| 5 | 128762098 | AX-170184797 | 0 | G | A | 0.149 | -8.68E-02 | 8.93E-02 | 2.27E+00 | 2.35E+00 | 3.68E-01 | no | Rimbp2 | intron_variant | MODIFIER |

| | | | | | | | | | | | | | | | |
|---|---|---|---|---|---|---|---|---|---|---|---|---|---|---|---|
| 5 | 128762177 | AX-97810069 | 0 | A | G | 0.149 | -8.68E-02 | 8.93E-02 | 2.27E+00 | 2.35E+00 | 3.68E-01 | no | Rimbp2 | intron_variant | MODIFIER |
| 5 | 128773750 | AX-170522843 | 0 | T | C | 0.149 | -8.68E-02 | 8.93E-02 | 2.27E+00 | 2.35E+00 | 3.68E-01 | no | Rimbp2 | intron_variant | MODIFIER |
| 5 | 128787760 | AX-170026143 | 0 | C | T | 0.149 | -8.68E-02 | 8.93E-02 | 2.27E+00 | 2.35E+00 | 3.68E-01 | no | Rimbp2 | intron_variant | MODIFIER |
| 5 | 128795932 | AX-170026147 | 0 | T | C | 0.149 | -8.68E-02 | 8.93E-02 | 2.27E+00 | 2.35E+00 | 3.68E-01 | no | Rimbp2 | intron_variant | MODIFIER |
| 5 | 128798225 | AX-170125900 | 0 | C | T | 0.149 | -8.68E-02 | 8.93E-02 | 2.27E+00 | 2.35E+00 | 3.68E-01 | no | Rimbp2 | intron_variant | MODIFIER |
| 5 | 128801216 | AX-170123738 | 0 | T | G | 0.149 | -8.68E-02 | 8.93E-02 | 2.27E+00 | 2.35E+00 | 3.68E-01 | no | Rimbp2 | intron_variant | MODIFIER |
| 5 | 128801245 | AX-170337874 | 0 | G | A | 0.149 | -8.68E-02 | 8.93E-02 | 2.27E+00 | 2.35E+00 | 3.68E-01 | no | Rimbp2 | intron_variant | MODIFIER |
| 5 | 128802461 | AX-170099013 | 0 | A | T | 0.149 | -8.68E-02 | 8.93E-02 | 2.27E+00 | 2.35E+00 | 3.68E-01 | no | Rimbp2 | intron_variant | MODIFIER |
| 5 | 128802991 | AX-170259829 | 0 | T | C | 0.149 | -8.68E-02 | 8.93E-02 | 2.27E+00 | 2.35E+00 | 3.68E-01 | no | Rimbp2 | intron_variant | MODIFIER |
| 5 | 128805611 | AX-97833846 | 0 | A | G | 0.149 | -8.68E-02 | 8.93E-02 | 2.27E+00 | 2.35E+00 | 3.68E-01 | no | Rimbp2 | intron_variant | MODIFIER |
| 5 | 128829911 | AX-170356170 | 0 | T | C | 0.149 | -8.68E-02 | 8.93E-02 | 2.27E+00 | 2.35E+00 | 3.68E-01 | no | Rimbp2 | intron_variant | MODIFIER |
| 5 | 128830030 | AX-170372528 | 0 | C | T | 0.149 | -8.68E-02 | 8.93E-02 | 2.27E+00 | 2.35E+00 | 3.68E-01 | no | Rimbp2 | intron_variant | MODIFIER |
| 5 | 128832988 | AX-170095466 | 0 | G | A | 0.149 | -8.68E-02 | 8.93E-02 | 2.27E+00 | 2.35E+00 | 3.68E-01 | no | Rimbp2 | intron_variant | MODIFIER |
| 5 | 128834898 | AX-170236410 | 0 | G | A | 0.149 | -8.68E-02 | 8.93E-02 | 2.27E+00 | 2.35E+00 | 3.68E-01 | no | Rimbp2 | intron_variant | MODIFIER |
| 1 | 96252562 | AX-170131429 | 0 | T | A | 0.167 | 1.56E-01 | 1.75E-01 | 2.64E+00 | 2.74E+00 | 3.69E-01 | Yes | | intergenic_variant | MODIFIER |
| 13 | 38263822 | AX-170114899 | 0 | A | G | 0.193 | 8.98E-02 | 9.87E-02 | 3.02E+00 | 3.16E+00 | 3.69E-01 | no | | intergenic_variant | MODIFIER |
| 7 | 144066850 | AX-97837107 | 1 | C | T | 0.205 | -6.70E-02 | 7.54E-02 | 2.63E+00 | 2.76E+00 | 3.69E-01 | no | Shank2 | intron_variant | MODIFIER |
| 4 | 96832540 | AX-97843506 | 0 | T | C | 0.237 | 1.05E-01 | 1.17E-01 | 2.58E+00 | 2.63E+00 | 3.69E-01 | no | | intergenic_variant | MODIFIER |
| 4 | 96856223 | AX-170206299 | 0 | G | A | 0.237 | 1.05E-01 | 1.17E-01 | 2.58E+00 | 2.63E+00 | 3.69E-01 | no | | intergenic_variant | MODIFIER |
| 4 | 96871257 | AX-97787432 | 0 | A | G | 0.237 | 1.05E-01 | 1.17E-01 | 2.58E+00 | 2.63E+00 | 3.69E-01 | no | | intergenic_variant | MODIFIER |
| 4 | 96873812 | AX-170416228 | 0 | A | T | 0.237 | 1.05E-01 | 1.17E-01 | 2.58E+00 | 2.63E+00 | 3.69E-01 | no | | intergenic_variant | MODIFIER |
| 4 | 96876075 | AX-170172907 | 0 | A | C | 0.237 | 1.05E-01 | 1.17E-01 | 2.58E+00 | 2.63E+00 | 3.69E-01 | no | | intergenic_variant | MODIFIER |
| 4 | 96877942 | AX-170435041 | 0 | C | T | 0.237 | 1.05E-01 | 1.17E-01 | 2.58E+00 | 2.63E+00 | 3.69E-01 | no | | intergenic_variant | MODIFIER |
| 4 | 96878560 | AX-170512743 | 0 | T | C | 0.237 | 1.05E-01 | 1.17E-01 | 2.58E+00 | 2.63E+00 | 3.69E-01 | no | | intergenic_variant | MODIFIER |
| 4 | 96879393 | AX-97822150 | 0 | C | T | 0.237 | 1.05E-01 | 1.17E-01 | 2.58E+00 | 2.63E+00 | 3.69E-01 | no | | intergenic_variant | MODIFIER |
| 4 | 97128339 | AX-170106976 | 0 | G | A | 0.237 | 1.05E-01 | 1.17E-01 | 2.58E+00 | 2.63E+00 | 3.69E-01 | no | | intergenic_variant | MODIFIER |
| 4 | 97143141 | AX-97848242 | 0 | A | G | 0.237 | 1.05E-01 | 1.17E-01 | 2.58E+00 | 2.63E+00 | 3.69E-01 | no | | intergenic_variant | MODIFIER |
| 4 | 97551849 | AX-97811788 | 0 | A | G | 0.237 | 1.05E-01 | 1.17E-01 | 2.58E+00 | 2.63E+00 | 3.69E-01 | no | Gm12697 | downstream_gene_variant | MODIFIER |
| 4 | 97605804 | AX-97782552 | 0 | A | G | 0.237 | 1.05E-01 | 1.17E-01 | 2.58E+00 | 2.63E+00 | 3.69E-01 | no | E130114P18Rik | intron_variant | MODIFIER |
| 4 | 97614019 | AX-170314690 | 0 | G | A | 0.237 | 1.05E-01 | 1.17E-01 | 2.58E+00 | 2.63E+00 | 3.69E-01 | no | E130114P18Rik | intron_variant | MODIFIER |
| 4 | 97622855 | AX-170023470 | 0 | G | A | 0.237 | 1.05E-01 | 1.17E-01 | 2.58E+00 | 2.63E+00 | 3.69E-01 | no | E130114P18Rik | intron_variant | MODIFIER |
| 4 | 97626682 | AX-170341188 | 0 | T | G | 0.237 | 1.05E-01 | 1.17E-01 | 2.58E+00 | 2.63E+00 | 3.69E-01 | no | E130114P18Rik | intron_variant | MODIFIER |
| 4 | 97643378 | AX-97855134 | 0 | G | A | 0.237 | 1.05E-01 | 1.17E-01 | 2.58E+00 | 2.63E+00 | 3.69E-01 | no | E130114P18Rik | intron_variant | MODIFIER |
| 4 | 97652366 | AX-170439145 | 0 | A | G | 0.237 | 1.05E-01 | 1.17E-01 | 2.58E+00 | 2.63E+00 | 3.69E-01 | no | E130114P18Rik | intron_variant | MODIFIER |
| 4 | 97661013 | AX-97807121 | 0 | C | T | 0.237 | 1.05E-01 | 1.17E-01 | 2.58E+00 | 2.63E+00 | 3.69E-01 | no | E130114P18Rik | intron_variant | MODIFIER |
| 4 | 97672334 | AX-170319057 | 0 | A | G | 0.237 | 1.05E-01 | 1.17E-01 | 2.58E+00 | 2.63E+00 | 3.69E-01 | no | E130114P18Rik | intron_variant | MODIFIER |
| 4 | 97672478 | AX-170105297 | 0 | T | A | 0.237 | 1.05E-01 | 1.17E-01 | 2.58E+00 | 2.63E+00 | 3.69E-01 | no | E130114P18Rik | intron_variant | MODIFIER |
| 4 | 97689872 | AX-97798547 | 0 | A | G | 0.237 | 1.05E-01 | 1.17E-01 | 2.58E+00 | 2.63E+00 | 3.69E-01 | no | E130114P18Rik | intron_variant | MODIFIER |
| 4 | 97693989 | AX-170436103 | 0 | G | T | 0.237 | 1.05E-01 | 1.17E-01 | 2.58E+00 | 2.63E+00 | 3.69E-01 | no | E130114P18Rik | intron_variant | MODIFIER |
| 4 | 97700469 | AX-170532350 | 0 | G | A | 0.237 | 1.05E-01 | 1.17E-01 | 2.58E+00 | 2.63E+00 | 3.69E-01 | no | E130114P18Rik | intron_variant | MODIFIER |
| 4 | 97708822 | AX-170148942 | 0 | G | A | 0.237 | 1.05E-01 | 1.17E-01 | 2.58E+00 | 2.63E+00 | 3.69E-01 | no | E130114P18Rik | intron_variant | MODIFIER |
| 4 | 97708904 | AX-97798336 | 0 | A | G | 0.237 | 1.05E-01 | 1.17E-01 | 2.58E+00 | 2.63E+00 | 3.69E-01 | no | E130114P18Rik | intron_variant | MODIFIER |
| 4 | 97736774 | AX-97821259 | 0 | G | A | 0.237 | 1.05E-01 | 1.17E-01 | 2.58E+00 | 2.63E+00 | 3.69E-01 | no | E130114P18Rik | intron_variant | MODIFIER |
| 4 | 97754594 | AX-97852636 | 0 | A | C | 0.237 | 1.05E-01 | 1.17E-01 | 2.58E+00 | 2.63E+00 | 3.69E-01 | no | E130114P18Rik | intron_variant | MODIFIER |
| 4 | 97754930 | AX-170430559 | 0 | A | T | 0.237 | 1.05E-01 | 1.17E-01 | 2.58E+00 | 2.63E+00 | 3.69E-01 | no | E130114P18Rik | intron_variant | MODIFIER |
| 4 | 97790236 | AX-97809004 | 0 | T | C | 0.237 | 1.05E-01 | 1.17E-01 | 2.58E+00 | 2.63E+00 | 3.69E-01 | no | Nfia | intron_variant | MODIFIER |
| 4 | 97821242 | AX-97839330 | 0 | C | T | 0.237 | 1.05E-01 | 1.17E-01 | 2.58E+00 | 2.63E+00 | 3.69E-01 | no | Nfia | intron_variant | MODIFIER |
| 4 | 97886753 | AX-97839322 | 0 | G | A | 0.237 | 1.05E-01 | 1.17E-01 | 2.58E+00 | 2.63E+00 | 3.69E-01 | no | Nfia | intron_variant | MODIFIER |
| 4 | 97898379 | AX-170385084 | 0 | A | G | 0.237 | 1.05E-01 | 1.17E-01 | 2.58E+00 | 2.63E+00 | 3.69E-01 | no | Nfia | intron_variant | MODIFIER |
| 4 | 97923519 | AX-170145941 | 0 | G | A | 0.237 | 1.05E-01 | 1.17E-01 | 2.58E+00 | 2.63E+00 | 3.69E-01 | no | Nfia | intron_variant | MODIFIER |
| 4 | 97924065 | AX-97812293 | 0 | C | T | 0.237 | 1.05E-01 | 1.17E-01 | 2.58E+00 | 2.63E+00 | 3.69E-01 | no | Nfia | intron_variant | MODIFIER |
| 4 | 98092080 | AX-170023479 | 0 | A | G | 0.237 | 1.05E-01 | 1.17E-01 | 2.58E+00 | 2.63E+00 | 3.69E-01 | no | Nfia | intron_variant | MODIFIER |
| 4 | 98093785 | AX-170070115 | 0 | G | A | 0.237 | 1.05E-01 | 1.17E-01 | 2.58E+00 | 2.63E+00 | 3.69E-01 | no | Nfia | intron_variant | MODIFIER |
| 4 | 98120782 | AX-97813646 | 0 | G | A | 0.237 | 1.05E-01 | 1.17E-01 | 2.58E+00 | 2.63E+00 | 3.69E-01 | no | Nfia | downstream_gene_variant | MODIFIER |

| | | | | | | | | | | | | | | | |
|---|---|---|---|---|---|---|---|---|---|---|---|---|---|---|---|
| 4 | 98364742 | AX-97820890 | 0 | C | T | 0.237 | 1.05E-01 | 1.17E-01 | 2.58E+00 | 2.63E+00 | 3.69E-01 | no | Tm2d1 | intron_variant | MODIFIER |
| 4 | 98421909 | AX-97853393 | 0 | C | T | 0.237 | 1.05E-01 | 1.17E-01 | 2.58E+00 | 2.63E+00 | 3.69E-01 | no | Patj | intron_variant | MODIFIER |
| 4 | 98512159 | AX-170180308 | 0 | C | T | 0.237 | 1.05E-01 | 1.17E-01 | 2.58E+00 | 2.63E+00 | 3.69E-01 | no | Patj | intron_variant | MODIFIER |
| 4 | 98578951 | AX-170159586 | 0 | C | T | 0.237 | 1.05E-01 | 1.17E-01 | 2.58E+00 | 2.63E+00 | 3.69E-01 | no | Patj | intron_variant | MODIFIER |
| 1 | 115689665 | AX-168407126 | 0 | C | A | 0.167 | -1.50E-01 | 1.68E-01 | 2.89E+00 | 3.03E+00 | 3.69E-01 | no | Cntnap5a | intron_variant | MODIFIER |
| 15 | 68199403 | AX-170161752 | 0 | G | C | 0.351 | -6.97E-02 | 7.45E-02 | 2.38E+00 | 2.46E+00 | 3.69E-01 | no | Zfat | intron_variant,non_coding_transcript_variant | MODIFIER |
| 10 | 18524079 | AX-170002853 | 1 | G | A | 0.205 | 9.27E-02 | 1.03E-01 | 2.96E+00 | 3.06E+00 | 3.69E-01 | no | Nhsl1 | missense_variant | MODERATE |
| 8 | 65727652 | AX-170111817 | 1 | T | A | 0.143 | -1.27E-01 | 1.42E-01 | 2.57E+00 | 2.64E+00 | 3.69E-01 | no | MAR1 | intron_variant | MODIFIER |
| 18 | 58755039 | AX-170287279 | 0 | A | G | 0.395 | 8.63E-02 | 9.40E-02 | 2.45E+00 | 2.50E+00 | 3.69E-01 | no | | regulatory_region_variant | MODIFIER |
| 5 | 47689429 | AX-170208079 | 0 | C | T | 0.491 | 9.07E-02 | 9.92E-02 | 2.46E+00 | 2.53E+00 | 3.69E-01 | no | | intergenic_variant | MODIFIER |
| 8 | 88366042 | AX-97804054 | 1 | T | G | 0.17 | -1.17E-01 | 1.31E-01 | 2.72E+00 | 2.80E+00 | 3.69E-01 | no | Brd7 | upstream_gene_variant | MODIFIER |
| 1 | 191643370 | AX-170392994 | 0 | G | A | 0.263 | 8.93E-02 | 9.57E-02 | 2.38E+00 | 2.46E+00 | 3.69E-01 | no | | intergenic_variant | MODIFIER |
| 15 | 69260945 | AX-170260040 | 0 | A | G | 0.553 | 7.12E-02 | 7.96E-02 | 2.58E+00 | 2.66E+00 | 3.70E-01 | no | | intergenic_variant | MODIFIER |
| 15 | 75196936 | AX-170435885 | 0 | C | T | 0.553 | 7.12E-02 | 7.96E-02 | 2.58E+00 | 2.66E+00 | 3.70E-01 | no | Gm28117 | intron_variant,non_coding_transcript_variant | MODIFIER |
| 4 | 95752886 | AX-170465752 | 1 | A | G | 0.241 | 9.94E-02 | 1.07E-01 | 2.41E+00 | 2.48E+00 | 3.70E-01 | no | Fggy | intron_variant | MODIFIER |
| 16 | 76896853 | AX-170445266 | 0 | C | G | 0.474 | 6.58E-02 | 7.28E-02 | 2.51E+00 | 2.60E+00 | 3.70E-01 | no | 1700041M19Rik | intron_variant,non_coding_transcript_variant | MODIFIER |
| 15 | 6925469 | AX-170494215 | 1 | T | C | 0.241 | -9.50E-02 | 1.06E-01 | 2.61E+00 | 2.66E+00 | 3.70E-01 | no | | intergenic_variant | MODIFIER |
| 15 | 19318143 | AX-170419038 | 1 | G | A | 0.241 | -9.50E-02 | 1.06E-01 | 2.61E+00 | 2.66E+00 | 3.70E-01 | no | Gm49162 | intron_variant,non_coding_transcript_variant | MODIFIER |
| 15 | 20875375 | AX-170412776 | 1 | T | C | 0.241 | -9.50E-02 | 1.06E-01 | 2.61E+00 | 2.66E+00 | 3.70E-01 | no | Cdh12 | intron_variant,non_coding_transcript_variant | MODIFIER |
| 13 | 5682017 | AX-170520637 | 1 | C | T | 0.205 | -1.10E-01 | 1.23E-01 | 2.61E+00 | 2.67E+00 | 3.70E-01 | no | Gm35330 | intron_variant,non_coding_transcript_variant | MODIFIER |
| 1 | 20795287 | AX-170000046 | 0 | T | G | 0.412 | 7.45E-02 | 8.33E-02 | 2.61E+00 | 2.66E+00 | 3.70E-01 | no | Il17f | upstream_gene_variant | MODIFIER |
| 1 | 20912886 | AX-170518579 | 0 | G | A | 0.412 | 7.45E-02 | 8.33E-02 | 2.61E+00 | 2.66E+00 | 3.70E-01 | no | Paqr8 | upstream_gene_variant | MODIFIER |
| 9 | 16302825 | AX-170366370 | 0 | G | A | 0.482 | 7.69E-02 | 8.57E-02 | 2.55E+00 | 2.63E+00 | 3.70E-01 | no | Fat3 | intron_variant | MODIFIER |
| 8 | 60813760 | AX-170080205 | 0 | T | G | 0.149 | -1.24E-01 | 1.34E-01 | 2.42E+00 | 2.49E+00 | 3.70E-01 | no | | intergenic_variant | MODIFIER |
| 10 | 93956408 | AX-170357766 | 2 | A | G | 0.282 | -7.83E-02 | 8.65E-02 | 2.51E+00 | 2.56E+00 | 3.70E-01 | no | Vezt | intron_variant | MODIFIER |
| 10 | 19277194 | AX-170171488 | 1 | G | T | 0.205 | 8.69E-02 | 9.74E-02 | 2.89E+00 | 3.01E+00 | 3.70E-01 | no | | regulatory_region_variant | MODIFIER |
| 10 | 17970319 | AX-170123729 | 0 | T | A | 0.193 | 9.15E-02 | 1.02E-01 | 2.97E+00 | 3.07E+00 | 3.70E-01 | no | | intergenic_variant | MODIFIER |
| 10 | 17974418 | AX-170511415 | 0 | C | T | 0.193 | 9.15E-02 | 1.02E-01 | 2.97E+00 | 3.07E+00 | 3.70E-01 | no | | intergenic_variant | MODIFIER |
| 10 | 17985933 | AX-97828962 | 0 | G | A | 0.193 | 9.15E-02 | 1.02E-01 | 2.97E+00 | 3.07E+00 | 3.70E-01 | no | | intergenic_variant | MODIFIER |
| 10 | 18009166 | AX-170518264 | 0 | C | T | 0.193 | 9.15E-02 | 1.02E-01 | 2.97E+00 | 3.07E+00 | 3.70E-01 | no | Abracl | downstream_gene_variant | MODIFIER |
| 10 | 18010503 | AX-97853127 | 0 | C | T | 0.193 | 9.15E-02 | 1.02E-01 | 2.97E+00 | 3.07E+00 | 3.70E-01 | no | Abracl | downstream_gene_variant | MODIFIER |
| 10 | 18019258 | AX-170228224 | 0 | T | C | 0.193 | 9.15E-02 | 1.02E-01 | 2.97E+00 | 3.07E+00 | 3.70E-01 | no | Abracl | intron_variant | MODIFIER |
| 10 | 18019475 | AX-170421729 | 0 | G | A | 0.193 | 9.15E-02 | 1.02E-01 | 2.97E+00 | 3.07E+00 | 3.70E-01 | no | Abracl | intron_variant | MODIFIER |
| 10 | 18033745 | AX-170099235 | 0 | A | G | 0.193 | 9.15E-02 | 1.02E-01 | 2.97E+00 | 3.07E+00 | 3.70E-01 | no | | intergenic_variant | MODIFIER |
| 10 | 18036086 | AX-97815011 | 0 | C | T | 0.193 | 9.15E-02 | 1.02E-01 | 2.97E+00 | 3.07E+00 | 3.70E-01 | no | | intergenic_variant | MODIFIER |
| 10 | 18044075 | AX-170432900 | 0 | C | T | 0.193 | 9.15E-02 | 1.02E-01 | 2.97E+00 | 3.07E+00 | 3.70E-01 | no | | intergenic_variant | MODIFIER |
| 10 | 18271348 | AX-170305338 | 0 | G | C | 0.193 | 9.15E-02 | 1.02E-01 | 2.97E+00 | 3.07E+00 | 3.70E-01 | no | | intergenic_variant | MODIFIER |
| 10 | 18277942 | AX-97798348 | 0 | C | T | 0.193 | 9.15E-02 | 1.02E-01 | 2.97E+00 | 3.07E+00 | 3.70E-01 | no | | intergenic_variant | MODIFIER |
| 10 | 18285499 | AX-170184906 | 0 | A | G | 0.193 | 9.15E-02 | 1.02E-01 | 2.97E+00 | 3.07E+00 | 3.70E-01 | no | | intergenic_variant | MODIFIER |
| 10 | 18286269 | AX-170002855 | 0 | T | C | 0.193 | 9.15E-02 | 1.02E-01 | 2.97E+00 | 3.07E+00 | 3.70E-01 | no | | intergenic_variant | MODIFIER |
| 10 | 18309936 | AX-170267510 | 0 | T | C | 0.193 | 9.15E-02 | 1.02E-01 | 2.97E+00 | 3.07E+00 | 3.70E-01 | no | | intergenic_variant | MODIFIER |
| 10 | 18340848 | AX-97806299 | 0 | T | C | 0.193 | 9.15E-02 | 1.02E-01 | 2.97E+00 | 3.07E+00 | 3.70E-01 | no | Nhsl1 | intron_variant | MODIFIER |
| 10 | 18354367 | AX-97816871 | 0 | C | T | 0.193 | 9.15E-02 | 1.02E-01 | 2.97E+00 | 3.07E+00 | 3.70E-01 | no | Nhsl1 | intron_variant | MODIFIER |
| 10 | 18393812 | AX-170163706 | 0 | C | T | 0.193 | 9.15E-02 | 1.02E-01 | 2.97E+00 | 3.07E+00 | 3.70E-01 | no | Nhsl1 | intron_variant,non_coding_transcript_variant | MODIFIER |
| 10 | 18400707 | AX-97815012 | 0 | T | C | 0.193 | 9.15E-02 | 1.02E-01 | 2.97E+00 | 3.07E+00 | 3.70E-01 | no | Nhsl1 | intron_variant,non_coding_transcript_variant | MODIFIER |
| 10 | 18919977 | AX-170287257 | 0 | T | C | 0.193 | 9.15E-02 | 1.02E-01 | 2.97E+00 | 3.07E+00 | 3.70E-01 | no | | intergenic_variant | MODIFIER |
| 10 | 18928090 | AX-170447253 | 0 | C | T | 0.193 | 9.15E-02 | 1.02E-01 | 2.97E+00 | 3.07E+00 | 3.70E-01 | no | | intergenic_variant | MODIFIER |
| 10 | 19082240 | AX-170213391 | 0 | A | G | 0.193 | 9.15E-02 | 1.02E-01 | 2.97E+00 | 3.07E+00 | 3.70E-01 | no | 1700124M09Rik | downstream_gene_variant | MODIFIER |
| 10 | 19082846 | AX-170409791 | 0 | G | A | 0.193 | 9.15E-02 | 1.02E-01 | 2.97E+00 | 3.07E+00 | 3.70E-01 | no | 1700124M09Rik | downstream_gene_variant | MODIFIER |
| 10 | 19083223 | AX-170355848 | 0 | G | A | 0.193 | 9.15E-02 | 1.02E-01 | 2.97E+00 | 3.07E+00 | 3.70E-01 | no | 1700124M09Rik | downstream_gene_variant | MODIFIER |
| 10 | 19083523 | AX-170501836 | 0 | T | A | 0.193 | 9.15E-02 | 1.02E-01 | 2.97E+00 | 3.07E+00 | 3.70E-01 | no | 1700124M09Rik | downstream_gene_variant | MODIFIER |
| 10 | 19085778 | AX-170515414 | 0 | T | C | 0.193 | 9.15E-02 | 1.02E-01 | 2.97E+00 | 3.07E+00 | 3.70E-01 | no | 1700124M09Rik | downstream_gene_variant | MODIFIER |
| 10 | 19085886 | AX-170437614 | 0 | A | T | 0.193 | 9.15E-02 | 1.02E-01 | 2.97E+00 | 3.07E+00 | 3.70E-01 | no | 1700124M09Rik | downstream_gene_variant | MODIFIER |
| 10 | 19094659 | AX-170236924 | 0 | T | C | 0.193 | 9.15E-02 | 1.02E-01 | 2.97E+00 | 3.07E+00 | 3.70E-01 | no | Gm48545 | non_coding_transcript_exon_variant | MODIFIER |

| | | | | | | | | | | | | | | | |
|---|---|---|---|---|---|---|---|---|---|---|---|---|---|---|---|
| 10 | 19112905 | AX-170310536 | 0 | A | G | 0.193 | 9.15E-02 | 1.02E-01 | 2.97E+00 | 3.07E+00 | 3.70E-01 | no | | intergenic_variant | MODIFIER |
| 10 | 19119820 | AX-170213923 | 0 | T | C | 0.193 | 9.15E-02 | 1.02E-01 | 2.97E+00 | 3.07E+00 | 3.70E-01 | no | Gm48546 | upstream_gene_variant | MODIFIER |
| 10 | 19126424 | AX-170482666 | 0 | G | A | 0.193 | 9.15E-02 | 1.02E-01 | 2.97E+00 | 3.07E+00 | 3.70E-01 | no | Gm48546 | downstream_gene_variant | MODIFIER |
| 10 | 19127594 | AX-170265073 | 0 | T | C | 0.193 | 9.15E-02 | 1.02E-01 | 2.97E+00 | 3.07E+00 | 3.70E-01 | no | Gm48546 | downstream_gene_variant | MODIFIER |
| 10 | 19172262 | AX-170516053 | 0 | T | C | 0.193 | 9.15E-02 | 1.02E-01 | 2.97E+00 | 3.07E+00 | 3.70E-01 | no | Gm20139 | intron_variant,non_coding_transcript_variant | MODIFIER |
| 10 | 19172855 | AX-170201161 | 0 | G | A | 0.193 | 9.15E-02 | 1.02E-01 | 2.97E+00 | 3.07E+00 | 3.70E-01 | no | Gm20139 | intron_variant,non_coding_transcript_variant | MODIFIER |
| 10 | 19209589 | AX-170405751 | 0 | C | T | 0.193 | 9.15E-02 | 1.02E-01 | 2.97E+00 | 3.07E+00 | 3.70E-01 | no | Gm20139 | downstream_gene_variant | MODIFIER |
| 10 | 19252918 | AX-170526471 | 0 | T | C | 0.193 | 9.15E-02 | 1.02E-01 | 2.97E+00 | 3.07E+00 | 3.70E-01 | no | | intergenic_variant | MODIFIER |
| 10 | 19280609 | AX-170428620 | 0 | A | T | 0.193 | 9.15E-02 | 1.02E-01 | 2.97E+00 | 3.07E+00 | 3.70E-01 | no | | intergenic_variant | MODIFIER |
| 10 | 19281246 | AX-170349695 | 0 | T | C | 0.193 | 9.15E-02 | 1.02E-01 | 2.97E+00 | 3.07E+00 | 3.70E-01 | no | | intergenic_variant | MODIFIER |
| 10 | 19281302 | AX-170328671 | 0 | A | G | 0.193 | 9.15E-02 | 1.02E-01 | 2.97E+00 | 3.07E+00 | 3.70E-01 | no | | intergenic_variant | MODIFIER |
| 10 | 19296246 | AX-170203212 | 0 | A | C | 0.193 | 9.15E-02 | 1.02E-01 | 2.97E+00 | 3.07E+00 | 3.70E-01 | no | Gm33056 | intron_variant,non_coding_transcript_variant | MODIFIER |
| 10 | 19318141 | AX-170282411 | 0 | A | G | 0.193 | 9.15E-02 | 1.02E-01 | 2.97E+00 | 3.07E+00 | 3.70E-01 | no | Gm33056 | intron_variant,non_coding_transcript_variant | MODIFIER |
| 5 | 5065798 | AX-170134117 | 1 | G | A | 0.446 | -5.89E-02 | 6.52E-02 | 2.50E+00 | 2.61E+00 | 3.70E-01 | no | Cdk14 | intron_variant | MODIFIER |
| 5 | 3884799 | AX-170434380 | 1 | C | T | 0.429 | -5.93E-02 | 6.55E-02 | 2.49E+00 | 2.60E+00 | 3.70E-01 | no | | intergenic_variant | MODIFIER |
| 16 | 50705946 | AX-170452982 | 0 | G | C | 0.518 | 7.38E-02 | 8.04E-02 | 2.44E+00 | 2.49E+00 | 3.70E-01 | no | | intergenic_variant | MODIFIER |
| 7 | 118381454 | AX-97832597 | 0 | G | A | 0.395 | 7.93E-02 | 8.95E-02 | 2.71E+00 | 2.77E+00 | 3.70E-01 | no | Syt17 | 3_prime_UTR_variant | MODIFIER |
| 7 | 39738013 | AX-170028870 | 1 | C | T | 0.348 | 8.47E-02 | 9.46E-02 | 2.57E+00 | 2.64E+00 | 3.70E-01 | no | Gm9230 | downstream_gene_variant | MODIFIER |
| 2 | 165559702 | AX-170124008 | 1 | C | A | 0.161 | 1.72E-01 | 1.95E-01 | 2.75E+00 | 2.83E+00 | 3.71E-01 | Yes | | regulatory_region_variant | MODIFIER |
| 4 | 97963384 | AX-170466890 | 1 | G | C | 0.232 | 1.06E-01 | 1.18E-01 | 2.57E+00 | 2.63E+00 | 3.71E-01 | no | Nfia | intron_variant | MODIFIER |
| 16 | 78490638 | AX-170476371 | 1 | A | G | 0.446 | 6.58E-02 | 7.39E-02 | 2.60E+00 | 2.68E+00 | 3.71E-01 | no | Gm49604 | non_coding_transcript_exon_variant | MODIFIER |
| 4 | 78758695 | AX-170023180 | 1 | T | C | 0.321 | 1.00E-01 | 1.13E-01 | 2.68E+00 | 2.74E+00 | 3.71E-01 | no | | intergenic_variant | MODIFIER |
| 4 | 72723684 | AX-170391925 | 0 | T | C | 0.447 | -7.48E-02 | 8.37E-02 | 2.57E+00 | 2.64E+00 | 3.71E-01 | no | | intergenic_variant | MODIFIER |
| 2 | 122174711 | AX-170215849 | 0 | T | G | 0.149 | 1.34E-01 | 1.51E-01 | 2.62E+00 | 2.74E+00 | 3.71E-01 | no | Patl2 | intron_variant | MODIFIER |
| 3 | 35216983 | AX-170119950 | 0 | C | G | 0.193 | 8.41E-02 | 9.49E-02 | 2.84E+00 | 2.93E+00 | 3.71E-01 | no | | intergenic_variant | MODIFIER |
| 3 | 35984248 | AX-170220703 | 0 | A | G | 0.193 | 8.41E-02 | 9.49E-02 | 2.84E+00 | 2.93E+00 | 3.71E-01 | no | Mccc1 | intron_variant | MODIFIER |
| 3 | 35986729 | AX-170393005 | 0 | T | G | 0.193 | 8.41E-02 | 9.49E-02 | 2.84E+00 | 2.93E+00 | 3.71E-01 | no | Mccc1 | intron_variant | MODIFIER |
| 3 | 35989707 | AX-170295353 | 0 | C | T | 0.193 | 8.41E-02 | 9.49E-02 | 2.84E+00 | 2.93E+00 | 3.71E-01 | no | Mccc1 | intron_variant | MODIFIER |
| 3 | 35990427 | AX-170365970 | 0 | G | A | 0.193 | 8.41E-02 | 9.49E-02 | 2.84E+00 | 2.93E+00 | 3.71E-01 | no | Mccc1 | intron_variant | MODIFIER |
| 3 | 36086157 | AX-170164482 | 0 | T | C | 0.193 | 8.41E-02 | 9.49E-02 | 2.84E+00 | 2.93E+00 | 3.71E-01 | no | Acad9 | intron_variant | MODIFIER |
| 3 | 36449161 | AX-97821408 | 0 | T | C | 0.193 | 8.41E-02 | 9.49E-02 | 2.84E+00 | 2.93E+00 | 3.71E-01 | no | Anxa5 | 3_prime_UTR_variant | MODIFIER |
| 3 | 36458475 | AX-170404339 | 0 | C | T | 0.193 | 8.41E-02 | 9.49E-02 | 2.84E+00 | 2.93E+00 | 3.71E-01 | no | Anxa5 | intron_variant | MODIFIER |
| 3 | 36498406 | AX-170405097 | 0 | G | C | 0.193 | 8.41E-02 | 9.49E-02 | 2.84E+00 | 2.93E+00 | 3.71E-01 | no | Gm11548 | downstream_gene_variant | MODIFIER |
| 3 | 36503331 | AX-170246923 | 0 | T | C | 0.193 | 8.41E-02 | 9.49E-02 | 2.84E+00 | 2.93E+00 | 3.71E-01 | no | Gm11548 | intron_variant,non_coding_transcript_variant | MODIFIER |
| 3 | 36519770 | AX-170395069 | 0 | C | T | 0.193 | 8.41E-02 | 9.49E-02 | 2.84E+00 | 2.93E+00 | 3.71E-01 | no | Gm11549 | intron_variant | MODIFIER |
| 3 | 36548656 | AX-170149552 | 0 | C | T | 0.193 | 8.41E-02 | 9.49E-02 | 2.84E+00 | 2.93E+00 | 3.71E-01 | no | Exosc9 | upstream_gene_variant | MODIFIER |
| 3 | 36549366 | AX-170379562 | 0 | A | T | 0.193 | 8.41E-02 | 9.49E-02 | 2.84E+00 | 2.93E+00 | 3.71E-01 | no | Exosc9 | upstream_gene_variant | MODIFIER |
| 3 | 36553442 | AX-170404859 | 0 | G | A | 0.193 | 8.41E-02 | 9.49E-02 | 2.84E+00 | 2.93E+00 | 3.71E-01 | no | Exosc9 | intron_variant | MODIFIER |
| 3 | 36563735 | AX-170522946 | 0 | A | G | 0.193 | 8.41E-02 | 9.49E-02 | 2.84E+00 | 2.93E+00 | 3.71E-01 | no | Exosc9 | intron_variant | MODIFIER |
| 3 | 36566110 | AX-97843970 | 0 | A | G | 0.193 | 8.41E-02 | 9.49E-02 | 2.84E+00 | 2.93E+00 | 3.71E-01 | no | Exosc9 | downstream_gene_variant | MODIFIER |
| 3 | 36568129 | AX-170020309 | 0 | C | T | 0.193 | 8.41E-02 | 9.49E-02 | 2.84E+00 | 2.93E+00 | 3.71E-01 | no | Exosc9 | downstream_gene_variant | MODIFIER |
| 3 | 36593026 | AX-170445038 | 0 | G | C | 0.193 | 8.41E-02 | 9.49E-02 | 2.84E+00 | 2.93E+00 | 3.71E-01 | no | Bbs7 | intron_variant | MODIFIER |
| 3 | 36597588 | AX-170020310 | 0 | T | C | 0.193 | 8.41E-02 | 9.49E-02 | 2.84E+00 | 2.93E+00 | 3.71E-01 | no | Bbs7 | intron_variant | MODIFIER |
| 3 | 36599358 | AX-97792854 | 0 | A | G | 0.193 | 8.41E-02 | 9.49E-02 | 2.84E+00 | 2.93E+00 | 3.71E-01 | no | Bbs7 | synonymous_variant | LOW |
| 3 | 36601849 | AX-170312082 | 0 | C | G | 0.193 | 8.41E-02 | 9.49E-02 | 2.84E+00 | 2.93E+00 | 3.71E-01 | no | Bbs7 | intron_variant | MODIFIER |
| 3 | 36606535 | AX-170454651 | 0 | A | G | 0.193 | 8.41E-02 | 9.49E-02 | 2.84E+00 | 2.93E+00 | 3.71E-01 | no | Bbs7 | intron_variant | MODIFIER |
| 3 | 36612449 | AX-97825149 | 0 | C | T | 0.193 | 8.41E-02 | 9.49E-02 | 2.84E+00 | 2.93E+00 | 3.71E-01 | no | Bbs7 | intron_variant | MODIFIER |
| 3 | 36655450 | AX-170394451 | 0 | A | G | 0.193 | 8.41E-02 | 9.49E-02 | 2.84E+00 | 2.93E+00 | 3.71E-01 | no | Trpc3 | intron_variant | MODIFIER |
| 3 | 36666837 | AX-170428840 | 0 | A | G | 0.193 | 8.41E-02 | 9.49E-02 | 2.84E+00 | 2.93E+00 | 3.71E-01 | no | Trpc3 | intron_variant | MODIFIER |
| 3 | 36666894 | AX-170345782 | 0 | T | C | 0.193 | 8.41E-02 | 9.49E-02 | 2.84E+00 | 2.93E+00 | 3.71E-01 | no | Trpc3 | intron_variant | MODIFIER |
| 3 | 36667900 | AX-170266249 | 0 | C | A | 0.193 | 8.41E-02 | 9.49E-02 | 2.84E+00 | 2.93E+00 | 3.71E-01 | no | Trpc3 | intron_variant | MODIFIER |
| 3 | 36668232 | AX-170460270 | 0 | G | A | 0.193 | 8.41E-02 | 9.49E-02 | 2.84E+00 | 2.93E+00 | 3.71E-01 | no | Trpc3 | intron_variant | MODIFIER |
| 3 | 36670473 | AX-170486296 | 0 | C | T | 0.193 | 8.41E-02 | 9.49E-02 | 2.84E+00 | 2.93E+00 | 3.71E-01 | no | Trpc3 | intron_variant | MODIFIER |
| 3 | 36676064 | AX-97840439 | 0 | G | A | 0.193 | 8.41E-02 | 9.49E-02 | 2.84E+00 | 2.93E+00 | 3.71E-01 | no | Trpc3 | intron_variant | MODIFIER |

| | | | | | | | | | | | | | | | |
|---|---|---|---|---|---|---|---|---|---|---|---|---|---|---|---|
| 3 | 36677633 | AX-170124332 | 0 | C | T | 0.193 | 8.41E-02 | 9.49E-02 | 2.84E+00 | 2.93E+00 | 3.71E-01 | no | Trpc3 | intron_variant | MODIFIER |
| 3 | 36687513 | AX-170464194 | 0 | T | C | 0.193 | 8.41E-02 | 9.49E-02 | 2.84E+00 | 2.93E+00 | 3.71E-01 | no | Trpc3 | intron_variant | MODIFIER |
| 3 | 36709757 | AX-170453552 | 0 | C | A | 0.193 | 8.41E-02 | 9.49E-02 | 2.84E+00 | 2.93E+00 | 3.71E-01 | no | | intergenic_variant | MODIFIER |
| 3 | 36716027 | AX-170293574 | 0 | C | T | 0.193 | 8.41E-02 | 9.49E-02 | 2.84E+00 | 2.93E+00 | 3.71E-01 | no | | intergenic_variant | MODIFIER |
| 3 | 36716166 | AX-170428761 | 0 | A | G | 0.193 | 8.41E-02 | 9.49E-02 | 2.84E+00 | 2.93E+00 | 3.71E-01 | no | | intergenic_variant | MODIFIER |
| 3 | 36720869 | AX-170209653 | 0 | T | A | 0.193 | 8.41E-02 | 9.49E-02 | 2.84E+00 | 2.93E+00 | 3.71E-01 | no | | intergenic_variant | MODIFIER |
| 3 | 36723149 | AX-97823003 | 0 | T | C | 0.193 | 8.41E-02 | 9.49E-02 | 2.84E+00 | 2.93E+00 | 3.71E-01 | no | | intergenic_variant | MODIFIER |
| 3 | 36749537 | AX-170238247 | 0 | A | G | 0.193 | 8.41E-02 | 9.49E-02 | 2.84E+00 | 2.93E+00 | 3.71E-01 | no | | regulatory_region_variant | MODIFIER |
| 3 | 36751608 | AX-170243152 | 0 | T | A | 0.193 | 8.41E-02 | 9.49E-02 | 2.84E+00 | 2.93E+00 | 3.71E-01 | no | | intergenic_variant | MODIFIER |
| 3 | 36751889 | AX-170514482 | 0 | T | G | 0.193 | 8.41E-02 | 9.49E-02 | 2.84E+00 | 2.93E+00 | 3.71E-01 | no | | intergenic_variant | MODIFIER |
| 3 | 36764006 | AX-97789675 | 0 | T | C | 0.193 | 8.41E-02 | 9.49E-02 | 2.84E+00 | 2.93E+00 | 3.71E-01 | no | | intergenic_variant | MODIFIER |
| 3 | 36764538 | AX-170506482 | 0 | T | C | 0.193 | 8.41E-02 | 9.49E-02 | 2.84E+00 | 2.93E+00 | 3.71E-01 | no | | intergenic_variant | MODIFIER |
| 3 | 36771568 | AX-170250607 | 0 | T | C | 0.193 | 8.41E-02 | 9.49E-02 | 2.84E+00 | 2.93E+00 | 3.71E-01 | no | | intergenic_variant | MODIFIER |
| 3 | 36780138 | AX-170188731 | 0 | G | A | 0.193 | 8.41E-02 | 9.49E-02 | 2.84E+00 | 2.93E+00 | 3.71E-01 | no | | intergenic_variant | MODIFIER |
| 3 | 36785120 | AX-170414756 | 0 | G | A | 0.193 | 8.41E-02 | 9.49E-02 | 2.84E+00 | 2.93E+00 | 3.71E-01 | no | | intergenic_variant | MODIFIER |
| 3 | 36810340 | AX-170066366 | 0 | T | C | 0.193 | 8.41E-02 | 9.49E-02 | 2.84E+00 | 2.93E+00 | 3.71E-01 | no | | intergenic_variant | MODIFIER |
| 3 | 36810766 | AX-170066362 | 0 | T | C | 0.193 | 8.41E-02 | 9.49E-02 | 2.84E+00 | 2.93E+00 | 3.71E-01 | no | | regulatory_region_variant | MODIFIER |
| 3 | 36812413 | AX-170066361 | 0 | A | G | 0.193 | 8.41E-02 | 9.49E-02 | 2.84E+00 | 2.93E+00 | 3.71E-01 | no | | intergenic_variant | MODIFIER |
| 3 | 36812443 | AX-170066363 | 0 | C | G | 0.193 | 8.41E-02 | 9.49E-02 | 2.84E+00 | 2.93E+00 | 3.71E-01 | no | | intergenic_variant | MODIFIER |
| 3 | 36816341 | AX-170412137 | 0 | A | G | 0.193 | 8.41E-02 | 9.49E-02 | 2.84E+00 | 2.93E+00 | 3.71E-01 | no | | intergenic_variant | MODIFIER |
| 3 | 36826538 | AX-170275644 | 0 | G | A | 0.193 | 8.41E-02 | 9.49E-02 | 2.84E+00 | 2.93E+00 | 3.71E-01 | no | Gm12376 | upstream_gene_variant | MODIFIER |
| 3 | 36834801 | AX-97803504 | 0 | A | G | 0.193 | 8.41E-02 | 9.49E-02 | 2.84E+00 | 2.93E+00 | 3.71E-01 | no | | intergenic_variant | MODIFIER |
| 3 | 36854743 | AX-170167115 | 0 | G | T | 0.193 | 8.41E-02 | 9.49E-02 | 2.84E+00 | 2.93E+00 | 3.71E-01 | no | | regulatory_region_variant | MODIFIER |
| 3 | 36862638 | AX-170311844 | 0 | C | T | 0.193 | 8.41E-02 | 9.49E-02 | 2.84E+00 | 2.93E+00 | 3.71E-01 | no | 4932438A13Rik | upstream_gene_variant | MODIFIER |
| 3 | 36870706 | AX-170196157 | 0 | C | T | 0.193 | 8.41E-02 | 9.49E-02 | 2.84E+00 | 2.93E+00 | 3.71E-01 | no | 4932438A13Rik | intron_variant | MODIFIER |
| 3 | 36870734 | AX-170232060 | 0 | G | A | 0.193 | 8.41E-02 | 9.49E-02 | 2.84E+00 | 2.93E+00 | 3.71E-01 | no | 4932438A13Rik | intron_variant | MODIFIER |
| 3 | 36875045 | AX-97846058 | 0 | G | A | 0.193 | 8.41E-02 | 9.49E-02 | 2.84E+00 | 2.93E+00 | 3.71E-01 | no | 4932438A13Rik | intron_variant | MODIFIER |
| 3 | 36924548 | AX-97849196 | 0 | G | C | 0.193 | 8.41E-02 | 9.49E-02 | 2.84E+00 | 2.93E+00 | 3.71E-01 | no | 4932438A13Rik | intron_variant | MODIFIER |
| 3 | 36945384 | AX-170143852 | 0 | A | G | 0.193 | 8.41E-02 | 9.49E-02 | 2.84E+00 | 2.93E+00 | 3.71E-01 | no | 4932438A13Rik | intron_variant | MODIFIER |
| 3 | 36954451 | AX-170346447 | 0 | T | C | 0.193 | 8.41E-02 | 9.49E-02 | 2.84E+00 | 2.93E+00 | 3.71E-01 | no | 4932438A13Rik | intron_variant | MODIFIER |
| 3 | 36960506 | AX-170275331 | 0 | T | C | 0.193 | 8.41E-02 | 9.49E-02 | 2.84E+00 | 2.93E+00 | 3.71E-01 | no | 4932438A13Rik | intron_variant | MODIFIER |
| 3 | 36986020 | AX-170381312 | 0 | T | A | 0.193 | 8.41E-02 | 9.49E-02 | 2.84E+00 | 2.93E+00 | 3.71E-01 | no | 4932438A13Rik | intron_variant | MODIFIER |
| 3 | 36987082 | AX-170387985 | 0 | C | T | 0.193 | 8.41E-02 | 9.49E-02 | 2.84E+00 | 2.93E+00 | 3.71E-01 | no | 4932438A13Rik | synonymous_variant | LOW |
| 3 | 36987427 | AX-170117787 | 0 | G | A | 0.193 | 8.41E-02 | 9.49E-02 | 2.84E+00 | 2.93E+00 | 3.71E-01 | no | 4932438A13Rik | synonymous_variant | LOW |
| 3 | 36997342 | AX-97804733 | 0 | T | C | 0.193 | 8.41E-02 | 9.49E-02 | 2.84E+00 | 2.93E+00 | 3.71E-01 | no | 4932438A13Rik | intron_variant | MODIFIER |
| 3 | 37007395 | AX-170066370 | 0 | A | T | 0.193 | 8.41E-02 | 9.49E-02 | 2.84E+00 | 2.93E+00 | 3.71E-01 | no | 4932438A13Rik | intron_variant | MODIFIER |
| 3 | 37028789 | AX-170404902 | 0 | A | G | 0.193 | 8.41E-02 | 9.49E-02 | 2.84E+00 | 2.93E+00 | 3.71E-01 | no | 4932438A13Rik | intron_variant | MODIFIER |
| 3 | 37086371 | AX-170236098 | 0 | T | C | 0.193 | 8.41E-02 | 9.49E-02 | 2.84E+00 | 2.93E+00 | 3.71E-01 | no | Adad1 | intron_variant | MODIFIER |
| 3 | 37091184 | AX-170020312 | 0 | C | T | 0.193 | 8.41E-02 | 9.49E-02 | 2.84E+00 | 2.93E+00 | 3.71E-01 | no | Adad1 | intron_variant | MODIFIER |
| 3 | 37123410 | AX-170020317 | 0 | G | A | 0.193 | 8.41E-02 | 9.49E-02 | 2.84E+00 | 2.93E+00 | 3.71E-01 | no | Il2 | intron_variant | MODIFIER |
| 3 | 37124055 | AX-170100097 | 0 | T | C | 0.193 | 8.41E-02 | 9.49E-02 | 2.84E+00 | 2.93E+00 | 3.71E-01 | no | Il2 | intron_variant | MODIFIER |
| 3 | 37128270 | AX-170394153 | 0 | G | C | 0.193 | 8.41E-02 | 9.49E-02 | 2.84E+00 | 2.93E+00 | 3.71E-01 | no | Il2 | upstream_gene_variant | MODIFIER |
| 3 | 37128641 | AX-170020313 | 0 | T | C | 0.193 | 8.41E-02 | 9.49E-02 | 2.84E+00 | 2.93E+00 | 3.71E-01 | no | Il2 | upstream_gene_variant | MODIFIER |
| 3 | 37134808 | AX-97783001 | 0 | C | T | 0.193 | 8.41E-02 | 9.49E-02 | 2.84E+00 | 2.93E+00 | 3.71E-01 | no | | intergenic_variant | MODIFIER |
| 3 | 37135808 | AX-170143954 | 0 | G | C | 0.193 | 8.41E-02 | 9.49E-02 | 2.84E+00 | 2.93E+00 | 3.71E-01 | no | | intergenic_variant | MODIFIER |
| 3 | 37166120 | AX-170297479 | 0 | A | G | 0.193 | 8.41E-02 | 9.49E-02 | 2.84E+00 | 2.93E+00 | 3.71E-01 | no | | intergenic_variant | MODIFIER |
| 3 | 37174320 | AX-97798848 | 0 | T | C | 0.193 | 8.41E-02 | 9.49E-02 | 2.84E+00 | 2.93E+00 | 3.71E-01 | no | | intergenic_variant | MODIFIER |
| 3 | 37186642 | AX-97857340 | 0 | G | A | 0.193 | 8.41E-02 | 9.49E-02 | 2.84E+00 | 2.93E+00 | 3.71E-01 | no | Gm12532 | non_coding_transcript_exon_variant | MODIFIER |
| 3 | 37197438 | AX-170253833 | 0 | T | G | 0.193 | 8.41E-02 | 9.49E-02 | 2.84E+00 | 2.93E+00 | 3.71E-01 | no | | intergenic_variant | MODIFIER |
| 3 | 37197520 | AX-170425826 | 0 | C | T | 0.193 | 8.41E-02 | 9.49E-02 | 2.84E+00 | 2.93E+00 | 3.71E-01 | no | | intergenic_variant | MODIFIER |
| 3 | 37211153 | AX-170139431 | 0 | G | A | 0.193 | 8.41E-02 | 9.49E-02 | 2.84E+00 | 2.93E+00 | 3.71E-01 | no | | intergenic_variant | MODIFIER |
| 3 | 37213901 | AX-97850577 | 0 | T | A | 0.193 | 8.41E-02 | 9.49E-02 | 2.84E+00 | 2.93E+00 | 3.71E-01 | no | | intergenic_variant | MODIFIER |
| 3 | 37219664 | AX-170201583 | 0 | C | T | 0.193 | 8.41E-02 | 9.49E-02 | 2.84E+00 | 2.93E+00 | 3.71E-01 | no | Il21 | downstream_gene_variant | MODIFIER |
| 3 | 37273433 | AX-97811749 | 0 | T | C | 0.193 | 8.41E-02 | 9.49E-02 | 2.84E+00 | 2.93E+00 | 3.71E-01 | no | | intergenic_variant | MODIFIER |

| | | | | | | | | | | | | | | | |
|---|---|---|---|---|---|---|---|---|---|---|---|---|---|---|---|
| 3 | 37297508 | AX-170088909 | 0 | T | C | 0.193 | 8.41E-02 | 9.49E-02 | 2.84E+00 | 2.93E+00 | 3.71E-01 | no | | intergenic_variant | MODIFIER |
| 3 | 37305523 | AX-170100548 | 0 | G | A | 0.193 | 8.41E-02 | 9.49E-02 | 2.84E+00 | 2.93E+00 | 3.71E-01 | no | Cetn4 | downstream_gene_variant | MODIFIER |
| 3 | 37345589 | AX-97791434 | 0 | T | C | 0.193 | 8.41E-02 | 9.49E-02 | 2.84E+00 | 2.93E+00 | 3.71E-01 | no | Fgf2 | upstream_gene_variant | MODIFIER |
| 3 | 37365377 | AX-170413876 | 0 | C | G | 0.193 | 8.41E-02 | 9.49E-02 | 2.84E+00 | 2.93E+00 | 3.71E-01 | no | Fgf2 | intron_variant | MODIFIER |
| 3 | 37378983 | AX-97847627 | 0 | C | T | 0.193 | 8.41E-02 | 9.49E-02 | 2.84E+00 | 2.93E+00 | 3.71E-01 | no | Fgf2 | intron_variant | MODIFIER |
| 3 | 37380615 | AX-170235873 | 0 | C | T | 0.193 | 8.41E-02 | 9.49E-02 | 2.84E+00 | 2.93E+00 | 3.71E-01 | no | Fgf2 | intron_variant | MODIFIER |
| 3 | 37386340 | AX-97814874 | 0 | A | G | 0.193 | 8.41E-02 | 9.49E-02 | 2.84E+00 | 2.93E+00 | 3.71E-01 | no | Fgf2 | intron_variant | MODIFIER |
| 3 | 37388360 | AX-170200967 | 0 | T | C | 0.193 | 8.41E-02 | 9.49E-02 | 2.84E+00 | 2.93E+00 | 3.71E-01 | no | Fgf2 | intron_variant | MODIFIER |
| 3 | 37397108 | AX-170066384 | 0 | A | G | 0.193 | 8.41E-02 | 9.49E-02 | 2.84E+00 | 2.93E+00 | 3.71E-01 | no | Fgf2 | intron_variant | MODIFIER |
| 3 | 37425800 | AX-170363524 | 0 | C | T | 0.193 | 8.41E-02 | 9.49E-02 | 2.84E+00 | 2.93E+00 | 3.71E-01 | no | Spata5 | intron_variant | MODIFIER |
| 3 | 37430574 | AX-170488254 | 0 | A | G | 0.193 | 8.41E-02 | 9.49E-02 | 2.84E+00 | 2.93E+00 | 3.71E-01 | no | Spata5 | intron_variant | MODIFIER |
| 3 | 37430671 | AX-170121979 | 0 | G | A | 0.193 | 8.41E-02 | 9.49E-02 | 2.84E+00 | 2.93E+00 | 3.71E-01 | no | Spata5 | intron_variant | MODIFIER |
| 3 | 37453870 | AX-170364734 | 0 | T | C | 0.193 | 8.41E-02 | 9.49E-02 | 2.84E+00 | 2.93E+00 | 3.71E-01 | no | Spata5 | intron_variant | MODIFIER |
| 3 | 37471304 | AX-170305190 | 0 | T | C | 0.193 | 8.41E-02 | 9.49E-02 | 2.84E+00 | 2.93E+00 | 3.71E-01 | no | Spata5 | intron_variant | MODIFIER |
| 3 | 37479146 | AX-170126824 | 0 | C | T | 0.193 | 8.41E-02 | 9.49E-02 | 2.84E+00 | 2.93E+00 | 3.71E-01 | no | Spata5 | intron_variant | MODIFIER |
| 3 | 37512519 | AX-170340694 | 0 | T | C | 0.193 | 8.41E-02 | 9.49E-02 | 2.84E+00 | 2.93E+00 | 3.71E-01 | no | Spata5 | intron_variant | MODIFIER |
| 3 | 37528277 | AX-170248505 | 0 | C | T | 0.193 | 8.41E-02 | 9.49E-02 | 2.84E+00 | 2.93E+00 | 3.71E-01 | no | Spata5 | synonymous_variant | LOW |
| 3 | 37532093 | AX-170144510 | 0 | A | G | 0.193 | 8.41E-02 | 9.49E-02 | 2.84E+00 | 2.93E+00 | 3.71E-01 | no | Spata5 | intron_variant | MODIFIER |
| 3 | 37547831 | AX-170532065 | 0 | A | G | 0.193 | 8.41E-02 | 9.49E-02 | 2.84E+00 | 2.93E+00 | 3.71E-01 | no | Spata5 | intron_variant | MODIFIER |
| 3 | 37548516 | AX-170210980 | 0 | A | G | 0.193 | 8.41E-02 | 9.49E-02 | 2.84E+00 | 2.93E+00 | 3.71E-01 | no | Spata5 | intron_variant | MODIFIER |
| 3 | 37575128 | AX-170285715 | 0 | A | G | 0.193 | 8.41E-02 | 9.49E-02 | 2.84E+00 | 2.93E+00 | 3.71E-01 | no | Spata5 | intron_variant | MODIFIER |
| 3 | 37585320 | AX-97797152 | 0 | G | A | 0.193 | 8.41E-02 | 9.49E-02 | 2.84E+00 | 2.93E+00 | 3.71E-01 | no | | regulatory_region_variant | MODIFIER |
| 3 | 37586281 | AX-170224381 | 0 | G | A | 0.193 | 8.41E-02 | 9.49E-02 | 2.84E+00 | 2.93E+00 | 3.71E-01 | no | | regulatory_region_variant | MODIFIER |
| 3 | 37600849 | AX-170381067 | 0 | G | A | 0.193 | 8.41E-02 | 9.49E-02 | 2.84E+00 | 2.93E+00 | 3.71E-01 | no | Gm12565 | upstream_gene_variant | MODIFIER |
| 3 | 37600969 | AX-170455472 | 0 | A | G | 0.193 | 8.41E-02 | 9.49E-02 | 2.84E+00 | 2.93E+00 | 3.71E-01 | no | Gm12565 | upstream_gene_variant | MODIFIER |
| 3 | 37609381 | AX-170530955 | 0 | T | G | 0.193 | 8.41E-02 | 9.49E-02 | 2.84E+00 | 2.93E+00 | 3.71E-01 | no | 4930594O21Rik | upstream_gene_variant | MODIFIER |
| 3 | 37609420 | AX-170262419 | 0 | T | C | 0.193 | 8.41E-02 | 9.49E-02 | 2.84E+00 | 2.93E+00 | 3.71E-01 | no | 4930594O21Rik | upstream_gene_variant | MODIFIER |
| 3 | 37614610 | AX-170066388 | 0 | T | G | 0.193 | 8.41E-02 | 9.49E-02 | 2.84E+00 | 2.93E+00 | 3.71E-01 | no | 4930594O21Rik | non_coding_transcript_exon_variant | MODIFIER |
| 3 | 37615345 | AX-170066385 | 0 | C | T | 0.193 | 8.41E-02 | 9.49E-02 | 2.84E+00 | 2.93E+00 | 3.71E-01 | no | 4930594O21Rik | downstream_gene_variant | MODIFIER |
| 3 | 37620736 | AX-170094582 | 0 | C | T | 0.193 | 8.41E-02 | 9.49E-02 | 2.84E+00 | 2.93E+00 | 3.71E-01 | no | | intergenic_variant | MODIFIER |
| 3 | 37630666 | AX-170066390 | 0 | T | C | 0.193 | 8.41E-02 | 9.49E-02 | 2.84E+00 | 2.93E+00 | 3.71E-01 | no | | intergenic_variant | MODIFIER |
| 3 | 37643452 | AX-170351348 | 0 | C | T | 0.193 | 8.41E-02 | 9.49E-02 | 2.84E+00 | 2.93E+00 | 3.71E-01 | no | Spry1 | synonymous_variant | LOW |
| 3 | 37677542 | AX-170424094 | 0 | T | C | 0.193 | 8.41E-02 | 9.49E-02 | 2.84E+00 | 2.93E+00 | 3.71E-01 | no | | intergenic_variant | MODIFIER |
| 3 | 37695023 | AX-168411401 | 0 | G | A | 0.193 | 8.41E-02 | 9.49E-02 | 2.84E+00 | 2.93E+00 | 3.71E-01 | no | | intergenic_variant | MODIFIER |
| 3 | 37710717 | AX-170194786 | 0 | T | G | 0.193 | 8.41E-02 | 9.49E-02 | 2.84E+00 | 2.93E+00 | 3.71E-01 | no | Gm5148 | downstream_gene_variant | MODIFIER |
| 3 | 37710883 | AX-170020320 | 0 | T | C | 0.193 | 8.41E-02 | 9.49E-02 | 2.84E+00 | 2.93E+00 | 3.71E-01 | no | Gm5148 | downstream_gene_variant | MODIFIER |
| 3 | 37715917 | AX-170158904 | 0 | G | A | 0.193 | 8.41E-02 | 9.49E-02 | 2.84E+00 | 2.93E+00 | 3.71E-01 | no | Gm5148 | intron_variant | MODIFIER |
| 3 | 37722491 | AX-170094583 | 0 | C | T | 0.193 | 8.41E-02 | 9.49E-02 | 2.84E+00 | 2.93E+00 | 3.71E-01 | no | Gm5148 | intron_variant | MODIFIER |
| 3 | 37760259 | AX-170020318 | 0 | A | G | 0.193 | 8.41E-02 | 9.49E-02 | 2.84E+00 | 2.93E+00 | 3.71E-01 | no | Gm42921 | upstream_gene_variant | MODIFIER |
| 3 | 37764986 | AX-170020319 | 0 | A | G | 0.193 | 8.41E-02 | 9.49E-02 | 2.84E+00 | 2.93E+00 | 3.71E-01 | no | Gm42921 | intron_variant,non_coding_transcript_variant | MODIFIER |
| 3 | 37766782 | AX-170534388 | 0 | G | A | 0.193 | 8.41E-02 | 9.49E-02 | 2.84E+00 | 2.93E+00 | 3.71E-01 | no | Gm42921 | non_coding_transcript_exon_variant | MODIFIER |
| 3 | 37766999 | AX-170514270 | 0 | A | T | 0.193 | 8.41E-02 | 9.49E-02 | 2.84E+00 | 2.93E+00 | 3.71E-01 | no | Gm42921 | non_coding_transcript_exon_variant | MODIFIER |
| 3 | 37791670 | AX-97821399 | 0 | C | T | 0.193 | 8.41E-02 | 9.49E-02 | 2.84E+00 | 2.93E+00 | 3.71E-01 | no | | intergenic_variant | MODIFIER |
| 3 | 37797496 | AX-170173358 | 0 | T | C | 0.193 | 8.41E-02 | 9.49E-02 | 2.84E+00 | 2.93E+00 | 3.71E-01 | no | | intergenic_variant | MODIFIER |
| 3 | 37832242 | AX-170363569 | 0 | A | G | 0.193 | 8.41E-02 | 9.49E-02 | 2.84E+00 | 2.93E+00 | 3.71E-01 | no | | intergenic_variant | MODIFIER |
| 3 | 37839673 | AX-170202002 | 0 | T | C | 0.193 | 8.41E-02 | 9.49E-02 | 2.84E+00 | 2.93E+00 | 3.71E-01 | no | Gm42753 | upstream_gene_variant | MODIFIER |
| 3 | 37845809 | AX-97789958 | 0 | C | T | 0.193 | 8.41E-02 | 9.49E-02 | 2.84E+00 | 2.93E+00 | 3.71E-01 | no | Gm42753 | downstream_gene_variant | MODIFIER |
| 3 | 37846343 | AX-170348071 | 0 | T | G | 0.193 | 8.41E-02 | 9.49E-02 | 2.84E+00 | 2.93E+00 | 3.71E-01 | no | Gm42753 | downstream_gene_variant | MODIFIER |
| 3 | 37848212 | AX-170440032 | 0 | T | C | 0.193 | 8.41E-02 | 9.49E-02 | 2.84E+00 | 2.93E+00 | 3.71E-01 | no | | intergenic_variant | MODIFIER |
| 3 | 37866764 | AX-170285833 | 0 | G | C | 0.193 | 8.41E-02 | 9.49E-02 | 2.84E+00 | 2.93E+00 | 3.71E-01 | no | | intergenic_variant | MODIFIER |
| 3 | 37868700 | AX-170321503 | 0 | T | C | 0.193 | 8.41E-02 | 9.49E-02 | 2.84E+00 | 2.93E+00 | 3.71E-01 | no | | regulatory_region_variant | MODIFIER |
| 3 | 37868742 | AX-170106169 | 0 | C | T | 0.193 | 8.41E-02 | 9.49E-02 | 2.84E+00 | 2.93E+00 | 3.71E-01 | no | | regulatory_region_variant | MODIFIER |
| 3 | 37869264 | AX-97783644 | 0 | G | T | 0.193 | 8.41E-02 | 9.49E-02 | 2.84E+00 | 2.93E+00 | 3.71E-01 | no | | regulatory_region_variant | MODIFIER |
| 3 | 37869279 | AX-170200100 | 0 | C | A | 0.193 | 8.41E-02 | 9.49E-02 | 2.84E+00 | 2.93E+00 | 3.71E-01 | no | | regulatory_region_variant | MODIFIER |

| | | | | | | | | | | | | | | | |
|---|---|---|---|---|---|---|---|---|---|---|---|---|---|---|---|
| 3 | 37877041 | AX-170160691 | 0 | A | G | 0.193 | 8.41E-02 | 9.49E-02 | 2.84E+00 | 2.93E+00 | 3.71E-01 | no | | intergenic_variant | MODIFIER |
| 3 | 37903025 | AX-97801554 | 0 | A | G | 0.193 | 8.41E-02 | 9.49E-02 | 2.84E+00 | 2.93E+00 | 3.71E-01 | no | Gm20755 | intron_variant,non_coding_transcript_variant | MODIFIER |
| 3 | 37921548 | AX-170270973 | 0 | A | G | 0.193 | 8.41E-02 | 9.49E-02 | 2.84E+00 | 2.93E+00 | 3.71E-01 | no | Gm20755 | intron_variant,non_coding_transcript_variant | MODIFIER |
| 3 | 37928842 | AX-97811160 | 0 | C | A | 0.193 | 8.41E-02 | 9.49E-02 | 2.84E+00 | 2.93E+00 | 3.71E-01 | no | Gm20755 | intron_variant,non_coding_transcript_variant | MODIFIER |
| 3 | 37934225 | AX-170514641 | 0 | T | C | 0.193 | 8.41E-02 | 9.49E-02 | 2.84E+00 | 2.93E+00 | 3.71E-01 | no | Gm20755 | intron_variant,non_coding_transcript_variant | MODIFIER |
| 3 | 37938044 | AX-170499411 | 0 | A | G | 0.193 | 8.41E-02 | 9.49E-02 | 2.84E+00 | 2.93E+00 | 3.71E-01 | no | Gm20755 | intron_variant,non_coding_transcript_variant | MODIFIER |
| 3 | 37954944 | AX-97806203 | 0 | T | G | 0.193 | 8.41E-02 | 9.49E-02 | 2.84E+00 | 2.93E+00 | 3.71E-01 | no | Gm7799 | downstream_gene_variant | MODIFIER |
| 3 | 37956452 | AX-170169924 | 0 | T | C | 0.193 | 8.41E-02 | 9.49E-02 | 2.84E+00 | 2.93E+00 | 3.71E-01 | no | Gm7799 | downstream_gene_variant | MODIFIER |
| 3 | 37962271 | AX-170234340 | 0 | A | G | 0.193 | 8.41E-02 | 9.49E-02 | 2.84E+00 | 2.93E+00 | 3.71E-01 | no | Gm7799 | upstream_gene_variant | MODIFIER |
| 3 | 37967875 | AX-170455935 | 0 | C | T | 0.193 | 8.41E-02 | 9.49E-02 | 2.84E+00 | 2.93E+00 | 3.71E-01 | no | Gm20755 | intron_variant,non_coding_transcript_variant | MODIFIER |
| 3 | 37984377 | AX-170524788 | 0 | A | G | 0.193 | 8.41E-02 | 9.49E-02 | 2.84E+00 | 2.93E+00 | 3.71E-01 | no | Gm20755 | intron_variant,non_coding_transcript_variant | MODIFIER |
| 3 | 37992133 | AX-170173810 | 0 | A | C | 0.193 | 8.41E-02 | 9.49E-02 | 2.84E+00 | 2.93E+00 | 3.71E-01 | no | Gm22899 | downstream_gene_variant | MODIFIER |
| 3 | 37995480 | AX-170333045 | 0 | A | C | 0.193 | 8.41E-02 | 9.49E-02 | 2.84E+00 | 2.93E+00 | 3.71E-01 | no | Gm22899 | upstream_gene_variant | MODIFIER |
| 3 | 37997606 | AX-170066398 | 0 | C | T | 0.193 | 8.41E-02 | 9.49E-02 | 2.84E+00 | 2.93E+00 | 3.71E-01 | no | Gm22899 | upstream_gene_variant | MODIFIER |
| 3 | 38001967 | AX-170066402 | 0 | T | C | 0.193 | 8.41E-02 | 9.49E-02 | 2.84E+00 | 2.93E+00 | 3.71E-01 | no | | intergenic_variant | MODIFIER |
| 3 | 38041493 | AX-170338594 | 0 | A | G | 0.193 | 8.41E-02 | 9.49E-02 | 2.84E+00 | 2.93E+00 | 3.71E-01 | no | | intergenic_variant | MODIFIER |
| 3 | 38041533 | AX-170146552 | 0 | T | C | 0.193 | 8.41E-02 | 9.49E-02 | 2.84E+00 | 2.93E+00 | 3.71E-01 | no | | intergenic_variant | MODIFIER |
| 3 | 38043343 | AX-170066399 | 0 | G | A | 0.193 | 8.41E-02 | 9.49E-02 | 2.84E+00 | 2.93E+00 | 3.71E-01 | no | | regulatory_region_variant | MODIFIER |
| 3 | 38071290 | AX-170525883 | 0 | C | A | 0.193 | 8.41E-02 | 9.49E-02 | 2.84E+00 | 2.93E+00 | 3.71E-01 | no | | intergenic_variant | MODIFIER |
| 6 | 138490539 | AX-170379805 | 0 | T | G | 0.57 | 7.76E-02 | 8.44E-02 | 2.42E+00 | 2.49E+00 | 3.72E-01 | no | Lmo3 | intron_variant | MODIFIER |
| 1 | 187943387 | AX-170172403 | 1 | G | T | 0.25 | 9.25E-02 | 1.01E-01 | 2.42E+00 | 2.51E+00 | 3.72E-01 | no | Esrrg | intron_variant | MODIFIER |
| 4 | 79682044 | AX-170214075 | 0 | C | A | 0.447 | -7.56E-02 | 8.42E-02 | 2.53E+00 | 2.60E+00 | 3.72E-01 | no | | intergenic_variant | MODIFIER |
| 14 | 113297339 | AX-170436131 | 1 | T | C | 0.339 | 6.50E-02 | 7.34E-02 | 2.61E+00 | 2.71E+00 | 3.72E-01 | no | | intergenic_variant | MODIFIER |
| 18 | 59801147 | AX-170339827 | 1 | A | G | 0.402 | 8.62E-02 | 9.37E-02 | 2.42E+00 | 2.47E+00 | 3.72E-01 | no | | intergenic_variant | MODIFIER |
| 10 | 71592718 | AX-170003446 | 1 | T | G | 0.42 | -7.12E-02 | 8.05E-02 | 2.65E+00 | 2.72E+00 | 3.72E-01 | no | Gm34544 | intron_variant,non_coding_transcript_variant | MODIFIER |
| 15 | 6632827 | AX-170378690 | 0 | A | C | 0.237 | -9.38E-02 | 1.06E-01 | 2.63E+00 | 2.69E+00 | 3.72E-01 | no | Fyb | intron_variant | MODIFIER |
| 3 | 37817560 | AX-170268033 | 1 | G | A | 0.188 | 8.34E-02 | 9.44E-02 | 2.83E+00 | 2.93E+00 | 3.72E-01 | no | | regulatory_region_variant | MODIFIER |
| 5 | 118096041 | AX-170463325 | 0 | G | A | 0.465 | -9.71E-02 | 1.08E-01 | 2.98E+00 | 3.07E+00 | 3.72E-01 | no | Fbxw8 | intron_variant | MODIFIER |
| 7 | 24723006 | AX-170263403 | 1 | C | G | 0.402 | 8.02E-02 | 9.01E-02 | 2.56E+00 | 2.67E+00 | 3.72E-01 | no | Gm4763 | intron_variant | MODIFIER |
| 5 | 45384956 | AX-170167481 | 1 | A | G | 0.5 | 9.29E-02 | 1.02E-01 | 2.44E+00 | 2.49E+00 | 3.72E-01 | no | D5Ertd615e | intron_variant,non_coding_transcript_variant | MODIFIER |
| 18 | 57259548 | AX-97843368 | 0 | G | A | 0.184 | -1.07E-01 | 1.21E-01 | 2.69E+00 | 2.74E+00 | 3.73E-01 | no | Megf10 | intron_variant | MODIFIER |
| 18 | 57407671 | AX-170153043 | 0 | T | C | 0.184 | -1.07E-01 | 1.21E-01 | 2.69E+00 | 2.74E+00 | 3.73E-01 | no | 4930511M06Rik | intron_variant,non_coding_transcript_variant | MODIFIER |
| 18 | 57427434 | AX-170425600 | 0 | A | G | 0.184 | -1.07E-01 | 1.21E-01 | 2.69E+00 | 2.74E+00 | 3.73E-01 | no | 4930511M06Rik | intron_variant,non_coding_transcript_variant | MODIFIER |
| 18 | 57437476 | AX-170133383 | 0 | A | G | 0.184 | -1.07E-01 | 1.21E-01 | 2.69E+00 | 2.74E+00 | 3.73E-01 | no | 4930511M06Rik | intron_variant,non_coding_transcript_variant | MODIFIER |
| 18 | 57437939 | AX-170314249 | 0 | G | A | 0.184 | -1.07E-01 | 1.21E-01 | 2.69E+00 | 2.74E+00 | 3.73E-01 | no | 4930511M06Rik | intron_variant,non_coding_transcript_variant | MODIFIER |
| 18 | 57438680 | AX-170213350 | 0 | T | C | 0.184 | -1.07E-01 | 1.21E-01 | 2.69E+00 | 2.74E+00 | 3.73E-01 | no | 4930511M06Rik | intron_variant,non_coding_transcript_variant | MODIFIER |
| 18 | 57440304 | AX-170386649 | 0 | C | T | 0.184 | -1.07E-01 | 1.21E-01 | 2.69E+00 | 2.74E+00 | 3.73E-01 | no | 4930511M06Rik | intron_variant,non_coding_transcript_variant | MODIFIER |
| 18 | 57473125 | AX-170125867 | 0 | T | C | 0.184 | -1.07E-01 | 1.21E-01 | 2.69E+00 | 2.74E+00 | 3.73E-01 | no | Ctxn3 | intron_variant | MODIFIER |
| 18 | 57501037 | AX-170119079 | 0 | A | G | 0.184 | -1.07E-01 | 1.21E-01 | 2.69E+00 | 2.74E+00 | 3.73E-01 | no | 4930511M06Rik | intron_variant,non_coding_transcript_variant | MODIFIER |
| 18 | 57502392 | AX-170428220 | 0 | A | G | 0.184 | -1.07E-01 | 1.21E-01 | 2.69E+00 | 2.74E+00 | 3.73E-01 | no | 4930511M06Rik | intron_variant,non_coding_transcript_variant | MODIFIER |
| 18 | 57502589 | AX-170535688 | 0 | G | C | 0.184 | -1.07E-01 | 1.21E-01 | 2.69E+00 | 2.74E+00 | 3.73E-01 | no | 4930511M06Rik | intron_variant,non_coding_transcript_variant | MODIFIER |
| 18 | 57504748 | AX-170515803 | 0 | C | G | 0.184 | -1.07E-01 | 1.21E-01 | 2.69E+00 | 2.74E+00 | 3.73E-01 | no | 4930511M06Rik | intron_variant,non_coding_transcript_variant | MODIFIER |
| 18 | 57598914 | AX-170463244 | 0 | T | A | 0.184 | -1.07E-01 | 1.21E-01 | 2.69E+00 | 2.74E+00 | 3.73E-01 | no | Ccdc192 | intron_variant | MODIFIER |
| 18 | 57657266 | AX-170165014 | 0 | C | T | 0.184 | -1.07E-01 | 1.21E-01 | 2.69E+00 | 2.74E+00 | 3.73E-01 | no | Ccdc192 | intron_variant | MODIFIER |
| 18 | 57669815 | AX-170503914 | 0 | G | T | 0.184 | -1.07E-01 | 1.21E-01 | 2.69E+00 | 2.74E+00 | 3.73E-01 | no | Ccdc192 | intron_variant | MODIFIER |
| 18 | 57742794 | AX-170349902 | 0 | A | C | 0.184 | -1.07E-01 | 1.21E-01 | 2.69E+00 | 2.74E+00 | 3.73E-01 | no | AC102232.1 | intron_variant,non_coding_transcript_variant | MODIFIER |
| 18 | 57742981 | AX-170093898 | 0 | T | C | 0.184 | -1.07E-01 | 1.21E-01 | 2.69E+00 | 2.74E+00 | 3.73E-01 | no | AC102232.1 | intron_variant,non_coding_transcript_variant | MODIFIER |
| 18 | 57746362 | AX-170449627 | 0 | G | A | 0.184 | -1.07E-01 | 1.21E-01 | 2.69E+00 | 2.74E+00 | 3.73E-01 | no | AC102232.1 | intron_variant,non_coding_transcript_variant | MODIFIER |
| 18 | 57789020 | AX-170403227 | 0 | T | C | 0.184 | -1.07E-01 | 1.21E-01 | 2.69E+00 | 2.74E+00 | 3.73E-01 | no | AC102232.1 | intron_variant,non_coding_transcript_variant | MODIFIER |
| 18 | 57806837 | AX-170184694 | 0 | C | A | 0.184 | -1.07E-01 | 1.21E-01 | 2.69E+00 | 2.74E+00 | 3.73E-01 | no | | regulatory_region_variant | MODIFIER |
| 18 | 57806858 | AX-170485108 | 0 | A | G | 0.184 | -1.07E-01 | 1.21E-01 | 2.69E+00 | 2.74E+00 | 3.73E-01 | no | | regulatory_region_variant | MODIFIER |
| 18 | 57822885 | AX-97834926 | 0 | A | G | 0.184 | -1.07E-01 | 1.21E-01 | 2.69E+00 | 2.74E+00 | 3.73E-01 | no | | intergenic_variant | MODIFIER |
| 18 | 57852415 | AX-170201510 | 0 | T | C | 0.184 | -1.07E-01 | 1.21E-01 | 2.69E+00 | 2.74E+00 | 3.73E-01 | no | | intergenic_variant | MODIFIER |
| 18 | 57862469 | AX-170420449 | 0 | A | G | 0.184 | -1.07E-01 | 1.21E-01 | 2.69E+00 | 2.74E+00 | 3.73E-01 | no | | intergenic_variant | MODIFIER |

| | | | | | | | | | | | | | | | |
|---|---|---|---|---|---|---|---|---|---|---|---|---|---|---|---|
| 18 | 57899980 | AX-97789644 | 0 | A | G | 0.184 | -1.07E-01 | 1.21E-01 | 2.69E+00 | 2.74E+00 | 3.73E-01 | no | Slc12a2 | intron_variant | MODIFIER |
| 18 | 57906966 | AX-170146722 | 0 | A | G | 0.184 | -1.07E-01 | 1.21E-01 | 2.69E+00 | 2.74E+00 | 3.73E-01 | no | Slc12a2 | intron_variant | MODIFIER |
| 18 | 57940964 | AX-170236829 | 0 | T | C | 0.184 | -1.07E-01 | 1.21E-01 | 2.69E+00 | 2.74E+00 | 3.73E-01 | no | Slc12a2 | intron_variant | MODIFIER |
| 18 | 57941013 | AX-97795453 | 0 | A | G | 0.184 | -1.07E-01 | 1.21E-01 | 2.69E+00 | 2.74E+00 | 3.73E-01 | no | Slc12a2 | missense_variant | MODERATE |
| 18 | 57960961 | AX-170310486 | 0 | T | G | 0.184 | -1.07E-01 | 1.21E-01 | 2.69E+00 | 2.74E+00 | 3.73E-01 | no | | intergenic_variant | MODIFIER |
| 18 | 58046442 | AX-170060954 | 0 | G | T | 0.184 | -1.07E-01 | 1.21E-01 | 2.69E+00 | 2.74E+00 | 3.73E-01 | no | Fbn2 | intron_variant | MODIFIER |
| 18 | 58060456 | AX-170099123 | 0 | A | G | 0.184 | -1.07E-01 | 1.21E-01 | 2.69E+00 | 2.74E+00 | 3.73E-01 | no | Fbn2 | intron_variant | MODIFIER |
| 18 | 58104197 | AX-170264104 | 0 | A | C | 0.184 | -1.07E-01 | 1.21E-01 | 2.69E+00 | 2.74E+00 | 3.73E-01 | no | Fbn2 | intron_variant | MODIFIER |
| 18 | 58145575 | AX-170323266 | 0 | A | G | 0.184 | -1.07E-01 | 1.21E-01 | 2.69E+00 | 2.74E+00 | 3.73E-01 | no | Fbn2 | intron_variant | MODIFIER |
| 18 | 58161894 | AX-170015785 | 0 | T | C | 0.184 | -1.07E-01 | 1.21E-01 | 2.69E+00 | 2.74E+00 | 3.73E-01 | no | Fbn2 | intron_variant | MODIFIER |
| 18 | 58169216 | AX-97849020 | 0 | A | G | 0.184 | -1.07E-01 | 1.21E-01 | 2.69E+00 | 2.74E+00 | 3.73E-01 | no | Fbn2 | intron_variant | MODIFIER |
| 18 | 58197513 | AX-170098625 | 0 | T | G | 0.184 | -1.07E-01 | 1.21E-01 | 2.69E+00 | 2.74E+00 | 3.73E-01 | no | Fbn2 | intron_variant | MODIFIER |
| 18 | 58197894 | AX-170174687 | 0 | T | G | 0.184 | -1.07E-01 | 1.21E-01 | 2.69E+00 | 2.74E+00 | 3.73E-01 | no | Fbn2 | intron_variant | MODIFIER |
| 18 | 58227522 | AX-170404241 | 0 | G | A | 0.184 | -1.07E-01 | 1.21E-01 | 2.69E+00 | 2.74E+00 | 3.73E-01 | no | Gm25660 | downstream_gene_variant | MODIFIER |
| 18 | 58248489 | AX-170194645 | 0 | T | G | 0.184 | -1.07E-01 | 1.21E-01 | 2.69E+00 | 2.74E+00 | 3.73E-01 | no | | intergenic_variant | MODIFIER |
| 18 | 58248908 | AX-170088348 | 0 | T | C | 0.184 | -1.07E-01 | 1.21E-01 | 2.69E+00 | 2.74E+00 | 3.73E-01 | no | | intergenic_variant | MODIFIER |
| 18 | 58294936 | AX-170102249 | 0 | A | G | 0.184 | -1.07E-01 | 1.21E-01 | 2.69E+00 | 2.74E+00 | 3.73E-01 | no | | intergenic_variant | MODIFIER |
| 18 | 58302553 | AX-170275674 | 0 | C | T | 0.184 | -1.07E-01 | 1.21E-01 | 2.69E+00 | 2.74E+00 | 3.73E-01 | no | | regulatory_region_variant | MODIFIER |
| 18 | 58305102 | AX-170159968 | 0 | T | C | 0.184 | -1.07E-01 | 1.21E-01 | 2.69E+00 | 2.74E+00 | 3.73E-01 | no | | intergenic_variant | MODIFIER |
| 18 | 58315844 | AX-170388756 | 0 | A | G | 0.184 | -1.07E-01 | 1.21E-01 | 2.69E+00 | 2.74E+00 | 3.73E-01 | no | | intergenic_variant | MODIFIER |
| 18 | 58338039 | AX-170316742 | 0 | G | A | 0.184 | -1.07E-01 | 1.21E-01 | 2.69E+00 | 2.74E+00 | 3.73E-01 | no | | intergenic_variant | MODIFIER |
| 18 | 58347340 | AX-170344807 | 0 | G | A | 0.184 | -1.07E-01 | 1.21E-01 | 2.69E+00 | 2.74E+00 | 3.73E-01 | no | | intergenic_variant | MODIFIER |
| 18 | 58352882 | AX-170514189 | 0 | C | G | 0.184 | -1.07E-01 | 1.21E-01 | 2.69E+00 | 2.74E+00 | 3.73E-01 | no | | intergenic_variant | MODIFIER |
| 18 | 58375420 | AX-170196882 | 0 | A | G | 0.184 | -1.07E-01 | 1.21E-01 | 2.69E+00 | 2.74E+00 | 3.73E-01 | no | | intergenic_variant | MODIFIER |
| 18 | 58386482 | AX-170143356 | 0 | T | C | 0.184 | -1.07E-01 | 1.21E-01 | 2.69E+00 | 2.74E+00 | 3.73E-01 | no | | regulatory_region_variant | MODIFIER |
| 18 | 58413984 | AX-170203048 | 0 | A | G | 0.184 | -1.07E-01 | 1.21E-01 | 2.69E+00 | 2.74E+00 | 3.73E-01 | no | | intergenic_variant | MODIFIER |
| 18 | 58608980 | AX-170523685 | 0 | T | C | 0.184 | -1.07E-01 | 1.21E-01 | 2.69E+00 | 2.74E+00 | 3.73E-01 | no | Slc27a6 | intron_variant | MODIFIER |
| 18 | 58660189 | AX-97823449 | 0 | C | T | 0.184 | -1.07E-01 | 1.21E-01 | 2.69E+00 | 2.74E+00 | 3.73E-01 | no | Isoc1 | intron_variant | MODIFIER |
| 18 | 58668333 | AX-170015792 | 0 | C | G | 0.184 | -1.07E-01 | 1.21E-01 | 2.69E+00 | 2.74E+00 | 3.73E-01 | no | Isoc1 | intron_variant | MODIFIER |
| 18 | 58673576 | AX-170409158 | 0 | T | C | 0.184 | -1.07E-01 | 1.21E-01 | 2.69E+00 | 2.74E+00 | 3.73E-01 | no | Isoc1 | intron_variant | MODIFIER |
| 18 | 58673614 | AX-170156810 | 0 | T | C | 0.184 | -1.07E-01 | 1.21E-01 | 2.69E+00 | 2.74E+00 | 3.73E-01 | no | Isoc1 | intron_variant | MODIFIER |
| 18 | 58678678 | AX-170015790 | 0 | T | A | 0.184 | -1.07E-01 | 1.21E-01 | 2.69E+00 | 2.74E+00 | 3.73E-01 | no | Isoc1 | 3_prime_UTR_variant | MODIFIER |
| 18 | 58860393 | AX-170088351 | 0 | T | G | 0.184 | -1.07E-01 | 1.21E-01 | 2.69E+00 | 2.74E+00 | 3.73E-01 | no | Adamts19 | intron_variant | MODIFIER |
| 18 | 58863009 | AX-170185002 | 0 | C | T | 0.184 | -1.07E-01 | 1.21E-01 | 2.69E+00 | 2.74E+00 | 3.73E-01 | no | Adamts19 | intron_variant | MODIFIER |
| 18 | 58867778 | AX-170434280 | 0 | C | T | 0.184 | -1.07E-01 | 1.21E-01 | 2.69E+00 | 2.74E+00 | 3.73E-01 | no | Adamts19 | intron_variant | MODIFIER |
| 18 | 58871643 | AX-170491082 | 0 | T | C | 0.184 | -1.07E-01 | 1.21E-01 | 2.69E+00 | 2.74E+00 | 3.73E-01 | no | Adamts19 | intron_variant | MODIFIER |
| 18 | 58873372 | AX-170164019 | 0 | T | C | 0.184 | -1.07E-01 | 1.21E-01 | 2.69E+00 | 2.74E+00 | 3.73E-01 | no | Adamts19 | intron_variant | MODIFIER |
| 18 | 58878949 | AX-170317761 | 0 | T | C | 0.184 | -1.07E-01 | 1.21E-01 | 2.69E+00 | 2.74E+00 | 3.73E-01 | no | Adamts19 | intron_variant | MODIFIER |
| 18 | 58878988 | AX-170412890 | 0 | G | A | 0.184 | -1.07E-01 | 1.21E-01 | 2.69E+00 | 2.74E+00 | 3.73E-01 | no | Adamts19 | intron_variant | MODIFIER |
| 18 | 58880064 | AX-168384569 | 0 | T | C | 0.184 | -1.07E-01 | 1.21E-01 | 2.69E+00 | 2.74E+00 | 3.73E-01 | no | Adamts19 | intron_variant | MODIFIER |
| 18 | 58948198 | AX-170098466 | 0 | T | A | 0.184 | -1.07E-01 | 1.21E-01 | 2.69E+00 | 2.74E+00 | 3.73E-01 | no | Adamts19 | intron_variant | MODIFIER |
| 18 | 58978444 | AX-170149059 | 0 | T | C | 0.184 | -1.07E-01 | 1.21E-01 | 2.69E+00 | 2.74E+00 | 3.73E-01 | no | Adamts19 | intron_variant | MODIFIER |
| 18 | 58992616 | AX-170060980 | 0 | G | A | 0.184 | -1.07E-01 | 1.21E-01 | 2.69E+00 | 2.74E+00 | 3.73E-01 | no | Adamts19 | intron_variant | MODIFIER |
| 18 | 59107697 | AX-170375754 | 0 | G | A | 0.184 | -1.07E-01 | 1.21E-01 | 2.69E+00 | 2.74E+00 | 3.73E-01 | no | | intergenic_variant | MODIFIER |
| 18 | 59140860 | AX-170240602 | 0 | T | G | 0.184 | -1.07E-01 | 1.21E-01 | 2.69E+00 | 2.74E+00 | 3.73E-01 | no | | intergenic_variant | MODIFIER |
| 18 | 59181846 | AX-170015800 | 0 | A | G | 0.184 | -1.07E-01 | 1.21E-01 | 2.69E+00 | 2.74E+00 | 3.73E-01 | no | Chsy3 | intron_variant | MODIFIER |
| 18 | 59205844 | AX-170323476 | 0 | T | C | 0.184 | -1.07E-01 | 1.21E-01 | 2.69E+00 | 2.74E+00 | 3.73E-01 | no | Chsy3 | intron_variant | MODIFIER |
| 18 | 59208057 | AX-170015795 | 0 | A | C | 0.184 | -1.07E-01 | 1.21E-01 | 2.69E+00 | 2.74E+00 | 3.73E-01 | no | Chsy3 | intron_variant | MODIFIER |
| 18 | 59243135 | AX-170129918 | 0 | T | A | 0.184 | -1.07E-01 | 1.21E-01 | 2.69E+00 | 2.74E+00 | 3.73E-01 | no | Chsy3 | intron_variant | MODIFIER |
| 18 | 59258386 | AX-170015798 | 0 | C | T | 0.184 | -1.07E-01 | 1.21E-01 | 2.69E+00 | 2.74E+00 | 3.73E-01 | no | Chsy3 | intron_variant | MODIFIER |
| 18 | 59261883 | AX-170435240 | 0 | C | G | 0.184 | -1.07E-01 | 1.21E-01 | 2.69E+00 | 2.74E+00 | 3.73E-01 | no | Chsy3 | intron_variant | MODIFIER |
| 18 | 59293908 | AX-170015797 | 0 | A | T | 0.184 | -1.07E-01 | 1.21E-01 | 2.69E+00 | 2.74E+00 | 3.73E-01 | no | Chsy3 | intron_variant | MODIFIER |
| 18 | 59328047 | AX-170060986 | 0 | G | T | 0.184 | -1.07E-01 | 1.21E-01 | 2.69E+00 | 2.74E+00 | 3.73E-01 | no | Chsy3 | intron_variant | MODIFIER |
| 18 | 59331052 | AX-170015796 | 0 | T | C | 0.184 | -1.07E-01 | 1.21E-01 | 2.69E+00 | 2.74E+00 | 3.73E-01 | no | Chsy3 | intron_variant | MODIFIER |

| | | | | | | | | | | | | | | | |
|---|---|---|---|---|---|---|---|---|---|---|---|---|---|---|---|
| 18 | 59342234 | AX-170226203 | 0 | C | T | 0.184 | -1.07E-01 | 1.21E-01 | 2.69E+00 | 2.74E+00 | 3.73E-01 | no | Chsy3 | intron_variant | MODIFIER |
| 18 | 59355917 | AX-170498422 | 0 | A | G | 0.184 | -1.07E-01 | 1.21E-01 | 2.69E+00 | 2.74E+00 | 3.73E-01 | no | Chsy3 | intron_variant | MODIFIER |
| 18 | 59375182 | AX-170392467 | 0 | T | A | 0.184 | -1.07E-01 | 1.21E-01 | 2.69E+00 | 2.74E+00 | 3.73E-01 | no | Chsy3 | intron_variant | MODIFIER |
| 18 | 59383394 | AX-170060990 | 0 | T | C | 0.184 | -1.07E-01 | 1.21E-01 | 2.69E+00 | 2.74E+00 | 3.73E-01 | no | Chsy3 | intron_variant | MODIFIER |
| 18 | 59405129 | AX-170455783 | 0 | T | C | 0.184 | -1.07E-01 | 1.21E-01 | 2.69E+00 | 2.74E+00 | 3.73E-01 | no | Chsy3 | intron_variant | MODIFIER |
| 18 | 59610730 | AX-170315104 | 0 | G | A | 0.184 | -1.07E-01 | 1.21E-01 | 2.69E+00 | 2.74E+00 | 3.73E-01 | no | | intergenic_variant | MODIFIER |
| 18 | 59821202 | AX-170015802 | 0 | A | G | 0.184 | -1.07E-01 | 1.21E-01 | 2.69E+00 | 2.74E+00 | 3.73E-01 | no | AC138134.1 | downstream_gene_variant | MODIFIER |
| 18 | 59828826 | AX-170015804 | 0 | G | A | 0.184 | -1.07E-01 | 1.21E-01 | 2.69E+00 | 2.74E+00 | 3.73E-01 | no | AC138134.1 | intron_variant,non_coding_transcript_variant | MODIFIER |
| 18 | 60123815 | AX-170509676 | 0 | G | T | 0.184 | -1.07E-01 | 1.21E-01 | 2.69E+00 | 2.74E+00 | 3.73E-01 | no | | intergenic_variant | MODIFIER |
| 18 | 60124259 | AX-170395028 | 0 | A | T | 0.184 | -1.07E-01 | 1.21E-01 | 2.69E+00 | 2.74E+00 | 3.73E-01 | no | | intergenic_variant | MODIFIER |
| 18 | 60125038 | AX-97857280 | 0 | C | T | 0.184 | -1.07E-01 | 1.21E-01 | 2.69E+00 | 2.74E+00 | 3.73E-01 | no | | intergenic_variant | MODIFIER |
| 18 | 60129043 | AX-170156714 | 0 | A | G | 0.184 | -1.07E-01 | 1.21E-01 | 2.69E+00 | 2.74E+00 | 3.73E-01 | no | | intergenic_variant | MODIFIER |
| 18 | 60137663 | AX-170402722 | 0 | G | A | 0.184 | -1.07E-01 | 1.21E-01 | 2.69E+00 | 2.74E+00 | 3.73E-01 | no | | intergenic_variant | MODIFIER |
| 18 | 60525050 | AX-97834928 | 0 | G | A | 0.184 | -1.07E-01 | 1.21E-01 | 2.69E+00 | 2.74E+00 | 3.73E-01 | no | Dctn4 | upstream_gene_variant | MODIFIER |
| 18 | 60707868 | AX-170474488 | 0 | T | C | 0.184 | -1.07E-01 | 1.21E-01 | 2.69E+00 | 2.74E+00 | 3.73E-01 | no | Ndst1 | synonymous_variant | LOW |
| 18 | 61345703 | AX-170385504 | 0 | T | C | 0.184 | -1.07E-01 | 1.21E-01 | 2.69E+00 | 2.74E+00 | 3.73E-01 | no | Ppargc1b | intron_variant | MODIFIER |
| 18 | 61350000 | AX-170345311 | 0 | T | C | 0.184 | -1.07E-01 | 1.21E-01 | 2.69E+00 | 2.74E+00 | 3.73E-01 | no | Ppargc1b | intron_variant | MODIFIER |
| 18 | 61351541 | AX-170499437 | 0 | A | G | 0.184 | -1.07E-01 | 1.21E-01 | 2.69E+00 | 2.74E+00 | 3.73E-01 | no | Ppargc1b | intron_variant | MODIFIER |
| 18 | 61384983 | AX-170514081 | 0 | G | T | 0.184 | -1.07E-01 | 1.21E-01 | 2.69E+00 | 2.74E+00 | 3.73E-01 | no | Ppargc1b | intron_variant | MODIFIER |
| 18 | 61385220 | AX-170487736 | 0 | T | C | 0.184 | -1.07E-01 | 1.21E-01 | 2.69E+00 | 2.74E+00 | 3.73E-01 | no | Ppargc1b | intron_variant | MODIFIER |
| 18 | 61385463 | AX-170104703 | 0 | A | G | 0.184 | -1.07E-01 | 1.21E-01 | 2.69E+00 | 2.74E+00 | 3.73E-01 | no | Ppargc1b | intron_variant | MODIFIER |
| 18 | 61479825 | AX-170313466 | 0 | A | G | 0.184 | -1.07E-01 | 1.21E-01 | 2.69E+00 | 2.74E+00 | 3.73E-01 | no | AC153155.2 | intron_variant,non_coding_transcript_variant | MODIFIER |
| 18 | 61495651 | AX-170373917 | 0 | A | C | 0.184 | -1.07E-01 | 1.21E-01 | 2.69E+00 | 2.74E+00 | 3.73E-01 | no | Arhgef37 | intron_variant | MODIFIER |
| 18 | 61500800 | AX-170061033 | 0 | T | C | 0.184 | -1.07E-01 | 1.21E-01 | 2.69E+00 | 2.74E+00 | 3.73E-01 | no | Arhgef37 | intron_variant | MODIFIER |
| 18 | 61502806 | AX-170457960 | 0 | G | A | 0.184 | -1.07E-01 | 1.21E-01 | 2.69E+00 | 2.74E+00 | 3.73E-01 | no | Arhgef37 | intron_variant | MODIFIER |
| 18 | 61578326 | AX-170015828 | 0 | C | T | 0.184 | -1.07E-01 | 1.21E-01 | 2.69E+00 | 2.74E+00 | 3.73E-01 | no | Csnk1a1 | intron_variant | MODIFIER |
| 18 | 61587335 | AX-170313925 | 0 | A | G | 0.184 | -1.07E-01 | 1.21E-01 | 2.69E+00 | 2.74E+00 | 3.73E-01 | no | Csnk1a1 | intron_variant | MODIFIER |
| 18 | 61635121 | AX-170015830 | 0 | G | C | 0.184 | -1.07E-01 | 1.21E-01 | 2.69E+00 | 2.74E+00 | 3.73E-01 | no | Bvht | upstream_gene_variant | MODIFIER |
| 18 | 61738800 | AX-97853419 | 0 | T | C | 0.184 | -1.07E-01 | 1.21E-01 | 2.69E+00 | 2.74E+00 | 3.73E-01 | no | Afap1l1 | intron_variant | MODIFIER |
| 18 | 61745732 | AX-170396041 | 0 | T | C | 0.184 | -1.07E-01 | 1.21E-01 | 2.69E+00 | 2.74E+00 | 3.73E-01 | no | Afap1l1 | intron_variant | MODIFIER |
| 18 | 61905123 | AX-170361493 | 0 | A | T | 0.184 | -1.07E-01 | 1.21E-01 | 2.69E+00 | 2.74E+00 | 3.73E-01 | no | Ablim3 | intron_variant | MODIFIER |
| 18 | 61922129 | AX-170505289 | 0 | A | G | 0.184 | -1.07E-01 | 1.21E-01 | 2.69E+00 | 2.74E+00 | 3.73E-01 | no | | intergenic_variant | MODIFIER |
| 18 | 61925948 | AX-170198929 | 0 | C | A | 0.184 | -1.07E-01 | 1.21E-01 | 2.69E+00 | 2.74E+00 | 3.73E-01 | no | | regulatory_region_variant | MODIFIER |
| 18 | 61947201 | AX-170207416 | 0 | T | G | 0.184 | -1.07E-01 | 1.21E-01 | 2.69E+00 | 2.74E+00 | 3.73E-01 | no | | intergenic_variant | MODIFIER |
| 18 | 61947313 | AX-170347851 | 0 | T | A | 0.184 | -1.07E-01 | 1.21E-01 | 2.69E+00 | 2.74E+00 | 3.73E-01 | no | | intergenic_variant | MODIFIER |
| 18 | 61975534 | AX-170444384 | 0 | C | T | 0.184 | -1.07E-01 | 1.21E-01 | 2.69E+00 | 2.74E+00 | 3.73E-01 | no | Sh3tc2 | intron_variant | MODIFIER |
| 18 | 62032622 | AX-170500906 | 0 | A | T | 0.184 | -1.07E-01 | 1.21E-01 | 2.69E+00 | 2.74E+00 | 3.73E-01 | no | | intergenic_variant | MODIFIER |
| 18 | 62036107 | AX-170093908 | 0 | A | G | 0.184 | -1.07E-01 | 1.21E-01 | 2.69E+00 | 2.74E+00 | 3.73E-01 | no | | intergenic_variant | MODIFIER |
| 18 | 62043720 | AX-170253463 | 0 | A | G | 0.184 | -1.07E-01 | 1.21E-01 | 2.69E+00 | 2.74E+00 | 3.73E-01 | no | | regulatory_region_variant | MODIFIER |
| 18 | 62044980 | AX-170206538 | 0 | C | T | 0.184 | -1.07E-01 | 1.21E-01 | 2.69E+00 | 2.74E+00 | 3.73E-01 | no | | regulatory_region_variant | MODIFIER |
| 18 | 62045088 | AX-170213447 | 0 | C | G | 0.184 | -1.07E-01 | 1.21E-01 | 2.69E+00 | 2.74E+00 | 3.73E-01 | no | | regulatory_region_variant | MODIFIER |
| 18 | 62050821 | AX-170015831 | 0 | A | C | 0.184 | -1.07E-01 | 1.21E-01 | 2.69E+00 | 2.74E+00 | 3.73E-01 | no | | intergenic_variant | MODIFIER |
| 18 | 62051455 | AX-170451627 | 0 | G | C | 0.184 | -1.07E-01 | 1.21E-01 | 2.69E+00 | 2.74E+00 | 3.73E-01 | no | | intergenic_variant | MODIFIER |
| 18 | 62094684 | AX-170015837 | 0 | T | G | 0.184 | -1.07E-01 | 1.21E-01 | 2.69E+00 | 2.74E+00 | 3.73E-01 | no | | intergenic_variant | MODIFIER |
| 18 | 62105645 | AX-97786745 | 0 | A | G | 0.184 | -1.07E-01 | 1.21E-01 | 2.69E+00 | 2.74E+00 | 3.73E-01 | no | | intergenic_variant | MODIFIER |
| 18 | 62117948 | AX-170111749 | 0 | A | G | 0.184 | -1.07E-01 | 1.21E-01 | 2.69E+00 | 2.74E+00 | 3.73E-01 | no | | regulatory_region_variant | MODIFIER |
| 18 | 62118442 | AX-170524254 | 0 | T | A | 0.184 | -1.07E-01 | 1.21E-01 | 2.69E+00 | 2.74E+00 | 3.73E-01 | no | | intergenic_variant | MODIFIER |
| 18 | 62121358 | AX-170122953 | 0 | A | G | 0.184 | -1.07E-01 | 1.21E-01 | 2.69E+00 | 2.74E+00 | 3.73E-01 | no | | intergenic_variant | MODIFIER |
| 18 | 62121448 | AX-170455764 | 0 | A | G | 0.184 | -1.07E-01 | 1.21E-01 | 2.69E+00 | 2.74E+00 | 3.73E-01 | no | | intergenic_variant | MODIFIER |
| 18 | 62124547 | AX-170199669 | 0 | G | C | 0.184 | -1.07E-01 | 1.21E-01 | 2.69E+00 | 2.74E+00 | 3.73E-01 | no | | intergenic_variant | MODIFIER |
| 18 | 62124792 | AX-170015832 | 0 | A | G | 0.184 | -1.07E-01 | 1.21E-01 | 2.69E+00 | 2.74E+00 | 3.73E-01 | no | | intergenic_variant | MODIFIER |
| 18 | 62125548 | AX-170240399 | 0 | G | C | 0.184 | -1.07E-01 | 1.21E-01 | 2.69E+00 | 2.74E+00 | 3.73E-01 | no | | intergenic_variant | MODIFIER |
| 18 | 62125663 | AX-97823311 | 0 | C | T | 0.184 | -1.07E-01 | 1.21E-01 | 2.69E+00 | 2.74E+00 | 3.73E-01 | no | | intergenic_variant | MODIFIER |
| 18 | 62180796 | AX-97835630 | 0 | C | T | 0.184 | -1.07E-01 | 1.21E-01 | 2.69E+00 | 2.74E+00 | 3.73E-01 | no | Adrb2 | upstream_gene_variant | MODIFIER |

| | | | | | | | | | | | | | | | |
|---|---|---|---|---|---|---|---|---|---|---|---|---|---|---|---|
| 18 | 62185164 | AX-170015836 | 0 | A | G | 0.184 | -1.07E-01 | 1.21E-01 | 2.69E+00 | 2.74E+00 | 3.73E-01 | no | Gm9949 | downstream_gene_variant | MODIFIER |
| 18 | 62230437 | AX-97807392 | 0 | T | C | 0.184 | -1.07E-01 | 1.21E-01 | 2.69E+00 | 2.74E+00 | 3.73E-01 | no | | regulatory_region_variant | MODIFIER |
| 18 | 62231800 | AX-170383831 | 0 | C | T | 0.184 | -1.07E-01 | 1.21E-01 | 2.69E+00 | 2.74E+00 | 3.73E-01 | no | | regulatory_region_variant | MODIFIER |
| 18 | 62235064 | AX-170390943 | 0 | T | G | 0.184 | -1.07E-01 | 1.21E-01 | 2.69E+00 | 2.74E+00 | 3.73E-01 | no | | intergenic_variant | MODIFIER |
| 18 | 62256502 | AX-97798073 | 0 | G | T | 0.184 | -1.07E-01 | 1.21E-01 | 2.69E+00 | 2.74E+00 | 3.73E-01 | no | | intergenic_variant | MODIFIER |
| 18 | 62256635 | AX-170195911 | 0 | A | G | 0.184 | -1.07E-01 | 1.21E-01 | 2.69E+00 | 2.74E+00 | 3.73E-01 | no | | intergenic_variant | MODIFIER |
| 18 | 62257558 | AX-170367664 | 0 | G | C | 0.184 | -1.07E-01 | 1.21E-01 | 2.69E+00 | 2.74E+00 | 3.73E-01 | no | | intergenic_variant | MODIFIER |
| 18 | 62504913 | AX-170061056 | 0 | C | T | 0.184 | -1.07E-01 | 1.21E-01 | 2.69E+00 | 2.74E+00 | 3.73E-01 | no | Fbxo38 | intron_variant | MODIFIER |
| 18 | 62505503 | AX-170417900 | 0 | G | T | 0.184 | -1.07E-01 | 1.21E-01 | 2.69E+00 | 2.74E+00 | 3.73E-01 | no | Fbxo38 | synonymous_variant | LOW |
| 18 | 62514485 | AX-170270034 | 0 | G | A | 0.184 | -1.07E-01 | 1.21E-01 | 2.69E+00 | 2.74E+00 | 3.73E-01 | no | Fbxo38 | intron_variant | MODIFIER |
| 18 | 62522999 | AX-170140546 | 0 | C | T | 0.184 | -1.07E-01 | 1.21E-01 | 2.69E+00 | 2.74E+00 | 3.73E-01 | no | Fbxo38 | intron_variant | MODIFIER |
| 18 | 62523358 | AX-168406704 | 0 | C | T | 0.184 | -1.07E-01 | 1.21E-01 | 2.69E+00 | 2.74E+00 | 3.73E-01 | no | Fbxo38 | intron_variant | MODIFIER |
| 18 | 62526910 | AX-170374895 | 0 | G | A | 0.184 | -1.07E-01 | 1.21E-01 | 2.69E+00 | 2.74E+00 | 3.73E-01 | no | Fbxo38 | intron_variant | MODIFIER |
| 18 | 62527075 | AX-170405918 | 0 | T | C | 0.184 | -1.07E-01 | 1.21E-01 | 2.69E+00 | 2.74E+00 | 3.73E-01 | no | Fbxo38 | intron_variant | MODIFIER |
| 18 | 62528984 | AX-97844974 | 0 | C | T | 0.184 | -1.07E-01 | 1.21E-01 | 2.69E+00 | 2.74E+00 | 3.73E-01 | no | Fbxo38 | intron_variant | MODIFIER |
| 18 | 62529433 | AX-170369472 | 0 | A | G | 0.184 | -1.07E-01 | 1.21E-01 | 2.69E+00 | 2.74E+00 | 3.73E-01 | no | Fbxo38 | intron_variant | MODIFIER |
| 18 | 62542663 | AX-170492191 | 0 | C | A | 0.184 | -1.07E-01 | 1.21E-01 | 2.69E+00 | 2.74E+00 | 3.73E-01 | no | Fbxo38 | intron_variant | MODIFIER |
| 18 | 62544213 | AX-170300082 | 0 | G | A | 0.184 | -1.07E-01 | 1.21E-01 | 2.69E+00 | 2.74E+00 | 3.73E-01 | no | Fbxo38 | intron_variant | MODIFIER |
| 18 | 62544485 | AX-170129912 | 0 | A | G | 0.184 | -1.07E-01 | 1.21E-01 | 2.69E+00 | 2.74E+00 | 3.73E-01 | no | Fbxo38 | intron_variant | MODIFIER |
| 18 | 62547866 | AX-170141560 | 0 | G | T | 0.184 | -1.07E-01 | 1.21E-01 | 2.69E+00 | 2.74E+00 | 3.73E-01 | no | Fbxo38 | intron_variant | MODIFIER |
| 18 | 62730776 | AX-170323827 | 0 | T | C | 0.184 | -1.07E-01 | 1.21E-01 | 2.69E+00 | 2.74E+00 | 3.73E-01 | no | Spink13 | intron_variant | MODIFIER |
| 18 | 62732643 | AX-170133703 | 0 | T | C | 0.184 | -1.07E-01 | 1.21E-01 | 2.69E+00 | 2.74E+00 | 3.73E-01 | no | Spink13 | intron_variant | MODIFIER |
| 18 | 62769557 | AX-97846297 | 0 | A | G | 0.184 | -1.07E-01 | 1.21E-01 | 2.69E+00 | 2.74E+00 | 3.73E-01 | no | | regulatory_region_variant | MODIFIER |
| 18 | 62769617 | AX-170191341 | 0 | C | G | 0.184 | -1.07E-01 | 1.21E-01 | 2.69E+00 | 2.74E+00 | 3.73E-01 | no | | intergenic_variant | MODIFIER |
| 18 | 62770259 | AX-170533511 | 0 | G | A | 0.184 | -1.07E-01 | 1.21E-01 | 2.69E+00 | 2.74E+00 | 3.73E-01 | no | | regulatory_region_variant | MODIFIER |
| 18 | 62774351 | AX-170122081 | 0 | C | T | 0.184 | -1.07E-01 | 1.21E-01 | 2.69E+00 | 2.74E+00 | 3.73E-01 | no | | regulatory_region_variant | MODIFIER |
| 18 | 62776819 | AX-170151827 | 0 | G | A | 0.184 | -1.07E-01 | 1.21E-01 | 2.69E+00 | 2.74E+00 | 3.73E-01 | no | | intergenic_variant | MODIFIER |
| 18 | 62820772 | AX-170378634 | 0 | T | A | 0.184 | -1.07E-01 | 1.21E-01 | 2.69E+00 | 2.74E+00 | 3.73E-01 | no | | intergenic_variant | MODIFIER |
| 18 | 62834791 | AX-97809793 | 0 | T | C | 0.184 | -1.07E-01 | 1.21E-01 | 2.69E+00 | 2.74E+00 | 3.73E-01 | no | | intergenic_variant | MODIFIER |
| 18 | 62840704 | AX-170170201 | 0 | T | C | 0.184 | -1.07E-01 | 1.21E-01 | 2.69E+00 | 2.74E+00 | 3.73E-01 | no | | intergenic_variant | MODIFIER |
| 18 | 62845346 | AX-170358273 | 0 | C | T | 0.184 | -1.07E-01 | 1.21E-01 | 2.69E+00 | 2.74E+00 | 3.73E-01 | no | | intergenic_variant | MODIFIER |
| 18 | 62846900 | AX-170155964 | 0 | G | A | 0.184 | -1.07E-01 | 1.21E-01 | 2.69E+00 | 2.74E+00 | 3.73E-01 | no | | intergenic_variant | MODIFIER |
| 18 | 62847429 | AX-170061065 | 0 | C | T | 0.184 | -1.07E-01 | 1.21E-01 | 2.69E+00 | 2.74E+00 | 3.73E-01 | no | | regulatory_region_variant | MODIFIER |
| 18 | 62848050 | AX-170093910 | 0 | G | A | 0.184 | -1.07E-01 | 1.21E-01 | 2.69E+00 | 2.74E+00 | 3.73E-01 | no | | regulatory_region_variant | MODIFIER |
| 18 | 62848724 | AX-170061063 | 0 | G | A | 0.184 | -1.07E-01 | 1.21E-01 | 2.69E+00 | 2.74E+00 | 3.73E-01 | no | | regulatory_region_variant | MODIFIER |
| 18 | 62853082 | AX-170321426 | 0 | C | A | 0.184 | -1.07E-01 | 1.21E-01 | 2.69E+00 | 2.74E+00 | 3.73E-01 | no | | intergenic_variant | MODIFIER |
| 18 | 62874835 | AX-170351437 | 0 | C | A | 0.184 | -1.07E-01 | 1.21E-01 | 2.69E+00 | 2.74E+00 | 3.73E-01 | no | | intergenic_variant | MODIFIER |
| 18 | 62882478 | AX-97834764 | 0 | T | C | 0.184 | -1.07E-01 | 1.21E-01 | 2.69E+00 | 2.74E+00 | 3.73E-01 | no | | intergenic_variant | MODIFIER |
| 18 | 62885851 | AX-170499187 | 0 | G | T | 0.184 | -1.07E-01 | 1.21E-01 | 2.69E+00 | 2.74E+00 | 3.73E-01 | no | | regulatory_region_variant | MODIFIER |
| 18 | 62960371 | AX-170187525 | 0 | A | G | 0.184 | -1.07E-01 | 1.21E-01 | 2.69E+00 | 2.74E+00 | 3.73E-01 | no | Apcdd1 | 3_prime_UTR_variant | MODIFIER |
| 18 | 62976980 | AX-170280337 | 0 | T | C | 0.184 | -1.07E-01 | 1.21E-01 | 2.69E+00 | 2.74E+00 | 3.73E-01 | no | Napg | upstream_gene_variant | MODIFIER |
| 18 | 63222871 | AX-170015845 | 0 | T | G | 0.184 | -1.07E-01 | 1.21E-01 | 2.69E+00 | 2.74E+00 | 3.73E-01 | no | Piezo2 | intron_variant | MODIFIER |
| 18 | 63314884 | AX-170225582 | 0 | T | A | 0.184 | -1.07E-01 | 1.21E-01 | 2.69E+00 | 2.74E+00 | 3.73E-01 | no | Piezo2 | intron_variant | MODIFIER |
| 18 | 63314915 | AX-170361560 | 0 | A | G | 0.184 | -1.07E-01 | 1.21E-01 | 2.69E+00 | 2.74E+00 | 3.73E-01 | no | Piezo2 | intron_variant | MODIFIER |
| 18 | 63329548 | AX-97833231 | 0 | A | G | 0.184 | -1.07E-01 | 1.21E-01 | 2.69E+00 | 2.74E+00 | 3.73E-01 | no | Piezo2 | upstream_gene_variant | MODIFIER |
| 18 | 63340841 | AX-170211478 | 0 | T | C | 0.184 | -1.07E-01 | 1.21E-01 | 2.69E+00 | 2.74E+00 | 3.73E-01 | no | Piezo2 | intron_variant | MODIFIER |
| 18 | 63341302 | AX-170362874 | 0 | A | G | 0.184 | -1.07E-01 | 1.21E-01 | 2.69E+00 | 2.74E+00 | 3.73E-01 | no | Piezo2 | intron_variant | MODIFIER |
| 18 | 63346585 | AX-170516556 | 0 | G | A | 0.184 | -1.07E-01 | 1.21E-01 | 2.69E+00 | 2.74E+00 | 3.73E-01 | no | Piezo2 | intron_variant | MODIFIER |
| 18 | 63348286 | AX-170335688 | 0 | C | G | 0.184 | -1.07E-01 | 1.21E-01 | 2.69E+00 | 2.74E+00 | 3.73E-01 | no | Piezo2 | intron_variant | MODIFIER |
| 18 | 63349042 | AX-170111421 | 0 | T | C | 0.184 | -1.07E-01 | 1.21E-01 | 2.69E+00 | 2.74E+00 | 3.73E-01 | no | Piezo2 | intron_variant | MODIFIER |
| 18 | 63350895 | AX-170315777 | 0 | T | A | 0.184 | -1.07E-01 | 1.21E-01 | 2.69E+00 | 2.74E+00 | 3.73E-01 | no | Piezo2 | intron_variant | MODIFIER |
| 18 | 63350946 | AX-170151616 | 0 | T | A | 0.184 | -1.07E-01 | 1.21E-01 | 2.69E+00 | 2.74E+00 | 3.73E-01 | no | Piezo2 | intron_variant | MODIFIER |
| 18 | 63363058 | AX-170354901 | 0 | A | G | 0.184 | -1.07E-01 | 1.21E-01 | 2.69E+00 | 2.74E+00 | 3.73E-01 | no | Piezo2 | intron_variant | MODIFIER |
| 18 | 63368737 | AX-170392240 | 0 | G | A | 0.184 | -1.07E-01 | 1.21E-01 | 2.69E+00 | 2.74E+00 | 3.73E-01 | no | Piezo2 | intron_variant | MODIFIER |

| | | | | | | | | | | | | | | | |
|---|---|---|---|---|---|---|---|---|---|---|---|---|---|---|---|
| 18 | 63411334 | AX-170061079 | 0 | A | G | 0.184 | -1.07E-01 | 1.21E-01 | 2.69E+00 | 2.74E+00 | 3.73E-01 | no | Gm26972 | intron_variant,non_coding_transcript_variant | MODIFIER |
| 18 | 63420471 | AX-170251434 | 0 | A | G | 0.184 | -1.07E-01 | 1.21E-01 | 2.69E+00 | 2.74E+00 | 3.73E-01 | no | Gm26972 | intron_variant,non_coding_transcript_variant | MODIFIER |
| 18 | 63420512 | AX-170061080 | 0 | G | A | 0.184 | -1.07E-01 | 1.21E-01 | 2.69E+00 | 2.74E+00 | 3.73E-01 | no | Gm26972 | intron_variant,non_coding_transcript_variant | MODIFIER |
| 18 | 63421900 | AX-170061082 | 0 | A | G | 0.184 | -1.07E-01 | 1.21E-01 | 2.69E+00 | 2.74E+00 | 3.73E-01 | no | Gm26972 | downstream_gene_variant | MODIFIER |
| 18 | 63421938 | AX-170061083 | 0 | G | A | 0.184 | -1.07E-01 | 1.21E-01 | 2.69E+00 | 2.74E+00 | 3.73E-01 | no | Gm26972 | downstream_gene_variant | MODIFIER |
| 18 | 63431601 | AX-170258691 | 0 | T | C | 0.184 | -1.07E-01 | 1.21E-01 | 2.69E+00 | 2.74E+00 | 3.73E-01 | no | | intergenic_variant | MODIFIER |
| 18 | 63440866 | AX-170157386 | 0 | A | G | 0.184 | -1.07E-01 | 1.21E-01 | 2.69E+00 | 2.74E+00 | 3.73E-01 | no | | intergenic_variant | MODIFIER |
| 18 | 63481628 | AX-170454097 | 0 | T | C | 0.184 | -1.07E-01 | 1.21E-01 | 2.69E+00 | 2.74E+00 | 3.73E-01 | no | | intergenic_variant | MODIFIER |
| 18 | 63483906 | AX-170465648 | 0 | T | C | 0.184 | -1.07E-01 | 1.21E-01 | 2.69E+00 | 2.74E+00 | 3.73E-01 | no | | intergenic_variant | MODIFIER |
| 18 | 63484034 | AX-170225773 | 0 | G | C | 0.184 | -1.07E-01 | 1.21E-01 | 2.69E+00 | 2.74E+00 | 3.73E-01 | no | | intergenic_variant | MODIFIER |
| 18 | 63573263 | AX-170256124 | 0 | G | A | 0.184 | -1.07E-01 | 1.21E-01 | 2.69E+00 | 2.74E+00 | 3.73E-01 | no | | intergenic_variant | MODIFIER |
| 18 | 63574070 | AX-170425873 | 0 | C | A | 0.184 | -1.07E-01 | 1.21E-01 | 2.69E+00 | 2.74E+00 | 3.73E-01 | no | | intergenic_variant | MODIFIER |
| 18 | 63574089 | AX-170135169 | 0 | T | C | 0.184 | -1.07E-01 | 1.21E-01 | 2.69E+00 | 2.74E+00 | 3.73E-01 | no | | intergenic_variant | MODIFIER |
| 18 | 63593940 | AX-170425366 | 0 | G | T | 0.184 | -1.07E-01 | 1.21E-01 | 2.69E+00 | 2.74E+00 | 3.73E-01 | no | | intergenic_variant | MODIFIER |
| 18 | 63599189 | AX-170169480 | 0 | A | G | 0.184 | -1.07E-01 | 1.21E-01 | 2.69E+00 | 2.74E+00 | 3.73E-01 | no | | intergenic_variant | MODIFIER |
| 18 | 63619132 | AX-170110098 | 0 | G | C | 0.184 | -1.07E-01 | 1.21E-01 | 2.69E+00 | 2.74E+00 | 3.73E-01 | no | | intergenic_variant | MODIFIER |
| 18 | 63619165 | AX-170088358 | 0 | C | T | 0.184 | -1.07E-01 | 1.21E-01 | 2.69E+00 | 2.74E+00 | 3.73E-01 | no | | intergenic_variant | MODIFIER |
| 18 | 63633614 | AX-170344490 | 0 | A | G | 0.184 | -1.07E-01 | 1.21E-01 | 2.69E+00 | 2.74E+00 | 3.73E-01 | no | | intergenic_variant | MODIFIER |
| 18 | 63900693 | AX-170482012 | 0 | T | C | 0.184 | -1.07E-01 | 1.21E-01 | 2.69E+00 | 2.74E+00 | 3.73E-01 | no | Wdr7 | intron_variant | MODIFIER |
| 18 | 63970963 | AX-170235281 | 0 | A | G | 0.184 | -1.07E-01 | 1.21E-01 | 2.69E+00 | 2.74E+00 | 3.73E-01 | no | Wdr7 | intron_variant | MODIFIER |
| 18 | 63973241 | AX-170290297 | 0 | T | C | 0.184 | -1.07E-01 | 1.21E-01 | 2.69E+00 | 2.74E+00 | 3.73E-01 | no | Wdr7 | intron_variant | MODIFIER |
| 18 | 63973364 | AX-170495970 | 0 | A | G | 0.184 | -1.07E-01 | 1.21E-01 | 2.69E+00 | 2.74E+00 | 3.73E-01 | no | Wdr7 | intron_variant | MODIFIER |
| 18 | 63973834 | AX-170362624 | 0 | A | G | 0.184 | -1.07E-01 | 1.21E-01 | 2.69E+00 | 2.74E+00 | 3.73E-01 | no | Wdr7 | intron_variant | MODIFIER |
| 18 | 63985392 | AX-170522771 | 0 | T | C | 0.184 | -1.07E-01 | 1.21E-01 | 2.69E+00 | 2.74E+00 | 3.73E-01 | no | Wdr7 | intron_variant | MODIFIER |
| 18 | 63985452 | AX-170335509 | 0 | T | C | 0.184 | -1.07E-01 | 1.21E-01 | 2.69E+00 | 2.74E+00 | 3.73E-01 | no | Wdr7 | intron_variant | MODIFIER |
| 18 | 63991505 | AX-170482541 | 0 | T | C | 0.184 | -1.07E-01 | 1.21E-01 | 2.69E+00 | 2.74E+00 | 3.73E-01 | no | Wdr7 | downstream_gene_variant | MODIFIER |
| 18 | 64000247 | AX-170061098 | 0 | A | G | 0.184 | -1.07E-01 | 1.21E-01 | 2.69E+00 | 2.74E+00 | 3.73E-01 | no | | intergenic_variant | MODIFIER |
| 18 | 64004339 | AX-170088359 | 0 | T | C | 0.184 | -1.07E-01 | 1.21E-01 | 2.69E+00 | 2.74E+00 | 3.73E-01 | no | | intergenic_variant | MODIFIER |
| 18 | 64008135 | AX-170458009 | 0 | C | T | 0.184 | -1.07E-01 | 1.21E-01 | 2.69E+00 | 2.74E+00 | 3.73E-01 | no | | intergenic_variant | MODIFIER |
| 18 | 64425180 | AX-170215047 | 0 | T | C | 0.184 | -1.07E-01 | 1.21E-01 | 2.69E+00 | 2.74E+00 | 3.73E-01 | no | AC102106.2 | downstream_gene_variant | MODIFIER |
| 18 | 64481040 | AX-170515531 | 0 | A | G | 0.184 | -1.07E-01 | 1.21E-01 | 2.69E+00 | 2.74E+00 | 3.73E-01 | no | Fech | intron_variant | MODIFIER |
| 18 | 64506747 | AX-170474530 | 0 | A | G | 0.184 | -1.07E-01 | 1.21E-01 | 2.69E+00 | 2.74E+00 | 3.73E-01 | no | Nars | intron_variant | MODIFIER |
| 18 | 64513912 | AX-170061107 | 0 | A | T | 0.184 | -1.07E-01 | 1.21E-01 | 2.69E+00 | 2.74E+00 | 3.73E-01 | no | Nars | intron_variant | MODIFIER |
| 18 | 64523384 | AX-170161024 | 0 | A | C | 0.184 | -1.07E-01 | 1.21E-01 | 2.69E+00 | 2.74E+00 | 3.73E-01 | no | | intergenic_variant | MODIFIER |
| 18 | 64644330 | AX-170469643 | 0 | C | A | 0.184 | -1.07E-01 | 1.21E-01 | 2.69E+00 | 2.74E+00 | 3.73E-01 | no | Atp8b1 | intron_variant | MODIFIER |
| 18 | 64663758 | AX-170412002 | 0 | T | C | 0.184 | -1.07E-01 | 1.21E-01 | 2.69E+00 | 2.74E+00 | 3.73E-01 | no | Atp8b1 | upstream_gene_variant | MODIFIER |
| 18 | 64663954 | AX-170223577 | 0 | T | A | 0.184 | -1.07E-01 | 1.21E-01 | 2.69E+00 | 2.74E+00 | 3.73E-01 | no | Atp8b1 | upstream_gene_variant | MODIFIER |
| 18 | 64674214 | AX-170362344 | 0 | C | T | 0.184 | -1.07E-01 | 1.21E-01 | 2.69E+00 | 2.74E+00 | 3.73E-01 | no | | intergenic_variant | MODIFIER |
| 18 | 64674746 | AX-97826264 | 0 | C | T | 0.184 | -1.07E-01 | 1.21E-01 | 2.69E+00 | 2.74E+00 | 3.73E-01 | no | | intergenic_variant | MODIFIER |
| 18 | 64837330 | AX-170276373 | 0 | T | C | 0.184 | -1.07E-01 | 1.21E-01 | 2.69E+00 | 2.74E+00 | 3.73E-01 | no | | intergenic_variant | MODIFIER |
| 18 | 64838285 | AX-170375160 | 0 | C | T | 0.184 | -1.07E-01 | 1.21E-01 | 2.69E+00 | 2.74E+00 | 3.73E-01 | no | | intergenic_variant | MODIFIER |
| 18 | 64845985 | AX-170363624 | 0 | G | A | 0.184 | -1.07E-01 | 1.21E-01 | 2.69E+00 | 2.74E+00 | 3.73E-01 | no | | intergenic_variant | MODIFIER |
| 18 | 65081860 | AX-170344011 | 0 | C | T | 0.184 | -1.07E-01 | 1.21E-01 | 2.69E+00 | 2.74E+00 | 3.73E-01 | no | Nedd4l | intron_variant | MODIFIER |
| 18 | 65088549 | AX-170334054 | 0 | C | T | 0.184 | -1.07E-01 | 1.21E-01 | 2.69E+00 | 2.74E+00 | 3.73E-01 | no | Nedd4l | intron_variant | MODIFIER |
| 18 | 65093285 | AX-170488180 | 0 | G | A | 0.184 | -1.07E-01 | 1.21E-01 | 2.69E+00 | 2.74E+00 | 3.73E-01 | no | Nedd4l | intron_variant | MODIFIER |
| 18 | 65095241 | AX-170246836 | 0 | A | G | 0.184 | -1.07E-01 | 1.21E-01 | 2.69E+00 | 2.74E+00 | 3.73E-01 | no | Nedd4l | intron_variant | MODIFIER |
| 18 | 65096902 | AX-170061121 | 0 | T | G | 0.184 | -1.07E-01 | 1.21E-01 | 2.69E+00 | 2.74E+00 | 3.73E-01 | no | Nedd4l | intron_variant | MODIFIER |
| 18 | 65097672 | AX-170517975 | 0 | G | A | 0.184 | -1.07E-01 | 1.21E-01 | 2.69E+00 | 2.74E+00 | 3.73E-01 | no | Nedd4l | intron_variant | MODIFIER |
| 18 | 65099587 | AX-170165651 | 0 | T | C | 0.184 | -1.07E-01 | 1.21E-01 | 2.69E+00 | 2.74E+00 | 3.73E-01 | no | Nedd4l | intron_variant | MODIFIER |
| 18 | 65105281 | AX-170442058 | 0 | T | C | 0.184 | -1.07E-01 | 1.21E-01 | 2.69E+00 | 2.74E+00 | 3.73E-01 | no | Nedd4l | intron_variant | MODIFIER |
| 18 | 65106819 | AX-170144810 | 0 | G | A | 0.184 | -1.07E-01 | 1.21E-01 | 2.69E+00 | 2.74E+00 | 3.73E-01 | no | Nedd4l | intron_variant | MODIFIER |
| 18 | 65107683 | AX-170149974 | 0 | A | G | 0.184 | -1.07E-01 | 1.21E-01 | 2.69E+00 | 2.74E+00 | 3.73E-01 | no | Nedd4l | intron_variant | MODIFIER |
| 18 | 65109498 | AX-170469537 | 0 | A | G | 0.184 | -1.07E-01 | 1.21E-01 | 2.69E+00 | 2.74E+00 | 3.73E-01 | no | Nedd4l | intron_variant | MODIFIER |
| 18 | 65109701 | AX-170397397 | 0 | G | A | 0.184 | -1.07E-01 | 1.21E-01 | 2.69E+00 | 2.74E+00 | 3.73E-01 | no | Nedd4l | intron_variant | MODIFIER |

| | | | | | | | | | | | | | | | |
|---|---|---|---|---|---|---|---|---|---|---|---|---|---|---|---|
| 18 | 65110304 | AX-170443345 | 0 | A | T | 0.184 | -1.07E-01 | 1.21E-01 | 2.69E+00 | 2.74E+00 | 3.73E-01 | no | Nedd4l | intron_variant | MODIFIER |
| 18 | 65110665 | AX-170240964 | 0 | G | A | 0.184 | -1.07E-01 | 1.21E-01 | 2.69E+00 | 2.74E+00 | 3.73E-01 | no | Nedd4l | intron_variant | MODIFIER |
| 18 | 65158249 | AX-170502036 | 0 | C | T | 0.184 | -1.07E-01 | 1.21E-01 | 2.69E+00 | 2.74E+00 | 3.73E-01 | no | Nedd4l | intron_variant | MODIFIER |
| 18 | 65174344 | AX-170271771 | 0 | T | C | 0.184 | -1.07E-01 | 1.21E-01 | 2.69E+00 | 2.74E+00 | 3.73E-01 | no | Nedd4l | intron_variant | MODIFIER |
| 18 | 65178489 | AX-170376286 | 0 | A | G | 0.184 | -1.07E-01 | 1.21E-01 | 2.69E+00 | 2.74E+00 | 3.73E-01 | no | Nedd4l | intron_variant | MODIFIER |
| 18 | 65179876 | AX-170374997 | 0 | G | C | 0.184 | -1.07E-01 | 1.21E-01 | 2.69E+00 | 2.74E+00 | 3.73E-01 | no | Nedd4l | intron_variant | MODIFIER |
| 18 | 65182420 | AX-170252594 | 0 | C | T | 0.184 | -1.07E-01 | 1.21E-01 | 2.69E+00 | 2.74E+00 | 3.73E-01 | no | Nedd4l | intron_variant | MODIFIER |
| 18 | 65191735 | AX-170149261 | 0 | A | G | 0.184 | -1.07E-01 | 1.21E-01 | 2.69E+00 | 2.74E+00 | 3.73E-01 | no | Nedd4l | intron_variant | MODIFIER |
| 18 | 65200505 | AX-170061124 | 0 | T | C | 0.184 | -1.07E-01 | 1.21E-01 | 2.69E+00 | 2.74E+00 | 3.73E-01 | no | Nedd4l | intron_variant | MODIFIER |
| 18 | 65200797 | AX-170384161 | 0 | G | C | 0.184 | -1.07E-01 | 1.21E-01 | 2.69E+00 | 2.74E+00 | 3.73E-01 | no | Nedd4l | intron_variant | MODIFIER |
| 18 | 65201453 | AX-170226358 | 0 | C | G | 0.184 | -1.07E-01 | 1.21E-01 | 2.69E+00 | 2.74E+00 | 3.73E-01 | no | Nedd4l | intron_variant | MODIFIER |
| 18 | 65201474 | AX-170417637 | 0 | C | T | 0.184 | -1.07E-01 | 1.21E-01 | 2.69E+00 | 2.74E+00 | 3.73E-01 | no | Nedd4l | intron_variant | MODIFIER |
| 18 | 65201573 | AX-170253423 | 0 | T | G | 0.184 | -1.07E-01 | 1.21E-01 | 2.69E+00 | 2.74E+00 | 3.73E-01 | no | Nedd4l | intron_variant | MODIFIER |
| 18 | 65216086 | AX-170146952 | 0 | G | A | 0.184 | -1.07E-01 | 1.21E-01 | 2.69E+00 | 2.74E+00 | 3.73E-01 | no | Nedd4l | downstream_gene_variant | MODIFIER |
| 18 | 65217331 | AX-170165500 | 0 | A | G | 0.184 | -1.07E-01 | 1.21E-01 | 2.69E+00 | 2.74E+00 | 3.73E-01 | no | Nedd4l | downstream_gene_variant | MODIFIER |
| 18 | 65219425 | AX-170135135 | 0 | G | A | 0.184 | -1.07E-01 | 1.21E-01 | 2.69E+00 | 2.74E+00 | 3.73E-01 | no | Nedd4l | downstream_gene_variant | MODIFIER |
| 18 | 65224771 | AX-170131648 | 0 | T | C | 0.184 | -1.07E-01 | 1.21E-01 | 2.69E+00 | 2.74E+00 | 3.73E-01 | no | | intergenic_variant | MODIFIER |
| 18 | 65224830 | AX-170520102 | 0 | G | C | 0.184 | -1.07E-01 | 1.21E-01 | 2.69E+00 | 2.74E+00 | 3.73E-01 | no | | intergenic_variant | MODIFIER |
| 18 | 65228055 | AX-97792094 | 0 | G | A | 0.184 | -1.07E-01 | 1.21E-01 | 2.69E+00 | 2.74E+00 | 3.73E-01 | no | | regulatory_region_variant | MODIFIER |
| 18 | 65235063 | AX-170142272 | 0 | A | G | 0.184 | -1.07E-01 | 1.21E-01 | 2.69E+00 | 2.74E+00 | 3.73E-01 | no | | intergenic_variant | MODIFIER |
| 18 | 65236401 | AX-170263229 | 0 | A | G | 0.184 | -1.07E-01 | 1.21E-01 | 2.69E+00 | 2.74E+00 | 3.73E-01 | no | | regulatory_region_variant | MODIFIER |
| 18 | 65261822 | AX-170213803 | 0 | T | C | 0.184 | -1.07E-01 | 1.21E-01 | 2.69E+00 | 2.74E+00 | 3.73E-01 | no | Alpk2 | downstream_gene_variant | MODIFIER |
| 18 | 65293882 | AX-170232068 | 0 | T | C | 0.184 | -1.07E-01 | 1.21E-01 | 2.69E+00 | 2.74E+00 | 3.73E-01 | no | Alpk2 | intron_variant | MODIFIER |
| 18 | 65313415 | AX-170061128 | 0 | C | T | 0.184 | -1.07E-01 | 1.21E-01 | 2.69E+00 | 2.74E+00 | 3.73E-01 | no | Alpk2 | intron_variant | MODIFIER |
| 18 | 65314843 | AX-170061129 | 0 | T | G | 0.184 | -1.07E-01 | 1.21E-01 | 2.69E+00 | 2.74E+00 | 3.73E-01 | no | Alpk2 | intron_variant | MODIFIER |
| 18 | 65316391 | AX-170309095 | 0 | C | T | 0.184 | -1.07E-01 | 1.21E-01 | 2.69E+00 | 2.74E+00 | 3.73E-01 | no | Alpk2 | intron_variant | MODIFIER |
| 18 | 65317035 | AX-170232497 | 0 | A | C | 0.184 | -1.07E-01 | 1.21E-01 | 2.69E+00 | 2.74E+00 | 3.73E-01 | no | Alpk2 | intron_variant | MODIFIER |
| 18 | 65350228 | AX-97854948 | 0 | G | A | 0.184 | -1.07E-01 | 1.21E-01 | 2.69E+00 | 2.74E+00 | 3.73E-01 | no | Alpk2 | synonymous_variant | LOW |
| 18 | 65465025 | AX-170133259 | 0 | C | G | 0.184 | -1.07E-01 | 1.21E-01 | 2.69E+00 | 2.74E+00 | 3.73E-01 | no | Malt1 | intron_variant | MODIFIER |
| 18 | 65465380 | AX-170130787 | 0 | A | G | 0.184 | -1.07E-01 | 1.21E-01 | 2.69E+00 | 2.74E+00 | 3.73E-01 | no | Malt1 | intron_variant | MODIFIER |
| 18 | 65476865 | AX-170435810 | 0 | G | T | 0.184 | -1.07E-01 | 1.21E-01 | 2.69E+00 | 2.74E+00 | 3.73E-01 | no | Malt1 | 3_prime_UTR_variant | MODIFIER |
| 18 | 65780689 | AX-97834977 | 0 | C | T | 0.184 | -1.07E-01 | 1.21E-01 | 2.69E+00 | 2.74E+00 | 3.73E-01 | no | | regulatory_region_variant | MODIFIER |
| 18 | 65799991 | AX-170466658 | 0 | C | T | 0.184 | -1.07E-01 | 1.21E-01 | 2.69E+00 | 2.74E+00 | 3.73E-01 | no | Sec11c | upstream_gene_variant | MODIFIER |
| 18 | 65811348 | AX-97835702 | 0 | G | A | 0.184 | -1.07E-01 | 1.21E-01 | 2.69E+00 | 2.74E+00 | 3.73E-01 | no | Sec11c | intron_variant | MODIFIER |
| 18 | 65822455 | AX-170220848 | 0 | T | C | 0.184 | -1.07E-01 | 1.21E-01 | 2.69E+00 | 2.74E+00 | 3.73E-01 | no | Sec11c | downstream_gene_variant | MODIFIER |
| 18 | 65825184 | AX-97855840 | 0 | T | G | 0.184 | -1.07E-01 | 1.21E-01 | 2.69E+00 | 2.74E+00 | 3.73E-01 | no | | intergenic_variant | MODIFIER |
| 18 | 65829654 | AX-170349639 | 0 | A | G | 0.184 | -1.07E-01 | 1.21E-01 | 2.69E+00 | 2.74E+00 | 3.73E-01 | no | | intergenic_variant | MODIFIER |
| 18 | 65841867 | AX-170346413 | 0 | G | A | 0.184 | -1.07E-01 | 1.21E-01 | 2.69E+00 | 2.74E+00 | 3.73E-01 | no | AC124578.3 | downstream_gene_variant | MODIFIER |
| 18 | 65842100 | AX-170173569 | 0 | T | C | 0.184 | -1.07E-01 | 1.21E-01 | 2.69E+00 | 2.74E+00 | 3.73E-01 | no | AC124578.3 | downstream_gene_variant | MODIFIER |
| 18 | 65842200 | AX-170429937 | 0 | T | C | 0.184 | -1.07E-01 | 1.21E-01 | 2.69E+00 | 2.74E+00 | 3.73E-01 | no | AC124578.3 | downstream_gene_variant | MODIFIER |
| 18 | 65850957 | AX-97834999 | 0 | T | C | 0.184 | -1.07E-01 | 1.21E-01 | 2.69E+00 | 2.74E+00 | 3.73E-01 | no | | regulatory_region_variant | MODIFIER |
| 18 | 65859569 | AX-170015880 | 0 | A | G | 0.184 | -1.07E-01 | 1.21E-01 | 2.69E+00 | 2.74E+00 | 3.73E-01 | no | AC152076.4 | downstream_gene_variant | MODIFIER |
| 18 | 65866728 | AX-170504107 | 0 | C | T | 0.184 | -1.07E-01 | 1.21E-01 | 2.69E+00 | 2.74E+00 | 3.73E-01 | no | AC152076.4 | intron_variant,non_coding_transcript_variant | MODIFIER |
| 18 | 65869864 | AX-170436961 | 0 | T | A | 0.184 | -1.07E-01 | 1.21E-01 | 2.69E+00 | 2.74E+00 | 3.73E-01 | no | Grp | upstream_gene_variant | MODIFIER |
| 18 | 66123291 | AX-170154713 | 0 | G | C | 0.184 | -1.07E-01 | 1.21E-01 | 2.69E+00 | 2.74E+00 | 3.73E-01 | no | Ccbe1 | intron_variant,NMD_transcript_variant | MODIFIER |
| 18 | 66132520 | AX-170364010 | 0 | C | T | 0.184 | -1.07E-01 | 1.21E-01 | 2.69E+00 | 2.74E+00 | 3.73E-01 | no | Ccbe1 | intron_variant,NMD_transcript_variant | MODIFIER |
| 18 | 66160313 | AX-170393672 | 0 | T | C | 0.184 | -1.07E-01 | 1.21E-01 | 2.69E+00 | 2.74E+00 | 3.73E-01 | no | Ccbe1 | intron_variant,NMD_transcript_variant | MODIFIER |
| 18 | 66160555 | AX-170123220 | 0 | C | T | 0.184 | -1.07E-01 | 1.21E-01 | 2.69E+00 | 2.74E+00 | 3.73E-01 | no | Ccbe1 | intron_variant,NMD_transcript_variant | MODIFIER |
| 18 | 66173254 | AX-97780931 | 0 | C | T | 0.184 | -1.07E-01 | 1.21E-01 | 2.69E+00 | 2.74E+00 | 3.73E-01 | no | Ccbe1 | intron_variant,NMD_transcript_variant | MODIFIER |
| 18 | 66173304 | AX-170197828 | 0 | T | A | 0.184 | -1.07E-01 | 1.21E-01 | 2.69E+00 | 2.74E+00 | 3.73E-01 | no | Ccbe1 | intron_variant,NMD_transcript_variant | MODIFIER |
| 18 | 66202616 | AX-97833079 | 0 | T | G | 0.184 | -1.07E-01 | 1.21E-01 | 2.69E+00 | 2.74E+00 | 3.73E-01 | no | Ccbe1 | intron_variant,NMD_transcript_variant | MODIFIER |
| 18 | 66202642 | AX-170144903 | 0 | C | T | 0.184 | -1.07E-01 | 1.21E-01 | 2.69E+00 | 2.74E+00 | 3.73E-01 | no | Ccbe1 | intron_variant,NMD_transcript_variant | MODIFIER |
| 18 | 66258382 | AX-170455303 | 0 | G | T | 0.184 | -1.07E-01 | 1.21E-01 | 2.69E+00 | 2.74E+00 | 3.73E-01 | no | Ccbe1 | intron_variant,NMD_transcript_variant | MODIFIER |
| 18 | 66277577 | AX-170460021 | 0 | A | G | 0.184 | -1.07E-01 | 1.21E-01 | 2.69E+00 | 2.74E+00 | 3.73E-01 | no | Ccbe1 | intron_variant,NMD_transcript_variant | MODIFIER |

| | | | | | | | | | | | | | | |
|---|---|---|---|---|---|---|---|---|---|---|---|---|---|---|
| 18 | 66328515 | AX-170503092 | 0 | T | C | 0.184 | -1.07E-01 | 1.21E-01 | 2.69E+00 | 2.74E+00 | 3.73E-01 | no | | intergenic_variant | MODIFIER |
| 18 | 66844696 | AX-170270855 | 0 | A | G | 0.184 | -1.07E-01 | 1.21E-01 | 2.69E+00 | 2.74E+00 | 3.73E-01 | no | | intergenic_variant | MODIFIER |
| 18 | 66868945 | AX-170425369 | 0 | G | A | 0.184 | -1.07E-01 | 1.21E-01 | 2.69E+00 | 2.74E+00 | 3.73E-01 | no | | intergenic_variant | MODIFIER |
| 18 | 66869139 | AX-170418075 | 0 | G | A | 0.184 | -1.07E-01 | 1.21E-01 | 2.69E+00 | 2.74E+00 | 3.73E-01 | no | | intergenic_variant | MODIFIER |
| 18 | 67063134 | AX-170223535 | 0 | A | G | 0.184 | -1.07E-01 | 1.21E-01 | 2.69E+00 | 2.74E+00 | 3.73E-01 | no | AC140207.2 | upstream_gene_variant | MODIFIER |
| 18 | 67138752 | AX-170440488 | 0 | C | A | 0.184 | -1.07E-01 | 1.21E-01 | 2.69E+00 | 2.74E+00 | 3.73E-01 | no | Gnal | intron_variant | MODIFIER |
| 18 | 67143844 | AX-170205603 | 0 | T | C | 0.184 | -1.07E-01 | 1.21E-01 | 2.69E+00 | 2.74E+00 | 3.73E-01 | no | Gnal | intron_variant | MODIFIER |
| 18 | 67144109 | AX-170137603 | 0 | A | T | 0.184 | -1.07E-01 | 1.21E-01 | 2.69E+00 | 2.74E+00 | 3.73E-01 | no | Gnal | intron_variant | MODIFIER |
| 18 | 67144583 | AX-170453570 | 0 | C | A | 0.184 | -1.07E-01 | 1.21E-01 | 2.69E+00 | 2.74E+00 | 3.73E-01 | no | Gnal | intron_variant | MODIFIER |
| 18 | 67177537 | AX-170362469 | 0 | G | A | 0.184 | -1.07E-01 | 1.21E-01 | 2.69E+00 | 2.74E+00 | 3.73E-01 | no | Gnal | intron_variant | MODIFIER |
| 18 | 67180458 | AX-170305322 | 0 | T | G | 0.184 | -1.07E-01 | 1.21E-01 | 2.69E+00 | 2.74E+00 | 3.73E-01 | no | Gnal | intron_variant | MODIFIER |
| 18 | 67181315 | AX-170467156 | 0 | G | A | 0.184 | -1.07E-01 | 1.21E-01 | 2.69E+00 | 2.74E+00 | 3.73E-01 | no | Gnal | intron_variant | MODIFIER |
| 18 | 67237538 | AX-170108292 | 0 | A | G | 0.184 | -1.07E-01 | 1.21E-01 | 2.69E+00 | 2.74E+00 | 3.73E-01 | no | Mppe1 | synonymous_variant | LOW |
| 18 | 67630065 | AX-97800631 | 0 | T | G | 0.184 | -1.07E-01 | 1.21E-01 | 2.69E+00 | 2.74E+00 | 3.73E-01 | no | Cep76 | intron_variant | MODIFIER |
| 18 | 67656368 | AX-170114494 | 0 | A | G | 0.184 | -1.07E-01 | 1.21E-01 | 2.69E+00 | 2.74E+00 | 3.73E-01 | no | Psmg2 | downstream_gene_variant | MODIFIER |
| 18 | 67685513 | AX-170136054 | 0 | G | T | 0.184 | -1.07E-01 | 1.21E-01 | 2.69E+00 | 2.74E+00 | 3.73E-01 | no | Ptpn2 | intron_variant | MODIFIER |
| 18 | 67696471 | AX-170138500 | 0 | A | G | 0.184 | -1.07E-01 | 1.21E-01 | 2.69E+00 | 2.74E+00 | 3.73E-01 | no | Ptpn2 | intron_variant | MODIFIER |
| 18 | 67731136 | AX-170305624 | 0 | G | A | 0.184 | -1.07E-01 | 1.21E-01 | 2.69E+00 | 2.74E+00 | 3.73E-01 | no | | regulatory_region_variant | MODIFIER |
| 18 | 67906556 | AX-170408287 | 0 | A | G | 0.184 | -1.07E-01 | 1.21E-01 | 2.69E+00 | 2.74E+00 | 3.73E-01 | no | AC127236.1 | upstream_gene_variant | MODIFIER |
| 18 | 67907978 | AX-170323830 | 0 | C | A | 0.184 | -1.07E-01 | 1.21E-01 | 2.69E+00 | 2.74E+00 | 3.73E-01 | no | AC127236.1 | non_coding_transcript_exon_variant | MODIFIER |
| 18 | 67908142 | AX-170518161 | 0 | C | T | 0.184 | -1.07E-01 | 1.21E-01 | 2.69E+00 | 2.74E+00 | 3.73E-01 | no | AC127236.1 | non_coding_transcript_exon_variant | MODIFIER |
| 18 | 67949311 | AX-170015908 | 0 | C | T | 0.184 | -1.07E-01 | 1.21E-01 | 2.69E+00 | 2.74E+00 | 3.73E-01 | no | Ldlrad4 | intron_variant | MODIFIER |
| 15 | 80075803 | AX-170164382 | 1 | A | G | 0.545 | 7.14E-02 | 8.00E-02 | 2.56E+00 | 2.65E+00 | 3.73E-01 | no | Rpl3 | downstream_gene_variant | MODIFIER |
| 9 | 91372969 | AX-97786063 | 0 | T | G | 0.14 | -1.03E-01 | 1.17E-01 | 2.64E+00 | 2.73E+00 | 3.73E-01 | no | Zic4 | intron_variant,non_coding_transcript_variant | MODIFIER |
| 9 | 95451164 | AX-170395579 | 1 | T | C | 0.143 | -1.01E-01 | 1.14E-01 | 2.67E+00 | 2.76E+00 | 3.73E-01 | no | | intergenic_variant | MODIFIER |
| 9 | 103748483 | AX-170484522 | 0 | C | T | 0.123 | -1.02E-01 | 1.14E-01 | 2.53E+00 | 2.61E+00 | 3.73E-01 | no | Tmem108 | downstream_gene_variant | MODIFIER |
| 13 | 6952729 | AX-170199525 | 1 | A | G | 0.196 | -1.09E-01 | 1.22E-01 | 2.56E+00 | 2.62E+00 | 3.74E-01 | no | | intergenic_variant | MODIFIER |
| 8 | 62550803 | AX-170370616 | 0 | T | G | 0.14 | -1.30E-01 | 1.46E-01 | 2.58E+00 | 2.65E+00 | 3.74E-01 | no | Gm7561 | upstream_gene_variant | MODIFIER |
| 8 | 62750013 | AX-170415079 | 0 | T | C | 0.14 | -1.30E-01 | 1.46E-01 | 2.58E+00 | 2.65E+00 | 3.74E-01 | no | Gm45459 | intron_variant,non_coding_transcript_variant | MODIFIER |
| 8 | 62756863 | AX-170203749 | 0 | T | C | 0.14 | -1.30E-01 | 1.46E-01 | 2.58E+00 | 2.65E+00 | 3.74E-01 | no | Gm45459 | intron_variant,non_coding_transcript_variant | MODIFIER |
| 8 | 62757160 | AX-170105646 | 0 | T | C | 0.14 | -1.30E-01 | 1.46E-01 | 2.58E+00 | 2.65E+00 | 3.74E-01 | no | Gm45459 | intron_variant,non_coding_transcript_variant | MODIFIER |
| 8 | 62757582 | AX-97830447 | 0 | A | G | 0.14 | -1.30E-01 | 1.46E-01 | 2.58E+00 | 2.65E+00 | 3.74E-01 | no | Gm45459 | intron_variant,non_coding_transcript_variant | MODIFIER |
| 8 | 62778169 | AX-170367257 | 0 | T | C | 0.14 | -1.30E-01 | 1.46E-01 | 2.58E+00 | 2.65E+00 | 3.74E-01 | no | Gm45459 | intron_variant,non_coding_transcript_variant | MODIFIER |
| 8 | 62792894 | AX-97780883 | 0 | G | A | 0.14 | -1.30E-01 | 1.46E-01 | 2.58E+00 | 2.65E+00 | 3.74E-01 | no | Gm45459 | intron_variant,non_coding_transcript_variant | MODIFIER |
| 8 | 62793994 | AX-170277116 | 0 | A | G | 0.14 | -1.30E-01 | 1.46E-01 | 2.58E+00 | 2.65E+00 | 3.74E-01 | no | Gm45459 | intron_variant,non_coding_transcript_variant | MODIFIER |
| 8 | 62813184 | AX-170336354 | 0 | G | A | 0.14 | -1.30E-01 | 1.46E-01 | 2.58E+00 | 2.65E+00 | 3.74E-01 | no | Gm45459 | intron_variant,non_coding_transcript_variant | MODIFIER |
| 8 | 63349126 | AX-170508498 | 0 | C | T | 0.14 | -1.30E-01 | 1.46E-01 | 2.58E+00 | 2.65E+00 | 3.74E-01 | no | Spock3 | intron_variant | MODIFIER |
| 8 | 63381173 | AX-97797977 | 0 | T | C | 0.14 | -1.30E-01 | 1.46E-01 | 2.58E+00 | 2.65E+00 | 3.74E-01 | no | | intergenic_variant | MODIFIER |
| 8 | 63396597 | AX-170136242 | 0 | G | A | 0.14 | -1.30E-01 | 1.46E-01 | 2.58E+00 | 2.65E+00 | 3.74E-01 | no | | intergenic_variant | MODIFIER |
| 8 | 63438757 | AX-170382594 | 0 | G | T | 0.14 | -1.30E-01 | 1.46E-01 | 2.58E+00 | 2.65E+00 | 3.74E-01 | no | | intergenic_variant | MODIFIER |
| 8 | 63438998 | AX-170161110 | 0 | T | C | 0.14 | -1.30E-01 | 1.46E-01 | 2.58E+00 | 2.65E+00 | 3.74E-01 | no | | intergenic_variant | MODIFIER |
| 8 | 63452481 | AX-170120785 | 0 | T | A | 0.14 | -1.30E-01 | 1.46E-01 | 2.58E+00 | 2.65E+00 | 3.74E-01 | no | | intergenic_variant | MODIFIER |
| 8 | 63560847 | AX-170030883 | 0 | A | T | 0.14 | -1.30E-01 | 1.46E-01 | 2.58E+00 | 2.65E+00 | 3.74E-01 | no | | intergenic_variant | MODIFIER |
| 8 | 63563996 | AX-97796774 | 0 | A | G | 0.14 | -1.30E-01 | 1.46E-01 | 2.58E+00 | 2.65E+00 | 3.74E-01 | no | | intergenic_variant | MODIFIER |
| 8 | 64160507 | AX-170080260 | 0 | T | G | 0.14 | -1.30E-01 | 1.46E-01 | 2.58E+00 | 2.65E+00 | 3.74E-01 | no | Tll1 | intron_variant | MODIFIER |
| 8 | 64165087 | AX-170096294 | 0 | A | G | 0.14 | -1.30E-01 | 1.46E-01 | 2.58E+00 | 2.65E+00 | 3.74E-01 | no | Tll1 | intron_variant | MODIFIER |
| 8 | 64167263 | AX-170080262 | 0 | G | A | 0.14 | -1.30E-01 | 1.46E-01 | 2.58E+00 | 2.65E+00 | 3.74E-01 | no | Tll1 | intron_variant | MODIFIER |
| 8 | 64167317 | AX-170080259 | 0 | C | T | 0.14 | -1.30E-01 | 1.46E-01 | 2.58E+00 | 2.65E+00 | 3.74E-01 | no | Tll1 | intron_variant | MODIFIER |
| 8 | 64167881 | AX-170317101 | 0 | C | A | 0.14 | -1.30E-01 | 1.46E-01 | 2.58E+00 | 2.65E+00 | 3.74E-01 | no | Tll1 | intron_variant | MODIFIER |
| 8 | 64170090 | AX-170030900 | 0 | T | C | 0.14 | -1.30E-01 | 1.46E-01 | 2.58E+00 | 2.65E+00 | 3.74E-01 | no | Tll1 | intron_variant | MODIFIER |
| 8 | 64170214 | AX-170030899 | 0 | C | T | 0.14 | -1.30E-01 | 1.46E-01 | 2.58E+00 | 2.65E+00 | 3.74E-01 | no | Tll1 | intron_variant | MODIFIER |
| 8 | 64178860 | AX-170348268 | 0 | A | G | 0.14 | -1.30E-01 | 1.46E-01 | 2.58E+00 | 2.65E+00 | 3.74E-01 | no | Tll1 | intron_variant | MODIFIER |
| 8 | 64220620 | AX-170497841 | 0 | G | C | 0.14 | -1.30E-01 | 1.46E-01 | 2.58E+00 | 2.65E+00 | 3.74E-01 | no | | intergenic_variant | MODIFIER |
| 8 | 64283666 | AX-170295487 | 0 | C | G | 0.14 | -1.30E-01 | 1.46E-01 | 2.58E+00 | 2.65E+00 | 3.74E-01 | no | | intergenic_variant | MODIFIER |

| | | | | | | | | | | | | | | | |
|---|---|---|---|---|---|---|---|---|---|---|---|---|---|---|---|
| 8 | 64323041 | AX-170096296 | 0 | T | C | 0.14 | -1.30E-01 | 1.46E-01 | 2.58E+00 | 2.65E+00 | 3.74E-01 | no | | intergenic_variant | MODIFIER |
| 8 | 64337721 | AX-170370173 | 0 | G | A | 0.14 | -1.30E-01 | 1.46E-01 | 2.58E+00 | 2.65E+00 | 3.74E-01 | no | | intergenic_variant | MODIFIER |
| 8 | 64338432 | AX-170313211 | 0 | T | C | 0.14 | -1.30E-01 | 1.46E-01 | 2.58E+00 | 2.65E+00 | 3.74E-01 | no | | intergenic_variant | MODIFIER |
| 8 | 64341947 | AX-170345517 | 0 | G | A | 0.14 | -1.30E-01 | 1.46E-01 | 2.58E+00 | 2.65E+00 | 3.74E-01 | no | | intergenic_variant | MODIFIER |
| 8 | 64362535 | AX-170525352 | 0 | C | T | 0.14 | -1.30E-01 | 1.46E-01 | 2.58E+00 | 2.65E+00 | 3.74E-01 | no | | regulatory_region_variant | MODIFIER |
| 8 | 64363378 | AX-170292716 | 0 | A | G | 0.14 | -1.30E-01 | 1.46E-01 | 2.58E+00 | 2.65E+00 | 3.74E-01 | no | | intergenic_variant | MODIFIER |
| 8 | 64364054 | AX-170428290 | 0 | G | C | 0.14 | -1.30E-01 | 1.46E-01 | 2.58E+00 | 2.65E+00 | 3.74E-01 | no | | intergenic_variant | MODIFIER |
| 8 | 64367592 | AX-170452751 | 0 | C | T | 0.14 | -1.30E-01 | 1.46E-01 | 2.58E+00 | 2.65E+00 | 3.74E-01 | no | | intergenic_variant | MODIFIER |
| 8 | 64383626 | AX-170507801 | 0 | A | G | 0.14 | -1.30E-01 | 1.46E-01 | 2.58E+00 | 2.65E+00 | 3.74E-01 | no | | intergenic_variant | MODIFIER |
| 8 | 64384859 | AX-170330234 | 0 | G | C | 0.14 | -1.30E-01 | 1.46E-01 | 2.58E+00 | 2.65E+00 | 3.74E-01 | no | | intergenic_variant | MODIFIER |
| 8 | 64389506 | AX-170372337 | 0 | C | T | 0.14 | -1.30E-01 | 1.46E-01 | 2.58E+00 | 2.65E+00 | 3.74E-01 | no | | intergenic_variant | MODIFIER |
| 8 | 64403284 | AX-170382349 | 0 | G | T | 0.14 | -1.30E-01 | 1.46E-01 | 2.58E+00 | 2.65E+00 | 3.74E-01 | no | | intergenic_variant | MODIFIER |
| 8 | 64486276 | AX-97848224 | 0 | A | G | 0.14 | -1.30E-01 | 1.46E-01 | 2.58E+00 | 2.65E+00 | 3.74E-01 | no | | intergenic_variant | MODIFIER |
| 8 | 64529414 | AX-170219066 | 0 | A | G | 0.14 | -1.30E-01 | 1.46E-01 | 2.58E+00 | 2.65E+00 | 3.74E-01 | no | | intergenic_variant | MODIFIER |
| 8 | 64533295 | AX-170268267 | 0 | C | G | 0.14 | -1.30E-01 | 1.46E-01 | 2.58E+00 | 2.65E+00 | 3.74E-01 | no | | intergenic_variant | MODIFIER |
| 8 | 64547819 | AX-170449590 | 0 | A | G | 0.14 | -1.30E-01 | 1.46E-01 | 2.58E+00 | 2.65E+00 | 3.74E-01 | no | | intergenic_variant | MODIFIER |
| 8 | 64548011 | AX-170030909 | 0 | A | G | 0.14 | -1.30E-01 | 1.46E-01 | 2.58E+00 | 2.65E+00 | 3.74E-01 | no | | regulatory_region_variant | MODIFIER |
| 8 | 64551872 | AX-97798762 | 0 | T | C | 0.14 | -1.30E-01 | 1.46E-01 | 2.58E+00 | 2.65E+00 | 3.74E-01 | no | | intergenic_variant | MODIFIER |
| 8 | 64569210 | AX-97838657 | 0 | A | G | 0.14 | -1.30E-01 | 1.46E-01 | 2.58E+00 | 2.65E+00 | 3.74E-01 | no | | intergenic_variant | MODIFIER |
| 8 | 64584670 | AX-97824344 | 0 | G | A | 0.14 | -1.30E-01 | 1.46E-01 | 2.58E+00 | 2.65E+00 | 3.74E-01 | no | | intergenic_variant | MODIFIER |
| 8 | 64611040 | AX-170030910 | 0 | C | A | 0.14 | -1.30E-01 | 1.46E-01 | 2.58E+00 | 2.65E+00 | 3.74E-01 | no | Cpe | intron_variant | MODIFIER |
| 8 | 64615998 | AX-170467808 | 0 | G | A | 0.14 | -1.30E-01 | 1.46E-01 | 2.58E+00 | 2.65E+00 | 3.74E-01 | no | Cpe | intron_variant | MODIFIER |
| 8 | 64624606 | AX-170116085 | 0 | C | G | 0.14 | -1.30E-01 | 1.46E-01 | 2.58E+00 | 2.65E+00 | 3.74E-01 | no | Cpe | intron_variant | MODIFIER |
| 8 | 64644161 | AX-170030908 | 0 | C | A | 0.14 | -1.30E-01 | 1.46E-01 | 2.58E+00 | 2.65E+00 | 3.74E-01 | no | Cpe | intron_variant | MODIFIER |
| 8 | 64647898 | AX-170030904 | 0 | C | G | 0.14 | -1.30E-01 | 1.46E-01 | 2.58E+00 | 2.65E+00 | 3.74E-01 | no | Cpe | intron_variant | MODIFIER |
| 8 | 64648490 | AX-97854256 | 0 | G | A | 0.14 | -1.30E-01 | 1.46E-01 | 2.58E+00 | 2.65E+00 | 3.74E-01 | no | Cpe | intron_variant | MODIFIER |
| 8 | 64680340 | AX-97804410 | 0 | A | C | 0.14 | -1.30E-01 | 1.46E-01 | 2.58E+00 | 2.65E+00 | 3.74E-01 | no | Cpe | intron_variant | MODIFIER |
| 8 | 64687687 | AX-170030907 | 0 | T | C | 0.14 | -1.30E-01 | 1.46E-01 | 2.58E+00 | 2.65E+00 | 3.74E-01 | no | Cpe | intron_variant | MODIFIER |
| 8 | 64696546 | AX-97846329 | 0 | G | A | 0.14 | -1.30E-01 | 1.46E-01 | 2.58E+00 | 2.65E+00 | 3.74E-01 | no | Cpe | upstream_gene_variant | MODIFIER |
| 8 | 64722557 | AX-170030906 | 0 | T | C | 0.14 | -1.30E-01 | 1.46E-01 | 2.58E+00 | 2.65E+00 | 3.74E-01 | no | Msmo1 | synonymous_variant | LOW |
| 8 | 64862530 | AX-97803817 | 0 | G | A | 0.14 | -1.30E-01 | 1.46E-01 | 2.58E+00 | 2.65E+00 | 3.74E-01 | no | | intergenic_variant | MODIFIER |
| 8 | 65213916 | AX-170090138 | 0 | T | C | 0.14 | -1.30E-01 | 1.46E-01 | 2.58E+00 | 2.65E+00 | 3.74E-01 | no | | intergenic_variant | MODIFIER |
| 8 | 65230399 | AX-170030911 | 0 | G | A | 0.14 | -1.30E-01 | 1.46E-01 | 2.58E+00 | 2.65E+00 | 3.74E-01 | no | | intergenic_variant | MODIFIER |
| 8 | 65237856 | AX-97852309 | 0 | A | C | 0.14 | -1.30E-01 | 1.46E-01 | 2.58E+00 | 2.65E+00 | 3.74E-01 | no | | intergenic_variant | MODIFIER |
| 8 | 65256894 | AX-170528733 | 0 | T | G | 0.14 | -1.30E-01 | 1.46E-01 | 2.58E+00 | 2.65E+00 | 3.74E-01 | no | | intergenic_variant | MODIFIER |
| 8 | 65264507 | AX-170030917 | 0 | T | C | 0.14 | -1.30E-01 | 1.46E-01 | 2.58E+00 | 2.65E+00 | 3.74E-01 | no | | intergenic_variant | MODIFIER |
| 8 | 65271084 | AX-97803816 | 0 | T | G | 0.14 | -1.30E-01 | 1.46E-01 | 2.58E+00 | 2.65E+00 | 3.74E-01 | no | | intergenic_variant | MODIFIER |
| 8 | 65339371 | AX-170320451 | 0 | G | A | 0.14 | -1.30E-01 | 1.46E-01 | 2.58E+00 | 2.65E+00 | 3.74E-01 | no | | intergenic_variant | MODIFIER |
| 8 | 65346034 | AX-170096298 | 0 | C | T | 0.14 | -1.30E-01 | 1.46E-01 | 2.58E+00 | 2.65E+00 | 3.74E-01 | no | | intergenic_variant | MODIFIER |
| 8 | 65353283 | AX-170080275 | 0 | G | A | 0.14 | -1.30E-01 | 1.46E-01 | 2.58E+00 | 2.65E+00 | 3.74E-01 | no | | intergenic_variant | MODIFIER |
| 8 | 65353318 | AX-170333449 | 0 | G | A | 0.14 | -1.30E-01 | 1.46E-01 | 2.58E+00 | 2.65E+00 | 3.74E-01 | no | | intergenic_variant | MODIFIER |
| 8 | 65368082 | AX-170434493 | 0 | A | T | 0.14 | -1.30E-01 | 1.46E-01 | 2.58E+00 | 2.65E+00 | 3.74E-01 | no | Gm25403 | downstream_gene_variant | MODIFIER |
| 8 | 65378949 | AX-170402655 | 0 | T | G | 0.14 | -1.30E-01 | 1.46E-01 | 2.58E+00 | 2.65E+00 | 3.74E-01 | no | | intergenic_variant | MODIFIER |
| 8 | 65383697 | AX-170380881 | 0 | G | T | 0.14 | -1.30E-01 | 1.46E-01 | 2.58E+00 | 2.65E+00 | 3.74E-01 | no | | intergenic_variant | MODIFIER |
| 8 | 65383797 | AX-170456190 | 0 | C | A | 0.14 | -1.30E-01 | 1.46E-01 | 2.58E+00 | 2.65E+00 | 3.74E-01 | no | | intergenic_variant | MODIFIER |
| 8 | 65384893 | AX-170485676 | 0 | C | T | 0.14 | -1.30E-01 | 1.46E-01 | 2.58E+00 | 2.65E+00 | 3.74E-01 | no | | intergenic_variant | MODIFIER |
| 8 | 65401567 | AX-170433524 | 0 | C | T | 0.14 | -1.30E-01 | 1.46E-01 | 2.58E+00 | 2.65E+00 | 3.74E-01 | no | | intergenic_variant | MODIFIER |
| 8 | 65472391 | AX-170096300 | 0 | T | G | 0.14 | -1.30E-01 | 1.46E-01 | 2.58E+00 | 2.65E+00 | 3.74E-01 | no | | intergenic_variant | MODIFIER |
| 8 | 65497766 | AX-170418775 | 0 | T | C | 0.14 | -1.30E-01 | 1.46E-01 | 2.58E+00 | 2.65E+00 | 3.74E-01 | no | | intergenic_variant | MODIFIER |
| 8 | 65498160 | AX-170171601 | 0 | T | C | 0.14 | -1.30E-01 | 1.46E-01 | 2.58E+00 | 2.65E+00 | 3.74E-01 | no | | intergenic_variant | MODIFIER |
| 8 | 65587495 | AX-170381212 | 0 | C | T | 0.14 | -1.30E-01 | 1.46E-01 | 2.58E+00 | 2.65E+00 | 3.74E-01 | no | | intergenic_variant | MODIFIER |
| 8 | 65588496 | AX-170406725 | 0 | T | C | 0.14 | -1.30E-01 | 1.46E-01 | 2.58E+00 | 2.65E+00 | 3.74E-01 | no | | intergenic_variant | MODIFIER |
| 8 | 65616001 | AX-170128957 | 0 | T | A | 0.14 | -1.30E-01 | 1.46E-01 | 2.58E+00 | 2.65E+00 | 3.74E-01 | no | MAR1 | upstream_gene_variant | MODIFIER |
| 8 | 65617851 | AX-97814086 | 0 | T | C | 0.14 | -1.30E-01 | 1.46E-01 | 2.58E+00 | 2.65E+00 | 3.74E-01 | no | MAR1 | upstream_gene_variant | MODIFIER |

| | | | | | | | | | | | | | | | |
|---|---|---|---|---|---|---|---|---|---|---|---|---|---|---|---|
| 8 | 65676034 | AX-170030929 | 0 | A | C | 0.14 | -1.30E-01 | 1.46E-01 | 2.58E+00 | 2.65E+00 | 3.74E-01 | no | MAR1 | intron_variant | MODIFIER |
| 8 | 65683375 | AX-97824202 | 0 | G | A | 0.14 | -1.30E-01 | 1.46E-01 | 2.58E+00 | 2.65E+00 | 3.74E-01 | no | MAR1 | intron_variant | MODIFIER |
| 8 | 65693043 | AX-97785451 | 0 | T | C | 0.14 | -1.30E-01 | 1.46E-01 | 2.58E+00 | 2.65E+00 | 3.74E-01 | no | MAR1 | intron_variant | MODIFIER |
| 8 | 65865015 | AX-170443620 | 0 | C | T | 0.14 | -1.30E-01 | 1.46E-01 | 2.58E+00 | 2.65E+00 | 3.74E-01 | no | MAR1 | intron_variant | MODIFIER |
| 8 | 65873170 | AX-170030924 | 0 | G | A | 0.14 | -1.30E-01 | 1.46E-01 | 2.58E+00 | 2.65E+00 | 3.74E-01 | no | MAR1 | intron_variant | MODIFIER |
| 8 | 65918470 | AX-170105788 | 0 | C | G | 0.14 | -1.30E-01 | 1.46E-01 | 2.58E+00 | 2.65E+00 | 3.74E-01 | no | MAR1 | intron_variant | MODIFIER |
| 8 | 65931286 | AX-170466725 | 0 | A | G | 0.14 | -1.30E-01 | 1.46E-01 | 2.58E+00 | 2.65E+00 | 3.74E-01 | no | MAR1 | intron_variant | MODIFIER |
| 8 | 65935758 | AX-97841597 | 0 | C | T | 0.14 | -1.30E-01 | 1.46E-01 | 2.58E+00 | 2.65E+00 | 3.74E-01 | no | MAR1 | intron_variant | MODIFIER |
| 8 | 65969815 | AX-170406630 | 0 | A | G | 0.14 | -1.30E-01 | 1.46E-01 | 2.58E+00 | 2.65E+00 | 3.74E-01 | no | MAR1 | intron_variant | MODIFIER |
| 8 | 65980530 | AX-170417679 | 0 | T | G | 0.14 | -1.30E-01 | 1.46E-01 | 2.58E+00 | 2.65E+00 | 3.74E-01 | no | MAR1 | intron_variant | MODIFIER |
| 8 | 65980903 | AX-170417276 | 0 | A | G | 0.14 | -1.30E-01 | 1.46E-01 | 2.58E+00 | 2.65E+00 | 3.74E-01 | no | MAR1 | intron_variant | MODIFIER |
| 8 | 65983113 | AX-170304335 | 0 | C | A | 0.14 | -1.30E-01 | 1.46E-01 | 2.58E+00 | 2.65E+00 | 3.74E-01 | no | MAR1 | intron_variant | MODIFIER |
| 8 | 66004343 | AX-170372379 | 0 | C | G | 0.14 | -1.30E-01 | 1.46E-01 | 2.58E+00 | 2.65E+00 | 3.74E-01 | no | MAR1 | intron_variant | MODIFIER |
| 8 | 66016529 | AX-170118515 | 0 | T | G | 0.14 | -1.30E-01 | 1.46E-01 | 2.58E+00 | 2.65E+00 | 3.74E-01 | no | MAR1 | intron_variant | MODIFIER |
| 8 | 66023495 | AX-170484832 | 0 | C | T | 0.14 | -1.30E-01 | 1.46E-01 | 2.58E+00 | 2.65E+00 | 3.74E-01 | no | MAR1 | intron_variant | MODIFIER |
| 8 | 66044105 | AX-170195374 | 0 | C | T | 0.14 | -1.30E-01 | 1.46E-01 | 2.58E+00 | 2.65E+00 | 3.74E-01 | no | MAR1 | intron_variant | MODIFIER |
| 8 | 66086649 | AX-170499452 | 0 | A | G | 0.14 | -1.30E-01 | 1.46E-01 | 2.58E+00 | 2.65E+00 | 3.74E-01 | no | MAR1 | intron_variant | MODIFIER |
| 8 | 66092603 | AX-97856989 | 0 | T | C | 0.14 | -1.30E-01 | 1.46E-01 | 2.58E+00 | 2.65E+00 | 3.74E-01 | no | MAR1 | intron_variant | MODIFIER |
| 8 | 66103661 | AX-170258805 | 0 | A | G | 0.14 | -1.30E-01 | 1.46E-01 | 2.58E+00 | 2.65E+00 | 3.74E-01 | no | MAR1 | intron_variant | MODIFIER |
| 8 | 66104587 | AX-170418361 | 0 | A | G | 0.14 | -1.30E-01 | 1.46E-01 | 2.58E+00 | 2.65E+00 | 3.74E-01 | no | MAR1 | intron_variant | MODIFIER |
| 8 | 66141399 | AX-170145280 | 0 | T | A | 0.14 | -1.30E-01 | 1.46E-01 | 2.58E+00 | 2.65E+00 | 3.74E-01 | no | MAR1 | intron_variant | MODIFIER |
| 8 | 66165285 | AX-97804441 | 0 | A | G | 0.14 | -1.30E-01 | 1.46E-01 | 2.58E+00 | 2.65E+00 | 3.74E-01 | no | MAR1 | intron_variant | MODIFIER |
| 8 | 66177576 | AX-170327851 | 0 | T | A | 0.14 | -1.30E-01 | 1.46E-01 | 2.58E+00 | 2.65E+00 | 3.74E-01 | no | MAR1 | intron_variant | MODIFIER |
| 8 | 66178480 | AX-170344885 | 0 | T | G | 0.14 | -1.30E-01 | 1.46E-01 | 2.58E+00 | 2.65E+00 | 3.74E-01 | no | MAR1 | intron_variant | MODIFIER |
| 8 | 66179912 | AX-170030934 | 0 | A | G | 0.14 | -1.30E-01 | 1.46E-01 | 2.58E+00 | 2.65E+00 | 3.74E-01 | no | MAR1 | intron_variant | MODIFIER |
| 8 | 66183603 | AX-170136992 | 0 | C | T | 0.14 | -1.30E-01 | 1.46E-01 | 2.58E+00 | 2.65E+00 | 3.74E-01 | no | MAR1 | intron_variant | MODIFIER |
| 8 | 66215359 | AX-170354827 | 0 | C | T | 0.14 | -1.30E-01 | 1.46E-01 | 2.58E+00 | 2.65E+00 | 3.74E-01 | no | MAR1 | intron_variant | MODIFIER |
| 8 | 66216251 | AX-170266283 | 0 | T | G | 0.14 | -1.30E-01 | 1.46E-01 | 2.58E+00 | 2.65E+00 | 3.74E-01 | no | MAR1 | intron_variant | MODIFIER |
| 8 | 66263674 | AX-170030935 | 0 | C | T | 0.14 | -1.30E-01 | 1.46E-01 | 2.58E+00 | 2.65E+00 | 3.74E-01 | no | MAR1 | intron_variant | MODIFIER |
| 8 | 66268397 | AX-170080290 | 0 | G | T | 0.14 | -1.30E-01 | 1.46E-01 | 2.58E+00 | 2.65E+00 | 3.74E-01 | no | MAR1 | intron_variant | MODIFIER |
| 8 | 66285135 | AX-170352806 | 0 | T | C | 0.14 | -1.30E-01 | 1.46E-01 | 2.58E+00 | 2.65E+00 | 3.74E-01 | no | MAR1 | intron_variant | MODIFIER |
| 8 | 66290327 | AX-97789187 | 0 | A | G | 0.14 | -1.30E-01 | 1.46E-01 | 2.58E+00 | 2.65E+00 | 3.74E-01 | no | MAR1 | intron_variant | MODIFIER |
| 8 | 66290732 | AX-170376896 | 0 | T | G | 0.14 | -1.30E-01 | 1.46E-01 | 2.58E+00 | 2.65E+00 | 3.74E-01 | no | MAR1 | intron_variant | MODIFIER |
| 8 | 66294857 | AX-170178912 | 0 | T | C | 0.14 | -1.30E-01 | 1.46E-01 | 2.58E+00 | 2.65E+00 | 3.74E-01 | no | MAR1 | intron_variant | MODIFIER |
| 8 | 66323824 | AX-170464046 | 0 | G | A | 0.14 | -1.30E-01 | 1.46E-01 | 2.58E+00 | 2.65E+00 | 3.74E-01 | no | MAR1 | intron_variant | MODIFIER |
| 8 | 66334194 | AX-170145951 | 0 | G | A | 0.14 | -1.30E-01 | 1.46E-01 | 2.58E+00 | 2.65E+00 | 3.74E-01 | no | MAR1 | intron_variant | MODIFIER |
| 8 | 66354671 | AX-170450886 | 0 | A | G | 0.14 | -1.30E-01 | 1.46E-01 | 2.58E+00 | 2.65E+00 | 3.74E-01 | no | MAR1 | intron_variant | MODIFIER |
| 8 | 66363643 | AX-170293160 | 0 | G | C | 0.14 | -1.30E-01 | 1.46E-01 | 2.58E+00 | 2.65E+00 | 3.74E-01 | no | MAR1 | intron_variant | MODIFIER |
| 8 | 66363903 | AX-170501820 | 0 | T | C | 0.14 | -1.30E-01 | 1.46E-01 | 2.58E+00 | 2.65E+00 | 3.74E-01 | no | MAR1 | intron_variant | MODIFIER |
| 8 | 66392123 | AX-170080293 | 0 | C | T | 0.14 | -1.30E-01 | 1.46E-01 | 2.58E+00 | 2.65E+00 | 3.74E-01 | no | MAR1 | intron_variant | MODIFIER |
| 8 | 66409798 | AX-170193054 | 0 | A | T | 0.14 | -1.30E-01 | 1.46E-01 | 2.58E+00 | 2.65E+00 | 3.74E-01 | no | MAR1 | intron_variant | MODIFIER |
| 8 | 66409985 | AX-170138174 | 0 | G | T | 0.14 | -1.30E-01 | 1.46E-01 | 2.58E+00 | 2.65E+00 | 3.74E-01 | no | MAR1 | intron_variant | MODIFIER |
| 8 | 66410380 | AX-170174070 | 0 | A | C | 0.14 | -1.30E-01 | 1.46E-01 | 2.58E+00 | 2.65E+00 | 3.74E-01 | no | MAR1 | intron_variant | MODIFIER |
| 8 | 66410480 | AX-170229885 | 0 | G | A | 0.14 | -1.30E-01 | 1.46E-01 | 2.58E+00 | 2.65E+00 | 3.74E-01 | no | MAR1 | intron_variant | MODIFIER |
| 8 | 66429389 | AX-170244469 | 0 | A | G | 0.14 | -1.30E-01 | 1.46E-01 | 2.58E+00 | 2.65E+00 | 3.74E-01 | no | MAR1 | intron_variant | MODIFIER |
| 8 | 66429907 | AX-170216104 | 0 | G | T | 0.14 | -1.30E-01 | 1.46E-01 | 2.58E+00 | 2.65E+00 | 3.74E-01 | no | MAR1 | intron_variant | MODIFIER |
| 8 | 66429998 | AX-170521749 | 0 | A | G | 0.14 | -1.30E-01 | 1.46E-01 | 2.58E+00 | 2.65E+00 | 3.74E-01 | no | MAR1 | intron_variant | MODIFIER |
| 8 | 66430131 | AX-97834812 | 0 | A | G | 0.14 | -1.30E-01 | 1.46E-01 | 2.58E+00 | 2.65E+00 | 3.74E-01 | no | MAR1 | intron_variant | MODIFIER |
| 8 | 66430177 | AX-170405559 | 0 | A | C | 0.14 | -1.30E-01 | 1.46E-01 | 2.58E+00 | 2.65E+00 | 3.74E-01 | no | MAR1 | intron_variant | MODIFIER |
| 8 | 66431486 | AX-170396692 | 0 | C | T | 0.14 | -1.30E-01 | 1.46E-01 | 2.58E+00 | 2.65E+00 | 3.74E-01 | no | MAR1 | intron_variant | MODIFIER |
| 8 | 66438954 | AX-170521293 | 0 | T | C | 0.14 | -1.30E-01 | 1.46E-01 | 2.58E+00 | 2.65E+00 | 3.74E-01 | no | MAR1 | intron_variant | MODIFIER |
| 8 | 66440939 | AX-170380797 | 0 | A | G | 0.14 | -1.30E-01 | 1.46E-01 | 2.58E+00 | 2.65E+00 | 3.74E-01 | no | MAR1 | intron_variant | MODIFIER |
| 8 | 66467544 | AX-170534979 | 0 | T | C | 0.14 | -1.30E-01 | 1.46E-01 | 2.58E+00 | 2.65E+00 | 3.74E-01 | no | MAR1 | intron_variant | MODIFIER |
| 8 | 66468893 | AX-97836915 | 0 | C | T | 0.14 | -1.30E-01 | 1.46E-01 | 2.58E+00 | 2.65E+00 | 3.74E-01 | no | Tma16 | downstream_gene_variant | MODIFIER |

| | | | | | | | | | | | | | | | |
|---|---|---|---|---|---|---|---|---|---|---|---|---|---|---|---|
| 8 | 66494031 | AX-170366528 | 0 | T | A | 0.14 | -1.30E-01 | 1.46E-01 | 2.58E+00 | 2.65E+00 | 3.74E-01 | no | Gm32568 | upstream_gene_variant | MODIFIER |
| 8 | 66502845 | AX-170276264 | 0 | C | A | 0.14 | -1.30E-01 | 1.46E-01 | 2.58E+00 | 2.65E+00 | 3.74E-01 | no | | intergenic_variant | MODIFIER |
| 8 | 66513225 | AX-170080298 | 0 | T | C | 0.14 | -1.30E-01 | 1.46E-01 | 2.58E+00 | 2.65E+00 | 3.74E-01 | no | Tktl2 | synonymous_variant | LOW |
| 8 | 66516292 | AX-97813388 | 0 | A | G | 0.14 | -1.30E-01 | 1.46E-01 | 2.58E+00 | 2.65E+00 | 3.74E-01 | no | Tktl2 | downstream_gene_variant | MODIFIER |
| 8 | 66516864 | AX-170080300 | 0 | T | C | 0.14 | -1.30E-01 | 1.46E-01 | 2.58E+00 | 2.65E+00 | 3.74E-01 | no | Tktl2 | downstream_gene_variant | MODIFIER |
| 8 | 66521153 | AX-170441503 | 0 | G | A | 0.14 | -1.30E-01 | 1.46E-01 | 2.58E+00 | 2.65E+00 | 3.74E-01 | no | Tktl2 | downstream_gene_variant | MODIFIER |
| 8 | 66610647 | AX-170234804 | 0 | G | A | 0.14 | -1.30E-01 | 1.46E-01 | 2.58E+00 | 2.65E+00 | 3.74E-01 | no | | intergenic_variant | MODIFIER |
| 8 | 66617767 | AX-170234129 | 0 | T | A | 0.14 | -1.30E-01 | 1.46E-01 | 2.58E+00 | 2.65E+00 | 3.74E-01 | no | | intergenic_variant | MODIFIER |
| 8 | 66619357 | AX-170304099 | 0 | A | G | 0.14 | -1.30E-01 | 1.46E-01 | 2.58E+00 | 2.65E+00 | 3.74E-01 | no | | intergenic_variant | MODIFIER |
| 8 | 66626403 | AX-170463557 | 0 | T | G | 0.14 | -1.30E-01 | 1.46E-01 | 2.58E+00 | 2.65E+00 | 3.74E-01 | no | | intergenic_variant | MODIFIER |
| 8 | 66674231 | AX-170356796 | 0 | G | A | 0.14 | -1.30E-01 | 1.46E-01 | 2.58E+00 | 2.65E+00 | 3.74E-01 | no | | intergenic_variant | MODIFIER |
| 8 | 66718237 | AX-170030944 | 0 | G | T | 0.14 | -1.30E-01 | 1.46E-01 | 2.58E+00 | 2.65E+00 | 3.74E-01 | no | | intergenic_variant | MODIFIER |
| 8 | 66724538 | AX-170030946 | 0 | G | A | 0.14 | -1.30E-01 | 1.46E-01 | 2.58E+00 | 2.65E+00 | 3.74E-01 | no | | intergenic_variant | MODIFIER |
| 8 | 66733971 | AX-170030945 | 0 | C | T | 0.14 | -1.30E-01 | 1.46E-01 | 2.58E+00 | 2.65E+00 | 3.74E-01 | no | | intergenic_variant | MODIFIER |
| 8 | 66756476 | AX-97804439 | 0 | C | T | 0.14 | -1.30E-01 | 1.46E-01 | 2.58E+00 | 2.65E+00 | 3.74E-01 | no | | intergenic_variant | MODIFIER |
| 8 | 66790973 | AX-170030947 | 0 | A | C | 0.14 | -1.30E-01 | 1.46E-01 | 2.58E+00 | 2.65E+00 | 3.74E-01 | no | Gm45911 | intron_variant,non_coding_transcript_variant | MODIFIER |
| 8 | 66817209 | AX-170030949 | 0 | A | G | 0.14 | -1.30E-01 | 1.46E-01 | 2.58E+00 | 2.65E+00 | 3.74E-01 | no | | intergenic_variant | MODIFIER |
| 8 | 66853629 | AX-97826367 | 0 | T | C | 0.14 | -1.30E-01 | 1.46E-01 | 2.58E+00 | 2.65E+00 | 3.74E-01 | no | | intergenic_variant | MODIFIER |
| 8 | 66863522 | AX-170030948 | 0 | C | T | 0.14 | -1.30E-01 | 1.46E-01 | 2.58E+00 | 2.65E+00 | 3.74E-01 | no | Naf1 | intron_variant | MODIFIER |
| 8 | 66876153 | AX-170283652 | 0 | G | A | 0.14 | -1.30E-01 | 1.46E-01 | 2.58E+00 | 2.65E+00 | 3.74E-01 | no | Naf1 | intron_variant | MODIFIER |
| 8 | 66879911 | AX-170030950 | 0 | G | C | 0.14 | -1.30E-01 | 1.46E-01 | 2.58E+00 | 2.65E+00 | 3.74E-01 | no | Naf1 | intron_variant | MODIFIER |
| 8 | 66879975 | AX-97794074 | 0 | A | G | 0.14 | -1.30E-01 | 1.46E-01 | 2.58E+00 | 2.65E+00 | 3.74E-01 | no | Naf1 | intron_variant | MODIFIER |
| 8 | 66898454 | AX-170173364 | 0 | G | C | 0.14 | -1.30E-01 | 1.46E-01 | 2.58E+00 | 2.65E+00 | 3.74E-01 | no | | intergenic_variant | MODIFIER |
| 8 | 66905885 | AX-97825443 | 0 | T | C | 0.14 | -1.30E-01 | 1.46E-01 | 2.58E+00 | 2.65E+00 | 3.74E-01 | no | Gm39185 | upstream_gene_variant | MODIFIER |
| 8 | 66922528 | AX-170030942 | 0 | C | A | 0.14 | -1.30E-01 | 1.46E-01 | 2.58E+00 | 2.65E+00 | 3.74E-01 | no | Gm25275 | upstream_gene_variant | MODIFIER |
| 8 | 66933788 | AX-170090139 | 0 | C | T | 0.14 | -1.30E-01 | 1.46E-01 | 2.58E+00 | 2.65E+00 | 3.74E-01 | no | Gm39185 | intron_variant,non_coding_transcript_variant | MODIFIER |
| 8 | 67021086 | AX-97814718 | 0 | C | G | 0.14 | -1.30E-01 | 1.46E-01 | 2.58E+00 | 2.65E+00 | 3.74E-01 | no | | intergenic_variant | MODIFIER |
| 8 | 67054656 | AX-97786938 | 0 | C | T | 0.14 | -1.30E-01 | 1.46E-01 | 2.58E+00 | 2.65E+00 | 3.74E-01 | no | Gm7639 | upstream_gene_variant | MODIFIER |
| 8 | 67070102 | AX-97837297 | 0 | G | T | 0.14 | -1.30E-01 | 1.46E-01 | 2.58E+00 | 2.65E+00 | 3.74E-01 | no | | intergenic_variant | MODIFIER |
| 8 | 67088335 | AX-170424344 | 0 | G | C | 0.14 | -1.30E-01 | 1.46E-01 | 2.58E+00 | 2.65E+00 | 3.74E-01 | no | | intergenic_variant | MODIFIER |
| 8 | 67330292 | AX-170090141 | 0 | T | G | 0.14 | -1.30E-01 | 1.46E-01 | 2.58E+00 | 2.65E+00 | 3.74E-01 | no | | intergenic_variant | MODIFIER |
| 8 | 67445822 | AX-97854823 | 0 | A | G | 0.14 | -1.30E-01 | 1.46E-01 | 2.58E+00 | 2.65E+00 | 3.74E-01 | no | | intergenic_variant | MODIFIER |
| 7 | 144089919 | AX-97843414 | 1 | T | C | 0.205 | -6.64E-02 | 7.55E-02 | 2.63E+00 | 2.76E+00 | 3.74E-01 | no | Shank2 | intron_variant | MODIFIER |
| 14 | 92001408 | AX-170503153 | 1 | A | C | 0.036 | 1.39E-01 | 1.57E-01 | 2.53E+00 | 2.70E+00 | 3.74E-01 | no | | intergenic_variant | MODIFIER |
| 1 | 169437577 | AX-170489618 | 0 | G | A | 0.421 | -6.18E-02 | 6.73E-02 | 3.12E+00 | 3.23E+00 | 3.74E-01 | no | | intergenic_variant | MODIFIER |
| 8 | 65866173 | AX-97843611 | 1 | G | A | 0.143 | -1.27E-01 | 1.43E-01 | 2.57E+00 | 2.65E+00 | 3.74E-01 | no | MAR1 | intron_variant | MODIFIER |
| 16 | 75708715 | AX-170413357 | 0 | A | G | 0.491 | 7.05E-02 | 7.76E-02 | 2.44E+00 | 2.51E+00 | 3.74E-01 | no | Gm30162 | downstream_gene_variant | MODIFIER |
| 4 | 72741808 | AX-170276445 | 0 | T | C | 0.456 | -7.41E-02 | 8.25E-02 | 2.51E+00 | 2.56E+00 | 3.74E-01 | no | | intergenic_variant | MODIFIER |
| 4 | 134581402 | AX-170278210 | 0 | C | G | 0.167 | 9.43E-02 | 1.05E-01 | 2.52E+00 | 2.58E+00 | 3.74E-01 | no | Man1c1 | intron_variant | MODIFIER |
| 2 | 169054183 | AX-170019655 | 1 | G | A | 0.161 | 1.53E-01 | 1.75E-01 | 2.68E+00 | 2.79E+00 | 3.74E-01 | Yes | | intergenic_variant | MODIFIER |
| 9 | 17081226 | AX-170031860 | 1 | A | G | 0.482 | 7.86E-02 | 8.81E-02 | 2.54E+00 | 2.60E+00 | 3.74E-01 | no | | intergenic_variant | MODIFIER |
| 8 | 90086894 | AX-170146375 | 0 | T | C | 0.193 | -9.75E-02 | 1.10E-01 | 2.59E+00 | 2.69E+00 | 3.75E-01 | no | | intergenic_variant | MODIFIER |
| 15 | 8629319 | AX-170087759 | 1 | G | T | 0.241 | -9.37E-02 | 1.06E-01 | 2.65E+00 | 2.70E+00 | 3.75E-01 | no | Slc1a3 | downstream_gene_variant | MODIFIER |
| 6 | 127846384 | AX-170244167 | 1 | G | T | 0.536 | 7.41E-02 | 8.23E-02 | 2.48E+00 | 2.55E+00 | 3.75E-01 | no | Gm23098 | upstream_gene_variant | MODIFIER |
| 10 | 99759076 | AX-170097017 | 0 | A | G | 0.272 | -7.62E-02 | 8.65E-02 | 2.62E+00 | 2.70E+00 | 3.75E-01 | no | Csl | synonymous_variant | LOW |
| 10 | 99987242 | AX-170124206 | 0 | A | T | 0.272 | -7.62E-02 | 8.65E-02 | 2.62E+00 | 2.70E+00 | 3.75E-01 | no | Gm17870 | upstream_gene_variant | MODIFIER |
| 6 | 130476518 | AX-97839263 | 0 | G | A | 0.561 | 7.43E-02 | 8.24E-02 | 2.48E+00 | 2.54E+00 | 3.75E-01 | no | Gm6590 | intron_variant,non_coding_transcript_variant | MODIFIER |
| 2 | 168855624 | AX-170116511 | 1 | A | T | 0.17 | 1.57E-01 | 1.78E-01 | 2.64E+00 | 2.73E+00 | 3.75E-01 | Yes | | intergenic_variant | MODIFIER |
| 18 | 80438734 | AX-170156993 | 1 | A | G | 0.375 | -7.47E-02 | 8.51E-02 | 2.66E+00 | 2.75E+00 | 3.76E-01 | no | Ctdp1 | intron_variant | MODIFIER |
| 16 | 49873433 | AX-97785565 | 1 | T | C | 0.473 | -7.03E-02 | 7.94E-02 | 2.57E+00 | 2.64E+00 | 3.76E-01 | no | Cd47 | intron_variant | MODIFIER |
| 4 | 137191393 | AX-97798747 | 0 | T | C | 0.175 | 9.58E-02 | 1.09E-01 | 2.62E+00 | 2.67E+00 | 3.76E-01 | no | | intergenic_variant | MODIFIER |
| 4 | 137868962 | AX-170444332 | 0 | A | G | 0.175 | 9.58E-02 | 1.09E-01 | 2.62E+00 | 2.67E+00 | 3.76E-01 | no | Ece1 | intron_variant | MODIFIER |
| 4 | 137888592 | AX-170481458 | 0 | C | T | 0.175 | 9.58E-02 | 1.09E-01 | 2.62E+00 | 2.67E+00 | 3.76E-01 | no | Ece1 | intron_variant | MODIFIER |
| 4 | 137895761 | AX-170506918 | 0 | C | A | 0.175 | 9.58E-02 | 1.09E-01 | 2.62E+00 | 2.67E+00 | 3.76E-01 | no | Ece1 | intron_variant | MODIFIER |

| | | | | | | | | | | | | | | | |
|---|---|---|---|---|---|---|---|---|---|---|---|---|---|---|---|
| 4 | 138187485 | AX-170171736 | 0 | G | A | 0.175 | 9.58E-02 | 1.09E-01 | 2.62E+00 | 2.67E+00 | 3.76E-01 | no | Eif4g3 | intron_variant | MODIFIER |
| 4 | 138284419 | AX-170274394 | 0 | G | A | 0.175 | 9.58E-02 | 1.09E-01 | 2.62E+00 | 2.67E+00 | 3.76E-01 | no | Kif17 | intron_variant | MODIFIER |
| 4 | 138293585 | AX-170070839 | 0 | G | A | 0.175 | 9.58E-02 | 1.09E-01 | 2.62E+00 | 2.67E+00 | 3.76E-01 | no | Kif17 | intron_variant | MODIFIER |
| 4 | 138340666 | AX-170205424 | 0 | C | G | 0.175 | 9.58E-02 | 1.09E-01 | 2.62E+00 | 2.67E+00 | 3.76E-01 | no | Cda | intron_variant | MODIFIER |
| 4 | 138382790 | AX-170297414 | 0 | A | G | 0.175 | 9.58E-02 | 1.09E-01 | 2.62E+00 | 2.67E+00 | 3.76E-01 | no | | regulatory_region_variant | MODIFIER |
| 4 | 138389187 | AX-170469044 | 0 | C | T | 0.175 | 9.58E-02 | 1.09E-01 | 2.62E+00 | 2.67E+00 | 3.76E-01 | no | Fam43b | downstream_gene_variant | MODIFIER |
| 4 | 138418743 | AX-170269172 | 0 | C | T | 0.175 | 9.58E-02 | 1.09E-01 | 2.62E+00 | 2.67E+00 | 3.76E-01 | no | | regulatory_region_variant | MODIFIER |
| 4 | 138512352 | AX-170501743 | 0 | C | T | 0.175 | 9.58E-02 | 1.09E-01 | 2.62E+00 | 2.67E+00 | 3.76E-01 | no | | intergenic_variant | MODIFIER |
| 4 | 138513106 | AX-170456428 | 0 | A | C | 0.175 | 9.58E-02 | 1.09E-01 | 2.62E+00 | 2.67E+00 | 3.76E-01 | no | | intergenic_variant | MODIFIER |
| 4 | 138544544 | AX-170125480 | 0 | T | C | 0.175 | 9.58E-02 | 1.09E-01 | 2.62E+00 | 2.67E+00 | 3.76E-01 | no | | intergenic_variant | MODIFIER |
| 4 | 138654448 | AX-170292237 | 0 | A | G | 0.175 | 9.58E-02 | 1.09E-01 | 2.62E+00 | 2.67E+00 | 3.76E-01 | no | | intergenic_variant | MODIFIER |
| 4 | 138994665 | AX-170197450 | 0 | C | T | 0.175 | 9.58E-02 | 1.09E-01 | 2.62E+00 | 2.67E+00 | 3.76E-01 | no | Tmco4 | intron_variant | MODIFIER |
| 4 | 139117665 | AX-170455238 | 0 | A | G | 0.175 | 9.58E-02 | 1.09E-01 | 2.62E+00 | 2.67E+00 | 3.76E-01 | no | Minos1 | intron_variant | MODIFIER |
| 4 | 139139041 | AX-97829008 | 0 | A | C | 0.175 | 9.58E-02 | 1.09E-01 | 2.62E+00 | 2.67E+00 | 3.76E-01 | no | | regulatory_region_variant | MODIFIER |
| 4 | 139174809 | AX-170322771 | 0 | C | T | 0.175 | 9.58E-02 | 1.09E-01 | 2.62E+00 | 2.67E+00 | 3.76E-01 | no | Gm16287 | downstream_gene_variant | MODIFIER |
| 4 | 139201281 | AX-170487710 | 0 | C | T | 0.175 | 9.58E-02 | 1.09E-01 | 2.62E+00 | 2.67E+00 | 3.76E-01 | no | Capzb | intron_variant | MODIFIER |
| 4 | 139204329 | AX-170024096 | 0 | G | A | 0.175 | 9.58E-02 | 1.09E-01 | 2.62E+00 | 2.67E+00 | 3.76E-01 | no | Capzb | intron_variant | MODIFIER |
| 4 | 139220748 | AX-170484276 | 0 | A | G | 0.175 | 9.58E-02 | 1.09E-01 | 2.62E+00 | 2.67E+00 | 3.76E-01 | no | Capzb | intron_variant | MODIFIER |
| 4 | 139224628 | AX-170215528 | 0 | C | A | 0.175 | 9.58E-02 | 1.09E-01 | 2.62E+00 | 2.67E+00 | 3.76E-01 | no | Capzb | intron_variant | MODIFIER |
| 4 | 139235440 | AX-170188489 | 0 | C | T | 0.175 | 9.58E-02 | 1.09E-01 | 2.62E+00 | 2.67E+00 | 3.76E-01 | no | Capzb | intron_variant | MODIFIER |
| 4 | 139253369 | AX-170430534 | 0 | T | G | 0.175 | 9.58E-02 | 1.09E-01 | 2.62E+00 | 2.67E+00 | 3.76E-01 | no | Capzb | intron_variant | MODIFIER |
| 4 | 139263601 | AX-170238884 | 0 | A | C | 0.175 | 9.58E-02 | 1.09E-01 | 2.62E+00 | 2.67E+00 | 3.76E-01 | no | Capzb | intron_variant | MODIFIER |
| 4 | 139302188 | AX-170358746 | 0 | G | A | 0.175 | 9.58E-02 | 1.09E-01 | 2.62E+00 | 2.67E+00 | 3.76E-01 | no | Pqlc2 | intron_variant | MODIFIER |
| 4 | 139318210 | AX-170277211 | 0 | C | G | 0.175 | 9.58E-02 | 1.09E-01 | 2.62E+00 | 2.67E+00 | 3.76E-01 | no | Akr7a5 | missense_variant | MODERATE |
| 4 | 139366339 | AX-170511280 | 0 | G | A | 0.175 | 9.58E-02 | 1.09E-01 | 2.62E+00 | 2.67E+00 | 3.76E-01 | no | Emc1 | intron_variant | MODIFIER |
| 4 | 139366568 | AX-170229558 | 0 | G | A | 0.175 | 9.58E-02 | 1.09E-01 | 2.62E+00 | 2.67E+00 | 3.76E-01 | no | Emc1 | intron_variant | MODIFIER |
| 4 | 139438981 | AX-170536106 | 0 | C | T | 0.175 | 9.58E-02 | 1.09E-01 | 2.62E+00 | 2.67E+00 | 3.76E-01 | no | Ubr4 | intron_variant | MODIFIER |
| 4 | 139451130 | AX-170190439 | 0 | C | T | 0.175 | 9.58E-02 | 1.09E-01 | 2.62E+00 | 2.67E+00 | 3.76E-01 | no | Ubr4 | intron_variant | MODIFIER |
| 4 | 139454022 | AX-170239760 | 0 | T | C | 0.175 | 9.58E-02 | 1.09E-01 | 2.62E+00 | 2.67E+00 | 3.76E-01 | no | Ubr4 | synonymous_variant | LOW |
| 4 | 139465569 | AX-170371330 | 0 | G | A | 0.175 | 9.58E-02 | 1.09E-01 | 2.62E+00 | 2.67E+00 | 3.76E-01 | no | Ubr4 | intron_variant | MODIFIER |
| 4 | 139489058 | AX-170232898 | 0 | C | A | 0.175 | 9.58E-02 | 1.09E-01 | 2.62E+00 | 2.67E+00 | 3.76E-01 | no | Ubr4 | intron_variant | MODIFIER |
| 4 | 139491584 | AX-170156949 | 0 | C | G | 0.175 | 9.58E-02 | 1.09E-01 | 2.62E+00 | 2.67E+00 | 3.76E-01 | no | Ubr4 | downstream_gene_variant | MODIFIER |
| 4 | 139492523 | AX-170112465 | 0 | G | A | 0.175 | 9.58E-02 | 1.09E-01 | 2.62E+00 | 2.67E+00 | 3.76E-01 | no | Ubr4 | downstream_gene_variant | MODIFIER |
| 4 | 139530859 | AX-170413798 | 0 | A | T | 0.175 | 9.58E-02 | 1.09E-01 | 2.62E+00 | 2.67E+00 | 3.76E-01 | no | Iffo2 | upstream_gene_variant | MODIFIER |
| 4 | 139542822 | AX-170510933 | 0 | A | G | 0.175 | 9.58E-02 | 1.09E-01 | 2.62E+00 | 2.67E+00 | 3.76E-01 | no | Iffo2 | intron_variant | MODIFIER |
| 4 | 139545181 | AX-170324048 | 0 | G | A | 0.175 | 9.58E-02 | 1.09E-01 | 2.62E+00 | 2.67E+00 | 3.76E-01 | no | Iffo2 | intron_variant | MODIFIER |
| 4 | 139550362 | AX-170327168 | 0 | C | T | 0.175 | 9.58E-02 | 1.09E-01 | 2.62E+00 | 2.67E+00 | 3.76E-01 | no | Iffo2 | intron_variant | MODIFIER |
| 4 | 139576298 | AX-170178119 | 0 | C | T | 0.175 | 9.58E-02 | 1.09E-01 | 2.62E+00 | 2.67E+00 | 3.76E-01 | no | Iffo2 | intron_variant | MODIFIER |
| 4 | 139579525 | AX-170070856 | 0 | G | T | 0.175 | 9.58E-02 | 1.09E-01 | 2.62E+00 | 2.67E+00 | 3.76E-01 | no | Iffo2 | intron_variant | MODIFIER |
| 4 | 139616304 | AX-97827859 | 0 | G | A | 0.175 | 9.58E-02 | 1.09E-01 | 2.62E+00 | 2.67E+00 | 3.76E-01 | no | Iffo2 | intron_variant | MODIFIER |
| 4 | 139660101 | AX-170512880 | 0 | C | T | 0.175 | 9.58E-02 | 1.09E-01 | 2.62E+00 | 2.67E+00 | 3.76E-01 | no | Tas1r2 | synonymous_variant | LOW |
| 4 | 139688265 | AX-170233507 | 0 | G | A | 0.175 | 9.58E-02 | 1.09E-01 | 2.62E+00 | 2.67E+00 | 3.76E-01 | no | | regulatory_region_variant | MODIFIER |
| 4 | 139688481 | AX-170503541 | 0 | G | C | 0.175 | 9.58E-02 | 1.09E-01 | 2.62E+00 | 2.67E+00 | 3.76E-01 | no | | regulatory_region_variant | MODIFIER |
| 4 | 140011101 | AX-170292977 | 0 | T | C | 0.175 | 9.58E-02 | 1.09E-01 | 2.62E+00 | 2.67E+00 | 3.76E-01 | no | | regulatory_region_variant | MODIFIER |
| 4 | 140712339 | AX-170488098 | 0 | G | A | 0.175 | 9.58E-02 | 1.09E-01 | 2.62E+00 | 2.67E+00 | 3.76E-01 | no | Rcc2 | intron_variant | MODIFIER |
| 4 | 140875300 | AX-170268070 | 0 | G | A | 0.175 | 9.58E-02 | 1.09E-01 | 2.62E+00 | 2.67E+00 | 3.76E-01 | no | 4930515B02Rik | intron_variant,non_coding_transcript_variant | MODIFIER |
| 4 | 156010585 | AX-97801862 | 1 | T | C | 0.098 | 1.38E-01 | 1.58E-01 | 2.70E+00 | 2.81E+00 | 3.76E-01 | no | Tnfrsf4 | upstream_gene_variant | MODIFIER |
| 5 | 35443316 | AX-170148851 | 0 | A | G | 0.342 | 8.65E-02 | 9.89E-02 | 2.75E+00 | 2.82E+00 | 3.76E-01 | no | | intergenic_variant | MODIFIER |
| 16 | 79133019 | AX-170531685 | 0 | G | A | 0.474 | 6.53E-02 | 7.36E-02 | 2.55E+00 | 2.63E+00 | 3.76E-01 | no | | intergenic_variant | MODIFIER |
| 5 | 121084061 | AX-97786813 | 0 | C | T | 0.456 | 9.10E-02 | 1.04E-01 | 2.70E+00 | 2.77E+00 | 3.76E-01 | no | | intergenic_variant | MODIFIER |
| 4 | 90564995 | AX-170107081 | 0 | T | C | 0.254 | 9.96E-02 | 1.11E-01 | 2.48E+00 | 2.53E+00 | 3.76E-01 | no | | intergenic_variant | MODIFIER |
| 8 | 16433527 | AX-170148574 | 1 | C | A | 0.241 | -6.57E-02 | 7.48E-02 | 2.60E+00 | 2.70E+00 | 3.76E-01 | no | Csmd1 | intron_variant | MODIFIER |
| 5 | 38515235 | AX-97781630 | 0 | A | G | 0.342 | 9.76E-02 | 1.11E-01 | 2.78E+00 | 2.85E+00 | 3.77E-01 | no | | intergenic_variant | MODIFIER |
| 4 | 69663176 | AX-170069614 | 0 | G | A | 0.316 | 9.84E-02 | 1.13E-01 | 2.72E+00 | 2.80E+00 | 3.77E-01 | no | | intergenic_variant | MODIFIER |

| | | | | | | | | | | | | | | | |
|---|---|---|---|---|---|---|---|---|---|---|---|---|---|---|---|
| 4 | 133589227 | AX-170141848 | 1 | A | G | 0.152 | 1.02E-01 | 1.14E-01 | 2.50E+00 | 2.56E+00 | 3.77E-01 | no | Gpn2 | intron_variant | MODIFIER |
| 4 | 134442165 | AX-97824193 | 0 | T | C | 0.167 | 9.45E-02 | 1.06E-01 | 2.53E+00 | 2.59E+00 | 3.77E-01 | no | | intergenic_variant | MODIFIER |
| 7 | 66390302 | AX-170029182 | 1 | T | A | 0.393 | -8.47E-02 | 9.65E-02 | 2.86E+00 | 2.96E+00 | 3.77E-01 | no | Lrrk1 | upstream_gene_variant | MODIFIER |
| 5 | 128543312 | AX-170357422 | 1 | C | T | 0.152 | -8.41E-02 | 8.86E-02 | 2.28E+00 | 2.38E+00 | 3.77E-01 | no | | intergenic_variant | MODIFIER |
| 15 | 84306843 | AX-170303763 | 0 | T | C | 0.088 | -1.13E-01 | 1.06E-01 | 2.04E+00 | 2.13E+00 | 3.77E-01 | no | Parvb | intron_variant | MODIFIER |
| 1 | 141977019 | AX-170476250 | 2 | C | T | 0.218 | 1.12E-01 | 1.29E-01 | 2.72E+00 | 2.80E+00 | 3.77E-01 | no | | intergenic_variant | MODIFIER |
| 8 | 65495531 | AX-170117112 | 0 | G | C | 0.149 | -1.17E-01 | 1.33E-01 | 2.56E+00 | 2.64E+00 | 3.77E-01 | no | | intergenic_variant | MODIFIER |
| 6 | 126352695 | AX-97851771 | 0 | A | C | 0.553 | 7.41E-02 | 8.29E-02 | 2.50E+00 | 2.57E+00 | 3.77E-01 | no | | intergenic_variant | MODIFIER |
| 6 | 126357516 | AX-170028194 | 0 | C | G | 0.553 | 7.41E-02 | 8.29E-02 | 2.50E+00 | 2.57E+00 | 3.77E-01 | no | | intergenic_variant | MODIFIER |
| 6 | 126385953 | AX-97807420 | 0 | C | T | 0.553 | 7.41E-02 | 8.29E-02 | 2.50E+00 | 2.57E+00 | 3.77E-01 | no | | intergenic_variant | MODIFIER |
| 6 | 126386319 | AX-170028192 | 0 | T | C | 0.553 | 7.41E-02 | 8.29E-02 | 2.50E+00 | 2.57E+00 | 3.77E-01 | no | | intergenic_variant | MODIFIER |
| 6 | 126410188 | AX-97804685 | 0 | A | G | 0.553 | 7.41E-02 | 8.29E-02 | 2.50E+00 | 2.57E+00 | 3.77E-01 | no | | intergenic_variant | MODIFIER |
| 6 | 126451396 | AX-170028190 | 0 | C | G | 0.553 | 7.41E-02 | 8.29E-02 | 2.50E+00 | 2.57E+00 | 3.77E-01 | no | | intergenic_variant | MODIFIER |
| 6 | 126484657 | AX-170028191 | 0 | C | T | 0.553 | 7.41E-02 | 8.29E-02 | 2.50E+00 | 2.57E+00 | 3.77E-01 | no | | intergenic_variant | MODIFIER |
| 6 | 126522104 | AX-170519225 | 0 | G | A | 0.553 | 7.41E-02 | 8.29E-02 | 2.50E+00 | 2.57E+00 | 3.77E-01 | no | | intergenic_variant | MODIFIER |
| 6 | 126522402 | AX-97821185 | 0 | T | G | 0.553 | 7.41E-02 | 8.29E-02 | 2.50E+00 | 2.57E+00 | 3.77E-01 | no | | intergenic_variant | MODIFIER |
| 6 | 126544503 | AX-170174755 | 0 | T | A | 0.553 | 7.41E-02 | 8.29E-02 | 2.50E+00 | 2.57E+00 | 3.77E-01 | no | | regulatory_region_variant | MODIFIER |
| 6 | 126555381 | AX-170421111 | 0 | G | T | 0.553 | 7.41E-02 | 8.29E-02 | 2.50E+00 | 2.57E+00 | 3.77E-01 | no | Gm33320 | upstream_gene_variant | MODIFIER |
| 6 | 126556449 | AX-97842420 | 0 | A | G | 0.553 | 7.41E-02 | 8.29E-02 | 2.50E+00 | 2.57E+00 | 3.77E-01 | no | Gm33320 | upstream_gene_variant | MODIFIER |
| 6 | 126570853 | AX-97847870 | 0 | G | T | 0.553 | 7.41E-02 | 8.29E-02 | 2.50E+00 | 2.57E+00 | 3.77E-01 | no | | intergenic_variant | MODIFIER |
| 6 | 126573752 | AX-170153260 | 0 | A | G | 0.553 | 7.41E-02 | 8.29E-02 | 2.50E+00 | 2.57E+00 | 3.77E-01 | no | | intergenic_variant | MODIFIER |
| 6 | 126596179 | AX-170167564 | 0 | A | G | 0.553 | 7.41E-02 | 8.29E-02 | 2.50E+00 | 2.57E+00 | 3.77E-01 | no | | intergenic_variant | MODIFIER |
| 6 | 126606438 | AX-97801829 | 0 | G | A | 0.553 | 7.41E-02 | 8.29E-02 | 2.50E+00 | 2.57E+00 | 3.77E-01 | no | | intergenic_variant | MODIFIER |
| 6 | 126654352 | AX-97856419 | 0 | C | T | 0.553 | 7.41E-02 | 8.29E-02 | 2.50E+00 | 2.57E+00 | 3.77E-01 | no | | intergenic_variant | MODIFIER |
| 6 | 126724691 | AX-97816640 | 0 | T | G | 0.553 | 7.41E-02 | 8.29E-02 | 2.50E+00 | 2.57E+00 | 3.77E-01 | no | Kcna6 | intron_variant,NMD_transcript_variant | MODIFIER |
| 6 | 126799800 | AX-170281647 | 0 | T | A | 0.553 | 7.41E-02 | 8.29E-02 | 2.50E+00 | 2.57E+00 | 3.77E-01 | no | | intergenic_variant | MODIFIER |
| 6 | 128163001 | AX-170460255 | 0 | C | T | 0.553 | 7.41E-02 | 8.29E-02 | 2.50E+00 | 2.57E+00 | 3.77E-01 | no | Gm44056 | downstream_gene_variant | MODIFIER |
| 6 | 128176312 | AX-170296771 | 0 | G | A | 0.553 | 7.41E-02 | 8.29E-02 | 2.50E+00 | 2.57E+00 | 3.77E-01 | no | 9330102E08Rik | intron_variant,non_coding_transcript_variant | MODIFIER |
| 6 | 128208655 | AX-170420130 | 0 | C | T | 0.553 | 7.41E-02 | 8.29E-02 | 2.50E+00 | 2.57E+00 | 3.77E-01 | no | Gm19065 | upstream_gene_variant | MODIFIER |
| 6 | 128220814 | AX-170454180 | 0 | C | T | 0.553 | 7.41E-02 | 8.29E-02 | 2.50E+00 | 2.57E+00 | 3.77E-01 | no | Tead4 | downstream_gene_variant | MODIFIER |
| 6 | 128221019 | AX-170478991 | 0 | C | T | 0.553 | 7.41E-02 | 8.29E-02 | 2.50E+00 | 2.57E+00 | 3.77E-01 | no | Tead4 | downstream_gene_variant | MODIFIER |
| 6 | 128234408 | AX-170120975 | 0 | C | A | 0.553 | 7.41E-02 | 8.29E-02 | 2.50E+00 | 2.57E+00 | 3.77E-01 | no | Tead4 | intron_variant | MODIFIER |
| 6 | 128234661 | AX-170535749 | 0 | A | G | 0.553 | 7.41E-02 | 8.29E-02 | 2.50E+00 | 2.57E+00 | 3.77E-01 | no | Tead4 | intron_variant | MODIFIER |
| 6 | 128235059 | AX-170525719 | 0 | G | C | 0.553 | 7.41E-02 | 8.29E-02 | 2.50E+00 | 2.57E+00 | 3.77E-01 | no | Tead4 | intron_variant | MODIFIER |
| 6 | 128241135 | AX-170148634 | 0 | C | T | 0.553 | 7.41E-02 | 8.29E-02 | 2.50E+00 | 2.57E+00 | 3.77E-01 | no | Tead4 | intron_variant | MODIFIER |
| 6 | 128245099 | AX-170367578 | 0 | G | A | 0.553 | 7.41E-02 | 8.29E-02 | 2.50E+00 | 2.57E+00 | 3.77E-01 | no | Tead4 | intron_variant | MODIFIER |
| 6 | 128247419 | AX-170255625 | 0 | C | T | 0.553 | 7.41E-02 | 8.29E-02 | 2.50E+00 | 2.57E+00 | 3.77E-01 | no | Tead4 | intron_variant | MODIFIER |
| 6 | 128478682 | AX-170428980 | 0 | G | A | 0.553 | 7.41E-02 | 8.29E-02 | 2.50E+00 | 2.57E+00 | 3.77E-01 | no | Pzp | downstream_gene_variant | MODIFIER |
| 6 | 128481655 | AX-97791517 | 0 | G | A | 0.553 | 7.41E-02 | 8.29E-02 | 2.50E+00 | 2.57E+00 | 3.77E-01 | no | Pzp | downstream_gene_variant | MODIFIER |
| 6 | 128518308 | AX-170129894 | 0 | T | C | 0.553 | 7.41E-02 | 8.29E-02 | 2.50E+00 | 2.57E+00 | 3.77E-01 | no | Pzp | intron_variant | MODIFIER |
| 6 | 128518680 | AX-170201868 | 0 | T | C | 0.553 | 7.41E-02 | 8.29E-02 | 2.50E+00 | 2.57E+00 | 3.77E-01 | no | Pzp | intron_variant | MODIFIER |
| 6 | 128518766 | AX-170095802 | 0 | A | C | 0.553 | 7.41E-02 | 8.29E-02 | 2.50E+00 | 2.57E+00 | 3.77E-01 | no | Pzp | intron_variant | MODIFIER |
| 6 | 128528739 | AX-170228836 | 0 | C | T | 0.553 | 7.41E-02 | 8.29E-02 | 2.50E+00 | 2.57E+00 | 3.77E-01 | no | Gm8724 | downstream_gene_variant | MODIFIER |
| 6 | 128529460 | AX-170184217 | 0 | C | T | 0.553 | 7.41E-02 | 8.29E-02 | 2.50E+00 | 2.57E+00 | 3.77E-01 | no | Gm8724 | downstream_gene_variant | MODIFIER |
| 6 | 128674180 | AX-170243388 | 0 | G | A | 0.553 | 7.41E-02 | 8.29E-02 | 2.50E+00 | 2.57E+00 | 3.77E-01 | no | Clec2h | intron_variant | MODIFIER |
| 6 | 128707350 | AX-170506724 | 0 | G | A | 0.553 | 7.41E-02 | 8.29E-02 | 2.50E+00 | 2.57E+00 | 3.77E-01 | no | Klrb1 | intron_variant | MODIFIER |
| 6 | 128713569 | AX-170417801 | 0 | C | G | 0.553 | 7.41E-02 | 8.29E-02 | 2.50E+00 | 2.57E+00 | 3.77E-01 | no | Klrb1 | intron_variant | MODIFIER |
| 6 | 128713707 | AX-170454354 | 0 | T | C | 0.553 | 7.41E-02 | 8.29E-02 | 2.50E+00 | 2.57E+00 | 3.77E-01 | no | Klrb1 | intron_variant | MODIFIER |
| 6 | 129002976 | AX-170364593 | 0 | T | A | 0.553 | 7.41E-02 | 8.29E-02 | 2.50E+00 | 2.57E+00 | 3.77E-01 | no | BC064078 | intron_variant,non_coding_transcript_variant | MODIFIER |
| 6 | 129006732 | AX-170485515 | 0 | A | G | 0.553 | 7.41E-02 | 8.29E-02 | 2.50E+00 | 2.57E+00 | 3.77E-01 | no | BC064078 | intron_variant,non_coding_transcript_variant | MODIFIER |
| 6 | 129006844 | AX-170344555 | 0 | T | C | 0.553 | 7.41E-02 | 8.29E-02 | 2.50E+00 | 2.57E+00 | 3.77E-01 | no | BC064078 | intron_variant,non_coding_transcript_variant | MODIFIER |
| 6 | 129007865 | AX-170391438 | 0 | T | C | 0.553 | 7.41E-02 | 8.29E-02 | 2.50E+00 | 2.57E+00 | 3.77E-01 | no | BC064078 | splice_acceptor_variant,non_coding_transcript_variant | HIGH |
| 6 | 129007956 | AX-170369638 | 0 | C | A | 0.553 | 7.41E-02 | 8.29E-02 | 2.50E+00 | 2.57E+00 | 3.77E-01 | no | BC064078 | non_coding_transcript_exon_variant | MODIFIER |
| 6 | 129013712 | AX-170501983 | 0 | A | G | 0.553 | 7.41E-02 | 8.29E-02 | 2.50E+00 | 2.57E+00 | 3.77E-01 | no | Clec2f | upstream_gene_variant | MODIFIER |

| | | | | | | | | | | | | | | | |
|---|---|---|---|---|---|---|---|---|---|---|---|---|---|---|---|
| 6 | 129013754 | AX-97801853 | 0 | C | A | 0.553 | 7.41E-02 | 8.29E-02 | 2.50E+00 | 2.57E+00 | 3.77E-01 | no | Clec2f | upstream_gene_variant | MODIFIER |
| 6 | 129014192 | AX-170374189 | 0 | T | C | 0.553 | 7.41E-02 | 8.29E-02 | 2.50E+00 | 2.57E+00 | 3.77E-01 | no | Clec2f | upstream_gene_variant | MODIFIER |
| 6 | 129015806 | AX-170422598 | 0 | G | T | 0.553 | 7.41E-02 | 8.29E-02 | 2.50E+00 | 2.57E+00 | 3.77E-01 | no | Clec2f | upstream_gene_variant | MODIFIER |
| 6 | 129018948 | AX-170168473 | 0 | G | A | 0.553 | 7.41E-02 | 8.29E-02 | 2.50E+00 | 2.57E+00 | 3.77E-01 | no | Clec2f | intron_variant,non_coding_transcript_variant | MODIFIER |
| 6 | 129020152 | AX-170280045 | 0 | A | T | 0.553 | 7.41E-02 | 8.29E-02 | 2.50E+00 | 2.57E+00 | 3.77E-01 | no | Clec2f | intron_variant,non_coding_transcript_variant | MODIFIER |
| 6 | 129029011 | AX-97855848 | 0 | T | C | 0.553 | 7.41E-02 | 8.29E-02 | 2.50E+00 | 2.57E+00 | 3.77E-01 | no | Gm16186 | downstream_gene_variant | MODIFIER |
| 6 | 129033324 | AX-170285382 | 0 | G | C | 0.553 | 7.41E-02 | 8.29E-02 | 2.50E+00 | 2.57E+00 | 3.77E-01 | no | Gm16186 | upstream_gene_variant | MODIFIER |
| 6 | 129053155 | AX-97785529 | 0 | T | C | 0.553 | 7.41E-02 | 8.29E-02 | 2.50E+00 | 2.57E+00 | 3.77E-01 | no | Klrb1f | intron_variant | MODIFIER |
| 6 | 129073128 | AX-170028237 | 0 | T | C | 0.553 | 7.41E-02 | 8.29E-02 | 2.50E+00 | 2.57E+00 | 3.77E-01 | no | | intergenic_variant | MODIFIER |
| 6 | 129073375 | AX-97786498 | 0 | C | T | 0.553 | 7.41E-02 | 8.29E-02 | 2.50E+00 | 2.57E+00 | 3.77E-01 | no | | intergenic_variant | MODIFIER |
| 6 | 129073453 | AX-170267634 | 0 | C | G | 0.553 | 7.41E-02 | 8.29E-02 | 2.50E+00 | 2.57E+00 | 3.77E-01 | no | | intergenic_variant | MODIFIER |
| 6 | 129087092 | AX-170476362 | 0 | T | G | 0.553 | 7.41E-02 | 8.29E-02 | 2.50E+00 | 2.57E+00 | 3.77E-01 | no | Clec2e | downstream_gene_variant | MODIFIER |
| 6 | 129101445 | AX-170532209 | 0 | C | T | 0.553 | 7.41E-02 | 8.29E-02 | 2.50E+00 | 2.57E+00 | 3.77E-01 | no | Clec2e | upstream_gene_variant | MODIFIER |
| 6 | 129105859 | AX-170271590 | 0 | T | G | 0.553 | 7.41E-02 | 8.29E-02 | 2.50E+00 | 2.57E+00 | 3.77E-01 | no | Clec2e | upstream_gene_variant | MODIFIER |
| 6 | 129105915 | AX-97788019 | 0 | T | C | 0.553 | 7.41E-02 | 8.29E-02 | 2.50E+00 | 2.57E+00 | 3.77E-01 | no | | intergenic_variant | MODIFIER |
| 6 | 129306422 | AX-170450088 | 0 | G | A | 0.553 | 7.41E-02 | 8.29E-02 | 2.50E+00 | 2.57E+00 | 3.77E-01 | no | | intergenic_variant | MODIFIER |
| 6 | 129376344 | AX-170369753 | 0 | A | G | 0.553 | 7.41E-02 | 8.29E-02 | 2.50E+00 | 2.57E+00 | 3.77E-01 | no | Clec12b | missense_variant,splice_region_variant | MODERATE |
| 6 | 129376940 | AX-170317420 | 0 | G | A | 0.553 | 7.41E-02 | 8.29E-02 | 2.50E+00 | 2.57E+00 | 3.77E-01 | no | Clec12b | intron_variant | MODIFIER |
| 6 | 129379349 | AX-170076212 | 0 | A | T | 0.553 | 7.41E-02 | 8.29E-02 | 2.50E+00 | 2.57E+00 | 3.77E-01 | no | Clec12b | intron_variant | MODIFIER |
| 6 | 129379363 | AX-170076213 | 0 | C | G | 0.553 | 7.41E-02 | 8.29E-02 | 2.50E+00 | 2.57E+00 | 3.77E-01 | no | Clec12b | intron_variant | MODIFIER |
| 6 | 129379501 | AX-170076214 | 0 | T | C | 0.553 | 7.41E-02 | 8.29E-02 | 2.50E+00 | 2.57E+00 | 3.77E-01 | no | Clec12b | intron_variant | MODIFIER |
| 6 | 129382663 | AX-97849034 | 0 | A | G | 0.553 | 7.41E-02 | 8.29E-02 | 2.50E+00 | 2.57E+00 | 3.77E-01 | no | Clec12b | intron_variant | MODIFIER |
| 6 | 129384558 | AX-170170363 | 0 | C | A | 0.553 | 7.41E-02 | 8.29E-02 | 2.50E+00 | 2.57E+00 | 3.77E-01 | no | Clec12b | intron_variant | MODIFIER |
| 6 | 129401547 | AX-170457758 | 0 | C | T | 0.553 | 7.41E-02 | 8.29E-02 | 2.50E+00 | 2.57E+00 | 3.77E-01 | no | Clec1b | missense_variant | MODERATE |
| 6 | 129403232 | AX-170149258 | 0 | T | C | 0.553 | 7.41E-02 | 8.29E-02 | 2.50E+00 | 2.57E+00 | 3.77E-01 | no | Clec1b | intron_variant | MODIFIER |
| 6 | 129405007 | AX-170166202 | 0 | C | A | 0.553 | 7.41E-02 | 8.29E-02 | 2.50E+00 | 2.57E+00 | 3.77E-01 | no | Clec1b | intron_variant | MODIFIER |
| 6 | 129406942 | AX-170436221 | 0 | G | A | 0.553 | 7.41E-02 | 8.29E-02 | 2.50E+00 | 2.57E+00 | 3.77E-01 | no | Clec1b | downstream_gene_variant | MODIFIER |
| 6 | 129411846 | AX-170208784 | 0 | C | T | 0.553 | 7.41E-02 | 8.29E-02 | 2.50E+00 | 2.57E+00 | 3.77E-01 | no | Clec9a | intron_variant | MODIFIER |
| 6 | 129418195 | AX-170249074 | 0 | C | T | 0.553 | 7.41E-02 | 8.29E-02 | 2.50E+00 | 2.57E+00 | 3.77E-01 | no | Clec9a | intron_variant | MODIFIER |
| 6 | 129420157 | AX-170136456 | 0 | C | T | 0.553 | 7.41E-02 | 8.29E-02 | 2.50E+00 | 2.57E+00 | 3.77E-01 | no | Clec9a | intron_variant | MODIFIER |
| 6 | 129468995 | AX-170396470 | 0 | G | A | 0.553 | 7.41E-02 | 8.29E-02 | 2.50E+00 | 2.57E+00 | 3.77E-01 | no | Clec7a | splice_acceptor_variant | HIGH |
| 6 | 129524704 | AX-170373597 | 0 | C | A | 0.553 | 7.41E-02 | 8.29E-02 | 2.50E+00 | 2.57E+00 | 3.77E-01 | no | Gm44120 | non_coding_transcript_exon_variant | MODIFIER |
| 6 | 129525765 | AX-170115578 | 0 | G | T | 0.553 | 7.41E-02 | 8.29E-02 | 2.50E+00 | 2.57E+00 | 3.77E-01 | no | Gm44120 | downstream_gene_variant | MODIFIER |
| 6 | 129537777 | AX-170307864 | 0 | A | C | 0.553 | 7.41E-02 | 8.29E-02 | 2.50E+00 | 2.57E+00 | 3.77E-01 | no | Gabarapl1 | intron_variant | MODIFIER |
| 6 | 129544091 | AX-170440830 | 0 | G | T | 0.553 | 7.41E-02 | 8.29E-02 | 2.50E+00 | 2.57E+00 | 3.77E-01 | no | Gabarapl1 | downstream_gene_variant | MODIFIER |
| 6 | 129545212 | AX-97807137 | 0 | C | T | 0.553 | 7.41E-02 | 8.29E-02 | 2.50E+00 | 2.57E+00 | 3.77E-01 | no | Gabarapl1 | downstream_gene_variant | MODIFIER |
| 6 | 129553710 | AX-170076217 | 0 | C | T | 0.553 | 7.41E-02 | 8.29E-02 | 2.50E+00 | 2.57E+00 | 3.77E-01 | no | Gm44251 | upstream_gene_variant | MODIFIER |
| 6 | 129589477 | AX-97852261 | 0 | G | T | 0.553 | 7.41E-02 | 8.29E-02 | 2.50E+00 | 2.57E+00 | 3.77E-01 | no | Klrd1 | upstream_gene_variant | MODIFIER |
| 6 | 129626237 | AX-170158177 | 0 | C | T | 0.553 | 7.41E-02 | 8.29E-02 | 2.50E+00 | 2.57E+00 | 3.77E-01 | no | Klrk1 | upstream_gene_variant | MODIFIER |
| 6 | 129637433 | AX-97802811 | 0 | G | A | 0.553 | 7.41E-02 | 8.29E-02 | 2.50E+00 | 2.57E+00 | 3.77E-01 | no | Klrc3 | downstream_gene_variant | MODIFIER |
| 6 | 129638796 | AX-170512050 | 0 | G | A | 0.553 | 7.41E-02 | 8.29E-02 | 2.50E+00 | 2.57E+00 | 3.77E-01 | no | Klrc3 | downstream_gene_variant | MODIFIER |
| 6 | 129691205 | AX-170319179 | 0 | C | T | 0.553 | 7.41E-02 | 8.29E-02 | 2.50E+00 | 2.57E+00 | 3.77E-01 | no | Mir680-1 | upstream_gene_variant | MODIFIER |
| 6 | 129882949 | AX-170193521 | 0 | C | G | 0.553 | 7.41E-02 | 8.29E-02 | 2.50E+00 | 2.57E+00 | 3.77E-01 | no | | intergenic_variant | MODIFIER |
| 6 | 130534541 | AX-170449972 | 0 | G | T | 0.553 | 7.41E-02 | 8.29E-02 | 2.50E+00 | 2.57E+00 | 3.77E-01 | no | | intergenic_variant | MODIFIER |
| 6 | 130554855 | AX-97824545 | 0 | T | G | 0.553 | 7.41E-02 | 8.29E-02 | 2.50E+00 | 2.57E+00 | 3.77E-01 | no | | intergenic_variant | MODIFIER |
| 6 | 130580629 | AX-170287779 | 0 | C | T | 0.553 | 7.41E-02 | 8.29E-02 | 2.50E+00 | 2.57E+00 | 3.77E-01 | no | Gm26996 | non_coding_transcript_exon_variant | MODIFIER |
| 6 | 130650464 | AX-97834245 | 0 | G | A | 0.553 | 7.41E-02 | 8.29E-02 | 2.50E+00 | 2.57E+00 | 3.77E-01 | no | | intergenic_variant | MODIFIER |
| 6 | 130774501 | AX-170455718 | 0 | C | A | 0.553 | 7.41E-02 | 8.29E-02 | 2.50E+00 | 2.57E+00 | 3.77E-01 | no | Gm18423 | upstream_gene_variant | MODIFIER |
| 6 | 130785871 | AX-97808901 | 0 | T | C | 0.553 | 7.41E-02 | 8.29E-02 | 2.50E+00 | 2.57E+00 | 3.77E-01 | no | Gm44525 | downstream_gene_variant | MODIFIER |
| 6 | 131126917 | AX-170185543 | 0 | A | G | 0.553 | 7.41E-02 | 8.29E-02 | 2.50E+00 | 2.57E+00 | 3.77E-01 | no | Gm8837 | upstream_gene_variant | MODIFIER |
| 6 | 131170261 | AX-170345954 | 0 | G | T | 0.553 | 7.41E-02 | 8.29E-02 | 2.50E+00 | 2.57E+00 | 3.77E-01 | no | Gm5581 | intron_variant,non_coding_transcript_variant | MODIFIER |
| 6 | 131217005 | AX-170028260 | 0 | G | A | 0.553 | 7.41E-02 | 8.29E-02 | 2.50E+00 | 2.57E+00 | 3.77E-01 | no | Klra2 | downstream_gene_variant | MODIFIER |
| 6 | 131231086 | AX-97798790 | 0 | T | G | 0.553 | 7.41E-02 | 8.29E-02 | 2.50E+00 | 2.57E+00 | 3.77E-01 | no | Klra2 | intron_variant | MODIFIER |
| 6 | 131375501 | AX-170266606 | 0 | C | G | 0.553 | 7.41E-02 | 8.29E-02 | 2.50E+00 | 2.57E+00 | 3.77E-01 | no | Ybx3 | intron_variant | MODIFIER |

| | | | | | | | | | | | | | | | |
|---|---|---|---|---|---|---|---|---|---|---|---|---|---|---|---|
| 6 | 131402016 | AX-170407135 | 0 | T | G | 0.553 | 7.41E-02 | 8.29E-02 | 2.50E+00 | 2.57E+00 | 3.77E-01 | no | | intergenic_variant | MODIFIER |
| 6 | 131406584 | AX-170224049 | 0 | A | G | 0.553 | 7.41E-02 | 8.29E-02 | 2.50E+00 | 2.57E+00 | 3.77E-01 | no | | intergenic_variant | MODIFIER |
| 6 | 131406639 | AX-97780773 | 0 | A | G | 0.553 | 7.41E-02 | 8.29E-02 | 2.50E+00 | 2.57E+00 | 3.77E-01 | no | | intergenic_variant | MODIFIER |
| 6 | 131408880 | AX-170076252 | 0 | A | C | 0.553 | 7.41E-02 | 8.29E-02 | 2.50E+00 | 2.57E+00 | 3.77E-01 | no | | intergenic_variant | MODIFIER |
| 6 | 131410721 | AX-170076251 | 0 | A | C | 0.553 | 7.41E-02 | 8.29E-02 | 2.50E+00 | 2.57E+00 | 3.77E-01 | no | | intergenic_variant | MODIFIER |
| 1 | 187677409 | AX-170286053 | 1 | C | T | 0.241 | 9.69E-02 | 1.05E-01 | 2.37E+00 | 2.45E+00 | 3.77E-01 | no | Esrrg | intron_variant | MODIFIER |
| 10 | 93582363 | AX-170003672 | 1 | A | T | 0.286 | -7.32E-02 | 8.26E-02 | 2.55E+00 | 2.61E+00 | 3.77E-01 | no | Snrpf | downstream_gene_variant | MODIFIER |
| 8 | 15549341 | AX-170509298 | 1 | C | T | 0.232 | -6.72E-02 | 7.62E-02 | 2.55E+00 | 2.65E+00 | 3.77E-01 | no | | intergenic_variant | MODIFIER |
| 8 | 78906452 | AX-170523064 | 0 | T | C | 0.219 | 9.07E-02 | 1.04E-01 | 2.72E+00 | 2.82E+00 | 3.78E-01 | no | | intergenic_variant | MODIFIER |
| 15 | 75422928 | AX-97832431 | 0 | G | A | 0.561 | 6.84E-02 | 7.78E-02 | 2.58E+00 | 2.66E+00 | 3.78E-01 | no | 9030619P08Rik | downstream_gene_variant | MODIFIER |
| 10 | 17971375 | AX-170236617 | 1 | C | T | 0.196 | 8.83E-02 | 1.00E-01 | 2.93E+00 | 3.03E+00 | 3.78E-01 | no | | intergenic_variant | MODIFIER |
| 5 | 79855803 | AX-170072583 | 0 | A | T | 0.447 | 8.81E-02 | 1.01E-01 | 2.70E+00 | 2.78E+00 | 3.78E-01 | no | | intergenic_variant | MODIFIER |
| 1 | 128788437 | AX-170001675 | 0 | C | T | 0.535 | 7.11E-02 | 8.12E-02 | 2.63E+00 | 2.70E+00 | 3.78E-01 | no | | regulatory_region_variant | MODIFIER |
| 9 | 101455221 | AX-170033150 | 1 | C | G | 0.134 | -1.03E-01 | 1.18E-01 | 2.63E+00 | 2.71E+00 | 3.78E-01 | no | | intergenic_variant | MODIFIER |
| 18 | 66132049 | AX-97820215 | 1 | G | A | 0.188 | -1.05E-01 | 1.20E-01 | 2.70E+00 | 2.76E+00 | 3.78E-01 | no | Ccbe1 | intron_variant,NMD_transcript_variant | MODIFIER |
| 18 | 66284995 | AX-170180541 | 1 | T | C | 0.188 | -1.05E-01 | 1.20E-01 | 2.70E+00 | 2.76E+00 | 3.78E-01 | no | Ccbe1 | intron_variant,NMD_transcript_variant | MODIFIER |
| 16 | 78226822 | AX-170458729 | 2 | C | T | 0.455 | 6.66E-02 | 7.58E-02 | 2.59E+00 | 2.66E+00 | 3.78E-01 | no | | intergenic_variant | MODIFIER |
| 3 | 26297042 | AX-170395474 | 0 | T | C | 0.14 | -8.99E-02 | 1.02E-01 | 2.55E+00 | 2.63E+00 | 3.78E-01 | no | Nlgn1 | intron_variant,non_coding_transcript_variant | MODIFIER |
| 15 | 8097396 | AX-170414178 | 0 | T | C | 0.246 | -8.79E-02 | 1.01E-01 | 2.67E+00 | 2.73E+00 | 3.78E-01 | no | Wdr70 | intron_variant | MODIFIER |
| 6 | 134887971 | AX-170400371 | 1 | C | T | 0.562 | 7.74E-02 | 8.68E-02 | 2.50E+00 | 2.55E+00 | 3.78E-01 | no | Gpr19 | intron_variant | MODIFIER |
| 6 | 129420508 | AX-97812285 | 1 | A | G | 0.554 | 7.37E-02 | 8.28E-02 | 2.51E+00 | 2.57E+00 | 3.78E-01 | no | Clec9a | intron_variant | MODIFIER |
| 9 | 16950857 | AX-170031859 | 0 | T | A | 0.482 | 7.61E-02 | 8.65E-02 | 2.57E+00 | 2.65E+00 | 3.78E-01 | no | | intergenic_variant | MODIFIER |
| 13 | 24169577 | AX-170155391 | 1 | A | T | 0.17 | -1.11E-01 | 1.20E-01 | 2.36E+00 | 2.41E+00 | 3.78E-01 | no | Carmil1 | intron_variant | MODIFIER |
| 18 | 69507582 | AX-170242774 | 1 | A | T | 0.259 | -8.66E-02 | 9.85E-02 | 2.59E+00 | 2.64E+00 | 3.79E-01 | no | Tcf4 | intron_variant | MODIFIER |
| 1 | 151257072 | AX-170043129 | 1 | G | A | 0.214 | 1.17E-01 | 1.34E-01 | 2.69E+00 | 2.74E+00 | 3.79E-01 | no | Gm24402 | downstream_gene_variant | MODIFIER |
| 5 | 48799985 | AX-97781059 | 1 | G | C | 0.473 | 9.20E-02 | 9.99E-02 | 2.37E+00 | 2.43E+00 | 3.79E-01 | no | Kcnip4 | intron_variant | MODIFIER |
| 5 | 50652530 | AX-170267989 | 1 | T | C | 0.473 | 9.20E-02 | 9.99E-02 | 2.37E+00 | 2.43E+00 | 3.79E-01 | no | Gm43016 | intron_variant,non_coding_transcript_variant | MODIFIER |
| 4 | 149607718 | AX-170225362 | 1 | T | G | 0.098 | 1.41E-01 | 1.62E-01 | 2.75E+00 | 2.87E+00 | 3.79E-01 | no | Clstn1 | intron_variant | MODIFIER |
| 8 | 16086219 | AX-170187271 | 1 | C | T | 0.232 | -6.81E-02 | 7.73E-02 | 2.56E+00 | 2.65E+00 | 3.79E-01 | no | Csmd1 | intron_variant | MODIFIER |
| 10 | 3126270 | AX-170267700 | 1 | G | A | 0.196 | -7.71E-02 | 8.83E-02 | 2.59E+00 | 2.71E+00 | 3.79E-01 | no | 9230019H11Rik | intron_variant,non_coding_transcript_variant | MODIFIER |
| 9 | 91957163 | AX-170254751 | 1 | G | A | 0.134 | -1.08E-01 | 1.24E-01 | 2.61E+00 | 2.69E+00 | 3.79E-01 | no | Gm31409 | intron_variant,non_coding_transcript_variant | MODIFIER |
| 9 | 94936475 | AX-97838734 | 0 | A | G | 0.132 | -1.06E-01 | 1.21E-01 | 2.67E+00 | 2.75E+00 | 3.79E-01 | no | Slc9a9 | intron_variant | MODIFIER |
| 9 | 95168766 | AX-170165764 | 0 | T | C | 0.132 | -1.06E-01 | 1.21E-01 | 2.67E+00 | 2.75E+00 | 3.79E-01 | no | Slc9a9 | intron_variant | MODIFIER |
| 9 | 95509660 | AX-168383916 | 0 | G | A | 0.132 | -1.06E-01 | 1.21E-01 | 2.67E+00 | 2.75E+00 | 3.79E-01 | no | U2surp | intron_variant | MODIFIER |
| 9 | 95964643 | AX-170397378 | 0 | A | G | 0.132 | -1.06E-01 | 1.21E-01 | 2.67E+00 | 2.75E+00 | 3.79E-01 | no | Xrn1 | intron_variant | MODIFIER |
| 9 | 96025993 | AX-97780972 | 0 | G | A | 0.132 | -1.06E-01 | 1.21E-01 | 2.67E+00 | 2.75E+00 | 3.79E-01 | no | Xrn1 | intron_variant | MODIFIER |
| 9 | 96031278 | AX-170195412 | 0 | A | G | 0.132 | -1.06E-01 | 1.21E-01 | 2.67E+00 | 2.75E+00 | 3.79E-01 | no | Xrn1 | intron_variant | MODIFIER |
| 9 | 96264404 | AX-170425003 | 0 | G | C | 0.132 | -1.06E-01 | 1.21E-01 | 2.67E+00 | 2.75E+00 | 3.79E-01 | no | Tfdp2 | intron_variant | MODIFIER |
| 9 | 96264626 | AX-170309689 | 0 | A | T | 0.132 | -1.06E-01 | 1.21E-01 | 2.67E+00 | 2.75E+00 | 3.79E-01 | no | Tfdp2 | intron_variant | MODIFIER |
| 9 | 96304215 | AX-170321926 | 0 | T | C | 0.132 | -1.06E-01 | 1.21E-01 | 2.67E+00 | 2.75E+00 | 3.79E-01 | no | Tfdp2 | intron_variant | MODIFIER |
| 9 | 96316913 | AX-170120179 | 0 | T | C | 0.132 | -1.06E-01 | 1.21E-01 | 2.67E+00 | 2.75E+00 | 3.79E-01 | no | Tfdp2 | intron_variant | MODIFIER |
| 9 | 96318011 | AX-170178866 | 0 | A | G | 0.132 | -1.06E-01 | 1.21E-01 | 2.67E+00 | 2.75E+00 | 3.79E-01 | no | Tfdp2 | 3_prime_UTR_variant | MODIFIER |
| 10 | 69566206 | AX-97822728 | 0 | G | A | 0.412 | -6.99E-02 | 8.02E-02 | 2.66E+00 | 2.73E+00 | 3.79E-01 | no | Ank3 | intron_variant | MODIFIER |
| 10 | 73115278 | AX-97785829 | 0 | G | A | 0.412 | -6.99E-02 | 8.02E-02 | 2.66E+00 | 2.73E+00 | 3.79E-01 | no | Pcdh15 | intron_variant,non_coding_transcript_variant | MODIFIER |
| 6 | 134397159 | AX-170122983 | 1 | A | G | 0.562 | 7.71E-02 | 8.66E-02 | 2.50E+00 | 2.56E+00 | 3.79E-01 | no | Bcl2l14 | intron_variant | MODIFIER |
| 6 | 135016159 | AX-170349160 | 1 | G | A | 0.562 | 7.71E-02 | 8.66E-02 | 2.50E+00 | 2.56E+00 | 3.79E-01 | no | Ddx47 | intron_variant | MODIFIER |
| 2 | 162810648 | AX-170503812 | 1 | T | A | 0.152 | 1.58E-01 | 1.82E-01 | 2.79E+00 | 2.89E+00 | 3.79E-01 | Yes | Gm11450 | downstream_gene_variant | MODIFIER |
| 1 | 174431570 | AX-170101721 | 1 | C | T | 0.232 | 1.03E-01 | 1.16E-01 | 2.51E+00 | 2.58E+00 | 3.79E-01 | no | Olfr414 | downstream_gene_variant | MODIFIER |
| 9 | 61495510 | AX-170149264 | 0 | T | C | 0.193 | -1.00E-01 | 1.16E-01 | 2.73E+00 | 2.83E+00 | 3.79E-01 | no | | intergenic_variant | MODIFIER |
| 9 | 67178207 | AX-170107300 | 0 | A | T | 0.193 | -1.00E-01 | 1.16E-01 | 2.73E+00 | 2.83E+00 | 3.79E-01 | no | Gm19299 | intron_variant,non_coding_transcript_variant | MODIFIER |
| 9 | 69753062 | AX-170101021 | 0 | C | T | 0.193 | -1.00E-01 | 1.16E-01 | 2.73E+00 | 2.83E+00 | 3.79E-01 | no | Foxb1 | downstream_gene_variant | MODIFIER |
| 9 | 74113208 | AX-170216565 | 0 | G | A | 0.193 | -1.00E-01 | 1.16E-01 | 2.73E+00 | 2.83E+00 | 3.79E-01 | no | Wdr72 | intron_variant | MODIFIER |
| 18 | 84244926 | AX-170061543 | 1 | C | T | 0.161 | -9.62E-02 | 1.07E-01 | 2.47E+00 | 2.55E+00 | 3.79E-01 | no | Zfp407 | intron_variant | MODIFIER |
| 12 | 60891587 | AX-97840914 | 0 | G | A | 0.035 | -1.87E-01 | 2.15E-01 | 2.61E+00 | 2.72E+00 | 3.79E-01 | Yes | | intergenic_variant | MODIFIER |

| | | | | | | | | | | | | | | | |
|---|---|---|---|---|---|---|---|---|---|---|---|---|---|---|---|
| 1 | 15794245 | AX-170294737 | 0 | C | T | 0.386 | 6.97E-02 | 7.92E-02 | 2.56E+00 | 2.62E+00 | 3.79E-01 | no | | intergenic_variant | MODIFIER |
| 8 | 37592696 | AX-170285977 | 0 | G | T | 0.158 | -1.11E-01 | 1.28E-01 | 2.78E+00 | 2.89E+00 | 3.80E-01 | no | Sgcz | intron_variant | MODIFIER |
| 12 | 101412671 | AX-170315625 | 0 | C | T | 0.096 | 1.45E-01 | 1.67E-01 | 2.77E+00 | 2.87E+00 | 3.80E-01 | Yes | Catsperb | intron_variant | MODIFIER |
| 18 | 88709107 | AX-170230407 | 0 | T | C | 0.158 | -9.23E-02 | 1.05E-01 | 2.53E+00 | 2.63E+00 | 3.80E-01 | no | Socs6 | intron_variant,non_coding_transcript_variant | MODIFIER |
| 1 | 13855369 | AX-170183105 | 0 | T | C | 0.377 | 6.98E-02 | 7.98E-02 | 2.60E+00 | 2.65E+00 | 3.80E-01 | no | | intergenic_variant | MODIFIER |
| 1 | 13860807 | AX-170140915 | 0 | G | A | 0.377 | 6.98E-02 | 7.98E-02 | 2.60E+00 | 2.65E+00 | 3.80E-01 | no | | intergenic_variant | MODIFIER |
| 1 | 13862613 | AX-170242769 | 0 | C | T | 0.377 | 6.98E-02 | 7.98E-02 | 2.60E+00 | 2.65E+00 | 3.80E-01 | no | | intergenic_variant | MODIFIER |
| 1 | 14252657 | AX-170200552 | 0 | A | C | 0.377 | 6.98E-02 | 7.98E-02 | 2.60E+00 | 2.65E+00 | 3.80E-01 | no | Eya1 | intron_variant | MODIFIER |
| 1 | 14275331 | AX-170133013 | 0 | A | G | 0.377 | 6.98E-02 | 7.98E-02 | 2.60E+00 | 2.65E+00 | 3.80E-01 | no | Eya1 | intron_variant | MODIFIER |
| 1 | 14333368 | AX-97791019 | 0 | T | C | 0.377 | 6.98E-02 | 7.98E-02 | 2.60E+00 | 2.65E+00 | 3.80E-01 | no | | intergenic_variant | MODIFIER |
| 1 | 14349168 | AX-170175992 | 0 | T | C | 0.377 | 6.98E-02 | 7.98E-02 | 2.60E+00 | 2.65E+00 | 3.80E-01 | no | | intergenic_variant | MODIFIER |
| 1 | 14349200 | AX-170461966 | 0 | G | C | 0.377 | 6.98E-02 | 7.98E-02 | 2.60E+00 | 2.65E+00 | 3.80E-01 | no | | intergenic_variant | MODIFIER |
| 1 | 14349537 | AX-170178848 | 0 | A | G | 0.377 | 6.98E-02 | 7.98E-02 | 2.60E+00 | 2.65E+00 | 3.80E-01 | no | | intergenic_variant | MODIFIER |
| 1 | 14373678 | AX-170481602 | 0 | T | G | 0.377 | 6.98E-02 | 7.98E-02 | 2.60E+00 | 2.65E+00 | 3.80E-01 | no | | intergenic_variant | MODIFIER |
| 1 | 14427436 | AX-170239905 | 0 | A | G | 0.377 | 6.98E-02 | 7.98E-02 | 2.60E+00 | 2.65E+00 | 3.80E-01 | no | | intergenic_variant | MODIFIER |
| 1 | 14457545 | AX-170120789 | 0 | A | G | 0.377 | 6.98E-02 | 7.98E-02 | 2.60E+00 | 2.65E+00 | 3.80E-01 | no | | intergenic_variant | MODIFIER |
| 1 | 14460254 | AX-97844884 | 0 | A | G | 0.377 | 6.98E-02 | 7.98E-02 | 2.60E+00 | 2.65E+00 | 3.80E-01 | no | | intergenic_variant | MODIFIER |
| 1 | 14474864 | AX-170235159 | 0 | C | G | 0.377 | 6.98E-02 | 7.98E-02 | 2.60E+00 | 2.65E+00 | 3.80E-01 | no | | intergenic_variant | MODIFIER |
| 1 | 14485626 | AX-170363492 | 0 | G | A | 0.377 | 6.98E-02 | 7.98E-02 | 2.60E+00 | 2.65E+00 | 3.80E-01 | no | | intergenic_variant | MODIFIER |
| 1 | 14504086 | AX-170238808 | 0 | A | C | 0.377 | 6.98E-02 | 7.98E-02 | 2.60E+00 | 2.65E+00 | 3.80E-01 | no | | regulatory_region_variant | MODIFIER |
| 1 | 14517993 | AX-97821656 | 0 | A | G | 0.377 | 6.98E-02 | 7.98E-02 | 2.60E+00 | 2.65E+00 | 3.80E-01 | no | | intergenic_variant | MODIFIER |
| 1 | 14567972 | AX-170086411 | 0 | C | T | 0.377 | 6.98E-02 | 7.98E-02 | 2.60E+00 | 2.65E+00 | 3.80E-01 | no | | intergenic_variant | MODIFIER |
| 1 | 14573595 | AX-170349973 | 0 | C | T | 0.377 | 6.98E-02 | 7.98E-02 | 2.60E+00 | 2.65E+00 | 3.80E-01 | no | | intergenic_variant | MODIFIER |
| 1 | 14576635 | AX-97810864 | 0 | G | C | 0.377 | 6.98E-02 | 7.98E-02 | 2.60E+00 | 2.65E+00 | 3.80E-01 | no | | intergenic_variant | MODIFIER |
| 1 | 14576853 | AX-170425904 | 0 | G | A | 0.377 | 6.98E-02 | 7.98E-02 | 2.60E+00 | 2.65E+00 | 3.80E-01 | no | | intergenic_variant | MODIFIER |
| 1 | 14579332 | AX-169999985 | 0 | G | A | 0.377 | 6.98E-02 | 7.98E-02 | 2.60E+00 | 2.65E+00 | 3.80E-01 | no | | intergenic_variant | MODIFIER |
| 1 | 14625759 | AX-169999984 | 0 | T | G | 0.377 | 6.98E-02 | 7.98E-02 | 2.60E+00 | 2.65E+00 | 3.80E-01 | no | | intergenic_variant | MODIFIER |
| 1 | 14677156 | AX-170197637 | 0 | T | C | 0.377 | 6.98E-02 | 7.98E-02 | 2.60E+00 | 2.65E+00 | 3.80E-01 | no | | intergenic_variant | MODIFIER |
| 1 | 14715817 | AX-170423348 | 0 | C | G | 0.377 | 6.98E-02 | 7.98E-02 | 2.60E+00 | 2.65E+00 | 3.80E-01 | no | | intergenic_variant | MODIFIER |
| 1 | 14717878 | AX-170493622 | 0 | G | A | 0.377 | 6.98E-02 | 7.98E-02 | 2.60E+00 | 2.65E+00 | 3.80E-01 | no | | intergenic_variant | MODIFIER |
| 1 | 14735091 | AX-97822109 | 0 | A | G | 0.377 | 6.98E-02 | 7.98E-02 | 2.60E+00 | 2.65E+00 | 3.80E-01 | no | | intergenic_variant | MODIFIER |
| 1 | 15654236 | AX-169999993 | 0 | A | G | 0.377 | 6.98E-02 | 7.98E-02 | 2.60E+00 | 2.65E+00 | 3.80E-01 | no | Kcnb2 | intron_variant | MODIFIER |
| 1 | 15680157 | AX-169999996 | 0 | G | A | 0.377 | 6.98E-02 | 7.98E-02 | 2.60E+00 | 2.65E+00 | 3.80E-01 | no | Kcnb2 | intron_variant | MODIFIER |
| 1 | 15682719 | AX-97844647 | 0 | G | A | 0.377 | 6.98E-02 | 7.98E-02 | 2.60E+00 | 2.65E+00 | 3.80E-01 | no | Gm25227 | downstream_gene_variant | MODIFIER |
| 1 | 15684051 | AX-169999994 | 0 | T | A | 0.377 | 6.98E-02 | 7.98E-02 | 2.60E+00 | 2.65E+00 | 3.80E-01 | no | Gm25227 | downstream_gene_variant | MODIFIER |
| 1 | 15702880 | AX-170525526 | 0 | C | T | 0.377 | 6.98E-02 | 7.98E-02 | 2.60E+00 | 2.65E+00 | 3.80E-01 | no | Kcnb2 | intron_variant | MODIFIER |
| 1 | 15708900 | AX-170462326 | 0 | T | C | 0.377 | 6.98E-02 | 7.98E-02 | 2.60E+00 | 2.65E+00 | 3.80E-01 | no | Kcnb2 | intron_variant | MODIFIER |
| 1 | 15711848 | AX-170193394 | 0 | G | T | 0.377 | 6.98E-02 | 7.98E-02 | 2.60E+00 | 2.65E+00 | 3.80E-01 | no | Kcnb2 | 3_prime_UTR_variant | MODIFIER |
| 1 | 15770142 | AX-170040467 | 0 | A | G | 0.377 | 6.98E-02 | 7.98E-02 | 2.60E+00 | 2.65E+00 | 3.80E-01 | no | | intergenic_variant | MODIFIER |
| 1 | 15798600 | AX-170338656 | 0 | T | C | 0.377 | 6.98E-02 | 7.98E-02 | 2.60E+00 | 2.65E+00 | 3.80E-01 | no | | intergenic_variant | MODIFIER |
| 1 | 15986759 | AX-170035602 | 0 | A | C | 0.377 | 6.98E-02 | 7.98E-02 | 2.60E+00 | 2.65E+00 | 3.80E-01 | no | | intergenic_variant | MODIFIER |
| 1 | 15988241 | AX-170192073 | 0 | G | A | 0.377 | 6.98E-02 | 7.98E-02 | 2.60E+00 | 2.65E+00 | 3.80E-01 | no | | intergenic_variant | MODIFIER |
| 1 | 15990044 | AX-170205850 | 0 | T | C | 0.377 | 6.98E-02 | 7.98E-02 | 2.60E+00 | 2.65E+00 | 3.80E-01 | no | | intergenic_variant | MODIFIER |
| 1 | 16048916 | AX-170370068 | 0 | T | A | 0.377 | 6.98E-02 | 7.98E-02 | 2.60E+00 | 2.65E+00 | 3.80E-01 | no | Gm7634 | upstream_gene_variant | MODIFIER |
| 1 | 16120607 | AX-170415553 | 0 | A | C | 0.377 | 6.98E-02 | 7.98E-02 | 2.60E+00 | 2.65E+00 | 3.80E-01 | no | Rdh10 | intron_variant | MODIFIER |
| 1 | 16130757 | AX-97787607 | 0 | A | G | 0.377 | 6.98E-02 | 7.98E-02 | 2.60E+00 | 2.65E+00 | 3.80E-01 | no | Rdh10 | intron_variant | MODIFIER |
| 1 | 16157468 | AX-170107361 | 0 | T | C | 0.377 | 6.98E-02 | 7.98E-02 | 2.60E+00 | 2.65E+00 | 3.80E-01 | no | | intergenic_variant | MODIFIER |
| 1 | 16159090 | AX-170040476 | 0 | G | A | 0.377 | 6.98E-02 | 7.98E-02 | 2.60E+00 | 2.65E+00 | 3.80E-01 | no | | intergenic_variant | MODIFIER |
| 1 | 16173414 | AX-170040474 | 0 | T | C | 0.377 | 6.98E-02 | 7.98E-02 | 2.60E+00 | 2.65E+00 | 3.80E-01 | no | | intergenic_variant | MODIFIER |
| 1 | 16174778 | AX-170232684 | 0 | G | A | 0.377 | 6.98E-02 | 7.98E-02 | 2.60E+00 | 2.65E+00 | 3.80E-01 | no | | intergenic_variant | MODIFIER |
| 1 | 16188253 | AX-169999998 | 0 | G | A | 0.377 | 6.98E-02 | 7.98E-02 | 2.60E+00 | 2.65E+00 | 3.80E-01 | no | | intergenic_variant | MODIFIER |
| 1 | 16239287 | AX-170417353 | 0 | G | A | 0.377 | 6.98E-02 | 7.98E-02 | 2.60E+00 | 2.65E+00 | 3.80E-01 | no | Stau2 | intron_variant | MODIFIER |
| 1 | 16241899 | AX-170467718 | 0 | T | C | 0.377 | 6.98E-02 | 7.98E-02 | 2.60E+00 | 2.65E+00 | 3.80E-01 | no | Stau2 | intron_variant | MODIFIER |
| 1 | 16245875 | AX-170531208 | 0 | T | C | 0.377 | 6.98E-02 | 7.98E-02 | 2.60E+00 | 2.65E+00 | 3.80E-01 | no | Stau2 | intron_variant | MODIFIER |

| | | | | | | | | | | | | | | | |
|---|---|---|---|---|---|---|---|---|---|---|---|---|---|---|---|
| 1 | 16247185 | AX-97852813 | 0 | A | G | 0.377 | 6.98E-02 | 7.98E-02 | 2.60E+00 | 2.65E+00 | 3.80E-01 | no | Stau2 | intron_variant | MODIFIER |
| 1 | 16278559 | AX-170262204 | 0 | C | T | 0.377 | 6.98E-02 | 7.98E-02 | 2.60E+00 | 2.65E+00 | 3.80E-01 | no | Stau2 | intron_variant | MODIFIER |
| 1 | 16282025 | AX-170392489 | 0 | T | G | 0.377 | 6.98E-02 | 7.98E-02 | 2.60E+00 | 2.65E+00 | 3.80E-01 | no | Stau2 | intron_variant | MODIFIER |
| 1 | 16626226 | AX-170390142 | 0 | T | A | 0.377 | 6.98E-02 | 7.98E-02 | 2.60E+00 | 2.65E+00 | 3.80E-01 | no | Gm38342 | downstream_gene_variant | MODIFIER |
| 1 | 16629027 | AX-170487246 | 0 | T | C | 0.377 | 6.98E-02 | 7.98E-02 | 2.60E+00 | 2.65E+00 | 3.80E-01 | no | | intergenic_variant | MODIFIER |
| 1 | 16668928 | AX-170150560 | 0 | A | G | 0.377 | 6.98E-02 | 7.98E-02 | 2.60E+00 | 2.65E+00 | 3.80E-01 | no | Tmem70 | intron_variant | MODIFIER |
| 1 | 16679742 | AX-170097243 | 0 | T | C | 0.377 | 6.98E-02 | 7.98E-02 | 2.60E+00 | 2.65E+00 | 3.80E-01 | no | Tmem70 | downstream_gene_variant | MODIFIER |
| 1 | 16698873 | AX-170484514 | 0 | C | T | 0.377 | 6.98E-02 | 7.98E-02 | 2.60E+00 | 2.65E+00 | 3.80E-01 | no | Ly96 | intron_variant | MODIFIER |
| 1 | 16698919 | AX-170465246 | 0 | T | C | 0.377 | 6.98E-02 | 7.98E-02 | 2.60E+00 | 2.65E+00 | 3.80E-01 | no | Ly96 | intron_variant | MODIFIER |
| 1 | 16700669 | AX-170206147 | 0 | A | G | 0.377 | 6.98E-02 | 7.98E-02 | 2.60E+00 | 2.65E+00 | 3.80E-01 | no | Ly96 | intron_variant | MODIFIER |
| 1 | 16769945 | AX-170000000 | 0 | T | C | 0.377 | 6.98E-02 | 7.98E-02 | 2.60E+00 | 2.65E+00 | 3.80E-01 | no | Gm5828 | non_coding_transcript_exon_variant | MODIFIER |
| 13 | 6457062 | AX-170156358 | 0 | C | T | 0.211 | -1.06E-01 | 1.21E-01 | 2.60E+00 | 2.67E+00 | 3.80E-01 | no | | intergenic_variant | MODIFIER |
| 12 | 114538058 | AX-170491921 | 2 | T | A | 0.136 | -1.40E-01 | 1.61E-01 | 2.67E+00 | 2.76E+00 | 3.80E-01 | no | Ighv1-7 | downstream_gene_variant | MODIFIER |
| 16 | 31508145 | AX-170264016 | 1 | G | A | 0.5 | 7.05E-02 | 8.02E-02 | 2.56E+00 | 2.63E+00 | 3.80E-01 | no | Gm46560 | upstream_gene_variant | MODIFIER |
| 9 | 50397360 | AX-170213406 | 0 | A | G | 0.211 | -8.97E-02 | 1.04E-01 | 2.71E+00 | 2.82E+00 | 3.80E-01 | no | | intergenic_variant | MODIFIER |
| 3 | 150458504 | AX-170099788 | 1 | C | T | 0.33 | 6.49E-02 | 7.48E-02 | 2.79E+00 | 2.88E+00 | 3.80E-01 | no | | intergenic_variant | MODIFIER |
| 5 | 32114310 | AX-170252872 | 0 | C | G | 0.368 | 8.99E-02 | 1.04E-01 | 2.76E+00 | 2.83E+00 | 3.80E-01 | no | | intergenic_variant | MODIFIER |
| 4 | 140459237 | AX-170528672 | 0 | G | A | 0.149 | -9.62E-02 | 1.08E-01 | 3.05E+00 | 3.24E+00 | 3.80E-01 | no | Gm13026 | intron_variant,non_coding_transcript_variant | MODIFIER |
| 5 | 37731912 | AX-170071831 | 1 | C | T | 0.348 | 9.35E-02 | 1.08E-01 | 2.79E+00 | 2.87E+00 | 3.80E-01 | no | Cytl1 | upstream_gene_variant | MODIFIER |
| 9 | 69075515 | AX-170332538 | 2 | C | T | 0.209 | 1.12E-01 | 1.28E-01 | 2.92E+00 | 2.99E+00 | 3.80E-01 | no | Rora | intron_variant | MODIFIER |
| 5 | 93022621 | AX-170449373 | 0 | C | T | 0.421 | -8.39E-02 | 9.69E-02 | 2.73E+00 | 2.82E+00 | 3.80E-01 | no | | regulatory_region_variant | MODIFIER |
| 1 | 147205914 | AX-170036048 | 0 | C | G | 0.404 | -6.02E-02 | 6.70E-02 | 3.08E+00 | 3.19E+00 | 3.80E-01 | no | | intergenic_variant | MODIFIER |
| 1 | 150178132 | AX-170356258 | 0 | G | A | 0.202 | -1.20E-01 | 1.34E-01 | 3.08E+00 | 3.19E+00 | 3.80E-01 | no | | intergenic_variant | MODIFIER |
| 1 | 152250800 | AX-170129841 | 0 | T | A | 0.202 | -1.20E-01 | 1.34E-01 | 3.08E+00 | 3.19E+00 | 3.80E-01 | no | | intergenic_variant | MODIFIER |
| 6 | 129096668 | AX-170028238 | 1 | T | C | 0.554 | 7.35E-02 | 8.29E-02 | 2.51E+00 | 2.57E+00 | 3.81E-01 | no | Clec2e | intron_variant | MODIFIER |
| 1 | 146810711 | AX-170001937 | 0 | C | T | 0.211 | 1.10E-01 | 1.27E-01 | 2.62E+00 | 2.69E+00 | 3.81E-01 | no | Brinp3 | intron_variant | MODIFIER |
| 1 | 149907194 | AX-170470305 | 0 | G | A | 0.211 | 1.10E-01 | 1.27E-01 | 2.62E+00 | 2.69E+00 | 3.81E-01 | no | Pla2g4a | intron_variant | MODIFIER |
| 5 | 120991478 | AX-170026021 | 0 | T | C | 0.447 | 9.39E-02 | 1.08E-01 | 2.75E+00 | 2.81E+00 | 3.81E-01 | no | Rph3a | intron_variant | MODIFIER |
| 5 | 122923359 | AX-170454400 | 0 | T | C | 0.447 | 9.39E-02 | 1.08E-01 | 2.75E+00 | 2.81E+00 | 3.81E-01 | no | Kdm2b | intron_variant | MODIFIER |
| 1 | 156312958 | AX-170298904 | 0 | T | C | 0.219 | -1.03E-01 | 1.17E-01 | 3.00E+00 | 3.16E+00 | 3.81E-01 | no | Nphs2 | intron_variant | MODIFIER |
| 13 | 19110681 | AX-170135061 | 1 | A | G | 0.205 | -9.51E-02 | 1.09E-01 | 2.59E+00 | 2.69E+00 | 3.81E-01 | no | Amph | intron_variant | MODIFIER |
| 1 | 192931898 | AX-170002595 | 0 | T | C | 0.272 | 8.31E-02 | 9.05E-02 | 2.36E+00 | 2.45E+00 | 3.81E-01 | no | Syt14 | intron_variant | MODIFIER |
| 5 | 49475877 | AX-170465310 | 0 | T | C | 0.5 | 9.34E-02 | 1.04E-01 | 2.45E+00 | 2.50E+00 | 3.81E-01 | no | Kcnip4 | intron_variant | MODIFIER |
| 9 | 108685351 | AX-97853725 | 0 | T | C | 0.123 | -1.02E-01 | 1.17E-01 | 2.61E+00 | 2.68E+00 | 3.81E-01 | no | Slc25a20 | downstream_gene_variant | MODIFIER |
| 18 | 82346173 | AX-168373241 | 0 | T | C | 0.149 | -9.97E-02 | 1.13E-01 | 2.53E+00 | 2.60E+00 | 3.81E-01 | no | AC157548.1 | intron_variant,non_coding_transcript_variant | MODIFIER |
| 6 | 128238233 | AX-170396180 | 1 | C | T | 0.554 | 7.34E-02 | 8.29E-02 | 2.51E+00 | 2.57E+00 | 3.81E-01 | no | Tead4 | intron_variant | MODIFIER |
| 4 | 76670717 | AX-170228133 | 0 | A | C | 0.316 | 9.64E-02 | 1.11E-01 | 2.72E+00 | 2.78E+00 | 3.81E-01 | no | Ptprd | intron_variant,non_coding_transcript_variant | MODIFIER |
| 18 | 69016021 | AX-170098117 | 0 | G | A | 0.175 | -1.01E-01 | 1.16E-01 | 2.62E+00 | 2.69E+00 | 3.81E-01 | no | | regulatory_region_variant | MODIFIER |
| 3 | 149192648 | AX-170472632 | 0 | T | C | 0.333 | 6.46E-02 | 7.46E-02 | 2.78E+00 | 2.87E+00 | 3.81E-01 | no | | intergenic_variant | MODIFIER |
| 3 | 149323918 | AX-170216506 | 0 | T | C | 0.333 | 6.46E-02 | 7.46E-02 | 2.78E+00 | 2.87E+00 | 3.81E-01 | no | Gm30382 | upstream_gene_variant | MODIFIER |
| 3 | 149432191 | AX-170364869 | 0 | A | G | 0.333 | 6.46E-02 | 7.46E-02 | 2.78E+00 | 2.87E+00 | 3.81E-01 | no | Gm30382 | intron_variant,non_coding_transcript_variant | MODIFIER |
| 3 | 149434040 | AX-170448204 | 0 | C | T | 0.333 | 6.46E-02 | 7.46E-02 | 2.78E+00 | 2.87E+00 | 3.81E-01 | no | Gm30382 | intron_variant,non_coding_transcript_variant | MODIFIER |
| 3 | 149438546 | AX-170374516 | 0 | T | C | 0.333 | 6.46E-02 | 7.46E-02 | 2.78E+00 | 2.87E+00 | 3.81E-01 | no | Gm30382 | intron_variant,non_coding_transcript_variant | MODIFIER |
| 3 | 149464791 | AX-170221698 | 0 | G | T | 0.333 | 6.46E-02 | 7.46E-02 | 2.78E+00 | 2.87E+00 | 3.81E-01 | no | | intergenic_variant | MODIFIER |
| 3 | 149675769 | AX-170193021 | 0 | G | A | 0.333 | 6.46E-02 | 7.46E-02 | 2.78E+00 | 2.87E+00 | 3.81E-01 | no | | intergenic_variant | MODIFIER |
| 3 | 149691491 | AX-97829154 | 0 | A | G | 0.333 | 6.46E-02 | 7.46E-02 | 2.78E+00 | 2.87E+00 | 3.81E-01 | no | | intergenic_variant | MODIFIER |
| 3 | 149724922 | AX-97854431 | 0 | A | C | 0.333 | 6.46E-02 | 7.46E-02 | 2.78E+00 | 2.87E+00 | 3.81E-01 | no | | intergenic_variant | MODIFIER |
| 3 | 149725056 | AX-170021900 | 0 | T | A | 0.333 | 6.46E-02 | 7.46E-02 | 2.78E+00 | 2.87E+00 | 3.81E-01 | no | | intergenic_variant | MODIFIER |
| 3 | 149781764 | AX-170021904 | 0 | T | C | 0.333 | 6.46E-02 | 7.46E-02 | 2.78E+00 | 2.87E+00 | 3.81E-01 | no | | intergenic_variant | MODIFIER |
| 3 | 149813653 | AX-170367284 | 0 | A | C | 0.333 | 6.46E-02 | 7.46E-02 | 2.78E+00 | 2.87E+00 | 3.81E-01 | no | | intergenic_variant | MODIFIER |
| 3 | 149820052 | AX-170169119 | 0 | A | G | 0.333 | 6.46E-02 | 7.46E-02 | 2.78E+00 | 2.87E+00 | 3.81E-01 | no | | regulatory_region_variant | MODIFIER |
| 3 | 149829853 | AX-170103906 | 0 | A | C | 0.333 | 6.46E-02 | 7.46E-02 | 2.78E+00 | 2.87E+00 | 3.81E-01 | no | | intergenic_variant | MODIFIER |
| 3 | 149834490 | AX-170199459 | 0 | G | A | 0.333 | 6.46E-02 | 7.46E-02 | 2.78E+00 | 2.87E+00 | 3.81E-01 | no | | intergenic_variant | MODIFIER |
| 3 | 149836761 | AX-170532750 | 0 | A | G | 0.333 | 6.46E-02 | 7.46E-02 | 2.78E+00 | 2.87E+00 | 3.81E-01 | no | | regulatory_region_variant | MODIFIER |

| | | | | | | | | | | | | | | | |
|---|---|---|---|---|---|---|---|---|---|---|---|---|---|---|---|
| 3 | 149837638 | AX-170165498 | 0 | A | G | 0.333 | 6.46E-02 | 7.46E-02 | 2.78E+00 | 2.87E+00 | 3.81E-01 | no | | intergenic_variant | MODIFIER |
| 3 | 149839266 | AX-97818171 | 0 | A | G | 0.333 | 6.46E-02 | 7.46E-02 | 2.78E+00 | 2.87E+00 | 3.81E-01 | no | | intergenic_variant | MODIFIER |
| 3 | 149839420 | AX-170021899 | 0 | T | G | 0.333 | 6.46E-02 | 7.46E-02 | 2.78E+00 | 2.87E+00 | 3.81E-01 | no | | intergenic_variant | MODIFIER |
| 3 | 149847492 | AX-170165200 | 0 | A | C | 0.333 | 6.46E-02 | 7.46E-02 | 2.78E+00 | 2.87E+00 | 3.81E-01 | no | | intergenic_variant | MODIFIER |
| 3 | 149848923 | AX-170198589 | 0 | T | C | 0.333 | 6.46E-02 | 7.46E-02 | 2.78E+00 | 2.87E+00 | 3.81E-01 | no | | intergenic_variant | MODIFIER |
| 3 | 149906538 | AX-97791112 | 0 | C | T | 0.333 | 6.46E-02 | 7.46E-02 | 2.78E+00 | 2.87E+00 | 3.81E-01 | no | | intergenic_variant | MODIFIER |
| 3 | 149909347 | AX-170021902 | 0 | A | C | 0.333 | 6.46E-02 | 7.46E-02 | 2.78E+00 | 2.87E+00 | 3.81E-01 | no | | intergenic_variant | MODIFIER |
| 3 | 149937987 | AX-170094852 | 0 | T | C | 0.333 | 6.46E-02 | 7.46E-02 | 2.78E+00 | 2.87E+00 | 3.81E-01 | no | | intergenic_variant | MODIFIER |
| 3 | 149942160 | AX-170228475 | 0 | T | G | 0.333 | 6.46E-02 | 7.46E-02 | 2.78E+00 | 2.87E+00 | 3.81E-01 | no | | intergenic_variant | MODIFIER |
| 3 | 149947330 | AX-170382700 | 0 | C | T | 0.333 | 6.46E-02 | 7.46E-02 | 2.78E+00 | 2.87E+00 | 3.81E-01 | no | | intergenic_variant | MODIFIER |
| 3 | 149954759 | AX-170021905 | 0 | G | A | 0.333 | 6.46E-02 | 7.46E-02 | 2.78E+00 | 2.87E+00 | 3.81E-01 | no | | intergenic_variant | MODIFIER |
| 3 | 149954873 | AX-170425192 | 0 | G | A | 0.333 | 6.46E-02 | 7.46E-02 | 2.78E+00 | 2.87E+00 | 3.81E-01 | no | | intergenic_variant | MODIFIER |
| 3 | 149979593 | AX-97827457 | 0 | A | C | 0.333 | 6.46E-02 | 7.46E-02 | 2.78E+00 | 2.87E+00 | 3.81E-01 | no | | intergenic_variant | MODIFIER |
| 3 | 149984139 | AX-170528456 | 0 | C | T | 0.333 | 6.46E-02 | 7.46E-02 | 2.78E+00 | 2.87E+00 | 3.81E-01 | no | | intergenic_variant | MODIFIER |
| 3 | 149990388 | AX-97839024 | 0 | C | T | 0.333 | 6.46E-02 | 7.46E-02 | 2.78E+00 | 2.87E+00 | 3.81E-01 | no | | intergenic_variant | MODIFIER |
| 3 | 150040147 | AX-170439482 | 0 | G | C | 0.333 | 6.46E-02 | 7.46E-02 | 2.78E+00 | 2.87E+00 | 3.81E-01 | no | | intergenic_variant | MODIFIER |
| 3 | 150051855 | AX-170089111 | 0 | T | G | 0.333 | 6.46E-02 | 7.46E-02 | 2.78E+00 | 2.87E+00 | 3.81E-01 | no | | intergenic_variant | MODIFIER |
| 3 | 150053955 | AX-170496464 | 0 | A | C | 0.333 | 6.46E-02 | 7.46E-02 | 2.78E+00 | 2.87E+00 | 3.81E-01 | no | | intergenic_variant | MODIFIER |
| 3 | 150433484 | AX-170021919 | 0 | A | T | 0.333 | 6.46E-02 | 7.46E-02 | 2.78E+00 | 2.87E+00 | 3.81E-01 | no | | intergenic_variant | MODIFIER |
| 3 | 150469067 | AX-170234051 | 0 | C | T | 0.333 | 6.46E-02 | 7.46E-02 | 2.78E+00 | 2.87E+00 | 3.81E-01 | no | | intergenic_variant | MODIFIER |
| 3 | 150469842 | AX-170382973 | 0 | T | C | 0.333 | 6.46E-02 | 7.46E-02 | 2.78E+00 | 2.87E+00 | 3.81E-01 | no | | intergenic_variant | MODIFIER |
| 3 | 150469988 | AX-170499634 | 0 | A | T | 0.333 | 6.46E-02 | 7.46E-02 | 2.78E+00 | 2.87E+00 | 3.81E-01 | no | | intergenic_variant | MODIFIER |
| 3 | 150486093 | AX-170021914 | 0 | A | G | 0.333 | 6.46E-02 | 7.46E-02 | 2.78E+00 | 2.87E+00 | 3.81E-01 | no | | intergenic_variant | MODIFIER |
| 3 | 150486314 | AX-170089113 | 0 | T | G | 0.333 | 6.46E-02 | 7.46E-02 | 2.78E+00 | 2.87E+00 | 3.81E-01 | no | | intergenic_variant | MODIFIER |
| 3 | 150604724 | AX-170160539 | 0 | A | G | 0.333 | 6.46E-02 | 7.46E-02 | 2.78E+00 | 2.87E+00 | 3.81E-01 | no | Gm43470 | intron_variant,non_coding_transcript_variant | MODIFIER |
| 3 | 150612392 | AX-170277964 | 0 | C | T | 0.333 | 6.46E-02 | 7.46E-02 | 2.78E+00 | 2.87E+00 | 3.81E-01 | no | Gm43470 | downstream_gene_variant | MODIFIER |
| 3 | 150616689 | AX-97830096 | 0 | T | C | 0.333 | 6.46E-02 | 7.46E-02 | 2.78E+00 | 2.87E+00 | 3.81E-01 | no | Gm43470 | downstream_gene_variant | MODIFIER |
| 3 | 150624465 | AX-170425499 | 0 | T | A | 0.333 | 6.46E-02 | 7.46E-02 | 2.78E+00 | 2.87E+00 | 3.81E-01 | no | | intergenic_variant | MODIFIER |
| 3 | 150640464 | AX-170211229 | 0 | T | C | 0.333 | 6.46E-02 | 7.46E-02 | 2.78E+00 | 2.87E+00 | 3.81E-01 | no | | intergenic_variant | MODIFIER |
| 3 | 151097749 | AX-170068446 | 0 | A | G | 0.333 | 6.46E-02 | 7.46E-02 | 2.78E+00 | 2.87E+00 | 3.81E-01 | no | Gm43469 | upstream_gene_variant | MODIFIER |
| 3 | 151098362 | AX-170068449 | 0 | C | T | 0.333 | 6.46E-02 | 7.46E-02 | 2.78E+00 | 2.87E+00 | 3.81E-01 | no | Gm43469 | upstream_gene_variant | MODIFIER |
| 3 | 151098390 | AX-170382544 | 0 | A | T | 0.333 | 6.46E-02 | 7.46E-02 | 2.78E+00 | 2.87E+00 | 3.81E-01 | no | Gm43469 | upstream_gene_variant | MODIFIER |
| 3 | 151124451 | AX-170517008 | 0 | T | C | 0.333 | 6.46E-02 | 7.46E-02 | 2.78E+00 | 2.87E+00 | 3.81E-01 | no | Gm43473 | upstream_gene_variant | MODIFIER |
| 3 | 151136280 | AX-97819849 | 0 | C | T | 0.333 | 6.46E-02 | 7.46E-02 | 2.78E+00 | 2.87E+00 | 3.81E-01 | no | | intergenic_variant | MODIFIER |
| 3 | 151498154 | AX-170189633 | 0 | T | C | 0.333 | 6.46E-02 | 7.46E-02 | 2.78E+00 | 2.87E+00 | 3.81E-01 | no | Adgrl4 | intron_variant | MODIFIER |
| 3 | 151615229 | AX-170433329 | 0 | C | T | 0.333 | 6.46E-02 | 7.46E-02 | 2.78E+00 | 2.87E+00 | 3.81E-01 | no | | intergenic_variant | MODIFIER |
| 3 | 151648127 | AX-170021933 | 0 | G | A | 0.333 | 6.46E-02 | 7.46E-02 | 2.78E+00 | 2.87E+00 | 3.81E-01 | no | | intergenic_variant | MODIFIER |
| 3 | 151662591 | AX-170021935 | 0 | A | G | 0.333 | 6.46E-02 | 7.46E-02 | 2.78E+00 | 2.87E+00 | 3.81E-01 | no | | intergenic_variant | MODIFIER |
| 3 | 151694695 | AX-170021930 | 0 | A | T | 0.333 | 6.46E-02 | 7.46E-02 | 2.78E+00 | 2.87E+00 | 3.81E-01 | no | | intergenic_variant | MODIFIER |
| 3 | 151707153 | AX-170131238 | 0 | G | A | 0.333 | 6.46E-02 | 7.46E-02 | 2.78E+00 | 2.87E+00 | 3.81E-01 | no | | intergenic_variant | MODIFIER |
| 3 | 151719094 | AX-170482832 | 0 | A | G | 0.333 | 6.46E-02 | 7.46E-02 | 2.78E+00 | 2.87E+00 | 3.81E-01 | no | | regulatory_region_variant | MODIFIER |
| 3 | 151737591 | AX-170201904 | 0 | A | C | 0.333 | 6.46E-02 | 7.46E-02 | 2.78E+00 | 2.87E+00 | 3.81E-01 | no | Ifi44 | intron_variant | MODIFIER |
| 3 | 151737870 | AX-170514272 | 0 | G | T | 0.333 | 6.46E-02 | 7.46E-02 | 2.78E+00 | 2.87E+00 | 3.81E-01 | no | Ifi44 | intron_variant | MODIFIER |
| 3 | 151748409 | AX-170318897 | 0 | A | G | 0.333 | 6.46E-02 | 7.46E-02 | 2.78E+00 | 2.87E+00 | 3.81E-01 | no | Ifi44 | intron_variant | MODIFIER |
| 3 | 151770547 | AX-170089116 | 0 | C | T | 0.333 | 6.46E-02 | 7.46E-02 | 2.78E+00 | 2.87E+00 | 3.81E-01 | no | | intergenic_variant | MODIFIER |
| 3 | 151773989 | AX-170402995 | 0 | G | A | 0.333 | 6.46E-02 | 7.46E-02 | 2.78E+00 | 2.87E+00 | 3.81E-01 | no | | intergenic_variant | MODIFIER |
| 3 | 151796156 | AX-170325891 | 0 | G | A | 0.333 | 6.46E-02 | 7.46E-02 | 2.78E+00 | 2.87E+00 | 3.81E-01 | no | Ptgfr | downstream_gene_variant | MODIFIER |
| 3 | 151800616 | AX-170443033 | 0 | A | C | 0.333 | 6.46E-02 | 7.46E-02 | 2.78E+00 | 2.87E+00 | 3.81E-01 | no | Ptgfr | 3_prime_UTR_variant | MODIFIER |
| 3 | 151811300 | AX-170068459 | 0 | C | T | 0.333 | 6.46E-02 | 7.46E-02 | 2.78E+00 | 2.87E+00 | 3.81E-01 | no | Ptgfr | intron_variant | MODIFIER |
| 3 | 151817526 | AX-170161920 | 0 | G | T | 0.333 | 6.46E-02 | 7.46E-02 | 2.78E+00 | 2.87E+00 | 3.81E-01 | no | Ptgfr | intron_variant | MODIFIER |
| 3 | 151828412 | AX-170257669 | 0 | A | T | 0.333 | 6.46E-02 | 7.46E-02 | 2.78E+00 | 2.87E+00 | 3.81E-01 | no | Ptgfr | intron_variant | MODIFIER |
| 3 | 151830174 | AX-170455540 | 0 | T | G | 0.333 | 6.46E-02 | 7.46E-02 | 2.78E+00 | 2.87E+00 | 3.81E-01 | no | Ptgfr | intron_variant | MODIFIER |
| 3 | 151842965 | AX-170323210 | 0 | A | C | 0.333 | 6.46E-02 | 7.46E-02 | 2.78E+00 | 2.87E+00 | 3.81E-01 | no | Gm43618 | downstream_gene_variant | MODIFIER |
| 3 | 151856204 | AX-170423870 | 0 | C | G | 0.333 | 6.46E-02 | 7.46E-02 | 2.78E+00 | 2.87E+00 | 3.81E-01 | no | | intergenic_variant | MODIFIER |

| | | | | | | | | | | | | | | | |
|---|---|---|---|---|---|---|---|---|---|---|---|---|---|---|---|
| 3 | 151886249 | AX-170258456 | 0 | G | A | 0.333 | 6.46E-02 | 7.46E-02 | 2.78E+00 | 2.87E+00 | 3.81E-01 | no | Gm43617 | upstream_gene_variant | MODIFIER |
| 3 | 151887965 | AX-170482252 | 0 | T | A | 0.333 | 6.46E-02 | 7.46E-02 | 2.78E+00 | 2.87E+00 | 3.81E-01 | no | Gm43617 | intron_variant,non_coding_transcript_variant | MODIFIER |
| 9 | 98534597 | AX-170253004 | 1 | G | A | 0.143 | -9.85E-02 | 1.12E-01 | 2.57E+00 | 2.64E+00 | 3.81E-01 | no | | intergenic_variant | MODIFIER |
| 8 | 111836828 | AX-170455661 | 0 | A | T | 0.228 | -8.95E-02 | 1.04E-01 | 2.76E+00 | 2.87E+00 | 3.81E-01 | no | Cfdp1 | intron_variant | MODIFIER |
| 1 | 145469258 | AX-170001914 | 0 | G | A | 0.228 | 1.01E-01 | 1.16E-01 | 2.68E+00 | 2.76E+00 | 3.81E-01 | no | | intergenic_variant | MODIFIER |
| 1 | 145048877 | AX-170158490 | 1 | G | A | 0.205 | 1.21E-01 | 1.39E-01 | 2.70E+00 | 2.75E+00 | 3.82E-01 | no | | intergenic_variant | MODIFIER |
| 1 | 149093474 | AX-170290726 | 1 | C | G | 0.205 | 1.21E-01 | 1.39E-01 | 2.70E+00 | 2.75E+00 | 3.82E-01 | no | | intergenic_variant | MODIFIER |
| 1 | 150139850 | AX-170422737 | 1 | C | T | 0.205 | 1.21E-01 | 1.39E-01 | 2.70E+00 | 2.75E+00 | 3.82E-01 | no | | intergenic_variant | MODIFIER |
| 1 | 153217729 | AX-97842902 | 1 | C | T | 0.205 | 1.21E-01 | 1.39E-01 | 2.70E+00 | 2.75E+00 | 3.82E-01 | no | Lamc1 | downstream_gene_variant | MODIFIER |
| 1 | 126619319 | AX-170042619 | 1 | G | A | 0.58 | 7.07E-02 | 7.97E-02 | 2.49E+00 | 2.56E+00 | 3.82E-01 | no | Nckap5 | intron_variant | MODIFIER |
| 18 | 60023858 | AX-97839765 | 0 | A | T | 0.404 | 8.48E-02 | 9.51E-02 | 2.47E+00 | 2.51E+00 | 3.82E-01 | no | | intergenic_variant | MODIFIER |
| 18 | 60525110 | AX-170015808 | 0 | A | G | 0.404 | 8.48E-02 | 9.51E-02 | 2.47E+00 | 2.51E+00 | 3.82E-01 | no | Dctn4 | upstream_gene_variant | MODIFIER |
| 5 | 85165355 | AX-97819646 | 0 | G | T | 0.439 | 9.22E-02 | 1.07E-01 | 2.79E+00 | 2.86E+00 | 3.82E-01 | no | | intergenic_variant | MODIFIER |
| 4 | 154980239 | AX-170478796 | 0 | A | G | 0.123 | 1.40E-01 | 1.62E-01 | 2.74E+00 | 2.85E+00 | 3.82E-01 | no | Pank4 | intron_variant | MODIFIER |
| 4 | 154991070 | AX-97804547 | 0 | G | A | 0.123 | 1.40E-01 | 1.62E-01 | 2.74E+00 | 2.85E+00 | 3.82E-01 | no | Plch2 | synonymous_variant | LOW |
| 4 | 154992163 | AX-170399618 | 0 | A | G | 0.123 | 1.40E-01 | 1.62E-01 | 2.74E+00 | 2.85E+00 | 3.82E-01 | no | Plch2 | intron_variant | MODIFIER |
| 4 | 154999940 | AX-170437456 | 0 | T | C | 0.123 | 1.40E-01 | 1.62E-01 | 2.74E+00 | 2.85E+00 | 3.82E-01 | no | Plch2 | intron_variant | MODIFIER |
| 4 | 155015439 | AX-170301691 | 0 | A | G | 0.123 | 1.40E-01 | 1.62E-01 | 2.74E+00 | 2.85E+00 | 3.82E-01 | no | Plch2 | upstream_gene_variant | MODIFIER |
| 4 | 155015810 | AX-170225889 | 0 | T | C | 0.123 | 1.40E-01 | 1.62E-01 | 2.74E+00 | 2.85E+00 | 3.82E-01 | no | Plch2 | upstream_gene_variant | MODIFIER |
| 4 | 155016835 | AX-170277708 | 0 | A | T | 0.123 | 1.40E-01 | 1.62E-01 | 2.74E+00 | 2.85E+00 | 3.82E-01 | no | Plch2 | intron_variant | MODIFIER |
| 4 | 155019836 | AX-170373332 | 0 | C | T | 0.123 | 1.40E-01 | 1.62E-01 | 2.74E+00 | 2.85E+00 | 3.82E-01 | no | Plch2 | upstream_gene_variant | MODIFIER |
| 4 | 155021102 | AX-170398277 | 0 | C | G | 0.123 | 1.40E-01 | 1.62E-01 | 2.74E+00 | 2.85E+00 | 3.82E-01 | no | Plch2 | upstream_gene_variant | MODIFIER |
| 4 | 155037655 | AX-170295192 | 0 | T | C | 0.123 | 1.40E-01 | 1.62E-01 | 2.74E+00 | 2.85E+00 | 3.82E-01 | no | Gm10564 | upstream_gene_variant | MODIFIER |
| 4 | 155070695 | AX-170354253 | 0 | G | A | 0.123 | 1.40E-01 | 1.62E-01 | 2.74E+00 | 2.85E+00 | 3.82E-01 | no | Rer1 | downstream_gene_variant | MODIFIER |
| 4 | 155174761 | AX-170229097 | 0 | C | T | 0.123 | 1.40E-01 | 1.62E-01 | 2.74E+00 | 2.85E+00 | 3.82E-01 | no | Ski | intron_variant | MODIFIER |
| 4 | 155210788 | AX-97793290 | 0 | C | T | 0.123 | 1.40E-01 | 1.62E-01 | 2.74E+00 | 2.85E+00 | 3.82E-01 | no | Ski | intron_variant | MODIFIER |
| 4 | 155211578 | AX-170400363 | 0 | T | C | 0.123 | 1.40E-01 | 1.62E-01 | 2.74E+00 | 2.85E+00 | 3.82E-01 | no | Ski | intron_variant | MODIFIER |
| 4 | 155228643 | AX-170024304 | 0 | G | C | 0.123 | 1.40E-01 | 1.62E-01 | 2.74E+00 | 2.85E+00 | 3.82E-01 | no | | intergenic_variant | MODIFIER |
| 4 | 155234223 | AX-97815787 | 0 | C | A | 0.123 | 1.40E-01 | 1.62E-01 | 2.74E+00 | 2.85E+00 | 3.82E-01 | no | | regulatory_region_variant | MODIFIER |
| 4 | 155454211 | AX-170421073 | 0 | C | A | 0.123 | 1.40E-01 | 1.62E-01 | 2.74E+00 | 2.85E+00 | 3.82E-01 | no | Cfap74 | intron_variant | MODIFIER |
| 4 | 155456179 | AX-170387781 | 0 | C | T | 0.123 | 1.40E-01 | 1.62E-01 | 2.74E+00 | 2.85E+00 | 3.82E-01 | no | Cfap74 | intron_variant | MODIFIER |
| 4 | 155513202 | AX-170292278 | 0 | C | A | 0.123 | 1.40E-01 | 1.62E-01 | 2.74E+00 | 2.85E+00 | 3.82E-01 | no | Gnb1 | intron_variant | MODIFIER |
| 4 | 155528514 | AX-170400000 | 0 | A | G | 0.123 | 1.40E-01 | 1.62E-01 | 2.74E+00 | 2.85E+00 | 3.82E-01 | no | Gnb1 | intron_variant | MODIFIER |
| 4 | 155533783 | AX-170446847 | 0 | G | A | 0.123 | 1.40E-01 | 1.62E-01 | 2.74E+00 | 2.85E+00 | 3.82E-01 | no | Gnb1 | intron_variant | MODIFIER |
| 4 | 155537910 | AX-170294628 | 0 | G | T | 0.123 | 1.40E-01 | 1.62E-01 | 2.74E+00 | 2.85E+00 | 3.82E-01 | no | Gnb1 | intron_variant | MODIFIER |
| 4 | 155540431 | AX-97795368 | 0 | A | G | 0.123 | 1.40E-01 | 1.62E-01 | 2.74E+00 | 2.85E+00 | 3.82E-01 | no | Gnb1 | intron_variant | MODIFIER |
| 4 | 155540688 | AX-170247538 | 0 | T | C | 0.123 | 1.40E-01 | 1.62E-01 | 2.74E+00 | 2.85E+00 | 3.82E-01 | no | Gnb1 | intron_variant | MODIFIER |
| 4 | 155666328 | AX-170089402 | 0 | T | C | 0.123 | 1.40E-01 | 1.62E-01 | 2.74E+00 | 2.85E+00 | 3.82E-01 | no | Mib2 | intron_variant | MODIFIER |
| 4 | 155741797 | AX-170089403 | 0 | C | T | 0.123 | 1.40E-01 | 1.62E-01 | 2.74E+00 | 2.85E+00 | 3.82E-01 | no | Atad3a | intron_variant | MODIFIER |
| 4 | 155745230 | AX-170320308 | 0 | G | A | 0.123 | 1.40E-01 | 1.62E-01 | 2.74E+00 | 2.85E+00 | 3.82E-01 | no | Atad3a | intron_variant | MODIFIER |
| 4 | 156035928 | AX-170035700 | 0 | T | C | 0.123 | 1.40E-01 | 1.62E-01 | 2.74E+00 | 2.85E+00 | 3.82E-01 | no | Ttll10 | intron_variant | MODIFIER |
| 2 | 168956149 | AX-170354957 | 0 | T | A | 0.175 | 1.34E-01 | 1.54E-01 | 2.56E+00 | 2.67E+00 | 3.82E-01 | no | Zfp64 | upstream_gene_variant | MODIFIER |
| 18 | 69965913 | AX-170188564 | 0 | C | G | 0.263 | -7.83E-02 | 9.07E-02 | 2.69E+00 | 2.78E+00 | 3.82E-01 | no | Ccdc68 | intron_variant | MODIFIER |
| 7 | 37753030 | AX-170268579 | 0 | C | A | 0.351 | 8.33E-02 | 9.51E-02 | 2.56E+00 | 2.63E+00 | 3.82E-01 | no | Zfp536 | intron_variant | MODIFIER |
| 7 | 37755198 | AX-170156327 | 0 | A | G | 0.351 | 8.33E-02 | 9.51E-02 | 2.56E+00 | 2.63E+00 | 3.82E-01 | no | Zfp536 | intron_variant | MODIFIER |
| 7 | 37755376 | AX-170028847 | 0 | A | G | 0.351 | 8.33E-02 | 9.51E-02 | 2.56E+00 | 2.63E+00 | 3.82E-01 | no | Zfp536 | intron_variant | MODIFIER |
| 7 | 37759608 | AX-170028841 | 0 | A | G | 0.351 | 8.33E-02 | 9.51E-02 | 2.56E+00 | 2.63E+00 | 3.82E-01 | no | Zfp536 | intron_variant | MODIFIER |
| 7 | 37765243 | AX-170028844 | 0 | T | C | 0.351 | 8.33E-02 | 9.51E-02 | 2.56E+00 | 2.63E+00 | 3.82E-01 | no | Zfp536 | intron_variant | MODIFIER |
| 7 | 37793730 | AX-170028846 | 0 | A | T | 0.351 | 8.33E-02 | 9.51E-02 | 2.56E+00 | 2.63E+00 | 3.82E-01 | no | | intergenic_variant | MODIFIER |
| 7 | 37825804 | AX-170275780 | 0 | C | T | 0.351 | 8.33E-02 | 9.51E-02 | 2.56E+00 | 2.63E+00 | 3.82E-01 | no | Gm22302 | upstream_gene_variant | MODIFIER |
| 7 | 37835804 | AX-170028843 | 0 | C | T | 0.351 | 8.33E-02 | 9.51E-02 | 2.56E+00 | 2.63E+00 | 3.82E-01 | no | | intergenic_variant | MODIFIER |
| 7 | 37836085 | AX-170440642 | 0 | C | A | 0.351 | 8.33E-02 | 9.51E-02 | 2.56E+00 | 2.63E+00 | 3.82E-01 | no | | intergenic_variant | MODIFIER |
| 7 | 37836507 | AX-170077102 | 0 | C | T | 0.351 | 8.33E-02 | 9.51E-02 | 2.56E+00 | 2.63E+00 | 3.82E-01 | no | | intergenic_variant | MODIFIER |
| 7 | 37839442 | AX-170028849 | 0 | C | T | 0.351 | 8.33E-02 | 9.51E-02 | 2.56E+00 | 2.63E+00 | 3.82E-01 | no | | intergenic_variant | MODIFIER |

| | | | | | | | | | | | | | | | |
|---|---|---|---|---|---|---|---|---|---|---|---|---|---|---|---|
| 7 | 37844790 | AX-170314474 | 0 | C | T | 0.351 | 8.33E-02 | 9.51E-02 | 2.56E+00 | 2.63E+00 | 3.82E-01 | no | | intergenic_variant | MODIFIER |
| 7 | 37849805 | AX-170512658 | 0 | C | T | 0.351 | 8.33E-02 | 9.51E-02 | 2.56E+00 | 2.63E+00 | 3.82E-01 | no | | intergenic_variant | MODIFIER |
| 7 | 37853982 | AX-170290713 | 0 | A | G | 0.351 | 8.33E-02 | 9.51E-02 | 2.56E+00 | 2.63E+00 | 3.82E-01 | no | | intergenic_variant | MODIFIER |
| 7 | 37888275 | AX-170228683 | 0 | A | G | 0.351 | 8.33E-02 | 9.51E-02 | 2.56E+00 | 2.63E+00 | 3.82E-01 | no | | intergenic_variant | MODIFIER |
| 7 | 37895586 | AX-97804301 | 0 | A | G | 0.351 | 8.33E-02 | 9.51E-02 | 2.56E+00 | 2.63E+00 | 3.82E-01 | no | | intergenic_variant | MODIFIER |
| 7 | 37918215 | AX-170399925 | 0 | T | C | 0.351 | 8.33E-02 | 9.51E-02 | 2.56E+00 | 2.63E+00 | 3.82E-01 | no | | intergenic_variant | MODIFIER |
| 7 | 37920698 | AX-170348685 | 0 | G | A | 0.351 | 8.33E-02 | 9.51E-02 | 2.56E+00 | 2.63E+00 | 3.82E-01 | no | | intergenic_variant | MODIFIER |
| 7 | 37923841 | AX-170100422 | 0 | A | G | 0.351 | 8.33E-02 | 9.51E-02 | 2.56E+00 | 2.63E+00 | 3.82E-01 | no | | intergenic_variant | MODIFIER |
| 7 | 37956263 | AX-170077105 | 0 | G | C | 0.351 | 8.33E-02 | 9.51E-02 | 2.56E+00 | 2.63E+00 | 3.82E-01 | no | Uri1 | downstream_gene_variant | MODIFIER |
| 7 | 37957683 | AX-170121204 | 0 | A | G | 0.351 | 8.33E-02 | 9.51E-02 | 2.56E+00 | 2.63E+00 | 3.82E-01 | no | Uri1 | downstream_gene_variant | MODIFIER |
| 7 | 37960294 | AX-170348186 | 0 | C | T | 0.351 | 8.33E-02 | 9.51E-02 | 2.56E+00 | 2.63E+00 | 3.82E-01 | no | Uri1 | 3_prime_UTR_variant | MODIFIER |
| 7 | 37976783 | AX-170143462 | 0 | C | T | 0.351 | 8.33E-02 | 9.51E-02 | 2.56E+00 | 2.63E+00 | 3.82E-01 | no | Uri1 | intron_variant | MODIFIER |
| 7 | 37976817 | AX-97795623 | 0 | T | G | 0.351 | 8.33E-02 | 9.51E-02 | 2.56E+00 | 2.63E+00 | 3.82E-01 | no | Uri1 | intron_variant | MODIFIER |
| 7 | 37999496 | AX-170323035 | 0 | G | C | 0.351 | 8.33E-02 | 9.51E-02 | 2.56E+00 | 2.63E+00 | 3.82E-01 | no | Uri1 | intron_variant | MODIFIER |
| 7 | 38003581 | AX-170439722 | 0 | G | T | 0.351 | 8.33E-02 | 9.51E-02 | 2.56E+00 | 2.63E+00 | 3.82E-01 | no | Uri1 | intron_variant | MODIFIER |
| 7 | 38043812 | AX-170089902 | 0 | A | G | 0.351 | 8.33E-02 | 9.51E-02 | 2.56E+00 | 2.63E+00 | 3.82E-01 | no | | intergenic_variant | MODIFIER |
| 7 | 38054365 | AX-170356827 | 0 | C | T | 0.351 | 8.33E-02 | 9.51E-02 | 2.56E+00 | 2.63E+00 | 3.82E-01 | no | Gm30684 | intron_variant,non_coding_transcript_variant | MODIFIER |
| 7 | 38071309 | AX-97856874 | 0 | G | A | 0.351 | 8.33E-02 | 9.51E-02 | 2.56E+00 | 2.63E+00 | 3.82E-01 | no | Gm30771 | intron_variant,non_coding_transcript_variant | MODIFIER |
| 7 | 38072023 | AX-170485003 | 0 | T | A | 0.351 | 8.33E-02 | 9.51E-02 | 2.56E+00 | 2.63E+00 | 3.82E-01 | no | Gm30771 | intron_variant,non_coding_transcript_variant | MODIFIER |
| 7 | 38074693 | AX-170449280 | 0 | C | G | 0.351 | 8.33E-02 | 9.51E-02 | 2.56E+00 | 2.63E+00 | 3.82E-01 | no | Gm30771 | intron_variant,non_coding_transcript_variant | MODIFIER |
| 7 | 38081874 | AX-97780240 | 0 | T | C | 0.351 | 8.33E-02 | 9.51E-02 | 2.56E+00 | 2.63E+00 | 3.82E-01 | no | | intergenic_variant | MODIFIER |
| 7 | 38104652 | AX-170227825 | 0 | C | A | 0.351 | 8.33E-02 | 9.51E-02 | 2.56E+00 | 2.63E+00 | 3.82E-01 | no | Ccne1 | intron_variant | MODIFIER |
| 7 | 38110473 | AX-97835236 | 0 | A | G | 0.351 | 8.33E-02 | 9.51E-02 | 2.56E+00 | 2.63E+00 | 3.82E-01 | no | Ccne1 | upstream_gene_variant | MODIFIER |
| 7 | 38117729 | AX-170226041 | 0 | T | C | 0.351 | 8.33E-02 | 9.51E-02 | 2.56E+00 | 2.63E+00 | 3.82E-01 | no | | intergenic_variant | MODIFIER |
| 7 | 38150011 | AX-170149627 | 0 | T | C | 0.351 | 8.33E-02 | 9.51E-02 | 2.56E+00 | 2.63E+00 | 3.82E-01 | no | | intergenic_variant | MODIFIER |
| 7 | 38151751 | AX-170152243 | 0 | G | A | 0.351 | 8.33E-02 | 9.51E-02 | 2.56E+00 | 2.63E+00 | 3.82E-01 | no | | intergenic_variant | MODIFIER |
| 7 | 38151922 | AX-170181069 | 0 | C | G | 0.351 | 8.33E-02 | 9.51E-02 | 2.56E+00 | 2.63E+00 | 3.82E-01 | no | | intergenic_variant | MODIFIER |
| 7 | 38175784 | AX-170271595 | 0 | T | C | 0.351 | 8.33E-02 | 9.51E-02 | 2.56E+00 | 2.63E+00 | 3.82E-01 | no | Gm22203 | downstream_gene_variant | MODIFIER |
| 7 | 38183268 | AX-170275563 | 0 | C | T | 0.351 | 8.33E-02 | 9.51E-02 | 2.56E+00 | 2.63E+00 | 3.82E-01 | no | 1600014C10Rik | upstream_gene_variant | MODIFIER |
| 7 | 38208863 | AX-97851817 | 0 | G | A | 0.351 | 8.33E-02 | 9.51E-02 | 2.56E+00 | 2.63E+00 | 3.82E-01 | no | | regulatory_region_variant | MODIFIER |
| 7 | 38211587 | AX-170028855 | 0 | G | C | 0.351 | 8.33E-02 | 9.51E-02 | 2.56E+00 | 2.63E+00 | 3.82E-01 | no | | intergenic_variant | MODIFIER |
| 7 | 38215331 | AX-170171598 | 0 | A | G | 0.351 | 8.33E-02 | 9.51E-02 | 2.56E+00 | 2.63E+00 | 3.82E-01 | no | Plekhf1 | downstream_gene_variant | MODIFIER |
| 7 | 38227151 | AX-170028856 | 0 | T | A | 0.351 | 8.33E-02 | 9.51E-02 | 2.56E+00 | 2.63E+00 | 3.82E-01 | no | Plekhf1 | intron_variant | MODIFIER |
| 7 | 38249208 | AX-170028852 | 0 | T | C | 0.351 | 8.33E-02 | 9.51E-02 | 2.56E+00 | 2.63E+00 | 3.82E-01 | no | | intergenic_variant | MODIFIER |
| 7 | 38266541 | AX-170201610 | 0 | C | T | 0.351 | 8.33E-02 | 9.51E-02 | 2.56E+00 | 2.63E+00 | 3.82E-01 | no | Pop4 | intron_variant | MODIFIER |
| 7 | 38269206 | AX-97839591 | 0 | A | T | 0.351 | 8.33E-02 | 9.51E-02 | 2.56E+00 | 2.63E+00 | 3.82E-01 | no | Pop4 | intron_variant | MODIFIER |
| 7 | 38366251 | AX-97847635 | 0 | G | C | 0.351 | 8.33E-02 | 9.51E-02 | 2.56E+00 | 2.63E+00 | 3.82E-01 | no | Gm4451 | non_coding_transcript_exon_variant | MODIFIER |
| 7 | 38407298 | AX-170461434 | 0 | C | T | 0.351 | 8.33E-02 | 9.51E-02 | 2.56E+00 | 2.63E+00 | 3.82E-01 | no | Gm28453 | intron_variant,non_coding_transcript_variant | MODIFIER |
| 7 | 38558210 | AX-170095897 | 0 | C | G | 0.351 | 8.33E-02 | 9.51E-02 | 2.56E+00 | 2.63E+00 | 3.82E-01 | no | | intergenic_variant | MODIFIER |
| 7 | 39274617 | AX-97849471 | 0 | T | C | 0.351 | 8.33E-02 | 9.51E-02 | 2.56E+00 | 2.63E+00 | 3.82E-01 | no | Gm9226 | non_coding_transcript_exon_variant | MODIFIER |
| 7 | 39445510 | AX-170184700 | 0 | G | C | 0.351 | 8.33E-02 | 9.51E-02 | 2.56E+00 | 2.63E+00 | 3.82E-01 | no | Gm5590 | downstream_gene_variant | MODIFIER |
| 7 | 39448687 | AX-97802677 | 0 | G | C | 0.351 | 8.33E-02 | 9.51E-02 | 2.56E+00 | 2.63E+00 | 3.82E-01 | no | Gm5590 | upstream_gene_variant | MODIFIER |
| 7 | 39457025 | AX-170313171 | 0 | A | G | 0.351 | 8.33E-02 | 9.51E-02 | 2.56E+00 | 2.63E+00 | 3.82E-01 | no | Zfp939 | upstream_gene_variant | MODIFIER |
| 7 | 39485947 | AX-170355001 | 0 | C | T | 0.351 | 8.33E-02 | 9.51E-02 | 2.56E+00 | 2.63E+00 | 3.82E-01 | no | Gm28455 | upstream_gene_variant | MODIFIER |
| 7 | 39518835 | AX-170089904 | 0 | T | C | 0.351 | 8.33E-02 | 9.51E-02 | 2.56E+00 | 2.63E+00 | 3.82E-01 | no | Zfp619 | intron_variant | MODIFIER |
| 7 | 39539208 | AX-168382893 | 0 | T | C | 0.351 | 8.33E-02 | 9.51E-02 | 2.56E+00 | 2.63E+00 | 3.82E-01 | no | Zfp619 | 3_prime_UTR_variant | MODIFIER |
| 7 | 39541616 | AX-170406517 | 0 | G | A | 0.351 | 8.33E-02 | 9.51E-02 | 2.56E+00 | 2.63E+00 | 3.82E-01 | no | Zfp619 | downstream_gene_variant | MODIFIER |
| 7 | 39548559 | AX-170368289 | 0 | G | T | 0.351 | 8.33E-02 | 9.51E-02 | 2.56E+00 | 2.63E+00 | 3.82E-01 | no | Gm37494 | upstream_gene_variant | MODIFIER |
| 7 | 39552531 | AX-170517685 | 0 | G | C | 0.351 | 8.33E-02 | 9.51E-02 | 2.56E+00 | 2.63E+00 | 3.82E-01 | no | Gm37494 | non_coding_transcript_exon_variant | MODIFIER |
| 7 | 39565380 | AX-170509634 | 0 | C | G | 0.351 | 8.33E-02 | 9.51E-02 | 2.56E+00 | 2.63E+00 | 3.82E-01 | no | Gm37494 | downstream_gene_variant | MODIFIER |
| 7 | 39605416 | AX-97850666 | 0 | A | G | 0.351 | 8.33E-02 | 9.51E-02 | 2.56E+00 | 2.63E+00 | 3.82E-01 | no | | intergenic_variant | MODIFIER |
| 7 | 39617055 | AX-170232421 | 0 | G | A | 0.351 | 8.33E-02 | 9.51E-02 | 2.56E+00 | 2.63E+00 | 3.82E-01 | no | | intergenic_variant | MODIFIER |
| 7 | 39620201 | AX-170364079 | 0 | T | C | 0.351 | 8.33E-02 | 9.51E-02 | 2.56E+00 | 2.63E+00 | 3.82E-01 | no | | intergenic_variant | MODIFIER |
| 7 | 39623437 | AX-170190427 | 0 | A | C | 0.351 | 8.33E-02 | 9.51E-02 | 2.56E+00 | 2.63E+00 | 3.82E-01 | no | | intergenic_variant | MODIFIER |

| | | | | | | | | | | | | | | | |
|---|---|---|---|---|---|---|---|---|---|---|---|---|---|---|---|
| 7 | 39639703 | AX-170352150 | 0 | T | C | 0.351 | 8.33E-02 | 9.51E-02 | 2.56E+00 | 2.63E+00 | 3.82E-01 | no | | intergenic_variant | MODIFIER |
| 7 | 39682564 | AX-170390709 | 0 | G | T | 0.351 | 8.33E-02 | 9.51E-02 | 2.56E+00 | 2.63E+00 | 3.82E-01 | no | | intergenic_variant | MODIFIER |
| 9 | 105463781 | AX-170454089 | 0 | C | A | 0.123 | -8.92E-02 | 1.02E-01 | 2.55E+00 | 2.66E+00 | 3.82E-01 | no | Atp2c1 | intron_variant | MODIFIER |
| 9 | 107842860 | AX-170097033 | 0 | A | C | 0.123 | -8.92E-02 | 1.02E-01 | 2.55E+00 | 2.66E+00 | 3.82E-01 | no | Rbm6 | intron_variant | MODIFIER |
| 18 | 86950121 | AX-170355266 | 1 | T | C | 0.17 | -9.32E-02 | 1.06E-01 | 2.50E+00 | 2.58E+00 | 3.82E-01 | no | | intergenic_variant | MODIFIER |
| 7 | 103469652 | AX-170029670 | 0 | A | T | 0.456 | 8.15E-02 | 9.39E-02 | 2.63E+00 | 2.69E+00 | 3.82E-01 | no | Olfr608 | upstream_gene_variant | MODIFIER |
| 2 | 129973580 | AX-170165118 | 0 | T | C | 0.14 | -1.55E-01 | 1.80E-01 | 2.77E+00 | 2.86E+00 | 3.83E-01 | no | | intergenic_variant | MODIFIER |
| 2 | 156405311 | AX-97856972 | 0 | C | T | 0.14 | -1.55E-01 | 1.80E-01 | 2.77E+00 | 2.86E+00 | 3.83E-01 | no | | intergenic_variant | MODIFIER |
| 5 | 45288127 | AX-170197867 | 0 | T | G | 0.184 | -9.20E-02 | 1.04E-01 | 2.50E+00 | 2.59E+00 | 3.83E-01 | no | D5Ertd615e | intron_variant,non_coding_transcript_variant | MODIFIER |
| 4 | 132947092 | AX-170403029 | 0 | C | A | 0.158 | 1.03E-01 | 1.18E-01 | 2.56E+00 | 2.61E+00 | 3.83E-01 | no | | regulatory_region_variant | MODIFIER |
| 4 | 133485447 | AX-170097532 | 0 | C | T | 0.158 | 1.03E-01 | 1.18E-01 | 2.56E+00 | 2.61E+00 | 3.83E-01 | no | Tent5b | intron_variant | MODIFIER |
| 13 | 23186000 | AX-170397606 | 1 | G | A | 0.161 | -9.53E-02 | 1.06E-01 | 2.42E+00 | 2.51E+00 | 3.83E-01 | no | Vmn1r220 | upstream_gene_variant | MODIFIER |
| 7 | 37821501 | AX-170477859 | 0 | T | C | 0.342 | 8.24E-02 | 9.43E-02 | 2.55E+00 | 2.62E+00 | 3.83E-01 | no | Gm22302 | downstream_gene_variant | MODIFIER |
| 7 | 37841255 | AX-170323906 | 0 | A | C | 0.342 | 8.24E-02 | 9.43E-02 | 2.55E+00 | 2.62E+00 | 3.83E-01 | no | | intergenic_variant | MODIFIER |
| 7 | 37848248 | AX-170420776 | 0 | A | C | 0.342 | 8.24E-02 | 9.43E-02 | 2.55E+00 | 2.62E+00 | 3.83E-01 | no | | intergenic_variant | MODIFIER |
| 7 | 37853900 | AX-97789262 | 0 | C | T | 0.342 | 8.24E-02 | 9.43E-02 | 2.55E+00 | 2.62E+00 | 3.83E-01 | no | | intergenic_variant | MODIFIER |
| 7 | 38133308 | AX-97856345 | 0 | T | C | 0.342 | 8.24E-02 | 9.43E-02 | 2.55E+00 | 2.62E+00 | 3.83E-01 | no | | intergenic_variant | MODIFIER |
| 7 | 136457538 | AX-170283249 | 1 | G | C | 0.304 | -6.70E-02 | 7.77E-02 | 2.65E+00 | 2.75E+00 | 3.83E-01 | no | | intergenic_variant | MODIFIER |
| 1 | 131758590 | AX-170361572 | 1 | T | C | 0.536 | 7.49E-02 | 8.66E-02 | 2.64E+00 | 2.71E+00 | 3.83E-01 | no | Slc26a9 | splice_region_variant,intron_variant | LOW |
| 4 | 116063298 | AX-170089355 | 1 | T | C | 0.214 | 1.01E-01 | 1.16E-01 | 2.60E+00 | 2.67E+00 | 3.83E-01 | no | Uqcrh | downstream_gene_variant | MODIFIER |
| 4 | 117391836 | AX-170415424 | 1 | G | A | 0.214 | 1.01E-01 | 1.16E-01 | 2.60E+00 | 2.67E+00 | 3.83E-01 | no | Rnf220 | intron_variant | MODIFIER |
| 15 | 74346842 | AX-170268412 | 0 | G | A | 0.553 | 6.96E-02 | 7.99E-02 | 2.58E+00 | 2.66E+00 | 3.83E-01 | no | | intergenic_variant | MODIFIER |
| 15 | 74545941 | AX-170372519 | 0 | C | T | 0.553 | 6.96E-02 | 7.99E-02 | 2.58E+00 | 2.66E+00 | 3.83E-01 | no | Adgrb1 | intron_variant | MODIFIER |
| 9 | 95520755 | AX-170033043 | 1 | G | A | 0.143 | -9.92E-02 | 1.15E-01 | 2.66E+00 | 2.74E+00 | 3.83E-01 | no | | regulatory_region_variant | MODIFIER |
| 4 | 63346567 | AX-170022948 | 0 | G | T | 0.342 | 9.59E-02 | 1.11E-01 | 2.83E+00 | 2.90E+00 | 3.83E-01 | no | Orm1 | intron_variant | MODIFIER |
| 1 | 169873813 | AX-170043528 | 0 | G | A | 0.219 | 1.09E-01 | 1.26E-01 | 2.59E+00 | 2.67E+00 | 3.83E-01 | no | | intergenic_variant | MODIFIER |
| 7 | 17976387 | AX-170417927 | 0 | G | A | 0.36 | 7.18E-02 | 8.33E-02 | 2.64E+00 | 2.76E+00 | 3.84E-01 | no | Ceacam11 | intron_variant | MODIFIER |
| 12 | 116453263 | AX-170228125 | 1 | A | G | 0.143 | -1.25E-01 | 1.45E-01 | 2.64E+00 | 2.75E+00 | 3.84E-01 | no | Ncapg2 | intron_variant | MODIFIER |
| 7 | 114099195 | AX-170262517 | 0 | G | T | 0.447 | 7.94E-02 | 9.21E-02 | 2.74E+00 | 2.80E+00 | 3.84E-01 | no | Rras2 | intron_variant | MODIFIER |
| 1 | 174854175 | AX-170325988 | 1 | T | C | 0.232 | 1.03E-01 | 1.17E-01 | 2.48E+00 | 2.55E+00 | 3.84E-01 | no | Grem2 | intron_variant | MODIFIER |
| 10 | 27681404 | AX-170044438 | 2 | C | G | 0.182 | 8.25E-02 | 9.58E-02 | 2.82E+00 | 2.94E+00 | 3.84E-01 | no | Gm30165 | intron_variant,non_coding_transcript_variant | MODIFIER |
| 2 | 38571049 | AX-170428758 | 1 | C | T | 0.045 | -1.30E-01 | 1.50E-01 | 2.57E+00 | 2.70E+00 | 3.84E-01 | no | Nek6 | intron_variant | MODIFIER |
| 8 | 30494273 | AX-170030435 | 0 | A | G | 0.167 | -8.62E-02 | 1.00E-01 | 2.71E+00 | 2.81E+00 | 3.84E-01 | no | Gm45489 | intron_variant,non_coding_transcript_variant | MODIFIER |
| 18 | 71029117 | AX-97791267 | 1 | C | T | 0.179 | -1.02E-01 | 1.17E-01 | 2.57E+00 | 2.63E+00 | 3.84E-01 | no | | intergenic_variant | MODIFIER |
| 7 | 70847942 | AX-170476178 | 0 | T | C | 0.36 | -8.75E-02 | 1.01E-01 | 2.87E+00 | 2.96E+00 | 3.84E-01 | no | | regulatory_region_variant | MODIFIER |
| 7 | 70871508 | AX-170184359 | 0 | C | G | 0.36 | -8.75E-02 | 1.01E-01 | 2.87E+00 | 2.96E+00 | 3.84E-01 | no | Gm42397 | intron_variant,non_coding_transcript_variant | MODIFIER |
| 7 | 70871798 | AX-170281986 | 0 | C | T | 0.36 | -8.75E-02 | 1.01E-01 | 2.87E+00 | 2.96E+00 | 3.84E-01 | no | Gm42397 | intron_variant,non_coding_transcript_variant | MODIFIER |
| 7 | 70874695 | AX-170460959 | 0 | T | C | 0.36 | -8.75E-02 | 1.01E-01 | 2.87E+00 | 2.96E+00 | 3.84E-01 | no | Gm42397 | intron_variant,non_coding_transcript_variant | MODIFIER |
| 7 | 70891257 | AX-170533579 | 0 | T | C | 0.36 | -8.75E-02 | 1.01E-01 | 2.87E+00 | 2.96E+00 | 3.84E-01 | no | Gm42397 | intron_variant,non_coding_transcript_variant | MODIFIER |
| 7 | 70946758 | AX-170089952 | 0 | A | G | 0.36 | -8.75E-02 | 1.01E-01 | 2.87E+00 | 2.96E+00 | 3.84E-01 | no | Gm42397 | intron_variant,non_coding_transcript_variant | MODIFIER |
| 7 | 71034524 | AX-170164885 | 0 | C | T | 0.36 | -8.75E-02 | 1.01E-01 | 2.87E+00 | 2.96E+00 | 3.84E-01 | no | Gm42397 | intron_variant,non_coding_transcript_variant | MODIFIER |
| 7 | 71034849 | AX-97809170 | 0 | G | A | 0.36 | -8.75E-02 | 1.01E-01 | 2.87E+00 | 2.96E+00 | 3.84E-01 | no | Gm42397 | intron_variant,non_coding_transcript_variant | MODIFIER |
| 7 | 71035528 | AX-170153511 | 0 | G | T | 0.36 | -8.75E-02 | 1.01E-01 | 2.87E+00 | 2.96E+00 | 3.84E-01 | no | Gm42397 | intron_variant,non_coding_transcript_variant | MODIFIER |
| 7 | 71047955 | AX-170257430 | 0 | C | T | 0.36 | -8.75E-02 | 1.01E-01 | 2.87E+00 | 2.96E+00 | 3.84E-01 | no | Gm42397 | intron_variant,non_coding_transcript_variant | MODIFIER |
| 7 | 71051964 | AX-170523386 | 0 | A | G | 0.36 | -8.75E-02 | 1.01E-01 | 2.87E+00 | 2.96E+00 | 3.84E-01 | no | Gm42397 | intron_variant,non_coding_transcript_variant | MODIFIER |
| 7 | 71052919 | AX-170289652 | 0 | A | G | 0.36 | -8.75E-02 | 1.01E-01 | 2.87E+00 | 2.96E+00 | 3.84E-01 | no | Gm42397 | intron_variant,non_coding_transcript_variant | MODIFIER |
| 7 | 71071342 | AX-168410424 | 0 | C | T | 0.36 | -8.75E-02 | 1.01E-01 | 2.87E+00 | 2.96E+00 | 3.84E-01 | no | Gm42397 | intron_variant,non_coding_transcript_variant | MODIFIER |
| 7 | 71299076 | AX-170029253 | 0 | A | G | 0.36 | -8.75E-02 | 1.01E-01 | 2.87E+00 | 2.96E+00 | 3.84E-01 | no | Gm29328 | intron_variant,non_coding_transcript_variant | MODIFIER |
| 7 | 71312444 | AX-170029254 | 0 | A | G | 0.36 | -8.75E-02 | 1.01E-01 | 2.87E+00 | 2.96E+00 | 3.84E-01 | no | Gm29328 | intron_variant,non_coding_transcript_variant | MODIFIER |
| 7 | 71330656 | AX-170200956 | 0 | T | C | 0.36 | -8.75E-02 | 1.01E-01 | 2.87E+00 | 2.96E+00 | 3.84E-01 | no | Gm10172 | downstream_gene_variant | MODIFIER |
| 7 | 71330829 | AX-170360511 | 0 | A | G | 0.36 | -8.75E-02 | 1.01E-01 | 2.87E+00 | 2.96E+00 | 3.84E-01 | no | Gm10172 | downstream_gene_variant | MODIFIER |
| 7 | 71333671 | AX-97821129 | 0 | A | G | 0.36 | -8.75E-02 | 1.01E-01 | 2.87E+00 | 2.96E+00 | 3.84E-01 | no | Gm10172 | upstream_gene_variant | MODIFIER |
| 7 | 71475824 | AX-170195262 | 0 | C | T | 0.36 | -8.75E-02 | 1.01E-01 | 2.87E+00 | 2.96E+00 | 3.84E-01 | no | | intergenic_variant | MODIFIER |
| 7 | 71479082 | AX-170183048 | 0 | C | T | 0.36 | -8.75E-02 | 1.01E-01 | 2.87E+00 | 2.96E+00 | 3.84E-01 | no | | regulatory_region_variant | MODIFIER |

| | | | | | | | | | | | | | | | |
|---|---|---|---|---|---|---|---|---|---|---|---|---|---|---|---|
| 7 | 71507142 | AX-170421295 | 0 | G | A | 0.36 | -8.75E-02 | 1.01E-01 | 2.87E+00 | 2.96E+00 | 3.84E-01 | no | | intergenic_variant | MODIFIER |
| 7 | 71538996 | AX-170424024 | 0 | A | G | 0.36 | -8.75E-02 | 1.01E-01 | 2.87E+00 | 2.96E+00 | 3.84E-01 | no | | intergenic_variant | MODIFIER |
| 1 | 170107103 | AX-170002266 | 0 | T | C | 0.211 | -1.18E-01 | 1.33E-01 | 3.06E+00 | 3.17E+00 | 3.84E-01 | no | Ddr2 | intron_variant | MODIFIER |
| 5 | 99691298 | AX-170236291 | 0 | G | A | 0.43 | -8.67E-02 | 1.01E-01 | 2.83E+00 | 2.92E+00 | 3.84E-01 | no | Rasgef1b | intron_variant,non_coding_transcript_variant | MODIFIER |
| 7 | 100925629 | AX-170441565 | 1 | T | A | 0.366 | -8.86E-02 | 1.03E-01 | 2.79E+00 | 2.88E+00 | 3.84E-01 | no | Arhgef17 | intron_variant | MODIFIER |
| 1 | 115157926 | AX-170351392 | 0 | A | T | 0.175 | 1.49E-01 | 1.71E-01 | 2.57E+00 | 2.67E+00 | 3.84E-01 | Yes | | intergenic_variant | MODIFIER |
| 8 | 61471583 | AX-170080224 | 1 | T | A | 0.152 | -1.19E-01 | 1.34E-01 | 2.48E+00 | 2.56E+00 | 3.84E-01 | no | | intergenic_variant | MODIFIER |
| 4 | 137226693 | AX-170338882 | 0 | A | G | 0.167 | 1.01E-01 | 1.16E-01 | 2.64E+00 | 2.69E+00 | 3.84E-01 | no | Gm13002 | upstream_gene_variant | MODIFIER |
| 10 | 67835938 | AX-170410411 | 1 | T | C | 0.411 | -6.89E-02 | 8.00E-02 | 2.65E+00 | 2.73E+00 | 3.85E-01 | no | Gm49766 | intron_variant,non_coding_transcript_variant | MODIFIER |
| 3 | 23750649 | AX-170228232 | 0 | G | A | 0.167 | -8.70E-02 | 9.98E-02 | 2.55E+00 | 2.63E+00 | 3.85E-01 | no | | intergenic_variant | MODIFIER |
| 5 | 118797611 | AX-170484940 | 0 | G | A | 0.456 | 9.42E-02 | 1.09E-01 | 2.75E+00 | 2.82E+00 | 3.85E-01 | no | | intergenic_variant | MODIFIER |
| 1 | 76400382 | AX-170041713 | 0 | C | A | 0.14 | -1.57E-01 | 1.83E-01 | 2.67E+00 | 2.78E+00 | 3.85E-01 | no | | intergenic_variant | MODIFIER |
| 9 | 118895932 | AX-170460625 | 1 | C | A | 0.125 | -7.76E-02 | 8.94E-02 | 2.56E+00 | 2.68E+00 | 3.85E-01 | no | Itga9 | intron_variant | MODIFIER |
| 4 | 80273493 | AX-97848897 | 2 | A | G | 0.227 | 8.34E-02 | 9.54E-02 | 2.52E+00 | 2.62E+00 | 3.85E-01 | no | | intergenic_variant | MODIFIER |
| 4 | 86887268 | AX-170476129 | 0 | T | G | 0.342 | 8.47E-02 | 9.85E-02 | 2.65E+00 | 2.74E+00 | 3.85E-01 | no | Acer2 | intron_variant | MODIFIER |
| 13 | 3417712 | AX-97828578 | 0 | T | G | 0.202 | -1.06E-01 | 1.23E-01 | 2.67E+00 | 2.73E+00 | 3.85E-01 | no | | intergenic_variant | MODIFIER |
| 7 | 72831059 | AX-170077706 | 0 | A | G | 0.368 | -8.76E-02 | 1.02E-01 | 2.79E+00 | 2.88E+00 | 3.85E-01 | no | | intergenic_variant | MODIFIER |
| 4 | 80102358 | AX-170023205 | 0 | T | G | 0.447 | -7.38E-02 | 8.41E-02 | 2.51E+00 | 2.57E+00 | 3.85E-01 | no | | intergenic_variant | MODIFIER |
| 5 | 50592952 | AX-170024918 | 0 | C | T | 0.482 | 8.64E-02 | 9.74E-02 | 2.46E+00 | 2.53E+00 | 3.85E-01 | no | Gm43016 | intron_variant,non_coding_transcript_variant | MODIFIER |
| 4 | 133105986 | AX-170023986 | 0 | C | T | 0.158 | 1.03E-01 | 1.19E-01 | 2.58E+00 | 2.62E+00 | 3.85E-01 | no | | intergenic_variant | MODIFIER |
| 16 | 50059504 | AX-97841423 | 0 | G | A | 0.518 | 7.07E-02 | 8.06E-02 | 2.51E+00 | 2.56E+00 | 3.85E-01 | no | | intergenic_variant | MODIFIER |
| 9 | 118595270 | AX-97843998 | 0 | C | T | 0.114 | -8.19E-02 | 9.45E-02 | 2.56E+00 | 2.67E+00 | 3.85E-01 | no | | intergenic_variant | MODIFIER |
| 2 | 162467362 | AX-170294745 | 0 | T | A | 0.149 | 1.49E-01 | 1.74E-01 | 2.79E+00 | 2.89E+00 | 3.85E-01 | Yes | Ptprt | intron_variant | MODIFIER |
| 2 | 162469197 | AX-170219532 | 0 | T | A | 0.149 | 1.49E-01 | 1.74E-01 | 2.79E+00 | 2.89E+00 | 3.85E-01 | Yes | Ptprt | intron_variant | MODIFIER |
| 5 | 48507252 | AX-97780940 | 0 | T | G | 0.491 | 9.14E-02 | 1.01E-01 | 2.40E+00 | 2.46E+00 | 3.86E-01 | no | Kcnip4 | intron_variant | MODIFIER |
| 5 | 51623221 | AX-170466518 | 0 | A | C | 0.491 | 9.14E-02 | 1.01E-01 | 2.40E+00 | 2.46E+00 | 3.86E-01 | no | | intergenic_variant | MODIFIER |
| 7 | 144898444 | AX-97849207 | 0 | C | T | 0.211 | -6.39E-02 | 7.45E-02 | 2.62E+00 | 2.75E+00 | 3.86E-01 | no | Fgf15 | intron_variant | MODIFIER |
| 7 | 72787490 | AX-170379732 | 1 | T | C | 0.348 | -8.57E-02 | 9.99E-02 | 2.80E+00 | 2.89E+00 | 3.86E-01 | no | | intergenic_variant | MODIFIER |
| 8 | 83046687 | AX-170277438 | 0 | T | C | 0.202 | -1.05E-01 | 1.21E-01 | 2.90E+00 | 2.98E+00 | 3.86E-01 | no | Rnf150 | intron_variant | MODIFIER |
| 1 | 128553940 | AX-97826432 | 0 | A | G | 0.535 | 7.35E-02 | 8.54E-02 | 2.64E+00 | 2.70E+00 | 3.86E-01 | no | | intergenic_variant | MODIFIER |
| 8 | 34194378 | AX-170030484 | 0 | G | A | 0.14 | -1.15E-01 | 1.33E-01 | 2.62E+00 | 2.69E+00 | 3.86E-01 | no | | intergenic_variant | MODIFIER |
| 8 | 62977998 | AX-170030881 | 1 | T | G | 0.143 | -1.24E-01 | 1.43E-01 | 2.59E+00 | 2.67E+00 | 3.86E-01 | no | Spock3 | intron_variant | MODIFIER |
| 16 | 39838628 | AX-97857226 | 0 | A | C | 0.5 | 7.00E-02 | 8.06E-02 | 2.56E+00 | 2.62E+00 | 3.86E-01 | no | | intergenic_variant | MODIFIER |
| 18 | 65236189 | AX-170229448 | 0 | G | A | 0.175 | -1.01E-01 | 1.18E-01 | 2.63E+00 | 2.69E+00 | 3.86E-01 | no | | intergenic_variant | MODIFIER |
| 16 | 53873617 | AX-170219635 | 1 | T | G | 0.518 | 7.29E-02 | 8.25E-02 | 2.47E+00 | 2.52E+00 | 3.86E-01 | no | | intergenic_variant | MODIFIER |
| 17 | 32298448 | AX-170203779 | 1 | C | T | 0.054 | 1.70E-01 | 1.99E-01 | 2.73E+00 | 2.86E+00 | 3.86E-01 | Yes | Gm26549 | intron_variant,non_coding_transcript_variant | MODIFIER |
| 8 | 16967023 | AX-97800089 | 0 | T | C | 0.219 | -6.62E-02 | 7.60E-02 | 2.53E+00 | 2.62E+00 | 3.86E-01 | no | Csmd1 | intron_variant | MODIFIER |
| 7 | 71853306 | AX-170156677 | 1 | G | C | 0.357 | -8.17E-02 | 9.54E-02 | 2.80E+00 | 2.89E+00 | 3.86E-01 | no | | intergenic_variant | MODIFIER |
| 10 | 70053051 | AX-170434130 | 0 | C | T | 0.421 | -6.88E-02 | 8.01E-02 | 2.65E+00 | 2.72E+00 | 3.86E-01 | no | | intergenic_variant | MODIFIER |
| 5 | 46044665 | AX-170490789 | 1 | C | T | 0.5 | 9.13E-02 | 1.02E-01 | 2.43E+00 | 2.48E+00 | 3.86E-01 | no | | intergenic_variant | MODIFIER |
| 15 | 71853677 | AX-170122496 | 0 | C | T | 0.491 | 2.15E-01 | 2.53E-01 | 2.56E+00 | 2.80E+00 | 3.86E-01 | Yes | Col22a1 | intron_variant | MODIFIER |
| 6 | 93650304 | AX-170475602 | 0 | C | T | 0.439 | -5.28E-02 | 6.05E-02 | 2.51E+00 | 2.62E+00 | 3.86E-01 | no | Gm7812 | upstream_gene_variant | MODIFIER |
| 6 | 93735609 | AX-170113340 | 0 | C | T | 0.439 | -5.28E-02 | 6.05E-02 | 2.51E+00 | 2.62E+00 | 3.86E-01 | no | Magi1 | intron_variant | MODIFIER |
| 1 | 20805148 | AX-97787576 | 0 | C | T | 0.421 | 7.40E-02 | 8.53E-02 | 2.56E+00 | 2.61E+00 | 3.86E-01 | no | Mcm3 | intron_variant | MODIFIER |
| 1 | 100568994 | AX-97822127 | 0 | G | T | 0.167 | 1.48E-01 | 1.70E-01 | 2.53E+00 | 2.63E+00 | 3.87E-01 | Yes | | intergenic_variant | MODIFIER |
| 1 | 131230981 | AX-170001714 | 0 | A | G | 0.535 | 7.32E-02 | 8.52E-02 | 2.64E+00 | 2.71E+00 | 3.87E-01 | no | Rassf5 | intron_variant | MODIFIER |
| 1 | 131679894 | AX-97813279 | 0 | T | C | 0.535 | 7.32E-02 | 8.52E-02 | 2.64E+00 | 2.71E+00 | 3.87E-01 | no | Ctse | downstream_gene_variant | MODIFIER |
| 3 | 149235555 | AX-170299058 | 1 | T | C | 0.339 | 6.24E-02 | 7.29E-02 | 2.77E+00 | 2.86E+00 | 3.87E-01 | no | | intergenic_variant | MODIFIER |
| 9 | 117903559 | AX-170396414 | 0 | G | T | 0.14 | -7.30E-02 | 8.57E-02 | 2.67E+00 | 2.81E+00 | 3.87E-01 | no | Zcwpw2 | intron_variant,non_coding_transcript_variant | MODIFIER |
| 1 | 90202173 | AX-170200887 | 1 | G | C | 0.161 | 1.44E-01 | 1.67E-01 | 2.60E+00 | 2.71E+00 | 3.87E-01 | no | Ackr3 | upstream_gene_variant | MODIFIER |
| 8 | 95889382 | AX-170536160 | 0 | A | G | 0.237 | 9.09E-02 | 1.06E-01 | 2.75E+00 | 2.83E+00 | 3.87E-01 | no | Got2 | upstream_gene_variant | MODIFIER |
| 9 | 22538126 | AX-170193382 | 0 | C | T | 0.465 | 7.85E-02 | 9.02E-02 | 2.53E+00 | 2.60E+00 | 3.87E-01 | no | Bbs9 | intron_variant | MODIFIER |
| 1 | 150820033 | AX-170043120 | 0 | A | T | 0.211 | -1.16E-01 | 1.31E-01 | 3.06E+00 | 3.17E+00 | 3.87E-01 | no | Hmcn1 | intron_variant | MODIFIER |
| 1 | 150947777 | AX-170473733 | 0 | T | C | 0.211 | -1.16E-01 | 1.31E-01 | 3.06E+00 | 3.17E+00 | 3.87E-01 | no | Hmcn1 | intron_variant | MODIFIER |

| | | | | | | | | | | | | | | | |
|---|---|---|---|---|---|---|---|---|---|---|---|---|---|---|---|
| 1 | 151636154 | AX-170371559 | 0 | T | A | 0.421 | -5.80E-02 | 6.56E-02 | 3.06E+00 | 3.17E+00 | 3.87E-01 | no | Fam129a | intron_variant,NMD_transcript_variant | MODIFIER |
| 2 | 169368149 | AX-170116020 | 2 | C | T | 0.145 | 1.50E-01 | 1.76E-01 | 2.65E+00 | 2.78E+00 | 3.87E-01 | Yes | | intergenic_variant | MODIFIER |
| 9 | 80254010 | AX-170082918 | 0 | A | T | 0.202 | -9.46E-02 | 1.11E-01 | 2.71E+00 | 2.82E+00 | 3.87E-01 | no | Myo6 | intron_variant | MODIFIER |
| 1 | 135629078 | AX-170192653 | 0 | C | T | 0.544 | 7.17E-02 | 8.30E-02 | 2.57E+00 | 2.63E+00 | 3.88E-01 | no | Gm38399 | intron_variant,non_coding_transcript_variant | MODIFIER |
| 7 | 142834927 | AX-170316977 | 1 | G | A | 0.214 | -6.40E-02 | 7.49E-02 | 2.63E+00 | 2.76E+00 | 3.88E-01 | no | Gm6471 | intron_variant,non_coding_transcript_variant | MODIFIER |
| 16 | 39508523 | AX-97831547 | 1 | T | A | 0.482 | -6.14E-02 | 7.14E-02 | 2.60E+00 | 2.68E+00 | 3.88E-01 | no | | intergenic_variant | MODIFIER |
| 12 | 69455399 | AX-170241715 | 0 | G | T | 0.07 | 1.79E-01 | 2.10E-01 | 2.76E+00 | 2.87E+00 | 3.88E-01 | Yes | | regulatory_region_variant | MODIFIER |
| 16 | 74227781 | AX-170534406 | 0 | A | C | 0.491 | 6.87E-02 | 7.77E-02 | 2.45E+00 | 2.51E+00 | 3.88E-01 | no | Robo2 | intron_variant | MODIFIER |
| 16 | 74233845 | AX-170246030 | 0 | A | G | 0.491 | 6.87E-02 | 7.77E-02 | 2.45E+00 | 2.51E+00 | 3.88E-01 | no | Robo2 | intron_variant | MODIFIER |
| 16 | 74237668 | AX-170375071 | 0 | A | G | 0.491 | 6.87E-02 | 7.77E-02 | 2.45E+00 | 2.51E+00 | 3.88E-01 | no | Robo2 | intron_variant | MODIFIER |
| 16 | 74237699 | AX-170178979 | 0 | C | T | 0.491 | 6.87E-02 | 7.77E-02 | 2.45E+00 | 2.51E+00 | 3.88E-01 | no | Robo2 | intron_variant | MODIFIER |
| 16 | 74237764 | AX-170436430 | 0 | C | G | 0.491 | 6.87E-02 | 7.77E-02 | 2.45E+00 | 2.51E+00 | 3.88E-01 | no | Robo2 | intron_variant | MODIFIER |
| 16 | 74246008 | AX-170146388 | 0 | G | A | 0.491 | 6.87E-02 | 7.77E-02 | 2.45E+00 | 2.51E+00 | 3.88E-01 | no | Robo2 | intron_variant | MODIFIER |
| 16 | 74246680 | AX-170341502 | 0 | G | A | 0.491 | 6.87E-02 | 7.77E-02 | 2.45E+00 | 2.51E+00 | 3.88E-01 | no | Robo2 | intron_variant | MODIFIER |
| 16 | 74285991 | AX-170139817 | 0 | G | T | 0.491 | 6.87E-02 | 7.77E-02 | 2.45E+00 | 2.51E+00 | 3.88E-01 | no | Robo2 | intron_variant | MODIFIER |
| 16 | 74286008 | AX-170430283 | 0 | C | A | 0.491 | 6.87E-02 | 7.77E-02 | 2.45E+00 | 2.51E+00 | 3.88E-01 | no | Robo2 | intron_variant | MODIFIER |
| 16 | 74899092 | AX-97831808 | 0 | A | C | 0.491 | 6.87E-02 | 7.77E-02 | 2.45E+00 | 2.51E+00 | 3.88E-01 | no | | intergenic_variant | MODIFIER |
| 16 | 75001490 | AX-97853921 | 0 | G | A | 0.491 | 6.87E-02 | 7.77E-02 | 2.45E+00 | 2.51E+00 | 3.88E-01 | no | | intergenic_variant | MODIFIER |
| 16 | 75001520 | AX-170057825 | 0 | G | A | 0.491 | 6.87E-02 | 7.77E-02 | 2.45E+00 | 2.51E+00 | 3.88E-01 | no | | intergenic_variant | MODIFIER |
| 16 | 75001554 | AX-170057822 | 0 | A | T | 0.491 | 6.87E-02 | 7.77E-02 | 2.45E+00 | 2.51E+00 | 3.88E-01 | no | | intergenic_variant | MODIFIER |
| 16 | 75063169 | AX-170111558 | 0 | C | T | 0.491 | 6.87E-02 | 7.77E-02 | 2.45E+00 | 2.51E+00 | 3.88E-01 | no | | intergenic_variant | MODIFIER |
| 13 | 25148964 | AX-170216703 | 0 | T | C | 0.167 | -1.02E-01 | 1.14E-01 | 2.41E+00 | 2.49E+00 | 3.88E-01 | no | Dcdc2a | intron_variant | MODIFIER |
| 1 | 21096083 | AX-170208965 | 0 | G | A | 0.412 | 7.38E-02 | 8.58E-02 | 2.60E+00 | 2.65E+00 | 3.88E-01 | no | | intergenic_variant | MODIFIER |
| 18 | 83749998 | AX-170302138 | 0 | T | C | 0.14 | -9.72E-02 | 1.11E-01 | 2.48E+00 | 2.56E+00 | 3.88E-01 | no | | intergenic_variant | MODIFIER |
| 18 | 84516845 | AX-170507950 | 0 | T | C | 0.14 | -9.72E-02 | 1.11E-01 | 2.48E+00 | 2.56E+00 | 3.88E-01 | no | Zfp407 | intron_variant | MODIFIER |
| 18 | 84611782 | AX-97808767 | 0 | T | C | 0.14 | -9.72E-02 | 1.11E-01 | 2.48E+00 | 2.56E+00 | 3.88E-01 | no | Cndp1 | intron_variant | MODIFIER |
| 10 | 92146818 | AX-170186968 | 1 | T | C | 0.295 | -7.31E-02 | 8.42E-02 | 2.54E+00 | 2.60E+00 | 3.88E-01 | no | Rmst | intron_variant,non_coding_transcript_variant | MODIFIER |
| 15 | 5175154 | AX-170409802 | 1 | C | G | 0.241 | -8.94E-02 | 1.04E-01 | 2.62E+00 | 2.68E+00 | 3.88E-01 | no | Prkaa1 | intron_variant | MODIFIER |
| 20 | 134245639 | AX-170085290 | 0 | A | G | 0.14 | 9.31E-02 | 1.10E-01 | 2.71E+00 | 2.82E+00 | 3.89E-01 | no | Trmt2b | intron_variant | MODIFIER |
| 16 | 50700493 | AX-170224874 | 2 | A | G | 0.445 | -6.68E-02 | 7.50E-02 | 2.41E+00 | 2.48E+00 | 3.89E-01 | no | | regulatory_region_variant | MODIFIER |
| 1 | 76998307 | AX-170337648 | 1 | G | A | 0.134 | -1.49E-01 | 1.75E-01 | 2.68E+00 | 2.80E+00 | 3.89E-01 | no | Gm816 | intron_variant,non_coding_transcript_variant | MODIFIER |
| 14 | 116724861 | AX-97824134 | 0 | C | T | 0.316 | 6.44E-02 | 7.51E-02 | 2.60E+00 | 2.69E+00 | 3.89E-01 | no | | intergenic_variant | MODIFIER |
| 3 | 38422783 | AX-170315084 | 0 | A | C | 0.202 | 7.75E-02 | 9.11E-02 | 2.78E+00 | 2.89E+00 | 3.89E-01 | no | Gm36823 | downstream_gene_variant | MODIFIER |
| 9 | 54327193 | AX-170302743 | 1 | C | G | 0.205 | -8.82E-02 | 1.04E-01 | 2.79E+00 | 2.91E+00 | 3.89E-01 | no | Gldn | intron_variant | MODIFIER |
| 5 | 34438863 | AX-97781181 | 1 | A | C | 0.321 | 9.36E-02 | 1.10E-01 | 2.78E+00 | 2.85E+00 | 3.89E-01 | no | Fam193a | intron_variant | MODIFIER |
| 5 | 35318640 | AX-170496714 | 1 | T | C | 0.321 | 9.36E-02 | 1.10E-01 | 2.78E+00 | 2.85E+00 | 3.89E-01 | no | | intergenic_variant | MODIFIER |
| 4 | 49994735 | AX-170435039 | 1 | A | G | 0.357 | -7.55E-02 | 8.88E-02 | 2.82E+00 | 2.94E+00 | 3.89E-01 | no | | intergenic_variant | MODIFIER |
| 3 | 20735950 | AX-170493378 | 1 | A | G | 0.143 | -8.86E-02 | 1.03E-01 | 2.57E+00 | 2.65E+00 | 3.89E-01 | no | | intergenic_variant | MODIFIER |
| 6 | 93951578 | AX-170268258 | 0 | T | A | 0.43 | -5.12E-02 | 5.94E-02 | 2.55E+00 | 2.67E+00 | 3.89E-01 | no | Magi1 | intron_variant | MODIFIER |
| 16 | 78858164 | AX-170436894 | 1 | C | T | 0.464 | 6.65E-02 | 7.70E-02 | 2.55E+00 | 2.61E+00 | 3.89E-01 | no | | intergenic_variant | MODIFIER |
| 5 | 45283183 | AX-170509722 | 1 | C | T | 0.179 | -8.99E-02 | 1.03E-01 | 2.50E+00 | 2.60E+00 | 3.89E-01 | no | D5Ertd615e | intron_variant,non_coding_transcript_variant | MODIFIER |
| 7 | 36966003 | AX-170180127 | 1 | T | C | 0.348 | 7.97E-02 | 9.27E-02 | 2.58E+00 | 2.66E+00 | 3.89E-01 | no | | intergenic_variant | MODIFIER |
| 7 | 37091139 | AX-170424406 | 1 | G | A | 0.348 | 7.97E-02 | 9.27E-02 | 2.58E+00 | 2.66E+00 | 3.89E-01 | no | | intergenic_variant | MODIFIER |
| 15 | 10549158 | AX-170332632 | 2 | C | T | 0.236 | -9.11E-02 | 1.06E-01 | 2.59E+00 | 2.65E+00 | 3.89E-01 | no | Ttc23l | intron_variant | MODIFIER |
| 3 | 36964769 | AX-170120479 | 0 | G | A | 0.202 | 7.76E-02 | 9.13E-02 | 2.78E+00 | 2.87E+00 | 3.89E-01 | no | 4932438A13Rik | intron_variant | MODIFIER |
| 9 | 98489462 | AX-170396407 | 1 | G | C | 0.143 | -9.71E-02 | 1.13E-01 | 2.58E+00 | 2.66E+00 | 3.89E-01 | no | Rbp2 | upstream_gene_variant | MODIFIER |
| 1 | 65762067 | AX-170000701 | 1 | T | C | 0.125 | -1.27E-01 | 1.49E-01 | 2.65E+00 | 2.78E+00 | 3.90E-01 | no | | intergenic_variant | MODIFIER |
| 13 | 5426500 | AX-170190883 | 0 | C | T | 0.202 | -9.86E-02 | 1.15E-01 | 2.57E+00 | 2.65E+00 | 3.90E-01 | no | | intergenic_variant | MODIFIER |
| 3 | 151117228 | AX-170409500 | 1 | G | A | 0.375 | 5.98E-02 | 7.05E-02 | 2.74E+00 | 2.84E+00 | 3.90E-01 | no | Gm43473 | downstream_gene_variant | MODIFIER |
| 3 | 37581790 | AX-170401115 | 1 | C | G | 0.196 | 8.08E-02 | 9.49E-02 | 2.82E+00 | 2.91E+00 | 3.90E-01 | no | Spata5 | downstream_gene_variant | MODIFIER |
| 1 | 156466246 | AX-170226968 | 1 | G | T | 0.223 | -1.12E-01 | 1.29E-01 | 3.00E+00 | 3.11E+00 | 3.90E-01 | no | Soat1 | intron_variant | MODIFIER |
| 5 | 50321585 | AX-170185001 | 0 | G | A | 0.482 | 9.12E-02 | 1.02E-01 | 2.41E+00 | 2.46E+00 | 3.90E-01 | no | Gm7205 | intron_variant,non_coding_transcript_variant | MODIFIER |
| 16 | 19705871 | AX-170432790 | 1 | A | G | 0.125 | 8.26E-02 | 9.75E-02 | 2.79E+00 | 2.93E+00 | 3.90E-01 | no | Lamp3 | intron_variant | MODIFIER |
| 11 | 35297452 | AX-170396680 | 0 | G | A | 0.105 | 1.33E-01 | 1.57E-01 | 2.67E+00 | 2.78E+00 | 3.90E-01 | no | Slit3 | intron_variant | MODIFIER |

| | | | | | | | | | | | | | | | |
|---|---|---|---|---|---|---|---|---|---|---|---|---|---|---|---|
| 1 | 135841997 | AX-170458863 | 0 | A | G | 0.219 | 9.78E-02 | 1.15E-01 | 2.63E+00 | 2.72E+00 | 3.90E-01 | no | Tnnt2 | intron_variant | MODIFIER |
| 18 | 80178649 | AX-170447647 | 0 | C | G | 0.395 | -7.21E-02 | 8.47E-02 | 2.64E+00 | 2.74E+00 | 3.90E-01 | no | | regulatory_region_variant | MODIFIER |
| 18 | 80262490 | AX-170259831 | 0 | A | T | 0.395 | -7.21E-02 | 8.47E-02 | 2.64E+00 | 2.74E+00 | 3.90E-01 | no | Pqlc1 | intron_variant | MODIFIER |
| 4 | 97298940 | AX-97798335 | 0 | A | G | 0.43 | -6.65E-02 | 7.71E-02 | 2.54E+00 | 2.62E+00 | 3.90E-01 | no | | intergenic_variant | MODIFIER |
| 4 | 97300056 | AX-170337471 | 0 | A | G | 0.43 | -6.65E-02 | 7.71E-02 | 2.54E+00 | 2.62E+00 | 3.90E-01 | no | | intergenic_variant | MODIFIER |
| 7 | 70578777 | AX-170264301 | 0 | A | G | 0.36 | -8.59E-02 | 1.01E-01 | 2.87E+00 | 2.97E+00 | 3.90E-01 | no | Gm44812 | intron_variant,non_coding_transcript_variant | MODIFIER |
| 16 | 77615176 | AX-97820747 | 1 | C | T | 0.464 | 6.65E-02 | 7.70E-02 | 2.54E+00 | 2.61E+00 | 3.90E-01 | no | Mir99ahg | intron_variant,non_coding_transcript_variant | MODIFIER |
| 8 | 97896129 | AX-170031345 | 0 | G | T | 0.219 | -9.69E-02 | 1.13E-01 | 2.88E+00 | 2.99E+00 | 3.90E-01 | no | | intergenic_variant | MODIFIER |
| 16 | 29536473 | AX-168392444 | 0 | C | T | 0.509 | 6.95E-02 | 8.06E-02 | 2.55E+00 | 2.59E+00 | 3.91E-01 | no | Atp13a4 | intron_variant | MODIFIER |
| 8 | 109292162 | AX-170278537 | 1 | C | T | 0.518 | 4.41E-02 | 5.20E-02 | 2.82E+00 | 2.94E+00 | 3.91E-01 | no | | intergenic_variant | MODIFIER |
| 11 | 6480924 | AX-170294034 | 0 | C | T | 0.044 | -1.57E-01 | 1.84E-01 | 2.56E+00 | 2.70E+00 | 3.91E-01 | no | Gm11973 | intron_variant,non_coding_transcript_variant | MODIFIER |
| 16 | 29759073 | AX-170126105 | 0 | C | T | 0.5 | 6.82E-02 | 7.95E-02 | 2.57E+00 | 2.63E+00 | 3.91E-01 | no | Gm49636 | upstream_gene_variant | MODIFIER |
| 9 | 71568398 | AX-170368178 | 1 | G | A | 0.196 | -1.11E-01 | 1.31E-01 | 2.78E+00 | 2.83E+00 | 3.91E-01 | no | Myzap | intron_variant | MODIFIER |
| 1 | 14904427 | AX-170506558 | 0 | A | G | 0.395 | 6.73E-02 | 7.88E-02 | 2.61E+00 | 2.66E+00 | 3.91E-01 | no | Trpa1 | intron_variant | MODIFIER |
| 7 | 3713237 | AX-170028531 | 2 | G | A | 0.2 | 6.46E-02 | 7.67E-02 | 2.73E+00 | 2.87E+00 | 3.91E-01 | no | Pirb | intron_variant | MODIFIER |
| 1 | 189711364 | AX-97852508 | 0 | T | C | 0.263 | 8.04E-02 | 9.05E-02 | 2.40E+00 | 2.50E+00 | 3.91E-01 | no | | intergenic_variant | MODIFIER |
| 18 | 85277488 | AX-97818741 | 1 | G | A | 0.161 | -8.97E-02 | 1.03E-01 | 2.48E+00 | 2.57E+00 | 3.91E-01 | no | | intergenic_variant | MODIFIER |
| 4 | 128760759 | AX-170267711 | 0 | T | A | 0.167 | 8.52E-02 | 9.91E-02 | 2.55E+00 | 2.62E+00 | 3.91E-01 | no | A3galt2 | intron_variant | MODIFIER |
| 5 | 84150078 | AX-97786427 | 0 | C | T | 0.439 | 9.16E-02 | 1.08E-01 | 2.73E+00 | 2.79E+00 | 3.91E-01 | no | Epha5 | intron_variant | MODIFIER |
| 1 | 169831658 | AX-170132644 | 0 | G | A | 0.219 | 1.14E-01 | 1.34E-01 | 2.67E+00 | 2.72E+00 | 3.91E-01 | no | | regulatory_region_variant | MODIFIER |
| 4 | 128540710 | AX-170269594 | 1 | C | T | 0.161 | 9.65E-02 | 1.13E-01 | 2.62E+00 | 2.68E+00 | 3.91E-01 | no | Csmd2 | intron_variant | MODIFIER |
| 2 | 169010847 | AX-97828510 | 0 | G | T | 0.167 | 1.52E-01 | 1.80E-01 | 2.66E+00 | 2.74E+00 | 3.91E-01 | Yes | | regulatory_region_variant | MODIFIER |
| 2 | 169065457 | AX-170340784 | 0 | C | T | 0.167 | 1.52E-01 | 1.80E-01 | 2.66E+00 | 2.74E+00 | 3.91E-01 | Yes | Gm26469 | downstream_gene_variant | MODIFIER |
| 4 | 140873166 | AX-170024126 | 0 | T | C | 0.167 | 9.03E-02 | 1.06E-01 | 2.63E+00 | 2.70E+00 | 3.91E-01 | no | 4930515B02Rik | intron_variant,non_coding_transcript_variant | MODIFIER |
| 7 | 100456693 | AX-97796267 | 0 | A | C | 0.36 | -8.81E-02 | 1.04E-01 | 2.78E+00 | 2.86E+00 | 3.91E-01 | no | C2cd3 | intron_variant | MODIFIER |
| 7 | 123905959 | AX-170518052 | 1 | G | A | 0.411 | 7.12E-02 | 8.40E-02 | 2.79E+00 | 2.88E+00 | 3.92E-01 | no | | intergenic_variant | MODIFIER |
| 16 | 70745811 | AX-170013348 | 1 | C | T | 0.509 | 6.43E-02 | 7.40E-02 | 2.49E+00 | 2.57E+00 | 3.92E-01 | no | Gm49665 | downstream_gene_variant | MODIFIER |
| 1 | 122937893 | AX-170261955 | 1 | C | T | 0.554 | 7.10E-02 | 8.27E-02 | 2.56E+00 | 2.62E+00 | 3.92E-01 | no | | intergenic_variant | MODIFIER |
| 6 | 136019426 | AX-170445039 | 1 | T | C | 0.571 | 7.61E-02 | 8.69E-02 | 2.46E+00 | 2.53E+00 | 3.92E-01 | no | Grin2b | intron_variant | MODIFIER |
| 4 | 134568933 | AX-170334662 | 0 | C | T | 0.167 | 1.00E-01 | 1.17E-01 | 2.57E+00 | 2.62E+00 | 3.92E-01 | no | Man1c1 | synonymous_variant | LOW |
| 1 | 99081660 | AX-170001182 | 1 | T | A | 0.152 | 1.66E-01 | 1.96E-01 | 2.70E+00 | 2.78E+00 | 3.92E-01 | Yes | | intergenic_variant | MODIFIER |
| 11 | 33491398 | AX-170237565 | 0 | A | G | 0.088 | -1.33E-01 | 1.57E-01 | 2.63E+00 | 2.74E+00 | 3.92E-01 | no | Ranbp17 | splice_region_variant,intron_variant | LOW |
| 7 | 73337030 | AX-170226695 | 0 | A | C | 0.377 | -8.41E-02 | 9.97E-02 | 2.73E+00 | 2.83E+00 | 3.92E-01 | no | A730056A06Rik | intron_variant,non_coding_transcript_variant | MODIFIER |
| 1 | 191559595 | AX-170167796 | 0 | G | A | 0.263 | 8.49E-02 | 9.51E-02 | 2.38E+00 | 2.46E+00 | 3.92E-01 | no | Dtl | intron_variant | MODIFIER |
| 20 | 50423972 | AX-170344392 | 1 | A | G | 0.241 | 4.65E-02 | 5.50E-02 | 2.61E+00 | 2.74E+00 | 3.93E-01 | no | Olfr1324 | downstream_gene_variant | MODIFIER |
| 7 | 4556686 | AX-170293206 | 0 | T | A | 0.175 | 6.65E-02 | 7.89E-02 | 2.81E+00 | 2.96E+00 | 3.93E-01 | no | Ptprh | intron_variant | MODIFIER |
| 7 | 4556992 | AX-170124391 | 0 | G | C | 0.175 | 6.65E-02 | 7.89E-02 | 2.81E+00 | 2.96E+00 | 3.93E-01 | no | Ptprh | intron_variant | MODIFIER |
| 7 | 4560270 | AX-170095854 | 0 | G | A | 0.175 | 6.65E-02 | 7.89E-02 | 2.81E+00 | 2.96E+00 | 3.93E-01 | no | Ptprh | intron_variant | MODIFIER |
| 7 | 4708598 | AX-170391304 | 0 | T | C | 0.175 | 6.65E-02 | 7.89E-02 | 2.81E+00 | 2.96E+00 | 3.93E-01 | no | Brsk1 | intron_variant | MODIFIER |
| 7 | 4736312 | AX-170521931 | 0 | C | T | 0.175 | 6.65E-02 | 7.89E-02 | 2.81E+00 | 2.96E+00 | 3.93E-01 | no | Kmt5c | upstream_gene_variant | MODIFIER |
| 7 | 4738813 | AX-170344400 | 0 | C | T | 0.175 | 6.65E-02 | 7.89E-02 | 2.81E+00 | 2.96E+00 | 3.93E-01 | no | Kmt5c | upstream_gene_variant | MODIFIER |
| 7 | 4743585 | AX-97853770 | 0 | A | G | 0.175 | 6.65E-02 | 7.89E-02 | 2.81E+00 | 2.96E+00 | 3.93E-01 | no | Kmt5c | intron_variant | MODIFIER |
| 7 | 4782747 | AX-170353923 | 0 | C | T | 0.175 | 6.65E-02 | 7.89E-02 | 2.81E+00 | 2.96E+00 | 3.93E-01 | no | Tmem190 | upstream_gene_variant | MODIFIER |
| 7 | 4784882 | AX-170217364 | 0 | T | C | 0.175 | 6.65E-02 | 7.89E-02 | 2.81E+00 | 2.96E+00 | 3.93E-01 | no | Tmem190 | downstream_gene_variant | MODIFIER |
| 7 | 4796049 | AX-170154431 | 0 | A | G | 0.175 | 6.65E-02 | 7.89E-02 | 2.81E+00 | 2.96E+00 | 3.93E-01 | no | Rpl28 | downstream_gene_variant | MODIFIER |
| 7 | 4822450 | AX-170533202 | 0 | G | A | 0.175 | 6.65E-02 | 7.89E-02 | 2.81E+00 | 2.96E+00 | 3.93E-01 | no | Shisa7 | downstream_gene_variant | MODIFIER |
| 7 | 4874185 | AX-170322931 | 0 | T | C | 0.175 | 6.65E-02 | 7.89E-02 | 2.81E+00 | 2.96E+00 | 3.93E-01 | no | Isoc2a | upstream_gene_variant | MODIFIER |
| 7 | 4874247 | AX-170140218 | 0 | C | T | 0.175 | 6.65E-02 | 7.89E-02 | 2.81E+00 | 2.96E+00 | 3.93E-01 | no | Isoc2a | upstream_gene_variant | MODIFIER |
| 7 | 4898707 | AX-170318059 | 0 | A | C | 0.175 | 6.65E-02 | 7.89E-02 | 2.81E+00 | 2.96E+00 | 3.93E-01 | no | Isoc2a | downstream_gene_variant | MODIFIER |
| 7 | 4898777 | AX-170380012 | 0 | T | G | 0.175 | 6.65E-02 | 7.89E-02 | 2.81E+00 | 2.96E+00 | 3.93E-01 | no | Isoc2a | downstream_gene_variant | MODIFIER |
| 7 | 4948555 | AX-170460791 | 0 | C | A | 0.175 | 6.65E-02 | 7.89E-02 | 2.81E+00 | 2.96E+00 | 3.93E-01 | no | Ssc5d | downstream_gene_variant | MODIFIER |
| 7 | 4952108 | AX-170285406 | 0 | G | C | 0.175 | 6.65E-02 | 7.89E-02 | 2.81E+00 | 2.96E+00 | 3.93E-01 | no | Sbk2 | downstream_gene_variant | MODIFIER |
| 7 | 4953986 | AX-170514785 | 0 | C | T | 0.175 | 6.65E-02 | 7.89E-02 | 2.81E+00 | 2.96E+00 | 3.93E-01 | no | Sbk2 | downstream_gene_variant | MODIFIER |
| 7 | 4955874 | AX-170186475 | 0 | A | G | 0.175 | 6.65E-02 | 7.89E-02 | 2.81E+00 | 2.96E+00 | 3.93E-01 | no | Sbk2 | downstream_gene_variant | MODIFIER |

| | | | | | | | | | | | | | | | |
|---|---|---|---|---|---|---|---|---|---|---|---|---|---|---|---|
| 7 | 4958074 | AX-170529515 | 0 | A | C | 0.175 | 6.65E-02 | 7.89E-02 | 2.81E+00 | 2.96E+00 | 3.93E-01 | no | Sbk2 | intron_variant | MODIFIER |
| 7 | 4965890 | AX-170076658 | 0 | G | A | 0.175 | 6.65E-02 | 7.89E-02 | 2.81E+00 | 2.96E+00 | 3.93E-01 | no | Sbk2 | upstream_gene_variant | MODIFIER |
| 7 | 4980256 | AX-170448933 | 0 | A | T | 0.175 | 6.65E-02 | 7.89E-02 | 2.81E+00 | 2.96E+00 | 3.93E-01 | no | Zfp579 | downstream_gene_variant | MODIFIER |
| 7 | 4980331 | AX-170371085 | 0 | G | A | 0.175 | 6.65E-02 | 7.89E-02 | 2.81E+00 | 2.96E+00 | 3.93E-01 | no | Zfp579 | downstream_gene_variant | MODIFIER |
| 7 | 5036123 | AX-170337056 | 0 | C | A | 0.175 | 6.65E-02 | 7.89E-02 | 2.81E+00 | 2.96E+00 | 3.93E-01 | no | Zfp784 | synonymous_variant | LOW |
| 7 | 5037831 | AX-170414874 | 0 | C | T | 0.175 | 6.65E-02 | 7.89E-02 | 2.81E+00 | 2.96E+00 | 3.93E-01 | no | Zfp784 | intron_variant | MODIFIER |
| 7 | 5042066 | AX-170492051 | 0 | A | G | 0.175 | 6.65E-02 | 7.89E-02 | 2.81E+00 | 2.96E+00 | 3.93E-01 | no | Zfp784 | upstream_gene_variant | MODIFIER |
| 7 | 5089499 | AX-170500771 | 0 | C | T | 0.175 | 6.65E-02 | 7.89E-02 | 2.81E+00 | 2.96E+00 | 3.93E-01 | no | Epn1 | intron_variant | MODIFIER |
| 7 | 5098684 | AX-170028546 | 0 | C | T | 0.175 | 6.65E-02 | 7.89E-02 | 2.81E+00 | 2.96E+00 | 3.93E-01 | no | Epn1 | downstream_gene_variant | MODIFIER |
| 7 | 5101199 | AX-170483487 | 0 | A | G | 0.175 | 6.65E-02 | 7.89E-02 | 2.81E+00 | 2.96E+00 | 3.93E-01 | no | Epn1 | downstream_gene_variant | MODIFIER |
| 7 | 5102761 | AX-170352187 | 0 | A | G | 0.175 | 6.65E-02 | 7.89E-02 | 2.81E+00 | 2.96E+00 | 3.93E-01 | no | Epn1 | downstream_gene_variant | MODIFIER |
| 7 | 5105662 | AX-170076662 | 0 | T | C | 0.175 | 6.65E-02 | 7.89E-02 | 2.81E+00 | 2.96E+00 | 3.93E-01 | no | Rfpl4 | downstream_gene_variant | MODIFIER |
| 7 | 5362021 | AX-170227610 | 0 | A | G | 0.175 | 6.65E-02 | 7.89E-02 | 2.81E+00 | 2.96E+00 | 3.93E-01 | no | Gm5065 | downstream_gene_variant | MODIFIER |
| 7 | 6011490 | AX-170460675 | 0 | G | A | 0.175 | 6.65E-02 | 7.89E-02 | 2.81E+00 | 2.96E+00 | 3.93E-01 | no | Vmn1r65 | upstream_gene_variant | MODIFIER |
| 1 | 135985522 | AX-170164531 | 0 | C | T | 0.219 | 9.72E-02 | 1.15E-01 | 2.68E+00 | 2.77E+00 | 3.93E-01 | no | Igfn1 | upstream_gene_variant | MODIFIER |
| 1 | 169583290 | AX-170398067 | 1 | G | A | 0.214 | 1.18E-01 | 1.39E-01 | 2.67E+00 | 2.71E+00 | 3.93E-01 | no | | intergenic_variant | MODIFIER |
| 1 | 14825850 | AX-170496432 | 0 | G | A | 0.404 | 6.57E-02 | 7.68E-02 | 2.57E+00 | 2.62E+00 | 3.93E-01 | no | | intergenic_variant | MODIFIER |
| 1 | 172894774 | AX-170221235 | 1 | C | T | 0.232 | 1.06E-01 | 1.23E-01 | 2.53E+00 | 2.59E+00 | 3.93E-01 | no | Apcs | intron_variant | MODIFIER |
| 9 | 52669516 | AX-170259753 | 0 | A | G | 0.219 | -9.43E-02 | 1.12E-01 | 2.77E+00 | 2.85E+00 | 3.93E-01 | no | AI593442 | downstream_gene_variant | MODIFIER |
| 7 | 95470399 | AX-170029564 | 0 | C | A | 0.14 | 1.46E-01 | 1.73E-01 | 2.74E+00 | 2.86E+00 | 3.93E-01 | Yes | Gm44603 | upstream_gene_variant | MODIFIER |
| 1 | 101233050 | AX-170329888 | 0 | T | C | 0.158 | 1.43E-01 | 1.68E-01 | 2.57E+00 | 2.67E+00 | 3.93E-01 | no | | intergenic_variant | MODIFIER |
| 10 | 19085398 | AX-97852875 | 0 | T | G | 0.202 | 8.28E-02 | 9.78E-02 | 2.87E+00 | 2.98E+00 | 3.93E-01 | no | 1700124M09Rik | downstream_gene_variant | MODIFIER |
| 7 | 3124122 | AX-170381708 | 0 | A | C | 0.193 | 6.43E-02 | 7.67E-02 | 2.73E+00 | 2.87E+00 | 3.93E-01 | no | Speer9-ps1 | upstream_gene_variant | MODIFIER |
| 7 | 3130432 | AX-170190105 | 0 | G | A | 0.193 | 6.43E-02 | 7.67E-02 | 2.73E+00 | 2.87E+00 | 3.93E-01 | no | Speer9-ps1 | intron_variant,non_coding_transcript_variant | MODIFIER |
| 7 | 3696957 | AX-170028529 | 0 | G | C | 0.193 | 6.43E-02 | 7.67E-02 | 2.73E+00 | 2.87E+00 | 3.93E-01 | no | Tsen34 | intron_variant | MODIFIER |
| 7 | 3697099 | AX-170028526 | 0 | T | C | 0.193 | 6.43E-02 | 7.67E-02 | 2.73E+00 | 2.87E+00 | 3.93E-01 | no | Tsen34 | intron_variant | MODIFIER |
| 7 | 3939315 | AX-170028530 | 0 | A | G | 0.193 | 6.43E-02 | 7.67E-02 | 2.73E+00 | 2.87E+00 | 3.93E-01 | no | Gm44580 | upstream_gene_variant | MODIFIER |
| 7 | 3981027 | AX-170028525 | 0 | A | C | 0.193 | 6.43E-02 | 7.67E-02 | 2.73E+00 | 2.87E+00 | 3.93E-01 | no | Gm7363 | downstream_gene_variant | MODIFIER |
| 7 | 4097342 | AX-170534866 | 0 | G | C | 0.193 | 6.43E-02 | 7.67E-02 | 2.73E+00 | 2.87E+00 | 3.93E-01 | no | Gm23741 | upstream_gene_variant | MODIFIER |
| 7 | 4098029 | AX-170076640 | 0 | G | A | 0.193 | 6.43E-02 | 7.67E-02 | 2.73E+00 | 2.87E+00 | 3.93E-01 | no | Gm23741 | upstream_gene_variant | MODIFIER |
| 7 | 4098057 | AX-170183976 | 0 | C | T | 0.193 | 6.43E-02 | 7.67E-02 | 2.73E+00 | 2.87E+00 | 3.93E-01 | no | Gm23741 | upstream_gene_variant | MODIFIER |
| 10 | 68888136 | AX-170510209 | 0 | A | T | 0.421 | -6.71E-02 | 7.95E-02 | 2.69E+00 | 2.77E+00 | 3.93E-01 | no | | intergenic_variant | MODIFIER |
| 3 | 37065405 | AX-97796482 | 0 | G | A | 0.211 | 7.51E-02 | 8.92E-02 | 2.77E+00 | 2.88E+00 | 3.93E-01 | no | Adad1 | intron_variant | MODIFIER |
| 4 | 138951829 | AX-170191009 | 0 | G | A | 0.167 | 9.39E-02 | 1.11E-01 | 2.61E+00 | 2.67E+00 | 3.94E-01 | no | | intergenic_variant | MODIFIER |
| 4 | 139119338 | AX-170278481 | 0 | G | A | 0.167 | 9.39E-02 | 1.11E-01 | 2.61E+00 | 2.67E+00 | 3.94E-01 | no | Minos1 | intron_variant | MODIFIER |
| 5 | 36626600 | AX-97819647 | 0 | C | G | 0.351 | 9.50E-02 | 1.13E-01 | 2.79E+00 | 2.85E+00 | 3.94E-01 | no | D5Ertd579e | intron_variant | MODIFIER |
| 1 | 53145810 | AX-170317394 | 1 | A | G | 0.098 | -1.49E-01 | 1.77E-01 | 2.75E+00 | 2.85E+00 | 3.94E-01 | no | | intergenic_variant | MODIFIER |
| 20 | 68043193 | AX-97850645 | 1 | A | G | 0.25 | 5.23E-02 | 6.21E-02 | 2.67E+00 | 2.76E+00 | 3.94E-01 | no | | intergenic_variant | MODIFIER |
| 7 | 36435246 | AX-170077066 | 0 | C | A | 0.342 | 7.85E-02 | 9.21E-02 | 2.57E+00 | 2.64E+00 | 3.94E-01 | no | Gm25247 | downstream_gene_variant | MODIFIER |
| 18 | 58862938 | AX-170516843 | 0 | C | T | 0.395 | 8.25E-02 | 9.41E-02 | 2.44E+00 | 2.49E+00 | 3.94E-01 | no | Adamts19 | intron_variant | MODIFIER |
| 2 | 50905238 | AX-97837051 | 1 | G | T | 0.045 | 1.70E-01 | 2.02E-01 | 2.85E+00 | 2.96E+00 | 3.95E-01 | Yes | Gm13498 | upstream_gene_variant | MODIFIER |
| 4 | 71645545 | AX-170095007 | 0 | G | A | 0.307 | 9.66E-02 | 1.15E-01 | 2.71E+00 | 2.76E+00 | 3.95E-01 | no | | intergenic_variant | MODIFIER |
| 9 | 52514798 | AX-97804872 | 0 | A | C | 0.228 | -8.57E-02 | 1.02E-01 | 2.76E+00 | 2.87E+00 | 3.95E-01 | no | | intergenic_variant | MODIFIER |
| 7 | 97959136 | AX-170277278 | 0 | A | G | 0.132 | 1.47E-01 | 1.75E-01 | 2.74E+00 | 2.87E+00 | 3.95E-01 | Yes | Gdpd4 | intron_variant | MODIFIER |
| 18 | 69488418 | AX-170402035 | 1 | T | G | 0.161 | -1.05E-01 | 1.23E-01 | 2.59E+00 | 2.64E+00 | 3.95E-01 | no | Tcf4 | intron_variant | MODIFIER |
| 16 | 76936096 | AX-170276062 | 0 | T | C | 0.465 | 6.57E-02 | 7.69E-02 | 2.54E+00 | 2.61E+00 | 3.95E-01 | no | 1700041M19Rik | intron_variant,non_coding_transcript_variant | MODIFIER |
| 16 | 77620745 | AX-170013434 | 0 | C | T | 0.465 | 6.57E-02 | 7.69E-02 | 2.54E+00 | 2.61E+00 | 3.95E-01 | no | Mir99ahg | intron_variant,non_coding_transcript_variant | MODIFIER |
| 16 | 77626522 | AX-170013432 | 0 | G | A | 0.465 | 6.57E-02 | 7.69E-02 | 2.54E+00 | 2.61E+00 | 3.95E-01 | no | Mir99ahg | intron_variant,non_coding_transcript_variant | MODIFIER |
| 16 | 77631174 | AX-170150477 | 0 | C | T | 0.465 | 6.57E-02 | 7.69E-02 | 2.54E+00 | 2.61E+00 | 3.95E-01 | no | Mir99ahg | intron_variant,non_coding_transcript_variant | MODIFIER |
| 16 | 77646231 | AX-97785199 | 0 | T | G | 0.465 | 6.57E-02 | 7.69E-02 | 2.54E+00 | 2.61E+00 | 3.95E-01 | no | Mir125b-2 | upstream_gene_variant | MODIFIER |
| 16 | 77663092 | AX-170443824 | 0 | T | G | 0.465 | 6.57E-02 | 7.69E-02 | 2.54E+00 | 2.61E+00 | 3.95E-01 | no | Mir99ahg | intron_variant,non_coding_transcript_variant | MODIFIER |
| 16 | 77695079 | AX-170198723 | 0 | A | G | 0.465 | 6.57E-02 | 7.69E-02 | 2.54E+00 | 2.61E+00 | 3.95E-01 | no | Mir99ahg | downstream_gene_variant | MODIFIER |
| 16 | 77738835 | AX-170220885 | 0 | T | C | 0.465 | 6.57E-02 | 7.69E-02 | 2.54E+00 | 2.61E+00 | 3.95E-01 | no | Mir99ahg | intron_variant,non_coding_transcript_variant | MODIFIER |
| 16 | 77739452 | AX-97811597 | 0 | A | G | 0.465 | 6.57E-02 | 7.69E-02 | 2.54E+00 | 2.61E+00 | 3.95E-01 | no | Mir99ahg | intron_variant,non_coding_transcript_variant | MODIFIER |

| | | | | | | | | | | | | | | | |
|---|---|---|---|---|---|---|---|---|---|---|---|---|---|---|---|
| 16 | 77749862 | AX-170057875 | 0 | T | C | 0.465 | 6.57E-02 | 7.69E-02 | 2.54E+00 | 2.61E+00 | 3.95E-01 | no | Mir99ahg | intron_variant,non_coding_transcript_variant | MODIFIER |
| 16 | 77773830 | AX-168419966 | 0 | C | T | 0.465 | 6.57E-02 | 7.69E-02 | 2.54E+00 | 2.61E+00 | 3.95E-01 | no | Mir99ahg | downstream_gene_variant | MODIFIER |
| 16 | 77774195 | AX-170104041 | 0 | C | T | 0.465 | 6.57E-02 | 7.69E-02 | 2.54E+00 | 2.61E+00 | 3.95E-01 | no | Mir99ahg | downstream_gene_variant | MODIFIER |
| 16 | 77795529 | AX-97842296 | 0 | C | T | 0.465 | 6.57E-02 | 7.69E-02 | 2.54E+00 | 2.61E+00 | 3.95E-01 | no | | intergenic_variant | MODIFIER |
| 16 | 77821575 | AX-170108202 | 0 | T | C | 0.465 | 6.57E-02 | 7.69E-02 | 2.54E+00 | 2.61E+00 | 3.95E-01 | no | | intergenic_variant | MODIFIER |
| 16 | 77823818 | AX-170330246 | 0 | A | G | 0.465 | 6.57E-02 | 7.69E-02 | 2.54E+00 | 2.61E+00 | 3.95E-01 | no | | intergenic_variant | MODIFIER |
| 16 | 77824526 | AX-170140850 | 0 | T | G | 0.465 | 6.57E-02 | 7.69E-02 | 2.54E+00 | 2.61E+00 | 3.95E-01 | no | | intergenic_variant | MODIFIER |
| 16 | 77876056 | AX-170088070 | 0 | C | G | 0.465 | 6.57E-02 | 7.69E-02 | 2.54E+00 | 2.61E+00 | 3.95E-01 | no | | intergenic_variant | MODIFIER |
| 16 | 77896744 | AX-170227669 | 0 | T | C | 0.465 | 6.57E-02 | 7.69E-02 | 2.54E+00 | 2.61E+00 | 3.95E-01 | no | | intergenic_variant | MODIFIER |
| 16 | 77898232 | AX-97853745 | 0 | G | A | 0.465 | 6.57E-02 | 7.69E-02 | 2.54E+00 | 2.61E+00 | 3.95E-01 | no | | intergenic_variant | MODIFIER |
| 16 | 77930434 | AX-170291511 | 0 | T | C | 0.465 | 6.57E-02 | 7.69E-02 | 2.54E+00 | 2.61E+00 | 3.95E-01 | no | | intergenic_variant | MODIFIER |
| 16 | 77932232 | AX-170202082 | 0 | A | G | 0.465 | 6.57E-02 | 7.69E-02 | 2.54E+00 | 2.61E+00 | 3.95E-01 | no | | intergenic_variant | MODIFIER |
| 16 | 77972753 | AX-97843267 | 0 | A | G | 0.465 | 6.57E-02 | 7.69E-02 | 2.54E+00 | 2.61E+00 | 3.95E-01 | no | | intergenic_variant | MODIFIER |
| 16 | 77987578 | AX-170525163 | 0 | T | C | 0.465 | 6.57E-02 | 7.69E-02 | 2.54E+00 | 2.61E+00 | 3.95E-01 | no | | regulatory_region_variant | MODIFIER |
| 16 | 77990298 | AX-170296592 | 0 | C | T | 0.465 | 6.57E-02 | 7.69E-02 | 2.54E+00 | 2.61E+00 | 3.95E-01 | no | Gm49589 | upstream_gene_variant | MODIFIER |
| 16 | 77992622 | AX-170335738 | 0 | G | A | 0.465 | 6.57E-02 | 7.69E-02 | 2.54E+00 | 2.61E+00 | 3.95E-01 | no | Gm49589 | upstream_gene_variant | MODIFIER |
| 16 | 78071664 | AX-170536139 | 0 | T | G | 0.465 | 6.57E-02 | 7.69E-02 | 2.54E+00 | 2.61E+00 | 3.95E-01 | no | | intergenic_variant | MODIFIER |
| 2 | 169530414 | AX-170065601 | 1 | A | G | 0.17 | 1.52E-01 | 1.80E-01 | 2.61E+00 | 2.70E+00 | 3.95E-01 | Yes | | regulatory_region_variant | MODIFIER |
| 1 | 132211653 | AX-170394302 | 1 | A | T | 0.527 | 7.15E-02 | 8.48E-02 | 2.63E+00 | 2.70E+00 | 3.95E-01 | no | Lemd1 | intron_variant | MODIFIER |
| 15 | 82396676 | AX-170264850 | 0 | A | C | 0.561 | 7.16E-02 | 8.34E-02 | 2.51E+00 | 2.58E+00 | 3.95E-01 | no | Cyp2d11 | upstream_gene_variant | MODIFIER |
| 12 | 41328983 | AX-170136225 | 0 | G | A | 0.035 | 1.22E-01 | 1.45E-01 | 2.57E+00 | 2.76E+00 | 3.95E-01 | no | Immp2l | intron_variant | MODIFIER |
| 1 | 128543327 | AX-170316526 | 2 | A | G | 0.536 | 6.97E-02 | 8.25E-02 | 2.62E+00 | 2.70E+00 | 3.96E-01 | no | | intergenic_variant | MODIFIER |
| 3 | 154523213 | AX-97815980 | 0 | T | C | 0.351 | 5.93E-02 | 7.06E-02 | 2.80E+00 | 2.90E+00 | 3.96E-01 | no | | intergenic_variant | MODIFIER |
| 16 | 77634923 | AX-170282475 | 1 | A | C | 0.455 | 6.51E-02 | 7.62E-02 | 2.54E+00 | 2.60E+00 | 3.96E-01 | no | Mir99ahg | intron_variant,non_coding_transcript_variant | MODIFIER |
| 4 | 134052772 | AX-170261761 | 0 | C | T | 0.184 | 8.89E-02 | 1.04E-01 | 2.56E+00 | 2.62E+00 | 3.96E-01 | no | Zfp683 | upstream_gene_variant | MODIFIER |
| 6 | 134343979 | AX-170132516 | 1 | T | G | 0.562 | 7.35E-02 | 8.56E-02 | 2.51E+00 | 2.57E+00 | 3.96E-01 | no | 4933406J09Rik | intron_variant,non_coding_transcript_variant | MODIFIER |
| 6 | 134397647 | AX-170207480 | 1 | A | G | 0.562 | 7.35E-02 | 8.56E-02 | 2.51E+00 | 2.57E+00 | 3.96E-01 | no | Bcl2l14 | intron_variant | MODIFIER |
| 18 | 84942394 | AX-170480784 | 1 | G | A | 0.161 | -9.37E-02 | 1.08E-01 | 2.47E+00 | 2.55E+00 | 3.96E-01 | no | Timm21 | downstream_gene_variant | MODIFIER |
| 15 | 101872103 | AX-170482969 | 0 | T | G | 0.132 | -1.24E-01 | 1.47E-01 | 2.89E+00 | 2.99E+00 | 3.96E-01 | no | Krt77 | upstream_gene_variant | MODIFIER |
| 4 | 116565159 | AX-97851127 | 1 | T | C | 0.205 | 9.93E-02 | 1.17E-01 | 2.60E+00 | 2.67E+00 | 3.96E-01 | no | Gpbp1l1 | intron_variant | MODIFIER |
| 8 | 95248300 | AX-170389394 | 0 | G | A | 0.544 | 4.69E-02 | 5.56E-02 | 2.87E+00 | 2.97E+00 | 3.96E-01 | no | Cngb1 | intron_variant | MODIFIER |
| 8 | 95540965 | AX-170251728 | 0 | T | C | 0.544 | 4.69E-02 | 5.56E-02 | 2.87E+00 | 2.97E+00 | 3.96E-01 | no | Ccdc113 | intron_variant | MODIFIER |
| 13 | 24850542 | AX-170007938 | 1 | C | T | 0.17 | -1.06E-01 | 1.18E-01 | 2.35E+00 | 2.40E+00 | 3.96E-01 | no | D130043K22Rik | intron_variant | MODIFIER |
| 6 | 133613679 | AX-170028287 | 2 | A | G | 0.564 | 7.46E-02 | 8.68E-02 | 2.50E+00 | 2.56E+00 | 3.96E-01 | no | Gm34314 | downstream_gene_variant | MODIFIER |
| 4 | 73944477 | AX-170340091 | 1 | T | C | 0.455 | -7.05E-02 | 8.22E-02 | 2.51E+00 | 2.58E+00 | 3.96E-01 | no | 2310002L09Rik | intron_variant | MODIFIER |
| 6 | 130317433 | AX-170439928 | 1 | A | T | 0.554 | 7.12E-02 | 8.30E-02 | 2.51E+00 | 2.57E+00 | 3.96E-01 | no | Gm24072 | upstream_gene_variant | MODIFIER |
| 1 | 132474867 | AX-170243295 | 1 | G | A | 0.536 | 7.14E-02 | 8.49E-02 | 2.64E+00 | 2.71E+00 | 3.97E-01 | no | Rbbp5 | upstream_gene_variant | MODIFIER |
| 9 | 98548158 | AX-170109277 | 0 | T | C | 0.14 | -9.41E-02 | 1.12E-01 | 2.61E+00 | 2.68E+00 | 3.97E-01 | no | | intergenic_variant | MODIFIER |
| 9 | 98557833 | AX-170355933 | 0 | T | C | 0.14 | -9.41E-02 | 1.12E-01 | 2.61E+00 | 2.68E+00 | 3.97E-01 | no | 4930579K19Rik | downstream_gene_variant | MODIFIER |
| 9 | 98561871 | AX-170478726 | 0 | G | A | 0.14 | -9.41E-02 | 1.12E-01 | 2.61E+00 | 2.68E+00 | 3.97E-01 | no | Copb2 | upstream_gene_variant | MODIFIER |
| 9 | 98562263 | AX-97791465 | 0 | A | G | 0.14 | -9.41E-02 | 1.12E-01 | 2.61E+00 | 2.68E+00 | 3.97E-01 | no | Copb2 | upstream_gene_variant | MODIFIER |
| 9 | 98570651 | AX-170149020 | 0 | G | A | 0.14 | -9.41E-02 | 1.12E-01 | 2.61E+00 | 2.68E+00 | 3.97E-01 | no | Copb2 | synonymous_variant | LOW |
| 9 | 98571323 | AX-170263967 | 0 | G | A | 0.14 | -9.41E-02 | 1.12E-01 | 2.61E+00 | 2.68E+00 | 3.97E-01 | no | Copb2 | intron_variant | MODIFIER |
| 9 | 98572536 | AX-170414003 | 0 | T | C | 0.14 | -9.41E-02 | 1.12E-01 | 2.61E+00 | 2.68E+00 | 3.97E-01 | no | Copb2 | intron_variant | MODIFIER |
| 9 | 98611492 | AX-170386883 | 0 | A | G | 0.14 | -9.41E-02 | 1.12E-01 | 2.61E+00 | 2.68E+00 | 3.97E-01 | no | | intergenic_variant | MODIFIER |
| 9 | 98611519 | AX-170386181 | 0 | C | T | 0.14 | -9.41E-02 | 1.12E-01 | 2.61E+00 | 2.68E+00 | 3.97E-01 | no | | intergenic_variant | MODIFIER |
| 9 | 98640793 | AX-170243061 | 0 | G | T | 0.14 | -9.41E-02 | 1.12E-01 | 2.61E+00 | 2.68E+00 | 3.97E-01 | no | | intergenic_variant | MODIFIER |
| 9 | 98645440 | AX-170262697 | 0 | T | C | 0.14 | -9.41E-02 | 1.12E-01 | 2.61E+00 | 2.68E+00 | 3.97E-01 | no | | intergenic_variant | MODIFIER |
| 9 | 98656014 | AX-170083209 | 0 | T | G | 0.14 | -9.41E-02 | 1.12E-01 | 2.61E+00 | 2.68E+00 | 3.97E-01 | no | | intergenic_variant | MODIFIER |
| 9 | 98657648 | AX-170083207 | 0 | T | G | 0.14 | -9.41E-02 | 1.12E-01 | 2.61E+00 | 2.68E+00 | 3.97E-01 | no | | intergenic_variant | MODIFIER |
| 9 | 98713870 | AX-170514774 | 0 | A | T | 0.14 | -9.41E-02 | 1.12E-01 | 2.61E+00 | 2.68E+00 | 3.97E-01 | no | | intergenic_variant | MODIFIER |
| 9 | 98736366 | AX-170303330 | 0 | C | T | 0.14 | -9.41E-02 | 1.12E-01 | 2.61E+00 | 2.68E+00 | 3.97E-01 | no | | intergenic_variant | MODIFIER |
| 9 | 100948644 | AX-97841797 | 0 | G | A | 0.14 | -9.41E-02 | 1.12E-01 | 2.61E+00 | 2.68E+00 | 3.97E-01 | no | Stag1 | intron_variant | MODIFIER |
| 9 | 100950781 | AX-170149871 | 0 | G | A | 0.14 | -9.41E-02 | 1.12E-01 | 2.61E+00 | 2.68E+00 | 3.97E-01 | no | Stag1 | intron_variant | MODIFIER |

| | | | | | | | | | | | | | | | |
|---|---|---|---|---|---|---|---|---|---|---|---|---|---|---|---|
| 9 | 101002196 | AX-170458573 | 0 | A | G | 0.14 | -9.41E-02 | 1.12E-01 | 2.61E+00 | 2.68E+00 | 3.97E-01 | no | Pccb | intron_variant | MODIFIER |
| 9 | 101192301 | AX-170302745 | 0 | A | G | 0.14 | -9.41E-02 | 1.12E-01 | 2.61E+00 | 2.68E+00 | 3.97E-01 | no | Ppp2r3a | intron_variant | MODIFIER |
| 9 | 101192651 | AX-97849444 | 0 | G | A | 0.14 | -9.41E-02 | 1.12E-01 | 2.61E+00 | 2.68E+00 | 3.97E-01 | no | Ppp2r3a | intron_variant | MODIFIER |
| 9 | 101214009 | AX-97781067 | 0 | T | C | 0.14 | -9.41E-02 | 1.12E-01 | 2.61E+00 | 2.68E+00 | 3.97E-01 | no | Ppp2r3a | intron_variant | MODIFIER |
| 9 | 101232059 | AX-170159898 | 0 | C | T | 0.14 | -9.41E-02 | 1.12E-01 | 2.61E+00 | 2.68E+00 | 3.97E-01 | no | Ppp2r3a | intron_variant | MODIFIER |
| 9 | 101256439 | AX-170090396 | 0 | T | C | 0.14 | -9.41E-02 | 1.12E-01 | 2.61E+00 | 2.68E+00 | 3.97E-01 | no | Ppp2r3a | upstream_gene_variant | MODIFIER |
| 9 | 101281482 | AX-170280500 | 0 | T | C | 0.14 | -9.41E-02 | 1.12E-01 | 2.61E+00 | 2.68E+00 | 3.97E-01 | no | Gm3121 | non_coding_transcript_exon_variant | MODIFIER |
| 9 | 101281605 | AX-170362519 | 0 | C | T | 0.14 | -9.41E-02 | 1.12E-01 | 2.61E+00 | 2.68E+00 | 3.97E-01 | no | Gm3121 | non_coding_transcript_exon_variant | MODIFIER |
| 9 | 101282314 | AX-170256791 | 0 | T | C | 0.14 | -9.41E-02 | 1.12E-01 | 2.61E+00 | 2.68E+00 | 3.97E-01 | no | Gm3121 | upstream_gene_variant | MODIFIER |
| 9 | 101287191 | AX-170502917 | 0 | T | C | 0.14 | -9.41E-02 | 1.12E-01 | 2.61E+00 | 2.68E+00 | 3.97E-01 | no | | intergenic_variant | MODIFIER |
| 9 | 101312666 | AX-170083267 | 0 | C | T | 0.14 | -9.41E-02 | 1.12E-01 | 2.61E+00 | 2.68E+00 | 3.97E-01 | no | | intergenic_variant | MODIFIER |
| 9 | 101315351 | AX-170124371 | 0 | C | T | 0.14 | -9.41E-02 | 1.12E-01 | 2.61E+00 | 2.68E+00 | 3.97E-01 | no | | intergenic_variant | MODIFIER |
| 9 | 101315400 | AX-170241686 | 0 | C | T | 0.14 | -9.41E-02 | 1.12E-01 | 2.61E+00 | 2.68E+00 | 3.97E-01 | no | | intergenic_variant | MODIFIER |
| 9 | 101565027 | AX-170033146 | 0 | C | T | 0.14 | -9.41E-02 | 1.12E-01 | 2.61E+00 | 2.68E+00 | 3.97E-01 | no | | intergenic_variant | MODIFIER |
| 9 | 101589905 | AX-97788922 | 0 | C | T | 0.14 | -9.41E-02 | 1.12E-01 | 2.61E+00 | 2.68E+00 | 3.97E-01 | no | | intergenic_variant | MODIFIER |
| 9 | 101624930 | AX-170394688 | 0 | T | A | 0.14 | -9.41E-02 | 1.12E-01 | 2.61E+00 | 2.68E+00 | 3.97E-01 | no | | intergenic_variant | MODIFIER |
| 9 | 101675735 | AX-170350623 | 0 | A | G | 0.14 | -9.41E-02 | 1.12E-01 | 2.61E+00 | 2.68E+00 | 3.97E-01 | no | | intergenic_variant | MODIFIER |
| 9 | 101676422 | AX-170344831 | 0 | G | A | 0.14 | -9.41E-02 | 1.12E-01 | 2.61E+00 | 2.68E+00 | 3.97E-01 | no | | intergenic_variant | MODIFIER |
| 9 | 101816773 | AX-97822513 | 0 | C | T | 0.14 | -9.41E-02 | 1.12E-01 | 2.61E+00 | 2.68E+00 | 3.97E-01 | no | Gm29521 | intron_variant,non_coding_transcript_variant | MODIFIER |
| 9 | 101880427 | AX-170534305 | 0 | A | C | 0.14 | -9.41E-02 | 1.12E-01 | 2.61E+00 | 2.68E+00 | 3.97E-01 | no | 9630041A04Rik | intron_variant,non_coding_transcript_variant | MODIFIER |
| 9 | 101918374 | AX-170090398 | 0 | A | G | 0.14 | -9.41E-02 | 1.12E-01 | 2.61E+00 | 2.68E+00 | 3.97E-01 | no | Ephb1 | downstream_gene_variant | MODIFIER |
| 9 | 101956881 | AX-170279854 | 0 | A | C | 0.14 | -9.41E-02 | 1.12E-01 | 2.61E+00 | 2.68E+00 | 3.97E-01 | no | Ephb1 | intron_variant | MODIFIER |
| 9 | 101956910 | AX-170458956 | 0 | A | G | 0.14 | -9.41E-02 | 1.12E-01 | 2.61E+00 | 2.68E+00 | 3.97E-01 | no | Ephb1 | intron_variant | MODIFIER |
| 9 | 101963533 | AX-170342619 | 0 | T | C | 0.14 | -9.41E-02 | 1.12E-01 | 2.61E+00 | 2.68E+00 | 3.97E-01 | no | Ephb1 | intron_variant | MODIFIER |
| 9 | 101974675 | AX-170418336 | 0 | C | G | 0.14 | -9.41E-02 | 1.12E-01 | 2.61E+00 | 2.68E+00 | 3.97E-01 | no | Ephb1 | intron_variant | MODIFIER |
| 9 | 101978444 | AX-170134911 | 0 | A | C | 0.14 | -9.41E-02 | 1.12E-01 | 2.61E+00 | 2.68E+00 | 3.97E-01 | no | Ephb1 | intron_variant | MODIFIER |
| 9 | 101994555 | AX-170224280 | 0 | T | G | 0.14 | -9.41E-02 | 1.12E-01 | 2.61E+00 | 2.68E+00 | 3.97E-01 | no | Ephb1 | intron_variant | MODIFIER |
| 9 | 101994819 | AX-97822292 | 0 | G | A | 0.14 | -9.41E-02 | 1.12E-01 | 2.61E+00 | 2.68E+00 | 3.97E-01 | no | Ephb1 | intron_variant | MODIFIER |
| 9 | 101996726 | AX-170374710 | 0 | C | T | 0.14 | -9.41E-02 | 1.12E-01 | 2.61E+00 | 2.68E+00 | 3.97E-01 | no | Ephb1 | intron_variant | MODIFIER |
| 9 | 102002183 | AX-170507698 | 0 | T | C | 0.14 | -9.41E-02 | 1.12E-01 | 2.61E+00 | 2.68E+00 | 3.97E-01 | no | Ephb1 | intron_variant | MODIFIER |
| 9 | 102003115 | AX-170419996 | 0 | G | A | 0.14 | -9.41E-02 | 1.12E-01 | 2.61E+00 | 2.68E+00 | 3.97E-01 | no | Ephb1 | intron_variant | MODIFIER |
| 9 | 102025319 | AX-97805249 | 0 | C | T | 0.14 | -9.41E-02 | 1.12E-01 | 2.61E+00 | 2.68E+00 | 3.97E-01 | no | Ephb1 | intron_variant | MODIFIER |
| 9 | 102035527 | AX-170512583 | 0 | C | G | 0.14 | -9.41E-02 | 1.12E-01 | 2.61E+00 | 2.68E+00 | 3.97E-01 | no | Ephb1 | intron_variant | MODIFIER |
| 9 | 102039824 | AX-170033155 | 0 | A | G | 0.14 | -9.41E-02 | 1.12E-01 | 2.61E+00 | 2.68E+00 | 3.97E-01 | no | Ephb1 | intron_variant | MODIFIER |
| 9 | 102044986 | AX-170161951 | 0 | C | T | 0.14 | -9.41E-02 | 1.12E-01 | 2.61E+00 | 2.68E+00 | 3.97E-01 | no | Ephb1 | intron_variant | MODIFIER |
| 9 | 102045514 | AX-97788993 | 0 | G | A | 0.14 | -9.41E-02 | 1.12E-01 | 2.61E+00 | 2.68E+00 | 3.97E-01 | no | Ephb1 | intron_variant | MODIFIER |
| 9 | 102075569 | AX-170083288 | 0 | T | A | 0.14 | -9.41E-02 | 1.12E-01 | 2.61E+00 | 2.68E+00 | 3.97E-01 | no | Ephb1 | intron_variant | MODIFIER |
| 1 | 149281962 | AX-170463854 | 0 | T | C | 0.202 | 1.14E-01 | 1.36E-01 | 2.67E+00 | 2.72E+00 | 3.97E-01 | no | | intergenic_variant | MODIFIER |
| 20 | 67721612 | AX-170034073 | 1 | A | G | 0.241 | 5.24E-02 | 6.27E-02 | 2.72E+00 | 2.81E+00 | 3.97E-01 | no | Rpl7a-ps13 | upstream_gene_variant | MODIFIER |
| 4 | 85678634 | AX-97850006 | 0 | G | A | 0.342 | 9.17E-02 | 1.10E-01 | 2.76E+00 | 2.85E+00 | 3.97E-01 | no | Adamtsl1 | intron_variant,non_coding_transcript_variant | MODIFIER |
| 4 | 58735380 | AX-170022891 | 0 | C | T | 0.368 | 9.47E-02 | 1.13E-01 | 2.67E+00 | 2.73E+00 | 3.97E-01 | no | | intergenic_variant | MODIFIER |
| 4 | 58764292 | AX-170022886 | 0 | T | A | 0.368 | 9.47E-02 | 1.13E-01 | 2.67E+00 | 2.73E+00 | 3.97E-01 | no | | intergenic_variant | MODIFIER |
| 4 | 58797948 | AX-97819814 | 0 | A | G | 0.368 | 9.47E-02 | 1.13E-01 | 2.67E+00 | 2.73E+00 | 3.97E-01 | no | Ecpas | downstream_gene_variant | MODIFIER |
| 4 | 58804557 | AX-97803228 | 0 | C | T | 0.368 | 9.47E-02 | 1.13E-01 | 2.67E+00 | 2.73E+00 | 3.97E-01 | no | Ecpas | intron_variant | MODIFIER |
| 4 | 58818951 | AX-170022887 | 0 | T | C | 0.368 | 9.47E-02 | 1.13E-01 | 2.67E+00 | 2.73E+00 | 3.97E-01 | no | Ecpas | intron_variant | MODIFIER |
| 4 | 58829926 | AX-170457720 | 0 | T | C | 0.368 | 9.47E-02 | 1.13E-01 | 2.67E+00 | 2.73E+00 | 3.97E-01 | no | Ecpas | intron_variant | MODIFIER |
| 4 | 58830776 | AX-170412108 | 0 | C | T | 0.368 | 9.47E-02 | 1.13E-01 | 2.67E+00 | 2.73E+00 | 3.97E-01 | no | Ecpas | intron_variant | MODIFIER |
| 4 | 58831312 | AX-170163108 | 0 | C | A | 0.368 | 9.47E-02 | 1.13E-01 | 2.67E+00 | 2.73E+00 | 3.97E-01 | no | Ecpas | intron_variant | MODIFIER |
| 4 | 58842420 | AX-170022892 | 0 | T | G | 0.368 | 9.47E-02 | 1.13E-01 | 2.67E+00 | 2.73E+00 | 3.97E-01 | no | Ecpas | intron_variant | MODIFIER |
| 4 | 58854673 | AX-97805746 | 0 | C | T | 0.368 | 9.47E-02 | 1.13E-01 | 2.67E+00 | 2.73E+00 | 3.97E-01 | no | Ecpas | intron_variant | MODIFIER |
| 4 | 58877792 | AX-170022893 | 0 | C | G | 0.368 | 9.47E-02 | 1.13E-01 | 2.67E+00 | 2.73E+00 | 3.97E-01 | no | Ecpas | intron_variant | MODIFIER |
| 4 | 58881309 | AX-170218231 | 0 | G | A | 0.368 | 9.47E-02 | 1.13E-01 | 2.67E+00 | 2.73E+00 | 3.97E-01 | no | Ecpas | intron_variant | MODIFIER |
| 4 | 58916769 | AX-170458746 | 0 | C | T | 0.368 | 9.47E-02 | 1.13E-01 | 2.67E+00 | 2.73E+00 | 3.97E-01 | no | Ecpas | upstream_gene_variant | MODIFIER |
| 4 | 58922534 | AX-170280986 | 0 | A | G | 0.368 | 9.47E-02 | 1.13E-01 | 2.67E+00 | 2.73E+00 | 3.97E-01 | no | | intergenic_variant | MODIFIER |

| | | | | | | | | | | | | | | | |
|---|---|---|---|---|---|---|---|---|---|---|---|---|---|---|---|
| 4 | 58922582 | AX-170451928 | 0 | G | A | 0.368 | 9.47E-02 | 1.13E-01 | 2.67E+00 | 2.73E+00 | 3.97E-01 | no | | intergenic_variant | MODIFIER |
| 4 | 58945000 | AX-170535480 | 0 | G | A | 0.368 | 9.47E-02 | 1.13E-01 | 2.67E+00 | 2.73E+00 | 3.97E-01 | no | Zkscan16 | intron_variant | MODIFIER |
| 4 | 58960573 | AX-170069474 | 0 | G | C | 0.368 | 9.47E-02 | 1.13E-01 | 2.67E+00 | 2.73E+00 | 3.97E-01 | no | Ptgr1 | downstream_gene_variant | MODIFIER |
| 4 | 58961056 | AX-170069475 | 0 | C | T | 0.368 | 9.47E-02 | 1.13E-01 | 2.67E+00 | 2.73E+00 | 3.97E-01 | no | Ptgr1 | downstream_gene_variant | MODIFIER |
| 4 | 58963397 | AX-170188982 | 0 | T | C | 0.368 | 9.47E-02 | 1.13E-01 | 2.67E+00 | 2.73E+00 | 3.97E-01 | no | Ptgr1 | downstream_gene_variant | MODIFIER |
| 4 | 58964767 | AX-170484543 | 0 | T | C | 0.368 | 9.47E-02 | 1.13E-01 | 2.67E+00 | 2.73E+00 | 3.97E-01 | no | Ptgr1 | downstream_gene_variant | MODIFIER |
| 4 | 58966440 | AX-170100907 | 0 | A | C | 0.368 | 9.47E-02 | 1.13E-01 | 2.67E+00 | 2.73E+00 | 3.97E-01 | no | Ptgr1 | 3_prime_UTR_variant | MODIFIER |
| 4 | 58967070 | AX-170261980 | 0 | A | T | 0.368 | 9.47E-02 | 1.13E-01 | 2.67E+00 | 2.73E+00 | 3.97E-01 | no | Ptgr1 | intron_variant | MODIFIER |
| 4 | 58968777 | AX-170477038 | 0 | C | T | 0.368 | 9.47E-02 | 1.13E-01 | 2.67E+00 | 2.73E+00 | 3.97E-01 | no | Ptgr1 | synonymous_variant | LOW |
| 4 | 58970159 | AX-170212647 | 0 | T | C | 0.368 | 9.47E-02 | 1.13E-01 | 2.67E+00 | 2.73E+00 | 3.97E-01 | no | Ptgr1 | intron_variant | MODIFIER |
| 4 | 58970482 | AX-170217127 | 0 | A | G | 0.368 | 9.47E-02 | 1.13E-01 | 2.67E+00 | 2.73E+00 | 3.97E-01 | no | Ptgr1 | intron_variant | MODIFIER |
| 4 | 58997329 | AX-170480826 | 0 | C | T | 0.368 | 9.47E-02 | 1.13E-01 | 2.67E+00 | 2.73E+00 | 3.97E-01 | no | Dnajc25 | intron_variant | MODIFIER |
| 4 | 59013917 | AX-170524905 | 0 | G | T | 0.368 | 9.47E-02 | 1.13E-01 | 2.67E+00 | 2.73E+00 | 3.97E-01 | no | Dnajc25 | intron_variant | MODIFIER |
| 4 | 59019003 | AX-170142416 | 0 | A | G | 0.368 | 9.47E-02 | 1.13E-01 | 2.67E+00 | 2.73E+00 | 3.97E-01 | no | Dnajc25 | intron_variant | MODIFIER |
| 4 | 59036258 | AX-170251213 | 0 | A | G | 0.368 | 9.47E-02 | 1.13E-01 | 2.67E+00 | 2.73E+00 | 3.97E-01 | no | Gng10 | intron_variant | MODIFIER |
| 4 | 59042176 | AX-170441691 | 0 | C | A | 0.368 | 9.47E-02 | 1.13E-01 | 2.67E+00 | 2.73E+00 | 3.97E-01 | no | Gng10 | downstream_gene_variant | MODIFIER |
| 4 | 59079686 | AX-170164116 | 0 | A | C | 0.368 | 9.47E-02 | 1.13E-01 | 2.67E+00 | 2.73E+00 | 3.97E-01 | no | AI481877 | intron_variant | MODIFIER |
| 4 | 59108035 | AX-97855936 | 0 | C | T | 0.368 | 9.47E-02 | 1.13E-01 | 2.67E+00 | 2.73E+00 | 3.97E-01 | no | AI481877 | intron_variant | MODIFIER |
| 4 | 59135384 | AX-170264605 | 0 | T | C | 0.368 | 9.47E-02 | 1.13E-01 | 2.67E+00 | 2.73E+00 | 3.97E-01 | no | AI481877 | intron_variant | MODIFIER |
| 4 | 59160837 | AX-170069477 | 0 | C | A | 0.368 | 9.47E-02 | 1.13E-01 | 2.67E+00 | 2.73E+00 | 3.97E-01 | no | Rpsa-ps11 | downstream_gene_variant | MODIFIER |
| 4 | 59161327 | AX-170431995 | 0 | C | T | 0.368 | 9.47E-02 | 1.13E-01 | 2.67E+00 | 2.73E+00 | 3.97E-01 | no | Rpsa-ps11 | downstream_gene_variant | MODIFIER |
| 4 | 59161731 | AX-97852111 | 0 | G | A | 0.368 | 9.47E-02 | 1.13E-01 | 2.67E+00 | 2.73E+00 | 3.97E-01 | no | Rpsa-ps11 | downstream_gene_variant | MODIFIER |
| 4 | 59213250 | AX-170406875 | 0 | C | T | 0.368 | 9.47E-02 | 1.13E-01 | 2.67E+00 | 2.73E+00 | 3.97E-01 | no | Ugcg | synonymous_variant | LOW |
| 4 | 59224764 | AX-170246197 | 0 | G | A | 0.368 | 9.47E-02 | 1.13E-01 | 2.67E+00 | 2.73E+00 | 3.97E-01 | no | Ugcg | downstream_gene_variant | MODIFIER |
| 4 | 59237284 | AX-170146199 | 0 | G | A | 0.368 | 9.47E-02 | 1.13E-01 | 2.67E+00 | 2.73E+00 | 3.97E-01 | no | | regulatory_region_variant | MODIFIER |
| 4 | 59238849 | AX-97802608 | 0 | T | G | 0.368 | 9.47E-02 | 1.13E-01 | 2.67E+00 | 2.73E+00 | 3.97E-01 | no | | regulatory_region_variant | MODIFIER |
| 4 | 59269462 | AX-170444816 | 0 | A | G | 0.368 | 9.47E-02 | 1.13E-01 | 2.67E+00 | 2.73E+00 | 3.97E-01 | no | | intergenic_variant | MODIFIER |
| 4 | 59293985 | AX-97792975 | 0 | T | C | 0.368 | 9.47E-02 | 1.13E-01 | 2.67E+00 | 2.73E+00 | 3.97E-01 | no | | intergenic_variant | MODIFIER |
| 4 | 59334362 | AX-170022894 | 0 | T | C | 0.368 | 9.47E-02 | 1.13E-01 | 2.67E+00 | 2.73E+00 | 3.97E-01 | no | Susd1 | intron_variant | MODIFIER |
| 4 | 59338704 | AX-97787425 | 0 | G | A | 0.368 | 9.47E-02 | 1.13E-01 | 2.67E+00 | 2.73E+00 | 3.97E-01 | no | Susd1 | intron_variant | MODIFIER |
| 4 | 59362763 | AX-170395421 | 0 | C | T | 0.368 | 9.47E-02 | 1.13E-01 | 2.67E+00 | 2.73E+00 | 3.97E-01 | no | Susd1 | intron_variant | MODIFIER |
| 4 | 59367164 | AX-97810803 | 0 | G | A | 0.368 | 9.47E-02 | 1.13E-01 | 2.67E+00 | 2.73E+00 | 3.97E-01 | no | Susd1 | intron_variant | MODIFIER |
| 4 | 59379708 | AX-170405911 | 0 | C | T | 0.368 | 9.47E-02 | 1.13E-01 | 2.67E+00 | 2.73E+00 | 3.97E-01 | no | Susd1 | synonymous_variant | LOW |
| 4 | 59382299 | AX-170022896 | 0 | C | A | 0.368 | 9.47E-02 | 1.13E-01 | 2.67E+00 | 2.73E+00 | 3.97E-01 | no | Susd1 | intron_variant | MODIFIER |
| 4 | 59385550 | AX-170137304 | 0 | C | A | 0.368 | 9.47E-02 | 1.13E-01 | 2.67E+00 | 2.73E+00 | 3.97E-01 | no | Susd1 | intron_variant | MODIFIER |
| 4 | 59418346 | AX-170534436 | 0 | G | A | 0.368 | 9.47E-02 | 1.13E-01 | 2.67E+00 | 2.73E+00 | 3.97E-01 | no | Susd1 | intron_variant | MODIFIER |
| 4 | 59427348 | AX-170022895 | 0 | G | A | 0.368 | 9.47E-02 | 1.13E-01 | 2.67E+00 | 2.73E+00 | 3.97E-01 | no | Susd1 | intron_variant | MODIFIER |
| 4 | 59440145 | AX-97811947 | 0 | G | C | 0.368 | 9.47E-02 | 1.13E-01 | 2.67E+00 | 2.73E+00 | 3.97E-01 | no | Susd1 | upstream_gene_variant | MODIFIER |
| 4 | 59443996 | AX-170022898 | 0 | A | G | 0.368 | 9.47E-02 | 1.13E-01 | 2.67E+00 | 2.73E+00 | 3.97E-01 | no | | regulatory_region_variant | MODIFIER |
| 4 | 59448999 | AX-170381101 | 0 | T | C | 0.368 | 9.47E-02 | 1.13E-01 | 2.67E+00 | 2.73E+00 | 3.97E-01 | no | | intergenic_variant | MODIFIER |
| 4 | 59700598 | AX-170335722 | 0 | T | C | 0.368 | 9.47E-02 | 1.13E-01 | 2.67E+00 | 2.73E+00 | 3.97E-01 | no | E130308A19Rik | intron_variant | MODIFIER |
| 11 | 19529614 | AX-170035490 | 1 | C | A | 0.054 | 1.40E-01 | 1.68E-01 | 2.65E+00 | 2.81E+00 | 3.97E-01 | no | | regulatory_region_variant | MODIFIER |
| 8 | 84969441 | AX-170345075 | 0 | T | C | 0.193 | -9.40E-02 | 1.13E-01 | 2.78E+00 | 2.90E+00 | 3.97E-01 | no | Prdx2 | upstream_gene_variant | MODIFIER |
| 7 | 41241543 | AX-170234739 | 0 | C | A | 0.342 | 7.78E-02 | 9.19E-02 | 2.56E+00 | 2.65E+00 | 3.97E-01 | no | Gm44748 | intron_variant,non_coding_transcript_variant | MODIFIER |
| 18 | 13792876 | AX-170462800 | 0 | G | T | 0.061 | 1.25E-01 | 1.50E-01 | 2.70E+00 | 2.84E+00 | 3.97E-01 | no | Zfp521 | intron_variant | MODIFIER |
| 15 | 4579177 | AX-97795299 | 0 | G | A | 0.246 | -8.76E-02 | 1.04E-01 | 2.61E+00 | 2.66E+00 | 3.97E-01 | no | Plcxd3 | downstream_gene_variant | MODIFIER |
| 2 | 164000472 | AX-170143751 | 0 | A | T | 0.149 | 1.47E-01 | 1.76E-01 | 2.72E+00 | 2.81E+00 | 3.97E-01 | Yes | Ywhab | intron_variant | MODIFIER |
| 15 | 7034179 | AX-170167854 | 0 | T | A | 0.237 | -9.04E-02 | 1.07E-01 | 2.62E+00 | 2.67E+00 | 3.97E-01 | no | | intergenic_variant | MODIFIER |
| 15 | 8152737 | AX-97828355 | 0 | A | C | 0.237 | -9.04E-02 | 1.07E-01 | 2.62E+00 | 2.67E+00 | 3.97E-01 | no | Nup155 | intron_variant | MODIFIER |
| 9 | 40747337 | AX-170384107 | 0 | A | T | 0.43 | -6.30E-02 | 7.54E-02 | 2.76E+00 | 2.85E+00 | 3.98E-01 | no | Clmp | intron_variant | MODIFIER |
| 9 | 21472429 | AX-170293358 | 1 | C | A | 0.473 | 7.63E-02 | 8.93E-02 | 2.51E+00 | 2.58E+00 | 3.98E-01 | no | Dnm2 | intron_variant | MODIFIER |
| 5 | 43809706 | AX-170089464 | 0 | A | G | 0.184 | -9.34E-02 | 1.11E-01 | 2.59E+00 | 2.68E+00 | 3.98E-01 | no | Fbxl5 | intron_variant | MODIFIER |
| 1 | 152534572 | AX-170245459 | 1 | A | G | 0.205 | 1.16E-01 | 1.38E-01 | 2.64E+00 | 2.69E+00 | 3.98E-01 | no | Rgl1 | intron_variant | MODIFIER |
| 15 | 79382629 | AX-170323612 | 1 | C | T | 0.571 | 6.86E-02 | 8.14E-02 | 2.58E+00 | 2.67E+00 | 3.98E-01 | no | Tmem184b | intron_variant | MODIFIER |

| | | | | | | | | | | | | | | | |
|---|---|---|---|---|---|---|---|---|---|---|---|---|---|---|---|
| 16 | 74217293 | AX-97807911 | 2 | T | C | 0.491 | 6.63E-02 | 7.69E-02 | 2.46E+00 | 2.54E+00 | 3.98E-01 | no | Robo2 | intron_variant | MODIFIER |
| 18 | 61955825 | AX-170061043 | 0 | A | T | 0.412 | 7.61E-02 | 8.75E-02 | 2.43E+00 | 2.50E+00 | 3.98E-01 | no | Sh3tc2 | intron_variant | MODIFIER |
| 19 | 20918369 | AX-170116030 | 1 | C | T | 0.036 | 1.21E-01 | 1.45E-01 | 2.58E+00 | 2.74E+00 | 3.98E-01 | no | Tmc1 | intron_variant | MODIFIER |
| 7 | 37307281 | AX-97818162 | 0 | T | C | 0.342 | 7.72E-02 | 9.14E-02 | 2.56E+00 | 2.64E+00 | 3.99E-01 | no | Gm28077 | downstream_gene_variant | MODIFIER |
| 6 | 134098652 | AX-170028298 | 1 | T | C | 0.545 | 7.47E-02 | 8.64E-02 | 2.46E+00 | 2.52E+00 | 3.99E-01 | no | Etv6 | intron_variant | MODIFIER |
| 8 | 88259970 | AX-170342776 | 0 | C | T | 0.228 | -1.03E-01 | 1.23E-01 | 2.82E+00 | 2.91E+00 | 3.99E-01 | no | Tent4b | downstream_gene_variant | MODIFIER |
| 2 | 104099075 | AX-170313422 | 0 | T | A | 0.061 | 1.82E-01 | 2.19E-01 | 2.78E+00 | 2.88E+00 | 3.99E-01 | Yes | Cd59a | intron_variant | MODIFIER |
| 5 | 34582143 | AX-170228385 | 1 | C | A | 0.339 | 8.85E-02 | 1.06E-01 | 2.73E+00 | 2.80E+00 | 3.99E-01 | no | Add1 | intron_variant | MODIFIER |
| 4 | 135009990 | AX-170109235 | 1 | G | T | 0.152 | 9.76E-02 | 1.16E-01 | 2.58E+00 | 2.63E+00 | 3.99E-01 | no | | intergenic_variant | MODIFIER |
| 9 | 105143835 | AX-97805524 | 1 | G | A | 0.125 | -8.59E-02 | 1.02E-01 | 2.57E+00 | 2.68E+00 | 3.99E-01 | no | 1700080E11Rik | intron_variant | MODIFIER |
| 5 | 83705935 | AX-170430913 | 1 | A | C | 0.438 | 9.15E-02 | 1.10E-01 | 2.75E+00 | 2.82E+00 | 3.99E-01 | no | | intergenic_variant | MODIFIER |
| 9 | 78649285 | AX-170456923 | 1 | A | T | 0.196 | -1.04E-01 | 1.25E-01 | 2.75E+00 | 2.81E+00 | 3.99E-01 | no | Cd109 | intron_variant | MODIFIER |
| 5 | 67395668 | AX-170072367 | 1 | T | C | 0.455 | 8.01E-02 | 9.30E-02 | 2.46E+00 | 2.55E+00 | 3.99E-01 | no | Bend4 | 3_prime_UTR_variant | MODIFIER |
| 4 | 72238309 | AX-97790585 | 0 | G | A | 0.456 | -7.08E-02 | 8.34E-02 | 2.54E+00 | 2.60E+00 | 3.99E-01 | no | | intergenic_variant | MODIFIER |
| 4 | 73349732 | AX-97848399 | 0 | T | C | 0.456 | -7.08E-02 | 8.34E-02 | 2.54E+00 | 2.60E+00 | 3.99E-01 | no | | intergenic_variant | MODIFIER |
| 8 | 38224848 | AX-170166358 | 0 | A | C | 0.149 | -9.80E-02 | 1.18E-01 | 2.71E+00 | 2.84E+00 | 3.99E-01 | no | Sgcz | intron_variant | MODIFIER |
| 9 | 102097561 | AX-170083285 | 1 | C | T | 0.134 | -9.68E-02 | 1.16E-01 | 2.63E+00 | 2.71E+00 | 3.99E-01 | no | Ephb1 | intron_variant | MODIFIER |
| 18 | 64670177 | AX-170100054 | 1 | A | T | 0.188 | -1.00E-01 | 1.20E-01 | 2.71E+00 | 2.77E+00 | 3.99E-01 | no | | intergenic_variant | MODIFIER |
| 18 | 64690680 | AX-170204339 | 1 | G | C | 0.188 | -1.00E-01 | 1.20E-01 | 2.71E+00 | 2.77E+00 | 3.99E-01 | no | AC137585.3 | downstream_gene_variant | MODIFIER |
| 18 | 64844855 | AX-170180066 | 1 | G | A | 0.188 | -1.00E-01 | 1.20E-01 | 2.71E+00 | 2.77E+00 | 3.99E-01 | no | | regulatory_region_variant | MODIFIER |
| 18 | 65478702 | AX-170461173 | 1 | T | C | 0.188 | -1.00E-01 | 1.20E-01 | 2.71E+00 | 2.77E+00 | 3.99E-01 | no | Malt1 | 3_prime_UTR_variant | MODIFIER |
| 4 | 89738155 | AX-170414796 | 0 | T | C | 0.316 | 8.33E-02 | 9.97E-02 | 2.63E+00 | 2.73E+00 | 3.99E-01 | no | | intergenic_variant | MODIFIER |
| 1 | 128723345 | AX-170394430 | 0 | C | T | 0.535 | 6.94E-02 | 8.28E-02 | 2.62E+00 | 2.68E+00 | 4.00E-01 | no | | regulatory_region_variant | MODIFIER |
| 5 | 83052955 | AX-170343089 | 1 | A | G | 0.438 | 8.72E-02 | 1.04E-01 | 2.62E+00 | 2.69E+00 | 4.00E-01 | no | | intergenic_variant | MODIFIER |
| 8 | 83490411 | AX-170204706 | 0 | T | C | 0.202 | -9.31E-02 | 1.12E-01 | 2.78E+00 | 2.90E+00 | 4.00E-01 | no | Gm33242 | non_coding_transcript_exon_variant | MODIFIER |
| 6 | 127753789 | AX-170422808 | 1 | A | G | 0.527 | 6.94E-02 | 8.09E-02 | 2.47E+00 | 2.55E+00 | 4.00E-01 | no | Prmt8 | intron_variant | MODIFIER |
| 5 | 42740858 | AX-170358167 | 1 | A | G | 0.196 | -8.89E-02 | 1.06E-01 | 2.59E+00 | 2.67E+00 | 4.00E-01 | no | | intergenic_variant | MODIFIER |
| 10 | 70028351 | AX-97839450 | 0 | C | T | 0.412 | -6.57E-02 | 7.85E-02 | 2.61E+00 | 2.69E+00 | 4.00E-01 | no | Ank3 | downstream_gene_variant | MODIFIER |
| 13 | 24169605 | AX-170313930 | 0 | G | A | 0.167 | -1.07E-01 | 1.20E-01 | 2.35E+00 | 2.40E+00 | 4.00E-01 | no | Carmil1 | intron_variant | MODIFIER |
| 13 | 24174427 | AX-170130312 | 0 | C | T | 0.167 | -1.07E-01 | 1.20E-01 | 2.35E+00 | 2.40E+00 | 4.00E-01 | no | Carmil1 | intron_variant | MODIFIER |
| 13 | 24174912 | AX-170233543 | 0 | G | A | 0.167 | -1.07E-01 | 1.20E-01 | 2.35E+00 | 2.40E+00 | 4.00E-01 | no | Carmil1 | intron_variant | MODIFIER |
| 13 | 24178717 | AX-170238592 | 0 | G | A | 0.167 | -1.07E-01 | 1.20E-01 | 2.35E+00 | 2.40E+00 | 4.00E-01 | no | Carmil1 | intron_variant | MODIFIER |
| 13 | 24178745 | AX-170050762 | 0 | C | T | 0.167 | -1.07E-01 | 1.20E-01 | 2.35E+00 | 2.40E+00 | 4.00E-01 | no | Carmil1 | intron_variant | MODIFIER |
| 13 | 24178838 | AX-170050764 | 0 | C | T | 0.167 | -1.07E-01 | 1.20E-01 | 2.35E+00 | 2.40E+00 | 4.00E-01 | no | Carmil1 | intron_variant | MODIFIER |
| 13 | 24199333 | AX-97846240 | 0 | T | C | 0.167 | -1.07E-01 | 1.20E-01 | 2.35E+00 | 2.40E+00 | 4.00E-01 | no | Carmil1 | intron_variant | MODIFIER |
| 13 | 24211376 | AX-97823123 | 0 | G | A | 0.167 | -1.07E-01 | 1.20E-01 | 2.35E+00 | 2.40E+00 | 4.00E-01 | no | Carmil1 | intron_variant | MODIFIER |
| 13 | 24211800 | AX-170301368 | 0 | A | T | 0.167 | -1.07E-01 | 1.20E-01 | 2.35E+00 | 2.40E+00 | 4.00E-01 | no | Carmil1 | intron_variant | MODIFIER |
| 13 | 24332602 | AX-170007936 | 0 | G | T | 0.167 | -1.07E-01 | 1.20E-01 | 2.35E+00 | 2.40E+00 | 4.00E-01 | no | Cmah | intron_variant | MODIFIER |
| 13 | 24339071 | AX-170092688 | 0 | A | C | 0.167 | -1.07E-01 | 1.20E-01 | 2.35E+00 | 2.40E+00 | 4.00E-01 | no | Cmah | intron_variant | MODIFIER |
| 13 | 24355357 | AX-170187906 | 0 | A | G | 0.167 | -1.07E-01 | 1.20E-01 | 2.35E+00 | 2.40E+00 | 4.00E-01 | no | Cmah | intron_variant | MODIFIER |
| 13 | 24371463 | AX-97818506 | 0 | A | C | 0.167 | -1.07E-01 | 1.20E-01 | 2.35E+00 | 2.40E+00 | 4.00E-01 | no | Cmah | intron_variant | MODIFIER |
| 13 | 24371649 | AX-170007932 | 0 | A | G | 0.167 | -1.07E-01 | 1.20E-01 | 2.35E+00 | 2.40E+00 | 4.00E-01 | no | Cmah | intron_variant | MODIFIER |
| 13 | 24387778 | AX-170210717 | 0 | T | A | 0.167 | -1.07E-01 | 1.20E-01 | 2.35E+00 | 2.40E+00 | 4.00E-01 | no | Cmah | intron_variant | MODIFIER |
| 13 | 24398227 | AX-97808617 | 0 | T | C | 0.167 | -1.07E-01 | 1.20E-01 | 2.35E+00 | 2.40E+00 | 4.00E-01 | no | Cmah | intron_variant | MODIFIER |
| 13 | 24398432 | AX-170494687 | 0 | G | T | 0.167 | -1.07E-01 | 1.20E-01 | 2.35E+00 | 2.40E+00 | 4.00E-01 | no | Cmah | intron_variant | MODIFIER |
| 13 | 24419593 | AX-170197833 | 0 | C | A | 0.167 | -1.07E-01 | 1.20E-01 | 2.35E+00 | 2.40E+00 | 4.00E-01 | no | Cmah | intron_variant | MODIFIER |
| 13 | 24442607 | AX-170386021 | 0 | C | T | 0.167 | -1.07E-01 | 1.20E-01 | 2.35E+00 | 2.40E+00 | 4.00E-01 | no | Cmah | intron_variant | MODIFIER |
| 13 | 24457623 | AX-170137947 | 0 | C | T | 0.167 | -1.07E-01 | 1.20E-01 | 2.35E+00 | 2.40E+00 | 4.00E-01 | no | Cmah | intron_variant | MODIFIER |
| 13 | 24465661 | AX-170424105 | 0 | C | G | 0.167 | -1.07E-01 | 1.20E-01 | 2.35E+00 | 2.40E+00 | 4.00E-01 | no | Cmah | intron_variant | MODIFIER |
| 13 | 24465812 | AX-97804036 | 0 | G | A | 0.167 | -1.07E-01 | 1.20E-01 | 2.35E+00 | 2.40E+00 | 4.00E-01 | no | Cmah | intron_variant | MODIFIER |
| 13 | 24482133 | AX-170134679 | 0 | T | C | 0.167 | -1.07E-01 | 1.20E-01 | 2.35E+00 | 2.40E+00 | 4.00E-01 | no | Cmah | downstream_gene_variant | MODIFIER |
| 13 | 24489310 | AX-170314740 | 0 | G | A | 0.167 | -1.07E-01 | 1.20E-01 | 2.35E+00 | 2.40E+00 | 4.00E-01 | no | Gm22013 | upstream_gene_variant | MODIFIER |
| 13 | 24504293 | AX-97794562 | 0 | T | C | 0.167 | -1.07E-01 | 1.20E-01 | 2.35E+00 | 2.40E+00 | 4.00E-01 | no | | intergenic_variant | MODIFIER |
| 13 | 24506277 | AX-170121296 | 0 | G | A | 0.167 | -1.07E-01 | 1.20E-01 | 2.35E+00 | 2.40E+00 | 4.00E-01 | no | | intergenic_variant | MODIFIER |

| | | | | | | | | | | | | | | | |
|---|---|---|---|---|---|---|---|---|---|---|---|---|---|---|---|
| 13 | 24510338 | AX-170050768 | 0 | G | A | 0.167 | -1.07E-01 | 1.20E-01 | 2.35E+00 | 2.40E+00 | 4.00E-01 | no | | intergenic_variant | MODIFIER |
| 13 | 24516518 | AX-97818366 | 0 | T | C | 0.167 | -1.07E-01 | 1.20E-01 | 2.35E+00 | 2.40E+00 | 4.00E-01 | no | | intergenic_variant | MODIFIER |
| 13 | 24521227 | AX-170422339 | 0 | G | A | 0.167 | -1.07E-01 | 1.20E-01 | 2.35E+00 | 2.40E+00 | 4.00E-01 | no | | regulatory_region_variant | MODIFIER |
| 13 | 24548534 | AX-170459670 | 0 | T | G | 0.167 | -1.07E-01 | 1.20E-01 | 2.35E+00 | 2.40E+00 | 4.00E-01 | no | | regulatory_region_variant | MODIFIER |
| 13 | 24576147 | AX-97827759 | 0 | C | T | 0.167 | -1.07E-01 | 1.20E-01 | 2.35E+00 | 2.40E+00 | 4.00E-01 | no | | regulatory_region_variant | MODIFIER |
| 13 | 24610748 | AX-170463910 | 0 | G | A | 0.167 | -1.07E-01 | 1.20E-01 | 2.35E+00 | 2.40E+00 | 4.00E-01 | no | Ripor2 | upstream_gene_variant | MODIFIER |
| 13 | 24611632 | AX-170175007 | 0 | A | G | 0.167 | -1.07E-01 | 1.20E-01 | 2.35E+00 | 2.40E+00 | 4.00E-01 | no | Ripor2 | upstream_gene_variant | MODIFIER |
| 13 | 24624983 | AX-97809099 | 0 | A | G | 0.167 | -1.07E-01 | 1.20E-01 | 2.35E+00 | 2.40E+00 | 4.00E-01 | no | Ripor2 | intron_variant | MODIFIER |
| 13 | 24629623 | AX-170376032 | 0 | T | C | 0.167 | -1.07E-01 | 1.20E-01 | 2.35E+00 | 2.40E+00 | 4.00E-01 | no | Ripor2 | intron_variant | MODIFIER |
| 13 | 24637339 | AX-170503945 | 0 | T | G | 0.167 | -1.07E-01 | 1.20E-01 | 2.35E+00 | 2.40E+00 | 4.00E-01 | no | Ripor2 | upstream_gene_variant | MODIFIER |
| 13 | 24651313 | AX-97829097 | 0 | T | C | 0.167 | -1.07E-01 | 1.20E-01 | 2.35E+00 | 2.40E+00 | 4.00E-01 | no | Ripor2 | intron_variant | MODIFIER |
| 13 | 24652310 | AX-170127145 | 0 | G | A | 0.167 | -1.07E-01 | 1.20E-01 | 2.35E+00 | 2.40E+00 | 4.00E-01 | no | Ripor2 | intron_variant | MODIFIER |
| 13 | 24661206 | AX-170472936 | 0 | T | G | 0.167 | -1.07E-01 | 1.20E-01 | 2.35E+00 | 2.40E+00 | 4.00E-01 | no | Ripor2 | intron_variant | MODIFIER |
| 13 | 24661330 | AX-170486884 | 0 | A | G | 0.167 | -1.07E-01 | 1.20E-01 | 2.35E+00 | 2.40E+00 | 4.00E-01 | no | Ripor2 | intron_variant | MODIFIER |
| 13 | 24661935 | AX-170414224 | 0 | G | A | 0.167 | -1.07E-01 | 1.20E-01 | 2.35E+00 | 2.40E+00 | 4.00E-01 | no | Ripor2 | intron_variant | MODIFIER |
| 13 | 24664484 | AX-170177105 | 0 | G | A | 0.167 | -1.07E-01 | 1.20E-01 | 2.35E+00 | 2.40E+00 | 4.00E-01 | no | Ripor2 | intron_variant | MODIFIER |
| 13 | 24664925 | AX-170474300 | 0 | C | T | 0.167 | -1.07E-01 | 1.20E-01 | 2.35E+00 | 2.40E+00 | 4.00E-01 | no | Ripor2 | intron_variant | MODIFIER |
| 13 | 24674246 | AX-170087388 | 0 | C | T | 0.167 | -1.07E-01 | 1.20E-01 | 2.35E+00 | 2.40E+00 | 4.00E-01 | no | Ripor2 | intron_variant | MODIFIER |
| 13 | 24687197 | AX-170466142 | 0 | C | A | 0.167 | -1.07E-01 | 1.20E-01 | 2.35E+00 | 2.40E+00 | 4.00E-01 | no | Ripor2 | intron_variant | MODIFIER |
| 13 | 24693611 | AX-97796534 | 0 | C | T | 0.167 | -1.07E-01 | 1.20E-01 | 2.35E+00 | 2.40E+00 | 4.00E-01 | no | Ripor2 | intron_variant | MODIFIER |
| 13 | 24710670 | AX-170433654 | 0 | G | C | 0.167 | -1.07E-01 | 1.20E-01 | 2.35E+00 | 2.40E+00 | 4.00E-01 | no | Ripor2 | downstream_gene_variant | MODIFIER |
| 13 | 24729231 | AX-170419092 | 0 | T | C | 0.167 | -1.07E-01 | 1.20E-01 | 2.35E+00 | 2.40E+00 | 4.00E-01 | no | Ripor2 | intron_variant | MODIFIER |
| 13 | 24731044 | AX-97823126 | 0 | G | T | 0.167 | -1.07E-01 | 1.20E-01 | 2.35E+00 | 2.40E+00 | 4.00E-01 | no | Ripor2 | intron_variant | MODIFIER |
| 13 | 24745423 | AX-97840698 | 0 | C | T | 0.167 | -1.07E-01 | 1.20E-01 | 2.35E+00 | 2.40E+00 | 4.00E-01 | no | Armh2 | intron_variant | MODIFIER |
| 13 | 24775415 | AX-97840848 | 0 | A | G | 0.167 | -1.07E-01 | 1.20E-01 | 2.35E+00 | 2.40E+00 | 4.00E-01 | no | | intergenic_variant | MODIFIER |
| 13 | 24781083 | AX-170275608 | 0 | A | G | 0.167 | -1.07E-01 | 1.20E-01 | 2.35E+00 | 2.40E+00 | 4.00E-01 | no | | intergenic_variant | MODIFIER |
| 13 | 24785612 | AX-170350463 | 0 | T | C | 0.167 | -1.07E-01 | 1.20E-01 | 2.35E+00 | 2.40E+00 | 4.00E-01 | no | | regulatory_region_variant | MODIFIER |
| 13 | 24794257 | AX-97807935 | 0 | T | C | 0.167 | -1.07E-01 | 1.20E-01 | 2.35E+00 | 2.40E+00 | 4.00E-01 | no | | intergenic_variant | MODIFIER |
| 13 | 24816068 | AX-97822993 | 0 | A | G | 0.167 | -1.07E-01 | 1.20E-01 | 2.35E+00 | 2.40E+00 | 4.00E-01 | no | Acot13 | downstream_gene_variant | MODIFIER |
| 13 | 24818894 | AX-170267597 | 0 | T | C | 0.167 | -1.07E-01 | 1.20E-01 | 2.35E+00 | 2.40E+00 | 4.00E-01 | no | Acot13 | intron_variant | MODIFIER |
| 13 | 24818981 | AX-170495046 | 0 | G | A | 0.167 | -1.07E-01 | 1.20E-01 | 2.35E+00 | 2.40E+00 | 4.00E-01 | no | Acot13 | intron_variant | MODIFIER |
| 13 | 24819279 | AX-170352005 | 0 | T | C | 0.167 | -1.07E-01 | 1.20E-01 | 2.35E+00 | 2.40E+00 | 4.00E-01 | no | Acot13 | intron_variant | MODIFIER |
| 13 | 24819460 | AX-170146523 | 0 | C | T | 0.167 | -1.07E-01 | 1.20E-01 | 2.35E+00 | 2.40E+00 | 4.00E-01 | no | Acot13 | intron_variant | MODIFIER |
| 13 | 24821643 | AX-170429590 | 0 | A | G | 0.167 | -1.07E-01 | 1.20E-01 | 2.35E+00 | 2.40E+00 | 4.00E-01 | no | Acot13 | intron_variant | MODIFIER |
| 13 | 24829143 | AX-170240010 | 0 | T | C | 0.167 | -1.07E-01 | 1.20E-01 | 2.35E+00 | 2.40E+00 | 4.00E-01 | no | Acot13 | intron_variant | MODIFIER |
| 13 | 24839467 | AX-170529884 | 0 | G | A | 0.167 | -1.07E-01 | 1.20E-01 | 2.35E+00 | 2.40E+00 | 4.00E-01 | no | Tdp2 | intron_variant | MODIFIER |
| 13 | 24844873 | AX-97847207 | 0 | G | A | 0.167 | -1.07E-01 | 1.20E-01 | 2.35E+00 | 2.40E+00 | 4.00E-01 | no | D130043K22Rik | upstream_gene_variant | MODIFIER |
| 13 | 24863714 | AX-170050773 | 0 | C | A | 0.167 | -1.07E-01 | 1.20E-01 | 2.35E+00 | 2.40E+00 | 4.00E-01 | no | D130043K22Rik | intron_variant | MODIFIER |
| 13 | 24864379 | AX-170270787 | 0 | G | C | 0.167 | -1.07E-01 | 1.20E-01 | 2.35E+00 | 2.40E+00 | 4.00E-01 | no | D130043K22Rik | intron_variant | MODIFIER |
| 13 | 24872098 | AX-97780628 | 0 | T | C | 0.167 | -1.07E-01 | 1.20E-01 | 2.35E+00 | 2.40E+00 | 4.00E-01 | no | D130043K22Rik | intron_variant | MODIFIER |
| 13 | 24876284 | AX-170207588 | 0 | T | C | 0.167 | -1.07E-01 | 1.20E-01 | 2.35E+00 | 2.40E+00 | 4.00E-01 | no | D130043K22Rik | intron_variant | MODIFIER |
| 13 | 24880683 | AX-170316082 | 0 | G | A | 0.167 | -1.07E-01 | 1.20E-01 | 2.35E+00 | 2.40E+00 | 4.00E-01 | no | D130043K22Rik | intron_variant | MODIFIER |
| 13 | 24880695 | AX-170410683 | 0 | T | C | 0.167 | -1.07E-01 | 1.20E-01 | 2.35E+00 | 2.40E+00 | 4.00E-01 | no | D130043K22Rik | intron_variant | MODIFIER |
| 13 | 24883973 | AX-170369458 | 0 | T | A | 0.167 | -1.07E-01 | 1.20E-01 | 2.35E+00 | 2.40E+00 | 4.00E-01 | no | D130043K22Rik | intron_variant | MODIFIER |
| 13 | 24883988 | AX-170427927 | 0 | C | T | 0.167 | -1.07E-01 | 1.20E-01 | 2.35E+00 | 2.40E+00 | 4.00E-01 | no | D130043K22Rik | intron_variant | MODIFIER |
| 13 | 24894264 | AX-97839224 | 0 | T | C | 0.167 | -1.07E-01 | 1.20E-01 | 2.35E+00 | 2.40E+00 | 4.00E-01 | no | D130043K22Rik | intron_variant | MODIFIER |
| 13 | 24898564 | AX-170314214 | 0 | T | G | 0.167 | -1.07E-01 | 1.20E-01 | 2.35E+00 | 2.40E+00 | 4.00E-01 | no | D130043K22Rik | intron_variant | MODIFIER |
| 1 | 120274670 | AX-170425147 | 1 | T | A | 0.589 | 6.97E-02 | 8.08E-02 | 2.45E+00 | 2.52E+00 | 4.00E-01 | no | Steap3 | upstream_gene_variant | MODIFIER |
| 1 | 120612817 | AX-170388811 | 1 | C | T | 0.589 | 6.97E-02 | 8.08E-02 | 2.45E+00 | 2.52E+00 | 4.00E-01 | no | En1 | downstream_gene_variant | MODIFIER |
| 4 | 89038771 | AX-170288874 | 1 | G | A | 0.438 | -6.94E-02 | 8.07E-02 | 2.46E+00 | 2.52E+00 | 4.00E-01 | no | AL805899.2 | intron_variant,non_coding_transcript_variant | MODIFIER |
| 3 | 151774035 | AX-170377028 | 0 | T | G | 0.342 | 5.96E-02 | 7.18E-02 | 2.74E+00 | 2.83E+00 | 4.00E-01 | no | | intergenic_variant | MODIFIER |
| 18 | 60038758 | AX-170061001 | 0 | T | A | 0.404 | 8.09E-02 | 9.43E-02 | 2.47E+00 | 2.52E+00 | 4.00E-01 | no | | intergenic_variant | MODIFIER |
| 7 | 37870647 | AX-170457328 | 0 | G | A | 0.342 | 7.82E-02 | 9.31E-02 | 2.58E+00 | 2.66E+00 | 4.00E-01 | no | | regulatory_region_variant | MODIFIER |
| 5 | 127034966 | AX-170387769 | 0 | T | G | 0.439 | 6.89E-02 | 8.21E-02 | 2.56E+00 | 2.68E+00 | 4.00E-01 | no | 4933438B17Rik | intron_variant,non_coding_transcript_variant | MODIFIER |

| | | | | | | | | | | | | | | | |
|---|---|---|---|---|---|---|---|---|---|---|---|---|---|---|---|
| 18 | 59974143 | AX-97846455 | 2 | A | G | 0.391 | 8.08E-02 | 9.30E-02 | 2.42E+00 | 2.48E+00 | 4.01E-01 | no | | intergenic_variant | MODIFIER |
| 2 | 169026398 | AX-170019654 | 0 | G | T | 0.167 | 1.51E-01 | 1.81E-01 | 2.64E+00 | 2.73E+00 | 4.01E-01 | Yes | | intergenic_variant | MODIFIER |
| 7 | 40999239 | AX-170147743 | 1 | T | C | 0.312 | 8.11E-02 | 9.68E-02 | 2.58E+00 | 2.67E+00 | 4.01E-01 | no | 4930433I11Rik | downstream_gene_variant | MODIFIER |
| 1 | 147404557 | AX-170397864 | 0 | G | T | 0.202 | 1.15E-01 | 1.38E-01 | 2.65E+00 | 2.70E+00 | 4.01E-01 | no | | intergenic_variant | MODIFIER |
| 5 | 46125000 | AX-170024850 | 0 | C | T | 0.5 | 8.59E-02 | 1.00E-01 | 2.47E+00 | 2.53E+00 | 4.01E-01 | no | | intergenic_variant | MODIFIER |
| 1 | 152494407 | AX-97829918 | 1 | G | A | 0.205 | 1.13E-01 | 1.36E-01 | 2.68E+00 | 2.73E+00 | 4.01E-01 | no | Colgalt2 | intron_variant | MODIFIER |
| 19 | 32310291 | AX-170498419 | 1 | A | C | 0.045 | -8.69E-02 | 1.04E-01 | 2.52E+00 | 2.68E+00 | 4.01E-01 | no | Sgms1 | intron_variant | MODIFIER |
| 18 | 62812655 | AX-170139339 | 1 | G | A | 0.188 | -9.83E-02 | 1.18E-01 | 2.66E+00 | 2.71E+00 | 4.01E-01 | no | | intergenic_variant | MODIFIER |
| 18 | 65442315 | AX-170494840 | 1 | C | T | 0.42 | 7.46E-02 | 8.64E-02 | 2.44E+00 | 2.50E+00 | 4.01E-01 | no | Malt1 | intron_variant | MODIFIER |
| 11 | 34820567 | AX-170004675 | 0 | C | G | 0.096 | -1.32E-01 | 1.59E-01 | 2.78E+00 | 2.91E+00 | 4.01E-01 | no | Spdl1 | intron_variant | MODIFIER |
| 18 | 21355153 | AX-97843670 | 0 | C | T | 0.061 | 1.14E-01 | 1.38E-01 | 2.72E+00 | 2.88E+00 | 4.01E-01 | no | | intergenic_variant | MODIFIER |
| 1 | 96521935 | AX-170124751 | 0 | A | T | 0.158 | 1.40E-01 | 1.69E-01 | 2.65E+00 | 2.75E+00 | 4.02E-01 | no | | intergenic_variant | MODIFIER |
| 8 | 90495636 | AX-170477703 | 0 | G | A | 0.193 | -9.21E-02 | 1.11E-01 | 2.65E+00 | 2.74E+00 | 4.02E-01 | no | | regulatory_region_variant | MODIFIER |
| 8 | 90822101 | AX-170179056 | 0 | A | G | 0.193 | -9.21E-02 | 1.11E-01 | 2.65E+00 | 2.74E+00 | 4.02E-01 | no | Gm19935 | upstream_gene_variant | MODIFIER |
| 1 | 172898634 | AX-170160135 | 1 | C | A | 0.232 | 9.80E-02 | 1.15E-01 | 2.47E+00 | 2.55E+00 | 4.02E-01 | no | Apcs | upstream_gene_variant | MODIFIER |
| 7 | 124718784 | AX-168408085 | 0 | G | A | 0.377 | 7.01E-02 | 8.46E-02 | 2.77E+00 | 2.85E+00 | 4.02E-01 | no | | intergenic_variant | MODIFIER |
| 2 | 161379375 | AX-170240401 | 0 | G | A | 0.158 | 1.63E-01 | 1.97E-01 | 2.76E+00 | 2.84E+00 | 4.02E-01 | Yes | | intergenic_variant | MODIFIER |
| 2 | 161380047 | AX-170476108 | 0 | T | C | 0.158 | 1.63E-01 | 1.97E-01 | 2.76E+00 | 2.84E+00 | 4.02E-01 | Yes | | intergenic_variant | MODIFIER |
| 2 | 161380222 | AX-170088821 | 0 | G | T | 0.158 | 1.63E-01 | 1.97E-01 | 2.76E+00 | 2.84E+00 | 4.02E-01 | Yes | | intergenic_variant | MODIFIER |
| 2 | 161391616 | AX-170283987 | 0 | C | T | 0.158 | 1.63E-01 | 1.97E-01 | 2.76E+00 | 2.84E+00 | 4.02E-01 | Yes | | regulatory_region_variant | MODIFIER |
| 2 | 161408011 | AX-170494899 | 0 | T | C | 0.158 | 1.63E-01 | 1.97E-01 | 2.76E+00 | 2.84E+00 | 4.02E-01 | Yes | | intergenic_variant | MODIFIER |
| 2 | 161417699 | AX-170378361 | 0 | C | T | 0.158 | 1.63E-01 | 1.97E-01 | 2.76E+00 | 2.84E+00 | 4.02E-01 | Yes | | intergenic_variant | MODIFIER |
| 2 | 161418497 | AX-170181869 | 0 | C | T | 0.158 | 1.63E-01 | 1.97E-01 | 2.76E+00 | 2.84E+00 | 4.02E-01 | Yes | | intergenic_variant | MODIFIER |
| 2 | 161419773 | AX-170340823 | 0 | A | G | 0.158 | 1.63E-01 | 1.97E-01 | 2.76E+00 | 2.84E+00 | 4.02E-01 | Yes | | intergenic_variant | MODIFIER |
| 2 | 161422205 | AX-170188640 | 0 | G | T | 0.158 | 1.63E-01 | 1.97E-01 | 2.76E+00 | 2.84E+00 | 4.02E-01 | Yes | | intergenic_variant | MODIFIER |
| 2 | 161426780 | AX-97823369 | 0 | C | T | 0.158 | 1.63E-01 | 1.97E-01 | 2.76E+00 | 2.84E+00 | 4.02E-01 | Yes | | intergenic_variant | MODIFIER |
| 2 | 161443315 | AX-170019538 | 0 | A | G | 0.158 | 1.63E-01 | 1.97E-01 | 2.76E+00 | 2.84E+00 | 4.02E-01 | Yes | | intergenic_variant | MODIFIER |
| 2 | 161443645 | AX-170186598 | 0 | T | C | 0.158 | 1.63E-01 | 1.97E-01 | 2.76E+00 | 2.84E+00 | 4.02E-01 | Yes | | intergenic_variant | MODIFIER |
| 2 | 161497016 | AX-97839748 | 0 | T | C | 0.158 | 1.63E-01 | 1.97E-01 | 2.76E+00 | 2.84E+00 | 4.02E-01 | Yes | | intergenic_variant | MODIFIER |
| 2 | 161690930 | AX-170422531 | 0 | T | C | 0.158 | 1.63E-01 | 1.97E-01 | 2.76E+00 | 2.84E+00 | 4.02E-01 | Yes | Ptprt | intron_variant | MODIFIER |
| 2 | 161693127 | AX-97855721 | 0 | G | C | 0.158 | 1.63E-01 | 1.97E-01 | 2.76E+00 | 2.84E+00 | 4.02E-01 | Yes | Ptprt | intron_variant | MODIFIER |
| 2 | 161698862 | AX-170215348 | 0 | C | A | 0.158 | 1.63E-01 | 1.97E-01 | 2.76E+00 | 2.84E+00 | 4.02E-01 | Yes | Ptprt | intron_variant | MODIFIER |
| 2 | 161704551 | AX-170468446 | 0 | A | G | 0.158 | 1.63E-01 | 1.97E-01 | 2.76E+00 | 2.84E+00 | 4.02E-01 | Yes | Ptprt | intron_variant | MODIFIER |
| 2 | 161705218 | AX-170527875 | 0 | C | G | 0.158 | 1.63E-01 | 1.97E-01 | 2.76E+00 | 2.84E+00 | 4.02E-01 | Yes | Ptprt | intron_variant | MODIFIER |
| 2 | 161705736 | AX-170322031 | 0 | G | A | 0.158 | 1.63E-01 | 1.97E-01 | 2.76E+00 | 2.84E+00 | 4.02E-01 | Yes | Ptprt | intron_variant | MODIFIER |
| 2 | 161706642 | AX-170275270 | 0 | C | G | 0.158 | 1.63E-01 | 1.97E-01 | 2.76E+00 | 2.84E+00 | 4.02E-01 | Yes | Ptprt | intron_variant | MODIFIER |
| 2 | 161710463 | AX-170182040 | 0 | A | G | 0.158 | 1.63E-01 | 1.97E-01 | 2.76E+00 | 2.84E+00 | 4.02E-01 | Yes | Ptprt | intron_variant | MODIFIER |
| 2 | 161720057 | AX-170100964 | 0 | T | C | 0.158 | 1.63E-01 | 1.97E-01 | 2.76E+00 | 2.84E+00 | 4.02E-01 | Yes | Ptprt | intron_variant | MODIFIER |
| 2 | 161731064 | AX-97792586 | 0 | G | A | 0.158 | 1.63E-01 | 1.97E-01 | 2.76E+00 | 2.84E+00 | 4.02E-01 | Yes | Ptprt | intron_variant | MODIFIER |
| 2 | 161736519 | AX-170418753 | 0 | C | A | 0.158 | 1.63E-01 | 1.97E-01 | 2.76E+00 | 2.84E+00 | 4.02E-01 | Yes | Ptprt | intron_variant | MODIFIER |
| 2 | 161746136 | AX-170112275 | 0 | A | G | 0.158 | 1.63E-01 | 1.97E-01 | 2.76E+00 | 2.84E+00 | 4.02E-01 | Yes | Ptprt | intron_variant | MODIFIER |
| 2 | 161761947 | AX-97812188 | 0 | C | T | 0.158 | 1.63E-01 | 1.97E-01 | 2.76E+00 | 2.84E+00 | 4.02E-01 | Yes | Ptprt | intron_variant | MODIFIER |
| 2 | 161763129 | AX-170186626 | 0 | A | C | 0.158 | 1.63E-01 | 1.97E-01 | 2.76E+00 | 2.84E+00 | 4.02E-01 | Yes | Ptprt | intron_variant | MODIFIER |
| 2 | 161763538 | AX-170179016 | 0 | A | G | 0.158 | 1.63E-01 | 1.97E-01 | 2.76E+00 | 2.84E+00 | 4.02E-01 | Yes | Ptprt | intron_variant | MODIFIER |
| 2 | 161763922 | AX-170486407 | 0 | C | T | 0.158 | 1.63E-01 | 1.97E-01 | 2.76E+00 | 2.84E+00 | 4.02E-01 | Yes | Ptprt | intron_variant | MODIFIER |
| 2 | 161767709 | AX-170065465 | 0 | C | T | 0.158 | 1.63E-01 | 1.97E-01 | 2.76E+00 | 2.84E+00 | 4.02E-01 | Yes | Ptprt | intron_variant | MODIFIER |
| 2 | 161770625 | AX-170311841 | 0 | G | C | 0.158 | 1.63E-01 | 1.97E-01 | 2.76E+00 | 2.84E+00 | 4.02E-01 | Yes | Ptprt | intron_variant | MODIFIER |
| 2 | 161795159 | AX-170166089 | 0 | C | T | 0.158 | 1.63E-01 | 1.97E-01 | 2.76E+00 | 2.84E+00 | 4.02E-01 | Yes | Ptprt | intron_variant | MODIFIER |
| 2 | 161795240 | AX-170460751 | 0 | C | T | 0.158 | 1.63E-01 | 1.97E-01 | 2.76E+00 | 2.84E+00 | 4.02E-01 | Yes | Ptprt | intron_variant | MODIFIER |
| 2 | 161815449 | AX-170251903 | 0 | C | A | 0.158 | 1.63E-01 | 1.97E-01 | 2.76E+00 | 2.84E+00 | 4.02E-01 | Yes | Ptprt | intron_variant | MODIFIER |
| 2 | 161815865 | AX-170164034 | 0 | T | C | 0.158 | 1.63E-01 | 1.97E-01 | 2.76E+00 | 2.84E+00 | 4.02E-01 | Yes | Ptprt | intron_variant | MODIFIER |
| 2 | 161823388 | AX-97833995 | 0 | T | C | 0.158 | 1.63E-01 | 1.97E-01 | 2.76E+00 | 2.84E+00 | 4.02E-01 | Yes | Ptprt | intron_variant | MODIFIER |
| 2 | 161825106 | AX-170156469 | 0 | G | C | 0.158 | 1.63E-01 | 1.97E-01 | 2.76E+00 | 2.84E+00 | 4.02E-01 | Yes | Ptprt | intron_variant | MODIFIER |
| 2 | 161825532 | AX-170449347 | 0 | T | A | 0.158 | 1.63E-01 | 1.97E-01 | 2.76E+00 | 2.84E+00 | 4.02E-01 | Yes | Ptprt | intron_variant | MODIFIER |

| | | | | | | | | | | | | | | | |
|---|---|---|---|---|---|---|---|---|---|---|---|---|---|---|---|
| 2 | 161876357 | AX-170265428 | 0 | A | G | 0.158 | 1.63E-01 | 1.97E-01 | 2.76E+00 | 2.84E+00 | 4.02E-01 | Yes | Ptprt | intron_variant | MODIFIER |
| 2 | 161877422 | AX-170382207 | 0 | A | G | 0.158 | 1.63E-01 | 1.97E-01 | 2.76E+00 | 2.84E+00 | 4.02E-01 | Yes | Ptprt | intron_variant | MODIFIER |
| 2 | 161881154 | AX-170376927 | 0 | A | G | 0.158 | 1.63E-01 | 1.97E-01 | 2.76E+00 | 2.84E+00 | 4.02E-01 | Yes | Ptprt | intron_variant | MODIFIER |
| 2 | 161886393 | AX-170367947 | 0 | A | G | 0.158 | 1.63E-01 | 1.97E-01 | 2.76E+00 | 2.84E+00 | 4.02E-01 | Yes | Ptprt | intron_variant | MODIFIER |
| 2 | 161895302 | AX-170099286 | 0 | T | C | 0.158 | 1.63E-01 | 1.97E-01 | 2.76E+00 | 2.84E+00 | 4.02E-01 | Yes | Ptprt | intron_variant | MODIFIER |
| 2 | 161898199 | AX-170065466 | 0 | A | G | 0.158 | 1.63E-01 | 1.97E-01 | 2.76E+00 | 2.84E+00 | 4.02E-01 | Yes | Ptprt | intron_variant | MODIFIER |
| 2 | 161900549 | AX-170498778 | 0 | T | C | 0.158 | 1.63E-01 | 1.97E-01 | 2.76E+00 | 2.84E+00 | 4.02E-01 | Yes | Ptprt | intron_variant | MODIFIER |
| 2 | 161900833 | AX-170382032 | 0 | C | T | 0.158 | 1.63E-01 | 1.97E-01 | 2.76E+00 | 2.84E+00 | 4.02E-01 | Yes | Ptprt | intron_variant | MODIFIER |
| 2 | 161915269 | AX-170164160 | 0 | A | G | 0.158 | 1.63E-01 | 1.97E-01 | 2.76E+00 | 2.84E+00 | 4.02E-01 | Yes | Ptprt | intron_variant | MODIFIER |
| 2 | 161917449 | AX-170357726 | 0 | T | C | 0.158 | 1.63E-01 | 1.97E-01 | 2.76E+00 | 2.84E+00 | 4.02E-01 | Yes | Ptprt | intron_variant | MODIFIER |
| 2 | 161955109 | AX-170430189 | 0 | C | T | 0.158 | 1.63E-01 | 1.97E-01 | 2.76E+00 | 2.84E+00 | 4.02E-01 | Yes | Ptprt | intron_variant | MODIFIER |
| 2 | 161956273 | AX-170217531 | 0 | T | C | 0.158 | 1.63E-01 | 1.97E-01 | 2.76E+00 | 2.84E+00 | 4.02E-01 | Yes | Ptprt | intron_variant | MODIFIER |
| 2 | 161958779 | AX-170019551 | 0 | A | C | 0.158 | 1.63E-01 | 1.97E-01 | 2.76E+00 | 2.84E+00 | 4.02E-01 | Yes | Ptprt | intron_variant | MODIFIER |
| 2 | 161959108 | AX-170226438 | 0 | T | C | 0.158 | 1.63E-01 | 1.97E-01 | 2.76E+00 | 2.84E+00 | 4.02E-01 | Yes | Ptprt | intron_variant | MODIFIER |
| 2 | 161959982 | AX-170019548 | 0 | G | A | 0.158 | 1.63E-01 | 1.97E-01 | 2.76E+00 | 2.84E+00 | 4.02E-01 | Yes | Ptprt | intron_variant | MODIFIER |
| 2 | 161960822 | AX-170124687 | 0 | C | T | 0.158 | 1.63E-01 | 1.97E-01 | 2.76E+00 | 2.84E+00 | 4.02E-01 | Yes | Ptprt | intron_variant | MODIFIER |
| 2 | 162003676 | AX-170267841 | 0 | A | G | 0.158 | 1.63E-01 | 1.97E-01 | 2.76E+00 | 2.84E+00 | 4.02E-01 | Yes | Ptprt | intron_variant | MODIFIER |
| 2 | 162015721 | AX-170019544 | 0 | T | C | 0.158 | 1.63E-01 | 1.97E-01 | 2.76E+00 | 2.84E+00 | 4.02E-01 | Yes | Ptprt | intron_variant | MODIFIER |
| 2 | 162016179 | AX-170065469 | 0 | A | G | 0.158 | 1.63E-01 | 1.97E-01 | 2.76E+00 | 2.84E+00 | 4.02E-01 | Yes | Ptprt | intron_variant | MODIFIER |
| 2 | 162024066 | AX-170201331 | 0 | T | C | 0.158 | 1.63E-01 | 1.97E-01 | 2.76E+00 | 2.84E+00 | 4.02E-01 | Yes | Ptprt | intron_variant | MODIFIER |
| 2 | 162032042 | AX-170019550 | 0 | G | A | 0.158 | 1.63E-01 | 1.97E-01 | 2.76E+00 | 2.84E+00 | 4.02E-01 | Yes | Ptprt | intron_variant | MODIFIER |
| 2 | 162032114 | AX-170019549 | 0 | A | G | 0.158 | 1.63E-01 | 1.97E-01 | 2.76E+00 | 2.84E+00 | 4.02E-01 | Yes | Ptprt | intron_variant | MODIFIER |
| 2 | 162041762 | AX-170019552 | 0 | A | G | 0.158 | 1.63E-01 | 1.97E-01 | 2.76E+00 | 2.84E+00 | 4.02E-01 | Yes | Ptprt | intron_variant | MODIFIER |
| 2 | 162045659 | AX-170134357 | 0 | A | G | 0.158 | 1.63E-01 | 1.97E-01 | 2.76E+00 | 2.84E+00 | 4.02E-01 | Yes | Ptprt | intron_variant | MODIFIER |
| 2 | 162070724 | AX-170316859 | 0 | G | A | 0.158 | 1.63E-01 | 1.97E-01 | 2.76E+00 | 2.84E+00 | 4.02E-01 | Yes | Ptprt | intron_variant | MODIFIER |
| 2 | 162071209 | AX-170301939 | 0 | A | G | 0.158 | 1.63E-01 | 1.97E-01 | 2.76E+00 | 2.84E+00 | 4.02E-01 | Yes | Ptprt | intron_variant | MODIFIER |
| 2 | 162086895 | AX-97850152 | 0 | T | G | 0.158 | 1.63E-01 | 1.97E-01 | 2.76E+00 | 2.84E+00 | 4.02E-01 | Yes | Ptprt | intron_variant | MODIFIER |
| 2 | 162087037 | AX-170166937 | 0 | A | G | 0.158 | 1.63E-01 | 1.97E-01 | 2.76E+00 | 2.84E+00 | 4.02E-01 | Yes | Ptprt | intron_variant | MODIFIER |
| 2 | 162087101 | AX-170200060 | 0 | T | C | 0.158 | 1.63E-01 | 1.97E-01 | 2.76E+00 | 2.84E+00 | 4.02E-01 | Yes | Ptprt | intron_variant | MODIFIER |
| 2 | 162088224 | AX-170447358 | 0 | C | T | 0.158 | 1.63E-01 | 1.97E-01 | 2.76E+00 | 2.84E+00 | 4.02E-01 | Yes | Ptprt | intron_variant | MODIFIER |
| 2 | 162101616 | AX-97815517 | 0 | G | A | 0.158 | 1.63E-01 | 1.97E-01 | 2.76E+00 | 2.84E+00 | 4.02E-01 | Yes | Ptprt | intron_variant | MODIFIER |
| 2 | 162131739 | AX-170223417 | 0 | G | C | 0.158 | 1.63E-01 | 1.97E-01 | 2.76E+00 | 2.84E+00 | 4.02E-01 | Yes | Ptprt | intron_variant | MODIFIER |
| 2 | 162133576 | AX-170345864 | 0 | G | A | 0.158 | 1.63E-01 | 1.97E-01 | 2.76E+00 | 2.84E+00 | 4.02E-01 | Yes | Ptprt | intron_variant | MODIFIER |
| 2 | 162134406 | AX-170516632 | 0 | T | A | 0.158 | 1.63E-01 | 1.97E-01 | 2.76E+00 | 2.84E+00 | 4.02E-01 | Yes | Ptprt | intron_variant | MODIFIER |
| 2 | 162134592 | AX-170116468 | 0 | T | C | 0.158 | 1.63E-01 | 1.97E-01 | 2.76E+00 | 2.84E+00 | 4.02E-01 | Yes | Ptprt | intron_variant | MODIFIER |
| 2 | 162137058 | AX-170460212 | 0 | T | A | 0.158 | 1.63E-01 | 1.97E-01 | 2.76E+00 | 2.84E+00 | 4.02E-01 | Yes | Ptprt | intron_variant | MODIFIER |
| 2 | 162138744 | AX-97806899 | 0 | G | A | 0.158 | 1.63E-01 | 1.97E-01 | 2.76E+00 | 2.84E+00 | 4.02E-01 | Yes | Ptprt | intron_variant | MODIFIER |
| 2 | 162139286 | AX-170255541 | 0 | T | G | 0.158 | 1.63E-01 | 1.97E-01 | 2.76E+00 | 2.84E+00 | 4.02E-01 | Yes | Ptprt | intron_variant | MODIFIER |
| 2 | 162139552 | AX-170251990 | 0 | C | T | 0.158 | 1.63E-01 | 1.97E-01 | 2.76E+00 | 2.84E+00 | 4.02E-01 | Yes | Ptprt | intron_variant | MODIFIER |
| 2 | 162203991 | AX-170505959 | 0 | T | A | 0.158 | 1.63E-01 | 1.97E-01 | 2.76E+00 | 2.84E+00 | 4.02E-01 | Yes | Ptprt | intron_variant | MODIFIER |
| 2 | 162205745 | AX-170259764 | 0 | A | G | 0.158 | 1.63E-01 | 1.97E-01 | 2.76E+00 | 2.84E+00 | 4.02E-01 | Yes | Ptprt | intron_variant | MODIFIER |
| 2 | 162214434 | AX-170157456 | 0 | G | C | 0.158 | 1.63E-01 | 1.97E-01 | 2.76E+00 | 2.84E+00 | 4.02E-01 | Yes | Ptprt | intron_variant | MODIFIER |
| 2 | 162217639 | AX-170276180 | 0 | C | T | 0.158 | 1.63E-01 | 1.97E-01 | 2.76E+00 | 2.84E+00 | 4.02E-01 | Yes | Ptprt | intron_variant | MODIFIER |
| 2 | 162223841 | AX-170296686 | 0 | A | G | 0.158 | 1.63E-01 | 1.97E-01 | 2.76E+00 | 2.84E+00 | 4.02E-01 | Yes | Ptprt | intron_variant | MODIFIER |
| 2 | 162224335 | AX-170153329 | 0 | T | C | 0.158 | 1.63E-01 | 1.97E-01 | 2.76E+00 | 2.84E+00 | 4.02E-01 | Yes | Ptprt | intron_variant | MODIFIER |
| 2 | 162232635 | AX-170184882 | 0 | C | T | 0.158 | 1.63E-01 | 1.97E-01 | 2.76E+00 | 2.84E+00 | 4.02E-01 | Yes | Ptprt | intron_variant | MODIFIER |
| 2 | 162233452 | AX-170278522 | 0 | A | G | 0.158 | 1.63E-01 | 1.97E-01 | 2.76E+00 | 2.84E+00 | 4.02E-01 | Yes | Ptprt | intron_variant | MODIFIER |
| 2 | 162233839 | AX-170492981 | 0 | A | G | 0.158 | 1.63E-01 | 1.97E-01 | 2.76E+00 | 2.84E+00 | 4.02E-01 | Yes | Ptprt | intron_variant | MODIFIER |
| 2 | 162234599 | AX-97794446 | 0 | A | G | 0.158 | 1.63E-01 | 1.97E-01 | 2.76E+00 | 2.84E+00 | 4.02E-01 | Yes | Ptprt | intron_variant | MODIFIER |
| 2 | 162279299 | AX-170118371 | 0 | T | C | 0.158 | 1.63E-01 | 1.97E-01 | 2.76E+00 | 2.84E+00 | 4.02E-01 | Yes | Ptprt | intron_variant | MODIFIER |
| 2 | 162350691 | AX-170245891 | 0 | G | A | 0.158 | 1.63E-01 | 1.97E-01 | 2.76E+00 | 2.84E+00 | 4.02E-01 | Yes | Ptprt | intron_variant | MODIFIER |
| 2 | 162353151 | AX-97797756 | 0 | C | T | 0.158 | 1.63E-01 | 1.97E-01 | 2.76E+00 | 2.84E+00 | 4.02E-01 | Yes | Ptprt | intron_variant | MODIFIER |
| 2 | 162353777 | AX-170128832 | 0 | G | A | 0.158 | 1.63E-01 | 1.97E-01 | 2.76E+00 | 2.84E+00 | 4.02E-01 | Yes | Ptprt | intron_variant | MODIFIER |
| 2 | 162353867 | AX-170391234 | 0 | T | G | 0.158 | 1.63E-01 | 1.97E-01 | 2.76E+00 | 2.84E+00 | 4.02E-01 | Yes | Ptprt | intron_variant | MODIFIER |

| | | | | | | | | | | | | | | | |
|---|---|---|---|---|---|---|---|---|---|---|---|---|---|---|---|
| 2 | 162355739 | AX-170173859 | 0 | C | A | 0.158 | 1.63E-01 | 1.97E-01 | 2.76E+00 | 2.84E+00 | 4.02E-01 | Yes | Ptprt | intron_variant | MODIFIER |
| 2 | 162365414 | AX-170019546 | 0 | A | G | 0.158 | 1.63E-01 | 1.97E-01 | 2.76E+00 | 2.84E+00 | 4.02E-01 | Yes | Ptprt | intron_variant | MODIFIER |
| 2 | 162381975 | AX-170372932 | 0 | T | C | 0.158 | 1.63E-01 | 1.97E-01 | 2.76E+00 | 2.84E+00 | 4.02E-01 | Yes | Ptprt | intron_variant | MODIFIER |
| 2 | 162382052 | AX-170260247 | 0 | T | C | 0.158 | 1.63E-01 | 1.97E-01 | 2.76E+00 | 2.84E+00 | 4.02E-01 | Yes | Ptprt | intron_variant | MODIFIER |
| 2 | 162383434 | AX-170157856 | 0 | A | G | 0.158 | 1.63E-01 | 1.97E-01 | 2.76E+00 | 2.84E+00 | 4.02E-01 | Yes | Ptprt | intron_variant | MODIFIER |
| 2 | 162385872 | AX-170440749 | 0 | G | T | 0.158 | 1.63E-01 | 1.97E-01 | 2.76E+00 | 2.84E+00 | 4.02E-01 | Yes | Ptprt | intron_variant | MODIFIER |
| 2 | 162391430 | AX-170466345 | 0 | G | C | 0.158 | 1.63E-01 | 1.97E-01 | 2.76E+00 | 2.84E+00 | 4.02E-01 | Yes | Ptprt | intron_variant | MODIFIER |
| 2 | 162411072 | AX-170243585 | 0 | C | G | 0.158 | 1.63E-01 | 1.97E-01 | 2.76E+00 | 2.84E+00 | 4.02E-01 | Yes | Ptprt | intron_variant | MODIFIER |
| 2 | 162418546 | AX-97829289 | 0 | C | T | 0.158 | 1.63E-01 | 1.97E-01 | 2.76E+00 | 2.84E+00 | 4.02E-01 | Yes | Ptprt | intron_variant | MODIFIER |
| 2 | 162421060 | AX-170471445 | 0 | G | A | 0.158 | 1.63E-01 | 1.97E-01 | 2.76E+00 | 2.84E+00 | 4.02E-01 | Yes | Ptprt | intron_variant | MODIFIER |
| 2 | 162444126 | AX-170368821 | 0 | C | T | 0.158 | 1.63E-01 | 1.97E-01 | 2.76E+00 | 2.84E+00 | 4.02E-01 | Yes | Ptprt | intron_variant | MODIFIER |
| 2 | 162454315 | AX-170192978 | 0 | C | T | 0.158 | 1.63E-01 | 1.97E-01 | 2.76E+00 | 2.84E+00 | 4.02E-01 | Yes | Ptprt | intron_variant | MODIFIER |
| 2 | 162458918 | AX-170226327 | 0 | T | C | 0.158 | 1.63E-01 | 1.97E-01 | 2.76E+00 | 2.84E+00 | 4.02E-01 | Yes | Ptprt | intron_variant | MODIFIER |
| 2 | 162460784 | AX-170326504 | 0 | C | G | 0.158 | 1.63E-01 | 1.97E-01 | 2.76E+00 | 2.84E+00 | 4.02E-01 | Yes | Ptprt | intron_variant | MODIFIER |
| 2 | 162462520 | AX-170494023 | 0 | G | A | 0.158 | 1.63E-01 | 1.97E-01 | 2.76E+00 | 2.84E+00 | 4.02E-01 | Yes | Ptprt | intron_variant | MODIFIER |
| 2 | 162485071 | AX-170088826 | 0 | C | T | 0.158 | 1.63E-01 | 1.97E-01 | 2.76E+00 | 2.84E+00 | 4.02E-01 | Yes | Ptprt | intron_variant | MODIFIER |
| 2 | 162495375 | AX-97812752 | 0 | G | A | 0.158 | 1.63E-01 | 1.97E-01 | 2.76E+00 | 2.84E+00 | 4.02E-01 | Yes | Ptprt | intron_variant | MODIFIER |
| 2 | 162498074 | AX-170441941 | 0 | A | G | 0.158 | 1.63E-01 | 1.97E-01 | 2.76E+00 | 2.84E+00 | 4.02E-01 | Yes | Ptprt | intron_variant | MODIFIER |
| 2 | 162498457 | AX-170347057 | 0 | C | T | 0.158 | 1.63E-01 | 1.97E-01 | 2.76E+00 | 2.84E+00 | 4.02E-01 | Yes | Ptprt | intron_variant | MODIFIER |
| 2 | 162500475 | AX-170388597 | 0 | A | G | 0.158 | 1.63E-01 | 1.97E-01 | 2.76E+00 | 2.84E+00 | 4.02E-01 | Yes | Ptprt | intron_variant | MODIFIER |
| 2 | 162500615 | AX-170175516 | 0 | C | G | 0.158 | 1.63E-01 | 1.97E-01 | 2.76E+00 | 2.84E+00 | 4.02E-01 | Yes | Ptprt | intron_variant | MODIFIER |
| 2 | 162501117 | AX-170385833 | 0 | A | G | 0.158 | 1.63E-01 | 1.97E-01 | 2.76E+00 | 2.84E+00 | 4.02E-01 | Yes | Ptprt | intron_variant | MODIFIER |
| 2 | 162503147 | AX-170449697 | 0 | A | G | 0.158 | 1.63E-01 | 1.97E-01 | 2.76E+00 | 2.84E+00 | 4.02E-01 | Yes | Ptprt | intron_variant | MODIFIER |
| 2 | 162524209 | AX-97793045 | 0 | T | G | 0.158 | 1.63E-01 | 1.97E-01 | 2.76E+00 | 2.84E+00 | 4.02E-01 | Yes | Ptprt | intron_variant | MODIFIER |
| 2 | 162528003 | AX-170019555 | 0 | A | G | 0.158 | 1.63E-01 | 1.97E-01 | 2.76E+00 | 2.84E+00 | 4.02E-01 | Yes | Ptprt | intron_variant | MODIFIER |
| 2 | 162604572 | AX-97791951 | 0 | T | C | 0.158 | 1.63E-01 | 1.97E-01 | 2.76E+00 | 2.84E+00 | 4.02E-01 | Yes | Ptprt | intron_variant | MODIFIER |
| 2 | 162615159 | AX-170019556 | 0 | A | G | 0.158 | 1.63E-01 | 1.97E-01 | 2.76E+00 | 2.84E+00 | 4.02E-01 | Yes | Ptprt | intron_variant | MODIFIER |
| 2 | 162616748 | AX-170019554 | 0 | A | G | 0.158 | 1.63E-01 | 1.97E-01 | 2.76E+00 | 2.84E+00 | 4.02E-01 | Yes | Ptprt | intron_variant | MODIFIER |
| 2 | 162636556 | AX-170065483 | 0 | G | C | 0.158 | 1.63E-01 | 1.97E-01 | 2.76E+00 | 2.84E+00 | 4.02E-01 | Yes | Ptprt | intron_variant | MODIFIER |
| 2 | 162637796 | AX-170458829 | 0 | T | C | 0.158 | 1.63E-01 | 1.97E-01 | 2.76E+00 | 2.84E+00 | 4.02E-01 | Yes | Ptprt | intron_variant | MODIFIER |
| 2 | 162671127 | AX-170019560 | 0 | C | T | 0.158 | 1.63E-01 | 1.97E-01 | 2.76E+00 | 2.84E+00 | 4.02E-01 | Yes | 9430021M05Rik | downstream_gene_variant | MODIFIER |
| 2 | 162672453 | AX-170378573 | 0 | T | C | 0.158 | 1.63E-01 | 1.97E-01 | 2.76E+00 | 2.84E+00 | 4.02E-01 | Yes | | intergenic_variant | MODIFIER |
| 2 | 162713966 | AX-170019561 | 0 | C | T | 0.158 | 1.63E-01 | 1.97E-01 | 2.76E+00 | 2.84E+00 | 4.02E-01 | Yes | | intergenic_variant | MODIFIER |
| 2 | 162741060 | AX-170429771 | 0 | C | T | 0.158 | 1.63E-01 | 1.97E-01 | 2.76E+00 | 2.84E+00 | 4.02E-01 | Yes | | intergenic_variant | MODIFIER |
| 2 | 162741072 | AX-170357641 | 0 | G | A | 0.158 | 1.63E-01 | 1.97E-01 | 2.76E+00 | 2.84E+00 | 4.02E-01 | Yes | | intergenic_variant | MODIFIER |
| 2 | 162742529 | AX-170512519 | 0 | C | T | 0.158 | 1.63E-01 | 1.97E-01 | 2.76E+00 | 2.84E+00 | 4.02E-01 | Yes | | intergenic_variant | MODIFIER |
| 2 | 162746990 | AX-170250448 | 0 | T | A | 0.158 | 1.63E-01 | 1.97E-01 | 2.76E+00 | 2.84E+00 | 4.02E-01 | Yes | | intergenic_variant | MODIFIER |
| 2 | 162779257 | AX-170151262 | 0 | A | G | 0.158 | 1.63E-01 | 1.97E-01 | 2.76E+00 | 2.84E+00 | 4.02E-01 | Yes | | intergenic_variant | MODIFIER |
| 2 | 162782152 | AX-97789330 | 0 | A | G | 0.158 | 1.63E-01 | 1.97E-01 | 2.76E+00 | 2.84E+00 | 4.02E-01 | Yes | | intergenic_variant | MODIFIER |
| 2 | 162793122 | AX-170118913 | 0 | T | C | 0.158 | 1.63E-01 | 1.97E-01 | 2.76E+00 | 2.84E+00 | 4.02E-01 | Yes | | intergenic_variant | MODIFIER |
| 2 | 162793195 | AX-170531377 | 0 | T | C | 0.158 | 1.63E-01 | 1.97E-01 | 2.76E+00 | 2.84E+00 | 4.02E-01 | Yes | | intergenic_variant | MODIFIER |
| 2 | 162794246 | AX-170307051 | 0 | G | A | 0.158 | 1.63E-01 | 1.97E-01 | 2.76E+00 | 2.84E+00 | 4.02E-01 | Yes | | intergenic_variant | MODIFIER |
| 2 | 162814184 | AX-170229103 | 0 | T | C | 0.158 | 1.63E-01 | 1.97E-01 | 2.76E+00 | 2.84E+00 | 4.02E-01 | Yes | Gm24478 | downstream_gene_variant | MODIFIER |
| 2 | 162818077 | AX-170519030 | 0 | G | T | 0.158 | 1.63E-01 | 1.97E-01 | 2.76E+00 | 2.84E+00 | 4.02E-01 | Yes | Gm24478 | upstream_gene_variant | MODIFIER |
| 2 | 162829245 | AX-170223720 | 0 | C | A | 0.158 | 1.63E-01 | 1.97E-01 | 2.76E+00 | 2.84E+00 | 4.02E-01 | Yes | | intergenic_variant | MODIFIER |
| 2 | 162830391 | AX-170104591 | 0 | C | T | 0.158 | 1.63E-01 | 1.97E-01 | 2.76E+00 | 2.84E+00 | 4.02E-01 | Yes | | intergenic_variant | MODIFIER |
| 2 | 162831543 | AX-170519787 | 0 | G | C | 0.158 | 1.63E-01 | 1.97E-01 | 2.76E+00 | 2.84E+00 | 4.02E-01 | Yes | | intergenic_variant | MODIFIER |
| 2 | 162833643 | AX-97790231 | 0 | C | T | 0.158 | 1.63E-01 | 1.97E-01 | 2.76E+00 | 2.84E+00 | 4.02E-01 | Yes | | intergenic_variant | MODIFIER |
| 2 | 162837849 | AX-170301517 | 0 | A | G | 0.158 | 1.63E-01 | 1.97E-01 | 2.76E+00 | 2.84E+00 | 4.02E-01 | Yes | | intergenic_variant | MODIFIER |
| 2 | 162849731 | AX-170304512 | 0 | A | G | 0.158 | 1.63E-01 | 1.97E-01 | 2.76E+00 | 2.84E+00 | 4.02E-01 | Yes | | intergenic_variant | MODIFIER |
| 2 | 162890805 | AX-170289136 | 0 | C | G | 0.158 | 1.63E-01 | 1.97E-01 | 2.76E+00 | 2.84E+00 | 4.02E-01 | Yes | | intergenic_variant | MODIFIER |
| 2 | 162909952 | AX-170384556 | 0 | G | A | 0.158 | 1.63E-01 | 1.97E-01 | 2.76E+00 | 2.84E+00 | 4.02E-01 | Yes | Gm11452 | downstream_gene_variant | MODIFIER |
| 2 | 162939859 | AX-170105898 | 0 | A | G | 0.158 | 1.63E-01 | 1.97E-01 | 2.76E+00 | 2.84E+00 | 4.02E-01 | Yes | Srsf6 | downstream_gene_variant | MODIFIER |
| 2 | 162941051 | AX-170249142 | 0 | A | T | 0.158 | 1.63E-01 | 1.97E-01 | 2.76E+00 | 2.84E+00 | 4.02E-01 | Yes | L3mbtl1 | upstream_gene_variant | MODIFIER |

| | | | | | | | | | | | | | | | |
|---|---|---|---|---|---|---|---|---|---|---|---|---|---|---|---|
| 2 | 162948169 | AX-97850688 | 0 | C | T | 0.158 | 1.63E-01 | 1.97E-01 | 2.76E+00 | 2.84E+00 | 4.02E-01 | Yes | L3mbtl1 | synonymous_variant | LOW |
| 2 | 162948476 | AX-170455847 | 0 | T | C | 0.158 | 1.63E-01 | 1.97E-01 | 2.76E+00 | 2.84E+00 | 4.02E-01 | Yes | L3mbtl1 | splice_region_variant,intron_variant | LOW |
| 2 | 162948518 | AX-170405271 | 0 | G | A | 0.158 | 1.63E-01 | 1.97E-01 | 2.76E+00 | 2.84E+00 | 4.02E-01 | Yes | L3mbtl1 | synonymous_variant | LOW |
| 2 | 162957294 | AX-170366807 | 0 | T | A | 0.158 | 1.63E-01 | 1.97E-01 | 2.76E+00 | 2.84E+00 | 4.02E-01 | Yes | L3mbtl1 | intron_variant | MODIFIER |
| 2 | 163141342 | AX-170019567 | 0 | T | C | 0.158 | 1.63E-01 | 1.97E-01 | 2.76E+00 | 2.84E+00 | 4.02E-01 | Yes | Gm11454 | intron_variant,non_coding_transcript_variant | MODIFIER |
| 2 | 163152539 | AX-97849791 | 0 | G | C | 0.158 | 1.63E-01 | 1.97E-01 | 2.76E+00 | 2.84E+00 | 4.02E-01 | Yes | Gm14254 | downstream_gene_variant | MODIFIER |
| 2 | 163156387 | AX-170019566 | 0 | T | C | 0.158 | 1.63E-01 | 1.97E-01 | 2.76E+00 | 2.84E+00 | 4.02E-01 | Yes | Gm14254 | upstream_gene_variant | MODIFIER |
| 2 | 163167366 | AX-170488532 | 0 | G | A | 0.158 | 1.63E-01 | 1.97E-01 | 2.76E+00 | 2.84E+00 | 4.02E-01 | Yes | | intergenic_variant | MODIFIER |
| 2 | 163167781 | AX-170372803 | 0 | G | A | 0.158 | 1.63E-01 | 1.97E-01 | 2.76E+00 | 2.84E+00 | 4.02E-01 | Yes | | intergenic_variant | MODIFIER |
| 2 | 163167833 | AX-170209470 | 0 | G | A | 0.158 | 1.63E-01 | 1.97E-01 | 2.76E+00 | 2.84E+00 | 4.02E-01 | Yes | | intergenic_variant | MODIFIER |
| 2 | 163211947 | AX-170460420 | 0 | C | T | 0.158 | 1.63E-01 | 1.97E-01 | 2.76E+00 | 2.84E+00 | 4.02E-01 | Yes | Tox2 | intron_variant | MODIFIER |
| 2 | 163224738 | AX-170065491 | 0 | C | T | 0.158 | 1.63E-01 | 1.97E-01 | 2.76E+00 | 2.84E+00 | 4.02E-01 | Yes | Tox2 | upstream_gene_variant | MODIFIER |
| 2 | 163235072 | AX-170308988 | 0 | T | C | 0.158 | 1.63E-01 | 1.97E-01 | 2.76E+00 | 2.84E+00 | 4.02E-01 | Yes | Tox2 | intron_variant | MODIFIER |
| 2 | 163237570 | AX-170276344 | 0 | G | A | 0.158 | 1.63E-01 | 1.97E-01 | 2.76E+00 | 2.84E+00 | 4.02E-01 | Yes | Tox2 | intron_variant | MODIFIER |
| 2 | 163246932 | AX-170299519 | 0 | A | G | 0.158 | 1.63E-01 | 1.97E-01 | 2.76E+00 | 2.84E+00 | 4.02E-01 | Yes | Tox2 | intron_variant | MODIFIER |
| 2 | 163247441 | AX-170303593 | 0 | T | C | 0.158 | 1.63E-01 | 1.97E-01 | 2.76E+00 | 2.84E+00 | 4.02E-01 | Yes | Tox2 | intron_variant | MODIFIER |
| 2 | 163254744 | AX-170420968 | 0 | T | C | 0.158 | 1.63E-01 | 1.97E-01 | 2.76E+00 | 2.84E+00 | 4.02E-01 | Yes | Tox2 | intron_variant | MODIFIER |
| 2 | 163254882 | AX-170246759 | 0 | A | G | 0.158 | 1.63E-01 | 1.97E-01 | 2.76E+00 | 2.84E+00 | 4.02E-01 | Yes | Tox2 | intron_variant | MODIFIER |
| 2 | 163258201 | AX-170239002 | 0 | C | T | 0.158 | 1.63E-01 | 1.97E-01 | 2.76E+00 | 2.84E+00 | 4.02E-01 | Yes | Tox2 | intron_variant | MODIFIER |
| 2 | 163260253 | AX-170184339 | 0 | T | C | 0.158 | 1.63E-01 | 1.97E-01 | 2.76E+00 | 2.84E+00 | 4.02E-01 | Yes | Tox2 | intron_variant | MODIFIER |
| 2 | 163272854 | AX-170189330 | 0 | A | G | 0.158 | 1.63E-01 | 1.97E-01 | 2.76E+00 | 2.84E+00 | 4.02E-01 | Yes | Tox2 | intron_variant | MODIFIER |
| 2 | 163272899 | AX-97839773 | 0 | A | G | 0.158 | 1.63E-01 | 1.97E-01 | 2.76E+00 | 2.84E+00 | 4.02E-01 | Yes | Tox2 | intron_variant | MODIFIER |
| 2 | 163370306 | AX-170257867 | 0 | T | C | 0.158 | 1.63E-01 | 1.97E-01 | 2.76E+00 | 2.84E+00 | 4.02E-01 | Yes | Jph2 | intron_variant | MODIFIER |
| 2 | 163468072 | AX-170373690 | 0 | G | T | 0.158 | 1.63E-01 | 1.97E-01 | 2.76E+00 | 2.84E+00 | 4.02E-01 | Yes | 2310001K24Rik | upstream_gene_variant | MODIFIER |
| 2 | 163755991 | AX-170393472 | 0 | C | T | 0.158 | 1.63E-01 | 1.97E-01 | 2.76E+00 | 2.84E+00 | 4.02E-01 | Yes | | intergenic_variant | MODIFIER |
| 2 | 163757065 | AX-170458066 | 0 | A | G | 0.158 | 1.63E-01 | 1.97E-01 | 2.76E+00 | 2.84E+00 | 4.02E-01 | Yes | | regulatory_region_variant | MODIFIER |
| 2 | 163757251 | AX-170412273 | 0 | C | T | 0.158 | 1.63E-01 | 1.97E-01 | 2.76E+00 | 2.84E+00 | 4.02E-01 | Yes | | regulatory_region_variant | MODIFIER |
| 2 | 163760372 | AX-170368639 | 0 | G | A | 0.158 | 1.63E-01 | 1.97E-01 | 2.76E+00 | 2.84E+00 | 4.02E-01 | Yes | | intergenic_variant | MODIFIER |
| 2 | 163764383 | AX-170260736 | 0 | T | C | 0.158 | 1.63E-01 | 1.97E-01 | 2.76E+00 | 2.84E+00 | 4.02E-01 | Yes | | intergenic_variant | MODIFIER |
| 2 | 163766371 | AX-170497114 | 0 | T | C | 0.158 | 1.63E-01 | 1.97E-01 | 2.76E+00 | 2.84E+00 | 4.02E-01 | Yes | | intergenic_variant | MODIFIER |
| 2 | 163766393 | AX-170499947 | 0 | T | C | 0.158 | 1.63E-01 | 1.97E-01 | 2.76E+00 | 2.84E+00 | 4.02E-01 | Yes | | intergenic_variant | MODIFIER |
| 2 | 163770387 | AX-170267692 | 0 | T | C | 0.158 | 1.63E-01 | 1.97E-01 | 2.76E+00 | 2.84E+00 | 4.02E-01 | Yes | | intergenic_variant | MODIFIER |
| 2 | 163786628 | AX-97789634 | 0 | T | C | 0.158 | 1.63E-01 | 1.97E-01 | 2.76E+00 | 2.84E+00 | 4.02E-01 | Yes | | intergenic_variant | MODIFIER |
| 2 | 163787139 | AX-170330294 | 0 | T | C | 0.158 | 1.63E-01 | 1.97E-01 | 2.76E+00 | 2.84E+00 | 4.02E-01 | Yes | | intergenic_variant | MODIFIER |
| 2 | 163809186 | AX-170476859 | 0 | G | A | 0.158 | 1.63E-01 | 1.97E-01 | 2.76E+00 | 2.84E+00 | 4.02E-01 | Yes | | intergenic_variant | MODIFIER |
| 2 | 163810743 | AX-170209325 | 0 | A | G | 0.158 | 1.63E-01 | 1.97E-01 | 2.76E+00 | 2.84E+00 | 4.02E-01 | Yes | | regulatory_region_variant | MODIFIER |
| 2 | 163827372 | AX-170450497 | 0 | G | A | 0.158 | 1.63E-01 | 1.97E-01 | 2.76E+00 | 2.84E+00 | 4.02E-01 | Yes | Ccn5 | intron_variant | MODIFIER |
| 2 | 163828401 | AX-170399667 | 0 | T | C | 0.158 | 1.63E-01 | 1.97E-01 | 2.76E+00 | 2.84E+00 | 4.02E-01 | Yes | Ccn5 | intron_variant | MODIFIER |
| 2 | 163849701 | AX-97856573 | 0 | T | C | 0.158 | 1.63E-01 | 1.97E-01 | 2.76E+00 | 2.84E+00 | 4.02E-01 | Yes | Kcnk15 | upstream_gene_variant | MODIFIER |
| 2 | 163875409 | AX-170291501 | 0 | C | T | 0.158 | 1.63E-01 | 1.97E-01 | 2.76E+00 | 2.84E+00 | 4.02E-01 | Yes | Rims4 | intron_variant | MODIFIER |
| 2 | 163876798 | AX-170382725 | 0 | C | A | 0.158 | 1.63E-01 | 1.97E-01 | 2.76E+00 | 2.84E+00 | 4.02E-01 | Yes | Rims4 | intron_variant | MODIFIER |
| 2 | 163877010 | AX-170269253 | 0 | A | G | 0.158 | 1.63E-01 | 1.97E-01 | 2.76E+00 | 2.84E+00 | 4.02E-01 | Yes | Rims4 | intron_variant | MODIFIER |
| 2 | 163877668 | AX-170372561 | 0 | C | G | 0.158 | 1.63E-01 | 1.97E-01 | 2.76E+00 | 2.84E+00 | 4.02E-01 | Yes | Rims4 | intron_variant | MODIFIER |
| 2 | 163896982 | AX-170257517 | 0 | T | C | 0.158 | 1.63E-01 | 1.97E-01 | 2.76E+00 | 2.84E+00 | 4.02E-01 | Yes | Rims4 | intron_variant | MODIFIER |
| 2 | 163925374 | AX-170342349 | 0 | C | T | 0.158 | 1.63E-01 | 1.97E-01 | 2.76E+00 | 2.84E+00 | 4.02E-01 | Yes | | intergenic_variant | MODIFIER |
| 2 | 163927722 | AX-170387955 | 0 | T | A | 0.158 | 1.63E-01 | 1.97E-01 | 2.76E+00 | 2.84E+00 | 4.02E-01 | Yes | | intergenic_variant | MODIFIER |
| 2 | 163928695 | AX-170193920 | 0 | C | G | 0.158 | 1.63E-01 | 1.97E-01 | 2.76E+00 | 2.84E+00 | 4.02E-01 | Yes | | intergenic_variant | MODIFIER |
| 2 | 163928737 | AX-170268419 | 0 | A | G | 0.158 | 1.63E-01 | 1.97E-01 | 2.76E+00 | 2.84E+00 | 4.02E-01 | Yes | | intergenic_variant | MODIFIER |
| 2 | 163928764 | AX-170094473 | 0 | T | C | 0.158 | 1.63E-01 | 1.97E-01 | 2.76E+00 | 2.84E+00 | 4.02E-01 | Yes | | intergenic_variant | MODIFIER |
| 2 | 163935711 | AX-170366616 | 0 | T | C | 0.158 | 1.63E-01 | 1.97E-01 | 2.76E+00 | 2.84E+00 | 4.02E-01 | Yes | | intergenic_variant | MODIFIER |
| 2 | 163941816 | AX-170065504 | 0 | A | G | 0.158 | 1.63E-01 | 1.97E-01 | 2.76E+00 | 2.84E+00 | 4.02E-01 | Yes | | intergenic_variant | MODIFIER |
| 2 | 163942799 | AX-170490284 | 0 | C | T | 0.158 | 1.63E-01 | 1.97E-01 | 2.76E+00 | 2.84E+00 | 4.02E-01 | Yes | | intergenic_variant | MODIFIER |
| 2 | 163954478 | AX-97830239 | 0 | T | C | 0.158 | 1.63E-01 | 1.97E-01 | 2.76E+00 | 2.84E+00 | 4.02E-01 | Yes | | intergenic_variant | MODIFIER |
| 2 | 164003122 | AX-170522911 | 0 | C | T | 0.158 | 1.63E-01 | 1.97E-01 | 2.76E+00 | 2.84E+00 | 4.02E-01 | Yes | Ywhab | intron_variant | MODIFIER |

| | | | | | | | | | | | | | | | |
|---|---|---|---|---|---|---|---|---|---|---|---|---|---|---|---|
| 2 | 164014231 | AX-170094474 | 0 | G | A | 0.158 | 1.63E-01 | 1.97E-01 | 2.76E+00 | 2.84E+00 | 4.02E-01 | Yes | Ywhab | intron_variant | MODIFIER |
| 2 | 164037080 | AX-97839826 | 0 | C | A | 0.158 | 1.63E-01 | 1.97E-01 | 2.76E+00 | 2.84E+00 | 4.02E-01 | Yes | Pabpc1l | intron_variant | MODIFIER |
| 2 | 164119319 | AX-97800580 | 0 | T | C | 0.158 | 1.63E-01 | 1.97E-01 | 2.76E+00 | 2.84E+00 | 4.02E-01 | Yes | Stk4 | intron_variant | MODIFIER |
| 2 | 164190920 | AX-170019579 | 0 | C | T | 0.158 | 1.63E-01 | 1.97E-01 | 2.76E+00 | 2.84E+00 | 4.02E-01 | Yes | Wfdc12 | upstream_gene_variant | MODIFIER |
| 2 | 164215949 | AX-170102534 | 0 | A | G | 0.158 | 1.63E-01 | 1.97E-01 | 2.76E+00 | 2.84E+00 | 4.02E-01 | Yes | Wfdc15b | intron_variant | MODIFIER |
| 2 | 164247427 | AX-97834210 | 0 | G | A | 0.158 | 1.63E-01 | 1.97E-01 | 2.76E+00 | 2.84E+00 | 4.02E-01 | Yes | | intergenic_variant | MODIFIER |
| 2 | 164390162 | AX-170286894 | 0 | A | G | 0.158 | 1.63E-01 | 1.97E-01 | 2.76E+00 | 2.84E+00 | 4.02E-01 | Yes | Matn4 | intron_variant | MODIFIER |
| 2 | 164414770 | AX-170291538 | 0 | A | T | 0.158 | 1.63E-01 | 1.97E-01 | 2.76E+00 | 2.84E+00 | 4.02E-01 | Yes | Rbpjl | 3_prime_UTR_variant | MODIFIER |
| 2 | 164417094 | AX-170344009 | 0 | T | C | 0.158 | 1.63E-01 | 1.97E-01 | 2.76E+00 | 2.84E+00 | 4.02E-01 | Yes | Rbpjl | downstream_gene_variant | MODIFIER |
| 2 | 164422917 | AX-170182015 | 0 | T | C | 0.158 | 1.63E-01 | 1.97E-01 | 2.76E+00 | 2.84E+00 | 4.02E-01 | Yes | Sdc4 | downstream_gene_variant | MODIFIER |
| 2 | 164422958 | AX-97799507 | 0 | T | C | 0.158 | 1.63E-01 | 1.97E-01 | 2.76E+00 | 2.84E+00 | 4.02E-01 | Yes | Sdc4 | downstream_gene_variant | MODIFIER |
| 2 | 164449766 | AX-170258440 | 0 | T | C | 0.158 | 1.63E-01 | 1.97E-01 | 2.76E+00 | 2.84E+00 | 4.02E-01 | Yes | | intergenic_variant | MODIFIER |
| 2 | 164452673 | AX-170456643 | 0 | C | T | 0.158 | 1.63E-01 | 1.97E-01 | 2.76E+00 | 2.84E+00 | 4.02E-01 | Yes | Sys1 | upstream_gene_variant | MODIFIER |
| 2 | 164460831 | AX-97845244 | 0 | C | T | 0.158 | 1.63E-01 | 1.97E-01 | 2.76E+00 | 2.84E+00 | 4.02E-01 | Yes | Sys1 | upstream_gene_variant | MODIFIER |
| 2 | 164472506 | AX-170208183 | 0 | A | G | 0.158 | 1.63E-01 | 1.97E-01 | 2.76E+00 | 2.84E+00 | 4.02E-01 | Yes | Trp53tg5 | intron_variant | MODIFIER |
| 2 | 164475512 | AX-170263513 | 0 | C | T | 0.158 | 1.63E-01 | 1.97E-01 | 2.76E+00 | 2.84E+00 | 4.02E-01 | Yes | Trp53tg5 | upstream_gene_variant | MODIFIER |
| 2 | 164475813 | AX-170512449 | 0 | A | G | 0.158 | 1.63E-01 | 1.97E-01 | 2.76E+00 | 2.84E+00 | 4.02E-01 | Yes | Trp53tg5 | upstream_gene_variant | MODIFIER |
| 2 | 164476284 | AX-97797018 | 0 | G | A | 0.158 | 1.63E-01 | 1.97E-01 | 2.76E+00 | 2.84E+00 | 4.02E-01 | Yes | Trp53tg5 | upstream_gene_variant | MODIFIER |
| 2 | 164508087 | AX-170509622 | 0 | T | A | 0.158 | 1.63E-01 | 1.97E-01 | 2.76E+00 | 2.84E+00 | 4.02E-01 | Yes | Pigt | 3_prime_UTR_variant | MODIFIER |
| 2 | 164512904 | AX-170311304 | 0 | G | A | 0.158 | 1.63E-01 | 1.97E-01 | 2.76E+00 | 2.84E+00 | 4.02E-01 | Yes | Pigt | downstream_gene_variant | MODIFIER |
| 2 | 164514315 | AX-170276758 | 0 | T | C | 0.158 | 1.63E-01 | 1.97E-01 | 2.76E+00 | 2.84E+00 | 4.02E-01 | Yes | | regulatory_region_variant | MODIFIER |
| 2 | 164521880 | AX-170235357 | 0 | A | G | 0.158 | 1.63E-01 | 1.97E-01 | 2.76E+00 | 2.84E+00 | 4.02E-01 | Yes | | regulatory_region_variant | MODIFIER |
| 2 | 164527738 | AX-97820773 | 0 | T | G | 0.158 | 1.63E-01 | 1.97E-01 | 2.76E+00 | 2.84E+00 | 4.02E-01 | Yes | | intergenic_variant | MODIFIER |
| 2 | 164530538 | AX-170527855 | 0 | C | T | 0.158 | 1.63E-01 | 1.97E-01 | 2.76E+00 | 2.84E+00 | 4.02E-01 | Yes | | intergenic_variant | MODIFIER |
| 2 | 164533948 | AX-170469661 | 0 | T | C | 0.158 | 1.63E-01 | 1.97E-01 | 2.76E+00 | 2.84E+00 | 4.02E-01 | Yes | | intergenic_variant | MODIFIER |
| 2 | 164546863 | AX-97793269 | 0 | G | T | 0.158 | 1.63E-01 | 1.97E-01 | 2.76E+00 | 2.84E+00 | 4.02E-01 | Yes | Gm14317 | upstream_gene_variant | MODIFIER |
| 2 | 164578276 | AX-170198049 | 0 | T | C | 0.158 | 1.63E-01 | 1.97E-01 | 2.76E+00 | 2.84E+00 | 4.02E-01 | Yes | Wfdc6a | downstream_gene_variant | MODIFIER |
| 2 | 164582079 | AX-97848395 | 0 | C | A | 0.158 | 1.63E-01 | 1.97E-01 | 2.76E+00 | 2.84E+00 | 4.02E-01 | Yes | Wfdc6a | intron_variant | MODIFIER |
| 2 | 164584733 | AX-170365040 | 0 | C | G | 0.158 | 1.63E-01 | 1.97E-01 | 2.76E+00 | 2.84E+00 | 4.02E-01 | Yes | Wfdc6a | intron_variant | MODIFIER |
| 2 | 164586457 | AX-170318242 | 0 | A | G | 0.158 | 1.63E-01 | 1.97E-01 | 2.76E+00 | 2.84E+00 | 4.02E-01 | Yes | Wfdc6a | upstream_gene_variant | MODIFIER |
| 2 | 164595006 | AX-170426016 | 0 | A | G | 0.158 | 1.63E-01 | 1.97E-01 | 2.76E+00 | 2.84E+00 | 4.02E-01 | Yes | Wfdc8 | downstream_gene_variant | MODIFIER |
| 2 | 164602407 | AX-97839804 | 0 | G | A | 0.158 | 1.63E-01 | 1.97E-01 | 2.76E+00 | 2.84E+00 | 4.02E-01 | Yes | Gm25512 | downstream_gene_variant | MODIFIER |
| 2 | 164602725 | AX-170368774 | 0 | T | C | 0.158 | 1.63E-01 | 1.97E-01 | 2.76E+00 | 2.84E+00 | 4.02E-01 | Yes | Gm25512 | downstream_gene_variant | MODIFIER |
| 2 | 164626091 | AX-170455556 | 0 | G | A | 0.158 | 1.63E-01 | 1.97E-01 | 2.76E+00 | 2.84E+00 | 4.02E-01 | Yes | Gm24473 | downstream_gene_variant | MODIFIER |
| 2 | 164641700 | AX-170521714 | 0 | A | G | 0.158 | 1.63E-01 | 1.97E-01 | 2.76E+00 | 2.84E+00 | 4.02E-01 | Yes | Wfdc16 | upstream_gene_variant | MODIFIER |
| 2 | 164642803 | AX-97839777 | 0 | G | A | 0.158 | 1.63E-01 | 1.97E-01 | 2.76E+00 | 2.84E+00 | 4.02E-01 | Yes | Wfdc16 | upstream_gene_variant | MODIFIER |
| 2 | 164644677 | AX-170321999 | 0 | C | T | 0.158 | 1.63E-01 | 1.97E-01 | 2.76E+00 | 2.84E+00 | 4.02E-01 | Yes | Wfdc9 | downstream_gene_variant | MODIFIER |
| 2 | 164846648 | AX-170125094 | 0 | T | C | 0.158 | 1.63E-01 | 1.97E-01 | 2.76E+00 | 2.84E+00 | 4.02E-01 | Yes | Pltp | synonymous_variant | LOW |
| 2 | 164851814 | AX-97823614 | 0 | T | C | 0.158 | 1.63E-01 | 1.97E-01 | 2.76E+00 | 2.84E+00 | 4.02E-01 | Yes | Pltp | intron_variant | MODIFIER |
| 2 | 164852656 | AX-170487869 | 0 | T | C | 0.158 | 1.63E-01 | 1.97E-01 | 2.76E+00 | 2.84E+00 | 4.02E-01 | Yes | Pltp | synonymous_variant | LOW |
| 2 | 164852739 | AX-170499469 | 0 | C | A | 0.158 | 1.63E-01 | 1.97E-01 | 2.76E+00 | 2.84E+00 | 4.02E-01 | Yes | Pltp | intron_variant | MODIFIER |
| 2 | 164868082 | AX-170483986 | 0 | G | A | 0.158 | 1.63E-01 | 1.97E-01 | 2.76E+00 | 2.84E+00 | 4.02E-01 | Yes | | intergenic_variant | MODIFIER |
| 2 | 164874058 | AX-170297062 | 0 | A | T | 0.158 | 1.63E-01 | 1.97E-01 | 2.76E+00 | 2.84E+00 | 4.02E-01 | Yes | | intergenic_variant | MODIFIER |
| 2 | 164874826 | AX-170312986 | 0 | A | C | 0.158 | 1.63E-01 | 1.97E-01 | 2.76E+00 | 2.84E+00 | 4.02E-01 | Yes | Pcif1 | upstream_gene_variant | MODIFIER |
| 2 | 164901666 | AX-170262969 | 0 | C | T | 0.158 | 1.63E-01 | 1.97E-01 | 2.76E+00 | 2.84E+00 | 4.02E-01 | Yes | Zfp335 | intron_variant | MODIFIER |
| 2 | 164921161 | AX-170279927 | 0 | G | T | 0.158 | 1.63E-01 | 1.97E-01 | 2.76E+00 | 2.84E+00 | 4.02E-01 | Yes | Zfp335os | downstream_gene_variant | MODIFIER |
| 2 | 164926755 | AX-170379477 | 0 | C | G | 0.158 | 1.63E-01 | 1.97E-01 | 2.76E+00 | 2.84E+00 | 4.02E-01 | Yes | | intergenic_variant | MODIFIER |
| 2 | 164947462 | AX-170298953 | 0 | A | G | 0.158 | 1.63E-01 | 1.97E-01 | 2.76E+00 | 2.84E+00 | 4.02E-01 | Yes | Mmp9 | upstream_gene_variant | MODIFIER |
| 2 | 164954845 | AX-170492898 | 0 | C | A | 0.158 | 1.63E-01 | 1.97E-01 | 2.76E+00 | 2.84E+00 | 4.02E-01 | Yes | Mmp9 | missense_variant | MODERATE |
| 2 | 164963580 | AX-170269454 | 0 | G | A | 0.158 | 1.63E-01 | 1.97E-01 | 2.76E+00 | 2.84E+00 | 4.02E-01 | Yes | Slc12a5 | upstream_gene_variant | MODIFIER |
| 2 | 164966344 | AX-170391053 | 0 | G | T | 0.158 | 1.63E-01 | 1.97E-01 | 2.76E+00 | 2.84E+00 | 4.02E-01 | Yes | Slc12a5 | upstream_gene_variant | MODIFIER |
| 2 | 164966553 | AX-170407290 | 0 | T | G | 0.158 | 1.63E-01 | 1.97E-01 | 2.76E+00 | 2.84E+00 | 4.02E-01 | Yes | Slc12a5 | upstream_gene_variant | MODIFIER |
| 2 | 165004951 | AX-170390009 | 0 | C | T | 0.158 | 1.63E-01 | 1.97E-01 | 2.76E+00 | 2.84E+00 | 4.02E-01 | Yes | Ncoa5 | intron_variant | MODIFIER |
| 2 | 165022725 | AX-170142338 | 0 | G | A | 0.158 | 1.63E-01 | 1.97E-01 | 2.76E+00 | 2.84E+00 | 4.02E-01 | Yes | Ncoa5 | intron_variant | MODIFIER |

| | | | | | | | | | | | | | | | |
|---|---|---|---|---|---|---|---|---|---|---|---|---|---|---|---|
| 2 | 165048168 | AX-97845627 | 0 | G | A | 0.158 | 1.63E-01 | 1.97E-01 | 2.76E+00 | 2.84E+00 | 4.02E-01 | Yes | | intergenic_variant | MODIFIER |
| 2 | 165049389 | AX-170326189 | 0 | T | C | 0.158 | 1.63E-01 | 1.97E-01 | 2.76E+00 | 2.84E+00 | 4.02E-01 | Yes | | intergenic_variant | MODIFIER |
| 2 | 165062095 | AX-170171560 | 0 | T | A | 0.158 | 1.63E-01 | 1.97E-01 | 2.76E+00 | 2.84E+00 | 4.02E-01 | Yes | Cd40 | intron_variant | MODIFIER |
| 2 | 165066963 | AX-170201005 | 0 | C | T | 0.158 | 1.63E-01 | 1.97E-01 | 2.76E+00 | 2.84E+00 | 4.02E-01 | Yes | Cd40 | intron_variant | MODIFIER |
| 2 | 165094269 | AX-97835167 | 0 | C | T | 0.158 | 1.63E-01 | 1.97E-01 | 2.76E+00 | 2.84E+00 | 4.02E-01 | Yes | 1700025C18Rik | upstream_gene_variant | MODIFIER |
| 2 | 165100418 | AX-170395807 | 0 | G | A | 0.158 | 1.63E-01 | 1.97E-01 | 2.76E+00 | 2.84E+00 | 4.02E-01 | Yes | | intergenic_variant | MODIFIER |
| 2 | 165100688 | AX-170334674 | 0 | G | A | 0.158 | 1.63E-01 | 1.97E-01 | 2.76E+00 | 2.84E+00 | 4.02E-01 | Yes | | intergenic_variant | MODIFIER |
| 2 | 165101295 | AX-170261748 | 0 | T | G | 0.158 | 1.63E-01 | 1.97E-01 | 2.76E+00 | 2.84E+00 | 4.02E-01 | Yes | | intergenic_variant | MODIFIER |
| 2 | 165153905 | AX-97839806 | 0 | C | T | 0.158 | 1.63E-01 | 1.97E-01 | 2.76E+00 | 2.84E+00 | 4.02E-01 | Yes | Cdh22 | intron_variant | MODIFIER |
| 2 | 165163765 | AX-170335172 | 0 | C | T | 0.158 | 1.63E-01 | 1.97E-01 | 2.76E+00 | 2.84E+00 | 4.02E-01 | Yes | Cdh22 | intron_variant | MODIFIER |
| 2 | 165167407 | AX-170311486 | 0 | T | C | 0.158 | 1.63E-01 | 1.97E-01 | 2.76E+00 | 2.84E+00 | 4.02E-01 | Yes | Cdh22 | intron_variant | MODIFIER |
| 2 | 165169199 | AX-168404781 | 0 | G | A | 0.158 | 1.63E-01 | 1.97E-01 | 2.76E+00 | 2.84E+00 | 4.02E-01 | Yes | Cdh22 | intron_variant | MODIFIER |
| 2 | 165169735 | AX-170448583 | 0 | T | C | 0.158 | 1.63E-01 | 1.97E-01 | 2.76E+00 | 2.84E+00 | 4.02E-01 | Yes | Cdh22 | intron_variant | MODIFIER |
| 2 | 165170083 | AX-170122292 | 0 | T | C | 0.158 | 1.63E-01 | 1.97E-01 | 2.76E+00 | 2.84E+00 | 4.02E-01 | Yes | Cdh22 | intron_variant | MODIFIER |
| 2 | 165171074 | AX-97791398 | 0 | T | C | 0.158 | 1.63E-01 | 1.97E-01 | 2.76E+00 | 2.84E+00 | 4.02E-01 | Yes | Cdh22 | intron_variant | MODIFIER |
| 2 | 165349674 | AX-170315006 | 0 | T | G | 0.158 | 1.63E-01 | 1.97E-01 | 2.76E+00 | 2.84E+00 | 4.02E-01 | Yes | Zfp663 | downstream_gene_variant | MODIFIER |
| 2 | 165349731 | AX-170216658 | 0 | T | G | 0.158 | 1.63E-01 | 1.97E-01 | 2.76E+00 | 2.84E+00 | 4.02E-01 | Yes | Zfp663 | downstream_gene_variant | MODIFIER |
| 2 | 165351245 | AX-170267324 | 0 | T | C | 0.158 | 1.63E-01 | 1.97E-01 | 2.76E+00 | 2.84E+00 | 4.02E-01 | Yes | Zfp663 | downstream_gene_variant | MODIFIER |
| 2 | 165351808 | AX-170152384 | 0 | T | A | 0.158 | 1.63E-01 | 1.97E-01 | 2.76E+00 | 2.84E+00 | 4.02E-01 | Yes | Zfp663 | 3_prime_UTR_variant | MODIFIER |
| 2 | 165353705 | AX-170135779 | 0 | A | G | 0.158 | 1.63E-01 | 1.97E-01 | 2.76E+00 | 2.84E+00 | 4.02E-01 | Yes | Zfp663 | missense_variant | MODERATE |
| 2 | 165368895 | AX-97800908 | 0 | C | T | 0.158 | 1.63E-01 | 1.97E-01 | 2.76E+00 | 2.84E+00 | 4.02E-01 | Yes | Zfp663 | upstream_gene_variant | MODIFIER |
| 2 | 165382352 | AX-97854310 | 0 | G | A | 0.158 | 1.63E-01 | 1.97E-01 | 2.76E+00 | 2.84E+00 | 4.02E-01 | Yes | Zfp334 | intron_variant | MODIFIER |
| 2 | 165549244 | AX-170130838 | 0 | T | C | 0.158 | 1.63E-01 | 1.97E-01 | 2.76E+00 | 2.84E+00 | 4.02E-01 | Yes | | intergenic_variant | MODIFIER |
| 2 | 165564544 | AX-170269711 | 0 | T | C | 0.158 | 1.63E-01 | 1.97E-01 | 2.76E+00 | 2.84E+00 | 4.02E-01 | Yes | | regulatory_region_variant | MODIFIER |
| 2 | 165659803 | AX-170019600 | 0 | C | G | 0.158 | 1.63E-01 | 1.97E-01 | 2.76E+00 | 2.84E+00 | 4.02E-01 | Yes | Eya2 | intron_variant | MODIFIER |
| 2 | 165685118 | AX-168419152 | 0 | T | G | 0.158 | 1.63E-01 | 1.97E-01 | 2.76E+00 | 2.84E+00 | 4.02E-01 | Yes | Eya2 | intron_variant | MODIFIER |
| 2 | 165686283 | AX-170296034 | 0 | T | C | 0.158 | 1.63E-01 | 1.97E-01 | 2.76E+00 | 2.84E+00 | 4.02E-01 | Yes | Eya2 | intron_variant | MODIFIER |
| 2 | 165703341 | AX-97834864 | 0 | G | T | 0.158 | 1.63E-01 | 1.97E-01 | 2.76E+00 | 2.84E+00 | 4.02E-01 | Yes | Eya2 | intron_variant | MODIFIER |
| 2 | 166129118 | AX-170292246 | 0 | T | C | 0.158 | 1.63E-01 | 1.97E-01 | 2.76E+00 | 2.84E+00 | 4.02E-01 | Yes | Sulf2 | intron_variant | MODIFIER |
| 2 | 166129149 | AX-97807674 | 0 | C | T | 0.158 | 1.63E-01 | 1.97E-01 | 2.76E+00 | 2.84E+00 | 4.02E-01 | Yes | Sulf2 | intron_variant | MODIFIER |
| 2 | 166129208 | AX-170227148 | 0 | G | A | 0.158 | 1.63E-01 | 1.97E-01 | 2.76E+00 | 2.84E+00 | 4.02E-01 | Yes | Sulf2 | intron_variant | MODIFIER |
| 2 | 166311392 | AX-170019607 | 0 | T | C | 0.158 | 1.63E-01 | 1.97E-01 | 2.76E+00 | 2.84E+00 | 4.02E-01 | Yes | | intergenic_variant | MODIFIER |
| 2 | 166314836 | AX-97828367 | 0 | T | C | 0.158 | 1.63E-01 | 1.97E-01 | 2.76E+00 | 2.84E+00 | 4.02E-01 | Yes | | intergenic_variant | MODIFIER |
| 2 | 166317951 | AX-170483844 | 0 | G | A | 0.158 | 1.63E-01 | 1.97E-01 | 2.76E+00 | 2.84E+00 | 4.02E-01 | Yes | | intergenic_variant | MODIFIER |
| 2 | 166330740 | AX-170329078 | 0 | C | A | 0.158 | 1.63E-01 | 1.97E-01 | 2.76E+00 | 2.84E+00 | 4.02E-01 | Yes | | regulatory_region_variant | MODIFIER |
| 2 | 166331529 | AX-170305367 | 0 | G | C | 0.158 | 1.63E-01 | 1.97E-01 | 2.76E+00 | 2.84E+00 | 4.02E-01 | Yes | | intergenic_variant | MODIFIER |
| 2 | 166331567 | AX-170144396 | 0 | A | G | 0.158 | 1.63E-01 | 1.97E-01 | 2.76E+00 | 2.84E+00 | 4.02E-01 | Yes | | intergenic_variant | MODIFIER |
| 2 | 166331658 | AX-170443364 | 0 | C | T | 0.158 | 1.63E-01 | 1.97E-01 | 2.76E+00 | 2.84E+00 | 4.02E-01 | Yes | | intergenic_variant | MODIFIER |
| 2 | 166332125 | AX-170201188 | 0 | A | G | 0.158 | 1.63E-01 | 1.97E-01 | 2.76E+00 | 2.84E+00 | 4.02E-01 | Yes | | regulatory_region_variant | MODIFIER |
| 2 | 166375236 | AX-170065543 | 0 | A | G | 0.158 | 1.63E-01 | 1.97E-01 | 2.76E+00 | 2.84E+00 | 4.02E-01 | Yes | | intergenic_variant | MODIFIER |
| 2 | 166378956 | AX-170019614 | 0 | G | A | 0.158 | 1.63E-01 | 1.97E-01 | 2.76E+00 | 2.84E+00 | 4.02E-01 | Yes | Gm14268 | downstream_gene_variant | MODIFIER |
| 2 | 166381199 | AX-170303186 | 0 | A | G | 0.158 | 1.63E-01 | 1.97E-01 | 2.76E+00 | 2.84E+00 | 4.02E-01 | Yes | Gm14268 | non_coding_transcript_exon_variant | MODIFIER |
| 2 | 166383541 | AX-170167488 | 0 | C | T | 0.158 | 1.63E-01 | 1.97E-01 | 2.76E+00 | 2.84E+00 | 4.02E-01 | Yes | Gm14268 | intron_variant,non_coding_transcript_variant | MODIFIER |
| 2 | 166394607 | AX-170094478 | 0 | T | C | 0.158 | 1.63E-01 | 1.97E-01 | 2.76E+00 | 2.84E+00 | 4.02E-01 | Yes | Gm14268 | intron_variant,non_coding_transcript_variant | MODIFIER |
| 2 | 166415629 | AX-170352163 | 0 | G | A | 0.158 | 1.63E-01 | 1.97E-01 | 2.76E+00 | 2.84E+00 | 4.02E-01 | Yes | | regulatory_region_variant | MODIFIER |
| 2 | 166458584 | AX-170428453 | 0 | T | C | 0.158 | 1.63E-01 | 1.97E-01 | 2.76E+00 | 2.84E+00 | 4.02E-01 | Yes | 5031425F14Rik | non_coding_transcript_exon_variant | MODIFIER |
| 2 | 166460668 | AX-170184255 | 0 | A | G | 0.158 | 1.63E-01 | 1.97E-01 | 2.76E+00 | 2.84E+00 | 4.02E-01 | Yes | 5031425F14Rik | downstream_gene_variant | MODIFIER |
| 2 | 166473927 | AX-97780515 | 0 | T | C | 0.158 | 1.63E-01 | 1.97E-01 | 2.76E+00 | 2.84E+00 | 4.02E-01 | Yes | | intergenic_variant | MODIFIER |
| 2 | 166475290 | AX-170242765 | 0 | G | C | 0.158 | 1.63E-01 | 1.97E-01 | 2.76E+00 | 2.84E+00 | 4.02E-01 | Yes | | regulatory_region_variant | MODIFIER |
| 2 | 166580926 | AX-97839821 | 0 | G | T | 0.158 | 1.63E-01 | 1.97E-01 | 2.76E+00 | 2.84E+00 | 4.02E-01 | Yes | Prex1 | intron_variant | MODIFIER |
| 2 | 166582231 | AX-170381031 | 0 | G | C | 0.158 | 1.63E-01 | 1.97E-01 | 2.76E+00 | 2.84E+00 | 4.02E-01 | Yes | Prex1 | intron_variant | MODIFIER |
| 2 | 166616342 | AX-97826573 | 0 | C | T | 0.158 | 1.63E-01 | 1.97E-01 | 2.76E+00 | 2.84E+00 | 4.02E-01 | Yes | Prex1 | intron_variant | MODIFIER |
| 2 | 166619064 | AX-170395954 | 0 | G | A | 0.158 | 1.63E-01 | 1.97E-01 | 2.76E+00 | 2.84E+00 | 4.02E-01 | Yes | Prex1 | intron_variant | MODIFIER |
| 2 | 166619706 | AX-170279737 | 0 | A | G | 0.158 | 1.63E-01 | 1.97E-01 | 2.76E+00 | 2.84E+00 | 4.02E-01 | Yes | Prex1 | intron_variant | MODIFIER |

| | | | | | | | | | | | | | | | |
|---|---|---|---|---|---|---|---|---|---|---|---|---|---|---|---|
| 2 | 166628728 | AX-97811415 | 0 | A | G | 0.158 | 1.63E-01 | 1.97E-01 | 2.76E+00 | 2.84E+00 | 4.02E-01 | Yes | Prex1 | intron_variant | MODIFIER |
| 2 | 166629260 | AX-170237524 | 0 | A | G | 0.158 | 1.63E-01 | 1.97E-01 | 2.76E+00 | 2.84E+00 | 4.02E-01 | Yes | Prex1 | intron_variant | MODIFIER |
| 2 | 166637522 | AX-170277518 | 0 | A | G | 0.158 | 1.63E-01 | 1.97E-01 | 2.76E+00 | 2.84E+00 | 4.02E-01 | Yes | Prex1 | intron_variant | MODIFIER |
| 2 | 166655031 | AX-97805420 | 0 | T | C | 0.158 | 1.63E-01 | 1.97E-01 | 2.76E+00 | 2.84E+00 | 4.02E-01 | Yes | Prex1 | intron_variant | MODIFIER |
| 2 | 166696155 | AX-170161522 | 0 | T | C | 0.158 | 1.63E-01 | 1.97E-01 | 2.76E+00 | 2.84E+00 | 4.02E-01 | Yes | Prex1 | intron_variant | MODIFIER |
| 2 | 166699884 | AX-170397724 | 0 | A | T | 0.158 | 1.63E-01 | 1.97E-01 | 2.76E+00 | 2.84E+00 | 4.02E-01 | Yes | Prex1 | intron_variant | MODIFIER |
| 2 | 166704392 | AX-97839822 | 0 | G | A | 0.158 | 1.63E-01 | 1.97E-01 | 2.76E+00 | 2.84E+00 | 4.02E-01 | Yes | Prex1 | intron_variant | MODIFIER |
| 2 | 166734320 | AX-170494738 | 0 | T | A | 0.158 | 1.63E-01 | 1.97E-01 | 2.76E+00 | 2.84E+00 | 4.02E-01 | Yes | | intergenic_variant | MODIFIER |
| 2 | 166852083 | AX-170019621 | 0 | T | C | 0.158 | 1.63E-01 | 1.97E-01 | 2.76E+00 | 2.84E+00 | 4.02E-01 | Yes | Arfgef2 | intron_variant | MODIFIER |
| 2 | 166857525 | AX-170401721 | 0 | A | C | 0.158 | 1.63E-01 | 1.97E-01 | 2.76E+00 | 2.84E+00 | 4.02E-01 | Yes | Arfgef2 | intron_variant | MODIFIER |
| 2 | 166858081 | AX-170272205 | 0 | G | A | 0.158 | 1.63E-01 | 1.97E-01 | 2.76E+00 | 2.84E+00 | 4.02E-01 | Yes | Arfgef2 | intron_variant | MODIFIER |
| 2 | 166858719 | AX-170019616 | 0 | C | T | 0.158 | 1.63E-01 | 1.97E-01 | 2.76E+00 | 2.84E+00 | 4.02E-01 | Yes | Arfgef2 | intron_variant | MODIFIER |
| 2 | 166862328 | AX-97787183 | 0 | A | G | 0.158 | 1.63E-01 | 1.97E-01 | 2.76E+00 | 2.84E+00 | 4.02E-01 | Yes | Arfgef2 | intron_variant | MODIFIER |
| 2 | 166866631 | AX-170403271 | 0 | T | C | 0.158 | 1.63E-01 | 1.97E-01 | 2.76E+00 | 2.84E+00 | 4.02E-01 | Yes | Arfgef2 | intron_variant | MODIFIER |
| 2 | 166869349 | AX-170504077 | 0 | T | C | 0.158 | 1.63E-01 | 1.97E-01 | 2.76E+00 | 2.84E+00 | 4.02E-01 | Yes | Arfgef2 | intron_variant | MODIFIER |
| 2 | 166871175 | AX-170360169 | 0 | C | T | 0.158 | 1.63E-01 | 1.97E-01 | 2.76E+00 | 2.84E+00 | 4.02E-01 | Yes | Arfgef2 | intron_variant | MODIFIER |
| 2 | 166871568 | AX-170355141 | 0 | A | G | 0.158 | 1.63E-01 | 1.97E-01 | 2.76E+00 | 2.84E+00 | 4.02E-01 | Yes | Arfgef2 | synonymous_variant | LOW |
| 2 | 166874133 | AX-170153788 | 0 | T | C | 0.158 | 1.63E-01 | 1.97E-01 | 2.76E+00 | 2.84E+00 | 4.02E-01 | Yes | Arfgef2 | intron_variant | MODIFIER |
| 2 | 166885749 | AX-170267287 | 0 | A | G | 0.158 | 1.63E-01 | 1.97E-01 | 2.76E+00 | 2.84E+00 | 4.02E-01 | Yes | Arfgef2 | intron_variant | MODIFIER |
| 2 | 166887254 | AX-170380708 | 0 | C | T | 0.158 | 1.63E-01 | 1.97E-01 | 2.76E+00 | 2.84E+00 | 4.02E-01 | Yes | Arfgef2 | intron_variant | MODIFIER |
| 2 | 166925540 | AX-170229757 | 0 | A | G | 0.158 | 1.63E-01 | 1.97E-01 | 2.76E+00 | 2.84E+00 | 4.02E-01 | Yes | Cse1l | intron_variant | MODIFIER |
| 2 | 166935535 | AX-170520076 | 0 | T | C | 0.158 | 1.63E-01 | 1.97E-01 | 2.76E+00 | 2.84E+00 | 4.02E-01 | Yes | Cse1l | intron_variant | MODIFIER |
| 2 | 166940076 | AX-170276966 | 0 | G | A | 0.158 | 1.63E-01 | 1.97E-01 | 2.76E+00 | 2.84E+00 | 4.02E-01 | Yes | Cse1l | synonymous_variant | LOW |
| 2 | 166976257 | AX-170293350 | 0 | A | C | 0.158 | 1.63E-01 | 1.97E-01 | 2.76E+00 | 2.84E+00 | 4.02E-01 | Yes | Stau1 | intron_variant | MODIFIER |
| 2 | 166985221 | AX-97802905 | 0 | C | T | 0.158 | 1.63E-01 | 1.97E-01 | 2.76E+00 | 2.84E+00 | 4.02E-01 | Yes | Stau1 | intron_variant | MODIFIER |
| 2 | 167027832 | AX-170065554 | 0 | C | T | 0.158 | 1.63E-01 | 1.97E-01 | 2.76E+00 | 2.84E+00 | 4.02E-01 | Yes | Ddx27 | intron_variant | MODIFIER |
| 2 | 167036734 | AX-170065553 | 0 | T | C | 0.158 | 1.63E-01 | 1.97E-01 | 2.76E+00 | 2.84E+00 | 4.02E-01 | Yes | Ddx27 | downstream_gene_variant | MODIFIER |
| 2 | 167109875 | AX-97801019 | 0 | C | T | 0.158 | 1.63E-01 | 1.97E-01 | 2.76E+00 | 2.84E+00 | 4.02E-01 | Yes | Kcnb1 | intron_variant | MODIFIER |
| 2 | 167110968 | AX-170284291 | 0 | G | A | 0.158 | 1.63E-01 | 1.97E-01 | 2.76E+00 | 2.84E+00 | 4.02E-01 | Yes | Kcnb1 | intron_variant | MODIFIER |
| 2 | 167124897 | AX-170362581 | 0 | A | G | 0.158 | 1.63E-01 | 1.97E-01 | 2.76E+00 | 2.84E+00 | 4.02E-01 | Yes | Kcnb1 | intron_variant | MODIFIER |
| 2 | 167132168 | AX-97850692 | 0 | C | T | 0.158 | 1.63E-01 | 1.97E-01 | 2.76E+00 | 2.84E+00 | 4.02E-01 | Yes | Kcnb1 | intron_variant | MODIFIER |
| 2 | 167134950 | AX-170351767 | 0 | C | A | 0.158 | 1.63E-01 | 1.97E-01 | 2.76E+00 | 2.84E+00 | 4.02E-01 | Yes | Kcnb1 | intron_variant | MODIFIER |
| 2 | 167163816 | AX-170389479 | 0 | T | C | 0.158 | 1.63E-01 | 1.97E-01 | 2.76E+00 | 2.84E+00 | 4.02E-01 | Yes | Kcnb1 | intron_variant | MODIFIER |
| 2 | 167213726 | AX-170174618 | 0 | G | A | 0.158 | 1.63E-01 | 1.97E-01 | 2.76E+00 | 2.84E+00 | 4.02E-01 | Yes | Ptgis | intron_variant | MODIFIER |
| 1 | 149316411 | AX-170197041 | 1 | C | A | 0.205 | 1.16E-01 | 1.39E-01 | 2.64E+00 | 2.69E+00 | 4.02E-01 | no | | intergenic_variant | MODIFIER |
| 3 | 38652992 | AX-97791183 | 0 | G | A | 0.202 | 7.05E-02 | 8.56E-02 | 2.72E+00 | 2.84E+00 | 4.03E-01 | no | | regulatory_region_variant | MODIFIER |
| 7 | 78220921 | AX-170077802 | 0 | G | A | 0.36 | -7.70E-02 | 9.33E-02 | 2.75E+00 | 2.85E+00 | 4.03E-01 | no | Ntrk3 | intron_variant | MODIFIER |
| 4 | 88199410 | AX-97801395 | 0 | T | C | 0.211 | -7.78E-02 | 9.24E-02 | 3.01E+00 | 3.22E+00 | 4.03E-01 | no | Focad | intron_variant | MODIFIER |
| 10 | 92996399 | AX-170156370 | 1 | T | G | 0.295 | -7.10E-02 | 8.41E-02 | 2.53E+00 | 2.58E+00 | 4.03E-01 | no | Cfap54 | intron_variant | MODIFIER |
| 2 | 163211469 | AX-170019564 | 1 | A | G | 0.152 | 1.62E-01 | 1.96E-01 | 2.77E+00 | 2.88E+00 | 4.03E-01 | Yes | Tox2 | intron_variant | MODIFIER |
| 8 | 29602401 | AX-170079622 | 0 | G | A | 0.158 | -8.34E-02 | 1.01E-01 | 2.73E+00 | 2.87E+00 | 4.03E-01 | no | | intergenic_variant | MODIFIER |
| 8 | 26402502 | AX-170134337 | 0 | A | T | 0.342 | -5.40E-02 | 6.40E-02 | 2.51E+00 | 2.60E+00 | 4.03E-01 | no | | intergenic_variant | MODIFIER |
| 18 | 54812185 | AX-170015729 | 0 | G | T | 0.447 | 8.32E-02 | 9.42E-02 | 2.36E+00 | 2.41E+00 | 4.03E-01 | no | AC129179.2 | intron_variant,non_coding_transcript_variant | MODIFIER |
| 13 | 34052620 | AX-170374993 | 0 | C | G | 0.175 | -1.02E-01 | 1.22E-01 | 2.54E+00 | 2.61E+00 | 4.03E-01 | no | Bphl | intron_variant | MODIFIER |
| 18 | 62530034 | AX-170252126 | 0 | G | C | 0.193 | -9.47E-02 | 1.15E-01 | 2.69E+00 | 2.76E+00 | 4.03E-01 | no | Fbxo38 | intron_variant | MODIFIER |
| 18 | 67697659 | AX-170344699 | 0 | G | T | 0.193 | -9.47E-02 | 1.15E-01 | 2.69E+00 | 2.76E+00 | 4.03E-01 | no | Ptpn2 | intron_variant | MODIFIER |
| 1 | 20654602 | AX-170363430 | 1 | C | T | 0.411 | 7.16E-02 | 8.59E-02 | 2.60E+00 | 2.64E+00 | 4.03E-01 | no | | regulatory_region_variant | MODIFIER |
| 1 | 20784893 | AX-97786384 | 1 | G | A | 0.411 | 7.16E-02 | 8.59E-02 | 2.60E+00 | 2.64E+00 | 4.03E-01 | no | Il17f | upstream_gene_variant | MODIFIER |
| 15 | 102424615 | AX-170228298 | 1 | G | T | 0.107 | 1.07E-01 | 1.30E-01 | 2.68E+00 | 2.79E+00 | 4.03E-01 | no | Sp1 | intron_variant | MODIFIER |
| 4 | 137891974 | AX-170453044 | 0 | T | C | 0.167 | 9.61E-02 | 1.15E-01 | 2.60E+00 | 2.65E+00 | 4.04E-01 | no | Ece1 | intron_variant | MODIFIER |
| 2 | 159635404 | AX-170143400 | 0 | A | G | 0.158 | 1.44E-01 | 1.75E-01 | 2.74E+00 | 2.86E+00 | 4.04E-01 | no | | intergenic_variant | MODIFIER |
| 5 | 104211705 | AX-97852427 | 0 | T | C | 0.456 | 8.99E-02 | 1.08E-01 | 2.60E+00 | 2.66E+00 | 4.04E-01 | no | Dmp1 | synonymous_variant | LOW |
| 5 | 114071860 | AX-170257357 | 0 | C | T | 0.456 | 8.99E-02 | 1.08E-01 | 2.60E+00 | 2.66E+00 | 4.04E-01 | no | Svop | intron_variant | MODIFIER |
| 5 | 115993517 | AX-170483635 | 0 | T | C | 0.456 | 8.99E-02 | 1.08E-01 | 2.60E+00 | 2.66E+00 | 4.04E-01 | no | Cit | intron_variant | MODIFIER |

| | | | | | | | | | | | | | | | |
|---|---|---|---|---|---|---|---|---|---|---|---|---|---|---|---|
| 5 | 99875475 | AX-170025711 | 0 | C | T | 0.43 | -8.55E-02 | 1.02E-01 | 2.94E+00 | 3.04E+00 | 4.04E-01 | no | | intergenic_variant | MODIFIER |
| 10 | 7258363 | AX-170002625 | 0 | A | C | 0.202 | -7.63E-02 | 9.27E-02 | 2.66E+00 | 2.78E+00 | 4.04E-01 | no | | regulatory_region_variant | MODIFIER |
| 5 | 110617276 | AX-170297268 | 0 | T | C | 0.465 | 9.11E-02 | 1.10E-01 | 2.68E+00 | 2.73E+00 | 4.04E-01 | no | Galnt9 | intron_variant | MODIFIER |
| 5 | 114386932 | AX-170073178 | 0 | T | C | 0.465 | 9.11E-02 | 1.10E-01 | 2.68E+00 | 2.73E+00 | 4.04E-01 | no | Ube3b | intron_variant | MODIFIER |
| 9 | 98687469 | AX-170215678 | 1 | C | T | 0.134 | -9.54E-02 | 1.15E-01 | 2.61E+00 | 2.69E+00 | 4.04E-01 | no | | intergenic_variant | MODIFIER |
| 6 | 135561991 | AX-170164634 | 1 | G | T | 0.58 | 7.35E-02 | 8.70E-02 | 2.51E+00 | 2.57E+00 | 4.04E-01 | no | | intergenic_variant | MODIFIER |
| 2 | 162485603 | AX-170019562 | 1 | A | G | 0.161 | 1.61E-01 | 1.96E-01 | 2.74E+00 | 2.82E+00 | 4.04E-01 | Yes | Ptprt | intron_variant | MODIFIER |
| 18 | 50018265 | AX-170088334 | 1 | A | G | 0.188 | -8.72E-02 | 1.04E-01 | 2.53E+00 | 2.62E+00 | 4.04E-01 | no | Tnfaip8 | intron_variant,non_coding_transcript_variant | MODIFIER |
| 5 | 85070838 | AX-170202566 | 0 | C | T | 0.447 | 8.32E-02 | 1.00E-01 | 2.89E+00 | 3.00E+00 | 4.05E-01 | no | | intergenic_variant | MODIFIER |
| 8 | 62814479 | AX-170382103 | 0 | G | T | 0.14 | -1.23E-01 | 1.49E-01 | 2.61E+00 | 2.68E+00 | 4.05E-01 | no | Gm45459 | intron_variant,non_coding_transcript_variant | MODIFIER |
| 11 | 61651159 | AX-170506006 | 0 | T | C | 0.053 | -1.27E-01 | 1.55E-01 | 2.65E+00 | 2.82E+00 | 4.05E-01 | no | Grap | upstream_gene_variant | MODIFIER |
| 8 | 17254859 | AX-170108769 | 0 | A | G | 0.228 | -6.48E-02 | 7.76E-02 | 2.56E+00 | 2.65E+00 | 4.05E-01 | no | Csmd1 | intron_variant | MODIFIER |
| 4 | 80491319 | AX-170212540 | 0 | G | A | 0.237 | 9.13E-02 | 1.08E-01 | 2.51E+00 | 2.57E+00 | 4.05E-01 | no | | intergenic_variant | MODIFIER |
| 18 | 61738669 | AX-170151723 | 1 | C | T | 0.188 | -9.89E-02 | 1.20E-01 | 2.67E+00 | 2.72E+00 | 4.05E-01 | no | Afap1l1 | intron_variant | MODIFIER |
| 7 | 37870308 | AX-170326948 | 0 | T | C | 0.351 | 7.67E-02 | 9.25E-02 | 2.60E+00 | 2.68E+00 | 4.05E-01 | no | | intergenic_variant | MODIFIER |
| 9 | 90683200 | AX-170416877 | 1 | G | A | 0.125 | -1.07E-01 | 1.29E-01 | 2.63E+00 | 2.71E+00 | 4.05E-01 | no | | intergenic_variant | MODIFIER |
| 4 | 134243555 | AX-170070754 | 1 | T | C | 0.17 | 8.96E-02 | 1.08E-01 | 2.58E+00 | 2.64E+00 | 4.05E-01 | no | Zfp593 | 3_prime_UTR_variant | MODIFIER |
| 16 | 76762263 | AX-170387869 | 0 | C | T | 0.456 | 6.13E-02 | 7.29E-02 | 2.51E+00 | 2.59E+00 | 4.05E-01 | no | Gm30790 | intron_variant,non_coding_transcript_variant | MODIFIER |
| 6 | 134270299 | AX-170305203 | 0 | T | C | 0.553 | 7.37E-02 | 8.67E-02 | 2.47E+00 | 2.52E+00 | 4.05E-01 | no | Etv6 | downstream_gene_variant | MODIFIER |
| 1 | 90816474 | AX-97816364 | 0 | C | T | 0.158 | 1.70E-01 | 2.06E-01 | 2.67E+00 | 2.74E+00 | 4.05E-01 | Yes | Col6a3 | synonymous_variant | LOW |
| 6 | 135304686 | AX-170296757 | 0 | C | T | 0.579 | 7.33E-02 | 8.70E-02 | 2.51E+00 | 2.57E+00 | 4.05E-01 | no | Pbp2 | downstream_gene_variant | MODIFIER |
| 6 | 135482510 | AX-170426050 | 0 | A | G | 0.579 | 7.33E-02 | 8.70E-02 | 2.51E+00 | 2.57E+00 | 4.05E-01 | no | | intergenic_variant | MODIFIER |
| 6 | 137531065 | AX-170182549 | 0 | G | A | 0.579 | 7.33E-02 | 8.70E-02 | 2.51E+00 | 2.57E+00 | 4.05E-01 | no | Eps8 | intron_variant | MODIFIER |
| 1 | 90182741 | AX-170163607 | 1 | C | T | 0.161 | 1.69E-01 | 2.05E-01 | 2.67E+00 | 2.73E+00 | 4.05E-01 | Yes | | intergenic_variant | MODIFIER |
| 7 | 46285699 | AX-97803530 | 0 | A | G | 0.333 | 7.82E-02 | 9.45E-02 | 2.60E+00 | 2.69E+00 | 4.05E-01 | no | Otog | intron_variant | MODIFIER |
| 2 | 162670010 | AX-170496988 | 1 | C | T | 0.161 | 1.61E-01 | 1.96E-01 | 2.74E+00 | 2.82E+00 | 4.05E-01 | Yes | 9430021M05Rik | downstream_gene_variant | MODIFIER |
| 5 | 48851041 | AX-170097652 | 0 | C | T | 0.5 | 8.82E-02 | 1.03E-01 | 2.43E+00 | 2.48E+00 | 4.05E-01 | no | Kcnip4 | intron_variant | MODIFIER |
| 5 | 51492947 | AX-170024937 | 0 | G | A | 0.5 | 8.82E-02 | 1.03E-01 | 2.43E+00 | 2.48E+00 | 4.05E-01 | no | Ppargc1a | intron_variant,NMD_transcript_variant | MODIFIER |
| 7 | 144705477 | AX-170398285 | 1 | C | A | 0.205 | -6.17E-02 | 7.50E-02 | 2.62E+00 | 2.75E+00 | 4.06E-01 | no | Ano1 | intron_variant | MODIFIER |
| 16 | 19006425 | AX-170037389 | 1 | C | T | 0.134 | -7.90E-02 | 9.67E-02 | 2.72E+00 | 2.87E+00 | 4.06E-01 | no | | intergenic_variant | MODIFIER |
| 4 | 139794939 | AX-170024109 | 1 | C | T | 0.161 | 9.76E-02 | 1.17E-01 | 2.59E+00 | 2.64E+00 | 4.06E-01 | no | Pax7 | intron_variant | MODIFIER |
| 6 | 137431738 | AX-170182241 | 0 | A | T | 0.57 | 7.09E-02 | 8.37E-02 | 2.48E+00 | 2.54E+00 | 4.06E-01 | no | Ptpro | intron_variant | MODIFIER |
| 18 | 84765953 | AX-170350479 | 1 | A | G | 0.161 | -9.36E-02 | 1.10E-01 | 2.47E+00 | 2.54E+00 | 4.06E-01 | no | | regulatory_region_variant | MODIFIER |
| 1 | 151215702 | AX-170001984 | 1 | T | C | 0.214 | 1.03E-01 | 1.24E-01 | 2.60E+00 | 2.65E+00 | 4.06E-01 | no | Gm47996 | downstream_gene_variant | MODIFIER |
| 4 | 136887586 | AX-97795432 | 1 | G | T | 0.152 | 9.91E-02 | 1.19E-01 | 2.60E+00 | 2.64E+00 | 4.06E-01 | no | C1qc | downstream_gene_variant | MODIFIER |
| 1 | 91145009 | AX-170449500 | 0 | T | A | 0.158 | -1.46E-01 | 1.75E-01 | 2.96E+00 | 3.08E+00 | 4.06E-01 | no | Rbm44 | upstream_gene_variant | MODIFIER |
| 15 | 66869886 | AX-97847412 | 0 | C | T | 0.342 | -6.82E-02 | 7.75E-02 | 2.35E+00 | 2.43E+00 | 4.06E-01 | no | | intergenic_variant | MODIFIER |
| 5 | 105533338 | AX-97845057 | 1 | G | A | 0.446 | 9.39E-02 | 1.13E-01 | 2.61E+00 | 2.65E+00 | 4.06E-01 | no | Lrrc8c | intron_variant | MODIFIER |
| 16 | 56176490 | AX-97831957 | 2 | A | G | 0.518 | 6.98E-02 | 8.34E-02 | 2.53E+00 | 2.58E+00 | 4.06E-01 | no | Senp7 | intron_variant | MODIFIER |
| 5 | 80784731 | AX-170072606 | 1 | T | C | 0.438 | 8.93E-02 | 1.09E-01 | 2.76E+00 | 2.81E+00 | 4.06E-01 | no | | intergenic_variant | MODIFIER |
| 7 | 96931642 | AX-97798939 | 0 | C | T | 0.14 | 1.20E-01 | 1.47E-01 | 2.68E+00 | 2.83E+00 | 4.06E-01 | no | C230038L03Rik | intron_variant,non_coding_transcript_variant | MODIFIER |
| 18 | 55670464 | AX-170015750 | 0 | A | T | 0.456 | 8.42E-02 | 9.41E-02 | 2.31E+00 | 2.36E+00 | 4.06E-01 | no | | intergenic_variant | MODIFIER |
| 15 | 6435326 | AX-170520363 | 1 | C | T | 0.241 | -8.79E-02 | 1.06E-01 | 2.61E+00 | 2.66E+00 | 4.06E-01 | no | Dab2 | missense_variant | MODERATE |
| 5 | 43374977 | AX-97804126 | 0 | C | T | 0.193 | -8.91E-02 | 1.08E-01 | 2.62E+00 | 2.71E+00 | 4.07E-01 | no | Gm43699 | upstream_gene_variant | MODIFIER |
| 4 | 50082417 | AX-170382291 | 0 | T | A | 0.36 | -7.71E-02 | 9.40E-02 | 2.81E+00 | 2.92E+00 | 4.07E-01 | no | | intergenic_variant | MODIFIER |
| 2 | 107769506 | AX-170372734 | 0 | A | G | 0.053 | 1.77E-01 | 2.17E-01 | 2.75E+00 | 2.88E+00 | 4.07E-01 | Yes | | intergenic_variant | MODIFIER |
| 4 | 136880948 | AX-170131261 | 0 | A | G | 0.158 | -9.54E-02 | 1.10E-01 | 3.17E+00 | 3.34E+00 | 4.07E-01 | no | C1qb | intron_variant | MODIFIER |
| 18 | 67146291 | AX-97841980 | 1 | A | G | 0.179 | -1.01E-01 | 1.23E-01 | 2.66E+00 | 2.71E+00 | 4.07E-01 | no | Gnal | intron_variant | MODIFIER |
| 5 | 88944719 | AX-170072722 | 1 | C | T | 0.438 | 8.81E-02 | 1.07E-01 | 2.81E+00 | 2.87E+00 | 4.07E-01 | no | Slc4a4 | intron_variant | MODIFIER |
| 7 | 17798440 | AX-170396861 | 0 | A | G | 0.307 | 7.05E-02 | 8.60E-02 | 2.60E+00 | 2.74E+00 | 4.07E-01 | no | Gm45149 | non_coding_transcript_exon_variant | MODIFIER |
| 9 | 110870420 | AX-170254814 | 0 | T | C | 0.096 | -9.26E-02 | 1.11E-01 | 2.54E+00 | 2.64E+00 | 4.08E-01 | no | Tmie | intron_variant | MODIFIER |
| 4 | 133055164 | AX-170329579 | 0 | C | A | 0.158 | 9.79E-02 | 1.18E-01 | 2.56E+00 | 2.61E+00 | 4.08E-01 | no | Ahdc1 | 5_prime_UTR_variant | MODIFIER |
| 10 | 91704714 | AX-97802446 | 1 | T | C | 0.304 | -7.01E-02 | 8.40E-02 | 2.53E+00 | 2.59E+00 | 4.08E-01 | no | | intergenic_variant | MODIFIER |
| 10 | 92352069 | AX-97830613 | 1 | G | A | 0.304 | -7.01E-02 | 8.40E-02 | 2.53E+00 | 2.59E+00 | 4.08E-01 | no | Gm20757 | intron_variant,non_coding_transcript_variant | MODIFIER |

| | | | | | | | | | | | | | | | |
|---|---|---|---|---|---|---|---|---|---|---|---|---|---|---|---|
| 8 | 40576415 | AX-170159263 | 1 | C | A | 0.152 | -1.18E-01 | 1.44E-01 | 2.63E+00 | 2.68E+00 | 4.08E-01 | no | Mtmr7 | intron_variant,NMD_transcript_variant | MODIFIER |
| 10 | 70463454 | AX-170419862 | 2 | C | G | 0.418 | -6.24E-02 | 7.60E-02 | 2.64E+00 | 2.74E+00 | 4.08E-01 | no | Fam13c | intron_variant | MODIFIER |
| 15 | 71439656 | AX-170334258 | 1 | G | A | 0.366 | 5.83E-02 | 6.78E-02 | 2.41E+00 | 2.50E+00 | 4.08E-01 | no | Fam135b | 3_prime_UTR_variant | MODIFIER |
| 18 | 65658848 | AX-170093918 | 1 | C | T | 0.42 | 7.32E-02 | 8.53E-02 | 2.42E+00 | 2.48E+00 | 4.08E-01 | no | Zfp532 | intron_variant | MODIFIER |
| 1 | 89320381 | AX-170281265 | 1 | G | A | 0.152 | 1.56E-01 | 1.90E-01 | 2.64E+00 | 2.73E+00 | 4.08E-01 | Yes | Gm28342 | intron_variant,non_coding_transcript_variant | MODIFIER |
| 1 | 14857531 | AX-170208133 | 0 | A | T | 0.395 | 6.51E-02 | 7.89E-02 | 2.60E+00 | 2.65E+00 | 4.08E-01 | no | | intergenic_variant | MODIFIER |
| 18 | 73131653 | AX-170403316 | 1 | A | C | 0.179 | -9.77E-02 | 1.18E-01 | 2.57E+00 | 2.62E+00 | 4.08E-01 | no | | intergenic_variant | MODIFIER |
| 4 | 116154454 | AX-170177800 | 0 | A | G | 0.219 | 9.54E-02 | 1.15E-01 | 2.57E+00 | 2.64E+00 | 4.08E-01 | no | Pomgnt1 | intron_variant | MODIFIER |
| 4 | 74075827 | AX-170023115 | 1 | C | T | 0.455 | -7.13E-02 | 8.53E-02 | 2.51E+00 | 2.57E+00 | 4.08E-01 | no | Frmd3 | intron_variant | MODIFIER |
| 10 | 99992727 | AX-170470188 | 0 | T | A | 0.272 | -7.04E-02 | 8.49E-02 | 2.56E+00 | 2.63E+00 | 4.08E-01 | no | Gm17870 | downstream_gene_variant | MODIFIER |
| 7 | 46235298 | AX-97791366 | 1 | T | C | 0.339 | 7.72E-02 | 9.40E-02 | 2.61E+00 | 2.70E+00 | 4.08E-01 | no | Ush1c | intron_variant | MODIFIER |
| 1 | 115693844 | AX-168403165 | 0 | C | T | 0.175 | 1.43E-01 | 1.74E-01 | 2.62E+00 | 2.72E+00 | 4.09E-01 | no | Cntnap5a | intron_variant | MODIFIER |
| 5 | 42593714 | AX-170349327 | 1 | G | A | 0.205 | -8.86E-02 | 1.08E-01 | 2.61E+00 | 2.69E+00 | 4.09E-01 | no | | intergenic_variant | MODIFIER |
| 7 | 91603352 | AX-170487203 | 0 | A | G | 0.14 | 1.45E-01 | 1.79E-01 | 2.74E+00 | 2.86E+00 | 4.09E-01 | Yes | Dlg2 | intron_variant | MODIFIER |
| 18 | 65223471 | AX-170328106 | 0 | T | A | 0.175 | -9.30E-02 | 1.13E-01 | 2.62E+00 | 2.69E+00 | 4.09E-01 | no | | regulatory_region_variant | MODIFIER |
| 8 | 88689813 | AX-97826919 | 1 | C | T | 0.179 | -1.02E-01 | 1.24E-01 | 2.65E+00 | 2.74E+00 | 4.09E-01 | no | Nod2 | downstream_gene_variant | MODIFIER |
| 20 | 121699886 | AX-170135526 | 1 | T | A | 0.045 | -8.58E-02 | 1.07E-01 | 2.69E+00 | 2.91E+00 | 4.09E-01 | no | | intergenic_variant | MODIFIER |
| 12 | 89603573 | AX-170187258 | 0 | G | T | 0.096 | -1.22E-01 | 1.49E-01 | 2.61E+00 | 2.72E+00 | 4.09E-01 | no | Nrxn3 | intron_variant | MODIFIER |
| 10 | 93818221 | AX-170045615 | 0 | T | G | 0.289 | -7.23E-02 | 8.69E-02 | 2.54E+00 | 2.59E+00 | 4.09E-01 | no | | intergenic_variant | MODIFIER |
| 10 | 93845744 | AX-170516521 | 0 | C | T | 0.289 | -7.23E-02 | 8.69E-02 | 2.54E+00 | 2.59E+00 | 4.09E-01 | no | Usp44 | non_coding_transcript_exon_variant | MODIFIER |
| 10 | 93846720 | AX-97824910 | 0 | C | T | 0.289 | -7.23E-02 | 8.69E-02 | 2.54E+00 | 2.59E+00 | 4.09E-01 | no | Usp44 | non_coding_transcript_exon_variant | MODIFIER |
| 10 | 93850026 | AX-170405591 | 0 | G | A | 0.289 | -7.23E-02 | 8.69E-02 | 2.54E+00 | 2.59E+00 | 4.09E-01 | no | Usp44 | downstream_gene_variant | MODIFIER |
| 10 | 93855494 | AX-170297906 | 0 | G | A | 0.289 | -7.23E-02 | 8.69E-02 | 2.54E+00 | 2.59E+00 | 4.09E-01 | no | Metap2 | downstream_gene_variant | MODIFIER |
| 10 | 93865154 | AX-170412067 | 0 | G | A | 0.289 | -7.23E-02 | 8.69E-02 | 2.54E+00 | 2.59E+00 | 4.09E-01 | no | Metap2 | intron_variant | MODIFIER |
| 10 | 93870386 | AX-170513794 | 0 | A | G | 0.289 | -7.23E-02 | 8.69E-02 | 2.54E+00 | 2.59E+00 | 4.09E-01 | no | Metap2 | intron_variant | MODIFIER |
| 10 | 93876513 | AX-170145186 | 0 | A | T | 0.289 | -7.23E-02 | 8.69E-02 | 2.54E+00 | 2.59E+00 | 4.09E-01 | no | Metap2 | intron_variant | MODIFIER |
| 10 | 93877196 | AX-170153095 | 0 | C | T | 0.289 | -7.23E-02 | 8.69E-02 | 2.54E+00 | 2.59E+00 | 4.09E-01 | no | Metap2 | intron_variant | MODIFIER |
| 10 | 93878210 | AX-170288524 | 0 | T | C | 0.289 | -7.23E-02 | 8.69E-02 | 2.54E+00 | 2.59E+00 | 4.09E-01 | no | Metap2 | intron_variant | MODIFIER |
| 10 | 93883356 | AX-170473920 | 0 | G | T | 0.289 | -7.23E-02 | 8.69E-02 | 2.54E+00 | 2.59E+00 | 4.09E-01 | no | Metap2 | intron_variant | MODIFIER |
| 10 | 93888594 | AX-170045618 | 0 | A | G | 0.289 | -7.23E-02 | 8.69E-02 | 2.54E+00 | 2.59E+00 | 4.09E-01 | no | Metap2 | intron_variant | MODIFIER |
| 10 | 93906639 | AX-170104423 | 0 | G | A | 0.289 | -7.23E-02 | 8.69E-02 | 2.54E+00 | 2.59E+00 | 4.09E-01 | no | 4930471D02Rik | downstream_gene_variant | MODIFIER |
| 10 | 93907302 | AX-170508149 | 0 | G | A | 0.289 | -7.23E-02 | 8.69E-02 | 2.54E+00 | 2.59E+00 | 4.09E-01 | no | | intergenic_variant | MODIFIER |
| 10 | 93910982 | AX-97816420 | 0 | C | T | 0.289 | -7.23E-02 | 8.69E-02 | 2.54E+00 | 2.59E+00 | 4.09E-01 | no | | regulatory_region_variant | MODIFIER |
| 10 | 93953204 | AX-170397315 | 0 | T | C | 0.289 | -7.23E-02 | 8.69E-02 | 2.54E+00 | 2.59E+00 | 4.09E-01 | no | Vezt | intron_variant | MODIFIER |
| 10 | 93959688 | AX-170457499 | 0 | A | T | 0.289 | -7.23E-02 | 8.69E-02 | 2.54E+00 | 2.59E+00 | 4.09E-01 | no | Mir331 | downstream_gene_variant | MODIFIER |
| 10 | 93963965 | AX-170442844 | 0 | A | G | 0.289 | -7.23E-02 | 8.69E-02 | 2.54E+00 | 2.59E+00 | 4.09E-01 | no | Vezt | downstream_gene_variant | MODIFIER |
| 10 | 93965640 | AX-170444199 | 0 | G | A | 0.289 | -7.23E-02 | 8.69E-02 | 2.54E+00 | 2.59E+00 | 4.09E-01 | no | Vezt | downstream_gene_variant | MODIFIER |
| 10 | 93971585 | AX-170383539 | 0 | C | T | 0.289 | -7.23E-02 | 8.69E-02 | 2.54E+00 | 2.59E+00 | 4.09E-01 | no | Vezt | intron_variant | MODIFIER |
| 10 | 93987242 | AX-170119008 | 0 | A | C | 0.289 | -7.23E-02 | 8.69E-02 | 2.54E+00 | 2.59E+00 | 4.09E-01 | no | Vezt | intron_variant | MODIFIER |
| 10 | 93992963 | AX-170333541 | 0 | G | A | 0.289 | -7.23E-02 | 8.69E-02 | 2.54E+00 | 2.59E+00 | 4.09E-01 | no | Vezt | intron_variant | MODIFIER |
| 10 | 93992992 | AX-97816417 | 0 | T | C | 0.289 | -7.23E-02 | 8.69E-02 | 2.54E+00 | 2.59E+00 | 4.09E-01 | no | Vezt | intron_variant | MODIFIER |
| 10 | 93994048 | AX-170003678 | 0 | T | C | 0.289 | -7.23E-02 | 8.69E-02 | 2.54E+00 | 2.59E+00 | 4.09E-01 | no | Vezt | intron_variant | MODIFIER |
| 10 | 93994302 | AX-170106066 | 0 | G | A | 0.289 | -7.23E-02 | 8.69E-02 | 2.54E+00 | 2.59E+00 | 4.09E-01 | no | Vezt | intron_variant | MODIFIER |
| 10 | 94017816 | AX-170003684 | 0 | A | G | 0.289 | -7.23E-02 | 8.69E-02 | 2.54E+00 | 2.59E+00 | 4.09E-01 | no | Vezt | intron_variant | MODIFIER |
| 10 | 94022270 | AX-97823569 | 0 | A | T | 0.289 | -7.23E-02 | 8.69E-02 | 2.54E+00 | 2.59E+00 | 4.09E-01 | no | Vezt | intron_variant | MODIFIER |
| 10 | 94067987 | AX-170003677 | 0 | T | G | 0.289 | -7.23E-02 | 8.69E-02 | 2.54E+00 | 2.59E+00 | 4.09E-01 | no | Fgd6 | intron_variant | MODIFIER |
| 10 | 94068691 | AX-170324777 | 0 | G | T | 0.289 | -7.23E-02 | 8.69E-02 | 2.54E+00 | 2.59E+00 | 4.09E-01 | no | Fgd6 | intron_variant | MODIFIER |
| 10 | 94094245 | AX-170472512 | 0 | T | C | 0.289 | -7.23E-02 | 8.69E-02 | 2.54E+00 | 2.59E+00 | 4.09E-01 | no | Fgd6 | intron_variant | MODIFIER |
| 10 | 94111174 | AX-170003683 | 0 | A | G | 0.289 | -7.23E-02 | 8.69E-02 | 2.54E+00 | 2.59E+00 | 4.09E-01 | no | Fgd6 | intron_variant | MODIFIER |
| 10 | 94124760 | AX-170108739 | 0 | G | A | 0.289 | -7.23E-02 | 8.69E-02 | 2.54E+00 | 2.59E+00 | 4.09E-01 | no | Fgd6 | intron_variant | MODIFIER |
| 10 | 94133965 | AX-97816444 | 0 | A | G | 0.289 | -7.23E-02 | 8.69E-02 | 2.54E+00 | 2.59E+00 | 4.09E-01 | no | Fgd6 | intron_variant | MODIFIER |
| 10 | 94148274 | AX-170085877 | 0 | A | G | 0.289 | -7.23E-02 | 8.69E-02 | 2.54E+00 | 2.59E+00 | 4.09E-01 | no | Fgd6 | downstream_gene_variant | MODIFIER |
| 10 | 94154831 | AX-170045619 | 0 | G | A | 0.289 | -7.23E-02 | 8.69E-02 | 2.54E+00 | 2.59E+00 | 4.09E-01 | no | Nr2c1 | intron_variant | MODIFIER |
| 10 | 94155178 | AX-170045620 | 0 | G | C | 0.289 | -7.23E-02 | 8.69E-02 | 2.54E+00 | 2.59E+00 | 4.09E-01 | no | Nr2c1 | intron_variant | MODIFIER |

| | | | | | | | | | | | | | | | |
|---|---|---|---|---|---|---|---|---|---|---|---|---|---|---|---|
| 10 | 94156010 | AX-170284574 | 0 | A | G | 0.289 | -7.23E-02 | 8.69E-02 | 2.54E+00 | 2.59E+00 | 4.09E-01 | no | Nr2c1 | intron_variant | MODIFIER |
| 10 | 94185823 | AX-170499831 | 0 | C | G | 0.289 | -7.23E-02 | 8.69E-02 | 2.54E+00 | 2.59E+00 | 4.09E-01 | no | Gm26122 | upstream_gene_variant | MODIFIER |
| 10 | 94188499 | AX-170003681 | 0 | G | A | 0.289 | -7.23E-02 | 8.69E-02 | 2.54E+00 | 2.59E+00 | 4.09E-01 | no | Gm26122 | upstream_gene_variant | MODIFIER |
| 10 | 94233465 | AX-170098764 | 0 | G | C | 0.289 | -7.23E-02 | 8.69E-02 | 2.54E+00 | 2.59E+00 | 4.09E-01 | no | | intergenic_variant | MODIFIER |
| 10 | 94233766 | AX-170003675 | 0 | G | C | 0.289 | -7.23E-02 | 8.69E-02 | 2.54E+00 | 2.59E+00 | 4.09E-01 | no | | intergenic_variant | MODIFIER |
| 10 | 94240606 | AX-170163414 | 0 | G | A | 0.289 | -7.23E-02 | 8.69E-02 | 2.54E+00 | 2.59E+00 | 4.09E-01 | no | | intergenic_variant | MODIFIER |
| 10 | 94245633 | AX-170138143 | 0 | G | A | 0.289 | -7.23E-02 | 8.69E-02 | 2.54E+00 | 2.59E+00 | 4.09E-01 | no | | intergenic_variant | MODIFIER |
| 10 | 94282375 | AX-170475450 | 0 | G | A | 0.289 | -7.23E-02 | 8.69E-02 | 2.54E+00 | 2.59E+00 | 4.09E-01 | no | | intergenic_variant | MODIFIER |
| 10 | 94294807 | AX-170163692 | 0 | A | G | 0.289 | -7.23E-02 | 8.69E-02 | 2.54E+00 | 2.59E+00 | 4.09E-01 | no | Gm4792 | intron_variant | MODIFIER |
| 10 | 94299095 | AX-170143232 | 0 | C | T | 0.289 | -7.23E-02 | 8.69E-02 | 2.54E+00 | 2.59E+00 | 4.09E-01 | no | Gm4792 | upstream_gene_variant | MODIFIER |
| 10 | 94299419 | AX-97829740 | 0 | T | C | 0.289 | -7.23E-02 | 8.69E-02 | 2.54E+00 | 2.59E+00 | 4.09E-01 | no | Gm4792 | upstream_gene_variant | MODIFIER |
| 10 | 94299813 | AX-170120503 | 0 | T | G | 0.289 | -7.23E-02 | 8.69E-02 | 2.54E+00 | 2.59E+00 | 4.09E-01 | no | Gm4792 | upstream_gene_variant | MODIFIER |
| 10 | 94301724 | AX-170466320 | 0 | A | G | 0.289 | -7.23E-02 | 8.69E-02 | 2.54E+00 | 2.59E+00 | 4.09E-01 | no | Gm4792 | upstream_gene_variant | MODIFIER |
| 10 | 94303673 | AX-170329928 | 0 | T | C | 0.289 | -7.23E-02 | 8.69E-02 | 2.54E+00 | 2.59E+00 | 4.09E-01 | no | Gm4792 | upstream_gene_variant | MODIFIER |
| 10 | 94306086 | AX-170109632 | 0 | G | A | 0.289 | -7.23E-02 | 8.69E-02 | 2.54E+00 | 2.59E+00 | 4.09E-01 | no | | intergenic_variant | MODIFIER |
| 10 | 94322584 | AX-170473472 | 0 | A | G | 0.289 | -7.23E-02 | 8.69E-02 | 2.54E+00 | 2.59E+00 | 4.09E-01 | no | Tmcc3 | intron_variant | MODIFIER |
| 10 | 94348968 | AX-170460539 | 0 | T | C | 0.289 | -7.23E-02 | 8.69E-02 | 2.54E+00 | 2.59E+00 | 4.09E-01 | no | Tmcc3 | intron_variant | MODIFIER |
| 10 | 95092693 | AX-170366388 | 0 | G | A | 0.289 | -7.23E-02 | 8.69E-02 | 2.54E+00 | 2.59E+00 | 4.09E-01 | no | | intergenic_variant | MODIFIER |
| 1 | 189793641 | AX-170148271 | 0 | G | C | 0.254 | 8.62E-02 | 1.02E-01 | 2.44E+00 | 2.53E+00 | 4.09E-01 | no | Ptpn14 | intron_variant | MODIFIER |
| 1 | 131384667 | AX-97813277 | 1 | G | T | 0.536 | 7.04E-02 | 8.62E-02 | 2.67E+00 | 2.74E+00 | 4.09E-01 | no | Srgap2 | intron_variant | MODIFIER |
| 9 | 80826948 | AX-170198278 | 1 | C | A | 0.196 | -9.04E-02 | 1.11E-01 | 2.65E+00 | 2.76E+00 | 4.09E-01 | no | Gm27224 | intron_variant,non_coding_transcript_variant | MODIFIER |
| 1 | 100526797 | AX-170322687 | 0 | T | C | 0.149 | 1.58E-01 | 1.93E-01 | 2.65E+00 | 2.74E+00 | 4.09E-01 | Yes | | intergenic_variant | MODIFIER |
| 2 | 161945628 | AX-168377355 | 1 | C | T | 0.161 | 1.61E-01 | 1.98E-01 | 2.76E+00 | 2.84E+00 | 4.09E-01 | Yes | Ptprt | intron_variant | MODIFIER |
| 1 | 188878726 | AX-170105350 | 1 | C | T | 0.241 | 8.75E-02 | 1.04E-01 | 2.45E+00 | 2.54E+00 | 4.09E-01 | no | Ush2a | intron_variant | MODIFIER |
| 18 | 53498213 | AX-170307718 | 0 | G | T | 0.237 | -8.66E-02 | 1.04E-01 | 2.53E+00 | 2.60E+00 | 4.09E-01 | no | Prdm6 | intron_variant | MODIFIER |
| 1 | 148028918 | AX-170218928 | 1 | A | G | 0.205 | 1.13E-01 | 1.39E-01 | 2.69E+00 | 2.73E+00 | 4.09E-01 | no | | intergenic_variant | MODIFIER |
| 1 | 148583498 | AX-170229597 | 1 | G | A | 0.205 | 1.13E-01 | 1.39E-01 | 2.69E+00 | 2.73E+00 | 4.09E-01 | no | | intergenic_variant | MODIFIER |
| 13 | 24918742 | AX-170007946 | 0 | A | G | 0.175 | -1.03E-01 | 1.20E-01 | 2.41E+00 | 2.46E+00 | 4.09E-01 | no | Aldh5a1 | intron_variant | MODIFIER |
| 16 | 41126314 | AX-170430403 | 0 | T | A | 0.491 | 6.82E-02 | 8.25E-02 | 2.56E+00 | 2.61E+00 | 4.09E-01 | no | Lsamp | intron_variant | MODIFIER |
| 5 | 104270563 | AX-170151019 | 0 | A | G | 0.439 | 9.46E-02 | 1.16E-01 | 2.66E+00 | 2.69E+00 | 4.09E-01 | no | | intergenic_variant | MODIFIER |
| 5 | 105708353 | AX-170115804 | 0 | C | T | 0.439 | 9.46E-02 | 1.16E-01 | 2.66E+00 | 2.69E+00 | 4.09E-01 | no | Lrrc8d | intron_variant | MODIFIER |
| 1 | 99016281 | AX-170437131 | 0 | G | A | 0.167 | 1.39E-01 | 1.69E-01 | 2.57E+00 | 2.67E+00 | 4.10E-01 | no | Gm23434 | upstream_gene_variant | MODIFIER |
| 16 | 52732736 | AX-170346097 | 0 | C | T | 0.518 | 6.74E-02 | 8.05E-02 | 2.49E+00 | 2.54E+00 | 4.10E-01 | no | | intergenic_variant | MODIFIER |
| 13 | 25325446 | AX-170007955 | 1 | C | T | 0.17 | -1.06E-01 | 1.25E-01 | 2.44E+00 | 2.49E+00 | 4.10E-01 | no | | intergenic_variant | MODIFIER |
| 9 | 17528295 | AX-170292248 | 1 | T | C | 0.473 | 7.17E-02 | 8.63E-02 | 2.52E+00 | 2.60E+00 | 4.10E-01 | no | Gm48846 | downstream_gene_variant | MODIFIER |
| 1 | 169182097 | AX-170527022 | 1 | C | T | 0.214 | 1.13E-01 | 1.39E-01 | 2.64E+00 | 2.69E+00 | 4.10E-01 | no | | intergenic_variant | MODIFIER |
| 1 | 170507692 | AX-170236211 | 1 | G | C | 0.214 | 1.13E-01 | 1.39E-01 | 2.64E+00 | 2.69E+00 | 4.10E-01 | no | Nos1ap | intron_variant,non_coding_transcript_variant | MODIFIER |
| 1 | 136304870 | AX-170447002 | 0 | T | C | 0.211 | 9.99E-02 | 1.22E-01 | 2.64E+00 | 2.72E+00 | 4.10E-01 | no | Camsap2 | intron_variant | MODIFIER |
| 16 | 49885695 | AX-170127609 | 0 | A | T | 0.5 | 6.87E-02 | 8.31E-02 | 2.56E+00 | 2.60E+00 | 4.10E-01 | no | Cd47 | intron_variant | MODIFIER |
| 15 | 86004920 | AX-170482636 | 0 | C | G | 0.026 | 1.03E-01 | 1.28E-01 | 2.78E+00 | 2.97E+00 | 4.11E-01 | no | Celsr1 | intron_variant | MODIFIER |
| 4 | 139298825 | AX-170447454 | 0 | A | G | 0.167 | 8.97E-02 | 1.09E-01 | 2.59E+00 | 2.65E+00 | 4.11E-01 | no | Pqlc2 | 3_prime_UTR_variant | MODIFIER |
| 4 | 139688279 | AX-170225286 | 0 | A | C | 0.167 | 8.97E-02 | 1.09E-01 | 2.59E+00 | 2.65E+00 | 4.11E-01 | no | | regulatory_region_variant | MODIFIER |
| 4 | 132614787 | AX-170304028 | 0 | G | A | 0.184 | 8.23E-02 | 9.96E-02 | 2.55E+00 | 2.61E+00 | 4.11E-01 | no | | intergenic_variant | MODIFIER |
| 4 | 124307519 | AX-170253869 | 0 | C | T | 0.228 | 8.72E-02 | 1.06E-01 | 2.56E+00 | 2.64E+00 | 4.11E-01 | no | 1700021L23Rik | intron_variant,non_coding_transcript_variant | MODIFIER |
| 6 | 130338028 | AX-170324622 | 1 | G | A | 0.545 | 6.80E-02 | 8.24E-02 | 2.55E+00 | 2.63E+00 | 4.11E-01 | no | Klra3 | upstream_gene_variant | MODIFIER |
| 8 | 15558646 | AX-170368469 | 0 | T | C | 0.228 | -6.32E-02 | 7.67E-02 | 2.55E+00 | 2.65E+00 | 4.11E-01 | no | | intergenic_variant | MODIFIER |
| 5 | 50375357 | AX-170499716 | 1 | A | G | 0.473 | 8.52E-02 | 9.87E-02 | 2.38E+00 | 2.45E+00 | 4.11E-01 | no | Gm7205 | intron_variant,non_coding_transcript_variant | MODIFIER |
| 3 | 151115692 | AX-170504890 | 1 | T | C | 0.33 | 6.09E-02 | 7.50E-02 | 2.76E+00 | 2.85E+00 | 4.11E-01 | no | Gm43473 | downstream_gene_variant | MODIFIER |
| 3 | 37380991 | AX-170181688 | 1 | G | A | 0.205 | 7.30E-02 | 9.01E-02 | 2.76E+00 | 2.87E+00 | 4.11E-01 | no | Fgf2 | intron_variant | MODIFIER |
| 20 | 156643526 | AX-170195636 | 0 | G | A | 0.14 | 9.21E-02 | 1.14E-01 | 2.71E+00 | 2.80E+00 | 4.11E-01 | no | | intergenic_variant | MODIFIER |
| 1 | 149144563 | AX-97844058 | 1 | G | C | 0.205 | 1.13E-01 | 1.38E-01 | 2.66E+00 | 2.71E+00 | 4.11E-01 | no | | intergenic_variant | MODIFIER |
| 1 | 149542209 | AX-170001965 | 1 | A | G | 0.205 | 1.13E-01 | 1.38E-01 | 2.66E+00 | 2.71E+00 | 4.11E-01 | no | | intergenic_variant | MODIFIER |
| 1 | 149845432 | AX-170359161 | 1 | A | G | 0.205 | 1.13E-01 | 1.38E-01 | 2.66E+00 | 2.71E+00 | 4.11E-01 | no | Pla2g4a | intron_variant | MODIFIER |
| 1 | 150619773 | AX-170473542 | 1 | C | T | 0.205 | 1.13E-01 | 1.38E-01 | 2.66E+00 | 2.71E+00 | 4.11E-01 | no | Hmcn1 | intron_variant | MODIFIER |

| | | | | | | | | | | | | | | | |
|---|---|---|---|---|---|---|---|---|---|---|---|---|---|---|---|
| 18 | 29850020 | AX-170123148 | 0 | G | T | 0.053 | 1.83E-01 | 2.27E-01 | 2.75E+00 | 2.89E+00 | 4.11E-01 | Yes | | intergenic_variant | MODIFIER |
| 3 | 152182395 | AX-170414722 | 0 | T | C | 0.333 | 5.91E-02 | 7.29E-02 | 2.79E+00 | 2.89E+00 | 4.11E-01 | no | Dnajb4 | downstream_gene_variant | MODIFIER |
| 3 | 152901329 | AX-170346739 | 0 | G | A | 0.333 | 5.91E-02 | 7.29E-02 | 2.79E+00 | 2.89E+00 | 4.11E-01 | no | St6galnac5 | intron_variant | MODIFIER |
| 3 | 153005173 | AX-170089119 | 0 | G | A | 0.333 | 5.91E-02 | 7.29E-02 | 2.79E+00 | 2.89E+00 | 4.11E-01 | no | | intergenic_variant | MODIFIER |
| 3 | 153019180 | AX-170454906 | 0 | C | T | 0.333 | 5.91E-02 | 7.29E-02 | 2.79E+00 | 2.89E+00 | 4.11E-01 | no | | intergenic_variant | MODIFIER |
| 9 | 23175297 | AX-97822904 | 1 | T | C | 0.455 | 7.45E-02 | 9.06E-02 | 2.57E+00 | 2.63E+00 | 4.11E-01 | no | | intergenic_variant | MODIFIER |
| 4 | 138516183 | AX-170488140 | 1 | T | G | 0.152 | -9.26E-02 | 1.12E-01 | 3.00E+00 | 3.17E+00 | 4.11E-01 | no | | intergenic_variant | MODIFIER |
| 1 | 123732097 | AX-170492302 | 0 | A | G | 0.561 | 6.75E-02 | 8.20E-02 | 2.57E+00 | 2.63E+00 | 4.11E-01 | no | Dpp10 | intron_variant | MODIFIER |
| 8 | 36140106 | AX-170351737 | 0 | T | C | 0.149 | -1.11E-01 | 1.36E-01 | 2.68E+00 | 2.75E+00 | 4.12E-01 | no | Prag1 | synonymous_variant | LOW |
| 14 | 64108767 | AX-170518762 | 0 | C | T | 0.053 | 1.53E-01 | 1.88E-01 | 2.81E+00 | 2.94E+00 | 4.12E-01 | Yes | Prss52 | intron_variant | MODIFIER |
| 14 | 64656398 | AX-170436098 | 0 | T | G | 0.053 | 1.53E-01 | 1.88E-01 | 2.81E+00 | 2.94E+00 | 4.12E-01 | Yes | Kif13b | intron_variant | MODIFIER |
| 16 | 92130928 | AX-97807706 | 0 | C | T | 0.465 | -6.47E-02 | 7.98E-02 | 2.77E+00 | 2.85E+00 | 4.12E-01 | no | Gm41496 | intron_variant,non_coding_transcript_variant | MODIFIER |
| 3 | 37135898 | AX-170193236 | 0 | T | C | 0.193 | 7.73E-02 | 9.50E-02 | 2.84E+00 | 2.94E+00 | 4.12E-01 | no | | intergenic_variant | MODIFIER |
| 8 | 29854553 | AX-170197152 | 0 | C | G | 0.175 | 9.17E-02 | 1.13E-01 | 2.81E+00 | 2.89E+00 | 4.12E-01 | no | | intergenic_variant | MODIFIER |
| 9 | 85670214 | AX-97788697 | 0 | T | C | 0.14 | -9.64E-02 | 1.18E-01 | 2.59E+00 | 2.68E+00 | 4.12E-01 | no | | intergenic_variant | MODIFIER |
| 6 | 136897946 | AX-170320732 | 0 | A | G | 0.57 | 7.32E-02 | 8.70E-02 | 2.46E+00 | 2.52E+00 | 4.12E-01 | no | | regulatory_region_variant | MODIFIER |
| 18 | 78339853 | AX-170498919 | 0 | A | G | 0.158 | -9.61E-02 | 1.17E-01 | 2.56E+00 | 2.62E+00 | 4.12E-01 | no | Slc14a2 | intron_variant | MODIFIER |
| 18 | 78469441 | AX-170404114 | 0 | T | G | 0.158 | -9.61E-02 | 1.17E-01 | 2.56E+00 | 2.62E+00 | 4.12E-01 | no | Slc14a2 | intron_variant | MODIFIER |
| 18 | 78547457 | AX-97802482 | 0 | T | C | 0.158 | -9.61E-02 | 1.17E-01 | 2.56E+00 | 2.62E+00 | 4.12E-01 | no | Slc14a2 | intron_variant | MODIFIER |
| 18 | 78576663 | AX-170248231 | 0 | T | G | 0.158 | -9.61E-02 | 1.17E-01 | 2.56E+00 | 2.62E+00 | 4.12E-01 | no | Slc14a2 | intron_variant | MODIFIER |
| 18 | 78583613 | AX-170519835 | 0 | C | T | 0.158 | -9.61E-02 | 1.17E-01 | 2.56E+00 | 2.62E+00 | 4.12E-01 | no | Slc14a2 | intron_variant | MODIFIER |
| 10 | 68260530 | AX-170264930 | 2 | A | G | 0.409 | -6.50E-02 | 7.99E-02 | 2.64E+00 | 2.71E+00 | 4.12E-01 | no | Arid5b | intron_variant,NMD_transcript_variant | MODIFIER |
| 13 | 6322879 | AX-170050451 | 0 | A | T | 0.211 | -9.81E-02 | 1.20E-01 | 2.58E+00 | 2.65E+00 | 4.12E-01 | no | | intergenic_variant | MODIFIER |
| 7 | 115023606 | AX-168396943 | 1 | C | T | 0.429 | 7.74E-02 | 9.52E-02 | 2.67E+00 | 2.72E+00 | 4.12E-01 | no | | intergenic_variant | MODIFIER |
| 16 | 96354085 | AX-170244753 | 2 | T | C | 0.455 | 6.32E-02 | 7.70E-02 | 2.95E+00 | 3.10E+00 | 4.12E-01 | no | | regulatory_region_variant | MODIFIER |
| 1 | 55660432 | AX-170477889 | 0 | A | G | 0.114 | 1.47E-01 | 1.82E-01 | 2.75E+00 | 2.87E+00 | 4.12E-01 | Yes | Plcl1 | intron_variant | MODIFIER |
| 8 | 68589935 | AX-170142083 | 1 | A | G | 0.205 | -9.23E-02 | 1.14E-01 | 2.68E+00 | 2.80E+00 | 4.12E-01 | no | Csgalnact1 | intron_variant | MODIFIER |
| 9 | 119259456 | AX-170033402 | 0 | A | G | 0.202 | -6.26E-02 | 7.76E-02 | 2.66E+00 | 2.80E+00 | 4.12E-01 | no | Oxsr1 | intron_variant | MODIFIER |
| 9 | 120382394 | AX-170358804 | 0 | A | G | 0.202 | -6.26E-02 | 7.76E-02 | 2.66E+00 | 2.80E+00 | 4.12E-01 | no | Myrip | intron_variant | MODIFIER |
| 9 | 105945247 | AX-170033206 | 0 | G | A | 0.123 | -8.99E-02 | 1.10E-01 | 2.61E+00 | 2.70E+00 | 4.13E-01 | no | Col6a5 | intron_variant | MODIFIER |
| 7 | 144693834 | AX-170440799 | 0 | A | C | 0.202 | -6.02E-02 | 7.46E-02 | 2.65E+00 | 2.78E+00 | 4.13E-01 | no | Ano1 | intron_variant | MODIFIER |
| 4 | 139506373 | AX-170113483 | 1 | T | C | 0.152 | 9.52E-02 | 1.17E-01 | 2.60E+00 | 2.65E+00 | 4.13E-01 | no | | regulatory_region_variant | MODIFIER |
| 8 | 94550826 | AX-170080861 | 2 | A | G | 0.209 | -7.62E-02 | 9.44E-02 | 2.71E+00 | 2.81E+00 | 4.13E-01 | no | Cpne2 | intron_variant | MODIFIER |
| 10 | 66106237 | AX-170134324 | 1 | A | T | 0.411 | -6.50E-02 | 8.01E-02 | 2.64E+00 | 2.72E+00 | 4.13E-01 | no | | intergenic_variant | MODIFIER |
| 18 | 85334427 | AX-170165329 | 0 | T | C | 0.167 | -7.85E-02 | 9.36E-02 | 2.45E+00 | 2.54E+00 | 4.13E-01 | no | | intergenic_variant | MODIFIER |
| 4 | 129791349 | AX-170296004 | 0 | T | C | 0.175 | 8.96E-02 | 1.10E-01 | 2.61E+00 | 2.66E+00 | 4.13E-01 | no | | regulatory_region_variant | MODIFIER |
| 1 | 14196251 | AX-170438049 | 0 | G | A | 0.395 | 6.43E-02 | 7.90E-02 | 2.62E+00 | 2.66E+00 | 4.14E-01 | no | Eya1 | intron_variant | MODIFIER |
| 1 | 14238688 | AX-170267704 | 0 | A | G | 0.395 | 6.43E-02 | 7.90E-02 | 2.62E+00 | 2.66E+00 | 4.14E-01 | no | Eya1 | intron_variant | MODIFIER |
| 1 | 14252750 | AX-170526778 | 0 | C | T | 0.395 | 6.43E-02 | 7.90E-02 | 2.62E+00 | 2.66E+00 | 4.14E-01 | no | Eya1 | intron_variant | MODIFIER |
| 1 | 14318787 | AX-170230320 | 0 | A | G | 0.395 | 6.43E-02 | 7.90E-02 | 2.62E+00 | 2.66E+00 | 4.14E-01 | no | | intergenic_variant | MODIFIER |
| 1 | 14349416 | AX-170211023 | 0 | G | A | 0.395 | 6.43E-02 | 7.90E-02 | 2.62E+00 | 2.66E+00 | 4.14E-01 | no | | intergenic_variant | MODIFIER |
| 1 | 14660935 | AX-170427297 | 0 | T | C | 0.395 | 6.43E-02 | 7.90E-02 | 2.62E+00 | 2.66E+00 | 4.14E-01 | no | | intergenic_variant | MODIFIER |
| 1 | 14668922 | AX-170264159 | 0 | G | C | 0.395 | 6.43E-02 | 7.90E-02 | 2.62E+00 | 2.66E+00 | 4.14E-01 | no | | intergenic_variant | MODIFIER |
| 1 | 14675443 | AX-170511208 | 0 | A | G | 0.395 | 6.43E-02 | 7.90E-02 | 2.62E+00 | 2.66E+00 | 4.14E-01 | no | | intergenic_variant | MODIFIER |
| 1 | 14753754 | AX-97851102 | 0 | C | T | 0.395 | 6.43E-02 | 7.90E-02 | 2.62E+00 | 2.66E+00 | 4.14E-01 | no | Msc | 3_prime_UTR_variant | MODIFIER |
| 1 | 14754121 | AX-170346671 | 0 | C | A | 0.395 | 6.43E-02 | 7.90E-02 | 2.62E+00 | 2.66E+00 | 4.14E-01 | no | Msc | 3_prime_UTR_variant | MODIFIER |
| 1 | 14754151 | AX-170224802 | 0 | C | T | 0.395 | 6.43E-02 | 7.90E-02 | 2.62E+00 | 2.66E+00 | 4.14E-01 | no | Msc | 3_prime_UTR_variant | MODIFIER |
| 1 | 14763878 | AX-170510380 | 0 | T | C | 0.395 | 6.43E-02 | 7.90E-02 | 2.62E+00 | 2.66E+00 | 4.14E-01 | no | Gm9947 | intron_variant,non_coding_transcript_variant | MODIFIER |
| 1 | 14779667 | AX-97798159 | 0 | C | T | 0.395 | 6.43E-02 | 7.90E-02 | 2.62E+00 | 2.66E+00 | 4.14E-01 | no | Gm9947 | downstream_gene_variant | MODIFIER |
| 1 | 14794379 | AX-170162803 | 0 | T | G | 0.395 | 6.43E-02 | 7.90E-02 | 2.62E+00 | 2.66E+00 | 4.14E-01 | no | Gm37412 | downstream_gene_variant | MODIFIER |
| 1 | 14825116 | AX-170521506 | 0 | C | T | 0.395 | 6.43E-02 | 7.90E-02 | 2.62E+00 | 2.66E+00 | 4.14E-01 | no | | intergenic_variant | MODIFIER |
| 1 | 14826114 | AX-97854011 | 0 | A | G | 0.395 | 6.43E-02 | 7.90E-02 | 2.62E+00 | 2.66E+00 | 4.14E-01 | no | | intergenic_variant | MODIFIER |
| 1 | 14842965 | AX-170293412 | 0 | G | A | 0.395 | 6.43E-02 | 7.90E-02 | 2.62E+00 | 2.66E+00 | 4.14E-01 | no | Smt3h2-ps4 | upstream_gene_variant | MODIFIER |
| 1 | 14857641 | AX-170456055 | 0 | G | T | 0.395 | 6.43E-02 | 7.90E-02 | 2.62E+00 | 2.66E+00 | 4.14E-01 | no | | regulatory_region_variant | MODIFIER |

| | | | | | | | | | | | | | | | |
|---|---|---|---|---|---|---|---|---|---|---|---|---|---|---|---|
| 1 | 14858509 | AX-170215762 | 0 | G | C | 0.395 | 6.43E-02 | 7.90E-02 | 2.62E+00 | 2.66E+00 | 4.14E-01 | no | | intergenic_variant | MODIFIER |
| 1 | 14890752 | AX-170230744 | 0 | T | C | 0.395 | 6.43E-02 | 7.90E-02 | 2.62E+00 | 2.66E+00 | 4.14E-01 | no | Trpa1 | intron_variant | MODIFIER |
| 1 | 14890841 | AX-170193291 | 0 | A | G | 0.395 | 6.43E-02 | 7.90E-02 | 2.62E+00 | 2.66E+00 | 4.14E-01 | no | Trpa1 | intron_variant | MODIFIER |
| 1 | 14907151 | AX-170403017 | 0 | T | C | 0.395 | 6.43E-02 | 7.90E-02 | 2.62E+00 | 2.66E+00 | 4.14E-01 | no | Trpa1 | intron_variant | MODIFIER |
| 1 | 14908332 | AX-170350359 | 0 | T | C | 0.395 | 6.43E-02 | 7.90E-02 | 2.62E+00 | 2.66E+00 | 4.14E-01 | no | Trpa1 | intron_variant | MODIFIER |
| 1 | 14909034 | AX-170191078 | 0 | G | T | 0.395 | 6.43E-02 | 7.90E-02 | 2.62E+00 | 2.66E+00 | 4.14E-01 | no | Trpa1 | intron_variant | MODIFIER |
| 1 | 15138185 | AX-169999988 | 0 | T | A | 0.395 | 6.43E-02 | 7.90E-02 | 2.62E+00 | 2.66E+00 | 4.14E-01 | no | | intergenic_variant | MODIFIER |
| 1 | 15359916 | AX-170280655 | 0 | C | G | 0.395 | 6.43E-02 | 7.90E-02 | 2.62E+00 | 2.66E+00 | 4.14E-01 | no | Kcnb2 | intron_variant | MODIFIER |
| 1 | 15389263 | AX-170329158 | 0 | T | C | 0.395 | 6.43E-02 | 7.90E-02 | 2.62E+00 | 2.66E+00 | 4.14E-01 | no | Kcnb2 | intron_variant | MODIFIER |
| 1 | 15454759 | AX-97810944 | 0 | T | C | 0.395 | 6.43E-02 | 7.90E-02 | 2.62E+00 | 2.66E+00 | 4.14E-01 | no | Kcnb2 | intron_variant | MODIFIER |
| 1 | 16997421 | AX-97841537 | 0 | C | T | 0.395 | 6.43E-02 | 7.90E-02 | 2.62E+00 | 2.66E+00 | 4.14E-01 | no | Jph1 | 3_prime_UTR_variant | MODIFIER |
| 16 | 76467005 | AX-97825199 | 0 | C | A | 0.474 | 6.27E-02 | 7.62E-02 | 2.53E+00 | 2.61E+00 | 4.14E-01 | no | Gm45030 | intron_variant,non_coding_transcript_variant | MODIFIER |
| 8 | 55726551 | AX-170030759 | 0 | C | T | 0.149 | -1.22E-01 | 1.49E-01 | 2.54E+00 | 2.60E+00 | 4.14E-01 | no | | intergenic_variant | MODIFIER |
| 1 | 176536871 | AX-170268944 | 0 | T | C | 0.228 | 9.29E-02 | 1.13E-01 | 2.56E+00 | 2.64E+00 | 4.14E-01 | no | | intergenic_variant | MODIFIER |
| 1 | 176873277 | AX-170457472 | 0 | T | C | 0.228 | 9.29E-02 | 1.13E-01 | 2.56E+00 | 2.64E+00 | 4.14E-01 | no | Sdccag8 | intron_variant | MODIFIER |
| 18 | 65691384 | AX-170271014 | 0 | C | G | 0.404 | 7.15E-02 | 8.52E-02 | 2.45E+00 | 2.51E+00 | 4.14E-01 | no | Zfp532 | downstream_gene_variant | MODIFIER |
| 8 | 68683901 | AX-170204225 | 0 | G | A | 0.193 | -9.16E-02 | 1.13E-01 | 2.59E+00 | 2.69E+00 | 4.14E-01 | no | Csgalnact1 | intron_variant | MODIFIER |
| 10 | 93755708 | AX-170205609 | 1 | T | C | 0.286 | -7.08E-02 | 8.57E-02 | 2.52E+00 | 2.57E+00 | 4.14E-01 | no | | intergenic_variant | MODIFIER |
| 8 | 73401887 | AX-170292252 | 0 | A | G | 0.193 | -1.01E-01 | 1.25E-01 | 2.72E+00 | 2.81E+00 | 4.14E-01 | no | | intergenic_variant | MODIFIER |
| 8 | 73804365 | AX-170080423 | 0 | A | G | 0.193 | -1.01E-01 | 1.25E-01 | 2.72E+00 | 2.81E+00 | 4.14E-01 | no | | intergenic_variant | MODIFIER |
| 10 | 8330801 | AX-170152781 | 0 | G | A | 0.219 | -7.54E-02 | 9.37E-02 | 2.68E+00 | 2.80E+00 | 4.14E-01 | no | Ust | intron_variant | MODIFIER |
| 10 | 93236381 | AX-170383738 | 1 | G | T | 0.304 | -7.10E-02 | 8.56E-02 | 2.50E+00 | 2.55E+00 | 4.14E-01 | no | Cdk17 | intron_variant | MODIFIER |
| 18 | 59074211 | AX-170330074 | 1 | G | A | 0.402 | 8.10E-02 | 9.53E-02 | 2.41E+00 | 2.46E+00 | 4.14E-01 | no | Minar2 | intron_variant | MODIFIER |
| 4 | 134893443 | AX-170262380 | 1 | C | T | 0.17 | 9.08E-02 | 1.12E-01 | 2.61E+00 | 2.65E+00 | 4.14E-01 | no | Tmem50a | downstream_gene_variant | MODIFIER |
| 16 | 77930554 | AX-97790994 | 1 | A | G | 0.473 | 6.23E-02 | 7.59E-02 | 2.54E+00 | 2.61E+00 | 4.14E-01 | no | | intergenic_variant | MODIFIER |
| 5 | 37201101 | AX-170497097 | 1 | A | G | 0.348 | 9.20E-02 | 1.14E-01 | 2.78E+00 | 2.85E+00 | 4.14E-01 | no | Gm1043 | intron_variant | MODIFIER |
| 9 | 96257737 | AX-97806043 | 2 | C | T | 0.127 | -8.74E-02 | 1.09E-01 | 2.72E+00 | 2.85E+00 | 4.14E-01 | no | Tfdp2 | intron_variant | MODIFIER |
| 6 | 131195959 | AX-170028261 | 0 | T | A | 0.544 | 6.38E-02 | 7.75E-02 | 2.52E+00 | 2.61E+00 | 4.14E-01 | no | | intergenic_variant | MODIFIER |
| 18 | 63503374 | AX-170511657 | 0 | A | G | 0.184 | -9.61E-02 | 1.19E-01 | 2.72E+00 | 2.77E+00 | 4.14E-01 | no | Gm23581 | upstream_gene_variant | MODIFIER |
| 1 | 14292496 | AX-170040438 | 0 | C | T | 0.404 | 6.38E-02 | 7.82E-02 | 2.58E+00 | 2.63E+00 | 4.15E-01 | no | Eya1 | intron_variant | MODIFIER |
| 4 | 63019978 | AX-97839767 | 0 | C | T | 0.377 | 9.50E-02 | 1.18E-01 | 2.69E+00 | 2.74E+00 | 4.15E-01 | no | Zfp618 | intron_variant | MODIFIER |
| 18 | 60316631 | AX-170507971 | 1 | G | A | 0.393 | 7.82E-02 | 9.27E-02 | 2.43E+00 | 2.49E+00 | 4.15E-01 | no | AC132320.1 | intron_variant,non_coding_transcript_variant | MODIFIER |
| 12 | 72930474 | AX-170124872 | 0 | C | T | 0.07 | 1.54E-01 | 1.93E-01 | 2.85E+00 | 3.05E+00 | 4.15E-01 | Yes | 4930447C04Rik | intron_variant | MODIFIER |
| 15 | 97840549 | AX-170253796 | 1 | G | T | 0.134 | 1.07E-01 | 1.34E-01 | 2.70E+00 | 2.85E+00 | 4.15E-01 | no | Hdac7 | intron_variant | MODIFIER |
| 8 | 60745107 | AX-170401277 | 0 | A | T | 0.149 | -1.14E-01 | 1.37E-01 | 2.48E+00 | 2.55E+00 | 4.15E-01 | no | | intergenic_variant | MODIFIER |
| 15 | 6002677 | AX-97799722 | 1 | A | C | 0.446 | -4.53E-02 | 5.62E-02 | 2.81E+00 | 2.92E+00 | 4.15E-01 | no | | intergenic_variant | MODIFIER |
| 9 | 92686323 | AX-170412267 | 1 | A | G | 0.143 | -1.01E-01 | 1.25E-01 | 2.62E+00 | 2.69E+00 | 4.15E-01 | no | | intergenic_variant | MODIFIER |
| 4 | 93729406 | AX-170215777 | 0 | T | G | 0.342 | 8.26E-02 | 1.02E-01 | 2.65E+00 | 2.73E+00 | 4.15E-01 | no | | intergenic_variant | MODIFIER |
| 9 | 80396675 | AX-170387914 | 0 | A | G | 0.211 | 9.93E-02 | 1.22E-01 | 2.91E+00 | 3.00E+00 | 4.15E-01 | no | Impg1 | intron_variant | MODIFIER |
| 7 | 37844227 | AX-170250658 | 0 | G | T | 0.351 | 7.63E-02 | 9.36E-02 | 2.57E+00 | 2.65E+00 | 4.15E-01 | no | | intergenic_variant | MODIFIER |
| 7 | 39445218 | AX-97795030 | 0 | A | G | 0.351 | 7.63E-02 | 9.36E-02 | 2.57E+00 | 2.65E+00 | 4.15E-01 | no | Gm5590 | downstream_gene_variant | MODIFIER |
| 1 | 189899338 | AX-170451040 | 1 | T | C | 0.25 | 8.41E-02 | 9.98E-02 | 2.43E+00 | 2.51E+00 | 4.15E-01 | no | Smyd2 | intron_variant | MODIFIER |
| 9 | 50383604 | AX-170505717 | 0 | T | G | 0.211 | -8.68E-02 | 1.08E-01 | 2.70E+00 | 2.79E+00 | 4.16E-01 | no | | intergenic_variant | MODIFIER |
| 1 | 149476545 | AX-170111704 | 0 | T | C | 0.193 | 1.07E-01 | 1.33E-01 | 2.69E+00 | 2.75E+00 | 4.16E-01 | no | | intergenic_variant | MODIFIER |
| 10 | 93348394 | AX-170375102 | 0 | A | G | 0.307 | -6.77E-02 | 8.23E-02 | 2.52E+00 | 2.58E+00 | 4.16E-01 | no | Gm17745 | non_coding_transcript_exon_variant | MODIFIER |
| 16 | 45184415 | AX-170154281 | 0 | A | G | 0.474 | -6.70E-02 | 8.18E-02 | 2.55E+00 | 2.60E+00 | 4.16E-01 | no | Atg3 | intron_variant | MODIFIER |
| 16 | 45510146 | AX-97802496 | 0 | T | G | 0.474 | -6.70E-02 | 8.18E-02 | 2.55E+00 | 2.60E+00 | 4.16E-01 | no | Gm17783 | downstream_gene_variant | MODIFIER |
| 1 | 53640077 | AX-170528500 | 0 | A | G | 0.114 | 1.49E-01 | 1.85E-01 | 2.67E+00 | 2.78E+00 | 4.16E-01 | Yes | Dnah7a | intron_variant | MODIFIER |
| 1 | 55473196 | AX-97808990 | 0 | C | T | 0.114 | 1.49E-01 | 1.85E-01 | 2.67E+00 | 2.78E+00 | 4.16E-01 | Yes | Plcl1 | intron_variant | MODIFIER |
| 9 | 117897841 | AX-170442030 | 0 | G | A | 0.123 | -7.85E-02 | 9.72E-02 | 2.60E+00 | 2.72E+00 | 4.16E-01 | no | Zcwpw2 | intron_variant,non_coding_transcript_variant | MODIFIER |
| 6 | 137420906 | AX-170028340 | 0 | G | A | 0.579 | 7.32E-02 | 8.81E-02 | 2.48E+00 | 2.53E+00 | 4.16E-01 | no | Ptpro | intron_variant | MODIFIER |
| 7 | 10277503 | AX-170467743 | 1 | G | C | 0.179 | 6.33E-02 | 7.86E-02 | 2.88E+00 | 3.04E+00 | 4.16E-01 | no | Vmn1r66 | upstream_gene_variant | MODIFIER |
| 6 | 134049834 | AX-170278669 | 0 | A | G | 0.553 | 7.20E-02 | 8.67E-02 | 2.48E+00 | 2.53E+00 | 4.16E-01 | no | Etv6 | intron_variant | MODIFIER |
| 6 | 134121403 | AX-170345683 | 0 | C | T | 0.553 | 7.20E-02 | 8.67E-02 | 2.48E+00 | 2.53E+00 | 4.16E-01 | no | Etv6 | intron_variant | MODIFIER |

| | | | | | | | | | | | | | | | |
|---|---|---|---|---|---|---|---|---|---|---|---|---|---|---|---|
| 6 | 134138033 | AX-170292969 | 0 | T | C | 0.553 | 7.20E-02 | 8.67E-02 | 2.48E+00 | 2.53E+00 | 4.16E-01 | no | Etv6 | intron_variant | MODIFIER |
| 6 | 134140533 | AX-97843661 | 0 | A | G | 0.553 | 7.20E-02 | 8.67E-02 | 2.48E+00 | 2.53E+00 | 4.16E-01 | no | Etv6 | intron_variant | MODIFIER |
| 6 | 134142963 | AX-170076295 | 0 | A | G | 0.553 | 7.20E-02 | 8.67E-02 | 2.48E+00 | 2.53E+00 | 4.16E-01 | no | Etv6 | intron_variant | MODIFIER |
| 6 | 134155361 | AX-97806442 | 0 | G | A | 0.553 | 7.20E-02 | 8.67E-02 | 2.48E+00 | 2.53E+00 | 4.16E-01 | no | Etv6 | intron_variant | MODIFIER |
| 6 | 134233270 | AX-170189257 | 0 | T | G | 0.553 | 7.20E-02 | 8.67E-02 | 2.48E+00 | 2.53E+00 | 4.16E-01 | no | Etv6 | intron_variant | MODIFIER |
| 6 | 134233523 | AX-170297687 | 0 | T | C | 0.553 | 7.20E-02 | 8.67E-02 | 2.48E+00 | 2.53E+00 | 4.16E-01 | no | Etv6 | synonymous_variant | LOW |
| 6 | 134253713 | AX-97798323 | 0 | G | A | 0.553 | 7.20E-02 | 8.67E-02 | 2.48E+00 | 2.53E+00 | 4.16E-01 | no | Etv6 | intron_variant | MODIFIER |
| 6 | 134257611 | AX-170455080 | 0 | T | C | 0.553 | 7.20E-02 | 8.67E-02 | 2.48E+00 | 2.53E+00 | 4.16E-01 | no | Etv6 | intron_variant | MODIFIER |
| 6 | 134269479 | AX-170238467 | 0 | A | G | 0.553 | 7.20E-02 | 8.67E-02 | 2.48E+00 | 2.53E+00 | 4.16E-01 | no | Etv6 | 3_prime_UTR_variant | MODIFIER |
| 6 | 134380769 | AX-97788411 | 0 | T | C | 0.553 | 7.20E-02 | 8.67E-02 | 2.48E+00 | 2.53E+00 | 4.16E-01 | no | 4933406J09Rik | intron_variant,non_coding_transcript_variant | MODIFIER |
| 6 | 134495284 | AX-170164263 | 0 | A | T | 0.553 | 7.20E-02 | 8.67E-02 | 2.48E+00 | 2.53E+00 | 4.16E-01 | no | Lrp6 | intron_variant | MODIFIER |
| 6 | 134584311 | AX-97845417 | 0 | A | G | 0.553 | 7.20E-02 | 8.67E-02 | 2.48E+00 | 2.53E+00 | 4.16E-01 | no | | intergenic_variant | MODIFIER |
| 6 | 134613744 | AX-170028303 | 0 | G | A | 0.553 | 7.20E-02 | 8.67E-02 | 2.48E+00 | 2.53E+00 | 4.16E-01 | no | Mansc1 | intron_variant | MODIFIER |
| 6 | 134655465 | AX-170076309 | 0 | C | T | 0.553 | 7.20E-02 | 8.67E-02 | 2.48E+00 | 2.53E+00 | 4.16E-01 | no | Borcs5 | intron_variant | MODIFIER |
| 6 | 134675400 | AX-170028306 | 0 | G | C | 0.553 | 7.20E-02 | 8.67E-02 | 2.48E+00 | 2.53E+00 | 4.16E-01 | no | Borcs5 | intron_variant | MODIFIER |
| 6 | 134696460 | AX-97856136 | 0 | C | T | 0.553 | 7.20E-02 | 8.67E-02 | 2.48E+00 | 2.53E+00 | 4.16E-01 | no | Borcs5 | intron_variant | MODIFIER |
| 6 | 134739370 | AX-97794976 | 0 | A | G | 0.553 | 7.20E-02 | 8.67E-02 | 2.48E+00 | 2.53E+00 | 4.16E-01 | no | Dusp16 | intron_variant | MODIFIER |
| 6 | 134981568 | AX-170225504 | 0 | T | C | 0.553 | 7.20E-02 | 8.67E-02 | 2.48E+00 | 2.53E+00 | 4.16E-01 | no | Apold1 | upstream_gene_variant | MODIFIER |
| 4 | 74849248 | AX-170287581 | 0 | T | A | 0.044 | 1.06E-01 | 1.33E-01 | 2.65E+00 | 2.85E+00 | 4.16E-01 | no | | intergenic_variant | MODIFIER |
| 13 | 23114850 | AX-170050739 | 0 | A | C | 0.158 | -9.55E-02 | 1.13E-01 | 2.43E+00 | 2.50E+00 | 4.16E-01 | no | Vmn1r217 | upstream_gene_variant | MODIFIER |
| 10 | 22891663 | AX-170517021 | 2 | T | A | 0.555 | -5.48E-02 | 6.77E-02 | 2.93E+00 | 3.11E+00 | 4.16E-01 | no | | intergenic_variant | MODIFIER |
| 9 | 17359887 | AX-97818035 | 1 | A | G | 0.482 | 7.26E-02 | 8.85E-02 | 2.53E+00 | 2.60E+00 | 4.16E-01 | no | | intergenic_variant | MODIFIER |
| 15 | 3749453 | AX-170460272 | 1 | T | G | 0.241 | -8.50E-02 | 1.05E-01 | 2.61E+00 | 2.66E+00 | 4.16E-01 | no | Gm4823 | upstream_gene_variant | MODIFIER |
| 15 | 7337015 | AX-97834215 | 1 | A | G | 0.241 | -8.50E-02 | 1.05E-01 | 2.61E+00 | 2.66E+00 | 4.16E-01 | no | Egflam | intron_variant | MODIFIER |
| 15 | 9336993 | AX-168375086 | 1 | G | A | 0.241 | -8.50E-02 | 1.05E-01 | 2.61E+00 | 2.66E+00 | 4.16E-01 | no | Ugt3a2 | intron_variant | MODIFIER |
| 15 | 94208000 | AX-170056349 | 0 | C | T | 0.158 | 1.23E-01 | 1.52E-01 | 2.59E+00 | 2.69E+00 | 4.16E-01 | no | | intergenic_variant | MODIFIER |
| 5 | 31038205 | AX-170433050 | 2 | C | A | 0.373 | 7.92E-02 | 9.87E-02 | 2.74E+00 | 2.82E+00 | 4.16E-01 | no | Slc5a6 | intron_variant | MODIFIER |
| 9 | 98615010 | AX-97815426 | 0 | A | G | 0.132 | -9.35E-02 | 1.16E-01 | 2.61E+00 | 2.69E+00 | 4.16E-01 | no | | intergenic_variant | MODIFIER |
| 9 | 98637738 | AX-170114427 | 0 | A | T | 0.132 | -9.35E-02 | 1.16E-01 | 2.61E+00 | 2.69E+00 | 4.16E-01 | no | | intergenic_variant | MODIFIER |
| 9 | 98647474 | AX-170283782 | 0 | G | C | 0.132 | -9.35E-02 | 1.16E-01 | 2.61E+00 | 2.69E+00 | 4.16E-01 | no | | regulatory_region_variant | MODIFIER |
| 9 | 98648943 | AX-170083208 | 0 | T | C | 0.132 | -9.35E-02 | 1.16E-01 | 2.61E+00 | 2.69E+00 | 4.16E-01 | no | | intergenic_variant | MODIFIER |
| 9 | 98659768 | AX-170272518 | 0 | T | C | 0.132 | -9.35E-02 | 1.16E-01 | 2.61E+00 | 2.69E+00 | 4.16E-01 | no | | intergenic_variant | MODIFIER |
| 9 | 98663516 | AX-170466499 | 0 | A | G | 0.132 | -9.35E-02 | 1.16E-01 | 2.61E+00 | 2.69E+00 | 4.16E-01 | no | | intergenic_variant | MODIFIER |
| 9 | 98705163 | AX-170394975 | 0 | A | G | 0.132 | -9.35E-02 | 1.16E-01 | 2.61E+00 | 2.69E+00 | 4.16E-01 | no | Gm27919 | downstream_gene_variant | MODIFIER |
| 9 | 98765402 | AX-170033102 | 0 | T | C | 0.132 | -9.35E-02 | 1.16E-01 | 2.61E+00 | 2.69E+00 | 4.16E-01 | no | E330023G01Rik | intron_variant,non_coding_transcript_variant | MODIFIER |
| 9 | 98765420 | AX-170033105 | 0 | A | G | 0.132 | -9.35E-02 | 1.16E-01 | 2.61E+00 | 2.69E+00 | 4.16E-01 | no | E330023G01Rik | intron_variant,non_coding_transcript_variant | MODIFIER |
| 9 | 98772009 | AX-97835838 | 0 | C | T | 0.132 | -9.35E-02 | 1.16E-01 | 2.61E+00 | 2.69E+00 | 4.16E-01 | no | E330023G01Rik | intron_variant,non_coding_transcript_variant | MODIFIER |
| 9 | 100927424 | AX-170033136 | 0 | T | C | 0.132 | -9.35E-02 | 1.16E-01 | 2.61E+00 | 2.69E+00 | 4.16E-01 | no | Stag1 | intron_variant | MODIFIER |
| 9 | 100954387 | AX-170033140 | 0 | A | G | 0.132 | -9.35E-02 | 1.16E-01 | 2.61E+00 | 2.69E+00 | 4.16E-01 | no | Stag1 | intron_variant | MODIFIER |
| 9 | 100980000 | AX-97817076 | 0 | T | G | 0.132 | -9.35E-02 | 1.16E-01 | 2.61E+00 | 2.69E+00 | 4.16E-01 | no | Pccb | downstream_gene_variant | MODIFIER |
| 9 | 100980146 | AX-170277580 | 0 | A | C | 0.132 | -9.35E-02 | 1.16E-01 | 2.61E+00 | 2.69E+00 | 4.16E-01 | no | Pccb | downstream_gene_variant | MODIFIER |
| 9 | 100985923 | AX-170418610 | 0 | C | G | 0.132 | -9.35E-02 | 1.16E-01 | 2.61E+00 | 2.69E+00 | 4.16E-01 | no | Pccb | intron_variant | MODIFIER |
| 9 | 100987583 | AX-97808416 | 0 | A | G | 0.132 | -9.35E-02 | 1.16E-01 | 2.61E+00 | 2.69E+00 | 4.16E-01 | no | Pccb | intron_variant | MODIFIER |
| 9 | 101004594 | AX-170474349 | 0 | C | T | 0.132 | -9.35E-02 | 1.16E-01 | 2.61E+00 | 2.69E+00 | 4.16E-01 | no | Pccb | intron_variant | MODIFIER |
| 9 | 101005540 | AX-170238163 | 0 | G | A | 0.132 | -9.35E-02 | 1.16E-01 | 2.61E+00 | 2.69E+00 | 4.16E-01 | no | Pccb | intron_variant | MODIFIER |
| 9 | 101006178 | AX-170373057 | 0 | A | T | 0.132 | -9.35E-02 | 1.16E-01 | 2.61E+00 | 2.69E+00 | 4.16E-01 | no | Pccb | intron_variant | MODIFIER |
| 9 | 101006716 | AX-170033138 | 0 | A | G | 0.132 | -9.35E-02 | 1.16E-01 | 2.61E+00 | 2.69E+00 | 4.16E-01 | no | Pccb | intron_variant | MODIFIER |
| 9 | 101115149 | AX-97857721 | 0 | A | C | 0.132 | -9.35E-02 | 1.16E-01 | 2.61E+00 | 2.69E+00 | 4.16E-01 | no | Ppp2r3a | intron_variant | MODIFIER |
| 9 | 101182391 | AX-170478667 | 0 | T | C | 0.132 | -9.35E-02 | 1.16E-01 | 2.61E+00 | 2.69E+00 | 4.16E-01 | no | Ppp2r3a | intron_variant | MODIFIER |
| 9 | 101399453 | AX-97803776 | 0 | T | C | 0.132 | -9.35E-02 | 1.16E-01 | 2.61E+00 | 2.69E+00 | 4.16E-01 | no | | intergenic_variant | MODIFIER |
| 9 | 101426084 | AX-97842502 | 0 | T | C | 0.132 | -9.35E-02 | 1.16E-01 | 2.61E+00 | 2.69E+00 | 4.16E-01 | no | | regulatory_region_variant | MODIFIER |
| 9 | 101436547 | AX-170135387 | 0 | T | C | 0.132 | -9.35E-02 | 1.16E-01 | 2.61E+00 | 2.69E+00 | 4.16E-01 | no | Gm29385 | upstream_gene_variant | MODIFIER |
| 9 | 101455610 | AX-170033141 | 0 | T | C | 0.132 | -9.35E-02 | 1.16E-01 | 2.61E+00 | 2.69E+00 | 4.16E-01 | no | | intergenic_variant | MODIFIER |
| 9 | 101457786 | AX-170161495 | 0 | T | C | 0.132 | -9.35E-02 | 1.16E-01 | 2.61E+00 | 2.69E+00 | 4.16E-01 | no | | intergenic_variant | MODIFIER |

| | | | | | | | | | | | | | | | |
|---|---|---|---|---|---|---|---|---|---|---|---|---|---|---|---|
| 9 | 101503563 | AX-170273016 | 0 | A | G | 0.132 | -9.35E-02 | 1.16E-01 | 2.61E+00 | 2.69E+00 | 4.16E-01 | no | | intergenic_variant | MODIFIER |
| 9 | 101565164 | AX-97788501 | 0 | T | C | 0.132 | -9.35E-02 | 1.16E-01 | 2.61E+00 | 2.69E+00 | 4.16E-01 | no | | intergenic_variant | MODIFIER |
| 9 | 101565654 | AX-170171789 | 0 | T | C | 0.132 | -9.35E-02 | 1.16E-01 | 2.61E+00 | 2.69E+00 | 4.16E-01 | no | | intergenic_variant | MODIFIER |
| 9 | 101568060 | AX-170191334 | 0 | A | T | 0.132 | -9.35E-02 | 1.16E-01 | 2.61E+00 | 2.69E+00 | 4.16E-01 | no | | intergenic_variant | MODIFIER |
| 9 | 101569933 | AX-170283495 | 0 | A | G | 0.132 | -9.35E-02 | 1.16E-01 | 2.61E+00 | 2.69E+00 | 4.16E-01 | no | | intergenic_variant | MODIFIER |
| 9 | 101619673 | AX-97811824 | 0 | T | C | 0.132 | -9.35E-02 | 1.16E-01 | 2.61E+00 | 2.69E+00 | 4.16E-01 | no | | intergenic_variant | MODIFIER |
| 9 | 101654026 | AX-170033142 | 0 | T | C | 0.132 | -9.35E-02 | 1.16E-01 | 2.61E+00 | 2.69E+00 | 4.16E-01 | no | | intergenic_variant | MODIFIER |
| 9 | 101657607 | AX-170427995 | 0 | C | T | 0.132 | -9.35E-02 | 1.16E-01 | 2.61E+00 | 2.69E+00 | 4.16E-01 | no | | intergenic_variant | MODIFIER |
| 9 | 101668615 | AX-168404569 | 0 | T | C | 0.132 | -9.35E-02 | 1.16E-01 | 2.61E+00 | 2.69E+00 | 4.16E-01 | no | | intergenic_variant | MODIFIER |
| 9 | 101674212 | AX-170173579 | 0 | T | C | 0.132 | -9.35E-02 | 1.16E-01 | 2.61E+00 | 2.69E+00 | 4.16E-01 | no | | intergenic_variant | MODIFIER |
| 9 | 102107013 | AX-170096651 | 0 | C | T | 0.132 | -9.35E-02 | 1.16E-01 | 2.61E+00 | 2.69E+00 | 4.16E-01 | no | Ephb1 | intron_variant | MODIFIER |
| 9 | 102124601 | AX-170170383 | 0 | A | T | 0.132 | -9.35E-02 | 1.16E-01 | 2.61E+00 | 2.69E+00 | 4.16E-01 | no | Ephb1 | intron_variant | MODIFIER |
| 9 | 102199286 | AX-170276650 | 0 | C | T | 0.132 | -9.35E-02 | 1.16E-01 | 2.61E+00 | 2.69E+00 | 4.16E-01 | no | Ephb1 | intron_variant | MODIFIER |
| 9 | 102244017 | AX-170287157 | 0 | T | C | 0.132 | -9.35E-02 | 1.16E-01 | 2.61E+00 | 2.69E+00 | 4.16E-01 | no | Ephb1 | intron_variant | MODIFIER |
| 9 | 102256863 | AX-170374308 | 0 | G | A | 0.132 | -9.35E-02 | 1.16E-01 | 2.61E+00 | 2.69E+00 | 4.16E-01 | no | Ephb1 | intron_variant | MODIFIER |
| 9 | 102256882 | AX-170145764 | 0 | A | T | 0.132 | -9.35E-02 | 1.16E-01 | 2.61E+00 | 2.69E+00 | 4.16E-01 | no | Ephb1 | intron_variant | MODIFIER |
| 9 | 102257583 | AX-170414147 | 0 | T | C | 0.132 | -9.35E-02 | 1.16E-01 | 2.61E+00 | 2.69E+00 | 4.16E-01 | no | Ephb1 | intron_variant | MODIFIER |
| 9 | 102262437 | AX-170339036 | 0 | G | C | 0.132 | -9.35E-02 | 1.16E-01 | 2.61E+00 | 2.69E+00 | 4.16E-01 | no | Ephb1 | intron_variant | MODIFIER |
| 9 | 102267369 | AX-170450690 | 0 | T | C | 0.132 | -9.35E-02 | 1.16E-01 | 2.61E+00 | 2.69E+00 | 4.16E-01 | no | Ephb1 | intron_variant | MODIFIER |
| 9 | 102286222 | AX-170320293 | 0 | A | G | 0.132 | -9.35E-02 | 1.16E-01 | 2.61E+00 | 2.69E+00 | 4.16E-01 | no | Ephb1 | intron_variant | MODIFIER |
| 9 | 102302379 | AX-97855851 | 0 | A | G | 0.132 | -9.35E-02 | 1.16E-01 | 2.61E+00 | 2.69E+00 | 4.16E-01 | no | Ephb1 | intron_variant | MODIFIER |
| 3 | 149088171 | AX-170221248 | 0 | T | A | 0.351 | 5.88E-02 | 7.32E-02 | 2.78E+00 | 2.87E+00 | 4.16E-01 | no | | regulatory_region_variant | MODIFIER |
| 3 | 149088473 | AX-170395002 | 0 | T | A | 0.351 | 5.88E-02 | 7.32E-02 | 2.78E+00 | 2.87E+00 | 4.16E-01 | no | | regulatory_region_variant | MODIFIER |
| 3 | 149102495 | AX-97827187 | 0 | T | C | 0.351 | 5.88E-02 | 7.32E-02 | 2.78E+00 | 2.87E+00 | 4.16E-01 | no | | intergenic_variant | MODIFIER |
| 3 | 149102950 | AX-170408786 | 0 | T | C | 0.351 | 5.88E-02 | 7.32E-02 | 2.78E+00 | 2.87E+00 | 4.16E-01 | no | | regulatory_region_variant | MODIFIER |
| 3 | 149105682 | AX-170115915 | 0 | C | T | 0.351 | 5.88E-02 | 7.32E-02 | 2.78E+00 | 2.87E+00 | 4.16E-01 | no | | regulatory_region_variant | MODIFIER |
| 3 | 149111186 | AX-170523380 | 0 | A | G | 0.351 | 5.88E-02 | 7.32E-02 | 2.78E+00 | 2.87E+00 | 4.16E-01 | no | | regulatory_region_variant | MODIFIER |
| 3 | 149114497 | AX-170021894 | 0 | T | C | 0.351 | 5.88E-02 | 7.32E-02 | 2.78E+00 | 2.87E+00 | 4.16E-01 | no | | intergenic_variant | MODIFIER |
| 3 | 149120009 | AX-170021895 | 0 | C | A | 0.351 | 5.88E-02 | 7.32E-02 | 2.78E+00 | 2.87E+00 | 4.16E-01 | no | | intergenic_variant | MODIFIER |
| 3 | 149120029 | AX-170021897 | 0 | T | G | 0.351 | 5.88E-02 | 7.32E-02 | 2.78E+00 | 2.87E+00 | 4.16E-01 | no | | intergenic_variant | MODIFIER |
| 3 | 149154729 | AX-170094849 | 0 | A | G | 0.351 | 5.88E-02 | 7.32E-02 | 2.78E+00 | 2.87E+00 | 4.16E-01 | no | | intergenic_variant | MODIFIER |
| 3 | 149159521 | AX-170021896 | 0 | T | C | 0.351 | 5.88E-02 | 7.32E-02 | 2.78E+00 | 2.87E+00 | 4.16E-01 | no | | intergenic_variant | MODIFIER |
| 3 | 149159613 | AX-170392230 | 0 | G | T | 0.351 | 5.88E-02 | 7.32E-02 | 2.78E+00 | 2.87E+00 | 4.16E-01 | no | | regulatory_region_variant | MODIFIER |
| 3 | 149160227 | AX-170160515 | 0 | T | C | 0.351 | 5.88E-02 | 7.32E-02 | 2.78E+00 | 2.87E+00 | 4.16E-01 | no | | intergenic_variant | MODIFIER |
| 3 | 149167260 | AX-170424871 | 0 | C | T | 0.351 | 5.88E-02 | 7.32E-02 | 2.78E+00 | 2.87E+00 | 4.16E-01 | no | | intergenic_variant | MODIFIER |
| 3 | 149170641 | AX-97844828 | 0 | C | T | 0.351 | 5.88E-02 | 7.32E-02 | 2.78E+00 | 2.87E+00 | 4.16E-01 | no | | intergenic_variant | MODIFIER |
| 3 | 149178217 | AX-170068417 | 0 | A | G | 0.351 | 5.88E-02 | 7.32E-02 | 2.78E+00 | 2.87E+00 | 4.16E-01 | no | | intergenic_variant | MODIFIER |
| 3 | 149185218 | AX-170300680 | 0 | G | A | 0.351 | 5.88E-02 | 7.32E-02 | 2.78E+00 | 2.87E+00 | 4.16E-01 | no | | regulatory_region_variant | MODIFIER |
| 3 | 149195817 | AX-170185261 | 0 | G | A | 0.351 | 5.88E-02 | 7.32E-02 | 2.78E+00 | 2.87E+00 | 4.16E-01 | no | | intergenic_variant | MODIFIER |
| 3 | 149226832 | AX-170240967 | 0 | C | T | 0.351 | 5.88E-02 | 7.32E-02 | 2.78E+00 | 2.87E+00 | 4.16E-01 | no | Gm10287 | upstream_gene_variant | MODIFIER |
| 3 | 149250362 | AX-170229300 | 0 | T | G | 0.351 | 5.88E-02 | 7.32E-02 | 2.78E+00 | 2.87E+00 | 4.16E-01 | no | | intergenic_variant | MODIFIER |
| 3 | 149252014 | AX-170317748 | 0 | T | C | 0.351 | 5.88E-02 | 7.32E-02 | 2.78E+00 | 2.87E+00 | 4.16E-01 | no | | intergenic_variant | MODIFIER |
| 3 | 149263624 | AX-97796165 | 0 | C | G | 0.351 | 5.88E-02 | 7.32E-02 | 2.78E+00 | 2.87E+00 | 4.16E-01 | no | | intergenic_variant | MODIFIER |
| 3 | 149263669 | AX-170187001 | 0 | C | T | 0.351 | 5.88E-02 | 7.32E-02 | 2.78E+00 | 2.87E+00 | 4.16E-01 | no | | intergenic_variant | MODIFIER |
| 3 | 149298187 | AX-170253608 | 0 | A | G | 0.351 | 5.88E-02 | 7.32E-02 | 2.78E+00 | 2.87E+00 | 4.16E-01 | no | Gm30382 | intron_variant,non_coding_transcript_variant | MODIFIER |
| 3 | 149300601 | AX-170414782 | 0 | C | T | 0.351 | 5.88E-02 | 7.32E-02 | 2.78E+00 | 2.87E+00 | 4.16E-01 | no | Gm30382 | intron_variant,non_coding_transcript_variant | MODIFIER |
| 3 | 149303562 | AX-170229450 | 0 | C | A | 0.351 | 5.88E-02 | 7.32E-02 | 2.78E+00 | 2.87E+00 | 4.16E-01 | no | Gm30382 | intron_variant,non_coding_transcript_variant | MODIFIER |
| 3 | 149318686 | AX-170483403 | 0 | T | C | 0.351 | 5.88E-02 | 7.32E-02 | 2.78E+00 | 2.87E+00 | 4.16E-01 | no | Gm30382 | intron_variant,non_coding_transcript_variant | MODIFIER |
| 3 | 149318932 | AX-170326409 | 0 | T | C | 0.351 | 5.88E-02 | 7.32E-02 | 2.78E+00 | 2.87E+00 | 4.16E-01 | no | Gm30382 | intron_variant,non_coding_transcript_variant | MODIFIER |
| 3 | 149319366 | AX-170267123 | 0 | A | C | 0.351 | 5.88E-02 | 7.32E-02 | 2.78E+00 | 2.87E+00 | 4.16E-01 | no | Gm30382 | intron_variant,non_coding_transcript_variant | MODIFIER |
| 3 | 149319629 | AX-170276278 | 0 | G | T | 0.351 | 5.88E-02 | 7.32E-02 | 2.78E+00 | 2.87E+00 | 4.16E-01 | no | Gm30382 | intron_variant,non_coding_transcript_variant | MODIFIER |
| 3 | 149347252 | AX-170192126 | 0 | C | T | 0.351 | 5.88E-02 | 7.32E-02 | 2.78E+00 | 2.87E+00 | 4.16E-01 | no | Gm30382 | intron_variant,non_coding_transcript_variant | MODIFIER |
| 3 | 149489486 | AX-170249682 | 0 | T | C | 0.351 | 5.88E-02 | 7.32E-02 | 2.78E+00 | 2.87E+00 | 4.16E-01 | no | | intergenic_variant | MODIFIER |
| 3 | 149493062 | AX-170505913 | 0 | A | C | 0.351 | 5.88E-02 | 7.32E-02 | 2.78E+00 | 2.87E+00 | 4.16E-01 | no | | intergenic_variant | MODIFIER |

| | | | | | | | | | | | | | | | |
|---|---|---|---|---|---|---|---|---|---|---|---|---|---|---|---|
| 3 | 149493467 | AX-170213571 | 0 | C | A | 0.351 | 5.88E-02 | 7.32E-02 | 2.78E+00 | 2.87E+00 | 4.16E-01 | no | | intergenic_variant | MODIFIER |
| 3 | 149493621 | AX-170213487 | 0 | A | G | 0.351 | 5.88E-02 | 7.32E-02 | 2.78E+00 | 2.87E+00 | 4.16E-01 | no | | intergenic_variant | MODIFIER |
| 3 | 149493818 | AX-97821526 | 0 | A | G | 0.351 | 5.88E-02 | 7.32E-02 | 2.78E+00 | 2.87E+00 | 4.16E-01 | no | | intergenic_variant | MODIFIER |
| 3 | 149494220 | AX-170068423 | 0 | C | G | 0.351 | 5.88E-02 | 7.32E-02 | 2.78E+00 | 2.87E+00 | 4.16E-01 | no | | intergenic_variant | MODIFIER |
| 3 | 149545886 | AX-97800131 | 0 | G | A | 0.351 | 5.88E-02 | 7.32E-02 | 2.78E+00 | 2.87E+00 | 4.16E-01 | no | | intergenic_variant | MODIFIER |
| 3 | 149675574 | AX-170393291 | 0 | C | T | 0.351 | 5.88E-02 | 7.32E-02 | 2.78E+00 | 2.87E+00 | 4.16E-01 | no | | intergenic_variant | MODIFIER |
| 3 | 149678380 | AX-170163411 | 0 | T | C | 0.351 | 5.88E-02 | 7.32E-02 | 2.78E+00 | 2.87E+00 | 4.16E-01 | no | | intergenic_variant | MODIFIER |
| 3 | 149749153 | AX-170177652 | 0 | G | A | 0.351 | 5.88E-02 | 7.32E-02 | 2.78E+00 | 2.87E+00 | 4.16E-01 | no | | intergenic_variant | MODIFIER |
| 3 | 150197686 | AX-97847074 | 0 | T | C | 0.351 | 5.88E-02 | 7.32E-02 | 2.78E+00 | 2.87E+00 | 4.16E-01 | no | | intergenic_variant | MODIFIER |
| 3 | 150207283 | AX-170459822 | 0 | A | C | 0.351 | 5.88E-02 | 7.32E-02 | 2.78E+00 | 2.87E+00 | 4.16E-01 | no | | intergenic_variant | MODIFIER |
| 3 | 150255774 | AX-97805827 | 0 | A | G | 0.351 | 5.88E-02 | 7.32E-02 | 2.78E+00 | 2.87E+00 | 4.16E-01 | no | | intergenic_variant | MODIFIER |
| 3 | 150277069 | AX-170021912 | 0 | C | A | 0.351 | 5.88E-02 | 7.32E-02 | 2.78E+00 | 2.87E+00 | 4.16E-01 | no | | intergenic_variant | MODIFIER |
| 3 | 150279042 | AX-170089112 | 0 | T | C | 0.351 | 5.88E-02 | 7.32E-02 | 2.78E+00 | 2.87E+00 | 4.16E-01 | no | | intergenic_variant | MODIFIER |
| 3 | 150286040 | AX-170068435 | 0 | G | A | 0.351 | 5.88E-02 | 7.32E-02 | 2.78E+00 | 2.87E+00 | 4.16E-01 | no | | intergenic_variant | MODIFIER |
| 3 | 150309609 | AX-170106688 | 0 | G | A | 0.351 | 5.88E-02 | 7.32E-02 | 2.78E+00 | 2.87E+00 | 4.16E-01 | no | | intergenic_variant | MODIFIER |
| 3 | 150312263 | AX-170192262 | 0 | T | C | 0.351 | 5.88E-02 | 7.32E-02 | 2.78E+00 | 2.87E+00 | 4.16E-01 | no | | intergenic_variant | MODIFIER |
| 3 | 150312284 | AX-170303132 | 0 | C | A | 0.351 | 5.88E-02 | 7.32E-02 | 2.78E+00 | 2.87E+00 | 4.16E-01 | no | | intergenic_variant | MODIFIER |
| 3 | 150525037 | AX-170426346 | 0 | C | A | 0.351 | 5.88E-02 | 7.32E-02 | 2.78E+00 | 2.87E+00 | 4.16E-01 | no | | intergenic_variant | MODIFIER |
| 3 | 150564037 | AX-170504978 | 0 | G | A | 0.351 | 5.88E-02 | 7.32E-02 | 2.78E+00 | 2.87E+00 | 4.16E-01 | no | | intergenic_variant | MODIFIER |
| 3 | 150609908 | AX-170216334 | 0 | T | C | 0.351 | 5.88E-02 | 7.32E-02 | 2.78E+00 | 2.87E+00 | 4.16E-01 | no | Gm43470 | intron_variant,non_coding_transcript_variant | MODIFIER |
| 3 | 150677426 | AX-170244589 | 0 | G | T | 0.351 | 5.88E-02 | 7.32E-02 | 2.78E+00 | 2.87E+00 | 4.16E-01 | no | | intergenic_variant | MODIFIER |
| 3 | 150681033 | AX-97844403 | 0 | T | C | 0.351 | 5.88E-02 | 7.32E-02 | 2.78E+00 | 2.87E+00 | 4.16E-01 | no | | intergenic_variant | MODIFIER |
| 3 | 150734642 | AX-170413687 | 0 | A | C | 0.351 | 5.88E-02 | 7.32E-02 | 2.78E+00 | 2.87E+00 | 4.16E-01 | no | | intergenic_variant | MODIFIER |
| 3 | 150815325 | AX-97831170 | 0 | A | C | 0.351 | 5.88E-02 | 7.32E-02 | 2.78E+00 | 2.87E+00 | 4.16E-01 | no | | intergenic_variant | MODIFIER |
| 3 | 150835973 | AX-170094855 | 0 | T | C | 0.351 | 5.88E-02 | 7.32E-02 | 2.78E+00 | 2.87E+00 | 4.16E-01 | no | | intergenic_variant | MODIFIER |
| 3 | 150843147 | AX-170021920 | 0 | A | G | 0.351 | 5.88E-02 | 7.32E-02 | 2.78E+00 | 2.87E+00 | 4.16E-01 | no | | intergenic_variant | MODIFIER |
| 3 | 150860244 | AX-170094856 | 0 | A | G | 0.351 | 5.88E-02 | 7.32E-02 | 2.78E+00 | 2.87E+00 | 4.16E-01 | no | | intergenic_variant | MODIFIER |
| 3 | 150865366 | AX-170177756 | 0 | A | C | 0.351 | 5.88E-02 | 7.32E-02 | 2.78E+00 | 2.87E+00 | 4.16E-01 | no | | intergenic_variant | MODIFIER |
| 3 | 150891795 | AX-170144319 | 0 | A | G | 0.351 | 5.88E-02 | 7.32E-02 | 2.78E+00 | 2.87E+00 | 4.16E-01 | no | | intergenic_variant | MODIFIER |
| 3 | 150896972 | AX-170194598 | 0 | G | A | 0.351 | 5.88E-02 | 7.32E-02 | 2.78E+00 | 2.87E+00 | 4.16E-01 | no | | intergenic_variant | MODIFIER |
| 3 | 150904836 | AX-170129312 | 0 | G | A | 0.351 | 5.88E-02 | 7.32E-02 | 2.78E+00 | 2.87E+00 | 4.16E-01 | no | | intergenic_variant | MODIFIER |
| 3 | 150917280 | AX-170520657 | 0 | A | G | 0.351 | 5.88E-02 | 7.32E-02 | 2.78E+00 | 2.87E+00 | 4.16E-01 | no | | intergenic_variant | MODIFIER |
| 3 | 150982467 | AX-170461113 | 0 | T | C | 0.351 | 5.88E-02 | 7.32E-02 | 2.78E+00 | 2.87E+00 | 4.16E-01 | no | | intergenic_variant | MODIFIER |
| 3 | 150992117 | AX-170515404 | 0 | C | T | 0.351 | 5.88E-02 | 7.32E-02 | 2.78E+00 | 2.87E+00 | 4.16E-01 | no | | intergenic_variant | MODIFIER |
| 3 | 151000486 | AX-170150037 | 0 | G | A | 0.351 | 5.88E-02 | 7.32E-02 | 2.78E+00 | 2.87E+00 | 4.16E-01 | no | | intergenic_variant | MODIFIER |
| 3 | 151000957 | AX-170456103 | 0 | C | T | 0.351 | 5.88E-02 | 7.32E-02 | 2.78E+00 | 2.87E+00 | 4.16E-01 | no | | intergenic_variant | MODIFIER |
| 3 | 151007388 | AX-170114488 | 0 | C | A | 0.351 | 5.88E-02 | 7.32E-02 | 2.78E+00 | 2.87E+00 | 4.16E-01 | no | | intergenic_variant | MODIFIER |
| 3 | 151023343 | AX-170424786 | 0 | A | C | 0.351 | 5.88E-02 | 7.32E-02 | 2.78E+00 | 2.87E+00 | 4.16E-01 | no | Mir3963 | downstream_gene_variant | MODIFIER |
| 3 | 151023748 | AX-97852599 | 0 | A | G | 0.351 | 5.88E-02 | 7.32E-02 | 2.78E+00 | 2.87E+00 | 4.16E-01 | no | Mir3963 | downstream_gene_variant | MODIFIER |
| 3 | 151042211 | AX-170068444 | 0 | G | T | 0.351 | 5.88E-02 | 7.32E-02 | 2.78E+00 | 2.87E+00 | 4.16E-01 | no | | intergenic_variant | MODIFIER |
| 3 | 151045322 | AX-97826892 | 0 | A | C | 0.351 | 5.88E-02 | 7.32E-02 | 2.78E+00 | 2.87E+00 | 4.16E-01 | no | | intergenic_variant | MODIFIER |
| 3 | 151046595 | AX-170068447 | 0 | T | C | 0.351 | 5.88E-02 | 7.32E-02 | 2.78E+00 | 2.87E+00 | 4.16E-01 | no | | intergenic_variant | MODIFIER |
| 3 | 151067844 | AX-170339656 | 0 | C | G | 0.351 | 5.88E-02 | 7.32E-02 | 2.78E+00 | 2.87E+00 | 4.16E-01 | no | Gm43468 | upstream_gene_variant | MODIFIER |
| 3 | 151071248 | AX-170400470 | 0 | T | C | 0.351 | 5.88E-02 | 7.32E-02 | 2.78E+00 | 2.87E+00 | 4.16E-01 | no | Gm43468 | upstream_gene_variant | MODIFIER |
| 3 | 151073393 | AX-170200994 | 0 | T | C | 0.351 | 5.88E-02 | 7.32E-02 | 2.78E+00 | 2.87E+00 | 4.16E-01 | no | Gm43468 | intron_variant,non_coding_transcript_variant | MODIFIER |
| 3 | 151226916 | AX-170279805 | 0 | A | G | 0.351 | 5.88E-02 | 7.32E-02 | 2.78E+00 | 2.87E+00 | 4.16E-01 | no | | intergenic_variant | MODIFIER |
| 3 | 151307200 | AX-170508761 | 0 | A | G | 0.351 | 5.88E-02 | 7.32E-02 | 2.78E+00 | 2.87E+00 | 4.16E-01 | no | | intergenic_variant | MODIFIER |
| 3 | 151334442 | AX-170523643 | 0 | T | C | 0.351 | 5.88E-02 | 7.32E-02 | 2.78E+00 | 2.87E+00 | 4.16E-01 | no | Gm42949 | intron_variant,non_coding_transcript_variant | MODIFIER |
| 3 | 151339091 | AX-170448858 | 0 | T | C | 0.351 | 5.88E-02 | 7.32E-02 | 2.78E+00 | 2.87E+00 | 4.16E-01 | no | Gm42949 | downstream_gene_variant | MODIFIER |
| 3 | 151350810 | AX-170368099 | 0 | T | C | 0.351 | 5.88E-02 | 7.32E-02 | 2.78E+00 | 2.87E+00 | 4.16E-01 | no | | intergenic_variant | MODIFIER |
| 3 | 151369235 | AX-97793960 | 0 | T | A | 0.351 | 5.88E-02 | 7.32E-02 | 2.78E+00 | 2.87E+00 | 4.16E-01 | no | | intergenic_variant | MODIFIER |
| 3 | 151382601 | AX-170499240 | 0 | C | T | 0.351 | 5.88E-02 | 7.32E-02 | 2.78E+00 | 2.87E+00 | 4.16E-01 | no | | intergenic_variant | MODIFIER |
| 3 | 151434416 | AX-170186760 | 0 | T | G | 0.351 | 5.88E-02 | 7.32E-02 | 2.78E+00 | 2.87E+00 | 4.16E-01 | no | Adgrl4 | upstream_gene_variant | MODIFIER |
| 3 | 151489527 | AX-170404681 | 0 | G | T | 0.351 | 5.88E-02 | 7.32E-02 | 2.78E+00 | 2.87E+00 | 4.16E-01 | no | Adgrl4 | intron_variant | MODIFIER |

| | | | | | | | | | | | | | | | |
|---|---|---|---|---|---|---|---|---|---|---|---|---|---|---|---|
| 3 | 151498026 | AX-97848382 | 0 | C | T | 0.351 | 5.88E-02 | 7.32E-02 | 2.78E+00 | 2.87E+00 | 4.16E-01 | no | Adgrl4 | intron_variant | MODIFIER |
| 3 | 151572255 | AX-170021937 | 0 | A | G | 0.351 | 5.88E-02 | 7.32E-02 | 2.78E+00 | 2.87E+00 | 4.16E-01 | no | | intergenic_variant | MODIFIER |
| 3 | 151576054 | AX-170094858 | 0 | G | A | 0.351 | 5.88E-02 | 7.32E-02 | 2.78E+00 | 2.87E+00 | 4.16E-01 | no | | intergenic_variant | MODIFIER |
| 3 | 151576234 | AX-170443718 | 0 | T | C | 0.351 | 5.88E-02 | 7.32E-02 | 2.78E+00 | 2.87E+00 | 4.16E-01 | no | | intergenic_variant | MODIFIER |
| 1 | 187433779 | AX-170418567 | 0 | G | A | 0.219 | 9.29E-02 | 1.12E-01 | 2.46E+00 | 2.54E+00 | 4.16E-01 | no | | intergenic_variant | MODIFIER |
| 3 | 151049660 | AX-170068448 | 1 | A | G | 0.348 | 5.88E-02 | 7.32E-02 | 2.79E+00 | 2.87E+00 | 4.17E-01 | no | | intergenic_variant | MODIFIER |
| 5 | 128218888 | AX-97789848 | 0 | G | A | 0.158 | -7.67E-02 | 8.88E-02 | 2.34E+00 | 2.43E+00 | 4.17E-01 | no | Tmem132d | intron_variant | MODIFIER |
| 3 | 31981544 | AX-170532474 | 1 | C | T | 0.161 | -7.44E-02 | 9.30E-02 | 2.78E+00 | 2.90E+00 | 4.17E-01 | no | Kcnmb2 | intron_variant | MODIFIER |
| 5 | 51042031 | AX-168388667 | 1 | C | T | 0.491 | 8.44E-02 | 9.97E-02 | 2.40E+00 | 2.47E+00 | 4.17E-01 | no | | intergenic_variant | MODIFIER |
| 3 | 149264035 | AX-170503002 | 1 | A | C | 0.339 | 5.82E-02 | 7.26E-02 | 2.77E+00 | 2.86E+00 | 4.17E-01 | no | | intergenic_variant | MODIFIER |
| 6 | 134735719 | AX-170245036 | 1 | C | T | 0.58 | 7.20E-02 | 8.58E-02 | 2.43E+00 | 2.49E+00 | 4.17E-01 | no | Dusp16 | intron_variant | MODIFIER |
| 16 | 49804384 | AX-170354779 | 0 | T | A | 0.509 | 6.92E-02 | 8.40E-02 | 2.51E+00 | 2.54E+00 | 4.17E-01 | no | Cd47 | intron_variant | MODIFIER |
| 16 | 49804469 | AX-170418309 | 0 | T | C | 0.509 | 6.92E-02 | 8.40E-02 | 2.51E+00 | 2.54E+00 | 4.17E-01 | no | Cd47 | intron_variant | MODIFIER |
| 16 | 49805451 | AX-170177352 | 0 | C | T | 0.509 | 6.92E-02 | 8.40E-02 | 2.51E+00 | 2.54E+00 | 4.17E-01 | no | Cd47 | intron_variant | MODIFIER |
| 16 | 49806373 | AX-97826334 | 0 | T | C | 0.509 | 6.92E-02 | 8.40E-02 | 2.51E+00 | 2.54E+00 | 4.17E-01 | no | Cd47 | intron_variant | MODIFIER |
| 16 | 49834716 | AX-170057396 | 0 | T | C | 0.509 | 6.92E-02 | 8.40E-02 | 2.51E+00 | 2.54E+00 | 4.17E-01 | no | Cd47 | intron_variant | MODIFIER |
| 16 | 97395508 | AX-170486030 | 1 | C | T | 0.473 | 6.16E-02 | 7.63E-02 | 2.89E+00 | 3.02E+00 | 4.17E-01 | no | Bace2 | intron_variant | MODIFIER |
| 18 | 64883630 | AX-170531298 | 0 | C | G | 0.193 | -8.90E-02 | 1.11E-01 | 2.71E+00 | 2.77E+00 | 4.17E-01 | no | A330084C13Rik | downstream_gene_variant | MODIFIER |
| 4 | 143440030 | AX-97830681 | 2 | C | T | 0.1 | 1.19E-01 | 1.48E-01 | 2.58E+00 | 2.70E+00 | 4.17E-01 | no | Oog4 | 5_prime_UTR_variant | MODIFIER |
| 16 | 73053593 | AX-170470142 | 0 | T | C | 0.491 | 6.23E-02 | 7.69E-02 | 2.59E+00 | 2.66E+00 | 4.17E-01 | no | | intergenic_variant | MODIFIER |
| 13 | 4979864 | AX-170320730 | 1 | C | A | 0.205 | -9.94E-02 | 1.21E-01 | 2.51E+00 | 2.57E+00 | 4.17E-01 | no | | intergenic_variant | MODIFIER |
| 13 | 6682102 | AX-170190556 | 1 | C | T | 0.205 | -9.94E-02 | 1.21E-01 | 2.51E+00 | 2.57E+00 | 4.17E-01 | no | | intergenic_variant | MODIFIER |
| 8 | 95316976 | AX-170357456 | 0 | T | C | 0.228 | 9.31E-02 | 1.16E-01 | 2.78E+00 | 2.85E+00 | 4.17E-01 | no | Tepp | intron_variant | MODIFIER |
| 9 | 17353813 | AX-170104020 | 0 | C | T | 0.482 | 6.89E-02 | 8.52E-02 | 2.59E+00 | 2.67E+00 | 4.17E-01 | no | | intergenic_variant | MODIFIER |
| 7 | 114286654 | AX-170363902 | 2 | A | G | 0.445 | 7.56E-02 | 9.42E-02 | 2.70E+00 | 2.76E+00 | 4.17E-01 | no | | intergenic_variant | MODIFIER |
| 8 | 35869708 | AX-170141152 | 0 | A | G | 0.158 | 9.64E-02 | 1.20E-01 | 2.83E+00 | 2.94E+00 | 4.17E-01 | no | 5430403N17Rik | intron_variant,non_coding_transcript_variant | MODIFIER |
| 1 | 6053972 | AX-97785436 | 0 | G | A | 0.114 | 1.02E-01 | 1.28E-01 | 2.64E+00 | 2.77E+00 | 4.17E-01 | no | Gm19214 | upstream_gene_variant | MODIFIER |
| 16 | 45710083 | AX-170507882 | 0 | A | G | 0.518 | 6.84E-02 | 8.36E-02 | 2.53E+00 | 2.57E+00 | 4.17E-01 | no | Tagln3 | downstream_gene_variant | MODIFIER |
| 1 | 149218253 | AX-170181200 | 1 | A | G | 0.205 | 1.10E-01 | 1.37E-01 | 2.65E+00 | 2.71E+00 | 4.17E-01 | no | | intergenic_variant | MODIFIER |
| 1 | 121826817 | AX-170421567 | 0 | C | A | 0.579 | 6.70E-02 | 8.20E-02 | 2.53E+00 | 2.60E+00 | 4.17E-01 | no | | intergenic_variant | MODIFIER |
| 4 | 118933871 | AX-170023803 | 1 | A | G | 0.214 | 9.42E-02 | 1.16E-01 | 2.59E+00 | 2.65E+00 | 4.17E-01 | no | Olfr1328 | downstream_gene_variant | MODIFIER |
| 1 | 190277411 | AX-97811853 | 1 | C | T | 0.25 | 8.22E-02 | 9.75E-02 | 2.41E+00 | 2.49E+00 | 4.17E-01 | no | | intergenic_variant | MODIFIER |
| 4 | 138032852 | AX-170438592 | 0 | T | C | 0.175 | 8.60E-02 | 1.05E-01 | 2.54E+00 | 2.60E+00 | 4.18E-01 | no | Eif4g3 | intron_variant | MODIFIER |
| 17 | 44297442 | AX-170480393 | 1 | T | C | 0.491 | -2.59E-01 | 3.25E-01 | 2.65E+00 | 2.80E+00 | 4.18E-01 | Yes | | intergenic_variant | MODIFIER |
| 7 | 70175937 | AX-170077661 | 0 | C | A | 0.36 | -8.22E-02 | 1.02E-01 | 2.89E+00 | 2.98E+00 | 4.18E-01 | no | Gm29683 | downstream_gene_variant | MODIFIER |
| 18 | 55561997 | AX-170516346 | 0 | A | G | 0.447 | 7.97E-02 | 9.16E-02 | 2.33E+00 | 2.39E+00 | 4.18E-01 | no | | intergenic_variant | MODIFIER |
| 1 | 170244759 | AX-97786501 | 0 | C | T | 0.219 | 1.04E-01 | 1.29E-01 | 2.61E+00 | 2.68E+00 | 4.18E-01 | no | Sh2d1b2 | intron_variant,NMD_transcript_variant | MODIFIER |
| 9 | 26722541 | AX-97833443 | 0 | A | G | 0.456 | 7.41E-02 | 9.12E-02 | 2.56E+00 | 2.63E+00 | 4.18E-01 | no | Gm48393 | non_coding_transcript_exon_variant | MODIFIER |
| 9 | 27187472 | AX-170280007 | 0 | C | T | 0.456 | 7.41E-02 | 9.12E-02 | 2.56E+00 | 2.63E+00 | 4.18E-01 | no | | intergenic_variant | MODIFIER |
| 9 | 27218098 | AX-170305335 | 0 | C | T | 0.456 | 7.41E-02 | 9.12E-02 | 2.56E+00 | 2.63E+00 | 4.18E-01 | no | | intergenic_variant | MODIFIER |
| 9 | 27219588 | AX-170505312 | 0 | A | C | 0.456 | 7.41E-02 | 9.12E-02 | 2.56E+00 | 2.63E+00 | 4.18E-01 | no | | intergenic_variant | MODIFIER |
| 9 | 27220944 | AX-170517176 | 0 | A | T | 0.456 | 7.41E-02 | 9.12E-02 | 2.56E+00 | 2.63E+00 | 4.18E-01 | no | | intergenic_variant | MODIFIER |
| 9 | 27221674 | AX-170447192 | 0 | G | T | 0.456 | 7.41E-02 | 9.12E-02 | 2.56E+00 | 2.63E+00 | 4.18E-01 | no | | intergenic_variant | MODIFIER |
| 9 | 27226097 | AX-97842801 | 0 | G | A | 0.456 | 7.41E-02 | 9.12E-02 | 2.56E+00 | 2.63E+00 | 4.18E-01 | no | | intergenic_variant | MODIFIER |
| 9 | 27226284 | AX-170495623 | 0 | T | C | 0.456 | 7.41E-02 | 9.12E-02 | 2.56E+00 | 2.63E+00 | 4.18E-01 | no | | intergenic_variant | MODIFIER |
| 9 | 27226474 | AX-170460323 | 0 | C | T | 0.456 | 7.41E-02 | 9.12E-02 | 2.56E+00 | 2.63E+00 | 4.18E-01 | no | | intergenic_variant | MODIFIER |
| 9 | 27226859 | AX-170502714 | 0 | T | C | 0.456 | 7.41E-02 | 9.12E-02 | 2.56E+00 | 2.63E+00 | 4.18E-01 | no | | intergenic_variant | MODIFIER |
| 20 | 168889374 | AX-170097006 | 0 | G | A | 0.193 | 8.86E-02 | 1.10E-01 | 2.60E+00 | 2.70E+00 | 4.18E-01 | no | Arhgap6 | intron_variant | MODIFIER |
| 9 | 54903703 | AX-170032395 | 0 | G | T | 0.219 | -8.41E-02 | 1.05E-01 | 2.80E+00 | 2.92E+00 | 4.18E-01 | no | Ireb2 | intron_variant | MODIFIER |
| 3 | 151811631 | AX-170408278 | 0 | C | T | 0.333 | 5.97E-02 | 7.46E-02 | 2.81E+00 | 2.91E+00 | 4.18E-01 | no | Ptgfr | intron_variant | MODIFIER |
| 4 | 103793605 | AX-170495431 | 0 | A | G | 0.237 | 9.02E-02 | 1.11E-01 | 2.55E+00 | 2.64E+00 | 4.18E-01 | no | Dab1 | intron_variant | MODIFIER |
| 15 | 4970018 | AX-170087752 | 0 | T | C | 0.079 | -9.84E-02 | 9.87E-02 | 2.05E+00 | 2.17E+00 | 4.18E-01 | no | | intergenic_variant | MODIFIER |
| 16 | 18427661 | AX-170012598 | 0 | C | T | 0.14 | 7.34E-02 | 9.22E-02 | 2.75E+00 | 2.88E+00 | 4.18E-01 | no | Comt | upstream_gene_variant | MODIFIER |
| 18 | 67521272 | AX-170222569 | 0 | A | G | 0.272 | -8.17E-02 | 1.02E-01 | 2.74E+00 | 2.81E+00 | 4.18E-01 | no | Spire1 | intron_variant | MODIFIER |

| | | | | | | | | | | | | | | | |
|---|---|---|---|---|---|---|---|---|---|---|---|---|---|---|---|
| 10 | 70904800 | AX-170003435 | 1 | C | T | 0.42 | -6.53E-02 | 8.11E-02 | 2.62E+00 | 2.70E+00 | 4.18E-01 | no | 4930533K18Rik | intron_variant,non_coding_transcript_variant | MODIFIER |
| 9 | 19135642 | AX-168391967 | 1 | C | A | 0.455 | 7.33E-02 | 9.03E-02 | 2.56E+00 | 2.63E+00 | 4.18E-01 | no | Olfr837 | upstream_gene_variant | MODIFIER |
| 9 | 21624351 | AX-170031939 | 1 | A | T | 0.455 | 7.33E-02 | 9.03E-02 | 2.56E+00 | 2.63E+00 | 4.18E-01 | no | Smarca4 | intron_variant | MODIFIER |
| 12 | 115812700 | AX-170050302 | 1 | G | A | 0.116 | 1.24E-01 | 1.56E-01 | 2.68E+00 | 2.78E+00 | 4.18E-01 | no | Ighv8-14 | upstream_gene_variant | MODIFIER |
| 9 | 101493178 | AX-97811219 | 1 | G | A | 0.143 | -8.93E-02 | 1.11E-01 | 2.62E+00 | 2.71E+00 | 4.18E-01 | no | | intergenic_variant | MODIFIER |
| 1 | 101914466 | AX-170342993 | 0 | T | G | 0.158 | 1.33E-01 | 1.64E-01 | 2.56E+00 | 2.67E+00 | 4.18E-01 | no | | intergenic_variant | MODIFIER |
| 4 | 137125232 | AX-97841038 | 0 | G | A | 0.167 | 9.49E-02 | 1.17E-01 | 2.57E+00 | 2.62E+00 | 4.18E-01 | no | | intergenic_variant | MODIFIER |
| 10 | 94054942 | AX-170242758 | 1 | T | A | 0.286 | -7.14E-02 | 8.70E-02 | 2.51E+00 | 2.56E+00 | 4.18E-01 | no | Fgd6 | intron_variant | MODIFIER |
| 18 | 88209539 | AX-170332622 | 1 | A | G | 0.161 | -8.56E-02 | 1.04E-01 | 2.48E+00 | 2.57E+00 | 4.18E-01 | no | | intergenic_variant | MODIFIER |
| 3 | 147425917 | AX-170288607 | 1 | C | T | 0.348 | 5.49E-02 | 6.83E-02 | 2.62E+00 | 2.71E+00 | 4.18E-01 | no | Gm37041 | upstream_gene_variant | MODIFIER |
| 7 | 87595813 | AX-170367535 | 0 | C | T | 0.114 | -1.41E-01 | 1.77E-01 | 2.73E+00 | 2.84E+00 | 4.19E-01 | no | Grm5 | intron_variant | MODIFIER |
| 16 | 78945891 | AX-170037516 | 0 | C | T | 0.465 | 6.24E-02 | 7.73E-02 | 2.59E+00 | 2.66E+00 | 4.19E-01 | no | Chodl | intron_variant | MODIFIER |
| 15 | 80074954 | AX-170236396 | 0 | C | T | 0.553 | 6.26E-02 | 7.75E-02 | 2.58E+00 | 2.67E+00 | 4.19E-01 | no | Rpl3 | downstream_gene_variant | MODIFIER |
| 4 | 127517708 | AX-170209593 | 0 | T | C | 0.14 | 9.87E-02 | 1.22E-01 | 2.59E+00 | 2.64E+00 | 4.19E-01 | no | | intergenic_variant | MODIFIER |
| 4 | 127674512 | AX-170226916 | 0 | T | C | 0.14 | 9.87E-02 | 1.22E-01 | 2.59E+00 | 2.64E+00 | 4.19E-01 | no | | intergenic_variant | MODIFIER |
| 4 | 127674650 | AX-170100226 | 0 | T | C | 0.14 | 9.87E-02 | 1.22E-01 | 2.59E+00 | 2.64E+00 | 4.19E-01 | no | | intergenic_variant | MODIFIER |
| 4 | 127678850 | AX-170527380 | 0 | T | C | 0.14 | 9.87E-02 | 1.22E-01 | 2.59E+00 | 2.64E+00 | 4.19E-01 | no | | intergenic_variant | MODIFIER |
| 4 | 127691257 | AX-170306936 | 0 | G | A | 0.14 | 9.87E-02 | 1.22E-01 | 2.59E+00 | 2.64E+00 | 4.19E-01 | no | | intergenic_variant | MODIFIER |
| 4 | 127692037 | AX-170442435 | 0 | T | C | 0.14 | 9.87E-02 | 1.22E-01 | 2.59E+00 | 2.64E+00 | 4.19E-01 | no | | intergenic_variant | MODIFIER |
| 4 | 127699576 | AX-170439987 | 0 | A | G | 0.14 | 9.87E-02 | 1.22E-01 | 2.59E+00 | 2.64E+00 | 4.19E-01 | no | | intergenic_variant | MODIFIER |
| 4 | 127702800 | AX-170256075 | 0 | G | A | 0.14 | 9.87E-02 | 1.22E-01 | 2.59E+00 | 2.64E+00 | 4.19E-01 | no | | intergenic_variant | MODIFIER |
| 4 | 127706931 | AX-170353227 | 0 | T | C | 0.14 | 9.87E-02 | 1.22E-01 | 2.59E+00 | 2.64E+00 | 4.19E-01 | no | | intergenic_variant | MODIFIER |
| 4 | 127711399 | AX-170483854 | 0 | T | C | 0.14 | 9.87E-02 | 1.22E-01 | 2.59E+00 | 2.64E+00 | 4.19E-01 | no | | regulatory_region_variant | MODIFIER |
| 4 | 127714501 | AX-170070622 | 0 | G | A | 0.14 | 9.87E-02 | 1.22E-01 | 2.59E+00 | 2.64E+00 | 4.19E-01 | no | | intergenic_variant | MODIFIER |
| 4 | 127715895 | AX-170070624 | 0 | A | G | 0.14 | 9.87E-02 | 1.22E-01 | 2.59E+00 | 2.64E+00 | 4.19E-01 | no | | intergenic_variant | MODIFIER |
| 4 | 127720319 | AX-170465228 | 0 | T | C | 0.14 | 9.87E-02 | 1.22E-01 | 2.59E+00 | 2.64E+00 | 4.19E-01 | no | | intergenic_variant | MODIFIER |
| 4 | 127720369 | AX-170492664 | 0 | G | A | 0.14 | 9.87E-02 | 1.22E-01 | 2.59E+00 | 2.64E+00 | 4.19E-01 | no | | intergenic_variant | MODIFIER |
| 4 | 127723873 | AX-97828979 | 0 | A | G | 0.14 | 9.87E-02 | 1.22E-01 | 2.59E+00 | 2.64E+00 | 4.19E-01 | no | | intergenic_variant | MODIFIER |
| 4 | 127745162 | AX-170395127 | 0 | G | A | 0.14 | 9.87E-02 | 1.22E-01 | 2.59E+00 | 2.64E+00 | 4.19E-01 | no | | intergenic_variant | MODIFIER |
| 4 | 127745457 | AX-170283809 | 0 | T | C | 0.14 | 9.87E-02 | 1.22E-01 | 2.59E+00 | 2.64E+00 | 4.19E-01 | no | | intergenic_variant | MODIFIER |
| 4 | 127746658 | AX-170123591 | 0 | T | G | 0.14 | 9.87E-02 | 1.22E-01 | 2.59E+00 | 2.64E+00 | 4.19E-01 | no | | intergenic_variant | MODIFIER |
| 4 | 127749363 | AX-170188987 | 0 | T | C | 0.14 | 9.87E-02 | 1.22E-01 | 2.59E+00 | 2.64E+00 | 4.19E-01 | no | | intergenic_variant | MODIFIER |
| 4 | 127749711 | AX-170415697 | 0 | A | G | 0.14 | 9.87E-02 | 1.22E-01 | 2.59E+00 | 2.64E+00 | 4.19E-01 | no | | intergenic_variant | MODIFIER |
| 4 | 127751192 | AX-170385335 | 0 | C | T | 0.14 | 9.87E-02 | 1.22E-01 | 2.59E+00 | 2.64E+00 | 4.19E-01 | no | | intergenic_variant | MODIFIER |
| 4 | 127755598 | AX-170275799 | 0 | T | C | 0.14 | 9.87E-02 | 1.22E-01 | 2.59E+00 | 2.64E+00 | 4.19E-01 | no | | intergenic_variant | MODIFIER |
| 4 | 127756591 | AX-170254388 | 0 | T | G | 0.14 | 9.87E-02 | 1.22E-01 | 2.59E+00 | 2.64E+00 | 4.19E-01 | no | | regulatory_region_variant | MODIFIER |
| 4 | 127759880 | AX-170134263 | 0 | A | G | 0.14 | 9.87E-02 | 1.22E-01 | 2.59E+00 | 2.64E+00 | 4.19E-01 | no | | intergenic_variant | MODIFIER |
| 4 | 127773698 | AX-170294199 | 0 | T | G | 0.14 | 9.87E-02 | 1.22E-01 | 2.59E+00 | 2.64E+00 | 4.19E-01 | no | | intergenic_variant | MODIFIER |
| 4 | 127773844 | AX-170122002 | 0 | A | C | 0.14 | 9.87E-02 | 1.22E-01 | 2.59E+00 | 2.64E+00 | 4.19E-01 | no | | intergenic_variant | MODIFIER |
| 4 | 127775536 | AX-170070625 | 0 | G | A | 0.14 | 9.87E-02 | 1.22E-01 | 2.59E+00 | 2.64E+00 | 4.19E-01 | no | | intergenic_variant | MODIFIER |
| 4 | 127775566 | AX-170233988 | 0 | T | C | 0.14 | 9.87E-02 | 1.22E-01 | 2.59E+00 | 2.64E+00 | 4.19E-01 | no | | intergenic_variant | MODIFIER |
| 4 | 127780339 | AX-170023905 | 0 | T | C | 0.14 | 9.87E-02 | 1.22E-01 | 2.59E+00 | 2.64E+00 | 4.19E-01 | no | | intergenic_variant | MODIFIER |
| 4 | 127839841 | AX-170285481 | 0 | A | G | 0.14 | 9.87E-02 | 1.22E-01 | 2.59E+00 | 2.64E+00 | 4.19E-01 | no | | intergenic_variant | MODIFIER |
| 4 | 127851148 | AX-170465703 | 0 | G | A | 0.14 | 9.87E-02 | 1.22E-01 | 2.59E+00 | 2.64E+00 | 4.19E-01 | no | | intergenic_variant | MODIFIER |
| 4 | 127852670 | AX-170261434 | 0 | G | A | 0.14 | 9.87E-02 | 1.22E-01 | 2.59E+00 | 2.64E+00 | 4.19E-01 | no | | intergenic_variant | MODIFIER |
| 4 | 127852979 | AX-170384777 | 0 | C | T | 0.14 | 9.87E-02 | 1.22E-01 | 2.59E+00 | 2.64E+00 | 4.19E-01 | no | | intergenic_variant | MODIFIER |
| 4 | 127868230 | AX-170223791 | 0 | A | G | 0.14 | 9.87E-02 | 1.22E-01 | 2.59E+00 | 2.64E+00 | 4.19E-01 | no | | intergenic_variant | MODIFIER |
| 4 | 127929098 | AX-170283402 | 0 | C | G | 0.14 | 9.87E-02 | 1.22E-01 | 2.59E+00 | 2.64E+00 | 4.19E-01 | no | CK137956 | intron_variant | MODIFIER |
| 4 | 127930609 | AX-170476494 | 0 | T | C | 0.14 | 9.87E-02 | 1.22E-01 | 2.59E+00 | 2.64E+00 | 4.19E-01 | no | CK137956 | intron_variant | MODIFIER |
| 4 | 128000547 | AX-170452264 | 0 | T | C | 0.14 | 9.87E-02 | 1.22E-01 | 2.59E+00 | 2.64E+00 | 4.19E-01 | no | Csmd2 | intron_variant,non_coding_transcript_variant | MODIFIER |
| 4 | 128003438 | AX-170346703 | 0 | G | A | 0.14 | 9.87E-02 | 1.22E-01 | 2.59E+00 | 2.64E+00 | 4.19E-01 | no | Csmd2 | intron_variant,non_coding_transcript_variant | MODIFIER |
| 4 | 128009282 | AX-170167982 | 0 | G | T | 0.14 | 9.87E-02 | 1.22E-01 | 2.59E+00 | 2.64E+00 | 4.19E-01 | no | Csmd2 | intron_variant,non_coding_transcript_variant | MODIFIER |
| 4 | 128019499 | AX-170260439 | 0 | A | C | 0.14 | 9.87E-02 | 1.22E-01 | 2.59E+00 | 2.64E+00 | 4.19E-01 | no | Csmd2 | intron_variant,non_coding_transcript_variant | MODIFIER |
| 4 | 128026696 | AX-97823536 | 0 | A | G | 0.14 | 9.87E-02 | 1.22E-01 | 2.59E+00 | 2.64E+00 | 4.19E-01 | no | Csmd2 | intron_variant,non_coding_transcript_variant | MODIFIER |

| | | | | | | | | | | | | | | | |
|---|---|---|---|---|---|---|---|---|---|---|---|---|---|---|---|
| 4 | 128032068 | AX-170245013 | 0 | G | A | 0.14 | 9.87E-02 | 1.22E-01 | 2.59E+00 | 2.64E+00 | 4.19E-01 | no | Csmd2 | intron_variant,non_coding_transcript_variant | MODIFIER |
| 4 | 128042169 | AX-170316855 | 0 | G | A | 0.14 | 9.87E-02 | 1.22E-01 | 2.59E+00 | 2.64E+00 | 4.19E-01 | no | Csmd2 | intron_variant,non_coding_transcript_variant | MODIFIER |
| 4 | 128043693 | AX-170225311 | 0 | T | C | 0.14 | 9.87E-02 | 1.22E-01 | 2.59E+00 | 2.64E+00 | 4.19E-01 | no | Csmd2 | intron_variant,non_coding_transcript_variant | MODIFIER |
| 4 | 128050832 | AX-170355672 | 0 | G | T | 0.14 | 9.87E-02 | 1.22E-01 | 2.59E+00 | 2.64E+00 | 4.19E-01 | no | Csmd2 | intron_variant,non_coding_transcript_variant | MODIFIER |
| 4 | 128078084 | AX-170303952 | 0 | T | C | 0.14 | 9.87E-02 | 1.22E-01 | 2.59E+00 | 2.64E+00 | 4.19E-01 | no | Csmd2 | intron_variant,non_coding_transcript_variant | MODIFIER |
| 4 | 128087462 | AX-170187378 | 0 | A | G | 0.14 | 9.87E-02 | 1.22E-01 | 2.59E+00 | 2.64E+00 | 4.19E-01 | no | Csmd2 | intron_variant,non_coding_transcript_variant | MODIFIER |
| 4 | 128105098 | AX-170070635 | 0 | T | A | 0.14 | 9.87E-02 | 1.22E-01 | 2.59E+00 | 2.64E+00 | 4.19E-01 | no | Csmd2 | intron_variant,non_coding_transcript_variant | MODIFIER |
| 4 | 128114628 | AX-170471190 | 0 | T | C | 0.14 | 9.87E-02 | 1.22E-01 | 2.59E+00 | 2.64E+00 | 4.19E-01 | no | Csmd2 | intron_variant,non_coding_transcript_variant | MODIFIER |
| 4 | 128114746 | AX-170373747 | 0 | A | G | 0.14 | 9.87E-02 | 1.22E-01 | 2.59E+00 | 2.64E+00 | 4.19E-01 | no | Csmd2 | intron_variant,non_coding_transcript_variant | MODIFIER |
| 4 | 128161202 | AX-170366449 | 0 | C | T | 0.14 | 9.87E-02 | 1.22E-01 | 2.59E+00 | 2.64E+00 | 4.19E-01 | no | Csmd2 | intron_variant,non_coding_transcript_variant | MODIFIER |
| 4 | 128161700 | AX-170276661 | 0 | T | C | 0.14 | 9.87E-02 | 1.22E-01 | 2.59E+00 | 2.64E+00 | 4.19E-01 | no | Csmd2 | intron_variant,non_coding_transcript_variant | MODIFIER |
| 4 | 128187074 | AX-170101105 | 0 | T | C | 0.14 | 9.87E-02 | 1.22E-01 | 2.59E+00 | 2.64E+00 | 4.19E-01 | no | Csmd2 | intron_variant,non_coding_transcript_variant | MODIFIER |
| 4 | 128205439 | AX-170515978 | 0 | C | T | 0.14 | 9.87E-02 | 1.22E-01 | 2.59E+00 | 2.64E+00 | 4.19E-01 | no | Csmd2 | intron_variant,non_coding_transcript_variant | MODIFIER |
| 4 | 128206737 | AX-170385929 | 0 | A | C | 0.14 | 9.87E-02 | 1.22E-01 | 2.59E+00 | 2.64E+00 | 4.19E-01 | no | Csmd2 | intron_variant,non_coding_transcript_variant | MODIFIER |
| 4 | 128210102 | AX-170203982 | 0 | T | A | 0.14 | 9.87E-02 | 1.22E-01 | 2.59E+00 | 2.64E+00 | 4.19E-01 | no | Csmd2 | intron_variant,non_coding_transcript_variant | MODIFIER |
| 4 | 128211382 | AX-97855191 | 0 | T | C | 0.14 | 9.87E-02 | 1.22E-01 | 2.59E+00 | 2.64E+00 | 4.19E-01 | no | Csmd2 | intron_variant,non_coding_transcript_variant | MODIFIER |
| 4 | 128244472 | AX-170269443 | 0 | G | A | 0.14 | 9.87E-02 | 1.22E-01 | 2.59E+00 | 2.64E+00 | 4.19E-01 | no | Csmd2 | intron_variant,non_coding_transcript_variant | MODIFIER |
| 4 | 128244904 | AX-97786141 | 0 | G | A | 0.14 | 9.87E-02 | 1.22E-01 | 2.59E+00 | 2.64E+00 | 4.19E-01 | no | Csmd2 | intron_variant,non_coding_transcript_variant | MODIFIER |
| 4 | 128258305 | AX-170339762 | 0 | G | A | 0.14 | 9.87E-02 | 1.22E-01 | 2.59E+00 | 2.64E+00 | 4.19E-01 | no | Hmgb4 | downstream_gene_variant | MODIFIER |
| 4 | 128263505 | AX-170198591 | 0 | A | G | 0.14 | 9.87E-02 | 1.22E-01 | 2.59E+00 | 2.64E+00 | 4.19E-01 | no | Hmgb4 | upstream_gene_variant | MODIFIER |
| 4 | 128265182 | AX-170336734 | 0 | G | A | 0.14 | 9.87E-02 | 1.22E-01 | 2.59E+00 | 2.64E+00 | 4.19E-01 | no | Hmgb4 | upstream_gene_variant | MODIFIER |
| 4 | 128269652 | AX-170446742 | 0 | A | G | 0.14 | 9.87E-02 | 1.22E-01 | 2.59E+00 | 2.64E+00 | 4.19E-01 | no | Csmd2 | intron_variant,non_coding_transcript_variant | MODIFIER |
| 4 | 128281285 | AX-170255592 | 0 | C | A | 0.14 | 9.87E-02 | 1.22E-01 | 2.59E+00 | 2.64E+00 | 4.19E-01 | no | Csmd2 | intron_variant,non_coding_transcript_variant | MODIFIER |
| 4 | 128303270 | AX-170399820 | 0 | A | G | 0.14 | 9.87E-02 | 1.22E-01 | 2.59E+00 | 2.64E+00 | 4.19E-01 | no | Csmd2 | intron_variant,non_coding_transcript_variant | MODIFIER |
| 4 | 128306659 | AX-170023912 | 0 | G | A | 0.14 | 9.87E-02 | 1.22E-01 | 2.59E+00 | 2.64E+00 | 4.19E-01 | no | Csmd2 | intron_variant,non_coding_transcript_variant | MODIFIER |
| 4 | 128324722 | AX-170185506 | 0 | A | G | 0.14 | 9.87E-02 | 1.22E-01 | 2.59E+00 | 2.64E+00 | 4.19E-01 | no | Csmd2 | intron_variant,non_coding_transcript_variant | MODIFIER |
| 4 | 128325961 | AX-170269730 | 0 | C | T | 0.14 | 9.87E-02 | 1.22E-01 | 2.59E+00 | 2.64E+00 | 4.19E-01 | no | Csmd2 | intron_variant,non_coding_transcript_variant | MODIFIER |
| 4 | 128326096 | AX-170140847 | 0 | C | T | 0.14 | 9.87E-02 | 1.22E-01 | 2.59E+00 | 2.64E+00 | 4.19E-01 | no | Csmd2 | intron_variant,non_coding_transcript_variant | MODIFIER |
| 4 | 128345111 | AX-170023915 | 0 | G | A | 0.14 | 9.87E-02 | 1.22E-01 | 2.59E+00 | 2.64E+00 | 4.19E-01 | no | Csmd2 | intron_variant,non_coding_transcript_variant | MODIFIER |
| 4 | 128350009 | AX-170023908 | 0 | G | A | 0.14 | 9.87E-02 | 1.22E-01 | 2.59E+00 | 2.64E+00 | 4.19E-01 | no | Csmd2 | intron_variant,non_coding_transcript_variant | MODIFIER |
| 4 | 128350410 | AX-170174982 | 0 | A | C | 0.14 | 9.87E-02 | 1.22E-01 | 2.59E+00 | 2.64E+00 | 4.19E-01 | no | Csmd2 | intron_variant,non_coding_transcript_variant | MODIFIER |
| 4 | 128396943 | AX-170023909 | 0 | T | G | 0.14 | 9.87E-02 | 1.22E-01 | 2.59E+00 | 2.64E+00 | 4.19E-01 | no | Csmd2 | intron_variant,non_coding_transcript_variant | MODIFIER |
| 4 | 128398302 | AX-170089367 | 0 | C | T | 0.14 | 9.87E-02 | 1.22E-01 | 2.59E+00 | 2.64E+00 | 4.19E-01 | no | Csmd2 | intron_variant,non_coding_transcript_variant | MODIFIER |
| 4 | 128410827 | AX-170193562 | 0 | A | G | 0.14 | 9.87E-02 | 1.22E-01 | 2.59E+00 | 2.64E+00 | 4.19E-01 | no | Csmd2 | intron_variant,non_coding_transcript_variant | MODIFIER |
| 4 | 128412477 | AX-170259950 | 0 | A | G | 0.14 | 9.87E-02 | 1.22E-01 | 2.59E+00 | 2.64E+00 | 4.19E-01 | no | Csmd2 | intron_variant,non_coding_transcript_variant | MODIFIER |
| 4 | 128416964 | AX-170101760 | 0 | A | G | 0.14 | 9.87E-02 | 1.22E-01 | 2.59E+00 | 2.64E+00 | 4.19E-01 | no | Csmd2 | intron_variant,non_coding_transcript_variant | MODIFIER |
| 4 | 128421222 | AX-170104421 | 0 | C | G | 0.14 | 9.87E-02 | 1.22E-01 | 2.59E+00 | 2.64E+00 | 4.19E-01 | no | Csmd2 | downstream_gene_variant | MODIFIER |
| 4 | 128421457 | AX-170191315 | 0 | G | A | 0.14 | 9.87E-02 | 1.22E-01 | 2.59E+00 | 2.64E+00 | 4.19E-01 | no | Csmd2 | downstream_gene_variant | MODIFIER |
| 4 | 128469760 | AX-170256559 | 0 | C | T | 0.14 | 9.87E-02 | 1.22E-01 | 2.59E+00 | 2.64E+00 | 4.19E-01 | no | Csmd2 | intron_variant | MODIFIER |
| 4 | 128469779 | AX-170484682 | 0 | C | G | 0.14 | 9.87E-02 | 1.22E-01 | 2.59E+00 | 2.64E+00 | 4.19E-01 | no | Csmd2 | intron_variant | MODIFIER |
| 4 | 128479258 | AX-170296229 | 0 | T | C | 0.14 | 9.87E-02 | 1.22E-01 | 2.59E+00 | 2.64E+00 | 4.19E-01 | no | Csmd2 | intron_variant,non_coding_transcript_variant | MODIFIER |
| 4 | 128480646 | AX-170421398 | 0 | T | C | 0.14 | 9.87E-02 | 1.22E-01 | 2.59E+00 | 2.64E+00 | 4.19E-01 | no | Csmd2 | intron_variant,non_coding_transcript_variant | MODIFIER |
| 4 | 128517738 | AX-170400658 | 0 | G | A | 0.14 | 9.87E-02 | 1.22E-01 | 2.59E+00 | 2.64E+00 | 4.19E-01 | no | Csmd2 | synonymous_variant | LOW |
| 4 | 128519444 | AX-97857520 | 0 | T | C | 0.14 | 9.87E-02 | 1.22E-01 | 2.59E+00 | 2.64E+00 | 4.19E-01 | no | Csmd2 | intron_variant | MODIFIER |
| 4 | 128545099 | AX-170329410 | 0 | A | G | 0.14 | 9.87E-02 | 1.22E-01 | 2.59E+00 | 2.64E+00 | 4.19E-01 | no | Csmd2 | upstream_gene_variant | MODIFIER |
| 4 | 128549273 | AX-170271951 | 0 | T | C | 0.14 | 9.87E-02 | 1.22E-01 | 2.59E+00 | 2.64E+00 | 4.19E-01 | no | Csmd2 | non_coding_transcript_exon_variant | MODIFIER |
| 4 | 128641312 | AX-97827884 | 0 | T | C | 0.14 | 9.87E-02 | 1.22E-01 | 2.59E+00 | 2.64E+00 | 4.19E-01 | no | Gm12958 | downstream_gene_variant | MODIFIER |
| 4 | 128641491 | AX-170282787 | 0 | A | C | 0.14 | 9.87E-02 | 1.22E-01 | 2.59E+00 | 2.64E+00 | 4.19E-01 | no | Gm12958 | downstream_gene_variant | MODIFIER |
| 4 | 128669474 | AX-97797575 | 0 | A | G | 0.14 | 9.87E-02 | 1.22E-01 | 2.59E+00 | 2.64E+00 | 4.19E-01 | no | Phc2 | intron_variant | MODIFIER |
| 4 | 128730568 | AX-170212367 | 0 | C | T | 0.14 | 9.87E-02 | 1.22E-01 | 2.59E+00 | 2.64E+00 | 4.19E-01 | no | Phc2 | intron_variant | MODIFIER |
| 8 | 61984765 | AX-170108252 | 0 | G | A | 0.149 | -1.16E-01 | 1.40E-01 | 2.47E+00 | 2.53E+00 | 4.19E-01 | no | Ddx60 | intron_variant | MODIFIER |
| 13 | 39372192 | AX-170321555 | 1 | T | C | 0.179 | -9.17E-02 | 1.09E-01 | 2.41E+00 | 2.49E+00 | 4.19E-01 | no | | intergenic_variant | MODIFIER |
| 2 | 162792871 | AX-170427607 | 1 | T | A | 0.161 | 1.57E-01 | 1.97E-01 | 2.74E+00 | 2.82E+00 | 4.19E-01 | Yes | | intergenic_variant | MODIFIER |
| 2 | 162965154 | AX-170302655 | 1 | G | A | 0.161 | 1.57E-01 | 1.97E-01 | 2.74E+00 | 2.82E+00 | 4.19E-01 | Yes | L3mbtl1 | intron_variant | MODIFIER |

| | | | | | | | | | | | | | | | |
|---|---|---|---|---|---|---|---|---|---|---|---|---|---|---|---|
| 9 | 26888792 | AX-97848652 | 0 | G | A | 0.465 | 7.42E-02 | 9.17E-02 | 2.57E+00 | 2.63E+00 | 4.19E-01 | no | Gm1110 | intron_variant | MODIFIER |
| 9 | 26939880 | AX-97818008 | 0 | G | A | 0.465 | 7.42E-02 | 9.17E-02 | 2.57E+00 | 2.63E+00 | 4.19E-01 | no | | intergenic_variant | MODIFIER |
| 9 | 26962662 | AX-170184436 | 0 | A | G | 0.465 | 7.42E-02 | 9.17E-02 | 2.57E+00 | 2.63E+00 | 4.19E-01 | no | Gm5120 | upstream_gene_variant | MODIFIER |
| 9 | 26991599 | AX-170283858 | 0 | T | C | 0.465 | 7.42E-02 | 9.17E-02 | 2.57E+00 | 2.63E+00 | 4.19E-01 | no | Acad8 | intron_variant | MODIFIER |
| 9 | 26992145 | AX-170445682 | 0 | C | T | 0.465 | 7.42E-02 | 9.17E-02 | 2.57E+00 | 2.63E+00 | 4.19E-01 | no | Acad8 | intron_variant | MODIFIER |
| 9 | 27001645 | AX-170410529 | 0 | C | T | 0.465 | 7.42E-02 | 9.17E-02 | 2.57E+00 | 2.63E+00 | 4.19E-01 | no | Thyn1 | intron_variant | MODIFIER |
| 9 | 27011483 | AX-170474616 | 0 | C | T | 0.465 | 7.42E-02 | 9.17E-02 | 2.57E+00 | 2.63E+00 | 4.19E-01 | no | Vps26b | intron_variant | MODIFIER |
| 9 | 27011601 | AX-170245654 | 0 | C | G | 0.465 | 7.42E-02 | 9.17E-02 | 2.57E+00 | 2.63E+00 | 4.19E-01 | no | Vps26b | intron_variant | MODIFIER |
| 9 | 27018752 | AX-170145704 | 0 | C | G | 0.465 | 7.42E-02 | 9.17E-02 | 2.57E+00 | 2.63E+00 | 4.19E-01 | no | Vps26b | intron_variant | MODIFIER |
| 9 | 27024614 | AX-170081922 | 0 | C | G | 0.465 | 7.42E-02 | 9.17E-02 | 2.57E+00 | 2.63E+00 | 4.19E-01 | no | Vps26b | intron_variant | MODIFIER |
| 9 | 27024840 | AX-97827336 | 0 | G | A | 0.465 | 7.42E-02 | 9.17E-02 | 2.57E+00 | 2.63E+00 | 4.19E-01 | no | Vps26b | intron_variant | MODIFIER |
| 9 | 27025001 | AX-170228048 | 0 | G | A | 0.465 | 7.42E-02 | 9.17E-02 | 2.57E+00 | 2.63E+00 | 4.19E-01 | no | Vps26b | intron_variant | MODIFIER |
| 9 | 27027386 | AX-170032018 | 0 | C | T | 0.465 | 7.42E-02 | 9.17E-02 | 2.57E+00 | 2.63E+00 | 4.19E-01 | no | Vps26b | intron_variant | MODIFIER |
| 4 | 54664462 | AX-170410790 | 1 | T | A | 0.348 | -7.70E-02 | 9.66E-02 | 2.71E+00 | 2.81E+00 | 4.19E-01 | no | Gm12478 | intron_variant,non_coding_transcript_variant | MODIFIER |
| 5 | 122127119 | AX-97832260 | 1 | G | A | 0.455 | -8.76E-02 | 1.09E-01 | 2.86E+00 | 2.94E+00 | 4.19E-01 | no | Ccdc63 | intron_variant | MODIFIER |
| 18 | 73632833 | AX-97835077 | 1 | A | T | 0.179 | -9.85E-02 | 1.22E-01 | 2.60E+00 | 2.65E+00 | 4.19E-01 | no | | intergenic_variant | MODIFIER |
| 1 | 187442505 | AX-170129112 | 0 | T | C | 0.246 | 8.87E-02 | 1.05E-01 | 2.39E+00 | 2.47E+00 | 4.19E-01 | no | | intergenic_variant | MODIFIER |
| 1 | 187444118 | AX-170477229 | 0 | C | T | 0.246 | 8.87E-02 | 1.05E-01 | 2.39E+00 | 2.47E+00 | 4.19E-01 | no | Gm37896 | upstream_gene_variant | MODIFIER |
| 1 | 187461286 | AX-170522496 | 0 | C | T | 0.246 | 8.87E-02 | 1.05E-01 | 2.39E+00 | 2.47E+00 | 4.19E-01 | no | Gm37896 | non_coding_transcript_exon_variant | MODIFIER |
| 1 | 187463600 | AX-170473485 | 0 | G | C | 0.246 | 8.87E-02 | 1.05E-01 | 2.39E+00 | 2.47E+00 | 4.19E-01 | no | Gm37896 | downstream_gene_variant | MODIFIER |
| 1 | 187477374 | AX-170162641 | 0 | G | A | 0.246 | 8.87E-02 | 1.05E-01 | 2.39E+00 | 2.47E+00 | 4.19E-01 | no | | intergenic_variant | MODIFIER |
| 1 | 187477527 | AX-170416734 | 0 | G | T | 0.246 | 8.87E-02 | 1.05E-01 | 2.39E+00 | 2.47E+00 | 4.19E-01 | no | | intergenic_variant | MODIFIER |
| 1 | 187478304 | AX-170505894 | 0 | A | G | 0.246 | 8.87E-02 | 1.05E-01 | 2.39E+00 | 2.47E+00 | 4.19E-01 | no | | intergenic_variant | MODIFIER |
| 1 | 187479809 | AX-170091828 | 0 | T | C | 0.246 | 8.87E-02 | 1.05E-01 | 2.39E+00 | 2.47E+00 | 4.19E-01 | no | | intergenic_variant | MODIFIER |
| 1 | 187485057 | AX-170002507 | 0 | G | T | 0.246 | 8.87E-02 | 1.05E-01 | 2.39E+00 | 2.47E+00 | 4.19E-01 | no | | regulatory_region_variant | MODIFIER |
| 1 | 187488865 | AX-170002511 | 0 | C | G | 0.246 | 8.87E-02 | 1.05E-01 | 2.39E+00 | 2.47E+00 | 4.19E-01 | no | | intergenic_variant | MODIFIER |
| 1 | 187517062 | AX-170043897 | 0 | T | C | 0.246 | 8.87E-02 | 1.05E-01 | 2.39E+00 | 2.47E+00 | 4.19E-01 | no | | intergenic_variant | MODIFIER |
| 1 | 187518427 | AX-170002510 | 0 | T | C | 0.246 | 8.87E-02 | 1.05E-01 | 2.39E+00 | 2.47E+00 | 4.19E-01 | no | | intergenic_variant | MODIFIER |
| 1 | 187521851 | AX-170501176 | 0 | A | T | 0.246 | 8.87E-02 | 1.05E-01 | 2.39E+00 | 2.47E+00 | 4.19E-01 | no | | intergenic_variant | MODIFIER |
| 1 | 187548524 | AX-170500884 | 0 | A | C | 0.246 | 8.87E-02 | 1.05E-01 | 2.39E+00 | 2.47E+00 | 4.19E-01 | no | | intergenic_variant | MODIFIER |
| 1 | 187563303 | AX-97814471 | 0 | A | T | 0.246 | 8.87E-02 | 1.05E-01 | 2.39E+00 | 2.47E+00 | 4.19E-01 | no | | intergenic_variant | MODIFIER |
| 1 | 187570577 | AX-170263660 | 0 | G | C | 0.246 | 8.87E-02 | 1.05E-01 | 2.39E+00 | 2.47E+00 | 4.19E-01 | no | | intergenic_variant | MODIFIER |
| 1 | 187571049 | AX-170309529 | 0 | T | C | 0.246 | 8.87E-02 | 1.05E-01 | 2.39E+00 | 2.47E+00 | 4.19E-01 | no | | intergenic_variant | MODIFIER |
| 1 | 187571357 | AX-97851830 | 0 | G | A | 0.246 | 8.87E-02 | 1.05E-01 | 2.39E+00 | 2.47E+00 | 4.19E-01 | no | | intergenic_variant | MODIFIER |
| 1 | 187598257 | AX-170002509 | 0 | T | C | 0.246 | 8.87E-02 | 1.05E-01 | 2.39E+00 | 2.47E+00 | 4.19E-01 | no | | intergenic_variant | MODIFIER |
| 1 | 187631128 | AX-170311979 | 0 | T | A | 0.246 | 8.87E-02 | 1.05E-01 | 2.39E+00 | 2.47E+00 | 4.19E-01 | no | Esrrg | intron_variant | MODIFIER |
| 1 | 187641177 | AX-170261098 | 0 | T | C | 0.246 | 8.87E-02 | 1.05E-01 | 2.39E+00 | 2.47E+00 | 4.19E-01 | no | Esrrg | intron_variant | MODIFIER |
| 1 | 187659770 | AX-170425866 | 0 | T | C | 0.246 | 8.87E-02 | 1.05E-01 | 2.39E+00 | 2.47E+00 | 4.19E-01 | no | Esrrg | intron_variant | MODIFIER |
| 1 | 187671005 | AX-170264713 | 0 | A | G | 0.246 | 8.87E-02 | 1.05E-01 | 2.39E+00 | 2.47E+00 | 4.19E-01 | no | Esrrg | intron_variant | MODIFIER |
| 1 | 187674267 | AX-170043900 | 0 | G | T | 0.246 | 8.87E-02 | 1.05E-01 | 2.39E+00 | 2.47E+00 | 4.19E-01 | no | Esrrg | intron_variant | MODIFIER |
| 1 | 187674377 | AX-170402691 | 0 | A | G | 0.246 | 8.87E-02 | 1.05E-01 | 2.39E+00 | 2.47E+00 | 4.19E-01 | no | Esrrg | intron_variant | MODIFIER |
| 1 | 187680446 | AX-97813631 | 0 | C | A | 0.246 | 8.87E-02 | 1.05E-01 | 2.39E+00 | 2.47E+00 | 4.19E-01 | no | Esrrg | intron_variant | MODIFIER |
| 1 | 187706396 | AX-170184274 | 0 | C | T | 0.246 | 8.87E-02 | 1.05E-01 | 2.39E+00 | 2.47E+00 | 4.19E-01 | no | Esrrg | intron_variant | MODIFIER |
| 1 | 187724437 | AX-170446136 | 0 | T | G | 0.246 | 8.87E-02 | 1.05E-01 | 2.39E+00 | 2.47E+00 | 4.19E-01 | no | Esrrg | intron_variant | MODIFIER |
| 1 | 187727948 | AX-170214796 | 0 | C | T | 0.246 | 8.87E-02 | 1.05E-01 | 2.39E+00 | 2.47E+00 | 4.19E-01 | no | Esrrg | intron_variant | MODIFIER |
| 1 | 187730446 | AX-170530401 | 0 | T | G | 0.246 | 8.87E-02 | 1.05E-01 | 2.39E+00 | 2.47E+00 | 4.19E-01 | no | Esrrg | intron_variant | MODIFIER |
| 1 | 187732746 | AX-97816147 | 0 | A | G | 0.246 | 8.87E-02 | 1.05E-01 | 2.39E+00 | 2.47E+00 | 4.19E-01 | no | Esrrg | intron_variant | MODIFIER |
| 1 | 187735271 | AX-170362045 | 0 | G | C | 0.246 | 8.87E-02 | 1.05E-01 | 2.39E+00 | 2.47E+00 | 4.19E-01 | no | Esrrg | intron_variant | MODIFIER |
| 1 | 187736484 | AX-170228841 | 0 | T | C | 0.246 | 8.87E-02 | 1.05E-01 | 2.39E+00 | 2.47E+00 | 4.19E-01 | no | Esrrg | intron_variant | MODIFIER |
| 1 | 187737438 | AX-170254887 | 0 | C | T | 0.246 | 8.87E-02 | 1.05E-01 | 2.39E+00 | 2.47E+00 | 4.19E-01 | no | Esrrg | intron_variant | MODIFIER |
| 1 | 187744313 | AX-170385535 | 0 | A | G | 0.246 | 8.87E-02 | 1.05E-01 | 2.39E+00 | 2.47E+00 | 4.19E-01 | no | Esrrg | intron_variant | MODIFIER |
| 1 | 187751766 | AX-170086091 | 0 | T | C | 0.246 | 8.87E-02 | 1.05E-01 | 2.39E+00 | 2.47E+00 | 4.19E-01 | no | Esrrg | intron_variant | MODIFIER |
| 1 | 187765926 | AX-170332869 | 0 | A | G | 0.246 | 8.87E-02 | 1.05E-01 | 2.39E+00 | 2.47E+00 | 4.19E-01 | no | Esrrg | intron_variant | MODIFIER |
| 1 | 187799707 | AX-170198065 | 0 | C | T | 0.246 | 8.87E-02 | 1.05E-01 | 2.39E+00 | 2.47E+00 | 4.19E-01 | no | Esrrg | intron_variant | MODIFIER |

| | | | | | | | | | | | | | | | |
|---|---|---|---|---|---|---|---|---|---|---|---|---|---|---|---|
| 7 | 122820181 | AX-170250747 | 1 | G | A | 0.411 | 6.70E-02 | 8.40E-02 | 2.76E+00 | 2.84E+00 | 4.19E-01 | no | | regulatory_region_variant | MODIFIER |
| 7 | 37115473 | AX-170354189 | 1 | C | T | 0.348 | 7.44E-02 | 9.20E-02 | 2.56E+00 | 2.64E+00 | 4.20E-01 | no | Gm28078 | downstream_gene_variant | MODIFIER |
| 10 | 107805046 | AX-170432377 | 1 | T | A | 0.321 | -6.61E-02 | 8.28E-02 | 2.68E+00 | 2.76E+00 | 4.20E-01 | no | Otogl | intron_variant | MODIFIER |
| 5 | 46649470 | AX-170205963 | 1 | A | C | 0.5 | 8.16E-02 | 9.72E-02 | 2.41E+00 | 2.48E+00 | 4.20E-01 | no | | intergenic_variant | MODIFIER |
| 18 | 58535402 | AX-170098691 | 1 | T | G | 0.411 | 7.66E-02 | 9.13E-02 | 2.41E+00 | 2.48E+00 | 4.20E-01 | no | | intergenic_variant | MODIFIER |
| 6 | 135729499 | AX-170076328 | 1 | A | G | 0.571 | 7.17E-02 | 8.67E-02 | 2.46E+00 | 2.53E+00 | 4.20E-01 | no | Grin2b | 3_prime_UTR_variant | MODIFIER |
| 6 | 137393315 | AX-170288701 | 1 | T | A | 0.571 | 7.17E-02 | 8.67E-02 | 2.46E+00 | 2.53E+00 | 4.20E-01 | no | Ptpro | intron_variant | MODIFIER |
| 1 | 16180172 | AX-169999997 | 1 | A | G | 0.375 | 6.44E-02 | 7.98E-02 | 2.58E+00 | 2.63E+00 | 4.20E-01 | no | | regulatory_region_variant | MODIFIER |
| 5 | 44739682 | AX-170186905 | 1 | T | C | 0.196 | -8.67E-02 | 1.08E-01 | 2.60E+00 | 2.68E+00 | 4.20E-01 | no | Ldb2 | intron_variant | MODIFIER |
| 9 | 103625468 | AX-170214346 | 0 | G | C | 0.114 | -9.23E-02 | 1.14E-01 | 2.53E+00 | 2.61E+00 | 4.20E-01 | no | Tmem108 | intron_variant | MODIFIER |
| 1 | 100145222 | AX-170291609 | 1 | A | C | 0.152 | 1.61E-01 | 2.02E-01 | 2.65E+00 | 2.73E+00 | 4.21E-01 | Yes | Cntnap5b | intron_variant | MODIFIER |
| 10 | 93719313 | AX-170272090 | 1 | C | T | 0.304 | -6.81E-02 | 8.37E-02 | 2.52E+00 | 2.58E+00 | 4.21E-01 | no | Ntn4 | intron_variant | MODIFIER |
| 1 | 65741912 | AX-170313784 | 2 | G | T | 0.136 | 1.18E-01 | 1.49E-01 | 2.76E+00 | 2.92E+00 | 4.21E-01 | no | | intergenic_variant | MODIFIER |
| 1 | 127223746 | AX-170440489 | 0 | A | C | 0.535 | 6.69E-02 | 8.34E-02 | 2.60E+00 | 2.66E+00 | 4.21E-01 | no | Mgat5 | intron_variant | MODIFIER |
| 1 | 66636909 | AX-97802537 | 0 | G | A | 0.114 | -1.51E-01 | 1.91E-01 | 2.69E+00 | 2.78E+00 | 4.21E-01 | no | Unc80 | intron_variant | MODIFIER |
| 10 | 93883419 | AX-97794309 | 0 | A | G | 0.289 | -7.02E-02 | 8.68E-02 | 2.55E+00 | 2.61E+00 | 4.21E-01 | no | Metap2 | intron_variant | MODIFIER |
| 20 | 67855642 | AX-97780932 | 1 | T | A | 0.25 | 5.01E-02 | 6.31E-02 | 2.69E+00 | 2.77E+00 | 4.21E-01 | no | | intergenic_variant | MODIFIER |
| 18 | 58212937 | AX-170015783 | 1 | T | C | 0.188 | -9.68E-02 | 1.21E-01 | 2.67E+00 | 2.72E+00 | 4.21E-01 | no | Fbn2 | upstream_gene_variant | MODIFIER |
| 5 | 45345758 | AX-170129360 | 0 | C | T | 0.491 | 7.54E-02 | 9.10E-02 | 2.44E+00 | 2.53E+00 | 4.21E-01 | no | D5Ertd615e | intron_variant,non_coding_transcript_variant | MODIFIER |
| 7 | 123269382 | AX-170029883 | 1 | G | A | 0.411 | 6.67E-02 | 8.40E-02 | 2.76E+00 | 2.85E+00 | 4.21E-01 | no | Slc5a11 | intron_variant | MODIFIER |
| 8 | 19425380 | AX-170086233 | 1 | T | G | 0.161 | -7.38E-02 | 9.37E-02 | 2.72E+00 | 2.88E+00 | 4.21E-01 | no | Defa-ps14 | downstream_gene_variant | MODIFIER |
| 13 | 6601436 | AX-170143595 | 0 | C | T | 0.211 | -9.52E-02 | 1.19E-01 | 2.58E+00 | 2.66E+00 | 4.21E-01 | no | Pfkp | intron_variant | MODIFIER |
| 13 | 6201500 | AX-170092641 | 1 | C | T | 0.214 | -9.43E-02 | 1.18E-01 | 2.59E+00 | 2.67E+00 | 4.22E-01 | no | | intergenic_variant | MODIFIER |
| 9 | 76920453 | AX-170238388 | 0 | A | G | 0.202 | 1.00E-01 | 1.25E-01 | 2.86E+00 | 2.91E+00 | 4.22E-01 | no | | intergenic_variant | MODIFIER |
| 9 | 19948891 | AX-97834747 | 0 | A | G | 0.456 | 7.36E-02 | 9.13E-02 | 2.56E+00 | 2.63E+00 | 4.22E-01 | no | | intergenic_variant | MODIFIER |
| 15 | 5937595 | AX-170370156 | 1 | G | A | 0.08 | -9.64E-02 | 9.88E-02 | 2.08E+00 | 2.19E+00 | 4.22E-01 | no | | intergenic_variant | MODIFIER |
| 2 | 169530388 | AX-170065600 | 0 | G | A | 0.167 | 1.45E-01 | 1.82E-01 | 2.64E+00 | 2.72E+00 | 4.22E-01 | Yes | | regulatory_region_variant | MODIFIER |
| 4 | 85476051 | AX-170467520 | 0 | T | A | 0.342 | 8.47E-02 | 1.07E-01 | 2.75E+00 | 2.84E+00 | 4.22E-01 | no | Sh3gl2 | intron_variant | MODIFIER |
| 7 | 137385632 | AX-97817421 | 1 | A | G | 0.268 | -6.26E-02 | 7.87E-02 | 2.62E+00 | 2.73E+00 | 4.22E-01 | no | 9430038I01Rik | intron_variant | MODIFIER |
| 4 | 133078197 | AX-170444428 | 0 | A | G | 0.167 | 9.25E-02 | 1.14E-01 | 2.54E+00 | 2.59E+00 | 4.22E-01 | no | Ahdc1 | downstream_gene_variant | MODIFIER |
| 8 | 34150018 | AX-170281157 | 1 | T | C | 0.143 | -1.07E-01 | 1.35E-01 | 2.70E+00 | 2.76E+00 | 4.22E-01 | no | Leprotl1 | upstream_gene_variant | MODIFIER |
| 1 | 144988851 | AX-170309537 | 1 | C | A | 0.214 | 1.08E-01 | 1.36E-01 | 2.69E+00 | 2.74E+00 | 4.23E-01 | no | | intergenic_variant | MODIFIER |
| 7 | 99198926 | AX-170203972 | 0 | G | A | 0.386 | -8.05E-02 | 1.02E-01 | 2.72E+00 | 2.81E+00 | 4.23E-01 | no | Gm45012 | upstream_gene_variant | MODIFIER |
| 15 | 74275508 | AX-97793934 | 0 | T | C | 0.553 | 6.78E-02 | 8.40E-02 | 2.54E+00 | 2.61E+00 | 4.23E-01 | no | | intergenic_variant | MODIFIER |
| 4 | 138064551 | AX-97841506 | 0 | G | T | 0.175 | 9.09E-02 | 1.13E-01 | 2.57E+00 | 2.63E+00 | 4.23E-01 | no | Eif4g3 | intron_variant | MODIFIER |
| 5 | 105788773 | AX-170491434 | 0 | T | A | 0.439 | 8.67E-02 | 1.08E-01 | 2.61E+00 | 2.65E+00 | 4.23E-01 | no | Lrrc8d | intron_variant | MODIFIER |
| 1 | 82305916 | AX-97815056 | 1 | T | G | 0.161 | 1.65E-01 | 2.07E-01 | 2.65E+00 | 2.71E+00 | 4.23E-01 | Yes | | intergenic_variant | MODIFIER |
| 5 | 77081528 | AX-170218463 | 0 | G | A | 0.439 | 8.02E-02 | 1.01E-01 | 2.67E+00 | 2.74E+00 | 4.23E-01 | no | | intergenic_variant | MODIFIER |
| 1 | 20029636 | AX-170040549 | 1 | A | T | 0.402 | 6.52E-02 | 8.16E-02 | 2.61E+00 | 2.66E+00 | 4.23E-01 | no | | intergenic_variant | MODIFIER |
| 6 | 129564332 | AX-170530115 | 0 | A | G | 0.561 | 6.58E-02 | 8.06E-02 | 2.49E+00 | 2.56E+00 | 4.23E-01 | no | | intergenic_variant | MODIFIER |
| 2 | 93795807 | AX-170294687 | 1 | G | A | 0.054 | -1.72E-01 | 2.17E-01 | 2.65E+00 | 2.76E+00 | 4.23E-01 | no | Ext2 | intron_variant | MODIFIER |
| 1 | 100424461 | AX-170449638 | 1 | T | A | 0.152 | 1.59E-01 | 2.01E-01 | 2.67E+00 | 2.75E+00 | 4.23E-01 | Yes | Cntnap5b | intron_variant | MODIFIER |
| 12 | 103666790 | AX-170515970 | 0 | C | T | 0.14 | -1.21E-01 | 1.53E-01 | 2.75E+00 | 2.86E+00 | 4.23E-01 | no | Serpina16 | downstream_gene_variant | MODIFIER |
| 18 | 66859495 | AX-170207101 | 1 | G | A | 0.188 | -9.71E-02 | 1.22E-01 | 2.69E+00 | 2.74E+00 | 4.23E-01 | no | Mc4r | synonymous_variant | LOW |
| 9 | 20513841 | AX-97804559 | 0 | A | G | 0.474 | 7.26E-02 | 8.92E-02 | 2.49E+00 | 2.56E+00 | 4.24E-01 | no | Zfp266 | intron_variant | MODIFIER |
| 6 | 134392966 | AX-170320874 | 1 | A | T | 0.58 | 6.79E-02 | 8.30E-02 | 2.47E+00 | 2.54E+00 | 4.24E-01 | no | Bcl2l14 | upstream_gene_variant | MODIFIER |
| 1 | 104795567 | AX-170193823 | 0 | T | A | 0.149 | 1.57E-01 | 1.98E-01 | 2.66E+00 | 2.72E+00 | 4.24E-01 | Yes | Cdh20 | intron_variant | MODIFIER |
| 6 | 137013809 | AX-97815143 | 1 | C | T | 0.562 | 6.94E-02 | 8.53E-02 | 2.49E+00 | 2.56E+00 | 4.24E-01 | no | | intergenic_variant | MODIFIER |
| 18 | 75880396 | AX-170518281 | 0 | C | T | 0.167 | -9.66E-02 | 1.20E-01 | 2.54E+00 | 2.60E+00 | 4.24E-01 | no | Zbtb7c | intron_variant | MODIFIER |
| 5 | 97189053 | AX-170025669 | 2 | G | A | 0.455 | 7.94E-02 | 1.01E-01 | 2.72E+00 | 2.81E+00 | 4.24E-01 | no | | intergenic_variant | MODIFIER |
| 8 | 38747476 | AX-97843903 | 0 | C | T | 0.158 | 1.05E-01 | 1.32E-01 | 2.86E+00 | 2.96E+00 | 4.24E-01 | no | | regulatory_region_variant | MODIFIER |
| 11 | 31803186 | AX-170392441 | 0 | A | G | 0.088 | 1.34E-01 | 1.69E-01 | 2.66E+00 | 2.77E+00 | 4.24E-01 | no | D630024D03Rik | intron_variant,non_coding_transcript_variant | MODIFIER |
| 15 | 5305788 | AX-170240802 | 2 | A | T | 0.245 | -8.39E-02 | 1.05E-01 | 2.60E+00 | 2.65E+00 | 4.24E-01 | no | Gm2157 | upstream_gene_variant | MODIFIER |
| 1 | 137322963 | AX-170509498 | 1 | A | G | 0.214 | 9.82E-02 | 1.24E-01 | 2.66E+00 | 2.73E+00 | 4.24E-01 | no | 1700006P03Rik | downstream_gene_variant | MODIFIER |

| | | | | | | | | | | | | | | | |
|---|---|---|---|---|---|---|---|---|---|---|---|---|---|---|---|
| 8 | 51676789 | AX-170090102 | 0 | T | C | 0.149 | -1.09E-01 | 1.36E-01 | 2.58E+00 | 2.65E+00 | 4.24E-01 | no | | intergenic_variant | MODIFIER |
| 8 | 51976733 | AX-170532914 | 0 | A | G | 0.149 | -1.09E-01 | 1.36E-01 | 2.58E+00 | 2.65E+00 | 4.24E-01 | no | | intergenic_variant | MODIFIER |
| 8 | 54045433 | AX-170222330 | 0 | A | G | 0.149 | -1.09E-01 | 1.36E-01 | 2.58E+00 | 2.65E+00 | 4.24E-01 | no | | intergenic_variant | MODIFIER |
| 8 | 55763315 | AX-170080122 | 0 | A | G | 0.149 | -1.09E-01 | 1.36E-01 | 2.58E+00 | 2.65E+00 | 4.24E-01 | no | | intergenic_variant | MODIFIER |
| 8 | 56914982 | AX-170030789 | 0 | A | G | 0.149 | -1.09E-01 | 1.36E-01 | 2.58E+00 | 2.65E+00 | 4.24E-01 | no | | intergenic_variant | MODIFIER |
| 6 | 134074543 | AX-170259141 | 1 | T | G | 0.545 | 6.99E-02 | 8.62E-02 | 2.51E+00 | 2.56E+00 | 4.24E-01 | no | Etv6 | intron_variant | MODIFIER |
| 1 | 144199481 | AX-170175612 | 0 | C | T | 0.228 | 9.24E-02 | 1.16E-01 | 2.57E+00 | 2.65E+00 | 4.24E-01 | no | | intergenic_variant | MODIFIER |
| 7 | 4825981 | AX-170153662 | 1 | A | T | 0.179 | 6.15E-02 | 7.84E-02 | 2.79E+00 | 2.93E+00 | 4.25E-01 | no | Shisa7 | 3_prime_UTR_variant | MODIFIER |
| 12 | 112218274 | AX-97790714 | 0 | A | G | 0.132 | 1.33E-01 | 1.70E-01 | 2.73E+00 | 2.83E+00 | 4.25E-01 | no | | intergenic_variant | MODIFIER |
| 16 | 46224715 | AX-170057323 | 0 | A | G | 0.465 | -6.38E-02 | 7.95E-02 | 2.55E+00 | 2.60E+00 | 4.25E-01 | no | Gm26297 | downstream_gene_variant | MODIFIER |
| 9 | 101322680 | AX-97831289 | 1 | G | A | 0.134 | -9.00E-02 | 1.14E-01 | 2.62E+00 | 2.71E+00 | 4.25E-01 | no | | intergenic_variant | MODIFIER |
| 10 | 107858565 | AX-97812099 | 0 | C | T | 0.333 | -6.55E-02 | 8.29E-02 | 2.66E+00 | 2.74E+00 | 4.25E-01 | no | Otogl | intron_variant | MODIFIER |
| 10 | 107881436 | AX-97856351 | 0 | A | G | 0.333 | -6.55E-02 | 8.29E-02 | 2.66E+00 | 2.74E+00 | 4.25E-01 | no | Otogl | synonymous_variant | LOW |
| 10 | 107892738 | AX-170532816 | 0 | T | C | 0.333 | -6.55E-02 | 8.29E-02 | 2.66E+00 | 2.74E+00 | 4.25E-01 | no | Otogl | intron_variant | MODIFIER |
| 10 | 107899210 | AX-170187243 | 0 | T | C | 0.333 | -6.55E-02 | 8.29E-02 | 2.66E+00 | 2.74E+00 | 4.25E-01 | no | Otogl | intron_variant | MODIFIER |
| 10 | 107905558 | AX-170253503 | 0 | G | A | 0.333 | -6.55E-02 | 8.29E-02 | 2.66E+00 | 2.74E+00 | 4.25E-01 | no | Otogl | intron_variant | MODIFIER |
| 10 | 107945837 | AX-97816596 | 0 | A | G | 0.333 | -6.55E-02 | 8.29E-02 | 2.66E+00 | 2.74E+00 | 4.25E-01 | no | Gm29685 | upstream_gene_variant | MODIFIER |
| 10 | 107953766 | AX-170361464 | 0 | C | T | 0.333 | -6.55E-02 | 8.29E-02 | 2.66E+00 | 2.74E+00 | 4.25E-01 | no | Gm29685 | intron_variant,non_coding_transcript_variant | MODIFIER |
| 10 | 107955369 | AX-170276484 | 0 | G | A | 0.333 | -6.55E-02 | 8.29E-02 | 2.66E+00 | 2.74E+00 | 4.25E-01 | no | Gm29685 | intron_variant,non_coding_transcript_variant | MODIFIER |
| 10 | 107963100 | AX-170447977 | 0 | T | C | 0.333 | -6.55E-02 | 8.29E-02 | 2.66E+00 | 2.74E+00 | 4.25E-01 | no | Gm29685 | intron_variant,non_coding_transcript_variant | MODIFIER |
| 10 | 107966531 | AX-170377769 | 0 | G | C | 0.333 | -6.55E-02 | 8.29E-02 | 2.66E+00 | 2.74E+00 | 4.25E-01 | no | Gm29685 | intron_variant,non_coding_transcript_variant | MODIFIER |
| 10 | 107974772 | AX-170186758 | 0 | C | A | 0.333 | -6.55E-02 | 8.29E-02 | 2.66E+00 | 2.74E+00 | 4.25E-01 | no | Gm29685 | intron_variant,non_coding_transcript_variant | MODIFIER |
| 10 | 107983758 | AX-170453701 | 0 | C | T | 0.333 | -6.55E-02 | 8.29E-02 | 2.66E+00 | 2.74E+00 | 4.25E-01 | no | | intergenic_variant | MODIFIER |
| 10 | 108007391 | AX-170108180 | 0 | C | T | 0.333 | -6.55E-02 | 8.29E-02 | 2.66E+00 | 2.74E+00 | 4.25E-01 | no | | intergenic_variant | MODIFIER |
| 10 | 108007607 | AX-170267844 | 0 | T | C | 0.333 | -6.55E-02 | 8.29E-02 | 2.66E+00 | 2.74E+00 | 4.25E-01 | no | | intergenic_variant | MODIFIER |
| 10 | 108010279 | AX-170045853 | 0 | A | C | 0.333 | -6.55E-02 | 8.29E-02 | 2.66E+00 | 2.74E+00 | 4.25E-01 | no | | regulatory_region_variant | MODIFIER |
| 10 | 108047687 | AX-170468954 | 0 | G | A | 0.333 | -6.55E-02 | 8.29E-02 | 2.66E+00 | 2.74E+00 | 4.25E-01 | no | | intergenic_variant | MODIFIER |
| 10 | 108066967 | AX-97816795 | 0 | G | A | 0.333 | -6.55E-02 | 8.29E-02 | 2.66E+00 | 2.74E+00 | 4.25E-01 | no | | regulatory_region_variant | MODIFIER |
| 10 | 108360797 | AX-170086897 | 0 | A | G | 0.333 | -6.55E-02 | 8.29E-02 | 2.66E+00 | 2.74E+00 | 4.25E-01 | no | Gm23105 | upstream_gene_variant | MODIFIER |
| 10 | 108415085 | AX-170444511 | 0 | G | A | 0.333 | -6.55E-02 | 8.29E-02 | 2.66E+00 | 2.74E+00 | 4.25E-01 | no | Pawr | downstream_gene_variant | MODIFIER |
| 10 | 108471411 | AX-170431310 | 0 | A | G | 0.333 | -6.55E-02 | 8.29E-02 | 2.66E+00 | 2.74E+00 | 4.25E-01 | no | | intergenic_variant | MODIFIER |
| 10 | 108473286 | AX-170309302 | 0 | C | T | 0.333 | -6.55E-02 | 8.29E-02 | 2.66E+00 | 2.74E+00 | 4.25E-01 | no | | intergenic_variant | MODIFIER |
| 10 | 108476466 | AX-170211765 | 0 | C | T | 0.333 | -6.55E-02 | 8.29E-02 | 2.66E+00 | 2.74E+00 | 4.25E-01 | no | | intergenic_variant | MODIFIER |
| 10 | 108495246 | AX-170276801 | 0 | T | C | 0.333 | -6.55E-02 | 8.29E-02 | 2.66E+00 | 2.74E+00 | 4.25E-01 | no | Syt1 | downstream_gene_variant | MODIFIER |
| 10 | 108496170 | AX-170250892 | 0 | T | C | 0.333 | -6.55E-02 | 8.29E-02 | 2.66E+00 | 2.74E+00 | 4.25E-01 | no | Syt1 | downstream_gene_variant | MODIFIER |
| 10 | 108497086 | AX-170230108 | 0 | T | C | 0.333 | -6.55E-02 | 8.29E-02 | 2.66E+00 | 2.74E+00 | 4.25E-01 | no | Syt1 | downstream_gene_variant | MODIFIER |
| 10 | 108521416 | AX-97850920 | 0 | G | A | 0.333 | -6.55E-02 | 8.29E-02 | 2.66E+00 | 2.74E+00 | 4.25E-01 | no | Syt1 | intron_variant | MODIFIER |
| 10 | 108525453 | AX-170147233 | 0 | C | T | 0.333 | -6.55E-02 | 8.29E-02 | 2.66E+00 | 2.74E+00 | 4.25E-01 | no | Syt1 | intron_variant | MODIFIER |
| 10 | 108534354 | AX-97832879 | 0 | A | C | 0.333 | -6.55E-02 | 8.29E-02 | 2.66E+00 | 2.74E+00 | 4.25E-01 | no | Syt1 | intron_variant | MODIFIER |
| 10 | 108539344 | AX-170112547 | 0 | G | A | 0.333 | -6.55E-02 | 8.29E-02 | 2.66E+00 | 2.74E+00 | 4.25E-01 | no | Syt1 | intron_variant | MODIFIER |
| 10 | 108558331 | AX-170414792 | 0 | G | C | 0.333 | -6.55E-02 | 8.29E-02 | 2.66E+00 | 2.74E+00 | 4.25E-01 | no | Syt1 | intron_variant | MODIFIER |
| 10 | 108576330 | AX-170324412 | 0 | A | G | 0.333 | -6.55E-02 | 8.29E-02 | 2.66E+00 | 2.74E+00 | 4.25E-01 | no | Syt1 | intron_variant | MODIFIER |
| 10 | 108576413 | AX-170045860 | 0 | T | C | 0.333 | -6.55E-02 | 8.29E-02 | 2.66E+00 | 2.74E+00 | 4.25E-01 | no | Syt1 | intron_variant | MODIFIER |
| 10 | 108577310 | AX-170402970 | 0 | A | G | 0.333 | -6.55E-02 | 8.29E-02 | 2.66E+00 | 2.74E+00 | 4.25E-01 | no | Syt1 | intron_variant | MODIFIER |
| 10 | 108595010 | AX-170045862 | 0 | C | T | 0.333 | -6.55E-02 | 8.29E-02 | 2.66E+00 | 2.74E+00 | 4.25E-01 | no | Syt1 | intron_variant | MODIFIER |
| 9 | 84228419 | AX-97781259 | 0 | G | A | 0.219 | 9.51E-02 | 1.20E-01 | 2.86E+00 | 2.94E+00 | 4.25E-01 | no | | intergenic_variant | MODIFIER |
| 5 | 36965755 | AX-170103050 | 0 | T | C | 0.333 | 8.30E-02 | 1.05E-01 | 2.71E+00 | 2.78E+00 | 4.25E-01 | no | Wfs1 | downstream_gene_variant | MODIFIER |
| 9 | 60109845 | AX-170101548 | 0 | T | C | 0.219 | -8.95E-02 | 1.13E-01 | 2.83E+00 | 2.93E+00 | 4.25E-01 | no | Thsd4 | intron_variant | MODIFIER |
| 1 | 96872795 | AX-170001148 | 1 | T | C | 0.152 | 1.58E-01 | 2.00E-01 | 2.65E+00 | 2.72E+00 | 4.25E-01 | Yes | Slco4c1 | upstream_gene_variant | MODIFIER |
| 1 | 101487289 | AX-170259658 | 1 | C | T | 0.152 | 1.58E-01 | 2.00E-01 | 2.65E+00 | 2.72E+00 | 4.25E-01 | Yes | | intergenic_variant | MODIFIER |
| 1 | 14124594 | AX-169999976 | 1 | A | G | 0.366 | 6.34E-02 | 7.94E-02 | 2.58E+00 | 2.64E+00 | 4.25E-01 | no | | intergenic_variant | MODIFIER |
| 3 | 36554023 | AX-170527686 | 1 | C | T | 0.205 | 7.15E-02 | 9.10E-02 | 2.75E+00 | 2.85E+00 | 4.25E-01 | no | Exosc9 | intron_variant | MODIFIER |
| 14 | 120239672 | AX-97828133 | 1 | C | T | 0.33 | 5.91E-02 | 7.47E-02 | 2.60E+00 | 2.71E+00 | 4.25E-01 | no | | intergenic_variant | MODIFIER |
| 5 | 128270043 | AX-170488850 | 0 | C | T | 0.149 | -7.69E-02 | 8.98E-02 | 2.32E+00 | 2.42E+00 | 4.26E-01 | no | Tmem132d | intron_variant | MODIFIER |

| | | | | | | | | | | | | | | | |
|---|---|---|---|---|---|---|---|---|---|---|---|---|---|---|---|
| 7 | 66353454 | AX-170095964 | 2 | A | C | 0.409 | -7.77E-02 | 9.82E-02 | 2.86E+00 | 2.95E+00 | 4.26E-01 | no | Lrrk1 | intron_variant | MODIFIER |
| 1 | 68293613 | AX-97850985 | 0 | C | T | 0.114 | -1.46E-01 | 1.86E-01 | 2.75E+00 | 2.85E+00 | 4.26E-01 | no | Gm15670 | non_coding_transcript_exon_variant | MODIFIER |
| 1 | 95056859 | AX-170517244 | 1 | C | T | 0.161 | 1.62E-01 | 2.05E-01 | 2.66E+00 | 2.73E+00 | 4.26E-01 | Yes | | intergenic_variant | MODIFIER |
| 6 | 94512692 | AX-170454480 | 1 | G | T | 0.446 | -4.80E-02 | 6.02E-02 | 2.54E+00 | 2.66E+00 | 4.26E-01 | no | Slc25a26 | intron_variant | MODIFIER |
| 8 | 40970263 | AX-170409348 | 0 | T | A | 0.149 | 1.07E-01 | 1.36E-01 | 2.83E+00 | 2.91E+00 | 4.26E-01 | no | Pdgfrl | intron_variant | MODIFIER |
| 5 | 121096852 | AX-170506735 | 0 | G | A | 0.447 | 8.50E-02 | 1.08E-01 | 2.67E+00 | 2.73E+00 | 4.26E-01 | no | | intergenic_variant | MODIFIER |
| 10 | 68547893 | AX-170410542 | 0 | G | A | 0.412 | -6.54E-02 | 8.30E-02 | 2.67E+00 | 2.74E+00 | 4.26E-01 | no | | intergenic_variant | MODIFIER |
| 10 | 68773027 | AX-170312537 | 0 | C | T | 0.412 | -6.54E-02 | 8.30E-02 | 2.67E+00 | 2.74E+00 | 4.26E-01 | no | Tmem26 | intron_variant | MODIFIER |
| 10 | 68931199 | AX-170484788 | 0 | T | C | 0.412 | -6.54E-02 | 8.30E-02 | 2.67E+00 | 2.74E+00 | 4.26E-01 | no | | intergenic_variant | MODIFIER |
| 9 | 92616846 | AX-170183524 | 0 | T | C | 0.149 | -9.03E-02 | 1.14E-01 | 2.61E+00 | 2.69E+00 | 4.26E-01 | no | Gm2495 | upstream_gene_variant | MODIFIER |
| 2 | 161905197 | AX-170410410 | 0 | G | T | 0.158 | 1.53E-01 | 1.95E-01 | 2.76E+00 | 2.84E+00 | 4.26E-01 | Yes | Ptprt | intron_variant | MODIFIER |
| 2 | 97275664 | AX-97855792 | 1 | C | A | 0.089 | -1.14E-01 | 1.47E-01 | 2.79E+00 | 2.94E+00 | 4.27E-01 | no | Lrrc4c | intron_variant | MODIFIER |
| 6 | 134659550 | AX-170136926 | 1 | G | A | 0.58 | 6.61E-02 | 8.29E-02 | 2.55E+00 | 2.63E+00 | 4.27E-01 | no | Borcs5 | intron_variant | MODIFIER |
| 7 | 144801249 | AX-97818233 | 0 | C | G | 0.211 | -5.88E-02 | 7.47E-02 | 2.62E+00 | 2.75E+00 | 4.27E-01 | no | | intergenic_variant | MODIFIER |
| 1 | 170306256 | AX-170166149 | 1 | C | T | 0.214 | 1.11E-01 | 1.40E-01 | 2.67E+00 | 2.71E+00 | 4.27E-01 | no | Spata46 | upstream_gene_variant | MODIFIER |
| 13 | 24695651 | AX-170111296 | 1 | G | A | 0.17 | -1.00E-01 | 1.19E-01 | 2.37E+00 | 2.42E+00 | 4.27E-01 | no | Ripor2 | missense_variant | MODERATE |
| 14 | 50374924 | AX-170297813 | 0 | T | C | 0.035 | -1.34E-01 | 1.68E-01 | 2.54E+00 | 2.66E+00 | 4.27E-01 | no | | intergenic_variant | MODIFIER |
| 18 | 84439379 | AX-168390722 | 0 | C | T | 0.167 | -8.30E-02 | 1.03E-01 | 2.51E+00 | 2.60E+00 | 4.27E-01 | no | Zfp407 | intron_variant | MODIFIER |
| 18 | 51132878 | AX-170497629 | 0 | C | T | 0.228 | -8.27E-02 | 1.04E-01 | 2.57E+00 | 2.64E+00 | 4.27E-01 | no | Prr16 | intron_variant | MODIFIER |
| 18 | 51139706 | AX-170195809 | 0 | A | G | 0.228 | -8.27E-02 | 1.04E-01 | 2.57E+00 | 2.64E+00 | 4.27E-01 | no | Prr16 | intron_variant | MODIFIER |
| 18 | 67437052 | AX-170015906 | 0 | C | A | 0.281 | -7.79E-02 | 9.92E-02 | 2.71E+00 | 2.78E+00 | 4.27E-01 | no | Afg3l2 | intron_variant | MODIFIER |
| 7 | 6884708 | AX-97839593 | 1 | G | T | 0.188 | 6.05E-02 | 7.70E-02 | 2.87E+00 | 3.03E+00 | 4.27E-01 | no | Usp29 | intron_variant | MODIFIER |
| 2 | 166311226 | AX-170019610 | 1 | C | T | 0.161 | 1.57E-01 | 2.01E-01 | 2.74E+00 | 2.82E+00 | 4.27E-01 | Yes | | intergenic_variant | MODIFIER |
| 8 | 66879936 | AX-170030940 | 0 | C | T | 0.149 | -1.04E-01 | 1.32E-01 | 2.60E+00 | 2.69E+00 | 4.28E-01 | no | Naf1 | intron_variant | MODIFIER |
| 10 | 108527453 | AX-170119055 | 0 | C | T | 0.333 | -6.48E-02 | 8.23E-02 | 2.63E+00 | 2.71E+00 | 4.28E-01 | no | Syt1 | intron_variant | MODIFIER |
| 12 | 88943242 | AX-97852688 | 1 | G | A | 0.071 | 1.67E-01 | 2.13E-01 | 2.78E+00 | 2.86E+00 | 4.28E-01 | Yes | Nrxn3 | intron_variant | MODIFIER |
| 12 | 92496143 | AX-170266830 | 1 | C | T | 0.071 | 1.67E-01 | 2.13E-01 | 2.78E+00 | 2.86E+00 | 4.28E-01 | Yes | | intergenic_variant | MODIFIER |
| 18 | 70708371 | AX-170126669 | 0 | A | C | 0.263 | -7.25E-02 | 9.11E-02 | 2.54E+00 | 2.63E+00 | 4.28E-01 | no | AC166779.4 | upstream_gene_variant | MODIFIER |
| 4 | 128070344 | AX-170230615 | 0 | G | A | 0.149 | 9.57E-02 | 1.21E-01 | 2.60E+00 | 2.65E+00 | 4.28E-01 | no | Csmd2 | intron_variant,non_coding_transcript_variant | MODIFIER |
| 5 | 78395186 | AX-170072562 | 1 | T | A | 0.42 | 8.33E-02 | 1.06E-01 | 2.73E+00 | 2.78E+00 | 4.28E-01 | no | | intergenic_variant | MODIFIER |
| 3 | 149318528 | AX-170068421 | 1 | G | C | 0.348 | 5.69E-02 | 7.28E-02 | 2.79E+00 | 2.87E+00 | 4.28E-01 | no | Gm30382 | intron_variant,non_coding_transcript_variant | MODIFIER |
| 6 | 36271685 | AX-97783192 | 0 | A | G | 0.061 | 1.20E-01 | 1.55E-01 | 2.72E+00 | 2.87E+00 | 4.28E-01 | no | 9330158H04Rik | intron_variant,non_coding_transcript_variant | MODIFIER |
| 4 | 59171897 | AX-170217172 | 1 | C | T | 0.366 | 8.93E-02 | 1.14E-01 | 2.67E+00 | 2.74E+00 | 4.29E-01 | no | | intergenic_variant | MODIFIER |
| 7 | 5104583 | AX-170329301 | 0 | A | G | 0.184 | 6.12E-02 | 7.88E-02 | 2.78E+00 | 2.92E+00 | 4.29E-01 | no | | intergenic_variant | MODIFIER |
| 3 | 149435980 | AX-170458442 | 1 | G | A | 0.339 | 5.86E-02 | 7.51E-02 | 2.74E+00 | 2.82E+00 | 4.29E-01 | no | Gm30382 | intron_variant,non_coding_transcript_variant | MODIFIER |
| 3 | 151703433 | AX-97801708 | 1 | A | G | 0.33 | 5.77E-02 | 7.39E-02 | 2.75E+00 | 2.83E+00 | 4.29E-01 | no | | intergenic_variant | MODIFIER |
| 1 | 71112189 | AX-170138848 | 0 | G | A | 0.14 | -1.54E-01 | 1.97E-01 | 2.68E+00 | 2.77E+00 | 4.29E-01 | no | | intergenic_variant | MODIFIER |
| 2 | 165098048 | AX-170356213 | 2 | A | C | 0.145 | 1.29E-01 | 1.66E-01 | 2.72E+00 | 2.84E+00 | 4.29E-01 | no | | intergenic_variant | MODIFIER |
| 13 | 24143116 | AX-170193537 | 0 | T | C | 0.158 | -1.03E-01 | 1.23E-01 | 2.37E+00 | 2.42E+00 | 4.29E-01 | no | Carmil1 | intron_variant | MODIFIER |
| 13 | 24164718 | AX-170487447 | 0 | A | G | 0.158 | -1.03E-01 | 1.23E-01 | 2.37E+00 | 2.42E+00 | 4.29E-01 | no | Carmil1 | intron_variant | MODIFIER |
| 13 | 24167110 | AX-170369518 | 0 | A | G | 0.158 | -1.03E-01 | 1.23E-01 | 2.37E+00 | 2.42E+00 | 4.29E-01 | no | Carmil1 | intron_variant | MODIFIER |
| 13 | 24167630 | AX-97826816 | 0 | G | A | 0.158 | -1.03E-01 | 1.23E-01 | 2.37E+00 | 2.42E+00 | 4.29E-01 | no | Carmil1 | intron_variant | MODIFIER |
| 1 | 122302354 | AX-170531465 | 0 | G | T | 0.561 | 6.57E-02 | 8.14E-02 | 2.48E+00 | 2.54E+00 | 4.29E-01 | no | | intergenic_variant | MODIFIER |
| 7 | 66381701 | AX-170029183 | 2 | T | C | 0.4 | -7.65E-02 | 9.77E-02 | 2.81E+00 | 2.89E+00 | 4.29E-01 | no | Lrrk1 | intron_variant | MODIFIER |
| 1 | 150801510 | AX-170220869 | 1 | G | A | 0.214 | 1.06E-01 | 1.35E-01 | 2.66E+00 | 2.71E+00 | 4.29E-01 | no | Hmcn1 | intron_variant | MODIFIER |
| 4 | 89404004 | AX-170427490 | 1 | G | T | 0.321 | 7.78E-02 | 9.95E-02 | 2.67E+00 | 2.75E+00 | 4.29E-01 | no | Gm12609 | downstream_gene_variant | MODIFIER |
| 5 | 44780829 | AX-170071947 | 0 | C | T | 0.184 | -8.62E-02 | 1.09E-01 | 2.57E+00 | 2.66E+00 | 4.30E-01 | no | Ldb2 | intron_variant | MODIFIER |
| 2 | 162372615 | AX-170185725 | 0 | C | G | 0.149 | 1.43E-01 | 1.83E-01 | 2.72E+00 | 2.81E+00 | 4.30E-01 | no | Ptprt | intron_variant | MODIFIER |
| 1 | 149975652 | AX-170383808 | 0 | G | A | 0.202 | 1.07E-01 | 1.37E-01 | 2.69E+00 | 2.75E+00 | 4.30E-01 | no | | regulatory_region_variant | MODIFIER |
| 7 | 24499318 | AX-97819108 | 0 | C | T | 0.404 | 6.99E-02 | 8.85E-02 | 2.56E+00 | 2.67E+00 | 4.30E-01 | no | Cadm4 | intron_variant | MODIFIER |
| 10 | 94295679 | AX-170165017 | 0 | G | A | 0.298 | -6.67E-02 | 8.40E-02 | 2.55E+00 | 2.62E+00 | 4.30E-01 | no | Gm4792 | intron_variant | MODIFIER |
| 9 | 68447956 | AX-97806713 | 1 | G | T | 0.205 | -9.06E-02 | 1.16E-01 | 2.74E+00 | 2.84E+00 | 4.30E-01 | no | | intergenic_variant | MODIFIER |
| 7 | 123069997 | AX-97857698 | 1 | T | C | 0.411 | 6.47E-02 | 8.31E-02 | 2.75E+00 | 2.84E+00 | 4.30E-01 | no | Gm45847 | downstream_gene_variant | MODIFIER |
| 7 | 99202326 | AX-170078205 | 0 | A | G | 0.377 | -7.95E-02 | 1.02E-01 | 2.72E+00 | 2.81E+00 | 4.30E-01 | no | Gm45012 | upstream_gene_variant | MODIFIER |

| | | | | | | | | | | | | | | | |
|---|---|---|---|---|---|---|---|---|---|---|---|---|---|---|---|
| 9 | 95157437 | AX-170199285 | 0 | G | C | 0.149 | -8.65E-02 | 1.11E-01 | 2.70E+00 | 2.79E+00 | 4.30E-01 | no | Slc9a9 | intron_variant | MODIFIER |
| 8 | 16859147 | AX-170375818 | 0 | C | T | 0.237 | -5.95E-02 | 7.50E-02 | 2.53E+00 | 2.63E+00 | 4.30E-01 | no | Csmd1 | intron_variant | MODIFIER |
| 6 | 138691373 | AX-170095827 | 0 | T | G | 0.553 | 6.84E-02 | 8.39E-02 | 2.44E+00 | 2.51E+00 | 4.30E-01 | no | | intergenic_variant | MODIFIER |
| 7 | 70409742 | AX-170234262 | 1 | T | G | 0.384 | -8.43E-02 | 1.07E-01 | 2.89E+00 | 2.99E+00 | 4.30E-01 | no | B130024G19Rik | intron_variant,non_coding_transcript_variant | MODIFIER |
| 3 | 23749857 | AX-170407599 | 0 | T | G | 0.167 | -7.81E-02 | 9.86E-02 | 2.55E+00 | 2.63E+00 | 4.30E-01 | no | | intergenic_variant | MODIFIER |
| 1 | 101806273 | AX-170314422 | 0 | C | A | 0.158 | 1.32E-01 | 1.67E-01 | 2.55E+00 | 2.64E+00 | 4.30E-01 | no | | intergenic_variant | MODIFIER |
| 12 | 107838528 | AX-97849505 | 0 | C | T | 0.114 | -1.33E-01 | 1.71E-01 | 2.75E+00 | 2.86E+00 | 4.30E-01 | no | | intergenic_variant | MODIFIER |
| 3 | 150315160 | AX-170193822 | 0 | T | G | 0.36 | 5.56E-02 | 7.15E-02 | 2.74E+00 | 2.83E+00 | 4.31E-01 | no | | intergenic_variant | MODIFIER |
| 10 | 28144055 | AX-170044448 | 1 | G | A | 0.036 | 1.17E-01 | 1.48E-01 | 2.50E+00 | 2.66E+00 | 4.31E-01 | no | Ptprk | intron_variant | MODIFIER |
| 13 | 23651851 | AX-170007919 | 1 | C | G | 0.17 | -8.98E-02 | 1.11E-01 | 2.46E+00 | 2.53E+00 | 4.31E-01 | no | | intergenic_variant | MODIFIER |
| 6 | 134273037 | AX-170076298 | 0 | C | T | 0.57 | 6.99E-02 | 8.71E-02 | 2.50E+00 | 2.55E+00 | 4.31E-01 | no | Etv6 | downstream_gene_variant | MODIFIER |
| 8 | 97975219 | AX-170440653 | 0 | C | T | 0.219 | 8.52E-02 | 1.10E-01 | 2.72E+00 | 2.80E+00 | 4.31E-01 | no | | intergenic_variant | MODIFIER |
| 5 | 104512357 | AX-97816882 | 1 | T | C | 0.464 | 8.27E-02 | 1.06E-01 | 2.67E+00 | 2.74E+00 | 4.31E-01 | no | BC005561 | intron_variant | MODIFIER |
| 2 | 169918146 | AX-170118055 | 0 | G | A | 0.175 | 1.24E-01 | 1.57E-01 | 2.55E+00 | 2.66E+00 | 4.31E-01 | no | Tshz2 | intron_variant | MODIFIER |
| 6 | 135158801 | AX-170509449 | 0 | C | T | 0.57 | 6.88E-02 | 8.60E-02 | 2.50E+00 | 2.56E+00 | 4.31E-01 | no | Hebp1 | intron_variant | MODIFIER |
| 1 | 135955967 | AX-170042836 | 0 | C | T | 0.219 | 9.04E-02 | 1.16E-01 | 2.61E+00 | 2.70E+00 | 4.31E-01 | no | Igfn1 | intron_variant | MODIFIER |
| 1 | 82143801 | AX-170342475 | 1 | C | T | 0.161 | 1.61E-01 | 2.07E-01 | 2.66E+00 | 2.73E+00 | 4.31E-01 | Yes | | intergenic_variant | MODIFIER |
| 18 | 55213986 | AX-170015742 | 0 | T | G | 0.447 | 7.82E-02 | 9.32E-02 | 2.35E+00 | 2.41E+00 | 4.31E-01 | no | | intergenic_variant | MODIFIER |
| 4 | 83587885 | AX-97812850 | 0 | T | G | 0.456 | -4.34E-02 | 5.36E-02 | 3.08E+00 | 3.23E+00 | 4.31E-01 | no | Ccdc171 | intron_variant | MODIFIER |
| 20 | 62396060 | AX-170186488 | 1 | G | T | 0.054 | -8.51E-02 | 1.10E-01 | 2.62E+00 | 2.79E+00 | 4.31E-01 | no | | intergenic_variant | MODIFIER |
| 6 | 128185043 | AX-170251708 | 0 | G | A | 0.553 | 6.45E-02 | 8.18E-02 | 2.56E+00 | 2.64E+00 | 4.31E-01 | no | Gm10010 | downstream_gene_variant | MODIFIER |
| 12 | 90628270 | AX-97790890 | 1 | T | A | 0.071 | 1.34E-01 | 1.73E-01 | 2.68E+00 | 2.79E+00 | 4.31E-01 | no | | intergenic_variant | MODIFIER |
| 6 | 138478284 | AX-170224031 | 1 | A | C | 0.554 | 6.87E-02 | 8.48E-02 | 2.45E+00 | 2.53E+00 | 4.32E-01 | no | Lmo3 | intron_variant | MODIFIER |
| 18 | 76072093 | AX-97783923 | 1 | C | T | 0.402 | -6.85E-02 | 8.78E-02 | 2.64E+00 | 2.71E+00 | 4.32E-01 | no | Zbtb7c | intron_variant | MODIFIER |
| 2 | 72598611 | AX-170320181 | 0 | A | G | 0.044 | 1.49E-01 | 1.93E-01 | 2.78E+00 | 2.94E+00 | 4.32E-01 | Yes | | intergenic_variant | MODIFIER |
| 1 | 149862969 | AX-170182840 | 1 | C | G | 0.205 | 1.04E-01 | 1.34E-01 | 2.64E+00 | 2.70E+00 | 4.32E-01 | no | Pla2g4a | intron_variant | MODIFIER |
| 1 | 143719261 | AX-170042978 | 0 | C | A | 0.202 | 9.48E-02 | 1.22E-01 | 2.70E+00 | 2.79E+00 | 4.32E-01 | no | Glrx2 | intron_variant,non_coding_transcript_variant | MODIFIER |
| 5 | 128566997 | AX-170122809 | 0 | A | G | 0.149 | -7.65E-02 | 9.00E-02 | 2.31E+00 | 2.40E+00 | 4.32E-01 | no | | intergenic_variant | MODIFIER |
| 9 | 26936645 | AX-170422292 | 2 | T | G | 0.464 | 7.31E-02 | 9.29E-02 | 2.57E+00 | 2.64E+00 | 4.32E-01 | no | | intergenic_variant | MODIFIER |
| 4 | 151288683 | AX-170190037 | 0 | G | A | 0.114 | 1.42E-01 | 1.83E-01 | 2.74E+00 | 2.83E+00 | 4.32E-01 | no | Camta1 | intron_variant | MODIFIER |
| 1 | 16177805 | AX-169999999 | 1 | G | T | 0.366 | 6.10E-02 | 7.75E-02 | 2.58E+00 | 2.63E+00 | 4.32E-01 | no | | regulatory_region_variant | MODIFIER |
| 18 | 74764691 | AX-170228364 | 1 | G | T | 0.438 | 6.94E-02 | 8.69E-02 | 2.50E+00 | 2.56E+00 | 4.32E-01 | no | Myo5b | intron_variant | MODIFIER |
| 18 | 74203765 | AX-170136732 | 0 | T | C | 0.184 | -8.69E-02 | 1.11E-01 | 2.59E+00 | 2.68E+00 | 4.32E-01 | no | Ska1 | intron_variant | MODIFIER |
| 10 | 70343750 | AX-170470535 | 0 | C | T | 0.421 | -6.32E-02 | 8.11E-02 | 2.63E+00 | 2.71E+00 | 4.32E-01 | no | | intergenic_variant | MODIFIER |
| 13 | 23236171 | AX-97797995 | 1 | A | G | 0.161 | -9.78E-02 | 1.19E-01 | 2.41E+00 | 2.46E+00 | 4.33E-01 | no | Vmn1r222 | upstream_gene_variant | MODIFIER |
| 18 | 80644851 | AX-170154663 | 1 | T | A | 0.384 | -6.67E-02 | 8.55E-02 | 2.61E+00 | 2.70E+00 | 4.33E-01 | no | Nfatc1 | downstream_gene_variant | MODIFIER |
| 6 | 134275055 | AX-170233361 | 0 | A | G | 0.57 | 6.96E-02 | 8.71E-02 | 2.50E+00 | 2.55E+00 | 4.33E-01 | no | Etv6 | downstream_gene_variant | MODIFIER |
| 2 | 144603634 | AX-97839303 | 0 | G | T | 0.149 | 1.38E-01 | 1.77E-01 | 2.65E+00 | 2.74E+00 | 4.33E-01 | no | Dtd1 | intron_variant | MODIFIER |
| 1 | 161352440 | AX-170415590 | 1 | C | A | 0.223 | 9.92E-02 | 1.27E-01 | 2.60E+00 | 2.67E+00 | 4.33E-01 | no | | intergenic_variant | MODIFIER |
| 1 | 21669118 | AX-170158147 | 0 | A | G | 0.412 | 6.59E-02 | 8.37E-02 | 2.56E+00 | 2.61E+00 | 4.33E-01 | no | Kcnq5 | intron_variant | MODIFIER |
| 5 | 4185228 | AX-170277758 | 1 | A | T | 0.429 | -5.06E-02 | 6.39E-02 | 2.50E+00 | 2.62E+00 | 4.33E-01 | no | | intergenic_variant | MODIFIER |
| 5 | 105682284 | AX-170352116 | 0 | G | C | 0.439 | 8.41E-02 | 1.08E-01 | 2.69E+00 | 2.74E+00 | 4.33E-01 | no | | intergenic_variant | MODIFIER |
| 18 | 55678057 | AX-170015754 | 1 | T | A | 0.446 | 7.85E-02 | 9.47E-02 | 2.38E+00 | 2.43E+00 | 4.33E-01 | no | | regulatory_region_variant | MODIFIER |
| 7 | 113915134 | AX-97790085 | 0 | C | T | 0.421 | 7.54E-02 | 9.73E-02 | 2.71E+00 | 2.75E+00 | 4.33E-01 | no | Spon1 | intron_variant | MODIFIER |
| 7 | 113946865 | AX-170514685 | 0 | A | C | 0.421 | 7.54E-02 | 9.73E-02 | 2.71E+00 | 2.75E+00 | 4.33E-01 | no | Spon1 | intron_variant | MODIFIER |
| 7 | 113946929 | AX-170434173 | 0 | C | G | 0.421 | 7.54E-02 | 9.73E-02 | 2.71E+00 | 2.75E+00 | 4.33E-01 | no | Spon1 | intron_variant | MODIFIER |
| 7 | 113951532 | AX-170029798 | 0 | C | T | 0.421 | 7.54E-02 | 9.73E-02 | 2.71E+00 | 2.75E+00 | 4.33E-01 | no | Spon1 | intron_variant | MODIFIER |
| 7 | 113959058 | AX-97840644 | 0 | T | C | 0.421 | 7.54E-02 | 9.73E-02 | 2.71E+00 | 2.75E+00 | 4.33E-01 | no | Spon1 | intron_variant | MODIFIER |
| 7 | 113966812 | AX-170287496 | 0 | T | A | 0.421 | 7.54E-02 | 9.73E-02 | 2.71E+00 | 2.75E+00 | 4.33E-01 | no | Spon1 | intron_variant | MODIFIER |
| 7 | 113972755 | AX-170299498 | 0 | A | T | 0.421 | 7.54E-02 | 9.73E-02 | 2.71E+00 | 2.75E+00 | 4.33E-01 | no | Spon1 | intron_variant | MODIFIER |
| 7 | 113976099 | AX-170237671 | 0 | T | C | 0.421 | 7.54E-02 | 9.73E-02 | 2.71E+00 | 2.75E+00 | 4.33E-01 | no | Spon1 | intron_variant | MODIFIER |
| 7 | 113993187 | AX-170458119 | 0 | T | C | 0.421 | 7.54E-02 | 9.73E-02 | 2.71E+00 | 2.75E+00 | 4.33E-01 | no | Spon1 | intron_variant | MODIFIER |
| 7 | 114707495 | AX-170203613 | 0 | G | A | 0.421 | 7.54E-02 | 9.73E-02 | 2.71E+00 | 2.75E+00 | 4.33E-01 | no | Gm15500 | upstream_gene_variant | MODIFIER |
| 7 | 114750026 | AX-170146382 | 0 | T | C | 0.421 | 7.54E-02 | 9.73E-02 | 2.71E+00 | 2.75E+00 | 4.33E-01 | no | Insc | intron_variant | MODIFIER |

| | | | | | | | | | | | | | | | |
|---|---|---|---|---|---|---|---|---|---|---|---|---|---|---|---|
| 7 | 114771781 | AX-170202151 | 0 | T | G | 0.421 | 7.54E-02 | 9.73E-02 | 2.71E+00 | 2.75E+00 | 4.33E-01 | no | Insc | intron_variant | MODIFIER |
| 7 | 114777363 | AX-97802652 | 0 | A | G | 0.421 | 7.54E-02 | 9.73E-02 | 2.71E+00 | 2.75E+00 | 4.33E-01 | no | Insc | intron_variant | MODIFIER |
| 7 | 114857545 | AX-170328381 | 0 | G | A | 0.421 | 7.54E-02 | 9.73E-02 | 2.71E+00 | 2.75E+00 | 4.33E-01 | no | Gm45017 | upstream_gene_variant | MODIFIER |
| 7 | 114880441 | AX-170150928 | 0 | T | C | 0.421 | 7.54E-02 | 9.73E-02 | 2.71E+00 | 2.75E+00 | 4.33E-01 | no | | intergenic_variant | MODIFIER |
| 7 | 114884939 | AX-170246050 | 0 | T | C | 0.421 | 7.54E-02 | 9.73E-02 | 2.71E+00 | 2.75E+00 | 4.33E-01 | no | | intergenic_variant | MODIFIER |
| 7 | 114892382 | AX-170096084 | 0 | T | C | 0.421 | 7.54E-02 | 9.73E-02 | 2.71E+00 | 2.75E+00 | 4.33E-01 | no | | intergenic_variant | MODIFIER |
| 7 | 114894169 | AX-170216841 | 0 | C | T | 0.421 | 7.54E-02 | 9.73E-02 | 2.71E+00 | 2.75E+00 | 4.33E-01 | no | | intergenic_variant | MODIFIER |
| 7 | 114913884 | AX-170206171 | 0 | G | A | 0.421 | 7.54E-02 | 9.73E-02 | 2.71E+00 | 2.75E+00 | 4.33E-01 | no | | intergenic_variant | MODIFIER |
| 7 | 114924675 | AX-170128067 | 0 | A | G | 0.421 | 7.54E-02 | 9.73E-02 | 2.71E+00 | 2.75E+00 | 4.33E-01 | no | Rps4l-ps | downstream_gene_variant | MODIFIER |
| 7 | 114933240 | AX-170517882 | 0 | G | T | 0.421 | 7.54E-02 | 9.73E-02 | 2.71E+00 | 2.75E+00 | 4.33E-01 | no | Gm22669 | upstream_gene_variant | MODIFIER |
| 7 | 114956082 | AX-170328700 | 0 | T | C | 0.421 | 7.54E-02 | 9.73E-02 | 2.71E+00 | 2.75E+00 | 4.33E-01 | no | A730082K24Rik | intron_variant,non_coding_transcript_variant | MODIFIER |
| 7 | 114956251 | AX-170090012 | 0 | T | C | 0.421 | 7.54E-02 | 9.73E-02 | 2.71E+00 | 2.75E+00 | 4.33E-01 | no | A730082K24Rik | intron_variant,non_coding_transcript_variant | MODIFIER |
| 7 | 114984653 | AX-97802999 | 0 | T | C | 0.421 | 7.54E-02 | 9.73E-02 | 2.71E+00 | 2.75E+00 | 4.33E-01 | no | A730082K24Rik | downstream_gene_variant | MODIFIER |
| 7 | 114984943 | AX-170029810 | 0 | A | G | 0.421 | 7.54E-02 | 9.73E-02 | 2.71E+00 | 2.75E+00 | 4.33E-01 | no | A730082K24Rik | downstream_gene_variant | MODIFIER |
| 7 | 114988281 | AX-170029809 | 0 | C | T | 0.421 | 7.54E-02 | 9.73E-02 | 2.71E+00 | 2.75E+00 | 4.33E-01 | no | A730082K24Rik | downstream_gene_variant | MODIFIER |
| 7 | 115001478 | AX-170338188 | 0 | T | C | 0.421 | 7.54E-02 | 9.73E-02 | 2.71E+00 | 2.75E+00 | 4.33E-01 | no | | regulatory_region_variant | MODIFIER |
| 7 | 115001510 | AX-170121873 | 0 | C | A | 0.421 | 7.54E-02 | 9.73E-02 | 2.71E+00 | 2.75E+00 | 4.33E-01 | no | | regulatory_region_variant | MODIFIER |
| 7 | 115001627 | AX-170341109 | 0 | C | T | 0.421 | 7.54E-02 | 9.73E-02 | 2.71E+00 | 2.75E+00 | 4.33E-01 | no | | intergenic_variant | MODIFIER |
| 7 | 115001654 | AX-170326622 | 0 | A | T | 0.421 | 7.54E-02 | 9.73E-02 | 2.71E+00 | 2.75E+00 | 4.33E-01 | no | | intergenic_variant | MODIFIER |
| 7 | 115002351 | AX-170338803 | 0 | G | A | 0.421 | 7.54E-02 | 9.73E-02 | 2.71E+00 | 2.75E+00 | 4.33E-01 | no | | intergenic_variant | MODIFIER |
| 7 | 115026880 | AX-170029811 | 0 | T | C | 0.421 | 7.54E-02 | 9.73E-02 | 2.71E+00 | 2.75E+00 | 4.33E-01 | no | | intergenic_variant | MODIFIER |
| 7 | 115046251 | AX-170124252 | 0 | A | G | 0.421 | 7.54E-02 | 9.73E-02 | 2.71E+00 | 2.75E+00 | 4.33E-01 | no | Gm34225 | intron_variant,non_coding_transcript_variant | MODIFIER |
| 7 | 115048592 | AX-170525226 | 0 | G | C | 0.421 | 7.54E-02 | 9.73E-02 | 2.71E+00 | 2.75E+00 | 4.33E-01 | no | Gm34225 | intron_variant,non_coding_transcript_variant | MODIFIER |
| 7 | 115064472 | AX-97844515 | 0 | A | G | 0.421 | 7.54E-02 | 9.73E-02 | 2.71E+00 | 2.75E+00 | 4.33E-01 | no | Gm34225 | intron_variant,non_coding_transcript_variant | MODIFIER |
| 7 | 115067635 | AX-170447095 | 0 | T | A | 0.421 | 7.54E-02 | 9.73E-02 | 2.71E+00 | 2.75E+00 | 4.33E-01 | no | Gm34225 | downstream_gene_variant | MODIFIER |
| 7 | 115069532 | AX-170274247 | 0 | G | A | 0.421 | 7.54E-02 | 9.73E-02 | 2.71E+00 | 2.75E+00 | 4.33E-01 | no | Gm34225 | downstream_gene_variant | MODIFIER |
| 7 | 115070069 | AX-170454473 | 0 | C | T | 0.421 | 7.54E-02 | 9.73E-02 | 2.71E+00 | 2.75E+00 | 4.33E-01 | no | Gm34225 | downstream_gene_variant | MODIFIER |
| 7 | 115084347 | AX-170390595 | 0 | C | T | 0.421 | 7.54E-02 | 9.73E-02 | 2.71E+00 | 2.75E+00 | 4.33E-01 | no | | intergenic_variant | MODIFIER |
| 7 | 115085765 | AX-170391453 | 0 | G | A | 0.421 | 7.54E-02 | 9.73E-02 | 2.71E+00 | 2.75E+00 | 4.33E-01 | no | | intergenic_variant | MODIFIER |
| 7 | 115087242 | AX-170192524 | 0 | A | G | 0.421 | 7.54E-02 | 9.73E-02 | 2.71E+00 | 2.75E+00 | 4.33E-01 | no | | intergenic_variant | MODIFIER |
| 7 | 115090364 | AX-170078504 | 0 | G | A | 0.421 | 7.54E-02 | 9.73E-02 | 2.71E+00 | 2.75E+00 | 4.33E-01 | no | | intergenic_variant | MODIFIER |
| 7 | 115095380 | AX-170096086 | 0 | G | A | 0.421 | 7.54E-02 | 9.73E-02 | 2.71E+00 | 2.75E+00 | 4.33E-01 | no | | intergenic_variant | MODIFIER |
| 7 | 115112848 | AX-170359744 | 0 | T | A | 0.421 | 7.54E-02 | 9.73E-02 | 2.71E+00 | 2.75E+00 | 4.33E-01 | no | | intergenic_variant | MODIFIER |
| 7 | 115117817 | AX-170493718 | 0 | C | G | 0.421 | 7.54E-02 | 9.73E-02 | 2.71E+00 | 2.75E+00 | 4.33E-01 | no | | intergenic_variant | MODIFIER |
| 7 | 115121366 | AX-170310994 | 0 | T | C | 0.421 | 7.54E-02 | 9.73E-02 | 2.71E+00 | 2.75E+00 | 4.33E-01 | no | | intergenic_variant | MODIFIER |
| 7 | 115132673 | AX-170510706 | 0 | C | T | 0.421 | 7.54E-02 | 9.73E-02 | 2.71E+00 | 2.75E+00 | 4.33E-01 | no | | regulatory_region_variant | MODIFIER |
| 7 | 115136349 | AX-170196720 | 0 | A | G | 0.421 | 7.54E-02 | 9.73E-02 | 2.71E+00 | 2.75E+00 | 4.33E-01 | no | | intergenic_variant | MODIFIER |
| 7 | 115144230 | AX-170414155 | 0 | A | G | 0.421 | 7.54E-02 | 9.73E-02 | 2.71E+00 | 2.75E+00 | 4.33E-01 | no | | intergenic_variant | MODIFIER |
| 7 | 115171546 | AX-170116729 | 0 | A | G | 0.421 | 7.54E-02 | 9.73E-02 | 2.71E+00 | 2.75E+00 | 4.33E-01 | no | Gm27684 | upstream_gene_variant | MODIFIER |
| 7 | 115204442 | AX-170502633 | 0 | C | G | 0.421 | 7.54E-02 | 9.73E-02 | 2.71E+00 | 2.75E+00 | 4.33E-01 | no | | intergenic_variant | MODIFIER |
| 7 | 115245416 | AX-170078507 | 0 | C | A | 0.421 | 7.54E-02 | 9.73E-02 | 2.71E+00 | 2.75E+00 | 4.33E-01 | no | | intergenic_variant | MODIFIER |
| 7 | 115248858 | AX-170078508 | 0 | A | T | 0.421 | 7.54E-02 | 9.73E-02 | 2.71E+00 | 2.75E+00 | 4.33E-01 | no | | intergenic_variant | MODIFIER |
| 7 | 115250092 | AX-170400436 | 0 | T | C | 0.421 | 7.54E-02 | 9.73E-02 | 2.71E+00 | 2.75E+00 | 4.33E-01 | no | | intergenic_variant | MODIFIER |
| 7 | 115250463 | AX-97806417 | 0 | C | T | 0.421 | 7.54E-02 | 9.73E-02 | 2.71E+00 | 2.75E+00 | 4.33E-01 | no | | intergenic_variant | MODIFIER |
| 7 | 115266931 | AX-170469180 | 0 | C | T | 0.421 | 7.54E-02 | 9.73E-02 | 2.71E+00 | 2.75E+00 | 4.33E-01 | no | | intergenic_variant | MODIFIER |
| 7 | 115324681 | AX-170125945 | 0 | A | G | 0.421 | 7.54E-02 | 9.73E-02 | 2.71E+00 | 2.75E+00 | 4.33E-01 | no | | intergenic_variant | MODIFIER |
| 7 | 115326140 | AX-170169045 | 0 | G | A | 0.421 | 7.54E-02 | 9.73E-02 | 2.71E+00 | 2.75E+00 | 4.33E-01 | no | | intergenic_variant | MODIFIER |
| 7 | 115332299 | AX-170439143 | 0 | A | G | 0.421 | 7.54E-02 | 9.73E-02 | 2.71E+00 | 2.75E+00 | 4.33E-01 | no | | intergenic_variant | MODIFIER |
| 7 | 115333269 | AX-170332652 | 0 | C | T | 0.421 | 7.54E-02 | 9.73E-02 | 2.71E+00 | 2.75E+00 | 4.33E-01 | no | | intergenic_variant | MODIFIER |
| 7 | 115333739 | AX-97791646 | 0 | T | C | 0.421 | 7.54E-02 | 9.73E-02 | 2.71E+00 | 2.75E+00 | 4.33E-01 | no | | intergenic_variant | MODIFIER |
| 7 | 115383572 | AX-170108416 | 0 | G | T | 0.421 | 7.54E-02 | 9.73E-02 | 2.71E+00 | 2.75E+00 | 4.33E-01 | no | | intergenic_variant | MODIFIER |
| 7 | 115401838 | AX-170483964 | 0 | C | G | 0.421 | 7.54E-02 | 9.73E-02 | 2.71E+00 | 2.75E+00 | 4.33E-01 | no | | intergenic_variant | MODIFIER |
| 18 | 56958677 | AX-170475362 | 1 | C | T | 0.411 | 8.36E-02 | 9.81E-02 | 2.31E+00 | 2.36E+00 | 4.33E-01 | no | C330018D20Rik | intron_variant | MODIFIER |
| 1 | 19976000 | AX-170040554 | 0 | C | T | 0.395 | 6.53E-02 | 8.35E-02 | 2.60E+00 | 2.65E+00 | 4.33E-01 | no | | intergenic_variant | MODIFIER |

| | | | | | | | | | | | | | | | |
|---|---|---|---|---|---|---|---|---|---|---|---|---|---|---|---|
| 10 | 108476003 | AX-97816594 | 1 | G | A | 0.33 | -6.45E-02 | 8.31E-02 | 2.64E+00 | 2.72E+00 | 4.33E-01 | no | | intergenic_variant | MODIFIER |
| 12 | 69389909 | AX-170154900 | 0 | C | T | 0.088 | 1.13E-01 | 1.48E-01 | 2.69E+00 | 2.85E+00 | 4.33E-01 | no | | regulatory_region_variant | MODIFIER |
| 9 | 81061058 | AX-170492417 | 1 | T | A | 0.214 | -8.42E-02 | 1.09E-01 | 2.68E+00 | 2.80E+00 | 4.34E-01 | no | Gm26713 | intron_variant,non_coding_transcript_variant | MODIFIER |
| 1 | 128150433 | AX-97846550 | 0 | C | T | 0.535 | 6.67E-02 | 8.56E-02 | 2.62E+00 | 2.68E+00 | 4.34E-01 | no | R3hdm1 | intron_variant | MODIFIER |
| 3 | 37041418 | AX-97797635 | 0 | G | A | 0.193 | 7.08E-02 | 9.13E-02 | 2.84E+00 | 2.96E+00 | 4.34E-01 | no | 4932438A13Rik | synonymous_variant | LOW |
| 5 | 4675473 | AX-170098748 | 1 | A | T | 0.402 | -5.30E-02 | 6.72E-02 | 2.51E+00 | 2.63E+00 | 4.34E-01 | no | | regulatory_region_variant | MODIFIER |
| 12 | 109771580 | AX-170092614 | 1 | G | A | 0.125 | -1.27E-01 | 1.64E-01 | 2.69E+00 | 2.80E+00 | 4.34E-01 | no | | intergenic_variant | MODIFIER |
| 2 | 102177795 | AX-170147756 | 0 | T | G | 0.044 | -1.57E-01 | 2.03E-01 | 2.62E+00 | 2.73E+00 | 4.34E-01 | no | Ldlrad3 | intron_variant | MODIFIER |
| 1 | 137070702 | AX-170135989 | 1 | A | G | 0.214 | 9.40E-02 | 1.21E-01 | 2.67E+00 | 2.75E+00 | 4.34E-01 | no | | intergenic_variant | MODIFIER |
| 10 | 66444139 | AX-170135825 | 0 | T | C | 0.404 | -6.32E-02 | 8.13E-02 | 2.62E+00 | 2.69E+00 | 4.34E-01 | no | | intergenic_variant | MODIFIER |
| 5 | 85624908 | AX-170212850 | 0 | G | A | 0.43 | 7.94E-02 | 1.03E-01 | 2.81E+00 | 2.89E+00 | 4.34E-01 | no | | intergenic_variant | MODIFIER |
| 1 | 148783142 | AX-170396424 | 0 | C | T | 0.219 | 8.99E-02 | 1.16E-01 | 2.61E+00 | 2.70E+00 | 4.34E-01 | no | | intergenic_variant | MODIFIER |
| 6 | 127438459 | AX-170329989 | 0 | G | A | 0.535 | 6.42E-02 | 8.15E-02 | 2.54E+00 | 2.61E+00 | 4.34E-01 | no | | intergenic_variant | MODIFIER |
| 7 | 44899432 | AX-170372314 | 0 | G | C | 0.342 | 7.38E-02 | 9.46E-02 | 2.59E+00 | 2.66E+00 | 4.34E-01 | no | Fuz | intron_variant | MODIFIER |
| 7 | 44913139 | AX-170105627 | 0 | A | G | 0.342 | 7.38E-02 | 9.46E-02 | 2.59E+00 | 2.66E+00 | 4.34E-01 | no | Ap2a1 | intron_variant | MODIFIER |
| 7 | 44919501 | AX-170534494 | 0 | A | G | 0.342 | 7.38E-02 | 9.46E-02 | 2.59E+00 | 2.66E+00 | 4.34E-01 | no | Ap2a1 | intron_variant | MODIFIER |
| 7 | 44919934 | AX-170340226 | 0 | A | T | 0.342 | 7.38E-02 | 9.46E-02 | 2.59E+00 | 2.66E+00 | 4.34E-01 | no | Ap2a1 | intron_variant | MODIFIER |
| 7 | 44958074 | AX-170467205 | 0 | A | G | 0.342 | 7.38E-02 | 9.46E-02 | 2.59E+00 | 2.66E+00 | 4.34E-01 | no | Cpt1c | downstream_gene_variant | MODIFIER |
| 7 | 44991837 | AX-170089912 | 0 | T | C | 0.342 | 7.38E-02 | 9.46E-02 | 2.59E+00 | 2.66E+00 | 4.34E-01 | no | Bcl2l12 | intron_variant | MODIFIER |
| 7 | 45005922 | AX-97802360 | 0 | T | C | 0.342 | 7.38E-02 | 9.46E-02 | 2.59E+00 | 2.66E+00 | 4.34E-01 | no | Irf3 | downstream_gene_variant | MODIFIER |
| 7 | 45008528 | AX-170028930 | 0 | T | C | 0.342 | 7.38E-02 | 9.46E-02 | 2.59E+00 | 2.66E+00 | 4.34E-01 | no | Scaf1 | missense_variant | MODERATE |
| 7 | 45025333 | AX-170472964 | 0 | C | T | 0.342 | 7.38E-02 | 9.46E-02 | 2.59E+00 | 2.66E+00 | 4.34E-01 | no | Rras | downstream_gene_variant | MODIFIER |
| 7 | 45032709 | AX-170028931 | 0 | A | G | 0.342 | 7.38E-02 | 9.46E-02 | 2.59E+00 | 2.66E+00 | 4.34E-01 | no | Prr12 | intron_variant | MODIFIER |
| 7 | 45055465 | AX-170028929 | 0 | C | T | 0.342 | 7.38E-02 | 9.46E-02 | 2.59E+00 | 2.66E+00 | 4.34E-01 | no | Prrg2 | intron_variant | MODIFIER |
| 7 | 45064425 | AX-170028932 | 0 | G | A | 0.342 | 7.38E-02 | 9.46E-02 | 2.59E+00 | 2.66E+00 | 4.34E-01 | no | Nosip | intron_variant | MODIFIER |
| 7 | 45073040 | AX-97802361 | 0 | G | A | 0.342 | 7.38E-02 | 9.46E-02 | 2.59E+00 | 2.66E+00 | 4.34E-01 | no | Nosip | intron_variant | MODIFIER |
| 7 | 45080165 | AX-170315674 | 0 | A | C | 0.342 | 7.38E-02 | 9.46E-02 | 2.59E+00 | 2.66E+00 | 4.34E-01 | no | Nosip | downstream_gene_variant | MODIFIER |
| 7 | 45083666 | AX-170347943 | 0 | T | C | 0.342 | 7.38E-02 | 9.46E-02 | 2.59E+00 | 2.66E+00 | 4.34E-01 | no | Rcn3 | missense_variant | MODERATE |
| 7 | 45084764 | AX-97838206 | 0 | C | T | 0.342 | 7.38E-02 | 9.46E-02 | 2.59E+00 | 2.66E+00 | 4.34E-01 | no | Rcn3 | intron_variant | MODIFIER |
| 7 | 45116978 | AX-170115096 | 0 | T | C | 0.342 | 7.38E-02 | 9.46E-02 | 2.59E+00 | 2.66E+00 | 4.34E-01 | no | Mir150 | upstream_gene_variant | MODIFIER |
| 2 | 163167075 | AX-170019568 | 1 | G | T | 0.152 | 1.34E-01 | 1.74E-01 | 2.69E+00 | 2.80E+00 | 4.34E-01 | no | | intergenic_variant | MODIFIER |
| 2 | 163829068 | AX-170352155 | 1 | G | A | 0.152 | 1.34E-01 | 1.74E-01 | 2.69E+00 | 2.80E+00 | 4.34E-01 | no | Ccn5 | missense_variant | MODERATE |
| 9 | 37256992 | AX-168381223 | 0 | G | A | 0.491 | -5.95E-02 | 7.71E-02 | 2.77E+00 | 2.88E+00 | 4.34E-01 | no | Slc37a2 | upstream_gene_variant | MODIFIER |
| 1 | 67659687 | AX-170086489 | 1 | C | T | 0.125 | 1.44E-01 | 1.86E-01 | 2.67E+00 | 2.76E+00 | 4.34E-01 | no | | intergenic_variant | MODIFIER |
| 18 | 65442571 | AX-97843492 | 0 | A | G | 0.421 | 7.04E-02 | 8.59E-02 | 2.40E+00 | 2.45E+00 | 4.35E-01 | no | Malt1 | intron_variant | MODIFIER |
| 1 | 101397231 | AX-170490611 | 1 | T | C | 0.143 | 1.49E-01 | 1.92E-01 | 2.64E+00 | 2.72E+00 | 4.35E-01 | Yes | | intergenic_variant | MODIFIER |
| 3 | 37506180 | AX-170184892 | 0 | T | G | 0.202 | 6.96E-02 | 9.04E-02 | 2.78E+00 | 2.88E+00 | 4.35E-01 | no | Spata5 | intron_variant | MODIFIER |
| 3 | 36574840 | AX-170020311 | 1 | A | G | 0.188 | 7.32E-02 | 9.47E-02 | 2.83E+00 | 2.92E+00 | 4.35E-01 | no | Ccna2 | upstream_gene_variant | MODIFIER |
| 7 | 20882842 | AX-170028676 | 0 | T | C | 0.053 | -1.38E-01 | 1.79E-01 | 2.63E+00 | 2.78E+00 | 4.35E-01 | no | Vmn1r117 | downstream_gene_variant | MODIFIER |
| 9 | 117119444 | AX-170471178 | 0 | A | G | 0.123 | -7.55E-02 | 9.73E-02 | 2.60E+00 | 2.71E+00 | 4.35E-01 | no | Rbms3 | intron_variant | MODIFIER |
| 2 | 125108819 | AX-170498543 | 0 | T | A | 0.158 | 9.46E-02 | 1.23E-01 | 2.64E+00 | 2.78E+00 | 4.35E-01 | no | Myef2 | intron_variant | MODIFIER |
| 4 | 128071508 | AX-170510376 | 1 | C | T | 0.161 | 8.66E-02 | 1.11E-01 | 2.61E+00 | 2.67E+00 | 4.35E-01 | no | Csmd2 | intron_variant,non_coding_transcript_variant | MODIFIER |
| 19 | 51892643 | AX-170016818 | 0 | G | C | 0.482 | 5.08E-02 | 6.54E-02 | 2.60E+00 | 2.69E+00 | 4.35E-01 | no | AC122763.1 | intron_variant,non_coding_transcript_variant | MODIFIER |
| 16 | 86115866 | AX-170088086 | 1 | C | T | 0.464 | 6.02E-02 | 7.79E-02 | 2.65E+00 | 2.73E+00 | 4.35E-01 | no | | intergenic_variant | MODIFIER |
| 16 | 80448828 | AX-170093539 | 1 | A | G | 0.455 | 5.87E-02 | 7.55E-02 | 2.60E+00 | 2.67E+00 | 4.35E-01 | no | | intergenic_variant | MODIFIER |
| 1 | 140354419 | AX-170467290 | 0 | G | C | 0.202 | 9.28E-02 | 1.20E-01 | 2.65E+00 | 2.72E+00 | 4.36E-01 | no | Kcnt2 | intron_variant | MODIFIER |
| 4 | 128231051 | AX-170499998 | 1 | T | C | 0.143 | 9.52E-02 | 1.22E-01 | 2.57E+00 | 2.63E+00 | 4.36E-01 | no | Csmd2 | intron_variant,non_coding_transcript_variant | MODIFIER |
| 18 | 62045862 | AX-170447366 | 1 | A | G | 0.188 | -9.48E-02 | 1.23E-01 | 2.68E+00 | 2.73E+00 | 4.36E-01 | no | | intergenic_variant | MODIFIER |
| 18 | 66792716 | AX-170061158 | 1 | C | T | 0.188 | -9.48E-02 | 1.23E-01 | 2.68E+00 | 2.73E+00 | 4.36E-01 | no | AC133607.3 | intron_variant,non_coding_transcript_variant | MODIFIER |
| 18 | 67683691 | AX-170099780 | 1 | T | C | 0.188 | -9.48E-02 | 1.23E-01 | 2.68E+00 | 2.73E+00 | 4.36E-01 | no | Ptpn2 | intron_variant | MODIFIER |
| 7 | 76712388 | AX-170089958 | 1 | C | A | 0.357 | -7.52E-02 | 9.78E-02 | 2.79E+00 | 2.89E+00 | 4.36E-01 | no | Agbl1 | intron_variant | MODIFIER |
| 7 | 36661732 | AX-170297498 | 0 | C | G | 0.342 | 7.18E-02 | 9.16E-02 | 2.55E+00 | 2.62E+00 | 4.36E-01 | no | | intergenic_variant | MODIFIER |
| 8 | 36627657 | AX-97807531 | 0 | G | A | 0.158 | 1.05E-01 | 1.36E-01 | 2.85E+00 | 2.93E+00 | 4.36E-01 | no | Dlc1 | intron_variant | MODIFIER |
| 8 | 40797486 | AX-170137879 | 0 | G | C | 0.158 | 1.05E-01 | 1.36E-01 | 2.85E+00 | 2.93E+00 | 4.36E-01 | no | Adam20 | downstream_gene_variant | MODIFIER |

| | | | | | | | | | | | | | | | |
|---|---|---|---|---|---|---|---|---|---|---|---|---|---|---|---|
| 8 | 42935017 | AX-170079904 | 0 | T | A | 0.158 | 1.05E-01 | 1.36E-01 | 2.85E+00 | 2.93E+00 | 4.36E-01 | no | | intergenic_variant | MODIFIER |
| 8 | 45563909 | AX-170492694 | 0 | T | C | 0.158 | 1.05E-01 | 1.36E-01 | 2.85E+00 | 2.93E+00 | 4.36E-01 | no | Sorbs2 | intron_variant | MODIFIER |
| 1 | 124855797 | AX-170434311 | 2 | A | T | 0.555 | 6.31E-02 | 8.07E-02 | 2.56E+00 | 2.63E+00 | 4.36E-01 | no | | intergenic_variant | MODIFIER |
| 6 | 37709328 | AX-170508055 | 0 | A | C | 0.035 | -1.59E-01 | 2.04E-01 | 2.58E+00 | 2.68E+00 | 4.36E-01 | no | | intergenic_variant | MODIFIER |
| 8 | 25983097 | AX-170164192 | 0 | T | G | 0.351 | -5.16E-02 | 6.59E-02 | 2.54E+00 | 2.63E+00 | 4.36E-01 | no | Pomk | missense_variant | MODERATE |
| 14 | 111242953 | AX-170054459 | 2 | A | G | 0.309 | 5.66E-02 | 7.30E-02 | 2.59E+00 | 2.70E+00 | 4.37E-01 | no | | intergenic_variant | MODIFIER |
| 7 | 73946665 | AX-170077728 | 0 | C | G | 0.377 | -7.65E-02 | 9.98E-02 | 2.72E+00 | 2.81E+00 | 4.37E-01 | no | St8sia2 | intron_variant | MODIFIER |
| 10 | 68340621 | AX-170391752 | 0 | T | C | 0.404 | -6.29E-02 | 8.17E-02 | 2.67E+00 | 2.74E+00 | 4.37E-01 | no | | intergenic_variant | MODIFIER |
| 5 | 114046183 | AX-170146567 | 1 | T | A | 0.464 | 9.18E-02 | 1.19E-01 | 2.63E+00 | 2.65E+00 | 4.37E-01 | no | Svop | intron_variant | MODIFIER |
| 5 | 117391643 | AX-97788564 | 1 | A | G | 0.464 | 9.18E-02 | 1.19E-01 | 2.63E+00 | 2.65E+00 | 4.37E-01 | no | Rfc5 | upstream_gene_variant | MODIFIER |
| 8 | 65992768 | AX-170387102 | 1 | A | G | 0.143 | -1.12E-01 | 1.44E-01 | 2.57E+00 | 2.64E+00 | 4.37E-01 | no | MAR1 | intron_variant | MODIFIER |
| 6 | 134086984 | AX-170191550 | 1 | T | C | 0.554 | 6.91E-02 | 8.69E-02 | 2.48E+00 | 2.53E+00 | 4.37E-01 | no | Etv6 | intron_variant | MODIFIER |
| 16 | 38018456 | AX-170412590 | 0 | G | C | 0.5 | 6.27E-02 | 8.02E-02 | 2.55E+00 | 2.61E+00 | 4.37E-01 | no | | regulatory_region_variant | MODIFIER |
| 3 | 151557651 | AX-170367635 | 2 | C | T | 0.355 | 5.52E-02 | 7.21E-02 | 2.75E+00 | 2.84E+00 | 4.37E-01 | no | | regulatory_region_variant | MODIFIER |
| 11 | 14022364 | AX-170359858 | 0 | T | C | 0.044 | -1.19E-01 | 1.55E-01 | 2.57E+00 | 2.73E+00 | 4.37E-01 | no | | intergenic_variant | MODIFIER |
| 5 | 113971947 | AX-170233646 | 1 | A | G | 0.446 | 7.76E-02 | 1.01E-01 | 2.62E+00 | 2.70E+00 | 4.37E-01 | no | Ssh1 | intron_variant | MODIFIER |
| 4 | 137922701 | AX-170476737 | 1 | A | G | 0.152 | 8.57E-02 | 1.11E-01 | 2.60E+00 | 2.69E+00 | 4.37E-01 | no | Ece1 | intron_variant | MODIFIER |
| 10 | 100224511 | AX-170003762 | 0 | C | G | 0.281 | -6.70E-02 | 8.58E-02 | 2.55E+00 | 2.62E+00 | 4.38E-01 | no | | intergenic_variant | MODIFIER |
| 5 | 37521529 | AX-170215606 | 0 | T | C | 0.333 | 7.89E-02 | 1.03E-01 | 2.69E+00 | 2.75E+00 | 4.38E-01 | no | Stk32b | intron_variant | MODIFIER |
| 7 | 123777537 | AX-97795649 | 0 | T | C | 0.404 | 6.54E-02 | 8.53E-02 | 2.77E+00 | 2.85E+00 | 4.38E-01 | no | | intergenic_variant | MODIFIER |
| 1 | 124327172 | AX-170086602 | 0 | T | C | 0.561 | 6.48E-02 | 8.31E-02 | 2.55E+00 | 2.62E+00 | 4.38E-01 | no | | intergenic_variant | MODIFIER |
| 3 | 149225160 | AX-97845740 | 0 | C | T | 0.351 | 5.65E-02 | 7.37E-02 | 2.80E+00 | 2.89E+00 | 4.38E-01 | no | Gm10287 | non_coding_transcript_exon_variant | MODIFIER |
| 4 | 59048168 | AX-97823697 | 1 | A | G | 0.366 | 8.55E-02 | 1.11E-01 | 2.66E+00 | 2.72E+00 | 4.38E-01 | no | AI481877 | synonymous_variant | LOW |
| 5 | 47774233 | AX-170199182 | 1 | G | C | 0.491 | 7.97E-02 | 9.97E-02 | 2.45E+00 | 2.52E+00 | 4.38E-01 | no | | intergenic_variant | MODIFIER |
| 18 | 83897140 | AX-170234424 | 1 | C | T | 0.161 | -8.55E-02 | 1.08E-01 | 2.47E+00 | 2.55E+00 | 4.38E-01 | no | | intergenic_variant | MODIFIER |
| 18 | 84161995 | AX-170445736 | 1 | C | T | 0.161 | -8.55E-02 | 1.08E-01 | 2.47E+00 | 2.55E+00 | 4.38E-01 | no | Zfp407 | intron_variant | MODIFIER |
| 4 | 148700998 | AX-170142909 | 0 | G | A | 0.096 | -1.31E-01 | 1.71E-01 | 2.70E+00 | 2.79E+00 | 4.38E-01 | no | | intergenic_variant | MODIFIER |
| 10 | 108749055 | AX-170164368 | 2 | G | T | 0.327 | -6.17E-02 | 8.05E-02 | 2.68E+00 | 2.76E+00 | 4.38E-01 | no | Syt1 | intron_variant | MODIFIER |
| 13 | 24963050 | AX-170139855 | 0 | A | G | 0.167 | -9.94E-02 | 1.23E-01 | 2.43E+00 | 2.48E+00 | 4.38E-01 | no | Gpld1 | intron_variant | MODIFIER |
| 13 | 24969218 | AX-170007940 | 0 | G | A | 0.167 | -9.94E-02 | 1.23E-01 | 2.43E+00 | 2.48E+00 | 4.38E-01 | no | Gpld1 | intron_variant | MODIFIER |
| 13 | 24969528 | AX-97809249 | 0 | G | T | 0.167 | -9.94E-02 | 1.23E-01 | 2.43E+00 | 2.48E+00 | 4.38E-01 | no | Gpld1 | intron_variant | MODIFIER |
| 13 | 24976543 | AX-170295991 | 0 | C | T | 0.167 | -9.94E-02 | 1.23E-01 | 2.43E+00 | 2.48E+00 | 4.38E-01 | no | Gpld1 | intron_variant | MODIFIER |
| 13 | 24981114 | AX-170050774 | 0 | G | A | 0.167 | -9.94E-02 | 1.23E-01 | 2.43E+00 | 2.48E+00 | 4.38E-01 | no | Gpld1 | intron_variant | MODIFIER |
| 13 | 24992415 | AX-170294783 | 0 | A | T | 0.167 | -9.94E-02 | 1.23E-01 | 2.43E+00 | 2.48E+00 | 4.38E-01 | no | Mrs2 | 3_prime_UTR_variant | MODIFIER |
| 13 | 24992690 | AX-170426633 | 0 | G | A | 0.167 | -9.94E-02 | 1.23E-01 | 2.43E+00 | 2.48E+00 | 4.38E-01 | no | Mrs2 | 3_prime_UTR_variant | MODIFIER |
| 13 | 24992772 | AX-170254106 | 0 | G | A | 0.167 | -9.94E-02 | 1.23E-01 | 2.43E+00 | 2.48E+00 | 4.38E-01 | no | Mrs2 | 3_prime_UTR_variant | MODIFIER |
| 13 | 24993541 | AX-97844294 | 0 | C | A | 0.167 | -9.94E-02 | 1.23E-01 | 2.43E+00 | 2.48E+00 | 4.38E-01 | no | Mrs2 | intron_variant | MODIFIER |
| 13 | 25007330 | AX-170007941 | 0 | C | T | 0.167 | -9.94E-02 | 1.23E-01 | 2.43E+00 | 2.48E+00 | 4.38E-01 | no | Mrs2 | intron_variant | MODIFIER |
| 13 | 25025685 | AX-170090926 | 0 | C | T | 0.167 | -9.94E-02 | 1.23E-01 | 2.43E+00 | 2.48E+00 | 4.38E-01 | no | | intergenic_variant | MODIFIER |
| 13 | 25038497 | AX-97855525 | 0 | A | G | 0.167 | -9.94E-02 | 1.23E-01 | 2.43E+00 | 2.48E+00 | 4.38E-01 | no | Gm47605 | downstream_gene_variant | MODIFIER |
| 13 | 25058616 | AX-97794308 | 0 | T | G | 0.167 | -9.94E-02 | 1.23E-01 | 2.43E+00 | 2.48E+00 | 4.38E-01 | no | Dcdc2a | intron_variant | MODIFIER |
| 13 | 25058892 | AX-170525432 | 0 | A | G | 0.167 | -9.94E-02 | 1.23E-01 | 2.43E+00 | 2.48E+00 | 4.38E-01 | no | Dcdc2a | intron_variant | MODIFIER |
| 13 | 25073281 | AX-170338073 | 0 | T | C | 0.167 | -9.94E-02 | 1.23E-01 | 2.43E+00 | 2.48E+00 | 4.38E-01 | no | Dcdc2a | intron_variant | MODIFIER |
| 13 | 25090613 | AX-170406362 | 0 | T | C | 0.167 | -9.94E-02 | 1.23E-01 | 2.43E+00 | 2.48E+00 | 4.38E-01 | no | Dcdc2a | intron_variant | MODIFIER |
| 13 | 25093569 | AX-97855756 | 0 | C | A | 0.167 | -9.94E-02 | 1.23E-01 | 2.43E+00 | 2.48E+00 | 4.38E-01 | no | Dcdc2a | intron_variant | MODIFIER |
| 13 | 25102362 | AX-170515705 | 0 | C | T | 0.167 | -9.94E-02 | 1.23E-01 | 2.43E+00 | 2.48E+00 | 4.38E-01 | no | Dcdc2a | synonymous_variant | LOW |
| 13 | 25121207 | AX-170322470 | 0 | G | A | 0.167 | -9.94E-02 | 1.23E-01 | 2.43E+00 | 2.48E+00 | 4.38E-01 | no | Dcdc2a | 3_prime_UTR_variant | MODIFIER |
| 13 | 25130600 | AX-97823149 | 0 | G | A | 0.167 | -9.94E-02 | 1.23E-01 | 2.43E+00 | 2.48E+00 | 4.38E-01 | no | Dcdc2a | intron_variant | MODIFIER |
| 13 | 25131524 | AX-170389744 | 0 | C | T | 0.167 | -9.94E-02 | 1.23E-01 | 2.43E+00 | 2.48E+00 | 4.38E-01 | no | Dcdc2a | intron_variant | MODIFIER |
| 13 | 25135879 | AX-170421301 | 0 | G | C | 0.167 | -9.94E-02 | 1.23E-01 | 2.43E+00 | 2.48E+00 | 4.38E-01 | no | Dcdc2a | intron_variant | MODIFIER |
| 13 | 25143950 | AX-170307154 | 0 | C | T | 0.167 | -9.94E-02 | 1.23E-01 | 2.43E+00 | 2.48E+00 | 4.38E-01 | no | Dcdc2a | intron_variant | MODIFIER |
| 13 | 25145308 | AX-97786173 | 0 | A | G | 0.167 | -9.94E-02 | 1.23E-01 | 2.43E+00 | 2.48E+00 | 4.38E-01 | no | Dcdc2a | intron_variant | MODIFIER |
| 13 | 25147535 | AX-170389743 | 0 | T | A | 0.167 | -9.94E-02 | 1.23E-01 | 2.43E+00 | 2.48E+00 | 4.38E-01 | no | Dcdc2a | intron_variant | MODIFIER |
| 13 | 25159985 | AX-170123886 | 0 | C | T | 0.167 | -9.94E-02 | 1.23E-01 | 2.43E+00 | 2.48E+00 | 4.38E-01 | no | Dcdc2a | intron_variant | MODIFIER |

| | | | | | | | | | | | | | | | |
|---|---|---|---|---|---|---|---|---|---|---|---|---|---|---|---|
| 13 | 25171008 | AX-97833965 | 0 | G | A | 0.167 | -9.94E-02 | 1.23E-01 | 2.43E+00 | 2.48E+00 | 4.38E-01 | no | Dcdc2a | intron_variant | MODIFIER |
| 13 | 25171361 | AX-170315796 | 0 | T | C | 0.167 | -9.94E-02 | 1.23E-01 | 2.43E+00 | 2.48E+00 | 4.38E-01 | no | Dcdc2a | intron_variant | MODIFIER |
| 13 | 25217370 | AX-97822994 | 0 | C | T | 0.167 | -9.94E-02 | 1.23E-01 | 2.43E+00 | 2.48E+00 | 4.38E-01 | no | | intergenic_variant | MODIFIER |
| 13 | 25251405 | AX-170224792 | 0 | C | A | 0.167 | -9.94E-02 | 1.23E-01 | 2.43E+00 | 2.48E+00 | 4.38E-01 | no | Nrsn1 | downstream_gene_variant | MODIFIER |
| 13 | 25252168 | AX-170334711 | 0 | T | C | 0.167 | -9.94E-02 | 1.23E-01 | 2.43E+00 | 2.48E+00 | 4.38E-01 | no | Nrsn1 | 3_prime_UTR_variant | MODIFIER |
| 13 | 25258177 | AX-170036880 | 0 | G | A | 0.167 | -9.94E-02 | 1.23E-01 | 2.43E+00 | 2.48E+00 | 4.38E-01 | no | Nrsn1 | intron_variant | MODIFIER |
| 13 | 25262592 | AX-170420841 | 0 | C | T | 0.167 | -9.94E-02 | 1.23E-01 | 2.43E+00 | 2.48E+00 | 4.38E-01 | no | Nrsn1 | intron_variant | MODIFIER |
| 13 | 25265705 | AX-170500602 | 0 | G | A | 0.167 | -9.94E-02 | 1.23E-01 | 2.43E+00 | 2.48E+00 | 4.38E-01 | no | Nrsn1 | intron_variant | MODIFIER |
| 13 | 25275792 | AX-97836685 | 0 | C | T | 0.167 | -9.94E-02 | 1.23E-01 | 2.43E+00 | 2.48E+00 | 4.38E-01 | no | | intergenic_variant | MODIFIER |
| 13 | 25275891 | AX-170358033 | 0 | T | C | 0.167 | -9.94E-02 | 1.23E-01 | 2.43E+00 | 2.48E+00 | 4.38E-01 | no | | intergenic_variant | MODIFIER |
| 13 | 25290387 | AX-170107984 | 0 | G | A | 0.167 | -9.94E-02 | 1.23E-01 | 2.43E+00 | 2.48E+00 | 4.38E-01 | no | | intergenic_variant | MODIFIER |
| 13 | 25290589 | AX-170502904 | 0 | T | C | 0.167 | -9.94E-02 | 1.23E-01 | 2.43E+00 | 2.48E+00 | 4.38E-01 | no | | intergenic_variant | MODIFIER |
| 13 | 25291486 | AX-170240467 | 0 | A | C | 0.167 | -9.94E-02 | 1.23E-01 | 2.43E+00 | 2.48E+00 | 4.38E-01 | no | | intergenic_variant | MODIFIER |
| 13 | 25291767 | AX-97820013 | 0 | G | A | 0.167 | -9.94E-02 | 1.23E-01 | 2.43E+00 | 2.48E+00 | 4.38E-01 | no | | intergenic_variant | MODIFIER |
| 13 | 25324613 | AX-97849529 | 0 | G | T | 0.167 | -9.94E-02 | 1.23E-01 | 2.43E+00 | 2.48E+00 | 4.38E-01 | no | | intergenic_variant | MODIFIER |
| 13 | 25341574 | AX-97847268 | 0 | T | C | 0.167 | -9.94E-02 | 1.23E-01 | 2.43E+00 | 2.48E+00 | 4.38E-01 | no | | intergenic_variant | MODIFIER |
| 13 | 25368650 | AX-97798081 | 0 | A | G | 0.167 | -9.94E-02 | 1.23E-01 | 2.43E+00 | 2.48E+00 | 4.38E-01 | no | | intergenic_variant | MODIFIER |
| 13 | 25375000 | AX-170389733 | 0 | A | G | 0.167 | -9.94E-02 | 1.23E-01 | 2.43E+00 | 2.48E+00 | 4.38E-01 | no | | intergenic_variant | MODIFIER |
| 13 | 25386410 | AX-170493095 | 0 | T | G | 0.167 | -9.94E-02 | 1.23E-01 | 2.43E+00 | 2.48E+00 | 4.38E-01 | no | | intergenic_variant | MODIFIER |
| 13 | 25386432 | AX-170289243 | 0 | A | G | 0.167 | -9.94E-02 | 1.23E-01 | 2.43E+00 | 2.48E+00 | 4.38E-01 | no | | intergenic_variant | MODIFIER |
| 13 | 25395155 | AX-97791076 | 0 | C | A | 0.167 | -9.94E-02 | 1.23E-01 | 2.43E+00 | 2.48E+00 | 4.38E-01 | no | | intergenic_variant | MODIFIER |
| 13 | 25404549 | AX-170007950 | 0 | G | C | 0.167 | -9.94E-02 | 1.23E-01 | 2.43E+00 | 2.48E+00 | 4.38E-01 | no | | intergenic_variant | MODIFIER |
| 13 | 25491227 | AX-170007951 | 0 | T | C | 0.167 | -9.94E-02 | 1.23E-01 | 2.43E+00 | 2.48E+00 | 4.38E-01 | no | | intergenic_variant | MODIFIER |
| 13 | 25491560 | AX-170050785 | 0 | T | C | 0.167 | -9.94E-02 | 1.23E-01 | 2.43E+00 | 2.48E+00 | 4.38E-01 | no | | intergenic_variant | MODIFIER |
| 13 | 25492803 | AX-170050786 | 0 | A | C | 0.167 | -9.94E-02 | 1.23E-01 | 2.43E+00 | 2.48E+00 | 4.38E-01 | no | | intergenic_variant | MODIFIER |
| 13 | 25494244 | AX-97781428 | 0 | G | A | 0.167 | -9.94E-02 | 1.23E-01 | 2.43E+00 | 2.48E+00 | 4.38E-01 | no | | intergenic_variant | MODIFIER |
| 13 | 25498578 | AX-170240000 | 0 | A | G | 0.167 | -9.94E-02 | 1.23E-01 | 2.43E+00 | 2.48E+00 | 4.38E-01 | no | | intergenic_variant | MODIFIER |
| 13 | 25498982 | AX-170236464 | 0 | T | C | 0.167 | -9.94E-02 | 1.23E-01 | 2.43E+00 | 2.48E+00 | 4.38E-01 | no | | intergenic_variant | MODIFIER |
| 13 | 25551173 | AX-170224293 | 0 | C | T | 0.167 | -9.94E-02 | 1.23E-01 | 2.43E+00 | 2.48E+00 | 4.38E-01 | no | | intergenic_variant | MODIFIER |
| 13 | 25556456 | AX-170458452 | 0 | T | C | 0.167 | -9.94E-02 | 1.23E-01 | 2.43E+00 | 2.48E+00 | 4.38E-01 | no | | intergenic_variant | MODIFIER |
| 7 | 113988586 | AX-170029797 | 0 | G | A | 0.421 | 7.38E-02 | 9.63E-02 | 2.72E+00 | 2.77E+00 | 4.38E-01 | no | Spon1 | intron_variant | MODIFIER |
| 3 | 36503638 | AX-97816683 | 1 | A | G | 0.188 | 7.15E-02 | 9.35E-02 | 2.80E+00 | 2.89E+00 | 4.39E-01 | no | Gm11548 | intron_variant,non_coding_transcript_variant | MODIFIER |
| 3 | 37740359 | AX-170020321 | 1 | T | C | 0.188 | 7.15E-02 | 9.35E-02 | 2.80E+00 | 2.89E+00 | 4.39E-01 | no | | intergenic_variant | MODIFIER |
| 7 | 46265462 | AX-170077224 | 0 | A | G | 0.325 | 7.34E-02 | 9.57E-02 | 2.64E+00 | 2.73E+00 | 4.39E-01 | no | Otog | intron_variant | MODIFIER |
| 2 | 168635317 | AX-170019645 | 0 | G | A | 0.167 | 1.20E-01 | 1.57E-01 | 2.66E+00 | 2.77E+00 | 4.39E-01 | no | Atp9a | intron_variant | MODIFIER |
| 16 | 91987269 | AX-170137305 | 2 | T | A | 0.445 | -5.92E-02 | 7.75E-02 | 2.73E+00 | 2.81E+00 | 4.39E-01 | no | | intergenic_variant | MODIFIER |
| 2 | 122089137 | AX-170260006 | 1 | C | T | 0.152 | 1.26E-01 | 1.66E-01 | 2.71E+00 | 2.86E+00 | 4.39E-01 | no | Spg11 | intron_variant | MODIFIER |
| 9 | 117904426 | AX-170254198 | 1 | C | T | 0.125 | -7.51E-02 | 9.80E-02 | 2.62E+00 | 2.74E+00 | 4.39E-01 | no | Zcwpw2 | intron_variant,non_coding_transcript_variant | MODIFIER |
| 18 | 53799716 | AX-97786679 | 0 | T | C | 0.395 | 7.52E-02 | 9.20E-02 | 2.38E+00 | 2.45E+00 | 4.39E-01 | no | | intergenic_variant | MODIFIER |
| 6 | 136229658 | AX-170333093 | 1 | A | T | 0.571 | 6.95E-02 | 8.82E-02 | 2.50E+00 | 2.56E+00 | 4.39E-01 | no | | intergenic_variant | MODIFIER |
| 12 | 57394578 | AX-97789685 | 0 | T | C | 0.07 | 1.70E-01 | 2.23E-01 | 2.68E+00 | 2.78E+00 | 4.39E-01 | Yes | Mipol1 | intron_variant | MODIFIER |
| 12 | 66660284 | AX-170399487 | 0 | G | T | 0.07 | 1.70E-01 | 2.23E-01 | 2.68E+00 | 2.78E+00 | 4.39E-01 | Yes | Mdga2 | intron_variant | MODIFIER |
| 12 | 72380775 | AX-170329041 | 0 | T | C | 0.07 | 1.70E-01 | 2.23E-01 | 2.68E+00 | 2.78E+00 | 4.39E-01 | Yes | Rtn1 | intron_variant | MODIFIER |
| 18 | 59565044 | AX-170473590 | 0 | G | A | 0.412 | -6.39E-02 | 8.28E-02 | 2.61E+00 | 2.67E+00 | 4.39E-01 | no | | intergenic_variant | MODIFIER |
| 18 | 87406017 | AX-170182685 | 0 | A | C | 0.149 | -8.14E-02 | 1.03E-01 | 2.49E+00 | 2.59E+00 | 4.39E-01 | no | | intergenic_variant | MODIFIER |
| 18 | 59256178 | AX-170501384 | 0 | C | T | 0.404 | 6.97E-02 | 8.48E-02 | 2.36E+00 | 2.44E+00 | 4.39E-01 | no | Chsy3 | intron_variant | MODIFIER |
| 5 | 77530655 | AX-170484998 | 0 | T | G | 0.421 | 8.15E-02 | 1.07E-01 | 2.72E+00 | 2.77E+00 | 4.39E-01 | no | | intergenic_variant | MODIFIER |
| 5 | 77530739 | AX-170464477 | 0 | A | G | 0.421 | 8.15E-02 | 1.07E-01 | 2.72E+00 | 2.77E+00 | 4.39E-01 | no | | intergenic_variant | MODIFIER |
| 5 | 77644616 | AX-170337596 | 0 | T | C | 0.421 | 8.15E-02 | 1.07E-01 | 2.72E+00 | 2.77E+00 | 4.39E-01 | no | | intergenic_variant | MODIFIER |
| 5 | 77644769 | AX-170307983 | 0 | A | C | 0.421 | 8.15E-02 | 1.07E-01 | 2.72E+00 | 2.77E+00 | 4.39E-01 | no | | intergenic_variant | MODIFIER |
| 5 | 77675761 | AX-170317492 | 0 | G | A | 0.421 | 8.15E-02 | 1.07E-01 | 2.72E+00 | 2.77E+00 | 4.39E-01 | no | | intergenic_variant | MODIFIER |
| 5 | 77792553 | AX-170246290 | 0 | T | C | 0.421 | 8.15E-02 | 1.07E-01 | 2.72E+00 | 2.77E+00 | 4.39E-01 | no | | intergenic_variant | MODIFIER |
| 5 | 77794106 | AX-170463059 | 0 | C | T | 0.421 | 8.15E-02 | 1.07E-01 | 2.72E+00 | 2.77E+00 | 4.39E-01 | no | | intergenic_variant | MODIFIER |
| 5 | 77801818 | AX-170095366 | 0 | G | A | 0.421 | 8.15E-02 | 1.07E-01 | 2.72E+00 | 2.77E+00 | 4.39E-01 | no | | intergenic_variant | MODIFIER |

| | | | | | | | | | | | | | | | |
|---|---|---|---|---|---|---|---|---|---|---|---|---|---|---|---|
| 5 | 77841836 | AX-170144982 | 0 | C | T | 0.421 | 8.15E-02 | 1.07E-01 | 2.72E+00 | 2.77E+00 | 4.39E-01 | no | | intergenic_variant | MODIFIER |
| 5 | 77842036 | AX-97783805 | 0 | G | A | 0.421 | 8.15E-02 | 1.07E-01 | 2.72E+00 | 2.77E+00 | 4.39E-01 | no | | intergenic_variant | MODIFIER |
| 5 | 77863329 | AX-170152476 | 0 | T | C | 0.421 | 8.15E-02 | 1.07E-01 | 2.72E+00 | 2.77E+00 | 4.39E-01 | no | | intergenic_variant | MODIFIER |
| 5 | 77894974 | AX-170371031 | 0 | A | T | 0.421 | 8.15E-02 | 1.07E-01 | 2.72E+00 | 2.77E+00 | 4.39E-01 | no | | intergenic_variant | MODIFIER |
| 5 | 78390834 | AX-170418585 | 0 | C | T | 0.421 | 8.15E-02 | 1.07E-01 | 2.72E+00 | 2.77E+00 | 4.39E-01 | no | | intergenic_variant | MODIFIER |
| 5 | 78393227 | AX-170222731 | 0 | T | C | 0.421 | 8.15E-02 | 1.07E-01 | 2.72E+00 | 2.77E+00 | 4.39E-01 | no | | intergenic_variant | MODIFIER |
| 5 | 78394426 | AX-170336395 | 0 | A | T | 0.421 | 8.15E-02 | 1.07E-01 | 2.72E+00 | 2.77E+00 | 4.39E-01 | no | | intergenic_variant | MODIFIER |
| 5 | 78394936 | AX-170072566 | 0 | T | C | 0.421 | 8.15E-02 | 1.07E-01 | 2.72E+00 | 2.77E+00 | 4.39E-01 | no | | intergenic_variant | MODIFIER |
| 5 | 78395210 | AX-170127198 | 0 | C | T | 0.421 | 8.15E-02 | 1.07E-01 | 2.72E+00 | 2.77E+00 | 4.39E-01 | no | | intergenic_variant | MODIFIER |
| 5 | 78410704 | AX-170148991 | 0 | C | T | 0.421 | 8.15E-02 | 1.07E-01 | 2.72E+00 | 2.77E+00 | 4.39E-01 | no | | intergenic_variant | MODIFIER |
| 5 | 78429956 | AX-170095368 | 0 | C | T | 0.421 | 8.15E-02 | 1.07E-01 | 2.72E+00 | 2.77E+00 | 4.39E-01 | no | | intergenic_variant | MODIFIER |
| 5 | 78485044 | AX-97824562 | 0 | C | T | 0.421 | 8.15E-02 | 1.07E-01 | 2.72E+00 | 2.77E+00 | 4.39E-01 | no | | intergenic_variant | MODIFIER |
| 5 | 78485104 | AX-170025359 | 0 | A | C | 0.421 | 8.15E-02 | 1.07E-01 | 2.72E+00 | 2.77E+00 | 4.39E-01 | no | | intergenic_variant | MODIFIER |
| 5 | 78506435 | AX-170025353 | 0 | G | C | 0.421 | 8.15E-02 | 1.07E-01 | 2.72E+00 | 2.77E+00 | 4.39E-01 | no | | intergenic_variant | MODIFIER |
| 5 | 78509976 | AX-97848019 | 0 | A | C | 0.421 | 8.15E-02 | 1.07E-01 | 2.72E+00 | 2.77E+00 | 4.39E-01 | no | | intergenic_variant | MODIFIER |
| 5 | 78519576 | AX-170025355 | 0 | A | C | 0.421 | 8.15E-02 | 1.07E-01 | 2.72E+00 | 2.77E+00 | 4.39E-01 | no | | intergenic_variant | MODIFIER |
| 5 | 78539158 | AX-97788938 | 0 | G | A | 0.421 | 8.15E-02 | 1.07E-01 | 2.72E+00 | 2.77E+00 | 4.39E-01 | no | Gm43232 | upstream_gene_variant | MODIFIER |
| 5 | 78547006 | AX-170196166 | 0 | G | A | 0.421 | 8.15E-02 | 1.07E-01 | 2.72E+00 | 2.77E+00 | 4.39E-01 | no | | intergenic_variant | MODIFIER |
| 5 | 78609007 | AX-170072569 | 0 | G | T | 0.421 | 8.15E-02 | 1.07E-01 | 2.72E+00 | 2.77E+00 | 4.39E-01 | no | | intergenic_variant | MODIFIER |
| 5 | 78628077 | AX-170025350 | 0 | C | A | 0.421 | 8.15E-02 | 1.07E-01 | 2.72E+00 | 2.77E+00 | 4.39E-01 | no | | intergenic_variant | MODIFIER |
| 5 | 78644746 | AX-170072568 | 0 | T | C | 0.421 | 8.15E-02 | 1.07E-01 | 2.72E+00 | 2.77E+00 | 4.39E-01 | no | | intergenic_variant | MODIFIER |
| 5 | 78663273 | AX-97854856 | 0 | C | T | 0.421 | 8.15E-02 | 1.07E-01 | 2.72E+00 | 2.77E+00 | 4.39E-01 | no | | intergenic_variant | MODIFIER |
| 5 | 78663316 | AX-170207935 | 0 | T | C | 0.421 | 8.15E-02 | 1.07E-01 | 2.72E+00 | 2.77E+00 | 4.39E-01 | no | | intergenic_variant | MODIFIER |
| 5 | 78712256 | AX-170355458 | 0 | C | A | 0.421 | 8.15E-02 | 1.07E-01 | 2.72E+00 | 2.77E+00 | 4.39E-01 | no | | intergenic_variant | MODIFIER |
| 5 | 78712708 | AX-170536129 | 0 | G | C | 0.421 | 8.15E-02 | 1.07E-01 | 2.72E+00 | 2.77E+00 | 4.39E-01 | no | | intergenic_variant | MODIFIER |
| 5 | 78713129 | AX-170508161 | 0 | T | C | 0.421 | 8.15E-02 | 1.07E-01 | 2.72E+00 | 2.77E+00 | 4.39E-01 | no | | intergenic_variant | MODIFIER |
| 5 | 78713205 | AX-170272936 | 0 | G | A | 0.421 | 8.15E-02 | 1.07E-01 | 2.72E+00 | 2.77E+00 | 4.39E-01 | no | | intergenic_variant | MODIFIER |
| 5 | 78713662 | AX-97796673 | 0 | G | A | 0.421 | 8.15E-02 | 1.07E-01 | 2.72E+00 | 2.77E+00 | 4.39E-01 | no | | intergenic_variant | MODIFIER |
| 5 | 78754526 | AX-170313281 | 0 | T | A | 0.421 | 8.15E-02 | 1.07E-01 | 2.72E+00 | 2.77E+00 | 4.39E-01 | no | | intergenic_variant | MODIFIER |
| 5 | 78760659 | AX-97801360 | 0 | T | C | 0.421 | 8.15E-02 | 1.07E-01 | 2.72E+00 | 2.77E+00 | 4.39E-01 | no | | intergenic_variant | MODIFIER |
| 5 | 78784560 | AX-97828599 | 0 | G | T | 0.421 | 8.15E-02 | 1.07E-01 | 2.72E+00 | 2.77E+00 | 4.39E-01 | no | Gm25756 | downstream_gene_variant | MODIFIER |
| 5 | 78810247 | AX-170324195 | 0 | C | T | 0.421 | 8.15E-02 | 1.07E-01 | 2.72E+00 | 2.77E+00 | 4.39E-01 | no | | intergenic_variant | MODIFIER |
| 5 | 78842952 | AX-97821616 | 0 | G | T | 0.421 | 8.15E-02 | 1.07E-01 | 2.72E+00 | 2.77E+00 | 4.39E-01 | no | | intergenic_variant | MODIFIER |
| 5 | 78844758 | AX-97784470 | 0 | G | A | 0.421 | 8.15E-02 | 1.07E-01 | 2.72E+00 | 2.77E+00 | 4.39E-01 | no | | intergenic_variant | MODIFIER |
| 5 | 78890837 | AX-170134104 | 0 | T | C | 0.421 | 8.15E-02 | 1.07E-01 | 2.72E+00 | 2.77E+00 | 4.39E-01 | no | | intergenic_variant | MODIFIER |
| 5 | 78968553 | AX-170025369 | 0 | C | A | 0.421 | 8.15E-02 | 1.07E-01 | 2.72E+00 | 2.77E+00 | 4.39E-01 | no | | intergenic_variant | MODIFIER |
| 5 | 79581718 | AX-170025383 | 0 | G | C | 0.421 | 8.15E-02 | 1.07E-01 | 2.72E+00 | 2.77E+00 | 4.39E-01 | no | | intergenic_variant | MODIFIER |
| 5 | 79596360 | AX-170025376 | 0 | A | G | 0.421 | 8.15E-02 | 1.07E-01 | 2.72E+00 | 2.77E+00 | 4.39E-01 | no | | intergenic_variant | MODIFIER |
| 5 | 79623083 | AX-170025380 | 0 | T | C | 0.421 | 8.15E-02 | 1.07E-01 | 2.72E+00 | 2.77E+00 | 4.39E-01 | no | | intergenic_variant | MODIFIER |
| 5 | 79626891 | AX-97832324 | 0 | G | A | 0.421 | 8.15E-02 | 1.07E-01 | 2.72E+00 | 2.77E+00 | 4.39E-01 | no | | intergenic_variant | MODIFIER |
| 5 | 79777395 | AX-170279343 | 0 | A | C | 0.421 | 8.15E-02 | 1.07E-01 | 2.72E+00 | 2.77E+00 | 4.39E-01 | no | | intergenic_variant | MODIFIER |
| 5 | 79777670 | AX-170348700 | 0 | A | G | 0.421 | 8.15E-02 | 1.07E-01 | 2.72E+00 | 2.77E+00 | 4.39E-01 | no | | intergenic_variant | MODIFIER |
| 5 | 79783571 | AX-170209955 | 0 | G | A | 0.421 | 8.15E-02 | 1.07E-01 | 2.72E+00 | 2.77E+00 | 4.39E-01 | no | | intergenic_variant | MODIFIER |
| 5 | 79791879 | AX-97836761 | 0 | G | A | 0.421 | 8.15E-02 | 1.07E-01 | 2.72E+00 | 2.77E+00 | 4.39E-01 | no | | intergenic_variant | MODIFIER |
| 5 | 79811079 | AX-97801931 | 0 | A | G | 0.421 | 8.15E-02 | 1.07E-01 | 2.72E+00 | 2.77E+00 | 4.39E-01 | no | | intergenic_variant | MODIFIER |
| 5 | 79871630 | AX-97790156 | 0 | C | T | 0.421 | 8.15E-02 | 1.07E-01 | 2.72E+00 | 2.77E+00 | 4.39E-01 | no | | intergenic_variant | MODIFIER |
| 5 | 79874796 | AX-170324413 | 0 | A | G | 0.421 | 8.15E-02 | 1.07E-01 | 2.72E+00 | 2.77E+00 | 4.39E-01 | no | | intergenic_variant | MODIFIER |
| 5 | 80069687 | AX-170436431 | 0 | T | C | 0.421 | 8.15E-02 | 1.07E-01 | 2.72E+00 | 2.77E+00 | 4.39E-01 | no | | intergenic_variant | MODIFIER |
| 5 | 80702550 | AX-170201797 | 0 | C | T | 0.421 | 8.15E-02 | 1.07E-01 | 2.72E+00 | 2.77E+00 | 4.39E-01 | no | | intergenic_variant | MODIFIER |
| 5 | 80712798 | AX-170379102 | 0 | G | A | 0.421 | 8.15E-02 | 1.07E-01 | 2.72E+00 | 2.77E+00 | 4.39E-01 | no | | intergenic_variant | MODIFIER |
| 5 | 81384867 | AX-170309522 | 0 | C | A | 0.421 | 8.15E-02 | 1.07E-01 | 2.72E+00 | 2.77E+00 | 4.39E-01 | no | Adgrl3 | intron_variant | MODIFIER |
| 5 | 81386875 | AX-97801414 | 0 | A | G | 0.421 | 8.15E-02 | 1.07E-01 | 2.72E+00 | 2.77E+00 | 4.39E-01 | no | Adgrl3 | intron_variant | MODIFIER |
| 5 | 81939714 | AX-170098701 | 0 | A | T | 0.421 | 8.15E-02 | 1.07E-01 | 2.72E+00 | 2.77E+00 | 4.39E-01 | no | | intergenic_variant | MODIFIER |
| 5 | 81940474 | AX-170115878 | 0 | A | G | 0.421 | 8.15E-02 | 1.07E-01 | 2.72E+00 | 2.77E+00 | 4.39E-01 | no | | intergenic_variant | MODIFIER |

| | | | | | | | | | | | | | | | |
|---|---|---|---|---|---|---|---|---|---|---|---|---|---|---|---|
| 5 | 81940533 | AX-170119779 | 0 | A | C | 0.421 | 8.15E-02 | 1.07E-01 | 2.72E+00 | 2.77E+00 | 4.39E-01 | no | | intergenic_variant | MODIFIER |
| 5 | 81967848 | AX-97847573 | 0 | T | C | 0.421 | 8.15E-02 | 1.07E-01 | 2.72E+00 | 2.77E+00 | 4.39E-01 | no | | intergenic_variant | MODIFIER |
| 5 | 82009348 | AX-170089534 | 0 | A | G | 0.421 | 8.15E-02 | 1.07E-01 | 2.72E+00 | 2.77E+00 | 4.39E-01 | no | | intergenic_variant | MODIFIER |
| 5 | 82019559 | AX-170089535 | 0 | G | A | 0.421 | 8.15E-02 | 1.07E-01 | 2.72E+00 | 2.77E+00 | 4.39E-01 | no | | intergenic_variant | MODIFIER |
| 5 | 82019605 | AX-170412403 | 0 | C | A | 0.421 | 8.15E-02 | 1.07E-01 | 2.72E+00 | 2.77E+00 | 4.39E-01 | no | | intergenic_variant | MODIFIER |
| 5 | 82044037 | AX-170141872 | 0 | C | A | 0.421 | 8.15E-02 | 1.07E-01 | 2.72E+00 | 2.77E+00 | 4.39E-01 | no | | intergenic_variant | MODIFIER |
| 5 | 82243731 | AX-170136482 | 0 | T | G | 0.421 | 8.15E-02 | 1.07E-01 | 2.72E+00 | 2.77E+00 | 4.39E-01 | no | | intergenic_variant | MODIFIER |
| 5 | 82281107 | AX-170189357 | 0 | T | G | 0.421 | 8.15E-02 | 1.07E-01 | 2.72E+00 | 2.77E+00 | 4.39E-01 | no | | intergenic_variant | MODIFIER |
| 5 | 82357322 | AX-170221058 | 0 | A | C | 0.421 | 8.15E-02 | 1.07E-01 | 2.72E+00 | 2.77E+00 | 4.39E-01 | no | | intergenic_variant | MODIFIER |
| 5 | 83065036 | AX-97827093 | 0 | T | G | 0.421 | 8.15E-02 | 1.07E-01 | 2.72E+00 | 2.77E+00 | 4.39E-01 | no | | intergenic_variant | MODIFIER |
| 16 | 29180486 | AX-170382336 | 0 | A | G | 0.491 | 6.17E-02 | 7.95E-02 | 2.57E+00 | 2.63E+00 | 4.39E-01 | no | | intergenic_variant | MODIFIER |
| 5 | 116964264 | AX-170292626 | 1 | G | A | 0.464 | 8.84E-02 | 1.15E-01 | 2.62E+00 | 2.66E+00 | 4.39E-01 | no | | intergenic_variant | MODIFIER |
| 4 | 90576216 | AX-170395867 | 0 | C | G | 0.254 | 8.85E-02 | 1.12E-01 | 2.50E+00 | 2.55E+00 | 4.40E-01 | no | | intergenic_variant | MODIFIER |
| 1 | 187921305 | AX-170159099 | 1 | G | A | 0.241 | 8.04E-02 | 1.01E-01 | 2.45E+00 | 2.54E+00 | 4.40E-01 | no | Esrrg | intron_variant | MODIFIER |
| 1 | 97004507 | AX-170124602 | 1 | A | G | 0.152 | 1.33E-01 | 1.74E-01 | 2.64E+00 | 2.75E+00 | 4.40E-01 | no | Gm9499 | upstream_gene_variant | MODIFIER |
| 10 | 94584550 | AX-170193825 | 0 | T | A | 0.272 | -6.80E-02 | 8.74E-02 | 2.55E+00 | 2.60E+00 | 4.40E-01 | no | Tmcc3 | intron_variant | MODIFIER |
| 8 | 88596819 | AX-170265862 | 0 | T | A | 0.175 | -9.85E-02 | 1.29E-01 | 2.65E+00 | 2.73E+00 | 4.40E-01 | no | Nkd1 | downstream_gene_variant | MODIFIER |
| 4 | 115888824 | AX-170471941 | 0 | G | A | 0.211 | 9.05E-02 | 1.17E-01 | 2.59E+00 | 2.65E+00 | 4.40E-01 | no | Kncn | downstream_gene_variant | MODIFIER |
| 4 | 115889444 | AX-97857138 | 0 | A | G | 0.211 | 9.05E-02 | 1.17E-01 | 2.59E+00 | 2.65E+00 | 4.40E-01 | no | Kncn | downstream_gene_variant | MODIFIER |
| 4 | 115909972 | AX-170367581 | 0 | A | G | 0.211 | 9.05E-02 | 1.17E-01 | 2.59E+00 | 2.65E+00 | 4.40E-01 | no | | intergenic_variant | MODIFIER |
| 4 | 115922287 | AX-170287113 | 0 | A | G | 0.211 | 9.05E-02 | 1.17E-01 | 2.59E+00 | 2.65E+00 | 4.40E-01 | no | Dmbx1 | intron_variant | MODIFIER |
| 4 | 115922509 | AX-170304801 | 0 | A | G | 0.211 | 9.05E-02 | 1.17E-01 | 2.59E+00 | 2.65E+00 | 4.40E-01 | no | Dmbx1 | intron_variant | MODIFIER |
| 4 | 115925225 | AX-170249439 | 0 | T | C | 0.211 | 9.05E-02 | 1.17E-01 | 2.59E+00 | 2.65E+00 | 4.40E-01 | no | Dmbx1 | 5_prime_UTR_variant | MODIFIER |
| 4 | 115938218 | AX-170351214 | 0 | G | A | 0.211 | 9.05E-02 | 1.17E-01 | 2.59E+00 | 2.65E+00 | 4.40E-01 | no | Dmbx1 | intron_variant | MODIFIER |
| 4 | 116015507 | AX-170477504 | 0 | G | A | 0.211 | 9.05E-02 | 1.17E-01 | 2.59E+00 | 2.65E+00 | 4.40E-01 | no | Faah | intron_variant | MODIFIER |
| 4 | 116019406 | AX-97828677 | 0 | T | C | 0.211 | 9.05E-02 | 1.17E-01 | 2.59E+00 | 2.65E+00 | 4.40E-01 | no | Faah | upstream_gene_variant | MODIFIER |
| 4 | 116036044 | AX-170023753 | 0 | C | T | 0.211 | 9.05E-02 | 1.17E-01 | 2.59E+00 | 2.65E+00 | 4.40E-01 | no | Nsun4 | intron_variant,NMD_transcript_variant | MODIFIER |
| 4 | 116063013 | AX-170023755 | 0 | G | A | 0.211 | 9.05E-02 | 1.17E-01 | 2.59E+00 | 2.65E+00 | 4.40E-01 | no | Uqcrh | downstream_gene_variant | MODIFIER |
| 4 | 116095556 | AX-170023752 | 0 | A | G | 0.211 | 9.05E-02 | 1.17E-01 | 2.59E+00 | 2.65E+00 | 4.40E-01 | no | Lrrc41 | intron_variant | MODIFIER |
| 4 | 116105462 | AX-97855834 | 0 | C | T | 0.211 | 9.05E-02 | 1.17E-01 | 2.59E+00 | 2.65E+00 | 4.40E-01 | no | Rad54l | intron_variant | MODIFIER |
| 4 | 116166578 | AX-170352082 | 0 | A | T | 0.211 | 9.05E-02 | 1.17E-01 | 2.59E+00 | 2.65E+00 | 4.40E-01 | no | Tspan1 | intron_variant | MODIFIER |
| 4 | 116166843 | AX-170265902 | 0 | A | G | 0.211 | 9.05E-02 | 1.17E-01 | 2.59E+00 | 2.65E+00 | 4.40E-01 | no | Tspan1 | intron_variant | MODIFIER |
| 4 | 116591936 | AX-170186374 | 0 | T | C | 0.211 | 9.05E-02 | 1.17E-01 | 2.59E+00 | 2.65E+00 | 4.40E-01 | no | Gpbp1l1 | intron_variant | MODIFIER |
| 4 | 116603158 | AX-170023766 | 0 | T | G | 0.211 | 9.05E-02 | 1.17E-01 | 2.59E+00 | 2.65E+00 | 4.40E-01 | no | Nasp | intron_variant | MODIFIER |
| 4 | 116604182 | AX-97801128 | 0 | A | G | 0.211 | 9.05E-02 | 1.17E-01 | 2.59E+00 | 2.65E+00 | 4.40E-01 | no | Nasp | synonymous_variant | LOW |
| 4 | 116626944 | AX-170536047 | 0 | C | G | 0.211 | 9.05E-02 | 1.17E-01 | 2.59E+00 | 2.65E+00 | 4.40E-01 | no | Nasp | intron_variant | MODIFIER |
| 4 | 116631133 | AX-170524689 | 0 | A | G | 0.211 | 9.05E-02 | 1.17E-01 | 2.59E+00 | 2.65E+00 | 4.40E-01 | no | Nasp | upstream_gene_variant | MODIFIER |
| 4 | 116684148 | AX-170023759 | 0 | A | C | 0.211 | 9.05E-02 | 1.17E-01 | 2.59E+00 | 2.65E+00 | 4.40E-01 | no | Prdx1 | upstream_gene_variant | MODIFIER |
| 4 | 116686877 | AX-170023761 | 0 | A | C | 0.211 | 9.05E-02 | 1.17E-01 | 2.59E+00 | 2.65E+00 | 4.40E-01 | no | Prdx1 | upstream_gene_variant | MODIFIER |
| 4 | 116698168 | AX-170488146 | 0 | G | T | 0.211 | 9.05E-02 | 1.17E-01 | 2.59E+00 | 2.65E+00 | 4.40E-01 | no | Mmachc | downstream_gene_variant | MODIFIER |
| 4 | 116803523 | AX-170420778 | 0 | C | A | 0.211 | 9.05E-02 | 1.17E-01 | 2.59E+00 | 2.65E+00 | 4.40E-01 | no | Toe1 | downstream_gene_variant | MODIFIER |
| 4 | 116803724 | AX-170457696 | 0 | A | G | 0.211 | 9.05E-02 | 1.17E-01 | 2.59E+00 | 2.65E+00 | 4.40E-01 | no | Toe1 | downstream_gene_variant | MODIFIER |
| 4 | 116811011 | AX-170492217 | 0 | A | T | 0.211 | 9.05E-02 | 1.17E-01 | 2.59E+00 | 2.65E+00 | 4.40E-01 | no | Toe1 | upstream_gene_variant | MODIFIER |
| 4 | 117183645 | AX-170492321 | 0 | A | G | 0.211 | 9.05E-02 | 1.17E-01 | 2.59E+00 | 2.65E+00 | 4.40E-01 | no | Kif2c | upstream_gene_variant | MODIFIER |
| 4 | 117320283 | AX-170246687 | 0 | T | A | 0.211 | 9.05E-02 | 1.17E-01 | 2.59E+00 | 2.65E+00 | 4.40E-01 | no | Rnf220 | intron_variant | MODIFIER |
| 4 | 117320625 | AX-170129884 | 0 | G | A | 0.211 | 9.05E-02 | 1.17E-01 | 2.59E+00 | 2.65E+00 | 4.40E-01 | no | Rnf220 | intron_variant | MODIFIER |
| 4 | 117333076 | AX-97787012 | 0 | C | T | 0.211 | 9.05E-02 | 1.17E-01 | 2.59E+00 | 2.65E+00 | 4.40E-01 | no | Rnf220 | intron_variant | MODIFIER |
| 4 | 117345229 | AX-170519230 | 0 | A | C | 0.211 | 9.05E-02 | 1.17E-01 | 2.59E+00 | 2.65E+00 | 4.40E-01 | no | Rnf220 | intron_variant | MODIFIER |
| 4 | 117358165 | AX-170130589 | 0 | T | C | 0.211 | 9.05E-02 | 1.17E-01 | 2.59E+00 | 2.65E+00 | 4.40E-01 | no | Rnf220 | intron_variant | MODIFIER |
| 4 | 117362980 | AX-170305460 | 0 | A | G | 0.211 | 9.05E-02 | 1.17E-01 | 2.59E+00 | 2.65E+00 | 4.40E-01 | no | Rnf220 | intron_variant | MODIFIER |
| 4 | 117390330 | AX-170120446 | 0 | T | C | 0.211 | 9.05E-02 | 1.17E-01 | 2.59E+00 | 2.65E+00 | 4.40E-01 | no | Rnf220 | intron_variant | MODIFIER |
| 4 | 117392520 | AX-170263288 | 0 | A | G | 0.211 | 9.05E-02 | 1.17E-01 | 2.59E+00 | 2.65E+00 | 4.40E-01 | no | Rnf220 | intron_variant | MODIFIER |
| 4 | 117407390 | AX-170282448 | 0 | T | C | 0.211 | 9.05E-02 | 1.17E-01 | 2.59E+00 | 2.65E+00 | 4.40E-01 | no | Rnf220 | intron_variant | MODIFIER |
| 4 | 117464970 | AX-97853913 | 0 | A | G | 0.211 | 9.05E-02 | 1.17E-01 | 2.59E+00 | 2.65E+00 | 4.40E-01 | no | Rnf220 | intron_variant | MODIFIER |

| | | | | | | | | | | | | | | | |
|---|---|---|---|---|---|---|---|---|---|---|---|---|---|---|---|
| 4 | 117466045 | AX-170100069 | 0 | T | G | 0.211 | 9.05E-02 | 1.17E-01 | 2.59E+00 | 2.65E+00 | 4.40E-01 | no | Rnf220 | intron_variant | MODIFIER |
| 4 | 117474398 | AX-168382364 | 0 | T | C | 0.211 | 9.05E-02 | 1.17E-01 | 2.59E+00 | 2.65E+00 | 4.40E-01 | no | Rnf220 | intron_variant | MODIFIER |
| 4 | 117497242 | AX-97819075 | 0 | T | C | 0.211 | 9.05E-02 | 1.17E-01 | 2.59E+00 | 2.65E+00 | 4.40E-01 | no | Rnf220 | upstream_gene_variant | MODIFIER |
| 4 | 117829818 | AX-170339918 | 0 | G | A | 0.211 | 9.05E-02 | 1.17E-01 | 2.59E+00 | 2.65E+00 | 4.40E-01 | no | Slc6a9 | upstream_gene_variant | MODIFIER |
| 4 | 117915377 | AX-170381291 | 0 | G | A | 0.211 | 9.05E-02 | 1.17E-01 | 2.59E+00 | 2.65E+00 | 4.40E-01 | no | Ipo13 | upstream_gene_variant | MODIFIER |
| 4 | 118017990 | AX-170148530 | 0 | A | G | 0.211 | 9.05E-02 | 1.17E-01 | 2.59E+00 | 2.65E+00 | 4.40E-01 | no | St3gal3 | intron_variant | MODIFIER |
| 4 | 118055082 | AX-170160645 | 0 | A | G | 0.211 | 9.05E-02 | 1.17E-01 | 2.59E+00 | 2.65E+00 | 4.40E-01 | no | St3gal3 | intron_variant | MODIFIER |
| 4 | 118056119 | AX-170333963 | 0 | T | G | 0.211 | 9.05E-02 | 1.17E-01 | 2.59E+00 | 2.65E+00 | 4.40E-01 | no | St3gal3 | intron_variant | MODIFIER |
| 4 | 118058222 | AX-170170719 | 0 | A | G | 0.211 | 9.05E-02 | 1.17E-01 | 2.59E+00 | 2.65E+00 | 4.40E-01 | no | St3gal3 | intron_variant | MODIFIER |
| 4 | 118058781 | AX-170432111 | 0 | G | A | 0.211 | 9.05E-02 | 1.17E-01 | 2.59E+00 | 2.65E+00 | 4.40E-01 | no | St3gal3 | intron_variant | MODIFIER |
| 4 | 118060106 | AX-170379293 | 0 | A | C | 0.211 | 9.05E-02 | 1.17E-01 | 2.59E+00 | 2.65E+00 | 4.40E-01 | no | St3gal3 | intron_variant | MODIFIER |
| 4 | 118064817 | AX-170322319 | 0 | G | A | 0.211 | 9.05E-02 | 1.17E-01 | 2.59E+00 | 2.65E+00 | 4.40E-01 | no | St3gal3 | intron_variant | MODIFIER |
| 4 | 118069218 | AX-168396391 | 0 | A | G | 0.211 | 9.05E-02 | 1.17E-01 | 2.59E+00 | 2.65E+00 | 4.40E-01 | no | St3gal3 | intron_variant | MODIFIER |
| 4 | 118069658 | AX-170126497 | 0 | A | G | 0.211 | 9.05E-02 | 1.17E-01 | 2.59E+00 | 2.65E+00 | 4.40E-01 | no | St3gal3 | intron_variant | MODIFIER |
| 4 | 118074220 | AX-170484890 | 0 | T | C | 0.211 | 9.05E-02 | 1.17E-01 | 2.59E+00 | 2.65E+00 | 4.40E-01 | no | St3gal3 | intron_variant | MODIFIER |
| 4 | 118074246 | AX-170524345 | 0 | C | T | 0.211 | 9.05E-02 | 1.17E-01 | 2.59E+00 | 2.65E+00 | 4.40E-01 | no | St3gal3 | intron_variant | MODIFIER |
| 4 | 118074282 | AX-170149173 | 0 | T | C | 0.211 | 9.05E-02 | 1.17E-01 | 2.59E+00 | 2.65E+00 | 4.40E-01 | no | St3gal3 | intron_variant | MODIFIER |
| 4 | 118098518 | AX-170443324 | 0 | T | A | 0.211 | 9.05E-02 | 1.17E-01 | 2.59E+00 | 2.65E+00 | 4.40E-01 | no | St3gal3 | intron_variant | MODIFIER |
| 4 | 118193446 | AX-168416112 | 0 | A | G | 0.211 | 9.05E-02 | 1.17E-01 | 2.59E+00 | 2.65E+00 | 4.40E-01 | no | | intergenic_variant | MODIFIER |
| 4 | 118222801 | AX-97787764 | 0 | G | A | 0.211 | 9.05E-02 | 1.17E-01 | 2.59E+00 | 2.65E+00 | 4.40E-01 | no | Ptprf | intron_variant | MODIFIER |
| 4 | 118226998 | AX-170365982 | 0 | C | T | 0.211 | 9.05E-02 | 1.17E-01 | 2.59E+00 | 2.65E+00 | 4.40E-01 | no | Ptprf | intron_variant | MODIFIER |
| 4 | 118297444 | AX-170023789 | 0 | T | C | 0.211 | 9.05E-02 | 1.17E-01 | 2.59E+00 | 2.65E+00 | 4.40E-01 | no | | regulatory_region_variant | MODIFIER |
| 4 | 118314817 | AX-170023787 | 0 | T | C | 0.211 | 9.05E-02 | 1.17E-01 | 2.59E+00 | 2.65E+00 | 4.40E-01 | no | | intergenic_variant | MODIFIER |
| 4 | 118320229 | AX-97848901 | 0 | G | A | 0.211 | 9.05E-02 | 1.17E-01 | 2.59E+00 | 2.65E+00 | 4.40E-01 | no | | regulatory_region_variant | MODIFIER |
| 4 | 118320307 | AX-170023788 | 0 | T | C | 0.211 | 9.05E-02 | 1.17E-01 | 2.59E+00 | 2.65E+00 | 4.40E-01 | no | | regulatory_region_variant | MODIFIER |
| 4 | 118350076 | AX-170252430 | 0 | T | G | 0.211 | 9.05E-02 | 1.17E-01 | 2.59E+00 | 2.65E+00 | 4.40E-01 | no | | intergenic_variant | MODIFIER |
| 4 | 118364287 | AX-170421158 | 0 | A | G | 0.211 | 9.05E-02 | 1.17E-01 | 2.59E+00 | 2.65E+00 | 4.40E-01 | no | Hyi | downstream_gene_variant | MODIFIER |
| 4 | 118370911 | AX-170436479 | 0 | T | G | 0.211 | 9.05E-02 | 1.17E-01 | 2.59E+00 | 2.65E+00 | 4.40E-01 | no | Szt2 | intron_variant | MODIFIER |
| 4 | 118372602 | AX-170142301 | 0 | A | G | 0.211 | 9.05E-02 | 1.17E-01 | 2.59E+00 | 2.65E+00 | 4.40E-01 | no | Szt2 | synonymous_variant | LOW |
| 4 | 118377869 | AX-170376737 | 0 | C | T | 0.211 | 9.05E-02 | 1.17E-01 | 2.59E+00 | 2.65E+00 | 4.40E-01 | no | Szt2 | missense_variant,splice_region_variant | MODERATE |
| 4 | 118381171 | AX-170023786 | 0 | A | C | 0.211 | 9.05E-02 | 1.17E-01 | 2.59E+00 | 2.65E+00 | 4.40E-01 | no | Szt2 | intron_variant | MODIFIER |
| 4 | 118381479 | AX-97789548 | 0 | A | G | 0.211 | 9.05E-02 | 1.17E-01 | 2.59E+00 | 2.65E+00 | 4.40E-01 | no | Szt2 | intron_variant | MODIFIER |
| 4 | 118396529 | AX-170023792 | 0 | T | C | 0.211 | 9.05E-02 | 1.17E-01 | 2.59E+00 | 2.65E+00 | 4.40E-01 | no | Szt2 | intron_variant | MODIFIER |
| 4 | 118397536 | AX-170293905 | 0 | C | T | 0.211 | 9.05E-02 | 1.17E-01 | 2.59E+00 | 2.65E+00 | 4.40E-01 | no | Szt2 | intron_variant | MODIFIER |
| 4 | 118406020 | AX-97803213 | 0 | G | A | 0.211 | 9.05E-02 | 1.17E-01 | 2.59E+00 | 2.65E+00 | 4.40E-01 | no | Med8 | upstream_gene_variant | MODIFIER |
| 4 | 118406706 | AX-170023793 | 0 | T | C | 0.211 | 9.05E-02 | 1.17E-01 | 2.59E+00 | 2.65E+00 | 4.40E-01 | no | Med8 | upstream_gene_variant | MODIFIER |
| 4 | 118425717 | AX-170493720 | 0 | G | A | 0.211 | 9.05E-02 | 1.17E-01 | 2.59E+00 | 2.65E+00 | 4.40E-01 | no | Elovl1 | upstream_gene_variant | MODIFIER |
| 4 | 118426291 | AX-170176681 | 0 | T | C | 0.211 | 9.05E-02 | 1.17E-01 | 2.59E+00 | 2.65E+00 | 4.40E-01 | no | Elovl1 | upstream_gene_variant | MODIFIER |
| 4 | 118430800 | AX-170447919 | 0 | G | A | 0.211 | 9.05E-02 | 1.17E-01 | 2.59E+00 | 2.65E+00 | 4.40E-01 | no | Elovl1 | synonymous_variant | LOW |
| 4 | 118443615 | AX-170336362 | 0 | C | T | 0.211 | 9.05E-02 | 1.17E-01 | 2.59E+00 | 2.65E+00 | 4.40E-01 | no | Mpl | synonymous_variant | LOW |
| 4 | 118443924 | AX-97782510 | 0 | G | A | 0.211 | 9.05E-02 | 1.17E-01 | 2.59E+00 | 2.65E+00 | 4.40E-01 | no | Mpl | intron_variant | MODIFIER |
| 4 | 118448697 | AX-170189288 | 0 | G | A | 0.211 | 9.05E-02 | 1.17E-01 | 2.59E+00 | 2.65E+00 | 4.40E-01 | no | Mpl | intron_variant | MODIFIER |
| 4 | 118455433 | AX-170425184 | 0 | A | G | 0.211 | 9.05E-02 | 1.17E-01 | 2.59E+00 | 2.65E+00 | 4.40E-01 | no | Mpl | intron_variant | MODIFIER |
| 4 | 118456124 | AX-97823441 | 0 | G | A | 0.211 | 9.05E-02 | 1.17E-01 | 2.59E+00 | 2.65E+00 | 4.40E-01 | no | Mpl | intron_variant | MODIFIER |
| 4 | 118477105 | AX-170070450 | 0 | A | G | 0.211 | 9.05E-02 | 1.17E-01 | 2.59E+00 | 2.65E+00 | 4.40E-01 | no | Tie1 | intron_variant | MODIFIER |
| 4 | 118478764 | AX-170365299 | 0 | T | C | 0.211 | 9.05E-02 | 1.17E-01 | 2.59E+00 | 2.65E+00 | 4.40E-01 | no | Tie1 | intron_variant | MODIFIER |
| 4 | 118478822 | AX-170520164 | 0 | A | G | 0.211 | 9.05E-02 | 1.17E-01 | 2.59E+00 | 2.65E+00 | 4.40E-01 | no | Tie1 | intron_variant | MODIFIER |
| 4 | 118480679 | AX-170205920 | 0 | C | G | 0.211 | 9.05E-02 | 1.17E-01 | 2.59E+00 | 2.65E+00 | 4.40E-01 | no | Tie1 | intron_variant | MODIFIER |
| 4 | 118483890 | AX-170224429 | 0 | A | G | 0.211 | 9.05E-02 | 1.17E-01 | 2.59E+00 | 2.65E+00 | 4.40E-01 | no | Tie1 | intron_variant | MODIFIER |
| 4 | 118487892 | AX-170513437 | 0 | A | G | 0.211 | 9.05E-02 | 1.17E-01 | 2.59E+00 | 2.65E+00 | 4.40E-01 | no | Tie1 | intron_variant | MODIFIER |
| 4 | 118499422 | AX-170098553 | 0 | G | A | 0.211 | 9.05E-02 | 1.17E-01 | 2.59E+00 | 2.65E+00 | 4.40E-01 | no | Gm12859 | upstream_gene_variant | MODIFIER |
| 4 | 118528528 | AX-170514537 | 0 | T | C | 0.211 | 9.05E-02 | 1.17E-01 | 2.59E+00 | 2.65E+00 | 4.40E-01 | no | 2610528J11Rik | intron_variant | MODIFIER |
| 4 | 118541357 | AX-170240751 | 0 | G | A | 0.211 | 9.05E-02 | 1.17E-01 | 2.59E+00 | 2.65E+00 | 4.40E-01 | no | Tmem125 | 3_prime_UTR_variant | MODIFIER |
| 4 | 118587462 | AX-170474954 | 0 | T | G | 0.211 | 9.05E-02 | 1.17E-01 | 2.59E+00 | 2.65E+00 | 4.40E-01 | no | Cfap57 | intron_variant | MODIFIER |

| | | | | | | | | | | | | | | | |
|---|---|---|---|---|---|---|---|---|---|---|---|---|---|---|---|
| 4 | 118587952 | AX-97814079 | 0 | T | A | 0.211 | 9.05E-02 | 1.17E-01 | 2.59E+00 | 2.65E+00 | 4.40E-01 | no | Cfap57 | intron_variant | MODIFIER |
| 4 | 118592680 | AX-170520195 | 0 | G | A | 0.211 | 9.05E-02 | 1.17E-01 | 2.59E+00 | 2.65E+00 | 4.40E-01 | no | Cfap57 | intron_variant | MODIFIER |
| 4 | 118598524 | AX-170210099 | 0 | A | G | 0.211 | 9.05E-02 | 1.17E-01 | 2.59E+00 | 2.65E+00 | 4.40E-01 | no | Cfap57 | intron_variant | MODIFIER |
| 4 | 118612003 | AX-97829118 | 0 | A | G | 0.211 | 9.05E-02 | 1.17E-01 | 2.59E+00 | 2.65E+00 | 4.40E-01 | no | Cfap57 | intron_variant | MODIFIER |
| 4 | 118700866 | AX-170439833 | 0 | A | G | 0.211 | 9.05E-02 | 1.17E-01 | 2.59E+00 | 2.65E+00 | 4.40E-01 | no | Olfr1341 | upstream_gene_variant | MODIFIER |
| 4 | 118702156 | AX-170456679 | 0 | C | T | 0.211 | 9.05E-02 | 1.17E-01 | 2.59E+00 | 2.65E+00 | 4.40E-01 | no | Olfr1341 | upstream_gene_variant | MODIFIER |
| 4 | 118704019 | AX-170485749 | 0 | T | C | 0.211 | 9.05E-02 | 1.17E-01 | 2.59E+00 | 2.65E+00 | 4.40E-01 | no | Olfr1341 | intron_variant | MODIFIER |
| 4 | 118705610 | AX-170250430 | 0 | A | G | 0.211 | 9.05E-02 | 1.17E-01 | 2.59E+00 | 2.65E+00 | 4.40E-01 | no | Olfr1341 | upstream_gene_variant | MODIFIER |
| 4 | 118709548 | AX-170314169 | 0 | T | C | 0.211 | 9.05E-02 | 1.17E-01 | 2.59E+00 | 2.65E+00 | 4.40E-01 | no | Olfr1341 | missense_variant | MODERATE |
| 4 | 118710224 | AX-168380660 | 0 | G | A | 0.211 | 9.05E-02 | 1.17E-01 | 2.59E+00 | 2.65E+00 | 4.40E-01 | no | Olfr1341 | synonymous_variant | LOW |
| 4 | 118731233 | AX-97798684 | 0 | C | A | 0.211 | 9.05E-02 | 1.17E-01 | 2.59E+00 | 2.65E+00 | 4.40E-01 | no | Olfr1339 | upstream_gene_variant | MODIFIER |
| 4 | 118786110 | AX-97806936 | 0 | T | C | 0.211 | 9.05E-02 | 1.17E-01 | 2.59E+00 | 2.65E+00 | 4.40E-01 | no | Olfr1337 | upstream_gene_variant | MODIFIER |
| 4 | 118792350 | AX-170462666 | 0 | A | G | 0.211 | 9.05E-02 | 1.17E-01 | 2.59E+00 | 2.65E+00 | 4.40E-01 | no | Gm12855 | upstream_gene_variant | MODIFIER |
| 4 | 118802967 | AX-97803370 | 0 | T | G | 0.211 | 9.05E-02 | 1.17E-01 | 2.59E+00 | 2.65E+00 | 4.40E-01 | no | | intergenic_variant | MODIFIER |
| 4 | 118803294 | AX-170023795 | 0 | T | C | 0.211 | 9.05E-02 | 1.17E-01 | 2.59E+00 | 2.65E+00 | 4.40E-01 | no | | intergenic_variant | MODIFIER |
| 4 | 118811028 | AX-170447424 | 0 | A | T | 0.211 | 9.05E-02 | 1.17E-01 | 2.59E+00 | 2.65E+00 | 4.40E-01 | no | Olfr1335 | upstream_gene_variant | MODIFIER |
| 4 | 118831186 | AX-97836154 | 0 | A | C | 0.211 | 9.05E-02 | 1.17E-01 | 2.59E+00 | 2.65E+00 | 4.40E-01 | no | Olfr1333 | upstream_gene_variant | MODIFIER |
| 4 | 118889335 | AX-170089359 | 0 | A | G | 0.211 | 9.05E-02 | 1.17E-01 | 2.59E+00 | 2.65E+00 | 4.40E-01 | no | Olfr1330 | upstream_gene_variant | MODIFIER |
| 4 | 118894290 | AX-170023801 | 0 | A | C | 0.211 | 9.05E-02 | 1.17E-01 | 2.59E+00 | 2.65E+00 | 4.40E-01 | no | Olfr1330 | downstream_gene_variant | MODIFIER |
| 4 | 118899073 | AX-170509192 | 0 | G | A | 0.211 | 9.05E-02 | 1.17E-01 | 2.59E+00 | 2.65E+00 | 4.40E-01 | no | Olfr1330 | downstream_gene_variant | MODIFIER |
| 4 | 118933155 | AX-170163447 | 0 | A | G | 0.211 | 9.05E-02 | 1.17E-01 | 2.59E+00 | 2.65E+00 | 4.40E-01 | no | Olfr1328 | downstream_gene_variant | MODIFIER |
| 4 | 118935155 | AX-170117973 | 0 | T | C | 0.211 | 9.05E-02 | 1.17E-01 | 2.59E+00 | 2.65E+00 | 4.40E-01 | no | Olfr1328 | upstream_gene_variant | MODIFIER |
| 4 | 118936627 | AX-170502732 | 0 | T | C | 0.211 | 9.05E-02 | 1.17E-01 | 2.59E+00 | 2.65E+00 | 4.40E-01 | no | Olfr1328 | upstream_gene_variant | MODIFIER |
| 4 | 118937606 | AX-170164573 | 0 | T | C | 0.211 | 9.05E-02 | 1.17E-01 | 2.59E+00 | 2.65E+00 | 4.40E-01 | no | Olfr1328 | upstream_gene_variant | MODIFIER |
| 4 | 118938519 | AX-170177387 | 0 | G | A | 0.211 | 9.05E-02 | 1.17E-01 | 2.59E+00 | 2.65E+00 | 4.40E-01 | no | Olfr1328 | upstream_gene_variant | MODIFIER |
| 4 | 118941024 | AX-170268138 | 0 | A | G | 0.211 | 9.05E-02 | 1.17E-01 | 2.59E+00 | 2.65E+00 | 4.40E-01 | no | Gm12856 | downstream_gene_variant | MODIFIER |
| 4 | 118941350 | AX-170341812 | 0 | C | T | 0.211 | 9.05E-02 | 1.17E-01 | 2.59E+00 | 2.65E+00 | 4.40E-01 | no | Gm12856 | downstream_gene_variant | MODIFIER |
| 4 | 118941383 | AX-170392366 | 0 | C | T | 0.211 | 9.05E-02 | 1.17E-01 | 2.59E+00 | 2.65E+00 | 4.40E-01 | no | Gm12856 | downstream_gene_variant | MODIFIER |
| 4 | 119018227 | AX-170023807 | 0 | C | T | 0.211 | 9.05E-02 | 1.17E-01 | 2.59E+00 | 2.65E+00 | 4.40E-01 | no | | regulatory_region_variant | MODIFIER |
| 4 | 119053497 | AX-170236225 | 0 | C | T | 0.211 | 9.05E-02 | 1.17E-01 | 2.59E+00 | 2.65E+00 | 4.40E-01 | no | Gm12866 | downstream_gene_variant | MODIFIER |
| 4 | 119054866 | AX-170198729 | 0 | A | C | 0.211 | 9.05E-02 | 1.17E-01 | 2.59E+00 | 2.65E+00 | 4.40E-01 | no | Gm12866 | non_coding_transcript_exon_variant | MODIFIER |
| 4 | 119055375 | AX-170295705 | 0 | A | G | 0.211 | 9.05E-02 | 1.17E-01 | 2.59E+00 | 2.65E+00 | 4.40E-01 | no | Gm12866 | non_coding_transcript_exon_variant | MODIFIER |
| 4 | 119172866 | AX-97787334 | 0 | T | C | 0.211 | 9.05E-02 | 1.17E-01 | 2.59E+00 | 2.65E+00 | 4.40E-01 | no | Ermap | downstream_gene_variant | MODIFIER |
| 4 | 119174294 | AX-170070465 | 0 | T | C | 0.211 | 9.05E-02 | 1.17E-01 | 2.59E+00 | 2.65E+00 | 4.40E-01 | no | Ermap | downstream_gene_variant | MODIFIER |
| 4 | 119181107 | AX-170496403 | 0 | A | G | 0.211 | 9.05E-02 | 1.17E-01 | 2.59E+00 | 2.65E+00 | 4.40E-01 | no | Ermap | intron_variant,NMD_transcript_variant | MODIFIER |
| 4 | 119185966 | AX-170210555 | 0 | A | C | 0.211 | 9.05E-02 | 1.17E-01 | 2.59E+00 | 2.65E+00 | 4.40E-01 | no | Ermap | intron_variant,NMD_transcript_variant | MODIFIER |
| 4 | 119188136 | AX-170214465 | 0 | A | G | 0.211 | 9.05E-02 | 1.17E-01 | 2.59E+00 | 2.65E+00 | 4.40E-01 | no | Ermap | intron_variant,NMD_transcript_variant | MODIFIER |
| 4 | 119191623 | AX-170414045 | 0 | T | G | 0.211 | 9.05E-02 | 1.17E-01 | 2.59E+00 | 2.65E+00 | 4.40E-01 | no | Svbp | upstream_gene_variant | MODIFIER |
| 4 | 119207798 | AX-170483098 | 0 | A | G | 0.211 | 9.05E-02 | 1.17E-01 | 2.59E+00 | 2.65E+00 | 4.40E-01 | no | Tmem269 | intron_variant | MODIFIER |
| 4 | 119222746 | AX-170384286 | 0 | A | G | 0.211 | 9.05E-02 | 1.17E-01 | 2.59E+00 | 2.65E+00 | 4.40E-01 | no | Tmem269 | upstream_gene_variant | MODIFIER |
| 4 | 119245028 | AX-170281936 | 0 | C | T | 0.211 | 9.05E-02 | 1.17E-01 | 2.59E+00 | 2.65E+00 | 4.40E-01 | no | P3h1 | synonymous_variant | LOW |
| 4 | 119246142 | AX-170468574 | 0 | A | G | 0.211 | 9.05E-02 | 1.17E-01 | 2.59E+00 | 2.65E+00 | 4.40E-01 | no | P3h1 | intron_variant | MODIFIER |
| 4 | 119253063 | AX-170100265 | 0 | G | A | 0.211 | 9.05E-02 | 1.17E-01 | 2.59E+00 | 2.65E+00 | 4.40E-01 | no | P3h1 | downstream_gene_variant | MODIFIER |
| 4 | 119258430 | AX-170388487 | 0 | A | G | 0.211 | 9.05E-02 | 1.17E-01 | 2.59E+00 | 2.65E+00 | 4.40E-01 | no | Cldn19 | intron_variant | MODIFIER |
| 4 | 119258704 | AX-170378999 | 0 | A | C | 0.211 | 9.05E-02 | 1.17E-01 | 2.59E+00 | 2.65E+00 | 4.40E-01 | no | Cldn19 | intron_variant | MODIFIER |
| 16 | 50165768 | AX-170037456 | 1 | A | G | 0.5 | 6.42E-02 | 8.22E-02 | 2.54E+00 | 2.57E+00 | 4.40E-01 | no | | intergenic_variant | MODIFIER |
| 17 | 70551791 | AX-170489428 | 1 | G | A | 0.054 | -1.39E-01 | 1.83E-01 | 2.73E+00 | 2.84E+00 | 4.40E-01 | no | Dlgap1 | intron_variant | MODIFIER |
| 5 | 46971821 | AX-170113901 | 0 | A | G | 0.474 | 8.00E-02 | 9.90E-02 | 2.40E+00 | 2.47E+00 | 4.40E-01 | no | | intergenic_variant | MODIFIER |
| 16 | 50219353 | AX-170088013 | 0 | A | C | 0.5 | 6.42E-02 | 8.24E-02 | 2.54E+00 | 2.57E+00 | 4.41E-01 | no | Bbx | intron_variant | MODIFIER |
| 18 | 68731233 | AX-170218302 | 1 | A | G | 0.277 | -7.70E-02 | 1.01E-01 | 2.73E+00 | 2.79E+00 | 4.41E-01 | no | | intergenic_variant | MODIFIER |
| 18 | 55308370 | AX-170015744 | 1 | C | T | 0.455 | 7.86E-02 | 9.54E-02 | 2.35E+00 | 2.41E+00 | 4.41E-01 | no | | intergenic_variant | MODIFIER |
| 10 | 91914208 | AX-170140501 | 0 | T | C | 0.298 | -6.57E-02 | 8.43E-02 | 2.53E+00 | 2.58E+00 | 4.41E-01 | no | Gm31592 | intron_variant,non_coding_transcript_variant | MODIFIER |
| 10 | 91914874 | AX-170238893 | 0 | A | C | 0.298 | -6.57E-02 | 8.43E-02 | 2.53E+00 | 2.58E+00 | 4.41E-01 | no | Gm31592 | intron_variant,non_coding_transcript_variant | MODIFIER |
| 10 | 91925343 | AX-170003662 | 0 | T | A | 0.298 | -6.57E-02 | 8.43E-02 | 2.53E+00 | 2.58E+00 | 4.41E-01 | no | Gm31592 | intron_variant,non_coding_transcript_variant | MODIFIER |

| | | | | | | | | | | | | | | | |
|---|---|---|---|---|---|---|---|---|---|---|---|---|---|---|---|
| 10 | 91925843 | AX-170045569 | 0 | T | C | 0.298 | -6.57E-02 | 8.43E-02 | 2.53E+00 | 2.58E+00 | 4.41E-01 | no | Gm31592 | intron_variant,non_coding_transcript_variant | MODIFIER |
| 10 | 91926331 | AX-170227318 | 0 | A | G | 0.298 | -6.57E-02 | 8.43E-02 | 2.53E+00 | 2.58E+00 | 4.41E-01 | no | Gm31592 | intron_variant,non_coding_transcript_variant | MODIFIER |
| 10 | 91926447 | AX-170160276 | 0 | C | G | 0.298 | -6.57E-02 | 8.43E-02 | 2.53E+00 | 2.58E+00 | 4.41E-01 | no | Gm31592 | intron_variant,non_coding_transcript_variant | MODIFIER |
| 10 | 91940504 | AX-170045568 | 0 | C | T | 0.298 | -6.57E-02 | 8.43E-02 | 2.53E+00 | 2.58E+00 | 4.41E-01 | no | Gm31592 | intron_variant,non_coding_transcript_variant | MODIFIER |
| 10 | 91950806 | AX-170231674 | 0 | T | C | 0.298 | -6.57E-02 | 8.43E-02 | 2.53E+00 | 2.58E+00 | 4.41E-01 | no | Gm31592 | intron_variant,non_coding_transcript_variant | MODIFIER |
| 10 | 91963135 | AX-170086863 | 0 | C | A | 0.298 | -6.57E-02 | 8.43E-02 | 2.53E+00 | 2.58E+00 | 4.41E-01 | no | | intergenic_variant | MODIFIER |
| 10 | 91978928 | AX-170035486 | 0 | A | G | 0.298 | -6.57E-02 | 8.43E-02 | 2.53E+00 | 2.58E+00 | 4.41E-01 | no | | intergenic_variant | MODIFIER |
| 10 | 91988423 | AX-170275528 | 0 | T | A | 0.298 | -6.57E-02 | 8.43E-02 | 2.53E+00 | 2.58E+00 | 4.41E-01 | no | | regulatory_region_variant | MODIFIER |
| 10 | 92009945 | AX-170102241 | 0 | A | G | 0.298 | -6.57E-02 | 8.43E-02 | 2.53E+00 | 2.58E+00 | 4.41E-01 | no | | regulatory_region_variant | MODIFIER |
| 10 | 92010118 | AX-170354042 | 0 | T | C | 0.298 | -6.57E-02 | 8.43E-02 | 2.53E+00 | 2.58E+00 | 4.41E-01 | no | | regulatory_region_variant | MODIFIER |
| 10 | 92013955 | AX-170225584 | 0 | A | T | 0.298 | -6.57E-02 | 8.43E-02 | 2.53E+00 | 2.58E+00 | 4.41E-01 | no | | intergenic_variant | MODIFIER |
| 10 | 92014321 | AX-170472586 | 0 | A | G | 0.298 | -6.57E-02 | 8.43E-02 | 2.53E+00 | 2.58E+00 | 4.41E-01 | no | | intergenic_variant | MODIFIER |
| 10 | 92016384 | AX-170504672 | 0 | C | T | 0.298 | -6.57E-02 | 8.43E-02 | 2.53E+00 | 2.58E+00 | 4.41E-01 | no | | intergenic_variant | MODIFIER |
| 10 | 92017887 | AX-170319205 | 0 | T | C | 0.298 | -6.57E-02 | 8.43E-02 | 2.53E+00 | 2.58E+00 | 4.41E-01 | no | | intergenic_variant | MODIFIER |
| 10 | 92021302 | AX-170490395 | 0 | A | C | 0.298 | -6.57E-02 | 8.43E-02 | 2.53E+00 | 2.58E+00 | 4.41E-01 | no | | intergenic_variant | MODIFIER |
| 10 | 92022079 | AX-170092063 | 0 | A | G | 0.298 | -6.57E-02 | 8.43E-02 | 2.53E+00 | 2.58E+00 | 4.41E-01 | no | | intergenic_variant | MODIFIER |
| 10 | 92026254 | AX-170409248 | 0 | G | A | 0.298 | -6.57E-02 | 8.43E-02 | 2.53E+00 | 2.58E+00 | 4.41E-01 | no | | intergenic_variant | MODIFIER |
| 10 | 92031781 | AX-170442903 | 0 | A | C | 0.298 | -6.57E-02 | 8.43E-02 | 2.53E+00 | 2.58E+00 | 4.41E-01 | no | | intergenic_variant | MODIFIER |
| 10 | 92048923 | AX-170232215 | 0 | G | A | 0.298 | -6.57E-02 | 8.43E-02 | 2.53E+00 | 2.58E+00 | 4.41E-01 | no | | intergenic_variant | MODIFIER |
| 10 | 92058429 | AX-97780438 | 0 | A | C | 0.298 | -6.57E-02 | 8.43E-02 | 2.53E+00 | 2.58E+00 | 4.41E-01 | no | | intergenic_variant | MODIFIER |
| 10 | 92081579 | AX-170045573 | 0 | G | A | 0.298 | -6.57E-02 | 8.43E-02 | 2.53E+00 | 2.58E+00 | 4.41E-01 | no | Rmst | intron_variant,non_coding_transcript_variant | MODIFIER |
| 10 | 92091568 | AX-170035483 | 0 | T | C | 0.298 | -6.57E-02 | 8.43E-02 | 2.53E+00 | 2.58E+00 | 4.41E-01 | no | Rmst | intron_variant,non_coding_transcript_variant | MODIFIER |
| 10 | 92095542 | AX-170338967 | 0 | A | G | 0.298 | -6.57E-02 | 8.43E-02 | 2.53E+00 | 2.58E+00 | 4.41E-01 | no | Rmst | intron_variant,non_coding_transcript_variant | MODIFIER |
| 10 | 92113677 | AX-170316357 | 0 | G | A | 0.298 | -6.57E-02 | 8.43E-02 | 2.53E+00 | 2.58E+00 | 4.41E-01 | no | Rmst | intron_variant,non_coding_transcript_variant | MODIFIER |
| 10 | 92144377 | AX-170045571 | 0 | G | C | 0.298 | -6.57E-02 | 8.43E-02 | 2.53E+00 | 2.58E+00 | 4.41E-01 | no | Rmst | intron_variant,non_coding_transcript_variant | MODIFIER |
| 10 | 92145354 | AX-170528631 | 0 | G | A | 0.298 | -6.57E-02 | 8.43E-02 | 2.53E+00 | 2.58E+00 | 4.41E-01 | no | Rmst | intron_variant,non_coding_transcript_variant | MODIFIER |
| 10 | 92145474 | AX-97827004 | 0 | C | T | 0.298 | -6.57E-02 | 8.43E-02 | 2.53E+00 | 2.58E+00 | 4.41E-01 | no | Rmst | intron_variant,non_coding_transcript_variant | MODIFIER |
| 10 | 92146528 | AX-170309935 | 0 | T | G | 0.298 | -6.57E-02 | 8.43E-02 | 2.53E+00 | 2.58E+00 | 4.41E-01 | no | Rmst | intron_variant,non_coding_transcript_variant | MODIFIER |
| 10 | 92147858 | AX-170391138 | 0 | A | T | 0.298 | -6.57E-02 | 8.43E-02 | 2.53E+00 | 2.58E+00 | 4.41E-01 | no | Rmst | intron_variant,non_coding_transcript_variant | MODIFIER |
| 10 | 92166806 | AX-170175636 | 0 | G | A | 0.298 | -6.57E-02 | 8.43E-02 | 2.53E+00 | 2.58E+00 | 4.41E-01 | no | Gm26700 | downstream_gene_variant | MODIFIER |
| 10 | 92192394 | AX-170104643 | 0 | A | C | 0.298 | -6.57E-02 | 8.43E-02 | 2.53E+00 | 2.58E+00 | 4.41E-01 | no | Gm20757 | intron_variant,non_coding_transcript_variant | MODIFIER |
| 10 | 92192446 | AX-170274730 | 0 | T | G | 0.298 | -6.57E-02 | 8.43E-02 | 2.53E+00 | 2.58E+00 | 4.41E-01 | no | Gm20757 | intron_variant,non_coding_transcript_variant | MODIFIER |
| 10 | 92192907 | AX-170482723 | 0 | G | A | 0.298 | -6.57E-02 | 8.43E-02 | 2.53E+00 | 2.58E+00 | 4.41E-01 | no | Gm20757 | intron_variant,non_coding_transcript_variant | MODIFIER |
| 10 | 92203129 | AX-170324062 | 0 | A | G | 0.298 | -6.57E-02 | 8.43E-02 | 2.53E+00 | 2.58E+00 | 4.41E-01 | no | Gm20757 | intron_variant,non_coding_transcript_variant | MODIFIER |
| 10 | 92223621 | AX-170115178 | 0 | T | C | 0.298 | -6.57E-02 | 8.43E-02 | 2.53E+00 | 2.58E+00 | 4.41E-01 | no | Gm20757 | intron_variant,non_coding_transcript_variant | MODIFIER |
| 10 | 92224019 | AX-170332646 | 0 | C | T | 0.298 | -6.57E-02 | 8.43E-02 | 2.53E+00 | 2.58E+00 | 4.41E-01 | no | Gm20757 | intron_variant,non_coding_transcript_variant | MODIFIER |
| 10 | 92232457 | AX-170494229 | 0 | C | G | 0.298 | -6.57E-02 | 8.43E-02 | 2.53E+00 | 2.58E+00 | 4.41E-01 | no | Gm20757 | intron_variant,non_coding_transcript_variant | MODIFIER |
| 10 | 92232558 | AX-170148730 | 0 | A | G | 0.298 | -6.57E-02 | 8.43E-02 | 2.53E+00 | 2.58E+00 | 4.41E-01 | no | Gm20757 | intron_variant,non_coding_transcript_variant | MODIFIER |
| 10 | 92239755 | AX-97812763 | 0 | A | G | 0.298 | -6.57E-02 | 8.43E-02 | 2.53E+00 | 2.58E+00 | 4.41E-01 | no | Gm20757 | intron_variant,non_coding_transcript_variant | MODIFIER |
| 10 | 92241525 | AX-170338364 | 0 | C | T | 0.298 | -6.57E-02 | 8.43E-02 | 2.53E+00 | 2.58E+00 | 4.41E-01 | no | Gm20757 | intron_variant,non_coding_transcript_variant | MODIFIER |
| 10 | 92252828 | AX-170467741 | 0 | G | A | 0.298 | -6.57E-02 | 8.43E-02 | 2.53E+00 | 2.58E+00 | 4.41E-01 | no | Gm20757 | intron_variant,non_coding_transcript_variant | MODIFIER |
| 10 | 92274533 | AX-170117998 | 0 | A | G | 0.298 | -6.57E-02 | 8.43E-02 | 2.53E+00 | 2.58E+00 | 4.41E-01 | no | Gm20757 | intron_variant,non_coding_transcript_variant | MODIFIER |
| 10 | 92275827 | AX-97825277 | 0 | C | T | 0.298 | -6.57E-02 | 8.43E-02 | 2.53E+00 | 2.58E+00 | 4.41E-01 | no | Gm20757 | intron_variant,non_coding_transcript_variant | MODIFIER |
| 10 | 92283435 | AX-170344294 | 0 | G | C | 0.298 | -6.57E-02 | 8.43E-02 | 2.53E+00 | 2.58E+00 | 4.41E-01 | no | Gm20757 | intron_variant,non_coding_transcript_variant | MODIFIER |
| 10 | 92291209 | AX-170163238 | 0 | A | T | 0.298 | -6.57E-02 | 8.43E-02 | 2.53E+00 | 2.58E+00 | 4.41E-01 | no | Gm20757 | intron_variant,non_coding_transcript_variant | MODIFIER |
| 10 | 92292028 | AX-170418959 | 0 | G | A | 0.298 | -6.57E-02 | 8.43E-02 | 2.53E+00 | 2.58E+00 | 4.41E-01 | no | Gm20757 | intron_variant,non_coding_transcript_variant | MODIFIER |
| 10 | 92292471 | AX-170340848 | 0 | C | G | 0.298 | -6.57E-02 | 8.43E-02 | 2.53E+00 | 2.58E+00 | 4.41E-01 | no | Gm20757 | intron_variant,non_coding_transcript_variant | MODIFIER |
| 10 | 92294391 | AX-170045577 | 0 | C | T | 0.298 | -6.57E-02 | 8.43E-02 | 2.53E+00 | 2.58E+00 | 4.41E-01 | no | Gm20757 | intron_variant,non_coding_transcript_variant | MODIFIER |
| 10 | 92302043 | AX-170279369 | 0 | G | C | 0.298 | -6.57E-02 | 8.43E-02 | 2.53E+00 | 2.58E+00 | 4.41E-01 | no | Gm20757 | intron_variant,non_coding_transcript_variant | MODIFIER |
| 10 | 92302072 | AX-170239939 | 0 | C | T | 0.298 | -6.57E-02 | 8.43E-02 | 2.53E+00 | 2.58E+00 | 4.41E-01 | no | Gm20757 | intron_variant,non_coding_transcript_variant | MODIFIER |
| 10 | 92310451 | AX-170105737 | 0 | C | T | 0.298 | -6.57E-02 | 8.43E-02 | 2.53E+00 | 2.58E+00 | 4.41E-01 | no | Gm20757 | intron_variant,non_coding_transcript_variant | MODIFIER |
| 10 | 92311019 | AX-97818785 | 0 | A | G | 0.298 | -6.57E-02 | 8.43E-02 | 2.53E+00 | 2.58E+00 | 4.41E-01 | no | Gm20757 | intron_variant,non_coding_transcript_variant | MODIFIER |
| 10 | 92314786 | AX-170379552 | 0 | G | T | 0.298 | -6.57E-02 | 8.43E-02 | 2.53E+00 | 2.58E+00 | 4.41E-01 | no | Gm20757 | intron_variant,non_coding_transcript_variant | MODIFIER |
| 10 | 92317622 | AX-170003663 | 0 | T | C | 0.298 | -6.57E-02 | 8.43E-02 | 2.53E+00 | 2.58E+00 | 4.41E-01 | no | Gm20757 | intron_variant,non_coding_transcript_variant | MODIFIER |

| | | | | | | | | | | | | | | | |
|---|---|---|---|---|---|---|---|---|---|---|---|---|---|---|---|
| 10 | 92318308 | AX-170517184 | 0 | T | A | 0.298 | -6.57E-02 | 8.43E-02 | 2.53E+00 | 2.58E+00 | 4.41E-01 | no | Gm20757 | intron_variant,non_coding_transcript_variant | MODIFIER |
| 10 | 92321283 | AX-170465451 | 0 | A | G | 0.298 | -6.57E-02 | 8.43E-02 | 2.53E+00 | 2.58E+00 | 4.41E-01 | no | Gm20757 | intron_variant,non_coding_transcript_variant | MODIFIER |
| 10 | 92379333 | AX-170455598 | 0 | G | A | 0.298 | -6.57E-02 | 8.43E-02 | 2.53E+00 | 2.58E+00 | 4.41E-01 | no | Gm20757 | upstream_gene_variant | MODIFIER |
| 10 | 92380486 | AX-170101430 | 0 | A | G | 0.298 | -6.57E-02 | 8.43E-02 | 2.53E+00 | 2.58E+00 | 4.41E-01 | no | | intergenic_variant | MODIFIER |
| 10 | 92389112 | AX-170427762 | 0 | C | T | 0.298 | -6.57E-02 | 8.43E-02 | 2.53E+00 | 2.58E+00 | 4.41E-01 | no | | intergenic_variant | MODIFIER |
| 10 | 92389203 | AX-170441475 | 0 | G | T | 0.298 | -6.57E-02 | 8.43E-02 | 2.53E+00 | 2.58E+00 | 4.41E-01 | no | | regulatory_region_variant | MODIFIER |
| 10 | 92395572 | AX-170180331 | 0 | C | A | 0.298 | -6.57E-02 | 8.43E-02 | 2.53E+00 | 2.58E+00 | 4.41E-01 | no | | intergenic_variant | MODIFIER |
| 10 | 92411785 | AX-170150898 | 0 | G | A | 0.298 | -6.57E-02 | 8.43E-02 | 2.53E+00 | 2.58E+00 | 4.41E-01 | no | 4930401A07Rik | intron_variant,non_coding_transcript_variant | MODIFIER |
| 10 | 92431630 | AX-170196857 | 0 | T | A | 0.298 | -6.57E-02 | 8.43E-02 | 2.53E+00 | 2.58E+00 | 4.41E-01 | no | | intergenic_variant | MODIFIER |
| 10 | 92448079 | AX-170045582 | 0 | A | G | 0.298 | -6.57E-02 | 8.43E-02 | 2.53E+00 | 2.58E+00 | 4.41E-01 | no | | intergenic_variant | MODIFIER |
| 10 | 92448780 | AX-170045583 | 0 | C | A | 0.298 | -6.57E-02 | 8.43E-02 | 2.53E+00 | 2.58E+00 | 4.41E-01 | no | | intergenic_variant | MODIFIER |
| 10 | 92482856 | AX-170206246 | 0 | A | G | 0.298 | -6.57E-02 | 8.43E-02 | 2.53E+00 | 2.58E+00 | 4.41E-01 | no | Gm25884 | upstream_gene_variant | MODIFIER |
| 10 | 92492391 | AX-170113739 | 0 | G | T | 0.298 | -6.57E-02 | 8.43E-02 | 2.53E+00 | 2.58E+00 | 4.41E-01 | no | | intergenic_variant | MODIFIER |
| 10 | 92492969 | AX-170187842 | 0 | A | G | 0.298 | -6.57E-02 | 8.43E-02 | 2.53E+00 | 2.58E+00 | 4.41E-01 | no | | intergenic_variant | MODIFIER |
| 10 | 92495630 | AX-170288500 | 0 | C | T | 0.298 | -6.57E-02 | 8.43E-02 | 2.53E+00 | 2.58E+00 | 4.41E-01 | no | | intergenic_variant | MODIFIER |
| 10 | 92528807 | AX-170160182 | 0 | G | T | 0.298 | -6.57E-02 | 8.43E-02 | 2.53E+00 | 2.58E+00 | 4.41E-01 | no | | intergenic_variant | MODIFIER |
| 10 | 92533430 | AX-170152092 | 0 | A | G | 0.298 | -6.57E-02 | 8.43E-02 | 2.53E+00 | 2.58E+00 | 4.41E-01 | no | | intergenic_variant | MODIFIER |
| 10 | 92534790 | AX-170248391 | 0 | C | G | 0.298 | -6.57E-02 | 8.43E-02 | 2.53E+00 | 2.58E+00 | 4.41E-01 | no | | intergenic_variant | MODIFIER |
| 10 | 92537736 | AX-170356892 | 0 | A | G | 0.298 | -6.57E-02 | 8.43E-02 | 2.53E+00 | 2.58E+00 | 4.41E-01 | no | Gm4800 | downstream_gene_variant | MODIFIER |
| 10 | 92568349 | AX-170311447 | 0 | C | T | 0.298 | -6.57E-02 | 8.43E-02 | 2.53E+00 | 2.58E+00 | 4.41E-01 | no | | intergenic_variant | MODIFIER |
| 10 | 92568754 | AX-170251665 | 0 | G | T | 0.298 | -6.57E-02 | 8.43E-02 | 2.53E+00 | 2.58E+00 | 4.41E-01 | no | | intergenic_variant | MODIFIER |
| 10 | 92570287 | AX-170253306 | 0 | A | G | 0.298 | -6.57E-02 | 8.43E-02 | 2.53E+00 | 2.58E+00 | 4.41E-01 | no | | intergenic_variant | MODIFIER |
| 10 | 92573634 | AX-170410399 | 0 | A | G | 0.298 | -6.57E-02 | 8.43E-02 | 2.53E+00 | 2.58E+00 | 4.41E-01 | no | | intergenic_variant | MODIFIER |
| 10 | 92583576 | AX-170289789 | 0 | A | G | 0.298 | -6.57E-02 | 8.43E-02 | 2.53E+00 | 2.58E+00 | 4.41E-01 | no | Gm22582 | upstream_gene_variant | MODIFIER |
| 10 | 92595794 | AX-97833439 | 0 | C | A | 0.298 | -6.57E-02 | 8.43E-02 | 2.53E+00 | 2.58E+00 | 4.41E-01 | no | | regulatory_region_variant | MODIFIER |
| 10 | 92596377 | AX-170154675 | 0 | G | T | 0.298 | -6.57E-02 | 8.43E-02 | 2.53E+00 | 2.58E+00 | 4.41E-01 | no | | regulatory_region_variant | MODIFIER |
| 10 | 92613524 | AX-170086864 | 0 | G | A | 0.298 | -6.57E-02 | 8.43E-02 | 2.53E+00 | 2.58E+00 | 4.41E-01 | no | | intergenic_variant | MODIFIER |
| 10 | 92631434 | AX-170162978 | 0 | A | G | 0.298 | -6.57E-02 | 8.43E-02 | 2.53E+00 | 2.58E+00 | 4.41E-01 | no | | intergenic_variant | MODIFIER |
| 10 | 92631675 | AX-170309509 | 0 | A | G | 0.298 | -6.57E-02 | 8.43E-02 | 2.53E+00 | 2.58E+00 | 4.41E-01 | no | | intergenic_variant | MODIFIER |
| 10 | 92655940 | AX-170118708 | 0 | C | G | 0.298 | -6.57E-02 | 8.43E-02 | 2.53E+00 | 2.58E+00 | 4.41E-01 | no | Gm31822 | downstream_gene_variant | MODIFIER |
| 10 | 92657184 | AX-170422256 | 0 | T | C | 0.298 | -6.57E-02 | 8.43E-02 | 2.53E+00 | 2.58E+00 | 4.41E-01 | no | | intergenic_variant | MODIFIER |
| 10 | 92673320 | AX-170506712 | 0 | T | C | 0.298 | -6.57E-02 | 8.43E-02 | 2.53E+00 | 2.58E+00 | 4.41E-01 | no | | intergenic_variant | MODIFIER |
| 10 | 92682455 | AX-170368324 | 0 | A | G | 0.298 | -6.57E-02 | 8.43E-02 | 2.53E+00 | 2.58E+00 | 4.41E-01 | no | Nedd1 | downstream_gene_variant | MODIFIER |
| 10 | 92692200 | AX-170502206 | 0 | G | A | 0.298 | -6.57E-02 | 8.43E-02 | 2.53E+00 | 2.58E+00 | 4.41E-01 | no | Nedd1 | intron_variant | MODIFIER |
| 10 | 92723902 | AX-170252638 | 0 | A | C | 0.298 | -6.57E-02 | 8.43E-02 | 2.53E+00 | 2.58E+00 | 4.41E-01 | no | Nedd1 | upstream_gene_variant | MODIFIER |
| 10 | 92732908 | AX-170100922 | 0 | T | C | 0.298 | -6.57E-02 | 8.43E-02 | 2.53E+00 | 2.58E+00 | 4.41E-01 | no | | intergenic_variant | MODIFIER |
| 10 | 92745802 | AX-170347775 | 0 | C | T | 0.298 | -6.57E-02 | 8.43E-02 | 2.53E+00 | 2.58E+00 | 4.41E-01 | no | | intergenic_variant | MODIFIER |
| 10 | 92746017 | AX-170391462 | 0 | G | C | 0.298 | -6.57E-02 | 8.43E-02 | 2.53E+00 | 2.58E+00 | 4.41E-01 | no | | intergenic_variant | MODIFIER |
| 10 | 92785514 | AX-97787179 | 0 | C | T | 0.298 | -6.57E-02 | 8.43E-02 | 2.53E+00 | 2.58E+00 | 4.41E-01 | no | Cfap54 | intron_variant | MODIFIER |
| 10 | 92790187 | AX-170245892 | 0 | A | G | 0.298 | -6.57E-02 | 8.43E-02 | 2.53E+00 | 2.58E+00 | 4.41E-01 | no | Cfap54 | intron_variant | MODIFIER |
| 10 | 92796565 | AX-170273722 | 0 | A | G | 0.298 | -6.57E-02 | 8.43E-02 | 2.53E+00 | 2.58E+00 | 4.41E-01 | no | Cfap54 | downstream_gene_variant | MODIFIER |
| 10 | 92796794 | AX-170407855 | 0 | T | C | 0.298 | -6.57E-02 | 8.43E-02 | 2.53E+00 | 2.58E+00 | 4.41E-01 | no | Cfap54 | 3_prime_UTR_variant,NMD_transcript_variant | MODIFIER |
| 10 | 92797214 | AX-170248073 | 0 | C | T | 0.298 | -6.57E-02 | 8.43E-02 | 2.53E+00 | 2.58E+00 | 4.41E-01 | no | Cfap54 | 3_prime_UTR_variant,NMD_transcript_variant | MODIFIER |
| 10 | 92817801 | AX-170271133 | 0 | C | T | 0.298 | -6.57E-02 | 8.43E-02 | 2.53E+00 | 2.58E+00 | 4.41E-01 | no | Cfap54 | intron_variant,NMD_transcript_variant | MODIFIER |
| 10 | 92818195 | AX-170393495 | 0 | C | T | 0.298 | -6.57E-02 | 8.43E-02 | 2.53E+00 | 2.58E+00 | 4.41E-01 | no | Cfap54 | intron_variant,NMD_transcript_variant | MODIFIER |
| 10 | 92829261 | AX-170045594 | 0 | C | T | 0.298 | -6.57E-02 | 8.43E-02 | 2.53E+00 | 2.58E+00 | 4.41E-01 | no | Cfap54 | intron_variant,NMD_transcript_variant | MODIFIER |
| 10 | 92830914 | AX-170155551 | 0 | A | C | 0.298 | -6.57E-02 | 8.43E-02 | 2.53E+00 | 2.58E+00 | 4.41E-01 | no | Cfap54 | intron_variant,NMD_transcript_variant | MODIFIER |
| 10 | 92841273 | AX-170274725 | 0 | G | A | 0.298 | -6.57E-02 | 8.43E-02 | 2.53E+00 | 2.58E+00 | 4.41E-01 | no | Cfap54 | upstream_gene_variant | MODIFIER |
| 10 | 92847125 | AX-170293441 | 0 | T | C | 0.298 | -6.57E-02 | 8.43E-02 | 2.53E+00 | 2.58E+00 | 4.41E-01 | no | Cfap54 | intron_variant | MODIFIER |
| 10 | 92855679 | AX-170138854 | 0 | G | T | 0.298 | -6.57E-02 | 8.43E-02 | 2.53E+00 | 2.58E+00 | 4.41E-01 | no | Cfap54 | intron_variant | MODIFIER |
| 10 | 92856084 | AX-170224800 | 0 | A | G | 0.298 | -6.57E-02 | 8.43E-02 | 2.53E+00 | 2.58E+00 | 4.41E-01 | no | Cfap54 | intron_variant | MODIFIER |
| 10 | 92859871 | AX-170291958 | 0 | C | T | 0.298 | -6.57E-02 | 8.43E-02 | 2.53E+00 | 2.58E+00 | 4.41E-01 | no | Cfap54 | intron_variant | MODIFIER |
| 10 | 92872723 | AX-170424403 | 0 | G | A | 0.298 | -6.57E-02 | 8.43E-02 | 2.53E+00 | 2.58E+00 | 4.41E-01 | no | Cfap54 | intron_variant | MODIFIER |
| 10 | 92882565 | AX-170317515 | 0 | C | A | 0.298 | -6.57E-02 | 8.43E-02 | 2.53E+00 | 2.58E+00 | 4.41E-01 | no | Cfap54 | intron_variant | MODIFIER |

| | | | | | | | | | | | | | | | |
|---|---|---|---|---|---|---|---|---|---|---|---|---|---|---|---|
| 10 | 92892100 | AX-170003666 | 0 | C | T | 0.298 | -6.57E-02 | 8.43E-02 | 2.53E+00 | 2.58E+00 | 4.41E-01 | no | Cfap54 | intron_variant | MODIFIER |
| 10 | 92903686 | AX-97817644 | 0 | A | G | 0.298 | -6.57E-02 | 8.43E-02 | 2.53E+00 | 2.58E+00 | 4.41E-01 | no | Cfap54 | intron_variant,NMD_transcript_variant | MODIFIER |
| 10 | 92926909 | AX-170430121 | 0 | A | G | 0.298 | -6.57E-02 | 8.43E-02 | 2.53E+00 | 2.58E+00 | 4.41E-01 | no | Cfap54 | intron_variant,NMD_transcript_variant | MODIFIER |
| 10 | 92943102 | AX-170131558 | 0 | G | A | 0.298 | -6.57E-02 | 8.43E-02 | 2.53E+00 | 2.58E+00 | 4.41E-01 | no | Cfap54 | upstream_gene_variant | MODIFIER |
| 10 | 92943133 | AX-170278752 | 0 | T | C | 0.298 | -6.57E-02 | 8.43E-02 | 2.53E+00 | 2.58E+00 | 4.41E-01 | no | Cfap54 | upstream_gene_variant | MODIFIER |
| 10 | 92944007 | AX-97834966 | 0 | T | G | 0.298 | -6.57E-02 | 8.43E-02 | 2.53E+00 | 2.58E+00 | 4.41E-01 | no | Cfap54 | upstream_gene_variant | MODIFIER |
| 10 | 92986620 | AX-170045596 | 0 | A | G | 0.298 | -6.57E-02 | 8.43E-02 | 2.53E+00 | 2.58E+00 | 4.41E-01 | no | Cfap54 | intron_variant | MODIFIER |
| 10 | 92988096 | AX-97783261 | 0 | A | G | 0.298 | -6.57E-02 | 8.43E-02 | 2.53E+00 | 2.58E+00 | 4.41E-01 | no | Cfap54 | intron_variant | MODIFIER |
| 10 | 92996794 | AX-170381275 | 0 | G | A | 0.298 | -6.57E-02 | 8.43E-02 | 2.53E+00 | 2.58E+00 | 4.41E-01 | no | Cfap54 | intron_variant | MODIFIER |
| 10 | 93032191 | AX-170167077 | 0 | G | C | 0.298 | -6.57E-02 | 8.43E-02 | 2.53E+00 | 2.58E+00 | 4.41E-01 | no | Cfap54 | intron_variant | MODIFIER |
| 10 | 93037888 | AX-170228936 | 0 | C | T | 0.298 | -6.57E-02 | 8.43E-02 | 2.53E+00 | 2.58E+00 | 4.41E-01 | no | Cfap54 | missense_variant | MODERATE |
| 10 | 93038024 | AX-170342757 | 0 | C | T | 0.298 | -6.57E-02 | 8.43E-02 | 2.53E+00 | 2.58E+00 | 4.41E-01 | no | Cfap54 | intron_variant | MODIFIER |
| 10 | 93046682 | AX-170326486 | 0 | T | C | 0.298 | -6.57E-02 | 8.43E-02 | 2.53E+00 | 2.58E+00 | 4.41E-01 | no | Cfap54 | intron_variant | MODIFIER |
| 10 | 93061346 | AX-97809596 | 0 | G | A | 0.298 | -6.57E-02 | 8.43E-02 | 2.53E+00 | 2.58E+00 | 4.41E-01 | no | Cfap54 | intron_variant | MODIFIER |
| 10 | 93067738 | AX-170406836 | 0 | A | G | 0.298 | -6.57E-02 | 8.43E-02 | 2.53E+00 | 2.58E+00 | 4.41E-01 | no | Cfap54 | intron_variant | MODIFIER |
| 10 | 93087102 | AX-170420974 | 0 | T | G | 0.298 | -6.57E-02 | 8.43E-02 | 2.53E+00 | 2.58E+00 | 4.41E-01 | no | Gm32468 | intron_variant,non_coding_transcript_variant | MODIFIER |
| 10 | 93087500 | AX-170193452 | 0 | T | G | 0.298 | -6.57E-02 | 8.43E-02 | 2.53E+00 | 2.58E+00 | 4.41E-01 | no | Gm32468 | intron_variant,non_coding_transcript_variant | MODIFIER |
| 10 | 93100615 | AX-170407543 | 0 | C | G | 0.298 | -6.57E-02 | 8.43E-02 | 2.53E+00 | 2.58E+00 | 4.41E-01 | no | | intergenic_variant | MODIFIER |
| 10 | 93116287 | AX-170223918 | 0 | T | C | 0.298 | -6.57E-02 | 8.43E-02 | 2.53E+00 | 2.58E+00 | 4.41E-01 | no | | intergenic_variant | MODIFIER |
| 10 | 93120633 | AX-170284884 | 0 | G | A | 0.298 | -6.57E-02 | 8.43E-02 | 2.53E+00 | 2.58E+00 | 4.41E-01 | no | | intergenic_variant | MODIFIER |
| 10 | 93145216 | AX-170045598 | 0 | T | G | 0.298 | -6.57E-02 | 8.43E-02 | 2.53E+00 | 2.58E+00 | 4.41E-01 | no | | regulatory_region_variant | MODIFIER |
| 10 | 93163915 | AX-170495572 | 0 | T | C | 0.298 | -6.57E-02 | 8.43E-02 | 2.53E+00 | 2.58E+00 | 4.41E-01 | no | Cdk17 | intron_variant | MODIFIER |
| 10 | 93164148 | AX-170086865 | 0 | C | T | 0.298 | -6.57E-02 | 8.43E-02 | 2.53E+00 | 2.58E+00 | 4.41E-01 | no | Cdk17 | intron_variant | MODIFIER |
| 10 | 93195160 | AX-170493836 | 0 | T | C | 0.298 | -6.57E-02 | 8.43E-02 | 2.53E+00 | 2.58E+00 | 4.41E-01 | no | Cdk17 | intron_variant | MODIFIER |
| 10 | 93199654 | AX-170098699 | 0 | C | A | 0.298 | -6.57E-02 | 8.43E-02 | 2.53E+00 | 2.58E+00 | 4.41E-01 | no | Cdk17 | intron_variant | MODIFIER |
| 10 | 93203201 | AX-170455010 | 0 | C | T | 0.298 | -6.57E-02 | 8.43E-02 | 2.53E+00 | 2.58E+00 | 4.41E-01 | no | Cdk17 | intron_variant | MODIFIER |
| 10 | 93236499 | AX-170521255 | 0 | T | C | 0.298 | -6.57E-02 | 8.43E-02 | 2.53E+00 | 2.58E+00 | 4.41E-01 | no | Cdk17 | intron_variant | MODIFIER |
| 10 | 93240027 | AX-97815182 | 0 | A | G | 0.298 | -6.57E-02 | 8.43E-02 | 2.53E+00 | 2.58E+00 | 4.41E-01 | no | Cdk17 | 3_prime_UTR_variant | MODIFIER |
| 10 | 93252661 | AX-170515270 | 0 | C | A | 0.298 | -6.57E-02 | 8.43E-02 | 2.53E+00 | 2.58E+00 | 4.41E-01 | no | Elk3 | intron_variant | MODIFIER |
| 10 | 93266058 | AX-97849742 | 0 | A | G | 0.298 | -6.57E-02 | 8.43E-02 | 2.53E+00 | 2.58E+00 | 4.41E-01 | no | Elk3 | intron_variant | MODIFIER |
| 10 | 93269645 | AX-170311039 | 0 | C | A | 0.298 | -6.57E-02 | 8.43E-02 | 2.53E+00 | 2.58E+00 | 4.41E-01 | no | Elk3 | intron_variant | MODIFIER |
| 10 | 93298485 | AX-170045605 | 0 | A | C | 0.298 | -6.57E-02 | 8.43E-02 | 2.53E+00 | 2.58E+00 | 4.41E-01 | no | Elk3 | intron_variant | MODIFIER |
| 10 | 93298897 | AX-97844732 | 0 | G | A | 0.298 | -6.57E-02 | 8.43E-02 | 2.53E+00 | 2.58E+00 | 4.41E-01 | no | Elk3 | intron_variant | MODIFIER |
| 10 | 93301917 | AX-170045604 | 0 | A | G | 0.298 | -6.57E-02 | 8.43E-02 | 2.53E+00 | 2.58E+00 | 4.41E-01 | no | Elk3 | intron_variant | MODIFIER |
| 10 | 93303038 | AX-97784459 | 0 | G | A | 0.298 | -6.57E-02 | 8.43E-02 | 2.53E+00 | 2.58E+00 | 4.41E-01 | no | Elk3 | intron_variant | MODIFIER |
| 10 | 93303140 | AX-170045607 | 0 | G | A | 0.298 | -6.57E-02 | 8.43E-02 | 2.53E+00 | 2.58E+00 | 4.41E-01 | no | Elk3 | intron_variant | MODIFIER |
| 10 | 93303977 | AX-170369939 | 0 | G | A | 0.298 | -6.57E-02 | 8.43E-02 | 2.53E+00 | 2.58E+00 | 4.41E-01 | no | Elk3 | intron_variant | MODIFIER |
| 10 | 93304196 | AX-170045609 | 0 | C | T | 0.298 | -6.57E-02 | 8.43E-02 | 2.53E+00 | 2.58E+00 | 4.41E-01 | no | Elk3 | intron_variant | MODIFIER |
| 10 | 93314551 | AX-170491581 | 0 | C | T | 0.298 | -6.57E-02 | 8.43E-02 | 2.53E+00 | 2.58E+00 | 4.41E-01 | no | Elk3 | upstream_gene_variant | MODIFIER |
| 10 | 93318990 | AX-170505888 | 0 | C | T | 0.298 | -6.57E-02 | 8.43E-02 | 2.53E+00 | 2.58E+00 | 4.41E-01 | no | Gm32688 | intron_variant,non_coding_transcript_variant | MODIFIER |
| 10 | 93343882 | AX-97787608 | 0 | A | G | 0.298 | -6.57E-02 | 8.43E-02 | 2.53E+00 | 2.58E+00 | 4.41E-01 | no | Gm17745 | intron_variant,non_coding_transcript_variant | MODIFIER |
| 10 | 93357443 | AX-170329883 | 0 | C | A | 0.298 | -6.57E-02 | 8.43E-02 | 2.53E+00 | 2.58E+00 | 4.41E-01 | no | | intergenic_variant | MODIFIER |
| 10 | 93358081 | AX-170368612 | 0 | C | G | 0.298 | -6.57E-02 | 8.43E-02 | 2.53E+00 | 2.58E+00 | 4.41E-01 | no | | intergenic_variant | MODIFIER |
| 10 | 93370778 | AX-170229362 | 0 | A | G | 0.298 | -6.57E-02 | 8.43E-02 | 2.53E+00 | 2.58E+00 | 4.41E-01 | no | | regulatory_region_variant | MODIFIER |
| 10 | 93375516 | AX-170378330 | 0 | C | T | 0.298 | -6.57E-02 | 8.43E-02 | 2.53E+00 | 2.58E+00 | 4.41E-01 | no | | regulatory_region_variant | MODIFIER |
| 10 | 93375585 | AX-170354241 | 0 | T | C | 0.298 | -6.57E-02 | 8.43E-02 | 2.53E+00 | 2.58E+00 | 4.41E-01 | no | | regulatory_region_variant | MODIFIER |
| 10 | 93399730 | AX-170164829 | 0 | A | C | 0.298 | -6.57E-02 | 8.43E-02 | 2.53E+00 | 2.58E+00 | 4.41E-01 | no | Gm47344 | upstream_gene_variant | MODIFIER |
| 10 | 93400364 | AX-170502393 | 0 | G | T | 0.298 | -6.57E-02 | 8.43E-02 | 2.53E+00 | 2.58E+00 | 4.41E-01 | no | Gm47344 | upstream_gene_variant | MODIFIER |
| 10 | 93443407 | AX-97792141 | 0 | C | T | 0.298 | -6.57E-02 | 8.43E-02 | 2.53E+00 | 2.58E+00 | 4.41E-01 | no | | regulatory_region_variant | MODIFIER |
| 10 | 93444145 | AX-170470429 | 0 | A | T | 0.298 | -6.57E-02 | 8.43E-02 | 2.53E+00 | 2.58E+00 | 4.41E-01 | no | | regulatory_region_variant | MODIFIER |
| 10 | 93448537 | AX-170120253 | 0 | A | C | 0.298 | -6.57E-02 | 8.43E-02 | 2.53E+00 | 2.58E+00 | 4.41E-01 | no | Lta4h | upstream_gene_variant | MODIFIER |
| 10 | 93448595 | AX-170420928 | 0 | C | T | 0.298 | -6.57E-02 | 8.43E-02 | 2.53E+00 | 2.58E+00 | 4.41E-01 | no | Lta4h | upstream_gene_variant | MODIFIER |
| 10 | 93469915 | AX-170003669 | 0 | A | G | 0.298 | -6.57E-02 | 8.43E-02 | 2.53E+00 | 2.58E+00 | 4.41E-01 | no | Lta4h | intron_variant | MODIFIER |
| 10 | 93479244 | AX-170003673 | 0 | A | G | 0.298 | -6.57E-02 | 8.43E-02 | 2.53E+00 | 2.58E+00 | 4.41E-01 | no | Lta4h | intron_variant | MODIFIER |

| | | | | | | | | | | | | | | | |
|---|---|---|---|---|---|---|---|---|---|---|---|---|---|---|---|
| 10 | 93491106 | AX-170395996 | 0 | C | T | 0.298 | -6.57E-02 | 8.43E-02 | 2.53E+00 | 2.58E+00 | 4.41E-01 | no | Hal | intron_variant,NMD_transcript_variant | MODIFIER |
| 10 | 93496661 | AX-170003668 | 0 | C | A | 0.298 | -6.57E-02 | 8.43E-02 | 2.53E+00 | 2.58E+00 | 4.41E-01 | no | Hal | intron_variant,NMD_transcript_variant | MODIFIER |
| 10 | 93516235 | AX-170003671 | 0 | A | G | 0.298 | -6.57E-02 | 8.43E-02 | 2.53E+00 | 2.58E+00 | 4.41E-01 | no | Hal | 3_prime_UTR_variant,NMD_transcript_variant | MODIFIER |
| 10 | 93582684 | AX-170240393 | 0 | G | A | 0.298 | -6.57E-02 | 8.43E-02 | 2.53E+00 | 2.58E+00 | 4.41E-01 | no | Snrpf | downstream_gene_variant | MODIFIER |
| 10 | 93586683 | AX-170003670 | 0 | C | T | 0.298 | -6.57E-02 | 8.43E-02 | 2.53E+00 | 2.58E+00 | 4.41E-01 | no | Snrpf | intron_variant | MODIFIER |
| 10 | 93618025 | AX-170191330 | 0 | T | C | 0.298 | -6.57E-02 | 8.43E-02 | 2.53E+00 | 2.58E+00 | 4.41E-01 | no | Gm47156 | upstream_gene_variant | MODIFIER |
| 10 | 93638706 | AX-97816350 | 0 | G | A | 0.298 | -6.57E-02 | 8.43E-02 | 2.53E+00 | 2.58E+00 | 4.41E-01 | no | Ntn4 | upstream_gene_variant | MODIFIER |
| 10 | 93639685 | AX-170514787 | 0 | T | C | 0.298 | -6.57E-02 | 8.43E-02 | 2.53E+00 | 2.58E+00 | 4.41E-01 | no | Ntn4 | upstream_gene_variant | MODIFIER |
| 10 | 93660463 | AX-97816421 | 0 | C | T | 0.298 | -6.57E-02 | 8.43E-02 | 2.53E+00 | 2.58E+00 | 4.41E-01 | no | Ntn4 | intron_variant | MODIFIER |
| 10 | 93662028 | AX-170281883 | 0 | T | C | 0.298 | -6.57E-02 | 8.43E-02 | 2.53E+00 | 2.58E+00 | 4.41E-01 | no | Ntn4 | intron_variant | MODIFIER |
| 10 | 93687113 | AX-170346521 | 0 | T | C | 0.298 | -6.57E-02 | 8.43E-02 | 2.53E+00 | 2.58E+00 | 4.41E-01 | no | Ntn4 | intron_variant | MODIFIER |
| 10 | 93702143 | AX-170275914 | 0 | C | T | 0.298 | -6.57E-02 | 8.43E-02 | 2.53E+00 | 2.58E+00 | 4.41E-01 | no | Ntn4 | intron_variant | MODIFIER |
| 10 | 93706286 | AX-170117053 | 0 | A | G | 0.298 | -6.57E-02 | 8.43E-02 | 2.53E+00 | 2.58E+00 | 4.41E-01 | no | Ntn4 | intron_variant | MODIFIER |
| 10 | 93706875 | AX-170286663 | 0 | G | A | 0.298 | -6.57E-02 | 8.43E-02 | 2.53E+00 | 2.58E+00 | 4.41E-01 | no | Ntn4 | intron_variant | MODIFIER |
| 10 | 93717178 | AX-170103040 | 0 | T | C | 0.298 | -6.57E-02 | 8.43E-02 | 2.53E+00 | 2.58E+00 | 4.41E-01 | no | Ntn4 | intron_variant | MODIFIER |
| 10 | 93721684 | AX-170163388 | 0 | C | T | 0.298 | -6.57E-02 | 8.43E-02 | 2.53E+00 | 2.58E+00 | 4.41E-01 | no | Ntn4 | intron_variant | MODIFIER |
| 10 | 93737439 | AX-170235486 | 0 | G | T | 0.298 | -6.57E-02 | 8.43E-02 | 2.53E+00 | 2.58E+00 | 4.41E-01 | no | Ntn4 | intron_variant | MODIFIER |
| 1 | 101534844 | AX-170042199 | 2 | A | T | 0.145 | 1.17E-01 | 1.51E-01 | 2.54E+00 | 2.65E+00 | 4.41E-01 | no | | intergenic_variant | MODIFIER |
| 1 | 96157052 | AX-97825716 | 1 | A | C | 0.152 | 1.52E-01 | 1.99E-01 | 2.64E+00 | 2.70E+00 | 4.41E-01 | Yes | | intergenic_variant | MODIFIER |
| 4 | 107785041 | AX-170070289 | 0 | G | A | 0.263 | 8.31E-02 | 1.08E-01 | 2.59E+00 | 2.67E+00 | 4.41E-01 | no | 4933424M12Rik | upstream_gene_variant | MODIFIER |
| 9 | 21127901 | AX-168410507 | 1 | A | C | 0.473 | 6.61E-02 | 8.52E-02 | 2.54E+00 | 2.61E+00 | 4.41E-01 | no | Tyk2 | upstream_gene_variant | MODIFIER |
| 7 | 116411166 | AX-170098179 | 1 | C | G | 0.42 | 6.88E-02 | 9.05E-02 | 2.71E+00 | 2.77E+00 | 4.41E-01 | no | Pik3c2a | intron_variant | MODIFIER |
| 8 | 31944425 | AX-97836900 | 0 | T | C | 0.14 | -1.03E-01 | 1.35E-01 | 2.66E+00 | 2.72E+00 | 4.41E-01 | no | Nrg1 | intron_variant | MODIFIER |
| 8 | 33857151 | AX-170162926 | 0 | G | A | 0.14 | -1.03E-01 | 1.35E-01 | 2.66E+00 | 2.72E+00 | 4.41E-01 | no | Rbpms | intron_variant | MODIFIER |
| 8 | 33858988 | AX-170090080 | 0 | C | A | 0.14 | -1.03E-01 | 1.35E-01 | 2.66E+00 | 2.72E+00 | 4.41E-01 | no | Rbpms | intron_variant | MODIFIER |
| 8 | 33886013 | AX-170402229 | 0 | A | G | 0.14 | -1.03E-01 | 1.35E-01 | 2.66E+00 | 2.72E+00 | 4.41E-01 | no | Rbpms | intron_variant | MODIFIER |
| 8 | 33886328 | AX-170519013 | 0 | C | T | 0.14 | -1.03E-01 | 1.35E-01 | 2.66E+00 | 2.72E+00 | 4.41E-01 | no | Rbpms | intron_variant | MODIFIER |
| 8 | 33886945 | AX-170286160 | 0 | G | A | 0.14 | -1.03E-01 | 1.35E-01 | 2.66E+00 | 2.72E+00 | 4.41E-01 | no | Rbpms | intron_variant | MODIFIER |
| 8 | 33907203 | AX-170264086 | 0 | A | G | 0.14 | -1.03E-01 | 1.35E-01 | 2.66E+00 | 2.72E+00 | 4.41E-01 | no | Rbpms | intron_variant | MODIFIER |
| 8 | 33907348 | AX-170224332 | 0 | T | A | 0.14 | -1.03E-01 | 1.35E-01 | 2.66E+00 | 2.72E+00 | 4.41E-01 | no | Rbpms | intron_variant | MODIFIER |
| 8 | 33955746 | AX-170316062 | 0 | A | T | 0.14 | -1.03E-01 | 1.35E-01 | 2.66E+00 | 2.72E+00 | 4.41E-01 | no | | intergenic_variant | MODIFIER |
| 8 | 33957464 | AX-170101072 | 0 | A | G | 0.14 | -1.03E-01 | 1.35E-01 | 2.66E+00 | 2.72E+00 | 4.41E-01 | no | | regulatory_region_variant | MODIFIER |
| 8 | 33994927 | AX-97803513 | 0 | C | T | 0.14 | -1.03E-01 | 1.35E-01 | 2.66E+00 | 2.72E+00 | 4.41E-01 | no | Gm39157 | downstream_gene_variant | MODIFIER |
| 8 | 33995066 | AX-170330139 | 0 | C | G | 0.14 | -1.03E-01 | 1.35E-01 | 2.66E+00 | 2.72E+00 | 4.41E-01 | no | Gm39157 | downstream_gene_variant | MODIFIER |
| 8 | 34054743 | AX-170109313 | 0 | T | A | 0.14 | -1.03E-01 | 1.35E-01 | 2.66E+00 | 2.72E+00 | 4.41E-01 | no | Gm9951 | intron_variant,non_coding_transcript_variant | MODIFIER |
| 8 | 34064439 | AX-170274751 | 0 | C | T | 0.14 | -1.03E-01 | 1.35E-01 | 2.66E+00 | 2.72E+00 | 4.41E-01 | no | | regulatory_region_variant | MODIFIER |
| 8 | 34065807 | AX-170504336 | 0 | T | C | 0.14 | -1.03E-01 | 1.35E-01 | 2.66E+00 | 2.72E+00 | 4.41E-01 | no | | regulatory_region_variant | MODIFIER |
| 8 | 34068176 | AX-170241285 | 0 | A | G | 0.14 | -1.03E-01 | 1.35E-01 | 2.66E+00 | 2.72E+00 | 4.41E-01 | no | | intergenic_variant | MODIFIER |
| 8 | 34104645 | AX-170079705 | 0 | G | C | 0.14 | -1.03E-01 | 1.35E-01 | 2.66E+00 | 2.72E+00 | 4.41E-01 | no | Dctn6 | intron_variant | MODIFIER |
| 8 | 34105177 | AX-97790494 | 0 | T | C | 0.14 | -1.03E-01 | 1.35E-01 | 2.66E+00 | 2.72E+00 | 4.41E-01 | no | Dctn6 | intron_variant | MODIFIER |
| 8 | 34107708 | AX-170135369 | 0 | C | G | 0.14 | -1.03E-01 | 1.35E-01 | 2.66E+00 | 2.72E+00 | 4.41E-01 | no | Dctn6 | intron_variant | MODIFIER |
| 8 | 34115184 | AX-170255844 | 0 | A | G | 0.14 | -1.03E-01 | 1.35E-01 | 2.66E+00 | 2.72E+00 | 4.41E-01 | no | Gm10131 | upstream_gene_variant | MODIFIER |
| 8 | 34118306 | AX-170254823 | 0 | T | C | 0.14 | -1.03E-01 | 1.35E-01 | 2.66E+00 | 2.72E+00 | 4.41E-01 | no | Mboat4 | intron_variant | MODIFIER |
| 8 | 34120771 | AX-170240711 | 0 | G | A | 0.14 | -1.03E-01 | 1.35E-01 | 2.66E+00 | 2.72E+00 | 4.41E-01 | no | Mboat4 | intron_variant | MODIFIER |
| 8 | 34120794 | AX-170400743 | 0 | G | T | 0.14 | -1.03E-01 | 1.35E-01 | 2.66E+00 | 2.72E+00 | 4.41E-01 | no | Mboat4 | intron_variant | MODIFIER |
| 8 | 34123629 | AX-170488141 | 0 | A | G | 0.14 | -1.03E-01 | 1.35E-01 | 2.66E+00 | 2.72E+00 | 4.41E-01 | no | Mboat4 | intron_variant | MODIFIER |
| 8 | 34136327 | AX-170224390 | 0 | C | T | 0.14 | -1.03E-01 | 1.35E-01 | 2.66E+00 | 2.72E+00 | 4.41E-01 | no | Leprotl1 | 3_prime_UTR_variant | MODIFIER |
| 8 | 34189212 | AX-170030486 | 0 | G | A | 0.14 | -1.03E-01 | 1.35E-01 | 2.66E+00 | 2.72E+00 | 4.41E-01 | no | | regulatory_region_variant | MODIFIER |
| 8 | 34189276 | AX-170481196 | 0 | G | A | 0.14 | -1.03E-01 | 1.35E-01 | 2.66E+00 | 2.72E+00 | 4.41E-01 | no | | regulatory_region_variant | MODIFIER |
| 8 | 34211963 | AX-170030485 | 0 | C | T | 0.14 | -1.03E-01 | 1.35E-01 | 2.66E+00 | 2.72E+00 | 4.41E-01 | no | Gm29243 | upstream_gene_variant | MODIFIER |
| 8 | 34213194 | AX-97851512 | 0 | G | A | 0.14 | -1.03E-01 | 1.35E-01 | 2.66E+00 | 2.72E+00 | 4.41E-01 | no | Gm29243 | upstream_gene_variant | MODIFIER |
| 8 | 34270973 | AX-97840652 | 0 | T | C | 0.14 | -1.03E-01 | 1.35E-01 | 2.66E+00 | 2.72E+00 | 4.41E-01 | no | | intergenic_variant | MODIFIER |
| 8 | 34284624 | AX-170363158 | 0 | C | T | 0.14 | -1.03E-01 | 1.35E-01 | 2.66E+00 | 2.72E+00 | 4.41E-01 | no | | intergenic_variant | MODIFIER |
| 8 | 34296691 | AX-97802512 | 0 | T | C | 0.14 | -1.03E-01 | 1.35E-01 | 2.66E+00 | 2.72E+00 | 4.41E-01 | no | Gm23177 | upstream_gene_variant | MODIFIER |

| | | | | | | | | | | | | | | | |
|---|---|---|---|---|---|---|---|---|---|---|---|---|---|---|---|
| 8 | 34297235 | AX-170079708 | 0 | G | A | 0.14 | -1.03E-01 | 1.35E-01 | 2.66E+00 | 2.72E+00 | 4.41E-01 | no | Gm23177 | upstream_gene_variant | MODIFIER |
| 8 | 34297291 | AX-170079707 | 0 | A | T | 0.14 | -1.03E-01 | 1.35E-01 | 2.66E+00 | 2.72E+00 | 4.41E-01 | no | Gm23177 | upstream_gene_variant | MODIFIER |
| 8 | 34300690 | AX-170479383 | 0 | A | G | 0.14 | -1.03E-01 | 1.35E-01 | 2.66E+00 | 2.72E+00 | 4.41E-01 | no | Gm23177 | upstream_gene_variant | MODIFIER |
| 8 | 34300905 | AX-170143345 | 0 | T | C | 0.14 | -1.03E-01 | 1.35E-01 | 2.66E+00 | 2.72E+00 | 4.41E-01 | no | Gm23177 | upstream_gene_variant | MODIFIER |
| 8 | 34301507 | AX-170259107 | 0 | G | A | 0.14 | -1.03E-01 | 1.35E-01 | 2.66E+00 | 2.72E+00 | 4.41E-01 | no | | intergenic_variant | MODIFIER |
| 8 | 34301535 | AX-170030489 | 0 | A | G | 0.14 | -1.03E-01 | 1.35E-01 | 2.66E+00 | 2.72E+00 | 4.41E-01 | no | | intergenic_variant | MODIFIER |
| 8 | 34301993 | AX-170312407 | 0 | A | G | 0.14 | -1.03E-01 | 1.35E-01 | 2.66E+00 | 2.72E+00 | 4.41E-01 | no | | intergenic_variant | MODIFIER |
| 8 | 34302497 | AX-170297710 | 0 | C | A | 0.14 | -1.03E-01 | 1.35E-01 | 2.66E+00 | 2.72E+00 | 4.41E-01 | no | Gm6100 | downstream_gene_variant | MODIFIER |
| 8 | 34302950 | AX-170352666 | 0 | G | A | 0.14 | -1.03E-01 | 1.35E-01 | 2.66E+00 | 2.72E+00 | 4.41E-01 | no | Gm6100 | downstream_gene_variant | MODIFIER |
| 8 | 34372230 | AX-170258257 | 0 | T | C | 0.14 | -1.03E-01 | 1.35E-01 | 2.66E+00 | 2.72E+00 | 4.41E-01 | no | | regulatory_region_variant | MODIFIER |
| 8 | 34374370 | AX-170246919 | 0 | G | A | 0.14 | -1.03E-01 | 1.35E-01 | 2.66E+00 | 2.72E+00 | 4.41E-01 | no | | intergenic_variant | MODIFIER |
| 8 | 34386254 | AX-170467892 | 0 | C | A | 0.14 | -1.03E-01 | 1.35E-01 | 2.66E+00 | 2.72E+00 | 4.41E-01 | no | Gm33831 | upstream_gene_variant | MODIFIER |
| 8 | 34386304 | AX-170360759 | 0 | C | T | 0.14 | -1.03E-01 | 1.35E-01 | 2.66E+00 | 2.72E+00 | 4.41E-01 | no | Gm33831 | upstream_gene_variant | MODIFIER |
| 8 | 34388168 | AX-170403873 | 0 | C | A | 0.14 | -1.03E-01 | 1.35E-01 | 2.66E+00 | 2.72E+00 | 4.41E-01 | no | Gm33831 | upstream_gene_variant | MODIFIER |
| 8 | 34388337 | AX-170139464 | 0 | A | G | 0.14 | -1.03E-01 | 1.35E-01 | 2.66E+00 | 2.72E+00 | 4.41E-01 | no | Gm33831 | upstream_gene_variant | MODIFIER |
| 8 | 34392958 | AX-170030490 | 0 | C | T | 0.14 | -1.03E-01 | 1.35E-01 | 2.66E+00 | 2.72E+00 | 4.41E-01 | no | | regulatory_region_variant | MODIFIER |
| 8 | 34396600 | AX-170477035 | 0 | G | A | 0.14 | -1.03E-01 | 1.35E-01 | 2.66E+00 | 2.72E+00 | 4.41E-01 | no | | regulatory_region_variant | MODIFIER |
| 8 | 34397566 | AX-170243849 | 0 | A | T | 0.14 | -1.03E-01 | 1.35E-01 | 2.66E+00 | 2.72E+00 | 4.41E-01 | no | | regulatory_region_variant | MODIFIER |
| 8 | 34404652 | AX-170079711 | 0 | G | T | 0.14 | -1.03E-01 | 1.35E-01 | 2.66E+00 | 2.72E+00 | 4.41E-01 | no | | intergenic_variant | MODIFIER |
| 8 | 34428842 | AX-170227777 | 0 | A | C | 0.14 | -1.03E-01 | 1.35E-01 | 2.66E+00 | 2.72E+00 | 4.41E-01 | no | Gm45349 | intron_variant,non_coding_transcript_variant | MODIFIER |
| 8 | 34432737 | AX-170148384 | 0 | C | T | 0.14 | -1.03E-01 | 1.35E-01 | 2.66E+00 | 2.72E+00 | 4.41E-01 | no | Gm45349 | intron_variant,non_coding_transcript_variant | MODIFIER |
| 8 | 34436415 | AX-170248524 | 0 | G | A | 0.14 | -1.03E-01 | 1.35E-01 | 2.66E+00 | 2.72E+00 | 4.41E-01 | no | Gm45349 | intron_variant,non_coding_transcript_variant | MODIFIER |
| 8 | 34437586 | AX-170376344 | 0 | C | T | 0.14 | -1.03E-01 | 1.35E-01 | 2.66E+00 | 2.72E+00 | 4.41E-01 | no | Gm45349 | intron_variant,non_coding_transcript_variant | MODIFIER |
| 8 | 34440305 | AX-170347489 | 0 | C | A | 0.14 | -1.03E-01 | 1.35E-01 | 2.66E+00 | 2.72E+00 | 4.41E-01 | no | Gm45349 | intron_variant,non_coding_transcript_variant | MODIFIER |
| 8 | 34450540 | AX-170446319 | 0 | C | T | 0.14 | -1.03E-01 | 1.35E-01 | 2.66E+00 | 2.72E+00 | 4.41E-01 | no | Gm45349 | intron_variant,non_coding_transcript_variant | MODIFIER |
| 8 | 34453678 | AX-170390505 | 0 | A | G | 0.14 | -1.03E-01 | 1.35E-01 | 2.66E+00 | 2.72E+00 | 4.41E-01 | no | Gm45349 | intron_variant,non_coding_transcript_variant | MODIFIER |
| 8 | 34456749 | AX-170286916 | 0 | A | G | 0.14 | -1.03E-01 | 1.35E-01 | 2.66E+00 | 2.72E+00 | 4.41E-01 | no | Gm45349 | intron_variant,non_coding_transcript_variant | MODIFIER |
| 8 | 34462376 | AX-170508756 | 0 | A | C | 0.14 | -1.03E-01 | 1.35E-01 | 2.66E+00 | 2.72E+00 | 4.41E-01 | no | Gm45349 | intron_variant,non_coding_transcript_variant | MODIFIER |
| 8 | 34475354 | AX-170193863 | 0 | A | C | 0.14 | -1.03E-01 | 1.35E-01 | 2.66E+00 | 2.72E+00 | 4.41E-01 | no | Gm45349 | upstream_gene_variant | MODIFIER |
| 8 | 34503023 | AX-97793172 | 0 | G | A | 0.14 | -1.03E-01 | 1.35E-01 | 2.66E+00 | 2.72E+00 | 4.41E-01 | no | Gm8268 | upstream_gene_variant | MODIFIER |
| 8 | 34503466 | AX-170524232 | 0 | C | T | 0.14 | -1.03E-01 | 1.35E-01 | 2.66E+00 | 2.72E+00 | 4.41E-01 | no | Gm8268 | upstream_gene_variant | MODIFIER |
| 8 | 34508627 | AX-170355781 | 0 | C | T | 0.14 | -1.03E-01 | 1.35E-01 | 2.66E+00 | 2.72E+00 | 4.41E-01 | no | Gm8268 | downstream_gene_variant | MODIFIER |
| 8 | 34508953 | AX-170270385 | 0 | G | A | 0.14 | -1.03E-01 | 1.35E-01 | 2.66E+00 | 2.72E+00 | 4.41E-01 | no | Gm8268 | downstream_gene_variant | MODIFIER |
| 8 | 34533679 | AX-170362485 | 0 | T | C | 0.14 | -1.03E-01 | 1.35E-01 | 2.66E+00 | 2.72E+00 | 4.41E-01 | no | | intergenic_variant | MODIFIER |
| 8 | 34533841 | AX-170079714 | 0 | A | G | 0.14 | -1.03E-01 | 1.35E-01 | 2.66E+00 | 2.72E+00 | 4.41E-01 | no | | regulatory_region_variant | MODIFIER |
| 8 | 34539127 | AX-170079713 | 0 | C | G | 0.14 | -1.03E-01 | 1.35E-01 | 2.66E+00 | 2.72E+00 | 4.41E-01 | no | | intergenic_variant | MODIFIER |
| 8 | 34554046 | AX-170079715 | 0 | G | A | 0.14 | -1.03E-01 | 1.35E-01 | 2.66E+00 | 2.72E+00 | 4.41E-01 | no | | regulatory_region_variant | MODIFIER |
| 8 | 34555313 | AX-170501997 | 0 | C | T | 0.14 | -1.03E-01 | 1.35E-01 | 2.66E+00 | 2.72E+00 | 4.41E-01 | no | | intergenic_variant | MODIFIER |
| 8 | 34556330 | AX-170432061 | 0 | G | A | 0.14 | -1.03E-01 | 1.35E-01 | 2.66E+00 | 2.72E+00 | 4.41E-01 | no | | intergenic_variant | MODIFIER |
| 8 | 34568400 | AX-170515669 | 0 | G | A | 0.14 | -1.03E-01 | 1.35E-01 | 2.66E+00 | 2.72E+00 | 4.41E-01 | no | | intergenic_variant | MODIFIER |
| 8 | 34573531 | AX-170270566 | 0 | T | C | 0.14 | -1.03E-01 | 1.35E-01 | 2.66E+00 | 2.72E+00 | 4.41E-01 | no | | regulatory_region_variant | MODIFIER |
| 8 | 34575536 | AX-170212638 | 0 | T | C | 0.14 | -1.03E-01 | 1.35E-01 | 2.66E+00 | 2.72E+00 | 4.41E-01 | no | | intergenic_variant | MODIFIER |
| 8 | 34578728 | AX-170486729 | 0 | G | A | 0.14 | -1.03E-01 | 1.35E-01 | 2.66E+00 | 2.72E+00 | 4.41E-01 | no | | intergenic_variant | MODIFIER |
| 8 | 34582786 | AX-170168429 | 0 | G | A | 0.14 | -1.03E-01 | 1.35E-01 | 2.66E+00 | 2.72E+00 | 4.41E-01 | no | | intergenic_variant | MODIFIER |
| 8 | 34587283 | AX-170175169 | 0 | C | T | 0.14 | -1.03E-01 | 1.35E-01 | 2.66E+00 | 2.72E+00 | 4.41E-01 | no | B930018H19Rik | downstream_gene_variant | MODIFIER |
| 8 | 34587344 | AX-170245093 | 0 | A | G | 0.14 | -1.03E-01 | 1.35E-01 | 2.66E+00 | 2.72E+00 | 4.41E-01 | no | B930018H19Rik | downstream_gene_variant | MODIFIER |
| 8 | 34588989 | AX-170264175 | 0 | T | C | 0.14 | -1.03E-01 | 1.35E-01 | 2.66E+00 | 2.72E+00 | 4.41E-01 | no | B930018H19Rik | downstream_gene_variant | MODIFIER |
| 8 | 34589281 | AX-170268928 | 0 | T | G | 0.14 | -1.03E-01 | 1.35E-01 | 2.66E+00 | 2.72E+00 | 4.41E-01 | no | B930018H19Rik | downstream_gene_variant | MODIFIER |
| 8 | 34600843 | AX-97821630 | 0 | G | A | 0.14 | -1.03E-01 | 1.35E-01 | 2.66E+00 | 2.72E+00 | 4.41E-01 | no | B930018H19Rik | intron_variant,non_coding_transcript_variant | MODIFIER |
| 8 | 34610247 | AX-97793526 | 0 | A | G | 0.14 | -1.03E-01 | 1.35E-01 | 2.66E+00 | 2.72E+00 | 4.41E-01 | no | Gm34096 | intron_variant,non_coding_transcript_variant | MODIFIER |
| 8 | 34614148 | AX-170267099 | 0 | G | A | 0.14 | -1.03E-01 | 1.35E-01 | 2.66E+00 | 2.72E+00 | 4.41E-01 | no | Gm34096 | intron_variant,non_coding_transcript_variant | MODIFIER |
| 8 | 34626521 | AX-170378718 | 0 | G | A | 0.14 | -1.03E-01 | 1.35E-01 | 2.66E+00 | 2.72E+00 | 4.41E-01 | no | B930018H19Rik | intron_variant,non_coding_transcript_variant | MODIFIER |
| 8 | 34642261 | AX-170079718 | 0 | G | T | 0.14 | -1.03E-01 | 1.35E-01 | 2.66E+00 | 2.72E+00 | 4.41E-01 | no | B930018H19Rik | upstream_gene_variant | MODIFIER |
| 8 | 34682644 | AX-97813636 | 0 | C | T | 0.14 | -1.03E-01 | 1.35E-01 | 2.66E+00 | 2.72E+00 | 4.41E-01 | no | | intergenic_variant | MODIFIER |

| | | | | | | | | | | | | | | | |
|---|---|---|---|---|---|---|---|---|---|---|---|---|---|---|---|
| 8 | 34695237 | AX-97824565 | 0 | T | C | 0.14 | -1.03E-01 | 1.35E-01 | 2.66E+00 | 2.72E+00 | 4.41E-01 | no | | intergenic_variant | MODIFIER |
| 8 | 34712322 | AX-170478720 | 0 | T | C | 0.14 | -1.03E-01 | 1.35E-01 | 2.66E+00 | 2.72E+00 | 4.41E-01 | no | | intergenic_variant | MODIFIER |
| 8 | 34716776 | AX-170096220 | 0 | T | G | 0.14 | -1.03E-01 | 1.35E-01 | 2.66E+00 | 2.72E+00 | 4.41E-01 | no | | intergenic_variant | MODIFIER |
| 8 | 34721639 | AX-170280667 | 0 | T | C | 0.14 | -1.03E-01 | 1.35E-01 | 2.66E+00 | 2.72E+00 | 4.41E-01 | no | | intergenic_variant | MODIFIER |
| 8 | 34721774 | AX-170164243 | 0 | A | G | 0.14 | -1.03E-01 | 1.35E-01 | 2.66E+00 | 2.72E+00 | 4.41E-01 | no | | intergenic_variant | MODIFIER |
| 8 | 34727127 | AX-170115631 | 0 | C | T | 0.14 | -1.03E-01 | 1.35E-01 | 2.66E+00 | 2.72E+00 | 4.41E-01 | no | | intergenic_variant | MODIFIER |
| 8 | 34727823 | AX-170335146 | 0 | T | G | 0.14 | -1.03E-01 | 1.35E-01 | 2.66E+00 | 2.72E+00 | 4.41E-01 | no | | intergenic_variant | MODIFIER |
| 8 | 34729712 | AX-170297582 | 0 | G | A | 0.14 | -1.03E-01 | 1.35E-01 | 2.66E+00 | 2.72E+00 | 4.41E-01 | no | | regulatory_region_variant | MODIFIER |
| 8 | 34731708 | AX-170282818 | 0 | C | T | 0.14 | -1.03E-01 | 1.35E-01 | 2.66E+00 | 2.72E+00 | 4.41E-01 | no | | regulatory_region_variant | MODIFIER |
| 8 | 34768784 | AX-170530561 | 0 | A | T | 0.14 | -1.03E-01 | 1.35E-01 | 2.66E+00 | 2.72E+00 | 4.41E-01 | no | | intergenic_variant | MODIFIER |
| 8 | 34768925 | AX-170030499 | 0 | T | C | 0.14 | -1.03E-01 | 1.35E-01 | 2.66E+00 | 2.72E+00 | 4.41E-01 | no | | intergenic_variant | MODIFIER |
| 8 | 34797760 | AX-170486541 | 0 | A | G | 0.14 | -1.03E-01 | 1.35E-01 | 2.66E+00 | 2.72E+00 | 4.41E-01 | no | | intergenic_variant | MODIFIER |
| 8 | 34798879 | AX-170248458 | 0 | T | A | 0.14 | -1.03E-01 | 1.35E-01 | 2.66E+00 | 2.72E+00 | 4.41E-01 | no | | intergenic_variant | MODIFIER |
| 8 | 34800054 | AX-97786805 | 0 | T | C | 0.14 | -1.03E-01 | 1.35E-01 | 2.66E+00 | 2.72E+00 | 4.41E-01 | no | | intergenic_variant | MODIFIER |
| 8 | 34801509 | AX-170189787 | 0 | T | C | 0.14 | -1.03E-01 | 1.35E-01 | 2.66E+00 | 2.72E+00 | 4.41E-01 | no | | intergenic_variant | MODIFIER |
| 8 | 34804350 | AX-170117481 | 0 | A | G | 0.14 | -1.03E-01 | 1.35E-01 | 2.66E+00 | 2.72E+00 | 4.41E-01 | no | Dusp4 | upstream_gene_variant | MODIFIER |
| 8 | 34810150 | AX-170030498 | 0 | G | A | 0.14 | -1.03E-01 | 1.35E-01 | 2.66E+00 | 2.72E+00 | 4.41E-01 | no | Dusp4 | intron_variant | MODIFIER |
| 1 | 99767782 | AX-170133192 | 1 | G | A | 0.152 | 1.51E-01 | 1.98E-01 | 2.65E+00 | 2.72E+00 | 4.42E-01 | Yes | Cntnap5b | upstream_gene_variant | MODIFIER |
| 4 | 88045738 | AX-170307195 | 1 | T | C | 0.25 | 8.79E-02 | 1.11E-01 | 2.46E+00 | 2.52E+00 | 4.42E-01 | no | Gm12646 | downstream_gene_variant | MODIFIER |
| 9 | 90792805 | AX-170103129 | 1 | C | A | 0.125 | -9.89E-02 | 1.29E-01 | 2.61E+00 | 2.69E+00 | 4.42E-01 | no | | intergenic_variant | MODIFIER |
| 8 | 65992524 | AX-170201859 | 1 | G | A | 0.143 | -1.15E-01 | 1.49E-01 | 2.57E+00 | 2.63E+00 | 4.42E-01 | no | MAR1 | intron_variant | MODIFIER |
| 18 | 65103093 | AX-170331887 | 2 | G | A | 0.191 | -9.28E-02 | 1.22E-01 | 2.69E+00 | 2.75E+00 | 4.42E-01 | no | Nedd4l | intron_variant | MODIFIER |
| 18 | 55451426 | AX-170308385 | 1 | T | A | 0.446 | 7.62E-02 | 9.38E-02 | 2.38E+00 | 2.44E+00 | 4.42E-01 | no | | intergenic_variant | MODIFIER |
| 1 | 91022033 | AX-170169465 | 1 | C | T | 0.161 | 1.55E-01 | 2.03E-01 | 2.66E+00 | 2.73E+00 | 4.42E-01 | Yes | Lrrfip1 | intron_variant | MODIFIER |
| 5 | 103340490 | AX-170365712 | 0 | G | A | 0.456 | 8.88E-02 | 1.16E-01 | 2.62E+00 | 2.65E+00 | 4.42E-01 | no | Gm42619 | upstream_gene_variant | MODIFIER |
| 5 | 103341530 | AX-170509913 | 0 | T | C | 0.456 | 8.88E-02 | 1.16E-01 | 2.62E+00 | 2.65E+00 | 4.42E-01 | no | Gm42619 | upstream_gene_variant | MODIFIER |
| 5 | 103354482 | AX-170111682 | 0 | G | A | 0.456 | 8.88E-02 | 1.16E-01 | 2.62E+00 | 2.65E+00 | 4.42E-01 | no | 5430427N15Rik | non_coding_transcript_exon_variant | MODIFIER |
| 5 | 103381525 | AX-168373113 | 0 | T | C | 0.456 | 8.88E-02 | 1.16E-01 | 2.62E+00 | 2.65E+00 | 4.42E-01 | no | | regulatory_region_variant | MODIFIER |
| 5 | 103381639 | AX-97810478 | 0 | T | C | 0.456 | 8.88E-02 | 1.16E-01 | 2.62E+00 | 2.65E+00 | 4.42E-01 | no | | intergenic_variant | MODIFIER |
| 7 | 72543740 | AX-97802620 | 1 | C | T | 0.357 | -7.45E-02 | 9.81E-02 | 2.80E+00 | 2.89E+00 | 4.42E-01 | no | | regulatory_region_variant | MODIFIER |
| 18 | 69409303 | AX-170127406 | 0 | G | T | 0.167 | -9.39E-02 | 1.22E-01 | 2.60E+00 | 2.65E+00 | 4.42E-01 | no | Tcf4 | intron_variant | MODIFIER |
| 6 | 134265844 | AX-170178313 | 1 | C | G | 0.58 | 6.92E-02 | 8.74E-02 | 2.46E+00 | 2.51E+00 | 4.42E-01 | no | Etv6 | intron_variant | MODIFIER |
| 1 | 9722387 | AX-170146829 | 0 | G | A | 0.105 | 1.02E-01 | 1.34E-01 | 2.59E+00 | 2.73E+00 | 4.42E-01 | no | Vcpip1 | 3_prime_UTR_variant | MODIFIER |
| 18 | 80053698 | AX-170169686 | 0 | G | A | 0.386 | -6.53E-02 | 8.55E-02 | 2.62E+00 | 2.71E+00 | 4.42E-01 | no | Pard6g | intron_variant | MODIFIER |
| 18 | 80061917 | AX-170406142 | 0 | A | G | 0.386 | -6.53E-02 | 8.55E-02 | 2.62E+00 | 2.71E+00 | 4.42E-01 | no | Pard6g | intron_variant | MODIFIER |
| 18 | 80086100 | AX-170299462 | 0 | G | A | 0.386 | -6.53E-02 | 8.55E-02 | 2.62E+00 | 2.71E+00 | 4.42E-01 | no | Pard6g | intron_variant | MODIFIER |
| 18 | 80088117 | AX-170203303 | 0 | G | C | 0.386 | -6.53E-02 | 8.55E-02 | 2.62E+00 | 2.71E+00 | 4.42E-01 | no | Pard6g | intron_variant | MODIFIER |
| 18 | 80116260 | AX-170468719 | 0 | A | G | 0.386 | -6.53E-02 | 8.55E-02 | 2.62E+00 | 2.71E+00 | 4.42E-01 | no | Pard6g | intron_variant | MODIFIER |
| 18 | 80118962 | AX-170125827 | 0 | G | A | 0.386 | -6.53E-02 | 8.55E-02 | 2.62E+00 | 2.71E+00 | 4.42E-01 | no | Pard6g | 3_prime_UTR_variant | MODIFIER |
| 18 | 80124830 | AX-170099006 | 0 | C | T | 0.386 | -6.53E-02 | 8.55E-02 | 2.62E+00 | 2.71E+00 | 4.42E-01 | no | Adnp2 | downstream_gene_variant | MODIFIER |
| 18 | 80133503 | AX-170356108 | 0 | C | T | 0.386 | -6.53E-02 | 8.55E-02 | 2.62E+00 | 2.71E+00 | 4.42E-01 | no | Adnp2 | intron_variant | MODIFIER |
| 18 | 80135725 | AX-170283958 | 0 | C | A | 0.386 | -6.53E-02 | 8.55E-02 | 2.62E+00 | 2.71E+00 | 4.42E-01 | no | Adnp2 | intron_variant | MODIFIER |
| 18 | 80148655 | AX-170272819 | 0 | C | A | 0.386 | -6.53E-02 | 8.55E-02 | 2.62E+00 | 2.71E+00 | 4.42E-01 | no | Adnp2 | intron_variant | MODIFIER |
| 18 | 80150200 | AX-170276048 | 0 | G | C | 0.386 | -6.53E-02 | 8.55E-02 | 2.62E+00 | 2.71E+00 | 4.42E-01 | no | Adnp2 | intron_variant | MODIFIER |
| 18 | 80175622 | AX-170061446 | 0 | A | G | 0.386 | -6.53E-02 | 8.55E-02 | 2.62E+00 | 2.71E+00 | 4.42E-01 | no | | intergenic_variant | MODIFIER |
| 18 | 80177551 | AX-170340549 | 0 | C | T | 0.386 | -6.53E-02 | 8.55E-02 | 2.62E+00 | 2.71E+00 | 4.42E-01 | no | | intergenic_variant | MODIFIER |
| 18 | 80188058 | AX-97835140 | 0 | A | G | 0.386 | -6.53E-02 | 8.55E-02 | 2.62E+00 | 2.71E+00 | 4.42E-01 | no | Rbfa | downstream_gene_variant | MODIFIER |
| 18 | 80195544 | AX-97785982 | 0 | G | A | 0.386 | -6.53E-02 | 8.55E-02 | 2.62E+00 | 2.71E+00 | 4.42E-01 | no | Rbfa | intron_variant | MODIFIER |
| 18 | 80197024 | AX-170208994 | 0 | A | G | 0.386 | -6.53E-02 | 8.55E-02 | 2.62E+00 | 2.71E+00 | 4.42E-01 | no | Rbfa | intron_variant | MODIFIER |
| 18 | 80220188 | AX-170460134 | 0 | C | G | 0.386 | -6.53E-02 | 8.55E-02 | 2.62E+00 | 2.71E+00 | 4.42E-01 | no | Txnl4a | intron_variant | MODIFIER |
| 18 | 80220795 | AX-170249977 | 0 | T | C | 0.386 | -6.53E-02 | 8.55E-02 | 2.62E+00 | 2.71E+00 | 4.42E-01 | no | Txnl4a | intron_variant | MODIFIER |
| 18 | 80221585 | AX-170187806 | 0 | T | C | 0.386 | -6.53E-02 | 8.55E-02 | 2.62E+00 | 2.71E+00 | 4.42E-01 | no | Txnl4a | intron_variant | MODIFIER |
| 18 | 80223492 | AX-170155770 | 0 | C | T | 0.386 | -6.53E-02 | 8.55E-02 | 2.62E+00 | 2.71E+00 | 4.42E-01 | no | Txnl4a | downstream_gene_variant | MODIFIER |
| 18 | 80226799 | AX-170242570 | 0 | C | G | 0.386 | -6.53E-02 | 8.55E-02 | 2.62E+00 | 2.71E+00 | 4.42E-01 | no | Txnl4a | downstream_gene_variant | MODIFIER |

| | | | | | | | | | | | | | | | |
|---|---|---|---|---|---|---|---|---|---|---|---|---|---|---|---|
| 18 | 80261843 | AX-170061447 | 0 | G | C | 0.386 | -6.53E-02 | 8.55E-02 | 2.62E+00 | 2.71E+00 | 4.42E-01 | no | Pqlc1 | intron_variant | MODIFIER |
| 18 | 80263001 | AX-170312917 | 0 | C | G | 0.386 | -6.53E-02 | 8.55E-02 | 2.62E+00 | 2.71E+00 | 4.42E-01 | no | Pqlc1 | intron_variant | MODIFIER |
| 18 | 80274903 | AX-170172496 | 0 | A | G | 0.386 | -6.53E-02 | 8.55E-02 | 2.62E+00 | 2.71E+00 | 4.42E-01 | no | Pqlc1 | intron_variant | MODIFIER |
| 18 | 80279895 | AX-97813833 | 0 | A | G | 0.386 | -6.53E-02 | 8.55E-02 | 2.62E+00 | 2.71E+00 | 4.42E-01 | no | Pqlc1 | intron_variant | MODIFIER |
| 18 | 80281384 | AX-170362748 | 0 | A | G | 0.386 | -6.53E-02 | 8.55E-02 | 2.62E+00 | 2.71E+00 | 4.42E-01 | no | Pqlc1 | intron_variant | MODIFIER |
| 18 | 80315164 | AX-170372611 | 0 | A | C | 0.386 | -6.53E-02 | 8.55E-02 | 2.62E+00 | 2.71E+00 | 4.42E-01 | no | Kcng2 | intron_variant | MODIFIER |
| 18 | 80339311 | AX-170453280 | 0 | G | A | 0.386 | -6.53E-02 | 8.55E-02 | 2.62E+00 | 2.71E+00 | 4.42E-01 | no | Kcng2 | intron_variant | MODIFIER |
| 18 | 80358565 | AX-170451304 | 0 | C | T | 0.386 | -6.53E-02 | 8.55E-02 | 2.62E+00 | 2.71E+00 | 4.42E-01 | no | Kcng2 | intron_variant | MODIFIER |
| 18 | 80359351 | AX-170284754 | 0 | A | G | 0.386 | -6.53E-02 | 8.55E-02 | 2.62E+00 | 2.71E+00 | 4.42E-01 | no | Kcng2 | intron_variant | MODIFIER |
| 18 | 80382621 | AX-170248755 | 0 | C | T | 0.386 | -6.53E-02 | 8.55E-02 | 2.62E+00 | 2.71E+00 | 4.42E-01 | no | | regulatory_region_variant | MODIFIER |
| 18 | 80387797 | AX-170106303 | 0 | T | C | 0.386 | -6.53E-02 | 8.55E-02 | 2.62E+00 | 2.71E+00 | 4.42E-01 | no | | intergenic_variant | MODIFIER |
| 18 | 80428618 | AX-170336932 | 0 | C | T | 0.386 | -6.53E-02 | 8.55E-02 | 2.62E+00 | 2.71E+00 | 4.42E-01 | no | Ctdp1 | intron_variant | MODIFIER |
| 18 | 80432326 | AX-170184247 | 0 | C | T | 0.386 | -6.53E-02 | 8.55E-02 | 2.62E+00 | 2.71E+00 | 4.42E-01 | no | Ctdp1 | intron_variant | MODIFIER |
| 18 | 80434436 | AX-170468713 | 0 | G | T | 0.386 | -6.53E-02 | 8.55E-02 | 2.62E+00 | 2.71E+00 | 4.42E-01 | no | Ctdp1 | intron_variant | MODIFIER |
| 18 | 80437177 | AX-170260696 | 0 | T | G | 0.386 | -6.53E-02 | 8.55E-02 | 2.62E+00 | 2.71E+00 | 4.42E-01 | no | Ctdp1 | intron_variant | MODIFIER |
| 18 | 80438832 | AX-170193952 | 0 | A | G | 0.386 | -6.53E-02 | 8.55E-02 | 2.62E+00 | 2.71E+00 | 4.42E-01 | no | Ctdp1 | intron_variant | MODIFIER |
| 18 | 80471796 | AX-170492682 | 0 | A | G | 0.386 | -6.53E-02 | 8.55E-02 | 2.62E+00 | 2.71E+00 | 4.42E-01 | no | Ctdp1 | upstream_gene_variant | MODIFIER |
| 18 | 80481062 | AX-170016081 | 0 | T | C | 0.386 | -6.53E-02 | 8.55E-02 | 2.62E+00 | 2.71E+00 | 4.42E-01 | no | AC127256.1 | downstream_gene_variant | MODIFIER |
| 18 | 80482026 | AX-170201284 | 0 | T | A | 0.386 | -6.53E-02 | 8.55E-02 | 2.62E+00 | 2.71E+00 | 4.42E-01 | no | | intergenic_variant | MODIFIER |
| 18 | 80482336 | AX-97816682 | 0 | T | C | 0.386 | -6.53E-02 | 8.55E-02 | 2.62E+00 | 2.71E+00 | 4.42E-01 | no | | intergenic_variant | MODIFIER |
| 18 | 80489802 | AX-170061453 | 0 | A | G | 0.386 | -6.53E-02 | 8.55E-02 | 2.62E+00 | 2.71E+00 | 4.42E-01 | no | | intergenic_variant | MODIFIER |
| 18 | 80519949 | AX-170016079 | 0 | T | C | 0.386 | -6.53E-02 | 8.55E-02 | 2.62E+00 | 2.71E+00 | 4.42E-01 | no | | intergenic_variant | MODIFIER |
| 18 | 80522157 | AX-170527869 | 0 | G | A | 0.386 | -6.53E-02 | 8.55E-02 | 2.62E+00 | 2.71E+00 | 4.42E-01 | no | | intergenic_variant | MODIFIER |
| 18 | 80524059 | AX-170339352 | 0 | T | C | 0.386 | -6.53E-02 | 8.55E-02 | 2.62E+00 | 2.71E+00 | 4.42E-01 | no | | intergenic_variant | MODIFIER |
| 18 | 80524950 | AX-170453349 | 0 | T | C | 0.386 | -6.53E-02 | 8.55E-02 | 2.62E+00 | 2.71E+00 | 4.42E-01 | no | | intergenic_variant | MODIFIER |
| 18 | 80539325 | AX-170519785 | 0 | A | G | 0.386 | -6.53E-02 | 8.55E-02 | 2.62E+00 | 2.71E+00 | 4.42E-01 | no | AC127256.2 | downstream_gene_variant | MODIFIER |
| 18 | 80550809 | AX-170363232 | 0 | G | A | 0.386 | -6.53E-02 | 8.55E-02 | 2.62E+00 | 2.71E+00 | 4.42E-01 | no | | intergenic_variant | MODIFIER |
| 18 | 80550931 | AX-170282714 | 0 | T | C | 0.386 | -6.53E-02 | 8.55E-02 | 2.62E+00 | 2.71E+00 | 4.42E-01 | no | | intergenic_variant | MODIFIER |
| 18 | 80558188 | AX-170500447 | 0 | T | G | 0.386 | -6.53E-02 | 8.55E-02 | 2.62E+00 | 2.71E+00 | 4.42E-01 | no | | intergenic_variant | MODIFIER |
| 18 | 80562491 | AX-170176161 | 0 | T | C | 0.386 | -6.53E-02 | 8.55E-02 | 2.62E+00 | 2.71E+00 | 4.42E-01 | no | | intergenic_variant | MODIFIER |
| 18 | 80586486 | AX-170480924 | 0 | C | T | 0.386 | -6.53E-02 | 8.55E-02 | 2.62E+00 | 2.71E+00 | 4.42E-01 | no | AC149054.1 | intron_variant,non_coding_transcript_variant | MODIFIER |
| 18 | 80594794 | AX-170181491 | 0 | G | A | 0.386 | -6.53E-02 | 8.55E-02 | 2.62E+00 | 2.71E+00 | 4.42E-01 | no | AC149054.1 | intron_variant,non_coding_transcript_variant | MODIFIER |
| 18 | 80603710 | AX-170291410 | 0 | T | C | 0.386 | -6.53E-02 | 8.55E-02 | 2.62E+00 | 2.71E+00 | 4.42E-01 | no | Nfatc1 | downstream_gene_variant | MODIFIER |
| 18 | 80614665 | AX-170211558 | 0 | A | G | 0.386 | -6.53E-02 | 8.55E-02 | 2.62E+00 | 2.71E+00 | 4.42E-01 | no | Nfatc1 | intron_variant | MODIFIER |
| 18 | 80615104 | AX-170147952 | 0 | G | A | 0.386 | -6.53E-02 | 8.55E-02 | 2.62E+00 | 2.71E+00 | 4.42E-01 | no | Nfatc1 | intron_variant | MODIFIER |
| 18 | 80619039 | AX-170276532 | 0 | A | G | 0.386 | -6.53E-02 | 8.55E-02 | 2.62E+00 | 2.71E+00 | 4.42E-01 | no | Nfatc1 | intron_variant | MODIFIER |
| 18 | 80619698 | AX-170166267 | 0 | A | G | 0.386 | -6.53E-02 | 8.55E-02 | 2.62E+00 | 2.71E+00 | 4.42E-01 | no | Nfatc1 | intron_variant | MODIFIER |
| 18 | 80619792 | AX-170485512 | 0 | T | A | 0.386 | -6.53E-02 | 8.55E-02 | 2.62E+00 | 2.71E+00 | 4.42E-01 | no | Nfatc1 | intron_variant | MODIFIER |
| 18 | 80626611 | AX-170250551 | 0 | T | A | 0.386 | -6.53E-02 | 8.55E-02 | 2.62E+00 | 2.71E+00 | 4.42E-01 | no | Nfatc1 | intron_variant | MODIFIER |
| 18 | 80639652 | AX-170324131 | 0 | C | T | 0.386 | -6.53E-02 | 8.55E-02 | 2.62E+00 | 2.71E+00 | 4.42E-01 | no | Nfatc1 | intron_variant | MODIFIER |
| 18 | 80641338 | AX-170263262 | 0 | G | A | 0.386 | -6.53E-02 | 8.55E-02 | 2.62E+00 | 2.71E+00 | 4.42E-01 | no | Nfatc1 | intron_variant | MODIFIER |
| 18 | 80641496 | AX-170306747 | 0 | G | T | 0.386 | -6.53E-02 | 8.55E-02 | 2.62E+00 | 2.71E+00 | 4.42E-01 | no | Nfatc1 | intron_variant | MODIFIER |
| 18 | 80644787 | AX-170374854 | 0 | T | C | 0.386 | -6.53E-02 | 8.55E-02 | 2.62E+00 | 2.71E+00 | 4.42E-01 | no | Nfatc1 | downstream_gene_variant | MODIFIER |
| 18 | 80655642 | AX-170294488 | 0 | A | G | 0.386 | -6.53E-02 | 8.55E-02 | 2.62E+00 | 2.71E+00 | 4.42E-01 | no | Nfatc1 | intron_variant | MODIFIER |
| 18 | 80656448 | AX-97832320 | 0 | T | C | 0.386 | -6.53E-02 | 8.55E-02 | 2.62E+00 | 2.71E+00 | 4.42E-01 | no | Nfatc1 | intron_variant | MODIFIER |
| 18 | 80657337 | AX-170061459 | 0 | T | C | 0.386 | -6.53E-02 | 8.55E-02 | 2.62E+00 | 2.71E+00 | 4.42E-01 | no | Nfatc1 | intron_variant | MODIFIER |
| 18 | 80660025 | AX-170061460 | 0 | A | G | 0.386 | -6.53E-02 | 8.55E-02 | 2.62E+00 | 2.71E+00 | 4.42E-01 | no | Nfatc1 | intron_variant | MODIFIER |
| 18 | 80663274 | AX-170243787 | 0 | A | G | 0.386 | -6.53E-02 | 8.55E-02 | 2.62E+00 | 2.71E+00 | 4.42E-01 | no | Nfatc1 | intron_variant | MODIFIER |
| 19 | 50763533 | AX-170016808 | 1 | A | G | 0.473 | 4.97E-02 | 6.52E-02 | 2.63E+00 | 2.72E+00 | 4.42E-01 | no | Gm26629 | intron_variant,non_coding_transcript_variant | MODIFIER |
| 18 | 59341184 | AX-170219242 | 0 | G | A | 0.404 | 7.36E-02 | 9.32E-02 | 2.47E+00 | 2.52E+00 | 4.42E-01 | no | Chsy3 | intron_variant | MODIFIER |
| 18 | 84561256 | AX-170122454 | 0 | G | T | 0.158 | -8.34E-02 | 1.07E-01 | 2.49E+00 | 2.58E+00 | 4.42E-01 | no | Zfp407 | upstream_gene_variant | MODIFIER |
| 5 | 50320170 | AX-170171319 | 0 | C | T | 0.482 | 8.44E-02 | 1.05E-01 | 2.42E+00 | 2.47E+00 | 4.42E-01 | no | Gm7205 | intron_variant,non_coding_transcript_variant | MODIFIER |
| 16 | 46109203 | AX-170364482 | 1 | G | A | 0.473 | -6.17E-02 | 7.97E-02 | 2.54E+00 | 2.60E+00 | 4.42E-01 | no | Cd96 | intron_variant | MODIFIER |
| 14 | 82760392 | AX-97826809 | 0 | T | C | 0.044 | -1.21E-01 | 1.61E-01 | 2.69E+00 | 2.88E+00 | 4.43E-01 | no | | intergenic_variant | MODIFIER |

| | | | | | | | | | | | | | | | |
|---|---|---|---|---|---|---|---|---|---|---|---|---|---|---|---|
| 1 | 151737008 | AX-170001986 | 1 | C | T | 0.205 | 1.05E-01 | 1.38E-01 | 2.67E+00 | 2.73E+00 | 4.43E-01 | no | | intergenic_variant | MODIFIER |
| 1 | 23835264 | AX-170040621 | 1 | C | A | 0.438 | 6.33E-02 | 8.31E-02 | 2.64E+00 | 2.71E+00 | 4.43E-01 | no | B3gat2 | intron_variant | MODIFIER |
| 18 | 54797694 | AX-170325343 | 1 | A | G | 0.446 | 7.68E-02 | 9.45E-02 | 2.38E+00 | 2.43E+00 | 4.43E-01 | no | AC129179.2 | intron_variant,non_coding_transcript_variant | MODIFIER |
| 10 | 92394010 | AX-97826514 | 1 | G | A | 0.304 | -6.49E-02 | 8.37E-02 | 2.53E+00 | 2.59E+00 | 4.43E-01 | no | | intergenic_variant | MODIFIER |
| 10 | 92493043 | AX-170513826 | 1 | G | A | 0.304 | -6.49E-02 | 8.37E-02 | 2.53E+00 | 2.59E+00 | 4.43E-01 | no | | intergenic_variant | MODIFIER |
| 13 | 23566906 | AX-97851911 | 1 | C | A | 0.17 | -9.70E-02 | 1.21E-01 | 2.42E+00 | 2.47E+00 | 4.43E-01 | no | Hist1h2bg | upstream_gene_variant | MODIFIER |
| 1 | 127968272 | AX-170345480 | 1 | C | A | 0.536 | 6.51E-02 | 8.53E-02 | 2.62E+00 | 2.68E+00 | 4.43E-01 | no | Zranb3 | intron_variant | MODIFIER |
| 10 | 68041647 | AX-170045116 | 2 | C | A | 0.427 | -5.84E-02 | 7.68E-02 | 2.62E+00 | 2.71E+00 | 4.43E-01 | no | Rtkn2 | missense_variant,NMD_transcript_variant | MODERATE |
| 8 | 15891799 | AX-170495988 | 1 | A | G | 0.232 | -5.93E-02 | 7.69E-02 | 2.55E+00 | 2.64E+00 | 4.43E-01 | no | Csmd1 | downstream_gene_variant | MODIFIER |
| 10 | 68080812 | AX-170265600 | 1 | C | T | 0.411 | -6.06E-02 | 7.96E-02 | 2.64E+00 | 2.71E+00 | 4.43E-01 | no | | regulatory_region_variant | MODIFIER |
| 1 | 71645499 | AX-170191084 | 0 | T | G | 0.114 | 1.17E-01 | 1.56E-01 | 2.79E+00 | 2.94E+00 | 4.43E-01 | no | Fn1 | intron_variant | MODIFIER |
| 18 | 65124201 | AX-170215506 | 1 | A | C | 0.188 | -9.15E-02 | 1.21E-01 | 2.70E+00 | 2.76E+00 | 4.43E-01 | no | Nedd4l | intron_variant | MODIFIER |
| 1 | 24575168 | AX-97788099 | 0 | T | C | 0.439 | 6.22E-02 | 8.10E-02 | 2.57E+00 | 2.64E+00 | 4.43E-01 | no | Col19a1 | intron_variant | MODIFIER |
| 1 | 177314812 | AX-170430089 | 0 | T | C | 0.228 | 8.79E-02 | 1.15E-01 | 2.60E+00 | 2.68E+00 | 4.43E-01 | no | | intergenic_variant | MODIFIER |
| 6 | 137385029 | AX-170028343 | 0 | C | A | 0.579 | 6.77E-02 | 8.63E-02 | 2.48E+00 | 2.54E+00 | 4.44E-01 | no | Ptpro | intron_variant | MODIFIER |
| 1 | 187677019 | AX-170265667 | 0 | C | T | 0.254 | 8.03E-02 | 9.84E-02 | 2.36E+00 | 2.45E+00 | 4.44E-01 | no | Esrrg | intron_variant | MODIFIER |
| 10 | 66003095 | AX-170003369 | 0 | A | C | 0.412 | -6.06E-02 | 7.97E-02 | 2.64E+00 | 2.71E+00 | 4.44E-01 | no | | intergenic_variant | MODIFIER |
| 10 | 66003836 | AX-170428902 | 0 | C | A | 0.412 | -6.06E-02 | 7.97E-02 | 2.64E+00 | 2.71E+00 | 4.44E-01 | no | | intergenic_variant | MODIFIER |
| 10 | 66005376 | AX-170120581 | 0 | G | T | 0.412 | -6.06E-02 | 7.97E-02 | 2.64E+00 | 2.71E+00 | 4.44E-01 | no | | intergenic_variant | MODIFIER |
| 10 | 66019392 | AX-170475008 | 0 | A | G | 0.412 | -6.06E-02 | 7.97E-02 | 2.64E+00 | 2.71E+00 | 4.44E-01 | no | | intergenic_variant | MODIFIER |
| 10 | 66046683 | AX-170003371 | 0 | G | A | 0.412 | -6.06E-02 | 7.97E-02 | 2.64E+00 | 2.71E+00 | 4.44E-01 | no | | intergenic_variant | MODIFIER |
| 10 | 66105411 | AX-170245663 | 0 | G | A | 0.412 | -6.06E-02 | 7.97E-02 | 2.64E+00 | 2.71E+00 | 4.44E-01 | no | | intergenic_variant | MODIFIER |
| 10 | 66113468 | AX-170424245 | 0 | G | A | 0.412 | -6.06E-02 | 7.97E-02 | 2.64E+00 | 2.71E+00 | 4.44E-01 | no | | intergenic_variant | MODIFIER |
| 10 | 66114507 | AX-170224595 | 0 | A | G | 0.412 | -6.06E-02 | 7.97E-02 | 2.64E+00 | 2.71E+00 | 4.44E-01 | no | | intergenic_variant | MODIFIER |
| 10 | 66114571 | AX-97781495 | 0 | G | A | 0.412 | -6.06E-02 | 7.97E-02 | 2.64E+00 | 2.71E+00 | 4.44E-01 | no | | intergenic_variant | MODIFIER |
| 10 | 66181594 | AX-170476925 | 0 | G | C | 0.412 | -6.06E-02 | 7.97E-02 | 2.64E+00 | 2.71E+00 | 4.44E-01 | no | 1700023F02Rik | intron_variant,non_coding_transcript_variant | MODIFIER |
| 10 | 66187092 | AX-97824446 | 0 | G | A | 0.412 | -6.06E-02 | 7.97E-02 | 2.64E+00 | 2.71E+00 | 4.44E-01 | no | 1700023F02Rik | intron_variant,non_coding_transcript_variant | MODIFIER |
| 10 | 66257850 | AX-97815836 | 0 | T | C | 0.412 | -6.06E-02 | 7.97E-02 | 2.64E+00 | 2.71E+00 | 4.44E-01 | no | 1700023F02Rik | upstream_gene_variant | MODIFIER |
| 10 | 66387150 | AX-97815780 | 0 | C | T | 0.412 | -6.06E-02 | 7.97E-02 | 2.64E+00 | 2.71E+00 | 4.44E-01 | no | | intergenic_variant | MODIFIER |
| 10 | 66444477 | AX-170086830 | 0 | T | G | 0.412 | -6.06E-02 | 7.97E-02 | 2.64E+00 | 2.71E+00 | 4.44E-01 | no | | intergenic_variant | MODIFIER |
| 10 | 66480387 | AX-170323469 | 0 | G | A | 0.412 | -6.06E-02 | 7.97E-02 | 2.64E+00 | 2.71E+00 | 4.44E-01 | no | Gm22594 | upstream_gene_variant | MODIFIER |
| 10 | 66491441 | AX-170107779 | 0 | T | C | 0.412 | -6.06E-02 | 7.97E-02 | 2.64E+00 | 2.71E+00 | 4.44E-01 | no | | intergenic_variant | MODIFIER |
| 10 | 66491583 | AX-170003381 | 0 | T | C | 0.412 | -6.06E-02 | 7.97E-02 | 2.64E+00 | 2.71E+00 | 4.44E-01 | no | | intergenic_variant | MODIFIER |
| 10 | 66498284 | AX-170284188 | 0 | T | C | 0.412 | -6.06E-02 | 7.97E-02 | 2.64E+00 | 2.71E+00 | 4.44E-01 | no | | intergenic_variant | MODIFIER |
| 10 | 66549084 | AX-97813963 | 0 | T | C | 0.412 | -6.06E-02 | 7.97E-02 | 2.64E+00 | 2.71E+00 | 4.44E-01 | no | | intergenic_variant | MODIFIER |
| 10 | 67365314 | AX-170086834 | 0 | A | G | 0.412 | -6.06E-02 | 7.97E-02 | 2.64E+00 | 2.71E+00 | 4.44E-01 | no | Gm49454 | intron_variant,non_coding_transcript_variant | MODIFIER |
| 10 | 67453110 | AX-170458789 | 0 | C | T | 0.412 | -6.06E-02 | 7.97E-02 | 2.64E+00 | 2.71E+00 | 4.44E-01 | no | | intergenic_variant | MODIFIER |
| 10 | 67464839 | AX-97832725 | 0 | A | C | 0.412 | -6.06E-02 | 7.97E-02 | 2.64E+00 | 2.71E+00 | 4.44E-01 | no | | intergenic_variant | MODIFIER |
| 10 | 67464912 | AX-170242972 | 0 | A | G | 0.412 | -6.06E-02 | 7.97E-02 | 2.64E+00 | 2.71E+00 | 4.44E-01 | no | | intergenic_variant | MODIFIER |
| 10 | 67688580 | AX-170142440 | 0 | C | T | 0.412 | -6.06E-02 | 7.97E-02 | 2.64E+00 | 2.71E+00 | 4.44E-01 | no | | intergenic_variant | MODIFIER |
| 10 | 67699961 | AX-170131205 | 0 | A | C | 0.412 | -6.06E-02 | 7.97E-02 | 2.64E+00 | 2.71E+00 | 4.44E-01 | no | | intergenic_variant | MODIFIER |
| 10 | 67702687 | AX-170003395 | 0 | A | C | 0.412 | -6.06E-02 | 7.97E-02 | 2.64E+00 | 2.71E+00 | 4.44E-01 | no | | intergenic_variant | MODIFIER |
| 10 | 67705591 | AX-170476039 | 0 | A | G | 0.412 | -6.06E-02 | 7.97E-02 | 2.64E+00 | 2.71E+00 | 4.44E-01 | no | 1700030E10Rik | downstream_gene_variant | MODIFIER |
| 10 | 67707720 | AX-170256074 | 0 | T | G | 0.412 | -6.06E-02 | 7.97E-02 | 2.64E+00 | 2.71E+00 | 4.44E-01 | no | 1700030E10Rik | downstream_gene_variant | MODIFIER |
| 10 | 67723768 | AX-170045108 | 0 | G | A | 0.412 | -6.06E-02 | 7.97E-02 | 2.64E+00 | 2.71E+00 | 4.44E-01 | no | | intergenic_variant | MODIFIER |
| 10 | 67726795 | AX-170293264 | 0 | G | A | 0.412 | -6.06E-02 | 7.97E-02 | 2.64E+00 | 2.71E+00 | 4.44E-01 | no | | intergenic_variant | MODIFIER |
| 10 | 67749400 | AX-170300186 | 0 | C | G | 0.412 | -6.06E-02 | 7.97E-02 | 2.64E+00 | 2.71E+00 | 4.44E-01 | no | | intergenic_variant | MODIFIER |
| 10 | 67751398 | AX-170234039 | 0 | C | T | 0.412 | -6.06E-02 | 7.97E-02 | 2.64E+00 | 2.71E+00 | 4.44E-01 | no | | intergenic_variant | MODIFIER |
| 10 | 67782796 | AX-170409863 | 0 | G | A | 0.412 | -6.06E-02 | 7.97E-02 | 2.64E+00 | 2.71E+00 | 4.44E-01 | no | 4930543I11Rik | upstream_gene_variant | MODIFIER |
| 10 | 67785680 | AX-97829371 | 0 | T | G | 0.412 | -6.06E-02 | 7.97E-02 | 2.64E+00 | 2.71E+00 | 4.44E-01 | no | 4930543I11Rik | upstream_gene_variant | MODIFIER |
| 10 | 67816638 | AX-170123579 | 0 | A | G | 0.412 | -6.06E-02 | 7.97E-02 | 2.64E+00 | 2.71E+00 | 4.44E-01 | no | Gm49766 | intron_variant,non_coding_transcript_variant | MODIFIER |
| 10 | 67829377 | AX-170496799 | 0 | A | G | 0.412 | -6.06E-02 | 7.97E-02 | 2.64E+00 | 2.71E+00 | 4.44E-01 | no | Gm49766 | intron_variant,non_coding_transcript_variant | MODIFIER |
| 10 | 67859584 | AX-170235645 | 0 | A | G | 0.412 | -6.06E-02 | 7.97E-02 | 2.64E+00 | 2.71E+00 | 4.44E-01 | no | | intergenic_variant | MODIFIER |
| 10 | 67859621 | AX-170098419 | 0 | C | G | 0.412 | -6.06E-02 | 7.97E-02 | 2.64E+00 | 2.71E+00 | 4.44E-01 | no | | intergenic_variant | MODIFIER |

| | | | | | | | | | | | | | | | |
|---|---|---|---|---|---|---|---|---|---|---|---|---|---|---|---|
| 10 | 67859707 | AX-97794402 | 0 | A | G | 0.412 | -6.06E-02 | 7.97E-02 | 2.64E+00 | 2.71E+00 | 4.44E-01 | no | | intergenic_variant | MODIFIER |
| 10 | 67885752 | AX-170474330 | 0 | T | C | 0.412 | -6.06E-02 | 7.97E-02 | 2.64E+00 | 2.71E+00 | 4.44E-01 | no | Zfp365 | downstream_gene_variant | MODIFIER |
| 10 | 67888989 | AX-170354210 | 0 | A | G | 0.412 | -6.06E-02 | 7.97E-02 | 2.64E+00 | 2.71E+00 | 4.44E-01 | no | Zfp365 | synonymous_variant | LOW |
| 10 | 67890113 | AX-170336440 | 0 | G | A | 0.412 | -6.06E-02 | 7.97E-02 | 2.64E+00 | 2.71E+00 | 4.44E-01 | no | Zfp365 | intron_variant | MODIFIER |
| 10 | 67891190 | AX-170379646 | 0 | A | G | 0.412 | -6.06E-02 | 7.97E-02 | 2.64E+00 | 2.71E+00 | 4.44E-01 | no | Zfp365 | intron_variant | MODIFIER |
| 10 | 67893185 | AX-170457271 | 0 | T | C | 0.412 | -6.06E-02 | 7.97E-02 | 2.64E+00 | 2.71E+00 | 4.44E-01 | no | Zfp365 | intron_variant | MODIFIER |
| 10 | 67897804 | AX-170212428 | 0 | A | G | 0.412 | -6.06E-02 | 7.97E-02 | 2.64E+00 | 2.71E+00 | 4.44E-01 | no | Zfp365 | intron_variant | MODIFIER |
| 10 | 67931240 | AX-170315535 | 0 | C | A | 0.412 | -6.06E-02 | 7.97E-02 | 2.64E+00 | 2.71E+00 | 4.44E-01 | no | | intergenic_variant | MODIFIER |
| 10 | 67945573 | AX-97843255 | 0 | C | A | 0.412 | -6.06E-02 | 7.97E-02 | 2.64E+00 | 2.71E+00 | 4.44E-01 | no | | intergenic_variant | MODIFIER |
| 10 | 67950293 | AX-170177808 | 0 | A | G | 0.412 | -6.06E-02 | 7.97E-02 | 2.64E+00 | 2.71E+00 | 4.44E-01 | no | | intergenic_variant | MODIFIER |
| 10 | 67953109 | AX-170377724 | 0 | T | C | 0.412 | -6.06E-02 | 7.97E-02 | 2.64E+00 | 2.71E+00 | 4.44E-01 | no | | intergenic_variant | MODIFIER |
| 10 | 67958032 | AX-170214884 | 0 | G | C | 0.412 | -6.06E-02 | 7.97E-02 | 2.64E+00 | 2.71E+00 | 4.44E-01 | no | 4930521O17Rik | downstream_gene_variant | MODIFIER |
| 10 | 67960380 | AX-170352590 | 0 | T | C | 0.412 | -6.06E-02 | 7.97E-02 | 2.64E+00 | 2.71E+00 | 4.44E-01 | no | 4930521O17Rik | downstream_gene_variant | MODIFIER |
| 10 | 67966320 | AX-170045112 | 0 | C | G | 0.412 | -6.06E-02 | 7.97E-02 | 2.64E+00 | 2.71E+00 | 4.44E-01 | no | 4930521O17Rik | intron_variant,non_coding_transcript_variant | MODIFIER |
| 10 | 67966796 | AX-170045113 | 0 | G | A | 0.412 | -6.06E-02 | 7.97E-02 | 2.64E+00 | 2.71E+00 | 4.44E-01 | no | 4930521O17Rik | intron_variant,non_coding_transcript_variant | MODIFIER |
| 10 | 67972264 | AX-170416942 | 0 | A | C | 0.412 | -6.06E-02 | 7.97E-02 | 2.64E+00 | 2.71E+00 | 4.44E-01 | no | 4930521O17Rik | intron_variant,non_coding_transcript_variant | MODIFIER |
| 10 | 67977938 | AX-170403763 | 0 | C | T | 0.412 | -6.06E-02 | 7.97E-02 | 2.64E+00 | 2.71E+00 | 4.44E-01 | no | Rtkn2 | upstream_gene_variant | MODIFIER |
| 10 | 68003849 | AX-170437371 | 0 | T | C | 0.412 | -6.06E-02 | 7.97E-02 | 2.64E+00 | 2.71E+00 | 4.44E-01 | no | Rtkn2 | intron_variant,NMD_transcript_variant | MODIFIER |
| 10 | 68004565 | AX-170353433 | 0 | C | T | 0.412 | -6.06E-02 | 7.97E-02 | 2.64E+00 | 2.71E+00 | 4.44E-01 | no | Rtkn2 | intron_variant,NMD_transcript_variant | MODIFIER |
| 10 | 68006408 | AX-170454285 | 0 | A | G | 0.412 | -6.06E-02 | 7.97E-02 | 2.64E+00 | 2.71E+00 | 4.44E-01 | no | Rtkn2 | intron_variant,NMD_transcript_variant | MODIFIER |
| 10 | 68006847 | AX-170497990 | 0 | C | A | 0.412 | -6.06E-02 | 7.97E-02 | 2.64E+00 | 2.71E+00 | 4.44E-01 | no | Rtkn2 | intron_variant,NMD_transcript_variant | MODIFIER |
| 10 | 68010894 | AX-170463216 | 0 | G | A | 0.412 | -6.06E-02 | 7.97E-02 | 2.64E+00 | 2.71E+00 | 4.44E-01 | no | Rtkn2 | intron_variant,NMD_transcript_variant | MODIFIER |
| 10 | 68017798 | AX-170419243 | 0 | A | G | 0.412 | -6.06E-02 | 7.97E-02 | 2.64E+00 | 2.71E+00 | 4.44E-01 | no | Rtkn2 | splice_region_variant,intron_variant,NMD_transcript_variant | LOW |
| 10 | 68023473 | AX-170360612 | 0 | C | T | 0.412 | -6.06E-02 | 7.97E-02 | 2.64E+00 | 2.71E+00 | 4.44E-01 | no | Rtkn2 | intron_variant,NMD_transcript_variant | MODIFIER |
| 10 | 68025358 | AX-97850802 | 0 | G | A | 0.412 | -6.06E-02 | 7.97E-02 | 2.64E+00 | 2.71E+00 | 4.44E-01 | no | Rtkn2 | intron_variant,NMD_transcript_variant | MODIFIER |
| 10 | 68026880 | AX-170373971 | 0 | A | G | 0.412 | -6.06E-02 | 7.97E-02 | 2.64E+00 | 2.71E+00 | 4.44E-01 | no | Rtkn2 | intron_variant,NMD_transcript_variant | MODIFIER |
| 10 | 68030418 | AX-170258122 | 0 | G | A | 0.412 | -6.06E-02 | 7.97E-02 | 2.64E+00 | 2.71E+00 | 4.44E-01 | no | Rtkn2 | intron_variant,NMD_transcript_variant | MODIFIER |
| 10 | 68031028 | AX-170342603 | 0 | C | T | 0.412 | -6.06E-02 | 7.97E-02 | 2.64E+00 | 2.71E+00 | 4.44E-01 | no | Rtkn2 | intron_variant,NMD_transcript_variant | MODIFIER |
| 10 | 68032848 | AX-170305209 | 0 | A | G | 0.412 | -6.06E-02 | 7.97E-02 | 2.64E+00 | 2.71E+00 | 4.44E-01 | no | Rtkn2 | intron_variant,NMD_transcript_variant | MODIFIER |
| 10 | 68069751 | AX-170509922 | 0 | C | T | 0.412 | -6.06E-02 | 7.97E-02 | 2.64E+00 | 2.71E+00 | 4.44E-01 | no | | intergenic_variant | MODIFIER |
| 10 | 68075678 | AX-170259397 | 0 | G | A | 0.412 | -6.06E-02 | 7.97E-02 | 2.64E+00 | 2.71E+00 | 4.44E-01 | no | | intergenic_variant | MODIFIER |
| 10 | 68079996 | AX-170265116 | 0 | C | T | 0.412 | -6.06E-02 | 7.97E-02 | 2.64E+00 | 2.71E+00 | 4.44E-01 | no | | regulatory_region_variant | MODIFIER |
| 10 | 68081531 | AX-170248507 | 0 | C | T | 0.412 | -6.06E-02 | 7.97E-02 | 2.64E+00 | 2.71E+00 | 4.44E-01 | no | | regulatory_region_variant | MODIFIER |
| 10 | 68089475 | AX-170130072 | 0 | T | A | 0.412 | -6.06E-02 | 7.97E-02 | 2.64E+00 | 2.71E+00 | 4.44E-01 | no | Arid5b | downstream_gene_variant | MODIFIER |
| 10 | 68089719 | AX-170360364 | 0 | A | G | 0.412 | -6.06E-02 | 7.97E-02 | 2.64E+00 | 2.71E+00 | 4.44E-01 | no | Arid5b | downstream_gene_variant | MODIFIER |
| 10 | 68114373 | AX-97815807 | 0 | A | G | 0.412 | -6.06E-02 | 7.97E-02 | 2.64E+00 | 2.71E+00 | 4.44E-01 | no | Arid5b | intron_variant,NMD_transcript_variant | MODIFIER |
| 10 | 68133106 | AX-170003404 | 0 | G | A | 0.412 | -6.06E-02 | 7.97E-02 | 2.64E+00 | 2.71E+00 | 4.44E-01 | no | Arid5b | intron_variant,NMD_transcript_variant | MODIFIER |
| 10 | 68134225 | AX-170003403 | 0 | T | G | 0.412 | -6.06E-02 | 7.97E-02 | 2.64E+00 | 2.71E+00 | 4.44E-01 | no | Arid5b | intron_variant,NMD_transcript_variant | MODIFIER |
| 10 | 68141440 | AX-97825128 | 0 | C | T | 0.412 | -6.06E-02 | 7.97E-02 | 2.64E+00 | 2.71E+00 | 4.44E-01 | no | Arid5b | intron_variant,NMD_transcript_variant | MODIFIER |
| 10 | 68145282 | AX-170388026 | 0 | C | T | 0.412 | -6.06E-02 | 7.97E-02 | 2.64E+00 | 2.71E+00 | 4.44E-01 | no | Arid5b | intron_variant,NMD_transcript_variant | MODIFIER |
| 10 | 68169377 | AX-170003405 | 0 | A | G | 0.412 | -6.06E-02 | 7.97E-02 | 2.64E+00 | 2.71E+00 | 4.44E-01 | no | Arid5b | intron_variant,NMD_transcript_variant | MODIFIER |
| 10 | 68173789 | AX-170187292 | 0 | G | T | 0.412 | -6.06E-02 | 7.97E-02 | 2.64E+00 | 2.71E+00 | 4.44E-01 | no | Arid5b | intron_variant,NMD_transcript_variant | MODIFIER |
| 10 | 68178116 | AX-170455446 | 0 | C | T | 0.412 | -6.06E-02 | 7.97E-02 | 2.64E+00 | 2.71E+00 | 4.44E-01 | no | Arid5b | intron_variant,NMD_transcript_variant | MODIFIER |
| 10 | 68181434 | AX-170189641 | 0 | T | G | 0.412 | -6.06E-02 | 7.97E-02 | 2.64E+00 | 2.71E+00 | 4.44E-01 | no | Arid5b | intron_variant,NMD_transcript_variant | MODIFIER |
| 10 | 68190605 | AX-170003406 | 0 | C | T | 0.412 | -6.06E-02 | 7.97E-02 | 2.64E+00 | 2.71E+00 | 4.44E-01 | no | Arid5b | intron_variant,NMD_transcript_variant | MODIFIER |
| 10 | 68199008 | AX-170303956 | 0 | G | T | 0.412 | -6.06E-02 | 7.97E-02 | 2.64E+00 | 2.71E+00 | 4.44E-01 | no | Arid5b | intron_variant,NMD_transcript_variant | MODIFIER |
| 10 | 68199100 | AX-170197955 | 0 | T | C | 0.412 | -6.06E-02 | 7.97E-02 | 2.64E+00 | 2.71E+00 | 4.44E-01 | no | Arid5b | intron_variant,NMD_transcript_variant | MODIFIER |
| 10 | 68201167 | AX-170411688 | 0 | C | A | 0.412 | -6.06E-02 | 7.97E-02 | 2.64E+00 | 2.71E+00 | 4.44E-01 | no | Arid5b | intron_variant,NMD_transcript_variant | MODIFIER |
| 10 | 68202519 | AX-170267290 | 0 | C | T | 0.412 | -6.06E-02 | 7.97E-02 | 2.64E+00 | 2.71E+00 | 4.44E-01 | no | Arid5b | intron_variant,NMD_transcript_variant | MODIFIER |
| 10 | 68224590 | AX-170194442 | 0 | A | G | 0.412 | -6.06E-02 | 7.97E-02 | 2.64E+00 | 2.71E+00 | 4.44E-01 | no | Arid5b | intron_variant,NMD_transcript_variant | MODIFIER |
| 10 | 68226448 | AX-97852561 | 0 | C | T | 0.412 | -6.06E-02 | 7.97E-02 | 2.64E+00 | 2.71E+00 | 4.44E-01 | no | Arid5b | intron_variant,NMD_transcript_variant | MODIFIER |
| 10 | 68230456 | AX-170045119 | 0 | A | G | 0.412 | -6.06E-02 | 7.97E-02 | 2.64E+00 | 2.71E+00 | 4.44E-01 | no | Arid5b | intron_variant,NMD_transcript_variant | MODIFIER |
| 10 | 68260750 | AX-170133323 | 0 | C | T | 0.412 | -6.06E-02 | 7.97E-02 | 2.64E+00 | 2.71E+00 | 4.44E-01 | no | Arid5b | intron_variant,NMD_transcript_variant | MODIFIER |
| 10 | 68285235 | AX-170177816 | 0 | A | G | 0.412 | -6.06E-02 | 7.97E-02 | 2.64E+00 | 2.71E+00 | 4.44E-01 | no | | intergenic_variant | MODIFIER |

| | | | | | | | | | | | | | | | |
|---|---|---|---|---|---|---|---|---|---|---|---|---|---|---|---|
| 13 | 35893825 | AX-170008110 | 1 | A | G | 0.179 | -7.55E-02 | 9.92E-02 | 2.60E+00 | 2.70E+00 | 4.44E-01 | no | Rpp40 | 3_prime_UTR_variant | MODIFIER |
| 4 | 58842616 | AX-97829507 | 0 | G | A | 0.368 | 8.35E-02 | 1.10E-01 | 2.70E+00 | 2.77E+00 | 4.44E-01 | no | Ecpas | synonymous_variant | LOW |
| 10 | 57048758 | AX-170435112 | 0 | C | T | 0.026 | -9.82E-02 | 1.30E-01 | 2.56E+00 | 2.73E+00 | 4.44E-01 | no | | intergenic_variant | MODIFIER |
| 9 | 27218436 | AX-170290959 | 1 | C | G | 0.455 | 7.03E-02 | 9.16E-02 | 2.57E+00 | 2.63E+00 | 4.44E-01 | no | | intergenic_variant | MODIFIER |
| 18 | 80382690 | AX-170474365 | 1 | C | T | 0.384 | -6.47E-02 | 8.52E-02 | 2.61E+00 | 2.70E+00 | 4.44E-01 | no | | regulatory_region_variant | MODIFIER |
| 1 | 191246437 | AX-170423304 | 0 | C | T | 0.263 | 7.57E-02 | 9.54E-02 | 2.43E+00 | 2.51E+00 | 4.44E-01 | no | Gm37074 | upstream_gene_variant | MODIFIER |
| 1 | 124500241 | AX-170252622 | 0 | G | A | 0.57 | 6.48E-02 | 8.45E-02 | 2.57E+00 | 2.63E+00 | 4.44E-01 | no | | intergenic_variant | MODIFIER |
| 18 | 80214917 | AX-170398729 | 1 | C | T | 0.384 | -6.47E-02 | 8.52E-02 | 2.62E+00 | 2.71E+00 | 4.44E-01 | no | Gm16286 | downstream_gene_variant | MODIFIER |
| 13 | 4597344 | AX-170404000 | 1 | G | A | 0.205 | -9.60E-02 | 1.25E-01 | 2.55E+00 | 2.61E+00 | 4.44E-01 | no | Akr1e1 | intron_variant | MODIFIER |
| 1 | 128031992 | AX-97851719 | 0 | T | C | 0.535 | 6.42E-02 | 8.43E-02 | 2.61E+00 | 2.68E+00 | 4.45E-01 | no | Zranb3 | intron_variant | MODIFIER |
| 7 | 74077517 | AX-97815876 | 2 | A | G | 0.373 | -7.37E-02 | 9.78E-02 | 2.75E+00 | 2.85E+00 | 4.45E-01 | no | | intergenic_variant | MODIFIER |
| 10 | 26558101 | AX-170272917 | 0 | G | A | 0.202 | 7.28E-02 | 9.66E-02 | 2.80E+00 | 2.92E+00 | 4.45E-01 | no | | intergenic_variant | MODIFIER |
| 5 | 128798256 | AX-170358831 | 1 | C | T | 0.161 | -7.13E-02 | 8.71E-02 | 2.34E+00 | 2.44E+00 | 4.45E-01 | no | Rimbp2 | intron_variant | MODIFIER |
| 1 | 170650928 | AX-170106780 | 0 | C | T | 0.246 | 9.23E-02 | 1.22E-01 | 2.65E+00 | 2.71E+00 | 4.45E-01 | no | Olfml2b | intron_variant | MODIFIER |
| 7 | 74851864 | AX-170391502 | 1 | A | T | 0.348 | -7.17E-02 | 9.53E-02 | 2.74E+00 | 2.85E+00 | 4.45E-01 | no | 0610006L08Rik | intron_variant,non_coding_transcript_variant | MODIFIER |
| 14 | 119918846 | AX-170010823 | 0 | T | C | 0.333 | 5.74E-02 | 7.57E-02 | 2.62E+00 | 2.73E+00 | 4.45E-01 | no | | regulatory_region_variant | MODIFIER |
| 5 | 35483650 | AX-170303111 | 0 | A | G | 0.325 | 8.44E-02 | 1.12E-01 | 2.75E+00 | 2.81E+00 | 4.45E-01 | no | | regulatory_region_variant | MODIFIER |
| 5 | 35941202 | AX-170209716 | 0 | A | G | 0.325 | 8.44E-02 | 1.12E-01 | 2.75E+00 | 2.81E+00 | 4.45E-01 | no | Afap1 | intron_variant | MODIFIER |
| 5 | 35984096 | AX-170519668 | 0 | C | T | 0.325 | 8.44E-02 | 1.12E-01 | 2.75E+00 | 2.81E+00 | 4.45E-01 | no | Afap1 | intron_variant | MODIFIER |
| 5 | 35985230 | AX-170302014 | 0 | T | C | 0.325 | 8.44E-02 | 1.12E-01 | 2.75E+00 | 2.81E+00 | 4.45E-01 | no | Afap1 | intron_variant | MODIFIER |
| 5 | 36009431 | AX-170142092 | 0 | T | C | 0.325 | 8.44E-02 | 1.12E-01 | 2.75E+00 | 2.81E+00 | 4.45E-01 | no | | intergenic_variant | MODIFIER |
| 5 | 36010254 | AX-170395661 | 0 | T | C | 0.325 | 8.44E-02 | 1.12E-01 | 2.75E+00 | 2.81E+00 | 4.45E-01 | no | | intergenic_variant | MODIFIER |
| 5 | 36021710 | AX-170525533 | 0 | A | C | 0.325 | 8.44E-02 | 1.12E-01 | 2.75E+00 | 2.81E+00 | 4.45E-01 | no | Sorcs2 | intron_variant | MODIFIER |
| 5 | 36030335 | AX-97804553 | 0 | C | A | 0.325 | 8.44E-02 | 1.12E-01 | 2.75E+00 | 2.81E+00 | 4.45E-01 | no | Sorcs2 | intron_variant | MODIFIER |
| 5 | 36050538 | AX-170448830 | 0 | G | T | 0.325 | 8.44E-02 | 1.12E-01 | 2.75E+00 | 2.81E+00 | 4.45E-01 | no | Sorcs2 | synonymous_variant | LOW |
| 5 | 36068266 | AX-97852067 | 0 | T | C | 0.325 | 8.44E-02 | 1.12E-01 | 2.75E+00 | 2.81E+00 | 4.45E-01 | no | Sorcs2 | intron_variant | MODIFIER |
| 5 | 36072441 | AX-170153366 | 0 | A | G | 0.325 | 8.44E-02 | 1.12E-01 | 2.75E+00 | 2.81E+00 | 4.45E-01 | no | Sorcs2 | intron_variant | MODIFIER |
| 5 | 36103219 | AX-170518654 | 0 | T | C | 0.325 | 8.44E-02 | 1.12E-01 | 2.75E+00 | 2.81E+00 | 4.45E-01 | no | Sorcs2 | intron_variant | MODIFIER |
| 5 | 36110353 | AX-170024684 | 0 | A | G | 0.325 | 8.44E-02 | 1.12E-01 | 2.75E+00 | 2.81E+00 | 4.45E-01 | no | Sorcs2 | intron_variant | MODIFIER |
| 5 | 36118144 | AX-170024683 | 0 | C | T | 0.325 | 8.44E-02 | 1.12E-01 | 2.75E+00 | 2.81E+00 | 4.45E-01 | no | Sorcs2 | intron_variant | MODIFIER |
| 5 | 36127636 | AX-170357148 | 0 | A | G | 0.325 | 8.44E-02 | 1.12E-01 | 2.75E+00 | 2.81E+00 | 4.45E-01 | no | Sorcs2 | intron_variant | MODIFIER |
| 5 | 36128287 | AX-170169431 | 0 | A | G | 0.325 | 8.44E-02 | 1.12E-01 | 2.75E+00 | 2.81E+00 | 4.45E-01 | no | Sorcs2 | intron_variant | MODIFIER |
| 5 | 36147066 | AX-170148109 | 0 | C | T | 0.325 | 8.44E-02 | 1.12E-01 | 2.75E+00 | 2.81E+00 | 4.45E-01 | no | Sorcs2 | intron_variant | MODIFIER |
| 5 | 36180578 | AX-97840901 | 0 | T | C | 0.325 | 8.44E-02 | 1.12E-01 | 2.75E+00 | 2.81E+00 | 4.45E-01 | no | Sorcs2 | intron_variant | MODIFIER |
| 5 | 36180876 | AX-170237395 | 0 | T | C | 0.325 | 8.44E-02 | 1.12E-01 | 2.75E+00 | 2.81E+00 | 4.45E-01 | no | Sorcs2 | intron_variant | MODIFIER |
| 5 | 36193458 | AX-170260030 | 0 | T | G | 0.325 | 8.44E-02 | 1.12E-01 | 2.75E+00 | 2.81E+00 | 4.45E-01 | no | Sorcs2 | intron_variant | MODIFIER |
| 5 | 36194054 | AX-170421464 | 0 | T | C | 0.325 | 8.44E-02 | 1.12E-01 | 2.75E+00 | 2.81E+00 | 4.45E-01 | no | Sorcs2 | intron_variant | MODIFIER |
| 5 | 36198424 | AX-170266460 | 0 | A | G | 0.325 | 8.44E-02 | 1.12E-01 | 2.75E+00 | 2.81E+00 | 4.45E-01 | no | Sorcs2 | intron_variant | MODIFIER |
| 5 | 36221334 | AX-170301927 | 0 | A | G | 0.325 | 8.44E-02 | 1.12E-01 | 2.75E+00 | 2.81E+00 | 4.45E-01 | no | Sorcs2 | intron_variant | MODIFIER |
| 5 | 36226159 | AX-97791523 | 0 | A | G | 0.325 | 8.44E-02 | 1.12E-01 | 2.75E+00 | 2.81E+00 | 4.45E-01 | no | Sorcs2 | intron_variant | MODIFIER |
| 5 | 36306721 | AX-170168614 | 0 | C | T | 0.325 | 8.44E-02 | 1.12E-01 | 2.75E+00 | 2.81E+00 | 4.45E-01 | no | Sorcs2 | intron_variant | MODIFIER |
| 5 | 36359370 | AX-170394184 | 0 | A | T | 0.325 | 8.44E-02 | 1.12E-01 | 2.75E+00 | 2.81E+00 | 4.45E-01 | no | Sorcs2 | intron_variant | MODIFIER |
| 5 | 36386421 | AX-170108831 | 0 | G | A | 0.325 | 8.44E-02 | 1.12E-01 | 2.75E+00 | 2.81E+00 | 4.45E-01 | no | Sorcs2 | intron_variant | MODIFIER |
| 5 | 36388228 | AX-170138089 | 0 | G | A | 0.325 | 8.44E-02 | 1.12E-01 | 2.75E+00 | 2.81E+00 | 4.45E-01 | no | Sorcs2 | intron_variant | MODIFIER |
| 5 | 36389204 | AX-170437459 | 0 | A | G | 0.325 | 8.44E-02 | 1.12E-01 | 2.75E+00 | 2.81E+00 | 4.45E-01 | no | Sorcs2 | intron_variant | MODIFIER |
| 5 | 36389755 | AX-170467644 | 0 | G | T | 0.325 | 8.44E-02 | 1.12E-01 | 2.75E+00 | 2.81E+00 | 4.45E-01 | no | Sorcs2 | intron_variant | MODIFIER |
| 5 | 36406602 | AX-170321931 | 0 | C | G | 0.325 | 8.44E-02 | 1.12E-01 | 2.75E+00 | 2.81E+00 | 4.45E-01 | no | | intergenic_variant | MODIFIER |
| 5 | 36413705 | AX-170071808 | 0 | C | T | 0.325 | 8.44E-02 | 1.12E-01 | 2.75E+00 | 2.81E+00 | 4.45E-01 | no | | intergenic_variant | MODIFIER |
| 5 | 36418831 | AX-170329276 | 0 | A | G | 0.325 | 8.44E-02 | 1.12E-01 | 2.75E+00 | 2.81E+00 | 4.45E-01 | no | | intergenic_variant | MODIFIER |
| 5 | 36474424 | AX-170114319 | 0 | T | C | 0.325 | 8.44E-02 | 1.12E-01 | 2.75E+00 | 2.81E+00 | 4.45E-01 | no | Tada2b | 3_prime_UTR_variant | MODIFIER |
| 5 | 36491610 | AX-170332802 | 0 | C | T | 0.325 | 8.44E-02 | 1.12E-01 | 2.75E+00 | 2.81E+00 | 4.45E-01 | no | Tbc1d14 | 3_prime_UTR_variant | MODIFIER |
| 5 | 36516016 | AX-170226885 | 0 | T | C | 0.325 | 8.44E-02 | 1.12E-01 | 2.75E+00 | 2.81E+00 | 4.45E-01 | no | Tbc1d14 | intron_variant | MODIFIER |
| 5 | 36529635 | AX-170183616 | 0 | A | G | 0.325 | 8.44E-02 | 1.12E-01 | 2.75E+00 | 2.81E+00 | 4.45E-01 | no | Tbc1d14 | intron_variant | MODIFIER |
| 5 | 36556460 | AX-170439572 | 0 | T | C | 0.325 | 8.44E-02 | 1.12E-01 | 2.75E+00 | 2.81E+00 | 4.45E-01 | no | Tbc1d14 | intron_variant | MODIFIER |

| | | | | | | | | | | | | | | | |
|---|---|---|---|---|---|---|---|---|---|---|---|---|---|---|---|
| 5 | 36563046 | AX-170349403 | 0 | T | C | 0.325 | 8.44E-02 | 1.12E-01 | 2.75E+00 | 2.81E+00 | 4.45E-01 | no | Tbc1d14 | intron_variant | MODIFIER |
| 5 | 36599811 | AX-170132922 | 0 | A | G | 0.325 | 8.44E-02 | 1.12E-01 | 2.75E+00 | 2.81E+00 | 4.45E-01 | no | D5Ertd579e | downstream_gene_variant | MODIFIER |
| 5 | 36607660 | AX-170437284 | 0 | T | C | 0.325 | 8.44E-02 | 1.12E-01 | 2.75E+00 | 2.81E+00 | 4.45E-01 | no | D5Ertd579e | intron_variant | MODIFIER |
| 5 | 36668821 | AX-170302658 | 0 | C | T | 0.325 | 8.44E-02 | 1.12E-01 | 2.75E+00 | 2.81E+00 | 4.45E-01 | no | D5Ertd579e | intron_variant | MODIFIER |
| 5 | 36691803 | AX-170024691 | 0 | T | C | 0.325 | 8.44E-02 | 1.12E-01 | 2.75E+00 | 2.81E+00 | 4.45E-01 | no | D5Ertd579e | intron_variant | MODIFIER |
| 5 | 36707101 | AX-170024686 | 0 | T | G | 0.325 | 8.44E-02 | 1.12E-01 | 2.75E+00 | 2.81E+00 | 4.45E-01 | no | | intergenic_variant | MODIFIER |
| 5 | 36713588 | AX-170024688 | 0 | C | T | 0.325 | 8.44E-02 | 1.12E-01 | 2.75E+00 | 2.81E+00 | 4.45E-01 | no | | regulatory_region_variant | MODIFIER |
| 5 | 36714039 | AX-97814646 | 0 | T | C | 0.325 | 8.44E-02 | 1.12E-01 | 2.75E+00 | 2.81E+00 | 4.45E-01 | no | | regulatory_region_variant | MODIFIER |
| 5 | 36731245 | AX-170400702 | 0 | C | T | 0.325 | 8.44E-02 | 1.12E-01 | 2.75E+00 | 2.81E+00 | 4.45E-01 | no | | regulatory_region_variant | MODIFIER |
| 5 | 36762984 | AX-170515846 | 0 | T | C | 0.325 | 8.44E-02 | 1.12E-01 | 2.75E+00 | 2.81E+00 | 4.45E-01 | no | Gm40318 | intron_variant,non_coding_transcript_variant | MODIFIER |
| 5 | 36891576 | AX-97793165 | 0 | A | C | 0.325 | 8.44E-02 | 1.12E-01 | 2.75E+00 | 2.81E+00 | 4.45E-01 | no | Ppp2r2c | intron_variant | MODIFIER |
| 5 | 36898142 | AX-170024695 | 0 | A | C | 0.325 | 8.44E-02 | 1.12E-01 | 2.75E+00 | 2.81E+00 | 4.45E-01 | no | Ppp2r2c | intron_variant | MODIFIER |
| 5 | 36899099 | AX-170071817 | 0 | A | G | 0.325 | 8.44E-02 | 1.12E-01 | 2.75E+00 | 2.81E+00 | 4.45E-01 | no | Ppp2r2c | intron_variant | MODIFIER |
| 5 | 36899999 | AX-170071815 | 0 | A | G | 0.325 | 8.44E-02 | 1.12E-01 | 2.75E+00 | 2.81E+00 | 4.45E-01 | no | Ppp2r2c | intron_variant | MODIFIER |
| 5 | 36905570 | AX-170334348 | 0 | C | G | 0.325 | 8.44E-02 | 1.12E-01 | 2.75E+00 | 2.81E+00 | 4.45E-01 | no | Ppp2r2c | intron_variant | MODIFIER |
| 5 | 36905600 | AX-170266064 | 0 | G | T | 0.325 | 8.44E-02 | 1.12E-01 | 2.75E+00 | 2.81E+00 | 4.45E-01 | no | Ppp2r2c | intron_variant | MODIFIER |
| 5 | 36951144 | AX-97781120 | 0 | G | A | 0.325 | 8.44E-02 | 1.12E-01 | 2.75E+00 | 2.81E+00 | 4.45E-01 | no | Ppp2r2c | intron_variant | MODIFIER |
| 5 | 37009887 | AX-97813851 | 0 | T | C | 0.325 | 8.44E-02 | 1.12E-01 | 2.75E+00 | 2.81E+00 | 4.45E-01 | no | | regulatory_region_variant | MODIFIER |
| 5 | 37035636 | AX-170400738 | 0 | C | T | 0.325 | 8.44E-02 | 1.12E-01 | 2.75E+00 | 2.81E+00 | 4.45E-01 | no | Jakmip1 | intron_variant | MODIFIER |
| 5 | 37048428 | AX-170305041 | 0 | A | G | 0.325 | 8.44E-02 | 1.12E-01 | 2.75E+00 | 2.81E+00 | 4.45E-01 | no | Jakmip1 | upstream_gene_variant | MODIFIER |
| 5 | 37049911 | AX-170071819 | 0 | C | T | 0.325 | 8.44E-02 | 1.12E-01 | 2.75E+00 | 2.81E+00 | 4.45E-01 | no | Jakmip1 | upstream_gene_variant | MODIFIER |
| 5 | 37065494 | AX-97830576 | 0 | C | T | 0.325 | 8.44E-02 | 1.12E-01 | 2.75E+00 | 2.81E+00 | 4.45E-01 | no | Jakmip1 | intron_variant | MODIFIER |
| 5 | 37097871 | AX-170394226 | 0 | A | G | 0.325 | 8.44E-02 | 1.12E-01 | 2.75E+00 | 2.81E+00 | 4.45E-01 | no | Jakmip1 | intron_variant | MODIFIER |
| 5 | 37100138 | AX-97833202 | 0 | G | A | 0.325 | 8.44E-02 | 1.12E-01 | 2.75E+00 | 2.81E+00 | 4.45E-01 | no | Jakmip1 | intron_variant | MODIFIER |
| 5 | 37100271 | AX-170329999 | 0 | A | T | 0.325 | 8.44E-02 | 1.12E-01 | 2.75E+00 | 2.81E+00 | 4.45E-01 | no | Jakmip1 | intron_variant | MODIFIER |
| 5 | 37126172 | AX-170336939 | 0 | G | C | 0.325 | 8.44E-02 | 1.12E-01 | 2.75E+00 | 2.81E+00 | 4.45E-01 | no | Jakmip1 | downstream_gene_variant | MODIFIER |
| 5 | 37194572 | AX-170114180 | 0 | G | T | 0.325 | 8.44E-02 | 1.12E-01 | 2.75E+00 | 2.81E+00 | 4.45E-01 | no | Gm1043 | downstream_gene_variant | MODIFIER |
| 5 | 37197367 | AX-170152080 | 0 | T | C | 0.325 | 8.44E-02 | 1.12E-01 | 2.75E+00 | 2.81E+00 | 4.45E-01 | no | Gm1043 | downstream_gene_variant | MODIFIER |
| 5 | 37336300 | AX-170371803 | 0 | T | C | 0.325 | 8.44E-02 | 1.12E-01 | 2.75E+00 | 2.81E+00 | 4.45E-01 | no | Evc | intron_variant | MODIFIER |
| 5 | 37380713 | AX-97814268 | 0 | T | G | 0.325 | 8.44E-02 | 1.12E-01 | 2.75E+00 | 2.81E+00 | 4.45E-01 | no | Evc2 | intron_variant | MODIFIER |
| 5 | 37384169 | AX-170505806 | 0 | G | A | 0.325 | 8.44E-02 | 1.12E-01 | 2.75E+00 | 2.81E+00 | 4.45E-01 | no | Evc2 | intron_variant | MODIFIER |
| 5 | 37388484 | AX-170401068 | 0 | C | T | 0.325 | 8.44E-02 | 1.12E-01 | 2.75E+00 | 2.81E+00 | 4.45E-01 | no | Evc2 | intron_variant | MODIFIER |
| 5 | 37448652 | AX-170401962 | 0 | T | C | 0.325 | 8.44E-02 | 1.12E-01 | 2.75E+00 | 2.81E+00 | 4.45E-01 | no | Stk32b | 3_prime_UTR_variant | MODIFIER |
| 5 | 37457191 | AX-97813364 | 0 | T | C | 0.325 | 8.44E-02 | 1.12E-01 | 2.75E+00 | 2.81E+00 | 4.45E-01 | no | Stk32b | synonymous_variant | LOW |
| 5 | 37463384 | AX-170115457 | 0 | A | G | 0.325 | 8.44E-02 | 1.12E-01 | 2.75E+00 | 2.81E+00 | 4.45E-01 | no | Stk32b | intron_variant | MODIFIER |
| 5 | 37463866 | AX-170071824 | 0 | A | T | 0.325 | 8.44E-02 | 1.12E-01 | 2.75E+00 | 2.81E+00 | 4.45E-01 | no | Stk32b | intron_variant | MODIFIER |
| 5 | 37496757 | AX-170348033 | 0 | T | C | 0.325 | 8.44E-02 | 1.12E-01 | 2.75E+00 | 2.81E+00 | 4.45E-01 | no | Stk32b | intron_variant | MODIFIER |
| 5 | 37498718 | AX-170311710 | 0 | A | C | 0.325 | 8.44E-02 | 1.12E-01 | 2.75E+00 | 2.81E+00 | 4.45E-01 | no | Stk32b | intron_variant | MODIFIER |
| 5 | 37520292 | AX-170169449 | 0 | G | A | 0.325 | 8.44E-02 | 1.12E-01 | 2.75E+00 | 2.81E+00 | 4.45E-01 | no | Stk32b | intron_variant | MODIFIER |
| 5 | 37520564 | AX-170210523 | 0 | T | C | 0.325 | 8.44E-02 | 1.12E-01 | 2.75E+00 | 2.81E+00 | 4.45E-01 | no | Stk32b | intron_variant | MODIFIER |
| 5 | 37550315 | AX-170413628 | 0 | C | G | 0.325 | 8.44E-02 | 1.12E-01 | 2.75E+00 | 2.81E+00 | 4.45E-01 | no | Stk32b | intron_variant | MODIFIER |
| 5 | 37550928 | AX-97853912 | 0 | C | T | 0.325 | 8.44E-02 | 1.12E-01 | 2.75E+00 | 2.81E+00 | 4.45E-01 | no | Stk32b | intron_variant | MODIFIER |
| 5 | 37570441 | AX-170392385 | 0 | A | G | 0.325 | 8.44E-02 | 1.12E-01 | 2.75E+00 | 2.81E+00 | 4.45E-01 | no | Stk32b | intron_variant | MODIFIER |
| 5 | 37576190 | AX-97843868 | 0 | C | T | 0.325 | 8.44E-02 | 1.12E-01 | 2.75E+00 | 2.81E+00 | 4.45E-01 | no | Stk32b | intron_variant | MODIFIER |
| 5 | 37602613 | AX-97826068 | 0 | T | G | 0.325 | 8.44E-02 | 1.12E-01 | 2.75E+00 | 2.81E+00 | 4.45E-01 | no | Stk32b | intron_variant | MODIFIER |
| 5 | 37723855 | AX-97821892 | 0 | T | C | 0.325 | 8.44E-02 | 1.12E-01 | 2.75E+00 | 2.81E+00 | 4.45E-01 | no | | intergenic_variant | MODIFIER |
| 5 | 37734663 | AX-170175548 | 0 | G | A | 0.325 | 8.44E-02 | 1.12E-01 | 2.75E+00 | 2.81E+00 | 4.45E-01 | no | Cytl1 | upstream_gene_variant | MODIFIER |
| 5 | 37770061 | AX-170024707 | 0 | C | T | 0.325 | 8.44E-02 | 1.12E-01 | 2.75E+00 | 2.81E+00 | 4.45E-01 | no | | intergenic_variant | MODIFIER |
| 5 | 37796338 | AX-170311048 | 0 | A | G | 0.325 | 8.44E-02 | 1.12E-01 | 2.75E+00 | 2.81E+00 | 4.45E-01 | no | | intergenic_variant | MODIFIER |
| 5 | 37796407 | AX-170203283 | 0 | G | T | 0.325 | 8.44E-02 | 1.12E-01 | 2.75E+00 | 2.81E+00 | 4.45E-01 | no | | intergenic_variant | MODIFIER |
| 5 | 37796461 | AX-97842413 | 0 | C | A | 0.325 | 8.44E-02 | 1.12E-01 | 2.75E+00 | 2.81E+00 | 4.45E-01 | no | | intergenic_variant | MODIFIER |
| 5 | 37832340 | AX-97817632 | 0 | A | G | 0.325 | 8.44E-02 | 1.12E-01 | 2.75E+00 | 2.81E+00 | 4.45E-01 | no | | intergenic_variant | MODIFIER |
| 5 | 37848265 | AX-170422807 | 0 | A | G | 0.325 | 8.44E-02 | 1.12E-01 | 2.75E+00 | 2.81E+00 | 4.45E-01 | no | | intergenic_variant | MODIFIER |
| 5 | 37872928 | AX-170325424 | 0 | T | C | 0.325 | 8.44E-02 | 1.12E-01 | 2.75E+00 | 2.81E+00 | 4.45E-01 | no | | intergenic_variant | MODIFIER |

| | | | | | | | | | | | | | | | |
|---|---|---|---|---|---|---|---|---|---|---|---|---|---|---|---|
| 5 | 37898348 | AX-97836659 | 0 | G | T | 0.325 | 8.44E-02 | 1.12E-01 | 2.75E+00 | 2.81E+00 | 4.45E-01 | no | Gm20052 | downstream_gene_variant | MODIFIER |
| 5 | 37899354 | AX-170331477 | 0 | A | G | 0.325 | 8.44E-02 | 1.12E-01 | 2.75E+00 | 2.81E+00 | 4.45E-01 | no | Gm20052 | downstream_gene_variant | MODIFIER |
| 5 | 37943324 | AX-97792689 | 0 | G | C | 0.325 | 8.44E-02 | 1.12E-01 | 2.75E+00 | 2.81E+00 | 4.45E-01 | no | | intergenic_variant | MODIFIER |
| 5 | 37949090 | AX-97845200 | 0 | A | G | 0.325 | 8.44E-02 | 1.12E-01 | 2.75E+00 | 2.81E+00 | 4.45E-01 | no | | intergenic_variant | MODIFIER |
| 5 | 37970054 | AX-97830467 | 0 | C | T | 0.325 | 8.44E-02 | 1.12E-01 | 2.75E+00 | 2.81E+00 | 4.45E-01 | no | | intergenic_variant | MODIFIER |
| 5 | 38018097 | AX-170500012 | 0 | T | C | 0.325 | 8.44E-02 | 1.12E-01 | 2.75E+00 | 2.81E+00 | 4.45E-01 | no | | intergenic_variant | MODIFIER |
| 5 | 38054455 | AX-97818139 | 0 | A | G | 0.325 | 8.44E-02 | 1.12E-01 | 2.75E+00 | 2.81E+00 | 4.45E-01 | no | Stx18 | intron_variant | MODIFIER |
| 5 | 38088758 | AX-170218939 | 0 | A | G | 0.325 | 8.44E-02 | 1.12E-01 | 2.75E+00 | 2.81E+00 | 4.45E-01 | no | Stx18 | intron_variant | MODIFIER |
| 5 | 38130131 | AX-97830945 | 0 | G | T | 0.325 | 8.44E-02 | 1.12E-01 | 2.75E+00 | 2.81E+00 | 4.45E-01 | no | Stx18 | intron_variant | MODIFIER |
| 5 | 38136349 | AX-170214252 | 0 | T | C | 0.325 | 8.44E-02 | 1.12E-01 | 2.75E+00 | 2.81E+00 | 4.45E-01 | no | Stx18 | splice_region_variant,intron_variant | LOW |
| 5 | 38157323 | AX-97784896 | 0 | T | C | 0.325 | 8.44E-02 | 1.12E-01 | 2.75E+00 | 2.81E+00 | 4.45E-01 | no | Nsg1 | intron_variant | MODIFIER |
| 5 | 38176634 | AX-97833938 | 0 | C | A | 0.325 | 8.44E-02 | 1.12E-01 | 2.75E+00 | 2.81E+00 | 4.45E-01 | no | | intergenic_variant | MODIFIER |
| 5 | 38202251 | AX-170024720 | 0 | T | C | 0.325 | 8.44E-02 | 1.12E-01 | 2.75E+00 | 2.81E+00 | 4.45E-01 | no | Zbtb49 | intron_variant | MODIFIER |
| 5 | 38202653 | AX-170024724 | 0 | A | C | 0.325 | 8.44E-02 | 1.12E-01 | 2.75E+00 | 2.81E+00 | 4.45E-01 | no | Zbtb49 | intron_variant | MODIFIER |
| 5 | 38204006 | AX-97801252 | 0 | A | C | 0.325 | 8.44E-02 | 1.12E-01 | 2.75E+00 | 2.81E+00 | 4.45E-01 | no | Zbtb49 | intron_variant | MODIFIER |
| 5 | 38291610 | AX-97835287 | 0 | A | C | 0.325 | 8.44E-02 | 1.12E-01 | 2.75E+00 | 2.81E+00 | 4.45E-01 | no | Otop1 | intron_variant | MODIFIER |
| 5 | 38296945 | AX-170275037 | 0 | T | G | 0.325 | 8.44E-02 | 1.12E-01 | 2.75E+00 | 2.81E+00 | 4.45E-01 | no | Otop1 | intron_variant | MODIFIER |
| 5 | 38298326 | AX-170334840 | 0 | A | G | 0.325 | 8.44E-02 | 1.12E-01 | 2.75E+00 | 2.81E+00 | 4.45E-01 | no | Otop1 | intron_variant | MODIFIER |
| 5 | 38302218 | AX-97802907 | 0 | A | G | 0.325 | 8.44E-02 | 1.12E-01 | 2.75E+00 | 2.81E+00 | 4.45E-01 | no | Otop1 | intron_variant | MODIFIER |
| 5 | 38344793 | AX-97832338 | 0 | C | T | 0.325 | 8.44E-02 | 1.12E-01 | 2.75E+00 | 2.81E+00 | 4.45E-01 | no | Slc2a9 | downstream_gene_variant | MODIFIER |
| 5 | 38345418 | AX-170024716 | 0 | A | G | 0.325 | 8.44E-02 | 1.12E-01 | 2.75E+00 | 2.81E+00 | 4.45E-01 | no | Slc2a9 | downstream_gene_variant | MODIFIER |
| 5 | 38361716 | AX-170253618 | 0 | G | A | 0.325 | 8.44E-02 | 1.12E-01 | 2.75E+00 | 2.81E+00 | 4.45E-01 | no | Slc2a9 | intron_variant | MODIFIER |
| 5 | 38397352 | AX-170507164 | 0 | G | C | 0.325 | 8.44E-02 | 1.12E-01 | 2.75E+00 | 2.81E+00 | 4.45E-01 | no | Slc2a9 | intron_variant | MODIFIER |
| 5 | 38424303 | AX-170071843 | 0 | T | C | 0.325 | 8.44E-02 | 1.12E-01 | 2.75E+00 | 2.81E+00 | 4.45E-01 | no | Slc2a9 | intron_variant | MODIFIER |
| 5 | 38424512 | AX-170024717 | 0 | A | G | 0.325 | 8.44E-02 | 1.12E-01 | 2.75E+00 | 2.81E+00 | 4.45E-01 | no | Slc2a9 | intron_variant | MODIFIER |
| 5 | 38490800 | AX-97788444 | 0 | A | G | 0.325 | 8.44E-02 | 1.12E-01 | 2.75E+00 | 2.81E+00 | 4.45E-01 | no | Slc2a9 | intron_variant | MODIFIER |
| 5 | 38491837 | AX-170437757 | 0 | A | G | 0.325 | 8.44E-02 | 1.12E-01 | 2.75E+00 | 2.81E+00 | 4.45E-01 | no | Slc2a9 | intron_variant | MODIFIER |
| 5 | 38527166 | AX-170182683 | 0 | T | C | 0.325 | 8.44E-02 | 1.12E-01 | 2.75E+00 | 2.81E+00 | 4.45E-01 | no | Wdr1 | 3_prime_UTR_variant | MODIFIER |
| 5 | 38546988 | AX-170279171 | 0 | C | G | 0.325 | 8.44E-02 | 1.12E-01 | 2.75E+00 | 2.81E+00 | 4.45E-01 | no | Wdr1 | synonymous_variant | LOW |
| 5 | 38550032 | AX-170310997 | 0 | A | G | 0.325 | 8.44E-02 | 1.12E-01 | 2.75E+00 | 2.81E+00 | 4.45E-01 | no | Wdr1 | intron_variant | MODIFIER |
| 5 | 38553646 | AX-170403450 | 0 | T | G | 0.325 | 8.44E-02 | 1.12E-01 | 2.75E+00 | 2.81E+00 | 4.45E-01 | no | Wdr1 | intron_variant | MODIFIER |
| 5 | 38571072 | AX-170228736 | 0 | T | G | 0.325 | 8.44E-02 | 1.12E-01 | 2.75E+00 | 2.81E+00 | 4.45E-01 | no | 4930421P07Rik | upstream_gene_variant | MODIFIER |
| 5 | 38604801 | AX-170071844 | 0 | A | G | 0.325 | 8.44E-02 | 1.12E-01 | 2.75E+00 | 2.81E+00 | 4.45E-01 | no | | intergenic_variant | MODIFIER |
| 5 | 38605813 | AX-97826789 | 0 | T | G | 0.325 | 8.44E-02 | 1.12E-01 | 2.75E+00 | 2.81E+00 | 4.45E-01 | no | | intergenic_variant | MODIFIER |
| 5 | 38609570 | AX-170471466 | 0 | C | T | 0.325 | 8.44E-02 | 1.12E-01 | 2.75E+00 | 2.81E+00 | 4.45E-01 | no | | intergenic_variant | MODIFIER |
| 5 | 38648772 | AX-170329098 | 0 | T | C | 0.325 | 8.44E-02 | 1.12E-01 | 2.75E+00 | 2.81E+00 | 4.45E-01 | no | | intergenic_variant | MODIFIER |
| 5 | 38651126 | AX-97784921 | 0 | C | T | 0.325 | 8.44E-02 | 1.12E-01 | 2.75E+00 | 2.81E+00 | 4.45E-01 | no | | intergenic_variant | MODIFIER |
| 5 | 38677159 | AX-170089453 | 0 | A | G | 0.325 | 8.44E-02 | 1.12E-01 | 2.75E+00 | 2.81E+00 | 4.45E-01 | no | Zfp518b | intron_variant | MODIFIER |
| 5 | 38710354 | AX-97818115 | 0 | C | T | 0.325 | 8.44E-02 | 1.12E-01 | 2.75E+00 | 2.81E+00 | 4.45E-01 | no | Clnk | intron_variant | MODIFIER |
| 18 | 66197150 | AX-170418419 | 0 | A | G | 0.114 | 9.49E-02 | 1.27E-01 | 2.71E+00 | 2.85E+00 | 4.45E-01 | no | Ccbe1 | intron_variant,NMD_transcript_variant | MODIFIER |
| 5 | 108242004 | AX-170147467 | 1 | C | T | 0.473 | 8.18E-02 | 1.08E-01 | 2.62E+00 | 2.68E+00 | 4.45E-01 | no | | intergenic_variant | MODIFIER |
| 12 | 69904586 | AX-97838162 | 0 | T | A | 0.061 | -1.67E-01 | 2.21E-01 | 2.64E+00 | 2.72E+00 | 4.45E-01 | no | Atl1 | intron_variant | MODIFIER |
| 12 | 71139137 | AX-170360604 | 0 | A | C | 0.061 | -1.67E-01 | 2.21E-01 | 2.64E+00 | 2.72E+00 | 4.45E-01 | no | Timm9 | upstream_gene_variant | MODIFIER |
| 13 | 25254043 | AX-97853762 | 1 | G | A | 0.17 | -9.71E-02 | 1.23E-01 | 2.45E+00 | 2.50E+00 | 4.45E-01 | no | Nrsn1 | intron_variant | MODIFIER |
| 11 | 32388628 | AX-97817988 | 0 | A | G | 0.088 | 1.25E-01 | 1.65E-01 | 2.58E+00 | 2.70E+00 | 4.45E-01 | no | Sh3pxd2b | intron_variant | MODIFIER |
| 9 | 79708848 | AX-170371184 | 1 | T | C | 0.214 | -7.83E-02 | 1.04E-01 | 2.71E+00 | 2.83E+00 | 4.45E-01 | no | Col12a1 | intron_variant | MODIFIER |
| 4 | 151640239 | AX-170382790 | 0 | G | A | 0.114 | 1.38E-01 | 1.83E-01 | 2.73E+00 | 2.81E+00 | 4.45E-01 | no | Camta1 | intron_variant | MODIFIER |
| 8 | 64339810 | AX-170286072 | 1 | G | A | 0.143 | -1.13E-01 | 1.48E-01 | 2.59E+00 | 2.65E+00 | 4.45E-01 | no | | intergenic_variant | MODIFIER |
| 5 | 43396588 | AX-170207523 | 2 | T | G | 0.191 | -7.96E-02 | 1.05E-01 | 2.60E+00 | 2.70E+00 | 4.45E-01 | no | | intergenic_variant | MODIFIER |
| 1 | 136553770 | AX-170277783 | 2 | G | A | 0.227 | 8.57E-02 | 1.13E-01 | 2.63E+00 | 2.72E+00 | 4.45E-01 | no | Gm37799 | intron_variant,non_coding_transcript_variant | MODIFIER |
| 13 | 5681784 | AX-170517239 | 1 | T | G | 0.214 | -8.48E-02 | 1.11E-01 | 2.59E+00 | 2.65E+00 | 4.45E-01 | no | Gm35330 | intron_variant,non_coding_transcript_variant | MODIFIER |
| 5 | 49719412 | AX-170421016 | 1 | A | G | 0.473 | 7.79E-02 | 9.66E-02 | 2.38E+00 | 2.45E+00 | 4.45E-01 | no | | intergenic_variant | MODIFIER |
| 10 | 6096233 | AX-170315998 | 0 | T | C | 0.211 | -7.43E-02 | 9.87E-02 | 2.67E+00 | 2.78E+00 | 4.45E-01 | no | | intergenic_variant | MODIFIER |
| 11 | 70852757 | AX-170267761 | 0 | A | G | 0.07 | 1.04E-01 | 1.37E-01 | 2.61E+00 | 2.74E+00 | 4.45E-01 | no | Rabep1 | intron_variant | MODIFIER |

| | | | | | | | | | | | | | | | |
|---|---|---|---|---|---|---|---|---|---|---|---|---|---|---|---|
| 2 | 124808694 | AX-170018923 | 0 | G | A | 0.149 | 1.13E-01 | 1.50E-01 | 2.65E+00 | 2.77E+00 | 4.46E-01 | no | | intergenic_variant | MODIFIER |
| 5 | 42926782 | AX-170119034 | 0 | C | A | 0.211 | -7.61E-02 | 1.00E-01 | 2.62E+00 | 2.71E+00 | 4.46E-01 | no | | intergenic_variant | MODIFIER |
| 20 | 12671555 | AX-170033547 | 0 | A | G | 0.202 | 4.48E-02 | 5.96E-02 | 2.63E+00 | 2.77E+00 | 4.46E-01 | no | 1810030O07Rik | intron_variant | MODIFIER |
| 7 | 124167423 | AX-170078725 | 2 | A | G | 0.382 | 6.42E-02 | 8.53E-02 | 2.75E+00 | 2.83E+00 | 4.46E-01 | no | Hs3st4 | intron_variant | MODIFIER |
| 12 | 88481814 | AX-170112999 | 0 | C | T | 0.035 | -1.03E-01 | 1.36E-01 | 2.56E+00 | 2.72E+00 | 4.46E-01 | no | | intergenic_variant | MODIFIER |
| 1 | 88127454 | AX-170317131 | 0 | A | G | 0.167 | 1.34E-01 | 1.76E-01 | 2.57E+00 | 2.66E+00 | 4.46E-01 | no | Ugt1a7c | intron_variant | MODIFIER |
| 18 | 88915812 | AX-170105549 | 0 | A | T | 0.325 | 4.59E-02 | 6.12E-02 | 2.64E+00 | 2.78E+00 | 4.46E-01 | no | Socs6 | intron_variant | MODIFIER |
| 9 | 18972141 | AX-97814768 | 1 | A | G | 0.464 | 6.87E-02 | 8.95E-02 | 2.55E+00 | 2.61E+00 | 4.46E-01 | no | Olfr833-ps1 | upstream_gene_variant | MODIFIER |
| 1 | 168166042 | AX-170243508 | 1 | A | C | 0.214 | 1.06E-01 | 1.40E-01 | 2.65E+00 | 2.70E+00 | 4.46E-01 | no | Pbx1 | intron_variant | MODIFIER |
| 1 | 168501421 | AX-170002256 | 1 | C | T | 0.214 | 1.06E-01 | 1.40E-01 | 2.65E+00 | 2.70E+00 | 4.46E-01 | no | | intergenic_variant | MODIFIER |
| 2 | 123982470 | AX-170192629 | 0 | G | A | 0.14 | 1.22E-01 | 1.63E-01 | 2.68E+00 | 2.79E+00 | 4.46E-01 | no | | intergenic_variant | MODIFIER |
| 10 | 19001055 | AX-170226529 | 0 | A | G | 0.193 | 7.02E-02 | 9.36E-02 | 2.79E+00 | 2.91E+00 | 4.46E-01 | no | Tnfaip3 | 3_prime_UTR_variant | MODIFIER |
| 18 | 77936117 | AX-170455271 | 2 | G | A | 0.445 | 6.56E-02 | 8.51E-02 | 2.52E+00 | 2.60E+00 | 4.46E-01 | no | Epg5 | upstream_gene_variant | MODIFIER |
| 9 | 99745698 | AX-170169581 | 1 | T | A | 0.143 | -8.45E-02 | 1.12E-01 | 2.67E+00 | 2.76E+00 | 4.46E-01 | no | Gm23949 | downstream_gene_variant | MODIFIER |
| 8 | 96131286 | AX-170429278 | 0 | G | C | 0.228 | 8.22E-02 | 1.09E-01 | 2.75E+00 | 2.83E+00 | 4.46E-01 | no | | intergenic_variant | MODIFIER |
| 5 | 106490053 | AX-97814791 | 0 | T | G | 0.447 | 8.04E-02 | 1.06E-01 | 2.61E+00 | 2.66E+00 | 4.46E-01 | no | | intergenic_variant | MODIFIER |
| 8 | 94843727 | AX-170431797 | 0 | C | A | 0.211 | -8.58E-02 | 1.14E-01 | 2.70E+00 | 2.78E+00 | 4.46E-01 | no | Coq9 | intron_variant | MODIFIER |
| 3 | 152144649 | AX-170089117 | 0 | G | T | 0.351 | 5.37E-02 | 7.15E-02 | 2.79E+00 | 2.88E+00 | 4.46E-01 | no | Gipc2 | intron_variant | MODIFIER |
| 3 | 152179248 | AX-97814945 | 0 | A | G | 0.351 | 5.37E-02 | 7.15E-02 | 2.79E+00 | 2.88E+00 | 4.46E-01 | no | Dnajb4 | downstream_gene_variant | MODIFIER |
| 3 | 152219576 | AX-97835585 | 0 | C | A | 0.351 | 5.37E-02 | 7.15E-02 | 2.79E+00 | 2.88E+00 | 4.46E-01 | no | Fubp1 | intron_variant | MODIFIER |
| 3 | 152225517 | AX-170272113 | 0 | A | G | 0.351 | 5.37E-02 | 7.15E-02 | 2.79E+00 | 2.88E+00 | 4.46E-01 | no | Fubp1 | intron_variant | MODIFIER |
| 3 | 152392550 | AX-97786271 | 0 | A | G | 0.351 | 5.37E-02 | 7.15E-02 | 2.79E+00 | 2.88E+00 | 4.46E-01 | no | Usp33 | 3_prime_UTR_variant | MODIFIER |
| 3 | 152488796 | AX-170137521 | 0 | G | A | 0.351 | 5.37E-02 | 7.15E-02 | 2.79E+00 | 2.88E+00 | 4.46E-01 | no | Ak5 | intron_variant | MODIFIER |
| 3 | 152584263 | AX-170021948 | 0 | C | T | 0.351 | 5.37E-02 | 7.15E-02 | 2.79E+00 | 2.88E+00 | 4.46E-01 | no | Ak5 | intron_variant | MODIFIER |
| 3 | 152624358 | AX-97797704 | 0 | C | T | 0.351 | 5.37E-02 | 7.15E-02 | 2.79E+00 | 2.88E+00 | 4.46E-01 | no | Ak5 | intron_variant | MODIFIER |
| 3 | 152728396 | AX-97809988 | 0 | G | A | 0.351 | 5.37E-02 | 7.15E-02 | 2.79E+00 | 2.88E+00 | 4.46E-01 | no | Pigk | intron_variant | MODIFIER |
| 3 | 152997493 | AX-170440518 | 0 | C | G | 0.351 | 5.37E-02 | 7.15E-02 | 2.79E+00 | 2.88E+00 | 4.46E-01 | no | | intergenic_variant | MODIFIER |
| 3 | 154401443 | AX-170021976 | 0 | G | A | 0.351 | 5.37E-02 | 7.15E-02 | 2.79E+00 | 2.88E+00 | 4.46E-01 | no | | intergenic_variant | MODIFIER |
| 3 | 154462184 | AX-97853209 | 0 | C | T | 0.351 | 5.37E-02 | 7.15E-02 | 2.79E+00 | 2.88E+00 | 4.46E-01 | no | | intergenic_variant | MODIFIER |
| 3 | 154511346 | AX-170291426 | 0 | A | G | 0.351 | 5.37E-02 | 7.15E-02 | 2.79E+00 | 2.88E+00 | 4.46E-01 | no | | intergenic_variant | MODIFIER |
| 3 | 154512120 | AX-170294584 | 0 | A | G | 0.351 | 5.37E-02 | 7.15E-02 | 2.79E+00 | 2.88E+00 | 4.46E-01 | no | | intergenic_variant | MODIFIER |
| 3 | 154515270 | AX-170200785 | 0 | G | A | 0.351 | 5.37E-02 | 7.15E-02 | 2.79E+00 | 2.88E+00 | 4.46E-01 | no | | intergenic_variant | MODIFIER |
| 3 | 154515877 | AX-170297357 | 0 | A | G | 0.351 | 5.37E-02 | 7.15E-02 | 2.79E+00 | 2.88E+00 | 4.46E-01 | no | | intergenic_variant | MODIFIER |
| 3 | 154517603 | AX-170407665 | 0 | G | A | 0.351 | 5.37E-02 | 7.15E-02 | 2.79E+00 | 2.88E+00 | 4.46E-01 | no | | intergenic_variant | MODIFIER |
| 3 | 154518909 | AX-170021969 | 0 | G | A | 0.351 | 5.37E-02 | 7.15E-02 | 2.79E+00 | 2.88E+00 | 4.46E-01 | no | | intergenic_variant | MODIFIER |
| 3 | 154520555 | AX-170212406 | 0 | T | G | 0.351 | 5.37E-02 | 7.15E-02 | 2.79E+00 | 2.88E+00 | 4.46E-01 | no | | intergenic_variant | MODIFIER |
| 3 | 154531639 | AX-170355084 | 0 | G | A | 0.351 | 5.37E-02 | 7.15E-02 | 2.79E+00 | 2.88E+00 | 4.46E-01 | no | | intergenic_variant | MODIFIER |
| 3 | 154571607 | AX-97848603 | 0 | C | T | 0.351 | 5.37E-02 | 7.15E-02 | 2.79E+00 | 2.88E+00 | 4.46E-01 | no | Tyw3 | downstream_gene_variant | MODIFIER |
| 3 | 154571766 | AX-170443968 | 0 | C | T | 0.351 | 5.37E-02 | 7.15E-02 | 2.79E+00 | 2.88E+00 | 4.46E-01 | no | Tyw3 | downstream_gene_variant | MODIFIER |
| 3 | 154587634 | AX-170148631 | 0 | A | G | 0.351 | 5.37E-02 | 7.15E-02 | 2.79E+00 | 2.88E+00 | 4.46E-01 | no | Tyw3 | intron_variant | MODIFIER |
| 3 | 154591219 | AX-170487551 | 0 | T | C | 0.351 | 5.37E-02 | 7.15E-02 | 2.79E+00 | 2.88E+00 | 4.46E-01 | no | Tyw3 | intron_variant | MODIFIER |
| 3 | 154592246 | AX-97829203 | 0 | A | C | 0.351 | 5.37E-02 | 7.15E-02 | 2.79E+00 | 2.88E+00 | 4.46E-01 | no | Cryz | upstream_gene_variant | MODIFIER |
| 3 | 154619677 | AX-97850285 | 0 | A | C | 0.351 | 5.37E-02 | 7.15E-02 | 2.79E+00 | 2.88E+00 | 4.46E-01 | no | Cryz | intron_variant | MODIFIER |
| 3 | 154737899 | AX-170206865 | 0 | C | T | 0.351 | 5.37E-02 | 7.15E-02 | 2.79E+00 | 2.88E+00 | 4.46E-01 | no | Erich3 | intron_variant | MODIFIER |
| 3 | 154738004 | AX-170299508 | 0 | C | T | 0.351 | 5.37E-02 | 7.15E-02 | 2.79E+00 | 2.88E+00 | 4.46E-01 | no | Erich3 | intron_variant | MODIFIER |
| 3 | 154738032 | AX-170313614 | 0 | C | A | 0.351 | 5.37E-02 | 7.15E-02 | 2.79E+00 | 2.88E+00 | 4.46E-01 | no | Erich3 | intron_variant | MODIFIER |
| 3 | 154758278 | AX-170068507 | 0 | G | A | 0.351 | 5.37E-02 | 7.15E-02 | 2.79E+00 | 2.88E+00 | 4.46E-01 | no | Erich3 | intron_variant | MODIFIER |
| 3 | 154758758 | AX-170477529 | 0 | T | C | 0.351 | 5.37E-02 | 7.15E-02 | 2.79E+00 | 2.88E+00 | 4.46E-01 | no | Erich3 | intron_variant | MODIFIER |
| 3 | 154763824 | AX-170284063 | 0 | C | A | 0.351 | 5.37E-02 | 7.15E-02 | 2.79E+00 | 2.88E+00 | 4.46E-01 | no | Erich3 | missense_variant | MODERATE |
| 3 | 154801523 | AX-170268027 | 0 | C | G | 0.351 | 5.37E-02 | 7.15E-02 | 2.79E+00 | 2.88E+00 | 4.46E-01 | no | Tnni3k | intron_variant | MODIFIER |
| 3 | 154809225 | AX-97782806 | 0 | T | C | 0.351 | 5.37E-02 | 7.15E-02 | 2.79E+00 | 2.88E+00 | 4.46E-01 | no | Tnni3k | intron_variant | MODIFIER |
| 3 | 154834121 | AX-170068508 | 0 | C | T | 0.351 | 5.37E-02 | 7.15E-02 | 2.79E+00 | 2.88E+00 | 4.46E-01 | no | Tnni3k | intron_variant | MODIFIER |
| 3 | 154839656 | AX-170021984 | 0 | G | A | 0.351 | 5.37E-02 | 7.15E-02 | 2.79E+00 | 2.88E+00 | 4.46E-01 | no | Tnni3k | intron_variant | MODIFIER |
| 15 | 6088776 | AX-170216052 | 1 | A | T | 0.089 | -7.02E-02 | 7.98E-02 | 2.19E+00 | 2.32E+00 | 4.46E-01 | no | | intergenic_variant | MODIFIER |

| | | | | | | | | | | | | | | | |
|---|---|---|---|---|---|---|---|---|---|---|---|---|---|---|---|
| 15 | 9228371 | AX-97818023 | 0 | T | C | 0.237 | -8.10E-02 | 1.07E-01 | 2.61E+00 | 2.67E+00 | 4.46E-01 | no | | intergenic_variant | MODIFIER |
| 15 | 10953278 | AX-170087768 | 0 | A | G | 0.237 | -8.10E-02 | 1.07E-01 | 2.61E+00 | 2.67E+00 | 4.46E-01 | no | C1qtnf3 | intron_variant | MODIFIER |
| 15 | 10685409 | AX-97851456 | 1 | C | A | 0.241 | -8.07E-02 | 1.06E-01 | 2.62E+00 | 2.67E+00 | 4.47E-01 | no | Rai14 | intron_variant | MODIFIER |
| 15 | 19409701 | AX-170152620 | 1 | C | T | 0.241 | -8.07E-02 | 1.06E-01 | 2.62E+00 | 2.67E+00 | 4.47E-01 | no | | intergenic_variant | MODIFIER |
| 15 | 22313304 | AX-170011216 | 1 | T | C | 0.241 | -8.07E-02 | 1.06E-01 | 2.62E+00 | 2.67E+00 | 4.47E-01 | no | | intergenic_variant | MODIFIER |
| 1 | 168186482 | AX-170302447 | 0 | T | A | 0.219 | 1.08E-01 | 1.43E-01 | 2.69E+00 | 2.73E+00 | 4.47E-01 | no | Pbx1 | intron_variant | MODIFIER |
| 5 | 79040365 | AX-97813459 | 1 | C | T | 0.438 | 7.28E-02 | 9.59E-02 | 2.59E+00 | 2.68E+00 | 4.47E-01 | no | | intergenic_variant | MODIFIER |
| 1 | 51797714 | AX-97786774 | 0 | T | C | 0.114 | 1.39E-01 | 1.85E-01 | 2.72E+00 | 2.83E+00 | 4.47E-01 | no | Myo1b | intron_variant | MODIFIER |
| 7 | 36386849 | AX-170235318 | 0 | G | C | 0.316 | 6.93E-02 | 9.07E-02 | 2.56E+00 | 2.63E+00 | 4.47E-01 | no | | regulatory_region_variant | MODIFIER |
| 7 | 36387082 | AX-170319603 | 0 | A | G | 0.316 | 6.93E-02 | 9.07E-02 | 2.56E+00 | 2.63E+00 | 4.47E-01 | no | | intergenic_variant | MODIFIER |
| 7 | 36393122 | AX-170529158 | 0 | T | C | 0.316 | 6.93E-02 | 9.07E-02 | 2.56E+00 | 2.63E+00 | 4.47E-01 | no | | intergenic_variant | MODIFIER |
| 7 | 36394426 | AX-170410257 | 0 | A | G | 0.316 | 6.93E-02 | 9.07E-02 | 2.56E+00 | 2.63E+00 | 4.47E-01 | no | | intergenic_variant | MODIFIER |
| 7 | 36395276 | AX-170516167 | 0 | C | T | 0.316 | 6.93E-02 | 9.07E-02 | 2.56E+00 | 2.63E+00 | 4.47E-01 | no | | intergenic_variant | MODIFIER |
| 12 | 106575166 | AX-97822337 | 0 | A | G | 0.123 | 1.19E-01 | 1.59E-01 | 2.73E+00 | 2.85E+00 | 4.47E-01 | no | Gm17033 | downstream_gene_variant | MODIFIER |
| 1 | 101021068 | AX-170157159 | 1 | C | A | 0.152 | -1.29E-01 | 1.72E-01 | 2.82E+00 | 2.94E+00 | 4.47E-01 | no | | intergenic_variant | MODIFIER |
| 9 | 67213871 | AX-170458002 | 0 | T | C | 0.193 | -8.54E-02 | 1.14E-01 | 2.74E+00 | 2.84E+00 | 4.47E-01 | no | Tln2 | downstream_gene_variant | MODIFIER |
| 7 | 36559783 | AX-97833527 | 0 | A | C | 0.351 | 7.02E-02 | 9.21E-02 | 2.56E+00 | 2.64E+00 | 4.47E-01 | no | | intergenic_variant | MODIFIER |
| 1 | 32627417 | AX-170149453 | 1 | A | G | 0.402 | 5.90E-02 | 7.89E-02 | 2.73E+00 | 2.83E+00 | 4.47E-01 | no | Khdrbs2 | intron_variant | MODIFIER |
| 4 | 140611129 | AX-170070873 | 1 | C | T | 0.152 | 8.69E-02 | 1.15E-01 | 2.63E+00 | 2.70E+00 | 4.47E-01 | no | Arhgef10l | intron_variant | MODIFIER |
| 5 | 105775924 | AX-170072982 | 1 | G | A | 0.455 | 8.04E-02 | 1.06E-01 | 2.60E+00 | 2.66E+00 | 4.48E-01 | no | Lrrc8d | intron_variant | MODIFIER |
| 2 | 169270036 | AX-170065594 | 0 | A | G | 0.167 | 1.36E-01 | 1.80E-01 | 2.64E+00 | 2.72E+00 | 4.48E-01 | no | | intergenic_variant | MODIFIER |
| 15 | 86068657 | AX-170172993 | 2 | T | A | 0.518 | 5.68E-02 | 7.28E-02 | 2.46E+00 | 2.55E+00 | 4.48E-01 | no | Gramd4 | intron_variant | MODIFIER |
| 5 | 88382213 | AX-170025537 | 0 | G | C | 0.439 | 7.81E-02 | 1.04E-01 | 2.80E+00 | 2.88E+00 | 4.48E-01 | no | Amtn | intron_variant | MODIFIER |
| 9 | 118715049 | AX-170096685 | 1 | G | T | 0.125 | -7.32E-02 | 9.72E-02 | 2.60E+00 | 2.71E+00 | 4.48E-01 | no | Itga9 | intron_variant | MODIFIER |
| 2 | 126037213 | AX-170358341 | 0 | T | C | 0.14 | -1.36E-01 | 1.82E-01 | 2.80E+00 | 2.89E+00 | 4.48E-01 | no | Fgf7 | intron_variant | MODIFIER |
| 1 | 172742225 | AX-170423802 | 0 | C | T | 0.237 | 8.88E-02 | 1.14E-01 | 2.47E+00 | 2.55E+00 | 4.48E-01 | no | Crp | intron_variant,non_coding_transcript_variant | MODIFIER |
| 6 | 132204039 | AX-170466848 | 0 | C | G | 0.57 | 6.73E-02 | 8.71E-02 | 2.50E+00 | 2.55E+00 | 4.48E-01 | no | Prb1 | downstream_gene_variant | MODIFIER |
| 6 | 132548420 | AX-170204458 | 0 | C | T | 0.57 | 6.73E-02 | 8.71E-02 | 2.50E+00 | 2.55E+00 | 4.48E-01 | no | | intergenic_variant | MODIFIER |
| 6 | 132957500 | AX-170076276 | 0 | C | A | 0.57 | 6.73E-02 | 8.71E-02 | 2.50E+00 | 2.55E+00 | 4.48E-01 | no | Tas2r131 | missense_variant | MODERATE |
| 6 | 133110126 | AX-170198727 | 0 | T | C | 0.57 | 6.73E-02 | 8.71E-02 | 2.50E+00 | 2.55E+00 | 4.48E-01 | no | Smim10l1 | downstream_gene_variant | MODIFIER |
| 6 | 133111963 | AX-170173770 | 0 | G | T | 0.57 | 6.73E-02 | 8.71E-02 | 2.50E+00 | 2.55E+00 | 4.48E-01 | no | Smim10l1 | downstream_gene_variant | MODIFIER |
| 6 | 133206083 | AX-170108132 | 0 | C | T | 0.57 | 6.73E-02 | 8.71E-02 | 2.50E+00 | 2.55E+00 | 4.48E-01 | no | | regulatory_region_variant | MODIFIER |
| 6 | 133303055 | AX-97787142 | 0 | A | C | 0.57 | 6.73E-02 | 8.71E-02 | 2.50E+00 | 2.55E+00 | 4.48E-01 | no | | intergenic_variant | MODIFIER |
| 6 | 133324623 | AX-170473556 | 0 | T | C | 0.57 | 6.73E-02 | 8.71E-02 | 2.50E+00 | 2.55E+00 | 4.48E-01 | no | | intergenic_variant | MODIFIER |
| 6 | 133328548 | AX-170475327 | 0 | G | A | 0.57 | 6.73E-02 | 8.71E-02 | 2.50E+00 | 2.55E+00 | 4.48E-01 | no | | intergenic_variant | MODIFIER |
| 6 | 133331982 | AX-170524389 | 0 | C | G | 0.57 | 6.73E-02 | 8.71E-02 | 2.50E+00 | 2.55E+00 | 4.48E-01 | no | | intergenic_variant | MODIFIER |
| 6 | 133348638 | AX-97816830 | 0 | T | C | 0.57 | 6.73E-02 | 8.71E-02 | 2.50E+00 | 2.55E+00 | 4.48E-01 | no | | intergenic_variant | MODIFIER |
| 6 | 133365853 | AX-170457264 | 0 | G | A | 0.57 | 6.73E-02 | 8.71E-02 | 2.50E+00 | 2.55E+00 | 4.48E-01 | no | | intergenic_variant | MODIFIER |
| 6 | 133373042 | AX-170116845 | 0 | T | A | 0.57 | 6.73E-02 | 8.71E-02 | 2.50E+00 | 2.55E+00 | 4.48E-01 | no | | intergenic_variant | MODIFIER |
| 6 | 133391279 | AX-170154500 | 0 | C | G | 0.57 | 6.73E-02 | 8.71E-02 | 2.50E+00 | 2.55E+00 | 4.48E-01 | no | | intergenic_variant | MODIFIER |
| 6 | 133516570 | AX-170494450 | 0 | G | C | 0.57 | 6.73E-02 | 8.71E-02 | 2.50E+00 | 2.55E+00 | 4.48E-01 | no | | intergenic_variant | MODIFIER |
| 6 | 133586948 | AX-97802356 | 0 | C | T | 0.57 | 6.73E-02 | 8.71E-02 | 2.50E+00 | 2.55E+00 | 4.48E-01 | no | | intergenic_variant | MODIFIER |
| 6 | 133607861 | AX-170095812 | 0 | G | A | 0.57 | 6.73E-02 | 8.71E-02 | 2.50E+00 | 2.55E+00 | 4.48E-01 | no | | intergenic_variant | MODIFIER |
| 6 | 133613560 | AX-170028294 | 0 | T | G | 0.57 | 6.73E-02 | 8.71E-02 | 2.50E+00 | 2.55E+00 | 4.48E-01 | no | Gm34314 | downstream_gene_variant | MODIFIER |
| 6 | 133616619 | AX-170502704 | 0 | G | A | 0.57 | 6.73E-02 | 8.71E-02 | 2.50E+00 | 2.55E+00 | 4.48E-01 | no | Gm34314 | downstream_gene_variant | MODIFIER |
| 6 | 133630055 | AX-97788785 | 0 | C | T | 0.57 | 6.73E-02 | 8.71E-02 | 2.50E+00 | 2.55E+00 | 4.48E-01 | no | | intergenic_variant | MODIFIER |
| 6 | 133634596 | AX-170322307 | 0 | G | T | 0.57 | 6.73E-02 | 8.71E-02 | 2.50E+00 | 2.55E+00 | 4.48E-01 | no | | intergenic_variant | MODIFIER |
| 6 | 133637503 | AX-170076284 | 0 | G | A | 0.57 | 6.73E-02 | 8.71E-02 | 2.50E+00 | 2.55E+00 | 4.48E-01 | no | | intergenic_variant | MODIFIER |
| 6 | 133653639 | AX-170028292 | 0 | C | G | 0.57 | 6.73E-02 | 8.71E-02 | 2.50E+00 | 2.55E+00 | 4.48E-01 | no | | intergenic_variant | MODIFIER |
| 6 | 133653721 | AX-170028291 | 0 | A | G | 0.57 | 6.73E-02 | 8.71E-02 | 2.50E+00 | 2.55E+00 | 4.48E-01 | no | | intergenic_variant | MODIFIER |
| 6 | 133654175 | AX-170028290 | 0 | G | A | 0.57 | 6.73E-02 | 8.71E-02 | 2.50E+00 | 2.55E+00 | 4.48E-01 | no | | intergenic_variant | MODIFIER |
| 6 | 133854950 | AX-97834347 | 0 | C | T | 0.57 | 6.73E-02 | 8.71E-02 | 2.50E+00 | 2.55E+00 | 4.48E-01 | no | Kap | upstream_gene_variant | MODIFIER |
| 6 | 133867514 | AX-97789564 | 0 | T | C | 0.57 | 6.73E-02 | 8.71E-02 | 2.50E+00 | 2.55E+00 | 4.48E-01 | no | Gm43968 | downstream_gene_variant | MODIFIER |
| 6 | 133877968 | AX-170516182 | 0 | T | C | 0.57 | 6.73E-02 | 8.71E-02 | 2.50E+00 | 2.55E+00 | 4.48E-01 | no | | intergenic_variant | MODIFIER |

| | | | | | | | | | | | | | | | |
|---|---|---|---|---|---|---|---|---|---|---|---|---|---|---|---|
| 6 | 133883226 | AX-170121110 | 0 | T | G | 0.57 | 6.73E-02 | 8.71E-02 | 2.50E+00 | 2.55E+00 | 4.48E-01 | no | | intergenic_variant | MODIFIER |
| 6 | 133886192 | AX-170161720 | 0 | T | C | 0.57 | 6.73E-02 | 8.71E-02 | 2.50E+00 | 2.55E+00 | 4.48E-01 | no | | regulatory_region_variant | MODIFIER |
| 6 | 133908477 | AX-170120256 | 0 | G | T | 0.57 | 6.73E-02 | 8.71E-02 | 2.50E+00 | 2.55E+00 | 4.48E-01 | no | | intergenic_variant | MODIFIER |
| 6 | 133916991 | AX-97815698 | 0 | C | T | 0.57 | 6.73E-02 | 8.71E-02 | 2.50E+00 | 2.55E+00 | 4.48E-01 | no | | intergenic_variant | MODIFIER |
| 6 | 133919443 | AX-170273971 | 0 | T | C | 0.57 | 6.73E-02 | 8.71E-02 | 2.50E+00 | 2.55E+00 | 4.48E-01 | no | | intergenic_variant | MODIFIER |
| 6 | 133928961 | AX-170028293 | 0 | A | G | 0.57 | 6.73E-02 | 8.71E-02 | 2.50E+00 | 2.55E+00 | 4.48E-01 | no | Gm8956 | downstream_gene_variant | MODIFIER |
| 6 | 133930475 | AX-170241376 | 0 | A | T | 0.57 | 6.73E-02 | 8.71E-02 | 2.50E+00 | 2.55E+00 | 4.48E-01 | no | Gm8956 | downstream_gene_variant | MODIFIER |
| 6 | 133960972 | AX-170440972 | 0 | G | A | 0.57 | 6.73E-02 | 8.71E-02 | 2.50E+00 | 2.55E+00 | 4.48E-01 | no | | regulatory_region_variant | MODIFIER |
| 6 | 133969389 | AX-97783391 | 0 | G | A | 0.57 | 6.73E-02 | 8.71E-02 | 2.50E+00 | 2.55E+00 | 4.48E-01 | no | | intergenic_variant | MODIFIER |
| 6 | 133977322 | AX-170424482 | 0 | G | A | 0.57 | 6.73E-02 | 8.71E-02 | 2.50E+00 | 2.55E+00 | 4.48E-01 | no | | intergenic_variant | MODIFIER |
| 6 | 133983908 | AX-170228219 | 0 | G | A | 0.57 | 6.73E-02 | 8.71E-02 | 2.50E+00 | 2.55E+00 | 4.48E-01 | no | | intergenic_variant | MODIFIER |
| 6 | 133984409 | AX-170204709 | 0 | C | T | 0.57 | 6.73E-02 | 8.71E-02 | 2.50E+00 | 2.55E+00 | 4.48E-01 | no | | regulatory_region_variant | MODIFIER |
| 6 | 133989858 | AX-170442584 | 0 | C | G | 0.57 | 6.73E-02 | 8.71E-02 | 2.50E+00 | 2.55E+00 | 4.48E-01 | no | | intergenic_variant | MODIFIER |
| 6 | 134000395 | AX-170076293 | 0 | G | C | 0.57 | 6.73E-02 | 8.71E-02 | 2.50E+00 | 2.55E+00 | 4.48E-01 | no | | intergenic_variant | MODIFIER |
| 6 | 134001502 | AX-97801160 | 0 | G | A | 0.57 | 6.73E-02 | 8.71E-02 | 2.50E+00 | 2.55E+00 | 4.48E-01 | no | | intergenic_variant | MODIFIER |
| 6 | 134096182 | AX-170265141 | 0 | C | T | 0.57 | 6.73E-02 | 8.71E-02 | 2.50E+00 | 2.55E+00 | 4.48E-01 | no | Etv6 | intron_variant | MODIFIER |
| 6 | 134133081 | AX-170028301 | 0 | G | T | 0.57 | 6.73E-02 | 8.71E-02 | 2.50E+00 | 2.55E+00 | 4.48E-01 | no | Etv6 | intron_variant | MODIFIER |
| 6 | 134141690 | AX-170076294 | 0 | G | A | 0.57 | 6.73E-02 | 8.71E-02 | 2.50E+00 | 2.55E+00 | 4.48E-01 | no | Etv6 | intron_variant | MODIFIER |
| 6 | 134145026 | AX-170512986 | 0 | A | G | 0.57 | 6.73E-02 | 8.71E-02 | 2.50E+00 | 2.55E+00 | 4.48E-01 | no | Etv6 | intron_variant | MODIFIER |
| 6 | 134145313 | AX-170312388 | 0 | C | A | 0.57 | 6.73E-02 | 8.71E-02 | 2.50E+00 | 2.55E+00 | 4.48E-01 | no | Etv6 | intron_variant | MODIFIER |
| 6 | 134155477 | AX-170028300 | 0 | T | C | 0.57 | 6.73E-02 | 8.71E-02 | 2.50E+00 | 2.55E+00 | 4.48E-01 | no | Etv6 | intron_variant | MODIFIER |
| 6 | 134163607 | AX-170347047 | 0 | C | T | 0.57 | 6.73E-02 | 8.71E-02 | 2.50E+00 | 2.55E+00 | 4.48E-01 | no | Etv6 | intron_variant | MODIFIER |
| 6 | 134164371 | AX-170195844 | 0 | C | T | 0.57 | 6.73E-02 | 8.71E-02 | 2.50E+00 | 2.55E+00 | 4.48E-01 | no | Etv6 | intron_variant | MODIFIER |
| 6 | 134170435 | AX-170169913 | 0 | T | A | 0.57 | 6.73E-02 | 8.71E-02 | 2.50E+00 | 2.55E+00 | 4.48E-01 | no | Etv6 | intron_variant | MODIFIER |
| 6 | 134177731 | AX-170525409 | 0 | C | T | 0.57 | 6.73E-02 | 8.71E-02 | 2.50E+00 | 2.55E+00 | 4.48E-01 | no | Etv6 | intron_variant | MODIFIER |
| 6 | 134230501 | AX-97820275 | 0 | A | C | 0.57 | 6.73E-02 | 8.71E-02 | 2.50E+00 | 2.55E+00 | 4.48E-01 | no | Etv6 | intron_variant | MODIFIER |
| 6 | 134240788 | AX-170139622 | 0 | T | C | 0.57 | 6.73E-02 | 8.71E-02 | 2.50E+00 | 2.55E+00 | 4.48E-01 | no | Etv6 | intron_variant | MODIFIER |
| 6 | 134263444 | AX-170282641 | 0 | C | T | 0.57 | 6.73E-02 | 8.71E-02 | 2.50E+00 | 2.55E+00 | 4.48E-01 | no | Etv6 | intron_variant | MODIFIER |
| 6 | 134266149 | AX-170125640 | 0 | A | G | 0.57 | 6.73E-02 | 8.71E-02 | 2.50E+00 | 2.55E+00 | 4.48E-01 | no | Etv6 | intron_variant | MODIFIER |
| 6 | 134269617 | AX-170505786 | 0 | A | G | 0.57 | 6.73E-02 | 8.71E-02 | 2.50E+00 | 2.55E+00 | 4.48E-01 | no | Etv6 | 3_prime_UTR_variant | MODIFIER |
| 6 | 134273008 | AX-170166919 | 0 | G | A | 0.57 | 6.73E-02 | 8.71E-02 | 2.50E+00 | 2.55E+00 | 4.48E-01 | no | Etv6 | downstream_gene_variant | MODIFIER |
| 6 | 134273673 | AX-170076299 | 0 | A | G | 0.57 | 6.73E-02 | 8.71E-02 | 2.50E+00 | 2.55E+00 | 4.48E-01 | no | Etv6 | downstream_gene_variant | MODIFIER |
| 6 | 134283405 | AX-170130278 | 0 | A | C | 0.57 | 6.73E-02 | 8.71E-02 | 2.50E+00 | 2.55E+00 | 4.48E-01 | no | | intergenic_variant | MODIFIER |
| 6 | 134291413 | AX-97823848 | 0 | T | C | 0.57 | 6.73E-02 | 8.71E-02 | 2.50E+00 | 2.55E+00 | 4.48E-01 | no | | intergenic_variant | MODIFIER |
| 6 | 134294340 | AX-170166861 | 0 | A | G | 0.57 | 6.73E-02 | 8.71E-02 | 2.50E+00 | 2.55E+00 | 4.48E-01 | no | | regulatory_region_variant | MODIFIER |
| 6 | 134324595 | AX-97833140 | 0 | A | G | 0.57 | 6.73E-02 | 8.71E-02 | 2.50E+00 | 2.55E+00 | 4.48E-01 | no | 4933406J09Rik | intron_variant,non_coding_transcript_variant | MODIFIER |
| 6 | 134342879 | AX-170139504 | 0 | A | G | 0.57 | 6.73E-02 | 8.71E-02 | 2.50E+00 | 2.55E+00 | 4.48E-01 | no | 4933406J09Rik | intron_variant,non_coding_transcript_variant | MODIFIER |
| 6 | 134343127 | AX-170535641 | 0 | G | A | 0.57 | 6.73E-02 | 8.71E-02 | 2.50E+00 | 2.55E+00 | 4.48E-01 | no | 4933406J09Rik | intron_variant,non_coding_transcript_variant | MODIFIER |
| 6 | 134346082 | AX-170329496 | 0 | A | T | 0.57 | 6.73E-02 | 8.71E-02 | 2.50E+00 | 2.55E+00 | 4.48E-01 | no | 4933406J09Rik | intron_variant,non_coding_transcript_variant | MODIFIER |
| 6 | 134393556 | AX-170354719 | 0 | T | C | 0.57 | 6.73E-02 | 8.71E-02 | 2.50E+00 | 2.55E+00 | 4.48E-01 | no | Bcl2l14 | upstream_gene_variant | MODIFIER |
| 6 | 134397401 | AX-170343225 | 0 | T | C | 0.57 | 6.73E-02 | 8.71E-02 | 2.50E+00 | 2.55E+00 | 4.48E-01 | no | Bcl2l14 | intron_variant | MODIFIER |
| 6 | 134401101 | AX-170227279 | 0 | C | T | 0.57 | 6.73E-02 | 8.71E-02 | 2.50E+00 | 2.55E+00 | 4.48E-01 | no | Bcl2l14 | intron_variant | MODIFIER |
| 6 | 134402445 | AX-170460411 | 0 | G | A | 0.57 | 6.73E-02 | 8.71E-02 | 2.50E+00 | 2.55E+00 | 4.48E-01 | no | Bcl2l14 | intron_variant | MODIFIER |
| 6 | 134405582 | AX-170076301 | 0 | T | C | 0.57 | 6.73E-02 | 8.71E-02 | 2.50E+00 | 2.55E+00 | 4.48E-01 | no | Bcl2l14 | intron_variant | MODIFIER |
| 6 | 134421566 | AX-170517482 | 0 | T | C | 0.57 | 6.73E-02 | 8.71E-02 | 2.50E+00 | 2.55E+00 | 4.48E-01 | no | Bcl2l14 | intron_variant | MODIFIER |
| 6 | 134421679 | AX-170390426 | 0 | T | C | 0.57 | 6.73E-02 | 8.71E-02 | 2.50E+00 | 2.55E+00 | 4.48E-01 | no | Bcl2l14 | intron_variant | MODIFIER |
| 6 | 134458599 | AX-170456311 | 0 | C | T | 0.57 | 6.73E-02 | 8.71E-02 | 2.50E+00 | 2.55E+00 | 4.48E-01 | no | Lrp6 | intron_variant | MODIFIER |
| 6 | 134460418 | AX-170374349 | 0 | T | G | 0.57 | 6.73E-02 | 8.71E-02 | 2.50E+00 | 2.55E+00 | 4.48E-01 | no | Lrp6 | intron_variant | MODIFIER |
| 6 | 134462709 | AX-170343095 | 0 | C | A | 0.57 | 6.73E-02 | 8.71E-02 | 2.50E+00 | 2.55E+00 | 4.48E-01 | no | Lrp6 | intron_variant | MODIFIER |
| 6 | 134464634 | AX-170482423 | 0 | A | G | 0.57 | 6.73E-02 | 8.71E-02 | 2.50E+00 | 2.55E+00 | 4.48E-01 | no | Lrp6 | intron_variant | MODIFIER |
| 6 | 134466255 | AX-170292997 | 0 | C | A | 0.57 | 6.73E-02 | 8.71E-02 | 2.50E+00 | 2.55E+00 | 4.48E-01 | no | Lrp6 | intron_variant | MODIFIER |
| 6 | 134466753 | AX-170496728 | 0 | A | G | 0.57 | 6.73E-02 | 8.71E-02 | 2.50E+00 | 2.55E+00 | 4.48E-01 | no | Lrp6 | intron_variant | MODIFIER |
| 6 | 134466885 | AX-170338298 | 0 | G | A | 0.57 | 6.73E-02 | 8.71E-02 | 2.50E+00 | 2.55E+00 | 4.48E-01 | no | Lrp6 | intron_variant | MODIFIER |
| 6 | 134469796 | AX-170099810 | 0 | T | C | 0.57 | 6.73E-02 | 8.71E-02 | 2.50E+00 | 2.55E+00 | 4.48E-01 | no | Lrp6 | intron_variant | MODIFIER |

| | | | | | | | | | | | | | | | |
|---|---|---|---|---|---|---|---|---|---|---|---|---|---|---|---|
| 6 | 134473556 | AX-170507156 | 0 | C | T | 0.57 | 6.73E-02 | 8.71E-02 | 2.50E+00 | 2.55E+00 | 4.48E-01 | no | Lrp6 | intron_variant | MODIFIER |
| 6 | 134474471 | AX-170499302 | 0 | G | A | 0.57 | 6.73E-02 | 8.71E-02 | 2.50E+00 | 2.55E+00 | 4.48E-01 | no | Lrp6 | intron_variant | MODIFIER |
| 6 | 134474917 | AX-170126383 | 0 | G | A | 0.57 | 6.73E-02 | 8.71E-02 | 2.50E+00 | 2.55E+00 | 4.48E-01 | no | Lrp6 | intron_variant | MODIFIER |
| 6 | 134487167 | AX-170114564 | 0 | G | A | 0.57 | 6.73E-02 | 8.71E-02 | 2.50E+00 | 2.55E+00 | 4.48E-01 | no | Lrp6 | intron_variant | MODIFIER |
| 6 | 134493223 | AX-170242622 | 0 | C | T | 0.57 | 6.73E-02 | 8.71E-02 | 2.50E+00 | 2.55E+00 | 4.48E-01 | no | Lrp6 | intron_variant | MODIFIER |
| 6 | 134507245 | AX-170446306 | 0 | A | G | 0.57 | 6.73E-02 | 8.71E-02 | 2.50E+00 | 2.55E+00 | 4.48E-01 | no | Lrp6 | intron_variant | MODIFIER |
| 6 | 134507850 | AX-170153120 | 0 | G | T | 0.57 | 6.73E-02 | 8.71E-02 | 2.50E+00 | 2.55E+00 | 4.48E-01 | no | Lrp6 | intron_variant | MODIFIER |
| 6 | 134513651 | AX-170276883 | 0 | G | A | 0.57 | 6.73E-02 | 8.71E-02 | 2.50E+00 | 2.55E+00 | 4.48E-01 | no | Lrp6 | intron_variant | MODIFIER |
| 6 | 134514180 | AX-170185519 | 0 | A | T | 0.57 | 6.73E-02 | 8.71E-02 | 2.50E+00 | 2.55E+00 | 4.48E-01 | no | Lrp6 | intron_variant | MODIFIER |
| 6 | 134526564 | AX-97812502 | 0 | A | G | 0.57 | 6.73E-02 | 8.71E-02 | 2.50E+00 | 2.55E+00 | 4.48E-01 | no | Lrp6 | intron_variant | MODIFIER |
| 6 | 134531076 | AX-170157124 | 0 | G | A | 0.57 | 6.73E-02 | 8.71E-02 | 2.50E+00 | 2.55E+00 | 4.48E-01 | no | Lrp6 | intron_variant | MODIFIER |
| 6 | 134540600 | AX-170076304 | 0 | A | G | 0.57 | 6.73E-02 | 8.71E-02 | 2.50E+00 | 2.55E+00 | 4.48E-01 | no | Lrp6 | intron_variant | MODIFIER |
| 6 | 134541853 | AX-170194671 | 0 | C | G | 0.57 | 6.73E-02 | 8.71E-02 | 2.50E+00 | 2.55E+00 | 4.48E-01 | no | Lrp6 | missense_variant | MODERATE |
| 6 | 134565277 | AX-170414295 | 0 | C | T | 0.57 | 6.73E-02 | 8.71E-02 | 2.50E+00 | 2.55E+00 | 4.48E-01 | no | Lrp6 | intron_variant | MODIFIER |
| 6 | 134570841 | AX-170196347 | 0 | G | A | 0.57 | 6.73E-02 | 8.71E-02 | 2.50E+00 | 2.55E+00 | 4.48E-01 | no | Lrp6 | upstream_gene_variant | MODIFIER |
| 6 | 134572720 | AX-170104900 | 0 | T | C | 0.57 | 6.73E-02 | 8.71E-02 | 2.50E+00 | 2.55E+00 | 4.48E-01 | no | | intergenic_variant | MODIFIER |
| 6 | 134580460 | AX-170292642 | 0 | A | G | 0.57 | 6.73E-02 | 8.71E-02 | 2.50E+00 | 2.55E+00 | 4.48E-01 | no | | intergenic_variant | MODIFIER |
| 6 | 134592708 | AX-170348491 | 0 | G | A | 0.57 | 6.73E-02 | 8.71E-02 | 2.50E+00 | 2.55E+00 | 4.48E-01 | no | | regulatory_region_variant | MODIFIER |
| 6 | 134595459 | AX-170275051 | 0 | G | A | 0.57 | 6.73E-02 | 8.71E-02 | 2.50E+00 | 2.55E+00 | 4.48E-01 | no | | regulatory_region_variant | MODIFIER |
| 6 | 134613543 | AX-170209377 | 0 | A | G | 0.57 | 6.73E-02 | 8.71E-02 | 2.50E+00 | 2.55E+00 | 4.48E-01 | no | Mansc1 | intron_variant | MODIFIER |
| 6 | 134646273 | AX-170028307 | 0 | C | T | 0.57 | 6.73E-02 | 8.71E-02 | 2.50E+00 | 2.55E+00 | 4.48E-01 | no | Borcs5 | intron_variant | MODIFIER |
| 6 | 134656557 | AX-170332112 | 0 | T | C | 0.57 | 6.73E-02 | 8.71E-02 | 2.50E+00 | 2.55E+00 | 4.48E-01 | no | Borcs5 | intron_variant | MODIFIER |
| 6 | 134688208 | AX-170286039 | 0 | C | T | 0.57 | 6.73E-02 | 8.71E-02 | 2.50E+00 | 2.55E+00 | 4.48E-01 | no | Borcs5 | intron_variant | MODIFIER |
| 6 | 134701950 | AX-170369207 | 0 | G | A | 0.57 | 6.73E-02 | 8.71E-02 | 2.50E+00 | 2.55E+00 | 4.48E-01 | no | Borcs5 | intron_variant | MODIFIER |
| 6 | 134711193 | AX-170466768 | 0 | C | T | 0.57 | 6.73E-02 | 8.71E-02 | 2.50E+00 | 2.55E+00 | 4.48E-01 | no | Borcs5 | downstream_gene_variant | MODIFIER |
| 6 | 134720118 | AX-170273517 | 0 | T | C | 0.57 | 6.73E-02 | 8.71E-02 | 2.50E+00 | 2.55E+00 | 4.48E-01 | no | Dusp16 | intron_variant | MODIFIER |
| 6 | 134726799 | AX-170194683 | 0 | A | G | 0.57 | 6.73E-02 | 8.71E-02 | 2.50E+00 | 2.55E+00 | 4.48E-01 | no | Dusp16 | intron_variant | MODIFIER |
| 6 | 134738586 | AX-170126646 | 0 | G | C | 0.57 | 6.73E-02 | 8.71E-02 | 2.50E+00 | 2.55E+00 | 4.48E-01 | no | Dusp16 | intron_variant | MODIFIER |
| 6 | 134745491 | AX-170130713 | 0 | T | C | 0.57 | 6.73E-02 | 8.71E-02 | 2.50E+00 | 2.55E+00 | 4.48E-01 | no | Dusp16 | intron_variant | MODIFIER |
| 6 | 134751606 | AX-170437508 | 0 | C | T | 0.57 | 6.73E-02 | 8.71E-02 | 2.50E+00 | 2.55E+00 | 4.48E-01 | no | Dusp16 | intron_variant | MODIFIER |
| 6 | 134752425 | AX-170525638 | 0 | A | G | 0.57 | 6.73E-02 | 8.71E-02 | 2.50E+00 | 2.55E+00 | 4.48E-01 | no | Dusp16 | intron_variant | MODIFIER |
| 6 | 134754406 | AX-170177798 | 0 | T | C | 0.57 | 6.73E-02 | 8.71E-02 | 2.50E+00 | 2.55E+00 | 4.48E-01 | no | Dusp16 | intron_variant | MODIFIER |
| 6 | 134806312 | AX-170204010 | 0 | T | C | 0.57 | 6.73E-02 | 8.71E-02 | 2.50E+00 | 2.55E+00 | 4.48E-01 | no | | regulatory_region_variant | MODIFIER |
| 6 | 134806608 | AX-170242533 | 0 | A | G | 0.57 | 6.73E-02 | 8.71E-02 | 2.50E+00 | 2.55E+00 | 4.48E-01 | no | | intergenic_variant | MODIFIER |
| 6 | 134811105 | AX-170434919 | 0 | G | A | 0.57 | 6.73E-02 | 8.71E-02 | 2.50E+00 | 2.55E+00 | 4.48E-01 | no | | intergenic_variant | MODIFIER |
| 6 | 134846192 | AX-170361949 | 0 | G | A | 0.57 | 6.73E-02 | 8.71E-02 | 2.50E+00 | 2.55E+00 | 4.48E-01 | no | Crebl2 | intron_variant | MODIFIER |
| 6 | 134852467 | AX-170436633 | 0 | C | T | 0.57 | 6.73E-02 | 8.71E-02 | 2.50E+00 | 2.55E+00 | 4.48E-01 | no | Crebl2 | intron_variant | MODIFIER |
| 6 | 134866115 | AX-170343759 | 0 | G | A | 0.57 | 6.73E-02 | 8.71E-02 | 2.50E+00 | 2.55E+00 | 4.48E-01 | no | Gpr19 | downstream_gene_variant | MODIFIER |
| 6 | 134866643 | AX-170396954 | 0 | G | T | 0.57 | 6.73E-02 | 8.71E-02 | 2.50E+00 | 2.55E+00 | 4.48E-01 | no | Gpr19 | downstream_gene_variant | MODIFIER |
| 6 | 134871170 | AX-170271534 | 0 | A | T | 0.57 | 6.73E-02 | 8.71E-02 | 2.50E+00 | 2.55E+00 | 4.48E-01 | no | Gpr19 | intron_variant | MODIFIER |
| 6 | 134872551 | AX-170183001 | 0 | G | A | 0.57 | 6.73E-02 | 8.71E-02 | 2.50E+00 | 2.55E+00 | 4.48E-01 | no | Gpr19 | intron_variant | MODIFIER |
| 6 | 134872583 | AX-170375681 | 0 | A | G | 0.57 | 6.73E-02 | 8.71E-02 | 2.50E+00 | 2.55E+00 | 4.48E-01 | no | Gpr19 | intron_variant | MODIFIER |
| 6 | 134875141 | AX-170440295 | 0 | T | C | 0.57 | 6.73E-02 | 8.71E-02 | 2.50E+00 | 2.55E+00 | 4.48E-01 | no | Gpr19 | intron_variant | MODIFIER |
| 6 | 134950958 | AX-170132165 | 0 | A | G | 0.57 | 6.73E-02 | 8.71E-02 | 2.50E+00 | 2.55E+00 | 4.48E-01 | no | Lockd | intron_variant,non_coding_transcript_variant | MODIFIER |
| 6 | 134961893 | AX-170110970 | 0 | C | A | 0.57 | 6.73E-02 | 8.71E-02 | 2.50E+00 | 2.55E+00 | 4.48E-01 | no | | regulatory_region_variant | MODIFIER |
| 6 | 134962529 | AX-170076314 | 0 | C | T | 0.57 | 6.73E-02 | 8.71E-02 | 2.50E+00 | 2.55E+00 | 4.48E-01 | no | | regulatory_region_variant | MODIFIER |
| 6 | 134968031 | AX-97788664 | 0 | T | C | 0.57 | 6.73E-02 | 8.71E-02 | 2.50E+00 | 2.55E+00 | 4.48E-01 | no | | intergenic_variant | MODIFIER |
| 6 | 134973053 | AX-170099142 | 0 | G | C | 0.57 | 6.73E-02 | 8.71E-02 | 2.50E+00 | 2.55E+00 | 4.48E-01 | no | | intergenic_variant | MODIFIER |
| 6 | 134981993 | AX-170294779 | 0 | G | A | 0.57 | 6.73E-02 | 8.71E-02 | 2.50E+00 | 2.55E+00 | 4.48E-01 | no | Apold1 | 5_prime_UTR_variant | MODIFIER |
| 6 | 135014591 | AX-170348406 | 0 | C | T | 0.57 | 6.73E-02 | 8.71E-02 | 2.50E+00 | 2.55E+00 | 4.48E-01 | no | Ddx47 | intron_variant | MODIFIER |
| 6 | 135019475 | AX-170294409 | 0 | T | C | 0.57 | 6.73E-02 | 8.71E-02 | 2.50E+00 | 2.55E+00 | 4.48E-01 | no | Ddx47 | intron_variant | MODIFIER |
| 6 | 135024459 | AX-170454288 | 0 | G | A | 0.57 | 6.73E-02 | 8.71E-02 | 2.50E+00 | 2.55E+00 | 4.48E-01 | no | Ddx47 | downstream_gene_variant | MODIFIER |
| 6 | 135025160 | AX-170482349 | 0 | C | T | 0.57 | 6.73E-02 | 8.71E-02 | 2.50E+00 | 2.55E+00 | 4.48E-01 | no | Ddx47 | downstream_gene_variant | MODIFIER |
| 2 | 164019139 | AX-170257705 | 0 | G | A | 0.158 | 1.47E-01 | 1.96E-01 | 2.76E+00 | 2.85E+00 | 4.48E-01 | Yes | Ywhab | downstream_gene_variant | MODIFIER |

| | | | | | | | | | | | | | | | |
|---|---|---|---|---|---|---|---|---|---|---|---|---|---|---|---|
| 2 | 167106476 | AX-170172197 | 0 | C | T | 0.158 | 1.47E-01 | 1.96E-01 | 2.76E+00 | 2.85E+00 | 4.48E-01 | Yes | Kcnb1 | intron_variant | MODIFIER |
| 8 | 97966969 | AX-170328113 | 1 | G | A | 0.223 | 8.68E-02 | 1.16E-01 | 2.76E+00 | 2.83E+00 | 4.48E-01 | no | | intergenic_variant | MODIFIER |
| 5 | 38239587 | AX-97818810 | 0 | G | A | 0.333 | 8.25E-02 | 1.10E-01 | 2.69E+00 | 2.76E+00 | 4.48E-01 | no | | intergenic_variant | MODIFIER |
| 5 | 38511799 | AX-170106892 | 0 | G | A | 0.333 | 8.25E-02 | 1.10E-01 | 2.69E+00 | 2.76E+00 | 4.48E-01 | no | Gm15796 | downstream_gene_variant | MODIFIER |
| 8 | 32591282 | AX-97854308 | 0 | C | T | 0.14 | -9.56E-02 | 1.28E-01 | 2.68E+00 | 2.75E+00 | 4.48E-01 | no | Nrg1 | intron_variant | MODIFIER |
| 8 | 23623585 | AX-170030350 | 1 | T | A | 0.161 | -7.44E-02 | 9.94E-02 | 2.65E+00 | 2.77E+00 | 4.48E-01 | no | | intergenic_variant | MODIFIER |
| 15 | 5673776 | AX-170360117 | 1 | T | A | 0.143 | -5.16E-02 | 5.60E-02 | 2.11E+00 | 2.23E+00 | 4.49E-01 | no | | intergenic_variant | MODIFIER |
| 1 | 131300577 | AX-170001711 | 0 | A | G | 0.535 | 6.44E-02 | 8.54E-02 | 2.62E+00 | 2.69E+00 | 4.49E-01 | no | Srgap2 | intron_variant | MODIFIER |
| 10 | 92628902 | AX-97810687 | 1 | T | C | 0.304 | -6.38E-02 | 8.34E-02 | 2.54E+00 | 2.60E+00 | 4.49E-01 | no | | intergenic_variant | MODIFIER |
| 15 | 85684580 | AX-170196039 | 1 | T | C | 0.509 | 5.55E-02 | 7.15E-02 | 2.47E+00 | 2.55E+00 | 4.49E-01 | no | Lncppara | intron_variant,non_coding_transcript_variant | MODIFIER |
| 7 | 75009137 | AX-170077742 | 0 | C | T | 0.368 | -6.88E-02 | 9.22E-02 | 2.78E+00 | 2.88E+00 | 4.49E-01 | no | | intergenic_variant | MODIFIER |
| 16 | 38652273 | AX-170181080 | 1 | T | C | 0.5 | 6.11E-02 | 8.02E-02 | 2.55E+00 | 2.60E+00 | 4.49E-01 | no | Arhgap31 | intron_variant | MODIFIER |
| 13 | 23370114 | AX-170036876 | 1 | A | G | 0.161 | -9.33E-02 | 1.20E-01 | 2.48E+00 | 2.54E+00 | 4.49E-01 | no | Zfp322a | upstream_gene_variant | MODIFIER |
| 1 | 153172146 | AX-170188918 | 0 | G | C | 0.211 | 9.81E-02 | 1.31E-01 | 2.65E+00 | 2.70E+00 | 4.49E-01 | no | Lamc2 | intron_variant | MODIFIER |
| 16 | 29175917 | AX-170119548 | 1 | G | A | 0.491 | 6.01E-02 | 7.92E-02 | 2.57E+00 | 2.63E+00 | 4.49E-01 | no | | intergenic_variant | MODIFIER |
| 15 | 3389348 | AX-170387492 | 0 | A | C | 0.088 | -8.59E-02 | 9.58E-02 | 2.15E+00 | 2.26E+00 | 4.49E-01 | no | Ghr | intron_variant | MODIFIER |
| 15 | 3570104 | AX-170248183 | 0 | C | G | 0.088 | -8.59E-02 | 9.58E-02 | 2.15E+00 | 2.26E+00 | 4.49E-01 | no | Ghr | intron_variant | MODIFIER |
| 15 | 3572979 | AX-170440034 | 0 | T | C | 0.088 | -8.59E-02 | 9.58E-02 | 2.15E+00 | 2.26E+00 | 4.49E-01 | no | Ghr | intron_variant | MODIFIER |
| 15 | 3612981 | AX-170357184 | 0 | A | G | 0.088 | -8.59E-02 | 9.58E-02 | 2.15E+00 | 2.26E+00 | 4.49E-01 | no | | intergenic_variant | MODIFIER |
| 15 | 3669113 | AX-170474847 | 0 | A | G | 0.088 | -8.59E-02 | 9.58E-02 | 2.15E+00 | 2.26E+00 | 4.49E-01 | no | | intergenic_variant | MODIFIER |
| 15 | 3669416 | AX-97848166 | 0 | A | G | 0.088 | -8.59E-02 | 9.58E-02 | 2.15E+00 | 2.26E+00 | 4.49E-01 | no | | intergenic_variant | MODIFIER |
| 15 | 3680551 | AX-170238697 | 0 | A | G | 0.088 | -8.59E-02 | 9.58E-02 | 2.15E+00 | 2.26E+00 | 4.49E-01 | no | | intergenic_variant | MODIFIER |
| 15 | 3693258 | AX-97806402 | 0 | A | G | 0.088 | -8.59E-02 | 9.58E-02 | 2.15E+00 | 2.26E+00 | 4.49E-01 | no | | intergenic_variant | MODIFIER |
| 15 | 3693286 | AX-170257286 | 0 | T | C | 0.088 | -8.59E-02 | 9.58E-02 | 2.15E+00 | 2.26E+00 | 4.49E-01 | no | | intergenic_variant | MODIFIER |
| 15 | 3704587 | AX-170149770 | 0 | T | C | 0.088 | -8.59E-02 | 9.58E-02 | 2.15E+00 | 2.26E+00 | 4.49E-01 | no | | intergenic_variant | MODIFIER |
| 15 | 3705081 | AX-170162840 | 0 | C | T | 0.088 | -8.59E-02 | 9.58E-02 | 2.15E+00 | 2.26E+00 | 4.49E-01 | no | | intergenic_variant | MODIFIER |
| 15 | 3711204 | AX-97834546 | 0 | T | C | 0.088 | -8.59E-02 | 9.58E-02 | 2.15E+00 | 2.26E+00 | 4.49E-01 | no | | intergenic_variant | MODIFIER |
| 15 | 3767744 | AX-97846369 | 0 | A | G | 0.088 | -8.59E-02 | 9.58E-02 | 2.15E+00 | 2.26E+00 | 4.49E-01 | no | | intergenic_variant | MODIFIER |
| 15 | 4004044 | AX-170010926 | 0 | T | C | 0.088 | -8.59E-02 | 9.58E-02 | 2.15E+00 | 2.26E+00 | 4.49E-01 | no | Gm2093 | intron_variant,non_coding_transcript_variant | MODIFIER |
| 15 | 4914244 | AX-170228950 | 0 | A | G | 0.088 | -8.59E-02 | 9.58E-02 | 2.15E+00 | 2.26E+00 | 4.49E-01 | no | Mroh2b | intron_variant | MODIFIER |
| 15 | 4914462 | AX-170378084 | 0 | A | G | 0.088 | -8.59E-02 | 9.58E-02 | 2.15E+00 | 2.26E+00 | 4.49E-01 | no | Mroh2b | intron_variant | MODIFIER |
| 15 | 4937338 | AX-170134176 | 0 | A | G | 0.088 | -8.59E-02 | 9.58E-02 | 2.15E+00 | 2.26E+00 | 4.49E-01 | no | Mroh2b | intron_variant | MODIFIER |
| 15 | 5306971 | AX-97829111 | 0 | A | G | 0.088 | -8.59E-02 | 9.58E-02 | 2.15E+00 | 2.26E+00 | 4.49E-01 | no | Gm2157 | upstream_gene_variant | MODIFIER |
| 15 | 5492872 | AX-97789067 | 0 | C | T | 0.088 | -8.59E-02 | 9.58E-02 | 2.15E+00 | 2.26E+00 | 4.49E-01 | no | 5430437J10Rik | upstream_gene_variant | MODIFIER |
| 15 | 6060843 | AX-170438468 | 0 | T | C | 0.088 | -8.59E-02 | 9.58E-02 | 2.15E+00 | 2.26E+00 | 4.49E-01 | no | | regulatory_region_variant | MODIFIER |
| 15 | 6082102 | AX-170202774 | 0 | C | T | 0.088 | -8.59E-02 | 9.58E-02 | 2.15E+00 | 2.26E+00 | 4.49E-01 | no | | intergenic_variant | MODIFIER |
| 15 | 6099610 | AX-170167110 | 0 | T | C | 0.088 | -8.59E-02 | 9.58E-02 | 2.15E+00 | 2.26E+00 | 4.49E-01 | no | | intergenic_variant | MODIFIER |
| 15 | 6123370 | AX-170130944 | 0 | C | A | 0.088 | -8.59E-02 | 9.58E-02 | 2.15E+00 | 2.26E+00 | 4.49E-01 | no | | intergenic_variant | MODIFIER |
| 1 | 15702698 | AX-170289993 | 2 | C | G | 0.382 | 6.00E-02 | 7.92E-02 | 2.58E+00 | 2.63E+00 | 4.49E-01 | no | Kcnb2 | intron_variant | MODIFIER |
| 11 | 33548367 | AX-97807855 | 0 | G | A | 0.105 | -1.23E-01 | 1.65E-01 | 2.72E+00 | 2.83E+00 | 4.50E-01 | no | Gabrp | downstream_gene_variant | MODIFIER |
| 7 | 74419878 | AX-97804087 | 1 | C | T | 0.384 | -7.56E-02 | 1.01E-01 | 2.78E+00 | 2.87E+00 | 4.50E-01 | no | Slco3a1 | intron_variant | MODIFIER |
| 7 | 75207039 | AX-170529343 | 1 | C | T | 0.384 | -7.56E-02 | 1.01E-01 | 2.78E+00 | 2.87E+00 | 4.50E-01 | no | Sv2b | intron_variant | MODIFIER |
| 6 | 136305567 | AX-170076345 | 0 | C | T | 0.579 | 6.66E-02 | 8.61E-02 | 2.48E+00 | 2.54E+00 | 4.50E-01 | no | | intergenic_variant | MODIFIER |
| 8 | 84407477 | AX-170154063 | 0 | T | G | 0.193 | -8.67E-02 | 1.16E-01 | 2.68E+00 | 2.79E+00 | 4.50E-01 | no | Cacna1a | intron_variant,non_coding_transcript_variant | MODIFIER |
| 9 | 27471836 | AX-168419957 | 0 | C | A | 0.456 | 6.93E-02 | 9.17E-02 | 2.59E+00 | 2.65E+00 | 4.50E-01 | no | | intergenic_variant | MODIFIER |
| 10 | 68119594 | AX-170362054 | 1 | G | A | 0.42 | -5.92E-02 | 7.91E-02 | 2.65E+00 | 2.73E+00 | 4.50E-01 | no | Arid5b | intron_variant,NMD_transcript_variant | MODIFIER |
| 10 | 68173466 | AX-170319269 | 1 | G | A | 0.42 | -5.92E-02 | 7.91E-02 | 2.65E+00 | 2.73E+00 | 4.50E-01 | no | Arid5b | intron_variant,NMD_transcript_variant | MODIFIER |
| 8 | 116564564 | AX-170355236 | 0 | C | T | 0.491 | 3.71E-02 | 5.00E-02 | 2.71E+00 | 2.83E+00 | 4.50E-01 | no | | intergenic_variant | MODIFIER |
| 1 | 19337113 | AX-170417159 | 1 | G | A | 0.411 | 6.24E-02 | 8.26E-02 | 2.59E+00 | 2.64E+00 | 4.50E-01 | no | Gm4849 | downstream_gene_variant | MODIFIER |
| 1 | 19961925 | AX-170315272 | 1 | C | T | 0.411 | 6.24E-02 | 8.26E-02 | 2.59E+00 | 2.64E+00 | 4.50E-01 | no | | intergenic_variant | MODIFIER |
| 7 | 45998089 | AX-170506413 | 1 | C | A | 0.33 | 6.85E-02 | 9.12E-02 | 2.62E+00 | 2.71E+00 | 4.50E-01 | no | Abcc6 | intron_variant | MODIFIER |
| 4 | 89141061 | AX-170360820 | 1 | A | G | 0.339 | 8.10E-02 | 1.08E-01 | 2.67E+00 | 2.75E+00 | 4.50E-01 | no | Mtap | intron_variant | MODIFIER |
| 7 | 37958519 | AX-170077106 | 0 | G | A | 0.351 | 6.85E-02 | 9.07E-02 | 2.58E+00 | 2.66E+00 | 4.50E-01 | no | Uri1 | downstream_gene_variant | MODIFIER |
| 1 | 13414624 | AX-97818400 | 0 | A | G | 0.377 | 5.76E-02 | 7.66E-02 | 2.62E+00 | 2.67E+00 | 4.50E-01 | no | | intergenic_variant | MODIFIER |

| | | | | | | | | | | | | | | | |
|---|---|---|---|---|---|---|---|---|---|---|---|---|---|---|---|
| 7 | 124708035 | AX-170321725 | 1 | C | T | 0.366 | 6.07E-02 | 8.16E-02 | 2.74E+00 | 2.84E+00 | 4.50E-01 | no | 3100003L05Rik | intron_variant,non_coding_transcript_variant | MODIFIER |
| 16 | 19501433 | AX-170037388 | 1 | A | T | 0.125 | 7.16E-02 | 9.67E-02 | 2.77E+00 | 2.90E+00 | 4.50E-01 | no | | intergenic_variant | MODIFIER |
| 7 | 77130538 | AX-170077788 | 1 | C | T | 0.375 | -6.89E-02 | 9.27E-02 | 2.69E+00 | 2.80E+00 | 4.50E-01 | no | | intergenic_variant | MODIFIER |
| 8 | 16657677 | AX-97854997 | 0 | C | A | 0.237 | -5.84E-02 | 7.68E-02 | 2.53E+00 | 2.63E+00 | 4.50E-01 | no | Csmd1 | intron_variant | MODIFIER |
| 8 | 17295085 | AX-170201686 | 0 | C | T | 0.237 | -5.84E-02 | 7.68E-02 | 2.53E+00 | 2.63E+00 | 4.50E-01 | no | Csmd1 | intron_variant | MODIFIER |
| 15 | 15139252 | AX-97797006 | 0 | C | T | 0.246 | -7.73E-02 | 1.03E-01 | 2.65E+00 | 2.71E+00 | 4.51E-01 | no | Gm49118 | downstream_gene_variant | MODIFIER |
| 8 | 102725096 | AX-170437934 | 1 | T | G | 0.214 | -7.62E-02 | 1.03E-01 | 2.72E+00 | 2.83E+00 | 4.51E-01 | no | Cdh11 | intron_variant | MODIFIER |
| 5 | 85215512 | AX-170227123 | 2 | G | A | 0.464 | 7.64E-02 | 1.03E-01 | 2.76E+00 | 2.84E+00 | 4.51E-01 | no | | intergenic_variant | MODIFIER |
| 6 | 102640825 | AX-170394336 | 0 | C | T | 0.061 | 1.62E-01 | 2.18E-01 | 2.69E+00 | 2.81E+00 | 4.51E-01 | Yes | | intergenic_variant | MODIFIER |
| 8 | 96117368 | AX-170080890 | 0 | T | G | 0.246 | 6.78E-02 | 9.12E-02 | 2.67E+00 | 2.76E+00 | 4.51E-01 | no | | intergenic_variant | MODIFIER |
| 19 | 50713104 | AX-170112419 | 1 | A | G | 0.482 | 4.76E-02 | 6.37E-02 | 2.62E+00 | 2.71E+00 | 4.51E-01 | no | Gm26629 | intron_variant,non_coding_transcript_variant | MODIFIER |
| 5 | 78891259 | AX-170259780 | 0 | A | G | 0.412 | 7.66E-02 | 1.03E-01 | 2.73E+00 | 2.79E+00 | 4.51E-01 | no | | intergenic_variant | MODIFIER |
| 10 | 107789347 | AX-170003860 | 0 | G | C | 0.316 | -6.24E-02 | 8.38E-02 | 2.67E+00 | 2.74E+00 | 4.52E-01 | no | Otogl | intron_variant | MODIFIER |
| 10 | 107824379 | AX-170430150 | 0 | C | T | 0.316 | -6.24E-02 | 8.38E-02 | 2.67E+00 | 2.74E+00 | 4.52E-01 | no | Otogl | intron_variant | MODIFIER |
| 10 | 107837764 | AX-170316708 | 0 | A | G | 0.316 | -6.24E-02 | 8.38E-02 | 2.67E+00 | 2.74E+00 | 4.52E-01 | no | Otogl | intron_variant | MODIFIER |
| 10 | 108041841 | AX-170045850 | 0 | G | T | 0.316 | -6.24E-02 | 8.38E-02 | 2.67E+00 | 2.74E+00 | 4.52E-01 | no | | intergenic_variant | MODIFIER |
| 10 | 108358395 | AX-170003869 | 0 | C | T | 0.316 | -6.24E-02 | 8.38E-02 | 2.67E+00 | 2.74E+00 | 4.52E-01 | no | Pawr | intron_variant | MODIFIER |
| 10 | 108367502 | AX-170003870 | 0 | A | T | 0.316 | -6.24E-02 | 8.38E-02 | 2.67E+00 | 2.74E+00 | 4.52E-01 | no | Gm23105 | downstream_gene_variant | MODIFIER |
| 10 | 108416536 | AX-170317670 | 0 | G | A | 0.316 | -6.24E-02 | 8.38E-02 | 2.67E+00 | 2.74E+00 | 4.52E-01 | no | Pawr | downstream_gene_variant | MODIFIER |
| 10 | 108417314 | AX-170237613 | 0 | C | G | 0.316 | -6.24E-02 | 8.38E-02 | 2.67E+00 | 2.74E+00 | 4.52E-01 | no | Pawr | downstream_gene_variant | MODIFIER |
| 10 | 108703679 | AX-170086899 | 0 | T | A | 0.316 | -6.24E-02 | 8.38E-02 | 2.67E+00 | 2.74E+00 | 4.52E-01 | no | Syt1 | intron_variant | MODIFIER |
| 10 | 108732590 | AX-170487669 | 0 | A | G | 0.316 | -6.24E-02 | 8.38E-02 | 2.67E+00 | 2.74E+00 | 4.52E-01 | no | Syt1 | intron_variant | MODIFIER |
| 10 | 108737793 | AX-170086900 | 0 | C | T | 0.316 | -6.24E-02 | 8.38E-02 | 2.67E+00 | 2.74E+00 | 4.52E-01 | no | Syt1 | intron_variant | MODIFIER |
| 10 | 108744465 | AX-170196985 | 0 | A | T | 0.316 | -6.24E-02 | 8.38E-02 | 2.67E+00 | 2.74E+00 | 4.52E-01 | no | Syt1 | intron_variant | MODIFIER |
| 10 | 108751940 | AX-170442128 | 0 | C | T | 0.316 | -6.24E-02 | 8.38E-02 | 2.67E+00 | 2.74E+00 | 4.52E-01 | no | Syt1 | intron_variant | MODIFIER |
| 10 | 108757075 | AX-170003875 | 0 | T | C | 0.316 | -6.24E-02 | 8.38E-02 | 2.67E+00 | 2.74E+00 | 4.52E-01 | no | Syt1 | intron_variant | MODIFIER |
| 13 | 7061503 | AX-170525347 | 0 | A | G | 0.211 | -9.06E-02 | 1.21E-01 | 2.60E+00 | 2.67E+00 | 4.52E-01 | no | Gm40662 | intron_variant,non_coding_transcript_variant | MODIFIER |
| 4 | 59423324 | AX-170022901 | 1 | G | A | 0.366 | 8.38E-02 | 1.12E-01 | 2.65E+00 | 2.71E+00 | 4.52E-01 | no | Susd1 | intron_variant | MODIFIER |
| 18 | 87775144 | AX-170229424 | 0 | C | A | 0.175 | -7.73E-02 | 1.01E-01 | 2.49E+00 | 2.59E+00 | 4.52E-01 | no | | intergenic_variant | MODIFIER |
| 5 | 78944021 | AX-97814182 | 0 | C | T | 0.43 | 7.71E-02 | 1.04E-01 | 2.68E+00 | 2.74E+00 | 4.52E-01 | no | | intergenic_variant | MODIFIER |
| 6 | 102776797 | AX-170323515 | 0 | T | C | 0.061 | 1.55E-01 | 2.09E-01 | 2.66E+00 | 2.78E+00 | 4.52E-01 | Yes | | intergenic_variant | MODIFIER |
| 8 | 74783778 | AX-170481960 | 0 | G | A | 0.193 | -9.79E-02 | 1.32E-01 | 2.70E+00 | 2.78E+00 | 4.52E-01 | no | Gm11033 | intron_variant,non_coding_transcript_variant | MODIFIER |
| 1 | 14732270 | AX-170513914 | 0 | C | T | 0.377 | 5.95E-02 | 7.91E-02 | 2.60E+00 | 2.65E+00 | 4.52E-01 | no | | intergenic_variant | MODIFIER |
| 13 | 25389564 | AX-170150979 | 1 | G | A | 0.17 | -9.40E-02 | 1.21E-01 | 2.45E+00 | 2.51E+00 | 4.52E-01 | no | | intergenic_variant | MODIFIER |
| 18 | 69078623 | AX-170377278 | 0 | T | C | 0.158 | -9.02E-02 | 1.20E-01 | 2.58E+00 | 2.63E+00 | 4.52E-01 | no | | intergenic_variant | MODIFIER |
| 4 | 128334250 | AX-170397075 | 1 | G | A | 0.152 | 8.70E-02 | 1.16E-01 | 2.57E+00 | 2.65E+00 | 4.52E-01 | no | Csmd2 | intron_variant,non_coding_transcript_variant | MODIFIER |
| 5 | 42320741 | AX-170377404 | 1 | A | G | 0.205 | -7.91E-02 | 1.05E-01 | 2.59E+00 | 2.67E+00 | 4.52E-01 | no | | intergenic_variant | MODIFIER |
| 14 | 112414678 | AX-170198777 | 0 | G | A | 0.307 | 5.34E-02 | 7.13E-02 | 2.58E+00 | 2.69E+00 | 4.52E-01 | no | | intergenic_variant | MODIFIER |
| 1 | 88155935 | AX-170086528 | 1 | C | T | 0.161 | 1.55E-01 | 2.07E-01 | 2.65E+00 | 2.71E+00 | 4.52E-01 | Yes | Ugt1a6a | intron_variant | MODIFIER |
| 6 | 134963158 | AX-170152395 | 0 | A | G | 0.561 | 6.65E-02 | 8.73E-02 | 2.52E+00 | 2.58E+00 | 4.52E-01 | no | | regulatory_region_variant | MODIFIER |
| 4 | 146532274 | AX-170454436 | 1 | T | C | 0.107 | 1.20E-01 | 1.63E-01 | 2.68E+00 | 2.79E+00 | 4.53E-01 | no | Zfp981 | intron_variant | MODIFIER |
| 15 | 86076815 | AX-170230008 | 0 | A | G | 0.518 | 5.42E-02 | 7.07E-02 | 2.48E+00 | 2.58E+00 | 4.53E-01 | no | Gramd4 | intron_variant | MODIFIER |
| 15 | 94569328 | AX-170137426 | 0 | T | C | 0.132 | -1.05E-01 | 1.42E-01 | 2.82E+00 | 2.91E+00 | 4.53E-01 | no | Irak4 | downstream_gene_variant | MODIFIER |
| 4 | 128078975 | AX-170434199 | 1 | G | A | 0.161 | 8.35E-02 | 1.11E-01 | 2.59E+00 | 2.65E+00 | 4.53E-01 | no | Csmd2 | intron_variant,non_coding_transcript_variant | MODIFIER |
| 5 | 36086699 | AX-170071798 | 0 | T | C | 0.333 | 8.21E-02 | 1.11E-01 | 2.75E+00 | 2.81E+00 | 4.53E-01 | no | Sorcs2 | intron_variant | MODIFIER |
| 5 | 36147502 | AX-170357199 | 0 | C | T | 0.333 | 8.21E-02 | 1.11E-01 | 2.75E+00 | 2.81E+00 | 4.53E-01 | no | Sorcs2 | intron_variant | MODIFIER |
| 6 | 133999064 | AX-170076290 | 1 | A | G | 0.571 | 6.65E-02 | 8.70E-02 | 2.50E+00 | 2.55E+00 | 4.53E-01 | no | | intergenic_variant | MODIFIER |
| 7 | 10373709 | AX-170432360 | 0 | T | C | 0.167 | 5.71E-02 | 7.66E-02 | 2.91E+00 | 3.07E+00 | 4.53E-01 | no | Nlrp4d | intron_variant,non_coding_transcript_variant | MODIFIER |
| 8 | 38297157 | AX-97791424 | 1 | T | A | 0.152 | 1.12E-01 | 1.49E-01 | 2.91E+00 | 2.97E+00 | 4.53E-01 | no | Sgcz | intron_variant | MODIFIER |
| 8 | 42560287 | AX-170254157 | 1 | C | T | 0.152 | 1.12E-01 | 1.49E-01 | 2.91E+00 | 2.97E+00 | 4.53E-01 | no | | intergenic_variant | MODIFIER |
| 15 | 71517331 | AX-97814690 | 1 | C | T | 0.375 | 5.65E-02 | 7.45E-02 | 2.52E+00 | 2.60E+00 | 4.53E-01 | no | Fam135b | intron_variant | MODIFIER |
| 10 | 108741880 | AX-170324443 | 0 | C | T | 0.316 | -6.19E-02 | 8.33E-02 | 2.64E+00 | 2.72E+00 | 4.54E-01 | no | Syt1 | intron_variant | MODIFIER |
| 6 | 124131162 | AX-170028158 | 1 | C | G | 0.179 | -6.10E-02 | 8.29E-02 | 2.72E+00 | 2.85E+00 | 4.54E-01 | no | Vmn2r-ps34 | non_coding_transcript_exon_variant | MODIFIER |
| 18 | 62504115 | AX-97801491 | 0 | C | A | 0.193 | -8.51E-02 | 1.14E-01 | 2.65E+00 | 2.71E+00 | 4.54E-01 | no | Fbxo38 | downstream_gene_variant | MODIFIER |

| | | | | | | | | | | | | | | | |
|---|---|---|---|---|---|---|---|---|---|---|---|---|---|---|---|
| 5 | 101619395 | AX-170440908 | 1 | G | A | 0.446 | 7.83E-02 | 1.06E-01 | 2.69E+00 | 2.76E+00 | 4.54E-01 | no | | intergenic_variant | MODIFIER |
| 5 | 41927913 | AX-170095298 | 0 | C | T | 0.202 | -8.04E-02 | 1.08E-01 | 2.60E+00 | 2.68E+00 | 4.54E-01 | no | | intergenic_variant | MODIFIER |
| 5 | 41940307 | AX-170230780 | 0 | T | A | 0.202 | -8.04E-02 | 1.08E-01 | 2.60E+00 | 2.68E+00 | 4.54E-01 | no | Gm20156 | upstream_gene_variant | MODIFIER |
| 5 | 41945350 | AX-97846406 | 0 | A | G | 0.202 | -8.04E-02 | 1.08E-01 | 2.60E+00 | 2.68E+00 | 4.54E-01 | no | | intergenic_variant | MODIFIER |
| 5 | 41948190 | AX-170169083 | 0 | G | C | 0.202 | -8.04E-02 | 1.08E-01 | 2.60E+00 | 2.68E+00 | 4.54E-01 | no | | intergenic_variant | MODIFIER |
| 5 | 41964168 | AX-170477811 | 0 | G | C | 0.202 | -8.04E-02 | 1.08E-01 | 2.60E+00 | 2.68E+00 | 4.54E-01 | no | | intergenic_variant | MODIFIER |
| 5 | 41994360 | AX-97793724 | 0 | T | C | 0.202 | -8.04E-02 | 1.08E-01 | 2.60E+00 | 2.68E+00 | 4.54E-01 | no | | intergenic_variant | MODIFIER |
| 5 | 42020358 | AX-170024788 | 0 | T | C | 0.202 | -8.04E-02 | 1.08E-01 | 2.60E+00 | 2.68E+00 | 4.54E-01 | no | | intergenic_variant | MODIFIER |
| 5 | 42020988 | AX-170426254 | 0 | C | T | 0.202 | -8.04E-02 | 1.08E-01 | 2.60E+00 | 2.68E+00 | 4.54E-01 | no | | intergenic_variant | MODIFIER |
| 5 | 42021853 | AX-170024785 | 0 | G | A | 0.202 | -8.04E-02 | 1.08E-01 | 2.60E+00 | 2.68E+00 | 4.54E-01 | no | | intergenic_variant | MODIFIER |
| 5 | 42049559 | AX-170294427 | 0 | T | C | 0.202 | -8.04E-02 | 1.08E-01 | 2.60E+00 | 2.68E+00 | 4.54E-01 | no | | intergenic_variant | MODIFIER |
| 5 | 42062601 | AX-170329826 | 0 | G | A | 0.202 | -8.04E-02 | 1.08E-01 | 2.60E+00 | 2.68E+00 | 4.54E-01 | no | | intergenic_variant | MODIFIER |
| 5 | 42063503 | AX-170201732 | 0 | C | T | 0.202 | -8.04E-02 | 1.08E-01 | 2.60E+00 | 2.68E+00 | 4.54E-01 | no | Gm16223 | upstream_gene_variant | MODIFIER |
| 5 | 42066831 | AX-170174383 | 0 | C | T | 0.202 | -8.04E-02 | 1.08E-01 | 2.60E+00 | 2.68E+00 | 4.54E-01 | no | Gm16223 | upstream_gene_variant | MODIFIER |
| 5 | 42068068 | AX-170407575 | 0 | G | A | 0.202 | -8.04E-02 | 1.08E-01 | 2.60E+00 | 2.68E+00 | 4.54E-01 | no | Gm16223 | missense_variant | MODERATE |
| 5 | 42071585 | AX-170451155 | 0 | C | T | 0.202 | -8.04E-02 | 1.08E-01 | 2.60E+00 | 2.68E+00 | 4.54E-01 | no | Gm16223 | intron_variant | MODIFIER |
| 5 | 42072460 | AX-97830060 | 0 | A | G | 0.202 | -8.04E-02 | 1.08E-01 | 2.60E+00 | 2.68E+00 | 4.54E-01 | no | Gm16223 | intron_variant | MODIFIER |
| 5 | 42111994 | AX-97817938 | 0 | A | G | 0.202 | -8.04E-02 | 1.08E-01 | 2.60E+00 | 2.68E+00 | 4.54E-01 | no | Gm16223 | intron_variant | MODIFIER |
| 5 | 42151275 | AX-170251513 | 0 | G | A | 0.202 | -8.04E-02 | 1.08E-01 | 2.60E+00 | 2.68E+00 | 4.54E-01 | no | Gm16223 | intron_variant | MODIFIER |
| 5 | 42175301 | AX-170289816 | 0 | A | G | 0.202 | -8.04E-02 | 1.08E-01 | 2.60E+00 | 2.68E+00 | 4.54E-01 | no | Gm16223 | intron_variant | MODIFIER |
| 5 | 42197224 | AX-170229842 | 0 | T | C | 0.202 | -8.04E-02 | 1.08E-01 | 2.60E+00 | 2.68E+00 | 4.54E-01 | no | Gm16223 | intron_variant | MODIFIER |
| 5 | 42236748 | AX-170291152 | 0 | A | G | 0.202 | -8.04E-02 | 1.08E-01 | 2.60E+00 | 2.68E+00 | 4.54E-01 | no | | intergenic_variant | MODIFIER |
| 5 | 42239803 | AX-97812226 | 0 | C | T | 0.202 | -8.04E-02 | 1.08E-01 | 2.60E+00 | 2.68E+00 | 4.54E-01 | no | | intergenic_variant | MODIFIER |
| 5 | 42240772 | AX-170249462 | 0 | C | T | 0.202 | -8.04E-02 | 1.08E-01 | 2.60E+00 | 2.68E+00 | 4.54E-01 | no | | intergenic_variant | MODIFIER |
| 5 | 42241190 | AX-170265129 | 0 | A | G | 0.202 | -8.04E-02 | 1.08E-01 | 2.60E+00 | 2.68E+00 | 4.54E-01 | no | | intergenic_variant | MODIFIER |
| 5 | 42297589 | AX-170324698 | 0 | T | C | 0.202 | -8.04E-02 | 1.08E-01 | 2.60E+00 | 2.68E+00 | 4.54E-01 | no | | intergenic_variant | MODIFIER |
| 5 | 42321055 | AX-170108314 | 0 | C | T | 0.202 | -8.04E-02 | 1.08E-01 | 2.60E+00 | 2.68E+00 | 4.54E-01 | no | | intergenic_variant | MODIFIER |
| 5 | 42426893 | AX-170130800 | 0 | A | G | 0.202 | -8.04E-02 | 1.08E-01 | 2.60E+00 | 2.68E+00 | 4.54E-01 | no | | intergenic_variant | MODIFIER |
| 5 | 42431524 | AX-97821875 | 0 | T | C | 0.202 | -8.04E-02 | 1.08E-01 | 2.60E+00 | 2.68E+00 | 4.54E-01 | no | | intergenic_variant | MODIFIER |
| 5 | 42433252 | AX-170490019 | 0 | A | C | 0.202 | -8.04E-02 | 1.08E-01 | 2.60E+00 | 2.68E+00 | 4.54E-01 | no | | intergenic_variant | MODIFIER |
| 5 | 42456761 | AX-170200167 | 0 | C | T | 0.202 | -8.04E-02 | 1.08E-01 | 2.60E+00 | 2.68E+00 | 4.54E-01 | no | | intergenic_variant | MODIFIER |
| 5 | 42456959 | AX-170151467 | 0 | G | C | 0.202 | -8.04E-02 | 1.08E-01 | 2.60E+00 | 2.68E+00 | 4.54E-01 | no | | intergenic_variant | MODIFIER |
| 5 | 42478380 | AX-170507524 | 0 | A | G | 0.202 | -8.04E-02 | 1.08E-01 | 2.60E+00 | 2.68E+00 | 4.54E-01 | no | | intergenic_variant | MODIFIER |
| 5 | 42480266 | AX-170024804 | 0 | G | A | 0.202 | -8.04E-02 | 1.08E-01 | 2.60E+00 | 2.68E+00 | 4.54E-01 | no | | intergenic_variant | MODIFIER |
| 5 | 42480738 | AX-170342410 | 0 | A | G | 0.202 | -8.04E-02 | 1.08E-01 | 2.60E+00 | 2.68E+00 | 4.54E-01 | no | | intergenic_variant | MODIFIER |
| 5 | 42480851 | AX-170186319 | 0 | C | T | 0.202 | -8.04E-02 | 1.08E-01 | 2.60E+00 | 2.68E+00 | 4.54E-01 | no | | intergenic_variant | MODIFIER |
| 5 | 42481637 | AX-170445458 | 0 | G | A | 0.202 | -8.04E-02 | 1.08E-01 | 2.60E+00 | 2.68E+00 | 4.54E-01 | no | | intergenic_variant | MODIFIER |
| 5 | 42488086 | AX-97799306 | 0 | A | G | 0.202 | -8.04E-02 | 1.08E-01 | 2.60E+00 | 2.68E+00 | 4.54E-01 | no | | intergenic_variant | MODIFIER |
| 5 | 42488181 | AX-170024798 | 0 | A | G | 0.202 | -8.04E-02 | 1.08E-01 | 2.60E+00 | 2.68E+00 | 4.54E-01 | no | | intergenic_variant | MODIFIER |
| 5 | 42554305 | AX-97785057 | 0 | A | G | 0.202 | -8.04E-02 | 1.08E-01 | 2.60E+00 | 2.68E+00 | 4.54E-01 | no | | intergenic_variant | MODIFIER |
| 5 | 42579990 | AX-170024797 | 0 | T | C | 0.202 | -8.04E-02 | 1.08E-01 | 2.60E+00 | 2.68E+00 | 4.54E-01 | no | | intergenic_variant | MODIFIER |
| 5 | 42580343 | AX-97848662 | 0 | T | C | 0.202 | -8.04E-02 | 1.08E-01 | 2.60E+00 | 2.68E+00 | 4.54E-01 | no | | intergenic_variant | MODIFIER |
| 5 | 42593681 | AX-170249315 | 0 | A | G | 0.202 | -8.04E-02 | 1.08E-01 | 2.60E+00 | 2.68E+00 | 4.54E-01 | no | | intergenic_variant | MODIFIER |
| 5 | 42596456 | AX-170337854 | 0 | T | G | 0.202 | -8.04E-02 | 1.08E-01 | 2.60E+00 | 2.68E+00 | 4.54E-01 | no | | intergenic_variant | MODIFIER |
| 5 | 42610375 | AX-170024801 | 0 | T | A | 0.202 | -8.04E-02 | 1.08E-01 | 2.60E+00 | 2.68E+00 | 4.54E-01 | no | | intergenic_variant | MODIFIER |
| 5 | 42637235 | AX-97799272 | 0 | T | C | 0.202 | -8.04E-02 | 1.08E-01 | 2.60E+00 | 2.68E+00 | 4.54E-01 | no | | intergenic_variant | MODIFIER |
| 5 | 42637524 | AX-170389548 | 0 | C | T | 0.202 | -8.04E-02 | 1.08E-01 | 2.60E+00 | 2.68E+00 | 4.54E-01 | no | | intergenic_variant | MODIFIER |
| 5 | 42644628 | AX-170374444 | 0 | T | G | 0.202 | -8.04E-02 | 1.08E-01 | 2.60E+00 | 2.68E+00 | 4.54E-01 | no | | intergenic_variant | MODIFIER |
| 5 | 42671709 | AX-170071907 | 0 | C | T | 0.202 | -8.04E-02 | 1.08E-01 | 2.60E+00 | 2.68E+00 | 4.54E-01 | no | | intergenic_variant | MODIFIER |
| 5 | 42672082 | AX-170071908 | 0 | G | T | 0.202 | -8.04E-02 | 1.08E-01 | 2.60E+00 | 2.68E+00 | 4.54E-01 | no | | intergenic_variant | MODIFIER |
| 5 | 42677711 | AX-97832233 | 0 | A | G | 0.202 | -8.04E-02 | 1.08E-01 | 2.60E+00 | 2.68E+00 | 4.54E-01 | no | | intergenic_variant | MODIFIER |
| 5 | 42677857 | AX-170469449 | 0 | T | C | 0.202 | -8.04E-02 | 1.08E-01 | 2.60E+00 | 2.68E+00 | 4.54E-01 | no | | intergenic_variant | MODIFIER |
| 5 | 42679382 | AX-170229921 | 0 | T | G | 0.202 | -8.04E-02 | 1.08E-01 | 2.60E+00 | 2.68E+00 | 4.54E-01 | no | | intergenic_variant | MODIFIER |
| 5 | 42697314 | AX-170484773 | 0 | G | A | 0.202 | -8.04E-02 | 1.08E-01 | 2.60E+00 | 2.68E+00 | 4.54E-01 | no | | intergenic_variant | MODIFIER |

| | | | | | | | | | | | | | | | |
|---|---|---|---|---|---|---|---|---|---|---|---|---|---|---|---|
| 5 | 42703317 | AX-170182951 | 0 | G | T | 0.202 | -8.04E-02 | 1.08E-01 | 2.60E+00 | 2.68E+00 | 4.54E-01 | no | | intergenic_variant | MODIFIER |
| 5 | 42719044 | AX-170211299 | 0 | C | T | 0.202 | -8.04E-02 | 1.08E-01 | 2.60E+00 | 2.68E+00 | 4.54E-01 | no | | intergenic_variant | MODIFIER |
| 5 | 42722118 | AX-97825196 | 0 | C | G | 0.202 | -8.04E-02 | 1.08E-01 | 2.60E+00 | 2.68E+00 | 4.54E-01 | no | | intergenic_variant | MODIFIER |
| 5 | 42727808 | AX-170126056 | 0 | A | G | 0.202 | -8.04E-02 | 1.08E-01 | 2.60E+00 | 2.68E+00 | 4.54E-01 | no | | intergenic_variant | MODIFIER |
| 5 | 42742940 | AX-170364947 | 0 | G | A | 0.202 | -8.04E-02 | 1.08E-01 | 2.60E+00 | 2.68E+00 | 4.54E-01 | no | | intergenic_variant | MODIFIER |
| 5 | 42762770 | AX-170038983 | 0 | A | G | 0.202 | -8.04E-02 | 1.08E-01 | 2.60E+00 | 2.68E+00 | 4.54E-01 | no | | intergenic_variant | MODIFIER |
| 5 | 42788295 | AX-170106021 | 0 | T | C | 0.202 | -8.04E-02 | 1.08E-01 | 2.60E+00 | 2.68E+00 | 4.54E-01 | no | | intergenic_variant | MODIFIER |
| 5 | 42790803 | AX-170472681 | 0 | G | C | 0.202 | -8.04E-02 | 1.08E-01 | 2.60E+00 | 2.68E+00 | 4.54E-01 | no | | intergenic_variant | MODIFIER |
| 5 | 42801078 | AX-170305587 | 0 | T | G | 0.202 | -8.04E-02 | 1.08E-01 | 2.60E+00 | 2.68E+00 | 4.54E-01 | no | | intergenic_variant | MODIFIER |
| 5 | 42801208 | AX-170505810 | 0 | T | C | 0.202 | -8.04E-02 | 1.08E-01 | 2.60E+00 | 2.68E+00 | 4.54E-01 | no | | intergenic_variant | MODIFIER |
| 5 | 42804617 | AX-170350783 | 0 | G | C | 0.202 | -8.04E-02 | 1.08E-01 | 2.60E+00 | 2.68E+00 | 4.54E-01 | no | | intergenic_variant | MODIFIER |
| 5 | 42845131 | AX-170135090 | 0 | G | A | 0.202 | -8.04E-02 | 1.08E-01 | 2.60E+00 | 2.68E+00 | 4.54E-01 | no | | intergenic_variant | MODIFIER |
| 5 | 42849691 | AX-170523179 | 0 | T | A | 0.202 | -8.04E-02 | 1.08E-01 | 2.60E+00 | 2.68E+00 | 4.54E-01 | no | | intergenic_variant | MODIFIER |
| 5 | 42849719 | AX-170535123 | 0 | G | T | 0.202 | -8.04E-02 | 1.08E-01 | 2.60E+00 | 2.68E+00 | 4.54E-01 | no | | intergenic_variant | MODIFIER |
| 5 | 42857965 | AX-170383367 | 0 | A | G | 0.202 | -8.04E-02 | 1.08E-01 | 2.60E+00 | 2.68E+00 | 4.54E-01 | no | | intergenic_variant | MODIFIER |
| 5 | 42884611 | AX-170165527 | 0 | T | C | 0.202 | -8.04E-02 | 1.08E-01 | 2.60E+00 | 2.68E+00 | 4.54E-01 | no | | regulatory_region_variant | MODIFIER |
| 5 | 42886086 | AX-168417006 | 0 | G | A | 0.202 | -8.04E-02 | 1.08E-01 | 2.60E+00 | 2.68E+00 | 4.54E-01 | no | | intergenic_variant | MODIFIER |
| 5 | 42890385 | AX-170071909 | 0 | A | G | 0.202 | -8.04E-02 | 1.08E-01 | 2.60E+00 | 2.68E+00 | 4.54E-01 | no | | intergenic_variant | MODIFIER |
| 5 | 42890962 | AX-170535023 | 0 | A | G | 0.202 | -8.04E-02 | 1.08E-01 | 2.60E+00 | 2.68E+00 | 4.54E-01 | no | | intergenic_variant | MODIFIER |
| 5 | 42903405 | AX-170288226 | 0 | T | G | 0.202 | -8.04E-02 | 1.08E-01 | 2.60E+00 | 2.68E+00 | 4.54E-01 | no | | intergenic_variant | MODIFIER |
| 5 | 42904203 | AX-97792942 | 0 | A | G | 0.202 | -8.04E-02 | 1.08E-01 | 2.60E+00 | 2.68E+00 | 4.54E-01 | no | | intergenic_variant | MODIFIER |
| 5 | 42924005 | AX-170462672 | 0 | G | A | 0.202 | -8.04E-02 | 1.08E-01 | 2.60E+00 | 2.68E+00 | 4.54E-01 | no | | intergenic_variant | MODIFIER |
| 5 | 42924036 | AX-170137707 | 0 | A | G | 0.202 | -8.04E-02 | 1.08E-01 | 2.60E+00 | 2.68E+00 | 4.54E-01 | no | | intergenic_variant | MODIFIER |
| 5 | 42951233 | AX-97819898 | 0 | G | T | 0.202 | -8.04E-02 | 1.08E-01 | 2.60E+00 | 2.68E+00 | 4.54E-01 | no | | intergenic_variant | MODIFIER |
| 5 | 42986254 | AX-170404403 | 0 | C | A | 0.202 | -8.04E-02 | 1.08E-01 | 2.60E+00 | 2.68E+00 | 4.54E-01 | no | | regulatory_region_variant | MODIFIER |
| 5 | 42991958 | AX-170024805 | 0 | G | C | 0.202 | -8.04E-02 | 1.08E-01 | 2.60E+00 | 2.68E+00 | 4.54E-01 | no | | intergenic_variant | MODIFIER |
| 5 | 42991999 | AX-170428284 | 0 | G | C | 0.202 | -8.04E-02 | 1.08E-01 | 2.60E+00 | 2.68E+00 | 4.54E-01 | no | | intergenic_variant | MODIFIER |
| 5 | 42995321 | AX-97815005 | 0 | A | G | 0.202 | -8.04E-02 | 1.08E-01 | 2.60E+00 | 2.68E+00 | 4.54E-01 | no | | intergenic_variant | MODIFIER |
| 5 | 42995896 | AX-170391701 | 0 | T | C | 0.202 | -8.04E-02 | 1.08E-01 | 2.60E+00 | 2.68E+00 | 4.54E-01 | no | | intergenic_variant | MODIFIER |
| 5 | 43016946 | AX-170410935 | 0 | G | A | 0.202 | -8.04E-02 | 1.08E-01 | 2.60E+00 | 2.68E+00 | 4.54E-01 | no | | intergenic_variant | MODIFIER |
| 5 | 43024856 | AX-170071911 | 0 | C | G | 0.202 | -8.04E-02 | 1.08E-01 | 2.60E+00 | 2.68E+00 | 4.54E-01 | no | Gm43700 | downstream_gene_variant | MODIFIER |
| 5 | 43026217 | AX-170071912 | 0 | C | T | 0.202 | -8.04E-02 | 1.08E-01 | 2.60E+00 | 2.68E+00 | 4.54E-01 | no | Gm43700 | downstream_gene_variant | MODIFIER |
| 5 | 43032525 | AX-170038980 | 0 | C | T | 0.202 | -8.04E-02 | 1.08E-01 | 2.60E+00 | 2.68E+00 | 4.54E-01 | no | Gm43700 | intron_variant,non_coding_transcript_variant | MODIFIER |
| 5 | 43053207 | AX-170024806 | 0 | G | A | 0.202 | -8.04E-02 | 1.08E-01 | 2.60E+00 | 2.68E+00 | 4.54E-01 | no | | intergenic_variant | MODIFIER |
| 5 | 43082237 | AX-170259158 | 0 | A | G | 0.202 | -8.04E-02 | 1.08E-01 | 2.60E+00 | 2.68E+00 | 4.54E-01 | no | | regulatory_region_variant | MODIFIER |
| 5 | 43082563 | AX-170038979 | 0 | G | T | 0.202 | -8.04E-02 | 1.08E-01 | 2.60E+00 | 2.68E+00 | 4.54E-01 | no | | regulatory_region_variant | MODIFIER |
| 5 | 43082625 | AX-97829473 | 0 | G | A | 0.202 | -8.04E-02 | 1.08E-01 | 2.60E+00 | 2.68E+00 | 4.54E-01 | no | | regulatory_region_variant | MODIFIER |
| 5 | 43107585 | AX-170363161 | 0 | T | G | 0.202 | -8.04E-02 | 1.08E-01 | 2.60E+00 | 2.68E+00 | 4.54E-01 | no | | intergenic_variant | MODIFIER |
| 5 | 43109761 | AX-97847352 | 0 | A | T | 0.202 | -8.04E-02 | 1.08E-01 | 2.60E+00 | 2.68E+00 | 4.54E-01 | no | | intergenic_variant | MODIFIER |
| 5 | 43117097 | AX-170242509 | 0 | C | G | 0.202 | -8.04E-02 | 1.08E-01 | 2.60E+00 | 2.68E+00 | 4.54E-01 | no | | regulatory_region_variant | MODIFIER |
| 5 | 43130223 | AX-97838180 | 0 | T | C | 0.202 | -8.04E-02 | 1.08E-01 | 2.60E+00 | 2.68E+00 | 4.54E-01 | no | | intergenic_variant | MODIFIER |
| 5 | 43133525 | AX-170171384 | 0 | A | G | 0.202 | -8.04E-02 | 1.08E-01 | 2.60E+00 | 2.68E+00 | 4.54E-01 | no | | intergenic_variant | MODIFIER |
| 5 | 43134634 | AX-170533931 | 0 | G | A | 0.202 | -8.04E-02 | 1.08E-01 | 2.60E+00 | 2.68E+00 | 4.54E-01 | no | | intergenic_variant | MODIFIER |
| 5 | 43138668 | AX-170399603 | 0 | G | A | 0.202 | -8.04E-02 | 1.08E-01 | 2.60E+00 | 2.68E+00 | 4.54E-01 | no | | intergenic_variant | MODIFIER |
| 5 | 43149036 | AX-97831451 | 0 | A | G | 0.202 | -8.04E-02 | 1.08E-01 | 2.60E+00 | 2.68E+00 | 4.54E-01 | no | Gm7854 | downstream_gene_variant | MODIFIER |
| 5 | 43166518 | AX-170138249 | 0 | G | C | 0.202 | -8.04E-02 | 1.08E-01 | 2.60E+00 | 2.68E+00 | 4.54E-01 | no | Gm7854 | intron_variant,non_coding_transcript_variant | MODIFIER |
| 5 | 43170461 | AX-170338351 | 0 | C | T | 0.202 | -8.04E-02 | 1.08E-01 | 2.60E+00 | 2.68E+00 | 4.54E-01 | no | Gm7854 | intron_variant,non_coding_transcript_variant | MODIFIER |
| 5 | 43191167 | AX-170445066 | 0 | G | T | 0.202 | -8.04E-02 | 1.08E-01 | 2.60E+00 | 2.68E+00 | 4.54E-01 | no | Gm7854 | intron_variant,non_coding_transcript_variant | MODIFIER |
| 5 | 43227907 | AX-170335742 | 0 | A | G | 0.202 | -8.04E-02 | 1.08E-01 | 2.60E+00 | 2.68E+00 | 4.54E-01 | no | Gm7854 | intron_variant,non_coding_transcript_variant | MODIFIER |
| 5 | 43979478 | AX-170089465 | 0 | C | T | 0.202 | -8.04E-02 | 1.08E-01 | 2.60E+00 | 2.68E+00 | 4.54E-01 | no | Fgfbp1 | synonymous_variant | LOW |
| 5 | 44014669 | AX-170410386 | 0 | T | C | 0.202 | -8.04E-02 | 1.08E-01 | 2.60E+00 | 2.68E+00 | 4.54E-01 | no | Prom1 | intron_variant | MODIFIER |
| 5 | 44032808 | AX-170455619 | 0 | G | C | 0.202 | -8.04E-02 | 1.08E-01 | 2.60E+00 | 2.68E+00 | 4.54E-01 | no | Prom1 | intron_variant | MODIFIER |
| 5 | 44058970 | AX-97788120 | 0 | A | G | 0.202 | -8.04E-02 | 1.08E-01 | 2.60E+00 | 2.68E+00 | 4.54E-01 | no | Prom1 | intron_variant | MODIFIER |
| 5 | 44068421 | AX-170278744 | 0 | T | G | 0.202 | -8.04E-02 | 1.08E-01 | 2.60E+00 | 2.68E+00 | 4.54E-01 | no | Prom1 | intron_variant | MODIFIER |

| | | | | | | | | | | | | | | | |
|---|---|---|---|---|---|---|---|---|---|---|---|---|---|---|---|
| 5 | 44079577 | AX-97850012 | 0 | T | C | 0.202 | -8.04E-02 | 1.08E-01 | 2.60E+00 | 2.68E+00 | 4.54E-01 | no | Prom1 | intron_variant | MODIFIER |
| 5 | 44084323 | AX-170024825 | 0 | G | A | 0.202 | -8.04E-02 | 1.08E-01 | 2.60E+00 | 2.68E+00 | 4.54E-01 | no | Prom1 | intron_variant | MODIFIER |
| 5 | 44089637 | AX-170024823 | 0 | T | C | 0.202 | -8.04E-02 | 1.08E-01 | 2.60E+00 | 2.68E+00 | 4.54E-01 | no | Prom1 | intron_variant | MODIFIER |
| 5 | 44095947 | AX-170024826 | 0 | G | A | 0.202 | -8.04E-02 | 1.08E-01 | 2.60E+00 | 2.68E+00 | 4.54E-01 | no | Prom1 | upstream_gene_variant | MODIFIER |
| 5 | 44111804 | AX-170024824 | 0 | C | T | 0.202 | -8.04E-02 | 1.08E-01 | 2.60E+00 | 2.68E+00 | 4.54E-01 | no | Gm42966 | downstream_gene_variant | MODIFIER |
| 5 | 44130411 | AX-170102913 | 0 | A | G | 0.202 | -8.04E-02 | 1.08E-01 | 2.60E+00 | 2.68E+00 | 4.54E-01 | no | Gm42429 | upstream_gene_variant | MODIFIER |
| 5 | 44157053 | AX-170231214 | 0 | G | A | 0.202 | -8.04E-02 | 1.08E-01 | 2.60E+00 | 2.68E+00 | 4.54E-01 | no | | intergenic_variant | MODIFIER |
| 5 | 44167099 | AX-170238537 | 0 | C | T | 0.202 | -8.04E-02 | 1.08E-01 | 2.60E+00 | 2.68E+00 | 4.54E-01 | no | | intergenic_variant | MODIFIER |
| 5 | 44168523 | AX-170214303 | 0 | A | T | 0.202 | -8.04E-02 | 1.08E-01 | 2.60E+00 | 2.68E+00 | 4.54E-01 | no | | intergenic_variant | MODIFIER |
| 5 | 44251942 | AX-170514778 | 0 | A | T | 0.202 | -8.04E-02 | 1.08E-01 | 2.60E+00 | 2.68E+00 | 4.54E-01 | no | | intergenic_variant | MODIFIER |
| 5 | 44254619 | AX-170477469 | 0 | G | C | 0.202 | -8.04E-02 | 1.08E-01 | 2.60E+00 | 2.68E+00 | 4.54E-01 | no | | intergenic_variant | MODIFIER |
| 5 | 44254908 | AX-170486732 | 0 | A | G | 0.202 | -8.04E-02 | 1.08E-01 | 2.60E+00 | 2.68E+00 | 4.54E-01 | no | | intergenic_variant | MODIFIER |
| 5 | 44267802 | AX-97856140 | 0 | C | T | 0.202 | -8.04E-02 | 1.08E-01 | 2.60E+00 | 2.68E+00 | 4.54E-01 | no | | intergenic_variant | MODIFIER |
| 5 | 44374321 | AX-170071939 | 0 | G | A | 0.202 | -8.04E-02 | 1.08E-01 | 2.60E+00 | 2.68E+00 | 4.54E-01 | no | | intergenic_variant | MODIFIER |
| 5 | 44377810 | AX-170442944 | 0 | G | T | 0.202 | -8.04E-02 | 1.08E-01 | 2.60E+00 | 2.68E+00 | 4.54E-01 | no | | intergenic_variant | MODIFIER |
| 5 | 44398917 | AX-170182937 | 0 | G | A | 0.202 | -8.04E-02 | 1.08E-01 | 2.60E+00 | 2.68E+00 | 4.54E-01 | no | | intergenic_variant | MODIFIER |
| 5 | 44425627 | AX-170515901 | 0 | G | A | 0.202 | -8.04E-02 | 1.08E-01 | 2.60E+00 | 2.68E+00 | 4.54E-01 | no | Gm36017 | intron_variant,non_coding_transcript_variant | MODIFIER |
| 5 | 44429104 | AX-170267747 | 0 | A | C | 0.202 | -8.04E-02 | 1.08E-01 | 2.60E+00 | 2.68E+00 | 4.54E-01 | no | Gm36017 | upstream_gene_variant | MODIFIER |
| 5 | 44446071 | AX-170237728 | 0 | C | T | 0.202 | -8.04E-02 | 1.08E-01 | 2.60E+00 | 2.68E+00 | 4.54E-01 | no | | intergenic_variant | MODIFIER |
| 5 | 44639993 | AX-170089466 | 0 | A | G | 0.202 | -8.04E-02 | 1.08E-01 | 2.60E+00 | 2.68E+00 | 4.54E-01 | no | Ldb2 | intron_variant | MODIFIER |
| 5 | 44657197 | AX-170205051 | 0 | G | A | 0.202 | -8.04E-02 | 1.08E-01 | 2.60E+00 | 2.68E+00 | 4.54E-01 | no | Ldb2 | intron_variant | MODIFIER |
| 5 | 44691624 | AX-97840075 | 0 | G | A | 0.202 | -8.04E-02 | 1.08E-01 | 2.60E+00 | 2.68E+00 | 4.54E-01 | no | Ldb2 | intron_variant | MODIFIER |
| 5 | 44730663 | AX-170283814 | 0 | C | A | 0.202 | -8.04E-02 | 1.08E-01 | 2.60E+00 | 2.68E+00 | 4.54E-01 | no | Ldb2 | intron_variant | MODIFIER |
| 5 | 44750337 | AX-170427014 | 0 | C | T | 0.202 | -8.04E-02 | 1.08E-01 | 2.60E+00 | 2.68E+00 | 4.54E-01 | no | Ldb2 | intron_variant | MODIFIER |
| 5 | 44844301 | AX-170526533 | 0 | C | T | 0.202 | -8.04E-02 | 1.08E-01 | 2.60E+00 | 2.68E+00 | 4.54E-01 | no | | intergenic_variant | MODIFIER |
| 5 | 44949431 | AX-170095304 | 0 | A | G | 0.202 | -8.04E-02 | 1.08E-01 | 2.60E+00 | 2.68E+00 | 4.54E-01 | no | | intergenic_variant | MODIFIER |
| 5 | 45003267 | AX-170222894 | 0 | A | G | 0.202 | -8.04E-02 | 1.08E-01 | 2.60E+00 | 2.68E+00 | 4.54E-01 | no | | intergenic_variant | MODIFIER |
| 7 | 24652899 | AX-170076859 | 0 | G | A | 0.395 | 6.50E-02 | 8.73E-02 | 2.59E+00 | 2.71E+00 | 4.54E-01 | no | | intergenic_variant | MODIFIER |
| 3 | 31930140 | AX-97838982 | 0 | A | G | 0.158 | -7.42E-02 | 1.00E-01 | 2.74E+00 | 2.82E+00 | 4.54E-01 | no | Kcnmb2 | intron_variant | MODIFIER |
| 9 | 92063498 | AX-170419413 | 0 | A | G | 0.14 | -8.86E-02 | 1.19E-01 | 2.59E+00 | 2.68E+00 | 4.54E-01 | no | | intergenic_variant | MODIFIER |
| 15 | 80249964 | AX-170336225 | 1 | C | G | 0.562 | 5.84E-02 | 7.73E-02 | 2.54E+00 | 2.62E+00 | 4.54E-01 | no | Mief1 | synonymous_variant | LOW |
| 6 | 134636475 | AX-170186738 | 1 | G | C | 0.562 | 6.60E-02 | 8.70E-02 | 2.53E+00 | 2.58E+00 | 4.54E-01 | no | Mansc1 | upstream_gene_variant | MODIFIER |
| 5 | 93732348 | AX-170089558 | 0 | T | C | 0.43 | 6.89E-02 | 9.34E-02 | 2.70E+00 | 2.80E+00 | 4.54E-01 | no | | intergenic_variant | MODIFIER |
| 1 | 192252318 | AX-170393566 | 1 | A | G | 0.268 | 7.13E-02 | 9.24E-02 | 2.45E+00 | 2.54E+00 | 4.54E-01 | no | Kcnh1 | intron_variant | MODIFIER |
| 10 | 66004097 | AX-170237136 | 0 | G | A | 0.421 | -5.81E-02 | 7.84E-02 | 2.65E+00 | 2.73E+00 | 4.54E-01 | no | | intergenic_variant | MODIFIER |
| 1 | 192237050 | AX-170217155 | 0 | A | T | 0.272 | 6.84E-02 | 8.85E-02 | 2.45E+00 | 2.54E+00 | 4.54E-01 | no | Kcnh1 | intron_variant | MODIFIER |
| 9 | 97893954 | AX-97830305 | 1 | C | T | 0.143 | -8.63E-02 | 1.15E-01 | 2.57E+00 | 2.64E+00 | 4.55E-01 | no | Clstn2 | intron_variant | MODIFIER |
| 7 | 135498768 | AX-170266431 | 0 | C | A | 0.254 | -5.99E-02 | 7.96E-02 | 2.54E+00 | 2.64E+00 | 4.55E-01 | no | | regulatory_region_variant | MODIFIER |
| 18 | 53776084 | AX-170099736 | 0 | A | G | 0.447 | 7.50E-02 | 9.45E-02 | 2.38E+00 | 2.44E+00 | 4.55E-01 | no | | intergenic_variant | MODIFIER |
| 18 | 53792742 | AX-170146127 | 0 | A | G | 0.447 | 7.50E-02 | 9.45E-02 | 2.38E+00 | 2.44E+00 | 4.55E-01 | no | | intergenic_variant | MODIFIER |
| 18 | 53792841 | AX-170123491 | 0 | C | G | 0.447 | 7.50E-02 | 9.45E-02 | 2.38E+00 | 2.44E+00 | 4.55E-01 | no | | intergenic_variant | MODIFIER |
| 18 | 53799912 | AX-170133857 | 0 | C | T | 0.447 | 7.50E-02 | 9.45E-02 | 2.38E+00 | 2.44E+00 | 4.55E-01 | no | | intergenic_variant | MODIFIER |
| 18 | 53981436 | AX-170102840 | 0 | A | G | 0.447 | 7.50E-02 | 9.45E-02 | 2.38E+00 | 2.44E+00 | 4.55E-01 | no | Gm5507 | upstream_gene_variant | MODIFIER |
| 18 | 54033812 | AX-170015725 | 0 | G | A | 0.447 | 7.50E-02 | 9.45E-02 | 2.38E+00 | 2.44E+00 | 4.55E-01 | no | AC148983.1 | upstream_gene_variant | MODIFIER |
| 18 | 54040590 | AX-170340344 | 0 | T | G | 0.447 | 7.50E-02 | 9.45E-02 | 2.38E+00 | 2.44E+00 | 4.55E-01 | no | | intergenic_variant | MODIFIER |
| 18 | 54054007 | AX-170015723 | 0 | C | T | 0.447 | 7.50E-02 | 9.45E-02 | 2.38E+00 | 2.44E+00 | 4.55E-01 | no | | intergenic_variant | MODIFIER |
| 18 | 54054641 | AX-170015727 | 0 | G | A | 0.447 | 7.50E-02 | 9.45E-02 | 2.38E+00 | 2.44E+00 | 4.55E-01 | no | | regulatory_region_variant | MODIFIER |
| 18 | 54054724 | AX-170129104 | 0 | C | T | 0.447 | 7.50E-02 | 9.45E-02 | 2.38E+00 | 2.44E+00 | 4.55E-01 | no | | regulatory_region_variant | MODIFIER |
| 18 | 54072098 | AX-170015724 | 0 | C | G | 0.447 | 7.50E-02 | 9.45E-02 | 2.38E+00 | 2.44E+00 | 4.55E-01 | no | | intergenic_variant | MODIFIER |
| 18 | 54089495 | AX-170449265 | 0 | C | G | 0.447 | 7.50E-02 | 9.45E-02 | 2.38E+00 | 2.44E+00 | 4.55E-01 | no | | intergenic_variant | MODIFIER |
| 18 | 54093437 | AX-97793175 | 0 | A | G | 0.447 | 7.50E-02 | 9.45E-02 | 2.38E+00 | 2.44E+00 | 4.55E-01 | no | | regulatory_region_variant | MODIFIER |
| 18 | 54098394 | AX-170015726 | 0 | G | C | 0.447 | 7.50E-02 | 9.45E-02 | 2.38E+00 | 2.44E+00 | 4.55E-01 | no | | intergenic_variant | MODIFIER |
| 18 | 54146708 | AX-170331761 | 0 | C | T | 0.447 | 7.50E-02 | 9.45E-02 | 2.38E+00 | 2.44E+00 | 4.55E-01 | no | | intergenic_variant | MODIFIER |
| 18 | 54172522 | AX-170392123 | 0 | G | A | 0.447 | 7.50E-02 | 9.45E-02 | 2.38E+00 | 2.44E+00 | 4.55E-01 | no | | regulatory_region_variant | MODIFIER |

| | | | | | | | | | | | | | | | |
|---|---|---|---|---|---|---|---|---|---|---|---|---|---|---|---|
| 18 | 54175783 | AX-170251404 | 0 | T | G | 0.447 | 7.50E-02 | 9.45E-02 | 2.38E+00 | 2.44E+00 | 4.55E-01 | no | | regulatory_region_variant | MODIFIER |
| 18 | 54235397 | AX-170158486 | 0 | T | C | 0.447 | 7.50E-02 | 9.45E-02 | 2.38E+00 | 2.44E+00 | 4.55E-01 | no | | regulatory_region_variant | MODIFIER |
| 18 | 54408559 | AX-170262609 | 0 | C | A | 0.447 | 7.50E-02 | 9.45E-02 | 2.38E+00 | 2.44E+00 | 4.55E-01 | no | | intergenic_variant | MODIFIER |
| 18 | 54517926 | AX-170440789 | 0 | T | A | 0.447 | 7.50E-02 | 9.45E-02 | 2.38E+00 | 2.44E+00 | 4.55E-01 | no | | intergenic_variant | MODIFIER |
| 18 | 54561888 | AX-97838800 | 0 | C | T | 0.447 | 7.50E-02 | 9.45E-02 | 2.38E+00 | 2.44E+00 | 4.55E-01 | no | | intergenic_variant | MODIFIER |
| 18 | 54614841 | AX-170490852 | 0 | C | A | 0.447 | 7.50E-02 | 9.45E-02 | 2.38E+00 | 2.44E+00 | 4.55E-01 | no | AC115119.1 | intron_variant,non_coding_transcript_variant | MODIFIER |
| 18 | 54614884 | AX-170015732 | 0 | T | C | 0.447 | 7.50E-02 | 9.45E-02 | 2.38E+00 | 2.44E+00 | 4.55E-01 | no | AC115119.1 | intron_variant,non_coding_transcript_variant | MODIFIER |
| 18 | 54639635 | AX-170093891 | 0 | T | C | 0.447 | 7.50E-02 | 9.45E-02 | 2.38E+00 | 2.44E+00 | 4.55E-01 | no | AC115119.1 | non_coding_transcript_exon_variant | MODIFIER |
| 18 | 54666381 | AX-170015735 | 0 | G | A | 0.447 | 7.50E-02 | 9.45E-02 | 2.38E+00 | 2.44E+00 | 4.55E-01 | no | | intergenic_variant | MODIFIER |
| 18 | 54666608 | AX-170088344 | 0 | C | T | 0.447 | 7.50E-02 | 9.45E-02 | 2.38E+00 | 2.44E+00 | 4.55E-01 | no | | intergenic_variant | MODIFIER |
| 18 | 54686251 | AX-170015731 | 0 | G | A | 0.447 | 7.50E-02 | 9.45E-02 | 2.38E+00 | 2.44E+00 | 4.55E-01 | no | | regulatory_region_variant | MODIFIER |
| 18 | 54688848 | AX-170149689 | 0 | G | C | 0.447 | 7.50E-02 | 9.45E-02 | 2.38E+00 | 2.44E+00 | 4.55E-01 | no | | intergenic_variant | MODIFIER |
| 18 | 54690698 | AX-97851569 | 0 | T | C | 0.447 | 7.50E-02 | 9.45E-02 | 2.38E+00 | 2.44E+00 | 4.55E-01 | no | | intergenic_variant | MODIFIER |
| 18 | 54722464 | AX-170257090 | 0 | C | A | 0.447 | 7.50E-02 | 9.45E-02 | 2.38E+00 | 2.44E+00 | 4.55E-01 | no | | regulatory_region_variant | MODIFIER |
| 18 | 54744985 | AX-170015730 | 0 | G | T | 0.447 | 7.50E-02 | 9.45E-02 | 2.38E+00 | 2.44E+00 | 4.55E-01 | no | | intergenic_variant | MODIFIER |
| 18 | 54746135 | AX-170015734 | 0 | A | G | 0.447 | 7.50E-02 | 9.45E-02 | 2.38E+00 | 2.44E+00 | 4.55E-01 | no | | intergenic_variant | MODIFIER |
| 18 | 54754214 | AX-97835523 | 0 | G | A | 0.447 | 7.50E-02 | 9.45E-02 | 2.38E+00 | 2.44E+00 | 4.55E-01 | no | | regulatory_region_variant | MODIFIER |
| 18 | 54765288 | AX-170015728 | 0 | A | G | 0.447 | 7.50E-02 | 9.45E-02 | 2.38E+00 | 2.44E+00 | 4.55E-01 | no | AC129179.3 | upstream_gene_variant | MODIFIER |
| 18 | 54768414 | AX-170129260 | 0 | G | A | 0.447 | 7.50E-02 | 9.45E-02 | 2.38E+00 | 2.44E+00 | 4.55E-01 | no | AC129179.3 | downstream_gene_variant | MODIFIER |
| 18 | 54781010 | AX-97803298 | 0 | G | A | 0.447 | 7.50E-02 | 9.45E-02 | 2.38E+00 | 2.44E+00 | 4.55E-01 | no | AC129179.2 | intron_variant,non_coding_transcript_variant | MODIFIER |
| 18 | 54796884 | AX-170310243 | 0 | C | A | 0.447 | 7.50E-02 | 9.45E-02 | 2.38E+00 | 2.44E+00 | 4.55E-01 | no | AC129179.2 | intron_variant,non_coding_transcript_variant | MODIFIER |
| 18 | 54799700 | AX-170407587 | 0 | T | C | 0.447 | 7.50E-02 | 9.45E-02 | 2.38E+00 | 2.44E+00 | 4.55E-01 | no | AC129179.2 | intron_variant,non_coding_transcript_variant | MODIFIER |
| 18 | 54800315 | AX-97849378 | 0 | G | A | 0.447 | 7.50E-02 | 9.45E-02 | 2.38E+00 | 2.44E+00 | 4.55E-01 | no | AC129179.2 | intron_variant,non_coding_transcript_variant | MODIFIER |
| 18 | 54804891 | AX-170060894 | 0 | A | C | 0.447 | 7.50E-02 | 9.45E-02 | 2.38E+00 | 2.44E+00 | 4.55E-01 | no | AC129179.2 | intron_variant,non_coding_transcript_variant | MODIFIER |
| 18 | 54807066 | AX-170247082 | 0 | A | G | 0.447 | 7.50E-02 | 9.45E-02 | 2.38E+00 | 2.44E+00 | 4.55E-01 | no | AC129179.2 | intron_variant,non_coding_transcript_variant | MODIFIER |
| 18 | 54816749 | AX-170250090 | 0 | A | G | 0.447 | 7.50E-02 | 9.45E-02 | 2.38E+00 | 2.44E+00 | 4.55E-01 | no | AC129179.2 | intron_variant,non_coding_transcript_variant | MODIFIER |
| 18 | 54822853 | AX-170212913 | 0 | A | G | 0.447 | 7.50E-02 | 9.45E-02 | 2.38E+00 | 2.44E+00 | 4.55E-01 | no | AC129179.2 | non_coding_transcript_exon_variant | MODIFIER |
| 18 | 54841121 | AX-170369954 | 0 | C | T | 0.447 | 7.50E-02 | 9.45E-02 | 2.38E+00 | 2.44E+00 | 4.55E-01 | no | AC129179.1 | downstream_gene_variant | MODIFIER |
| 18 | 54841258 | AX-170217168 | 0 | G | A | 0.447 | 7.50E-02 | 9.45E-02 | 2.38E+00 | 2.44E+00 | 4.55E-01 | no | AC129179.1 | downstream_gene_variant | MODIFIER |
| 18 | 54841416 | AX-97803434 | 0 | A | G | 0.447 | 7.50E-02 | 9.45E-02 | 2.38E+00 | 2.44E+00 | 4.55E-01 | no | AC129179.1 | downstream_gene_variant | MODIFIER |
| 18 | 54841771 | AX-170415592 | 0 | G | A | 0.447 | 7.50E-02 | 9.45E-02 | 2.38E+00 | 2.44E+00 | 4.55E-01 | no | AC129179.1 | downstream_gene_variant | MODIFIER |
| 18 | 54893483 | AX-170116054 | 0 | G | A | 0.447 | 7.50E-02 | 9.45E-02 | 2.38E+00 | 2.44E+00 | 4.55E-01 | no | Zfp608 | intron_variant | MODIFIER |
| 18 | 55018004 | AX-170245117 | 0 | A | C | 0.447 | 7.50E-02 | 9.45E-02 | 2.38E+00 | 2.44E+00 | 4.55E-01 | no | Gm4221 | intron_variant,non_coding_transcript_variant | MODIFIER |
| 18 | 55097846 | AX-170015743 | 0 | G | A | 0.447 | 7.50E-02 | 9.45E-02 | 2.38E+00 | 2.44E+00 | 4.55E-01 | no | AC163347.1 | intron_variant,non_coding_transcript_variant | MODIFIER |
| 18 | 55183937 | AX-170088346 | 0 | T | C | 0.447 | 7.50E-02 | 9.45E-02 | 2.38E+00 | 2.44E+00 | 4.55E-01 | no | Gm22597 | downstream_gene_variant | MODIFIER |
| 18 | 55243391 | AX-97839340 | 0 | A | G | 0.447 | 7.50E-02 | 9.45E-02 | 2.38E+00 | 2.44E+00 | 4.55E-01 | no | | intergenic_variant | MODIFIER |
| 18 | 55243626 | AX-170159520 | 0 | G | A | 0.447 | 7.50E-02 | 9.45E-02 | 2.38E+00 | 2.44E+00 | 4.55E-01 | no | | intergenic_variant | MODIFIER |
| 18 | 55273373 | AX-170329085 | 0 | C | T | 0.447 | 7.50E-02 | 9.45E-02 | 2.38E+00 | 2.44E+00 | 4.55E-01 | no | | intergenic_variant | MODIFIER |
| 18 | 55285184 | AX-97834869 | 0 | G | A | 0.447 | 7.50E-02 | 9.45E-02 | 2.38E+00 | 2.44E+00 | 4.55E-01 | no | | intergenic_variant | MODIFIER |
| 18 | 55285866 | AX-170015738 | 0 | C | T | 0.447 | 7.50E-02 | 9.45E-02 | 2.38E+00 | 2.44E+00 | 4.55E-01 | no | | intergenic_variant | MODIFIER |
| 18 | 55288721 | AX-170117538 | 0 | C | T | 0.447 | 7.50E-02 | 9.45E-02 | 2.38E+00 | 2.44E+00 | 4.55E-01 | no | | regulatory_region_variant | MODIFIER |
| 18 | 55298023 | AX-170060901 | 0 | G | A | 0.447 | 7.50E-02 | 9.45E-02 | 2.38E+00 | 2.44E+00 | 4.55E-01 | no | | intergenic_variant | MODIFIER |
| 18 | 55325243 | AX-170060902 | 0 | A | G | 0.447 | 7.50E-02 | 9.45E-02 | 2.38E+00 | 2.44E+00 | 4.55E-01 | no | | intergenic_variant | MODIFIER |
| 18 | 55325724 | AX-97853481 | 0 | G | T | 0.447 | 7.50E-02 | 9.45E-02 | 2.38E+00 | 2.44E+00 | 4.55E-01 | no | | intergenic_variant | MODIFIER |
| 18 | 55338683 | AX-170015745 | 0 | G | C | 0.447 | 7.50E-02 | 9.45E-02 | 2.38E+00 | 2.44E+00 | 4.55E-01 | no | Gm37828 | upstream_gene_variant | MODIFIER |
| 18 | 55339513 | AX-170243273 | 0 | C | A | 0.447 | 7.50E-02 | 9.45E-02 | 2.38E+00 | 2.44E+00 | 4.55E-01 | no | Gm37828 | non_coding_transcript_exon_variant | MODIFIER |
| 18 | 55341890 | AX-97846046 | 0 | C | T | 0.447 | 7.50E-02 | 9.45E-02 | 2.38E+00 | 2.44E+00 | 4.55E-01 | no | Gm37828 | downstream_gene_variant | MODIFIER |
| 18 | 55344205 | AX-170257947 | 0 | G | A | 0.447 | 7.50E-02 | 9.45E-02 | 2.38E+00 | 2.44E+00 | 4.55E-01 | no | Gm37828 | downstream_gene_variant | MODIFIER |
| 18 | 55409332 | AX-170015740 | 0 | G | T | 0.447 | 7.50E-02 | 9.45E-02 | 2.38E+00 | 2.44E+00 | 4.55E-01 | no | | regulatory_region_variant | MODIFIER |
| 18 | 55444380 | AX-170291115 | 0 | G | T | 0.447 | 7.50E-02 | 9.45E-02 | 2.38E+00 | 2.44E+00 | 4.55E-01 | no | | intergenic_variant | MODIFIER |
| 18 | 55446106 | AX-170273011 | 0 | C | T | 0.447 | 7.50E-02 | 9.45E-02 | 2.38E+00 | 2.44E+00 | 4.55E-01 | no | | intergenic_variant | MODIFIER |
| 18 | 55448762 | AX-97797363 | 0 | C | T | 0.447 | 7.50E-02 | 9.45E-02 | 2.38E+00 | 2.44E+00 | 4.55E-01 | no | | intergenic_variant | MODIFIER |
| 18 | 55451469 | AX-170310146 | 0 | G | A | 0.447 | 7.50E-02 | 9.45E-02 | 2.38E+00 | 2.44E+00 | 4.55E-01 | no | | intergenic_variant | MODIFIER |
| 18 | 55488881 | AX-170477306 | 0 | A | G | 0.447 | 7.50E-02 | 9.45E-02 | 2.38E+00 | 2.44E+00 | 4.55E-01 | no | | intergenic_variant | MODIFIER |

| | | | | | | | | | | | | | | | |
|---|---|---|---|---|---|---|---|---|---|---|---|---|---|---|---|
| 18 | 55508939 | AX-170437839 | 0 | T | G | 0.447 | 7.50E-02 | 9.45E-02 | 2.38E+00 | 2.44E+00 | 4.55E-01 | no | | regulatory_region_variant | MODIFIER |
| 18 | 55532500 | AX-168395042 | 0 | A | G | 0.447 | 7.50E-02 | 9.45E-02 | 2.38E+00 | 2.44E+00 | 4.55E-01 | no | | intergenic_variant | MODIFIER |
| 18 | 55533587 | AX-170450600 | 0 | C | G | 0.447 | 7.50E-02 | 9.45E-02 | 2.38E+00 | 2.44E+00 | 4.55E-01 | no | | intergenic_variant | MODIFIER |
| 18 | 55562344 | AX-97846868 | 0 | G | A | 0.447 | 7.50E-02 | 9.45E-02 | 2.38E+00 | 2.44E+00 | 4.55E-01 | no | | intergenic_variant | MODIFIER |
| 18 | 55582913 | AX-170093894 | 0 | G | A | 0.447 | 7.50E-02 | 9.45E-02 | 2.38E+00 | 2.44E+00 | 4.55E-01 | no | | regulatory_region_variant | MODIFIER |
| 18 | 55588337 | AX-170345689 | 0 | G | A | 0.447 | 7.50E-02 | 9.45E-02 | 2.38E+00 | 2.44E+00 | 4.55E-01 | no | | intergenic_variant | MODIFIER |
| 18 | 55594192 | AX-170309578 | 0 | T | C | 0.447 | 7.50E-02 | 9.45E-02 | 2.38E+00 | 2.44E+00 | 4.55E-01 | no | | intergenic_variant | MODIFIER |
| 18 | 55602128 | AX-97856983 | 0 | T | C | 0.447 | 7.50E-02 | 9.45E-02 | 2.38E+00 | 2.44E+00 | 4.55E-01 | no | | intergenic_variant | MODIFIER |
| 18 | 55604598 | AX-170133126 | 0 | A | C | 0.447 | 7.50E-02 | 9.45E-02 | 2.38E+00 | 2.44E+00 | 4.55E-01 | no | | regulatory_region_variant | MODIFIER |
| 18 | 55617933 | AX-170315544 | 0 | C | T | 0.447 | 7.50E-02 | 9.45E-02 | 2.38E+00 | 2.44E+00 | 4.55E-01 | no | Gm24514 | downstream_gene_variant | MODIFIER |
| 18 | 55617982 | AX-170060907 | 0 | A | C | 0.447 | 7.50E-02 | 9.45E-02 | 2.38E+00 | 2.44E+00 | 4.55E-01 | no | Gm24514 | downstream_gene_variant | MODIFIER |
| 18 | 55620073 | AX-170246366 | 0 | A | G | 0.447 | 7.50E-02 | 9.45E-02 | 2.38E+00 | 2.44E+00 | 4.55E-01 | no | Gm24514 | downstream_gene_variant | MODIFIER |
| 18 | 55622626 | AX-170186633 | 0 | G | A | 0.447 | 7.50E-02 | 9.45E-02 | 2.38E+00 | 2.44E+00 | 4.55E-01 | no | Gm24514 | downstream_gene_variant | MODIFIER |
| 18 | 55626782 | AX-97848887 | 0 | C | A | 0.447 | 7.50E-02 | 9.45E-02 | 2.38E+00 | 2.44E+00 | 4.55E-01 | no | Gm24514 | upstream_gene_variant | MODIFIER |
| 18 | 55645120 | AX-170530353 | 0 | A | G | 0.447 | 7.50E-02 | 9.45E-02 | 2.38E+00 | 2.44E+00 | 4.55E-01 | no | | intergenic_variant | MODIFIER |
| 18 | 55653649 | AX-170015749 | 0 | G | A | 0.447 | 7.50E-02 | 9.45E-02 | 2.38E+00 | 2.44E+00 | 4.55E-01 | no | | intergenic_variant | MODIFIER |
| 18 | 55669231 | AX-170252733 | 0 | T | C | 0.447 | 7.50E-02 | 9.45E-02 | 2.38E+00 | 2.44E+00 | 4.55E-01 | no | | intergenic_variant | MODIFIER |
| 18 | 55676118 | AX-170359602 | 0 | G | A | 0.447 | 7.50E-02 | 9.45E-02 | 2.38E+00 | 2.44E+00 | 4.55E-01 | no | | regulatory_region_variant | MODIFIER |
| 18 | 55700784 | AX-97824863 | 0 | T | C | 0.447 | 7.50E-02 | 9.45E-02 | 2.38E+00 | 2.44E+00 | 4.55E-01 | no | | intergenic_variant | MODIFIER |
| 6 | 134402675 | AX-170274139 | 1 | G | A | 0.571 | 6.64E-02 | 8.71E-02 | 2.50E+00 | 2.55E+00 | 4.55E-01 | no | Bcl2l14 | intron_variant | MODIFIER |
| 7 | 115326081 | AX-170191621 | 1 | A | G | 0.42 | 7.21E-02 | 9.74E-02 | 2.70E+00 | 2.74E+00 | 4.55E-01 | no | | intergenic_variant | MODIFIER |
| 1 | 176788582 | AX-170421692 | 0 | C | T | 0.228 | 8.48E-02 | 1.13E-01 | 2.58E+00 | 2.66E+00 | 4.55E-01 | no | Cep170 | intron_variant | MODIFIER |
| 18 | 84872456 | AX-170287487 | 1 | T | C | 0.17 | -7.58E-02 | 9.97E-02 | 2.50E+00 | 2.59E+00 | 4.55E-01 | no | Cyb5a | intron_variant,NMD_transcript_variant | MODIFIER |
| 16 | 9212865 | AX-170484335 | 0 | C | G | 0.061 | 1.15E-01 | 1.58E-01 | 2.73E+00 | 2.89E+00 | 4.55E-01 | no | | intergenic_variant | MODIFIER |
| 4 | 114649999 | AX-170120303 | 1 | G | A | 0.232 | 8.12E-02 | 1.09E-01 | 2.61E+00 | 2.67E+00 | 4.55E-01 | no | | intergenic_variant | MODIFIER |
| 9 | 103634525 | AX-170033174 | 0 | T | C | 0.123 | -8.64E-02 | 1.15E-01 | 2.54E+00 | 2.62E+00 | 4.55E-01 | no | Tmem108 | intron_variant | MODIFIER |
| 9 | 103660428 | AX-170083318 | 0 | G | T | 0.123 | -8.64E-02 | 1.15E-01 | 2.54E+00 | 2.62E+00 | 4.55E-01 | no | Tmem108 | intron_variant | MODIFIER |
| 9 | 103677877 | AX-170083319 | 0 | A | G | 0.123 | -8.64E-02 | 1.15E-01 | 2.54E+00 | 2.62E+00 | 4.55E-01 | no | Tmem108 | intron_variant | MODIFIER |
| 9 | 103677923 | AX-170083320 | 0 | G | T | 0.123 | -8.64E-02 | 1.15E-01 | 2.54E+00 | 2.62E+00 | 4.55E-01 | no | Tmem108 | intron_variant | MODIFIER |
| 9 | 103686134 | AX-170331003 | 0 | A | G | 0.123 | -8.64E-02 | 1.15E-01 | 2.54E+00 | 2.62E+00 | 4.55E-01 | no | Tmem108 | intron_variant | MODIFIER |
| 9 | 103708701 | AX-170483093 | 0 | G | A | 0.123 | -8.64E-02 | 1.15E-01 | 2.54E+00 | 2.62E+00 | 4.55E-01 | no | Tmem108 | intron_variant | MODIFIER |
| 9 | 103715124 | AX-170394424 | 0 | A | G | 0.123 | -8.64E-02 | 1.15E-01 | 2.54E+00 | 2.62E+00 | 4.55E-01 | no | Tmem108 | intron_variant | MODIFIER |
| 9 | 103715158 | AX-97804230 | 0 | C | T | 0.123 | -8.64E-02 | 1.15E-01 | 2.54E+00 | 2.62E+00 | 4.55E-01 | no | Tmem108 | intron_variant | MODIFIER |
| 9 | 103715209 | AX-170328037 | 0 | C | G | 0.123 | -8.64E-02 | 1.15E-01 | 2.54E+00 | 2.62E+00 | 4.55E-01 | no | Tmem108 | intron_variant | MODIFIER |
| 9 | 103718499 | AX-170225610 | 0 | C | T | 0.123 | -8.64E-02 | 1.15E-01 | 2.54E+00 | 2.62E+00 | 4.55E-01 | no | Tmem108 | intron_variant | MODIFIER |
| 9 | 103721315 | AX-170493354 | 0 | A | G | 0.123 | -8.64E-02 | 1.15E-01 | 2.54E+00 | 2.62E+00 | 4.55E-01 | no | Tmem108 | intron_variant | MODIFIER |
| 9 | 103722021 | AX-170485040 | 0 | A | G | 0.123 | -8.64E-02 | 1.15E-01 | 2.54E+00 | 2.62E+00 | 4.55E-01 | no | Tmem108 | intron_variant | MODIFIER |
| 9 | 103723786 | AX-170352099 | 0 | C | T | 0.123 | -8.64E-02 | 1.15E-01 | 2.54E+00 | 2.62E+00 | 4.55E-01 | no | Tmem108 | intron_variant | MODIFIER |
| 9 | 103732775 | AX-170425544 | 0 | G | A | 0.123 | -8.64E-02 | 1.15E-01 | 2.54E+00 | 2.62E+00 | 4.55E-01 | no | Tmem108 | intron_variant | MODIFIER |
| 9 | 103732807 | AX-170329311 | 0 | G | T | 0.123 | -8.64E-02 | 1.15E-01 | 2.54E+00 | 2.62E+00 | 4.55E-01 | no | Tmem108 | intron_variant | MODIFIER |
| 9 | 103737226 | AX-170387237 | 0 | T | C | 0.123 | -8.64E-02 | 1.15E-01 | 2.54E+00 | 2.62E+00 | 4.55E-01 | no | Tmem108 | intron_variant | MODIFIER |
| 9 | 103868113 | AX-170282357 | 0 | C | T | 0.123 | -8.64E-02 | 1.15E-01 | 2.54E+00 | 2.62E+00 | 4.55E-01 | no | | intergenic_variant | MODIFIER |
| 9 | 103868499 | AX-170402640 | 0 | G | A | 0.123 | -8.64E-02 | 1.15E-01 | 2.54E+00 | 2.62E+00 | 4.55E-01 | no | | intergenic_variant | MODIFIER |
| 9 | 103883199 | AX-170431873 | 0 | C | T | 0.123 | -8.64E-02 | 1.15E-01 | 2.54E+00 | 2.62E+00 | 4.55E-01 | no | Gm5372 | upstream_gene_variant | MODIFIER |
| 9 | 103903884 | AX-170179601 | 0 | A | G | 0.123 | -8.64E-02 | 1.15E-01 | 2.54E+00 | 2.62E+00 | 4.55E-01 | no | Gm29154 | intron_variant,non_coding_transcript_variant | MODIFIER |
| 9 | 103930101 | AX-170169036 | 0 | C | A | 0.123 | -8.64E-02 | 1.15E-01 | 2.54E+00 | 2.62E+00 | 4.55E-01 | no | Gm29154 | intron_variant,non_coding_transcript_variant | MODIFIER |
| 2 | 166606647 | AX-170393767 | 1 | C | G | 0.152 | 1.32E-01 | 1.80E-01 | 2.75E+00 | 2.84E+00 | 4.55E-01 | no | Prex1 | intron_variant | MODIFIER |
| 15 | 9197358 | AX-170087761 | 0 | T | C | 0.237 | -7.56E-02 | 1.02E-01 | 2.62E+00 | 2.69E+00 | 4.55E-01 | no | Lmbrd2 | 3_prime_UTR_variant | MODIFIER |
| 6 | 134469982 | AX-170095814 | 1 | G | A | 0.571 | 6.61E-02 | 8.70E-02 | 2.50E+00 | 2.56E+00 | 4.56E-01 | no | Lrp6 | intron_variant | MODIFIER |
| 10 | 67850083 | AX-170333324 | 1 | G | C | 0.411 | -5.94E-02 | 8.01E-02 | 2.64E+00 | 2.71E+00 | 4.56E-01 | no | | intergenic_variant | MODIFIER |
| 18 | 39678326 | AX-97848348 | 1 | G | A | 0.062 | -1.55E-01 | 2.09E-01 | 2.62E+00 | 2.73E+00 | 4.56E-01 | no | AC124518.1 | intron_variant,non_coding_transcript_variant | MODIFIER |
| 7 | 74050971 | AX-170456676 | 0 | G | A | 0.386 | -7.36E-02 | 1.00E-01 | 2.78E+00 | 2.88E+00 | 4.56E-01 | no | | intergenic_variant | MODIFIER |
| 7 | 78111302 | AX-170029365 | 0 | T | C | 0.386 | -7.36E-02 | 1.00E-01 | 2.78E+00 | 2.88E+00 | 4.56E-01 | no | | intergenic_variant | MODIFIER |
| 5 | 93732650 | AX-170025627 | 0 | C | G | 0.43 | 9.47E-02 | 1.26E-01 | 2.51E+00 | 2.66E+00 | 4.56E-01 | no | | intergenic_variant | MODIFIER |

| | | | | | | | | | | | | | | | |
|---|---|---|---|---|---|---|---|---|---|---|---|---|---|---|---|
| 9 | 66210551 | AX-170330439 | 0 | C | T | 0.202 | 9.63E-02 | 1.29E-01 | 2.88E+00 | 2.94E+00 | 4.56E-01 | no | Dapk2 | intron_variant | MODIFIER |
| 9 | 17179401 | AX-170090250 | 0 | T | C | 0.482 | 6.61E-02 | 8.81E-02 | 2.55E+00 | 2.62E+00 | 4.56E-01 | no | | intergenic_variant | MODIFIER |
| 10 | 92739412 | AX-170520266 | 1 | G | A | 0.295 | -6.36E-02 | 8.40E-02 | 2.52E+00 | 2.58E+00 | 4.56E-01 | no | | intergenic_variant | MODIFIER |
| 10 | 93383395 | AX-170224859 | 1 | C | T | 0.295 | -6.36E-02 | 8.40E-02 | 2.52E+00 | 2.58E+00 | 4.56E-01 | no | | intergenic_variant | MODIFIER |
| 10 | 93543246 | AX-170086867 | 1 | A | G | 0.295 | -6.36E-02 | 8.40E-02 | 2.52E+00 | 2.58E+00 | 4.56E-01 | no | Amdhd1 | upstream_gene_variant | MODIFIER |
| 1 | 144227929 | AX-170266430 | 1 | T | C | 0.214 | 9.00E-02 | 1.21E-01 | 2.61E+00 | 2.67E+00 | 4.56E-01 | no | Rbm6-ps2 | upstream_gene_variant | MODIFIER |
| 10 | 93734307 | AX-170476274 | 1 | A | C | 0.286 | -6.70E-02 | 8.78E-02 | 2.49E+00 | 2.55E+00 | 4.56E-01 | no | Ntn4 | intron_variant | MODIFIER |
| 2 | 37749258 | AX-170299503 | 0 | A | G | 0.035 | -1.50E-01 | 2.03E-01 | 2.63E+00 | 2.73E+00 | 4.56E-01 | no | | regulatory_region_variant | MODIFIER |
| 5 | 145218118 | AX-170140126 | 0 | T | C | 0.035 | -1.50E-01 | 2.03E-01 | 2.63E+00 | 2.73E+00 | 4.56E-01 | no | Zkscan5 | intron_variant | MODIFIER |
| 11 | 99676093 | AX-170411259 | 0 | C | T | 0.035 | -1.50E-01 | 2.03E-01 | 2.63E+00 | 2.73E+00 | 4.56E-01 | no | Gm11557 | upstream_gene_variant | MODIFIER |
| 1 | 101346647 | AX-170253370 | 2 | A | C | 0.145 | 1.55E-01 | 2.10E-01 | 2.72E+00 | 2.80E+00 | 4.56E-01 | Yes | | intergenic_variant | MODIFIER |
| 13 | 3801748 | AX-97856281 | 1 | A | G | 0.205 | -9.30E-02 | 1.24E-01 | 2.56E+00 | 2.62E+00 | 4.56E-01 | no | Calml3 | downstream_gene_variant | MODIFIER |
| 7 | 98566259 | AX-170404115 | 1 | A | G | 0.125 | -1.19E-01 | 1.61E-01 | 2.62E+00 | 2.72E+00 | 4.56E-01 | no | A630091E08Rik | upstream_gene_variant | MODIFIER |
| 18 | 55279153 | AX-170122622 | 1 | A | G | 0.446 | 7.46E-02 | 9.46E-02 | 2.39E+00 | 2.45E+00 | 4.56E-01 | no | | intergenic_variant | MODIFIER |
| 6 | 134002475 | AX-170482932 | 1 | A | G | 0.571 | 6.62E-02 | 8.72E-02 | 2.50E+00 | 2.55E+00 | 4.56E-01 | no | | intergenic_variant | MODIFIER |
| 4 | 129826611 | AX-170503675 | 0 | T | G | 0.167 | 8.56E-02 | 1.13E-01 | 2.52E+00 | 2.57E+00 | 4.56E-01 | no | Ptp4a2 | intron_variant | MODIFIER |
| 15 | 83233939 | AX-97837672 | 1 | A | G | 0.098 | -9.81E-02 | 1.04E-01 | 2.06E+00 | 2.17E+00 | 4.56E-01 | no | A4galt | intron_variant | MODIFIER |
| 7 | 73304020 | AX-170276448 | 0 | A | C | 0.377 | -7.32E-02 | 9.97E-02 | 2.75E+00 | 2.85E+00 | 4.56E-01 | no | | regulatory_region_variant | MODIFIER |
| 7 | 73310865 | AX-170384441 | 0 | C | T | 0.377 | -7.32E-02 | 9.97E-02 | 2.75E+00 | 2.85E+00 | 4.56E-01 | no | A730056A06Rik | downstream_gene_variant | MODIFIER |
| 7 | 73326892 | AX-170282231 | 0 | C | T | 0.377 | -7.32E-02 | 9.97E-02 | 2.75E+00 | 2.85E+00 | 4.56E-01 | no | A730056A06Rik | intron_variant,non_coding_transcript_variant | MODIFIER |
| 7 | 73343171 | AX-170496031 | 0 | G | A | 0.377 | -7.32E-02 | 9.97E-02 | 2.75E+00 | 2.85E+00 | 4.56E-01 | no | A730056A06Rik | intron_variant,non_coding_transcript_variant | MODIFIER |
| 7 | 73361372 | AX-170337843 | 0 | T | G | 0.377 | -7.32E-02 | 9.97E-02 | 2.75E+00 | 2.85E+00 | 4.56E-01 | no | A730056A06Rik | intron_variant,non_coding_transcript_variant | MODIFIER |
| 7 | 73507603 | AX-170465974 | 0 | G | A | 0.377 | -7.32E-02 | 9.97E-02 | 2.75E+00 | 2.85E+00 | 4.56E-01 | no | Chd2 | intron_variant | MODIFIER |
| 7 | 73746984 | AX-170029296 | 0 | A | G | 0.377 | -7.32E-02 | 9.97E-02 | 2.75E+00 | 2.85E+00 | 4.56E-01 | no | Fam174b | intron_variant | MODIFIER |
| 7 | 73747368 | AX-97811919 | 0 | G | A | 0.377 | -7.32E-02 | 9.97E-02 | 2.75E+00 | 2.85E+00 | 4.56E-01 | no | Fam174b | intron_variant | MODIFIER |
| 7 | 73747397 | AX-170029290 | 0 | A | C | 0.377 | -7.32E-02 | 9.97E-02 | 2.75E+00 | 2.85E+00 | 4.56E-01 | no | Fam174b | intron_variant | MODIFIER |
| 7 | 73829194 | AX-170077726 | 0 | C | T | 0.377 | -7.32E-02 | 9.97E-02 | 2.75E+00 | 2.85E+00 | 4.56E-01 | no | Gm21269 | intron_variant,non_coding_transcript_variant | MODIFIER |
| 7 | 73833935 | AX-170281910 | 0 | C | T | 0.377 | -7.32E-02 | 9.97E-02 | 2.75E+00 | 2.85E+00 | 4.56E-01 | no | Gm21269 | intron_variant,non_coding_transcript_variant | MODIFIER |
| 7 | 73886706 | AX-97853076 | 0 | A | G | 0.377 | -7.32E-02 | 9.97E-02 | 2.75E+00 | 2.85E+00 | 4.56E-01 | no | | intergenic_variant | MODIFIER |
| 7 | 73942263 | AX-170425244 | 0 | T | C | 0.377 | -7.32E-02 | 9.97E-02 | 2.75E+00 | 2.85E+00 | 4.56E-01 | no | St8sia2 | 3_prime_UTR_variant | MODIFIER |
| 7 | 73942631 | AX-170346982 | 0 | A | G | 0.377 | -7.32E-02 | 9.97E-02 | 2.75E+00 | 2.85E+00 | 4.56E-01 | no | St8sia2 | 3_prime_UTR_variant | MODIFIER |
| 7 | 73942680 | AX-170479533 | 0 | C | T | 0.377 | -7.32E-02 | 9.97E-02 | 2.75E+00 | 2.85E+00 | 4.56E-01 | no | St8sia2 | 3_prime_UTR_variant | MODIFIER |
| 7 | 73945762 | AX-170265310 | 0 | C | A | 0.377 | -7.32E-02 | 9.97E-02 | 2.75E+00 | 2.85E+00 | 4.56E-01 | no | St8sia2 | intron_variant | MODIFIER |
| 7 | 74029104 | AX-170401096 | 0 | G | A | 0.377 | -7.32E-02 | 9.97E-02 | 2.75E+00 | 2.85E+00 | 4.56E-01 | no | | intergenic_variant | MODIFIER |
| 7 | 74029545 | AX-170386596 | 0 | G | A | 0.377 | -7.32E-02 | 9.97E-02 | 2.75E+00 | 2.85E+00 | 4.56E-01 | no | | intergenic_variant | MODIFIER |
| 7 | 74046134 | AX-170212427 | 0 | C | T | 0.377 | -7.32E-02 | 9.97E-02 | 2.75E+00 | 2.85E+00 | 4.56E-01 | no | Gm45004 | upstream_gene_variant | MODIFIER |
| 7 | 74070996 | AX-170382563 | 0 | G | A | 0.377 | -7.32E-02 | 9.97E-02 | 2.75E+00 | 2.85E+00 | 4.56E-01 | no | | intergenic_variant | MODIFIER |
| 7 | 74072707 | AX-170465975 | 0 | G | A | 0.377 | -7.32E-02 | 9.97E-02 | 2.75E+00 | 2.85E+00 | 4.56E-01 | no | | intergenic_variant | MODIFIER |
| 7 | 74076763 | AX-170448331 | 0 | G | A | 0.377 | -7.32E-02 | 9.97E-02 | 2.75E+00 | 2.85E+00 | 4.56E-01 | no | | intergenic_variant | MODIFIER |
| 7 | 74077503 | AX-170283339 | 0 | G | T | 0.377 | -7.32E-02 | 9.97E-02 | 2.75E+00 | 2.85E+00 | 4.56E-01 | no | | intergenic_variant | MODIFIER |
| 7 | 74111251 | AX-170107719 | 0 | G | A | 0.377 | -7.32E-02 | 9.97E-02 | 2.75E+00 | 2.85E+00 | 4.56E-01 | no | | intergenic_variant | MODIFIER |
| 7 | 74114669 | AX-170504289 | 0 | C | T | 0.377 | -7.32E-02 | 9.97E-02 | 2.75E+00 | 2.85E+00 | 4.56E-01 | no | | intergenic_variant | MODIFIER |
| 7 | 74148002 | AX-170410562 | 0 | A | C | 0.377 | -7.32E-02 | 9.97E-02 | 2.75E+00 | 2.85E+00 | 4.56E-01 | no | | intergenic_variant | MODIFIER |
| 7 | 74157699 | AX-170116691 | 0 | C | T | 0.377 | -7.32E-02 | 9.97E-02 | 2.75E+00 | 2.85E+00 | 4.56E-01 | no | | intergenic_variant | MODIFIER |
| 7 | 74226542 | AX-97785740 | 0 | T | G | 0.377 | -7.32E-02 | 9.97E-02 | 2.75E+00 | 2.85E+00 | 4.56E-01 | no | | intergenic_variant | MODIFIER |
| 7 | 74248842 | AX-170187192 | 0 | C | G | 0.377 | -7.32E-02 | 9.97E-02 | 2.75E+00 | 2.85E+00 | 4.56E-01 | no | | intergenic_variant | MODIFIER |
| 7 | 74250452 | AX-170423095 | 0 | A | G | 0.377 | -7.32E-02 | 9.97E-02 | 2.75E+00 | 2.85E+00 | 4.56E-01 | no | | intergenic_variant | MODIFIER |
| 7 | 74250616 | AX-170363996 | 0 | T | C | 0.377 | -7.32E-02 | 9.97E-02 | 2.75E+00 | 2.85E+00 | 4.56E-01 | no | | intergenic_variant | MODIFIER |
| 7 | 74256428 | AX-170356224 | 0 | A | C | 0.377 | -7.32E-02 | 9.97E-02 | 2.75E+00 | 2.85E+00 | 4.56E-01 | no | | intergenic_variant | MODIFIER |
| 7 | 74267766 | AX-97827199 | 0 | T | C | 0.377 | -7.32E-02 | 9.97E-02 | 2.75E+00 | 2.85E+00 | 4.56E-01 | no | | intergenic_variant | MODIFIER |
| 7 | 74421458 | AX-170354077 | 0 | A | G | 0.377 | -7.32E-02 | 9.97E-02 | 2.75E+00 | 2.85E+00 | 4.56E-01 | no | Slco3a1 | intron_variant | MODIFIER |
| 7 | 74432364 | AX-170029300 | 0 | T | C | 0.377 | -7.32E-02 | 9.97E-02 | 2.75E+00 | 2.85E+00 | 4.56E-01 | no | Slco3a1 | intron_variant | MODIFIER |
| 7 | 74444454 | AX-170128085 | 0 | A | G | 0.377 | -7.32E-02 | 9.97E-02 | 2.75E+00 | 2.85E+00 | 4.56E-01 | no | Slco3a1 | intron_variant | MODIFIER |
| 7 | 74458399 | AX-170468958 | 0 | A | C | 0.377 | -7.32E-02 | 9.97E-02 | 2.75E+00 | 2.85E+00 | 4.56E-01 | no | Slco3a1 | intron_variant | MODIFIER |

| | | | | | | | | | | | | | | | |
|---|---|---|---|---|---|---|---|---|---|---|---|---|---|---|---|
| 7 | 74524456 | AX-170397785 | 0 | C | A | 0.377 | -7.32E-02 | 9.97E-02 | 2.75E+00 | 2.85E+00 | 4.56E-01 | no | Slco3a1 | intron_variant | MODIFIER |
| 7 | 74528120 | AX-170388624 | 0 | G | A | 0.377 | -7.32E-02 | 9.97E-02 | 2.75E+00 | 2.85E+00 | 4.56E-01 | no | Slco3a1 | intron_variant | MODIFIER |
| 7 | 74534452 | AX-170251078 | 0 | C | T | 0.377 | -7.32E-02 | 9.97E-02 | 2.75E+00 | 2.85E+00 | 4.56E-01 | no | Slco3a1 | intron_variant | MODIFIER |
| 7 | 74538683 | AX-170417020 | 0 | C | A | 0.377 | -7.32E-02 | 9.97E-02 | 2.75E+00 | 2.85E+00 | 4.56E-01 | no | Slco3a1 | intron_variant | MODIFIER |
| 7 | 74788642 | AX-170136711 | 0 | A | T | 0.377 | -7.32E-02 | 9.97E-02 | 2.75E+00 | 2.85E+00 | 4.56E-01 | no | | intergenic_variant | MODIFIER |
| 7 | 74986014 | AX-97856108 | 0 | G | A | 0.377 | -7.32E-02 | 9.97E-02 | 2.75E+00 | 2.85E+00 | 4.56E-01 | no | | intergenic_variant | MODIFIER |
| 7 | 75215247 | AX-170463539 | 0 | C | T | 0.377 | -7.32E-02 | 9.97E-02 | 2.75E+00 | 2.85E+00 | 4.56E-01 | no | Sv2b | intron_variant | MODIFIER |
| 7 | 75301035 | AX-170029321 | 0 | G | A | 0.377 | -7.32E-02 | 9.97E-02 | 2.75E+00 | 2.85E+00 | 4.56E-01 | no | Sv2b | intron_variant | MODIFIER |
| 7 | 75442499 | AX-170514024 | 0 | G | A | 0.377 | -7.32E-02 | 9.97E-02 | 2.75E+00 | 2.85E+00 | 4.56E-01 | no | | intergenic_variant | MODIFIER |
| 7 | 75855213 | AX-170366445 | 0 | T | C | 0.377 | -7.32E-02 | 9.97E-02 | 2.75E+00 | 2.85E+00 | 4.56E-01 | no | Klhl25 | intron_variant | MODIFIER |
| 7 | 76294233 | AX-170424505 | 0 | T | C | 0.377 | -7.32E-02 | 9.97E-02 | 2.75E+00 | 2.85E+00 | 4.56E-01 | no | Agbl1 | intron_variant | MODIFIER |
| 7 | 76294276 | AX-170413662 | 0 | G | A | 0.377 | -7.32E-02 | 9.97E-02 | 2.75E+00 | 2.85E+00 | 4.56E-01 | no | Agbl1 | intron_variant | MODIFIER |
| 7 | 76381786 | AX-170238810 | 0 | T | A | 0.377 | -7.32E-02 | 9.97E-02 | 2.75E+00 | 2.85E+00 | 4.56E-01 | no | Agbl1 | intron_variant | MODIFIER |
| 7 | 76383548 | AX-170372106 | 0 | G | A | 0.377 | -7.32E-02 | 9.97E-02 | 2.75E+00 | 2.85E+00 | 4.56E-01 | no | Agbl1 | intron_variant | MODIFIER |
| 7 | 76385665 | AX-170509757 | 0 | C | T | 0.377 | -7.32E-02 | 9.97E-02 | 2.75E+00 | 2.85E+00 | 4.56E-01 | no | Agbl1 | intron_variant | MODIFIER |
| 7 | 76385682 | AX-170329091 | 0 | T | G | 0.377 | -7.32E-02 | 9.97E-02 | 2.75E+00 | 2.85E+00 | 4.56E-01 | no | Agbl1 | intron_variant | MODIFIER |
| 7 | 76386930 | AX-170470297 | 0 | T | C | 0.377 | -7.32E-02 | 9.97E-02 | 2.75E+00 | 2.85E+00 | 4.56E-01 | no | Agbl1 | intron_variant | MODIFIER |
| 7 | 76390729 | AX-170447423 | 0 | T | C | 0.377 | -7.32E-02 | 9.97E-02 | 2.75E+00 | 2.85E+00 | 4.56E-01 | no | Agbl1 | intron_variant | MODIFIER |
| 7 | 76420066 | AX-170466690 | 0 | A | G | 0.377 | -7.32E-02 | 9.97E-02 | 2.75E+00 | 2.85E+00 | 4.56E-01 | no | Agbl1 | missense_variant | MODERATE |
| 7 | 76440298 | AX-170403799 | 0 | C | T | 0.377 | -7.32E-02 | 9.97E-02 | 2.75E+00 | 2.85E+00 | 4.56E-01 | no | Agbl1 | intron_variant | MODIFIER |
| 7 | 77129120 | AX-170077786 | 0 | A | G | 0.377 | -7.32E-02 | 9.97E-02 | 2.75E+00 | 2.85E+00 | 4.56E-01 | no | Agbl1 | downstream_gene_variant | MODIFIER |
| 7 | 77154204 | AX-170315124 | 0 | T | C | 0.377 | -7.32E-02 | 9.97E-02 | 2.75E+00 | 2.85E+00 | 4.56E-01 | no | | intergenic_variant | MODIFIER |
| 7 | 77183569 | AX-170108011 | 0 | G | A | 0.377 | -7.32E-02 | 9.97E-02 | 2.75E+00 | 2.85E+00 | 4.56E-01 | no | | intergenic_variant | MODIFIER |
| 7 | 77241895 | AX-170109595 | 0 | A | G | 0.377 | -7.32E-02 | 9.97E-02 | 2.75E+00 | 2.85E+00 | 4.56E-01 | no | | regulatory_region_variant | MODIFIER |
| 7 | 77272137 | AX-170507596 | 0 | C | G | 0.377 | -7.32E-02 | 9.97E-02 | 2.75E+00 | 2.85E+00 | 4.56E-01 | no | | intergenic_variant | MODIFIER |
| 7 | 77280721 | AX-170418517 | 0 | G | A | 0.377 | -7.32E-02 | 9.97E-02 | 2.75E+00 | 2.85E+00 | 4.56E-01 | no | | intergenic_variant | MODIFIER |
| 7 | 77298997 | AX-170164666 | 0 | G | C | 0.377 | -7.32E-02 | 9.97E-02 | 2.75E+00 | 2.85E+00 | 4.56E-01 | no | Gm44879 | upstream_gene_variant | MODIFIER |
| 7 | 77622079 | AX-97783451 | 0 | C | A | 0.377 | -7.32E-02 | 9.97E-02 | 2.75E+00 | 2.85E+00 | 4.56E-01 | no | | intergenic_variant | MODIFIER |
| 7 | 77844319 | AX-97798907 | 0 | G | A | 0.377 | -7.32E-02 | 9.97E-02 | 2.75E+00 | 2.85E+00 | 4.56E-01 | no | | intergenic_variant | MODIFIER |
| 7 | 77861494 | AX-170089961 | 0 | A | G | 0.377 | -7.32E-02 | 9.97E-02 | 2.75E+00 | 2.85E+00 | 4.56E-01 | no | | intergenic_variant | MODIFIER |
| 7 | 77861612 | AX-170271526 | 0 | A | G | 0.377 | -7.32E-02 | 9.97E-02 | 2.75E+00 | 2.85E+00 | 4.56E-01 | no | | intergenic_variant | MODIFIER |
| 7 | 77896794 | AX-170236281 | 0 | G | A | 0.377 | -7.32E-02 | 9.97E-02 | 2.75E+00 | 2.85E+00 | 4.56E-01 | no | | intergenic_variant | MODIFIER |
| 7 | 77897164 | AX-97812020 | 0 | A | C | 0.377 | -7.32E-02 | 9.97E-02 | 2.75E+00 | 2.85E+00 | 4.56E-01 | no | | intergenic_variant | MODIFIER |
| 7 | 77905744 | AX-97856024 | 0 | G | A | 0.377 | -7.32E-02 | 9.97E-02 | 2.75E+00 | 2.85E+00 | 4.56E-01 | no | Gm23239 | downstream_gene_variant | MODIFIER |
| 7 | 77921809 | AX-170219402 | 0 | A | T | 0.377 | -7.32E-02 | 9.97E-02 | 2.75E+00 | 2.85E+00 | 4.56E-01 | no | | intergenic_variant | MODIFIER |
| 7 | 77922778 | AX-170435011 | 0 | G | T | 0.377 | -7.32E-02 | 9.97E-02 | 2.75E+00 | 2.85E+00 | 4.56E-01 | no | | intergenic_variant | MODIFIER |
| 7 | 77946384 | AX-97817591 | 0 | C | A | 0.377 | -7.32E-02 | 9.97E-02 | 2.75E+00 | 2.85E+00 | 4.56E-01 | no | | intergenic_variant | MODIFIER |
| 7 | 77971659 | AX-170349266 | 0 | T | G | 0.377 | -7.32E-02 | 9.97E-02 | 2.75E+00 | 2.85E+00 | 4.56E-01 | no | | intergenic_variant | MODIFIER |
| 7 | 77978793 | AX-170448618 | 0 | G | T | 0.377 | -7.32E-02 | 9.97E-02 | 2.75E+00 | 2.85E+00 | 4.56E-01 | no | | intergenic_variant | MODIFIER |
| 7 | 77979001 | AX-170424522 | 0 | T | C | 0.377 | -7.32E-02 | 9.97E-02 | 2.75E+00 | 2.85E+00 | 4.56E-01 | no | | intergenic_variant | MODIFIER |
| 7 | 78037677 | AX-97813256 | 0 | T | C | 0.377 | -7.32E-02 | 9.97E-02 | 2.75E+00 | 2.85E+00 | 4.56E-01 | no | | intergenic_variant | MODIFIER |
| 7 | 78039088 | AX-170029362 | 0 | G | A | 0.377 | -7.32E-02 | 9.97E-02 | 2.75E+00 | 2.85E+00 | 4.56E-01 | no | | intergenic_variant | MODIFIER |
| 7 | 78076013 | AX-170029368 | 0 | G | A | 0.377 | -7.32E-02 | 9.97E-02 | 2.75E+00 | 2.85E+00 | 4.56E-01 | no | | intergenic_variant | MODIFIER |
| 7 | 78085803 | AX-97793162 | 0 | A | G | 0.377 | -7.32E-02 | 9.97E-02 | 2.75E+00 | 2.85E+00 | 4.56E-01 | no | | regulatory_region_variant | MODIFIER |
| 7 | 78109286 | AX-170210553 | 0 | T | G | 0.377 | -7.32E-02 | 9.97E-02 | 2.75E+00 | 2.85E+00 | 4.56E-01 | no | | intergenic_variant | MODIFIER |
| 7 | 78112052 | AX-170223415 | 0 | T | C | 0.377 | -7.32E-02 | 9.97E-02 | 2.75E+00 | 2.85E+00 | 4.56E-01 | no | | intergenic_variant | MODIFIER |
| 7 | 78147661 | AX-170029361 | 0 | C | A | 0.377 | -7.32E-02 | 9.97E-02 | 2.75E+00 | 2.85E+00 | 4.56E-01 | no | | intergenic_variant | MODIFIER |
| 7 | 78149173 | AX-170505319 | 0 | C | A | 0.377 | -7.32E-02 | 9.97E-02 | 2.75E+00 | 2.85E+00 | 4.56E-01 | no | | intergenic_variant | MODIFIER |
| 7 | 78149530 | AX-170317829 | 0 | G | A | 0.377 | -7.32E-02 | 9.97E-02 | 2.75E+00 | 2.85E+00 | 4.56E-01 | no | | intergenic_variant | MODIFIER |
| 7 | 78149798 | AX-170335598 | 0 | T | C | 0.377 | -7.32E-02 | 9.97E-02 | 2.75E+00 | 2.85E+00 | 4.56E-01 | no | | intergenic_variant | MODIFIER |
| 7 | 78150191 | AX-170254091 | 0 | A | G | 0.377 | -7.32E-02 | 9.97E-02 | 2.75E+00 | 2.85E+00 | 4.56E-01 | no | | intergenic_variant | MODIFIER |
| 7 | 78151207 | AX-170431505 | 0 | G | A | 0.377 | -7.32E-02 | 9.97E-02 | 2.75E+00 | 2.85E+00 | 4.56E-01 | no | | intergenic_variant | MODIFIER |
| 7 | 78151414 | AX-170393407 | 0 | G | C | 0.377 | -7.32E-02 | 9.97E-02 | 2.75E+00 | 2.85E+00 | 4.56E-01 | no | | intergenic_variant | MODIFIER |
| 7 | 78152107 | AX-170029366 | 0 | C | T | 0.377 | -7.32E-02 | 9.97E-02 | 2.75E+00 | 2.85E+00 | 4.56E-01 | no | | intergenic_variant | MODIFIER |

| | | | | | | | | | | | | | | | |
|---|---|---|---|---|---|---|---|---|---|---|---|---|---|---|---|
| 7 | 78162338 | AX-170089962 | 0 | G | T | 0.377 | -7.32E-02 | 9.97E-02 | 2.75E+00 | 2.85E+00 | 4.56E-01 | no | | intergenic_variant | MODIFIER |
| 7 | 78288165 | AX-170422550 | 0 | G | A | 0.377 | -7.32E-02 | 9.97E-02 | 2.75E+00 | 2.85E+00 | 4.56E-01 | no | Ntrk3 | intron_variant | MODIFIER |
| 20 | 49434852 | AX-170033815 | 1 | G | A | 0.241 | 4.29E-02 | 5.79E-02 | 2.58E+00 | 2.70E+00 | 4.56E-01 | no | Gm14697 | intron_variant,non_coding_transcript_variant | MODIFIER |
| 18 | 86640695 | AX-170475946 | 1 | G | C | 0.161 | -8.10E-02 | 1.07E-01 | 2.51E+00 | 2.60E+00 | 4.57E-01 | no | | intergenic_variant | MODIFIER |
| 7 | 61201077 | AX-97789424 | 0 | G | T | 0.044 | -8.18E-02 | 1.12E-01 | 2.58E+00 | 2.76E+00 | 4.57E-01 | no | A230006K03Rik | intron_variant,non_coding_transcript_variant | MODIFIER |
| 12 | 27565282 | AX-170123230 | 1 | G | T | 0.045 | 1.19E-01 | 1.63E-01 | 2.66E+00 | 2.81E+00 | 4.57E-01 | no | | regulatory_region_variant | MODIFIER |
| 18 | 88827027 | AX-170324116 | 0 | A | G | 0.325 | 4.50E-02 | 6.15E-02 | 2.65E+00 | 2.78E+00 | 4.57E-01 | no | Socs6 | intron_variant,non_coding_transcript_variant | MODIFIER |
| 18 | 88852223 | AX-170120900 | 0 | A | T | 0.325 | 4.50E-02 | 6.15E-02 | 2.65E+00 | 2.78E+00 | 4.57E-01 | no | Socs6 | intron_variant,non_coding_transcript_variant | MODIFIER |
| 18 | 88857590 | AX-170340063 | 0 | G | T | 0.325 | 4.50E-02 | 6.15E-02 | 2.65E+00 | 2.78E+00 | 4.57E-01 | no | Socs6 | intron_variant,non_coding_transcript_variant | MODIFIER |
| 18 | 88867531 | AX-170016165 | 0 | T | C | 0.325 | 4.50E-02 | 6.15E-02 | 2.65E+00 | 2.78E+00 | 4.57E-01 | no | Socs6 | 3_prime_UTR_variant | MODIFIER |
| 18 | 88905003 | AX-170016164 | 0 | A | G | 0.325 | 4.50E-02 | 6.15E-02 | 2.65E+00 | 2.78E+00 | 4.57E-01 | no | Socs6 | intron_variant | MODIFIER |
| 18 | 88905169 | AX-170366976 | 0 | G | A | 0.325 | 4.50E-02 | 6.15E-02 | 2.65E+00 | 2.78E+00 | 4.57E-01 | no | Socs6 | intron_variant | MODIFIER |
| 18 | 88906443 | AX-170151357 | 0 | G | T | 0.325 | 4.50E-02 | 6.15E-02 | 2.65E+00 | 2.78E+00 | 4.57E-01 | no | Socs6 | intron_variant | MODIFIER |
| 18 | 88908048 | AX-170260054 | 0 | G | A | 0.325 | 4.50E-02 | 6.15E-02 | 2.65E+00 | 2.78E+00 | 4.57E-01 | no | Socs6 | intron_variant | MODIFIER |
| 18 | 88911714 | AX-170270631 | 0 | T | G | 0.325 | 4.50E-02 | 6.15E-02 | 2.65E+00 | 2.78E+00 | 4.57E-01 | no | Socs6 | intron_variant | MODIFIER |
| 18 | 88913855 | AX-170191789 | 0 | C | T | 0.325 | 4.50E-02 | 6.15E-02 | 2.65E+00 | 2.78E+00 | 4.57E-01 | no | Socs6 | intron_variant | MODIFIER |
| 18 | 88922564 | AX-170061643 | 0 | T | C | 0.325 | 4.50E-02 | 6.15E-02 | 2.65E+00 | 2.78E+00 | 4.57E-01 | no | Socs6 | intron_variant | MODIFIER |
| 18 | 88922687 | AX-170061639 | 0 | T | A | 0.325 | 4.50E-02 | 6.15E-02 | 2.65E+00 | 2.78E+00 | 4.57E-01 | no | Socs6 | intron_variant | MODIFIER |
| 9 | 22694531 | AX-97816661 | 0 | G | A | 0.465 | 6.41E-02 | 8.66E-02 | 2.61E+00 | 2.70E+00 | 4.57E-01 | no | Bbs9 | intron_variant | MODIFIER |
| 2 | 168819310 | AX-97827186 | 1 | A | G | 0.143 | 1.26E-01 | 1.69E-01 | 2.57E+00 | 2.67E+00 | 4.57E-01 | no | | intergenic_variant | MODIFIER |
| 7 | 70894427 | AX-97808709 | 1 | A | G | 0.348 | -7.26E-02 | 9.90E-02 | 2.79E+00 | 2.88E+00 | 4.57E-01 | no | Gm42397 | intron_variant,non_coding_transcript_variant | MODIFIER |
| 9 | 16995911 | AX-97805731 | 0 | T | C | 0.474 | 6.58E-02 | 8.74E-02 | 2.52E+00 | 2.59E+00 | 4.57E-01 | no | | intergenic_variant | MODIFIER |
| 14 | 91549224 | AX-97795520 | 1 | G | T | 0.045 | 1.31E-01 | 1.81E-01 | 2.72E+00 | 2.87E+00 | 4.58E-01 | no | | intergenic_variant | MODIFIER |
| 1 | 169537726 | AX-97824087 | 1 | C | T | 0.214 | 1.02E-01 | 1.39E-01 | 2.65E+00 | 2.69E+00 | 4.58E-01 | no | | intergenic_variant | MODIFIER |
| 7 | 124154189 | AX-168387317 | 0 | G | C | 0.377 | 6.18E-02 | 8.44E-02 | 2.75E+00 | 2.83E+00 | 4.58E-01 | no | Hs3st4 | intron_variant | MODIFIER |
| 7 | 124154699 | AX-170444400 | 0 | G | C | 0.377 | 6.18E-02 | 8.44E-02 | 2.75E+00 | 2.83E+00 | 4.58E-01 | no | Hs3st4 | intron_variant | MODIFIER |
| 7 | 124167777 | AX-97853916 | 0 | T | G | 0.377 | 6.18E-02 | 8.44E-02 | 2.75E+00 | 2.83E+00 | 4.58E-01 | no | Hs3st4 | intron_variant | MODIFIER |
| 7 | 124176195 | AX-170115321 | 0 | A | G | 0.377 | 6.18E-02 | 8.44E-02 | 2.75E+00 | 2.83E+00 | 4.58E-01 | no | Hs3st4 | intron_variant | MODIFIER |
| 7 | 124179379 | AX-97820632 | 0 | T | C | 0.377 | 6.18E-02 | 8.44E-02 | 2.75E+00 | 2.83E+00 | 4.58E-01 | no | Hs3st4 | intron_variant | MODIFIER |
| 7 | 124669523 | AX-170449457 | 0 | G | T | 0.377 | 6.18E-02 | 8.44E-02 | 2.75E+00 | 2.83E+00 | 4.58E-01 | no | 3100003L05Rik | intron_variant,non_coding_transcript_variant | MODIFIER |
| 7 | 124708143 | AX-170513670 | 0 | C | T | 0.377 | 6.18E-02 | 8.44E-02 | 2.75E+00 | 2.83E+00 | 4.58E-01 | no | 3100003L05Rik | intron_variant,non_coding_transcript_variant | MODIFIER |
| 7 | 124710851 | AX-97854589 | 0 | A | G | 0.377 | 6.18E-02 | 8.44E-02 | 2.75E+00 | 2.83E+00 | 4.58E-01 | no | 3100003L05Rik | upstream_gene_variant | MODIFIER |
| 9 | 96322082 | AX-170425964 | 0 | C | T | 0.158 | -7.59E-02 | 1.04E-01 | 2.70E+00 | 2.81E+00 | 4.58E-01 | no | Tfdp2 | 3_prime_UTR_variant | MODIFIER |
| 5 | 36671754 | AX-170332905 | 1 | A | C | 0.321 | 8.25E-02 | 1.12E-01 | 2.77E+00 | 2.82E+00 | 4.58E-01 | no | D5Ertd579e | intron_variant | MODIFIER |
| 5 | 38082471 | AX-170166366 | 1 | G | C | 0.321 | 8.25E-02 | 1.12E-01 | 2.77E+00 | 2.82E+00 | 4.58E-01 | no | Stx18 | intron_variant | MODIFIER |
| 18 | 89047455 | AX-170488537 | 0 | A | G | 0.333 | 4.50E-02 | 6.17E-02 | 2.65E+00 | 2.78E+00 | 4.58E-01 | no | Rttn | intron_variant | MODIFIER |
| 4 | 85302179 | AX-170412048 | 1 | T | A | 0.339 | 7.71E-02 | 1.05E-01 | 2.73E+00 | 2.82E+00 | 4.58E-01 | no | Sh3gl2 | intron_variant | MODIFIER |
| 3 | 36997242 | AX-170066368 | 0 | A | T | 0.193 | 6.98E-02 | 9.52E-02 | 2.79E+00 | 2.89E+00 | 4.58E-01 | no | 4932438A13Rik | intron_variant | MODIFIER |
| 5 | 109548677 | AX-170025864 | 1 | G | C | 0.348 | 7.86E-02 | 1.06E-01 | 2.54E+00 | 2.68E+00 | 4.58E-01 | no | Gm8493 | downstream_gene_variant | MODIFIER |
| 3 | 36611913 | AX-170374346 | 1 | T | C | 0.188 | 6.96E-02 | 9.50E-02 | 2.78E+00 | 2.87E+00 | 4.58E-01 | no | Bbs7 | intron_variant | MODIFIER |
| 7 | 36965796 | AX-170160185 | 0 | G | T | 0.333 | 6.56E-02 | 8.85E-02 | 2.59E+00 | 2.66E+00 | 4.58E-01 | no | | intergenic_variant | MODIFIER |
| 12 | 91989594 | AX-170324352 | 1 | T | C | 0.062 | 1.39E-01 | 1.91E-01 | 2.69E+00 | 2.81E+00 | 4.58E-01 | no | | intergenic_variant | MODIFIER |
| 16 | 38894024 | AX-168420014 | 0 | A | G | 0.491 | 5.99E-02 | 7.99E-02 | 2.54E+00 | 2.60E+00 | 4.58E-01 | no | Gm15802 | intron_variant,non_coding_transcript_variant | MODIFIER |
| 4 | 101445306 | AX-170438442 | 1 | T | G | 0.348 | 6.66E-02 | 9.07E-02 | 2.64E+00 | 2.74E+00 | 4.58E-01 | no | Ak4 | intron_variant | MODIFIER |
| 16 | 49929977 | AX-170357041 | 1 | A | C | 0.509 | 6.27E-02 | 8.39E-02 | 2.55E+00 | 2.58E+00 | 4.59E-01 | no | | intergenic_variant | MODIFIER |
| 4 | 127325758 | AX-97786677 | 1 | C | T | 0.143 | 8.86E-02 | 1.19E-01 | 2.56E+00 | 2.63E+00 | 4.59E-01 | no | Gjb3 | 3_prime_UTR_variant | MODIFIER |
| 7 | 124191337 | AX-170029894 | 0 | T | C | 0.395 | 6.16E-02 | 8.43E-02 | 2.77E+00 | 2.86E+00 | 4.59E-01 | no | Hs3st4 | intron_variant | MODIFIER |
| 18 | 59380475 | AX-170439748 | 0 | A | C | 0.184 | -8.74E-02 | 1.19E-01 | 2.68E+00 | 2.73E+00 | 4.59E-01 | no | Chsy3 | intron_variant | MODIFIER |
| 1 | 149790314 | AX-170169747 | 0 | C | T | 0.211 | 9.47E-02 | 1.29E-01 | 2.67E+00 | 2.73E+00 | 4.59E-01 | no | | intergenic_variant | MODIFIER |
| 4 | 125626713 | AX-170354208 | 0 | T | C | 0.219 | 8.25E-02 | 1.09E-01 | 2.50E+00 | 2.58E+00 | 4.59E-01 | no | Grik3 | intron_variant | MODIFIER |
| 3 | 37826000 | AX-170189374 | 0 | A | G | 0.202 | 6.74E-02 | 9.23E-02 | 2.77E+00 | 2.86E+00 | 4.59E-01 | no | | intergenic_variant | MODIFIER |
| 16 | 45103194 | AX-170012950 | 0 | T | G | 0.474 | -5.98E-02 | 8.01E-02 | 2.55E+00 | 2.60E+00 | 4.59E-01 | no | Ccdc80 | intron_variant | MODIFIER |
| 7 | 45679463 | AX-170282103 | 1 | G | A | 0.348 | 6.92E-02 | 9.41E-02 | 2.62E+00 | 2.71E+00 | 4.59E-01 | no | Sec1 | synonymous_variant | LOW |
| 15 | 23208795 | AX-170221860 | 0 | C | T | 0.246 | -7.70E-02 | 1.04E-01 | 2.62E+00 | 2.67E+00 | 4.59E-01 | no | Cdh18 | intron_variant,non_coding_transcript_variant | MODIFIER |

| | | | | | | | | | | | | | | | |
|---|---|---|---|---|---|---|---|---|---|---|---|---|---|---|---|
| 15 | 23209903 | AX-170117465 | 0 | G | A | 0.246 | -7.70E-02 | 1.04E-01 | 2.62E+00 | 2.67E+00 | 4.59E-01 | no | Cdh18 | intron_variant,non_coding_transcript_variant | MODIFIER |
| 15 | 23265252 | AX-170055025 | 0 | A | G | 0.246 | -7.70E-02 | 1.04E-01 | 2.62E+00 | 2.67E+00 | 4.59E-01 | no | Cdh18 | intron_variant,non_coding_transcript_variant | MODIFIER |
| 15 | 23272004 | AX-170120449 | 0 | G | A | 0.246 | -7.70E-02 | 1.04E-01 | 2.62E+00 | 2.67E+00 | 4.59E-01 | no | Cdh18 | intron_variant,non_coding_transcript_variant | MODIFIER |
| 15 | 23278474 | AX-170222090 | 0 | G | A | 0.246 | -7.70E-02 | 1.04E-01 | 2.62E+00 | 2.67E+00 | 4.59E-01 | no | Cdh18 | intron_variant,non_coding_transcript_variant | MODIFIER |
| 15 | 23313567 | AX-170494784 | 0 | T | C | 0.246 | -7.70E-02 | 1.04E-01 | 2.62E+00 | 2.67E+00 | 4.59E-01 | no | Cdh18 | intron_variant,non_coding_transcript_variant | MODIFIER |
| 15 | 23390382 | AX-170011234 | 0 | G | T | 0.246 | -7.70E-02 | 1.04E-01 | 2.62E+00 | 2.67E+00 | 4.59E-01 | no | Cdh18 | intron_variant,non_coding_transcript_variant | MODIFIER |
| 15 | 23425254 | AX-170141937 | 0 | A | G | 0.246 | -7.70E-02 | 1.04E-01 | 2.62E+00 | 2.67E+00 | 4.59E-01 | no | Cdh18 | intron_variant,non_coding_transcript_variant | MODIFIER |
| 15 | 23433145 | AX-170011235 | 0 | C | A | 0.246 | -7.70E-02 | 1.04E-01 | 2.62E+00 | 2.67E+00 | 4.59E-01 | no | Cdh18 | intron_variant,non_coding_transcript_variant | MODIFIER |
| 15 | 23437997 | AX-170179179 | 0 | G | C | 0.246 | -7.70E-02 | 1.04E-01 | 2.62E+00 | 2.67E+00 | 4.59E-01 | no | Cdh18 | intron_variant,non_coding_transcript_variant | MODIFIER |
| 15 | 23441481 | AX-170151674 | 0 | C | T | 0.246 | -7.70E-02 | 1.04E-01 | 2.62E+00 | 2.67E+00 | 4.59E-01 | no | Cdh18 | intron_variant,non_coding_transcript_variant | MODIFIER |
| 15 | 23441554 | AX-170499467 | 0 | A | T | 0.246 | -7.70E-02 | 1.04E-01 | 2.62E+00 | 2.67E+00 | 4.59E-01 | no | Cdh18 | intron_variant,non_coding_transcript_variant | MODIFIER |
| 15 | 23449285 | AX-170441084 | 0 | A | C | 0.246 | -7.70E-02 | 1.04E-01 | 2.62E+00 | 2.67E+00 | 4.59E-01 | no | Cdh18 | downstream_gene_variant | MODIFIER |
| 15 | 23609943 | AX-170520758 | 0 | G | T | 0.246 | -7.70E-02 | 1.04E-01 | 2.62E+00 | 2.67E+00 | 4.59E-01 | no | | intergenic_variant | MODIFIER |
| 15 | 23664931 | AX-170011240 | 0 | T | C | 0.246 | -7.70E-02 | 1.04E-01 | 2.62E+00 | 2.67E+00 | 4.59E-01 | no | | intergenic_variant | MODIFIER |
| 15 | 24185076 | AX-170495211 | 0 | A | T | 0.246 | -7.70E-02 | 1.04E-01 | 2.62E+00 | 2.67E+00 | 4.59E-01 | no | | intergenic_variant | MODIFIER |
| 15 | 24188785 | AX-170348370 | 0 | G | C | 0.246 | -7.70E-02 | 1.04E-01 | 2.62E+00 | 2.67E+00 | 4.59E-01 | no | | intergenic_variant | MODIFIER |
| 15 | 24257319 | AX-170220980 | 0 | A | G | 0.246 | -7.70E-02 | 1.04E-01 | 2.62E+00 | 2.67E+00 | 4.59E-01 | no | Gm49729 | intron_variant,non_coding_transcript_variant | MODIFIER |
| 15 | 24532469 | AX-170011261 | 0 | T | G | 0.246 | -7.70E-02 | 1.04E-01 | 2.62E+00 | 2.67E+00 | 4.59E-01 | no | | intergenic_variant | MODIFIER |
| 15 | 24899974 | AX-170344751 | 0 | T | C | 0.246 | -7.70E-02 | 1.04E-01 | 2.62E+00 | 2.67E+00 | 4.59E-01 | no | Gm35769 | intron_variant,non_coding_transcript_variant | MODIFIER |
| 15 | 25322575 | AX-170331309 | 0 | T | C | 0.246 | -7.70E-02 | 1.04E-01 | 2.62E+00 | 2.67E+00 | 4.59E-01 | no | 4930445E18Rik | intron_variant,non_coding_transcript_variant | MODIFIER |
| 15 | 25342336 | AX-170340318 | 0 | A | C | 0.246 | -7.70E-02 | 1.04E-01 | 2.62E+00 | 2.67E+00 | 4.59E-01 | no | | intergenic_variant | MODIFIER |
| 15 | 25351808 | AX-170509577 | 0 | C | A | 0.246 | -7.70E-02 | 1.04E-01 | 2.62E+00 | 2.67E+00 | 4.59E-01 | no | | intergenic_variant | MODIFIER |
| 15 | 25352973 | AX-170263920 | 0 | A | G | 0.246 | -7.70E-02 | 1.04E-01 | 2.62E+00 | 2.67E+00 | 4.59E-01 | no | | regulatory_region_variant | MODIFIER |
| 15 | 25385464 | AX-170219231 | 0 | T | A | 0.246 | -7.70E-02 | 1.04E-01 | 2.62E+00 | 2.67E+00 | 4.59E-01 | no | Basp1 | intron_variant | MODIFIER |
| 15 | 25386287 | AX-170237602 | 0 | C | A | 0.246 | -7.70E-02 | 1.04E-01 | 2.62E+00 | 2.67E+00 | 4.59E-01 | no | Basp1 | intron_variant | MODIFIER |
| 15 | 25387762 | AX-170370856 | 0 | C | A | 0.246 | -7.70E-02 | 1.04E-01 | 2.62E+00 | 2.67E+00 | 4.59E-01 | no | Basp1 | intron_variant | MODIFIER |
| 15 | 25387806 | AX-170112599 | 0 | G | A | 0.246 | -7.70E-02 | 1.04E-01 | 2.62E+00 | 2.67E+00 | 4.59E-01 | no | Basp1 | intron_variant | MODIFIER |
| 15 | 25389920 | AX-170318700 | 0 | A | C | 0.246 | -7.70E-02 | 1.04E-01 | 2.62E+00 | 2.67E+00 | 4.59E-01 | no | Basp1 | intron_variant | MODIFIER |
| 15 | 25392459 | AX-170488855 | 0 | C | T | 0.246 | -7.70E-02 | 1.04E-01 | 2.62E+00 | 2.67E+00 | 4.59E-01 | no | Basp1 | intron_variant | MODIFIER |
| 15 | 25407107 | AX-170334831 | 0 | A | G | 0.246 | -7.70E-02 | 1.04E-01 | 2.62E+00 | 2.67E+00 | 4.59E-01 | no | Basp1 | intron_variant | MODIFIER |
| 15 | 25439314 | AX-97823519 | 0 | T | C | 0.246 | -7.70E-02 | 1.04E-01 | 2.62E+00 | 2.67E+00 | 4.59E-01 | no | Gm48956 | intron_variant,non_coding_transcript_variant | MODIFIER |
| 15 | 25440745 | AX-170379019 | 0 | C | A | 0.246 | -7.70E-02 | 1.04E-01 | 2.62E+00 | 2.67E+00 | 4.59E-01 | no | Gm48956 | intron_variant,non_coding_transcript_variant | MODIFIER |
| 15 | 25498520 | AX-170165607 | 0 | G | C | 0.246 | -7.70E-02 | 1.04E-01 | 2.62E+00 | 2.67E+00 | 4.59E-01 | no | | intergenic_variant | MODIFIER |
| 15 | 25515097 | AX-170420539 | 0 | G | A | 0.246 | -7.70E-02 | 1.04E-01 | 2.62E+00 | 2.67E+00 | 4.59E-01 | no | | intergenic_variant | MODIFIER |
| 15 | 25519332 | AX-170184076 | 0 | A | C | 0.246 | -7.70E-02 | 1.04E-01 | 2.62E+00 | 2.67E+00 | 4.59E-01 | no | | intergenic_variant | MODIFIER |
| 15 | 25519371 | AX-170412704 | 0 | A | G | 0.246 | -7.70E-02 | 1.04E-01 | 2.62E+00 | 2.67E+00 | 4.59E-01 | no | | intergenic_variant | MODIFIER |
| 15 | 25534023 | AX-168414785 | 0 | A | G | 0.246 | -7.70E-02 | 1.04E-01 | 2.62E+00 | 2.67E+00 | 4.59E-01 | no | Gm36642 | intron_variant,non_coding_transcript_variant | MODIFIER |
| 15 | 25538901 | AX-170292781 | 0 | T | C | 0.246 | -7.70E-02 | 1.04E-01 | 2.62E+00 | 2.67E+00 | 4.59E-01 | no | Gm36642 | intron_variant,non_coding_transcript_variant | MODIFIER |
| 15 | 25543865 | AX-170203883 | 0 | A | C | 0.246 | -7.70E-02 | 1.04E-01 | 2.62E+00 | 2.67E+00 | 4.59E-01 | no | Gm36642 | intron_variant,non_coding_transcript_variant | MODIFIER |
| 15 | 25544044 | AX-170388287 | 0 | G | C | 0.246 | -7.70E-02 | 1.04E-01 | 2.62E+00 | 2.67E+00 | 4.59E-01 | no | Gm36642 | intron_variant,non_coding_transcript_variant | MODIFIER |
| 15 | 25550478 | AX-170310759 | 0 | G | C | 0.246 | -7.70E-02 | 1.04E-01 | 2.62E+00 | 2.67E+00 | 4.59E-01 | no | Gm36642 | intron_variant,non_coding_transcript_variant | MODIFIER |
| 15 | 25577816 | AX-170093185 | 0 | A | G | 0.246 | -7.70E-02 | 1.04E-01 | 2.62E+00 | 2.67E+00 | 4.59E-01 | no | Gm36642 | intron_variant,non_coding_transcript_variant | MODIFIER |
| 15 | 25592211 | AX-170253663 | 0 | C | T | 0.246 | -7.70E-02 | 1.04E-01 | 2.62E+00 | 2.67E+00 | 4.59E-01 | no | Gm36642 | intron_variant,non_coding_transcript_variant | MODIFIER |
| 15 | 25708008 | AX-170055058 | 0 | T | C | 0.246 | -7.70E-02 | 1.04E-01 | 2.62E+00 | 2.67E+00 | 4.59E-01 | no | Myo10 | intron_variant | MODIFIER |
| 1 | 175196012 | AX-170499647 | 0 | T | C | 0.228 | 9.30E-02 | 1.23E-01 | 2.51E+00 | 2.56E+00 | 4.59E-01 | no | Rgs7 | intron_variant | MODIFIER |
| 20 | 120098852 | AX-170230813 | 0 | G | T | 0.035 | 9.68E-02 | 1.34E-01 | 2.70E+00 | 2.88E+00 | 4.59E-01 | no | | intergenic_variant | MODIFIER |
| 11 | 31916131 | AX-97817956 | 0 | T | C | 0.079 | -1.25E-01 | 1.71E-01 | 2.77E+00 | 2.89E+00 | 4.60E-01 | no | Cpeb4 | intron_variant | MODIFIER |
| 8 | 88306862 | AX-170090174 | 0 | T | C | 0.175 | -9.38E-02 | 1.28E-01 | 2.68E+00 | 2.76E+00 | 4.60E-01 | no | Adcy7 | intron_variant | MODIFIER |
| 8 | 88316972 | AX-170274282 | 0 | C | T | 0.175 | -9.38E-02 | 1.28E-01 | 2.68E+00 | 2.76E+00 | 4.60E-01 | no | Adcy7 | intron_variant | MODIFIER |
| 8 | 88317695 | AX-170201631 | 0 | T | C | 0.175 | -9.38E-02 | 1.28E-01 | 2.68E+00 | 2.76E+00 | 4.60E-01 | no | Adcy7 | intron_variant | MODIFIER |
| 8 | 88325675 | AX-170317920 | 0 | G | A | 0.175 | -9.38E-02 | 1.28E-01 | 2.68E+00 | 2.76E+00 | 4.60E-01 | no | Adcy7 | intron_variant | MODIFIER |
| 8 | 88333412 | AX-170210610 | 0 | G | A | 0.175 | -9.38E-02 | 1.28E-01 | 2.68E+00 | 2.76E+00 | 4.60E-01 | no | Brd7 | intron_variant | MODIFIER |
| 8 | 88340683 | AX-168404926 | 0 | T | C | 0.175 | -9.38E-02 | 1.28E-01 | 2.68E+00 | 2.76E+00 | 4.60E-01 | no | Brd7 | intron_variant | MODIFIER |
| 8 | 88350372 | AX-170255984 | 0 | A | G | 0.175 | -9.38E-02 | 1.28E-01 | 2.68E+00 | 2.76E+00 | 4.60E-01 | no | Brd7 | intron_variant | MODIFIER |

| | | | | | | | | | | | | | | | |
|---|---|---|---|---|---|---|---|---|---|---|---|---|---|---|---|
| 8 | 88393925 | AX-170528671 | 0 | G | C | 0.175 | -9.38E-02 | 1.28E-01 | 2.68E+00 | 2.76E+00 | 4.60E-01 | no | Gm45497 | downstream_gene_variant | MODIFIER |
| 8 | 88396574 | AX-170160997 | 0 | A | G | 0.175 | -9.38E-02 | 1.28E-01 | 2.68E+00 | 2.76E+00 | 4.60E-01 | no | Gm45497 | downstream_gene_variant | MODIFIER |
| 8 | 88417379 | AX-170080700 | 0 | G | A | 0.175 | -9.38E-02 | 1.28E-01 | 2.68E+00 | 2.76E+00 | 4.60E-01 | no | | regulatory_region_variant | MODIFIER |
| 8 | 88418728 | AX-170080702 | 0 | C | G | 0.175 | -9.38E-02 | 1.28E-01 | 2.68E+00 | 2.76E+00 | 4.60E-01 | no | | intergenic_variant | MODIFIER |
| 8 | 88418809 | AX-170432674 | 0 | T | C | 0.175 | -9.38E-02 | 1.28E-01 | 2.68E+00 | 2.76E+00 | 4.60E-01 | no | | intergenic_variant | MODIFIER |
| 8 | 88421317 | AX-97804050 | 0 | T | C | 0.175 | -9.38E-02 | 1.28E-01 | 2.68E+00 | 2.76E+00 | 4.60E-01 | no | | intergenic_variant | MODIFIER |
| 8 | 88448081 | AX-170031216 | 0 | G | A | 0.175 | -9.38E-02 | 1.28E-01 | 2.68E+00 | 2.76E+00 | 4.60E-01 | no | | regulatory_region_variant | MODIFIER |
| 8 | 88448317 | AX-170031224 | 0 | T | C | 0.175 | -9.38E-02 | 1.28E-01 | 2.68E+00 | 2.76E+00 | 4.60E-01 | no | | regulatory_region_variant | MODIFIER |
| 8 | 88448558 | AX-170090175 | 0 | T | C | 0.175 | -9.38E-02 | 1.28E-01 | 2.68E+00 | 2.76E+00 | 4.60E-01 | no | | regulatory_region_variant | MODIFIER |
| 8 | 88479041 | AX-170212363 | 0 | T | C | 0.175 | -9.38E-02 | 1.28E-01 | 2.68E+00 | 2.76E+00 | 4.60E-01 | no | | intergenic_variant | MODIFIER |
| 8 | 88482166 | AX-170031218 | 0 | T | C | 0.175 | -9.38E-02 | 1.28E-01 | 2.68E+00 | 2.76E+00 | 4.60E-01 | no | | intergenic_variant | MODIFIER |
| 8 | 88482302 | AX-170031221 | 0 | G | T | 0.175 | -9.38E-02 | 1.28E-01 | 2.68E+00 | 2.76E+00 | 4.60E-01 | no | | intergenic_variant | MODIFIER |
| 8 | 88482587 | AX-170031222 | 0 | C | T | 0.175 | -9.38E-02 | 1.28E-01 | 2.68E+00 | 2.76E+00 | 4.60E-01 | no | | intergenic_variant | MODIFIER |
| 8 | 88486821 | AX-170296822 | 0 | C | G | 0.175 | -9.38E-02 | 1.28E-01 | 2.68E+00 | 2.76E+00 | 4.60E-01 | no | | intergenic_variant | MODIFIER |
| 8 | 88506754 | AX-170303640 | 0 | A | G | 0.175 | -9.38E-02 | 1.28E-01 | 2.68E+00 | 2.76E+00 | 4.60E-01 | no | | intergenic_variant | MODIFIER |
| 8 | 88512604 | AX-97814430 | 0 | C | T | 0.175 | -9.38E-02 | 1.28E-01 | 2.68E+00 | 2.76E+00 | 4.60E-01 | no | | regulatory_region_variant | MODIFIER |
| 8 | 88514213 | AX-170363609 | 0 | A | G | 0.175 | -9.38E-02 | 1.28E-01 | 2.68E+00 | 2.76E+00 | 4.60E-01 | no | | intergenic_variant | MODIFIER |
| 8 | 88529963 | AX-170031225 | 0 | A | G | 0.175 | -9.38E-02 | 1.28E-01 | 2.68E+00 | 2.76E+00 | 4.60E-01 | no | Nkd1 | intron_variant | MODIFIER |
| 8 | 88530819 | AX-170031223 | 0 | A | G | 0.175 | -9.38E-02 | 1.28E-01 | 2.68E+00 | 2.76E+00 | 4.60E-01 | no | Nkd1 | intron_variant | MODIFIER |
| 8 | 88579813 | AX-170203990 | 0 | G | A | 0.175 | -9.38E-02 | 1.28E-01 | 2.68E+00 | 2.76E+00 | 4.60E-01 | no | Nkd1 | intron_variant | MODIFIER |
| 8 | 88592054 | AX-97857474 | 0 | G | A | 0.175 | -9.38E-02 | 1.28E-01 | 2.68E+00 | 2.76E+00 | 4.60E-01 | no | Nkd1 | missense_variant | MODERATE |
| 8 | 88611043 | AX-170440269 | 0 | T | C | 0.175 | -9.38E-02 | 1.28E-01 | 2.68E+00 | 2.76E+00 | 4.60E-01 | no | | intergenic_variant | MODIFIER |
| 8 | 88615208 | AX-170080705 | 0 | A | T | 0.175 | -9.38E-02 | 1.28E-01 | 2.68E+00 | 2.76E+00 | 4.60E-01 | no | | regulatory_region_variant | MODIFIER |
| 8 | 88615830 | AX-170080704 | 0 | A | G | 0.175 | -9.38E-02 | 1.28E-01 | 2.68E+00 | 2.76E+00 | 4.60E-01 | no | | intergenic_variant | MODIFIER |
| 8 | 88616098 | AX-170080706 | 0 | C | A | 0.175 | -9.38E-02 | 1.28E-01 | 2.68E+00 | 2.76E+00 | 4.60E-01 | no | | intergenic_variant | MODIFIER |
| 8 | 88623059 | AX-170339693 | 0 | G | A | 0.175 | -9.38E-02 | 1.28E-01 | 2.68E+00 | 2.76E+00 | 4.60E-01 | no | Snx20 | downstream_gene_variant | MODIFIER |
| 8 | 88624738 | AX-97854105 | 0 | G | A | 0.175 | -9.38E-02 | 1.28E-01 | 2.68E+00 | 2.76E+00 | 4.60E-01 | no | Snx20 | downstream_gene_variant | MODIFIER |
| 8 | 88624904 | AX-170125018 | 0 | T | C | 0.175 | -9.38E-02 | 1.28E-01 | 2.68E+00 | 2.76E+00 | 4.60E-01 | no | Snx20 | downstream_gene_variant | MODIFIER |
| 8 | 88625509 | AX-170494649 | 0 | T | G | 0.175 | -9.38E-02 | 1.28E-01 | 2.68E+00 | 2.76E+00 | 4.60E-01 | no | Snx20 | downstream_gene_variant | MODIFIER |
| 8 | 88652250 | AX-170327806 | 0 | A | C | 0.175 | -9.38E-02 | 1.28E-01 | 2.68E+00 | 2.76E+00 | 4.60E-01 | no | Nod2 | intron_variant | MODIFIER |
| 8 | 88654653 | AX-170287754 | 0 | C | T | 0.175 | -9.38E-02 | 1.28E-01 | 2.68E+00 | 2.76E+00 | 4.60E-01 | no | Nod2 | intron_variant | MODIFIER |
| 8 | 88656475 | AX-170204415 | 0 | T | C | 0.175 | -9.38E-02 | 1.28E-01 | 2.68E+00 | 2.76E+00 | 4.60E-01 | no | Nod2 | intron_variant | MODIFIER |
| 8 | 88657573 | AX-170327013 | 0 | C | G | 0.175 | -9.38E-02 | 1.28E-01 | 2.68E+00 | 2.76E+00 | 4.60E-01 | no | Nod2 | intron_variant | MODIFIER |
| 8 | 88659380 | AX-170285141 | 0 | G | A | 0.175 | -9.38E-02 | 1.28E-01 | 2.68E+00 | 2.76E+00 | 4.60E-01 | no | Nod2 | intron_variant | MODIFIER |
| 8 | 88660402 | AX-170223008 | 0 | T | G | 0.175 | -9.38E-02 | 1.28E-01 | 2.68E+00 | 2.76E+00 | 4.60E-01 | no | Nod2 | intron_variant | MODIFIER |
| 8 | 88668263 | AX-170269486 | 0 | C | T | 0.175 | -9.38E-02 | 1.28E-01 | 2.68E+00 | 2.76E+00 | 4.60E-01 | no | Nod2 | intron_variant | MODIFIER |
| 8 | 88670769 | AX-170463400 | 0 | T | C | 0.175 | -9.38E-02 | 1.28E-01 | 2.68E+00 | 2.76E+00 | 4.60E-01 | no | Nod2 | intron_variant | MODIFIER |
| 8 | 88674992 | AX-170489408 | 0 | A | G | 0.175 | -9.38E-02 | 1.28E-01 | 2.68E+00 | 2.76E+00 | 4.60E-01 | no | Nod2 | splice_region_variant,intron_variant | LOW |
| 8 | 88680202 | AX-170080708 | 0 | C | T | 0.175 | -9.38E-02 | 1.28E-01 | 2.68E+00 | 2.76E+00 | 4.60E-01 | no | Nod2 | intron_variant | MODIFIER |
| 8 | 88683261 | AX-170145203 | 0 | G | A | 0.175 | -9.38E-02 | 1.28E-01 | 2.68E+00 | 2.76E+00 | 4.60E-01 | no | Nod2 | intron_variant | MODIFIER |
| 8 | 88688728 | AX-170511216 | 0 | G | A | 0.175 | -9.38E-02 | 1.28E-01 | 2.68E+00 | 2.76E+00 | 4.60E-01 | no | Nod2 | downstream_gene_variant | MODIFIER |
| 8 | 88709727 | AX-170193785 | 0 | A | T | 0.175 | -9.38E-02 | 1.28E-01 | 2.68E+00 | 2.76E+00 | 4.60E-01 | no | Cyld | intron_variant | MODIFIER |
| 8 | 88732747 | AX-170167451 | 0 | T | A | 0.175 | -9.38E-02 | 1.28E-01 | 2.68E+00 | 2.76E+00 | 4.60E-01 | no | Cyld | intron_variant | MODIFIER |
| 8 | 88746337 | AX-170534284 | 0 | G | A | 0.175 | -9.38E-02 | 1.28E-01 | 2.68E+00 | 2.76E+00 | 4.60E-01 | no | Cyld | intron_variant | MODIFIER |
| 8 | 88755197 | AX-170486457 | 0 | A | G | 0.175 | -9.38E-02 | 1.28E-01 | 2.68E+00 | 2.76E+00 | 4.60E-01 | no | Cyld | downstream_gene_variant | MODIFIER |
| 8 | 88756334 | AX-170503471 | 0 | A | T | 0.175 | -9.38E-02 | 1.28E-01 | 2.68E+00 | 2.76E+00 | 4.60E-01 | no | Cyld | downstream_gene_variant | MODIFIER |
| 8 | 88757051 | AX-170359407 | 0 | C | A | 0.175 | -9.38E-02 | 1.28E-01 | 2.68E+00 | 2.76E+00 | 4.60E-01 | no | | intergenic_variant | MODIFIER |
| 8 | 88759899 | AX-170376104 | 0 | G | A | 0.175 | -9.38E-02 | 1.28E-01 | 2.68E+00 | 2.76E+00 | 4.60E-01 | no | | regulatory_region_variant | MODIFIER |
| 8 | 88772929 | AX-170436830 | 0 | G | T | 0.175 | -9.38E-02 | 1.28E-01 | 2.68E+00 | 2.76E+00 | 4.60E-01 | no | | intergenic_variant | MODIFIER |
| 8 | 88773292 | AX-170363261 | 0 | T | C | 0.175 | -9.38E-02 | 1.28E-01 | 2.68E+00 | 2.76E+00 | 4.60E-01 | no | | intergenic_variant | MODIFIER |
| 8 | 88780403 | AX-170257922 | 0 | C | T | 0.175 | -9.38E-02 | 1.28E-01 | 2.68E+00 | 2.76E+00 | 4.60E-01 | no | | intergenic_variant | MODIFIER |
| 8 | 88780433 | AX-170457979 | 0 | G | A | 0.175 | -9.38E-02 | 1.28E-01 | 2.68E+00 | 2.76E+00 | 4.60E-01 | no | | intergenic_variant | MODIFIER |
| 8 | 88780446 | AX-97798856 | 0 | A | G | 0.175 | -9.38E-02 | 1.28E-01 | 2.68E+00 | 2.76E+00 | 4.60E-01 | no | | intergenic_variant | MODIFIER |
| 8 | 88790591 | AX-170245638 | 0 | G | C | 0.175 | -9.38E-02 | 1.28E-01 | 2.68E+00 | 2.76E+00 | 4.60E-01 | no | | intergenic_variant | MODIFIER |

| | | | | | | | | | | | | | | | |
|---|---|---|---|---|---|---|---|---|---|---|---|---|---|---|---|
| 12 | 89934645 | AX-97852961 | 1 | C | T | 0.098 | 1.40E-01 | 1.92E-01 | 2.78E+00 | 2.85E+00 | 4.60E-01 | no | Nrxn3 | intron_variant | MODIFIER |
| 12 | 90085012 | AX-170049746 | 1 | A | G | 0.098 | 1.40E-01 | 1.92E-01 | 2.78E+00 | 2.85E+00 | 4.60E-01 | no | Nrxn3 | intron_variant | MODIFIER |
| 1 | 105696964 | AX-170434212 | 1 | G | C | 0.143 | 1.12E-01 | 1.52E-01 | 2.54E+00 | 2.67E+00 | 4.60E-01 | no | Relch | intron_variant | MODIFIER |
| 18 | 64932289 | AX-170477121 | 2 | C | A | 0.418 | 7.03E-02 | 9.21E-02 | 2.46E+00 | 2.51E+00 | 4.60E-01 | no | Nedd4l | intron_variant | MODIFIER |
| 5 | 112826493 | AX-97792641 | 0 | T | C | 0.456 | 8.60E-02 | 1.17E-01 | 2.67E+00 | 2.70E+00 | 4.60E-01 | no | Myo18b | intron_variant | MODIFIER |
| 5 | 114159810 | AX-170126322 | 0 | G | A | 0.456 | 7.89E-02 | 1.07E-01 | 2.59E+00 | 2.66E+00 | 4.60E-01 | no | Acacb | intron_variant | MODIFIER |
| 1 | 91244410 | AX-97824634 | 1 | C | T | 0.161 | 1.52E-01 | 2.08E-01 | 2.66E+00 | 2.72E+00 | 4.60E-01 | Yes | | intergenic_variant | MODIFIER |
| 18 | 81091838 | AX-170016083 | 0 | G | A | 0.298 | -6.29E-02 | 8.50E-02 | 2.55E+00 | 2.65E+00 | 4.60E-01 | no | AC105065.1 | intron_variant,non_coding_transcript_variant | MODIFIER |
| 16 | 41045355 | AX-97841799 | 1 | C | T | 0.491 | 5.92E-02 | 7.93E-02 | 2.53E+00 | 2.60E+00 | 4.60E-01 | no | Lsamp | intron_variant | MODIFIER |
| 11 | 107774415 | AX-97816054 | 0 | G | A | 0.044 | -1.45E-01 | 1.96E-01 | 2.55E+00 | 2.65E+00 | 4.60E-01 | no | Cacng4 | intron_variant | MODIFIER |
| 9 | 79269996 | AX-170155426 | 0 | G | A | 0.202 | -8.16E-02 | 1.12E-01 | 2.65E+00 | 2.75E+00 | 4.60E-01 | no | Gm47501 | upstream_gene_variant | MODIFIER |
| 9 | 79632102 | AX-170534768 | 0 | C | T | 0.202 | -8.16E-02 | 1.12E-01 | 2.65E+00 | 2.75E+00 | 4.60E-01 | no | Col12a1 | intron_variant | MODIFIER |
| 9 | 118219538 | AX-170201604 | 1 | C | T | 0.125 | -6.98E-02 | 9.56E-02 | 2.62E+00 | 2.74E+00 | 4.60E-01 | no | | intergenic_variant | MODIFIER |
| 1 | 147527883 | AX-170098667 | 1 | T | C | 0.196 | 1.02E-01 | 1.39E-01 | 2.68E+00 | 2.73E+00 | 4.61E-01 | no | | intergenic_variant | MODIFIER |
| 18 | 68234342 | AX-170088369 | 0 | A | C | 0.272 | -7.41E-02 | 1.02E-01 | 2.69E+00 | 2.75E+00 | 4.61E-01 | no | Ldlrad4 | intron_variant | MODIFIER |
| 18 | 68241894 | AX-97793226 | 0 | A | G | 0.272 | -7.41E-02 | 1.02E-01 | 2.69E+00 | 2.75E+00 | 4.61E-01 | no | Ldlrad4 | intron_variant | MODIFIER |
| 18 | 68262924 | AX-170535395 | 0 | T | C | 0.272 | -7.41E-02 | 1.02E-01 | 2.69E+00 | 2.75E+00 | 4.61E-01 | no | Fam210a | 3_prime_UTR_variant | MODIFIER |
| 18 | 68263267 | AX-170155922 | 0 | T | C | 0.272 | -7.41E-02 | 1.02E-01 | 2.69E+00 | 2.75E+00 | 4.61E-01 | no | Fam210a | 3_prime_UTR_variant | MODIFIER |
| 18 | 68269916 | AX-170139064 | 0 | C | G | 0.272 | -7.41E-02 | 1.02E-01 | 2.69E+00 | 2.75E+00 | 4.61E-01 | no | Fam210a | intron_variant | MODIFIER |
| 18 | 68278830 | AX-97782119 | 0 | A | G | 0.272 | -7.41E-02 | 1.02E-01 | 2.69E+00 | 2.75E+00 | 4.61E-01 | no | Fam210a | intron_variant | MODIFIER |
| 18 | 68309753 | AX-170498970 | 0 | G | A | 0.272 | -7.41E-02 | 1.02E-01 | 2.69E+00 | 2.75E+00 | 4.61E-01 | no | Rnmt | intron_variant | MODIFIER |
| 18 | 68311573 | AX-97835048 | 0 | G | T | 0.272 | -7.41E-02 | 1.02E-01 | 2.69E+00 | 2.75E+00 | 4.61E-01 | no | Rnmt | intron_variant | MODIFIER |
| 18 | 68313817 | AX-97828109 | 0 | T | C | 0.272 | -7.41E-02 | 1.02E-01 | 2.69E+00 | 2.75E+00 | 4.61E-01 | no | Rnmt | synonymous_variant | LOW |
| 18 | 68321100 | AX-170398057 | 0 | A | T | 0.272 | -7.41E-02 | 1.02E-01 | 2.69E+00 | 2.75E+00 | 4.61E-01 | no | Rnmt | intron_variant | MODIFIER |
| 18 | 68339640 | AX-170348057 | 0 | G | A | 0.272 | -7.41E-02 | 1.02E-01 | 2.69E+00 | 2.75E+00 | 4.61E-01 | no | Mc5r | missense_variant | MODERATE |
| 18 | 68339716 | AX-97811913 | 0 | G | A | 0.272 | -7.41E-02 | 1.02E-01 | 2.69E+00 | 2.75E+00 | 4.61E-01 | no | Mc5r | 3_prime_UTR_variant | MODIFIER |
| 18 | 68355581 | AX-170214519 | 0 | C | T | 0.272 | -7.41E-02 | 1.02E-01 | 2.69E+00 | 2.75E+00 | 4.61E-01 | no | | intergenic_variant | MODIFIER |
| 18 | 68399036 | AX-170458435 | 0 | G | A | 0.272 | -7.41E-02 | 1.02E-01 | 2.69E+00 | 2.75E+00 | 4.61E-01 | no | | regulatory_region_variant | MODIFIER |
| 18 | 68407144 | AX-170477759 | 0 | C | T | 0.272 | -7.41E-02 | 1.02E-01 | 2.69E+00 | 2.75E+00 | 4.61E-01 | no | Mc2r | 3_prime_UTR_variant | MODIFIER |
| 18 | 68407262 | AX-97837751 | 0 | T | G | 0.272 | -7.41E-02 | 1.02E-01 | 2.69E+00 | 2.75E+00 | 4.61E-01 | no | Mc2r | 3_prime_UTR_variant | MODIFIER |
| 18 | 68409783 | AX-170343906 | 0 | C | A | 0.272 | -7.41E-02 | 1.02E-01 | 2.69E+00 | 2.75E+00 | 4.61E-01 | no | Mc2r | intron_variant | MODIFIER |
| 18 | 68423681 | AX-97824430 | 0 | C | A | 0.272 | -7.41E-02 | 1.02E-01 | 2.69E+00 | 2.75E+00 | 4.61E-01 | no | Mc2r | intron_variant | MODIFIER |
| 18 | 68431044 | AX-170299611 | 0 | C | G | 0.272 | -7.41E-02 | 1.02E-01 | 2.69E+00 | 2.75E+00 | 4.61E-01 | no | Mc2r | upstream_gene_variant | MODIFIER |
| 18 | 68468343 | AX-170162481 | 0 | G | C | 0.272 | -7.41E-02 | 1.02E-01 | 2.69E+00 | 2.75E+00 | 4.61E-01 | no | | intergenic_variant | MODIFIER |
| 18 | 68471999 | AX-170491231 | 0 | A | C | 0.272 | -7.41E-02 | 1.02E-01 | 2.69E+00 | 2.75E+00 | 4.61E-01 | no | | intergenic_variant | MODIFIER |
| 18 | 68491817 | AX-170015914 | 0 | A | G | 0.272 | -7.41E-02 | 1.02E-01 | 2.69E+00 | 2.75E+00 | 4.61E-01 | no | | regulatory_region_variant | MODIFIER |
| 18 | 68586402 | AX-170476465 | 0 | T | C | 0.272 | -7.41E-02 | 1.02E-01 | 2.69E+00 | 2.75E+00 | 4.61E-01 | no | AC147020.3 | downstream_gene_variant | MODIFIER |
| 18 | 68726767 | AX-170514120 | 0 | G | A | 0.272 | -7.41E-02 | 1.02E-01 | 2.69E+00 | 2.75E+00 | 4.61E-01 | no | | intergenic_variant | MODIFIER |
| 18 | 68732590 | AX-97855669 | 0 | C | A | 0.272 | -7.41E-02 | 1.02E-01 | 2.69E+00 | 2.75E+00 | 4.61E-01 | no | | intergenic_variant | MODIFIER |
| 18 | 68732951 | AX-170384781 | 0 | A | G | 0.272 | -7.41E-02 | 1.02E-01 | 2.69E+00 | 2.75E+00 | 4.61E-01 | no | | intergenic_variant | MODIFIER |
| 18 | 68739096 | AX-170188656 | 0 | A | G | 0.272 | -7.41E-02 | 1.02E-01 | 2.69E+00 | 2.75E+00 | 4.61E-01 | no | Mir6356 | upstream_gene_variant | MODIFIER |
| 18 | 68739252 | AX-170464562 | 0 | C | T | 0.272 | -7.41E-02 | 1.02E-01 | 2.69E+00 | 2.75E+00 | 4.61E-01 | no | Mir6356 | upstream_gene_variant | MODIFIER |
| 18 | 68825480 | AX-170191020 | 0 | A | G | 0.272 | -7.41E-02 | 1.02E-01 | 2.69E+00 | 2.75E+00 | 4.61E-01 | no | | intergenic_variant | MODIFIER |
| 16 | 45598943 | AX-170012956 | 1 | G | A | 0.473 | -5.97E-02 | 8.02E-02 | 2.55E+00 | 2.60E+00 | 4.61E-01 | no | Slc9c1 | intron_variant | MODIFIER |
| 7 | 73760114 | AX-97857069 | 1 | C | T | 0.366 | -7.08E-02 | 9.74E-02 | 2.78E+00 | 2.88E+00 | 4.61E-01 | no | Fam174b | intron_variant | MODIFIER |
| 18 | 80840826 | AX-170319532 | 1 | T | G | 0.295 | -6.35E-02 | 8.59E-02 | 2.56E+00 | 2.66E+00 | 4.61E-01 | no | Atp9b | intron_variant | MODIFIER |
| 6 | 94059804 | AX-97807522 | 0 | C | A | 0.43 | -4.50E-02 | 6.04E-02 | 2.51E+00 | 2.63E+00 | 4.61E-01 | no | Magi1 | intron_variant | MODIFIER |
| 15 | 3319911 | AX-170489614 | 1 | C | T | 0.241 | -7.63E-02 | 1.03E-01 | 2.58E+00 | 2.64E+00 | 4.61E-01 | no | Ghr | missense_variant | MODERATE |
| 6 | 94409408 | AX-170499868 | 1 | T | C | 0.429 | -4.50E-02 | 6.03E-02 | 2.51E+00 | 2.62E+00 | 4.61E-01 | no | | intergenic_variant | MODIFIER |
| 12 | 91809260 | AX-170518350 | 1 | T | C | 0.071 | 1.55E-01 | 2.14E-01 | 2.76E+00 | 2.84E+00 | 4.61E-01 | Yes | Sel1l | 3_prime_UTR_variant | MODIFIER |
| 16 | 49846156 | AX-170330026 | 0 | A | G | 0.509 | 6.24E-02 | 8.40E-02 | 2.55E+00 | 2.58E+00 | 4.61E-01 | no | Cd47 | intron_variant | MODIFIER |
| 16 | 49849657 | AX-170228142 | 0 | C | T | 0.509 | 6.24E-02 | 8.40E-02 | 2.55E+00 | 2.58E+00 | 4.61E-01 | no | Cd47 | intron_variant | MODIFIER |
| 16 | 49893491 | AX-170236262 | 0 | T | C | 0.509 | 6.24E-02 | 8.40E-02 | 2.55E+00 | 2.58E+00 | 4.61E-01 | no | Cd47 | intron_variant | MODIFIER |
| 16 | 49906317 | AX-170379559 | 0 | A | G | 0.509 | 6.24E-02 | 8.40E-02 | 2.55E+00 | 2.58E+00 | 4.61E-01 | no | Cd47 | intron_variant | MODIFIER |

| | | | | | | | | | | | | | | | |
|---|---|---|---|---|---|---|---|---|---|---|---|---|---|---|---|
| 16 | 49911541 | AX-170192612 | 0 | C | T | 0.509 | 6.24E-02 | 8.40E-02 | 2.55E+00 | 2.58E+00 | 4.61E-01 | no | Cd47 | 3_prime_UTR_variant | MODIFIER |
| 16 | 49915974 | AX-170013018 | 0 | T | C | 0.509 | 6.24E-02 | 8.40E-02 | 2.55E+00 | 2.58E+00 | 4.61E-01 | no | Cd47 | downstream_gene_variant | MODIFIER |
| 16 | 49924842 | AX-97790664 | 0 | T | C | 0.509 | 6.24E-02 | 8.40E-02 | 2.55E+00 | 2.58E+00 | 4.61E-01 | no | | intergenic_variant | MODIFIER |
| 16 | 49925034 | AX-170201658 | 0 | G | A | 0.509 | 6.24E-02 | 8.40E-02 | 2.55E+00 | 2.58E+00 | 4.61E-01 | no | | intergenic_variant | MODIFIER |
| 16 | 49933461 | AX-170368779 | 0 | A | G | 0.509 | 6.24E-02 | 8.40E-02 | 2.55E+00 | 2.58E+00 | 4.61E-01 | no | | intergenic_variant | MODIFIER |
| 16 | 49935615 | AX-170439647 | 0 | C | T | 0.509 | 6.24E-02 | 8.40E-02 | 2.55E+00 | 2.58E+00 | 4.61E-01 | no | | intergenic_variant | MODIFIER |
| 16 | 49938804 | AX-170253748 | 0 | A | G | 0.509 | 6.24E-02 | 8.40E-02 | 2.55E+00 | 2.58E+00 | 4.61E-01 | no | | intergenic_variant | MODIFIER |
| 16 | 49952120 | AX-170105663 | 0 | A | G | 0.509 | 6.24E-02 | 8.40E-02 | 2.55E+00 | 2.58E+00 | 4.61E-01 | no | | regulatory_region_variant | MODIFIER |
| 16 | 49952485 | AX-170442771 | 0 | T | C | 0.509 | 6.24E-02 | 8.40E-02 | 2.55E+00 | 2.58E+00 | 4.61E-01 | no | | regulatory_region_variant | MODIFIER |
| 16 | 49953412 | AX-170057397 | 0 | T | C | 0.509 | 6.24E-02 | 8.40E-02 | 2.55E+00 | 2.58E+00 | 4.61E-01 | no | | regulatory_region_variant | MODIFIER |
| 16 | 49960742 | AX-170453178 | 0 | C | T | 0.509 | 6.24E-02 | 8.40E-02 | 2.55E+00 | 2.58E+00 | 4.61E-01 | no | | intergenic_variant | MODIFIER |
| 16 | 49990808 | AX-170013013 | 0 | G | A | 0.509 | 6.24E-02 | 8.40E-02 | 2.55E+00 | 2.58E+00 | 4.61E-01 | no | | intergenic_variant | MODIFIER |
| 16 | 49991728 | AX-97831726 | 0 | C | T | 0.509 | 6.24E-02 | 8.40E-02 | 2.55E+00 | 2.58E+00 | 4.61E-01 | no | | intergenic_variant | MODIFIER |
| 16 | 50004757 | AX-170013015 | 0 | C | T | 0.509 | 6.24E-02 | 8.40E-02 | 2.55E+00 | 2.58E+00 | 4.61E-01 | no | | intergenic_variant | MODIFIER |
| 16 | 50010515 | AX-170013011 | 0 | G | C | 0.509 | 6.24E-02 | 8.40E-02 | 2.55E+00 | 2.58E+00 | 4.61E-01 | no | | regulatory_region_variant | MODIFIER |
| 16 | 50014183 | AX-170383636 | 0 | C | G | 0.509 | 6.24E-02 | 8.40E-02 | 2.55E+00 | 2.58E+00 | 4.61E-01 | no | | intergenic_variant | MODIFIER |
| 16 | 50014281 | AX-170504535 | 0 | A | T | 0.509 | 6.24E-02 | 8.40E-02 | 2.55E+00 | 2.58E+00 | 4.61E-01 | no | | intergenic_variant | MODIFIER |
| 16 | 50017225 | AX-97841077 | 0 | C | T | 0.509 | 6.24E-02 | 8.40E-02 | 2.55E+00 | 2.58E+00 | 4.61E-01 | no | | intergenic_variant | MODIFIER |
| 16 | 50017690 | AX-170136008 | 0 | C | T | 0.509 | 6.24E-02 | 8.40E-02 | 2.55E+00 | 2.58E+00 | 4.61E-01 | no | | intergenic_variant | MODIFIER |
| 16 | 50017779 | AX-170232831 | 0 | T | A | 0.509 | 6.24E-02 | 8.40E-02 | 2.55E+00 | 2.58E+00 | 4.61E-01 | no | | intergenic_variant | MODIFIER |
| 16 | 50018168 | AX-170279177 | 0 | A | G | 0.509 | 6.24E-02 | 8.40E-02 | 2.55E+00 | 2.58E+00 | 4.61E-01 | no | | intergenic_variant | MODIFIER |
| 16 | 50048489 | AX-170088012 | 0 | A | G | 0.509 | 6.24E-02 | 8.40E-02 | 2.55E+00 | 2.58E+00 | 4.61E-01 | no | | intergenic_variant | MODIFIER |
| 16 | 50053609 | AX-170299932 | 0 | T | C | 0.509 | 6.24E-02 | 8.40E-02 | 2.55E+00 | 2.58E+00 | 4.61E-01 | no | | intergenic_variant | MODIFIER |
| 16 | 50074028 | AX-170013004 | 0 | C | T | 0.509 | 6.24E-02 | 8.40E-02 | 2.55E+00 | 2.58E+00 | 4.61E-01 | no | | regulatory_region_variant | MODIFIER |
| 16 | 50082612 | AX-97850531 | 0 | T | G | 0.509 | 6.24E-02 | 8.40E-02 | 2.55E+00 | 2.58E+00 | 4.61E-01 | no | | intergenic_variant | MODIFIER |
| 16 | 50106834 | AX-170013012 | 0 | G | A | 0.509 | 6.24E-02 | 8.40E-02 | 2.55E+00 | 2.58E+00 | 4.61E-01 | no | Gm8824 | upstream_gene_variant | MODIFIER |
| 16 | 50112553 | AX-170517606 | 0 | A | C | 0.509 | 6.24E-02 | 8.40E-02 | 2.55E+00 | 2.58E+00 | 4.61E-01 | no | Gm8824 | downstream_gene_variant | MODIFIER |
| 16 | 50113738 | AX-170107590 | 0 | A | G | 0.509 | 6.24E-02 | 8.40E-02 | 2.55E+00 | 2.58E+00 | 4.61E-01 | no | Gm8824 | downstream_gene_variant | MODIFIER |
| 16 | 50147656 | AX-170013014 | 0 | C | G | 0.509 | 6.24E-02 | 8.40E-02 | 2.55E+00 | 2.58E+00 | 4.61E-01 | no | | intergenic_variant | MODIFIER |
| 16 | 50147941 | AX-97811143 | 0 | C | T | 0.509 | 6.24E-02 | 8.40E-02 | 2.55E+00 | 2.58E+00 | 4.61E-01 | no | | regulatory_region_variant | MODIFIER |
| 16 | 50178846 | AX-97847490 | 0 | C | A | 0.509 | 6.24E-02 | 8.40E-02 | 2.55E+00 | 2.58E+00 | 4.61E-01 | no | | intergenic_variant | MODIFIER |
| 16 | 50185717 | AX-170396163 | 0 | G | A | 0.509 | 6.24E-02 | 8.40E-02 | 2.55E+00 | 2.58E+00 | 4.61E-01 | no | | regulatory_region_variant | MODIFIER |
| 16 | 50187861 | AX-170529977 | 0 | A | C | 0.509 | 6.24E-02 | 8.40E-02 | 2.55E+00 | 2.58E+00 | 4.61E-01 | no | Bbx | downstream_gene_variant | MODIFIER |
| 16 | 50191724 | AX-170184429 | 0 | G | A | 0.509 | 6.24E-02 | 8.40E-02 | 2.55E+00 | 2.58E+00 | 4.61E-01 | no | Bbx | downstream_gene_variant | MODIFIER |
| 16 | 50205834 | AX-170381744 | 0 | T | C | 0.509 | 6.24E-02 | 8.40E-02 | 2.55E+00 | 2.58E+00 | 4.61E-01 | no | Bbx | intron_variant | MODIFIER |
| 16 | 50207781 | AX-170157737 | 0 | A | G | 0.509 | 6.24E-02 | 8.40E-02 | 2.55E+00 | 2.58E+00 | 4.61E-01 | no | Bbx | intron_variant | MODIFIER |
| 16 | 50208177 | AX-170421956 | 0 | T | G | 0.509 | 6.24E-02 | 8.40E-02 | 2.55E+00 | 2.58E+00 | 4.61E-01 | no | Bbx | intron_variant | MODIFIER |
| 16 | 50209376 | AX-170057399 | 0 | G | A | 0.509 | 6.24E-02 | 8.40E-02 | 2.55E+00 | 2.58E+00 | 4.61E-01 | no | Bbx | intron_variant | MODIFIER |
| 16 | 50222999 | AX-170107302 | 0 | T | G | 0.509 | 6.24E-02 | 8.40E-02 | 2.55E+00 | 2.58E+00 | 4.61E-01 | no | Bbx | intron_variant | MODIFIER |
| 16 | 50236256 | AX-170295901 | 0 | T | C | 0.509 | 6.24E-02 | 8.40E-02 | 2.55E+00 | 2.58E+00 | 4.61E-01 | no | Bbx | intron_variant | MODIFIER |
| 16 | 50255575 | AX-170366257 | 0 | A | G | 0.509 | 6.24E-02 | 8.40E-02 | 2.55E+00 | 2.58E+00 | 4.61E-01 | no | Bbx | intron_variant | MODIFIER |
| 16 | 50257079 | AX-170276994 | 0 | A | C | 0.509 | 6.24E-02 | 8.40E-02 | 2.55E+00 | 2.58E+00 | 4.61E-01 | no | Bbx | intron_variant | MODIFIER |
| 16 | 50274488 | AX-97854026 | 0 | T | C | 0.509 | 6.24E-02 | 8.40E-02 | 2.55E+00 | 2.58E+00 | 4.61E-01 | no | Bbx | intron_variant | MODIFIER |
| 16 | 50305404 | AX-170321419 | 0 | T | C | 0.509 | 6.24E-02 | 8.40E-02 | 2.55E+00 | 2.58E+00 | 4.61E-01 | no | Bbx | intron_variant | MODIFIER |
| 16 | 50323107 | AX-170206249 | 0 | G | A | 0.509 | 6.24E-02 | 8.40E-02 | 2.55E+00 | 2.58E+00 | 4.61E-01 | no | Bbx | intron_variant | MODIFIER |
| 16 | 50343981 | AX-97831753 | 0 | A | G | 0.509 | 6.24E-02 | 8.40E-02 | 2.55E+00 | 2.58E+00 | 4.61E-01 | no | Bbx | intron_variant | MODIFIER |
| 16 | 50348462 | AX-170013019 | 0 | C | G | 0.509 | 6.24E-02 | 8.40E-02 | 2.55E+00 | 2.58E+00 | 4.61E-01 | no | Bbx | intron_variant | MODIFIER |
| 16 | 50348493 | AX-170013008 | 0 | A | C | 0.509 | 6.24E-02 | 8.40E-02 | 2.55E+00 | 2.58E+00 | 4.61E-01 | no | Bbx | intron_variant | MODIFIER |
| 1 | 110788681 | AX-170276446 | 0 | A | T | 0.175 | 1.21E-01 | 1.65E-01 | 2.59E+00 | 2.70E+00 | 4.61E-01 | no | | intergenic_variant | MODIFIER |
| 1 | 115591695 | AX-170001471 | 0 | C | A | 0.175 | 1.21E-01 | 1.65E-01 | 2.59E+00 | 2.70E+00 | 4.61E-01 | no | | intergenic_variant | MODIFIER |
| 9 | 20698085 | AX-170443451 | 0 | C | A | 0.474 | 6.74E-02 | 9.08E-02 | 2.55E+00 | 2.61E+00 | 4.62E-01 | no | Olfm2 | intron_variant | MODIFIER |
| 9 | 20702036 | AX-170031928 | 0 | G | T | 0.474 | 6.74E-02 | 9.08E-02 | 2.55E+00 | 2.61E+00 | 4.62E-01 | no | Olfm2 | intron_variant | MODIFIER |
| 9 | 20703416 | AX-170471016 | 0 | G | T | 0.474 | 6.74E-02 | 9.08E-02 | 2.55E+00 | 2.61E+00 | 4.62E-01 | no | Olfm2 | intron_variant | MODIFIER |
| 9 | 20799977 | AX-97835728 | 0 | T | C | 0.474 | 6.74E-02 | 9.08E-02 | 2.55E+00 | 2.61E+00 | 4.62E-01 | no | Col5a3 | intron_variant | MODIFIER |

| | | | | | | | | | | | | | | | |
|---|---|---|---|---|---|---|---|---|---|---|---|---|---|---|---|
| 9 | 20805571 | AX-168405450 | 0 | C | T | 0.474 | 6.74E-02 | 9.08E-02 | 2.55E+00 | 2.61E+00 | 4.62E-01 | no | Col5a3 | intron_variant | MODIFIER |
| 9 | 20812798 | AX-170401249 | 0 | A | G | 0.474 | 6.74E-02 | 9.08E-02 | 2.55E+00 | 2.61E+00 | 4.62E-01 | no | Col5a3 | intron_variant | MODIFIER |
| 9 | 20829387 | AX-170031919 | 0 | C | T | 0.474 | 6.74E-02 | 9.08E-02 | 2.55E+00 | 2.61E+00 | 4.62E-01 | no | Rdh8 | downstream_gene_variant | MODIFIER |
| 9 | 20831391 | AX-170487938 | 0 | C | T | 0.474 | 6.74E-02 | 9.08E-02 | 2.55E+00 | 2.61E+00 | 4.62E-01 | no | | regulatory_region_variant | MODIFIER |
| 9 | 20859749 | AX-97811795 | 0 | T | C | 0.474 | 6.74E-02 | 9.08E-02 | 2.55E+00 | 2.61E+00 | 4.62E-01 | no | | intergenic_variant | MODIFIER |
| 9 | 20886999 | AX-170317329 | 0 | T | C | 0.474 | 6.74E-02 | 9.08E-02 | 2.55E+00 | 2.61E+00 | 4.62E-01 | no | Ppan | upstream_gene_variant | MODIFIER |
| 9 | 20910718 | AX-170193170 | 0 | A | T | 0.474 | 6.74E-02 | 9.08E-02 | 2.55E+00 | 2.61E+00 | 4.62E-01 | no | Dnmt1 | intron_variant | MODIFIER |
| 9 | 20911777 | AX-170332177 | 0 | A | G | 0.474 | 6.74E-02 | 9.08E-02 | 2.55E+00 | 2.61E+00 | 4.62E-01 | no | Dnmt1 | synonymous_variant | LOW |
| 9 | 20912020 | AX-170204841 | 0 | C | T | 0.474 | 6.74E-02 | 9.08E-02 | 2.55E+00 | 2.61E+00 | 4.62E-01 | no | Dnmt1 | splice_region_variant,intron_variant | LOW |
| 9 | 20916570 | AX-170183952 | 0 | A | C | 0.474 | 6.74E-02 | 9.08E-02 | 2.55E+00 | 2.61E+00 | 4.62E-01 | no | Dnmt1 | intron_variant | MODIFIER |
| 9 | 20920223 | AX-97789549 | 0 | G | A | 0.474 | 6.74E-02 | 9.08E-02 | 2.55E+00 | 2.61E+00 | 4.62E-01 | no | Dnmt1 | intron_variant | MODIFIER |
| 9 | 20943893 | AX-170255936 | 0 | G | A | 0.474 | 6.74E-02 | 9.08E-02 | 2.55E+00 | 2.61E+00 | 4.62E-01 | no | Dnmt1 | intron_variant | MODIFIER |
| 9 | 20944088 | AX-170137632 | 0 | G | A | 0.474 | 6.74E-02 | 9.08E-02 | 2.55E+00 | 2.61E+00 | 4.62E-01 | no | Dnmt1 | intron_variant | MODIFIER |
| 9 | 20947433 | AX-97787668 | 0 | C | T | 0.474 | 6.74E-02 | 9.08E-02 | 2.55E+00 | 2.61E+00 | 4.62E-01 | no | Dnmt1 | intron_variant | MODIFIER |
| 9 | 21137074 | AX-97810218 | 0 | A | G | 0.474 | 6.74E-02 | 9.08E-02 | 2.55E+00 | 2.61E+00 | 4.62E-01 | no | Cdc37 | 3_prime_UTR_variant | MODIFIER |
| 9 | 22854061 | AX-97831288 | 0 | G | A | 0.474 | 6.74E-02 | 9.08E-02 | 2.55E+00 | 2.61E+00 | 4.62E-01 | no | Bbs9 | intron_variant | MODIFIER |
| 9 | 24789420 | AX-170254409 | 0 | C | T | 0.474 | 6.74E-02 | 9.08E-02 | 2.55E+00 | 2.61E+00 | 4.62E-01 | no | Gm29824 | intron_variant,non_coding_transcript_variant | MODIFIER |
| 9 | 52665510 | AX-170499680 | 0 | T | C | 0.219 | -8.03E-02 | 1.11E-01 | 2.78E+00 | 2.87E+00 | 4.62E-01 | no | | intergenic_variant | MODIFIER |
| 13 | 37751202 | AX-170234506 | 0 | T | G | 0.175 | -7.98E-02 | 1.08E-01 | 2.57E+00 | 2.64E+00 | 4.62E-01 | no | | regulatory_region_variant | MODIFIER |
| 1 | 116681087 | AX-170411416 | 2 | T | G | 0.164 | 1.35E-01 | 1.84E-01 | 2.61E+00 | 2.70E+00 | 4.62E-01 | no | | intergenic_variant | MODIFIER |
| 15 | 4702309 | AX-170225236 | 0 | A | T | 0.246 | -7.43E-02 | 1.02E-01 | 2.65E+00 | 2.72E+00 | 4.62E-01 | no | | intergenic_variant | MODIFIER |
| 8 | 66312135 | AX-97826791 | 0 | G | A | 0.158 | -8.98E-02 | 1.23E-01 | 2.62E+00 | 2.73E+00 | 4.62E-01 | no | MAR1 | intron_variant | MODIFIER |
| 15 | 95365684 | AX-170133260 | 0 | G | T | 0.149 | -1.16E-01 | 1.60E-01 | 2.78E+00 | 2.87E+00 | 4.62E-01 | no | Nell2 | intron_variant | MODIFIER |
| 4 | 136472183 | AX-170024034 | 1 | G | A | 0.161 | 8.15E-02 | 1.10E-01 | 2.55E+00 | 2.62E+00 | 4.62E-01 | no | Luzp1 | intron_variant | MODIFIER |
| 5 | 36722252 | AX-170390362 | 0 | A | G | 0.325 | 8.26E-02 | 1.14E-01 | 2.76E+00 | 2.82E+00 | 4.62E-01 | no | | regulatory_region_variant | MODIFIER |
| 3 | 26341277 | AX-168415878 | 2 | G | A | 0.136 | -7.77E-02 | 1.05E-01 | 2.56E+00 | 2.64E+00 | 4.62E-01 | no | | intergenic_variant | MODIFIER |
| 9 | 18819793 | AX-97853197 | 0 | T | C | 0.465 | 6.67E-02 | 8.99E-02 | 2.55E+00 | 2.61E+00 | 4.62E-01 | no | Olfr828 | upstream_gene_variant | MODIFIER |
| 9 | 18827417 | AX-170031888 | 0 | A | T | 0.465 | 6.67E-02 | 8.99E-02 | 2.55E+00 | 2.61E+00 | 4.62E-01 | no | | intergenic_variant | MODIFIER |
| 9 | 18928501 | AX-97786993 | 0 | C | T | 0.465 | 6.67E-02 | 8.99E-02 | 2.55E+00 | 2.61E+00 | 4.62E-01 | no | Olfr831-ps1 | downstream_gene_variant | MODIFIER |
| 9 | 19031578 | AX-170031892 | 0 | G | C | 0.465 | 6.67E-02 | 8.99E-02 | 2.55E+00 | 2.61E+00 | 4.62E-01 | no | Olfr835 | upstream_gene_variant | MODIFIER |
| 9 | 19039320 | AX-97801687 | 0 | C | A | 0.465 | 6.67E-02 | 8.99E-02 | 2.55E+00 | 2.61E+00 | 4.62E-01 | no | Olfr835 | downstream_gene_variant | MODIFIER |
| 9 | 19039363 | AX-170031889 | 0 | G | A | 0.465 | 6.67E-02 | 8.99E-02 | 2.55E+00 | 2.61E+00 | 4.62E-01 | no | Olfr835 | downstream_gene_variant | MODIFIER |
| 9 | 19075470 | AX-170031890 | 0 | C | G | 0.465 | 6.67E-02 | 8.99E-02 | 2.55E+00 | 2.61E+00 | 4.62E-01 | no | | intergenic_variant | MODIFIER |
| 9 | 19083783 | AX-97798551 | 0 | C | T | 0.465 | 6.67E-02 | 8.99E-02 | 2.55E+00 | 2.61E+00 | 4.62E-01 | no | | intergenic_variant | MODIFIER |
| 9 | 19102908 | AX-97795907 | 0 | C | T | 0.465 | 6.67E-02 | 8.99E-02 | 2.55E+00 | 2.61E+00 | 4.62E-01 | no | | intergenic_variant | MODIFIER |
| 9 | 19135464 | AX-168384982 | 0 | T | G | 0.465 | 6.67E-02 | 8.99E-02 | 2.55E+00 | 2.61E+00 | 4.62E-01 | no | Olfr837 | upstream_gene_variant | MODIFIER |
| 9 | 19137146 | AX-97818843 | 0 | T | G | 0.465 | 6.67E-02 | 8.99E-02 | 2.55E+00 | 2.61E+00 | 4.62E-01 | no | Olfr837 | missense_variant | MODERATE |
| 9 | 19204979 | AX-170209645 | 0 | A | T | 0.465 | 6.67E-02 | 8.99E-02 | 2.55E+00 | 2.61E+00 | 4.62E-01 | no | Olfr840-ps1 | non_coding_transcript_exon_variant | MODIFIER |
| 9 | 19232868 | AX-97782272 | 0 | G | A | 0.465 | 6.67E-02 | 8.99E-02 | 2.55E+00 | 2.61E+00 | 4.62E-01 | no | AC119987.1 | upstream_gene_variant | MODIFIER |
| 9 | 19240716 | AX-170381871 | 0 | T | G | 0.465 | 6.67E-02 | 8.99E-02 | 2.55E+00 | 2.61E+00 | 4.62E-01 | no | Olfr843 | downstream_gene_variant | MODIFIER |
| 9 | 19250951 | AX-170031905 | 0 | G | A | 0.465 | 6.67E-02 | 8.99E-02 | 2.55E+00 | 2.61E+00 | 4.62E-01 | no | Olfr843 | upstream_gene_variant | MODIFIER |
| 9 | 19341392 | AX-97790769 | 0 | A | G | 0.465 | 6.67E-02 | 8.99E-02 | 2.55E+00 | 2.61E+00 | 4.62E-01 | no | Olfr845 | downstream_gene_variant | MODIFIER |
| 9 | 19343954 | AX-170031894 | 0 | C | T | 0.465 | 6.67E-02 | 8.99E-02 | 2.55E+00 | 2.61E+00 | 4.62E-01 | no | Olfr845 | downstream_gene_variant | MODIFIER |
| 9 | 19372352 | AX-97789637 | 0 | A | G | 0.465 | 6.67E-02 | 8.99E-02 | 2.55E+00 | 2.61E+00 | 4.62E-01 | no | Olfr847 | downstream_gene_variant | MODIFIER |
| 9 | 19437927 | AX-170031903 | 0 | T | C | 0.465 | 6.67E-02 | 8.99E-02 | 2.55E+00 | 2.61E+00 | 4.62E-01 | no | Olfr849 | upstream_gene_variant | MODIFIER |
| 9 | 19438540 | AX-97786964 | 0 | T | C | 0.465 | 6.67E-02 | 8.99E-02 | 2.55E+00 | 2.61E+00 | 4.62E-01 | no | Olfr849 | upstream_gene_variant | MODIFIER |
| 9 | 19465572 | AX-97836969 | 0 | A | G | 0.465 | 6.67E-02 | 8.99E-02 | 2.55E+00 | 2.61E+00 | 4.62E-01 | no | | intergenic_variant | MODIFIER |
| 9 | 19693918 | AX-97788764 | 0 | T | C | 0.465 | 6.67E-02 | 8.99E-02 | 2.55E+00 | 2.61E+00 | 4.62E-01 | no | Gm7769 | downstream_gene_variant | MODIFIER |
| 9 | 19770172 | AX-97822362 | 0 | T | C | 0.465 | 6.67E-02 | 8.99E-02 | 2.55E+00 | 2.61E+00 | 4.62E-01 | no | | regulatory_region_variant | MODIFIER |
| 9 | 19873465 | AX-97806790 | 0 | G | T | 0.465 | 6.67E-02 | 8.99E-02 | 2.55E+00 | 2.61E+00 | 4.62E-01 | no | Olfr861-ps1 | upstream_gene_variant | MODIFIER |
| 9 | 20468068 | AX-170510367 | 0 | C | G | 0.465 | 6.67E-02 | 8.99E-02 | 2.55E+00 | 2.61E+00 | 4.62E-01 | no | Zfp426 | downstream_gene_variant | MODIFIER |
| 9 | 20469776 | AX-170284024 | 0 | C | T | 0.465 | 6.67E-02 | 8.99E-02 | 2.55E+00 | 2.61E+00 | 4.62E-01 | no | Zfp426 | 3_prime_UTR_variant | MODIFIER |
| 9 | 20469983 | AX-170231927 | 0 | C | T | 0.465 | 6.67E-02 | 8.99E-02 | 2.55E+00 | 2.61E+00 | 4.62E-01 | no | Zfp426 | 3_prime_UTR_variant | MODIFIER |
| 9 | 20471213 | AX-170333146 | 0 | G | A | 0.465 | 6.67E-02 | 8.99E-02 | 2.55E+00 | 2.61E+00 | 4.62E-01 | no | Zfp426 | synonymous_variant | LOW |

| | | | | | | | | | | | | | | | |
|---|---|---|---|---|---|---|---|---|---|---|---|---|---|---|---|
| 9 | 20471398 | AX-97816898 | 0 | C | T | 0.465 | 6.67E-02 | 8.99E-02 | 2.55E+00 | 2.61E+00 | 4.62E-01 | no | Zfp426 | intron_variant | MODIFIER |
| 9 | 20472398 | AX-170105189 | 0 | G | C | 0.465 | 6.67E-02 | 8.99E-02 | 2.55E+00 | 2.61E+00 | 4.62E-01 | no | Zfp426 | intron_variant | MODIFIER |
| 9 | 20476257 | AX-170351043 | 0 | C | T | 0.465 | 6.67E-02 | 8.99E-02 | 2.55E+00 | 2.61E+00 | 4.62E-01 | no | Zfp426 | intron_variant | MODIFIER |
| 9 | 20478169 | AX-170211485 | 0 | C | G | 0.465 | 6.67E-02 | 8.99E-02 | 2.55E+00 | 2.61E+00 | 4.62E-01 | no | Zfp426 | 5_prime_UTR_variant | MODIFIER |
| 9 | 20489742 | AX-170153920 | 0 | A | G | 0.465 | 6.67E-02 | 8.99E-02 | 2.55E+00 | 2.61E+00 | 4.62E-01 | no | Zfp426 | intron_variant | MODIFIER |
| 9 | 20534366 | AX-170090260 | 0 | A | G | 0.465 | 6.67E-02 | 8.99E-02 | 2.55E+00 | 2.61E+00 | 4.62E-01 | no | Gm26733 | intron_variant,non_coding_transcript_variant | MODIFIER |
| 9 | 20583493 | AX-97825148 | 0 | T | C | 0.465 | 6.67E-02 | 8.99E-02 | 2.55E+00 | 2.61E+00 | 4.62E-01 | no | Zfp846 | upstream_gene_variant | MODIFIER |
| 9 | 20583543 | AX-170081821 | 0 | A | G | 0.465 | 6.67E-02 | 8.99E-02 | 2.55E+00 | 2.61E+00 | 4.62E-01 | no | Zfp846 | upstream_gene_variant | MODIFIER |
| 9 | 20612307 | AX-97805211 | 0 | C | T | 0.465 | 6.67E-02 | 8.99E-02 | 2.55E+00 | 2.61E+00 | 4.62E-01 | no | Fbxl12os | upstream_gene_variant | MODIFIER |
| 9 | 20618570 | AX-170031923 | 0 | G | C | 0.465 | 6.67E-02 | 8.99E-02 | 2.55E+00 | 2.61E+00 | 4.62E-01 | no | Fbxl12os | downstream_gene_variant | MODIFIER |
| 9 | 20655634 | AX-97833992 | 0 | A | G | 0.465 | 6.67E-02 | 8.99E-02 | 2.55E+00 | 2.61E+00 | 4.62E-01 | no | Pin1 | intron_variant | MODIFIER |
| 9 | 20660445 | AX-170233385 | 0 | C | G | 0.465 | 6.67E-02 | 8.99E-02 | 2.55E+00 | 2.61E+00 | 4.62E-01 | no | Pin1 | intron_variant | MODIFIER |
| 9 | 20672808 | AX-170031918 | 0 | T | C | 0.465 | 6.67E-02 | 8.99E-02 | 2.55E+00 | 2.61E+00 | 4.62E-01 | no | Olfm2 | intron_variant | MODIFIER |
| 9 | 20672890 | AX-97815778 | 0 | A | C | 0.465 | 6.67E-02 | 8.99E-02 | 2.55E+00 | 2.61E+00 | 4.62E-01 | no | Olfm2 | intron_variant | MODIFIER |
| 9 | 20673726 | AX-170488221 | 0 | A | G | 0.465 | 6.67E-02 | 8.99E-02 | 2.55E+00 | 2.61E+00 | 4.62E-01 | no | Olfm2 | intron_variant | MODIFIER |
| 9 | 20673902 | AX-170427218 | 0 | C | T | 0.465 | 6.67E-02 | 8.99E-02 | 2.55E+00 | 2.61E+00 | 4.62E-01 | no | Olfm2 | intron_variant | MODIFIER |
| 9 | 20695794 | AX-97817660 | 0 | G | A | 0.465 | 6.67E-02 | 8.99E-02 | 2.55E+00 | 2.61E+00 | 4.62E-01 | no | Olfm2 | intron_variant | MODIFIER |
| 9 | 20707927 | AX-170031920 | 0 | C | G | 0.465 | 6.67E-02 | 8.99E-02 | 2.55E+00 | 2.61E+00 | 4.62E-01 | no | Olfm2 | intron_variant | MODIFIER |
| 9 | 20814085 | AX-97803893 | 0 | T | C | 0.465 | 6.67E-02 | 8.99E-02 | 2.55E+00 | 2.61E+00 | 4.62E-01 | no | Col5a3 | intron_variant | MODIFIER |
| 9 | 21192633 | AX-170031935 | 0 | A | G | 0.465 | 6.67E-02 | 8.99E-02 | 2.55E+00 | 2.61E+00 | 4.62E-01 | no | Pde4a | upstream_gene_variant | MODIFIER |
| 9 | 21205350 | AX-170234564 | 0 | G | A | 0.465 | 6.67E-02 | 8.99E-02 | 2.55E+00 | 2.61E+00 | 4.62E-01 | no | Pde4a | synonymous_variant | LOW |
| 9 | 21213241 | AX-170181529 | 0 | C | T | 0.465 | 6.67E-02 | 8.99E-02 | 2.55E+00 | 2.61E+00 | 4.62E-01 | no | Pde4a | downstream_gene_variant | MODIFIER |
| 9 | 21227875 | AX-97843784 | 0 | T | C | 0.465 | 6.67E-02 | 8.99E-02 | 2.55E+00 | 2.61E+00 | 4.62E-01 | no | Keap1 | downstream_gene_variant | MODIFIER |
| 9 | 21227911 | AX-170031934 | 0 | A | G | 0.465 | 6.67E-02 | 8.99E-02 | 2.55E+00 | 2.61E+00 | 4.62E-01 | no | Keap1 | downstream_gene_variant | MODIFIER |
| 9 | 21253447 | AX-170031933 | 0 | A | T | 0.465 | 6.67E-02 | 8.99E-02 | 2.55E+00 | 2.61E+00 | 4.62E-01 | no | Gm48303 | downstream_gene_variant | MODIFIER |
| 9 | 21266919 | AX-170374572 | 0 | A | G | 0.465 | 6.67E-02 | 8.99E-02 | 2.55E+00 | 2.61E+00 | 4.62E-01 | no | Atg4d | missense_variant | MODERATE |
| 9 | 21287144 | AX-170211444 | 0 | A | G | 0.465 | 6.67E-02 | 8.99E-02 | 2.55E+00 | 2.61E+00 | 4.62E-01 | no | Kri1 | intron_variant | MODIFIER |
| 9 | 21302577 | AX-170123467 | 0 | T | G | 0.465 | 6.67E-02 | 8.99E-02 | 2.55E+00 | 2.61E+00 | 4.62E-01 | no | Ap1m2 | intron_variant | MODIFIER |
| 9 | 21306882 | AX-170112139 | 0 | T | C | 0.465 | 6.67E-02 | 8.99E-02 | 2.55E+00 | 2.61E+00 | 4.62E-01 | no | Ap1m2 | intron_variant | MODIFIER |
| 9 | 21312237 | AX-170291401 | 0 | G | C | 0.465 | 6.67E-02 | 8.99E-02 | 2.55E+00 | 2.61E+00 | 4.62E-01 | no | Ap1m2 | synonymous_variant | LOW |
| 9 | 21352526 | AX-170203970 | 0 | T | C | 0.465 | 6.67E-02 | 8.99E-02 | 2.55E+00 | 2.61E+00 | 4.62E-01 | no | Slc44a2 | synonymous_variant | LOW |
| 9 | 21391433 | AX-170394632 | 0 | A | G | 0.465 | 6.67E-02 | 8.99E-02 | 2.55E+00 | 2.61E+00 | 4.62E-01 | no | Ilf3 | intron_variant | MODIFIER |
| 9 | 21409716 | AX-170236606 | 0 | A | G | 0.465 | 6.67E-02 | 8.99E-02 | 2.55E+00 | 2.61E+00 | 4.62E-01 | no | Qtrt1 | upstream_gene_variant | MODIFIER |
| 9 | 21435613 | AX-170371912 | 0 | A | G | 0.465 | 6.67E-02 | 8.99E-02 | 2.55E+00 | 2.61E+00 | 4.62E-01 | no | Dnm2 | intron_variant | MODIFIER |
| 9 | 21476641 | AX-170124271 | 0 | C | T | 0.465 | 6.67E-02 | 8.99E-02 | 2.55E+00 | 2.61E+00 | 4.62E-01 | no | Dnm2 | intron_variant | MODIFIER |
| 9 | 21491381 | AX-170533310 | 0 | G | A | 0.465 | 6.67E-02 | 8.99E-02 | 2.55E+00 | 2.61E+00 | 4.62E-01 | no | Dnm2 | intron_variant | MODIFIER |
| 9 | 21492060 | AX-170383262 | 0 | T | C | 0.465 | 6.67E-02 | 8.99E-02 | 2.55E+00 | 2.61E+00 | 4.62E-01 | no | Dnm2 | intron_variant | MODIFIER |
| 9 | 21529123 | AX-170330504 | 0 | T | C | 0.465 | 6.67E-02 | 8.99E-02 | 2.55E+00 | 2.61E+00 | 4.62E-01 | no | AB124611 | intron_variant,NMD_transcript_variant | MODIFIER |
| 9 | 21553777 | AX-170357240 | 0 | T | C | 0.465 | 6.67E-02 | 8.99E-02 | 2.55E+00 | 2.61E+00 | 4.62E-01 | no | Carm1 | intron_variant | MODIFIER |
| 9 | 21558675 | AX-97855069 | 0 | A | G | 0.465 | 6.67E-02 | 8.99E-02 | 2.55E+00 | 2.61E+00 | 4.62E-01 | no | Carm1 | intron_variant | MODIFIER |
| 9 | 21559299 | AX-170366341 | 0 | C | G | 0.465 | 6.67E-02 | 8.99E-02 | 2.55E+00 | 2.61E+00 | 4.62E-01 | no | Carm1 | intron_variant | MODIFIER |
| 9 | 21596937 | AX-97781802 | 0 | C | T | 0.465 | 6.67E-02 | 8.99E-02 | 2.55E+00 | 2.61E+00 | 4.62E-01 | no | Yipf2 | upstream_gene_variant | MODIFIER |
| 9 | 21606110 | AX-170254721 | 0 | A | G | 0.465 | 6.67E-02 | 8.99E-02 | 2.55E+00 | 2.61E+00 | 4.62E-01 | no | Gm7904 | downstream_gene_variant | MODIFIER |
| 9 | 21606909 | AX-170511436 | 0 | G | A | 0.465 | 6.67E-02 | 8.99E-02 | 2.55E+00 | 2.61E+00 | 4.62E-01 | no | Gm7904 | downstream_gene_variant | MODIFIER |
| 9 | 21636986 | AX-168410185 | 0 | A | G | 0.465 | 6.67E-02 | 8.99E-02 | 2.55E+00 | 2.61E+00 | 4.62E-01 | no | Smarca4 | intron_variant | MODIFIER |
| 9 | 21643266 | AX-170420482 | 0 | A | G | 0.465 | 6.67E-02 | 8.99E-02 | 2.55E+00 | 2.61E+00 | 4.62E-01 | no | Smarca4 | intron_variant | MODIFIER |
| 9 | 21649772 | AX-170031937 | 0 | C | T | 0.465 | 6.67E-02 | 8.99E-02 | 2.55E+00 | 2.61E+00 | 4.62E-01 | no | Smarca4 | intron_variant | MODIFIER |
| 9 | 21670111 | AX-170031938 | 0 | C | T | 0.465 | 6.67E-02 | 8.99E-02 | 2.55E+00 | 2.61E+00 | 4.62E-01 | no | Smarca4 | intron_variant | MODIFIER |
| 9 | 21675571 | AX-170435860 | 0 | G | A | 0.465 | 6.67E-02 | 8.99E-02 | 2.55E+00 | 2.61E+00 | 4.62E-01 | no | Smarca4 | intron_variant | MODIFIER |
| 9 | 21693547 | AX-170522317 | 0 | C | G | 0.465 | 6.67E-02 | 8.99E-02 | 2.55E+00 | 2.61E+00 | 4.62E-01 | no | Smarca4 | intron_variant | MODIFIER |
| 9 | 21713698 | AX-170240604 | 0 | T | G | 0.465 | 6.67E-02 | 8.99E-02 | 2.55E+00 | 2.61E+00 | 4.62E-01 | no | Gm26511 | downstream_gene_variant | MODIFIER |
| 9 | 21746409 | AX-170483686 | 0 | G | A | 0.465 | 6.67E-02 | 8.99E-02 | 2.55E+00 | 2.61E+00 | 4.62E-01 | no | Ldlr | missense_variant | MODERATE |
| 9 | 21747261 | AX-170529370 | 0 | T | C | 0.465 | 6.67E-02 | 8.99E-02 | 2.55E+00 | 2.61E+00 | 4.62E-01 | no | Ldlr | synonymous_variant | LOW |
| 9 | 21747288 | AX-170141043 | 0 | C | T | 0.465 | 6.67E-02 | 8.99E-02 | 2.55E+00 | 2.61E+00 | 4.62E-01 | no | Ldlr | synonymous_variant | LOW |

| | | | | | | | | | | | | | | | |
|---|---|---|---|---|---|---|---|---|---|---|---|---|---|---|---|
| 9 | 21756171 | AX-170452821 | 0 | G | A | 0.465 | 6.67E-02 | 8.99E-02 | 2.55E+00 | 2.61E+00 | 4.62E-01 | no | Spc24 | 3_prime_UTR_variant | MODIFIER |
| 9 | 21784749 | AX-170260866 | 0 | C | T | 0.465 | 6.67E-02 | 8.99E-02 | 2.55E+00 | 2.61E+00 | 4.62E-01 | no | Kank2 | intron_variant | MODIFIER |
| 9 | 21835040 | AX-170248973 | 0 | G | A | 0.465 | 6.67E-02 | 8.99E-02 | 2.55E+00 | 2.61E+00 | 4.62E-01 | no | Dock6 | intron_variant | MODIFIER |
| 9 | 21835572 | AX-170354820 | 0 | A | G | 0.465 | 6.67E-02 | 8.99E-02 | 2.55E+00 | 2.61E+00 | 4.62E-01 | no | Dock6 | intron_variant | MODIFIER |
| 9 | 21873319 | AX-170367628 | 0 | T | C | 0.465 | 6.67E-02 | 8.99E-02 | 2.55E+00 | 2.61E+00 | 4.62E-01 | no | Gm47873 | intron_variant,non_coding_transcript_variant | MODIFIER |
| 9 | 21901836 | AX-170509980 | 0 | T | C | 0.465 | 6.67E-02 | 8.99E-02 | 2.55E+00 | 2.61E+00 | 4.62E-01 | no | | regulatory_region_variant | MODIFIER |
| 9 | 21929467 | AX-170161463 | 0 | T | G | 0.465 | 6.67E-02 | 8.99E-02 | 2.55E+00 | 2.61E+00 | 4.62E-01 | no | Ccdc159 | missense_variant | MODERATE |
| 9 | 21959719 | AX-170150404 | 0 | G | A | 0.465 | 6.67E-02 | 8.99E-02 | 2.55E+00 | 2.61E+00 | 4.62E-01 | no | Epor | intron_variant | MODIFIER |
| 9 | 21963417 | AX-170163898 | 0 | T | C | 0.465 | 6.67E-02 | 8.99E-02 | 2.55E+00 | 2.61E+00 | 4.62E-01 | no | Epor | synonymous_variant | LOW |
| 9 | 21970852 | AX-170351846 | 0 | C | G | 0.465 | 6.67E-02 | 8.99E-02 | 2.55E+00 | 2.61E+00 | 4.62E-01 | no | Rgl3 | downstream_gene_variant | MODIFIER |
| 9 | 22027767 | AX-170380449 | 0 | C | G | 0.465 | 6.67E-02 | 8.99E-02 | 2.55E+00 | 2.61E+00 | 4.62E-01 | no | Elavl3 | intron_variant | MODIFIER |
| 9 | 22043173 | AX-170172561 | 0 | G | C | 0.465 | 6.67E-02 | 8.99E-02 | 2.55E+00 | 2.61E+00 | 4.62E-01 | no | Elavl3 | intron_variant | MODIFIER |
| 9 | 22063122 | AX-97823128 | 0 | G | A | 0.465 | 6.67E-02 | 8.99E-02 | 2.55E+00 | 2.61E+00 | 4.62E-01 | no | Zfp653 | intron_variant | MODIFIER |
| 9 | 22063851 | AX-170311448 | 0 | A | G | 0.465 | 6.67E-02 | 8.99E-02 | 2.55E+00 | 2.61E+00 | 4.62E-01 | no | Zfp653 | intron_variant | MODIFIER |
| 9 | 22064251 | AX-170121380 | 0 | A | G | 0.465 | 6.67E-02 | 8.99E-02 | 2.55E+00 | 2.61E+00 | 4.62E-01 | no | Zfp653 | intron_variant | MODIFIER |
| 9 | 22107795 | AX-170111442 | 0 | T | C | 0.465 | 6.67E-02 | 8.99E-02 | 2.55E+00 | 2.61E+00 | 4.62E-01 | no | Cnn1 | synonymous_variant | LOW |
| 9 | 22142383 | AX-97811153 | 0 | A | G | 0.465 | 6.67E-02 | 8.99E-02 | 2.55E+00 | 2.61E+00 | 4.62E-01 | no | 6530413G14Rik | downstream_gene_variant | MODIFIER |
| 9 | 22153508 | AX-170231120 | 0 | A | G | 0.465 | 6.67E-02 | 8.99E-02 | 2.55E+00 | 2.61E+00 | 4.62E-01 | no | Pigyl | upstream_gene_variant | MODIFIER |
| 9 | 22208826 | AX-170456926 | 0 | A | C | 0.465 | 6.67E-02 | 8.99E-02 | 2.55E+00 | 2.61E+00 | 4.62E-01 | no | 1810064F22Rik | upstream_gene_variant | MODIFIER |
| 9 | 22216412 | AX-170031948 | 0 | G | A | 0.465 | 6.67E-02 | 8.99E-02 | 2.55E+00 | 2.61E+00 | 4.62E-01 | no | 1810064F22Rik | upstream_gene_variant | MODIFIER |
| 9 | 22219006 | AX-97802643 | 0 | G | A | 0.465 | 6.67E-02 | 8.99E-02 | 2.55E+00 | 2.61E+00 | 4.62E-01 | no | | regulatory_region_variant | MODIFIER |
| 9 | 22233977 | AX-170031949 | 0 | T | C | 0.465 | 6.67E-02 | 8.99E-02 | 2.55E+00 | 2.61E+00 | 4.62E-01 | no | Zfp809 | intron_variant | MODIFIER |
| 9 | 22244783 | AX-170081844 | 0 | T | C | 0.465 | 6.67E-02 | 8.99E-02 | 2.55E+00 | 2.61E+00 | 4.62E-01 | no | Zfp809 | downstream_gene_variant | MODIFIER |
| 9 | 22253898 | AX-170081847 | 0 | A | G | 0.465 | 6.67E-02 | 8.99E-02 | 2.55E+00 | 2.61E+00 | 4.62E-01 | no | Zfp599 | synonymous_variant | LOW |
| 9 | 22254025 | AX-170081846 | 0 | T | C | 0.465 | 6.67E-02 | 8.99E-02 | 2.55E+00 | 2.61E+00 | 4.62E-01 | no | Zfp599 | intron_variant | MODIFIER |
| 9 | 22257547 | AX-170296233 | 0 | A | G | 0.465 | 6.67E-02 | 8.99E-02 | 2.55E+00 | 2.61E+00 | 4.62E-01 | no | Zfp599 | intron_variant | MODIFIER |
| 9 | 22258019 | AX-170031946 | 0 | C | T | 0.465 | 6.67E-02 | 8.99E-02 | 2.55E+00 | 2.61E+00 | 4.62E-01 | no | Zfp599 | intron_variant | MODIFIER |
| 9 | 22258151 | AX-170463101 | 0 | A | G | 0.465 | 6.67E-02 | 8.99E-02 | 2.55E+00 | 2.61E+00 | 4.62E-01 | no | Zfp599 | synonymous_variant | LOW |
| 9 | 22258320 | AX-170031950 | 0 | A | G | 0.465 | 6.67E-02 | 8.99E-02 | 2.55E+00 | 2.61E+00 | 4.62E-01 | no | Zfp599 | intron_variant | MODIFIER |
| 9 | 22291403 | AX-170031944 | 0 | G | T | 0.465 | 6.67E-02 | 8.99E-02 | 2.55E+00 | 2.61E+00 | 4.62E-01 | no | Zfp810 | intron_variant | MODIFIER |
| 9 | 22291946 | AX-170031940 | 0 | C | T | 0.465 | 6.67E-02 | 8.99E-02 | 2.55E+00 | 2.61E+00 | 4.62E-01 | no | Zfp810 | intron_variant | MODIFIER |
| 9 | 22311111 | AX-170031951 | 0 | G | A | 0.465 | 6.67E-02 | 8.99E-02 | 2.55E+00 | 2.61E+00 | 4.62E-01 | no | Zfp810 | upstream_gene_variant | MODIFIER |
| 9 | 22380982 | AX-97819359 | 0 | G | A | 0.465 | 6.67E-02 | 8.99E-02 | 2.55E+00 | 2.61E+00 | 4.62E-01 | no | Anln | intron_variant | MODIFIER |
| 9 | 22405074 | AX-97807206 | 0 | A | G | 0.465 | 6.67E-02 | 8.99E-02 | 2.55E+00 | 2.61E+00 | 4.62E-01 | no | Gm48652 | downstream_gene_variant | MODIFIER |
| 9 | 22405185 | AX-170031947 | 0 | G | T | 0.465 | 6.67E-02 | 8.99E-02 | 2.55E+00 | 2.61E+00 | 4.62E-01 | no | Gm48652 | downstream_gene_variant | MODIFIER |
| 9 | 22411254 | AX-170503201 | 0 | A | T | 0.465 | 6.67E-02 | 8.99E-02 | 2.55E+00 | 2.61E+00 | 4.62E-01 | no | 9530077C05Rik | upstream_gene_variant | MODIFIER |
| 9 | 22411395 | AX-170174451 | 0 | C | T | 0.465 | 6.67E-02 | 8.99E-02 | 2.55E+00 | 2.61E+00 | 4.62E-01 | no | 9530077C05Rik | upstream_gene_variant | MODIFIER |
| 9 | 22417789 | AX-170031941 | 0 | A | C | 0.465 | 6.67E-02 | 8.99E-02 | 2.55E+00 | 2.61E+00 | 4.62E-01 | no | 9530077C05Rik | intron_variant | MODIFIER |
| 9 | 22417877 | AX-97841146 | 0 | A | G | 0.465 | 6.67E-02 | 8.99E-02 | 2.55E+00 | 2.61E+00 | 4.62E-01 | no | 9530077C05Rik | intron_variant | MODIFIER |
| 9 | 22422610 | AX-170031945 | 0 | T | A | 0.465 | 6.67E-02 | 8.99E-02 | 2.55E+00 | 2.61E+00 | 4.62E-01 | no | 9530077C05Rik | intron_variant | MODIFIER |
| 9 | 22456995 | AX-170090261 | 0 | A | G | 0.465 | 6.67E-02 | 8.99E-02 | 2.55E+00 | 2.61E+00 | 4.62E-01 | no | Rp9 | intron_variant | MODIFIER |
| 9 | 22477339 | AX-97825921 | 0 | T | C | 0.465 | 6.67E-02 | 8.99E-02 | 2.55E+00 | 2.61E+00 | 4.62E-01 | no | Bbs9 | intron_variant | MODIFIER |
| 9 | 22509650 | AX-170266179 | 0 | A | G | 0.465 | 6.67E-02 | 8.99E-02 | 2.55E+00 | 2.61E+00 | 4.62E-01 | no | Bbs9 | intron_variant | MODIFIER |
| 9 | 22510141 | AX-170397713 | 0 | G | A | 0.465 | 6.67E-02 | 8.99E-02 | 2.55E+00 | 2.61E+00 | 4.62E-01 | no | Bbs9 | intron_variant | MODIFIER |
| 9 | 22550751 | AX-170234170 | 0 | C | G | 0.465 | 6.67E-02 | 8.99E-02 | 2.55E+00 | 2.61E+00 | 4.62E-01 | no | Bbs9 | intron_variant | MODIFIER |
| 9 | 22609564 | AX-97804938 | 0 | A | G | 0.465 | 6.67E-02 | 8.99E-02 | 2.55E+00 | 2.61E+00 | 4.62E-01 | no | Bbs9 | intron_variant | MODIFIER |
| 9 | 22615066 | AX-170370193 | 0 | A | G | 0.465 | 6.67E-02 | 8.99E-02 | 2.55E+00 | 2.61E+00 | 4.62E-01 | no | Bbs9 | intron_variant | MODIFIER |
| 9 | 22638592 | AX-97803260 | 0 | C | A | 0.465 | 6.67E-02 | 8.99E-02 | 2.55E+00 | 2.61E+00 | 4.62E-01 | no | Bbs9 | intron_variant | MODIFIER |
| 9 | 22663476 | AX-97857431 | 0 | T | G | 0.465 | 6.67E-02 | 8.99E-02 | 2.55E+00 | 2.61E+00 | 4.62E-01 | no | Bbs9 | intron_variant | MODIFIER |
| 9 | 22680223 | AX-170207607 | 0 | C | T | 0.465 | 6.67E-02 | 8.99E-02 | 2.55E+00 | 2.61E+00 | 4.62E-01 | no | Bbs9 | intron_variant | MODIFIER |
| 9 | 22690412 | AX-170332274 | 0 | C | T | 0.465 | 6.67E-02 | 8.99E-02 | 2.55E+00 | 2.61E+00 | 4.62E-01 | no | Bbs9 | intron_variant | MODIFIER |
| 9 | 22716417 | AX-97805239 | 0 | A | G | 0.465 | 6.67E-02 | 8.99E-02 | 2.55E+00 | 2.61E+00 | 4.62E-01 | no | Bbs9 | intron_variant | MODIFIER |
| 9 | 22780335 | AX-97812963 | 0 | C | T | 0.465 | 6.67E-02 | 8.99E-02 | 2.55E+00 | 2.61E+00 | 4.62E-01 | no | Bbs9 | intron_variant | MODIFIER |
| 9 | 22780442 | AX-170498111 | 0 | C | T | 0.465 | 6.67E-02 | 8.99E-02 | 2.55E+00 | 2.61E+00 | 4.62E-01 | no | Bbs9 | intron_variant | MODIFIER |

| | | | | | | | | | | | | | | | |
|---|---|---|---|---|---|---|---|---|---|---|---|---|---|---|---|
| 9 | 22787075 | AX-170031955 | 0 | C | T | 0.465 | 6.67E-02 | 8.99E-02 | 2.55E+00 | 2.61E+00 | 4.62E-01 | no | Bbs9 | intron_variant | MODIFIER |
| 9 | 22826202 | AX-170031956 | 0 | G | C | 0.465 | 6.67E-02 | 8.99E-02 | 2.55E+00 | 2.61E+00 | 4.62E-01 | no | Bbs9 | intron_variant | MODIFIER |
| 9 | 22826345 | AX-170031959 | 0 | C | G | 0.465 | 6.67E-02 | 8.99E-02 | 2.55E+00 | 2.61E+00 | 4.62E-01 | no | Bbs9 | intron_variant | MODIFIER |
| 9 | 22827016 | AX-97812977 | 0 | A | G | 0.465 | 6.67E-02 | 8.99E-02 | 2.55E+00 | 2.61E+00 | 4.62E-01 | no | Bbs9 | intron_variant | MODIFIER |
| 9 | 22865326 | AX-170243627 | 0 | A | G | 0.465 | 6.67E-02 | 8.99E-02 | 2.55E+00 | 2.61E+00 | 4.62E-01 | no | Bbs9 | intron_variant | MODIFIER |
| 9 | 22865360 | AX-170284469 | 0 | T | C | 0.465 | 6.67E-02 | 8.99E-02 | 2.55E+00 | 2.61E+00 | 4.62E-01 | no | Bbs9 | intron_variant | MODIFIER |
| 9 | 22875808 | AX-170359488 | 0 | C | A | 0.465 | 6.67E-02 | 8.99E-02 | 2.55E+00 | 2.61E+00 | 4.62E-01 | no | Bbs9 | intron_variant | MODIFIER |
| 9 | 22887944 | AX-170081853 | 0 | A | C | 0.465 | 6.67E-02 | 8.99E-02 | 2.55E+00 | 2.61E+00 | 4.62E-01 | no | Bbs9 | 3_prime_UTR_variant | MODIFIER |
| 9 | 22906981 | AX-97815932 | 0 | G | A | 0.465 | 6.67E-02 | 8.99E-02 | 2.55E+00 | 2.61E+00 | 4.62E-01 | no | | intergenic_variant | MODIFIER |
| 9 | 23203380 | AX-97823275 | 0 | A | G | 0.465 | 6.67E-02 | 8.99E-02 | 2.55E+00 | 2.61E+00 | 4.62E-01 | no | | intergenic_variant | MODIFIER |
| 9 | 23209093 | AX-170179779 | 0 | C | A | 0.465 | 6.67E-02 | 8.99E-02 | 2.55E+00 | 2.61E+00 | 4.62E-01 | no | | intergenic_variant | MODIFIER |
| 9 | 23209176 | AX-170427341 | 0 | C | T | 0.465 | 6.67E-02 | 8.99E-02 | 2.55E+00 | 2.61E+00 | 4.62E-01 | no | | intergenic_variant | MODIFIER |
| 9 | 23209356 | AX-170443693 | 0 | T | A | 0.465 | 6.67E-02 | 8.99E-02 | 2.55E+00 | 2.61E+00 | 4.62E-01 | no | | intergenic_variant | MODIFIER |
| 9 | 23215849 | AX-170096491 | 0 | A | G | 0.465 | 6.67E-02 | 8.99E-02 | 2.55E+00 | 2.61E+00 | 4.62E-01 | no | | intergenic_variant | MODIFIER |
| 9 | 23217151 | AX-170316435 | 0 | T | C | 0.465 | 6.67E-02 | 8.99E-02 | 2.55E+00 | 2.61E+00 | 4.62E-01 | no | | intergenic_variant | MODIFIER |
| 9 | 23231117 | AX-170382260 | 0 | G | A | 0.465 | 6.67E-02 | 8.99E-02 | 2.55E+00 | 2.61E+00 | 4.62E-01 | no | Bmper | intron_variant | MODIFIER |
| 9 | 23240166 | AX-170490837 | 0 | C | G | 0.465 | 6.67E-02 | 8.99E-02 | 2.55E+00 | 2.61E+00 | 4.62E-01 | no | Bmper | intron_variant | MODIFIER |
| 9 | 23240824 | AX-170382969 | 0 | C | T | 0.465 | 6.67E-02 | 8.99E-02 | 2.55E+00 | 2.61E+00 | 4.62E-01 | no | Bmper | intron_variant | MODIFIER |
| 9 | 23245258 | AX-170168109 | 0 | A | G | 0.465 | 6.67E-02 | 8.99E-02 | 2.55E+00 | 2.61E+00 | 4.62E-01 | no | Bmper | intron_variant | MODIFIER |
| 9 | 23245298 | AX-97855326 | 0 | C | T | 0.465 | 6.67E-02 | 8.99E-02 | 2.55E+00 | 2.61E+00 | 4.62E-01 | no | Bmper | intron_variant | MODIFIER |
| 9 | 23300649 | AX-97850340 | 0 | C | T | 0.465 | 6.67E-02 | 8.99E-02 | 2.55E+00 | 2.61E+00 | 4.62E-01 | no | Bmper | intron_variant | MODIFIER |
| 9 | 23313103 | AX-97855490 | 0 | C | T | 0.465 | 6.67E-02 | 8.99E-02 | 2.55E+00 | 2.61E+00 | 4.62E-01 | no | Bmper | intron_variant | MODIFIER |
| 9 | 23336434 | AX-170233094 | 0 | C | T | 0.465 | 6.67E-02 | 8.99E-02 | 2.55E+00 | 2.61E+00 | 4.62E-01 | no | Bmper | intron_variant | MODIFIER |
| 9 | 23339915 | AX-170266779 | 0 | C | T | 0.465 | 6.67E-02 | 8.99E-02 | 2.55E+00 | 2.61E+00 | 4.62E-01 | no | Bmper | intron_variant | MODIFIER |
| 9 | 23347604 | AX-170031961 | 0 | C | T | 0.465 | 6.67E-02 | 8.99E-02 | 2.55E+00 | 2.61E+00 | 4.62E-01 | no | Bmper | intron_variant | MODIFIER |
| 9 | 23354609 | AX-170424352 | 0 | T | C | 0.465 | 6.67E-02 | 8.99E-02 | 2.55E+00 | 2.61E+00 | 4.62E-01 | no | Bmper | intron_variant | MODIFIER |
| 9 | 23358209 | AX-170198759 | 0 | A | G | 0.465 | 6.67E-02 | 8.99E-02 | 2.55E+00 | 2.61E+00 | 4.62E-01 | no | Bmper | intron_variant | MODIFIER |
| 9 | 23379015 | AX-97823485 | 0 | T | C | 0.465 | 6.67E-02 | 8.99E-02 | 2.55E+00 | 2.61E+00 | 4.62E-01 | no | Bmper | intron_variant | MODIFIER |
| 16 | 79149346 | AX-97802099 | 0 | C | T | 0.465 | 5.57E-02 | 7.59E-02 | 2.59E+00 | 2.66E+00 | 4.62E-01 | no | | intergenic_variant | MODIFIER |
| 12 | 109488650 | AX-170007503 | 0 | C | T | 0.14 | -1.20E-01 | 1.66E-01 | 2.70E+00 | 2.80E+00 | 4.62E-01 | no | Gm34081 | downstream_gene_variant | MODIFIER |
| 6 | 32811092 | AX-170520744 | 1 | C | T | 0.027 | -9.20E-02 | 1.26E-01 | 2.53E+00 | 2.72E+00 | 4.62E-01 | no | Chchd3 | intron_variant | MODIFIER |
| 6 | 93771234 | AX-170202459 | 1 | T | A | 0.438 | -4.41E-02 | 5.97E-02 | 2.53E+00 | 2.65E+00 | 4.62E-01 | no | Magi1 | intron_variant | MODIFIER |
| 16 | 50831169 | AX-170362138 | 0 | A | G | 0.518 | 5.93E-02 | 7.83E-02 | 2.47E+00 | 2.52E+00 | 4.63E-01 | no | | intergenic_variant | MODIFIER |
| 4 | 154705329 | AX-97805071 | 0 | C | G | 0.096 | 1.34E-01 | 1.84E-01 | 2.67E+00 | 2.76E+00 | 4.63E-01 | no | | intergenic_variant | MODIFIER |
| 4 | 154705523 | AX-170024295 | 0 | A | C | 0.096 | 1.34E-01 | 1.84E-01 | 2.67E+00 | 2.76E+00 | 4.63E-01 | no | | intergenic_variant | MODIFIER |
| 4 | 154709482 | AX-170024296 | 0 | A | C | 0.096 | 1.34E-01 | 1.84E-01 | 2.67E+00 | 2.76E+00 | 4.63E-01 | no | | regulatory_region_variant | MODIFIER |
| 4 | 154751560 | AX-170467534 | 0 | A | G | 0.096 | 1.34E-01 | 1.84E-01 | 2.67E+00 | 2.76E+00 | 4.63E-01 | no | | regulatory_region_variant | MODIFIER |
| 4 | 154755483 | AX-170332974 | 0 | C | T | 0.096 | 1.34E-01 | 1.84E-01 | 2.67E+00 | 2.76E+00 | 4.63E-01 | no | | intergenic_variant | MODIFIER |
| 4 | 154758394 | AX-170178814 | 0 | C | T | 0.096 | 1.34E-01 | 1.84E-01 | 2.67E+00 | 2.76E+00 | 4.63E-01 | no | | intergenic_variant | MODIFIER |
| 4 | 154774965 | AX-170332566 | 0 | A | G | 0.096 | 1.34E-01 | 1.84E-01 | 2.67E+00 | 2.76E+00 | 4.63E-01 | no | | intergenic_variant | MODIFIER |
| 4 | 154778399 | AX-170438490 | 0 | C | T | 0.096 | 1.34E-01 | 1.84E-01 | 2.67E+00 | 2.76E+00 | 4.63E-01 | no | | intergenic_variant | MODIFIER |
| 4 | 154782576 | AX-170240661 | 0 | C | T | 0.096 | 1.34E-01 | 1.84E-01 | 2.67E+00 | 2.76E+00 | 4.63E-01 | no | | regulatory_region_variant | MODIFIER |
| 4 | 154783994 | AX-97808325 | 0 | G | A | 0.096 | 1.34E-01 | 1.84E-01 | 2.67E+00 | 2.76E+00 | 4.63E-01 | no | | intergenic_variant | MODIFIER |
| 4 | 154784147 | AX-170254104 | 0 | C | G | 0.096 | 1.34E-01 | 1.84E-01 | 2.67E+00 | 2.76E+00 | 4.63E-01 | no | | intergenic_variant | MODIFIER |
| 5 | 3363243 | AX-170470608 | 0 | T | C | 0.43 | -4.91E-02 | 6.64E-02 | 2.53E+00 | 2.64E+00 | 4.63E-01 | no | Cdk6 | intron_variant | MODIFIER |
| 2 | 130058542 | AX-170251688 | 0 | C | A | 0.149 | 1.29E-01 | 1.76E-01 | 2.59E+00 | 2.68E+00 | 4.63E-01 | no | | regulatory_region_variant | MODIFIER |
| 8 | 108555088 | AX-170258192 | 1 | G | A | 0.223 | 7.88E-02 | 1.09E-01 | 2.74E+00 | 2.82E+00 | 4.63E-01 | no | Lncbate1 | intron_variant,non_coding_transcript_variant | MODIFIER |
| 8 | 17313271 | AX-170393149 | 0 | G | A | 0.228 | -5.64E-02 | 7.62E-02 | 2.53E+00 | 2.62E+00 | 4.63E-01 | no | Csmd1 | intron_variant | MODIFIER |
| 5 | 50724129 | AX-170072055 | 0 | G | A | 0.5 | 7.69E-02 | 1.03E-01 | 2.50E+00 | 2.56E+00 | 4.63E-01 | no | | intergenic_variant | MODIFIER |
| 5 | 42120332 | AX-170071899 | 0 | T | C | 0.202 | -7.24E-02 | 9.83E-02 | 2.55E+00 | 2.64E+00 | 4.64E-01 | no | Gm16223 | intron_variant | MODIFIER |
| 18 | 65463333 | AX-170217084 | 0 | T | A | 0.412 | 6.56E-02 | 8.61E-02 | 2.44E+00 | 2.50E+00 | 4.64E-01 | no | Malt1 | intron_variant | MODIFIER |
| 18 | 72095594 | AX-97853091 | 1 | C | T | 0.188 | -8.48E-02 | 1.17E-01 | 2.67E+00 | 2.73E+00 | 4.64E-01 | no | Dcc | intron_variant | MODIFIER |
| 1 | 176847300 | AX-170349832 | 1 | A | C | 0.223 | 8.82E-02 | 1.20E-01 | 2.59E+00 | 2.65E+00 | 4.64E-01 | no | Sdccag8 | intron_variant | MODIFIER |
| 1 | 176901839 | AX-170144652 | 1 | T | C | 0.223 | 8.82E-02 | 1.20E-01 | 2.59E+00 | 2.65E+00 | 4.64E-01 | no | Sdccag8 | intron_variant | MODIFIER |

| | | | | | | | | | | | | | | | |
|---|---|---|---|---|---|---|---|---|---|---|---|---|---|---|---|
| 14 | 112294882 | AX-170390384 | 0 | T | C | 0.316 | 5.30E-02 | 7.22E-02 | 2.55E+00 | 2.66E+00 | 4.64E-01 | no | | intergenic_variant | MODIFIER |
| 6 | 135254976 | AX-170028310 | 1 | C | G | 0.571 | 6.65E-02 | 8.76E-02 | 2.45E+00 | 2.51E+00 | 4.64E-01 | no | Gsg1 | upstream_gene_variant | MODIFIER |
| 6 | 137053944 | AX-170028331 | 1 | A | G | 0.571 | 6.65E-02 | 8.76E-02 | 2.45E+00 | 2.51E+00 | 4.64E-01 | no | Rerg | downstream_gene_variant | MODIFIER |
| 6 | 94439710 | AX-97801216 | 1 | G | A | 0.438 | -4.40E-02 | 5.97E-02 | 2.53E+00 | 2.65E+00 | 4.64E-01 | no | | intergenic_variant | MODIFIER |
| 10 | 9431055 | AX-170125111 | 0 | T | C | 0.202 | 7.08E-02 | 9.81E-02 | 2.81E+00 | 2.92E+00 | 4.64E-01 | no | | intergenic_variant | MODIFIER |
| 2 | 166456574 | AX-97855353 | 0 | C | A | 0.158 | 1.42E-01 | 1.97E-01 | 2.74E+00 | 2.82E+00 | 4.64E-01 | no | 5031425F14Rik | intron_variant,non_coding_transcript_variant | MODIFIER |
| 6 | 130826337 | AX-170255239 | 1 | A | G | 0.527 | 6.04E-02 | 8.10E-02 | 2.50E+00 | 2.58E+00 | 4.64E-01 | no | Gm44118 | downstream_gene_variant | MODIFIER |
| 4 | 116023690 | AX-170023757 | 0 | C | T | 0.202 | 8.19E-02 | 1.12E-01 | 2.60E+00 | 2.69E+00 | 4.64E-01 | no | | intergenic_variant | MODIFIER |
| 4 | 116586646 | AX-97845017 | 0 | A | G | 0.202 | 8.19E-02 | 1.12E-01 | 2.60E+00 | 2.69E+00 | 4.64E-01 | no | Gpbp1l1 | intron_variant | MODIFIER |
| 4 | 118357751 | AX-97792686 | 0 | A | G | 0.202 | 8.19E-02 | 1.12E-01 | 2.60E+00 | 2.69E+00 | 4.64E-01 | no | Hyi | upstream_gene_variant | MODIFIER |
| 4 | 128202197 | AX-170373439 | 0 | A | G | 0.14 | 8.92E-02 | 1.22E-01 | 2.61E+00 | 2.67E+00 | 4.65E-01 | no | Csmd2 | intron_variant,non_coding_transcript_variant | MODIFIER |
| 8 | 37832586 | AX-170379951 | 1 | A | G | 0.161 | -8.58E-02 | 1.19E-01 | 2.64E+00 | 2.75E+00 | 4.65E-01 | no | Sgcz | intron_variant | MODIFIER |
| 4 | 115933641 | AX-170265515 | 1 | G | A | 0.214 | 8.45E-02 | 1.16E-01 | 2.59E+00 | 2.66E+00 | 4.65E-01 | no | Dmbx1 | intron_variant | MODIFIER |
| 7 | 37468664 | AX-170077096 | 0 | G | A | 0.342 | 6.97E-02 | 9.48E-02 | 2.56E+00 | 2.62E+00 | 4.65E-01 | no | Zfp536 | downstream_gene_variant | MODIFIER |
| 7 | 37474352 | AX-170527144 | 0 | G | T | 0.342 | 6.97E-02 | 9.48E-02 | 2.56E+00 | 2.62E+00 | 4.65E-01 | no | Zfp536 | downstream_gene_variant | MODIFIER |
| 7 | 37487379 | AX-170519140 | 0 | T | G | 0.342 | 6.97E-02 | 9.48E-02 | 2.56E+00 | 2.62E+00 | 4.65E-01 | no | Zfp536 | intron_variant | MODIFIER |
| 7 | 37496906 | AX-170167724 | 0 | C | T | 0.342 | 6.97E-02 | 9.48E-02 | 2.56E+00 | 2.62E+00 | 4.65E-01 | no | Zfp536 | intron_variant | MODIFIER |
| 7 | 37497158 | AX-170322683 | 0 | G | A | 0.342 | 6.97E-02 | 9.48E-02 | 2.56E+00 | 2.62E+00 | 4.65E-01 | no | Zfp536 | intron_variant | MODIFIER |
| 7 | 37498520 | AX-170374089 | 0 | T | G | 0.342 | 6.97E-02 | 9.48E-02 | 2.56E+00 | 2.62E+00 | 4.65E-01 | no | Zfp536 | intron_variant | MODIFIER |
| 7 | 37498670 | AX-170427365 | 0 | T | C | 0.342 | 6.97E-02 | 9.48E-02 | 2.56E+00 | 2.62E+00 | 4.65E-01 | no | Zfp536 | intron_variant | MODIFIER |
| 7 | 37500082 | AX-170357268 | 0 | A | G | 0.342 | 6.97E-02 | 9.48E-02 | 2.56E+00 | 2.62E+00 | 4.65E-01 | no | Zfp536 | intron_variant | MODIFIER |
| 7 | 37504696 | AX-170520224 | 0 | T | G | 0.342 | 6.97E-02 | 9.48E-02 | 2.56E+00 | 2.62E+00 | 4.65E-01 | no | Zfp536 | intron_variant | MODIFIER |
| 7 | 37507845 | AX-169999809 | 0 | G | A | 0.342 | 6.97E-02 | 9.48E-02 | 2.56E+00 | 2.62E+00 | 4.65E-01 | no | Zfp536 | intron_variant | MODIFIER |
| 7 | 37511760 | AX-170343313 | 0 | G | A | 0.342 | 6.97E-02 | 9.48E-02 | 2.56E+00 | 2.62E+00 | 4.65E-01 | no | Zfp536 | intron_variant | MODIFIER |
| 7 | 37531133 | AX-170122116 | 0 | C | T | 0.342 | 6.97E-02 | 9.48E-02 | 2.56E+00 | 2.62E+00 | 4.65E-01 | no | Zfp536 | intron_variant | MODIFIER |
| 7 | 37544549 | AX-170158403 | 0 | A | G | 0.342 | 6.97E-02 | 9.48E-02 | 2.56E+00 | 2.62E+00 | 4.65E-01 | no | Zfp536 | intron_variant | MODIFIER |
| 7 | 37550079 | AX-170288392 | 0 | A | T | 0.342 | 6.97E-02 | 9.48E-02 | 2.56E+00 | 2.62E+00 | 4.65E-01 | no | Zfp536 | intron_variant | MODIFIER |
| 7 | 37559200 | AX-170325576 | 0 | G | A | 0.342 | 6.97E-02 | 9.48E-02 | 2.56E+00 | 2.62E+00 | 4.65E-01 | no | Zfp536 | intron_variant | MODIFIER |
| 7 | 37562172 | AX-170453262 | 0 | G | A | 0.342 | 6.97E-02 | 9.48E-02 | 2.56E+00 | 2.62E+00 | 4.65E-01 | no | Zfp536 | intron_variant | MODIFIER |
| 7 | 37562446 | AX-170360478 | 0 | A | G | 0.342 | 6.97E-02 | 9.48E-02 | 2.56E+00 | 2.62E+00 | 4.65E-01 | no | Zfp536 | intron_variant | MODIFIER |
| 7 | 37572987 | AX-97855699 | 0 | G | A | 0.342 | 6.97E-02 | 9.48E-02 | 2.56E+00 | 2.62E+00 | 4.65E-01 | no | Zfp536 | intron_variant | MODIFIER |
| 7 | 37613871 | AX-170291055 | 0 | G | A | 0.342 | 6.97E-02 | 9.48E-02 | 2.56E+00 | 2.62E+00 | 4.65E-01 | no | Zfp536 | intron_variant | MODIFIER |
| 7 | 37615345 | AX-170335536 | 0 | A | G | 0.342 | 6.97E-02 | 9.48E-02 | 2.56E+00 | 2.62E+00 | 4.65E-01 | no | Zfp536 | intron_variant | MODIFIER |
| 7 | 37619420 | AX-170400976 | 0 | G | A | 0.342 | 6.97E-02 | 9.48E-02 | 2.56E+00 | 2.62E+00 | 4.65E-01 | no | Zfp536 | intron_variant | MODIFIER |
| 7 | 37644783 | AX-97817597 | 0 | C | T | 0.342 | 6.97E-02 | 9.48E-02 | 2.56E+00 | 2.62E+00 | 4.65E-01 | no | Zfp536 | intron_variant | MODIFIER |
| 7 | 37644934 | AX-170474038 | 0 | G | A | 0.342 | 6.97E-02 | 9.48E-02 | 2.56E+00 | 2.62E+00 | 4.65E-01 | no | Zfp536 | intron_variant | MODIFIER |
| 7 | 37653591 | AX-97837930 | 0 | T | C | 0.342 | 6.97E-02 | 9.48E-02 | 2.56E+00 | 2.62E+00 | 4.65E-01 | no | Zfp536 | intron_variant | MODIFIER |
| 7 | 37662122 | AX-170355759 | 0 | A | C | 0.342 | 6.97E-02 | 9.48E-02 | 2.56E+00 | 2.62E+00 | 4.65E-01 | no | Zfp536 | intron_variant | MODIFIER |
| 7 | 37667057 | AX-170272532 | 0 | C | T | 0.342 | 6.97E-02 | 9.48E-02 | 2.56E+00 | 2.62E+00 | 4.65E-01 | no | Zfp536 | intron_variant | MODIFIER |
| 7 | 37667854 | AX-170259066 | 0 | T | G | 0.342 | 6.97E-02 | 9.48E-02 | 2.56E+00 | 2.62E+00 | 4.65E-01 | no | Zfp536 | intron_variant | MODIFIER |
| 7 | 37668995 | AX-170028842 | 0 | C | A | 0.342 | 6.97E-02 | 9.48E-02 | 2.56E+00 | 2.62E+00 | 4.65E-01 | no | Zfp536 | intron_variant | MODIFIER |
| 7 | 37669477 | AX-170333513 | 0 | T | C | 0.342 | 6.97E-02 | 9.48E-02 | 2.56E+00 | 2.62E+00 | 4.65E-01 | no | Zfp536 | intron_variant | MODIFIER |
| 7 | 37683330 | AX-170348934 | 0 | G | C | 0.342 | 6.97E-02 | 9.48E-02 | 2.56E+00 | 2.62E+00 | 4.65E-01 | no | Zfp536 | intron_variant | MODIFIER |
| 7 | 37695478 | AX-170077099 | 0 | G | C | 0.342 | 6.97E-02 | 9.48E-02 | 2.56E+00 | 2.62E+00 | 4.65E-01 | no | Zfp536 | intron_variant | MODIFIER |
| 7 | 37696452 | AX-170077100 | 0 | G | A | 0.342 | 6.97E-02 | 9.48E-02 | 2.56E+00 | 2.62E+00 | 4.65E-01 | no | Zfp536 | intron_variant | MODIFIER |
| 7 | 37713862 | AX-170028845 | 0 | T | C | 0.342 | 6.97E-02 | 9.48E-02 | 2.56E+00 | 2.62E+00 | 4.65E-01 | no | Zfp536 | intron_variant | MODIFIER |
| 7 | 37730146 | AX-97796321 | 0 | A | G | 0.342 | 6.97E-02 | 9.48E-02 | 2.56E+00 | 2.62E+00 | 4.65E-01 | no | Zfp536 | intron_variant | MODIFIER |
| 7 | 37746015 | AX-170343989 | 0 | C | T | 0.342 | 6.97E-02 | 9.48E-02 | 2.56E+00 | 2.62E+00 | 4.65E-01 | no | Zfp536 | intron_variant | MODIFIER |
| 7 | 37751183 | AX-97823545 | 0 | A | G | 0.342 | 6.97E-02 | 9.48E-02 | 2.56E+00 | 2.62E+00 | 4.65E-01 | no | Zfp536 | intron_variant | MODIFIER |
| 3 | 147380197 | AX-170161882 | 0 | A | G | 0.342 | 5.57E-02 | 7.70E-02 | 2.65E+00 | 2.72E+00 | 4.65E-01 | no | | intergenic_variant | MODIFIER |
| 10 | 107819273 | AX-97816796 | 2 | A | G | 0.3 | -5.94E-02 | 8.15E-02 | 2.58E+00 | 2.68E+00 | 4.65E-01 | no | Otogl | intron_variant | MODIFIER |
| 7 | 37512824 | AX-170396719 | 0 | G | T | 0.333 | 6.90E-02 | 9.38E-02 | 2.55E+00 | 2.62E+00 | 4.65E-01 | no | Zfp536 | intron_variant | MODIFIER |
| 7 | 37516498 | AX-170115980 | 0 | T | C | 0.333 | 6.90E-02 | 9.38E-02 | 2.55E+00 | 2.62E+00 | 4.65E-01 | no | Zfp536 | intron_variant | MODIFIER |
| 7 | 37530047 | AX-170097854 | 0 | T | C | 0.333 | 6.90E-02 | 9.38E-02 | 2.55E+00 | 2.62E+00 | 4.65E-01 | no | Zfp536 | intron_variant | MODIFIER |

| | | | | | | | | | | | | | | | |
|---|---|---|---|---|---|---|---|---|---|---|---|---|---|---|---|
| 7 | 37578274 | AX-97791884 | 0 | A | T | 0.333 | 6.90E-02 | 9.38E-02 | 2.55E+00 | 2.62E+00 | 4.65E-01 | no | Zfp536 | intron_variant | MODIFIER |
| 7 | 37684832 | AX-170308037 | 0 | T | G | 0.333 | 6.90E-02 | 9.38E-02 | 2.55E+00 | 2.62E+00 | 4.65E-01 | no | Zfp536 | intron_variant | MODIFIER |
| 7 | 37696635 | AX-170254663 | 0 | T | C | 0.333 | 6.90E-02 | 9.38E-02 | 2.55E+00 | 2.62E+00 | 4.65E-01 | no | Zfp536 | intron_variant | MODIFIER |
| 1 | 185162914 | AX-97814473 | 0 | G | A | 0.237 | 8.68E-02 | 1.15E-01 | 2.45E+00 | 2.52E+00 | 4.65E-01 | no | | intergenic_variant | MODIFIER |
| 10 | 12821051 | AX-170438477 | 0 | A | G | 0.202 | -6.95E-02 | 9.69E-02 | 2.69E+00 | 2.81E+00 | 4.65E-01 | no | Utrn | intron_variant | MODIFIER |
| 10 | 92487731 | AX-170374115 | 1 | C | T | 0.304 | -6.18E-02 | 8.40E-02 | 2.55E+00 | 2.61E+00 | 4.66E-01 | no | Gm25884 | downstream_gene_variant | MODIFIER |
| 3 | 151816533 | AX-170225925 | 1 | C | T | 0.339 | 5.36E-02 | 7.46E-02 | 2.72E+00 | 2.81E+00 | 4.66E-01 | no | Ptgfr | intron_variant | MODIFIER |
| 5 | 39984432 | AX-97788502 | 2 | C | T | 0.209 | -7.48E-02 | 1.02E-01 | 2.53E+00 | 2.62E+00 | 4.66E-01 | no | Gm35191 | intron_variant,non_coding_transcript_variant | MODIFIER |
| 15 | 10895202 | AX-97854715 | 1 | C | T | 0.241 | -7.77E-02 | 1.07E-01 | 2.59E+00 | 2.64E+00 | 4.66E-01 | no | | intergenic_variant | MODIFIER |
| 20 | 68968373 | AX-170034094 | 1 | C | T | 0.25 | 4.54E-02 | 6.31E-02 | 2.68E+00 | 2.76E+00 | 4.66E-01 | no | | intergenic_variant | MODIFIER |
| 4 | 128068314 | AX-170242163 | 0 | C | T | 0.158 | 8.19E-02 | 1.13E-01 | 2.61E+00 | 2.67E+00 | 4.66E-01 | no | Csmd2 | intron_variant,non_coding_transcript_variant | MODIFIER |
| 4 | 128069692 | AX-170181967 | 0 | G | A | 0.158 | 8.19E-02 | 1.13E-01 | 2.61E+00 | 2.67E+00 | 4.66E-01 | no | Csmd2 | intron_variant,non_coding_transcript_variant | MODIFIER |
| 4 | 128069845 | AX-170419310 | 0 | T | C | 0.158 | 8.19E-02 | 1.13E-01 | 2.61E+00 | 2.67E+00 | 4.66E-01 | no | Csmd2 | intron_variant,non_coding_transcript_variant | MODIFIER |
| 4 | 128077322 | AX-170172030 | 0 | G | T | 0.158 | 8.19E-02 | 1.13E-01 | 2.61E+00 | 2.67E+00 | 4.66E-01 | no | Csmd2 | intron_variant,non_coding_transcript_variant | MODIFIER |
| 4 | 128081339 | AX-170451076 | 0 | A | C | 0.158 | 8.19E-02 | 1.13E-01 | 2.61E+00 | 2.67E+00 | 4.66E-01 | no | Csmd2 | intron_variant,non_coding_transcript_variant | MODIFIER |
| 4 | 128082188 | AX-170157244 | 0 | T | C | 0.158 | 8.19E-02 | 1.13E-01 | 2.61E+00 | 2.67E+00 | 4.66E-01 | no | Csmd2 | intron_variant,non_coding_transcript_variant | MODIFIER |
| 4 | 128085527 | AX-97784515 | 0 | C | T | 0.158 | 8.19E-02 | 1.13E-01 | 2.61E+00 | 2.67E+00 | 4.66E-01 | no | Csmd2 | intron_variant,non_coding_transcript_variant | MODIFIER |
| 4 | 128105230 | AX-170070636 | 0 | C | T | 0.158 | 8.19E-02 | 1.13E-01 | 2.61E+00 | 2.67E+00 | 4.66E-01 | no | Csmd2 | intron_variant,non_coding_transcript_variant | MODIFIER |
| 4 | 128106988 | AX-170487491 | 0 | T | C | 0.158 | 8.19E-02 | 1.13E-01 | 2.61E+00 | 2.67E+00 | 4.66E-01 | no | Csmd2 | intron_variant,non_coding_transcript_variant | MODIFIER |
| 4 | 128179201 | AX-170119430 | 0 | G | A | 0.158 | 8.19E-02 | 1.13E-01 | 2.61E+00 | 2.67E+00 | 4.66E-01 | no | Csmd2 | intron_variant,non_coding_transcript_variant | MODIFIER |
| 4 | 128188795 | AX-170291053 | 0 | A | G | 0.158 | 8.19E-02 | 1.13E-01 | 2.61E+00 | 2.67E+00 | 4.66E-01 | no | Csmd2 | intron_variant,non_coding_transcript_variant | MODIFIER |
| 4 | 128195474 | AX-97807014 | 0 | C | T | 0.158 | 8.19E-02 | 1.13E-01 | 2.61E+00 | 2.67E+00 | 4.66E-01 | no | Csmd2 | intron_variant,non_coding_transcript_variant | MODIFIER |
| 4 | 128416038 | AX-97822171 | 0 | T | C | 0.158 | 8.19E-02 | 1.13E-01 | 2.61E+00 | 2.67E+00 | 4.66E-01 | no | Csmd2 | intron_variant,non_coding_transcript_variant | MODIFIER |
| 4 | 128416696 | AX-170270395 | 0 | G | A | 0.158 | 8.19E-02 | 1.13E-01 | 2.61E+00 | 2.67E+00 | 4.66E-01 | no | Csmd2 | intron_variant,non_coding_transcript_variant | MODIFIER |
| 4 | 128417677 | AX-170364972 | 0 | G | A | 0.158 | 8.19E-02 | 1.13E-01 | 2.61E+00 | 2.67E+00 | 4.66E-01 | no | Csmd2 | intron_variant,non_coding_transcript_variant | MODIFIER |
| 4 | 128484239 | AX-170099594 | 0 | G | A | 0.158 | 8.19E-02 | 1.13E-01 | 2.61E+00 | 2.67E+00 | 4.66E-01 | no | Csmd2 | intron_variant,non_coding_transcript_variant | MODIFIER |
| 4 | 128537449 | AX-170341957 | 0 | A | G | 0.158 | 8.19E-02 | 1.13E-01 | 2.61E+00 | 2.67E+00 | 4.66E-01 | no | Csmd2 | intron_variant | MODIFIER |
| 4 | 128539821 | AX-170391424 | 0 | C | T | 0.158 | 8.19E-02 | 1.13E-01 | 2.61E+00 | 2.67E+00 | 4.66E-01 | no | Csmd2 | intron_variant | MODIFIER |
| 4 | 128540030 | AX-170463818 | 0 | A | G | 0.158 | 8.19E-02 | 1.13E-01 | 2.61E+00 | 2.67E+00 | 4.66E-01 | no | Csmd2 | intron_variant | MODIFIER |
| 4 | 128541471 | AX-170232852 | 0 | G | A | 0.158 | 8.19E-02 | 1.13E-01 | 2.61E+00 | 2.67E+00 | 4.66E-01 | no | Csmd2 | intron_variant | MODIFIER |
| 4 | 128591097 | AX-170426218 | 0 | A | G | 0.158 | 8.19E-02 | 1.13E-01 | 2.61E+00 | 2.67E+00 | 4.66E-01 | no | Zscan20 | upstream_gene_variant | MODIFIER |
| 4 | 128604995 | AX-170258507 | 0 | T | G | 0.158 | 8.19E-02 | 1.13E-01 | 2.61E+00 | 2.67E+00 | 4.66E-01 | no | Zscan20 | intron_variant | MODIFIER |
| 4 | 128611405 | AX-170333540 | 0 | C | G | 0.158 | 8.19E-02 | 1.13E-01 | 2.61E+00 | 2.67E+00 | 4.66E-01 | no | Tlr12 | downstream_gene_variant | MODIFIER |
| 4 | 128613603 | AX-170194685 | 0 | A | G | 0.158 | 8.19E-02 | 1.13E-01 | 2.61E+00 | 2.67E+00 | 4.66E-01 | no | Tlr12 | downstream_gene_variant | MODIFIER |
| 4 | 128613659 | AX-170373727 | 0 | C | A | 0.158 | 8.19E-02 | 1.13E-01 | 2.61E+00 | 2.67E+00 | 4.66E-01 | no | Tlr12 | downstream_gene_variant | MODIFIER |
| 4 | 128635786 | AX-170138486 | 0 | G | A | 0.158 | 8.19E-02 | 1.13E-01 | 2.61E+00 | 2.67E+00 | 4.66E-01 | no | Gm12958 | upstream_gene_variant | MODIFIER |
| 4 | 128656373 | AX-170272468 | 0 | G | A | 0.158 | 8.19E-02 | 1.13E-01 | 2.61E+00 | 2.67E+00 | 4.66E-01 | no | Phc2 | intron_variant | MODIFIER |
| 4 | 128699508 | AX-170070648 | 0 | C | T | 0.158 | 8.19E-02 | 1.13E-01 | 2.61E+00 | 2.67E+00 | 4.66E-01 | no | Phc2 | intron_variant | MODIFIER |
| 4 | 128703905 | AX-170139859 | 0 | G | A | 0.158 | 8.19E-02 | 1.13E-01 | 2.61E+00 | 2.67E+00 | 4.66E-01 | no | Phc2 | intron_variant | MODIFIER |
| 4 | 128744213 | AX-97803201 | 0 | A | G | 0.158 | 8.19E-02 | 1.13E-01 | 2.61E+00 | 2.67E+00 | 4.66E-01 | no | Phc2 | intron_variant | MODIFIER |
| 4 | 128747265 | AX-170235018 | 0 | C | T | 0.158 | 8.19E-02 | 1.13E-01 | 2.61E+00 | 2.67E+00 | 4.66E-01 | no | Phc2 | intron_variant | MODIFIER |
| 4 | 128752489 | AX-170439639 | 0 | A | G | 0.158 | 8.19E-02 | 1.13E-01 | 2.61E+00 | 2.67E+00 | 4.66E-01 | no | Phc2 | 3_prime_UTR_variant | MODIFIER |
| 4 | 128759626 | AX-97791630 | 0 | C | A | 0.158 | 8.19E-02 | 1.13E-01 | 2.61E+00 | 2.67E+00 | 4.66E-01 | no | A3galt2 | 5_prime_UTR_variant | MODIFIER |
| 4 | 128823487 | AX-170285059 | 0 | G | A | 0.158 | 8.19E-02 | 1.13E-01 | 2.61E+00 | 2.67E+00 | 4.66E-01 | no | | intergenic_variant | MODIFIER |
| 4 | 128845802 | AX-97805176 | 0 | A | G | 0.158 | 8.19E-02 | 1.13E-01 | 2.61E+00 | 2.67E+00 | 4.66E-01 | no | Gm12968 | upstream_gene_variant | MODIFIER |
| 4 | 128853212 | AX-170446807 | 0 | C | T | 0.158 | 8.19E-02 | 1.13E-01 | 2.61E+00 | 2.67E+00 | 4.66E-01 | no | Gm12968 | intron_variant,non_coding_transcript_variant | MODIFIER |
| 6 | 130476592 | AX-170028248 | 0 | G | A | 0.535 | 6.02E-02 | 8.14E-02 | 2.52E+00 | 2.59E+00 | 4.66E-01 | no | Gm6590 | intron_variant,non_coding_transcript_variant | MODIFIER |
| 5 | 73406948 | AX-170498532 | 1 | T | A | 0.438 | 6.39E-02 | 8.88E-02 | 2.64E+00 | 2.75E+00 | 4.66E-01 | no | Cwh43 | intron_variant | MODIFIER |
| 7 | 70216744 | AX-170099607 | 1 | C | T | 0.357 | -7.28E-02 | 1.01E-01 | 2.83E+00 | 2.92E+00 | 4.67E-01 | no | Gm29683 | intron_variant,non_coding_transcript_variant | MODIFIER |
| 1 | 99883325 | AX-97832408 | 1 | G | T | 0.161 | 1.49E-01 | 2.06E-01 | 2.65E+00 | 2.71E+00 | 4.67E-01 | Yes | Cntnap5b | intron_variant | MODIFIER |
| 1 | 20802781 | AX-170138226 | 0 | G | A | 0.404 | 6.12E-02 | 8.41E-02 | 2.59E+00 | 2.63E+00 | 4.67E-01 | no | Mcm3 | downstream_gene_variant | MODIFIER |
| 7 | 14676220 | AX-170479670 | 0 | G | T | 0.316 | 6.21E-02 | 8.59E-02 | 2.58E+00 | 2.71E+00 | 4.67E-01 | no | | intergenic_variant | MODIFIER |
| 6 | 93835389 | AX-170204624 | 1 | A | G | 0.438 | -4.37E-02 | 5.97E-02 | 2.53E+00 | 2.65E+00 | 4.67E-01 | no | Magi1 | intron_variant | MODIFIER |
| 2 | 162046409 | AX-170409186 | 1 | C | T | 0.161 | 1.40E-01 | 1.95E-01 | 2.72E+00 | 2.80E+00 | 4.67E-01 | no | Ptprt | intron_variant | MODIFIER |

| | | | | | | | | | | | | | | | |
|---|---|---|---|---|---|---|---|---|---|---|---|---|---|---|---|
| 2 | 148931222 | AX-170019346 | 1 | C | T | 0.143 | -1.31E-01 | 1.83E-01 | 2.76E+00 | 2.84E+00 | 4.67E-01 | no | Gm14129 | intron_variant,non_coding_transcript_variant | MODIFIER |
| 5 | 107914107 | AX-97805828 | 1 | T | C | 0.027 | -1.03E-01 | 1.43E-01 | 2.59E+00 | 2.77E+00 | 4.67E-01 | no | Fam69a | intron_variant | MODIFIER |
| 5 | 80074072 | AX-170072593 | 0 | C | A | 0.412 | 7.33E-02 | 1.02E-01 | 2.73E+00 | 2.79E+00 | 4.67E-01 | no | | intergenic_variant | MODIFIER |
| 1 | 101744511 | AX-97844875 | 1 | G | A | 0.152 | 1.44E-01 | 1.99E-01 | 2.64E+00 | 2.70E+00 | 4.67E-01 | no | | intergenic_variant | MODIFIER |
| 5 | 112313817 | AX-170483860 | 0 | G | A | 0.465 | 7.94E-02 | 1.10E-01 | 2.63E+00 | 2.68E+00 | 4.67E-01 | no | Tpst2 | intron_variant | MODIFIER |
| 7 | 114031595 | AX-170161423 | 0 | A | G | 0.439 | 7.04E-02 | 9.79E-02 | 2.71E+00 | 2.76E+00 | 4.67E-01 | no | Spon1 | intron_variant | MODIFIER |
| 7 | 114049727 | AX-170299329 | 0 | C | T | 0.439 | 7.04E-02 | 9.79E-02 | 2.71E+00 | 2.76E+00 | 4.67E-01 | no | Rras2 | intron_variant | MODIFIER |
| 7 | 114050470 | AX-170349259 | 0 | T | C | 0.439 | 7.04E-02 | 9.79E-02 | 2.71E+00 | 2.76E+00 | 4.67E-01 | no | Rras2 | intron_variant | MODIFIER |
| 7 | 114055691 | AX-170306619 | 0 | A | G | 0.439 | 7.04E-02 | 9.79E-02 | 2.71E+00 | 2.76E+00 | 4.67E-01 | no | Rras2 | intron_variant | MODIFIER |
| 7 | 114056539 | AX-170102162 | 0 | G | A | 0.439 | 7.04E-02 | 9.79E-02 | 2.71E+00 | 2.76E+00 | 4.67E-01 | no | Rras2 | intron_variant | MODIFIER |
| 7 | 114056619 | AX-97781611 | 0 | G | T | 0.439 | 7.04E-02 | 9.79E-02 | 2.71E+00 | 2.76E+00 | 4.67E-01 | no | Rras2 | intron_variant | MODIFIER |
| 7 | 114093436 | AX-97794941 | 0 | C | T | 0.439 | 7.04E-02 | 9.79E-02 | 2.71E+00 | 2.76E+00 | 4.67E-01 | no | Rras2 | intron_variant | MODIFIER |
| 7 | 114096554 | AX-170284612 | 0 | T | G | 0.439 | 7.04E-02 | 9.79E-02 | 2.71E+00 | 2.76E+00 | 4.67E-01 | no | Rras2 | intron_variant | MODIFIER |
| 7 | 114101279 | AX-170320454 | 0 | A | G | 0.439 | 7.04E-02 | 9.79E-02 | 2.71E+00 | 2.76E+00 | 4.67E-01 | no | Rras2 | intron_variant | MODIFIER |
| 7 | 114155023 | AX-170320848 | 0 | T | A | 0.439 | 7.04E-02 | 9.79E-02 | 2.71E+00 | 2.76E+00 | 4.67E-01 | no | | intergenic_variant | MODIFIER |
| 7 | 114158859 | AX-170279305 | 0 | A | C | 0.439 | 7.04E-02 | 9.79E-02 | 2.71E+00 | 2.76E+00 | 4.67E-01 | no | | intergenic_variant | MODIFIER |
| 7 | 114162301 | AX-97791405 | 0 | C | T | 0.439 | 7.04E-02 | 9.79E-02 | 2.71E+00 | 2.76E+00 | 4.67E-01 | no | | intergenic_variant | MODIFIER |
| 7 | 114175426 | AX-170482107 | 0 | G | A | 0.439 | 7.04E-02 | 9.79E-02 | 2.71E+00 | 2.76E+00 | 4.67E-01 | no | | intergenic_variant | MODIFIER |
| 7 | 114185799 | AX-97828981 | 0 | T | G | 0.439 | 7.04E-02 | 9.79E-02 | 2.71E+00 | 2.76E+00 | 4.67E-01 | no | | intergenic_variant | MODIFIER |
| 7 | 114210406 | AX-170107633 | 0 | A | G | 0.439 | 7.04E-02 | 9.79E-02 | 2.71E+00 | 2.76E+00 | 4.67E-01 | no | Gm45454 | upstream_gene_variant | MODIFIER |
| 7 | 114257099 | AX-170190145 | 0 | G | A | 0.439 | 7.04E-02 | 9.79E-02 | 2.71E+00 | 2.76E+00 | 4.67E-01 | no | Copb1 | upstream_gene_variant | MODIFIER |
| 7 | 114291766 | AX-170278077 | 0 | C | G | 0.439 | 7.04E-02 | 9.79E-02 | 2.71E+00 | 2.76E+00 | 4.67E-01 | no | | regulatory_region_variant | MODIFIER |
| 7 | 114297244 | AX-170314451 | 0 | G | A | 0.439 | 7.04E-02 | 9.79E-02 | 2.71E+00 | 2.76E+00 | 4.67E-01 | no | | intergenic_variant | MODIFIER |
| 7 | 114297821 | AX-170481471 | 0 | A | G | 0.439 | 7.04E-02 | 9.79E-02 | 2.71E+00 | 2.76E+00 | 4.67E-01 | no | | intergenic_variant | MODIFIER |
| 7 | 114299309 | AX-170144442 | 0 | T | G | 0.439 | 7.04E-02 | 9.79E-02 | 2.71E+00 | 2.76E+00 | 4.67E-01 | no | | intergenic_variant | MODIFIER |
| 7 | 114361222 | AX-170456581 | 0 | A | G | 0.439 | 7.04E-02 | 9.79E-02 | 2.71E+00 | 2.76E+00 | 4.67E-01 | no | 4933406I18Rik | intron_variant,non_coding_transcript_variant | MODIFIER |
| 7 | 114363384 | AX-170414421 | 0 | T | C | 0.439 | 7.04E-02 | 9.79E-02 | 2.71E+00 | 2.76E+00 | 4.67E-01 | no | 4933406I18Rik | intron_variant,non_coding_transcript_variant | MODIFIER |
| 7 | 114427472 | AX-170029808 | 0 | G | A | 0.439 | 7.04E-02 | 9.79E-02 | 2.71E+00 | 2.76E+00 | 4.67E-01 | no | Pde3b | intron_variant | MODIFIER |
| 7 | 114452893 | AX-170344147 | 0 | A | G | 0.439 | 7.04E-02 | 9.79E-02 | 2.71E+00 | 2.76E+00 | 4.67E-01 | no | Pde3b | intron_variant | MODIFIER |
| 7 | 114464353 | AX-170029805 | 0 | A | T | 0.439 | 7.04E-02 | 9.79E-02 | 2.71E+00 | 2.76E+00 | 4.67E-01 | no | Pde3b | intron_variant | MODIFIER |
| 7 | 114470170 | AX-170029804 | 0 | C | T | 0.439 | 7.04E-02 | 9.79E-02 | 2.71E+00 | 2.76E+00 | 4.67E-01 | no | Pde3b | intron_variant | MODIFIER |
| 7 | 114476748 | AX-97854535 | 0 | T | C | 0.439 | 7.04E-02 | 9.79E-02 | 2.71E+00 | 2.76E+00 | 4.67E-01 | no | Pde3b | intron_variant | MODIFIER |
| 7 | 114511774 | AX-170029806 | 0 | A | C | 0.439 | 7.04E-02 | 9.79E-02 | 2.71E+00 | 2.76E+00 | 4.67E-01 | no | Pde3b | intron_variant | MODIFIER |
| 7 | 114554917 | AX-170029803 | 0 | C | T | 0.439 | 7.04E-02 | 9.79E-02 | 2.71E+00 | 2.76E+00 | 4.67E-01 | no | Cyp2r1 | intron_variant | MODIFIER |
| 7 | 114605031 | AX-170029802 | 0 | T | G | 0.439 | 7.04E-02 | 9.79E-02 | 2.71E+00 | 2.76E+00 | 4.67E-01 | no | | intergenic_variant | MODIFIER |
| 7 | 114606529 | AX-170485246 | 0 | T | C | 0.439 | 7.04E-02 | 9.79E-02 | 2.71E+00 | 2.76E+00 | 4.67E-01 | no | | intergenic_variant | MODIFIER |
| 7 | 114617255 | AX-170256888 | 0 | C | T | 0.439 | 7.04E-02 | 9.79E-02 | 2.71E+00 | 2.76E+00 | 4.67E-01 | no | | intergenic_variant | MODIFIER |
| 5 | 82282899 | AX-170518203 | 0 | A | T | 0.421 | 7.94E-02 | 1.10E-01 | 2.79E+00 | 2.83E+00 | 4.68E-01 | no | | intergenic_variant | MODIFIER |
| 8 | 46473258 | AX-170399026 | 0 | T | A | 0.149 | 9.65E-02 | 1.35E-01 | 2.77E+00 | 2.85E+00 | 4.68E-01 | no | Acsl1 | intron_variant | MODIFIER |
| 17 | 52125646 | AX-170516078 | 0 | G | A | 0.053 | 1.10E-01 | 1.55E-01 | 2.74E+00 | 2.89E+00 | 4.68E-01 | no | Gm19585 | intron_variant,non_coding_transcript_variant | MODIFIER |
| 20 | 66564549 | AX-170034040 | 0 | A | G | 0.246 | 4.50E-02 | 6.28E-02 | 2.68E+00 | 2.77E+00 | 4.68E-01 | no | | intergenic_variant | MODIFIER |
| 20 | 66629112 | AX-97785846 | 0 | T | G | 0.246 | 4.50E-02 | 6.28E-02 | 2.68E+00 | 2.77E+00 | 4.68E-01 | no | | intergenic_variant | MODIFIER |
| 20 | 66654921 | AX-170202203 | 0 | C | T | 0.246 | 4.50E-02 | 6.28E-02 | 2.68E+00 | 2.77E+00 | 4.68E-01 | no | Slitrk2 | synonymous_variant | LOW |
| 20 | 66674144 | AX-170398410 | 0 | T | C | 0.246 | 4.50E-02 | 6.28E-02 | 2.68E+00 | 2.77E+00 | 4.68E-01 | no | | intergenic_variant | MODIFIER |
| 20 | 66710515 | AX-97794520 | 0 | C | T | 0.246 | 4.50E-02 | 6.28E-02 | 2.68E+00 | 2.77E+00 | 4.68E-01 | no | | intergenic_variant | MODIFIER |
| 20 | 66736337 | AX-97803030 | 0 | A | C | 0.246 | 4.50E-02 | 6.28E-02 | 2.68E+00 | 2.77E+00 | 4.68E-01 | no | | intergenic_variant | MODIFIER |
| 20 | 66748971 | AX-170084404 | 0 | C | T | 0.246 | 4.50E-02 | 6.28E-02 | 2.68E+00 | 2.77E+00 | 4.68E-01 | no | | intergenic_variant | MODIFIER |
| 20 | 66754982 | AX-170319835 | 0 | T | C | 0.246 | 4.50E-02 | 6.28E-02 | 2.68E+00 | 2.77E+00 | 4.68E-01 | no | | intergenic_variant | MODIFIER |
| 20 | 66796838 | AX-170278594 | 0 | C | T | 0.246 | 4.50E-02 | 6.28E-02 | 2.68E+00 | 2.77E+00 | 4.68E-01 | no | Mir463 | downstream_gene_variant | MODIFIER |
| 20 | 66840671 | AX-170159992 | 0 | A | C | 0.246 | 4.50E-02 | 6.28E-02 | 2.68E+00 | 2.77E+00 | 4.68E-01 | no | Mir465 | upstream_gene_variant | MODIFIER |
| 20 | 66884994 | AX-170308981 | 0 | T | C | 0.246 | 4.50E-02 | 6.28E-02 | 2.68E+00 | 2.77E+00 | 4.68E-01 | no | | intergenic_variant | MODIFIER |
| 20 | 66964948 | AX-170034057 | 0 | T | C | 0.246 | 4.50E-02 | 6.28E-02 | 2.68E+00 | 2.77E+00 | 4.68E-01 | no | Gm6783 | downstream_gene_variant | MODIFIER |
| 20 | 66969877 | AX-170034060 | 0 | T | C | 0.246 | 4.50E-02 | 6.28E-02 | 2.68E+00 | 2.77E+00 | 4.68E-01 | no | Gm14682 | upstream_gene_variant | MODIFIER |
| 20 | 67070365 | AX-97814579 | 0 | A | G | 0.246 | 4.50E-02 | 6.28E-02 | 2.68E+00 | 2.77E+00 | 4.68E-01 | no | | intergenic_variant | MODIFIER |

| | | | | | | | | | | | | | | | |
|---|---|---|---|---|---|---|---|---|---|---|---|---|---|---|---|
| 20 | 67124928 | AX-170034054 | 0 | T | C | 0.246 | 4.50E-02 | 6.28E-02 | 2.68E+00 | 2.77E+00 | 4.68E-01 | no | | intergenic_variant | MODIFIER |
| 20 | 67156722 | AX-170034056 | 0 | A | G | 0.246 | 4.50E-02 | 6.28E-02 | 2.68E+00 | 2.77E+00 | 4.68E-01 | no | | intergenic_variant | MODIFIER |
| 20 | 67157128 | AX-170090530 | 0 | G | A | 0.246 | 4.50E-02 | 6.28E-02 | 2.68E+00 | 2.77E+00 | 4.68E-01 | no | | intergenic_variant | MODIFIER |
| 20 | 67178965 | AX-170034053 | 0 | G | A | 0.246 | 4.50E-02 | 6.28E-02 | 2.68E+00 | 2.77E+00 | 4.68E-01 | no | | intergenic_variant | MODIFIER |
| 20 | 67185913 | AX-170034051 | 0 | A | G | 0.246 | 4.50E-02 | 6.28E-02 | 2.68E+00 | 2.77E+00 | 4.68E-01 | no | | intergenic_variant | MODIFIER |
| 20 | 67287616 | AX-170497389 | 0 | G | A | 0.246 | 4.50E-02 | 6.28E-02 | 2.68E+00 | 2.77E+00 | 4.68E-01 | no | | intergenic_variant | MODIFIER |
| 20 | 67289436 | AX-97838390 | 0 | G | A | 0.246 | 4.50E-02 | 6.28E-02 | 2.68E+00 | 2.77E+00 | 4.68E-01 | no | | intergenic_variant | MODIFIER |
| 20 | 67329420 | AX-170224442 | 0 | T | A | 0.246 | 4.50E-02 | 6.28E-02 | 2.68E+00 | 2.77E+00 | 4.68E-01 | no | | intergenic_variant | MODIFIER |
| 20 | 67329987 | AX-170090531 | 0 | T | C | 0.246 | 4.50E-02 | 6.28E-02 | 2.68E+00 | 2.77E+00 | 4.68E-01 | no | | intergenic_variant | MODIFIER |
| 20 | 67360784 | AX-97798548 | 0 | A | G | 0.246 | 4.50E-02 | 6.28E-02 | 2.68E+00 | 2.77E+00 | 4.68E-01 | no | | intergenic_variant | MODIFIER |
| 20 | 67417287 | AX-97838745 | 0 | G | A | 0.246 | 4.50E-02 | 6.28E-02 | 2.68E+00 | 2.77E+00 | 4.68E-01 | no | | intergenic_variant | MODIFIER |
| 20 | 67457878 | AX-97818598 | 0 | T | A | 0.246 | 4.50E-02 | 6.28E-02 | 2.68E+00 | 2.77E+00 | 4.68E-01 | no | 1700036O09Rik | upstream_gene_variant | MODIFIER |
| 20 | 67591890 | AX-170034066 | 0 | C | T | 0.246 | 4.50E-02 | 6.28E-02 | 2.68E+00 | 2.77E+00 | 4.68E-01 | no | Gm26276 | downstream_gene_variant | MODIFIER |
| 20 | 67720451 | AX-170034077 | 0 | G | A | 0.246 | 4.50E-02 | 6.28E-02 | 2.68E+00 | 2.77E+00 | 4.68E-01 | no | | intergenic_variant | MODIFIER |
| 20 | 67727233 | AX-170034061 | 0 | A | G | 0.246 | 4.50E-02 | 6.28E-02 | 2.68E+00 | 2.77E+00 | 4.68E-01 | no | Rpl7a-ps13 | downstream_gene_variant | MODIFIER |
| 20 | 67743055 | AX-97798851 | 0 | G | A | 0.246 | 4.50E-02 | 6.28E-02 | 2.68E+00 | 2.77E+00 | 4.68E-01 | no | | intergenic_variant | MODIFIER |
| 20 | 67780985 | AX-97786311 | 0 | A | G | 0.246 | 4.50E-02 | 6.28E-02 | 2.68E+00 | 2.77E+00 | 4.68E-01 | no | | intergenic_variant | MODIFIER |
| 20 | 67825443 | AX-97785147 | 0 | A | G | 0.246 | 4.50E-02 | 6.28E-02 | 2.68E+00 | 2.77E+00 | 4.68E-01 | no | | intergenic_variant | MODIFIER |
| 20 | 67885957 | AX-168388314 | 0 | A | G | 0.246 | 4.50E-02 | 6.28E-02 | 2.68E+00 | 2.77E+00 | 4.68E-01 | no | | intergenic_variant | MODIFIER |
| 20 | 67976881 | AX-170090533 | 0 | G | A | 0.246 | 4.50E-02 | 6.28E-02 | 2.68E+00 | 2.77E+00 | 4.68E-01 | no | | intergenic_variant | MODIFIER |
| 20 | 68089265 | AX-97798898 | 0 | A | C | 0.246 | 4.50E-02 | 6.28E-02 | 2.68E+00 | 2.77E+00 | 4.68E-01 | no | | intergenic_variant | MODIFIER |
| 20 | 68126383 | AX-170034075 | 0 | G | T | 0.246 | 4.50E-02 | 6.28E-02 | 2.68E+00 | 2.77E+00 | 4.68E-01 | no | | intergenic_variant | MODIFIER |
| 20 | 68331448 | AX-170034082 | 0 | A | G | 0.246 | 4.50E-02 | 6.28E-02 | 2.68E+00 | 2.77E+00 | 4.68E-01 | no | | intergenic_variant | MODIFIER |
| 20 | 68483760 | AX-170433206 | 0 | T | A | 0.246 | 4.50E-02 | 6.28E-02 | 2.68E+00 | 2.77E+00 | 4.68E-01 | no | | intergenic_variant | MODIFIER |
| 20 | 68544764 | AX-170409614 | 0 | G | A | 0.246 | 4.50E-02 | 6.28E-02 | 2.68E+00 | 2.77E+00 | 4.68E-01 | no | | intergenic_variant | MODIFIER |
| 20 | 68554782 | AX-97819562 | 0 | C | T | 0.246 | 4.50E-02 | 6.28E-02 | 2.68E+00 | 2.77E+00 | 4.68E-01 | no | | intergenic_variant | MODIFIER |
| 20 | 68576734 | AX-170347676 | 0 | A | G | 0.246 | 4.50E-02 | 6.28E-02 | 2.68E+00 | 2.77E+00 | 4.68E-01 | no | | intergenic_variant | MODIFIER |
| 20 | 68577587 | AX-170482861 | 0 | A | G | 0.246 | 4.50E-02 | 6.28E-02 | 2.68E+00 | 2.77E+00 | 4.68E-01 | no | | intergenic_variant | MODIFIER |
| 20 | 68658349 | AX-170436913 | 0 | G | A | 0.246 | 4.50E-02 | 6.28E-02 | 2.68E+00 | 2.77E+00 | 4.68E-01 | no | | intergenic_variant | MODIFIER |
| 20 | 68720300 | AX-170411982 | 0 | A | G | 0.246 | 4.50E-02 | 6.28E-02 | 2.68E+00 | 2.77E+00 | 4.68E-01 | no | Fmr1 | downstream_gene_variant | MODIFIER |
| 20 | 68720539 | AX-170364283 | 0 | T | C | 0.246 | 4.50E-02 | 6.28E-02 | 2.68E+00 | 2.77E+00 | 4.68E-01 | no | Fmr1 | downstream_gene_variant | MODIFIER |
| 20 | 68728490 | AX-170438852 | 0 | A | G | 0.246 | 4.50E-02 | 6.28E-02 | 2.68E+00 | 2.77E+00 | 4.68E-01 | no | | intergenic_variant | MODIFIER |
| 20 | 68850873 | AX-170329336 | 0 | A | G | 0.246 | 4.50E-02 | 6.28E-02 | 2.68E+00 | 2.77E+00 | 4.68E-01 | no | | intergenic_variant | MODIFIER |
| 20 | 68850910 | AX-170310226 | 0 | C | T | 0.246 | 4.50E-02 | 6.28E-02 | 2.68E+00 | 2.77E+00 | 4.68E-01 | no | | intergenic_variant | MODIFIER |
| 20 | 68851627 | AX-97836806 | 0 | G | A | 0.246 | 4.50E-02 | 6.28E-02 | 2.68E+00 | 2.77E+00 | 4.68E-01 | no | | intergenic_variant | MODIFIER |
| 20 | 68870975 | AX-170120930 | 0 | T | A | 0.246 | 4.50E-02 | 6.28E-02 | 2.68E+00 | 2.77E+00 | 4.68E-01 | no | | intergenic_variant | MODIFIER |
| 20 | 68921264 | AX-170034093 | 0 | G | A | 0.246 | 4.50E-02 | 6.28E-02 | 2.68E+00 | 2.77E+00 | 4.68E-01 | no | Gm8344 | non_coding_transcript_exon_variant | MODIFIER |
| 20 | 68988294 | AX-170034088 | 0 | G | A | 0.246 | 4.50E-02 | 6.28E-02 | 2.68E+00 | 2.77E+00 | 4.68E-01 | no | | intergenic_variant | MODIFIER |
| 20 | 69026685 | AX-170208442 | 0 | T | C | 0.246 | 4.50E-02 | 6.28E-02 | 2.68E+00 | 2.77E+00 | 4.68E-01 | no | | intergenic_variant | MODIFIER |
| 20 | 69043929 | AX-97853401 | 0 | G | A | 0.246 | 4.50E-02 | 6.28E-02 | 2.68E+00 | 2.77E+00 | 4.68E-01 | no | | intergenic_variant | MODIFIER |
| 20 | 69044565 | AX-170034091 | 0 | A | C | 0.246 | 4.50E-02 | 6.28E-02 | 2.68E+00 | 2.77E+00 | 4.68E-01 | no | | intergenic_variant | MODIFIER |
| 20 | 69103084 | AX-170034095 | 0 | G | A | 0.246 | 4.50E-02 | 6.28E-02 | 2.68E+00 | 2.77E+00 | 4.68E-01 | no | | intergenic_variant | MODIFIER |
| 20 | 69160214 | AX-97820885 | 0 | A | G | 0.246 | 4.50E-02 | 6.28E-02 | 2.68E+00 | 2.77E+00 | 4.68E-01 | no | | intergenic_variant | MODIFIER |
| 20 | 69162369 | AX-170034090 | 0 | T | G | 0.246 | 4.50E-02 | 6.28E-02 | 2.68E+00 | 2.77E+00 | 4.68E-01 | no | | intergenic_variant | MODIFIER |
| 20 | 69174264 | AX-170341031 | 0 | T | C | 0.246 | 4.50E-02 | 6.28E-02 | 2.68E+00 | 2.77E+00 | 4.68E-01 | no | | intergenic_variant | MODIFIER |
| 20 | 69213997 | AX-170447676 | 0 | A | C | 0.246 | 4.50E-02 | 6.28E-02 | 2.68E+00 | 2.77E+00 | 4.68E-01 | no | | intergenic_variant | MODIFIER |
| 16 | 85957178 | AX-170313020 | 1 | T | C | 0.438 | 5.69E-02 | 7.91E-02 | 2.64E+00 | 2.72E+00 | 4.68E-01 | no | | intergenic_variant | MODIFIER |
| 1 | 187919469 | AX-170450348 | 1 | G | C | 0.25 | 7.62E-02 | 1.01E-01 | 2.44E+00 | 2.53E+00 | 4.68E-01 | no | Esrrg | intron_variant | MODIFIER |
| 12 | 108429018 | AX-170050138 | 0 | T | C | 0.114 | -1.17E-01 | 1.63E-01 | 2.73E+00 | 2.84E+00 | 4.68E-01 | no | Eml1 | intron_variant | MODIFIER |
| 3 | 37799968 | AX-168390585 | 1 | A | G | 0.188 | 6.87E-02 | 9.59E-02 | 2.79E+00 | 2.88E+00 | 4.68E-01 | no | Gm26404 | downstream_gene_variant | MODIFIER |
| 1 | 156444981 | AX-170394261 | 0 | A | G | 0.219 | -8.50E-02 | 1.18E-01 | 2.88E+00 | 3.01E+00 | 4.68E-01 | no | Soat1 | intron_variant | MODIFIER |
| 6 | 137117434 | AX-170347854 | 1 | A | C | 0.571 | 6.64E-02 | 8.76E-02 | 2.43E+00 | 2.49E+00 | 4.68E-01 | no | Rerg | intron_variant | MODIFIER |
| 6 | 93615891 | AX-170027742 | 0 | A | C | 0.439 | -4.36E-02 | 5.97E-02 | 2.53E+00 | 2.65E+00 | 4.69E-01 | no | | intergenic_variant | MODIFIER |
| 6 | 93616211 | AX-97842137 | 0 | T | C | 0.439 | -4.36E-02 | 5.97E-02 | 2.53E+00 | 2.65E+00 | 4.69E-01 | no | | intergenic_variant | MODIFIER |

| | | | | | | | | | | | | | | | |
|---|---|---|---|---|---|---|---|---|---|---|---|---|---|---|---|
| 6 | 93643694 | AX-170429860 | 0 | T | C | 0.439 | -4.36E-02 | 5.97E-02 | 2.53E+00 | 2.65E+00 | 4.69E-01 | no | | intergenic_variant | MODIFIER |
| 6 | 93646617 | AX-170198672 | 0 | G | T | 0.439 | -4.36E-02 | 5.97E-02 | 2.53E+00 | 2.65E+00 | 4.69E-01 | no | Gm7812 | downstream_gene_variant | MODIFIER |
| 6 | 93649894 | AX-170027738 | 0 | A | G | 0.439 | -4.36E-02 | 5.97E-02 | 2.53E+00 | 2.65E+00 | 4.69E-01 | no | Gm7812 | non_coding_transcript_exon_variant | MODIFIER |
| 6 | 93650814 | AX-170430282 | 0 | A | G | 0.439 | -4.36E-02 | 5.97E-02 | 2.53E+00 | 2.65E+00 | 4.69E-01 | no | Gm7812 | upstream_gene_variant | MODIFIER |
| 6 | 93655349 | AX-97798319 | 0 | C | T | 0.439 | -4.36E-02 | 5.97E-02 | 2.53E+00 | 2.65E+00 | 4.69E-01 | no | | intergenic_variant | MODIFIER |
| 6 | 93700706 | AX-170311631 | 0 | T | C | 0.439 | -4.36E-02 | 5.97E-02 | 2.53E+00 | 2.65E+00 | 4.69E-01 | no | Magi1 | intron_variant | MODIFIER |
| 6 | 93706156 | AX-170089789 | 0 | C | T | 0.439 | -4.36E-02 | 5.97E-02 | 2.53E+00 | 2.65E+00 | 4.69E-01 | no | Magi1 | intron_variant | MODIFIER |
| 6 | 93706362 | AX-170165460 | 0 | C | T | 0.439 | -4.36E-02 | 5.97E-02 | 2.53E+00 | 2.65E+00 | 4.69E-01 | no | Magi1 | intron_variant | MODIFIER |
| 6 | 93710185 | AX-170323927 | 0 | G | A | 0.439 | -4.36E-02 | 5.97E-02 | 2.53E+00 | 2.65E+00 | 4.69E-01 | no | Magi1 | intron_variant | MODIFIER |
| 6 | 93713663 | AX-170222015 | 0 | A | G | 0.439 | -4.36E-02 | 5.97E-02 | 2.53E+00 | 2.65E+00 | 4.69E-01 | no | Magi1 | intron_variant | MODIFIER |
| 6 | 93714083 | AX-170027740 | 0 | T | C | 0.439 | -4.36E-02 | 5.97E-02 | 2.53E+00 | 2.65E+00 | 4.69E-01 | no | Magi1 | intron_variant | MODIFIER |
| 6 | 93723312 | AX-170296739 | 0 | T | C | 0.439 | -4.36E-02 | 5.97E-02 | 2.53E+00 | 2.65E+00 | 4.69E-01 | no | Magi1 | intron_variant | MODIFIER |
| 6 | 93729996 | AX-170244942 | 0 | T | G | 0.439 | -4.36E-02 | 5.97E-02 | 2.53E+00 | 2.65E+00 | 4.69E-01 | no | Magi1 | intron_variant | MODIFIER |
| 6 | 93743168 | AX-170259338 | 0 | A | T | 0.439 | -4.36E-02 | 5.97E-02 | 2.53E+00 | 2.65E+00 | 4.69E-01 | no | Magi1 | intron_variant | MODIFIER |
| 6 | 93745729 | AX-170357920 | 0 | G | A | 0.439 | -4.36E-02 | 5.97E-02 | 2.53E+00 | 2.65E+00 | 4.69E-01 | no | Magi1 | synonymous_variant | LOW |
| 6 | 93747468 | AX-170179775 | 0 | A | T | 0.439 | -4.36E-02 | 5.97E-02 | 2.53E+00 | 2.65E+00 | 4.69E-01 | no | Magi1 | intron_variant | MODIFIER |
| 6 | 93748071 | AX-170204664 | 0 | C | T | 0.439 | -4.36E-02 | 5.97E-02 | 2.53E+00 | 2.65E+00 | 4.69E-01 | no | Magi1 | intron_variant | MODIFIER |
| 6 | 93771484 | AX-170267191 | 0 | C | T | 0.439 | -4.36E-02 | 5.97E-02 | 2.53E+00 | 2.65E+00 | 4.69E-01 | no | Magi1 | intron_variant | MODIFIER |
| 6 | 93771889 | AX-170517647 | 0 | C | T | 0.439 | -4.36E-02 | 5.97E-02 | 2.53E+00 | 2.65E+00 | 4.69E-01 | no | Magi1 | intron_variant | MODIFIER |
| 6 | 93800702 | AX-170360261 | 0 | G | A | 0.439 | -4.36E-02 | 5.97E-02 | 2.53E+00 | 2.65E+00 | 4.69E-01 | no | Magi1 | intron_variant | MODIFIER |
| 6 | 93801444 | AX-170186226 | 0 | G | A | 0.439 | -4.36E-02 | 5.97E-02 | 2.53E+00 | 2.65E+00 | 4.69E-01 | no | Magi1 | intron_variant | MODIFIER |
| 6 | 93821500 | AX-170378666 | 0 | C | T | 0.439 | -4.36E-02 | 5.97E-02 | 2.53E+00 | 2.65E+00 | 4.69E-01 | no | Magi1 | intron_variant | MODIFIER |
| 6 | 93835196 | AX-170267725 | 0 | C | T | 0.439 | -4.36E-02 | 5.97E-02 | 2.53E+00 | 2.65E+00 | 4.69E-01 | no | Magi1 | intron_variant | MODIFIER |
| 6 | 93835693 | AX-170466109 | 0 | C | A | 0.439 | -4.36E-02 | 5.97E-02 | 2.53E+00 | 2.65E+00 | 4.69E-01 | no | Magi1 | intron_variant | MODIFIER |
| 6 | 93843110 | AX-170214490 | 0 | G | C | 0.439 | -4.36E-02 | 5.97E-02 | 2.53E+00 | 2.65E+00 | 4.69E-01 | no | Magi1 | intron_variant | MODIFIER |
| 6 | 93863961 | AX-97811686 | 0 | T | C | 0.439 | -4.36E-02 | 5.97E-02 | 2.53E+00 | 2.65E+00 | 4.69E-01 | no | Magi1 | intron_variant | MODIFIER |
| 6 | 93865652 | AX-170163118 | 0 | C | T | 0.439 | -4.36E-02 | 5.97E-02 | 2.53E+00 | 2.65E+00 | 4.69E-01 | no | Magi1 | intron_variant | MODIFIER |
| 6 | 93903855 | AX-170265946 | 0 | C | T | 0.439 | -4.36E-02 | 5.97E-02 | 2.53E+00 | 2.65E+00 | 4.69E-01 | no | Magi1 | intron_variant | MODIFIER |
| 6 | 93904001 | AX-170184080 | 0 | A | T | 0.439 | -4.36E-02 | 5.97E-02 | 2.53E+00 | 2.65E+00 | 4.69E-01 | no | Magi1 | intron_variant | MODIFIER |
| 6 | 93904215 | AX-170496739 | 0 | C | T | 0.439 | -4.36E-02 | 5.97E-02 | 2.53E+00 | 2.65E+00 | 4.69E-01 | no | Magi1 | intron_variant | MODIFIER |
| 6 | 93909191 | AX-170239345 | 0 | A | T | 0.439 | -4.36E-02 | 5.97E-02 | 2.53E+00 | 2.65E+00 | 4.69E-01 | no | Magi1 | intron_variant | MODIFIER |
| 6 | 93914713 | AX-170221515 | 0 | C | T | 0.439 | -4.36E-02 | 5.97E-02 | 2.53E+00 | 2.65E+00 | 4.69E-01 | no | Magi1 | intron_variant | MODIFIER |
| 6 | 93915240 | AX-170356993 | 0 | G | A | 0.439 | -4.36E-02 | 5.97E-02 | 2.53E+00 | 2.65E+00 | 4.69E-01 | no | Magi1 | intron_variant | MODIFIER |
| 6 | 94019680 | AX-170488449 | 0 | A | C | 0.439 | -4.36E-02 | 5.97E-02 | 2.53E+00 | 2.65E+00 | 4.69E-01 | no | Magi1 | intron_variant | MODIFIER |
| 6 | 94051325 | AX-170352508 | 0 | C | T | 0.439 | -4.36E-02 | 5.97E-02 | 2.53E+00 | 2.65E+00 | 4.69E-01 | no | Magi1 | intron_variant | MODIFIER |
| 6 | 94079849 | AX-170380221 | 0 | A | C | 0.439 | -4.36E-02 | 5.97E-02 | 2.53E+00 | 2.65E+00 | 4.69E-01 | no | Magi1 | intron_variant | MODIFIER |
| 6 | 94087618 | AX-170027748 | 0 | A | T | 0.439 | -4.36E-02 | 5.97E-02 | 2.53E+00 | 2.65E+00 | 4.69E-01 | no | Magi1 | intron_variant | MODIFIER |
| 6 | 94095264 | AX-170027753 | 0 | G | A | 0.439 | -4.36E-02 | 5.97E-02 | 2.53E+00 | 2.65E+00 | 4.69E-01 | no | Magi1 | intron_variant | MODIFIER |
| 6 | 94115577 | AX-97826282 | 0 | A | T | 0.439 | -4.36E-02 | 5.97E-02 | 2.53E+00 | 2.65E+00 | 4.69E-01 | no | Magi1 | intron_variant | MODIFIER |
| 6 | 94119513 | AX-170329327 | 0 | C | T | 0.439 | -4.36E-02 | 5.97E-02 | 2.53E+00 | 2.65E+00 | 4.69E-01 | no | Magi1 | intron_variant | MODIFIER |
| 6 | 94131578 | AX-170075536 | 0 | T | G | 0.439 | -4.36E-02 | 5.97E-02 | 2.53E+00 | 2.65E+00 | 4.69E-01 | no | Magi1 | intron_variant | MODIFIER |
| 6 | 94141688 | AX-170027751 | 0 | C | T | 0.439 | -4.36E-02 | 5.97E-02 | 2.53E+00 | 2.65E+00 | 4.69E-01 | no | Magi1 | intron_variant | MODIFIER |
| 6 | 94146610 | AX-170208852 | 0 | G | A | 0.439 | -4.36E-02 | 5.97E-02 | 2.53E+00 | 2.65E+00 | 4.69E-01 | no | Magi1 | intron_variant | MODIFIER |
| 6 | 94147808 | AX-170027747 | 0 | C | T | 0.439 | -4.36E-02 | 5.97E-02 | 2.53E+00 | 2.65E+00 | 4.69E-01 | no | Magi1 | intron_variant | MODIFIER |
| 6 | 94150670 | AX-170217735 | 0 | A | G | 0.439 | -4.36E-02 | 5.97E-02 | 2.53E+00 | 2.65E+00 | 4.69E-01 | no | Magi1 | intron_variant | MODIFIER |
| 6 | 94151696 | AX-170200037 | 0 | C | T | 0.439 | -4.36E-02 | 5.97E-02 | 2.53E+00 | 2.65E+00 | 4.69E-01 | no | Magi1 | intron_variant | MODIFIER |
| 6 | 94155283 | AX-170027749 | 0 | T | C | 0.439 | -4.36E-02 | 5.97E-02 | 2.53E+00 | 2.65E+00 | 4.69E-01 | no | Magi1 | intron_variant | MODIFIER |
| 6 | 94193763 | AX-170027746 | 0 | A | C | 0.439 | -4.36E-02 | 5.97E-02 | 2.53E+00 | 2.65E+00 | 4.69E-01 | no | Magi1 | intron_variant | MODIFIER |
| 6 | 94224152 | AX-97822173 | 0 | A | G | 0.439 | -4.36E-02 | 5.97E-02 | 2.53E+00 | 2.65E+00 | 4.69E-01 | no | Magi1 | intron_variant | MODIFIER |
| 6 | 94224471 | AX-170027750 | 0 | C | T | 0.439 | -4.36E-02 | 5.97E-02 | 2.53E+00 | 2.65E+00 | 4.69E-01 | no | Magi1 | intron_variant | MODIFIER |
| 6 | 94248175 | AX-170366522 | 0 | A | T | 0.439 | -4.36E-02 | 5.97E-02 | 2.53E+00 | 2.65E+00 | 4.69E-01 | no | Magi1 | intron_variant | MODIFIER |
| 6 | 94263839 | AX-170224690 | 0 | C | T | 0.439 | -4.36E-02 | 5.97E-02 | 2.53E+00 | 2.65E+00 | 4.69E-01 | no | Magi1 | intron_variant | MODIFIER |
| 6 | 94296445 | AX-170355995 | 0 | T | C | 0.439 | -4.36E-02 | 5.97E-02 | 2.53E+00 | 2.65E+00 | 4.69E-01 | no | | intergenic_variant | MODIFIER |
| 6 | 94305488 | AX-97854300 | 0 | A | G | 0.439 | -4.36E-02 | 5.97E-02 | 2.53E+00 | 2.65E+00 | 4.69E-01 | no | | intergenic_variant | MODIFIER |

| | | | | | | | | | | | | | | | |
|---|---|---|---|---|---|---|---|---|---|---|---|---|---|---|---|
| 6 | 94326846 | AX-170184787 | 0 | A | T | 0.439 | -4.36E-02 | 5.97E-02 | 2.53E+00 | 2.65E+00 | 4.69E-01 | no | Gm44307 | upstream_gene_variant | MODIFIER |
| 6 | 94332756 | AX-170350144 | 0 | G | A | 0.439 | -4.36E-02 | 5.97E-02 | 2.53E+00 | 2.65E+00 | 4.69E-01 | no | | intergenic_variant | MODIFIER |
| 6 | 94374384 | AX-97852039 | 0 | C | T | 0.439 | -4.36E-02 | 5.97E-02 | 2.53E+00 | 2.65E+00 | 4.69E-01 | no | | intergenic_variant | MODIFIER |
| 6 | 94385341 | AX-170391489 | 0 | C | T | 0.439 | -4.36E-02 | 5.97E-02 | 2.53E+00 | 2.65E+00 | 4.69E-01 | no | | regulatory_region_variant | MODIFIER |
| 6 | 94386037 | AX-170208038 | 0 | C | T | 0.439 | -4.36E-02 | 5.97E-02 | 2.53E+00 | 2.65E+00 | 4.69E-01 | no | | intergenic_variant | MODIFIER |
| 6 | 94393634 | AX-170125833 | 0 | G | A | 0.439 | -4.36E-02 | 5.97E-02 | 2.53E+00 | 2.65E+00 | 4.69E-01 | no | | intergenic_variant | MODIFIER |
| 6 | 94404849 | AX-97793461 | 0 | T | C | 0.439 | -4.36E-02 | 5.97E-02 | 2.53E+00 | 2.65E+00 | 4.69E-01 | no | | intergenic_variant | MODIFIER |
| 6 | 94405526 | AX-170178029 | 0 | A | G | 0.439 | -4.36E-02 | 5.97E-02 | 2.53E+00 | 2.65E+00 | 4.69E-01 | no | | intergenic_variant | MODIFIER |
| 6 | 94410995 | AX-170517816 | 0 | A | G | 0.439 | -4.36E-02 | 5.97E-02 | 2.53E+00 | 2.65E+00 | 4.69E-01 | no | | regulatory_region_variant | MODIFIER |
| 6 | 94432748 | AX-170411477 | 0 | C | A | 0.439 | -4.36E-02 | 5.97E-02 | 2.53E+00 | 2.65E+00 | 4.69E-01 | no | | intergenic_variant | MODIFIER |
| 6 | 94495065 | AX-170460074 | 0 | A | T | 0.439 | -4.36E-02 | 5.97E-02 | 2.53E+00 | 2.65E+00 | 4.69E-01 | no | | intergenic_variant | MODIFIER |
| 6 | 94503144 | AX-97851493 | 0 | C | T | 0.439 | -4.36E-02 | 5.97E-02 | 2.53E+00 | 2.65E+00 | 4.69E-01 | no | Slc25a26 | intron_variant | MODIFIER |
| 6 | 94526098 | AX-170530977 | 0 | T | G | 0.439 | -4.36E-02 | 5.97E-02 | 2.53E+00 | 2.65E+00 | 4.69E-01 | no | Slc25a26 | intron_variant | MODIFIER |
| 6 | 94539011 | AX-170089792 | 0 | T | C | 0.439 | -4.36E-02 | 5.97E-02 | 2.53E+00 | 2.65E+00 | 4.69E-01 | no | Slc25a26 | intron_variant | MODIFIER |
| 6 | 94542074 | AX-170508220 | 0 | T | C | 0.439 | -4.36E-02 | 5.97E-02 | 2.53E+00 | 2.65E+00 | 4.69E-01 | no | Slc25a26 | intron_variant | MODIFIER |
| 6 | 94554434 | AX-97813521 | 0 | T | C | 0.439 | -4.36E-02 | 5.97E-02 | 2.53E+00 | 2.65E+00 | 4.69E-01 | no | Slc25a26 | intron_variant | MODIFIER |
| 6 | 94576561 | AX-170326898 | 0 | T | C | 0.439 | -4.36E-02 | 5.97E-02 | 2.53E+00 | 2.65E+00 | 4.69E-01 | no | Slc25a26 | intron_variant | MODIFIER |
| 6 | 94577824 | AX-170174516 | 0 | G | A | 0.439 | -4.36E-02 | 5.97E-02 | 2.53E+00 | 2.65E+00 | 4.69E-01 | no | Slc25a26 | intron_variant | MODIFIER |
| 6 | 94581318 | AX-170492150 | 0 | G | A | 0.439 | -4.36E-02 | 5.97E-02 | 2.53E+00 | 2.65E+00 | 4.69E-01 | no | Slc25a26 | intron_variant | MODIFIER |
| 6 | 94581773 | AX-170372087 | 0 | T | C | 0.439 | -4.36E-02 | 5.97E-02 | 2.53E+00 | 2.65E+00 | 4.69E-01 | no | Slc25a26 | intron_variant | MODIFIER |
| 6 | 94589553 | AX-170075543 | 0 | A | C | 0.439 | -4.36E-02 | 5.97E-02 | 2.53E+00 | 2.65E+00 | 4.69E-01 | no | Slc25a26 | intron_variant | MODIFIER |
| 6 | 94597892 | AX-170075544 | 0 | T | A | 0.439 | -4.36E-02 | 5.97E-02 | 2.53E+00 | 2.65E+00 | 4.69E-01 | no | Slc25a26 | intron_variant | MODIFIER |
| 6 | 94603041 | AX-170513366 | 0 | C | T | 0.439 | -4.36E-02 | 5.97E-02 | 2.53E+00 | 2.65E+00 | 4.69E-01 | no | Lrig1 | downstream_gene_variant | MODIFIER |
| 6 | 94609550 | AX-170327172 | 0 | C | T | 0.439 | -4.36E-02 | 5.97E-02 | 2.53E+00 | 2.65E+00 | 4.69E-01 | no | Lrig1 | intron_variant | MODIFIER |
| 6 | 94609925 | AX-170210950 | 0 | T | C | 0.439 | -4.36E-02 | 5.97E-02 | 2.53E+00 | 2.65E+00 | 4.69E-01 | no | Lrig1 | intron_variant | MODIFIER |
| 6 | 94656295 | AX-170089793 | 0 | T | C | 0.439 | -4.36E-02 | 5.97E-02 | 2.53E+00 | 2.65E+00 | 4.69E-01 | no | Lrig1 | intron_variant | MODIFIER |
| 6 | 94668372 | AX-170095707 | 0 | G | A | 0.439 | -4.36E-02 | 5.97E-02 | 2.53E+00 | 2.65E+00 | 4.69E-01 | no | Lrig1 | intron_variant | MODIFIER |
| 6 | 94794096 | AX-170165168 | 0 | C | T | 0.439 | -4.36E-02 | 5.97E-02 | 2.53E+00 | 2.65E+00 | 4.69E-01 | no | | intergenic_variant | MODIFIER |
| 1 | 170838396 | AX-170423206 | 1 | C | T | 0.205 | 9.93E-02 | 1.38E-01 | 2.69E+00 | 2.73E+00 | 4.69E-01 | no | Atf6 | intron_variant | MODIFIER |
| 16 | 46421484 | AX-170316388 | 0 | C | T | 0.518 | 6.01E-02 | 8.18E-02 | 2.53E+00 | 2.58E+00 | 4.69E-01 | no | Nectin3 | intron_variant | MODIFIER |
| 1 | 22290408 | AX-170512858 | 0 | A | G | 0.421 | 6.02E-02 | 8.34E-02 | 2.61E+00 | 2.67E+00 | 4.69E-01 | no | Rims1 | intron_variant | MODIFIER |
| 5 | 46061646 | AX-170517216 | 0 | T | C | 0.474 | 7.17E-02 | 9.64E-02 | 2.48E+00 | 2.55E+00 | 4.69E-01 | no | | intergenic_variant | MODIFIER |
| 5 | 106450137 | AX-170491518 | 1 | T | C | 0.446 | 8.16E-02 | 1.13E-01 | 2.65E+00 | 2.69E+00 | 4.69E-01 | no | Barhl2 | downstream_gene_variant | MODIFIER |
| 8 | 87120460 | AX-97850929 | 1 | A | G | 0.196 | -9.35E-02 | 1.31E-01 | 2.74E+00 | 2.80E+00 | 4.69E-01 | no | Gm27246 | intron_variant,non_coding_transcript_variant | MODIFIER |
| 1 | 105245784 | AX-97803837 | 1 | G | A | 0.161 | 1.48E-01 | 2.07E-01 | 2.66E+00 | 2.72E+00 | 4.69E-01 | Yes | Gm29013 | downstream_gene_variant | MODIFIER |
| 4 | 128499669 | AX-170251838 | 1 | G | A | 0.161 | 8.06E-02 | 1.12E-01 | 2.59E+00 | 2.65E+00 | 4.70E-01 | no | Csmd2 | non_coding_transcript_exon_variant | MODIFIER |
| 6 | 93496574 | AX-170521957 | 0 | C | T | 0.421 | -4.21E-02 | 5.80E-02 | 2.54E+00 | 2.67E+00 | 4.70E-01 | no | | intergenic_variant | MODIFIER |
| 6 | 93535816 | AX-170155105 | 0 | C | T | 0.421 | -4.21E-02 | 5.80E-02 | 2.54E+00 | 2.67E+00 | 4.70E-01 | no | | intergenic_variant | MODIFIER |
| 6 | 93576042 | AX-170075524 | 0 | T | G | 0.421 | -4.21E-02 | 5.80E-02 | 2.54E+00 | 2.67E+00 | 4.70E-01 | no | | intergenic_variant | MODIFIER |
| 6 | 93577355 | AX-97818859 | 0 | A | G | 0.421 | -4.21E-02 | 5.80E-02 | 2.54E+00 | 2.67E+00 | 4.70E-01 | no | | intergenic_variant | MODIFIER |
| 6 | 93577537 | AX-170075523 | 0 | G | A | 0.421 | -4.21E-02 | 5.80E-02 | 2.54E+00 | 2.67E+00 | 4.70E-01 | no | | intergenic_variant | MODIFIER |
| 6 | 93582168 | AX-170027741 | 0 | T | C | 0.421 | -4.21E-02 | 5.80E-02 | 2.54E+00 | 2.67E+00 | 4.70E-01 | no | | intergenic_variant | MODIFIER |
| 6 | 93589607 | AX-170027744 | 0 | G | A | 0.421 | -4.21E-02 | 5.80E-02 | 2.54E+00 | 2.67E+00 | 4.70E-01 | no | | intergenic_variant | MODIFIER |
| 6 | 93589708 | AX-170027739 | 0 | G | A | 0.421 | -4.21E-02 | 5.80E-02 | 2.54E+00 | 2.67E+00 | 4.70E-01 | no | | intergenic_variant | MODIFIER |
| 12 | 61046684 | AX-170501513 | 0 | C | A | 0.061 | -1.50E-01 | 2.09E-01 | 2.62E+00 | 2.72E+00 | 4.70E-01 | no | | intergenic_variant | MODIFIER |
| 12 | 63955565 | AX-170006910 | 0 | G | A | 0.061 | -1.50E-01 | 2.09E-01 | 2.62E+00 | 2.72E+00 | 4.70E-01 | no | | intergenic_variant | MODIFIER |
| 18 | 41271065 | AX-170251209 | 0 | A | G | 0.061 | -1.50E-01 | 2.09E-01 | 2.62E+00 | 2.72E+00 | 4.70E-01 | no | | intergenic_variant | MODIFIER |
| 18 | 41968629 | AX-170015599 | 0 | C | A | 0.061 | -1.50E-01 | 2.09E-01 | 2.62E+00 | 2.72E+00 | 4.70E-01 | no | | intergenic_variant | MODIFIER |
| 15 | 75741689 | AX-170248190 | 0 | G | A | 0.561 | 5.96E-02 | 8.24E-02 | 2.58E+00 | 2.65E+00 | 4.70E-01 | no | Mafa | downstream_gene_variant | MODIFIER |
| 9 | 90732175 | AX-170083079 | 0 | C | T | 0.14 | -8.15E-02 | 1.12E-01 | 2.53E+00 | 2.63E+00 | 4.70E-01 | no | Gm2497 | downstream_gene_variant | MODIFIER |
| 10 | 66011411 | AX-170142291 | 0 | T | C | 0.412 | -5.71E-02 | 7.97E-02 | 2.64E+00 | 2.71E+00 | 4.70E-01 | no | | intergenic_variant | MODIFIER |
| 16 | 81175920 | AX-170118155 | 0 | T | C | 0.465 | 5.47E-02 | 7.57E-02 | 2.59E+00 | 2.66E+00 | 4.70E-01 | no | | regulatory_region_variant | MODIFIER |
| 16 | 50215296 | AX-170013009 | 1 | G | C | 0.5 | 5.98E-02 | 8.21E-02 | 2.56E+00 | 2.60E+00 | 4.70E-01 | no | Bbx | intron_variant | MODIFIER |
| 18 | 57781742 | AX-170263583 | 0 | A | G | 0.412 | 6.18E-02 | 8.42E-02 | 2.99E+00 | 3.09E+00 | 4.70E-01 | no | AC102232.1 | upstream_gene_variant | MODIFIER |

| | | | | | | | | | | | | | | | |
|---|---|---|---|---|---|---|---|---|---|---|---|---|---|---|---|
| 9 | 76290614 | AX-170191451 | 0 | A | T | 0.211 | 9.10E-02 | 1.27E-01 | 2.86E+00 | 2.93E+00 | 4.70E-01 | no | Hcrtr2 | intron_variant | MODIFIER |
| 9 | 84146909 | AX-170341833 | 0 | G | T | 0.211 | -7.86E-02 | 1.10E-01 | 2.67E+00 | 2.78E+00 | 4.71E-01 | no | | intergenic_variant | MODIFIER |
| 9 | 84437564 | AX-170032856 | 0 | A | G | 0.211 | -7.86E-02 | 1.10E-01 | 2.67E+00 | 2.78E+00 | 4.71E-01 | no | | intergenic_variant | MODIFIER |
| 3 | 14871980 | AX-170483441 | 1 | A | T | 0.429 | -4.56E-02 | 6.46E-02 | 2.73E+00 | 2.88E+00 | 4.71E-01 | no | Car3 | 3_prime_UTR_variant | MODIFIER |
| 4 | 64969028 | AX-170410948 | 0 | G | T | 0.325 | 8.53E-02 | 1.19E-01 | 2.69E+00 | 2.74E+00 | 4.71E-01 | no | | intergenic_variant | MODIFIER |
| 4 | 69118626 | AX-170097995 | 0 | A | G | 0.325 | 8.53E-02 | 1.19E-01 | 2.69E+00 | 2.74E+00 | 4.71E-01 | no | | intergenic_variant | MODIFIER |
| 4 | 75821150 | AX-170520420 | 0 | G | A | 0.325 | 8.53E-02 | 1.19E-01 | 2.69E+00 | 2.74E+00 | 4.71E-01 | no | | intergenic_variant | MODIFIER |
| 5 | 47016603 | AX-170254796 | 1 | T | C | 0.473 | 7.46E-02 | 9.87E-02 | 2.42E+00 | 2.49E+00 | 4.71E-01 | no | | intergenic_variant | MODIFIER |
| 18 | 55491869 | AX-170399920 | 0 | C | T | 0.456 | 7.05E-02 | 9.16E-02 | 2.38E+00 | 2.45E+00 | 4.71E-01 | no | | regulatory_region_variant | MODIFIER |
| 20 | 68577518 | AX-170403599 | 0 | A | G | 0.246 | 4.44E-02 | 6.24E-02 | 2.69E+00 | 2.78E+00 | 4.71E-01 | no | | intergenic_variant | MODIFIER |
| 4 | 132433173 | AX-170356483 | 1 | G | A | 0.152 | 8.54E-02 | 1.18E-01 | 2.59E+00 | 2.64E+00 | 4.71E-01 | no | | regulatory_region_variant | MODIFIER |
| 1 | 187901038 | AX-170230024 | 0 | T | C | 0.246 | 7.58E-02 | 1.02E-01 | 2.45E+00 | 2.54E+00 | 4.71E-01 | no | Esrrg | intron_variant | MODIFIER |
| 1 | 187901304 | AX-170502255 | 0 | A | G | 0.246 | 7.58E-02 | 1.02E-01 | 2.45E+00 | 2.54E+00 | 4.71E-01 | no | Esrrg | intron_variant | MODIFIER |
| 1 | 187901546 | AX-170272777 | 0 | A | G | 0.246 | 7.58E-02 | 1.02E-01 | 2.45E+00 | 2.54E+00 | 4.71E-01 | no | Esrrg | intron_variant | MODIFIER |
| 1 | 187907710 | AX-170360276 | 0 | C | T | 0.246 | 7.58E-02 | 1.02E-01 | 2.45E+00 | 2.54E+00 | 4.71E-01 | no | Esrrg | intron_variant | MODIFIER |
| 1 | 187916279 | AX-170323369 | 0 | T | G | 0.246 | 7.58E-02 | 1.02E-01 | 2.45E+00 | 2.54E+00 | 4.71E-01 | no | Esrrg | intron_variant | MODIFIER |
| 1 | 187919330 | AX-170347657 | 0 | G | A | 0.246 | 7.58E-02 | 1.02E-01 | 2.45E+00 | 2.54E+00 | 4.71E-01 | no | Esrrg | intron_variant | MODIFIER |
| 1 | 187919386 | AX-170153363 | 0 | T | C | 0.246 | 7.58E-02 | 1.02E-01 | 2.45E+00 | 2.54E+00 | 4.71E-01 | no | Esrrg | intron_variant | MODIFIER |
| 1 | 187923944 | AX-170505321 | 0 | C | T | 0.246 | 7.58E-02 | 1.02E-01 | 2.45E+00 | 2.54E+00 | 4.71E-01 | no | Esrrg | intron_variant | MODIFIER |
| 1 | 187924827 | AX-170296752 | 0 | C | G | 0.246 | 7.58E-02 | 1.02E-01 | 2.45E+00 | 2.54E+00 | 4.71E-01 | no | Esrrg | intron_variant | MODIFIER |
| 1 | 187926495 | AX-170269381 | 0 | A | G | 0.246 | 7.58E-02 | 1.02E-01 | 2.45E+00 | 2.54E+00 | 4.71E-01 | no | Esrrg | intron_variant | MODIFIER |
| 1 | 187926786 | AX-170043906 | 0 | T | C | 0.246 | 7.58E-02 | 1.02E-01 | 2.45E+00 | 2.54E+00 | 4.71E-01 | no | Esrrg | intron_variant | MODIFIER |
| 1 | 187926919 | AX-170043908 | 0 | A | G | 0.246 | 7.58E-02 | 1.02E-01 | 2.45E+00 | 2.54E+00 | 4.71E-01 | no | Esrrg | intron_variant | MODIFIER |
| 1 | 187927195 | AX-170043905 | 0 | A | G | 0.246 | 7.58E-02 | 1.02E-01 | 2.45E+00 | 2.54E+00 | 4.71E-01 | no | Esrrg | intron_variant | MODIFIER |
| 1 | 187927478 | AX-170043909 | 0 | G | A | 0.246 | 7.58E-02 | 1.02E-01 | 2.45E+00 | 2.54E+00 | 4.71E-01 | no | Esrrg | intron_variant | MODIFIER |
| 1 | 187928180 | AX-97831900 | 0 | G | A | 0.246 | 7.58E-02 | 1.02E-01 | 2.45E+00 | 2.54E+00 | 4.71E-01 | no | Esrrg | intron_variant | MODIFIER |
| 1 | 187931168 | AX-170043907 | 0 | A | T | 0.246 | 7.58E-02 | 1.02E-01 | 2.45E+00 | 2.54E+00 | 4.71E-01 | no | Esrrg | intron_variant | MODIFIER |
| 1 | 187936464 | AX-170469309 | 0 | C | T | 0.246 | 7.58E-02 | 1.02E-01 | 2.45E+00 | 2.54E+00 | 4.71E-01 | no | Esrrg | intron_variant | MODIFIER |
| 1 | 187948894 | AX-170307153 | 0 | G | A | 0.246 | 7.58E-02 | 1.02E-01 | 2.45E+00 | 2.54E+00 | 4.71E-01 | no | Esrrg | intron_variant | MODIFIER |
| 1 | 187972316 | AX-97815623 | 0 | T | C | 0.246 | 7.58E-02 | 1.02E-01 | 2.45E+00 | 2.54E+00 | 4.71E-01 | no | Esrrg | intron_variant | MODIFIER |
| 1 | 188006398 | AX-170002516 | 0 | C | T | 0.246 | 7.58E-02 | 1.02E-01 | 2.45E+00 | 2.54E+00 | 4.71E-01 | no | Esrrg | intron_variant | MODIFIER |
| 1 | 188010190 | AX-170002514 | 0 | C | T | 0.246 | 7.58E-02 | 1.02E-01 | 2.45E+00 | 2.54E+00 | 4.71E-01 | no | Esrrg | intron_variant | MODIFIER |
| 1 | 188015764 | AX-170195984 | 0 | C | T | 0.246 | 7.58E-02 | 1.02E-01 | 2.45E+00 | 2.54E+00 | 4.71E-01 | no | Esrrg | intron_variant | MODIFIER |
| 1 | 188016130 | AX-170002515 | 0 | C | T | 0.246 | 7.58E-02 | 1.02E-01 | 2.45E+00 | 2.54E+00 | 4.71E-01 | no | Esrrg | intron_variant | MODIFIER |
| 1 | 188016248 | AX-170306593 | 0 | C | T | 0.246 | 7.58E-02 | 1.02E-01 | 2.45E+00 | 2.54E+00 | 4.71E-01 | no | Esrrg | intron_variant | MODIFIER |
| 1 | 188021979 | AX-170002513 | 0 | A | G | 0.246 | 7.58E-02 | 1.02E-01 | 2.45E+00 | 2.54E+00 | 4.71E-01 | no | Esrrg | intron_variant | MODIFIER |
| 1 | 188035807 | AX-170119686 | 0 | T | C | 0.246 | 7.58E-02 | 1.02E-01 | 2.45E+00 | 2.54E+00 | 4.71E-01 | no | Esrrg | intron_variant | MODIFIER |
| 1 | 188059357 | AX-170091830 | 0 | G | C | 0.246 | 7.58E-02 | 1.02E-01 | 2.45E+00 | 2.54E+00 | 4.71E-01 | no | Esrrg | intron_variant | MODIFIER |
| 1 | 188062764 | AX-170162024 | 0 | G | A | 0.246 | 7.58E-02 | 1.02E-01 | 2.45E+00 | 2.54E+00 | 4.71E-01 | no | Esrrg | intron_variant | MODIFIER |
| 1 | 188063344 | AX-170132402 | 0 | C | T | 0.246 | 7.58E-02 | 1.02E-01 | 2.45E+00 | 2.54E+00 | 4.71E-01 | no | Esrrg | intron_variant | MODIFIER |
| 1 | 188065848 | AX-170252008 | 0 | A | G | 0.246 | 7.58E-02 | 1.02E-01 | 2.45E+00 | 2.54E+00 | 4.71E-01 | no | Esrrg | intron_variant | MODIFIER |
| 1 | 188081035 | AX-170275408 | 0 | A | C | 0.246 | 7.58E-02 | 1.02E-01 | 2.45E+00 | 2.54E+00 | 4.71E-01 | no | Esrrg | intron_variant | MODIFIER |
| 1 | 188635448 | AX-170187481 | 0 | A | C | 0.246 | 7.58E-02 | 1.02E-01 | 2.45E+00 | 2.54E+00 | 4.71E-01 | no | Ush2a | intron_variant | MODIFIER |
| 1 | 188638865 | AX-170489394 | 0 | C | T | 0.246 | 7.58E-02 | 1.02E-01 | 2.45E+00 | 2.54E+00 | 4.71E-01 | no | Ush2a | intron_variant | MODIFIER |
| 1 | 188638962 | AX-170259679 | 0 | C | T | 0.246 | 7.58E-02 | 1.02E-01 | 2.45E+00 | 2.54E+00 | 4.71E-01 | no | Ush2a | intron_variant | MODIFIER |
| 1 | 188643243 | AX-170463996 | 0 | G | T | 0.246 | 7.58E-02 | 1.02E-01 | 2.45E+00 | 2.54E+00 | 4.71E-01 | no | Ush2a | intron_variant | MODIFIER |
| 1 | 188659784 | AX-97847814 | 0 | G | A | 0.246 | 7.58E-02 | 1.02E-01 | 2.45E+00 | 2.54E+00 | 4.71E-01 | no | Ush2a | intron_variant | MODIFIER |
| 1 | 188676091 | AX-170531252 | 0 | G | A | 0.246 | 7.58E-02 | 1.02E-01 | 2.45E+00 | 2.54E+00 | 4.71E-01 | no | Ush2a | intron_variant | MODIFIER |
| 1 | 188687756 | AX-170091834 | 0 | A | C | 0.246 | 7.58E-02 | 1.02E-01 | 2.45E+00 | 2.54E+00 | 4.71E-01 | no | Ush2a | intron_variant | MODIFIER |
| 1 | 188689458 | AX-170476170 | 0 | C | A | 0.246 | 7.58E-02 | 1.02E-01 | 2.45E+00 | 2.54E+00 | 4.71E-01 | no | Ush2a | intron_variant | MODIFIER |
| 1 | 188692034 | AX-170429732 | 0 | C | T | 0.246 | 7.58E-02 | 1.02E-01 | 2.45E+00 | 2.54E+00 | 4.71E-01 | no | Ush2a | intron_variant | MODIFIER |
| 1 | 188699450 | AX-170520165 | 0 | C | T | 0.246 | 7.58E-02 | 1.02E-01 | 2.45E+00 | 2.54E+00 | 4.71E-01 | no | Ush2a | intron_variant | MODIFIER |
| 1 | 188702509 | AX-97810739 | 0 | A | G | 0.246 | 7.58E-02 | 1.02E-01 | 2.45E+00 | 2.54E+00 | 4.71E-01 | no | Ush2a | intron_variant | MODIFIER |
| 1 | 188705209 | AX-170158146 | 0 | T | C | 0.246 | 7.58E-02 | 1.02E-01 | 2.45E+00 | 2.54E+00 | 4.71E-01 | no | Ush2a | intron_variant | MODIFIER |

| | | | | | | | | | | | | | | | |
|---|---|---|---|---|---|---|---|---|---|---|---|---|---|---|---|
| 1 | 188711672 | AX-170410479 | 0 | G | A | 0.246 | 7.58E-02 | 1.02E-01 | 2.45E+00 | 2.54E+00 | 4.71E-01 | no | Ush2a | intron_variant | MODIFIER |
| 1 | 188722008 | AX-170123801 | 0 | C | T | 0.246 | 7.58E-02 | 1.02E-01 | 2.45E+00 | 2.54E+00 | 4.71E-01 | no | Ush2a | intron_variant | MODIFIER |
| 1 | 188722765 | AX-170043927 | 0 | G | A | 0.246 | 7.58E-02 | 1.02E-01 | 2.45E+00 | 2.54E+00 | 4.71E-01 | no | Ush2a | intron_variant | MODIFIER |
| 1 | 188727827 | AX-170313708 | 0 | G | A | 0.246 | 7.58E-02 | 1.02E-01 | 2.45E+00 | 2.54E+00 | 4.71E-01 | no | Ush2a | intron_variant | MODIFIER |
| 1 | 188734688 | AX-170479612 | 0 | T | C | 0.246 | 7.58E-02 | 1.02E-01 | 2.45E+00 | 2.54E+00 | 4.71E-01 | no | Ush2a | missense_variant | MODERATE |
| 1 | 188734987 | AX-170351334 | 0 | A | G | 0.246 | 7.58E-02 | 1.02E-01 | 2.45E+00 | 2.54E+00 | 4.71E-01 | no | Ush2a | intron_variant | MODIFIER |
| 1 | 188864187 | AX-170516289 | 0 | G | A | 0.246 | 7.58E-02 | 1.02E-01 | 2.45E+00 | 2.54E+00 | 4.71E-01 | no | Ush2a | intron_variant | MODIFIER |
| 1 | 188866029 | AX-170376681 | 0 | G | T | 0.246 | 7.58E-02 | 1.02E-01 | 2.45E+00 | 2.54E+00 | 4.71E-01 | no | Ush2a | intron_variant | MODIFIER |
| 1 | 188866494 | AX-170150388 | 0 | G | A | 0.246 | 7.58E-02 | 1.02E-01 | 2.45E+00 | 2.54E+00 | 4.71E-01 | no | Ush2a | intron_variant | MODIFIER |
| 1 | 188867121 | AX-170424225 | 0 | T | G | 0.246 | 7.58E-02 | 1.02E-01 | 2.45E+00 | 2.54E+00 | 4.71E-01 | no | Ush2a | intron_variant | MODIFIER |
| 1 | 188868729 | AX-170186401 | 0 | C | G | 0.246 | 7.58E-02 | 1.02E-01 | 2.45E+00 | 2.54E+00 | 4.71E-01 | no | Ush2a | intron_variant | MODIFIER |
| 1 | 188877656 | AX-170309441 | 0 | T | C | 0.246 | 7.58E-02 | 1.02E-01 | 2.45E+00 | 2.54E+00 | 4.71E-01 | no | Ush2a | intron_variant | MODIFIER |
| 1 | 188877806 | AX-170424761 | 0 | T | C | 0.246 | 7.58E-02 | 1.02E-01 | 2.45E+00 | 2.54E+00 | 4.71E-01 | no | Ush2a | intron_variant | MODIFIER |
| 1 | 188879292 | AX-97822823 | 0 | G | A | 0.246 | 7.58E-02 | 1.02E-01 | 2.45E+00 | 2.54E+00 | 4.71E-01 | no | Ush2a | intron_variant | MODIFIER |
| 1 | 189042748 | AX-170002529 | 0 | C | T | 0.246 | 7.58E-02 | 1.02E-01 | 2.45E+00 | 2.54E+00 | 4.71E-01 | no | | intergenic_variant | MODIFIER |
| 1 | 189101206 | AX-170002525 | 0 | A | G | 0.246 | 7.58E-02 | 1.02E-01 | 2.45E+00 | 2.54E+00 | 4.71E-01 | no | | intergenic_variant | MODIFIER |
| 1 | 189120865 | AX-170439268 | 0 | C | A | 0.246 | 7.58E-02 | 1.02E-01 | 2.45E+00 | 2.54E+00 | 4.71E-01 | no | | intergenic_variant | MODIFIER |
| 1 | 189126550 | AX-170220213 | 0 | A | T | 0.246 | 7.58E-02 | 1.02E-01 | 2.45E+00 | 2.54E+00 | 4.71E-01 | no | | intergenic_variant | MODIFIER |
| 8 | 118612864 | AX-170187885 | 1 | A | C | 0.25 | 6.24E-02 | 8.83E-02 | 2.74E+00 | 2.86E+00 | 4.71E-01 | no | Cdh13 | intron_variant | MODIFIER |
| 5 | 77534031 | AX-170528028 | 0 | A | T | 0.439 | 7.74E-02 | 1.09E-01 | 2.73E+00 | 2.78E+00 | 4.71E-01 | no | | intergenic_variant | MODIFIER |
| 5 | 77544867 | AX-170371946 | 0 | G | A | 0.439 | 7.74E-02 | 1.09E-01 | 2.73E+00 | 2.78E+00 | 4.71E-01 | no | | intergenic_variant | MODIFIER |
| 5 | 77545091 | AX-170248872 | 0 | G | A | 0.439 | 7.74E-02 | 1.09E-01 | 2.73E+00 | 2.78E+00 | 4.71E-01 | no | | intergenic_variant | MODIFIER |
| 5 | 77545180 | AX-170072552 | 0 | A | G | 0.439 | 7.74E-02 | 1.09E-01 | 2.73E+00 | 2.78E+00 | 4.71E-01 | no | | intergenic_variant | MODIFIER |
| 5 | 77853360 | AX-97854267 | 0 | G | T | 0.439 | 7.74E-02 | 1.09E-01 | 2.73E+00 | 2.78E+00 | 4.71E-01 | no | | intergenic_variant | MODIFIER |
| 5 | 78266860 | AX-97840710 | 0 | C | T | 0.439 | 7.74E-02 | 1.09E-01 | 2.73E+00 | 2.78E+00 | 4.71E-01 | no | | intergenic_variant | MODIFIER |
| 5 | 78305064 | AX-170266087 | 0 | C | T | 0.439 | 7.74E-02 | 1.09E-01 | 2.73E+00 | 2.78E+00 | 4.71E-01 | no | | intergenic_variant | MODIFIER |
| 5 | 78319250 | AX-170474359 | 0 | C | G | 0.439 | 7.74E-02 | 1.09E-01 | 2.73E+00 | 2.78E+00 | 4.71E-01 | no | | intergenic_variant | MODIFIER |
| 5 | 78342026 | AX-97805025 | 0 | C | T | 0.439 | 7.74E-02 | 1.09E-01 | 2.73E+00 | 2.78E+00 | 4.71E-01 | no | | intergenic_variant | MODIFIER |
| 5 | 78375229 | AX-170095367 | 0 | A | G | 0.439 | 7.74E-02 | 1.09E-01 | 2.73E+00 | 2.78E+00 | 4.71E-01 | no | | intergenic_variant | MODIFIER |
| 5 | 78403584 | AX-97837601 | 0 | C | T | 0.439 | 7.74E-02 | 1.09E-01 | 2.73E+00 | 2.78E+00 | 4.71E-01 | no | | intergenic_variant | MODIFIER |
| 5 | 78453464 | AX-97815977 | 0 | A | T | 0.439 | 7.74E-02 | 1.09E-01 | 2.73E+00 | 2.78E+00 | 4.71E-01 | no | | intergenic_variant | MODIFIER |
| 5 | 78568560 | AX-170025354 | 0 | G | C | 0.439 | 7.74E-02 | 1.09E-01 | 2.73E+00 | 2.78E+00 | 4.71E-01 | no | | intergenic_variant | MODIFIER |
| 5 | 78568846 | AX-170025352 | 0 | T | C | 0.439 | 7.74E-02 | 1.09E-01 | 2.73E+00 | 2.78E+00 | 4.71E-01 | no | | intergenic_variant | MODIFIER |
| 5 | 78578406 | AX-170025357 | 0 | A | T | 0.439 | 7.74E-02 | 1.09E-01 | 2.73E+00 | 2.78E+00 | 4.71E-01 | no | | regulatory_region_variant | MODIFIER |
| 5 | 78985796 | AX-97819888 | 0 | G | C | 0.439 | 7.74E-02 | 1.09E-01 | 2.73E+00 | 2.78E+00 | 4.71E-01 | no | | intergenic_variant | MODIFIER |
| 5 | 79003292 | AX-170025371 | 0 | A | C | 0.439 | 7.74E-02 | 1.09E-01 | 2.73E+00 | 2.78E+00 | 4.71E-01 | no | | intergenic_variant | MODIFIER |
| 5 | 79017762 | AX-170025368 | 0 | A | G | 0.439 | 7.74E-02 | 1.09E-01 | 2.73E+00 | 2.78E+00 | 4.71E-01 | no | | intergenic_variant | MODIFIER |
| 5 | 79017837 | AX-170025372 | 0 | A | G | 0.439 | 7.74E-02 | 1.09E-01 | 2.73E+00 | 2.78E+00 | 4.71E-01 | no | | intergenic_variant | MODIFIER |
| 5 | 79086494 | AX-97796783 | 0 | C | A | 0.439 | 7.74E-02 | 1.09E-01 | 2.73E+00 | 2.78E+00 | 4.71E-01 | no | | intergenic_variant | MODIFIER |
| 5 | 79086562 | AX-170025362 | 0 | G | T | 0.439 | 7.74E-02 | 1.09E-01 | 2.73E+00 | 2.78E+00 | 4.71E-01 | no | | intergenic_variant | MODIFIER |
| 5 | 79128943 | AX-170072576 | 0 | C | T | 0.439 | 7.74E-02 | 1.09E-01 | 2.73E+00 | 2.78E+00 | 4.71E-01 | no | | intergenic_variant | MODIFIER |
| 5 | 79143285 | AX-170025367 | 0 | T | G | 0.439 | 7.74E-02 | 1.09E-01 | 2.73E+00 | 2.78E+00 | 4.71E-01 | no | | intergenic_variant | MODIFIER |
| 5 | 79156615 | AX-97822895 | 0 | T | G | 0.439 | 7.74E-02 | 1.09E-01 | 2.73E+00 | 2.78E+00 | 4.71E-01 | no | | intergenic_variant | MODIFIER |
| 5 | 79188345 | AX-97851798 | 0 | T | C | 0.439 | 7.74E-02 | 1.09E-01 | 2.73E+00 | 2.78E+00 | 4.71E-01 | no | | intergenic_variant | MODIFIER |
| 5 | 79194246 | AX-170145208 | 0 | C | T | 0.439 | 7.74E-02 | 1.09E-01 | 2.73E+00 | 2.78E+00 | 4.71E-01 | no | | intergenic_variant | MODIFIER |
| 5 | 79199207 | AX-170025364 | 0 | T | A | 0.439 | 7.74E-02 | 1.09E-01 | 2.73E+00 | 2.78E+00 | 4.71E-01 | no | | intergenic_variant | MODIFIER |
| 5 | 79244256 | AX-170328681 | 0 | A | C | 0.439 | 7.74E-02 | 1.09E-01 | 2.73E+00 | 2.78E+00 | 4.71E-01 | no | | intergenic_variant | MODIFIER |
| 5 | 79431331 | AX-170350884 | 0 | G | A | 0.439 | 7.74E-02 | 1.09E-01 | 2.73E+00 | 2.78E+00 | 4.71E-01 | no | | intergenic_variant | MODIFIER |
| 5 | 79431705 | AX-97800325 | 0 | T | C | 0.439 | 7.74E-02 | 1.09E-01 | 2.73E+00 | 2.78E+00 | 4.71E-01 | no | | intergenic_variant | MODIFIER |
| 5 | 79463549 | AX-97790862 | 0 | T | C | 0.439 | 7.74E-02 | 1.09E-01 | 2.73E+00 | 2.78E+00 | 4.71E-01 | no | | intergenic_variant | MODIFIER |
| 5 | 79471600 | AX-97799757 | 0 | C | T | 0.439 | 7.74E-02 | 1.09E-01 | 2.73E+00 | 2.78E+00 | 4.71E-01 | no | | intergenic_variant | MODIFIER |
| 5 | 79665787 | AX-170141789 | 0 | C | A | 0.439 | 7.74E-02 | 1.09E-01 | 2.73E+00 | 2.78E+00 | 4.71E-01 | no | | intergenic_variant | MODIFIER |
| 5 | 79682189 | AX-97795959 | 0 | G | A | 0.439 | 7.74E-02 | 1.09E-01 | 2.73E+00 | 2.78E+00 | 4.71E-01 | no | | intergenic_variant | MODIFIER |
| 5 | 80735890 | AX-97820819 | 0 | G | A | 0.439 | 7.74E-02 | 1.09E-01 | 2.73E+00 | 2.78E+00 | 4.71E-01 | no | | intergenic_variant | MODIFIER |

| | | | | | | | | | | | | | | | |
|---|---|---|---|---|---|---|---|---|---|---|---|---|---|---|---|
| 5 | 80742157 | AX-170355676 | 0 | A | C | 0.439 | 7.74E-02 | 1.09E-01 | 2.73E+00 | 2.78E+00 | 4.71E-01 | no | | intergenic_variant | MODIFIER |
| 5 | 80743073 | AX-170314072 | 0 | G | T | 0.439 | 7.74E-02 | 1.09E-01 | 2.73E+00 | 2.78E+00 | 4.71E-01 | no | | intergenic_variant | MODIFIER |
| 5 | 80746325 | AX-97805236 | 0 | T | G | 0.439 | 7.74E-02 | 1.09E-01 | 2.73E+00 | 2.78E+00 | 4.71E-01 | no | | intergenic_variant | MODIFIER |
| 5 | 80810168 | AX-170494173 | 0 | G | C | 0.439 | 7.74E-02 | 1.09E-01 | 2.73E+00 | 2.78E+00 | 4.71E-01 | no | | intergenic_variant | MODIFIER |
| 5 | 80820030 | AX-170209428 | 0 | G | A | 0.439 | 7.74E-02 | 1.09E-01 | 2.73E+00 | 2.78E+00 | 4.71E-01 | no | | intergenic_variant | MODIFIER |
| 5 | 80825289 | AX-170438097 | 0 | A | C | 0.439 | 7.74E-02 | 1.09E-01 | 2.73E+00 | 2.78E+00 | 4.71E-01 | no | | intergenic_variant | MODIFIER |
| 5 | 80828256 | AX-170331207 | 0 | T | G | 0.439 | 7.74E-02 | 1.09E-01 | 2.73E+00 | 2.78E+00 | 4.71E-01 | no | | intergenic_variant | MODIFIER |
| 5 | 80836593 | AX-97801858 | 0 | G | A | 0.439 | 7.74E-02 | 1.09E-01 | 2.73E+00 | 2.78E+00 | 4.71E-01 | no | | intergenic_variant | MODIFIER |
| 5 | 80869984 | AX-170118225 | 0 | A | G | 0.439 | 7.74E-02 | 1.09E-01 | 2.73E+00 | 2.78E+00 | 4.71E-01 | no | | intergenic_variant | MODIFIER |
| 5 | 80871722 | AX-170519105 | 0 | A | G | 0.439 | 7.74E-02 | 1.09E-01 | 2.73E+00 | 2.78E+00 | 4.71E-01 | no | | intergenic_variant | MODIFIER |
| 5 | 80876927 | AX-170482208 | 0 | C | T | 0.439 | 7.74E-02 | 1.09E-01 | 2.73E+00 | 2.78E+00 | 4.71E-01 | no | | intergenic_variant | MODIFIER |
| 5 | 80880833 | AX-170311117 | 0 | A | C | 0.439 | 7.74E-02 | 1.09E-01 | 2.73E+00 | 2.78E+00 | 4.71E-01 | no | | intergenic_variant | MODIFIER |
| 5 | 81023521 | AX-97799758 | 0 | T | G | 0.439 | 7.74E-02 | 1.09E-01 | 2.73E+00 | 2.78E+00 | 4.71E-01 | no | Adgrl3 | intron_variant | MODIFIER |
| 5 | 81042530 | AX-170072610 | 0 | A | T | 0.439 | 7.74E-02 | 1.09E-01 | 2.73E+00 | 2.78E+00 | 4.71E-01 | no | Adgrl3 | intron_variant | MODIFIER |
| 5 | 81061329 | AX-97800267 | 0 | C | T | 0.439 | 7.74E-02 | 1.09E-01 | 2.73E+00 | 2.78E+00 | 4.71E-01 | no | Adgrl3 | intron_variant | MODIFIER |
| 5 | 81071324 | AX-170025404 | 0 | T | C | 0.439 | 7.74E-02 | 1.09E-01 | 2.73E+00 | 2.78E+00 | 4.71E-01 | no | Adgrl3 | intron_variant | MODIFIER |
| 5 | 81071818 | AX-170025396 | 0 | A | C | 0.439 | 7.74E-02 | 1.09E-01 | 2.73E+00 | 2.78E+00 | 4.71E-01 | no | Adgrl3 | intron_variant | MODIFIER |
| 5 | 81090971 | AX-170284853 | 0 | C | T | 0.439 | 7.74E-02 | 1.09E-01 | 2.73E+00 | 2.78E+00 | 4.71E-01 | no | Adgrl3 | intron_variant | MODIFIER |
| 5 | 81096703 | AX-170025399 | 0 | G | A | 0.439 | 7.74E-02 | 1.09E-01 | 2.73E+00 | 2.78E+00 | 4.71E-01 | no | Adgrl3 | intron_variant | MODIFIER |
| 5 | 81099059 | AX-170025402 | 0 | C | T | 0.439 | 7.74E-02 | 1.09E-01 | 2.73E+00 | 2.78E+00 | 4.71E-01 | no | Adgrl3 | intron_variant | MODIFIER |
| 5 | 81146857 | AX-170025397 | 0 | G | C | 0.439 | 7.74E-02 | 1.09E-01 | 2.73E+00 | 2.78E+00 | 4.71E-01 | no | Adgrl3 | intron_variant | MODIFIER |
| 5 | 81148572 | AX-170025400 | 0 | G | T | 0.439 | 7.74E-02 | 1.09E-01 | 2.73E+00 | 2.78E+00 | 4.71E-01 | no | Adgrl3 | intron_variant | MODIFIER |
| 5 | 81151327 | AX-97806416 | 0 | T | G | 0.439 | 7.74E-02 | 1.09E-01 | 2.73E+00 | 2.78E+00 | 4.71E-01 | no | Adgrl3 | intron_variant | MODIFIER |
| 5 | 81174264 | AX-170025398 | 0 | C | T | 0.439 | 7.74E-02 | 1.09E-01 | 2.73E+00 | 2.78E+00 | 4.71E-01 | no | Adgrl3 | intron_variant | MODIFIER |
| 5 | 81180214 | AX-170025403 | 0 | C | T | 0.439 | 7.74E-02 | 1.09E-01 | 2.73E+00 | 2.78E+00 | 4.71E-01 | no | Adgrl3 | intron_variant | MODIFIER |
| 5 | 81256634 | AX-97799010 | 0 | C | T | 0.439 | 7.74E-02 | 1.09E-01 | 2.73E+00 | 2.78E+00 | 4.71E-01 | no | Adgrl3 | intron_variant | MODIFIER |
| 5 | 81290472 | AX-170523494 | 0 | G | A | 0.439 | 7.74E-02 | 1.09E-01 | 2.73E+00 | 2.78E+00 | 4.71E-01 | no | Adgrl3 | intron_variant | MODIFIER |
| 5 | 81415921 | AX-97781792 | 0 | T | C | 0.439 | 7.74E-02 | 1.09E-01 | 2.73E+00 | 2.78E+00 | 4.71E-01 | no | Adgrl3 | intron_variant | MODIFIER |
| 5 | 81481954 | AX-170467680 | 0 | G | A | 0.439 | 7.74E-02 | 1.09E-01 | 2.73E+00 | 2.78E+00 | 4.71E-01 | no | Adgrl3 | intron_variant | MODIFIER |
| 5 | 81489189 | AX-170483068 | 0 | G | A | 0.439 | 7.74E-02 | 1.09E-01 | 2.73E+00 | 2.78E+00 | 4.71E-01 | no | Adgrl3 | intron_variant | MODIFIER |
| 5 | 81500382 | AX-170131326 | 0 | T | C | 0.439 | 7.74E-02 | 1.09E-01 | 2.73E+00 | 2.78E+00 | 4.71E-01 | no | Adgrl3 | intron_variant | MODIFIER |
| 5 | 81509665 | AX-170095377 | 0 | T | C | 0.439 | 7.74E-02 | 1.09E-01 | 2.73E+00 | 2.78E+00 | 4.71E-01 | no | Adgrl3 | intron_variant | MODIFIER |
| 5 | 81549017 | AX-170112505 | 0 | C | T | 0.439 | 7.74E-02 | 1.09E-01 | 2.73E+00 | 2.78E+00 | 4.71E-01 | no | Adgrl3 | intron_variant | MODIFIER |
| 5 | 81549085 | AX-170519812 | 0 | G | A | 0.439 | 7.74E-02 | 1.09E-01 | 2.73E+00 | 2.78E+00 | 4.71E-01 | no | Adgrl3 | intron_variant | MODIFIER |
| 5 | 81549126 | AX-97856982 | 0 | G | T | 0.439 | 7.74E-02 | 1.09E-01 | 2.73E+00 | 2.78E+00 | 4.71E-01 | no | Adgrl3 | intron_variant | MODIFIER |
| 5 | 81581039 | AX-170217124 | 0 | G | A | 0.439 | 7.74E-02 | 1.09E-01 | 2.73E+00 | 2.78E+00 | 4.71E-01 | no | Adgrl3 | intron_variant | MODIFIER |
| 5 | 81581864 | AX-97856614 | 0 | C | A | 0.439 | 7.74E-02 | 1.09E-01 | 2.73E+00 | 2.78E+00 | 4.71E-01 | no | Adgrl3 | intron_variant | MODIFIER |
| 5 | 81597768 | AX-170025410 | 0 | G | A | 0.439 | 7.74E-02 | 1.09E-01 | 2.73E+00 | 2.78E+00 | 4.71E-01 | no | Adgrl3 | intron_variant | MODIFIER |
| 5 | 81598137 | AX-97803911 | 0 | G | A | 0.439 | 7.74E-02 | 1.09E-01 | 2.73E+00 | 2.78E+00 | 4.71E-01 | no | Adgrl3 | intron_variant | MODIFIER |
| 5 | 81616770 | AX-170025412 | 0 | G | A | 0.439 | 7.74E-02 | 1.09E-01 | 2.73E+00 | 2.78E+00 | 4.71E-01 | no | Adgrl3 | intron_variant | MODIFIER |
| 5 | 81639729 | AX-170025406 | 0 | T | A | 0.439 | 7.74E-02 | 1.09E-01 | 2.73E+00 | 2.78E+00 | 4.71E-01 | no | Adgrl3 | intron_variant | MODIFIER |
| 5 | 81640142 | AX-170025409 | 0 | C | T | 0.439 | 7.74E-02 | 1.09E-01 | 2.73E+00 | 2.78E+00 | 4.71E-01 | no | Adgrl3 | intron_variant | MODIFIER |
| 5 | 81655626 | AX-170072618 | 0 | A | G | 0.439 | 7.74E-02 | 1.09E-01 | 2.73E+00 | 2.78E+00 | 4.71E-01 | no | Adgrl3 | intron_variant | MODIFIER |
| 5 | 81659332 | AX-170072617 | 0 | G | T | 0.439 | 7.74E-02 | 1.09E-01 | 2.73E+00 | 2.78E+00 | 4.71E-01 | no | Adgrl3 | intron_variant | MODIFIER |
| 5 | 81659488 | AX-170399563 | 0 | T | C | 0.439 | 7.74E-02 | 1.09E-01 | 2.73E+00 | 2.78E+00 | 4.71E-01 | no | Adgrl3 | intron_variant | MODIFIER |
| 5 | 81665229 | AX-170025414 | 0 | A | G | 0.439 | 7.74E-02 | 1.09E-01 | 2.73E+00 | 2.78E+00 | 4.71E-01 | no | Adgrl3 | intron_variant | MODIFIER |
| 5 | 81671864 | AX-97781574 | 0 | T | G | 0.439 | 7.74E-02 | 1.09E-01 | 2.73E+00 | 2.78E+00 | 4.71E-01 | no | Adgrl3 | intron_variant | MODIFIER |
| 5 | 81676647 | AX-170245754 | 0 | G | A | 0.439 | 7.74E-02 | 1.09E-01 | 2.73E+00 | 2.78E+00 | 4.71E-01 | no | Adgrl3 | intron_variant | MODIFIER |
| 5 | 81694967 | AX-170025405 | 0 | G | A | 0.439 | 7.74E-02 | 1.09E-01 | 2.73E+00 | 2.78E+00 | 4.71E-01 | no | Adgrl3 | intron_variant | MODIFIER |
| 5 | 81716258 | AX-170270257 | 0 | G | A | 0.439 | 7.74E-02 | 1.09E-01 | 2.73E+00 | 2.78E+00 | 4.71E-01 | no | Adgrl3 | intron_variant | MODIFIER |
| 5 | 81738763 | AX-170025415 | 0 | T | C | 0.439 | 7.74E-02 | 1.09E-01 | 2.73E+00 | 2.78E+00 | 4.71E-01 | no | Adgrl3 | intron_variant | MODIFIER |
| 5 | 81776185 | AX-170170155 | 0 | T | G | 0.439 | 7.74E-02 | 1.09E-01 | 2.73E+00 | 2.78E+00 | 4.71E-01 | no | Adgrl3 | intron_variant | MODIFIER |
| 5 | 81781591 | AX-170089532 | 0 | A | C | 0.439 | 7.74E-02 | 1.09E-01 | 2.73E+00 | 2.78E+00 | 4.71E-01 | no | Adgrl3 | intron_variant | MODIFIER |
| 5 | 81794537 | AX-170025411 | 0 | A | G | 0.439 | 7.74E-02 | 1.09E-01 | 2.73E+00 | 2.78E+00 | 4.71E-01 | no | Adgrl3 | missense_variant | MODERATE |

| | | | | | | | | | | | | | | | |
|---|---|---|---|---|---|---|---|---|---|---|---|---|---|---|---|
| 5 | 81848593 | AX-170437021 | 0 | T | C | 0.439 | 7.74E-02 | 1.09E-01 | 2.73E+00 | 2.78E+00 | 4.71E-01 | no | | intergenic_variant | MODIFIER |
| 5 | 81868895 | AX-170317169 | 0 | T | C | 0.439 | 7.74E-02 | 1.09E-01 | 2.73E+00 | 2.78E+00 | 4.71E-01 | no | | intergenic_variant | MODIFIER |
| 5 | 81871822 | AX-170324609 | 0 | A | G | 0.439 | 7.74E-02 | 1.09E-01 | 2.73E+00 | 2.78E+00 | 4.71E-01 | no | | intergenic_variant | MODIFIER |
| 5 | 81892841 | AX-170072623 | 0 | G | A | 0.439 | 7.74E-02 | 1.09E-01 | 2.73E+00 | 2.78E+00 | 4.71E-01 | no | | intergenic_variant | MODIFIER |
| 5 | 81893846 | AX-170199044 | 0 | A | G | 0.439 | 7.74E-02 | 1.09E-01 | 2.73E+00 | 2.78E+00 | 4.71E-01 | no | | intergenic_variant | MODIFIER |
| 5 | 81893959 | AX-170380875 | 0 | G | A | 0.439 | 7.74E-02 | 1.09E-01 | 2.73E+00 | 2.78E+00 | 4.71E-01 | no | | intergenic_variant | MODIFIER |
| 5 | 81894063 | AX-97831889 | 0 | T | C | 0.439 | 7.74E-02 | 1.09E-01 | 2.73E+00 | 2.78E+00 | 4.71E-01 | no | | intergenic_variant | MODIFIER |
| 5 | 81904086 | AX-97798642 | 0 | A | G | 0.439 | 7.74E-02 | 1.09E-01 | 2.73E+00 | 2.78E+00 | 4.71E-01 | no | | intergenic_variant | MODIFIER |
| 5 | 81905673 | AX-170258045 | 0 | A | G | 0.439 | 7.74E-02 | 1.09E-01 | 2.73E+00 | 2.78E+00 | 4.71E-01 | no | | regulatory_region_variant | MODIFIER |
| 5 | 81933241 | AX-170355682 | 0 | C | T | 0.439 | 7.74E-02 | 1.09E-01 | 2.73E+00 | 2.78E+00 | 4.71E-01 | no | | intergenic_variant | MODIFIER |
| 5 | 82095925 | AX-170089536 | 0 | G | A | 0.439 | 7.74E-02 | 1.09E-01 | 2.73E+00 | 2.78E+00 | 4.71E-01 | no | | intergenic_variant | MODIFIER |
| 5 | 82102565 | AX-170025422 | 0 | C | A | 0.439 | 7.74E-02 | 1.09E-01 | 2.73E+00 | 2.78E+00 | 4.71E-01 | no | | intergenic_variant | MODIFIER |
| 5 | 82210734 | AX-97801037 | 0 | A | G | 0.439 | 7.74E-02 | 1.09E-01 | 2.73E+00 | 2.78E+00 | 4.71E-01 | no | | intergenic_variant | MODIFIER |
| 5 | 82231460 | AX-97800353 | 0 | G | A | 0.439 | 7.74E-02 | 1.09E-01 | 2.73E+00 | 2.78E+00 | 4.71E-01 | no | | intergenic_variant | MODIFIER |
| 5 | 82296225 | AX-170025420 | 0 | C | T | 0.439 | 7.74E-02 | 1.09E-01 | 2.73E+00 | 2.78E+00 | 4.71E-01 | no | | intergenic_variant | MODIFIER |
| 5 | 82932063 | AX-170170578 | 0 | T | G | 0.439 | 7.74E-02 | 1.09E-01 | 2.73E+00 | 2.78E+00 | 4.71E-01 | no | | intergenic_variant | MODIFIER |
| 5 | 83064448 | AX-170506750 | 0 | G | A | 0.439 | 7.74E-02 | 1.09E-01 | 2.73E+00 | 2.78E+00 | 4.71E-01 | no | | intergenic_variant | MODIFIER |
| 5 | 83069893 | AX-170333639 | 0 | G | A | 0.439 | 7.74E-02 | 1.09E-01 | 2.73E+00 | 2.78E+00 | 4.71E-01 | no | | intergenic_variant | MODIFIER |
| 5 | 83108615 | AX-170491155 | 0 | T | C | 0.439 | 7.74E-02 | 1.09E-01 | 2.73E+00 | 2.78E+00 | 4.71E-01 | no | | intergenic_variant | MODIFIER |
| 5 | 83167550 | AX-97832886 | 0 | C | T | 0.439 | 7.74E-02 | 1.09E-01 | 2.73E+00 | 2.78E+00 | 4.71E-01 | no | | intergenic_variant | MODIFIER |
| 5 | 83189617 | AX-97846197 | 0 | C | T | 0.439 | 7.74E-02 | 1.09E-01 | 2.73E+00 | 2.78E+00 | 4.71E-01 | no | | intergenic_variant | MODIFIER |
| 5 | 83189685 | AX-170072642 | 0 | A | T | 0.439 | 7.74E-02 | 1.09E-01 | 2.73E+00 | 2.78E+00 | 4.71E-01 | no | | intergenic_variant | MODIFIER |
| 5 | 83267553 | AX-170025443 | 0 | G | T | 0.439 | 7.74E-02 | 1.09E-01 | 2.73E+00 | 2.78E+00 | 4.71E-01 | no | | regulatory_region_variant | MODIFIER |
| 5 | 83284423 | AX-170496725 | 0 | A | G | 0.439 | 7.74E-02 | 1.09E-01 | 2.73E+00 | 2.78E+00 | 4.71E-01 | no | Tecrl | intron_variant | MODIFIER |
| 5 | 83300623 | AX-97803055 | 0 | T | C | 0.439 | 7.74E-02 | 1.09E-01 | 2.73E+00 | 2.78E+00 | 4.71E-01 | no | Tecrl | intron_variant | MODIFIER |
| 5 | 83325887 | AX-170135442 | 0 | C | T | 0.439 | 7.74E-02 | 1.09E-01 | 2.73E+00 | 2.78E+00 | 4.71E-01 | no | Tecrl | intron_variant | MODIFIER |
| 5 | 83336803 | AX-97796734 | 0 | A | G | 0.439 | 7.74E-02 | 1.09E-01 | 2.73E+00 | 2.78E+00 | 4.71E-01 | no | Tecrl | intron_variant | MODIFIER |
| 5 | 83341725 | AX-170282240 | 0 | A | G | 0.439 | 7.74E-02 | 1.09E-01 | 2.73E+00 | 2.78E+00 | 4.71E-01 | no | Tecrl | intron_variant | MODIFIER |
| 5 | 83344905 | AX-170025444 | 0 | G | T | 0.439 | 7.74E-02 | 1.09E-01 | 2.73E+00 | 2.78E+00 | 4.71E-01 | no | Tecrl | intron_variant | MODIFIER |
| 5 | 83346512 | AX-97834884 | 0 | T | C | 0.439 | 7.74E-02 | 1.09E-01 | 2.73E+00 | 2.78E+00 | 4.71E-01 | no | Tecrl | intron_variant | MODIFIER |
| 5 | 83492959 | AX-170137941 | 0 | C | T | 0.439 | 7.74E-02 | 1.09E-01 | 2.73E+00 | 2.78E+00 | 4.71E-01 | no | | intergenic_variant | MODIFIER |
| 5 | 83534177 | AX-170454008 | 0 | C | T | 0.439 | 7.74E-02 | 1.09E-01 | 2.73E+00 | 2.78E+00 | 4.71E-01 | no | | intergenic_variant | MODIFIER |
| 5 | 83534278 | AX-170393394 | 0 | A | C | 0.439 | 7.74E-02 | 1.09E-01 | 2.73E+00 | 2.78E+00 | 4.71E-01 | no | | intergenic_variant | MODIFIER |
| 5 | 83554431 | AX-97784506 | 0 | A | G | 0.439 | 7.74E-02 | 1.09E-01 | 2.73E+00 | 2.78E+00 | 4.71E-01 | no | | intergenic_variant | MODIFIER |
| 5 | 83618530 | AX-170072652 | 0 | T | C | 0.439 | 7.74E-02 | 1.09E-01 | 2.73E+00 | 2.78E+00 | 4.71E-01 | no | | intergenic_variant | MODIFIER |
| 5 | 83622384 | AX-97783075 | 0 | A | G | 0.439 | 7.74E-02 | 1.09E-01 | 2.73E+00 | 2.78E+00 | 4.71E-01 | no | | intergenic_variant | MODIFIER |
| 2 | 141047845 | AX-170435189 | 0 | C | T | 0.149 | 1.30E-01 | 1.82E-01 | 2.67E+00 | 2.75E+00 | 4.71E-01 | no | Macrod2 | intron_variant | MODIFIER |
| 2 | 148295488 | AX-97840564 | 0 | G | A | 0.149 | 1.30E-01 | 1.82E-01 | 2.67E+00 | 2.75E+00 | 4.71E-01 | no | | intergenic_variant | MODIFIER |
| 5 | 35977784 | AX-170332176 | 1 | T | A | 0.339 | 8.02E-02 | 1.13E-01 | 2.76E+00 | 2.82E+00 | 4.71E-01 | no | Afap1 | intron_variant | MODIFIER |
| 18 | 74160512 | AX-97835096 | 1 | G | A | 0.179 | -8.98E-02 | 1.25E-01 | 2.61E+00 | 2.66E+00 | 4.72E-01 | no | | regulatory_region_variant | MODIFIER |
| 1 | 189455418 | AX-97831715 | 2 | A | G | 0.264 | 7.31E-02 | 9.84E-02 | 2.46E+00 | 2.55E+00 | 4.72E-01 | no | | intergenic_variant | MODIFIER |
| 16 | 51052708 | AX-170441047 | 1 | T | C | 0.509 | 5.97E-02 | 8.13E-02 | 2.51E+00 | 2.56E+00 | 4.72E-01 | no | Gm32431 | intron_variant,non_coding_transcript_variant | MODIFIER |
| 15 | 9466812 | AX-97822499 | 2 | C | T | 0.245 | -7.57E-02 | 1.05E-01 | 2.58E+00 | 2.64E+00 | 4.72E-01 | no | Capsl | downstream_gene_variant | MODIFIER |
| 8 | 52473731 | AX-170090108 | 1 | C | A | 0.152 | -1.04E-01 | 1.44E-01 | 2.58E+00 | 2.65E+00 | 4.72E-01 | no | | intergenic_variant | MODIFIER |
| 3 | 149193649 | AX-170164963 | 1 | G | A | 0.348 | 5.16E-02 | 7.28E-02 | 2.76E+00 | 2.85E+00 | 4.72E-01 | no | | intergenic_variant | MODIFIER |
| 5 | 82285770 | AX-97831906 | 1 | C | A | 0.429 | 7.50E-02 | 1.05E-01 | 2.66E+00 | 2.72E+00 | 4.72E-01 | no | | intergenic_variant | MODIFIER |
| 4 | 95913538 | AX-97813608 | 0 | G | A | 0.254 | 7.95E-02 | 1.08E-01 | 2.50E+00 | 2.57E+00 | 4.72E-01 | no | Fggy | intron_variant | MODIFIER |
| 7 | 73170738 | AX-170263608 | 0 | C | T | 0.36 | -6.67E-02 | 9.42E-02 | 2.72E+00 | 2.82E+00 | 4.72E-01 | no | | intergenic_variant | MODIFIER |
| 18 | 77568731 | AX-170016032 | 1 | G | C | 0.438 | 6.40E-02 | 8.67E-02 | 2.48E+00 | 2.55E+00 | 4.72E-01 | no | Rnf165 | upstream_gene_variant | MODIFIER |
| 7 | 92642977 | AX-170516923 | 1 | G | A | 0.125 | 1.23E-01 | 1.72E-01 | 2.61E+00 | 2.73E+00 | 4.72E-01 | no | Pcf11 | downstream_gene_variant | MODIFIER |
| 17 | 22340001 | AX-170140613 | 1 | G | A | 0.036 | -1.22E-01 | 1.70E-01 | 2.57E+00 | 2.70E+00 | 4.72E-01 | no | Zfp944 | synonymous_variant | LOW |
| 1 | 147386436 | AX-170448527 | 1 | T | A | 0.205 | 9.78E-02 | 1.37E-01 | 2.66E+00 | 2.71E+00 | 4.72E-01 | no | | intergenic_variant | MODIFIER |
| 1 | 149941794 | AX-170140738 | 1 | T | C | 0.205 | 9.78E-02 | 1.37E-01 | 2.66E+00 | 2.71E+00 | 4.72E-01 | no | Pla2g4a | intron_variant | MODIFIER |
| 8 | 26499432 | AX-170090067 | 0 | A | C | 0.351 | -4.77E-02 | 6.59E-02 | 2.54E+00 | 2.63E+00 | 4.73E-01 | no | | intergenic_variant | MODIFIER |

| | | | | | | | | | | | | | | | |
|---|---|---|---|---|---|---|---|---|---|---|---|---|---|---|---|
| 8 | 28359838 | AX-170309950 | 0 | T | C | 0.167 | -7.36E-02 | 1.04E-01 | 2.72E+00 | 2.83E+00 | 4.73E-01 | no | | intergenic_variant | MODIFIER |
| 8 | 30941039 | AX-170420648 | 0 | C | T | 0.167 | -7.36E-02 | 1.04E-01 | 2.72E+00 | 2.83E+00 | 4.73E-01 | no | | intergenic_variant | MODIFIER |
| 1 | 105987800 | AX-170432618 | 1 | T | C | 0.161 | 1.47E-01 | 2.06E-01 | 2.64E+00 | 2.70E+00 | 4.73E-01 | Yes | Zcchc2 | upstream_gene_variant | MODIFIER |
| 16 | 50403592 | AX-170369613 | 1 | T | C | 0.464 | -5.67E-02 | 7.65E-02 | 2.46E+00 | 2.53E+00 | 4.73E-01 | no | Bbx | intron_variant | MODIFIER |
| 18 | 55974695 | AX-97801537 | 0 | G | A | 0.404 | 7.55E-02 | 9.78E-02 | 2.37E+00 | 2.42E+00 | 4.73E-01 | no | Gm6944 | downstream_gene_variant | MODIFIER |
| 18 | 55977431 | AX-170269832 | 0 | C | G | 0.404 | 7.55E-02 | 9.78E-02 | 2.37E+00 | 2.42E+00 | 4.73E-01 | no | Gm6944 | downstream_gene_variant | MODIFIER |
| 18 | 55982964 | AX-170385441 | 0 | G | A | 0.404 | 7.55E-02 | 9.78E-02 | 2.37E+00 | 2.42E+00 | 4.73E-01 | no | Gm22484 | downstream_gene_variant | MODIFIER |
| 18 | 55990308 | AX-170370803 | 0 | G | T | 0.404 | 7.55E-02 | 9.78E-02 | 2.37E+00 | 2.42E+00 | 4.73E-01 | no | Gm22484 | upstream_gene_variant | MODIFIER |
| 18 | 55996971 | AX-170251654 | 0 | T | C | 0.404 | 7.55E-02 | 9.78E-02 | 2.37E+00 | 2.42E+00 | 4.73E-01 | no | | regulatory_region_variant | MODIFIER |
| 18 | 56002689 | AX-170422781 | 0 | A | G | 0.404 | 7.55E-02 | 9.78E-02 | 2.37E+00 | 2.42E+00 | 4.73E-01 | no | | intergenic_variant | MODIFIER |
| 18 | 56012349 | AX-170296041 | 0 | A | G | 0.404 | 7.55E-02 | 9.78E-02 | 2.37E+00 | 2.42E+00 | 4.73E-01 | no | | intergenic_variant | MODIFIER |
| 18 | 56022235 | AX-170378168 | 0 | T | C | 0.404 | 7.55E-02 | 9.78E-02 | 2.37E+00 | 2.42E+00 | 4.73E-01 | no | | regulatory_region_variant | MODIFIER |
| 18 | 56032607 | AX-170350793 | 0 | A | G | 0.404 | 7.55E-02 | 9.78E-02 | 2.37E+00 | 2.42E+00 | 4.73E-01 | no | | intergenic_variant | MODIFIER |
| 18 | 56033003 | AX-170244193 | 0 | C | T | 0.404 | 7.55E-02 | 9.78E-02 | 2.37E+00 | 2.42E+00 | 4.73E-01 | no | | intergenic_variant | MODIFIER |
| 18 | 56033226 | AX-170127483 | 0 | C | T | 0.404 | 7.55E-02 | 9.78E-02 | 2.37E+00 | 2.42E+00 | 4.73E-01 | no | | intergenic_variant | MODIFIER |
| 18 | 56085428 | AX-97837754 | 0 | C | T | 0.404 | 7.55E-02 | 9.78E-02 | 2.37E+00 | 2.42E+00 | 4.73E-01 | no | | intergenic_variant | MODIFIER |
| 18 | 56087157 | AX-170453258 | 0 | G | A | 0.404 | 7.55E-02 | 9.78E-02 | 2.37E+00 | 2.42E+00 | 4.73E-01 | no | | intergenic_variant | MODIFIER |
| 18 | 56117990 | AX-170015763 | 0 | C | T | 0.404 | 7.55E-02 | 9.78E-02 | 2.37E+00 | 2.42E+00 | 4.73E-01 | no | | intergenic_variant | MODIFIER |
| 18 | 56138758 | AX-170534119 | 0 | G | A | 0.404 | 7.55E-02 | 9.78E-02 | 2.37E+00 | 2.42E+00 | 4.73E-01 | no | | regulatory_region_variant | MODIFIER |
| 18 | 56603036 | AX-97834900 | 0 | G | A | 0.404 | 7.55E-02 | 9.78E-02 | 2.37E+00 | 2.42E+00 | 4.73E-01 | no | AC162035.1 | upstream_gene_variant | MODIFIER |
| 18 | 56603471 | AX-170015769 | 0 | A | C | 0.404 | 7.55E-02 | 9.78E-02 | 2.37E+00 | 2.42E+00 | 4.73E-01 | no | AC162035.1 | upstream_gene_variant | MODIFIER |
| 18 | 56763284 | AX-170338356 | 0 | T | C | 0.404 | 7.55E-02 | 9.78E-02 | 2.37E+00 | 2.42E+00 | 4.73E-01 | no | MAR3 | intron_variant | MODIFIER |
| 18 | 56945639 | AX-97854809 | 0 | T | C | 0.404 | 7.55E-02 | 9.78E-02 | 2.37E+00 | 2.42E+00 | 4.73E-01 | no | | intergenic_variant | MODIFIER |
| 18 | 57001525 | AX-97847377 | 0 | A | G | 0.404 | 7.55E-02 | 9.78E-02 | 2.37E+00 | 2.42E+00 | 4.73E-01 | no | | intergenic_variant | MODIFIER |
| 18 | 57007152 | AX-170295216 | 0 | T | C | 0.404 | 7.55E-02 | 9.78E-02 | 2.37E+00 | 2.42E+00 | 4.73E-01 | no | | intergenic_variant | MODIFIER |
| 18 | 57065565 | AX-97782588 | 0 | T | G | 0.404 | 7.55E-02 | 9.78E-02 | 2.37E+00 | 2.42E+00 | 4.73E-01 | no | | intergenic_variant | MODIFIER |
| 18 | 57083049 | AX-170260564 | 0 | G | A | 0.404 | 7.55E-02 | 9.78E-02 | 2.37E+00 | 2.42E+00 | 4.73E-01 | no | | intergenic_variant | MODIFIER |
| 4 | 55970495 | AX-170491010 | 0 | A | T | 0.36 | -7.16E-02 | 1.01E-01 | 2.72E+00 | 2.81E+00 | 4.73E-01 | no | Gm12519 | intron_variant,non_coding_transcript_variant | MODIFIER |
| 1 | 19906477 | AX-170131920 | 0 | C | A | 0.421 | 5.84E-02 | 8.01E-02 | 2.52E+00 | 2.58E+00 | 4.73E-01 | no | | intergenic_variant | MODIFIER |
| 6 | 127384105 | AX-170095800 | 0 | G | A | 0.535 | 5.90E-02 | 8.10E-02 | 2.52E+00 | 2.60E+00 | 4.73E-01 | no | | intergenic_variant | MODIFIER |
| 4 | 98578756 | AX-170267627 | 1 | T | A | 0.223 | 8.40E-02 | 1.16E-01 | 2.55E+00 | 2.61E+00 | 4.73E-01 | no | Patj | intron_variant | MODIFIER |
| 1 | 26095788 | AX-170198662 | 0 | C | G | 0.395 | 6.04E-02 | 8.48E-02 | 2.63E+00 | 2.71E+00 | 4.73E-01 | no | | intergenic_variant | MODIFIER |
| 18 | 59064744 | AX-170194838 | 0 | C | T | 0.404 | 7.19E-02 | 9.64E-02 | 2.45E+00 | 2.50E+00 | 4.73E-01 | no | Minar2 | intron_variant | MODIFIER |
| 7 | 114055186 | AX-170383081 | 1 | A | G | 0.429 | 6.77E-02 | 9.55E-02 | 2.70E+00 | 2.75E+00 | 4.73E-01 | no | Rras2 | intron_variant | MODIFIER |
| 18 | 38934047 | AX-170102754 | 0 | G | T | 0.07 | 1.53E-01 | 2.17E-01 | 2.76E+00 | 2.86E+00 | 4.73E-01 | Yes | AC110730.1 | intron_variant,non_coding_transcript_variant | MODIFIER |
| 10 | 16061835 | AX-97814880 | 0 | A | C | 0.202 | 7.31E-02 | 1.02E-01 | 2.89E+00 | 2.99E+00 | 4.73E-01 | no | | intergenic_variant | MODIFIER |
| 15 | 25495241 | AX-170158279 | 0 | T | C | 0.246 | -7.24E-02 | 1.01E-01 | 2.57E+00 | 2.64E+00 | 4.73E-01 | no | | intergenic_variant | MODIFIER |
| 5 | 37013228 | AX-170263936 | 1 | T | C | 0.339 | 7.87E-02 | 1.11E-01 | 2.75E+00 | 2.82E+00 | 4.73E-01 | no | | intergenic_variant | MODIFIER |
| 6 | 137215634 | AX-170523112 | 1 | A | G | 0.554 | 6.40E-02 | 8.66E-02 | 2.47E+00 | 2.55E+00 | 4.73E-01 | no | | intergenic_variant | MODIFIER |
| 1 | 89707503 | AX-170374251 | 0 | A | T | 0.14 | 1.44E-01 | 2.02E-01 | 2.61E+00 | 2.68E+00 | 4.74E-01 | Yes | Agap1 | intron_variant | MODIFIER |
| 1 | 93673177 | AX-170273442 | 0 | A | T | 0.14 | 1.44E-01 | 2.02E-01 | 2.61E+00 | 2.68E+00 | 4.74E-01 | Yes | | intergenic_variant | MODIFIER |
| 1 | 94623271 | AX-168405748 | 0 | A | G | 0.14 | 1.44E-01 | 2.02E-01 | 2.61E+00 | 2.68E+00 | 4.74E-01 | Yes | | intergenic_variant | MODIFIER |
| 1 | 99803298 | AX-97807497 | 0 | C | T | 0.14 | 1.44E-01 | 2.02E-01 | 2.61E+00 | 2.68E+00 | 4.74E-01 | Yes | Cntnap5b | intron_variant | MODIFIER |
| 1 | 101153473 | AX-170380793 | 0 | A | C | 0.14 | 1.44E-01 | 2.02E-01 | 2.61E+00 | 2.68E+00 | 4.74E-01 | Yes | Gm20268 | upstream_gene_variant | MODIFIER |
| 1 | 101560572 | AX-170127060 | 0 | C | T | 0.14 | 1.44E-01 | 2.02E-01 | 2.61E+00 | 2.68E+00 | 4.74E-01 | Yes | | intergenic_variant | MODIFIER |
| 1 | 104551934 | AX-97812542 | 0 | A | C | 0.14 | 1.44E-01 | 2.02E-01 | 2.61E+00 | 2.68E+00 | 4.74E-01 | Yes | | intergenic_variant | MODIFIER |
| 1 | 141977398 | AX-170406167 | 1 | C | A | 0.241 | 7.72E-02 | 1.08E-01 | 2.57E+00 | 2.67E+00 | 4.74E-01 | no | | intergenic_variant | MODIFIER |
| 15 | 5243740 | AX-170010944 | 1 | C | T | 0.241 | -7.64E-02 | 1.07E-01 | 2.60E+00 | 2.65E+00 | 4.74E-01 | no | Ptger4 | 5_prime_UTR_variant | MODIFIER |
| 15 | 19422177 | AX-170054954 | 1 | C | G | 0.241 | -7.64E-02 | 1.07E-01 | 2.60E+00 | 2.65E+00 | 4.74E-01 | no | | intergenic_variant | MODIFIER |
| 8 | 65924138 | AX-170140178 | 0 | C | T | 0.149 | -9.75E-02 | 1.36E-01 | 2.57E+00 | 2.65E+00 | 4.74E-01 | no | MAR1 | intron_variant | MODIFIER |
| 1 | 135838679 | AX-170494358 | 0 | G | A | 0.579 | 4.29E-02 | 6.01E-02 | 2.89E+00 | 3.01E+00 | 4.74E-01 | no | Tnnt2 | upstream_gene_variant | MODIFIER |
| 18 | 80908617 | AX-170161756 | 0 | T | A | 0.298 | -6.00E-02 | 8.34E-02 | 2.55E+00 | 2.66E+00 | 4.74E-01 | no | Atp9b | intron_variant | MODIFIER |
| 6 | 129035693 | AX-170134615 | 0 | C | T | 0.553 | 6.01E-02 | 8.30E-02 | 2.53E+00 | 2.60E+00 | 4.74E-01 | no | Gm16186 | upstream_gene_variant | MODIFIER |
| 6 | 129626171 | AX-170255551 | 0 | C | G | 0.553 | 6.01E-02 | 8.30E-02 | 2.53E+00 | 2.60E+00 | 4.74E-01 | no | Klrk1 | upstream_gene_variant | MODIFIER |

| | | | | | | | | | | | | | | | |
|---|---|---|---|---|---|---|---|---|---|---|---|---|---|---|---|
| 6 | 130911320 | AX-170100748 | 0 | G | T | 0.553 | 6.01E-02 | 8.30E-02 | 2.53E+00 | 2.60E+00 | 4.74E-01 | no | Gm44128 | non_coding_transcript_exon_variant | MODIFIER |
| 5 | 42269300 | AX-170346076 | 1 | C | T | 0.205 | -7.75E-02 | 1.09E-01 | 2.62E+00 | 2.70E+00 | 4.74E-01 | no | | intergenic_variant | MODIFIER |
| 5 | 42317437 | AX-170102415 | 1 | C | T | 0.205 | -7.75E-02 | 1.09E-01 | 2.62E+00 | 2.70E+00 | 4.74E-01 | no | | intergenic_variant | MODIFIER |
| 5 | 42884536 | AX-170318847 | 1 | C | T | 0.205 | -7.75E-02 | 1.09E-01 | 2.62E+00 | 2.70E+00 | 4.74E-01 | no | | regulatory_region_variant | MODIFIER |
| 7 | 78110002 | AX-170099186 | 1 | T | A | 0.384 | -7.17E-02 | 1.02E-01 | 2.78E+00 | 2.88E+00 | 4.74E-01 | no | | intergenic_variant | MODIFIER |
| 15 | 4019146 | AX-170244168 | 2 | A | C | 0.082 | -8.63E-02 | 9.87E-02 | 2.13E+00 | 2.25E+00 | 4.74E-01 | no | BC037032 | downstream_gene_variant | MODIFIER |
| 3 | 149328569 | AX-170136834 | 0 | G | A | 0.351 | 5.22E-02 | 7.39E-02 | 2.77E+00 | 2.85E+00 | 4.74E-01 | no | Gm30382 | upstream_gene_variant | MODIFIER |
| 18 | 75946512 | AX-170149426 | 0 | G | A | 0.395 | -6.07E-02 | 8.54E-02 | 2.62E+00 | 2.70E+00 | 4.74E-01 | no | Zbtb7c | intron_variant | MODIFIER |
| 18 | 76154473 | AX-170367502 | 0 | T | C | 0.395 | -6.07E-02 | 8.54E-02 | 2.62E+00 | 2.70E+00 | 4.74E-01 | no | | intergenic_variant | MODIFIER |
| 5 | 44118933 | AX-170484320 | 1 | A | G | 0.188 | -7.71E-02 | 1.08E-01 | 2.60E+00 | 2.69E+00 | 4.75E-01 | no | | intergenic_variant | MODIFIER |
| 4 | 151475555 | AX-170380904 | 1 | G | T | 0.116 | 1.29E-01 | 1.83E-01 | 2.70E+00 | 2.79E+00 | 4.75E-01 | no | Camta1 | intron_variant | MODIFIER |
| 10 | 12560165 | AX-170326342 | 1 | A | G | 0.205 | -7.01E-02 | 9.93E-02 | 2.66E+00 | 2.76E+00 | 4.75E-01 | no | Utrn | intron_variant | MODIFIER |
| 2 | 71036294 | AX-170173791 | 0 | C | T | 0.061 | -1.01E-01 | 1.41E-01 | 2.54E+00 | 2.66E+00 | 4.75E-01 | no | Mettl8 | intron_variant | MODIFIER |
| 18 | 58355219 | AX-170464838 | 0 | G | A | 0.412 | 7.22E-02 | 9.62E-02 | 2.42E+00 | 2.48E+00 | 4.75E-01 | no | | intergenic_variant | MODIFIER |
| 11 | 8071845 | AX-97783052 | 0 | T | C | 0.053 | 1.40E-01 | 2.01E-01 | 2.71E+00 | 2.85E+00 | 4.75E-01 | no | | intergenic_variant | MODIFIER |
| 8 | 118655439 | AX-170388434 | 1 | G | T | 0.25 | 6.71E-02 | 9.58E-02 | 2.75E+00 | 2.88E+00 | 4.75E-01 | no | Cdh13 | intron_variant | MODIFIER |
| 1 | 149311529 | AX-170086644 | 0 | C | A | 0.202 | 9.80E-02 | 1.38E-01 | 2.65E+00 | 2.71E+00 | 4.75E-01 | no | | intergenic_variant | MODIFIER |
| 7 | 36636826 | AX-170077070 | 1 | G | A | 0.339 | 6.54E-02 | 9.14E-02 | 2.58E+00 | 2.65E+00 | 4.75E-01 | no | | intergenic_variant | MODIFIER |
| 5 | 38034988 | AX-170402825 | 1 | T | C | 0.33 | 7.80E-02 | 1.11E-01 | 2.75E+00 | 2.81E+00 | 4.75E-01 | no | Stx18 | upstream_gene_variant | MODIFIER |
| 15 | 6581161 | AX-170122599 | 0 | A | T | 0.237 | -7.55E-02 | 1.06E-01 | 2.59E+00 | 2.65E+00 | 4.75E-01 | no | Fyb | intron_variant | MODIFIER |
| 5 | 13970794 | AX-170261072 | 0 | C | A | 0.342 | 6.08E-02 | 8.65E-02 | 2.68E+00 | 2.79E+00 | 4.75E-01 | no | | intergenic_variant | MODIFIER |
| 14 | 124700477 | AX-97787324 | 0 | G | A | 0.325 | 5.51E-02 | 7.75E-02 | 2.57E+00 | 2.69E+00 | 4.76E-01 | no | | intergenic_variant | MODIFIER |
| 4 | 139631903 | AX-170260158 | 0 | C | T | 0.184 | 7.49E-02 | 1.05E-01 | 2.59E+00 | 2.65E+00 | 4.76E-01 | no | Aldh4a1 | intron_variant | MODIFIER |
| 4 | 97947101 | AX-170145404 | 0 | G | C | 0.228 | 8.44E-02 | 1.18E-01 | 2.58E+00 | 2.64E+00 | 4.76E-01 | no | Nfia | intron_variant | MODIFIER |
| 7 | 52415988 | AX-170380754 | 0 | C | T | 0.439 | 6.85E-02 | 9.71E-02 | 2.65E+00 | 2.73E+00 | 4.76E-01 | no | Gm20074 | downstream_gene_variant | MODIFIER |
| 7 | 57155574 | AX-170510734 | 0 | C | T | 0.439 | 6.85E-02 | 9.71E-02 | 2.65E+00 | 2.73E+00 | 4.76E-01 | no | Gabrg3 | intron_variant | MODIFIER |
| 18 | 65558714 | AX-170517813 | 1 | T | C | 0.42 | 6.40E-02 | 8.61E-02 | 2.44E+00 | 2.50E+00 | 4.76E-01 | no | | regulatory_region_variant | MODIFIER |
| 12 | 104296953 | AX-170098551 | 0 | C | A | 0.132 | -1.16E-01 | 1.65E-01 | 2.69E+00 | 2.79E+00 | 4.76E-01 | no | | intergenic_variant | MODIFIER |
| 1 | 135839082 | AX-170373927 | 1 | T | G | 0.214 | 8.64E-02 | 1.22E-01 | 2.65E+00 | 2.73E+00 | 4.76E-01 | no | Tnnt2 | upstream_gene_variant | MODIFIER |
| 9 | 17554432 | AX-170224251 | 0 | T | C | 0.474 | 6.13E-02 | 8.59E-02 | 2.58E+00 | 2.65E+00 | 4.76E-01 | no | | intergenic_variant | MODIFIER |
| 2 | 108018544 | AX-170018614 | 0 | A | G | 0.07 | -1.13E-01 | 1.61E-01 | 2.66E+00 | 2.79E+00 | 4.76E-01 | no | | intergenic_variant | MODIFIER |
| 5 | 128910129 | AX-170386033 | 1 | A | G | 0.17 | -6.49E-02 | 8.39E-02 | 2.34E+00 | 2.45E+00 | 4.76E-01 | no | Rimbp2 | intron_variant | MODIFIER |
| 8 | 41917277 | AX-170520970 | 2 | G | A | 0.155 | 1.03E-01 | 1.45E-01 | 2.91E+00 | 2.97E+00 | 4.76E-01 | no | 2810404M03Rik | intron_variant,non_coding_transcript_variant | MODIFIER |
| 8 | 65487952 | AX-170422617 | 0 | A | G | 0.14 | -1.03E-01 | 1.45E-01 | 2.60E+00 | 2.68E+00 | 4.76E-01 | no | | intergenic_variant | MODIFIER |
| 8 | 65613192 | AX-170536022 | 0 | T | C | 0.14 | -1.03E-01 | 1.45E-01 | 2.60E+00 | 2.68E+00 | 4.76E-01 | no | MAR1 | upstream_gene_variant | MODIFIER |
| 18 | 74715106 | AX-170242460 | 0 | G | A | 0.43 | 6.21E-02 | 8.62E-02 | 2.54E+00 | 2.60E+00 | 4.76E-01 | no | Myo5b | intron_variant | MODIFIER |
| 1 | 19979973 | AX-170149008 | 2 | T | A | 0.418 | 5.83E-02 | 8.09E-02 | 2.54E+00 | 2.60E+00 | 4.76E-01 | no | | intergenic_variant | MODIFIER |
| 18 | 58011227 | AX-170060958 | 1 | G | A | 0.411 | 7.16E-02 | 9.66E-02 | 2.45E+00 | 2.50E+00 | 4.76E-01 | no | Fbn2 | intron_variant | MODIFIER |
| 3 | 29774335 | AX-170204828 | 0 | A | G | 0.167 | -6.41E-02 | 9.14E-02 | 2.70E+00 | 2.80E+00 | 4.76E-01 | no | Gm37557 | downstream_gene_variant | MODIFIER |
| 3 | 31841896 | AX-170359188 | 0 | T | C | 0.167 | -6.41E-02 | 9.14E-02 | 2.70E+00 | 2.80E+00 | 4.76E-01 | no | | intergenic_variant | MODIFIER |
| 6 | 94419422 | AX-170519282 | 2 | A | C | 0.391 | -6.37E-02 | 9.14E-02 | 2.79E+00 | 2.96E+00 | 4.76E-01 | no | Gm7825 | downstream_gene_variant | MODIFIER |
| 5 | 43953351 | AX-170105091 | 1 | C | A | 0.205 | -7.56E-02 | 1.06E-01 | 2.58E+00 | 2.66E+00 | 4.76E-01 | no | Gm43184 | upstream_gene_variant | MODIFIER |
| 5 | 82208684 | AX-170072626 | 1 | C | G | 0.446 | 7.55E-02 | 1.07E-01 | 2.70E+00 | 2.76E+00 | 4.77E-01 | no | | intergenic_variant | MODIFIER |
| 15 | 24432862 | AX-97852776 | 1 | T | C | 0.241 | -7.42E-02 | 1.04E-01 | 2.58E+00 | 2.64E+00 | 4.77E-01 | no | | intergenic_variant | MODIFIER |
| 9 | 16390559 | AX-170031850 | 0 | G | T | 0.482 | 6.45E-02 | 8.98E-02 | 2.55E+00 | 2.61E+00 | 4.77E-01 | no | Fat3 | intron_variant | MODIFIER |
| 18 | 65135571 | AX-170179716 | 0 | C | T | 0.412 | 6.37E-02 | 8.64E-02 | 2.46E+00 | 2.51E+00 | 4.77E-01 | no | Nedd4l | intron_variant | MODIFIER |
| 18 | 65140117 | AX-170421297 | 0 | T | C | 0.412 | 6.37E-02 | 8.64E-02 | 2.46E+00 | 2.51E+00 | 4.77E-01 | no | Nedd4l | intron_variant | MODIFIER |
| 18 | 65140446 | AX-97852783 | 0 | G | A | 0.412 | 6.37E-02 | 8.64E-02 | 2.46E+00 | 2.51E+00 | 4.77E-01 | no | Nedd4l | intron_variant | MODIFIER |
| 18 | 65140983 | AX-170438931 | 0 | G | A | 0.412 | 6.37E-02 | 8.64E-02 | 2.46E+00 | 2.51E+00 | 4.77E-01 | no | Nedd4l | intron_variant | MODIFIER |
| 18 | 65141178 | AX-170310430 | 0 | T | C | 0.412 | 6.37E-02 | 8.64E-02 | 2.46E+00 | 2.51E+00 | 4.77E-01 | no | Nedd4l | intron_variant | MODIFIER |
| 18 | 65305748 | AX-170293510 | 0 | G | A | 0.412 | 6.37E-02 | 8.64E-02 | 2.46E+00 | 2.51E+00 | 4.77E-01 | no | Alpk2 | missense_variant | MODERATE |
| 18 | 65307880 | AX-170405847 | 0 | A | G | 0.412 | 6.37E-02 | 8.64E-02 | 2.46E+00 | 2.51E+00 | 4.77E-01 | no | Alpk2 | intron_variant | MODIFIER |
| 18 | 65309314 | AX-170298521 | 0 | T | C | 0.412 | 6.37E-02 | 8.64E-02 | 2.46E+00 | 2.51E+00 | 4.77E-01 | no | Alpk2 | intron_variant | MODIFIER |
| 18 | 65309643 | AX-170061130 | 0 | G | C | 0.412 | 6.37E-02 | 8.64E-02 | 2.46E+00 | 2.51E+00 | 4.77E-01 | no | Alpk2 | intron_variant | MODIFIER |

| | | | | | | | | | | | | | | | |
|---|---|---|---|---|---|---|---|---|---|---|---|---|---|---|---|
| 18 | 65393282 | AX-170015872 | 0 | G | C | 0.412 | 6.37E-02 | 8.64E-02 | 2.46E+00 | 2.51E+00 | 4.77E-01 | no | Alpk2 | intron_variant | MODIFIER |
| 18 | 65393490 | AX-170515968 | 0 | G | A | 0.412 | 6.37E-02 | 8.64E-02 | 2.46E+00 | 2.51E+00 | 4.77E-01 | no | Alpk2 | intron_variant | MODIFIER |
| 18 | 65428854 | AX-170323114 | 0 | A | G | 0.412 | 6.37E-02 | 8.64E-02 | 2.46E+00 | 2.51E+00 | 4.77E-01 | no | Malt1 | upstream_gene_variant | MODIFIER |
| 18 | 65442288 | AX-170323543 | 0 | C | T | 0.412 | 6.37E-02 | 8.64E-02 | 2.46E+00 | 2.51E+00 | 4.77E-01 | no | Malt1 | intron_variant | MODIFIER |
| 18 | 65449703 | AX-170367830 | 0 | A | G | 0.412 | 6.37E-02 | 8.64E-02 | 2.46E+00 | 2.51E+00 | 4.77E-01 | no | Malt1 | intron_variant | MODIFIER |
| 18 | 65477820 | AX-170372489 | 0 | G | C | 0.412 | 6.37E-02 | 8.64E-02 | 2.46E+00 | 2.51E+00 | 4.77E-01 | no | Malt1 | 3_prime_UTR_variant | MODIFIER |
| 18 | 65545388 | AX-170281496 | 0 | A | G | 0.412 | 6.37E-02 | 8.64E-02 | 2.46E+00 | 2.51E+00 | 4.77E-01 | no | AC132307.4 | upstream_gene_variant | MODIFIER |
| 18 | 65545451 | AX-170496481 | 0 | G | A | 0.412 | 6.37E-02 | 8.64E-02 | 2.46E+00 | 2.51E+00 | 4.77E-01 | no | AC132307.4 | upstream_gene_variant | MODIFIER |
| 18 | 65556457 | AX-97801937 | 0 | T | C | 0.412 | 6.37E-02 | 8.64E-02 | 2.46E+00 | 2.51E+00 | 4.77E-01 | no | | intergenic_variant | MODIFIER |
| 18 | 65571472 | AX-170357427 | 0 | A | G | 0.412 | 6.37E-02 | 8.64E-02 | 2.46E+00 | 2.51E+00 | 4.77E-01 | no | | intergenic_variant | MODIFIER |
| 18 | 65600328 | AX-170402325 | 0 | C | G | 0.412 | 6.37E-02 | 8.64E-02 | 2.46E+00 | 2.51E+00 | 4.77E-01 | no | Zfp532 | intron_variant | MODIFIER |
| 18 | 65602348 | AX-170352833 | 0 | T | C | 0.412 | 6.37E-02 | 8.64E-02 | 2.46E+00 | 2.51E+00 | 4.77E-01 | no | Zfp532 | intron_variant | MODIFIER |
| 18 | 65603380 | AX-170475889 | 0 | C | A | 0.412 | 6.37E-02 | 8.64E-02 | 2.46E+00 | 2.51E+00 | 4.77E-01 | no | Zfp532 | intron_variant | MODIFIER |
| 18 | 65613054 | AX-97855588 | 0 | C | T | 0.412 | 6.37E-02 | 8.64E-02 | 2.46E+00 | 2.51E+00 | 4.77E-01 | no | Zfp532 | intron_variant | MODIFIER |
| 18 | 65613188 | AX-170521865 | 0 | G | A | 0.412 | 6.37E-02 | 8.64E-02 | 2.46E+00 | 2.51E+00 | 4.77E-01 | no | Zfp532 | intron_variant | MODIFIER |
| 18 | 65627838 | AX-97857570 | 0 | G | A | 0.412 | 6.37E-02 | 8.64E-02 | 2.46E+00 | 2.51E+00 | 4.77E-01 | no | Zfp532 | intron_variant | MODIFIER |
| 18 | 65658961 | AX-170483939 | 0 | A | G | 0.412 | 6.37E-02 | 8.64E-02 | 2.46E+00 | 2.51E+00 | 4.77E-01 | no | Zfp532 | intron_variant | MODIFIER |
| 18 | 65659162 | AX-170526677 | 0 | T | C | 0.412 | 6.37E-02 | 8.64E-02 | 2.46E+00 | 2.51E+00 | 4.77E-01 | no | Zfp532 | intron_variant | MODIFIER |
| 18 | 65669513 | AX-170242350 | 0 | T | A | 0.412 | 6.37E-02 | 8.64E-02 | 2.46E+00 | 2.51E+00 | 4.77E-01 | no | Zfp532 | intron_variant | MODIFIER |
| 18 | 65670431 | AX-170235812 | 0 | G | A | 0.412 | 6.37E-02 | 8.64E-02 | 2.46E+00 | 2.51E+00 | 4.77E-01 | no | Zfp532 | intron_variant | MODIFIER |
| 18 | 65671270 | AX-170394271 | 0 | A | G | 0.412 | 6.37E-02 | 8.64E-02 | 2.46E+00 | 2.51E+00 | 4.77E-01 | no | Zfp532 | intron_variant | MODIFIER |
| 18 | 65674672 | AX-170513956 | 0 | C | A | 0.412 | 6.37E-02 | 8.64E-02 | 2.46E+00 | 2.51E+00 | 4.77E-01 | no | Zfp532 | intron_variant | MODIFIER |
| 18 | 65678257 | AX-170061136 | 0 | A | G | 0.412 | 6.37E-02 | 8.64E-02 | 2.46E+00 | 2.51E+00 | 4.77E-01 | no | Zfp532 | intron_variant | MODIFIER |
| 18 | 65681254 | AX-97795585 | 0 | G | A | 0.412 | 6.37E-02 | 8.64E-02 | 2.46E+00 | 2.51E+00 | 4.77E-01 | no | Zfp532 | intron_variant | MODIFIER |
| 18 | 65681743 | AX-170299663 | 0 | T | C | 0.412 | 6.37E-02 | 8.64E-02 | 2.46E+00 | 2.51E+00 | 4.77E-01 | no | Zfp532 | intron_variant | MODIFIER |
| 18 | 66093919 | AX-97829057 | 0 | C | T | 0.412 | 6.37E-02 | 8.64E-02 | 2.46E+00 | 2.51E+00 | 4.77E-01 | no | Ccbe1 | intron_variant,NMD_transcript_variant | MODIFIER |
| 18 | 66105938 | AX-170061141 | 0 | T | A | 0.412 | 6.37E-02 | 8.64E-02 | 2.46E+00 | 2.51E+00 | 4.77E-01 | no | Ccbe1 | intron_variant,NMD_transcript_variant | MODIFIER |
| 18 | 66106532 | AX-170504115 | 0 | A | G | 0.412 | 6.37E-02 | 8.64E-02 | 2.46E+00 | 2.51E+00 | 4.77E-01 | no | Ccbe1 | intron_variant,NMD_transcript_variant | MODIFIER |
| 18 | 66109953 | AX-170208549 | 0 | C | T | 0.412 | 6.37E-02 | 8.64E-02 | 2.46E+00 | 2.51E+00 | 4.77E-01 | no | Ccbe1 | intron_variant,NMD_transcript_variant | MODIFIER |
| 18 | 66110910 | AX-170183120 | 0 | G | A | 0.412 | 6.37E-02 | 8.64E-02 | 2.46E+00 | 2.51E+00 | 4.77E-01 | no | Ccbe1 | intron_variant,NMD_transcript_variant | MODIFIER |
| 20 | 61590480 | AX-170198654 | 1 | G | A | 0.027 | -9.32E-02 | 1.32E-01 | 2.56E+00 | 2.74E+00 | 4.77E-01 | no | | intergenic_variant | MODIFIER |
| 8 | 85990631 | AX-170096344 | 1 | C | T | 0.196 | -9.30E-02 | 1.32E-01 | 2.77E+00 | 2.84E+00 | 4.77E-01 | no | Phkb | intron_variant | MODIFIER |
| 7 | 74014903 | AX-170394850 | 1 | T | C | 0.366 | -6.75E-02 | 9.64E-02 | 2.75E+00 | 2.85E+00 | 4.77E-01 | no | St8sia2 | upstream_gene_variant | MODIFIER |
| 7 | 75217074 | AX-170225383 | 1 | A | T | 0.366 | -6.75E-02 | 9.64E-02 | 2.75E+00 | 2.85E+00 | 4.77E-01 | no | Sv2b | intron_variant | MODIFIER |
| 7 | 76383828 | AX-170165152 | 1 | C | T | 0.375 | -6.94E-02 | 9.91E-02 | 2.74E+00 | 2.84E+00 | 4.77E-01 | no | Agbl1 | intron_variant | MODIFIER |
| 9 | 19495927 | AX-97792209 | 1 | C | T | 0.464 | 6.51E-02 | 9.07E-02 | 2.55E+00 | 2.61E+00 | 4.77E-01 | no | Olfr851 | upstream_gene_variant | MODIFIER |
| 9 | 22551428 | AX-97806260 | 1 | A | G | 0.464 | 6.51E-02 | 9.07E-02 | 2.55E+00 | 2.61E+00 | 4.77E-01 | no | Bbs9 | intron_variant | MODIFIER |
| 6 | 148074858 | AX-97791669 | 0 | T | A | 0.053 | 1.25E-01 | 1.79E-01 | 2.76E+00 | 2.88E+00 | 4.77E-01 | no | Far2 | intron_variant | MODIFIER |
| 18 | 70754141 | AX-170015940 | 2 | C | A | 0.182 | -8.77E-02 | 1.24E-01 | 2.67E+00 | 2.72E+00 | 4.77E-01 | no | | intergenic_variant | MODIFIER |
| 10 | 127439170 | AX-170465684 | 0 | G | A | 0.342 | -5.02E-02 | 7.12E-02 | 2.61E+00 | 2.72E+00 | 4.77E-01 | no | R3hdm2 | intron_variant | MODIFIER |
| 2 | 162799564 | AX-170290446 | 0 | C | T | 0.167 | 1.17E-01 | 1.66E-01 | 2.62E+00 | 2.74E+00 | 4.77E-01 | no | | intergenic_variant | MODIFIER |
| 10 | 115637595 | AX-170418119 | 1 | C | T | 0.393 | 5.47E-02 | 7.81E-02 | 2.78E+00 | 2.87E+00 | 4.77E-01 | no | | intergenic_variant | MODIFIER |
| 2 | 167225641 | AX-97845730 | 1 | A | T | 0.143 | 1.28E-01 | 1.82E-01 | 2.67E+00 | 2.76E+00 | 4.77E-01 | no | Ptgis | intron_variant | MODIFIER |
| 7 | 123219643 | AX-170453528 | 2 | T | G | 0.4 | 5.87E-02 | 8.39E-02 | 2.76E+00 | 2.86E+00 | 4.77E-01 | no | Slc5a11 | intron_variant | MODIFIER |
| 9 | 102134269 | AX-170083282 | 1 | T | C | 0.134 | -8.06E-02 | 1.14E-01 | 2.61E+00 | 2.70E+00 | 4.78E-01 | no | Ephb1 | intron_variant | MODIFIER |
| 7 | 82010915 | AX-170491676 | 0 | G | A | 0.105 | -1.11E-01 | 1.57E-01 | 2.61E+00 | 2.73E+00 | 4.78E-01 | no | | intergenic_variant | MODIFIER |
| 9 | 98715880 | AX-170170686 | 1 | T | A | 0.134 | -8.09E-02 | 1.14E-01 | 2.60E+00 | 2.68E+00 | 4.78E-01 | no | | intergenic_variant | MODIFIER |
| 9 | 102134334 | AX-170083287 | 1 | G | C | 0.134 | -8.09E-02 | 1.14E-01 | 2.60E+00 | 2.68E+00 | 4.78E-01 | no | Ephb1 | intron_variant | MODIFIER |
| 5 | 89824568 | AX-170101859 | 0 | A | C | 0.43 | 7.67E-02 | 1.09E-01 | 2.70E+00 | 2.76E+00 | 4.78E-01 | no | Adamts3 | intron_variant | MODIFIER |
| 1 | 126699260 | AX-170261531 | 0 | G | A | 0.544 | 5.78E-02 | 8.18E-02 | 2.61E+00 | 2.68E+00 | 4.78E-01 | no | Nckap5 | intron_variant | MODIFIER |
| 1 | 127366182 | AX-170396324 | 0 | G | T | 0.544 | 5.78E-02 | 8.18E-02 | 2.61E+00 | 2.68E+00 | 4.78E-01 | no | Mgat5 | intron_variant | MODIFIER |
| 1 | 128461902 | AX-170140692 | 0 | C | T | 0.544 | 5.78E-02 | 8.18E-02 | 2.61E+00 | 2.68E+00 | 4.78E-01 | no | | intergenic_variant | MODIFIER |
| 5 | 69890254 | AX-170244657 | 1 | G | A | 0.455 | 6.45E-02 | 9.20E-02 | 2.66E+00 | 2.76E+00 | 4.78E-01 | no | | intergenic_variant | MODIFIER |
| 15 | 82141147 | AX-170157665 | 0 | C | T | 0.561 | 5.65E-02 | 7.90E-02 | 2.55E+00 | 2.62E+00 | 4.78E-01 | no | Ccdc134 | 3_prime_UTR_variant | MODIFIER |

| | | | | | | | | | | | | | | | |
|---|---|---|---|---|---|---|---|---|---|---|---|---|---|---|---|
| 4 | 97796134 | AX-170393867 | 0 | T | C | 0.237 | 7.93E-02 | 1.11E-01 | 2.56E+00 | 2.62E+00 | 4.78E-01 | no | Nfia | intron_variant | MODIFIER |
| 1 | 187720614 | AX-97846607 | 1 | T | C | 0.241 | 7.98E-02 | 1.06E-01 | 2.40E+00 | 2.48E+00 | 4.78E-01 | no | Esrrg | intron_variant | MODIFIER |
| 9 | 92339401 | AX-170032996 | 1 | A | G | 0.134 | -8.91E-02 | 1.26E-01 | 2.60E+00 | 2.67E+00 | 4.78E-01 | no | 1700057G04Rik | intron_variant | MODIFIER |
| 15 | 4374899 | AX-170470972 | 0 | C | T | 0.237 | -7.50E-02 | 1.06E-01 | 2.60E+00 | 2.65E+00 | 4.78E-01 | no | Plcxd3 | upstream_gene_variant | MODIFIER |
| 2 | 169476857 | AX-170333829 | 0 | T | C | 0.158 | 1.30E-01 | 1.84E-01 | 2.60E+00 | 2.67E+00 | 4.79E-01 | no | Gm14249 | upstream_gene_variant | MODIFIER |
| 8 | 64615570 | AX-170267539 | 1 | A | T | 0.152 | -9.54E-02 | 1.35E-01 | 2.58E+00 | 2.66E+00 | 4.79E-01 | no | Cpe | intron_variant | MODIFIER |
| 9 | 20507943 | AX-170031926 | 1 | A | C | 0.455 | 5.92E-02 | 8.25E-02 | 2.52E+00 | 2.61E+00 | 4.79E-01 | no | Zfp266 | intron_variant | MODIFIER |
| 9 | 23212413 | AX-170118078 | 1 | T | A | 0.455 | 5.92E-02 | 8.25E-02 | 2.52E+00 | 2.61E+00 | 4.79E-01 | no | | regulatory_region_variant | MODIFIER |
| 8 | 22310845 | AX-170223785 | 1 | T | A | 0.188 | 6.88E-02 | 9.88E-02 | 2.69E+00 | 2.80E+00 | 4.79E-01 | no | Tpte | intron_variant | MODIFIER |
| 7 | 115145980 | AX-170472932 | 0 | G | A | 0.421 | 6.72E-02 | 9.61E-02 | 2.74E+00 | 2.79E+00 | 4.79E-01 | no | | intergenic_variant | MODIFIER |
| 9 | 76861202 | AX-97821556 | 0 | C | T | 0.193 | -8.91E-02 | 1.27E-01 | 2.76E+00 | 2.83E+00 | 4.79E-01 | no | | intergenic_variant | MODIFIER |
| 10 | 69906221 | AX-170457320 | 1 | A | C | 0.42 | -5.64E-02 | 8.00E-02 | 2.62E+00 | 2.70E+00 | 4.79E-01 | no | Ank3 | intron_variant | MODIFIER |
| 1 | 182221906 | AX-170002433 | 0 | A | T | 0.228 | 7.88E-02 | 1.08E-01 | 2.47E+00 | 2.55E+00 | 4.79E-01 | no | | intergenic_variant | MODIFIER |
| 8 | 23071957 | AX-170444267 | 2 | C | A | 0.164 | -7.62E-02 | 1.09E-01 | 2.65E+00 | 2.74E+00 | 4.79E-01 | no | Ank1 | intron_variant | MODIFIER |
| 16 | 27911662 | AX-97854245 | 1 | G | A | 0.25 | 6.21E-02 | 8.88E-02 | 2.62E+00 | 2.74E+00 | 4.79E-01 | no | Gm10823 | non_coding_transcript_exon_variant | MODIFIER |
| 9 | 94936364 | AX-170266192 | 0 | G | T | 0.132 | -8.32E-02 | 1.19E-01 | 2.66E+00 | 2.75E+00 | 4.79E-01 | no | Slc9a9 | synonymous_variant | LOW |
| 8 | 21947940 | AX-170102927 | 0 | T | C | 0.167 | 6.95E-02 | 9.99E-02 | 2.70E+00 | 2.82E+00 | 4.79E-01 | no | Defb13 | intron_variant | MODIFIER |
| 3 | 149834439 | AX-170470924 | 0 | G | A | 0.333 | 5.20E-02 | 7.45E-02 | 2.74E+00 | 2.83E+00 | 4.79E-01 | no | | intergenic_variant | MODIFIER |
| 7 | 73326538 | AX-170163686 | 0 | C | A | 0.36 | -6.69E-02 | 9.60E-02 | 2.72E+00 | 2.81E+00 | 4.79E-01 | no | A730056A06Rik | intron_variant,non_coding_transcript_variant | MODIFIER |
| 18 | 59381407 | AX-170508979 | 1 | T | A | 0.411 | 6.50E-02 | 8.78E-02 | 2.43E+00 | 2.50E+00 | 4.79E-01 | no | Chsy3 | intron_variant | MODIFIER |
| 1 | 70029188 | AX-170091548 | 1 | C | T | 0.143 | 1.14E-01 | 1.65E-01 | 2.72E+00 | 2.86E+00 | 4.80E-01 | no | Spag16 | intron_variant | MODIFIER |
| 3 | 37132751 | AX-97848185 | 1 | G | A | 0.188 | 6.59E-02 | 9.45E-02 | 2.79E+00 | 2.89E+00 | 4.80E-01 | no | | intergenic_variant | MODIFIER |
| 13 | 25200438 | AX-97793067 | 1 | G | A | 0.17 | -8.92E-02 | 1.21E-01 | 2.45E+00 | 2.51E+00 | 4.80E-01 | no | Dcdc2a | intron_variant | MODIFIER |
| 19 | 50629822 | AX-170314975 | 1 | T | C | 0.473 | 4.51E-02 | 6.43E-02 | 2.61E+00 | 2.71E+00 | 4.80E-01 | no | Sorcs1 | intron_variant | MODIFIER |
| 18 | 65626514 | AX-170473758 | 1 | A | G | 0.411 | 6.29E-02 | 8.61E-02 | 2.47E+00 | 2.52E+00 | 4.80E-01 | no | Zfp532 | intron_variant | MODIFIER |
| 15 | 68072141 | AX-170093291 | 1 | A | G | 0.339 | -5.69E-02 | 7.57E-02 | 2.39E+00 | 2.47E+00 | 4.80E-01 | no | | regulatory_region_variant | MODIFIER |
| 4 | 114630203 | AX-97831469 | 0 | T | C | 0.228 | 7.72E-02 | 1.10E-01 | 2.61E+00 | 2.68E+00 | 4.80E-01 | no | | intergenic_variant | MODIFIER |
| 4 | 114631638 | AX-170267493 | 0 | A | C | 0.228 | 7.72E-02 | 1.10E-01 | 2.61E+00 | 2.68E+00 | 4.80E-01 | no | | intergenic_variant | MODIFIER |
| 4 | 114650713 | AX-170481719 | 0 | G | A | 0.228 | 7.72E-02 | 1.10E-01 | 2.61E+00 | 2.68E+00 | 4.80E-01 | no | | intergenic_variant | MODIFIER |
| 4 | 114661159 | AX-170229582 | 0 | T | G | 0.228 | 7.72E-02 | 1.10E-01 | 2.61E+00 | 2.68E+00 | 4.80E-01 | no | | intergenic_variant | MODIFIER |
| 4 | 114666373 | AX-97849425 | 0 | G | A | 0.228 | 7.72E-02 | 1.10E-01 | 2.61E+00 | 2.68E+00 | 4.80E-01 | no | | regulatory_region_variant | MODIFIER |
| 4 | 114666397 | AX-170294293 | 0 | C | G | 0.228 | 7.72E-02 | 1.10E-01 | 2.61E+00 | 2.68E+00 | 4.80E-01 | no | | regulatory_region_variant | MODIFIER |
| 4 | 114666908 | AX-170267393 | 0 | A | G | 0.228 | 7.72E-02 | 1.10E-01 | 2.61E+00 | 2.68E+00 | 4.80E-01 | no | | regulatory_region_variant | MODIFIER |
| 4 | 114768655 | AX-170519468 | 0 | T | G | 0.228 | 7.72E-02 | 1.10E-01 | 2.61E+00 | 2.68E+00 | 4.80E-01 | no | | intergenic_variant | MODIFIER |
| 4 | 114773171 | AX-97851541 | 0 | A | G | 0.228 | 7.72E-02 | 1.10E-01 | 2.61E+00 | 2.68E+00 | 4.80E-01 | no | | intergenic_variant | MODIFIER |
| 4 | 114773986 | AX-170211898 | 0 | C | G | 0.228 | 7.72E-02 | 1.10E-01 | 2.61E+00 | 2.68E+00 | 4.80E-01 | no | | intergenic_variant | MODIFIER |
| 4 | 117930173 | AX-170180149 | 0 | G | A | 0.228 | 7.72E-02 | 1.10E-01 | 2.61E+00 | 2.68E+00 | 4.80E-01 | no | St3gal3 | downstream_gene_variant | MODIFIER |
| 2 | 159844041 | AX-170356932 | 1 | G | A | 0.152 | 9.59E-02 | 1.38E-01 | 2.67E+00 | 2.80E+00 | 4.80E-01 | no | | intergenic_variant | MODIFIER |
| 5 | 28134431 | AX-170441262 | 1 | G | A | 0.366 | 7.23E-02 | 1.04E-01 | 2.73E+00 | 2.79E+00 | 4.80E-01 | no | | intergenic_variant | MODIFIER |
| 18 | 70856968 | AX-97799485 | 0 | C | T | 0.184 | -8.24E-02 | 1.18E-01 | 2.64E+00 | 2.69E+00 | 4.80E-01 | no | | intergenic_variant | MODIFIER |
| 18 | 70885153 | AX-170180920 | 0 | A | G | 0.184 | -8.24E-02 | 1.18E-01 | 2.64E+00 | 2.69E+00 | 4.80E-01 | no | | intergenic_variant | MODIFIER |
| 18 | 70890307 | AX-170318648 | 0 | T | C | 0.184 | -8.24E-02 | 1.18E-01 | 2.64E+00 | 2.69E+00 | 4.80E-01 | no | | intergenic_variant | MODIFIER |
| 18 | 70905853 | AX-170310359 | 0 | T | G | 0.184 | -8.24E-02 | 1.18E-01 | 2.64E+00 | 2.69E+00 | 4.80E-01 | no | AC103658.1 | downstream_gene_variant | MODIFIER |
| 18 | 70919434 | AX-170250955 | 0 | G | A | 0.184 | -8.24E-02 | 1.18E-01 | 2.64E+00 | 2.69E+00 | 4.80E-01 | no | | intergenic_variant | MODIFIER |
| 18 | 71102108 | AX-97852556 | 0 | G | A | 0.184 | -8.24E-02 | 1.18E-01 | 2.64E+00 | 2.69E+00 | 4.80E-01 | no | | intergenic_variant | MODIFIER |
| 18 | 71262972 | AX-170365425 | 0 | A | G | 0.184 | -8.24E-02 | 1.18E-01 | 2.64E+00 | 2.69E+00 | 4.80E-01 | no | Dcc | intron_variant | MODIFIER |
| 18 | 71817252 | AX-97844995 | 0 | T | C | 0.184 | -8.24E-02 | 1.18E-01 | 2.64E+00 | 2.69E+00 | 4.80E-01 | no | Dcc | intron_variant | MODIFIER |
| 18 | 71924019 | AX-97786914 | 0 | G | A | 0.184 | -8.24E-02 | 1.18E-01 | 2.64E+00 | 2.69E+00 | 4.80E-01 | no | Dcc | intron_variant | MODIFIER |
| 18 | 71949089 | AX-170313654 | 0 | A | T | 0.184 | -8.24E-02 | 1.18E-01 | 2.64E+00 | 2.69E+00 | 4.80E-01 | no | Dcc | intron_variant | MODIFIER |
| 18 | 71975839 | AX-170015960 | 0 | G | T | 0.184 | -8.24E-02 | 1.18E-01 | 2.64E+00 | 2.69E+00 | 4.80E-01 | no | Dcc | intron_variant | MODIFIER |
| 18 | 71975888 | AX-170061279 | 0 | G | A | 0.184 | -8.24E-02 | 1.18E-01 | 2.64E+00 | 2.69E+00 | 4.80E-01 | no | Dcc | intron_variant | MODIFIER |
| 18 | 71975911 | AX-97828503 | 0 | G | A | 0.184 | -8.24E-02 | 1.18E-01 | 2.64E+00 | 2.69E+00 | 4.80E-01 | no | Dcc | intron_variant | MODIFIER |
| 18 | 72009090 | AX-97835790 | 0 | G | A | 0.184 | -8.24E-02 | 1.18E-01 | 2.64E+00 | 2.69E+00 | 4.80E-01 | no | Dcc | intron_variant | MODIFIER |
| 18 | 72046242 | AX-170088373 | 0 | C | T | 0.184 | -8.24E-02 | 1.18E-01 | 2.64E+00 | 2.69E+00 | 4.80E-01 | no | Dcc | intron_variant | MODIFIER |

| | | | | | | | | | | | | | | | |
|---|---|---|---|---|---|---|---|---|---|---|---|---|---|---|---|
| 18 | 72093900 | AX-170295728 | 0 | C | A | 0.184 | -8.24E-02 | 1.18E-01 | 2.64E+00 | 2.69E+00 | 4.80E-01 | no | Dcc | intron_variant | MODIFIER |
| 18 | 72096932 | AX-170393746 | 0 | A | G | 0.184 | -8.24E-02 | 1.18E-01 | 2.64E+00 | 2.69E+00 | 4.80E-01 | no | Dcc | intron_variant | MODIFIER |
| 18 | 72097054 | AX-170521101 | 0 | T | G | 0.184 | -8.24E-02 | 1.18E-01 | 2.64E+00 | 2.69E+00 | 4.80E-01 | no | Dcc | intron_variant | MODIFIER |
| 18 | 72097471 | AX-170420696 | 0 | A | G | 0.184 | -8.24E-02 | 1.18E-01 | 2.64E+00 | 2.69E+00 | 4.80E-01 | no | Dcc | intron_variant | MODIFIER |
| 18 | 72097743 | AX-170405550 | 0 | C | T | 0.184 | -8.24E-02 | 1.18E-01 | 2.64E+00 | 2.69E+00 | 4.80E-01 | no | Dcc | intron_variant | MODIFIER |
| 18 | 72100506 | AX-170402454 | 0 | C | T | 0.184 | -8.24E-02 | 1.18E-01 | 2.64E+00 | 2.69E+00 | 4.80E-01 | no | Dcc | intron_variant | MODIFIER |
| 18 | 72100665 | AX-170300596 | 0 | A | G | 0.184 | -8.24E-02 | 1.18E-01 | 2.64E+00 | 2.69E+00 | 4.80E-01 | no | Dcc | intron_variant | MODIFIER |
| 18 | 72135611 | AX-97835786 | 0 | A | C | 0.184 | -8.24E-02 | 1.18E-01 | 2.64E+00 | 2.69E+00 | 4.80E-01 | no | Dcc | intron_variant | MODIFIER |
| 18 | 72147517 | AX-170447686 | 0 | T | A | 0.184 | -8.24E-02 | 1.18E-01 | 2.64E+00 | 2.69E+00 | 4.80E-01 | no | Dcc | intron_variant | MODIFIER |
| 18 | 72152640 | AX-97840120 | 0 | T | C | 0.184 | -8.24E-02 | 1.18E-01 | 2.64E+00 | 2.69E+00 | 4.80E-01 | no | Dcc | intron_variant | MODIFIER |
| 18 | 72152668 | AX-170035588 | 0 | A | G | 0.184 | -8.24E-02 | 1.18E-01 | 2.64E+00 | 2.69E+00 | 4.80E-01 | no | Dcc | intron_variant | MODIFIER |
| 18 | 72154642 | AX-170147084 | 0 | C | G | 0.184 | -8.24E-02 | 1.18E-01 | 2.64E+00 | 2.69E+00 | 4.80E-01 | no | Dcc | intron_variant | MODIFIER |
| 18 | 72173859 | AX-170269610 | 0 | G | A | 0.184 | -8.24E-02 | 1.18E-01 | 2.64E+00 | 2.69E+00 | 4.80E-01 | no | Dcc | intron_variant | MODIFIER |
| 18 | 72176557 | AX-97806064 | 0 | T | G | 0.184 | -8.24E-02 | 1.18E-01 | 2.64E+00 | 2.69E+00 | 4.80E-01 | no | Dcc | intron_variant | MODIFIER |
| 18 | 72182654 | AX-170061283 | 0 | T | C | 0.184 | -8.24E-02 | 1.18E-01 | 2.64E+00 | 2.69E+00 | 4.80E-01 | no | Dcc | intron_variant | MODIFIER |
| 18 | 72189070 | AX-170121397 | 0 | A | G | 0.184 | -8.24E-02 | 1.18E-01 | 2.64E+00 | 2.69E+00 | 4.80E-01 | no | Dcc | intron_variant | MODIFIER |
| 18 | 72191482 | AX-97790534 | 0 | C | T | 0.184 | -8.24E-02 | 1.18E-01 | 2.64E+00 | 2.69E+00 | 4.80E-01 | no | Dcc | intron_variant | MODIFIER |
| 18 | 72191639 | AX-170149098 | 0 | T | A | 0.184 | -8.24E-02 | 1.18E-01 | 2.64E+00 | 2.69E+00 | 4.80E-01 | no | Dcc | intron_variant | MODIFIER |
| 18 | 72194894 | AX-170467089 | 0 | C | T | 0.184 | -8.24E-02 | 1.18E-01 | 2.64E+00 | 2.69E+00 | 4.80E-01 | no | Dcc | intron_variant | MODIFIER |
| 18 | 72196738 | AX-170517359 | 0 | C | T | 0.184 | -8.24E-02 | 1.18E-01 | 2.64E+00 | 2.69E+00 | 4.80E-01 | no | Dcc | intron_variant | MODIFIER |
| 18 | 72196899 | AX-170343586 | 0 | A | G | 0.184 | -8.24E-02 | 1.18E-01 | 2.64E+00 | 2.69E+00 | 4.80E-01 | no | Dcc | intron_variant | MODIFIER |
| 18 | 72197944 | AX-170104165 | 0 | T | G | 0.184 | -8.24E-02 | 1.18E-01 | 2.64E+00 | 2.69E+00 | 4.80E-01 | no | Dcc | intron_variant | MODIFIER |
| 18 | 72198428 | AX-170403855 | 0 | A | C | 0.184 | -8.24E-02 | 1.18E-01 | 2.64E+00 | 2.69E+00 | 4.80E-01 | no | Dcc | intron_variant | MODIFIER |
| 18 | 72240364 | AX-170271262 | 0 | A | C | 0.184 | -8.24E-02 | 1.18E-01 | 2.64E+00 | 2.69E+00 | 4.80E-01 | no | Dcc | intron_variant | MODIFIER |
| 18 | 72325113 | AX-170035587 | 0 | T | C | 0.184 | -8.24E-02 | 1.18E-01 | 2.64E+00 | 2.69E+00 | 4.80E-01 | no | Dcc | intron_variant | MODIFIER |
| 18 | 72327417 | AX-170457532 | 0 | C | T | 0.184 | -8.24E-02 | 1.18E-01 | 2.64E+00 | 2.69E+00 | 4.80E-01 | no | Dcc | intron_variant | MODIFIER |
| 18 | 72364892 | AX-168374727 | 0 | C | A | 0.184 | -8.24E-02 | 1.18E-01 | 2.64E+00 | 2.69E+00 | 4.80E-01 | no | | intergenic_variant | MODIFIER |
| 18 | 72409211 | AX-97857251 | 0 | T | G | 0.184 | -8.24E-02 | 1.18E-01 | 2.64E+00 | 2.69E+00 | 4.80E-01 | no | | intergenic_variant | MODIFIER |
| 18 | 72471200 | AX-170015964 | 0 | C | T | 0.184 | -8.24E-02 | 1.18E-01 | 2.64E+00 | 2.69E+00 | 4.80E-01 | no | | intergenic_variant | MODIFIER |
| 7 | 78103495 | AX-170029363 | 0 | G | A | 0.368 | -6.47E-02 | 9.31E-02 | 2.72E+00 | 2.82E+00 | 4.81E-01 | no | | regulatory_region_variant | MODIFIER |
| 7 | 70114210 | AX-97806069 | 0 | C | G | 0.377 | -7.49E-02 | 1.07E-01 | 2.85E+00 | 2.94E+00 | 4.81E-01 | no | | intergenic_variant | MODIFIER |
| 7 | 70371483 | AX-170512980 | 0 | C | A | 0.377 | -7.49E-02 | 1.07E-01 | 2.85E+00 | 2.94E+00 | 4.81E-01 | no | Nr2f2 | upstream_gene_variant | MODIFIER |
| 7 | 70403960 | AX-97835763 | 0 | C | T | 0.377 | -7.49E-02 | 1.07E-01 | 2.85E+00 | 2.94E+00 | 4.81E-01 | no | B130024G19Rik | intron_variant,non_coding_transcript_variant | MODIFIER |
| 7 | 70408711 | AX-170405742 | 0 | G | C | 0.377 | -7.49E-02 | 1.07E-01 | 2.85E+00 | 2.94E+00 | 4.81E-01 | no | B130024G19Rik | intron_variant,non_coding_transcript_variant | MODIFIER |
| 7 | 70489073 | AX-97851468 | 0 | A | G | 0.377 | -7.49E-02 | 1.07E-01 | 2.85E+00 | 2.94E+00 | 4.81E-01 | no | Gm7656 | upstream_gene_variant | MODIFIER |
| 7 | 70490322 | AX-170244484 | 0 | T | A | 0.377 | -7.49E-02 | 1.07E-01 | 2.85E+00 | 2.94E+00 | 4.81E-01 | no | Gm7656 | upstream_gene_variant | MODIFIER |
| 7 | 70531355 | AX-170243806 | 0 | C | T | 0.377 | -7.49E-02 | 1.07E-01 | 2.85E+00 | 2.94E+00 | 4.81E-01 | no | Gm44811 | non_coding_transcript_exon_variant | MODIFIER |
| 7 | 70532764 | AX-170236199 | 0 | A | T | 0.377 | -7.49E-02 | 1.07E-01 | 2.85E+00 | 2.94E+00 | 4.81E-01 | no | Gm44811 | downstream_gene_variant | MODIFIER |
| 7 | 70553117 | AX-170273451 | 0 | A | G | 0.377 | -7.49E-02 | 1.07E-01 | 2.85E+00 | 2.94E+00 | 4.81E-01 | no | Gm35842 | non_coding_transcript_exon_variant | MODIFIER |
| 7 | 70593328 | AX-170529294 | 0 | T | A | 0.377 | -7.49E-02 | 1.07E-01 | 2.85E+00 | 2.94E+00 | 4.81E-01 | no | Gm44809 | non_coding_transcript_exon_variant | MODIFIER |
| 7 | 70604367 | AX-170369631 | 0 | A | C | 0.377 | -7.49E-02 | 1.07E-01 | 2.85E+00 | 2.94E+00 | 4.81E-01 | no | | intergenic_variant | MODIFIER |
| 7 | 70616980 | AX-97801925 | 0 | A | G | 0.377 | -7.49E-02 | 1.07E-01 | 2.85E+00 | 2.94E+00 | 4.81E-01 | no | | intergenic_variant | MODIFIER |
| 7 | 70632869 | AX-97853704 | 0 | T | C | 0.377 | -7.49E-02 | 1.07E-01 | 2.85E+00 | 2.94E+00 | 4.81E-01 | no | | intergenic_variant | MODIFIER |
| 7 | 70657830 | AX-170029248 | 0 | T | A | 0.377 | -7.49E-02 | 1.07E-01 | 2.85E+00 | 2.94E+00 | 4.81E-01 | no | | intergenic_variant | MODIFIER |
| 7 | 70658163 | AX-170029247 | 0 | G | A | 0.377 | -7.49E-02 | 1.07E-01 | 2.85E+00 | 2.94E+00 | 4.81E-01 | no | | intergenic_variant | MODIFIER |
| 7 | 70658473 | AX-170029250 | 0 | C | G | 0.377 | -7.49E-02 | 1.07E-01 | 2.85E+00 | 2.94E+00 | 4.81E-01 | no | | intergenic_variant | MODIFIER |
| 13 | 6470502 | AX-170092644 | 0 | T | C | 0.202 | -8.86E-02 | 1.25E-01 | 2.57E+00 | 2.63E+00 | 4.81E-01 | no | | regulatory_region_variant | MODIFIER |
| 7 | 36856189 | AX-170522157 | 0 | T | C | 0.325 | 5.93E-02 | 8.45E-02 | 2.61E+00 | 2.70E+00 | 4.81E-01 | no | | intergenic_variant | MODIFIER |
| 5 | 51088071 | AX-170072069 | 1 | G | T | 0.491 | 7.20E-02 | 9.69E-02 | 2.41E+00 | 2.49E+00 | 4.81E-01 | no | | intergenic_variant | MODIFIER |
| 4 | 139420227 | AX-170024091 | 0 | G | A | 0.158 | 8.22E-02 | 1.17E-01 | 2.60E+00 | 2.65E+00 | 4.81E-01 | no | Ubr4 | intron_variant | MODIFIER |
| 16 | 46224587 | AX-170358911 | 2 | T | A | 0.491 | 5.54E-02 | 7.88E-02 | 2.60E+00 | 2.67E+00 | 4.81E-01 | no | Gm26297 | downstream_gene_variant | MODIFIER |
| 9 | 78789130 | AX-97835475 | 1 | G | A | 0.196 | 8.97E-02 | 1.28E-01 | 2.83E+00 | 2.89E+00 | 4.81E-01 | no | | intergenic_variant | MODIFIER |
| 20 | 71167599 | AX-170424207 | 1 | A | T | 0.268 | 4.28E-02 | 6.12E-02 | 2.62E+00 | 2.71E+00 | 4.81E-01 | no | | intergenic_variant | MODIFIER |
| 16 | 27855921 | AX-170379780 | 0 | A | G | 0.246 | 6.17E-02 | 8.86E-02 | 2.63E+00 | 2.76E+00 | 4.81E-01 | no | Gm10823 | intron_variant,non_coding_transcript_variant | MODIFIER |

| | | | | | | | | | | | | | | | |
|---|---|---|---|---|---|---|---|---|---|---|---|---|---|---|---|
| 16 | 27872934 | AX-170290225 | 0 | A | T | 0.246 | 6.17E-02 | 8.86E-02 | 2.63E+00 | 2.76E+00 | 4.81E-01 | no | Gm10823 | intron_variant,non_coding_transcript_variant | MODIFIER |
| 16 | 27878323 | AX-97788986 | 0 | G | A | 0.246 | 6.17E-02 | 8.86E-02 | 2.63E+00 | 2.76E+00 | 4.81E-01 | no | Gm10823 | intron_variant,non_coding_transcript_variant | MODIFIER |
| 16 | 27879135 | AX-170488171 | 0 | C | G | 0.246 | 6.17E-02 | 8.86E-02 | 2.63E+00 | 2.76E+00 | 4.81E-01 | no | Gm10823 | intron_variant,non_coding_transcript_variant | MODIFIER |
| 16 | 27880848 | AX-97844575 | 0 | A | G | 0.246 | 6.17E-02 | 8.86E-02 | 2.63E+00 | 2.76E+00 | 4.81E-01 | no | Gm10823 | intron_variant,non_coding_transcript_variant | MODIFIER |
| 16 | 27937057 | AX-170508744 | 0 | T | G | 0.246 | 6.17E-02 | 8.86E-02 | 2.63E+00 | 2.76E+00 | 4.81E-01 | no | | intergenic_variant | MODIFIER |
| 16 | 27973698 | AX-97794540 | 0 | T | C | 0.246 | 6.17E-02 | 8.86E-02 | 2.63E+00 | 2.76E+00 | 4.81E-01 | no | Gm49765 | upstream_gene_variant | MODIFIER |
| 16 | 27974490 | AX-170148242 | 0 | T | C | 0.246 | 6.17E-02 | 8.86E-02 | 2.63E+00 | 2.76E+00 | 4.81E-01 | no | Gm49765 | upstream_gene_variant | MODIFIER |
| 16 | 27976824 | AX-170447818 | 0 | A | G | 0.246 | 6.17E-02 | 8.86E-02 | 2.63E+00 | 2.76E+00 | 4.81E-01 | no | | intergenic_variant | MODIFIER |
| 16 | 28042169 | AX-97785368 | 0 | C | T | 0.246 | 6.17E-02 | 8.86E-02 | 2.63E+00 | 2.76E+00 | 4.81E-01 | no | | intergenic_variant | MODIFIER |
| 16 | 28051149 | AX-170223059 | 0 | T | C | 0.246 | 6.17E-02 | 8.86E-02 | 2.63E+00 | 2.76E+00 | 4.81E-01 | no | | intergenic_variant | MODIFIER |
| 16 | 28083467 | AX-170211091 | 0 | A | G | 0.246 | 6.17E-02 | 8.86E-02 | 2.63E+00 | 2.76E+00 | 4.81E-01 | no | | intergenic_variant | MODIFIER |
| 16 | 28108620 | AX-97784657 | 0 | A | G | 0.246 | 6.17E-02 | 8.86E-02 | 2.63E+00 | 2.76E+00 | 4.81E-01 | no | | intergenic_variant | MODIFIER |
| 16 | 28203428 | AX-97781383 | 0 | A | G | 0.246 | 6.17E-02 | 8.86E-02 | 2.63E+00 | 2.76E+00 | 4.81E-01 | no | Fgf12 | intron_variant | MODIFIER |
| 16 | 28206886 | AX-170112763 | 0 | G | A | 0.246 | 6.17E-02 | 8.86E-02 | 2.63E+00 | 2.76E+00 | 4.81E-01 | no | Fgf12 | intron_variant | MODIFIER |
| 16 | 28232451 | AX-97825332 | 0 | C | T | 0.246 | 6.17E-02 | 8.86E-02 | 2.63E+00 | 2.76E+00 | 4.81E-01 | no | Fgf12 | intron_variant | MODIFIER |
| 16 | 28278388 | AX-170294072 | 0 | C | G | 0.246 | 6.17E-02 | 8.86E-02 | 2.63E+00 | 2.76E+00 | 4.81E-01 | no | Fgf12 | intron_variant | MODIFIER |
| 16 | 28280056 | AX-170056991 | 0 | T | G | 0.246 | 6.17E-02 | 8.86E-02 | 2.63E+00 | 2.76E+00 | 4.81E-01 | no | Fgf12 | intron_variant | MODIFIER |
| 16 | 28330209 | AX-170511801 | 0 | T | C | 0.246 | 6.17E-02 | 8.86E-02 | 2.63E+00 | 2.76E+00 | 4.81E-01 | no | Fgf12 | intron_variant | MODIFIER |
| 16 | 28339961 | AX-170413807 | 0 | A | G | 0.246 | 6.17E-02 | 8.86E-02 | 2.63E+00 | 2.76E+00 | 4.81E-01 | no | Fgf12 | intron_variant | MODIFIER |
| 16 | 28345671 | AX-97830734 | 0 | G | T | 0.246 | 6.17E-02 | 8.86E-02 | 2.63E+00 | 2.76E+00 | 4.81E-01 | no | Fgf12 | intron_variant | MODIFIER |
| 16 | 28387902 | AX-97800596 | 0 | T | C | 0.246 | 6.17E-02 | 8.86E-02 | 2.63E+00 | 2.76E+00 | 4.81E-01 | no | Fgf12 | intron_variant | MODIFIER |
| 16 | 28388598 | AX-170109243 | 0 | A | G | 0.246 | 6.17E-02 | 8.86E-02 | 2.63E+00 | 2.76E+00 | 4.81E-01 | no | Fgf12 | intron_variant | MODIFIER |
| 16 | 28404039 | AX-170179661 | 0 | A | C | 0.246 | 6.17E-02 | 8.86E-02 | 2.63E+00 | 2.76E+00 | 4.81E-01 | no | Fgf12 | intron_variant | MODIFIER |
| 16 | 28524086 | AX-168415666 | 0 | A | T | 0.246 | 6.17E-02 | 8.86E-02 | 2.63E+00 | 2.76E+00 | 4.81E-01 | no | Fgf12 | intron_variant | MODIFIER |
| 8 | 37903097 | AX-170433760 | 0 | G | A | 0.149 | -8.33E-02 | 1.20E-01 | 2.65E+00 | 2.76E+00 | 4.81E-01 | no | Sgcz | intron_variant | MODIFIER |
| 8 | 43101531 | AX-170315501 | 0 | A | G | 0.149 | -8.33E-02 | 1.20E-01 | 2.65E+00 | 2.76E+00 | 4.81E-01 | no | | intergenic_variant | MODIFIER |
| 8 | 43335908 | AX-170344022 | 0 | G | A | 0.149 | -8.33E-02 | 1.20E-01 | 2.65E+00 | 2.76E+00 | 4.81E-01 | no | | intergenic_variant | MODIFIER |
| 13 | 25150604 | AX-170509970 | 0 | A | C | 0.175 | -9.14E-02 | 1.24E-01 | 2.44E+00 | 2.50E+00 | 4.81E-01 | no | Dcdc2a | intron_variant | MODIFIER |
| 5 | 93025229 | AX-170244069 | 0 | G | A | 0.43 | 7.24E-02 | 1.04E-01 | 2.70E+00 | 2.77E+00 | 4.81E-01 | no | | regulatory_region_variant | MODIFIER |
| 1 | 21651855 | AX-170146152 | 0 | A | G | 0.404 | 5.95E-02 | 8.42E-02 | 2.58E+00 | 2.63E+00 | 4.81E-01 | no | Kcnq5 | intron_variant | MODIFIER |
| 15 | 86612999 | AX-170308465 | 0 | G | A | 0.518 | 5.12E-02 | 7.07E-02 | 2.47E+00 | 2.57E+00 | 4.81E-01 | no | | intergenic_variant | MODIFIER |
| 4 | 133078454 | AX-170509537 | 1 | C | T | 0.161 | 8.37E-02 | 1.18E-01 | 2.56E+00 | 2.60E+00 | 4.82E-01 | no | Ahdc1 | downstream_gene_variant | MODIFIER |
| 7 | 36582163 | AX-97848729 | 0 | A | G | 0.333 | 6.42E-02 | 9.10E-02 | 2.57E+00 | 2.65E+00 | 4.82E-01 | no | | intergenic_variant | MODIFIER |
| 7 | 36600208 | AX-97786130 | 0 | C | T | 0.333 | 6.42E-02 | 9.10E-02 | 2.57E+00 | 2.65E+00 | 4.82E-01 | no | | intergenic_variant | MODIFIER |
| 7 | 36627260 | AX-170282760 | 0 | T | A | 0.333 | 6.42E-02 | 9.10E-02 | 2.57E+00 | 2.65E+00 | 4.82E-01 | no | | intergenic_variant | MODIFIER |
| 7 | 36628852 | AX-97851216 | 0 | T | C | 0.333 | 6.42E-02 | 9.10E-02 | 2.57E+00 | 2.65E+00 | 4.82E-01 | no | | intergenic_variant | MODIFIER |
| 7 | 36631463 | AX-170028833 | 0 | G | A | 0.333 | 6.42E-02 | 9.10E-02 | 2.57E+00 | 2.65E+00 | 4.82E-01 | no | | intergenic_variant | MODIFIER |
| 7 | 36639199 | AX-170374406 | 0 | T | C | 0.333 | 6.42E-02 | 9.10E-02 | 2.57E+00 | 2.65E+00 | 4.82E-01 | no | | intergenic_variant | MODIFIER |
| 7 | 36651437 | AX-97848295 | 0 | A | C | 0.333 | 6.42E-02 | 9.10E-02 | 2.57E+00 | 2.65E+00 | 4.82E-01 | no | | intergenic_variant | MODIFIER |
| 7 | 36665310 | AX-170446113 | 0 | A | G | 0.333 | 6.42E-02 | 9.10E-02 | 2.57E+00 | 2.65E+00 | 4.82E-01 | no | | intergenic_variant | MODIFIER |
| 7 | 36675886 | AX-97788649 | 0 | G | A | 0.333 | 6.42E-02 | 9.10E-02 | 2.57E+00 | 2.65E+00 | 4.82E-01 | no | | intergenic_variant | MODIFIER |
| 7 | 36712182 | AX-170483681 | 0 | C | A | 0.333 | 6.42E-02 | 9.10E-02 | 2.57E+00 | 2.65E+00 | 4.82E-01 | no | Tshz3 | intron_variant | MODIFIER |
| 7 | 36713745 | AX-170240471 | 0 | C | G | 0.333 | 6.42E-02 | 9.10E-02 | 2.57E+00 | 2.65E+00 | 4.82E-01 | no | Tshz3 | intron_variant | MODIFIER |
| 7 | 36714159 | AX-170320310 | 0 | T | C | 0.333 | 6.42E-02 | 9.10E-02 | 2.57E+00 | 2.65E+00 | 4.82E-01 | no | Tshz3 | intron_variant | MODIFIER |
| 7 | 36720188 | AX-170292748 | 0 | T | C | 0.333 | 6.42E-02 | 9.10E-02 | 2.57E+00 | 2.65E+00 | 4.82E-01 | no | Tshz3 | intron_variant | MODIFIER |
| 7 | 36725097 | AX-170239878 | 0 | T | C | 0.333 | 6.42E-02 | 9.10E-02 | 2.57E+00 | 2.65E+00 | 4.82E-01 | no | Tshz3 | intron_variant | MODIFIER |
| 7 | 36732721 | AX-170077075 | 0 | C | T | 0.333 | 6.42E-02 | 9.10E-02 | 2.57E+00 | 2.65E+00 | 4.82E-01 | no | Tshz3 | intron_variant | MODIFIER |
| 7 | 36738967 | AX-170268961 | 0 | T | C | 0.333 | 6.42E-02 | 9.10E-02 | 2.57E+00 | 2.65E+00 | 4.82E-01 | no | Tshz3 | intron_variant | MODIFIER |
| 7 | 36741608 | AX-170493559 | 0 | C | G | 0.333 | 6.42E-02 | 9.10E-02 | 2.57E+00 | 2.65E+00 | 4.82E-01 | no | Tshz3 | intron_variant | MODIFIER |
| 7 | 36744756 | AX-170493749 | 0 | C | A | 0.333 | 6.42E-02 | 9.10E-02 | 2.57E+00 | 2.65E+00 | 4.82E-01 | no | Tshz3 | intron_variant | MODIFIER |
| 7 | 36758570 | AX-170247115 | 0 | G | A | 0.333 | 6.42E-02 | 9.10E-02 | 2.57E+00 | 2.65E+00 | 4.82E-01 | no | Tshz3 | intron_variant | MODIFIER |
| 7 | 36758604 | AX-170112415 | 0 | T | C | 0.333 | 6.42E-02 | 9.10E-02 | 2.57E+00 | 2.65E+00 | 4.82E-01 | no | Tshz3 | intron_variant | MODIFIER |
| 7 | 36766884 | AX-170175942 | 0 | G | A | 0.333 | 6.42E-02 | 9.10E-02 | 2.57E+00 | 2.65E+00 | 4.82E-01 | no | Tshz3 | intron_variant | MODIFIER |
| 7 | 36780100 | AX-170335661 | 0 | G | C | 0.333 | 6.42E-02 | 9.10E-02 | 2.57E+00 | 2.65E+00 | 4.82E-01 | no | | intergenic_variant | MODIFIER |

| | | | | | | | | | | | | | | | |
|---|---|---|---|---|---|---|---|---|---|---|---|---|---|---|---|
| 7 | 36783540 | AX-170389093 | 0 | C | T | 0.333 | 6.42E-02 | 9.10E-02 | 2.57E+00 | 2.65E+00 | 4.82E-01 | no | | intergenic_variant | MODIFIER |
| 7 | 36791859 | AX-170290198 | 0 | C | T | 0.333 | 6.42E-02 | 9.10E-02 | 2.57E+00 | 2.65E+00 | 4.82E-01 | no | | intergenic_variant | MODIFIER |
| 7 | 36792667 | AX-170353867 | 0 | G | A | 0.333 | 6.42E-02 | 9.10E-02 | 2.57E+00 | 2.65E+00 | 4.82E-01 | no | | intergenic_variant | MODIFIER |
| 7 | 36808329 | AX-170232092 | 0 | G | A | 0.333 | 6.42E-02 | 9.10E-02 | 2.57E+00 | 2.65E+00 | 4.82E-01 | no | | intergenic_variant | MODIFIER |
| 7 | 36811787 | AX-170323532 | 0 | G | A | 0.333 | 6.42E-02 | 9.10E-02 | 2.57E+00 | 2.65E+00 | 4.82E-01 | no | | intergenic_variant | MODIFIER |
| 7 | 36815259 | AX-170095893 | 0 | T | G | 0.333 | 6.42E-02 | 9.10E-02 | 2.57E+00 | 2.65E+00 | 4.82E-01 | no | | intergenic_variant | MODIFIER |
| 7 | 36829466 | AX-170343876 | 0 | T | C | 0.333 | 6.42E-02 | 9.10E-02 | 2.57E+00 | 2.65E+00 | 4.82E-01 | no | | intergenic_variant | MODIFIER |
| 7 | 36393646 | AX-170140707 | 0 | T | C | 0.342 | 6.47E-02 | 9.18E-02 | 2.58E+00 | 2.65E+00 | 4.82E-01 | no | | intergenic_variant | MODIFIER |
| 7 | 36396465 | AX-170266802 | 0 | A | G | 0.342 | 6.47E-02 | 9.18E-02 | 2.58E+00 | 2.65E+00 | 4.82E-01 | no | | intergenic_variant | MODIFIER |
| 7 | 36398785 | AX-170462273 | 0 | T | C | 0.342 | 6.47E-02 | 9.18E-02 | 2.58E+00 | 2.65E+00 | 4.82E-01 | no | | intergenic_variant | MODIFIER |
| 7 | 36399004 | AX-170255695 | 0 | C | G | 0.342 | 6.47E-02 | 9.18E-02 | 2.58E+00 | 2.65E+00 | 4.82E-01 | no | | intergenic_variant | MODIFIER |
| 7 | 36400045 | AX-170503574 | 0 | T | G | 0.342 | 6.47E-02 | 9.18E-02 | 2.58E+00 | 2.65E+00 | 4.82E-01 | no | | intergenic_variant | MODIFIER |
| 7 | 36406254 | AX-170469957 | 0 | A | G | 0.342 | 6.47E-02 | 9.18E-02 | 2.58E+00 | 2.65E+00 | 4.82E-01 | no | | intergenic_variant | MODIFIER |
| 7 | 36410683 | AX-170474231 | 0 | T | G | 0.342 | 6.47E-02 | 9.18E-02 | 2.58E+00 | 2.65E+00 | 4.82E-01 | no | | intergenic_variant | MODIFIER |
| 7 | 36417247 | AX-170386983 | 0 | T | G | 0.342 | 6.47E-02 | 9.18E-02 | 2.58E+00 | 2.65E+00 | 4.82E-01 | no | | intergenic_variant | MODIFIER |
| 7 | 36417579 | AX-170447326 | 0 | T | C | 0.342 | 6.47E-02 | 9.18E-02 | 2.58E+00 | 2.65E+00 | 4.82E-01 | no | | intergenic_variant | MODIFIER |
| 7 | 36418012 | AX-170486422 | 0 | G | A | 0.342 | 6.47E-02 | 9.18E-02 | 2.58E+00 | 2.65E+00 | 4.82E-01 | no | | intergenic_variant | MODIFIER |
| 7 | 36424453 | AX-97844227 | 0 | G | A | 0.342 | 6.47E-02 | 9.18E-02 | 2.58E+00 | 2.65E+00 | 4.82E-01 | no | | intergenic_variant | MODIFIER |
| 7 | 36429237 | AX-170316537 | 0 | A | G | 0.342 | 6.47E-02 | 9.18E-02 | 2.58E+00 | 2.65E+00 | 4.82E-01 | no | Gm25247 | upstream_gene_variant | MODIFIER |
| 7 | 36430441 | AX-170077063 | 0 | T | C | 0.342 | 6.47E-02 | 9.18E-02 | 2.58E+00 | 2.65E+00 | 4.82E-01 | no | Gm25247 | upstream_gene_variant | MODIFIER |
| 7 | 36437223 | AX-170235998 | 0 | G | C | 0.342 | 6.47E-02 | 9.18E-02 | 2.58E+00 | 2.65E+00 | 4.82E-01 | no | | intergenic_variant | MODIFIER |
| 7 | 36442219 | AX-170400782 | 0 | C | A | 0.342 | 6.47E-02 | 9.18E-02 | 2.58E+00 | 2.65E+00 | 4.82E-01 | no | | intergenic_variant | MODIFIER |
| 7 | 36444153 | AX-170393709 | 0 | C | T | 0.342 | 6.47E-02 | 9.18E-02 | 2.58E+00 | 2.65E+00 | 4.82E-01 | no | | intergenic_variant | MODIFIER |
| 7 | 36468116 | AX-170160305 | 0 | G | A | 0.342 | 6.47E-02 | 9.18E-02 | 2.58E+00 | 2.65E+00 | 4.82E-01 | no | | intergenic_variant | MODIFIER |
| 7 | 36468213 | AX-170363663 | 0 | T | C | 0.342 | 6.47E-02 | 9.18E-02 | 2.58E+00 | 2.65E+00 | 4.82E-01 | no | | intergenic_variant | MODIFIER |
| 7 | 36474596 | AX-170086030 | 0 | G | T | 0.342 | 6.47E-02 | 9.18E-02 | 2.58E+00 | 2.65E+00 | 4.82E-01 | no | | intergenic_variant | MODIFIER |
| 7 | 36474658 | AX-170166651 | 0 | C | T | 0.342 | 6.47E-02 | 9.18E-02 | 2.58E+00 | 2.65E+00 | 4.82E-01 | no | | intergenic_variant | MODIFIER |
| 7 | 36504836 | AX-170193529 | 0 | T | C | 0.342 | 6.47E-02 | 9.18E-02 | 2.58E+00 | 2.65E+00 | 4.82E-01 | no | | regulatory_region_variant | MODIFIER |
| 7 | 36505160 | AX-170288360 | 0 | T | C | 0.342 | 6.47E-02 | 9.18E-02 | 2.58E+00 | 2.65E+00 | 4.82E-01 | no | | regulatory_region_variant | MODIFIER |
| 7 | 36508410 | AX-170143899 | 0 | G | A | 0.342 | 6.47E-02 | 9.18E-02 | 2.58E+00 | 2.65E+00 | 4.82E-01 | no | | regulatory_region_variant | MODIFIER |
| 7 | 36515744 | AX-170353448 | 0 | G | A | 0.342 | 6.47E-02 | 9.18E-02 | 2.58E+00 | 2.65E+00 | 4.82E-01 | no | | intergenic_variant | MODIFIER |
| 7 | 36520277 | AX-97822624 | 0 | C | T | 0.342 | 6.47E-02 | 9.18E-02 | 2.58E+00 | 2.65E+00 | 4.82E-01 | no | | intergenic_variant | MODIFIER |
| 7 | 36527535 | AX-170185218 | 0 | C | T | 0.342 | 6.47E-02 | 9.18E-02 | 2.58E+00 | 2.65E+00 | 4.82E-01 | no | | intergenic_variant | MODIFIER |
| 7 | 36536778 | AX-170165499 | 0 | A | C | 0.342 | 6.47E-02 | 9.18E-02 | 2.58E+00 | 2.65E+00 | 4.82E-01 | no | | intergenic_variant | MODIFIER |
| 7 | 36549159 | AX-170534806 | 0 | T | C | 0.342 | 6.47E-02 | 9.18E-02 | 2.58E+00 | 2.65E+00 | 4.82E-01 | no | | intergenic_variant | MODIFIER |
| 7 | 36549376 | AX-170111278 | 0 | C | T | 0.342 | 6.47E-02 | 9.18E-02 | 2.58E+00 | 2.65E+00 | 4.82E-01 | no | | intergenic_variant | MODIFIER |
| 7 | 36559663 | AX-170378638 | 0 | C | T | 0.342 | 6.47E-02 | 9.18E-02 | 2.58E+00 | 2.65E+00 | 4.82E-01 | no | | intergenic_variant | MODIFIER |
| 7 | 36561522 | AX-170289991 | 0 | C | T | 0.342 | 6.47E-02 | 9.18E-02 | 2.58E+00 | 2.65E+00 | 4.82E-01 | no | | intergenic_variant | MODIFIER |
| 7 | 36565614 | AX-170326102 | 0 | C | T | 0.342 | 6.47E-02 | 9.18E-02 | 2.58E+00 | 2.65E+00 | 4.82E-01 | no | Gm29129 | upstream_gene_variant | MODIFIER |
| 7 | 36578347 | AX-170294402 | 0 | T | G | 0.342 | 6.47E-02 | 9.18E-02 | 2.58E+00 | 2.65E+00 | 4.82E-01 | no | | intergenic_variant | MODIFIER |
| 7 | 36580305 | AX-170408092 | 0 | G | T | 0.342 | 6.47E-02 | 9.18E-02 | 2.58E+00 | 2.65E+00 | 4.82E-01 | no | | intergenic_variant | MODIFIER |
| 7 | 36599017 | AX-170286485 | 0 | T | C | 0.342 | 6.47E-02 | 9.18E-02 | 2.58E+00 | 2.65E+00 | 4.82E-01 | no | | intergenic_variant | MODIFIER |
| 7 | 36631543 | AX-170077071 | 0 | A | G | 0.342 | 6.47E-02 | 9.18E-02 | 2.58E+00 | 2.65E+00 | 4.82E-01 | no | | intergenic_variant | MODIFIER |
| 7 | 36662607 | AX-170348359 | 0 | A | G | 0.342 | 6.47E-02 | 9.18E-02 | 2.58E+00 | 2.65E+00 | 4.82E-01 | no | | intergenic_variant | MODIFIER |
| 7 | 36663359 | AX-170383201 | 0 | A | C | 0.342 | 6.47E-02 | 9.18E-02 | 2.58E+00 | 2.65E+00 | 4.82E-01 | no | | intergenic_variant | MODIFIER |
| 7 | 36664027 | AX-170235035 | 0 | A | G | 0.342 | 6.47E-02 | 9.18E-02 | 2.58E+00 | 2.65E+00 | 4.82E-01 | no | | intergenic_variant | MODIFIER |
| 7 | 36667403 | AX-170357591 | 0 | T | C | 0.342 | 6.47E-02 | 9.18E-02 | 2.58E+00 | 2.65E+00 | 4.82E-01 | no | | intergenic_variant | MODIFIER |
| 7 | 36678008 | AX-170449810 | 0 | C | A | 0.342 | 6.47E-02 | 9.18E-02 | 2.58E+00 | 2.65E+00 | 4.82E-01 | no | | intergenic_variant | MODIFIER |
| 7 | 36688320 | AX-170028836 | 0 | T | C | 0.342 | 6.47E-02 | 9.18E-02 | 2.58E+00 | 2.65E+00 | 4.82E-01 | no | | intergenic_variant | MODIFIER |
| 7 | 36688951 | AX-170286541 | 0 | A | G | 0.342 | 6.47E-02 | 9.18E-02 | 2.58E+00 | 2.65E+00 | 4.82E-01 | no | | intergenic_variant | MODIFIER |
| 7 | 36694138 | AX-170308283 | 0 | T | G | 0.342 | 6.47E-02 | 9.18E-02 | 2.58E+00 | 2.65E+00 | 4.82E-01 | no | Tshz3 | upstream_gene_variant | MODIFIER |
| 7 | 36844276 | AX-170077081 | 0 | T | A | 0.342 | 6.47E-02 | 9.18E-02 | 2.58E+00 | 2.65E+00 | 4.82E-01 | no | | intergenic_variant | MODIFIER |
| 7 | 36847704 | AX-170077080 | 0 | C | T | 0.342 | 6.47E-02 | 9.18E-02 | 2.58E+00 | 2.65E+00 | 4.82E-01 | no | | intergenic_variant | MODIFIER |
| 7 | 36848387 | AX-97857309 | 0 | T | G | 0.342 | 6.47E-02 | 9.18E-02 | 2.58E+00 | 2.65E+00 | 4.82E-01 | no | | intergenic_variant | MODIFIER |

| | | | | | | | | | | | | | | | |
|---|---|---|---|---|---|---|---|---|---|---|---|---|---|---|---|
| 7 | 36864341 | AX-170124317 | 0 | G | A | 0.342 | 6.47E-02 | 9.18E-02 | 2.58E+00 | 2.65E+00 | 4.82E-01 | no | | intergenic_variant | MODIFIER |
| 7 | 36873009 | AX-170181992 | 0 | C | T | 0.342 | 6.47E-02 | 9.18E-02 | 2.58E+00 | 2.65E+00 | 4.82E-01 | no | | intergenic_variant | MODIFIER |
| 7 | 36888054 | AX-170129627 | 0 | T | C | 0.342 | 6.47E-02 | 9.18E-02 | 2.58E+00 | 2.65E+00 | 4.82E-01 | no | | intergenic_variant | MODIFIER |
| 7 | 36900194 | AX-170115193 | 0 | A | G | 0.342 | 6.47E-02 | 9.18E-02 | 2.58E+00 | 2.65E+00 | 4.82E-01 | no | | intergenic_variant | MODIFIER |
| 7 | 36910456 | AX-170269295 | 0 | T | C | 0.342 | 6.47E-02 | 9.18E-02 | 2.58E+00 | 2.65E+00 | 4.82E-01 | no | | intergenic_variant | MODIFIER |
| 7 | 36931893 | AX-170422575 | 0 | C | T | 0.342 | 6.47E-02 | 9.18E-02 | 2.58E+00 | 2.65E+00 | 4.82E-01 | no | | intergenic_variant | MODIFIER |
| 7 | 36945764 | AX-170464061 | 0 | T | C | 0.342 | 6.47E-02 | 9.18E-02 | 2.58E+00 | 2.65E+00 | 4.82E-01 | no | | regulatory_region_variant | MODIFIER |
| 7 | 36951841 | AX-170190638 | 0 | T | A | 0.342 | 6.47E-02 | 9.18E-02 | 2.58E+00 | 2.65E+00 | 4.82E-01 | no | | intergenic_variant | MODIFIER |
| 7 | 36955819 | AX-170115235 | 0 | C | T | 0.342 | 6.47E-02 | 9.18E-02 | 2.58E+00 | 2.65E+00 | 4.82E-01 | no | | intergenic_variant | MODIFIER |
| 7 | 36961325 | AX-170264306 | 0 | T | C | 0.342 | 6.47E-02 | 9.18E-02 | 2.58E+00 | 2.65E+00 | 4.82E-01 | no | | intergenic_variant | MODIFIER |
| 7 | 36963214 | AX-170120031 | 0 | C | G | 0.342 | 6.47E-02 | 9.18E-02 | 2.58E+00 | 2.65E+00 | 4.82E-01 | no | | intergenic_variant | MODIFIER |
| 7 | 36963346 | AX-170305113 | 0 | G | A | 0.342 | 6.47E-02 | 9.18E-02 | 2.58E+00 | 2.65E+00 | 4.82E-01 | no | | intergenic_variant | MODIFIER |
| 7 | 36966730 | AX-170527516 | 0 | C | T | 0.342 | 6.47E-02 | 9.18E-02 | 2.58E+00 | 2.65E+00 | 4.82E-01 | no | | intergenic_variant | MODIFIER |
| 7 | 36976875 | AX-170138709 | 0 | T | C | 0.342 | 6.47E-02 | 9.18E-02 | 2.58E+00 | 2.65E+00 | 4.82E-01 | no | | intergenic_variant | MODIFIER |
| 7 | 36977239 | AX-170279685 | 0 | A | G | 0.342 | 6.47E-02 | 9.18E-02 | 2.58E+00 | 2.65E+00 | 4.82E-01 | no | | intergenic_variant | MODIFIER |
| 7 | 36987707 | AX-170418508 | 0 | T | C | 0.342 | 6.47E-02 | 9.18E-02 | 2.58E+00 | 2.65E+00 | 4.82E-01 | no | | intergenic_variant | MODIFIER |
| 7 | 36987960 | AX-170405261 | 0 | G | T | 0.342 | 6.47E-02 | 9.18E-02 | 2.58E+00 | 2.65E+00 | 4.82E-01 | no | | regulatory_region_variant | MODIFIER |
| 7 | 36991759 | AX-97832161 | 0 | T | C | 0.342 | 6.47E-02 | 9.18E-02 | 2.58E+00 | 2.65E+00 | 4.82E-01 | no | | intergenic_variant | MODIFIER |
| 7 | 37005262 | AX-97847056 | 0 | A | G | 0.342 | 6.47E-02 | 9.18E-02 | 2.58E+00 | 2.65E+00 | 4.82E-01 | no | | intergenic_variant | MODIFIER |
| 7 | 37020607 | AX-170387433 | 0 | T | C | 0.342 | 6.47E-02 | 9.18E-02 | 2.58E+00 | 2.65E+00 | 4.82E-01 | no | | intergenic_variant | MODIFIER |
| 7 | 37035537 | AX-170296225 | 0 | T | C | 0.342 | 6.47E-02 | 9.18E-02 | 2.58E+00 | 2.65E+00 | 4.82E-01 | no | | intergenic_variant | MODIFIER |
| 7 | 37044163 | AX-170260899 | 0 | A | C | 0.342 | 6.47E-02 | 9.18E-02 | 2.58E+00 | 2.65E+00 | 4.82E-01 | no | | intergenic_variant | MODIFIER |
| 7 | 37044300 | AX-170379665 | 0 | G | A | 0.342 | 6.47E-02 | 9.18E-02 | 2.58E+00 | 2.65E+00 | 4.82E-01 | no | | intergenic_variant | MODIFIER |
| 7 | 37070353 | AX-170388668 | 0 | T | C | 0.342 | 6.47E-02 | 9.18E-02 | 2.58E+00 | 2.65E+00 | 4.82E-01 | no | | intergenic_variant | MODIFIER |
| 7 | 37077114 | AX-170278811 | 0 | G | A | 0.342 | 6.47E-02 | 9.18E-02 | 2.58E+00 | 2.65E+00 | 4.82E-01 | no | | intergenic_variant | MODIFIER |
| 7 | 37095455 | AX-170259755 | 0 | A | G | 0.342 | 6.47E-02 | 9.18E-02 | 2.58E+00 | 2.65E+00 | 4.82E-01 | no | | intergenic_variant | MODIFIER |
| 7 | 37095745 | AX-170223285 | 0 | A | G | 0.342 | 6.47E-02 | 9.18E-02 | 2.58E+00 | 2.65E+00 | 4.82E-01 | no | | intergenic_variant | MODIFIER |
| 7 | 37100342 | AX-170095894 | 0 | A | G | 0.342 | 6.47E-02 | 9.18E-02 | 2.58E+00 | 2.65E+00 | 4.82E-01 | no | | intergenic_variant | MODIFIER |
| 7 | 37101560 | AX-170521466 | 0 | T | C | 0.342 | 6.47E-02 | 9.18E-02 | 2.58E+00 | 2.65E+00 | 4.82E-01 | no | | intergenic_variant | MODIFIER |
| 7 | 37105698 | AX-170181468 | 0 | T | G | 0.342 | 6.47E-02 | 9.18E-02 | 2.58E+00 | 2.65E+00 | 4.82E-01 | no | | intergenic_variant | MODIFIER |
| 7 | 37115685 | AX-170464697 | 0 | C | T | 0.342 | 6.47E-02 | 9.18E-02 | 2.58E+00 | 2.65E+00 | 4.82E-01 | no | Gm28078 | downstream_gene_variant | MODIFIER |
| 7 | 37128938 | AX-170290965 | 0 | T | C | 0.342 | 6.47E-02 | 9.18E-02 | 2.58E+00 | 2.65E+00 | 4.82E-01 | no | Gm28078 | upstream_gene_variant | MODIFIER |
| 7 | 37131261 | AX-170315976 | 0 | G | C | 0.342 | 6.47E-02 | 9.18E-02 | 2.58E+00 | 2.65E+00 | 4.82E-01 | no | Gm28078 | upstream_gene_variant | MODIFIER |
| 7 | 37132062 | AX-170077085 | 0 | T | C | 0.342 | 6.47E-02 | 9.18E-02 | 2.58E+00 | 2.65E+00 | 4.82E-01 | no | Gm28076 | downstream_gene_variant | MODIFIER |
| 7 | 37133634 | AX-170077084 | 0 | C | A | 0.342 | 6.47E-02 | 9.18E-02 | 2.58E+00 | 2.65E+00 | 4.82E-01 | no | Gm28076 | downstream_gene_variant | MODIFIER |
| 7 | 37139216 | AX-170174329 | 0 | T | C | 0.342 | 6.47E-02 | 9.18E-02 | 2.58E+00 | 2.65E+00 | 4.82E-01 | no | Gm28076 | intron_variant,non_coding_transcript_variant | MODIFIER |
| 7 | 37139353 | AX-170254717 | 0 | C | A | 0.342 | 6.47E-02 | 9.18E-02 | 2.58E+00 | 2.65E+00 | 4.82E-01 | no | Gm28076 | intron_variant,non_coding_transcript_variant | MODIFIER |
| 7 | 37147049 | AX-170028838 | 0 | C | A | 0.342 | 6.47E-02 | 9.18E-02 | 2.58E+00 | 2.65E+00 | 4.82E-01 | no | | regulatory_region_variant | MODIFIER |
| 7 | 37148662 | AX-170162037 | 0 | A | G | 0.342 | 6.47E-02 | 9.18E-02 | 2.58E+00 | 2.65E+00 | 4.82E-01 | no | | intergenic_variant | MODIFIER |
| 7 | 37154897 | AX-170028839 | 0 | A | G | 0.342 | 6.47E-02 | 9.18E-02 | 2.58E+00 | 2.65E+00 | 4.82E-01 | no | | intergenic_variant | MODIFIER |
| 7 | 37168234 | AX-170295590 | 0 | C | A | 0.342 | 6.47E-02 | 9.18E-02 | 2.58E+00 | 2.65E+00 | 4.82E-01 | no | | intergenic_variant | MODIFIER |
| 7 | 37184866 | AX-170395442 | 0 | T | A | 0.342 | 6.47E-02 | 9.18E-02 | 2.58E+00 | 2.65E+00 | 4.82E-01 | no | | intergenic_variant | MODIFIER |
| 7 | 37186345 | AX-170473566 | 0 | A | C | 0.342 | 6.47E-02 | 9.18E-02 | 2.58E+00 | 2.65E+00 | 4.82E-01 | no | | intergenic_variant | MODIFIER |
| 7 | 37190472 | AX-170501282 | 0 | G | T | 0.342 | 6.47E-02 | 9.18E-02 | 2.58E+00 | 2.65E+00 | 4.82E-01 | no | | intergenic_variant | MODIFIER |
| 7 | 37196313 | AX-170109784 | 0 | C | G | 0.342 | 6.47E-02 | 9.18E-02 | 2.58E+00 | 2.65E+00 | 4.82E-01 | no | | intergenic_variant | MODIFIER |
| 7 | 37198559 | AX-170348228 | 0 | T | C | 0.342 | 6.47E-02 | 9.18E-02 | 2.58E+00 | 2.65E+00 | 4.82E-01 | no | | intergenic_variant | MODIFIER |
| 7 | 37201532 | AX-170323010 | 0 | C | T | 0.342 | 6.47E-02 | 9.18E-02 | 2.58E+00 | 2.65E+00 | 4.82E-01 | no | | intergenic_variant | MODIFIER |
| 7 | 37230078 | AX-170378709 | 0 | A | G | 0.342 | 6.47E-02 | 9.18E-02 | 2.58E+00 | 2.65E+00 | 4.82E-01 | no | | intergenic_variant | MODIFIER |
| 7 | 37236786 | AX-170153698 | 0 | T | C | 0.342 | 6.47E-02 | 9.18E-02 | 2.58E+00 | 2.65E+00 | 4.82E-01 | no | | intergenic_variant | MODIFIER |
| 7 | 37240179 | AX-170221369 | 0 | A | G | 0.342 | 6.47E-02 | 9.18E-02 | 2.58E+00 | 2.65E+00 | 4.82E-01 | no | | intergenic_variant | MODIFIER |
| 7 | 37240310 | AX-170362382 | 0 | T | C | 0.342 | 6.47E-02 | 9.18E-02 | 2.58E+00 | 2.65E+00 | 4.82E-01 | no | | intergenic_variant | MODIFIER |
| 7 | 37240420 | AX-170219587 | 0 | G | A | 0.342 | 6.47E-02 | 9.18E-02 | 2.58E+00 | 2.65E+00 | 4.82E-01 | no | | intergenic_variant | MODIFIER |
| 7 | 37244696 | AX-170338521 | 0 | T | C | 0.342 | 6.47E-02 | 9.18E-02 | 2.58E+00 | 2.65E+00 | 4.82E-01 | no | | intergenic_variant | MODIFIER |
| 7 | 37245002 | AX-170318585 | 0 | A | G | 0.342 | 6.47E-02 | 9.18E-02 | 2.58E+00 | 2.65E+00 | 4.82E-01 | no | | intergenic_variant | MODIFIER |

| | | | | | | | | | | | | | | | |
|---|---|---|---|---|---|---|---|---|---|---|---|---|---|---|---|
| 7 | 37254299 | AX-170354058 | 0 | T | C | 0.342 | 6.47E-02 | 9.18E-02 | 2.58E+00 | 2.65E+00 | 4.82E-01 | no | | regulatory_region_variant | MODIFIER |
| 7 | 37256940 | AX-170299986 | 0 | A | G | 0.342 | 6.47E-02 | 9.18E-02 | 2.58E+00 | 2.65E+00 | 4.82E-01 | no | | intergenic_variant | MODIFIER |
| 7 | 37259855 | AX-170241276 | 0 | T | A | 0.342 | 6.47E-02 | 9.18E-02 | 2.58E+00 | 2.65E+00 | 4.82E-01 | no | | intergenic_variant | MODIFIER |
| 7 | 37261826 | AX-170357623 | 0 | A | G | 0.342 | 6.47E-02 | 9.18E-02 | 2.58E+00 | 2.65E+00 | 4.82E-01 | no | | intergenic_variant | MODIFIER |
| 7 | 37275656 | AX-170315499 | 0 | C | T | 0.342 | 6.47E-02 | 9.18E-02 | 2.58E+00 | 2.65E+00 | 4.82E-01 | no | | intergenic_variant | MODIFIER |
| 7 | 37300108 | AX-170100324 | 0 | C | T | 0.342 | 6.47E-02 | 9.18E-02 | 2.58E+00 | 2.65E+00 | 4.82E-01 | no | Gm28077 | intron_variant,non_coding_transcript_variant | MODIFIER |
| 7 | 37303798 | AX-170366461 | 0 | T | C | 0.342 | 6.47E-02 | 9.18E-02 | 2.58E+00 | 2.65E+00 | 4.82E-01 | no | Gm28077 | non_coding_transcript_exon_variant | MODIFIER |
| 7 | 37312064 | AX-170170610 | 0 | G | A | 0.342 | 6.47E-02 | 9.18E-02 | 2.58E+00 | 2.65E+00 | 4.82E-01 | no | | intergenic_variant | MODIFIER |
| 7 | 37314139 | AX-170373730 | 0 | T | G | 0.342 | 6.47E-02 | 9.18E-02 | 2.58E+00 | 2.65E+00 | 4.82E-01 | no | Gm45768 | downstream_gene_variant | MODIFIER |
| 7 | 37324256 | AX-170357660 | 0 | A | C | 0.342 | 6.47E-02 | 9.18E-02 | 2.58E+00 | 2.65E+00 | 4.82E-01 | no | Gm45768 | upstream_gene_variant | MODIFIER |
| 7 | 37342409 | AX-170527521 | 0 | C | T | 0.342 | 6.47E-02 | 9.18E-02 | 2.58E+00 | 2.65E+00 | 4.82E-01 | no | | regulatory_region_variant | MODIFIER |
| 7 | 37346084 | AX-97809129 | 0 | T | A | 0.342 | 6.47E-02 | 9.18E-02 | 2.58E+00 | 2.65E+00 | 4.82E-01 | no | | regulatory_region_variant | MODIFIER |
| 7 | 37372476 | AX-170393400 | 0 | C | T | 0.342 | 6.47E-02 | 9.18E-02 | 2.58E+00 | 2.65E+00 | 4.82E-01 | no | | intergenic_variant | MODIFIER |
| 7 | 37372501 | AX-97820612 | 0 | T | C | 0.342 | 6.47E-02 | 9.18E-02 | 2.58E+00 | 2.65E+00 | 4.82E-01 | no | | intergenic_variant | MODIFIER |
| 7 | 37372974 | AX-170440583 | 0 | T | C | 0.342 | 6.47E-02 | 9.18E-02 | 2.58E+00 | 2.65E+00 | 4.82E-01 | no | | intergenic_variant | MODIFIER |
| 7 | 37373472 | AX-170324569 | 0 | T | A | 0.342 | 6.47E-02 | 9.18E-02 | 2.58E+00 | 2.65E+00 | 4.82E-01 | no | | intergenic_variant | MODIFIER |
| 7 | 37387387 | AX-170077093 | 0 | T | C | 0.342 | 6.47E-02 | 9.18E-02 | 2.58E+00 | 2.65E+00 | 4.82E-01 | no | | intergenic_variant | MODIFIER |
| 7 | 37389183 | AX-170077091 | 0 | C | T | 0.342 | 6.47E-02 | 9.18E-02 | 2.58E+00 | 2.65E+00 | 4.82E-01 | no | | regulatory_region_variant | MODIFIER |
| 7 | 37391975 | AX-170358785 | 0 | A | G | 0.342 | 6.47E-02 | 9.18E-02 | 2.58E+00 | 2.65E+00 | 4.82E-01 | no | | intergenic_variant | MODIFIER |
| 7 | 37392508 | AX-170273537 | 0 | A | G | 0.342 | 6.47E-02 | 9.18E-02 | 2.58E+00 | 2.65E+00 | 4.82E-01 | no | | intergenic_variant | MODIFIER |
| 7 | 37394318 | AX-170457397 | 0 | T | G | 0.342 | 6.47E-02 | 9.18E-02 | 2.58E+00 | 2.65E+00 | 4.82E-01 | no | | intergenic_variant | MODIFIER |
| 7 | 37400353 | AX-170373701 | 0 | G | A | 0.342 | 6.47E-02 | 9.18E-02 | 2.58E+00 | 2.65E+00 | 4.82E-01 | no | | intergenic_variant | MODIFIER |
| 7 | 37416554 | AX-170423325 | 0 | T | C | 0.342 | 6.47E-02 | 9.18E-02 | 2.58E+00 | 2.65E+00 | 4.82E-01 | no | | intergenic_variant | MODIFIER |
| 7 | 37426672 | AX-170485752 | 0 | T | C | 0.342 | 6.47E-02 | 9.18E-02 | 2.58E+00 | 2.65E+00 | 4.82E-01 | no | | intergenic_variant | MODIFIER |
| 7 | 37431938 | AX-170275590 | 0 | C | A | 0.342 | 6.47E-02 | 9.18E-02 | 2.58E+00 | 2.65E+00 | 4.82E-01 | no | | regulatory_region_variant | MODIFIER |
| 7 | 37444858 | AX-170170137 | 0 | G | T | 0.342 | 6.47E-02 | 9.18E-02 | 2.58E+00 | 2.65E+00 | 4.82E-01 | no | | intergenic_variant | MODIFIER |
| 7 | 37447263 | AX-170115440 | 0 | G | A | 0.342 | 6.47E-02 | 9.18E-02 | 2.58E+00 | 2.65E+00 | 4.82E-01 | no | | intergenic_variant | MODIFIER |
| 7 | 37450705 | AX-170373686 | 0 | C | A | 0.342 | 6.47E-02 | 9.18E-02 | 2.58E+00 | 2.65E+00 | 4.82E-01 | no | | intergenic_variant | MODIFIER |
| 7 | 37452210 | AX-170387421 | 0 | A | G | 0.342 | 6.47E-02 | 9.18E-02 | 2.58E+00 | 2.65E+00 | 4.82E-01 | no | | intergenic_variant | MODIFIER |
| 7 | 37459335 | AX-170350324 | 0 | A | G | 0.342 | 6.47E-02 | 9.18E-02 | 2.58E+00 | 2.65E+00 | 4.82E-01 | no | | intergenic_variant | MODIFIER |
| 7 | 37459538 | AX-170398990 | 0 | G | A | 0.342 | 6.47E-02 | 9.18E-02 | 2.58E+00 | 2.65E+00 | 4.82E-01 | no | | intergenic_variant | MODIFIER |
| 7 | 37464856 | AX-170262413 | 0 | A | G | 0.342 | 6.47E-02 | 9.18E-02 | 2.58E+00 | 2.65E+00 | 4.82E-01 | no | Gm44806 | downstream_gene_variant | MODIFIER |
| 1 | 170774149 | AX-170138855 | 1 | G | T | 0.214 | 9.98E-02 | 1.43E-01 | 2.64E+00 | 2.68E+00 | 4.82E-01 | no | Atf6 | intron_variant | MODIFIER |
| 13 | 25550920 | AX-170417095 | 1 | C | T | 0.161 | -9.13E-02 | 1.23E-01 | 2.42E+00 | 2.48E+00 | 4.82E-01 | no | | intergenic_variant | MODIFIER |
| 5 | 43553082 | AX-170526852 | 1 | C | T | 0.188 | -7.75E-02 | 1.11E-01 | 2.61E+00 | 2.69E+00 | 4.82E-01 | no | C1qtnf7 | intron_variant | MODIFIER |
| 5 | 80821529 | AX-170269125 | 1 | C | T | 0.446 | 7.34E-02 | 1.06E-01 | 2.69E+00 | 2.75E+00 | 4.82E-01 | no | | intergenic_variant | MODIFIER |
| 5 | 81042663 | AX-170072611 | 1 | C | T | 0.446 | 7.34E-02 | 1.06E-01 | 2.69E+00 | 2.75E+00 | 4.82E-01 | no | Adgrl3 | intron_variant | MODIFIER |
| 15 | 68382994 | AX-170104241 | 1 | G | T | 0.339 | -5.68E-02 | 7.58E-02 | 2.38E+00 | 2.47E+00 | 4.82E-01 | no | | intergenic_variant | MODIFIER |
| 1 | 101973166 | AX-170193491 | 1 | A | T | 0.134 | 1.23E-01 | 1.75E-01 | 2.60E+00 | 2.67E+00 | 4.82E-01 | no | | intergenic_variant | MODIFIER |
| 14 | 86065494 | AX-97821235 | 1 | G | A | 0.036 | -1.28E-01 | 1.72E-01 | 2.40E+00 | 2.51E+00 | 4.82E-01 | no | | intergenic_variant | MODIFIER |
| 8 | 96973624 | AX-97798860 | 0 | G | A | 0.228 | 7.84E-02 | 1.13E-01 | 2.73E+00 | 2.81E+00 | 4.82E-01 | no | | intergenic_variant | MODIFIER |
| 1 | 151217528 | AX-170001981 | 0 | T | C | 0.211 | 8.74E-02 | 1.24E-01 | 2.59E+00 | 2.65E+00 | 4.82E-01 | no | | regulatory_region_variant | MODIFIER |
| 5 | 38540889 | AX-170234197 | 1 | G | A | 0.348 | 7.81E-02 | 1.13E-01 | 2.72E+00 | 2.79E+00 | 4.82E-01 | no | Wdr1 | intron_variant | MODIFIER |
| 8 | 33886900 | AX-97809350 | 1 | G | A | 0.143 | -9.35E-02 | 1.34E-01 | 2.64E+00 | 2.70E+00 | 4.83E-01 | no | Rbpms | intron_variant | MODIFIER |
| 16 | 73977527 | AX-170421322 | 0 | T | C | 0.482 | 5.59E-02 | 7.73E-02 | 2.48E+00 | 2.55E+00 | 4.83E-01 | no | Gm24272 | downstream_gene_variant | MODIFIER |
| 16 | 73990790 | AX-170521027 | 0 | T | G | 0.482 | 5.59E-02 | 7.73E-02 | 2.48E+00 | 2.55E+00 | 4.83E-01 | no | Robo2 | intron_variant | MODIFIER |
| 16 | 74059455 | AX-170013380 | 0 | A | G | 0.482 | 5.59E-02 | 7.73E-02 | 2.48E+00 | 2.55E+00 | 4.83E-01 | no | Robo2 | intron_variant | MODIFIER |
| 16 | 74063489 | AX-170013383 | 0 | T | G | 0.482 | 5.59E-02 | 7.73E-02 | 2.48E+00 | 2.55E+00 | 4.83E-01 | no | Robo2 | intron_variant | MODIFIER |
| 16 | 74081392 | AX-170272650 | 0 | T | C | 0.482 | 5.59E-02 | 7.73E-02 | 2.48E+00 | 2.55E+00 | 4.83E-01 | no | Robo2 | intron_variant | MODIFIER |
| 16 | 74084707 | AX-170013379 | 0 | A | G | 0.482 | 5.59E-02 | 7.73E-02 | 2.48E+00 | 2.55E+00 | 4.83E-01 | no | Robo2 | intron_variant | MODIFIER |
| 16 | 74099769 | AX-170013378 | 0 | T | C | 0.482 | 5.59E-02 | 7.73E-02 | 2.48E+00 | 2.55E+00 | 4.83E-01 | no | Robo2 | intron_variant | MODIFIER |
| 16 | 74113478 | AX-97842836 | 0 | G | A | 0.482 | 5.59E-02 | 7.73E-02 | 2.48E+00 | 2.55E+00 | 4.83E-01 | no | Robo2 | intron_variant | MODIFIER |
| 16 | 74123743 | AX-170013381 | 0 | A | T | 0.482 | 5.59E-02 | 7.73E-02 | 2.48E+00 | 2.55E+00 | 4.83E-01 | no | Robo2 | intron_variant | MODIFIER |
| 16 | 74139825 | AX-170013385 | 0 | C | A | 0.482 | 5.59E-02 | 7.73E-02 | 2.48E+00 | 2.55E+00 | 4.83E-01 | no | Robo2 | intron_variant | MODIFIER |

| | | | | | | | | | | | | | | | |
|---|---|---|---|---|---|---|---|---|---|---|---|---|---|---|---|
| 16 | 74148498 | AX-170192400 | 0 | C | G | 0.482 | 5.59E-02 | 7.73E-02 | 2.48E+00 | 2.55E+00 | 4.83E-01 | no | Robo2 | intron_variant | MODIFIER |
| 16 | 74149339 | AX-97813497 | 0 | C | A | 0.482 | 5.59E-02 | 7.73E-02 | 2.48E+00 | 2.55E+00 | 4.83E-01 | no | Robo2 | intron_variant | MODIFIER |
| 16 | 74168064 | AX-170477458 | 0 | C | G | 0.482 | 5.59E-02 | 7.73E-02 | 2.48E+00 | 2.55E+00 | 4.83E-01 | no | Robo2 | intron_variant | MODIFIER |
| 16 | 74168111 | AX-170269374 | 0 | G | A | 0.482 | 5.59E-02 | 7.73E-02 | 2.48E+00 | 2.55E+00 | 4.83E-01 | no | Robo2 | intron_variant | MODIFIER |
| 16 | 74168560 | AX-170518965 | 0 | T | G | 0.482 | 5.59E-02 | 7.73E-02 | 2.48E+00 | 2.55E+00 | 4.83E-01 | no | Robo2 | intron_variant | MODIFIER |
| 16 | 74173364 | AX-170013382 | 0 | C | T | 0.482 | 5.59E-02 | 7.73E-02 | 2.48E+00 | 2.55E+00 | 4.83E-01 | no | Robo2 | intron_variant | MODIFIER |
| 16 | 74178191 | AX-170395475 | 0 | C | T | 0.482 | 5.59E-02 | 7.73E-02 | 2.48E+00 | 2.55E+00 | 4.83E-01 | no | Robo2 | intron_variant | MODIFIER |
| 15 | 75314108 | AX-170410427 | 1 | G | A | 0.562 | 6.00E-02 | 8.42E-02 | 2.53E+00 | 2.60E+00 | 4.83E-01 | no | Gm3454 | intron_variant,non_coding_transcript_variant | MODIFIER |
| 1 | 136942816 | AX-170042856 | 1 | A | C | 0.214 | 8.14E-02 | 1.17E-01 | 2.62E+00 | 2.71E+00 | 4.83E-01 | no | Nr5a2 | intron_variant | MODIFIER |
| 5 | 40128004 | AX-97838892 | 1 | G | T | 0.205 | -7.46E-02 | 1.06E-01 | 2.58E+00 | 2.66E+00 | 4.83E-01 | no | | intergenic_variant | MODIFIER |
| 1 | 189126296 | AX-170107032 | 1 | C | A | 0.25 | 7.38E-02 | 1.02E-01 | 2.45E+00 | 2.54E+00 | 4.83E-01 | no | | intergenic_variant | MODIFIER |
| 4 | 71617348 | AX-170189737 | 2 | C | A | 0.236 | 7.33E-02 | 1.03E-01 | 2.51E+00 | 2.59E+00 | 4.83E-01 | no | | intergenic_variant | MODIFIER |
| 1 | 39301899 | AX-170283836 | 0 | T | C | 0.404 | 5.33E-02 | 7.72E-02 | 2.78E+00 | 2.88E+00 | 4.83E-01 | no | Npas2 | intron_variant | MODIFIER |
| 1 | 190007213 | AX-170282961 | 0 | A | G | 0.246 | 7.09E-02 | 9.74E-02 | 2.45E+00 | 2.53E+00 | 4.83E-01 | no | | regulatory_region_variant | MODIFIER |
| 1 | 190011355 | AX-170113188 | 0 | G | T | 0.246 | 7.09E-02 | 9.74E-02 | 2.45E+00 | 2.53E+00 | 4.83E-01 | no | | intergenic_variant | MODIFIER |
| 1 | 190019140 | AX-170118462 | 0 | C | A | 0.246 | 7.09E-02 | 9.74E-02 | 2.45E+00 | 2.53E+00 | 4.83E-01 | no | | intergenic_variant | MODIFIER |
| 1 | 190040562 | AX-97814549 | 0 | T | G | 0.246 | 7.09E-02 | 9.74E-02 | 2.45E+00 | 2.53E+00 | 4.83E-01 | no | | intergenic_variant | MODIFIER |
| 1 | 190040696 | AX-170002546 | 0 | T | C | 0.246 | 7.09E-02 | 9.74E-02 | 2.45E+00 | 2.53E+00 | 4.83E-01 | no | | regulatory_region_variant | MODIFIER |
| 1 | 190043257 | AX-170002553 | 0 | C | A | 0.246 | 7.09E-02 | 9.74E-02 | 2.45E+00 | 2.53E+00 | 4.83E-01 | no | | intergenic_variant | MODIFIER |
| 15 | 4016885 | AX-97828272 | 0 | G | A | 0.096 | -7.39E-02 | 9.06E-02 | 2.21E+00 | 2.34E+00 | 4.83E-01 | no | BC037032 | downstream_gene_variant | MODIFIER |
| 6 | 131053189 | AX-170192344 | 1 | T | C | 0.536 | 5.82E-02 | 8.19E-02 | 2.52E+00 | 2.60E+00 | 4.83E-01 | no | Gm44134 | upstream_gene_variant | MODIFIER |
| 9 | 22591970 | AX-97788574 | 0 | G | T | 0.456 | 6.16E-02 | 8.74E-02 | 2.56E+00 | 2.63E+00 | 4.83E-01 | no | Bbs9 | intron_variant | MODIFIER |
| 7 | 144407532 | AX-170030170 | 2 | T | A | 0.218 | -5.22E-02 | 7.54E-02 | 2.62E+00 | 2.75E+00 | 4.84E-01 | no | Shank2 | intron_variant | MODIFIER |
| 5 | 110581780 | AX-170323573 | 1 | C | T | 0.464 | 7.81E-02 | 1.12E-01 | 2.64E+00 | 2.68E+00 | 4.84E-01 | no | Galnt9 | intron_variant | MODIFIER |
| 5 | 112796080 | AX-170138818 | 1 | C | T | 0.464 | 7.81E-02 | 1.12E-01 | 2.64E+00 | 2.68E+00 | 4.84E-01 | no | Myo18b | intron_variant | MODIFIER |
| 17 | 36291030 | AX-170204155 | 0 | T | G | 0.044 | -1.20E-01 | 1.74E-01 | 2.63E+00 | 2.74E+00 | 4.84E-01 | no | H2-M10.2 | upstream_gene_variant | MODIFIER |
| 1 | 169074553 | AX-170002260 | 0 | G | A | 0.211 | 9.82E-02 | 1.41E-01 | 2.66E+00 | 2.69E+00 | 4.84E-01 | no | | intergenic_variant | MODIFIER |
| 1 | 169077767 | AX-170448833 | 0 | C | T | 0.211 | 9.82E-02 | 1.41E-01 | 2.66E+00 | 2.69E+00 | 4.84E-01 | no | | intergenic_variant | MODIFIER |
| 1 | 169094484 | AX-170213522 | 0 | G | A | 0.211 | 9.82E-02 | 1.41E-01 | 2.66E+00 | 2.69E+00 | 4.84E-01 | no | | intergenic_variant | MODIFIER |
| 1 | 169097616 | AX-170302051 | 0 | C | A | 0.211 | 9.82E-02 | 1.41E-01 | 2.66E+00 | 2.69E+00 | 4.84E-01 | no | | intergenic_variant | MODIFIER |
| 1 | 169115473 | AX-170002261 | 0 | C | T | 0.211 | 9.82E-02 | 1.41E-01 | 2.66E+00 | 2.69E+00 | 4.84E-01 | no | | intergenic_variant | MODIFIER |
| 1 | 169115528 | AX-170391403 | 0 | C | T | 0.211 | 9.82E-02 | 1.41E-01 | 2.66E+00 | 2.69E+00 | 4.84E-01 | no | | intergenic_variant | MODIFIER |
| 1 | 169118157 | AX-170169284 | 0 | A | T | 0.211 | 9.82E-02 | 1.41E-01 | 2.66E+00 | 2.69E+00 | 4.84E-01 | no | | intergenic_variant | MODIFIER |
| 1 | 169127224 | AX-170348746 | 0 | T | C | 0.211 | 9.82E-02 | 1.41E-01 | 2.66E+00 | 2.69E+00 | 4.84E-01 | no | | intergenic_variant | MODIFIER |
| 1 | 169145562 | AX-170374068 | 0 | A | C | 0.211 | 9.82E-02 | 1.41E-01 | 2.66E+00 | 2.69E+00 | 4.84E-01 | no | | intergenic_variant | MODIFIER |
| 1 | 169182054 | AX-170498165 | 0 | C | T | 0.211 | 9.82E-02 | 1.41E-01 | 2.66E+00 | 2.69E+00 | 4.84E-01 | no | | intergenic_variant | MODIFIER |
| 1 | 169211115 | AX-170394971 | 0 | A | G | 0.211 | 9.82E-02 | 1.41E-01 | 2.66E+00 | 2.69E+00 | 4.84E-01 | no | | intergenic_variant | MODIFIER |
| 1 | 169227328 | AX-170534741 | 0 | T | C | 0.211 | 9.82E-02 | 1.41E-01 | 2.66E+00 | 2.69E+00 | 4.84E-01 | no | | intergenic_variant | MODIFIER |
| 1 | 169241850 | AX-170116979 | 0 | C | T | 0.211 | 9.82E-02 | 1.41E-01 | 2.66E+00 | 2.69E+00 | 4.84E-01 | no | | intergenic_variant | MODIFIER |
| 1 | 169241886 | AX-170043507 | 0 | G | A | 0.211 | 9.82E-02 | 1.41E-01 | 2.66E+00 | 2.69E+00 | 4.84E-01 | no | | intergenic_variant | MODIFIER |
| 1 | 169242289 | AX-170043504 | 0 | A | G | 0.211 | 9.82E-02 | 1.41E-01 | 2.66E+00 | 2.69E+00 | 4.84E-01 | no | | intergenic_variant | MODIFIER |
| 1 | 169246356 | AX-170043506 | 0 | T | C | 0.211 | 9.82E-02 | 1.41E-01 | 2.66E+00 | 2.69E+00 | 4.84E-01 | no | | intergenic_variant | MODIFIER |
| 1 | 169247225 | AX-170259022 | 0 | G | A | 0.211 | 9.82E-02 | 1.41E-01 | 2.66E+00 | 2.69E+00 | 4.84E-01 | no | | regulatory_region_variant | MODIFIER |
| 1 | 169250939 | AX-170279286 | 0 | C | T | 0.211 | 9.82E-02 | 1.41E-01 | 2.66E+00 | 2.69E+00 | 4.84E-01 | no | | intergenic_variant | MODIFIER |
| 1 | 169252756 | AX-170167661 | 0 | T | G | 0.211 | 9.82E-02 | 1.41E-01 | 2.66E+00 | 2.69E+00 | 4.84E-01 | no | | intergenic_variant | MODIFIER |
| 1 | 169278986 | AX-170185210 | 0 | C | T | 0.211 | 9.82E-02 | 1.41E-01 | 2.66E+00 | 2.69E+00 | 4.84E-01 | no | | intergenic_variant | MODIFIER |
| 1 | 169284168 | AX-170161390 | 0 | A | G | 0.211 | 9.82E-02 | 1.41E-01 | 2.66E+00 | 2.69E+00 | 4.84E-01 | no | | intergenic_variant | MODIFIER |
| 1 | 169299570 | AX-170176436 | 0 | C | T | 0.211 | 9.82E-02 | 1.41E-01 | 2.66E+00 | 2.69E+00 | 4.84E-01 | no | | intergenic_variant | MODIFIER |
| 1 | 169305175 | AX-170379523 | 0 | G | A | 0.211 | 9.82E-02 | 1.41E-01 | 2.66E+00 | 2.69E+00 | 4.84E-01 | no | | intergenic_variant | MODIFIER |
| 1 | 169348604 | AX-97848363 | 0 | C | A | 0.211 | 9.82E-02 | 1.41E-01 | 2.66E+00 | 2.69E+00 | 4.84E-01 | no | Gm37839 | upstream_gene_variant | MODIFIER |
| 1 | 169352900 | AX-170219830 | 0 | T | G | 0.211 | 9.82E-02 | 1.41E-01 | 2.66E+00 | 2.69E+00 | 4.84E-01 | no | | intergenic_variant | MODIFIER |
| 1 | 169358271 | AX-170328656 | 0 | T | C | 0.211 | 9.82E-02 | 1.41E-01 | 2.66E+00 | 2.69E+00 | 4.84E-01 | no | | regulatory_region_variant | MODIFIER |
| 1 | 169358550 | AX-170513074 | 0 | G | A | 0.211 | 9.82E-02 | 1.41E-01 | 2.66E+00 | 2.69E+00 | 4.84E-01 | no | | intergenic_variant | MODIFIER |
| 1 | 169360266 | AX-170286480 | 0 | C | G | 0.211 | 9.82E-02 | 1.41E-01 | 2.66E+00 | 2.69E+00 | 4.84E-01 | no | | intergenic_variant | MODIFIER |

| | | | | | | | | | | | | | | | |
|---|---|---|---|---|---|---|---|---|---|---|---|---|---|---|---|
| 1 | 169360937 | AX-170043509 | 0 | T | C | 0.211 | 9.82E-02 | 1.41E-01 | 2.66E+00 | 2.69E+00 | 4.84E-01 | no | | intergenic_variant | MODIFIER |
| 1 | 169362113 | AX-170354855 | 0 | A | C | 0.211 | 9.82E-02 | 1.41E-01 | 2.66E+00 | 2.69E+00 | 4.84E-01 | no | | intergenic_variant | MODIFIER |
| 1 | 169371536 | AX-170436563 | 0 | C | T | 0.211 | 9.82E-02 | 1.41E-01 | 2.66E+00 | 2.69E+00 | 4.84E-01 | no | | intergenic_variant | MODIFIER |
| 1 | 169372182 | AX-170527995 | 0 | A | G | 0.211 | 9.82E-02 | 1.41E-01 | 2.66E+00 | 2.69E+00 | 4.84E-01 | no | | intergenic_variant | MODIFIER |
| 1 | 169373474 | AX-170252506 | 0 | C | T | 0.211 | 9.82E-02 | 1.41E-01 | 2.66E+00 | 2.69E+00 | 4.84E-01 | no | | intergenic_variant | MODIFIER |
| 1 | 169392512 | AX-170231519 | 0 | C | G | 0.211 | 9.82E-02 | 1.41E-01 | 2.66E+00 | 2.69E+00 | 4.84E-01 | no | | intergenic_variant | MODIFIER |
| 1 | 169440274 | AX-170332270 | 0 | A | G | 0.211 | 9.82E-02 | 1.41E-01 | 2.66E+00 | 2.69E+00 | 4.84E-01 | no | | regulatory_region_variant | MODIFIER |
| 1 | 169440707 | AX-170173768 | 0 | A | G | 0.211 | 9.82E-02 | 1.41E-01 | 2.66E+00 | 2.69E+00 | 4.84E-01 | no | | intergenic_variant | MODIFIER |
| 1 | 169448158 | AX-170153110 | 0 | G | C | 0.211 | 9.82E-02 | 1.41E-01 | 2.66E+00 | 2.69E+00 | 4.84E-01 | no | | intergenic_variant | MODIFIER |
| 1 | 169460775 | AX-170154340 | 0 | A | G | 0.211 | 9.82E-02 | 1.41E-01 | 2.66E+00 | 2.69E+00 | 4.84E-01 | no | | intergenic_variant | MODIFIER |
| 1 | 169467373 | AX-170201280 | 0 | G | A | 0.211 | 9.82E-02 | 1.41E-01 | 2.66E+00 | 2.69E+00 | 4.84E-01 | no | | intergenic_variant | MODIFIER |
| 1 | 169469811 | AX-170043511 | 0 | C | T | 0.211 | 9.82E-02 | 1.41E-01 | 2.66E+00 | 2.69E+00 | 4.84E-01 | no | | intergenic_variant | MODIFIER |
| 1 | 169475182 | AX-170366759 | 0 | A | G | 0.211 | 9.82E-02 | 1.41E-01 | 2.66E+00 | 2.69E+00 | 4.84E-01 | no | | intergenic_variant | MODIFIER |
| 1 | 169485268 | AX-170180368 | 0 | T | C | 0.211 | 9.82E-02 | 1.41E-01 | 2.66E+00 | 2.69E+00 | 4.84E-01 | no | | regulatory_region_variant | MODIFIER |
| 1 | 169485389 | AX-170241397 | 0 | T | C | 0.211 | 9.82E-02 | 1.41E-01 | 2.66E+00 | 2.69E+00 | 4.84E-01 | no | | regulatory_region_variant | MODIFIER |
| 1 | 169485489 | AX-170366930 | 0 | C | T | 0.211 | 9.82E-02 | 1.41E-01 | 2.66E+00 | 2.69E+00 | 4.84E-01 | no | | regulatory_region_variant | MODIFIER |
| 1 | 169491612 | AX-170446768 | 0 | G | A | 0.211 | 9.82E-02 | 1.41E-01 | 2.66E+00 | 2.69E+00 | 4.84E-01 | no | | intergenic_variant | MODIFIER |
| 1 | 169492280 | AX-170289372 | 0 | C | T | 0.211 | 9.82E-02 | 1.41E-01 | 2.66E+00 | 2.69E+00 | 4.84E-01 | no | | intergenic_variant | MODIFIER |
| 1 | 169497022 | AX-170358160 | 0 | C | T | 0.211 | 9.82E-02 | 1.41E-01 | 2.66E+00 | 2.69E+00 | 4.84E-01 | no | Nuf2 | downstream_gene_variant | MODIFIER |
| 1 | 169528459 | AX-170394853 | 0 | T | C | 0.211 | 9.82E-02 | 1.41E-01 | 2.66E+00 | 2.69E+00 | 4.84E-01 | no | Nuf2 | intron_variant | MODIFIER |
| 1 | 169537908 | AX-170002263 | 0 | C | G | 0.211 | 9.82E-02 | 1.41E-01 | 2.66E+00 | 2.69E+00 | 4.84E-01 | no | | intergenic_variant | MODIFIER |
| 1 | 169573892 | AX-170342044 | 0 | A | G | 0.211 | 9.82E-02 | 1.41E-01 | 2.66E+00 | 2.69E+00 | 4.84E-01 | no | | intergenic_variant | MODIFIER |
| 1 | 169574673 | AX-170471516 | 0 | C | A | 0.211 | 9.82E-02 | 1.41E-01 | 2.66E+00 | 2.69E+00 | 4.84E-01 | no | | intergenic_variant | MODIFIER |
| 1 | 169574811 | AX-170043515 | 0 | C | T | 0.211 | 9.82E-02 | 1.41E-01 | 2.66E+00 | 2.69E+00 | 4.84E-01 | no | | intergenic_variant | MODIFIER |
| 1 | 169574868 | AX-170043514 | 0 | C | G | 0.211 | 9.82E-02 | 1.41E-01 | 2.66E+00 | 2.69E+00 | 4.84E-01 | no | | intergenic_variant | MODIFIER |
| 1 | 169577160 | AX-170043516 | 0 | T | C | 0.211 | 9.82E-02 | 1.41E-01 | 2.66E+00 | 2.69E+00 | 4.84E-01 | no | | intergenic_variant | MODIFIER |
| 1 | 169577184 | AX-170043517 | 0 | A | G | 0.211 | 9.82E-02 | 1.41E-01 | 2.66E+00 | 2.69E+00 | 4.84E-01 | no | | intergenic_variant | MODIFIER |
| 1 | 169582904 | AX-170462820 | 0 | G | C | 0.211 | 9.82E-02 | 1.41E-01 | 2.66E+00 | 2.69E+00 | 4.84E-01 | no | | intergenic_variant | MODIFIER |
| 1 | 169585991 | AX-170333791 | 0 | G | A | 0.211 | 9.82E-02 | 1.41E-01 | 2.66E+00 | 2.69E+00 | 4.84E-01 | no | | intergenic_variant | MODIFIER |
| 1 | 169586241 | AX-170284490 | 0 | G | A | 0.211 | 9.82E-02 | 1.41E-01 | 2.66E+00 | 2.69E+00 | 4.84E-01 | no | | intergenic_variant | MODIFIER |
| 1 | 169619920 | AX-170190986 | 0 | G | T | 0.211 | 9.82E-02 | 1.41E-01 | 2.66E+00 | 2.69E+00 | 4.84E-01 | no | | intergenic_variant | MODIFIER |
| 1 | 169644680 | AX-170520247 | 0 | A | G | 0.211 | 9.82E-02 | 1.41E-01 | 2.66E+00 | 2.69E+00 | 4.84E-01 | no | | intergenic_variant | MODIFIER |
| 1 | 169660811 | AX-170495394 | 0 | T | C | 0.211 | 9.82E-02 | 1.41E-01 | 2.66E+00 | 2.69E+00 | 4.84E-01 | no | Rgs5 | intron_variant | MODIFIER |
| 1 | 169671015 | AX-170157488 | 0 | T | A | 0.211 | 9.82E-02 | 1.41E-01 | 2.66E+00 | 2.69E+00 | 4.84E-01 | no | Rgs5 | intron_variant | MODIFIER |
| 1 | 169681202 | AX-170430171 | 0 | A | G | 0.211 | 9.82E-02 | 1.41E-01 | 2.66E+00 | 2.69E+00 | 4.84E-01 | no | Rgs5 | intron_variant | MODIFIER |
| 1 | 169687540 | AX-170394143 | 0 | C | T | 0.211 | 9.82E-02 | 1.41E-01 | 2.66E+00 | 2.69E+00 | 4.84E-01 | no | Rgs5 | intron_variant | MODIFIER |
| 1 | 169689523 | AX-170453045 | 0 | A | G | 0.211 | 9.82E-02 | 1.41E-01 | 2.66E+00 | 2.69E+00 | 4.84E-01 | no | Rgs5 | intron_variant | MODIFIER |
| 1 | 169702964 | AX-170311807 | 0 | C | T | 0.211 | 9.82E-02 | 1.41E-01 | 2.66E+00 | 2.69E+00 | 4.84E-01 | no | | intergenic_variant | MODIFIER |
| 1 | 169703017 | AX-170043520 | 0 | A | G | 0.211 | 9.82E-02 | 1.41E-01 | 2.66E+00 | 2.69E+00 | 4.84E-01 | no | | intergenic_variant | MODIFIER |
| 1 | 169703278 | AX-170043519 | 0 | A | G | 0.211 | 9.82E-02 | 1.41E-01 | 2.66E+00 | 2.69E+00 | 4.84E-01 | no | | regulatory_region_variant | MODIFIER |
| 1 | 169717355 | AX-170155230 | 0 | T | C | 0.211 | 9.82E-02 | 1.41E-01 | 2.66E+00 | 2.69E+00 | 4.84E-01 | no | | intergenic_variant | MODIFIER |
| 1 | 169735589 | AX-170205929 | 0 | T | C | 0.211 | 9.82E-02 | 1.41E-01 | 2.66E+00 | 2.69E+00 | 4.84E-01 | no | | intergenic_variant | MODIFIER |
| 1 | 169741731 | AX-170456708 | 0 | A | G | 0.211 | 9.82E-02 | 1.41E-01 | 2.66E+00 | 2.69E+00 | 4.84E-01 | no | Rgs4 | 3_prime_UTR_variant | MODIFIER |
| 1 | 169747820 | AX-170201754 | 0 | A | G | 0.211 | 9.82E-02 | 1.41E-01 | 2.66E+00 | 2.69E+00 | 4.84E-01 | no | Rgs4 | upstream_gene_variant | MODIFIER |
| 1 | 169782994 | AX-170452720 | 0 | A | G | 0.211 | 9.82E-02 | 1.41E-01 | 2.66E+00 | 2.69E+00 | 4.84E-01 | no | | intergenic_variant | MODIFIER |
| 1 | 169802466 | AX-97812959 | 0 | C | A | 0.211 | 9.82E-02 | 1.41E-01 | 2.66E+00 | 2.69E+00 | 4.84E-01 | no | Gm22114 | upstream_gene_variant | MODIFIER |
| 1 | 169842909 | AX-170043522 | 0 | T | C | 0.211 | 9.82E-02 | 1.41E-01 | 2.66E+00 | 2.69E+00 | 4.84E-01 | no | | intergenic_variant | MODIFIER |
| 1 | 169847098 | AX-170143313 | 0 | G | A | 0.211 | 9.82E-02 | 1.41E-01 | 2.66E+00 | 2.69E+00 | 4.84E-01 | no | | intergenic_variant | MODIFIER |
| 1 | 169882577 | AX-170276508 | 0 | T | C | 0.211 | 9.82E-02 | 1.41E-01 | 2.66E+00 | 2.69E+00 | 4.84E-01 | no | | intergenic_variant | MODIFIER |
| 1 | 169898306 | AX-170153352 | 0 | A | C | 0.211 | 9.82E-02 | 1.41E-01 | 2.66E+00 | 2.69E+00 | 4.84E-01 | no | | intergenic_variant | MODIFIER |
| 1 | 169909144 | AX-170323685 | 0 | C | T | 0.211 | 9.82E-02 | 1.41E-01 | 2.66E+00 | 2.69E+00 | 4.84E-01 | no | | intergenic_variant | MODIFIER |
| 1 | 169909328 | AX-170439627 | 0 | T | C | 0.211 | 9.82E-02 | 1.41E-01 | 2.66E+00 | 2.69E+00 | 4.84E-01 | no | | intergenic_variant | MODIFIER |
| 1 | 169920176 | AX-170113100 | 0 | T | C | 0.211 | 9.82E-02 | 1.41E-01 | 2.66E+00 | 2.69E+00 | 4.84E-01 | no | | intergenic_variant | MODIFIER |
| 1 | 169920877 | AX-170531919 | 0 | A | G | 0.211 | 9.82E-02 | 1.41E-01 | 2.66E+00 | 2.69E+00 | 4.84E-01 | no | | intergenic_variant | MODIFIER |

| | | | | | | | | | | | | | | | |
|---|---|---|---|---|---|---|---|---|---|---|---|---|---|---|---|
| 1 | 169927061 | AX-170140023 | 0 | C | A | 0.211 | 9.82E-02 | 1.41E-01 | 2.66E+00 | 2.69E+00 | 4.84E-01 | no | Ccdc190 | upstream_gene_variant | MODIFIER |
| 1 | 169970578 | AX-170241850 | 0 | G | A | 0.211 | 9.82E-02 | 1.41E-01 | 2.66E+00 | 2.69E+00 | 4.84E-01 | no | Hsd17b7 | upstream_gene_variant | MODIFIER |
| 1 | 169975556 | AX-170153408 | 0 | T | C | 0.211 | 9.82E-02 | 1.41E-01 | 2.66E+00 | 2.69E+00 | 4.84E-01 | no | Ddr2 | downstream_gene_variant | MODIFIER |
| 1 | 169975619 | AX-170043529 | 0 | G | A | 0.211 | 9.82E-02 | 1.41E-01 | 2.66E+00 | 2.69E+00 | 4.84E-01 | no | Ddr2 | downstream_gene_variant | MODIFIER |
| 1 | 169975814 | AX-170469385 | 0 | C | G | 0.211 | 9.82E-02 | 1.41E-01 | 2.66E+00 | 2.69E+00 | 4.84E-01 | no | Ddr2 | downstream_gene_variant | MODIFIER |
| 1 | 169990266 | AX-170270500 | 0 | A | G | 0.211 | 9.82E-02 | 1.41E-01 | 2.66E+00 | 2.69E+00 | 4.84E-01 | no | Ddr2 | intron_variant | MODIFIER |
| 1 | 169995053 | AX-97853521 | 0 | T | C | 0.211 | 9.82E-02 | 1.41E-01 | 2.66E+00 | 2.69E+00 | 4.84E-01 | no | Ddr2 | intron_variant | MODIFIER |
| 1 | 170004432 | AX-170230811 | 0 | T | G | 0.211 | 9.82E-02 | 1.41E-01 | 2.66E+00 | 2.69E+00 | 4.84E-01 | no | Ddr2 | intron_variant | MODIFIER |
| 1 | 170014642 | AX-170200925 | 0 | G | A | 0.211 | 9.82E-02 | 1.41E-01 | 2.66E+00 | 2.69E+00 | 4.84E-01 | no | Ddr2 | intron_variant | MODIFIER |
| 1 | 170032969 | AX-170425505 | 0 | C | T | 0.211 | 9.82E-02 | 1.41E-01 | 2.66E+00 | 2.69E+00 | 4.84E-01 | no | Ddr2 | intron_variant | MODIFIER |
| 1 | 170035236 | AX-170257797 | 0 | A | G | 0.211 | 9.82E-02 | 1.41E-01 | 2.66E+00 | 2.69E+00 | 4.84E-01 | no | Ddr2 | intron_variant | MODIFIER |
| 1 | 170061540 | AX-170198677 | 0 | A | G | 0.211 | 9.82E-02 | 1.41E-01 | 2.66E+00 | 2.69E+00 | 4.84E-01 | no | Ddr2 | intron_variant | MODIFIER |
| 1 | 170061797 | AX-170361307 | 0 | T | C | 0.211 | 9.82E-02 | 1.41E-01 | 2.66E+00 | 2.69E+00 | 4.84E-01 | no | Ddr2 | intron_variant | MODIFIER |
| 1 | 170079677 | AX-170293442 | 0 | C | T | 0.211 | 9.82E-02 | 1.41E-01 | 2.66E+00 | 2.69E+00 | 4.84E-01 | no | Ddr2 | intron_variant | MODIFIER |
| 1 | 170079985 | AX-170179088 | 0 | G | A | 0.211 | 9.82E-02 | 1.41E-01 | 2.66E+00 | 2.69E+00 | 4.84E-01 | no | Ddr2 | intron_variant | MODIFIER |
| 1 | 170106714 | AX-170002264 | 0 | T | C | 0.211 | 9.82E-02 | 1.41E-01 | 2.66E+00 | 2.69E+00 | 4.84E-01 | no | Ddr2 | intron_variant | MODIFIER |
| 1 | 170107062 | AX-97850541 | 0 | G | A | 0.211 | 9.82E-02 | 1.41E-01 | 2.66E+00 | 2.69E+00 | 4.84E-01 | no | Ddr2 | intron_variant | MODIFIER |
| 1 | 170109618 | AX-170258797 | 0 | T | C | 0.211 | 9.82E-02 | 1.41E-01 | 2.66E+00 | 2.69E+00 | 4.84E-01 | no | Ddr2 | intron_variant | MODIFIER |
| 1 | 170123339 | AX-170358878 | 0 | G | A | 0.211 | 9.82E-02 | 1.41E-01 | 2.66E+00 | 2.69E+00 | 4.84E-01 | no | | intergenic_variant | MODIFIER |
| 1 | 170124060 | AX-170235221 | 0 | A | C | 0.211 | 9.82E-02 | 1.41E-01 | 2.66E+00 | 2.69E+00 | 4.84E-01 | no | | intergenic_variant | MODIFIER |
| 1 | 170124316 | AX-170465965 | 0 | C | T | 0.211 | 9.82E-02 | 1.41E-01 | 2.66E+00 | 2.69E+00 | 4.84E-01 | no | | intergenic_variant | MODIFIER |
| 1 | 170127297 | AX-170249192 | 0 | T | C | 0.211 | 9.82E-02 | 1.41E-01 | 2.66E+00 | 2.69E+00 | 4.84E-01 | no | | intergenic_variant | MODIFIER |
| 1 | 170128044 | AX-170002267 | 0 | T | C | 0.211 | 9.82E-02 | 1.41E-01 | 2.66E+00 | 2.69E+00 | 4.84E-01 | no | | intergenic_variant | MODIFIER |
| 1 | 170128129 | AX-170289318 | 0 | T | C | 0.211 | 9.82E-02 | 1.41E-01 | 2.66E+00 | 2.69E+00 | 4.84E-01 | no | | intergenic_variant | MODIFIER |
| 1 | 170136223 | AX-170043538 | 0 | G | A | 0.211 | 9.82E-02 | 1.41E-01 | 2.66E+00 | 2.69E+00 | 4.84E-01 | no | | regulatory_region_variant | MODIFIER |
| 1 | 170172020 | AX-170043539 | 0 | G | A | 0.211 | 9.82E-02 | 1.41E-01 | 2.66E+00 | 2.69E+00 | 4.84E-01 | no | Uap1 | intron_variant | MODIFIER |
| 1 | 170188716 | AX-170356200 | 0 | C | T | 0.211 | 9.82E-02 | 1.41E-01 | 2.66E+00 | 2.69E+00 | 4.84E-01 | no | Uhmk1 | downstream_gene_variant | MODIFIER |
| 1 | 170190320 | AX-170433096 | 0 | T | C | 0.211 | 9.82E-02 | 1.41E-01 | 2.66E+00 | 2.69E+00 | 4.84E-01 | no | Uhmk1 | downstream_gene_variant | MODIFIER |
| 1 | 170195712 | AX-170128104 | 0 | A | G | 0.211 | 9.82E-02 | 1.41E-01 | 2.66E+00 | 2.69E+00 | 4.84E-01 | no | Uhmk1 | 3_prime_UTR_variant | MODIFIER |
| 1 | 170202882 | AX-170111192 | 0 | C | G | 0.211 | 9.82E-02 | 1.41E-01 | 2.66E+00 | 2.69E+00 | 4.84E-01 | no | Uhmk1 | intron_variant | MODIFIER |
| 1 | 170208544 | AX-170351722 | 0 | A | G | 0.211 | 9.82E-02 | 1.41E-01 | 2.66E+00 | 2.69E+00 | 4.84E-01 | no | Uhmk1 | intron_variant | MODIFIER |
| 1 | 170208614 | AX-170387839 | 0 | G | C | 0.211 | 9.82E-02 | 1.41E-01 | 2.66E+00 | 2.69E+00 | 4.84E-01 | no | Uhmk1 | intron_variant | MODIFIER |
| 1 | 170252747 | AX-170419454 | 0 | T | C | 0.211 | 9.82E-02 | 1.41E-01 | 2.66E+00 | 2.69E+00 | 4.84E-01 | no | Sh2d1b2 | downstream_gene_variant | MODIFIER |
| 1 | 170252930 | AX-170367242 | 0 | T | C | 0.211 | 9.82E-02 | 1.41E-01 | 2.66E+00 | 2.69E+00 | 4.84E-01 | no | Sh2d1b2 | downstream_gene_variant | MODIFIER |
| 1 | 170283815 | AX-170501245 | 0 | T | C | 0.211 | 9.82E-02 | 1.41E-01 | 2.66E+00 | 2.69E+00 | 4.84E-01 | no | Sh2d1b1 | intron_variant | MODIFIER |
| 1 | 170293148 | AX-170203163 | 0 | G | A | 0.211 | 9.82E-02 | 1.41E-01 | 2.66E+00 | 2.69E+00 | 4.84E-01 | no | | regulatory_region_variant | MODIFIER |
| 1 | 170293517 | AX-170139426 | 0 | T | G | 0.211 | 9.82E-02 | 1.41E-01 | 2.66E+00 | 2.69E+00 | 4.84E-01 | no | Gm7694 | downstream_gene_variant | MODIFIER |
| 1 | 170294864 | AX-170444538 | 0 | T | G | 0.211 | 9.82E-02 | 1.41E-01 | 2.66E+00 | 2.69E+00 | 4.84E-01 | no | Gm7694 | downstream_gene_variant | MODIFIER |
| 1 | 170300332 | AX-170274927 | 0 | A | G | 0.211 | 9.82E-02 | 1.41E-01 | 2.66E+00 | 2.69E+00 | 4.84E-01 | no | Nos1ap | downstream_gene_variant | MODIFIER |
| 1 | 170300785 | AX-170376994 | 0 | C | T | 0.211 | 9.82E-02 | 1.41E-01 | 2.66E+00 | 2.69E+00 | 4.84E-01 | no | Nos1ap | downstream_gene_variant | MODIFIER |
| 1 | 170302591 | AX-170371826 | 0 | T | C | 0.211 | 9.82E-02 | 1.41E-01 | 2.66E+00 | 2.69E+00 | 4.84E-01 | no | Nos1ap | downstream_gene_variant | MODIFIER |
| 1 | 170306377 | AX-170333744 | 0 | G | A | 0.211 | 9.82E-02 | 1.41E-01 | 2.66E+00 | 2.69E+00 | 4.84E-01 | no | Spata46 | upstream_gene_variant | MODIFIER |
| 1 | 170306485 | AX-170116048 | 0 | C | T | 0.211 | 9.82E-02 | 1.41E-01 | 2.66E+00 | 2.69E+00 | 4.84E-01 | no | Spata46 | upstream_gene_variant | MODIFIER |
| 1 | 170311169 | AX-170514008 | 0 | C | G | 0.211 | 9.82E-02 | 1.41E-01 | 2.66E+00 | 2.69E+00 | 4.84E-01 | no | Spata46 | intron_variant | MODIFIER |
| 1 | 170312693 | AX-170318922 | 0 | T | A | 0.211 | 9.82E-02 | 1.41E-01 | 2.66E+00 | 2.69E+00 | 4.84E-01 | no | Spata46 | downstream_gene_variant | MODIFIER |
| 1 | 170313069 | AX-170097373 | 0 | A | G | 0.211 | 9.82E-02 | 1.41E-01 | 2.66E+00 | 2.69E+00 | 4.84E-01 | no | Spata46 | downstream_gene_variant | MODIFIER |
| 1 | 170320723 | AX-170168488 | 0 | A | G | 0.211 | 9.82E-02 | 1.41E-01 | 2.66E+00 | 2.69E+00 | 4.84E-01 | no | Nos1ap | intron_variant | MODIFIER |
| 1 | 170320753 | AX-170321728 | 0 | T | C | 0.211 | 9.82E-02 | 1.41E-01 | 2.66E+00 | 2.69E+00 | 4.84E-01 | no | Nos1ap | intron_variant | MODIFIER |
| 1 | 170326142 | AX-170276623 | 0 | T | G | 0.211 | 9.82E-02 | 1.41E-01 | 2.66E+00 | 2.69E+00 | 4.84E-01 | no | Nos1ap | intron_variant | MODIFIER |
| 1 | 170330465 | AX-170268561 | 0 | A | G | 0.211 | 9.82E-02 | 1.41E-01 | 2.66E+00 | 2.69E+00 | 4.84E-01 | no | Nos1ap | intron_variant | MODIFIER |
| 1 | 170331374 | AX-170369854 | 0 | T | C | 0.211 | 9.82E-02 | 1.41E-01 | 2.66E+00 | 2.69E+00 | 4.84E-01 | no | Nos1ap | intron_variant | MODIFIER |
| 1 | 170343465 | AX-170483533 | 0 | G | C | 0.211 | 9.82E-02 | 1.41E-01 | 2.66E+00 | 2.69E+00 | 4.84E-01 | no | Nos1ap | intron_variant | MODIFIER |
| 1 | 170343770 | AX-170163476 | 0 | C | T | 0.211 | 9.82E-02 | 1.41E-01 | 2.66E+00 | 2.69E+00 | 4.84E-01 | no | Nos1ap | intron_variant | MODIFIER |
| 1 | 170368853 | AX-170149738 | 0 | C | T | 0.211 | 9.82E-02 | 1.41E-01 | 2.66E+00 | 2.69E+00 | 4.84E-01 | no | Nos1ap | downstream_gene_variant | MODIFIER |

| | | | | | | | | | | | | | | | |
|---|---|---|---|---|---|---|---|---|---|---|---|---|---|---|---|
| 1 | 170383905 | AX-170383525 | 0 | G | A | 0.211 | 9.82E-02 | 1.41E-01 | 2.66E+00 | 2.69E+00 | 4.84E-01 | no | Nos1ap | intron_variant,non_coding_transcript_variant | MODIFIER |
| 1 | 170385353 | AX-170351739 | 0 | G | A | 0.211 | 9.82E-02 | 1.41E-01 | 2.66E+00 | 2.69E+00 | 4.84E-01 | no | Nos1ap | intron_variant,non_coding_transcript_variant | MODIFIER |
| 1 | 170385673 | AX-170508840 | 0 | G | A | 0.211 | 9.82E-02 | 1.41E-01 | 2.66E+00 | 2.69E+00 | 4.84E-01 | no | Nos1ap | intron_variant,non_coding_transcript_variant | MODIFIER |
| 1 | 170386576 | AX-170232457 | 0 | T | C | 0.211 | 9.82E-02 | 1.41E-01 | 2.66E+00 | 2.69E+00 | 4.84E-01 | no | Nos1ap | intron_variant,non_coding_transcript_variant | MODIFIER |
| 1 | 170388830 | AX-97847971 | 0 | G | A | 0.211 | 9.82E-02 | 1.41E-01 | 2.66E+00 | 2.69E+00 | 4.84E-01 | no | Nos1ap | intron_variant,non_coding_transcript_variant | MODIFIER |
| 1 | 170389054 | AX-170453547 | 0 | T | G | 0.211 | 9.82E-02 | 1.41E-01 | 2.66E+00 | 2.69E+00 | 4.84E-01 | no | Nos1ap | intron_variant,non_coding_transcript_variant | MODIFIER |
| 1 | 170411977 | AX-97838582 | 0 | A | G | 0.211 | 9.82E-02 | 1.41E-01 | 2.66E+00 | 2.69E+00 | 4.84E-01 | no | Nos1ap | intron_variant,non_coding_transcript_variant | MODIFIER |
| 1 | 170416731 | AX-170132501 | 0 | G | A | 0.211 | 9.82E-02 | 1.41E-01 | 2.66E+00 | 2.69E+00 | 4.84E-01 | no | Nos1ap | intron_variant,non_coding_transcript_variant | MODIFIER |
| 1 | 170416768 | AX-170526936 | 0 | T | A | 0.211 | 9.82E-02 | 1.41E-01 | 2.66E+00 | 2.69E+00 | 4.84E-01 | no | Nos1ap | intron_variant,non_coding_transcript_variant | MODIFIER |
| 1 | 170423925 | AX-170126592 | 0 | T | G | 0.211 | 9.82E-02 | 1.41E-01 | 2.66E+00 | 2.69E+00 | 4.84E-01 | no | Nos1ap | intron_variant,non_coding_transcript_variant | MODIFIER |
| 1 | 170424321 | AX-170207801 | 0 | C | T | 0.211 | 9.82E-02 | 1.41E-01 | 2.66E+00 | 2.69E+00 | 4.84E-01 | no | Nos1ap | intron_variant,non_coding_transcript_variant | MODIFIER |
| 1 | 170439243 | AX-170163415 | 0 | T | G | 0.211 | 9.82E-02 | 1.41E-01 | 2.66E+00 | 2.69E+00 | 4.84E-01 | no | Nos1ap | intron_variant,non_coding_transcript_variant | MODIFIER |
| 1 | 170439281 | AX-170307317 | 0 | A | G | 0.211 | 9.82E-02 | 1.41E-01 | 2.66E+00 | 2.69E+00 | 4.84E-01 | no | Nos1ap | intron_variant,non_coding_transcript_variant | MODIFIER |
| 1 | 170439484 | AX-170305743 | 0 | C | A | 0.211 | 9.82E-02 | 1.41E-01 | 2.66E+00 | 2.69E+00 | 4.84E-01 | no | Nos1ap | intron_variant,non_coding_transcript_variant | MODIFIER |
| 1 | 170442548 | AX-170439016 | 0 | A | G | 0.211 | 9.82E-02 | 1.41E-01 | 2.66E+00 | 2.69E+00 | 4.84E-01 | no | Nos1ap | intron_variant,non_coding_transcript_variant | MODIFIER |
| 1 | 170458499 | AX-170233429 | 0 | A | T | 0.211 | 9.82E-02 | 1.41E-01 | 2.66E+00 | 2.69E+00 | 4.84E-01 | no | Nos1ap | intron_variant,non_coding_transcript_variant | MODIFIER |
| 1 | 170460913 | AX-97784852 | 0 | T | C | 0.211 | 9.82E-02 | 1.41E-01 | 2.66E+00 | 2.69E+00 | 4.84E-01 | no | Nos1ap | intron_variant,non_coding_transcript_variant | MODIFIER |
| 1 | 170462835 | AX-170489699 | 0 | T | C | 0.211 | 9.82E-02 | 1.41E-01 | 2.66E+00 | 2.69E+00 | 4.84E-01 | no | Nos1ap | intron_variant,non_coding_transcript_variant | MODIFIER |
| 1 | 170464581 | AX-170263622 | 0 | G | T | 0.211 | 9.82E-02 | 1.41E-01 | 2.66E+00 | 2.69E+00 | 4.84E-01 | no | Nos1ap | intron_variant,non_coding_transcript_variant | MODIFIER |
| 1 | 170489572 | AX-170404028 | 0 | G | A | 0.211 | 9.82E-02 | 1.41E-01 | 2.66E+00 | 2.69E+00 | 4.84E-01 | no | Nos1ap | intron_variant,non_coding_transcript_variant | MODIFIER |
| 1 | 170489653 | AX-170322548 | 0 | C | T | 0.211 | 9.82E-02 | 1.41E-01 | 2.66E+00 | 2.69E+00 | 4.84E-01 | no | Nos1ap | intron_variant,non_coding_transcript_variant | MODIFIER |
| 1 | 170490687 | AX-170373749 | 0 | G | A | 0.211 | 9.82E-02 | 1.41E-01 | 2.66E+00 | 2.69E+00 | 4.84E-01 | no | Nos1ap | intron_variant,non_coding_transcript_variant | MODIFIER |
| 1 | 170512101 | AX-170333736 | 0 | A | C | 0.211 | 9.82E-02 | 1.41E-01 | 2.66E+00 | 2.69E+00 | 4.84E-01 | no | Nos1ap | intron_variant,non_coding_transcript_variant | MODIFIER |
| 1 | 170535929 | AX-170176610 | 0 | C | T | 0.211 | 9.82E-02 | 1.41E-01 | 2.66E+00 | 2.69E+00 | 4.84E-01 | no | Nos1ap | intron_variant,non_coding_transcript_variant | MODIFIER |
| 1 | 170542504 | AX-170303198 | 0 | G | C | 0.211 | 9.82E-02 | 1.41E-01 | 2.66E+00 | 2.69E+00 | 4.84E-01 | no | Nos1ap | intron_variant,non_coding_transcript_variant | MODIFIER |
| 1 | 170542583 | AX-170431857 | 0 | G | A | 0.211 | 9.82E-02 | 1.41E-01 | 2.66E+00 | 2.69E+00 | 4.84E-01 | no | Nos1ap | intron_variant,non_coding_transcript_variant | MODIFIER |
| 1 | 170546667 | AX-170291552 | 0 | A | T | 0.211 | 9.82E-02 | 1.41E-01 | 2.66E+00 | 2.69E+00 | 4.84E-01 | no | Nos1ap | intron_variant,non_coding_transcript_variant | MODIFIER |
| 1 | 170546727 | AX-170397929 | 0 | A | G | 0.211 | 9.82E-02 | 1.41E-01 | 2.66E+00 | 2.69E+00 | 4.84E-01 | no | Nos1ap | intron_variant,non_coding_transcript_variant | MODIFIER |
| 1 | 170554836 | AX-170310693 | 0 | T | C | 0.211 | 9.82E-02 | 1.41E-01 | 2.66E+00 | 2.69E+00 | 4.84E-01 | no | Nos1ap | intron_variant,non_coding_transcript_variant | MODIFIER |
| 1 | 170557897 | AX-170043551 | 0 | A | C | 0.211 | 9.82E-02 | 1.41E-01 | 2.66E+00 | 2.69E+00 | 4.84E-01 | no | Nos1ap | intron_variant,non_coding_transcript_variant | MODIFIER |
| 1 | 170574196 | AX-170154444 | 0 | A | G | 0.211 | 9.82E-02 | 1.41E-01 | 2.66E+00 | 2.69E+00 | 4.84E-01 | no | Nos1ap | intron_variant,non_coding_transcript_variant | MODIFIER |
| 1 | 170574769 | AX-170002271 | 0 | C | T | 0.211 | 9.82E-02 | 1.41E-01 | 2.66E+00 | 2.69E+00 | 4.84E-01 | no | Nos1ap | intron_variant,non_coding_transcript_variant | MODIFIER |
| 1 | 170577758 | AX-170043550 | 0 | G | A | 0.211 | 9.82E-02 | 1.41E-01 | 2.66E+00 | 2.69E+00 | 4.84E-01 | no | Nos1ap | intron_variant,non_coding_transcript_variant | MODIFIER |
| 1 | 170579687 | AX-170257928 | 0 | T | C | 0.211 | 9.82E-02 | 1.41E-01 | 2.66E+00 | 2.69E+00 | 4.84E-01 | no | Nos1ap | intron_variant,non_coding_transcript_variant | MODIFIER |
| 1 | 170583752 | AX-170344680 | 0 | A | T | 0.211 | 9.82E-02 | 1.41E-01 | 2.66E+00 | 2.69E+00 | 4.84E-01 | no | Nos1ap | intron_variant,non_coding_transcript_variant | MODIFIER |
| 1 | 170583897 | AX-170345961 | 0 | T | A | 0.211 | 9.82E-02 | 1.41E-01 | 2.66E+00 | 2.69E+00 | 4.84E-01 | no | Nos1ap | intron_variant,non_coding_transcript_variant | MODIFIER |
| 1 | 170587384 | AX-170002268 | 0 | C | T | 0.211 | 9.82E-02 | 1.41E-01 | 2.66E+00 | 2.69E+00 | 4.84E-01 | no | Nos1ap | intron_variant,non_coding_transcript_variant | MODIFIER |
| 1 | 170603085 | AX-170125520 | 0 | A | G | 0.211 | 9.82E-02 | 1.41E-01 | 2.66E+00 | 2.69E+00 | 4.84E-01 | no | | intergenic_variant | MODIFIER |
| 1 | 170610417 | AX-170002269 | 0 | T | C | 0.211 | 9.82E-02 | 1.41E-01 | 2.66E+00 | 2.69E+00 | 4.84E-01 | no | | regulatory_region_variant | MODIFIER |
| 1 | 170641290 | AX-170128007 | 0 | T | C | 0.211 | 9.82E-02 | 1.41E-01 | 2.66E+00 | 2.69E+00 | 4.84E-01 | no | Olfml2b | upstream_gene_variant | MODIFIER |
| 1 | 170649399 | AX-170433886 | 0 | A | G | 0.211 | 9.82E-02 | 1.41E-01 | 2.66E+00 | 2.69E+00 | 4.84E-01 | no | Olfml2b | intron_variant | MODIFIER |
| 1 | 170653813 | AX-170290445 | 0 | G | A | 0.211 | 9.82E-02 | 1.41E-01 | 2.66E+00 | 2.69E+00 | 4.84E-01 | no | Olfml2b | intron_variant | MODIFIER |
| 1 | 170654729 | AX-170284082 | 0 | C | T | 0.211 | 9.82E-02 | 1.41E-01 | 2.66E+00 | 2.69E+00 | 4.84E-01 | no | Olfml2b | intron_variant | MODIFIER |
| 1 | 170659871 | AX-170139393 | 0 | A | G | 0.211 | 9.82E-02 | 1.41E-01 | 2.66E+00 | 2.69E+00 | 4.84E-01 | no | Olfml2b | intron_variant | MODIFIER |
| 1 | 170670515 | AX-170043555 | 0 | T | A | 0.211 | 9.82E-02 | 1.41E-01 | 2.66E+00 | 2.69E+00 | 4.84E-01 | no | Olfml2b | intron_variant | MODIFIER |
| 1 | 170674204 | AX-170091785 | 0 | G | A | 0.211 | 9.82E-02 | 1.41E-01 | 2.66E+00 | 2.69E+00 | 4.84E-01 | no | Olfml2b | intron_variant | MODIFIER |
| 1 | 170704305 | AX-170182903 | 0 | A | C | 0.211 | 9.82E-02 | 1.41E-01 | 2.66E+00 | 2.69E+00 | 4.84E-01 | no | Atf6 | downstream_gene_variant | MODIFIER |
| 1 | 170706023 | AX-170289806 | 0 | A | G | 0.211 | 9.82E-02 | 1.41E-01 | 2.66E+00 | 2.69E+00 | 4.84E-01 | no | Atf6 | 3_prime_UTR_variant | MODIFIER |
| 1 | 170716705 | AX-170375924 | 0 | A | G | 0.211 | 9.82E-02 | 1.41E-01 | 2.66E+00 | 2.69E+00 | 4.84E-01 | no | Atf6 | intron_variant | MODIFIER |
| 1 | 170782073 | AX-170350912 | 0 | A | T | 0.211 | 9.82E-02 | 1.41E-01 | 2.66E+00 | 2.69E+00 | 4.84E-01 | no | Atf6 | intron_variant | MODIFIER |
| 1 | 170782557 | AX-170149574 | 0 | A | C | 0.211 | 9.82E-02 | 1.41E-01 | 2.66E+00 | 2.69E+00 | 4.84E-01 | no | Atf6 | intron_variant | MODIFIER |
| 1 | 170784975 | AX-170128417 | 0 | T | C | 0.211 | 9.82E-02 | 1.41E-01 | 2.66E+00 | 2.69E+00 | 4.84E-01 | no | Atf6 | intron_variant | MODIFIER |
| 1 | 170787361 | AX-170479706 | 0 | A | C | 0.211 | 9.82E-02 | 1.41E-01 | 2.66E+00 | 2.69E+00 | 4.84E-01 | no | Atf6 | synonymous_variant | LOW |
| 1 | 170801746 | AX-170217760 | 0 | T | C | 0.211 | 9.82E-02 | 1.41E-01 | 2.66E+00 | 2.69E+00 | 4.84E-01 | no | Atf6 | intron_variant | MODIFIER |

| | | | | | | | | | | | | | | | |
|---|---|---|---|---|---|---|---|---|---|---|---|---|---|---|---|
| 1 | 170806454 | AX-170387965 | 0 | A | G | 0.211 | 9.82E-02 | 1.41E-01 | 2.66E+00 | 2.69E+00 | 4.84E-01 | no | Atf6 | intron_variant | MODIFIER |
| 1 | 170812141 | AX-97821201 | 0 | G | T | 0.211 | 9.82E-02 | 1.41E-01 | 2.66E+00 | 2.69E+00 | 4.84E-01 | no | Atf6 | intron_variant | MODIFIER |
| 1 | 170818789 | AX-170330589 | 0 | A | C | 0.211 | 9.82E-02 | 1.41E-01 | 2.66E+00 | 2.69E+00 | 4.84E-01 | no | Atf6 | intron_variant | MODIFIER |
| 1 | 170819926 | AX-170043560 | 0 | G | A | 0.211 | 9.82E-02 | 1.41E-01 | 2.66E+00 | 2.69E+00 | 4.84E-01 | no | Atf6 | intron_variant | MODIFIER |
| 1 | 170831537 | AX-170043558 | 0 | G | A | 0.211 | 9.82E-02 | 1.41E-01 | 2.66E+00 | 2.69E+00 | 4.84E-01 | no | Atf6 | intron_variant | MODIFIER |
| 1 | 170850827 | AX-170282896 | 0 | A | C | 0.211 | 9.82E-02 | 1.41E-01 | 2.66E+00 | 2.69E+00 | 4.84E-01 | no | Atf6 | intron_variant | MODIFIER |
| 1 | 170861273 | AX-170366481 | 0 | A | G | 0.211 | 9.82E-02 | 1.41E-01 | 2.66E+00 | 2.69E+00 | 4.84E-01 | no | Atf6 | intron_variant | MODIFIER |
| 1 | 170861378 | AX-170423065 | 0 | T | C | 0.211 | 9.82E-02 | 1.41E-01 | 2.66E+00 | 2.69E+00 | 4.84E-01 | no | Atf6 | intron_variant | MODIFIER |
| 1 | 170868582 | AX-170403598 | 0 | A | G | 0.211 | 9.82E-02 | 1.41E-01 | 2.66E+00 | 2.69E+00 | 4.84E-01 | no | Atf6 | upstream_gene_variant | MODIFIER |
| 1 | 170878562 | AX-170318534 | 0 | G | C | 0.211 | 9.82E-02 | 1.41E-01 | 2.66E+00 | 2.69E+00 | 4.84E-01 | no | Dusp12 | intron_variant | MODIFIER |
| 1 | 170878847 | AX-170493992 | 0 | A | G | 0.211 | 9.82E-02 | 1.41E-01 | 2.66E+00 | 2.69E+00 | 4.84E-01 | no | Dusp12 | intron_variant | MODIFIER |
| 1 | 170884677 | AX-170290533 | 0 | T | A | 0.211 | 9.82E-02 | 1.41E-01 | 2.66E+00 | 2.69E+00 | 4.84E-01 | no | Dusp12 | intron_variant | MODIFIER |
| 1 | 170884742 | AX-170201278 | 0 | T | C | 0.211 | 9.82E-02 | 1.41E-01 | 2.66E+00 | 2.69E+00 | 4.84E-01 | no | Dusp12 | intron_variant | MODIFIER |
| 1 | 170885169 | AX-169999711 | 0 | G | T | 0.211 | 9.82E-02 | 1.41E-01 | 2.66E+00 | 2.69E+00 | 4.84E-01 | no | Dusp12 | missense_variant | MODERATE |
| 1 | 170893051 | AX-170460616 | 0 | G | A | 0.211 | 9.82E-02 | 1.41E-01 | 2.66E+00 | 2.69E+00 | 4.84E-01 | no | | intergenic_variant | MODIFIER |
| 1 | 170893246 | AX-170155515 | 0 | C | T | 0.211 | 9.82E-02 | 1.41E-01 | 2.66E+00 | 2.69E+00 | 4.84E-01 | no | | intergenic_variant | MODIFIER |
| 1 | 170898368 | AX-170293762 | 0 | T | C | 0.211 | 9.82E-02 | 1.41E-01 | 2.66E+00 | 2.69E+00 | 4.84E-01 | no | | regulatory_region_variant | MODIFIER |
| 1 | 170920883 | AX-170447715 | 0 | C | T | 0.211 | 9.82E-02 | 1.41E-01 | 2.66E+00 | 2.69E+00 | 4.84E-01 | no | Fcrla | synonymous_variant | LOW |
| 1 | 170930841 | AX-170119710 | 0 | G | T | 0.211 | 9.82E-02 | 1.41E-01 | 2.66E+00 | 2.69E+00 | 4.84E-01 | no | Fcrla | upstream_gene_variant | MODIFIER |
| 1 | 170938264 | AX-170268952 | 0 | A | T | 0.211 | 9.82E-02 | 1.41E-01 | 2.66E+00 | 2.69E+00 | 4.84E-01 | no | Gm37030 | downstream_gene_variant | MODIFIER |
| 1 | 170952269 | AX-170131812 | 0 | A | G | 0.211 | 9.82E-02 | 1.41E-01 | 2.66E+00 | 2.69E+00 | 4.84E-01 | no | | intergenic_variant | MODIFIER |
| 1 | 170960884 | AX-97837911 | 0 | G | T | 0.211 | 9.82E-02 | 1.41E-01 | 2.66E+00 | 2.69E+00 | 4.84E-01 | no | Fcgr2b | 3_prime_UTR_variant | MODIFIER |
| 1 | 170971667 | AX-170494779 | 0 | G | A | 0.211 | 9.82E-02 | 1.41E-01 | 2.66E+00 | 2.69E+00 | 4.84E-01 | no | Fcgr2b | intron_variant | MODIFIER |
| 1 | 170988866 | AX-170043563 | 0 | T | C | 0.211 | 9.82E-02 | 1.41E-01 | 2.66E+00 | 2.69E+00 | 4.84E-01 | no | Gm10522 | downstream_gene_variant | MODIFIER |
| 1 | 170991807 | AX-170043562 | 0 | T | C | 0.211 | 9.82E-02 | 1.41E-01 | 2.66E+00 | 2.69E+00 | 4.84E-01 | no | | intergenic_variant | MODIFIER |
| 1 | 170997961 | AX-170043564 | 0 | G | A | 0.211 | 9.82E-02 | 1.41E-01 | 2.66E+00 | 2.69E+00 | 4.84E-01 | no | | regulatory_region_variant | MODIFIER |
| 1 | 171000013 | AX-170297405 | 0 | T | C | 0.211 | 9.82E-02 | 1.41E-01 | 2.66E+00 | 2.69E+00 | 4.84E-01 | no | | intergenic_variant | MODIFIER |
| 1 | 171004566 | AX-170194376 | 0 | G | A | 0.211 | 9.82E-02 | 1.41E-01 | 2.66E+00 | 2.69E+00 | 4.84E-01 | no | | intergenic_variant | MODIFIER |
| 1 | 171007804 | AX-170388475 | 0 | T | C | 0.211 | 9.82E-02 | 1.41E-01 | 2.66E+00 | 2.69E+00 | 4.84E-01 | no | | intergenic_variant | MODIFIER |
| 1 | 171020937 | AX-170457699 | 0 | G | A | 0.211 | 9.82E-02 | 1.41E-01 | 2.66E+00 | 2.69E+00 | 4.84E-01 | no | Fcgr4 | intron_variant | MODIFIER |
| 1 | 171021474 | AX-170459372 | 0 | T | C | 0.211 | 9.82E-02 | 1.41E-01 | 2.66E+00 | 2.69E+00 | 4.84E-01 | no | Fcgr4 | intron_variant | MODIFIER |
| 1 | 171023589 | AX-97796209 | 0 | G | A | 0.211 | 9.82E-02 | 1.41E-01 | 2.66E+00 | 2.69E+00 | 4.84E-01 | no | Fcgr4 | intron_variant | MODIFIER |
| 1 | 171025842 | AX-170462246 | 0 | G | A | 0.211 | 9.82E-02 | 1.41E-01 | 2.66E+00 | 2.69E+00 | 4.84E-01 | no | Fcgr4 | intron_variant | MODIFIER |
| 1 | 171027241 | AX-170373405 | 0 | C | T | 0.211 | 9.82E-02 | 1.41E-01 | 2.66E+00 | 2.69E+00 | 4.84E-01 | no | Fcgr4 | intron_variant | MODIFIER |
| 5 | 81665450 | AX-97780946 | 1 | C | T | 0.446 | 7.01E-02 | 1.02E-01 | 2.73E+00 | 2.81E+00 | 4.84E-01 | no | Adgrl3 | intron_variant | MODIFIER |
| 1 | 24357425 | AX-97832686 | 0 | T | C | 0.43 | 5.86E-02 | 8.42E-02 | 2.62E+00 | 2.69E+00 | 4.84E-01 | no | Col19a1 | intron_variant | MODIFIER |
| 5 | 89340139 | AX-170266306 | 1 | G | T | 0.438 | 7.70E-02 | 1.11E-01 | 2.77E+00 | 2.82E+00 | 4.84E-01 | no | | intergenic_variant | MODIFIER |
| 6 | 139616949 | AX-97821982 | 1 | C | T | 0.054 | 1.41E-01 | 2.05E-01 | 2.65E+00 | 2.77E+00 | 4.84E-01 | no | Pik3c2g | upstream_gene_variant | MODIFIER |
| 18 | 81038092 | AX-170436909 | 0 | C | T | 0.298 | -5.99E-02 | 8.50E-02 | 2.54E+00 | 2.65E+00 | 4.85E-01 | no | | intergenic_variant | MODIFIER |
| 17 | 24307644 | AX-170120825 | 0 | A | G | 0.044 | -1.17E-01 | 1.66E-01 | 2.54E+00 | 2.66E+00 | 4.85E-01 | no | Abca17 | intron_variant | MODIFIER |
| 9 | 87809236 | AX-170325025 | 1 | C | T | 0.143 | -8.92E-02 | 1.28E-01 | 2.60E+00 | 2.66E+00 | 4.85E-01 | no | | intergenic_variant | MODIFIER |
| 4 | 89401847 | AX-170497005 | 1 | T | A | 0.33 | 7.32E-02 | 1.06E-01 | 2.66E+00 | 2.74E+00 | 4.85E-01 | no | Gm12609 | downstream_gene_variant | MODIFIER |
| 5 | 78393461 | AX-170072564 | 0 | T | G | 0.421 | 7.61E-02 | 1.10E-01 | 2.80E+00 | 2.84E+00 | 4.85E-01 | no | | intergenic_variant | MODIFIER |
| 8 | 115898906 | AX-170339274 | 0 | T | C | 0.254 | 6.87E-02 | 1.00E-01 | 2.75E+00 | 2.86E+00 | 4.85E-01 | no | | intergenic_variant | MODIFIER |
| 6 | 131368984 | AX-170177947 | 1 | G | A | 0.554 | 5.78E-02 | 8.19E-02 | 2.54E+00 | 2.62E+00 | 4.85E-01 | no | Ybx3 | intron_variant | MODIFIER |
| 6 | 128518724 | AX-170299306 | 0 | C | T | 0.553 | 5.85E-02 | 8.27E-02 | 2.54E+00 | 2.61E+00 | 4.85E-01 | no | Pzp | intron_variant | MODIFIER |
| 10 | 66137207 | AX-97800169 | 1 | A | G | 0.402 | -5.44E-02 | 7.85E-02 | 2.65E+00 | 2.72E+00 | 4.85E-01 | no | 1700023F02Rik | intron_variant,non_coding_transcript_variant | MODIFIER |
| 5 | 35972491 | AX-170071796 | 0 | T | C | 0.342 | 7.76E-02 | 1.12E-01 | 2.76E+00 | 2.81E+00 | 4.85E-01 | no | Afap1 | intron_variant | MODIFIER |
| 5 | 35974575 | AX-170454387 | 0 | C | T | 0.342 | 7.76E-02 | 1.12E-01 | 2.76E+00 | 2.81E+00 | 4.85E-01 | no | Afap1 | synonymous_variant | LOW |
| 5 | 35984377 | AX-170520765 | 0 | C | T | 0.342 | 7.76E-02 | 1.12E-01 | 2.76E+00 | 2.81E+00 | 4.85E-01 | no | Afap1 | intron_variant | MODIFIER |
| 5 | 35985371 | AX-170130330 | 0 | C | T | 0.342 | 7.76E-02 | 1.12E-01 | 2.76E+00 | 2.81E+00 | 4.85E-01 | no | Afap1 | intron_variant | MODIFIER |
| 5 | 35992075 | AX-170157561 | 0 | C | T | 0.342 | 7.76E-02 | 1.12E-01 | 2.76E+00 | 2.81E+00 | 4.85E-01 | no | Afap1 | intron_variant | MODIFIER |
| 5 | 35994641 | AX-97831386 | 0 | C | T | 0.342 | 7.76E-02 | 1.12E-01 | 2.76E+00 | 2.81E+00 | 4.85E-01 | no | Afap1 | intron_variant | MODIFIER |
| 5 | 36009984 | AX-170173965 | 0 | C | T | 0.342 | 7.76E-02 | 1.12E-01 | 2.76E+00 | 2.81E+00 | 4.85E-01 | no | | intergenic_variant | MODIFIER |

| | | | | | | | | | | | | | | | |
|---|---|---|---|---|---|---|---|---|---|---|---|---|---|---|---|
| 5 | 36023159 | AX-170345353 | 0 | C | T | 0.342 | 7.76E-02 | 1.12E-01 | 2.76E+00 | 2.81E+00 | 4.85E-01 | no | Sorcs2 | intron_variant | MODIFIER |
| 5 | 36048621 | AX-170250459 | 0 | G | A | 0.342 | 7.76E-02 | 1.12E-01 | 2.76E+00 | 2.81E+00 | 4.85E-01 | no | Sorcs2 | intron_variant | MODIFIER |
| 5 | 36072354 | AX-170298038 | 0 | A | C | 0.342 | 7.76E-02 | 1.12E-01 | 2.76E+00 | 2.81E+00 | 4.85E-01 | no | Sorcs2 | intron_variant | MODIFIER |
| 5 | 36085904 | AX-97841659 | 0 | G | T | 0.342 | 7.76E-02 | 1.12E-01 | 2.76E+00 | 2.81E+00 | 4.85E-01 | no | Sorcs2 | intron_variant | MODIFIER |
| 5 | 36134691 | AX-170491344 | 0 | C | T | 0.342 | 7.76E-02 | 1.12E-01 | 2.76E+00 | 2.81E+00 | 4.85E-01 | no | Sorcs2 | intron_variant | MODIFIER |
| 5 | 36181253 | AX-170393015 | 0 | T | G | 0.342 | 7.76E-02 | 1.12E-01 | 2.76E+00 | 2.81E+00 | 4.85E-01 | no | Sorcs2 | intron_variant | MODIFIER |
| 5 | 36181893 | AX-170243490 | 0 | G | A | 0.342 | 7.76E-02 | 1.12E-01 | 2.76E+00 | 2.81E+00 | 4.85E-01 | no | Sorcs2 | intron_variant | MODIFIER |
| 5 | 36191613 | AX-170147537 | 0 | C | T | 0.342 | 7.76E-02 | 1.12E-01 | 2.76E+00 | 2.81E+00 | 4.85E-01 | no | Sorcs2 | intron_variant | MODIFIER |
| 5 | 36222102 | AX-170200773 | 0 | A | G | 0.342 | 7.76E-02 | 1.12E-01 | 2.76E+00 | 2.81E+00 | 4.85E-01 | no | Sorcs2 | intron_variant | MODIFIER |
| 5 | 36222149 | AX-170260149 | 0 | C | T | 0.342 | 7.76E-02 | 1.12E-01 | 2.76E+00 | 2.81E+00 | 4.85E-01 | no | Sorcs2 | intron_variant | MODIFIER |
| 5 | 36334052 | AX-97806998 | 0 | G | A | 0.342 | 7.76E-02 | 1.12E-01 | 2.76E+00 | 2.81E+00 | 4.85E-01 | no | Sorcs2 | intron_variant | MODIFIER |
| 5 | 36344790 | AX-170382126 | 0 | G | A | 0.342 | 7.76E-02 | 1.12E-01 | 2.76E+00 | 2.81E+00 | 4.85E-01 | no | Sorcs2 | intron_variant | MODIFIER |
| 5 | 36361412 | AX-170326271 | 0 | G | A | 0.342 | 7.76E-02 | 1.12E-01 | 2.76E+00 | 2.81E+00 | 4.85E-01 | no | Sorcs2 | intron_variant | MODIFIER |
| 5 | 36381130 | AX-170137817 | 0 | G | A | 0.342 | 7.76E-02 | 1.12E-01 | 2.76E+00 | 2.81E+00 | 4.85E-01 | no | Sorcs2 | intron_variant | MODIFIER |
| 5 | 36433053 | AX-170364174 | 0 | G | A | 0.342 | 7.76E-02 | 1.12E-01 | 2.76E+00 | 2.81E+00 | 4.85E-01 | no | | intergenic_variant | MODIFIER |
| 5 | 36545077 | AX-170507565 | 0 | C | T | 0.342 | 7.76E-02 | 1.12E-01 | 2.76E+00 | 2.81E+00 | 4.85E-01 | no | Tbc1d14 | intron_variant | MODIFIER |
| 5 | 36553074 | AX-170377926 | 0 | G | A | 0.342 | 7.76E-02 | 1.12E-01 | 2.76E+00 | 2.81E+00 | 4.85E-01 | no | Tbc1d14 | intron_variant | MODIFIER |
| 5 | 36562177 | AX-170146862 | 0 | C | T | 0.342 | 7.76E-02 | 1.12E-01 | 2.76E+00 | 2.81E+00 | 4.85E-01 | no | Tbc1d14 | intron_variant | MODIFIER |
| 5 | 36565400 | AX-170330222 | 0 | T | C | 0.342 | 7.76E-02 | 1.12E-01 | 2.76E+00 | 2.81E+00 | 4.85E-01 | no | Tbc1d14 | intron_variant | MODIFIER |
| 5 | 36567809 | AX-170071810 | 0 | A | G | 0.342 | 7.76E-02 | 1.12E-01 | 2.76E+00 | 2.81E+00 | 4.85E-01 | no | Tbc1d14 | intron_variant | MODIFIER |
| 5 | 36570206 | AX-170164425 | 0 | G | A | 0.342 | 7.76E-02 | 1.12E-01 | 2.76E+00 | 2.81E+00 | 4.85E-01 | no | Tbc1d14 | intron_variant | MODIFIER |
| 5 | 36621864 | AX-170114520 | 0 | G | A | 0.342 | 7.76E-02 | 1.12E-01 | 2.76E+00 | 2.81E+00 | 4.85E-01 | no | D5Ertd579e | intron_variant | MODIFIER |
| 5 | 36622662 | AX-170024696 | 0 | T | A | 0.342 | 7.76E-02 | 1.12E-01 | 2.76E+00 | 2.81E+00 | 4.85E-01 | no | D5Ertd579e | intron_variant | MODIFIER |
| 5 | 36664395 | AX-170024687 | 0 | G | A | 0.342 | 7.76E-02 | 1.12E-01 | 2.76E+00 | 2.81E+00 | 4.85E-01 | no | D5Ertd579e | intron_variant | MODIFIER |
| 5 | 36713032 | AX-170311016 | 0 | G | T | 0.342 | 7.76E-02 | 1.12E-01 | 2.76E+00 | 2.81E+00 | 4.85E-01 | no | | regulatory_region_variant | MODIFIER |
| 5 | 36718380 | AX-170024690 | 0 | T | C | 0.342 | 7.76E-02 | 1.12E-01 | 2.76E+00 | 2.81E+00 | 4.85E-01 | no | | intergenic_variant | MODIFIER |
| 5 | 36761924 | AX-170275909 | 0 | A | T | 0.342 | 7.76E-02 | 1.12E-01 | 2.76E+00 | 2.81E+00 | 4.85E-01 | no | Gm40318 | downstream_gene_variant | MODIFIER |
| 5 | 36786015 | AX-170425875 | 0 | A | C | 0.342 | 7.76E-02 | 1.12E-01 | 2.76E+00 | 2.81E+00 | 4.85E-01 | no | | intergenic_variant | MODIFIER |
| 5 | 36898249 | AX-170223115 | 0 | G | A | 0.342 | 7.76E-02 | 1.12E-01 | 2.76E+00 | 2.81E+00 | 4.85E-01 | no | Ppp2r2c | intron_variant | MODIFIER |
| 5 | 36905056 | AX-170237736 | 0 | T | C | 0.342 | 7.76E-02 | 1.12E-01 | 2.76E+00 | 2.81E+00 | 4.85E-01 | no | Ppp2r2c | intron_variant | MODIFIER |
| 5 | 37009762 | AX-170225601 | 0 | G | A | 0.342 | 7.76E-02 | 1.12E-01 | 2.76E+00 | 2.81E+00 | 4.85E-01 | no | | intergenic_variant | MODIFIER |
| 5 | 37050516 | AX-170071820 | 0 | C | T | 0.342 | 7.76E-02 | 1.12E-01 | 2.76E+00 | 2.81E+00 | 4.85E-01 | no | Jakmip1 | upstream_gene_variant | MODIFIER |
| 5 | 37066375 | AX-170188824 | 0 | A | C | 0.342 | 7.76E-02 | 1.12E-01 | 2.76E+00 | 2.81E+00 | 4.85E-01 | no | Jakmip1 | intron_variant | MODIFIER |
| 5 | 37098513 | AX-170325231 | 0 | A | G | 0.342 | 7.76E-02 | 1.12E-01 | 2.76E+00 | 2.81E+00 | 4.85E-01 | no | Jakmip1 | intron_variant | MODIFIER |
| 5 | 37115941 | AX-170371180 | 0 | C | T | 0.342 | 7.76E-02 | 1.12E-01 | 2.76E+00 | 2.81E+00 | 4.85E-01 | no | Jakmip1 | intron_variant | MODIFIER |
| 5 | 37189667 | AX-170024701 | 0 | T | C | 0.342 | 7.76E-02 | 1.12E-01 | 2.76E+00 | 2.81E+00 | 4.85E-01 | no | Gm1043 | intron_variant,non_coding_transcript_variant | MODIFIER |
| 5 | 37197590 | AX-170146126 | 0 | G | C | 0.342 | 7.76E-02 | 1.12E-01 | 2.76E+00 | 2.81E+00 | 4.85E-01 | no | Gm1043 | downstream_gene_variant | MODIFIER |
| 5 | 37197656 | AX-170407749 | 0 | T | G | 0.342 | 7.76E-02 | 1.12E-01 | 2.76E+00 | 2.81E+00 | 4.85E-01 | no | Gm1043 | downstream_gene_variant | MODIFIER |
| 5 | 37220927 | AX-170276139 | 0 | G | T | 0.342 | 7.76E-02 | 1.12E-01 | 2.76E+00 | 2.81E+00 | 4.85E-01 | no | Gm1043 | intron_variant | MODIFIER |
| 5 | 37222226 | AX-170024704 | 0 | A | G | 0.342 | 7.76E-02 | 1.12E-01 | 2.76E+00 | 2.81E+00 | 4.85E-01 | no | Gm1043 | intron_variant | MODIFIER |
| 5 | 37324779 | AX-170148869 | 0 | A | G | 0.342 | 7.76E-02 | 1.12E-01 | 2.76E+00 | 2.81E+00 | 4.85E-01 | no | Evc | intron_variant | MODIFIER |
| 5 | 37392955 | AX-170275659 | 0 | C | G | 0.342 | 7.76E-02 | 1.12E-01 | 2.76E+00 | 2.81E+00 | 4.85E-01 | no | Evc2 | intron_variant | MODIFIER |
| 5 | 37394281 | AX-170534574 | 0 | G | A | 0.342 | 7.76E-02 | 1.12E-01 | 2.76E+00 | 2.81E+00 | 4.85E-01 | no | Evc2 | intron_variant | MODIFIER |
| 5 | 37395315 | AX-170474935 | 0 | C | T | 0.342 | 7.76E-02 | 1.12E-01 | 2.76E+00 | 2.81E+00 | 4.85E-01 | no | Evc2 | intron_variant | MODIFIER |
| 5 | 37408264 | AX-170089450 | 0 | C | T | 0.342 | 7.76E-02 | 1.12E-01 | 2.76E+00 | 2.81E+00 | 4.85E-01 | no | Evc2 | intron_variant | MODIFIER |
| 5 | 37423176 | AX-170328356 | 0 | T | C | 0.342 | 7.76E-02 | 1.12E-01 | 2.76E+00 | 2.81E+00 | 4.85E-01 | no | Evc2 | intron_variant | MODIFIER |
| 5 | 37423375 | AX-170172635 | 0 | C | T | 0.342 | 7.76E-02 | 1.12E-01 | 2.76E+00 | 2.81E+00 | 4.85E-01 | no | Evc2 | intron_variant | MODIFIER |
| 5 | 37451343 | AX-170293226 | 0 | G | A | 0.342 | 7.76E-02 | 1.12E-01 | 2.76E+00 | 2.81E+00 | 4.85E-01 | no | Stk32b | intron_variant | MODIFIER |
| 5 | 37451623 | AX-170481179 | 0 | T | G | 0.342 | 7.76E-02 | 1.12E-01 | 2.76E+00 | 2.81E+00 | 4.85E-01 | no | Stk32b | intron_variant | MODIFIER |
| 5 | 37459470 | AX-170156148 | 0 | T | C | 0.342 | 7.76E-02 | 1.12E-01 | 2.76E+00 | 2.81E+00 | 4.85E-01 | no | Stk32b | intron_variant | MODIFIER |
| 5 | 37490287 | AX-97788905 | 0 | C | T | 0.342 | 7.76E-02 | 1.12E-01 | 2.76E+00 | 2.81E+00 | 4.85E-01 | no | Stk32b | intron_variant | MODIFIER |
| 5 | 37490794 | AX-170216714 | 0 | G | A | 0.342 | 7.76E-02 | 1.12E-01 | 2.76E+00 | 2.81E+00 | 4.85E-01 | no | Stk32b | intron_variant | MODIFIER |
| 5 | 37495638 | AX-97832862 | 0 | T | C | 0.342 | 7.76E-02 | 1.12E-01 | 2.76E+00 | 2.81E+00 | 4.85E-01 | no | Stk32b | intron_variant | MODIFIER |
| 5 | 37538113 | AX-170318302 | 0 | T | C | 0.342 | 7.76E-02 | 1.12E-01 | 2.76E+00 | 2.81E+00 | 4.85E-01 | no | Stk32b | intron_variant | MODIFIER |

| | | | | | | | | | | | | | | | |
|---|---|---|---|---|---|---|---|---|---|---|---|---|---|---|---|
| 5 | 37558726 | AX-170071830 | 0 | C | T | 0.342 | 7.76E-02 | 1.12E-01 | 2.76E+00 | 2.81E+00 | 4.85E-01 | no | Stk32b | intron_variant | MODIFIER |
| 5 | 37561011 | AX-170071827 | 0 | C | T | 0.342 | 7.76E-02 | 1.12E-01 | 2.76E+00 | 2.81E+00 | 4.85E-01 | no | Stk32b | intron_variant | MODIFIER |
| 5 | 37715178 | AX-170289436 | 0 | G | A | 0.342 | 7.76E-02 | 1.12E-01 | 2.76E+00 | 2.81E+00 | 4.85E-01 | no | Stk32b | intron_variant | MODIFIER |
| 5 | 37771413 | AX-97856921 | 0 | A | G | 0.342 | 7.76E-02 | 1.12E-01 | 2.76E+00 | 2.81E+00 | 4.85E-01 | no | | intergenic_variant | MODIFIER |
| 5 | 37845537 | AX-97799245 | 0 | T | G | 0.342 | 7.76E-02 | 1.12E-01 | 2.76E+00 | 2.81E+00 | 4.85E-01 | no | | intergenic_variant | MODIFIER |
| 5 | 37894741 | AX-170486332 | 0 | T | A | 0.342 | 7.76E-02 | 1.12E-01 | 2.76E+00 | 2.81E+00 | 4.85E-01 | no | Gm20052 | intron_variant,non_coding_transcript_variant | MODIFIER |
| 5 | 37899434 | AX-170277759 | 0 | C | A | 0.342 | 7.76E-02 | 1.12E-01 | 2.76E+00 | 2.81E+00 | 4.85E-01 | no | Gm20052 | downstream_gene_variant | MODIFIER |
| 5 | 37950495 | AX-170071835 | 0 | C | T | 0.342 | 7.76E-02 | 1.12E-01 | 2.76E+00 | 2.81E+00 | 4.85E-01 | no | | intergenic_variant | MODIFIER |
| 5 | 38020913 | AX-170397229 | 0 | C | T | 0.342 | 7.76E-02 | 1.12E-01 | 2.76E+00 | 2.81E+00 | 4.85E-01 | no | | intergenic_variant | MODIFIER |
| 5 | 38054662 | AX-170352713 | 0 | G | A | 0.342 | 7.76E-02 | 1.12E-01 | 2.76E+00 | 2.81E+00 | 4.85E-01 | no | Stx18 | intron_variant | MODIFIER |
| 5 | 38080935 | AX-170095291 | 0 | G | A | 0.342 | 7.76E-02 | 1.12E-01 | 2.76E+00 | 2.81E+00 | 4.85E-01 | no | Stx18 | intron_variant | MODIFIER |
| 5 | 38086864 | AX-170390958 | 0 | C | T | 0.342 | 7.76E-02 | 1.12E-01 | 2.76E+00 | 2.81E+00 | 4.85E-01 | no | Stx18 | intron_variant | MODIFIER |
| 5 | 38096208 | AX-170142951 | 0 | G | A | 0.342 | 7.76E-02 | 1.12E-01 | 2.76E+00 | 2.81E+00 | 4.85E-01 | no | Stx18 | intron_variant | MODIFIER |
| 5 | 38113510 | AX-97856619 | 0 | C | T | 0.342 | 7.76E-02 | 1.12E-01 | 2.76E+00 | 2.81E+00 | 4.85E-01 | no | Stx18 | intron_variant | MODIFIER |
| 5 | 38188823 | AX-170024722 | 0 | G | A | 0.342 | 7.76E-02 | 1.12E-01 | 2.76E+00 | 2.81E+00 | 4.85E-01 | no | Zbtb49 | downstream_gene_variant | MODIFIER |
| 5 | 38246530 | AX-97838092 | 0 | C | T | 0.342 | 7.76E-02 | 1.12E-01 | 2.76E+00 | 2.81E+00 | 4.85E-01 | no | | intergenic_variant | MODIFIER |
| 5 | 38291486 | AX-170024721 | 0 | T | A | 0.342 | 7.76E-02 | 1.12E-01 | 2.76E+00 | 2.81E+00 | 4.85E-01 | no | Otop1 | intron_variant | MODIFIER |
| 5 | 38293528 | AX-170024719 | 0 | G | A | 0.342 | 7.76E-02 | 1.12E-01 | 2.76E+00 | 2.81E+00 | 4.85E-01 | no | Otop1 | intron_variant | MODIFIER |
| 5 | 38345519 | AX-170024723 | 0 | C | T | 0.342 | 7.76E-02 | 1.12E-01 | 2.76E+00 | 2.81E+00 | 4.85E-01 | no | Slc2a9 | downstream_gene_variant | MODIFIER |
| 5 | 38367849 | AX-170024714 | 0 | G | A | 0.342 | 7.76E-02 | 1.12E-01 | 2.76E+00 | 2.81E+00 | 4.85E-01 | no | Slc2a9 | intron_variant | MODIFIER |
| 5 | 38369551 | AX-170089452 | 0 | C | T | 0.342 | 7.76E-02 | 1.12E-01 | 2.76E+00 | 2.81E+00 | 4.85E-01 | no | Slc2a9 | intron_variant | MODIFIER |
| 5 | 38385455 | AX-170492861 | 0 | G | A | 0.342 | 7.76E-02 | 1.12E-01 | 2.76E+00 | 2.81E+00 | 4.85E-01 | no | Slc2a9 | intron_variant | MODIFIER |
| 5 | 38405176 | AX-97812035 | 0 | C | T | 0.342 | 7.76E-02 | 1.12E-01 | 2.76E+00 | 2.81E+00 | 4.85E-01 | no | Slc2a9 | intron_variant | MODIFIER |
| 5 | 38449216 | AX-170147144 | 0 | C | T | 0.342 | 7.76E-02 | 1.12E-01 | 2.76E+00 | 2.81E+00 | 4.85E-01 | no | Slc2a9 | intron_variant | MODIFIER |
| 5 | 38449630 | AX-170128240 | 0 | G | C | 0.342 | 7.76E-02 | 1.12E-01 | 2.76E+00 | 2.81E+00 | 4.85E-01 | no | Slc2a9 | intron_variant | MODIFIER |
| 5 | 38450571 | AX-97793896 | 0 | C | A | 0.342 | 7.76E-02 | 1.12E-01 | 2.76E+00 | 2.81E+00 | 4.85E-01 | no | Slc2a9 | intron_variant | MODIFIER |
| 5 | 38542114 | AX-170270792 | 0 | G | C | 0.342 | 7.76E-02 | 1.12E-01 | 2.76E+00 | 2.81E+00 | 4.85E-01 | no | Wdr1 | intron_variant | MODIFIER |
| 5 | 38550178 | AX-97808742 | 0 | G | A | 0.342 | 7.76E-02 | 1.12E-01 | 2.76E+00 | 2.81E+00 | 4.85E-01 | no | Wdr1 | intron_variant | MODIFIER |
| 5 | 38691183 | AX-170024730 | 0 | C | T | 0.342 | 7.76E-02 | 1.12E-01 | 2.76E+00 | 2.81E+00 | 4.85E-01 | no | | intergenic_variant | MODIFIER |
| 7 | 65516165 | AX-170504202 | 0 | C | T | 0.421 | -6.90E-02 | 1.00E-01 | 2.79E+00 | 2.87E+00 | 4.85E-01 | no | Tjp1 | intron_variant | MODIFIER |
| 7 | 65534881 | AX-170164575 | 0 | C | A | 0.421 | -6.90E-02 | 1.00E-01 | 2.79E+00 | 2.87E+00 | 4.85E-01 | no | | intergenic_variant | MODIFIER |
| 7 | 65534899 | AX-170267776 | 0 | T | C | 0.421 | -6.90E-02 | 1.00E-01 | 2.79E+00 | 2.87E+00 | 4.85E-01 | no | | intergenic_variant | MODIFIER |
| 7 | 65669678 | AX-170240228 | 0 | T | C | 0.421 | -6.90E-02 | 1.00E-01 | 2.79E+00 | 2.87E+00 | 4.85E-01 | no | Tarsl2 | intron_variant | MODIFIER |
| 7 | 65669934 | AX-170303798 | 0 | T | C | 0.421 | -6.90E-02 | 1.00E-01 | 2.79E+00 | 2.87E+00 | 4.85E-01 | no | Tarsl2 | intron_variant | MODIFIER |
| 7 | 65679485 | AX-170230975 | 0 | G | A | 0.421 | -6.90E-02 | 1.00E-01 | 2.79E+00 | 2.87E+00 | 4.85E-01 | no | Tarsl2 | intron_variant | MODIFIER |
| 7 | 65679534 | AX-97840704 | 0 | A | G | 0.421 | -6.90E-02 | 1.00E-01 | 2.79E+00 | 2.87E+00 | 4.85E-01 | no | Tarsl2 | intron_variant | MODIFIER |
| 7 | 65679712 | AX-170281866 | 0 | T | C | 0.421 | -6.90E-02 | 1.00E-01 | 2.79E+00 | 2.87E+00 | 4.85E-01 | no | Tarsl2 | intron_variant | MODIFIER |
| 7 | 65682742 | AX-170412573 | 0 | T | A | 0.421 | -6.90E-02 | 1.00E-01 | 2.79E+00 | 2.87E+00 | 4.85E-01 | no | Tarsl2 | intron_variant | MODIFIER |
| 7 | 65684556 | AX-170496473 | 0 | A | G | 0.421 | -6.90E-02 | 1.00E-01 | 2.79E+00 | 2.87E+00 | 4.85E-01 | no | Tarsl2 | intron_variant | MODIFIER |
| 7 | 65699317 | AX-170169871 | 0 | G | T | 0.421 | -6.90E-02 | 1.00E-01 | 2.79E+00 | 2.87E+00 | 4.85E-01 | no | Tm2d3 | intron_variant | MODIFIER |
| 7 | 65717775 | AX-170077566 | 0 | T | G | 0.421 | -6.90E-02 | 1.00E-01 | 2.79E+00 | 2.87E+00 | 4.85E-01 | no | | intergenic_variant | MODIFIER |
| 7 | 65737127 | AX-170304015 | 0 | T | C | 0.421 | -6.90E-02 | 1.00E-01 | 2.79E+00 | 2.87E+00 | 4.85E-01 | no | | regulatory_region_variant | MODIFIER |
| 7 | 65781440 | AX-170410006 | 0 | G | A | 0.421 | -6.90E-02 | 1.00E-01 | 2.79E+00 | 2.87E+00 | 4.85E-01 | no | Gm7551 | upstream_gene_variant | MODIFIER |
| 7 | 65881156 | AX-97820540 | 0 | A | G | 0.421 | -6.90E-02 | 1.00E-01 | 2.79E+00 | 2.87E+00 | 4.85E-01 | no | Pcsk6 | intron_variant | MODIFIER |
| 7 | 65900652 | AX-170089940 | 0 | C | T | 0.421 | -6.90E-02 | 1.00E-01 | 2.79E+00 | 2.87E+00 | 4.85E-01 | no | Pcsk6 | intron_variant | MODIFIER |
| 7 | 65980459 | AX-97814697 | 0 | G | A | 0.421 | -6.90E-02 | 1.00E-01 | 2.79E+00 | 2.87E+00 | 4.85E-01 | no | Pcsk6 | intron_variant | MODIFIER |
| 7 | 65992200 | AX-170323846 | 0 | T | C | 0.421 | -6.90E-02 | 1.00E-01 | 2.79E+00 | 2.87E+00 | 4.85E-01 | no | Pcsk6 | intron_variant | MODIFIER |
| 7 | 66043088 | AX-170421947 | 0 | C | T | 0.421 | -6.90E-02 | 1.00E-01 | 2.79E+00 | 2.87E+00 | 4.85E-01 | no | Pcsk6 | intron_variant | MODIFIER |
| 7 | 66046478 | AX-97817194 | 0 | C | T | 0.421 | -6.90E-02 | 1.00E-01 | 2.79E+00 | 2.87E+00 | 4.85E-01 | no | Pcsk6 | intron_variant | MODIFIER |
| 7 | 66047617 | AX-170336553 | 0 | T | C | 0.421 | -6.90E-02 | 1.00E-01 | 2.79E+00 | 2.87E+00 | 4.85E-01 | no | Pcsk6 | synonymous_variant | LOW |
| 7 | 66120123 | AX-170357269 | 0 | T | C | 0.421 | -6.90E-02 | 1.00E-01 | 2.79E+00 | 2.87E+00 | 4.85E-01 | no | Chsy1 | intron_variant | MODIFIER |
| 7 | 66159524 | AX-97847355 | 0 | G | A | 0.421 | -6.90E-02 | 1.00E-01 | 2.79E+00 | 2.87E+00 | 4.85E-01 | no | Chsy1 | intron_variant | MODIFIER |
| 7 | 66172651 | AX-170489368 | 0 | C | T | 0.421 | -6.90E-02 | 1.00E-01 | 2.79E+00 | 2.87E+00 | 4.85E-01 | no | Chsy1 | 3_prime_UTR_variant | MODIFIER |
| 7 | 66188674 | AX-170266348 | 0 | T | C | 0.421 | -6.90E-02 | 1.00E-01 | 2.79E+00 | 2.87E+00 | 4.85E-01 | no | Gm45080 | upstream_gene_variant | MODIFIER |

| | | | | | | | | | | | | | | | |
|---|---|---|---|---|---|---|---|---|---|---|---|---|---|---|---|
| 7 | 66192313 | AX-170305062 | 0 | C | T | 0.421 | -6.90E-02 | 1.00E-01 | 2.79E+00 | 2.87E+00 | 4.85E-01 | no | Gm45080 | downstream_gene_variant | MODIFIER |
| 7 | 66192404 | AX-170303373 | 0 | A | G | 0.421 | -6.90E-02 | 1.00E-01 | 2.79E+00 | 2.87E+00 | 4.85E-01 | no | Gm45080 | downstream_gene_variant | MODIFIER |
| 7 | 66193716 | AX-170278380 | 0 | T | C | 0.421 | -6.90E-02 | 1.00E-01 | 2.79E+00 | 2.87E+00 | 4.85E-01 | no | Gm45080 | downstream_gene_variant | MODIFIER |
| 7 | 66194327 | AX-170515532 | 0 | T | C | 0.421 | -6.90E-02 | 1.00E-01 | 2.79E+00 | 2.87E+00 | 4.85E-01 | no | Gm45080 | downstream_gene_variant | MODIFIER |
| 7 | 66194660 | AX-170176426 | 0 | G | C | 0.421 | -6.90E-02 | 1.00E-01 | 2.79E+00 | 2.87E+00 | 4.85E-01 | no | Gm45080 | downstream_gene_variant | MODIFIER |
| 7 | 66212493 | AX-170495098 | 0 | T | C | 0.421 | -6.90E-02 | 1.00E-01 | 2.79E+00 | 2.87E+00 | 4.85E-01 | no | | regulatory_region_variant | MODIFIER |
| 7 | 66233770 | AX-170483239 | 0 | G | A | 0.421 | -6.90E-02 | 1.00E-01 | 2.79E+00 | 2.87E+00 | 4.85E-01 | no | Lrrk1 | intron_variant,non_coding_transcript_variant | MODIFIER |
| 7 | 66254382 | AX-170329153 | 0 | G | A | 0.421 | -6.90E-02 | 1.00E-01 | 2.79E+00 | 2.87E+00 | 4.85E-01 | no | Lrrk1 | downstream_gene_variant | MODIFIER |
| 7 | 66287554 | AX-170306027 | 0 | C | T | 0.421 | -6.90E-02 | 1.00E-01 | 2.79E+00 | 2.87E+00 | 4.85E-01 | no | Lrrk1 | missense_variant | MODERATE |
| 7 | 66297369 | AX-170180071 | 0 | G | C | 0.421 | -6.90E-02 | 1.00E-01 | 2.79E+00 | 2.87E+00 | 4.85E-01 | no | Lrrk1 | intron_variant | MODIFIER |
| 7 | 66298190 | AX-170452947 | 0 | C | T | 0.421 | -6.90E-02 | 1.00E-01 | 2.79E+00 | 2.87E+00 | 4.85E-01 | no | Lrrk1 | intron_variant | MODIFIER |
| 7 | 66304384 | AX-170373546 | 0 | G | T | 0.421 | -6.90E-02 | 1.00E-01 | 2.79E+00 | 2.87E+00 | 4.85E-01 | no | Lrrk1 | intron_variant | MODIFIER |
| 7 | 66472412 | AX-170248498 | 0 | C | T | 0.421 | -6.90E-02 | 1.00E-01 | 2.79E+00 | 2.87E+00 | 4.85E-01 | no | | intergenic_variant | MODIFIER |
| 7 | 66525320 | AX-170123042 | 0 | A | G | 0.421 | -6.90E-02 | 1.00E-01 | 2.79E+00 | 2.87E+00 | 4.85E-01 | no | | intergenic_variant | MODIFIER |
| 7 | 66529326 | AX-170363174 | 0 | A | G | 0.421 | -6.90E-02 | 1.00E-01 | 2.79E+00 | 2.87E+00 | 4.85E-01 | no | | regulatory_region_variant | MODIFIER |
| 7 | 66573637 | AX-97856934 | 0 | A | C | 0.421 | -6.90E-02 | 1.00E-01 | 2.79E+00 | 2.87E+00 | 4.85E-01 | no | | intergenic_variant | MODIFIER |
| 10 | 71592974 | AX-170003445 | 0 | T | C | 0.412 | -5.59E-02 | 8.07E-02 | 2.64E+00 | 2.72E+00 | 4.85E-01 | no | Gm34544 | intron_variant,non_coding_transcript_variant | MODIFIER |
| 1 | 143354418 | AX-170321247 | 0 | A | C | 0.202 | 9.76E-02 | 1.41E-01 | 2.67E+00 | 2.71E+00 | 4.85E-01 | no | | intergenic_variant | MODIFIER |
| 1 | 145023724 | AX-170507114 | 0 | G | A | 0.202 | 9.76E-02 | 1.41E-01 | 2.67E+00 | 2.71E+00 | 4.85E-01 | no | | intergenic_variant | MODIFIER |
| 1 | 145481731 | AX-170001915 | 0 | T | C | 0.202 | 9.76E-02 | 1.41E-01 | 2.67E+00 | 2.71E+00 | 4.85E-01 | no | | intergenic_variant | MODIFIER |
| 1 | 145521884 | AX-170001912 | 0 | C | A | 0.202 | 9.76E-02 | 1.41E-01 | 2.67E+00 | 2.71E+00 | 4.85E-01 | no | | intergenic_variant | MODIFIER |
| 1 | 145599206 | AX-170448898 | 0 | G | C | 0.202 | 9.76E-02 | 1.41E-01 | 2.67E+00 | 2.71E+00 | 4.85E-01 | no | | regulatory_region_variant | MODIFIER |
| 1 | 146473882 | AX-97800537 | 0 | T | C | 0.202 | 9.76E-02 | 1.41E-01 | 2.67E+00 | 2.71E+00 | 4.85E-01 | no | Gm29514 | intron_variant,non_coding_transcript_variant | MODIFIER |
| 1 | 146486066 | AX-170450558 | 0 | A | T | 0.202 | 9.76E-02 | 1.41E-01 | 2.67E+00 | 2.71E+00 | 4.85E-01 | no | Gm29514 | intron_variant,non_coding_transcript_variant | MODIFIER |
| 1 | 146503968 | AX-97855479 | 0 | T | C | 0.202 | 9.76E-02 | 1.41E-01 | 2.67E+00 | 2.71E+00 | 4.85E-01 | no | Brinp3 | intron_variant | MODIFIER |
| 1 | 146632122 | AX-170001938 | 0 | A | T | 0.202 | 9.76E-02 | 1.41E-01 | 2.67E+00 | 2.71E+00 | 4.85E-01 | no | Brinp3 | intron_variant | MODIFIER |
| 1 | 146656910 | AX-170001927 | 0 | C | A | 0.202 | 9.76E-02 | 1.41E-01 | 2.67E+00 | 2.71E+00 | 4.85E-01 | no | Brinp3 | intron_variant | MODIFIER |
| 1 | 146693265 | AX-170482335 | 0 | G | A | 0.202 | 9.76E-02 | 1.41E-01 | 2.67E+00 | 2.71E+00 | 4.85E-01 | no | Brinp3 | intron_variant | MODIFIER |
| 1 | 146827194 | AX-97802587 | 0 | A | G | 0.202 | 9.76E-02 | 1.41E-01 | 2.67E+00 | 2.71E+00 | 4.85E-01 | no | Brinp3 | intron_variant | MODIFIER |
| 1 | 146839357 | AX-170001939 | 0 | A | G | 0.202 | 9.76E-02 | 1.41E-01 | 2.67E+00 | 2.71E+00 | 4.85E-01 | no | Brinp3 | intron_variant | MODIFIER |
| 1 | 146844427 | AX-170001928 | 0 | A | C | 0.202 | 9.76E-02 | 1.41E-01 | 2.67E+00 | 2.71E+00 | 4.85E-01 | no | Brinp3 | intron_variant | MODIFIER |
| 1 | 147035175 | AX-97832522 | 0 | G | A | 0.202 | 9.76E-02 | 1.41E-01 | 2.67E+00 | 2.71E+00 | 4.85E-01 | no | | intergenic_variant | MODIFIER |
| 1 | 147127220 | AX-170001941 | 0 | A | G | 0.202 | 9.76E-02 | 1.41E-01 | 2.67E+00 | 2.71E+00 | 4.85E-01 | no | | intergenic_variant | MODIFIER |
| 1 | 147127426 | AX-170189040 | 0 | G | T | 0.202 | 9.76E-02 | 1.41E-01 | 2.67E+00 | 2.71E+00 | 4.85E-01 | no | | intergenic_variant | MODIFIER |
| 1 | 147129805 | AX-170289708 | 0 | T | C | 0.202 | 9.76E-02 | 1.41E-01 | 2.67E+00 | 2.71E+00 | 4.85E-01 | no | | intergenic_variant | MODIFIER |
| 1 | 147130116 | AX-170127815 | 0 | T | G | 0.202 | 9.76E-02 | 1.41E-01 | 2.67E+00 | 2.71E+00 | 4.85E-01 | no | | intergenic_variant | MODIFIER |
| 1 | 147130437 | AX-170502363 | 0 | A | C | 0.202 | 9.76E-02 | 1.41E-01 | 2.67E+00 | 2.71E+00 | 4.85E-01 | no | | intergenic_variant | MODIFIER |
| 1 | 147130682 | AX-170117554 | 0 | G | A | 0.202 | 9.76E-02 | 1.41E-01 | 2.67E+00 | 2.71E+00 | 4.85E-01 | no | | intergenic_variant | MODIFIER |
| 1 | 147149602 | AX-97844678 | 0 | A | G | 0.202 | 9.76E-02 | 1.41E-01 | 2.67E+00 | 2.71E+00 | 4.85E-01 | no | | intergenic_variant | MODIFIER |
| 1 | 147174772 | AX-170193450 | 0 | C | T | 0.202 | 9.76E-02 | 1.41E-01 | 2.67E+00 | 2.71E+00 | 4.85E-01 | no | | intergenic_variant | MODIFIER |
| 1 | 147176540 | AX-170001940 | 0 | C | T | 0.202 | 9.76E-02 | 1.41E-01 | 2.67E+00 | 2.71E+00 | 4.85E-01 | no | | intergenic_variant | MODIFIER |
| 1 | 147187376 | AX-170001942 | 0 | A | G | 0.202 | 9.76E-02 | 1.41E-01 | 2.67E+00 | 2.71E+00 | 4.85E-01 | no | | intergenic_variant | MODIFIER |
| 1 | 147188548 | AX-170275995 | 0 | C | A | 0.202 | 9.76E-02 | 1.41E-01 | 2.67E+00 | 2.71E+00 | 4.85E-01 | no | | regulatory_region_variant | MODIFIER |
| 1 | 147188779 | AX-170407976 | 0 | A | G | 0.202 | 9.76E-02 | 1.41E-01 | 2.67E+00 | 2.71E+00 | 4.85E-01 | no | | intergenic_variant | MODIFIER |
| 1 | 147210117 | AX-170471688 | 0 | T | C | 0.202 | 9.76E-02 | 1.41E-01 | 2.67E+00 | 2.71E+00 | 4.85E-01 | no | | intergenic_variant | MODIFIER |
| 1 | 147246915 | AX-170143881 | 0 | C | G | 0.202 | 9.76E-02 | 1.41E-01 | 2.67E+00 | 2.71E+00 | 4.85E-01 | no | | intergenic_variant | MODIFIER |
| 1 | 147248781 | AX-170001943 | 0 | T | C | 0.202 | 9.76E-02 | 1.41E-01 | 2.67E+00 | 2.71E+00 | 4.85E-01 | no | | intergenic_variant | MODIFIER |
| 1 | 147253185 | AX-97800273 | 0 | T | C | 0.202 | 9.76E-02 | 1.41E-01 | 2.67E+00 | 2.71E+00 | 4.85E-01 | no | | intergenic_variant | MODIFIER |
| 1 | 147280575 | AX-97806057 | 0 | A | G | 0.202 | 9.76E-02 | 1.41E-01 | 2.67E+00 | 2.71E+00 | 4.85E-01 | no | Gm9931 | downstream_gene_variant | MODIFIER |
| 1 | 147281852 | AX-170534575 | 0 | G | A | 0.202 | 9.76E-02 | 1.41E-01 | 2.67E+00 | 2.71E+00 | 4.85E-01 | no | Gm9931 | non_coding_transcript_exon_variant | MODIFIER |
| 1 | 147298797 | AX-170091734 | 0 | G | A | 0.202 | 9.76E-02 | 1.41E-01 | 2.67E+00 | 2.71E+00 | 4.85E-01 | no | | intergenic_variant | MODIFIER |
| 1 | 147321795 | AX-170533051 | 0 | G | A | 0.202 | 9.76E-02 | 1.41E-01 | 2.67E+00 | 2.71E+00 | 4.85E-01 | no | | intergenic_variant | MODIFIER |
| 1 | 147357896 | AX-170043049 | 0 | A | G | 0.202 | 9.76E-02 | 1.41E-01 | 2.67E+00 | 2.71E+00 | 4.85E-01 | no | | intergenic_variant | MODIFIER |
| 1 | 147404581 | AX-97801367 | 0 | A | C | 0.202 | 9.76E-02 | 1.41E-01 | 2.67E+00 | 2.71E+00 | 4.85E-01 | no | | intergenic_variant | MODIFIER |

| | | | | | | | | | | | | | | | |
|---|---|---|---|---|---|---|---|---|---|---|---|---|---|---|---|
| 1 | 147422233 | AX-170110784 | 0 | T | C | 0.202 | 9.76E-02 | 1.41E-01 | 2.67E+00 | 2.71E+00 | 4.85E-01 | no | | intergenic_variant | MODIFIER |
| 1 | 147432170 | AX-97825075 | 0 | T | G | 0.202 | 9.76E-02 | 1.41E-01 | 2.67E+00 | 2.71E+00 | 4.85E-01 | no | | intergenic_variant | MODIFIER |
| 1 | 147434793 | AX-170533120 | 0 | C | T | 0.202 | 9.76E-02 | 1.41E-01 | 2.67E+00 | 2.71E+00 | 4.85E-01 | no | | intergenic_variant | MODIFIER |
| 1 | 147514996 | AX-97813640 | 0 | A | G | 0.202 | 9.76E-02 | 1.41E-01 | 2.67E+00 | 2.71E+00 | 4.85E-01 | no | Gm38284 | upstream_gene_variant | MODIFIER |
| 1 | 147515265 | AX-170227047 | 0 | C | G | 0.202 | 9.76E-02 | 1.41E-01 | 2.67E+00 | 2.71E+00 | 4.85E-01 | no | Gm38284 | upstream_gene_variant | MODIFIER |
| 1 | 147532966 | AX-170412310 | 0 | C | G | 0.202 | 9.76E-02 | 1.41E-01 | 2.67E+00 | 2.71E+00 | 4.85E-01 | no | | intergenic_variant | MODIFIER |
| 1 | 147536523 | AX-170488611 | 0 | C | T | 0.202 | 9.76E-02 | 1.41E-01 | 2.67E+00 | 2.71E+00 | 4.85E-01 | no | | intergenic_variant | MODIFIER |
| 1 | 147542366 | AX-170317971 | 0 | T | C | 0.202 | 9.76E-02 | 1.41E-01 | 2.67E+00 | 2.71E+00 | 4.85E-01 | no | | intergenic_variant | MODIFIER |
| 1 | 147591351 | AX-170097726 | 0 | A | C | 0.202 | 9.76E-02 | 1.41E-01 | 2.67E+00 | 2.71E+00 | 4.85E-01 | no | | intergenic_variant | MODIFIER |
| 1 | 147613737 | AX-97798316 | 0 | C | T | 0.202 | 9.76E-02 | 1.41E-01 | 2.67E+00 | 2.71E+00 | 4.85E-01 | no | | intergenic_variant | MODIFIER |
| 1 | 147631553 | AX-170043050 | 0 | G | A | 0.202 | 9.76E-02 | 1.41E-01 | 2.67E+00 | 2.71E+00 | 4.85E-01 | no | | intergenic_variant | MODIFIER |
| 1 | 147658748 | AX-97813671 | 0 | A | G | 0.202 | 9.76E-02 | 1.41E-01 | 2.67E+00 | 2.71E+00 | 4.85E-01 | no | | intergenic_variant | MODIFIER |
| 1 | 147658798 | AX-170001948 | 0 | T | C | 0.202 | 9.76E-02 | 1.41E-01 | 2.67E+00 | 2.71E+00 | 4.85E-01 | no | | intergenic_variant | MODIFIER |
| 1 | 147682189 | AX-170043057 | 0 | T | C | 0.202 | 9.76E-02 | 1.41E-01 | 2.67E+00 | 2.71E+00 | 4.85E-01 | no | | intergenic_variant | MODIFIER |
| 1 | 147690296 | AX-170043055 | 0 | A | G | 0.202 | 9.76E-02 | 1.41E-01 | 2.67E+00 | 2.71E+00 | 4.85E-01 | no | | intergenic_variant | MODIFIER |
| 1 | 147690753 | AX-97796602 | 0 | A | G | 0.202 | 9.76E-02 | 1.41E-01 | 2.67E+00 | 2.71E+00 | 4.85E-01 | no | | intergenic_variant | MODIFIER |
| 1 | 147690905 | AX-170522541 | 0 | A | G | 0.202 | 9.76E-02 | 1.41E-01 | 2.67E+00 | 2.71E+00 | 4.85E-01 | no | | intergenic_variant | MODIFIER |
| 1 | 147722074 | AX-170434562 | 0 | C | A | 0.202 | 9.76E-02 | 1.41E-01 | 2.67E+00 | 2.71E+00 | 4.85E-01 | no | | intergenic_variant | MODIFIER |
| 1 | 147744255 | AX-97783704 | 0 | T | C | 0.202 | 9.76E-02 | 1.41E-01 | 2.67E+00 | 2.71E+00 | 4.85E-01 | no | | intergenic_variant | MODIFIER |
| 1 | 147779617 | AX-170310349 | 0 | G | T | 0.202 | 9.76E-02 | 1.41E-01 | 2.67E+00 | 2.71E+00 | 4.85E-01 | no | | intergenic_variant | MODIFIER |
| 1 | 147787230 | AX-170429059 | 0 | T | C | 0.202 | 9.76E-02 | 1.41E-01 | 2.67E+00 | 2.71E+00 | 4.85E-01 | no | | intergenic_variant | MODIFIER |
| 1 | 147787364 | AX-170328494 | 0 | T | G | 0.202 | 9.76E-02 | 1.41E-01 | 2.67E+00 | 2.71E+00 | 4.85E-01 | no | | intergenic_variant | MODIFIER |
| 1 | 147943076 | AX-97805573 | 0 | T | C | 0.202 | 9.76E-02 | 1.41E-01 | 2.67E+00 | 2.71E+00 | 4.85E-01 | no | | intergenic_variant | MODIFIER |
| 1 | 147959222 | AX-170153448 | 0 | G | A | 0.202 | 9.76E-02 | 1.41E-01 | 2.67E+00 | 2.71E+00 | 4.85E-01 | no | | intergenic_variant | MODIFIER |
| 1 | 147984690 | AX-170043064 | 0 | G | T | 0.202 | 9.76E-02 | 1.41E-01 | 2.67E+00 | 2.71E+00 | 4.85E-01 | no | | intergenic_variant | MODIFIER |
| 1 | 148017672 | AX-170043063 | 0 | T | G | 0.202 | 9.76E-02 | 1.41E-01 | 2.67E+00 | 2.71E+00 | 4.85E-01 | no | | intergenic_variant | MODIFIER |
| 1 | 148033034 | AX-170387561 | 0 | T | C | 0.202 | 9.76E-02 | 1.41E-01 | 2.67E+00 | 2.71E+00 | 4.85E-01 | no | | intergenic_variant | MODIFIER |
| 1 | 148037146 | AX-170289650 | 0 | A | G | 0.202 | 9.76E-02 | 1.41E-01 | 2.67E+00 | 2.71E+00 | 4.85E-01 | no | | intergenic_variant | MODIFIER |
| 1 | 148038550 | AX-170261420 | 0 | A | T | 0.202 | 9.76E-02 | 1.41E-01 | 2.67E+00 | 2.71E+00 | 4.85E-01 | no | | intergenic_variant | MODIFIER |
| 1 | 148056689 | AX-97850122 | 0 | G | T | 0.202 | 9.76E-02 | 1.41E-01 | 2.67E+00 | 2.71E+00 | 4.85E-01 | no | | intergenic_variant | MODIFIER |
| 1 | 148228103 | AX-170155880 | 0 | C | T | 0.202 | 9.76E-02 | 1.41E-01 | 2.67E+00 | 2.71E+00 | 4.85E-01 | no | | intergenic_variant | MODIFIER |
| 1 | 148246163 | AX-170111323 | 0 | T | G | 0.202 | 9.76E-02 | 1.41E-01 | 2.67E+00 | 2.71E+00 | 4.85E-01 | no | Gm29515 | intron_variant,non_coding_transcript_variant | MODIFIER |
| 1 | 148262393 | AX-170120328 | 0 | G | A | 0.202 | 9.76E-02 | 1.41E-01 | 2.67E+00 | 2.71E+00 | 4.85E-01 | no | | intergenic_variant | MODIFIER |
| 1 | 148280485 | AX-170487119 | 0 | A | G | 0.202 | 9.76E-02 | 1.41E-01 | 2.67E+00 | 2.71E+00 | 4.85E-01 | no | | intergenic_variant | MODIFIER |
| 1 | 148286615 | AX-170043070 | 0 | A | G | 0.202 | 9.76E-02 | 1.41E-01 | 2.67E+00 | 2.71E+00 | 4.85E-01 | no | | intergenic_variant | MODIFIER |
| 1 | 148288151 | AX-170043066 | 0 | A | T | 0.202 | 9.76E-02 | 1.41E-01 | 2.67E+00 | 2.71E+00 | 4.85E-01 | no | | intergenic_variant | MODIFIER |
| 1 | 148313904 | AX-97818872 | 0 | C | A | 0.202 | 9.76E-02 | 1.41E-01 | 2.67E+00 | 2.71E+00 | 4.85E-01 | no | | intergenic_variant | MODIFIER |
| 1 | 148354285 | AX-97812874 | 0 | A | C | 0.202 | 9.76E-02 | 1.41E-01 | 2.67E+00 | 2.71E+00 | 4.85E-01 | no | | intergenic_variant | MODIFIER |
| 1 | 148393348 | AX-97816559 | 0 | A | G | 0.202 | 9.76E-02 | 1.41E-01 | 2.67E+00 | 2.71E+00 | 4.85E-01 | no | | intergenic_variant | MODIFIER |
| 1 | 148409011 | AX-97797823 | 0 | A | G | 0.202 | 9.76E-02 | 1.41E-01 | 2.67E+00 | 2.71E+00 | 4.85E-01 | no | | intergenic_variant | MODIFIER |
| 1 | 148409076 | AX-170388765 | 0 | A | G | 0.202 | 9.76E-02 | 1.41E-01 | 2.67E+00 | 2.71E+00 | 4.85E-01 | no | | intergenic_variant | MODIFIER |
| 1 | 148420988 | AX-170001952 | 0 | A | G | 0.202 | 9.76E-02 | 1.41E-01 | 2.67E+00 | 2.71E+00 | 4.85E-01 | no | | intergenic_variant | MODIFIER |
| 1 | 148455535 | AX-97856991 | 0 | A | G | 0.202 | 9.76E-02 | 1.41E-01 | 2.67E+00 | 2.71E+00 | 4.85E-01 | no | | intergenic_variant | MODIFIER |
| 1 | 148455657 | AX-170001956 | 0 | T | A | 0.202 | 9.76E-02 | 1.41E-01 | 2.67E+00 | 2.71E+00 | 4.85E-01 | no | | intergenic_variant | MODIFIER |
| 1 | 148484164 | AX-97794871 | 0 | C | T | 0.202 | 9.76E-02 | 1.41E-01 | 2.67E+00 | 2.71E+00 | 4.85E-01 | no | | intergenic_variant | MODIFIER |
| 1 | 148495574 | AX-170126865 | 0 | G | A | 0.202 | 9.76E-02 | 1.41E-01 | 2.67E+00 | 2.71E+00 | 4.85E-01 | no | Gm19891 | upstream_gene_variant | MODIFIER |
| 1 | 148577565 | AX-97844011 | 0 | G | T | 0.202 | 9.76E-02 | 1.41E-01 | 2.67E+00 | 2.71E+00 | 4.85E-01 | no | | intergenic_variant | MODIFIER |
| 1 | 148585983 | AX-170351560 | 0 | C | G | 0.202 | 9.76E-02 | 1.41E-01 | 2.67E+00 | 2.71E+00 | 4.85E-01 | no | | intergenic_variant | MODIFIER |
| 1 | 148621346 | AX-170152097 | 0 | T | G | 0.202 | 9.76E-02 | 1.41E-01 | 2.67E+00 | 2.71E+00 | 4.85E-01 | no | | intergenic_variant | MODIFIER |
| 1 | 148648623 | AX-170310178 | 0 | C | T | 0.202 | 9.76E-02 | 1.41E-01 | 2.67E+00 | 2.71E+00 | 4.85E-01 | no | Gm18505 | downstream_gene_variant | MODIFIER |
| 1 | 148650872 | AX-170151737 | 0 | T | C | 0.202 | 9.76E-02 | 1.41E-01 | 2.67E+00 | 2.71E+00 | 4.85E-01 | no | Gm18505 | non_coding_transcript_exon_variant | MODIFIER |
| 1 | 148669370 | AX-170331928 | 0 | C | G | 0.202 | 9.76E-02 | 1.41E-01 | 2.67E+00 | 2.71E+00 | 4.85E-01 | no | | intergenic_variant | MODIFIER |
| 1 | 148696532 | AX-170309756 | 0 | A | G | 0.202 | 9.76E-02 | 1.41E-01 | 2.67E+00 | 2.71E+00 | 4.85E-01 | no | | intergenic_variant | MODIFIER |
| 1 | 148704221 | AX-170531289 | 0 | C | T | 0.202 | 9.76E-02 | 1.41E-01 | 2.67E+00 | 2.71E+00 | 4.85E-01 | no | | intergenic_variant | MODIFIER |

| | | | | | | | | | | | | | | | |
|---|---|---|---|---|---|---|---|---|---|---|---|---|---|---|---|
| 1 | 148711069 | AX-97820484 | 0 | A | G | 0.202 | 9.76E-02 | 1.41E-01 | 2.67E+00 | 2.71E+00 | 4.85E-01 | no | | intergenic_variant | MODIFIER |
| 1 | 148755769 | AX-170287610 | 0 | T | G | 0.202 | 9.76E-02 | 1.41E-01 | 2.67E+00 | 2.71E+00 | 4.85E-01 | no | | intergenic_variant | MODIFIER |
| 1 | 148775095 | AX-170043073 | 0 | A | G | 0.202 | 9.76E-02 | 1.41E-01 | 2.67E+00 | 2.71E+00 | 4.85E-01 | no | | intergenic_variant | MODIFIER |
| 1 | 148775122 | AX-170506161 | 0 | A | C | 0.202 | 9.76E-02 | 1.41E-01 | 2.67E+00 | 2.71E+00 | 4.85E-01 | no | | intergenic_variant | MODIFIER |
| 1 | 148783014 | AX-170192468 | 0 | A | C | 0.202 | 9.76E-02 | 1.41E-01 | 2.67E+00 | 2.71E+00 | 4.85E-01 | no | | intergenic_variant | MODIFIER |
| 1 | 148792672 | AX-170144150 | 0 | A | G | 0.202 | 9.76E-02 | 1.41E-01 | 2.67E+00 | 2.71E+00 | 4.85E-01 | no | | intergenic_variant | MODIFIER |
| 1 | 148814893 | AX-170164035 | 0 | G | C | 0.202 | 9.76E-02 | 1.41E-01 | 2.67E+00 | 2.71E+00 | 4.85E-01 | no | | intergenic_variant | MODIFIER |
| 1 | 148827960 | AX-170344621 | 0 | T | C | 0.202 | 9.76E-02 | 1.41E-01 | 2.67E+00 | 2.71E+00 | 4.85E-01 | no | | intergenic_variant | MODIFIER |
| 1 | 148827984 | AX-170293822 | 0 | T | G | 0.202 | 9.76E-02 | 1.41E-01 | 2.67E+00 | 2.71E+00 | 4.85E-01 | no | | intergenic_variant | MODIFIER |
| 1 | 148829936 | AX-170205185 | 0 | A | G | 0.202 | 9.76E-02 | 1.41E-01 | 2.67E+00 | 2.71E+00 | 4.85E-01 | no | | intergenic_variant | MODIFIER |
| 1 | 148832774 | AX-170417941 | 0 | G | C | 0.202 | 9.76E-02 | 1.41E-01 | 2.67E+00 | 2.71E+00 | 4.85E-01 | no | | intergenic_variant | MODIFIER |
| 1 | 148851991 | AX-97850100 | 0 | T | C | 0.202 | 9.76E-02 | 1.41E-01 | 2.67E+00 | 2.71E+00 | 4.85E-01 | no | | intergenic_variant | MODIFIER |
| 1 | 148907084 | AX-170349534 | 0 | A | G | 0.202 | 9.76E-02 | 1.41E-01 | 2.67E+00 | 2.71E+00 | 4.85E-01 | no | | intergenic_variant | MODIFIER |
| 1 | 148914149 | AX-97842809 | 0 | C | T | 0.202 | 9.76E-02 | 1.41E-01 | 2.67E+00 | 2.71E+00 | 4.85E-01 | no | | intergenic_variant | MODIFIER |
| 1 | 148920775 | AX-170001961 | 0 | T | G | 0.202 | 9.76E-02 | 1.41E-01 | 2.67E+00 | 2.71E+00 | 4.85E-01 | no | | intergenic_variant | MODIFIER |
| 1 | 148931381 | AX-170500366 | 0 | T | C | 0.202 | 9.76E-02 | 1.41E-01 | 2.67E+00 | 2.71E+00 | 4.85E-01 | no | | intergenic_variant | MODIFIER |
| 1 | 148950988 | AX-170227815 | 0 | C | T | 0.202 | 9.76E-02 | 1.41E-01 | 2.67E+00 | 2.71E+00 | 4.85E-01 | no | | intergenic_variant | MODIFIER |
| 1 | 148963600 | AX-170001960 | 0 | A | C | 0.202 | 9.76E-02 | 1.41E-01 | 2.67E+00 | 2.71E+00 | 4.85E-01 | no | | intergenic_variant | MODIFIER |
| 1 | 148978382 | AX-170043078 | 0 | C | A | 0.202 | 9.76E-02 | 1.41E-01 | 2.67E+00 | 2.71E+00 | 4.85E-01 | no | Gm5703 | non_coding_transcript_exon_variant | MODIFIER |
| 1 | 148978796 | AX-170043076 | 0 | A | G | 0.202 | 9.76E-02 | 1.41E-01 | 2.67E+00 | 2.71E+00 | 4.85E-01 | no | Gm5703 | non_coding_transcript_exon_variant | MODIFIER |
| 1 | 148982538 | AX-97802241 | 0 | T | C | 0.202 | 9.76E-02 | 1.41E-01 | 2.67E+00 | 2.71E+00 | 4.85E-01 | no | Gm5703 | downstream_gene_variant | MODIFIER |
| 1 | 149020613 | AX-170509776 | 0 | G | A | 0.202 | 9.76E-02 | 1.41E-01 | 2.67E+00 | 2.71E+00 | 4.85E-01 | no | Gm15443 | downstream_gene_variant | MODIFIER |
| 1 | 149020789 | AX-170256558 | 0 | A | G | 0.202 | 9.76E-02 | 1.41E-01 | 2.67E+00 | 2.71E+00 | 4.85E-01 | no | Gm15443 | downstream_gene_variant | MODIFIER |
| 1 | 149088812 | AX-97792950 | 0 | T | C | 0.202 | 9.76E-02 | 1.41E-01 | 2.67E+00 | 2.71E+00 | 4.85E-01 | no | | intergenic_variant | MODIFIER |
| 1 | 149089076 | AX-170464092 | 0 | C | G | 0.202 | 9.76E-02 | 1.41E-01 | 2.67E+00 | 2.71E+00 | 4.85E-01 | no | | intergenic_variant | MODIFIER |
| 1 | 149106421 | AX-170190695 | 0 | A | C | 0.202 | 9.76E-02 | 1.41E-01 | 2.67E+00 | 2.71E+00 | 4.85E-01 | no | Gm29020 | intron_variant,non_coding_transcript_variant | MODIFIER |
| 1 | 149107463 | AX-170370053 | 0 | A | G | 0.202 | 9.76E-02 | 1.41E-01 | 2.67E+00 | 2.71E+00 | 4.85E-01 | no | Gm29020 | intron_variant,non_coding_transcript_variant | MODIFIER |
| 1 | 149107860 | AX-170097595 | 0 | G | A | 0.202 | 9.76E-02 | 1.41E-01 | 2.67E+00 | 2.71E+00 | 4.85E-01 | no | Gm29020 | non_coding_transcript_exon_variant | MODIFIER |
| 1 | 149113793 | AX-170091740 | 0 | G | A | 0.202 | 9.76E-02 | 1.41E-01 | 2.67E+00 | 2.71E+00 | 4.85E-01 | no | Gm29020 | intron_variant,non_coding_transcript_variant | MODIFIER |
| 1 | 149165052 | AX-170196573 | 0 | A | G | 0.202 | 9.76E-02 | 1.41E-01 | 2.67E+00 | 2.71E+00 | 4.85E-01 | no | | intergenic_variant | MODIFIER |
| 1 | 149173671 | AX-97820834 | 0 | C | A | 0.202 | 9.76E-02 | 1.41E-01 | 2.67E+00 | 2.71E+00 | 4.85E-01 | no | Gm18701 | non_coding_transcript_exon_variant | MODIFIER |
| 1 | 149200690 | AX-170311262 | 0 | T | C | 0.202 | 9.76E-02 | 1.41E-01 | 2.67E+00 | 2.71E+00 | 4.85E-01 | no | | intergenic_variant | MODIFIER |
| 1 | 149201796 | AX-170260711 | 0 | C | T | 0.202 | 9.76E-02 | 1.41E-01 | 2.67E+00 | 2.71E+00 | 4.85E-01 | no | | intergenic_variant | MODIFIER |
| 1 | 149205362 | AX-170440624 | 0 | G | A | 0.202 | 9.76E-02 | 1.41E-01 | 2.67E+00 | 2.71E+00 | 4.85E-01 | no | | intergenic_variant | MODIFIER |
| 1 | 149216080 | AX-170220393 | 0 | C | G | 0.202 | 9.76E-02 | 1.41E-01 | 2.67E+00 | 2.71E+00 | 4.85E-01 | no | | intergenic_variant | MODIFIER |
| 1 | 149218515 | AX-170036052 | 0 | A | C | 0.202 | 9.76E-02 | 1.41E-01 | 2.67E+00 | 2.71E+00 | 4.85E-01 | no | | intergenic_variant | MODIFIER |
| 1 | 149237202 | AX-170131439 | 0 | C | A | 0.202 | 9.76E-02 | 1.41E-01 | 2.67E+00 | 2.71E+00 | 4.85E-01 | no | | intergenic_variant | MODIFIER |
| 1 | 149290666 | AX-97813128 | 0 | T | C | 0.202 | 9.76E-02 | 1.41E-01 | 2.67E+00 | 2.71E+00 | 4.85E-01 | no | | intergenic_variant | MODIFIER |
| 1 | 149307620 | AX-170469007 | 0 | T | G | 0.202 | 9.76E-02 | 1.41E-01 | 2.67E+00 | 2.71E+00 | 4.85E-01 | no | | intergenic_variant | MODIFIER |
| 1 | 149328744 | AX-170286168 | 0 | G | A | 0.202 | 9.76E-02 | 1.41E-01 | 2.67E+00 | 2.71E+00 | 4.85E-01 | no | | intergenic_variant | MODIFIER |
| 1 | 149328969 | AX-97841479 | 0 | A | T | 0.202 | 9.76E-02 | 1.41E-01 | 2.67E+00 | 2.71E+00 | 4.85E-01 | no | | intergenic_variant | MODIFIER |
| 1 | 149333309 | AX-170036053 | 0 | C | T | 0.202 | 9.76E-02 | 1.41E-01 | 2.67E+00 | 2.71E+00 | 4.85E-01 | no | | intergenic_variant | MODIFIER |
| 1 | 149333411 | AX-170036054 | 0 | T | C | 0.202 | 9.76E-02 | 1.41E-01 | 2.67E+00 | 2.71E+00 | 4.85E-01 | no | | intergenic_variant | MODIFIER |
| 1 | 149353390 | AX-170043087 | 0 | A | G | 0.202 | 9.76E-02 | 1.41E-01 | 2.67E+00 | 2.71E+00 | 4.85E-01 | no | | intergenic_variant | MODIFIER |
| 1 | 149366790 | AX-170298431 | 0 | G | A | 0.202 | 9.76E-02 | 1.41E-01 | 2.67E+00 | 2.71E+00 | 4.85E-01 | no | | intergenic_variant | MODIFIER |
| 1 | 149381852 | AX-170036056 | 0 | G | A | 0.202 | 9.76E-02 | 1.41E-01 | 2.67E+00 | 2.71E+00 | 4.85E-01 | no | | intergenic_variant | MODIFIER |
| 1 | 149390512 | AX-170282012 | 0 | C | A | 0.202 | 9.76E-02 | 1.41E-01 | 2.67E+00 | 2.71E+00 | 4.85E-01 | no | | intergenic_variant | MODIFIER |
| 1 | 149391029 | AX-97789819 | 0 | T | C | 0.202 | 9.76E-02 | 1.41E-01 | 2.67E+00 | 2.71E+00 | 4.85E-01 | no | | intergenic_variant | MODIFIER |
| 1 | 149410769 | AX-170403632 | 0 | G | A | 0.202 | 9.76E-02 | 1.41E-01 | 2.67E+00 | 2.71E+00 | 4.85E-01 | no | Gm29398 | downstream_gene_variant | MODIFIER |
| 1 | 149436097 | AX-170280395 | 0 | C | T | 0.202 | 9.76E-02 | 1.41E-01 | 2.67E+00 | 2.71E+00 | 4.85E-01 | no | Gm29398 | intron_variant,non_coding_transcript_variant | MODIFIER |
| 1 | 149460506 | AX-97795752 | 0 | A | G | 0.202 | 9.76E-02 | 1.41E-01 | 2.67E+00 | 2.71E+00 | 4.85E-01 | no | | intergenic_variant | MODIFIER |
| 1 | 149462361 | AX-170402977 | 0 | T | C | 0.202 | 9.76E-02 | 1.41E-01 | 2.67E+00 | 2.71E+00 | 4.85E-01 | no | | intergenic_variant | MODIFIER |
| 1 | 149525441 | AX-170115688 | 0 | A | T | 0.202 | 9.76E-02 | 1.41E-01 | 2.67E+00 | 2.71E+00 | 4.85E-01 | no | | intergenic_variant | MODIFIER |
| 1 | 149531086 | AX-170515019 | 0 | G | A | 0.202 | 9.76E-02 | 1.41E-01 | 2.67E+00 | 2.71E+00 | 4.85E-01 | no | | intergenic_variant | MODIFIER |

| | | | | | | | | | | | | | | | |
|---|---|---|---|---|---|---|---|---|---|---|---|---|---|---|---|
| 1 | 149533115 | AX-170043091 | 0 | A | G | 0.202 | 9.76E-02 | 1.41E-01 | 2.67E+00 | 2.71E+00 | 4.85E-01 | no | | intergenic_variant | MODIFIER |
| 1 | 149542559 | AX-170515502 | 0 | T | C | 0.202 | 9.76E-02 | 1.41E-01 | 2.67E+00 | 2.71E+00 | 4.85E-01 | no | | intergenic_variant | MODIFIER |
| 1 | 149543007 | AX-170266495 | 0 | C | G | 0.202 | 9.76E-02 | 1.41E-01 | 2.67E+00 | 2.71E+00 | 4.85E-01 | no | | intergenic_variant | MODIFIER |
| 1 | 149546564 | AX-170001968 | 0 | G | T | 0.202 | 9.76E-02 | 1.41E-01 | 2.67E+00 | 2.71E+00 | 4.85E-01 | no | | intergenic_variant | MODIFIER |
| 1 | 149599336 | AX-170091744 | 0 | C | T | 0.202 | 9.76E-02 | 1.41E-01 | 2.67E+00 | 2.71E+00 | 4.85E-01 | no | | intergenic_variant | MODIFIER |
| 1 | 149647854 | AX-170197436 | 0 | A | G | 0.202 | 9.76E-02 | 1.41E-01 | 2.67E+00 | 2.71E+00 | 4.85E-01 | no | | intergenic_variant | MODIFIER |
| 1 | 149669735 | AX-170134273 | 0 | A | C | 0.202 | 9.76E-02 | 1.41E-01 | 2.67E+00 | 2.71E+00 | 4.85E-01 | no | | intergenic_variant | MODIFIER |
| 1 | 149694699 | AX-170043094 | 0 | C | T | 0.202 | 9.76E-02 | 1.41E-01 | 2.67E+00 | 2.71E+00 | 4.85E-01 | no | | intergenic_variant | MODIFIER |
| 1 | 149715455 | AX-170333333 | 0 | A | C | 0.202 | 9.76E-02 | 1.41E-01 | 2.67E+00 | 2.71E+00 | 4.85E-01 | no | | intergenic_variant | MODIFIER |
| 1 | 149718771 | AX-170423837 | 0 | G | A | 0.202 | 9.76E-02 | 1.41E-01 | 2.67E+00 | 2.71E+00 | 4.85E-01 | no | | intergenic_variant | MODIFIER |
| 1 | 149720104 | AX-97819701 | 0 | A | G | 0.202 | 9.76E-02 | 1.41E-01 | 2.67E+00 | 2.71E+00 | 4.85E-01 | no | | intergenic_variant | MODIFIER |
| 1 | 149720379 | AX-170119973 | 0 | A | G | 0.202 | 9.76E-02 | 1.41E-01 | 2.67E+00 | 2.71E+00 | 4.85E-01 | no | | intergenic_variant | MODIFIER |
| 1 | 149726795 | AX-170473976 | 0 | C | T | 0.202 | 9.76E-02 | 1.41E-01 | 2.67E+00 | 2.71E+00 | 4.85E-01 | no | | intergenic_variant | MODIFIER |
| 1 | 149730371 | AX-170170001 | 0 | G | C | 0.202 | 9.76E-02 | 1.41E-01 | 2.67E+00 | 2.71E+00 | 4.85E-01 | no | n-R5s218 | downstream_gene_variant | MODIFIER |
| 1 | 149738604 | AX-170435595 | 0 | A | G | 0.202 | 9.76E-02 | 1.41E-01 | 2.67E+00 | 2.71E+00 | 4.85E-01 | no | n-R5s218 | upstream_gene_variant | MODIFIER |
| 1 | 149764712 | AX-170285817 | 0 | C | A | 0.202 | 9.76E-02 | 1.41E-01 | 2.67E+00 | 2.71E+00 | 4.85E-01 | no | | intergenic_variant | MODIFIER |
| 1 | 149764939 | AX-170453651 | 0 | T | C | 0.202 | 9.76E-02 | 1.41E-01 | 2.67E+00 | 2.71E+00 | 4.85E-01 | no | | intergenic_variant | MODIFIER |
| 1 | 149764962 | AX-170043095 | 0 | A | G | 0.202 | 9.76E-02 | 1.41E-01 | 2.67E+00 | 2.71E+00 | 4.85E-01 | no | | intergenic_variant | MODIFIER |
| 1 | 149765488 | AX-170179073 | 0 | T | C | 0.202 | 9.76E-02 | 1.41E-01 | 2.67E+00 | 2.71E+00 | 4.85E-01 | no | | intergenic_variant | MODIFIER |
| 1 | 149765573 | AX-170115603 | 0 | A | G | 0.202 | 9.76E-02 | 1.41E-01 | 2.67E+00 | 2.71E+00 | 4.85E-01 | no | | intergenic_variant | MODIFIER |
| 1 | 149765653 | AX-170492911 | 0 | G | A | 0.202 | 9.76E-02 | 1.41E-01 | 2.67E+00 | 2.71E+00 | 4.85E-01 | no | | intergenic_variant | MODIFIER |
| 1 | 149766118 | AX-170412825 | 0 | T | A | 0.202 | 9.76E-02 | 1.41E-01 | 2.67E+00 | 2.71E+00 | 4.85E-01 | no | | intergenic_variant | MODIFIER |
| 1 | 149799698 | AX-170207151 | 0 | G | A | 0.202 | 9.76E-02 | 1.41E-01 | 2.67E+00 | 2.71E+00 | 4.85E-01 | no | | intergenic_variant | MODIFIER |
| 1 | 149799807 | AX-170350082 | 0 | C | T | 0.202 | 9.76E-02 | 1.41E-01 | 2.67E+00 | 2.71E+00 | 4.85E-01 | no | | intergenic_variant | MODIFIER |
| 1 | 149803229 | AX-170269097 | 0 | C | T | 0.202 | 9.76E-02 | 1.41E-01 | 2.67E+00 | 2.71E+00 | 4.85E-01 | no | | intergenic_variant | MODIFIER |
| 1 | 149819244 | AX-170172431 | 0 | C | A | 0.202 | 9.76E-02 | 1.41E-01 | 2.67E+00 | 2.71E+00 | 4.85E-01 | no | | intergenic_variant | MODIFIER |
| 1 | 149829493 | AX-170226610 | 0 | T | A | 0.202 | 9.76E-02 | 1.41E-01 | 2.67E+00 | 2.71E+00 | 4.85E-01 | no | Pla2g4a | downstream_gene_variant | MODIFIER |
| 1 | 149829751 | AX-97840993 | 0 | C | T | 0.202 | 9.76E-02 | 1.41E-01 | 2.67E+00 | 2.71E+00 | 4.85E-01 | no | Pla2g4a | 3_prime_UTR_variant | MODIFIER |
| 1 | 149835974 | AX-170226082 | 0 | A | G | 0.202 | 9.76E-02 | 1.41E-01 | 2.67E+00 | 2.71E+00 | 4.85E-01 | no | Pla2g4a | intron_variant | MODIFIER |
| 1 | 149836002 | AX-170506533 | 0 | C | T | 0.202 | 9.76E-02 | 1.41E-01 | 2.67E+00 | 2.71E+00 | 4.85E-01 | no | Pla2g4a | intron_variant | MODIFIER |
| 1 | 149838770 | AX-170108531 | 0 | T | C | 0.202 | 9.76E-02 | 1.41E-01 | 2.67E+00 | 2.71E+00 | 4.85E-01 | no | Pla2g4a | intron_variant | MODIFIER |
| 1 | 149845511 | AX-170107057 | 0 | C | T | 0.202 | 9.76E-02 | 1.41E-01 | 2.67E+00 | 2.71E+00 | 4.85E-01 | no | Pla2g4a | intron_variant | MODIFIER |
| 1 | 149848746 | AX-170395362 | 0 | A | G | 0.202 | 9.76E-02 | 1.41E-01 | 2.67E+00 | 2.71E+00 | 4.85E-01 | no | Pla2g4a | intron_variant | MODIFIER |
| 1 | 149876043 | AX-170194242 | 0 | A | G | 0.202 | 9.76E-02 | 1.41E-01 | 2.67E+00 | 2.71E+00 | 4.85E-01 | no | Pla2g4a | intron_variant | MODIFIER |
| 1 | 149897688 | AX-170333871 | 0 | A | G | 0.202 | 9.76E-02 | 1.41E-01 | 2.67E+00 | 2.71E+00 | 4.85E-01 | no | Pla2g4a | intron_variant | MODIFIER |
| 1 | 149908921 | AX-170118810 | 0 | T | C | 0.202 | 9.76E-02 | 1.41E-01 | 2.67E+00 | 2.71E+00 | 4.85E-01 | no | Pla2g4a | intron_variant | MODIFIER |
| 1 | 149909670 | AX-170254399 | 0 | A | G | 0.202 | 9.76E-02 | 1.41E-01 | 2.67E+00 | 2.71E+00 | 4.85E-01 | no | Pla2g4a | intron_variant | MODIFIER |
| 1 | 149910478 | AX-170205487 | 0 | T | C | 0.202 | 9.76E-02 | 1.41E-01 | 2.67E+00 | 2.71E+00 | 4.85E-01 | no | Pla2g4a | intron_variant | MODIFIER |
| 1 | 149911017 | AX-170420198 | 0 | T | C | 0.202 | 9.76E-02 | 1.41E-01 | 2.67E+00 | 2.71E+00 | 4.85E-01 | no | Pla2g4a | intron_variant | MODIFIER |
| 1 | 149915194 | AX-170363219 | 0 | G | C | 0.202 | 9.76E-02 | 1.41E-01 | 2.67E+00 | 2.71E+00 | 4.85E-01 | no | Pla2g4a | intron_variant | MODIFIER |
| 1 | 149919858 | AX-170263522 | 0 | T | C | 0.202 | 9.76E-02 | 1.41E-01 | 2.67E+00 | 2.71E+00 | 4.85E-01 | no | Pla2g4a | intron_variant | MODIFIER |
| 1 | 149928656 | AX-170308630 | 0 | A | C | 0.202 | 9.76E-02 | 1.41E-01 | 2.67E+00 | 2.71E+00 | 4.85E-01 | no | Pla2g4a | intron_variant | MODIFIER |
| 1 | 149937955 | AX-170455170 | 0 | G | A | 0.202 | 9.76E-02 | 1.41E-01 | 2.67E+00 | 2.71E+00 | 4.85E-01 | no | Pla2g4a | intron_variant | MODIFIER |
| 1 | 149939893 | AX-170126998 | 0 | A | G | 0.202 | 9.76E-02 | 1.41E-01 | 2.67E+00 | 2.71E+00 | 4.85E-01 | no | Pla2g4a | intron_variant | MODIFIER |
| 1 | 149943418 | AX-170152979 | 0 | A | C | 0.202 | 9.76E-02 | 1.41E-01 | 2.67E+00 | 2.71E+00 | 4.85E-01 | no | Pla2g4a | intron_variant | MODIFIER |
| 1 | 149948277 | AX-170349564 | 0 | G | A | 0.202 | 9.76E-02 | 1.41E-01 | 2.67E+00 | 2.71E+00 | 4.85E-01 | no | Pla2g4a | intron_variant | MODIFIER |
| 1 | 149959632 | AX-97787486 | 0 | T | G | 0.202 | 9.76E-02 | 1.41E-01 | 2.67E+00 | 2.71E+00 | 4.85E-01 | no | Pla2g4a | intron_variant | MODIFIER |
| 1 | 149983530 | AX-170470255 | 0 | T | A | 0.202 | 9.76E-02 | 1.41E-01 | 2.67E+00 | 2.71E+00 | 4.85E-01 | no | | intergenic_variant | MODIFIER |
| 1 | 149988638 | AX-170253664 | 0 | C | T | 0.202 | 9.76E-02 | 1.41E-01 | 2.67E+00 | 2.71E+00 | 4.85E-01 | no | | intergenic_variant | MODIFIER |
| 1 | 149989698 | AX-97806509 | 0 | T | C | 0.202 | 9.76E-02 | 1.41E-01 | 2.67E+00 | 2.71E+00 | 4.85E-01 | no | | intergenic_variant | MODIFIER |
| 1 | 150024010 | AX-170249653 | 0 | G | A | 0.202 | 9.76E-02 | 1.41E-01 | 2.67E+00 | 2.71E+00 | 4.85E-01 | no | Gm19136 | non_coding_transcript_exon_variant | MODIFIER |
| 1 | 150030135 | AX-170412761 | 0 | G | A | 0.202 | 9.76E-02 | 1.41E-01 | 2.67E+00 | 2.71E+00 | 4.85E-01 | no | | intergenic_variant | MODIFIER |
| 1 | 150054160 | AX-170001972 | 0 | A | G | 0.202 | 9.76E-02 | 1.41E-01 | 2.67E+00 | 2.71E+00 | 4.85E-01 | no | | intergenic_variant | MODIFIER |
| 1 | 150054306 | AX-170001969 | 0 | A | G | 0.202 | 9.76E-02 | 1.41E-01 | 2.67E+00 | 2.71E+00 | 4.85E-01 | no | | intergenic_variant | MODIFIER |

| | | | | | | | | | | | | | | | |
|---|---|---|---|---|---|---|---|---|---|---|---|---|---|---|---|
| 1 | 150058998 | AX-170001971 | 0 | A | T | 0.202 | 9.76E-02 | 1.41E-01 | 2.67E+00 | 2.71E+00 | 4.85E-01 | no | Gm44329 | downstream_gene_variant | MODIFIER |
| 1 | 150059440 | AX-170515927 | 0 | G | C | 0.202 | 9.76E-02 | 1.41E-01 | 2.67E+00 | 2.71E+00 | 4.85E-01 | no | Gm44329 | downstream_gene_variant | MODIFIER |
| 1 | 150068008 | AX-170043107 | 0 | T | C | 0.202 | 9.76E-02 | 1.41E-01 | 2.67E+00 | 2.71E+00 | 4.85E-01 | no | Gm44329 | upstream_gene_variant | MODIFIER |
| 1 | 150101693 | AX-170288583 | 0 | G | A | 0.202 | 9.76E-02 | 1.41E-01 | 2.67E+00 | 2.71E+00 | 4.85E-01 | no | Ptgs2 | intron_variant | MODIFIER |
| 1 | 150102097 | AX-170446219 | 0 | C | A | 0.202 | 9.76E-02 | 1.41E-01 | 2.67E+00 | 2.71E+00 | 4.85E-01 | no | Ptgs2 | synonymous_variant | LOW |
| 1 | 150102492 | AX-170112676 | 0 | G | A | 0.202 | 9.76E-02 | 1.41E-01 | 2.67E+00 | 2.71E+00 | 4.85E-01 | no | Ptgs2 | intron_variant | MODIFIER |
| 1 | 150111769 | AX-170201464 | 0 | C | T | 0.202 | 9.76E-02 | 1.41E-01 | 2.67E+00 | 2.71E+00 | 4.85E-01 | no | Ptgs2 | downstream_gene_variant | MODIFIER |
| 1 | 150111792 | AX-97808979 | 0 | C | A | 0.202 | 9.76E-02 | 1.41E-01 | 2.67E+00 | 2.71E+00 | 4.85E-01 | no | Ptgs2 | downstream_gene_variant | MODIFIER |
| 1 | 150111820 | AX-170401576 | 0 | C | T | 0.202 | 9.76E-02 | 1.41E-01 | 2.67E+00 | 2.71E+00 | 4.85E-01 | no | Ptgs2 | downstream_gene_variant | MODIFIER |
| 1 | 150118210 | AX-170001975 | 0 | G | C | 0.202 | 9.76E-02 | 1.41E-01 | 2.67E+00 | 2.71E+00 | 4.85E-01 | no | | intergenic_variant | MODIFIER |
| 1 | 150166717 | AX-97840753 | 0 | A | G | 0.202 | 9.76E-02 | 1.41E-01 | 2.67E+00 | 2.71E+00 | 4.85E-01 | no | Ptgs2os2 | upstream_gene_variant | MODIFIER |
| 1 | 150166984 | AX-170001970 | 0 | A | G | 0.202 | 9.76E-02 | 1.41E-01 | 2.67E+00 | 2.71E+00 | 4.85E-01 | no | Ptgs2os2 | upstream_gene_variant | MODIFIER |
| 1 | 150167086 | AX-170166713 | 0 | G | A | 0.202 | 9.76E-02 | 1.41E-01 | 2.67E+00 | 2.71E+00 | 4.85E-01 | no | Ptgs2os2 | upstream_gene_variant | MODIFIER |
| 1 | 150177837 | AX-170324113 | 0 | G | C | 0.202 | 9.76E-02 | 1.41E-01 | 2.67E+00 | 2.71E+00 | 4.85E-01 | no | | intergenic_variant | MODIFIER |
| 1 | 150203204 | AX-170349460 | 0 | T | C | 0.202 | 9.76E-02 | 1.41E-01 | 2.67E+00 | 2.71E+00 | 4.85E-01 | no | | intergenic_variant | MODIFIER |
| 1 | 150225930 | AX-170189767 | 0 | G | A | 0.202 | 9.76E-02 | 1.41E-01 | 2.67E+00 | 2.71E+00 | 4.85E-01 | no | | regulatory_region_variant | MODIFIER |
| 1 | 150226401 | AX-97784503 | 0 | A | G | 0.202 | 9.76E-02 | 1.41E-01 | 2.67E+00 | 2.71E+00 | 4.85E-01 | no | | intergenic_variant | MODIFIER |
| 1 | 150241285 | AX-170161066 | 0 | A | G | 0.202 | 9.76E-02 | 1.41E-01 | 2.67E+00 | 2.71E+00 | 4.85E-01 | no | | intergenic_variant | MODIFIER |
| 1 | 150256227 | AX-170369675 | 0 | T | A | 0.202 | 9.76E-02 | 1.41E-01 | 2.67E+00 | 2.71E+00 | 4.85E-01 | no | | intergenic_variant | MODIFIER |
| 1 | 150256762 | AX-170216999 | 0 | T | C | 0.202 | 9.76E-02 | 1.41E-01 | 2.67E+00 | 2.71E+00 | 4.85E-01 | no | | intergenic_variant | MODIFIER |
| 1 | 150258217 | AX-170302878 | 0 | G | A | 0.202 | 9.76E-02 | 1.41E-01 | 2.67E+00 | 2.71E+00 | 4.85E-01 | no | | intergenic_variant | MODIFIER |
| 1 | 150258259 | AX-170341114 | 0 | A | G | 0.202 | 9.76E-02 | 1.41E-01 | 2.67E+00 | 2.71E+00 | 4.85E-01 | no | | intergenic_variant | MODIFIER |
| 1 | 150258317 | AX-170514925 | 0 | T | G | 0.202 | 9.76E-02 | 1.41E-01 | 2.67E+00 | 2.71E+00 | 4.85E-01 | no | | intergenic_variant | MODIFIER |
| 1 | 150260582 | AX-170367518 | 0 | A | G | 0.202 | 9.76E-02 | 1.41E-01 | 2.67E+00 | 2.71E+00 | 4.85E-01 | no | Rbm3-ps | downstream_gene_variant | MODIFIER |
| 1 | 150322255 | AX-170186526 | 0 | A | G | 0.202 | 9.76E-02 | 1.41E-01 | 2.67E+00 | 2.71E+00 | 4.85E-01 | no | Pdc | intron_variant | MODIFIER |
| 1 | 150322814 | AX-170356156 | 0 | T | C | 0.202 | 9.76E-02 | 1.41E-01 | 2.67E+00 | 2.71E+00 | 4.85E-01 | no | Pdc | intron_variant | MODIFIER |
| 1 | 150346013 | AX-170043108 | 0 | C | T | 0.202 | 9.76E-02 | 1.41E-01 | 2.67E+00 | 2.71E+00 | 4.85E-01 | no | 2310030A07Rik | intron_variant,non_coding_transcript_variant | MODIFIER |
| 1 | 150360911 | AX-170437928 | 0 | G | A | 0.202 | 9.76E-02 | 1.41E-01 | 2.67E+00 | 2.71E+00 | 4.85E-01 | no | Odr4 | downstream_gene_variant | MODIFIER |
| 1 | 150387065 | AX-170255651 | 0 | A | G | 0.202 | 9.76E-02 | 1.41E-01 | 2.67E+00 | 2.71E+00 | 4.85E-01 | no | Odr4 | intron_variant | MODIFIER |
| 1 | 150387104 | AX-170461736 | 0 | A | G | 0.202 | 9.76E-02 | 1.41E-01 | 2.67E+00 | 2.71E+00 | 4.85E-01 | no | Odr4 | intron_variant | MODIFIER |
| 1 | 150387273 | AX-170480148 | 0 | C | T | 0.202 | 9.76E-02 | 1.41E-01 | 2.67E+00 | 2.71E+00 | 4.85E-01 | no | Odr4 | intron_variant | MODIFIER |
| 1 | 150403644 | AX-170283857 | 0 | G | A | 0.202 | 9.76E-02 | 1.41E-01 | 2.67E+00 | 2.71E+00 | 4.85E-01 | no | Tpr | synonymous_variant | LOW |
| 1 | 150421860 | AX-170500656 | 0 | T | C | 0.202 | 9.76E-02 | 1.41E-01 | 2.67E+00 | 2.71E+00 | 4.85E-01 | no | Tpr | intron_variant | MODIFIER |
| 1 | 150429151 | AX-170312461 | 0 | A | G | 0.202 | 9.76E-02 | 1.41E-01 | 2.67E+00 | 2.71E+00 | 4.85E-01 | no | Tpr | intron_variant | MODIFIER |
| 1 | 150449795 | AX-170043109 | 0 | C | A | 0.202 | 9.76E-02 | 1.41E-01 | 2.67E+00 | 2.71E+00 | 4.85E-01 | no | Prg4 | 3_prime_UTR_variant | MODIFIER |
| 1 | 150450406 | AX-170197529 | 0 | A | G | 0.202 | 9.76E-02 | 1.41E-01 | 2.67E+00 | 2.71E+00 | 4.85E-01 | no | Prg4 | intron_variant | MODIFIER |
| 1 | 150468749 | AX-170194658 | 0 | T | C | 0.202 | 9.76E-02 | 1.41E-01 | 2.67E+00 | 2.71E+00 | 4.85E-01 | no | Prg4 | upstream_gene_variant | MODIFIER |
| 1 | 150479213 | AX-170427566 | 0 | T | C | 0.202 | 9.76E-02 | 1.41E-01 | 2.67E+00 | 2.71E+00 | 4.85E-01 | no | | intergenic_variant | MODIFIER |
| 1 | 150480098 | AX-170098793 | 0 | T | A | 0.202 | 9.76E-02 | 1.41E-01 | 2.67E+00 | 2.71E+00 | 4.85E-01 | no | | intergenic_variant | MODIFIER |
| 1 | 150485231 | AX-170318587 | 0 | T | C | 0.202 | 9.76E-02 | 1.41E-01 | 2.67E+00 | 2.71E+00 | 4.85E-01 | no | | intergenic_variant | MODIFIER |
| 1 | 150487173 | AX-170154303 | 0 | T | C | 0.202 | 9.76E-02 | 1.41E-01 | 2.67E+00 | 2.71E+00 | 4.85E-01 | no | | intergenic_variant | MODIFIER |
| 1 | 150487705 | AX-170470626 | 0 | A | T | 0.202 | 9.76E-02 | 1.41E-01 | 2.67E+00 | 2.71E+00 | 4.85E-01 | no | | intergenic_variant | MODIFIER |
| 1 | 150488172 | AX-170512711 | 0 | G | A | 0.202 | 9.76E-02 | 1.41E-01 | 2.67E+00 | 2.71E+00 | 4.85E-01 | no | | regulatory_region_variant | MODIFIER |
| 1 | 150494384 | AX-170290250 | 0 | A | C | 0.202 | 9.76E-02 | 1.41E-01 | 2.67E+00 | 2.71E+00 | 4.85E-01 | no | | intergenic_variant | MODIFIER |
| 1 | 150494814 | AX-170219579 | 0 | A | G | 0.202 | 9.76E-02 | 1.41E-01 | 2.67E+00 | 2.71E+00 | 4.85E-01 | no | | intergenic_variant | MODIFIER |
| 1 | 150496227 | AX-170505779 | 0 | A | G | 0.202 | 9.76E-02 | 1.41E-01 | 2.67E+00 | 2.71E+00 | 4.85E-01 | no | | intergenic_variant | MODIFIER |
| 1 | 150503234 | AX-170223983 | 0 | T | C | 0.202 | 9.76E-02 | 1.41E-01 | 2.67E+00 | 2.71E+00 | 4.85E-01 | no | | intergenic_variant | MODIFIER |
| 1 | 150507414 | AX-170124009 | 0 | C | T | 0.202 | 9.76E-02 | 1.41E-01 | 2.67E+00 | 2.71E+00 | 4.85E-01 | no | | intergenic_variant | MODIFIER |
| 1 | 150529091 | AX-170416663 | 0 | T | G | 0.202 | 9.76E-02 | 1.41E-01 | 2.67E+00 | 2.71E+00 | 4.85E-01 | no | | intergenic_variant | MODIFIER |
| 1 | 150529256 | AX-170480600 | 0 | A | T | 0.202 | 9.76E-02 | 1.41E-01 | 2.67E+00 | 2.71E+00 | 4.85E-01 | no | | intergenic_variant | MODIFIER |
| 1 | 150554068 | AX-170224235 | 0 | G | A | 0.202 | 9.76E-02 | 1.41E-01 | 2.67E+00 | 2.71E+00 | 4.85E-01 | no | Ska2l-ps | downstream_gene_variant | MODIFIER |
| 1 | 150577348 | AX-170409792 | 0 | T | C | 0.202 | 9.76E-02 | 1.41E-01 | 2.67E+00 | 2.71E+00 | 4.85E-01 | no | Hmcn1 | synonymous_variant | LOW |
| 1 | 150577781 | AX-170001977 | 0 | T | C | 0.202 | 9.76E-02 | 1.41E-01 | 2.67E+00 | 2.71E+00 | 4.85E-01 | no | Hmcn1 | intron_variant | MODIFIER |
| 1 | 150578115 | AX-170491514 | 0 | G | A | 0.202 | 9.76E-02 | 1.41E-01 | 2.67E+00 | 2.71E+00 | 4.85E-01 | no | Hmcn1 | intron_variant | MODIFIER |

| | | | | | | | | | | | | | | | |
|---|---|---|---|---|---|---|---|---|---|---|---|---|---|---|---|
| 1 | 150598296 | AX-170234716 | 0 | T | G | 0.202 | 9.76E-02 | 1.41E-01 | 2.67E+00 | 2.71E+00 | 4.85E-01 | no | Hmcn1 | synonymous_variant | LOW |
| 1 | 150612880 | AX-170200638 | 0 | A | C | 0.202 | 9.76E-02 | 1.41E-01 | 2.67E+00 | 2.71E+00 | 4.85E-01 | no | Hmcn1 | intron_variant | MODIFIER |
| 1 | 150619147 | AX-170001980 | 0 | T | C | 0.202 | 9.76E-02 | 1.41E-01 | 2.67E+00 | 2.71E+00 | 4.85E-01 | no | Hmcn1 | intron_variant | MODIFIER |
| 1 | 150619336 | AX-170001978 | 0 | C | T | 0.202 | 9.76E-02 | 1.41E-01 | 2.67E+00 | 2.71E+00 | 4.85E-01 | no | Hmcn1 | intron_variant | MODIFIER |
| 1 | 150619425 | AX-170101282 | 0 | T | C | 0.202 | 9.76E-02 | 1.41E-01 | 2.67E+00 | 2.71E+00 | 4.85E-01 | no | Hmcn1 | intron_variant | MODIFIER |
| 1 | 150657806 | AX-170188530 | 0 | G | T | 0.202 | 9.76E-02 | 1.41E-01 | 2.67E+00 | 2.71E+00 | 4.85E-01 | no | Hmcn1 | intron_variant | MODIFIER |
| 1 | 150658169 | AX-97813158 | 0 | G | A | 0.202 | 9.76E-02 | 1.41E-01 | 2.67E+00 | 2.71E+00 | 4.85E-01 | no | Hmcn1 | intron_variant | MODIFIER |
| 1 | 150658463 | AX-170497197 | 0 | T | C | 0.202 | 9.76E-02 | 1.41E-01 | 2.67E+00 | 2.71E+00 | 4.85E-01 | no | Hmcn1 | intron_variant | MODIFIER |
| 1 | 150663845 | AX-170477112 | 0 | C | T | 0.202 | 9.76E-02 | 1.41E-01 | 2.67E+00 | 2.71E+00 | 4.85E-01 | no | Hmcn1 | intron_variant | MODIFIER |
| 1 | 150676821 | AX-170525062 | 0 | T | G | 0.202 | 9.76E-02 | 1.41E-01 | 2.67E+00 | 2.71E+00 | 4.85E-01 | no | Hmcn1 | intron_variant | MODIFIER |
| 1 | 150676875 | AX-170362634 | 0 | C | A | 0.202 | 9.76E-02 | 1.41E-01 | 2.67E+00 | 2.71E+00 | 4.85E-01 | no | Hmcn1 | intron_variant | MODIFIER |
| 1 | 150681165 | AX-170043118 | 0 | C | G | 0.202 | 9.76E-02 | 1.41E-01 | 2.67E+00 | 2.71E+00 | 4.85E-01 | no | Hmcn1 | intron_variant | MODIFIER |
| 1 | 150735325 | AX-170190742 | 0 | A | G | 0.202 | 9.76E-02 | 1.41E-01 | 2.67E+00 | 2.71E+00 | 4.85E-01 | no | Hmcn1 | intron_variant | MODIFIER |
| 1 | 150755421 | AX-170144096 | 0 | A | G | 0.202 | 9.76E-02 | 1.41E-01 | 2.67E+00 | 2.71E+00 | 4.85E-01 | no | Hmcn1 | intron_variant | MODIFIER |
| 1 | 150791354 | AX-97814924 | 0 | T | C | 0.202 | 9.76E-02 | 1.41E-01 | 2.67E+00 | 2.71E+00 | 4.85E-01 | no | Hmcn1 | intron_variant | MODIFIER |
| 1 | 150807920 | AX-170262685 | 0 | G | A | 0.202 | 9.76E-02 | 1.41E-01 | 2.67E+00 | 2.71E+00 | 4.85E-01 | no | Hmcn1 | intron_variant | MODIFIER |
| 1 | 150820574 | AX-170278549 | 0 | A | G | 0.202 | 9.76E-02 | 1.41E-01 | 2.67E+00 | 2.71E+00 | 4.85E-01 | no | Hmcn1 | intron_variant | MODIFIER |
| 1 | 150896283 | AX-170162775 | 0 | A | G | 0.202 | 9.76E-02 | 1.41E-01 | 2.67E+00 | 2.71E+00 | 4.85E-01 | no | Hmcn1 | intron_variant | MODIFIER |
| 1 | 150950221 | AX-170244632 | 0 | C | T | 0.202 | 9.76E-02 | 1.41E-01 | 2.67E+00 | 2.71E+00 | 4.85E-01 | no | Hmcn1 | intron_variant | MODIFIER |
| 1 | 150951818 | AX-170246550 | 0 | T | C | 0.202 | 9.76E-02 | 1.41E-01 | 2.67E+00 | 2.71E+00 | 4.85E-01 | no | Hmcn1 | intron_variant | MODIFIER |
| 1 | 150955028 | AX-97845758 | 0 | G | A | 0.202 | 9.76E-02 | 1.41E-01 | 2.67E+00 | 2.71E+00 | 4.85E-01 | no | Hmcn1 | intron_variant | MODIFIER |
| 1 | 150983693 | AX-97799940 | 0 | G | C | 0.202 | 9.76E-02 | 1.41E-01 | 2.67E+00 | 2.71E+00 | 4.85E-01 | no | Hmcn1 | intron_variant | MODIFIER |
| 1 | 150985433 | AX-170521711 | 0 | G | A | 0.202 | 9.76E-02 | 1.41E-01 | 2.67E+00 | 2.71E+00 | 4.85E-01 | no | Hmcn1 | intron_variant | MODIFIER |
| 1 | 151007466 | AX-97839570 | 0 | T | C | 0.202 | 9.76E-02 | 1.41E-01 | 2.67E+00 | 2.71E+00 | 4.85E-01 | no | Gm29188 | downstream_gene_variant | MODIFIER |
| 1 | 151011437 | AX-170313580 | 0 | A | C | 0.202 | 9.76E-02 | 1.41E-01 | 2.67E+00 | 2.71E+00 | 4.85E-01 | no | | intergenic_variant | MODIFIER |
| 1 | 151056542 | AX-97785162 | 0 | A | G | 0.202 | 9.76E-02 | 1.41E-01 | 2.67E+00 | 2.71E+00 | 4.85E-01 | no | | intergenic_variant | MODIFIER |
| 1 | 151064247 | AX-170098752 | 0 | A | G | 0.202 | 9.76E-02 | 1.41E-01 | 2.67E+00 | 2.71E+00 | 4.85E-01 | no | Gm22756 | upstream_gene_variant | MODIFIER |
| 1 | 151074132 | AX-170395232 | 0 | T | C | 0.202 | 9.76E-02 | 1.41E-01 | 2.67E+00 | 2.71E+00 | 4.85E-01 | no | | intergenic_variant | MODIFIER |
| 1 | 151099179 | AX-97813200 | 0 | A | G | 0.202 | 9.76E-02 | 1.41E-01 | 2.67E+00 | 2.71E+00 | 4.85E-01 | no | | intergenic_variant | MODIFIER |
| 1 | 151142747 | AX-97804802 | 0 | A | G | 0.202 | 9.76E-02 | 1.41E-01 | 2.67E+00 | 2.71E+00 | 4.85E-01 | no | Gm8941 | downstream_gene_variant | MODIFIER |
| 1 | 151177868 | AX-170001985 | 0 | A | G | 0.202 | 9.76E-02 | 1.41E-01 | 2.67E+00 | 2.71E+00 | 4.85E-01 | no | Gm47985 | upstream_gene_variant | MODIFIER |
| 1 | 151214895 | AX-170154617 | 0 | A | G | 0.202 | 9.76E-02 | 1.41E-01 | 2.67E+00 | 2.71E+00 | 4.85E-01 | no | Gm47996 | downstream_gene_variant | MODIFIER |
| 1 | 151238409 | AX-170271439 | 0 | A | C | 0.202 | 9.76E-02 | 1.41E-01 | 2.67E+00 | 2.71E+00 | 4.85E-01 | no | | regulatory_region_variant | MODIFIER |
| 1 | 151244670 | AX-170318476 | 0 | T | C | 0.202 | 9.76E-02 | 1.41E-01 | 2.67E+00 | 2.71E+00 | 4.85E-01 | no | | intergenic_variant | MODIFIER |
| 1 | 151340019 | AX-170296798 | 0 | T | C | 0.202 | 9.76E-02 | 1.41E-01 | 2.67E+00 | 2.71E+00 | 4.85E-01 | no | Ivns1abp | upstream_gene_variant | MODIFIER |
| 1 | 151356839 | AX-170283911 | 0 | T | C | 0.202 | 9.76E-02 | 1.41E-01 | 2.67E+00 | 2.71E+00 | 4.85E-01 | no | Ivns1abp | intron_variant | MODIFIER |
| 1 | 151726479 | AX-170152930 | 0 | G | A | 0.202 | 9.76E-02 | 1.41E-01 | 2.67E+00 | 2.71E+00 | 4.85E-01 | no | Fam129a | downstream_gene_variant | MODIFIER |
| 1 | 151736978 | AX-170099430 | 0 | G | T | 0.202 | 9.76E-02 | 1.41E-01 | 2.67E+00 | 2.71E+00 | 4.85E-01 | no | | intergenic_variant | MODIFIER |
| 1 | 151752098 | AX-170247444 | 0 | C | T | 0.202 | 9.76E-02 | 1.41E-01 | 2.67E+00 | 2.71E+00 | 4.85E-01 | no | Edem3 | upstream_gene_variant | MODIFIER |
| 1 | 151763543 | AX-97846902 | 0 | A | G | 0.202 | 9.76E-02 | 1.41E-01 | 2.67E+00 | 2.71E+00 | 4.85E-01 | no | Edem3 | intron_variant | MODIFIER |
| 1 | 151773691 | AX-170413149 | 0 | T | C | 0.202 | 9.76E-02 | 1.41E-01 | 2.67E+00 | 2.71E+00 | 4.85E-01 | no | Edem3 | intron_variant | MODIFIER |
| 1 | 151777225 | AX-170269080 | 0 | T | C | 0.202 | 9.76E-02 | 1.41E-01 | 2.67E+00 | 2.71E+00 | 4.85E-01 | no | Edem3 | intron_variant | MODIFIER |
| 1 | 151777309 | AX-97813740 | 0 | A | G | 0.202 | 9.76E-02 | 1.41E-01 | 2.67E+00 | 2.71E+00 | 4.85E-01 | no | Edem3 | intron_variant | MODIFIER |
| 1 | 151792887 | AX-170480786 | 0 | T | C | 0.202 | 9.76E-02 | 1.41E-01 | 2.67E+00 | 2.71E+00 | 4.85E-01 | no | Edem3 | intron_variant | MODIFIER |
| 1 | 151815159 | AX-170337936 | 0 | T | C | 0.202 | 9.76E-02 | 1.41E-01 | 2.67E+00 | 2.71E+00 | 4.85E-01 | no | Edem3 | intron_variant | MODIFIER |
| 1 | 151816650 | AX-170099718 | 0 | C | T | 0.202 | 9.76E-02 | 1.41E-01 | 2.67E+00 | 2.71E+00 | 4.85E-01 | no | Edem3 | intron_variant | MODIFIER |
| 1 | 151847672 | AX-170243285 | 0 | A | T | 0.202 | 9.76E-02 | 1.41E-01 | 2.67E+00 | 2.71E+00 | 4.85E-01 | no | 1700025G04Rik | downstream_gene_variant | MODIFIER |
| 1 | 151862153 | AX-170336604 | 0 | C | T | 0.202 | 9.76E-02 | 1.41E-01 | 2.67E+00 | 2.71E+00 | 4.85E-01 | no | 1700025G04Rik | intron_variant,NMD_transcript_variant | MODIFIER |
| 1 | 151865218 | AX-97838592 | 0 | A | G | 0.202 | 9.76E-02 | 1.41E-01 | 2.67E+00 | 2.71E+00 | 4.85E-01 | no | 1700025G04Rik | intron_variant,NMD_transcript_variant | MODIFIER |
| 1 | 151873920 | AX-97857090 | 0 | T | C | 0.202 | 9.76E-02 | 1.41E-01 | 2.67E+00 | 2.71E+00 | 4.85E-01 | no | 1700025G04Rik | intron_variant,NMD_transcript_variant | MODIFIER |
| 1 | 151884314 | AX-170370740 | 0 | G | A | 0.202 | 9.76E-02 | 1.41E-01 | 2.67E+00 | 2.71E+00 | 4.85E-01 | no | 1700025G04Rik | downstream_gene_variant | MODIFIER |
| 1 | 151884348 | AX-170281250 | 0 | C | G | 0.202 | 9.76E-02 | 1.41E-01 | 2.67E+00 | 2.71E+00 | 4.85E-01 | no | 1700025G04Rik | downstream_gene_variant | MODIFIER |
| 1 | 151891286 | AX-170403989 | 0 | T | C | 0.202 | 9.76E-02 | 1.41E-01 | 2.67E+00 | 2.71E+00 | 4.85E-01 | no | 1700025G04Rik | 3_prime_UTR_variant | MODIFIER |
| 1 | 151899731 | AX-170504214 | 0 | T | C | 0.202 | 9.76E-02 | 1.41E-01 | 2.67E+00 | 2.71E+00 | 4.85E-01 | no | 1700025G04Rik | intron_variant | MODIFIER |

| | | | | | | | | | | | | | | | |
|---|---|---|---|---|---|---|---|---|---|---|---|---|---|---|---|
| 1 | 151913471 | AX-170173421 | 0 | C | T | 0.202 | 9.76E-02 | 1.41E-01 | 2.67E+00 | 2.71E+00 | 4.85E-01 | no | 1700025G04Rik | intron_variant | MODIFIER |
| 1 | 151915736 | AX-170482453 | 0 | G | C | 0.202 | 9.76E-02 | 1.41E-01 | 2.67E+00 | 2.71E+00 | 4.85E-01 | no | 1700025G04Rik | intron_variant | MODIFIER |
| 1 | 151917608 | AX-170409435 | 0 | A | C | 0.202 | 9.76E-02 | 1.41E-01 | 2.67E+00 | 2.71E+00 | 4.85E-01 | no | 1700025G04Rik | intron_variant | MODIFIER |
| 1 | 151921167 | AX-170228659 | 0 | A | G | 0.202 | 9.76E-02 | 1.41E-01 | 2.67E+00 | 2.71E+00 | 4.85E-01 | no | 1700025G04Rik | intron_variant | MODIFIER |
| 1 | 151929907 | AX-170043153 | 0 | C | T | 0.202 | 9.76E-02 | 1.41E-01 | 2.67E+00 | 2.71E+00 | 4.85E-01 | no | 1700025G04Rik | intron_variant | MODIFIER |
| 1 | 151938447 | AX-170491004 | 0 | A | G | 0.202 | 9.76E-02 | 1.41E-01 | 2.67E+00 | 2.71E+00 | 4.85E-01 | no | 1700025G04Rik | intron_variant | MODIFIER |
| 1 | 151962262 | AX-170155630 | 0 | A | G | 0.202 | 9.76E-02 | 1.41E-01 | 2.67E+00 | 2.71E+00 | 4.85E-01 | no | 1700025G04Rik | intron_variant | MODIFIER |
| 1 | 151980923 | AX-170213770 | 0 | G | C | 0.202 | 9.76E-02 | 1.41E-01 | 2.67E+00 | 2.71E+00 | 4.85E-01 | no | 1700025G04Rik | intron_variant | MODIFIER |
| 1 | 151991206 | AX-97843645 | 0 | T | C | 0.202 | 9.76E-02 | 1.41E-01 | 2.67E+00 | 2.71E+00 | 4.85E-01 | no | 1700025G04Rik | intron_variant | MODIFIER |
| 1 | 151992672 | AX-170099765 | 0 | G | A | 0.202 | 9.76E-02 | 1.41E-01 | 2.67E+00 | 2.71E+00 | 4.85E-01 | no | 1700025G04Rik | intron_variant | MODIFIER |
| 1 | 151999548 | AX-170253787 | 0 | G | T | 0.202 | 9.76E-02 | 1.41E-01 | 2.67E+00 | 2.71E+00 | 4.85E-01 | no | 1700025G04Rik | intron_variant | MODIFIER |
| 1 | 152002042 | AX-170452552 | 0 | T | G | 0.202 | 9.76E-02 | 1.41E-01 | 2.67E+00 | 2.71E+00 | 4.85E-01 | no | 1700025G04Rik | intron_variant | MODIFIER |
| 1 | 152015244 | AX-97800229 | 0 | A | G | 0.202 | 9.76E-02 | 1.41E-01 | 2.67E+00 | 2.71E+00 | 4.85E-01 | no | 1700025G04Rik | intron_variant | MODIFIER |
| 1 | 152015402 | AX-170158237 | 0 | C | T | 0.202 | 9.76E-02 | 1.41E-01 | 2.67E+00 | 2.71E+00 | 4.85E-01 | no | 1700025G04Rik | intron_variant | MODIFIER |
| 1 | 152056012 | AX-170234283 | 0 | C | T | 0.202 | 9.76E-02 | 1.41E-01 | 2.67E+00 | 2.71E+00 | 4.85E-01 | no | 1700025G04Rik | intron_variant | MODIFIER |
| 1 | 152056280 | AX-170422593 | 0 | A | T | 0.202 | 9.76E-02 | 1.41E-01 | 2.67E+00 | 2.71E+00 | 4.85E-01 | no | 1700025G04Rik | intron_variant | MODIFIER |
| 1 | 152056312 | AX-170208193 | 0 | C | T | 0.202 | 9.76E-02 | 1.41E-01 | 2.67E+00 | 2.71E+00 | 4.85E-01 | no | 1700025G04Rik | intron_variant | MODIFIER |
| 1 | 152064793 | AX-170349212 | 0 | T | C | 0.202 | 9.76E-02 | 1.41E-01 | 2.67E+00 | 2.71E+00 | 4.85E-01 | no | 1700025G04Rik | intron_variant | MODIFIER |
| 1 | 152111301 | AX-97850110 | 0 | C | G | 0.202 | 9.76E-02 | 1.41E-01 | 2.67E+00 | 2.71E+00 | 4.85E-01 | no | | intergenic_variant | MODIFIER |
| 1 | 152113824 | AX-170423061 | 0 | T | C | 0.202 | 9.76E-02 | 1.41E-01 | 2.67E+00 | 2.71E+00 | 4.85E-01 | no | | intergenic_variant | MODIFIER |
| 1 | 152121940 | AX-170355816 | 0 | A | G | 0.202 | 9.76E-02 | 1.41E-01 | 2.67E+00 | 2.71E+00 | 4.85E-01 | no | | intergenic_variant | MODIFIER |
| 1 | 152136092 | AX-170486090 | 0 | T | C | 0.202 | 9.76E-02 | 1.41E-01 | 2.67E+00 | 2.71E+00 | 4.85E-01 | no | | intergenic_variant | MODIFIER |
| 1 | 152143818 | AX-170043158 | 0 | T | C | 0.202 | 9.76E-02 | 1.41E-01 | 2.67E+00 | 2.71E+00 | 4.85E-01 | no | | intergenic_variant | MODIFIER |
| 1 | 152164845 | AX-170043157 | 0 | A | G | 0.202 | 9.76E-02 | 1.41E-01 | 2.67E+00 | 2.71E+00 | 4.85E-01 | no | | regulatory_region_variant | MODIFIER |
| 1 | 152170286 | AX-170043156 | 0 | A | G | 0.202 | 9.76E-02 | 1.41E-01 | 2.67E+00 | 2.71E+00 | 4.85E-01 | no | | intergenic_variant | MODIFIER |
| 1 | 152172889 | AX-170365376 | 0 | G | A | 0.202 | 9.76E-02 | 1.41E-01 | 2.67E+00 | 2.71E+00 | 4.85E-01 | no | | intergenic_variant | MODIFIER |
| 1 | 152177083 | AX-170504979 | 0 | G | A | 0.202 | 9.76E-02 | 1.41E-01 | 2.67E+00 | 2.71E+00 | 4.85E-01 | no | | intergenic_variant | MODIFIER |
| 1 | 152177275 | AX-170377401 | 0 | G | C | 0.202 | 9.76E-02 | 1.41E-01 | 2.67E+00 | 2.71E+00 | 4.85E-01 | no | | intergenic_variant | MODIFIER |
| 1 | 152201334 | AX-170144599 | 0 | T | C | 0.202 | 9.76E-02 | 1.41E-01 | 2.67E+00 | 2.71E+00 | 4.85E-01 | no | | intergenic_variant | MODIFIER |
| 1 | 152212189 | AX-170295407 | 0 | T | C | 0.202 | 9.76E-02 | 1.41E-01 | 2.67E+00 | 2.71E+00 | 4.85E-01 | no | | intergenic_variant | MODIFIER |
| 1 | 152214580 | AX-170371138 | 0 | A | G | 0.202 | 9.76E-02 | 1.41E-01 | 2.67E+00 | 2.71E+00 | 4.85E-01 | no | | regulatory_region_variant | MODIFIER |
| 1 | 152218411 | AX-170238333 | 0 | A | G | 0.202 | 9.76E-02 | 1.41E-01 | 2.67E+00 | 2.71E+00 | 4.85E-01 | no | | intergenic_variant | MODIFIER |
| 1 | 152228683 | AX-170220155 | 0 | T | C | 0.202 | 9.76E-02 | 1.41E-01 | 2.67E+00 | 2.71E+00 | 4.85E-01 | no | | intergenic_variant | MODIFIER |
| 1 | 152234775 | AX-170220477 | 0 | A | G | 0.202 | 9.76E-02 | 1.41E-01 | 2.67E+00 | 2.71E+00 | 4.85E-01 | no | | intergenic_variant | MODIFIER |
| 1 | 152253278 | AX-170147425 | 0 | A | G | 0.202 | 9.76E-02 | 1.41E-01 | 2.67E+00 | 2.71E+00 | 4.85E-01 | no | | regulatory_region_variant | MODIFIER |
| 1 | 152284975 | AX-97828113 | 0 | T | G | 0.202 | 9.76E-02 | 1.41E-01 | 2.67E+00 | 2.71E+00 | 4.85E-01 | no | | regulatory_region_variant | MODIFIER |
| 1 | 152293952 | AX-170001995 | 0 | A | G | 0.202 | 9.76E-02 | 1.41E-01 | 2.67E+00 | 2.71E+00 | 4.85E-01 | no | Gm38238 | upstream_gene_variant | MODIFIER |
| 1 | 152293983 | AX-170001996 | 0 | A | G | 0.202 | 9.76E-02 | 1.41E-01 | 2.67E+00 | 2.71E+00 | 4.85E-01 | no | Gm38238 | upstream_gene_variant | MODIFIER |
| 1 | 152300732 | AX-170408902 | 0 | T | C | 0.202 | 9.76E-02 | 1.41E-01 | 2.67E+00 | 2.71E+00 | 4.85E-01 | no | | intergenic_variant | MODIFIER |
| 1 | 152301152 | AX-170408018 | 0 | T | C | 0.202 | 9.76E-02 | 1.41E-01 | 2.67E+00 | 2.71E+00 | 4.85E-01 | no | | intergenic_variant | MODIFIER |
| 1 | 152305245 | AX-170043163 | 0 | T | C | 0.202 | 9.76E-02 | 1.41E-01 | 2.67E+00 | 2.71E+00 | 4.85E-01 | no | | regulatory_region_variant | MODIFIER |
| 1 | 152306573 | AX-170182494 | 0 | A | G | 0.202 | 9.76E-02 | 1.41E-01 | 2.67E+00 | 2.71E+00 | 4.85E-01 | no | | intergenic_variant | MODIFIER |
| 1 | 152310201 | AX-170361835 | 0 | T | C | 0.202 | 9.76E-02 | 1.41E-01 | 2.67E+00 | 2.71E+00 | 4.85E-01 | no | | intergenic_variant | MODIFIER |
| 1 | 152310644 | AX-170001997 | 0 | T | G | 0.202 | 9.76E-02 | 1.41E-01 | 2.67E+00 | 2.71E+00 | 4.85E-01 | no | | intergenic_variant | MODIFIER |
| 1 | 152327757 | AX-170257572 | 0 | A | C | 0.202 | 9.76E-02 | 1.41E-01 | 2.67E+00 | 2.71E+00 | 4.85E-01 | no | | intergenic_variant | MODIFIER |
| 1 | 152331044 | AX-97814575 | 0 | G | A | 0.202 | 9.76E-02 | 1.41E-01 | 2.67E+00 | 2.71E+00 | 4.85E-01 | no | | intergenic_variant | MODIFIER |
| 1 | 152332914 | AX-170001991 | 0 | G | A | 0.202 | 9.76E-02 | 1.41E-01 | 2.67E+00 | 2.71E+00 | 4.85E-01 | no | | intergenic_variant | MODIFIER |
| 1 | 152332979 | AX-170001994 | 0 | T | C | 0.202 | 9.76E-02 | 1.41E-01 | 2.67E+00 | 2.71E+00 | 4.85E-01 | no | | intergenic_variant | MODIFIER |
| 1 | 152350190 | AX-170362700 | 0 | G | T | 0.202 | 9.76E-02 | 1.41E-01 | 2.67E+00 | 2.71E+00 | 4.85E-01 | no | | intergenic_variant | MODIFIER |
| 1 | 152350686 | AX-170455985 | 0 | G | A | 0.202 | 9.76E-02 | 1.41E-01 | 2.67E+00 | 2.71E+00 | 4.85E-01 | no | | intergenic_variant | MODIFIER |
| 1 | 152351574 | AX-97854570 | 0 | A | G | 0.202 | 9.76E-02 | 1.41E-01 | 2.67E+00 | 2.71E+00 | 4.85E-01 | no | | intergenic_variant | MODIFIER |
| 1 | 152355246 | AX-170380334 | 0 | T | G | 0.202 | 9.76E-02 | 1.41E-01 | 2.67E+00 | 2.71E+00 | 4.85E-01 | no | | intergenic_variant | MODIFIER |
| 1 | 152363857 | AX-170322144 | 0 | C | G | 0.202 | 9.76E-02 | 1.41E-01 | 2.67E+00 | 2.71E+00 | 4.85E-01 | no | | intergenic_variant | MODIFIER |
| 1 | 152369668 | AX-170216075 | 0 | T | G | 0.202 | 9.76E-02 | 1.41E-01 | 2.67E+00 | 2.71E+00 | 4.85E-01 | no | Tsen15 | downstream_gene_variant | MODIFIER |

| | | | | | | | | | | | | | | | |
|---|---|---|---|---|---|---|---|---|---|---|---|---|---|---|---|
| 1 | 152370545 | AX-170478832 | 0 | G | A | 0.202 | 9.76E-02 | 1.41E-01 | 2.67E+00 | 2.71E+00 | 4.85E-01 | no | Tsen15 | downstream_gene_variant | MODIFIER |
| 1 | 152383233 | AX-170160257 | 0 | T | G | 0.202 | 9.76E-02 | 1.41E-01 | 2.67E+00 | 2.71E+00 | 4.85E-01 | no | Tsen15 | intron_variant | MODIFIER |
| 1 | 152409319 | AX-170105550 | 0 | T | G | 0.202 | 9.76E-02 | 1.41E-01 | 2.67E+00 | 2.71E+00 | 4.85E-01 | no | Colgalt2 | intron_variant | MODIFIER |
| 1 | 152409790 | AX-170043166 | 0 | G | A | 0.202 | 9.76E-02 | 1.41E-01 | 2.67E+00 | 2.71E+00 | 4.85E-01 | no | Colgalt2 | intron_variant | MODIFIER |
| 1 | 152417081 | AX-170119542 | 0 | T | C | 0.202 | 9.76E-02 | 1.41E-01 | 2.67E+00 | 2.71E+00 | 4.85E-01 | no | Colgalt2 | intron_variant | MODIFIER |
| 1 | 152418185 | AX-97790387 | 0 | A | G | 0.202 | 9.76E-02 | 1.41E-01 | 2.67E+00 | 2.71E+00 | 4.85E-01 | no | Colgalt2 | intron_variant | MODIFIER |
| 1 | 152423756 | AX-170501752 | 0 | T | C | 0.202 | 9.76E-02 | 1.41E-01 | 2.67E+00 | 2.71E+00 | 4.85E-01 | no | Colgalt2 | intron_variant | MODIFIER |
| 1 | 152434237 | AX-170377490 | 0 | G | C | 0.202 | 9.76E-02 | 1.41E-01 | 2.67E+00 | 2.71E+00 | 4.85E-01 | no | Colgalt2 | intron_variant | MODIFIER |
| 1 | 152446015 | AX-170164110 | 0 | A | G | 0.202 | 9.76E-02 | 1.41E-01 | 2.67E+00 | 2.71E+00 | 4.85E-01 | no | Colgalt2 | intron_variant | MODIFIER |
| 1 | 152447620 | AX-170411325 | 0 | A | G | 0.202 | 9.76E-02 | 1.41E-01 | 2.67E+00 | 2.71E+00 | 4.85E-01 | no | Colgalt2 | intron_variant | MODIFIER |
| 1 | 152451573 | AX-170495775 | 0 | A | G | 0.202 | 9.76E-02 | 1.41E-01 | 2.67E+00 | 2.71E+00 | 4.85E-01 | no | Colgalt2 | intron_variant | MODIFIER |
| 1 | 152453540 | AX-170139710 | 0 | G | A | 0.202 | 9.76E-02 | 1.41E-01 | 2.67E+00 | 2.71E+00 | 4.85E-01 | no | Colgalt2 | intron_variant | MODIFIER |
| 1 | 152458296 | AX-170145665 | 0 | C | T | 0.202 | 9.76E-02 | 1.41E-01 | 2.67E+00 | 2.71E+00 | 4.85E-01 | no | Colgalt2 | intron_variant | MODIFIER |
| 1 | 152466717 | AX-170532798 | 0 | G | A | 0.202 | 9.76E-02 | 1.41E-01 | 2.67E+00 | 2.71E+00 | 4.85E-01 | no | Colgalt2 | intron_variant | MODIFIER |
| 1 | 152470228 | AX-170120354 | 0 | A | G | 0.202 | 9.76E-02 | 1.41E-01 | 2.67E+00 | 2.71E+00 | 4.85E-01 | no | Colgalt2 | intron_variant | MODIFIER |
| 1 | 152470244 | AX-170325064 | 0 | A | C | 0.202 | 9.76E-02 | 1.41E-01 | 2.67E+00 | 2.71E+00 | 4.85E-01 | no | Colgalt2 | intron_variant | MODIFIER |
| 1 | 152470422 | AX-97797456 | 0 | A | C | 0.202 | 9.76E-02 | 1.41E-01 | 2.67E+00 | 2.71E+00 | 4.85E-01 | no | Colgalt2 | intron_variant | MODIFIER |
| 1 | 152472075 | AX-170359285 | 0 | C | A | 0.202 | 9.76E-02 | 1.41E-01 | 2.67E+00 | 2.71E+00 | 4.85E-01 | no | Colgalt2 | intron_variant | MODIFIER |
| 1 | 152495833 | AX-170443651 | 0 | C | T | 0.202 | 9.76E-02 | 1.41E-01 | 2.67E+00 | 2.71E+00 | 4.85E-01 | no | Colgalt2 | synonymous_variant | LOW |
| 1 | 152497241 | AX-170468717 | 0 | G | A | 0.202 | 9.76E-02 | 1.41E-01 | 2.67E+00 | 2.71E+00 | 4.85E-01 | no | Colgalt2 | intron_variant | MODIFIER |
| 1 | 152503145 | AX-170140614 | 0 | T | C | 0.202 | 9.76E-02 | 1.41E-01 | 2.67E+00 | 2.71E+00 | 4.85E-01 | no | Colgalt2 | intron_variant | MODIFIER |
| 1 | 152504566 | AX-170241079 | 0 | C | G | 0.202 | 9.76E-02 | 1.41E-01 | 2.67E+00 | 2.71E+00 | 4.85E-01 | no | Colgalt2 | intron_variant | MODIFIER |
| 1 | 152506397 | AX-170401119 | 0 | G | A | 0.202 | 9.76E-02 | 1.41E-01 | 2.67E+00 | 2.71E+00 | 4.85E-01 | no | Colgalt2 | intron_variant | MODIFIER |
| 1 | 152506582 | AX-97781555 | 0 | C | T | 0.202 | 9.76E-02 | 1.41E-01 | 2.67E+00 | 2.71E+00 | 4.85E-01 | no | Colgalt2 | intron_variant | MODIFIER |
| 1 | 152524473 | AX-170318336 | 0 | A | G | 0.202 | 9.76E-02 | 1.41E-01 | 2.67E+00 | 2.71E+00 | 4.85E-01 | no | Rgl1 | intron_variant | MODIFIER |
| 1 | 152524944 | AX-170441150 | 0 | T | C | 0.202 | 9.76E-02 | 1.41E-01 | 2.67E+00 | 2.71E+00 | 4.85E-01 | no | Rgl1 | intron_variant | MODIFIER |
| 1 | 152528104 | AX-170502790 | 0 | T | C | 0.202 | 9.76E-02 | 1.41E-01 | 2.67E+00 | 2.71E+00 | 4.85E-01 | no | Rgl1 | intron_variant | MODIFIER |
| 1 | 152533704 | AX-170127278 | 0 | A | T | 0.202 | 9.76E-02 | 1.41E-01 | 2.67E+00 | 2.71E+00 | 4.85E-01 | no | Rgl1 | intron_variant | MODIFIER |
| 1 | 152534466 | AX-170383146 | 0 | C | T | 0.202 | 9.76E-02 | 1.41E-01 | 2.67E+00 | 2.71E+00 | 4.85E-01 | no | Rgl1 | intron_variant | MODIFIER |
| 1 | 152534644 | AX-170415026 | 0 | A | G | 0.202 | 9.76E-02 | 1.41E-01 | 2.67E+00 | 2.71E+00 | 4.85E-01 | no | Rgl1 | intron_variant | MODIFIER |
| 1 | 152548529 | AX-170387949 | 0 | T | A | 0.202 | 9.76E-02 | 1.41E-01 | 2.67E+00 | 2.71E+00 | 4.85E-01 | no | Rgl1 | intron_variant | MODIFIER |
| 1 | 152548575 | AX-170328215 | 0 | A | T | 0.202 | 9.76E-02 | 1.41E-01 | 2.67E+00 | 2.71E+00 | 4.85E-01 | no | Rgl1 | intron_variant | MODIFIER |
| 1 | 152548837 | AX-97813936 | 0 | T | C | 0.202 | 9.76E-02 | 1.41E-01 | 2.67E+00 | 2.71E+00 | 4.85E-01 | no | Rgl1 | intron_variant | MODIFIER |
| 1 | 152577027 | AX-170192053 | 0 | A | G | 0.202 | 9.76E-02 | 1.41E-01 | 2.67E+00 | 2.71E+00 | 4.85E-01 | no | Rgl1 | intron_variant | MODIFIER |
| 1 | 152580443 | AX-97799531 | 0 | C | T | 0.202 | 9.76E-02 | 1.41E-01 | 2.67E+00 | 2.71E+00 | 4.85E-01 | no | Rgl1 | intron_variant | MODIFIER |
| 1 | 152587770 | AX-170249877 | 0 | A | G | 0.202 | 9.76E-02 | 1.41E-01 | 2.67E+00 | 2.71E+00 | 4.85E-01 | no | Rgl1 | intron_variant | MODIFIER |
| 1 | 152587869 | AX-170143580 | 0 | C | T | 0.202 | 9.76E-02 | 1.41E-01 | 2.67E+00 | 2.71E+00 | 4.85E-01 | no | Rgl1 | intron_variant | MODIFIER |
| 1 | 152589205 | AX-170475476 | 0 | A | G | 0.202 | 9.76E-02 | 1.41E-01 | 2.67E+00 | 2.71E+00 | 4.85E-01 | no | Rgl1 | intron_variant | MODIFIER |
| 1 | 152591658 | AX-170260671 | 0 | C | T | 0.202 | 9.76E-02 | 1.41E-01 | 2.67E+00 | 2.71E+00 | 4.85E-01 | no | Rgl1 | intron_variant | MODIFIER |
| 1 | 152599690 | AX-170412939 | 0 | A | G | 0.202 | 9.76E-02 | 1.41E-01 | 2.67E+00 | 2.71E+00 | 4.85E-01 | no | Rgl1 | intron_variant | MODIFIER |
| 1 | 152604489 | AX-170169921 | 0 | C | T | 0.202 | 9.76E-02 | 1.41E-01 | 2.67E+00 | 2.71E+00 | 4.85E-01 | no | Rgl1 | intron_variant | MODIFIER |
| 1 | 152604692 | AX-170246965 | 0 | G | C | 0.202 | 9.76E-02 | 1.41E-01 | 2.67E+00 | 2.71E+00 | 4.85E-01 | no | Rgl1 | intron_variant | MODIFIER |
| 1 | 152606846 | AX-170111054 | 0 | C | T | 0.202 | 9.76E-02 | 1.41E-01 | 2.67E+00 | 2.71E+00 | 4.85E-01 | no | Rgl1 | intron_variant | MODIFIER |
| 1 | 152607210 | AX-170502841 | 0 | C | A | 0.202 | 9.76E-02 | 1.41E-01 | 2.67E+00 | 2.71E+00 | 4.85E-01 | no | Rgl1 | intron_variant | MODIFIER |
| 1 | 152612049 | AX-170397521 | 0 | A | G | 0.202 | 9.76E-02 | 1.41E-01 | 2.67E+00 | 2.71E+00 | 4.85E-01 | no | Rgl1 | intron_variant | MODIFIER |
| 1 | 152627093 | AX-170086648 | 0 | A | G | 0.202 | 9.76E-02 | 1.41E-01 | 2.67E+00 | 2.71E+00 | 4.85E-01 | no | Rgl1 | upstream_gene_variant | MODIFIER |
| 1 | 152631813 | AX-170422555 | 0 | G | A | 0.202 | 9.76E-02 | 1.41E-01 | 2.67E+00 | 2.71E+00 | 4.85E-01 | no | Rgl1 | intron_variant | MODIFIER |
| 1 | 152632845 | AX-170487857 | 0 | A | T | 0.202 | 9.76E-02 | 1.41E-01 | 2.67E+00 | 2.71E+00 | 4.85E-01 | no | Rgl1 | intron_variant | MODIFIER |
| 1 | 152633307 | AX-170285012 | 0 | T | C | 0.202 | 9.76E-02 | 1.41E-01 | 2.67E+00 | 2.71E+00 | 4.85E-01 | no | Rgl1 | intron_variant | MODIFIER |
| 1 | 152641373 | AX-170477817 | 0 | C | T | 0.202 | 9.76E-02 | 1.41E-01 | 2.67E+00 | 2.71E+00 | 4.85E-01 | no | Rgl1 | intron_variant | MODIFIER |
| 1 | 152656290 | AX-97808806 | 0 | A | G | 0.202 | 9.76E-02 | 1.41E-01 | 2.67E+00 | 2.71E+00 | 4.85E-01 | no | Rgl1 | intron_variant | MODIFIER |
| 1 | 152662844 | AX-170106448 | 0 | T | C | 0.202 | 9.76E-02 | 1.41E-01 | 2.67E+00 | 2.71E+00 | 4.85E-01 | no | Rgl1 | intron_variant | MODIFIER |
| 1 | 152700775 | AX-170379278 | 0 | T | C | 0.202 | 9.76E-02 | 1.41E-01 | 2.67E+00 | 2.71E+00 | 4.85E-01 | no | Rgl1 | intron_variant | MODIFIER |
| 1 | 152709697 | AX-170043173 | 0 | C | A | 0.202 | 9.76E-02 | 1.41E-01 | 2.67E+00 | 2.71E+00 | 4.85E-01 | no | Rgl1 | intron_variant | MODIFIER |

| | | | | | | | | | | | | | | | |
|---|---|---|---|---|---|---|---|---|---|---|---|---|---|---|---|
| 1 | 152732575 | AX-170346978 | 0 | A | C | 0.202 | 9.76E-02 | 1.41E-01 | 2.67E+00 | 2.71E+00 | 4.85E-01 | no | Rgl1 | intron_variant | MODIFIER |
| 1 | 152733342 | AX-170203191 | 0 | C | T | 0.202 | 9.76E-02 | 1.41E-01 | 2.67E+00 | 2.71E+00 | 4.85E-01 | no | Rgl1 | intron_variant | MODIFIER |
| 1 | 152733382 | AX-97842998 | 0 | C | T | 0.202 | 9.76E-02 | 1.41E-01 | 2.67E+00 | 2.71E+00 | 4.85E-01 | no | Rgl1 | intron_variant | MODIFIER |
| 1 | 152736706 | AX-170404038 | 0 | A | G | 0.202 | 9.76E-02 | 1.41E-01 | 2.67E+00 | 2.71E+00 | 4.85E-01 | no | Rgl1 | intron_variant | MODIFIER |
| 1 | 152738340 | AX-170390056 | 0 | G | A | 0.202 | 9.76E-02 | 1.41E-01 | 2.67E+00 | 2.71E+00 | 4.85E-01 | no | Rgl1 | intron_variant | MODIFIER |
| 1 | 152743360 | AX-170284759 | 0 | T | A | 0.202 | 9.76E-02 | 1.41E-01 | 2.67E+00 | 2.71E+00 | 4.85E-01 | no | Rgl1 | intron_variant | MODIFIER |
| 1 | 152743685 | AX-170381073 | 0 | G | T | 0.202 | 9.76E-02 | 1.41E-01 | 2.67E+00 | 2.71E+00 | 4.85E-01 | no | Rgl1 | intron_variant | MODIFIER |
| 1 | 152750573 | AX-97782989 | 0 | A | G | 0.202 | 9.76E-02 | 1.41E-01 | 2.67E+00 | 2.71E+00 | 4.85E-01 | no | Apobec4 | 5_prime_UTR_variant | MODIFIER |
| 1 | 152776089 | AX-97846109 | 0 | T | G | 0.202 | 9.76E-02 | 1.41E-01 | 2.67E+00 | 2.71E+00 | 4.85E-01 | no | Arpc5 | downstream_gene_variant | MODIFIER |
| 1 | 152776693 | AX-170228060 | 0 | A | G | 0.202 | 9.76E-02 | 1.41E-01 | 2.67E+00 | 2.71E+00 | 4.85E-01 | no | Arpc5 | downstream_gene_variant | MODIFIER |
| 1 | 152776868 | AX-170347956 | 0 | T | G | 0.202 | 9.76E-02 | 1.41E-01 | 2.67E+00 | 2.71E+00 | 4.85E-01 | no | Arpc5 | downstream_gene_variant | MODIFIER |
| 1 | 152780223 | AX-170143628 | 0 | G | A | 0.202 | 9.76E-02 | 1.41E-01 | 2.67E+00 | 2.71E+00 | 4.85E-01 | no | Arpc5 | downstream_gene_variant | MODIFIER |
| 1 | 152794000 | AX-170308627 | 0 | C | T | 0.202 | 9.76E-02 | 1.41E-01 | 2.67E+00 | 2.71E+00 | 4.85E-01 | no | | intergenic_variant | MODIFIER |
| 1 | 152817795 | AX-170184442 | 0 | A | G | 0.202 | 9.76E-02 | 1.41E-01 | 2.67E+00 | 2.71E+00 | 4.85E-01 | no | Ncf2 | intron_variant | MODIFIER |
| 1 | 152818210 | AX-170002000 | 0 | A | G | 0.202 | 9.76E-02 | 1.41E-01 | 2.67E+00 | 2.71E+00 | 4.85E-01 | no | Ncf2 | intron_variant | MODIFIER |
| 1 | 152818652 | AX-170002004 | 0 | G | A | 0.202 | 9.76E-02 | 1.41E-01 | 2.67E+00 | 2.71E+00 | 4.85E-01 | no | Ncf2 | intron_variant | MODIFIER |
| 1 | 152832690 | AX-170340449 | 0 | T | C | 0.202 | 9.76E-02 | 1.41E-01 | 2.67E+00 | 2.71E+00 | 4.85E-01 | no | Ncf2 | intron_variant | MODIFIER |
| 1 | 152832924 | AX-170502890 | 0 | A | T | 0.202 | 9.76E-02 | 1.41E-01 | 2.67E+00 | 2.71E+00 | 4.85E-01 | no | Ncf2 | intron_variant | MODIFIER |
| 1 | 152834295 | AX-170400549 | 0 | A | G | 0.202 | 9.76E-02 | 1.41E-01 | 2.67E+00 | 2.71E+00 | 4.85E-01 | no | Ncf2 | intron_variant | MODIFIER |
| 1 | 152834949 | AX-170043178 | 0 | A | C | 0.202 | 9.76E-02 | 1.41E-01 | 2.67E+00 | 2.71E+00 | 4.85E-01 | no | Ncf2 | intron_variant | MODIFIER |
| 1 | 152835438 | AX-170043176 | 0 | C | A | 0.202 | 9.76E-02 | 1.41E-01 | 2.67E+00 | 2.71E+00 | 4.85E-01 | no | Ncf2 | intron_variant | MODIFIER |
| 1 | 152842119 | AX-170001999 | 0 | C | G | 0.202 | 9.76E-02 | 1.41E-01 | 2.67E+00 | 2.71E+00 | 4.85E-01 | no | Smg7 | intron_variant | MODIFIER |
| 1 | 152844587 | AX-97803081 | 0 | G | A | 0.202 | 9.76E-02 | 1.41E-01 | 2.67E+00 | 2.71E+00 | 4.85E-01 | no | Smg7 | intron_variant | MODIFIER |
| 1 | 152849932 | AX-170043174 | 0 | C | T | 0.202 | 9.76E-02 | 1.41E-01 | 2.67E+00 | 2.71E+00 | 4.85E-01 | no | Smg7 | missense_variant | MODERATE |
| 1 | 152853840 | AX-170512324 | 0 | A | T | 0.202 | 9.76E-02 | 1.41E-01 | 2.67E+00 | 2.71E+00 | 4.85E-01 | no | Smg7 | intron_variant | MODIFIER |
| 1 | 152855992 | AX-170002001 | 0 | G | A | 0.202 | 9.76E-02 | 1.41E-01 | 2.67E+00 | 2.71E+00 | 4.85E-01 | no | Smg7 | synonymous_variant | LOW |
| 1 | 152856055 | AX-170002002 | 0 | A | G | 0.202 | 9.76E-02 | 1.41E-01 | 2.67E+00 | 2.71E+00 | 4.85E-01 | no | Smg7 | intron_variant | MODIFIER |
| 1 | 152856770 | AX-170157239 | 0 | T | C | 0.202 | 9.76E-02 | 1.41E-01 | 2.67E+00 | 2.71E+00 | 4.85E-01 | no | Smg7 | intron_variant | MODIFIER |
| 1 | 152889995 | AX-97792116 | 0 | A | G | 0.202 | 9.76E-02 | 1.41E-01 | 2.67E+00 | 2.71E+00 | 4.85E-01 | no | Smg7 | intron_variant | MODIFIER |
| 1 | 152894569 | AX-170382444 | 0 | T | C | 0.202 | 9.76E-02 | 1.41E-01 | 2.67E+00 | 2.71E+00 | 4.85E-01 | no | Smg7 | intron_variant | MODIFIER |
| 1 | 152896619 | AX-170001998 | 0 | T | C | 0.202 | 9.76E-02 | 1.41E-01 | 2.67E+00 | 2.71E+00 | 4.85E-01 | no | Smg7 | intron_variant | MODIFIER |
| 1 | 152911670 | AX-170393661 | 0 | C | T | 0.202 | 9.76E-02 | 1.41E-01 | 2.67E+00 | 2.71E+00 | 4.85E-01 | no | | intergenic_variant | MODIFIER |
| 1 | 152912341 | AX-97844573 | 0 | T | C | 0.202 | 9.76E-02 | 1.41E-01 | 2.67E+00 | 2.71E+00 | 4.85E-01 | no | | intergenic_variant | MODIFIER |
| 1 | 152914150 | AX-170002005 | 0 | C | T | 0.202 | 9.76E-02 | 1.41E-01 | 2.67E+00 | 2.71E+00 | 4.85E-01 | no | | regulatory_region_variant | MODIFIER |
| 1 | 152923140 | AX-170091753 | 0 | T | C | 0.202 | 9.76E-02 | 1.41E-01 | 2.67E+00 | 2.71E+00 | 4.85E-01 | no | | intergenic_variant | MODIFIER |
| 1 | 152929561 | AX-170316921 | 0 | C | T | 0.202 | 9.76E-02 | 1.41E-01 | 2.67E+00 | 2.71E+00 | 4.85E-01 | no | | intergenic_variant | MODIFIER |
| 1 | 152952403 | AX-170511542 | 0 | C | T | 0.202 | 9.76E-02 | 1.41E-01 | 2.67E+00 | 2.71E+00 | 4.85E-01 | no | Nmnat2 | upstream_gene_variant | MODIFIER |
| 1 | 152956095 | AX-170397556 | 0 | G | A | 0.202 | 9.76E-02 | 1.41E-01 | 2.67E+00 | 2.71E+00 | 4.85E-01 | no | Nmnat2 | intron_variant | MODIFIER |
| 1 | 152956129 | AX-170474747 | 0 | A | G | 0.202 | 9.76E-02 | 1.41E-01 | 2.67E+00 | 2.71E+00 | 4.85E-01 | no | Nmnat2 | intron_variant | MODIFIER |
| 1 | 152956171 | AX-170236602 | 0 | G | C | 0.202 | 9.76E-02 | 1.41E-01 | 2.67E+00 | 2.71E+00 | 4.85E-01 | no | Nmnat2 | intron_variant | MODIFIER |
| 1 | 152961649 | AX-170383989 | 0 | C | G | 0.202 | 9.76E-02 | 1.41E-01 | 2.67E+00 | 2.71E+00 | 4.85E-01 | no | Nmnat2 | intron_variant | MODIFIER |
| 1 | 152970258 | AX-170180034 | 0 | G | T | 0.202 | 9.76E-02 | 1.41E-01 | 2.67E+00 | 2.71E+00 | 4.85E-01 | no | Nmnat2 | intron_variant | MODIFIER |
| 1 | 152970284 | AX-170394369 | 0 | T | C | 0.202 | 9.76E-02 | 1.41E-01 | 2.67E+00 | 2.71E+00 | 4.85E-01 | no | Nmnat2 | intron_variant | MODIFIER |
| 1 | 152971292 | AX-170043179 | 0 | G | A | 0.202 | 9.76E-02 | 1.41E-01 | 2.67E+00 | 2.71E+00 | 4.85E-01 | no | Nmnat2 | intron_variant | MODIFIER |
| 1 | 152971679 | AX-97813772 | 0 | G | A | 0.202 | 9.76E-02 | 1.41E-01 | 2.67E+00 | 2.71E+00 | 4.85E-01 | no | Nmnat2 | intron_variant | MODIFIER |
| 1 | 153001864 | AX-170462416 | 0 | A | G | 0.202 | 9.76E-02 | 1.41E-01 | 2.67E+00 | 2.71E+00 | 4.85E-01 | no | Nmnat2 | intron_variant | MODIFIER |
| 1 | 153006430 | AX-170409829 | 0 | G | A | 0.202 | 9.76E-02 | 1.41E-01 | 2.67E+00 | 2.71E+00 | 4.85E-01 | no | Nmnat2 | intron_variant | MODIFIER |
| 1 | 153009090 | AX-97814440 | 0 | C | T | 0.202 | 9.76E-02 | 1.41E-01 | 2.67E+00 | 2.71E+00 | 4.85E-01 | no | Nmnat2 | intron_variant | MODIFIER |
| 1 | 153013668 | AX-170291216 | 0 | C | T | 0.202 | 9.76E-02 | 1.41E-01 | 2.67E+00 | 2.71E+00 | 4.85E-01 | no | Nmnat2 | intron_variant | MODIFIER |
| 1 | 153015588 | AX-170356048 | 0 | C | T | 0.202 | 9.76E-02 | 1.41E-01 | 2.67E+00 | 2.71E+00 | 4.85E-01 | no | Nmnat2 | intron_variant | MODIFIER |
| 1 | 153015716 | AX-170268788 | 0 | T | C | 0.202 | 9.76E-02 | 1.41E-01 | 2.67E+00 | 2.71E+00 | 4.85E-01 | no | Nmnat2 | intron_variant | MODIFIER |
| 1 | 153018357 | AX-170225985 | 0 | G | A | 0.202 | 9.76E-02 | 1.41E-01 | 2.67E+00 | 2.71E+00 | 4.85E-01 | no | Nmnat2 | intron_variant | MODIFIER |
| 1 | 153040063 | AX-170268143 | 0 | A | C | 0.202 | 9.76E-02 | 1.41E-01 | 2.67E+00 | 2.71E+00 | 4.85E-01 | no | Nmnat2 | intron_variant | MODIFIER |
| 1 | 153049642 | AX-170320133 | 0 | C | T | 0.202 | 9.76E-02 | 1.41E-01 | 2.67E+00 | 2.71E+00 | 4.85E-01 | no | Nmnat2 | intron_variant | MODIFIER |

| | | | | | | | | | | | | | | | |
|---|---|---|---|---|---|---|---|---|---|---|---|---|---|---|---|
| 1 | 153055850 | AX-170189939 | 0 | C | A | 0.202 | 9.76E-02 | 1.41E-01 | 2.67E+00 | 2.71E+00 | 4.85E-01 | no | Nmnat2 | intron_variant | MODIFIER |
| 1 | 153059692 | AX-170445151 | 0 | A | G | 0.202 | 9.76E-02 | 1.41E-01 | 2.67E+00 | 2.71E+00 | 4.85E-01 | no | Nmnat2 | intron_variant | MODIFIER |
| 1 | 153060249 | AX-170043181 | 0 | T | A | 0.202 | 9.76E-02 | 1.41E-01 | 2.67E+00 | 2.71E+00 | 4.85E-01 | no | Nmnat2 | intron_variant | MODIFIER |
| 1 | 153060320 | AX-170410072 | 0 | G | C | 0.202 | 9.76E-02 | 1.41E-01 | 2.67E+00 | 2.71E+00 | 4.85E-01 | no | Nmnat2 | intron_variant | MODIFIER |
| 1 | 153065998 | AX-97838483 | 0 | C | T | 0.202 | 9.76E-02 | 1.41E-01 | 2.67E+00 | 2.71E+00 | 4.85E-01 | no | Nmnat2 | intron_variant | MODIFIER |
| 1 | 153068250 | AX-170362853 | 0 | C | T | 0.202 | 9.76E-02 | 1.41E-01 | 2.67E+00 | 2.71E+00 | 4.85E-01 | no | Nmnat2 | intron_variant | MODIFIER |
| 1 | 153069504 | AX-170262034 | 0 | C | A | 0.202 | 9.76E-02 | 1.41E-01 | 2.67E+00 | 2.71E+00 | 4.85E-01 | no | Nmnat2 | intron_variant | MODIFIER |
| 1 | 153071968 | AX-170385202 | 0 | G | A | 0.202 | 9.76E-02 | 1.41E-01 | 2.67E+00 | 2.71E+00 | 4.85E-01 | no | Nmnat2 | intron_variant | MODIFIER |
| 1 | 153072472 | AX-170420414 | 0 | G | A | 0.202 | 9.76E-02 | 1.41E-01 | 2.67E+00 | 2.71E+00 | 4.85E-01 | no | Nmnat2 | intron_variant | MODIFIER |
| 1 | 153087436 | AX-168411674 | 0 | T | C | 0.202 | 9.76E-02 | 1.41E-01 | 2.67E+00 | 2.71E+00 | 4.85E-01 | no | Nmnat2 | intron_variant | MODIFIER |
| 1 | 153087660 | AX-170169406 | 0 | C | G | 0.202 | 9.76E-02 | 1.41E-01 | 2.67E+00 | 2.71E+00 | 4.85E-01 | no | Nmnat2 | intron_variant | MODIFIER |
| 1 | 153094271 | AX-97848111 | 0 | C | T | 0.202 | 9.76E-02 | 1.41E-01 | 2.67E+00 | 2.71E+00 | 4.85E-01 | no | Nmnat2 | intron_variant | MODIFIER |
| 1 | 153104273 | AX-97809929 | 0 | C | T | 0.202 | 9.76E-02 | 1.41E-01 | 2.67E+00 | 2.71E+00 | 4.85E-01 | no | Nmnat2 | intron_variant | MODIFIER |
| 1 | 153126179 | AX-170485725 | 0 | A | G | 0.202 | 9.76E-02 | 1.41E-01 | 2.67E+00 | 2.71E+00 | 4.85E-01 | no | Lamc2 | intron_variant | MODIFIER |
| 1 | 153126619 | AX-170263570 | 0 | A | C | 0.202 | 9.76E-02 | 1.41E-01 | 2.67E+00 | 2.71E+00 | 4.85E-01 | no | Lamc2 | intron_variant | MODIFIER |
| 1 | 153126949 | AX-97788955 | 0 | T | G | 0.202 | 9.76E-02 | 1.41E-01 | 2.67E+00 | 2.71E+00 | 4.85E-01 | no | Lamc2 | intron_variant | MODIFIER |
| 1 | 153135976 | AX-170296675 | 0 | C | T | 0.202 | 9.76E-02 | 1.41E-01 | 2.67E+00 | 2.71E+00 | 4.85E-01 | no | Lamc2 | intron_variant | MODIFIER |
| 1 | 153144507 | AX-170275351 | 0 | C | T | 0.202 | 9.76E-02 | 1.41E-01 | 2.67E+00 | 2.71E+00 | 4.85E-01 | no | Lamc2 | splice_region_variant,intron_variant | LOW |
| 1 | 153149437 | AX-170295282 | 0 | C | A | 0.202 | 9.76E-02 | 1.41E-01 | 2.67E+00 | 2.71E+00 | 4.85E-01 | no | Lamc2 | intron_variant | MODIFIER |
| 1 | 153154547 | AX-97819714 | 0 | A | G | 0.202 | 9.76E-02 | 1.41E-01 | 2.67E+00 | 2.71E+00 | 4.85E-01 | no | Lamc2 | intron_variant | MODIFIER |
| 1 | 153159350 | AX-170486241 | 0 | C | T | 0.202 | 9.76E-02 | 1.41E-01 | 2.67E+00 | 2.71E+00 | 4.85E-01 | no | Lamc2 | intron_variant | MODIFIER |
| 1 | 153172174 | AX-170159978 | 0 | G | A | 0.202 | 9.76E-02 | 1.41E-01 | 2.67E+00 | 2.71E+00 | 4.85E-01 | no | Lamc2 | intron_variant | MODIFIER |
| 1 | 153180350 | AX-97856141 | 0 | C | A | 0.202 | 9.76E-02 | 1.41E-01 | 2.67E+00 | 2.71E+00 | 4.85E-01 | no | Lamc2 | intron_variant | MODIFIER |
| 1 | 153185208 | AX-170126733 | 0 | A | T | 0.202 | 9.76E-02 | 1.41E-01 | 2.67E+00 | 2.71E+00 | 4.85E-01 | no | Lamc2 | intron_variant | MODIFIER |
| 1 | 153200313 | AX-170430444 | 0 | A | T | 0.202 | 9.76E-02 | 1.41E-01 | 2.67E+00 | 2.71E+00 | 4.85E-01 | no | | intergenic_variant | MODIFIER |
| 1 | 153218525 | AX-170147493 | 0 | T | C | 0.202 | 9.76E-02 | 1.41E-01 | 2.67E+00 | 2.71E+00 | 4.85E-01 | no | Lamc1 | downstream_gene_variant | MODIFIER |
| 1 | 153232496 | AX-170404328 | 0 | A | T | 0.202 | 9.76E-02 | 1.41E-01 | 2.67E+00 | 2.71E+00 | 4.85E-01 | no | Lamc1 | intron_variant | MODIFIER |
| 1 | 153235430 | AX-170043184 | 0 | C | A | 0.202 | 9.76E-02 | 1.41E-01 | 2.67E+00 | 2.71E+00 | 4.85E-01 | no | Lamc1 | intron_variant | MODIFIER |
| 1 | 153241759 | AX-170200665 | 0 | C | T | 0.202 | 9.76E-02 | 1.41E-01 | 2.67E+00 | 2.71E+00 | 4.85E-01 | no | Lamc1 | intron_variant | MODIFIER |
| 1 | 153241935 | AX-170257963 | 0 | G | A | 0.202 | 9.76E-02 | 1.41E-01 | 2.67E+00 | 2.71E+00 | 4.85E-01 | no | Lamc1 | synonymous_variant | LOW |
| 1 | 153242095 | AX-170333125 | 0 | G | T | 0.202 | 9.76E-02 | 1.41E-01 | 2.67E+00 | 2.71E+00 | 4.85E-01 | no | Lamc1 | intron_variant | MODIFIER |
| 16 | 93921301 | AX-170399791 | 1 | G | A | 0.411 | 5.46E-02 | 7.83E-02 | 2.92E+00 | 3.05E+00 | 4.85E-01 | no | Cldn14 | intron_variant | MODIFIER |
| 7 | 89679181 | AX-170520270 | 0 | C | A | 0.14 | 1.19E-01 | 1.72E-01 | 2.65E+00 | 2.77E+00 | 4.85E-01 | no | Me3 | intron_variant | MODIFIER |
| 5 | 40761789 | AX-170152501 | 0 | C | T | 0.184 | -7.68E-02 | 1.10E-01 | 2.61E+00 | 2.69E+00 | 4.85E-01 | no | | intergenic_variant | MODIFIER |
| 5 | 41273802 | AX-170401936 | 0 | A | G | 0.184 | -7.68E-02 | 1.10E-01 | 2.61E+00 | 2.69E+00 | 4.85E-01 | no | | intergenic_variant | MODIFIER |
| 5 | 43188950 | AX-170251104 | 0 | A | G | 0.184 | -7.68E-02 | 1.10E-01 | 2.61E+00 | 2.69E+00 | 4.85E-01 | no | Gm7854 | splice_region_variant,intron_variant,non_coding_transcript_variant | LOW |
| 5 | 43205710 | AX-170321818 | 0 | C | T | 0.184 | -7.68E-02 | 1.10E-01 | 2.61E+00 | 2.69E+00 | 4.85E-01 | no | Gm7854 | intron_variant,non_coding_transcript_variant | MODIFIER |
| 5 | 43228533 | AX-170217176 | 0 | C | A | 0.184 | -7.68E-02 | 1.10E-01 | 2.61E+00 | 2.69E+00 | 4.85E-01 | no | Cpeb2 | upstream_gene_variant | MODIFIER |
| 5 | 43228685 | AX-97800547 | 0 | A | G | 0.184 | -7.68E-02 | 1.10E-01 | 2.61E+00 | 2.69E+00 | 4.85E-01 | no | Cpeb2 | upstream_gene_variant | MODIFIER |
| 5 | 43334285 | AX-97806403 | 0 | G | A | 0.184 | -7.68E-02 | 1.10E-01 | 2.61E+00 | 2.69E+00 | 4.85E-01 | no | | intergenic_variant | MODIFIER |
| 5 | 43338650 | AX-170376808 | 0 | G | A | 0.184 | -7.68E-02 | 1.10E-01 | 2.61E+00 | 2.69E+00 | 4.85E-01 | no | | intergenic_variant | MODIFIER |
| 5 | 43343258 | AX-170071919 | 0 | C | G | 0.184 | -7.68E-02 | 1.10E-01 | 2.61E+00 | 2.69E+00 | 4.85E-01 | no | | intergenic_variant | MODIFIER |
| 5 | 43374909 | AX-170342961 | 0 | T | C | 0.184 | -7.68E-02 | 1.10E-01 | 2.61E+00 | 2.69E+00 | 4.85E-01 | no | Gm43699 | upstream_gene_variant | MODIFIER |
| 5 | 43376792 | AX-170264067 | 0 | C | T | 0.184 | -7.68E-02 | 1.10E-01 | 2.61E+00 | 2.69E+00 | 4.85E-01 | no | Gm43699 | intron_variant,non_coding_transcript_variant | MODIFIER |
| 5 | 43396531 | AX-170391454 | 0 | A | G | 0.184 | -7.68E-02 | 1.10E-01 | 2.61E+00 | 2.69E+00 | 4.85E-01 | no | | intergenic_variant | MODIFIER |
| 5 | 43396820 | AX-170100923 | 0 | G | A | 0.184 | -7.68E-02 | 1.10E-01 | 2.61E+00 | 2.69E+00 | 4.85E-01 | no | | intergenic_variant | MODIFIER |
| 5 | 43403564 | AX-170104705 | 0 | C | A | 0.184 | -7.68E-02 | 1.10E-01 | 2.61E+00 | 2.69E+00 | 4.85E-01 | no | | intergenic_variant | MODIFIER |
| 5 | 43415731 | AX-170157964 | 0 | A | G | 0.184 | -7.68E-02 | 1.10E-01 | 2.61E+00 | 2.69E+00 | 4.85E-01 | no | | intergenic_variant | MODIFIER |
| 5 | 43415770 | AX-170309599 | 0 | C | G | 0.184 | -7.68E-02 | 1.10E-01 | 2.61E+00 | 2.69E+00 | 4.85E-01 | no | | intergenic_variant | MODIFIER |
| 5 | 43416014 | AX-170402527 | 0 | C | G | 0.184 | -7.68E-02 | 1.10E-01 | 2.61E+00 | 2.69E+00 | 4.85E-01 | no | | intergenic_variant | MODIFIER |
| 5 | 43437953 | AX-170528382 | 0 | C | T | 0.184 | -7.68E-02 | 1.10E-01 | 2.61E+00 | 2.69E+00 | 4.85E-01 | no | | intergenic_variant | MODIFIER |
| 5 | 43438079 | AX-170229374 | 0 | C | A | 0.184 | -7.68E-02 | 1.10E-01 | 2.61E+00 | 2.69E+00 | 4.85E-01 | no | | intergenic_variant | MODIFIER |
| 5 | 43443105 | AX-170237267 | 0 | C | G | 0.184 | -7.68E-02 | 1.10E-01 | 2.61E+00 | 2.69E+00 | 4.85E-01 | no | | intergenic_variant | MODIFIER |
| 5 | 43464371 | AX-97784699 | 0 | T | C | 0.184 | -7.68E-02 | 1.10E-01 | 2.61E+00 | 2.69E+00 | 4.85E-01 | no | | intergenic_variant | MODIFIER |

| | | | | | | | | | | | | | | | |
|---|---|---|---|---|---|---|---|---|---|---|---|---|---|---|---|
| 5 | 43479229 | AX-97789709 | 0 | G | A | 0.184 | -7.68E-02 | 1.10E-01 | 2.61E+00 | 2.69E+00 | 4.85E-01 | no | Gm43021 | downstream_gene_variant | MODIFIER |
| 5 | 43483176 | AX-170502125 | 0 | A | T | 0.184 | -7.68E-02 | 1.10E-01 | 2.61E+00 | 2.69E+00 | 4.85E-01 | no | | intergenic_variant | MODIFIER |
| 5 | 43498566 | AX-170160782 | 0 | T | C | 0.184 | -7.68E-02 | 1.10E-01 | 2.61E+00 | 2.69E+00 | 4.85E-01 | no | | intergenic_variant | MODIFIER |
| 5 | 43502723 | AX-170232889 | 0 | A | G | 0.184 | -7.68E-02 | 1.10E-01 | 2.61E+00 | 2.69E+00 | 4.85E-01 | no | | intergenic_variant | MODIFIER |
| 5 | 43520268 | AX-170024814 | 0 | A | G | 0.184 | -7.68E-02 | 1.10E-01 | 2.61E+00 | 2.69E+00 | 4.85E-01 | no | C1qtnf7 | intron_variant | MODIFIER |
| 5 | 43531107 | AX-170487755 | 0 | T | G | 0.184 | -7.68E-02 | 1.10E-01 | 2.61E+00 | 2.69E+00 | 4.85E-01 | no | C1qtnf7 | intron_variant | MODIFIER |
| 5 | 43538501 | AX-170258041 | 0 | T | A | 0.184 | -7.68E-02 | 1.10E-01 | 2.61E+00 | 2.69E+00 | 4.85E-01 | no | C1qtnf7 | intron_variant | MODIFIER |
| 5 | 43539781 | AX-170370522 | 0 | T | C | 0.184 | -7.68E-02 | 1.10E-01 | 2.61E+00 | 2.69E+00 | 4.85E-01 | no | C1qtnf7 | intron_variant | MODIFIER |
| 5 | 43549032 | AX-170199513 | 0 | A | G | 0.184 | -7.68E-02 | 1.10E-01 | 2.61E+00 | 2.69E+00 | 4.85E-01 | no | C1qtnf7 | intron_variant | MODIFIER |
| 5 | 43550565 | AX-170150005 | 0 | G | C | 0.184 | -7.68E-02 | 1.10E-01 | 2.61E+00 | 2.69E+00 | 4.85E-01 | no | C1qtnf7 | intron_variant | MODIFIER |
| 5 | 43566382 | AX-170311659 | 0 | G | C | 0.184 | -7.68E-02 | 1.10E-01 | 2.61E+00 | 2.69E+00 | 4.85E-01 | no | C1qtnf7 | intron_variant | MODIFIER |
| 5 | 43567715 | AX-170024820 | 0 | G | C | 0.184 | -7.68E-02 | 1.10E-01 | 2.61E+00 | 2.69E+00 | 4.85E-01 | no | C1qtnf7 | intron_variant | MODIFIER |
| 5 | 43568310 | AX-170024821 | 0 | C | G | 0.184 | -7.68E-02 | 1.10E-01 | 2.61E+00 | 2.69E+00 | 4.85E-01 | no | C1qtnf7 | intron_variant | MODIFIER |
| 5 | 43569153 | AX-97824266 | 0 | G | A | 0.184 | -7.68E-02 | 1.10E-01 | 2.61E+00 | 2.69E+00 | 4.85E-01 | no | C1qtnf7 | intron_variant | MODIFIER |
| 5 | 43569366 | AX-97854298 | 0 | A | G | 0.184 | -7.68E-02 | 1.10E-01 | 2.61E+00 | 2.69E+00 | 4.85E-01 | no | C1qtnf7 | intron_variant | MODIFIER |
| 5 | 43569755 | AX-170138130 | 0 | A | G | 0.184 | -7.68E-02 | 1.10E-01 | 2.61E+00 | 2.69E+00 | 4.85E-01 | no | C1qtnf7 | intron_variant | MODIFIER |
| 5 | 43573112 | AX-170071925 | 0 | A | T | 0.184 | -7.68E-02 | 1.10E-01 | 2.61E+00 | 2.69E+00 | 4.85E-01 | no | C1qtnf7 | intron_variant | MODIFIER |
| 5 | 43574580 | AX-170464258 | 0 | T | A | 0.184 | -7.68E-02 | 1.10E-01 | 2.61E+00 | 2.69E+00 | 4.85E-01 | no | C1qtnf7 | intron_variant | MODIFIER |
| 5 | 43575636 | AX-170284237 | 0 | A | G | 0.184 | -7.68E-02 | 1.10E-01 | 2.61E+00 | 2.69E+00 | 4.85E-01 | no | C1qtnf7 | intron_variant | MODIFIER |
| 5 | 43577766 | AX-170223771 | 0 | G | C | 0.184 | -7.68E-02 | 1.10E-01 | 2.61E+00 | 2.69E+00 | 4.85E-01 | no | C1qtnf7 | intron_variant | MODIFIER |
| 5 | 43583393 | AX-170453560 | 0 | A | G | 0.184 | -7.68E-02 | 1.10E-01 | 2.61E+00 | 2.69E+00 | 4.85E-01 | no | C1qtnf7 | intron_variant | MODIFIER |
| 5 | 43591167 | AX-170152207 | 0 | C | T | 0.184 | -7.68E-02 | 1.10E-01 | 2.61E+00 | 2.69E+00 | 4.85E-01 | no | C1qtnf7 | intron_variant | MODIFIER |
| 5 | 43591633 | AX-170472341 | 0 | A | G | 0.184 | -7.68E-02 | 1.10E-01 | 2.61E+00 | 2.69E+00 | 4.85E-01 | no | C1qtnf7 | intron_variant | MODIFIER |
| 5 | 43596124 | AX-97798058 | 0 | T | G | 0.184 | -7.68E-02 | 1.10E-01 | 2.61E+00 | 2.69E+00 | 4.85E-01 | no | C1qtnf7 | intron_variant | MODIFIER |
| 5 | 43599409 | AX-170390635 | 0 | C | T | 0.184 | -7.68E-02 | 1.10E-01 | 2.61E+00 | 2.69E+00 | 4.85E-01 | no | C1qtnf7 | intron_variant | MODIFIER |
| 5 | 43617794 | AX-170306973 | 0 | G | A | 0.184 | -7.68E-02 | 1.10E-01 | 2.61E+00 | 2.69E+00 | 4.85E-01 | no | C1qtnf7 | 3_prime_UTR_variant | MODIFIER |
| 5 | 43634971 | AX-97831735 | 0 | G | A | 0.184 | -7.68E-02 | 1.10E-01 | 2.61E+00 | 2.69E+00 | 4.85E-01 | no | | regulatory_region_variant | MODIFIER |
| 5 | 43643394 | AX-170498702 | 0 | T | C | 0.184 | -7.68E-02 | 1.10E-01 | 2.61E+00 | 2.69E+00 | 4.85E-01 | no | | intergenic_variant | MODIFIER |
| 5 | 43643916 | AX-170160682 | 0 | T | C | 0.184 | -7.68E-02 | 1.10E-01 | 2.61E+00 | 2.69E+00 | 4.85E-01 | no | | intergenic_variant | MODIFIER |
| 5 | 43668357 | AX-170482449 | 0 | G | A | 0.184 | -7.68E-02 | 1.10E-01 | 2.61E+00 | 2.69E+00 | 4.85E-01 | no | Cc2d2a | intron_variant | MODIFIER |
| 5 | 43689927 | AX-97788079 | 0 | A | C | 0.184 | -7.68E-02 | 1.10E-01 | 2.61E+00 | 2.69E+00 | 4.85E-01 | no | Cc2d2a | intron_variant | MODIFIER |
| 5 | 43693179 | AX-170024815 | 0 | C | T | 0.184 | -7.68E-02 | 1.10E-01 | 2.61E+00 | 2.69E+00 | 4.85E-01 | no | Cc2d2a | intron_variant | MODIFIER |
| 5 | 43706543 | AX-170361674 | 0 | C | T | 0.184 | -7.68E-02 | 1.10E-01 | 2.61E+00 | 2.69E+00 | 4.85E-01 | no | Cc2d2a | intron_variant | MODIFIER |
| 5 | 43827918 | AX-97784696 | 0 | A | G | 0.184 | -7.68E-02 | 1.10E-01 | 2.61E+00 | 2.69E+00 | 4.85E-01 | no | Bst1 | intron_variant | MODIFIER |
| 5 | 43859356 | AX-97813571 | 0 | A | G | 0.184 | -7.68E-02 | 1.10E-01 | 2.61E+00 | 2.69E+00 | 4.85E-01 | no | | intergenic_variant | MODIFIER |
| 5 | 43872419 | AX-97852867 | 0 | A | G | 0.184 | -7.68E-02 | 1.10E-01 | 2.61E+00 | 2.69E+00 | 4.85E-01 | no | Cd38 | intron_variant | MODIFIER |
| 5 | 43894721 | AX-170305588 | 0 | C | T | 0.184 | -7.68E-02 | 1.10E-01 | 2.61E+00 | 2.69E+00 | 4.85E-01 | no | Cd38 | intron_variant | MODIFIER |
| 5 | 43895785 | AX-97792399 | 0 | T | C | 0.184 | -7.68E-02 | 1.10E-01 | 2.61E+00 | 2.69E+00 | 4.85E-01 | no | Cd38 | intron_variant | MODIFIER |
| 5 | 43918506 | AX-170467945 | 0 | C | G | 0.184 | -7.68E-02 | 1.10E-01 | 2.61E+00 | 2.69E+00 | 4.85E-01 | no | Gm7879 | downstream_gene_variant | MODIFIER |
| 5 | 43975863 | AX-170211767 | 0 | A | G | 0.184 | -7.68E-02 | 1.10E-01 | 2.61E+00 | 2.69E+00 | 4.85E-01 | no | Fgfbp1 | downstream_gene_variant | MODIFIER |
| 5 | 44010024 | AX-97857521 | 0 | T | G | 0.184 | -7.68E-02 | 1.10E-01 | 2.61E+00 | 2.69E+00 | 4.85E-01 | no | Prom1 | intron_variant | MODIFIER |
| 5 | 44011374 | AX-170458795 | 0 | G | C | 0.184 | -7.68E-02 | 1.10E-01 | 2.61E+00 | 2.69E+00 | 4.85E-01 | no | Prom1 | intron_variant | MODIFIER |
| 5 | 44014058 | AX-170375446 | 0 | T | C | 0.184 | -7.68E-02 | 1.10E-01 | 2.61E+00 | 2.69E+00 | 4.85E-01 | no | Prom1 | intron_variant | MODIFIER |
| 5 | 44041919 | AX-97794618 | 0 | C | T | 0.184 | -7.68E-02 | 1.10E-01 | 2.61E+00 | 2.69E+00 | 4.85E-01 | no | Prom1 | intron_variant | MODIFIER |
| 5 | 44068083 | AX-170218429 | 0 | A | G | 0.184 | -7.68E-02 | 1.10E-01 | 2.61E+00 | 2.69E+00 | 4.85E-01 | no | Prom1 | intron_variant | MODIFIER |
| 5 | 44099142 | AX-170373015 | 0 | A | G | 0.184 | -7.68E-02 | 1.10E-01 | 2.61E+00 | 2.69E+00 | 4.85E-01 | no | Prom1 | upstream_gene_variant | MODIFIER |
| 5 | 44240319 | AX-170240021 | 0 | T | C | 0.184 | -7.68E-02 | 1.10E-01 | 2.61E+00 | 2.69E+00 | 4.85E-01 | no | Gm42984 | intron_variant,non_coding_transcript_variant | MODIFIER |
| 5 | 44240993 | AX-170463844 | 0 | G | A | 0.184 | -7.68E-02 | 1.10E-01 | 2.61E+00 | 2.69E+00 | 4.85E-01 | no | Gm42984 | intron_variant,non_coding_transcript_variant | MODIFIER |
| 5 | 44266454 | AX-170164861 | 0 | T | G | 0.184 | -7.68E-02 | 1.10E-01 | 2.61E+00 | 2.69E+00 | 4.85E-01 | no | | intergenic_variant | MODIFIER |
| 5 | 44303618 | AX-170318441 | 0 | C | T | 0.184 | -7.68E-02 | 1.10E-01 | 2.61E+00 | 2.69E+00 | 4.85E-01 | no | | intergenic_variant | MODIFIER |
| 5 | 44338923 | AX-170323826 | 0 | C | G | 0.184 | -7.68E-02 | 1.10E-01 | 2.61E+00 | 2.69E+00 | 4.85E-01 | no | | intergenic_variant | MODIFIER |
| 5 | 44360882 | AX-170277703 | 0 | A | G | 0.184 | -7.68E-02 | 1.10E-01 | 2.61E+00 | 2.69E+00 | 4.85E-01 | no | Gm42698 | upstream_gene_variant | MODIFIER |
| 5 | 44369274 | AX-170071937 | 0 | T | C | 0.184 | -7.68E-02 | 1.10E-01 | 2.61E+00 | 2.69E+00 | 4.85E-01 | no | | intergenic_variant | MODIFIER |
| 5 | 44383220 | AX-97857559 | 0 | G | A | 0.184 | -7.68E-02 | 1.10E-01 | 2.61E+00 | 2.69E+00 | 4.85E-01 | no | | intergenic_variant | MODIFIER |

| | | | | | | | | | | | | | | | |
|---|---|---|---|---|---|---|---|---|---|---|---|---|---|---|---|
| 5 | 44427209 | AX-170125355 | 0 | G | T | 0.184 | -7.68E-02 | 1.10E-01 | 2.61E+00 | 2.69E+00 | 4.85E-01 | no | Gm36017 | intron_variant,non_coding_transcript_variant | MODIFIER |
| 5 | 44435748 | AX-170416201 | 0 | T | C | 0.184 | -7.68E-02 | 1.10E-01 | 2.61E+00 | 2.69E+00 | 4.85E-01 | no | | intergenic_variant | MODIFIER |
| 5 | 44445626 | AX-170460778 | 0 | A | T | 0.184 | -7.68E-02 | 1.10E-01 | 2.61E+00 | 2.69E+00 | 4.85E-01 | no | | intergenic_variant | MODIFIER |
| 5 | 44481694 | AX-170354789 | 0 | A | G | 0.184 | -7.68E-02 | 1.10E-01 | 2.61E+00 | 2.69E+00 | 4.85E-01 | no | Ldb2 | intron_variant | MODIFIER |
| 5 | 44491091 | AX-97785889 | 0 | C | A | 0.184 | -7.68E-02 | 1.10E-01 | 2.61E+00 | 2.69E+00 | 4.85E-01 | no | Ldb2 | intron_variant | MODIFIER |
| 5 | 44494277 | AX-170305204 | 0 | A | G | 0.184 | -7.68E-02 | 1.10E-01 | 2.61E+00 | 2.69E+00 | 4.85E-01 | no | Ldb2 | intron_variant | MODIFIER |
| 5 | 44501815 | AX-170205764 | 0 | A | G | 0.184 | -7.68E-02 | 1.10E-01 | 2.61E+00 | 2.69E+00 | 4.85E-01 | no | Ldb2 | intron_variant | MODIFIER |
| 5 | 44623664 | AX-170024828 | 0 | A | T | 0.184 | -7.68E-02 | 1.10E-01 | 2.61E+00 | 2.69E+00 | 4.85E-01 | no | Ldb2 | intron_variant | MODIFIER |
| 5 | 44652384 | AX-170024831 | 0 | A | C | 0.184 | -7.68E-02 | 1.10E-01 | 2.61E+00 | 2.69E+00 | 4.85E-01 | no | Ldb2 | intron_variant | MODIFIER |
| 5 | 44698540 | AX-170359007 | 0 | A | C | 0.184 | -7.68E-02 | 1.10E-01 | 2.61E+00 | 2.69E+00 | 4.85E-01 | no | Ldb2 | intron_variant | MODIFIER |
| 5 | 44735862 | AX-170499460 | 0 | T | C | 0.184 | -7.68E-02 | 1.10E-01 | 2.61E+00 | 2.69E+00 | 4.85E-01 | no | Ldb2 | intron_variant | MODIFIER |
| 5 | 44772300 | AX-170089467 | 0 | A | G | 0.184 | -7.68E-02 | 1.10E-01 | 2.61E+00 | 2.69E+00 | 4.85E-01 | no | Ldb2 | intron_variant | MODIFIER |
| 5 | 44971175 | AX-170525287 | 0 | T | C | 0.184 | -7.68E-02 | 1.10E-01 | 2.61E+00 | 2.69E+00 | 4.85E-01 | no | 4930431F12Rik | downstream_gene_variant | MODIFIER |
| 5 | 44974378 | AX-170422006 | 0 | C | T | 0.184 | -7.68E-02 | 1.10E-01 | 2.61E+00 | 2.69E+00 | 4.85E-01 | no | 4930431F12Rik | intron_variant,non_coding_transcript_variant | MODIFIER |
| 5 | 45045100 | AX-170479039 | 0 | T | C | 0.184 | -7.68E-02 | 1.10E-01 | 2.61E+00 | 2.69E+00 | 4.85E-01 | no | D5Ertd615e | intron_variant,non_coding_transcript_variant | MODIFIER |
| 5 | 45208374 | AX-170488958 | 0 | T | C | 0.184 | -7.68E-02 | 1.10E-01 | 2.61E+00 | 2.69E+00 | 4.85E-01 | no | D5Ertd615e | intron_variant,non_coding_transcript_variant | MODIFIER |
| 5 | 51840323 | AX-170307196 | 0 | C | T | 0.5 | 7.22E-02 | 9.96E-02 | 2.45E+00 | 2.52E+00 | 4.85E-01 | no | | intergenic_variant | MODIFIER |
| 8 | 64568924 | AX-170220542 | 0 | T | G | 0.14 | -1.00E-01 | 1.43E-01 | 2.60E+00 | 2.67E+00 | 4.85E-01 | no | | intergenic_variant | MODIFIER |
| 4 | 75062048 | AX-170035687 | 0 | T | A | 0.228 | -6.76E-02 | 9.64E-02 | 2.96E+00 | 3.12E+00 | 4.85E-01 | no | | intergenic_variant | MODIFIER |
| 5 | 38541956 | AX-97810949 | 1 | G | A | 0.339 | 7.72E-02 | 1.12E-01 | 2.75E+00 | 2.81E+00 | 4.85E-01 | no | Wdr1 | intron_variant | MODIFIER |
| 1 | 174344758 | AX-170509273 | 1 | C | G | 0.232 | 8.73E-02 | 1.23E-01 | 2.52E+00 | 2.57E+00 | 4.85E-01 | no | | intergenic_variant | MODIFIER |
| 1 | 174695014 | AX-170002325 | 1 | C | T | 0.232 | 8.73E-02 | 1.23E-01 | 2.52E+00 | 2.57E+00 | 4.85E-01 | no | Fmn2 | intron_variant | MODIFIER |
| 4 | 69495556 | AX-170267654 | 1 | G | A | 0.348 | 7.64E-02 | 1.11E-01 | 2.70E+00 | 2.77E+00 | 4.85E-01 | no | | intergenic_variant | MODIFIER |
| 9 | 22175622 | AX-97830700 | 1 | T | C | 0.473 | 6.38E-02 | 8.95E-02 | 2.51E+00 | 2.57E+00 | 4.85E-01 | no | Gm18227 | downstream_gene_variant | MODIFIER |
| 9 | 15696346 | AX-170336853 | 0 | C | T | 0.474 | 6.20E-02 | 8.78E-02 | 2.54E+00 | 2.61E+00 | 4.85E-01 | no | | intergenic_variant | MODIFIER |
| 18 | 74695140 | AX-170015996 | 1 | T | A | 0.42 | 5.93E-02 | 8.28E-02 | 2.48E+00 | 2.56E+00 | 4.85E-01 | no | Myo5b | intron_variant | MODIFIER |
| 2 | 162872714 | AX-170260910 | 0 | A | G | 0.158 | 1.12E-01 | 1.64E-01 | 2.72E+00 | 2.84E+00 | 4.86E-01 | no | Gm14255 | upstream_gene_variant | MODIFIER |
| 1 | 170447611 | AX-170260148 | 0 | T | A | 0.237 | 8.86E-02 | 1.28E-01 | 2.68E+00 | 2.73E+00 | 4.86E-01 | no | Nos1ap | intron_variant,non_coding_transcript_variant | MODIFIER |
| 18 | 71655738 | AX-170458158 | 1 | A | T | 0.179 | -8.45E-02 | 1.22E-01 | 2.65E+00 | 2.70E+00 | 4.86E-01 | no | Dcc | intron_variant | MODIFIER |
| 9 | 38619059 | AX-170356776 | 0 | C | T | 0.5 | -5.32E-02 | 7.75E-02 | 2.75E+00 | 2.84E+00 | 4.86E-01 | no | | intergenic_variant | MODIFIER |
| 4 | 151783878 | AX-97840074 | 0 | C | A | 0.105 | 9.43E-02 | 1.37E-01 | 2.61E+00 | 2.73E+00 | 4.86E-01 | no | Camta1 | intron_variant | MODIFIER |
| 1 | 48458141 | AX-170132457 | 0 | T | C | 0.114 | 1.26E-01 | 1.85E-01 | 2.71E+00 | 2.82E+00 | 4.86E-01 | no | Gm23240 | downstream_gene_variant | MODIFIER |
| 11 | 62911531 | AX-170387575 | 0 | T | C | 0.035 | -1.08E-01 | 1.56E-01 | 2.56E+00 | 2.71E+00 | 4.86E-01 | no | Tvp23b | intron_variant,non_coding_transcript_variant | MODIFIER |
| 4 | 95298198 | AX-170310931 | 0 | T | C | 0.333 | 7.31E-02 | 1.06E-01 | 2.67E+00 | 2.74E+00 | 4.86E-01 | no | | intergenic_variant | MODIFIER |
| 1 | 175129374 | AX-170481182 | 1 | G | A | 0.232 | 8.68E-02 | 1.23E-01 | 2.53E+00 | 2.59E+00 | 4.86E-01 | no | Rgs7 | intron_variant | MODIFIER |
| 5 | 32642623 | AX-170213550 | 2 | G | A | 0.364 | 7.10E-02 | 1.03E-01 | 2.73E+00 | 2.80E+00 | 4.86E-01 | no | Yes1 | intron_variant | MODIFIER |
| 15 | 83523314 | AX-170467234 | 0 | G | A | 0.096 | -9.26E-02 | 1.05E-01 | 2.10E+00 | 2.20E+00 | 4.87E-01 | no | Bik | upstream_gene_variant | MODIFIER |
| 3 | 147422119 | AX-170236864 | 0 | T | C | 0.342 | 5.15E-02 | 7.49E-02 | 2.70E+00 | 2.77E+00 | 4.87E-01 | no | | intergenic_variant | MODIFIER |
| 5 | 40432948 | AX-170203349 | 1 | T | C | 0.214 | -7.51E-02 | 1.08E-01 | 2.59E+00 | 2.68E+00 | 4.87E-01 | no | | intergenic_variant | MODIFIER |
| 2 | 72209636 | AX-170018030 | 2 | C | T | 0.055 | 9.72E-02 | 1.44E-01 | 2.77E+00 | 2.95E+00 | 4.87E-01 | no | Rapgef4 | intron_variant | MODIFIER |
| 5 | 117379940 | AX-170150513 | 1 | G | A | 0.464 | 7.65E-02 | 1.11E-01 | 2.63E+00 | 2.68E+00 | 4.87E-01 | no | Wsb2 | downstream_gene_variant | MODIFIER |
| 1 | 11058927 | AX-170098172 | 0 | G | A | 0.096 | -9.59E-02 | 1.41E-01 | 2.72E+00 | 2.87E+00 | 4.87E-01 | no | Prex2 | intron_variant | MODIFIER |
| 1 | 21690793 | AX-170389280 | 1 | A | G | 0.402 | 5.87E-02 | 8.44E-02 | 2.60E+00 | 2.64E+00 | 4.87E-01 | no | Kcnq5 | intron_variant | MODIFIER |
| 1 | 136374307 | AX-170267161 | 1 | T | C | 0.214 | 8.40E-02 | 1.22E-01 | 2.66E+00 | 2.73E+00 | 4.87E-01 | no | | intergenic_variant | MODIFIER |
| 1 | 77398746 | AX-170370901 | 0 | A | G | 0.132 | -1.14E-01 | 1.66E-01 | 2.67E+00 | 2.76E+00 | 4.87E-01 | no | Epha4 | intron_variant | MODIFIER |
| 4 | 83899726 | AX-97802172 | 1 | A | G | 0.33 | 7.32E-02 | 1.07E-01 | 2.70E+00 | 2.77E+00 | 4.87E-01 | no | | intergenic_variant | MODIFIER |
| 7 | 37608988 | AX-97821532 | 1 | C | T | 0.348 | 6.67E-02 | 9.51E-02 | 2.55E+00 | 2.62E+00 | 4.87E-01 | no | Zfp536 | intron_variant | MODIFIER |
| 1 | 146924939 | AX-97852464 | 1 | G | T | 0.214 | 9.06E-02 | 1.31E-01 | 2.62E+00 | 2.68E+00 | 4.87E-01 | no | | intergenic_variant | MODIFIER |
| 18 | 65142575 | AX-170454027 | 0 | G | C | 0.412 | 6.09E-02 | 8.50E-02 | 2.48E+00 | 2.54E+00 | 4.87E-01 | no | Nedd4l | intron_variant | MODIFIER |
| 13 | 6434194 | AX-170527102 | 0 | G | A | 0.219 | -7.87E-02 | 1.14E-01 | 2.61E+00 | 2.68E+00 | 4.87E-01 | no | | intergenic_variant | MODIFIER |
| 4 | 154828856 | AX-170337674 | 1 | G | T | 0.116 | 1.26E-01 | 1.84E-01 | 2.72E+00 | 2.80E+00 | 4.87E-01 | no | | intergenic_variant | MODIFIER |
| 7 | 36670539 | AX-170101714 | 1 | T | C | 0.33 | 6.23E-02 | 8.95E-02 | 2.57E+00 | 2.65E+00 | 4.87E-01 | no | | intergenic_variant | MODIFIER |
| 9 | 9378324 | AX-170465808 | 0 | G | T | 0.474 | 5.86E-02 | 8.17E-02 | 2.46E+00 | 2.56E+00 | 4.87E-01 | no | Gm46102 | intron_variant,non_coding_transcript_variant | MODIFIER |
| 1 | 36323387 | AX-170000284 | 1 | G | A | 0.464 | 5.09E-02 | 7.45E-02 | 2.74E+00 | 2.85E+00 | 4.87E-01 | no | Arid5a | downstream_gene_variant | MODIFIER |

| | | | | | | | | | | | | | | | |
|---|---|---|---|---|---|---|---|---|---|---|---|---|---|---|---|
| 8 | 93169029 | AX-170498409 | 0 | T | A | 0.211 | -7.62E-02 | 1.11E-01 | 2.72E+00 | 2.81E+00 | 4.87E-01 | no | Ces1d | intron_variant | MODIFIER |
| 9 | 84495853 | AX-170032852 | 0 | C | T | 0.219 | -7.84E-02 | 1.15E-01 | 2.71E+00 | 2.81E+00 | 4.87E-01 | no | | intergenic_variant | MODIFIER |
| 1 | 103350337 | AX-170001232 | 1 | A | T | 0.143 | 1.18E-01 | 1.72E-01 | 2.65E+00 | 2.72E+00 | 4.88E-01 | no | | intergenic_variant | MODIFIER |
| 1 | 128499206 | AX-170455436 | 0 | A | G | 0.544 | 5.63E-02 | 8.14E-02 | 2.61E+00 | 2.69E+00 | 4.88E-01 | no | Gm29598 | upstream_gene_variant | MODIFIER |
| 8 | 85639508 | AX-170031178 | 1 | A | G | 0.196 | -8.19E-02 | 1.20E-01 | 2.74E+00 | 2.84E+00 | 4.88E-01 | no | Neto2 | 3_prime_UTR_variant | MODIFIER |
| 9 | 20891716 | AX-97805095 | 1 | T | C | 0.473 | 6.38E-02 | 9.11E-02 | 2.55E+00 | 2.62E+00 | 4.88E-01 | no | Ppan | synonymous_variant | LOW |
| 14 | 95627355 | AX-170267735 | 1 | C | T | 0.045 | 1.06E-01 | 1.56E-01 | 2.65E+00 | 2.80E+00 | 4.88E-01 | no | | intergenic_variant | MODIFIER |
| 8 | 34214969 | AX-170030487 | 0 | T | C | 0.14 | -9.18E-02 | 1.34E-01 | 2.70E+00 | 2.77E+00 | 4.88E-01 | no | Gm29243 | upstream_gene_variant | MODIFIER |
| 2 | 161452964 | AX-170019540 | 1 | C | T | 0.161 | 1.34E-01 | 1.97E-01 | 2.73E+00 | 2.81E+00 | 4.88E-01 | no | | intergenic_variant | MODIFIER |
| 3 | 151226783 | AX-170386663 | 2 | A | T | 0.355 | 4.86E-02 | 7.13E-02 | 2.73E+00 | 2.83E+00 | 4.88E-01 | no | | intergenic_variant | MODIFIER |
| 3 | 29043336 | AX-170356912 | 1 | A | C | 0.152 | -6.85E-02 | 1.00E-01 | 2.68E+00 | 2.77E+00 | 4.88E-01 | no | | intergenic_variant | MODIFIER |
| 1 | 91192560 | AX-97838099 | 1 | G | A | 0.161 | 1.42E-01 | 2.07E-01 | 2.64E+00 | 2.70E+00 | 4.88E-01 | no | Ramp1 | intron_variant | MODIFIER |
| 1 | 99305492 | AX-170267228 | 1 | T | C | 0.161 | 1.42E-01 | 2.07E-01 | 2.64E+00 | 2.70E+00 | 4.88E-01 | no | | intergenic_variant | MODIFIER |
| 5 | 108151725 | AX-170419966 | 1 | G | A | 0.455 | 7.60E-02 | 1.10E-01 | 2.63E+00 | 2.68E+00 | 4.89E-01 | no | Ccdc18 | intron_variant | MODIFIER |
| 18 | 88736329 | AX-170255497 | 0 | T | C | 0.167 | -6.94E-02 | 9.99E-02 | 2.56E+00 | 2.66E+00 | 4.89E-01 | no | Socs6 | intron_variant,non_coding_transcript_variant | MODIFIER |
| 1 | 170383504 | AX-170043544 | 1 | G | A | 0.214 | 9.68E-02 | 1.41E-01 | 2.66E+00 | 2.70E+00 | 4.89E-01 | no | Nos1ap | intron_variant,non_coding_transcript_variant | MODIFIER |
| 8 | 35087107 | AX-97804052 | 1 | G | A | 0.143 | -9.76E-02 | 1.41E-01 | 2.61E+00 | 2.66E+00 | 4.89E-01 | no | | regulatory_region_variant | MODIFIER |
| 8 | 35181569 | AX-97780605 | 1 | T | C | 0.143 | -9.76E-02 | 1.41E-01 | 2.61E+00 | 2.66E+00 | 4.89E-01 | no | Gm45625 | intron_variant,non_coding_transcript_variant | MODIFIER |
| 9 | 98316651 | AX-170276844 | 0 | G | C | 0.149 | -7.27E-02 | 1.05E-01 | 2.59E+00 | 2.67E+00 | 4.89E-01 | no | Nmnat3 | intron_variant | MODIFIER |
| 1 | 138780330 | AX-97837101 | 0 | G | A | 0.211 | 8.52E-02 | 1.24E-01 | 2.65E+00 | 2.72E+00 | 4.89E-01 | no | | intergenic_variant | MODIFIER |
| 1 | 170322124 | AX-170280811 | 0 | G | T | 0.211 | 9.62E-02 | 1.40E-01 | 2.66E+00 | 2.71E+00 | 4.89E-01 | no | Nos1ap | intron_variant | MODIFIER |
| 13 | 19883791 | AX-170401453 | 0 | C | T | 0.184 | -8.57E-02 | 1.20E-01 | 2.47E+00 | 2.53E+00 | 4.89E-01 | no | | intergenic_variant | MODIFIER |
| 1 | 170466172 | AX-170238775 | 1 | G | T | 0.214 | 9.49E-02 | 1.38E-01 | 2.64E+00 | 2.68E+00 | 4.89E-01 | no | Nos1ap | intron_variant,non_coding_transcript_variant | MODIFIER |
| 4 | 139361509 | AX-170414385 | 0 | G | C | 0.298 | -3.99E-02 | 5.76E-02 | 2.93E+00 | 3.07E+00 | 4.89E-01 | no | Emc1 | intron_variant | MODIFIER |
| 7 | 73164130 | AX-170351471 | 1 | T | A | 0.366 | -6.87E-02 | 1.01E-01 | 2.79E+00 | 2.88E+00 | 4.89E-01 | no | | regulatory_region_variant | MODIFIER |
| 1 | 107843485 | AX-170292862 | 0 | A | G | 0.175 | 1.16E-01 | 1.67E-01 | 2.54E+00 | 2.65E+00 | 4.90E-01 | no | | intergenic_variant | MODIFIER |
| 15 | 97745249 | AX-170346859 | 2 | C | T | 0.1 | 9.56E-02 | 1.39E-01 | 2.58E+00 | 2.70E+00 | 4.90E-01 | no | Rapgef3 | downstream_gene_variant | MODIFIER |
| 4 | 6231035 | AX-170447285 | 2 | C | T | 0.045 | -1.22E-01 | 1.79E-01 | 2.61E+00 | 2.72E+00 | 4.90E-01 | no | | intergenic_variant | MODIFIER |
| 1 | 152200773 | AX-170438724 | 1 | C | T | 0.205 | 9.43E-02 | 1.38E-01 | 2.65E+00 | 2.70E+00 | 4.90E-01 | no | | intergenic_variant | MODIFIER |
| 18 | 58132368 | AX-170367606 | 0 | C | G | 0.193 | -8.06E-02 | 1.18E-01 | 2.65E+00 | 2.70E+00 | 4.90E-01 | no | Fbn2 | intron_variant | MODIFIER |
| 12 | 102286559 | AX-170489170 | 0 | G | C | 0.096 | -1.05E-01 | 1.55E-01 | 2.73E+00 | 2.88E+00 | 4.90E-01 | no | Rin3 | intron_variant | MODIFIER |
| 1 | 95669819 | AX-170086543 | 1 | C | T | 0.152 | 1.40E-01 | 2.04E-01 | 2.65E+00 | 2.71E+00 | 4.90E-01 | no | St8sia4 | upstream_gene_variant | MODIFIER |
| 1 | 136283698 | AX-170397086 | 0 | T | C | 0.202 | 8.63E-02 | 1.26E-01 | 2.66E+00 | 2.73E+00 | 4.90E-01 | no | Camsap2 | intron_variant | MODIFIER |
| 4 | 88439207 | AX-170273721 | 1 | G | T | 0.339 | 7.33E-02 | 1.06E-01 | 2.57E+00 | 2.64E+00 | 4.90E-01 | no | Hacd4 | upstream_gene_variant | MODIFIER |
| 3 | 31602038 | AX-170275474 | 1 | G | C | 0.152 | -6.88E-02 | 1.01E-01 | 2.75E+00 | 2.84E+00 | 4.90E-01 | no | | intergenic_variant | MODIFIER |
| 8 | 37170002 | AX-170291657 | 1 | C | T | 0.152 | 1.02E-01 | 1.48E-01 | 2.89E+00 | 2.95E+00 | 4.90E-01 | no | Gm45524 | intron_variant,non_coding_transcript_variant | MODIFIER |
| 7 | 122675420 | AX-170029877 | 0 | A | C | 0.404 | 5.79E-02 | 8.51E-02 | 2.75E+00 | 2.83E+00 | 4.90E-01 | no | Cacng3 | intron_variant | MODIFIER |
| 7 | 122679834 | AX-170029872 | 0 | A | G | 0.404 | 5.79E-02 | 8.51E-02 | 2.75E+00 | 2.83E+00 | 4.90E-01 | no | Cacng3 | intron_variant | MODIFIER |
| 7 | 122688915 | AX-97825846 | 0 | A | G | 0.404 | 5.79E-02 | 8.51E-02 | 2.75E+00 | 2.83E+00 | 4.90E-01 | no | Cacng3 | intron_variant | MODIFIER |
| 7 | 122695133 | AX-168403540 | 0 | A | G | 0.404 | 5.79E-02 | 8.51E-02 | 2.75E+00 | 2.83E+00 | 4.90E-01 | no | Cacng3 | intron_variant | MODIFIER |
| 7 | 122695433 | AX-170180207 | 0 | C | T | 0.404 | 5.79E-02 | 8.51E-02 | 2.75E+00 | 2.83E+00 | 4.90E-01 | no | Cacng3 | intron_variant | MODIFIER |
| 7 | 122717161 | AX-97848524 | 0 | A | G | 0.404 | 5.79E-02 | 8.51E-02 | 2.75E+00 | 2.83E+00 | 4.90E-01 | no | Cacng3 | intron_variant | MODIFIER |
| 7 | 122739042 | AX-170393598 | 0 | C | T | 0.404 | 5.79E-02 | 8.51E-02 | 2.75E+00 | 2.83E+00 | 4.90E-01 | no | Cacng3 | intron_variant | MODIFIER |
| 7 | 122742955 | AX-170029876 | 0 | G | A | 0.404 | 5.79E-02 | 8.51E-02 | 2.75E+00 | 2.83E+00 | 4.90E-01 | no | Cacng3 | intron_variant | MODIFIER |
| 7 | 122743042 | AX-170290268 | 0 | T | C | 0.404 | 5.79E-02 | 8.51E-02 | 2.75E+00 | 2.83E+00 | 4.90E-01 | no | Cacng3 | intron_variant | MODIFIER |
| 7 | 122750785 | AX-170223896 | 0 | G | A | 0.404 | 5.79E-02 | 8.51E-02 | 2.75E+00 | 2.83E+00 | 4.90E-01 | no | Cacng3 | intron_variant | MODIFIER |
| 7 | 122756947 | AX-170118983 | 0 | A | G | 0.404 | 5.79E-02 | 8.51E-02 | 2.75E+00 | 2.83E+00 | 4.90E-01 | no | Cacng3 | intron_variant | MODIFIER |
| 7 | 122758709 | AX-97841739 | 0 | C | T | 0.404 | 5.79E-02 | 8.51E-02 | 2.75E+00 | 2.83E+00 | 4.90E-01 | no | Cacng3 | intron_variant | MODIFIER |
| 7 | 122776489 | AX-97803059 | 0 | C | T | 0.404 | 5.79E-02 | 8.51E-02 | 2.75E+00 | 2.83E+00 | 4.90E-01 | no | | intergenic_variant | MODIFIER |
| 7 | 122776530 | AX-170229284 | 0 | G | A | 0.404 | 5.79E-02 | 8.51E-02 | 2.75E+00 | 2.83E+00 | 4.90E-01 | no | | intergenic_variant | MODIFIER |
| 7 | 122820268 | AX-170239833 | 0 | G | C | 0.404 | 5.79E-02 | 8.51E-02 | 2.75E+00 | 2.83E+00 | 4.90E-01 | no | | regulatory_region_variant | MODIFIER |
| 7 | 122821082 | AX-170113837 | 0 | T | C | 0.404 | 5.79E-02 | 8.51E-02 | 2.75E+00 | 2.83E+00 | 4.90E-01 | no | | regulatory_region_variant | MODIFIER |
| 7 | 122821096 | AX-170153915 | 0 | T | C | 0.404 | 5.79E-02 | 8.51E-02 | 2.75E+00 | 2.83E+00 | 4.90E-01 | no | | regulatory_region_variant | MODIFIER |
| 7 | 122822548 | AX-170078690 | 0 | C | T | 0.404 | 5.79E-02 | 8.51E-02 | 2.75E+00 | 2.83E+00 | 4.90E-01 | no | | intergenic_variant | MODIFIER |

| | | | | | | | | | | | | | | | |
|---|---|---|---|---|---|---|---|---|---|---|---|---|---|---|---|
| 7 | 122823119 | AX-170321118 | 0 | C | T | 0.404 | 5.79E-02 | 8.51E-02 | 2.75E+00 | 2.83E+00 | 4.90E-01 | no | | intergenic_variant | MODIFIER |
| 7 | 122877467 | AX-170145101 | 0 | G | A | 0.404 | 5.79E-02 | 8.51E-02 | 2.75E+00 | 2.83E+00 | 4.90E-01 | no | | intergenic_variant | MODIFIER |
| 7 | 123063563 | AX-170153902 | 0 | A | G | 0.404 | 5.79E-02 | 8.51E-02 | 2.75E+00 | 2.83E+00 | 4.90E-01 | no | Gm45847 | intron_variant,non_coding_transcript_variant | MODIFIER |
| 7 | 123129405 | AX-170106797 | 0 | T | G | 0.404 | 5.79E-02 | 8.51E-02 | 2.75E+00 | 2.83E+00 | 4.90E-01 | no | Tnrc6a | intron_variant | MODIFIER |
| 7 | 123137713 | AX-170115543 | 0 | C | T | 0.404 | 5.79E-02 | 8.51E-02 | 2.75E+00 | 2.83E+00 | 4.90E-01 | no | Tnrc6a | intron_variant | MODIFIER |
| 7 | 123139688 | AX-170218743 | 0 | T | C | 0.404 | 5.79E-02 | 8.51E-02 | 2.75E+00 | 2.83E+00 | 4.90E-01 | no | Tnrc6a | intron_variant | MODIFIER |
| 7 | 123142675 | AX-170515157 | 0 | A | G | 0.404 | 5.79E-02 | 8.51E-02 | 2.75E+00 | 2.83E+00 | 4.90E-01 | no | Tnrc6a | intron_variant | MODIFIER |
| 7 | 123154499 | AX-170078700 | 0 | T | C | 0.404 | 5.79E-02 | 8.51E-02 | 2.75E+00 | 2.83E+00 | 4.90E-01 | no | Tnrc6a | intron_variant | MODIFIER |
| 7 | 123177798 | AX-170029881 | 0 | T | C | 0.404 | 5.79E-02 | 8.51E-02 | 2.75E+00 | 2.83E+00 | 4.90E-01 | no | Tnrc6a | intron_variant | MODIFIER |
| 7 | 123200916 | AX-170029879 | 0 | G | A | 0.404 | 5.79E-02 | 8.51E-02 | 2.75E+00 | 2.83E+00 | 4.90E-01 | no | | regulatory_region_variant | MODIFIER |
| 7 | 123201888 | AX-170218966 | 0 | A | G | 0.404 | 5.79E-02 | 8.51E-02 | 2.75E+00 | 2.83E+00 | 4.90E-01 | no | | intergenic_variant | MODIFIER |
| 7 | 123207749 | AX-97804278 | 0 | T | C | 0.404 | 5.79E-02 | 8.51E-02 | 2.75E+00 | 2.83E+00 | 4.90E-01 | no | | regulatory_region_variant | MODIFIER |
| 7 | 123207783 | AX-170356130 | 0 | C | G | 0.404 | 5.79E-02 | 8.51E-02 | 2.75E+00 | 2.83E+00 | 4.90E-01 | no | | regulatory_region_variant | MODIFIER |
| 7 | 123219913 | AX-170243814 | 0 | A | G | 0.404 | 5.79E-02 | 8.51E-02 | 2.75E+00 | 2.83E+00 | 4.90E-01 | no | Slc5a11 | intron_variant | MODIFIER |
| 7 | 123246153 | AX-170029880 | 0 | T | C | 0.404 | 5.79E-02 | 8.51E-02 | 2.75E+00 | 2.83E+00 | 4.90E-01 | no | Slc5a11 | intron_variant | MODIFIER |
| 7 | 123255615 | AX-170433620 | 0 | C | G | 0.404 | 5.79E-02 | 8.51E-02 | 2.75E+00 | 2.83E+00 | 4.90E-01 | no | Slc5a11 | intron_variant | MODIFIER |
| 7 | 123255649 | AX-97854447 | 0 | T | C | 0.404 | 5.79E-02 | 8.51E-02 | 2.75E+00 | 2.83E+00 | 4.90E-01 | no | Slc5a11 | intron_variant | MODIFIER |
| 7 | 123260399 | AX-170078702 | 0 | G | C | 0.404 | 5.79E-02 | 8.51E-02 | 2.75E+00 | 2.83E+00 | 4.90E-01 | no | Slc5a11 | intron_variant | MODIFIER |
| 7 | 123279561 | AX-170513513 | 0 | C | T | 0.404 | 5.79E-02 | 8.51E-02 | 2.75E+00 | 2.83E+00 | 4.90E-01 | no | Arhgap17 | 3_prime_UTR_variant | MODIFIER |
| 7 | 123282088 | AX-97839118 | 0 | C | T | 0.404 | 5.79E-02 | 8.51E-02 | 2.75E+00 | 2.83E+00 | 4.90E-01 | no | Arhgap17 | intron_variant | MODIFIER |
| 7 | 123285138 | AX-170156844 | 0 | G | A | 0.404 | 5.79E-02 | 8.51E-02 | 2.75E+00 | 2.83E+00 | 4.90E-01 | no | Arhgap17 | intron_variant | MODIFIER |
| 7 | 123290576 | AX-170097426 | 0 | C | T | 0.404 | 5.79E-02 | 8.51E-02 | 2.75E+00 | 2.83E+00 | 4.90E-01 | no | Arhgap17 | intron_variant | MODIFIER |
| 7 | 123294790 | AX-170327056 | 0 | G | T | 0.404 | 5.79E-02 | 8.51E-02 | 2.75E+00 | 2.83E+00 | 4.90E-01 | no | Arhgap17 | intron_variant | MODIFIER |
| 7 | 123301574 | AX-170346564 | 0 | A | T | 0.404 | 5.79E-02 | 8.51E-02 | 2.75E+00 | 2.83E+00 | 4.90E-01 | no | Arhgap17 | intron_variant | MODIFIER |
| 7 | 123309416 | AX-170151780 | 0 | G | A | 0.404 | 5.79E-02 | 8.51E-02 | 2.75E+00 | 2.83E+00 | 4.90E-01 | no | Arhgap17 | intron_variant | MODIFIER |
| 7 | 123309443 | AX-97817319 | 0 | G | T | 0.404 | 5.79E-02 | 8.51E-02 | 2.75E+00 | 2.83E+00 | 4.90E-01 | no | Arhgap17 | intron_variant | MODIFIER |
| 7 | 123310455 | AX-170410012 | 0 | T | C | 0.404 | 5.79E-02 | 8.51E-02 | 2.75E+00 | 2.83E+00 | 4.90E-01 | no | Arhgap17 | intron_variant | MODIFIER |
| 7 | 123315155 | AX-170235813 | 0 | A | C | 0.404 | 5.79E-02 | 8.51E-02 | 2.75E+00 | 2.83E+00 | 4.90E-01 | no | Arhgap17 | intron_variant | MODIFIER |
| 7 | 123317697 | AX-170441709 | 0 | T | C | 0.404 | 5.79E-02 | 8.51E-02 | 2.75E+00 | 2.83E+00 | 4.90E-01 | no | Arhgap17 | intron_variant | MODIFIER |
| 7 | 123325244 | AX-170507518 | 0 | T | C | 0.404 | 5.79E-02 | 8.51E-02 | 2.75E+00 | 2.83E+00 | 4.90E-01 | no | Arhgap17 | intron_variant | MODIFIER |
| 7 | 123326587 | AX-170156726 | 0 | T | C | 0.404 | 5.79E-02 | 8.51E-02 | 2.75E+00 | 2.83E+00 | 4.90E-01 | no | Arhgap17 | intron_variant | MODIFIER |
| 7 | 123328355 | AX-170376751 | 0 | C | G | 0.404 | 5.79E-02 | 8.51E-02 | 2.75E+00 | 2.83E+00 | 4.90E-01 | no | Arhgap17 | intron_variant | MODIFIER |
| 7 | 123330733 | AX-170334236 | 0 | A | G | 0.404 | 5.79E-02 | 8.51E-02 | 2.75E+00 | 2.83E+00 | 4.90E-01 | no | Arhgap17 | intron_variant | MODIFIER |
| 7 | 123338879 | AX-170318567 | 0 | C | T | 0.404 | 5.79E-02 | 8.51E-02 | 2.75E+00 | 2.83E+00 | 4.90E-01 | no | Arhgap17 | intron_variant | MODIFIER |
| 7 | 123340686 | AX-97828312 | 0 | A | G | 0.404 | 5.79E-02 | 8.51E-02 | 2.75E+00 | 2.83E+00 | 4.90E-01 | no | Arhgap17 | intron_variant | MODIFIER |
| 7 | 123340765 | AX-170078703 | 0 | A | G | 0.404 | 5.79E-02 | 8.51E-02 | 2.75E+00 | 2.83E+00 | 4.90E-01 | no | Arhgap17 | intron_variant | MODIFIER |
| 7 | 123340903 | AX-170078704 | 0 | A | T | 0.404 | 5.79E-02 | 8.51E-02 | 2.75E+00 | 2.83E+00 | 4.90E-01 | no | Arhgap17 | intron_variant | MODIFIER |
| 7 | 123344062 | AX-170434963 | 0 | C | T | 0.404 | 5.79E-02 | 8.51E-02 | 2.75E+00 | 2.83E+00 | 4.90E-01 | no | Arhgap17 | intron_variant | MODIFIER |
| 7 | 123348478 | AX-170228345 | 0 | A | G | 0.404 | 5.79E-02 | 8.51E-02 | 2.75E+00 | 2.83E+00 | 4.90E-01 | no | Arhgap17 | intron_variant | MODIFIER |
| 7 | 123353246 | AX-170478495 | 0 | A | G | 0.404 | 5.79E-02 | 8.51E-02 | 2.75E+00 | 2.83E+00 | 4.90E-01 | no | Arhgap17 | intron_variant | MODIFIER |
| 7 | 123356359 | AX-170420897 | 0 | T | G | 0.404 | 5.79E-02 | 8.51E-02 | 2.75E+00 | 2.83E+00 | 4.90E-01 | no | Arhgap17 | intron_variant | MODIFIER |
| 7 | 123364400 | AX-170194222 | 0 | A | C | 0.404 | 5.79E-02 | 8.51E-02 | 2.75E+00 | 2.83E+00 | 4.90E-01 | no | Arhgap17 | intron_variant | MODIFIER |
| 7 | 123389851 | AX-170494146 | 0 | A | T | 0.404 | 5.79E-02 | 8.51E-02 | 2.75E+00 | 2.83E+00 | 4.90E-01 | no | Lcmt1 | intron_variant | MODIFIER |
| 7 | 123399792 | AX-170138179 | 0 | C | T | 0.404 | 5.79E-02 | 8.51E-02 | 2.75E+00 | 2.83E+00 | 4.90E-01 | no | Lcmt1 | intron_variant | MODIFIER |
| 7 | 123404824 | AX-170448294 | 0 | T | C | 0.404 | 5.79E-02 | 8.51E-02 | 2.75E+00 | 2.83E+00 | 4.90E-01 | no | Lcmt1 | intron_variant | MODIFIER |
| 7 | 123407717 | AX-170376916 | 0 | C | T | 0.404 | 5.79E-02 | 8.51E-02 | 2.75E+00 | 2.83E+00 | 4.90E-01 | no | Lcmt1 | intron_variant | MODIFIER |
| 7 | 123409036 | AX-170441719 | 0 | C | T | 0.404 | 5.79E-02 | 8.51E-02 | 2.75E+00 | 2.83E+00 | 4.90E-01 | no | Lcmt1 | intron_variant | MODIFIER |
| 7 | 123409446 | AX-97807600 | 0 | A | G | 0.404 | 5.79E-02 | 8.51E-02 | 2.75E+00 | 2.83E+00 | 4.90E-01 | no | Lcmt1 | intron_variant | MODIFIER |
| 7 | 123414243 | AX-170426547 | 0 | G | A | 0.404 | 5.79E-02 | 8.51E-02 | 2.75E+00 | 2.83E+00 | 4.90E-01 | no | Lcmt1 | intron_variant | MODIFIER |
| 7 | 123419961 | AX-170244182 | 0 | A | G | 0.404 | 5.79E-02 | 8.51E-02 | 2.75E+00 | 2.83E+00 | 4.90E-01 | no | Lcmt1 | intron_variant | MODIFIER |
| 7 | 123421471 | AX-170452279 | 0 | A | G | 0.404 | 5.79E-02 | 8.51E-02 | 2.75E+00 | 2.83E+00 | 4.90E-01 | no | Lcmt1 | intron_variant | MODIFIER |
| 7 | 123425806 | AX-170506310 | 0 | T | C | 0.404 | 5.79E-02 | 8.51E-02 | 2.75E+00 | 2.83E+00 | 4.90E-01 | no | Lcmt1 | intron_variant | MODIFIER |
| 7 | 123515461 | AX-168377854 | 0 | C | A | 0.404 | 5.79E-02 | 8.51E-02 | 2.75E+00 | 2.83E+00 | 4.90E-01 | no | | intergenic_variant | MODIFIER |
| 7 | 123516786 | AX-170329216 | 0 | T | G | 0.404 | 5.79E-02 | 8.51E-02 | 2.75E+00 | 2.83E+00 | 4.90E-01 | no | | intergenic_variant | MODIFIER |

| | | | | | | | | | | | | | | | |
|---|---|---|---|---|---|---|---|---|---|---|---|---|---|---|---|
| 7 | 123685876 | AX-170029884 | 0 | A | T | 0.404 | 5.79E-02 | 8.51E-02 | 2.75E+00 | 2.83E+00 | 4.90E-01 | no | | intergenic_variant | MODIFIER |
| 7 | 123735520 | AX-170096115 | 0 | G | A | 0.404 | 5.79E-02 | 8.51E-02 | 2.75E+00 | 2.83E+00 | 4.90E-01 | no | | intergenic_variant | MODIFIER |
| 7 | 123751812 | AX-170478915 | 0 | T | C | 0.404 | 5.79E-02 | 8.51E-02 | 2.75E+00 | 2.83E+00 | 4.90E-01 | no | | intergenic_variant | MODIFIER |
| 7 | 123758298 | AX-97827434 | 0 | C | T | 0.404 | 5.79E-02 | 8.51E-02 | 2.75E+00 | 2.83E+00 | 4.90E-01 | no | | intergenic_variant | MODIFIER |
| 7 | 123775080 | AX-170475264 | 0 | A | G | 0.404 | 5.79E-02 | 8.51E-02 | 2.75E+00 | 2.83E+00 | 4.90E-01 | no | | intergenic_variant | MODIFIER |
| 7 | 123784646 | AX-170214149 | 0 | A | G | 0.404 | 5.79E-02 | 8.51E-02 | 2.75E+00 | 2.83E+00 | 4.90E-01 | no | | intergenic_variant | MODIFIER |
| 7 | 123800641 | AX-170029888 | 0 | T | C | 0.404 | 5.79E-02 | 8.51E-02 | 2.75E+00 | 2.83E+00 | 4.90E-01 | no | | intergenic_variant | MODIFIER |
| 7 | 123812027 | AX-97786075 | 0 | A | G | 0.404 | 5.79E-02 | 8.51E-02 | 2.75E+00 | 2.83E+00 | 4.90E-01 | no | | intergenic_variant | MODIFIER |
| 7 | 123824536 | AX-97786162 | 0 | G | A | 0.404 | 5.79E-02 | 8.51E-02 | 2.75E+00 | 2.83E+00 | 4.90E-01 | no | | intergenic_variant | MODIFIER |
| 7 | 123834537 | AX-170078718 | 0 | C | G | 0.404 | 5.79E-02 | 8.51E-02 | 2.75E+00 | 2.83E+00 | 4.90E-01 | no | | intergenic_variant | MODIFIER |
| 7 | 123851006 | AX-170443375 | 0 | T | G | 0.404 | 5.79E-02 | 8.51E-02 | 2.75E+00 | 2.83E+00 | 4.90E-01 | no | | intergenic_variant | MODIFIER |
| 7 | 123851187 | AX-97834018 | 0 | A | C | 0.404 | 5.79E-02 | 8.51E-02 | 2.75E+00 | 2.83E+00 | 4.90E-01 | no | | intergenic_variant | MODIFIER |
| 7 | 123870700 | AX-170224070 | 0 | A | C | 0.404 | 5.79E-02 | 8.51E-02 | 2.75E+00 | 2.83E+00 | 4.90E-01 | no | | intergenic_variant | MODIFIER |
| 7 | 123878313 | AX-170437309 | 0 | T | C | 0.404 | 5.79E-02 | 8.51E-02 | 2.75E+00 | 2.83E+00 | 4.90E-01 | no | | intergenic_variant | MODIFIER |
| 7 | 123901917 | AX-170124769 | 0 | C | T | 0.404 | 5.79E-02 | 8.51E-02 | 2.75E+00 | 2.83E+00 | 4.90E-01 | no | | intergenic_variant | MODIFIER |
| 7 | 123901967 | AX-170512044 | 0 | T | C | 0.404 | 5.79E-02 | 8.51E-02 | 2.75E+00 | 2.83E+00 | 4.90E-01 | no | | intergenic_variant | MODIFIER |
| 7 | 123902337 | AX-170367452 | 0 | A | G | 0.404 | 5.79E-02 | 8.51E-02 | 2.75E+00 | 2.83E+00 | 4.90E-01 | no | | intergenic_variant | MODIFIER |
| 7 | 123905550 | AX-170136996 | 0 | T | C | 0.404 | 5.79E-02 | 8.51E-02 | 2.75E+00 | 2.83E+00 | 4.90E-01 | no | | intergenic_variant | MODIFIER |
| 7 | 123907887 | AX-170524079 | 0 | T | C | 0.404 | 5.79E-02 | 8.51E-02 | 2.75E+00 | 2.83E+00 | 4.90E-01 | no | | intergenic_variant | MODIFIER |
| 7 | 123909208 | AX-97837691 | 0 | A | C | 0.404 | 5.79E-02 | 8.51E-02 | 2.75E+00 | 2.83E+00 | 4.90E-01 | no | | intergenic_variant | MODIFIER |
| 7 | 123916130 | AX-170331864 | 0 | A | G | 0.404 | 5.79E-02 | 8.51E-02 | 2.75E+00 | 2.83E+00 | 4.90E-01 | no | | intergenic_variant | MODIFIER |
| 7 | 123916302 | AX-170426820 | 0 | A | G | 0.404 | 5.79E-02 | 8.51E-02 | 2.75E+00 | 2.83E+00 | 4.90E-01 | no | | intergenic_variant | MODIFIER |
| 7 | 123921506 | AX-170459342 | 0 | A | G | 0.404 | 5.79E-02 | 8.51E-02 | 2.75E+00 | 2.83E+00 | 4.90E-01 | no | | intergenic_variant | MODIFIER |
| 7 | 123959465 | AX-97807764 | 0 | C | A | 0.404 | 5.79E-02 | 8.51E-02 | 2.75E+00 | 2.83E+00 | 4.90E-01 | no | Gm26974 | intron_variant,non_coding_transcript_variant | MODIFIER |
| 7 | 123963053 | AX-170078720 | 0 | G | A | 0.404 | 5.79E-02 | 8.51E-02 | 2.75E+00 | 2.83E+00 | 4.90E-01 | no | Gm26974 | downstream_gene_variant | MODIFIER |
| 7 | 124077014 | AX-170090018 | 0 | T | A | 0.404 | 5.79E-02 | 8.51E-02 | 2.75E+00 | 2.83E+00 | 4.90E-01 | no | Hs3st4 | intron_variant | MODIFIER |
| 7 | 124349506 | AX-170219636 | 0 | G | A | 0.404 | 5.79E-02 | 8.51E-02 | 2.75E+00 | 2.83E+00 | 4.90E-01 | no | Hs3st4 | intron_variant | MODIFIER |
| 7 | 124619491 | AX-97849316 | 0 | C | T | 0.404 | 5.79E-02 | 8.51E-02 | 2.75E+00 | 2.83E+00 | 4.90E-01 | no | | intergenic_variant | MODIFIER |
| 7 | 124663199 | AX-97798605 | 0 | G | A | 0.404 | 5.79E-02 | 8.51E-02 | 2.75E+00 | 2.83E+00 | 4.90E-01 | no | 3100003L05Rik | intron_variant,non_coding_transcript_variant | MODIFIER |
| 7 | 124709857 | AX-170473330 | 0 | T | A | 0.404 | 5.79E-02 | 8.51E-02 | 2.75E+00 | 2.83E+00 | 4.90E-01 | no | 3100003L05Rik | upstream_gene_variant | MODIFIER |
| 7 | 124710882 | AX-170078738 | 0 | G | A | 0.404 | 5.79E-02 | 8.51E-02 | 2.75E+00 | 2.83E+00 | 4.90E-01 | no | 3100003L05Rik | upstream_gene_variant | MODIFIER |
| 17 | 14534899 | AX-170058477 | 0 | T | C | 0.018 | 1.27E-01 | 1.87E-01 | 2.59E+00 | 2.77E+00 | 4.90E-01 | no | | intergenic_variant | MODIFIER |
| 18 | 74273513 | AX-170015987 | 0 | G | T | 0.202 | -7.04E-02 | 1.01E-01 | 2.54E+00 | 2.63E+00 | 4.90E-01 | no | Mbd1 | missense_variant | MODERATE |
| 5 | 93061954 | AX-170134469 | 1 | C | T | 0.429 | 7.14E-02 | 1.05E-01 | 2.71E+00 | 2.78E+00 | 4.90E-01 | no | | intergenic_variant | MODIFIER |
| 9 | 16340355 | AX-170311378 | 0 | T | C | 0.482 | 6.06E-02 | 8.72E-02 | 2.56E+00 | 2.63E+00 | 4.90E-01 | no | Fat3 | intron_variant | MODIFIER |
| 8 | 95309844 | AX-170360548 | 1 | A | G | 0.223 | 7.64E-02 | 1.12E-01 | 2.73E+00 | 2.81E+00 | 4.90E-01 | no | Cngb1 | upstream_gene_variant | MODIFIER |
| 1 | 151920841 | AX-170294522 | 1 | T | C | 0.205 | 9.51E-02 | 1.39E-01 | 2.65E+00 | 2.70E+00 | 4.90E-01 | no | 1700025G04Rik | intron_variant | MODIFIER |
| 1 | 152200917 | AX-170189639 | 1 | G | A | 0.205 | 9.51E-02 | 1.39E-01 | 2.65E+00 | 2.70E+00 | 4.90E-01 | no | | intergenic_variant | MODIFIER |
| 1 | 152461687 | AX-170432701 | 1 | C | A | 0.205 | 9.51E-02 | 1.39E-01 | 2.65E+00 | 2.70E+00 | 4.90E-01 | no | Colgalt2 | intron_variant | MODIFIER |
| 1 | 152919938 | AX-170335100 | 1 | G | A | 0.205 | 9.51E-02 | 1.39E-01 | 2.65E+00 | 2.70E+00 | 4.90E-01 | no | | intergenic_variant | MODIFIER |
| 6 | 136591215 | AX-170499955 | 1 | A | G | 0.571 | 6.27E-02 | 8.76E-02 | 2.46E+00 | 2.52E+00 | 4.90E-01 | no | Atf7ip | intron_variant | MODIFIER |
| 6 | 137030452 | AX-170211005 | 1 | C | T | 0.571 | 6.27E-02 | 8.76E-02 | 2.46E+00 | 2.52E+00 | 4.90E-01 | no | | regulatory_region_variant | MODIFIER |
| 8 | 40361268 | AX-170030543 | 0 | G | A | 0.158 | 9.15E-02 | 1.34E-01 | 2.84E+00 | 2.93E+00 | 4.91E-01 | no | Micu3 | intron_variant | MODIFIER |
| 4 | 86553480 | AX-170407341 | 0 | A | G | 0.342 | 7.20E-02 | 1.05E-01 | 2.62E+00 | 2.71E+00 | 4.91E-01 | no | Saxo1 | intron_variant | MODIFIER |
| 9 | 33896837 | AX-170327667 | 0 | G | C | 0.447 | 5.88E-02 | 8.59E-02 | 2.63E+00 | 2.70E+00 | 4.91E-01 | no | | intergenic_variant | MODIFIER |
| 15 | 67969514 | AX-170430662 | 0 | A | G | 0.342 | -5.56E-02 | 7.57E-02 | 2.39E+00 | 2.47E+00 | 4.91E-01 | no | | intergenic_variant | MODIFIER |
| 15 | 68002314 | AX-97781961 | 0 | T | G | 0.342 | -5.56E-02 | 7.57E-02 | 2.39E+00 | 2.47E+00 | 4.91E-01 | no | | intergenic_variant | MODIFIER |
| 15 | 68004313 | AX-170124611 | 0 | A | G | 0.342 | -5.56E-02 | 7.57E-02 | 2.39E+00 | 2.47E+00 | 4.91E-01 | no | | intergenic_variant | MODIFIER |
| 15 | 68004413 | AX-170131327 | 0 | G | A | 0.342 | -5.56E-02 | 7.57E-02 | 2.39E+00 | 2.47E+00 | 4.91E-01 | no | | intergenic_variant | MODIFIER |
| 15 | 68072174 | AX-170366537 | 0 | G | T | 0.342 | -5.56E-02 | 7.57E-02 | 2.39E+00 | 2.47E+00 | 4.91E-01 | no | | regulatory_region_variant | MODIFIER |
| 15 | 68087015 | AX-170500462 | 0 | T | C | 0.342 | -5.56E-02 | 7.57E-02 | 2.39E+00 | 2.47E+00 | 4.91E-01 | no | Zfat | intron_variant | MODIFIER |
| 15 | 68129427 | AX-170291908 | 0 | C | T | 0.342 | -5.56E-02 | 7.57E-02 | 2.39E+00 | 2.47E+00 | 4.91E-01 | no | Zfat | intron_variant | MODIFIER |
| 15 | 68130960 | AX-170303894 | 0 | T | C | 0.342 | -5.56E-02 | 7.57E-02 | 2.39E+00 | 2.47E+00 | 4.91E-01 | no | Zfat | intron_variant | MODIFIER |
| 15 | 68131903 | AX-170162270 | 0 | G | C | 0.342 | -5.56E-02 | 7.57E-02 | 2.39E+00 | 2.47E+00 | 4.91E-01 | no | Zfat | intron_variant | MODIFIER |

| | | | | | | | | | | | | | | | |
|---|---|---|---|---|---|---|---|---|---|---|---|---|---|---|---|
| 15 | 68158182 | AX-170168176 | 0 | G | T | 0.342 | -5.56E-02 | 7.57E-02 | 2.39E+00 | 2.47E+00 | 4.91E-01 | no | Zfat | intron_variant | MODIFIER |
| 15 | 68158377 | AX-170319592 | 0 | G | A | 0.342 | -5.56E-02 | 7.57E-02 | 2.39E+00 | 2.47E+00 | 4.91E-01 | no | Zfat | intron_variant | MODIFIER |
| 15 | 68158549 | AX-170504220 | 0 | T | G | 0.342 | -5.56E-02 | 7.57E-02 | 2.39E+00 | 2.47E+00 | 4.91E-01 | no | Zfat | intron_variant | MODIFIER |
| 15 | 68162728 | AX-170141760 | 0 | G | T | 0.342 | -5.56E-02 | 7.57E-02 | 2.39E+00 | 2.47E+00 | 4.91E-01 | no | Zfat | intron_variant | MODIFIER |
| 15 | 68181475 | AX-170106138 | 0 | A | G | 0.342 | -5.56E-02 | 7.57E-02 | 2.39E+00 | 2.47E+00 | 4.91E-01 | no | Zfat | intron_variant,non_coding_transcript_variant | MODIFIER |
| 15 | 68181811 | AX-170317550 | 0 | T | C | 0.342 | -5.56E-02 | 7.57E-02 | 2.39E+00 | 2.47E+00 | 4.91E-01 | no | Zfat | intron_variant,non_coding_transcript_variant | MODIFIER |
| 15 | 68188496 | AX-170330397 | 0 | T | C | 0.342 | -5.56E-02 | 7.57E-02 | 2.39E+00 | 2.47E+00 | 4.91E-01 | no | Zfat | intron_variant,non_coding_transcript_variant | MODIFIER |
| 15 | 68188621 | AX-170178254 | 0 | A | T | 0.342 | -5.56E-02 | 7.57E-02 | 2.39E+00 | 2.47E+00 | 4.91E-01 | no | Zfat | intron_variant,non_coding_transcript_variant | MODIFIER |
| 15 | 68189468 | AX-170055849 | 0 | G | A | 0.342 | -5.56E-02 | 7.57E-02 | 2.39E+00 | 2.47E+00 | 4.91E-01 | no | Zfat | intron_variant,non_coding_transcript_variant | MODIFIER |
| 15 | 68195554 | AX-97832086 | 0 | A | G | 0.342 | -5.56E-02 | 7.57E-02 | 2.39E+00 | 2.47E+00 | 4.91E-01 | no | Zfat | intron_variant,non_coding_transcript_variant | MODIFIER |
| 15 | 68200255 | AX-170270543 | 0 | G | A | 0.342 | -5.56E-02 | 7.57E-02 | 2.39E+00 | 2.47E+00 | 4.91E-01 | no | Zfat | intron_variant,non_coding_transcript_variant | MODIFIER |
| 15 | 68211567 | AX-170373122 | 0 | G | A | 0.342 | -5.56E-02 | 7.57E-02 | 2.39E+00 | 2.47E+00 | 4.91E-01 | no | Zfat | intron_variant,non_coding_transcript_variant | MODIFIER |
| 15 | 68227830 | AX-170177705 | 0 | T | C | 0.342 | -5.56E-02 | 7.57E-02 | 2.39E+00 | 2.47E+00 | 4.91E-01 | no | Zfat | intron_variant,non_coding_transcript_variant | MODIFIER |
| 15 | 68228648 | AX-170331376 | 0 | T | C | 0.342 | -5.56E-02 | 7.57E-02 | 2.39E+00 | 2.47E+00 | 4.91E-01 | no | Zfat | intron_variant,non_coding_transcript_variant | MODIFIER |
| 15 | 68509759 | AX-170011916 | 0 | G | A | 0.342 | -5.56E-02 | 7.57E-02 | 2.39E+00 | 2.47E+00 | 4.91E-01 | no | | intergenic_variant | MODIFIER |
| 15 | 68513898 | AX-170508351 | 0 | T | A | 0.342 | -5.56E-02 | 7.57E-02 | 2.39E+00 | 2.47E+00 | 4.91E-01 | no | | intergenic_variant | MODIFIER |
| 15 | 68521131 | AX-170383509 | 0 | G | A | 0.342 | -5.56E-02 | 7.57E-02 | 2.39E+00 | 2.47E+00 | 4.91E-01 | no | | intergenic_variant | MODIFIER |
| 15 | 68525280 | AX-170481620 | 0 | C | T | 0.342 | -5.56E-02 | 7.57E-02 | 2.39E+00 | 2.47E+00 | 4.91E-01 | no | | regulatory_region_variant | MODIFIER |
| 15 | 68525458 | AX-170295059 | 0 | A | T | 0.342 | -5.56E-02 | 7.57E-02 | 2.39E+00 | 2.47E+00 | 4.91E-01 | no | | regulatory_region_variant | MODIFIER |
| 15 | 68530907 | AX-168378815 | 0 | T | C | 0.342 | -5.56E-02 | 7.57E-02 | 2.39E+00 | 2.47E+00 | 4.91E-01 | no | | intergenic_variant | MODIFIER |
| 15 | 68557192 | AX-170235750 | 0 | A | G | 0.342 | -5.56E-02 | 7.57E-02 | 2.39E+00 | 2.47E+00 | 4.91E-01 | no | | intergenic_variant | MODIFIER |
| 15 | 68567708 | AX-170197804 | 0 | C | A | 0.342 | -5.56E-02 | 7.57E-02 | 2.39E+00 | 2.47E+00 | 4.91E-01 | no | | intergenic_variant | MODIFIER |
| 15 | 68574295 | AX-97810686 | 0 | G | A | 0.342 | -5.56E-02 | 7.57E-02 | 2.39E+00 | 2.47E+00 | 4.91E-01 | no | | intergenic_variant | MODIFIER |
| 15 | 68576685 | AX-170394511 | 0 | T | C | 0.342 | -5.56E-02 | 7.57E-02 | 2.39E+00 | 2.47E+00 | 4.91E-01 | no | | intergenic_variant | MODIFIER |
| 15 | 68583055 | AX-170179144 | 0 | C | T | 0.342 | -5.56E-02 | 7.57E-02 | 2.39E+00 | 2.47E+00 | 4.91E-01 | no | | intergenic_variant | MODIFIER |
| 15 | 68583080 | AX-170055859 | 0 | T | C | 0.342 | -5.56E-02 | 7.57E-02 | 2.39E+00 | 2.47E+00 | 4.91E-01 | no | | intergenic_variant | MODIFIER |
| 15 | 68590052 | AX-170167347 | 0 | G | A | 0.342 | -5.56E-02 | 7.57E-02 | 2.39E+00 | 2.47E+00 | 4.91E-01 | no | | intergenic_variant | MODIFIER |
| 5 | 36407033 | AX-170225512 | 0 | G | A | 0.351 | 7.50E-02 | 1.10E-01 | 2.70E+00 | 2.77E+00 | 4.91E-01 | no | | intergenic_variant | MODIFIER |
| 5 | 37694674 | AX-170024708 | 0 | G | A | 0.351 | 7.50E-02 | 1.10E-01 | 2.70E+00 | 2.77E+00 | 4.91E-01 | no | Stk32b | intron_variant | MODIFIER |
| 10 | 6408238 | AX-170411145 | 0 | C | A | 0.211 | -6.75E-02 | 9.91E-02 | 2.64E+00 | 2.75E+00 | 4.91E-01 | no | | intergenic_variant | MODIFIER |
| 5 | 78517249 | AX-170025358 | 0 | G | T | 0.447 | 7.03E-02 | 1.03E-01 | 2.68E+00 | 2.74E+00 | 4.91E-01 | no | | intergenic_variant | MODIFIER |
| 5 | 80830186 | AX-170387976 | 0 | T | C | 0.447 | 7.03E-02 | 1.03E-01 | 2.68E+00 | 2.74E+00 | 4.91E-01 | no | | intergenic_variant | MODIFIER |
| 1 | 146793528 | AX-170001931 | 1 | T | C | 0.223 | 8.59E-02 | 1.25E-01 | 2.59E+00 | 2.66E+00 | 4.91E-01 | no | Brinp3 | intron_variant | MODIFIER |
| 6 | 147307965 | AX-97838111 | 0 | C | T | 0.079 | 1.37E-01 | 2.02E-01 | 2.70E+00 | 2.80E+00 | 4.91E-01 | no | | intergenic_variant | MODIFIER |
| 5 | 37555411 | AX-170527039 | 1 | G | C | 0.321 | 7.60E-02 | 1.12E-01 | 2.73E+00 | 2.79E+00 | 4.91E-01 | no | Stk32b | intron_variant | MODIFIER |
| 10 | 14098614 | AX-170002778 | 2 | C | T | 0.209 | -7.01E-02 | 9.99E-02 | 2.50E+00 | 2.60E+00 | 4.91E-01 | no | Hivep2 | intron_variant | MODIFIER |
| 9 | 8995815 | AX-170031776 | 0 | C | T | 0.465 | 5.70E-02 | 8.04E-02 | 2.47E+00 | 2.56E+00 | 4.91E-01 | no | Arhgap42 | 3_prime_UTR_variant | MODIFIER |
| 10 | 18989275 | AX-170395758 | 1 | A | G | 0.196 | 7.15E-02 | 1.04E-01 | 2.87E+00 | 2.97E+00 | 4.91E-01 | no | | intergenic_variant | MODIFIER |
| 12 | 89582533 | AX-97821380 | 0 | T | C | 0.079 | 1.26E-01 | 1.86E-01 | 2.75E+00 | 2.84E+00 | 4.91E-01 | no | Nrxn3 | intron_variant | MODIFIER |
| 1 | 99581076 | AX-170001187 | 1 | C | G | 0.161 | 1.41E-01 | 2.06E-01 | 2.64E+00 | 2.70E+00 | 4.92E-01 | no | | intergenic_variant | MODIFIER |
| 12 | 27183165 | AX-97799772 | 1 | A | G | 0.027 | -1.12E-01 | 1.62E-01 | 2.50E+00 | 2.67E+00 | 4.92E-01 | no | | regulatory_region_variant | MODIFIER |
| 5 | 122984283 | AX-170073350 | 0 | A | T | 0.439 | -7.39E-02 | 1.07E-01 | 2.89E+00 | 2.96E+00 | 4.92E-01 | no | Kdm2b | intron_variant | MODIFIER |
| 12 | 90642654 | AX-97794831 | 1 | G | A | 0.098 | 1.31E-01 | 1.93E-01 | 2.77E+00 | 2.84E+00 | 4.92E-01 | no | | intergenic_variant | MODIFIER |
| 7 | 77183407 | AX-170255232 | 0 | T | C | 0.386 | -6.84E-02 | 1.01E-01 | 2.78E+00 | 2.89E+00 | 4.92E-01 | no | | intergenic_variant | MODIFIER |
| 4 | 124418697 | AX-170351848 | 0 | T | C | 0.237 | 7.23E-02 | 1.04E-01 | 2.53E+00 | 2.61E+00 | 4.92E-01 | no | 1700057H15Rik | non_coding_transcript_exon_variant | MODIFIER |
| 12 | 103832572 | AX-97848251 | 2 | T | C | 0.1 | 9.55E-02 | 1.42E-01 | 2.68E+00 | 2.84E+00 | 4.92E-01 | no | Serpina1b | upstream_gene_variant | MODIFIER |
| 5 | 4354204 | AX-170071181 | 0 | G | A | 0.439 | -4.38E-02 | 6.28E-02 | 2.50E+00 | 2.62E+00 | 4.92E-01 | no | | intergenic_variant | MODIFIER |
| 1 | 170557714 | AX-170241604 | 1 | T | C | 0.214 | 9.46E-02 | 1.38E-01 | 2.63E+00 | 2.68E+00 | 4.92E-01 | no | Nos1ap | intron_variant,non_coding_transcript_variant | MODIFIER |
| 8 | 36786684 | AX-170030515 | 1 | C | T | 0.152 | 1.02E-01 | 1.49E-01 | 2.89E+00 | 2.94E+00 | 4.92E-01 | no | Dlc1 | intron_variant | MODIFIER |
| 8 | 40625120 | AX-170141057 | 1 | G | A | 0.152 | 1.02E-01 | 1.49E-01 | 2.89E+00 | 2.94E+00 | 4.92E-01 | no | Mtmr7 | intron_variant,NMD_transcript_variant | MODIFIER |
| 8 | 41376660 | AX-170287087 | 1 | T | C | 0.152 | 1.02E-01 | 1.49E-01 | 2.89E+00 | 2.94E+00 | 4.92E-01 | no | Asah1 | upstream_gene_variant | MODIFIER |
| 8 | 42388251 | AX-170400819 | 1 | C | T | 0.152 | 1.02E-01 | 1.49E-01 | 2.89E+00 | 2.94E+00 | 4.92E-01 | no | | intergenic_variant | MODIFIER |
| 1 | 106555536 | AX-97812652 | 0 | A | G | 0.167 | 1.13E-01 | 1.63E-01 | 2.54E+00 | 2.65E+00 | 4.93E-01 | no | Bcl2 | intron_variant | MODIFIER |
| 4 | 73864976 | AX-170491920 | 1 | G | T | 0.25 | 7.08E-02 | 1.01E-01 | 2.50E+00 | 2.58E+00 | 4.93E-01 | no | Gm13867 | non_coding_transcript_exon_variant | MODIFIER |

| | | | | | | | | | | | | | | | |
|---|---|---|---|---|---|---|---|---|---|---|---|---|---|---|---|
| 1 | 96238350 | AX-97788455 | 1 | T | C | 0.134 | 1.30E-01 | 1.90E-01 | 2.62E+00 | 2.68E+00 | 4.93E-01 | no | | intergenic_variant | MODIFIER |
| 9 | 15786015 | AX-170031841 | 0 | C | T | 0.474 | 6.03E-02 | 8.70E-02 | 2.55E+00 | 2.62E+00 | 4.93E-01 | no | | intergenic_variant | MODIFIER |
| 2 | 104128837 | AX-97807622 | 0 | C | T | 0.061 | -1.58E-01 | 2.33E-01 | 2.77E+00 | 2.87E+00 | 4.93E-01 | no | A930018P22Rik | downstream_gene_variant | MODIFIER |
| 1 | 151135704 | AX-170142764 | 1 | G | A | 0.214 | 8.48E-02 | 1.24E-01 | 2.62E+00 | 2.69E+00 | 4.93E-01 | no | Gm8941 | upstream_gene_variant | MODIFIER |
| 6 | 138409144 | AX-170089853 | 0 | C | T | 0.561 | 6.08E-02 | 8.51E-02 | 2.45E+00 | 2.52E+00 | 4.93E-01 | no | Lmo3 | intron_variant,NMD_transcript_variant | MODIFIER |
| 6 | 138413506 | AX-170460761 | 0 | A | G | 0.561 | 6.08E-02 | 8.51E-02 | 2.45E+00 | 2.52E+00 | 4.93E-01 | no | Lmo3 | intron_variant,NMD_transcript_variant | MODIFIER |
| 6 | 138421068 | AX-170076390 | 0 | G | C | 0.561 | 6.08E-02 | 8.51E-02 | 2.45E+00 | 2.52E+00 | 4.93E-01 | no | A830011K09Rik | upstream_gene_variant | MODIFIER |
| 6 | 138429407 | AX-97813980 | 0 | T | C | 0.561 | 6.08E-02 | 8.51E-02 | 2.45E+00 | 2.52E+00 | 4.93E-01 | no | A830011K09Rik | downstream_gene_variant | MODIFIER |
| 6 | 138470701 | AX-170202444 | 0 | C | T | 0.561 | 6.08E-02 | 8.51E-02 | 2.45E+00 | 2.52E+00 | 4.93E-01 | no | Lmo3 | intron_variant | MODIFIER |
| 6 | 138474133 | AX-170343015 | 0 | G | A | 0.561 | 6.08E-02 | 8.51E-02 | 2.45E+00 | 2.52E+00 | 4.93E-01 | no | Lmo3 | intron_variant | MODIFIER |
| 6 | 138477905 | AX-170482127 | 0 | A | T | 0.561 | 6.08E-02 | 8.51E-02 | 2.45E+00 | 2.52E+00 | 4.93E-01 | no | Lmo3 | intron_variant | MODIFIER |
| 6 | 138482962 | AX-170412789 | 0 | G | A | 0.561 | 6.08E-02 | 8.51E-02 | 2.45E+00 | 2.52E+00 | 4.93E-01 | no | Lmo3 | intron_variant | MODIFIER |
| 6 | 138487938 | AX-170108526 | 0 | C | A | 0.561 | 6.08E-02 | 8.51E-02 | 2.45E+00 | 2.52E+00 | 4.93E-01 | no | Lmo3 | intron_variant | MODIFIER |
| 6 | 138496388 | AX-170028359 | 0 | C | A | 0.561 | 6.08E-02 | 8.51E-02 | 2.45E+00 | 2.52E+00 | 4.93E-01 | no | Lmo3 | intron_variant | MODIFIER |
| 6 | 138496494 | AX-170028358 | 0 | A | G | 0.561 | 6.08E-02 | 8.51E-02 | 2.45E+00 | 2.52E+00 | 4.93E-01 | no | Lmo3 | intron_variant | MODIFIER |
| 6 | 138507320 | AX-97816335 | 0 | G | T | 0.561 | 6.08E-02 | 8.51E-02 | 2.45E+00 | 2.52E+00 | 4.93E-01 | no | Lmo3 | intron_variant | MODIFIER |
| 6 | 138547682 | AX-97825691 | 0 | C | T | 0.561 | 6.08E-02 | 8.51E-02 | 2.45E+00 | 2.52E+00 | 4.93E-01 | no | Lmo3 | intron_variant | MODIFIER |
| 6 | 138555945 | AX-170517069 | 0 | G | T | 0.561 | 6.08E-02 | 8.51E-02 | 2.45E+00 | 2.52E+00 | 4.93E-01 | no | Lmo3 | intron_variant | MODIFIER |
| 6 | 138565899 | AX-170028356 | 0 | T | C | 0.561 | 6.08E-02 | 8.51E-02 | 2.45E+00 | 2.52E+00 | 4.93E-01 | no | Lmo3 | intron_variant | MODIFIER |
| 6 | 138629286 | AX-170191289 | 0 | C | A | 0.561 | 6.08E-02 | 8.51E-02 | 2.45E+00 | 2.52E+00 | 4.93E-01 | no | | intergenic_variant | MODIFIER |
| 6 | 138633544 | AX-97814134 | 0 | A | C | 0.561 | 6.08E-02 | 8.51E-02 | 2.45E+00 | 2.52E+00 | 4.93E-01 | no | | intergenic_variant | MODIFIER |
| 6 | 138657268 | AX-97807113 | 0 | G | A | 0.561 | 6.08E-02 | 8.51E-02 | 2.45E+00 | 2.52E+00 | 4.93E-01 | no | Igbp1b | 3_prime_UTR_variant | MODIFIER |
| 6 | 138658431 | AX-170141962 | 0 | G | A | 0.561 | 6.08E-02 | 8.51E-02 | 2.45E+00 | 2.52E+00 | 4.93E-01 | no | Igbp1b | missense_variant | MODERATE |
| 6 | 138668127 | AX-170100287 | 0 | T | C | 0.561 | 6.08E-02 | 8.51E-02 | 2.45E+00 | 2.52E+00 | 4.93E-01 | no | | intergenic_variant | MODIFIER |
| 6 | 138668201 | AX-170438993 | 0 | G | A | 0.561 | 6.08E-02 | 8.51E-02 | 2.45E+00 | 2.52E+00 | 4.93E-01 | no | | intergenic_variant | MODIFIER |
| 6 | 138668468 | AX-170417675 | 0 | T | C | 0.561 | 6.08E-02 | 8.51E-02 | 2.45E+00 | 2.52E+00 | 4.93E-01 | no | | intergenic_variant | MODIFIER |
| 6 | 138670066 | AX-170488685 | 0 | G | A | 0.561 | 6.08E-02 | 8.51E-02 | 2.45E+00 | 2.52E+00 | 4.93E-01 | no | | intergenic_variant | MODIFIER |
| 6 | 138681615 | AX-97823733 | 0 | C | T | 0.561 | 6.08E-02 | 8.51E-02 | 2.45E+00 | 2.52E+00 | 4.93E-01 | no | | intergenic_variant | MODIFIER |
| 15 | 25596303 | AX-170164866 | 1 | G | A | 0.25 | -6.99E-02 | 1.03E-01 | 2.63E+00 | 2.70E+00 | 4.93E-01 | no | Gm36642 | upstream_gene_variant | MODIFIER |
| 1 | 187674004 | AX-170165852 | 0 | T | C | 0.254 | 7.19E-02 | 9.93E-02 | 2.41E+00 | 2.50E+00 | 4.93E-01 | no | Esrrg | intron_variant | MODIFIER |
| 2 | 137230708 | AX-170019153 | 0 | T | A | 0.149 | 1.20E-01 | 1.78E-01 | 2.68E+00 | 2.77E+00 | 4.93E-01 | no | | intergenic_variant | MODIFIER |
| 6 | 125550114 | AX-97814988 | 0 | C | T | 0.167 | -5.60E-02 | 8.34E-02 | 2.69E+00 | 2.82E+00 | 4.93E-01 | no | Vwf | intron_variant | MODIFIER |
| 18 | 59379001 | AX-97857695 | 0 | G | T | 0.404 | 6.65E-02 | 9.43E-02 | 2.49E+00 | 2.54E+00 | 4.93E-01 | no | Chsy3 | intron_variant | MODIFIER |
| 12 | 89604345 | AX-170135147 | 0 | C | T | 0.07 | 1.44E-01 | 2.13E-01 | 2.75E+00 | 2.83E+00 | 4.93E-01 | Yes | Nrxn3 | intron_variant | MODIFIER |
| 12 | 91534868 | AX-170376766 | 0 | T | A | 0.07 | 1.44E-01 | 2.13E-01 | 2.75E+00 | 2.83E+00 | 4.93E-01 | Yes | Tshr | intron_variant | MODIFIER |
| 12 | 91782905 | AX-170116704 | 0 | A | C | 0.07 | 1.44E-01 | 2.13E-01 | 2.75E+00 | 2.83E+00 | 4.93E-01 | Yes | Ston2 | intron_variant | MODIFIER |
| 12 | 91788985 | AX-170116685 | 0 | G | C | 0.07 | 1.44E-01 | 2.13E-01 | 2.75E+00 | 2.83E+00 | 4.93E-01 | Yes | Ston2 | upstream_gene_variant | MODIFIER |
| 12 | 91805961 | AX-97855574 | 0 | G | A | 0.07 | 1.44E-01 | 2.13E-01 | 2.75E+00 | 2.83E+00 | 4.93E-01 | Yes | Sel1l | downstream_gene_variant | MODIFIER |
| 12 | 91814089 | AX-97821988 | 0 | A | G | 0.07 | 1.44E-01 | 2.13E-01 | 2.75E+00 | 2.83E+00 | 4.93E-01 | Yes | Sel1l | intron_variant | MODIFIER |
| 12 | 91817627 | AX-170349702 | 0 | A | G | 0.07 | 1.44E-01 | 2.13E-01 | 2.75E+00 | 2.83E+00 | 4.93E-01 | Yes | Sel1l | intron_variant | MODIFIER |
| 12 | 91836360 | AX-170362872 | 0 | T | C | 0.14 | 7.21E-02 | 1.07E-01 | 2.75E+00 | 2.83E+00 | 4.93E-01 | no | Sel1l | intron_variant | MODIFIER |
| 12 | 91836491 | AX-97810775 | 0 | G | T | 0.07 | 1.44E-01 | 2.13E-01 | 2.75E+00 | 2.83E+00 | 4.93E-01 | Yes | Sel1l | intron_variant | MODIFIER |
| 12 | 91836782 | AX-170382936 | 0 | C | T | 0.07 | 1.44E-01 | 2.13E-01 | 2.75E+00 | 2.83E+00 | 4.93E-01 | Yes | Sel1l | intron_variant | MODIFIER |
| 12 | 91847386 | AX-170498475 | 0 | T | C | 0.07 | 1.44E-01 | 2.13E-01 | 2.75E+00 | 2.83E+00 | 4.93E-01 | Yes | Sel1l | intron_variant | MODIFIER |
| 12 | 91847600 | AX-170387768 | 0 | A | C | 0.07 | 1.44E-01 | 2.13E-01 | 2.75E+00 | 2.83E+00 | 4.93E-01 | Yes | Sel1l | intron_variant | MODIFIER |
| 12 | 91850850 | AX-170483738 | 0 | T | G | 0.07 | 1.44E-01 | 2.13E-01 | 2.75E+00 | 2.83E+00 | 4.93E-01 | Yes | Sel1l | upstream_gene_variant | MODIFIER |
| 12 | 91856897 | AX-97822015 | 0 | C | T | 0.07 | 1.44E-01 | 2.13E-01 | 2.75E+00 | 2.83E+00 | 4.93E-01 | Yes | | intergenic_variant | MODIFIER |
| 12 | 91914049 | AX-170111263 | 0 | C | T | 0.07 | 1.44E-01 | 2.13E-01 | 2.75E+00 | 2.83E+00 | 4.93E-01 | Yes | | intergenic_variant | MODIFIER |
| 12 | 91931939 | AX-170197633 | 0 | T | G | 0.07 | 1.44E-01 | 2.13E-01 | 2.75E+00 | 2.83E+00 | 4.93E-01 | Yes | | intergenic_variant | MODIFIER |
| 12 | 91945251 | AX-170186228 | 0 | G | C | 0.07 | 1.44E-01 | 2.13E-01 | 2.75E+00 | 2.83E+00 | 4.93E-01 | Yes | | intergenic_variant | MODIFIER |
| 12 | 91946219 | AX-170526731 | 0 | T | C | 0.07 | 1.44E-01 | 2.13E-01 | 2.75E+00 | 2.83E+00 | 4.93E-01 | Yes | | intergenic_variant | MODIFIER |
| 12 | 91948760 | AX-170357584 | 0 | T | C | 0.07 | 1.44E-01 | 2.13E-01 | 2.75E+00 | 2.83E+00 | 4.93E-01 | Yes | | regulatory_region_variant | MODIFIER |
| 12 | 91952676 | AX-170258003 | 0 | A | C | 0.07 | 1.44E-01 | 2.13E-01 | 2.75E+00 | 2.83E+00 | 4.93E-01 | Yes | | intergenic_variant | MODIFIER |
| 12 | 91953241 | AX-170524192 | 0 | G | A | 0.07 | 1.44E-01 | 2.13E-01 | 2.75E+00 | 2.83E+00 | 4.93E-01 | Yes | | intergenic_variant | MODIFIER |

| | | | | | | | | | | | | | | | |
|---|---|---|---|---|---|---|---|---|---|---|---|---|---|---|---|
| 12 | 91955127 | AX-170049784 | 0 | A | G | 0.07 | 1.44E-01 | 2.13E-01 | 2.75E+00 | 2.83E+00 | 4.93E-01 | Yes | | intergenic_variant | MODIFIER |
| 12 | 91956660 | AX-170379877 | 0 | C | T | 0.07 | 1.44E-01 | 2.13E-01 | 2.75E+00 | 2.83E+00 | 4.93E-01 | Yes | | intergenic_variant | MODIFIER |
| 12 | 91956833 | AX-97824107 | 0 | T | C | 0.07 | 1.44E-01 | 2.13E-01 | 2.75E+00 | 2.83E+00 | 4.93E-01 | Yes | | intergenic_variant | MODIFIER |
| 12 | 91958755 | AX-170123741 | 0 | A | T | 0.07 | 1.44E-01 | 2.13E-01 | 2.75E+00 | 2.83E+00 | 4.93E-01 | Yes | | intergenic_variant | MODIFIER |
| 12 | 91964272 | AX-170098558 | 0 | A | T | 0.07 | 1.44E-01 | 2.13E-01 | 2.75E+00 | 2.83E+00 | 4.93E-01 | Yes | Gm21614 | downstream_gene_variant | MODIFIER |
| 12 | 91968729 | AX-170420439 | 0 | T | C | 0.07 | 1.44E-01 | 2.13E-01 | 2.75E+00 | 2.83E+00 | 4.93E-01 | Yes | Gm21614 | non_coding_transcript_exon_variant | MODIFIER |
| 12 | 91984351 | AX-170144827 | 0 | A | G | 0.07 | 1.44E-01 | 2.13E-01 | 2.75E+00 | 2.83E+00 | 4.93E-01 | Yes | | intergenic_variant | MODIFIER |
| 12 | 91984558 | AX-170422284 | 0 | A | C | 0.07 | 1.44E-01 | 2.13E-01 | 2.75E+00 | 2.83E+00 | 4.93E-01 | Yes | | intergenic_variant | MODIFIER |
| 12 | 91987631 | AX-170196208 | 0 | C | T | 0.07 | 1.44E-01 | 2.13E-01 | 2.75E+00 | 2.83E+00 | 4.93E-01 | Yes | | intergenic_variant | MODIFIER |
| 12 | 91990025 | AX-170253819 | 0 | C | T | 0.07 | 1.44E-01 | 2.13E-01 | 2.75E+00 | 2.83E+00 | 4.93E-01 | Yes | | intergenic_variant | MODIFIER |
| 12 | 91990911 | AX-97789276 | 0 | C | T | 0.07 | 1.44E-01 | 2.13E-01 | 2.75E+00 | 2.83E+00 | 4.93E-01 | Yes | | intergenic_variant | MODIFIER |
| 12 | 92001550 | AX-170514697 | 0 | A | G | 0.07 | 1.44E-01 | 2.13E-01 | 2.75E+00 | 2.83E+00 | 4.93E-01 | Yes | | intergenic_variant | MODIFIER |
| 12 | 92005052 | AX-170188232 | 0 | A | G | 0.07 | 1.44E-01 | 2.13E-01 | 2.75E+00 | 2.83E+00 | 4.93E-01 | Yes | Gm47984 | downstream_gene_variant | MODIFIER |
| 12 | 92008729 | AX-170449101 | 0 | G | A | 0.07 | 1.44E-01 | 2.13E-01 | 2.75E+00 | 2.83E+00 | 4.93E-01 | Yes | Gm47984 | non_coding_transcript_exon_variant | MODIFIER |
| 12 | 92021742 | AX-97804670 | 0 | G | A | 0.07 | 1.44E-01 | 2.13E-01 | 2.75E+00 | 2.83E+00 | 4.93E-01 | Yes | | intergenic_variant | MODIFIER |
| 12 | 92022250 | AX-170244813 | 0 | A | C | 0.07 | 1.44E-01 | 2.13E-01 | 2.75E+00 | 2.83E+00 | 4.93E-01 | Yes | | intergenic_variant | MODIFIER |
| 12 | 92023020 | AX-170184370 | 0 | C | G | 0.07 | 1.44E-01 | 2.13E-01 | 2.75E+00 | 2.83E+00 | 4.93E-01 | Yes | | intergenic_variant | MODIFIER |
| 12 | 92028930 | AX-170522938 | 0 | C | T | 0.07 | 1.44E-01 | 2.13E-01 | 2.75E+00 | 2.83E+00 | 4.93E-01 | Yes | | intergenic_variant | MODIFIER |
| 12 | 92031786 | AX-170156057 | 0 | G | A | 0.07 | 1.44E-01 | 2.13E-01 | 2.75E+00 | 2.83E+00 | 4.93E-01 | Yes | | intergenic_variant | MODIFIER |
| 12 | 92033323 | AX-170373391 | 0 | C | T | 0.07 | 1.44E-01 | 2.13E-01 | 2.75E+00 | 2.83E+00 | 4.93E-01 | Yes | | intergenic_variant | MODIFIER |
| 12 | 92037601 | AX-170523955 | 0 | A | G | 0.07 | 1.44E-01 | 2.13E-01 | 2.75E+00 | 2.83E+00 | 4.93E-01 | Yes | | intergenic_variant | MODIFIER |
| 12 | 92047029 | AX-170120863 | 0 | T | G | 0.07 | 1.44E-01 | 2.13E-01 | 2.75E+00 | 2.83E+00 | 4.93E-01 | Yes | | intergenic_variant | MODIFIER |
| 12 | 92053218 | AX-170108175 | 0 | T | A | 0.07 | 1.44E-01 | 2.13E-01 | 2.75E+00 | 2.83E+00 | 4.93E-01 | Yes | | intergenic_variant | MODIFIER |
| 12 | 92053416 | AX-97804514 | 0 | A | G | 0.07 | 1.44E-01 | 2.13E-01 | 2.75E+00 | 2.83E+00 | 4.93E-01 | Yes | | intergenic_variant | MODIFIER |
| 12 | 92123949 | AX-97814196 | 0 | T | C | 0.07 | 1.44E-01 | 2.13E-01 | 2.75E+00 | 2.83E+00 | 4.93E-01 | Yes | | intergenic_variant | MODIFIER |
| 12 | 92182330 | AX-170462291 | 0 | A | G | 0.07 | 1.44E-01 | 2.13E-01 | 2.75E+00 | 2.83E+00 | 4.93E-01 | Yes | | intergenic_variant | MODIFIER |
| 12 | 92186468 | AX-168419910 | 0 | A | C | 0.07 | 1.44E-01 | 2.13E-01 | 2.75E+00 | 2.83E+00 | 4.93E-01 | Yes | | regulatory_region_variant | MODIFIER |
| 12 | 92198699 | AX-170324272 | 0 | G | C | 0.07 | 1.44E-01 | 2.13E-01 | 2.75E+00 | 2.83E+00 | 4.93E-01 | Yes | | intergenic_variant | MODIFIER |
| 12 | 92308104 | AX-170524707 | 0 | A | G | 0.07 | 1.44E-01 | 2.13E-01 | 2.75E+00 | 2.83E+00 | 4.93E-01 | Yes | | intergenic_variant | MODIFIER |
| 12 | 92390825 | AX-170332339 | 0 | C | T | 0.07 | 1.44E-01 | 2.13E-01 | 2.75E+00 | 2.83E+00 | 4.93E-01 | Yes | | intergenic_variant | MODIFIER |
| 12 | 92397359 | AX-97839646 | 0 | T | C | 0.07 | 1.44E-01 | 2.13E-01 | 2.75E+00 | 2.83E+00 | 4.93E-01 | Yes | | intergenic_variant | MODIFIER |
| 12 | 92473151 | AX-97839051 | 0 | T | C | 0.07 | 1.44E-01 | 2.13E-01 | 2.75E+00 | 2.83E+00 | 4.93E-01 | Yes | Gm18500 | non_coding_transcript_exon_variant | MODIFIER |
| 12 | 92473197 | AX-170103375 | 0 | C | T | 0.07 | 1.44E-01 | 2.13E-01 | 2.75E+00 | 2.83E+00 | 4.93E-01 | Yes | Gm18500 | non_coding_transcript_exon_variant | MODIFIER |
| 12 | 92475181 | AX-170232435 | 0 | T | C | 0.07 | 1.44E-01 | 2.13E-01 | 2.75E+00 | 2.83E+00 | 4.93E-01 | Yes | Gm18500 | upstream_gene_variant | MODIFIER |
| 12 | 92508913 | AX-170333989 | 0 | G | C | 0.07 | 1.44E-01 | 2.13E-01 | 2.75E+00 | 2.83E+00 | 4.93E-01 | Yes | | regulatory_region_variant | MODIFIER |
| 12 | 92509122 | AX-170344126 | 0 | C | A | 0.07 | 1.44E-01 | 2.13E-01 | 2.75E+00 | 2.83E+00 | 4.93E-01 | Yes | | intergenic_variant | MODIFIER |
| 12 | 92509376 | AX-170441315 | 0 | G | A | 0.07 | 1.44E-01 | 2.13E-01 | 2.75E+00 | 2.83E+00 | 4.93E-01 | Yes | | intergenic_variant | MODIFIER |
| 12 | 92520206 | AX-97782628 | 0 | G | A | 0.07 | 1.44E-01 | 2.13E-01 | 2.75E+00 | 2.83E+00 | 4.93E-01 | Yes | | intergenic_variant | MODIFIER |
| 6 | 130408970 | AX-170195205 | 0 | G | A | 0.535 | 5.76E-02 | 8.23E-02 | 2.51E+00 | 2.58E+00 | 4.93E-01 | no | Gm6584 | upstream_gene_variant | MODIFIER |
| 4 | 90746040 | AX-170423885 | 0 | G | A | 0.325 | 7.07E-02 | 1.04E-01 | 2.67E+00 | 2.74E+00 | 4.93E-01 | no | | intergenic_variant | MODIFIER |
| 4 | 80207878 | AX-170023206 | 0 | A | G | 0.237 | 7.24E-02 | 1.04E-01 | 2.53E+00 | 2.60E+00 | 4.93E-01 | no | | intergenic_variant | MODIFIER |
| 6 | 110489486 | AX-170027966 | 1 | G | T | 0.08 | 1.35E-01 | 2.01E-01 | 2.69E+00 | 2.79E+00 | 4.94E-01 | no | | intergenic_variant | MODIFIER |
| 7 | 113995973 | AX-170397921 | 0 | T | C | 0.412 | 6.41E-02 | 9.48E-02 | 2.70E+00 | 2.75E+00 | 4.94E-01 | no | Spon1 | intron_variant | MODIFIER |
| 1 | 151891455 | AX-170181825 | 1 | C | T | 0.205 | 9.34E-02 | 1.37E-01 | 2.64E+00 | 2.69E+00 | 4.94E-01 | no | 1700025G04Rik | 3_prime_UTR_variant | MODIFIER |
| 1 | 153018280 | AX-170197980 | 1 | G | A | 0.205 | 9.34E-02 | 1.37E-01 | 2.64E+00 | 2.69E+00 | 4.94E-01 | no | Nmnat2 | intron_variant | MODIFIER |
| 9 | 62185967 | AX-170082609 | 0 | C | T | 0.193 | -8.32E-02 | 1.23E-01 | 2.68E+00 | 2.74E+00 | 4.94E-01 | no | | intergenic_variant | MODIFIER |
| 7 | 40437881 | AX-97798637 | 0 | A | C | 0.333 | 6.19E-02 | 9.10E-02 | 2.62E+00 | 2.70E+00 | 4.94E-01 | no | Gm28807 | intron_variant,non_coding_transcript_variant | MODIFIER |
| 9 | 37869343 | AX-170082148 | 0 | C | G | 0.5 | -5.12E-02 | 7.61E-02 | 2.74E+00 | 2.84E+00 | 4.94E-01 | no | | intergenic_variant | MODIFIER |
| 17 | 55704132 | AX-170331032 | 0 | C | T | 0.053 | -1.09E-01 | 1.63E-01 | 2.71E+00 | 2.84E+00 | 4.94E-01 | no | Pot1b | intron_variant | MODIFIER |
| 10 | 92058379 | AX-170169587 | 1 | T | G | 0.295 | -5.70E-02 | 8.23E-02 | 2.54E+00 | 2.61E+00 | 4.94E-01 | no | | intergenic_variant | MODIFIER |
| 11 | 22344184 | AX-97802331 | 1 | G | A | 0.045 | 1.27E-01 | 1.89E-01 | 2.69E+00 | 2.83E+00 | 4.94E-01 | no | Ehbp1 | upstream_gene_variant | MODIFIER |
| 7 | 36744669 | AX-97784103 | 0 | C | A | 0.342 | 6.07E-02 | 8.77E-02 | 2.53E+00 | 2.62E+00 | 4.94E-01 | no | Tshz3 | intron_variant | MODIFIER |
| 7 | 36833259 | AX-170172821 | 0 | G | A | 0.342 | 6.07E-02 | 8.77E-02 | 2.53E+00 | 2.62E+00 | 4.94E-01 | no | | intergenic_variant | MODIFIER |
| 1 | 174687861 | AX-170526198 | 1 | C | T | 0.232 | 8.46E-02 | 1.21E-01 | 2.51E+00 | 2.57E+00 | 4.94E-01 | no | Fmn2 | intron_variant | MODIFIER |

| | | | | | | | | | | | | | | | |
|---|---|---|---|---|---|---|---|---|---|---|---|---|---|---|---|
| 2 | 179104545 | AX-170476712 | 0 | T | C | 0.053 | 1.12E-01 | 1.67E-01 | 2.65E+00 | 2.79E+00 | 4.94E-01 | no | | intergenic_variant | MODIFIER |
| 1 | 44412034 | AX-97828007 | 1 | C | T | 0.098 | -1.42E-01 | 2.11E-01 | 2.75E+00 | 2.84E+00 | 4.94E-01 | no | | intergenic_variant | MODIFIER |
| 10 | 93416548 | AX-170142946 | 1 | A | G | 0.286 | -5.69E-02 | 8.22E-02 | 2.54E+00 | 2.60E+00 | 4.94E-01 | no | | intergenic_variant | MODIFIER |
| 5 | 78343139 | AX-170134037 | 2 | C | A | 0.445 | 7.15E-02 | 1.06E-01 | 2.71E+00 | 2.77E+00 | 4.95E-01 | no | | intergenic_variant | MODIFIER |
| 5 | 37383695 | AX-170446459 | 0 | G | T | 0.333 | 7.34E-02 | 1.09E-01 | 2.74E+00 | 2.80E+00 | 4.95E-01 | no | Evc2 | intron_variant | MODIFIER |
| 13 | 25362261 | AX-170461398 | 0 | T | A | 0.167 | -8.43E-02 | 1.20E-01 | 2.49E+00 | 2.56E+00 | 4.95E-01 | no | | regulatory_region_variant | MODIFIER |
| 7 | 36470535 | AX-170143662 | 0 | G | A | 0.351 | 6.10E-02 | 8.84E-02 | 2.54E+00 | 2.63E+00 | 4.95E-01 | no | | intergenic_variant | MODIFIER |
| 9 | 57484305 | AX-170323151 | 1 | C | G | 0.214 | -7.76E-02 | 1.15E-01 | 2.79E+00 | 2.88E+00 | 4.95E-01 | no | Gm6166 | non_coding_transcript_exon_variant | MODIFIER |
| 4 | 153222148 | AX-170394384 | 0 | A | G | 0.096 | 1.24E-01 | 1.84E-01 | 2.67E+00 | 2.76E+00 | 4.95E-01 | no | Gm13174 | downstream_gene_variant | MODIFIER |
| 15 | 85364746 | AX-97830664 | 0 | G | T | 0.5 | 4.81E-02 | 6.86E-02 | 2.49E+00 | 2.57E+00 | 4.95E-01 | no | Atxn10 | intron_variant | MODIFIER |
| 1 | 20509040 | AX-170395339 | 1 | C | T | 0.411 | 5.79E-02 | 8.42E-02 | 2.57E+00 | 2.61E+00 | 4.95E-01 | no | Pkhd1 | missense_variant | MODERATE |
| 1 | 21599835 | AX-97790072 | 1 | C | T | 0.411 | 5.79E-02 | 8.42E-02 | 2.57E+00 | 2.61E+00 | 4.95E-01 | no | Kcnq5 | intron_variant | MODIFIER |
| 8 | 14721328 | AX-97794004 | 1 | C | T | 0.241 | -5.22E-02 | 7.60E-02 | 2.55E+00 | 2.64E+00 | 4.95E-01 | no | Dlgap2 | intron_variant | MODIFIER |
| 10 | 123620019 | AX-170519153 | 1 | G | A | 0.045 | 8.37E-02 | 1.24E-01 | 2.58E+00 | 2.74E+00 | 4.95E-01 | no | Fam19a2 | intron_variant | MODIFIER |
| 18 | 81019539 | AX-170302460 | 1 | G | A | 0.295 | -5.63E-02 | 8.26E-02 | 2.58E+00 | 2.69E+00 | 4.95E-01 | no | | intergenic_variant | MODIFIER |
| 11 | 18841664 | AX-97814384 | 0 | G | A | 0.053 | 1.19E-01 | 1.79E-01 | 2.67E+00 | 2.83E+00 | 4.95E-01 | no | 8430419K02Rik | downstream_gene_variant | MODIFIER |
| 12 | 24393225 | AX-170369290 | 0 | A | T | 0.044 | -1.15E-01 | 1.72E-01 | 2.67E+00 | 2.79E+00 | 4.95E-01 | no | | intergenic_variant | MODIFIER |
| 18 | 62812980 | AX-170160199 | 1 | A | T | 0.196 | -7.51E-02 | 1.11E-01 | 2.64E+00 | 2.72E+00 | 4.95E-01 | no | | intergenic_variant | MODIFIER |
| 8 | 30859839 | AX-170431718 | 1 | A | G | 0.17 | 6.70E-02 | 9.99E-02 | 2.75E+00 | 2.85E+00 | 4.95E-01 | no | | intergenic_variant | MODIFIER |
| 5 | 28519615 | AX-170158362 | 1 | C | A | 0.384 | 6.57E-02 | 9.73E-02 | 2.64E+00 | 2.73E+00 | 4.96E-01 | no | 9530036O11Rik | intron_variant,non_coding_transcript_variant | MODIFIER |
| 7 | 112415473 | AX-170437329 | 0 | C | T | 0.456 | -6.60E-02 | 9.75E-02 | 2.85E+00 | 2.92E+00 | 4.96E-01 | no | Mical2 | downstream_gene_variant | MODIFIER |
| 1 | 44566761 | AX-97811477 | 0 | G | T | 0.096 | -1.42E-01 | 2.11E-01 | 2.73E+00 | 2.82E+00 | 4.96E-01 | no | Gulp1 | intron_variant | MODIFIER |
| 9 | 117830889 | AX-170441417 | 1 | A | T | 0.125 | -6.44E-02 | 9.53E-02 | 2.60E+00 | 2.72E+00 | 4.96E-01 | no | Gm45881 | intron_variant,non_coding_transcript_variant | MODIFIER |
| 8 | 73766158 | AX-170320073 | 0 | C | T | 0.202 | -7.95E-02 | 1.19E-01 | 2.67E+00 | 2.77E+00 | 4.96E-01 | no | | intergenic_variant | MODIFIER |
| 11 | 29673434 | AX-170187671 | 1 | G | C | 0.071 | 1.45E-01 | 2.15E-01 | 2.71E+00 | 2.78E+00 | 4.96E-01 | Yes | | intergenic_variant | MODIFIER |
| 1 | 30863698 | AX-97834872 | 1 | T | A | 0.411 | 5.65E-02 | 8.42E-02 | 2.72E+00 | 2.80E+00 | 4.96E-01 | no | Phf3 | upstream_gene_variant | MODIFIER |
| 5 | 31058720 | AX-170024613 | 1 | G | A | 0.366 | 7.07E-02 | 1.05E-01 | 2.73E+00 | 2.79E+00 | 4.96E-01 | no | Atraid | downstream_gene_variant | MODIFIER |
| 1 | 19473729 | AX-170327558 | 1 | T | C | 0.411 | 5.70E-02 | 8.33E-02 | 2.57E+00 | 2.62E+00 | 4.96E-01 | no | | regulatory_region_variant | MODIFIER |
| 1 | 123430360 | AX-170489739 | 0 | C | T | 0.57 | 5.51E-02 | 7.92E-02 | 2.50E+00 | 2.57E+00 | 4.97E-01 | no | Dpp10 | intron_variant | MODIFIER |
| 1 | 191280708 | AX-170321964 | 0 | T | C | 0.263 | 6.73E-02 | 9.42E-02 | 2.43E+00 | 2.52E+00 | 4.97E-01 | no | Gm37168 | intron_variant,non_coding_transcript_variant | MODIFIER |
| 10 | 28651381 | AX-170298261 | 0 | C | G | 0.202 | 6.50E-02 | 9.73E-02 | 2.77E+00 | 2.87E+00 | 4.97E-01 | no | | intergenic_variant | MODIFIER |
| 8 | 78906251 | AX-170192955 | 0 | A | G | 0.202 | -8.13E-02 | 1.22E-01 | 2.72E+00 | 2.81E+00 | 4.97E-01 | no | | intergenic_variant | MODIFIER |
| 9 | 19120099 | AX-97805180 | 0 | T | C | 0.474 | 6.17E-02 | 8.85E-02 | 2.49E+00 | 2.57E+00 | 4.97E-01 | no | Olfr836 | upstream_gene_variant | MODIFIER |
| 18 | 81015065 | AX-170319775 | 1 | A | T | 0.286 | -5.90E-02 | 8.62E-02 | 2.54E+00 | 2.65E+00 | 4.97E-01 | no | | intergenic_variant | MODIFIER |
| 9 | 91856356 | AX-170180824 | 0 | A | G | 0.132 | -8.29E-02 | 1.22E-01 | 2.60E+00 | 2.69E+00 | 4.97E-01 | no | | intergenic_variant | MODIFIER |
| 1 | 16737386 | AX-170000005 | 0 | A | C | 0.377 | 5.57E-02 | 8.21E-02 | 2.61E+00 | 2.65E+00 | 4.97E-01 | no | | intergenic_variant | MODIFIER |
| 16 | 29499460 | AX-170012706 | 0 | T | C | 0.491 | 5.50E-02 | 8.04E-02 | 2.56E+00 | 2.62E+00 | 4.97E-01 | no | Atp13a4 | intron_variant | MODIFIER |
| 1 | 19864482 | AX-170317542 | 0 | G | A | 0.412 | 5.73E-02 | 8.37E-02 | 2.56E+00 | 2.61E+00 | 4.97E-01 | no | | intergenic_variant | MODIFIER |
| 1 | 20066181 | AX-170224454 | 0 | C | T | 0.412 | 5.73E-02 | 8.37E-02 | 2.56E+00 | 2.61E+00 | 4.97E-01 | no | Pkhd1 | intron_variant | MODIFIER |
| 16 | 45853750 | AX-97814253 | 0 | A | G | 0.509 | 5.56E-02 | 8.12E-02 | 2.57E+00 | 2.61E+00 | 4.97E-01 | no | Phldb2 | intron_variant | MODIFIER |
| 3 | 32410105 | AX-97834541 | 1 | T | C | 0.179 | 6.31E-02 | 9.45E-02 | 2.75E+00 | 2.84E+00 | 4.97E-01 | no | Pik3ca | intron_variant | MODIFIER |
| 1 | 192534110 | AX-170402253 | 0 | A | G | 0.272 | 6.68E-02 | 9.19E-02 | 2.38E+00 | 2.47E+00 | 4.97E-01 | no | Hhat | intron_variant | MODIFIER |
| 7 | 24877464 | AX-170462969 | 0 | G | A | 0.404 | 6.41E-02 | 9.37E-02 | 2.54E+00 | 2.64E+00 | 4.97E-01 | no | Dmrtc2 | 3_prime_UTR_variant | MODIFIER |
| 7 | 24878158 | AX-170454269 | 0 | T | C | 0.404 | 6.41E-02 | 9.37E-02 | 2.54E+00 | 2.64E+00 | 4.97E-01 | no | Dmrtc2 | downstream_gene_variant | MODIFIER |
| 7 | 24879966 | AX-170393906 | 0 | A | G | 0.404 | 6.41E-02 | 9.37E-02 | 2.54E+00 | 2.64E+00 | 4.97E-01 | no | Dmrtc2 | downstream_gene_variant | MODIFIER |
| 7 | 24881446 | AX-97788678 | 0 | C | T | 0.404 | 6.41E-02 | 9.37E-02 | 2.54E+00 | 2.64E+00 | 4.97E-01 | no | Dmrtc2 | downstream_gene_variant | MODIFIER |
| 7 | 24912751 | AX-97809725 | 0 | G | C | 0.404 | 6.41E-02 | 9.37E-02 | 2.54E+00 | 2.64E+00 | 4.97E-01 | no | Arhgef1 | synonymous_variant | LOW |
| 7 | 24937916 | AX-170268456 | 0 | C | T | 0.404 | 6.41E-02 | 9.37E-02 | 2.54E+00 | 2.64E+00 | 4.97E-01 | no | Gm4881 | intron_variant | MODIFIER |
| 7 | 25026268 | AX-170244796 | 0 | A | G | 0.404 | 6.41E-02 | 9.37E-02 | 2.54E+00 | 2.64E+00 | 4.97E-01 | no | Grik5 | intron_variant | MODIFIER |
| 7 | 25031225 | AX-170467847 | 0 | T | C | 0.404 | 6.41E-02 | 9.37E-02 | 2.54E+00 | 2.64E+00 | 4.97E-01 | no | Grik5 | intron_variant | MODIFIER |
| 7 | 25031275 | AX-97840436 | 0 | T | C | 0.404 | 6.41E-02 | 9.37E-02 | 2.54E+00 | 2.64E+00 | 4.97E-01 | no | Grik5 | intron_variant | MODIFIER |
| 7 | 25138715 | AX-97797495 | 0 | G | A | 0.404 | 6.41E-02 | 9.37E-02 | 2.54E+00 | 2.64E+00 | 4.97E-01 | no | Pou2f2 | intron_variant,non_coding_transcript_variant | MODIFIER |
| 7 | 25284796 | AX-170311191 | 0 | A | T | 0.404 | 6.41E-02 | 9.37E-02 | 2.54E+00 | 2.64E+00 | 4.97E-01 | no | Cic | intron_variant | MODIFIER |
| 7 | 25341373 | AX-97835566 | 0 | G | A | 0.404 | 6.41E-02 | 9.37E-02 | 2.54E+00 | 2.64E+00 | 4.97E-01 | no | Megf8 | intron_variant | MODIFIER |

| | | | | | | | | | | | | | | | |
|---|---|---|---|---|---|---|---|---|---|---|---|---|---|---|---|
| 7 | 25344211 | AX-170184808 | 0 | G | A | 0.404 | 6.41E-02 | 9.37E-02 | 2.54E+00 | 2.64E+00 | 4.97E-01 | no | Megf8 | intron_variant | MODIFIER |
| 7 | 25349028 | AX-170076873 | 0 | T | C | 0.404 | 6.41E-02 | 9.37E-02 | 2.54E+00 | 2.64E+00 | 4.97E-01 | no | Megf8 | missense_variant | MODERATE |
| 7 | 25394498 | AX-97781030 | 0 | G | T | 0.404 | 6.41E-02 | 9.37E-02 | 2.54E+00 | 2.64E+00 | 4.97E-01 | no | Lipe | intron_variant | MODIFIER |
| 7 | 25400007 | AX-97821444 | 0 | T | C | 0.404 | 6.41E-02 | 9.37E-02 | 2.54E+00 | 2.64E+00 | 4.97E-01 | no | Lipe | upstream_gene_variant | MODIFIER |
| 7 | 25451361 | AX-170436729 | 0 | G | A | 0.404 | 6.41E-02 | 9.37E-02 | 2.54E+00 | 2.64E+00 | 4.97E-01 | no | 4732471J01Rik | intron_variant,non_coding_transcript_variant | MODIFIER |
| 7 | 25459491 | AX-97828842 | 0 | C | T | 0.404 | 6.41E-02 | 9.37E-02 | 2.54E+00 | 2.64E+00 | 4.97E-01 | no | Ceacam1 | downstream_gene_variant | MODIFIER |
| 7 | 25465095 | AX-170524455 | 0 | T | C | 0.404 | 6.41E-02 | 9.37E-02 | 2.54E+00 | 2.64E+00 | 4.97E-01 | no | Ceacam1 | intron_variant | MODIFIER |
| 7 | 25487604 | AX-97789614 | 0 | A | T | 0.404 | 6.41E-02 | 9.37E-02 | 2.54E+00 | 2.64E+00 | 4.97E-01 | no | 4732471J01Rik | intron_variant,non_coding_transcript_variant | MODIFIER |
| 7 | 25513512 | AX-170245210 | 0 | C | A | 0.404 | 6.41E-02 | 9.37E-02 | 2.54E+00 | 2.64E+00 | 4.97E-01 | no | Ceacam2 | downstream_gene_variant | MODIFIER |
| 7 | 25577912 | AX-170076879 | 0 | C | T | 0.404 | 6.41E-02 | 9.37E-02 | 2.54E+00 | 2.64E+00 | 4.97E-01 | no | Gm29918 | upstream_gene_variant | MODIFIER |
| 7 | 25635329 | AX-97830256 | 0 | T | C | 0.404 | 6.41E-02 | 9.37E-02 | 2.54E+00 | 2.64E+00 | 4.97E-01 | no | Bckdha | intron_variant | MODIFIER |
| 7 | 25636953 | AX-170301066 | 0 | A | C | 0.404 | 6.41E-02 | 9.37E-02 | 2.54E+00 | 2.64E+00 | 4.97E-01 | no | Bckdha | intron_variant | MODIFIER |
| 7 | 25678857 | AX-97850821 | 0 | C | T | 0.404 | 6.41E-02 | 9.37E-02 | 2.54E+00 | 2.64E+00 | 4.97E-01 | no | Tmem91 | upstream_gene_variant | MODIFIER |
| 7 | 25851998 | AX-170028704 | 0 | G | A | 0.404 | 6.41E-02 | 9.37E-02 | 2.54E+00 | 2.64E+00 | 4.97E-01 | no | | intergenic_variant | MODIFIER |
| 7 | 25924091 | AX-97815668 | 0 | A | G | 0.404 | 6.41E-02 | 9.37E-02 | 2.54E+00 | 2.64E+00 | 4.97E-01 | no | Cyp2b10 | intron_variant | MODIFIER |
| 7 | 26018705 | AX-170296208 | 0 | T | C | 0.404 | 6.41E-02 | 9.37E-02 | 2.54E+00 | 2.64E+00 | 4.97E-01 | no | Cyp2b27-ps | intron_variant,non_coding_transcript_variant | MODIFIER |
| 11 | 47552296 | AX-97819711 | 0 | T | C | 0.096 | -1.01E-01 | 1.51E-01 | 2.63E+00 | 2.77E+00 | 4.97E-01 | no | Sgcd | intron_variant | MODIFIER |
| 8 | 15459335 | AX-170377471 | 1 | A | G | 0.232 | -5.19E-02 | 7.58E-02 | 2.54E+00 | 2.63E+00 | 4.97E-01 | no | | intergenic_variant | MODIFIER |
| 18 | 64986216 | AX-170287066 | 1 | G | T | 0.411 | 6.36E-02 | 9.11E-02 | 2.49E+00 | 2.54E+00 | 4.98E-01 | no | Nedd4l | intron_variant | MODIFIER |
| 10 | 93675559 | AX-170157712 | 0 | T | C | 0.289 | -5.81E-02 | 8.31E-02 | 2.48E+00 | 2.55E+00 | 4.98E-01 | no | Ntn4 | intron_variant | MODIFIER |
| 10 | 93281582 | AX-170140639 | 1 | A | T | 0.295 | -5.78E-02 | 8.38E-02 | 2.53E+00 | 2.59E+00 | 4.98E-01 | no | Elk3 | intron_variant | MODIFIER |
| 18 | 43409707 | AX-170394902 | 0 | A | C | 0.07 | 1.41E-01 | 2.11E-01 | 2.76E+00 | 2.88E+00 | 4.98E-01 | no | Dpysl3 | intron_variant | MODIFIER |
| 20 | 67272269 | AX-97783592 | 1 | G | T | 0.25 | 4.20E-02 | 6.28E-02 | 2.67E+00 | 2.76E+00 | 4.98E-01 | no | | intergenic_variant | MODIFIER |
| 16 | 52136475 | AX-170288750 | 0 | T | C | 0.474 | -5.77E-02 | 8.31E-02 | 2.51E+00 | 2.55E+00 | 4.98E-01 | no | Cblb | intron_variant | MODIFIER |
| 1 | 68202080 | AX-170504008 | 1 | C | T | 0.116 | 1.19E-01 | 1.77E-01 | 2.61E+00 | 2.73E+00 | 4.98E-01 | no | Erbb4 | intron_variant | MODIFIER |
| 12 | 91914385 | AX-97826605 | 2 | T | G | 0.073 | 1.43E-01 | 2.15E-01 | 2.74E+00 | 2.81E+00 | 4.98E-01 | no | | intergenic_variant | MODIFIER |
| 18 | 57936287 | AX-170099741 | 0 | T | A | 0.412 | 5.57E-02 | 8.17E-02 | 2.93E+00 | 3.03E+00 | 4.98E-01 | no | Slc12a2 | intron_variant | MODIFIER |
| 7 | 37427821 | AX-170176359 | 1 | G | T | 0.339 | 6.24E-02 | 9.24E-02 | 2.59E+00 | 2.67E+00 | 4.99E-01 | no | | intergenic_variant | MODIFIER |
| 4 | 85588989 | AX-170320050 | 0 | G | A | 0.333 | 7.56E-02 | 1.13E-01 | 2.69E+00 | 2.76E+00 | 4.99E-01 | no | Sh3gl2 | intron_variant | MODIFIER |
| 4 | 73471568 | AX-168401955 | 0 | G | A | 0.456 | -5.68E-02 | 8.29E-02 | 2.54E+00 | 2.61E+00 | 4.99E-01 | no | | intergenic_variant | MODIFIER |
| 2 | 77755214 | AX-170018107 | 1 | G | T | 0.071 | 1.19E-01 | 1.79E-01 | 2.67E+00 | 2.80E+00 | 4.99E-01 | no | Zfp385b | intron_variant | MODIFIER |
| 1 | 69087443 | AX-170041557 | 0 | G | A | 0.149 | 1.21E-01 | 1.82E-01 | 2.68E+00 | 2.80E+00 | 4.99E-01 | no | Erbb4 | intron_variant | MODIFIER |
| 6 | 146770846 | AX-170028479 | 0 | C | T | 0.07 | -1.36E-01 | 2.03E-01 | 2.65E+00 | 2.76E+00 | 4.99E-01 | no | Stk38l | intron_variant | MODIFIER |
| 1 | 19331210 | AX-170317360 | 0 | G | A | 0.404 | 5.70E-02 | 8.41E-02 | 2.59E+00 | 2.64E+00 | 4.99E-01 | no | Gm4849 | non_coding_transcript_exon_variant | MODIFIER |
| 1 | 19473322 | AX-170221945 | 0 | T | C | 0.404 | 5.70E-02 | 8.41E-02 | 2.59E+00 | 2.64E+00 | 4.99E-01 | no | | intergenic_variant | MODIFIER |
| 1 | 19473397 | AX-170185791 | 0 | G | A | 0.404 | 5.70E-02 | 8.41E-02 | 2.59E+00 | 2.64E+00 | 4.99E-01 | no | | intergenic_variant | MODIFIER |
| 1 | 19615328 | AX-170512480 | 0 | G | A | 0.404 | 5.70E-02 | 8.41E-02 | 2.59E+00 | 2.64E+00 | 4.99E-01 | no | Gm38382 | non_coding_transcript_exon_variant | MODIFIER |
| 1 | 19633600 | AX-170000035 | 0 | G | A | 0.404 | 5.70E-02 | 8.41E-02 | 2.59E+00 | 2.64E+00 | 4.99E-01 | no | | intergenic_variant | MODIFIER |
| 1 | 19638906 | AX-97832487 | 0 | G | A | 0.404 | 5.70E-02 | 8.41E-02 | 2.59E+00 | 2.64E+00 | 4.99E-01 | no | | intergenic_variant | MODIFIER |
| 1 | 19680313 | AX-97846315 | 0 | T | C | 0.404 | 5.70E-02 | 8.41E-02 | 2.59E+00 | 2.64E+00 | 4.99E-01 | no | | intergenic_variant | MODIFIER |
| 1 | 19705932 | AX-170109453 | 0 | T | C | 0.404 | 5.70E-02 | 8.41E-02 | 2.59E+00 | 2.64E+00 | 4.99E-01 | no | | intergenic_variant | MODIFIER |
| 1 | 19706707 | AX-170192804 | 0 | T | C | 0.404 | 5.70E-02 | 8.41E-02 | 2.59E+00 | 2.64E+00 | 4.99E-01 | no | | intergenic_variant | MODIFIER |
| 1 | 19715283 | AX-170405240 | 0 | C | T | 0.404 | 5.70E-02 | 8.41E-02 | 2.59E+00 | 2.64E+00 | 4.99E-01 | no | | intergenic_variant | MODIFIER |
| 1 | 19715318 | AX-170261047 | 0 | A | T | 0.404 | 5.70E-02 | 8.41E-02 | 2.59E+00 | 2.64E+00 | 4.99E-01 | no | | intergenic_variant | MODIFIER |
| 1 | 19715907 | AX-170434383 | 0 | G | A | 0.404 | 5.70E-02 | 8.41E-02 | 2.59E+00 | 2.64E+00 | 4.99E-01 | no | | intergenic_variant | MODIFIER |
| 1 | 19732512 | AX-170234614 | 0 | A | G | 0.404 | 5.70E-02 | 8.41E-02 | 2.59E+00 | 2.64E+00 | 4.99E-01 | no | | intergenic_variant | MODIFIER |
| 1 | 19752395 | AX-97787411 | 0 | T | C | 0.404 | 5.70E-02 | 8.41E-02 | 2.59E+00 | 2.64E+00 | 4.99E-01 | no | | intergenic_variant | MODIFIER |
| 1 | 19759789 | AX-170085966 | 0 | G | A | 0.404 | 5.70E-02 | 8.41E-02 | 2.59E+00 | 2.64E+00 | 4.99E-01 | no | | intergenic_variant | MODIFIER |
| 1 | 19761898 | AX-170040545 | 0 | A | G | 0.404 | 5.70E-02 | 8.41E-02 | 2.59E+00 | 2.64E+00 | 4.99E-01 | no | | intergenic_variant | MODIFIER |
| 1 | 19765671 | AX-170281535 | 0 | C | G | 0.404 | 5.70E-02 | 8.41E-02 | 2.59E+00 | 2.64E+00 | 4.99E-01 | no | | intergenic_variant | MODIFIER |
| 1 | 19774749 | AX-170091423 | 0 | T | C | 0.404 | 5.70E-02 | 8.41E-02 | 2.59E+00 | 2.64E+00 | 4.99E-01 | no | | intergenic_variant | MODIFIER |
| 1 | 19777242 | AX-170354033 | 0 | G | A | 0.404 | 5.70E-02 | 8.41E-02 | 2.59E+00 | 2.64E+00 | 4.99E-01 | no | | intergenic_variant | MODIFIER |
| 1 | 19783423 | AX-170210316 | 0 | T | C | 0.404 | 5.70E-02 | 8.41E-02 | 2.59E+00 | 2.64E+00 | 4.99E-01 | no | | intergenic_variant | MODIFIER |
| 1 | 19795922 | AX-170398306 | 0 | G | T | 0.404 | 5.70E-02 | 8.41E-02 | 2.59E+00 | 2.64E+00 | 4.99E-01 | no | | intergenic_variant | MODIFIER |

| | | | | | | | | | | | | | | | |
|---|---|---|---|---|---|---|---|---|---|---|---|---|---|---|---|
| 1 | 19824304 | AX-170416003 | 0 | A | G | 0.404 | 5.70E-02 | 8.41E-02 | 2.59E+00 | 2.64E+00 | 4.99E-01 | no | | intergenic_variant | MODIFIER |
| 1 | 19832993 | AX-170195427 | 0 | C | A | 0.404 | 5.70E-02 | 8.41E-02 | 2.59E+00 | 2.64E+00 | 4.99E-01 | no | | intergenic_variant | MODIFIER |
| 1 | 19846802 | AX-170421603 | 0 | G | A | 0.404 | 5.70E-02 | 8.41E-02 | 2.59E+00 | 2.64E+00 | 4.99E-01 | no | | intergenic_variant | MODIFIER |
| 1 | 19847117 | AX-170423966 | 0 | T | C | 0.404 | 5.70E-02 | 8.41E-02 | 2.59E+00 | 2.64E+00 | 4.99E-01 | no | | intergenic_variant | MODIFIER |
| 1 | 19851427 | AX-170529791 | 0 | G | A | 0.404 | 5.70E-02 | 8.41E-02 | 2.59E+00 | 2.64E+00 | 4.99E-01 | no | | intergenic_variant | MODIFIER |
| 1 | 19861107 | AX-97798505 | 0 | T | C | 0.404 | 5.70E-02 | 8.41E-02 | 2.59E+00 | 2.64E+00 | 4.99E-01 | no | | intergenic_variant | MODIFIER |
| 1 | 19862013 | AX-170109373 | 0 | T | C | 0.404 | 5.70E-02 | 8.41E-02 | 2.59E+00 | 2.64E+00 | 4.99E-01 | no | | intergenic_variant | MODIFIER |
| 1 | 19877796 | AX-170155749 | 0 | C | T | 0.404 | 5.70E-02 | 8.41E-02 | 2.59E+00 | 2.64E+00 | 4.99E-01 | no | | intergenic_variant | MODIFIER |
| 1 | 19878145 | AX-170445365 | 0 | T | C | 0.404 | 5.70E-02 | 8.41E-02 | 2.59E+00 | 2.64E+00 | 4.99E-01 | no | | intergenic_variant | MODIFIER |
| 1 | 19887304 | AX-170290784 | 0 | C | T | 0.404 | 5.70E-02 | 8.41E-02 | 2.59E+00 | 2.64E+00 | 4.99E-01 | no | | intergenic_variant | MODIFIER |
| 1 | 19906494 | AX-170406082 | 0 | C | T | 0.404 | 5.70E-02 | 8.41E-02 | 2.59E+00 | 2.64E+00 | 4.99E-01 | no | | intergenic_variant | MODIFIER |
| 1 | 19906532 | AX-97846571 | 0 | A | G | 0.404 | 5.70E-02 | 8.41E-02 | 2.59E+00 | 2.64E+00 | 4.99E-01 | no | | intergenic_variant | MODIFIER |
| 1 | 19931415 | AX-170355778 | 0 | G | C | 0.404 | 5.70E-02 | 8.41E-02 | 2.59E+00 | 2.64E+00 | 4.99E-01 | no | | intergenic_variant | MODIFIER |
| 1 | 19939334 | AX-170228729 | 0 | A | T | 0.404 | 5.70E-02 | 8.41E-02 | 2.59E+00 | 2.64E+00 | 4.99E-01 | no | | intergenic_variant | MODIFIER |
| 1 | 19942467 | AX-170198960 | 0 | A | G | 0.404 | 5.70E-02 | 8.41E-02 | 2.59E+00 | 2.64E+00 | 4.99E-01 | no | | intergenic_variant | MODIFIER |
| 1 | 19945577 | AX-170257707 | 0 | C | G | 0.404 | 5.70E-02 | 8.41E-02 | 2.59E+00 | 2.64E+00 | 4.99E-01 | no | | intergenic_variant | MODIFIER |
| 1 | 19955143 | AX-170371278 | 0 | C | T | 0.404 | 5.70E-02 | 8.41E-02 | 2.59E+00 | 2.64E+00 | 4.99E-01 | no | | intergenic_variant | MODIFIER |
| 1 | 19963706 | AX-170424073 | 0 | C | T | 0.404 | 5.70E-02 | 8.41E-02 | 2.59E+00 | 2.64E+00 | 4.99E-01 | no | | intergenic_variant | MODIFIER |
| 1 | 19964836 | AX-170284567 | 0 | C | A | 0.404 | 5.70E-02 | 8.41E-02 | 2.59E+00 | 2.64E+00 | 4.99E-01 | no | | intergenic_variant | MODIFIER |
| 1 | 19965820 | AX-170487694 | 0 | T | G | 0.404 | 5.70E-02 | 8.41E-02 | 2.59E+00 | 2.64E+00 | 4.99E-01 | no | | intergenic_variant | MODIFIER |
| 1 | 19969889 | AX-170451026 | 0 | G | A | 0.404 | 5.70E-02 | 8.41E-02 | 2.59E+00 | 2.64E+00 | 4.99E-01 | no | | intergenic_variant | MODIFIER |
| 1 | 19979128 | AX-170471502 | 0 | A | G | 0.404 | 5.70E-02 | 8.41E-02 | 2.59E+00 | 2.64E+00 | 4.99E-01 | no | | intergenic_variant | MODIFIER |
| 1 | 20016491 | AX-170214371 | 0 | A | C | 0.404 | 5.70E-02 | 8.41E-02 | 2.59E+00 | 2.64E+00 | 4.99E-01 | no | | intergenic_variant | MODIFIER |
| 1 | 20016527 | AX-170248865 | 0 | A | C | 0.404 | 5.70E-02 | 8.41E-02 | 2.59E+00 | 2.64E+00 | 4.99E-01 | no | | intergenic_variant | MODIFIER |
| 1 | 20035303 | AX-97797430 | 0 | C | T | 0.404 | 5.70E-02 | 8.41E-02 | 2.59E+00 | 2.64E+00 | 4.99E-01 | no | | intergenic_variant | MODIFIER |
| 1 | 20035602 | AX-170439376 | 0 | G | T | 0.404 | 5.70E-02 | 8.41E-02 | 2.59E+00 | 2.64E+00 | 4.99E-01 | no | | intergenic_variant | MODIFIER |
| 1 | 20050469 | AX-170091425 | 0 | G | A | 0.404 | 5.70E-02 | 8.41E-02 | 2.59E+00 | 2.64E+00 | 4.99E-01 | no | | intergenic_variant | MODIFIER |
| 1 | 20072148 | AX-170405345 | 0 | C | G | 0.404 | 5.70E-02 | 8.41E-02 | 2.59E+00 | 2.64E+00 | 4.99E-01 | no | Pkhd1 | intron_variant | MODIFIER |
| 1 | 20073948 | AX-97854920 | 0 | C | T | 0.404 | 5.70E-02 | 8.41E-02 | 2.59E+00 | 2.64E+00 | 4.99E-01 | no | Pkhd1 | intron_variant | MODIFIER |
| 1 | 20111580 | AX-170204194 | 0 | C | T | 0.404 | 5.70E-02 | 8.41E-02 | 2.59E+00 | 2.64E+00 | 4.99E-01 | no | Pkhd1 | intron_variant | MODIFIER |
| 1 | 20137532 | AX-97822973 | 0 | G | A | 0.404 | 5.70E-02 | 8.41E-02 | 2.59E+00 | 2.64E+00 | 4.99E-01 | no | Pkhd1 | intron_variant | MODIFIER |
| 1 | 20149687 | AX-170000040 | 0 | T | C | 0.404 | 5.70E-02 | 8.41E-02 | 2.59E+00 | 2.64E+00 | 4.99E-01 | no | Pkhd1 | intron_variant | MODIFIER |
| 1 | 20158019 | AX-170454829 | 0 | A | T | 0.404 | 5.70E-02 | 8.41E-02 | 2.59E+00 | 2.64E+00 | 4.99E-01 | no | Pkhd1 | intron_variant | MODIFIER |
| 1 | 20163903 | AX-170000036 | 0 | C | A | 0.404 | 5.70E-02 | 8.41E-02 | 2.59E+00 | 2.64E+00 | 4.99E-01 | no | Pkhd1 | intron_variant | MODIFIER |
| 1 | 20197671 | AX-97844482 | 0 | A | T | 0.404 | 5.70E-02 | 8.41E-02 | 2.59E+00 | 2.64E+00 | 4.99E-01 | no | Pkhd1 | intron_variant | MODIFIER |
| 1 | 20201277 | AX-170000038 | 0 | A | G | 0.404 | 5.70E-02 | 8.41E-02 | 2.59E+00 | 2.64E+00 | 4.99E-01 | no | Pkhd1 | synonymous_variant | LOW |
| 1 | 20203398 | AX-170040558 | 0 | G | A | 0.404 | 5.70E-02 | 8.41E-02 | 2.59E+00 | 2.64E+00 | 4.99E-01 | no | Pkhd1 | intron_variant | MODIFIER |
| 1 | 20228307 | AX-97830108 | 0 | A | G | 0.404 | 5.70E-02 | 8.41E-02 | 2.59E+00 | 2.64E+00 | 4.99E-01 | no | Pkhd1 | intron_variant | MODIFIER |
| 1 | 20288127 | AX-170166862 | 0 | A | G | 0.404 | 5.70E-02 | 8.41E-02 | 2.59E+00 | 2.64E+00 | 4.99E-01 | no | Pkhd1 | intron_variant | MODIFIER |
| 1 | 20333064 | AX-170167028 | 0 | T | A | 0.404 | 5.70E-02 | 8.41E-02 | 2.59E+00 | 2.64E+00 | 4.99E-01 | no | Pkhd1 | intron_variant | MODIFIER |
| 1 | 20357813 | AX-170136356 | 0 | C | A | 0.404 | 5.70E-02 | 8.41E-02 | 2.59E+00 | 2.64E+00 | 4.99E-01 | no | Pkhd1 | intron_variant | MODIFIER |
| 1 | 20392340 | AX-170117823 | 0 | G | A | 0.404 | 5.70E-02 | 8.41E-02 | 2.59E+00 | 2.64E+00 | 4.99E-01 | no | Pkhd1 | intron_variant | MODIFIER |
| 1 | 20395737 | AX-170124863 | 0 | G | A | 0.404 | 5.70E-02 | 8.41E-02 | 2.59E+00 | 2.64E+00 | 4.99E-01 | no | Pkhd1 | intron_variant | MODIFIER |
| 1 | 20397625 | AX-170411450 | 0 | G | A | 0.404 | 5.70E-02 | 8.41E-02 | 2.59E+00 | 2.64E+00 | 4.99E-01 | no | Pkhd1 | intron_variant | MODIFIER |
| 1 | 20442341 | AX-97791476 | 0 | G | A | 0.404 | 5.70E-02 | 8.41E-02 | 2.59E+00 | 2.64E+00 | 4.99E-01 | no | Pkhd1 | intron_variant | MODIFIER |
| 1 | 20513599 | AX-170086418 | 0 | C | T | 0.404 | 5.70E-02 | 8.41E-02 | 2.59E+00 | 2.64E+00 | 4.99E-01 | no | Pkhd1 | intron_variant | MODIFIER |
| 1 | 20522938 | AX-170294318 | 0 | A | G | 0.404 | 5.70E-02 | 8.41E-02 | 2.59E+00 | 2.64E+00 | 4.99E-01 | no | Pkhd1 | synonymous_variant | LOW |
| 1 | 20522979 | AX-170035604 | 0 | T | C | 0.404 | 5.70E-02 | 8.41E-02 | 2.59E+00 | 2.64E+00 | 4.99E-01 | no | Pkhd1 | missense_variant | MODERATE |
| 1 | 20523125 | AX-170426629 | 0 | G | A | 0.404 | 5.70E-02 | 8.41E-02 | 2.59E+00 | 2.64E+00 | 4.99E-01 | no | Pkhd1 | missense_variant | MODERATE |
| 1 | 20538400 | AX-170100529 | 0 | G | C | 0.404 | 5.70E-02 | 8.41E-02 | 2.59E+00 | 2.64E+00 | 4.99E-01 | no | Pkhd1 | intron_variant | MODIFIER |
| 1 | 20551885 | AX-170511963 | 0 | C | G | 0.404 | 5.70E-02 | 8.41E-02 | 2.59E+00 | 2.64E+00 | 4.99E-01 | no | Pkhd1 | intron_variant | MODIFIER |
| 1 | 20571515 | AX-97799472 | 0 | C | T | 0.404 | 5.70E-02 | 8.41E-02 | 2.59E+00 | 2.64E+00 | 4.99E-01 | no | Pkhd1 | intron_variant | MODIFIER |
| 1 | 20582342 | AX-170358083 | 0 | A | G | 0.404 | 5.70E-02 | 8.41E-02 | 2.59E+00 | 2.64E+00 | 4.99E-01 | no | Pkhd1 | intron_variant | MODIFIER |
| 1 | 20613073 | AX-170354337 | 0 | A | C | 0.404 | 5.70E-02 | 8.41E-02 | 2.59E+00 | 2.64E+00 | 4.99E-01 | no | Pkhd1 | intron_variant | MODIFIER |

| | | | | | | | | | | | | | | | |
|---|---|---|---|---|---|---|---|---|---|---|---|---|---|---|---|
| 1 | 20617669 | AX-170304986 | 0 | G | A | 0.404 | 5.70E-02 | 8.41E-02 | 2.59E+00 | 2.64E+00 | 4.99E-01 | no | Pkhd1 | intron_variant | MODIFIER |
| 1 | 20621995 | AX-170459217 | 0 | G | A | 0.404 | 5.70E-02 | 8.41E-02 | 2.59E+00 | 2.64E+00 | 4.99E-01 | no | Pkhd1 | upstream_gene_variant | MODIFIER |
| 1 | 20633818 | AX-170367463 | 0 | G | A | 0.404 | 5.70E-02 | 8.41E-02 | 2.59E+00 | 2.64E+00 | 4.99E-01 | no | | intergenic_variant | MODIFIER |
| 1 | 20653063 | AX-170437300 | 0 | A | G | 0.404 | 5.70E-02 | 8.41E-02 | 2.59E+00 | 2.64E+00 | 4.99E-01 | no | | intergenic_variant | MODIFIER |
| 1 | 20682536 | AX-170000043 | 0 | T | C | 0.404 | 5.70E-02 | 8.41E-02 | 2.59E+00 | 2.64E+00 | 4.99E-01 | no | Mir133b | upstream_gene_variant | MODIFIER |
| 1 | 20745888 | AX-97791508 | 0 | G | A | 0.404 | 5.70E-02 | 8.41E-02 | 2.59E+00 | 2.64E+00 | 4.99E-01 | no | | intergenic_variant | MODIFIER |
| 1 | 20775218 | AX-170040569 | 0 | G | A | 0.404 | 5.70E-02 | 8.41E-02 | 2.59E+00 | 2.64E+00 | 4.99E-01 | no | Il17f | downstream_gene_variant | MODIFIER |
| 1 | 20780177 | AX-170040568 | 0 | G | A | 0.404 | 5.70E-02 | 8.41E-02 | 2.59E+00 | 2.64E+00 | 4.99E-01 | no | Il17f | intron_variant | MODIFIER |
| 1 | 20801833 | AX-170201588 | 0 | A | G | 0.404 | 5.70E-02 | 8.41E-02 | 2.59E+00 | 2.64E+00 | 4.99E-01 | no | Mcm3 | downstream_gene_variant | MODIFIER |
| 1 | 20810101 | AX-170451860 | 0 | C | A | 0.404 | 5.70E-02 | 8.41E-02 | 2.59E+00 | 2.64E+00 | 4.99E-01 | no | Mcm3 | synonymous_variant | LOW |
| 1 | 20816043 | AX-170489363 | 0 | G | A | 0.404 | 5.70E-02 | 8.41E-02 | 2.59E+00 | 2.64E+00 | 4.99E-01 | no | Mcm3 | intron_variant | MODIFIER |
| 1 | 20871006 | AX-170202909 | 0 | T | C | 0.404 | 5.70E-02 | 8.41E-02 | 2.59E+00 | 2.64E+00 | 4.99E-01 | no | | regulatory_region_variant | MODIFIER |
| 1 | 20894616 | AX-170320453 | 0 | C | T | 0.404 | 5.70E-02 | 8.41E-02 | 2.59E+00 | 2.64E+00 | 4.99E-01 | no | 6720483E21Rik | upstream_gene_variant | MODIFIER |
| 1 | 20897541 | AX-170534609 | 0 | C | T | 0.404 | 5.70E-02 | 8.41E-02 | 2.59E+00 | 2.64E+00 | 4.99E-01 | no | Paqr8 | intron_variant | MODIFIER |
| 1 | 20897664 | AX-170313836 | 0 | G | A | 0.404 | 5.70E-02 | 8.41E-02 | 2.59E+00 | 2.64E+00 | 4.99E-01 | no | Paqr8 | intron_variant | MODIFIER |
| 1 | 20907943 | AX-170530254 | 0 | C | A | 0.404 | 5.70E-02 | 8.41E-02 | 2.59E+00 | 2.64E+00 | 4.99E-01 | no | Paqr8 | intron_variant | MODIFIER |
| 1 | 20915583 | AX-170506107 | 0 | G | A | 0.404 | 5.70E-02 | 8.41E-02 | 2.59E+00 | 2.64E+00 | 4.99E-01 | no | Paqr8 | upstream_gene_variant | MODIFIER |
| 1 | 20952903 | AX-97829807 | 0 | C | T | 0.404 | 5.70E-02 | 8.41E-02 | 2.59E+00 | 2.64E+00 | 4.99E-01 | no | Efhc1 | intron_variant | MODIFIER |
| 1 | 20967981 | AX-170303765 | 0 | C | T | 0.404 | 5.70E-02 | 8.41E-02 | 2.59E+00 | 2.64E+00 | 4.99E-01 | no | Efhc1 | intron_variant | MODIFIER |
| 1 | 20969965 | AX-170185814 | 0 | C | T | 0.404 | 5.70E-02 | 8.41E-02 | 2.59E+00 | 2.64E+00 | 4.99E-01 | no | Efhc1 | intron_variant | MODIFIER |
| 1 | 20978558 | AX-170355445 | 0 | A | C | 0.404 | 5.70E-02 | 8.41E-02 | 2.59E+00 | 2.64E+00 | 4.99E-01 | no | Efhc1 | intron_variant | MODIFIER |
| 1 | 20992425 | AX-170333028 | 0 | C | T | 0.404 | 5.70E-02 | 8.41E-02 | 2.59E+00 | 2.64E+00 | 4.99E-01 | no | Tram2 | downstream_gene_variant | MODIFIER |
| 1 | 21001912 | AX-97793865 | 0 | G | A | 0.404 | 5.70E-02 | 8.41E-02 | 2.59E+00 | 2.64E+00 | 4.99E-01 | no | Tram2 | intron_variant | MODIFIER |
| 1 | 21006782 | AX-170440492 | 0 | C | A | 0.404 | 5.70E-02 | 8.41E-02 | 2.59E+00 | 2.64E+00 | 4.99E-01 | no | Tram2 | intron_variant | MODIFIER |
| 1 | 21012383 | AX-170040572 | 0 | C | G | 0.404 | 5.70E-02 | 8.41E-02 | 2.59E+00 | 2.64E+00 | 4.99E-01 | no | Tram2 | intron_variant | MODIFIER |
| 1 | 21023242 | AX-97805055 | 0 | T | C | 0.404 | 5.70E-02 | 8.41E-02 | 2.59E+00 | 2.64E+00 | 4.99E-01 | no | Tram2 | intron_variant | MODIFIER |
| 1 | 21023322 | AX-170040576 | 0 | G | A | 0.404 | 5.70E-02 | 8.41E-02 | 2.59E+00 | 2.64E+00 | 4.99E-01 | no | Tram2 | intron_variant | MODIFIER |
| 1 | 21036484 | AX-170040574 | 0 | T | C | 0.404 | 5.70E-02 | 8.41E-02 | 2.59E+00 | 2.64E+00 | 4.99E-01 | no | Tram2 | intron_variant | MODIFIER |
| 1 | 21036504 | AX-170527673 | 0 | A | G | 0.404 | 5.70E-02 | 8.41E-02 | 2.59E+00 | 2.64E+00 | 4.99E-01 | no | Tram2 | intron_variant | MODIFIER |
| 1 | 21040992 | AX-170226032 | 0 | T | C | 0.404 | 5.70E-02 | 8.41E-02 | 2.59E+00 | 2.64E+00 | 4.99E-01 | no | Tram2 | intron_variant | MODIFIER |
| 1 | 21041383 | AX-170085967 | 0 | T | C | 0.404 | 5.70E-02 | 8.41E-02 | 2.59E+00 | 2.64E+00 | 4.99E-01 | no | Tram2 | intron_variant | MODIFIER |
| 1 | 21061199 | AX-97811616 | 0 | G | A | 0.404 | 5.70E-02 | 8.41E-02 | 2.59E+00 | 2.64E+00 | 4.99E-01 | no | Tram2 | intron_variant | MODIFIER |
| 1 | 21080639 | AX-170476204 | 0 | G | A | 0.404 | 5.70E-02 | 8.41E-02 | 2.59E+00 | 2.64E+00 | 4.99E-01 | no | Tram2 | upstream_gene_variant | MODIFIER |
| 1 | 21080784 | AX-170171466 | 0 | G | C | 0.404 | 5.70E-02 | 8.41E-02 | 2.59E+00 | 2.64E+00 | 4.99E-01 | no | Tram2 | upstream_gene_variant | MODIFIER |
| 1 | 21095763 | AX-97840313 | 0 | C | T | 0.404 | 5.70E-02 | 8.41E-02 | 2.59E+00 | 2.64E+00 | 4.99E-01 | no | | intergenic_variant | MODIFIER |
| 1 | 21101610 | AX-97839049 | 0 | T | C | 0.404 | 5.70E-02 | 8.41E-02 | 2.59E+00 | 2.64E+00 | 4.99E-01 | no | | regulatory_region_variant | MODIFIER |
| 1 | 21127252 | AX-170489238 | 0 | A | G | 0.404 | 5.70E-02 | 8.41E-02 | 2.59E+00 | 2.64E+00 | 4.99E-01 | no | | intergenic_variant | MODIFIER |
| 1 | 21214358 | AX-170263787 | 0 | T | C | 0.404 | 5.70E-02 | 8.41E-02 | 2.59E+00 | 2.64E+00 | 4.99E-01 | no | Tmem14a | upstream_gene_variant | MODIFIER |
| 1 | 21284643 | AX-170000053 | 0 | C | G | 0.404 | 5.70E-02 | 8.41E-02 | 2.59E+00 | 2.64E+00 | 4.99E-01 | no | Gm4956 | downstream_gene_variant | MODIFIER |
| 1 | 21292126 | AX-97856685 | 0 | T | C | 0.404 | 5.70E-02 | 8.41E-02 | 2.59E+00 | 2.64E+00 | 4.99E-01 | no | Gm4956 | intron_variant,non_coding_transcript_variant | MODIFIER |
| 1 | 21292145 | AX-170000055 | 0 | T | G | 0.404 | 5.70E-02 | 8.41E-02 | 2.59E+00 | 2.64E+00 | 4.99E-01 | no | Gm4956 | intron_variant,non_coding_transcript_variant | MODIFIER |
| 1 | 21304039 | AX-170483769 | 0 | C | G | 0.404 | 5.70E-02 | 8.41E-02 | 2.59E+00 | 2.64E+00 | 4.99E-01 | no | | intergenic_variant | MODIFIER |
| 1 | 21324240 | AX-97814640 | 0 | A | G | 0.404 | 5.70E-02 | 8.41E-02 | 2.59E+00 | 2.64E+00 | 4.99E-01 | no | | regulatory_region_variant | MODIFIER |
| 1 | 21387490 | AX-97799561 | 0 | T | C | 0.404 | 5.70E-02 | 8.41E-02 | 2.59E+00 | 2.64E+00 | 4.99E-01 | no | Gm16328 | upstream_gene_variant | MODIFIER |
| 1 | 21388582 | AX-170302598 | 0 | G | A | 0.404 | 5.70E-02 | 8.41E-02 | 2.59E+00 | 2.64E+00 | 4.99E-01 | no | Gm16328 | upstream_gene_variant | MODIFIER |
| 1 | 21399811 | AX-170000054 | 0 | G | A | 0.404 | 5.70E-02 | 8.41E-02 | 2.59E+00 | 2.64E+00 | 4.99E-01 | no | Kcnq5 | 3_prime_UTR_variant | MODIFIER |
| 1 | 21409090 | AX-170040583 | 0 | A | G | 0.404 | 5.70E-02 | 8.41E-02 | 2.59E+00 | 2.64E+00 | 4.99E-01 | no | Kcnq5 | intron_variant | MODIFIER |
| 1 | 21413011 | AX-170197519 | 0 | G | T | 0.404 | 5.70E-02 | 8.41E-02 | 2.59E+00 | 2.64E+00 | 4.99E-01 | no | Kcnq5 | intron_variant | MODIFIER |
| 1 | 21415602 | AX-170447013 | 0 | C | T | 0.404 | 5.70E-02 | 8.41E-02 | 2.59E+00 | 2.64E+00 | 4.99E-01 | no | Kcnq5 | intron_variant | MODIFIER |
| 1 | 21419775 | AX-170000057 | 0 | C | A | 0.404 | 5.70E-02 | 8.41E-02 | 2.59E+00 | 2.64E+00 | 4.99E-01 | no | Kcnq5 | intron_variant | MODIFIER |
| 1 | 21428139 | AX-170000051 | 0 | G | A | 0.404 | 5.70E-02 | 8.41E-02 | 2.59E+00 | 2.64E+00 | 4.99E-01 | no | Kcnq5 | intron_variant | MODIFIER |
| 1 | 21564901 | AX-97810886 | 0 | A | G | 0.404 | 5.70E-02 | 8.41E-02 | 2.59E+00 | 2.64E+00 | 4.99E-01 | no | Kcnq5 | intron_variant | MODIFIER |
| 1 | 21598764 | AX-170116733 | 0 | C | T | 0.404 | 5.70E-02 | 8.41E-02 | 2.59E+00 | 2.64E+00 | 4.99E-01 | no | Kcnq5 | intron_variant | MODIFIER |
| 1 | 21599176 | AX-170468249 | 0 | A | G | 0.404 | 5.70E-02 | 8.41E-02 | 2.59E+00 | 2.64E+00 | 4.99E-01 | no | Kcnq5 | intron_variant | MODIFIER |

| | | | | | | | | | | | | | | | |
|---|---|---|---|---|---|---|---|---|---|---|---|---|---|---|---|
| 1 | 21622710 | AX-170086419 | 0 | T | G | 0.404 | 5.70E-02 | 8.41E-02 | 2.59E+00 | 2.64E+00 | 4.99E-01 | no | Kcnq5 | intron_variant | MODIFIER |
| 1 | 21635936 | AX-170250653 | 0 | T | C | 0.404 | 5.70E-02 | 8.41E-02 | 2.59E+00 | 2.64E+00 | 4.99E-01 | no | Kcnq5 | intron_variant | MODIFIER |
| 1 | 21641040 | AX-170165177 | 0 | C | T | 0.404 | 5.70E-02 | 8.41E-02 | 2.59E+00 | 2.64E+00 | 4.99E-01 | no | Kcnq5 | intron_variant | MODIFIER |
| 1 | 21649184 | AX-170252191 | 0 | G | A | 0.404 | 5.70E-02 | 8.41E-02 | 2.59E+00 | 2.64E+00 | 4.99E-01 | no | Kcnq5 | intron_variant | MODIFIER |
| 1 | 21650918 | AX-170194407 | 0 | C | T | 0.404 | 5.70E-02 | 8.41E-02 | 2.59E+00 | 2.64E+00 | 4.99E-01 | no | Kcnq5 | intron_variant | MODIFIER |
| 1 | 21651072 | AX-170262205 | 0 | C | T | 0.404 | 5.70E-02 | 8.41E-02 | 2.59E+00 | 2.64E+00 | 4.99E-01 | no | Kcnq5 | intron_variant | MODIFIER |
| 1 | 21651288 | AX-170525477 | 0 | A | G | 0.404 | 5.70E-02 | 8.41E-02 | 2.59E+00 | 2.64E+00 | 4.99E-01 | no | Kcnq5 | intron_variant | MODIFIER |
| 1 | 21652493 | AX-170395519 | 0 | A | G | 0.404 | 5.70E-02 | 8.41E-02 | 2.59E+00 | 2.64E+00 | 4.99E-01 | no | Kcnq5 | intron_variant | MODIFIER |
| 1 | 21656617 | AX-170332485 | 0 | A | G | 0.404 | 5.70E-02 | 8.41E-02 | 2.59E+00 | 2.64E+00 | 4.99E-01 | no | Kcnq5 | intron_variant | MODIFIER |
| 1 | 21662482 | AX-170301189 | 0 | G | C | 0.404 | 5.70E-02 | 8.41E-02 | 2.59E+00 | 2.64E+00 | 4.99E-01 | no | Kcnq5 | intron_variant | MODIFIER |
| 1 | 21664453 | AX-170214874 | 0 | T | A | 0.404 | 5.70E-02 | 8.41E-02 | 2.59E+00 | 2.64E+00 | 4.99E-01 | no | Kcnq5 | intron_variant | MODIFIER |
| 1 | 21671046 | AX-170411126 | 0 | G | A | 0.404 | 5.70E-02 | 8.41E-02 | 2.59E+00 | 2.64E+00 | 4.99E-01 | no | Kcnq5 | intron_variant | MODIFIER |
| 1 | 21684162 | AX-170168900 | 0 | C | A | 0.404 | 5.70E-02 | 8.41E-02 | 2.59E+00 | 2.64E+00 | 4.99E-01 | no | Kcnq5 | intron_variant | MODIFIER |
| 1 | 21686268 | AX-170144950 | 0 | C | G | 0.404 | 5.70E-02 | 8.41E-02 | 2.59E+00 | 2.64E+00 | 4.99E-01 | no | Kcnq5 | intron_variant | MODIFIER |
| 1 | 21690888 | AX-170474583 | 0 | T | C | 0.404 | 5.70E-02 | 8.41E-02 | 2.59E+00 | 2.64E+00 | 4.99E-01 | no | Kcnq5 | intron_variant | MODIFIER |
| 1 | 21713949 | AX-170386697 | 0 | T | C | 0.404 | 5.70E-02 | 8.41E-02 | 2.59E+00 | 2.64E+00 | 4.99E-01 | no | Kcnq5 | intron_variant | MODIFIER |
| 1 | 21714324 | AX-170134020 | 0 | T | C | 0.404 | 5.70E-02 | 8.41E-02 | 2.59E+00 | 2.64E+00 | 4.99E-01 | no | Kcnq5 | intron_variant | MODIFIER |
| 1 | 21730383 | AX-170131251 | 0 | G | A | 0.404 | 5.70E-02 | 8.41E-02 | 2.59E+00 | 2.64E+00 | 4.99E-01 | no | Kcnq5 | intron_variant | MODIFIER |
| 1 | 21734712 | AX-170335892 | 0 | G | T | 0.404 | 5.70E-02 | 8.41E-02 | 2.59E+00 | 2.64E+00 | 4.99E-01 | no | Kcnq5 | intron_variant | MODIFIER |
| 1 | 21735943 | AX-170204694 | 0 | T | C | 0.404 | 5.70E-02 | 8.41E-02 | 2.59E+00 | 2.64E+00 | 4.99E-01 | no | Kcnq5 | intron_variant | MODIFIER |
| 1 | 21815784 | AX-97810603 | 0 | A | C | 0.404 | 5.70E-02 | 8.41E-02 | 2.59E+00 | 2.64E+00 | 4.99E-01 | no | Kcnq5 | intron_variant | MODIFIER |
| 1 | 21852313 | AX-170282283 | 0 | A | G | 0.404 | 5.70E-02 | 8.41E-02 | 2.59E+00 | 2.64E+00 | 4.99E-01 | no | Kcnq5 | intron_variant | MODIFIER |
| 1 | 21867986 | AX-170241589 | 0 | T | C | 0.404 | 5.70E-02 | 8.41E-02 | 2.59E+00 | 2.64E+00 | 4.99E-01 | no | Kcnq5 | intron_variant | MODIFIER |
| 1 | 96936362 | AX-97798623 | 1 | C | T | 0.152 | 1.33E-01 | 1.98E-01 | 2.62E+00 | 2.68E+00 | 4.99E-01 | no | Slco6b1 | intron_variant,non_coding_transcript_variant | MODIFIER |
| 1 | 189996770 | AX-170102139 | 0 | G | A | 0.246 | 6.61E-02 | 9.49E-02 | 2.48E+00 | 2.57E+00 | 4.99E-01 | no | | intergenic_variant | MODIFIER |
| 1 | 143275656 | AX-97833255 | 0 | C | A | 0.211 | 9.26E-02 | 1.38E-01 | 2.68E+00 | 2.72E+00 | 4.99E-01 | no | | intergenic_variant | MODIFIER |
| 1 | 143313305 | AX-97812842 | 0 | A | G | 0.211 | 9.26E-02 | 1.38E-01 | 2.68E+00 | 2.72E+00 | 4.99E-01 | no | | regulatory_region_variant | MODIFIER |
| 1 | 143339245 | AX-170086632 | 0 | G | A | 0.211 | 9.26E-02 | 1.38E-01 | 2.68E+00 | 2.72E+00 | 4.99E-01 | no | | intergenic_variant | MODIFIER |
| 1 | 143349596 | AX-170001877 | 0 | A | G | 0.211 | 9.26E-02 | 1.38E-01 | 2.68E+00 | 2.72E+00 | 4.99E-01 | no | | intergenic_variant | MODIFIER |
| 1 | 143719386 | AX-170091726 | 0 | G | A | 0.211 | 9.26E-02 | 1.38E-01 | 2.68E+00 | 2.72E+00 | 4.99E-01 | no | Glrx2 | non_coding_transcript_exon_variant | MODIFIER |
| 1 | 143720758 | AX-170387023 | 0 | T | C | 0.211 | 9.26E-02 | 1.38E-01 | 2.68E+00 | 2.72E+00 | 4.99E-01 | no | Glrx2 | intron_variant,non_coding_transcript_variant | MODIFIER |
| 1 | 143726206 | AX-170179214 | 0 | A | C | 0.211 | 9.26E-02 | 1.38E-01 | 2.68E+00 | 2.72E+00 | 4.99E-01 | no | Glrx2 | intron_variant,non_coding_transcript_variant | MODIFIER |
| 1 | 143745345 | AX-170228904 | 0 | G | A | 0.211 | 9.26E-02 | 1.38E-01 | 2.68E+00 | 2.72E+00 | 4.99E-01 | no | Glrx2 | intron_variant,NMD_transcript_variant | MODIFIER |
| 1 | 143745922 | AX-170169025 | 0 | C | T | 0.211 | 9.26E-02 | 1.38E-01 | 2.68E+00 | 2.72E+00 | 4.99E-01 | no | Glrx2 | intron_variant,NMD_transcript_variant | MODIFIER |
| 1 | 144118457 | AX-97815190 | 0 | G | A | 0.211 | 9.26E-02 | 1.38E-01 | 2.68E+00 | 2.72E+00 | 4.99E-01 | no | | intergenic_variant | MODIFIER |
| 1 | 144124126 | AX-97785995 | 0 | G | A | 0.211 | 9.26E-02 | 1.38E-01 | 2.68E+00 | 2.72E+00 | 4.99E-01 | no | | intergenic_variant | MODIFIER |
| 1 | 144176600 | AX-170189994 | 0 | A | C | 0.211 | 9.26E-02 | 1.38E-01 | 2.68E+00 | 2.72E+00 | 4.99E-01 | no | Rgs13 | intron_variant | MODIFIER |
| 1 | 144179223 | AX-97815890 | 0 | G | A | 0.211 | 9.26E-02 | 1.38E-01 | 2.68E+00 | 2.72E+00 | 4.99E-01 | no | Rgs13 | upstream_gene_variant | MODIFIER |
| 1 | 144198084 | AX-170155425 | 0 | A | G | 0.211 | 9.26E-02 | 1.38E-01 | 2.68E+00 | 2.72E+00 | 4.99E-01 | no | | regulatory_region_variant | MODIFIER |
| 1 | 144198949 | AX-170042986 | 0 | T | C | 0.211 | 9.26E-02 | 1.38E-01 | 2.68E+00 | 2.72E+00 | 4.99E-01 | no | | intergenic_variant | MODIFIER |
| 1 | 144204890 | AX-170190099 | 0 | T | C | 0.211 | 9.26E-02 | 1.38E-01 | 2.68E+00 | 2.72E+00 | 4.99E-01 | no | | regulatory_region_variant | MODIFIER |
| 1 | 144206447 | AX-170291006 | 0 | T | C | 0.211 | 9.26E-02 | 1.38E-01 | 2.68E+00 | 2.72E+00 | 4.99E-01 | no | | intergenic_variant | MODIFIER |
| 1 | 144229404 | AX-170459341 | 0 | T | C | 0.211 | 9.26E-02 | 1.38E-01 | 2.68E+00 | 2.72E+00 | 4.99E-01 | no | | intergenic_variant | MODIFIER |
| 1 | 144237666 | AX-170207599 | 0 | G | A | 0.211 | 9.26E-02 | 1.38E-01 | 2.68E+00 | 2.72E+00 | 4.99E-01 | no | Rgs1 | downstream_gene_variant | MODIFIER |
| 1 | 144242893 | AX-170447653 | 0 | C | G | 0.211 | 9.26E-02 | 1.38E-01 | 2.68E+00 | 2.72E+00 | 4.99E-01 | no | Rgs1 | downstream_gene_variant | MODIFIER |
| 1 | 144246000 | AX-170509814 | 0 | T | C | 0.211 | 9.26E-02 | 1.38E-01 | 2.68E+00 | 2.72E+00 | 4.99E-01 | no | Rgs1 | intron_variant,NMD_transcript_variant | MODIFIER |
| 1 | 144246889 | AX-170291988 | 0 | A | G | 0.211 | 9.26E-02 | 1.38E-01 | 2.68E+00 | 2.72E+00 | 4.99E-01 | no | Rgs1 | intron_variant,NMD_transcript_variant | MODIFIER |
| 1 | 144250561 | AX-170042988 | 0 | A | G | 0.211 | 9.26E-02 | 1.38E-01 | 2.68E+00 | 2.72E+00 | 4.99E-01 | no | Rgs1 | upstream_gene_variant | MODIFIER |
| 1 | 144257621 | AX-170042990 | 0 | C | T | 0.211 | 9.26E-02 | 1.38E-01 | 2.68E+00 | 2.72E+00 | 4.99E-01 | no | | intergenic_variant | MODIFIER |
| 1 | 144283509 | AX-170405168 | 0 | A | G | 0.211 | 9.26E-02 | 1.38E-01 | 2.68E+00 | 2.72E+00 | 4.99E-01 | no | | regulatory_region_variant | MODIFIER |
| 1 | 144558625 | AX-97786691 | 0 | T | C | 0.211 | 9.26E-02 | 1.38E-01 | 2.68E+00 | 2.72E+00 | 4.99E-01 | no | Rgs21 | intron_variant,non_coding_transcript_variant | MODIFIER |
| 1 | 144574781 | AX-170001899 | 0 | A | T | 0.211 | 9.26E-02 | 1.38E-01 | 2.68E+00 | 2.72E+00 | 4.99E-01 | no | | intergenic_variant | MODIFIER |
| 1 | 144576596 | AX-97834074 | 0 | A | C | 0.211 | 9.26E-02 | 1.38E-01 | 2.68E+00 | 2.72E+00 | 4.99E-01 | no | | intergenic_variant | MODIFIER |
| 1 | 144604808 | AX-97849676 | 0 | G | C | 0.211 | 9.26E-02 | 1.38E-01 | 2.68E+00 | 2.72E+00 | 4.99E-01 | no | Gm23672 | downstream_gene_variant | MODIFIER |

| | | | | | | | | | | | | | | | |
|---|---|---|---|---|---|---|---|---|---|---|---|---|---|---|---|
| 1 | 144629587 | AX-170001900 | 0 | C | T | 0.211 | 9.26E-02 | 1.38E-01 | 2.68E+00 | 2.72E+00 | 4.99E-01 | no | | intergenic_variant | MODIFIER |
| 1 | 144644820 | AX-170385134 | 0 | A | C | 0.211 | 9.26E-02 | 1.38E-01 | 2.68E+00 | 2.72E+00 | 4.99E-01 | no | | intergenic_variant | MODIFIER |
| 1 | 144651240 | AX-170510967 | 0 | T | C | 0.211 | 9.26E-02 | 1.38E-01 | 2.68E+00 | 2.72E+00 | 4.99E-01 | no | | intergenic_variant | MODIFIER |
| 1 | 144657314 | AX-170416862 | 0 | G | T | 0.211 | 9.26E-02 | 1.38E-01 | 2.68E+00 | 2.72E+00 | 4.99E-01 | no | | intergenic_variant | MODIFIER |
| 1 | 144681676 | AX-170105201 | 0 | A | G | 0.211 | 9.26E-02 | 1.38E-01 | 2.68E+00 | 2.72E+00 | 4.99E-01 | no | | intergenic_variant | MODIFIER |
| 1 | 144681704 | AX-97791777 | 0 | A | G | 0.211 | 9.26E-02 | 1.38E-01 | 2.68E+00 | 2.72E+00 | 4.99E-01 | no | | intergenic_variant | MODIFIER |
| 1 | 144698565 | AX-170086636 | 0 | C | G | 0.211 | 9.26E-02 | 1.38E-01 | 2.68E+00 | 2.72E+00 | 4.99E-01 | no | | intergenic_variant | MODIFIER |
| 1 | 144713107 | AX-170001901 | 0 | T | C | 0.211 | 9.26E-02 | 1.38E-01 | 2.68E+00 | 2.72E+00 | 4.99E-01 | no | | intergenic_variant | MODIFIER |
| 1 | 144754715 | AX-170212483 | 0 | G | C | 0.211 | 9.26E-02 | 1.38E-01 | 2.68E+00 | 2.72E+00 | 4.99E-01 | no | Rgs18 | intron_variant | MODIFIER |
| 1 | 144755783 | AX-97813562 | 0 | T | C | 0.211 | 9.26E-02 | 1.38E-01 | 2.68E+00 | 2.72E+00 | 4.99E-01 | no | Rgs18 | intron_variant | MODIFIER |
| 1 | 144772617 | AX-97836270 | 0 | C | T | 0.211 | 9.26E-02 | 1.38E-01 | 2.68E+00 | 2.72E+00 | 4.99E-01 | no | Rgs18 | intron_variant | MODIFIER |
| 1 | 144817818 | AX-170356783 | 0 | G | C | 0.211 | 9.26E-02 | 1.38E-01 | 2.68E+00 | 2.72E+00 | 4.99E-01 | no | | intergenic_variant | MODIFIER |
| 1 | 145037751 | AX-170465851 | 0 | G | A | 0.211 | 9.26E-02 | 1.38E-01 | 2.68E+00 | 2.72E+00 | 4.99E-01 | no | | intergenic_variant | MODIFIER |
| 1 | 145041544 | AX-170202674 | 0 | T | C | 0.211 | 9.26E-02 | 1.38E-01 | 2.68E+00 | 2.72E+00 | 4.99E-01 | no | | intergenic_variant | MODIFIER |
| 1 | 145063318 | AX-170186650 | 0 | G | T | 0.211 | 9.26E-02 | 1.38E-01 | 2.68E+00 | 2.72E+00 | 4.99E-01 | no | | intergenic_variant | MODIFIER |
| 1 | 145069312 | AX-170352756 | 0 | C | A | 0.211 | 9.26E-02 | 1.38E-01 | 2.68E+00 | 2.72E+00 | 4.99E-01 | no | | intergenic_variant | MODIFIER |
| 1 | 145145582 | AX-97813618 | 0 | C | G | 0.211 | 9.26E-02 | 1.38E-01 | 2.68E+00 | 2.72E+00 | 4.99E-01 | no | | intergenic_variant | MODIFIER |
| 1 | 145463714 | AX-170001916 | 0 | C | A | 0.211 | 9.26E-02 | 1.38E-01 | 2.68E+00 | 2.72E+00 | 4.99E-01 | no | | intergenic_variant | MODIFIER |
| 1 | 145526095 | AX-170413602 | 0 | G | C | 0.211 | 9.26E-02 | 1.38E-01 | 2.68E+00 | 2.72E+00 | 4.99E-01 | no | | intergenic_variant | MODIFIER |
| 1 | 145597328 | AX-170256858 | 0 | G | A | 0.211 | 9.26E-02 | 1.38E-01 | 2.68E+00 | 2.72E+00 | 4.99E-01 | no | | intergenic_variant | MODIFIER |
| 1 | 145605124 | AX-170481760 | 0 | T | A | 0.211 | 9.26E-02 | 1.38E-01 | 2.68E+00 | 2.72E+00 | 4.99E-01 | no | | intergenic_variant | MODIFIER |
| 1 | 146455453 | AX-170086638 | 0 | C | T | 0.211 | 9.26E-02 | 1.38E-01 | 2.68E+00 | 2.72E+00 | 4.99E-01 | no | Gm29514 | intron_variant,non_coding_transcript_variant | MODIFIER |
| 1 | 146498415 | AX-170237468 | 0 | A | T | 0.211 | 9.26E-02 | 1.38E-01 | 2.68E+00 | 2.72E+00 | 4.99E-01 | no | Brinp3 | intron_variant | MODIFIER |
| 1 | 146507856 | AX-170198354 | 0 | C | T | 0.211 | 9.26E-02 | 1.38E-01 | 2.68E+00 | 2.72E+00 | 4.99E-01 | no | Brinp3 | intron_variant | MODIFIER |
| 1 | 146538273 | AX-170314098 | 0 | G | T | 0.211 | 9.26E-02 | 1.38E-01 | 2.68E+00 | 2.72E+00 | 4.99E-01 | no | Brinp3 | intron_variant | MODIFIER |
| 1 | 146549371 | AX-170186494 | 0 | A | T | 0.211 | 9.26E-02 | 1.38E-01 | 2.68E+00 | 2.72E+00 | 4.99E-01 | no | Brinp3 | intron_variant | MODIFIER |
| 1 | 146550812 | AX-97826564 | 0 | T | C | 0.211 | 9.26E-02 | 1.38E-01 | 2.68E+00 | 2.72E+00 | 4.99E-01 | no | Brinp3 | intron_variant | MODIFIER |
| 1 | 146642081 | AX-170284981 | 0 | T | A | 0.211 | 9.26E-02 | 1.38E-01 | 2.68E+00 | 2.72E+00 | 4.99E-01 | no | Brinp3 | intron_variant | MODIFIER |
| 1 | 146656876 | AX-97806681 | 0 | T | C | 0.211 | 9.26E-02 | 1.38E-01 | 2.68E+00 | 2.72E+00 | 4.99E-01 | no | Brinp3 | intron_variant | MODIFIER |
| 1 | 146715323 | AX-170494286 | 0 | C | A | 0.211 | 9.26E-02 | 1.38E-01 | 2.68E+00 | 2.72E+00 | 4.99E-01 | no | Brinp3 | intron_variant | MODIFIER |
| 1 | 146782668 | AX-170001930 | 0 | G | A | 0.211 | 9.26E-02 | 1.38E-01 | 2.68E+00 | 2.72E+00 | 4.99E-01 | no | Brinp3 | intron_variant | MODIFIER |
| 1 | 146806692 | AX-97813366 | 0 | G | T | 0.211 | 9.26E-02 | 1.38E-01 | 2.68E+00 | 2.72E+00 | 4.99E-01 | no | Brinp3 | intron_variant | MODIFIER |
| 1 | 146825048 | AX-170001934 | 0 | C | T | 0.211 | 9.26E-02 | 1.38E-01 | 2.68E+00 | 2.72E+00 | 4.99E-01 | no | Brinp3 | intron_variant | MODIFIER |
| 1 | 146861920 | AX-97794073 | 0 | C | T | 0.211 | 9.26E-02 | 1.38E-01 | 2.68E+00 | 2.72E+00 | 4.99E-01 | no | Brinp3 | intron_variant | MODIFIER |
| 1 | 150710820 | AX-170389378 | 0 | T | C | 0.211 | 9.26E-02 | 1.38E-01 | 2.68E+00 | 2.72E+00 | 4.99E-01 | no | Hmcn1 | intron_variant | MODIFIER |
| 1 | 150716507 | AX-170380980 | 0 | C | G | 0.211 | 9.26E-02 | 1.38E-01 | 2.68E+00 | 2.72E+00 | 4.99E-01 | no | Hmcn1 | intron_variant | MODIFIER |
| 1 | 150717282 | AX-170420034 | 0 | C | A | 0.211 | 9.26E-02 | 1.38E-01 | 2.68E+00 | 2.72E+00 | 4.99E-01 | no | Hmcn1 | intron_variant | MODIFIER |
| 1 | 150721169 | AX-170159627 | 0 | A | G | 0.211 | 9.26E-02 | 1.38E-01 | 2.68E+00 | 2.72E+00 | 4.99E-01 | no | Hmcn1 | intron_variant | MODIFIER |
| 1 | 150752224 | AX-170453727 | 0 | A | G | 0.211 | 9.26E-02 | 1.38E-01 | 2.68E+00 | 2.72E+00 | 4.99E-01 | no | Hmcn1 | intron_variant | MODIFIER |
| 1 | 150752276 | AX-170277415 | 0 | C | T | 0.211 | 9.26E-02 | 1.38E-01 | 2.68E+00 | 2.72E+00 | 4.99E-01 | no | Hmcn1 | intron_variant | MODIFIER |
| 1 | 150759136 | AX-170164541 | 0 | T | G | 0.211 | 9.26E-02 | 1.38E-01 | 2.68E+00 | 2.72E+00 | 4.99E-01 | no | Hmcn1 | intron_variant | MODIFIER |
| 1 | 150759411 | AX-170225938 | 0 | C | T | 0.211 | 9.26E-02 | 1.38E-01 | 2.68E+00 | 2.72E+00 | 4.99E-01 | no | Hmcn1 | intron_variant | MODIFIER |
| 1 | 150763728 | AX-170224133 | 0 | G | A | 0.211 | 9.26E-02 | 1.38E-01 | 2.68E+00 | 2.72E+00 | 4.99E-01 | no | Hmcn1 | intron_variant | MODIFIER |
| 1 | 150823541 | AX-170277144 | 0 | G | A | 0.211 | 9.26E-02 | 1.38E-01 | 2.68E+00 | 2.72E+00 | 4.99E-01 | no | Hmcn1 | intron_variant | MODIFIER |
| 1 | 150826731 | AX-97803254 | 0 | G | T | 0.211 | 9.26E-02 | 1.38E-01 | 2.68E+00 | 2.72E+00 | 4.99E-01 | no | Hmcn1 | intron_variant | MODIFIER |
| 1 | 150886077 | AX-170451209 | 0 | G | A | 0.211 | 9.26E-02 | 1.38E-01 | 2.68E+00 | 2.72E+00 | 4.99E-01 | no | Hmcn1 | intron_variant | MODIFIER |
| 1 | 150901227 | AX-170256310 | 0 | C | T | 0.211 | 9.26E-02 | 1.38E-01 | 2.68E+00 | 2.72E+00 | 4.99E-01 | no | Hmcn1 | intron_variant | MODIFIER |
| 1 | 150907731 | AX-170312866 | 0 | T | G | 0.211 | 9.26E-02 | 1.38E-01 | 2.68E+00 | 2.72E+00 | 4.99E-01 | no | Hmcn1 | intron_variant | MODIFIER |
| 1 | 150913507 | AX-170411617 | 0 | T | A | 0.211 | 9.26E-02 | 1.38E-01 | 2.68E+00 | 2.72E+00 | 4.99E-01 | no | Hmcn1 | intron_variant | MODIFIER |
| 1 | 150914584 | AX-97837481 | 0 | A | G | 0.211 | 9.26E-02 | 1.38E-01 | 2.68E+00 | 2.72E+00 | 4.99E-01 | no | Hmcn1 | intron_variant | MODIFIER |
| 1 | 150927862 | AX-170476312 | 0 | C | T | 0.211 | 9.26E-02 | 1.38E-01 | 2.68E+00 | 2.72E+00 | 4.99E-01 | no | Hmcn1 | intron_variant | MODIFIER |
| 1 | 150952760 | AX-170131525 | 0 | G | A | 0.211 | 9.26E-02 | 1.38E-01 | 2.68E+00 | 2.72E+00 | 4.99E-01 | no | Hmcn1 | intron_variant | MODIFIER |
| 1 | 150956326 | AX-170524424 | 0 | A | G | 0.211 | 9.26E-02 | 1.38E-01 | 2.68E+00 | 2.72E+00 | 4.99E-01 | no | Hmcn1 | intron_variant | MODIFIER |
| 1 | 150956357 | AX-170469540 | 0 | T | C | 0.211 | 9.26E-02 | 1.38E-01 | 2.68E+00 | 2.72E+00 | 4.99E-01 | no | Hmcn1 | intron_variant | MODIFIER |

| | | | | | | | | | | | | | | | |
|---|---|---|---|---|---|---|---|---|---|---|---|---|---|---|---|
| 1 | 151013329 | AX-170360458 | 0 | G | A | 0.211 | 9.26E-02 | 1.38E-01 | 2.68E+00 | 2.72E+00 | 4.99E-01 | no | | intergenic_variant | MODIFIER |
| 1 | 151013358 | AX-170462594 | 0 | T | C | 0.211 | 9.26E-02 | 1.38E-01 | 2.68E+00 | 2.72E+00 | 4.99E-01 | no | | intergenic_variant | MODIFIER |
| 1 | 151022652 | AX-170199396 | 0 | T | C | 0.211 | 9.26E-02 | 1.38E-01 | 2.68E+00 | 2.72E+00 | 4.99E-01 | no | | intergenic_variant | MODIFIER |
| 1 | 151023369 | AX-170493538 | 0 | C | T | 0.211 | 9.26E-02 | 1.38E-01 | 2.68E+00 | 2.72E+00 | 4.99E-01 | no | | intergenic_variant | MODIFIER |
| 1 | 151025150 | AX-170291625 | 0 | C | G | 0.211 | 9.26E-02 | 1.38E-01 | 2.68E+00 | 2.72E+00 | 4.99E-01 | no | | regulatory_region_variant | MODIFIER |
| 1 | 151046496 | AX-170473180 | 0 | G | A | 0.211 | 9.26E-02 | 1.38E-01 | 2.68E+00 | 2.72E+00 | 4.99E-01 | no | | regulatory_region_variant | MODIFIER |
| 1 | 151059870 | AX-170331121 | 0 | T | C | 0.211 | 9.26E-02 | 1.38E-01 | 2.68E+00 | 2.72E+00 | 4.99E-01 | no | Gm22756 | downstream_gene_variant | MODIFIER |
| 1 | 151070558 | AX-170393706 | 0 | C | A | 0.211 | 9.26E-02 | 1.38E-01 | 2.68E+00 | 2.72E+00 | 4.99E-01 | no | | intergenic_variant | MODIFIER |
| 1 | 151071437 | AX-97788591 | 0 | C | T | 0.211 | 9.26E-02 | 1.38E-01 | 2.68E+00 | 2.72E+00 | 4.99E-01 | no | | intergenic_variant | MODIFIER |
| 1 | 151094025 | AX-170128201 | 0 | T | G | 0.211 | 9.26E-02 | 1.38E-01 | 2.68E+00 | 2.72E+00 | 4.99E-01 | no | | intergenic_variant | MODIFIER |
| 1 | 151103040 | AX-170043128 | 0 | G | A | 0.211 | 9.26E-02 | 1.38E-01 | 2.68E+00 | 2.72E+00 | 4.99E-01 | no | Gm19087 | upstream_gene_variant | MODIFIER |
| 1 | 151136402 | AX-170240714 | 0 | A | G | 0.211 | 9.26E-02 | 1.38E-01 | 2.68E+00 | 2.72E+00 | 4.99E-01 | no | Gm8941 | upstream_gene_variant | MODIFIER |
| 1 | 151139385 | AX-170505183 | 0 | T | C | 0.211 | 9.26E-02 | 1.38E-01 | 2.68E+00 | 2.72E+00 | 4.99E-01 | no | Gm8941 | downstream_gene_variant | MODIFIER |
| 1 | 151139616 | AX-170339966 | 0 | C | A | 0.211 | 9.26E-02 | 1.38E-01 | 2.68E+00 | 2.72E+00 | 4.99E-01 | no | Gm8941 | downstream_gene_variant | MODIFIER |
| 1 | 151141793 | AX-170261491 | 0 | G | A | 0.211 | 9.26E-02 | 1.38E-01 | 2.68E+00 | 2.72E+00 | 4.99E-01 | no | Gm8941 | downstream_gene_variant | MODIFIER |
| 1 | 151142527 | AX-170382607 | 0 | C | T | 0.211 | 9.26E-02 | 1.38E-01 | 2.68E+00 | 2.72E+00 | 4.99E-01 | no | Gm8941 | downstream_gene_variant | MODIFIER |
| 1 | 151142951 | AX-170101031 | 0 | T | C | 0.211 | 9.26E-02 | 1.38E-01 | 2.68E+00 | 2.72E+00 | 4.99E-01 | no | C730036E19Rik | downstream_gene_variant | MODIFIER |
| 1 | 151216661 | AX-170435728 | 0 | G | A | 0.211 | 9.26E-02 | 1.38E-01 | 2.68E+00 | 2.72E+00 | 4.99E-01 | no | | regulatory_region_variant | MODIFIER |
| 1 | 151229525 | AX-170432604 | 0 | A | G | 0.211 | 9.26E-02 | 1.38E-01 | 2.68E+00 | 2.72E+00 | 4.99E-01 | no | | intergenic_variant | MODIFIER |
| 1 | 151265846 | AX-170043130 | 0 | C | T | 0.211 | 9.26E-02 | 1.38E-01 | 2.68E+00 | 2.72E+00 | 4.99E-01 | no | Gm24402 | upstream_gene_variant | MODIFIER |
| 1 | 151299791 | AX-170482952 | 0 | G | T | 0.211 | 9.26E-02 | 1.38E-01 | 2.68E+00 | 2.72E+00 | 4.99E-01 | no | | intergenic_variant | MODIFIER |
| 1 | 151310097 | AX-170276330 | 0 | A | G | 0.211 | 9.26E-02 | 1.38E-01 | 2.68E+00 | 2.72E+00 | 4.99E-01 | no | | intergenic_variant | MODIFIER |
| 1 | 151325090 | AX-170374065 | 0 | A | T | 0.211 | 9.26E-02 | 1.38E-01 | 2.68E+00 | 2.72E+00 | 4.99E-01 | no | | regulatory_region_variant | MODIFIER |
| 1 | 151325551 | AX-170440764 | 0 | C | G | 0.211 | 9.26E-02 | 1.38E-01 | 2.68E+00 | 2.72E+00 | 4.99E-01 | no | | regulatory_region_variant | MODIFIER |
| 1 | 151329566 | AX-170189761 | 0 | A | C | 0.211 | 9.26E-02 | 1.38E-01 | 2.68E+00 | 2.72E+00 | 4.99E-01 | no | | regulatory_region_variant | MODIFIER |
| 1 | 151337341 | AX-170190158 | 0 | G | C | 0.211 | 9.26E-02 | 1.38E-01 | 2.68E+00 | 2.72E+00 | 4.99E-01 | no | Gm10138 | downstream_gene_variant | MODIFIER |
| 1 | 151357111 | AX-170513160 | 0 | C | T | 0.211 | 9.26E-02 | 1.38E-01 | 2.68E+00 | 2.72E+00 | 4.99E-01 | no | Ivns1abp | intron_variant | MODIFIER |
| 1 | 151358315 | AX-170119946 | 0 | G | T | 0.211 | 9.26E-02 | 1.38E-01 | 2.68E+00 | 2.72E+00 | 4.99E-01 | no | Ivns1abp | intron_variant | MODIFIER |
| 1 | 151371177 | AX-170198626 | 0 | T | C | 0.211 | 9.26E-02 | 1.38E-01 | 2.68E+00 | 2.72E+00 | 4.99E-01 | no | Swt1 | intron_variant | MODIFIER |
| 1 | 151371289 | AX-170036059 | 0 | C | T | 0.211 | 9.26E-02 | 1.38E-01 | 2.68E+00 | 2.72E+00 | 4.99E-01 | no | Swt1 | intron_variant | MODIFIER |
| 1 | 151391702 | AX-97796661 | 0 | G | A | 0.211 | 9.26E-02 | 1.38E-01 | 2.68E+00 | 2.72E+00 | 4.99E-01 | no | Swt1 | intron_variant | MODIFIER |
| 1 | 151402908 | AX-170043138 | 0 | C | T | 0.211 | 9.26E-02 | 1.38E-01 | 2.68E+00 | 2.72E+00 | 4.99E-01 | no | Swt1 | intron_variant | MODIFIER |
| 1 | 151409587 | AX-170043133 | 0 | T | C | 0.211 | 9.26E-02 | 1.38E-01 | 2.68E+00 | 2.72E+00 | 4.99E-01 | no | Swt1 | intron_variant | MODIFIER |
| 1 | 151409633 | AX-170043134 | 0 | T | G | 0.211 | 9.26E-02 | 1.38E-01 | 2.68E+00 | 2.72E+00 | 4.99E-01 | no | Swt1 | intron_variant | MODIFIER |
| 1 | 151476777 | AX-170099319 | 0 | G | A | 0.211 | 9.26E-02 | 1.38E-01 | 2.68E+00 | 2.72E+00 | 4.99E-01 | no | Rnf2 | intron_variant | MODIFIER |
| 1 | 151491200 | AX-170370783 | 0 | G | C | 0.211 | 9.26E-02 | 1.38E-01 | 2.68E+00 | 2.72E+00 | 4.99E-01 | no | Rnf2 | intron_variant | MODIFIER |
| 1 | 151509305 | AX-170286212 | 0 | C | T | 0.211 | 9.26E-02 | 1.38E-01 | 2.68E+00 | 2.72E+00 | 4.99E-01 | no | Gm29072 | non_coding_transcript_exon_variant | MODIFIER |
| 1 | 151509500 | AX-170320744 | 0 | T | C | 0.211 | 9.26E-02 | 1.38E-01 | 2.68E+00 | 2.72E+00 | 4.99E-01 | no | Gm29072 | non_coding_transcript_exon_variant | MODIFIER |
| 1 | 151528184 | AX-97818553 | 0 | T | C | 0.211 | 9.26E-02 | 1.38E-01 | 2.68E+00 | 2.72E+00 | 4.99E-01 | no | | intergenic_variant | MODIFIER |
| 1 | 151550577 | AX-170148540 | 0 | A | G | 0.211 | 9.26E-02 | 1.38E-01 | 2.68E+00 | 2.72E+00 | 4.99E-01 | no | Gm28181 | intron_variant,non_coding_transcript_variant | MODIFIER |
| 1 | 151578210 | AX-170451579 | 0 | C | T | 0.211 | 9.26E-02 | 1.38E-01 | 2.68E+00 | 2.72E+00 | 4.99E-01 | no | Fam129a | intron_variant,NMD_transcript_variant | MODIFIER |
| 1 | 151593080 | AX-170260663 | 0 | T | C | 0.211 | 9.26E-02 | 1.38E-01 | 2.68E+00 | 2.72E+00 | 4.99E-01 | no | Fam129a | intron_variant,NMD_transcript_variant | MODIFIER |
| 1 | 151593215 | AX-170091750 | 0 | A | G | 0.211 | 9.26E-02 | 1.38E-01 | 2.68E+00 | 2.72E+00 | 4.99E-01 | no | Fam129a | intron_variant,NMD_transcript_variant | MODIFIER |
| 1 | 151593442 | AX-170386523 | 0 | C | T | 0.211 | 9.26E-02 | 1.38E-01 | 2.68E+00 | 2.72E+00 | 4.99E-01 | no | Fam129a | intron_variant,NMD_transcript_variant | MODIFIER |
| 1 | 151593558 | AX-170337429 | 0 | A | T | 0.211 | 9.26E-02 | 1.38E-01 | 2.68E+00 | 2.72E+00 | 4.99E-01 | no | Fam129a | intron_variant,NMD_transcript_variant | MODIFIER |
| 1 | 151598278 | AX-97841823 | 0 | A | T | 0.211 | 9.26E-02 | 1.38E-01 | 2.68E+00 | 2.72E+00 | 4.99E-01 | no | Fam129a | intron_variant,NMD_transcript_variant | MODIFIER |
| 1 | 151612000 | AX-170357301 | 0 | G | A | 0.211 | 9.26E-02 | 1.38E-01 | 2.68E+00 | 2.72E+00 | 4.99E-01 | no | Fam129a | intron_variant,NMD_transcript_variant | MODIFIER |
| 1 | 151633481 | AX-170043139 | 0 | A | G | 0.211 | 9.26E-02 | 1.38E-01 | 2.68E+00 | 2.72E+00 | 4.99E-01 | no | Fam129a | intron_variant,NMD_transcript_variant | MODIFIER |
| 1 | 151633537 | AX-97847165 | 0 | C | A | 0.211 | 9.26E-02 | 1.38E-01 | 2.68E+00 | 2.72E+00 | 4.99E-01 | no | Fam129a | intron_variant,NMD_transcript_variant | MODIFIER |
| 1 | 151633804 | AX-170036060 | 0 | C | G | 0.211 | 9.26E-02 | 1.38E-01 | 2.68E+00 | 2.72E+00 | 4.99E-01 | no | Fam129a | intron_variant,NMD_transcript_variant | MODIFIER |
| 1 | 151633873 | AX-170276436 | 0 | A | G | 0.211 | 9.26E-02 | 1.38E-01 | 2.68E+00 | 2.72E+00 | 4.99E-01 | no | Fam129a | intron_variant,NMD_transcript_variant | MODIFIER |
| 1 | 151636250 | AX-170175040 | 0 | A | C | 0.211 | 9.26E-02 | 1.38E-01 | 2.68E+00 | 2.72E+00 | 4.99E-01 | no | Fam129a | intron_variant,NMD_transcript_variant | MODIFIER |
| 1 | 151637268 | AX-170178448 | 0 | G | A | 0.211 | 9.26E-02 | 1.38E-01 | 2.68E+00 | 2.72E+00 | 4.99E-01 | no | Fam129a | intron_variant,NMD_transcript_variant | MODIFIER |
| 1 | 151638785 | AX-170383162 | 0 | A | G | 0.211 | 9.26E-02 | 1.38E-01 | 2.68E+00 | 2.72E+00 | 4.99E-01 | no | Fam129a | intron_variant,NMD_transcript_variant | MODIFIER |

| | | | | | | | | | | | | | | | |
|---|---|---|---|---|---|---|---|---|---|---|---|---|---|---|---|
| 1 | 151638928 | AX-170107407 | 0 | G | T | 0.211 | 9.26E-02 | 1.38E-01 | 2.68E+00 | 2.72E+00 | 4.99E-01 | no | Fam129a | intron_variant,NMD_transcript_variant | MODIFIER |
| 1 | 151640138 | AX-170354492 | 0 | A | G | 0.211 | 9.26E-02 | 1.38E-01 | 2.68E+00 | 2.72E+00 | 4.99E-01 | no | Fam129a | intron_variant,NMD_transcript_variant | MODIFIER |
| 1 | 151642739 | AX-170379789 | 0 | G | A | 0.211 | 9.26E-02 | 1.38E-01 | 2.68E+00 | 2.72E+00 | 4.99E-01 | no | Fam129a | intron_variant,NMD_transcript_variant | MODIFIER |
| 1 | 151687800 | AX-170495201 | 0 | A | G | 0.211 | 9.26E-02 | 1.38E-01 | 2.68E+00 | 2.72E+00 | 4.99E-01 | no | Fam129a | intron_variant,non_coding_transcript_variant | MODIFIER |
| 1 | 151695529 | AX-170177486 | 0 | A | G | 0.211 | 9.26E-02 | 1.38E-01 | 2.68E+00 | 2.72E+00 | 4.99E-01 | no | Fam129a | intron_variant,non_coding_transcript_variant | MODIFIER |
| 1 | 151696008 | AX-170204635 | 0 | G | A | 0.211 | 9.26E-02 | 1.38E-01 | 2.68E+00 | 2.72E+00 | 4.99E-01 | no | Fam129a | intron_variant,non_coding_transcript_variant | MODIFIER |
| 1 | 151704337 | AX-170001989 | 0 | T | C | 0.211 | 9.26E-02 | 1.38E-01 | 2.68E+00 | 2.72E+00 | 4.99E-01 | no | Fam129a | intron_variant,non_coding_transcript_variant | MODIFIER |
| 1 | 151708588 | AX-170214883 | 0 | A | G | 0.211 | 9.26E-02 | 1.38E-01 | 2.68E+00 | 2.72E+00 | 4.99E-01 | no | Fam129a | intron_variant,non_coding_transcript_variant | MODIFIER |
| 1 | 151711740 | AX-97783658 | 0 | A | G | 0.211 | 9.26E-02 | 1.38E-01 | 2.68E+00 | 2.72E+00 | 4.99E-01 | no | Fam129a | intron_variant,non_coding_transcript_variant | MODIFIER |
| 1 | 151717678 | AX-170288188 | 0 | A | G | 0.211 | 9.26E-02 | 1.38E-01 | 2.68E+00 | 2.72E+00 | 4.99E-01 | no | Fam129a | non_coding_transcript_exon_variant | MODIFIER |
| 1 | 151720790 | AX-170372525 | 0 | T | C | 0.211 | 9.26E-02 | 1.38E-01 | 2.68E+00 | 2.72E+00 | 4.99E-01 | no | Fam129a | downstream_gene_variant | MODIFIER |
| 5 | 38431412 | AX-97838325 | 0 | C | T | 0.325 | 7.63E-02 | 1.14E-01 | 2.73E+00 | 2.79E+00 | 4.99E-01 | no | Slc2a9 | intron_variant | MODIFIER |
| 9 | 116844412 | AX-170282685 | 0 | T | C | 0.123 | -6.46E-02 | 9.63E-02 | 2.61E+00 | 2.72E+00 | 4.99E-01 | no | Rbms3 | intron_variant | MODIFIER |
| 9 | 116844491 | AX-97813208 | 0 | T | C | 0.123 | -6.46E-02 | 9.63E-02 | 2.61E+00 | 2.72E+00 | 4.99E-01 | no | Rbms3 | intron_variant | MODIFIER |
| 9 | 116847329 | AX-170238724 | 0 | A | G | 0.123 | -6.46E-02 | 9.63E-02 | 2.61E+00 | 2.72E+00 | 4.99E-01 | no | Rbms3 | intron_variant | MODIFIER |
| 9 | 116848404 | AX-170261595 | 0 | G | T | 0.123 | -6.46E-02 | 9.63E-02 | 2.61E+00 | 2.72E+00 | 4.99E-01 | no | Rbms3 | intron_variant | MODIFIER |
| 9 | 116857940 | AX-170155874 | 0 | C | T | 0.123 | -6.46E-02 | 9.63E-02 | 2.61E+00 | 2.72E+00 | 4.99E-01 | no | Rbms3 | intron_variant | MODIFIER |
| 9 | 116896226 | AX-97840142 | 0 | T | C | 0.123 | -6.46E-02 | 9.63E-02 | 2.61E+00 | 2.72E+00 | 4.99E-01 | no | Rbms3 | intron_variant | MODIFIER |
| 9 | 116899950 | AX-170033363 | 0 | G | T | 0.123 | -6.46E-02 | 9.63E-02 | 2.61E+00 | 2.72E+00 | 4.99E-01 | no | Rbms3 | intron_variant | MODIFIER |
| 9 | 116903793 | AX-170033362 | 0 | C | T | 0.123 | -6.46E-02 | 9.63E-02 | 2.61E+00 | 2.72E+00 | 4.99E-01 | no | Rbms3 | intron_variant | MODIFIER |
| 9 | 116925895 | AX-170083575 | 0 | T | C | 0.123 | -6.46E-02 | 9.63E-02 | 2.61E+00 | 2.72E+00 | 4.99E-01 | no | Rbms3 | intron_variant | MODIFIER |
| 9 | 116926279 | AX-170083573 | 0 | A | C | 0.123 | -6.46E-02 | 9.63E-02 | 2.61E+00 | 2.72E+00 | 4.99E-01 | no | Rbms3 | intron_variant | MODIFIER |
| 9 | 116927151 | AX-170304575 | 0 | C | T | 0.123 | -6.46E-02 | 9.63E-02 | 2.61E+00 | 2.72E+00 | 4.99E-01 | no | Rbms3 | intron_variant | MODIFIER |
| 9 | 116927234 | AX-170279793 | 0 | T | C | 0.123 | -6.46E-02 | 9.63E-02 | 2.61E+00 | 2.72E+00 | 4.99E-01 | no | Rbms3 | intron_variant | MODIFIER |
| 9 | 116935343 | AX-170417890 | 0 | G | A | 0.123 | -6.46E-02 | 9.63E-02 | 2.61E+00 | 2.72E+00 | 4.99E-01 | no | Rbms3 | intron_variant | MODIFIER |
| 9 | 116935459 | AX-170489670 | 0 | C | T | 0.123 | -6.46E-02 | 9.63E-02 | 2.61E+00 | 2.72E+00 | 4.99E-01 | no | Rbms3 | intron_variant | MODIFIER |
| 9 | 116943157 | AX-170033365 | 0 | T | A | 0.123 | -6.46E-02 | 9.63E-02 | 2.61E+00 | 2.72E+00 | 4.99E-01 | no | Rbms3 | intron_variant | MODIFIER |
| 9 | 116951518 | AX-170477947 | 0 | A | G | 0.123 | -6.46E-02 | 9.63E-02 | 2.61E+00 | 2.72E+00 | 4.99E-01 | no | Rbms3 | intron_variant | MODIFIER |
| 9 | 116951773 | AX-170350140 | 0 | A | G | 0.123 | -6.46E-02 | 9.63E-02 | 2.61E+00 | 2.72E+00 | 4.99E-01 | no | Rbms3 | intron_variant | MODIFIER |
| 9 | 116952510 | AX-170119379 | 0 | C | T | 0.123 | -6.46E-02 | 9.63E-02 | 2.61E+00 | 2.72E+00 | 4.99E-01 | no | Rbms3 | intron_variant | MODIFIER |
| 9 | 116956965 | AX-170033364 | 0 | A | C | 0.123 | -6.46E-02 | 9.63E-02 | 2.61E+00 | 2.72E+00 | 4.99E-01 | no | Rbms3 | intron_variant | MODIFIER |
| 9 | 116961694 | AX-170033359 | 0 | G | C | 0.123 | -6.46E-02 | 9.63E-02 | 2.61E+00 | 2.72E+00 | 4.99E-01 | no | Rbms3 | intron_variant | MODIFIER |
| 9 | 116990051 | AX-97840757 | 0 | T | C | 0.123 | -6.46E-02 | 9.63E-02 | 2.61E+00 | 2.72E+00 | 4.99E-01 | no | Rbms3 | intron_variant | MODIFIER |
| 9 | 117018045 | AX-97823417 | 0 | T | C | 0.123 | -6.46E-02 | 9.63E-02 | 2.61E+00 | 2.72E+00 | 4.99E-01 | no | Rbms3 | intron_variant | MODIFIER |
| 9 | 117051240 | AX-170489095 | 0 | A | G | 0.123 | -6.46E-02 | 9.63E-02 | 2.61E+00 | 2.72E+00 | 4.99E-01 | no | Rbms3 | intron_variant | MODIFIER |
| 9 | 117053138 | AX-170090427 | 0 | A | G | 0.123 | -6.46E-02 | 9.63E-02 | 2.61E+00 | 2.72E+00 | 4.99E-01 | no | Rbms3 | intron_variant | MODIFIER |
| 9 | 117063935 | AX-97820524 | 0 | T | C | 0.123 | -6.46E-02 | 9.63E-02 | 2.61E+00 | 2.72E+00 | 4.99E-01 | no | Rbms3 | intron_variant | MODIFIER |
| 9 | 117084039 | AX-170121357 | 0 | C | T | 0.123 | -6.46E-02 | 9.63E-02 | 2.61E+00 | 2.72E+00 | 4.99E-01 | no | Rbms3 | intron_variant | MODIFIER |
| 9 | 117099192 | AX-97794671 | 0 | A | G | 0.123 | -6.46E-02 | 9.63E-02 | 2.61E+00 | 2.72E+00 | 4.99E-01 | no | Rbms3 | intron_variant | MODIFIER |
| 9 | 117105938 | AX-170346577 | 0 | T | C | 0.123 | -6.46E-02 | 9.63E-02 | 2.61E+00 | 2.72E+00 | 4.99E-01 | no | Rbms3 | intron_variant | MODIFIER |
| 9 | 117110892 | AX-170465215 | 0 | A | T | 0.123 | -6.46E-02 | 9.63E-02 | 2.61E+00 | 2.72E+00 | 4.99E-01 | no | Rbms3 | intron_variant | MODIFIER |
| 9 | 117113707 | AX-170491028 | 0 | G | A | 0.123 | -6.46E-02 | 9.63E-02 | 2.61E+00 | 2.72E+00 | 4.99E-01 | no | Rbms3 | intron_variant | MODIFIER |
| 9 | 117114185 | AX-170366421 | 0 | A | G | 0.123 | -6.46E-02 | 9.63E-02 | 2.61E+00 | 2.72E+00 | 4.99E-01 | no | Rbms3 | intron_variant | MODIFIER |
| 9 | 117117696 | AX-170394766 | 0 | T | A | 0.123 | -6.46E-02 | 9.63E-02 | 2.61E+00 | 2.72E+00 | 4.99E-01 | no | Rbms3 | intron_variant | MODIFIER |
| 9 | 117148783 | AX-170276082 | 0 | T | C | 0.123 | -6.46E-02 | 9.63E-02 | 2.61E+00 | 2.72E+00 | 4.99E-01 | no | Rbms3 | intron_variant | MODIFIER |
| 9 | 117154973 | AX-170232335 | 0 | G | C | 0.123 | -6.46E-02 | 9.63E-02 | 2.61E+00 | 2.72E+00 | 4.99E-01 | no | Rbms3 | intron_variant | MODIFIER |
| 9 | 117383524 | AX-170279267 | 0 | C | T | 0.123 | -6.46E-02 | 9.63E-02 | 2.61E+00 | 2.72E+00 | 4.99E-01 | no | Rbms3 | intron_variant | MODIFIER |
| 9 | 117383580 | AX-170193259 | 0 | C | A | 0.123 | -6.46E-02 | 9.63E-02 | 2.61E+00 | 2.72E+00 | 4.99E-01 | no | Rbms3 | intron_variant | MODIFIER |
| 9 | 117385056 | AX-170307266 | 0 | T | C | 0.123 | -6.46E-02 | 9.63E-02 | 2.61E+00 | 2.72E+00 | 4.99E-01 | no | Rbms3 | intron_variant | MODIFIER |
| 9 | 117388805 | AX-97820867 | 0 | G | C | 0.123 | -6.46E-02 | 9.63E-02 | 2.61E+00 | 2.72E+00 | 4.99E-01 | no | Rbms3 | intron_variant | MODIFIER |
| 9 | 117409996 | AX-170300165 | 0 | C | T | 0.123 | -6.46E-02 | 9.63E-02 | 2.61E+00 | 2.72E+00 | 4.99E-01 | no | Rbms3 | intron_variant | MODIFIER |
| 9 | 117410138 | AX-170185869 | 0 | T | C | 0.123 | -6.46E-02 | 9.63E-02 | 2.61E+00 | 2.72E+00 | 4.99E-01 | no | Rbms3 | intron_variant | MODIFIER |
| 9 | 117445565 | AX-170201678 | 0 | T | C | 0.123 | -6.46E-02 | 9.63E-02 | 2.61E+00 | 2.72E+00 | 4.99E-01 | no | Rbms3 | intron_variant | MODIFIER |
| 9 | 117445604 | AX-97786694 | 0 | T | C | 0.123 | -6.46E-02 | 9.63E-02 | 2.61E+00 | 2.72E+00 | 4.99E-01 | no | Rbms3 | intron_variant | MODIFIER |

| | | | | | | | | | | | | | | | |
|---|---|---|---|---|---|---|---|---|---|---|---|---|---|---|---|
| 9 | 117447920 | AX-170291575 | 0 | T | G | 0.123 | -6.46E-02 | 9.63E-02 | 2.61E+00 | 2.72E+00 | 4.99E-01 | no | Rbms3 | intron_variant | MODIFIER |
| 9 | 117448069 | AX-170448021 | 0 | T | G | 0.123 | -6.46E-02 | 9.63E-02 | 2.61E+00 | 2.72E+00 | 4.99E-01 | no | Rbms3 | intron_variant | MODIFIER |
| 9 | 117453271 | AX-170033371 | 0 | A | G | 0.123 | -6.46E-02 | 9.63E-02 | 2.61E+00 | 2.72E+00 | 4.99E-01 | no | Rbms3 | intron_variant | MODIFIER |
| 9 | 117454582 | AX-170083584 | 0 | G | T | 0.123 | -6.46E-02 | 9.63E-02 | 2.61E+00 | 2.72E+00 | 4.99E-01 | no | Rbms3 | intron_variant | MODIFIER |
| 9 | 117455014 | AX-170083585 | 0 | T | C | 0.123 | -6.46E-02 | 9.63E-02 | 2.61E+00 | 2.72E+00 | 4.99E-01 | no | Rbms3 | intron_variant | MODIFIER |
| 9 | 117476273 | AX-170335466 | 0 | T | C | 0.123 | -6.46E-02 | 9.63E-02 | 2.61E+00 | 2.72E+00 | 4.99E-01 | no | Rbms3 | intron_variant | MODIFIER |
| 9 | 117491466 | AX-97806741 | 0 | A | G | 0.123 | -6.46E-02 | 9.63E-02 | 2.61E+00 | 2.72E+00 | 4.99E-01 | no | Rbms3 | intron_variant | MODIFIER |
| 9 | 117519731 | AX-97798002 | 0 | T | C | 0.123 | -6.46E-02 | 9.63E-02 | 2.61E+00 | 2.72E+00 | 4.99E-01 | no | Rbms3 | intron_variant | MODIFIER |
| 9 | 117519941 | AX-170101331 | 0 | C | T | 0.123 | -6.46E-02 | 9.63E-02 | 2.61E+00 | 2.72E+00 | 4.99E-01 | no | Rbms3 | intron_variant | MODIFIER |
| 9 | 117571114 | AX-97812341 | 0 | T | C | 0.123 | -6.46E-02 | 9.63E-02 | 2.61E+00 | 2.72E+00 | 4.99E-01 | no | Rbms3 | intron_variant | MODIFIER |
| 9 | 117604214 | AX-170311404 | 0 | G | A | 0.123 | -6.46E-02 | 9.63E-02 | 2.61E+00 | 2.72E+00 | 4.99E-01 | no | Rbms3 | intron_variant | MODIFIER |
| 9 | 117610615 | AX-170033368 | 0 | C | T | 0.123 | -6.46E-02 | 9.63E-02 | 2.61E+00 | 2.72E+00 | 4.99E-01 | no | Rbms3 | intron_variant | MODIFIER |
| 9 | 117614001 | AX-170283084 | 0 | A | G | 0.123 | -6.46E-02 | 9.63E-02 | 2.61E+00 | 2.72E+00 | 4.99E-01 | no | Rbms3 | intron_variant | MODIFIER |
| 9 | 117728628 | AX-97828372 | 0 | T | C | 0.123 | -6.46E-02 | 9.63E-02 | 2.61E+00 | 2.72E+00 | 4.99E-01 | no | Gm45881 | intron_variant,non_coding_transcript_variant | MODIFIER |
| 9 | 117729336 | AX-170520447 | 0 | C | G | 0.123 | -6.46E-02 | 9.63E-02 | 2.61E+00 | 2.72E+00 | 4.99E-01 | no | Gm45881 | intron_variant,non_coding_transcript_variant | MODIFIER |
| 9 | 117750011 | AX-170248301 | 0 | A | G | 0.123 | -6.46E-02 | 9.63E-02 | 2.61E+00 | 2.72E+00 | 4.99E-01 | no | Gm45881 | intron_variant,non_coding_transcript_variant | MODIFIER |
| 9 | 117750124 | AX-170364918 | 0 | C | G | 0.123 | -6.46E-02 | 9.63E-02 | 2.61E+00 | 2.72E+00 | 4.99E-01 | no | Gm45881 | intron_variant,non_coding_transcript_variant | MODIFIER |
| 9 | 117750864 | AX-170118015 | 0 | A | G | 0.123 | -6.46E-02 | 9.63E-02 | 2.61E+00 | 2.72E+00 | 4.99E-01 | no | Gm45881 | intron_variant,non_coding_transcript_variant | MODIFIER |
| 9 | 117754940 | AX-170352032 | 0 | C | T | 0.123 | -6.46E-02 | 9.63E-02 | 2.61E+00 | 2.72E+00 | 4.99E-01 | no | Gm45881 | intron_variant,non_coding_transcript_variant | MODIFIER |
| 9 | 117755665 | AX-170112539 | 0 | A | C | 0.123 | -6.46E-02 | 9.63E-02 | 2.61E+00 | 2.72E+00 | 4.99E-01 | no | Gm45881 | intron_variant,non_coding_transcript_variant | MODIFIER |
| 9 | 117755687 | AX-170243847 | 0 | G | T | 0.123 | -6.46E-02 | 9.63E-02 | 2.61E+00 | 2.72E+00 | 4.99E-01 | no | Gm45881 | intron_variant,non_coding_transcript_variant | MODIFIER |
| 9 | 117756982 | AX-170105256 | 0 | A | G | 0.123 | -6.46E-02 | 9.63E-02 | 2.61E+00 | 2.72E+00 | 4.99E-01 | no | Gm45881 | intron_variant,non_coding_transcript_variant | MODIFIER |
| 9 | 117772006 | AX-97809970 | 0 | C | T | 0.123 | -6.46E-02 | 9.63E-02 | 2.61E+00 | 2.72E+00 | 4.99E-01 | no | Gm45881 | intron_variant,non_coding_transcript_variant | MODIFIER |
| 9 | 117772722 | AX-170310308 | 0 | A | G | 0.123 | -6.46E-02 | 9.63E-02 | 2.61E+00 | 2.72E+00 | 4.99E-01 | no | Gm45881 | intron_variant,non_coding_transcript_variant | MODIFIER |
| 9 | 117772792 | AX-170083590 | 0 | G | A | 0.123 | -6.46E-02 | 9.63E-02 | 2.61E+00 | 2.72E+00 | 4.99E-01 | no | Gm45881 | intron_variant,non_coding_transcript_variant | MODIFIER |
| 9 | 117777011 | AX-170473346 | 0 | G | A | 0.123 | -6.46E-02 | 9.63E-02 | 2.61E+00 | 2.72E+00 | 4.99E-01 | no | Gm45881 | intron_variant,non_coding_transcript_variant | MODIFIER |
| 9 | 117796787 | AX-97829343 | 0 | T | C | 0.123 | -6.46E-02 | 9.63E-02 | 2.61E+00 | 2.72E+00 | 4.99E-01 | no | Gm45881 | intron_variant,non_coding_transcript_variant | MODIFIER |
| 9 | 117798155 | AX-170293427 | 0 | G | A | 0.123 | -6.46E-02 | 9.63E-02 | 2.61E+00 | 2.72E+00 | 4.99E-01 | no | Gm45881 | intron_variant,non_coding_transcript_variant | MODIFIER |
| 9 | 117801969 | AX-170505034 | 0 | T | C | 0.123 | -6.46E-02 | 9.63E-02 | 2.61E+00 | 2.72E+00 | 4.99E-01 | no | Gm45881 | intron_variant,non_coding_transcript_variant | MODIFIER |
| 9 | 117803672 | AX-170246424 | 0 | T | G | 0.123 | -6.46E-02 | 9.63E-02 | 2.61E+00 | 2.72E+00 | 4.99E-01 | no | Gm45881 | intron_variant,non_coding_transcript_variant | MODIFIER |
| 9 | 117811165 | AX-170477982 | 0 | C | T | 0.123 | -6.46E-02 | 9.63E-02 | 2.61E+00 | 2.72E+00 | 4.99E-01 | no | Gm45881 | intron_variant,non_coding_transcript_variant | MODIFIER |
| 9 | 117829323 | AX-170367376 | 0 | G | A | 0.123 | -6.46E-02 | 9.63E-02 | 2.61E+00 | 2.72E+00 | 4.99E-01 | no | Gm45881 | intron_variant,non_coding_transcript_variant | MODIFIER |
| 9 | 117830915 | AX-170110596 | 0 | A | C | 0.123 | -6.46E-02 | 9.63E-02 | 2.61E+00 | 2.72E+00 | 4.99E-01 | no | Gm45881 | intron_variant,non_coding_transcript_variant | MODIFIER |
| 9 | 117867289 | AX-97830616 | 0 | A | G | 0.123 | -6.46E-02 | 9.63E-02 | 2.61E+00 | 2.72E+00 | 4.99E-01 | no | Gm45881 | intron_variant,non_coding_transcript_variant | MODIFIER |
| 9 | 117876939 | AX-170287746 | 0 | G | C | 0.123 | -6.46E-02 | 9.63E-02 | 2.61E+00 | 2.72E+00 | 4.99E-01 | no | Zcwpw2 | intron_variant,non_coding_transcript_variant | MODIFIER |
| 9 | 117895563 | AX-97846801 | 0 | A | G | 0.123 | -6.46E-02 | 9.63E-02 | 2.61E+00 | 2.72E+00 | 4.99E-01 | no | Zcwpw2 | intron_variant,non_coding_transcript_variant | MODIFIER |
| 9 | 117898004 | AX-170173751 | 0 | A | G | 0.123 | -6.46E-02 | 9.63E-02 | 2.61E+00 | 2.72E+00 | 4.99E-01 | no | Zcwpw2 | intron_variant,non_coding_transcript_variant | MODIFIER |
| 9 | 117899717 | AX-170462848 | 0 | A | G | 0.123 | -6.46E-02 | 9.63E-02 | 2.61E+00 | 2.72E+00 | 4.99E-01 | no | Zcwpw2 | intron_variant,non_coding_transcript_variant | MODIFIER |
| 9 | 117899737 | AX-170300500 | 0 | T | C | 0.123 | -6.46E-02 | 9.63E-02 | 2.61E+00 | 2.72E+00 | 4.99E-01 | no | Zcwpw2 | intron_variant,non_coding_transcript_variant | MODIFIER |
| 9 | 117902571 | AX-170083595 | 0 | T | C | 0.123 | -6.46E-02 | 9.63E-02 | 2.61E+00 | 2.72E+00 | 4.99E-01 | no | Zcwpw2 | intron_variant,non_coding_transcript_variant | MODIFIER |
| 9 | 117903684 | AX-170158236 | 0 | G | A | 0.123 | -6.46E-02 | 9.63E-02 | 2.61E+00 | 2.72E+00 | 4.99E-01 | no | Zcwpw2 | intron_variant,non_coding_transcript_variant | MODIFIER |
| 9 | 117905046 | AX-170484906 | 0 | C | T | 0.123 | -6.46E-02 | 9.63E-02 | 2.61E+00 | 2.72E+00 | 4.99E-01 | no | Zcwpw2 | intron_variant,non_coding_transcript_variant | MODIFIER |
| 9 | 117905395 | AX-170261645 | 0 | C | G | 0.123 | -6.46E-02 | 9.63E-02 | 2.61E+00 | 2.72E+00 | 4.99E-01 | no | Zcwpw2 | intron_variant,non_coding_transcript_variant | MODIFIER |
| 9 | 117906265 | AX-170317120 | 0 | T | C | 0.123 | -6.46E-02 | 9.63E-02 | 2.61E+00 | 2.72E+00 | 4.99E-01 | no | Zcwpw2 | intron_variant,non_coding_transcript_variant | MODIFIER |
| 9 | 117908600 | AX-170478671 | 0 | A | T | 0.123 | -6.46E-02 | 9.63E-02 | 2.61E+00 | 2.72E+00 | 4.99E-01 | no | Zcwpw2 | intron_variant,non_coding_transcript_variant | MODIFIER |
| 9 | 117908650 | AX-170521946 | 0 | A | G | 0.123 | -6.46E-02 | 9.63E-02 | 2.61E+00 | 2.72E+00 | 4.99E-01 | no | Zcwpw2 | intron_variant,non_coding_transcript_variant | MODIFIER |
| 9 | 117909189 | AX-97808905 | 0 | T | G | 0.123 | -6.46E-02 | 9.63E-02 | 2.61E+00 | 2.72E+00 | 4.99E-01 | no | Zcwpw2 | intron_variant,non_coding_transcript_variant | MODIFIER |
| 9 | 117909901 | AX-170461699 | 0 | C | A | 0.123 | -6.46E-02 | 9.63E-02 | 2.61E+00 | 2.72E+00 | 4.99E-01 | no | Zcwpw2 | intron_variant,non_coding_transcript_variant | MODIFIER |
| 9 | 117916720 | AX-170354524 | 0 | T | C | 0.123 | -6.46E-02 | 9.63E-02 | 2.61E+00 | 2.72E+00 | 4.99E-01 | no | Zcwpw2 | intron_variant,non_coding_transcript_variant | MODIFIER |
| 9 | 117960836 | AX-170033376 | 0 | T | C | 0.123 | -6.46E-02 | 9.63E-02 | 2.61E+00 | 2.72E+00 | 4.99E-01 | no | Zcwpw2 | intron_variant,non_coding_transcript_variant | MODIFIER |
| 9 | 117963035 | AX-170033382 | 0 | T | A | 0.123 | -6.46E-02 | 9.63E-02 | 2.61E+00 | 2.72E+00 | 4.99E-01 | no | Zcwpw2 | intron_variant,non_coding_transcript_variant | MODIFIER |
| 9 | 117973441 | AX-170215848 | 0 | T | C | 0.123 | -6.46E-02 | 9.63E-02 | 2.61E+00 | 2.72E+00 | 4.99E-01 | no | Zcwpw2 | intron_variant,non_coding_transcript_variant | MODIFIER |
| 9 | 118042384 | AX-97805660 | 0 | T | C | 0.123 | -6.46E-02 | 9.63E-02 | 2.61E+00 | 2.72E+00 | 4.99E-01 | no | Azi2 | intron_variant | MODIFIER |
| 9 | 118078114 | AX-170033378 | 0 | A | G | 0.123 | -6.46E-02 | 9.63E-02 | 2.61E+00 | 2.72E+00 | 4.99E-01 | no | Cmc1 | intron_variant | MODIFIER |

| | | | | | | | | | | | | | | | |
|---|---|---|---|---|---|---|---|---|---|---|---|---|---|---|---|
| 9 | 118078590 | AX-170357529 | 0 | T | C | 0.123 | -6.46E-02 | 9.63E-02 | 2.61E+00 | 2.72E+00 | 4.99E-01 | no | Cmc1 | intron_variant | MODIFIER |
| 9 | 118081820 | AX-170033377 | 0 | G | C | 0.123 | -6.46E-02 | 9.63E-02 | 2.61E+00 | 2.72E+00 | 4.99E-01 | no | Cmc1 | intron_variant | MODIFIER |
| 9 | 118115380 | AX-97790333 | 0 | T | C | 0.123 | -6.46E-02 | 9.63E-02 | 2.61E+00 | 2.72E+00 | 4.99E-01 | no | Cmc1 | missense_variant | MODERATE |
| 9 | 118185855 | AX-170260217 | 0 | C | T | 0.123 | -6.46E-02 | 9.63E-02 | 2.61E+00 | 2.72E+00 | 4.99E-01 | no | | intergenic_variant | MODIFIER |
| 9 | 118217189 | AX-170382227 | 0 | T | C | 0.123 | -6.46E-02 | 9.63E-02 | 2.61E+00 | 2.72E+00 | 4.99E-01 | no | | intergenic_variant | MODIFIER |
| 9 | 118219612 | AX-97805661 | 0 | A | G | 0.123 | -6.46E-02 | 9.63E-02 | 2.61E+00 | 2.72E+00 | 4.99E-01 | no | | intergenic_variant | MODIFIER |
| 9 | 118252695 | AX-170090428 | 0 | T | G | 0.123 | -6.46E-02 | 9.63E-02 | 2.61E+00 | 2.72E+00 | 4.99E-01 | no | | intergenic_variant | MODIFIER |
| 9 | 118256142 | AX-170033381 | 0 | C | T | 0.123 | -6.46E-02 | 9.63E-02 | 2.61E+00 | 2.72E+00 | 4.99E-01 | no | | regulatory_region_variant | MODIFIER |
| 9 | 118267969 | AX-170370563 | 0 | A | G | 0.123 | -6.46E-02 | 9.63E-02 | 2.61E+00 | 2.72E+00 | 4.99E-01 | no | | intergenic_variant | MODIFIER |
| 9 | 118280680 | AX-170534456 | 0 | C | T | 0.123 | -6.46E-02 | 9.63E-02 | 2.61E+00 | 2.72E+00 | 4.99E-01 | no | | intergenic_variant | MODIFIER |
| 9 | 118297113 | AX-97802551 | 0 | G | T | 0.123 | -6.46E-02 | 9.63E-02 | 2.61E+00 | 2.72E+00 | 4.99E-01 | no | | intergenic_variant | MODIFIER |
| 9 | 118299940 | AX-170328324 | 0 | T | C | 0.123 | -6.46E-02 | 9.63E-02 | 2.61E+00 | 2.72E+00 | 4.99E-01 | no | | intergenic_variant | MODIFIER |
| 9 | 118309819 | AX-97841314 | 0 | T | C | 0.123 | -6.46E-02 | 9.63E-02 | 2.61E+00 | 2.72E+00 | 4.99E-01 | no | | intergenic_variant | MODIFIER |
| 9 | 118318178 | AX-170376284 | 0 | G | A | 0.123 | -6.46E-02 | 9.63E-02 | 2.61E+00 | 2.72E+00 | 4.99E-01 | no | | intergenic_variant | MODIFIER |
| 9 | 118336403 | AX-170518596 | 0 | C | T | 0.123 | -6.46E-02 | 9.63E-02 | 2.61E+00 | 2.72E+00 | 4.99E-01 | no | | intergenic_variant | MODIFIER |
| 9 | 118337342 | AX-170359477 | 0 | A | G | 0.123 | -6.46E-02 | 9.63E-02 | 2.61E+00 | 2.72E+00 | 4.99E-01 | no | | intergenic_variant | MODIFIER |
| 9 | 118337771 | AX-170377193 | 0 | C | T | 0.123 | -6.46E-02 | 9.63E-02 | 2.61E+00 | 2.72E+00 | 4.99E-01 | no | | intergenic_variant | MODIFIER |
| 9 | 118341310 | AX-170146077 | 0 | T | G | 0.123 | -6.46E-02 | 9.63E-02 | 2.61E+00 | 2.72E+00 | 4.99E-01 | no | | regulatory_region_variant | MODIFIER |
| 9 | 118485763 | AX-97837246 | 0 | T | A | 0.123 | -6.46E-02 | 9.63E-02 | 2.61E+00 | 2.72E+00 | 4.99E-01 | no | Eomes | 3_prime_UTR_variant | MODIFIER |
| 9 | 118590243 | AX-170033389 | 0 | T | C | 0.123 | -6.46E-02 | 9.63E-02 | 2.61E+00 | 2.72E+00 | 4.99E-01 | no | Gm45897 | upstream_gene_variant | MODIFIER |
| 9 | 118649608 | AX-97799603 | 0 | C | T | 0.123 | -6.46E-02 | 9.63E-02 | 2.61E+00 | 2.72E+00 | 4.99E-01 | no | Itga9 | intron_variant | MODIFIER |
| 9 | 118654687 | AX-170033390 | 0 | C | T | 0.123 | -6.46E-02 | 9.63E-02 | 2.61E+00 | 2.72E+00 | 4.99E-01 | no | Itga9 | intron_variant | MODIFIER |
| 9 | 118672684 | AX-170174774 | 0 | G | C | 0.123 | -6.46E-02 | 9.63E-02 | 2.61E+00 | 2.72E+00 | 4.99E-01 | no | Itga9 | intron_variant | MODIFIER |
| 9 | 118676285 | AX-168418097 | 0 | C | T | 0.123 | -6.46E-02 | 9.63E-02 | 2.61E+00 | 2.72E+00 | 4.99E-01 | no | Itga9 | intron_variant | MODIFIER |
| 9 | 118681061 | AX-170381662 | 0 | A | T | 0.123 | -6.46E-02 | 9.63E-02 | 2.61E+00 | 2.72E+00 | 4.99E-01 | no | Itga9 | intron_variant | MODIFIER |
| 9 | 118690679 | AX-170033387 | 0 | G | A | 0.123 | -6.46E-02 | 9.63E-02 | 2.61E+00 | 2.72E+00 | 4.99E-01 | no | Itga9 | intron_variant | MODIFIER |
| 9 | 118690910 | AX-170033391 | 0 | T | C | 0.123 | -6.46E-02 | 9.63E-02 | 2.61E+00 | 2.72E+00 | 4.99E-01 | no | Itga9 | intron_variant | MODIFIER |
| 9 | 118694192 | AX-170090429 | 0 | A | G | 0.123 | -6.46E-02 | 9.63E-02 | 2.61E+00 | 2.72E+00 | 4.99E-01 | no | Itga9 | intron_variant | MODIFIER |
| 9 | 118702603 | AX-170288024 | 0 | G | A | 0.123 | -6.46E-02 | 9.63E-02 | 2.61E+00 | 2.72E+00 | 4.99E-01 | no | Itga9 | intron_variant | MODIFIER |
| 9 | 118711088 | AX-170154336 | 0 | T | C | 0.123 | -6.46E-02 | 9.63E-02 | 2.61E+00 | 2.72E+00 | 4.99E-01 | no | Itga9 | intron_variant | MODIFIER |
| 9 | 118714095 | AX-170033393 | 0 | C | A | 0.123 | -6.46E-02 | 9.63E-02 | 2.61E+00 | 2.72E+00 | 4.99E-01 | no | Itga9 | intron_variant | MODIFIER |
| 9 | 118719802 | AX-170033394 | 0 | G | T | 0.123 | -6.46E-02 | 9.63E-02 | 2.61E+00 | 2.72E+00 | 4.99E-01 | no | Itga9 | intron_variant | MODIFIER |
| 9 | 118739387 | AX-97809402 | 0 | C | T | 0.123 | -6.46E-02 | 9.63E-02 | 2.61E+00 | 2.72E+00 | 4.99E-01 | no | Itga9 | intron_variant | MODIFIER |
| 9 | 118766528 | AX-170465193 | 0 | A | G | 0.123 | -6.46E-02 | 9.63E-02 | 2.61E+00 | 2.72E+00 | 4.99E-01 | no | Itga9 | intron_variant | MODIFIER |
| 9 | 118767028 | AX-170219309 | 0 | T | C | 0.123 | -6.46E-02 | 9.63E-02 | 2.61E+00 | 2.72E+00 | 4.99E-01 | no | Itga9 | intron_variant | MODIFIER |
| 9 | 118779602 | AX-97780738 | 0 | G | A | 0.123 | -6.46E-02 | 9.63E-02 | 2.61E+00 | 2.72E+00 | 4.99E-01 | no | Itga9 | intron_variant | MODIFIER |
| 9 | 118807365 | AX-97809173 | 0 | A | C | 0.123 | -6.46E-02 | 9.63E-02 | 2.61E+00 | 2.72E+00 | 4.99E-01 | no | Itga9 | intron_variant | MODIFIER |
| 9 | 118811652 | AX-97790615 | 0 | A | G | 0.123 | -6.46E-02 | 9.63E-02 | 2.61E+00 | 2.72E+00 | 4.99E-01 | no | Itga9 | intron_variant | MODIFIER |
| 9 | 118847462 | AX-97851058 | 0 | A | G | 0.123 | -6.46E-02 | 9.63E-02 | 2.61E+00 | 2.72E+00 | 4.99E-01 | no | Itga9 | intron_variant | MODIFIER |
| 9 | 118847623 | AX-170290569 | 0 | T | C | 0.123 | -6.46E-02 | 9.63E-02 | 2.61E+00 | 2.72E+00 | 4.99E-01 | no | Itga9 | intron_variant | MODIFIER |
| 9 | 118847813 | AX-170408757 | 0 | C | T | 0.123 | -6.46E-02 | 9.63E-02 | 2.61E+00 | 2.72E+00 | 4.99E-01 | no | Itga9 | intron_variant | MODIFIER |
| 9 | 118863722 | AX-170238831 | 0 | C | T | 0.123 | -6.46E-02 | 9.63E-02 | 2.61E+00 | 2.72E+00 | 4.99E-01 | no | Itga9 | intron_variant | MODIFIER |
| 9 | 118864089 | AX-170107210 | 0 | G | A | 0.123 | -6.46E-02 | 9.63E-02 | 2.61E+00 | 2.72E+00 | 4.99E-01 | no | Itga9 | intron_variant | MODIFIER |
| 9 | 118866374 | AX-170490875 | 0 | G | C | 0.123 | -6.46E-02 | 9.63E-02 | 2.61E+00 | 2.72E+00 | 4.99E-01 | no | Itga9 | intron_variant | MODIFIER |
| 9 | 118867833 | AX-170149050 | 0 | A | G | 0.123 | -6.46E-02 | 9.63E-02 | 2.61E+00 | 2.72E+00 | 4.99E-01 | no | Itga9 | intron_variant | MODIFIER |
| 9 | 118869489 | AX-170354620 | 0 | A | G | 0.123 | -6.46E-02 | 9.63E-02 | 2.61E+00 | 2.72E+00 | 4.99E-01 | no | Itga9 | intron_variant | MODIFIER |
| 9 | 118872301 | AX-170180330 | 0 | T | G | 0.123 | -6.46E-02 | 9.63E-02 | 2.61E+00 | 2.72E+00 | 4.99E-01 | no | Itga9 | intron_variant | MODIFIER |
| 9 | 118873887 | AX-170526136 | 0 | C | T | 0.123 | -6.46E-02 | 9.63E-02 | 2.61E+00 | 2.72E+00 | 4.99E-01 | no | Itga9 | intron_variant | MODIFIER |
| 9 | 118893324 | AX-170303555 | 0 | T | C | 0.123 | -6.46E-02 | 9.63E-02 | 2.61E+00 | 2.72E+00 | 4.99E-01 | no | Itga9 | intron_variant | MODIFIER |
| 9 | 118951052 | AX-170413859 | 0 | G | A | 0.123 | -6.46E-02 | 9.63E-02 | 2.61E+00 | 2.72E+00 | 4.99E-01 | no | Ctdspl | intron_variant | MODIFIER |
| 9 | 119081634 | AX-170359406 | 0 | T | G | 0.123 | -6.46E-02 | 9.63E-02 | 2.61E+00 | 2.72E+00 | 4.99E-01 | no | Plcd1 | intron_variant | MODIFIER |
| 9 | 119084693 | AX-97793089 | 0 | C | T | 0.123 | -6.46E-02 | 9.63E-02 | 2.61E+00 | 2.72E+00 | 4.99E-01 | no | Plcd1 | synonymous_variant | LOW |
| 9 | 119119559 | AX-170033404 | 0 | C | T | 0.123 | -6.46E-02 | 9.63E-02 | 2.61E+00 | 2.72E+00 | 4.99E-01 | no | Dlec1 | intron_variant,NMD_transcript_variant | MODIFIER |
| 9 | 119206137 | AX-170033405 | 0 | T | C | 0.123 | -6.46E-02 | 9.63E-02 | 2.61E+00 | 2.72E+00 | 4.99E-01 | no | Slc22a13 | intron_variant | MODIFIER |

| | | | | | | | | | | | | | | | |
|---|---|---|---|---|---|---|---|---|---|---|---|---|---|---|---|
| 9 | 119208638 | AX-170336713 | 0 | C | T | 0.123 | -6.46E-02 | 9.63E-02 | 2.61E+00 | 2.72E+00 | 4.99E-01 | no | Slc22a13 | intron_variant | MODIFIER |
| 9 | 119218777 | AX-170033398 | 0 | T | C | 0.123 | -6.46E-02 | 9.63E-02 | 2.61E+00 | 2.72E+00 | 4.99E-01 | no | Slc22a14 | intron_variant | MODIFIER |
| 14 | 91579147 | AX-97827556 | 0 | T | C | 0.053 | 1.02E-01 | 1.54E-01 | 2.67E+00 | 2.82E+00 | 4.99E-01 | no | | intergenic_variant | MODIFIER |
| 18 | 63100428 | AX-170197279 | 1 | A | G | 0.188 | -7.49E-02 | 1.12E-01 | 2.70E+00 | 2.75E+00 | 4.99E-01 | no | Piezo2 | intron_variant | MODIFIER |
| 8 | 34710445 | AX-170097217 | 1 | C | A | 0.143 | -8.85E-02 | 1.32E-01 | 2.65E+00 | 2.73E+00 | 4.99E-01 | no | | regulatory_region_variant | MODIFIER |
| 1 | 175187012 | AX-170002326 | 0 | A | C | 0.237 | 7.74E-02 | 1.13E-01 | 2.54E+00 | 2.60E+00 | 5.00E-01 | no | Rgs7 | intron_variant | MODIFIER |
| 18 | 57019005 | AX-170252462 | 1 | C | T | 0.411 | 6.76E-02 | 9.32E-02 | 2.38E+00 | 2.46E+00 | 5.00E-01 | no | | intergenic_variant | MODIFIER |
| 8 | 82991088 | AX-97804672 | 0 | T | C | 0.211 | 7.87E-02 | 1.18E-01 | 2.74E+00 | 2.81E+00 | 5.00E-01 | no | Rnf150 | intron_variant | MODIFIER |
| 18 | 69868374 | AX-97781249 | 0 | T | C | 0.175 | -8.31E-02 | 1.24E-01 | 2.65E+00 | 2.70E+00 | 5.00E-01 | no | | intergenic_variant | MODIFIER |
| 18 | 69893717 | AX-170061236 | 0 | T | C | 0.175 | -8.31E-02 | 1.24E-01 | 2.65E+00 | 2.70E+00 | 5.00E-01 | no | AC108433.1 | intron_variant,non_coding_transcript_variant | MODIFIER |
| 18 | 69895322 | AX-170534526 | 0 | T | C | 0.175 | -8.31E-02 | 1.24E-01 | 2.65E+00 | 2.70E+00 | 5.00E-01 | no | AC108433.1 | intron_variant,non_coding_transcript_variant | MODIFIER |
| 18 | 70052507 | AX-97835756 | 0 | A | G | 0.175 | -8.31E-02 | 1.24E-01 | 2.65E+00 | 2.70E+00 | 5.00E-01 | no | Rab27b | intron_variant | MODIFIER |
| 18 | 70785271 | AX-170174844 | 0 | C | T | 0.175 | -8.31E-02 | 1.24E-01 | 2.65E+00 | 2.70E+00 | 5.00E-01 | no | AC166779.3 | downstream_gene_variant | MODIFIER |
| 18 | 70814978 | AX-97790940 | 0 | G | A | 0.175 | -8.31E-02 | 1.24E-01 | 2.65E+00 | 2.70E+00 | 5.00E-01 | no | | intergenic_variant | MODIFIER |
| 18 | 70871899 | AX-170397717 | 0 | T | C | 0.175 | -8.31E-02 | 1.24E-01 | 2.65E+00 | 2.70E+00 | 5.00E-01 | no | | intergenic_variant | MODIFIER |
| 18 | 70872252 | AX-170396000 | 0 | T | C | 0.175 | -8.31E-02 | 1.24E-01 | 2.65E+00 | 2.70E+00 | 5.00E-01 | no | | intergenic_variant | MODIFIER |
| 18 | 70877284 | AX-170520209 | 0 | G | C | 0.175 | -8.31E-02 | 1.24E-01 | 2.65E+00 | 2.70E+00 | 5.00E-01 | no | | intergenic_variant | MODIFIER |
| 18 | 70932782 | AX-170522195 | 0 | A | G | 0.175 | -8.31E-02 | 1.24E-01 | 2.65E+00 | 2.70E+00 | 5.00E-01 | no | | intergenic_variant | MODIFIER |
| 18 | 70992572 | AX-97815253 | 0 | C | T | 0.175 | -8.31E-02 | 1.24E-01 | 2.65E+00 | 2.70E+00 | 5.00E-01 | no | | intergenic_variant | MODIFIER |
| 18 | 71015149 | AX-170162743 | 0 | T | C | 0.175 | -8.31E-02 | 1.24E-01 | 2.65E+00 | 2.70E+00 | 5.00E-01 | no | | intergenic_variant | MODIFIER |
| 18 | 71015897 | AX-170475089 | 0 | A | G | 0.175 | -8.31E-02 | 1.24E-01 | 2.65E+00 | 2.70E+00 | 5.00E-01 | no | | intergenic_variant | MODIFIER |
| 18 | 71016369 | AX-170123160 | 0 | T | C | 0.175 | -8.31E-02 | 1.24E-01 | 2.65E+00 | 2.70E+00 | 5.00E-01 | no | | intergenic_variant | MODIFIER |
| 18 | 71016411 | AX-170218015 | 0 | G | A | 0.175 | -8.31E-02 | 1.24E-01 | 2.65E+00 | 2.70E+00 | 5.00E-01 | no | | intergenic_variant | MODIFIER |
| 18 | 71040156 | AX-170105639 | 0 | T | C | 0.175 | -8.31E-02 | 1.24E-01 | 2.65E+00 | 2.70E+00 | 5.00E-01 | no | | intergenic_variant | MODIFIER |
| 18 | 71696821 | AX-170230346 | 0 | A | G | 0.175 | -8.31E-02 | 1.24E-01 | 2.65E+00 | 2.70E+00 | 5.00E-01 | no | Dcc | intron_variant | MODIFIER |
| 18 | 71744132 | AX-170035586 | 0 | T | G | 0.175 | -8.31E-02 | 1.24E-01 | 2.65E+00 | 2.70E+00 | 5.00E-01 | no | Dcc | intron_variant | MODIFIER |
| 18 | 71817352 | AX-170457487 | 0 | T | C | 0.175 | -8.31E-02 | 1.24E-01 | 2.65E+00 | 2.70E+00 | 5.00E-01 | no | Dcc | intron_variant | MODIFIER |
| 18 | 71929204 | AX-170015957 | 0 | T | A | 0.175 | -8.31E-02 | 1.24E-01 | 2.65E+00 | 2.70E+00 | 5.00E-01 | no | Dcc | intron_variant | MODIFIER |
| 18 | 71947707 | AX-170242751 | 0 | C | T | 0.175 | -8.31E-02 | 1.24E-01 | 2.65E+00 | 2.70E+00 | 5.00E-01 | no | Dcc | intron_variant | MODIFIER |
| 4 | 88922862 | AX-97798252 | 0 | G | T | 0.254 | 7.43E-02 | 1.07E-01 | 2.49E+00 | 2.56E+00 | 5.00E-01 | no | AL805899.1 | intron_variant,non_coding_transcript_variant | MODIFIER |
| 18 | 43643551 | AX-170267195 | 0 | C | T | 0.079 | 1.18E-01 | 1.77E-01 | 2.65E+00 | 2.78E+00 | 5.00E-01 | no | Jakmip2 | intron_variant | MODIFIER |
| 10 | 28597891 | AX-170002967 | 1 | T | A | 0.196 | 6.54E-02 | 9.84E-02 | 2.81E+00 | 2.91E+00 | 5.00E-01 | no | Ptprk | downstream_gene_variant | MODIFIER |
| 7 | 36498569 | AX-170241675 | 0 | G | A | 0.351 | 6.00E-02 | 8.97E-02 | 2.63E+00 | 2.72E+00 | 5.00E-01 | no | | intergenic_variant | MODIFIER |
| 9 | 40673497 | AX-170082213 | 0 | C | T | 0.43 | -4.97E-02 | 7.48E-02 | 2.70E+00 | 2.80E+00 | 5.00E-01 | no | Gm21915 | downstream_gene_variant | MODIFIER |
| 6 | 134786652 | AX-170076311 | 0 | A | G | 0.579 | 5.85E-02 | 8.43E-02 | 2.49E+00 | 2.55E+00 | 5.00E-01 | no | Dusp16 | intron_variant | MODIFIER |
| 15 | 79377174 | AX-170012069 | 0 | T | C | 0.57 | 5.39E-02 | 7.84E-02 | 2.52E+00 | 2.59E+00 | 5.00E-01 | no | Tmem184b | synonymous_variant | LOW |
| 4 | 128703836 | AX-170235536 | 1 | C | T | 0.161 | 7.48E-02 | 1.11E-01 | 2.61E+00 | 2.67E+00 | 5.00E-01 | no | Phc2 | intron_variant | MODIFIER |
| 1 | 144249217 | AX-170283982 | 0 | G | T | 0.211 | 9.22E-02 | 1.38E-01 | 2.69E+00 | 2.74E+00 | 5.00E-01 | no | Rgs1 | upstream_gene_variant | MODIFIER |
| 8 | 100647610 | AX-97838902 | 1 | G | A | 0.241 | 7.50E-02 | 1.13E-01 | 2.73E+00 | 2.80E+00 | 5.01E-01 | no | | intergenic_variant | MODIFIER |
| 18 | 49697566 | AX-170250673 | 0 | A | G | 0.088 | 9.77E-02 | 1.43E-01 | 2.50E+00 | 2.64E+00 | 5.01E-01 | no | Dtwd2 | downstream_gene_variant | MODIFIER |
| 18 | 65572378 | AX-97834973 | 0 | G | A | 0.404 | 5.77E-02 | 8.31E-02 | 2.48E+00 | 2.55E+00 | 5.01E-01 | no | | regulatory_region_variant | MODIFIER |
| 1 | 173674084 | AX-170043628 | 1 | C | G | 0.223 | 8.02E-02 | 1.17E-01 | 2.53E+00 | 2.59E+00 | 5.01E-01 | no | Ifi208 | intron_variant | MODIFIER |
| 11 | 74260023 | AX-170276870 | 0 | C | A | 0.281 | 4.76E-02 | 7.21E-02 | 2.66E+00 | 2.80E+00 | 5.01E-01 | no | Olfr405-ps1 | upstream_gene_variant | MODIFIER |
| 10 | 107985652 | AX-170367707 | 0 | A | G | 0.333 | -5.60E-02 | 8.40E-02 | 2.65E+00 | 2.73E+00 | 5.01E-01 | no | | intergenic_variant | MODIFIER |
| 5 | 84071223 | AX-170230773 | 1 | C | T | 0.438 | 7.47E-02 | 1.12E-01 | 2.81E+00 | 2.86E+00 | 5.01E-01 | no | Epha5 | synonymous_variant | LOW |
| 16 | 50114070 | AX-97803169 | 0 | G | A | 0.509 | 5.57E-02 | 8.27E-02 | 2.59E+00 | 2.64E+00 | 5.01E-01 | no | Gm8824 | downstream_gene_variant | MODIFIER |
| 1 | 93347168 | AX-170341053 | 1 | A | G | 0.152 | 1.33E-01 | 1.98E-01 | 2.63E+00 | 2.70E+00 | 5.01E-01 | no | Pask | upstream_gene_variant | MODIFIER |
| 3 | 23003226 | AX-97795546 | 1 | G | A | 0.045 | -7.38E-02 | 1.10E-01 | 2.54E+00 | 2.72E+00 | 5.01E-01 | no | | intergenic_variant | MODIFIER |
| 1 | 14624812 | AX-97810992 | 1 | T | C | 0.366 | 5.29E-02 | 7.86E-02 | 2.60E+00 | 2.66E+00 | 5.01E-01 | no | | intergenic_variant | MODIFIER |
| 1 | 135835456 | AX-170086616 | 0 | A | C | 0.211 | 8.29E-02 | 1.24E-01 | 2.66E+00 | 2.73E+00 | 5.01E-01 | no | Lad1 | downstream_gene_variant | MODIFIER |
| 1 | 135837872 | AX-170478082 | 0 | C | T | 0.211 | 8.29E-02 | 1.24E-01 | 2.66E+00 | 2.73E+00 | 5.01E-01 | no | Tnnt2 | upstream_gene_variant | MODIFIER |
| 1 | 135838160 | AX-170180145 | 0 | C | T | 0.211 | 8.29E-02 | 1.24E-01 | 2.66E+00 | 2.73E+00 | 5.01E-01 | no | Tnnt2 | upstream_gene_variant | MODIFIER |
| 1 | 135838648 | AX-170248279 | 0 | T | C | 0.211 | 8.29E-02 | 1.24E-01 | 2.66E+00 | 2.73E+00 | 5.01E-01 | no | Tnnt2 | upstream_gene_variant | MODIFIER |
| 1 | 135844368 | AX-170353270 | 0 | G | A | 0.211 | 8.29E-02 | 1.24E-01 | 2.66E+00 | 2.73E+00 | 5.01E-01 | no | Tnnt2 | intron_variant | MODIFIER |

| | | | | | | | | | | | | | | | |
|---|---|---|---|---|---|---|---|---|---|---|---|---|---|---|---|
| 1 | 135862578 | AX-170154623 | 0 | T | C | 0.211 | 8.29E-02 | 1.24E-01 | 2.66E+00 | 2.73E+00 | 5.01E-01 | no | | regulatory_region_variant | MODIFIER |
| 1 | 135862614 | AX-170042834 | 0 | G | A | 0.211 | 8.29E-02 | 1.24E-01 | 2.66E+00 | 2.73E+00 | 5.01E-01 | no | | regulatory_region_variant | MODIFIER |
| 1 | 135863244 | AX-170265676 | 0 | G | A | 0.211 | 8.29E-02 | 1.24E-01 | 2.66E+00 | 2.73E+00 | 5.01E-01 | no | | intergenic_variant | MODIFIER |
| 1 | 135866357 | AX-97851339 | 0 | A | G | 0.211 | 8.29E-02 | 1.24E-01 | 2.66E+00 | 2.73E+00 | 5.01E-01 | no | Gm28883 | upstream_gene_variant | MODIFIER |
| 1 | 135883165 | AX-170086617 | 0 | T | C | 0.211 | 8.29E-02 | 1.24E-01 | 2.66E+00 | 2.73E+00 | 5.01E-01 | no | Pkp1 | intron_variant | MODIFIER |
| 1 | 135902617 | AX-170450742 | 0 | C | T | 0.211 | 8.29E-02 | 1.24E-01 | 2.66E+00 | 2.73E+00 | 5.01E-01 | no | Pkp1 | intron_variant | MODIFIER |
| 1 | 135903758 | AX-170367371 | 0 | T | C | 0.211 | 8.29E-02 | 1.24E-01 | 2.66E+00 | 2.73E+00 | 5.01E-01 | no | Pkp1 | intron_variant | MODIFIER |
| 1 | 135907260 | AX-170086618 | 0 | T | C | 0.211 | 8.29E-02 | 1.24E-01 | 2.66E+00 | 2.73E+00 | 5.01E-01 | no | Pkp1 | intron_variant | MODIFIER |
| 1 | 135911278 | AX-170355668 | 0 | T | A | 0.211 | 8.29E-02 | 1.24E-01 | 2.66E+00 | 2.73E+00 | 5.01E-01 | no | Pkp1 | intron_variant | MODIFIER |
| 1 | 135951762 | AX-170370491 | 0 | G | A | 0.211 | 8.29E-02 | 1.24E-01 | 2.66E+00 | 2.73E+00 | 5.01E-01 | no | Igfn1 | downstream_gene_variant | MODIFIER |
| 1 | 135952005 | AX-170162677 | 0 | T | C | 0.211 | 8.29E-02 | 1.24E-01 | 2.66E+00 | 2.73E+00 | 5.01E-01 | no | Igfn1 | downstream_gene_variant | MODIFIER |
| 1 | 135954235 | AX-170259306 | 0 | T | C | 0.211 | 8.29E-02 | 1.24E-01 | 2.66E+00 | 2.73E+00 | 5.01E-01 | no | Igfn1 | downstream_gene_variant | MODIFIER |
| 1 | 135955688 | AX-170042835 | 0 | A | G | 0.211 | 8.29E-02 | 1.24E-01 | 2.66E+00 | 2.73E+00 | 5.01E-01 | no | Igfn1 | synonymous_variant | LOW |
| 1 | 135957598 | AX-97829369 | 0 | A | G | 0.211 | 8.29E-02 | 1.24E-01 | 2.66E+00 | 2.73E+00 | 5.01E-01 | no | Igfn1 | intron_variant | MODIFIER |
| 1 | 135978841 | AX-170138306 | 0 | T | C | 0.211 | 8.29E-02 | 1.24E-01 | 2.66E+00 | 2.73E+00 | 5.01E-01 | no | Igfn1 | intron_variant | MODIFIER |
| 1 | 135980102 | AX-170420063 | 0 | C | G | 0.211 | 8.29E-02 | 1.24E-01 | 2.66E+00 | 2.73E+00 | 5.01E-01 | no | Igfn1 | intron_variant | MODIFIER |
| 1 | 135989704 | AX-170206003 | 0 | C | T | 0.211 | 8.29E-02 | 1.24E-01 | 2.66E+00 | 2.73E+00 | 5.01E-01 | no | Igfn1 | intron_variant | MODIFIER |
| 1 | 135989754 | AX-97839770 | 0 | A | G | 0.211 | 8.29E-02 | 1.24E-01 | 2.66E+00 | 2.73E+00 | 5.01E-01 | no | Igfn1 | intron_variant | MODIFIER |
| 1 | 135990012 | AX-170312297 | 0 | T | C | 0.211 | 8.29E-02 | 1.24E-01 | 2.66E+00 | 2.73E+00 | 5.01E-01 | no | Igfn1 | intron_variant | MODIFIER |
| 1 | 136001694 | AX-170277409 | 0 | G | A | 0.211 | 8.29E-02 | 1.24E-01 | 2.66E+00 | 2.73E+00 | 5.01E-01 | no | Igfn1 | intron_variant | MODIFIER |
| 1 | 136020793 | AX-170351836 | 0 | T | C | 0.211 | 8.29E-02 | 1.24E-01 | 2.66E+00 | 2.73E+00 | 5.01E-01 | no | Tmem9 | intron_variant | MODIFIER |
| 1 | 136028093 | AX-170337880 | 0 | G | A | 0.211 | 8.29E-02 | 1.24E-01 | 2.66E+00 | 2.73E+00 | 5.01E-01 | no | Tmem9 | intron_variant | MODIFIER |
| 1 | 136039900 | AX-170001774 | 0 | T | A | 0.211 | 8.29E-02 | 1.24E-01 | 2.66E+00 | 2.73E+00 | 5.01E-01 | no | Tmem9 | downstream_gene_variant | MODIFIER |
| 1 | 136040194 | AX-170001775 | 0 | A | G | 0.211 | 8.29E-02 | 1.24E-01 | 2.66E+00 | 2.73E+00 | 5.01E-01 | no | Tmem9 | downstream_gene_variant | MODIFIER |
| 1 | 136040214 | AX-170142926 | 0 | T | C | 0.211 | 8.29E-02 | 1.24E-01 | 2.66E+00 | 2.73E+00 | 5.01E-01 | no | Tmem9 | downstream_gene_variant | MODIFIER |
| 1 | 136040387 | AX-170146893 | 0 | A | G | 0.211 | 8.29E-02 | 1.24E-01 | 2.66E+00 | 2.73E+00 | 5.01E-01 | no | | regulatory_region_variant | MODIFIER |
| 1 | 136042714 | AX-170447053 | 0 | A | T | 0.211 | 8.29E-02 | 1.24E-01 | 2.66E+00 | 2.73E+00 | 5.01E-01 | no | | intergenic_variant | MODIFIER |
| 1 | 136043054 | AX-170272306 | 0 | A | G | 0.211 | 8.29E-02 | 1.24E-01 | 2.66E+00 | 2.73E+00 | 5.01E-01 | no | | intergenic_variant | MODIFIER |
| 1 | 136047542 | AX-170042838 | 0 | G | C | 0.211 | 8.29E-02 | 1.24E-01 | 2.66E+00 | 2.73E+00 | 5.01E-01 | no | Ascl5 | upstream_gene_variant | MODIFIER |
| 1 | 136069821 | AX-170434435 | 0 | C | T | 0.211 | 8.29E-02 | 1.24E-01 | 2.66E+00 | 2.73E+00 | 5.01E-01 | no | Cacna1s | intron_variant | MODIFIER |
| 1 | 136069968 | AX-170507181 | 0 | A | C | 0.211 | 8.29E-02 | 1.24E-01 | 2.66E+00 | 2.73E+00 | 5.01E-01 | no | Cacna1s | intron_variant | MODIFIER |
| 1 | 136070498 | AX-170519690 | 0 | A | C | 0.211 | 8.29E-02 | 1.24E-01 | 2.66E+00 | 2.73E+00 | 5.01E-01 | no | Cacna1s | intron_variant | MODIFIER |
| 1 | 136074903 | AX-170159524 | 0 | C | T | 0.211 | 8.29E-02 | 1.24E-01 | 2.66E+00 | 2.73E+00 | 5.01E-01 | no | Cacna1s | intron_variant | MODIFIER |
| 1 | 136085196 | AX-170284070 | 0 | C | G | 0.211 | 8.29E-02 | 1.24E-01 | 2.66E+00 | 2.73E+00 | 5.01E-01 | no | Cacna1s | intron_variant | MODIFIER |
| 1 | 136106722 | AX-170405136 | 0 | A | C | 0.211 | 8.29E-02 | 1.24E-01 | 2.66E+00 | 2.73E+00 | 5.01E-01 | no | Cacna1s | intron_variant | MODIFIER |
| 1 | 136113171 | AX-170454257 | 0 | A | C | 0.211 | 8.29E-02 | 1.24E-01 | 2.66E+00 | 2.73E+00 | 5.01E-01 | no | Cacna1s | intron_variant | MODIFIER |
| 1 | 136116992 | AX-170292856 | 0 | A | G | 0.211 | 8.29E-02 | 1.24E-01 | 2.66E+00 | 2.73E+00 | 5.01E-01 | no | Cacna1s | intron_variant | MODIFIER |
| 1 | 136126896 | AX-170336857 | 0 | G | A | 0.211 | 8.29E-02 | 1.24E-01 | 2.66E+00 | 2.73E+00 | 5.01E-01 | no | Kif21b | upstream_gene_variant | MODIFIER |
| 1 | 136134074 | AX-97850853 | 0 | T | C | 0.211 | 8.29E-02 | 1.24E-01 | 2.66E+00 | 2.73E+00 | 5.01E-01 | no | Kif21b | intron_variant | MODIFIER |
| 1 | 136138959 | AX-170121611 | 0 | C | T | 0.211 | 8.29E-02 | 1.24E-01 | 2.66E+00 | 2.73E+00 | 5.01E-01 | no | Kif21b | intron_variant | MODIFIER |
| 1 | 136138995 | AX-170383351 | 0 | C | T | 0.211 | 8.29E-02 | 1.24E-01 | 2.66E+00 | 2.73E+00 | 5.01E-01 | no | Kif21b | intron_variant | MODIFIER |
| 1 | 136148275 | AX-170354120 | 0 | T | G | 0.211 | 8.29E-02 | 1.24E-01 | 2.66E+00 | 2.73E+00 | 5.01E-01 | no | Kif21b | synonymous_variant | LOW |
| 1 | 136151190 | AX-97834562 | 0 | C | T | 0.211 | 8.29E-02 | 1.24E-01 | 2.66E+00 | 2.73E+00 | 5.01E-01 | no | Kif21b | synonymous_variant | LOW |
| 1 | 136163059 | AX-170184733 | 0 | A | G | 0.211 | 8.29E-02 | 1.24E-01 | 2.66E+00 | 2.73E+00 | 5.01E-01 | no | Kif21b | intron_variant | MODIFIER |
| 1 | 136163156 | AX-170323915 | 0 | T | A | 0.211 | 8.29E-02 | 1.24E-01 | 2.66E+00 | 2.73E+00 | 5.01E-01 | no | Kif21b | splice_region_variant,intron_variant | LOW |
| 1 | 136163185 | AX-170381696 | 0 | C | T | 0.211 | 8.29E-02 | 1.24E-01 | 2.66E+00 | 2.73E+00 | 5.01E-01 | no | Kif21b | intron_variant | MODIFIER |
| 1 | 136190925 | AX-170366427 | 0 | A | G | 0.211 | 8.29E-02 | 1.24E-01 | 2.66E+00 | 2.73E+00 | 5.01E-01 | no | Mroh3 | upstream_gene_variant | MODIFIER |
| 1 | 136199130 | AX-170320487 | 0 | A | G | 0.211 | 8.29E-02 | 1.24E-01 | 2.66E+00 | 2.73E+00 | 5.01E-01 | no | Mroh3 | intron_variant,non_coding_transcript_variant | MODIFIER |
| 1 | 136241629 | AX-170136408 | 0 | T | G | 0.211 | 8.29E-02 | 1.24E-01 | 2.66E+00 | 2.73E+00 | 5.01E-01 | no | | intergenic_variant | MODIFIER |
| 1 | 136249116 | AX-170463465 | 0 | A | G | 0.211 | 8.29E-02 | 1.24E-01 | 2.66E+00 | 2.73E+00 | 5.01E-01 | no | Gm37552 | downstream_gene_variant | MODIFIER |
| 1 | 136249464 | AX-97831143 | 0 | C | T | 0.211 | 8.29E-02 | 1.24E-01 | 2.66E+00 | 2.73E+00 | 5.01E-01 | no | Gm37552 | downstream_gene_variant | MODIFIER |
| 1 | 136249856 | AX-170326248 | 0 | C | A | 0.211 | 8.29E-02 | 1.24E-01 | 2.66E+00 | 2.73E+00 | 5.01E-01 | no | Gm37552 | downstream_gene_variant | MODIFIER |
| 1 | 136251431 | AX-170292282 | 0 | T | G | 0.211 | 8.29E-02 | 1.24E-01 | 2.66E+00 | 2.73E+00 | 5.01E-01 | no | Gm37552 | downstream_gene_variant | MODIFIER |
| 1 | 136251877 | AX-170180471 | 0 | A | G | 0.211 | 8.29E-02 | 1.24E-01 | 2.66E+00 | 2.73E+00 | 5.01E-01 | no | Gm37552 | downstream_gene_variant | MODIFIER |

| | | | | | | | | | | | | | | | |
|---|---|---|---|---|---|---|---|---|---|---|---|---|---|---|---|
| 1 | 136252264 | AX-170098571 | 0 | A | G | 0.211 | 8.29E-02 | 1.24E-01 | 2.66E+00 | 2.73E+00 | 5.01E-01 | no | Gm37552 | downstream_gene_variant | MODIFIER |
| 1 | 136277955 | AX-170449019 | 0 | A | G | 0.211 | 8.29E-02 | 1.24E-01 | 2.66E+00 | 2.73E+00 | 5.01E-01 | no | Camsap2 | intron_variant | MODIFIER |
| 1 | 136279906 | AX-97841292 | 0 | A | C | 0.211 | 8.29E-02 | 1.24E-01 | 2.66E+00 | 2.73E+00 | 5.01E-01 | no | Camsap2 | intron_variant | MODIFIER |
| 1 | 136284241 | AX-170459890 | 0 | G | C | 0.211 | 8.29E-02 | 1.24E-01 | 2.66E+00 | 2.73E+00 | 5.01E-01 | no | Camsap2 | intron_variant | MODIFIER |
| 1 | 136284278 | AX-170369556 | 0 | T | C | 0.211 | 8.29E-02 | 1.24E-01 | 2.66E+00 | 2.73E+00 | 5.01E-01 | no | Camsap2 | intron_variant | MODIFIER |
| 1 | 136295203 | AX-170430959 | 0 | A | G | 0.211 | 8.29E-02 | 1.24E-01 | 2.66E+00 | 2.73E+00 | 5.01E-01 | no | Camsap2 | intron_variant | MODIFIER |
| 1 | 136304895 | AX-170042845 | 0 | A | G | 0.211 | 8.29E-02 | 1.24E-01 | 2.66E+00 | 2.73E+00 | 5.01E-01 | no | Camsap2 | intron_variant | MODIFIER |
| 1 | 136322794 | AX-170416241 | 0 | G | A | 0.211 | 8.29E-02 | 1.24E-01 | 2.66E+00 | 2.73E+00 | 5.01E-01 | no | Camsap2 | intron_variant | MODIFIER |
| 1 | 136323658 | AX-97855463 | 0 | G | A | 0.211 | 8.29E-02 | 1.24E-01 | 2.66E+00 | 2.73E+00 | 5.01E-01 | no | Camsap2 | intron_variant | MODIFIER |
| 1 | 136352118 | AX-170233565 | 0 | T | A | 0.211 | 8.29E-02 | 1.24E-01 | 2.66E+00 | 2.73E+00 | 5.01E-01 | no | | intergenic_variant | MODIFIER |
| 1 | 136362087 | AX-170300722 | 0 | T | C | 0.211 | 8.29E-02 | 1.24E-01 | 2.66E+00 | 2.73E+00 | 5.01E-01 | no | | regulatory_region_variant | MODIFIER |
| 1 | 136362126 | AX-97813324 | 0 | C | T | 0.211 | 8.29E-02 | 1.24E-01 | 2.66E+00 | 2.73E+00 | 5.01E-01 | no | | regulatory_region_variant | MODIFIER |
| 1 | 136373284 | AX-170499514 | 0 | T | C | 0.211 | 8.29E-02 | 1.24E-01 | 2.66E+00 | 2.73E+00 | 5.01E-01 | no | | intergenic_variant | MODIFIER |
| 1 | 136410637 | AX-170472008 | 0 | C | T | 0.211 | 8.29E-02 | 1.24E-01 | 2.66E+00 | 2.73E+00 | 5.01E-01 | no | Ddx59 | upstream_gene_variant | MODIFIER |
| 1 | 136431800 | AX-170195241 | 0 | G | A | 0.211 | 8.29E-02 | 1.24E-01 | 2.66E+00 | 2.73E+00 | 5.01E-01 | no | Ddx59 | intron_variant | MODIFIER |
| 1 | 136432051 | AX-170166947 | 0 | A | C | 0.211 | 8.29E-02 | 1.24E-01 | 2.66E+00 | 2.73E+00 | 5.01E-01 | no | Ddx59 | intron_variant | MODIFIER |
| 1 | 136486216 | AX-170290271 | 0 | G | C | 0.211 | 8.29E-02 | 1.24E-01 | 2.66E+00 | 2.73E+00 | 5.01E-01 | no | Kif14 | intron_variant | MODIFIER |
| 1 | 136500221 | AX-170444665 | 0 | G | A | 0.211 | 8.29E-02 | 1.24E-01 | 2.66E+00 | 2.73E+00 | 5.01E-01 | no | Kif14 | intron_variant | MODIFIER |
| 1 | 136506707 | AX-170395399 | 0 | A | T | 0.211 | 8.29E-02 | 1.24E-01 | 2.66E+00 | 2.73E+00 | 5.01E-01 | no | Kif14 | intron_variant | MODIFIER |
| 1 | 136537577 | AX-170443355 | 0 | A | G | 0.211 | 8.29E-02 | 1.24E-01 | 2.66E+00 | 2.73E+00 | 5.01E-01 | no | Gm37799 | upstream_gene_variant | MODIFIER |
| 1 | 136552477 | AX-170404584 | 0 | A | C | 0.211 | 8.29E-02 | 1.24E-01 | 2.66E+00 | 2.73E+00 | 5.01E-01 | no | Gm37799 | intron_variant,non_coding_transcript_variant | MODIFIER |
| 1 | 136553642 | AX-170346819 | 0 | C | T | 0.211 | 8.29E-02 | 1.24E-01 | 2.66E+00 | 2.73E+00 | 5.01E-01 | no | Gm37799 | intron_variant,non_coding_transcript_variant | MODIFIER |
| 1 | 136553892 | AX-170234938 | 0 | C | T | 0.211 | 8.29E-02 | 1.24E-01 | 2.66E+00 | 2.73E+00 | 5.01E-01 | no | Gm37799 | intron_variant,non_coding_transcript_variant | MODIFIER |
| 1 | 136554165 | AX-170001786 | 0 | G | A | 0.211 | 8.29E-02 | 1.24E-01 | 2.66E+00 | 2.73E+00 | 5.01E-01 | no | Gm37799 | intron_variant,non_coding_transcript_variant | MODIFIER |
| 1 | 136554384 | AX-170151885 | 0 | A | G | 0.211 | 8.29E-02 | 1.24E-01 | 2.66E+00 | 2.73E+00 | 5.01E-01 | no | Gm37799 | intron_variant,non_coding_transcript_variant | MODIFIER |
| 1 | 136561318 | AX-170522327 | 0 | C | G | 0.211 | 8.29E-02 | 1.24E-01 | 2.66E+00 | 2.73E+00 | 5.01E-01 | no | | intergenic_variant | MODIFIER |
| 1 | 136566409 | AX-97795885 | 0 | T | C | 0.211 | 8.29E-02 | 1.24E-01 | 2.66E+00 | 2.73E+00 | 5.01E-01 | no | | intergenic_variant | MODIFIER |
| 1 | 136588512 | AX-170401376 | 0 | A | T | 0.211 | 8.29E-02 | 1.24E-01 | 2.66E+00 | 2.73E+00 | 5.01E-01 | no | | regulatory_region_variant | MODIFIER |
| 1 | 136595236 | AX-170432409 | 0 | G | A | 0.211 | 8.29E-02 | 1.24E-01 | 2.66E+00 | 2.73E+00 | 5.01E-01 | no | | intergenic_variant | MODIFIER |
| 1 | 136595255 | AX-97853769 | 0 | A | G | 0.211 | 8.29E-02 | 1.24E-01 | 2.66E+00 | 2.73E+00 | 5.01E-01 | no | | intergenic_variant | MODIFIER |
| 1 | 136596414 | AX-170282469 | 0 | C | A | 0.211 | 8.29E-02 | 1.24E-01 | 2.66E+00 | 2.73E+00 | 5.01E-01 | no | | intergenic_variant | MODIFIER |
| 1 | 136612790 | AX-97819537 | 0 | T | C | 0.211 | 8.29E-02 | 1.24E-01 | 2.66E+00 | 2.73E+00 | 5.01E-01 | no | | intergenic_variant | MODIFIER |
| 1 | 136638105 | AX-170185864 | 0 | G | A | 0.211 | 8.29E-02 | 1.24E-01 | 2.66E+00 | 2.73E+00 | 5.01E-01 | no | | regulatory_region_variant | MODIFIER |
| 1 | 136682065 | AX-170001783 | 0 | A | G | 0.211 | 8.29E-02 | 1.24E-01 | 2.66E+00 | 2.73E+00 | 5.01E-01 | no | Gm19705 | intron_variant,non_coding_transcript_variant | MODIFIER |
| 1 | 136691925 | AX-170001785 | 0 | G | A | 0.211 | 8.29E-02 | 1.24E-01 | 2.66E+00 | 2.73E+00 | 5.01E-01 | no | Gm19705 | downstream_gene_variant | MODIFIER |
| 1 | 136717077 | AX-170373339 | 0 | G | A | 0.211 | 8.29E-02 | 1.24E-01 | 2.66E+00 | 2.73E+00 | 5.01E-01 | no | Platr22 | intron_variant,non_coding_transcript_variant | MODIFIER |
| 1 | 136719375 | AX-170260146 | 0 | A | T | 0.211 | 8.29E-02 | 1.24E-01 | 2.66E+00 | 2.73E+00 | 5.01E-01 | no | Platr22 | intron_variant,non_coding_transcript_variant | MODIFIER |
| 1 | 136749531 | AX-170343226 | 0 | C | G | 0.211 | 8.29E-02 | 1.24E-01 | 2.66E+00 | 2.73E+00 | 5.01E-01 | no | | regulatory_region_variant | MODIFIER |
| 1 | 136796684 | AX-170001778 | 0 | C | T | 0.211 | 8.29E-02 | 1.24E-01 | 2.66E+00 | 2.73E+00 | 5.01E-01 | no | Gm29718 | downstream_gene_variant | MODIFIER |
| 1 | 136803866 | AX-170001787 | 0 | G | A | 0.211 | 8.29E-02 | 1.24E-01 | 2.66E+00 | 2.73E+00 | 5.01E-01 | no | Gm17781 | upstream_gene_variant | MODIFIER |
| 1 | 136812624 | AX-97838463 | 0 | T | C | 0.211 | 8.29E-02 | 1.24E-01 | 2.66E+00 | 2.73E+00 | 5.01E-01 | no | | intergenic_variant | MODIFIER |
| 1 | 136827050 | AX-170281230 | 0 | G | T | 0.211 | 8.29E-02 | 1.24E-01 | 2.66E+00 | 2.73E+00 | 5.01E-01 | no | | intergenic_variant | MODIFIER |
| 1 | 136827108 | AX-170457855 | 0 | T | C | 0.211 | 8.29E-02 | 1.24E-01 | 2.66E+00 | 2.73E+00 | 5.01E-01 | no | | intergenic_variant | MODIFIER |
| 1 | 136827267 | AX-170086620 | 0 | G | T | 0.211 | 8.29E-02 | 1.24E-01 | 2.66E+00 | 2.73E+00 | 5.01E-01 | no | | intergenic_variant | MODIFIER |
| 1 | 136847914 | AX-170252801 | 0 | A | G | 0.211 | 8.29E-02 | 1.24E-01 | 2.66E+00 | 2.73E+00 | 5.01E-01 | no | Nr5a2 | intron_variant | MODIFIER |
| 1 | 136847981 | AX-170490440 | 0 | C | G | 0.211 | 8.29E-02 | 1.24E-01 | 2.66E+00 | 2.73E+00 | 5.01E-01 | no | Nr5a2 | intron_variant | MODIFIER |
| 1 | 136848684 | AX-170498273 | 0 | A | T | 0.211 | 8.29E-02 | 1.24E-01 | 2.66E+00 | 2.73E+00 | 5.01E-01 | no | Nr5a2 | intron_variant | MODIFIER |
| 1 | 136849281 | AX-170440693 | 0 | G | A | 0.211 | 8.29E-02 | 1.24E-01 | 2.66E+00 | 2.73E+00 | 5.01E-01 | no | Nr5a2 | intron_variant | MODIFIER |
| 1 | 136853881 | AX-170042854 | 0 | C | G | 0.211 | 8.29E-02 | 1.24E-01 | 2.66E+00 | 2.73E+00 | 5.01E-01 | no | Nr5a2 | intron_variant | MODIFIER |
| 1 | 136856450 | AX-97784607 | 0 | T | C | 0.211 | 8.29E-02 | 1.24E-01 | 2.66E+00 | 2.73E+00 | 5.01E-01 | no | Nr5a2 | intron_variant | MODIFIER |
| 1 | 136861306 | AX-170499075 | 0 | T | C | 0.211 | 8.29E-02 | 1.24E-01 | 2.66E+00 | 2.73E+00 | 5.01E-01 | no | Nr5a2 | intron_variant | MODIFIER |
| 1 | 136861707 | AX-170287063 | 0 | C | T | 0.211 | 8.29E-02 | 1.24E-01 | 2.66E+00 | 2.73E+00 | 5.01E-01 | no | Nr5a2 | intron_variant | MODIFIER |
| 1 | 136873521 | AX-170394872 | 0 | A | G | 0.211 | 8.29E-02 | 1.24E-01 | 2.66E+00 | 2.73E+00 | 5.01E-01 | no | Nr5a2 | intron_variant | MODIFIER |
| 1 | 136873626 | AX-170299354 | 0 | G | A | 0.211 | 8.29E-02 | 1.24E-01 | 2.66E+00 | 2.73E+00 | 5.01E-01 | no | Nr5a2 | intron_variant | MODIFIER |

| | | | | | | | | | | | | | | | |
|---|---|---|---|---|---|---|---|---|---|---|---|---|---|---|---|
| 1 | 136875035 | AX-170472399 | 0 | T | G | 0.211 | 8.29E-02 | 1.24E-01 | 2.66E+00 | 2.73E+00 | 5.01E-01 | no | Nr5a2 | intron_variant | MODIFIER |
| 1 | 136875167 | AX-170188757 | 0 | A | G | 0.211 | 8.29E-02 | 1.24E-01 | 2.66E+00 | 2.73E+00 | 5.01E-01 | no | Nr5a2 | intron_variant | MODIFIER |
| 1 | 136884921 | AX-170319883 | 0 | G | A | 0.211 | 8.29E-02 | 1.24E-01 | 2.66E+00 | 2.73E+00 | 5.01E-01 | no | Nr5a2 | intron_variant | MODIFIER |
| 1 | 136885021 | AX-97813326 | 0 | T | C | 0.211 | 8.29E-02 | 1.24E-01 | 2.66E+00 | 2.73E+00 | 5.01E-01 | no | Nr5a2 | intron_variant | MODIFIER |
| 1 | 136885316 | AX-170126897 | 0 | T | C | 0.211 | 8.29E-02 | 1.24E-01 | 2.66E+00 | 2.73E+00 | 5.01E-01 | no | Nr5a2 | intron_variant | MODIFIER |
| 1 | 136897684 | AX-170518462 | 0 | C | T | 0.211 | 8.29E-02 | 1.24E-01 | 2.66E+00 | 2.73E+00 | 5.01E-01 | no | Nr5a2 | intron_variant | MODIFIER |
| 1 | 136906662 | AX-97815663 | 0 | G | A | 0.211 | 8.29E-02 | 1.24E-01 | 2.66E+00 | 2.73E+00 | 5.01E-01 | no | Nr5a2 | intron_variant | MODIFIER |
| 1 | 136913460 | AX-170291829 | 0 | T | C | 0.211 | 8.29E-02 | 1.24E-01 | 2.66E+00 | 2.73E+00 | 5.01E-01 | no | Nr5a2 | intron_variant | MODIFIER |
| 1 | 136918756 | AX-170228661 | 0 | C | T | 0.211 | 8.29E-02 | 1.24E-01 | 2.66E+00 | 2.73E+00 | 5.01E-01 | no | Nr5a2 | intron_variant | MODIFIER |
| 1 | 136923091 | AX-170370692 | 0 | T | G | 0.211 | 8.29E-02 | 1.24E-01 | 2.66E+00 | 2.73E+00 | 5.01E-01 | no | Nr5a2 | intron_variant | MODIFIER |
| 1 | 136928071 | AX-170308201 | 0 | T | C | 0.211 | 8.29E-02 | 1.24E-01 | 2.66E+00 | 2.73E+00 | 5.01E-01 | no | Nr5a2 | intron_variant | MODIFIER |
| 1 | 136956786 | AX-170249093 | 0 | C | A | 0.211 | 8.29E-02 | 1.24E-01 | 2.66E+00 | 2.73E+00 | 5.01E-01 | no | Nr5a2 | intron_variant | MODIFIER |
| 1 | 136961248 | AX-170278314 | 0 | A | C | 0.211 | 8.29E-02 | 1.24E-01 | 2.66E+00 | 2.73E+00 | 5.01E-01 | no | Nr5a2 | upstream_gene_variant | MODIFIER |
| 1 | 136973169 | AX-170340059 | 0 | C | G | 0.211 | 8.29E-02 | 1.24E-01 | 2.66E+00 | 2.73E+00 | 5.01E-01 | no | | intergenic_variant | MODIFIER |
| 1 | 137044532 | AX-170196056 | 0 | G | A | 0.211 | 8.29E-02 | 1.24E-01 | 2.66E+00 | 2.73E+00 | 5.01E-01 | no | | intergenic_variant | MODIFIER |
| 1 | 137053174 | AX-97811193 | 0 | C | T | 0.211 | 8.29E-02 | 1.24E-01 | 2.66E+00 | 2.73E+00 | 5.01E-01 | no | | intergenic_variant | MODIFIER |
| 1 | 137058658 | AX-170107833 | 0 | A | G | 0.211 | 8.29E-02 | 1.24E-01 | 2.66E+00 | 2.73E+00 | 5.01E-01 | no | | intergenic_variant | MODIFIER |
| 1 | 137070192 | AX-170305201 | 0 | A | G | 0.211 | 8.29E-02 | 1.24E-01 | 2.66E+00 | 2.73E+00 | 5.01E-01 | no | | intergenic_variant | MODIFIER |
| 1 | 137073897 | AX-170184655 | 0 | C | G | 0.211 | 8.29E-02 | 1.24E-01 | 2.66E+00 | 2.73E+00 | 5.01E-01 | no | | regulatory_region_variant | MODIFIER |
| 1 | 137074347 | AX-170375219 | 0 | G | A | 0.211 | 8.29E-02 | 1.24E-01 | 2.66E+00 | 2.73E+00 | 5.01E-01 | no | | regulatory_region_variant | MODIFIER |
| 1 | 137074928 | AX-170273813 | 0 | C | T | 0.211 | 8.29E-02 | 1.24E-01 | 2.66E+00 | 2.73E+00 | 5.01E-01 | no | | regulatory_region_variant | MODIFIER |
| 1 | 137083023 | AX-97813349 | 0 | T | C | 0.211 | 8.29E-02 | 1.24E-01 | 2.66E+00 | 2.73E+00 | 5.01E-01 | no | | intergenic_variant | MODIFIER |
| 1 | 137088537 | AX-170360888 | 0 | G | T | 0.211 | 8.29E-02 | 1.24E-01 | 2.66E+00 | 2.73E+00 | 5.01E-01 | no | | regulatory_region_variant | MODIFIER |
| 1 | 137107283 | AX-170412960 | 0 | A | G | 0.211 | 8.29E-02 | 1.24E-01 | 2.66E+00 | 2.73E+00 | 5.01E-01 | no | | intergenic_variant | MODIFIER |
| 1 | 137111116 | AX-170091707 | 0 | T | C | 0.211 | 8.29E-02 | 1.24E-01 | 2.66E+00 | 2.73E+00 | 5.01E-01 | no | | intergenic_variant | MODIFIER |
| 1 | 137115162 | AX-170506263 | 0 | G | A | 0.211 | 8.29E-02 | 1.24E-01 | 2.66E+00 | 2.73E+00 | 5.01E-01 | no | | intergenic_variant | MODIFIER |
| 1 | 137141201 | AX-97841174 | 0 | T | G | 0.211 | 8.29E-02 | 1.24E-01 | 2.66E+00 | 2.73E+00 | 5.01E-01 | no | | intergenic_variant | MODIFIER |
| 1 | 137194563 | AX-170301512 | 0 | G | A | 0.211 | 8.29E-02 | 1.24E-01 | 2.66E+00 | 2.73E+00 | 5.01E-01 | no | | intergenic_variant | MODIFIER |
| 1 | 137205554 | AX-170001796 | 0 | T | C | 0.211 | 8.29E-02 | 1.24E-01 | 2.66E+00 | 2.73E+00 | 5.01E-01 | no | | intergenic_variant | MODIFIER |
| 1 | 137207411 | AX-170001793 | 0 | A | C | 0.211 | 8.29E-02 | 1.24E-01 | 2.66E+00 | 2.73E+00 | 5.01E-01 | no | | intergenic_variant | MODIFIER |
| 1 | 137236675 | AX-170224311 | 0 | A | G | 0.211 | 8.29E-02 | 1.24E-01 | 2.66E+00 | 2.73E+00 | 5.01E-01 | no | | intergenic_variant | MODIFIER |
| 1 | 137245747 | AX-170357472 | 0 | T | C | 0.211 | 8.29E-02 | 1.24E-01 | 2.66E+00 | 2.73E+00 | 5.01E-01 | no | | intergenic_variant | MODIFIER |
| 1 | 137301227 | AX-97810966 | 0 | A | G | 0.211 | 8.29E-02 | 1.24E-01 | 2.66E+00 | 2.73E+00 | 5.01E-01 | no | | intergenic_variant | MODIFIER |
| 1 | 137301600 | AX-170001789 | 0 | G | A | 0.211 | 8.29E-02 | 1.24E-01 | 2.66E+00 | 2.73E+00 | 5.01E-01 | no | | intergenic_variant | MODIFIER |
| 1 | 137315049 | AX-170204423 | 0 | T | C | 0.211 | 8.29E-02 | 1.24E-01 | 2.66E+00 | 2.73E+00 | 5.01E-01 | no | | intergenic_variant | MODIFIER |
| 1 | 137325622 | AX-170522856 | 0 | A | G | 0.211 | 8.29E-02 | 1.24E-01 | 2.66E+00 | 2.73E+00 | 5.01E-01 | no | 1700006P03Rik | non_coding_transcript_exon_variant | MODIFIER |
| 1 | 137325773 | AX-170240159 | 0 | G | A | 0.211 | 8.29E-02 | 1.24E-01 | 2.66E+00 | 2.73E+00 | 5.01E-01 | no | 1700006P03Rik | non_coding_transcript_exon_variant | MODIFIER |
| 1 | 137335848 | AX-170440132 | 0 | T | C | 0.211 | 8.29E-02 | 1.24E-01 | 2.66E+00 | 2.73E+00 | 5.01E-01 | no | | intergenic_variant | MODIFIER |
| 1 | 137359412 | AX-97813351 | 0 | T | C | 0.211 | 8.29E-02 | 1.24E-01 | 2.66E+00 | 2.73E+00 | 5.01E-01 | no | | intergenic_variant | MODIFIER |
| 1 | 137376446 | AX-170386129 | 0 | T | C | 0.211 | 8.29E-02 | 1.24E-01 | 2.66E+00 | 2.73E+00 | 5.01E-01 | no | | regulatory_region_variant | MODIFIER |
| 1 | 137442280 | AX-170465572 | 0 | T | C | 0.211 | 8.29E-02 | 1.24E-01 | 2.66E+00 | 2.73E+00 | 5.01E-01 | no | | intergenic_variant | MODIFIER |
| 1 | 137445698 | AX-170418087 | 0 | T | C | 0.211 | 8.29E-02 | 1.24E-01 | 2.66E+00 | 2.73E+00 | 5.01E-01 | no | | intergenic_variant | MODIFIER |
| 1 | 137466813 | AX-170500224 | 0 | T | C | 0.211 | 8.29E-02 | 1.24E-01 | 2.66E+00 | 2.73E+00 | 5.01E-01 | no | | intergenic_variant | MODIFIER |
| 1 | 137483701 | AX-170217863 | 0 | G | C | 0.211 | 8.29E-02 | 1.24E-01 | 2.66E+00 | 2.73E+00 | 5.01E-01 | no | | intergenic_variant | MODIFIER |
| 1 | 137484254 | AX-170521334 | 0 | G | T | 0.211 | 8.29E-02 | 1.24E-01 | 2.66E+00 | 2.73E+00 | 5.01E-01 | no | | intergenic_variant | MODIFIER |
| 1 | 137493904 | AX-170196828 | 0 | T | G | 0.211 | 8.29E-02 | 1.24E-01 | 2.66E+00 | 2.73E+00 | 5.01E-01 | no | | intergenic_variant | MODIFIER |
| 1 | 137523071 | AX-97834310 | 0 | C | T | 0.211 | 8.29E-02 | 1.24E-01 | 2.66E+00 | 2.73E+00 | 5.01E-01 | no | | intergenic_variant | MODIFIER |
| 1 | 137539926 | AX-170409266 | 0 | A | T | 0.211 | 8.29E-02 | 1.24E-01 | 2.66E+00 | 2.73E+00 | 5.01E-01 | no | | regulatory_region_variant | MODIFIER |
| 1 | 137546039 | AX-170382156 | 0 | T | C | 0.211 | 8.29E-02 | 1.24E-01 | 2.66E+00 | 2.73E+00 | 5.01E-01 | no | | regulatory_region_variant | MODIFIER |
| 1 | 137546297 | AX-170510224 | 0 | T | C | 0.211 | 8.29E-02 | 1.24E-01 | 2.66E+00 | 2.73E+00 | 5.01E-01 | no | | regulatory_region_variant | MODIFIER |
| 1 | 137546389 | AX-170371646 | 0 | A | C | 0.211 | 8.29E-02 | 1.24E-01 | 2.66E+00 | 2.73E+00 | 5.01E-01 | no | | regulatory_region_variant | MODIFIER |
| 1 | 137554617 | AX-170308438 | 0 | C | T | 0.211 | 8.29E-02 | 1.24E-01 | 2.66E+00 | 2.73E+00 | 5.01E-01 | no | | intergenic_variant | MODIFIER |
| 1 | 137568267 | AX-97852890 | 0 | G | A | 0.211 | 8.29E-02 | 1.24E-01 | 2.66E+00 | 2.73E+00 | 5.01E-01 | no | | intergenic_variant | MODIFIER |
| 1 | 137875043 | AX-170371707 | 0 | A | G | 0.211 | 8.29E-02 | 1.24E-01 | 2.66E+00 | 2.73E+00 | 5.01E-01 | no | | intergenic_variant | MODIFIER |

| | | | | | | | | | | | | | | | |
|---|---|---|---|---|---|---|---|---|---|---|---|---|---|---|---|
| 1 | 137886138 | AX-170294869 | 0 | C | T | 0.211 | 8.29E-02 | 1.24E-01 | 2.66E+00 | 2.73E+00 | 5.01E-01 | no | | intergenic_variant | MODIFIER |
| 1 | 137891142 | AX-170153811 | 0 | G | A | 0.211 | 8.29E-02 | 1.24E-01 | 2.66E+00 | 2.73E+00 | 5.01E-01 | no | | intergenic_variant | MODIFIER |
| 1 | 137893922 | AX-170191630 | 0 | G | A | 0.211 | 8.29E-02 | 1.24E-01 | 2.66E+00 | 2.73E+00 | 5.01E-01 | no | Gm4258 | upstream_gene_variant | MODIFIER |
| 1 | 137923006 | AX-170134099 | 0 | G | T | 0.211 | 8.29E-02 | 1.24E-01 | 2.66E+00 | 2.73E+00 | 5.01E-01 | no | Gm4258 | intron_variant,non_coding_transcript_variant | MODIFIER |
| 1 | 137925259 | AX-97813762 | 0 | G | T | 0.211 | 8.29E-02 | 1.24E-01 | 2.66E+00 | 2.73E+00 | 5.01E-01 | no | Gm4258 | intron_variant,non_coding_transcript_variant | MODIFIER |
| 1 | 137927151 | AX-170427445 | 0 | C | G | 0.211 | 8.29E-02 | 1.24E-01 | 2.66E+00 | 2.73E+00 | 5.01E-01 | no | Gm4258 | intron_variant,non_coding_transcript_variant | MODIFIER |
| 1 | 137927771 | AX-170187120 | 0 | G | C | 0.211 | 8.29E-02 | 1.24E-01 | 2.66E+00 | 2.73E+00 | 5.01E-01 | no | Gm4258 | intron_variant,non_coding_transcript_variant | MODIFIER |
| 1 | 137932744 | AX-170342286 | 0 | T | A | 0.211 | 8.29E-02 | 1.24E-01 | 2.66E+00 | 2.73E+00 | 5.01E-01 | no | Gm4258 | intron_variant,non_coding_transcript_variant | MODIFIER |
| 1 | 137932986 | AX-170042874 | 0 | C | T | 0.211 | 8.29E-02 | 1.24E-01 | 2.66E+00 | 2.73E+00 | 5.01E-01 | no | Gm4258 | intron_variant,non_coding_transcript_variant | MODIFIER |
| 1 | 138042917 | AX-97834988 | 0 | G | A | 0.211 | 8.29E-02 | 1.24E-01 | 2.66E+00 | 2.73E+00 | 5.01E-01 | no | | intergenic_variant | MODIFIER |
| 18 | 54838314 | AX-170256405 | 0 | C | T | 0.447 | 6.47E-02 | 9.19E-02 | 2.44E+00 | 2.51E+00 | 5.01E-01 | no | AC129179.1 | non_coding_transcript_exon_variant | MODIFIER |
| 1 | 16247316 | AX-170526325 | 0 | A | G | 0.377 | 5.38E-02 | 7.99E-02 | 2.60E+00 | 2.65E+00 | 5.02E-01 | no | Stau2 | intron_variant | MODIFIER |
| 18 | 88915723 | AX-170223332 | 0 | C | A | 0.333 | 4.07E-02 | 6.15E-02 | 2.63E+00 | 2.76E+00 | 5.02E-01 | no | Socs6 | intron_variant | MODIFIER |
| 5 | 42645024 | AX-170302756 | 1 | C | T | 0.205 | -7.24E-02 | 1.08E-01 | 2.61E+00 | 2.69E+00 | 5.02E-01 | no | | intergenic_variant | MODIFIER |
| 9 | 119213185 | AX-170256309 | 1 | G | A | 0.125 | -6.38E-02 | 9.58E-02 | 2.61E+00 | 2.73E+00 | 5.02E-01 | no | Slc22a13 | upstream_gene_variant | MODIFIER |
| 2 | 138310185 | AX-170188965 | 0 | A | T | 0.149 | 1.20E-01 | 1.80E-01 | 2.65E+00 | 2.73E+00 | 5.02E-01 | no | Btbd3 | intron_variant,non_coding_transcript_variant | MODIFIER |
| 9 | 118865641 | AX-170530856 | 1 | A | T | 0.125 | -6.39E-02 | 9.60E-02 | 2.62E+00 | 2.73E+00 | 5.02E-01 | no | Itga9 | intron_variant | MODIFIER |
| 16 | 29482079 | AX-170173718 | 1 | A | G | 0.491 | 5.35E-02 | 7.86E-02 | 2.54E+00 | 2.60E+00 | 5.02E-01 | no | Atp13a4 | intron_variant | MODIFIER |
| 18 | 80593685 | AX-170314142 | 0 | C | T | 0.395 | -5.87E-02 | 8.77E-02 | 2.61E+00 | 2.70E+00 | 5.02E-01 | no | AC149054.1 | intron_variant,non_coding_transcript_variant | MODIFIER |
| 1 | 53952180 | AX-170201176 | 2 | T | A | 0.118 | -1.11E-01 | 1.69E-01 | 2.74E+00 | 2.89E+00 | 5.02E-01 | no | Hecw2 | intron_variant | MODIFIER |
| 8 | 42021185 | AX-97817195 | 0 | A | G | 0.149 | -8.74E-02 | 1.32E-01 | 2.68E+00 | 2.75E+00 | 5.02E-01 | no | 2810404M03Rik | intron_variant,non_coding_transcript_variant | MODIFIER |
| 6 | 138593429 | AX-170076391 | 0 | T | C | 0.553 | 5.87E-02 | 8.45E-02 | 2.47E+00 | 2.55E+00 | 5.02E-01 | no | B230110G15Rik | intron_variant,non_coding_transcript_variant | MODIFIER |
| 2 | 110200721 | AX-97786123 | 0 | A | G | 0.061 | 1.35E-01 | 2.05E-01 | 2.65E+00 | 2.77E+00 | 5.02E-01 | no | Gm13936 | intron_variant,non_coding_transcript_variant | MODIFIER |
| 2 | 51614337 | AX-170063654 | 0 | T | C | 0.061 | 8.57E-02 | 1.31E-01 | 2.71E+00 | 2.88E+00 | 5.02E-01 | no | | intergenic_variant | MODIFIER |
| 8 | 86535858 | AX-170290881 | 1 | C | G | 0.205 | -8.74E-02 | 1.32E-01 | 2.76E+00 | 2.83E+00 | 5.02E-01 | no | Abcc12 | intron_variant | MODIFIER |
| 4 | 154195507 | AX-170503501 | 1 | A | T | 0.107 | 1.20E-01 | 1.81E-01 | 2.70E+00 | 2.78E+00 | 5.02E-01 | no | Megf6 | intron_variant | MODIFIER |
| 8 | 75376927 | AX-170229651 | 0 | A | C | 0.193 | -8.08E-02 | 1.22E-01 | 2.70E+00 | 2.80E+00 | 5.03E-01 | no | | intergenic_variant | MODIFIER |
| 14 | 95311912 | AX-170383616 | 0 | C | T | 0.035 | -1.33E-01 | 1.99E-01 | 2.58E+00 | 2.68E+00 | 5.03E-01 | no | | intergenic_variant | MODIFIER |
| 2 | 117366257 | AX-170426888 | 0 | T | A | 0.14 | 1.12E-01 | 1.68E-01 | 2.65E+00 | 2.73E+00 | 5.03E-01 | no | | regulatory_region_variant | MODIFIER |
| 4 | 89658336 | AX-170218067 | 0 | A | T | 0.439 | -5.48E-02 | 7.95E-02 | 2.49E+00 | 2.56E+00 | 5.03E-01 | no | | intergenic_variant | MODIFIER |
| 7 | 72923384 | AX-170128918 | 1 | G | A | 0.357 | -6.55E-02 | 9.93E-02 | 2.74E+00 | 2.83E+00 | 5.03E-01 | no | | intergenic_variant | MODIFIER |
| 9 | 117833599 | AX-170322213 | 0 | T | C | 0.123 | -6.35E-02 | 9.57E-02 | 2.62E+00 | 2.73E+00 | 5.03E-01 | no | Gm45881 | intron_variant,non_coding_transcript_variant | MODIFIER |
| 7 | 129035694 | AX-170278030 | 0 | G | A | 0.237 | -6.38E-02 | 9.45E-02 | 2.55E+00 | 2.63E+00 | 5.03E-01 | no | | intergenic_variant | MODIFIER |
| 9 | 74945699 | AX-170426842 | 0 | T | A | 0.193 | 8.46E-02 | 1.27E-01 | 2.82E+00 | 2.87E+00 | 5.03E-01 | no | | intergenic_variant | MODIFIER |
| 8 | 62200171 | AX-170389166 | 1 | C | T | 0.152 | -9.68E-02 | 1.41E-01 | 2.49E+00 | 2.56E+00 | 5.03E-01 | no | | intergenic_variant | MODIFIER |
| 17 | 48343545 | AX-168396660 | 0 | A | G | 0.053 | 1.11E-01 | 1.70E-01 | 2.72E+00 | 2.88E+00 | 5.03E-01 | no | Trem2 | upstream_gene_variant | MODIFIER |
| 1 | 169074732 | AX-170043501 | 1 | C | A | 0.241 | 8.45E-02 | 1.27E-01 | 2.66E+00 | 2.71E+00 | 5.03E-01 | no | | intergenic_variant | MODIFIER |
| 15 | 3751346 | AX-170144175 | 0 | A | T | 0.096 | -7.37E-02 | 9.35E-02 | 2.22E+00 | 2.33E+00 | 5.03E-01 | no | Gm4823 | upstream_gene_variant | MODIFIER |
| 8 | 66098117 | AX-170193637 | 2 | T | C | 0.136 | -9.63E-02 | 1.43E-01 | 2.56E+00 | 2.65E+00 | 5.03E-01 | no | MAR1 | intron_variant | MODIFIER |
| 18 | 63338002 | AX-170428952 | 1 | T | C | 0.188 | -7.86E-02 | 1.19E-01 | 2.67E+00 | 2.73E+00 | 5.03E-01 | no | Piezo2 | intron_variant | MODIFIER |
| 1 | 173057517 | AX-170242707 | 0 | C | T | 0.237 | 7.60E-02 | 1.11E-01 | 2.50E+00 | 2.59E+00 | 5.03E-01 | no | Gm42472 | upstream_gene_variant | MODIFIER |
| 5 | 104098345 | AX-170429090 | 1 | T | A | 0.446 | 7.26E-02 | 1.09E-01 | 2.65E+00 | 2.71E+00 | 5.03E-01 | no | Sparcl1 | intron_variant | MODIFIER |
| 16 | 37212868 | AX-170350279 | 0 | T | A | 0.482 | 5.52E-02 | 8.16E-02 | 2.55E+00 | 2.60E+00 | 5.03E-01 | no | Stxbp5l | intron_variant,non_coding_transcript_variant | MODIFIER |
| 8 | 26625065 | AX-97822892 | 1 | A | T | 0.179 | 6.90E-02 | 1.05E-01 | 2.71E+00 | 2.82E+00 | 5.03E-01 | no | Gm32050 | downstream_gene_variant | MODIFIER |
| 5 | 37464114 | AX-170524240 | 1 | G | A | 0.339 | 7.37E-02 | 1.12E-01 | 2.75E+00 | 2.82E+00 | 5.04E-01 | no | Stk32b | intron_variant | MODIFIER |
| 16 | 31334698 | AX-170396703 | 0 | G | A | 0.5 | 5.32E-02 | 7.90E-02 | 2.56E+00 | 2.63E+00 | 5.04E-01 | no | | intergenic_variant | MODIFIER |
| 7 | 73157186 | AX-170455276 | 0 | A | G | 0.36 | -6.42E-02 | 9.77E-02 | 2.75E+00 | 2.85E+00 | 5.04E-01 | no | | intergenic_variant | MODIFIER |
| 7 | 73157349 | AX-170029285 | 0 | A | G | 0.36 | -6.42E-02 | 9.77E-02 | 2.75E+00 | 2.85E+00 | 5.04E-01 | no | | intergenic_variant | MODIFIER |
| 7 | 73161383 | AX-170095982 | 0 | T | C | 0.36 | -6.42E-02 | 9.77E-02 | 2.75E+00 | 2.85E+00 | 5.04E-01 | no | | regulatory_region_variant | MODIFIER |
| 7 | 73162958 | AX-170356131 | 0 | G | A | 0.36 | -6.42E-02 | 9.77E-02 | 2.75E+00 | 2.85E+00 | 5.04E-01 | no | | intergenic_variant | MODIFIER |
| 7 | 73163049 | AX-170226750 | 0 | G | A | 0.36 | -6.42E-02 | 9.77E-02 | 2.75E+00 | 2.85E+00 | 5.04E-01 | no | | intergenic_variant | MODIFIER |
| 7 | 73172621 | AX-170029287 | 0 | A | G | 0.36 | -6.42E-02 | 9.77E-02 | 2.75E+00 | 2.85E+00 | 5.04E-01 | no | | intergenic_variant | MODIFIER |
| 7 | 73187931 | AX-170029284 | 0 | C | T | 0.36 | -6.42E-02 | 9.77E-02 | 2.75E+00 | 2.85E+00 | 5.04E-01 | no | Gm28746 | upstream_gene_variant | MODIFIER |
| 7 | 73189547 | AX-170508354 | 0 | G | A | 0.36 | -6.42E-02 | 9.77E-02 | 2.75E+00 | 2.85E+00 | 5.04E-01 | no | Gm28746 | upstream_gene_variant | MODIFIER |

| | | | | | | | | | | | | | | | |
|---|---|---|---|---|---|---|---|---|---|---|---|---|---|---|---|
| 7 | 73325652 | AX-170464496 | 0 | G | T | 0.36 | -6.42E-02 | 9.77E-02 | 2.75E+00 | 2.85E+00 | 5.04E-01 | no | A730056A06Rik | intron_variant,non_coding_transcript_variant | MODIFIER |
| 7 | 73326720 | AX-170376226 | 0 | C | T | 0.36 | -6.42E-02 | 9.77E-02 | 2.75E+00 | 2.85E+00 | 5.04E-01 | no | A730056A06Rik | intron_variant,non_coding_transcript_variant | MODIFIER |
| 7 | 73334576 | AX-170384458 | 0 | A | G | 0.36 | -6.42E-02 | 9.77E-02 | 2.75E+00 | 2.85E+00 | 5.04E-01 | no | A730056A06Rik | intron_variant,non_coding_transcript_variant | MODIFIER |
| 7 | 73336961 | AX-97825619 | 0 | A | G | 0.36 | -6.42E-02 | 9.77E-02 | 2.75E+00 | 2.85E+00 | 5.04E-01 | no | A730056A06Rik | intron_variant,non_coding_transcript_variant | MODIFIER |
| 7 | 73337478 | AX-170142501 | 0 | T | C | 0.36 | -6.42E-02 | 9.77E-02 | 2.75E+00 | 2.85E+00 | 5.04E-01 | no | A730056A06Rik | intron_variant,non_coding_transcript_variant | MODIFIER |
| 7 | 73343761 | AX-170170628 | 0 | A | G | 0.36 | -6.42E-02 | 9.77E-02 | 2.75E+00 | 2.85E+00 | 5.04E-01 | no | A730056A06Rik | intron_variant,non_coding_transcript_variant | MODIFIER |
| 7 | 73344187 | AX-170386166 | 0 | T | C | 0.36 | -6.42E-02 | 9.77E-02 | 2.75E+00 | 2.85E+00 | 5.04E-01 | no | A730056A06Rik | intron_variant,non_coding_transcript_variant | MODIFIER |
| 7 | 73819333 | AX-170308145 | 0 | T | C | 0.36 | -6.42E-02 | 9.77E-02 | 2.75E+00 | 2.85E+00 | 5.04E-01 | no | Gm10619 | upstream_gene_variant | MODIFIER |
| 7 | 73878935 | AX-170180821 | 0 | T | C | 0.36 | -6.42E-02 | 9.77E-02 | 2.75E+00 | 2.85E+00 | 5.04E-01 | no | | intergenic_variant | MODIFIER |
| 7 | 73890123 | AX-170139783 | 0 | T | C | 0.36 | -6.42E-02 | 9.77E-02 | 2.75E+00 | 2.85E+00 | 5.04E-01 | no | | intergenic_variant | MODIFIER |
| 7 | 73895307 | AX-170300907 | 0 | G | A | 0.36 | -6.42E-02 | 9.77E-02 | 2.75E+00 | 2.85E+00 | 5.04E-01 | no | | intergenic_variant | MODIFIER |
| 7 | 73898048 | AX-170386530 | 0 | A | G | 0.36 | -6.42E-02 | 9.77E-02 | 2.75E+00 | 2.85E+00 | 5.04E-01 | no | | intergenic_variant | MODIFIER |
| 7 | 73938604 | AX-97848332 | 0 | T | C | 0.36 | -6.42E-02 | 9.77E-02 | 2.75E+00 | 2.85E+00 | 5.04E-01 | no | St8sia2 | downstream_gene_variant | MODIFIER |
| 7 | 74032045 | AX-170326073 | 0 | C | T | 0.36 | -6.42E-02 | 9.77E-02 | 2.75E+00 | 2.85E+00 | 5.04E-01 | no | Gm45004 | downstream_gene_variant | MODIFIER |
| 7 | 74037164 | AX-170095984 | 0 | T | C | 0.36 | -6.42E-02 | 9.77E-02 | 2.75E+00 | 2.85E+00 | 5.04E-01 | no | Gm45004 | intron_variant,non_coding_transcript_variant | MODIFIER |
| 7 | 74055451 | AX-170217817 | 0 | G | A | 0.36 | -6.42E-02 | 9.77E-02 | 2.75E+00 | 2.85E+00 | 5.04E-01 | no | | intergenic_variant | MODIFIER |
| 7 | 74058986 | AX-97833033 | 0 | A | G | 0.36 | -6.42E-02 | 9.77E-02 | 2.75E+00 | 2.85E+00 | 5.04E-01 | no | | intergenic_variant | MODIFIER |
| 7 | 74270053 | AX-170274144 | 0 | A | G | 0.36 | -6.42E-02 | 9.77E-02 | 2.75E+00 | 2.85E+00 | 5.04E-01 | no | | regulatory_region_variant | MODIFIER |
| 7 | 74459961 | AX-170294911 | 0 | C | G | 0.36 | -6.42E-02 | 9.77E-02 | 2.75E+00 | 2.85E+00 | 5.04E-01 | no | Slco3a1 | intron_variant | MODIFIER |
| 7 | 74512886 | AX-170119252 | 0 | G | A | 0.36 | -6.42E-02 | 9.77E-02 | 2.75E+00 | 2.85E+00 | 5.04E-01 | no | Slco3a1 | intron_variant | MODIFIER |
| 7 | 74748799 | AX-170448513 | 0 | G | A | 0.36 | -6.42E-02 | 9.77E-02 | 2.75E+00 | 2.85E+00 | 5.04E-01 | no | | intergenic_variant | MODIFIER |
| 7 | 74749844 | AX-170203850 | 0 | T | C | 0.36 | -6.42E-02 | 9.77E-02 | 2.75E+00 | 2.85E+00 | 5.04E-01 | no | | intergenic_variant | MODIFIER |
| 7 | 74768350 | AX-170029311 | 0 | T | G | 0.36 | -6.42E-02 | 9.77E-02 | 2.75E+00 | 2.85E+00 | 5.04E-01 | no | Gm30459 | intron_variant,non_coding_transcript_variant | MODIFIER |
| 7 | 74851294 | AX-170456669 | 0 | C | A | 0.36 | -6.42E-02 | 9.77E-02 | 2.75E+00 | 2.85E+00 | 5.04E-01 | no | 0610006L08Rik | intron_variant,non_coding_transcript_variant | MODIFIER |
| 7 | 74987821 | AX-170337794 | 0 | A | G | 0.36 | -6.42E-02 | 9.77E-02 | 2.75E+00 | 2.85E+00 | 5.04E-01 | no | | intergenic_variant | MODIFIER |
| 7 | 74990745 | AX-170204393 | 0 | T | C | 0.36 | -6.42E-02 | 9.77E-02 | 2.75E+00 | 2.85E+00 | 5.04E-01 | no | | intergenic_variant | MODIFIER |
| 7 | 74990787 | AX-170304964 | 0 | G | T | 0.36 | -6.42E-02 | 9.77E-02 | 2.75E+00 | 2.85E+00 | 5.04E-01 | no | | intergenic_variant | MODIFIER |
| 7 | 74993450 | AX-170138164 | 0 | T | C | 0.36 | -6.42E-02 | 9.77E-02 | 2.75E+00 | 2.85E+00 | 5.04E-01 | no | | intergenic_variant | MODIFIER |
| 7 | 75000575 | AX-170172541 | 0 | A | G | 0.36 | -6.42E-02 | 9.77E-02 | 2.75E+00 | 2.85E+00 | 5.04E-01 | no | | intergenic_variant | MODIFIER |
| 7 | 75000958 | AX-170500802 | 0 | T | G | 0.36 | -6.42E-02 | 9.77E-02 | 2.75E+00 | 2.85E+00 | 5.04E-01 | no | | intergenic_variant | MODIFIER |
| 7 | 75025712 | AX-170077745 | 0 | C | G | 0.36 | -6.42E-02 | 9.77E-02 | 2.75E+00 | 2.85E+00 | 5.04E-01 | no | 4930533N22Rik | upstream_gene_variant | MODIFIER |
| 7 | 75043812 | AX-170458831 | 0 | C | T | 0.36 | -6.42E-02 | 9.77E-02 | 2.75E+00 | 2.85E+00 | 5.04E-01 | no | 4930533N22Rik | intron_variant,non_coding_transcript_variant | MODIFIER |
| 7 | 75043877 | AX-97782767 | 0 | A | G | 0.36 | -6.42E-02 | 9.77E-02 | 2.75E+00 | 2.85E+00 | 5.04E-01 | no | 4930533N22Rik | intron_variant,non_coding_transcript_variant | MODIFIER |
| 7 | 75044011 | AX-170490402 | 0 | A | G | 0.36 | -6.42E-02 | 9.77E-02 | 2.75E+00 | 2.85E+00 | 5.04E-01 | no | 4930533N22Rik | intron_variant,non_coding_transcript_variant | MODIFIER |
| 7 | 75098698 | AX-168418451 | 0 | T | C | 0.36 | -6.42E-02 | 9.77E-02 | 2.75E+00 | 2.85E+00 | 5.04E-01 | no | | intergenic_variant | MODIFIER |
| 7 | 75101636 | AX-170386614 | 0 | A | G | 0.36 | -6.42E-02 | 9.77E-02 | 2.75E+00 | 2.85E+00 | 5.04E-01 | no | | intergenic_variant | MODIFIER |
| 7 | 75241508 | AX-170146313 | 0 | A | G | 0.36 | -6.42E-02 | 9.77E-02 | 2.75E+00 | 2.85E+00 | 5.04E-01 | no | Sv2b | intron_variant | MODIFIER |
| 7 | 75242691 | AX-170353965 | 0 | T | C | 0.36 | -6.42E-02 | 9.77E-02 | 2.75E+00 | 2.85E+00 | 5.04E-01 | no | Sv2b | intron_variant | MODIFIER |
| 7 | 75258422 | AX-97799781 | 0 | G | T | 0.36 | -6.42E-02 | 9.77E-02 | 2.75E+00 | 2.85E+00 | 5.04E-01 | no | Sv2b | intron_variant | MODIFIER |
| 7 | 75373629 | AX-97839465 | 0 | C | T | 0.36 | -6.42E-02 | 9.77E-02 | 2.75E+00 | 2.85E+00 | 5.04E-01 | no | | intergenic_variant | MODIFIER |
| 7 | 75455096 | AX-170512791 | 0 | G | A | 0.36 | -6.42E-02 | 9.77E-02 | 2.75E+00 | 2.85E+00 | 5.04E-01 | no | Akap13 | upstream_gene_variant | MODIFIER |
| 7 | 75587847 | AX-170444661 | 0 | A | G | 0.36 | -6.42E-02 | 9.77E-02 | 2.75E+00 | 2.85E+00 | 5.04E-01 | no | Akap13 | intron_variant | MODIFIER |
| 7 | 75622227 | AX-97851075 | 0 | C | A | 0.36 | -6.42E-02 | 9.77E-02 | 2.75E+00 | 2.85E+00 | 5.04E-01 | no | Akap13 | intron_variant | MODIFIER |
| 7 | 75622556 | AX-170133727 | 0 | G | C | 0.36 | -6.42E-02 | 9.77E-02 | 2.75E+00 | 2.85E+00 | 5.04E-01 | no | Akap13 | intron_variant | MODIFIER |
| 7 | 75623184 | AX-170162050 | 0 | C | T | 0.36 | -6.42E-02 | 9.77E-02 | 2.75E+00 | 2.85E+00 | 5.04E-01 | no | Akap13 | intron_variant | MODIFIER |
| 7 | 75628053 | AX-97797694 | 0 | A | G | 0.36 | -6.42E-02 | 9.77E-02 | 2.75E+00 | 2.85E+00 | 5.04E-01 | no | Akap13 | intron_variant | MODIFIER |
| 7 | 75645929 | AX-170230341 | 0 | T | C | 0.36 | -6.42E-02 | 9.77E-02 | 2.75E+00 | 2.85E+00 | 5.04E-01 | no | Akap13 | intron_variant | MODIFIER |
| 7 | 76334659 | AX-97803832 | 0 | A | G | 0.36 | -6.42E-02 | 9.77E-02 | 2.75E+00 | 2.85E+00 | 5.04E-01 | no | Agbl1 | intron_variant | MODIFIER |
| 7 | 76341783 | AX-170294177 | 0 | T | C | 0.36 | -6.42E-02 | 9.77E-02 | 2.75E+00 | 2.85E+00 | 5.04E-01 | no | Agbl1 | intron_variant | MODIFIER |
| 7 | 76391486 | AX-170418240 | 0 | G | A | 0.36 | -6.42E-02 | 9.77E-02 | 2.75E+00 | 2.85E+00 | 5.04E-01 | no | Agbl1 | intron_variant | MODIFIER |
| 7 | 76401957 | AX-170148692 | 0 | T | G | 0.36 | -6.42E-02 | 9.77E-02 | 2.75E+00 | 2.85E+00 | 5.04E-01 | no | Agbl1 | intron_variant | MODIFIER |
| 7 | 76420092 | AX-170519281 | 0 | A | C | 0.36 | -6.42E-02 | 9.77E-02 | 2.75E+00 | 2.85E+00 | 5.04E-01 | no | Agbl1 | missense_variant | MODERATE |
| 7 | 76422656 | AX-170029341 | 0 | A | G | 0.36 | -6.42E-02 | 9.77E-02 | 2.75E+00 | 2.85E+00 | 5.04E-01 | no | Agbl1 | intron_variant | MODIFIER |
| 7 | 76436967 | AX-97810352 | 0 | C | T | 0.36 | -6.42E-02 | 9.77E-02 | 2.75E+00 | 2.85E+00 | 5.04E-01 | no | Agbl1 | intron_variant | MODIFIER |

| | | | | | | | | | | | | | | | |
|---|---|---|---|---|---|---|---|---|---|---|---|---|---|---|---|
| 7 | 76462343 | AX-170029337 | 0 | A | G | 0.36 | -6.42E-02 | 9.77E-02 | 2.75E+00 | 2.85E+00 | 5.04E-01 | no | Agbl1 | intron_variant | MODIFIER |
| 7 | 76468082 | AX-97830351 | 0 | T | C | 0.36 | -6.42E-02 | 9.77E-02 | 2.75E+00 | 2.85E+00 | 5.04E-01 | no | Agbl1 | intron_variant | MODIFIER |
| 7 | 76472191 | AX-170416928 | 0 | T | C | 0.36 | -6.42E-02 | 9.77E-02 | 2.75E+00 | 2.85E+00 | 5.04E-01 | no | Agbl1 | intron_variant | MODIFIER |
| 7 | 76499176 | AX-97802679 | 0 | A | G | 0.36 | -6.42E-02 | 9.77E-02 | 2.75E+00 | 2.85E+00 | 5.04E-01 | no | Agbl1 | intron_variant | MODIFIER |
| 7 | 76523764 | AX-170029336 | 0 | A | G | 0.36 | -6.42E-02 | 9.77E-02 | 2.75E+00 | 2.85E+00 | 5.04E-01 | no | Agbl1 | intron_variant | MODIFIER |
| 7 | 76544974 | AX-170029339 | 0 | C | A | 0.36 | -6.42E-02 | 9.77E-02 | 2.75E+00 | 2.85E+00 | 5.04E-01 | no | Agbl1 | intron_variant | MODIFIER |
| 7 | 76546780 | AX-170029340 | 0 | G | T | 0.36 | -6.42E-02 | 9.77E-02 | 2.75E+00 | 2.85E+00 | 5.04E-01 | no | Agbl1 | intron_variant | MODIFIER |
| 7 | 76848144 | AX-97803371 | 0 | A | G | 0.36 | -6.42E-02 | 9.77E-02 | 2.75E+00 | 2.85E+00 | 5.04E-01 | no | Agbl1 | intron_variant | MODIFIER |
| 7 | 76857273 | AX-97782050 | 0 | A | G | 0.36 | -6.42E-02 | 9.77E-02 | 2.75E+00 | 2.85E+00 | 5.04E-01 | no | Agbl1 | intron_variant | MODIFIER |
| 7 | 76873861 | AX-170367251 | 0 | C | G | 0.36 | -6.42E-02 | 9.77E-02 | 2.75E+00 | 2.85E+00 | 5.04E-01 | no | Agbl1 | intron_variant | MODIFIER |
| 7 | 76874934 | AX-170518324 | 0 | G | A | 0.36 | -6.42E-02 | 9.77E-02 | 2.75E+00 | 2.85E+00 | 5.04E-01 | no | Agbl1 | intron_variant | MODIFIER |
| 7 | 78195657 | AX-97835593 | 0 | C | T | 0.36 | -6.42E-02 | 9.77E-02 | 2.75E+00 | 2.85E+00 | 5.04E-01 | no | Ntrk3 | intron_variant | MODIFIER |
| 7 | 78220834 | AX-170077804 | 0 | A | G | 0.36 | -6.42E-02 | 9.77E-02 | 2.75E+00 | 2.85E+00 | 5.04E-01 | no | Ntrk3 | intron_variant | MODIFIER |
| 7 | 78221674 | AX-170175749 | 0 | C | T | 0.36 | -6.42E-02 | 9.77E-02 | 2.75E+00 | 2.85E+00 | 5.04E-01 | no | Ntrk3 | intron_variant | MODIFIER |
| 7 | 78225707 | AX-170427747 | 0 | C | G | 0.36 | -6.42E-02 | 9.77E-02 | 2.75E+00 | 2.85E+00 | 5.04E-01 | no | Ntrk3 | intron_variant | MODIFIER |
| 7 | 78227342 | AX-170299522 | 0 | C | T | 0.36 | -6.42E-02 | 9.77E-02 | 2.75E+00 | 2.85E+00 | 5.04E-01 | no | Ntrk3 | intron_variant | MODIFIER |
| 7 | 78244670 | AX-170309827 | 0 | A | G | 0.36 | -6.42E-02 | 9.77E-02 | 2.75E+00 | 2.85E+00 | 5.04E-01 | no | Ntrk3 | intron_variant | MODIFIER |
| 7 | 78255128 | AX-97853724 | 0 | C | T | 0.36 | -6.42E-02 | 9.77E-02 | 2.75E+00 | 2.85E+00 | 5.04E-01 | no | Ntrk3 | intron_variant | MODIFIER |
| 7 | 78281763 | AX-170350511 | 0 | A | G | 0.36 | -6.42E-02 | 9.77E-02 | 2.75E+00 | 2.85E+00 | 5.04E-01 | no | Ntrk3 | intron_variant | MODIFIER |
| 7 | 78298435 | AX-170177867 | 0 | A | G | 0.36 | -6.42E-02 | 9.77E-02 | 2.75E+00 | 2.85E+00 | 5.04E-01 | no | Ntrk3 | intron_variant | MODIFIER |
| 7 | 78301226 | AX-170387139 | 0 | G | A | 0.36 | -6.42E-02 | 9.77E-02 | 2.75E+00 | 2.85E+00 | 5.04E-01 | no | Ntrk3 | intron_variant | MODIFIER |
| 7 | 78301852 | AX-170439697 | 0 | T | G | 0.36 | -6.42E-02 | 9.77E-02 | 2.75E+00 | 2.85E+00 | 5.04E-01 | no | Ntrk3 | intron_variant | MODIFIER |
| 11 | 31849725 | AX-170046712 | 0 | A | C | 0.079 | 1.32E-01 | 2.00E-01 | 2.66E+00 | 2.74E+00 | 5.04E-01 | no | | intergenic_variant | MODIFIER |
| 4 | 154753736 | AX-170142046 | 1 | G | A | 0.107 | 1.21E-01 | 1.84E-01 | 2.69E+00 | 2.78E+00 | 5.04E-01 | no | | intergenic_variant | MODIFIER |
| 5 | 45288181 | AX-170240479 | 1 | G | A | 0.17 | -7.09E-02 | 1.04E-01 | 2.52E+00 | 2.61E+00 | 5.04E-01 | no | D5Ertd615e | intron_variant,non_coding_transcript_variant | MODIFIER |
| 7 | 37619189 | AX-170175519 | 1 | G | A | 0.339 | 6.38E-02 | 9.49E-02 | 2.57E+00 | 2.64E+00 | 5.04E-01 | no | Zfp536 | intron_variant | MODIFIER |
| 16 | 45375398 | AX-170159404 | 0 | G | A | 0.518 | 5.53E-02 | 8.04E-02 | 2.49E+00 | 2.54E+00 | 5.04E-01 | no | | intergenic_variant | MODIFIER |
| 16 | 46090747 | AX-170012964 | 0 | A | G | 0.518 | 5.53E-02 | 8.04E-02 | 2.49E+00 | 2.54E+00 | 5.04E-01 | no | Cd96 | intron_variant | MODIFIER |
| 14 | 93682571 | AX-170421387 | 0 | G | T | 0.044 | -1.12E-01 | 1.71E-01 | 2.68E+00 | 2.80E+00 | 5.04E-01 | no | Pcdh9 | intron_variant | MODIFIER |
| 6 | 94458423 | AX-170216010 | 1 | C | T | 0.446 | -4.09E-02 | 6.13E-02 | 2.57E+00 | 2.68E+00 | 5.04E-01 | no | | regulatory_region_variant | MODIFIER |
| 7 | 37614969 | AX-170338671 | 2 | A | G | 0.345 | 6.42E-02 | 9.57E-02 | 2.57E+00 | 2.64E+00 | 5.04E-01 | no | Zfp536 | intron_variant | MODIFIER |
| 3 | 76429579 | AX-170205645 | 0 | C | T | 0.044 | -1.14E-01 | 1.70E-01 | 2.56E+00 | 2.67E+00 | 5.04E-01 | no | Fstl5 | intron_variant | MODIFIER |
| 16 | 79467406 | AX-170103356 | 0 | G | C | 0.456 | 4.89E-02 | 7.35E-02 | 2.60E+00 | 2.68E+00 | 5.05E-01 | no | | intergenic_variant | MODIFIER |
| 15 | 84746536 | AX-97830051 | 0 | C | T | 0.509 | 4.66E-02 | 6.81E-02 | 2.49E+00 | 2.58E+00 | 5.05E-01 | no | Arhgap8 | intron_variant | MODIFIER |
| 1 | 20072127 | AX-170119987 | 1 | G | A | 0.411 | 5.59E-02 | 8.37E-02 | 2.60E+00 | 2.64E+00 | 5.05E-01 | no | Pkhd1 | intron_variant | MODIFIER |
| 3 | 29762232 | AX-170180062 | 0 | A | G | 0.158 | -6.25E-02 | 9.53E-02 | 2.72E+00 | 2.82E+00 | 5.05E-01 | no | | intergenic_variant | MODIFIER |
| 2 | 72276593 | AX-170063966 | 0 | A | G | 0.061 | 1.08E-01 | 1.65E-01 | 2.71E+00 | 2.86E+00 | 5.05E-01 | no | | intergenic_variant | MODIFIER |
| 2 | 124393980 | AX-170094400 | 2 | T | G | 0.136 | -1.12E-01 | 1.70E-01 | 2.76E+00 | 2.86E+00 | 5.05E-01 | no | Sema6d | intron_variant | MODIFIER |
| 16 | 54013771 | AX-97780821 | 0 | T | A | 0.474 | -5.68E-02 | 8.31E-02 | 2.51E+00 | 2.55E+00 | 5.05E-01 | no | | intergenic_variant | MODIFIER |
| 3 | 23694674 | AX-170160662 | 0 | C | T | 0.158 | -6.95E-02 | 1.03E-01 | 2.56E+00 | 2.63E+00 | 5.05E-01 | no | | intergenic_variant | MODIFIER |
| 10 | 94241699 | AX-170461723 | 0 | G | A | 0.307 | -5.55E-02 | 8.15E-02 | 2.51E+00 | 2.59E+00 | 5.05E-01 | no | | intergenic_variant | MODIFIER |
| 14 | 122885185 | AX-97827844 | 0 | T | C | 0.298 | 5.17E-02 | 7.70E-02 | 2.53E+00 | 2.65E+00 | 5.06E-01 | no | Pcca | intron_variant | MODIFIER |
| 5 | 77499905 | AX-97797492 | 1 | A | T | 0.438 | 7.15E-02 | 1.09E-01 | 2.74E+00 | 2.79E+00 | 5.06E-01 | no | | intergenic_variant | MODIFIER |
| 8 | 67088670 | AX-97847470 | 1 | C | T | 0.143 | -9.72E-02 | 1.46E-01 | 2.58E+00 | 2.65E+00 | 5.06E-01 | no | | intergenic_variant | MODIFIER |
| 7 | 25031394 | AX-170272203 | 0 | C | T | 0.412 | 6.18E-02 | 9.11E-02 | 2.50E+00 | 2.61E+00 | 5.06E-01 | no | Grik5 | intron_variant | MODIFIER |
| 15 | 68090540 | AX-97826467 | 1 | G | A | 0.348 | -5.23E-02 | 7.41E-02 | 2.41E+00 | 2.50E+00 | 5.06E-01 | no | Zfat | intron_variant | MODIFIER |
| 7 | 75442127 | AX-170029316 | 1 | G | C | 0.348 | -6.14E-02 | 9.41E-02 | 2.74E+00 | 2.85E+00 | 5.06E-01 | no | | intergenic_variant | MODIFIER |
| 18 | 64483193 | AX-170523656 | 0 | A | G | 0.395 | 6.41E-02 | 9.40E-02 | 2.51E+00 | 2.55E+00 | 5.06E-01 | no | Fech | intron_variant | MODIFIER |
| 9 | 104483742 | AX-170368171 | 0 | A | G | 0.123 | -7.41E-02 | 1.11E-01 | 2.59E+00 | 2.67E+00 | 5.06E-01 | no | Gm25904 | upstream_gene_variant | MODIFIER |
| 13 | 32372695 | AX-170276554 | 0 | G | A | 0.184 | -7.61E-02 | 1.14E-01 | 2.56E+00 | 2.63E+00 | 5.06E-01 | no | | regulatory_region_variant | MODIFIER |
| 9 | 95148195 | AX-170237761 | 0 | G | C | 0.14 | -7.61E-02 | 1.15E-01 | 2.64E+00 | 2.72E+00 | 5.06E-01 | no | Slc9a9 | intron_variant | MODIFIER |
| 18 | 69895826 | AX-170249508 | 0 | C | T | 0.184 | -7.97E-02 | 1.21E-01 | 2.65E+00 | 2.70E+00 | 5.06E-01 | no | AC108433.1 | intron_variant,non_coding_transcript_variant | MODIFIER |
| 13 | 47098448 | AX-170534001 | 0 | G | A | 0.482 | -4.13E-02 | 6.32E-02 | 2.77E+00 | 2.88E+00 | 5.06E-01 | no | Dek | intron_variant | MODIFIER |
| 6 | 94767639 | AX-97830618 | 1 | T | C | 0.438 | -3.95E-02 | 5.95E-02 | 2.57E+00 | 2.69E+00 | 5.06E-01 | no | | intergenic_variant | MODIFIER |

| | | | | | | | | | | | | | | | |
|---|---|---|---|---|---|---|---|---|---|---|---|---|---|---|---|
| 18 | 58877330 | AX-170259186 | 1 | C | T | 0.402 | 6.78E-02 | 9.70E-02 | 2.44E+00 | 2.49E+00 | 5.06E-01 | no | Adamts19 | intron_variant | MODIFIER |
| 18 | 58970640 | AX-170401151 | 1 | A | G | 0.402 | 6.78E-02 | 9.70E-02 | 2.44E+00 | 2.49E+00 | 5.06E-01 | no | Adamts19 | intron_variant | MODIFIER |
| 15 | 25363047 | AX-170220625 | 1 | C | A | 0.25 | -6.95E-02 | 1.05E-01 | 2.62E+00 | 2.69E+00 | 5.07E-01 | no | Basp1 | downstream_gene_variant | MODIFIER |
| 6 | 138680833 | AX-170231817 | 1 | C | T | 0.562 | 5.89E-02 | 8.51E-02 | 2.45E+00 | 2.53E+00 | 5.07E-01 | no | | intergenic_variant | MODIFIER |
| 10 | 31340531 | AX-170247609 | 1 | A | T | 0.402 | -4.91E-02 | 7.49E-02 | 2.64E+00 | 2.76E+00 | 5.07E-01 | no | Tpd52l1 | intron_variant | MODIFIER |
| 5 | 104287918 | AX-170025787 | 1 | G | C | 0.429 | 7.26E-02 | 1.10E-01 | 2.66E+00 | 2.71E+00 | 5.07E-01 | no | | intergenic_variant | MODIFIER |
| 11 | 34378684 | AX-170160553 | 0 | A | T | 0.105 | 1.06E-01 | 1.62E-01 | 2.65E+00 | 2.76E+00 | 5.07E-01 | no | Fam196b | intron_variant | MODIFIER |
| 9 | 72548762 | AX-170353076 | 0 | A | T | 0.219 | -7.38E-02 | 1.13E-01 | 2.70E+00 | 2.81E+00 | 5.07E-01 | no | Rfx7 | intron_variant | MODIFIER |
| 10 | 23627133 | AX-170352928 | 1 | A | G | 0.562 | -4.67E-02 | 7.10E-02 | 2.90E+00 | 3.07E+00 | 5.07E-01 | no | 4930520K02Rik | upstream_gene_variant | MODIFIER |
| 16 | 88900440 | AX-170221594 | 2 | T | C | 0.473 | 4.71E-02 | 7.23E-02 | 2.72E+00 | 2.82E+00 | 5.07E-01 | no | Krtap19-5 | upstream_gene_variant | MODIFIER |
| 18 | 61971817 | AX-170384109 | 0 | G | A | 0.412 | -5.64E-02 | 8.43E-02 | 2.56E+00 | 2.62E+00 | 5.07E-01 | no | Sh3tc2 | intron_variant | MODIFIER |
| 18 | 74374050 | AX-170093952 | 0 | C | T | 0.184 | -7.46E-02 | 1.12E-01 | 2.59E+00 | 2.65E+00 | 5.07E-01 | no | | intergenic_variant | MODIFIER |
| 5 | 93021506 | AX-170124136 | 1 | T | A | 0.429 | 6.98E-02 | 1.07E-01 | 2.70E+00 | 2.75E+00 | 5.07E-01 | no | | intergenic_variant | MODIFIER |
| 6 | 93914330 | AX-170513935 | 2 | C | T | 0.464 | -3.92E-02 | 5.88E-02 | 2.54E+00 | 2.67E+00 | 5.07E-01 | no | Magi1 | intron_variant | MODIFIER |
| 5 | 27906611 | AX-170071624 | 1 | T | G | 0.321 | 6.91E-02 | 1.06E-01 | 2.72E+00 | 2.79E+00 | 5.07E-01 | no | | intergenic_variant | MODIFIER |
| 5 | 34975553 | AX-170024669 | 1 | T | C | 0.321 | 6.91E-02 | 1.06E-01 | 2.72E+00 | 2.79E+00 | 5.07E-01 | no | Rgs12 | intron_variant | MODIFIER |
| 5 | 128800285 | AX-170480809 | 1 | C | T | 0.152 | -6.64E-02 | 9.02E-02 | 2.32E+00 | 2.42E+00 | 5.07E-01 | no | Rimbp2 | intron_variant | MODIFIER |
| 16 | 78349576 | AX-170263986 | 0 | C | T | 0.456 | 5.03E-02 | 7.58E-02 | 2.59E+00 | 2.66E+00 | 5.07E-01 | no | Cxadr | intron_variant | MODIFIER |
| 4 | 108678076 | AX-170447025 | 0 | C | G | 0.254 | 7.71E-02 | 1.15E-01 | 2.56E+00 | 2.62E+00 | 5.07E-01 | no | Zfyve9 | intron_variant | MODIFIER |
| 3 | 149318555 | AX-170396666 | 1 | G | A | 0.357 | 4.77E-02 | 7.30E-02 | 2.72E+00 | 2.81E+00 | 5.07E-01 | no | Gm30382 | intron_variant,non_coding_transcript_variant | MODIFIER |
| 10 | 30916795 | AX-170416454 | 0 | C | G | 0.184 | 6.14E-02 | 9.45E-02 | 2.75E+00 | 2.86E+00 | 5.07E-01 | no | | intergenic_variant | MODIFIER |
| 8 | 19062628 | AX-97830941 | 0 | T | C | 0.184 | 6.15E-02 | 9.42E-02 | 2.64E+00 | 2.76E+00 | 5.08E-01 | no | Defb39 | intron_variant | MODIFIER |
| 4 | 81273677 | AX-170471411 | 0 | G | T | 0.254 | 6.41E-02 | 9.49E-02 | 2.51E+00 | 2.60E+00 | 5.08E-01 | no | Mpdz | downstream_gene_variant | MODIFIER |
| 1 | 41412360 | AX-170287214 | 0 | C | T | 0.412 | -4.76E-02 | 7.29E-02 | 2.69E+00 | 2.78E+00 | 5.08E-01 | no | | intergenic_variant | MODIFIER |
| 18 | 55668841 | AX-170512070 | 1 | T | A | 0.438 | 6.50E-02 | 9.01E-02 | 2.36E+00 | 2.44E+00 | 5.08E-01 | no | | intergenic_variant | MODIFIER |
| 7 | 52200827 | AX-170077343 | 0 | G | T | 0.43 | 6.40E-02 | 9.66E-02 | 2.59E+00 | 2.66E+00 | 5.08E-01 | no | | intergenic_variant | MODIFIER |
| 5 | 119253116 | AX-97853383 | 1 | C | T | 0.446 | -7.24E-02 | 1.10E-01 | 2.85E+00 | 2.92E+00 | 5.08E-01 | no | | intergenic_variant | MODIFIER |
| 18 | 81031045 | AX-170471018 | 0 | A | G | 0.298 | -5.72E-02 | 8.55E-02 | 2.53E+00 | 2.64E+00 | 5.08E-01 | no | Gm25718 | downstream_gene_variant | MODIFIER |
| 1 | 149901211 | AX-170408528 | 1 | G | A | 0.205 | 9.06E-02 | 1.38E-01 | 2.67E+00 | 2.71E+00 | 5.08E-01 | no | Pla2g4a | splice_region_variant,synonymous_variant | LOW |
| 2 | 164488890 | AX-170445755 | 0 | A | G | 0.167 | 1.10E-01 | 1.69E-01 | 2.70E+00 | 2.80E+00 | 5.08E-01 | no | Dbndd2 | intron_variant | MODIFIER |
| 20 | 78085049 | AX-170497898 | 1 | G | A | 0.045 | 7.27E-02 | 1.08E-01 | 2.48E+00 | 2.66E+00 | 5.08E-01 | no | Obp1a | downstream_gene_variant | MODIFIER |
| 10 | 92654005 | AX-170227874 | 1 | G | A | 0.304 | -5.65E-02 | 8.41E-02 | 2.54E+00 | 2.60E+00 | 5.08E-01 | no | Gm31822 | downstream_gene_variant | MODIFIER |
| 15 | 6311007 | AX-170276173 | 2 | C | A | 0.245 | -6.76E-02 | 1.02E-01 | 2.60E+00 | 2.67E+00 | 5.08E-01 | no | Dab2 | intron_variant | MODIFIER |
| 9 | 34153107 | AX-170220631 | 0 | G | A | 0.439 | 5.46E-02 | 8.31E-02 | 2.63E+00 | 2.70E+00 | 5.08E-01 | no | | intergenic_variant | MODIFIER |
| 6 | 145448902 | AX-170341113 | 2 | C | T | 0.064 | -1.31E-01 | 2.02E-01 | 2.72E+00 | 2.83E+00 | 5.08E-01 | no | Lmntd1 | intron_variant | MODIFIER |
| 11 | 49215451 | AX-170103374 | 0 | C | T | 0.096 | 9.91E-02 | 1.53E-01 | 2.70E+00 | 2.84E+00 | 5.08E-01 | no | Zfp62 | missense_variant | MODERATE |
| 4 | 133209280 | AX-170023987 | 0 | A | T | 0.149 | 7.42E-02 | 1.12E-01 | 2.60E+00 | 2.66E+00 | 5.08E-01 | no | Gpr3 | downstream_gene_variant | MODIFIER |
| 12 | 91818784 | AX-170510500 | 1 | G | A | 0.071 | 1.40E-01 | 2.15E-01 | 2.74E+00 | 2.82E+00 | 5.09E-01 | no | Sel1l | intron_variant | MODIFIER |
| 11 | 102264959 | AX-170362745 | 0 | C | T | 0.053 | -1.37E-01 | 2.11E-01 | 2.71E+00 | 2.81E+00 | 5.09E-01 | no | Asb16 | upstream_gene_variant | MODIFIER |
| 11 | 116544707 | AX-170478093 | 0 | T | G | 0.053 | -1.37E-01 | 2.11E-01 | 2.71E+00 | 2.81E+00 | 5.09E-01 | no | Ube2o | synonymous_variant | LOW |
| 1 | 174621771 | AX-170523145 | 1 | G | A | 0.232 | 8.31E-02 | 1.23E-01 | 2.51E+00 | 2.57E+00 | 5.09E-01 | no | Fmn2 | intron_variant | MODIFIER |
| 8 | 49352949 | AX-170030647 | 0 | G | A | 0.158 | 9.13E-02 | 1.37E-01 | 2.94E+00 | 3.03E+00 | 5.09E-01 | no | | intergenic_variant | MODIFIER |
| 18 | 57806496 | AX-170240310 | 1 | C | T | 0.411 | 6.75E-02 | 9.78E-02 | 2.46E+00 | 2.51E+00 | 5.09E-01 | no | | intergenic_variant | MODIFIER |
| 18 | 58235032 | AX-170220272 | 1 | G | T | 0.411 | 6.75E-02 | 9.78E-02 | 2.46E+00 | 2.51E+00 | 5.09E-01 | no | Gm25660 | upstream_gene_variant | MODIFIER |
| 4 | 72903950 | AX-168383789 | 1 | T | C | 0.223 | 6.98E-02 | 1.05E-01 | 2.55E+00 | 2.62E+00 | 5.09E-01 | no | Gm25653 | downstream_gene_variant | MODIFIER |
| 16 | 28280162 | AX-170101260 | 0 | G | T | 0.246 | 5.74E-02 | 8.83E-02 | 2.65E+00 | 2.78E+00 | 5.09E-01 | no | Fgf12 | intron_variant | MODIFIER |
| 4 | 144436919 | AX-170024178 | 0 | C | T | 0.114 | 1.11E-01 | 1.70E-01 | 2.69E+00 | 2.80E+00 | 5.09E-01 | no | 1700012P22Rik | intron_variant | MODIFIER |
| 9 | 123278190 | AX-170527652 | 0 | T | A | 0.14 | -5.85E-02 | 9.02E-02 | 2.69E+00 | 2.81E+00 | 5.09E-01 | no | | intergenic_variant | MODIFIER |
| 9 | 94551001 | AX-97821083 | 0 | G | A | 0.14 | -7.82E-02 | 1.19E-01 | 2.64E+00 | 2.72E+00 | 5.09E-01 | no | | intergenic_variant | MODIFIER |
| 10 | 93888864 | AX-170451328 | 1 | T | C | 0.295 | -5.75E-02 | 8.57E-02 | 2.54E+00 | 2.59E+00 | 5.09E-01 | no | Metap2 | intron_variant | MODIFIER |
| 1 | 136252098 | AX-170217750 | 1 | C | T | 0.214 | 8.00E-02 | 1.22E-01 | 2.64E+00 | 2.72E+00 | 5.09E-01 | no | Gm37552 | downstream_gene_variant | MODIFIER |
| 12 | 116863876 | AX-170349445 | 0 | A | G | 0.132 | 1.07E-01 | 1.63E-01 | 2.61E+00 | 2.73E+00 | 5.09E-01 | no | Ptprn2 | intron_variant | MODIFIER |
| 15 | 19751115 | AX-97792165 | 0 | G | A | 0.246 | -6.73E-02 | 1.01E-01 | 2.58E+00 | 2.64E+00 | 5.09E-01 | no | Gm18741 | downstream_gene_variant | MODIFIER |
| 15 | 79610768 | AX-170360171 | 0 | G | A | 0.57 | 5.34E-02 | 8.11E-02 | 2.61E+00 | 2.69E+00 | 5.09E-01 | no | Fam227a | downstream_gene_variant | MODIFIER |

| | | | | | | | | | | | | | | | |
|---|---|---|---|---|---|---|---|---|---|---|---|---|---|---|---|
| 7 | 66405003 | AX-170346137 | 1 | A | G | 0.402 | -6.31E-02 | 9.68E-02 | 2.79E+00 | 2.87E+00 | 5.10E-01 | no | Aldh1a3 | intron_variant | MODIFIER |
| 2 | 127004630 | AX-170281413 | 0 | T | C | 0.184 | -6.02E-02 | 9.33E-02 | 2.64E+00 | 2.80E+00 | 5.10E-01 | no | Ap4e1 | upstream_gene_variant | MODIFIER |
| 4 | 137957616 | AX-170355365 | 0 | G | T | 0.167 | 6.10E-02 | 9.29E-02 | 2.59E+00 | 2.69E+00 | 5.10E-01 | no | Ece1 | intron_variant | MODIFIER |
| 5 | 44011133 | AX-170314830 | 1 | C | T | 0.196 | -7.15E-02 | 1.08E-01 | 2.58E+00 | 2.66E+00 | 5.10E-01 | no | Prom1 | intron_variant | MODIFIER |
| 4 | 118381226 | AX-170335069 | 0 | A | G | 0.211 | 7.66E-02 | 1.16E-01 | 2.60E+00 | 2.67E+00 | 5.10E-01 | no | Szt2 | intron_variant | MODIFIER |
| 2 | 61523439 | AX-97837349 | 0 | A | C | 0.053 | 1.11E-01 | 1.70E-01 | 2.62E+00 | 2.75E+00 | 5.10E-01 | no | | intergenic_variant | MODIFIER |
| 5 | 50006032 | AX-97801341 | 0 | A | G | 0.482 | 7.20E-02 | 1.04E-01 | 2.45E+00 | 2.50E+00 | 5.10E-01 | no | Adgra3 | intron_variant | MODIFIER |
| 5 | 51980950 | AX-170481846 | 0 | C | T | 0.482 | 7.20E-02 | 1.04E-01 | 2.45E+00 | 2.50E+00 | 5.10E-01 | no | | intergenic_variant | MODIFIER |
| 1 | 121324701 | AX-170130603 | 0 | G | A | 0.588 | 5.67E-02 | 8.24E-02 | 2.45E+00 | 2.52E+00 | 5.10E-01 | no | Insig2 | intron_variant | MODIFIER |
| 5 | 102801101 | AX-170403215 | 0 | T | C | 0.43 | 7.06E-02 | 1.08E-01 | 2.67E+00 | 2.73E+00 | 5.10E-01 | no | Arhgap24 | intron_variant | MODIFIER |
| 6 | 139311939 | AX-170397753 | 0 | G | T | 0.061 | 1.39E-01 | 2.11E-01 | 2.57E+00 | 2.67E+00 | 5.10E-01 | no | 4922502N22Rik | downstream_gene_variant | MODIFIER |
| 10 | 17900285 | AX-170202006 | 1 | A | C | 0.196 | 6.55E-02 | 1.01E-01 | 2.84E+00 | 2.95E+00 | 5.11E-01 | no | Heca | 3_prime_UTR_variant | MODIFIER |
| 1 | 172687457 | AX-170532036 | 1 | A | C | 0.232 | 8.16E-02 | 1.22E-01 | 2.54E+00 | 2.59E+00 | 5.11E-01 | no | Gm22383 | upstream_gene_variant | MODIFIER |
| 1 | 175222432 | AX-170384059 | 1 | A | G | 0.232 | 8.16E-02 | 1.22E-01 | 2.54E+00 | 2.59E+00 | 5.11E-01 | no | Rgs7 | intron_variant | MODIFIER |
| 1 | 175348645 | AX-170230767 | 1 | T | A | 0.232 | 8.16E-02 | 1.22E-01 | 2.54E+00 | 2.59E+00 | 5.11E-01 | no | Rgs7 | intron_variant | MODIFIER |
| 18 | 76149022 | AX-97853648 | 1 | G | A | 0.161 | -8.11E-02 | 1.21E-01 | 2.53E+00 | 2.58E+00 | 5.11E-01 | no | Zbtb7c | downstream_gene_variant | MODIFIER |
| 7 | 72787755 | AX-170184886 | 0 | A | G | 0.368 | -6.45E-02 | 9.97E-02 | 2.71E+00 | 2.81E+00 | 5.11E-01 | no | | intergenic_variant | MODIFIER |
| 5 | 119282640 | AX-170110522 | 1 | C | G | 0.446 | 7.01E-02 | 1.07E-01 | 2.65E+00 | 2.71E+00 | 5.11E-01 | no | | intergenic_variant | MODIFIER |
| 18 | 50107096 | AX-170237855 | 1 | G | A | 0.188 | -6.89E-02 | 1.04E-01 | 2.54E+00 | 2.63E+00 | 5.11E-01 | no | Tnfaip8 | non_coding_transcript_exon_variant | MODIFIER |
| 9 | 19308635 | AX-170090256 | 1 | C | T | 0.473 | 6.01E-02 | 8.99E-02 | 2.53E+00 | 2.60E+00 | 5.11E-01 | no | | intergenic_variant | MODIFIER |
| 4 | 74200741 | AX-170023119 | 0 | G | T | 0.263 | 6.26E-02 | 9.26E-02 | 2.48E+00 | 2.58E+00 | 5.11E-01 | no | Frmd3 | intron_variant | MODIFIER |
| 8 | 93552429 | AX-170504084 | 0 | T | A | 0.211 | -7.60E-02 | 1.17E-01 | 2.72E+00 | 2.79E+00 | 5.11E-01 | no | | intergenic_variant | MODIFIER |
| 1 | 25227218 | AX-170000126 | 0 | T | G | 0.421 | 5.59E-02 | 8.53E-02 | 2.62E+00 | 2.68E+00 | 5.11E-01 | no | Adgrb3 | intron_variant | MODIFIER |
| 8 | 31912888 | AX-170182792 | 1 | A | G | 0.134 | -8.32E-02 | 1.28E-01 | 2.65E+00 | 2.72E+00 | 5.11E-01 | no | Nrg1 | intron_variant | MODIFIER |
| 5 | 43416190 | AX-97783409 | 2 | G | A | 0.191 | -6.78E-02 | 1.03E-01 | 2.57E+00 | 2.67E+00 | 5.11E-01 | no | | intergenic_variant | MODIFIER |
| 10 | 67703815 | AX-170003401 | 0 | G | A | 0.421 | -5.18E-02 | 7.94E-02 | 2.63E+00 | 2.70E+00 | 5.12E-01 | no | 1700030E10Rik | downstream_gene_variant | MODIFIER |
| 20 | 87202785 | AX-97785200 | 0 | G | A | 0.263 | -3.87E-02 | 6.02E-02 | 2.74E+00 | 2.85E+00 | 5.12E-01 | no | Il1rapl1 | intron_variant | MODIFIER |
| 7 | 117861609 | AX-170322580 | 0 | C | T | 0.395 | 5.84E-02 | 9.01E-02 | 2.71E+00 | 2.76E+00 | 5.12E-01 | no | | intergenic_variant | MODIFIER |
| 5 | 96391020 | AX-97856913 | 1 | G | A | 0.429 | 6.88E-02 | 1.06E-01 | 2.70E+00 | 2.76E+00 | 5.12E-01 | no | Fras1 | intron_variant | MODIFIER |
| 1 | 78668915 | AX-170086507 | 0 | A | T | 0.132 | -1.04E-01 | 1.59E-01 | 2.59E+00 | 2.70E+00 | 5.12E-01 | no | Acsl3 | intron_variant | MODIFIER |
| 4 | 135209005 | AX-170070780 | 0 | G | A | 0.175 | 6.99E-02 | 1.07E-01 | 2.62E+00 | 2.67E+00 | 5.12E-01 | no | Clic4 | downstream_gene_variant | MODIFIER |
| 3 | 26230868 | AX-97796410 | 0 | A | G | 0.14 | -6.66E-02 | 1.01E-01 | 2.55E+00 | 2.63E+00 | 5.12E-01 | no | Nlgn1 | intron_variant,non_coding_transcript_variant | MODIFIER |
| 3 | 28806031 | AX-170393423 | 1 | G | A | 0.17 | -6.53E-02 | 1.01E-01 | 2.70E+00 | 2.77E+00 | 5.12E-01 | no | Rpl22l1 | intron_variant | MODIFIER |
| 12 | 89159407 | AX-170159810 | 0 | A | G | 0.096 | 1.24E-01 | 1.92E-01 | 2.76E+00 | 2.83E+00 | 5.12E-01 | no | Nrxn3 | intron_variant | MODIFIER |
| 12 | 89174765 | AX-170368976 | 0 | A | C | 0.096 | 1.24E-01 | 1.92E-01 | 2.76E+00 | 2.83E+00 | 5.12E-01 | no | Nrxn3 | intron_variant | MODIFIER |
| 12 | 89174856 | AX-170428018 | 0 | T | C | 0.096 | 1.24E-01 | 1.92E-01 | 2.76E+00 | 2.83E+00 | 5.12E-01 | no | Nrxn3 | intron_variant | MODIFIER |
| 12 | 89240307 | AX-170162782 | 0 | C | T | 0.096 | 1.24E-01 | 1.92E-01 | 2.76E+00 | 2.83E+00 | 5.12E-01 | no | Nrxn3 | intron_variant | MODIFIER |
| 12 | 89241649 | AX-170489286 | 0 | G | A | 0.096 | 1.24E-01 | 1.92E-01 | 2.76E+00 | 2.83E+00 | 5.12E-01 | no | Nrxn3 | intron_variant | MODIFIER |
| 12 | 89249416 | AX-170181808 | 0 | G | A | 0.096 | 1.24E-01 | 1.92E-01 | 2.76E+00 | 2.83E+00 | 5.12E-01 | no | Nrxn3 | intron_variant | MODIFIER |
| 12 | 89249522 | AX-170114166 | 0 | C | A | 0.096 | 1.24E-01 | 1.92E-01 | 2.76E+00 | 2.83E+00 | 5.12E-01 | no | Nrxn3 | intron_variant | MODIFIER |
| 12 | 89402493 | AX-97838102 | 0 | G | A | 0.096 | 1.24E-01 | 1.92E-01 | 2.76E+00 | 2.83E+00 | 5.12E-01 | no | Nrxn3 | intron_variant | MODIFIER |
| 12 | 89405608 | AX-170120557 | 0 | A | G | 0.096 | 1.24E-01 | 1.92E-01 | 2.76E+00 | 2.83E+00 | 5.12E-01 | no | Nrxn3 | intron_variant | MODIFIER |
| 12 | 89406352 | AX-170406665 | 0 | G | A | 0.096 | 1.24E-01 | 1.92E-01 | 2.76E+00 | 2.83E+00 | 5.12E-01 | no | Nrxn3 | intron_variant | MODIFIER |
| 12 | 89406890 | AX-170007225 | 0 | A | C | 0.096 | 1.24E-01 | 1.92E-01 | 2.76E+00 | 2.83E+00 | 5.12E-01 | no | Nrxn3 | intron_variant | MODIFIER |
| 12 | 89407312 | AX-97819402 | 0 | A | G | 0.096 | 1.24E-01 | 1.92E-01 | 2.76E+00 | 2.83E+00 | 5.12E-01 | no | Nrxn3 | intron_variant | MODIFIER |
| 12 | 89408684 | AX-170158979 | 0 | C | T | 0.096 | 1.24E-01 | 1.92E-01 | 2.76E+00 | 2.83E+00 | 5.12E-01 | no | Nrxn3 | intron_variant | MODIFIER |
| 12 | 89416724 | AX-170106402 | 0 | G | A | 0.096 | 1.24E-01 | 1.92E-01 | 2.76E+00 | 2.83E+00 | 5.12E-01 | no | Nrxn3 | intron_variant | MODIFIER |
| 12 | 90084828 | AX-170258538 | 0 | C | G | 0.096 | 1.24E-01 | 1.92E-01 | 2.76E+00 | 2.83E+00 | 5.12E-01 | no | Nrxn3 | intron_variant | MODIFIER |
| 12 | 90169955 | AX-97804406 | 0 | A | C | 0.096 | 1.24E-01 | 1.92E-01 | 2.76E+00 | 2.83E+00 | 5.12E-01 | no | Nrxn3 | intron_variant | MODIFIER |
| 12 | 90174060 | AX-170007230 | 0 | A | T | 0.096 | 1.24E-01 | 1.92E-01 | 2.76E+00 | 2.83E+00 | 5.12E-01 | no | Nrxn3 | intron_variant | MODIFIER |
| 12 | 90207301 | AX-170049747 | 0 | A | G | 0.096 | 1.24E-01 | 1.92E-01 | 2.76E+00 | 2.83E+00 | 5.12E-01 | no | Nrxn3 | intron_variant | MODIFIER |
| 12 | 90350885 | AX-97844438 | 0 | T | C | 0.096 | 1.24E-01 | 1.92E-01 | 2.76E+00 | 2.83E+00 | 5.12E-01 | no | | intergenic_variant | MODIFIER |
| 12 | 90641717 | AX-170165869 | 0 | T | C | 0.096 | 1.24E-01 | 1.92E-01 | 2.76E+00 | 2.83E+00 | 5.12E-01 | no | | intergenic_variant | MODIFIER |
| 12 | 90648246 | AX-170007246 | 0 | C | T | 0.096 | 1.24E-01 | 1.92E-01 | 2.76E+00 | 2.83E+00 | 5.12E-01 | no | | intergenic_variant | MODIFIER |

| | | | | | | | | | | | | | | | |
|---|---|---|---|---|---|---|---|---|---|---|---|---|---|---|---|
| 12 | 90651264 | AX-170007242 | 0 | G | A | 0.096 | 1.24E-01 | 1.92E-01 | 2.76E+00 | 2.83E+00 | 5.12E-01 | no | | intergenic_variant | MODIFIER |
| 12 | 90697871 | AX-170007247 | 0 | C | T | 0.096 | 1.24E-01 | 1.92E-01 | 2.76E+00 | 2.83E+00 | 5.12E-01 | no | | intergenic_variant | MODIFIER |
| 12 | 90705385 | AX-97823907 | 0 | G | A | 0.096 | 1.24E-01 | 1.92E-01 | 2.76E+00 | 2.83E+00 | 5.12E-01 | no | | intergenic_variant | MODIFIER |
| 12 | 92302146 | AX-97854226 | 0 | C | T | 0.096 | 1.24E-01 | 1.92E-01 | 2.76E+00 | 2.83E+00 | 5.12E-01 | no | | intergenic_variant | MODIFIER |
| 7 | 117245484 | AX-170197635 | 1 | T | G | 0.42 | 6.30E-02 | 9.71E-02 | 2.69E+00 | 2.73E+00 | 5.12E-01 | no | Gm39075 | intron_variant,non_coding_transcript_variant | MODIFIER |
| 1 | 148033718 | AX-170499784 | 0 | C | G | 0.202 | -8.53E-02 | 1.29E-01 | 2.93E+00 | 3.04E+00 | 5.12E-01 | no | | intergenic_variant | MODIFIER |
| 7 | 37480797 | AX-170449490 | 0 | T | C | 0.342 | 5.99E-02 | 9.12E-02 | 2.58E+00 | 2.67E+00 | 5.12E-01 | no | Zfp536 | synonymous_variant | LOW |
| 3 | 38340692 | AX-170334676 | 0 | T | C | 0.219 | 5.08E-02 | 7.88E-02 | 2.67E+00 | 2.79E+00 | 5.12E-01 | no | | intergenic_variant | MODIFIER |
| 7 | 17155365 | AX-170470398 | 0 | T | C | 0.36 | 5.52E-02 | 8.47E-02 | 2.59E+00 | 2.71E+00 | 5.12E-01 | no | Ceacam3 | intron_variant | MODIFIER |
| 7 | 17156564 | AX-170460051 | 0 | T | A | 0.36 | 5.52E-02 | 8.47E-02 | 2.59E+00 | 2.71E+00 | 5.12E-01 | no | Ceacam3 | intron_variant | MODIFIER |
| 7 | 17156976 | AX-170298816 | 0 | C | T | 0.36 | 5.52E-02 | 8.47E-02 | 2.59E+00 | 2.71E+00 | 5.12E-01 | no | Ceacam3 | intron_variant | MODIFIER |
| 7 | 17157035 | AX-170076787 | 0 | A | G | 0.36 | 5.52E-02 | 8.47E-02 | 2.59E+00 | 2.71E+00 | 5.12E-01 | no | Ceacam3 | intron_variant | MODIFIER |
| 7 | 17157256 | AX-170076789 | 0 | C | T | 0.36 | 5.52E-02 | 8.47E-02 | 2.59E+00 | 2.71E+00 | 5.12E-01 | no | Ceacam3 | intron_variant | MODIFIER |
| 7 | 17157962 | AX-170138964 | 0 | G | C | 0.36 | 5.52E-02 | 8.47E-02 | 2.59E+00 | 2.71E+00 | 5.12E-01 | no | Ceacam3 | intron_variant | MODIFIER |
| 7 | 17158231 | AX-170423315 | 0 | A | G | 0.36 | 5.52E-02 | 8.47E-02 | 2.59E+00 | 2.71E+00 | 5.12E-01 | no | Ceacam3 | missense_variant | MODERATE |
| 7 | 17161233 | AX-170327087 | 0 | A | G | 0.36 | 5.52E-02 | 8.47E-02 | 2.59E+00 | 2.71E+00 | 5.12E-01 | no | Ceacam3 | intron_variant | MODIFIER |
| 7 | 17767382 | AX-170028649 | 0 | G | T | 0.36 | 5.52E-02 | 8.47E-02 | 2.59E+00 | 2.71E+00 | 5.12E-01 | no | | intergenic_variant | MODIFIER |
| 7 | 17886968 | AX-170452633 | 0 | T | C | 0.36 | 5.52E-02 | 8.47E-02 | 2.59E+00 | 2.71E+00 | 5.12E-01 | no | Gm5155 | intron_variant | MODIFIER |
| 7 | 17888549 | AX-170316905 | 0 | G | A | 0.36 | 5.52E-02 | 8.47E-02 | 2.59E+00 | 2.71E+00 | 5.12E-01 | no | Gm5155 | intron_variant | MODIFIER |
| 7 | 17909452 | AX-170153681 | 0 | A | G | 0.36 | 5.52E-02 | 8.47E-02 | 2.59E+00 | 2.71E+00 | 5.12E-01 | no | Gm5155 | intron_variant | MODIFIER |
| 7 | 17910252 | AX-170335576 | 0 | A | G | 0.36 | 5.52E-02 | 8.47E-02 | 2.59E+00 | 2.71E+00 | 5.12E-01 | no | Gm5155 | intron_variant | MODIFIER |
| 7 | 17910818 | AX-97813409 | 0 | T | C | 0.36 | 5.52E-02 | 8.47E-02 | 2.59E+00 | 2.71E+00 | 5.12E-01 | no | Gm5155 | intron_variant | MODIFIER |
| 7 | 18001891 | AX-170516335 | 0 | C | T | 0.36 | 5.52E-02 | 8.47E-02 | 2.59E+00 | 2.71E+00 | 5.12E-01 | no | | intergenic_variant | MODIFIER |
| 7 | 18002393 | AX-170459946 | 0 | T | C | 0.36 | 5.52E-02 | 8.47E-02 | 2.59E+00 | 2.71E+00 | 5.12E-01 | no | | intergenic_variant | MODIFIER |
| 7 | 18002451 | AX-170162812 | 0 | C | T | 0.36 | 5.52E-02 | 8.47E-02 | 2.59E+00 | 2.71E+00 | 5.12E-01 | no | | intergenic_variant | MODIFIER |
| 7 | 18002487 | AX-170287882 | 0 | C | T | 0.36 | 5.52E-02 | 8.47E-02 | 2.59E+00 | 2.71E+00 | 5.12E-01 | no | | intergenic_variant | MODIFIER |
| 7 | 18003506 | AX-170274069 | 0 | T | G | 0.36 | 5.52E-02 | 8.47E-02 | 2.59E+00 | 2.71E+00 | 5.12E-01 | no | | intergenic_variant | MODIFIER |
| 7 | 18005612 | AX-170230055 | 0 | C | T | 0.36 | 5.52E-02 | 8.47E-02 | 2.59E+00 | 2.71E+00 | 5.12E-01 | no | Ceacam13 | upstream_gene_variant | MODIFIER |
| 7 | 18010796 | AX-170466038 | 0 | T | C | 0.36 | 5.52E-02 | 8.47E-02 | 2.59E+00 | 2.71E+00 | 5.12E-01 | no | Ceacam13 | intron_variant | MODIFIER |
| 7 | 18564040 | AX-97819334 | 0 | G | A | 0.36 | 5.52E-02 | 8.47E-02 | 2.59E+00 | 2.71E+00 | 5.12E-01 | no | Psg27 | intron_variant | MODIFIER |
| 7 | 18564288 | AX-170243420 | 0 | G | T | 0.36 | 5.52E-02 | 8.47E-02 | 2.59E+00 | 2.71E+00 | 5.12E-01 | no | Psg27 | intron_variant | MODIFIER |
| 7 | 18649601 | AX-170182003 | 0 | G | C | 0.36 | 5.52E-02 | 8.47E-02 | 2.59E+00 | 2.71E+00 | 5.12E-01 | no | Psg21 | intron_variant | MODIFIER |
| 7 | 18686869 | AX-170424189 | 0 | A | G | 0.36 | 5.52E-02 | 8.47E-02 | 2.59E+00 | 2.71E+00 | 5.12E-01 | no | Psg20 | upstream_gene_variant | MODIFIER |
| 7 | 18687473 | AX-170438584 | 0 | G | A | 0.36 | 5.52E-02 | 8.47E-02 | 2.59E+00 | 2.71E+00 | 5.12E-01 | no | Psg20 | upstream_gene_variant | MODIFIER |
| 7 | 18693604 | AX-170035747 | 0 | T | C | 0.36 | 5.52E-02 | 8.47E-02 | 2.59E+00 | 2.71E+00 | 5.12E-01 | no | Gm44655 | upstream_gene_variant | MODIFIER |
| 7 | 18713738 | AX-170103084 | 0 | G | A | 0.36 | 5.52E-02 | 8.47E-02 | 2.59E+00 | 2.71E+00 | 5.12E-01 | no | Psg22 | upstream_gene_variant | MODIFIER |
| 7 | 18726756 | AX-170174612 | 0 | G | A | 0.36 | 5.52E-02 | 8.47E-02 | 2.59E+00 | 2.71E+00 | 5.12E-01 | no | Psg22 | missense_variant | MODERATE |
| 7 | 18749397 | AX-170189301 | 0 | T | A | 0.36 | 5.52E-02 | 8.47E-02 | 2.59E+00 | 2.71E+00 | 5.12E-01 | no | Gm24464 | upstream_gene_variant | MODIFIER |
| 7 | 18750125 | AX-97804455 | 0 | A | G | 0.36 | 5.52E-02 | 8.47E-02 | 2.59E+00 | 2.71E+00 | 5.12E-01 | no | Gm24464 | upstream_gene_variant | MODIFIER |
| 7 | 18789230 | AX-97786348 | 0 | G | A | 0.36 | 5.52E-02 | 8.47E-02 | 2.59E+00 | 2.71E+00 | 5.12E-01 | no | Psg19 | downstream_gene_variant | MODIFIER |
| 7 | 18826426 | AX-170299434 | 0 | T | C | 0.36 | 5.52E-02 | 8.47E-02 | 2.59E+00 | 2.71E+00 | 5.12E-01 | no | Psg17 | upstream_gene_variant | MODIFIER |
| 7 | 18826989 | AX-170379606 | 0 | A | G | 0.36 | 5.52E-02 | 8.47E-02 | 2.59E+00 | 2.71E+00 | 5.12E-01 | no | Gm18855 | upstream_gene_variant | MODIFIER |
| 7 | 18842560 | AX-170109044 | 0 | A | G | 0.36 | 5.52E-02 | 8.47E-02 | 2.59E+00 | 2.71E+00 | 5.12E-01 | no | Mill2 | intron_variant | MODIFIER |
| 7 | 18868912 | AX-170508767 | 0 | T | C | 0.36 | 5.52E-02 | 8.47E-02 | 2.59E+00 | 2.71E+00 | 5.12E-01 | no | Pglyrp1 | upstream_gene_variant | MODIFIER |
| 7 | 18877400 | AX-169999743 | 0 | T | A | 0.36 | 5.52E-02 | 8.47E-02 | 2.59E+00 | 2.71E+00 | 5.12E-01 | no | Pglyrp1 | intron_variant,non_coding_transcript_variant | MODIFIER |
| 7 | 18882337 | AX-97844741 | 0 | C | T | 0.36 | 5.52E-02 | 8.47E-02 | 2.59E+00 | 2.71E+00 | 5.12E-01 | no | Pglyrp1 | upstream_gene_variant | MODIFIER |
| 7 | 18884677 | AX-170271293 | 0 | C | T | 0.36 | 5.52E-02 | 8.47E-02 | 2.59E+00 | 2.71E+00 | 5.12E-01 | no | Pglyrp1 | upstream_gene_variant | MODIFIER |
| 7 | 19031872 | AX-170219020 | 0 | C | T | 0.36 | 5.52E-02 | 8.47E-02 | 2.59E+00 | 2.71E+00 | 5.12E-01 | no | Sympk | intron_variant | MODIFIER |
| 7 | 19038252 | AX-97833474 | 0 | A | G | 0.36 | 5.52E-02 | 8.47E-02 | 2.59E+00 | 2.71E+00 | 5.12E-01 | no | Sympk | intron_variant | MODIFIER |
| 7 | 19222709 | AX-170510112 | 0 | A | G | 0.36 | 5.52E-02 | 8.47E-02 | 2.59E+00 | 2.71E+00 | 5.12E-01 | no | Gpr4 | synonymous_variant | LOW |
| 7 | 19247330 | AX-170338388 | 0 | T | C | 0.36 | 5.52E-02 | 8.47E-02 | 2.59E+00 | 2.71E+00 | 5.12E-01 | no | Opa3 | 3_prime_UTR_variant | MODIFIER |
| 7 | 19249618 | AX-170327815 | 0 | C | A | 0.36 | 5.52E-02 | 8.47E-02 | 2.59E+00 | 2.71E+00 | 5.12E-01 | no | Opa3 | downstream_gene_variant | MODIFIER |
| 7 | 19269220 | AX-170290767 | 0 | C | T | 0.36 | 5.52E-02 | 8.47E-02 | 2.59E+00 | 2.71E+00 | 5.12E-01 | no | Vasp | intron_variant | MODIFIER |
| 7 | 19343567 | AX-170156189 | 0 | C | T | 0.36 | 5.52E-02 | 8.47E-02 | 2.59E+00 | 2.71E+00 | 5.12E-01 | no | Ercc1 | upstream_gene_variant | MODIFIER |

| | | | | | | | | | | | | | | | |
|---|---|---|---|---|---|---|---|---|---|---|---|---|---|---|---|
| 7 | 19356696 | AX-170076823 | 0 | A | G | 0.36 | 5.52E-02 | 8.47E-02 | 2.59E+00 | 2.71E+00 | 5.12E-01 | no | Ercc1 | downstream_gene_variant | MODIFIER |
| 7 | 19360106 | AX-97820382 | 0 | C | G | 0.36 | 5.52E-02 | 8.47E-02 | 2.59E+00 | 2.71E+00 | 5.12E-01 | no | Ercc1 | downstream_gene_variant | MODIFIER |
| 7 | 19414284 | AX-97814483 | 0 | T | C | 0.36 | 5.52E-02 | 8.47E-02 | 2.59E+00 | 2.71E+00 | 5.12E-01 | no | Ckm | intron_variant | MODIFIER |
| 7 | 19414710 | AX-170341732 | 0 | T | C | 0.36 | 5.52E-02 | 8.47E-02 | 2.59E+00 | 2.71E+00 | 5.12E-01 | no | Ckm | intron_variant | MODIFIER |
| 7 | 19555902 | AX-170350120 | 0 | G | A | 0.36 | 5.52E-02 | 8.47E-02 | 2.59E+00 | 2.71E+00 | 5.12E-01 | no | Ppp1r37 | intron_variant | MODIFIER |
| 3 | 37249740 | AX-170020316 | 1 | T | A | 0.188 | 6.09E-02 | 9.44E-02 | 2.77E+00 | 2.86E+00 | 5.12E-01 | no | | intergenic_variant | MODIFIER |
| 7 | 115385604 | AX-97841464 | 1 | C | A | 0.42 | 6.31E-02 | 9.74E-02 | 2.69E+00 | 2.73E+00 | 5.12E-01 | no | | intergenic_variant | MODIFIER |
| 7 | 92081261 | AX-170078069 | 0 | C | A | 0.132 | -1.11E-01 | 1.72E-01 | 2.65E+00 | 2.77E+00 | 5.12E-01 | no | Dlg2 | intron_variant | MODIFIER |
| 8 | 34561976 | AX-170515259 | 1 | G | A | 0.143 | -8.79E-02 | 1.35E-01 | 2.64E+00 | 2.70E+00 | 5.13E-01 | no | | intergenic_variant | MODIFIER |
| 13 | 37393408 | AX-170223708 | 0 | T | C | 0.175 | -7.28E-02 | 1.10E-01 | 2.55E+00 | 2.64E+00 | 5.13E-01 | no | Ly86 | intron_variant | MODIFIER |
| 1 | 3887902 | AX-170192080 | 0 | A | G | 0.518 | 4.68E-02 | 6.94E-02 | 2.49E+00 | 2.56E+00 | 5.13E-01 | no | | intergenic_variant | MODIFIER |
| 1 | 4895450 | AX-170309252 | 0 | T | C | 0.518 | 4.68E-02 | 6.94E-02 | 2.49E+00 | 2.56E+00 | 5.13E-01 | no | Tcea1 | intron_variant | MODIFIER |
| 4 | 154190922 | AX-170150971 | 0 | G | C | 0.096 | 1.13E-01 | 1.75E-01 | 2.66E+00 | 2.76E+00 | 5.13E-01 | no | Megf6 | intron_variant | MODIFIER |
| 20 | 142015423 | AX-97781935 | 1 | G | A | 0.143 | 7.22E-02 | 1.11E-01 | 2.61E+00 | 2.70E+00 | 5.13E-01 | no | | intergenic_variant | MODIFIER |
| 9 | 118661256 | AX-97838071 | 1 | C | A | 0.125 | -6.25E-02 | 9.59E-02 | 2.60E+00 | 2.71E+00 | 5.13E-01 | no | Itga9 | intron_variant | MODIFIER |
| 16 | 9433132 | AX-170113110 | 0 | C | T | 0.053 | 1.13E-01 | 1.75E-01 | 2.59E+00 | 2.74E+00 | 5.13E-01 | no | | intergenic_variant | MODIFIER |
| 7 | 70296786 | AX-97789986 | 0 | A | C | 0.36 | -6.59E-02 | 1.02E-01 | 2.81E+00 | 2.91E+00 | 5.13E-01 | no | Gm29683 | intron_variant,non_coding_transcript_variant | MODIFIER |
| 4 | 135954618 | AX-170252963 | 0 | T | G | 0.167 | 6.90E-02 | 1.05E-01 | 2.59E+00 | 2.66E+00 | 5.13E-01 | no | Hmgcl | intron_variant | MODIFIER |
| 1 | 53230068 | AX-170524992 | 0 | G | A | 0.105 | -1.21E-01 | 1.88E-01 | 2.69E+00 | 2.79E+00 | 5.13E-01 | no | Pms1 | intron_variant | MODIFIER |
| 11 | 74734133 | AX-170454272 | 0 | A | G | 0.246 | 4.52E-02 | 7.05E-02 | 2.66E+00 | 2.81E+00 | 5.13E-01 | no | Gm16032 | upstream_gene_variant | MODIFIER |
| 9 | 55918199 | AX-170416695 | 0 | T | C | 0.202 | -7.75E-02 | 1.20E-01 | 2.78E+00 | 2.85E+00 | 5.13E-01 | no | Scaper | intron_variant | MODIFIER |
| 2 | 169602098 | AX-170437981 | 1 | C | T | 0.161 | 1.13E-01 | 1.74E-01 | 2.64E+00 | 2.73E+00 | 5.13E-01 | no | | intergenic_variant | MODIFIER |
| 9 | 96277643 | AX-97784538 | 1 | A | G | 0.143 | -7.56E-02 | 1.16E-01 | 2.63E+00 | 2.71E+00 | 5.14E-01 | no | Tfdp2 | intron_variant | MODIFIER |
| 5 | 32662821 | AX-170288853 | 2 | C | T | 0.373 | 6.68E-02 | 1.04E-01 | 2.69E+00 | 2.76E+00 | 5.14E-01 | no | Yes1 | intron_variant | MODIFIER |
| 1 | 20351137 | AX-170232075 | 0 | G | T | 0.395 | 5.29E-02 | 8.10E-02 | 2.60E+00 | 2.66E+00 | 5.14E-01 | no | Pkhd1 | intron_variant | MODIFIER |
| 18 | 58558868 | AX-170183744 | 1 | A | G | 0.393 | 6.63E-02 | 9.69E-02 | 2.45E+00 | 2.50E+00 | 5.14E-01 | no | Slc27a6 | intron_variant | MODIFIER |
| 9 | 97510974 | AX-170521879 | 0 | T | C | 0.14 | -7.35E-02 | 1.12E-01 | 2.58E+00 | 2.65E+00 | 5.14E-01 | no | Clstn2 | intron_variant | MODIFIER |
| 5 | 113304398 | AX-170025920 | 1 | A | T | 0.464 | 7.45E-02 | 1.15E-01 | 2.66E+00 | 2.69E+00 | 5.14E-01 | no | Sgsm1 | intron_variant | MODIFIER |
| 5 | 114265604 | AX-170228916 | 1 | C | T | 0.464 | 7.45E-02 | 1.15E-01 | 2.66E+00 | 2.69E+00 | 5.14E-01 | no | Foxn4 | intron_variant | MODIFIER |
| 18 | 55120942 | AX-97835527 | 1 | G | A | 0.455 | 6.48E-02 | 9.33E-02 | 2.41E+00 | 2.48E+00 | 5.14E-01 | no | | intergenic_variant | MODIFIER |
| 10 | 13295359 | AX-97818441 | 1 | A | C | 0.214 | 6.42E-02 | 9.93E-02 | 2.84E+00 | 2.95E+00 | 5.14E-01 | no | Phactr2 | intron_variant | MODIFIER |
| 18 | 80670972 | AX-170236222 | 0 | G | A | 0.298 | -5.67E-02 | 8.53E-02 | 2.51E+00 | 2.61E+00 | 5.14E-01 | no | Nfatc1 | intron_variant | MODIFIER |
| 13 | 48745258 | AX-97814662 | 0 | C | T | 0.491 | -4.26E-02 | 6.64E-02 | 2.81E+00 | 2.93E+00 | 5.14E-01 | no | | intergenic_variant | MODIFIER |
| 15 | 75213072 | AX-170510475 | 1 | A | G | 0.545 | 5.31E-02 | 8.15E-02 | 2.59E+00 | 2.67E+00 | 5.15E-01 | no | BC025446 | upstream_gene_variant | MODIFIER |
| 1 | 78827608 | AX-170456549 | 0 | G | A | 0.14 | 1.18E-01 | 1.85E-01 | 2.68E+00 | 2.79E+00 | 5.15E-01 | no | Gm29187 | downstream_gene_variant | MODIFIER |
| 13 | 38118787 | AX-170051009 | 1 | G | A | 0.179 | -7.21E-02 | 1.10E-01 | 2.56E+00 | 2.66E+00 | 5.15E-01 | no | | intergenic_variant | MODIFIER |
| 15 | 10637246 | AX-170414093 | 0 | A | G | 0.237 | -6.98E-02 | 1.07E-01 | 2.61E+00 | 2.66E+00 | 5.15E-01 | no | Rai14 | intron_variant | MODIFIER |
| 19 | 50629683 | AX-97836927 | 1 | T | G | 0.5 | -4.33E-02 | 6.74E-02 | 2.65E+00 | 2.75E+00 | 5.15E-01 | no | Sorcs1 | intron_variant | MODIFIER |
| 7 | 37941797 | AX-97828837 | 1 | A | C | 0.33 | 6.12E-02 | 9.21E-02 | 2.51E+00 | 2.59E+00 | 5.15E-01 | no | | intergenic_variant | MODIFIER |
| 4 | 155562069 | AX-170425482 | 0 | G | A | 0.123 | 9.04E-02 | 1.42E-01 | 2.71E+00 | 2.84E+00 | 5.15E-01 | no | Nadk | upstream_gene_variant | MODIFIER |
| 7 | 114921650 | AX-97807319 | 1 | T | C | 0.411 | 5.98E-02 | 9.30E-02 | 2.69E+00 | 2.75E+00 | 5.15E-01 | no | | intergenic_variant | MODIFIER |
| 1 | 47828524 | AX-170129648 | 0 | C | G | 0.105 | -1.20E-01 | 1.87E-01 | 2.69E+00 | 2.78E+00 | 5.16E-01 | no | | intergenic_variant | MODIFIER |
| 2 | 154707646 | AX-170274469 | 0 | G | T | 0.149 | 1.14E-01 | 1.79E-01 | 2.67E+00 | 2.77E+00 | 5.16E-01 | no | Rpl5-ps2 | downstream_gene_variant | MODIFIER |
| 9 | 38581159 | AX-97817350 | 1 | A | G | 0.482 | -4.88E-02 | 7.64E-02 | 2.73E+00 | 2.83E+00 | 5.16E-01 | no | Olfr912 | intron_variant | MODIFIER |
| 13 | 24155656 | AX-170330855 | 1 | A | T | 0.161 | -8.38E-02 | 1.21E-01 | 2.41E+00 | 2.48E+00 | 5.16E-01 | no | Carmil1 | intron_variant | MODIFIER |
| 2 | 70945036 | AX-170359725 | 0 | A | T | 0.053 | 1.25E-01 | 1.96E-01 | 2.73E+00 | 2.85E+00 | 5.16E-01 | no | | regulatory_region_variant | MODIFIER |
| 9 | 52463977 | AX-170311025 | 0 | C | T | 0.219 | -7.09E-02 | 1.11E-01 | 2.73E+00 | 2.82E+00 | 5.16E-01 | no | AC166902.1 | intron_variant,non_coding_transcript_variant | MODIFIER |
| 8 | 101912752 | AX-170112287 | 0 | A | T | 0.211 | 6.56E-02 | 1.03E-01 | 2.74E+00 | 2.85E+00 | 5.16E-01 | no | | intergenic_variant | MODIFIER |
| 11 | 29665753 | AX-170046676 | 1 | G | A | 0.08 | 1.29E-01 | 2.01E-01 | 2.65E+00 | 2.73E+00 | 5.16E-01 | no | | intergenic_variant | MODIFIER |
| 1 | 191452985 | AX-170503579 | 0 | G | T | 0.263 | 6.71E-02 | 9.63E-02 | 2.39E+00 | 2.48E+00 | 5.17E-01 | no | Gm32200 | intron_variant,non_coding_transcript_variant | MODIFIER |
| 4 | 154790281 | AX-170397760 | 0 | C | T | 0.105 | 1.04E-01 | 1.61E-01 | 2.59E+00 | 2.70E+00 | 5.17E-01 | no | | intergenic_variant | MODIFIER |
| 16 | 44299927 | AX-170446105 | 1 | A | G | 0.509 | 5.33E-02 | 8.15E-02 | 2.57E+00 | 2.61E+00 | 5.17E-01 | no | Sidt1 | synonymous_variant | LOW |
| 4 | 140371826 | AX-170478097 | 1 | G | T | 0.152 | 7.47E-02 | 1.14E-01 | 2.55E+00 | 2.61E+00 | 5.17E-01 | no | | intergenic_variant | MODIFIER |
| 1 | 99247124 | AX-170001183 | 1 | G | A | 0.161 | 1.31E-01 | 2.04E-01 | 2.65E+00 | 2.72E+00 | 5.17E-01 | no | | intergenic_variant | MODIFIER |

| | | | | | | | | | | | | | | | |
|---|---|---|---|---|---|---|---|---|---|---|---|---|---|---|---|
| 8 | 40487026 | AX-170103578 | 0 | T | C | 0.158 | 8.68E-02 | 1.36E-01 | 2.80E+00 | 2.87E+00 | 5.17E-01 | no | Zdhhc2 | downstream_gene_variant | MODIFIER |
| 6 | 136499391 | AX-170308524 | 0 | G | C | 0.579 | 5.48E-02 | 8.39E-02 | 2.56E+00 | 2.64E+00 | 5.17E-01 | no | | regulatory_region_variant | MODIFIER |
| 7 | 36547745 | AX-170340990 | 0 | C | T | 0.342 | 5.93E-02 | 9.15E-02 | 2.59E+00 | 2.67E+00 | 5.17E-01 | no | | regulatory_region_variant | MODIFIER |
| 1 | 95965917 | AX-170427767 | 0 | C | G | 0.158 | 1.09E-01 | 1.70E-01 | 2.62E+00 | 2.69E+00 | 5.17E-01 | no | | intergenic_variant | MODIFIER |
| 2 | 110215410 | AX-170018661 | 0 | C | T | 0.061 | -1.39E-01 | 2.18E-01 | 2.76E+00 | 2.87E+00 | 5.17E-01 | no | Gm22997 | upstream_gene_variant | MODIFIER |
| 7 | 39929079 | AX-170406385 | 0 | A | G | 0.351 | 6.19E-02 | 9.52E-02 | 2.58E+00 | 2.65E+00 | 5.17E-01 | no | Gm44992 | intron_variant,non_coding_transcript_variant | MODIFIER |
| 18 | 67820261 | AX-170398564 | 0 | A | T | 0.289 | -6.44E-02 | 1.01E-01 | 2.70E+00 | 2.76E+00 | 5.17E-01 | no | Cep192 | intron_variant | MODIFIER |
| 1 | 187269566 | AX-170099133 | 1 | C | T | 0.232 | 7.40E-02 | 1.10E-01 | 2.46E+00 | 2.54E+00 | 5.18E-01 | no | Gpatch2 | intron_variant | MODIFIER |
| 18 | 67868799 | AX-170313746 | 0 | G | T | 0.281 | -6.21E-02 | 9.70E-02 | 2.66E+00 | 2.73E+00 | 5.18E-01 | no | Cep192 | intron_variant | MODIFIER |
| 1 | 107059658 | AX-97795736 | 1 | G | T | 0.17 | 1.26E-01 | 1.94E-01 | 2.59E+00 | 2.66E+00 | 5.18E-01 | no | | intergenic_variant | MODIFIER |
| 1 | 131743383 | AX-97834875 | 0 | T | C | 0.544 | 5.42E-02 | 8.44E-02 | 2.64E+00 | 2.71E+00 | 5.18E-01 | no | Slc26a9 | upstream_gene_variant | MODIFIER |
| 11 | 23738558 | AX-170260527 | 0 | C | T | 0.053 | 1.25E-01 | 1.97E-01 | 2.66E+00 | 2.80E+00 | 5.18E-01 | no | Rel | 3_prime_UTR_variant | MODIFIER |
| 6 | 133695934 | AX-170028286 | 0 | T | A | 0.57 | 5.52E-02 | 8.41E-02 | 2.54E+00 | 2.61E+00 | 5.18E-01 | no | | intergenic_variant | MODIFIER |
| 1 | 149879803 | AX-170405459 | 1 | T | A | 0.205 | 8.87E-02 | 1.38E-01 | 2.67E+00 | 2.72E+00 | 5.18E-01 | no | Pla2g4a | intron_variant | MODIFIER |
| 1 | 152776254 | AX-170235628 | 1 | C | T | 0.205 | 8.87E-02 | 1.38E-01 | 2.67E+00 | 2.72E+00 | 5.18E-01 | no | Arpc5 | downstream_gene_variant | MODIFIER |
| 20 | 76126008 | AX-97802301 | 0 | C | T | 0.263 | 3.97E-02 | 6.20E-02 | 2.64E+00 | 2.73E+00 | 5.18E-01 | no | | intergenic_variant | MODIFIER |
| 1 | 148345065 | AX-170091738 | 0 | C | A | 0.219 | -6.90E-02 | 1.09E-01 | 2.78E+00 | 2.91E+00 | 5.18E-01 | no | | intergenic_variant | MODIFIER |
| 2 | 109229046 | AX-170018637 | 1 | C | T | 0.054 | 7.33E-02 | 1.17E-01 | 2.69E+00 | 2.87E+00 | 5.18E-01 | no | Mettl15 | intron_variant | MODIFIER |
| 1 | 170322393 | AX-170276276 | 0 | T | C | 0.211 | 9.11E-02 | 1.42E-01 | 2.67E+00 | 2.71E+00 | 5.18E-01 | no | Nos1ap | intron_variant | MODIFIER |
| 6 | 134606384 | AX-170028308 | 0 | A | T | 0.553 | 5.59E-02 | 8.52E-02 | 2.54E+00 | 2.60E+00 | 5.18E-01 | no | Mansc1 | downstream_gene_variant | MODIFIER |
| 10 | 43906118 | AX-170106291 | 0 | A | G | 0.368 | 4.85E-02 | 7.63E-02 | 2.78E+00 | 2.89E+00 | 5.18E-01 | no | Qrsl1 | upstream_gene_variant | MODIFIER |
| 16 | 73977996 | AX-170272786 | 0 | A | G | 0.491 | 5.17E-02 | 7.71E-02 | 2.47E+00 | 2.55E+00 | 5.18E-01 | no | Gm24272 | downstream_gene_variant | MODIFIER |
| 12 | 111856369 | AX-97802207 | 0 | T | C | 0.123 | -1.07E-01 | 1.69E-01 | 2.71E+00 | 2.81E+00 | 5.19E-01 | no | Ppp1r13b | intron_variant | MODIFIER |
| 11 | 74289187 | AX-170478179 | 1 | T | C | 0.232 | -5.11E-02 | 8.07E-02 | 2.64E+00 | 2.79E+00 | 5.19E-01 | no | Olfr59 | synonymous_variant | LOW |
| 1 | 24317131 | AX-170145287 | 1 | G | A | 0.446 | 5.22E-02 | 8.13E-02 | 2.62E+00 | 2.70E+00 | 5.19E-01 | no | Col19a1 | intron_variant | MODIFIER |
| 7 | 53113910 | AX-170525549 | 0 | C | T | 0.439 | 6.27E-02 | 9.78E-02 | 2.63E+00 | 2.71E+00 | 5.19E-01 | no | | intergenic_variant | MODIFIER |
| 16 | 76349542 | AX-170399877 | 1 | T | A | 0.473 | 4.77E-02 | 7.28E-02 | 2.53E+00 | 2.61E+00 | 5.19E-01 | no | Nrip1 | intron_variant | MODIFIER |
| 1 | 168371418 | AX-170250070 | 0 | A | C | 0.228 | 8.00E-02 | 1.24E-01 | 2.59E+00 | 2.66E+00 | 5.19E-01 | no | Pbx1 | intron_variant | MODIFIER |
| 1 | 45689273 | AX-97785877 | 1 | T | C | 0.089 | -1.06E-01 | 1.65E-01 | 2.59E+00 | 2.71E+00 | 5.19E-01 | no | Gm23216 | upstream_gene_variant | MODIFIER |
| 4 | 141245963 | AX-97857142 | 0 | T | C | 0.096 | -1.08E-01 | 1.70E-01 | 2.67E+00 | 2.76E+00 | 5.19E-01 | no | Arhgef19 | intron_variant | MODIFIER |
| 4 | 148703635 | AX-170139166 | 0 | A | G | 0.096 | -1.08E-01 | 1.70E-01 | 2.67E+00 | 2.76E+00 | 5.19E-01 | no | | intergenic_variant | MODIFIER |
| 4 | 150250892 | AX-170310727 | 0 | T | C | 0.096 | -1.08E-01 | 1.70E-01 | 2.67E+00 | 2.76E+00 | 5.19E-01 | no | Eno1 | downstream_gene_variant | MODIFIER |
| 5 | 83711836 | AX-170372550 | 0 | T | C | 0.439 | 7.18E-02 | 1.12E-01 | 2.80E+00 | 2.85E+00 | 5.19E-01 | no | | intergenic_variant | MODIFIER |
| 5 | 83746148 | AX-97819343 | 0 | C | T | 0.439 | 7.18E-02 | 1.12E-01 | 2.80E+00 | 2.85E+00 | 5.19E-01 | no | | intergenic_variant | MODIFIER |
| 5 | 83746207 | AX-170025454 | 0 | C | A | 0.439 | 7.18E-02 | 1.12E-01 | 2.80E+00 | 2.85E+00 | 5.19E-01 | no | | intergenic_variant | MODIFIER |
| 5 | 83768684 | AX-170486955 | 0 | C | G | 0.439 | 7.18E-02 | 1.12E-01 | 2.80E+00 | 2.85E+00 | 5.19E-01 | no | | intergenic_variant | MODIFIER |
| 5 | 83768719 | AX-170389427 | 0 | T | C | 0.439 | 7.18E-02 | 1.12E-01 | 2.80E+00 | 2.85E+00 | 5.19E-01 | no | | intergenic_variant | MODIFIER |
| 5 | 83827111 | AX-170025458 | 0 | G | A | 0.439 | 7.18E-02 | 1.12E-01 | 2.80E+00 | 2.85E+00 | 5.19E-01 | no | | intergenic_variant | MODIFIER |
| 5 | 83844237 | AX-97788178 | 0 | T | C | 0.439 | 7.18E-02 | 1.12E-01 | 2.80E+00 | 2.85E+00 | 5.19E-01 | no | | intergenic_variant | MODIFIER |
| 5 | 83867159 | AX-170039054 | 0 | C | T | 0.439 | 7.18E-02 | 1.12E-01 | 2.80E+00 | 2.85E+00 | 5.19E-01 | no | Gm4866 | downstream_gene_variant | MODIFIER |
| 5 | 83868036 | AX-97780679 | 0 | T | C | 0.439 | 7.18E-02 | 1.12E-01 | 2.80E+00 | 2.85E+00 | 5.19E-01 | no | Gm4866 | downstream_gene_variant | MODIFIER |
| 5 | 83877110 | AX-170089541 | 0 | G | A | 0.439 | 7.18E-02 | 1.12E-01 | 2.80E+00 | 2.85E+00 | 5.19E-01 | no | | intergenic_variant | MODIFIER |
| 5 | 83915165 | AX-170318121 | 0 | G | C | 0.439 | 7.18E-02 | 1.12E-01 | 2.80E+00 | 2.85E+00 | 5.19E-01 | no | | intergenic_variant | MODIFIER |
| 5 | 83930034 | AX-170170740 | 0 | T | C | 0.439 | 7.18E-02 | 1.12E-01 | 2.80E+00 | 2.85E+00 | 5.19E-01 | no | | intergenic_variant | MODIFIER |
| 5 | 83960701 | AX-97831909 | 0 | G | C | 0.439 | 7.18E-02 | 1.12E-01 | 2.80E+00 | 2.85E+00 | 5.19E-01 | no | | intergenic_variant | MODIFIER |
| 5 | 84034977 | AX-170369795 | 0 | G | A | 0.439 | 7.18E-02 | 1.12E-01 | 2.80E+00 | 2.85E+00 | 5.19E-01 | no | | intergenic_variant | MODIFIER |
| 5 | 84067332 | AX-97835319 | 0 | T | C | 0.439 | 7.18E-02 | 1.12E-01 | 2.80E+00 | 2.85E+00 | 5.19E-01 | no | Epha5 | intron_variant | MODIFIER |
| 5 | 84097548 | AX-97826747 | 0 | G | A | 0.439 | 7.18E-02 | 1.12E-01 | 2.80E+00 | 2.85E+00 | 5.19E-01 | no | Epha5 | intron_variant | MODIFIER |
| 5 | 84111154 | AX-170169034 | 0 | A | G | 0.439 | 7.18E-02 | 1.12E-01 | 2.80E+00 | 2.85E+00 | 5.19E-01 | no | Epha5 | intron_variant | MODIFIER |
| 5 | 84122505 | AX-170280927 | 0 | T | G | 0.439 | 7.18E-02 | 1.12E-01 | 2.80E+00 | 2.85E+00 | 5.19E-01 | no | Epha5 | intron_variant | MODIFIER |
| 5 | 84132178 | AX-170216711 | 0 | G | A | 0.439 | 7.18E-02 | 1.12E-01 | 2.80E+00 | 2.85E+00 | 5.19E-01 | no | Epha5 | intron_variant | MODIFIER |
| 5 | 84170418 | AX-97790037 | 0 | A | G | 0.439 | 7.18E-02 | 1.12E-01 | 2.80E+00 | 2.85E+00 | 5.19E-01 | no | Epha5 | intron_variant | MODIFIER |
| 5 | 84200795 | AX-97804342 | 0 | G | A | 0.439 | 7.18E-02 | 1.12E-01 | 2.80E+00 | 2.85E+00 | 5.19E-01 | no | Epha5 | intron_variant | MODIFIER |
| 5 | 84217576 | AX-170144803 | 0 | T | A | 0.439 | 7.18E-02 | 1.12E-01 | 2.80E+00 | 2.85E+00 | 5.19E-01 | no | Epha5 | intron_variant | MODIFIER |

| | | | | | | | | | | | | | | | |
|---|---|---|---|---|---|---|---|---|---|---|---|---|---|---|---|
| 5 | 84222236 | AX-170416513 | 0 | G | C | 0.439 | 7.18E-02 | 1.12E-01 | 2.80E+00 | 2.85E+00 | 5.19E-01 | no | Epha5 | intron_variant | MODIFIER |
| 5 | 84235637 | AX-170349743 | 0 | A | G | 0.439 | 7.18E-02 | 1.12E-01 | 2.80E+00 | 2.85E+00 | 5.19E-01 | no | Epha5 | intron_variant | MODIFIER |
| 5 | 84241143 | AX-170162706 | 0 | C | T | 0.439 | 7.18E-02 | 1.12E-01 | 2.80E+00 | 2.85E+00 | 5.19E-01 | no | Epha5 | intron_variant | MODIFIER |
| 5 | 84241766 | AX-97784897 | 0 | G | A | 0.439 | 7.18E-02 | 1.12E-01 | 2.80E+00 | 2.85E+00 | 5.19E-01 | no | Epha5 | intron_variant | MODIFIER |
| 5 | 84273365 | AX-97855447 | 0 | T | C | 0.439 | 7.18E-02 | 1.12E-01 | 2.80E+00 | 2.85E+00 | 5.19E-01 | no | Epha5 | intron_variant | MODIFIER |
| 5 | 84983164 | AX-170025478 | 0 | A | G | 0.439 | 7.18E-02 | 1.12E-01 | 2.80E+00 | 2.85E+00 | 5.19E-01 | no | | intergenic_variant | MODIFIER |
| 5 | 85015646 | AX-170095384 | 0 | C | T | 0.439 | 7.18E-02 | 1.12E-01 | 2.80E+00 | 2.85E+00 | 5.19E-01 | no | | intergenic_variant | MODIFIER |
| 5 | 85016011 | AX-170169277 | 0 | C | T | 0.439 | 7.18E-02 | 1.12E-01 | 2.80E+00 | 2.85E+00 | 5.19E-01 | no | | intergenic_variant | MODIFIER |
| 5 | 85050262 | AX-170238311 | 0 | T | G | 0.439 | 7.18E-02 | 1.12E-01 | 2.80E+00 | 2.85E+00 | 5.19E-01 | no | | intergenic_variant | MODIFIER |
| 5 | 85100340 | AX-97803653 | 0 | T | C | 0.439 | 7.18E-02 | 1.12E-01 | 2.80E+00 | 2.85E+00 | 5.19E-01 | no | | intergenic_variant | MODIFIER |
| 5 | 85100941 | AX-170388071 | 0 | T | C | 0.439 | 7.18E-02 | 1.12E-01 | 2.80E+00 | 2.85E+00 | 5.19E-01 | no | | intergenic_variant | MODIFIER |
| 5 | 85222156 | AX-170448029 | 0 | A | G | 0.439 | 7.18E-02 | 1.12E-01 | 2.80E+00 | 2.85E+00 | 5.19E-01 | no | | intergenic_variant | MODIFIER |
| 5 | 85229514 | AX-168419976 | 0 | C | T | 0.439 | 7.18E-02 | 1.12E-01 | 2.80E+00 | 2.85E+00 | 5.19E-01 | no | | intergenic_variant | MODIFIER |
| 1 | 95912494 | AX-170374903 | 1 | C | T | 0.143 | 1.15E-01 | 1.79E-01 | 2.59E+00 | 2.68E+00 | 5.19E-01 | no | | intergenic_variant | MODIFIER |
| 7 | 16999352 | AX-170190360 | 1 | T | C | 0.375 | 5.47E-02 | 8.48E-02 | 2.57E+00 | 2.68E+00 | 5.20E-01 | no | Ccdc8 | downstream_gene_variant | MODIFIER |
| 9 | 7926597 | AX-170090240 | 0 | T | A | 0.465 | 5.04E-02 | 7.53E-02 | 2.46E+00 | 2.56E+00 | 5.20E-01 | no | | intergenic_variant | MODIFIER |
| 9 | 113279505 | AX-170293819 | 0 | G | A | 0.123 | -6.83E-02 | 1.07E-01 | 2.66E+00 | 2.75E+00 | 5.20E-01 | no | | intergenic_variant | MODIFIER |
| 1 | 88127791 | AX-170243946 | 0 | T | C | 0.149 | 1.19E-01 | 1.85E-01 | 2.61E+00 | 2.69E+00 | 5.20E-01 | no | Ugt1a7c | intron_variant | MODIFIER |
| 1 | 100355216 | AX-170375454 | 0 | T | C | 0.149 | 1.19E-01 | 1.85E-01 | 2.61E+00 | 2.69E+00 | 5.20E-01 | no | Cntnap5b | intron_variant | MODIFIER |
| 1 | 100569111 | AX-170042179 | 0 | T | A | 0.149 | 1.19E-01 | 1.85E-01 | 2.61E+00 | 2.69E+00 | 5.20E-01 | no | | intergenic_variant | MODIFIER |
| 9 | 116950709 | AX-170128978 | 1 | G | A | 0.125 | -6.10E-02 | 9.56E-02 | 2.62E+00 | 2.73E+00 | 5.20E-01 | no | Rbms3 | intron_variant | MODIFIER |
| 18 | 37404735 | AX-170518810 | 0 | A | G | 0.07 | -1.36E-01 | 2.15E-01 | 2.73E+00 | 2.82E+00 | 5.20E-01 | no | Pcdhb9 | downstream_gene_variant | MODIFIER |
| 4 | 94781294 | AX-170458111 | 1 | T | A | 0.321 | 6.81E-02 | 1.06E-01 | 2.59E+00 | 2.68E+00 | 5.20E-01 | no | Tek | intron_variant | MODIFIER |
| 4 | 80726124 | AX-170424536 | 0 | C | T | 0.465 | -5.41E-02 | 8.22E-02 | 2.51E+00 | 2.58E+00 | 5.20E-01 | no | | intergenic_variant | MODIFIER |
| 20 | 84862453 | AX-170096835 | 0 | C | T | 0.044 | 7.86E-02 | 1.26E-01 | 2.68E+00 | 2.86E+00 | 5.20E-01 | no | Dmd | intron_variant | MODIFIER |
| 10 | 68241874 | AX-97828788 | 0 | C | G | 0.412 | -5.14E-02 | 8.04E-02 | 2.63E+00 | 2.70E+00 | 5.20E-01 | no | Arid5b | intron_variant,NMD_transcript_variant | MODIFIER |
| 7 | 113100417 | AX-170498591 | 0 | G | A | 0.456 | 5.45E-02 | 8.61E-02 | 2.69E+00 | 2.77E+00 | 5.20E-01 | no | 4930543E12Rik | non_coding_transcript_exon_variant | MODIFIER |
| 1 | 137371411 | AX-170525097 | 1 | G | A | 0.214 | 7.99E-02 | 1.26E-01 | 2.67E+00 | 2.74E+00 | 5.21E-01 | no | Gm23534 | upstream_gene_variant | MODIFIER |
| 13 | 95727347 | AX-170201165 | 1 | C | T | 0.036 | -1.11E-01 | 1.70E-01 | 2.52E+00 | 2.64E+00 | 5.21E-01 | no | Iqgap2 | intron_variant | MODIFIER |
| 8 | 87114285 | AX-170535478 | 0 | G | C | 0.211 | -7.96E-02 | 1.26E-01 | 2.76E+00 | 2.84E+00 | 5.21E-01 | no | Gm27246 | upstream_gene_variant | MODIFIER |
| 8 | 88260809 | AX-97832248 | 0 | G | A | 0.211 | -7.96E-02 | 1.26E-01 | 2.76E+00 | 2.84E+00 | 5.21E-01 | no | Tent4b | downstream_gene_variant | MODIFIER |
| 18 | 47677716 | AX-97834714 | 0 | T | A | 0.053 | 1.50E-01 | 2.35E-01 | 2.59E+00 | 2.70E+00 | 5.21E-01 | Yes | AC141639.4 | intron_variant,non_coding_transcript_variant | MODIFIER |
| 18 | 49282915 | AX-170263695 | 0 | T | C | 0.053 | 1.50E-01 | 2.35E-01 | 2.59E+00 | 2.70E+00 | 5.21E-01 | Yes | | intergenic_variant | MODIFIER |
| 5 | 28097222 | AX-170153188 | 0 | T | C | 0.368 | 6.59E-02 | 1.04E-01 | 2.71E+00 | 2.78E+00 | 5.21E-01 | no | | intergenic_variant | MODIFIER |
| 5 | 31815504 | AX-97781386 | 0 | G | A | 0.368 | 6.59E-02 | 1.04E-01 | 2.71E+00 | 2.78E+00 | 5.21E-01 | no | Babam2 | intron_variant | MODIFIER |
| 5 | 88331030 | AX-97786876 | 1 | C | T | 0.429 | 6.34E-02 | 1.00E-01 | 2.77E+00 | 2.86E+00 | 5.21E-01 | no | Prol1 | downstream_gene_variant | MODIFIER |
| 1 | 91073656 | AX-170434576 | 1 | A | G | 0.161 | 1.13E-01 | 1.78E-01 | 2.64E+00 | 2.72E+00 | 5.21E-01 | no | Lrrfip1 | intron_variant | MODIFIER |
| 9 | 98668535 | AX-170258928 | 1 | G | C | 0.143 | -7.16E-02 | 1.12E-01 | 2.60E+00 | 2.68E+00 | 5.21E-01 | no | | regulatory_region_variant | MODIFIER |
| 1 | 88562054 | AX-170381396 | 0 | A | G | 0.149 | 1.18E-01 | 1.85E-01 | 2.62E+00 | 2.70E+00 | 5.21E-01 | no | Gm19589 | non_coding_transcript_exon_variant | MODIFIER |
| 3 | 150205506 | AX-170527560 | 0 | G | A | 0.351 | 4.61E-02 | 7.30E-02 | 2.74E+00 | 2.83E+00 | 5.21E-01 | no | | intergenic_variant | MODIFIER |
| 14 | 112358777 | AX-97795261 | 0 | C | T | 0.316 | 4.43E-02 | 6.91E-02 | 2.57E+00 | 2.68E+00 | 5.21E-01 | no | | intergenic_variant | MODIFIER |
| 1 | 101949618 | AX-170246445 | 1 | C | G | 0.152 | 1.30E-01 | 2.04E-01 | 2.64E+00 | 2.70E+00 | 5.21E-01 | no | | intergenic_variant | MODIFIER |
| 9 | 116951543 | AX-170033361 | 1 | T | A | 0.125 | -6.09E-02 | 9.56E-02 | 2.60E+00 | 2.72E+00 | 5.22E-01 | no | Rbms3 | intron_variant | MODIFIER |
| 15 | 68943648 | AX-170087896 | 0 | T | C | 0.544 | 5.25E-02 | 8.18E-02 | 2.59E+00 | 2.67E+00 | 5.22E-01 | no | Khdrbs3 | intron_variant | MODIFIER |
| 15 | 69074052 | AX-170011920 | 0 | C | G | 0.544 | 5.25E-02 | 8.18E-02 | 2.59E+00 | 2.67E+00 | 5.22E-01 | no | Khdrbs3 | intron_variant | MODIFIER |
| 5 | 122999050 | AX-170381818 | 0 | G | A | 0.447 | -7.17E-02 | 1.12E-01 | 2.88E+00 | 2.94E+00 | 5.22E-01 | no | A930024E05Rik | downstream_gene_variant | MODIFIER |
| 6 | 135889742 | AX-170272438 | 0 | T | C | 0.579 | 5.66E-02 | 8.71E-02 | 2.55E+00 | 2.61E+00 | 5.22E-01 | no | Grin2b | intron_variant | MODIFIER |
| 1 | 170859661 | AX-170341902 | 0 | A | T | 0.219 | 8.21E-02 | 1.28E-01 | 2.59E+00 | 2.65E+00 | 5.22E-01 | no | Atf6 | intron_variant | MODIFIER |
| 9 | 15692199 | AX-170173578 | 0 | G | C | 0.447 | 5.29E-02 | 8.14E-02 | 2.54E+00 | 2.61E+00 | 5.22E-01 | no | | intergenic_variant | MODIFIER |
| 1 | 20201542 | AX-170000039 | 1 | G | A | 0.402 | 5.41E-02 | 8.43E-02 | 2.60E+00 | 2.64E+00 | 5.22E-01 | no | Pkhd1 | intron_variant | MODIFIER |
| 4 | 84004134 | AX-170402959 | 1 | A | G | 0.241 | 6.77E-02 | 1.03E-01 | 2.50E+00 | 2.57E+00 | 5.22E-01 | no | | intergenic_variant | MODIFIER |
| 1 | 20894744 | AX-170390296 | 0 | A | G | 0.395 | 5.43E-02 | 8.42E-02 | 2.57E+00 | 2.62E+00 | 5.22E-01 | no | 6720483E21Rik | upstream_gene_variant | MODIFIER |
| 7 | 14569252 | AX-97851726 | 2 | A | C | 0.355 | 5.14E-02 | 7.97E-02 | 2.54E+00 | 2.66E+00 | 5.22E-01 | no | Nlrp5-ps | intron_variant,non_coding_transcript_variant | MODIFIER |
| 18 | 57806585 | AX-170093899 | 1 | C | T | 0.429 | 5.34E-02 | 8.37E-02 | 2.88E+00 | 2.98E+00 | 5.22E-01 | no | | intergenic_variant | MODIFIER |

| | | | | | | | | | | | | | | | |
|---|---|---|---|---|---|---|---|---|---|---|---|---|---|---|---|
| 18 | 80226168 | AX-97790889 | 1 | A | C | 0.384 | -5.46E-02 | 8.60E-02 | 2.62E+00 | 2.71E+00 | 5.23E-01 | no | Txnl4a | downstream_gene_variant | MODIFIER |
| 7 | 138945283 | AX-97827189 | 1 | C | T | 0.295 | 4.93E-02 | 7.86E-02 | 2.77E+00 | 2.89E+00 | 5.23E-01 | no | Jakmip3 | intron_variant | MODIFIER |
| 18 | 80939526 | AX-170290893 | 2 | T | A | 0.291 | -5.49E-02 | 8.60E-02 | 2.58E+00 | 2.69E+00 | 5.23E-01 | no | | intergenic_variant | MODIFIER |
| 7 | 123391687 | AX-170090017 | 1 | G | A | 0.402 | 5.36E-02 | 8.51E-02 | 2.74E+00 | 2.82E+00 | 5.23E-01 | no | Lcmt1 | intron_variant | MODIFIER |
| 1 | 188677313 | AX-170159223 | 0 | C | T | 0.254 | 6.50E-02 | 9.85E-02 | 2.48E+00 | 2.57E+00 | 5.23E-01 | no | Ush2a | intron_variant | MODIFIER |
| 20 | 157374402 | AX-170035319 | 2 | G | A | 0.027 | -9.35E-02 | 1.46E-01 | 2.53E+00 | 2.69E+00 | 5.23E-01 | no | Phex | intron_variant | MODIFIER |
| 7 | 74161293 | AX-170433356 | 0 | T | C | 0.386 | -6.01E-02 | 9.59E-02 | 2.72E+00 | 2.82E+00 | 5.23E-01 | no | | intergenic_variant | MODIFIER |
| 1 | 151976140 | AX-170293006 | 1 | C | T | 0.205 | 8.79E-02 | 1.39E-01 | 2.65E+00 | 2.70E+00 | 5.23E-01 | no | 1700025G04Rik | intron_variant | MODIFIER |
| 2 | 162008338 | AX-170263340 | 1 | G | A | 0.152 | 1.12E-01 | 1.78E-01 | 2.71E+00 | 2.81E+00 | 5.23E-01 | no | Ptprt | intron_variant | MODIFIER |
| 11 | 17722418 | AX-97782397 | 0 | C | T | 0.044 | 1.00E-01 | 1.62E-01 | 2.68E+00 | 2.85E+00 | 5.24E-01 | no | | intergenic_variant | MODIFIER |
| 1 | 150329282 | AX-97826112 | 0 | G | A | 0.211 | 8.11E-02 | 1.27E-01 | 2.60E+00 | 2.67E+00 | 5.24E-01 | no | Pdc | intron_variant | MODIFIER |
| 1 | 151433555 | AX-170150294 | 0 | G | A | 0.211 | 8.11E-02 | 1.27E-01 | 2.60E+00 | 2.67E+00 | 5.24E-01 | no | Trmt1l | intron_variant | MODIFIER |
| 1 | 152409411 | AX-170271918 | 0 | C | T | 0.211 | 8.11E-02 | 1.27E-01 | 2.60E+00 | 2.67E+00 | 5.24E-01 | no | Colgalt2 | intron_variant | MODIFIER |
| 1 | 152588528 | AX-170383430 | 0 | C | T | 0.211 | 8.11E-02 | 1.27E-01 | 2.60E+00 | 2.67E+00 | 5.24E-01 | no | Rgl1 | intron_variant | MODIFIER |
| 2 | 138988510 | AX-97848252 | 0 | G | A | 0.149 | 1.14E-01 | 1.81E-01 | 2.67E+00 | 2.75E+00 | 5.24E-01 | no | | intergenic_variant | MODIFIER |
| 2 | 143699991 | AX-170125228 | 0 | A | G | 0.149 | 1.14E-01 | 1.81E-01 | 2.67E+00 | 2.75E+00 | 5.24E-01 | no | Pcsk2 | intron_variant | MODIFIER |
| 15 | 71477171 | AX-170111620 | 0 | A | C | 0.368 | 4.85E-02 | 7.46E-02 | 2.52E+00 | 2.60E+00 | 5.24E-01 | no | Fam135b | intron_variant | MODIFIER |
| 9 | 69025079 | AX-170090324 | 0 | G | T | 0.202 | 8.08E-02 | 1.27E-01 | 2.84E+00 | 2.91E+00 | 5.24E-01 | no | Rora | intron_variant | MODIFIER |
| 17 | 7750161 | AX-168417083 | 0 | T | C | 0.053 | 1.05E-01 | 1.69E-01 | 2.72E+00 | 2.85E+00 | 5.24E-01 | no | Fndc1 | intron_variant,non_coding_transcript_variant | MODIFIER |
| 13 | 6839599 | AX-170223218 | 2 | C | T | 0.218 | -7.50E-02 | 1.18E-01 | 2.60E+00 | 2.69E+00 | 5.24E-01 | no | | intergenic_variant | MODIFIER |
| 4 | 138984954 | AX-170398979 | 1 | C | A | 0.152 | 7.36E-02 | 1.15E-01 | 2.59E+00 | 2.65E+00 | 5.24E-01 | no | Tmco4 | intron_variant | MODIFIER |
| 4 | 92565814 | AX-170156147 | 0 | G | A | 0.342 | 6.39E-02 | 1.00E-01 | 2.59E+00 | 2.68E+00 | 5.24E-01 | no | | intergenic_variant | MODIFIER |
| 2 | 157433203 | AX-170405382 | 1 | C | A | 0.152 | 1.10E-01 | 1.73E-01 | 2.60E+00 | 2.70E+00 | 5.24E-01 | no | Src | intron_variant | MODIFIER |
| 20 | 49069245 | AX-170229797 | 0 | T | A | 0.237 | 3.66E-02 | 5.78E-02 | 2.59E+00 | 2.71E+00 | 5.24E-01 | no | Enox2 | intron_variant | MODIFIER |
| 20 | 49090291 | AX-97853727 | 0 | C | T | 0.237 | 3.66E-02 | 5.78E-02 | 2.59E+00 | 2.71E+00 | 5.24E-01 | no | Enox2 | intron_variant | MODIFIER |
| 20 | 49256289 | AX-170286045 | 0 | C | A | 0.237 | 3.66E-02 | 5.78E-02 | 2.59E+00 | 2.71E+00 | 5.24E-01 | no | Enox2 | intron_variant | MODIFIER |
| 20 | 49291542 | AX-97832436 | 0 | T | C | 0.237 | 3.66E-02 | 5.78E-02 | 2.59E+00 | 2.71E+00 | 5.24E-01 | no | Enox2 | upstream_gene_variant | MODIFIER |
| 20 | 49292960 | AX-170033818 | 0 | G | A | 0.237 | 3.66E-02 | 5.78E-02 | 2.59E+00 | 2.71E+00 | 5.24E-01 | no | Enox2 | upstream_gene_variant | MODIFIER |
| 20 | 49310179 | AX-170193057 | 0 | G | C | 0.237 | 3.66E-02 | 5.78E-02 | 2.59E+00 | 2.71E+00 | 5.24E-01 | no | Gm14696 | downstream_gene_variant | MODIFIER |
| 20 | 49334626 | AX-170033814 | 0 | T | C | 0.237 | 3.66E-02 | 5.78E-02 | 2.59E+00 | 2.71E+00 | 5.24E-01 | no | Gm14696 | intron_variant,non_coding_transcript_variant | MODIFIER |
| 20 | 49644788 | AX-170132822 | 0 | A | G | 0.237 | 3.66E-02 | 5.78E-02 | 2.59E+00 | 2.71E+00 | 5.24E-01 | no | | intergenic_variant | MODIFIER |
| 20 | 49671338 | AX-170296813 | 0 | G | A | 0.237 | 3.66E-02 | 5.78E-02 | 2.59E+00 | 2.71E+00 | 5.24E-01 | no | Olfr1320 | upstream_gene_variant | MODIFIER |
| 20 | 49711616 | AX-170443065 | 0 | A | G | 0.237 | 3.66E-02 | 5.78E-02 | 2.59E+00 | 2.71E+00 | 5.24E-01 | no | | intergenic_variant | MODIFIER |
| 20 | 49765807 | AX-170272127 | 0 | C | G | 0.237 | 3.66E-02 | 5.78E-02 | 2.59E+00 | 2.71E+00 | 5.24E-01 | no | | intergenic_variant | MODIFIER |
| 20 | 49850514 | AX-97853953 | 0 | T | C | 0.237 | 3.66E-02 | 5.78E-02 | 2.59E+00 | 2.71E+00 | 5.24E-01 | no | | intergenic_variant | MODIFIER |
| 20 | 49853958 | AX-170033823 | 0 | C | G | 0.237 | 3.66E-02 | 5.78E-02 | 2.59E+00 | 2.71E+00 | 5.24E-01 | no | | intergenic_variant | MODIFIER |
| 20 | 49860788 | AX-170033827 | 0 | T | G | 0.237 | 3.66E-02 | 5.78E-02 | 2.59E+00 | 2.71E+00 | 5.24E-01 | no | | intergenic_variant | MODIFIER |
| 20 | 50699853 | AX-170141940 | 0 | A | G | 0.237 | 3.66E-02 | 5.78E-02 | 2.59E+00 | 2.71E+00 | 5.24E-01 | no | | intergenic_variant | MODIFIER |
| 20 | 50730537 | AX-170116081 | 0 | C | A | 0.237 | 3.66E-02 | 5.78E-02 | 2.59E+00 | 2.71E+00 | 5.24E-01 | no | | intergenic_variant | MODIFIER |
| 5 | 105501957 | AX-170072978 | 1 | C | T | 0.438 | 7.35E-02 | 1.16E-01 | 2.65E+00 | 2.68E+00 | 5.24E-01 | no | Gm42141 | downstream_gene_variant | MODIFIER |
| 20 | 67607793 | AX-97797564 | 0 | G | A | 0.254 | 3.93E-02 | 6.24E-02 | 2.66E+00 | 2.75E+00 | 5.24E-01 | no | | intergenic_variant | MODIFIER |
| 4 | 154182335 | AX-170401933 | 2 | C | T | 0.109 | 1.13E-01 | 1.80E-01 | 2.68E+00 | 2.76E+00 | 5.24E-01 | no | Megf6 | intron_variant | MODIFIER |
| 5 | 79233855 | AX-97841454 | 1 | G | T | 0.438 | 6.85E-02 | 1.09E-01 | 2.72E+00 | 2.77E+00 | 5.25E-01 | no | | intergenic_variant | MODIFIER |
| 1 | 186978857 | AX-170002500 | 1 | A | G | 0.232 | 7.10E-02 | 1.07E-01 | 2.47E+00 | 2.54E+00 | 5.25E-01 | no | | intergenic_variant | MODIFIER |
| 1 | 146681987 | AX-170001936 | 1 | G | T | 0.205 | 8.75E-02 | 1.39E-01 | 2.66E+00 | 2.71E+00 | 5.25E-01 | no | Brinp3 | intron_variant | MODIFIER |
| 1 | 149975551 | AX-170155229 | 1 | G | A | 0.205 | 8.75E-02 | 1.39E-01 | 2.66E+00 | 2.71E+00 | 5.25E-01 | no | | intergenic_variant | MODIFIER |
| 5 | 83781314 | AX-170131166 | 1 | A | C | 0.438 | 6.23E-02 | 9.93E-02 | 2.76E+00 | 2.84E+00 | 5.25E-01 | no | | intergenic_variant | MODIFIER |
| 18 | 74189632 | AX-170216385 | 1 | T | C | 0.17 | -7.43E-02 | 1.15E-01 | 2.54E+00 | 2.62E+00 | 5.25E-01 | no | | intergenic_variant | MODIFIER |
| 4 | 136563281 | AX-97825283 | 0 | A | C | 0.167 | 7.24E-02 | 1.14E-01 | 2.61E+00 | 2.66E+00 | 5.25E-01 | no | Kdm1a | intron_variant | MODIFIER |
| 8 | 30737765 | AX-170238533 | 0 | C | G | 0.167 | -6.39E-02 | 1.03E-01 | 2.74E+00 | 2.87E+00 | 5.25E-01 | no | | intergenic_variant | MODIFIER |
| 4 | 154867619 | AX-170439289 | 0 | G | C | 0.14 | 7.04E-02 | 1.13E-01 | 2.66E+00 | 2.78E+00 | 5.25E-01 | no | Ttc34 | downstream_gene_variant | MODIFIER |
| 5 | 45551168 | AX-170227383 | 1 | C | T | 0.491 | 6.33E-02 | 9.54E-02 | 2.46E+00 | 2.54E+00 | 5.25E-01 | no | Fam184b | intron_variant | MODIFIER |
| 10 | 6824945 | AX-170160299 | 0 | T | C | 0.193 | -6.31E-02 | 1.01E-01 | 2.65E+00 | 2.75E+00 | 5.25E-01 | no | Oprm1 | intron_variant | MODIFIER |
| 1 | 91043909 | AX-97782941 | 1 | A | C | 0.161 | 1.31E-01 | 2.08E-01 | 2.63E+00 | 2.69E+00 | 5.25E-01 | no | Lrrfip1 | intron_variant | MODIFIER |

| | | | | | | | | | | | | | | | |
|---|---|---|---|---|---|---|---|---|---|---|---|---|---|---|---|
| 6 | 130504107 | AX-170095804 | 0 | A | G | 0.561 | 5.16E-02 | 7.81E-02 | 2.47E+00 | 2.55E+00 | 5.25E-01 | no | Gm38706 | upstream_gene_variant | MODIFIER |
| 1 | 185391825 | AX-170002475 | 0 | T | C | 0.456 | 3.78E-02 | 5.72E-02 | 2.47E+00 | 2.55E+00 | 5.25E-01 | no | Eprs | intron_variant | MODIFIER |
| 8 | 25491532 | AX-97813104 | 0 | T | G | 0.351 | -4.07E-02 | 6.42E-02 | 2.59E+00 | 2.69E+00 | 5.25E-01 | no | Gm31045 | upstream_gene_variant | MODIFIER |
| 8 | 30700043 | AX-170200291 | 0 | T | G | 0.167 | 7.19E-02 | 1.15E-01 | 2.76E+00 | 2.84E+00 | 5.26E-01 | no | | intergenic_variant | MODIFIER |
| 18 | 77645392 | AX-170128003 | 0 | A | G | 0.395 | -5.51E-02 | 8.66E-02 | 2.58E+00 | 2.67E+00 | 5.26E-01 | no | 8030462N17Rik | intron_variant | MODIFIER |
| 7 | 101375520 | AX-170078251 | 0 | T | C | 0.377 | -6.25E-02 | 1.00E-01 | 2.72E+00 | 2.83E+00 | 5.26E-01 | no | Arap1 | upstream_gene_variant | MODIFIER |
| 15 | 75215789 | AX-170264283 | 0 | T | C | 0.561 | 5.08E-02 | 7.92E-02 | 2.55E+00 | 2.63E+00 | 5.26E-01 | no | BC025446 | upstream_gene_variant | MODIFIER |
| 3 | 155497255 | AX-170295441 | 0 | A | C | 0.325 | 4.73E-02 | 7.56E-02 | 2.68E+00 | 2.76E+00 | 5.26E-01 | no | | intergenic_variant | MODIFIER |
| 8 | 40185239 | AX-170119242 | 0 | A | G | 0.149 | 9.15E-02 | 1.46E-01 | 2.82E+00 | 2.88E+00 | 5.26E-01 | no | | intergenic_variant | MODIFIER |
| 7 | 45851729 | AX-97790752 | 1 | G | A | 0.33 | 5.90E-02 | 9.38E-02 | 2.63E+00 | 2.71E+00 | 5.26E-01 | no | Grin2d | intron_variant | MODIFIER |
| 4 | 153321351 | AX-97821486 | 1 | T | C | 0.107 | 1.13E-01 | 1.80E-01 | 2.68E+00 | 2.76E+00 | 5.26E-01 | no | | regulatory_region_variant | MODIFIER |
| 1 | 51596973 | AX-170496693 | 0 | A | G | 0.105 | -1.20E-01 | 1.92E-01 | 2.70E+00 | 2.79E+00 | 5.26E-01 | no | Gm17767 | upstream_gene_variant | MODIFIER |
| 1 | 52750139 | AX-170000504 | 0 | T | C | 0.105 | -1.20E-01 | 1.92E-01 | 2.70E+00 | 2.79E+00 | 5.26E-01 | no | | regulatory_region_variant | MODIFIER |
| 18 | 53432149 | AX-170323777 | 1 | C | T | 0.232 | -6.92E-02 | 1.08E-01 | 2.58E+00 | 2.63E+00 | 5.26E-01 | no | | intergenic_variant | MODIFIER |
| 2 | 71397422 | AX-170018004 | 0 | A | G | 0.044 | 1.20E-01 | 1.95E-01 | 2.70E+00 | 2.84E+00 | 5.26E-01 | no | Hat1 | intron_variant | MODIFIER |
| 7 | 25638747 | AX-170372598 | 1 | A | G | 0.402 | 6.00E-02 | 9.39E-02 | 2.55E+00 | 2.65E+00 | 5.26E-01 | no | Bckdha | intron_variant | MODIFIER |
| 4 | 82326377 | AX-170069819 | 0 | A | G | 0.228 | 6.54E-02 | 1.02E-01 | 2.54E+00 | 2.62E+00 | 5.26E-01 | no | Nfib | intron_variant | MODIFIER |
| 9 | 67564269 | AX-170275646 | 0 | T | C | 0.193 | -7.86E-02 | 1.25E-01 | 2.66E+00 | 2.71E+00 | 5.27E-01 | no | Tln2 | upstream_gene_variant | MODIFIER |
| 9 | 72037909 | AX-170415702 | 0 | G | C | 0.193 | -7.86E-02 | 1.25E-01 | 2.66E+00 | 2.71E+00 | 5.27E-01 | no | Tcf12 | intron_variant | MODIFIER |
| 1 | 136872280 | AX-170222197 | 1 | G | A | 0.214 | 7.59E-02 | 1.21E-01 | 2.66E+00 | 2.73E+00 | 5.27E-01 | no | Nr5a2 | intron_variant | MODIFIER |
| 18 | 51609934 | AX-170266350 | 0 | C | T | 0.044 | -1.41E-01 | 2.24E-01 | 2.61E+00 | 2.74E+00 | 5.27E-01 | no | Gm25785 | upstream_gene_variant | MODIFIER |
| 8 | 100388535 | AX-170329618 | 0 | T | C | 0.228 | -6.36E-02 | 1.03E-01 | 2.77E+00 | 2.89E+00 | 5.27E-01 | no | | intergenic_variant | MODIFIER |
| 1 | 88152536 | AX-170041947 | 1 | C | T | 0.152 | 1.14E-01 | 1.82E-01 | 2.66E+00 | 2.74E+00 | 5.27E-01 | no | Ugt1a6a | intron_variant | MODIFIER |
| 1 | 89228300 | AX-170001028 | 1 | C | T | 0.152 | 1.14E-01 | 1.82E-01 | 2.66E+00 | 2.74E+00 | 5.27E-01 | no | | intergenic_variant | MODIFIER |
| 1 | 95245104 | AX-170532792 | 1 | G | T | 0.152 | 1.14E-01 | 1.82E-01 | 2.66E+00 | 2.74E+00 | 5.27E-01 | no | | intergenic_variant | MODIFIER |
| 7 | 16975401 | AX-170294464 | 2 | C | T | 0.364 | 5.44E-02 | 8.63E-02 | 2.59E+00 | 2.70E+00 | 5.27E-01 | no | | regulatory_region_variant | MODIFIER |
| 9 | 96367239 | AX-170160647 | 1 | A | T | 0.143 | -7.45E-02 | 1.18E-01 | 2.63E+00 | 2.70E+00 | 5.27E-01 | no | Atp1b3 | upstream_gene_variant | MODIFIER |
| 4 | 137888999 | AX-97792899 | 0 | G | A | 0.175 | 6.85E-02 | 1.09E-01 | 2.62E+00 | 2.68E+00 | 5.27E-01 | no | Ece1 | intron_variant | MODIFIER |
| 9 | 21381326 | AX-170137109 | 1 | T | C | 0.446 | 5.74E-02 | 8.98E-02 | 2.56E+00 | 2.63E+00 | 5.27E-01 | no | Ilf3 | intron_variant | MODIFIER |
| 12 | 61583705 | AX-168373232 | 0 | G | A | 0.07 | 1.31E-01 | 2.11E-01 | 2.76E+00 | 2.88E+00 | 5.27E-01 | no | Lrfn5 | intron_variant | MODIFIER |
| 4 | 54480517 | AX-170214794 | 0 | G | A | 0.377 | -6.23E-02 | 1.00E-01 | 2.69E+00 | 2.78E+00 | 5.27E-01 | no | | intergenic_variant | MODIFIER |
| 4 | 128115916 | AX-170438749 | 1 | C | T | 0.143 | 7.74E-02 | 1.22E-01 | 2.58E+00 | 2.64E+00 | 5.27E-01 | no | Csmd2 | intron_variant,non_coding_transcript_variant | MODIFIER |
| 8 | 68845743 | AX-170080335 | 1 | G | A | 0.188 | -8.16E-02 | 1.30E-01 | 2.64E+00 | 2.71E+00 | 5.27E-01 | no | | intergenic_variant | MODIFIER |
| 1 | 170782575 | AX-170389559 | 1 | A | T | 0.205 | 8.70E-02 | 1.39E-01 | 2.66E+00 | 2.71E+00 | 5.27E-01 | no | Atf6 | intron_variant | MODIFIER |
| 18 | 80872281 | AX-170352678 | 0 | C | A | 0.289 | -5.17E-02 | 8.11E-02 | 2.55E+00 | 2.66E+00 | 5.27E-01 | no | Atp9b | intron_variant | MODIFIER |
| 18 | 53218009 | AX-170381431 | 1 | C | G | 0.241 | -6.45E-02 | 1.02E-01 | 2.58E+00 | 2.66E+00 | 5.27E-01 | no | Snx2 | intron_variant | MODIFIER |
| 1 | 168428630 | AX-170378798 | 1 | G | A | 0.205 | 8.11E-02 | 1.29E-01 | 2.64E+00 | 2.71E+00 | 5.27E-01 | no | Pbx1 | intron_variant | MODIFIER |
| 5 | 108556418 | AX-168396540 | 1 | C | T | 0.473 | 6.97E-02 | 1.11E-01 | 2.61E+00 | 2.67E+00 | 5.28E-01 | no | Gm43364 | upstream_gene_variant | MODIFIER |
| 16 | 96474064 | AX-170013710 | 0 | A | G | 0.456 | 4.98E-02 | 7.98E-02 | 2.84E+00 | 2.96E+00 | 5.28E-01 | no | Pcp4 | intron_variant | MODIFIER |
| 8 | 64363980 | AX-97831591 | 0 | C | T | 0.149 | -8.58E-02 | 1.36E-01 | 2.59E+00 | 2.69E+00 | 5.28E-01 | no | | intergenic_variant | MODIFIER |
| 1 | 23675949 | AX-170086420 | 0 | A | C | 0.421 | 5.38E-02 | 8.55E-02 | 2.62E+00 | 2.68E+00 | 5.28E-01 | no | | intergenic_variant | MODIFIER |
| 1 | 23689383 | AX-170000092 | 0 | T | C | 0.421 | 5.38E-02 | 8.55E-02 | 2.62E+00 | 2.68E+00 | 5.28E-01 | no | | intergenic_variant | MODIFIER |
| 1 | 23756940 | AX-170000086 | 0 | G | A | 0.421 | 5.38E-02 | 8.55E-02 | 2.62E+00 | 2.68E+00 | 5.28E-01 | no | B3gat2 | upstream_gene_variant | MODIFIER |
| 1 | 24479240 | AX-170161451 | 0 | G | A | 0.421 | 5.38E-02 | 8.55E-02 | 2.62E+00 | 2.68E+00 | 5.28E-01 | no | Col19a1 | intron_variant | MODIFIER |
| 1 | 24797212 | AX-97815665 | 0 | T | C | 0.421 | 5.38E-02 | 8.55E-02 | 2.62E+00 | 2.68E+00 | 5.28E-01 | no | | intergenic_variant | MODIFIER |
| 1 | 24853901 | AX-97824277 | 0 | G | A | 0.421 | 5.38E-02 | 8.55E-02 | 2.62E+00 | 2.68E+00 | 5.28E-01 | no | | intergenic_variant | MODIFIER |
| 1 | 24866952 | AX-170040639 | 0 | A | G | 0.421 | 5.38E-02 | 8.55E-02 | 2.62E+00 | 2.68E+00 | 5.28E-01 | no | | intergenic_variant | MODIFIER |
| 1 | 24867795 | AX-170000117 | 0 | T | C | 0.421 | 5.38E-02 | 8.55E-02 | 2.62E+00 | 2.68E+00 | 5.28E-01 | no | | intergenic_variant | MODIFIER |
| 1 | 24868642 | AX-170437507 | 0 | T | A | 0.421 | 5.38E-02 | 8.55E-02 | 2.62E+00 | 2.68E+00 | 5.28E-01 | no | | intergenic_variant | MODIFIER |
| 1 | 24878019 | AX-170000120 | 0 | T | C | 0.421 | 5.38E-02 | 8.55E-02 | 2.62E+00 | 2.68E+00 | 5.28E-01 | no | | intergenic_variant | MODIFIER |
| 1 | 24885284 | AX-97783201 | 0 | G | T | 0.421 | 5.38E-02 | 8.55E-02 | 2.62E+00 | 2.68E+00 | 5.28E-01 | no | | intergenic_variant | MODIFIER |
| 1 | 24885350 | AX-170451168 | 0 | G | C | 0.421 | 5.38E-02 | 8.55E-02 | 2.62E+00 | 2.68E+00 | 5.28E-01 | no | | intergenic_variant | MODIFIER |
| 1 | 24898356 | AX-170098106 | 0 | C | T | 0.421 | 5.38E-02 | 8.55E-02 | 2.62E+00 | 2.68E+00 | 5.28E-01 | no | | intergenic_variant | MODIFIER |
| 1 | 24901796 | AX-170339791 | 0 | A | G | 0.421 | 5.38E-02 | 8.55E-02 | 2.62E+00 | 2.68E+00 | 5.28E-01 | no | | intergenic_variant | MODIFIER |

| | | | | | | | | | | | | | | | |
|---|---|---|---|---|---|---|---|---|---|---|---|---|---|---|---|
| 1 | 24901851 | AX-97827624 | 0 | A | G | 0.421 | 5.38E-02 | 8.55E-02 | 2.62E+00 | 2.68E+00 | 5.28E-01 | no | | intergenic_variant | MODIFIER |
| 1 | 25162868 | AX-97810998 | 0 | C | T | 0.421 | 5.38E-02 | 8.55E-02 | 2.62E+00 | 2.68E+00 | 5.28E-01 | no | Adgrb3 | intron_variant | MODIFIER |
| 1 | 25166118 | AX-170000121 | 0 | A | G | 0.421 | 5.38E-02 | 8.55E-02 | 2.62E+00 | 2.68E+00 | 5.28E-01 | no | Adgrb3 | intron_variant | MODIFIER |
| 1 | 25173722 | AX-170400923 | 0 | G | T | 0.421 | 5.38E-02 | 8.55E-02 | 2.62E+00 | 2.68E+00 | 5.28E-01 | no | Adgrb3 | intron_variant | MODIFIER |
| 1 | 25189544 | AX-170150217 | 0 | C | G | 0.421 | 5.38E-02 | 8.55E-02 | 2.62E+00 | 2.68E+00 | 5.28E-01 | no | Adgrb3 | intron_variant | MODIFIER |
| 1 | 25196153 | AX-97790088 | 0 | A | G | 0.421 | 5.38E-02 | 8.55E-02 | 2.62E+00 | 2.68E+00 | 5.28E-01 | no | Adgrb3 | intron_variant | MODIFIER |
| 1 | 25213091 | AX-170477807 | 0 | T | C | 0.421 | 5.38E-02 | 8.55E-02 | 2.62E+00 | 2.68E+00 | 5.28E-01 | no | Adgrb3 | intron_variant | MODIFIER |
| 1 | 21040395 | AX-170335002 | 1 | A | C | 0.402 | 5.31E-02 | 8.39E-02 | 2.59E+00 | 2.64E+00 | 5.28E-01 | no | Tram2 | intron_variant | MODIFIER |
| 16 | 46415403 | AX-170480275 | 1 | A | G | 0.509 | 5.15E-02 | 8.05E-02 | 2.55E+00 | 2.59E+00 | 5.29E-01 | no | Nectin3 | intron_variant | MODIFIER |
| 1 | 136249479 | AX-170311153 | 0 | G | A | 0.219 | 7.25E-02 | 1.16E-01 | 2.61E+00 | 2.70E+00 | 5.29E-01 | no | Gm37552 | downstream_gene_variant | MODIFIER |
| 1 | 21671029 | AX-170254491 | 0 | C | A | 0.404 | 5.22E-02 | 8.29E-02 | 2.61E+00 | 2.67E+00 | 5.29E-01 | no | Kcnq5 | intron_variant | MODIFIER |
| 18 | 61650990 | AX-170015825 | 0 | C | T | 0.404 | 6.08E-02 | 9.35E-02 | 2.50E+00 | 2.55E+00 | 5.29E-01 | no | Mir143 | upstream_gene_variant | MODIFIER |
| 1 | 177080853 | AX-170507987 | 2 | C | T | 0.236 | 6.78E-02 | 1.08E-01 | 2.58E+00 | 2.68E+00 | 5.29E-01 | no | Akt3 | intron_variant | MODIFIER |
| 10 | 93381983 | AX-97780973 | 0 | C | A | 0.307 | -5.28E-02 | 8.21E-02 | 2.52E+00 | 2.59E+00 | 5.29E-01 | no | | intergenic_variant | MODIFIER |
| 7 | 57700082 | AX-170496495 | 0 | G | T | 0.43 | 5.90E-02 | 9.46E-02 | 2.64E+00 | 2.73E+00 | 5.29E-01 | no | Gabrb3 | intron_variant | MODIFIER |
| 20 | 87288862 | AX-170034351 | 0 | T | A | 0.254 | -3.65E-02 | 5.91E-02 | 2.70E+00 | 2.82E+00 | 5.29E-01 | no | Il1rapl1 | intron_variant | MODIFIER |
| 18 | 62747084 | AX-97786674 | 1 | G | A | 0.188 | -7.39E-02 | 1.17E-01 | 2.61E+00 | 2.67E+00 | 5.29E-01 | no | 2700046A07Rik | downstream_gene_variant | MODIFIER |
| 9 | 101695938 | AX-97807579 | 1 | G | A | 0.143 | -6.95E-02 | 1.10E-01 | 2.58E+00 | 2.67E+00 | 5.29E-01 | no | | intergenic_variant | MODIFIER |
| 9 | 120327461 | AX-170216548 | 0 | A | G | 0.202 | -4.99E-02 | 8.06E-02 | 2.64E+00 | 2.77E+00 | 5.29E-01 | no | Myrip | intron_variant | MODIFIER |
| 4 | 149728518 | AX-170024234 | 0 | C | G | 0.105 | 1.06E-01 | 1.71E-01 | 2.69E+00 | 2.79E+00 | 5.29E-01 | no | Tmem201 | intron_variant | MODIFIER |
| 4 | 150243239 | AX-170484678 | 0 | C | T | 0.105 | 1.06E-01 | 1.71E-01 | 2.69E+00 | 2.79E+00 | 5.29E-01 | no | Eno1 | upstream_gene_variant | MODIFIER |
| 1 | 167304120 | AX-170086678 | 0 | T | C | 0.219 | -8.34E-02 | 1.33E-01 | 2.90E+00 | 3.00E+00 | 5.30E-01 | no | Tmco1 | upstream_gene_variant | MODIFIER |
| 9 | 22742437 | AX-97808786 | 2 | T | C | 0.464 | 5.78E-02 | 9.08E-02 | 2.55E+00 | 2.62E+00 | 5.30E-01 | no | Bbs9 | intron_variant | MODIFIER |
| 8 | 68812851 | AX-170347271 | 0 | G | A | 0.193 | -7.64E-02 | 1.22E-01 | 2.59E+00 | 2.69E+00 | 5.30E-01 | no | Ints10 | intron_variant | MODIFIER |
| 15 | 25515456 | AX-97820426 | 1 | C | T | 0.25 | -6.57E-02 | 1.05E-01 | 2.62E+00 | 2.68E+00 | 5.30E-01 | no | | intergenic_variant | MODIFIER |
| 1 | 24885493 | AX-170244641 | 1 | A | G | 0.411 | 5.33E-02 | 8.54E-02 | 2.63E+00 | 2.70E+00 | 5.30E-01 | no | | intergenic_variant | MODIFIER |
| 15 | 25391217 | AX-170333171 | 1 | G | A | 0.25 | -6.57E-02 | 1.05E-01 | 2.60E+00 | 2.67E+00 | 5.30E-01 | no | Basp1 | intron_variant | MODIFIER |
| 15 | 25494498 | AX-170302174 | 1 | C | T | 0.25 | -6.57E-02 | 1.05E-01 | 2.60E+00 | 2.67E+00 | 5.30E-01 | no | | intergenic_variant | MODIFIER |
| 16 | 46380069 | AX-170225403 | 1 | C | T | 0.518 | 5.07E-02 | 7.95E-02 | 2.54E+00 | 2.61E+00 | 5.30E-01 | no | | intergenic_variant | MODIFIER |
| 5 | 5167181 | AX-170115624 | 0 | G | A | 0.43 | -4.20E-02 | 6.52E-02 | 2.49E+00 | 2.61E+00 | 5.31E-01 | no | Cdk14 | intron_variant | MODIFIER |
| 1 | 82142302 | AX-170134979 | 0 | C | T | 0.167 | 1.11E-01 | 1.74E-01 | 2.54E+00 | 2.63E+00 | 5.31E-01 | no | | intergenic_variant | MODIFIER |
| 1 | 88585109 | AX-97834751 | 0 | G | T | 0.167 | 1.11E-01 | 1.74E-01 | 2.54E+00 | 2.63E+00 | 5.31E-01 | no | | intergenic_variant | MODIFIER |
| 18 | 58560005 | AX-170434415 | 0 | C | T | 0.395 | 6.38E-02 | 9.69E-02 | 2.46E+00 | 2.51E+00 | 5.31E-01 | no | Slc27a6 | intron_variant | MODIFIER |
| 4 | 88290608 | AX-170221403 | 0 | C | T | 0.342 | 6.90E-02 | 1.10E-01 | 2.61E+00 | 2.68E+00 | 5.31E-01 | no | Focad | intron_variant | MODIFIER |
| 4 | 88291689 | AX-97780411 | 0 | A | G | 0.342 | 6.90E-02 | 1.10E-01 | 2.61E+00 | 2.68E+00 | 5.31E-01 | no | Focad | intron_variant | MODIFIER |
| 4 | 88292819 | AX-170505686 | 0 | A | G | 0.342 | 6.90E-02 | 1.10E-01 | 2.61E+00 | 2.68E+00 | 5.31E-01 | no | Focad | intron_variant | MODIFIER |
| 4 | 88369620 | AX-97789597 | 0 | T | C | 0.342 | 6.90E-02 | 1.10E-01 | 2.61E+00 | 2.68E+00 | 5.31E-01 | no | Focad | intron_variant | MODIFIER |
| 4 | 88392215 | AX-97837555 | 0 | T | C | 0.342 | 6.90E-02 | 1.10E-01 | 2.61E+00 | 2.68E+00 | 5.31E-01 | no | Focad | intron_variant | MODIFIER |
| 9 | 111318081 | AX-170040043 | 1 | T | A | 0.107 | -6.41E-02 | 1.02E-01 | 2.57E+00 | 2.67E+00 | 5.31E-01 | no | Trank1 | intron_variant | MODIFIER |
| 6 | 110996101 | AX-170198598 | 0 | G | A | 0.088 | 1.12E-01 | 1.82E-01 | 2.66E+00 | 2.77E+00 | 5.31E-01 | no | Grm7 | intron_variant | MODIFIER |
| 1 | 28663121 | AX-97844765 | 0 | C | T | 0.421 | 5.01E-02 | 8.11E-02 | 2.73E+00 | 2.79E+00 | 5.31E-01 | no | | intergenic_variant | MODIFIER |
| 6 | 138474612 | AX-170189511 | 0 | T | A | 0.579 | 5.53E-02 | 8.50E-02 | 2.48E+00 | 2.55E+00 | 5.31E-01 | no | Lmo3 | intron_variant | MODIFIER |
| 6 | 138478674 | AX-170331815 | 0 | C | T | 0.579 | 5.53E-02 | 8.50E-02 | 2.48E+00 | 2.55E+00 | 5.31E-01 | no | Lmo3 | intron_variant | MODIFIER |
| 6 | 138483034 | AX-170405469 | 0 | G | A | 0.579 | 5.53E-02 | 8.50E-02 | 2.48E+00 | 2.55E+00 | 5.31E-01 | no | Lmo3 | intron_variant | MODIFIER |
| 6 | 138493913 | AX-170146931 | 0 | A | G | 0.579 | 5.53E-02 | 8.50E-02 | 2.48E+00 | 2.55E+00 | 5.31E-01 | no | Lmo3 | intron_variant | MODIFIER |
| 6 | 138503968 | AX-170411750 | 0 | C | T | 0.579 | 5.53E-02 | 8.50E-02 | 2.48E+00 | 2.55E+00 | 5.31E-01 | no | Lmo3 | intron_variant | MODIFIER |
| 6 | 138515260 | AX-170239181 | 0 | G | A | 0.579 | 5.53E-02 | 8.50E-02 | 2.48E+00 | 2.55E+00 | 5.31E-01 | no | Lmo3 | intron_variant | MODIFIER |
| 6 | 138611685 | AX-170228401 | 0 | C | T | 0.579 | 5.53E-02 | 8.50E-02 | 2.48E+00 | 2.55E+00 | 5.31E-01 | no | | intergenic_variant | MODIFIER |
| 1 | 90647086 | AX-97802562 | 1 | G | A | 0.152 | 1.11E-01 | 1.79E-01 | 2.61E+00 | 2.71E+00 | 5.31E-01 | no | | intergenic_variant | MODIFIER |
| 1 | 108655888 | AX-170420459 | 1 | T | C | 0.152 | 1.11E-01 | 1.79E-01 | 2.61E+00 | 2.71E+00 | 5.31E-01 | no | | intergenic_variant | MODIFIER |
| 7 | 92346459 | AX-170153421 | 0 | G | A | 0.132 | 9.57E-02 | 1.53E-01 | 2.58E+00 | 2.70E+00 | 5.31E-01 | no | Dlg2 | intron_variant | MODIFIER |
| 15 | 68075056 | AX-170055845 | 1 | C | T | 0.348 | -4.97E-02 | 7.51E-02 | 2.44E+00 | 2.52E+00 | 5.31E-01 | no | | intergenic_variant | MODIFIER |
| 1 | 33112783 | AX-97786192 | 0 | T | C | 0.421 | 4.94E-02 | 8.01E-02 | 2.69E+00 | 2.77E+00 | 5.31E-01 | no | Gm4785 | upstream_gene_variant | MODIFIER |
| 5 | 102306255 | AX-170424484 | 0 | A | T | 0.43 | 6.58E-02 | 1.07E-01 | 2.77E+00 | 2.84E+00 | 5.31E-01 | no | | intergenic_variant | MODIFIER |

| | | | | | | | | | | | | | | | |
|---|---|---|---|---|---|---|---|---|---|---|---|---|---|---|---|
| 9 | 40510503 | AX-170308092 | 0 | T | A | 0.439 | -4.48E-02 | 7.29E-02 | 2.70E+00 | 2.81E+00 | 5.31E-01 | no | Gramd1b | intron_variant | MODIFIER |
| 6 | 144317471 | AX-170105677 | 0 | G | A | 0.061 | 1.37E-01 | 2.18E-01 | 2.56E+00 | 2.67E+00 | 5.31E-01 | no | Sox5 | intron_variant,non_coding_transcript_variant | MODIFIER |
| 8 | 39008408 | AX-97795054 | 0 | G | A | 0.158 | 8.77E-02 | 1.41E-01 | 2.84E+00 | 2.91E+00 | 5.31E-01 | no | Tusc3 | intron_variant | MODIFIER |
| 8 | 39360180 | AX-170487241 | 0 | C | A | 0.158 | 8.77E-02 | 1.41E-01 | 2.84E+00 | 2.91E+00 | 5.31E-01 | no | Gm6213 | intron_variant,non_coding_transcript_variant | MODIFIER |
| 8 | 42964042 | AX-97851612 | 0 | T | C | 0.158 | 8.77E-02 | 1.41E-01 | 2.84E+00 | 2.91E+00 | 5.31E-01 | no | | intergenic_variant | MODIFIER |
| 15 | 5943299 | AX-170229686 | 1 | C | T | 0.107 | -6.48E-02 | 8.92E-02 | 2.27E+00 | 2.39E+00 | 5.31E-01 | no | | intergenic_variant | MODIFIER |
| 11 | 32111305 | AX-170325619 | 1 | G | A | 0.054 | 1.14E-01 | 1.85E-01 | 2.64E+00 | 2.75E+00 | 5.32E-01 | no | | regulatory_region_variant | MODIFIER |
| 9 | 69834411 | AX-170332400 | 1 | A | G | 0.214 | 6.94E-02 | 1.12E-01 | 2.78E+00 | 2.85E+00 | 5.32E-01 | no | B230323A14Rik | downstream_gene_variant | MODIFIER |
| 1 | 137207997 | AX-170086621 | 1 | G | A | 0.214 | 7.71E-02 | 1.24E-01 | 2.65E+00 | 2.72E+00 | 5.32E-01 | no | | intergenic_variant | MODIFIER |
| 11 | 74290280 | AX-170308085 | 1 | A | C | 0.25 | 4.56E-02 | 7.48E-02 | 2.70E+00 | 2.84E+00 | 5.32E-01 | no | Olfr59 | 3_prime_UTR_variant | MODIFIER |
| 19 | 50613198 | AX-170062653 | 0 | C | T | 0.482 | 4.08E-02 | 6.57E-02 | 2.62E+00 | 2.71E+00 | 5.32E-01 | no | Sorcs1 | intron_variant | MODIFIER |
| 9 | 54139135 | AX-168388370 | 0 | G | A | 0.202 | -7.10E-02 | 1.15E-01 | 2.73E+00 | 2.82E+00 | 5.32E-01 | no | | intergenic_variant | MODIFIER |
| 5 | 105714817 | AX-170473094 | 1 | A | G | 0.438 | 7.20E-02 | 1.16E-01 | 2.65E+00 | 2.68E+00 | 5.32E-01 | no | Lrrc8d | intron_variant | MODIFIER |
| 9 | 61581969 | AX-170348820 | 0 | T | C | 0.202 | 7.90E-02 | 1.28E-01 | 2.79E+00 | 2.85E+00 | 5.32E-01 | no | | intergenic_variant | MODIFIER |
| 18 | 55804918 | AX-170105574 | 0 | C | T | 0.053 | 7.45E-02 | 1.22E-01 | 2.59E+00 | 2.77E+00 | 5.32E-01 | no | | intergenic_variant | MODIFIER |
| 1 | 193329022 | AX-170091841 | 0 | G | A | 0.272 | 6.07E-02 | 9.09E-02 | 2.41E+00 | 2.51E+00 | 5.33E-01 | no | Lamb3 | intron_variant | MODIFIER |
| 10 | 92362340 | AX-170305314 | 1 | C | T | 0.295 | -5.37E-02 | 8.41E-02 | 2.52E+00 | 2.58E+00 | 5.33E-01 | no | Gm20757 | intron_variant,non_coding_transcript_variant | MODIFIER |
| 6 | 94415464 | AX-170514456 | 0 | T | C | 0.439 | -3.82E-02 | 6.12E-02 | 2.57E+00 | 2.69E+00 | 5.33E-01 | no | Gm7825 | upstream_gene_variant | MODIFIER |
| 1 | 20623542 | AX-97817725 | 1 | C | T | 0.411 | 5.17E-02 | 8.27E-02 | 2.60E+00 | 2.65E+00 | 5.33E-01 | no | | intergenic_variant | MODIFIER |
| 18 | 72932303 | AX-97826547 | 1 | T | C | 0.179 | -7.73E-02 | 1.23E-01 | 2.59E+00 | 2.64E+00 | 5.33E-01 | no | | intergenic_variant | MODIFIER |
| 13 | 11153719 | AX-170166032 | 1 | G | A | 0.188 | -7.06E-02 | 1.13E-01 | 2.59E+00 | 2.67E+00 | 5.33E-01 | no | | intergenic_variant | MODIFIER |
| 8 | 3112702 | AX-97852584 | 1 | G | A | 0.277 | 4.61E-02 | 7.54E-02 | 2.70E+00 | 2.81E+00 | 5.33E-01 | no | | intergenic_variant | MODIFIER |
| 4 | 88076606 | AX-170375448 | 0 | G | A | 0.333 | 6.75E-02 | 1.08E-01 | 2.61E+00 | 2.68E+00 | 5.33E-01 | no | | intergenic_variant | MODIFIER |
| 4 | 88280589 | AX-170378566 | 0 | T | C | 0.333 | 6.75E-02 | 1.08E-01 | 2.61E+00 | 2.68E+00 | 5.33E-01 | no | Focad | intron_variant | MODIFIER |
| 4 | 88434841 | AX-170224687 | 0 | T | C | 0.333 | 6.75E-02 | 1.08E-01 | 2.61E+00 | 2.68E+00 | 5.33E-01 | no | Hacd4 | intron_variant | MODIFIER |
| 5 | 51549031 | AX-170024941 | 1 | T | C | 0.473 | 6.22E-02 | 9.57E-02 | 2.47E+00 | 2.55E+00 | 5.33E-01 | no | Ppargc1a | intron_variant,NMD_transcript_variant | MODIFIER |
| 5 | 43703149 | AX-97808285 | 1 | G | A | 0.188 | -6.89E-02 | 1.11E-01 | 2.62E+00 | 2.70E+00 | 5.33E-01 | no | Cc2d2a | intron_variant | MODIFIER |
| 18 | 64844551 | AX-170200607 | 0 | G | A | 0.404 | 6.01E-02 | 9.27E-02 | 2.48E+00 | 2.53E+00 | 5.33E-01 | no | | intergenic_variant | MODIFIER |
| 12 | 91375779 | AX-170163849 | 1 | A | G | 0.08 | 1.14E-01 | 1.85E-01 | 2.74E+00 | 2.82E+00 | 5.33E-01 | no | Cep128 | intron_variant,non_coding_transcript_variant | MODIFIER |
| 7 | 10410291 | AX-170431640 | 0 | A | G | 0.167 | 4.88E-02 | 8.02E-02 | 2.80E+00 | 2.95E+00 | 5.33E-01 | no | Gm3726 | downstream_gene_variant | MODIFIER |
| 1 | 107314915 | AX-170001300 | 0 | A | T | 0.184 | 9.45E-02 | 1.52E-01 | 2.57E+00 | 2.69E+00 | 5.33E-01 | no | | intergenic_variant | MODIFIER |
| 1 | 45908458 | AX-170442823 | 0 | G | C | 0.096 | -1.29E-01 | 2.11E-01 | 2.73E+00 | 2.81E+00 | 5.34E-01 | no | Slc40a1 | 3_prime_UTR_variant | MODIFIER |
| 13 | 24133042 | AX-170201177 | 0 | G | A | 0.158 | -8.36E-02 | 1.27E-01 | 2.44E+00 | 2.49E+00 | 5.34E-01 | no | Carmil1 | intron_variant | MODIFIER |
| 10 | 65945100 | AX-170152953 | 0 | A | G | 0.412 | -4.94E-02 | 7.98E-02 | 2.63E+00 | 2.70E+00 | 5.34E-01 | no | | intergenic_variant | MODIFIER |
| 5 | 125830211 | AX-97845413 | 0 | G | A | 0.447 | -6.72E-02 | 1.09E-01 | 2.79E+00 | 2.85E+00 | 5.34E-01 | no | | intergenic_variant | MODIFIER |
| 11 | 31572746 | AX-97811335 | 0 | C | T | 0.07 | 1.12E-01 | 1.81E-01 | 2.61E+00 | 2.72E+00 | 5.34E-01 | no | | intergenic_variant | MODIFIER |
| 5 | 104255739 | AX-170396085 | 1 | C | A | 0.438 | 7.14E-02 | 1.15E-01 | 2.65E+00 | 2.68E+00 | 5.34E-01 | no | | intergenic_variant | MODIFIER |
| 7 | 145160759 | AX-170079134 | 0 | A | G | 0.211 | -4.41E-02 | 7.19E-02 | 2.62E+00 | 2.75E+00 | 5.34E-01 | no | Gm45181 | upstream_gene_variant | MODIFIER |
| 16 | 46407653 | AX-170452923 | 0 | T | G | 0.518 | 5.10E-02 | 7.99E-02 | 2.51E+00 | 2.57E+00 | 5.34E-01 | no | Nectin3 | intron_variant | MODIFIER |
| 5 | 36381684 | AX-170223973 | 0 | T | A | 0.325 | 6.76E-02 | 1.10E-01 | 2.76E+00 | 2.82E+00 | 5.34E-01 | no | Sorcs2 | intron_variant | MODIFIER |
| 12 | 89020826 | AX-97801490 | 2 | G | A | 0.082 | 1.15E-01 | 1.88E-01 | 2.73E+00 | 2.82E+00 | 5.34E-01 | no | Nrxn3 | intron_variant | MODIFIER |
| 20 | 11187790 | AX-170262238 | 0 | T | C | 0.254 | 3.34E-02 | 5.47E-02 | 2.62E+00 | 2.76E+00 | 5.34E-01 | no | | intergenic_variant | MODIFIER |
| 20 | 11188063 | AX-170195128 | 0 | G | A | 0.254 | 3.34E-02 | 5.47E-02 | 2.62E+00 | 2.76E+00 | 5.34E-01 | no | | intergenic_variant | MODIFIER |
| 20 | 11188403 | AX-170350704 | 0 | A | C | 0.254 | 3.34E-02 | 5.47E-02 | 2.62E+00 | 2.76E+00 | 5.34E-01 | no | | intergenic_variant | MODIFIER |
| 20 | 11206714 | AX-170174025 | 0 | A | G | 0.254 | 3.34E-02 | 5.47E-02 | 2.62E+00 | 2.76E+00 | 5.34E-01 | no | | intergenic_variant | MODIFIER |
| 8 | 34278162 | AX-170263066 | 1 | G | A | 0.134 | -7.96E-02 | 1.29E-01 | 2.62E+00 | 2.70E+00 | 5.34E-01 | no | | intergenic_variant | MODIFIER |
| 1 | 167820439 | AX-170504151 | 0 | T | C | 0.237 | 7.52E-02 | 1.21E-01 | 2.60E+00 | 2.67E+00 | 5.35E-01 | no | Lmx1a | intron_variant | MODIFIER |
| 1 | 173185197 | AX-170091789 | 0 | A | G | 0.228 | 7.72E-02 | 1.20E-01 | 2.49E+00 | 2.55E+00 | 5.35E-01 | no | Olfr1406 | upstream_gene_variant | MODIFIER |
| 2 | 102736124 | AX-170476214 | 0 | A | C | 0.061 | 1.26E-01 | 2.08E-01 | 2.70E+00 | 2.81E+00 | 5.35E-01 | no | Slc1a2 | intron_variant | MODIFIER |
| 1 | 191840656 | AX-170402861 | 0 | T | A | 0.272 | 5.82E-02 | 8.74E-02 | 2.40E+00 | 2.51E+00 | 5.35E-01 | no | | regulatory_region_variant | MODIFIER |
| 1 | 193179588 | AX-170519958 | 0 | G | C | 0.272 | 5.82E-02 | 8.74E-02 | 2.40E+00 | 2.51E+00 | 5.35E-01 | no | Traf3ip3 | intron_variant | MODIFIER |
| 18 | 84127612 | AX-170161784 | 2 | A | G | 0.164 | -6.92E-02 | 1.07E-01 | 2.48E+00 | 2.56E+00 | 5.35E-01 | no | Zfp407 | downstream_gene_variant | MODIFIER |
| 1 | 63465159 | AX-170225576 | 0 | G | A | 0.132 | -1.16E-01 | 1.90E-01 | 2.68E+00 | 2.78E+00 | 5.35E-01 | no | Adam23 | intron_variant | MODIFIER |
| 4 | 145340768 | AX-170527278 | 0 | G | A | 0.114 | 1.05E-01 | 1.72E-01 | 2.69E+00 | 2.80E+00 | 5.35E-01 | no | | intergenic_variant | MODIFIER |

| | | | | | | | | | | | | | | | |
|---|---|---|---|---|---|---|---|---|---|---|---|---|---|---|---|
| 10 | 11327005 | AX-170215847 | 0 | C | T | 0.044 | -8.97E-02 | 1.48E-01 | 2.67E+00 | 2.81E+00 | 5.35E-01 | no | Gm25979 | downstream_gene_variant | MODIFIER |
| 18 | 68999566 | AX-170378266 | 0 | C | T | 0.263 | -6.24E-02 | 1.01E-01 | 2.66E+00 | 2.72E+00 | 5.35E-01 | no | | intergenic_variant | MODIFIER |
| 18 | 69024647 | AX-170491806 | 0 | C | A | 0.263 | -6.24E-02 | 1.01E-01 | 2.66E+00 | 2.72E+00 | 5.35E-01 | no | Mir145b | upstream_gene_variant | MODIFIER |
| 18 | 69036496 | AX-170150062 | 0 | C | T | 0.263 | -6.24E-02 | 1.01E-01 | 2.66E+00 | 2.72E+00 | 5.35E-01 | no | | intergenic_variant | MODIFIER |
| 18 | 69060239 | AX-170528768 | 0 | G | T | 0.263 | -6.24E-02 | 1.01E-01 | 2.66E+00 | 2.72E+00 | 5.35E-01 | no | | intergenic_variant | MODIFIER |
| 18 | 69065061 | AX-170525385 | 0 | G | A | 0.263 | -6.24E-02 | 1.01E-01 | 2.66E+00 | 2.72E+00 | 5.35E-01 | no | | intergenic_variant | MODIFIER |
| 18 | 69066690 | AX-170524623 | 0 | T | C | 0.263 | -6.24E-02 | 1.01E-01 | 2.66E+00 | 2.72E+00 | 5.35E-01 | no | | intergenic_variant | MODIFIER |
| 18 | 69067090 | AX-170271484 | 0 | T | C | 0.263 | -6.24E-02 | 1.01E-01 | 2.66E+00 | 2.72E+00 | 5.35E-01 | no | | intergenic_variant | MODIFIER |
| 18 | 69067195 | AX-170532338 | 0 | G | C | 0.263 | -6.24E-02 | 1.01E-01 | 2.66E+00 | 2.72E+00 | 5.35E-01 | no | | intergenic_variant | MODIFIER |
| 18 | 69072478 | AX-170339877 | 0 | A | G | 0.263 | -6.24E-02 | 1.01E-01 | 2.66E+00 | 2.72E+00 | 5.35E-01 | no | | intergenic_variant | MODIFIER |
| 18 | 69072594 | AX-170425368 | 0 | C | A | 0.263 | -6.24E-02 | 1.01E-01 | 2.66E+00 | 2.72E+00 | 5.35E-01 | no | | intergenic_variant | MODIFIER |
| 18 | 69072756 | AX-170135906 | 0 | A | G | 0.263 | -6.24E-02 | 1.01E-01 | 2.66E+00 | 2.72E+00 | 5.35E-01 | no | | intergenic_variant | MODIFIER |
| 18 | 69183864 | AX-170061220 | 0 | C | G | 0.263 | -6.24E-02 | 1.01E-01 | 2.66E+00 | 2.72E+00 | 5.35E-01 | no | | regulatory_region_variant | MODIFIER |
| 18 | 69186712 | AX-170061218 | 0 | G | A | 0.263 | -6.24E-02 | 1.01E-01 | 2.66E+00 | 2.72E+00 | 5.35E-01 | no | | intergenic_variant | MODIFIER |
| 18 | 69202541 | AX-170450373 | 0 | G | T | 0.263 | -6.24E-02 | 1.01E-01 | 2.66E+00 | 2.72E+00 | 5.35E-01 | no | AC121263.1 | downstream_gene_variant | MODIFIER |
| 18 | 69203608 | AX-170204312 | 0 | G | A | 0.263 | -6.24E-02 | 1.01E-01 | 2.66E+00 | 2.72E+00 | 5.35E-01 | no | AC121263.1 | downstream_gene_variant | MODIFIER |
| 18 | 69234127 | AX-97798863 | 0 | A | G | 0.263 | -6.24E-02 | 1.01E-01 | 2.66E+00 | 2.72E+00 | 5.35E-01 | no | | regulatory_region_variant | MODIFIER |
| 18 | 69443996 | AX-170127754 | 0 | C | T | 0.263 | -6.24E-02 | 1.01E-01 | 2.66E+00 | 2.72E+00 | 5.35E-01 | no | Tcf4 | intron_variant | MODIFIER |
| 18 | 69449974 | AX-170515131 | 0 | C | A | 0.263 | -6.24E-02 | 1.01E-01 | 2.66E+00 | 2.72E+00 | 5.35E-01 | no | Tcf4 | intron_variant | MODIFIER |
| 18 | 69450139 | AX-170435879 | 0 | G | A | 0.263 | -6.24E-02 | 1.01E-01 | 2.66E+00 | 2.72E+00 | 5.35E-01 | no | Tcf4 | intron_variant | MODIFIER |
| 18 | 69477456 | AX-170204198 | 0 | T | G | 0.263 | -6.24E-02 | 1.01E-01 | 2.66E+00 | 2.72E+00 | 5.35E-01 | no | Tcf4 | intron_variant | MODIFIER |
| 18 | 69486583 | AX-170459837 | 0 | G | A | 0.263 | -6.24E-02 | 1.01E-01 | 2.66E+00 | 2.72E+00 | 5.35E-01 | no | Tcf4 | intron_variant | MODIFIER |
| 18 | 69490228 | AX-170505179 | 0 | C | A | 0.263 | -6.24E-02 | 1.01E-01 | 2.66E+00 | 2.72E+00 | 5.35E-01 | no | Tcf4 | intron_variant | MODIFIER |
| 18 | 69508151 | AX-97797075 | 0 | G | A | 0.263 | -6.24E-02 | 1.01E-01 | 2.66E+00 | 2.72E+00 | 5.35E-01 | no | Tcf4 | intron_variant | MODIFIER |
| 18 | 69518530 | AX-170481818 | 0 | G | C | 0.263 | -6.24E-02 | 1.01E-01 | 2.66E+00 | 2.72E+00 | 5.35E-01 | no | Tcf4 | intron_variant | MODIFIER |
| 18 | 69526325 | AX-170346709 | 0 | C | G | 0.263 | -6.24E-02 | 1.01E-01 | 2.66E+00 | 2.72E+00 | 5.35E-01 | no | Tcf4 | intron_variant | MODIFIER |
| 18 | 69544926 | AX-170320775 | 0 | A | G | 0.263 | -6.24E-02 | 1.01E-01 | 2.66E+00 | 2.72E+00 | 5.35E-01 | no | Tcf4 | intron_variant | MODIFIER |
| 18 | 69568610 | AX-170386238 | 0 | A | G | 0.263 | -6.24E-02 | 1.01E-01 | 2.66E+00 | 2.72E+00 | 5.35E-01 | no | Tcf4 | intron_variant | MODIFIER |
| 18 | 69582771 | AX-170061225 | 0 | A | G | 0.263 | -6.24E-02 | 1.01E-01 | 2.66E+00 | 2.72E+00 | 5.35E-01 | no | Tcf4 | intron_variant | MODIFIER |
| 18 | 69606794 | AX-170186233 | 0 | A | C | 0.263 | -6.24E-02 | 1.01E-01 | 2.66E+00 | 2.72E+00 | 5.35E-01 | no | Tcf4 | intron_variant | MODIFIER |
| 18 | 69618551 | AX-170516052 | 0 | T | C | 0.263 | -6.24E-02 | 1.01E-01 | 2.66E+00 | 2.72E+00 | 5.35E-01 | no | Tcf4 | intron_variant | MODIFIER |
| 18 | 69620013 | AX-170521490 | 0 | T | C | 0.263 | -6.24E-02 | 1.01E-01 | 2.66E+00 | 2.72E+00 | 5.35E-01 | no | Tcf4 | intron_variant | MODIFIER |
| 18 | 69622610 | AX-170276106 | 0 | T | C | 0.263 | -6.24E-02 | 1.01E-01 | 2.66E+00 | 2.72E+00 | 5.35E-01 | no | Tcf4 | intron_variant | MODIFIER |
| 18 | 69628582 | AX-97843330 | 0 | A | C | 0.263 | -6.24E-02 | 1.01E-01 | 2.66E+00 | 2.72E+00 | 5.35E-01 | no | Tcf4 | intron_variant | MODIFIER |
| 18 | 69636583 | AX-170015924 | 0 | C | T | 0.263 | -6.24E-02 | 1.01E-01 | 2.66E+00 | 2.72E+00 | 5.35E-01 | no | Tcf4 | intron_variant | MODIFIER |
| 18 | 69637514 | AX-170499259 | 0 | G | A | 0.263 | -6.24E-02 | 1.01E-01 | 2.66E+00 | 2.72E+00 | 5.35E-01 | no | Tcf4 | intron_variant | MODIFIER |
| 18 | 69641305 | AX-168416814 | 0 | A | G | 0.263 | -6.24E-02 | 1.01E-01 | 2.66E+00 | 2.72E+00 | 5.35E-01 | no | Tcf4 | intron_variant | MODIFIER |
| 18 | 69646460 | AX-170532163 | 0 | C | A | 0.263 | -6.24E-02 | 1.01E-01 | 2.66E+00 | 2.72E+00 | 5.35E-01 | no | Tcf4 | intron_variant | MODIFIER |
| 18 | 69664782 | AX-170015925 | 0 | G | A | 0.263 | -6.24E-02 | 1.01E-01 | 2.66E+00 | 2.72E+00 | 5.35E-01 | no | Tcf4 | intron_variant | MODIFIER |
| 18 | 69669621 | AX-170357111 | 0 | T | C | 0.263 | -6.24E-02 | 1.01E-01 | 2.66E+00 | 2.72E+00 | 5.35E-01 | no | Tcf4 | intron_variant | MODIFIER |
| 18 | 69669638 | AX-170408770 | 0 | T | G | 0.263 | -6.24E-02 | 1.01E-01 | 2.66E+00 | 2.72E+00 | 5.35E-01 | no | Tcf4 | intron_variant | MODIFIER |
| 18 | 69669932 | AX-170285354 | 0 | G | C | 0.263 | -6.24E-02 | 1.01E-01 | 2.66E+00 | 2.72E+00 | 5.35E-01 | no | Tcf4 | intron_variant | MODIFIER |
| 18 | 69676156 | AX-170061228 | 0 | C | T | 0.263 | -6.24E-02 | 1.01E-01 | 2.66E+00 | 2.72E+00 | 5.35E-01 | no | Tcf4 | intron_variant | MODIFIER |
| 18 | 69676633 | AX-170061231 | 0 | G | A | 0.263 | -6.24E-02 | 1.01E-01 | 2.66E+00 | 2.72E+00 | 5.35E-01 | no | Tcf4 | intron_variant | MODIFIER |
| 18 | 69689848 | AX-170098935 | 0 | G | T | 0.263 | -6.24E-02 | 1.01E-01 | 2.66E+00 | 2.72E+00 | 5.35E-01 | no | Tcf4 | downstream_gene_variant | MODIFIER |
| 18 | 69695476 | AX-170196286 | 0 | T | G | 0.263 | -6.24E-02 | 1.01E-01 | 2.66E+00 | 2.72E+00 | 5.35E-01 | no | | intergenic_variant | MODIFIER |
| 18 | 69695585 | AX-170015927 | 0 | C | A | 0.263 | -6.24E-02 | 1.01E-01 | 2.66E+00 | 2.72E+00 | 5.35E-01 | no | | intergenic_variant | MODIFIER |
| 18 | 69701460 | AX-170194219 | 0 | C | T | 0.263 | -6.24E-02 | 1.01E-01 | 2.66E+00 | 2.72E+00 | 5.35E-01 | no | | intergenic_variant | MODIFIER |
| 18 | 69704016 | AX-170423805 | 0 | C | T | 0.263 | -6.24E-02 | 1.01E-01 | 2.66E+00 | 2.72E+00 | 5.35E-01 | no | | intergenic_variant | MODIFIER |
| 18 | 69704107 | AX-170141240 | 0 | C | T | 0.263 | -6.24E-02 | 1.01E-01 | 2.66E+00 | 2.72E+00 | 5.35E-01 | no | | intergenic_variant | MODIFIER |
| 18 | 69709214 | AX-170446405 | 0 | G | A | 0.263 | -6.24E-02 | 1.01E-01 | 2.66E+00 | 2.72E+00 | 5.35E-01 | no | | intergenic_variant | MODIFIER |
| 18 | 69718869 | AX-170530613 | 0 | T | C | 0.263 | -6.24E-02 | 1.01E-01 | 2.66E+00 | 2.72E+00 | 5.35E-01 | no | | intergenic_variant | MODIFIER |
| 18 | 69719445 | AX-170275613 | 0 | T | C | 0.263 | -6.24E-02 | 1.01E-01 | 2.66E+00 | 2.72E+00 | 5.35E-01 | no | | intergenic_variant | MODIFIER |
| 18 | 69722036 | AX-170462980 | 0 | G | A | 0.263 | -6.24E-02 | 1.01E-01 | 2.66E+00 | 2.72E+00 | 5.35E-01 | no | | regulatory_region_variant | MODIFIER |

| | | | | | | | | | | | | | | | |
|---|---|---|---|---|---|---|---|---|---|---|---|---|---|---|---|
| 18 | 69741247 | AX-170334486 | 0 | A | C | 0.263 | -6.24E-02 | 1.01E-01 | 2.66E+00 | 2.72E+00 | 5.35E-01 | no | | intergenic_variant | MODIFIER |
| 18 | 69743630 | AX-170249852 | 0 | G | C | 0.263 | -6.24E-02 | 1.01E-01 | 2.66E+00 | 2.72E+00 | 5.35E-01 | no | | intergenic_variant | MODIFIER |
| 18 | 69745425 | AX-170201949 | 0 | T | C | 0.263 | -6.24E-02 | 1.01E-01 | 2.66E+00 | 2.72E+00 | 5.35E-01 | no | | intergenic_variant | MODIFIER |
| 18 | 69774543 | AX-170381622 | 0 | C | T | 0.263 | -6.24E-02 | 1.01E-01 | 2.66E+00 | 2.72E+00 | 5.35E-01 | no | Gm22508 | downstream_gene_variant | MODIFIER |
| 18 | 69775646 | AX-170061235 | 0 | G | T | 0.263 | -6.24E-02 | 1.01E-01 | 2.66E+00 | 2.72E+00 | 5.35E-01 | no | Gm22508 | downstream_gene_variant | MODIFIER |
| 18 | 69808020 | AX-97801501 | 0 | T | C | 0.263 | -6.24E-02 | 1.01E-01 | 2.66E+00 | 2.72E+00 | 5.35E-01 | no | | intergenic_variant | MODIFIER |
| 18 | 69817692 | AX-170136286 | 0 | C | T | 0.263 | -6.24E-02 | 1.01E-01 | 2.66E+00 | 2.72E+00 | 5.35E-01 | no | | intergenic_variant | MODIFIER |
| 18 | 69822115 | AX-170460022 | 0 | G | A | 0.263 | -6.24E-02 | 1.01E-01 | 2.66E+00 | 2.72E+00 | 5.35E-01 | no | | intergenic_variant | MODIFIER |
| 18 | 69831336 | AX-170152042 | 0 | T | C | 0.263 | -6.24E-02 | 1.01E-01 | 2.66E+00 | 2.72E+00 | 5.35E-01 | no | | intergenic_variant | MODIFIER |
| 18 | 69836392 | AX-170408890 | 0 | C | T | 0.263 | -6.24E-02 | 1.01E-01 | 2.66E+00 | 2.72E+00 | 5.35E-01 | no | | regulatory_region_variant | MODIFIER |
| 18 | 62812942 | AX-170227745 | 0 | T | A | 0.193 | -7.34E-02 | 1.20E-01 | 2.70E+00 | 2.76E+00 | 5.35E-01 | no | | intergenic_variant | MODIFIER |
| 15 | 9202158 | AX-170408305 | 0 | G | A | 0.237 | -6.63E-02 | 1.06E-01 | 2.59E+00 | 2.64E+00 | 5.35E-01 | no | Lmbrd2 | downstream_gene_variant | MODIFIER |
| 15 | 11027657 | AX-170450680 | 0 | G | C | 0.237 | -6.63E-02 | 1.06E-01 | 2.59E+00 | 2.64E+00 | 5.35E-01 | no | Slc45a2 | intron_variant | MODIFIER |
| 15 | 20449359 | AX-97828881 | 0 | G | C | 0.237 | -6.63E-02 | 1.06E-01 | 2.59E+00 | 2.64E+00 | 5.35E-01 | no | Cdh12 | non_coding_transcript_exon_variant | MODIFIER |
| 14 | 117040983 | AX-170342107 | 0 | T | G | 0.316 | 4.58E-02 | 7.32E-02 | 2.55E+00 | 2.65E+00 | 5.35E-01 | no | Gpc6 | intron_variant | MODIFIER |
| 14 | 117700339 | AX-170149516 | 0 | C | T | 0.316 | 4.58E-02 | 7.32E-02 | 2.55E+00 | 2.65E+00 | 5.35E-01 | no | Gpc6 | intron_variant | MODIFIER |
| 18 | 72097691 | AX-170325560 | 1 | T | A | 0.179 | -7.23E-02 | 1.17E-01 | 2.63E+00 | 2.68E+00 | 5.35E-01 | no | Dcc | intron_variant | MODIFIER |
| 9 | 103970242 | AX-97803054 | 0 | G | A | 0.018 | -7.80E-02 | 1.27E-01 | 2.55E+00 | 2.72E+00 | 5.35E-01 | no | | intergenic_variant | MODIFIER |
| 10 | 19126459 | AX-170326037 | 1 | T | A | 0.188 | 6.19E-02 | 1.01E-01 | 2.83E+00 | 2.93E+00 | 5.35E-01 | no | Gm48546 | downstream_gene_variant | MODIFIER |
| 7 | 70113923 | AX-170109421 | 0 | T | C | 0.36 | -6.39E-02 | 1.04E-01 | 2.83E+00 | 2.92E+00 | 5.35E-01 | no | | intergenic_variant | MODIFIER |
| 7 | 70132579 | AX-170442925 | 0 | G | A | 0.36 | -6.39E-02 | 1.04E-01 | 2.83E+00 | 2.92E+00 | 5.35E-01 | no | | intergenic_variant | MODIFIER |
| 7 | 70132644 | AX-170285613 | 0 | G | A | 0.36 | -6.39E-02 | 1.04E-01 | 2.83E+00 | 2.92E+00 | 5.35E-01 | no | | intergenic_variant | MODIFIER |
| 7 | 70153467 | AX-170104149 | 0 | T | C | 0.36 | -6.39E-02 | 1.04E-01 | 2.83E+00 | 2.92E+00 | 5.35E-01 | no | | intergenic_variant | MODIFIER |
| 7 | 70184886 | AX-170029237 | 0 | T | C | 0.36 | -6.39E-02 | 1.04E-01 | 2.83E+00 | 2.92E+00 | 5.35E-01 | no | Gm29683 | intron_variant,non_coding_transcript_variant | MODIFIER |
| 7 | 70220063 | AX-170111816 | 0 | C | T | 0.36 | -6.39E-02 | 1.04E-01 | 2.83E+00 | 2.92E+00 | 5.35E-01 | no | Gm29683 | intron_variant,non_coding_transcript_variant | MODIFIER |
| 7 | 70220227 | AX-170446930 | 0 | C | G | 0.36 | -6.39E-02 | 1.04E-01 | 2.83E+00 | 2.92E+00 | 5.35E-01 | no | Gm29683 | intron_variant,non_coding_transcript_variant | MODIFIER |
| 7 | 70225110 | AX-97824395 | 0 | A | G | 0.36 | -6.39E-02 | 1.04E-01 | 2.83E+00 | 2.92E+00 | 5.35E-01 | no | Gm29683 | intron_variant,non_coding_transcript_variant | MODIFIER |
| 7 | 70281433 | AX-170368238 | 0 | A | T | 0.36 | -6.39E-02 | 1.04E-01 | 2.83E+00 | 2.92E+00 | 5.35E-01 | no | Gm29683 | intron_variant,non_coding_transcript_variant | MODIFIER |
| 7 | 70282100 | AX-170509207 | 0 | G | A | 0.36 | -6.39E-02 | 1.04E-01 | 2.83E+00 | 2.92E+00 | 5.35E-01 | no | Gm29683 | intron_variant,non_coding_transcript_variant | MODIFIER |
| 7 | 70292848 | AX-170029240 | 0 | T | C | 0.36 | -6.39E-02 | 1.04E-01 | 2.83E+00 | 2.92E+00 | 5.35E-01 | no | Gm29683 | intron_variant,non_coding_transcript_variant | MODIFIER |
| 7 | 70292865 | AX-170431789 | 0 | T | C | 0.36 | -6.39E-02 | 1.04E-01 | 2.83E+00 | 2.92E+00 | 5.35E-01 | no | Gm29683 | intron_variant,non_coding_transcript_variant | MODIFIER |
| 7 | 70295956 | AX-170183847 | 0 | A | G | 0.36 | -6.39E-02 | 1.04E-01 | 2.83E+00 | 2.92E+00 | 5.35E-01 | no | Gm29683 | intron_variant,non_coding_transcript_variant | MODIFIER |
| 7 | 70303818 | AX-170128905 | 0 | A | G | 0.36 | -6.39E-02 | 1.04E-01 | 2.83E+00 | 2.92E+00 | 5.35E-01 | no | Gm29683 | intron_variant,non_coding_transcript_variant | MODIFIER |
| 7 | 70304458 | AX-170099675 | 0 | C | A | 0.36 | -6.39E-02 | 1.04E-01 | 2.83E+00 | 2.92E+00 | 5.35E-01 | no | Gm29683 | downstream_gene_variant | MODIFIER |
| 7 | 70313875 | AX-170372458 | 0 | T | G | 0.36 | -6.39E-02 | 1.04E-01 | 2.83E+00 | 2.92E+00 | 5.35E-01 | no | Gm29683 | intron_variant,non_coding_transcript_variant | MODIFIER |
| 7 | 70314450 | AX-170461686 | 0 | T | G | 0.36 | -6.39E-02 | 1.04E-01 | 2.83E+00 | 2.92E+00 | 5.35E-01 | no | Gm29683 | intron_variant,non_coding_transcript_variant | MODIFIER |
| 7 | 70316243 | AX-170341562 | 0 | C | T | 0.36 | -6.39E-02 | 1.04E-01 | 2.83E+00 | 2.92E+00 | 5.35E-01 | no | Gm29683 | intron_variant,non_coding_transcript_variant | MODIFIER |
| 7 | 70506465 | AX-170089951 | 0 | T | C | 0.36 | -6.39E-02 | 1.04E-01 | 2.83E+00 | 2.92E+00 | 5.35E-01 | no | | regulatory_region_variant | MODIFIER |
| 7 | 70507297 | AX-170104208 | 0 | G | T | 0.36 | -6.39E-02 | 1.04E-01 | 2.83E+00 | 2.92E+00 | 5.35E-01 | no | | intergenic_variant | MODIFIER |
| 7 | 70541651 | AX-97797998 | 0 | G | A | 0.36 | -6.39E-02 | 1.04E-01 | 2.83E+00 | 2.92E+00 | 5.35E-01 | no | | intergenic_variant | MODIFIER |
| 7 | 70541692 | AX-170327396 | 0 | A | G | 0.36 | -6.39E-02 | 1.04E-01 | 2.83E+00 | 2.92E+00 | 5.35E-01 | no | | intergenic_variant | MODIFIER |
| 7 | 70542233 | AX-170414850 | 0 | T | A | 0.36 | -6.39E-02 | 1.04E-01 | 2.83E+00 | 2.92E+00 | 5.35E-01 | no | | regulatory_region_variant | MODIFIER |
| 7 | 70542841 | AX-170219021 | 0 | T | C | 0.36 | -6.39E-02 | 1.04E-01 | 2.83E+00 | 2.92E+00 | 5.35E-01 | no | | regulatory_region_variant | MODIFIER |
| 7 | 70543408 | AX-170473554 | 0 | T | A | 0.36 | -6.39E-02 | 1.04E-01 | 2.83E+00 | 2.92E+00 | 5.35E-01 | no | | intergenic_variant | MODIFIER |
| 7 | 70546815 | AX-170409328 | 0 | G | A | 0.36 | -6.39E-02 | 1.04E-01 | 2.83E+00 | 2.92E+00 | 5.35E-01 | no | Gm35842 | upstream_gene_variant | MODIFIER |
| 7 | 70581173 | AX-170167199 | 0 | T | A | 0.36 | -6.39E-02 | 1.04E-01 | 2.83E+00 | 2.92E+00 | 5.35E-01 | no | Gm44812 | intron_variant,non_coding_transcript_variant | MODIFIER |
| 18 | 65119997 | AX-170240241 | 0 | T | C | 0.412 | 5.91E-02 | 9.12E-02 | 2.47E+00 | 2.52E+00 | 5.35E-01 | no | Nedd4l | intron_variant | MODIFIER |
| 1 | 143745130 | AX-170461812 | 0 | C | A | 0.211 | 8.49E-02 | 1.38E-01 | 2.69E+00 | 2.73E+00 | 5.35E-01 | no | Glrx2 | 3_prime_UTR_variant,NMD_transcript_variant | MODIFIER |
| 1 | 144536776 | AX-97845708 | 0 | A | C | 0.211 | 8.49E-02 | 1.38E-01 | 2.69E+00 | 2.73E+00 | 5.35E-01 | no | Rgs21 | intron_variant,non_coding_transcript_variant | MODIFIER |
| 5 | 114285126 | AX-170227058 | 0 | A | C | 0.456 | 6.95E-02 | 1.11E-01 | 2.58E+00 | 2.62E+00 | 5.35E-01 | no | Myo1h | upstream_gene_variant | MODIFIER |
| 18 | 80299415 | AX-170304091 | 0 | G | A | 0.404 | 3.15E-02 | 5.19E-02 | 2.67E+00 | 2.81E+00 | 5.36E-01 | no | Kcng2 | intron_variant | MODIFIER |
| 11 | 117384612 | AX-170453465 | 0 | A | T | 0.044 | -1.14E-01 | 1.86E-01 | 2.61E+00 | 2.74E+00 | 5.36E-01 | no | | regulatory_region_variant | MODIFIER |
| 4 | 80493547 | AX-170023218 | 1 | A | G | 0.232 | 6.64E-02 | 1.06E-01 | 2.54E+00 | 2.62E+00 | 5.36E-01 | no | | intergenic_variant | MODIFIER |
| 5 | 45896037 | AX-170163102 | 0 | A | G | 0.491 | 6.35E-02 | 9.72E-02 | 2.45E+00 | 2.52E+00 | 5.36E-01 | no | | intergenic_variant | MODIFIER |

| | | | | | | | | | | | | | | | |
|---|---|---|---|---|---|---|---|---|---|---|---|---|---|---|---|
| 1 | 176615330 | AX-170287248 | 1 | C | A | 0.223 | 7.40E-02 | 1.19E-01 | 2.59E+00 | 2.65E+00 | 5.36E-01 | no | | intergenic_variant | MODIFIER |
| 7 | 14607629 | AX-170089884 | 1 | C | A | 0.384 | 5.03E-02 | 8.00E-02 | 2.53E+00 | 2.64E+00 | 5.36E-01 | no | Nlrp5-ps | intron_variant,non_coding_transcript_variant | MODIFIER |
| 1 | 118723521 | AX-170283999 | 2 | T | A | 0.164 | 1.30E-01 | 2.10E-01 | 2.61E+00 | 2.67E+00 | 5.36E-01 | no | | intergenic_variant | MODIFIER |
| 7 | 90282520 | AX-170484531 | 1 | C | T | 0.116 | -1.09E-01 | 1.78E-01 | 2.64E+00 | 2.76E+00 | 5.36E-01 | no | | regulatory_region_variant | MODIFIER |
| 20 | 72467193 | AX-170196454 | 1 | G | T | 0.259 | 3.82E-02 | 6.26E-02 | 2.67E+00 | 2.76E+00 | 5.36E-01 | no | Gabra3 | intron_variant | MODIFIER |
| 4 | 149987194 | AX-170501903 | 0 | G | A | 0.105 | 1.10E-01 | 1.81E-01 | 2.69E+00 | 2.77E+00 | 5.36E-01 | no | H6pd | intron_variant | MODIFIER |
| 4 | 153028979 | AX-170484760 | 0 | A | T | 0.105 | 1.10E-01 | 1.81E-01 | 2.69E+00 | 2.77E+00 | 5.36E-01 | no | | intergenic_variant | MODIFIER |
| 4 | 153067782 | AX-97811893 | 0 | G | A | 0.105 | 1.10E-01 | 1.81E-01 | 2.69E+00 | 2.77E+00 | 5.36E-01 | no | | intergenic_variant | MODIFIER |
| 4 | 153198750 | AX-170071091 | 0 | C | A | 0.105 | 1.10E-01 | 1.81E-01 | 2.69E+00 | 2.77E+00 | 5.36E-01 | no | | intergenic_variant | MODIFIER |
| 4 | 153200621 | AX-170252451 | 0 | A | T | 0.105 | 1.10E-01 | 1.81E-01 | 2.69E+00 | 2.77E+00 | 5.36E-01 | no | | intergenic_variant | MODIFIER |
| 4 | 153224433 | AX-170239879 | 0 | A | C | 0.105 | 1.10E-01 | 1.81E-01 | 2.69E+00 | 2.77E+00 | 5.36E-01 | no | | regulatory_region_variant | MODIFIER |
| 4 | 153238878 | AX-170374604 | 0 | C | T | 0.105 | 1.10E-01 | 1.81E-01 | 2.69E+00 | 2.77E+00 | 5.36E-01 | no | | intergenic_variant | MODIFIER |
| 4 | 153321958 | AX-170373087 | 0 | G | A | 0.105 | 1.10E-01 | 1.81E-01 | 2.69E+00 | 2.77E+00 | 5.36E-01 | no | | intergenic_variant | MODIFIER |
| 4 | 153322516 | AX-170296429 | 0 | G | A | 0.105 | 1.10E-01 | 1.81E-01 | 2.69E+00 | 2.77E+00 | 5.36E-01 | no | | intergenic_variant | MODIFIER |
| 4 | 153322588 | AX-170464792 | 0 | C | T | 0.105 | 1.10E-01 | 1.81E-01 | 2.69E+00 | 2.77E+00 | 5.36E-01 | no | | intergenic_variant | MODIFIER |
| 4 | 153322779 | AX-170238208 | 0 | C | T | 0.105 | 1.10E-01 | 1.81E-01 | 2.69E+00 | 2.77E+00 | 5.36E-01 | no | | intergenic_variant | MODIFIER |
| 4 | 153324415 | AX-170394312 | 0 | C | G | 0.105 | 1.10E-01 | 1.81E-01 | 2.69E+00 | 2.77E+00 | 5.36E-01 | no | | intergenic_variant | MODIFIER |
| 4 | 153332439 | AX-170484348 | 0 | A | G | 0.105 | 1.10E-01 | 1.81E-01 | 2.69E+00 | 2.77E+00 | 5.36E-01 | no | | intergenic_variant | MODIFIER |
| 4 | 153340089 | AX-170304496 | 0 | A | G | 0.105 | 1.10E-01 | 1.81E-01 | 2.69E+00 | 2.77E+00 | 5.36E-01 | no | | intergenic_variant | MODIFIER |
| 4 | 153340481 | AX-97845066 | 0 | T | G | 0.105 | 1.10E-01 | 1.81E-01 | 2.69E+00 | 2.77E+00 | 5.36E-01 | no | | intergenic_variant | MODIFIER |
| 4 | 153414160 | AX-97844606 | 0 | T | C | 0.105 | 1.10E-01 | 1.81E-01 | 2.69E+00 | 2.77E+00 | 5.36E-01 | no | Ajap1 | intron_variant | MODIFIER |
| 4 | 153933969 | AX-170337191 | 0 | T | C | 0.105 | 1.10E-01 | 1.81E-01 | 2.69E+00 | 2.77E+00 | 5.36E-01 | no | | intergenic_variant | MODIFIER |
| 4 | 153939301 | AX-97834130 | 0 | G | A | 0.105 | 1.10E-01 | 1.81E-01 | 2.69E+00 | 2.77E+00 | 5.36E-01 | no | | intergenic_variant | MODIFIER |
| 4 | 153940375 | AX-170395175 | 0 | G | A | 0.105 | 1.10E-01 | 1.81E-01 | 2.69E+00 | 2.77E+00 | 5.36E-01 | no | | intergenic_variant | MODIFIER |
| 4 | 153947277 | AX-170104108 | 0 | G | A | 0.105 | 1.10E-01 | 1.81E-01 | 2.69E+00 | 2.77E+00 | 5.36E-01 | no | BC039966 | downstream_gene_variant | MODIFIER |
| 4 | 153987460 | AX-170212030 | 0 | G | A | 0.105 | 1.10E-01 | 1.81E-01 | 2.69E+00 | 2.77E+00 | 5.36E-01 | no | Cep104 | intron_variant | MODIFIER |
| 4 | 153988028 | AX-97819101 | 0 | T | C | 0.105 | 1.10E-01 | 1.81E-01 | 2.69E+00 | 2.77E+00 | 5.36E-01 | no | Cep104 | intron_variant | MODIFIER |
| 4 | 154019324 | AX-170346751 | 0 | G | C | 0.105 | 1.10E-01 | 1.81E-01 | 2.69E+00 | 2.77E+00 | 5.36E-01 | no | Lrrc47 | intron_variant | MODIFIER |
| 4 | 154019443 | AX-97815664 | 0 | A | G | 0.105 | 1.10E-01 | 1.81E-01 | 2.69E+00 | 2.77E+00 | 5.36E-01 | no | Lrrc47 | missense_variant | MODERATE |
| 4 | 154046273 | AX-170071111 | 0 | C | A | 0.105 | 1.10E-01 | 1.81E-01 | 2.69E+00 | 2.77E+00 | 5.36E-01 | no | Ccdc27 | upstream_gene_variant | MODIFIER |
| 4 | 154047827 | AX-170071109 | 0 | G | T | 0.105 | 1.10E-01 | 1.81E-01 | 2.69E+00 | 2.77E+00 | 5.36E-01 | no | Gm13114 | downstream_gene_variant | MODIFIER |
| 4 | 154054281 | AX-170337968 | 0 | T | C | 0.105 | 1.10E-01 | 1.81E-01 | 2.69E+00 | 2.77E+00 | 5.36E-01 | no | Trp73 | downstream_gene_variant | MODIFIER |
| 4 | 154056493 | AX-170473449 | 0 | C | T | 0.105 | 1.10E-01 | 1.81E-01 | 2.69E+00 | 2.77E+00 | 5.36E-01 | no | Trp73 | downstream_gene_variant | MODIFIER |
| 4 | 154069482 | AX-170182020 | 0 | A | G | 0.105 | 1.10E-01 | 1.81E-01 | 2.69E+00 | 2.77E+00 | 5.36E-01 | no | Trp73 | intron_variant | MODIFIER |
| 4 | 154072413 | AX-170024287 | 0 | A | T | 0.105 | 1.10E-01 | 1.81E-01 | 2.69E+00 | 2.77E+00 | 5.36E-01 | no | Trp73 | intron_variant | MODIFIER |
| 4 | 154119880 | AX-170390939 | 0 | A | G | 0.105 | 1.10E-01 | 1.81E-01 | 2.69E+00 | 2.77E+00 | 5.36E-01 | no | Trp73 | intron_variant | MODIFIER |
| 4 | 154125909 | AX-97794611 | 0 | A | G | 0.105 | 1.10E-01 | 1.81E-01 | 2.69E+00 | 2.77E+00 | 5.36E-01 | no | Trp73 | intron_variant | MODIFIER |
| 4 | 154178411 | AX-170024289 | 0 | T | C | 0.105 | 1.10E-01 | 1.81E-01 | 2.69E+00 | 2.77E+00 | 5.36E-01 | no | Megf6 | intron_variant | MODIFIER |
| 4 | 154183320 | AX-170376302 | 0 | C | T | 0.105 | 1.10E-01 | 1.81E-01 | 2.69E+00 | 2.77E+00 | 5.36E-01 | no | Megf6 | intron_variant | MODIFIER |
| 4 | 154188992 | AX-170380297 | 0 | C | T | 0.105 | 1.10E-01 | 1.81E-01 | 2.69E+00 | 2.77E+00 | 5.36E-01 | no | Megf6 | intron_variant | MODIFIER |
| 4 | 154196557 | AX-170280638 | 0 | A | G | 0.105 | 1.10E-01 | 1.81E-01 | 2.69E+00 | 2.77E+00 | 5.36E-01 | no | Megf6 | intron_variant | MODIFIER |
| 4 | 154197167 | AX-170369740 | 0 | A | G | 0.105 | 1.10E-01 | 1.81E-01 | 2.69E+00 | 2.77E+00 | 5.36E-01 | no | Megf6 | intron_variant | MODIFIER |
| 4 | 154207632 | AX-97803543 | 0 | T | C | 0.105 | 1.10E-01 | 1.81E-01 | 2.69E+00 | 2.77E+00 | 5.36E-01 | no | Megf6 | intron_variant | MODIFIER |
| 4 | 154225375 | AX-170487976 | 0 | C | T | 0.105 | 1.10E-01 | 1.81E-01 | 2.69E+00 | 2.77E+00 | 5.36E-01 | no | Megf6 | intron_variant | MODIFIER |
| 4 | 154326749 | AX-170303744 | 0 | T | G | 0.105 | 1.10E-01 | 1.81E-01 | 2.69E+00 | 2.77E+00 | 5.36E-01 | no | Prdm16 | intron_variant | MODIFIER |
| 4 | 154372450 | AX-170134387 | 0 | T | C | 0.105 | 1.10E-01 | 1.81E-01 | 2.69E+00 | 2.77E+00 | 5.36E-01 | no | Prdm16 | intron_variant | MODIFIER |
| 4 | 154430058 | AX-97848100 | 0 | T | C | 0.105 | 1.10E-01 | 1.81E-01 | 2.69E+00 | 2.77E+00 | 5.36E-01 | no | Prdm16 | intron_variant | MODIFIER |
| 4 | 154450781 | AX-170331901 | 0 | T | A | 0.105 | 1.10E-01 | 1.81E-01 | 2.69E+00 | 2.77E+00 | 5.36E-01 | no | Prdm16 | intron_variant | MODIFIER |
| 4 | 154526989 | AX-170148478 | 0 | T | C | 0.105 | 1.10E-01 | 1.81E-01 | 2.69E+00 | 2.77E+00 | 5.36E-01 | no | Prdm16 | intron_variant | MODIFIER |
| 4 | 154533779 | AX-170361534 | 0 | A | G | 0.105 | 1.10E-01 | 1.81E-01 | 2.69E+00 | 2.77E+00 | 5.36E-01 | no | Prdm16 | intron_variant | MODIFIER |
| 4 | 154539849 | AX-97852302 | 0 | A | G | 0.105 | 1.10E-01 | 1.81E-01 | 2.69E+00 | 2.77E+00 | 5.36E-01 | no | Prdm16 | intron_variant | MODIFIER |
| 4 | 154576933 | AX-97830992 | 0 | G | A | 0.105 | 1.10E-01 | 1.81E-01 | 2.69E+00 | 2.77E+00 | 5.36E-01 | no | Prdm16 | intron_variant | MODIFIER |
| 4 | 154589189 | AX-170180354 | 0 | A | G | 0.105 | 1.10E-01 | 1.81E-01 | 2.69E+00 | 2.77E+00 | 5.36E-01 | no | Prdm16 | intron_variant | MODIFIER |
| 4 | 154594213 | AX-170396685 | 0 | T | C | 0.105 | 1.10E-01 | 1.81E-01 | 2.69E+00 | 2.77E+00 | 5.36E-01 | no | Prdm16 | intron_variant | MODIFIER |

| | | | | | | | | | | | | | | | |
|---|---|---|---|---|---|---|---|---|---|---|---|---|---|---|---|
| 4 | 154597888 | AX-170071121 | 0 | A | G | 0.105 | 1.10E-01 | 1.81E-01 | 2.69E+00 | 2.77E+00 | 5.36E-01 | no | Prdm16 | intron_variant | MODIFIER |
| 7 | 66250564 | AX-170423277 | 0 | A | G | 0.404 | -5.93E-02 | 9.72E-02 | 2.78E+00 | 2.86E+00 | 5.36E-01 | no | Lrrk1 | intron_variant,non_coding_transcript_variant | MODIFIER |
| 7 | 66309541 | AX-97809109 | 0 | T | C | 0.404 | -5.93E-02 | 9.72E-02 | 2.78E+00 | 2.86E+00 | 5.36E-01 | no | Lrrk1 | intron_variant | MODIFIER |
| 7 | 66341907 | AX-170240413 | 0 | A | G | 0.404 | -5.93E-02 | 9.72E-02 | 2.78E+00 | 2.86E+00 | 5.36E-01 | no | Lrrk1 | intron_variant | MODIFIER |
| 7 | 66342748 | AX-170332591 | 0 | T | G | 0.404 | -5.93E-02 | 9.72E-02 | 2.78E+00 | 2.86E+00 | 5.36E-01 | no | Lrrk1 | intron_variant | MODIFIER |
| 7 | 66343308 | AX-170282290 | 0 | G | A | 0.404 | -5.93E-02 | 9.72E-02 | 2.78E+00 | 2.86E+00 | 5.36E-01 | no | Lrrk1 | intron_variant | MODIFIER |
| 7 | 66371162 | AX-170029186 | 0 | C | T | 0.404 | -5.93E-02 | 9.72E-02 | 2.78E+00 | 2.86E+00 | 5.36E-01 | no | Lrrk1 | intron_variant | MODIFIER |
| 7 | 66390229 | AX-170095965 | 0 | A | G | 0.404 | -5.93E-02 | 9.72E-02 | 2.78E+00 | 2.86E+00 | 5.36E-01 | no | Lrrk1 | upstream_gene_variant | MODIFIER |
| 7 | 66395813 | AX-170481776 | 0 | C | T | 0.404 | -5.93E-02 | 9.72E-02 | 2.78E+00 | 2.86E+00 | 5.36E-01 | no | Aldh1a3 | intron_variant | MODIFIER |
| 7 | 66397529 | AX-170147184 | 0 | A | G | 0.404 | -5.93E-02 | 9.72E-02 | 2.78E+00 | 2.86E+00 | 5.36E-01 | no | Aldh1a3 | intron_variant | MODIFIER |
| 7 | 66403052 | AX-170311834 | 0 | A | G | 0.404 | -5.93E-02 | 9.72E-02 | 2.78E+00 | 2.86E+00 | 5.36E-01 | no | Aldh1a3 | intron_variant | MODIFIER |
| 7 | 66403756 | AX-170197946 | 0 | A | G | 0.404 | -5.93E-02 | 9.72E-02 | 2.78E+00 | 2.86E+00 | 5.36E-01 | no | Aldh1a3 | intron_variant | MODIFIER |
| 7 | 66403827 | AX-97786193 | 0 | T | C | 0.404 | -5.93E-02 | 9.72E-02 | 2.78E+00 | 2.86E+00 | 5.36E-01 | no | Aldh1a3 | intron_variant | MODIFIER |
| 7 | 66409516 | AX-170493564 | 0 | G | C | 0.404 | -5.93E-02 | 9.72E-02 | 2.78E+00 | 2.86E+00 | 5.36E-01 | no | Aldh1a3 | intron_variant | MODIFIER |
| 7 | 66449992 | AX-170029184 | 0 | G | A | 0.404 | -5.93E-02 | 9.72E-02 | 2.78E+00 | 2.86E+00 | 5.36E-01 | no | | regulatory_region_variant | MODIFIER |
| 15 | 70606884 | AX-97852512 | 0 | C | T | 0.368 | 4.70E-02 | 7.46E-02 | 2.53E+00 | 2.61E+00 | 5.36E-01 | no | | intergenic_variant | MODIFIER |
| 15 | 70833234 | AX-170093297 | 0 | A | G | 0.368 | 4.70E-02 | 7.46E-02 | 2.53E+00 | 2.61E+00 | 5.36E-01 | no | | intergenic_variant | MODIFIER |
| 15 | 70834808 | AX-170093298 | 0 | A | G | 0.368 | 4.70E-02 | 7.46E-02 | 2.53E+00 | 2.61E+00 | 5.36E-01 | no | | intergenic_variant | MODIFIER |
| 15 | 70847768 | AX-170122962 | 0 | T | A | 0.368 | 4.70E-02 | 7.46E-02 | 2.53E+00 | 2.61E+00 | 5.36E-01 | no | | intergenic_variant | MODIFIER |
| 15 | 70920477 | AX-97830362 | 0 | C | T | 0.368 | 4.70E-02 | 7.46E-02 | 2.53E+00 | 2.61E+00 | 5.36E-01 | no | | regulatory_region_variant | MODIFIER |
| 15 | 71032501 | AX-170491905 | 0 | G | T | 0.368 | 4.70E-02 | 7.46E-02 | 2.53E+00 | 2.61E+00 | 5.36E-01 | no | | intergenic_variant | MODIFIER |
| 15 | 71259129 | AX-97847822 | 0 | A | G | 0.368 | 4.70E-02 | 7.46E-02 | 2.53E+00 | 2.61E+00 | 5.36E-01 | no | | intergenic_variant | MODIFIER |
| 15 | 71275663 | AX-170011965 | 0 | G | A | 0.368 | 4.70E-02 | 7.46E-02 | 2.53E+00 | 2.61E+00 | 5.36E-01 | no | | intergenic_variant | MODIFIER |
| 15 | 71289351 | AX-170011962 | 0 | C | A | 0.368 | 4.70E-02 | 7.46E-02 | 2.53E+00 | 2.61E+00 | 5.36E-01 | no | | intergenic_variant | MODIFIER |
| 15 | 71297130 | AX-97810448 | 0 | C | T | 0.368 | 4.70E-02 | 7.46E-02 | 2.53E+00 | 2.61E+00 | 5.36E-01 | no | | intergenic_variant | MODIFIER |
| 15 | 71311113 | AX-170011963 | 0 | C | T | 0.368 | 4.70E-02 | 7.46E-02 | 2.53E+00 | 2.61E+00 | 5.36E-01 | no | Gm5217 | non_coding_transcript_exon_variant | MODIFIER |
| 15 | 71312908 | AX-170502414 | 0 | G | A | 0.368 | 4.70E-02 | 7.46E-02 | 2.53E+00 | 2.61E+00 | 5.36E-01 | no | Gm5217 | downstream_gene_variant | MODIFIER |
| 15 | 71315723 | AX-170341998 | 0 | T | C | 0.368 | 4.70E-02 | 7.46E-02 | 2.53E+00 | 2.61E+00 | 5.36E-01 | no | Gm5217 | downstream_gene_variant | MODIFIER |
| 15 | 71318097 | AX-97832273 | 0 | T | C | 0.368 | 4.70E-02 | 7.46E-02 | 2.53E+00 | 2.61E+00 | 5.36E-01 | no | | intergenic_variant | MODIFIER |
| 15 | 71370122 | AX-97820979 | 0 | C | A | 0.368 | 4.70E-02 | 7.46E-02 | 2.53E+00 | 2.61E+00 | 5.36E-01 | no | | intergenic_variant | MODIFIER |
| 15 | 71442639 | AX-170331534 | 0 | C | T | 0.368 | 4.70E-02 | 7.46E-02 | 2.53E+00 | 2.61E+00 | 5.36E-01 | no | Fam135b | 3_prime_UTR_variant | MODIFIER |
| 15 | 71443197 | AX-170277721 | 0 | G | C | 0.368 | 4.70E-02 | 7.46E-02 | 2.53E+00 | 2.61E+00 | 5.36E-01 | no | Fam135b | 3_prime_UTR_variant | MODIFIER |
| 15 | 71481668 | AX-170525850 | 0 | T | C | 0.368 | 4.70E-02 | 7.46E-02 | 2.53E+00 | 2.61E+00 | 5.36E-01 | no | Fam135b | intron_variant | MODIFIER |
| 15 | 71485141 | AX-170273440 | 0 | C | A | 0.368 | 4.70E-02 | 7.46E-02 | 2.53E+00 | 2.61E+00 | 5.36E-01 | no | Fam135b | intron_variant | MODIFIER |
| 15 | 71504445 | AX-170055913 | 0 | A | G | 0.368 | 4.70E-02 | 7.46E-02 | 2.53E+00 | 2.61E+00 | 5.36E-01 | no | Fam135b | intron_variant | MODIFIER |
| 15 | 71527338 | AX-170118624 | 0 | A | C | 0.368 | 4.70E-02 | 7.46E-02 | 2.53E+00 | 2.61E+00 | 5.36E-01 | no | Fam135b | intron_variant | MODIFIER |
| 15 | 71540865 | AX-170471306 | 0 | C | T | 0.368 | 4.70E-02 | 7.46E-02 | 2.53E+00 | 2.61E+00 | 5.36E-01 | no | Fam135b | intron_variant | MODIFIER |
| 15 | 71541464 | AX-170430327 | 0 | T | G | 0.368 | 4.70E-02 | 7.46E-02 | 2.53E+00 | 2.61E+00 | 5.36E-01 | no | Fam135b | intron_variant | MODIFIER |
| 15 | 71542922 | AX-170429929 | 0 | C | T | 0.368 | 4.70E-02 | 7.46E-02 | 2.53E+00 | 2.61E+00 | 5.36E-01 | no | Fam135b | intron_variant | MODIFIER |
| 15 | 71544062 | AX-97832930 | 0 | A | G | 0.368 | 4.70E-02 | 7.46E-02 | 2.53E+00 | 2.61E+00 | 5.36E-01 | no | Fam135b | intron_variant | MODIFIER |
| 15 | 71544159 | AX-170214466 | 0 | C | T | 0.368 | 4.70E-02 | 7.46E-02 | 2.53E+00 | 2.61E+00 | 5.36E-01 | no | Fam135b | intron_variant | MODIFIER |
| 15 | 71552329 | AX-170411228 | 0 | T | G | 0.368 | 4.70E-02 | 7.46E-02 | 2.53E+00 | 2.61E+00 | 5.36E-01 | no | Fam135b | intron_variant | MODIFIER |
| 15 | 71554067 | AX-170340542 | 0 | T | C | 0.368 | 4.70E-02 | 7.46E-02 | 2.53E+00 | 2.61E+00 | 5.36E-01 | no | Fam135b | intron_variant | MODIFIER |
| 8 | 101671260 | AX-170031395 | 1 | G | A | 0.232 | 6.95E-02 | 1.14E-01 | 2.71E+00 | 2.78E+00 | 5.36E-01 | no | | intergenic_variant | MODIFIER |
| 1 | 21691129 | AX-170480513 | 1 | T | C | 0.393 | 5.06E-02 | 8.09E-02 | 2.57E+00 | 2.62E+00 | 5.36E-01 | no | Kcnq5 | intron_variant | MODIFIER |
| 10 | 30886592 | AX-170280584 | 1 | C | T | 0.179 | 5.97E-02 | 9.85E-02 | 2.77E+00 | 2.87E+00 | 5.36E-01 | no | | intergenic_variant | MODIFIER |
| 7 | 16109430 | AX-170350990 | 0 | G | T | 0.377 | 5.27E-02 | 8.45E-02 | 2.55E+00 | 2.66E+00 | 5.36E-01 | no | Napa | intron_variant | MODIFIER |
| 7 | 45946623 | AX-170464176 | 0 | C | T | 0.325 | 5.41E-02 | 8.82E-02 | 2.62E+00 | 2.71E+00 | 5.37E-01 | no | Ccdc114 | intron_variant | MODIFIER |
| 1 | 25792617 | AX-170000128 | 1 | G | A | 0.42 | 5.15E-02 | 8.43E-02 | 2.67E+00 | 2.74E+00 | 5.37E-01 | no | Adgrb3 | intron_variant | MODIFIER |
| 13 | 54715388 | AX-170253382 | 0 | T | A | 0.518 | 4.32E-02 | 6.87E-02 | 2.52E+00 | 2.63E+00 | 5.37E-01 | no | Cdhr2 | intron_variant | MODIFIER |
| 4 | 151284270 | AX-170071051 | 0 | A | G | 0.114 | 1.12E-01 | 1.84E-01 | 2.70E+00 | 2.78E+00 | 5.37E-01 | no | Camta1 | intron_variant | MODIFIER |
| 4 | 151286312 | AX-170071052 | 0 | A | G | 0.114 | 1.12E-01 | 1.84E-01 | 2.70E+00 | 2.78E+00 | 5.37E-01 | no | Camta1 | intron_variant | MODIFIER |
| 4 | 151286481 | AX-170336754 | 0 | G | A | 0.114 | 1.12E-01 | 1.84E-01 | 2.70E+00 | 2.78E+00 | 5.37E-01 | no | Camta1 | intron_variant | MODIFIER |
| 4 | 151286929 | AX-170483430 | 0 | C | T | 0.114 | 1.12E-01 | 1.84E-01 | 2.70E+00 | 2.78E+00 | 5.37E-01 | no | Camta1 | intron_variant | MODIFIER |

| | | | | | | | | | | | | | | | |
|---|---|---|---|---|---|---|---|---|---|---|---|---|---|---|---|
| 4 | 151362194 | AX-170412544 | 0 | C | T | 0.114 | 1.12E-01 | 1.84E-01 | 2.70E+00 | 2.78E+00 | 5.37E-01 | no | Camta1 | intron_variant | MODIFIER |
| 4 | 151371208 | AX-170024257 | 0 | G | T | 0.114 | 1.12E-01 | 1.84E-01 | 2.70E+00 | 2.78E+00 | 5.37E-01 | no | Camta1 | intron_variant | MODIFIER |
| 4 | 151371232 | AX-170024258 | 0 | C | G | 0.114 | 1.12E-01 | 1.84E-01 | 2.70E+00 | 2.78E+00 | 5.37E-01 | no | Camta1 | intron_variant | MODIFIER |
| 4 | 151382681 | AX-170203909 | 0 | C | T | 0.114 | 1.12E-01 | 1.84E-01 | 2.70E+00 | 2.78E+00 | 5.37E-01 | no | Camta1 | intron_variant | MODIFIER |
| 4 | 151475307 | AX-170448899 | 0 | T | C | 0.114 | 1.12E-01 | 1.84E-01 | 2.70E+00 | 2.78E+00 | 5.37E-01 | no | Camta1 | intron_variant | MODIFIER |
| 4 | 151476079 | AX-170291691 | 0 | C | G | 0.114 | 1.12E-01 | 1.84E-01 | 2.70E+00 | 2.78E+00 | 5.37E-01 | no | Camta1 | intron_variant | MODIFIER |
| 4 | 151481316 | AX-170431458 | 0 | T | G | 0.114 | 1.12E-01 | 1.84E-01 | 2.70E+00 | 2.78E+00 | 5.37E-01 | no | Camta1 | intron_variant | MODIFIER |
| 4 | 151485333 | AX-170381015 | 0 | G | A | 0.114 | 1.12E-01 | 1.84E-01 | 2.70E+00 | 2.78E+00 | 5.37E-01 | no | Camta1 | intron_variant | MODIFIER |
| 4 | 151487854 | AX-170144174 | 0 | G | A | 0.114 | 1.12E-01 | 1.84E-01 | 2.70E+00 | 2.78E+00 | 5.37E-01 | no | Camta1 | intron_variant | MODIFIER |
| 4 | 151498047 | AX-170258995 | 0 | C | T | 0.114 | 1.12E-01 | 1.84E-01 | 2.70E+00 | 2.78E+00 | 5.37E-01 | no | Camta1 | intron_variant | MODIFIER |
| 4 | 151505117 | AX-170212579 | 0 | G | A | 0.114 | 1.12E-01 | 1.84E-01 | 2.70E+00 | 2.78E+00 | 5.37E-01 | no | Camta1 | intron_variant | MODIFIER |
| 4 | 151506753 | AX-170336179 | 0 | A | G | 0.114 | 1.12E-01 | 1.84E-01 | 2.70E+00 | 2.78E+00 | 5.37E-01 | no | Camta1 | intron_variant | MODIFIER |
| 4 | 151538502 | AX-170478020 | 0 | C | T | 0.114 | 1.12E-01 | 1.84E-01 | 2.70E+00 | 2.78E+00 | 5.37E-01 | no | Camta1 | intron_variant | MODIFIER |
| 4 | 151564579 | AX-170267264 | 0 | C | T | 0.114 | 1.12E-01 | 1.84E-01 | 2.70E+00 | 2.78E+00 | 5.37E-01 | no | Camta1 | intron_variant | MODIFIER |
| 4 | 151569468 | AX-170380807 | 0 | G | A | 0.114 | 1.12E-01 | 1.84E-01 | 2.70E+00 | 2.78E+00 | 5.37E-01 | no | Camta1 | intron_variant | MODIFIER |
| 4 | 151585401 | AX-170511247 | 0 | G | T | 0.114 | 1.12E-01 | 1.84E-01 | 2.70E+00 | 2.78E+00 | 5.37E-01 | no | Camta1 | intron_variant | MODIFIER |
| 4 | 151589200 | AX-170317989 | 0 | A | C | 0.114 | 1.12E-01 | 1.84E-01 | 2.70E+00 | 2.78E+00 | 5.37E-01 | no | Camta1 | intron_variant | MODIFIER |
| 4 | 151590887 | AX-170462201 | 0 | G | C | 0.114 | 1.12E-01 | 1.84E-01 | 2.70E+00 | 2.78E+00 | 5.37E-01 | no | Camta1 | intron_variant | MODIFIER |
| 4 | 151608124 | AX-170271088 | 0 | C | T | 0.114 | 1.12E-01 | 1.84E-01 | 2.70E+00 | 2.78E+00 | 5.37E-01 | no | Camta1 | intron_variant | MODIFIER |
| 4 | 151628305 | AX-170474919 | 0 | C | T | 0.114 | 1.12E-01 | 1.84E-01 | 2.70E+00 | 2.78E+00 | 5.37E-01 | no | Camta1 | intron_variant | MODIFIER |
| 4 | 151639667 | AX-170164542 | 0 | C | T | 0.114 | 1.12E-01 | 1.84E-01 | 2.70E+00 | 2.78E+00 | 5.37E-01 | no | Camta1 | intron_variant | MODIFIER |
| 4 | 151648424 | AX-170224821 | 0 | A | T | 0.114 | 1.12E-01 | 1.84E-01 | 2.70E+00 | 2.78E+00 | 5.37E-01 | no | Camta1 | intron_variant | MODIFIER |
| 4 | 151665428 | AX-97852010 | 0 | G | A | 0.114 | 1.12E-01 | 1.84E-01 | 2.70E+00 | 2.78E+00 | 5.37E-01 | no | Camta1 | intron_variant | MODIFIER |
| 4 | 151673324 | AX-170394554 | 0 | C | T | 0.114 | 1.12E-01 | 1.84E-01 | 2.70E+00 | 2.78E+00 | 5.37E-01 | no | Camta1 | intron_variant | MODIFIER |
| 4 | 151680562 | AX-170173474 | 0 | T | C | 0.114 | 1.12E-01 | 1.84E-01 | 2.70E+00 | 2.78E+00 | 5.37E-01 | no | Camta1 | intron_variant | MODIFIER |
| 4 | 151808923 | AX-170269236 | 0 | T | A | 0.114 | 1.12E-01 | 1.84E-01 | 2.70E+00 | 2.78E+00 | 5.37E-01 | no | Camta1 | intron_variant | MODIFIER |
| 4 | 151968275 | AX-170518469 | 0 | G | C | 0.114 | 1.12E-01 | 1.84E-01 | 2.70E+00 | 2.78E+00 | 5.37E-01 | no | Dnajc11 | intron_variant | MODIFIER |
| 4 | 154100672 | AX-97853166 | 0 | T | C | 0.114 | 1.12E-01 | 1.84E-01 | 2.70E+00 | 2.78E+00 | 5.37E-01 | no | Trp73 | upstream_gene_variant | MODIFIER |
| 4 | 154101046 | AX-170024284 | 0 | A | G | 0.114 | 1.12E-01 | 1.84E-01 | 2.70E+00 | 2.78E+00 | 5.37E-01 | no | Trp73 | upstream_gene_variant | MODIFIER |
| 4 | 154101339 | AX-170299425 | 0 | G | T | 0.114 | 1.12E-01 | 1.84E-01 | 2.70E+00 | 2.78E+00 | 5.37E-01 | no | Trp73 | upstream_gene_variant | MODIFIER |
| 4 | 154111146 | AX-170487269 | 0 | G | A | 0.114 | 1.12E-01 | 1.84E-01 | 2.70E+00 | 2.78E+00 | 5.37E-01 | no | Trp73 | intron_variant | MODIFIER |
| 4 | 154112807 | AX-170180450 | 0 | C | T | 0.114 | 1.12E-01 | 1.84E-01 | 2.70E+00 | 2.78E+00 | 5.37E-01 | no | Trp73 | intron_variant | MODIFIER |
| 4 | 154197217 | AX-170440316 | 0 | G | A | 0.114 | 1.12E-01 | 1.84E-01 | 2.70E+00 | 2.78E+00 | 5.37E-01 | no | Megf6 | intron_variant | MODIFIER |
| 4 | 154197997 | AX-170271600 | 0 | T | C | 0.114 | 1.12E-01 | 1.84E-01 | 2.70E+00 | 2.78E+00 | 5.37E-01 | no | Megf6 | intron_variant | MODIFIER |
| 4 | 154204454 | AX-170281962 | 0 | G | A | 0.114 | 1.12E-01 | 1.84E-01 | 2.70E+00 | 2.78E+00 | 5.37E-01 | no | Megf6 | intron_variant | MODIFIER |
| 4 | 154257171 | AX-170177982 | 0 | G | A | 0.114 | 1.12E-01 | 1.84E-01 | 2.70E+00 | 2.78E+00 | 5.37E-01 | no | Megf6 | intron_variant | MODIFIER |
| 4 | 154269607 | AX-170307170 | 0 | C | T | 0.114 | 1.12E-01 | 1.84E-01 | 2.70E+00 | 2.78E+00 | 5.37E-01 | no | Megf6 | intron_variant | MODIFIER |
| 4 | 154322833 | AX-97804780 | 0 | G | A | 0.114 | 1.12E-01 | 1.84E-01 | 2.70E+00 | 2.78E+00 | 5.37E-01 | no | Prdm16 | intron_variant | MODIFIER |
| 4 | 154323585 | AX-170409122 | 0 | T | C | 0.114 | 1.12E-01 | 1.84E-01 | 2.70E+00 | 2.78E+00 | 5.37E-01 | no | Prdm16 | intron_variant | MODIFIER |
| 4 | 154323910 | AX-170360577 | 0 | G | A | 0.114 | 1.12E-01 | 1.84E-01 | 2.70E+00 | 2.78E+00 | 5.37E-01 | no | Prdm16 | intron_variant | MODIFIER |
| 4 | 154529786 | AX-170377091 | 0 | C | T | 0.114 | 1.12E-01 | 1.84E-01 | 2.70E+00 | 2.78E+00 | 5.37E-01 | no | Prdm16 | intron_variant | MODIFIER |
| 4 | 154540042 | AX-170197657 | 0 | G | C | 0.114 | 1.12E-01 | 1.84E-01 | 2.70E+00 | 2.78E+00 | 5.37E-01 | no | Prdm16 | intron_variant | MODIFIER |
| 4 | 154551639 | AX-170411493 | 0 | G | A | 0.114 | 1.12E-01 | 1.84E-01 | 2.70E+00 | 2.78E+00 | 5.37E-01 | no | Prdm16 | intron_variant | MODIFIER |
| 4 | 154575679 | AX-170512086 | 0 | C | T | 0.114 | 1.12E-01 | 1.84E-01 | 2.70E+00 | 2.78E+00 | 5.37E-01 | no | Prdm16 | intron_variant | MODIFIER |
| 4 | 154579329 | AX-170187388 | 0 | C | T | 0.114 | 1.12E-01 | 1.84E-01 | 2.70E+00 | 2.78E+00 | 5.37E-01 | no | Prdm16 | intron_variant | MODIFIER |
| 4 | 154594233 | AX-170071120 | 0 | G | C | 0.114 | 1.12E-01 | 1.84E-01 | 2.70E+00 | 2.78E+00 | 5.37E-01 | no | Prdm16 | intron_variant | MODIFIER |
| 4 | 154596368 | AX-170071119 | 0 | C | G | 0.114 | 1.12E-01 | 1.84E-01 | 2.70E+00 | 2.78E+00 | 5.37E-01 | no | Prdm16 | intron_variant | MODIFIER |
| 4 | 154707617 | AX-170253864 | 0 | C | A | 0.114 | 1.12E-01 | 1.84E-01 | 2.70E+00 | 2.78E+00 | 5.37E-01 | no | | regulatory_region_variant | MODIFIER |
| 4 | 154727748 | AX-170371115 | 0 | C | A | 0.114 | 1.12E-01 | 1.84E-01 | 2.70E+00 | 2.78E+00 | 5.37E-01 | no | | regulatory_region_variant | MODIFIER |
| 4 | 154750253 | AX-170351210 | 0 | T | C | 0.114 | 1.12E-01 | 1.84E-01 | 2.70E+00 | 2.78E+00 | 5.37E-01 | no | | regulatory_region_variant | MODIFIER |
| 4 | 154754521 | AX-170106612 | 0 | A | G | 0.114 | 1.12E-01 | 1.84E-01 | 2.70E+00 | 2.78E+00 | 5.37E-01 | no | | regulatory_region_variant | MODIFIER |
| 4 | 154755097 | AX-170282111 | 0 | G | A | 0.114 | 1.12E-01 | 1.84E-01 | 2.70E+00 | 2.78E+00 | 5.37E-01 | no | | regulatory_region_variant | MODIFIER |
| 4 | 154758258 | AX-97822544 | 0 | T | C | 0.114 | 1.12E-01 | 1.84E-01 | 2.70E+00 | 2.78E+00 | 5.37E-01 | no | | intergenic_variant | MODIFIER |
| 4 | 154799715 | AX-170071128 | 0 | T | C | 0.114 | 1.12E-01 | 1.84E-01 | 2.70E+00 | 2.78E+00 | 5.37E-01 | no | | intergenic_variant | MODIFIER |

| | | | | | | | | | | | | | | | |
|---|---|---|---|---|---|---|---|---|---|---|---|---|---|---|---|
| 4 | 154804358 | AX-170071127 | 0 | C | A | 0.114 | 1.12E-01 | 1.84E-01 | 2.70E+00 | 2.78E+00 | 5.37E-01 | no | | intergenic_variant | MODIFIER |
| 4 | 154817878 | AX-170095203 | 0 | T | C | 0.114 | 1.12E-01 | 1.84E-01 | 2.70E+00 | 2.78E+00 | 5.37E-01 | no | | intergenic_variant | MODIFIER |
| 4 | 154820856 | AX-170450071 | 0 | C | A | 0.114 | 1.12E-01 | 1.84E-01 | 2.70E+00 | 2.78E+00 | 5.37E-01 | no | | intergenic_variant | MODIFIER |
| 4 | 154823258 | AX-170192759 | 0 | T | A | 0.114 | 1.12E-01 | 1.84E-01 | 2.70E+00 | 2.78E+00 | 5.37E-01 | no | | intergenic_variant | MODIFIER |
| 4 | 154823678 | AX-170533969 | 0 | T | C | 0.114 | 1.12E-01 | 1.84E-01 | 2.70E+00 | 2.78E+00 | 5.37E-01 | no | | intergenic_variant | MODIFIER |
| 4 | 154824897 | AX-170531719 | 0 | A | C | 0.114 | 1.12E-01 | 1.84E-01 | 2.70E+00 | 2.78E+00 | 5.37E-01 | no | | intergenic_variant | MODIFIER |
| 4 | 154826257 | AX-170148768 | 0 | A | C | 0.114 | 1.12E-01 | 1.84E-01 | 2.70E+00 | 2.78E+00 | 5.37E-01 | no | | regulatory_region_variant | MODIFIER |
| 4 | 154827215 | AX-170175157 | 0 | T | C | 0.114 | 1.12E-01 | 1.84E-01 | 2.70E+00 | 2.78E+00 | 5.37E-01 | no | | regulatory_region_variant | MODIFIER |
| 4 | 154828174 | AX-170149003 | 0 | G | A | 0.114 | 1.12E-01 | 1.84E-01 | 2.70E+00 | 2.78E+00 | 5.37E-01 | no | | regulatory_region_variant | MODIFIER |
| 4 | 154867926 | AX-170450994 | 0 | G | A | 0.114 | 1.12E-01 | 1.84E-01 | 2.70E+00 | 2.78E+00 | 5.37E-01 | no | Ttc34 | downstream_gene_variant | MODIFIER |
| 4 | 154869706 | AX-170164171 | 0 | A | C | 0.114 | 1.12E-01 | 1.84E-01 | 2.70E+00 | 2.78E+00 | 5.37E-01 | no | Ttc34 | downstream_gene_variant | MODIFIER |
| 4 | 154871382 | AX-170354507 | 0 | A | G | 0.114 | 1.12E-01 | 1.84E-01 | 2.70E+00 | 2.78E+00 | 5.37E-01 | no | Ttc34 | downstream_gene_variant | MODIFIER |
| 4 | 154872258 | AX-170440340 | 0 | T | C | 0.114 | 1.12E-01 | 1.84E-01 | 2.70E+00 | 2.78E+00 | 5.37E-01 | no | Mmel1 | intron_variant | MODIFIER |
| 9 | 80036620 | AX-170307179 | 1 | T | A | 0.214 | 7.49E-02 | 1.22E-01 | 2.82E+00 | 2.89E+00 | 5.37E-01 | no | Gm25433 | upstream_gene_variant | MODIFIER |
| 1 | 173312471 | AX-170043617 | 1 | G | A | 0.214 | -8.20E-02 | 1.30E-01 | 2.99E+00 | 3.11E+00 | 5.37E-01 | no | | intergenic_variant | MODIFIER |
| 1 | 20197225 | AX-170040555 | 0 | G | A | 0.404 | 5.21E-02 | 8.42E-02 | 2.61E+00 | 2.66E+00 | 5.37E-01 | no | Pkhd1 | intron_variant | MODIFIER |
| 4 | 84046747 | AX-97800414 | 1 | T | C | 0.232 | 6.61E-02 | 1.05E-01 | 2.53E+00 | 2.60E+00 | 5.37E-01 | no | 6030471H07Rik | upstream_gene_variant | MODIFIER |
| 12 | 81688889 | AX-170007145 | 0 | A | G | 0.044 | -1.28E-01 | 2.08E-01 | 2.59E+00 | 2.70E+00 | 5.37E-01 | no | | regulatory_region_variant | MODIFIER |
| 10 | 68080321 | AX-170435973 | 1 | C | A | 0.42 | -4.97E-02 | 8.09E-02 | 2.62E+00 | 2.70E+00 | 5.37E-01 | no | | regulatory_region_variant | MODIFIER |
| 14 | 119175706 | AX-170133261 | 0 | T | A | 0.316 | 4.34E-02 | 7.05E-02 | 2.59E+00 | 2.70E+00 | 5.37E-01 | no | Hs6st3 | intron_variant | MODIFIER |
| 5 | 33441729 | AX-170122266 | 0 | T | C | 0.333 | 6.61E-02 | 1.08E-01 | 2.69E+00 | 2.74E+00 | 5.37E-01 | no | Gm43851 | upstream_gene_variant | MODIFIER |
| 8 | 124932729 | AX-170443616 | 0 | A | G | 0.053 | -1.19E-01 | 1.96E-01 | 2.66E+00 | 2.77E+00 | 5.38E-01 | no | Egln1 | intron_variant | MODIFIER |
| 15 | 82947008 | AX-170518057 | 0 | C | G | 0.553 | 5.24E-02 | 8.32E-02 | 2.53E+00 | 2.60E+00 | 5.38E-01 | no | Tcf20 | intron_variant | MODIFIER |
| 8 | 97983091 | AX-170035788 | 0 | C | A | 0.219 | 7.17E-02 | 1.18E-01 | 2.72E+00 | 2.79E+00 | 5.38E-01 | no | | intergenic_variant | MODIFIER |
| 7 | 37381896 | AX-97787141 | 1 | A | G | 0.339 | 5.66E-02 | 9.15E-02 | 2.58E+00 | 2.66E+00 | 5.38E-01 | no | | intergenic_variant | MODIFIER |
| 1 | 23257693 | AX-170000081 | 0 | A | G | 0.43 | 5.17E-02 | 8.43E-02 | 2.62E+00 | 2.68E+00 | 5.38E-01 | no | Gm6420 | upstream_gene_variant | MODIFIER |
| 1 | 23291334 | AX-168378924 | 0 | G | A | 0.43 | 5.17E-02 | 8.43E-02 | 2.62E+00 | 2.68E+00 | 5.38E-01 | no | Mir30c-2 | upstream_gene_variant | MODIFIER |
| 1 | 23344369 | AX-170141809 | 0 | T | A | 0.43 | 5.17E-02 | 8.43E-02 | 2.62E+00 | 2.68E+00 | 5.38E-01 | no | Gm20954 | downstream_gene_variant | MODIFIER |
| 1 | 23354566 | AX-97824233 | 0 | C | T | 0.43 | 5.17E-02 | 8.43E-02 | 2.62E+00 | 2.68E+00 | 5.38E-01 | no | | intergenic_variant | MODIFIER |
| 1 | 23378841 | AX-170477604 | 0 | A | G | 0.43 | 5.17E-02 | 8.43E-02 | 2.62E+00 | 2.68E+00 | 5.38E-01 | no | Ogfrl1 | intron_variant | MODIFIER |
| 1 | 23379335 | AX-97839459 | 0 | G | A | 0.43 | 5.17E-02 | 8.43E-02 | 2.62E+00 | 2.68E+00 | 5.38E-01 | no | Ogfrl1 | intron_variant | MODIFIER |
| 1 | 23498510 | AX-170266419 | 0 | G | A | 0.43 | 5.17E-02 | 8.43E-02 | 2.62E+00 | 2.68E+00 | 5.38E-01 | no | | intergenic_variant | MODIFIER |
| 1 | 23506980 | AX-170211206 | 0 | G | A | 0.43 | 5.17E-02 | 8.43E-02 | 2.62E+00 | 2.68E+00 | 5.38E-01 | no | | intergenic_variant | MODIFIER |
| 1 | 23589618 | AX-170000093 | 0 | A | G | 0.43 | 5.17E-02 | 8.43E-02 | 2.62E+00 | 2.68E+00 | 5.38E-01 | no | Gm28822 | intron_variant,non_coding_transcript_variant | MODIFIER |
| 1 | 23590127 | AX-170000088 | 0 | A | G | 0.43 | 5.17E-02 | 8.43E-02 | 2.62E+00 | 2.68E+00 | 5.38E-01 | no | Gm28822 | intron_variant,non_coding_transcript_variant | MODIFIER |
| 1 | 23683509 | AX-170000098 | 0 | A | G | 0.43 | 5.17E-02 | 8.43E-02 | 2.62E+00 | 2.68E+00 | 5.38E-01 | no | | intergenic_variant | MODIFIER |
| 1 | 23690398 | AX-170000087 | 0 | A | G | 0.43 | 5.17E-02 | 8.43E-02 | 2.62E+00 | 2.68E+00 | 5.38E-01 | no | | intergenic_variant | MODIFIER |
| 1 | 23692058 | AX-170000096 | 0 | T | C | 0.43 | 5.17E-02 | 8.43E-02 | 2.62E+00 | 2.68E+00 | 5.38E-01 | no | | intergenic_variant | MODIFIER |
| 1 | 23701270 | AX-170172784 | 0 | A | G | 0.43 | 5.17E-02 | 8.43E-02 | 2.62E+00 | 2.68E+00 | 5.38E-01 | no | | intergenic_variant | MODIFIER |
| 1 | 23730862 | AX-170091433 | 0 | T | C | 0.43 | 5.17E-02 | 8.43E-02 | 2.62E+00 | 2.68E+00 | 5.38E-01 | no | | intergenic_variant | MODIFIER |
| 1 | 23730960 | AX-170479041 | 0 | C | A | 0.43 | 5.17E-02 | 8.43E-02 | 2.62E+00 | 2.68E+00 | 5.38E-01 | no | | intergenic_variant | MODIFIER |
| 1 | 23742803 | AX-170321910 | 0 | G | A | 0.43 | 5.17E-02 | 8.43E-02 | 2.62E+00 | 2.68E+00 | 5.38E-01 | no | | intergenic_variant | MODIFIER |
| 1 | 23762255 | AX-97810999 | 0 | G | A | 0.43 | 5.17E-02 | 8.43E-02 | 2.62E+00 | 2.68E+00 | 5.38E-01 | no | B3gat2 | 5_prime_UTR_variant | MODIFIER |
| 1 | 23808343 | AX-97791817 | 0 | T | C | 0.43 | 5.17E-02 | 8.43E-02 | 2.62E+00 | 2.68E+00 | 5.38E-01 | no | B3gat2 | intron_variant | MODIFIER |
| 1 | 23824136 | AX-170000095 | 0 | G | C | 0.43 | 5.17E-02 | 8.43E-02 | 2.62E+00 | 2.68E+00 | 5.38E-01 | no | B3gat2 | intron_variant | MODIFIER |
| 1 | 23826371 | AX-170235693 | 0 | C | A | 0.43 | 5.17E-02 | 8.43E-02 | 2.62E+00 | 2.68E+00 | 5.38E-01 | no | B3gat2 | intron_variant | MODIFIER |
| 1 | 23826973 | AX-170296367 | 0 | A | G | 0.43 | 5.17E-02 | 8.43E-02 | 2.62E+00 | 2.68E+00 | 5.38E-01 | no | B3gat2 | intron_variant | MODIFIER |
| 1 | 23829185 | AX-170170858 | 0 | C | T | 0.43 | 5.17E-02 | 8.43E-02 | 2.62E+00 | 2.68E+00 | 5.38E-01 | no | B3gat2 | intron_variant | MODIFIER |
| 1 | 23832598 | AX-97793268 | 0 | A | G | 0.43 | 5.17E-02 | 8.43E-02 | 2.62E+00 | 2.68E+00 | 5.38E-01 | no | B3gat2 | intron_variant | MODIFIER |
| 1 | 23906982 | AX-97824258 | 0 | A | C | 0.43 | 5.17E-02 | 8.43E-02 | 2.62E+00 | 2.68E+00 | 5.38E-01 | no | Smap1 | intron_variant | MODIFIER |
| 1 | 24072208 | AX-170231855 | 0 | A | G | 0.43 | 5.17E-02 | 8.43E-02 | 2.62E+00 | 2.68E+00 | 5.38E-01 | no | Fam135a | intron_variant | MODIFIER |
| 1 | 24142352 | AX-170000103 | 0 | G | A | 0.43 | 5.17E-02 | 8.43E-02 | 2.62E+00 | 2.68E+00 | 5.38E-01 | no | | intergenic_variant | MODIFIER |
| 1 | 24240857 | AX-170000107 | 0 | G | C | 0.43 | 5.17E-02 | 8.43E-02 | 2.62E+00 | 2.68E+00 | 5.38E-01 | no | Col9a1 | intron_variant | MODIFIER |
| 1 | 24376303 | AX-170465818 | 0 | T | A | 0.43 | 5.17E-02 | 8.43E-02 | 2.62E+00 | 2.68E+00 | 5.38E-01 | no | Col19a1 | intron_variant | MODIFIER |

| | | | | | | | | | | | | | | | |
|---|---|---|---|---|---|---|---|---|---|---|---|---|---|---|---|
| 1 | 24384239 | AX-170260482 | 0 | T | C | 0.43 | 5.17E-02 | 8.43E-02 | 2.62E+00 | 2.68E+00 | 5.38E-01 | no | Col19a1 | intron_variant | MODIFIER |
| 8 | 95445150 | AX-170277207 | 0 | A | G | 0.237 | 6.56E-02 | 1.08E-01 | 2.74E+00 | 2.81E+00 | 5.38E-01 | no | Csnk2a2 | downstream_gene_variant | MODIFIER |
| 5 | 40745186 | AX-97850896 | 1 | A | G | 0.205 | -6.70E-02 | 1.09E-01 | 2.60E+00 | 2.67E+00 | 5.38E-01 | no | | intergenic_variant | MODIFIER |
| 1 | 46322485 | AX-170458780 | 1 | T | C | 0.054 | -1.08E-01 | 1.80E-01 | 2.69E+00 | 2.82E+00 | 5.38E-01 | no | Dnah7b | intron_variant | MODIFIER |
| 1 | 76137999 | AX-170086501 | 0 | A | G | 0.14 | -9.60E-02 | 1.59E-01 | 2.66E+00 | 2.78E+00 | 5.38E-01 | no | | regulatory_region_variant | MODIFIER |
| 3 | 7839161 | AX-170277695 | 2 | T | C | 0.409 | -3.79E-02 | 6.30E-02 | 2.66E+00 | 2.80E+00 | 5.38E-01 | no | | intergenic_variant | MODIFIER |
| 8 | 96035904 | AX-170339401 | 1 | A | G | 0.223 | 7.08E-02 | 1.17E-01 | 2.73E+00 | 2.80E+00 | 5.38E-01 | no | | intergenic_variant | MODIFIER |
| 8 | 96074583 | AX-170147879 | 1 | C | A | 0.223 | 7.08E-02 | 1.17E-01 | 2.73E+00 | 2.80E+00 | 5.38E-01 | no | | intergenic_variant | MODIFIER |
| 12 | 91989175 | AX-170408877 | 0 | G | T | 0.07 | 1.29E-01 | 2.14E-01 | 2.73E+00 | 2.81E+00 | 5.39E-01 | no | | intergenic_variant | MODIFIER |
| 12 | 92259388 | AX-170404900 | 0 | C | G | 0.07 | 1.29E-01 | 2.14E-01 | 2.73E+00 | 2.81E+00 | 5.39E-01 | no | | intergenic_variant | MODIFIER |
| 5 | 92892796 | AX-170025624 | 1 | G | A | 0.429 | 6.40E-02 | 1.05E-01 | 2.68E+00 | 2.74E+00 | 5.39E-01 | no | Shroom3 | intron_variant | MODIFIER |
| 5 | 93160765 | AX-170189370 | 1 | C | T | 0.429 | 6.40E-02 | 1.05E-01 | 2.68E+00 | 2.74E+00 | 5.39E-01 | no | SEP11 | intron_variant | MODIFIER |
| 1 | 36195580 | AX-170386156 | 1 | A | G | 0.446 | 4.82E-02 | 7.97E-02 | 2.73E+00 | 2.81E+00 | 5.39E-01 | no | Uggt1 | intron_variant | MODIFIER |
| 18 | 67482829 | AX-170268957 | 0 | T | C | 0.272 | -6.36E-02 | 1.05E-01 | 2.67E+00 | 2.74E+00 | 5.39E-01 | no | Prelid3a | downstream_gene_variant | MODIFIER |
| 15 | 78859410 | AX-170309363 | 1 | T | C | 0.571 | 5.07E-02 | 8.02E-02 | 2.51E+00 | 2.58E+00 | 5.39E-01 | no | Lgals2 | upstream_gene_variant | MODIFIER |
| 7 | 36503742 | AX-170353251 | 0 | G | C | 0.36 | 4.97E-02 | 7.91E-02 | 2.50E+00 | 2.62E+00 | 5.39E-01 | no | | regulatory_region_variant | MODIFIER |
| 6 | 110603979 | AX-97801675 | 0 | C | A | 0.079 | 1.19E-01 | 1.98E-01 | 2.70E+00 | 2.83E+00 | 5.39E-01 | no | | intergenic_variant | MODIFIER |
| 8 | 38257068 | AX-170096230 | 0 | T | G | 0.158 | 8.44E-02 | 1.39E-01 | 2.79E+00 | 2.86E+00 | 5.39E-01 | no | Mir383 | upstream_gene_variant | MODIFIER |
| 8 | 41631853 | AX-170409972 | 0 | G | A | 0.158 | 8.44E-02 | 1.39E-01 | 2.79E+00 | 2.86E+00 | 5.39E-01 | no | | intergenic_variant | MODIFIER |
| 1 | 169478713 | AX-97815284 | 0 | C | T | 0.211 | 8.67E-02 | 1.43E-01 | 2.69E+00 | 2.73E+00 | 5.39E-01 | no | | intergenic_variant | MODIFIER |
| 15 | 90920968 | AX-170124338 | 0 | A | C | 0.158 | 9.60E-02 | 1.58E-01 | 2.63E+00 | 2.73E+00 | 5.39E-01 | no | Gm36618 | intron_variant,non_coding_transcript_variant | MODIFIER |
| 5 | 110412569 | AX-170289878 | 0 | G | A | 0.456 | 6.45E-02 | 1.05E-01 | 2.59E+00 | 2.65E+00 | 5.39E-01 | no | Fbrsl1 | intron_variant | MODIFIER |
| 18 | 30657787 | AX-170375903 | 0 | C | G | 0.07 | 1.30E-01 | 2.17E-01 | 2.73E+00 | 2.84E+00 | 5.39E-01 | no | | intergenic_variant | MODIFIER |
| 1 | 188034862 | AX-170210486 | 0 | G | A | 0.246 | 6.38E-02 | 1.01E-01 | 2.49E+00 | 2.59E+00 | 5.40E-01 | no | Esrrg | intron_variant | MODIFIER |
| 5 | 111457375 | AX-170296652 | 1 | T | C | 0.438 | 6.99E-02 | 1.14E-01 | 2.64E+00 | 2.67E+00 | 5.40E-01 | no | Mn1 | downstream_gene_variant | MODIFIER |
| 4 | 58764884 | AX-170069468 | 0 | G | A | 0.368 | 6.83E-02 | 1.12E-01 | 2.65E+00 | 2.71E+00 | 5.40E-01 | no | | intergenic_variant | MODIFIER |
| 7 | 123225672 | AX-170254053 | 1 | C | T | 0.411 | 5.09E-02 | 8.44E-02 | 2.72E+00 | 2.80E+00 | 5.40E-01 | no | Slc5a11 | intron_variant | MODIFIER |
| 7 | 123348441 | AX-170412400 | 1 | G | A | 0.411 | 5.09E-02 | 8.44E-02 | 2.72E+00 | 2.80E+00 | 5.40E-01 | no | Arhgap17 | intron_variant | MODIFIER |
| 8 | 15995345 | AX-170503657 | 0 | T | C | 0.219 | -4.71E-02 | 7.58E-02 | 2.54E+00 | 2.64E+00 | 5.40E-01 | no | Csmd1 | intron_variant | MODIFIER |
| 11 | 93309637 | AX-170224524 | 1 | C | T | 0.071 | 8.65E-02 | 1.42E-01 | 2.59E+00 | 2.72E+00 | 5.40E-01 | no | Car10 | intron_variant | MODIFIER |
| 7 | 66392198 | AX-170383133 | 1 | A | C | 0.402 | -5.85E-02 | 9.67E-02 | 2.78E+00 | 2.86E+00 | 5.40E-01 | no | Lrrk1 | upstream_gene_variant | MODIFIER |
| 18 | 68307997 | AX-170147594 | 1 | G | T | 0.277 | -6.22E-02 | 1.03E-01 | 2.70E+00 | 2.76E+00 | 5.40E-01 | no | Rnmt | intron_variant | MODIFIER |
| 1 | 175129607 | AX-170149772 | 1 | G | T | 0.241 | 7.23E-02 | 1.15E-01 | 2.52E+00 | 2.59E+00 | 5.40E-01 | no | Rgs7 | intron_variant | MODIFIER |
| 16 | 76763480 | AX-170535821 | 1 | C | T | 0.464 | 4.75E-02 | 7.61E-02 | 2.54E+00 | 2.61E+00 | 5.40E-01 | no | Gm30790 | intron_variant,non_coding_transcript_variant | MODIFIER |
| 10 | 11302877 | AX-170429152 | 2 | T | A | 0.036 | 8.99E-02 | 1.50E-01 | 2.64E+00 | 2.80E+00 | 5.40E-01 | no | Fbxo30 | downstream_gene_variant | MODIFIER |
| 2 | 136885665 | AX-170147469 | 0 | G | A | 0.149 | 1.10E-01 | 1.82E-01 | 2.67E+00 | 2.76E+00 | 5.41E-01 | no | Mkks | 5_prime_UTR_variant | MODIFIER |
| 2 | 156420253 | AX-170512203 | 0 | G | C | 0.149 | 1.10E-01 | 1.82E-01 | 2.67E+00 | 2.76E+00 | 5.41E-01 | no | Epb41l1 | upstream_gene_variant | MODIFIER |
| 1 | 99792698 | AX-170498190 | 0 | A | C | 0.132 | 1.09E-01 | 1.80E-01 | 2.64E+00 | 2.72E+00 | 5.41E-01 | no | Cntnap5b | intron_variant | MODIFIER |
| 10 | 28060294 | AX-170531016 | 0 | G | C | 0.184 | 5.60E-02 | 9.32E-02 | 2.74E+00 | 2.85E+00 | 5.41E-01 | no | | intergenic_variant | MODIFIER |
| 18 | 41447189 | AX-170015594 | 0 | C | T | 0.07 | 1.28E-01 | 2.14E-01 | 2.74E+00 | 2.85E+00 | 5.41E-01 | no | | intergenic_variant | MODIFIER |
| 1 | 19775001 | AX-170040542 | 0 | G | A | 0.412 | 5.03E-02 | 8.21E-02 | 2.60E+00 | 2.65E+00 | 5.41E-01 | no | | intergenic_variant | MODIFIER |
| 1 | 20542562 | AX-170179595 | 0 | C | T | 0.412 | 5.03E-02 | 8.21E-02 | 2.60E+00 | 2.65E+00 | 5.41E-01 | no | Pkhd1 | intron_variant | MODIFIER |
| 5 | 83792791 | AX-170500832 | 0 | G | A | 0.447 | 6.40E-02 | 1.06E-01 | 2.73E+00 | 2.80E+00 | 5.41E-01 | no | | intergenic_variant | MODIFIER |
| 1 | 60670236 | AX-97810581 | 0 | C | T | 0.105 | -1.11E-01 | 1.85E-01 | 2.69E+00 | 2.78E+00 | 5.41E-01 | no | | regulatory_region_variant | MODIFIER |
| 7 | 90218723 | AX-170250134 | 1 | G | T | 0.125 | -8.08E-02 | 1.34E-01 | 2.60E+00 | 2.75E+00 | 5.41E-01 | no | | intergenic_variant | MODIFIER |
| 8 | 46360619 | AX-170491315 | 0 | C | T | 0.158 | -6.87E-02 | 1.13E-01 | 2.62E+00 | 2.73E+00 | 5.41E-01 | no | Gm30931 | upstream_gene_variant | MODIFIER |
| 1 | 44470167 | AX-170192522 | 0 | G | A | 0.105 | -1.03E-01 | 1.72E-01 | 2.72E+00 | 2.85E+00 | 5.41E-01 | no | | intergenic_variant | MODIFIER |
| 4 | 128312093 | AX-170023917 | 0 | T | C | 0.14 | 7.55E-02 | 1.23E-01 | 2.60E+00 | 2.66E+00 | 5.41E-01 | no | Csmd2 | intron_variant,non_coding_transcript_variant | MODIFIER |
| 5 | 10060660 | AX-170133676 | 1 | G | T | 0.045 | 8.14E-02 | 1.35E-01 | 2.58E+00 | 2.76E+00 | 5.41E-01 | no | | intergenic_variant | MODIFIER |
| 5 | 100943137 | AX-170072888 | 1 | T | A | 0.438 | 6.09E-02 | 9.98E-02 | 2.60E+00 | 2.68E+00 | 5.41E-01 | no | | intergenic_variant | MODIFIER |
| 16 | 27970928 | AX-170180624 | 2 | A | G | 0.236 | 5.40E-02 | 8.97E-02 | 2.63E+00 | 2.76E+00 | 5.42E-01 | no | Gm49765 | upstream_gene_variant | MODIFIER |
| 1 | 146677264 | AX-170516698 | 1 | G | A | 0.205 | 8.61E-02 | 1.42E-01 | 2.66E+00 | 2.70E+00 | 5.42E-01 | no | Brinp3 | intron_variant | MODIFIER |
| 9 | 61222828 | AX-170483510 | 0 | T | C | 0.193 | -7.71E-02 | 1.28E-01 | 2.71E+00 | 2.77E+00 | 5.42E-01 | no | | intergenic_variant | MODIFIER |
| 9 | 61773243 | AX-170082603 | 0 | G | A | 0.193 | -7.71E-02 | 1.28E-01 | 2.71E+00 | 2.77E+00 | 5.42E-01 | no | | intergenic_variant | MODIFIER |

| | | | | | | | | | | | | | | | |
|---|---|---|---|---|---|---|---|---|---|---|---|---|---|---|---|
| 9 | 64203257 | AX-170485746 | 0 | T | C | 0.193 | -7.71E-02 | 1.28E-01 | 2.71E+00 | 2.77E+00 | 5.42E-01 | no | Map2k1 | intron_variant | MODIFIER |
| 9 | 65265264 | AX-170195149 | 0 | A | C | 0.193 | -7.71E-02 | 1.28E-01 | 2.71E+00 | 2.77E+00 | 5.42E-01 | no | Cilp | intron_variant | MODIFIER |
| 9 | 66152448 | AX-170369023 | 0 | A | G | 0.193 | -7.71E-02 | 1.28E-01 | 2.71E+00 | 2.77E+00 | 5.42E-01 | no | | intergenic_variant | MODIFIER |
| 9 | 66152538 | AX-170086239 | 0 | G | A | 0.193 | -7.71E-02 | 1.28E-01 | 2.71E+00 | 2.77E+00 | 5.42E-01 | no | | intergenic_variant | MODIFIER |
| 9 | 67572883 | AX-170249625 | 0 | A | G | 0.193 | -7.71E-02 | 1.28E-01 | 2.71E+00 | 2.77E+00 | 5.42E-01 | no | | regulatory_region_variant | MODIFIER |
| 9 | 71840355 | AX-170370632 | 0 | T | C | 0.193 | -7.71E-02 | 1.28E-01 | 2.71E+00 | 2.77E+00 | 5.42E-01 | no | Tcf12 | downstream_gene_variant | MODIFIER |
| 9 | 75214567 | AX-170090336 | 0 | T | C | 0.193 | -7.71E-02 | 1.28E-01 | 2.71E+00 | 2.77E+00 | 5.42E-01 | no | Myo5a | intron_variant | MODIFIER |
| 11 | 22793728 | AX-170264895 | 1 | T | G | 0.054 | 1.11E-01 | 1.83E-01 | 2.59E+00 | 2.73E+00 | 5.42E-01 | no | B3gnt2 | intron_variant,non_coding_transcript_variant | MODIFIER |
| 9 | 62276465 | AX-170338753 | 0 | G | A | 0.193 | 6.95E-02 | 1.16E-01 | 2.78E+00 | 2.88E+00 | 5.42E-01 | no | Spesp1 | intron_variant | MODIFIER |
| 6 | 129319140 | AX-170335577 | 0 | G | C | 0.553 | 5.05E-02 | 8.14E-02 | 2.54E+00 | 2.62E+00 | 5.42E-01 | no | 4922502D21Rik | downstream_gene_variant | MODIFIER |
| 13 | 5485707 | AX-170117186 | 0 | A | T | 0.193 | -6.99E-02 | 1.13E-01 | 2.53E+00 | 2.62E+00 | 5.42E-01 | no | Gm40660 | intron_variant,non_coding_transcript_variant | MODIFIER |
| 4 | 154359380 | AX-170301122 | 1 | T | A | 0.089 | 1.02E-01 | 1.67E-01 | 2.58E+00 | 2.70E+00 | 5.42E-01 | no | Prdm16 | intron_variant | MODIFIER |
| 9 | 104961154 | AX-170250657 | 1 | G | A | 0.125 | -6.87E-02 | 1.12E-01 | 2.58E+00 | 2.66E+00 | 5.42E-01 | no | Cpne4 | intron_variant | MODIFIER |
| 1 | 98878963 | AX-170496820 | 1 | A | T | 0.161 | 1.27E-01 | 2.09E-01 | 2.63E+00 | 2.69E+00 | 5.42E-01 | no | | intergenic_variant | MODIFIER |
| 8 | 67059284 | AX-170458486 | 1 | G | C | 0.143 | -8.93E-02 | 1.45E-01 | 2.57E+00 | 2.64E+00 | 5.42E-01 | no | | intergenic_variant | MODIFIER |
| 6 | 94080222 | AX-170409895 | 0 | G | T | 0.447 | -3.72E-02 | 6.02E-02 | 2.53E+00 | 2.65E+00 | 5.42E-01 | no | Magi1 | intron_variant | MODIFIER |
| 2 | 118059668 | AX-97807455 | 0 | C | A | 0.14 | 1.08E-01 | 1.79E-01 | 2.72E+00 | 2.79E+00 | 5.42E-01 | no | Gm13986 | intron_variant,non_coding_transcript_variant | MODIFIER |
| 16 | 28392216 | AX-170522642 | 0 | C | T | 0.246 | 5.15E-02 | 8.64E-02 | 2.66E+00 | 2.80E+00 | 5.42E-01 | no | Fgf12 | intron_variant | MODIFIER |
| 1 | 63491717 | AX-170103868 | 1 | G | A | 0.134 | 1.16E-01 | 1.94E-01 | 2.70E+00 | 2.80E+00 | 5.42E-01 | no | Adam23 | intron_variant | MODIFIER |
| 15 | 67989428 | AX-170011907 | 0 | T | C | 0.342 | -4.92E-02 | 7.71E-02 | 2.46E+00 | 2.55E+00 | 5.43E-01 | no | | intergenic_variant | MODIFIER |
| 2 | 162456919 | AX-170228135 | 0 | G | A | 0.167 | 1.02E-01 | 1.70E-01 | 2.63E+00 | 2.74E+00 | 5.43E-01 | no | Ptprt | intron_variant | MODIFIER |
| 7 | 45950230 | AX-170516065 | 1 | C | T | 0.33 | 5.62E-02 | 9.30E-02 | 2.63E+00 | 2.71E+00 | 5.43E-01 | no | Ccdc114 | downstream_gene_variant | MODIFIER |
| 1 | 82066868 | AX-170409292 | 0 | T | C | 0.158 | 1.26E-01 | 2.09E-01 | 2.65E+00 | 2.71E+00 | 5.43E-01 | no | | intergenic_variant | MODIFIER |
| 1 | 82067984 | AX-170430300 | 0 | T | C | 0.158 | 1.26E-01 | 2.09E-01 | 2.65E+00 | 2.71E+00 | 5.43E-01 | no | | intergenic_variant | MODIFIER |
| 1 | 82073932 | AX-170222341 | 0 | G | A | 0.158 | 1.26E-01 | 2.09E-01 | 2.65E+00 | 2.71E+00 | 5.43E-01 | no | | intergenic_variant | MODIFIER |
| 1 | 82076785 | AX-170398859 | 0 | T | C | 0.158 | 1.26E-01 | 2.09E-01 | 2.65E+00 | 2.71E+00 | 5.43E-01 | no | | intergenic_variant | MODIFIER |
| 1 | 82079974 | AX-97796279 | 0 | A | C | 0.158 | 1.26E-01 | 2.09E-01 | 2.65E+00 | 2.71E+00 | 5.43E-01 | no | | intergenic_variant | MODIFIER |
| 1 | 82080577 | AX-170497946 | 0 | T | G | 0.158 | 1.26E-01 | 2.09E-01 | 2.65E+00 | 2.71E+00 | 5.43E-01 | no | | intergenic_variant | MODIFIER |
| 1 | 82086036 | AX-170256871 | 0 | T | G | 0.158 | 1.26E-01 | 2.09E-01 | 2.65E+00 | 2.71E+00 | 5.43E-01 | no | | intergenic_variant | MODIFIER |
| 1 | 82088936 | AX-170185676 | 0 | T | C | 0.158 | 1.26E-01 | 2.09E-01 | 2.65E+00 | 2.71E+00 | 5.43E-01 | no | | intergenic_variant | MODIFIER |
| 1 | 82088975 | AX-170238045 | 0 | T | C | 0.158 | 1.26E-01 | 2.09E-01 | 2.65E+00 | 2.71E+00 | 5.43E-01 | no | | intergenic_variant | MODIFIER |
| 1 | 82141120 | AX-170385855 | 0 | A | C | 0.158 | 1.26E-01 | 2.09E-01 | 2.65E+00 | 2.71E+00 | 5.43E-01 | no | | intergenic_variant | MODIFIER |
| 1 | 82143863 | AX-170240969 | 0 | C | T | 0.158 | 1.26E-01 | 2.09E-01 | 2.65E+00 | 2.71E+00 | 5.43E-01 | no | | intergenic_variant | MODIFIER |
| 1 | 82145082 | AX-170358924 | 0 | T | C | 0.158 | 1.26E-01 | 2.09E-01 | 2.65E+00 | 2.71E+00 | 5.43E-01 | no | | intergenic_variant | MODIFIER |
| 1 | 82146988 | AX-97783560 | 0 | A | G | 0.158 | 1.26E-01 | 2.09E-01 | 2.65E+00 | 2.71E+00 | 5.43E-01 | no | | intergenic_variant | MODIFIER |
| 1 | 82149705 | AX-170381754 | 0 | G | A | 0.158 | 1.26E-01 | 2.09E-01 | 2.65E+00 | 2.71E+00 | 5.43E-01 | no | | intergenic_variant | MODIFIER |
| 1 | 82183548 | AX-170322703 | 0 | T | C | 0.158 | 1.26E-01 | 2.09E-01 | 2.65E+00 | 2.71E+00 | 5.43E-01 | no | | intergenic_variant | MODIFIER |
| 1 | 82184344 | AX-170256271 | 0 | A | G | 0.158 | 1.26E-01 | 2.09E-01 | 2.65E+00 | 2.71E+00 | 5.43E-01 | no | | regulatory_region_variant | MODIFIER |
| 1 | 82184404 | AX-170271205 | 0 | A | G | 0.158 | 1.26E-01 | 2.09E-01 | 2.65E+00 | 2.71E+00 | 5.43E-01 | no | | regulatory_region_variant | MODIFIER |
| 1 | 82196450 | AX-170504988 | 0 | A | G | 0.158 | 1.26E-01 | 2.09E-01 | 2.65E+00 | 2.71E+00 | 5.43E-01 | no | | intergenic_variant | MODIFIER |
| 1 | 82215278 | AX-170242344 | 0 | A | G | 0.158 | 1.26E-01 | 2.09E-01 | 2.65E+00 | 2.71E+00 | 5.43E-01 | no | | intergenic_variant | MODIFIER |
| 1 | 82218390 | AX-170412901 | 0 | T | C | 0.158 | 1.26E-01 | 2.09E-01 | 2.65E+00 | 2.71E+00 | 5.43E-01 | no | | intergenic_variant | MODIFIER |
| 1 | 82258972 | AX-97851007 | 0 | T | G | 0.158 | 1.26E-01 | 2.09E-01 | 2.65E+00 | 2.71E+00 | 5.43E-01 | no | Irs1 | intron_variant | MODIFIER |
| 1 | 82268523 | AX-170211374 | 0 | A | C | 0.158 | 1.26E-01 | 2.09E-01 | 2.65E+00 | 2.71E+00 | 5.43E-01 | no | Irs1 | intron_variant | MODIFIER |
| 1 | 82268759 | AX-170264537 | 0 | A | G | 0.158 | 1.26E-01 | 2.09E-01 | 2.65E+00 | 2.71E+00 | 5.43E-01 | no | Irs1 | intron_variant | MODIFIER |
| 1 | 82268828 | AX-170404555 | 0 | G | A | 0.158 | 1.26E-01 | 2.09E-01 | 2.65E+00 | 2.71E+00 | 5.43E-01 | no | Irs1 | intron_variant | MODIFIER |
| 1 | 82282034 | AX-170152766 | 0 | G | A | 0.158 | 1.26E-01 | 2.09E-01 | 2.65E+00 | 2.71E+00 | 5.43E-01 | no | Irs1 | intron_variant | MODIFIER |
| 1 | 82298507 | AX-170179341 | 0 | A | G | 0.158 | 1.26E-01 | 2.09E-01 | 2.65E+00 | 2.71E+00 | 5.43E-01 | no | | intergenic_variant | MODIFIER |
| 1 | 82300779 | AX-170344878 | 0 | C | T | 0.158 | 1.26E-01 | 2.09E-01 | 2.65E+00 | 2.71E+00 | 5.43E-01 | no | | intergenic_variant | MODIFIER |
| 1 | 82303905 | AX-170497883 | 0 | T | A | 0.158 | 1.26E-01 | 2.09E-01 | 2.65E+00 | 2.71E+00 | 5.43E-01 | no | | intergenic_variant | MODIFIER |
| 1 | 82333737 | AX-170222256 | 0 | G | A | 0.158 | 1.26E-01 | 2.09E-01 | 2.65E+00 | 2.71E+00 | 5.43E-01 | no | Rhbdd1 | intron_variant | MODIFIER |
| 1 | 82640132 | AX-170098868 | 0 | C | T | 0.158 | 1.26E-01 | 2.09E-01 | 2.65E+00 | 2.71E+00 | 5.43E-01 | no | Col4a3 | intron_variant | MODIFIER |
| 1 | 82904440 | AX-170278856 | 0 | G | A | 0.158 | 1.26E-01 | 2.09E-01 | 2.65E+00 | 2.71E+00 | 5.43E-01 | no | Agfg1 | downstream_gene_variant | MODIFIER |
| 1 | 88085995 | AX-170259774 | 0 | T | C | 0.158 | 1.26E-01 | 2.09E-01 | 2.65E+00 | 2.71E+00 | 5.43E-01 | no | Ugt1a9 | intron_variant | MODIFIER |

| | | | | | | | | | | | | | | | |
|---|---|---|---|---|---|---|---|---|---|---|---|---|---|---|---|
| 1 | 88106342 | AX-170254900 | 0 | G | T | 0.158 | 1.26E-01 | 2.09E-01 | 2.65E+00 | 2.71E+00 | 5.43E-01 | no | Ugt1a7c | intron_variant | MODIFIER |
| 1 | 88154122 | AX-170514412 | 0 | T | C | 0.158 | 1.26E-01 | 2.09E-01 | 2.65E+00 | 2.71E+00 | 5.43E-01 | no | Ugt1a6a | intron_variant | MODIFIER |
| 1 | 88156770 | AX-170416507 | 0 | T | C | 0.158 | 1.26E-01 | 2.09E-01 | 2.65E+00 | 2.71E+00 | 5.43E-01 | no | Ugt1a6a | intron_variant | MODIFIER |
| 1 | 88162049 | AX-170184938 | 0 | T | C | 0.158 | 1.26E-01 | 2.09E-01 | 2.65E+00 | 2.71E+00 | 5.43E-01 | no | Ugt1a6a | intron_variant | MODIFIER |
| 1 | 88227935 | AX-97806539 | 0 | T | C | 0.158 | 1.26E-01 | 2.09E-01 | 2.65E+00 | 2.71E+00 | 5.43E-01 | no | Mroh2a | intron_variant | MODIFIER |
| 1 | 88347051 | AX-170332696 | 0 | G | A | 0.158 | 1.26E-01 | 2.09E-01 | 2.65E+00 | 2.71E+00 | 5.43E-01 | no | Trpm8 | intron_variant | MODIFIER |
| 1 | 88349770 | AX-170086529 | 0 | T | C | 0.158 | 1.26E-01 | 2.09E-01 | 2.65E+00 | 2.71E+00 | 5.43E-01 | no | Trpm8 | intron_variant | MODIFIER |
| 1 | 88349831 | AX-170106981 | 0 | C | T | 0.158 | 1.26E-01 | 2.09E-01 | 2.65E+00 | 2.71E+00 | 5.43E-01 | no | Trpm8 | intron_variant | MODIFIER |
| 1 | 88354486 | AX-170151223 | 0 | T | C | 0.158 | 1.26E-01 | 2.09E-01 | 2.65E+00 | 2.71E+00 | 5.43E-01 | no | Trpm8 | synonymous_variant | LOW |
| 1 | 88355543 | AX-170444714 | 0 | C | T | 0.158 | 1.26E-01 | 2.09E-01 | 2.65E+00 | 2.71E+00 | 5.43E-01 | no | Trpm8 | intron_variant | MODIFIER |
| 1 | 88443701 | AX-97795586 | 0 | G | T | 0.158 | 1.26E-01 | 2.09E-01 | 2.65E+00 | 2.71E+00 | 5.43E-01 | no | Gm29538 | downstream_gene_variant | MODIFIER |
| 1 | 88546806 | AX-170001021 | 0 | G | A | 0.158 | 1.26E-01 | 2.09E-01 | 2.65E+00 | 2.71E+00 | 5.43E-01 | no | Gm19589 | upstream_gene_variant | MODIFIER |
| 1 | 88558141 | AX-97849422 | 0 | T | C | 0.158 | 1.26E-01 | 2.09E-01 | 2.65E+00 | 2.71E+00 | 5.43E-01 | no | Gm19589 | intron_variant,non_coding_transcript_variant | MODIFIER |
| 1 | 88592751 | AX-170151584 | 0 | A | C | 0.158 | 1.26E-01 | 2.09E-01 | 2.65E+00 | 2.71E+00 | 5.43E-01 | no | | regulatory_region_variant | MODIFIER |
| 1 | 88597337 | AX-97835206 | 0 | G | A | 0.158 | 1.26E-01 | 2.09E-01 | 2.65E+00 | 2.71E+00 | 5.43E-01 | no | | intergenic_variant | MODIFIER |
| 1 | 88621187 | AX-170529102 | 0 | C | T | 0.158 | 1.26E-01 | 2.09E-01 | 2.65E+00 | 2.71E+00 | 5.43E-01 | no | | regulatory_region_variant | MODIFIER |
| 1 | 88634360 | AX-170125923 | 0 | T | G | 0.158 | 1.26E-01 | 2.09E-01 | 2.65E+00 | 2.71E+00 | 5.43E-01 | no | | intergenic_variant | MODIFIER |
| 1 | 88642053 | AX-170090771 | 0 | G | A | 0.158 | 1.26E-01 | 2.09E-01 | 2.65E+00 | 2.71E+00 | 5.43E-01 | no | | intergenic_variant | MODIFIER |
| 1 | 88655158 | AX-170465127 | 0 | G | A | 0.158 | 1.26E-01 | 2.09E-01 | 2.65E+00 | 2.71E+00 | 5.43E-01 | no | Gm29336 | upstream_gene_variant | MODIFIER |
| 1 | 88655350 | AX-170211224 | 0 | T | C | 0.158 | 1.26E-01 | 2.09E-01 | 2.65E+00 | 2.71E+00 | 5.43E-01 | no | Gm29336 | upstream_gene_variant | MODIFIER |
| 1 | 88656070 | AX-170451998 | 0 | C | T | 0.158 | 1.26E-01 | 2.09E-01 | 2.65E+00 | 2.71E+00 | 5.43E-01 | no | Gm29336 | upstream_gene_variant | MODIFIER |
| 1 | 88667441 | AX-170268118 | 0 | T | C | 0.158 | 1.26E-01 | 2.09E-01 | 2.65E+00 | 2.71E+00 | 5.43E-01 | no | | regulatory_region_variant | MODIFIER |
| 1 | 88667546 | AX-170001016 | 0 | C | A | 0.158 | 1.26E-01 | 2.09E-01 | 2.65E+00 | 2.71E+00 | 5.43E-01 | no | | regulatory_region_variant | MODIFIER |
| 1 | 88690851 | AX-170373567 | 0 | A | G | 0.158 | 1.26E-01 | 2.09E-01 | 2.65E+00 | 2.71E+00 | 5.43E-01 | no | Arl4c | intron_variant,NMD_transcript_variant | MODIFIER |
| 1 | 88729305 | AX-170260698 | 0 | C | A | 0.158 | 1.26E-01 | 2.09E-01 | 2.65E+00 | 2.71E+00 | 5.43E-01 | no | | intergenic_variant | MODIFIER |
| 1 | 88729506 | AX-97822740 | 0 | A | G | 0.158 | 1.26E-01 | 2.09E-01 | 2.65E+00 | 2.71E+00 | 5.43E-01 | no | | intergenic_variant | MODIFIER |
| 1 | 88738496 | AX-170172199 | 0 | C | T | 0.158 | 1.26E-01 | 2.09E-01 | 2.65E+00 | 2.71E+00 | 5.43E-01 | no | | intergenic_variant | MODIFIER |
| 1 | 88746016 | AX-170454618 | 0 | A | T | 0.158 | 1.26E-01 | 2.09E-01 | 2.65E+00 | 2.71E+00 | 5.43E-01 | no | | regulatory_region_variant | MODIFIER |
| 1 | 88746178 | AX-170162360 | 0 | A | G | 0.158 | 1.26E-01 | 2.09E-01 | 2.65E+00 | 2.71E+00 | 5.43E-01 | no | | regulatory_region_variant | MODIFIER |
| 1 | 88746364 | AX-170377813 | 0 | A | C | 0.158 | 1.26E-01 | 2.09E-01 | 2.65E+00 | 2.71E+00 | 5.43E-01 | no | | regulatory_region_variant | MODIFIER |
| 1 | 88822053 | AX-170231228 | 0 | T | C | 0.158 | 1.26E-01 | 2.09E-01 | 2.65E+00 | 2.71E+00 | 5.43E-01 | no | | intergenic_variant | MODIFIER |
| 1 | 88838535 | AX-170125773 | 0 | A | G | 0.158 | 1.26E-01 | 2.09E-01 | 2.65E+00 | 2.71E+00 | 5.43E-01 | no | | regulatory_region_variant | MODIFIER |
| 1 | 88859601 | AX-170421496 | 0 | C | T | 0.158 | 1.26E-01 | 2.09E-01 | 2.65E+00 | 2.71E+00 | 5.43E-01 | no | | intergenic_variant | MODIFIER |
| 1 | 88920703 | AX-170146738 | 0 | G | A | 0.158 | 1.26E-01 | 2.09E-01 | 2.65E+00 | 2.71E+00 | 5.43E-01 | no | | intergenic_variant | MODIFIER |
| 1 | 88935016 | AX-170233970 | 0 | G | C | 0.158 | 1.26E-01 | 2.09E-01 | 2.65E+00 | 2.71E+00 | 5.43E-01 | no | Gm4753 | downstream_gene_variant | MODIFIER |
| 1 | 88955071 | AX-170241004 | 0 | A | C | 0.158 | 1.26E-01 | 2.09E-01 | 2.65E+00 | 2.71E+00 | 5.43E-01 | no | | regulatory_region_variant | MODIFIER |
| 1 | 88980654 | AX-97816502 | 0 | G | A | 0.158 | 1.26E-01 | 2.09E-01 | 2.65E+00 | 2.71E+00 | 5.43E-01 | no | | intergenic_variant | MODIFIER |
| 1 | 88981830 | AX-170469353 | 0 | G | A | 0.158 | 1.26E-01 | 2.09E-01 | 2.65E+00 | 2.71E+00 | 5.43E-01 | no | | intergenic_variant | MODIFIER |
| 1 | 88983109 | AX-170001025 | 0 | G | A | 0.158 | 1.26E-01 | 2.09E-01 | 2.65E+00 | 2.71E+00 | 5.43E-01 | no | | intergenic_variant | MODIFIER |
| 1 | 89137283 | AX-97852930 | 0 | A | G | 0.158 | 1.26E-01 | 2.09E-01 | 2.65E+00 | 2.71E+00 | 5.43E-01 | no | Sh3bp4 | intron_variant | MODIFIER |
| 1 | 89146909 | AX-97812375 | 0 | G | A | 0.158 | 1.26E-01 | 2.09E-01 | 2.65E+00 | 2.71E+00 | 5.43E-01 | no | Sh3bp4 | intron_variant | MODIFIER |
| 1 | 89203820 | AX-170143601 | 0 | C | T | 0.158 | 1.26E-01 | 2.09E-01 | 2.65E+00 | 2.71E+00 | 5.43E-01 | no | | intergenic_variant | MODIFIER |
| 1 | 89204333 | AX-170329927 | 0 | T | C | 0.158 | 1.26E-01 | 2.09E-01 | 2.65E+00 | 2.71E+00 | 5.43E-01 | no | | intergenic_variant | MODIFIER |
| 1 | 89204789 | AX-170372512 | 0 | A | G | 0.158 | 1.26E-01 | 2.09E-01 | 2.65E+00 | 2.71E+00 | 5.43E-01 | no | | intergenic_variant | MODIFIER |
| 1 | 89208725 | AX-97828002 | 0 | A | G | 0.158 | 1.26E-01 | 2.09E-01 | 2.65E+00 | 2.71E+00 | 5.43E-01 | no | | intergenic_variant | MODIFIER |
| 1 | 89227793 | AX-170041964 | 0 | G | C | 0.158 | 1.26E-01 | 2.09E-01 | 2.65E+00 | 2.71E+00 | 5.43E-01 | no | | intergenic_variant | MODIFIER |
| 1 | 89311461 | AX-170425992 | 0 | G | C | 0.158 | 1.26E-01 | 2.09E-01 | 2.65E+00 | 2.71E+00 | 5.43E-01 | no | Gm28342 | intron_variant,non_coding_transcript_variant | MODIFIER |
| 1 | 89321405 | AX-170250522 | 0 | C | T | 0.158 | 1.26E-01 | 2.09E-01 | 2.65E+00 | 2.71E+00 | 5.43E-01 | no | Gm28342 | intron_variant,non_coding_transcript_variant | MODIFIER |
| 1 | 89342066 | AX-97788047 | 0 | T | C | 0.158 | 1.26E-01 | 2.09E-01 | 2.65E+00 | 2.71E+00 | 5.43E-01 | no | Gm28342 | intron_variant,non_coding_transcript_variant | MODIFIER |
| 1 | 89403876 | AX-170190375 | 0 | A | G | 0.158 | 1.26E-01 | 2.09E-01 | 2.65E+00 | 2.71E+00 | 5.43E-01 | no | C030007H22Rik | intron_variant,non_coding_transcript_variant | MODIFIER |
| 1 | 89406197 | AX-170402089 | 0 | G | A | 0.158 | 1.26E-01 | 2.09E-01 | 2.65E+00 | 2.71E+00 | 5.43E-01 | no | C030007H22Rik | upstream_gene_variant | MODIFIER |
| 1 | 89415190 | AX-170523730 | 0 | T | C | 0.158 | 1.26E-01 | 2.09E-01 | 2.65E+00 | 2.71E+00 | 5.43E-01 | no | | intergenic_variant | MODIFIER |
| 1 | 89416466 | AX-170524271 | 0 | A | G | 0.158 | 1.26E-01 | 2.09E-01 | 2.65E+00 | 2.71E+00 | 5.43E-01 | no | | intergenic_variant | MODIFIER |
| 1 | 89417970 | AX-170308748 | 0 | T | C | 0.158 | 1.26E-01 | 2.09E-01 | 2.65E+00 | 2.71E+00 | 5.43E-01 | no | | intergenic_variant | MODIFIER |

| | | | | | | | | | | | | | | | |
|---|---|---|---|---|---|---|---|---|---|---|---|---|---|---|---|
| 1 | 89419801 | AX-170041967 | 0 | T | C | 0.158 | 1.26E-01 | 2.09E-01 | 2.65E+00 | 2.71E+00 | 5.43E-01 | no | | intergenic_variant | MODIFIER |
| 1 | 89428398 | AX-170035621 | 0 | C | T | 0.158 | 1.26E-01 | 2.09E-01 | 2.65E+00 | 2.71E+00 | 5.43E-01 | no | | intergenic_variant | MODIFIER |
| 1 | 89432662 | AX-97829394 | 0 | T | C | 0.158 | 1.26E-01 | 2.09E-01 | 2.65E+00 | 2.71E+00 | 5.43E-01 | no | | intergenic_variant | MODIFIER |
| 1 | 89471051 | AX-170173221 | 0 | A | T | 0.158 | 1.26E-01 | 2.09E-01 | 2.65E+00 | 2.71E+00 | 5.43E-01 | no | Agap1 | intron_variant | MODIFIER |
| 1 | 89486316 | AX-170492471 | 0 | T | C | 0.158 | 1.26E-01 | 2.09E-01 | 2.65E+00 | 2.71E+00 | 5.43E-01 | no | Agap1 | intron_variant | MODIFIER |
| 1 | 89496983 | AX-170530153 | 0 | C | T | 0.158 | 1.26E-01 | 2.09E-01 | 2.65E+00 | 2.71E+00 | 5.43E-01 | no | Agap1 | intron_variant | MODIFIER |
| 1 | 89514615 | AX-97793802 | 0 | C | T | 0.158 | 1.26E-01 | 2.09E-01 | 2.65E+00 | 2.71E+00 | 5.43E-01 | no | Agap1 | intron_variant | MODIFIER |
| 1 | 89544887 | AX-170001035 | 0 | A | T | 0.158 | 1.26E-01 | 2.09E-01 | 2.65E+00 | 2.71E+00 | 5.43E-01 | no | Agap1 | intron_variant | MODIFIER |
| 1 | 89555734 | AX-170001039 | 0 | T | C | 0.158 | 1.26E-01 | 2.09E-01 | 2.65E+00 | 2.71E+00 | 5.43E-01 | no | Agap1 | intron_variant | MODIFIER |
| 1 | 89609878 | AX-170001042 | 0 | C | T | 0.158 | 1.26E-01 | 2.09E-01 | 2.65E+00 | 2.71E+00 | 5.43E-01 | no | Agap1 | intron_variant | MODIFIER |
| 1 | 89610894 | AX-170476758 | 0 | C | T | 0.158 | 1.26E-01 | 2.09E-01 | 2.65E+00 | 2.71E+00 | 5.43E-01 | no | Agap1 | intron_variant | MODIFIER |
| 1 | 89611213 | AX-170001037 | 0 | A | G | 0.158 | 1.26E-01 | 2.09E-01 | 2.65E+00 | 2.71E+00 | 5.43E-01 | no | Agap1 | intron_variant | MODIFIER |
| 1 | 89676373 | AX-170001038 | 0 | T | G | 0.158 | 1.26E-01 | 2.09E-01 | 2.65E+00 | 2.71E+00 | 5.43E-01 | no | Agap1 | intron_variant | MODIFIER |
| 1 | 89676568 | AX-170001041 | 0 | A | G | 0.158 | 1.26E-01 | 2.09E-01 | 2.65E+00 | 2.71E+00 | 5.43E-01 | no | Agap1 | intron_variant | MODIFIER |
| 1 | 89691106 | AX-97813954 | 0 | C | T | 0.158 | 1.26E-01 | 2.09E-01 | 2.65E+00 | 2.71E+00 | 5.43E-01 | no | Agap1 | intron_variant | MODIFIER |
| 1 | 89701618 | AX-170478987 | 0 | T | C | 0.158 | 1.26E-01 | 2.09E-01 | 2.65E+00 | 2.71E+00 | 5.43E-01 | no | Agap1 | intron_variant | MODIFIER |
| 1 | 89710203 | AX-97808968 | 0 | C | A | 0.158 | 1.26E-01 | 2.09E-01 | 2.65E+00 | 2.71E+00 | 5.43E-01 | no | Agap1 | intron_variant | MODIFIER |
| 1 | 89717706 | AX-170383117 | 0 | T | C | 0.158 | 1.26E-01 | 2.09E-01 | 2.65E+00 | 2.71E+00 | 5.43E-01 | no | Agap1 | intron_variant | MODIFIER |
| 1 | 89726831 | AX-97812376 | 0 | T | G | 0.158 | 1.26E-01 | 2.09E-01 | 2.65E+00 | 2.71E+00 | 5.43E-01 | no | Agap1 | intron_variant | MODIFIER |
| 1 | 89781691 | AX-97840346 | 0 | T | C | 0.158 | 1.26E-01 | 2.09E-01 | 2.65E+00 | 2.71E+00 | 5.43E-01 | no | Agap1 | intron_variant | MODIFIER |
| 1 | 89796001 | AX-170041973 | 0 | C | G | 0.158 | 1.26E-01 | 2.09E-01 | 2.65E+00 | 2.71E+00 | 5.43E-01 | no | Agap1 | intron_variant | MODIFIER |
| 1 | 89807365 | AX-170203034 | 0 | A | G | 0.158 | 1.26E-01 | 2.09E-01 | 2.65E+00 | 2.71E+00 | 5.43E-01 | no | Agap1 | intron_variant | MODIFIER |
| 1 | 89807401 | AX-170086530 | 0 | G | A | 0.158 | 1.26E-01 | 2.09E-01 | 2.65E+00 | 2.71E+00 | 5.43E-01 | no | Agap1 | intron_variant | MODIFIER |
| 1 | 89807894 | AX-170350579 | 0 | T | A | 0.158 | 1.26E-01 | 2.09E-01 | 2.65E+00 | 2.71E+00 | 5.43E-01 | no | Agap1 | intron_variant | MODIFIER |
| 1 | 89810876 | AX-170407505 | 0 | T | C | 0.158 | 1.26E-01 | 2.09E-01 | 2.65E+00 | 2.71E+00 | 5.43E-01 | no | Agap1 | intron_variant | MODIFIER |
| 1 | 89814529 | AX-170209336 | 0 | A | C | 0.158 | 1.26E-01 | 2.09E-01 | 2.65E+00 | 2.71E+00 | 5.43E-01 | no | Agap1 | intron_variant | MODIFIER |
| 1 | 89829745 | AX-170492585 | 0 | C | A | 0.158 | 1.26E-01 | 2.09E-01 | 2.65E+00 | 2.71E+00 | 5.43E-01 | no | Agap1 | intron_variant | MODIFIER |
| 1 | 89832514 | AX-97808836 | 0 | C | T | 0.158 | 1.26E-01 | 2.09E-01 | 2.65E+00 | 2.71E+00 | 5.43E-01 | no | Agap1 | intron_variant | MODIFIER |
| 1 | 89833237 | AX-170379859 | 0 | G | A | 0.158 | 1.26E-01 | 2.09E-01 | 2.65E+00 | 2.71E+00 | 5.43E-01 | no | Agap1 | intron_variant | MODIFIER |
| 1 | 90003047 | AX-97815170 | 0 | A | G | 0.158 | 1.26E-01 | 2.09E-01 | 2.65E+00 | 2.71E+00 | 5.43E-01 | no | Asb18 | intron_variant | MODIFIER |
| 1 | 90182876 | AX-170091594 | 0 | T | C | 0.158 | 1.26E-01 | 2.09E-01 | 2.65E+00 | 2.71E+00 | 5.43E-01 | no | | intergenic_variant | MODIFIER |
| 1 | 90189039 | AX-170289822 | 0 | G | A | 0.158 | 1.26E-01 | 2.09E-01 | 2.65E+00 | 2.71E+00 | 5.43E-01 | no | | regulatory_region_variant | MODIFIER |
| 1 | 90201884 | AX-170422539 | 0 | A | G | 0.158 | 1.26E-01 | 2.09E-01 | 2.65E+00 | 2.71E+00 | 5.43E-01 | no | Ackr3 | upstream_gene_variant | MODIFIER |
| 1 | 90207349 | AX-170407402 | 0 | A | G | 0.158 | 1.26E-01 | 2.09E-01 | 2.65E+00 | 2.71E+00 | 5.43E-01 | no | Ackr3 | intron_variant | MODIFIER |
| 1 | 90214765 | AX-170452844 | 0 | T | C | 0.158 | 1.26E-01 | 2.09E-01 | 2.65E+00 | 2.71E+00 | 5.43E-01 | no | Ackr3 | synonymous_variant | LOW |
| 1 | 90221460 | AX-170484397 | 0 | A | G | 0.158 | 1.26E-01 | 2.09E-01 | 2.65E+00 | 2.71E+00 | 5.43E-01 | no | Ackr3 | downstream_gene_variant | MODIFIER |
| 1 | 90221856 | AX-170041981 | 0 | C | A | 0.158 | 1.26E-01 | 2.09E-01 | 2.65E+00 | 2.71E+00 | 5.43E-01 | no | | regulatory_region_variant | MODIFIER |
| 1 | 90233605 | AX-170424364 | 0 | G | A | 0.158 | 1.26E-01 | 2.09E-01 | 2.65E+00 | 2.71E+00 | 5.43E-01 | no | | regulatory_region_variant | MODIFIER |
| 1 | 90236800 | AX-170398678 | 0 | C | T | 0.158 | 1.26E-01 | 2.09E-01 | 2.65E+00 | 2.71E+00 | 5.43E-01 | no | | regulatory_region_variant | MODIFIER |
| 1 | 90271240 | AX-170130293 | 0 | G | A | 0.158 | 1.26E-01 | 2.09E-01 | 2.65E+00 | 2.71E+00 | 5.43E-01 | no | | intergenic_variant | MODIFIER |
| 1 | 90275355 | AX-97852080 | 0 | C | T | 0.158 | 1.26E-01 | 2.09E-01 | 2.65E+00 | 2.71E+00 | 5.43E-01 | no | | regulatory_region_variant | MODIFIER |
| 1 | 90368792 | AX-97812373 | 0 | A | G | 0.158 | 1.26E-01 | 2.09E-01 | 2.65E+00 | 2.71E+00 | 5.43E-01 | no | | regulatory_region_variant | MODIFIER |
| 1 | 90385008 | AX-97826354 | 0 | A | G | 0.158 | 1.26E-01 | 2.09E-01 | 2.65E+00 | 2.71E+00 | 5.43E-01 | no | | intergenic_variant | MODIFIER |
| 1 | 90390150 | AX-170126555 | 0 | T | C | 0.158 | 1.26E-01 | 2.09E-01 | 2.65E+00 | 2.71E+00 | 5.43E-01 | no | | intergenic_variant | MODIFIER |
| 1 | 90398581 | AX-170237538 | 0 | G | A | 0.158 | 1.26E-01 | 2.09E-01 | 2.65E+00 | 2.71E+00 | 5.43E-01 | no | | regulatory_region_variant | MODIFIER |
| 1 | 90398863 | AX-170295263 | 0 | C | T | 0.158 | 1.26E-01 | 2.09E-01 | 2.65E+00 | 2.71E+00 | 5.43E-01 | no | | regulatory_region_variant | MODIFIER |
| 1 | 90407951 | AX-97839502 | 0 | C | T | 0.158 | 1.26E-01 | 2.09E-01 | 2.65E+00 | 2.71E+00 | 5.43E-01 | no | | intergenic_variant | MODIFIER |
| 1 | 90413212 | AX-170201408 | 0 | T | C | 0.158 | 1.26E-01 | 2.09E-01 | 2.65E+00 | 2.71E+00 | 5.43E-01 | no | | intergenic_variant | MODIFIER |
| 1 | 90413916 | AX-170458686 | 0 | T | A | 0.158 | 1.26E-01 | 2.09E-01 | 2.65E+00 | 2.71E+00 | 5.43E-01 | no | | intergenic_variant | MODIFIER |
| 1 | 90414392 | AX-170171370 | 0 | A | G | 0.158 | 1.26E-01 | 2.09E-01 | 2.65E+00 | 2.71E+00 | 5.43E-01 | no | | intergenic_variant | MODIFIER |
| 1 | 90415121 | AX-170530452 | 0 | A | G | 0.158 | 1.26E-01 | 2.09E-01 | 2.65E+00 | 2.71E+00 | 5.43E-01 | no | | intergenic_variant | MODIFIER |
| 1 | 90416736 | AX-170183188 | 0 | A | G | 0.158 | 1.26E-01 | 2.09E-01 | 2.65E+00 | 2.71E+00 | 5.43E-01 | no | | intergenic_variant | MODIFIER |
| 1 | 90417234 | AX-170407127 | 0 | G | A | 0.158 | 1.26E-01 | 2.09E-01 | 2.65E+00 | 2.71E+00 | 5.43E-01 | no | | intergenic_variant | MODIFIER |
| 1 | 90422306 | AX-170295959 | 0 | G | A | 0.158 | 1.26E-01 | 2.09E-01 | 2.65E+00 | 2.71E+00 | 5.43E-01 | no | | intergenic_variant | MODIFIER |

| | | | | | | | | | | | | | | | |
|---|---|---|---|---|---|---|---|---|---|---|---|---|---|---|---|
| 1 | 90443954 | AX-97816134 | 0 | T | C | 0.158 | 1.26E-01 | 2.09E-01 | 2.65E+00 | 2.71E+00 | 5.43E-01 | no | | regulatory_region_variant | MODIFIER |
| 1 | 90475829 | AX-97783349 | 0 | A | G | 0.158 | 1.26E-01 | 2.09E-01 | 2.65E+00 | 2.71E+00 | 5.43E-01 | no | | regulatory_region_variant | MODIFIER |
| 1 | 90483195 | AX-170231063 | 0 | A | G | 0.158 | 1.26E-01 | 2.09E-01 | 2.65E+00 | 2.71E+00 | 5.43E-01 | no | | intergenic_variant | MODIFIER |
| 1 | 90581952 | AX-97848026 | 0 | A | G | 0.158 | 1.26E-01 | 2.09E-01 | 2.65E+00 | 2.71E+00 | 5.43E-01 | no | | intergenic_variant | MODIFIER |
| 1 | 90583219 | AX-170477132 | 0 | G | A | 0.158 | 1.26E-01 | 2.09E-01 | 2.65E+00 | 2.71E+00 | 5.43E-01 | no | | intergenic_variant | MODIFIER |
| 1 | 90586706 | AX-170248236 | 0 | A | C | 0.158 | 1.26E-01 | 2.09E-01 | 2.65E+00 | 2.71E+00 | 5.43E-01 | no | | intergenic_variant | MODIFIER |
| 1 | 90587829 | AX-170340565 | 0 | T | G | 0.158 | 1.26E-01 | 2.09E-01 | 2.65E+00 | 2.71E+00 | 5.43E-01 | no | | intergenic_variant | MODIFIER |
| 1 | 90588419 | AX-170329302 | 0 | C | T | 0.158 | 1.26E-01 | 2.09E-01 | 2.65E+00 | 2.71E+00 | 5.43E-01 | no | | regulatory_region_variant | MODIFIER |
| 1 | 90596554 | AX-170001051 | 0 | C | G | 0.158 | 1.26E-01 | 2.09E-01 | 2.65E+00 | 2.71E+00 | 5.43E-01 | no | | intergenic_variant | MODIFIER |
| 1 | 90598041 | AX-170091595 | 0 | A | G | 0.158 | 1.26E-01 | 2.09E-01 | 2.65E+00 | 2.71E+00 | 5.43E-01 | no | Cops8 | upstream_gene_variant | MODIFIER |
| 1 | 90601263 | AX-170371273 | 0 | G | A | 0.158 | 1.26E-01 | 2.09E-01 | 2.65E+00 | 2.71E+00 | 5.43E-01 | no | Cops8 | upstream_gene_variant | MODIFIER |
| 1 | 90639505 | AX-97824635 | 0 | A | G | 0.158 | 1.26E-01 | 2.09E-01 | 2.65E+00 | 2.71E+00 | 5.43E-01 | no | | intergenic_variant | MODIFIER |
| 1 | 90647056 | AX-170001055 | 0 | A | G | 0.158 | 1.26E-01 | 2.09E-01 | 2.65E+00 | 2.71E+00 | 5.43E-01 | no | | intergenic_variant | MODIFIER |
| 1 | 90648790 | AX-170203529 | 0 | T | C | 0.158 | 1.26E-01 | 2.09E-01 | 2.65E+00 | 2.71E+00 | 5.43E-01 | no | | regulatory_region_variant | MODIFIER |
| 1 | 90650708 | AX-170184943 | 0 | C | G | 0.158 | 1.26E-01 | 2.09E-01 | 2.65E+00 | 2.71E+00 | 5.43E-01 | no | | regulatory_region_variant | MODIFIER |
| 1 | 90674953 | AX-170001049 | 0 | G | A | 0.158 | 1.26E-01 | 2.09E-01 | 2.65E+00 | 2.71E+00 | 5.43E-01 | no | Gm9991 | upstream_gene_variant | MODIFIER |
| 1 | 90674987 | AX-170001050 | 0 | A | G | 0.158 | 1.26E-01 | 2.09E-01 | 2.65E+00 | 2.71E+00 | 5.43E-01 | no | Gm9991 | upstream_gene_variant | MODIFIER |
| 1 | 90675939 | AX-97820812 | 0 | T | C | 0.158 | 1.26E-01 | 2.09E-01 | 2.65E+00 | 2.71E+00 | 5.43E-01 | no | Gm9991 | intron_variant,non_coding_transcript_variant | MODIFIER |
| 1 | 90677721 | AX-170420601 | 0 | G | T | 0.158 | 1.26E-01 | 2.09E-01 | 2.65E+00 | 2.71E+00 | 5.43E-01 | no | Gm9991 | intron_variant,non_coding_transcript_variant | MODIFIER |
| 1 | 90705647 | AX-170001054 | 0 | G | A | 0.158 | 1.26E-01 | 2.09E-01 | 2.65E+00 | 2.71E+00 | 5.43E-01 | no | | intergenic_variant | MODIFIER |
| 1 | 90706731 | AX-170001056 | 0 | G | T | 0.158 | 1.26E-01 | 2.09E-01 | 2.65E+00 | 2.71E+00 | 5.43E-01 | no | | intergenic_variant | MODIFIER |
| 1 | 90709985 | AX-97831309 | 0 | T | G | 0.158 | 1.26E-01 | 2.09E-01 | 2.65E+00 | 2.71E+00 | 5.43E-01 | no | | intergenic_variant | MODIFIER |
| 1 | 90747103 | AX-170001053 | 0 | T | A | 0.158 | 1.26E-01 | 2.09E-01 | 2.65E+00 | 2.71E+00 | 5.43E-01 | no | | intergenic_variant | MODIFIER |
| 1 | 90755964 | AX-97815945 | 0 | A | G | 0.158 | 1.26E-01 | 2.09E-01 | 2.65E+00 | 2.71E+00 | 5.43E-01 | no | | intergenic_variant | MODIFIER |
| 1 | 90769772 | AX-170001058 | 0 | T | C | 0.158 | 1.26E-01 | 2.09E-01 | 2.65E+00 | 2.71E+00 | 5.43E-01 | no | Col6a3 | intron_variant | MODIFIER |
| 1 | 90772572 | AX-170421605 | 0 | T | C | 0.158 | 1.26E-01 | 2.09E-01 | 2.65E+00 | 2.71E+00 | 5.43E-01 | no | Col6a3 | intron_variant | MODIFIER |
| 1 | 90773424 | AX-97788578 | 0 | C | T | 0.158 | 1.26E-01 | 2.09E-01 | 2.65E+00 | 2.71E+00 | 5.43E-01 | no | Col6a3 | intron_variant | MODIFIER |
| 1 | 90790760 | AX-170235802 | 0 | G | A | 0.158 | 1.26E-01 | 2.09E-01 | 2.65E+00 | 2.71E+00 | 5.43E-01 | no | Col6a3 | intron_variant | MODIFIER |
| 1 | 90794479 | AX-170001057 | 0 | T | C | 0.158 | 1.26E-01 | 2.09E-01 | 2.65E+00 | 2.71E+00 | 5.43E-01 | no | Col6a3 | intron_variant | MODIFIER |
| 1 | 90816806 | AX-170143811 | 0 | A | G | 0.158 | 1.26E-01 | 2.09E-01 | 2.65E+00 | 2.71E+00 | 5.43E-01 | no | Col6a3 | intron_variant | MODIFIER |
| 1 | 90818213 | AX-170126618 | 0 | G | A | 0.158 | 1.26E-01 | 2.09E-01 | 2.65E+00 | 2.71E+00 | 5.43E-01 | no | Col6a3 | intron_variant | MODIFIER |
| 1 | 90822849 | AX-170408248 | 0 | A | C | 0.158 | 1.26E-01 | 2.09E-01 | 2.65E+00 | 2.71E+00 | 5.43E-01 | no | Col6a3 | intron_variant | MODIFIER |
| 1 | 90838457 | AX-170456256 | 0 | C | G | 0.158 | 1.26E-01 | 2.09E-01 | 2.65E+00 | 2.71E+00 | 5.43E-01 | no | Col6a3 | intron_variant | MODIFIER |
| 1 | 90840028 | AX-170434539 | 0 | G | A | 0.158 | 1.26E-01 | 2.09E-01 | 2.65E+00 | 2.71E+00 | 5.43E-01 | no | Col6a3 | intron_variant | MODIFIER |
| 1 | 90846434 | AX-97834425 | 0 | A | G | 0.158 | 1.26E-01 | 2.09E-01 | 2.65E+00 | 2.71E+00 | 5.43E-01 | no | Col6a3 | upstream_gene_variant | MODIFIER |
| 1 | 90850372 | AX-170109827 | 0 | C | T | 0.158 | 1.26E-01 | 2.09E-01 | 2.65E+00 | 2.71E+00 | 5.43E-01 | no | | regulatory_region_variant | MODIFIER |
| 1 | 90871973 | AX-170506061 | 0 | A | T | 0.158 | 1.26E-01 | 2.09E-01 | 2.65E+00 | 2.71E+00 | 5.43E-01 | no | 4930474B08Rik | upstream_gene_variant | MODIFIER |
| 1 | 90878644 | AX-170041993 | 0 | A | G | 0.158 | 1.26E-01 | 2.09E-01 | 2.65E+00 | 2.71E+00 | 5.43E-01 | no | | intergenic_variant | MODIFIER |
| 1 | 90882519 | AX-97856544 | 0 | G | A | 0.158 | 1.26E-01 | 2.09E-01 | 2.65E+00 | 2.71E+00 | 5.43E-01 | no | | intergenic_variant | MODIFIER |
| 1 | 90883452 | AX-170104304 | 0 | G | A | 0.158 | 1.26E-01 | 2.09E-01 | 2.65E+00 | 2.71E+00 | 5.43E-01 | no | | intergenic_variant | MODIFIER |
| 1 | 90896422 | AX-97817464 | 0 | C | A | 0.158 | 1.26E-01 | 2.09E-01 | 2.65E+00 | 2.71E+00 | 5.43E-01 | no | | regulatory_region_variant | MODIFIER |
| 1 | 90904680 | AX-170341542 | 0 | T | C | 0.158 | 1.26E-01 | 2.09E-01 | 2.65E+00 | 2.71E+00 | 5.43E-01 | no | | regulatory_region_variant | MODIFIER |
| 1 | 90916311 | AX-170313066 | 0 | A | G | 0.158 | 1.26E-01 | 2.09E-01 | 2.65E+00 | 2.71E+00 | 5.43E-01 | no | Mlph | intron_variant | MODIFIER |
| 1 | 90933038 | AX-97797141 | 0 | T | C | 0.158 | 1.26E-01 | 2.09E-01 | 2.65E+00 | 2.71E+00 | 5.43E-01 | no | Mlph | intron_variant | MODIFIER |
| 1 | 90936514 | AX-170116673 | 0 | G | A | 0.158 | 1.26E-01 | 2.09E-01 | 2.65E+00 | 2.71E+00 | 5.43E-01 | no | Mlph | intron_variant | MODIFIER |
| 1 | 90938672 | AX-170105497 | 0 | T | C | 0.158 | 1.26E-01 | 2.09E-01 | 2.65E+00 | 2.71E+00 | 5.43E-01 | no | Mlph | intron_variant | MODIFIER |
| 1 | 90946931 | AX-170285145 | 0 | C | G | 0.158 | 1.26E-01 | 2.09E-01 | 2.65E+00 | 2.71E+00 | 5.43E-01 | no | Mlph | intron_variant | MODIFIER |
| 1 | 90960718 | AX-97811429 | 0 | T | C | 0.158 | 1.26E-01 | 2.09E-01 | 2.65E+00 | 2.71E+00 | 5.43E-01 | no | Rab17 | intron_variant | MODIFIER |
| 1 | 90976516 | AX-170197463 | 0 | T | C | 0.158 | 1.26E-01 | 2.09E-01 | 2.65E+00 | 2.71E+00 | 5.43E-01 | no | Gm28199 | upstream_gene_variant | MODIFIER |
| 1 | 90976600 | AX-170411813 | 0 | A | G | 0.158 | 1.26E-01 | 2.09E-01 | 2.65E+00 | 2.71E+00 | 5.43E-01 | no | Gm28199 | upstream_gene_variant | MODIFIER |
| 1 | 90983092 | AX-97818845 | 0 | C | T | 0.158 | 1.26E-01 | 2.09E-01 | 2.65E+00 | 2.71E+00 | 5.43E-01 | no | | intergenic_variant | MODIFIER |
| 1 | 90988311 | AX-170513734 | 0 | A | G | 0.158 | 1.26E-01 | 2.09E-01 | 2.65E+00 | 2.71E+00 | 5.43E-01 | no | | intergenic_variant | MODIFIER |
| 1 | 91043098 | AX-170453270 | 0 | T | C | 0.158 | 1.26E-01 | 2.09E-01 | 2.65E+00 | 2.71E+00 | 5.43E-01 | no | Lrrfip1 | intron_variant | MODIFIER |
| 1 | 91049321 | AX-170476097 | 0 | C | T | 0.158 | 1.26E-01 | 2.09E-01 | 2.65E+00 | 2.71E+00 | 5.43E-01 | no | Lrrfip1 | intron_variant | MODIFIER |

| | | | | | | | | | | | | | | | |
|---|---|---|---|---|---|---|---|---|---|---|---|---|---|---|---|
| 1 | 91061354 | AX-97825971 | 0 | A | G | 0.158 | 1.26E-01 | 2.09E-01 | 2.65E+00 | 2.71E+00 | 5.43E-01 | no | Lrrfip1 | intron_variant | MODIFIER |
| 1 | 91061389 | AX-170349284 | 0 | T | C | 0.158 | 1.26E-01 | 2.09E-01 | 2.65E+00 | 2.71E+00 | 5.43E-01 | no | Lrrfip1 | intron_variant | MODIFIER |
| 1 | 91077630 | AX-170399745 | 0 | C | T | 0.158 | 1.26E-01 | 2.09E-01 | 2.65E+00 | 2.71E+00 | 5.43E-01 | no | Lrrfip1 | intron_variant | MODIFIER |
| 1 | 91079994 | AX-170353358 | 0 | T | C | 0.158 | 1.26E-01 | 2.09E-01 | 2.65E+00 | 2.71E+00 | 5.43E-01 | no | Lrrfip1 | intron_variant | MODIFIER |
| 1 | 91084590 | AX-170451481 | 0 | G | A | 0.158 | 1.26E-01 | 2.09E-01 | 2.65E+00 | 2.71E+00 | 5.43E-01 | no | Lrrfip1 | intron_variant | MODIFIER |
| 1 | 91089939 | AX-170209458 | 0 | T | C | 0.158 | 1.26E-01 | 2.09E-01 | 2.65E+00 | 2.71E+00 | 5.43E-01 | no | Lrrfip1 | intron_variant | MODIFIER |
| 1 | 91093922 | AX-170126484 | 0 | A | G | 0.158 | 1.26E-01 | 2.09E-01 | 2.65E+00 | 2.71E+00 | 5.43E-01 | no | Lrrfip1 | intron_variant | MODIFIER |
| 1 | 91094925 | AX-170236254 | 0 | T | C | 0.158 | 1.26E-01 | 2.09E-01 | 2.65E+00 | 2.71E+00 | 5.43E-01 | no | Lrrfip1 | intron_variant | MODIFIER |
| 1 | 91096523 | AX-170386383 | 0 | G | T | 0.158 | 1.26E-01 | 2.09E-01 | 2.65E+00 | 2.71E+00 | 5.43E-01 | no | Lrrfip1 | intron_variant | MODIFIER |
| 1 | 91096711 | AX-170286425 | 0 | C | T | 0.158 | 1.26E-01 | 2.09E-01 | 2.65E+00 | 2.71E+00 | 5.43E-01 | no | Lrrfip1 | intron_variant | MODIFIER |
| 1 | 91101434 | AX-170035622 | 0 | G | T | 0.158 | 1.26E-01 | 2.09E-01 | 2.65E+00 | 2.71E+00 | 5.43E-01 | no | Lrrfip1 | intron_variant | MODIFIER |
| 1 | 91105085 | AX-170342117 | 0 | A | G | 0.158 | 1.26E-01 | 2.09E-01 | 2.65E+00 | 2.71E+00 | 5.43E-01 | no | Lrrfip1 | intron_variant | MODIFIER |
| 1 | 91108023 | AX-170246580 | 0 | C | T | 0.158 | 1.26E-01 | 2.09E-01 | 2.65E+00 | 2.71E+00 | 5.43E-01 | no | Lrrfip1 | intron_variant | MODIFIER |
| 1 | 91109253 | AX-97812326 | 0 | A | G | 0.158 | 1.26E-01 | 2.09E-01 | 2.65E+00 | 2.71E+00 | 5.43E-01 | no | Lrrfip1 | intron_variant | MODIFIER |
| 1 | 91114140 | AX-170400523 | 0 | A | G | 0.158 | 1.26E-01 | 2.09E-01 | 2.65E+00 | 2.71E+00 | 5.43E-01 | no | Lrrfip1 | intron_variant | MODIFIER |
| 1 | 91118075 | AX-170035623 | 0 | C | T | 0.158 | 1.26E-01 | 2.09E-01 | 2.65E+00 | 2.71E+00 | 5.43E-01 | no | Lrrfip1 | intron_variant | MODIFIER |
| 1 | 91136955 | AX-170041999 | 0 | A | G | 0.158 | 1.26E-01 | 2.09E-01 | 2.65E+00 | 2.71E+00 | 5.43E-01 | no | | intergenic_variant | MODIFIER |
| 1 | 91139124 | AX-170001059 | 0 | T | C | 0.158 | 1.26E-01 | 2.09E-01 | 2.65E+00 | 2.71E+00 | 5.43E-01 | no | | intergenic_variant | MODIFIER |
| 1 | 91154956 | AX-170303068 | 0 | A | G | 0.158 | 1.26E-01 | 2.09E-01 | 2.65E+00 | 2.71E+00 | 5.43E-01 | no | Rbm44 | intron_variant | MODIFIER |
| 1 | 91161030 | AX-170518917 | 0 | A | G | 0.158 | 1.26E-01 | 2.09E-01 | 2.65E+00 | 2.71E+00 | 5.43E-01 | no | Rbm44 | intron_variant | MODIFIER |
| 1 | 91161786 | AX-170342864 | 0 | T | C | 0.158 | 1.26E-01 | 2.09E-01 | 2.65E+00 | 2.71E+00 | 5.43E-01 | no | Rbm44 | intron_variant | MODIFIER |
| 1 | 91163276 | AX-170394940 | 0 | C | A | 0.158 | 1.26E-01 | 2.09E-01 | 2.65E+00 | 2.71E+00 | 5.43E-01 | no | Rbm44 | intron_variant | MODIFIER |
| 1 | 91163622 | AX-170431907 | 0 | G | A | 0.158 | 1.26E-01 | 2.09E-01 | 2.65E+00 | 2.71E+00 | 5.43E-01 | no | Rbm44 | intron_variant | MODIFIER |
| 1 | 91165314 | AX-170001060 | 0 | T | C | 0.158 | 1.26E-01 | 2.09E-01 | 2.65E+00 | 2.71E+00 | 5.43E-01 | no | Rbm44 | intron_variant | MODIFIER |
| 1 | 91173707 | AX-170158720 | 0 | C | T | 0.158 | 1.26E-01 | 2.09E-01 | 2.65E+00 | 2.71E+00 | 5.43E-01 | no | Rbm44 | downstream_gene_variant | MODIFIER |
| 1 | 91176389 | AX-170472689 | 0 | A | G | 0.158 | 1.26E-01 | 2.09E-01 | 2.65E+00 | 2.71E+00 | 5.43E-01 | no | Ramp1 | upstream_gene_variant | MODIFIER |
| 1 | 91182380 | AX-170001063 | 0 | G | A | 0.158 | 1.26E-01 | 2.09E-01 | 2.65E+00 | 2.71E+00 | 5.43E-01 | no | Ramp1 | intron_variant | MODIFIER |
| 1 | 91206616 | AX-170403411 | 0 | T | C | 0.158 | 1.26E-01 | 2.09E-01 | 2.65E+00 | 2.71E+00 | 5.43E-01 | no | Ramp1 | intron_variant | MODIFIER |
| 1 | 91207075 | AX-170290450 | 0 | G | T | 0.158 | 1.26E-01 | 2.09E-01 | 2.65E+00 | 2.71E+00 | 5.43E-01 | no | Ramp1 | intron_variant | MODIFIER |
| 1 | 91207410 | AX-170490436 | 0 | T | C | 0.158 | 1.26E-01 | 2.09E-01 | 2.65E+00 | 2.71E+00 | 5.43E-01 | no | Ramp1 | intron_variant | MODIFIER |
| 1 | 91209204 | AX-97783677 | 0 | T | C | 0.158 | 1.26E-01 | 2.09E-01 | 2.65E+00 | 2.71E+00 | 5.43E-01 | no | Ramp1 | intron_variant | MODIFIER |
| 1 | 91212126 | AX-170146671 | 0 | A | G | 0.158 | 1.26E-01 | 2.09E-01 | 2.65E+00 | 2.71E+00 | 5.43E-01 | no | Ramp1 | intron_variant | MODIFIER |
| 1 | 91249142 | AX-170469940 | 0 | G | A | 0.158 | 1.26E-01 | 2.09E-01 | 2.65E+00 | 2.71E+00 | 5.43E-01 | no | Ube2f | upstream_gene_variant | MODIFIER |
| 1 | 91262352 | AX-170108581 | 0 | C | T | 0.158 | 1.26E-01 | 2.09E-01 | 2.65E+00 | 2.71E+00 | 5.43E-01 | no | Ube2f | intron_variant | MODIFIER |
| 1 | 91263590 | AX-170386762 | 0 | A | G | 0.158 | 1.26E-01 | 2.09E-01 | 2.65E+00 | 2.71E+00 | 5.43E-01 | no | Ube2f | intron_variant | MODIFIER |
| 1 | 91272647 | AX-170035617 | 0 | G | A | 0.158 | 1.26E-01 | 2.09E-01 | 2.65E+00 | 2.71E+00 | 5.43E-01 | no | Ube2f | intron_variant | MODIFIER |
| 1 | 91277131 | AX-170243629 | 0 | C | T | 0.158 | 1.26E-01 | 2.09E-01 | 2.65E+00 | 2.71E+00 | 5.43E-01 | no | Ube2f | intron_variant | MODIFIER |
| 1 | 91289831 | AX-170035619 | 0 | C | A | 0.158 | 1.26E-01 | 2.09E-01 | 2.65E+00 | 2.71E+00 | 5.43E-01 | no | Ube2f | downstream_gene_variant | MODIFIER |
| 1 | 91308168 | AX-170212155 | 0 | C | T | 0.158 | 1.26E-01 | 2.09E-01 | 2.65E+00 | 2.71E+00 | 5.43E-01 | no | Scly | intron_variant | MODIFIER |
| 1 | 91308248 | AX-97851346 | 0 | A | G | 0.158 | 1.26E-01 | 2.09E-01 | 2.65E+00 | 2.71E+00 | 5.43E-01 | no | Scly | intron_variant | MODIFIER |
| 1 | 91314387 | AX-170298134 | 0 | A | G | 0.158 | 1.26E-01 | 2.09E-01 | 2.65E+00 | 2.71E+00 | 5.43E-01 | no | Scly | intron_variant | MODIFIER |
| 1 | 91348440 | AX-97811451 | 0 | A | G | 0.158 | 1.26E-01 | 2.09E-01 | 2.65E+00 | 2.71E+00 | 5.43E-01 | no | Klhl30 | upstream_gene_variant | MODIFIER |
| 1 | 91436960 | AX-170189777 | 0 | T | A | 0.158 | 1.26E-01 | 2.09E-01 | 2.65E+00 | 2.71E+00 | 5.43E-01 | no | Per2 | intron_variant | MODIFIER |
| 1 | 91437389 | AX-170346676 | 0 | T | C | 0.158 | 1.26E-01 | 2.09E-01 | 2.65E+00 | 2.71E+00 | 5.43E-01 | no | Per2 | intron_variant | MODIFIER |
| 1 | 91440634 | AX-170349055 | 0 | C | T | 0.158 | 1.26E-01 | 2.09E-01 | 2.65E+00 | 2.71E+00 | 5.43E-01 | no | Per2 | intron_variant | MODIFIER |
| 1 | 91451195 | AX-170486200 | 0 | C | T | 0.158 | 1.26E-01 | 2.09E-01 | 2.65E+00 | 2.71E+00 | 5.43E-01 | no | Per2 | intron_variant | MODIFIER |
| 1 | 91462064 | AX-97804790 | 0 | A | C | 0.158 | 1.26E-01 | 2.09E-01 | 2.65E+00 | 2.71E+00 | 5.43E-01 | no | Per2 | upstream_gene_variant | MODIFIER |
| 1 | 91466102 | AX-170229579 | 0 | G | A | 0.158 | 1.26E-01 | 2.09E-01 | 2.65E+00 | 2.71E+00 | 5.43E-01 | no | Gm28381 | downstream_gene_variant | MODIFIER |
| 1 | 91497858 | AX-97793377 | 0 | C | T | 0.158 | 1.26E-01 | 2.09E-01 | 2.65E+00 | 2.71E+00 | 5.43E-01 | no | Traf3ip1 | intron_variant | MODIFIER |
| 1 | 91503370 | AX-170494886 | 0 | C | T | 0.158 | 1.26E-01 | 2.09E-01 | 2.65E+00 | 2.71E+00 | 5.43E-01 | no | Traf3ip1 | intron_variant | MODIFIER |
| 1 | 91504975 | AX-170042011 | 0 | A | G | 0.158 | 1.26E-01 | 2.09E-01 | 2.65E+00 | 2.71E+00 | 5.43E-01 | no | Traf3ip1 | intron_variant | MODIFIER |
| 1 | 91521580 | AX-170234440 | 0 | A | T | 0.158 | 1.26E-01 | 2.09E-01 | 2.65E+00 | 2.71E+00 | 5.43E-01 | no | Traf3ip1 | intron_variant | MODIFIER |
| 1 | 91522237 | AX-170042009 | 0 | G | A | 0.158 | 1.26E-01 | 2.09E-01 | 2.65E+00 | 2.71E+00 | 5.43E-01 | no | Traf3ip1 | intron_variant | MODIFIER |
| 1 | 91530551 | AX-170042008 | 0 | T | C | 0.158 | 1.26E-01 | 2.09E-01 | 2.65E+00 | 2.71E+00 | 5.43E-01 | no | Traf3ip1 | downstream_gene_variant | MODIFIER |

| | | | | | | | | | | | | | | | |
|---|---|---|---|---|---|---|---|---|---|---|---|---|---|---|---|
| 1 | 91546181 | AX-170358973 | 0 | A | G | 0.158 | 1.26E-01 | 2.09E-01 | 2.65E+00 | 2.71E+00 | 5.43E-01 | no | Asb1 | intron_variant | MODIFIER |
| 1 | 91548423 | AX-170364264 | 0 | T | C | 0.158 | 1.26E-01 | 2.09E-01 | 2.65E+00 | 2.71E+00 | 5.43E-01 | no | Asb1 | intron_variant | MODIFIER |
| 1 | 91554703 | AX-97824641 | 0 | C | T | 0.158 | 1.26E-01 | 2.09E-01 | 2.65E+00 | 2.71E+00 | 5.43E-01 | no | Asb1 | intron_variant | MODIFIER |
| 1 | 91658160 | AX-97835273 | 0 | A | G | 0.158 | 1.26E-01 | 2.09E-01 | 2.65E+00 | 2.71E+00 | 5.43E-01 | no | | intergenic_variant | MODIFIER |
| 1 | 91667226 | AX-170001067 | 0 | C | T | 0.158 | 1.26E-01 | 2.09E-01 | 2.65E+00 | 2.71E+00 | 5.43E-01 | no | | intergenic_variant | MODIFIER |
| 1 | 93119799 | AX-170178903 | 0 | A | G | 0.158 | 1.26E-01 | 2.09E-01 | 2.65E+00 | 2.71E+00 | 5.43E-01 | no | Gm28086 | downstream_gene_variant | MODIFIER |
| 1 | 93206098 | AX-170001094 | 0 | G | A | 0.158 | 1.26E-01 | 2.09E-01 | 2.65E+00 | 2.71E+00 | 5.43E-01 | no | Crocc2 | intron_variant | MODIFIER |
| 1 | 93214589 | AX-170328707 | 0 | C | T | 0.158 | 1.26E-01 | 2.09E-01 | 2.65E+00 | 2.71E+00 | 5.43E-01 | no | Crocc2 | intron_variant | MODIFIER |
| 1 | 93225959 | AX-170001092 | 0 | C | T | 0.158 | 1.26E-01 | 2.09E-01 | 2.65E+00 | 2.71E+00 | 5.43E-01 | no | Crocc2 | intron_variant | MODIFIER |
| 1 | 93226912 | AX-97829557 | 0 | G | T | 0.158 | 1.26E-01 | 2.09E-01 | 2.65E+00 | 2.71E+00 | 5.43E-01 | no | Crocc2 | intron_variant | MODIFIER |
| 1 | 93230395 | AX-170042043 | 0 | C | A | 0.158 | 1.26E-01 | 2.09E-01 | 2.65E+00 | 2.71E+00 | 5.43E-01 | no | Crocc2 | 3_prime_UTR_variant | MODIFIER |
| 1 | 93261707 | AX-170001095 | 0 | T | C | 0.158 | 1.26E-01 | 2.09E-01 | 2.65E+00 | 2.71E+00 | 5.43E-01 | no | Sned1 | synonymous_variant | LOW |
| 1 | 93276794 | AX-170001093 | 0 | C | A | 0.158 | 1.26E-01 | 2.09E-01 | 2.65E+00 | 2.71E+00 | 5.43E-01 | no | Sned1 | intron_variant | MODIFIER |
| 1 | 93279208 | AX-170103563 | 0 | G | A | 0.158 | 1.26E-01 | 2.09E-01 | 2.65E+00 | 2.71E+00 | 5.43E-01 | no | Sned1 | intron_variant | MODIFIER |
| 1 | 93295909 | AX-170386139 | 0 | G | A | 0.158 | 1.26E-01 | 2.09E-01 | 2.65E+00 | 2.71E+00 | 5.43E-01 | no | Sned1 | intron_variant | MODIFIER |
| 1 | 93311869 | AX-170172700 | 0 | T | C | 0.158 | 1.26E-01 | 2.09E-01 | 2.65E+00 | 2.71E+00 | 5.43E-01 | no | Pask | intron_variant | MODIFIER |
| 1 | 93316866 | AX-170486649 | 0 | A | G | 0.158 | 1.26E-01 | 2.09E-01 | 2.65E+00 | 2.71E+00 | 5.43E-01 | no | Pask | synonymous_variant | LOW |
| 1 | 93348696 | AX-170217280 | 0 | G | C | 0.158 | 1.26E-01 | 2.09E-01 | 2.65E+00 | 2.71E+00 | 5.43E-01 | no | Ppp1r7 | intron_variant | MODIFIER |
| 1 | 93403982 | AX-170186713 | 0 | T | C | 0.158 | 1.26E-01 | 2.09E-01 | 2.65E+00 | 2.71E+00 | 5.43E-01 | no | Hdlbp | downstream_gene_variant | MODIFIER |
| 1 | 93404123 | AX-170383372 | 0 | C | G | 0.158 | 1.26E-01 | 2.09E-01 | 2.65E+00 | 2.71E+00 | 5.43E-01 | no | Hdlbp | downstream_gene_variant | MODIFIER |
| 1 | 93404204 | AX-170355205 | 0 | A | G | 0.158 | 1.26E-01 | 2.09E-01 | 2.65E+00 | 2.71E+00 | 5.43E-01 | no | Hdlbp | downstream_gene_variant | MODIFIER |
| 1 | 93426581 | AX-170431261 | 0 | T | A | 0.158 | 1.26E-01 | 2.09E-01 | 2.65E+00 | 2.71E+00 | 5.43E-01 | no | Hdlbp | intron_variant | MODIFIER |
| 1 | 93426683 | AX-170250259 | 0 | C | T | 0.158 | 1.26E-01 | 2.09E-01 | 2.65E+00 | 2.71E+00 | 5.43E-01 | no | Hdlbp | intron_variant | MODIFIER |
| 1 | 93469049 | AX-170447889 | 0 | G | A | 0.158 | 1.26E-01 | 2.09E-01 | 2.65E+00 | 2.71E+00 | 5.43E-01 | no | Hdlbp | intron_variant | MODIFIER |
| 1 | 93496124 | AX-170042048 | 0 | T | C | 0.158 | 1.26E-01 | 2.09E-01 | 2.65E+00 | 2.71E+00 | 5.43E-01 | no | SEP2 | intron_variant | MODIFIER |
| 1 | 93513540 | AX-170269734 | 0 | A | C | 0.158 | 1.26E-01 | 2.09E-01 | 2.65E+00 | 2.71E+00 | 5.43E-01 | no | SEP2 | downstream_gene_variant | MODIFIER |
| 1 | 93513613 | AX-170475540 | 0 | C | T | 0.158 | 1.26E-01 | 2.09E-01 | 2.65E+00 | 2.71E+00 | 5.43E-01 | no | SEP2 | downstream_gene_variant | MODIFIER |
| 1 | 93526021 | AX-97839085 | 0 | T | C | 0.158 | 1.26E-01 | 2.09E-01 | 2.65E+00 | 2.71E+00 | 5.43E-01 | no | Farp2 | intron_variant | MODIFIER |
| 1 | 93552893 | AX-170086535 | 0 | A | G | 0.158 | 1.26E-01 | 2.09E-01 | 2.65E+00 | 2.71E+00 | 5.43E-01 | no | Farp2 | intron_variant | MODIFIER |
| 1 | 93614692 | AX-97808272 | 0 | C | T | 0.158 | 1.26E-01 | 2.09E-01 | 2.65E+00 | 2.71E+00 | 5.43E-01 | no | Farp2 | intron_variant | MODIFIER |
| 1 | 93622090 | AX-170138871 | 0 | T | G | 0.158 | 1.26E-01 | 2.09E-01 | 2.65E+00 | 2.71E+00 | 5.43E-01 | no | Stk25 | 3_prime_UTR_variant | MODIFIER |
| 1 | 93624538 | AX-170117790 | 0 | C | T | 0.158 | 1.26E-01 | 2.09E-01 | 2.65E+00 | 2.71E+00 | 5.43E-01 | no | Stk25 | intron_variant | MODIFIER |
| 1 | 93625106 | AX-170086536 | 0 | T | C | 0.158 | 1.26E-01 | 2.09E-01 | 2.65E+00 | 2.71E+00 | 5.43E-01 | no | Stk25 | synonymous_variant | LOW |
| 1 | 93667326 | AX-97852988 | 0 | G | A | 0.158 | 1.26E-01 | 2.09E-01 | 2.65E+00 | 2.71E+00 | 5.43E-01 | no | | regulatory_region_variant | MODIFIER |
| 1 | 95110483 | AX-97799371 | 0 | C | T | 0.158 | 1.26E-01 | 2.09E-01 | 2.65E+00 | 2.71E+00 | 5.43E-01 | no | | intergenic_variant | MODIFIER |
| 1 | 95110541 | AX-170530911 | 0 | T | C | 0.158 | 1.26E-01 | 2.09E-01 | 2.65E+00 | 2.71E+00 | 5.43E-01 | no | | intergenic_variant | MODIFIER |
| 1 | 95138968 | AX-170522898 | 0 | G | A | 0.158 | 1.26E-01 | 2.09E-01 | 2.65E+00 | 2.71E+00 | 5.43E-01 | no | | intergenic_variant | MODIFIER |
| 1 | 95140249 | AX-170253385 | 0 | A | G | 0.158 | 1.26E-01 | 2.09E-01 | 2.65E+00 | 2.71E+00 | 5.43E-01 | no | | intergenic_variant | MODIFIER |
| 1 | 95190129 | AX-170257062 | 0 | G | A | 0.158 | 1.26E-01 | 2.09E-01 | 2.65E+00 | 2.71E+00 | 5.43E-01 | no | | intergenic_variant | MODIFIER |
| 1 | 95201644 | AX-170042074 | 0 | G | A | 0.158 | 1.26E-01 | 2.09E-01 | 2.65E+00 | 2.71E+00 | 5.43E-01 | no | | intergenic_variant | MODIFIER |
| 1 | 95242799 | AX-97789978 | 0 | G | A | 0.158 | 1.26E-01 | 2.09E-01 | 2.65E+00 | 2.71E+00 | 5.43E-01 | no | | intergenic_variant | MODIFIER |
| 1 | 95242863 | AX-170042073 | 0 | T | C | 0.158 | 1.26E-01 | 2.09E-01 | 2.65E+00 | 2.71E+00 | 5.43E-01 | no | | intergenic_variant | MODIFIER |
| 1 | 95245425 | AX-170122652 | 0 | G | A | 0.158 | 1.26E-01 | 2.09E-01 | 2.65E+00 | 2.71E+00 | 5.43E-01 | no | | intergenic_variant | MODIFIER |
| 1 | 95950220 | AX-97796447 | 0 | T | G | 0.158 | 1.26E-01 | 2.09E-01 | 2.65E+00 | 2.71E+00 | 5.43E-01 | no | | intergenic_variant | MODIFIER |
| 1 | 96262046 | AX-170466201 | 0 | T | C | 0.158 | 1.26E-01 | 2.09E-01 | 2.65E+00 | 2.71E+00 | 5.43E-01 | no | Gm24586 | downstream_gene_variant | MODIFIER |
| 1 | 96267116 | AX-170001141 | 0 | C | T | 0.158 | 1.26E-01 | 2.09E-01 | 2.65E+00 | 2.71E+00 | 5.43E-01 | no | Gm24586 | upstream_gene_variant | MODIFIER |
| 1 | 97772567 | AX-170446471 | 0 | C | T | 0.158 | 1.26E-01 | 2.09E-01 | 2.65E+00 | 2.71E+00 | 5.43E-01 | no | Gin1 | intron_variant | MODIFIER |
| 1 | 97773018 | AX-170445152 | 0 | C | A | 0.158 | 1.26E-01 | 2.09E-01 | 2.65E+00 | 2.71E+00 | 5.43E-01 | no | Gin1 | intron_variant | MODIFIER |
| 1 | 97775126 | AX-168395856 | 0 | C | T | 0.158 | 1.26E-01 | 2.09E-01 | 2.65E+00 | 2.71E+00 | 5.43E-01 | no | Gin1 | intron_variant | MODIFIER |
| 1 | 97926627 | AX-97821980 | 0 | C | T | 0.158 | 1.26E-01 | 2.09E-01 | 2.65E+00 | 2.71E+00 | 5.43E-01 | no | Pam | intron_variant | MODIFIER |
| 1 | 98062075 | AX-170086548 | 0 | G | A | 0.158 | 1.26E-01 | 2.09E-01 | 2.65E+00 | 2.71E+00 | 5.43E-01 | no | Pam | intron_variant | MODIFIER |
| 1 | 98504776 | AX-170086551 | 0 | C | G | 0.158 | 1.26E-01 | 2.09E-01 | 2.65E+00 | 2.71E+00 | 5.43E-01 | no | Slco6d1 | intron_variant | MODIFIER |
| 1 | 98587846 | AX-170455315 | 0 | C | G | 0.158 | 1.26E-01 | 2.09E-01 | 2.65E+00 | 2.71E+00 | 5.43E-01 | no | Gm7967 | upstream_gene_variant | MODIFIER |
| 1 | 98606933 | AX-168375031 | 0 | G | A | 0.158 | 1.26E-01 | 2.09E-01 | 2.65E+00 | 2.71E+00 | 5.43E-01 | no | Gm7967 | intron_variant,non_coding_transcript_variant | MODIFIER |

| | | | | | | | | | | | | | | | |
|---|---|---|---|---|---|---|---|---|---|---|---|---|---|---|---|
| 1 | 98637686 | AX-170124064 | 0 | A | G | 0.158 | 1.26E-01 | 2.09E-01 | 2.65E+00 | 2.71E+00 | 5.43E-01 | no | | intergenic_variant | MODIFIER |
| 1 | 98917850 | AX-170373368 | 0 | C | T | 0.158 | 1.26E-01 | 2.09E-01 | 2.65E+00 | 2.71E+00 | 5.43E-01 | no | | intergenic_variant | MODIFIER |
| 1 | 98932890 | AX-170265300 | 0 | C | G | 0.158 | 1.26E-01 | 2.09E-01 | 2.65E+00 | 2.71E+00 | 5.43E-01 | no | | intergenic_variant | MODIFIER |
| 1 | 98960097 | AX-170378730 | 0 | A | G | 0.158 | 1.26E-01 | 2.09E-01 | 2.65E+00 | 2.71E+00 | 5.43E-01 | no | | intergenic_variant | MODIFIER |
| 1 | 98999005 | AX-170001177 | 0 | C | T | 0.158 | 1.26E-01 | 2.09E-01 | 2.65E+00 | 2.71E+00 | 5.43E-01 | no | | intergenic_variant | MODIFIER |
| 1 | 99008975 | AX-170326942 | 0 | G | C | 0.158 | 1.26E-01 | 2.09E-01 | 2.65E+00 | 2.71E+00 | 5.43E-01 | no | | intergenic_variant | MODIFIER |
| 1 | 99016735 | AX-170464974 | 0 | A | G | 0.158 | 1.26E-01 | 2.09E-01 | 2.65E+00 | 2.71E+00 | 5.43E-01 | no | Gm23434 | upstream_gene_variant | MODIFIER |
| 1 | 99019406 | AX-170001175 | 0 | T | A | 0.158 | 1.26E-01 | 2.09E-01 | 2.65E+00 | 2.71E+00 | 5.43E-01 | no | Gm23434 | downstream_gene_variant | MODIFIER |
| 1 | 99027001 | AX-97842644 | 0 | A | G | 0.158 | 1.26E-01 | 2.09E-01 | 2.65E+00 | 2.71E+00 | 5.43E-01 | no | | intergenic_variant | MODIFIER |
| 1 | 99048846 | AX-170356444 | 0 | G | A | 0.158 | 1.26E-01 | 2.09E-01 | 2.65E+00 | 2.71E+00 | 5.43E-01 | no | | intergenic_variant | MODIFIER |
| 1 | 99067208 | AX-170382545 | 0 | T | C | 0.158 | 1.26E-01 | 2.09E-01 | 2.65E+00 | 2.71E+00 | 5.43E-01 | no | | intergenic_variant | MODIFIER |
| 1 | 99070410 | AX-170001181 | 0 | A | C | 0.158 | 1.26E-01 | 2.09E-01 | 2.65E+00 | 2.71E+00 | 5.43E-01 | no | | intergenic_variant | MODIFIER |
| 1 | 99124026 | AX-170001180 | 0 | A | G | 0.158 | 1.26E-01 | 2.09E-01 | 2.65E+00 | 2.71E+00 | 5.43E-01 | no | | intergenic_variant | MODIFIER |
| 1 | 99126891 | AX-170001184 | 0 | G | A | 0.158 | 1.26E-01 | 2.09E-01 | 2.65E+00 | 2.71E+00 | 5.43E-01 | no | | intergenic_variant | MODIFIER |
| 1 | 99140758 | AX-170001178 | 0 | C | G | 0.158 | 1.26E-01 | 2.09E-01 | 2.65E+00 | 2.71E+00 | 5.43E-01 | no | | intergenic_variant | MODIFIER |
| 1 | 99211464 | AX-170392091 | 0 | T | G | 0.158 | 1.26E-01 | 2.09E-01 | 2.65E+00 | 2.71E+00 | 5.43E-01 | no | | intergenic_variant | MODIFIER |
| 1 | 99317236 | AX-170229463 | 0 | G | C | 0.158 | 1.26E-01 | 2.09E-01 | 2.65E+00 | 2.71E+00 | 5.43E-01 | no | | intergenic_variant | MODIFIER |
| 1 | 99338614 | AX-97843754 | 0 | T | C | 0.158 | 1.26E-01 | 2.09E-01 | 2.65E+00 | 2.71E+00 | 5.43E-01 | no | | intergenic_variant | MODIFIER |
| 1 | 99344660 | AX-170217834 | 0 | T | C | 0.158 | 1.26E-01 | 2.09E-01 | 2.65E+00 | 2.71E+00 | 5.43E-01 | no | | intergenic_variant | MODIFIER |
| 1 | 99400091 | AX-170186965 | 0 | G | A | 0.158 | 1.26E-01 | 2.09E-01 | 2.65E+00 | 2.71E+00 | 5.43E-01 | no | | intergenic_variant | MODIFIER |
| 1 | 99411425 | AX-170108954 | 0 | T | C | 0.158 | 1.26E-01 | 2.09E-01 | 2.65E+00 | 2.71E+00 | 5.43E-01 | no | | intergenic_variant | MODIFIER |
| 1 | 99417823 | AX-170525006 | 0 | A | G | 0.158 | 1.26E-01 | 2.09E-01 | 2.65E+00 | 2.71E+00 | 5.43E-01 | no | | intergenic_variant | MODIFIER |
| 1 | 99447469 | AX-97809903 | 0 | A | G | 0.158 | 1.26E-01 | 2.09E-01 | 2.65E+00 | 2.71E+00 | 5.43E-01 | no | | intergenic_variant | MODIFIER |
| 1 | 99502622 | AX-170207331 | 0 | T | G | 0.158 | 1.26E-01 | 2.09E-01 | 2.65E+00 | 2.71E+00 | 5.43E-01 | no | | intergenic_variant | MODIFIER |
| 1 | 99522025 | AX-170001192 | 0 | A | G | 0.158 | 1.26E-01 | 2.09E-01 | 2.65E+00 | 2.71E+00 | 5.43E-01 | no | | intergenic_variant | MODIFIER |
| 1 | 99529904 | AX-97797648 | 0 | G | A | 0.158 | 1.26E-01 | 2.09E-01 | 2.65E+00 | 2.71E+00 | 5.43E-01 | no | | intergenic_variant | MODIFIER |
| 1 | 99568197 | AX-97829836 | 0 | T | A | 0.158 | 1.26E-01 | 2.09E-01 | 2.65E+00 | 2.71E+00 | 5.43E-01 | no | | intergenic_variant | MODIFIER |
| 1 | 99629263 | AX-170001188 | 0 | T | G | 0.158 | 1.26E-01 | 2.09E-01 | 2.65E+00 | 2.71E+00 | 5.43E-01 | no | | intergenic_variant | MODIFIER |
| 1 | 99629782 | AX-97807379 | 0 | G | A | 0.158 | 1.26E-01 | 2.09E-01 | 2.65E+00 | 2.71E+00 | 5.43E-01 | no | | intergenic_variant | MODIFIER |
| 1 | 99657653 | AX-97850145 | 0 | G | T | 0.158 | 1.26E-01 | 2.09E-01 | 2.65E+00 | 2.71E+00 | 5.43E-01 | no | | intergenic_variant | MODIFIER |
| 1 | 99690074 | AX-170235785 | 0 | G | A | 0.158 | 1.26E-01 | 2.09E-01 | 2.65E+00 | 2.71E+00 | 5.43E-01 | no | | intergenic_variant | MODIFIER |
| 1 | 99774594 | AX-170412844 | 0 | G | T | 0.158 | 1.26E-01 | 2.09E-01 | 2.65E+00 | 2.71E+00 | 5.43E-01 | no | Cntnap5b | intron_variant | MODIFIER |
| 1 | 99892814 | AX-170099989 | 0 | G | C | 0.158 | 1.26E-01 | 2.09E-01 | 2.65E+00 | 2.71E+00 | 5.43E-01 | no | Cntnap5b | intron_variant | MODIFIER |
| 1 | 99899093 | AX-97838864 | 0 | G | A | 0.158 | 1.26E-01 | 2.09E-01 | 2.65E+00 | 2.71E+00 | 5.43E-01 | no | Cntnap5b | intron_variant | MODIFIER |
| 1 | 99899645 | AX-170221436 | 0 | C | G | 0.158 | 1.26E-01 | 2.09E-01 | 2.65E+00 | 2.71E+00 | 5.43E-01 | no | Cntnap5b | intron_variant | MODIFIER |
| 1 | 99929758 | AX-170099551 | 0 | G | C | 0.158 | 1.26E-01 | 2.09E-01 | 2.65E+00 | 2.71E+00 | 5.43E-01 | no | Cntnap5b | intron_variant | MODIFIER |
| 1 | 99930096 | AX-170373470 | 0 | C | G | 0.158 | 1.26E-01 | 2.09E-01 | 2.65E+00 | 2.71E+00 | 5.43E-01 | no | Cntnap5b | intron_variant | MODIFIER |
| 1 | 99930227 | AX-170403813 | 0 | C | A | 0.158 | 1.26E-01 | 2.09E-01 | 2.65E+00 | 2.71E+00 | 5.43E-01 | no | Cntnap5b | intron_variant | MODIFIER |
| 1 | 99930556 | AX-170408033 | 0 | G | A | 0.158 | 1.26E-01 | 2.09E-01 | 2.65E+00 | 2.71E+00 | 5.43E-01 | no | Cntnap5b | intron_variant | MODIFIER |
| 1 | 100019942 | AX-170112520 | 0 | A | C | 0.158 | 1.26E-01 | 2.09E-01 | 2.65E+00 | 2.71E+00 | 5.43E-01 | no | Cntnap5b | intron_variant | MODIFIER |
| 1 | 100065779 | AX-170195576 | 0 | G | A | 0.158 | 1.26E-01 | 2.09E-01 | 2.65E+00 | 2.71E+00 | 5.43E-01 | no | Cntnap5b | intron_variant | MODIFIER |
| 1 | 100141387 | AX-170244394 | 0 | T | C | 0.158 | 1.26E-01 | 2.09E-01 | 2.65E+00 | 2.71E+00 | 5.43E-01 | no | Cntnap5b | synonymous_variant | LOW |
| 1 | 100141632 | AX-97840812 | 0 | T | C | 0.158 | 1.26E-01 | 2.09E-01 | 2.65E+00 | 2.71E+00 | 5.43E-01 | no | Cntnap5b | intron_variant | MODIFIER |
| 1 | 100141864 | AX-170001198 | 0 | G | A | 0.158 | 1.26E-01 | 2.09E-01 | 2.65E+00 | 2.71E+00 | 5.43E-01 | no | Cntnap5b | intron_variant | MODIFIER |
| 1 | 100150041 | AX-170103499 | 0 | T | C | 0.158 | 1.26E-01 | 2.09E-01 | 2.65E+00 | 2.71E+00 | 5.43E-01 | no | Cntnap5b | intron_variant | MODIFIER |
| 1 | 100170994 | AX-170216846 | 0 | G | A | 0.158 | 1.26E-01 | 2.09E-01 | 2.65E+00 | 2.71E+00 | 5.43E-01 | no | Cntnap5b | intron_variant | MODIFIER |
| 1 | 100220098 | AX-170471133 | 0 | T | G | 0.158 | 1.26E-01 | 2.09E-01 | 2.65E+00 | 2.71E+00 | 5.43E-01 | no | Cntnap5b | intron_variant | MODIFIER |
| 1 | 100221445 | AX-170426959 | 0 | T | G | 0.158 | 1.26E-01 | 2.09E-01 | 2.65E+00 | 2.71E+00 | 5.43E-01 | no | Cntnap5b | intron_variant | MODIFIER |
| 1 | 100264839 | AX-97824401 | 0 | G | T | 0.158 | 1.26E-01 | 2.09E-01 | 2.65E+00 | 2.71E+00 | 5.43E-01 | no | Cntnap5b | intron_variant | MODIFIER |
| 1 | 100270236 | AX-170163471 | 0 | A | G | 0.158 | 1.26E-01 | 2.09E-01 | 2.65E+00 | 2.71E+00 | 5.43E-01 | no | Cntnap5b | intron_variant | MODIFIER |
| 1 | 100280147 | AX-170417412 | 0 | A | G | 0.158 | 1.26E-01 | 2.09E-01 | 2.65E+00 | 2.71E+00 | 5.43E-01 | no | Cntnap5b | intron_variant | MODIFIER |
| 1 | 100285806 | AX-170265168 | 0 | G | A | 0.158 | 1.26E-01 | 2.09E-01 | 2.65E+00 | 2.71E+00 | 5.43E-01 | no | Cntnap5b | intron_variant | MODIFIER |
| 1 | 100295327 | AX-170252905 | 0 | G | A | 0.158 | 1.26E-01 | 2.09E-01 | 2.65E+00 | 2.71E+00 | 5.43E-01 | no | Cntnap5b | intron_variant | MODIFIER |
| 1 | 100295890 | AX-170409228 | 0 | T | C | 0.158 | 1.26E-01 | 2.09E-01 | 2.65E+00 | 2.71E+00 | 5.43E-01 | no | Cntnap5b | intron_variant | MODIFIER |

| | | | | | | | | | | | | | | | |
|---|---|---|---|---|---|---|---|---|---|---|---|---|---|---|---|
| 1 | 100330780 | AX-170231755 | 0 | C | T | 0.158 | 1.26E-01 | 2.09E-01 | 2.65E+00 | 2.71E+00 | 5.43E-01 | no | Cntnap5b | intron_variant | MODIFIER |
| 1 | 100345299 | AX-97841957 | 0 | T | C | 0.158 | 1.26E-01 | 2.09E-01 | 2.65E+00 | 2.71E+00 | 5.43E-01 | no | Cntnap5b | intron_variant | MODIFIER |
| 1 | 100348193 | AX-170135860 | 0 | C | T | 0.158 | 1.26E-01 | 2.09E-01 | 2.65E+00 | 2.71E+00 | 5.43E-01 | no | Cntnap5b | intron_variant | MODIFIER |
| 1 | 100356632 | AX-170307272 | 0 | A | G | 0.158 | 1.26E-01 | 2.09E-01 | 2.65E+00 | 2.71E+00 | 5.43E-01 | no | Cntnap5b | intron_variant | MODIFIER |
| 1 | 100356825 | AX-170309934 | 0 | C | A | 0.158 | 1.26E-01 | 2.09E-01 | 2.65E+00 | 2.71E+00 | 5.43E-01 | no | Cntnap5b | intron_variant | MODIFIER |
| 1 | 100383536 | AX-170246300 | 0 | C | A | 0.158 | 1.26E-01 | 2.09E-01 | 2.65E+00 | 2.71E+00 | 5.43E-01 | no | Cntnap5b | missense_variant | MODERATE |
| 1 | 100383950 | AX-170444244 | 0 | G | T | 0.158 | 1.26E-01 | 2.09E-01 | 2.65E+00 | 2.71E+00 | 5.43E-01 | no | Cntnap5b | intron_variant | MODIFIER |
| 1 | 100385999 | AX-97833808 | 0 | A | G | 0.158 | 1.26E-01 | 2.09E-01 | 2.65E+00 | 2.71E+00 | 5.43E-01 | no | Cntnap5b | intron_variant | MODIFIER |
| 1 | 100391124 | AX-170326708 | 0 | C | G | 0.158 | 1.26E-01 | 2.09E-01 | 2.65E+00 | 2.71E+00 | 5.43E-01 | no | Cntnap5b | intron_variant | MODIFIER |
| 1 | 100421150 | AX-170492641 | 0 | A | G | 0.158 | 1.26E-01 | 2.09E-01 | 2.65E+00 | 2.71E+00 | 5.43E-01 | no | Cntnap5b | intron_variant | MODIFIER |
| 1 | 100428435 | AX-170284832 | 0 | G | T | 0.158 | 1.26E-01 | 2.09E-01 | 2.65E+00 | 2.71E+00 | 5.43E-01 | no | Cntnap5b | intron_variant | MODIFIER |
| 1 | 100428474 | AX-170218488 | 0 | T | C | 0.158 | 1.26E-01 | 2.09E-01 | 2.65E+00 | 2.71E+00 | 5.43E-01 | no | Cntnap5b | intron_variant | MODIFIER |
| 1 | 100430328 | AX-170119174 | 0 | G | C | 0.158 | 1.26E-01 | 2.09E-01 | 2.65E+00 | 2.71E+00 | 5.43E-01 | no | Cntnap5b | intron_variant | MODIFIER |
| 1 | 100456654 | AX-170332406 | 0 | G | A | 0.158 | 1.26E-01 | 2.09E-01 | 2.65E+00 | 2.71E+00 | 5.43E-01 | no | Cntnap5b | intron_variant | MODIFIER |
| 1 | 100478831 | AX-170517708 | 0 | G | C | 0.158 | 1.26E-01 | 2.09E-01 | 2.65E+00 | 2.71E+00 | 5.43E-01 | no | Cntnap5b | intron_variant | MODIFIER |
| 1 | 100479864 | AX-170134376 | 0 | A | T | 0.158 | 1.26E-01 | 2.09E-01 | 2.65E+00 | 2.71E+00 | 5.43E-01 | no | Cntnap5b | intron_variant | MODIFIER |
| 1 | 100491515 | AX-170324960 | 0 | C | G | 0.158 | 1.26E-01 | 2.09E-01 | 2.65E+00 | 2.71E+00 | 5.43E-01 | no | | intergenic_variant | MODIFIER |
| 1 | 100497662 | AX-97781884 | 0 | T | C | 0.158 | 1.26E-01 | 2.09E-01 | 2.65E+00 | 2.71E+00 | 5.43E-01 | no | | intergenic_variant | MODIFIER |
| 1 | 100504430 | AX-170292729 | 0 | A | G | 0.158 | 1.26E-01 | 2.09E-01 | 2.65E+00 | 2.71E+00 | 5.43E-01 | no | | intergenic_variant | MODIFIER |
| 1 | 100526261 | AX-97806166 | 0 | T | A | 0.158 | 1.26E-01 | 2.09E-01 | 2.65E+00 | 2.71E+00 | 5.43E-01 | no | | intergenic_variant | MODIFIER |
| 1 | 100533427 | AX-170042180 | 0 | A | G | 0.158 | 1.26E-01 | 2.09E-01 | 2.65E+00 | 2.71E+00 | 5.43E-01 | no | | intergenic_variant | MODIFIER |
| 1 | 100553948 | AX-170163122 | 0 | A | G | 0.158 | 1.26E-01 | 2.09E-01 | 2.65E+00 | 2.71E+00 | 5.43E-01 | no | | intergenic_variant | MODIFIER |
| 1 | 100568878 | AX-170116783 | 0 | T | G | 0.158 | 1.26E-01 | 2.09E-01 | 2.65E+00 | 2.71E+00 | 5.43E-01 | no | | intergenic_variant | MODIFIER |
| 1 | 100570547 | AX-170486208 | 0 | C | T | 0.158 | 1.26E-01 | 2.09E-01 | 2.65E+00 | 2.71E+00 | 5.43E-01 | no | | intergenic_variant | MODIFIER |
| 1 | 100571374 | AX-170091611 | 0 | T | G | 0.158 | 1.26E-01 | 2.09E-01 | 2.65E+00 | 2.71E+00 | 5.43E-01 | no | | intergenic_variant | MODIFIER |
| 1 | 100600413 | AX-170001201 | 0 | A | T | 0.158 | 1.26E-01 | 2.09E-01 | 2.65E+00 | 2.71E+00 | 5.43E-01 | no | Gm18700 | downstream_gene_variant | MODIFIER |
| 1 | 100606632 | AX-97787219 | 0 | C | T | 0.158 | 1.26E-01 | 2.09E-01 | 2.65E+00 | 2.71E+00 | 5.43E-01 | no | | intergenic_variant | MODIFIER |
| 1 | 100615104 | AX-170001200 | 0 | G | C | 0.158 | 1.26E-01 | 2.09E-01 | 2.65E+00 | 2.71E+00 | 5.43E-01 | no | | intergenic_variant | MODIFIER |
| 1 | 100644501 | AX-97846756 | 0 | T | C | 0.158 | 1.26E-01 | 2.09E-01 | 2.65E+00 | 2.71E+00 | 5.43E-01 | no | Gm18185 | downstream_gene_variant | MODIFIER |
| 1 | 100650349 | AX-170370152 | 0 | A | T | 0.158 | 1.26E-01 | 2.09E-01 | 2.65E+00 | 2.71E+00 | 5.43E-01 | no | Gm29334 | upstream_gene_variant | MODIFIER |
| 1 | 100651046 | AX-170450214 | 0 | A | G | 0.158 | 1.26E-01 | 2.09E-01 | 2.65E+00 | 2.71E+00 | 5.43E-01 | no | Gm29334 | upstream_gene_variant | MODIFIER |
| 1 | 100651250 | AX-170132656 | 0 | G | A | 0.158 | 1.26E-01 | 2.09E-01 | 2.65E+00 | 2.71E+00 | 5.43E-01 | no | Gm29334 | upstream_gene_variant | MODIFIER |
| 1 | 100654978 | AX-97815717 | 0 | A | G | 0.158 | 1.26E-01 | 2.09E-01 | 2.65E+00 | 2.71E+00 | 5.43E-01 | no | Gm29334 | downstream_gene_variant | MODIFIER |
| 1 | 100666822 | AX-170402421 | 0 | G | A | 0.158 | 1.26E-01 | 2.09E-01 | 2.65E+00 | 2.71E+00 | 5.43E-01 | no | | intergenic_variant | MODIFIER |
| 1 | 100669650 | AX-170223725 | 0 | C | T | 0.158 | 1.26E-01 | 2.09E-01 | 2.65E+00 | 2.71E+00 | 5.43E-01 | no | | intergenic_variant | MODIFIER |
| 1 | 100685780 | AX-170154178 | 0 | T | C | 0.158 | 1.26E-01 | 2.09E-01 | 2.65E+00 | 2.71E+00 | 5.43E-01 | no | | intergenic_variant | MODIFIER |
| 1 | 100686023 | AX-170308776 | 0 | A | G | 0.158 | 1.26E-01 | 2.09E-01 | 2.65E+00 | 2.71E+00 | 5.43E-01 | no | | intergenic_variant | MODIFIER |
| 1 | 100698947 | AX-170452207 | 0 | A | T | 0.158 | 1.26E-01 | 2.09E-01 | 2.65E+00 | 2.71E+00 | 5.43E-01 | no | | intergenic_variant | MODIFIER |
| 1 | 100703015 | AX-170506504 | 0 | T | G | 0.158 | 1.26E-01 | 2.09E-01 | 2.65E+00 | 2.71E+00 | 5.43E-01 | no | | intergenic_variant | MODIFIER |
| 1 | 100725290 | AX-170495383 | 0 | G | T | 0.158 | 1.26E-01 | 2.09E-01 | 2.65E+00 | 2.71E+00 | 5.43E-01 | no | | intergenic_variant | MODIFIER |
| 1 | 100736174 | AX-97794241 | 0 | G | A | 0.158 | 1.26E-01 | 2.09E-01 | 2.65E+00 | 2.71E+00 | 5.43E-01 | no | | intergenic_variant | MODIFIER |
| 1 | 100736343 | AX-170350487 | 0 | G | T | 0.158 | 1.26E-01 | 2.09E-01 | 2.65E+00 | 2.71E+00 | 5.43E-01 | no | | intergenic_variant | MODIFIER |
| 1 | 100736374 | AX-170188178 | 0 | G | C | 0.158 | 1.26E-01 | 2.09E-01 | 2.65E+00 | 2.71E+00 | 5.43E-01 | no | | intergenic_variant | MODIFIER |
| 1 | 100739481 | AX-170042182 | 0 | A | G | 0.158 | 1.26E-01 | 2.09E-01 | 2.65E+00 | 2.71E+00 | 5.43E-01 | no | | intergenic_variant | MODIFIER |
| 1 | 100742558 | AX-170166781 | 0 | T | G | 0.158 | 1.26E-01 | 2.09E-01 | 2.65E+00 | 2.71E+00 | 5.43E-01 | no | | intergenic_variant | MODIFIER |
| 1 | 100745145 | AX-170042186 | 0 | C | T | 0.158 | 1.26E-01 | 2.09E-01 | 2.65E+00 | 2.71E+00 | 5.43E-01 | no | | intergenic_variant | MODIFIER |
| 1 | 100770343 | AX-170210824 | 0 | C | T | 0.158 | 1.26E-01 | 2.09E-01 | 2.65E+00 | 2.71E+00 | 5.43E-01 | no | | intergenic_variant | MODIFIER |
| 1 | 102068900 | AX-170153962 | 0 | T | C | 0.158 | 1.26E-01 | 2.09E-01 | 2.65E+00 | 2.71E+00 | 5.43E-01 | no | | intergenic_variant | MODIFIER |
| 1 | 102101054 | AX-170226431 | 0 | C | G | 0.158 | 1.26E-01 | 2.09E-01 | 2.65E+00 | 2.71E+00 | 5.43E-01 | no | | intergenic_variant | MODIFIER |
| 1 | 102197824 | AX-170001213 | 0 | T | C | 0.158 | 1.26E-01 | 2.09E-01 | 2.65E+00 | 2.71E+00 | 5.43E-01 | no | | intergenic_variant | MODIFIER |
| 1 | 102813199 | AX-170001223 | 0 | G | A | 0.158 | 1.26E-01 | 2.09E-01 | 2.65E+00 | 2.71E+00 | 5.43E-01 | no | | intergenic_variant | MODIFIER |
| 1 | 102951617 | AX-170382309 | 0 | A | T | 0.158 | 1.26E-01 | 2.09E-01 | 2.65E+00 | 2.71E+00 | 5.43E-01 | no | Gm22034 | downstream_gene_variant | MODIFIER |
| 1 | 103175675 | AX-170445223 | 0 | T | A | 0.158 | 1.26E-01 | 2.09E-01 | 2.65E+00 | 2.71E+00 | 5.43E-01 | no | Gm18728 | intron_variant,non_coding_transcript_variant | MODIFIER |
| 1 | 104589638 | AX-170256525 | 0 | A | G | 0.158 | 1.26E-01 | 2.09E-01 | 2.65E+00 | 2.71E+00 | 5.43E-01 | no | | intergenic_variant | MODIFIER |

| | | | | | | | | | | | | | | | |
|---|---|---|---|---|---|---|---|---|---|---|---|---|---|---|---|
| 1 | 104986195 | AX-170091624 | 0 | A | G | 0.158 | 1.26E-01 | 2.09E-01 | 2.65E+00 | 2.71E+00 | 5.43E-01 | no | Cdh20 | intron_variant | MODIFIER |
| 1 | 105245972 | AX-170338610 | 0 | T | A | 0.158 | 1.26E-01 | 2.09E-01 | 2.65E+00 | 2.71E+00 | 5.43E-01 | no | Gm29013 | downstream_gene_variant | MODIFIER |
| 1 | 105260632 | AX-170509793 | 0 | T | C | 0.158 | 1.26E-01 | 2.09E-01 | 2.65E+00 | 2.71E+00 | 5.43E-01 | no | | intergenic_variant | MODIFIER |
| 1 | 105264641 | AX-170433441 | 0 | C | A | 0.158 | 1.26E-01 | 2.09E-01 | 2.65E+00 | 2.71E+00 | 5.43E-01 | no | | intergenic_variant | MODIFIER |
| 1 | 105264829 | AX-170430587 | 0 | T | A | 0.158 | 1.26E-01 | 2.09E-01 | 2.65E+00 | 2.71E+00 | 5.43E-01 | no | | intergenic_variant | MODIFIER |
| 1 | 105290410 | AX-170042263 | 0 | C | T | 0.158 | 1.26E-01 | 2.09E-01 | 2.65E+00 | 2.71E+00 | 5.43E-01 | no | Rnf152 | intron_variant | MODIFIER |
| 1 | 105290585 | AX-170196832 | 0 | A | G | 0.158 | 1.26E-01 | 2.09E-01 | 2.65E+00 | 2.71E+00 | 5.43E-01 | no | Rnf152 | intron_variant | MODIFIER |
| 1 | 105293236 | AX-170474176 | 0 | A | G | 0.158 | 1.26E-01 | 2.09E-01 | 2.65E+00 | 2.71E+00 | 5.43E-01 | no | Rnf152 | intron_variant | MODIFIER |
| 1 | 105293385 | AX-170444426 | 0 | T | C | 0.158 | 1.26E-01 | 2.09E-01 | 2.65E+00 | 2.71E+00 | 5.43E-01 | no | Rnf152 | intron_variant | MODIFIER |
| 1 | 105294423 | AX-170401302 | 0 | A | G | 0.158 | 1.26E-01 | 2.09E-01 | 2.65E+00 | 2.71E+00 | 5.43E-01 | no | Rnf152 | intron_variant | MODIFIER |
| 1 | 105301217 | AX-170309468 | 0 | G | C | 0.158 | 1.26E-01 | 2.09E-01 | 2.65E+00 | 2.71E+00 | 5.43E-01 | no | Rnf152 | intron_variant | MODIFIER |
| 1 | 105302194 | AX-170459990 | 0 | G | A | 0.158 | 1.26E-01 | 2.09E-01 | 2.65E+00 | 2.71E+00 | 5.43E-01 | no | Rnf152 | intron_variant | MODIFIER |
| 1 | 105329258 | AX-170513138 | 0 | C | G | 0.158 | 1.26E-01 | 2.09E-01 | 2.65E+00 | 2.71E+00 | 5.43E-01 | no | Rnf152 | intron_variant | MODIFIER |
| 1 | 105331144 | AX-170263566 | 0 | T | G | 0.158 | 1.26E-01 | 2.09E-01 | 2.65E+00 | 2.71E+00 | 5.43E-01 | no | Rnf152 | intron_variant | MODIFIER |
| 1 | 105333250 | AX-170136871 | 0 | A | G | 0.158 | 1.26E-01 | 2.09E-01 | 2.65E+00 | 2.71E+00 | 5.43E-01 | no | Rnf152 | intron_variant | MODIFIER |
| 1 | 105334710 | AX-170458975 | 0 | C | G | 0.158 | 1.26E-01 | 2.09E-01 | 2.65E+00 | 2.71E+00 | 5.43E-01 | no | Rnf152 | intron_variant | MODIFIER |
| 1 | 105347291 | AX-170473332 | 0 | C | T | 0.158 | 1.26E-01 | 2.09E-01 | 2.65E+00 | 2.71E+00 | 5.43E-01 | no | Rnf152 | intron_variant | MODIFIER |
| 1 | 105348014 | AX-170152698 | 0 | G | A | 0.158 | 1.26E-01 | 2.09E-01 | 2.65E+00 | 2.71E+00 | 5.43E-01 | no | Rnf152 | intron_variant | MODIFIER |
| 1 | 105354474 | AX-170363966 | 0 | T | G | 0.158 | 1.26E-01 | 2.09E-01 | 2.65E+00 | 2.71E+00 | 5.43E-01 | no | Rnf152 | intron_variant | MODIFIER |
| 1 | 105423644 | AX-170042264 | 0 | T | A | 0.158 | 1.26E-01 | 2.09E-01 | 2.65E+00 | 2.71E+00 | 5.43E-01 | no | AC124553.1 | intron_variant,non_coding_transcript_variant | MODIFIER |
| 1 | 105425428 | AX-170174985 | 0 | T | C | 0.158 | 1.26E-01 | 2.09E-01 | 2.65E+00 | 2.71E+00 | 5.43E-01 | no | AC124553.1 | intron_variant,non_coding_transcript_variant | MODIFIER |
| 1 | 105425516 | AX-170042267 | 0 | C | T | 0.158 | 1.26E-01 | 2.09E-01 | 2.65E+00 | 2.71E+00 | 5.43E-01 | no | AC124553.1 | intron_variant,non_coding_transcript_variant | MODIFIER |
| 1 | 105427439 | AX-97826300 | 0 | C | T | 0.158 | 1.26E-01 | 2.09E-01 | 2.65E+00 | 2.71E+00 | 5.43E-01 | no | AC124553.1 | intron_variant,non_coding_transcript_variant | MODIFIER |
| 1 | 105462432 | AX-170298722 | 0 | C | T | 0.158 | 1.26E-01 | 2.09E-01 | 2.65E+00 | 2.71E+00 | 5.43E-01 | no | Gm3608 | downstream_gene_variant | MODIFIER |
| 1 | 105462722 | AX-97812630 | 0 | A | C | 0.158 | 1.26E-01 | 2.09E-01 | 2.65E+00 | 2.71E+00 | 5.43E-01 | no | Gm3608 | downstream_gene_variant | MODIFIER |
| 1 | 105462785 | AX-170121797 | 0 | G | A | 0.158 | 1.26E-01 | 2.09E-01 | 2.65E+00 | 2.71E+00 | 5.43E-01 | no | Gm3608 | downstream_gene_variant | MODIFIER |
| 1 | 105502964 | AX-170001270 | 0 | C | T | 0.158 | 1.26E-01 | 2.09E-01 | 2.65E+00 | 2.71E+00 | 5.43E-01 | no | AC124553.1 | intron_variant,non_coding_transcript_variant | MODIFIER |
| 1 | 105564058 | AX-170183720 | 0 | G | A | 0.158 | 1.26E-01 | 2.09E-01 | 2.65E+00 | 2.71E+00 | 5.43E-01 | no | Pign | intron_variant | MODIFIER |
| 1 | 105578737 | AX-97852734 | 0 | T | A | 0.158 | 1.26E-01 | 2.09E-01 | 2.65E+00 | 2.71E+00 | 5.43E-01 | no | Pign | intron_variant | MODIFIER |
| 1 | 105582623 | AX-170311424 | 0 | T | G | 0.158 | 1.26E-01 | 2.09E-01 | 2.65E+00 | 2.71E+00 | 5.43E-01 | no | Pign | intron_variant | MODIFIER |
| 1 | 105636786 | AX-170195986 | 0 | G | A | 0.158 | 1.26E-01 | 2.09E-01 | 2.65E+00 | 2.71E+00 | 5.43E-01 | no | Pign | intron_variant | MODIFIER |
| 1 | 105644528 | AX-170453090 | 0 | G | A | 0.158 | 1.26E-01 | 2.09E-01 | 2.65E+00 | 2.71E+00 | 5.43E-01 | no | Pign | intron_variant | MODIFIER |
| 1 | 105740650 | AX-170143160 | 0 | T | C | 0.158 | 1.26E-01 | 2.09E-01 | 2.65E+00 | 2.71E+00 | 5.43E-01 | no | Relch | intron_variant | MODIFIER |
| 1 | 105750845 | AX-170519547 | 0 | G | A | 0.158 | 1.26E-01 | 2.09E-01 | 2.65E+00 | 2.71E+00 | 5.43E-01 | no | Relch | intron_variant | MODIFIER |
| 1 | 105896692 | AX-170166676 | 0 | C | T | 0.158 | 1.26E-01 | 2.09E-01 | 2.65E+00 | 2.71E+00 | 5.43E-01 | no | | intergenic_variant | MODIFIER |
| 1 | 105986396 | AX-97842859 | 0 | A | G | 0.158 | 1.26E-01 | 2.09E-01 | 2.65E+00 | 2.71E+00 | 5.43E-01 | no | Zcchc2 | upstream_gene_variant | MODIFIER |
| 12 | 101636711 | AX-170007404 | 0 | A | T | 0.096 | 8.78E-02 | 1.47E-01 | 2.72E+00 | 2.84E+00 | 5.43E-01 | no | | intergenic_variant | MODIFIER |
| 9 | 89413600 | AX-170153251 | 2 | A | G | 0.118 | -7.90E-02 | 1.30E-01 | 2.60E+00 | 2.68E+00 | 5.43E-01 | no | | intergenic_variant | MODIFIER |
| 7 | 10328442 | AX-170095861 | 0 | T | A | 0.167 | 4.70E-02 | 7.93E-02 | 2.79E+00 | 2.94E+00 | 5.43E-01 | no | Gm45402 | upstream_gene_variant | MODIFIER |
| 12 | 104979444 | AX-168378863 | 0 | G | A | 0.132 | -1.16E-01 | 1.94E-01 | 2.77E+00 | 2.84E+00 | 5.43E-01 | no | Syne3 | intron_variant | MODIFIER |
| 5 | 32075141 | AX-97820106 | 1 | A | G | 0.357 | 6.33E-02 | 1.05E-01 | 2.71E+00 | 2.76E+00 | 5.43E-01 | no | Babam2 | intron_variant | MODIFIER |
| 1 | 100293308 | AX-170348413 | 1 | G | A | 0.161 | 1.24E-01 | 2.05E-01 | 2.66E+00 | 2.72E+00 | 5.43E-01 | no | Cntnap5b | intron_variant | MODIFIER |
| 1 | 20802709 | AX-170000044 | 0 | T | A | 0.412 | 5.09E-02 | 8.30E-02 | 2.58E+00 | 2.63E+00 | 5.43E-01 | no | Mcm3 | downstream_gene_variant | MODIFIER |
| 15 | 79794344 | AX-170452199 | 1 | A | G | 0.554 | 4.89E-02 | 7.97E-02 | 2.56E+00 | 2.64E+00 | 5.43E-01 | no | Nptxr | synonymous_variant | LOW |
| 4 | 69604034 | AX-170235638 | 0 | T | C | 0.325 | 6.74E-02 | 1.12E-01 | 2.68E+00 | 2.75E+00 | 5.44E-01 | no | | intergenic_variant | MODIFIER |
| 7 | 37927464 | AX-170295767 | 0 | C | T | 0.351 | 5.69E-02 | 9.35E-02 | 2.59E+00 | 2.67E+00 | 5.44E-01 | no | | intergenic_variant | MODIFIER |
| 12 | 111874082 | AX-97843952 | 1 | C | T | 0.143 | 9.57E-02 | 1.61E-01 | 2.69E+00 | 2.80E+00 | 5.44E-01 | no | Ppp1r13b | intron_variant | MODIFIER |
| 9 | 75826962 | AX-170096589 | 0 | C | A | 0.211 | 7.54E-02 | 1.26E-01 | 2.78E+00 | 2.85E+00 | 5.44E-01 | no | Bmp5 | intron_variant | MODIFIER |
| 2 | 114659791 | AX-170348815 | 2 | A | G | 0.1 | 1.02E-01 | 1.71E-01 | 2.66E+00 | 2.75E+00 | 5.45E-01 | no | Dph6 | upstream_gene_variant | MODIFIER |
| 15 | 74333912 | AX-170442730 | 1 | G | C | 0.545 | 4.92E-02 | 8.06E-02 | 2.58E+00 | 2.65E+00 | 5.45E-01 | no | | intergenic_variant | MODIFIER |
| 10 | 71614268 | AX-170492016 | 1 | T | C | 0.42 | -4.73E-02 | 7.85E-02 | 2.63E+00 | 2.71E+00 | 5.45E-01 | no | Gm34544 | intron_variant,non_coding_transcript_variant | MODIFIER |
| 11 | 60164603 | AX-97796899 | 0 | T | G | 0.044 | -9.99E-02 | 1.64E-01 | 2.55E+00 | 2.68E+00 | 5.45E-01 | no | Rai1 | intron_variant | MODIFIER |
| 6 | 129681856 | AX-170508281 | 0 | T | C | 0.544 | 5.16E-02 | 8.32E-02 | 2.52E+00 | 2.59E+00 | 5.45E-01 | no | Klrc1 | upstream_gene_variant | MODIFIER |
| 10 | 5636905 | AX-170344057 | 0 | A | T | 0.211 | -6.31E-02 | 1.05E-01 | 2.65E+00 | 2.74E+00 | 5.45E-01 | no | Vip | upstream_gene_variant | MODIFIER |

| | | | | | | | | | | | | | | |
|---|---|---|---|---|---|---|---|---|---|---|---|---|---|---|
| 4 | 97886259 | AX-170400180 | 1 | C | T | 0.223 | 7.24E-02 | 1.18E-01 | 2.56E+00 | 2.62E+00 | 5.45E-01 | no | Nfia | intron_variant | MODIFIER |
| 8 | 22120841 | AX-170421700 | 0 | T | C | 0.175 | -6.31E-02 | 1.06E-01 | 2.71E+00 | 2.82E+00 | 5.45E-01 | no | Nek5 | missense_variant | MODERATE |
| 1 | 20978171 | AX-170155217 | 0 | G | A | 0.404 | 5.06E-02 | 8.38E-02 | 2.63E+00 | 2.68E+00 | 5.45E-01 | no | Efhc1 | intron_variant | MODIFIER |
| 18 | 52630384 | AX-170093885 | 0 | T | C | 0.053 | 1.36E-01 | 2.30E-01 | 2.70E+00 | 2.82E+00 | 5.45E-01 | no | Zfp474 | intron_variant | MODIFIER |
| 5 | 79682028 | AX-168384443 | 0 | G | A | 0.439 | 6.20E-02 | 1.04E-01 | 2.72E+00 | 2.79E+00 | 5.45E-01 | no | | intergenic_variant | MODIFIER |
| 10 | 114763585 | AX-170278487 | 2 | G | A | 0.382 | 4.82E-02 | 8.10E-02 | 2.74E+00 | 2.83E+00 | 5.45E-01 | no | Trhde | intron_variant | MODIFIER |
| 13 | 31070618 | AX-170275404 | 1 | T | A | 0.17 | -7.33E-02 | 1.20E-01 | 2.58E+00 | 2.64E+00 | 5.45E-01 | no | Gm48662 | downstream_gene_variant | MODIFIER |
| 4 | 90545974 | AX-170394818 | 1 | C | T | 0.259 | 6.81E-02 | 1.06E-01 | 2.43E+00 | 2.51E+00 | 5.45E-01 | no | Gm12636 | upstream_gene_variant | MODIFIER |
| 5 | 5683034 | AX-97850283 | 1 | T | G | 0.411 | -4.01E-02 | 6.59E-02 | 2.55E+00 | 2.68E+00 | 5.45E-01 | no | Steap2 | 5_prime_UTR_variant | MODIFIER |
| 5 | 97142480 | AX-170025666 | 1 | T | A | 0.446 | 6.43E-02 | 1.08E-01 | 2.71E+00 | 2.77E+00 | 5.45E-01 | no | Gm33609 | upstream_gene_variant | MODIFIER |
| 5 | 35460434 | AX-170364951 | 0 | A | G | 0.333 | 6.42E-02 | 1.07E-01 | 2.68E+00 | 2.73E+00 | 5.45E-01 | no | | intergenic_variant | MODIFIER |
| 6 | 128703285 | AX-170215736 | 1 | A | G | 0.545 | 5.16E-02 | 8.26E-02 | 2.50E+00 | 2.58E+00 | 5.45E-01 | no | Klrb1 | downstream_gene_variant | MODIFIER |
| 5 | 103671180 | AX-170185303 | 0 | G | A | 0.439 | 6.96E-02 | 1.16E-01 | 2.65E+00 | 2.68E+00 | 5.45E-01 | no | | intergenic_variant | MODIFIER |
| 5 | 103671434 | AX-170483985 | 0 | G | A | 0.439 | 6.96E-02 | 1.16E-01 | 2.65E+00 | 2.68E+00 | 5.45E-01 | no | | intergenic_variant | MODIFIER |
| 5 | 103671541 | AX-170410299 | 0 | A | G | 0.439 | 6.96E-02 | 1.16E-01 | 2.65E+00 | 2.68E+00 | 5.45E-01 | no | | intergenic_variant | MODIFIER |
| 5 | 103674091 | AX-170221129 | 0 | G | C | 0.439 | 6.96E-02 | 1.16E-01 | 2.65E+00 | 2.68E+00 | 5.45E-01 | no | | regulatory_region_variant | MODIFIER |
| 5 | 103677400 | AX-170172650 | 0 | C | A | 0.439 | 6.96E-02 | 1.16E-01 | 2.65E+00 | 2.68E+00 | 5.45E-01 | no | | intergenic_variant | MODIFIER |
| 5 | 103875885 | AX-170335584 | 0 | G | A | 0.439 | 6.96E-02 | 1.16E-01 | 2.65E+00 | 2.68E+00 | 5.45E-01 | no | Klhl8 | synonymous_variant | LOW |
| 5 | 103901129 | AX-170475213 | 0 | A | C | 0.439 | 6.96E-02 | 1.16E-01 | 2.65E+00 | 2.68E+00 | 5.45E-01 | no | Klhl8 | intron_variant | MODIFIER |
| 5 | 103901305 | AX-97809666 | 0 | A | C | 0.439 | 6.96E-02 | 1.16E-01 | 2.65E+00 | 2.68E+00 | 5.45E-01 | no | Klhl8 | intron_variant | MODIFIER |
| 5 | 103962860 | AX-170524728 | 0 | C | T | 0.439 | 6.96E-02 | 1.16E-01 | 2.65E+00 | 2.68E+00 | 5.45E-01 | no | Hsd17b13 | intron_variant | MODIFIER |
| 5 | 103963801 | AX-170350108 | 0 | T | A | 0.439 | 6.96E-02 | 1.16E-01 | 2.65E+00 | 2.68E+00 | 5.45E-01 | no | Hsd17b13 | intron_variant | MODIFIER |
| 5 | 103967032 | AX-170186907 | 0 | T | G | 0.439 | 6.96E-02 | 1.16E-01 | 2.65E+00 | 2.68E+00 | 5.45E-01 | no | Hsd17b13 | missense_variant | MODERATE |
| 5 | 103967587 | AX-170192663 | 0 | T | C | 0.439 | 6.96E-02 | 1.16E-01 | 2.65E+00 | 2.68E+00 | 5.45E-01 | no | Hsd17b13 | intron_variant | MODIFIER |
| 5 | 103970828 | AX-170396354 | 0 | A | G | 0.439 | 6.96E-02 | 1.16E-01 | 2.65E+00 | 2.68E+00 | 5.45E-01 | no | Hsd17b13 | synonymous_variant | LOW |
| 5 | 104128672 | AX-170035714 | 0 | C | T | 0.439 | 6.96E-02 | 1.16E-01 | 2.65E+00 | 2.68E+00 | 5.45E-01 | no | | regulatory_region_variant | MODIFIER |
| 5 | 104129182 | AX-170072950 | 0 | G | A | 0.439 | 6.96E-02 | 1.16E-01 | 2.65E+00 | 2.68E+00 | 5.45E-01 | no | | regulatory_region_variant | MODIFIER |
| 5 | 104140588 | AX-170113138 | 0 | C | T | 0.439 | 6.96E-02 | 1.16E-01 | 2.65E+00 | 2.68E+00 | 5.45E-01 | no | | intergenic_variant | MODIFIER |
| 5 | 104153804 | AX-170025783 | 0 | T | C | 0.439 | 6.96E-02 | 1.16E-01 | 2.65E+00 | 2.68E+00 | 5.45E-01 | no | | intergenic_variant | MODIFIER |
| 5 | 104155090 | AX-97833093 | 0 | A | G | 0.439 | 6.96E-02 | 1.16E-01 | 2.65E+00 | 2.68E+00 | 5.45E-01 | no | | intergenic_variant | MODIFIER |
| 5 | 104193441 | AX-170288065 | 0 | A | G | 0.439 | 6.96E-02 | 1.16E-01 | 2.65E+00 | 2.68E+00 | 5.45E-01 | no | | intergenic_variant | MODIFIER |
| 5 | 104203088 | AX-170025788 | 0 | C | T | 0.439 | 6.96E-02 | 1.16E-01 | 2.65E+00 | 2.68E+00 | 5.45E-01 | no | Dmp1 | intron_variant | MODIFIER |
| 5 | 104203119 | AX-170106699 | 0 | C | T | 0.439 | 6.96E-02 | 1.16E-01 | 2.65E+00 | 2.68E+00 | 5.45E-01 | no | Dmp1 | intron_variant | MODIFIER |
| 5 | 104220469 | AX-170136989 | 0 | T | C | 0.439 | 6.96E-02 | 1.16E-01 | 2.65E+00 | 2.68E+00 | 5.45E-01 | no | | intergenic_variant | MODIFIER |
| 5 | 104224009 | AX-170488153 | 0 | C | T | 0.439 | 6.96E-02 | 1.16E-01 | 2.65E+00 | 2.68E+00 | 5.45E-01 | no | | intergenic_variant | MODIFIER |
| 5 | 104248869 | AX-170286323 | 0 | C | G | 0.439 | 6.96E-02 | 1.16E-01 | 2.65E+00 | 2.68E+00 | 5.45E-01 | no | Gm31048 | upstream_gene_variant | MODIFIER |
| 5 | 104249733 | AX-170454703 | 0 | C | T | 0.439 | 6.96E-02 | 1.16E-01 | 2.65E+00 | 2.68E+00 | 5.45E-01 | no | Gm31048 | upstream_gene_variant | MODIFIER |
| 5 | 104250394 | AX-170292247 | 0 | T | C | 0.439 | 6.96E-02 | 1.16E-01 | 2.65E+00 | 2.68E+00 | 5.45E-01 | no | Gm31048 | upstream_gene_variant | MODIFIER |
| 5 | 104257718 | AX-97832123 | 0 | T | C | 0.439 | 6.96E-02 | 1.16E-01 | 2.65E+00 | 2.68E+00 | 5.45E-01 | no | | intergenic_variant | MODIFIER |
| 5 | 104258457 | AX-170025784 | 0 | G | A | 0.439 | 6.96E-02 | 1.16E-01 | 2.65E+00 | 2.68E+00 | 5.45E-01 | no | | intergenic_variant | MODIFIER |
| 5 | 104267925 | AX-170130993 | 0 | T | G | 0.439 | 6.96E-02 | 1.16E-01 | 2.65E+00 | 2.68E+00 | 5.45E-01 | no | | intergenic_variant | MODIFIER |
| 5 | 104268304 | AX-170113419 | 0 | T | C | 0.439 | 6.96E-02 | 1.16E-01 | 2.65E+00 | 2.68E+00 | 5.45E-01 | no | | intergenic_variant | MODIFIER |
| 5 | 104268481 | AX-170505067 | 0 | A | G | 0.439 | 6.96E-02 | 1.16E-01 | 2.65E+00 | 2.68E+00 | 5.45E-01 | no | | intergenic_variant | MODIFIER |
| 5 | 104287774 | AX-97819720 | 0 | G | A | 0.439 | 6.96E-02 | 1.16E-01 | 2.65E+00 | 2.68E+00 | 5.45E-01 | no | | intergenic_variant | MODIFIER |
| 5 | 104287820 | AX-170025778 | 0 | C | T | 0.439 | 6.96E-02 | 1.16E-01 | 2.65E+00 | 2.68E+00 | 5.45E-01 | no | | intergenic_variant | MODIFIER |
| 5 | 104358335 | AX-97793446 | 0 | A | G | 0.439 | 6.96E-02 | 1.16E-01 | 2.65E+00 | 2.68E+00 | 5.45E-01 | no | | intergenic_variant | MODIFIER |
| 5 | 104384803 | AX-170025781 | 0 | A | C | 0.439 | 6.96E-02 | 1.16E-01 | 2.65E+00 | 2.68E+00 | 5.45E-01 | no | | intergenic_variant | MODIFIER |
| 5 | 104385696 | AX-170412741 | 0 | C | T | 0.439 | 6.96E-02 | 1.16E-01 | 2.65E+00 | 2.68E+00 | 5.45E-01 | no | | intergenic_variant | MODIFIER |
| 5 | 104948585 | AX-170416592 | 0 | G | A | 0.439 | 6.96E-02 | 1.16E-01 | 2.65E+00 | 2.68E+00 | 5.45E-01 | no | Abcg3 | intron_variant | MODIFIER |
| 5 | 104987437 | AX-170311653 | 0 | T | C | 0.439 | 6.96E-02 | 1.16E-01 | 2.65E+00 | 2.68E+00 | 5.45E-01 | no | Abcg3 | upstream_gene_variant | MODIFIER |
| 5 | 104987600 | AX-170231251 | 0 | T | G | 0.439 | 6.96E-02 | 1.16E-01 | 2.65E+00 | 2.68E+00 | 5.45E-01 | no | Abcg3 | upstream_gene_variant | MODIFIER |
| 5 | 105134985 | AX-170337932 | 0 | A | G | 0.439 | 6.96E-02 | 1.16E-01 | 2.65E+00 | 2.68E+00 | 5.45E-01 | no | Gbp8 | intron_variant | MODIFIER |
| 5 | 105135326 | AX-170377438 | 0 | G | A | 0.439 | 6.96E-02 | 1.16E-01 | 2.65E+00 | 2.68E+00 | 5.45E-01 | no | Gbp8 | intron_variant | MODIFIER |
| 5 | 105260059 | AX-97840291 | 0 | T | C | 0.439 | 6.96E-02 | 1.16E-01 | 2.65E+00 | 2.68E+00 | 5.45E-01 | no | Gm43302 | intron_variant | MODIFIER |

| | | | | | | | | | | | | | | | |
|---|---|---|---|---|---|---|---|---|---|---|---|---|---|---|---|
| 5 | 105268693 | AX-170354749 | 0 | G | A | 0.439 | 6.96E-02 | 1.16E-01 | 2.65E+00 | 2.68E+00 | 5.45E-01 | no | Gm43302 | intron_variant | MODIFIER |
| 5 | 105304706 | AX-170146171 | 0 | C | T | 0.439 | 6.96E-02 | 1.16E-01 | 2.65E+00 | 2.68E+00 | 5.45E-01 | no | | intergenic_variant | MODIFIER |
| 5 | 105311392 | AX-170349993 | 0 | G | A | 0.439 | 6.96E-02 | 1.16E-01 | 2.65E+00 | 2.68E+00 | 5.45E-01 | no | | regulatory_region_variant | MODIFIER |
| 5 | 105327032 | AX-170240752 | 0 | C | A | 0.439 | 6.96E-02 | 1.16E-01 | 2.65E+00 | 2.68E+00 | 5.45E-01 | no | Gbp11 | intron_variant | MODIFIER |
| 5 | 105400321 | AX-170072974 | 0 | C | A | 0.439 | 6.96E-02 | 1.16E-01 | 2.65E+00 | 2.68E+00 | 5.45E-01 | no | Gm26519 | upstream_gene_variant | MODIFIER |
| 5 | 105403990 | AX-170119293 | 0 | T | C | 0.439 | 6.96E-02 | 1.16E-01 | 2.65E+00 | 2.68E+00 | 5.45E-01 | no | Gm26519 | intron_variant,non_coding_transcript_variant | MODIFIER |
| 5 | 105407498 | AX-170383389 | 0 | A | C | 0.439 | 6.96E-02 | 1.16E-01 | 2.65E+00 | 2.68E+00 | 5.45E-01 | no | Gm26519 | downstream_gene_variant | MODIFIER |
| 5 | 105407895 | AX-170252048 | 0 | G | A | 0.439 | 6.96E-02 | 1.16E-01 | 2.65E+00 | 2.68E+00 | 5.45E-01 | no | Gm26519 | downstream_gene_variant | MODIFIER |
| 5 | 105408251 | AX-170025806 | 0 | C | T | 0.439 | 6.96E-02 | 1.16E-01 | 2.65E+00 | 2.68E+00 | 5.45E-01 | no | Gm26519 | downstream_gene_variant | MODIFIER |
| 5 | 105411088 | AX-170473745 | 0 | C | A | 0.439 | 6.96E-02 | 1.16E-01 | 2.65E+00 | 2.68E+00 | 5.45E-01 | no | Lrrc8b | upstream_gene_variant | MODIFIER |
| 5 | 105430341 | AX-170323896 | 0 | T | G | 0.439 | 6.96E-02 | 1.16E-01 | 2.65E+00 | 2.68E+00 | 5.45E-01 | no | Lrrc8b | intron_variant | MODIFIER |
| 5 | 105434355 | AX-97784803 | 0 | G | A | 0.439 | 6.96E-02 | 1.16E-01 | 2.65E+00 | 2.68E+00 | 5.45E-01 | no | Lrrc8b | intron_variant | MODIFIER |
| 5 | 105434377 | AX-170181218 | 0 | T | C | 0.439 | 6.96E-02 | 1.16E-01 | 2.65E+00 | 2.68E+00 | 5.45E-01 | no | Lrrc8b | intron_variant | MODIFIER |
| 5 | 105439725 | AX-170025805 | 0 | C | T | 0.439 | 6.96E-02 | 1.16E-01 | 2.65E+00 | 2.68E+00 | 5.45E-01 | no | Lrrc8b | intron_variant | MODIFIER |
| 5 | 105461957 | AX-170412311 | 0 | T | C | 0.439 | 6.96E-02 | 1.16E-01 | 2.65E+00 | 2.68E+00 | 5.45E-01 | no | Lrrc8b | intron_variant | MODIFIER |
| 5 | 105534906 | AX-170340712 | 0 | T | C | 0.439 | 6.96E-02 | 1.16E-01 | 2.65E+00 | 2.68E+00 | 5.45E-01 | no | Lrrc8c | intron_variant | MODIFIER |
| 5 | 105536587 | AX-170025802 | 0 | G | T | 0.439 | 6.96E-02 | 1.16E-01 | 2.65E+00 | 2.68E+00 | 5.45E-01 | no | Lrrc8c | intron_variant | MODIFIER |
| 5 | 105536910 | AX-170211675 | 0 | T | G | 0.439 | 6.96E-02 | 1.16E-01 | 2.65E+00 | 2.68E+00 | 5.45E-01 | no | Lrrc8c | intron_variant | MODIFIER |
| 5 | 105537470 | AX-170459679 | 0 | A | C | 0.439 | 6.96E-02 | 1.16E-01 | 2.65E+00 | 2.68E+00 | 5.45E-01 | no | Lrrc8c | intron_variant | MODIFIER |
| 5 | 105538597 | AX-170025804 | 0 | T | C | 0.439 | 6.96E-02 | 1.16E-01 | 2.65E+00 | 2.68E+00 | 5.45E-01 | no | Lrrc8c | intron_variant | MODIFIER |
| 5 | 105539458 | AX-170407616 | 0 | T | C | 0.439 | 6.96E-02 | 1.16E-01 | 2.65E+00 | 2.68E+00 | 5.45E-01 | no | Lrrc8c | intron_variant | MODIFIER |
| 5 | 105542648 | AX-170025799 | 0 | C | T | 0.439 | 6.96E-02 | 1.16E-01 | 2.65E+00 | 2.68E+00 | 5.45E-01 | no | Lrrc8c | intron_variant | MODIFIER |
| 5 | 105569995 | AX-170484327 | 0 | A | G | 0.439 | 6.96E-02 | 1.16E-01 | 2.65E+00 | 2.68E+00 | 5.45E-01 | no | Lrrc8c | intron_variant | MODIFIER |
| 5 | 105580350 | AX-97820225 | 0 | C | A | 0.439 | 6.96E-02 | 1.16E-01 | 2.65E+00 | 2.68E+00 | 5.45E-01 | no | Lrrc8c | intron_variant | MODIFIER |
| 5 | 105584223 | AX-170025798 | 0 | C | T | 0.439 | 6.96E-02 | 1.16E-01 | 2.65E+00 | 2.68E+00 | 5.45E-01 | no | Lrrc8c | intron_variant | MODIFIER |
| 5 | 105593642 | AX-170025801 | 0 | G | A | 0.439 | 6.96E-02 | 1.16E-01 | 2.65E+00 | 2.68E+00 | 5.45E-01 | no | Lrrc8c | intron_variant | MODIFIER |
| 5 | 105597818 | AX-170388791 | 0 | T | G | 0.439 | 6.96E-02 | 1.16E-01 | 2.65E+00 | 2.68E+00 | 5.45E-01 | no | Lrrc8c | intron_variant | MODIFIER |
| 5 | 105602514 | AX-97798069 | 0 | A | G | 0.439 | 6.96E-02 | 1.16E-01 | 2.65E+00 | 2.68E+00 | 5.45E-01 | no | Lrrc8c | intron_variant | MODIFIER |
| 5 | 105613209 | AX-170072981 | 0 | A | G | 0.439 | 6.96E-02 | 1.16E-01 | 2.65E+00 | 2.68E+00 | 5.45E-01 | no | Lrrc8c | downstream_gene_variant | MODIFIER |
| 5 | 105614015 | AX-170072979 | 0 | A | G | 0.439 | 6.96E-02 | 1.16E-01 | 2.65E+00 | 2.68E+00 | 5.45E-01 | no | Lrrc8c | downstream_gene_variant | MODIFIER |
| 5 | 105622381 | AX-97789598 | 0 | T | C | 0.439 | 6.96E-02 | 1.16E-01 | 2.65E+00 | 2.68E+00 | 5.45E-01 | no | | regulatory_region_variant | MODIFIER |
| 5 | 105622553 | AX-170131246 | 0 | G | C | 0.439 | 6.96E-02 | 1.16E-01 | 2.65E+00 | 2.68E+00 | 5.45E-01 | no | | regulatory_region_variant | MODIFIER |
| 5 | 105623239 | AX-170451894 | 0 | T | A | 0.439 | 6.96E-02 | 1.16E-01 | 2.65E+00 | 2.68E+00 | 5.45E-01 | no | | regulatory_region_variant | MODIFIER |
| 5 | 105646100 | AX-170208554 | 0 | C | T | 0.439 | 6.96E-02 | 1.16E-01 | 2.65E+00 | 2.68E+00 | 5.45E-01 | no | | intergenic_variant | MODIFIER |
| 5 | 105650195 | AX-170125379 | 0 | A | T | 0.439 | 6.96E-02 | 1.16E-01 | 2.65E+00 | 2.68E+00 | 5.45E-01 | no | | regulatory_region_variant | MODIFIER |
| 5 | 105650208 | AX-170484436 | 0 | A | G | 0.439 | 6.96E-02 | 1.16E-01 | 2.65E+00 | 2.68E+00 | 5.45E-01 | no | | regulatory_region_variant | MODIFIER |
| 5 | 105660472 | AX-97788218 | 0 | C | T | 0.439 | 6.96E-02 | 1.16E-01 | 2.65E+00 | 2.68E+00 | 5.45E-01 | no | | intergenic_variant | MODIFIER |
| 5 | 105664279 | AX-170416991 | 0 | G | A | 0.439 | 6.96E-02 | 1.16E-01 | 2.65E+00 | 2.68E+00 | 5.45E-01 | no | | intergenic_variant | MODIFIER |
| 5 | 105677964 | AX-97828288 | 0 | T | A | 0.439 | 6.96E-02 | 1.16E-01 | 2.65E+00 | 2.68E+00 | 5.45E-01 | no | | intergenic_variant | MODIFIER |
| 5 | 105682135 | AX-170528513 | 0 | T | G | 0.439 | 6.96E-02 | 1.16E-01 | 2.65E+00 | 2.68E+00 | 5.45E-01 | no | | regulatory_region_variant | MODIFIER |
| 5 | 105682469 | AX-170333413 | 0 | A | G | 0.439 | 6.96E-02 | 1.16E-01 | 2.65E+00 | 2.68E+00 | 5.45E-01 | no | | intergenic_variant | MODIFIER |
| 5 | 105705438 | AX-170256585 | 0 | C | T | 0.439 | 6.96E-02 | 1.16E-01 | 2.65E+00 | 2.68E+00 | 5.45E-01 | no | Lrrc8d | intron_variant | MODIFIER |
| 5 | 105708068 | AX-170471666 | 0 | T | C | 0.439 | 6.96E-02 | 1.16E-01 | 2.65E+00 | 2.68E+00 | 5.45E-01 | no | Lrrc8d | intron_variant | MODIFIER |
| 5 | 105743150 | AX-97803470 | 0 | C | T | 0.439 | 6.96E-02 | 1.16E-01 | 2.65E+00 | 2.68E+00 | 5.45E-01 | no | Lrrc8d | intron_variant | MODIFIER |
| 5 | 105743244 | AX-170302034 | 0 | G | A | 0.439 | 6.96E-02 | 1.16E-01 | 2.65E+00 | 2.68E+00 | 5.45E-01 | no | Lrrc8d | intron_variant | MODIFIER |
| 5 | 105746366 | AX-170095426 | 0 | G | C | 0.439 | 6.96E-02 | 1.16E-01 | 2.65E+00 | 2.68E+00 | 5.45E-01 | no | Lrrc8d | intron_variant | MODIFIER |
| 5 | 105772138 | AX-170409202 | 0 | G | A | 0.439 | 6.96E-02 | 1.16E-01 | 2.65E+00 | 2.68E+00 | 5.45E-01 | no | Lrrc8d | intron_variant | MODIFIER |
| 5 | 105778393 | AX-170395880 | 0 | G | A | 0.439 | 6.96E-02 | 1.16E-01 | 2.65E+00 | 2.68E+00 | 5.45E-01 | no | Lrrc8d | intron_variant | MODIFIER |
| 5 | 105784331 | AX-97853828 | 0 | G | A | 0.439 | 6.96E-02 | 1.16E-01 | 2.65E+00 | 2.68E+00 | 5.45E-01 | no | Lrrc8d | intron_variant | MODIFIER |
| 5 | 105807163 | AX-170191459 | 0 | A | C | 0.439 | 6.96E-02 | 1.16E-01 | 2.65E+00 | 2.68E+00 | 5.45E-01 | no | Lrrc8d | intron_variant | MODIFIER |
| 5 | 105811553 | AX-170488172 | 0 | G | A | 0.439 | 6.96E-02 | 1.16E-01 | 2.65E+00 | 2.68E+00 | 5.45E-01 | no | Lrrc8d | intron_variant | MODIFIER |
| 5 | 105811948 | AX-170216935 | 0 | G | A | 0.439 | 6.96E-02 | 1.16E-01 | 2.65E+00 | 2.68E+00 | 5.45E-01 | no | Lrrc8d | missense_variant | MODERATE |
| 5 | 105817791 | AX-97800263 | 0 | A | C | 0.439 | 6.96E-02 | 1.16E-01 | 2.65E+00 | 2.68E+00 | 5.45E-01 | no | Lrrc8d | downstream_gene_variant | MODIFIER |
| 5 | 105819417 | AX-170393942 | 0 | A | C | 0.439 | 6.96E-02 | 1.16E-01 | 2.65E+00 | 2.68E+00 | 5.45E-01 | no | Lrrc8d | downstream_gene_variant | MODIFIER |

| | | | | | | | | | | | | | | | |
|---|---|---|---|---|---|---|---|---|---|---|---|---|---|---|---|
| 5 | 105819553 | AX-170205206 | 0 | T | C | 0.439 | 6.96E-02 | 1.16E-01 | 2.65E+00 | 2.68E+00 | 5.45E-01 | no | Lrrc8d | downstream_gene_variant | MODIFIER |
| 5 | 106380472 | AX-170184846 | 0 | A | G | 0.439 | 6.96E-02 | 1.16E-01 | 2.65E+00 | 2.68E+00 | 5.45E-01 | no | | intergenic_variant | MODIFIER |
| 5 | 106399186 | AX-170442427 | 0 | A | T | 0.439 | 6.96E-02 | 1.16E-01 | 2.65E+00 | 2.68E+00 | 5.45E-01 | no | | intergenic_variant | MODIFIER |
| 5 | 106431903 | AX-170208657 | 0 | T | C | 0.439 | 6.96E-02 | 1.16E-01 | 2.65E+00 | 2.68E+00 | 5.45E-01 | no | | intergenic_variant | MODIFIER |
| 5 | 106433979 | AX-97829641 | 0 | C | T | 0.439 | 6.96E-02 | 1.16E-01 | 2.65E+00 | 2.68E+00 | 5.45E-01 | no | | intergenic_variant | MODIFIER |
| 5 | 106467839 | AX-97842509 | 0 | C | T | 0.439 | 6.96E-02 | 1.16E-01 | 2.65E+00 | 2.68E+00 | 5.45E-01 | no | Gm26872 | non_coding_transcript_exon_variant | MODIFIER |
| 5 | 106479700 | AX-170072995 | 0 | A | C | 0.439 | 6.96E-02 | 1.16E-01 | 2.65E+00 | 2.68E+00 | 5.45E-01 | no | | intergenic_variant | MODIFIER |
| 5 | 106481793 | AX-170072996 | 0 | A | G | 0.439 | 6.96E-02 | 1.16E-01 | 2.65E+00 | 2.68E+00 | 5.45E-01 | no | | intergenic_variant | MODIFIER |
| 5 | 106494185 | AX-170025823 | 0 | G | A | 0.439 | 6.96E-02 | 1.16E-01 | 2.65E+00 | 2.68E+00 | 5.45E-01 | no | | intergenic_variant | MODIFIER |
| 5 | 106494931 | AX-170493778 | 0 | C | A | 0.439 | 6.96E-02 | 1.16E-01 | 2.65E+00 | 2.68E+00 | 5.45E-01 | no | | intergenic_variant | MODIFIER |
| 5 | 106496648 | AX-170519142 | 0 | A | G | 0.439 | 6.96E-02 | 1.16E-01 | 2.65E+00 | 2.68E+00 | 5.45E-01 | no | Gm28050 | downstream_gene_variant | MODIFIER |
| 5 | 106497391 | AX-170110601 | 0 | T | C | 0.439 | 6.96E-02 | 1.16E-01 | 2.65E+00 | 2.68E+00 | 5.45E-01 | no | Gm28050 | downstream_gene_variant | MODIFIER |
| 5 | 106499967 | AX-170025822 | 0 | C | T | 0.439 | 6.96E-02 | 1.16E-01 | 2.65E+00 | 2.68E+00 | 5.45E-01 | no | Gm28050 | downstream_gene_variant | MODIFIER |
| 5 | 106563439 | AX-170025818 | 0 | T | C | 0.439 | 6.96E-02 | 1.16E-01 | 2.65E+00 | 2.68E+00 | 5.45E-01 | no | Gm28050 | intron_variant,non_coding_transcript_variant | MODIFIER |
| 5 | 106583019 | AX-97840238 | 0 | G | A | 0.439 | 6.96E-02 | 1.16E-01 | 2.65E+00 | 2.68E+00 | 5.45E-01 | no | Gm28039 | intron_variant,non_coding_transcript_variant | MODIFIER |
| 5 | 106595418 | AX-170391682 | 0 | T | C | 0.439 | 6.96E-02 | 1.16E-01 | 2.65E+00 | 2.68E+00 | 5.45E-01 | no | Gm28039 | intron_variant,non_coding_transcript_variant | MODIFIER |
| 5 | 106602210 | AX-170277537 | 0 | G | C | 0.439 | 6.96E-02 | 1.16E-01 | 2.65E+00 | 2.68E+00 | 5.45E-01 | no | Gm28039 | intron_variant,non_coding_transcript_variant | MODIFIER |
| 5 | 106603869 | AX-170146045 | 0 | A | G | 0.439 | 6.96E-02 | 1.16E-01 | 2.65E+00 | 2.68E+00 | 5.45E-01 | no | Gm28039 | intron_variant,non_coding_transcript_variant | MODIFIER |
| 5 | 106627912 | AX-170186202 | 0 | G | T | 0.439 | 6.96E-02 | 1.16E-01 | 2.65E+00 | 2.68E+00 | 5.45E-01 | no | Zfp644 | intron_variant | MODIFIER |
| 5 | 106685738 | AX-170514391 | 0 | G | A | 0.439 | 6.96E-02 | 1.16E-01 | 2.65E+00 | 2.68E+00 | 5.45E-01 | no | Zfp644 | intron_variant | MODIFIER |
| 5 | 106767334 | AX-97833396 | 0 | T | G | 0.439 | 6.96E-02 | 1.16E-01 | 2.65E+00 | 2.68E+00 | 5.45E-01 | no | | intergenic_variant | MODIFIER |
| 5 | 106769184 | AX-170267113 | 0 | A | T | 0.439 | 6.96E-02 | 1.16E-01 | 2.65E+00 | 2.68E+00 | 5.45E-01 | no | | intergenic_variant | MODIFIER |
| 5 | 106790493 | AX-170350894 | 0 | A | C | 0.439 | 6.96E-02 | 1.16E-01 | 2.65E+00 | 2.68E+00 | 5.45E-01 | no | | intergenic_variant | MODIFIER |
| 5 | 106794668 | AX-170348206 | 0 | A | G | 0.439 | 6.96E-02 | 1.16E-01 | 2.65E+00 | 2.68E+00 | 5.45E-01 | no | | intergenic_variant | MODIFIER |
| 5 | 106795083 | AX-170403282 | 0 | C | T | 0.439 | 6.96E-02 | 1.16E-01 | 2.65E+00 | 2.68E+00 | 5.45E-01 | no | | intergenic_variant | MODIFIER |
| 5 | 106810158 | AX-170101940 | 0 | A | G | 0.439 | 6.96E-02 | 1.16E-01 | 2.65E+00 | 2.68E+00 | 5.45E-01 | no | | intergenic_variant | MODIFIER |
| 5 | 106814151 | AX-97854339 | 0 | A | G | 0.439 | 6.96E-02 | 1.16E-01 | 2.65E+00 | 2.68E+00 | 5.45E-01 | no | | intergenic_variant | MODIFIER |
| 5 | 106818409 | AX-170277545 | 0 | C | G | 0.439 | 6.96E-02 | 1.16E-01 | 2.65E+00 | 2.68E+00 | 5.45E-01 | no | | intergenic_variant | MODIFIER |
| 5 | 106819088 | AX-170200834 | 0 | T | G | 0.439 | 6.96E-02 | 1.16E-01 | 2.65E+00 | 2.68E+00 | 5.45E-01 | no | | intergenic_variant | MODIFIER |
| 5 | 106831312 | AX-97819840 | 0 | T | C | 0.439 | 6.96E-02 | 1.16E-01 | 2.65E+00 | 2.68E+00 | 5.45E-01 | no | | intergenic_variant | MODIFIER |
| 5 | 106847033 | AX-168403115 | 0 | A | G | 0.439 | 6.96E-02 | 1.16E-01 | 2.65E+00 | 2.68E+00 | 5.45E-01 | no | Hfm1 | intron_variant | MODIFIER |
| 5 | 106851961 | AX-170073008 | 0 | C | T | 0.439 | 6.96E-02 | 1.16E-01 | 2.65E+00 | 2.68E+00 | 5.45E-01 | no | Hfm1 | intron_variant | MODIFIER |
| 5 | 106858719 | AX-97851012 | 0 | C | T | 0.439 | 6.96E-02 | 1.16E-01 | 2.65E+00 | 2.68E+00 | 5.45E-01 | no | Hfm1 | intron_variant | MODIFIER |
| 5 | 106870450 | AX-168394969 | 0 | A | G | 0.439 | 6.96E-02 | 1.16E-01 | 2.65E+00 | 2.68E+00 | 5.45E-01 | no | Hfm1 | intron_variant | MODIFIER |
| 5 | 106874237 | AX-97810743 | 0 | A | G | 0.439 | 6.96E-02 | 1.16E-01 | 2.65E+00 | 2.68E+00 | 5.45E-01 | no | Hfm1 | missense_variant | MODERATE |
| 5 | 106891547 | AX-170322078 | 0 | A | G | 0.439 | 6.96E-02 | 1.16E-01 | 2.65E+00 | 2.68E+00 | 5.45E-01 | no | Hfm1 | intron_variant | MODIFIER |
| 5 | 106891706 | AX-170327088 | 0 | C | A | 0.439 | 6.96E-02 | 1.16E-01 | 2.65E+00 | 2.68E+00 | 5.45E-01 | no | Hfm1 | intron_variant | MODIFIER |
| 5 | 106898124 | AX-170419368 | 0 | T | C | 0.439 | 6.96E-02 | 1.16E-01 | 2.65E+00 | 2.68E+00 | 5.45E-01 | no | Hfm1 | intron_variant | MODIFIER |
| 5 | 106904788 | AX-170123652 | 0 | C | T | 0.439 | 6.96E-02 | 1.16E-01 | 2.65E+00 | 2.68E+00 | 5.45E-01 | no | Hfm1 | synonymous_variant | LOW |
| 5 | 106905495 | AX-168419958 | 0 | G | A | 0.439 | 6.96E-02 | 1.16E-01 | 2.65E+00 | 2.68E+00 | 5.45E-01 | no | Hfm1 | intron_variant | MODIFIER |
| 5 | 106907906 | AX-170298926 | 0 | A | G | 0.439 | 6.96E-02 | 1.16E-01 | 2.65E+00 | 2.68E+00 | 5.45E-01 | no | Hfm1 | intron_variant | MODIFIER |
| 5 | 106917047 | AX-170532095 | 0 | A | G | 0.439 | 6.96E-02 | 1.16E-01 | 2.65E+00 | 2.68E+00 | 5.45E-01 | no | Hfm1 | intron_variant | MODIFIER |
| 5 | 106921129 | AX-97828409 | 0 | T | C | 0.439 | 6.96E-02 | 1.16E-01 | 2.65E+00 | 2.68E+00 | 5.45E-01 | no | Hfm1 | intron_variant | MODIFIER |
| 5 | 106993607 | AX-170227691 | 0 | T | G | 0.439 | 6.96E-02 | 1.16E-01 | 2.65E+00 | 2.68E+00 | 5.45E-01 | no | | intergenic_variant | MODIFIER |
| 5 | 107016950 | AX-170118345 | 0 | G | T | 0.439 | 6.96E-02 | 1.16E-01 | 2.65E+00 | 2.68E+00 | 5.45E-01 | no | | intergenic_variant | MODIFIER |
| 5 | 107017934 | AX-170459250 | 0 | C | T | 0.439 | 6.96E-02 | 1.16E-01 | 2.65E+00 | 2.68E+00 | 5.45E-01 | no | | regulatory_region_variant | MODIFIER |
| 5 | 107055427 | AX-97805848 | 0 | A | G | 0.439 | 6.96E-02 | 1.16E-01 | 2.65E+00 | 2.68E+00 | 5.45E-01 | no | Gm33474 | non_coding_transcript_exon_variant | MODIFIER |
| 5 | 107074921 | AX-170153295 | 0 | T | C | 0.439 | 6.96E-02 | 1.16E-01 | 2.65E+00 | 2.68E+00 | 5.45E-01 | no | | regulatory_region_variant | MODIFIER |
| 5 | 107104855 | AX-97851100 | 0 | A | G | 0.439 | 6.96E-02 | 1.16E-01 | 2.65E+00 | 2.68E+00 | 5.45E-01 | no | Tgfbr3 | downstream_gene_variant | MODIFIER |
| 5 | 107117519 | AX-170025831 | 0 | A | G | 0.439 | 6.96E-02 | 1.16E-01 | 2.65E+00 | 2.68E+00 | 5.45E-01 | no | Tgfbr3 | intron_variant | MODIFIER |
| 5 | 107119361 | AX-170330064 | 0 | G | A | 0.439 | 6.96E-02 | 1.16E-01 | 2.65E+00 | 2.68E+00 | 5.45E-01 | no | Tgfbr3 | intron_variant | MODIFIER |
| 5 | 107119862 | AX-170528890 | 0 | G | T | 0.439 | 6.96E-02 | 1.16E-01 | 2.65E+00 | 2.68E+00 | 5.45E-01 | no | Tgfbr3 | intron_variant | MODIFIER |
| 5 | 107121287 | AX-170025828 | 0 | G | A | 0.439 | 6.96E-02 | 1.16E-01 | 2.65E+00 | 2.68E+00 | 5.45E-01 | no | Tgfbr3 | intron_variant | MODIFIER |
| 5 | 107121740 | AX-170025827 | 0 | A | G | 0.439 | 6.96E-02 | 1.16E-01 | 2.65E+00 | 2.68E+00 | 5.45E-01 | no | Tgfbr3 | intron_variant | MODIFIER |

| | | | | | | | | | | | | | | | |
|---|---|---|---|---|---|---|---|---|---|---|---|---|---|---|---|
| 5 | 107121806 | AX-170086345 | 0 | T | C | 0.439 | 6.96E-02 | 1.16E-01 | 2.65E+00 | 2.68E+00 | 5.45E-01 | no | Tgfbr3 | intron_variant | MODIFIER |
| 5 | 107126013 | AX-170151863 | 0 | C | T | 0.439 | 6.96E-02 | 1.16E-01 | 2.65E+00 | 2.68E+00 | 5.45E-01 | no | Tgfbr3 | intron_variant | MODIFIER |
| 5 | 107129359 | AX-170297845 | 0 | G | A | 0.439 | 6.96E-02 | 1.16E-01 | 2.65E+00 | 2.68E+00 | 5.45E-01 | no | Tgfbr3 | intron_variant | MODIFIER |
| 5 | 107130564 | AX-170241728 | 0 | A | G | 0.439 | 6.96E-02 | 1.16E-01 | 2.65E+00 | 2.68E+00 | 5.45E-01 | no | Tgfbr3 | synonymous_variant | LOW |
| 5 | 107137879 | AX-170025825 | 0 | A | G | 0.439 | 6.96E-02 | 1.16E-01 | 2.65E+00 | 2.68E+00 | 5.45E-01 | no | Tgfbr3 | intron_variant | MODIFIER |
| 5 | 107301071 | AX-170512625 | 0 | G | A | 0.439 | 6.96E-02 | 1.16E-01 | 2.65E+00 | 2.68E+00 | 5.45E-01 | no | Gm22557 | upstream_gene_variant | MODIFIER |
| 5 | 107304434 | AX-170534394 | 0 | C | T | 0.439 | 6.96E-02 | 1.16E-01 | 2.65E+00 | 2.68E+00 | 5.45E-01 | no | Gm22557 | downstream_gene_variant | MODIFIER |
| 10 | 70231402 | AX-170287484 | 0 | G | T | 0.421 | -4.72E-02 | 7.86E-02 | 2.63E+00 | 2.71E+00 | 5.46E-01 | no | | intergenic_variant | MODIFIER |
| 7 | 36992583 | AX-170077082 | 1 | T | C | 0.339 | 5.61E-02 | 9.23E-02 | 2.58E+00 | 2.65E+00 | 5.46E-01 | no | | intergenic_variant | MODIFIER |
| 1 | 121104917 | AX-170205792 | 1 | A | C | 0.562 | 5.10E-02 | 8.19E-02 | 2.51E+00 | 2.58E+00 | 5.46E-01 | no | Celrr | intron_variant,non_coding_transcript_variant | MODIFIER |
| 1 | 32145252 | AX-97843915 | 1 | C | T | 0.402 | 4.60E-02 | 7.74E-02 | 2.68E+00 | 2.78E+00 | 5.46E-01 | no | | intergenic_variant | MODIFIER |
| 2 | 165084529 | AX-170245436 | 0 | T | C | 0.167 | 1.03E-01 | 1.72E-01 | 2.66E+00 | 2.75E+00 | 5.46E-01 | no | 1700025C18Rik | intron_variant | MODIFIER |
| 5 | 77133855 | AX-97784399 | 0 | A | G | 0.439 | 5.89E-02 | 9.76E-02 | 2.61E+00 | 2.68E+00 | 5.46E-01 | no | Chaer1 | downstream_gene_variant | MODIFIER |
| 5 | 39947147 | AX-170521099 | 1 | G | T | 0.205 | -6.50E-02 | 1.07E-01 | 2.58E+00 | 2.66E+00 | 5.46E-01 | no | | intergenic_variant | MODIFIER |
| 18 | 65138802 | AX-170403145 | 1 | A | G | 0.411 | 5.46E-02 | 8.60E-02 | 2.46E+00 | 2.52E+00 | 5.46E-01 | no | Nedd4l | intron_variant | MODIFIER |
| 1 | 188873711 | AX-170305136 | 1 | T | G | 0.241 | -5.76E-02 | 9.56E-02 | 2.93E+00 | 3.10E+00 | 5.46E-01 | no | Ush2a | intron_variant | MODIFIER |
| 1 | 105616253 | AX-97826030 | 1 | T | G | 0.143 | 9.24E-02 | 1.53E-01 | 2.60E+00 | 2.69E+00 | 5.46E-01 | no | Pign | intron_variant | MODIFIER |
| 9 | 103714518 | AX-170319692 | 0 | G | A | 0.123 | -6.83E-02 | 1.13E-01 | 2.58E+00 | 2.67E+00 | 5.46E-01 | no | Tmem108 | intron_variant | MODIFIER |
| 9 | 103723902 | AX-170188191 | 0 | T | G | 0.123 | -6.83E-02 | 1.13E-01 | 2.58E+00 | 2.67E+00 | 5.46E-01 | no | Tmem108 | intron_variant | MODIFIER |
| 18 | 74368379 | AX-170154011 | 0 | T | A | 0.184 | -7.23E-02 | 1.19E-01 | 2.59E+00 | 2.65E+00 | 5.46E-01 | no | | intergenic_variant | MODIFIER |
| 4 | 93051100 | AX-170369287 | 0 | T | C | 0.333 | 6.49E-02 | 1.08E-01 | 2.63E+00 | 2.71E+00 | 5.46E-01 | no | | intergenic_variant | MODIFIER |
| 10 | 18989645 | AX-170491528 | 0 | C | A | 0.202 | 5.94E-02 | 9.93E-02 | 2.86E+00 | 2.97E+00 | 5.47E-01 | no | | intergenic_variant | MODIFIER |
| 2 | 98155267 | AX-170018440 | 0 | C | A | 0.053 | -1.33E-01 | 2.18E-01 | 2.56E+00 | 2.66E+00 | 5.47E-01 | no | | intergenic_variant | MODIFIER |
| 5 | 46895135 | AX-170089469 | 0 | G | T | 0.5 | 6.19E-02 | 9.83E-02 | 2.48E+00 | 2.55E+00 | 5.47E-01 | no | | intergenic_variant | MODIFIER |
| 5 | 47732537 | AX-170071993 | 0 | C | G | 0.5 | 6.19E-02 | 9.83E-02 | 2.48E+00 | 2.55E+00 | 5.47E-01 | no | | intergenic_variant | MODIFIER |
| 5 | 27920243 | AX-97838310 | 1 | G | T | 0.339 | 6.24E-02 | 1.05E-01 | 2.69E+00 | 2.75E+00 | 5.47E-01 | no | | intergenic_variant | MODIFIER |
| 18 | 83997607 | AX-97811497 | 2 | C | T | 0.155 | -6.74E-02 | 1.08E-01 | 2.48E+00 | 2.56E+00 | 5.47E-01 | no | | intergenic_variant | MODIFIER |
| 3 | 33713226 | AX-170395447 | 0 | G | C | 0.491 | 1.93E-01 | 3.32E-01 | 2.69E+00 | 2.86E+00 | 5.47E-01 | Yes | Gm43666 | intron_variant,non_coding_transcript_variant | MODIFIER |
| 6 | 138533866 | AX-97801946 | 1 | T | C | 0.545 | 5.20E-02 | 8.44E-02 | 2.52E+00 | 2.61E+00 | 5.47E-01 | no | Lmo3 | intron_variant | MODIFIER |
| 5 | 88052585 | AX-170072705 | 0 | T | A | 0.421 | 6.37E-02 | 1.07E-01 | 2.77E+00 | 2.83E+00 | 5.47E-01 | no | | intergenic_variant | MODIFIER |
| 5 | 88383754 | AX-170025539 | 0 | C | T | 0.421 | 6.37E-02 | 1.07E-01 | 2.77E+00 | 2.83E+00 | 5.47E-01 | no | Amtn | intron_variant | MODIFIER |
| 6 | 102646174 | AX-170431012 | 0 | T | C | 0.061 | 1.31E-01 | 2.20E-01 | 2.63E+00 | 2.73E+00 | 5.47E-01 | no | | intergenic_variant | MODIFIER |
| 8 | 64523948 | AX-97806390 | 0 | G | A | 0.14 | -8.63E-02 | 1.43E-01 | 2.58E+00 | 2.66E+00 | 5.47E-01 | no | | intergenic_variant | MODIFIER |
| 1 | 20654352 | AX-170409520 | 0 | C | T | 0.404 | 5.04E-02 | 8.37E-02 | 2.61E+00 | 2.66E+00 | 5.47E-01 | no | | regulatory_region_variant | MODIFIER |
| 18 | 64945914 | AX-170342199 | 1 | T | A | 0.411 | 5.77E-02 | 9.19E-02 | 2.49E+00 | 2.53E+00 | 5.47E-01 | no | Nedd4l | intron_variant | MODIFIER |
| 3 | 145299452 | AX-170136004 | 0 | C | T | 0.36 | -4.10E-02 | 6.93E-02 | 2.76E+00 | 2.86E+00 | 5.47E-01 | no | Col24a1 | intron_variant | MODIFIER |
| 18 | 88502935 | AX-170234080 | 0 | C | G | 0.149 | -6.08E-02 | 9.83E-02 | 2.50E+00 | 2.61E+00 | 5.47E-01 | no | | intergenic_variant | MODIFIER |
| 9 | 85949780 | AX-170512134 | 0 | T | C | 0.132 | -7.92E-02 | 1.31E-01 | 2.59E+00 | 2.66E+00 | 5.48E-01 | no | | intergenic_variant | MODIFIER |
| 9 | 87409563 | AX-170216811 | 0 | A | C | 0.132 | -7.92E-02 | 1.31E-01 | 2.59E+00 | 2.66E+00 | 5.48E-01 | no | | intergenic_variant | MODIFIER |
| 16 | 75433114 | AX-97845993 | 1 | A | G | 0.5 | 4.66E-02 | 7.54E-02 | 2.51E+00 | 2.59E+00 | 5.48E-01 | no | Gm49678 | intron_variant,non_coding_transcript_variant | MODIFIER |
| 3 | 29767873 | AX-170252123 | 0 | G | T | 0.184 | -5.66E-02 | 9.55E-02 | 2.69E+00 | 2.77E+00 | 5.48E-01 | no | Gm37557 | upstream_gene_variant | MODIFIER |
| 3 | 30723326 | AX-170242341 | 0 | G | T | 0.175 | -5.94E-02 | 1.00E-01 | 2.69E+00 | 2.76E+00 | 5.48E-01 | no | | regulatory_region_variant | MODIFIER |
| 6 | 129714512 | AX-168388780 | 0 | T | C | 0.544 | 5.01E-02 | 8.16E-02 | 2.53E+00 | 2.60E+00 | 5.48E-01 | no | Klri1 | intron_variant | MODIFIER |
| 8 | 90184568 | AX-170505404 | 1 | G | A | 0.179 | -6.95E-02 | 1.16E-01 | 2.61E+00 | 2.71E+00 | 5.48E-01 | no | | intergenic_variant | MODIFIER |
| 4 | 153948392 | AX-170295235 | 1 | C | A | 0.107 | 1.07E-01 | 1.80E-01 | 2.68E+00 | 2.76E+00 | 5.48E-01 | no | BC039966 | non_coding_transcript_exon_variant | MODIFIER |
| 18 | 75864517 | AX-170016011 | 0 | C | T | 0.167 | -7.30E-02 | 1.20E-01 | 2.57E+00 | 2.62E+00 | 5.48E-01 | no | Zbtb7c | intron_variant | MODIFIER |
| 18 | 75876474 | AX-97857608 | 0 | A | G | 0.167 | -7.30E-02 | 1.20E-01 | 2.57E+00 | 2.62E+00 | 5.48E-01 | no | Zbtb7c | intron_variant | MODIFIER |
| 18 | 75877884 | AX-170061355 | 0 | T | C | 0.167 | -7.30E-02 | 1.20E-01 | 2.57E+00 | 2.62E+00 | 5.48E-01 | no | Zbtb7c | intron_variant | MODIFIER |
| 18 | 75877952 | AX-170353754 | 0 | G | A | 0.167 | -7.30E-02 | 1.20E-01 | 2.57E+00 | 2.62E+00 | 5.48E-01 | no | Zbtb7c | intron_variant | MODIFIER |
| 18 | 75879794 | AX-170294048 | 0 | C | T | 0.167 | -7.30E-02 | 1.20E-01 | 2.57E+00 | 2.62E+00 | 5.48E-01 | no | Zbtb7c | intron_variant | MODIFIER |
| 18 | 75879873 | AX-170244460 | 0 | T | C | 0.167 | -7.30E-02 | 1.20E-01 | 2.57E+00 | 2.62E+00 | 5.48E-01 | no | Zbtb7c | intron_variant | MODIFIER |
| 18 | 75880302 | AX-170226109 | 0 | T | G | 0.167 | -7.30E-02 | 1.20E-01 | 2.57E+00 | 2.62E+00 | 5.48E-01 | no | Zbtb7c | intron_variant | MODIFIER |
| 18 | 75880471 | AX-170375388 | 0 | C | T | 0.167 | -7.30E-02 | 1.20E-01 | 2.57E+00 | 2.62E+00 | 5.48E-01 | no | Zbtb7c | intron_variant | MODIFIER |
| 18 | 75894641 | AX-170016010 | 0 | T | A | 0.167 | -7.30E-02 | 1.20E-01 | 2.57E+00 | 2.62E+00 | 5.48E-01 | no | Zbtb7c | intron_variant | MODIFIER |

| | | | | | | | | | | | | | | | |
|---|---|---|---|---|---|---|---|---|---|---|---|---|---|---|---|
| 18 | 75898416 | AX-97837074 | 0 | G | A | 0.167 | -7.30E-02 | 1.20E-01 | 2.57E+00 | 2.62E+00 | 5.48E-01 | no | Zbtb7c | intron_variant | MODIFIER |
| 18 | 75900786 | AX-170488746 | 0 | C | A | 0.167 | -7.30E-02 | 1.20E-01 | 2.57E+00 | 2.62E+00 | 5.48E-01 | no | Zbtb7c | intron_variant | MODIFIER |
| 18 | 75909186 | AX-170348167 | 0 | G | C | 0.167 | -7.30E-02 | 1.20E-01 | 2.57E+00 | 2.62E+00 | 5.48E-01 | no | Zbtb7c | intron_variant | MODIFIER |
| 18 | 75910667 | AX-170282420 | 0 | T | C | 0.167 | -7.30E-02 | 1.20E-01 | 2.57E+00 | 2.62E+00 | 5.48E-01 | no | Zbtb7c | intron_variant | MODIFIER |
| 18 | 75950460 | AX-97807228 | 0 | A | G | 0.167 | -7.30E-02 | 1.20E-01 | 2.57E+00 | 2.62E+00 | 5.48E-01 | no | Zbtb7c | intron_variant | MODIFIER |
| 18 | 75972388 | AX-97791157 | 0 | T | C | 0.167 | -7.30E-02 | 1.20E-01 | 2.57E+00 | 2.62E+00 | 5.48E-01 | no | Zbtb7c | intron_variant | MODIFIER |
| 18 | 76024875 | AX-170371373 | 0 | T | C | 0.167 | -7.30E-02 | 1.20E-01 | 2.57E+00 | 2.62E+00 | 5.48E-01 | no | Zbtb7c | intron_variant | MODIFIER |
| 18 | 76034967 | AX-170240239 | 0 | A | G | 0.167 | -7.30E-02 | 1.20E-01 | 2.57E+00 | 2.62E+00 | 5.48E-01 | no | Zbtb7c | intron_variant | MODIFIER |
| 18 | 76109685 | AX-97835930 | 0 | T | C | 0.167 | -7.30E-02 | 1.20E-01 | 2.57E+00 | 2.62E+00 | 5.48E-01 | no | Zbtb7c | intron_variant | MODIFIER |
| 18 | 76186957 | AX-97781612 | 0 | A | G | 0.167 | -7.30E-02 | 1.20E-01 | 2.57E+00 | 2.62E+00 | 5.48E-01 | no | | intergenic_variant | MODIFIER |
| 18 | 76228994 | AX-170199509 | 0 | A | G | 0.167 | -7.30E-02 | 1.20E-01 | 2.57E+00 | 2.62E+00 | 5.48E-01 | no | AC107768.2 | upstream_gene_variant | MODIFIER |
| 18 | 76229018 | AX-170424862 | 0 | G | C | 0.167 | -7.30E-02 | 1.20E-01 | 2.57E+00 | 2.62E+00 | 5.48E-01 | no | AC107768.2 | upstream_gene_variant | MODIFIER |
| 18 | 76263179 | AX-170090769 | 0 | C | T | 0.167 | -7.30E-02 | 1.20E-01 | 2.57E+00 | 2.62E+00 | 5.48E-01 | no | Smad2 | intron_variant | MODIFIER |
| 18 | 76312999 | AX-170254722 | 0 | G | A | 0.167 | -7.30E-02 | 1.20E-01 | 2.57E+00 | 2.62E+00 | 5.48E-01 | no | Smad2 | downstream_gene_variant | MODIFIER |
| 18 | 76313729 | AX-170443419 | 0 | T | C | 0.167 | -7.30E-02 | 1.20E-01 | 2.57E+00 | 2.62E+00 | 5.48E-01 | no | Smad2 | downstream_gene_variant | MODIFIER |
| 18 | 76321422 | AX-170416277 | 0 | A | G | 0.167 | -7.30E-02 | 1.20E-01 | 2.57E+00 | 2.62E+00 | 5.48E-01 | no | AC164009.1 | upstream_gene_variant | MODIFIER |
| 18 | 76920151 | AX-170306110 | 0 | A | G | 0.167 | -7.30E-02 | 1.20E-01 | 2.57E+00 | 2.62E+00 | 5.48E-01 | no | | intergenic_variant | MODIFIER |
| 18 | 76922453 | AX-97827432 | 0 | A | G | 0.167 | -7.30E-02 | 1.20E-01 | 2.57E+00 | 2.62E+00 | 5.48E-01 | no | | intergenic_variant | MODIFIER |
| 18 | 76924001 | AX-170272487 | 0 | T | C | 0.167 | -7.30E-02 | 1.20E-01 | 2.57E+00 | 2.62E+00 | 5.48E-01 | no | | intergenic_variant | MODIFIER |
| 18 | 76974118 | AX-97835954 | 0 | A | G | 0.167 | -7.30E-02 | 1.20E-01 | 2.57E+00 | 2.62E+00 | 5.48E-01 | no | Hdhd2 | downstream_gene_variant | MODIFIER |
| 18 | 77128328 | AX-97814923 | 0 | T | C | 0.167 | -7.30E-02 | 1.20E-01 | 2.57E+00 | 2.62E+00 | 5.48E-01 | no | Pias2 | intron_variant | MODIFIER |
| 18 | 77727600 | AX-170235167 | 0 | T | C | 0.167 | -7.30E-02 | 1.20E-01 | 2.57E+00 | 2.62E+00 | 5.48E-01 | no | A330094K24Rik | downstream_gene_variant | MODIFIER |
| 9 | 33119771 | AX-97805333 | 0 | T | C | 0.43 | 4.86E-02 | 8.14E-02 | 2.63E+00 | 2.70E+00 | 5.48E-01 | no | | intergenic_variant | MODIFIER |
| 20 | 70877805 | AX-170413207 | 1 | C | T | 0.259 | 3.70E-02 | 6.24E-02 | 2.67E+00 | 2.76E+00 | 5.48E-01 | no | | intergenic_variant | MODIFIER |
| 7 | 16259553 | AX-170298645 | 0 | C | T | 0.368 | 4.98E-02 | 8.25E-02 | 2.55E+00 | 2.67E+00 | 5.48E-01 | no | C5ar1 | upstream_gene_variant | MODIFIER |
| 8 | 34639459 | AX-170079721 | 1 | C | T | 0.143 | -7.89E-02 | 1.32E-01 | 2.63E+00 | 2.70E+00 | 5.48E-01 | no | B930018H19Rik | intron_variant,non_coding_transcript_variant | MODIFIER |
| 1 | 121312058 | AX-170285720 | 0 | G | A | 0.579 | 5.20E-02 | 8.35E-02 | 2.49E+00 | 2.55E+00 | 5.48E-01 | no | Insig2 | intron_variant | MODIFIER |
| 1 | 121597564 | AX-170403448 | 0 | G | A | 0.579 | 5.20E-02 | 8.35E-02 | 2.49E+00 | 2.55E+00 | 5.48E-01 | no | | regulatory_region_variant | MODIFIER |
| 7 | 16201857 | AX-170487488 | 1 | A | G | 0.375 | 5.15E-02 | 8.53E-02 | 2.56E+00 | 2.67E+00 | 5.48E-01 | no | Dhx34 | intron_variant | MODIFIER |
| 7 | 109496239 | AX-170203502 | 0 | A | T | 0.474 | -5.59E-02 | 9.47E-02 | 2.77E+00 | 2.85E+00 | 5.48E-01 | no | Trim66 | intron_variant | MODIFIER |
| 5 | 5243496 | AX-170481021 | 0 | C | T | 0.439 | -4.03E-02 | 6.52E-02 | 2.49E+00 | 2.61E+00 | 5.49E-01 | no | Cdk14 | intron_variant | MODIFIER |
| 18 | 60919850 | AX-170143182 | 1 | G | A | 0.402 | -5.04E-02 | 8.29E-02 | 2.56E+00 | 2.62E+00 | 5.49E-01 | no | Arsi | downstream_gene_variant | MODIFIER |
| 10 | 28693961 | AX-170086772 | 0 | C | T | 0.202 | 5.50E-02 | 9.34E-02 | 2.77E+00 | 2.88E+00 | 5.49E-01 | no | Themis | intron_variant | MODIFIER |
| 4 | 63667478 | AX-170308226 | 0 | C | T | 0.342 | 6.82E-02 | 1.15E-01 | 2.68E+00 | 2.73E+00 | 5.49E-01 | no | Gm11214 | upstream_gene_variant | MODIFIER |
| 10 | 114132157 | AX-97816852 | 1 | T | C | 0.339 | -4.64E-02 | 7.86E-02 | 2.68E+00 | 2.77E+00 | 5.49E-01 | no | | intergenic_variant | MODIFIER |
| 1 | 98957724 | AX-170239073 | 2 | C | A | 0.164 | 8.47E-02 | 1.43E-01 | 2.61E+00 | 2.73E+00 | 5.49E-01 | no | | intergenic_variant | MODIFIER |
| 4 | 83187988 | AX-97794834 | 1 | G | A | 0.25 | 6.43E-02 | 1.05E-01 | 2.53E+00 | 2.60E+00 | 5.49E-01 | no | | intergenic_variant | MODIFIER |
| 5 | 82106091 | AX-170025427 | 1 | C | G | 0.411 | 6.10E-02 | 1.03E-01 | 2.66E+00 | 2.72E+00 | 5.49E-01 | no | | intergenic_variant | MODIFIER |
| 9 | 52785119 | AX-170295454 | 0 | T | G | 0.219 | 6.24E-02 | 1.06E-01 | 2.70E+00 | 2.78E+00 | 5.49E-01 | no | Gm1715 | intron_variant,non_coding_transcript_variant | MODIFIER |
| 16 | 75615488 | AX-170128789 | 2 | T | C | 0.491 | 4.69E-02 | 7.58E-02 | 2.50E+00 | 2.57E+00 | 5.49E-01 | no | | intergenic_variant | MODIFIER |
| 2 | 161839010 | AX-170443390 | 1 | T | G | 0.152 | 1.04E-01 | 1.77E-01 | 2.70E+00 | 2.80E+00 | 5.49E-01 | no | Ptprt | intron_variant | MODIFIER |
| 5 | 43583316 | AX-170431323 | 1 | G | A | 0.188 | -6.79E-02 | 1.14E-01 | 2.63E+00 | 2.71E+00 | 5.50E-01 | no | C1qtnf7 | intron_variant | MODIFIER |
| 5 | 43889750 | AX-170097357 | 1 | T | C | 0.188 | -6.79E-02 | 1.14E-01 | 2.63E+00 | 2.71E+00 | 5.50E-01 | no | Cd38 | intron_variant | MODIFIER |
| 9 | 78749090 | AX-97854264 | 0 | A | G | 0.184 | -7.80E-02 | 1.32E-01 | 2.72E+00 | 2.77E+00 | 5.50E-01 | no | | intergenic_variant | MODIFIER |
| 12 | 92337662 | AX-170049797 | 1 | T | C | 0.071 | 1.28E-01 | 2.17E-01 | 2.73E+00 | 2.81E+00 | 5.50E-01 | no | | intergenic_variant | MODIFIER |
| 20 | 72966830 | AX-97837633 | 0 | A | G | 0.263 | 3.66E-02 | 6.20E-02 | 2.67E+00 | 2.76E+00 | 5.50E-01 | no | | intergenic_variant | MODIFIER |
| 5 | 112828235 | AX-170143002 | 0 | T | C | 0.447 | 6.78E-02 | 1.14E-01 | 2.67E+00 | 2.72E+00 | 5.50E-01 | no | Myo18b | intron_variant | MODIFIER |
| 7 | 144703973 | AX-170383381 | 1 | G | A | 0.223 | -4.39E-02 | 7.43E-02 | 2.61E+00 | 2.74E+00 | 5.50E-01 | no | Ano1 | intron_variant | MODIFIER |
| 5 | 85573034 | AX-170414522 | 2 | A | G | 0.418 | 5.97E-02 | 1.01E-01 | 2.71E+00 | 2.79E+00 | 5.50E-01 | no | | intergenic_variant | MODIFIER |
| 1 | 96401621 | AX-170231531 | 1 | G | C | 0.152 | 1.19E-01 | 2.00E-01 | 2.62E+00 | 2.69E+00 | 5.50E-01 | no | | intergenic_variant | MODIFIER |
| 1 | 99756190 | AX-170001186 | 1 | C | T | 0.152 | 1.19E-01 | 2.00E-01 | 2.62E+00 | 2.69E+00 | 5.50E-01 | no | | intergenic_variant | MODIFIER |
| 10 | 19371019 | AX-170390853 | 0 | A | C | 0.412 | -4.04E-02 | 6.47E-02 | 2.45E+00 | 2.59E+00 | 5.50E-01 | no | | intergenic_variant | MODIFIER |
| 5 | 121083329 | AX-170486668 | 1 | G | T | 0.455 | 6.06E-02 | 1.02E-01 | 2.64E+00 | 2.71E+00 | 5.50E-01 | no | | intergenic_variant | MODIFIER |
| 5 | 121095938 | AX-170026020 | 1 | T | G | 0.455 | 6.06E-02 | 1.02E-01 | 2.64E+00 | 2.71E+00 | 5.50E-01 | no | | intergenic_variant | MODIFIER |

| | | | | | | | | | | | | | | | |
|---|---|---|---|---|---|---|---|---|---|---|---|---|---|---|---|
| 5 | 28745728 | AX-97783187 | 0 | T | G | 0.36 | -5.50E-02 | 9.35E-02 | 2.72E+00 | 2.81E+00 | 5.50E-01 | no | 9530036O11Rik | intron_variant,non_coding_transcript_variant | MODIFIER |
| 1 | 114861980 | AX-170086582 | 1 | A | C | 0.161 | -9.74E-02 | 1.66E-01 | 2.82E+00 | 2.95E+00 | 5.50E-01 | no | | intergenic_variant | MODIFIER |
| 7 | 36694979 | AX-170442168 | 0 | T | C | 0.342 | 5.31E-02 | 8.80E-02 | 2.56E+00 | 2.65E+00 | 5.50E-01 | no | Tshz3 | upstream_gene_variant | MODIFIER |
| 2 | 147521969 | AX-170019308 | 1 | A | T | 0.134 | 8.89E-02 | 1.51E-01 | 2.65E+00 | 2.75E+00 | 5.50E-01 | no | | intergenic_variant | MODIFIER |
| 8 | 75148123 | AX-170379187 | 0 | A | G | 0.193 | -7.43E-02 | 1.26E-01 | 2.66E+00 | 2.74E+00 | 5.50E-01 | no | | intergenic_variant | MODIFIER |
| 1 | 20257967 | AX-97851793 | 0 | T | C | 0.404 | 4.79E-02 | 8.06E-02 | 2.63E+00 | 2.69E+00 | 5.50E-01 | no | Pkhd1 | intron_variant | MODIFIER |
| 1 | 184779241 | AX-170146454 | 1 | G | A | 0.232 | 6.08E-02 | 9.89E-02 | 2.50E+00 | 2.61E+00 | 5.50E-01 | no | | intergenic_variant | MODIFIER |
| 9 | 118721011 | AX-170451698 | 1 | G | A | 0.125 | -5.71E-02 | 9.60E-02 | 2.60E+00 | 2.71E+00 | 5.50E-01 | no | Itga9 | intron_variant | MODIFIER |
| 16 | 45429271 | AX-97831079 | 1 | T | G | 0.518 | 4.97E-02 | 8.20E-02 | 2.56E+00 | 2.60E+00 | 5.50E-01 | no | Gm609 | intron_variant | MODIFIER |
| 1 | 88681612 | AX-170401884 | 0 | C | T | 0.149 | 1.19E-01 | 2.00E-01 | 2.64E+00 | 2.70E+00 | 5.51E-01 | no | Arl4c | intron_variant,NMD_transcript_variant | MODIFIER |
| 1 | 88821844 | AX-170159414 | 0 | A | C | 0.149 | 1.19E-01 | 2.00E-01 | 2.64E+00 | 2.70E+00 | 5.51E-01 | no | | intergenic_variant | MODIFIER |
| 1 | 88956288 | AX-170164879 | 0 | A | G | 0.149 | 1.19E-01 | 2.00E-01 | 2.64E+00 | 2.70E+00 | 5.51E-01 | no | | intergenic_variant | MODIFIER |
| 1 | 89142553 | AX-170232505 | 0 | A | G | 0.149 | 1.19E-01 | 2.00E-01 | 2.64E+00 | 2.70E+00 | 5.51E-01 | no | Sh3bp4 | intron_variant | MODIFIER |
| 1 | 89149908 | AX-170214131 | 0 | A | G | 0.149 | 1.19E-01 | 2.00E-01 | 2.64E+00 | 2.70E+00 | 5.51E-01 | no | Sh3bp4 | intron_variant | MODIFIER |
| 1 | 89319954 | AX-170131801 | 0 | C | T | 0.149 | 1.19E-01 | 2.00E-01 | 2.64E+00 | 2.70E+00 | 5.51E-01 | no | Gm28342 | intron_variant,non_coding_transcript_variant | MODIFIER |
| 1 | 89419875 | AX-170041968 | 0 | A | C | 0.149 | 1.19E-01 | 2.00E-01 | 2.64E+00 | 2.70E+00 | 5.51E-01 | no | | intergenic_variant | MODIFIER |
| 1 | 89419992 | AX-170172082 | 0 | C | A | 0.149 | 1.19E-01 | 2.00E-01 | 2.64E+00 | 2.70E+00 | 5.51E-01 | no | | intergenic_variant | MODIFIER |
| 1 | 89716793 | AX-170247868 | 0 | T | C | 0.149 | 1.19E-01 | 2.00E-01 | 2.64E+00 | 2.70E+00 | 5.51E-01 | no | Agap1 | intron_variant | MODIFIER |
| 1 | 94919277 | AX-170196511 | 0 | A | G | 0.149 | 1.19E-01 | 2.00E-01 | 2.64E+00 | 2.70E+00 | 5.51E-01 | no | | intergenic_variant | MODIFIER |
| 1 | 95592841 | AX-170001123 | 0 | C | G | 0.149 | 1.19E-01 | 2.00E-01 | 2.64E+00 | 2.70E+00 | 5.51E-01 | no | St8sia4 | intron_variant | MODIFIER |
| 1 | 95629491 | AX-97847895 | 0 | T | C | 0.149 | 1.19E-01 | 2.00E-01 | 2.64E+00 | 2.70E+00 | 5.51E-01 | no | St8sia4 | intron_variant | MODIFIER |
| 1 | 95639537 | AX-170001122 | 0 | A | C | 0.149 | 1.19E-01 | 2.00E-01 | 2.64E+00 | 2.70E+00 | 5.51E-01 | no | St8sia4 | intron_variant | MODIFIER |
| 1 | 95667642 | AX-170001132 | 0 | G | C | 0.149 | 1.19E-01 | 2.00E-01 | 2.64E+00 | 2.70E+00 | 5.51E-01 | no | St8sia4 | upstream_gene_variant | MODIFIER |
| 1 | 95707039 | AX-97811238 | 0 | C | T | 0.149 | 1.19E-01 | 2.00E-01 | 2.64E+00 | 2.70E+00 | 5.51E-01 | no | 4930598F16Rik | intron_variant,non_coding_transcript_variant | MODIFIER |
| 1 | 95761758 | AX-170326217 | 0 | T | C | 0.149 | 1.19E-01 | 2.00E-01 | 2.64E+00 | 2.70E+00 | 5.51E-01 | no | | intergenic_variant | MODIFIER |
| 1 | 95766566 | AX-170001127 | 0 | T | C | 0.149 | 1.19E-01 | 2.00E-01 | 2.64E+00 | 2.70E+00 | 5.51E-01 | no | | intergenic_variant | MODIFIER |
| 1 | 95801070 | AX-97847881 | 0 | C | T | 0.149 | 1.19E-01 | 2.00E-01 | 2.64E+00 | 2.70E+00 | 5.51E-01 | no | | intergenic_variant | MODIFIER |
| 1 | 95839923 | AX-170158789 | 0 | T | A | 0.149 | 1.19E-01 | 2.00E-01 | 2.64E+00 | 2.70E+00 | 5.51E-01 | no | | intergenic_variant | MODIFIER |
| 1 | 95869550 | AX-170170286 | 0 | A | G | 0.149 | 1.19E-01 | 2.00E-01 | 2.64E+00 | 2.70E+00 | 5.51E-01 | no | | intergenic_variant | MODIFIER |
| 1 | 95880870 | AX-170413050 | 0 | G | A | 0.149 | 1.19E-01 | 2.00E-01 | 2.64E+00 | 2.70E+00 | 5.51E-01 | no | | intergenic_variant | MODIFIER |
| 1 | 95905648 | AX-170114153 | 0 | A | G | 0.149 | 1.19E-01 | 2.00E-01 | 2.64E+00 | 2.70E+00 | 5.51E-01 | no | | intergenic_variant | MODIFIER |
| 1 | 95914096 | AX-170197236 | 0 | C | G | 0.149 | 1.19E-01 | 2.00E-01 | 2.64E+00 | 2.70E+00 | 5.51E-01 | no | | intergenic_variant | MODIFIER |
| 1 | 95914167 | AX-97856694 | 0 | G | A | 0.149 | 1.19E-01 | 2.00E-01 | 2.64E+00 | 2.70E+00 | 5.51E-01 | no | | intergenic_variant | MODIFIER |
| 1 | 95918270 | AX-170528340 | 0 | T | C | 0.149 | 1.19E-01 | 2.00E-01 | 2.64E+00 | 2.70E+00 | 5.51E-01 | no | | intergenic_variant | MODIFIER |
| 1 | 95933752 | AX-97850911 | 0 | C | T | 0.149 | 1.19E-01 | 2.00E-01 | 2.64E+00 | 2.70E+00 | 5.51E-01 | no | | intergenic_variant | MODIFIER |
| 1 | 95991872 | AX-97780465 | 0 | C | T | 0.149 | 1.19E-01 | 2.00E-01 | 2.64E+00 | 2.70E+00 | 5.51E-01 | no | 1700063A18Rik | intron_variant,non_coding_transcript_variant | MODIFIER |
| 1 | 96073965 | AX-170174751 | 0 | G | A | 0.149 | 1.19E-01 | 2.00E-01 | 2.64E+00 | 2.70E+00 | 5.51E-01 | no | | intergenic_variant | MODIFIER |
| 1 | 96074916 | AX-170001136 | 0 | A | G | 0.149 | 1.19E-01 | 2.00E-01 | 2.64E+00 | 2.70E+00 | 5.51E-01 | no | | intergenic_variant | MODIFIER |
| 1 | 96155460 | AX-170001134 | 0 | T | A | 0.149 | 1.19E-01 | 2.00E-01 | 2.64E+00 | 2.70E+00 | 5.51E-01 | no | | intergenic_variant | MODIFIER |
| 1 | 96167092 | AX-170001138 | 0 | A | C | 0.149 | 1.19E-01 | 2.00E-01 | 2.64E+00 | 2.70E+00 | 5.51E-01 | no | | intergenic_variant | MODIFIER |
| 1 | 96168485 | AX-170291190 | 0 | A | G | 0.149 | 1.19E-01 | 2.00E-01 | 2.64E+00 | 2.70E+00 | 5.51E-01 | no | | intergenic_variant | MODIFIER |
| 1 | 96289830 | AX-170001133 | 0 | T | C | 0.149 | 1.19E-01 | 2.00E-01 | 2.64E+00 | 2.70E+00 | 5.51E-01 | no | | intergenic_variant | MODIFIER |
| 1 | 96300522 | AX-97810221 | 0 | C | T | 0.149 | 1.19E-01 | 2.00E-01 | 2.64E+00 | 2.70E+00 | 5.51E-01 | no | Gm22917 | upstream_gene_variant | MODIFIER |
| 1 | 96310710 | AX-170001137 | 0 | G | A | 0.149 | 1.19E-01 | 2.00E-01 | 2.64E+00 | 2.70E+00 | 5.51E-01 | no | Gm37076 | upstream_gene_variant | MODIFIER |
| 1 | 96320286 | AX-170001140 | 0 | A | G | 0.149 | 1.19E-01 | 2.00E-01 | 2.64E+00 | 2.70E+00 | 5.51E-01 | no | Gm37076 | downstream_gene_variant | MODIFIER |
| 1 | 96352834 | AX-97843820 | 0 | T | C | 0.149 | 1.19E-01 | 2.00E-01 | 2.64E+00 | 2.70E+00 | 5.51E-01 | no | | intergenic_variant | MODIFIER |
| 1 | 96419244 | AX-170110560 | 0 | A | G | 0.149 | 1.19E-01 | 2.00E-01 | 2.64E+00 | 2.70E+00 | 5.51E-01 | no | | intergenic_variant | MODIFIER |
| 1 | 96425404 | AX-170480716 | 0 | A | G | 0.149 | 1.19E-01 | 2.00E-01 | 2.64E+00 | 2.70E+00 | 5.51E-01 | no | | intergenic_variant | MODIFIER |
| 1 | 96437174 | AX-170220390 | 0 | G | C | 0.149 | 1.19E-01 | 2.00E-01 | 2.64E+00 | 2.70E+00 | 5.51E-01 | no | | intergenic_variant | MODIFIER |
| 1 | 96437766 | AX-170426326 | 0 | C | T | 0.149 | 1.19E-01 | 2.00E-01 | 2.64E+00 | 2.70E+00 | 5.51E-01 | no | | intergenic_variant | MODIFIER |
| 1 | 96438166 | AX-170469196 | 0 | A | C | 0.149 | 1.19E-01 | 2.00E-01 | 2.64E+00 | 2.70E+00 | 5.51E-01 | no | | intergenic_variant | MODIFIER |
| 1 | 96473304 | AX-170327810 | 0 | T | G | 0.149 | 1.19E-01 | 2.00E-01 | 2.64E+00 | 2.70E+00 | 5.51E-01 | no | | intergenic_variant | MODIFIER |
| 1 | 96473485 | AX-170516244 | 0 | C | T | 0.149 | 1.19E-01 | 2.00E-01 | 2.64E+00 | 2.70E+00 | 5.51E-01 | no | | intergenic_variant | MODIFIER |
| 1 | 96482226 | AX-170042099 | 0 | G | T | 0.149 | 1.19E-01 | 2.00E-01 | 2.64E+00 | 2.70E+00 | 5.51E-01 | no | | intergenic_variant | MODIFIER |

| | | | | | | | | | | | | | | | |
|---|---|---|---|---|---|---|---|---|---|---|---|---|---|---|---|
| 1 | 96483056 | AX-170464847 | 0 | C | T | 0.149 | 1.19E-01 | 2.00E-01 | 2.64E+00 | 2.70E+00 | 5.51E-01 | no | | intergenic_variant | MODIFIER |
| 1 | 96489003 | AX-170042097 | 0 | C | T | 0.149 | 1.19E-01 | 2.00E-01 | 2.64E+00 | 2.70E+00 | 5.51E-01 | no | | intergenic_variant | MODIFIER |
| 1 | 96504141 | AX-170466154 | 0 | A | G | 0.149 | 1.19E-01 | 2.00E-01 | 2.64E+00 | 2.70E+00 | 5.51E-01 | no | | intergenic_variant | MODIFIER |
| 1 | 96513584 | AX-170254174 | 0 | G | T | 0.149 | 1.19E-01 | 2.00E-01 | 2.64E+00 | 2.70E+00 | 5.51E-01 | no | | intergenic_variant | MODIFIER |
| 1 | 96513969 | AX-97838865 | 0 | A | C | 0.149 | 1.19E-01 | 2.00E-01 | 2.64E+00 | 2.70E+00 | 5.51E-01 | no | | intergenic_variant | MODIFIER |
| 1 | 96514281 | AX-170155541 | 0 | T | C | 0.149 | 1.19E-01 | 2.00E-01 | 2.64E+00 | 2.70E+00 | 5.51E-01 | no | | intergenic_variant | MODIFIER |
| 1 | 96514298 | AX-170276487 | 0 | A | G | 0.149 | 1.19E-01 | 2.00E-01 | 2.64E+00 | 2.70E+00 | 5.51E-01 | no | | intergenic_variant | MODIFIER |
| 1 | 96522003 | AX-97842587 | 0 | T | G | 0.149 | 1.19E-01 | 2.00E-01 | 2.64E+00 | 2.70E+00 | 5.51E-01 | no | | intergenic_variant | MODIFIER |
| 1 | 96525096 | AX-170276307 | 0 | A | G | 0.149 | 1.19E-01 | 2.00E-01 | 2.64E+00 | 2.70E+00 | 5.51E-01 | no | | intergenic_variant | MODIFIER |
| 1 | 96540227 | AX-170217165 | 0 | A | G | 0.149 | 1.19E-01 | 2.00E-01 | 2.64E+00 | 2.70E+00 | 5.51E-01 | no | Gm29601 | downstream_gene_variant | MODIFIER |
| 1 | 96558916 | AX-170135188 | 0 | G | C | 0.149 | 1.19E-01 | 2.00E-01 | 2.64E+00 | 2.70E+00 | 5.51E-01 | no | Gm29601 | upstream_gene_variant | MODIFIER |
| 1 | 96710277 | AX-170001150 | 0 | A | C | 0.149 | 1.19E-01 | 2.00E-01 | 2.64E+00 | 2.70E+00 | 5.51E-01 | no | | intergenic_variant | MODIFIER |
| 1 | 96714229 | AX-170042103 | 0 | C | A | 0.149 | 1.19E-01 | 2.00E-01 | 2.64E+00 | 2.70E+00 | 5.51E-01 | no | | intergenic_variant | MODIFIER |
| 1 | 96729350 | AX-97803386 | 0 | T | G | 0.149 | 1.19E-01 | 2.00E-01 | 2.64E+00 | 2.70E+00 | 5.51E-01 | no | | intergenic_variant | MODIFIER |
| 1 | 96772132 | AX-170001147 | 0 | T | C | 0.149 | 1.19E-01 | 2.00E-01 | 2.64E+00 | 2.70E+00 | 5.51E-01 | no | | intergenic_variant | MODIFIER |
| 1 | 96780119 | AX-97812466 | 0 | C | A | 0.149 | 1.19E-01 | 2.00E-01 | 2.64E+00 | 2.70E+00 | 5.51E-01 | no | | intergenic_variant | MODIFIER |
| 1 | 96799857 | AX-170535590 | 0 | A | C | 0.149 | 1.19E-01 | 2.00E-01 | 2.64E+00 | 2.70E+00 | 5.51E-01 | no | | intergenic_variant | MODIFIER |
| 1 | 96803484 | AX-170001151 | 0 | G | A | 0.149 | 1.19E-01 | 2.00E-01 | 2.64E+00 | 2.70E+00 | 5.51E-01 | no | | intergenic_variant | MODIFIER |
| 1 | 96803637 | AX-170172611 | 0 | T | C | 0.149 | 1.19E-01 | 2.00E-01 | 2.64E+00 | 2.70E+00 | 5.51E-01 | no | | intergenic_variant | MODIFIER |
| 1 | 96833804 | AX-170001142 | 0 | G | T | 0.149 | 1.19E-01 | 2.00E-01 | 2.64E+00 | 2.70E+00 | 5.51E-01 | no | Slco4c1 | intron_variant | MODIFIER |
| 1 | 96837300 | AX-170001152 | 0 | T | C | 0.149 | 1.19E-01 | 2.00E-01 | 2.64E+00 | 2.70E+00 | 5.51E-01 | no | Slco4c1 | intron_variant | MODIFIER |
| 1 | 96869845 | AX-97825758 | 0 | A | G | 0.149 | 1.19E-01 | 2.00E-01 | 2.64E+00 | 2.70E+00 | 5.51E-01 | no | Slco4c1 | intron_variant | MODIFIER |
| 1 | 96873541 | AX-170042107 | 0 | A | G | 0.149 | 1.19E-01 | 2.00E-01 | 2.64E+00 | 2.70E+00 | 5.51E-01 | no | Slco4c1 | upstream_gene_variant | MODIFIER |
| 1 | 96880423 | AX-170042105 | 0 | A | T | 0.149 | 1.19E-01 | 2.00E-01 | 2.64E+00 | 2.70E+00 | 5.51E-01 | no | Panct2 | non_coding_transcript_exon_variant | MODIFIER |
| 1 | 96894865 | AX-170425594 | 0 | A | G | 0.149 | 1.19E-01 | 2.00E-01 | 2.64E+00 | 2.70E+00 | 5.51E-01 | no | Panct2 | intron_variant,non_coding_transcript_variant | MODIFIER |
| 1 | 96937929 | AX-170186665 | 0 | C | T | 0.149 | 1.19E-01 | 2.00E-01 | 2.64E+00 | 2.70E+00 | 5.51E-01 | no | Slco6b1 | intron_variant,non_coding_transcript_variant | MODIFIER |
| 1 | 96948597 | AX-168390623 | 0 | T | C | 0.149 | 1.19E-01 | 2.00E-01 | 2.64E+00 | 2.70E+00 | 5.51E-01 | no | Slco6b1 | intron_variant,non_coding_transcript_variant | MODIFIER |
| 1 | 96948905 | AX-97812467 | 0 | A | G | 0.149 | 1.19E-01 | 2.00E-01 | 2.64E+00 | 2.70E+00 | 5.51E-01 | no | Slco6b1 | intron_variant,non_coding_transcript_variant | MODIFIER |
| 1 | 96988698 | AX-170086546 | 0 | T | C | 0.149 | 1.19E-01 | 2.00E-01 | 2.64E+00 | 2.70E+00 | 5.51E-01 | no | Slco6b1 | non_coding_transcript_exon_variant | MODIFIER |
| 1 | 96993164 | AX-170135488 | 0 | G | C | 0.149 | 1.19E-01 | 2.00E-01 | 2.64E+00 | 2.70E+00 | 5.51E-01 | no | Slco6b1 | intron_variant,non_coding_transcript_variant | MODIFIER |
| 1 | 96997354 | AX-170531199 | 0 | T | C | 0.149 | 1.19E-01 | 2.00E-01 | 2.64E+00 | 2.70E+00 | 5.51E-01 | no | Slco6b1 | non_coding_transcript_exon_variant | MODIFIER |
| 1 | 96997494 | AX-170284615 | 0 | A | T | 0.149 | 1.19E-01 | 2.00E-01 | 2.64E+00 | 2.70E+00 | 5.51E-01 | no | Slco6b1 | non_coding_transcript_exon_variant | MODIFIER |
| 1 | 97019052 | AX-170361752 | 0 | G | T | 0.149 | 1.19E-01 | 2.00E-01 | 2.64E+00 | 2.70E+00 | 5.51E-01 | no | | regulatory_region_variant | MODIFIER |
| 1 | 97042912 | AX-170377867 | 0 | A | G | 0.149 | 1.19E-01 | 2.00E-01 | 2.64E+00 | 2.70E+00 | 5.51E-01 | no | | intergenic_variant | MODIFIER |
| 1 | 97044070 | AX-170233024 | 0 | G | A | 0.149 | 1.19E-01 | 2.00E-01 | 2.64E+00 | 2.70E+00 | 5.51E-01 | no | | intergenic_variant | MODIFIER |
| 1 | 99398062 | AX-170386692 | 0 | A | G | 0.149 | 1.19E-01 | 2.00E-01 | 2.64E+00 | 2.70E+00 | 5.51E-01 | no | | intergenic_variant | MODIFIER |
| 1 | 99683394 | AX-170340971 | 0 | T | C | 0.149 | 1.19E-01 | 2.00E-01 | 2.64E+00 | 2.70E+00 | 5.51E-01 | no | | intergenic_variant | MODIFIER |
| 1 | 99687761 | AX-170091606 | 0 | G | C | 0.149 | 1.19E-01 | 2.00E-01 | 2.64E+00 | 2.70E+00 | 5.51E-01 | no | | intergenic_variant | MODIFIER |
| 1 | 99720250 | AX-170001190 | 0 | T | C | 0.149 | 1.19E-01 | 2.00E-01 | 2.64E+00 | 2.70E+00 | 5.51E-01 | no | | intergenic_variant | MODIFIER |
| 1 | 99741464 | AX-170431973 | 0 | T | G | 0.149 | 1.19E-01 | 2.00E-01 | 2.64E+00 | 2.70E+00 | 5.51E-01 | no | | intergenic_variant | MODIFIER |
| 1 | 99784114 | AX-170474659 | 0 | T | G | 0.149 | 1.19E-01 | 2.00E-01 | 2.64E+00 | 2.70E+00 | 5.51E-01 | no | Cntnap5b | intron_variant | MODIFIER |
| 1 | 99792341 | AX-170091608 | 0 | C | T | 0.149 | 1.19E-01 | 2.00E-01 | 2.64E+00 | 2.70E+00 | 5.51E-01 | no | Cntnap5b | intron_variant | MODIFIER |
| 1 | 99811668 | AX-170184448 | 0 | C | G | 0.149 | 1.19E-01 | 2.00E-01 | 2.64E+00 | 2.70E+00 | 5.51E-01 | no | Cntnap5b | intron_variant | MODIFIER |
| 1 | 99838605 | AX-170498993 | 0 | C | G | 0.149 | 1.19E-01 | 2.00E-01 | 2.64E+00 | 2.70E+00 | 5.51E-01 | no | Cntnap5b | intron_variant | MODIFIER |
| 1 | 99839152 | AX-170396296 | 0 | G | A | 0.149 | 1.19E-01 | 2.00E-01 | 2.64E+00 | 2.70E+00 | 5.51E-01 | no | Cntnap5b | intron_variant | MODIFIER |
| 1 | 99843579 | AX-170460930 | 0 | A | G | 0.149 | 1.19E-01 | 2.00E-01 | 2.64E+00 | 2.70E+00 | 5.51E-01 | no | Cntnap5b | intron_variant | MODIFIER |
| 1 | 99848776 | AX-97794137 | 0 | G | A | 0.149 | 1.19E-01 | 2.00E-01 | 2.64E+00 | 2.70E+00 | 5.51E-01 | no | Cntnap5b | intron_variant | MODIFIER |
| 1 | 99853192 | AX-170123286 | 0 | C | T | 0.149 | 1.19E-01 | 2.00E-01 | 2.64E+00 | 2.70E+00 | 5.51E-01 | no | Cntnap5b | intron_variant | MODIFIER |
| 1 | 101000818 | AX-170432579 | 0 | T | G | 0.149 | 1.19E-01 | 2.00E-01 | 2.64E+00 | 2.70E+00 | 5.51E-01 | no | | intergenic_variant | MODIFIER |
| 1 | 101005216 | AX-170220106 | 0 | G | A | 0.149 | 1.19E-01 | 2.00E-01 | 2.64E+00 | 2.70E+00 | 5.51E-01 | no | | intergenic_variant | MODIFIER |
| 1 | 101008104 | AX-170346093 | 0 | T | G | 0.149 | 1.19E-01 | 2.00E-01 | 2.64E+00 | 2.70E+00 | 5.51E-01 | no | | intergenic_variant | MODIFIER |
| 1 | 101012194 | AX-170042189 | 0 | A | G | 0.149 | 1.19E-01 | 2.00E-01 | 2.64E+00 | 2.70E+00 | 5.51E-01 | no | | intergenic_variant | MODIFIER |
| 1 | 101012816 | AX-97812490 | 0 | G | T | 0.149 | 1.19E-01 | 2.00E-01 | 2.64E+00 | 2.70E+00 | 5.51E-01 | no | | intergenic_variant | MODIFIER |
| 1 | 101019424 | AX-170042190 | 0 | A | G | 0.149 | 1.19E-01 | 2.00E-01 | 2.64E+00 | 2.70E+00 | 5.51E-01 | no | | intergenic_variant | MODIFIER |

| | | | | | | | | | | | | | | | |
|---|---|---|---|---|---|---|---|---|---|---|---|---|---|---|---|
| 1 | 101034318 | AX-170193569 | 0 | T | A | 0.149 | 1.19E-01 | 2.00E-01 | 2.64E+00 | 2.70E+00 | 5.51E-01 | no | | intergenic_variant | MODIFIER |
| 1 | 101060897 | AX-170521055 | 0 | A | G | 0.149 | 1.19E-01 | 2.00E-01 | 2.64E+00 | 2.70E+00 | 5.51E-01 | no | | intergenic_variant | MODIFIER |
| 1 | 101063259 | AX-170254070 | 0 | T | C | 0.149 | 1.19E-01 | 2.00E-01 | 2.64E+00 | 2.70E+00 | 5.51E-01 | no | | intergenic_variant | MODIFIER |
| 1 | 101074953 | AX-170001204 | 0 | C | T | 0.149 | 1.19E-01 | 2.00E-01 | 2.64E+00 | 2.70E+00 | 5.51E-01 | no | | intergenic_variant | MODIFIER |
| 1 | 101077564 | AX-170086554 | 0 | A | G | 0.149 | 1.19E-01 | 2.00E-01 | 2.64E+00 | 2.70E+00 | 5.51E-01 | no | | intergenic_variant | MODIFIER |
| 1 | 101102334 | AX-170219441 | 0 | T | C | 0.149 | 1.19E-01 | 2.00E-01 | 2.64E+00 | 2.70E+00 | 5.51E-01 | no | | intergenic_variant | MODIFIER |
| 1 | 101123452 | AX-170001206 | 0 | A | G | 0.149 | 1.19E-01 | 2.00E-01 | 2.64E+00 | 2.70E+00 | 5.51E-01 | no | | intergenic_variant | MODIFIER |
| 1 | 101123892 | AX-170255631 | 0 | C | T | 0.149 | 1.19E-01 | 2.00E-01 | 2.64E+00 | 2.70E+00 | 5.51E-01 | no | | intergenic_variant | MODIFIER |
| 1 | 101124995 | AX-170001203 | 0 | G | T | 0.149 | 1.19E-01 | 2.00E-01 | 2.64E+00 | 2.70E+00 | 5.51E-01 | no | | intergenic_variant | MODIFIER |
| 1 | 101125023 | AX-170407827 | 0 | C | G | 0.149 | 1.19E-01 | 2.00E-01 | 2.64E+00 | 2.70E+00 | 5.51E-01 | no | | intergenic_variant | MODIFIER |
| 1 | 101129614 | AX-170488852 | 0 | T | G | 0.149 | 1.19E-01 | 2.00E-01 | 2.64E+00 | 2.70E+00 | 5.51E-01 | no | | intergenic_variant | MODIFIER |
| 1 | 101130894 | AX-97821019 | 0 | T | C | 0.149 | 1.19E-01 | 2.00E-01 | 2.64E+00 | 2.70E+00 | 5.51E-01 | no | | intergenic_variant | MODIFIER |
| 1 | 101137501 | AX-170437636 | 0 | G | T | 0.149 | 1.19E-01 | 2.00E-01 | 2.64E+00 | 2.70E+00 | 5.51E-01 | no | | intergenic_variant | MODIFIER |
| 1 | 101137541 | AX-170159321 | 0 | C | G | 0.149 | 1.19E-01 | 2.00E-01 | 2.64E+00 | 2.70E+00 | 5.51E-01 | no | | intergenic_variant | MODIFIER |
| 1 | 101156467 | AX-170430009 | 0 | C | T | 0.149 | 1.19E-01 | 2.00E-01 | 2.64E+00 | 2.70E+00 | 5.51E-01 | no | Gm20268 | intron_variant,non_coding_transcript_variant | MODIFIER |
| 1 | 101156517 | AX-170412713 | 0 | T | A | 0.149 | 1.19E-01 | 2.00E-01 | 2.64E+00 | 2.70E+00 | 5.51E-01 | no | Gm20268 | intron_variant,non_coding_transcript_variant | MODIFIER |
| 1 | 101167095 | AX-170042191 | 0 | A | G | 0.149 | 1.19E-01 | 2.00E-01 | 2.64E+00 | 2.70E+00 | 5.51E-01 | no | Gm20268 | intron_variant,non_coding_transcript_variant | MODIFIER |
| 1 | 101260735 | AX-170452453 | 0 | A | T | 0.149 | 1.19E-01 | 2.00E-01 | 2.64E+00 | 2.70E+00 | 5.51E-01 | no | | intergenic_variant | MODIFIER |
| 1 | 101292851 | AX-170319662 | 0 | A | G | 0.149 | 1.19E-01 | 2.00E-01 | 2.64E+00 | 2.70E+00 | 5.51E-01 | no | | intergenic_variant | MODIFIER |
| 1 | 101324097 | AX-170379914 | 0 | G | A | 0.149 | 1.19E-01 | 2.00E-01 | 2.64E+00 | 2.70E+00 | 5.51E-01 | no | | intergenic_variant | MODIFIER |
| 1 | 101356892 | AX-97812610 | 0 | A | C | 0.149 | 1.19E-01 | 2.00E-01 | 2.64E+00 | 2.70E+00 | 5.51E-01 | no | | intergenic_variant | MODIFIER |
| 1 | 101370221 | AX-170042195 | 0 | T | C | 0.149 | 1.19E-01 | 2.00E-01 | 2.64E+00 | 2.70E+00 | 5.51E-01 | no | Gm28801 | upstream_gene_variant | MODIFIER |
| 1 | 101391184 | AX-170508448 | 0 | A | C | 0.149 | 1.19E-01 | 2.00E-01 | 2.64E+00 | 2.70E+00 | 5.51E-01 | no | Psmb6-ps2 | downstream_gene_variant | MODIFIER |
| 1 | 101407883 | AX-170325326 | 0 | C | G | 0.149 | 1.19E-01 | 2.00E-01 | 2.64E+00 | 2.70E+00 | 5.51E-01 | no | | intergenic_variant | MODIFIER |
| 1 | 101417327 | AX-97812511 | 0 | A | G | 0.149 | 1.19E-01 | 2.00E-01 | 2.64E+00 | 2.70E+00 | 5.51E-01 | no | | intergenic_variant | MODIFIER |
| 1 | 101469452 | AX-97812539 | 0 | T | C | 0.149 | 1.19E-01 | 2.00E-01 | 2.64E+00 | 2.70E+00 | 5.51E-01 | no | | intergenic_variant | MODIFIER |
| 1 | 101487768 | AX-170501685 | 0 | C | G | 0.149 | 1.19E-01 | 2.00E-01 | 2.64E+00 | 2.70E+00 | 5.51E-01 | no | | intergenic_variant | MODIFIER |
| 1 | 101511426 | AX-170110586 | 0 | T | C | 0.149 | 1.19E-01 | 2.00E-01 | 2.64E+00 | 2.70E+00 | 5.51E-01 | no | | intergenic_variant | MODIFIER |
| 1 | 101511606 | AX-170254078 | 0 | C | T | 0.149 | 1.19E-01 | 2.00E-01 | 2.64E+00 | 2.70E+00 | 5.51E-01 | no | | intergenic_variant | MODIFIER |
| 1 | 101516785 | AX-170415650 | 0 | A | C | 0.149 | 1.19E-01 | 2.00E-01 | 2.64E+00 | 2.70E+00 | 5.51E-01 | no | | intergenic_variant | MODIFIER |
| 1 | 101519257 | AX-170221454 | 0 | A | T | 0.149 | 1.19E-01 | 2.00E-01 | 2.64E+00 | 2.70E+00 | 5.51E-01 | no | | intergenic_variant | MODIFIER |
| 1 | 101542944 | AX-170336002 | 0 | A | G | 0.149 | 1.19E-01 | 2.00E-01 | 2.64E+00 | 2.70E+00 | 5.51E-01 | no | | intergenic_variant | MODIFIER |
| 1 | 101543608 | AX-170513836 | 0 | C | T | 0.149 | 1.19E-01 | 2.00E-01 | 2.64E+00 | 2.70E+00 | 5.51E-01 | no | | intergenic_variant | MODIFIER |
| 1 | 101552147 | AX-97832667 | 0 | T | C | 0.149 | 1.19E-01 | 2.00E-01 | 2.64E+00 | 2.70E+00 | 5.51E-01 | no | | intergenic_variant | MODIFIER |
| 1 | 101560647 | AX-170482719 | 0 | C | A | 0.149 | 1.19E-01 | 2.00E-01 | 2.64E+00 | 2.70E+00 | 5.51E-01 | no | | intergenic_variant | MODIFIER |
| 1 | 101660905 | AX-170120814 | 0 | G | A | 0.149 | 1.19E-01 | 2.00E-01 | 2.64E+00 | 2.70E+00 | 5.51E-01 | no | | intergenic_variant | MODIFIER |
| 1 | 101681078 | AX-170419599 | 0 | T | C | 0.149 | 1.19E-01 | 2.00E-01 | 2.64E+00 | 2.70E+00 | 5.51E-01 | no | | intergenic_variant | MODIFIER |
| 1 | 101682749 | AX-170258171 | 0 | G | A | 0.149 | 1.19E-01 | 2.00E-01 | 2.64E+00 | 2.70E+00 | 5.51E-01 | no | | intergenic_variant | MODIFIER |
| 1 | 101687382 | AX-170042202 | 0 | T | C | 0.149 | 1.19E-01 | 2.00E-01 | 2.64E+00 | 2.70E+00 | 5.51E-01 | no | | intergenic_variant | MODIFIER |
| 1 | 101687838 | AX-170042203 | 0 | A | T | 0.149 | 1.19E-01 | 2.00E-01 | 2.64E+00 | 2.70E+00 | 5.51E-01 | no | | intergenic_variant | MODIFIER |
| 1 | 101687957 | AX-170042200 | 0 | C | T | 0.149 | 1.19E-01 | 2.00E-01 | 2.64E+00 | 2.70E+00 | 5.51E-01 | no | | intergenic_variant | MODIFIER |
| 1 | 101721489 | AX-170001212 | 0 | T | A | 0.149 | 1.19E-01 | 2.00E-01 | 2.64E+00 | 2.70E+00 | 5.51E-01 | no | | intergenic_variant | MODIFIER |
| 1 | 101794660 | AX-170183592 | 0 | C | T | 0.149 | 1.19E-01 | 2.00E-01 | 2.64E+00 | 2.70E+00 | 5.51E-01 | no | | intergenic_variant | MODIFIER |
| 1 | 101801985 | AX-170184959 | 0 | T | G | 0.149 | 1.19E-01 | 2.00E-01 | 2.64E+00 | 2.70E+00 | 5.51E-01 | no | | intergenic_variant | MODIFIER |
| 1 | 101821744 | AX-170235110 | 0 | C | T | 0.149 | 1.19E-01 | 2.00E-01 | 2.64E+00 | 2.70E+00 | 5.51E-01 | no | | intergenic_variant | MODIFIER |
| 1 | 101838869 | AX-170426705 | 0 | T | C | 0.149 | 1.19E-01 | 2.00E-01 | 2.64E+00 | 2.70E+00 | 5.51E-01 | no | | intergenic_variant | MODIFIER |
| 1 | 101843404 | AX-170464479 | 0 | C | A | 0.149 | 1.19E-01 | 2.00E-01 | 2.64E+00 | 2.70E+00 | 5.51E-01 | no | | intergenic_variant | MODIFIER |
| 1 | 101872984 | AX-170219986 | 0 | A | G | 0.149 | 1.19E-01 | 2.00E-01 | 2.64E+00 | 2.70E+00 | 5.51E-01 | no | | intergenic_variant | MODIFIER |
| 1 | 101873080 | AX-170413247 | 0 | T | G | 0.149 | 1.19E-01 | 2.00E-01 | 2.64E+00 | 2.70E+00 | 5.51E-01 | no | | intergenic_variant | MODIFIER |
| 1 | 101876225 | AX-170443683 | 0 | G | C | 0.149 | 1.19E-01 | 2.00E-01 | 2.64E+00 | 2.70E+00 | 5.51E-01 | no | Gm28188 | downstream_gene_variant | MODIFIER |
| 1 | 101912981 | AX-170486010 | 0 | T | C | 0.149 | 1.19E-01 | 2.00E-01 | 2.64E+00 | 2.70E+00 | 5.51E-01 | no | | intergenic_variant | MODIFIER |
| 1 | 101913125 | AX-170287665 | 0 | T | G | 0.149 | 1.19E-01 | 2.00E-01 | 2.64E+00 | 2.70E+00 | 5.51E-01 | no | | intergenic_variant | MODIFIER |
| 1 | 101916043 | AX-170477101 | 0 | C | T | 0.149 | 1.19E-01 | 2.00E-01 | 2.64E+00 | 2.70E+00 | 5.51E-01 | no | | intergenic_variant | MODIFIER |
| 1 | 101943842 | AX-170164457 | 0 | T | A | 0.149 | 1.19E-01 | 2.00E-01 | 2.64E+00 | 2.70E+00 | 5.51E-01 | no | | intergenic_variant | MODIFIER |

| | | | | | | | | | | | | | | | |
|---|---|---|---|---|---|---|---|---|---|---|---|---|---|---|---|
| 1 | 101962766 | AX-170161124 | 0 | A | C | 0.149 | 1.19E-01 | 2.00E-01 | 2.64E+00 | 2.70E+00 | 5.51E-01 | no | | intergenic_variant | MODIFIER |
| 1 | 101964188 | AX-170533445 | 0 | A | T | 0.149 | 1.19E-01 | 2.00E-01 | 2.64E+00 | 2.70E+00 | 5.51E-01 | no | | intergenic_variant | MODIFIER |
| 1 | 102273047 | AX-170001220 | 0 | A | G | 0.149 | 1.19E-01 | 2.00E-01 | 2.64E+00 | 2.70E+00 | 5.51E-01 | no | | intergenic_variant | MODIFIER |
| 1 | 102436369 | AX-170249837 | 0 | T | G | 0.149 | 1.19E-01 | 2.00E-01 | 2.64E+00 | 2.70E+00 | 5.51E-01 | no | | intergenic_variant | MODIFIER |
| 1 | 102551572 | AX-97829095 | 0 | A | G | 0.149 | 1.19E-01 | 2.00E-01 | 2.64E+00 | 2.70E+00 | 5.51E-01 | no | | intergenic_variant | MODIFIER |
| 1 | 102554925 | AX-170456112 | 0 | G | A | 0.149 | 1.19E-01 | 2.00E-01 | 2.64E+00 | 2.70E+00 | 5.51E-01 | no | | intergenic_variant | MODIFIER |
| 1 | 103129820 | AX-97801932 | 0 | A | C | 0.149 | 1.19E-01 | 2.00E-01 | 2.64E+00 | 2.70E+00 | 5.51E-01 | no | | intergenic_variant | MODIFIER |
| 1 | 103348112 | AX-97848592 | 0 | T | G | 0.149 | 1.19E-01 | 2.00E-01 | 2.64E+00 | 2.70E+00 | 5.51E-01 | no | | intergenic_variant | MODIFIER |
| 1 | 103536820 | AX-97785595 | 0 | A | G | 0.149 | 1.19E-01 | 2.00E-01 | 2.64E+00 | 2.70E+00 | 5.51E-01 | no | | intergenic_variant | MODIFIER |
| 1 | 104799608 | AX-170042249 | 0 | A | G | 0.149 | 1.19E-01 | 2.00E-01 | 2.64E+00 | 2.70E+00 | 5.51E-01 | no | Cdh20 | intron_variant | MODIFIER |
| 1 | 104800339 | AX-170223450 | 0 | A | C | 0.149 | 1.19E-01 | 2.00E-01 | 2.64E+00 | 2.70E+00 | 5.51E-01 | no | Cdh20 | intron_variant | MODIFIER |
| 1 | 104800721 | AX-170491054 | 0 | C | T | 0.149 | 1.19E-01 | 2.00E-01 | 2.64E+00 | 2.70E+00 | 5.51E-01 | no | Cdh20 | intron_variant | MODIFIER |
| 1 | 104802977 | AX-170428298 | 0 | A | G | 0.149 | 1.19E-01 | 2.00E-01 | 2.64E+00 | 2.70E+00 | 5.51E-01 | no | Cdh20 | intron_variant | MODIFIER |
| 1 | 104803006 | AX-170303048 | 0 | T | G | 0.149 | 1.19E-01 | 2.00E-01 | 2.64E+00 | 2.70E+00 | 5.51E-01 | no | Cdh20 | intron_variant | MODIFIER |
| 1 | 104804094 | AX-170304932 | 0 | G | A | 0.149 | 1.19E-01 | 2.00E-01 | 2.64E+00 | 2.70E+00 | 5.51E-01 | no | Cdh20 | intron_variant | MODIFIER |
| 1 | 104820625 | AX-170366688 | 0 | G | A | 0.149 | 1.19E-01 | 2.00E-01 | 2.64E+00 | 2.70E+00 | 5.51E-01 | no | Cdh20 | intron_variant | MODIFIER |
| 1 | 104821703 | AX-170429026 | 0 | T | C | 0.149 | 1.19E-01 | 2.00E-01 | 2.64E+00 | 2.70E+00 | 5.51E-01 | no | Cdh20 | intron_variant | MODIFIER |
| 1 | 104823857 | AX-170468296 | 0 | T | C | 0.149 | 1.19E-01 | 2.00E-01 | 2.64E+00 | 2.70E+00 | 5.51E-01 | no | Cdh20 | intron_variant | MODIFIER |
| 1 | 104825091 | AX-170176726 | 0 | G | C | 0.149 | 1.19E-01 | 2.00E-01 | 2.64E+00 | 2.70E+00 | 5.51E-01 | no | Cdh20 | intron_variant | MODIFIER |
| 1 | 104838896 | AX-170182963 | 0 | A | G | 0.149 | 1.19E-01 | 2.00E-01 | 2.64E+00 | 2.70E+00 | 5.51E-01 | no | Cdh20 | intron_variant | MODIFIER |
| 1 | 104871023 | AX-170345970 | 0 | G | C | 0.149 | 1.19E-01 | 2.00E-01 | 2.64E+00 | 2.70E+00 | 5.51E-01 | no | Cdh20 | intron_variant | MODIFIER |
| 1 | 104887662 | AX-170467762 | 0 | C | T | 0.149 | 1.19E-01 | 2.00E-01 | 2.64E+00 | 2.70E+00 | 5.51E-01 | no | Cdh20 | intron_variant | MODIFIER |
| 1 | 104888201 | AX-170291264 | 0 | G | A | 0.149 | 1.19E-01 | 2.00E-01 | 2.64E+00 | 2.70E+00 | 5.51E-01 | no | Cdh20 | intron_variant | MODIFIER |
| 1 | 104909317 | AX-170375113 | 0 | G | A | 0.149 | 1.19E-01 | 2.00E-01 | 2.64E+00 | 2.70E+00 | 5.51E-01 | no | Cdh20 | intron_variant | MODIFIER |
| 1 | 104919238 | AX-170133321 | 0 | A | C | 0.149 | 1.19E-01 | 2.00E-01 | 2.64E+00 | 2.70E+00 | 5.51E-01 | no | Cdh20 | intron_variant | MODIFIER |
| 1 | 104919550 | AX-170437464 | 0 | A | G | 0.149 | 1.19E-01 | 2.00E-01 | 2.64E+00 | 2.70E+00 | 5.51E-01 | no | Cdh20 | intron_variant | MODIFIER |
| 1 | 104935024 | AX-170042256 | 0 | C | A | 0.149 | 1.19E-01 | 2.00E-01 | 2.64E+00 | 2.70E+00 | 5.51E-01 | no | Cdh20 | intron_variant | MODIFIER |
| 1 | 104976105 | AX-170217591 | 0 | A | T | 0.149 | 1.19E-01 | 2.00E-01 | 2.64E+00 | 2.70E+00 | 5.51E-01 | no | Cdh20 | intron_variant | MODIFIER |
| 1 | 105387848 | AX-97812104 | 0 | A | G | 0.149 | 1.19E-01 | 2.00E-01 | 2.64E+00 | 2.70E+00 | 5.51E-01 | no | AC124553.1 | intron_variant,non_coding_transcript_variant | MODIFIER |
| 1 | 105502590 | AX-170001271 | 0 | T | C | 0.149 | 1.19E-01 | 2.00E-01 | 2.64E+00 | 2.70E+00 | 5.51E-01 | no | AC124553.1 | intron_variant,non_coding_transcript_variant | MODIFIER |
| 1 | 105502624 | AX-170001269 | 0 | A | G | 0.149 | 1.19E-01 | 2.00E-01 | 2.64E+00 | 2.70E+00 | 5.51E-01 | no | AC124553.1 | intron_variant,non_coding_transcript_variant | MODIFIER |
| 1 | 105561039 | AX-170201645 | 0 | G | T | 0.149 | 1.19E-01 | 2.00E-01 | 2.64E+00 | 2.70E+00 | 5.51E-01 | no | Pign | intron_variant | MODIFIER |
| 1 | 105650281 | AX-170414405 | 0 | T | A | 0.149 | 1.19E-01 | 2.00E-01 | 2.64E+00 | 2.70E+00 | 5.51E-01 | no | Pign | intron_variant | MODIFIER |
| 1 | 105653281 | AX-97842298 | 0 | C | T | 0.149 | 1.19E-01 | 2.00E-01 | 2.64E+00 | 2.70E+00 | 5.51E-01 | no | Pign | intron_variant | MODIFIER |
| 1 | 105663160 | AX-170254175 | 0 | C | A | 0.149 | 1.19E-01 | 2.00E-01 | 2.64E+00 | 2.70E+00 | 5.51E-01 | no | Relch | upstream_gene_variant | MODIFIER |
| 1 | 105697860 | AX-170417942 | 0 | T | G | 0.149 | 1.19E-01 | 2.00E-01 | 2.64E+00 | 2.70E+00 | 5.51E-01 | no | Relch | intron_variant | MODIFIER |
| 1 | 105963964 | AX-170319215 | 0 | G | A | 0.149 | 1.19E-01 | 2.00E-01 | 2.64E+00 | 2.70E+00 | 5.51E-01 | no | | intergenic_variant | MODIFIER |
| 1 | 105964753 | AX-170197256 | 0 | G | A | 0.149 | 1.19E-01 | 2.00E-01 | 2.64E+00 | 2.70E+00 | 5.51E-01 | no | | intergenic_variant | MODIFIER |
| 1 | 105965469 | AX-170418523 | 0 | T | C | 0.149 | 1.19E-01 | 2.00E-01 | 2.64E+00 | 2.70E+00 | 5.51E-01 | no | | regulatory_region_variant | MODIFIER |
| 1 | 106259210 | AX-97857535 | 0 | T | C | 0.149 | 1.19E-01 | 2.00E-01 | 2.64E+00 | 2.70E+00 | 5.51E-01 | no | Phlpp1 | intron_variant | MODIFIER |
| 3 | 35955990 | AX-170270281 | 1 | C | A | 0.196 | 5.69E-02 | 9.69E-02 | 2.74E+00 | 2.81E+00 | 5.51E-01 | no | Mccc1 | downstream_gene_variant | MODIFIER |
| 5 | 100601455 | AX-170434537 | 1 | G | C | 0.429 | -5.98E-02 | 1.02E-01 | 2.76E+00 | 2.85E+00 | 5.51E-01 | no | | intergenic_variant | MODIFIER |
| 7 | 114175067 | AX-170078488 | 0 | G | A | 0.447 | 5.62E-02 | 9.51E-02 | 2.67E+00 | 2.72E+00 | 5.51E-01 | no | | intergenic_variant | MODIFIER |
| 5 | 115602998 | AX-97839378 | 1 | C | A | 0.464 | 6.76E-02 | 1.14E-01 | 2.65E+00 | 2.69E+00 | 5.51E-01 | no | Gcn1l1 | intron_variant | MODIFIER |
| 9 | 33645138 | AX-170372099 | 0 | C | A | 0.43 | 5.01E-02 | 8.45E-02 | 2.63E+00 | 2.70E+00 | 5.51E-01 | no | | intergenic_variant | MODIFIER |
| 8 | 32282684 | AX-170283322 | 0 | G | A | 0.167 | -5.99E-02 | 1.02E-01 | 2.67E+00 | 2.78E+00 | 5.51E-01 | no | Nrg1 | intron_variant | MODIFIER |
| 1 | 45285463 | AX-170448463 | 2 | G | A | 0.082 | -1.15E-01 | 1.96E-01 | 2.65E+00 | 2.77E+00 | 5.51E-01 | no | | intergenic_variant | MODIFIER |
| 7 | 37655163 | AX-170400203 | 1 | A | G | 0.339 | 5.70E-02 | 9.43E-02 | 2.55E+00 | 2.62E+00 | 5.51E-01 | no | Zfp536 | intron_variant | MODIFIER |
| 9 | 120697838 | AX-170137073 | 0 | C | T | 0.202 | -4.72E-02 | 8.07E-02 | 2.65E+00 | 2.78E+00 | 5.51E-01 | no | | regulatory_region_variant | MODIFIER |
| 5 | 108719433 | AX-170528108 | 1 | A | G | 0.455 | 6.67E-02 | 1.13E-01 | 2.65E+00 | 2.69E+00 | 5.52E-01 | no | | intergenic_variant | MODIFIER |
| 1 | 169114410 | AX-170320139 | 1 | T | C | 0.223 | 8.09E-02 | 1.37E-01 | 2.67E+00 | 2.71E+00 | 5.52E-01 | no | | intergenic_variant | MODIFIER |
| 7 | 66188441 | AX-170507081 | 0 | T | C | 0.43 | -5.50E-02 | 9.37E-02 | 2.79E+00 | 2.88E+00 | 5.52E-01 | no | Gm45080 | upstream_gene_variant | MODIFIER |
| 5 | 36470764 | AX-170492584 | 1 | C | T | 0.339 | 6.57E-02 | 1.12E-01 | 2.73E+00 | 2.79E+00 | 5.52E-01 | no | Tada2b | downstream_gene_variant | MODIFIER |
| 5 | 110150049 | AX-170141126 | 0 | A | G | 0.439 | 6.63E-02 | 1.12E-01 | 2.67E+00 | 2.71E+00 | 5.52E-01 | no | Chfr | intron_variant | MODIFIER |

| | | | | | | | | | | | | | | | |
|---|---|---|---|---|---|---|---|---|---|---|---|---|---|---|---|
| 12 | 104005248 | AX-97819375 | 0 | G | A | 0.14 | -1.11E-01 | 1.89E-01 | 2.75E+00 | 2.84E+00 | 5.52E-01 | no | Serpina9 | intron_variant | MODIFIER |
| 16 | 46413249 | AX-170357803 | 1 | A | T | 0.518 | 4.94E-02 | 8.19E-02 | 2.56E+00 | 2.60E+00 | 5.52E-01 | no | Nectin3 | intron_variant | MODIFIER |
| 1 | 173536812 | AX-170489336 | 1 | G | A | 0.232 | 7.33E-02 | 1.20E-01 | 2.53E+00 | 2.59E+00 | 5.52E-01 | no | Ifi214 | upstream_gene_variant | MODIFIER |
| 16 | 38567463 | AX-97793818 | 1 | G | A | 0.491 | 4.82E-02 | 7.96E-02 | 2.54E+00 | 2.60E+00 | 5.52E-01 | no | Tmem39a | intron_variant | MODIFIER |
| 1 | 169813885 | AX-170217647 | 0 | A | G | 0.228 | 8.12E-02 | 1.38E-01 | 2.67E+00 | 2.71E+00 | 5.52E-01 | no | | intergenic_variant | MODIFIER |
| 1 | 170465510 | AX-170308124 | 0 | C | G | 0.228 | 8.12E-02 | 1.38E-01 | 2.67E+00 | 2.71E+00 | 5.52E-01 | no | Nos1ap | intron_variant,non_coding_transcript_variant | MODIFIER |
| 7 | 106887136 | AX-170029723 | 1 | G | A | 0.491 | 5.35E-02 | 9.06E-02 | 2.62E+00 | 2.71E+00 | 5.53E-01 | no | Olfr707 | downstream_gene_variant | MODIFIER |
| 10 | 20650861 | AX-170405042 | 2 | G | A | 0.055 | 8.21E-02 | 1.35E-01 | 2.48E+00 | 2.64E+00 | 5.53E-01 | no | Pde7b | intron_variant | MODIFIER |
| 18 | 57297163 | AX-170181574 | 0 | G | A | 0.412 | 6.16E-02 | 9.67E-02 | 2.43E+00 | 2.48E+00 | 5.53E-01 | no | Megf10 | 3_prime_UTR_variant | MODIFIER |
| 4 | 132977681 | AX-170462799 | 1 | A | G | 0.17 | 6.48E-02 | 1.09E-01 | 2.59E+00 | 2.64E+00 | 5.53E-01 | no | Fgr | intron_variant | MODIFIER |
| 16 | 95415508 | AX-97853084 | 0 | A | G | 0.447 | 4.37E-02 | 7.49E-02 | 2.82E+00 | 2.95E+00 | 5.53E-01 | no | Erg | intron_variant | MODIFIER |
| 13 | 40084414 | AX-97786319 | 0 | T | C | 0.167 | -7.07E-02 | 1.12E-01 | 2.45E+00 | 2.53E+00 | 5.53E-01 | no | Ofcc1 | intron_variant | MODIFIER |
| 13 | 40911908 | AX-97824671 | 0 | A | G | 0.167 | -7.07E-02 | 1.12E-01 | 2.45E+00 | 2.53E+00 | 5.53E-01 | no | Gcnt2 | intron_variant | MODIFIER |
| 2 | 139984278 | AX-170065060 | 2 | T | A | 0.127 | -8.71E-02 | 1.49E-01 | 2.70E+00 | 2.80E+00 | 5.53E-01 | no | Tasp1 | intron_variant | MODIFIER |
| 18 | 66202218 | AX-170061145 | 0 | G | A | 0.096 | 9.92E-02 | 1.68E-01 | 2.60E+00 | 2.72E+00 | 5.53E-01 | no | Ccbe1 | intron_variant,NMD_transcript_variant | MODIFIER |
| 5 | 84466377 | AX-97796171 | 1 | G | A | 0.438 | 6.57E-02 | 1.12E-01 | 2.79E+00 | 2.84E+00 | 5.53E-01 | no | | intergenic_variant | MODIFIER |
| 12 | 109556397 | AX-170496582 | 0 | C | G | 0.123 | -9.44E-02 | 1.63E-01 | 2.70E+00 | 2.81E+00 | 5.54E-01 | no | Meg3 | intron_variant,non_coding_transcript_variant | MODIFIER |
| 8 | 20990630 | AX-170505922 | 0 | C | G | 0.184 | 5.52E-02 | 9.54E-02 | 2.70E+00 | 2.82E+00 | 5.54E-01 | no | | intergenic_variant | MODIFIER |
| 5 | 38127184 | AX-170347301 | 1 | C | T | 0.339 | 6.12E-02 | 1.05E-01 | 2.66E+00 | 2.74E+00 | 5.54E-01 | no | Stx18 | intron_variant | MODIFIER |
| 16 | 44226920 | AX-170483785 | 0 | G | A | 0.518 | 4.92E-02 | 8.20E-02 | 2.56E+00 | 2.60E+00 | 5.54E-01 | no | Usf3 | 3_prime_UTR_variant | MODIFIER |
| 16 | 44262142 | AX-170152249 | 0 | A | G | 0.518 | 4.92E-02 | 8.20E-02 | 2.56E+00 | 2.60E+00 | 5.54E-01 | no | Sidt1 | splice_region_variant,synonymous_variant | LOW |
| 16 | 44265090 | AX-97797619 | 0 | T | C | 0.518 | 4.92E-02 | 8.20E-02 | 2.56E+00 | 2.60E+00 | 5.54E-01 | no | Sidt1 | intron_variant | MODIFIER |
| 16 | 44270937 | AX-170413934 | 0 | A | G | 0.518 | 4.92E-02 | 8.20E-02 | 2.56E+00 | 2.60E+00 | 5.54E-01 | no | Sidt1 | intron_variant | MODIFIER |
| 16 | 44271198 | AX-170443641 | 0 | G | A | 0.518 | 4.92E-02 | 8.20E-02 | 2.56E+00 | 2.60E+00 | 5.54E-01 | no | Sidt1 | intron_variant | MODIFIER |
| 16 | 44288766 | AX-170115110 | 0 | C | G | 0.518 | 4.92E-02 | 8.20E-02 | 2.56E+00 | 2.60E+00 | 5.54E-01 | no | Sidt1 | intron_variant | MODIFIER |
| 16 | 44300187 | AX-170245664 | 0 | A | C | 0.518 | 4.92E-02 | 8.20E-02 | 2.56E+00 | 2.60E+00 | 5.54E-01 | no | Sidt1 | intron_variant | MODIFIER |
| 16 | 44300217 | AX-170186720 | 0 | A | G | 0.518 | 4.92E-02 | 8.20E-02 | 2.56E+00 | 2.60E+00 | 5.54E-01 | no | Sidt1 | intron_variant | MODIFIER |
| 16 | 44307372 | AX-170406924 | 0 | A | G | 0.518 | 4.92E-02 | 8.20E-02 | 2.56E+00 | 2.60E+00 | 5.54E-01 | no | Sidt1 | intron_variant | MODIFIER |
| 16 | 44354870 | AX-170133178 | 0 | T | C | 0.518 | 4.92E-02 | 8.20E-02 | 2.56E+00 | 2.60E+00 | 5.54E-01 | no | Spice1 | intron_variant | MODIFIER |
| 16 | 44392507 | AX-170012932 | 0 | G | A | 0.518 | 4.92E-02 | 8.20E-02 | 2.56E+00 | 2.60E+00 | 5.54E-01 | no | Spice1 | downstream_gene_variant | MODIFIER |
| 16 | 44430360 | AX-170012942 | 0 | T | C | 0.518 | 4.92E-02 | 8.20E-02 | 2.56E+00 | 2.60E+00 | 5.54E-01 | no | Cfap44 | intron_variant | MODIFIER |
| 16 | 44451020 | AX-170349416 | 0 | C | T | 0.518 | 4.92E-02 | 8.20E-02 | 2.56E+00 | 2.60E+00 | 5.54E-01 | no | Cfap44 | intron_variant | MODIFIER |
| 16 | 44460839 | AX-170088006 | 0 | A | T | 0.518 | 4.92E-02 | 8.20E-02 | 2.56E+00 | 2.60E+00 | 5.54E-01 | no | Cfap44 | intron_variant | MODIFIER |
| 16 | 44525399 | AX-170012936 | 0 | G | A | 0.518 | 4.92E-02 | 8.20E-02 | 2.56E+00 | 2.60E+00 | 5.54E-01 | no | Boc | intron_variant | MODIFIER |
| 16 | 44534503 | AX-170400838 | 0 | A | C | 0.518 | 4.92E-02 | 8.20E-02 | 2.56E+00 | 2.60E+00 | 5.54E-01 | no | Boc | intron_variant | MODIFIER |
| 16 | 44556465 | AX-170012933 | 0 | G | A | 0.518 | 4.92E-02 | 8.20E-02 | 2.56E+00 | 2.60E+00 | 5.54E-01 | no | Boc | intron_variant | MODIFIER |
| 16 | 44574414 | AX-170012941 | 0 | C | T | 0.518 | 4.92E-02 | 8.20E-02 | 2.56E+00 | 2.60E+00 | 5.54E-01 | no | | intergenic_variant | MODIFIER |
| 16 | 44574565 | AX-97841662 | 0 | C | T | 0.518 | 4.92E-02 | 8.20E-02 | 2.56E+00 | 2.60E+00 | 5.54E-01 | no | | intergenic_variant | MODIFIER |
| 16 | 44583970 | AX-170012937 | 0 | C | A | 0.518 | 4.92E-02 | 8.20E-02 | 2.56E+00 | 2.60E+00 | 5.54E-01 | no | | intergenic_variant | MODIFIER |
| 16 | 44618329 | AX-97841049 | 0 | C | T | 0.518 | 4.92E-02 | 8.20E-02 | 2.56E+00 | 2.60E+00 | 5.54E-01 | no | | regulatory_region_variant | MODIFIER |
| 16 | 44753065 | AX-97783028 | 0 | C | T | 0.518 | 4.92E-02 | 8.20E-02 | 2.56E+00 | 2.60E+00 | 5.54E-01 | no | | intergenic_variant | MODIFIER |
| 16 | 45031938 | AX-170483346 | 0 | A | G | 0.518 | 4.92E-02 | 8.20E-02 | 2.56E+00 | 2.60E+00 | 5.54E-01 | no | | intergenic_variant | MODIFIER |
| 16 | 45031970 | AX-97786284 | 0 | C | T | 0.518 | 4.92E-02 | 8.20E-02 | 2.56E+00 | 2.60E+00 | 5.54E-01 | no | | intergenic_variant | MODIFIER |
| 16 | 45033690 | AX-170320867 | 0 | G | A | 0.518 | 4.92E-02 | 8.20E-02 | 2.56E+00 | 2.60E+00 | 5.54E-01 | no | | intergenic_variant | MODIFIER |
| 16 | 45037051 | AX-170012944 | 0 | A | G | 0.518 | 4.92E-02 | 8.20E-02 | 2.56E+00 | 2.60E+00 | 5.54E-01 | no | | intergenic_variant | MODIFIER |
| 16 | 45037286 | AX-170388583 | 0 | G | C | 0.518 | 4.92E-02 | 8.20E-02 | 2.56E+00 | 2.60E+00 | 5.54E-01 | no | | intergenic_variant | MODIFIER |
| 16 | 45049751 | AX-170340345 | 0 | A | G | 0.518 | 4.92E-02 | 8.20E-02 | 2.56E+00 | 2.60E+00 | 5.54E-01 | no | | intergenic_variant | MODIFIER |
| 16 | 45097777 | AX-170531146 | 0 | A | C | 0.518 | 4.92E-02 | 8.20E-02 | 2.56E+00 | 2.60E+00 | 5.54E-01 | no | Ccdc80 | intron_variant | MODIFIER |
| 16 | 45100797 | AX-170184008 | 0 | G | A | 0.518 | 4.92E-02 | 8.20E-02 | 2.56E+00 | 2.60E+00 | 5.54E-01 | no | Ccdc80 | intron_variant | MODIFIER |
| 16 | 45102168 | AX-170106314 | 0 | C | T | 0.518 | 4.92E-02 | 8.20E-02 | 2.56E+00 | 2.60E+00 | 5.54E-01 | no | Ccdc80 | intron_variant | MODIFIER |
| 16 | 45103430 | AX-170012947 | 0 | C | T | 0.518 | 4.92E-02 | 8.20E-02 | 2.56E+00 | 2.60E+00 | 5.54E-01 | no | Ccdc80 | intron_variant | MODIFIER |
| 16 | 45142017 | AX-97788779 | 0 | C | T | 0.518 | 4.92E-02 | 8.20E-02 | 2.56E+00 | 2.60E+00 | 5.54E-01 | no | Slc35a5 | 3_prime_UTR_variant | MODIFIER |
| 16 | 45142760 | AX-170012951 | 0 | C | T | 0.518 | 4.92E-02 | 8.20E-02 | 2.56E+00 | 2.60E+00 | 5.54E-01 | no | Slc35a5 | intron_variant | MODIFIER |
| 16 | 45147563 | AX-170220075 | 0 | T | C | 0.518 | 4.92E-02 | 8.20E-02 | 2.56E+00 | 2.60E+00 | 5.54E-01 | no | Slc35a5 | intron_variant | MODIFIER |

| | | | | | | | | | | | | | | | |
|---|---|---|---|---|---|---|---|---|---|---|---|---|---|---|---|
| 16 | 45150755 | AX-170161936 | 0 | A | T | 0.518 | 4.92E-02 | 8.20E-02 | 2.56E+00 | 2.60E+00 | 5.54E-01 | no | Slc35a5 | intron_variant | MODIFIER |
| 16 | 45159770 | AX-97856489 | 0 | A | G | 0.518 | 4.92E-02 | 8.20E-02 | 2.56E+00 | 2.60E+00 | 5.54E-01 | no | Atg3 | intron_variant | MODIFIER |
| 16 | 45162082 | AX-170473092 | 0 | C | T | 0.518 | 4.92E-02 | 8.20E-02 | 2.56E+00 | 2.60E+00 | 5.54E-01 | no | Atg3 | intron_variant | MODIFIER |
| 16 | 45179324 | AX-170317750 | 0 | C | A | 0.518 | 4.92E-02 | 8.20E-02 | 2.56E+00 | 2.60E+00 | 5.54E-01 | no | Atg3 | intron_variant | MODIFIER |
| 16 | 45185332 | AX-97840047 | 0 | T | C | 0.518 | 4.92E-02 | 8.20E-02 | 2.56E+00 | 2.60E+00 | 5.54E-01 | no | Atg3 | intron_variant | MODIFIER |
| 16 | 45186787 | AX-170281454 | 0 | T | C | 0.518 | 4.92E-02 | 8.20E-02 | 2.56E+00 | 2.60E+00 | 5.54E-01 | no | Atg3 | intron_variant | MODIFIER |
| 16 | 45221359 | AX-170486816 | 0 | G | A | 0.518 | 4.92E-02 | 8.20E-02 | 2.56E+00 | 2.60E+00 | 5.54E-01 | no | Btla | upstream_gene_variant | MODIFIER |
| 16 | 45234746 | AX-170505392 | 0 | G | A | 0.518 | 4.92E-02 | 8.20E-02 | 2.56E+00 | 2.60E+00 | 5.54E-01 | no | Btla | intron_variant | MODIFIER |
| 16 | 45287318 | AX-97794658 | 0 | A | G | 0.518 | 4.92E-02 | 8.20E-02 | 2.56E+00 | 2.60E+00 | 5.54E-01 | no | | intergenic_variant | MODIFIER |
| 16 | 45288045 | AX-170415801 | 0 | G | C | 0.518 | 4.92E-02 | 8.20E-02 | 2.56E+00 | 2.60E+00 | 5.54E-01 | no | | intergenic_variant | MODIFIER |
| 16 | 45295490 | AX-170382422 | 0 | G | A | 0.518 | 4.92E-02 | 8.20E-02 | 2.56E+00 | 2.60E+00 | 5.54E-01 | no | | intergenic_variant | MODIFIER |
| 16 | 45295874 | AX-170057302 | 0 | C | T | 0.518 | 4.92E-02 | 8.20E-02 | 2.56E+00 | 2.60E+00 | 5.54E-01 | no | | intergenic_variant | MODIFIER |
| 16 | 45365492 | AX-170057301 | 0 | G | A | 0.518 | 4.92E-02 | 8.20E-02 | 2.56E+00 | 2.60E+00 | 5.54E-01 | no | | regulatory_region_variant | MODIFIER |
| 16 | 45366937 | AX-170259272 | 0 | A | G | 0.518 | 4.92E-02 | 8.20E-02 | 2.56E+00 | 2.60E+00 | 5.54E-01 | no | | regulatory_region_variant | MODIFIER |
| 16 | 45375358 | AX-170145225 | 0 | C | T | 0.518 | 4.92E-02 | 8.20E-02 | 2.56E+00 | 2.60E+00 | 5.54E-01 | no | | intergenic_variant | MODIFIER |
| 16 | 45398215 | AX-170333353 | 0 | C | A | 0.518 | 4.92E-02 | 8.20E-02 | 2.56E+00 | 2.60E+00 | 5.54E-01 | no | Cd200 | intron_variant | MODIFIER |
| 16 | 45411527 | AX-170133334 | 0 | A | G | 0.518 | 4.92E-02 | 8.20E-02 | 2.56E+00 | 2.60E+00 | 5.54E-01 | no | Cd200 | upstream_gene_variant | MODIFIER |
| 16 | 45428836 | AX-170150907 | 0 | T | C | 0.518 | 4.92E-02 | 8.20E-02 | 2.56E+00 | 2.60E+00 | 5.54E-01 | no | Gm609 | intron_variant | MODIFIER |
| 16 | 45471284 | AX-170133066 | 0 | C | T | 0.518 | 4.92E-02 | 8.20E-02 | 2.56E+00 | 2.60E+00 | 5.54E-01 | no | Gm609 | intron_variant | MODIFIER |
| 16 | 45520660 | AX-170530090 | 0 | G | A | 0.518 | 4.92E-02 | 8.20E-02 | 2.56E+00 | 2.60E+00 | 5.54E-01 | no | Gm17783 | intron_variant | MODIFIER |
| 16 | 45524061 | AX-170124399 | 0 | C | T | 0.518 | 4.92E-02 | 8.20E-02 | 2.56E+00 | 2.60E+00 | 5.54E-01 | no | Gm17783 | intron_variant | MODIFIER |
| 16 | 45559106 | AX-170012957 | 0 | G | T | 0.518 | 4.92E-02 | 8.20E-02 | 2.56E+00 | 2.60E+00 | 5.54E-01 | no | Slc9c1 | intron_variant | MODIFIER |
| 16 | 45559250 | AX-170264610 | 0 | C | T | 0.518 | 4.92E-02 | 8.20E-02 | 2.56E+00 | 2.60E+00 | 5.54E-01 | no | Slc9c1 | intron_variant | MODIFIER |
| 16 | 45559896 | AX-170380749 | 0 | G | A | 0.518 | 4.92E-02 | 8.20E-02 | 2.56E+00 | 2.60E+00 | 5.54E-01 | no | Slc9c1 | intron_variant | MODIFIER |
| 16 | 45573354 | AX-170515121 | 0 | C | T | 0.518 | 4.92E-02 | 8.20E-02 | 2.56E+00 | 2.60E+00 | 5.54E-01 | no | Slc9c1 | synonymous_variant | LOW |
| 16 | 45573621 | AX-170057305 | 0 | A | G | 0.518 | 4.92E-02 | 8.20E-02 | 2.56E+00 | 2.60E+00 | 5.54E-01 | no | Slc9c1 | intron_variant | MODIFIER |
| 16 | 45577540 | AX-170138519 | 0 | A | T | 0.518 | 4.92E-02 | 8.20E-02 | 2.56E+00 | 2.60E+00 | 5.54E-01 | no | Slc9c1 | intron_variant | MODIFIER |
| 16 | 45581418 | AX-170012958 | 0 | C | G | 0.518 | 4.92E-02 | 8.20E-02 | 2.56E+00 | 2.60E+00 | 5.54E-01 | no | Slc9c1 | intron_variant | MODIFIER |
| 16 | 45588892 | AX-170372773 | 0 | C | T | 0.518 | 4.92E-02 | 8.20E-02 | 2.56E+00 | 2.60E+00 | 5.54E-01 | no | Slc9c1 | intron_variant | MODIFIER |
| 16 | 45599193 | AX-97841584 | 0 | A | C | 0.518 | 4.92E-02 | 8.20E-02 | 2.56E+00 | 2.60E+00 | 5.54E-01 | no | Slc9c1 | intron_variant | MODIFIER |
| 16 | 45606988 | AX-170213823 | 0 | T | C | 0.518 | 4.92E-02 | 8.20E-02 | 2.56E+00 | 2.60E+00 | 5.54E-01 | no | Gcsam | upstream_gene_variant | MODIFIER |
| 16 | 45612703 | AX-170467112 | 0 | C | T | 0.518 | 4.92E-02 | 8.20E-02 | 2.56E+00 | 2.60E+00 | 5.54E-01 | no | Gcsam | intron_variant | MODIFIER |
| 16 | 45618970 | AX-170108144 | 0 | A | G | 0.518 | 4.92E-02 | 8.20E-02 | 2.56E+00 | 2.60E+00 | 5.54E-01 | no | Gcsam | intron_variant | MODIFIER |
| 16 | 45625918 | AX-170101428 | 0 | C | T | 0.518 | 4.92E-02 | 8.20E-02 | 2.56E+00 | 2.60E+00 | 5.54E-01 | no | Gcsam | downstream_gene_variant | MODIFIER |
| 16 | 45628754 | AX-170257800 | 0 | C | T | 0.518 | 4.92E-02 | 8.20E-02 | 2.56E+00 | 2.60E+00 | 5.54E-01 | no | BC016579 | intron_variant | MODIFIER |
| 16 | 45637334 | AX-170012953 | 0 | A | G | 0.518 | 4.92E-02 | 8.20E-02 | 2.56E+00 | 2.60E+00 | 5.54E-01 | no | BC016579 | intron_variant | MODIFIER |
| 16 | 45650049 | AX-170057309 | 0 | T | G | 0.518 | 4.92E-02 | 8.20E-02 | 2.56E+00 | 2.60E+00 | 5.54E-01 | no | BC016579 | intron_variant | MODIFIER |
| 16 | 45656552 | AX-170057308 | 0 | G | A | 0.518 | 4.92E-02 | 8.20E-02 | 2.56E+00 | 2.60E+00 | 5.54E-01 | no | BC016579 | upstream_gene_variant | MODIFIER |
| 16 | 45660952 | AX-170105904 | 0 | G | A | 0.518 | 4.92E-02 | 8.20E-02 | 2.56E+00 | 2.60E+00 | 5.54E-01 | no | Tmprss7 | intron_variant | MODIFIER |
| 16 | 45663040 | AX-170479684 | 0 | G | A | 0.518 | 4.92E-02 | 8.20E-02 | 2.56E+00 | 2.60E+00 | 5.54E-01 | no | Tmprss7 | intron_variant | MODIFIER |
| 16 | 45682055 | AX-170428288 | 0 | A | C | 0.518 | 4.92E-02 | 8.20E-02 | 2.56E+00 | 2.60E+00 | 5.54E-01 | no | Tmprss7 | intron_variant | MODIFIER |
| 16 | 45709849 | AX-170302636 | 0 | C | T | 0.518 | 4.92E-02 | 8.20E-02 | 2.56E+00 | 2.60E+00 | 5.54E-01 | no | Tagln3 | downstream_gene_variant | MODIFIER |
| 16 | 45709994 | AX-170249624 | 0 | G | A | 0.518 | 4.92E-02 | 8.20E-02 | 2.56E+00 | 2.60E+00 | 5.54E-01 | no | Tagln3 | downstream_gene_variant | MODIFIER |
| 16 | 45719304 | AX-170466888 | 0 | C | T | 0.518 | 4.92E-02 | 8.20E-02 | 2.56E+00 | 2.60E+00 | 5.54E-01 | no | Tagln3 | intron_variant | MODIFIER |
| 16 | 45730623 | AX-97853178 | 0 | C | T | 0.518 | 4.92E-02 | 8.20E-02 | 2.56E+00 | 2.60E+00 | 5.54E-01 | no | Abhd10 | 3_prime_UTR_variant | MODIFIER |
| 16 | 45737527 | AX-170225326 | 0 | T | C | 0.518 | 4.92E-02 | 8.20E-02 | 2.56E+00 | 2.60E+00 | 5.54E-01 | no | Abhd10 | synonymous_variant | LOW |
| 16 | 45738772 | AX-170486758 | 0 | C | T | 0.518 | 4.92E-02 | 8.20E-02 | 2.56E+00 | 2.60E+00 | 5.54E-01 | no | Abhd10 | intron_variant | MODIFIER |
| 16 | 45752494 | AX-170099270 | 0 | A | C | 0.482 | -4.92E-02 | 8.20E-02 | 2.56E+00 | 2.60E+00 | 5.54E-01 | no | Phldb2 | intron_variant | MODIFIER |
| 16 | 45777219 | AX-170503217 | 0 | C | T | 0.518 | 4.92E-02 | 8.20E-02 | 2.56E+00 | 2.60E+00 | 5.54E-01 | no | Phldb2 | intron_variant | MODIFIER |
| 16 | 45792886 | AX-97853997 | 0 | G | A | 0.518 | 4.92E-02 | 8.20E-02 | 2.56E+00 | 2.60E+00 | 5.54E-01 | no | Phldb2 | intron_variant | MODIFIER |
| 16 | 45815257 | AX-170357322 | 0 | T | C | 0.518 | 4.92E-02 | 8.20E-02 | 2.56E+00 | 2.60E+00 | 5.54E-01 | no | Phldb2 | intron_variant | MODIFIER |
| 16 | 45831057 | AX-170118076 | 0 | C | G | 0.518 | 4.92E-02 | 8.20E-02 | 2.56E+00 | 2.60E+00 | 5.54E-01 | no | Phldb2 | intron_variant | MODIFIER |
| 16 | 45847668 | AX-170354809 | 0 | T | C | 0.518 | 4.92E-02 | 8.20E-02 | 2.56E+00 | 2.60E+00 | 5.54E-01 | no | Phldb2 | upstream_gene_variant | MODIFIER |
| 16 | 45857355 | AX-170446022 | 0 | T | C | 0.518 | 4.92E-02 | 8.20E-02 | 2.56E+00 | 2.60E+00 | 5.54E-01 | no | Phldb2 | intron_variant | MODIFIER |

| | | | | | | | | | | | | | | | |
|---|---|---|---|---|---|---|---|---|---|---|---|---|---|---|---|
| 16 | 45857448 | AX-170110330 | 0 | C | T | 0.518 | 4.92E-02 | 8.20E-02 | 2.56E+00 | 2.60E+00 | 5.54E-01 | no | Phldb2 | intron_variant | MODIFIER |
| 16 | 45860533 | AX-170289654 | 0 | G | A | 0.518 | 4.92E-02 | 8.20E-02 | 2.56E+00 | 2.60E+00 | 5.54E-01 | no | Phldb2 | intron_variant | MODIFIER |
| 16 | 45871073 | AX-170500758 | 0 | A | T | 0.518 | 4.92E-02 | 8.20E-02 | 2.56E+00 | 2.60E+00 | 5.54E-01 | no | Phldb2 | intron_variant | MODIFIER |
| 16 | 45873239 | AX-97831666 | 0 | T | C | 0.518 | 4.92E-02 | 8.20E-02 | 2.56E+00 | 2.60E+00 | 5.54E-01 | no | Phldb2 | intron_variant | MODIFIER |
| 16 | 45890541 | AX-170165800 | 0 | T | C | 0.518 | 4.92E-02 | 8.20E-02 | 2.56E+00 | 2.60E+00 | 5.54E-01 | no | Phldb2 | intron_variant | MODIFIER |
| 16 | 45893525 | AX-97829070 | 0 | T | C | 0.518 | 4.92E-02 | 8.20E-02 | 2.56E+00 | 2.60E+00 | 5.54E-01 | no | Phldb2 | intron_variant | MODIFIER |
| 16 | 45893929 | AX-170182490 | 0 | A | C | 0.518 | 4.92E-02 | 8.20E-02 | 2.56E+00 | 2.60E+00 | 5.54E-01 | no | Phldb2 | intron_variant | MODIFIER |
| 16 | 45894178 | AX-170212000 | 0 | G | T | 0.518 | 4.92E-02 | 8.20E-02 | 2.56E+00 | 2.60E+00 | 5.54E-01 | no | Phldb2 | intron_variant | MODIFIER |
| 16 | 45894272 | AX-170235222 | 0 | G | A | 0.518 | 4.92E-02 | 8.20E-02 | 2.56E+00 | 2.60E+00 | 5.54E-01 | no | Phldb2 | intron_variant | MODIFIER |
| 16 | 45894902 | AX-170447882 | 0 | G | C | 0.518 | 4.92E-02 | 8.20E-02 | 2.56E+00 | 2.60E+00 | 5.54E-01 | no | Phldb2 | intron_variant | MODIFIER |
| 16 | 45947347 | AX-97852747 | 0 | T | C | 0.518 | 4.92E-02 | 8.20E-02 | 2.56E+00 | 2.60E+00 | 5.54E-01 | no | Gm15638 | intron_variant,non_coding_transcript_variant | MODIFIER |
| 16 | 45947367 | AX-170182198 | 0 | T | C | 0.518 | 4.92E-02 | 8.20E-02 | 2.56E+00 | 2.60E+00 | 5.54E-01 | no | Gm15638 | intron_variant,non_coding_transcript_variant | MODIFIER |
| 16 | 45947956 | AX-170247452 | 0 | C | T | 0.518 | 4.92E-02 | 8.20E-02 | 2.56E+00 | 2.60E+00 | 5.54E-01 | no | Gm15638 | non_coding_transcript_exon_variant | MODIFIER |
| 16 | 45952861 | AX-170329072 | 0 | G | C | 0.518 | 4.92E-02 | 8.20E-02 | 2.56E+00 | 2.60E+00 | 5.54E-01 | no | Gm15638 | intron_variant,non_coding_transcript_variant | MODIFIER |
| 16 | 45970582 | AX-97787215 | 0 | C | T | 0.518 | 4.92E-02 | 8.20E-02 | 2.56E+00 | 2.60E+00 | 5.54E-01 | no | Plcxd2 | intron_variant | MODIFIER |
| 16 | 45970656 | AX-170523249 | 0 | A | T | 0.518 | 4.92E-02 | 8.20E-02 | 2.56E+00 | 2.60E+00 | 5.54E-01 | no | Plcxd2 | intron_variant | MODIFIER |
| 16 | 45972844 | AX-170499693 | 0 | C | T | 0.518 | 4.92E-02 | 8.20E-02 | 2.56E+00 | 2.60E+00 | 5.54E-01 | no | Plcxd2 | intron_variant | MODIFIER |
| 16 | 45980323 | AX-170360996 | 0 | T | C | 0.518 | 4.92E-02 | 8.20E-02 | 2.56E+00 | 2.60E+00 | 5.54E-01 | no | Plcxd2 | synonymous_variant | LOW |
| 16 | 45984088 | AX-170522473 | 0 | C | T | 0.518 | 4.92E-02 | 8.20E-02 | 2.56E+00 | 2.60E+00 | 5.54E-01 | no | Plcxd2 | intron_variant | MODIFIER |
| 16 | 45986356 | AX-170229520 | 0 | G | A | 0.518 | 4.92E-02 | 8.20E-02 | 2.56E+00 | 2.60E+00 | 5.54E-01 | no | Plcxd2 | intron_variant | MODIFIER |
| 16 | 45993136 | AX-170088008 | 0 | A | C | 0.518 | 4.92E-02 | 8.20E-02 | 2.56E+00 | 2.60E+00 | 5.54E-01 | no | Plcxd2 | intron_variant | MODIFIER |
| 16 | 46036560 | AX-170093466 | 0 | A | C | 0.518 | 4.92E-02 | 8.20E-02 | 2.56E+00 | 2.60E+00 | 5.54E-01 | no | Cd96 | intron_variant | MODIFIER |
| 16 | 46050992 | AX-170012961 | 0 | T | C | 0.518 | 4.92E-02 | 8.20E-02 | 2.56E+00 | 2.60E+00 | 5.54E-01 | no | Cd96 | intron_variant | MODIFIER |
| 16 | 46062309 | AX-170012966 | 0 | C | T | 0.518 | 4.92E-02 | 8.20E-02 | 2.56E+00 | 2.60E+00 | 5.54E-01 | no | Cd96 | intron_variant | MODIFIER |
| 16 | 46063414 | AX-170468229 | 0 | A | T | 0.518 | 4.92E-02 | 8.20E-02 | 2.56E+00 | 2.60E+00 | 5.54E-01 | no | Cd96 | intron_variant | MODIFIER |
| 16 | 46068534 | AX-97849864 | 0 | A | G | 0.518 | 4.92E-02 | 8.20E-02 | 2.56E+00 | 2.60E+00 | 5.54E-01 | no | Cd96 | intron_variant | MODIFIER |
| 16 | 46068799 | AX-170372285 | 0 | T | G | 0.518 | 4.92E-02 | 8.20E-02 | 2.56E+00 | 2.60E+00 | 5.54E-01 | no | Cd96 | intron_variant | MODIFIER |
| 16 | 46076895 | AX-170457934 | 0 | A | G | 0.518 | 4.92E-02 | 8.20E-02 | 2.56E+00 | 2.60E+00 | 5.54E-01 | no | Cd96 | intron_variant | MODIFIER |
| 16 | 46089611 | AX-170131274 | 0 | A | G | 0.518 | 4.92E-02 | 8.20E-02 | 2.56E+00 | 2.60E+00 | 5.54E-01 | no | Cd96 | intron_variant | MODIFIER |
| 16 | 46090831 | AX-170012960 | 0 | G | C | 0.518 | 4.92E-02 | 8.20E-02 | 2.56E+00 | 2.60E+00 | 5.54E-01 | no | Cd96 | intron_variant | MODIFIER |
| 16 | 46091447 | AX-97791222 | 0 | A | G | 0.518 | 4.92E-02 | 8.20E-02 | 2.56E+00 | 2.60E+00 | 5.54E-01 | no | Cd96 | intron_variant | MODIFIER |
| 16 | 46112557 | AX-170284396 | 0 | C | T | 0.518 | 4.92E-02 | 8.20E-02 | 2.56E+00 | 2.60E+00 | 5.54E-01 | no | Cd96 | intron_variant | MODIFIER |
| 16 | 46117980 | AX-97820838 | 0 | C | T | 0.518 | 4.92E-02 | 8.20E-02 | 2.56E+00 | 2.60E+00 | 5.54E-01 | no | Cd96 | missense_variant | MODERATE |
| 16 | 46144834 | AX-170435034 | 0 | T | C | 0.518 | 4.92E-02 | 8.20E-02 | 2.56E+00 | 2.60E+00 | 5.54E-01 | no | | intergenic_variant | MODIFIER |
| 16 | 46152542 | AX-170485060 | 0 | C | T | 0.518 | 4.92E-02 | 8.20E-02 | 2.56E+00 | 2.60E+00 | 5.54E-01 | no | Gm4737 | downstream_gene_variant | MODIFIER |
| 16 | 46159664 | AX-170130521 | 0 | T | C | 0.518 | 4.92E-02 | 8.20E-02 | 2.56E+00 | 2.60E+00 | 5.54E-01 | no | Gm4737 | upstream_gene_variant | MODIFIER |
| 16 | 46179272 | AX-170176597 | 0 | T | C | 0.518 | 4.92E-02 | 8.20E-02 | 2.56E+00 | 2.60E+00 | 5.54E-01 | no | | intergenic_variant | MODIFIER |
| 16 | 46179369 | AX-170216861 | 0 | T | G | 0.518 | 4.92E-02 | 8.20E-02 | 2.56E+00 | 2.60E+00 | 5.54E-01 | no | | intergenic_variant | MODIFIER |
| 16 | 46196124 | AX-170125353 | 0 | G | A | 0.518 | 4.92E-02 | 8.20E-02 | 2.56E+00 | 2.60E+00 | 5.54E-01 | no | | intergenic_variant | MODIFIER |
| 16 | 46196215 | AX-170289943 | 0 | T | C | 0.518 | 4.92E-02 | 8.20E-02 | 2.56E+00 | 2.60E+00 | 5.54E-01 | no | | intergenic_variant | MODIFIER |
| 16 | 46196393 | AX-170236776 | 0 | A | G | 0.518 | 4.92E-02 | 8.20E-02 | 2.56E+00 | 2.60E+00 | 5.54E-01 | no | | intergenic_variant | MODIFIER |
| 16 | 46201224 | AX-170427136 | 0 | C | G | 0.518 | 4.92E-02 | 8.20E-02 | 2.56E+00 | 2.60E+00 | 5.54E-01 | no | | intergenic_variant | MODIFIER |
| 16 | 46203951 | AX-170325485 | 0 | C | T | 0.518 | 4.92E-02 | 8.20E-02 | 2.56E+00 | 2.60E+00 | 5.54E-01 | no | | intergenic_variant | MODIFIER |
| 16 | 46215188 | AX-97831109 | 0 | G | A | 0.518 | 4.92E-02 | 8.20E-02 | 2.56E+00 | 2.60E+00 | 5.54E-01 | no | Gm26297 | upstream_gene_variant | MODIFIER |
| 16 | 46218074 | AX-170511834 | 0 | G | A | 0.518 | 4.92E-02 | 8.20E-02 | 2.56E+00 | 2.60E+00 | 5.54E-01 | no | Gm26297 | upstream_gene_variant | MODIFIER |
| 16 | 46244562 | AX-170516015 | 0 | C | T | 0.518 | 4.92E-02 | 8.20E-02 | 2.56E+00 | 2.60E+00 | 5.54E-01 | no | | intergenic_variant | MODIFIER |
| 16 | 46250652 | AX-170302403 | 0 | C | T | 0.518 | 4.92E-02 | 8.20E-02 | 2.56E+00 | 2.60E+00 | 5.54E-01 | no | | intergenic_variant | MODIFIER |
| 16 | 46335205 | AX-170109488 | 0 | G | A | 0.518 | 4.92E-02 | 8.20E-02 | 2.56E+00 | 2.60E+00 | 5.54E-01 | no | | intergenic_variant | MODIFIER |
| 16 | 46353818 | AX-170267828 | 0 | C | G | 0.518 | 4.92E-02 | 8.20E-02 | 2.56E+00 | 2.60E+00 | 5.54E-01 | no | | intergenic_variant | MODIFIER |
| 16 | 46357304 | AX-170532806 | 0 | G | A | 0.518 | 4.92E-02 | 8.20E-02 | 2.56E+00 | 2.60E+00 | 5.54E-01 | no | | intergenic_variant | MODIFIER |
| 16 | 46375300 | AX-170482548 | 0 | G | A | 0.518 | 4.92E-02 | 8.20E-02 | 2.56E+00 | 2.60E+00 | 5.54E-01 | no | | intergenic_variant | MODIFIER |
| 16 | 46378604 | AX-170487100 | 0 | G | A | 0.518 | 4.92E-02 | 8.20E-02 | 2.56E+00 | 2.60E+00 | 5.54E-01 | no | | intergenic_variant | MODIFIER |
| 16 | 46398961 | AX-170518304 | 0 | G | A | 0.518 | 4.92E-02 | 8.20E-02 | 2.56E+00 | 2.60E+00 | 5.54E-01 | no | Nectin3 | intron_variant | MODIFIER |
| 16 | 46412138 | AX-170176376 | 0 | G | C | 0.518 | 4.92E-02 | 8.20E-02 | 2.56E+00 | 2.60E+00 | 5.54E-01 | no | Nectin3 | intron_variant | MODIFIER |

| | | | | | | | | | | | | | | | |
|---|---|---|---|---|---|---|---|---|---|---|---|---|---|---|---|
| 16 | 46412962 | AX-170206183 | 0 | T | A | 0.518 | 4.92E-02 | 8.20E-02 | 2.56E+00 | 2.60E+00 | 5.54E-01 | no | Nectin3 | intron_variant | MODIFIER |
| 16 | 45738241 | AX-170282418 | 1 | G | A | 0.518 | 4.92E-02 | 8.20E-02 | 2.56E+00 | 2.60E+00 | 5.54E-01 | no | Abhd10 | intron_variant | MODIFIER |
| 4 | 72871072 | AX-170106307 | 0 | T | A | 0.456 | -5.12E-02 | 8.33E-02 | 2.49E+00 | 2.56E+00 | 5.54E-01 | no | | intergenic_variant | MODIFIER |
| 9 | 118898170 | AX-170126010 | 0 | G | A | 0.132 | -5.42E-02 | 9.31E-02 | 2.63E+00 | 2.76E+00 | 5.55E-01 | no | Itga9 | 3_prime_UTR_variant | MODIFIER |
| 6 | 111027742 | AX-97818715 | 0 | A | G | 0.053 | -1.27E-01 | 2.19E-01 | 2.66E+00 | 2.77E+00 | 5.55E-01 | no | Grm7 | intron_variant | MODIFIER |
| 10 | 23296728 | AX-170411018 | 1 | C | A | 0.58 | -4.08E-02 | 6.98E-02 | 2.89E+00 | 3.06E+00 | 5.55E-01 | no | Eya4 | intron_variant | MODIFIER |
| 12 | 107898573 | AX-170518729 | 0 | A | T | 0.123 | -9.36E-02 | 1.61E-01 | 2.67E+00 | 2.77E+00 | 5.55E-01 | no | | intergenic_variant | MODIFIER |
| 4 | 82445664 | AX-170490839 | 1 | A | T | 0.232 | 6.02E-02 | 1.00E-01 | 2.53E+00 | 2.62E+00 | 5.55E-01 | no | Nfib | intron_variant | MODIFIER |
| 7 | 45531755 | AX-170245948 | 0 | T | C | 0.333 | 5.52E-02 | 9.38E-02 | 2.61E+00 | 2.69E+00 | 5.55E-01 | no | Plekha4 | intron_variant | MODIFIER |
| 7 | 45533978 | AX-97827414 | 0 | C | T | 0.333 | 5.52E-02 | 9.38E-02 | 2.61E+00 | 2.69E+00 | 5.55E-01 | no | Plekha4 | intron_variant | MODIFIER |
| 7 | 45640399 | AX-97838249 | 0 | G | A | 0.333 | 5.52E-02 | 9.38E-02 | 2.61E+00 | 2.69E+00 | 5.55E-01 | no | Rasip1 | downstream_gene_variant | MODIFIER |
| 7 | 45640461 | AX-170362331 | 0 | T | A | 0.333 | 5.52E-02 | 9.38E-02 | 2.61E+00 | 2.69E+00 | 5.55E-01 | no | Rasip1 | downstream_gene_variant | MODIFIER |
| 7 | 45674664 | AX-97826648 | 0 | T | C | 0.333 | 5.52E-02 | 9.38E-02 | 2.61E+00 | 2.69E+00 | 5.55E-01 | no | Sec1 | downstream_gene_variant | MODIFIER |
| 7 | 45869789 | AX-170328866 | 0 | C | T | 0.333 | 5.52E-02 | 9.38E-02 | 2.61E+00 | 2.69E+00 | 5.55E-01 | no | Grin2d | upstream_gene_variant | MODIFIER |
| 7 | 45871125 | AX-170148555 | 0 | T | C | 0.333 | 5.52E-02 | 9.38E-02 | 2.61E+00 | 2.69E+00 | 5.55E-01 | no | Grin2d | upstream_gene_variant | MODIFIER |
| 7 | 45874308 | AX-170310942 | 0 | T | C | 0.333 | 5.52E-02 | 9.38E-02 | 2.61E+00 | 2.69E+00 | 5.55E-01 | no | Kdelr1 | intron_variant | MODIFIER |
| 7 | 45880972 | AX-170330420 | 0 | T | G | 0.333 | 5.52E-02 | 9.38E-02 | 2.61E+00 | 2.69E+00 | 5.55E-01 | no | Kdelr1 | intron_variant | MODIFIER |
| 7 | 45893930 | AX-97782685 | 0 | G | A | 0.333 | 5.52E-02 | 9.38E-02 | 2.61E+00 | 2.69E+00 | 5.55E-01 | no | Syngr4 | intron_variant | MODIFIER |
| 7 | 45914234 | AX-97856342 | 0 | T | G | 0.333 | 5.52E-02 | 9.38E-02 | 2.61E+00 | 2.69E+00 | 5.55E-01 | no | Emp3 | downstream_gene_variant | MODIFIER |
| 7 | 45939168 | AX-170226088 | 0 | T | C | 0.333 | 5.52E-02 | 9.38E-02 | 2.61E+00 | 2.69E+00 | 5.55E-01 | no | Ccdc114 | intron_variant | MODIFIER |
| 7 | 45940370 | AX-170373672 | 0 | A | C | 0.333 | 5.52E-02 | 9.38E-02 | 2.61E+00 | 2.69E+00 | 5.55E-01 | no | Ccdc114 | intron_variant | MODIFIER |
| 7 | 45946334 | AX-97852000 | 0 | T | C | 0.333 | 5.52E-02 | 9.38E-02 | 2.61E+00 | 2.69E+00 | 5.55E-01 | no | Ccdc114 | intron_variant | MODIFIER |
| 7 | 45950818 | AX-170399955 | 0 | G | A | 0.333 | 5.52E-02 | 9.38E-02 | 2.61E+00 | 2.69E+00 | 5.55E-01 | no | Ccdc114 | downstream_gene_variant | MODIFIER |
| 7 | 45960188 | AX-170394593 | 0 | A | C | 0.333 | 5.52E-02 | 9.38E-02 | 2.61E+00 | 2.69E+00 | 5.55E-01 | no | | intergenic_variant | MODIFIER |
| 7 | 45969476 | AX-170252718 | 0 | G | A | 0.333 | 5.52E-02 | 9.38E-02 | 2.61E+00 | 2.69E+00 | 5.55E-01 | no | Gm45674 | upstream_gene_variant | MODIFIER |
| 7 | 45969585 | AX-170504963 | 0 | G | A | 0.333 | 5.52E-02 | 9.38E-02 | 2.61E+00 | 2.69E+00 | 5.55E-01 | no | Gm45674 | upstream_gene_variant | MODIFIER |
| 7 | 45970437 | AX-170258888 | 0 | G | A | 0.333 | 5.52E-02 | 9.38E-02 | 2.61E+00 | 2.69E+00 | 5.55E-01 | no | Gm45674 | upstream_gene_variant | MODIFIER |
| 7 | 45970615 | AX-170475186 | 0 | G | A | 0.333 | 5.52E-02 | 9.38E-02 | 2.61E+00 | 2.69E+00 | 5.55E-01 | no | Gm45674 | upstream_gene_variant | MODIFIER |
| 7 | 45973220 | AX-97834901 | 0 | C | T | 0.333 | 5.52E-02 | 9.38E-02 | 2.61E+00 | 2.69E+00 | 5.55E-01 | no | Abcc6 | downstream_gene_variant | MODIFIER |
| 7 | 45992069 | AX-97814350 | 0 | G | A | 0.333 | 5.52E-02 | 9.38E-02 | 2.61E+00 | 2.69E+00 | 5.55E-01 | no | Abcc6 | intron_variant | MODIFIER |
| 7 | 45992221 | AX-170403259 | 0 | T | C | 0.333 | 5.52E-02 | 9.38E-02 | 2.61E+00 | 2.69E+00 | 5.55E-01 | no | Abcc6 | intron_variant | MODIFIER |
| 7 | 45992812 | AX-170520048 | 0 | C | T | 0.333 | 5.52E-02 | 9.38E-02 | 2.61E+00 | 2.69E+00 | 5.55E-01 | no | Abcc6 | intron_variant | MODIFIER |
| 7 | 45996870 | AX-170376604 | 0 | T | C | 0.333 | 5.52E-02 | 9.38E-02 | 2.61E+00 | 2.69E+00 | 5.55E-01 | no | Abcc6 | intron_variant | MODIFIER |
| 7 | 45998774 | AX-170348800 | 0 | G | A | 0.333 | 5.52E-02 | 9.38E-02 | 2.61E+00 | 2.69E+00 | 5.55E-01 | no | Abcc6 | missense_variant | MODERATE |
| 7 | 46006150 | AX-97805377 | 0 | A | G | 0.333 | 5.52E-02 | 9.38E-02 | 2.61E+00 | 2.69E+00 | 5.55E-01 | no | Abcc6 | intron_variant | MODIFIER |
| 7 | 46059074 | AX-97855949 | 0 | T | G | 0.333 | 5.52E-02 | 9.38E-02 | 2.61E+00 | 2.69E+00 | 5.55E-01 | no | Nomo1 | intron_variant | MODIFIER |
| 7 | 46157774 | AX-170323328 | 0 | G | A | 0.333 | 5.52E-02 | 9.38E-02 | 2.61E+00 | 2.69E+00 | 5.55E-01 | no | Abcc8 | intron_variant | MODIFIER |
| 7 | 46167040 | AX-170206728 | 0 | T | C | 0.333 | 5.52E-02 | 9.38E-02 | 2.61E+00 | 2.69E+00 | 5.55E-01 | no | Abcc8 | synonymous_variant | LOW |
| 7 | 46174304 | AX-170115460 | 0 | T | C | 0.333 | 5.52E-02 | 9.38E-02 | 2.61E+00 | 2.69E+00 | 5.55E-01 | no | Abcc8 | intron_variant | MODIFIER |
| 7 | 46176862 | AX-170249868 | 0 | A | G | 0.333 | 5.52E-02 | 9.38E-02 | 2.61E+00 | 2.69E+00 | 5.55E-01 | no | Abcc8 | intron_variant | MODIFIER |
| 7 | 46189414 | AX-170491961 | 0 | A | C | 0.333 | 5.52E-02 | 9.38E-02 | 2.61E+00 | 2.69E+00 | 5.55E-01 | no | | intergenic_variant | MODIFIER |
| 7 | 46205197 | AX-170162081 | 0 | G | A | 0.333 | 5.52E-02 | 9.38E-02 | 2.61E+00 | 2.69E+00 | 5.55E-01 | no | Ush1c | synonymous_variant | LOW |
| 7 | 46205760 | AX-170159145 | 0 | T | C | 0.333 | 5.52E-02 | 9.38E-02 | 2.61E+00 | 2.69E+00 | 5.55E-01 | no | Ush1c | intron_variant | MODIFIER |
| 7 | 46205983 | AX-170265618 | 0 | A | G | 0.333 | 5.52E-02 | 9.38E-02 | 2.61E+00 | 2.69E+00 | 5.55E-01 | no | Ush1c | intron_variant | MODIFIER |
| 7 | 46209842 | AX-170186955 | 0 | T | C | 0.333 | 5.52E-02 | 9.38E-02 | 2.61E+00 | 2.69E+00 | 5.55E-01 | no | Ush1c | intron_variant | MODIFIER |
| 7 | 46209882 | AX-170167044 | 0 | G | T | 0.333 | 5.52E-02 | 9.38E-02 | 2.61E+00 | 2.69E+00 | 5.55E-01 | no | Ush1c | synonymous_variant | LOW |
| 7 | 46210478 | AX-170504184 | 0 | G | A | 0.333 | 5.52E-02 | 9.38E-02 | 2.61E+00 | 2.69E+00 | 5.55E-01 | no | Ush1c | intron_variant | MODIFIER |
| 7 | 46215885 | AX-170449036 | 0 | C | T | 0.333 | 5.52E-02 | 9.38E-02 | 2.61E+00 | 2.69E+00 | 5.55E-01 | no | Ush1c | intron_variant | MODIFIER |
| 7 | 46216161 | AX-170317838 | 0 | T | C | 0.333 | 5.52E-02 | 9.38E-02 | 2.61E+00 | 2.69E+00 | 5.55E-01 | no | Ush1c | intron_variant | MODIFIER |
| 7 | 46222754 | AX-170352104 | 0 | T | C | 0.333 | 5.52E-02 | 9.38E-02 | 2.61E+00 | 2.69E+00 | 5.55E-01 | no | Ush1c | intron_variant | MODIFIER |
| 7 | 46236637 | AX-170181458 | 0 | T | C | 0.333 | 5.52E-02 | 9.38E-02 | 2.61E+00 | 2.69E+00 | 5.55E-01 | no | Ush1c | intron_variant | MODIFIER |
| 7 | 46258515 | AX-170156531 | 0 | T | C | 0.333 | 5.52E-02 | 9.38E-02 | 2.61E+00 | 2.69E+00 | 5.55E-01 | no | Otog | intron_variant | MODIFIER |
| 7 | 46260305 | AX-170179977 | 0 | C | T | 0.333 | 5.52E-02 | 9.38E-02 | 2.61E+00 | 2.69E+00 | 5.55E-01 | no | Otog | intron_variant | MODIFIER |
| 7 | 46260592 | AX-170336427 | 0 | G | C | 0.333 | 5.52E-02 | 9.38E-02 | 2.61E+00 | 2.69E+00 | 5.55E-01 | no | Otog | intron_variant | MODIFIER |

| | | | | | | | | | | | | | | | |
|---|---|---|---|---|---|---|---|---|---|---|---|---|---|---|---|
| 7 | 46261609 | AX-170254922 | 0 | G | T | 0.333 | 5.52E-02 | 9.38E-02 | 2.61E+00 | 2.69E+00 | 5.55E-01 | no | Otog | intron_variant | MODIFIER |
| 7 | 46261912 | AX-170077223 | 0 | C | T | 0.333 | 5.52E-02 | 9.38E-02 | 2.61E+00 | 2.69E+00 | 5.55E-01 | no | Otog | intron_variant | MODIFIER |
| 7 | 46263380 | AX-170216285 | 0 | C | T | 0.333 | 5.52E-02 | 9.38E-02 | 2.61E+00 | 2.69E+00 | 5.55E-01 | no | Otog | intron_variant | MODIFIER |
| 7 | 46263573 | AX-170130365 | 0 | T | A | 0.333 | 5.52E-02 | 9.38E-02 | 2.61E+00 | 2.69E+00 | 5.55E-01 | no | Otog | intron_variant | MODIFIER |
| 7 | 46263852 | AX-97812346 | 0 | G | A | 0.333 | 5.52E-02 | 9.38E-02 | 2.61E+00 | 2.69E+00 | 5.55E-01 | no | Otog | intron_variant | MODIFIER |
| 7 | 46266073 | AX-170167950 | 0 | A | G | 0.333 | 5.52E-02 | 9.38E-02 | 2.61E+00 | 2.69E+00 | 5.55E-01 | no | Otog | intron_variant | MODIFIER |
| 7 | 46268240 | AX-170414800 | 0 | G | A | 0.333 | 5.52E-02 | 9.38E-02 | 2.61E+00 | 2.69E+00 | 5.55E-01 | no | Otog | intron_variant | MODIFIER |
| 7 | 46297780 | AX-170233370 | 0 | C | T | 0.333 | 5.52E-02 | 9.38E-02 | 2.61E+00 | 2.69E+00 | 5.55E-01 | no | Otog | intron_variant | MODIFIER |
| 2 | 165150853 | AX-170519424 | 1 | G | A | 0.152 | 1.04E-01 | 1.80E-01 | 2.71E+00 | 2.79E+00 | 5.55E-01 | no | Cdh22 | intron_variant | MODIFIER |
| 8 | 97847069 | AX-170031344 | 0 | T | G | 0.211 | 6.73E-02 | 1.16E-01 | 2.72E+00 | 2.79E+00 | 5.55E-01 | no | Gm39231 | intron_variant,non_coding_transcript_variant | MODIFIER |
| 18 | 50012683 | AX-170428596 | 1 | C | T | 0.196 | -6.21E-02 | 1.05E-01 | 2.58E+00 | 2.67E+00 | 5.55E-01 | no | Tnfaip8 | intron_variant,non_coding_transcript_variant | MODIFIER |
| 13 | 24039235 | AX-97819974 | 0 | C | T | 0.167 | -7.37E-02 | 1.20E-01 | 2.49E+00 | 2.55E+00 | 5.55E-01 | no | Carmil1 | intron_variant | MODIFIER |
| 6 | 134244764 | AX-170097238 | 0 | G | A | 0.553 | 5.17E-02 | 8.54E-02 | 2.53E+00 | 2.59E+00 | 5.55E-01 | no | Etv6 | intron_variant | MODIFIER |
| 1 | 23479711 | AX-97803867 | 2 | T | G | 0.427 | 4.94E-02 | 8.40E-02 | 2.62E+00 | 2.68E+00 | 5.56E-01 | no | Gm7784 | non_coding_transcript_exon_variant | MODIFIER |
| 7 | 104487112 | AX-170120663 | 0 | A | G | 0.447 | -5.16E-02 | 8.90E-02 | 2.77E+00 | 2.86E+00 | 5.56E-01 | no | Trim30d | intron_variant | MODIFIER |
| 16 | 19250462 | AX-170517661 | 1 | T | C | 0.125 | 5.69E-02 | 9.88E-02 | 2.70E+00 | 2.83E+00 | 5.56E-01 | no | | intergenic_variant | MODIFIER |
| 5 | 115921956 | AX-170351738 | 1 | G | A | 0.464 | 6.38E-02 | 1.09E-01 | 2.64E+00 | 2.70E+00 | 5.56E-01 | no | Cit | intron_variant | MODIFIER |
| 5 | 28524160 | AX-170514797 | 1 | G | A | 0.375 | 5.78E-02 | 9.93E-02 | 2.67E+00 | 2.75E+00 | 5.56E-01 | no | 9530036O11Rik | intron_variant,non_coding_transcript_variant | MODIFIER |
| 18 | 66158315 | AX-97832271 | 0 | T | G | 0.281 | -5.84E-02 | 1.00E-01 | 2.70E+00 | 2.76E+00 | 5.56E-01 | no | Ccbe1 | intron_variant,NMD_transcript_variant | MODIFIER |
| 18 | 66158340 | AX-170399090 | 0 | T | C | 0.281 | -5.84E-02 | 1.00E-01 | 2.70E+00 | 2.76E+00 | 5.56E-01 | no | Ccbe1 | intron_variant,NMD_transcript_variant | MODIFIER |
| 18 | 66158570 | AX-170124001 | 0 | A | G | 0.281 | -5.84E-02 | 1.00E-01 | 2.70E+00 | 2.76E+00 | 5.56E-01 | no | Ccbe1 | intron_variant,NMD_transcript_variant | MODIFIER |
| 18 | 66158820 | AX-170128115 | 0 | A | G | 0.281 | -5.84E-02 | 1.00E-01 | 2.70E+00 | 2.76E+00 | 5.56E-01 | no | Ccbe1 | intron_variant,NMD_transcript_variant | MODIFIER |
| 18 | 66160861 | AX-170480680 | 0 | C | T | 0.281 | -5.84E-02 | 1.00E-01 | 2.70E+00 | 2.76E+00 | 5.56E-01 | no | Ccbe1 | intron_variant,NMD_transcript_variant | MODIFIER |
| 18 | 66165047 | AX-170115492 | 0 | G | T | 0.281 | -5.84E-02 | 1.00E-01 | 2.70E+00 | 2.76E+00 | 5.56E-01 | no | Ccbe1 | intron_variant,NMD_transcript_variant | MODIFIER |
| 18 | 66171109 | AX-170346543 | 0 | G | A | 0.281 | -5.84E-02 | 1.00E-01 | 2.70E+00 | 2.76E+00 | 5.56E-01 | no | Ccbe1 | intron_variant,NMD_transcript_variant | MODIFIER |
| 18 | 66202179 | AX-170061143 | 0 | G | A | 0.281 | -5.84E-02 | 1.00E-01 | 2.70E+00 | 2.76E+00 | 5.56E-01 | no | Ccbe1 | intron_variant,NMD_transcript_variant | MODIFIER |
| 18 | 66244987 | AX-170173400 | 0 | A | G | 0.281 | -5.84E-02 | 1.00E-01 | 2.70E+00 | 2.76E+00 | 5.56E-01 | no | Ccbe1 | intron_variant,NMD_transcript_variant | MODIFIER |
| 18 | 66251178 | AX-170513509 | 0 | A | G | 0.281 | -5.84E-02 | 1.00E-01 | 2.70E+00 | 2.76E+00 | 5.56E-01 | no | Ccbe1 | intron_variant,NMD_transcript_variant | MODIFIER |
| 18 | 66251293 | AX-170358098 | 0 | A | T | 0.281 | -5.84E-02 | 1.00E-01 | 2.70E+00 | 2.76E+00 | 5.56E-01 | no | Ccbe1 | intron_variant,NMD_transcript_variant | MODIFIER |
| 18 | 66252590 | AX-170362808 | 0 | T | C | 0.281 | -5.84E-02 | 1.00E-01 | 2.70E+00 | 2.76E+00 | 5.56E-01 | no | Ccbe1 | intron_variant,NMD_transcript_variant | MODIFIER |
| 18 | 66256067 | AX-170511975 | 0 | A | G | 0.281 | -5.84E-02 | 1.00E-01 | 2.70E+00 | 2.76E+00 | 5.56E-01 | no | Ccbe1 | intron_variant,NMD_transcript_variant | MODIFIER |
| 18 | 66256834 | AX-97790760 | 0 | A | G | 0.281 | -5.84E-02 | 1.00E-01 | 2.70E+00 | 2.76E+00 | 5.56E-01 | no | Ccbe1 | intron_variant,NMD_transcript_variant | MODIFIER |
| 18 | 66271579 | AX-97816033 | 0 | G | A | 0.281 | -5.84E-02 | 1.00E-01 | 2.70E+00 | 2.76E+00 | 5.56E-01 | no | Ccbe1 | intron_variant,NMD_transcript_variant | MODIFIER |
| 18 | 66341919 | AX-170251003 | 0 | G | A | 0.281 | -5.84E-02 | 1.00E-01 | 2.70E+00 | 2.76E+00 | 5.56E-01 | no | | intergenic_variant | MODIFIER |
| 18 | 66341969 | AX-170259731 | 0 | A | G | 0.281 | -5.84E-02 | 1.00E-01 | 2.70E+00 | 2.76E+00 | 5.56E-01 | no | | intergenic_variant | MODIFIER |
| 18 | 66350139 | AX-170530684 | 0 | A | G | 0.281 | -5.84E-02 | 1.00E-01 | 2.70E+00 | 2.76E+00 | 5.56E-01 | no | | intergenic_variant | MODIFIER |
| 18 | 66379572 | AX-170148264 | 0 | G | A | 0.281 | -5.84E-02 | 1.00E-01 | 2.70E+00 | 2.76E+00 | 5.56E-01 | no | | intergenic_variant | MODIFIER |
| 18 | 66405329 | AX-170443800 | 0 | T | A | 0.281 | -5.84E-02 | 1.00E-01 | 2.70E+00 | 2.76E+00 | 5.56E-01 | no | AC161932.2 | downstream_gene_variant | MODIFIER |
| 18 | 66604880 | AX-170429674 | 0 | G | A | 0.281 | -5.84E-02 | 1.00E-01 | 2.70E+00 | 2.76E+00 | 5.56E-01 | no | AC100212.2 | downstream_gene_variant | MODIFIER |
| 18 | 66605730 | AX-170242917 | 0 | C | T | 0.281 | -5.84E-02 | 1.00E-01 | 2.70E+00 | 2.76E+00 | 5.56E-01 | no | AC100212.2 | downstream_gene_variant | MODIFIER |
| 18 | 66610376 | AX-170304268 | 0 | C | T | 0.281 | -5.84E-02 | 1.00E-01 | 2.70E+00 | 2.76E+00 | 5.56E-01 | no | AC100212.2 | upstream_gene_variant | MODIFIER |
| 18 | 66968639 | AX-97816744 | 0 | A | G | 0.281 | -5.84E-02 | 1.00E-01 | 2.70E+00 | 2.76E+00 | 5.56E-01 | no | | intergenic_variant | MODIFIER |
| 18 | 66969101 | AX-170061160 | 0 | C | T | 0.281 | -5.84E-02 | 1.00E-01 | 2.70E+00 | 2.76E+00 | 5.56E-01 | no | | intergenic_variant | MODIFIER |
| 18 | 66989157 | AX-170015898 | 0 | G | A | 0.281 | -5.84E-02 | 1.00E-01 | 2.70E+00 | 2.76E+00 | 5.56E-01 | no | | intergenic_variant | MODIFIER |
| 18 | 67003562 | AX-170158459 | 0 | G | A | 0.281 | -5.84E-02 | 1.00E-01 | 2.70E+00 | 2.76E+00 | 5.56E-01 | no | | intergenic_variant | MODIFIER |
| 18 | 67392835 | AX-97803926 | 0 | T | C | 0.281 | -5.84E-02 | 1.00E-01 | 2.70E+00 | 2.76E+00 | 5.56E-01 | no | Tubb6 | intron_variant | MODIFIER |
| 18 | 67427140 | AX-170460918 | 0 | G | A | 0.281 | -5.84E-02 | 1.00E-01 | 2.70E+00 | 2.76E+00 | 5.56E-01 | no | Afg3l2 | intron_variant | MODIFIER |
| 18 | 67440644 | AX-170527321 | 0 | A | C | 0.281 | -5.84E-02 | 1.00E-01 | 2.70E+00 | 2.76E+00 | 5.56E-01 | no | Afg3l2 | intron_variant | MODIFIER |
| 18 | 67441461 | AX-97833801 | 0 | A | G | 0.281 | -5.84E-02 | 1.00E-01 | 2.70E+00 | 2.76E+00 | 5.56E-01 | no | Afg3l2 | intron_variant | MODIFIER |
| 18 | 67461292 | AX-170015903 | 0 | T | C | 0.281 | -5.84E-02 | 1.00E-01 | 2.70E+00 | 2.76E+00 | 5.56E-01 | no | Prelid3a | upstream_gene_variant | MODIFIER |
| 18 | 67478797 | AX-97835208 | 0 | A | G | 0.281 | -5.84E-02 | 1.00E-01 | 2.70E+00 | 2.76E+00 | 5.56E-01 | no | Prelid3a | intron_variant | MODIFIER |
| 18 | 67482279 | AX-170490008 | 0 | A | G | 0.281 | -5.84E-02 | 1.00E-01 | 2.70E+00 | 2.76E+00 | 5.56E-01 | no | Prelid3a | downstream_gene_variant | MODIFIER |
| 18 | 67484591 | AX-170520036 | 0 | G | A | 0.281 | -5.84E-02 | 1.00E-01 | 2.70E+00 | 2.76E+00 | 5.56E-01 | no | Prelid3a | downstream_gene_variant | MODIFIER |
| 18 | 67495511 | AX-97827146 | 0 | A | G | 0.281 | -5.84E-02 | 1.00E-01 | 2.70E+00 | 2.76E+00 | 5.56E-01 | no | Spire1 | intron_variant | MODIFIER |

| | | | | | | | | | | | | | | | |
|---|---|---|---|---|---|---|---|---|---|---|---|---|---|---|---|
| 18 | 67536145 | AX-170256828 | 0 | C | T | 0.281 | -5.84E-02 | 1.00E-01 | 2.70E+00 | 2.76E+00 | 5.56E-01 | no | Spire1 | intron_variant | MODIFIER |
| 18 | 67537570 | AX-97842206 | 0 | A | C | 0.281 | -5.84E-02 | 1.00E-01 | 2.70E+00 | 2.76E+00 | 5.56E-01 | no | Spire1 | intron_variant | MODIFIER |
| 18 | 67561304 | AX-97811993 | 0 | A | G | 0.281 | -5.84E-02 | 1.00E-01 | 2.70E+00 | 2.76E+00 | 5.56E-01 | no | Spire1 | intron_variant | MODIFIER |
| 18 | 67568523 | AX-97831782 | 0 | T | C | 0.281 | -5.84E-02 | 1.00E-01 | 2.70E+00 | 2.76E+00 | 5.56E-01 | no | Spire1 | intron_variant | MODIFIER |
| 18 | 67597517 | AX-170517531 | 0 | C | T | 0.281 | -5.84E-02 | 1.00E-01 | 2.70E+00 | 2.76E+00 | 5.56E-01 | no | Spire1 | intron_variant | MODIFIER |
| 18 | 67597544 | AX-170061172 | 0 | A | G | 0.281 | -5.84E-02 | 1.00E-01 | 2.70E+00 | 2.76E+00 | 5.56E-01 | no | Spire1 | intron_variant | MODIFIER |
| 18 | 67598164 | AX-170104888 | 0 | A | G | 0.281 | -5.84E-02 | 1.00E-01 | 2.70E+00 | 2.76E+00 | 5.56E-01 | no | Spire1 | intron_variant | MODIFIER |
| 18 | 67655204 | AX-170351971 | 0 | G | T | 0.281 | -5.84E-02 | 1.00E-01 | 2.70E+00 | 2.76E+00 | 5.56E-01 | no | Psmg2 | downstream_gene_variant | MODIFIER |
| 18 | 67662727 | AX-170521273 | 0 | A | G | 0.281 | -5.84E-02 | 1.00E-01 | 2.70E+00 | 2.76E+00 | 5.56E-01 | no | Ptpn2 | downstream_gene_variant | MODIFIER |
| 18 | 67680165 | AX-170179385 | 0 | C | T | 0.281 | -5.84E-02 | 1.00E-01 | 2.70E+00 | 2.76E+00 | 5.56E-01 | no | Ptpn2 | intron_variant | MODIFIER |
| 18 | 67734998 | AX-170061174 | 0 | A | G | 0.281 | -5.84E-02 | 1.00E-01 | 2.70E+00 | 2.76E+00 | 5.56E-01 | no | | intergenic_variant | MODIFIER |
| 18 | 67738205 | AX-97785424 | 0 | T | C | 0.281 | -5.84E-02 | 1.00E-01 | 2.70E+00 | 2.76E+00 | 5.56E-01 | no | | regulatory_region_variant | MODIFIER |
| 18 | 67738650 | AX-170467095 | 0 | A | G | 0.281 | -5.84E-02 | 1.00E-01 | 2.70E+00 | 2.76E+00 | 5.56E-01 | no | | regulatory_region_variant | MODIFIER |
| 18 | 67744555 | AX-170391052 | 0 | G | T | 0.281 | -5.84E-02 | 1.00E-01 | 2.70E+00 | 2.76E+00 | 5.56E-01 | no | | intergenic_variant | MODIFIER |
| 18 | 67768440 | AX-170200848 | 0 | A | G | 0.281 | -5.84E-02 | 1.00E-01 | 2.70E+00 | 2.76E+00 | 5.56E-01 | no | 4930549G23Rik | intron_variant,non_coding_transcript_variant | MODIFIER |
| 18 | 67768455 | AX-170371505 | 0 | C | G | 0.281 | -5.84E-02 | 1.00E-01 | 2.70E+00 | 2.76E+00 | 5.56E-01 | no | 4930549G23Rik | intron_variant,non_coding_transcript_variant | MODIFIER |
| 18 | 67848305 | AX-97835736 | 0 | C | T | 0.281 | -5.84E-02 | 1.00E-01 | 2.70E+00 | 2.76E+00 | 5.56E-01 | no | Cep192 | intron_variant | MODIFIER |
| 18 | 67867113 | AX-170507363 | 0 | G | A | 0.281 | -5.84E-02 | 1.00E-01 | 2.70E+00 | 2.76E+00 | 5.56E-01 | no | Cep192 | intron_variant | MODIFIER |
| 18 | 67867146 | AX-170453464 | 0 | C | A | 0.281 | -5.84E-02 | 1.00E-01 | 2.70E+00 | 2.76E+00 | 5.56E-01 | no | Cep192 | intron_variant | MODIFIER |
| 18 | 67868202 | AX-170401974 | 0 | T | C | 0.281 | -5.84E-02 | 1.00E-01 | 2.70E+00 | 2.76E+00 | 5.56E-01 | no | Cep192 | intron_variant | MODIFIER |
| 18 | 67879118 | AX-170219120 | 0 | C | T | 0.281 | -5.84E-02 | 1.00E-01 | 2.70E+00 | 2.76E+00 | 5.56E-01 | no | Cep192 | intron_variant | MODIFIER |
| 18 | 67886186 | AX-170213621 | 0 | T | C | 0.281 | -5.84E-02 | 1.00E-01 | 2.70E+00 | 2.76E+00 | 5.56E-01 | no | Cep192 | downstream_gene_variant | MODIFIER |
| 18 | 67899508 | AX-97841880 | 0 | G | A | 0.281 | -5.84E-02 | 1.00E-01 | 2.70E+00 | 2.76E+00 | 5.56E-01 | no | | intergenic_variant | MODIFIER |
| 18 | 67908758 | AX-170238072 | 0 | T | C | 0.281 | -5.84E-02 | 1.00E-01 | 2.70E+00 | 2.76E+00 | 5.56E-01 | no | AC127236.1 | non_coding_transcript_exon_variant | MODIFIER |
| 18 | 67909560 | AX-170338221 | 0 | A | G | 0.281 | -5.84E-02 | 1.00E-01 | 2.70E+00 | 2.76E+00 | 5.56E-01 | no | AC127236.1 | downstream_gene_variant | MODIFIER |
| 18 | 67909717 | AX-170180834 | 0 | A | G | 0.281 | -5.84E-02 | 1.00E-01 | 2.70E+00 | 2.76E+00 | 5.56E-01 | no | AC127236.1 | downstream_gene_variant | MODIFIER |
| 7 | 37148576 | AX-170473734 | 0 | T | C | 0.342 | 5.39E-02 | 9.13E-02 | 2.59E+00 | 2.67E+00 | 5.56E-01 | no | | intergenic_variant | MODIFIER |
| 20 | 69726933 | AX-97786067 | 0 | C | T | 0.263 | 3.65E-02 | 6.27E-02 | 2.67E+00 | 2.75E+00 | 5.56E-01 | no | Aff2 | intron_variant | MODIFIER |
| 1 | 136383933 | AX-170369590 | 1 | C | T | 0.205 | 7.26E-02 | 1.24E-01 | 2.65E+00 | 2.72E+00 | 5.56E-01 | no | | intergenic_variant | MODIFIER |
| 1 | 137442572 | AX-170086622 | 1 | A | T | 0.205 | 7.26E-02 | 1.24E-01 | 2.65E+00 | 2.72E+00 | 5.56E-01 | no | | intergenic_variant | MODIFIER |
| 6 | 134239587 | AX-170289581 | 0 | G | C | 0.57 | 5.11E-02 | 8.55E-02 | 2.56E+00 | 2.62E+00 | 5.56E-01 | no | Etv6 | intron_variant | MODIFIER |
| 11 | 30286727 | AX-97790863 | 1 | C | T | 0.08 | 1.18E-01 | 2.02E-01 | 2.65E+00 | 2.72E+00 | 5.56E-01 | no | | intergenic_variant | MODIFIER |
| 8 | 34474715 | AX-170142735 | 1 | G | A | 0.143 | -7.78E-02 | 1.33E-01 | 2.63E+00 | 2.70E+00 | 5.56E-01 | no | Gm45349 | upstream_gene_variant | MODIFIER |
| 8 | 95215868 | AX-170391692 | 1 | A | G | 0.223 | 6.55E-02 | 1.13E-01 | 2.72E+00 | 2.80E+00 | 5.56E-01 | no | | regulatory_region_variant | MODIFIER |
| 13 | 25171174 | AX-170516315 | 2 | G | A | 0.173 | -7.57E-02 | 1.22E-01 | 2.47E+00 | 2.53E+00 | 5.56E-01 | no | Dcdc2a | intron_variant | MODIFIER |
| 10 | 26458465 | AX-170116642 | 0 | C | A | 0.184 | 5.86E-02 | 1.01E-01 | 2.81E+00 | 2.92E+00 | 5.56E-01 | no | | intergenic_variant | MODIFIER |
| 1 | 127900843 | AX-170156681 | 2 | T | C | 0.527 | 4.82E-02 | 8.17E-02 | 2.60E+00 | 2.67E+00 | 5.56E-01 | no | Rab3gap1 | intron_variant | MODIFIER |
| 2 | 168732639 | AX-97798326 | 0 | C | T | 0.167 | 1.06E-01 | 1.81E-01 | 2.61E+00 | 2.69E+00 | 5.57E-01 | no | Atp9a | intron_variant | MODIFIER |
| 8 | 91022740 | AX-170198315 | 1 | G | C | 0.179 | -7.00E-02 | 1.20E-01 | 2.63E+00 | 2.71E+00 | 5.57E-01 | no | Chd9 | intron_variant | MODIFIER |
| 5 | 118194695 | AX-170492776 | 1 | C | T | 0.446 | -6.42E-02 | 1.11E-01 | 2.78E+00 | 2.85E+00 | 5.57E-01 | no | Rnft2 | intron_variant | MODIFIER |
| 1 | 21679501 | AX-170193915 | 0 | G | A | 0.404 | 4.87E-02 | 8.32E-02 | 2.63E+00 | 2.69E+00 | 5.57E-01 | no | Kcnq5 | intron_variant | MODIFIER |
| 11 | 5926738 | AX-170402149 | 0 | G | A | 0.053 | 1.07E-01 | 1.84E-01 | 2.62E+00 | 2.76E+00 | 5.57E-01 | no | Gck | intron_variant | MODIFIER |
| 8 | 73762411 | AX-170510908 | 2 | A | G | 0.191 | 7.23E-02 | 1.25E-01 | 2.74E+00 | 2.82E+00 | 5.57E-01 | no | | intergenic_variant | MODIFIER |
| 8 | 81068708 | AX-170279450 | 0 | C | A | 0.202 | -7.07E-02 | 1.22E-01 | 2.74E+00 | 2.83E+00 | 5.57E-01 | no | Gm45538 | intron_variant,non_coding_transcript_variant | MODIFIER |
| 8 | 84089490 | AX-170280560 | 0 | A | C | 0.202 | -7.07E-02 | 1.22E-01 | 2.74E+00 | 2.83E+00 | 5.57E-01 | no | Rfx1 | intron_variant | MODIFIER |
| 8 | 85538166 | AX-170148070 | 0 | A | G | 0.202 | -7.07E-02 | 1.22E-01 | 2.74E+00 | 2.83E+00 | 5.57E-01 | no | Dnaja2 | 3_prime_UTR_variant | MODIFIER |
| 6 | 135832889 | AX-170485818 | 0 | C | T | 0.579 | 5.32E-02 | 8.79E-02 | 2.52E+00 | 2.57E+00 | 5.57E-01 | no | Grin2b | intron_variant | MODIFIER |
| 18 | 58990370 | AX-170111522 | 1 | C | T | 0.188 | -6.99E-02 | 1.20E-01 | 2.65E+00 | 2.70E+00 | 5.57E-01 | no | Adamts19 | intron_variant | MODIFIER |
| 7 | 40448264 | AX-170077127 | 0 | C | T | 0.333 | 5.35E-02 | 9.20E-02 | 2.65E+00 | 2.73E+00 | 5.57E-01 | no | Gm28807 | intron_variant,non_coding_transcript_variant | MODIFIER |
| 4 | 78224331 | AX-170109580 | 0 | G | T | 0.298 | 6.33E-02 | 1.09E-01 | 2.65E+00 | 2.72E+00 | 5.57E-01 | no | | intergenic_variant | MODIFIER |
| 20 | 70803170 | AX-170034111 | 1 | C | G | 0.259 | 3.62E-02 | 6.25E-02 | 2.67E+00 | 2.76E+00 | 5.57E-01 | no | | intergenic_variant | MODIFIER |
| 4 | 49300463 | AX-170089225 | 1 | G | C | 0.357 | -5.31E-02 | 9.21E-02 | 2.69E+00 | 2.79E+00 | 5.57E-01 | no | Plppr1 | intron_variant | MODIFIER |
| 7 | 45146346 | AX-170197019 | 0 | C | T | 0.342 | 5.51E-02 | 9.42E-02 | 2.62E+00 | 2.70E+00 | 5.57E-01 | no | Aldh16a1 | intron_variant | MODIFIER |
| 7 | 45148108 | AX-170233564 | 0 | A | G | 0.342 | 5.51E-02 | 9.42E-02 | 2.62E+00 | 2.70E+00 | 5.57E-01 | no | Aldh16a1 | intron_variant | MODIFIER |

| | | | | | | | | | | | | | | | |
|---|---|---|---|---|---|---|---|---|---|---|---|---|---|---|---|
| 7 | 45151778 | AX-97799648 | 0 | A | T | 0.342 | 5.51E-02 | 9.42E-02 | 2.62E+00 | 2.70E+00 | 5.57E-01 | no | Pih1d1 | upstream_gene_variant | MODIFIER |
| 7 | 45155616 | AX-170495340 | 0 | C | T | 0.342 | 5.51E-02 | 9.42E-02 | 2.62E+00 | 2.70E+00 | 5.57E-01 | no | Pih1d1 | intron_variant | MODIFIER |
| 7 | 45504843 | AX-97815593 | 0 | T | C | 0.342 | 5.51E-02 | 9.42E-02 | 2.62E+00 | 2.70E+00 | 5.57E-01 | no | Nucb1 | intron_variant | MODIFIER |
| 7 | 45505741 | AX-170358518 | 0 | C | T | 0.342 | 5.51E-02 | 9.42E-02 | 2.62E+00 | 2.70E+00 | 5.57E-01 | no | Nucb1 | intron_variant | MODIFIER |
| 7 | 45518959 | AX-170077206 | 0 | A | G | 0.342 | 5.51E-02 | 9.42E-02 | 2.62E+00 | 2.70E+00 | 5.57E-01 | no | Tulp2 | intron_variant | MODIFIER |
| 7 | 45552563 | AX-170095914 | 0 | C | A | 0.342 | 5.51E-02 | 9.42E-02 | 2.62E+00 | 2.70E+00 | 5.57E-01 | no | Plekha4 | intron_variant | MODIFIER |
| 7 | 45635347 | AX-170283362 | 0 | C | T | 0.342 | 5.51E-02 | 9.42E-02 | 2.62E+00 | 2.70E+00 | 5.57E-01 | no | Rasip1 | intron_variant | MODIFIER |
| 7 | 45723064 | AX-97830128 | 0 | C | A | 0.342 | 5.51E-02 | 9.42E-02 | 2.62E+00 | 2.70E+00 | 5.57E-01 | no | Rpl18 | downstream_gene_variant | MODIFIER |
| 7 | 45734081 | AX-97834902 | 0 | A | G | 0.342 | 5.51E-02 | 9.42E-02 | 2.62E+00 | 2.70E+00 | 5.57E-01 | no | Sult2b1 | synonymous_variant | LOW |
| 7 | 45771762 | AX-97829098 | 0 | A | G | 0.342 | 5.51E-02 | 9.42E-02 | 2.62E+00 | 2.70E+00 | 5.57E-01 | no | Sult2b1 | intron_variant | MODIFIER |
| 9 | 17311699 | AX-170525863 | 0 | G | A | 0.482 | 5.22E-02 | 8.76E-02 | 2.55E+00 | 2.62E+00 | 5.57E-01 | no | | intergenic_variant | MODIFIER |
| 1 | 119205129 | AX-170001530 | 1 | G | C | 0.152 | 1.05E-01 | 1.80E-01 | 2.61E+00 | 2.71E+00 | 5.58E-01 | no | | intergenic_variant | MODIFIER |
| 7 | 69871060 | AX-170098803 | 1 | A | C | 0.366 | -5.42E-02 | 9.27E-02 | 2.89E+00 | 3.02E+00 | 5.58E-01 | no | | intergenic_variant | MODIFIER |
| 1 | 170458748 | AX-170043546 | 1 | G | T | 0.241 | 7.56E-02 | 1.30E-01 | 2.69E+00 | 2.74E+00 | 5.58E-01 | no | Nos1ap | intron_variant,non_coding_transcript_variant | MODIFIER |
| 13 | 3649309 | AX-170139007 | 1 | G | C | 0.205 | -7.07E-02 | 1.19E-01 | 2.56E+00 | 2.64E+00 | 5.58E-01 | no | Asb13 | intron_variant | MODIFIER |
| 13 | 48990720 | AX-97848306 | 0 | C | T | 0.491 | -3.83E-02 | 6.66E-02 | 2.79E+00 | 2.90E+00 | 5.58E-01 | no | | intergenic_variant | MODIFIER |
| 18 | 57717244 | AX-170174031 | 1 | C | G | 0.196 | -4.27E-02 | 7.45E-02 | 2.79E+00 | 2.92E+00 | 5.58E-01 | no | Ccdc192 | intron_variant | MODIFIER |
| 18 | 63513573 | AX-170499683 | 1 | C | A | 0.188 | -7.02E-02 | 1.20E-01 | 2.64E+00 | 2.69E+00 | 5.58E-01 | no | Gm22560 | downstream_gene_variant | MODIFIER |
| 7 | 8290737 | AX-170039412 | 0 | T | G | 0.175 | 4.23E-02 | 7.44E-02 | 2.76E+00 | 2.91E+00 | 5.58E-01 | no | Gm3654 | upstream_gene_variant | MODIFIER |
| 1 | 14435701 | AX-170114874 | 2 | C | T | 0.4 | 4.45E-02 | 7.57E-02 | 2.59E+00 | 2.67E+00 | 5.58E-01 | no | | intergenic_variant | MODIFIER |
| 20 | 9158170 | AX-170409416 | 0 | G | A | 0.228 | 3.28E-02 | 5.73E-02 | 2.64E+00 | 2.79E+00 | 5.58E-01 | no | | intergenic_variant | MODIFIER |
| 18 | 55649406 | AX-97850353 | 0 | A | C | 0.447 | 5.91E-02 | 9.47E-02 | 2.44E+00 | 2.50E+00 | 5.59E-01 | no | | intergenic_variant | MODIFIER |
| 12 | 88936904 | AX-170294950 | 0 | T | A | 0.079 | 1.07E-01 | 1.86E-01 | 2.73E+00 | 2.81E+00 | 5.59E-01 | no | Nrxn3 | intron_variant | MODIFIER |
| 12 | 88998481 | AX-168374337 | 0 | A | G | 0.079 | 1.07E-01 | 1.86E-01 | 2.73E+00 | 2.81E+00 | 5.59E-01 | no | Nrxn3 | intron_variant | MODIFIER |
| 12 | 89001237 | AX-170007221 | 0 | C | T | 0.079 | 1.07E-01 | 1.86E-01 | 2.73E+00 | 2.81E+00 | 5.59E-01 | no | Nrxn3 | intron_variant | MODIFIER |
| 12 | 89001334 | AX-170007217 | 0 | C | T | 0.079 | 1.07E-01 | 1.86E-01 | 2.73E+00 | 2.81E+00 | 5.59E-01 | no | Nrxn3 | intron_variant | MODIFIER |
| 12 | 89002568 | AX-170007219 | 0 | C | T | 0.079 | 1.07E-01 | 1.86E-01 | 2.73E+00 | 2.81E+00 | 5.59E-01 | no | Nrxn3 | intron_variant | MODIFIER |
| 12 | 89003168 | AX-170337847 | 0 | T | C | 0.079 | 1.07E-01 | 1.86E-01 | 2.73E+00 | 2.81E+00 | 5.59E-01 | no | Nrxn3 | intron_variant | MODIFIER |
| 12 | 89011504 | AX-170501605 | 0 | G | A | 0.079 | 1.07E-01 | 1.86E-01 | 2.73E+00 | 2.81E+00 | 5.59E-01 | no | Nrxn3 | intron_variant | MODIFIER |
| 12 | 89012482 | AX-170007220 | 0 | G | A | 0.079 | 1.07E-01 | 1.86E-01 | 2.73E+00 | 2.81E+00 | 5.59E-01 | no | Nrxn3 | intron_variant | MODIFIER |
| 12 | 89013961 | AX-170505418 | 0 | C | G | 0.079 | 1.07E-01 | 1.86E-01 | 2.73E+00 | 2.81E+00 | 5.59E-01 | no | Nrxn3 | intron_variant | MODIFIER |
| 12 | 89020978 | AX-170049725 | 0 | G | A | 0.079 | 1.07E-01 | 1.86E-01 | 2.73E+00 | 2.81E+00 | 5.59E-01 | no | Nrxn3 | intron_variant | MODIFIER |
| 12 | 89036046 | AX-170497014 | 0 | A | G | 0.079 | 1.07E-01 | 1.86E-01 | 2.73E+00 | 2.81E+00 | 5.59E-01 | no | Nrxn3 | intron_variant | MODIFIER |
| 12 | 89041725 | AX-97801200 | 0 | T | C | 0.079 | 1.07E-01 | 1.86E-01 | 2.73E+00 | 2.81E+00 | 5.59E-01 | no | Nrxn3 | intron_variant | MODIFIER |
| 12 | 89064382 | AX-170492431 | 0 | A | C | 0.079 | 1.07E-01 | 1.86E-01 | 2.73E+00 | 2.81E+00 | 5.59E-01 | no | Nrxn3 | intron_variant | MODIFIER |
| 12 | 89065880 | AX-97826602 | 0 | G | A | 0.079 | 1.07E-01 | 1.86E-01 | 2.73E+00 | 2.81E+00 | 5.59E-01 | no | Nrxn3 | intron_variant | MODIFIER |
| 12 | 89067783 | AX-170235388 | 0 | C | T | 0.079 | 1.07E-01 | 1.86E-01 | 2.73E+00 | 2.81E+00 | 5.59E-01 | no | Nrxn3 | intron_variant | MODIFIER |
| 12 | 89085109 | AX-170376333 | 0 | T | C | 0.079 | 1.07E-01 | 1.86E-01 | 2.73E+00 | 2.81E+00 | 5.59E-01 | no | Nrxn3 | intron_variant | MODIFIER |
| 12 | 89091651 | AX-170099998 | 0 | T | C | 0.079 | 1.07E-01 | 1.86E-01 | 2.73E+00 | 2.81E+00 | 5.59E-01 | no | Nrxn3 | intron_variant | MODIFIER |
| 12 | 89099249 | AX-97822664 | 0 | T | C | 0.079 | 1.07E-01 | 1.86E-01 | 2.73E+00 | 2.81E+00 | 5.59E-01 | no | Nrxn3 | intron_variant | MODIFIER |
| 12 | 89105976 | AX-170164547 | 0 | T | G | 0.079 | 1.07E-01 | 1.86E-01 | 2.73E+00 | 2.81E+00 | 5.59E-01 | no | Nrxn3 | intron_variant | MODIFIER |
| 12 | 89130760 | AX-97831140 | 0 | C | T | 0.079 | 1.07E-01 | 1.86E-01 | 2.73E+00 | 2.81E+00 | 5.59E-01 | no | Nrxn3 | intron_variant | MODIFIER |
| 12 | 89164863 | AX-170527051 | 0 | A | G | 0.079 | 1.07E-01 | 1.86E-01 | 2.73E+00 | 2.81E+00 | 5.59E-01 | no | Nrxn3 | intron_variant | MODIFIER |
| 12 | 89164976 | AX-170406598 | 0 | G | T | 0.079 | 1.07E-01 | 1.86E-01 | 2.73E+00 | 2.81E+00 | 5.59E-01 | no | Nrxn3 | intron_variant | MODIFIER |
| 12 | 89166590 | AX-97819659 | 0 | T | C | 0.079 | 1.07E-01 | 1.86E-01 | 2.73E+00 | 2.81E+00 | 5.59E-01 | no | Nrxn3 | intron_variant | MODIFIER |
| 12 | 89167101 | AX-170141974 | 0 | T | C | 0.079 | 1.07E-01 | 1.86E-01 | 2.73E+00 | 2.81E+00 | 5.59E-01 | no | Nrxn3 | intron_variant | MODIFIER |
| 12 | 89172971 | AX-170049730 | 0 | G | T | 0.079 | 1.07E-01 | 1.86E-01 | 2.73E+00 | 2.81E+00 | 5.59E-01 | no | Nrxn3 | intron_variant | MODIFIER |
| 12 | 89173004 | AX-170049731 | 0 | A | G | 0.079 | 1.07E-01 | 1.86E-01 | 2.73E+00 | 2.81E+00 | 5.59E-01 | no | Nrxn3 | intron_variant | MODIFIER |
| 12 | 89173057 | AX-170049729 | 0 | T | C | 0.079 | 1.07E-01 | 1.86E-01 | 2.73E+00 | 2.81E+00 | 5.59E-01 | no | Nrxn3 | intron_variant | MODIFIER |
| 12 | 89210737 | AX-170420524 | 0 | C | T | 0.079 | 1.07E-01 | 1.86E-01 | 2.73E+00 | 2.81E+00 | 5.59E-01 | no | Nrxn3 | intron_variant | MODIFIER |
| 12 | 89217848 | AX-97821991 | 0 | C | T | 0.079 | 1.07E-01 | 1.86E-01 | 2.73E+00 | 2.81E+00 | 5.59E-01 | no | Nrxn3 | intron_variant | MODIFIER |
| 12 | 89225586 | AX-170395705 | 0 | G | A | 0.079 | 1.07E-01 | 1.86E-01 | 2.73E+00 | 2.81E+00 | 5.59E-01 | no | Nrxn3 | intron_variant | MODIFIER |
| 12 | 89228725 | AX-170285794 | 0 | C | T | 0.079 | 1.07E-01 | 1.86E-01 | 2.73E+00 | 2.81E+00 | 5.59E-01 | no | Nrxn3 | intron_variant | MODIFIER |
| 12 | 89236796 | AX-97815453 | 0 | T | C | 0.079 | 1.07E-01 | 1.86E-01 | 2.73E+00 | 2.81E+00 | 5.59E-01 | no | Nrxn3 | intron_variant | MODIFIER |

| | | | | | | | | | | | | | | | |
|---|---|---|---|---|---|---|---|---|---|---|---|---|---|---|---|
| 12 | 89532272 | AX-170049738 | 0 | T | G | 0.079 | 1.07E-01 | 1.86E-01 | 2.73E+00 | 2.81E+00 | 5.59E-01 | no | Nrxn3 | intron_variant | MODIFIER |
| 12 | 89580944 | AX-97813029 | 0 | G | A | 0.079 | 1.07E-01 | 1.86E-01 | 2.73E+00 | 2.81E+00 | 5.59E-01 | no | Nrxn3 | intron_variant | MODIFIER |
| 12 | 89581039 | AX-170173129 | 0 | A | G | 0.079 | 1.07E-01 | 1.86E-01 | 2.73E+00 | 2.81E+00 | 5.59E-01 | no | Nrxn3 | intron_variant | MODIFIER |
| 12 | 89581206 | AX-170107772 | 0 | T | C | 0.079 | 1.07E-01 | 1.86E-01 | 2.73E+00 | 2.81E+00 | 5.59E-01 | no | Nrxn3 | intron_variant | MODIFIER |
| 12 | 89582015 | AX-170499599 | 0 | A | G | 0.079 | 1.07E-01 | 1.86E-01 | 2.73E+00 | 2.81E+00 | 5.59E-01 | no | Nrxn3 | intron_variant | MODIFIER |
| 12 | 89623413 | AX-97831459 | 0 | G | A | 0.079 | 1.07E-01 | 1.86E-01 | 2.73E+00 | 2.81E+00 | 5.59E-01 | no | Nrxn3 | intron_variant | MODIFIER |
| 12 | 89640510 | AX-170451126 | 0 | T | G | 0.079 | 1.07E-01 | 1.86E-01 | 2.73E+00 | 2.81E+00 | 5.59E-01 | no | Nrxn3 | intron_variant | MODIFIER |
| 12 | 89653972 | AX-97815145 | 0 | G | A | 0.079 | 1.07E-01 | 1.86E-01 | 2.73E+00 | 2.81E+00 | 5.59E-01 | no | Nrxn3 | intron_variant | MODIFIER |
| 12 | 90226396 | AX-170216425 | 0 | C | A | 0.079 | 1.07E-01 | 1.86E-01 | 2.73E+00 | 2.81E+00 | 5.59E-01 | no | Nrxn3 | intron_variant | MODIFIER |
| 12 | 90229205 | AX-168400867 | 0 | C | G | 0.079 | 1.07E-01 | 1.86E-01 | 2.73E+00 | 2.81E+00 | 5.59E-01 | no | Nrxn3 | intron_variant | MODIFIER |
| 12 | 90232671 | AX-170291639 | 0 | C | T | 0.079 | 1.07E-01 | 1.86E-01 | 2.73E+00 | 2.81E+00 | 5.59E-01 | no | Nrxn3 | intron_variant | MODIFIER |
| 12 | 90234405 | AX-170240350 | 0 | T | A | 0.079 | 1.07E-01 | 1.86E-01 | 2.73E+00 | 2.81E+00 | 5.59E-01 | no | Nrxn3 | intron_variant | MODIFIER |
| 12 | 90234504 | AX-170106085 | 0 | A | G | 0.079 | 1.07E-01 | 1.86E-01 | 2.73E+00 | 2.81E+00 | 5.59E-01 | no | Nrxn3 | intron_variant | MODIFIER |
| 12 | 90234728 | AX-170338379 | 0 | T | A | 0.079 | 1.07E-01 | 1.86E-01 | 2.73E+00 | 2.81E+00 | 5.59E-01 | no | Nrxn3 | intron_variant | MODIFIER |
| 12 | 90246451 | AX-170148787 | 0 | C | T | 0.079 | 1.07E-01 | 1.86E-01 | 2.73E+00 | 2.81E+00 | 5.59E-01 | no | Nrxn3 | intron_variant | MODIFIER |
| 12 | 90268656 | AX-97793493 | 0 | T | C | 0.079 | 1.07E-01 | 1.86E-01 | 2.73E+00 | 2.81E+00 | 5.59E-01 | no | Nrxn3 | intron_variant | MODIFIER |
| 12 | 90422498 | AX-170112405 | 0 | G | T | 0.079 | 1.07E-01 | 1.86E-01 | 2.73E+00 | 2.81E+00 | 5.59E-01 | no | | intergenic_variant | MODIFIER |
| 12 | 90454088 | AX-97848646 | 0 | T | C | 0.079 | 1.07E-01 | 1.86E-01 | 2.73E+00 | 2.81E+00 | 5.59E-01 | no | | intergenic_variant | MODIFIER |
| 12 | 90461377 | AX-170049752 | 0 | G | A | 0.079 | 1.07E-01 | 1.86E-01 | 2.73E+00 | 2.81E+00 | 5.59E-01 | no | | intergenic_variant | MODIFIER |
| 12 | 90532563 | AX-170007248 | 0 | T | C | 0.079 | 1.07E-01 | 1.86E-01 | 2.73E+00 | 2.81E+00 | 5.59E-01 | no | | intergenic_variant | MODIFIER |
| 12 | 90590833 | AX-170511091 | 0 | C | T | 0.079 | 1.07E-01 | 1.86E-01 | 2.73E+00 | 2.81E+00 | 5.59E-01 | no | Gm2139 | upstream_gene_variant | MODIFIER |
| 12 | 90602259 | AX-97845258 | 0 | T | C | 0.079 | 1.07E-01 | 1.86E-01 | 2.73E+00 | 2.81E+00 | 5.59E-01 | no | | intergenic_variant | MODIFIER |
| 12 | 90644756 | AX-170518420 | 0 | A | G | 0.079 | 1.07E-01 | 1.86E-01 | 2.73E+00 | 2.81E+00 | 5.59E-01 | no | | intergenic_variant | MODIFIER |
| 12 | 90707015 | AX-97845336 | 0 | A | G | 0.079 | 1.07E-01 | 1.86E-01 | 2.73E+00 | 2.81E+00 | 5.59E-01 | no | | intergenic_variant | MODIFIER |
| 12 | 90772402 | AX-97802278 | 0 | G | A | 0.079 | 1.07E-01 | 1.86E-01 | 2.73E+00 | 2.81E+00 | 5.59E-01 | no | 4930544I03Rik | intron_variant,non_coding_transcript_variant | MODIFIER |
| 12 | 90780946 | AX-170214933 | 0 | C | T | 0.079 | 1.07E-01 | 1.86E-01 | 2.73E+00 | 2.81E+00 | 5.59E-01 | no | 4930544I03Rik | intron_variant,non_coding_transcript_variant | MODIFIER |
| 12 | 91211967 | AX-97848021 | 0 | C | T | 0.079 | 1.07E-01 | 1.86E-01 | 2.73E+00 | 2.81E+00 | 5.59E-01 | no | Cep128 | downstream_gene_variant | MODIFIER |
| 12 | 91340166 | AX-170174657 | 0 | T | C | 0.079 | 1.07E-01 | 1.86E-01 | 2.73E+00 | 2.81E+00 | 5.59E-01 | no | Cep128 | intron_variant,non_coding_transcript_variant | MODIFIER |
| 12 | 91340465 | AX-170177820 | 0 | C | G | 0.079 | 1.07E-01 | 1.86E-01 | 2.73E+00 | 2.81E+00 | 5.59E-01 | no | Cep128 | intron_variant,non_coding_transcript_variant | MODIFIER |
| 12 | 91340494 | AX-170341137 | 0 | C | T | 0.079 | 1.07E-01 | 1.86E-01 | 2.73E+00 | 2.81E+00 | 5.59E-01 | no | Cep128 | intron_variant,non_coding_transcript_variant | MODIFIER |
| 12 | 91340506 | AX-170104339 | 0 | T | C | 0.079 | 1.07E-01 | 1.86E-01 | 2.73E+00 | 2.81E+00 | 5.59E-01 | no | Cep128 | intron_variant,non_coding_transcript_variant | MODIFIER |
| 12 | 91355762 | AX-170181818 | 0 | A | G | 0.079 | 1.07E-01 | 1.86E-01 | 2.73E+00 | 2.81E+00 | 5.59E-01 | no | Cep128 | intron_variant,non_coding_transcript_variant | MODIFIER |
| 12 | 91375754 | AX-170347910 | 0 | T | C | 0.079 | 1.07E-01 | 1.86E-01 | 2.73E+00 | 2.81E+00 | 5.59E-01 | no | Cep128 | intron_variant,non_coding_transcript_variant | MODIFIER |
| 12 | 91387347 | AX-97826985 | 0 | T | G | 0.079 | 1.07E-01 | 1.86E-01 | 2.73E+00 | 2.81E+00 | 5.59E-01 | no | Cep128 | upstream_gene_variant | MODIFIER |
| 12 | 91403027 | AX-170257709 | 0 | A | G | 0.079 | 1.07E-01 | 1.86E-01 | 2.73E+00 | 2.81E+00 | 5.59E-01 | no | Tshr | intron_variant | MODIFIER |
| 12 | 91409988 | AX-170250902 | 0 | C | G | 0.079 | 1.07E-01 | 1.86E-01 | 2.73E+00 | 2.81E+00 | 5.59E-01 | no | Tshr | intron_variant | MODIFIER |
| 12 | 91410276 | AX-97786073 | 0 | G | A | 0.079 | 1.07E-01 | 1.86E-01 | 2.73E+00 | 2.81E+00 | 5.59E-01 | no | Tshr | intron_variant | MODIFIER |
| 12 | 91514095 | AX-170460228 | 0 | A | C | 0.079 | 1.07E-01 | 1.86E-01 | 2.73E+00 | 2.81E+00 | 5.59E-01 | no | Tshr | intron_variant | MODIFIER |
| 12 | 91514345 | AX-170173146 | 0 | G | A | 0.079 | 1.07E-01 | 1.86E-01 | 2.73E+00 | 2.81E+00 | 5.59E-01 | no | Tshr | intron_variant | MODIFIER |
| 12 | 91514454 | AX-170303340 | 0 | G | A | 0.079 | 1.07E-01 | 1.86E-01 | 2.73E+00 | 2.81E+00 | 5.59E-01 | no | Tshr | intron_variant | MODIFIER |
| 12 | 91516041 | AX-97830755 | 0 | G | A | 0.079 | 1.07E-01 | 1.86E-01 | 2.73E+00 | 2.81E+00 | 5.59E-01 | no | Tshr | intron_variant | MODIFIER |
| 12 | 91516844 | AX-170049772 | 0 | G | A | 0.079 | 1.07E-01 | 1.86E-01 | 2.73E+00 | 2.81E+00 | 5.59E-01 | no | Tshr | intron_variant | MODIFIER |
| 12 | 91528513 | AX-170151661 | 0 | G | T | 0.079 | 1.07E-01 | 1.86E-01 | 2.73E+00 | 2.81E+00 | 5.59E-01 | no | Tshr | intron_variant | MODIFIER |
| 12 | 91528824 | AX-170177773 | 0 | C | T | 0.079 | 1.07E-01 | 1.86E-01 | 2.73E+00 | 2.81E+00 | 5.59E-01 | no | Tshr | intron_variant | MODIFIER |
| 12 | 91530834 | AX-170163907 | 0 | T | A | 0.079 | 1.07E-01 | 1.86E-01 | 2.73E+00 | 2.81E+00 | 5.59E-01 | no | Tshr | intron_variant | MODIFIER |
| 12 | 91531378 | AX-170162631 | 0 | A | G | 0.079 | 1.07E-01 | 1.86E-01 | 2.73E+00 | 2.81E+00 | 5.59E-01 | no | Tshr | intron_variant | MODIFIER |
| 12 | 91533083 | AX-170118292 | 0 | T | C | 0.079 | 1.07E-01 | 1.86E-01 | 2.73E+00 | 2.81E+00 | 5.59E-01 | no | Tshr | intron_variant | MODIFIER |
| 12 | 91534351 | AX-170507515 | 0 | T | C | 0.079 | 1.07E-01 | 1.86E-01 | 2.73E+00 | 2.81E+00 | 5.59E-01 | no | Tshr | intron_variant | MODIFIER |
| 12 | 91534408 | AX-170459889 | 0 | C | T | 0.079 | 1.07E-01 | 1.86E-01 | 2.73E+00 | 2.81E+00 | 5.59E-01 | no | Tshr | intron_variant | MODIFIER |
| 12 | 91534492 | AX-97834050 | 0 | A | G | 0.079 | 1.07E-01 | 1.86E-01 | 2.73E+00 | 2.81E+00 | 5.59E-01 | no | Tshr | intron_variant | MODIFIER |
| 12 | 91690198 | AX-170007261 | 0 | G | A | 0.079 | 1.07E-01 | 1.86E-01 | 2.73E+00 | 2.81E+00 | 5.59E-01 | no | Ston2 | intron_variant | MODIFIER |
| 12 | 91731748 | AX-170264898 | 0 | T | C | 0.079 | 1.07E-01 | 1.86E-01 | 2.73E+00 | 2.81E+00 | 5.59E-01 | no | Ston2 | intron_variant | MODIFIER |
| 12 | 91732487 | AX-170457593 | 0 | A | G | 0.079 | 1.07E-01 | 1.86E-01 | 2.73E+00 | 2.81E+00 | 5.59E-01 | no | Ston2 | intron_variant | MODIFIER |
| 12 | 91732552 | AX-170248299 | 0 | T | G | 0.079 | 1.07E-01 | 1.86E-01 | 2.73E+00 | 2.81E+00 | 5.59E-01 | no | Ston2 | intron_variant | MODIFIER |

| | | | | | | | | | | | | | | | |
|---|---|---|---|---|---|---|---|---|---|---|---|---|---|---|---|
| 12 | 91734837 | AX-170209983 | 0 | G | A | 0.079 | 1.07E-01 | 1.86E-01 | 2.73E+00 | 2.81E+00 | 5.59E-01 | no | Ston2 | intron_variant | MODIFIER |
| 12 | 91735575 | AX-170475218 | 0 | G | A | 0.079 | 1.07E-01 | 1.86E-01 | 2.73E+00 | 2.81E+00 | 5.59E-01 | no | Ston2 | intron_variant | MODIFIER |
| 12 | 91737899 | AX-170334115 | 0 | A | G | 0.079 | 1.07E-01 | 1.86E-01 | 2.73E+00 | 2.81E+00 | 5.59E-01 | no | Ston2 | intron_variant | MODIFIER |
| 12 | 91739532 | AX-170217443 | 0 | G | A | 0.079 | 1.07E-01 | 1.86E-01 | 2.73E+00 | 2.81E+00 | 5.59E-01 | no | Ston2 | intron_variant | MODIFIER |
| 12 | 91739728 | AX-170287647 | 0 | G | C | 0.079 | 1.07E-01 | 1.86E-01 | 2.73E+00 | 2.81E+00 | 5.59E-01 | no | Ston2 | intron_variant | MODIFIER |
| 12 | 91746010 | AX-170340939 | 0 | A | G | 0.079 | 1.07E-01 | 1.86E-01 | 2.73E+00 | 2.81E+00 | 5.59E-01 | no | Ston2 | intron_variant | MODIFIER |
| 12 | 91748581 | AX-97817315 | 0 | T | C | 0.079 | 1.07E-01 | 1.86E-01 | 2.73E+00 | 2.81E+00 | 5.59E-01 | no | Ston2 | intron_variant | MODIFIER |
| 12 | 91753203 | AX-170446173 | 0 | T | G | 0.079 | 1.07E-01 | 1.86E-01 | 2.73E+00 | 2.81E+00 | 5.59E-01 | no | Ston2 | intron_variant | MODIFIER |
| 12 | 91755174 | AX-170525877 | 0 | T | G | 0.079 | 1.07E-01 | 1.86E-01 | 2.73E+00 | 2.81E+00 | 5.59E-01 | no | Ston2 | intron_variant | MODIFIER |
| 12 | 91755450 | AX-170257503 | 0 | A | G | 0.079 | 1.07E-01 | 1.86E-01 | 2.73E+00 | 2.81E+00 | 5.59E-01 | no | Ston2 | intron_variant | MODIFIER |
| 12 | 91755765 | AX-170379967 | 0 | A | C | 0.079 | 1.07E-01 | 1.86E-01 | 2.73E+00 | 2.81E+00 | 5.59E-01 | no | Ston2 | intron_variant | MODIFIER |
| 12 | 91761407 | AX-170092570 | 0 | T | C | 0.079 | 1.07E-01 | 1.86E-01 | 2.73E+00 | 2.81E+00 | 5.59E-01 | no | Ston2 | intron_variant | MODIFIER |
| 12 | 91764671 | AX-170243368 | 0 | T | C | 0.079 | 1.07E-01 | 1.86E-01 | 2.73E+00 | 2.81E+00 | 5.59E-01 | no | Ston2 | intron_variant | MODIFIER |
| 1 | 23291299 | AX-168385336 | 0 | A | G | 0.439 | 4.83E-02 | 8.29E-02 | 2.62E+00 | 2.70E+00 | 5.59E-01 | no | Mir30c-2 | upstream_gene_variant | MODIFIER |
| 2 | 161838224 | AX-170514843 | 1 | C | T | 0.161 | 1.16E-01 | 2.01E-01 | 2.72E+00 | 2.79E+00 | 5.59E-01 | no | Ptprt | intron_variant | MODIFIER |
| 7 | 116510659 | AX-170078541 | 1 | G | A | 0.411 | 5.03E-02 | 8.70E-02 | 2.66E+00 | 2.74E+00 | 5.59E-01 | no | Nucb2 | intron_variant | MODIFIER |
| 12 | 92000918 | AX-170277971 | 0 | C | T | 0.079 | 1.05E-01 | 1.83E-01 | 2.66E+00 | 2.76E+00 | 5.59E-01 | no | | intergenic_variant | MODIFIER |
| 10 | 70620802 | AX-170497966 | 0 | G | A | 0.412 | -4.64E-02 | 7.93E-02 | 2.60E+00 | 2.68E+00 | 5.59E-01 | no | Phyhipl | intron_variant,non_coding_transcript_variant | MODIFIER |
| 7 | 111781854 | AX-170130693 | 0 | T | G | 0.474 | 5.21E-02 | 9.01E-02 | 2.68E+00 | 2.75E+00 | 5.59E-01 | no | Galnt18 | upstream_gene_variant | MODIFIER |
| 8 | 66429246 | AX-170429976 | 1 | C | T | 0.161 | -7.55E-02 | 1.31E-01 | 2.65E+00 | 2.75E+00 | 5.59E-01 | no | MAR1 | intron_variant | MODIFIER |
| 8 | 101960463 | AX-97838431 | 0 | C | T | 0.219 | -5.83E-02 | 1.02E-01 | 2.69E+00 | 2.80E+00 | 5.59E-01 | no | | intergenic_variant | MODIFIER |
| 8 | 102057567 | AX-170525619 | 0 | T | A | 0.219 | -5.83E-02 | 1.02E-01 | 2.69E+00 | 2.80E+00 | 5.59E-01 | no | | intergenic_variant | MODIFIER |
| 8 | 85260543 | AX-170532570 | 0 | A | G | 0.193 | -7.08E-02 | 1.23E-01 | 2.74E+00 | 2.83E+00 | 5.59E-01 | no | Vps35 | 3_prime_UTR_variant | MODIFIER |
| 5 | 110415217 | AX-170288445 | 1 | T | C | 0.446 | 6.44E-02 | 1.11E-01 | 2.64E+00 | 2.68E+00 | 5.59E-01 | no | Fbrsl1 | intron_variant | MODIFIER |
| 7 | 123759478 | AX-170029887 | 1 | A | G | 0.411 | 4.59E-02 | 7.98E-02 | 2.67E+00 | 2.77E+00 | 5.59E-01 | no | | intergenic_variant | MODIFIER |
| 1 | 45912375 | AX-170446585 | 0 | G | A | 0.096 | -1.21E-01 | 2.10E-01 | 2.71E+00 | 2.80E+00 | 5.59E-01 | no | Slc40a1 | synonymous_variant | LOW |
| 1 | 171006300 | AX-170106701 | 0 | C | T | 0.219 | 7.73E-02 | 1.33E-01 | 2.66E+00 | 2.71E+00 | 5.59E-01 | no | | intergenic_variant | MODIFIER |
| 5 | 36390573 | AX-170477584 | 0 | G | A | 0.325 | 6.48E-02 | 1.13E-01 | 2.71E+00 | 2.78E+00 | 5.59E-01 | no | Sorcs2 | intron_variant | MODIFIER |
| 5 | 113052127 | AX-170147882 | 0 | G | A | 0.447 | 6.37E-02 | 1.10E-01 | 2.67E+00 | 2.71E+00 | 5.60E-01 | no | | intergenic_variant | MODIFIER |
| 4 | 72965650 | AX-170197687 | 1 | A | C | 0.223 | -5.88E-02 | 1.01E-01 | 2.89E+00 | 3.02E+00 | 5.60E-01 | no | | intergenic_variant | MODIFIER |
| 5 | 105538129 | AX-170404219 | 0 | G | A | 0.447 | 6.45E-02 | 1.10E-01 | 2.60E+00 | 2.65E+00 | 5.60E-01 | no | Lrrc8c | intron_variant | MODIFIER |
| 5 | 105759819 | AX-170176119 | 0 | G | A | 0.447 | 6.45E-02 | 1.10E-01 | 2.60E+00 | 2.65E+00 | 5.60E-01 | no | Lrrc8d | intron_variant | MODIFIER |
| 3 | 31724175 | AX-170020244 | 0 | C | G | 0.158 | -5.84E-02 | 1.01E-01 | 2.69E+00 | 2.76E+00 | 5.60E-01 | no | | intergenic_variant | MODIFIER |
| 8 | 67538804 | AX-170279750 | 0 | T | G | 0.149 | -8.48E-02 | 1.44E-01 | 2.57E+00 | 2.64E+00 | 5.60E-01 | no | Nat3 | intron_variant | MODIFIER |
| 8 | 67541743 | AX-170143611 | 0 | T | G | 0.149 | -8.48E-02 | 1.44E-01 | 2.57E+00 | 2.64E+00 | 5.60E-01 | no | Nat3 | intron_variant | MODIFIER |
| 8 | 67544265 | AX-170257726 | 0 | C | T | 0.149 | -8.48E-02 | 1.44E-01 | 2.57E+00 | 2.64E+00 | 5.60E-01 | no | Nat3 | intron_variant | MODIFIER |
| 8 | 67546176 | AX-170246192 | 0 | T | C | 0.149 | -8.48E-02 | 1.44E-01 | 2.57E+00 | 2.64E+00 | 5.60E-01 | no | Nat3 | intron_variant | MODIFIER |
| 8 | 67546690 | AX-97856786 | 0 | C | T | 0.149 | -8.48E-02 | 1.44E-01 | 2.57E+00 | 2.64E+00 | 5.60E-01 | no | Nat3 | intron_variant | MODIFIER |
| 8 | 67547442 | AX-170429799 | 0 | C | T | 0.149 | -8.48E-02 | 1.44E-01 | 2.57E+00 | 2.64E+00 | 5.60E-01 | no | Nat3 | intron_variant | MODIFIER |
| 8 | 67554510 | AX-170201330 | 0 | T | C | 0.149 | -8.48E-02 | 1.44E-01 | 2.57E+00 | 2.64E+00 | 5.60E-01 | no | | intergenic_variant | MODIFIER |
| 8 | 67593025 | AX-97849352 | 0 | G | A | 0.149 | -8.48E-02 | 1.44E-01 | 2.57E+00 | 2.64E+00 | 5.60E-01 | no | | intergenic_variant | MODIFIER |
| 8 | 67633483 | AX-97790035 | 0 | A | G | 0.149 | -8.48E-02 | 1.44E-01 | 2.57E+00 | 2.64E+00 | 5.60E-01 | no | Gm44391 | upstream_gene_variant | MODIFIER |
| 8 | 67634337 | AX-170311868 | 0 | A | G | 0.149 | -8.48E-02 | 1.44E-01 | 2.57E+00 | 2.64E+00 | 5.60E-01 | no | | intergenic_variant | MODIFIER |
| 8 | 67694496 | AX-170272287 | 0 | C | T | 0.149 | -8.48E-02 | 1.44E-01 | 2.57E+00 | 2.64E+00 | 5.60E-01 | no | Psd3 | 3_prime_UTR_variant | MODIFIER |
| 8 | 67696996 | AX-170356449 | 0 | C | T | 0.149 | -8.48E-02 | 1.44E-01 | 2.57E+00 | 2.64E+00 | 5.60E-01 | no | Psd3 | synonymous_variant | LOW |
| 8 | 67697710 | AX-170533521 | 0 | C | T | 0.149 | -8.48E-02 | 1.44E-01 | 2.57E+00 | 2.64E+00 | 5.60E-01 | no | Psd3 | intron_variant | MODIFIER |
| 8 | 67716848 | AX-170247174 | 0 | G | C | 0.149 | -8.48E-02 | 1.44E-01 | 2.57E+00 | 2.64E+00 | 5.60E-01 | no | Psd3 | intron_variant | MODIFIER |
| 8 | 67719752 | AX-170148639 | 0 | A | C | 0.149 | -8.48E-02 | 1.44E-01 | 2.57E+00 | 2.64E+00 | 5.60E-01 | no | Psd3 | intron_variant | MODIFIER |
| 8 | 67719787 | AX-170156154 | 0 | A | G | 0.149 | -8.48E-02 | 1.44E-01 | 2.57E+00 | 2.64E+00 | 5.60E-01 | no | Psd3 | intron_variant | MODIFIER |
| 8 | 67732364 | AX-170534035 | 0 | A | G | 0.149 | -8.48E-02 | 1.44E-01 | 2.57E+00 | 2.64E+00 | 5.60E-01 | no | Psd3 | intron_variant | MODIFIER |
| 8 | 67888434 | AX-97803845 | 0 | A | G | 0.149 | -8.48E-02 | 1.44E-01 | 2.57E+00 | 2.64E+00 | 5.60E-01 | no | Psd3 | intron_variant | MODIFIER |
| 8 | 67934975 | AX-168413170 | 0 | G | A | 0.149 | -8.48E-02 | 1.44E-01 | 2.57E+00 | 2.64E+00 | 5.60E-01 | no | Psd3 | intron_variant | MODIFIER |
| 8 | 68010668 | AX-170274190 | 0 | T | C | 0.149 | -8.48E-02 | 1.44E-01 | 2.57E+00 | 2.64E+00 | 5.60E-01 | no | Gm22018 | downstream_gene_variant | MODIFIER |
| 8 | 68057313 | AX-97781225 | 0 | A | T | 0.149 | -8.48E-02 | 1.44E-01 | 2.57E+00 | 2.64E+00 | 5.60E-01 | no | Psd3 | intron_variant,non_coding_transcript_variant | MODIFIER |

| | | | | | | | | | | | | | | | |
|---|---|---|---|---|---|---|---|---|---|---|---|---|---|---|---|
| 8 | 68083386 | AX-170113249 | 0 | C | T | 0.149 | -8.48E-02 | 1.44E-01 | 2.57E+00 | 2.64E+00 | 5.60E-01 | no | Psd3 | intron_variant | MODIFIER |
| 8 | 68083563 | AX-170030959 | 0 | C | G | 0.149 | -8.48E-02 | 1.44E-01 | 2.57E+00 | 2.64E+00 | 5.60E-01 | no | Psd3 | intron_variant | MODIFIER |
| 8 | 68086219 | AX-170399950 | 0 | C | G | 0.149 | -8.48E-02 | 1.44E-01 | 2.57E+00 | 2.64E+00 | 5.60E-01 | no | Psd3 | intron_variant | MODIFIER |
| 8 | 68098522 | AX-170391516 | 0 | A | G | 0.149 | -8.48E-02 | 1.44E-01 | 2.57E+00 | 2.64E+00 | 5.60E-01 | no | Psd3 | downstream_gene_variant | MODIFIER |
| 8 | 68100107 | AX-170270294 | 0 | A | G | 0.149 | -8.48E-02 | 1.44E-01 | 2.57E+00 | 2.64E+00 | 5.60E-01 | no | Psd3 | intron_variant,non_coding_transcript_variant | MODIFIER |
| 8 | 68105670 | AX-170080318 | 0 | C | T | 0.149 | -8.48E-02 | 1.44E-01 | 2.57E+00 | 2.64E+00 | 5.60E-01 | no | Psd3 | intron_variant,non_coding_transcript_variant | MODIFIER |
| 8 | 68126448 | AX-97830881 | 0 | C | T | 0.149 | -8.48E-02 | 1.44E-01 | 2.57E+00 | 2.64E+00 | 5.60E-01 | no | Psd3 | upstream_gene_variant | MODIFIER |
| 8 | 68131614 | AX-170109338 | 0 | A | G | 0.149 | -8.48E-02 | 1.44E-01 | 2.57E+00 | 2.64E+00 | 5.60E-01 | no | Psd3 | intron_variant | MODIFIER |
| 8 | 68131783 | AX-170148071 | 0 | C | T | 0.149 | -8.48E-02 | 1.44E-01 | 2.57E+00 | 2.64E+00 | 5.60E-01 | no | Psd3 | intron_variant | MODIFIER |
| 8 | 68132642 | AX-170438736 | 0 | T | C | 0.149 | -8.48E-02 | 1.44E-01 | 2.57E+00 | 2.64E+00 | 5.60E-01 | no | Psd3 | intron_variant | MODIFIER |
| 8 | 68134163 | AX-97851781 | 0 | C | T | 0.149 | -8.48E-02 | 1.44E-01 | 2.57E+00 | 2.64E+00 | 5.60E-01 | no | Psd3 | intron_variant | MODIFIER |
| 8 | 68182051 | AX-170438976 | 0 | C | T | 0.149 | -8.48E-02 | 1.44E-01 | 2.57E+00 | 2.64E+00 | 5.60E-01 | no | Psd3 | intron_variant | MODIFIER |
| 8 | 68200819 | AX-170416740 | 0 | T | A | 0.149 | -8.48E-02 | 1.44E-01 | 2.57E+00 | 2.64E+00 | 5.60E-01 | no | Psd3 | intron_variant | MODIFIER |
| 8 | 68207471 | AX-170357667 | 0 | A | C | 0.149 | -8.48E-02 | 1.44E-01 | 2.57E+00 | 2.64E+00 | 5.60E-01 | no | Psd3 | upstream_gene_variant | MODIFIER |
| 8 | 68209135 | AX-170309214 | 0 | T | G | 0.149 | -8.48E-02 | 1.44E-01 | 2.57E+00 | 2.64E+00 | 5.60E-01 | no | Psd3 | upstream_gene_variant | MODIFIER |
| 8 | 68215037 | AX-170154405 | 0 | A | G | 0.149 | -8.48E-02 | 1.44E-01 | 2.57E+00 | 2.64E+00 | 5.60E-01 | no | Gm26659 | downstream_gene_variant | MODIFIER |
| 8 | 68220604 | AX-170114048 | 0 | T | A | 0.149 | -8.48E-02 | 1.44E-01 | 2.57E+00 | 2.64E+00 | 5.60E-01 | no | Gm26659 | non_coding_transcript_exon_variant | MODIFIER |
| 8 | 68230005 | AX-170350227 | 0 | C | T | 0.149 | -8.48E-02 | 1.44E-01 | 2.57E+00 | 2.64E+00 | 5.60E-01 | no | Gm26659 | upstream_gene_variant | MODIFIER |
| 8 | 68247273 | AX-170433480 | 0 | T | C | 0.149 | -8.48E-02 | 1.44E-01 | 2.57E+00 | 2.64E+00 | 5.60E-01 | no | | regulatory_region_variant | MODIFIER |
| 8 | 68247615 | AX-97807106 | 0 | A | G | 0.149 | -8.48E-02 | 1.44E-01 | 2.57E+00 | 2.64E+00 | 5.60E-01 | no | | regulatory_region_variant | MODIFIER |
| 8 | 68295586 | AX-170245468 | 0 | A | G | 0.149 | -8.48E-02 | 1.44E-01 | 2.57E+00 | 2.64E+00 | 5.60E-01 | no | Sh2d4a | intron_variant | MODIFIER |
| 8 | 68307584 | AX-170362781 | 0 | T | C | 0.149 | -8.48E-02 | 1.44E-01 | 2.57E+00 | 2.64E+00 | 5.60E-01 | no | Sh2d4a | intron_variant | MODIFIER |
| 9 | 117520206 | AX-170342234 | 1 | C | T | 0.125 | -5.56E-02 | 9.60E-02 | 2.60E+00 | 2.72E+00 | 5.60E-01 | no | Rbms3 | intron_variant | MODIFIER |
| 11 | 31861617 | AX-170447458 | 1 | A | G | 0.08 | 1.16E-01 | 2.01E-01 | 2.64E+00 | 2.72E+00 | 5.60E-01 | no | | regulatory_region_variant | MODIFIER |
| 9 | 104597883 | AX-97806525 | 0 | A | G | 0.123 | -6.58E-02 | 1.12E-01 | 2.57E+00 | 2.65E+00 | 5.60E-01 | no | Cpne4 | intron_variant | MODIFIER |
| 9 | 105827485 | AX-170262904 | 0 | A | G | 0.123 | -6.58E-02 | 1.12E-01 | 2.57E+00 | 2.65E+00 | 5.60E-01 | no | Col6a6 | intron_variant | MODIFIER |
| 2 | 171456426 | AX-170065637 | 1 | C | T | 0.071 | 6.42E-02 | 1.13E-01 | 2.63E+00 | 2.78E+00 | 5.60E-01 | no | | intergenic_variant | MODIFIER |
| 18 | 63162328 | AX-170061072 | 1 | A | T | 0.411 | 5.58E-02 | 9.10E-02 | 2.47E+00 | 2.54E+00 | 5.60E-01 | no | Piezo2 | intron_variant | MODIFIER |
| 1 | 100445249 | AX-170265268 | 1 | T | C | 0.161 | 8.12E-02 | 1.38E-01 | 2.56E+00 | 2.67E+00 | 5.60E-01 | no | Cntnap5b | intron_variant | MODIFIER |
| 12 | 111858294 | AX-170050218 | 0 | G | T | 0.14 | -9.46E-02 | 1.65E-01 | 2.67E+00 | 2.76E+00 | 5.60E-01 | no | Ppp1r13b | splice_region_variant,intron_variant | LOW |
| 1 | 176839836 | AX-170431713 | 0 | T | A | 0.211 | -7.40E-02 | 1.27E-01 | 2.90E+00 | 3.00E+00 | 5.60E-01 | no | Sdccag8 | intron_variant | MODIFIER |
| 7 | 138919733 | AX-170030098 | 0 | A | C | 0.298 | -4.32E-02 | 7.48E-02 | 2.62E+00 | 2.73E+00 | 5.60E-01 | no | Gm9358 | downstream_gene_variant | MODIFIER |
| 16 | 19249549 | AX-170339075 | 1 | G | A | 0.134 | 5.55E-02 | 9.75E-02 | 2.70E+00 | 2.83E+00 | 5.60E-01 | no | A530030E21Rik | downstream_gene_variant | MODIFIER |
| 18 | 66604610 | AX-170441145 | 1 | G | A | 0.286 | -5.73E-02 | 9.98E-02 | 2.71E+00 | 2.77E+00 | 5.60E-01 | no | AC100212.2 | downstream_gene_variant | MODIFIER |
| 7 | 122626881 | AX-170090015 | 2 | T | C | 0.391 | 4.76E-02 | 8.31E-02 | 2.74E+00 | 2.83E+00 | 5.60E-01 | no | Prkcb | intron_variant | MODIFIER |
| 1 | 90904828 | AX-170514705 | 1 | A | G | 0.152 | 1.05E-01 | 1.82E-01 | 2.66E+00 | 2.74E+00 | 5.61E-01 | no | | regulatory_region_variant | MODIFIER |
| 4 | 128417712 | AX-170400025 | 0 | G | A | 0.158 | 6.53E-02 | 1.12E-01 | 2.61E+00 | 2.67E+00 | 5.61E-01 | no | Csmd2 | intron_variant,non_coding_transcript_variant | MODIFIER |
| 15 | 74581818 | AX-97818894 | 0 | A | G | 0.544 | 4.74E-02 | 8.09E-02 | 2.58E+00 | 2.66E+00 | 5.61E-01 | no | Adgrb1 | intron_variant | MODIFIER |
| 7 | 36444172 | AX-170231598 | 0 | C | T | 0.351 | 5.41E-02 | 9.21E-02 | 2.56E+00 | 2.65E+00 | 5.61E-01 | no | | intergenic_variant | MODIFIER |
| 7 | 36564912 | AX-170469372 | 0 | T | A | 0.351 | 5.41E-02 | 9.21E-02 | 2.56E+00 | 2.65E+00 | 5.61E-01 | no | Gm29129 | upstream_gene_variant | MODIFIER |
| 3 | 149431654 | AX-170136145 | 1 | C | T | 0.339 | 4.25E-02 | 7.43E-02 | 2.70E+00 | 2.79E+00 | 5.61E-01 | no | Gm30382 | intron_variant,non_coding_transcript_variant | MODIFIER |
| 1 | 173451285 | AX-170316780 | 0 | C | T | 0.228 | -6.85E-02 | 1.18E-01 | 2.87E+00 | 2.97E+00 | 5.61E-01 | no | Aim2 | upstream_gene_variant | MODIFIER |
| 6 | 122858517 | AX-97830422 | 1 | G | A | 0.188 | -4.67E-02 | 8.21E-02 | 2.68E+00 | 2.81E+00 | 5.61E-01 | no | C3ar1 | upstream_gene_variant | MODIFIER |
| 6 | 126290418 | AX-170521351 | 1 | T | C | 0.188 | -4.67E-02 | 8.21E-02 | 2.68E+00 | 2.81E+00 | 5.61E-01 | no | Gm10415 | intron_variant,non_coding_transcript_variant | MODIFIER |
| 7 | 122704036 | AX-170029874 | 1 | G | C | 0.393 | 4.73E-02 | 8.28E-02 | 2.73E+00 | 2.82E+00 | 5.61E-01 | no | Cacng3 | intron_variant | MODIFIER |
| 10 | 107788825 | AX-170003863 | 1 | G | T | 0.321 | -4.77E-02 | 8.27E-02 | 2.64E+00 | 2.72E+00 | 5.61E-01 | no | Otogl | intron_variant | MODIFIER |
| 3 | 36700612 | AX-170066359 | 1 | A | T | 0.188 | 5.42E-02 | 9.50E-02 | 2.73E+00 | 2.82E+00 | 5.62E-01 | no | | intergenic_variant | MODIFIER |
| 5 | 6504698 | AX-170035701 | 0 | T | A | 0.439 | 3.62E-02 | 6.41E-02 | 2.70E+00 | 2.84E+00 | 5.62E-01 | no | | intergenic_variant | MODIFIER |
| 15 | 4951897 | AX-170226938 | 0 | A | G | 0.237 | -6.27E-02 | 1.08E-01 | 2.61E+00 | 2.66E+00 | 5.62E-01 | no | Mroh2b | intron_variant | MODIFIER |
| 7 | 75725219 | AX-170191001 | 1 | C | T | 0.348 | -5.17E-02 | 9.12E-02 | 2.71E+00 | 2.82E+00 | 5.62E-01 | no | Akap13 | intron_variant | MODIFIER |
| 7 | 78221969 | AX-170141227 | 1 | T | G | 0.348 | -5.17E-02 | 9.12E-02 | 2.71E+00 | 2.82E+00 | 5.62E-01 | no | Ntrk3 | intron_variant | MODIFIER |
| 3 | 32225670 | AX-170020253 | 0 | T | C | 0.167 | -5.57E-02 | 9.75E-02 | 2.69E+00 | 2.77E+00 | 5.62E-01 | no | | intergenic_variant | MODIFIER |
| 9 | 35310896 | AX-170345672 | 1 | T | G | 0.429 | -4.80E-02 | 8.41E-02 | 2.80E+00 | 2.89E+00 | 5.62E-01 | no | Gm33838 | upstream_gene_variant | MODIFIER |
| 7 | 16264750 | AX-170366915 | 2 | C | T | 0.373 | 4.98E-02 | 8.57E-02 | 2.57E+00 | 2.69E+00 | 5.62E-01 | no | | intergenic_variant | MODIFIER |

| | | | | | | | | | | | | | | | |
|---|---|---|---|---|---|---|---|---|---|---|---|---|---|---|---|
| 16 | 46014258 | AX-97792738 | 1 | G | A | 0.518 | 4.83E-02 | 8.20E-02 | 2.56E+00 | 2.60E+00 | 5.62E-01 | no | Plcxd2 | upstream_gene_variant | MODIFIER |
| 16 | 73980885 | AX-170211649 | 1 | G | A | 0.473 | 4.63E-02 | 7.54E-02 | 2.45E+00 | 2.53E+00 | 5.62E-01 | no | Robo2 | intron_variant | MODIFIER |
| 1 | 131753251 | AX-97788483 | 1 | T | C | 0.536 | 4.96E-02 | 8.54E-02 | 2.60E+00 | 2.67E+00 | 5.62E-01 | no | Slc26a9 | intron_variant | MODIFIER |
| 9 | 53947119 | AX-97830368 | 0 | C | T | 0.202 | -6.30E-02 | 1.10E-01 | 2.69E+00 | 2.77E+00 | 5.63E-01 | no | Elmod1 | intron_variant | MODIFIER |
| 11 | 12757745 | AX-97821108 | 0 | C | T | 0.035 | 6.93E-02 | 1.23E-01 | 2.58E+00 | 2.77E+00 | 5.63E-01 | no | | intergenic_variant | MODIFIER |
| 1 | 90221484 | AX-170220574 | 0 | A | T | 0.158 | 1.19E-01 | 2.07E-01 | 2.66E+00 | 2.73E+00 | 5.63E-01 | no | Ackr3 | downstream_gene_variant | MODIFIER |
| 1 | 78232446 | AX-170330402 | 0 | C | T | 0.114 | 1.22E-01 | 2.14E-01 | 2.65E+00 | 2.75E+00 | 5.63E-01 | no | | regulatory_region_variant | MODIFIER |
| 20 | 50730575 | AX-170517908 | 1 | G | C | 0.25 | 3.37E-02 | 5.79E-02 | 2.56E+00 | 2.68E+00 | 5.63E-01 | no | | intergenic_variant | MODIFIER |
| 16 | 45635450 | AX-97812507 | 0 | T | C | 0.518 | 4.73E-02 | 7.97E-02 | 2.53E+00 | 2.59E+00 | 5.63E-01 | no | BC016579 | intron_variant | MODIFIER |
| 16 | 45695792 | AX-170370226 | 0 | T | C | 0.518 | 4.73E-02 | 7.97E-02 | 2.53E+00 | 2.59E+00 | 5.63E-01 | no | Tmprss7 | upstream_gene_variant | MODIFIER |
| 14 | 106504442 | AX-97827713 | 1 | T | C | 0.054 | 8.95E-02 | 1.59E-01 | 2.65E+00 | 2.81E+00 | 5.63E-01 | no | | intergenic_variant | MODIFIER |
| 1 | 151878765 | AX-170156961 | 0 | C | T | 0.193 | 7.61E-02 | 1.32E-01 | 2.65E+00 | 2.71E+00 | 5.63E-01 | no | 1700025G04Rik | intron_variant,NMD_transcript_variant | MODIFIER |
| 5 | 124305948 | AX-170131573 | 1 | T | C | 0.464 | 5.93E-02 | 1.03E-01 | 2.64E+00 | 2.71E+00 | 5.63E-01 | no | Mphosph9 | intron_variant | MODIFIER |
| 18 | 83946682 | AX-170432018 | 0 | G | A | 0.158 | -6.62E-02 | 1.11E-01 | 2.51E+00 | 2.59E+00 | 5.63E-01 | no | | intergenic_variant | MODIFIER |
| 11 | 33213360 | AX-97827431 | 1 | G | T | 0.098 | 9.03E-02 | 1.56E-01 | 2.58E+00 | 2.69E+00 | 5.63E-01 | no | Ranbp17 | 3_prime_UTR_variant | MODIFIER |
| 7 | 93064255 | AX-170078081 | 0 | T | A | 0.14 | -8.78E-02 | 1.55E-01 | 2.67E+00 | 2.79E+00 | 5.63E-01 | no | Gm9934 | intron_variant,non_coding_transcript_variant | MODIFIER |
| 1 | 22830804 | AX-170176006 | 1 | A | G | 0.42 | 4.80E-02 | 8.30E-02 | 2.61E+00 | 2.67E+00 | 5.64E-01 | no | | intergenic_variant | MODIFIER |
| 8 | 97616912 | AX-170239075 | 0 | T | C | 0.219 | 6.72E-02 | 1.18E-01 | 2.71E+00 | 2.77E+00 | 5.64E-01 | no | | intergenic_variant | MODIFIER |
| 5 | 88212279 | AX-170513677 | 1 | G | A | 0.438 | 6.16E-02 | 1.08E-01 | 2.74E+00 | 2.79E+00 | 5.64E-01 | no | Gm42164 | downstream_gene_variant | MODIFIER |
| 8 | 18665266 | AX-170174037 | 0 | C | T | 0.175 | -5.35E-02 | 9.49E-02 | 2.70E+00 | 2.83E+00 | 5.64E-01 | no | Mcph1 | intron_variant | MODIFIER |
| 8 | 21794745 | AX-170535531 | 0 | A | G | 0.175 | -5.35E-02 | 9.49E-02 | 2.70E+00 | 2.83E+00 | 5.64E-01 | no | Defb1 | 3_prime_UTR_variant | MODIFIER |
| 15 | 3292173 | AX-97825717 | 2 | C | T | 0.236 | -6.02E-02 | 1.04E-01 | 2.59E+00 | 2.66E+00 | 5.64E-01 | no | Ccdc152 | intron_variant | MODIFIER |
| 5 | 42061853 | AX-170342165 | 1 | G | T | 0.205 | -6.20E-02 | 1.07E-01 | 2.58E+00 | 2.67E+00 | 5.64E-01 | no | | intergenic_variant | MODIFIER |
| 16 | 93052819 | AX-170215367 | 0 | C | G | 0.386 | -4.50E-02 | 7.70E-02 | 2.55E+00 | 2.64E+00 | 5.64E-01 | no | | intergenic_variant | MODIFIER |
| 17 | 27336742 | AX-170514436 | 1 | T | C | 0.054 | -8.20E-02 | 1.44E-01 | 2.64E+00 | 2.76E+00 | 5.64E-01 | no | Gm10505 | intron_variant,non_coding_transcript_variant | MODIFIER |
| 8 | 74445654 | AX-170031015 | 0 | C | G | 0.193 | -7.68E-02 | 1.35E-01 | 2.68E+00 | 2.74E+00 | 5.64E-01 | no | | intergenic_variant | MODIFIER |
| 11 | 74825133 | AX-97818946 | 0 | G | T | 0.228 | 4.21E-02 | 7.48E-02 | 2.67E+00 | 2.80E+00 | 5.64E-01 | no | Mettl16 | 3_prime_UTR_variant | MODIFIER |
| 8 | 60842887 | AX-170296670 | 0 | C | T | 0.149 | -8.02E-02 | 1.33E-01 | 2.47E+00 | 2.56E+00 | 5.64E-01 | no | Gm45485 | upstream_gene_variant | MODIFIER |
| 18 | 66376237 | AX-170015884 | 0 | T | C | 0.289 | -5.30E-02 | 9.31E-02 | 2.68E+00 | 2.75E+00 | 5.64E-01 | no | | intergenic_variant | MODIFIER |
| 1 | 43202705 | AX-170396477 | 0 | G | A | 0.096 | -1.20E-01 | 2.11E-01 | 2.71E+00 | 2.79E+00 | 5.64E-01 | no | | regulatory_region_variant | MODIFIER |
| 16 | 72695144 | AX-170115635 | 0 | C | T | 0.491 | 4.47E-02 | 7.65E-02 | 2.55E+00 | 2.63E+00 | 5.65E-01 | no | Robo1 | intron_variant | MODIFIER |
| 1 | 35558389 | AX-170520974 | 1 | A | G | 0.446 | 4.46E-02 | 7.85E-02 | 2.79E+00 | 2.87E+00 | 5.65E-01 | no | | intergenic_variant | MODIFIER |
| 6 | 138491472 | AX-170206326 | 2 | T | C | 0.564 | 4.89E-02 | 8.29E-02 | 2.53E+00 | 2.62E+00 | 5.65E-01 | no | Lmo3 | intron_variant | MODIFIER |
| 10 | 88500625 | AX-170228596 | 0 | G | A | 0.053 | -1.02E-01 | 1.80E-01 | 2.66E+00 | 2.77E+00 | 5.65E-01 | no | Chpt1 | intron_variant | MODIFIER |
| 8 | 34388394 | AX-170143837 | 1 | G | T | 0.152 | -7.08E-02 | 1.23E-01 | 2.63E+00 | 2.70E+00 | 5.65E-01 | no | Gm33831 | upstream_gene_variant | MODIFIER |
| 8 | 100503914 | AX-170255528 | 0 | C | T | 0.237 | 5.51E-02 | 9.72E-02 | 2.64E+00 | 2.76E+00 | 5.65E-01 | no | | intergenic_variant | MODIFIER |
| 18 | 80816591 | AX-170420312 | 0 | A | C | 0.158 | -6.28E-02 | 1.08E-01 | 2.57E+00 | 2.64E+00 | 5.65E-01 | no | Atp9b | intron_variant | MODIFIER |
| 7 | 73845499 | AX-170174091 | 0 | G | A | 0.351 | -5.22E-02 | 9.24E-02 | 2.67E+00 | 2.78E+00 | 5.66E-01 | no | | intergenic_variant | MODIFIER |
| 18 | 51575213 | AX-170149171 | 0 | A | G | 0.193 | -6.62E-02 | 1.14E-01 | 2.58E+00 | 2.63E+00 | 5.66E-01 | no | | intergenic_variant | MODIFIER |
| 18 | 52142849 | AX-170521672 | 0 | A | C | 0.193 | -6.62E-02 | 1.14E-01 | 2.58E+00 | 2.63E+00 | 5.66E-01 | no | | intergenic_variant | MODIFIER |
| 18 | 52151868 | AX-170388094 | 0 | T | C | 0.193 | -6.62E-02 | 1.14E-01 | 2.58E+00 | 2.63E+00 | 5.66E-01 | no | Gm22259 | downstream_gene_variant | MODIFIER |
| 18 | 52152085 | AX-170254145 | 0 | A | G | 0.193 | -6.62E-02 | 1.14E-01 | 2.58E+00 | 2.63E+00 | 5.66E-01 | no | Gm22259 | downstream_gene_variant | MODIFIER |
| 18 | 52166050 | AX-170494933 | 0 | T | C | 0.193 | -6.62E-02 | 1.14E-01 | 2.58E+00 | 2.63E+00 | 5.66E-01 | no | | intergenic_variant | MODIFIER |
| 18 | 52173479 | AX-170162660 | 0 | A | G | 0.193 | -6.62E-02 | 1.14E-01 | 2.58E+00 | 2.63E+00 | 5.66E-01 | no | | intergenic_variant | MODIFIER |
| 18 | 52346064 | AX-170143869 | 0 | A | C | 0.193 | -6.62E-02 | 1.14E-01 | 2.58E+00 | 2.63E+00 | 5.66E-01 | no | | intergenic_variant | MODIFIER |
| 18 | 52346462 | AX-170293490 | 0 | T | C | 0.193 | -6.62E-02 | 1.14E-01 | 2.58E+00 | 2.63E+00 | 5.66E-01 | no | | intergenic_variant | MODIFIER |
| 18 | 52427426 | AX-170015707 | 0 | C | T | 0.193 | -6.62E-02 | 1.14E-01 | 2.58E+00 | 2.63E+00 | 5.66E-01 | no | | intergenic_variant | MODIFIER |
| 18 | 52458057 | AX-97843053 | 0 | T | C | 0.193 | -6.62E-02 | 1.14E-01 | 2.58E+00 | 2.63E+00 | 5.66E-01 | no | | regulatory_region_variant | MODIFIER |
| 18 | 52472971 | AX-170015705 | 0 | T | G | 0.193 | -6.62E-02 | 1.14E-01 | 2.58E+00 | 2.63E+00 | 5.66E-01 | no | Srfbp1 | intron_variant | MODIFIER |
| 18 | 52480612 | AX-97844981 | 0 | C | T | 0.193 | -6.62E-02 | 1.14E-01 | 2.58E+00 | 2.63E+00 | 5.66E-01 | no | Srfbp1 | intron_variant | MODIFIER |
| 18 | 52489328 | AX-170254307 | 0 | A | C | 0.193 | -6.62E-02 | 1.14E-01 | 2.58E+00 | 2.63E+00 | 5.66E-01 | no | Srfbp1 | intron_variant | MODIFIER |
| 18 | 52515561 | AX-170015709 | 0 | T | C | 0.193 | -6.62E-02 | 1.14E-01 | 2.58E+00 | 2.63E+00 | 5.66E-01 | no | Lox | downstream_gene_variant | MODIFIER |
| 18 | 52522058 | AX-170015704 | 0 | G | A | 0.193 | -6.62E-02 | 1.14E-01 | 2.58E+00 | 2.63E+00 | 5.66E-01 | no | Lox | intron_variant | MODIFIER |
| 18 | 52522090 | AX-170329353 | 0 | C | T | 0.193 | -6.62E-02 | 1.14E-01 | 2.58E+00 | 2.63E+00 | 5.66E-01 | no | Lox | intron_variant | MODIFIER |

| | | | | | | | | | | | | | | | |
|---|---|---|---|---|---|---|---|---|---|---|---|---|---|---|---|
| 18 | 52690307 | AX-170513556 | 0 | C | G | 0.193 | -6.62E-02 | 1.14E-01 | 2.58E+00 | 2.63E+00 | 5.66E-01 | no | Gykl1 | upstream_gene_variant | MODIFIER |
| 18 | 52727653 | AX-170405396 | 0 | A | G | 0.193 | -6.62E-02 | 1.14E-01 | 2.58E+00 | 2.63E+00 | 5.66E-01 | no | | intergenic_variant | MODIFIER |
| 18 | 52766301 | AX-97846142 | 0 | T | C | 0.193 | -6.62E-02 | 1.14E-01 | 2.58E+00 | 2.63E+00 | 5.66E-01 | no | Sncaip | upstream_gene_variant | MODIFIER |
| 18 | 52772548 | AX-170178052 | 0 | C | A | 0.193 | -6.62E-02 | 1.14E-01 | 2.58E+00 | 2.63E+00 | 5.66E-01 | no | Sncaip | intron_variant | MODIFIER |
| 18 | 52776035 | AX-170117256 | 0 | C | T | 0.193 | -6.62E-02 | 1.14E-01 | 2.58E+00 | 2.63E+00 | 5.66E-01 | no | Sncaip | intron_variant | MODIFIER |
| 18 | 52816544 | AX-97834795 | 0 | T | A | 0.193 | -6.62E-02 | 1.14E-01 | 2.58E+00 | 2.63E+00 | 5.66E-01 | no | Sncaip | intron_variant | MODIFIER |
| 18 | 52833315 | AX-170527391 | 0 | T | G | 0.193 | -6.62E-02 | 1.14E-01 | 2.58E+00 | 2.63E+00 | 5.66E-01 | no | Sncaip | intron_variant | MODIFIER |
| 18 | 53363694 | AX-170514591 | 0 | A | G | 0.193 | -6.62E-02 | 1.14E-01 | 2.58E+00 | 2.63E+00 | 5.66E-01 | no | Snx24 | intron_variant | MODIFIER |
| 18 | 53370155 | AX-170380862 | 0 | A | G | 0.193 | -6.62E-02 | 1.14E-01 | 2.58E+00 | 2.63E+00 | 5.66E-01 | no | Snx24 | intron_variant | MODIFIER |
| 18 | 53371117 | AX-170150519 | 0 | A | G | 0.193 | -6.62E-02 | 1.14E-01 | 2.58E+00 | 2.63E+00 | 5.66E-01 | no | Snx24 | intron_variant | MODIFIER |
| 18 | 53389929 | AX-170508107 | 0 | T | C | 0.193 | -6.62E-02 | 1.14E-01 | 2.58E+00 | 2.63E+00 | 5.66E-01 | no | Snx24 | intron_variant | MODIFIER |
| 18 | 53484665 | AX-170291269 | 0 | A | C | 0.193 | -6.62E-02 | 1.14E-01 | 2.58E+00 | 2.63E+00 | 5.66E-01 | no | Prdm6 | intron_variant | MODIFIER |
| 18 | 53484812 | AX-170483981 | 0 | T | C | 0.193 | -6.62E-02 | 1.14E-01 | 2.58E+00 | 2.63E+00 | 5.66E-01 | no | Prdm6 | intron_variant | MODIFIER |
| 18 | 53485059 | AX-170216070 | 0 | A | C | 0.193 | -6.62E-02 | 1.14E-01 | 2.58E+00 | 2.63E+00 | 5.66E-01 | no | Prdm6 | intron_variant | MODIFIER |
| 18 | 53581243 | AX-170343631 | 0 | T | C | 0.193 | -6.62E-02 | 1.14E-01 | 2.58E+00 | 2.63E+00 | 5.66E-01 | no | | intergenic_variant | MODIFIER |
| 18 | 53657366 | AX-170278901 | 0 | T | C | 0.193 | -6.62E-02 | 1.14E-01 | 2.58E+00 | 2.63E+00 | 5.66E-01 | no | AC116597.1 | intron_variant,non_coding_transcript_variant | MODIFIER |
| 18 | 53694397 | AX-170318340 | 0 | T | C | 0.193 | -6.62E-02 | 1.14E-01 | 2.58E+00 | 2.63E+00 | 5.66E-01 | no | Cep120 | intron_variant | MODIFIER |
| 18 | 53713296 | AX-170307953 | 0 | G | C | 0.193 | -6.62E-02 | 1.14E-01 | 2.58E+00 | 2.63E+00 | 5.66E-01 | no | Cep120 | intron_variant | MODIFIER |
| 18 | 53750302 | AX-170376042 | 0 | A | G | 0.193 | -6.62E-02 | 1.14E-01 | 2.58E+00 | 2.63E+00 | 5.66E-01 | no | | intergenic_variant | MODIFIER |
| 18 | 55789960 | AX-170015751 | 0 | C | G | 0.193 | -6.62E-02 | 1.14E-01 | 2.58E+00 | 2.63E+00 | 5.66E-01 | no | | intergenic_variant | MODIFIER |
| 18 | 55794484 | AX-170015752 | 0 | G | C | 0.193 | -6.62E-02 | 1.14E-01 | 2.58E+00 | 2.63E+00 | 5.66E-01 | no | | intergenic_variant | MODIFIER |
| 18 | 55820749 | AX-170508805 | 0 | T | C | 0.193 | -6.62E-02 | 1.14E-01 | 2.58E+00 | 2.63E+00 | 5.66E-01 | no | | intergenic_variant | MODIFIER |
| 18 | 55826440 | AX-97837058 | 0 | A | G | 0.193 | -6.62E-02 | 1.14E-01 | 2.58E+00 | 2.63E+00 | 5.66E-01 | no | | intergenic_variant | MODIFIER |
| 18 | 55826785 | AX-170278981 | 0 | T | A | 0.193 | -6.62E-02 | 1.14E-01 | 2.58E+00 | 2.63E+00 | 5.66E-01 | no | | intergenic_variant | MODIFIER |
| 18 | 55827493 | AX-170376477 | 0 | C | T | 0.193 | -6.62E-02 | 1.14E-01 | 2.58E+00 | 2.63E+00 | 5.66E-01 | no | | intergenic_variant | MODIFIER |
| 18 | 55833279 | AX-170309416 | 0 | A | G | 0.193 | -6.62E-02 | 1.14E-01 | 2.58E+00 | 2.63E+00 | 5.66E-01 | no | Gm8614 | downstream_gene_variant | MODIFIER |
| 18 | 55837901 | AX-170116212 | 0 | G | A | 0.193 | -6.62E-02 | 1.14E-01 | 2.58E+00 | 2.63E+00 | 5.66E-01 | no | Gm8614 | downstream_gene_variant | MODIFIER |
| 18 | 55849267 | AX-170419814 | 0 | C | T | 0.193 | -6.62E-02 | 1.14E-01 | 2.58E+00 | 2.63E+00 | 5.66E-01 | no | | intergenic_variant | MODIFIER |
| 18 | 56370127 | AX-170321940 | 0 | C | T | 0.193 | -6.62E-02 | 1.14E-01 | 2.58E+00 | 2.63E+00 | 5.66E-01 | no | | intergenic_variant | MODIFIER |
| 18 | 56509966 | AX-170318054 | 0 | C | A | 0.193 | -6.62E-02 | 1.14E-01 | 2.58E+00 | 2.63E+00 | 5.66E-01 | no | Aldh7a1 | non_coding_transcript_exon_variant | MODIFIER |
| 18 | 56534206 | AX-170106011 | 0 | A | G | 0.193 | -6.62E-02 | 1.14E-01 | 2.58E+00 | 2.63E+00 | 5.66E-01 | no | Aldh7a1 | synonymous_variant | LOW |
| 18 | 56538775 | AX-170450160 | 0 | C | T | 0.193 | -6.62E-02 | 1.14E-01 | 2.58E+00 | 2.63E+00 | 5.66E-01 | no | Aldh7a1 | intron_variant | MODIFIER |
| 18 | 56538866 | AX-170159853 | 0 | C | T | 0.193 | -6.62E-02 | 1.14E-01 | 2.58E+00 | 2.63E+00 | 5.66E-01 | no | Aldh7a1 | intron_variant | MODIFIER |
| 18 | 56544636 | AX-170310869 | 0 | A | C | 0.193 | -6.62E-02 | 1.14E-01 | 2.58E+00 | 2.63E+00 | 5.66E-01 | no | Aldh7a1 | intron_variant | MODIFIER |
| 18 | 56546219 | AX-170355423 | 0 | G | A | 0.193 | -6.62E-02 | 1.14E-01 | 2.58E+00 | 2.63E+00 | 5.66E-01 | no | Aldh7a1 | intron_variant | MODIFIER |
| 18 | 56554825 | AX-170501896 | 0 | A | G | 0.193 | -6.62E-02 | 1.14E-01 | 2.58E+00 | 2.63E+00 | 5.66E-01 | no | Aldh7a1 | intron_variant | MODIFIER |
| 18 | 56570839 | AX-170497383 | 0 | T | C | 0.193 | -6.62E-02 | 1.14E-01 | 2.58E+00 | 2.63E+00 | 5.66E-01 | no | Phax | upstream_gene_variant | MODIFIER |
| 18 | 56575308 | AX-170383432 | 0 | A | C | 0.193 | -6.62E-02 | 1.14E-01 | 2.58E+00 | 2.63E+00 | 5.66E-01 | no | Phax | intron_variant | MODIFIER |
| 18 | 56737246 | AX-170327632 | 0 | G | A | 0.193 | -6.62E-02 | 1.14E-01 | 2.58E+00 | 2.63E+00 | 5.66E-01 | no | Lmnb1 | intron_variant | MODIFIER |
| 18 | 56948156 | AX-170396939 | 0 | T | G | 0.193 | -6.62E-02 | 1.14E-01 | 2.58E+00 | 2.63E+00 | 5.66E-01 | no | | intergenic_variant | MODIFIER |
| 18 | 56958633 | AX-170404485 | 0 | T | C | 0.193 | -6.62E-02 | 1.14E-01 | 2.58E+00 | 2.63E+00 | 5.66E-01 | no | C330018D20Rik | intron_variant | MODIFIER |
| 18 | 57019347 | AX-170211616 | 0 | G | A | 0.193 | -6.62E-02 | 1.14E-01 | 2.58E+00 | 2.63E+00 | 5.66E-01 | no | | intergenic_variant | MODIFIER |
| 18 | 57027428 | AX-170112929 | 0 | A | T | 0.193 | -6.62E-02 | 1.14E-01 | 2.58E+00 | 2.63E+00 | 5.66E-01 | no | | intergenic_variant | MODIFIER |
| 18 | 57027905 | AX-97798383 | 0 | A | C | 0.193 | -6.62E-02 | 1.14E-01 | 2.58E+00 | 2.63E+00 | 5.66E-01 | no | | intergenic_variant | MODIFIER |
| 18 | 57160442 | AX-97841197 | 0 | T | C | 0.193 | -6.62E-02 | 1.14E-01 | 2.58E+00 | 2.63E+00 | 5.66E-01 | no | Megf10 | intron_variant | MODIFIER |
| 18 | 57221563 | AX-97825045 | 0 | A | C | 0.193 | -6.62E-02 | 1.14E-01 | 2.58E+00 | 2.63E+00 | 5.66E-01 | no | Megf10 | intron_variant | MODIFIER |
| 18 | 57239113 | AX-170463886 | 0 | A | T | 0.193 | -6.62E-02 | 1.14E-01 | 2.58E+00 | 2.63E+00 | 5.66E-01 | no | Megf10 | intron_variant | MODIFIER |
| 18 | 57245153 | AX-170146615 | 0 | A | C | 0.193 | -6.62E-02 | 1.14E-01 | 2.58E+00 | 2.63E+00 | 5.66E-01 | no | Megf10 | intron_variant | MODIFIER |
| 18 | 57249304 | AX-170060934 | 0 | A | G | 0.193 | -6.62E-02 | 1.14E-01 | 2.58E+00 | 2.63E+00 | 5.66E-01 | no | Megf10 | intron_variant | MODIFIER |
| 5 | 28003141 | AX-170360351 | 1 | T | C | 0.339 | 5.74E-02 | 1.01E-01 | 2.70E+00 | 2.76E+00 | 5.66E-01 | no | Gm4865 | upstream_gene_variant | MODIFIER |
| 8 | 32087217 | AX-170110466 | 2 | C | T | 0.145 | -6.86E-02 | 1.20E-01 | 2.62E+00 | 2.72E+00 | 5.66E-01 | no | Nrg1 | intron_variant | MODIFIER |
| 4 | 136561443 | AX-170175278 | 0 | A | C | 0.158 | 6.48E-02 | 1.09E-01 | 2.50E+00 | 2.58E+00 | 5.66E-01 | no | Kdm1a | intron_variant | MODIFIER |
| 12 | 73102265 | AX-170475092 | 0 | T | C | 0.061 | -1.28E-01 | 2.23E-01 | 2.59E+00 | 2.69E+00 | 5.66E-01 | no | Six4 | 3_prime_UTR_variant | MODIFIER |
| 7 | 73157137 | AX-170355169 | 0 | C | T | 0.36 | -5.10E-02 | 9.03E-02 | 2.66E+00 | 2.77E+00 | 5.66E-01 | no | | intergenic_variant | MODIFIER |

| | | | | | | | | | | | | | | | |
|---|---|---|---|---|---|---|---|---|---|---|---|---|---|---|---|
| 5 | 28405141 | AX-97855518 | 1 | A | G | 0.348 | 5.86E-02 | 1.04E-01 | 2.72E+00 | 2.79E+00 | 5.66E-01 | no | Rbm33 | intron_variant | MODIFIER |
| 1 | 105273045 | AX-170377647 | 1 | C | G | 0.161 | 1.18E-01 | 2.07E-01 | 2.63E+00 | 2.69E+00 | 5.66E-01 | no | Rnf152 | downstream_gene_variant | MODIFIER |
| 7 | 37276112 | AX-170473294 | 0 | T | G | 0.342 | 5.26E-02 | 9.16E-02 | 2.59E+00 | 2.67E+00 | 5.66E-01 | no | | intergenic_variant | MODIFIER |
| 18 | 85546549 | AX-170093984 | 0 | G | A | 0.149 | -6.44E-02 | 1.09E-01 | 2.52E+00 | 2.61E+00 | 5.66E-01 | no | | intergenic_variant | MODIFIER |
| 13 | 53819512 | AX-97791710 | 1 | G | A | 0.518 | 3.97E-02 | 6.80E-02 | 2.53E+00 | 2.64E+00 | 5.67E-01 | no | | intergenic_variant | MODIFIER |
| 1 | 172377316 | AX-97811857 | 0 | A | T | 0.228 | 7.22E-02 | 1.23E-01 | 2.54E+00 | 2.60E+00 | 5.67E-01 | no | Pigm | missense_variant | MODERATE |
| 1 | 172414490 | AX-170332927 | 0 | C | T | 0.228 | 7.22E-02 | 1.23E-01 | 2.54E+00 | 2.60E+00 | 5.67E-01 | no | | intergenic_variant | MODIFIER |
| 1 | 172443527 | AX-170533868 | 0 | T | C | 0.228 | 7.22E-02 | 1.23E-01 | 2.54E+00 | 2.60E+00 | 5.67E-01 | no | | intergenic_variant | MODIFIER |
| 1 | 172444585 | AX-170449821 | 0 | T | C | 0.228 | 7.22E-02 | 1.23E-01 | 2.54E+00 | 2.60E+00 | 5.67E-01 | no | | intergenic_variant | MODIFIER |
| 1 | 172448250 | AX-170274472 | 0 | C | T | 0.228 | 7.22E-02 | 1.23E-01 | 2.54E+00 | 2.60E+00 | 5.67E-01 | no | | intergenic_variant | MODIFIER |
| 1 | 172482778 | AX-170126213 | 0 | A | G | 0.228 | 7.22E-02 | 1.23E-01 | 2.54E+00 | 2.60E+00 | 5.67E-01 | no | Slamf9 | downstream_gene_variant | MODIFIER |
| 1 | 172487253 | AX-170301345 | 0 | A | G | 0.228 | 7.22E-02 | 1.23E-01 | 2.54E+00 | 2.60E+00 | 5.67E-01 | no | Igsf9 | intron_variant | MODIFIER |
| 1 | 172489036 | AX-170043599 | 0 | C | A | 0.228 | 7.22E-02 | 1.23E-01 | 2.54E+00 | 2.60E+00 | 5.67E-01 | no | Igsf9 | intron_variant | MODIFIER |
| 1 | 172489471 | AX-170043601 | 0 | C | T | 0.228 | 7.22E-02 | 1.23E-01 | 2.54E+00 | 2.60E+00 | 5.67E-01 | no | Igsf9 | intron_variant | MODIFIER |
| 1 | 172490225 | AX-170043598 | 0 | T | C | 0.228 | 7.22E-02 | 1.23E-01 | 2.54E+00 | 2.60E+00 | 5.67E-01 | no | Igsf9 | intron_variant | MODIFIER |
| 1 | 172493185 | AX-170091788 | 0 | G | T | 0.228 | 7.22E-02 | 1.23E-01 | 2.54E+00 | 2.60E+00 | 5.67E-01 | no | Igsf9 | intron_variant | MODIFIER |
| 1 | 172493461 | AX-170260452 | 0 | T | G | 0.228 | 7.22E-02 | 1.23E-01 | 2.54E+00 | 2.60E+00 | 5.67E-01 | no | Igsf9 | intron_variant | MODIFIER |
| 1 | 172495844 | AX-170181872 | 0 | C | G | 0.228 | 7.22E-02 | 1.23E-01 | 2.54E+00 | 2.60E+00 | 5.67E-01 | no | Igsf9 | intron_variant | MODIFIER |
| 1 | 172508661 | AX-170343462 | 0 | G | C | 0.228 | 7.22E-02 | 1.23E-01 | 2.54E+00 | 2.60E+00 | 5.67E-01 | no | Tagln2 | downstream_gene_variant | MODIFIER |
| 1 | 172511944 | AX-170487450 | 0 | C | G | 0.228 | 7.22E-02 | 1.23E-01 | 2.54E+00 | 2.60E+00 | 5.67E-01 | no | Tagln2 | downstream_gene_variant | MODIFIER |
| 1 | 172530124 | AX-170514212 | 0 | T | C | 0.228 | 7.22E-02 | 1.23E-01 | 2.54E+00 | 2.60E+00 | 5.67E-01 | no | Cfap45 | intron_variant | MODIFIER |
| 1 | 172538538 | AX-170103700 | 0 | T | C | 0.228 | 7.22E-02 | 1.23E-01 | 2.54E+00 | 2.60E+00 | 5.67E-01 | no | Cfap45 | synonymous_variant | LOW |
| 1 | 172538781 | AX-170303940 | 0 | C | T | 0.228 | 7.22E-02 | 1.23E-01 | 2.54E+00 | 2.60E+00 | 5.67E-01 | no | Cfap45 | intron_variant | MODIFIER |
| 1 | 172576522 | AX-170143220 | 0 | G | C | 0.228 | 7.22E-02 | 1.23E-01 | 2.54E+00 | 2.60E+00 | 5.67E-01 | no | | intergenic_variant | MODIFIER |
| 1 | 172576873 | AX-97810930 | 0 | C | T | 0.228 | 7.22E-02 | 1.23E-01 | 2.54E+00 | 2.60E+00 | 5.67E-01 | no | Slamf8 | downstream_gene_variant | MODIFIER |
| 1 | 172580596 | AX-170321138 | 0 | C | G | 0.228 | 7.22E-02 | 1.23E-01 | 2.54E+00 | 2.60E+00 | 5.67E-01 | no | Slamf8 | downstream_gene_variant | MODIFIER |
| 1 | 172580781 | AX-170433988 | 0 | A | G | 0.228 | 7.22E-02 | 1.23E-01 | 2.54E+00 | 2.60E+00 | 5.67E-01 | no | Slamf8 | downstream_gene_variant | MODIFIER |
| 1 | 172611708 | AX-170043602 | 0 | T | C | 0.228 | 7.22E-02 | 1.23E-01 | 2.54E+00 | 2.60E+00 | 5.67E-01 | no | | intergenic_variant | MODIFIER |
| 1 | 172623365 | AX-170097423 | 0 | G | A | 0.228 | 7.22E-02 | 1.23E-01 | 2.54E+00 | 2.60E+00 | 5.67E-01 | no | | intergenic_variant | MODIFIER |
| 1 | 172624575 | AX-170338600 | 0 | G | A | 0.228 | 7.22E-02 | 1.23E-01 | 2.54E+00 | 2.60E+00 | 5.67E-01 | no | | intergenic_variant | MODIFIER |
| 1 | 172637826 | AX-97845459 | 0 | T | C | 0.228 | 7.22E-02 | 1.23E-01 | 2.54E+00 | 2.60E+00 | 5.67E-01 | no | Dusp23 | upstream_gene_variant | MODIFIER |
| 1 | 172658134 | AX-170473855 | 0 | C | G | 0.228 | 7.22E-02 | 1.23E-01 | 2.54E+00 | 2.60E+00 | 5.67E-01 | no | | intergenic_variant | MODIFIER |
| 1 | 172663897 | AX-97815050 | 0 | C | T | 0.228 | 7.22E-02 | 1.23E-01 | 2.54E+00 | 2.60E+00 | 5.67E-01 | no | | intergenic_variant | MODIFIER |
| 1 | 172680721 | AX-97783800 | 0 | A | C | 0.228 | 7.22E-02 | 1.23E-01 | 2.54E+00 | 2.60E+00 | 5.67E-01 | no | Gm22383 | downstream_gene_variant | MODIFIER |
| 1 | 172681389 | AX-170268405 | 0 | T | A | 0.228 | 7.22E-02 | 1.23E-01 | 2.54E+00 | 2.60E+00 | 5.67E-01 | no | Gm22383 | downstream_gene_variant | MODIFIER |
| 1 | 172687914 | AX-170129093 | 0 | G | A | 0.228 | 7.22E-02 | 1.23E-01 | 2.54E+00 | 2.60E+00 | 5.67E-01 | no | Gm22383 | upstream_gene_variant | MODIFIER |
| 1 | 172706981 | AX-170155781 | 0 | T | G | 0.228 | 7.22E-02 | 1.23E-01 | 2.54E+00 | 2.60E+00 | 5.67E-01 | no | Crp | intron_variant,non_coding_transcript_variant | MODIFIER |
| 1 | 172707310 | AX-97787627 | 0 | A | C | 0.228 | 7.22E-02 | 1.23E-01 | 2.54E+00 | 2.60E+00 | 5.67E-01 | no | Crp | intron_variant,non_coding_transcript_variant | MODIFIER |
| 1 | 172738269 | AX-97824627 | 0 | G | A | 0.228 | 7.22E-02 | 1.23E-01 | 2.54E+00 | 2.60E+00 | 5.67E-01 | no | Crp | intron_variant,non_coding_transcript_variant | MODIFIER |
| 1 | 172739227 | AX-170470571 | 0 | T | C | 0.228 | 7.22E-02 | 1.23E-01 | 2.54E+00 | 2.60E+00 | 5.67E-01 | no | Crp | intron_variant,non_coding_transcript_variant | MODIFIER |
| 1 | 172741762 | AX-170277610 | 0 | A | G | 0.228 | 7.22E-02 | 1.23E-01 | 2.54E+00 | 2.60E+00 | 5.67E-01 | no | Crp | intron_variant,non_coding_transcript_variant | MODIFIER |
| 1 | 172742334 | AX-170152552 | 0 | C | A | 0.228 | 7.22E-02 | 1.23E-01 | 2.54E+00 | 2.60E+00 | 5.67E-01 | no | Crp | intron_variant,non_coding_transcript_variant | MODIFIER |
| 1 | 172742472 | AX-170316313 | 0 | A | G | 0.228 | 7.22E-02 | 1.23E-01 | 2.54E+00 | 2.60E+00 | 5.67E-01 | no | Crp | intron_variant,non_coding_transcript_variant | MODIFIER |
| 1 | 172748354 | AX-170330958 | 0 | C | G | 0.228 | 7.22E-02 | 1.23E-01 | 2.54E+00 | 2.60E+00 | 5.67E-01 | no | Crp | intron_variant,non_coding_transcript_variant | MODIFIER |
| 1 | 172751095 | AX-170236465 | 0 | G | A | 0.228 | 7.22E-02 | 1.23E-01 | 2.54E+00 | 2.60E+00 | 5.67E-01 | no | Crp | intron_variant,non_coding_transcript_variant | MODIFIER |
| 1 | 172751392 | AX-170418735 | 0 | C | G | 0.228 | 7.22E-02 | 1.23E-01 | 2.54E+00 | 2.60E+00 | 5.67E-01 | no | Crp | intron_variant,non_coding_transcript_variant | MODIFIER |
| 1 | 172753081 | AX-170348642 | 0 | G | A | 0.228 | 7.22E-02 | 1.23E-01 | 2.54E+00 | 2.60E+00 | 5.67E-01 | no | Crp | intron_variant,non_coding_transcript_variant | MODIFIER |
| 1 | 172753426 | AX-170152279 | 0 | G | A | 0.228 | 7.22E-02 | 1.23E-01 | 2.54E+00 | 2.60E+00 | 5.67E-01 | no | Crp | intron_variant,non_coding_transcript_variant | MODIFIER |
| 1 | 172753612 | AX-170196174 | 0 | G | A | 0.228 | 7.22E-02 | 1.23E-01 | 2.54E+00 | 2.60E+00 | 5.67E-01 | no | Crp | intron_variant,non_coding_transcript_variant | MODIFIER |
| 1 | 172766166 | AX-170002296 | 0 | T | C | 0.228 | 7.22E-02 | 1.23E-01 | 2.54E+00 | 2.60E+00 | 5.67E-01 | no | Crp | intron_variant,non_coding_transcript_variant | MODIFIER |
| 1 | 172778247 | AX-170002301 | 0 | T | G | 0.228 | 7.22E-02 | 1.23E-01 | 2.54E+00 | 2.60E+00 | 5.67E-01 | no | Gm38051 | upstream_gene_variant | MODIFIER |
| 1 | 172789545 | AX-97842415 | 0 | T | C | 0.228 | 7.22E-02 | 1.23E-01 | 2.54E+00 | 2.60E+00 | 5.67E-01 | no | Crp | intron_variant,non_coding_transcript_variant | MODIFIER |
| 1 | 172831769 | AX-170388551 | 0 | A | G | 0.228 | 7.22E-02 | 1.23E-01 | 2.54E+00 | 2.60E+00 | 5.67E-01 | no | Crp | non_coding_transcript_exon_variant | MODIFIER |
| 1 | 172832045 | AX-97855395 | 0 | C | T | 0.228 | 7.22E-02 | 1.23E-01 | 2.54E+00 | 2.60E+00 | 5.67E-01 | no | Crp | non_coding_transcript_exon_variant | MODIFIER |

| | | | | | | | | | | | | | | | |
|---|---|---|---|---|---|---|---|---|---|---|---|---|---|---|---|
| 1 | 172859925 | AX-170317532 | 0 | C | A | 0.228 | 7.22E-02 | 1.23E-01 | 2.54E+00 | 2.60E+00 | 5.67E-01 | no | | intergenic_variant | MODIFIER |
| 1 | 172860775 | AX-170191082 | 0 | A | C | 0.228 | 7.22E-02 | 1.23E-01 | 2.54E+00 | 2.60E+00 | 5.67E-01 | no | | intergenic_variant | MODIFIER |
| 1 | 172865246 | AX-170162996 | 0 | C | A | 0.228 | 7.22E-02 | 1.23E-01 | 2.54E+00 | 2.60E+00 | 5.67E-01 | no | | intergenic_variant | MODIFIER |
| 1 | 172865618 | AX-170237272 | 0 | A | G | 0.228 | 7.22E-02 | 1.23E-01 | 2.54E+00 | 2.60E+00 | 5.67E-01 | no | | intergenic_variant | MODIFIER |
| 1 | 172866389 | AX-170425324 | 0 | T | C | 0.228 | 7.22E-02 | 1.23E-01 | 2.54E+00 | 2.60E+00 | 5.67E-01 | no | | intergenic_variant | MODIFIER |
| 1 | 172866886 | AX-170258579 | 0 | C | T | 0.228 | 7.22E-02 | 1.23E-01 | 2.54E+00 | 2.60E+00 | 5.67E-01 | no | | intergenic_variant | MODIFIER |
| 1 | 172886032 | AX-170111462 | 0 | A | T | 0.228 | 7.22E-02 | 1.23E-01 | 2.54E+00 | 2.60E+00 | 5.67E-01 | no | | intergenic_variant | MODIFIER |
| 1 | 172886069 | AX-170043610 | 0 | T | C | 0.228 | 7.22E-02 | 1.23E-01 | 2.54E+00 | 2.60E+00 | 5.67E-01 | no | | intergenic_variant | MODIFIER |
| 1 | 172886138 | AX-170043609 | 0 | G | C | 0.228 | 7.22E-02 | 1.23E-01 | 2.54E+00 | 2.60E+00 | 5.67E-01 | no | | intergenic_variant | MODIFIER |
| 1 | 172886359 | AX-170245622 | 0 | G | C | 0.228 | 7.22E-02 | 1.23E-01 | 2.54E+00 | 2.60E+00 | 5.67E-01 | no | | intergenic_variant | MODIFIER |
| 1 | 172886937 | AX-170287198 | 0 | T | G | 0.228 | 7.22E-02 | 1.23E-01 | 2.54E+00 | 2.60E+00 | 5.67E-01 | no | | intergenic_variant | MODIFIER |
| 1 | 172894298 | AX-170272998 | 0 | G | A | 0.228 | 7.22E-02 | 1.23E-01 | 2.54E+00 | 2.60E+00 | 5.67E-01 | no | Apcs | synonymous_variant | LOW |
| 1 | 172897060 | AX-170326478 | 0 | A | T | 0.228 | 7.22E-02 | 1.23E-01 | 2.54E+00 | 2.60E+00 | 5.67E-01 | no | Apcs | upstream_gene_variant | MODIFIER |
| 1 | 172924358 | AX-97830593 | 0 | C | T | 0.228 | 7.22E-02 | 1.23E-01 | 2.54E+00 | 2.60E+00 | 5.67E-01 | no | | intergenic_variant | MODIFIER |
| 1 | 172937148 | AX-170002302 | 0 | T | C | 0.228 | 7.22E-02 | 1.23E-01 | 2.54E+00 | 2.60E+00 | 5.67E-01 | no | Gm26077 | upstream_gene_variant | MODIFIER |
| 1 | 172955306 | AX-170376733 | 0 | T | C | 0.228 | 7.22E-02 | 1.23E-01 | 2.54E+00 | 2.60E+00 | 5.67E-01 | no | Olfr16 | upstream_gene_variant | MODIFIER |
| 1 | 172956255 | AX-170382028 | 0 | G | A | 0.228 | 7.22E-02 | 1.23E-01 | 2.54E+00 | 2.60E+00 | 5.67E-01 | no | Olfr16 | upstream_gene_variant | MODIFIER |
| 1 | 172956602 | AX-170188955 | 0 | A | G | 0.228 | 7.22E-02 | 1.23E-01 | 2.54E+00 | 2.60E+00 | 5.67E-01 | no | Olfr16 | upstream_gene_variant | MODIFIER |
| 1 | 172957114 | AX-97826608 | 0 | C | T | 0.228 | 7.22E-02 | 1.23E-01 | 2.54E+00 | 2.60E+00 | 5.67E-01 | no | Olfr16 | synonymous_variant | LOW |
| 1 | 172957231 | AX-170202044 | 0 | T | C | 0.228 | 7.22E-02 | 1.23E-01 | 2.54E+00 | 2.60E+00 | 5.67E-01 | no | Olfr16 | synonymous_variant | LOW |
| 1 | 172957630 | AX-170389006 | 0 | C | T | 0.228 | 7.22E-02 | 1.23E-01 | 2.54E+00 | 2.60E+00 | 5.67E-01 | no | Olfr16 | synonymous_variant | LOW |
| 1 | 173005499 | AX-170527480 | 0 | C | T | 0.228 | 7.22E-02 | 1.23E-01 | 2.54E+00 | 2.60E+00 | 5.67E-01 | no | | intergenic_variant | MODIFIER |
| 1 | 173005759 | AX-170123609 | 0 | T | C | 0.228 | 7.22E-02 | 1.23E-01 | 2.54E+00 | 2.60E+00 | 5.67E-01 | no | | intergenic_variant | MODIFIER |
| 1 | 173006022 | AX-97827109 | 0 | T | C | 0.228 | 7.22E-02 | 1.23E-01 | 2.54E+00 | 2.60E+00 | 5.67E-01 | no | | intergenic_variant | MODIFIER |
| 1 | 173009377 | AX-170043613 | 0 | T | C | 0.228 | 7.22E-02 | 1.23E-01 | 2.54E+00 | 2.60E+00 | 5.67E-01 | no | | intergenic_variant | MODIFIER |
| 1 | 173011761 | AX-170462304 | 0 | C | T | 0.228 | 7.22E-02 | 1.23E-01 | 2.54E+00 | 2.60E+00 | 5.67E-01 | no | | intergenic_variant | MODIFIER |
| 1 | 173040888 | AX-170452227 | 0 | C | T | 0.228 | 7.22E-02 | 1.23E-01 | 2.54E+00 | 2.60E+00 | 5.67E-01 | no | | intergenic_variant | MODIFIER |
| 1 | 173050280 | AX-170497296 | 0 | T | C | 0.228 | 7.22E-02 | 1.23E-01 | 2.54E+00 | 2.60E+00 | 5.67E-01 | no | Gm42472 | downstream_gene_variant | MODIFIER |
| 1 | 173063851 | AX-170086689 | 0 | G | C | 0.228 | 7.22E-02 | 1.23E-01 | 2.54E+00 | 2.60E+00 | 5.67E-01 | no | | intergenic_variant | MODIFIER |
| 1 | 173072839 | AX-170172663 | 0 | C | T | 0.228 | 7.22E-02 | 1.23E-01 | 2.54E+00 | 2.60E+00 | 5.67E-01 | no | Gm42475 | upstream_gene_variant | MODIFIER |
| 1 | 173072867 | AX-170086690 | 0 | A | G | 0.228 | 7.22E-02 | 1.23E-01 | 2.54E+00 | 2.60E+00 | 5.67E-01 | no | Gm42475 | upstream_gene_variant | MODIFIER |
| 1 | 173119144 | AX-170344137 | 0 | A | G | 0.228 | 7.22E-02 | 1.23E-01 | 2.54E+00 | 2.60E+00 | 5.67E-01 | no | | intergenic_variant | MODIFIER |
| 1 | 173119191 | AX-170497418 | 0 | C | T | 0.228 | 7.22E-02 | 1.23E-01 | 2.54E+00 | 2.60E+00 | 5.67E-01 | no | | intergenic_variant | MODIFIER |
| 1 | 173137609 | AX-170432628 | 0 | C | G | 0.228 | 7.22E-02 | 1.23E-01 | 2.54E+00 | 2.60E+00 | 5.67E-01 | no | Olfr1408 | upstream_gene_variant | MODIFIER |
| 1 | 173140111 | AX-97800968 | 0 | A | C | 0.228 | 7.22E-02 | 1.23E-01 | 2.54E+00 | 2.60E+00 | 5.67E-01 | no | Olfr1408 | upstream_gene_variant | MODIFIER |
| 1 | 173183633 | AX-170248138 | 0 | C | T | 0.228 | 7.22E-02 | 1.23E-01 | 2.54E+00 | 2.60E+00 | 5.67E-01 | no | Olfr1406 | missense_variant | MODERATE |
| 1 | 173206689 | AX-170330232 | 0 | T | G | 0.228 | 7.22E-02 | 1.23E-01 | 2.54E+00 | 2.60E+00 | 5.67E-01 | no | Olfr218 | downstream_gene_variant | MODIFIER |
| 1 | 173213441 | AX-170317018 | 0 | A | C | 0.228 | 7.22E-02 | 1.23E-01 | 2.54E+00 | 2.60E+00 | 5.67E-01 | no | Olfr1404 | upstream_gene_variant | MODIFIER |
| 1 | 173213504 | AX-170275609 | 0 | A | C | 0.228 | 7.22E-02 | 1.23E-01 | 2.54E+00 | 2.60E+00 | 5.67E-01 | no | Olfr1404 | upstream_gene_variant | MODIFIER |
| 1 | 173221935 | AX-97814244 | 0 | C | T | 0.228 | 7.22E-02 | 1.23E-01 | 2.54E+00 | 2.60E+00 | 5.67E-01 | no | Fcer1a | intron_variant | MODIFIER |
| 1 | 173236138 | AX-170346159 | 0 | T | C | 0.228 | 7.22E-02 | 1.23E-01 | 2.54E+00 | 2.60E+00 | 5.67E-01 | no | | intergenic_variant | MODIFIER |
| 1 | 173281381 | AX-170091790 | 0 | A | G | 0.228 | 7.22E-02 | 1.23E-01 | 2.54E+00 | 2.60E+00 | 5.67E-01 | no | Mptx2 | upstream_gene_variant | MODIFIER |
| 1 | 173289519 | AX-170229954 | 0 | C | T | 0.228 | 7.22E-02 | 1.23E-01 | 2.54E+00 | 2.60E+00 | 5.67E-01 | no | | intergenic_variant | MODIFIER |
| 1 | 173289583 | AX-170043619 | 0 | C | T | 0.228 | 7.22E-02 | 1.23E-01 | 2.54E+00 | 2.60E+00 | 5.67E-01 | no | | intergenic_variant | MODIFIER |
| 1 | 173312762 | AX-170043616 | 0 | G | C | 0.228 | 7.22E-02 | 1.23E-01 | 2.54E+00 | 2.60E+00 | 5.67E-01 | no | | intergenic_variant | MODIFIER |
| 1 | 173317348 | AX-170247318 | 0 | T | A | 0.228 | 7.22E-02 | 1.23E-01 | 2.54E+00 | 2.60E+00 | 5.67E-01 | no | | intergenic_variant | MODIFIER |
| 1 | 173321686 | AX-170156308 | 0 | C | T | 0.228 | 7.22E-02 | 1.23E-01 | 2.54E+00 | 2.60E+00 | 5.67E-01 | no | | regulatory_region_variant | MODIFIER |
| 1 | 173338628 | AX-170508005 | 0 | C | T | 0.228 | 7.22E-02 | 1.23E-01 | 2.54E+00 | 2.60E+00 | 5.67E-01 | no | Cadm3 | intron_variant | MODIFIER |
| 1 | 173339174 | AX-170433123 | 0 | A | G | 0.228 | 7.22E-02 | 1.23E-01 | 2.54E+00 | 2.60E+00 | 5.67E-01 | no | Cadm3 | intron_variant | MODIFIER |
| 1 | 173340126 | AX-170441331 | 0 | C | A | 0.228 | 7.22E-02 | 1.23E-01 | 2.54E+00 | 2.60E+00 | 5.67E-01 | no | Cadm3 | intron_variant | MODIFIER |
| 1 | 173342829 | AX-170411948 | 0 | A | G | 0.228 | 7.22E-02 | 1.23E-01 | 2.54E+00 | 2.60E+00 | 5.67E-01 | no | Cadm3 | intron_variant | MODIFIER |
| 1 | 173346309 | AX-170111239 | 0 | A | G | 0.228 | 7.22E-02 | 1.23E-01 | 2.54E+00 | 2.60E+00 | 5.67E-01 | no | Cadm3 | intron_variant | MODIFIER |
| 1 | 173347893 | AX-170426531 | 0 | C | T | 0.228 | 7.22E-02 | 1.23E-01 | 2.54E+00 | 2.60E+00 | 5.67E-01 | no | Cadm3 | intron_variant | MODIFIER |
| 1 | 173351504 | AX-170388408 | 0 | T | A | 0.228 | 7.22E-02 | 1.23E-01 | 2.54E+00 | 2.60E+00 | 5.67E-01 | no | Cadm3 | intron_variant | MODIFIER |

| | | | | | | | | | | | | | | | |
|---|---|---|---|---|---|---|---|---|---|---|---|---|---|---|---|
| 1 | 173363930 | AX-170231852 | 0 | A | G | 0.228 | 7.22E-02 | 1.23E-01 | 2.54E+00 | 2.60E+00 | 5.67E-01 | no | Cadm3 | intron_variant | MODIFIER |
| 1 | 173364081 | AX-170491996 | 0 | G | A | 0.228 | 7.22E-02 | 1.23E-01 | 2.54E+00 | 2.60E+00 | 5.67E-01 | no | Cadm3 | intron_variant | MODIFIER |
| 1 | 173370140 | AX-170175552 | 0 | C | A | 0.228 | 7.22E-02 | 1.23E-01 | 2.54E+00 | 2.60E+00 | 5.67E-01 | no | Cadm3 | upstream_gene_variant | MODIFIER |
| 1 | 173370918 | AX-170433068 | 0 | T | C | 0.228 | 7.22E-02 | 1.23E-01 | 2.54E+00 | 2.60E+00 | 5.67E-01 | no | Cadm3 | upstream_gene_variant | MODIFIER |
| 1 | 173384889 | AX-97807827 | 0 | T | C | 0.228 | 7.22E-02 | 1.23E-01 | 2.54E+00 | 2.60E+00 | 5.67E-01 | no | Aim2 | intron_variant | MODIFIER |
| 1 | 173403855 | AX-170252393 | 0 | G | A | 0.228 | 7.22E-02 | 1.23E-01 | 2.54E+00 | 2.60E+00 | 5.67E-01 | no | Aim2 | intron_variant | MODIFIER |
| 1 | 173410907 | AX-170091791 | 0 | C | T | 0.228 | 7.22E-02 | 1.23E-01 | 2.54E+00 | 2.60E+00 | 5.67E-01 | no | Aim2 | intron_variant | MODIFIER |
| 1 | 173413497 | AX-170532531 | 0 | A | G | 0.228 | 7.22E-02 | 1.23E-01 | 2.54E+00 | 2.60E+00 | 5.67E-01 | no | Aim2 | intron_variant | MODIFIER |
| 1 | 173417785 | AX-170288713 | 0 | A | G | 0.228 | 7.22E-02 | 1.23E-01 | 2.54E+00 | 2.60E+00 | 5.67E-01 | no | Aim2 | intron_variant | MODIFIER |
| 1 | 173429208 | AX-170465266 | 0 | G | A | 0.228 | 7.22E-02 | 1.23E-01 | 2.54E+00 | 2.60E+00 | 5.67E-01 | no | Aim2 | intron_variant | MODIFIER |
| 1 | 173454097 | AX-170373824 | 0 | A | T | 0.228 | 7.22E-02 | 1.23E-01 | 2.54E+00 | 2.60E+00 | 5.67E-01 | no | Aim2 | upstream_gene_variant | MODIFIER |
| 1 | 173456096 | AX-170369324 | 0 | C | T | 0.228 | 7.22E-02 | 1.23E-01 | 2.54E+00 | 2.60E+00 | 5.67E-01 | no | Aim2 | non_coding_transcript_exon_variant | MODIFIER |
| 1 | 173457382 | AX-170140240 | 0 | T | C | 0.228 | 7.22E-02 | 1.23E-01 | 2.54E+00 | 2.60E+00 | 5.67E-01 | no | Aim2 | downstream_gene_variant | MODIFIER |
| 1 | 173462814 | AX-170462208 | 0 | C | G | 0.228 | 7.22E-02 | 1.23E-01 | 2.54E+00 | 2.60E+00 | 5.67E-01 | no | Aim2 | intron_variant | MODIFIER |
| 1 | 173486559 | AX-170309437 | 0 | T | G | 0.228 | 7.22E-02 | 1.23E-01 | 2.54E+00 | 2.60E+00 | 5.67E-01 | no | Ifi206 | intron_variant | MODIFIER |
| 1 | 173509116 | AX-170528915 | 0 | C | T | 0.228 | 7.22E-02 | 1.23E-01 | 2.54E+00 | 2.60E+00 | 5.67E-01 | no | | intergenic_variant | MODIFIER |
| 1 | 173582067 | AX-170202465 | 0 | G | A | 0.228 | 7.22E-02 | 1.23E-01 | 2.54E+00 | 2.60E+00 | 5.67E-01 | no | Ifi213 | intron_variant | MODIFIER |
| 1 | 173582101 | AX-170479141 | 0 | G | T | 0.228 | 7.22E-02 | 1.23E-01 | 2.54E+00 | 2.60E+00 | 5.67E-01 | no | Ifi213 | intron_variant | MODIFIER |
| 1 | 173608404 | AX-170438657 | 0 | T | C | 0.228 | 7.22E-02 | 1.23E-01 | 2.54E+00 | 2.60E+00 | 5.67E-01 | no | | intergenic_variant | MODIFIER |
| 1 | 173645553 | AX-170224918 | 0 | A | C | 0.228 | 7.22E-02 | 1.23E-01 | 2.54E+00 | 2.60E+00 | 5.67E-01 | no | Ifi209 | intron_variant | MODIFIER |
| 1 | 173646000 | AX-97812525 | 0 | G | A | 0.228 | 7.22E-02 | 1.23E-01 | 2.54E+00 | 2.60E+00 | 5.67E-01 | no | Ifi209 | missense_variant | MODERATE |
| 1 | 173722904 | AX-170222075 | 0 | G | C | 0.228 | 7.22E-02 | 1.23E-01 | 2.54E+00 | 2.60E+00 | 5.67E-01 | no | Ifi207 | downstream_gene_variant | MODIFIER |
| 1 | 173742993 | AX-170415134 | 0 | G | A | 0.228 | 7.22E-02 | 1.23E-01 | 2.54E+00 | 2.60E+00 | 5.67E-01 | no | Ifi207 | upstream_gene_variant | MODIFIER |
| 1 | 173781904 | AX-97839760 | 0 | A | C | 0.228 | 7.22E-02 | 1.23E-01 | 2.54E+00 | 2.60E+00 | 5.67E-01 | no | Ifi203-ps | intron_variant,non_coding_transcript_variant | MODIFIER |
| 1 | 173824798 | AX-170214782 | 0 | C | G | 0.228 | 7.22E-02 | 1.23E-01 | 2.54E+00 | 2.60E+00 | 5.67E-01 | no | | intergenic_variant | MODIFIER |
| 1 | 173840015 | AX-170134693 | 0 | T | A | 0.228 | 7.22E-02 | 1.23E-01 | 2.54E+00 | 2.60E+00 | 5.67E-01 | no | | intergenic_variant | MODIFIER |
| 1 | 173971432 | AX-170345555 | 0 | A | G | 0.228 | 7.22E-02 | 1.23E-01 | 2.54E+00 | 2.60E+00 | 5.67E-01 | no | Ifi202b | missense_variant | MODERATE |
| 1 | 174036719 | AX-170223464 | 0 | G | A | 0.228 | 7.22E-02 | 1.23E-01 | 2.54E+00 | 2.60E+00 | 5.67E-01 | no | Olfr433 | upstream_gene_variant | MODIFIER |
| 1 | 174037187 | AX-170129147 | 0 | C | T | 0.228 | 7.22E-02 | 1.23E-01 | 2.54E+00 | 2.60E+00 | 5.67E-01 | no | Olfr433 | upstream_gene_variant | MODIFIER |
| 1 | 174037671 | AX-170304812 | 0 | T | C | 0.228 | 7.22E-02 | 1.23E-01 | 2.54E+00 | 2.60E+00 | 5.67E-01 | no | Olfr433 | upstream_gene_variant | MODIFIER |
| 1 | 174047092 | AX-170423356 | 0 | G | A | 0.228 | 7.22E-02 | 1.23E-01 | 2.54E+00 | 2.60E+00 | 5.67E-01 | no | Olfr433 | downstream_gene_variant | MODIFIER |
| 1 | 174066622 | AX-97811058 | 0 | C | T | 0.228 | 7.22E-02 | 1.23E-01 | 2.54E+00 | 2.60E+00 | 5.67E-01 | no | Olfr430 | upstream_gene_variant | MODIFIER |
| 1 | 174066845 | AX-170002316 | 0 | G | A | 0.228 | 7.22E-02 | 1.23E-01 | 2.54E+00 | 2.60E+00 | 5.67E-01 | no | Olfr430 | upstream_gene_variant | MODIFIER |
| 1 | 174100264 | AX-170002307 | 0 | T | C | 0.228 | 7.22E-02 | 1.23E-01 | 2.54E+00 | 2.60E+00 | 5.67E-01 | no | Olfr427 | synonymous_variant | LOW |
| 1 | 174151673 | AX-97830327 | 0 | C | T | 0.228 | 7.22E-02 | 1.23E-01 | 2.54E+00 | 2.60E+00 | 5.67E-01 | no | Olfr421-ps1 | synonymous_variant | LOW |
| 1 | 174169336 | AX-170513627 | 0 | T | C | 0.228 | 7.22E-02 | 1.23E-01 | 2.54E+00 | 2.60E+00 | 5.67E-01 | no | Spta1 | upstream_gene_variant | MODIFIER |
| 1 | 174170805 | AX-170446545 | 0 | T | A | 0.228 | 7.22E-02 | 1.23E-01 | 2.54E+00 | 2.60E+00 | 5.67E-01 | no | Spta1 | upstream_gene_variant | MODIFIER |
| 1 | 174210967 | AX-170331523 | 0 | C | G | 0.228 | 7.22E-02 | 1.23E-01 | 2.54E+00 | 2.60E+00 | 5.67E-01 | no | Spta1 | intron_variant | MODIFIER |
| 1 | 174211021 | AX-170449242 | 0 | C | T | 0.228 | 7.22E-02 | 1.23E-01 | 2.54E+00 | 2.60E+00 | 5.67E-01 | no | Spta1 | intron_variant | MODIFIER |
| 1 | 174217144 | AX-170515130 | 0 | T | G | 0.228 | 7.22E-02 | 1.23E-01 | 2.54E+00 | 2.60E+00 | 5.67E-01 | no | Spta1 | missense_variant | MODERATE |
| 1 | 174217616 | AX-170299156 | 0 | C | T | 0.228 | 7.22E-02 | 1.23E-01 | 2.54E+00 | 2.60E+00 | 5.67E-01 | no | Spta1 | synonymous_variant | LOW |
| 1 | 174219588 | AX-170471973 | 0 | T | C | 0.228 | 7.22E-02 | 1.23E-01 | 2.54E+00 | 2.60E+00 | 5.67E-01 | no | Spta1 | intron_variant | MODIFIER |
| 1 | 174220172 | AX-170229946 | 0 | C | T | 0.228 | 7.22E-02 | 1.23E-01 | 2.54E+00 | 2.60E+00 | 5.67E-01 | no | Spta1 | intron_variant | MODIFIER |
| 1 | 174225758 | AX-170219197 | 0 | G | C | 0.228 | 7.22E-02 | 1.23E-01 | 2.54E+00 | 2.60E+00 | 5.67E-01 | no | Spta1 | intron_variant | MODIFIER |
| 1 | 174226439 | AX-97855926 | 0 | T | C | 0.228 | 7.22E-02 | 1.23E-01 | 2.54E+00 | 2.60E+00 | 5.67E-01 | no | Spta1 | intron_variant | MODIFIER |
| 1 | 174235643 | AX-170530015 | 0 | G | A | 0.228 | 7.22E-02 | 1.23E-01 | 2.54E+00 | 2.60E+00 | 5.67E-01 | no | Spta1 | intron_variant | MODIFIER |
| 1 | 174242078 | AX-170233236 | 0 | C | T | 0.228 | 7.22E-02 | 1.23E-01 | 2.54E+00 | 2.60E+00 | 5.67E-01 | no | Spta1 | intron_variant | MODIFIER |
| 1 | 174242862 | AX-170149965 | 0 | A | G | 0.228 | 7.22E-02 | 1.23E-01 | 2.54E+00 | 2.60E+00 | 5.67E-01 | no | Spta1 | intron_variant | MODIFIER |
| 1 | 174245339 | AX-170434136 | 0 | A | G | 0.228 | 7.22E-02 | 1.23E-01 | 2.54E+00 | 2.60E+00 | 5.67E-01 | no | Spta1 | intron_variant | MODIFIER |
| 1 | 174245966 | AX-170185241 | 0 | T | C | 0.228 | 7.22E-02 | 1.23E-01 | 2.54E+00 | 2.60E+00 | 5.67E-01 | no | Spta1 | intron_variant | MODIFIER |
| 1 | 174246095 | AX-170306527 | 0 | T | G | 0.228 | 7.22E-02 | 1.23E-01 | 2.54E+00 | 2.60E+00 | 5.67E-01 | no | Spta1 | intron_variant | MODIFIER |
| 1 | 174246720 | AX-170282321 | 0 | T | C | 0.228 | 7.22E-02 | 1.23E-01 | 2.54E+00 | 2.60E+00 | 5.67E-01 | no | Spta1 | intron_variant | MODIFIER |
| 1 | 174249092 | AX-170443691 | 0 | T | C | 0.228 | 7.22E-02 | 1.23E-01 | 2.54E+00 | 2.60E+00 | 5.67E-01 | no | Spta1 | downstream_gene_variant | MODIFIER |
| 1 | 174282475 | AX-97814845 | 0 | G | C | 0.228 | 7.22E-02 | 1.23E-01 | 2.54E+00 | 2.60E+00 | 5.67E-01 | no | | intergenic_variant | MODIFIER |

| | | | | | | | | | | | | | | | |
|---|---|---|---|---|---|---|---|---|---|---|---|---|---|---|---|
| 1 | 174301879 | AX-170497706 | 0 | C | T | 0.228 | 7.22E-02 | 1.23E-01 | 2.54E+00 | 2.60E+00 | 5.67E-01 | no | | intergenic_variant | MODIFIER |
| 1 | 174358309 | AX-170043645 | 0 | T | G | 0.228 | 7.22E-02 | 1.23E-01 | 2.54E+00 | 2.60E+00 | 5.67E-01 | no | | intergenic_variant | MODIFIER |
| 1 | 174374842 | AX-170232245 | 0 | G | A | 0.228 | 7.22E-02 | 1.23E-01 | 2.54E+00 | 2.60E+00 | 5.67E-01 | no | Olfr417 | downstream_gene_variant | MODIFIER |
| 1 | 174431497 | AX-170115164 | 0 | G | A | 0.228 | 7.22E-02 | 1.23E-01 | 2.54E+00 | 2.60E+00 | 5.67E-01 | no | Olfr414 | downstream_gene_variant | MODIFIER |
| 1 | 174431904 | AX-170099394 | 0 | A | G | 0.228 | 7.22E-02 | 1.23E-01 | 2.54E+00 | 2.60E+00 | 5.67E-01 | no | Olfr414 | downstream_gene_variant | MODIFIER |
| 1 | 174431943 | AX-170184002 | 0 | C | T | 0.228 | 7.22E-02 | 1.23E-01 | 2.54E+00 | 2.60E+00 | 5.67E-01 | no | Olfr414 | downstream_gene_variant | MODIFIER |
| 1 | 174537391 | AX-170464335 | 0 | A | G | 0.228 | 7.22E-02 | 1.23E-01 | 2.54E+00 | 2.60E+00 | 5.67E-01 | no | Fmn2 | intron_variant | MODIFIER |
| 1 | 174546004 | AX-97811059 | 0 | A | C | 0.228 | 7.22E-02 | 1.23E-01 | 2.54E+00 | 2.60E+00 | 5.67E-01 | no | Fmn2 | intron_variant | MODIFIER |
| 1 | 174546388 | AX-170199648 | 0 | C | G | 0.228 | 7.22E-02 | 1.23E-01 | 2.54E+00 | 2.60E+00 | 5.67E-01 | no | Fmn2 | intron_variant | MODIFIER |
| 1 | 174547758 | AX-170323381 | 0 | A | T | 0.228 | 7.22E-02 | 1.23E-01 | 2.54E+00 | 2.60E+00 | 5.67E-01 | no | Fmn2 | intron_variant | MODIFIER |
| 1 | 174557890 | AX-170457009 | 0 | T | C | 0.228 | 7.22E-02 | 1.23E-01 | 2.54E+00 | 2.60E+00 | 5.67E-01 | no | Fmn2 | intron_variant | MODIFIER |
| 1 | 174583205 | AX-170043650 | 0 | A | G | 0.228 | 7.22E-02 | 1.23E-01 | 2.54E+00 | 2.60E+00 | 5.67E-01 | no | Fmn2 | intron_variant | MODIFIER |
| 1 | 174633081 | AX-170433896 | 0 | A | T | 0.228 | 7.22E-02 | 1.23E-01 | 2.54E+00 | 2.60E+00 | 5.67E-01 | no | Fmn2 | intron_variant | MODIFIER |
| 1 | 174635145 | AX-170347158 | 0 | G | A | 0.228 | 7.22E-02 | 1.23E-01 | 2.54E+00 | 2.60E+00 | 5.67E-01 | no | Fmn2 | intron_variant | MODIFIER |
| 1 | 174639127 | AX-170155595 | 0 | C | T | 0.228 | 7.22E-02 | 1.23E-01 | 2.54E+00 | 2.60E+00 | 5.67E-01 | no | Fmn2 | intron_variant | MODIFIER |
| 1 | 174639647 | AX-170385122 | 0 | C | T | 0.228 | 7.22E-02 | 1.23E-01 | 2.54E+00 | 2.60E+00 | 5.67E-01 | no | Fmn2 | intron_variant | MODIFIER |
| 1 | 174662275 | AX-170091797 | 0 | G | T | 0.228 | 7.22E-02 | 1.23E-01 | 2.54E+00 | 2.60E+00 | 5.67E-01 | no | Fmn2 | intron_variant | MODIFIER |
| 1 | 174675038 | AX-170002318 | 0 | G | A | 0.228 | 7.22E-02 | 1.23E-01 | 2.54E+00 | 2.60E+00 | 5.67E-01 | no | Fmn2 | intron_variant | MODIFIER |
| 1 | 174676808 | AX-170169364 | 0 | C | G | 0.228 | 7.22E-02 | 1.23E-01 | 2.54E+00 | 2.60E+00 | 5.67E-01 | no | Fmn2 | intron_variant | MODIFIER |
| 1 | 174689368 | AX-170502137 | 0 | C | G | 0.228 | 7.22E-02 | 1.23E-01 | 2.54E+00 | 2.60E+00 | 5.67E-01 | no | Fmn2 | intron_variant | MODIFIER |
| 1 | 174757620 | AX-170149389 | 0 | A | T | 0.228 | 7.22E-02 | 1.23E-01 | 2.54E+00 | 2.60E+00 | 5.67E-01 | no | Fmn2 | intron_variant | MODIFIER |
| 1 | 174759347 | AX-170110064 | 0 | C | T | 0.228 | 7.22E-02 | 1.23E-01 | 2.54E+00 | 2.60E+00 | 5.67E-01 | no | Fmn2 | intron_variant | MODIFIER |
| 1 | 174759458 | AX-170294835 | 0 | T | G | 0.228 | 7.22E-02 | 1.23E-01 | 2.54E+00 | 2.60E+00 | 5.67E-01 | no | Fmn2 | intron_variant | MODIFIER |
| 1 | 174759487 | AX-170098672 | 0 | G | A | 0.228 | 7.22E-02 | 1.23E-01 | 2.54E+00 | 2.60E+00 | 5.67E-01 | no | Fmn2 | intron_variant | MODIFIER |
| 1 | 174794863 | AX-170503447 | 0 | G | A | 0.228 | 7.22E-02 | 1.23E-01 | 2.54E+00 | 2.60E+00 | 5.67E-01 | no | Fmn2 | intron_variant | MODIFIER |
| 1 | 174815734 | AX-170507294 | 0 | A | G | 0.228 | 7.22E-02 | 1.23E-01 | 2.54E+00 | 2.60E+00 | 5.67E-01 | no | Fmn2 | intron_variant | MODIFIER |
| 1 | 174840751 | AX-170181520 | 0 | T | C | 0.228 | 7.22E-02 | 1.23E-01 | 2.54E+00 | 2.60E+00 | 5.67E-01 | no | Grem2 | intron_variant | MODIFIER |
| 1 | 174840928 | AX-97813043 | 0 | G | A | 0.228 | 7.22E-02 | 1.23E-01 | 2.54E+00 | 2.60E+00 | 5.67E-01 | no | Grem2 | intron_variant | MODIFIER |
| 1 | 174843412 | AX-170334329 | 0 | G | A | 0.228 | 7.22E-02 | 1.23E-01 | 2.54E+00 | 2.60E+00 | 5.67E-01 | no | Grem2 | intron_variant | MODIFIER |
| 1 | 174844026 | AX-170436288 | 0 | A | G | 0.228 | 7.22E-02 | 1.23E-01 | 2.54E+00 | 2.60E+00 | 5.67E-01 | no | Grem2 | intron_variant | MODIFIER |
| 1 | 174854882 | AX-170520615 | 0 | G | A | 0.228 | 7.22E-02 | 1.23E-01 | 2.54E+00 | 2.60E+00 | 5.67E-01 | no | Grem2 | intron_variant | MODIFIER |
| 1 | 174861038 | AX-170450318 | 0 | A | G | 0.228 | 7.22E-02 | 1.23E-01 | 2.54E+00 | 2.60E+00 | 5.67E-01 | no | Grem2 | intron_variant | MODIFIER |
| 1 | 174902526 | AX-170233081 | 0 | T | G | 0.228 | 7.22E-02 | 1.23E-01 | 2.54E+00 | 2.60E+00 | 5.67E-01 | no | Grem2 | intron_variant | MODIFIER |
| 1 | 174912821 | AX-170394033 | 0 | G | A | 0.228 | 7.22E-02 | 1.23E-01 | 2.54E+00 | 2.60E+00 | 5.67E-01 | no | Grem2 | intron_variant | MODIFIER |
| 1 | 174918680 | AX-170392020 | 0 | G | A | 0.228 | 7.22E-02 | 1.23E-01 | 2.54E+00 | 2.60E+00 | 5.67E-01 | no | Grem2 | intron_variant | MODIFIER |
| 1 | 174922565 | AX-170383528 | 0 | C | T | 0.228 | 7.22E-02 | 1.23E-01 | 2.54E+00 | 2.60E+00 | 5.67E-01 | no | Grem2 | upstream_gene_variant | MODIFIER |
| 1 | 174941203 | AX-170086694 | 0 | T | C | 0.228 | 7.22E-02 | 1.23E-01 | 2.54E+00 | 2.60E+00 | 5.67E-01 | no | | intergenic_variant | MODIFIER |
| 1 | 174951303 | AX-170174926 | 0 | C | T | 0.228 | 7.22E-02 | 1.23E-01 | 2.54E+00 | 2.60E+00 | 5.67E-01 | no | | intergenic_variant | MODIFIER |
| 1 | 174951416 | AX-97814299 | 0 | A | G | 0.228 | 7.22E-02 | 1.23E-01 | 2.54E+00 | 2.60E+00 | 5.67E-01 | no | | intergenic_variant | MODIFIER |
| 1 | 174975782 | AX-170148843 | 0 | A | C | 0.228 | 7.22E-02 | 1.23E-01 | 2.54E+00 | 2.60E+00 | 5.67E-01 | no | | intergenic_variant | MODIFIER |
| 1 | 174976368 | AX-170398211 | 0 | G | A | 0.228 | 7.22E-02 | 1.23E-01 | 2.54E+00 | 2.60E+00 | 5.67E-01 | no | | intergenic_variant | MODIFIER |
| 1 | 174981277 | AX-170513661 | 0 | A | G | 0.228 | 7.22E-02 | 1.23E-01 | 2.54E+00 | 2.60E+00 | 5.67E-01 | no | | intergenic_variant | MODIFIER |
| 1 | 174981953 | AX-170145547 | 0 | T | C | 0.228 | 7.22E-02 | 1.23E-01 | 2.54E+00 | 2.60E+00 | 5.67E-01 | no | | intergenic_variant | MODIFIER |
| 1 | 175007523 | AX-170043657 | 0 | G | A | 0.228 | 7.22E-02 | 1.23E-01 | 2.54E+00 | 2.60E+00 | 5.67E-01 | no | | intergenic_variant | MODIFIER |
| 1 | 175007841 | AX-97782815 | 0 | C | A | 0.228 | 7.22E-02 | 1.23E-01 | 2.54E+00 | 2.60E+00 | 5.67E-01 | no | | intergenic_variant | MODIFIER |
| 1 | 175041914 | AX-170530205 | 0 | T | C | 0.228 | 7.22E-02 | 1.23E-01 | 2.54E+00 | 2.60E+00 | 5.67E-01 | no | | intergenic_variant | MODIFIER |
| 1 | 175041949 | AX-170419369 | 0 | T | C | 0.228 | 7.22E-02 | 1.23E-01 | 2.54E+00 | 2.60E+00 | 5.67E-01 | no | | intergenic_variant | MODIFIER |
| 1 | 175049618 | AX-170107309 | 0 | G | A | 0.228 | 7.22E-02 | 1.23E-01 | 2.54E+00 | 2.60E+00 | 5.67E-01 | no | | intergenic_variant | MODIFIER |
| 1 | 175054333 | AX-170002334 | 0 | A | G | 0.228 | 7.22E-02 | 1.23E-01 | 2.54E+00 | 2.60E+00 | 5.67E-01 | no | Rgs7 | downstream_gene_variant | MODIFIER |
| 1 | 175056753 | AX-170002330 | 0 | T | C | 0.228 | 7.22E-02 | 1.23E-01 | 2.54E+00 | 2.60E+00 | 5.67E-01 | no | Rgs7 | downstream_gene_variant | MODIFIER |
| 1 | 175082196 | AX-170451679 | 0 | A | G | 0.228 | 7.22E-02 | 1.23E-01 | 2.54E+00 | 2.60E+00 | 5.67E-01 | no | Rgs7 | intron_variant | MODIFIER |
| 1 | 175090891 | AX-97844993 | 0 | C | G | 0.228 | 7.22E-02 | 1.23E-01 | 2.54E+00 | 2.60E+00 | 5.67E-01 | no | Rgs7 | intron_variant | MODIFIER |
| 1 | 175097361 | AX-170222587 | 0 | A | T | 0.228 | 7.22E-02 | 1.23E-01 | 2.54E+00 | 2.60E+00 | 5.67E-01 | no | Rgs7 | intron_variant | MODIFIER |
| 1 | 175100020 | AX-170135514 | 0 | T | C | 0.228 | 7.22E-02 | 1.23E-01 | 2.54E+00 | 2.60E+00 | 5.67E-01 | no | Rgs7 | intron_variant | MODIFIER |

| | | | | | | | | | | | | | | | |
|---|---|---|---|---|---|---|---|---|---|---|---|---|---|---|---|
| 1 | 175100041 | AX-170191159 | 0 | G | A | 0.228 | 7.22E-02 | 1.23E-01 | 2.54E+00 | 2.60E+00 | 5.67E-01 | no | Rgs7 | intron_variant | MODIFIER |
| 1 | 175128987 | AX-170491971 | 0 | C | A | 0.228 | 7.22E-02 | 1.23E-01 | 2.54E+00 | 2.60E+00 | 5.67E-01 | no | Rgs7 | intron_variant | MODIFIER |
| 1 | 175144102 | AX-170002327 | 0 | T | G | 0.228 | 7.22E-02 | 1.23E-01 | 2.54E+00 | 2.60E+00 | 5.67E-01 | no | Rgs7 | intron_variant | MODIFIER |
| 1 | 175170188 | AX-170471732 | 0 | C | T | 0.228 | 7.22E-02 | 1.23E-01 | 2.54E+00 | 2.60E+00 | 5.67E-01 | no | Rgs7 | intron_variant | MODIFIER |
| 1 | 175187979 | AX-170002329 | 0 | C | T | 0.228 | 7.22E-02 | 1.23E-01 | 2.54E+00 | 2.60E+00 | 5.67E-01 | no | Rgs7 | intron_variant | MODIFIER |
| 1 | 175188025 | AX-170253770 | 0 | T | A | 0.228 | 7.22E-02 | 1.23E-01 | 2.54E+00 | 2.60E+00 | 5.67E-01 | no | Rgs7 | intron_variant | MODIFIER |
| 1 | 175245876 | AX-170212299 | 0 | A | G | 0.228 | 7.22E-02 | 1.23E-01 | 2.54E+00 | 2.60E+00 | 5.67E-01 | no | Rgs7 | intron_variant | MODIFIER |
| 1 | 175262198 | AX-170002328 | 0 | C | T | 0.228 | 7.22E-02 | 1.23E-01 | 2.54E+00 | 2.60E+00 | 5.67E-01 | no | Rgs7 | intron_variant | MODIFIER |
| 1 | 175262251 | AX-170107751 | 0 | G | A | 0.228 | 7.22E-02 | 1.23E-01 | 2.54E+00 | 2.60E+00 | 5.67E-01 | no | Rgs7 | intron_variant | MODIFIER |
| 1 | 175269981 | AX-170527545 | 0 | G | A | 0.228 | 7.22E-02 | 1.23E-01 | 2.54E+00 | 2.60E+00 | 5.67E-01 | no | Rgs7 | intron_variant | MODIFIER |
| 1 | 175276446 | AX-168403106 | 0 | C | A | 0.228 | 7.22E-02 | 1.23E-01 | 2.54E+00 | 2.60E+00 | 5.67E-01 | no | Rgs7 | intron_variant | MODIFIER |
| 1 | 175276551 | AX-170135414 | 0 | T | G | 0.228 | 7.22E-02 | 1.23E-01 | 2.54E+00 | 2.60E+00 | 5.67E-01 | no | Rgs7 | intron_variant | MODIFIER |
| 1 | 175276584 | AX-170492010 | 0 | C | T | 0.228 | 7.22E-02 | 1.23E-01 | 2.54E+00 | 2.60E+00 | 5.67E-01 | no | Rgs7 | intron_variant | MODIFIER |
| 1 | 175286465 | AX-170124230 | 0 | T | C | 0.228 | 7.22E-02 | 1.23E-01 | 2.54E+00 | 2.60E+00 | 5.67E-01 | no | Rgs7 | intron_variant | MODIFIER |
| 1 | 175292528 | AX-170292667 | 0 | A | T | 0.228 | 7.22E-02 | 1.23E-01 | 2.54E+00 | 2.60E+00 | 5.67E-01 | no | Rgs7 | intron_variant | MODIFIER |
| 1 | 175298245 | AX-170434789 | 0 | C | T | 0.228 | 7.22E-02 | 1.23E-01 | 2.54E+00 | 2.60E+00 | 5.67E-01 | no | Rgs7 | intron_variant | MODIFIER |
| 1 | 175318661 | AX-170187918 | 0 | C | T | 0.228 | 7.22E-02 | 1.23E-01 | 2.54E+00 | 2.60E+00 | 5.67E-01 | no | Rgs7 | intron_variant | MODIFIER |
| 1 | 175319069 | AX-170373981 | 0 | C | A | 0.228 | 7.22E-02 | 1.23E-01 | 2.54E+00 | 2.60E+00 | 5.67E-01 | no | Rgs7 | intron_variant | MODIFIER |
| 1 | 175331265 | AX-170205262 | 0 | A | G | 0.228 | 7.22E-02 | 1.23E-01 | 2.54E+00 | 2.60E+00 | 5.67E-01 | no | Rgs7 | intron_variant | MODIFIER |
| 1 | 175338950 | AX-170043662 | 0 | T | C | 0.228 | 7.22E-02 | 1.23E-01 | 2.54E+00 | 2.60E+00 | 5.67E-01 | no | Rgs7 | intron_variant | MODIFIER |
| 1 | 175348591 | AX-170191446 | 0 | T | G | 0.228 | 7.22E-02 | 1.23E-01 | 2.54E+00 | 2.60E+00 | 5.67E-01 | no | Rgs7 | intron_variant | MODIFIER |
| 1 | 175348667 | AX-97824857 | 0 | G | T | 0.228 | 7.22E-02 | 1.23E-01 | 2.54E+00 | 2.60E+00 | 5.67E-01 | no | Rgs7 | intron_variant | MODIFIER |
| 1 | 175349305 | AX-170280586 | 0 | T | C | 0.228 | 7.22E-02 | 1.23E-01 | 2.54E+00 | 2.60E+00 | 5.67E-01 | no | Rgs7 | intron_variant | MODIFIER |
| 1 | 175497347 | AX-170253001 | 0 | C | A | 0.228 | 7.22E-02 | 1.23E-01 | 2.54E+00 | 2.60E+00 | 5.67E-01 | no | Rgs7 | upstream_gene_variant | MODIFIER |
| 1 | 175505148 | AX-170528077 | 0 | T | A | 0.228 | 7.22E-02 | 1.23E-01 | 2.54E+00 | 2.60E+00 | 5.67E-01 | no | | intergenic_variant | MODIFIER |
| 1 | 175509366 | AX-170104177 | 0 | T | C | 0.228 | 7.22E-02 | 1.23E-01 | 2.54E+00 | 2.60E+00 | 5.67E-01 | no | | intergenic_variant | MODIFIER |
| 1 | 175510657 | AX-170316243 | 0 | G | A | 0.228 | 7.22E-02 | 1.23E-01 | 2.54E+00 | 2.60E+00 | 5.67E-01 | no | | intergenic_variant | MODIFIER |
| 1 | 175510999 | AX-170160169 | 0 | C | T | 0.228 | 7.22E-02 | 1.23E-01 | 2.54E+00 | 2.60E+00 | 5.67E-01 | no | | intergenic_variant | MODIFIER |
| 1 | 175511362 | AX-170440682 | 0 | T | C | 0.228 | 7.22E-02 | 1.23E-01 | 2.54E+00 | 2.60E+00 | 5.67E-01 | no | | intergenic_variant | MODIFIER |
| 1 | 175513768 | AX-170465588 | 0 | A | G | 0.228 | 7.22E-02 | 1.23E-01 | 2.54E+00 | 2.60E+00 | 5.67E-01 | no | | intergenic_variant | MODIFIER |
| 1 | 175523386 | AX-170433080 | 0 | T | G | 0.228 | 7.22E-02 | 1.23E-01 | 2.54E+00 | 2.60E+00 | 5.67E-01 | no | | intergenic_variant | MODIFIER |
| 1 | 175529006 | AX-170151842 | 0 | C | T | 0.228 | 7.22E-02 | 1.23E-01 | 2.54E+00 | 2.60E+00 | 5.67E-01 | no | | intergenic_variant | MODIFIER |
| 1 | 175533715 | AX-97840096 | 0 | G | A | 0.228 | 7.22E-02 | 1.23E-01 | 2.54E+00 | 2.60E+00 | 5.67E-01 | no | | intergenic_variant | MODIFIER |
| 1 | 175537265 | AX-170466021 | 0 | G | C | 0.228 | 7.22E-02 | 1.23E-01 | 2.54E+00 | 2.60E+00 | 5.67E-01 | no | | intergenic_variant | MODIFIER |
| 1 | 175539981 | AX-170107974 | 0 | T | C | 0.228 | 7.22E-02 | 1.23E-01 | 2.54E+00 | 2.60E+00 | 5.67E-01 | no | | intergenic_variant | MODIFIER |
| 1 | 175548756 | AX-170392180 | 0 | C | T | 0.228 | 7.22E-02 | 1.23E-01 | 2.54E+00 | 2.60E+00 | 5.67E-01 | no | | intergenic_variant | MODIFIER |
| 1 | 175565246 | AX-170109622 | 0 | A | G | 0.228 | 7.22E-02 | 1.23E-01 | 2.54E+00 | 2.60E+00 | 5.67E-01 | no | | intergenic_variant | MODIFIER |
| 1 | 175573816 | AX-170116595 | 0 | A | G | 0.228 | 7.22E-02 | 1.23E-01 | 2.54E+00 | 2.60E+00 | 5.67E-01 | no | Gm36307 | upstream_gene_variant | MODIFIER |
| 1 | 175574343 | AX-170245089 | 0 | A | C | 0.228 | 7.22E-02 | 1.23E-01 | 2.54E+00 | 2.60E+00 | 5.67E-01 | no | Gm36307 | non_coding_transcript_exon_variant | MODIFIER |
| 1 | 175588034 | AX-170309907 | 0 | A | C | 0.228 | 7.22E-02 | 1.23E-01 | 2.54E+00 | 2.60E+00 | 5.67E-01 | no | | intergenic_variant | MODIFIER |
| 1 | 175588323 | AX-97849559 | 0 | G | A | 0.228 | 7.22E-02 | 1.23E-01 | 2.54E+00 | 2.60E+00 | 5.67E-01 | no | | intergenic_variant | MODIFIER |
| 1 | 175604556 | AX-170160719 | 0 | G | A | 0.228 | 7.22E-02 | 1.23E-01 | 2.54E+00 | 2.60E+00 | 5.67E-01 | no | Fh1 | intron_variant | MODIFIER |
| 1 | 175609773 | AX-97818192 | 0 | C | T | 0.228 | 7.22E-02 | 1.23E-01 | 2.54E+00 | 2.60E+00 | 5.67E-01 | no | Fh1 | synonymous_variant | LOW |
| 1 | 175794941 | AX-97807391 | 0 | A | G | 0.228 | 7.22E-02 | 1.23E-01 | 2.54E+00 | 2.60E+00 | 5.67E-01 | no | Wdr64 | intron_variant | MODIFIER |
| 1 | 175862820 | AX-97795353 | 0 | G | A | 0.228 | 7.22E-02 | 1.23E-01 | 2.54E+00 | 2.60E+00 | 5.67E-01 | no | | intergenic_variant | MODIFIER |
| 1 | 175863165 | AX-170272887 | 0 | A | G | 0.228 | 7.22E-02 | 1.23E-01 | 2.54E+00 | 2.60E+00 | 5.67E-01 | no | | intergenic_variant | MODIFIER |
| 1 | 175871230 | AX-170199825 | 0 | A | G | 0.228 | 7.22E-02 | 1.23E-01 | 2.54E+00 | 2.60E+00 | 5.67E-01 | no | | intergenic_variant | MODIFIER |
| 1 | 175871578 | AX-97824885 | 0 | G | A | 0.228 | 7.22E-02 | 1.23E-01 | 2.54E+00 | 2.60E+00 | 5.67E-01 | no | | intergenic_variant | MODIFIER |
| 1 | 175872154 | AX-170326301 | 0 | T | A | 0.228 | 7.22E-02 | 1.23E-01 | 2.54E+00 | 2.60E+00 | 5.67E-01 | no | | regulatory_region_variant | MODIFIER |
| 16 | 75473434 | AX-170520493 | 0 | C | T | 0.491 | 4.51E-02 | 7.59E-02 | 2.50E+00 | 2.58E+00 | 5.67E-01 | no | | intergenic_variant | MODIFIER |
| 16 | 75474734 | AX-170109664 | 0 | G | A | 0.491 | 4.51E-02 | 7.59E-02 | 2.50E+00 | 2.58E+00 | 5.67E-01 | no | | intergenic_variant | MODIFIER |
| 16 | 75509185 | AX-170533715 | 0 | G | A | 0.491 | 4.51E-02 | 7.59E-02 | 2.50E+00 | 2.58E+00 | 5.67E-01 | no | | intergenic_variant | MODIFIER |
| 16 | 75539802 | AX-170057829 | 0 | T | C | 0.491 | 4.51E-02 | 7.59E-02 | 2.50E+00 | 2.58E+00 | 5.67E-01 | no | Lipi | downstream_gene_variant | MODIFIER |
| 16 | 75562752 | AX-170479855 | 0 | C | T | 0.491 | 4.51E-02 | 7.59E-02 | 2.50E+00 | 2.58E+00 | 5.67E-01 | no | Lipi | intron_variant | MODIFIER |

| | | | | | | | | | | | | | | | |
|---|---|---|---|---|---|---|---|---|---|---|---|---|---|---|---|
| 16 | 75574985 | AX-170013408 | 0 | G | A | 0.491 | 4.51E-02 | 7.59E-02 | 2.50E+00 | 2.58E+00 | 5.67E-01 | no | Lipi | intron_variant | MODIFIER |
| 16 | 75575432 | AX-170383885 | 0 | G | A | 0.491 | 4.51E-02 | 7.59E-02 | 2.50E+00 | 2.58E+00 | 5.67E-01 | no | Lipi | intron_variant | MODIFIER |
| 16 | 75602393 | AX-97832366 | 0 | A | G | 0.491 | 4.51E-02 | 7.59E-02 | 2.50E+00 | 2.58E+00 | 5.67E-01 | no | Rbm11 | downstream_gene_variant | MODIFIER |
| 16 | 75609727 | AX-170013409 | 0 | A | T | 0.491 | 4.51E-02 | 7.59E-02 | 2.50E+00 | 2.58E+00 | 5.67E-01 | no | | intergenic_variant | MODIFIER |
| 16 | 75609789 | AX-170013403 | 0 | T | C | 0.491 | 4.51E-02 | 7.59E-02 | 2.50E+00 | 2.58E+00 | 5.67E-01 | no | | intergenic_variant | MODIFIER |
| 16 | 75616174 | AX-170236144 | 0 | C | G | 0.491 | 4.51E-02 | 7.59E-02 | 2.50E+00 | 2.58E+00 | 5.67E-01 | no | | intergenic_variant | MODIFIER |
| 16 | 75616271 | AX-97848651 | 0 | T | C | 0.491 | 4.51E-02 | 7.59E-02 | 2.50E+00 | 2.58E+00 | 5.67E-01 | no | | intergenic_variant | MODIFIER |
| 16 | 75616493 | AX-170509748 | 0 | G | C | 0.491 | 4.51E-02 | 7.59E-02 | 2.50E+00 | 2.58E+00 | 5.67E-01 | no | | regulatory_region_variant | MODIFIER |
| 16 | 75618900 | AX-170397571 | 0 | G | A | 0.491 | 4.51E-02 | 7.59E-02 | 2.50E+00 | 2.58E+00 | 5.67E-01 | no | | intergenic_variant | MODIFIER |
| 16 | 75619142 | AX-170380674 | 0 | A | G | 0.491 | 4.51E-02 | 7.59E-02 | 2.50E+00 | 2.58E+00 | 5.67E-01 | no | | intergenic_variant | MODIFIER |
| 16 | 75648921 | AX-170013402 | 0 | C | T | 0.491 | 4.51E-02 | 7.59E-02 | 2.50E+00 | 2.58E+00 | 5.67E-01 | no | Gm30162 | intron_variant,non_coding_transcript_variant | MODIFIER |
| 16 | 75649318 | AX-170013410 | 0 | T | C | 0.491 | 4.51E-02 | 7.59E-02 | 2.50E+00 | 2.58E+00 | 5.67E-01 | no | Gm30162 | intron_variant,non_coding_transcript_variant | MODIFIER |
| 16 | 75665665 | AX-97793980 | 0 | T | C | 0.491 | 4.51E-02 | 7.59E-02 | 2.50E+00 | 2.58E+00 | 5.67E-01 | no | Gm30162 | intron_variant,non_coding_transcript_variant | MODIFIER |
| 16 | 75672068 | AX-170312961 | 0 | G | A | 0.491 | 4.51E-02 | 7.59E-02 | 2.50E+00 | 2.58E+00 | 5.67E-01 | no | Gm30162 | intron_variant,non_coding_transcript_variant | MODIFIER |
| 16 | 75680825 | AX-97791890 | 0 | C | T | 0.491 | 4.51E-02 | 7.59E-02 | 2.50E+00 | 2.58E+00 | 5.67E-01 | no | Gm30162 | downstream_gene_variant | MODIFIER |
| 16 | 75751836 | AX-97841364 | 0 | A | G | 0.491 | 4.51E-02 | 7.59E-02 | 2.50E+00 | 2.58E+00 | 5.67E-01 | no | Hspa13 | downstream_gene_variant | MODIFIER |
| 16 | 75756987 | AX-170136982 | 0 | T | C | 0.491 | 4.51E-02 | 7.59E-02 | 2.50E+00 | 2.58E+00 | 5.67E-01 | no | Hspa13 | 3_prime_UTR_variant | MODIFIER |
| 16 | 75761387 | AX-170430069 | 0 | G | A | 0.491 | 4.51E-02 | 7.59E-02 | 2.50E+00 | 2.58E+00 | 5.67E-01 | no | Hspa13 | intron_variant | MODIFIER |
| 16 | 75941367 | AX-97791296 | 0 | A | G | 0.491 | 4.51E-02 | 7.59E-02 | 2.50E+00 | 2.58E+00 | 5.67E-01 | no | Samsn1 | intron_variant,non_coding_transcript_variant | MODIFIER |
| 16 | 75951885 | AX-168388257 | 0 | T | C | 0.491 | 4.51E-02 | 7.59E-02 | 2.50E+00 | 2.58E+00 | 5.67E-01 | no | Samsn1 | intron_variant,non_coding_transcript_variant | MODIFIER |
| 16 | 75966198 | AX-97791295 | 0 | T | C | 0.491 | 4.51E-02 | 7.59E-02 | 2.50E+00 | 2.58E+00 | 5.67E-01 | no | Samsn1 | intron_variant,non_coding_transcript_variant | MODIFIER |
| 16 | 76250356 | AX-170300512 | 0 | C | T | 0.491 | 4.51E-02 | 7.59E-02 | 2.50E+00 | 2.58E+00 | 5.67E-01 | no | | intergenic_variant | MODIFIER |
| 16 | 76251180 | AX-97792869 | 0 | G | A | 0.491 | 4.51E-02 | 7.59E-02 | 2.50E+00 | 2.58E+00 | 5.67E-01 | no | | intergenic_variant | MODIFIER |
| 16 | 76276938 | AX-97802599 | 0 | C | T | 0.491 | 4.51E-02 | 7.59E-02 | 2.50E+00 | 2.58E+00 | 5.67E-01 | no | | intergenic_variant | MODIFIER |
| 13 | 3967430 | AX-170177247 | 0 | T | G | 0.211 | -7.01E-02 | 1.21E-01 | 2.56E+00 | 2.63E+00 | 5.67E-01 | no | | intergenic_variant | MODIFIER |
| 1 | 116309101 | AX-97809965 | 1 | G | A | 0.17 | 1.11E-01 | 1.94E-01 | 2.59E+00 | 2.67E+00 | 5.67E-01 | no | Cntnap5a | intron_variant | MODIFIER |
| 1 | 24113962 | AX-170533333 | 1 | C | T | 0.446 | 4.78E-02 | 8.34E-02 | 2.61E+00 | 2.68E+00 | 5.67E-01 | no | | intergenic_variant | MODIFIER |
| 1 | 173182497 | AX-170453104 | 1 | G | C | 0.232 | 7.12E-02 | 1.21E-01 | 2.53E+00 | 2.59E+00 | 5.67E-01 | no | Olfr1406 | downstream_gene_variant | MODIFIER |
| 1 | 150480218 | AX-170361865 | 0 | C | T | 0.202 | 7.96E-02 | 1.41E-01 | 2.69E+00 | 2.74E+00 | 5.67E-01 | no | | intergenic_variant | MODIFIER |
| 4 | 128543350 | AX-170452600 | 0 | A | G | 0.14 | 6.36E-02 | 1.12E-01 | 2.63E+00 | 2.71E+00 | 5.67E-01 | no | Csmd2 | upstream_gene_variant | MODIFIER |
| 1 | 106762087 | AX-170394360 | 0 | T | C | 0.167 | 1.11E-01 | 1.94E-01 | 2.60E+00 | 2.68E+00 | 5.67E-01 | no | Kdsr | upstream_gene_variant | MODIFIER |
| 1 | 113767387 | AX-170366306 | 0 | G | A | 0.167 | 1.11E-01 | 1.94E-01 | 2.60E+00 | 2.68E+00 | 5.67E-01 | no | | intergenic_variant | MODIFIER |
| 1 | 115707348 | AX-170303091 | 0 | C | T | 0.167 | 1.11E-01 | 1.94E-01 | 2.60E+00 | 2.68E+00 | 5.67E-01 | no | Cntnap5a | intron_variant | MODIFIER |
| 15 | 74217891 | AX-170438575 | 2 | C | T | 0.545 | 4.64E-02 | 8.06E-02 | 2.58E+00 | 2.66E+00 | 5.67E-01 | no | | regulatory_region_variant | MODIFIER |
| 5 | 42702550 | AX-170532815 | 0 | T | C | 0.202 | -6.21E-02 | 1.09E-01 | 2.60E+00 | 2.68E+00 | 5.67E-01 | no | | intergenic_variant | MODIFIER |
| 7 | 137171021 | AX-170370048 | 1 | T | C | 0.277 | -4.66E-02 | 8.19E-02 | 2.61E+00 | 2.71E+00 | 5.67E-01 | no | Gm45499 | downstream_gene_variant | MODIFIER |
| 9 | 104757619 | AX-170033189 | 0 | C | A | 0.114 | -6.50E-02 | 1.11E-01 | 2.53E+00 | 2.62E+00 | 5.67E-01 | no | Cpne4 | intron_variant | MODIFIER |
| 9 | 83096145 | AX-170398302 | 0 | C | T | 0.211 | -6.84E-02 | 1.19E-01 | 2.59E+00 | 2.66E+00 | 5.67E-01 | no | Gm38398 | upstream_gene_variant | MODIFIER |
| 4 | 98372836 | AX-170509596 | 0 | C | T | 0.237 | 6.73E-02 | 1.17E-01 | 2.59E+00 | 2.66E+00 | 5.67E-01 | no | Tm2d1 | intron_variant | MODIFIER |
| 4 | 107548713 | AX-170149151 | 0 | T | C | 0.254 | 6.34E-02 | 1.09E-01 | 2.54E+00 | 2.62E+00 | 5.67E-01 | no | Glis1 | intron_variant | MODIFIER |
| 12 | 73728588 | AX-97794732 | 0 | C | T | 0.035 | 9.49E-02 | 1.71E-01 | 2.68E+00 | 2.84E+00 | 5.67E-01 | no | Prkch | intron_variant | MODIFIER |
| 6 | 134259507 | AX-170488091 | 0 | G | T | 0.57 | 5.06E-02 | 8.66E-02 | 2.54E+00 | 2.60E+00 | 5.68E-01 | no | Etv6 | intron_variant | MODIFIER |
| 5 | 42025006 | AX-170455394 | 0 | G | A | 0.202 | -6.20E-02 | 1.08E-01 | 2.60E+00 | 2.68E+00 | 5.68E-01 | no | | intergenic_variant | MODIFIER |
| 1 | 169358581 | AX-170364974 | 1 | C | T | 0.214 | 7.99E-02 | 1.40E-01 | 2.63E+00 | 2.67E+00 | 5.68E-01 | no | | intergenic_variant | MODIFIER |
| 8 | 81005979 | AX-170193719 | 0 | C | T | 0.202 | -7.36E-02 | 1.31E-01 | 2.73E+00 | 2.80E+00 | 5.68E-01 | no | Usp38 | intron_variant | MODIFIER |
| 8 | 85704768 | AX-170031174 | 0 | C | G | 0.202 | -7.36E-02 | 1.31E-01 | 2.73E+00 | 2.80E+00 | 5.68E-01 | no | Mir8109 | downstream_gene_variant | MODIFIER |
| 8 | 85717522 | AX-170031170 | 0 | G | A | 0.202 | -7.36E-02 | 1.31E-01 | 2.73E+00 | 2.80E+00 | 5.68E-01 | no | Itfg1 | downstream_gene_variant | MODIFIER |
| 8 | 85718517 | AX-170475334 | 0 | A | G | 0.202 | -7.36E-02 | 1.31E-01 | 2.73E+00 | 2.80E+00 | 5.68E-01 | no | Itfg1 | 3_prime_UTR_variant | MODIFIER |
| 8 | 85722676 | AX-97837145 | 0 | G | A | 0.202 | -7.36E-02 | 1.31E-01 | 2.73E+00 | 2.80E+00 | 5.68E-01 | no | Itfg1 | synonymous_variant | LOW |
| 8 | 85726898 | AX-170031180 | 0 | G | T | 0.202 | -7.36E-02 | 1.31E-01 | 2.73E+00 | 2.80E+00 | 5.68E-01 | no | Itfg1 | intron_variant | MODIFIER |
| 8 | 85754332 | AX-97809413 | 0 | A | G | 0.202 | -7.36E-02 | 1.31E-01 | 2.73E+00 | 2.80E+00 | 5.68E-01 | no | Itfg1 | intron_variant | MODIFIER |
| 8 | 85779126 | AX-97827878 | 0 | C | T | 0.202 | -7.36E-02 | 1.31E-01 | 2.73E+00 | 2.80E+00 | 5.68E-01 | no | Itfg1 | intron_variant | MODIFIER |
| 8 | 85853407 | AX-97813674 | 0 | T | C | 0.202 | -7.36E-02 | 1.31E-01 | 2.73E+00 | 2.80E+00 | 5.68E-01 | no | Phkb | intron_variant | MODIFIER |
| 8 | 85861955 | AX-170460913 | 0 | T | C | 0.202 | -7.36E-02 | 1.31E-01 | 2.73E+00 | 2.80E+00 | 5.68E-01 | no | Phkb | intron_variant | MODIFIER |

| | | | | | | | | | | | | | | | |
|---|---|---|---|---|---|---|---|---|---|---|---|---|---|---|---|
| 8 | 85869995 | AX-170126154 | 0 | A | T | 0.202 | -7.36E-02 | 1.31E-01 | 2.73E+00 | 2.80E+00 | 5.68E-01 | no | Phkb | intron_variant | MODIFIER |
| 8 | 85904111 | AX-170385516 | 0 | G | A | 0.202 | -7.36E-02 | 1.31E-01 | 2.73E+00 | 2.80E+00 | 5.68E-01 | no | Phkb | intron_variant | MODIFIER |
| 8 | 85908359 | AX-170412036 | 0 | G | C | 0.202 | -7.36E-02 | 1.31E-01 | 2.73E+00 | 2.80E+00 | 5.68E-01 | no | Phkb | intron_variant | MODIFIER |
| 8 | 85908592 | AX-97824314 | 0 | A | G | 0.202 | -7.36E-02 | 1.31E-01 | 2.73E+00 | 2.80E+00 | 5.68E-01 | no | Phkb | intron_variant | MODIFIER |
| 8 | 85923514 | AX-170031169 | 0 | T | A | 0.202 | -7.36E-02 | 1.31E-01 | 2.73E+00 | 2.80E+00 | 5.68E-01 | no | Phkb | intron_variant | MODIFIER |
| 8 | 85939252 | AX-170375451 | 0 | A | G | 0.202 | -7.36E-02 | 1.31E-01 | 2.73E+00 | 2.80E+00 | 5.68E-01 | no | Phkb | intron_variant | MODIFIER |
| 8 | 85946542 | AX-97819853 | 0 | A | G | 0.202 | -7.36E-02 | 1.31E-01 | 2.73E+00 | 2.80E+00 | 5.68E-01 | no | Phkb | intron_variant | MODIFIER |
| 8 | 85971625 | AX-170424421 | 0 | A | C | 0.202 | -7.36E-02 | 1.31E-01 | 2.73E+00 | 2.80E+00 | 5.68E-01 | no | Phkb | intron_variant | MODIFIER |
| 8 | 86019688 | AX-170113395 | 0 | C | T | 0.202 | -7.36E-02 | 1.31E-01 | 2.73E+00 | 2.80E+00 | 5.68E-01 | no | Phkb | intron_variant | MODIFIER |
| 8 | 86035409 | AX-170101182 | 0 | A | G | 0.202 | -7.36E-02 | 1.31E-01 | 2.73E+00 | 2.80E+00 | 5.68E-01 | no | Phkb | intron_variant | MODIFIER |
| 8 | 86052001 | AX-97799249 | 0 | G | T | 0.202 | -7.36E-02 | 1.31E-01 | 2.73E+00 | 2.80E+00 | 5.68E-01 | no | Phkb | intron_variant | MODIFIER |
| 8 | 86056762 | AX-170349425 | 0 | G | A | 0.202 | -7.36E-02 | 1.31E-01 | 2.73E+00 | 2.80E+00 | 5.68E-01 | no | Phkb | intron_variant | MODIFIER |
| 8 | 86087106 | AX-97817894 | 0 | T | C | 0.202 | -7.36E-02 | 1.31E-01 | 2.73E+00 | 2.80E+00 | 5.68E-01 | no | | regulatory_region_variant | MODIFIER |
| 8 | 86087916 | AX-170524119 | 0 | A | T | 0.202 | -7.36E-02 | 1.31E-01 | 2.73E+00 | 2.80E+00 | 5.68E-01 | no | | regulatory_region_variant | MODIFIER |
| 8 | 86265554 | AX-97827952 | 0 | C | T | 0.202 | -7.36E-02 | 1.31E-01 | 2.73E+00 | 2.80E+00 | 5.68E-01 | no | | intergenic_variant | MODIFIER |
| 8 | 86279191 | AX-170171549 | 0 | C | T | 0.202 | -7.36E-02 | 1.31E-01 | 2.73E+00 | 2.80E+00 | 5.68E-01 | no | | intergenic_variant | MODIFIER |
| 8 | 86328013 | AX-97809895 | 0 | A | C | 0.202 | -7.36E-02 | 1.31E-01 | 2.73E+00 | 2.80E+00 | 5.68E-01 | no | | intergenic_variant | MODIFIER |
| 8 | 86360016 | AX-170090170 | 0 | G | A | 0.202 | -7.36E-02 | 1.31E-01 | 2.73E+00 | 2.80E+00 | 5.68E-01 | no | | regulatory_region_variant | MODIFIER |
| 8 | 86398398 | AX-170509044 | 0 | C | G | 0.202 | -7.36E-02 | 1.31E-01 | 2.73E+00 | 2.80E+00 | 5.68E-01 | no | | intergenic_variant | MODIFIER |
| 8 | 86404914 | AX-170363578 | 0 | T | C | 0.202 | -7.36E-02 | 1.31E-01 | 2.73E+00 | 2.80E+00 | 5.68E-01 | no | | intergenic_variant | MODIFIER |
| 8 | 86405747 | AX-170399168 | 0 | C | G | 0.202 | -7.36E-02 | 1.31E-01 | 2.73E+00 | 2.80E+00 | 5.68E-01 | no | | intergenic_variant | MODIFIER |
| 8 | 86409226 | AX-170215580 | 0 | T | C | 0.202 | -7.36E-02 | 1.31E-01 | 2.73E+00 | 2.80E+00 | 5.68E-01 | no | | intergenic_variant | MODIFIER |
| 8 | 86424194 | AX-97834649 | 0 | C | T | 0.202 | -7.36E-02 | 1.31E-01 | 2.73E+00 | 2.80E+00 | 5.68E-01 | no | | intergenic_variant | MODIFIER |
| 8 | 86430321 | AX-170178871 | 0 | T | A | 0.202 | -7.36E-02 | 1.31E-01 | 2.73E+00 | 2.80E+00 | 5.68E-01 | no | | intergenic_variant | MODIFIER |
| 8 | 86442923 | AX-170080658 | 0 | A | G | 0.202 | -7.36E-02 | 1.31E-01 | 2.73E+00 | 2.80E+00 | 5.68E-01 | no | | intergenic_variant | MODIFIER |
| 8 | 86448293 | AX-170080659 | 0 | T | C | 0.202 | -7.36E-02 | 1.31E-01 | 2.73E+00 | 2.80E+00 | 5.68E-01 | no | | intergenic_variant | MODIFIER |
| 8 | 86455336 | AX-97818252 | 0 | C | T | 0.202 | -7.36E-02 | 1.31E-01 | 2.73E+00 | 2.80E+00 | 5.68E-01 | no | | intergenic_variant | MODIFIER |
| 8 | 86465666 | AX-97830699 | 0 | T | G | 0.202 | -7.36E-02 | 1.31E-01 | 2.73E+00 | 2.80E+00 | 5.68E-01 | no | | intergenic_variant | MODIFIER |
| 8 | 86491596 | AX-170287119 | 0 | A | G | 0.202 | -7.36E-02 | 1.31E-01 | 2.73E+00 | 2.80E+00 | 5.68E-01 | no | Abcc12 | intron_variant,NMD_transcript_variant | MODIFIER |
| 8 | 86497755 | AX-97854449 | 0 | T | C | 0.202 | -7.36E-02 | 1.31E-01 | 2.73E+00 | 2.80E+00 | 5.68E-01 | no | Abcc12 | intron_variant,NMD_transcript_variant | MODIFIER |
| 8 | 86505263 | AX-170150151 | 0 | G | A | 0.202 | -7.36E-02 | 1.31E-01 | 2.73E+00 | 2.80E+00 | 5.68E-01 | no | Abcc12 | synonymous_variant | LOW |
| 8 | 86505404 | AX-170182895 | 0 | T | C | 0.202 | -7.36E-02 | 1.31E-01 | 2.73E+00 | 2.80E+00 | 5.68E-01 | no | Abcc12 | intron_variant | MODIFIER |
| 8 | 86517364 | AX-170121666 | 0 | A | C | 0.202 | -7.36E-02 | 1.31E-01 | 2.73E+00 | 2.80E+00 | 5.68E-01 | no | Abcc12 | synonymous_variant | LOW |
| 8 | 86519950 | AX-97851097 | 0 | T | C | 0.202 | -7.36E-02 | 1.31E-01 | 2.73E+00 | 2.80E+00 | 5.68E-01 | no | Abcc12 | intron_variant | MODIFIER |
| 8 | 86535400 | AX-170099475 | 0 | C | A | 0.202 | -7.36E-02 | 1.31E-01 | 2.73E+00 | 2.80E+00 | 5.68E-01 | no | Abcc12 | intron_variant | MODIFIER |
| 8 | 86535746 | AX-170168654 | 0 | A | G | 0.202 | -7.36E-02 | 1.31E-01 | 2.73E+00 | 2.80E+00 | 5.68E-01 | no | Abcc12 | intron_variant | MODIFIER |
| 8 | 86535832 | AX-170187758 | 0 | A | C | 0.202 | -7.36E-02 | 1.31E-01 | 2.73E+00 | 2.80E+00 | 5.68E-01 | no | Abcc12 | intron_variant | MODIFIER |
| 8 | 86551224 | AX-170090171 | 0 | A | C | 0.202 | -7.36E-02 | 1.31E-01 | 2.73E+00 | 2.80E+00 | 5.68E-01 | no | Abcc12 | intron_variant | MODIFIER |
| 8 | 86802908 | AX-97821329 | 0 | G | A | 0.202 | -7.36E-02 | 1.31E-01 | 2.73E+00 | 2.80E+00 | 5.68E-01 | no | | intergenic_variant | MODIFIER |
| 8 | 86841843 | AX-170376777 | 0 | G | A | 0.202 | -7.36E-02 | 1.31E-01 | 2.73E+00 | 2.80E+00 | 5.68E-01 | no | N4bp1 | 3_prime_UTR_variant | MODIFIER |
| 8 | 86847741 | AX-97783915 | 0 | C | T | 0.202 | -7.36E-02 | 1.31E-01 | 2.73E+00 | 2.80E+00 | 5.68E-01 | no | N4bp1 | intron_variant | MODIFIER |
| 8 | 86865439 | AX-97825721 | 0 | A | C | 0.202 | -7.36E-02 | 1.31E-01 | 2.73E+00 | 2.80E+00 | 5.68E-01 | no | N4bp1 | intron_variant | MODIFIER |
| 8 | 86895330 | AX-170338968 | 0 | T | C | 0.202 | -7.36E-02 | 1.31E-01 | 2.73E+00 | 2.80E+00 | 5.68E-01 | no | | intergenic_variant | MODIFIER |
| 8 | 86899831 | AX-97782855 | 0 | A | G | 0.202 | -7.36E-02 | 1.31E-01 | 2.73E+00 | 2.80E+00 | 5.68E-01 | no | Gm27167 | upstream_gene_variant | MODIFIER |
| 8 | 86922292 | AX-170495802 | 0 | A | T | 0.202 | -7.36E-02 | 1.31E-01 | 2.73E+00 | 2.80E+00 | 5.68E-01 | no | Gm27168 | upstream_gene_variant | MODIFIER |
| 8 | 86922342 | AX-97827881 | 0 | C | T | 0.202 | -7.36E-02 | 1.31E-01 | 2.73E+00 | 2.80E+00 | 5.68E-01 | no | Gm27168 | upstream_gene_variant | MODIFIER |
| 8 | 86923769 | AX-170350208 | 0 | A | G | 0.202 | -7.36E-02 | 1.31E-01 | 2.73E+00 | 2.80E+00 | 5.68E-01 | no | Gm27168 | upstream_gene_variant | MODIFIER |
| 8 | 86983269 | AX-97793873 | 0 | C | T | 0.202 | -7.36E-02 | 1.31E-01 | 2.73E+00 | 2.80E+00 | 5.68E-01 | no | | intergenic_variant | MODIFIER |
| 8 | 87009360 | AX-170103821 | 0 | C | A | 0.202 | -7.36E-02 | 1.31E-01 | 2.73E+00 | 2.80E+00 | 5.68E-01 | no | | intergenic_variant | MODIFIER |
| 8 | 87010033 | AX-170255428 | 0 | G | A | 0.202 | -7.36E-02 | 1.31E-01 | 2.73E+00 | 2.80E+00 | 5.68E-01 | no | | intergenic_variant | MODIFIER |
| 8 | 87010070 | AX-170086373 | 0 | C | T | 0.202 | -7.36E-02 | 1.31E-01 | 2.73E+00 | 2.80E+00 | 5.68E-01 | no | | intergenic_variant | MODIFIER |
| 8 | 87010268 | AX-97812747 | 0 | C | T | 0.202 | -7.36E-02 | 1.31E-01 | 2.73E+00 | 2.80E+00 | 5.68E-01 | no | | intergenic_variant | MODIFIER |
| 8 | 87010488 | AX-170430669 | 0 | G | A | 0.202 | -7.36E-02 | 1.31E-01 | 2.73E+00 | 2.80E+00 | 5.68E-01 | no | | intergenic_variant | MODIFIER |
| 8 | 87011495 | AX-170458859 | 0 | G | C | 0.202 | -7.36E-02 | 1.31E-01 | 2.73E+00 | 2.80E+00 | 5.68E-01 | no | | intergenic_variant | MODIFIER |

| | | | | | | | | | | | | | | | |
|---|---|---|---|---|---|---|---|---|---|---|---|---|---|---|---|
| 8 | 87013482 | AX-170191763 | 0 | A | C | 0.202 | -7.36E-02 | 1.31E-01 | 2.73E+00 | 2.80E+00 | 5.68E-01 | no | | intergenic_variant | MODIFIER |
| 8 | 87016068 | AX-170346454 | 0 | G | A | 0.202 | -7.36E-02 | 1.31E-01 | 2.73E+00 | 2.80E+00 | 5.68E-01 | no | | regulatory_region_variant | MODIFIER |
| 8 | 87086095 | AX-170180816 | 0 | C | G | 0.202 | -7.36E-02 | 1.31E-01 | 2.73E+00 | 2.80E+00 | 5.68E-01 | no | | intergenic_variant | MODIFIER |
| 8 | 87090649 | AX-170244899 | 0 | C | T | 0.202 | -7.36E-02 | 1.31E-01 | 2.73E+00 | 2.80E+00 | 5.68E-01 | no | Gm5118 | upstream_gene_variant | MODIFIER |
| 8 | 87092709 | AX-170256331 | 0 | G | A | 0.202 | -7.36E-02 | 1.31E-01 | 2.73E+00 | 2.80E+00 | 5.68E-01 | no | Gm5118 | upstream_gene_variant | MODIFIER |
| 8 | 87093324 | AX-170466763 | 0 | A | G | 0.202 | -7.36E-02 | 1.31E-01 | 2.73E+00 | 2.80E+00 | 5.68E-01 | no | Gm5118 | non_coding_transcript_exon_variant | MODIFIER |
| 8 | 87094070 | AX-170353818 | 0 | T | C | 0.202 | -7.36E-02 | 1.31E-01 | 2.73E+00 | 2.80E+00 | 5.68E-01 | no | Gm5118 | intron_variant,non_coding_transcript_variant | MODIFIER |
| 8 | 87096939 | AX-170080668 | 0 | T | C | 0.202 | -7.36E-02 | 1.31E-01 | 2.73E+00 | 2.80E+00 | 5.68E-01 | no | Gm5118 | intron_variant,non_coding_transcript_variant | MODIFIER |
| 8 | 87099118 | AX-170521970 | 0 | C | T | 0.202 | -7.36E-02 | 1.31E-01 | 2.73E+00 | 2.80E+00 | 5.68E-01 | no | Gm5118 | non_coding_transcript_exon_variant | MODIFIER |
| 8 | 87101152 | AX-170420540 | 0 | C | T | 0.202 | -7.36E-02 | 1.31E-01 | 2.73E+00 | 2.80E+00 | 5.68E-01 | no | Gm5118 | downstream_gene_variant | MODIFIER |
| 8 | 87102466 | AX-170484973 | 0 | C | T | 0.202 | -7.36E-02 | 1.31E-01 | 2.73E+00 | 2.80E+00 | 5.68E-01 | no | Gm5118 | downstream_gene_variant | MODIFIER |
| 8 | 87106939 | AX-97802508 | 0 | T | C | 0.202 | -7.36E-02 | 1.31E-01 | 2.73E+00 | 2.80E+00 | 5.68E-01 | no | | intergenic_variant | MODIFIER |
| 8 | 87119644 | AX-170196515 | 0 | A | G | 0.202 | -7.36E-02 | 1.31E-01 | 2.73E+00 | 2.80E+00 | 5.68E-01 | no | Gm27246 | intron_variant,non_coding_transcript_variant | MODIFIER |
| 8 | 87120235 | AX-170520509 | 0 | A | G | 0.202 | -7.36E-02 | 1.31E-01 | 2.73E+00 | 2.80E+00 | 5.68E-01 | no | Gm27246 | intron_variant,non_coding_transcript_variant | MODIFIER |
| 8 | 87122564 | AX-170361012 | 0 | T | C | 0.202 | -7.36E-02 | 1.31E-01 | 2.73E+00 | 2.80E+00 | 5.68E-01 | no | Gm27246 | intron_variant,non_coding_transcript_variant | MODIFIER |
| 8 | 87124842 | AX-170259879 | 0 | C | T | 0.202 | -7.36E-02 | 1.31E-01 | 2.73E+00 | 2.80E+00 | 5.68E-01 | no | Gm27246 | intron_variant,non_coding_transcript_variant | MODIFIER |
| 8 | 87198932 | AX-97811434 | 0 | C | T | 0.202 | -7.36E-02 | 1.31E-01 | 2.73E+00 | 2.80E+00 | 5.68E-01 | no | Gm27169 | upstream_gene_variant | MODIFIER |
| 18 | 77626765 | AX-97831200 | 0 | A | G | 0.395 | -4.93E-02 | 8.66E-02 | 2.62E+00 | 2.71E+00 | 5.68E-01 | no | 8030462N17Rik | downstream_gene_variant | MODIFIER |
| 18 | 77646481 | AX-170093963 | 0 | T | C | 0.395 | -4.93E-02 | 8.66E-02 | 2.62E+00 | 2.71E+00 | 5.68E-01 | no | 8030462N17Rik | intron_variant | MODIFIER |
| 18 | 77650354 | AX-170436266 | 0 | G | A | 0.395 | -4.93E-02 | 8.66E-02 | 2.62E+00 | 2.71E+00 | 5.68E-01 | no | 8030462N17Rik | intron_variant | MODIFIER |
| 18 | 77654902 | AX-170016038 | 0 | C | T | 0.395 | -4.93E-02 | 8.66E-02 | 2.62E+00 | 2.71E+00 | 5.68E-01 | no | 8030462N17Rik | intron_variant | MODIFIER |
| 18 | 77680318 | AX-170016035 | 0 | G | A | 0.395 | -4.93E-02 | 8.66E-02 | 2.62E+00 | 2.71E+00 | 5.68E-01 | no | 8030462N17Rik | intron_variant | MODIFIER |
| 18 | 78483316 | AX-170483991 | 0 | T | C | 0.395 | -4.93E-02 | 8.66E-02 | 2.62E+00 | 2.71E+00 | 5.68E-01 | no | Slc14a2 | intron_variant | MODIFIER |
| 18 | 78501466 | AX-97805642 | 0 | C | T | 0.395 | -4.93E-02 | 8.66E-02 | 2.62E+00 | 2.71E+00 | 5.68E-01 | no | Slc14a2 | intron_variant | MODIFIER |
| 18 | 78502258 | AX-170130901 | 0 | T | A | 0.395 | -4.93E-02 | 8.66E-02 | 2.62E+00 | 2.71E+00 | 5.68E-01 | no | Slc14a2 | intron_variant | MODIFIER |
| 18 | 78521676 | AX-97835115 | 0 | G | A | 0.395 | -4.93E-02 | 8.66E-02 | 2.62E+00 | 2.71E+00 | 5.68E-01 | no | Slc14a2 | intron_variant | MODIFIER |
| 18 | 78542246 | AX-170453858 | 0 | A | G | 0.395 | -4.93E-02 | 8.66E-02 | 2.62E+00 | 2.71E+00 | 5.68E-01 | no | Slc14a2 | intron_variant | MODIFIER |
| 18 | 78644037 | AX-170308740 | 0 | G | C | 0.395 | -4.93E-02 | 8.66E-02 | 2.62E+00 | 2.71E+00 | 5.68E-01 | no | Gm23895 | upstream_gene_variant | MODIFIER |
| 18 | 78646237 | AX-170130177 | 0 | G | C | 0.395 | -4.93E-02 | 8.66E-02 | 2.62E+00 | 2.71E+00 | 5.68E-01 | no | Gm23895 | upstream_gene_variant | MODIFIER |
| 18 | 78648762 | AX-170516426 | 0 | C | G | 0.395 | -4.93E-02 | 8.66E-02 | 2.62E+00 | 2.71E+00 | 5.68E-01 | no | Gm23895 | downstream_gene_variant | MODIFIER |
| 7 | 40101831 | AX-170408271 | 1 | G | A | 0.357 | 5.46E-02 | 9.53E-02 | 2.59E+00 | 2.67E+00 | 5.68E-01 | no | | intergenic_variant | MODIFIER |
| 5 | 106608845 | AX-170025816 | 1 | C | T | 0.455 | 6.66E-02 | 1.17E-01 | 2.66E+00 | 2.69E+00 | 5.68E-01 | no | Gm28039 | intron_variant,non_coding_transcript_variant | MODIFIER |
| 5 | 127842352 | AX-170227822 | 0 | C | G | 0.412 | 4.63E-02 | 8.17E-02 | 2.61E+00 | 2.72E+00 | 5.68E-01 | no | Tmem132d | intron_variant | MODIFIER |
| 2 | 51898908 | AX-170523538 | 1 | C | A | 0.036 | -1.20E-01 | 2.06E-01 | 2.54E+00 | 2.62E+00 | 5.68E-01 | no | | intergenic_variant | MODIFIER |
| 9 | 79319446 | AX-170231208 | 0 | G | C | 0.202 | -6.84E-02 | 1.21E-01 | 2.67E+00 | 2.75E+00 | 5.69E-01 | no | | intergenic_variant | MODIFIER |
| 16 | 79177290 | AX-97830603 | 0 | G | A | 0.465 | 4.24E-02 | 7.43E-02 | 2.60E+00 | 2.68E+00 | 5.69E-01 | no | | regulatory_region_variant | MODIFIER |
| 18 | 80665069 | AX-170104948 | 0 | A | G | 0.298 | -4.90E-02 | 8.55E-02 | 2.57E+00 | 2.67E+00 | 5.69E-01 | no | Nfatc1 | intron_variant | MODIFIER |
| 18 | 80679154 | AX-170306863 | 0 | A | G | 0.298 | -4.90E-02 | 8.55E-02 | 2.57E+00 | 2.67E+00 | 5.69E-01 | no | Nfatc1 | intron_variant | MODIFIER |
| 18 | 80679192 | AX-97854415 | 0 | G | A | 0.298 | -4.90E-02 | 8.55E-02 | 2.57E+00 | 2.67E+00 | 5.69E-01 | no | Nfatc1 | intron_variant | MODIFIER |
| 18 | 80690721 | AX-170357603 | 0 | C | T | 0.298 | -4.90E-02 | 8.55E-02 | 2.57E+00 | 2.67E+00 | 5.69E-01 | no | Nfatc1 | intron_variant | MODIFIER |
| 18 | 80694665 | AX-170498902 | 0 | A | G | 0.298 | -4.90E-02 | 8.55E-02 | 2.57E+00 | 2.67E+00 | 5.69E-01 | no | Nfatc1 | intron_variant | MODIFIER |
| 18 | 80696277 | AX-170348520 | 0 | T | C | 0.298 | -4.90E-02 | 8.55E-02 | 2.57E+00 | 2.67E+00 | 5.69E-01 | no | Nfatc1 | intron_variant | MODIFIER |
| 18 | 80696361 | AX-170271642 | 0 | A | G | 0.298 | -4.90E-02 | 8.55E-02 | 2.57E+00 | 2.67E+00 | 5.69E-01 | no | Nfatc1 | intron_variant | MODIFIER |
| 18 | 80699847 | AX-170102805 | 0 | G | T | 0.298 | -4.90E-02 | 8.55E-02 | 2.57E+00 | 2.67E+00 | 5.69E-01 | no | Nfatc1 | intron_variant | MODIFIER |
| 18 | 80702680 | AX-170169437 | 0 | C | T | 0.298 | -4.90E-02 | 8.55E-02 | 2.57E+00 | 2.67E+00 | 5.69E-01 | no | Nfatc1 | intron_variant | MODIFIER |
| 18 | 80704621 | AX-170379351 | 0 | C | T | 0.298 | -4.90E-02 | 8.55E-02 | 2.57E+00 | 2.67E+00 | 5.69E-01 | no | Nfatc1 | intron_variant | MODIFIER |
| 18 | 80731128 | AX-170174537 | 0 | C | A | 0.298 | -4.90E-02 | 8.55E-02 | 2.57E+00 | 2.67E+00 | 5.69E-01 | no | Atp9b | downstream_gene_variant | MODIFIER |
| 18 | 80733113 | AX-170518007 | 0 | T | C | 0.298 | -4.90E-02 | 8.55E-02 | 2.57E+00 | 2.67E+00 | 5.69E-01 | no | Atp9b | downstream_gene_variant | MODIFIER |
| 18 | 80733861 | AX-170260770 | 0 | T | A | 0.298 | -4.90E-02 | 8.55E-02 | 2.57E+00 | 2.67E+00 | 5.69E-01 | no | Atp9b | downstream_gene_variant | MODIFIER |
| 18 | 80787378 | AX-170269349 | 0 | C | T | 0.298 | -4.90E-02 | 8.55E-02 | 2.57E+00 | 2.67E+00 | 5.69E-01 | no | Atp9b | intron_variant | MODIFIER |
| 18 | 80805547 | AX-170499549 | 0 | A | G | 0.298 | -4.90E-02 | 8.55E-02 | 2.57E+00 | 2.67E+00 | 5.69E-01 | no | Atp9b | intron_variant | MODIFIER |
| 18 | 80858680 | AX-170382069 | 0 | C | T | 0.298 | -4.90E-02 | 8.55E-02 | 2.57E+00 | 2.67E+00 | 5.69E-01 | no | Atp9b | intron_variant | MODIFIER |
| 18 | 80905369 | AX-170404362 | 0 | G | A | 0.298 | -4.90E-02 | 8.55E-02 | 2.57E+00 | 2.67E+00 | 5.69E-01 | no | Atp9b | intron_variant | MODIFIER |
| 18 | 80908513 | AX-170061464 | 0 | T | C | 0.298 | -4.90E-02 | 8.55E-02 | 2.57E+00 | 2.67E+00 | 5.69E-01 | no | Atp9b | intron_variant | MODIFIER |

| | | | | | | | | | | | | | | | |
|---|---|---|---|---|---|---|---|---|---|---|---|---|---|---|---|
| 18 | 80939476 | AX-170203662 | 0 | G | T | 0.298 | -4.90E-02 | 8.55E-02 | 2.57E+00 | 2.67E+00 | 5.69E-01 | no | | intergenic_variant | MODIFIER |
| 18 | 81006742 | AX-170395606 | 0 | T | G | 0.298 | -4.90E-02 | 8.55E-02 | 2.57E+00 | 2.67E+00 | 5.69E-01 | no | | intergenic_variant | MODIFIER |
| 18 | 81007812 | AX-170154113 | 0 | G | A | 0.298 | -4.90E-02 | 8.55E-02 | 2.57E+00 | 2.67E+00 | 5.69E-01 | no | | intergenic_variant | MODIFIER |
| 18 | 81008047 | AX-170467897 | 0 | C | T | 0.298 | -4.90E-02 | 8.55E-02 | 2.57E+00 | 2.67E+00 | 5.69E-01 | no | | intergenic_variant | MODIFIER |
| 18 | 81008396 | AX-170346456 | 0 | T | G | 0.298 | -4.90E-02 | 8.55E-02 | 2.57E+00 | 2.67E+00 | 5.69E-01 | no | | intergenic_variant | MODIFIER |
| 18 | 81014984 | AX-170213915 | 0 | T | C | 0.298 | -4.90E-02 | 8.55E-02 | 2.57E+00 | 2.67E+00 | 5.69E-01 | no | | intergenic_variant | MODIFIER |
| 18 | 81026193 | AX-170350375 | 0 | G | A | 0.298 | -4.90E-02 | 8.55E-02 | 2.57E+00 | 2.67E+00 | 5.69E-01 | no | Gm25718 | upstream_gene_variant | MODIFIER |
| 18 | 81031015 | AX-170337842 | 0 | C | G | 0.298 | -4.90E-02 | 8.55E-02 | 2.57E+00 | 2.67E+00 | 5.69E-01 | no | Gm25718 | downstream_gene_variant | MODIFIER |
| 18 | 81033152 | AX-170432560 | 0 | G | C | 0.298 | -4.90E-02 | 8.55E-02 | 2.57E+00 | 2.67E+00 | 5.69E-01 | no | | intergenic_variant | MODIFIER |
| 18 | 81050641 | AX-170329326 | 0 | G | A | 0.298 | -4.90E-02 | 8.55E-02 | 2.57E+00 | 2.67E+00 | 5.69E-01 | no | AC125210.2 | downstream_gene_variant | MODIFIER |
| 18 | 81051843 | AX-170221060 | 0 | C | A | 0.298 | -4.90E-02 | 8.55E-02 | 2.57E+00 | 2.67E+00 | 5.69E-01 | no | AC125210.2 | downstream_gene_variant | MODIFIER |
| 18 | 81051860 | AX-170061467 | 0 | T | A | 0.298 | -4.90E-02 | 8.55E-02 | 2.57E+00 | 2.67E+00 | 5.69E-01 | no | AC125210.2 | downstream_gene_variant | MODIFIER |
| 18 | 81065559 | AX-170061468 | 0 | G | C | 0.298 | -4.90E-02 | 8.55E-02 | 2.57E+00 | 2.67E+00 | 5.69E-01 | no | | intergenic_variant | MODIFIER |
| 18 | 81067976 | AX-170523424 | 0 | T | C | 0.298 | -4.90E-02 | 8.55E-02 | 2.57E+00 | 2.67E+00 | 5.69E-01 | no | | intergenic_variant | MODIFIER |
| 18 | 81076735 | AX-170445744 | 0 | G | A | 0.298 | -4.90E-02 | 8.55E-02 | 2.57E+00 | 2.67E+00 | 5.69E-01 | no | | intergenic_variant | MODIFIER |
| 18 | 81083251 | AX-170016082 | 0 | G | A | 0.298 | -4.90E-02 | 8.55E-02 | 2.57E+00 | 2.67E+00 | 5.69E-01 | no | AC105065.1 | non_coding_transcript_exon_variant | MODIFIER |
| 18 | 81084747 | AX-97782563 | 0 | T | C | 0.298 | -4.90E-02 | 8.55E-02 | 2.57E+00 | 2.67E+00 | 5.69E-01 | no | AC105065.1 | intron_variant,non_coding_transcript_variant | MODIFIER |
| 18 | 81088479 | AX-170115784 | 0 | G | A | 0.298 | -4.90E-02 | 8.55E-02 | 2.57E+00 | 2.67E+00 | 5.69E-01 | no | AC105065.1 | intron_variant,non_coding_transcript_variant | MODIFIER |
| 18 | 81105415 | AX-97856985 | 0 | G | A | 0.298 | -4.90E-02 | 8.55E-02 | 2.57E+00 | 2.67E+00 | 5.69E-01 | no | | intergenic_variant | MODIFIER |
| 18 | 81110035 | AX-170477255 | 0 | T | G | 0.298 | -4.90E-02 | 8.55E-02 | 2.57E+00 | 2.67E+00 | 5.69E-01 | no | | intergenic_variant | MODIFIER |
| 18 | 81128854 | AX-170424832 | 0 | A | C | 0.298 | -4.90E-02 | 8.55E-02 | 2.57E+00 | 2.67E+00 | 5.69E-01 | no | | intergenic_variant | MODIFIER |
| 18 | 81129093 | AX-170490390 | 0 | C | T | 0.298 | -4.90E-02 | 8.55E-02 | 2.57E+00 | 2.67E+00 | 5.69E-01 | no | | intergenic_variant | MODIFIER |
| 18 | 81134420 | AX-170016086 | 0 | T | G | 0.298 | -4.90E-02 | 8.55E-02 | 2.57E+00 | 2.67E+00 | 5.69E-01 | no | | intergenic_variant | MODIFIER |
| 18 | 81137236 | AX-170428939 | 0 | C | T | 0.298 | -4.90E-02 | 8.55E-02 | 2.57E+00 | 2.67E+00 | 5.69E-01 | no | | intergenic_variant | MODIFIER |
| 18 | 81140539 | AX-170186253 | 0 | T | C | 0.298 | -4.90E-02 | 8.55E-02 | 2.57E+00 | 2.67E+00 | 5.69E-01 | no | | intergenic_variant | MODIFIER |
| 18 | 81141894 | AX-170263828 | 0 | A | G | 0.298 | -4.90E-02 | 8.55E-02 | 2.57E+00 | 2.67E+00 | 5.69E-01 | no | | intergenic_variant | MODIFIER |
| 18 | 81142040 | AX-170424975 | 0 | A | G | 0.298 | -4.90E-02 | 8.55E-02 | 2.57E+00 | 2.67E+00 | 5.69E-01 | no | | intergenic_variant | MODIFIER |
| 13 | 47279443 | AX-97808260 | 0 | G | A | 0.465 | -3.56E-02 | 6.38E-02 | 2.73E+00 | 2.84E+00 | 5.69E-01 | no | | regulatory_region_variant | MODIFIER |
| 1 | 175349177 | AX-170360267 | 1 | C | A | 0.232 | 7.12E-02 | 1.21E-01 | 2.52E+00 | 2.58E+00 | 5.69E-01 | no | Rgs7 | intron_variant | MODIFIER |
| 15 | 79844334 | AX-170121982 | 1 | A | G | 0.554 | 4.70E-02 | 8.12E-02 | 2.55E+00 | 2.62E+00 | 5.69E-01 | no | Gm10856 | non_coding_transcript_exon_variant | MODIFIER |
| 9 | 16574775 | AX-170081776 | 0 | G | A | 0.482 | 4.80E-02 | 8.36E-02 | 2.57E+00 | 2.65E+00 | 5.69E-01 | no | | intergenic_variant | MODIFIER |
| 11 | 48026653 | AX-170087016 | 0 | A | G | 0.088 | -9.11E-02 | 1.60E-01 | 2.58E+00 | 2.69E+00 | 5.69E-01 | no | | intergenic_variant | MODIFIER |
| 13 | 37250600 | AX-170050982 | 0 | G | A | 0.175 | -7.07E-02 | 1.23E-01 | 2.57E+00 | 2.63E+00 | 5.69E-01 | no | Gm47709 | intron_variant,non_coding_transcript_variant | MODIFIER |
| 1 | 128682844 | AX-170001677 | 0 | T | A | 0.544 | 4.84E-02 | 8.56E-02 | 2.64E+00 | 2.70E+00 | 5.69E-01 | no | | regulatory_region_variant | MODIFIER |
| 2 | 162096051 | AX-170506294 | 0 | G | A | 0.158 | 9.20E-02 | 1.65E-01 | 2.71E+00 | 2.82E+00 | 5.70E-01 | no | Ptprt | intron_variant | MODIFIER |
| 2 | 162552746 | AX-97810119 | 0 | C | T | 0.158 | 9.20E-02 | 1.65E-01 | 2.71E+00 | 2.82E+00 | 5.70E-01 | no | Ptprt | intron_variant | MODIFIER |
| 9 | 79453420 | AX-170194311 | 0 | C | T | 0.193 | -6.94E-02 | 1.23E-01 | 2.66E+00 | 2.73E+00 | 5.70E-01 | no | Gm10635 | intron_variant,non_coding_transcript_variant | MODIFIER |
| 8 | 106770464 | AX-97815245 | 1 | G | A | 0.232 | 5.99E-02 | 1.07E-01 | 2.71E+00 | 2.78E+00 | 5.70E-01 | no | Tango6 | intron_variant | MODIFIER |
| 3 | 38758073 | AX-170066423 | 0 | G | A | 0.202 | 4.84E-02 | 8.69E-02 | 2.69E+00 | 2.80E+00 | 5.70E-01 | no | | intergenic_variant | MODIFIER |
| 3 | 38906500 | AX-170478061 | 0 | T | C | 0.202 | 4.84E-02 | 8.69E-02 | 2.69E+00 | 2.80E+00 | 5.70E-01 | no | Fat4 | intron_variant | MODIFIER |
| 1 | 174343003 | AX-97830088 | 1 | G | A | 0.232 | 7.11E-02 | 1.22E-01 | 2.53E+00 | 2.58E+00 | 5.70E-01 | no | | intergenic_variant | MODIFIER |
| 1 | 169342908 | AX-170522007 | 0 | C | G | 0.211 | 7.86E-02 | 1.40E-01 | 2.67E+00 | 2.72E+00 | 5.70E-01 | no | Gm37839 | intron_variant,non_coding_transcript_variant | MODIFIER |
| 16 | 29274361 | AX-170293556 | 0 | T | G | 0.509 | 4.20E-02 | 7.36E-02 | 2.59E+00 | 2.66E+00 | 5.70E-01 | no | Atp13a5 | intron_variant | MODIFIER |
| 8 | 34635084 | AX-170402935 | 2 | T | G | 0.136 | -7.18E-02 | 1.27E-01 | 2.63E+00 | 2.71E+00 | 5.70E-01 | no | B930018H19Rik | intron_variant,non_coding_transcript_variant | MODIFIER |
| 7 | 37430599 | AX-170252621 | 0 | G | A | 0.342 | 5.23E-02 | 9.20E-02 | 2.60E+00 | 2.67E+00 | 5.70E-01 | no | | intergenic_variant | MODIFIER |
| 1 | 23041813 | AX-97829536 | 0 | A | C | 0.43 | 4.71E-02 | 8.34E-02 | 2.63E+00 | 2.69E+00 | 5.71E-01 | no | | intergenic_variant | MODIFIER |
| 1 | 23043337 | AX-170000077 | 0 | A | G | 0.43 | 4.71E-02 | 8.34E-02 | 2.63E+00 | 2.69E+00 | 5.71E-01 | no | Gm29506 | downstream_gene_variant | MODIFIER |
| 1 | 23626807 | AX-170000094 | 0 | A | G | 0.43 | 4.71E-02 | 8.34E-02 | 2.63E+00 | 2.69E+00 | 5.71E-01 | no | | intergenic_variant | MODIFIER |
| 1 | 23626952 | AX-170000097 | 0 | T | C | 0.43 | 4.71E-02 | 8.34E-02 | 2.63E+00 | 2.69E+00 | 5.71E-01 | no | | intergenic_variant | MODIFIER |
| 1 | 23627161 | AX-170000090 | 0 | T | G | 0.43 | 4.71E-02 | 8.34E-02 | 2.63E+00 | 2.69E+00 | 5.71E-01 | no | | intergenic_variant | MODIFIER |
| 1 | 23629446 | AX-170292804 | 0 | C | T | 0.43 | 4.71E-02 | 8.34E-02 | 2.63E+00 | 2.69E+00 | 5.71E-01 | no | | intergenic_variant | MODIFIER |
| 1 | 23651997 | AX-168380113 | 0 | C | T | 0.43 | 4.71E-02 | 8.34E-02 | 2.63E+00 | 2.69E+00 | 5.71E-01 | no | | intergenic_variant | MODIFIER |
| 1 | 24312463 | AX-170144666 | 0 | A | C | 0.43 | 4.71E-02 | 8.34E-02 | 2.63E+00 | 2.69E+00 | 5.71E-01 | no | Col19a1 | missense_variant | MODERATE |
| 1 | 24946778 | AX-168403696 | 0 | C | T | 0.43 | 4.71E-02 | 8.34E-02 | 2.63E+00 | 2.69E+00 | 5.71E-01 | no | | intergenic_variant | MODIFIER |

| | | | | | | | | | | | | | | | |
|---|---|---|---|---|---|---|---|---|---|---|---|---|---|---|---|
| 1 | 24227973 | AX-170270522 | 0 | T | G | 0.43 | 4.76E-02 | 8.44E-02 | 2.64E+00 | 2.71E+00 | 5.71E-01 | no | Col9a1 | intron_variant | MODIFIER |
| 5 | 103591577 | AX-170178199 | 1 | T | C | 0.455 | 6.62E-02 | 1.18E-01 | 2.66E+00 | 2.69E+00 | 5.71E-01 | no | Ptpn13 | intron_variant | MODIFIER |
| 5 | 116729707 | AX-170117277 | 1 | C | G | 0.455 | 6.62E-02 | 1.18E-01 | 2.66E+00 | 2.69E+00 | 5.71E-01 | no | Gm43122 | intron_variant,non_coding_transcript_variant | MODIFIER |
| 5 | 116762487 | AX-170317903 | 1 | G | A | 0.455 | 6.62E-02 | 1.18E-01 | 2.66E+00 | 2.69E+00 | 5.71E-01 | no | Gm43122 | intron_variant,non_coding_transcript_variant | MODIFIER |
| 4 | 89627944 | AX-97848103 | 0 | A | G | 0.439 | 4.36E-02 | 7.64E-02 | 2.93E+00 | 3.06E+00 | 5.71E-01 | no | | intergenic_variant | MODIFIER |
| 1 | 23063992 | AX-170231142 | 0 | T | C | 0.439 | 4.75E-02 | 8.42E-02 | 2.63E+00 | 2.69E+00 | 5.71E-01 | no | Gm29506 | intron_variant,non_coding_transcript_variant | MODIFIER |
| 1 | 23074216 | AX-170196683 | 0 | G | A | 0.439 | 4.75E-02 | 8.42E-02 | 2.63E+00 | 2.69E+00 | 5.71E-01 | no | Gm29506 | intron_variant,non_coding_transcript_variant | MODIFIER |
| 1 | 23074701 | AX-170204141 | 0 | T | C | 0.439 | 4.75E-02 | 8.42E-02 | 2.63E+00 | 2.69E+00 | 5.71E-01 | no | Gm29506 | intron_variant,non_coding_transcript_variant | MODIFIER |
| 1 | 23268315 | AX-170391859 | 0 | G | A | 0.439 | 4.75E-02 | 8.42E-02 | 2.63E+00 | 2.69E+00 | 5.71E-01 | no | Mir30a | upstream_gene_variant | MODIFIER |
| 1 | 23284883 | AX-170401138 | 0 | A | G | 0.439 | 4.75E-02 | 8.42E-02 | 2.63E+00 | 2.69E+00 | 5.71E-01 | no | | regulatory_region_variant | MODIFIER |
| 1 | 23337362 | AX-170224642 | 0 | G | A | 0.439 | 4.75E-02 | 8.42E-02 | 2.63E+00 | 2.69E+00 | 5.71E-01 | no | | regulatory_region_variant | MODIFIER |
| 1 | 23373596 | AX-170446141 | 0 | G | A | 0.439 | 4.75E-02 | 8.42E-02 | 2.63E+00 | 2.69E+00 | 5.71E-01 | no | Ogfrl1 | intron_variant | MODIFIER |
| 1 | 23397340 | AX-170098576 | 0 | A | G | 0.439 | 4.75E-02 | 8.42E-02 | 2.63E+00 | 2.69E+00 | 5.71E-01 | no | Ogfrl1 | intron_variant | MODIFIER |
| 1 | 23467141 | AX-170104301 | 0 | C | T | 0.439 | 4.75E-02 | 8.42E-02 | 2.63E+00 | 2.69E+00 | 5.71E-01 | no | | intergenic_variant | MODIFIER |
| 1 | 23588345 | AX-170241780 | 0 | T | C | 0.439 | 4.75E-02 | 8.42E-02 | 2.63E+00 | 2.69E+00 | 5.71E-01 | no | Gm28822 | intron_variant,non_coding_transcript_variant | MODIFIER |
| 1 | 24114061 | AX-170511132 | 0 | C | T | 0.439 | 4.75E-02 | 8.42E-02 | 2.63E+00 | 2.69E+00 | 5.71E-01 | no | | regulatory_region_variant | MODIFIER |
| 1 | 24124719 | AX-170305643 | 0 | T | C | 0.439 | 4.75E-02 | 8.42E-02 | 2.63E+00 | 2.69E+00 | 5.71E-01 | no | Gm26607 | upstream_gene_variant | MODIFIER |
| 1 | 24157041 | AX-170472739 | 0 | G | A | 0.439 | 4.75E-02 | 8.42E-02 | 2.63E+00 | 2.69E+00 | 5.71E-01 | no | | intergenic_variant | MODIFIER |
| 1 | 24159929 | AX-170131019 | 0 | G | C | 0.439 | 4.75E-02 | 8.42E-02 | 2.63E+00 | 2.69E+00 | 5.71E-01 | no | | intergenic_variant | MODIFIER |
| 1 | 24160223 | AX-170522014 | 0 | C | T | 0.439 | 4.75E-02 | 8.42E-02 | 2.63E+00 | 2.69E+00 | 5.71E-01 | no | | intergenic_variant | MODIFIER |
| 1 | 24231528 | AX-170406798 | 0 | A | C | 0.439 | 4.75E-02 | 8.42E-02 | 2.63E+00 | 2.69E+00 | 5.71E-01 | no | Col9a1 | intron_variant | MODIFIER |
| 1 | 24235210 | AX-170211078 | 0 | T | C | 0.439 | 4.75E-02 | 8.42E-02 | 2.63E+00 | 2.69E+00 | 5.71E-01 | no | Col9a1 | intron_variant | MODIFIER |
| 1 | 24246655 | AX-170243478 | 0 | T | C | 0.439 | 4.75E-02 | 8.42E-02 | 2.63E+00 | 2.69E+00 | 5.71E-01 | no | Col9a1 | intron_variant | MODIFIER |
| 1 | 24249626 | AX-170040630 | 0 | G | T | 0.439 | 4.75E-02 | 8.42E-02 | 2.63E+00 | 2.69E+00 | 5.71E-01 | no | Col9a1 | intron_variant | MODIFIER |
| 1 | 24264904 | AX-170000105 | 0 | C | A | 0.439 | 4.75E-02 | 8.42E-02 | 2.63E+00 | 2.69E+00 | 5.71E-01 | no | Col19a1 | intron_variant | MODIFIER |
| 1 | 24265652 | AX-170489041 | 0 | A | G | 0.439 | 4.75E-02 | 8.42E-02 | 2.63E+00 | 2.69E+00 | 5.71E-01 | no | Col19a1 | intron_variant | MODIFIER |
| 1 | 24299976 | AX-170530328 | 0 | A | G | 0.439 | 4.75E-02 | 8.42E-02 | 2.63E+00 | 2.69E+00 | 5.71E-01 | no | Col19a1 | intron_variant | MODIFIER |
| 1 | 24313138 | AX-170224468 | 0 | C | G | 0.439 | 4.75E-02 | 8.42E-02 | 2.63E+00 | 2.69E+00 | 5.71E-01 | no | Col19a1 | intron_variant | MODIFIER |
| 1 | 24313729 | AX-170203472 | 0 | A | G | 0.439 | 4.75E-02 | 8.42E-02 | 2.63E+00 | 2.69E+00 | 5.71E-01 | no | Col19a1 | intron_variant | MODIFIER |
| 1 | 24314887 | AX-170229535 | 0 | C | T | 0.439 | 4.75E-02 | 8.42E-02 | 2.63E+00 | 2.69E+00 | 5.71E-01 | no | Col19a1 | intron_variant | MODIFIER |
| 1 | 24315410 | AX-170529058 | 0 | T | G | 0.439 | 4.75E-02 | 8.42E-02 | 2.63E+00 | 2.69E+00 | 5.71E-01 | no | Col19a1 | synonymous_variant | LOW |
| 1 | 24319593 | AX-170317083 | 0 | A | G | 0.439 | 4.75E-02 | 8.42E-02 | 2.63E+00 | 2.69E+00 | 5.71E-01 | no | Col19a1 | intron_variant | MODIFIER |
| 1 | 24319747 | AX-170501676 | 0 | T | C | 0.439 | 4.75E-02 | 8.42E-02 | 2.63E+00 | 2.69E+00 | 5.71E-01 | no | Col19a1 | intron_variant | MODIFIER |
| 1 | 24349448 | AX-170110725 | 0 | A | G | 0.439 | 4.75E-02 | 8.42E-02 | 2.63E+00 | 2.69E+00 | 5.71E-01 | no | Col19a1 | intron_variant | MODIFIER |
| 1 | 24397722 | AX-170305064 | 0 | A | G | 0.439 | 4.75E-02 | 8.42E-02 | 2.63E+00 | 2.69E+00 | 5.71E-01 | no | Col19a1 | intron_variant | MODIFIER |
| 1 | 24422694 | AX-170184716 | 0 | G | C | 0.439 | 4.75E-02 | 8.42E-02 | 2.63E+00 | 2.69E+00 | 5.71E-01 | no | Col19a1 | intron_variant | MODIFIER |
| 2 | 92869272 | AX-170018343 | 0 | G | A | 0.061 | -1.27E-01 | 2.27E-01 | 2.72E+00 | 2.81E+00 | 5.71E-01 | no | Gm13792 | downstream_gene_variant | MODIFIER |
| 2 | 97506070 | AX-170226065 | 0 | G | T | 0.061 | -1.27E-01 | 2.27E-01 | 2.72E+00 | 2.81E+00 | 5.71E-01 | no | Lrrc4c | intron_variant | MODIFIER |
| 18 | 65124234 | AX-170103669 | 0 | T | A | 0.404 | 5.33E-02 | 9.08E-02 | 2.51E+00 | 2.56E+00 | 5.71E-01 | no | Nedd4l | intron_variant | MODIFIER |
| 1 | 172582700 | AX-170043603 | 1 | A | T | 0.223 | -7.56E-02 | 1.31E-01 | 2.96E+00 | 3.07E+00 | 5.71E-01 | no | Slamf8 | intron_variant | MODIFIER |
| 15 | 81323886 | AX-170012105 | 0 | A | G | 0.553 | 4.69E-02 | 8.14E-02 | 2.55E+00 | 2.62E+00 | 5.71E-01 | no | Slc25a17 | intron_variant | MODIFIER |
| 15 | 81336696 | AX-170270196 | 0 | C | T | 0.553 | 4.69E-02 | 8.14E-02 | 2.55E+00 | 2.62E+00 | 5.71E-01 | no | Slc25a17 | intron_variant | MODIFIER |
| 15 | 81338704 | AX-170294376 | 0 | C | G | 0.553 | 4.69E-02 | 8.14E-02 | 2.55E+00 | 2.62E+00 | 5.71E-01 | no | Slc25a17 | intron_variant | MODIFIER |
| 15 | 81342799 | AX-97781335 | 0 | T | G | 0.553 | 4.69E-02 | 8.14E-02 | 2.55E+00 | 2.62E+00 | 5.71E-01 | no | Slc25a17 | intron_variant | MODIFIER |
| 15 | 81398815 | AX-170535013 | 0 | C | T | 0.553 | 4.69E-02 | 8.14E-02 | 2.55E+00 | 2.62E+00 | 5.71E-01 | no | St13 | intron_variant | MODIFIER |
| 15 | 81541869 | AX-170372303 | 0 | G | A | 0.553 | 4.69E-02 | 8.14E-02 | 2.55E+00 | 2.62E+00 | 5.71E-01 | no | | intergenic_variant | MODIFIER |
| 15 | 81593668 | AX-97841405 | 0 | T | C | 0.553 | 4.69E-02 | 8.14E-02 | 2.55E+00 | 2.62E+00 | 5.71E-01 | no | Ep300 | intron_variant | MODIFIER |
| 15 | 81633607 | AX-97830587 | 0 | A | G | 0.553 | 4.69E-02 | 8.14E-02 | 2.55E+00 | 2.62E+00 | 5.71E-01 | no | Ep300 | intron_variant | MODIFIER |
| 15 | 81643506 | AX-170426253 | 0 | T | C | 0.553 | 4.69E-02 | 8.14E-02 | 2.55E+00 | 2.62E+00 | 5.71E-01 | no | Ep300 | intron_variant | MODIFIER |
| 15 | 81717045 | AX-170267039 | 0 | G | T | 0.553 | 4.69E-02 | 8.14E-02 | 2.55E+00 | 2.62E+00 | 5.71E-01 | no | Rangap1 | intron_variant | MODIFIER |
| 15 | 81780135 | AX-170236167 | 0 | T | C | 0.553 | 4.69E-02 | 8.14E-02 | 2.55E+00 | 2.62E+00 | 5.71E-01 | no | Zc3h7b | intron_variant | MODIFIER |
| 15 | 81792720 | AX-170171819 | 0 | G | T | 0.553 | 4.69E-02 | 8.14E-02 | 2.55E+00 | 2.62E+00 | 5.71E-01 | no | Zc3h7b | intron_variant | MODIFIER |
| 15 | 82106657 | AX-170458231 | 0 | G | A | 0.553 | 4.69E-02 | 8.14E-02 | 2.55E+00 | 2.62E+00 | 5.71E-01 | no | Mei1 | intron_variant | MODIFIER |
| 15 | 82107530 | AX-170277447 | 0 | A | G | 0.553 | 4.69E-02 | 8.14E-02 | 2.55E+00 | 2.62E+00 | 5.71E-01 | no | Mei1 | intron_variant | MODIFIER |

| | | | | | | | | | | | | | | | |
|---|---|---|---|---|---|---|---|---|---|---|---|---|---|---|---|
| 15 | 82130528 | AX-170056126 | 0 | C | T | 0.553 | 4.69E-02 | 8.14E-02 | 2.55E+00 | 2.62E+00 | 5.71E-01 | no | Ccdc134 | intron_variant | MODIFIER |
| 15 | 82137176 | AX-168413625 | 0 | C | T | 0.553 | 4.69E-02 | 8.14E-02 | 2.55E+00 | 2.62E+00 | 5.71E-01 | no | Ccdc134 | intron_variant | MODIFIER |
| 15 | 82141918 | AX-97831393 | 0 | T | G | 0.553 | 4.69E-02 | 8.14E-02 | 2.55E+00 | 2.62E+00 | 5.71E-01 | no | Ccdc134 | 3_prime_UTR_variant | MODIFIER |
| 15 | 82154518 | AX-170507513 | 0 | G | A | 0.553 | 4.69E-02 | 8.14E-02 | 2.55E+00 | 2.62E+00 | 5.71E-01 | no | Srebf2 | intron_variant | MODIFIER |
| 15 | 82233332 | AX-97857445 | 0 | A | C | 0.553 | 4.69E-02 | 8.14E-02 | 2.55E+00 | 2.62E+00 | 5.71E-01 | no | Cenpm | downstream_gene_variant | MODIFIER |
| 15 | 82233382 | AX-170298067 | 0 | C | T | 0.553 | 4.69E-02 | 8.14E-02 | 2.55E+00 | 2.62E+00 | 5.71E-01 | no | Cenpm | downstream_gene_variant | MODIFIER |
| 15 | 82235784 | AX-170388837 | 0 | C | G | 0.553 | 4.69E-02 | 8.14E-02 | 2.55E+00 | 2.62E+00 | 5.71E-01 | no | Cenpm | intron_variant | MODIFIER |
| 15 | 82264706 | AX-97855425 | 0 | G | A | 0.553 | 4.69E-02 | 8.14E-02 | 2.55E+00 | 2.62E+00 | 5.71E-01 | no | 1500009C09Rik | downstream_gene_variant | MODIFIER |
| 15 | 82287413 | AX-170098510 | 0 | C | T | 0.553 | 4.69E-02 | 8.14E-02 | 2.55E+00 | 2.62E+00 | 5.71E-01 | no | SEP3 | intron_variant | MODIFIER |
| 15 | 82288725 | AX-170411796 | 0 | A | T | 0.553 | 4.69E-02 | 8.14E-02 | 2.55E+00 | 2.62E+00 | 5.71E-01 | no | SEP3 | intron_variant | MODIFIER |
| 15 | 82288924 | AX-97835238 | 0 | T | C | 0.553 | 4.69E-02 | 8.14E-02 | 2.55E+00 | 2.62E+00 | 5.71E-01 | no | SEP3 | intron_variant | MODIFIER |
| 15 | 82339645 | AX-170266572 | 0 | C | T | 0.553 | 4.69E-02 | 8.14E-02 | 2.55E+00 | 2.62E+00 | 5.71E-01 | no | Naga | upstream_gene_variant | MODIFIER |
| 15 | 82421108 | AX-97821143 | 0 | G | A | 0.553 | 4.69E-02 | 8.14E-02 | 2.55E+00 | 2.62E+00 | 5.71E-01 | no | Gm19267 | downstream_gene_variant | MODIFIER |
| 15 | 82435020 | AX-170238143 | 0 | T | C | 0.553 | 4.69E-02 | 8.14E-02 | 2.55E+00 | 2.62E+00 | 5.71E-01 | no | Cyp2d9 | intron_variant | MODIFIER |
| 15 | 82437955 | AX-97823689 | 0 | T | C | 0.553 | 4.69E-02 | 8.14E-02 | 2.55E+00 | 2.62E+00 | 5.71E-01 | no | Cyp2d9 | intron_variant | MODIFIER |
| 15 | 82567433 | AX-97839963 | 0 | C | A | 0.553 | 4.69E-02 | 8.14E-02 | 2.55E+00 | 2.62E+00 | 5.71E-01 | no | Gm49429 | upstream_gene_variant | MODIFIER |
| 15 | 82612763 | AX-97789778 | 0 | T | C | 0.553 | 4.69E-02 | 8.14E-02 | 2.55E+00 | 2.62E+00 | 5.71E-01 | no | Cyp2d34 | downstream_gene_variant | MODIFIER |
| 15 | 82648218 | AX-170409079 | 0 | G | A | 0.553 | 4.69E-02 | 8.14E-02 | 2.55E+00 | 2.62E+00 | 5.71E-01 | no | Cyp2d35-ps | downstream_gene_variant | MODIFIER |
| 15 | 82652558 | AX-97798541 | 0 | G | A | 0.553 | 4.69E-02 | 8.14E-02 | 2.55E+00 | 2.62E+00 | 5.71E-01 | no | Cyp2d35-ps | intron_variant,non_coding_transcript_variant | MODIFIER |
| 15 | 82671742 | AX-97829903 | 0 | T | C | 0.553 | 4.69E-02 | 8.14E-02 | 2.55E+00 | 2.62E+00 | 5.71E-01 | no | Cyp2d36-ps | intron_variant,non_coding_transcript_variant | MODIFIER |
| 15 | 82702066 | AX-97845674 | 0 | T | C | 0.553 | 4.69E-02 | 8.14E-02 | 2.55E+00 | 2.62E+00 | 5.71E-01 | no | Cyp2d38-ps | downstream_gene_variant | MODIFIER |
| 15 | 82704269 | AX-170326958 | 0 | A | G | 0.553 | 4.69E-02 | 8.14E-02 | 2.55E+00 | 2.62E+00 | 5.71E-01 | no | Cyp2d38-ps | downstream_gene_variant | MODIFIER |
| 15 | 82714086 | AX-170099254 | 0 | G | C | 0.553 | 4.69E-02 | 8.14E-02 | 2.55E+00 | 2.62E+00 | 5.71E-01 | no | Cyp2d39-ps | downstream_gene_variant | MODIFIER |
| 15 | 82759008 | AX-170316822 | 0 | G | C | 0.553 | 4.69E-02 | 8.14E-02 | 2.55E+00 | 2.62E+00 | 5.71E-01 | no | Cyp2d40 | downstream_gene_variant | MODIFIER |
| 15 | 82765929 | AX-170093321 | 0 | T | C | 0.553 | 4.69E-02 | 8.14E-02 | 2.55E+00 | 2.62E+00 | 5.71E-01 | no | Cyp2d40 | upstream_gene_variant | MODIFIER |
| 15 | 82776463 | AX-170330821 | 0 | T | C | 0.553 | 4.69E-02 | 8.14E-02 | 2.55E+00 | 2.62E+00 | 5.71E-01 | no | Cyp2d41-ps | downstream_gene_variant | MODIFIER |
| 15 | 82779172 | AX-170085895 | 0 | A | C | 0.553 | 4.69E-02 | 8.14E-02 | 2.55E+00 | 2.62E+00 | 5.71E-01 | no | Cyp2d41-ps | intron_variant,non_coding_transcript_variant | MODIFIER |
| 15 | 82791722 | AX-97848133 | 0 | A | G | 0.553 | 4.69E-02 | 8.14E-02 | 2.55E+00 | 2.62E+00 | 5.71E-01 | no | Cyp2d26 | missense_variant | MODERATE |
| 15 | 82888930 | AX-97817703 | 0 | C | T | 0.553 | 4.69E-02 | 8.14E-02 | 2.55E+00 | 2.62E+00 | 5.71E-01 | no | Tcf20 | intron_variant | MODIFIER |
| 15 | 82988381 | AX-170481315 | 0 | G | A | 0.553 | 4.69E-02 | 8.14E-02 | 2.55E+00 | 2.62E+00 | 5.71E-01 | no | AW121686 | non_coding_transcript_exon_variant | MODIFIER |
| 15 | 82994403 | AX-170106596 | 0 | A | G | 0.553 | 4.69E-02 | 8.14E-02 | 2.55E+00 | 2.62E+00 | 5.71E-01 | no | Nfam1 | downstream_gene_variant | MODIFIER |
| 15 | 82995038 | AX-170260440 | 0 | A | C | 0.553 | 4.69E-02 | 8.14E-02 | 2.55E+00 | 2.62E+00 | 5.71E-01 | no | Nfam1 | downstream_gene_variant | MODIFIER |
| 15 | 82995749 | AX-97803463 | 0 | C | T | 0.553 | 4.69E-02 | 8.14E-02 | 2.55E+00 | 2.62E+00 | 5.71E-01 | no | Nfam1 | downstream_gene_variant | MODIFIER |
| 15 | 82998455 | AX-170295265 | 0 | A | G | 0.553 | 4.69E-02 | 8.14E-02 | 2.55E+00 | 2.62E+00 | 5.71E-01 | no | Nfam1 | 3_prime_UTR_variant | MODIFIER |
| 15 | 83037363 | AX-170012125 | 0 | C | T | 0.553 | 4.69E-02 | 8.14E-02 | 2.55E+00 | 2.62E+00 | 5.71E-01 | no | Nfam1 | upstream_gene_variant | MODIFIER |
| 15 | 83037547 | AX-170012126 | 0 | G | C | 0.553 | 4.69E-02 | 8.14E-02 | 2.55E+00 | 2.62E+00 | 5.71E-01 | no | Nfam1 | upstream_gene_variant | MODIFIER |
| 15 | 83044101 | AX-170056139 | 0 | T | C | 0.553 | 4.69E-02 | 8.14E-02 | 2.55E+00 | 2.62E+00 | 5.71E-01 | no | | regulatory_region_variant | MODIFIER |
| 15 | 83044360 | AX-170056140 | 0 | A | G | 0.553 | 4.69E-02 | 8.14E-02 | 2.55E+00 | 2.62E+00 | 5.71E-01 | no | | regulatory_region_variant | MODIFIER |
| 1 | 82307262 | AX-170147556 | 1 | A | T | 0.161 | 1.20E-01 | 2.12E-01 | 2.64E+00 | 2.69E+00 | 5.71E-01 | no | | intergenic_variant | MODIFIER |
| 1 | 88106882 | AX-170348668 | 1 | G | T | 0.161 | 1.20E-01 | 2.12E-01 | 2.64E+00 | 2.69E+00 | 5.71E-01 | no | Ugt1a7c | intron_variant | MODIFIER |
| 1 | 99500460 | AX-170129293 | 1 | C | A | 0.161 | 1.20E-01 | 2.12E-01 | 2.64E+00 | 2.69E+00 | 5.71E-01 | no | | intergenic_variant | MODIFIER |
| 9 | 113786625 | AX-170206791 | 0 | G | A | 0.132 | -5.76E-02 | 1.03E-01 | 2.63E+00 | 2.74E+00 | 5.71E-01 | no | Clasp2 | missense_variant | MODERATE |
| 5 | 32685726 | AX-97806410 | 0 | A | G | 0.36 | 4.94E-02 | 8.88E-02 | 2.69E+00 | 2.79E+00 | 5.71E-01 | no | Yes1 | 3_prime_UTR_variant | MODIFIER |
| 1 | 149723971 | AX-170335877 | 1 | A | T | 0.196 | 7.77E-02 | 1.38E-01 | 2.67E+00 | 2.72E+00 | 5.71E-01 | no | | intergenic_variant | MODIFIER |
| 8 | 86388770 | AX-170180015 | 0 | G | A | 0.193 | -7.33E-02 | 1.31E-01 | 2.73E+00 | 2.80E+00 | 5.71E-01 | no | | regulatory_region_variant | MODIFIER |
| 18 | 59140235 | AX-170060983 | 1 | C | G | 0.393 | 5.80E-02 | 9.74E-02 | 2.48E+00 | 2.53E+00 | 5.71E-01 | no | | intergenic_variant | MODIFIER |
| 7 | 75204181 | AX-170526745 | 0 | C | A | 0.377 | -5.57E-02 | 1.00E-01 | 2.72E+00 | 2.82E+00 | 5.71E-01 | no | Sv2b | intron_variant | MODIFIER |
| 7 | 78149395 | AX-170285711 | 0 | A | T | 0.377 | -5.57E-02 | 1.00E-01 | 2.72E+00 | 2.82E+00 | 5.71E-01 | no | | intergenic_variant | MODIFIER |
| 6 | 126143230 | AX-170195599 | 1 | A | C | 0.188 | -4.62E-02 | 8.36E-02 | 2.69E+00 | 2.81E+00 | 5.72E-01 | no | Ntf3 | intron_variant | MODIFIER |
| 8 | 3319022 | AX-170157583 | 0 | G | T | 0.272 | 4.25E-02 | 7.64E-02 | 2.67E+00 | 2.78E+00 | 5.72E-01 | no | | intergenic_variant | MODIFIER |
| 10 | 69334380 | AX-170315402 | 0 | C | A | 0.421 | -4.39E-02 | 7.82E-02 | 2.64E+00 | 2.72E+00 | 5.72E-01 | no | Cdk1 | downstream_gene_variant | MODIFIER |
| 5 | 18228582 | AX-97799248 | 1 | G | T | 0.036 | -7.90E-02 | 1.38E-01 | 2.52E+00 | 2.68E+00 | 5.72E-01 | no | | intergenic_variant | MODIFIER |
| 15 | 93884746 | AX-170404770 | 0 | G | A | 0.158 | 8.74E-02 | 1.56E-01 | 2.62E+00 | 2.72E+00 | 5.72E-01 | no | | intergenic_variant | MODIFIER |
| 12 | 68875038 | AX-170271034 | 0 | T | C | 0.07 | 1.18E-01 | 2.13E-01 | 2.73E+00 | 2.84E+00 | 5.72E-01 | no | | intergenic_variant | MODIFIER |

| | | | | | | | | | | | | | | | |
|---|---|---|---|---|---|---|---|---|---|---|---|---|---|---|---|
| 1 | 98861619 | AX-170391191 | 0 | C | A | 0.158 | 1.16E-01 | 2.06E-01 | 2.65E+00 | 2.72E+00 | 5.72E-01 | no | | intergenic_variant | MODIFIER |
| 1 | 46466558 | AX-170196547 | 1 | A | G | 0.098 | -1.18E-01 | 2.13E-01 | 2.72E+00 | 2.81E+00 | 5.72E-01 | no | Dnah7c | intron_variant | MODIFIER |
| 5 | 78328316 | AX-170307567 | 1 | C | T | 0.438 | 6.04E-02 | 1.08E-01 | 2.71E+00 | 2.76E+00 | 5.72E-01 | no | | intergenic_variant | MODIFIER |
| 4 | 82324337 | AX-170069822 | 2 | C | T | 0.227 | 5.76E-02 | 1.01E-01 | 2.58E+00 | 2.67E+00 | 5.72E-01 | no | Nfib | intron_variant | MODIFIER |
| 5 | 89659975 | AX-170345403 | 0 | C | T | 0.439 | 5.71E-02 | 1.03E-01 | 2.75E+00 | 2.82E+00 | 5.72E-01 | no | | intergenic_variant | MODIFIER |
| 10 | 18534551 | AX-170186985 | 0 | C | T | 0.202 | 5.17E-02 | 9.38E-02 | 2.74E+00 | 2.85E+00 | 5.73E-01 | no | Hebp2 | downstream_gene_variant | MODIFIER |
| 5 | 115168370 | AX-170418139 | 1 | A | T | 0.455 | 5.91E-02 | 1.04E-01 | 2.57E+00 | 2.64E+00 | 5.73E-01 | no | Cabp1 | downstream_gene_variant | MODIFIER |
| 4 | 56880802 | AX-168397556 | 0 | A | G | 0.368 | -5.77E-02 | 1.04E-01 | 2.71E+00 | 2.79E+00 | 5.73E-01 | no | Tmem245 | intron_variant | MODIFIER |
| 18 | 58882512 | AX-170487900 | 2 | C | G | 0.382 | 5.57E-02 | 9.48E-02 | 2.50E+00 | 2.56E+00 | 5.73E-01 | no | Adamts19 | intron_variant | MODIFIER |
| 8 | 127789443 | AX-170494002 | 2 | T | C | 0.036 | -7.31E-02 | 1.34E-01 | 2.67E+00 | 2.86E+00 | 5.73E-01 | no | | intergenic_variant | MODIFIER |
| 20 | 68870592 | AX-170191839 | 1 | T | A | 0.241 | 3.49E-02 | 6.31E-02 | 2.70E+00 | 2.80E+00 | 5.73E-01 | no | | intergenic_variant | MODIFIER |
| 2 | 102670955 | AX-170427696 | 0 | A | G | 0.061 | 1.02E-01 | 1.88E-01 | 2.70E+00 | 2.89E+00 | 5.73E-01 | no | Slc1a2 | intron_variant | MODIFIER |
| 3 | 7384249 | AX-170356948 | 0 | A | G | 0.386 | -3.72E-02 | 6.77E-02 | 2.69E+00 | 2.83E+00 | 5.73E-01 | no | Pkia | intron_variant | MODIFIER |
| 5 | 40298586 | AX-97831739 | 1 | C | T | 0.214 | -6.11E-02 | 1.08E-01 | 2.60E+00 | 2.68E+00 | 5.73E-01 | no | | intergenic_variant | MODIFIER |
| 10 | 19219241 | AX-170364687 | 1 | G | A | 0.196 | 5.61E-02 | 1.01E-01 | 2.82E+00 | 2.92E+00 | 5.73E-01 | no | | intergenic_variant | MODIFIER |
| 3 | 38905281 | AX-170424557 | 0 | C | G | 0.202 | 4.79E-02 | 8.67E-02 | 2.67E+00 | 2.79E+00 | 5.73E-01 | no | Fat4 | intron_variant | MODIFIER |
| 7 | 46182384 | AX-97826627 | 1 | G | A | 0.33 | 5.28E-02 | 9.44E-02 | 2.63E+00 | 2.71E+00 | 5.74E-01 | no | Abcc8 | upstream_gene_variant | MODIFIER |
| 1 | 38895147 | AX-170117643 | 0 | T | C | 0.456 | -4.34E-02 | 7.77E-02 | 2.64E+00 | 2.72E+00 | 5.74E-01 | no | Chst10 | intron_variant | MODIFIER |
| 7 | 37402261 | AX-170171405 | 1 | G | C | 0.348 | 5.15E-02 | 9.05E-02 | 2.57E+00 | 2.65E+00 | 5.74E-01 | no | | intergenic_variant | MODIFIER |
| 9 | 109236359 | AX-97781933 | 0 | C | T | 0.035 | -7.07E-02 | 1.29E-01 | 2.62E+00 | 2.79E+00 | 5.74E-01 | no | Fbxw20 | upstream_gene_variant | MODIFIER |
| 1 | 95599388 | AX-97821425 | 1 | G | T | 0.152 | 1.10E-01 | 1.97E-01 | 2.64E+00 | 2.70E+00 | 5.74E-01 | no | St8sia4 | intron_variant | MODIFIER |
| 9 | 33711622 | AX-97804680 | 0 | C | T | 0.43 | 4.78E-02 | 8.52E-02 | 2.63E+00 | 2.69E+00 | 5.74E-01 | no | | intergenic_variant | MODIFIER |
| 1 | 22290334 | AX-170103776 | 1 | T | C | 0.411 | 4.84E-02 | 8.61E-02 | 2.62E+00 | 2.67E+00 | 5.74E-01 | no | Rims1 | intron_variant | MODIFIER |
| 12 | 92033523 | AX-170358731 | 0 | C | T | 0.079 | 9.66E-02 | 1.74E-01 | 2.62E+00 | 2.74E+00 | 5.74E-01 | no | | intergenic_variant | MODIFIER |
| 11 | 3268999 | AX-97845342 | 0 | G | A | 0.07 | -9.74E-02 | 1.78E-01 | 2.63E+00 | 2.80E+00 | 5.74E-01 | no | Drg1 | upstream_gene_variant | MODIFIER |
| 5 | 35121467 | AX-170218529 | 0 | G | A | 0.342 | 6.01E-02 | 1.08E-01 | 2.68E+00 | 2.74E+00 | 5.74E-01 | no | | intergenic_variant | MODIFIER |
| 1 | 116285667 | AX-170042424 | 1 | G | A | 0.152 | 8.23E-02 | 1.49E-01 | 2.62E+00 | 2.77E+00 | 5.74E-01 | no | Cntnap5a | intron_variant | MODIFIER |
| 15 | 82237257 | AX-97840093 | 1 | T | C | 0.554 | 4.65E-02 | 8.14E-02 | 2.56E+00 | 2.62E+00 | 5.74E-01 | no | Cenpm | intron_variant | MODIFIER |
| 8 | 103866322 | AX-170404941 | 0 | G | A | 0.228 | 5.83E-02 | 1.05E-01 | 2.69E+00 | 2.78E+00 | 5.74E-01 | no | | intergenic_variant | MODIFIER |
| 1 | 101563368 | AX-170356615 | 0 | A | G | 0.14 | 9.53E-02 | 1.70E-01 | 2.60E+00 | 2.69E+00 | 5.74E-01 | no | | intergenic_variant | MODIFIER |
| 20 | 134134977 | AX-97857616 | 0 | T | C | 0.14 | 6.31E-02 | 1.14E-01 | 2.67E+00 | 2.76E+00 | 5.74E-01 | no | Nox1 | intron_variant | MODIFIER |
| 9 | 27882193 | AX-97846172 | 1 | A | G | 0.446 | 4.97E-02 | 8.89E-02 | 2.62E+00 | 2.70E+00 | 5.75E-01 | no | Opcml | intron_variant | MODIFIER |
| 8 | 38545719 | AX-170030528 | 1 | C | G | 0.152 | 8.18E-02 | 1.48E-01 | 2.77E+00 | 2.82E+00 | 5.75E-01 | no | Sgcz | intron_variant | MODIFIER |
| 16 | 75503003 | AX-97825571 | 1 | C | T | 0.5 | 4.35E-02 | 7.39E-02 | 2.48E+00 | 2.56E+00 | 5.75E-01 | no | | intergenic_variant | MODIFIER |
| 20 | 145657924 | AX-97848215 | 0 | C | T | 0.158 | 6.84E-02 | 1.24E-01 | 2.64E+00 | 2.76E+00 | 5.75E-01 | no | | intergenic_variant | MODIFIER |
| 5 | 89458340 | AX-170025555 | 1 | T | C | 0.446 | 5.69E-02 | 1.03E-01 | 2.74E+00 | 2.80E+00 | 5.75E-01 | no | Gc | upstream_gene_variant | MODIFIER |
| 10 | 14189614 | AX-170376844 | 0 | G | T | 0.211 | -5.42E-02 | 9.77E-02 | 2.64E+00 | 2.74E+00 | 5.75E-01 | no | | intergenic_variant | MODIFIER |
| 13 | 106381136 | AX-170147706 | 0 | A | C | 0.061 | 7.51E-02 | 1.37E-01 | 2.64E+00 | 2.78E+00 | 5.75E-01 | no | | intergenic_variant | MODIFIER |
| 15 | 82375773 | AX-97829927 | 1 | T | C | 0.554 | 4.45E-02 | 7.80E-02 | 2.55E+00 | 2.63E+00 | 5.75E-01 | no | Cyp2d22 | intron_variant | MODIFIER |
| 17 | 70023544 | AX-170532594 | 0 | A | G | 0.044 | -9.84E-02 | 1.78E-01 | 2.63E+00 | 2.74E+00 | 5.75E-01 | no | Dlgap1 | intron_variant | MODIFIER |
| 9 | 22257425 | AX-97781669 | 0 | A | T | 0.465 | 5.00E-02 | 8.88E-02 | 2.59E+00 | 2.67E+00 | 5.75E-01 | no | Zfp599 | intron_variant | MODIFIER |
| 16 | 51392427 | AX-97855542 | 0 | A | T | 0.518 | 4.78E-02 | 8.21E-02 | 2.51E+00 | 2.56E+00 | 5.75E-01 | no | | intergenic_variant | MODIFIER |
| 16 | 86205571 | AX-170058012 | 0 | G | A | 0.456 | 4.39E-02 | 7.87E-02 | 2.63E+00 | 2.70E+00 | 5.75E-01 | no | | intergenic_variant | MODIFIER |
| 7 | 98755107 | AX-170274298 | 1 | T | C | 0.143 | 1.06E-01 | 1.93E-01 | 2.70E+00 | 2.81E+00 | 5.75E-01 | no | | regulatory_region_variant | MODIFIER |
| 5 | 97054638 | AX-170193593 | 1 | G | A | 0.446 | 5.90E-02 | 1.06E-01 | 2.69E+00 | 2.75E+00 | 5.75E-01 | no | Bmp2k | intron_variant | MODIFIER |
| 1 | 96346122 | AX-97812423 | 1 | A | T | 0.152 | 8.00E-02 | 1.41E-01 | 2.56E+00 | 2.66E+00 | 5.75E-01 | no | | intergenic_variant | MODIFIER |
| 3 | 148029237 | AX-170485900 | 0 | A | G | 0.342 | 3.99E-02 | 7.21E-02 | 2.67E+00 | 2.75E+00 | 5.75E-01 | no | | intergenic_variant | MODIFIER |
| 6 | 128243993 | AX-170477088 | 0 | A | G | 0.561 | 4.55E-02 | 7.85E-02 | 2.51E+00 | 2.59E+00 | 5.75E-01 | no | Tead4 | intron_variant | MODIFIER |
| 18 | 28742093 | AX-97834324 | 0 | T | C | 0.053 | 1.22E-01 | 2.21E-01 | 2.61E+00 | 2.75E+00 | 5.76E-01 | no | | regulatory_region_variant | MODIFIER |
| 11 | 11412482 | AX-170279092 | 0 | A | T | 0.491 | -1.65E-01 | 2.93E-01 | 2.50E+00 | 2.70E+00 | 5.76E-01 | no | Zpbp | intron_variant | MODIFIER |
| 4 | 80248067 | AX-170023209 | 0 | A | G | 0.237 | -5.28E-02 | 9.57E-02 | 2.84E+00 | 2.98E+00 | 5.76E-01 | no | | intergenic_variant | MODIFIER |
| 3 | 143929516 | AX-170094839 | 2 | C | T | 0.382 | 3.70E-02 | 6.65E-02 | 2.61E+00 | 2.71E+00 | 5.76E-01 | no | Gm42707 | intron_variant,non_coding_transcript_variant | MODIFIER |
| 1 | 170308754 | AX-170507077 | 0 | C | A | 0.211 | 7.52E-02 | 1.36E-01 | 2.67E+00 | 2.72E+00 | 5.76E-01 | no | Spata46 | upstream_gene_variant | MODIFIER |
| 9 | 39853651 | AX-170032176 | 1 | C | T | 0.402 | -4.05E-02 | 7.35E-02 | 2.66E+00 | 2.76E+00 | 5.76E-01 | no | | intergenic_variant | MODIFIER |

| | | | | | | | | | | | | | | | |
|---|---|---|---|---|---|---|---|---|---|---|---|---|---|---|---|
| 4 | 118041202 | AX-170488268 | 0 | A | T | 0.211 | -6.39E-02 | 1.15E-01 | 2.88E+00 | 3.00E+00 | 5.76E-01 | no | St3gal3 | intron_variant | MODIFIER |
| 4 | 118196538 | AX-170495589 | 0 | C | T | 0.211 | -6.39E-02 | 1.15E-01 | 2.88E+00 | 3.00E+00 | 5.76E-01 | no | | regulatory_region_variant | MODIFIER |
| 4 | 118705726 | AX-170518930 | 0 | C | T | 0.211 | -6.39E-02 | 1.15E-01 | 2.88E+00 | 3.00E+00 | 5.76E-01 | no | Olfr1341 | upstream_gene_variant | MODIFIER |
| 7 | 14607574 | AX-170197370 | 0 | T | G | 0.368 | 4.76E-02 | 8.50E-02 | 2.58E+00 | 2.69E+00 | 5.76E-01 | no | Nlrp5-ps | intron_variant,non_coding_transcript_variant | MODIFIER |
| 7 | 14608546 | AX-170339538 | 0 | G | A | 0.368 | 4.76E-02 | 8.50E-02 | 2.58E+00 | 2.69E+00 | 5.76E-01 | no | Nlrp5-ps | intron_variant,non_coding_transcript_variant | MODIFIER |
| 7 | 14608856 | AX-170161388 | 0 | C | T | 0.368 | 4.76E-02 | 8.50E-02 | 2.58E+00 | 2.69E+00 | 5.76E-01 | no | Nlrp5-ps | intron_variant,non_coding_transcript_variant | MODIFIER |
| 7 | 14867888 | AX-170166617 | 0 | T | A | 0.368 | 4.76E-02 | 8.50E-02 | 2.58E+00 | 2.69E+00 | 5.76E-01 | no | Obox4-ps38 | non_coding_transcript_exon_variant | MODIFIER |
| 7 | 15006269 | AX-170085868 | 0 | G | A | 0.368 | 4.76E-02 | 8.50E-02 | 2.58E+00 | 2.69E+00 | 5.76E-01 | no | Obox4-ps8 | upstream_gene_variant | MODIFIER |
| 7 | 15892938 | AX-170160582 | 0 | G | A | 0.368 | 4.76E-02 | 8.50E-02 | 2.58E+00 | 2.69E+00 | 5.76E-01 | no | Crxos | upstream_gene_variant | MODIFIER |
| 7 | 15897374 | AX-97839131 | 0 | T | G | 0.368 | 4.76E-02 | 8.50E-02 | 2.58E+00 | 2.69E+00 | 5.76E-01 | no | Crxos | intron_variant | MODIFIER |
| 7 | 15901407 | AX-97830654 | 0 | G | A | 0.368 | 4.76E-02 | 8.50E-02 | 2.58E+00 | 2.69E+00 | 5.76E-01 | no | Crxos | intron_variant | MODIFIER |
| 7 | 16009846 | AX-97851870 | 0 | C | T | 0.368 | 4.76E-02 | 8.50E-02 | 2.58E+00 | 2.69E+00 | 5.76E-01 | no | Bicra | intron_variant | MODIFIER |
| 7 | 16036173 | AX-170285868 | 0 | T | C | 0.368 | 4.76E-02 | 8.50E-02 | 2.58E+00 | 2.69E+00 | 5.76E-01 | no | Bicra | intron_variant | MODIFIER |
| 7 | 16109085 | AX-170360585 | 0 | G | A | 0.368 | 4.76E-02 | 8.50E-02 | 2.58E+00 | 2.69E+00 | 5.76E-01 | no | Napa | synonymous_variant | LOW |
| 7 | 16128144 | AX-170125091 | 0 | A | G | 0.368 | 4.76E-02 | 8.50E-02 | 2.58E+00 | 2.69E+00 | 5.76E-01 | no | Kptn | downstream_gene_variant | MODIFIER |
| 7 | 16128746 | AX-170494789 | 0 | A | G | 0.368 | 4.76E-02 | 8.50E-02 | 2.58E+00 | 2.69E+00 | 5.76E-01 | no | Kptn | downstream_gene_variant | MODIFIER |
| 7 | 16135312 | AX-170330341 | 0 | T | C | 0.368 | 4.76E-02 | 8.50E-02 | 2.58E+00 | 2.69E+00 | 5.76E-01 | no | Slc8a2 | intron_variant | MODIFIER |
| 7 | 16138175 | AX-97856254 | 0 | C | T | 0.368 | 4.76E-02 | 8.50E-02 | 2.58E+00 | 2.69E+00 | 5.76E-01 | no | Slc8a2 | intron_variant | MODIFIER |
| 7 | 16138312 | AX-170374356 | 0 | T | C | 0.368 | 4.76E-02 | 8.50E-02 | 2.58E+00 | 2.69E+00 | 5.76E-01 | no | Slc8a2 | intron_variant | MODIFIER |
| 7 | 16153290 | AX-170481469 | 0 | T | C | 0.368 | 4.76E-02 | 8.50E-02 | 2.58E+00 | 2.69E+00 | 5.76E-01 | no | Slc8a2 | synonymous_variant | LOW |
| 7 | 16153977 | AX-170146404 | 0 | A | G | 0.368 | 4.76E-02 | 8.50E-02 | 2.58E+00 | 2.69E+00 | 5.76E-01 | no | Slc8a2 | intron_variant | MODIFIER |
| 7 | 16163638 | AX-97786251 | 0 | A | G | 0.368 | 4.76E-02 | 8.50E-02 | 2.58E+00 | 2.69E+00 | 5.76E-01 | no | Slc8a2 | downstream_gene_variant | MODIFIER |
| 7 | 16179622 | AX-170232214 | 0 | A | G | 0.368 | 4.76E-02 | 8.50E-02 | 2.58E+00 | 2.69E+00 | 5.76E-01 | no | Meis3 | intron_variant | MODIFIER |
| 7 | 16179921 | AX-168394569 | 0 | T | C | 0.368 | 4.76E-02 | 8.50E-02 | 2.58E+00 | 2.69E+00 | 5.76E-01 | no | Meis3 | intron_variant | MODIFIER |
| 7 | 16187521 | AX-170113534 | 0 | C | G | 0.368 | 4.76E-02 | 8.50E-02 | 2.58E+00 | 2.69E+00 | 5.76E-01 | no | Meis3 | downstream_gene_variant | MODIFIER |
| 7 | 16189936 | AX-97792703 | 0 | T | C | 0.368 | 4.76E-02 | 8.50E-02 | 2.58E+00 | 2.69E+00 | 5.76E-01 | no | Meis3 | downstream_gene_variant | MODIFIER |
| 7 | 16195027 | AX-170330881 | 0 | C | T | 0.368 | 4.76E-02 | 8.50E-02 | 2.58E+00 | 2.69E+00 | 5.76E-01 | no | Dhx34 | downstream_gene_variant | MODIFIER |
| 7 | 16195220 | AX-170333856 | 0 | G | T | 0.368 | 4.76E-02 | 8.50E-02 | 2.58E+00 | 2.69E+00 | 5.76E-01 | no | Dhx34 | downstream_gene_variant | MODIFIER |
| 7 | 16201947 | AX-170205156 | 0 | G | T | 0.368 | 4.76E-02 | 8.50E-02 | 2.58E+00 | 2.69E+00 | 5.76E-01 | no | Dhx34 | synonymous_variant | LOW |
| 7 | 16210434 | AX-170210096 | 0 | A | G | 0.368 | 4.76E-02 | 8.50E-02 | 2.58E+00 | 2.69E+00 | 5.76E-01 | no | Dhx34 | intron_variant | MODIFIER |
| 7 | 16214314 | AX-170454678 | 0 | T | C | 0.368 | 4.76E-02 | 8.50E-02 | 2.58E+00 | 2.69E+00 | 5.76E-01 | no | Dhx34 | intron_variant | MODIFIER |
| 7 | 16228312 | AX-170409619 | 0 | G | A | 0.368 | 4.76E-02 | 8.50E-02 | 2.58E+00 | 2.69E+00 | 5.76E-01 | no | Gm45409 | upstream_gene_variant | MODIFIER |
| 7 | 16236094 | AX-97844475 | 0 | A | G | 0.368 | 4.76E-02 | 8.50E-02 | 2.58E+00 | 2.69E+00 | 5.76E-01 | no | C5ar2 | 3_prime_UTR_variant | MODIFIER |
| 7 | 16236707 | AX-170035744 | 0 | C | G | 0.368 | 4.76E-02 | 8.50E-02 | 2.58E+00 | 2.69E+00 | 5.76E-01 | no | C5ar2 | 3_prime_UTR_variant | MODIFIER |
| 7 | 16236782 | AX-170086363 | 0 | C | G | 0.368 | 4.76E-02 | 8.50E-02 | 2.58E+00 | 2.69E+00 | 5.76E-01 | no | C5ar2 | 3_prime_UTR_variant | MODIFIER |
| 7 | 16262752 | AX-170260884 | 0 | A | G | 0.368 | 4.76E-02 | 8.50E-02 | 2.58E+00 | 2.69E+00 | 5.76E-01 | no | C5ar1 | upstream_gene_variant | MODIFIER |
| 7 | 16528344 | AX-97820365 | 0 | C | T | 0.368 | 4.76E-02 | 8.50E-02 | 2.58E+00 | 2.69E+00 | 5.76E-01 | no | Arhgap35 | intron_variant | MODIFIER |
| 7 | 16563519 | AX-97844051 | 0 | G | A | 0.368 | 4.76E-02 | 8.50E-02 | 2.58E+00 | 2.69E+00 | 5.76E-01 | no | Arhgap35 | synonymous_variant | LOW |
| 7 | 16919522 | AX-170035748 | 0 | G | A | 0.368 | 4.76E-02 | 8.50E-02 | 2.58E+00 | 2.69E+00 | 5.76E-01 | no | Calm3 | intron_variant | MODIFIER |
| 7 | 16964719 | AX-170350028 | 0 | T | C | 0.368 | 4.76E-02 | 8.50E-02 | 2.58E+00 | 2.69E+00 | 5.76E-01 | no | Pnmal1 | downstream_gene_variant | MODIFIER |
| 7 | 16985317 | AX-170435834 | 0 | G | C | 0.368 | 4.76E-02 | 8.50E-02 | 2.58E+00 | 2.69E+00 | 5.76E-01 | no | Gm42372 | 3_prime_UTR_variant | MODIFIER |
| 7 | 17005819 | AX-170368055 | 0 | G | A | 0.368 | 4.76E-02 | 8.50E-02 | 2.58E+00 | 2.69E+00 | 5.76E-01 | no | Ppp5c | intron_variant | MODIFIER |
| 7 | 17013354 | AX-170196598 | 0 | T | G | 0.368 | 4.76E-02 | 8.50E-02 | 2.58E+00 | 2.69E+00 | 5.76E-01 | no | Ppp5c | intron_variant | MODIFIER |
| 7 | 17014976 | AX-170186936 | 0 | T | C | 0.368 | 4.76E-02 | 8.50E-02 | 2.58E+00 | 2.69E+00 | 5.76E-01 | no | Ppp5c | intron_variant | MODIFIER |
| 7 | 17015351 | AX-170342730 | 0 | G | A | 0.368 | 4.76E-02 | 8.50E-02 | 2.58E+00 | 2.69E+00 | 5.76E-01 | no | Ppp5c | intron_variant | MODIFIER |
| 7 | 17022025 | AX-170199452 | 0 | C | T | 0.368 | 4.76E-02 | 8.50E-02 | 2.58E+00 | 2.69E+00 | 5.76E-01 | no | Ppp5c | intron_variant | MODIFIER |
| 7 | 17029623 | AX-97826692 | 0 | G | A | 0.368 | 4.76E-02 | 8.50E-02 | 2.58E+00 | 2.69E+00 | 5.76E-01 | no | Ppp5c | upstream_gene_variant | MODIFIER |
| 7 | 17076556 | AX-170484975 | 0 | A | T | 0.368 | 4.76E-02 | 8.50E-02 | 2.58E+00 | 2.69E+00 | 5.76E-01 | no | Psg16 | intron_variant | MODIFIER |
| 7 | 17088316 | AX-170228404 | 0 | G | A | 0.368 | 4.76E-02 | 8.50E-02 | 2.58E+00 | 2.69E+00 | 5.76E-01 | no | Psg16 | intron_variant | MODIFIER |
| 7 | 17092834 | AX-170328142 | 0 | G | T | 0.368 | 4.76E-02 | 8.50E-02 | 2.58E+00 | 2.69E+00 | 5.76E-01 | no | Psg16 | intron_variant | MODIFIER |
| 7 | 17094129 | AX-97846654 | 0 | T | C | 0.368 | 4.76E-02 | 8.50E-02 | 2.58E+00 | 2.69E+00 | 5.76E-01 | no | Psg16 | intron_variant | MODIFIER |
| 5 | 76866058 | AX-170237674 | 0 | T | C | 0.43 | 5.69E-02 | 1.02E-01 | 2.65E+00 | 2.71E+00 | 5.76E-01 | no | C530008M17Rik | synonymous_variant | LOW |
| 5 | 76965978 | AX-170466754 | 0 | G | A | 0.43 | 5.69E-02 | 1.02E-01 | 2.65E+00 | 2.71E+00 | 5.76E-01 | no | Paics | intron_variant | MODIFIER |
| 5 | 77057041 | AX-170319184 | 0 | C | T | 0.43 | 5.69E-02 | 1.02E-01 | 2.65E+00 | 2.71E+00 | 5.76E-01 | no | Thegl | intron_variant | MODIFIER |

| | | | | | | | | | | | | | | | |
|---|---|---|---|---|---|---|---|---|---|---|---|---|---|---|---|
| 5 | 77069389 | AX-170290358 | 0 | G | A | 0.43 | 5.69E-02 | 1.02E-01 | 2.65E+00 | 2.71E+00 | 5.76E-01 | no | | intergenic_variant | MODIFIER |
| 5 | 77073697 | AX-170099116 | 0 | C | T | 0.43 | 5.69E-02 | 1.02E-01 | 2.65E+00 | 2.71E+00 | 5.76E-01 | no | | intergenic_variant | MODIFIER |
| 5 | 77075555 | AX-170360039 | 0 | A | T | 0.43 | 5.69E-02 | 1.02E-01 | 2.65E+00 | 2.71E+00 | 5.76E-01 | no | | regulatory_region_variant | MODIFIER |
| 5 | 77076895 | AX-170167892 | 0 | A | G | 0.43 | 5.69E-02 | 1.02E-01 | 2.65E+00 | 2.71E+00 | 5.76E-01 | no | | intergenic_variant | MODIFIER |
| 5 | 77105323 | AX-170221896 | 0 | G | C | 0.43 | 5.69E-02 | 1.02E-01 | 2.65E+00 | 2.71E+00 | 5.76E-01 | no | Hopx | intron_variant | MODIFIER |
| 5 | 77107161 | AX-168415534 | 0 | C | A | 0.43 | 5.69E-02 | 1.02E-01 | 2.65E+00 | 2.71E+00 | 5.76E-01 | no | Hopx | intron_variant | MODIFIER |
| 5 | 77112262 | AX-97854384 | 0 | T | C | 0.43 | 5.69E-02 | 1.02E-01 | 2.65E+00 | 2.71E+00 | 5.76E-01 | no | Hopx | intron_variant | MODIFIER |
| 5 | 77121185 | AX-170336676 | 0 | C | T | 0.43 | 5.69E-02 | 1.02E-01 | 2.65E+00 | 2.71E+00 | 5.76E-01 | no | Gm15831 | intron_variant,non_coding_transcript_variant | MODIFIER |
| 5 | 77123038 | AX-170464904 | 0 | G | T | 0.43 | 5.69E-02 | 1.02E-01 | 2.65E+00 | 2.71E+00 | 5.76E-01 | no | Gm15831 | upstream_gene_variant | MODIFIER |
| 5 | 77125924 | AX-170193594 | 0 | T | C | 0.43 | 5.69E-02 | 1.02E-01 | 2.65E+00 | 2.71E+00 | 5.76E-01 | no | Gm15831 | upstream_gene_variant | MODIFIER |
| 5 | 77130822 | AX-170237824 | 0 | G | A | 0.43 | 5.69E-02 | 1.02E-01 | 2.65E+00 | 2.71E+00 | 5.76E-01 | no | | regulatory_region_variant | MODIFIER |
| 5 | 77130853 | AX-170298807 | 0 | A | G | 0.43 | 5.69E-02 | 1.02E-01 | 2.65E+00 | 2.71E+00 | 5.76E-01 | no | | regulatory_region_variant | MODIFIER |
| 5 | 77130867 | AX-170440495 | 0 | A | G | 0.43 | 5.69E-02 | 1.02E-01 | 2.65E+00 | 2.71E+00 | 5.76E-01 | no | | regulatory_region_variant | MODIFIER |
| 5 | 77154170 | AX-170143660 | 0 | T | C | 0.43 | 5.69E-02 | 1.02E-01 | 2.65E+00 | 2.71E+00 | 5.76E-01 | no | | regulatory_region_variant | MODIFIER |
| 6 | 136712941 | AX-170165104 | 0 | C | G | 0.57 | 4.73E-02 | 8.23E-02 | 2.53E+00 | 2.60E+00 | 5.76E-01 | no | Gucy2c | intron_variant | MODIFIER |
| 5 | 36411157 | AX-170370553 | 0 | C | A | 0.325 | 6.03E-02 | 1.10E-01 | 2.72E+00 | 2.78E+00 | 5.76E-01 | no | | intergenic_variant | MODIFIER |
| 9 | 95502683 | AX-170310128 | 0 | G | A | 0.14 | -6.23E-02 | 1.13E-01 | 2.64E+00 | 2.73E+00 | 5.76E-01 | no | U2surp | intron_variant | MODIFIER |
| 10 | 16081880 | AX-170420299 | 0 | T | C | 0.193 | -5.58E-02 | 1.00E-01 | 2.61E+00 | 2.70E+00 | 5.76E-01 | no | | intergenic_variant | MODIFIER |
| 16 | 35699627 | AX-170228109 | 0 | C | T | 0.491 | 4.56E-02 | 7.94E-02 | 2.53E+00 | 2.59E+00 | 5.76E-01 | no | Dirc2 | intron_variant | MODIFIER |
| 9 | 29719264 | AX-97810815 | 0 | G | A | 0.439 | 4.71E-02 | 8.47E-02 | 2.63E+00 | 2.70E+00 | 5.77E-01 | no | Ntm | intron_variant | MODIFIER |
| 9 | 108777544 | AX-170281064 | 0 | G | A | 0.123 | 6.30E-02 | 1.14E-01 | 2.82E+00 | 2.91E+00 | 5.77E-01 | no | | regulatory_region_variant | MODIFIER |
| 11 | 30245931 | AX-97803579 | 1 | A | G | 0.071 | 1.19E-01 | 2.16E-01 | 2.68E+00 | 2.76E+00 | 5.77E-01 | no | Sptbn1 | intron_variant | MODIFIER |
| 4 | 95197347 | AX-170070045 | 0 | T | C | 0.316 | 5.64E-02 | 1.02E-01 | 2.64E+00 | 2.72E+00 | 5.77E-01 | no | Gm12708 | intron_variant,non_coding_transcript_variant | MODIFIER |
| 13 | 31920218 | AX-170008030 | 0 | T | C | 0.175 | -6.91E-02 | 1.22E-01 | 2.56E+00 | 2.62E+00 | 5.77E-01 | no | Gmds | intron_variant | MODIFIER |
| 13 | 32228788 | AX-170391292 | 0 | C | T | 0.175 | -7.07E-02 | 1.24E-01 | 2.56E+00 | 2.61E+00 | 5.77E-01 | no | Gmds | intron_variant | MODIFIER |
| 18 | 49937953 | AX-170404606 | 0 | A | G | 0.193 | -5.89E-02 | 1.04E-01 | 2.56E+00 | 2.64E+00 | 5.77E-01 | no | Dmxl1 | intron_variant | MODIFIER |
| 18 | 49941981 | AX-97808174 | 0 | A | G | 0.193 | -5.89E-02 | 1.04E-01 | 2.56E+00 | 2.64E+00 | 5.77E-01 | no | Dmxl1 | intron_variant | MODIFIER |
| 18 | 49987238 | AX-97828472 | 0 | G | A | 0.193 | -5.89E-02 | 1.04E-01 | 2.56E+00 | 2.64E+00 | 5.77E-01 | no | Tnfaip8 | intron_variant,non_coding_transcript_variant | MODIFIER |
| 18 | 49994756 | AX-170169403 | 0 | A | G | 0.193 | -5.89E-02 | 1.04E-01 | 2.56E+00 | 2.64E+00 | 5.77E-01 | no | Tnfaip8 | intron_variant,non_coding_transcript_variant | MODIFIER |
| 18 | 49996721 | AX-170535059 | 0 | A | G | 0.193 | -5.89E-02 | 1.04E-01 | 2.56E+00 | 2.64E+00 | 5.77E-01 | no | Tnfaip8 | intron_variant,non_coding_transcript_variant | MODIFIER |
| 18 | 50003995 | AX-170163580 | 0 | C | T | 0.193 | -5.89E-02 | 1.04E-01 | 2.56E+00 | 2.64E+00 | 5.77E-01 | no | Tnfaip8 | intron_variant,non_coding_transcript_variant | MODIFIER |
| 18 | 50028057 | AX-170515598 | 0 | T | C | 0.193 | -5.89E-02 | 1.04E-01 | 2.56E+00 | 2.64E+00 | 5.77E-01 | no | Tnfaip8 | intron_variant,non_coding_transcript_variant | MODIFIER |
| 18 | 50042472 | AX-170354824 | 0 | T | G | 0.193 | -5.89E-02 | 1.04E-01 | 2.56E+00 | 2.64E+00 | 5.77E-01 | no | Tnfaip8 | intron_variant,non_coding_transcript_variant | MODIFIER |
| 18 | 50042681 | AX-170191642 | 0 | C | T | 0.193 | -5.89E-02 | 1.04E-01 | 2.56E+00 | 2.64E+00 | 5.77E-01 | no | Tnfaip8 | intron_variant,non_coding_transcript_variant | MODIFIER |
| 18 | 50047506 | AX-170015681 | 0 | G | A | 0.193 | -5.89E-02 | 1.04E-01 | 2.56E+00 | 2.64E+00 | 5.77E-01 | no | Tnfaip8 | non_coding_transcript_exon_variant | MODIFIER |
| 18 | 50048958 | AX-170015679 | 0 | T | C | 0.193 | -5.89E-02 | 1.04E-01 | 2.56E+00 | 2.64E+00 | 5.77E-01 | no | Tnfaip8 | non_coding_transcript_exon_variant | MODIFIER |
| 18 | 50058271 | AX-170393033 | 0 | A | G | 0.193 | -5.89E-02 | 1.04E-01 | 2.56E+00 | 2.64E+00 | 5.77E-01 | no | Tnfaip8 | intron_variant | MODIFIER |
| 18 | 50061673 | AX-170015680 | 0 | C | T | 0.193 | -5.89E-02 | 1.04E-01 | 2.56E+00 | 2.64E+00 | 5.77E-01 | no | Tnfaip8 | intron_variant | MODIFIER |
| 18 | 50061856 | AX-170281626 | 0 | A | C | 0.193 | -5.89E-02 | 1.04E-01 | 2.56E+00 | 2.64E+00 | 5.77E-01 | no | Tnfaip8 | intron_variant | MODIFIER |
| 18 | 50097940 | AX-170464420 | 0 | C | T | 0.193 | -5.89E-02 | 1.04E-01 | 2.56E+00 | 2.64E+00 | 5.77E-01 | no | Tnfaip8 | intron_variant,non_coding_transcript_variant | MODIFIER |
| 18 | 50107015 | AX-170337665 | 0 | C | T | 0.193 | -5.89E-02 | 1.04E-01 | 2.56E+00 | 2.64E+00 | 5.77E-01 | no | Tnfaip8 | non_coding_transcript_exon_variant | MODIFIER |
| 18 | 50107284 | AX-170212542 | 0 | G | A | 0.193 | -5.89E-02 | 1.04E-01 | 2.56E+00 | 2.64E+00 | 5.77E-01 | no | Tnfaip8 | downstream_gene_variant | MODIFIER |
| 18 | 50133625 | AX-170410027 | 0 | C | A | 0.193 | -5.89E-02 | 1.04E-01 | 2.56E+00 | 2.64E+00 | 5.77E-01 | no | Hsd17b4 | intron_variant | MODIFIER |
| 18 | 50179321 | AX-170282484 | 0 | C | A | 0.193 | -5.89E-02 | 1.04E-01 | 2.56E+00 | 2.64E+00 | 5.77E-01 | no | Hsd17b4 | intron_variant | MODIFIER |
| 1 | 180897615 | AX-170152100 | 0 | A | G | 0.228 | 6.72E-02 | 1.15E-01 | 2.47E+00 | 2.56E+00 | 5.77E-01 | no | Lefty2 | synonymous_variant | LOW |
| 6 | 126246278 | AX-97810781 | 0 | G | T | 0.167 | -4.60E-02 | 8.44E-02 | 2.66E+00 | 2.79E+00 | 5.77E-01 | no | | intergenic_variant | MODIFIER |
| 10 | 26678735 | AX-170243286 | 0 | G | A | 0.184 | 5.32E-02 | 9.76E-02 | 2.71E+00 | 2.82E+00 | 5.78E-01 | no | Gm8709 | upstream_gene_variant | MODIFIER |
| 9 | 27010243 | AX-170356544 | 0 | C | G | 0.465 | 5.08E-02 | 9.04E-02 | 2.58E+00 | 2.65E+00 | 5.78E-01 | no | Vps26b | intron_variant | MODIFIER |
| 1 | 140708372 | AX-97835008 | 0 | A | G | 0.202 | 7.41E-02 | 1.35E-01 | 2.68E+00 | 2.74E+00 | 5.78E-01 | no | | intergenic_variant | MODIFIER |
| 1 | 140765592 | AX-170001835 | 0 | A | T | 0.202 | 7.41E-02 | 1.35E-01 | 2.68E+00 | 2.74E+00 | 5.78E-01 | no | | intergenic_variant | MODIFIER |
| 1 | 140765800 | AX-170360563 | 0 | C | T | 0.202 | 7.41E-02 | 1.35E-01 | 2.68E+00 | 2.74E+00 | 5.78E-01 | no | | intergenic_variant | MODIFIER |
| 1 | 140767585 | AX-97835039 | 0 | C | T | 0.202 | 7.41E-02 | 1.35E-01 | 2.68E+00 | 2.74E+00 | 5.78E-01 | no | | intergenic_variant | MODIFIER |
| 1 | 140767676 | AX-170525364 | 0 | A | G | 0.202 | 7.41E-02 | 1.35E-01 | 2.68E+00 | 2.74E+00 | 5.78E-01 | no | | intergenic_variant | MODIFIER |
| 1 | 140773405 | AX-170042922 | 0 | C | T | 0.202 | 7.41E-02 | 1.35E-01 | 2.68E+00 | 2.74E+00 | 5.78E-01 | no | | intergenic_variant | MODIFIER |

| | | | | | | | | | | | | | | | |
|---|---|---|---|---|---|---|---|---|---|---|---|---|---|---|---|
| 1 | 140916270 | AX-97853145 | 0 | G | A | 0.202 | 7.41E-02 | 1.35E-01 | 2.68E+00 | 2.74E+00 | 5.78E-01 | no | | intergenic_variant | MODIFIER |
| 1 | 140995242 | AX-97824855 | 0 | G | A | 0.202 | 7.41E-02 | 1.35E-01 | 2.68E+00 | 2.74E+00 | 5.78E-01 | no | | intergenic_variant | MODIFIER |
| 1 | 141057407 | AX-170215919 | 0 | C | T | 0.202 | 7.41E-02 | 1.35E-01 | 2.68E+00 | 2.74E+00 | 5.78E-01 | no | | intergenic_variant | MODIFIER |
| 1 | 141059833 | AX-97799672 | 0 | A | G | 0.202 | 7.41E-02 | 1.35E-01 | 2.68E+00 | 2.74E+00 | 5.78E-01 | no | | intergenic_variant | MODIFIER |
| 1 | 141065397 | AX-168381950 | 0 | T | G | 0.202 | 7.41E-02 | 1.35E-01 | 2.68E+00 | 2.74E+00 | 5.78E-01 | no | | intergenic_variant | MODIFIER |
| 1 | 141077250 | AX-170438788 | 0 | C | G | 0.202 | 7.41E-02 | 1.35E-01 | 2.68E+00 | 2.74E+00 | 5.78E-01 | no | | intergenic_variant | MODIFIER |
| 1 | 141078348 | AX-170530395 | 0 | T | A | 0.202 | 7.41E-02 | 1.35E-01 | 2.68E+00 | 2.74E+00 | 5.78E-01 | no | | intergenic_variant | MODIFIER |
| 1 | 141339385 | AX-170435544 | 0 | A | G | 0.202 | 7.41E-02 | 1.35E-01 | 2.68E+00 | 2.74E+00 | 5.78E-01 | no | | intergenic_variant | MODIFIER |
| 1 | 141468687 | AX-97813531 | 0 | A | T | 0.202 | 7.41E-02 | 1.35E-01 | 2.68E+00 | 2.74E+00 | 5.78E-01 | no | | intergenic_variant | MODIFIER |
| 1 | 150763636 | AX-170414604 | 0 | G | C | 0.202 | 7.41E-02 | 1.35E-01 | 2.68E+00 | 2.74E+00 | 5.78E-01 | no | Hmcn1 | intron_variant | MODIFIER |
| 1 | 153039829 | AX-170406487 | 0 | G | A | 0.202 | 7.41E-02 | 1.35E-01 | 2.68E+00 | 2.74E+00 | 5.78E-01 | no | Nmnat2 | intron_variant | MODIFIER |
| 8 | 88324302 | AX-97852457 | 1 | C | T | 0.179 | -6.93E-02 | 1.26E-01 | 2.64E+00 | 2.73E+00 | 5.78E-01 | no | Adcy7 | intron_variant | MODIFIER |
| 1 | 88352332 | AX-170441030 | 0 | T | C | 0.158 | 1.14E-01 | 2.07E-01 | 2.65E+00 | 2.71E+00 | 5.78E-01 | no | Trpm8 | intron_variant | MODIFIER |
| 1 | 105733884 | AX-170441896 | 0 | C | G | 0.158 | 1.14E-01 | 2.07E-01 | 2.65E+00 | 2.71E+00 | 5.78E-01 | no | Relch | intron_variant | MODIFIER |
| 8 | 51467667 | AX-170238426 | 1 | C | A | 0.161 | 7.49E-02 | 1.35E-01 | 2.85E+00 | 2.95E+00 | 5.78E-01 | no | | intergenic_variant | MODIFIER |
| 11 | 47593798 | AX-97818406 | 0 | G | A | 0.088 | -8.72E-02 | 1.56E-01 | 2.57E+00 | 2.69E+00 | 5.78E-01 | no | Sgcd | intron_variant,non_coding_transcript_variant | MODIFIER |
| 4 | 151555375 | AX-170275758 | 1 | A | G | 0.098 | 8.79E-02 | 1.60E-01 | 2.68E+00 | 2.76E+00 | 5.78E-01 | no | Camta1 | intron_variant | MODIFIER |
| 6 | 147540746 | AX-170028502 | 0 | T | C | 0.053 | -1.03E-01 | 1.90E-01 | 2.70E+00 | 2.81E+00 | 5.78E-01 | no | Ccdc91 | intron_variant | MODIFIER |
| 6 | 147913255 | AX-97811319 | 0 | T | C | 0.053 | -1.03E-01 | 1.90E-01 | 2.70E+00 | 2.81E+00 | 5.78E-01 | no | | intergenic_variant | MODIFIER |
| 2 | 168566801 | AX-97782217 | 0 | G | T | 0.167 | 1.01E-01 | 1.82E-01 | 2.62E+00 | 2.70E+00 | 5.78E-01 | no | Nfatc2 | intron_variant | MODIFIER |
| 2 | 169068274 | AX-170177946 | 0 | C | T | 0.167 | 1.01E-01 | 1.82E-01 | 2.62E+00 | 2.70E+00 | 5.78E-01 | no | | intergenic_variant | MODIFIER |
| 1 | 128550330 | AX-170345606 | 0 | C | T | 0.544 | 4.46E-02 | 8.06E-02 | 2.63E+00 | 2.71E+00 | 5.78E-01 | no | | intergenic_variant | MODIFIER |
| 8 | 95869455 | AX-170459674 | 0 | C | G | 0.228 | 6.35E-02 | 1.16E-01 | 2.71E+00 | 2.77E+00 | 5.78E-01 | no | Got2 | intron_variant | MODIFIER |
| 16 | 27062881 | AX-97843333 | 1 | A | G | 0.25 | 4.92E-02 | 9.01E-02 | 2.65E+00 | 2.78E+00 | 5.78E-01 | no | | intergenic_variant | MODIFIER |
| 1 | 169089943 | AX-97799528 | 0 | G | A | 0.237 | 7.22E-02 | 1.31E-01 | 2.68E+00 | 2.72E+00 | 5.78E-01 | no | | intergenic_variant | MODIFIER |
| 1 | 169122348 | AX-170167917 | 0 | C | T | 0.237 | 7.22E-02 | 1.31E-01 | 2.68E+00 | 2.72E+00 | 5.78E-01 | no | | intergenic_variant | MODIFIER |
| 1 | 169126046 | AX-170238686 | 0 | G | C | 0.237 | 7.22E-02 | 1.31E-01 | 2.68E+00 | 2.72E+00 | 5.78E-01 | no | | intergenic_variant | MODIFIER |
| 1 | 169126170 | AX-168404437 | 0 | C | T | 0.237 | 7.22E-02 | 1.31E-01 | 2.68E+00 | 2.72E+00 | 5.78E-01 | no | | intergenic_variant | MODIFIER |
| 1 | 169135488 | AX-170388916 | 0 | G | C | 0.237 | 7.22E-02 | 1.31E-01 | 2.68E+00 | 2.72E+00 | 5.78E-01 | no | | intergenic_variant | MODIFIER |
| 1 | 169283579 | AX-97787262 | 0 | G | A | 0.237 | 7.22E-02 | 1.31E-01 | 2.68E+00 | 2.72E+00 | 5.78E-01 | no | | intergenic_variant | MODIFIER |
| 1 | 169283855 | AX-170318181 | 0 | C | G | 0.237 | 7.22E-02 | 1.31E-01 | 2.68E+00 | 2.72E+00 | 5.78E-01 | no | | intergenic_variant | MODIFIER |
| 1 | 169416162 | AX-170521686 | 0 | A | G | 0.237 | 7.22E-02 | 1.31E-01 | 2.68E+00 | 2.72E+00 | 5.78E-01 | no | | intergenic_variant | MODIFIER |
| 1 | 169416784 | AX-170387042 | 0 | T | C | 0.237 | 7.22E-02 | 1.31E-01 | 2.68E+00 | 2.72E+00 | 5.78E-01 | no | | intergenic_variant | MODIFIER |
| 1 | 169416934 | AX-170155618 | 0 | C | T | 0.237 | 7.22E-02 | 1.31E-01 | 2.68E+00 | 2.72E+00 | 5.78E-01 | no | | intergenic_variant | MODIFIER |
| 1 | 169439121 | AX-170426212 | 0 | T | G | 0.237 | 7.22E-02 | 1.31E-01 | 2.68E+00 | 2.72E+00 | 5.78E-01 | no | | intergenic_variant | MODIFIER |
| 1 | 169608568 | AX-97850711 | 0 | A | G | 0.237 | 7.22E-02 | 1.31E-01 | 2.68E+00 | 2.72E+00 | 5.78E-01 | no | | regulatory_region_variant | MODIFIER |
| 1 | 169626804 | AX-97820517 | 0 | G | A | 0.237 | 7.22E-02 | 1.31E-01 | 2.68E+00 | 2.72E+00 | 5.78E-01 | no | | intergenic_variant | MODIFIER |
| 1 | 169653200 | AX-170413992 | 0 | A | C | 0.237 | 7.22E-02 | 1.31E-01 | 2.68E+00 | 2.72E+00 | 5.78E-01 | no | Rgs5 | upstream_gene_variant | MODIFIER |
| 1 | 169653348 | AX-170273924 | 0 | T | A | 0.237 | 7.22E-02 | 1.31E-01 | 2.68E+00 | 2.72E+00 | 5.78E-01 | no | Rgs5 | upstream_gene_variant | MODIFIER |
| 1 | 169658730 | AX-170229648 | 0 | T | C | 0.237 | 7.22E-02 | 1.31E-01 | 2.68E+00 | 2.72E+00 | 5.78E-01 | no | Rgs5 | intron_variant | MODIFIER |
| 1 | 169718501 | AX-97829005 | 0 | G | A | 0.237 | 7.22E-02 | 1.31E-01 | 2.68E+00 | 2.72E+00 | 5.78E-01 | no | | intergenic_variant | MODIFIER |
| 1 | 169731290 | AX-170533301 | 0 | C | T | 0.237 | 7.22E-02 | 1.31E-01 | 2.68E+00 | 2.72E+00 | 5.78E-01 | no | | intergenic_variant | MODIFIER |
| 1 | 169737033 | AX-170307580 | 0 | G | A | 0.237 | 7.22E-02 | 1.31E-01 | 2.68E+00 | 2.72E+00 | 5.78E-01 | no | Rgs4 | downstream_gene_variant | MODIFIER |
| 1 | 169742828 | AX-170330607 | 0 | G | A | 0.237 | 7.22E-02 | 1.31E-01 | 2.68E+00 | 2.72E+00 | 5.78E-01 | no | Rgs4 | 3_prime_UTR_variant | MODIFIER |
| 1 | 169782968 | AX-170449879 | 0 | T | C | 0.237 | 7.22E-02 | 1.31E-01 | 2.68E+00 | 2.72E+00 | 5.78E-01 | no | | intergenic_variant | MODIFIER |
| 1 | 169809054 | AX-170432087 | 0 | G | A | 0.237 | 7.22E-02 | 1.31E-01 | 2.68E+00 | 2.72E+00 | 5.78E-01 | no | | intergenic_variant | MODIFIER |
| 1 | 169813674 | AX-170138653 | 0 | A | G | 0.237 | 7.22E-02 | 1.31E-01 | 2.68E+00 | 2.72E+00 | 5.78E-01 | no | | intergenic_variant | MODIFIER |
| 1 | 169877481 | AX-170253609 | 0 | C | T | 0.237 | 7.22E-02 | 1.31E-01 | 2.68E+00 | 2.72E+00 | 5.78E-01 | no | | intergenic_variant | MODIFIER |
| 1 | 169877586 | AX-168394622 | 0 | C | A | 0.237 | 7.22E-02 | 1.31E-01 | 2.68E+00 | 2.72E+00 | 5.78E-01 | no | | intergenic_variant | MODIFIER |
| 1 | 169877977 | AX-170404794 | 0 | G | A | 0.237 | 7.22E-02 | 1.31E-01 | 2.68E+00 | 2.72E+00 | 5.78E-01 | no | | intergenic_variant | MODIFIER |
| 1 | 169879143 | AX-168374819 | 0 | T | G | 0.237 | 7.22E-02 | 1.31E-01 | 2.68E+00 | 2.72E+00 | 5.78E-01 | no | | intergenic_variant | MODIFIER |
| 1 | 170060571 | AX-170111297 | 0 | C | T | 0.237 | 7.22E-02 | 1.31E-01 | 2.68E+00 | 2.72E+00 | 5.78E-01 | no | Ddr2 | intron_variant | MODIFIER |
| 1 | 170203955 | AX-168390153 | 0 | C | T | 0.237 | 7.22E-02 | 1.31E-01 | 2.68E+00 | 2.72E+00 | 5.78E-01 | no | Uhmk1 | intron_variant | MODIFIER |
| 1 | 170206428 | AX-170262959 | 0 | G | A | 0.237 | 7.22E-02 | 1.31E-01 | 2.68E+00 | 2.72E+00 | 5.78E-01 | no | Uhmk1 | intron_variant | MODIFIER |

| | | | | | | | | | | | | | | | |
|---|---|---|---|---|---|---|---|---|---|---|---|---|---|---|---|
| 1 | 170387137 | AX-170328624 | 0 | G | C | 0.237 | 7.22E-02 | 1.31E-01 | 2.68E+00 | 2.72E+00 | 5.78E-01 | no | Nos1ap | intron_variant,non_coding_transcript_variant | MODIFIER |
| 1 | 170451140 | AX-170476743 | 0 | T | G | 0.237 | 7.22E-02 | 1.31E-01 | 2.68E+00 | 2.72E+00 | 5.78E-01 | no | Nos1ap | intron_variant,non_coding_transcript_variant | MODIFIER |
| 1 | 170459563 | AX-168401667 | 0 | A | G | 0.237 | 7.22E-02 | 1.31E-01 | 2.68E+00 | 2.72E+00 | 5.78E-01 | no | Nos1ap | intron_variant,non_coding_transcript_variant | MODIFIER |
| 1 | 170464904 | AX-170100497 | 0 | G | C | 0.237 | 7.22E-02 | 1.31E-01 | 2.68E+00 | 2.72E+00 | 5.78E-01 | no | Nos1ap | intron_variant,non_coding_transcript_variant | MODIFIER |
| 1 | 170466113 | AX-170215541 | 0 | T | C | 0.237 | 7.22E-02 | 1.31E-01 | 2.68E+00 | 2.72E+00 | 5.78E-01 | no | Nos1ap | intron_variant,non_coding_transcript_variant | MODIFIER |
| 1 | 170578973 | AX-170495987 | 0 | T | C | 0.237 | 7.22E-02 | 1.31E-01 | 2.68E+00 | 2.72E+00 | 5.78E-01 | no | Nos1ap | intron_variant,non_coding_transcript_variant | MODIFIER |
| 1 | 170698538 | AX-97852642 | 0 | T | C | 0.237 | 7.22E-02 | 1.31E-01 | 2.68E+00 | 2.72E+00 | 5.78E-01 | no | | regulatory_region_variant | MODIFIER |
| 1 | 170772762 | AX-97780638 | 0 | T | G | 0.237 | 7.22E-02 | 1.31E-01 | 2.68E+00 | 2.72E+00 | 5.78E-01 | no | Atf6 | intron_variant | MODIFIER |
| 2 | 159842940 | AX-170213926 | 2 | T | C | 0.136 | 9.41E-02 | 1.73E-01 | 2.70E+00 | 2.81E+00 | 5.78E-01 | no | | intergenic_variant | MODIFIER |
| 7 | 78270260 | AX-170502368 | 1 | C | T | 0.357 | -5.39E-02 | 9.89E-02 | 2.72E+00 | 2.81E+00 | 5.79E-01 | no | Ntrk3 | intron_variant | MODIFIER |
| 9 | 40780950 | AX-170428383 | 1 | T | G | 0.42 | -4.01E-02 | 7.34E-02 | 2.69E+00 | 2.79E+00 | 5.79E-01 | no | Clmp | intron_variant | MODIFIER |
| 18 | 80737138 | AX-170356347 | 0 | C | G | 0.307 | -4.68E-02 | 8.41E-02 | 2.58E+00 | 2.69E+00 | 5.79E-01 | no | Atp9b | intron_variant | MODIFIER |
| 10 | 71173737 | AX-170172911 | 2 | A | G | 0.409 | -4.09E-02 | 7.26E-02 | 2.55E+00 | 2.65E+00 | 5.79E-01 | no | C730027H18Rik | intron_variant,non_coding_transcript_variant | MODIFIER |
| 15 | 53592928 | AX-170465536 | 1 | C | T | 0.036 | 8.60E-02 | 1.55E-01 | 2.56E+00 | 2.71E+00 | 5.79E-01 | no | Samd12 | intron_variant | MODIFIER |
| 15 | 83069702 | AX-97792283 | 0 | A | G | 0.544 | 4.53E-02 | 8.06E-02 | 2.56E+00 | 2.64E+00 | 5.79E-01 | no | | intergenic_variant | MODIFIER |
| 4 | 49583071 | AX-170225109 | 0 | A | G | 0.377 | -4.87E-02 | 8.95E-02 | 2.70E+00 | 2.80E+00 | 5.79E-01 | no | Tmem246 | downstream_gene_variant | MODIFIER |
| 5 | 7278310 | AX-97794398 | 0 | G | A | 0.439 | 3.36E-02 | 6.24E-02 | 2.72E+00 | 2.87E+00 | 5.79E-01 | no | Gm43841 | downstream_gene_variant | MODIFIER |
| 9 | 40142346 | AX-97843796 | 1 | A | C | 0.411 | -4.01E-02 | 7.36E-02 | 2.69E+00 | 2.79E+00 | 5.80E-01 | no | | intergenic_variant | MODIFIER |
| 5 | 83622908 | AX-170467746 | 0 | G | C | 0.439 | 5.96E-02 | 1.09E-01 | 2.73E+00 | 2.78E+00 | 5.80E-01 | no | | regulatory_region_variant | MODIFIER |
| 1 | 96409264 | AX-170271511 | 0 | T | C | 0.149 | 1.09E-01 | 1.98E-01 | 2.65E+00 | 2.72E+00 | 5.80E-01 | no | | intergenic_variant | MODIFIER |
| 8 | 14536658 | AX-170126556 | 0 | A | G | 0.237 | -4.28E-02 | 7.65E-02 | 2.56E+00 | 2.65E+00 | 5.80E-01 | no | Dlgap2 | intron_variant | MODIFIER |
| 18 | 63440840 | AX-170103744 | 0 | G | A | 0.193 | -6.25E-02 | 1.14E-01 | 2.64E+00 | 2.70E+00 | 5.80E-01 | no | | intergenic_variant | MODIFIER |
| 10 | 23572600 | AX-97794560 | 0 | G | A | 0.57 | -3.71E-02 | 6.90E-02 | 2.79E+00 | 2.95E+00 | 5.80E-01 | no | Gm47843 | downstream_gene_variant | MODIFIER |
| 7 | 57550655 | AX-170223433 | 1 | T | A | 0.438 | 5.44E-02 | 9.91E-02 | 2.64E+00 | 2.72E+00 | 5.80E-01 | no | Gabrb3 | intron_variant | MODIFIER |
| 15 | 81451381 | AX-97838923 | 1 | G | A | 0.554 | 4.58E-02 | 8.14E-02 | 2.55E+00 | 2.62E+00 | 5.80E-01 | no | Xpnpep3 | intron_variant | MODIFIER |
| 8 | 14476494 | AX-170164652 | 0 | C | T | 0.237 | -4.28E-02 | 7.63E-02 | 2.55E+00 | 2.65E+00 | 5.81E-01 | no | Dlgap2 | intron_variant | MODIFIER |
| 8 | 14481431 | AX-170335358 | 0 | T | C | 0.237 | -4.28E-02 | 7.63E-02 | 2.55E+00 | 2.65E+00 | 5.81E-01 | no | Dlgap2 | intron_variant | MODIFIER |
| 8 | 14494594 | AX-170096175 | 0 | G | A | 0.237 | -4.28E-02 | 7.63E-02 | 2.55E+00 | 2.65E+00 | 5.81E-01 | no | Dlgap2 | intron_variant | MODIFIER |
| 8 | 14496838 | AX-170218639 | 0 | C | T | 0.237 | -4.28E-02 | 7.63E-02 | 2.55E+00 | 2.65E+00 | 5.81E-01 | no | Dlgap2 | intron_variant | MODIFIER |
| 8 | 14514879 | AX-170395791 | 0 | A | G | 0.237 | -4.28E-02 | 7.63E-02 | 2.55E+00 | 2.65E+00 | 5.81E-01 | no | Dlgap2 | intron_variant | MODIFIER |
| 8 | 14528134 | AX-170477095 | 0 | T | C | 0.237 | -4.28E-02 | 7.63E-02 | 2.55E+00 | 2.65E+00 | 5.81E-01 | no | Dlgap2 | intron_variant | MODIFIER |
| 8 | 14532029 | AX-170258079 | 0 | G | A | 0.237 | -4.28E-02 | 7.63E-02 | 2.55E+00 | 2.65E+00 | 5.81E-01 | no | Dlgap2 | intron_variant | MODIFIER |
| 8 | 14539109 | AX-97817656 | 0 | T | C | 0.237 | -4.28E-02 | 7.63E-02 | 2.55E+00 | 2.65E+00 | 5.81E-01 | no | Dlgap2 | intron_variant | MODIFIER |
| 8 | 14574101 | AX-170402634 | 0 | T | C | 0.237 | -4.28E-02 | 7.63E-02 | 2.55E+00 | 2.65E+00 | 5.81E-01 | no | Dlgap2 | intron_variant | MODIFIER |
| 8 | 14574800 | AX-170468764 | 0 | G | A | 0.237 | -4.28E-02 | 7.63E-02 | 2.55E+00 | 2.65E+00 | 5.81E-01 | no | Dlgap2 | intron_variant | MODIFIER |
| 8 | 14580396 | AX-170108107 | 0 | A | G | 0.237 | -4.28E-02 | 7.63E-02 | 2.55E+00 | 2.65E+00 | 5.81E-01 | no | Dlgap2 | intron_variant | MODIFIER |
| 8 | 14608117 | AX-170327412 | 0 | G | A | 0.237 | -4.28E-02 | 7.63E-02 | 2.55E+00 | 2.65E+00 | 5.81E-01 | no | Dlgap2 | intron_variant | MODIFIER |
| 8 | 14636179 | AX-170460133 | 0 | G | A | 0.237 | -4.28E-02 | 7.63E-02 | 2.55E+00 | 2.65E+00 | 5.81E-01 | no | Dlgap2 | intron_variant | MODIFIER |
| 8 | 14652046 | AX-170151415 | 0 | C | T | 0.237 | -4.28E-02 | 7.63E-02 | 2.55E+00 | 2.65E+00 | 5.81E-01 | no | Dlgap2 | intron_variant | MODIFIER |
| 8 | 14652091 | AX-97825064 | 0 | A | C | 0.237 | -4.28E-02 | 7.63E-02 | 2.55E+00 | 2.65E+00 | 5.81E-01 | no | Dlgap2 | intron_variant | MODIFIER |
| 8 | 14652417 | AX-170422390 | 0 | A | C | 0.237 | -4.28E-02 | 7.63E-02 | 2.55E+00 | 2.65E+00 | 5.81E-01 | no | Dlgap2 | intron_variant | MODIFIER |
| 8 | 14652945 | AX-170367637 | 0 | C | T | 0.237 | -4.28E-02 | 7.63E-02 | 2.55E+00 | 2.65E+00 | 5.81E-01 | no | Dlgap2 | intron_variant | MODIFIER |
| 8 | 14656317 | AX-170215244 | 0 | G | A | 0.237 | -4.28E-02 | 7.63E-02 | 2.55E+00 | 2.65E+00 | 5.81E-01 | no | Dlgap2 | intron_variant | MODIFIER |
| 8 | 14656371 | AX-170530315 | 0 | G | A | 0.237 | -4.28E-02 | 7.63E-02 | 2.55E+00 | 2.65E+00 | 5.81E-01 | no | Dlgap2 | intron_variant | MODIFIER |
| 8 | 14669040 | AX-170187660 | 0 | T | C | 0.237 | -4.28E-02 | 7.63E-02 | 2.55E+00 | 2.65E+00 | 5.81E-01 | no | Dlgap2 | intron_variant | MODIFIER |
| 8 | 14670422 | AX-170328561 | 0 | A | C | 0.237 | -4.28E-02 | 7.63E-02 | 2.55E+00 | 2.65E+00 | 5.81E-01 | no | Dlgap2 | intron_variant | MODIFIER |
| 8 | 14672142 | AX-170320018 | 0 | T | C | 0.237 | -4.28E-02 | 7.63E-02 | 2.55E+00 | 2.65E+00 | 5.81E-01 | no | Dlgap2 | intron_variant | MODIFIER |
| 8 | 14673789 | AX-97847440 | 0 | A | G | 0.237 | -4.28E-02 | 7.63E-02 | 2.55E+00 | 2.65E+00 | 5.81E-01 | no | Dlgap2 | intron_variant | MODIFIER |
| 8 | 14725795 | AX-170372632 | 0 | C | T | 0.237 | -4.28E-02 | 7.63E-02 | 2.55E+00 | 2.65E+00 | 5.81E-01 | no | Dlgap2 | intron_variant | MODIFIER |
| 8 | 14726403 | AX-169999738 | 0 | G | A | 0.237 | -4.28E-02 | 7.63E-02 | 2.55E+00 | 2.65E+00 | 5.81E-01 | no | Dlgap2 | intron_variant | MODIFIER |
| 8 | 14779871 | AX-170030297 | 0 | G | A | 0.237 | -4.28E-02 | 7.63E-02 | 2.55E+00 | 2.65E+00 | 5.81E-01 | no | Dlgap2 | intron_variant | MODIFIER |
| 8 | 14783051 | AX-170221753 | 0 | T | C | 0.237 | -4.28E-02 | 7.63E-02 | 2.55E+00 | 2.65E+00 | 5.81E-01 | no | Dlgap2 | intron_variant | MODIFIER |
| 8 | 14838067 | AX-170441054 | 0 | G | A | 0.237 | -4.28E-02 | 7.63E-02 | 2.55E+00 | 2.65E+00 | 5.81E-01 | no | Dlgap2 | intron_variant | MODIFIER |
| 8 | 14881784 | AX-170454446 | 0 | G | C | 0.237 | -4.28E-02 | 7.63E-02 | 2.55E+00 | 2.65E+00 | 5.81E-01 | no | Cln8 | 5_prime_UTR_variant | MODIFIER |

| | | | | | | | | | | | | | | | |
|---|---|---|---|---|---|---|---|---|---|---|---|---|---|---|---|
| 8 | 14922194 | AX-170251928 | 0 | T | C | 0.237 | -4.28E-02 | 7.63E-02 | 2.55E+00 | 2.65E+00 | 5.81E-01 | no | Arhgef10 | intron_variant | MODIFIER |
| 8 | 14978445 | AX-170130724 | 0 | C | T | 0.237 | -4.28E-02 | 7.63E-02 | 2.55E+00 | 2.65E+00 | 5.81E-01 | no | Arhgef10 | synonymous_variant | LOW |
| 8 | 14979195 | AX-170144726 | 0 | A | G | 0.237 | -4.28E-02 | 7.63E-02 | 2.55E+00 | 2.65E+00 | 5.81E-01 | no | Arhgef10 | intron_variant | MODIFIER |
| 8 | 14980073 | AX-170165484 | 0 | T | G | 0.237 | -4.28E-02 | 7.63E-02 | 2.55E+00 | 2.65E+00 | 5.81E-01 | no | Arhgef10 | intron_variant | MODIFIER |
| 8 | 14995961 | AX-170180546 | 0 | A | G | 0.237 | -4.28E-02 | 7.63E-02 | 2.55E+00 | 2.65E+00 | 5.81E-01 | no | Arhgef10 | intron_variant | MODIFIER |
| 8 | 15002874 | AX-170449282 | 0 | T | G | 0.237 | -4.28E-02 | 7.63E-02 | 2.55E+00 | 2.65E+00 | 5.81E-01 | no | Arhgef10 | downstream_gene_variant | MODIFIER |
| 8 | 15039420 | AX-170079380 | 0 | G | A | 0.237 | -4.28E-02 | 7.63E-02 | 2.55E+00 | 2.65E+00 | 5.81E-01 | no | BB014433 | downstream_gene_variant | MODIFIER |
| 8 | 15056954 | AX-170226668 | 0 | A | G | 0.237 | -4.28E-02 | 7.63E-02 | 2.55E+00 | 2.65E+00 | 5.81E-01 | no | Myom2 | upstream_gene_variant | MODIFIER |
| 8 | 15058800 | AX-170243786 | 0 | C | T | 0.237 | -4.28E-02 | 7.63E-02 | 2.55E+00 | 2.65E+00 | 5.81E-01 | no | Myom2 | intron_variant | MODIFIER |
| 8 | 15064689 | AX-170153237 | 0 | T | C | 0.237 | -4.28E-02 | 7.63E-02 | 2.55E+00 | 2.65E+00 | 5.81E-01 | no | Myom2 | intron_variant | MODIFIER |
| 8 | 15158736 | AX-170265475 | 0 | C | T | 0.237 | -4.28E-02 | 7.63E-02 | 2.55E+00 | 2.65E+00 | 5.81E-01 | no | Gm7706 | intron_variant,non_coding_transcript_variant | MODIFIER |
| 8 | 15175120 | AX-170255970 | 0 | C | G | 0.237 | -4.28E-02 | 7.63E-02 | 2.55E+00 | 2.65E+00 | 5.81E-01 | no | Gm7706 | intron_variant,non_coding_transcript_variant | MODIFIER |
| 8 | 15210201 | AX-170499318 | 0 | G | A | 0.237 | -4.28E-02 | 7.63E-02 | 2.55E+00 | 2.65E+00 | 5.81E-01 | no | Gm7706 | intron_variant,non_coding_transcript_variant | MODIFIER |
| 8 | 15243539 | AX-170495054 | 0 | C | A | 0.237 | -4.28E-02 | 7.63E-02 | 2.55E+00 | 2.65E+00 | 5.81E-01 | no | Gm7706 | intron_variant,non_coding_transcript_variant | MODIFIER |
| 8 | 15264518 | AX-97837307 | 0 | T | C | 0.237 | -4.28E-02 | 7.63E-02 | 2.55E+00 | 2.65E+00 | 5.81E-01 | no | Gm7706 | intron_variant,non_coding_transcript_variant | MODIFIER |
| 8 | 15270431 | AX-170355727 | 0 | A | C | 0.237 | -4.28E-02 | 7.63E-02 | 2.55E+00 | 2.65E+00 | 5.81E-01 | no | Gm7706 | intron_variant,non_coding_transcript_variant | MODIFIER |
| 8 | 15287937 | AX-170125051 | 0 | G | T | 0.237 | -4.28E-02 | 7.63E-02 | 2.55E+00 | 2.65E+00 | 5.81E-01 | no | Gm7706 | intron_variant,non_coding_transcript_variant | MODIFIER |
| 8 | 15297126 | AX-170272764 | 0 | C | T | 0.237 | -4.28E-02 | 7.63E-02 | 2.55E+00 | 2.65E+00 | 5.81E-01 | no | Gm7706 | intron_variant,non_coding_transcript_variant | MODIFIER |
| 8 | 15300942 | AX-97798012 | 0 | T | C | 0.237 | -4.28E-02 | 7.63E-02 | 2.55E+00 | 2.65E+00 | 5.81E-01 | no | Gm7706 | intron_variant,non_coding_transcript_variant | MODIFIER |
| 8 | 15301284 | AX-170305354 | 0 | T | G | 0.237 | -4.28E-02 | 7.63E-02 | 2.55E+00 | 2.65E+00 | 5.81E-01 | no | Gm7706 | intron_variant,non_coding_transcript_variant | MODIFIER |
| 8 | 15344885 | AX-170079387 | 0 | T | C | 0.237 | -4.28E-02 | 7.63E-02 | 2.55E+00 | 2.65E+00 | 5.81E-01 | no | Gm7706 | intron_variant,non_coding_transcript_variant | MODIFIER |
| 8 | 15345582 | AX-97823822 | 0 | G | C | 0.237 | -4.28E-02 | 7.63E-02 | 2.55E+00 | 2.65E+00 | 5.81E-01 | no | Gm7706 | intron_variant,non_coding_transcript_variant | MODIFIER |
| 8 | 15355117 | AX-170030303 | 0 | C | G | 0.237 | -4.28E-02 | 7.63E-02 | 2.55E+00 | 2.65E+00 | 5.81E-01 | no | Gm7706 | intron_variant,non_coding_transcript_variant | MODIFIER |
| 8 | 15361363 | AX-97837904 | 0 | G | C | 0.237 | -4.28E-02 | 7.63E-02 | 2.55E+00 | 2.65E+00 | 5.81E-01 | no | Gm7706 | intron_variant,non_coding_transcript_variant | MODIFIER |
| 8 | 15403876 | AX-170148470 | 0 | G | A | 0.237 | -4.28E-02 | 7.63E-02 | 2.55E+00 | 2.65E+00 | 5.81E-01 | no | | intergenic_variant | MODIFIER |
| 8 | 15404558 | AX-170217999 | 0 | C | G | 0.237 | -4.28E-02 | 7.63E-02 | 2.55E+00 | 2.65E+00 | 5.81E-01 | no | | intergenic_variant | MODIFIER |
| 8 | 15458776 | AX-170141824 | 0 | T | C | 0.237 | -4.28E-02 | 7.63E-02 | 2.55E+00 | 2.65E+00 | 5.81E-01 | no | | intergenic_variant | MODIFIER |
| 8 | 15459017 | AX-170361844 | 0 | T | C | 0.237 | -4.28E-02 | 7.63E-02 | 2.55E+00 | 2.65E+00 | 5.81E-01 | no | | intergenic_variant | MODIFIER |
| 6 | 121624648 | AX-170263077 | 0 | C | T | 0.061 | 1.14E-01 | 2.08E-01 | 2.59E+00 | 2.71E+00 | 5.81E-01 | no | Gm43945 | upstream_gene_variant | MODIFIER |
| 8 | 83218451 | AX-170485610 | 0 | G | A | 0.211 | -6.27E-02 | 1.16E-01 | 2.74E+00 | 2.84E+00 | 5.81E-01 | no | Tbc1d9 | intron_variant | MODIFIER |
| 2 | 71859393 | AX-168392261 | 0 | C | T | 0.044 | -1.08E-01 | 1.91E-01 | 2.53E+00 | 2.65E+00 | 5.81E-01 | no | Itga6 | downstream_gene_variant | MODIFIER |
| 12 | 57762014 | AX-170155603 | 0 | C | T | 0.07 | 1.22E-01 | 2.25E-01 | 2.75E+00 | 2.84E+00 | 5.81E-01 | no | | intergenic_variant | MODIFIER |
| 18 | 34598140 | AX-170411179 | 0 | T | C | 0.07 | 1.22E-01 | 2.25E-01 | 2.75E+00 | 2.84E+00 | 5.81E-01 | no | Brd8 | downstream_gene_variant | MODIFIER |
| 16 | 77209387 | AX-97791294 | 0 | C | T | 0.456 | 4.30E-02 | 7.66E-02 | 2.56E+00 | 2.63E+00 | 5.81E-01 | no | | intergenic_variant | MODIFIER |
| 1 | 20535799 | AX-170035600 | 0 | T | C | 0.412 | 4.48E-02 | 8.02E-02 | 2.58E+00 | 2.63E+00 | 5.81E-01 | no | Pkhd1 | intron_variant | MODIFIER |
| 15 | 82701982 | AX-170439406 | 1 | C | G | 0.554 | 4.57E-02 | 8.13E-02 | 2.56E+00 | 2.62E+00 | 5.81E-01 | no | Cyp2d38-ps | downstream_gene_variant | MODIFIER |
| 8 | 36158527 | AX-97830189 | 1 | G | A | 0.152 | 8.14E-02 | 1.48E-01 | 2.83E+00 | 2.89E+00 | 5.81E-01 | no | Gm38414 | intron_variant,non_coding_transcript_variant | MODIFIER |
| 8 | 40888054 | AX-170368931 | 1 | T | C | 0.152 | 8.14E-02 | 1.48E-01 | 2.83E+00 | 2.89E+00 | 5.81E-01 | no | Slc7a2 | intron_variant | MODIFIER |
| 1 | 161212137 | AX-170002141 | 1 | A | G | 0.223 | 7.52E-02 | 1.37E-01 | 2.65E+00 | 2.71E+00 | 5.81E-01 | no | Gm6185 | non_coding_transcript_exon_variant | MODIFIER |
| 9 | 105956961 | AX-170205673 | 1 | C | A | 0.134 | 5.68E-02 | 1.05E-01 | 2.77E+00 | 2.88E+00 | 5.81E-01 | no | Col6a5 | intron_variant | MODIFIER |
| 13 | 32115537 | AX-170008025 | 0 | A | T | 0.167 | -6.65E-02 | 1.18E-01 | 2.54E+00 | 2.62E+00 | 5.81E-01 | no | Gmds | intron_variant | MODIFIER |
| 9 | 107854470 | AX-170143257 | 1 | A | G | 0.125 | -6.20E-02 | 1.12E-01 | 2.58E+00 | 2.67E+00 | 5.81E-01 | no | Rbm6 | intron_variant | MODIFIER |
| 2 | 76533561 | AX-97819214 | 0 | G | A | 0.044 | -1.02E-01 | 1.87E-01 | 2.61E+00 | 2.73E+00 | 5.81E-01 | no | Osbpl6 | intron_variant | MODIFIER |
| 5 | 105294995 | AX-170104427 | 1 | C | A | 0.429 | 5.81E-02 | 1.06E-01 | 2.65E+00 | 2.71E+00 | 5.81E-01 | no | Gm43302 | upstream_gene_variant | MODIFIER |
| 5 | 77123233 | AX-170269198 | 1 | C | T | 0.438 | 5.51E-02 | 1.00E-01 | 2.62E+00 | 2.69E+00 | 5.82E-01 | no | Gm15831 | upstream_gene_variant | MODIFIER |
| 1 | 69674285 | AX-97811976 | 2 | T | A | 0.127 | -9.39E-02 | 1.74E-01 | 2.65E+00 | 2.77E+00 | 5.82E-01 | no | Ikzf2 | intron_variant | MODIFIER |
| 1 | 106461837 | AX-170091627 | 0 | T | C | 0.158 | 1.03E-01 | 1.87E-01 | 2.59E+00 | 2.67E+00 | 5.82E-01 | no | | intergenic_variant | MODIFIER |
| 18 | 66681123 | AX-170452748 | 0 | A | G | 0.105 | 8.67E-02 | 1.59E-01 | 2.62E+00 | 2.73E+00 | 5.82E-01 | no | Gm22219 | downstream_gene_variant | MODIFIER |
| 9 | 20907726 | AX-170497566 | 0 | G | A | 0.474 | 5.03E-02 | 9.03E-02 | 2.57E+00 | 2.64E+00 | 5.82E-01 | no | Dnmt1 | intron_variant | MODIFIER |
| 1 | 187169478 | AX-97815150 | 0 | A | G | 0.237 | 6.36E-02 | 1.08E-01 | 2.44E+00 | 2.52E+00 | 5.82E-01 | no | Spata17 | intron_variant | MODIFIER |
| 1 | 11355221 | AX-169999946 | 0 | A | T | 0.386 | -3.97E-02 | 7.22E-02 | 2.88E+00 | 2.97E+00 | 5.82E-01 | no | | intergenic_variant | MODIFIER |
| 8 | 32514536 | AX-170366913 | 1 | A | G | 0.143 | -6.90E-02 | 1.27E-01 | 2.68E+00 | 2.75E+00 | 5.83E-01 | no | Nrg1 | intron_variant | MODIFIER |
| 4 | 92465850 | AX-170344481 | 1 | T | C | 0.339 | 5.86E-02 | 1.08E-01 | 2.65E+00 | 2.73E+00 | 5.83E-01 | no | | intergenic_variant | MODIFIER |
| 1 | 127396557 | AX-170443077 | 0 | G | A | 0.544 | 4.57E-02 | 8.34E-02 | 2.62E+00 | 2.69E+00 | 5.83E-01 | no | Mgat5 | intron_variant | MODIFIER |

| | | | | | | | | | | | | | | | |
|---|---|---|---|---|---|---|---|---|---|---|---|---|---|---|---|
| 1 | 36356467 | AX-97784646 | 0 | T | C | 0.465 | 4.14E-02 | 7.67E-02 | 2.71E+00 | 2.79E+00 | 5.83E-01 | no | Kansl3 | intron_variant | MODIFIER |
| 8 | 54199952 | AX-170422858 | 1 | G | T | 0.152 | -6.95E-02 | 1.24E-01 | 2.55E+00 | 2.64E+00 | 5.83E-01 | no | | intergenic_variant | MODIFIER |
| 18 | 69895101 | AX-97837823 | 0 | C | A | 0.088 | 9.16E-02 | 1.69E-01 | 2.65E+00 | 2.76E+00 | 5.83E-01 | no | AC108433.1 | intron_variant,non_coding_transcript_variant | MODIFIER |
| 18 | 69915708 | AX-170332129 | 0 | A | T | 0.088 | 9.16E-02 | 1.69E-01 | 2.65E+00 | 2.76E+00 | 5.83E-01 | no | AC108433.1 | upstream_gene_variant | MODIFIER |
| 18 | 69922317 | AX-170099076 | 0 | G | T | 0.088 | 9.16E-02 | 1.69E-01 | 2.65E+00 | 2.76E+00 | 5.83E-01 | no | Ccdc68 | upstream_gene_variant | MODIFIER |
| 18 | 70410888 | AX-97853907 | 0 | A | G | 0.088 | 9.16E-02 | 1.69E-01 | 2.65E+00 | 2.76E+00 | 5.83E-01 | no | | regulatory_region_variant | MODIFIER |
| 9 | 21623829 | AX-97844856 | 0 | G | A | 0.456 | 5.01E-02 | 8.93E-02 | 2.55E+00 | 2.62E+00 | 5.83E-01 | no | Smarca4 | intron_variant | MODIFIER |
| 9 | 23237931 | AX-170228163 | 0 | A | G | 0.456 | 5.01E-02 | 8.93E-02 | 2.55E+00 | 2.62E+00 | 5.83E-01 | no | Bmper | intron_variant | MODIFIER |
| 16 | 97197081 | AX-170172619 | 0 | A | G | 0.447 | 4.00E-02 | 7.45E-02 | 2.77E+00 | 2.89E+00 | 5.83E-01 | no | | intergenic_variant | MODIFIER |
| 7 | 73172217 | AX-170327690 | 1 | G | A | 0.366 | -5.30E-02 | 9.85E-02 | 2.73E+00 | 2.83E+00 | 5.83E-01 | no | | intergenic_variant | MODIFIER |
| 12 | 91768340 | AX-170148577 | 1 | C | T | 0.098 | 7.96E-02 | 1.46E-01 | 2.61E+00 | 2.72E+00 | 5.83E-01 | no | Ston2 | intron_variant | MODIFIER |
| 6 | 89274439 | AX-170027668 | 0 | A | G | 0.061 | 7.95E-02 | 1.48E-01 | 2.66E+00 | 2.79E+00 | 5.83E-01 | no | 4930512J16Rik | intron_variant,non_coding_transcript_variant | MODIFIER |
| 1 | 115955100 | AX-170295726 | 1 | T | A | 0.152 | 1.03E-01 | 1.86E-01 | 2.57E+00 | 2.65E+00 | 5.83E-01 | no | Cntnap5a | intron_variant | MODIFIER |
| 8 | 14596271 | AX-170266889 | 1 | C | T | 0.241 | -4.21E-02 | 7.58E-02 | 2.56E+00 | 2.65E+00 | 5.83E-01 | no | Dlgap2 | intron_variant | MODIFIER |
| 9 | 117777090 | AX-170364219 | 0 | G | C | 0.123 | -5.24E-02 | 9.63E-02 | 2.61E+00 | 2.73E+00 | 5.84E-01 | no | Gm45881 | intron_variant,non_coding_transcript_variant | MODIFIER |
| 2 | 145575653 | AX-170065168 | 2 | C | G | 0.136 | 1.11E-01 | 2.06E-01 | 2.70E+00 | 2.77E+00 | 5.84E-01 | no | Slc24a3 | intron_variant | MODIFIER |
| 10 | 59104737 | AX-170372529 | 1 | G | A | 0.295 | 4.29E-02 | 7.92E-02 | 2.86E+00 | 2.99E+00 | 5.84E-01 | no | Sh3rf3 | non_coding_transcript_exon_variant | MODIFIER |
| 9 | 111957839 | AX-97805580 | 0 | A | C | 0.105 | -5.99E-02 | 1.09E-01 | 2.60E+00 | 2.70E+00 | 5.84E-01 | no | | regulatory_region_variant | MODIFIER |
| 18 | 58863043 | AX-170149885 | 0 | C | A | 0.395 | 5.62E-02 | 9.68E-02 | 2.47E+00 | 2.52E+00 | 5.84E-01 | no | Adamts19 | intron_variant | MODIFIER |
| 18 | 66214824 | AX-170377942 | 1 | G | A | 0.286 | -5.10E-02 | 9.44E-02 | 2.69E+00 | 2.77E+00 | 5.84E-01 | no | Ccbe1 | intron_variant,NMD_transcript_variant | MODIFIER |
| 4 | 81273719 | AX-170110384 | 1 | G | T | 0.241 | 5.92E-02 | 1.05E-01 | 2.54E+00 | 2.61E+00 | 5.84E-01 | no | Mpdz | downstream_gene_variant | MODIFIER |
| 2 | 95127476 | AX-170251282 | 0 | C | T | 0.061 | 1.01E-01 | 1.89E-01 | 2.64E+00 | 2.77E+00 | 5.84E-01 | no | | intergenic_variant | MODIFIER |
| 1 | 24319399 | AX-170269795 | 1 | T | C | 0.438 | 4.59E-02 | 8.41E-02 | 2.64E+00 | 2.70E+00 | 5.84E-01 | no | Col19a1 | intron_variant | MODIFIER |
| 8 | 56479367 | AX-170363172 | 1 | A | G | 0.152 | -7.59E-02 | 1.37E-01 | 2.57E+00 | 2.65E+00 | 5.84E-01 | no | 4930518J21Rik | intron_variant,non_coding_transcript_variant | MODIFIER |
| 5 | 113942171 | AX-170377227 | 2 | T | C | 0.464 | 6.33E-02 | 1.16E-01 | 2.64E+00 | 2.68E+00 | 5.84E-01 | no | Ssh1 | downstream_gene_variant | MODIFIER |
| 1 | 23633138 | AX-170414269 | 1 | C | T | 0.429 | 4.55E-02 | 8.34E-02 | 2.63E+00 | 2.69E+00 | 5.84E-01 | no | | intergenic_variant | MODIFIER |
| 3 | 31836504 | AX-97835466 | 0 | C | A | 0.167 | -5.20E-02 | 9.65E-02 | 2.69E+00 | 2.77E+00 | 5.85E-01 | no | | intergenic_variant | MODIFIER |
| 7 | 93814029 | AX-97815617 | 1 | C | A | 0.116 | 1.08E-01 | 2.01E-01 | 2.66E+00 | 2.77E+00 | 5.85E-01 | no | | intergenic_variant | MODIFIER |
| 7 | 57422268 | AX-170328584 | 0 | G | A | 0.43 | 5.25E-02 | 9.70E-02 | 2.65E+00 | 2.73E+00 | 5.85E-01 | no | Gabra5 | intron_variant | MODIFIER |
| 5 | 113050894 | AX-170413304 | 1 | A | G | 0.455 | 6.37E-02 | 1.17E-01 | 2.66E+00 | 2.69E+00 | 5.85E-01 | no | | regulatory_region_variant | MODIFIER |
| 5 | 114352259 | AX-170479339 | 1 | A | G | 0.455 | 6.37E-02 | 1.17E-01 | 2.66E+00 | 2.69E+00 | 5.85E-01 | no | Myo1h | intron_variant | MODIFIER |
| 5 | 114642775 | AX-97822035 | 1 | C | T | 0.455 | 6.37E-02 | 1.17E-01 | 2.66E+00 | 2.69E+00 | 5.85E-01 | no | Trpv4 | intron_variant | MODIFIER |
| 16 | 19834906 | AX-97845926 | 1 | C | T | 0.134 | -5.04E-02 | 9.34E-02 | 2.60E+00 | 2.74E+00 | 5.85E-01 | no | | intergenic_variant | MODIFIER |
| 8 | 93244880 | AX-170232870 | 0 | G | A | 0.219 | 5.97E-02 | 1.11E-01 | 2.75E+00 | 2.83E+00 | 5.85E-01 | no | Gm45727 | intron_variant,non_coding_transcript_variant | MODIFIER |
| 3 | 32596324 | AX-170268491 | 1 | G | A | 0.17 | 5.29E-02 | 9.88E-02 | 2.73E+00 | 2.82E+00 | 5.85E-01 | no | Gnb4 | synonymous_variant | LOW |
| 18 | 81638677 | AX-170493070 | 0 | A | G | 0.421 | 3.33E-02 | 6.24E-02 | 2.70E+00 | 2.82E+00 | 5.85E-01 | no | AC102255.1 | upstream_gene_variant | MODIFIER |
| 14 | 52031205 | AX-170009794 | 0 | C | T | 0.044 | -1.13E-01 | 2.10E-01 | 2.66E+00 | 2.75E+00 | 5.85E-01 | no | Olfr221 | downstream_gene_variant | MODIFIER |
| 4 | 151528328 | AX-97818833 | 1 | G | A | 0.08 | -8.26E-02 | 1.54E-01 | 2.65E+00 | 2.76E+00 | 5.86E-01 | no | Camta1 | intron_variant | MODIFIER |
| 5 | 39886127 | AX-170024751 | 1 | G | A | 0.214 | -5.84E-02 | 1.07E-01 | 2.60E+00 | 2.68E+00 | 5.86E-01 | no | | intergenic_variant | MODIFIER |
| 16 | 74029221 | AX-170013386 | 0 | C | T | 0.482 | 4.51E-02 | 7.97E-02 | 2.51E+00 | 2.58E+00 | 5.86E-01 | no | Robo2 | intron_variant | MODIFIER |
| 4 | 151291687 | AX-170320045 | 0 | T | C | 0.096 | 8.41E-02 | 1.57E-01 | 2.67E+00 | 2.76E+00 | 5.86E-01 | no | Camta1 | intron_variant | MODIFIER |
| 4 | 151364852 | AX-97810922 | 0 | T | C | 0.096 | 8.41E-02 | 1.57E-01 | 2.67E+00 | 2.76E+00 | 5.86E-01 | no | Camta1 | intron_variant | MODIFIER |
| 4 | 151403941 | AX-170024254 | 0 | C | T | 0.096 | 8.41E-02 | 1.57E-01 | 2.67E+00 | 2.76E+00 | 5.86E-01 | no | Camta1 | intron_variant | MODIFIER |
| 4 | 151413660 | AX-170071056 | 0 | C | T | 0.096 | 8.41E-02 | 1.57E-01 | 2.67E+00 | 2.76E+00 | 5.86E-01 | no | Camta1 | intron_variant | MODIFIER |
| 4 | 151415495 | AX-170024255 | 0 | A | G | 0.096 | 8.41E-02 | 1.57E-01 | 2.67E+00 | 2.76E+00 | 5.86E-01 | no | Camta1 | intron_variant | MODIFIER |
| 4 | 151419589 | AX-97856094 | 0 | A | G | 0.096 | 8.41E-02 | 1.57E-01 | 2.67E+00 | 2.76E+00 | 5.86E-01 | no | Camta1 | intron_variant | MODIFIER |
| 4 | 151439676 | AX-97848504 | 0 | G | A | 0.096 | 8.41E-02 | 1.57E-01 | 2.67E+00 | 2.76E+00 | 5.86E-01 | no | Camta1 | intron_variant | MODIFIER |
| 4 | 151445537 | AX-170138493 | 0 | G | A | 0.096 | 8.41E-02 | 1.57E-01 | 2.67E+00 | 2.76E+00 | 5.86E-01 | no | Camta1 | intron_variant | MODIFIER |
| 4 | 151459952 | AX-97813749 | 0 | G | T | 0.096 | 8.41E-02 | 1.57E-01 | 2.67E+00 | 2.76E+00 | 5.86E-01 | no | Camta1 | intron_variant | MODIFIER |
| 4 | 151460398 | AX-170370467 | 0 | G | C | 0.096 | 8.41E-02 | 1.57E-01 | 2.67E+00 | 2.76E+00 | 5.86E-01 | no | Camta1 | intron_variant | MODIFIER |
| 4 | 151505212 | AX-170215630 | 0 | A | G | 0.096 | 8.41E-02 | 1.57E-01 | 2.67E+00 | 2.76E+00 | 5.86E-01 | no | Camta1 | intron_variant | MODIFIER |
| 4 | 151519182 | AX-170305411 | 0 | T | C | 0.096 | 8.41E-02 | 1.57E-01 | 2.67E+00 | 2.76E+00 | 5.86E-01 | no | Camta1 | intron_variant | MODIFIER |
| 4 | 151560410 | AX-170071058 | 0 | T | C | 0.096 | 8.41E-02 | 1.57E-01 | 2.67E+00 | 2.76E+00 | 5.86E-01 | no | Camta1 | intron_variant | MODIFIER |
| 4 | 151562276 | AX-170071059 | 0 | G | A | 0.096 | 8.41E-02 | 1.57E-01 | 2.67E+00 | 2.76E+00 | 5.86E-01 | no | Camta1 | intron_variant | MODIFIER |

| | | | | | | | | | | | | | | | |
|---|---|---|---|---|---|---|---|---|---|---|---|---|---|---|---|
| 4 | 151578635 | AX-170466102 | 0 | G | A | 0.096 | 8.41E-02 | 1.57E-01 | 2.67E+00 | 2.76E+00 | 5.86E-01 | no | Camta1 | intron_variant | MODIFIER |
| 4 | 151584346 | AX-170236893 | 0 | G | A | 0.096 | 8.41E-02 | 1.57E-01 | 2.67E+00 | 2.76E+00 | 5.86E-01 | no | Camta1 | intron_variant | MODIFIER |
| 4 | 151585819 | AX-97780958 | 0 | A | G | 0.096 | 8.41E-02 | 1.57E-01 | 2.67E+00 | 2.76E+00 | 5.86E-01 | no | Camta1 | intron_variant | MODIFIER |
| 4 | 151616369 | AX-97800536 | 0 | G | T | 0.096 | 8.41E-02 | 1.57E-01 | 2.67E+00 | 2.76E+00 | 5.86E-01 | no | Camta1 | intron_variant | MODIFIER |
| 4 | 151617223 | AX-170298585 | 0 | G | C | 0.096 | 8.41E-02 | 1.57E-01 | 2.67E+00 | 2.76E+00 | 5.86E-01 | no | Camta1 | intron_variant | MODIFIER |
| 6 | 93591353 | AX-170370961 | 1 | G | A | 0.42 | -3.24E-02 | 5.83E-02 | 2.52E+00 | 2.65E+00 | 5.86E-01 | no | | intergenic_variant | MODIFIER |
| 18 | 89663000 | AX-170491871 | 1 | T | C | 0.5 | -3.44E-02 | 6.49E-02 | 2.72E+00 | 2.86E+00 | 5.86E-01 | no | Dok6 | intron_variant | MODIFIER |
| 8 | 19408352 | AX-170112359 | 0 | T | C | 0.175 | 5.65E-02 | 1.05E-01 | 2.67E+00 | 2.76E+00 | 5.86E-01 | no | | intergenic_variant | MODIFIER |
| 18 | 64918175 | AX-170466895 | 0 | C | T | 0.404 | 5.13E-02 | 9.12E-02 | 2.53E+00 | 2.57E+00 | 5.86E-01 | no | Nedd4l | intron_variant | MODIFIER |
| 18 | 64918738 | AX-170257732 | 0 | T | G | 0.404 | 5.13E-02 | 9.12E-02 | 2.53E+00 | 2.57E+00 | 5.86E-01 | no | Nedd4l | intron_variant | MODIFIER |
| 18 | 64919336 | AX-170340917 | 0 | A | G | 0.404 | 5.13E-02 | 9.12E-02 | 2.53E+00 | 2.57E+00 | 5.86E-01 | no | Nedd4l | intron_variant | MODIFIER |
| 18 | 64935248 | AX-170119647 | 0 | C | T | 0.404 | 5.13E-02 | 9.12E-02 | 2.53E+00 | 2.57E+00 | 5.86E-01 | no | Nedd4l | intron_variant | MODIFIER |
| 18 | 64945835 | AX-97834727 | 0 | C | T | 0.404 | 5.13E-02 | 9.12E-02 | 2.53E+00 | 2.57E+00 | 5.86E-01 | no | Nedd4l | intron_variant | MODIFIER |
| 18 | 64946596 | AX-170381951 | 0 | T | G | 0.404 | 5.13E-02 | 9.12E-02 | 2.53E+00 | 2.57E+00 | 5.86E-01 | no | Nedd4l | intron_variant | MODIFIER |
| 18 | 64952710 | AX-170301321 | 0 | T | C | 0.404 | 5.13E-02 | 9.12E-02 | 2.53E+00 | 2.57E+00 | 5.86E-01 | no | Nedd4l | intron_variant | MODIFIER |
| 18 | 64976643 | AX-170282736 | 0 | T | A | 0.404 | 5.13E-02 | 9.12E-02 | 2.53E+00 | 2.57E+00 | 5.86E-01 | no | Nedd4l | intron_variant | MODIFIER |
| 18 | 64976692 | AX-170322253 | 0 | C | A | 0.404 | 5.13E-02 | 9.12E-02 | 2.53E+00 | 2.57E+00 | 5.86E-01 | no | Nedd4l | intron_variant | MODIFIER |
| 18 | 64981080 | AX-170221320 | 0 | C | A | 0.404 | 5.13E-02 | 9.12E-02 | 2.53E+00 | 2.57E+00 | 5.86E-01 | no | Nedd4l | intron_variant | MODIFIER |
| 18 | 64986171 | AX-97799468 | 0 | C | T | 0.404 | 5.13E-02 | 9.12E-02 | 2.53E+00 | 2.57E+00 | 5.86E-01 | no | Nedd4l | intron_variant | MODIFIER |
| 18 | 65008578 | AX-170343505 | 0 | C | A | 0.404 | 5.13E-02 | 9.12E-02 | 2.53E+00 | 2.57E+00 | 5.86E-01 | no | Nedd4l | intron_variant | MODIFIER |
| 18 | 65010609 | AX-170391898 | 0 | C | T | 0.404 | 5.13E-02 | 9.12E-02 | 2.53E+00 | 2.57E+00 | 5.86E-01 | no | Nedd4l | intron_variant | MODIFIER |
| 18 | 65091124 | AX-170403900 | 0 | A | G | 0.404 | 5.13E-02 | 9.12E-02 | 2.53E+00 | 2.57E+00 | 5.86E-01 | no | Nedd4l | intron_variant | MODIFIER |
| 18 | 65116888 | AX-170411790 | 0 | T | C | 0.404 | 5.13E-02 | 9.12E-02 | 2.53E+00 | 2.57E+00 | 5.86E-01 | no | Nedd4l | intron_variant | MODIFIER |
| 18 | 65120273 | AX-170164256 | 0 | C | T | 0.404 | 5.13E-02 | 9.12E-02 | 2.53E+00 | 2.57E+00 | 5.86E-01 | no | Nedd4l | intron_variant | MODIFIER |
| 6 | 131051147 | AX-170193305 | 0 | A | G | 0.535 | 4.47E-02 | 7.92E-02 | 2.51E+00 | 2.59E+00 | 5.86E-01 | no | Gm44134 | intron_variant,non_coding_transcript_variant | MODIFIER |
| 9 | 79045975 | AX-170505163 | 0 | A | C | 0.202 | 7.10E-02 | 1.31E-01 | 2.82E+00 | 2.87E+00 | 5.86E-01 | no | | intergenic_variant | MODIFIER |
| 15 | 92637581 | AX-170087939 | 2 | A | T | 0.127 | 9.85E-02 | 1.81E-01 | 2.61E+00 | 2.70E+00 | 5.86E-01 | no | Pdzrn4 | intron_variant | MODIFIER |
| 9 | 69029107 | AX-170114846 | 0 | T | C | 0.193 | -6.71E-02 | 1.25E-01 | 2.69E+00 | 2.74E+00 | 5.87E-01 | no | Rora | intron_variant | MODIFIER |
| 5 | 104085527 | AX-170485146 | 1 | G | C | 0.438 | 6.21E-02 | 1.15E-01 | 2.65E+00 | 2.68E+00 | 5.87E-01 | no | Sparcl1 | intron_variant | MODIFIER |
| 7 | 137348711 | AX-97822880 | 0 | G | A | 0.298 | -4.43E-02 | 8.14E-02 | 2.59E+00 | 2.69E+00 | 5.87E-01 | no | | intergenic_variant | MODIFIER |
| 11 | 22371807 | AX-170301432 | 0 | G | A | 0.053 | 1.03E-01 | 1.92E-01 | 2.62E+00 | 2.76E+00 | 5.87E-01 | no | | intergenic_variant | MODIFIER |
| 3 | 6594544 | AX-170019836 | 1 | C | T | 0.045 | -7.15E-02 | 1.34E-01 | 2.61E+00 | 2.77E+00 | 5.87E-01 | no | | intergenic_variant | MODIFIER |
| 12 | 89211216 | AX-170333793 | 1 | G | A | 0.071 | 8.52E-02 | 1.59E-01 | 2.68E+00 | 2.78E+00 | 5.87E-01 | no | Nrxn3 | intron_variant | MODIFIER |
| 9 | 40569615 | AX-170352284 | 2 | C | T | 0.436 | -4.05E-02 | 7.59E-02 | 2.70E+00 | 2.79E+00 | 5.87E-01 | no | | intergenic_variant | MODIFIER |
| 12 | 91987356 | AX-170522160 | 0 | G | T | 0.07 | 9.24E-02 | 1.74E-01 | 2.70E+00 | 2.82E+00 | 5.87E-01 | no | | intergenic_variant | MODIFIER |
| 1 | 173942037 | AX-170043637 | 2 | A | G | 0.218 | -6.79E-02 | 1.25E-01 | 2.88E+00 | 3.00E+00 | 5.87E-01 | no | Ifi203 | intron_variant | MODIFIER |
| 1 | 174873890 | AX-97824077 | 0 | G | A | 0.237 | 6.51E-02 | 1.15E-01 | 2.51E+00 | 2.58E+00 | 5.87E-01 | no | Grem2 | intron_variant | MODIFIER |
| 1 | 175392352 | AX-97818469 | 0 | T | G | 0.237 | 6.51E-02 | 1.15E-01 | 2.51E+00 | 2.58E+00 | 5.87E-01 | no | Rgs7 | intron_variant | MODIFIER |
| 11 | 29541766 | AX-170280494 | 0 | A | G | 0.079 | 1.09E-01 | 2.02E-01 | 2.64E+00 | 2.72E+00 | 5.87E-01 | no | Mtif2 | intron_variant | MODIFIER |
| 11 | 29556157 | AX-170199246 | 0 | A | G | 0.079 | 1.09E-01 | 2.02E-01 | 2.64E+00 | 2.72E+00 | 5.87E-01 | no | Clhc1 | missense_variant | MODERATE |
| 11 | 29560800 | AX-97818019 | 0 | G | A | 0.079 | 1.09E-01 | 2.02E-01 | 2.64E+00 | 2.72E+00 | 5.87E-01 | no | Clhc1 | synonymous_variant | LOW |
| 11 | 29563439 | AX-97818294 | 0 | T | G | 0.079 | 1.09E-01 | 2.02E-01 | 2.64E+00 | 2.72E+00 | 5.87E-01 | no | Clhc1 | intron_variant | MODIFIER |
| 11 | 29604352 | AX-97810090 | 0 | C | A | 0.079 | 1.09E-01 | 2.02E-01 | 2.64E+00 | 2.72E+00 | 5.87E-01 | no | Gm12091 | upstream_gene_variant | MODIFIER |
| 11 | 29608127 | AX-170109452 | 0 | T | C | 0.079 | 1.09E-01 | 2.02E-01 | 2.64E+00 | 2.72E+00 | 5.87E-01 | no | Gm12092 | downstream_gene_variant | MODIFIER |
| 11 | 29616552 | AX-170113887 | 0 | A | G | 0.079 | 1.09E-01 | 2.02E-01 | 2.64E+00 | 2.72E+00 | 5.87E-01 | no | Gm12092 | intron_variant,non_coding_transcript_variant | MODIFIER |
| 11 | 29617540 | AX-170410154 | 0 | A | G | 0.079 | 1.09E-01 | 2.02E-01 | 2.64E+00 | 2.72E+00 | 5.87E-01 | no | Gm12092 | intron_variant,non_coding_transcript_variant | MODIFIER |
| 11 | 29618682 | AX-170144953 | 0 | C | G | 0.079 | 1.09E-01 | 2.02E-01 | 2.64E+00 | 2.72E+00 | 5.87E-01 | no | Gm12092 | intron_variant,non_coding_transcript_variant | MODIFIER |
| 11 | 29619340 | AX-170232063 | 0 | T | C | 0.079 | 1.09E-01 | 2.02E-01 | 2.64E+00 | 2.72E+00 | 5.87E-01 | no | Gm12092 | intron_variant,non_coding_transcript_variant | MODIFIER |
| 11 | 29619486 | AX-170435382 | 0 | T | C | 0.079 | 1.09E-01 | 2.02E-01 | 2.64E+00 | 2.72E+00 | 5.87E-01 | no | Gm12092 | intron_variant,non_coding_transcript_variant | MODIFIER |
| 11 | 29623866 | AX-97853768 | 0 | T | G | 0.079 | 1.09E-01 | 2.02E-01 | 2.64E+00 | 2.72E+00 | 5.87E-01 | no | Gm12092 | intron_variant,non_coding_transcript_variant | MODIFIER |
| 11 | 29625205 | AX-170098191 | 0 | A | C | 0.079 | 1.09E-01 | 2.02E-01 | 2.64E+00 | 2.72E+00 | 5.87E-01 | no | Gm12092 | intron_variant,non_coding_transcript_variant | MODIFIER |
| 11 | 29625305 | AX-170512425 | 0 | G | T | 0.079 | 1.09E-01 | 2.02E-01 | 2.64E+00 | 2.72E+00 | 5.87E-01 | no | Gm12092 | intron_variant,non_coding_transcript_variant | MODIFIER |
| 11 | 29642167 | AX-97835453 | 0 | G | T | 0.079 | 1.09E-01 | 2.02E-01 | 2.64E+00 | 2.72E+00 | 5.87E-01 | no | Gm12092 | intron_variant,non_coding_transcript_variant | MODIFIER |
| 11 | 29661543 | AX-170046675 | 0 | C | A | 0.079 | 1.09E-01 | 2.02E-01 | 2.64E+00 | 2.72E+00 | 5.87E-01 | no | | intergenic_variant | MODIFIER |

| | | | | | | | | | | | | | | | |
|---|---|---|---|---|---|---|---|---|---|---|---|---|---|---|---|
| 11 | 29664111 | AX-170046672 | 0 | A | G | 0.079 | 1.09E-01 | 2.02E-01 | 2.64E+00 | 2.72E+00 | 5.87E-01 | no | | intergenic_variant | MODIFIER |
| 11 | 29664654 | AX-170214538 | 0 | C | T | 0.079 | 1.09E-01 | 2.02E-01 | 2.64E+00 | 2.72E+00 | 5.87E-01 | no | | intergenic_variant | MODIFIER |
| 11 | 29666592 | AX-170150183 | 0 | C | G | 0.079 | 1.09E-01 | 2.02E-01 | 2.64E+00 | 2.72E+00 | 5.87E-01 | no | | intergenic_variant | MODIFIER |
| 11 | 29669132 | AX-97815015 | 0 | G | A | 0.079 | 1.09E-01 | 2.02E-01 | 2.64E+00 | 2.72E+00 | 5.87E-01 | no | | intergenic_variant | MODIFIER |
| 11 | 29971748 | AX-170086985 | 0 | C | T | 0.079 | 1.09E-01 | 2.02E-01 | 2.64E+00 | 2.72E+00 | 5.87E-01 | no | Eml6 | intron_variant | MODIFIER |
| 11 | 30007135 | AX-170203642 | 0 | A | G | 0.079 | 1.09E-01 | 2.02E-01 | 2.64E+00 | 2.72E+00 | 5.87E-01 | no | Eml6 | intron_variant | MODIFIER |
| 11 | 30010017 | AX-170274105 | 0 | A | G | 0.079 | 1.09E-01 | 2.02E-01 | 2.64E+00 | 2.72E+00 | 5.87E-01 | no | Eml6 | intron_variant | MODIFIER |
| 11 | 30117927 | AX-97826463 | 0 | C | T | 0.079 | 1.09E-01 | 2.02E-01 | 2.64E+00 | 2.72E+00 | 5.87E-01 | no | Sptbn1 | synonymous_variant | LOW |
| 11 | 30117954 | AX-170191260 | 0 | T | C | 0.079 | 1.09E-01 | 2.02E-01 | 2.64E+00 | 2.72E+00 | 5.87E-01 | no | Sptbn1 | synonymous_variant | LOW |
| 11 | 30124371 | AX-170418090 | 0 | T | C | 0.079 | 1.09E-01 | 2.02E-01 | 2.64E+00 | 2.72E+00 | 5.87E-01 | no | Sptbn1 | intron_variant | MODIFIER |
| 11 | 30145785 | AX-170115212 | 0 | G | T | 0.079 | 1.09E-01 | 2.02E-01 | 2.64E+00 | 2.72E+00 | 5.87E-01 | no | Sptbn1 | intron_variant | MODIFIER |
| 11 | 30157049 | AX-170365267 | 0 | T | C | 0.079 | 1.09E-01 | 2.02E-01 | 2.64E+00 | 2.72E+00 | 5.87E-01 | no | Sptbn1 | intron_variant | MODIFIER |
| 11 | 30161786 | AX-170215662 | 0 | G | A | 0.079 | 1.09E-01 | 2.02E-01 | 2.64E+00 | 2.72E+00 | 5.87E-01 | no | Sptbn1 | intron_variant | MODIFIER |
| 11 | 30173553 | AX-170046684 | 0 | A | G | 0.079 | 1.09E-01 | 2.02E-01 | 2.64E+00 | 2.72E+00 | 5.87E-01 | no | Sptbn1 | intron_variant | MODIFIER |
| 11 | 30175897 | AX-170046682 | 0 | A | G | 0.079 | 1.09E-01 | 2.02E-01 | 2.64E+00 | 2.72E+00 | 5.87E-01 | no | Sptbn1 | intron_variant | MODIFIER |
| 11 | 30176643 | AX-170046685 | 0 | G | A | 0.079 | 1.09E-01 | 2.02E-01 | 2.64E+00 | 2.72E+00 | 5.87E-01 | no | Sptbn1 | intron_variant | MODIFIER |
| 11 | 30193308 | AX-170165213 | 0 | C | A | 0.079 | 1.09E-01 | 2.02E-01 | 2.64E+00 | 2.72E+00 | 5.87E-01 | no | Sptbn1 | intron_variant | MODIFIER |
| 11 | 30195755 | AX-170220442 | 0 | A | G | 0.079 | 1.09E-01 | 2.02E-01 | 2.64E+00 | 2.72E+00 | 5.87E-01 | no | Sptbn1 | intron_variant | MODIFIER |
| 11 | 30196778 | AX-170154714 | 0 | A | G | 0.079 | 1.09E-01 | 2.02E-01 | 2.64E+00 | 2.72E+00 | 5.87E-01 | no | Sptbn1 | intron_variant | MODIFIER |
| 11 | 30200714 | AX-170474139 | 0 | A | T | 0.079 | 1.09E-01 | 2.02E-01 | 2.64E+00 | 2.72E+00 | 5.87E-01 | no | Sptbn1 | intron_variant | MODIFIER |
| 11 | 30203022 | AX-97805924 | 0 | T | G | 0.079 | 1.09E-01 | 2.02E-01 | 2.64E+00 | 2.72E+00 | 5.87E-01 | no | Sptbn1 | intron_variant | MODIFIER |
| 11 | 30209082 | AX-170237419 | 0 | C | T | 0.079 | 1.09E-01 | 2.02E-01 | 2.64E+00 | 2.72E+00 | 5.87E-01 | no | Sptbn1 | intron_variant | MODIFIER |
| 11 | 30209146 | AX-170369615 | 0 | T | G | 0.079 | 1.09E-01 | 2.02E-01 | 2.64E+00 | 2.72E+00 | 5.87E-01 | no | Sptbn1 | intron_variant | MODIFIER |
| 11 | 30243601 | AX-170282825 | 0 | T | C | 0.079 | 1.09E-01 | 2.02E-01 | 2.64E+00 | 2.72E+00 | 5.87E-01 | no | Sptbn1 | intron_variant | MODIFIER |
| 11 | 30243736 | AX-170510133 | 0 | A | G | 0.079 | 1.09E-01 | 2.02E-01 | 2.64E+00 | 2.72E+00 | 5.87E-01 | no | Sptbn1 | intron_variant | MODIFIER |
| 11 | 30245897 | AX-170263663 | 0 | G | A | 0.079 | 1.09E-01 | 2.02E-01 | 2.64E+00 | 2.72E+00 | 5.87E-01 | no | Sptbn1 | intron_variant | MODIFIER |
| 11 | 30292849 | AX-170428190 | 0 | G | A | 0.079 | 1.09E-01 | 2.02E-01 | 2.64E+00 | 2.72E+00 | 5.87E-01 | no | | regulatory_region_variant | MODIFIER |
| 11 | 31364299 | AX-97783970 | 0 | T | C | 0.079 | 1.09E-01 | 2.02E-01 | 2.64E+00 | 2.72E+00 | 5.87E-01 | no | Stc2 | intron_variant | MODIFIER |
| 11 | 31376523 | AX-170502316 | 0 | T | C | 0.079 | 1.09E-01 | 2.02E-01 | 2.64E+00 | 2.72E+00 | 5.87E-01 | no | | intergenic_variant | MODIFIER |
| 11 | 31386883 | AX-170518902 | 0 | C | G | 0.079 | 1.09E-01 | 2.02E-01 | 2.64E+00 | 2.72E+00 | 5.87E-01 | no | | regulatory_region_variant | MODIFIER |
| 11 | 31396006 | AX-97803756 | 0 | A | G | 0.079 | 1.09E-01 | 2.02E-01 | 2.64E+00 | 2.72E+00 | 5.87E-01 | no | | intergenic_variant | MODIFIER |
| 11 | 31400335 | AX-170004632 | 0 | T | C | 0.079 | 1.09E-01 | 2.02E-01 | 2.64E+00 | 2.72E+00 | 5.87E-01 | no | | intergenic_variant | MODIFIER |
| 11 | 31401041 | AX-170004640 | 0 | C | T | 0.079 | 1.09E-01 | 2.02E-01 | 2.64E+00 | 2.72E+00 | 5.87E-01 | no | | intergenic_variant | MODIFIER |
| 11 | 31416625 | AX-170004638 | 0 | T | C | 0.079 | 1.09E-01 | 2.02E-01 | 2.64E+00 | 2.72E+00 | 5.87E-01 | no | | intergenic_variant | MODIFIER |
| 11 | 31429813 | AX-97854603 | 0 | G | A | 0.079 | 1.09E-01 | 2.02E-01 | 2.64E+00 | 2.72E+00 | 5.87E-01 | no | | intergenic_variant | MODIFIER |
| 11 | 31430151 | AX-170004633 | 0 | A | G | 0.079 | 1.09E-01 | 2.02E-01 | 2.64E+00 | 2.72E+00 | 5.87E-01 | no | | intergenic_variant | MODIFIER |
| 11 | 31443516 | AX-97826338 | 0 | A | G | 0.079 | 1.09E-01 | 2.02E-01 | 2.64E+00 | 2.72E+00 | 5.87E-01 | no | | intergenic_variant | MODIFIER |
| 11 | 31465897 | AX-170418621 | 0 | T | C | 0.079 | 1.09E-01 | 2.02E-01 | 2.64E+00 | 2.72E+00 | 5.87E-01 | no | | intergenic_variant | MODIFIER |
| 11 | 31478946 | AX-97818096 | 0 | T | C | 0.079 | 1.09E-01 | 2.02E-01 | 2.64E+00 | 2.72E+00 | 5.87E-01 | no | | intergenic_variant | MODIFIER |
| 11 | 31504234 | AX-170534772 | 0 | C | T | 0.079 | 1.09E-01 | 2.02E-01 | 2.64E+00 | 2.72E+00 | 5.87E-01 | no | | intergenic_variant | MODIFIER |
| 11 | 31523332 | AX-97787785 | 0 | A | C | 0.079 | 1.09E-01 | 2.02E-01 | 2.64E+00 | 2.72E+00 | 5.87E-01 | no | | intergenic_variant | MODIFIER |
| 11 | 31524109 | AX-170251082 | 0 | G | C | 0.079 | 1.09E-01 | 2.02E-01 | 2.64E+00 | 2.72E+00 | 5.87E-01 | no | | intergenic_variant | MODIFIER |
| 11 | 31532399 | AX-170004637 | 0 | C | T | 0.079 | 1.09E-01 | 2.02E-01 | 2.64E+00 | 2.72E+00 | 5.87E-01 | no | | intergenic_variant | MODIFIER |
| 11 | 31532438 | AX-170004635 | 0 | A | G | 0.079 | 1.09E-01 | 2.02E-01 | 2.64E+00 | 2.72E+00 | 5.87E-01 | no | | intergenic_variant | MODIFIER |
| 11 | 31563840 | AX-170004634 | 0 | G | A | 0.079 | 1.09E-01 | 2.02E-01 | 2.64E+00 | 2.72E+00 | 5.87E-01 | no | | regulatory_region_variant | MODIFIER |
| 11 | 31566004 | AX-170141591 | 0 | T | C | 0.079 | 1.09E-01 | 2.02E-01 | 2.64E+00 | 2.72E+00 | 5.87E-01 | no | | intergenic_variant | MODIFIER |
| 11 | 31571240 | AX-170004639 | 0 | T | A | 0.079 | 1.09E-01 | 2.02E-01 | 2.64E+00 | 2.72E+00 | 5.87E-01 | no | | intergenic_variant | MODIFIER |
| 11 | 31590963 | AX-97833107 | 0 | T | G | 0.079 | 1.09E-01 | 2.02E-01 | 2.64E+00 | 2.72E+00 | 5.87E-01 | no | | intergenic_variant | MODIFIER |
| 11 | 31596017 | AX-170004641 | 0 | A | G | 0.079 | 1.09E-01 | 2.02E-01 | 2.64E+00 | 2.72E+00 | 5.87E-01 | no | | intergenic_variant | MODIFIER |
| 11 | 31602692 | AX-170004631 | 0 | C | T | 0.079 | 1.09E-01 | 2.02E-01 | 2.64E+00 | 2.72E+00 | 5.87E-01 | no | | regulatory_region_variant | MODIFIER |
| 11 | 31619509 | AX-170373781 | 0 | A | G | 0.079 | 1.09E-01 | 2.02E-01 | 2.64E+00 | 2.72E+00 | 5.87E-01 | no | | intergenic_variant | MODIFIER |
| 11 | 31627247 | AX-170513800 | 0 | C | G | 0.079 | 1.09E-01 | 2.02E-01 | 2.64E+00 | 2.72E+00 | 5.87E-01 | no | | intergenic_variant | MODIFIER |
| 11 | 31627322 | AX-97815475 | 0 | C | T | 0.079 | 1.09E-01 | 2.02E-01 | 2.64E+00 | 2.72E+00 | 5.87E-01 | no | | intergenic_variant | MODIFIER |
| 11 | 31630924 | AX-170363829 | 0 | A | G | 0.079 | 1.09E-01 | 2.02E-01 | 2.64E+00 | 2.72E+00 | 5.87E-01 | no | | intergenic_variant | MODIFIER |

| | | | | | | | | | | | | | | | |
|---|---|---|---|---|---|---|---|---|---|---|---|---|---|---|---|
| 11 | 31639834 | AX-97817930 | 0 | T | C | 0.079 | 1.09E-01 | 2.02E-01 | 2.64E+00 | 2.72E+00 | 5.87E-01 | no | | intergenic_variant | MODIFIER |
| 11 | 31679414 | AX-170004636 | 0 | T | C | 0.079 | 1.09E-01 | 2.02E-01 | 2.64E+00 | 2.72E+00 | 5.87E-01 | no | | intergenic_variant | MODIFIER |
| 11 | 31679640 | AX-97825500 | 0 | G | A | 0.158 | 5.43E-02 | 1.01E-01 | 2.64E+00 | 2.72E+00 | 5.87E-01 | no | | intergenic_variant | MODIFIER |
| 11 | 31679682 | AX-170334343 | 0 | T | A | 0.079 | 1.09E-01 | 2.02E-01 | 2.64E+00 | 2.72E+00 | 5.87E-01 | no | | intergenic_variant | MODIFIER |
| 11 | 31720430 | AX-97817872 | 0 | A | G | 0.079 | 1.09E-01 | 2.02E-01 | 2.64E+00 | 2.72E+00 | 5.87E-01 | no | | intergenic_variant | MODIFIER |
| 11 | 31739897 | AX-170374376 | 0 | C | T | 0.079 | 1.09E-01 | 2.02E-01 | 2.64E+00 | 2.72E+00 | 5.87E-01 | no | | intergenic_variant | MODIFIER |
| 11 | 31741167 | AX-170391137 | 0 | C | A | 0.079 | 1.09E-01 | 2.02E-01 | 2.64E+00 | 2.72E+00 | 5.87E-01 | no | | regulatory_region_variant | MODIFIER |
| 11 | 31742817 | AX-170212278 | 0 | A | C | 0.079 | 1.09E-01 | 2.02E-01 | 2.64E+00 | 2.72E+00 | 5.87E-01 | no | | regulatory_region_variant | MODIFIER |
| 11 | 31765041 | AX-170171026 | 0 | T | G | 0.079 | 1.09E-01 | 2.02E-01 | 2.64E+00 | 2.72E+00 | 5.87E-01 | no | | intergenic_variant | MODIFIER |
| 11 | 31768562 | AX-170495172 | 0 | G | C | 0.079 | 1.09E-01 | 2.02E-01 | 2.64E+00 | 2.72E+00 | 5.87E-01 | no | | regulatory_region_variant | MODIFIER |
| 11 | 31768620 | AX-170213316 | 0 | T | A | 0.079 | 1.09E-01 | 2.02E-01 | 2.64E+00 | 2.72E+00 | 5.87E-01 | no | | regulatory_region_variant | MODIFIER |
| 11 | 31768842 | AX-97817875 | 0 | A | G | 0.079 | 1.09E-01 | 2.02E-01 | 2.64E+00 | 2.72E+00 | 5.87E-01 | no | | regulatory_region_variant | MODIFIER |
| 11 | 31807106 | AX-97842758 | 0 | C | T | 0.079 | 1.09E-01 | 2.02E-01 | 2.64E+00 | 2.72E+00 | 5.87E-01 | no | D630024D03Rik | intron_variant,non_coding_transcript_variant | MODIFIER |
| 11 | 31811537 | AX-170536053 | 0 | T | A | 0.079 | 1.09E-01 | 2.02E-01 | 2.64E+00 | 2.72E+00 | 5.87E-01 | no | D630024D03Rik | intron_variant,non_coding_transcript_variant | MODIFIER |
| 11 | 31839426 | AX-170485223 | 0 | G | A | 0.079 | 1.09E-01 | 2.02E-01 | 2.64E+00 | 2.72E+00 | 5.87E-01 | no | Gm12107 | downstream_gene_variant | MODIFIER |
| 11 | 31851516 | AX-97809979 | 0 | A | G | 0.079 | 1.09E-01 | 2.02E-01 | 2.64E+00 | 2.72E+00 | 5.87E-01 | no | | regulatory_region_variant | MODIFIER |
| 11 | 31851545 | AX-170046709 | 0 | C | A | 0.079 | 1.09E-01 | 2.02E-01 | 2.64E+00 | 2.72E+00 | 5.87E-01 | no | | regulatory_region_variant | MODIFIER |
| 11 | 31861410 | AX-170441878 | 0 | G | T | 0.079 | 1.09E-01 | 2.02E-01 | 2.64E+00 | 2.72E+00 | 5.87E-01 | no | | regulatory_region_variant | MODIFIER |
| 11 | 31861486 | AX-170213266 | 0 | T | C | 0.079 | 1.09E-01 | 2.02E-01 | 2.64E+00 | 2.72E+00 | 5.87E-01 | no | | regulatory_region_variant | MODIFIER |
| 11 | 31907677 | AX-170534466 | 0 | A | C | 0.079 | 1.09E-01 | 2.02E-01 | 2.64E+00 | 2.72E+00 | 5.87E-01 | no | Cpeb4 | intron_variant | MODIFIER |
| 11 | 31913206 | AX-170504938 | 0 | C | G | 0.079 | 1.09E-01 | 2.02E-01 | 2.64E+00 | 2.72E+00 | 5.87E-01 | no | Cpeb4 | intron_variant | MODIFIER |
| 11 | 31978296 | AX-97817958 | 0 | A | G | 0.079 | 1.09E-01 | 2.02E-01 | 2.64E+00 | 2.72E+00 | 5.87E-01 | no | 4930524B15Rik | 3_prime_UTR_variant | MODIFIER |
| 11 | 31978521 | AX-170004643 | 0 | G | T | 0.079 | 1.09E-01 | 2.02E-01 | 2.64E+00 | 2.72E+00 | 5.87E-01 | no | 4930524B15Rik | 3_prime_UTR_variant | MODIFIER |
| 11 | 31985010 | AX-170004649 | 0 | T | C | 0.079 | 1.09E-01 | 2.02E-01 | 2.64E+00 | 2.72E+00 | 5.87E-01 | no | | intergenic_variant | MODIFIER |
| 11 | 31995377 | AX-170275932 | 0 | G | A | 0.079 | 1.09E-01 | 2.02E-01 | 2.64E+00 | 2.72E+00 | 5.87E-01 | no | | intergenic_variant | MODIFIER |
| 11 | 32013378 | AX-170165496 | 0 | C | A | 0.079 | 1.09E-01 | 2.02E-01 | 2.64E+00 | 2.72E+00 | 5.87E-01 | no | Nsg2 | intron_variant | MODIFIER |
| 11 | 32014204 | AX-170004647 | 0 | G | A | 0.079 | 1.09E-01 | 2.02E-01 | 2.64E+00 | 2.72E+00 | 5.87E-01 | no | Nsg2 | intron_variant | MODIFIER |
| 11 | 32019456 | AX-97828474 | 0 | C | T | 0.079 | 1.09E-01 | 2.02E-01 | 2.64E+00 | 2.72E+00 | 5.87E-01 | no | Nsg2 | intron_variant | MODIFIER |
| 11 | 32048732 | AX-97794192 | 0 | C | T | 0.079 | 1.09E-01 | 2.02E-01 | 2.64E+00 | 2.72E+00 | 5.87E-01 | no | Nsg2 | intron_variant | MODIFIER |
| 11 | 32072458 | AX-170004645 | 0 | A | G | 0.079 | 1.09E-01 | 2.02E-01 | 2.64E+00 | 2.72E+00 | 5.87E-01 | no | | intergenic_variant | MODIFIER |
| 11 | 32072818 | AX-170364276 | 0 | G | A | 0.079 | 1.09E-01 | 2.02E-01 | 2.64E+00 | 2.72E+00 | 5.87E-01 | no | | intergenic_variant | MODIFIER |
| 11 | 32075414 | AX-97838569 | 0 | T | G | 0.079 | 1.09E-01 | 2.02E-01 | 2.64E+00 | 2.72E+00 | 5.87E-01 | no | | regulatory_region_variant | MODIFIER |
| 11 | 32111004 | AX-170086987 | 0 | G | A | 0.079 | 1.09E-01 | 2.02E-01 | 2.64E+00 | 2.72E+00 | 5.87E-01 | no | | intergenic_variant | MODIFIER |
| 11 | 32111281 | AX-170497724 | 0 | A | G | 0.079 | 1.09E-01 | 2.02E-01 | 2.64E+00 | 2.72E+00 | 5.87E-01 | no | | regulatory_region_variant | MODIFIER |
| 11 | 32131692 | AX-170046714 | 0 | A | G | 0.079 | 1.09E-01 | 2.02E-01 | 2.64E+00 | 2.72E+00 | 5.87E-01 | no | | intergenic_variant | MODIFIER |
| 11 | 32131782 | AX-170046713 | 0 | T | C | 0.079 | 1.09E-01 | 2.02E-01 | 2.64E+00 | 2.72E+00 | 5.87E-01 | no | | intergenic_variant | MODIFIER |
| 11 | 32132263 | AX-170323952 | 0 | T | C | 0.079 | 1.09E-01 | 2.02E-01 | 2.64E+00 | 2.72E+00 | 5.87E-01 | no | | regulatory_region_variant | MODIFIER |
| 11 | 32132330 | AX-170483462 | 0 | G | T | 0.079 | 1.09E-01 | 2.02E-01 | 2.64E+00 | 2.72E+00 | 5.87E-01 | no | | regulatory_region_variant | MODIFIER |
| 11 | 32140924 | AX-170098426 | 0 | C | T | 0.079 | 1.09E-01 | 2.02E-01 | 2.64E+00 | 2.72E+00 | 5.87E-01 | no | | regulatory_region_variant | MODIFIER |
| 11 | 32141103 | AX-97821917 | 0 | G | A | 0.079 | 1.09E-01 | 2.02E-01 | 2.64E+00 | 2.72E+00 | 5.87E-01 | no | | regulatory_region_variant | MODIFIER |
| 11 | 32162638 | AX-170122097 | 0 | T | C | 0.079 | 1.09E-01 | 2.02E-01 | 2.64E+00 | 2.72E+00 | 5.87E-01 | no | | regulatory_region_variant | MODIFIER |
| 11 | 32162706 | AX-97844456 | 0 | G | A | 0.079 | 1.09E-01 | 2.02E-01 | 2.64E+00 | 2.72E+00 | 5.87E-01 | no | | regulatory_region_variant | MODIFIER |
| 11 | 32190953 | AX-170233937 | 0 | T | C | 0.079 | 1.09E-01 | 2.02E-01 | 2.64E+00 | 2.72E+00 | 5.87E-01 | no | Il9r | 3_prime_UTR_variant,NMD_transcript_variant | MODIFIER |
| 11 | 32228975 | AX-170326397 | 0 | A | C | 0.079 | 1.09E-01 | 2.02E-01 | 2.64E+00 | 2.72E+00 | 5.87E-01 | no | Mpg | intron_variant | MODIFIER |
| 11 | 32236141 | AX-170313890 | 0 | T | C | 0.079 | 1.09E-01 | 2.02E-01 | 2.64E+00 | 2.72E+00 | 5.87E-01 | no | Mpg | downstream_gene_variant | MODIFIER |
| 11 | 32288750 | AX-170260105 | 0 | T | G | 0.079 | 1.09E-01 | 2.02E-01 | 2.64E+00 | 2.72E+00 | 5.87E-01 | no | Hba-a1 | downstream_gene_variant | MODIFIER |
| 11 | 32304476 | AX-170215722 | 0 | T | C | 0.079 | 1.09E-01 | 2.02E-01 | 2.64E+00 | 2.72E+00 | 5.87E-01 | no | Hbq1a | downstream_gene_variant | MODIFIER |
| 11 | 32318317 | AX-170346694 | 0 | T | C | 0.079 | 1.09E-01 | 2.02E-01 | 2.64E+00 | 2.72E+00 | 5.87E-01 | no | | intergenic_variant | MODIFIER |
| 11 | 32356887 | AX-97823396 | 0 | G | A | 0.079 | 1.09E-01 | 2.02E-01 | 2.64E+00 | 2.72E+00 | 5.87E-01 | no | Sh3pxd2b | intron_variant | MODIFIER |
| 11 | 32357728 | AX-170348859 | 0 | A | G | 0.079 | 1.09E-01 | 2.02E-01 | 2.64E+00 | 2.72E+00 | 5.87E-01 | no | Sh3pxd2b | intron_variant | MODIFIER |
| 11 | 32360071 | AX-170296613 | 0 | A | G | 0.079 | 1.09E-01 | 2.02E-01 | 2.64E+00 | 2.72E+00 | 5.87E-01 | no | Sh3pxd2b | intron_variant | MODIFIER |
| 11 | 32404463 | AX-170322088 | 0 | G | T | 0.079 | 1.09E-01 | 2.02E-01 | 2.64E+00 | 2.72E+00 | 5.87E-01 | no | Sh3pxd2b | intron_variant | MODIFIER |
| 11 | 32542069 | AX-97809242 | 0 | T | C | 0.079 | 1.09E-01 | 2.02E-01 | 2.64E+00 | 2.72E+00 | 5.87E-01 | no | Stk10 | intron_variant | MODIFIER |
| 11 | 32552886 | AX-170359639 | 0 | T | C | 0.079 | 1.09E-01 | 2.02E-01 | 2.64E+00 | 2.72E+00 | 5.87E-01 | no | Stk10 | intron_variant | MODIFIER |

| | | | | | | | | | | | | | | | |
|---|---|---|---|---|---|---|---|---|---|---|---|---|---|---|---|
| 11 | 32553820 | AX-170004652 | 0 | A | T | 0.079 | 1.09E-01 | 2.02E-01 | 2.64E+00 | 2.72E+00 | 5.87E-01 | no | Stk10 | intron_variant | MODIFIER |
| 11 | 32554864 | AX-170190762 | 0 | T | C | 0.079 | 1.09E-01 | 2.02E-01 | 2.64E+00 | 2.72E+00 | 5.87E-01 | no | Stk10 | intron_variant | MODIFIER |
| 11 | 32595591 | AX-170309431 | 0 | G | A | 0.079 | 1.09E-01 | 2.02E-01 | 2.64E+00 | 2.72E+00 | 5.87E-01 | no | Stk10 | intron_variant | MODIFIER |
| 11 | 32598783 | AX-170004653 | 0 | G | A | 0.079 | 1.09E-01 | 2.02E-01 | 2.64E+00 | 2.72E+00 | 5.87E-01 | no | Stk10 | synonymous_variant | LOW |
| 11 | 32600699 | AX-170189743 | 0 | A | G | 0.079 | 1.09E-01 | 2.02E-01 | 2.64E+00 | 2.72E+00 | 5.87E-01 | no | Stk10 | intron_variant | MODIFIER |
| 11 | 32670165 | AX-170402084 | 0 | G | A | 0.079 | 1.09E-01 | 2.02E-01 | 2.64E+00 | 2.72E+00 | 5.87E-01 | no | Fbxw11 | intron_variant | MODIFIER |
| 11 | 32675635 | AX-97792128 | 0 | T | C | 0.079 | 1.09E-01 | 2.02E-01 | 2.64E+00 | 2.72E+00 | 5.87E-01 | no | Fbxw11 | intron_variant | MODIFIER |
| 11 | 32681448 | AX-170046727 | 0 | T | G | 0.079 | 1.09E-01 | 2.02E-01 | 2.64E+00 | 2.72E+00 | 5.87E-01 | no | Fbxw11 | intron_variant | MODIFIER |
| 11 | 32689880 | AX-170136652 | 0 | T | C | 0.079 | 1.09E-01 | 2.02E-01 | 2.64E+00 | 2.72E+00 | 5.87E-01 | no | Fbxw11 | intron_variant | MODIFIER |
| 11 | 32691246 | AX-170238246 | 0 | G | A | 0.079 | 1.09E-01 | 2.02E-01 | 2.64E+00 | 2.72E+00 | 5.87E-01 | no | Fbxw11 | intron_variant | MODIFIER |
| 11 | 32701630 | AX-170442885 | 0 | G | A | 0.079 | 1.09E-01 | 2.02E-01 | 2.64E+00 | 2.72E+00 | 5.87E-01 | no | Fbxw11 | intron_variant | MODIFIER |
| 11 | 32703635 | AX-170308800 | 0 | T | A | 0.079 | 1.09E-01 | 2.02E-01 | 2.64E+00 | 2.72E+00 | 5.87E-01 | no | Fbxw11 | intron_variant | MODIFIER |
| 11 | 32727501 | AX-170105004 | 0 | T | C | 0.079 | 1.09E-01 | 2.02E-01 | 2.64E+00 | 2.72E+00 | 5.87E-01 | no | Fbxw11 | intron_variant | MODIFIER |
| 11 | 32732147 | AX-170191704 | 0 | A | G | 0.079 | 1.09E-01 | 2.02E-01 | 2.64E+00 | 2.72E+00 | 5.87E-01 | no | Fbxw11 | intron_variant | MODIFIER |
| 11 | 32762518 | AX-170401186 | 0 | A | G | 0.079 | 1.09E-01 | 2.02E-01 | 2.64E+00 | 2.72E+00 | 5.87E-01 | no | | intergenic_variant | MODIFIER |
| 11 | 32795292 | AX-170480526 | 0 | A | G | 0.079 | 1.09E-01 | 2.02E-01 | 2.64E+00 | 2.72E+00 | 5.87E-01 | no | 4930555O08Rik | upstream_gene_variant | MODIFIER |
| 6 | 107636635 | AX-170270805 | 0 | A | G | 0.053 | -1.14E-01 | 2.14E-01 | 2.66E+00 | 2.78E+00 | 5.87E-01 | no | 4933431M02Rik | downstream_gene_variant | MODIFIER |
| 2 | 162551841 | AX-168406990 | 0 | G | A | 0.167 | 1.01E-01 | 1.88E-01 | 2.67E+00 | 2.76E+00 | 5.87E-01 | no | Ptprt | intron_variant | MODIFIER |
| 5 | 125186322 | AX-170256003 | 0 | G | T | 0.456 | -5.76E-02 | 1.08E-01 | 2.77E+00 | 2.84E+00 | 5.87E-01 | no | | intergenic_variant | MODIFIER |
| 2 | 101708260 | AX-170523411 | 0 | G | A | 0.07 | -9.64E-02 | 1.83E-01 | 2.72E+00 | 2.86E+00 | 5.87E-01 | no | | intergenic_variant | MODIFIER |
| 15 | 81171552 | AX-170246378 | 0 | C | A | 0.561 | 4.51E-02 | 8.18E-02 | 2.56E+00 | 2.63E+00 | 5.88E-01 | no | Mrtfa | intron_variant,non_coding_transcript_variant | MODIFIER |
| 4 | 72808504 | AX-170023101 | 1 | G | C | 0.223 | 5.71E-02 | 1.04E-01 | 2.55E+00 | 2.63E+00 | 5.88E-01 | no | | intergenic_variant | MODIFIER |
| 13 | 36737583 | AX-170191416 | 1 | A | G | 0.188 | -6.61E-02 | 1.19E-01 | 2.55E+00 | 2.61E+00 | 5.88E-01 | no | Nrn1 | upstream_gene_variant | MODIFIER |
| 8 | 34427927 | AX-170039722 | 0 | G | A | 0.149 | -6.68E-02 | 1.24E-01 | 2.63E+00 | 2.71E+00 | 5.88E-01 | no | Gm45349 | intron_variant,non_coding_transcript_variant | MODIFIER |
| 11 | 29666609 | AX-170472310 | 1 | G | A | 0.08 | 1.08E-01 | 2.01E-01 | 2.64E+00 | 2.72E+00 | 5.88E-01 | no | | intergenic_variant | MODIFIER |
| 11 | 31741464 | AX-97826295 | 1 | G | A | 0.08 | 1.08E-01 | 2.01E-01 | 2.64E+00 | 2.72E+00 | 5.88E-01 | no | | intergenic_variant | MODIFIER |
| 18 | 58337446 | AX-170382982 | 0 | A | G | 0.412 | 4.44E-02 | 8.27E-02 | 2.84E+00 | 2.93E+00 | 5.88E-01 | no | | intergenic_variant | MODIFIER |
| 15 | 93302292 | AX-170012283 | 1 | C | T | 0.116 | 9.02E-02 | 1.68E-01 | 2.62E+00 | 2.72E+00 | 5.88E-01 | no | Yaf2 | intron_variant | MODIFIER |
| 6 | 103999239 | AX-170101076 | 1 | A | G | 0.062 | 1.19E-01 | 2.21E-01 | 2.63E+00 | 2.73E+00 | 5.88E-01 | no | | intergenic_variant | MODIFIER |
| 7 | 112284105 | AX-170519983 | 0 | C | A | 0.456 | -5.25E-02 | 9.80E-02 | 2.80E+00 | 2.86E+00 | 5.88E-01 | no | Mical2 | intron_variant | MODIFIER |
| 5 | 88893631 | AX-170453444 | 0 | T | C | 0.412 | 5.92E-02 | 1.11E-01 | 2.75E+00 | 2.80E+00 | 5.88E-01 | no | Slc4a4 | intron_variant | MODIFIER |
| 5 | 89135132 | AX-170072723 | 0 | T | C | 0.412 | 5.92E-02 | 1.11E-01 | 2.75E+00 | 2.80E+00 | 5.88E-01 | no | Slc4a4 | intron_variant | MODIFIER |
| 5 | 90054399 | AX-97808792 | 0 | A | G | 0.412 | 5.92E-02 | 1.11E-01 | 2.75E+00 | 2.80E+00 | 5.88E-01 | no | | intergenic_variant | MODIFIER |
| 5 | 90122366 | AX-97816196 | 0 | A | G | 0.412 | 5.92E-02 | 1.11E-01 | 2.75E+00 | 2.80E+00 | 5.88E-01 | no | | intergenic_variant | MODIFIER |
| 5 | 90155537 | AX-97831950 | 0 | A | G | 0.412 | 5.92E-02 | 1.11E-01 | 2.75E+00 | 2.80E+00 | 5.88E-01 | no | | intergenic_variant | MODIFIER |
| 5 | 90252935 | AX-170089555 | 0 | T | C | 0.412 | 5.92E-02 | 1.11E-01 | 2.75E+00 | 2.80E+00 | 5.88E-01 | no | Ankrd17 | synonymous_variant | LOW |
| 5 | 113436578 | AX-170108664 | 0 | A | G | 0.456 | 6.33E-02 | 1.18E-01 | 2.65E+00 | 2.68E+00 | 5.89E-01 | no | | intergenic_variant | MODIFIER |
| 5 | 101489590 | AX-170072901 | 0 | T | C | 0.412 | 5.49E-02 | 1.01E-01 | 2.61E+00 | 2.68E+00 | 5.89E-01 | no | | intergenic_variant | MODIFIER |
| 5 | 89633316 | AX-170197514 | 1 | C | T | 0.438 | 5.92E-02 | 1.11E-01 | 2.76E+00 | 2.81E+00 | 5.89E-01 | no | | intergenic_variant | MODIFIER |
| 1 | 189438852 | AX-170305493 | 1 | T | C | 0.259 | 5.49E-02 | 9.55E-02 | 2.45E+00 | 2.56E+00 | 5.89E-01 | no | | intergenic_variant | MODIFIER |
| 8 | 83262972 | AX-170450984 | 0 | A | G | 0.202 | 6.89E-02 | 1.29E-01 | 2.73E+00 | 2.80E+00 | 5.89E-01 | no | Tbc1d9 | intron_variant | MODIFIER |
| 4 | 95506137 | AX-97792315 | 0 | C | T | 0.333 | -5.54E-02 | 1.05E-01 | 2.74E+00 | 2.83E+00 | 5.89E-01 | no | | intergenic_variant | MODIFIER |
| 9 | 81097679 | AX-97850126 | 0 | T | C | 0.211 | -6.45E-02 | 1.20E-01 | 2.64E+00 | 2.71E+00 | 5.89E-01 | no | Gm26713 | intron_variant,non_coding_transcript_variant | MODIFIER |
| 9 | 35475540 | AX-170334991 | 0 | G | C | 0.421 | -4.68E-02 | 8.78E-02 | 2.81E+00 | 2.89E+00 | 5.89E-01 | no | Cdon | intron_variant | MODIFIER |
| 18 | 58391031 | AX-170183293 | 1 | A | G | 0.402 | 5.55E-02 | 9.76E-02 | 2.48E+00 | 2.54E+00 | 5.89E-01 | no | | intergenic_variant | MODIFIER |
| 4 | 154729326 | AX-170397805 | 0 | G | A | 0.123 | 8.52E-02 | 1.59E-01 | 2.62E+00 | 2.73E+00 | 5.89E-01 | no | | intergenic_variant | MODIFIER |
| 18 | 43396742 | AX-170109462 | 0 | G | T | 0.061 | -1.14E-01 | 2.10E-01 | 2.57E+00 | 2.67E+00 | 5.89E-01 | no | Dpysl3 | upstream_gene_variant | MODIFIER |
| 1 | 98478060 | AX-170001167 | 0 | C | T | 0.167 | 1.00E-01 | 1.84E-01 | 2.58E+00 | 2.67E+00 | 5.90E-01 | no | Slco6d1 | intron_variant | MODIFIER |
| 2 | 93755821 | AX-170456651 | 0 | C | A | 0.061 | -1.24E-01 | 2.34E-01 | 2.73E+00 | 2.82E+00 | 5.90E-01 | no | Ext2 | intron_variant | MODIFIER |
| 2 | 96728194 | AX-170111804 | 0 | C | T | 0.061 | -1.24E-01 | 2.34E-01 | 2.73E+00 | 2.82E+00 | 5.90E-01 | no | Lrrc4c | intron_variant | MODIFIER |
| 2 | 102651173 | AX-170492876 | 0 | C | T | 0.061 | -1.24E-01 | 2.34E-01 | 2.73E+00 | 2.82E+00 | 5.90E-01 | no | | intergenic_variant | MODIFIER |
| 5 | 23942234 | AX-97799103 | 0 | C | T | 0.061 | -1.24E-01 | 2.34E-01 | 2.73E+00 | 2.82E+00 | 5.90E-01 | no | Fam126a | intron_variant | MODIFIER |
| 16 | 44656977 | AX-170012939 | 1 | A | G | 0.509 | 4.56E-02 | 8.26E-02 | 2.55E+00 | 2.60E+00 | 5.90E-01 | no | | intergenic_variant | MODIFIER |
| 12 | 90074066 | AX-97815904 | 0 | G | A | 0.105 | 8.57E-02 | 1.61E-01 | 2.66E+00 | 2.76E+00 | 5.90E-01 | no | Nrxn3 | intron_variant | MODIFIER |

| | | | | | | | | | | | | | | | |
|---|---|---|---|---|---|---|---|---|---|---|---|---|---|---|---|
| 1 | 42080878 | AX-170171646 | 0 | C | T | 0.439 | -3.98E-02 | 7.45E-02 | 2.64E+00 | 2.73E+00 | 5.90E-01 | no | | intergenic_variant | MODIFIER |
| 2 | 107931070 | AX-97850761 | 1 | G | T | 0.062 | -1.24E-01 | 2.34E-01 | 2.74E+00 | 2.83E+00 | 5.90E-01 | no | | intergenic_variant | MODIFIER |
| 15 | 101026084 | AX-97845146 | 0 | G | A | 0.14 | 7.58E-02 | 1.43E-01 | 2.66E+00 | 2.77E+00 | 5.90E-01 | no | Scn8a | intron_variant | MODIFIER |
| 5 | 106496989 | AX-170355035 | 1 | T | C | 0.438 | 6.18E-02 | 1.15E-01 | 2.65E+00 | 2.68E+00 | 5.90E-01 | no | Gm28050 | downstream_gene_variant | MODIFIER |
| 5 | 103409378 | AX-97807908 | 0 | A | C | 0.456 | 6.29E-02 | 1.17E-01 | 2.66E+00 | 2.69E+00 | 5.90E-01 | no | | intergenic_variant | MODIFIER |
| 5 | 103409413 | AX-170257791 | 0 | A | G | 0.456 | 6.29E-02 | 1.17E-01 | 2.66E+00 | 2.69E+00 | 5.90E-01 | no | | intergenic_variant | MODIFIER |
| 5 | 103418911 | AX-170212978 | 0 | A | G | 0.456 | 6.29E-02 | 1.17E-01 | 2.66E+00 | 2.69E+00 | 5.90E-01 | no | | regulatory_region_variant | MODIFIER |
| 5 | 103599308 | AX-170025774 | 0 | G | C | 0.456 | 6.29E-02 | 1.17E-01 | 2.66E+00 | 2.69E+00 | 5.90E-01 | no | Ptpn13 | downstream_gene_variant | MODIFIER |
| 5 | 103600564 | AX-170305863 | 0 | C | T | 0.456 | 6.29E-02 | 1.17E-01 | 2.66E+00 | 2.69E+00 | 5.90E-01 | no | Ptpn13 | downstream_gene_variant | MODIFIER |
| 5 | 103602843 | AX-170377778 | 0 | G | C | 0.456 | 6.29E-02 | 1.17E-01 | 2.66E+00 | 2.69E+00 | 5.90E-01 | no | Slc10a6 | downstream_gene_variant | MODIFIER |
| 5 | 103615059 | AX-170025771 | 0 | G | A | 0.456 | 6.29E-02 | 1.17E-01 | 2.66E+00 | 2.69E+00 | 5.90E-01 | no | Slc10a6 | intron_variant | MODIFIER |
| 5 | 103617922 | AX-170025775 | 0 | C | G | 0.456 | 6.29E-02 | 1.17E-01 | 2.66E+00 | 2.69E+00 | 5.90E-01 | no | Slc10a6 | intron_variant | MODIFIER |
| 5 | 103626540 | AX-170363701 | 0 | C | A | 0.456 | 6.29E-02 | 1.17E-01 | 2.66E+00 | 2.69E+00 | 5.90E-01 | no | Slc10a6 | intron_variant | MODIFIER |
| 5 | 103761832 | AX-168414485 | 0 | G | A | 0.456 | 6.29E-02 | 1.17E-01 | 2.66E+00 | 2.69E+00 | 5.90E-01 | no | Aff1 | intron_variant | MODIFIER |
| 5 | 103763474 | AX-170447722 | 0 | G | A | 0.456 | 6.29E-02 | 1.17E-01 | 2.66E+00 | 2.69E+00 | 5.90E-01 | no | Aff1 | intron_variant | MODIFIER |
| 5 | 103775118 | AX-97833095 | 0 | C | A | 0.456 | 6.29E-02 | 1.17E-01 | 2.66E+00 | 2.69E+00 | 5.90E-01 | no | Aff1 | intron_variant | MODIFIER |
| 5 | 103863996 | AX-170310686 | 0 | G | T | 0.456 | 6.29E-02 | 1.17E-01 | 2.66E+00 | 2.69E+00 | 5.90E-01 | no | Klhl8 | intron_variant | MODIFIER |
| 5 | 103968727 | AX-170149588 | 0 | G | A | 0.456 | 6.29E-02 | 1.17E-01 | 2.66E+00 | 2.69E+00 | 5.90E-01 | no | Hsd17b13 | synonymous_variant | LOW |
| 5 | 103979469 | AX-170447075 | 0 | C | T | 0.456 | 6.29E-02 | 1.17E-01 | 2.66E+00 | 2.69E+00 | 5.90E-01 | no | Hsd17b13 | upstream_gene_variant | MODIFIER |
| 5 | 104154033 | AX-170524891 | 0 | A | G | 0.456 | 6.29E-02 | 1.17E-01 | 2.66E+00 | 2.69E+00 | 5.90E-01 | no | | intergenic_variant | MODIFIER |
| 5 | 104211903 | AX-170140928 | 0 | C | T | 0.456 | 6.29E-02 | 1.17E-01 | 2.66E+00 | 2.69E+00 | 5.90E-01 | no | Dmp1 | synonymous_variant | LOW |
| 5 | 104219968 | AX-170185144 | 0 | T | C | 0.456 | 6.29E-02 | 1.17E-01 | 2.66E+00 | 2.69E+00 | 5.90E-01 | no | | intergenic_variant | MODIFIER |
| 5 | 104222766 | AX-97834663 | 0 | A | G | 0.456 | 6.29E-02 | 1.17E-01 | 2.66E+00 | 2.69E+00 | 5.90E-01 | no | | intergenic_variant | MODIFIER |
| 5 | 104430384 | AX-170327647 | 0 | T | G | 0.456 | 6.29E-02 | 1.17E-01 | 2.66E+00 | 2.69E+00 | 5.90E-01 | no | Spp1 | upstream_gene_variant | MODIFIER |
| 5 | 104431451 | AX-97821833 | 0 | G | A | 0.456 | 6.29E-02 | 1.17E-01 | 2.66E+00 | 2.69E+00 | 5.90E-01 | no | Spp1 | upstream_gene_variant | MODIFIER |
| 5 | 104434007 | AX-170142055 | 0 | A | T | 0.456 | 6.29E-02 | 1.17E-01 | 2.66E+00 | 2.69E+00 | 5.90E-01 | no | Spp1 | upstream_gene_variant | MODIFIER |
| 5 | 104465361 | AX-170469892 | 0 | C | T | 0.456 | 6.29E-02 | 1.17E-01 | 2.66E+00 | 2.69E+00 | 5.90E-01 | no | Pkd2 | intron_variant | MODIFIER |
| 5 | 104471327 | AX-97804238 | 0 | T | C | 0.456 | 6.29E-02 | 1.17E-01 | 2.66E+00 | 2.69E+00 | 5.90E-01 | no | Pkd2 | intron_variant | MODIFIER |
| 5 | 104517635 | AX-170288504 | 0 | C | G | 0.456 | 6.29E-02 | 1.17E-01 | 2.66E+00 | 2.69E+00 | 5.90E-01 | no | BC005561 | missense_variant | MODERATE |
| 5 | 104517658 | AX-170150057 | 0 | T | C | 0.456 | 6.29E-02 | 1.17E-01 | 2.66E+00 | 2.69E+00 | 5.90E-01 | no | BC005561 | synonymous_variant | LOW |
| 5 | 104521066 | AX-170304603 | 0 | A | C | 0.456 | 6.29E-02 | 1.17E-01 | 2.66E+00 | 2.69E+00 | 5.90E-01 | no | BC005561 | synonymous_variant | LOW |
| 5 | 104521303 | AX-170089572 | 0 | G | A | 0.456 | 6.29E-02 | 1.17E-01 | 2.66E+00 | 2.69E+00 | 5.90E-01 | no | BC005561 | synonymous_variant | LOW |
| 5 | 104538029 | AX-170467862 | 0 | T | A | 0.456 | 6.29E-02 | 1.17E-01 | 2.66E+00 | 2.69E+00 | 5.90E-01 | no | D930016D06Rik | upstream_gene_variant | MODIFIER |
| 5 | 104605480 | AX-170089574 | 0 | T | G | 0.456 | 6.29E-02 | 1.17E-01 | 2.66E+00 | 2.69E+00 | 5.90E-01 | no | | intergenic_variant | MODIFIER |
| 5 | 104702188 | AX-97798269 | 0 | A | G | 0.456 | 6.29E-02 | 1.17E-01 | 2.66E+00 | 2.69E+00 | 5.90E-01 | no | | intergenic_variant | MODIFIER |
| 5 | 104918928 | AX-170025795 | 0 | C | A | 0.456 | 6.29E-02 | 1.17E-01 | 2.66E+00 | 2.69E+00 | 5.90E-01 | no | | intergenic_variant | MODIFIER |
| 5 | 104939711 | AX-97846567 | 0 | T | C | 0.456 | 6.29E-02 | 1.17E-01 | 2.66E+00 | 2.69E+00 | 5.90E-01 | no | Abcg3 | intron_variant | MODIFIER |
| 5 | 104945715 | AX-170025796 | 0 | G | A | 0.456 | 6.29E-02 | 1.17E-01 | 2.66E+00 | 2.69E+00 | 5.90E-01 | no | Abcg3 | intron_variant | MODIFIER |
| 5 | 104952725 | AX-170478642 | 0 | T | G | 0.456 | 6.29E-02 | 1.17E-01 | 2.66E+00 | 2.69E+00 | 5.90E-01 | no | Abcg3 | intron_variant | MODIFIER |
| 5 | 104952770 | AX-170257236 | 0 | G | T | 0.456 | 6.29E-02 | 1.17E-01 | 2.66E+00 | 2.69E+00 | 5.90E-01 | no | Abcg3 | intron_variant | MODIFIER |
| 5 | 104961454 | AX-170497324 | 0 | A | G | 0.456 | 6.29E-02 | 1.17E-01 | 2.66E+00 | 2.69E+00 | 5.90E-01 | no | Abcg3 | intron_variant | MODIFIER |
| 5 | 104962100 | AX-170072967 | 0 | C | T | 0.456 | 6.29E-02 | 1.17E-01 | 2.66E+00 | 2.69E+00 | 5.90E-01 | no | Abcg3 | intron_variant | MODIFIER |
| 5 | 104968559 | AX-170025790 | 0 | T | C | 0.456 | 6.29E-02 | 1.17E-01 | 2.66E+00 | 2.69E+00 | 5.90E-01 | no | Abcg3 | intron_variant | MODIFIER |
| 5 | 104978520 | AX-97854577 | 0 | G | A | 0.456 | 6.29E-02 | 1.17E-01 | 2.66E+00 | 2.69E+00 | 5.90E-01 | no | Abcg3 | intron_variant | MODIFIER |
| 5 | 106042839 | AX-97847759 | 0 | C | T | 0.456 | 6.29E-02 | 1.17E-01 | 2.66E+00 | 2.69E+00 | 5.90E-01 | no | | intergenic_variant | MODIFIER |
| 5 | 106053639 | AX-170035716 | 0 | T | C | 0.456 | 6.29E-02 | 1.17E-01 | 2.66E+00 | 2.69E+00 | 5.90E-01 | no | | intergenic_variant | MODIFIER |
| 5 | 106080867 | AX-170347130 | 0 | G | A | 0.456 | 6.29E-02 | 1.17E-01 | 2.66E+00 | 2.69E+00 | 5.90E-01 | no | | intergenic_variant | MODIFIER |
| 5 | 106335353 | AX-97800659 | 0 | C | T | 0.456 | 6.29E-02 | 1.17E-01 | 2.66E+00 | 2.69E+00 | 5.90E-01 | no | | intergenic_variant | MODIFIER |
| 5 | 106366345 | AX-170448755 | 0 | G | A | 0.456 | 6.29E-02 | 1.17E-01 | 2.66E+00 | 2.69E+00 | 5.90E-01 | no | Gm42916 | upstream_gene_variant | MODIFIER |
| 5 | 106366580 | AX-170341224 | 0 | C | A | 0.456 | 6.29E-02 | 1.17E-01 | 2.66E+00 | 2.69E+00 | 5.90E-01 | no | Gm42916 | upstream_gene_variant | MODIFIER |
| 5 | 106370152 | AX-170154203 | 0 | G | A | 0.456 | 6.29E-02 | 1.17E-01 | 2.66E+00 | 2.69E+00 | 5.90E-01 | no | Gm42916 | intron_variant,non_coding_transcript_variant | MODIFIER |
| 5 | 106378004 | AX-170415542 | 0 | A | T | 0.456 | 6.29E-02 | 1.17E-01 | 2.66E+00 | 2.69E+00 | 5.90E-01 | no | | intergenic_variant | MODIFIER |
| 5 | 106378792 | AX-97821361 | 0 | T | C | 0.456 | 6.29E-02 | 1.17E-01 | 2.66E+00 | 2.69E+00 | 5.90E-01 | no | | intergenic_variant | MODIFIER |
| 5 | 106393127 | AX-170114401 | 0 | C | T | 0.456 | 6.29E-02 | 1.17E-01 | 2.66E+00 | 2.69E+00 | 5.90E-01 | no | | intergenic_variant | MODIFIER |

| | | | | | | | | | | | | | | | |
|---|---|---|---|---|---|---|---|---|---|---|---|---|---|---|---|
| 5 | 106405634 | AX-97832131 | 0 | G | A | 0.456 | 6.29E-02 | 1.17E-01 | 2.66E+00 | 2.69E+00 | 5.90E-01 | no | | intergenic_variant | MODIFIER |
| 5 | 106434430 | AX-170345185 | 0 | G | A | 0.456 | 6.29E-02 | 1.17E-01 | 2.66E+00 | 2.69E+00 | 5.90E-01 | no | | intergenic_variant | MODIFIER |
| 5 | 106440852 | AX-170134652 | 0 | G | T | 0.456 | 6.29E-02 | 1.17E-01 | 2.66E+00 | 2.69E+00 | 5.90E-01 | no | | intergenic_variant | MODIFIER |
| 5 | 106475357 | AX-170072997 | 0 | C | T | 0.456 | 6.29E-02 | 1.17E-01 | 2.66E+00 | 2.69E+00 | 5.90E-01 | no | Gm26872 | upstream_gene_variant | MODIFIER |
| 5 | 106502160 | AX-170186126 | 0 | C | A | 0.456 | 6.29E-02 | 1.17E-01 | 2.66E+00 | 2.69E+00 | 5.90E-01 | no | Gm28050 | intron_variant,non_coding_transcript_variant | MODIFIER |
| 5 | 106505749 | AX-170453147 | 0 | A | G | 0.456 | 6.29E-02 | 1.17E-01 | 2.66E+00 | 2.69E+00 | 5.90E-01 | no | Gm28050 | intron_variant,non_coding_transcript_variant | MODIFIER |
| 5 | 106507798 | AX-170436715 | 0 | C | A | 0.456 | 6.29E-02 | 1.17E-01 | 2.66E+00 | 2.69E+00 | 5.90E-01 | no | Gm28050 | intron_variant,non_coding_transcript_variant | MODIFIER |
| 5 | 106510873 | AX-170025817 | 0 | T | C | 0.456 | 6.29E-02 | 1.17E-01 | 2.66E+00 | 2.69E+00 | 5.90E-01 | no | Gm28050 | intron_variant,non_coding_transcript_variant | MODIFIER |
| 5 | 106569439 | AX-170025819 | 0 | C | T | 0.456 | 6.29E-02 | 1.17E-01 | 2.66E+00 | 2.69E+00 | 5.90E-01 | no | Gm28050 | intron_variant,non_coding_transcript_variant | MODIFIER |
| 5 | 106638048 | AX-170302674 | 0 | C | T | 0.456 | 6.29E-02 | 1.17E-01 | 2.66E+00 | 2.69E+00 | 5.90E-01 | no | Zfp644 | missense_variant | MODERATE |
| 5 | 106683524 | AX-170395594 | 0 | A | G | 0.456 | 6.29E-02 | 1.17E-01 | 2.66E+00 | 2.69E+00 | 5.90E-01 | no | Zfp644 | intron_variant | MODIFIER |
| 5 | 106685166 | AX-97808776 | 0 | A | T | 0.456 | 6.29E-02 | 1.17E-01 | 2.66E+00 | 2.69E+00 | 5.90E-01 | no | Zfp644 | intron_variant | MODIFIER |
| 5 | 106717520 | AX-97781672 | 0 | T | C | 0.456 | 6.29E-02 | 1.17E-01 | 2.66E+00 | 2.69E+00 | 5.90E-01 | no | | intergenic_variant | MODIFIER |
| 5 | 106745364 | AX-170311460 | 0 | C | T | 0.456 | 6.29E-02 | 1.17E-01 | 2.66E+00 | 2.69E+00 | 5.90E-01 | no | | intergenic_variant | MODIFIER |
| 5 | 106822820 | AX-170175194 | 0 | C | T | 0.456 | 6.29E-02 | 1.17E-01 | 2.66E+00 | 2.69E+00 | 5.90E-01 | no | | intergenic_variant | MODIFIER |
| 5 | 107022390 | AX-170500555 | 0 | A | C | 0.456 | 6.29E-02 | 1.17E-01 | 2.66E+00 | 2.69E+00 | 5.90E-01 | no | | intergenic_variant | MODIFIER |
| 5 | 107028943 | AX-170418320 | 0 | A | G | 0.456 | 6.29E-02 | 1.17E-01 | 2.66E+00 | 2.69E+00 | 5.90E-01 | no | | regulatory_region_variant | MODIFIER |
| 5 | 107036014 | AX-170285764 | 0 | G | A | 0.456 | 6.29E-02 | 1.17E-01 | 2.66E+00 | 2.69E+00 | 5.90E-01 | no | | regulatory_region_variant | MODIFIER |
| 5 | 107042160 | AX-170095428 | 0 | A | G | 0.456 | 6.29E-02 | 1.17E-01 | 2.66E+00 | 2.69E+00 | 5.90E-01 | no | | intergenic_variant | MODIFIER |
| 5 | 107071219 | AX-170025826 | 0 | A | G | 0.456 | 6.29E-02 | 1.17E-01 | 2.66E+00 | 2.69E+00 | 5.90E-01 | no | | intergenic_variant | MODIFIER |
| 5 | 107183572 | AX-170349205 | 0 | G | A | 0.456 | 6.29E-02 | 1.17E-01 | 2.66E+00 | 2.69E+00 | 5.90E-01 | no | Tgfbr3 | intron_variant | MODIFIER |
| 5 | 107184309 | AX-170106626 | 0 | C | T | 0.456 | 6.29E-02 | 1.17E-01 | 2.66E+00 | 2.69E+00 | 5.90E-01 | no | Tgfbr3 | intron_variant | MODIFIER |
| 5 | 107185580 | AX-170025829 | 0 | T | C | 0.456 | 6.29E-02 | 1.17E-01 | 2.66E+00 | 2.69E+00 | 5.90E-01 | no | Tgfbr3 | intron_variant | MODIFIER |
| 5 | 107199598 | AX-170025830 | 0 | G | A | 0.456 | 6.29E-02 | 1.17E-01 | 2.66E+00 | 2.69E+00 | 5.90E-01 | no | Tgfbr3 | intron_variant | MODIFIER |
| 5 | 107205501 | AX-170415679 | 0 | G | C | 0.456 | 6.29E-02 | 1.17E-01 | 2.66E+00 | 2.69E+00 | 5.90E-01 | no | Tgfbr3 | intron_variant | MODIFIER |
| 5 | 107205546 | AX-170144072 | 0 | G | A | 0.456 | 6.29E-02 | 1.17E-01 | 2.66E+00 | 2.69E+00 | 5.90E-01 | no | Tgfbr3 | intron_variant | MODIFIER |
| 5 | 107205804 | AX-97787926 | 0 | T | C | 0.456 | 6.29E-02 | 1.17E-01 | 2.66E+00 | 2.69E+00 | 5.90E-01 | no | Tgfbr3 | intron_variant | MODIFIER |
| 5 | 107350502 | AX-170089577 | 0 | C | T | 0.456 | 6.29E-02 | 1.17E-01 | 2.66E+00 | 2.69E+00 | 5.90E-01 | no | Brdt | intron_variant | MODIFIER |
| 5 | 107372421 | AX-170173655 | 0 | T | C | 0.456 | 6.29E-02 | 1.17E-01 | 2.66E+00 | 2.69E+00 | 5.90E-01 | no | Brdt | intron_variant | MODIFIER |
| 5 | 107568702 | AX-97800081 | 0 | A | C | 0.456 | 6.29E-02 | 1.17E-01 | 2.66E+00 | 2.69E+00 | 5.90E-01 | no | Glmn | intron_variant | MODIFIER |
| 5 | 107600956 | AX-170290227 | 0 | G | A | 0.456 | 6.29E-02 | 1.17E-01 | 2.66E+00 | 2.69E+00 | 5.90E-01 | no | Rpap2 | intron_variant | MODIFIER |
| 5 | 107608141 | AX-97833146 | 0 | T | G | 0.456 | 6.29E-02 | 1.17E-01 | 2.66E+00 | 2.69E+00 | 5.90E-01 | no | Rpap2 | intron_variant | MODIFIER |
| 5 | 107625192 | AX-170338238 | 0 | G | A | 0.456 | 6.29E-02 | 1.17E-01 | 2.66E+00 | 2.69E+00 | 5.90E-01 | no | Rpap2 | intron_variant | MODIFIER |
| 5 | 107627125 | AX-97790219 | 0 | C | T | 0.456 | 6.29E-02 | 1.17E-01 | 2.66E+00 | 2.69E+00 | 5.90E-01 | no | Rpap2 | intron_variant | MODIFIER |
| 5 | 107637845 | AX-170457276 | 0 | G | A | 0.456 | 6.29E-02 | 1.17E-01 | 2.66E+00 | 2.69E+00 | 5.90E-01 | no | Rpap2 | intron_variant | MODIFIER |
| 5 | 107650256 | AX-170025838 | 0 | C | T | 0.456 | 6.29E-02 | 1.17E-01 | 2.66E+00 | 2.69E+00 | 5.90E-01 | no | Rpap2 | intron_variant | MODIFIER |
| 5 | 107655534 | AX-170365574 | 0 | G | A | 0.456 | 6.29E-02 | 1.17E-01 | 2.66E+00 | 2.69E+00 | 5.90E-01 | no | Rpap2 | intron_variant | MODIFIER |
| 5 | 107689628 | AX-170025835 | 0 | C | T | 0.456 | 6.29E-02 | 1.17E-01 | 2.66E+00 | 2.69E+00 | 5.90E-01 | no | Gm26692 | upstream_gene_variant | MODIFIER |
| 5 | 108115285 | AX-170320119 | 0 | C | T | 0.456 | 6.29E-02 | 1.17E-01 | 2.66E+00 | 2.69E+00 | 5.90E-01 | no | Tmed5 | downstream_gene_variant | MODIFIER |
| 5 | 108250474 | AX-170247879 | 0 | G | A | 0.456 | 6.29E-02 | 1.17E-01 | 2.66E+00 | 2.69E+00 | 5.90E-01 | no | | intergenic_variant | MODIFIER |
| 5 | 108278401 | AX-168419914 | 0 | G | T | 0.456 | 6.29E-02 | 1.17E-01 | 2.66E+00 | 2.69E+00 | 5.90E-01 | no | Dr1 | intron_variant | MODIFIER |
| 5 | 108378254 | AX-170025843 | 0 | T | C | 0.456 | 6.29E-02 | 1.17E-01 | 2.66E+00 | 2.69E+00 | 5.90E-01 | no | Gm10419 | downstream_gene_variant | MODIFIER |
| 5 | 108557232 | AX-170336476 | 0 | T | C | 0.456 | 6.29E-02 | 1.17E-01 | 2.66E+00 | 2.69E+00 | 5.90E-01 | no | Gm43364 | upstream_gene_variant | MODIFIER |
| 5 | 108557299 | AX-170103708 | 0 | A | G | 0.456 | 6.29E-02 | 1.17E-01 | 2.66E+00 | 2.69E+00 | 5.90E-01 | no | Gm43364 | upstream_gene_variant | MODIFIER |
| 5 | 108559501 | AX-170073044 | 0 | G | A | 0.456 | 6.29E-02 | 1.17E-01 | 2.66E+00 | 2.69E+00 | 5.90E-01 | no | | intergenic_variant | MODIFIER |
| 5 | 108565390 | AX-170427534 | 0 | C | G | 0.456 | 6.29E-02 | 1.17E-01 | 2.66E+00 | 2.69E+00 | 5.90E-01 | no | Gak | downstream_gene_variant | MODIFIER |
| 5 | 108565588 | AX-170326923 | 0 | C | T | 0.456 | 6.29E-02 | 1.17E-01 | 2.66E+00 | 2.69E+00 | 5.90E-01 | no | Gak | downstream_gene_variant | MODIFIER |
| 5 | 108566663 | AX-170531448 | 0 | C | T | 0.456 | 6.29E-02 | 1.17E-01 | 2.66E+00 | 2.69E+00 | 5.90E-01 | no | Gak | downstream_gene_variant | MODIFIER |
| 5 | 108567606 | AX-170371956 | 0 | G | A | 0.456 | 6.29E-02 | 1.17E-01 | 2.66E+00 | 2.69E+00 | 5.90E-01 | no | Gak | downstream_gene_variant | MODIFIER |
| 5 | 108567952 | AX-170293831 | 0 | T | A | 0.456 | 6.29E-02 | 1.17E-01 | 2.66E+00 | 2.69E+00 | 5.90E-01 | no | Gak | downstream_gene_variant | MODIFIER |
| 5 | 108568531 | AX-170348100 | 0 | C | T | 0.456 | 6.29E-02 | 1.17E-01 | 2.66E+00 | 2.69E+00 | 5.90E-01 | no | Gak | downstream_gene_variant | MODIFIER |
| 5 | 108570699 | AX-170299740 | 0 | A | G | 0.456 | 6.29E-02 | 1.17E-01 | 2.66E+00 | 2.69E+00 | 5.90E-01 | no | Gak | intron_variant | MODIFIER |
| 5 | 108571611 | AX-170326984 | 0 | T | C | 0.456 | 6.29E-02 | 1.17E-01 | 2.66E+00 | 2.69E+00 | 5.90E-01 | no | Gak | intron_variant | MODIFIER |
| 5 | 108572190 | AX-170369624 | 0 | C | A | 0.456 | 6.29E-02 | 1.17E-01 | 2.66E+00 | 2.69E+00 | 5.90E-01 | no | Gak | intron_variant | MODIFIER |

| | | | | | | | | | | | | | | | |
|---|---|---|---|---|---|---|---|---|---|---|---|---|---|---|---|
| 5 | 108586311 | AX-170511759 | 0 | T | C | 0.456 | 6.29E-02 | 1.17E-01 | 2.66E+00 | 2.69E+00 | 5.90E-01 | no | Gak | intron_variant | MODIFIER |
| 5 | 108586822 | AX-170298274 | 0 | G | A | 0.456 | 6.29E-02 | 1.17E-01 | 2.66E+00 | 2.69E+00 | 5.90E-01 | no | Gak | intron_variant | MODIFIER |
| 5 | 108608962 | AX-170177968 | 0 | A | G | 0.456 | 6.29E-02 | 1.17E-01 | 2.66E+00 | 2.69E+00 | 5.90E-01 | no | Gak | intron_variant | MODIFIER |
| 5 | 108609637 | AX-170238443 | 0 | C | G | 0.456 | 6.29E-02 | 1.17E-01 | 2.66E+00 | 2.69E+00 | 5.90E-01 | no | Gak | intron_variant | MODIFIER |
| 5 | 108609705 | AX-170457784 | 0 | G | C | 0.456 | 6.29E-02 | 1.17E-01 | 2.66E+00 | 2.69E+00 | 5.90E-01 | no | Gak | intron_variant | MODIFIER |
| 5 | 108611012 | AX-170441518 | 0 | A | G | 0.456 | 6.29E-02 | 1.17E-01 | 2.66E+00 | 2.69E+00 | 5.90E-01 | no | Gak | intron_variant | MODIFIER |
| 5 | 108612554 | AX-170352325 | 0 | A | G | 0.456 | 6.29E-02 | 1.17E-01 | 2.66E+00 | 2.69E+00 | 5.90E-01 | no | Gak | intron_variant | MODIFIER |
| 5 | 108619041 | AX-170411844 | 0 | A | G | 0.456 | 6.29E-02 | 1.17E-01 | 2.66E+00 | 2.69E+00 | 5.90E-01 | no | Gak | intron_variant | MODIFIER |
| 5 | 108621063 | AX-170429727 | 0 | G | T | 0.456 | 6.29E-02 | 1.17E-01 | 2.66E+00 | 2.69E+00 | 5.90E-01 | no | Gak | intron_variant | MODIFIER |
| 5 | 108622211 | AX-170256477 | 0 | C | T | 0.456 | 6.29E-02 | 1.17E-01 | 2.66E+00 | 2.69E+00 | 5.90E-01 | no | Gak | intron_variant | MODIFIER |
| 5 | 108653568 | AX-170073048 | 0 | G | C | 0.456 | 6.29E-02 | 1.17E-01 | 2.66E+00 | 2.69E+00 | 5.90E-01 | no | Dgkq | intron_variant | MODIFIER |
| 5 | 108657135 | AX-170322410 | 0 | C | T | 0.456 | 6.29E-02 | 1.17E-01 | 2.66E+00 | 2.69E+00 | 5.90E-01 | no | Dgkq | intron_variant | MODIFIER |
| 5 | 108723633 | AX-170106783 | 0 | T | C | 0.456 | 6.29E-02 | 1.17E-01 | 2.66E+00 | 2.69E+00 | 5.90E-01 | no | | intergenic_variant | MODIFIER |
| 5 | 108727614 | AX-170469492 | 0 | C | T | 0.456 | 6.29E-02 | 1.17E-01 | 2.66E+00 | 2.69E+00 | 5.90E-01 | no | Rnf212 | downstream_gene_variant | MODIFIER |
| 5 | 108744427 | AX-170246219 | 0 | A | G | 0.456 | 6.29E-02 | 1.17E-01 | 2.66E+00 | 2.69E+00 | 5.90E-01 | no | Rnf212 | intron_variant | MODIFIER |
| 5 | 109237625 | AX-97834336 | 0 | T | C | 0.456 | 6.29E-02 | 1.17E-01 | 2.66E+00 | 2.69E+00 | 5.90E-01 | no | Gm8473 | non_coding_transcript_exon_variant | MODIFIER |
| 5 | 109340301 | AX-170102025 | 0 | T | A | 0.456 | 6.29E-02 | 1.17E-01 | 2.66E+00 | 2.69E+00 | 5.90E-01 | no | Vmn2r16 | missense_variant | MODERATE |
| 5 | 109395596 | AX-170098547 | 0 | T | A | 0.456 | 6.29E-02 | 1.17E-01 | 2.66E+00 | 2.69E+00 | 5.90E-01 | no | Gm7171 | upstream_gene_variant | MODIFIER |
| 5 | 109419274 | AX-170466032 | 0 | A | G | 0.456 | 6.29E-02 | 1.17E-01 | 2.66E+00 | 2.69E+00 | 5.90E-01 | no | Vmn2r17 | upstream_gene_variant | MODIFIER |
| 5 | 109523535 | AX-170025869 | 0 | G | A | 0.456 | 6.29E-02 | 1.17E-01 | 2.66E+00 | 2.69E+00 | 5.90E-01 | no | Gm8488 | downstream_gene_variant | MODIFIER |
| 5 | 109550059 | AX-170171814 | 0 | T | C | 0.456 | 6.29E-02 | 1.17E-01 | 2.66E+00 | 2.69E+00 | 5.90E-01 | no | Crlf2 | downstream_gene_variant | MODIFIER |
| 5 | 109559328 | AX-170385161 | 0 | C | T | 0.456 | 6.29E-02 | 1.17E-01 | 2.66E+00 | 2.69E+00 | 5.90E-01 | no | Crlf2 | upstream_gene_variant | MODIFIER |
| 5 | 109566688 | AX-170343033 | 0 | C | T | 0.456 | 6.29E-02 | 1.17E-01 | 2.66E+00 | 2.69E+00 | 5.90E-01 | no | | intergenic_variant | MODIFIER |
| 5 | 109587641 | AX-170025866 | 0 | T | C | 0.456 | 6.29E-02 | 1.17E-01 | 2.66E+00 | 2.69E+00 | 5.90E-01 | no | | intergenic_variant | MODIFIER |
| 5 | 109720150 | AX-170304637 | 0 | G | A | 0.456 | 6.29E-02 | 1.17E-01 | 2.66E+00 | 2.69E+00 | 5.90E-01 | no | | intergenic_variant | MODIFIER |
| 5 | 110054911 | AX-170533862 | 0 | C | T | 0.456 | 6.29E-02 | 1.17E-01 | 2.66E+00 | 2.69E+00 | 5.90E-01 | no | Gm17655 | intron_variant | MODIFIER |
| 5 | 110075199 | AX-170447201 | 0 | T | C | 0.456 | 6.29E-02 | 1.17E-01 | 2.66E+00 | 2.69E+00 | 5.90E-01 | no | Gm35315 | downstream_gene_variant | MODIFIER |
| 5 | 110180308 | AX-170442265 | 0 | A | C | 0.456 | 6.29E-02 | 1.17E-01 | 2.66E+00 | 2.69E+00 | 5.90E-01 | no | Golga3 | intron_variant | MODIFIER |
| 5 | 110267804 | AX-97798379 | 0 | A | G | 0.456 | 6.29E-02 | 1.17E-01 | 2.66E+00 | 2.69E+00 | 5.90E-01 | no | Pgam5 | intron_variant | MODIFIER |
| 5 | 110273540 | AX-170295669 | 0 | C | T | 0.456 | 6.29E-02 | 1.17E-01 | 2.66E+00 | 2.69E+00 | 5.90E-01 | no | Pxmp2 | downstream_gene_variant | MODIFIER |
| 5 | 110297879 | AX-170341470 | 0 | T | C | 0.456 | 6.29E-02 | 1.17E-01 | 2.66E+00 | 2.69E+00 | 5.90E-01 | no | Pole | synonymous_variant | LOW |
| 5 | 110305342 | AX-170233610 | 0 | C | T | 0.456 | 6.29E-02 | 1.17E-01 | 2.66E+00 | 2.69E+00 | 5.90E-01 | no | Pole | intron_variant | MODIFIER |
| 5 | 110341706 | AX-170303949 | 0 | G | A | 0.456 | 6.29E-02 | 1.17E-01 | 2.66E+00 | 2.69E+00 | 5.90E-01 | no | Pole | downstream_gene_variant | MODIFIER |
| 5 | 110385941 | AX-170333911 | 0 | T | C | 0.456 | 6.29E-02 | 1.17E-01 | 2.66E+00 | 2.69E+00 | 5.90E-01 | no | Fbrsl1 | intron_variant | MODIFIER |
| 5 | 110412543 | AX-170431629 | 0 | C | T | 0.456 | 6.29E-02 | 1.17E-01 | 2.66E+00 | 2.69E+00 | 5.90E-01 | no | Fbrsl1 | intron_variant | MODIFIER |
| 5 | 110421322 | AX-170174571 | 0 | G | A | 0.456 | 6.29E-02 | 1.17E-01 | 2.66E+00 | 2.69E+00 | 5.90E-01 | no | Fbrsl1 | intron_variant | MODIFIER |
| 5 | 110422242 | AX-170377139 | 0 | T | C | 0.456 | 6.29E-02 | 1.17E-01 | 2.66E+00 | 2.69E+00 | 5.90E-01 | no | Fbrsl1 | intron_variant | MODIFIER |
| 5 | 110582845 | AX-170226240 | 0 | C | T | 0.456 | 6.29E-02 | 1.17E-01 | 2.66E+00 | 2.69E+00 | 5.90E-01 | no | Galnt9 | intron_variant | MODIFIER |
| 5 | 110609450 | AX-170114850 | 0 | A | G | 0.456 | 6.29E-02 | 1.17E-01 | 2.66E+00 | 2.69E+00 | 5.90E-01 | no | Galnt9 | intron_variant | MODIFIER |
| 5 | 110666462 | AX-170350605 | 0 | C | T | 0.456 | 6.29E-02 | 1.17E-01 | 2.66E+00 | 2.69E+00 | 5.90E-01 | no | Ep400 | intron_variant | MODIFIER |
| 5 | 110670487 | AX-170150677 | 0 | A | T | 0.456 | 6.29E-02 | 1.17E-01 | 2.66E+00 | 2.69E+00 | 5.90E-01 | no | Ep400 | intron_variant | MODIFIER |
| 5 | 110737486 | AX-170422841 | 0 | G | A | 0.456 | 6.29E-02 | 1.17E-01 | 2.66E+00 | 2.69E+00 | 5.90E-01 | no | Ep400 | intron_variant | MODIFIER |
| 5 | 110800328 | AX-170341499 | 0 | C | T | 0.456 | 6.29E-02 | 1.17E-01 | 2.66E+00 | 2.69E+00 | 5.90E-01 | no | Ulk1 | intron_variant | MODIFIER |
| 5 | 110823292 | AX-170466221 | 0 | T | C | 0.456 | 6.29E-02 | 1.17E-01 | 2.66E+00 | 2.69E+00 | 5.90E-01 | no | | intergenic_variant | MODIFIER |
| 5 | 110827450 | AX-170073089 | 0 | T | C | 0.456 | 6.29E-02 | 1.17E-01 | 2.66E+00 | 2.69E+00 | 5.90E-01 | no | Hscb | downstream_gene_variant | MODIFIER |
| 5 | 110905537 | AX-97830491 | 0 | C | T | 0.456 | 6.29E-02 | 1.17E-01 | 2.66E+00 | 2.69E+00 | 5.90E-01 | no | Ttc28 | intron_variant | MODIFIER |
| 5 | 110922101 | AX-97837932 | 0 | G | T | 0.456 | 6.29E-02 | 1.17E-01 | 2.66E+00 | 2.69E+00 | 5.90E-01 | no | Ttc28 | intron_variant | MODIFIER |
| 5 | 110993810 | AX-97835952 | 0 | G | C | 0.456 | 6.29E-02 | 1.17E-01 | 2.66E+00 | 2.69E+00 | 5.90E-01 | no | Ttc28 | intron_variant | MODIFIER |
| 5 | 110998251 | AX-170491714 | 0 | T | G | 0.456 | 6.29E-02 | 1.17E-01 | 2.66E+00 | 2.69E+00 | 5.90E-01 | no | Ttc28 | intron_variant | MODIFIER |
| 5 | 111003645 | AX-170519440 | 0 | C | T | 0.456 | 6.29E-02 | 1.17E-01 | 2.66E+00 | 2.69E+00 | 5.90E-01 | no | Ttc28 | intron_variant | MODIFIER |
| 5 | 111013722 | AX-170329337 | 0 | A | G | 0.456 | 6.29E-02 | 1.17E-01 | 2.66E+00 | 2.69E+00 | 5.90E-01 | no | Ttc28 | intron_variant | MODIFIER |
| 5 | 111014838 | AX-170402417 | 0 | C | T | 0.456 | 6.29E-02 | 1.17E-01 | 2.66E+00 | 2.69E+00 | 5.90E-01 | no | Ttc28 | intron_variant | MODIFIER |
| 5 | 111056742 | AX-170471985 | 0 | G | A | 0.456 | 6.29E-02 | 1.17E-01 | 2.66E+00 | 2.69E+00 | 5.90E-01 | no | Ttc28 | intron_variant | MODIFIER |
| 5 | 111113943 | AX-170158559 | 0 | C | A | 0.456 | 6.29E-02 | 1.17E-01 | 2.66E+00 | 2.69E+00 | 5.90E-01 | no | Ttc28 | intron_variant | MODIFIER |

| | | | | | | | | | | | | | | | |
|---|---|---|---|---|---|---|---|---|---|---|---|---|---|---|---|
| 5 | 111158220 | AX-170459313 | 0 | C | T | 0.456 | 6.29E-02 | 1.17E-01 | 2.66E+00 | 2.69E+00 | 5.90E-01 | no | Ttc28 | intron_variant | MODIFIER |
| 5 | 111198464 | AX-170483483 | 0 | T | C | 0.456 | 6.29E-02 | 1.17E-01 | 2.66E+00 | 2.69E+00 | 5.90E-01 | no | Ttc28 | intron_variant | MODIFIER |
| 5 | 111276554 | AX-170181811 | 0 | A | G | 0.456 | 6.29E-02 | 1.17E-01 | 2.66E+00 | 2.69E+00 | 5.90E-01 | no | Ttc28 | intron_variant | MODIFIER |
| 5 | 111282845 | AX-97820030 | 0 | A | C | 0.456 | 6.29E-02 | 1.17E-01 | 2.66E+00 | 2.69E+00 | 5.90E-01 | no | Ttc28 | synonymous_variant | LOW |
| 5 | 111286097 | AX-170125513 | 0 | A | G | 0.456 | 6.29E-02 | 1.17E-01 | 2.66E+00 | 2.69E+00 | 5.90E-01 | no | Ttc28 | synonymous_variant | LOW |
| 5 | 111327051 | AX-97835162 | 0 | C | T | 0.456 | 6.29E-02 | 1.17E-01 | 2.66E+00 | 2.69E+00 | 5.90E-01 | no | Pitpnb | upstream_gene_variant | MODIFIER |
| 5 | 111357323 | AX-170485878 | 0 | A | G | 0.456 | 6.29E-02 | 1.17E-01 | 2.66E+00 | 2.69E+00 | 5.90E-01 | no | Pitpnb | intron_variant | MODIFIER |
| 5 | 111884385 | AX-170376368 | 0 | G | A | 0.456 | 6.29E-02 | 1.17E-01 | 2.66E+00 | 2.69E+00 | 5.90E-01 | no | | intergenic_variant | MODIFIER |
| 5 | 111900964 | AX-170396494 | 0 | T | C | 0.456 | 6.29E-02 | 1.17E-01 | 2.66E+00 | 2.69E+00 | 5.90E-01 | no | | intergenic_variant | MODIFIER |
| 5 | 111902193 | AX-170458237 | 0 | C | T | 0.456 | 6.29E-02 | 1.17E-01 | 2.66E+00 | 2.69E+00 | 5.90E-01 | no | | intergenic_variant | MODIFIER |
| 5 | 111909690 | AX-170398468 | 0 | G | T | 0.456 | 6.29E-02 | 1.17E-01 | 2.66E+00 | 2.69E+00 | 5.90E-01 | no | | intergenic_variant | MODIFIER |
| 5 | 111945592 | AX-170025905 | 0 | T | C | 0.456 | 6.29E-02 | 1.17E-01 | 2.66E+00 | 2.69E+00 | 5.90E-01 | no | Gm42488 | upstream_gene_variant | MODIFIER |
| 5 | 111945605 | AX-97827502 | 0 | G | A | 0.456 | 6.29E-02 | 1.17E-01 | 2.66E+00 | 2.69E+00 | 5.90E-01 | no | Gm42488 | upstream_gene_variant | MODIFIER |
| 5 | 111970707 | AX-170249496 | 0 | T | G | 0.456 | 6.29E-02 | 1.17E-01 | 2.66E+00 | 2.69E+00 | 5.90E-01 | no | | intergenic_variant | MODIFIER |
| 5 | 111971082 | AX-170351295 | 0 | C | T | 0.456 | 6.29E-02 | 1.17E-01 | 2.66E+00 | 2.69E+00 | 5.90E-01 | no | | intergenic_variant | MODIFIER |
| 5 | 111988140 | AX-97781678 | 0 | T | C | 0.456 | 6.29E-02 | 1.17E-01 | 2.66E+00 | 2.69E+00 | 5.90E-01 | no | | intergenic_variant | MODIFIER |
| 5 | 112011087 | AX-97857492 | 0 | T | C | 0.456 | 6.29E-02 | 1.17E-01 | 2.66E+00 | 2.69E+00 | 5.90E-01 | no | | intergenic_variant | MODIFIER |
| 5 | 112013791 | AX-170073118 | 0 | C | T | 0.456 | 6.29E-02 | 1.17E-01 | 2.66E+00 | 2.69E+00 | 5.90E-01 | no | | intergenic_variant | MODIFIER |
| 5 | 112015096 | AX-170025899 | 0 | G | A | 0.456 | 6.29E-02 | 1.17E-01 | 2.66E+00 | 2.69E+00 | 5.90E-01 | no | | intergenic_variant | MODIFIER |
| 5 | 112029380 | AX-97802373 | 0 | C | T | 0.456 | 6.29E-02 | 1.17E-01 | 2.66E+00 | 2.69E+00 | 5.90E-01 | no | | regulatory_region_variant | MODIFIER |
| 5 | 112051303 | AX-170165932 | 0 | G | A | 0.456 | 6.29E-02 | 1.17E-01 | 2.66E+00 | 2.69E+00 | 5.90E-01 | no | | regulatory_region_variant | MODIFIER |
| 5 | 112094018 | AX-170293723 | 0 | A | G | 0.456 | 6.29E-02 | 1.17E-01 | 2.66E+00 | 2.69E+00 | 5.90E-01 | no | | regulatory_region_variant | MODIFIER |
| 5 | 112107851 | AX-170025898 | 0 | C | T | 0.456 | 6.29E-02 | 1.17E-01 | 2.66E+00 | 2.69E+00 | 5.90E-01 | no | | regulatory_region_variant | MODIFIER |
| 5 | 112134825 | AX-97833195 | 0 | G | A | 0.456 | 6.29E-02 | 1.17E-01 | 2.66E+00 | 2.69E+00 | 5.90E-01 | no | | regulatory_region_variant | MODIFIER |
| 5 | 112135329 | AX-170181240 | 0 | T | C | 0.456 | 6.29E-02 | 1.17E-01 | 2.66E+00 | 2.69E+00 | 5.90E-01 | no | | intergenic_variant | MODIFIER |
| 5 | 112138916 | AX-170025895 | 0 | G | A | 0.456 | 6.29E-02 | 1.17E-01 | 2.66E+00 | 2.69E+00 | 5.90E-01 | no | | intergenic_variant | MODIFIER |
| 5 | 112147668 | AX-97851404 | 0 | C | T | 0.456 | 6.29E-02 | 1.17E-01 | 2.66E+00 | 2.69E+00 | 5.90E-01 | no | | intergenic_variant | MODIFIER |
| 5 | 112162571 | AX-170025901 | 0 | T | A | 0.456 | 6.29E-02 | 1.17E-01 | 2.66E+00 | 2.69E+00 | 5.90E-01 | no | 1700016B01Rik | intron_variant,non_coding_transcript_variant | MODIFIER |
| 5 | 112178894 | AX-170025903 | 0 | G | T | 0.456 | 6.29E-02 | 1.17E-01 | 2.66E+00 | 2.69E+00 | 5.90E-01 | no | | regulatory_region_variant | MODIFIER |
| 5 | 112179146 | AX-170025896 | 0 | G | A | 0.456 | 6.29E-02 | 1.17E-01 | 2.66E+00 | 2.69E+00 | 5.90E-01 | no | | regulatory_region_variant | MODIFIER |
| 5 | 112182366 | AX-97850272 | 0 | T | C | 0.456 | 6.29E-02 | 1.17E-01 | 2.66E+00 | 2.69E+00 | 5.90E-01 | no | | intergenic_variant | MODIFIER |
| 5 | 112260273 | AX-97856961 | 0 | C | T | 0.456 | 6.29E-02 | 1.17E-01 | 2.66E+00 | 2.69E+00 | 5.90E-01 | no | Crybb1 | intron_variant | MODIFIER |
| 5 | 112294664 | AX-170485915 | 0 | C | T | 0.456 | 6.29E-02 | 1.17E-01 | 2.66E+00 | 2.69E+00 | 5.90E-01 | no | Tpst2 | intron_variant | MODIFIER |
| 5 | 112299966 | AX-170296537 | 0 | G | A | 0.456 | 6.29E-02 | 1.17E-01 | 2.66E+00 | 2.69E+00 | 5.90E-01 | no | Tpst2 | intron_variant | MODIFIER |
| 5 | 112318410 | AX-170347353 | 0 | A | G | 0.456 | 6.29E-02 | 1.17E-01 | 2.66E+00 | 2.69E+00 | 5.90E-01 | no | Tpst2 | downstream_gene_variant | MODIFIER |
| 5 | 112329733 | AX-97841353 | 0 | T | C | 0.456 | 6.29E-02 | 1.17E-01 | 2.66E+00 | 2.69E+00 | 5.90E-01 | no | Tfip11 | intron_variant | MODIFIER |
| 5 | 112331547 | AX-170300789 | 0 | A | T | 0.456 | 6.29E-02 | 1.17E-01 | 2.66E+00 | 2.69E+00 | 5.90E-01 | no | Tfip11 | intron_variant | MODIFIER |
| 5 | 112332352 | AX-170486019 | 0 | G | A | 0.456 | 6.29E-02 | 1.17E-01 | 2.66E+00 | 2.69E+00 | 5.90E-01 | no | Tfip11 | intron_variant | MODIFIER |
| 5 | 112373843 | AX-170399075 | 0 | A | G | 0.456 | 6.29E-02 | 1.17E-01 | 2.66E+00 | 2.69E+00 | 5.90E-01 | no | Hps4 | intron_variant | MODIFIER |
| 5 | 112382575 | AX-170289435 | 0 | C | G | 0.456 | 6.29E-02 | 1.17E-01 | 2.66E+00 | 2.69E+00 | 5.90E-01 | no | Asphd2 | downstream_gene_variant | MODIFIER |
| 5 | 112382719 | AX-97788467 | 0 | T | C | 0.456 | 6.29E-02 | 1.17E-01 | 2.66E+00 | 2.69E+00 | 5.90E-01 | no | Asphd2 | downstream_gene_variant | MODIFIER |
| 5 | 112384972 | AX-170208339 | 0 | C | T | 0.456 | 6.29E-02 | 1.17E-01 | 2.66E+00 | 2.69E+00 | 5.90E-01 | no | Asphd2 | 3_prime_UTR_variant | MODIFIER |
| 5 | 112385761 | AX-170073127 | 0 | T | C | 0.456 | 6.29E-02 | 1.17E-01 | 2.66E+00 | 2.69E+00 | 5.90E-01 | no | Asphd2 | synonymous_variant | LOW |
| 5 | 112390167 | AX-170195976 | 0 | T | G | 0.456 | 6.29E-02 | 1.17E-01 | 2.66E+00 | 2.69E+00 | 5.90E-01 | no | Asphd2 | intron_variant | MODIFIER |
| 5 | 112391067 | AX-170402410 | 0 | A | G | 0.456 | 6.29E-02 | 1.17E-01 | 2.66E+00 | 2.69E+00 | 5.90E-01 | no | Asphd2 | intron_variant | MODIFIER |
| 5 | 112796209 | AX-97800709 | 0 | G | A | 0.456 | 6.29E-02 | 1.17E-01 | 2.66E+00 | 2.69E+00 | 5.90E-01 | no | Myo18b | intron_variant | MODIFIER |
| 5 | 112796470 | AX-170492946 | 0 | C | T | 0.456 | 6.29E-02 | 1.17E-01 | 2.66E+00 | 2.69E+00 | 5.90E-01 | no | Myo18b | intron_variant | MODIFIER |
| 5 | 112822246 | AX-170163193 | 0 | C | A | 0.456 | 6.29E-02 | 1.17E-01 | 2.66E+00 | 2.69E+00 | 5.90E-01 | no | Myo18b | intron_variant | MODIFIER |
| 5 | 112822464 | AX-170099737 | 0 | C | T | 0.456 | 6.29E-02 | 1.17E-01 | 2.66E+00 | 2.69E+00 | 5.90E-01 | no | Myo18b | intron_variant | MODIFIER |
| 5 | 112824324 | AX-170187029 | 0 | T | C | 0.456 | 6.29E-02 | 1.17E-01 | 2.66E+00 | 2.69E+00 | 5.90E-01 | no | Myo18b | intron_variant | MODIFIER |
| 5 | 112824426 | AX-170121037 | 0 | T | C | 0.456 | 6.29E-02 | 1.17E-01 | 2.66E+00 | 2.69E+00 | 5.90E-01 | no | Myo18b | intron_variant | MODIFIER |
| 5 | 112827958 | AX-170266385 | 0 | T | C | 0.456 | 6.29E-02 | 1.17E-01 | 2.66E+00 | 2.69E+00 | 5.90E-01 | no | Myo18b | intron_variant | MODIFIER |
| 5 | 112830829 | AX-170189266 | 0 | A | T | 0.456 | 6.29E-02 | 1.17E-01 | 2.66E+00 | 2.69E+00 | 5.90E-01 | no | Myo18b | intron_variant | MODIFIER |
| 5 | 112831327 | AX-170484161 | 0 | T | C | 0.456 | 6.29E-02 | 1.17E-01 | 2.66E+00 | 2.69E+00 | 5.90E-01 | no | Myo18b | intron_variant | MODIFIER |

| | | | | | | | | | | | | | | | |
|---|---|---|---|---|---|---|---|---|---|---|---|---|---|---|---|
| 5 | 112936177 | AX-170298459 | 0 | G | T | 0.456 | 6.29E-02 | 1.17E-01 | 2.66E+00 | 2.69E+00 | 5.90E-01 | no | Grk3 | intron_variant | MODIFIER |
| 5 | 112947096 | AX-170239057 | 0 | G | A | 0.456 | 6.29E-02 | 1.17E-01 | 2.66E+00 | 2.69E+00 | 5.90E-01 | no | Grk3 | intron_variant | MODIFIER |
| 5 | 112948441 | AX-170233865 | 0 | A | C | 0.456 | 6.29E-02 | 1.17E-01 | 2.66E+00 | 2.69E+00 | 5.90E-01 | no | Grk3 | intron_variant | MODIFIER |
| 5 | 112949063 | AX-170175382 | 0 | T | C | 0.456 | 6.29E-02 | 1.17E-01 | 2.66E+00 | 2.69E+00 | 5.90E-01 | no | Grk3 | intron_variant | MODIFIER |
| 5 | 112953448 | AX-170383479 | 0 | C | A | 0.456 | 6.29E-02 | 1.17E-01 | 2.66E+00 | 2.69E+00 | 5.90E-01 | no | Grk3 | intron_variant | MODIFIER |
| 5 | 112957056 | AX-170385127 | 0 | T | C | 0.456 | 6.29E-02 | 1.17E-01 | 2.66E+00 | 2.69E+00 | 5.90E-01 | no | Grk3 | intron_variant | MODIFIER |
| 5 | 112959685 | AX-170265685 | 0 | A | G | 0.456 | 6.29E-02 | 1.17E-01 | 2.66E+00 | 2.69E+00 | 5.90E-01 | no | Grk3 | intron_variant | MODIFIER |
| 5 | 112962108 | AX-170212717 | 0 | A | G | 0.456 | 6.29E-02 | 1.17E-01 | 2.66E+00 | 2.69E+00 | 5.90E-01 | no | Grk3 | intron_variant | MODIFIER |
| 5 | 112962456 | AX-170471627 | 0 | G | A | 0.456 | 6.29E-02 | 1.17E-01 | 2.66E+00 | 2.69E+00 | 5.90E-01 | no | Grk3 | intron_variant | MODIFIER |
| 5 | 112963704 | AX-170455232 | 0 | G | A | 0.456 | 6.29E-02 | 1.17E-01 | 2.66E+00 | 2.69E+00 | 5.90E-01 | no | Grk3 | intron_variant | MODIFIER |
| 5 | 112974586 | AX-170490079 | 0 | G | A | 0.456 | 6.29E-02 | 1.17E-01 | 2.66E+00 | 2.69E+00 | 5.90E-01 | no | Grk3 | intron_variant | MODIFIER |
| 5 | 112974628 | AX-170282114 | 0 | C | A | 0.456 | 6.29E-02 | 1.17E-01 | 2.66E+00 | 2.69E+00 | 5.90E-01 | no | Grk3 | intron_variant | MODIFIER |
| 5 | 112981990 | AX-97822992 | 0 | A | G | 0.456 | 6.29E-02 | 1.17E-01 | 2.66E+00 | 2.69E+00 | 5.90E-01 | no | Grk3 | intron_variant | MODIFIER |
| 5 | 112984727 | AX-170169417 | 0 | A | G | 0.456 | 6.29E-02 | 1.17E-01 | 2.66E+00 | 2.69E+00 | 5.90E-01 | no | Grk3 | intron_variant | MODIFIER |
| 5 | 112987104 | AX-170447197 | 0 | C | T | 0.456 | 6.29E-02 | 1.17E-01 | 2.66E+00 | 2.69E+00 | 5.90E-01 | no | Grk3 | intron_variant | MODIFIER |
| 5 | 113017855 | AX-170377553 | 0 | T | C | 0.456 | 6.29E-02 | 1.17E-01 | 2.66E+00 | 2.69E+00 | 5.90E-01 | no | Grk3 | upstream_gene_variant | MODIFIER |
| 5 | 113042110 | AX-170441313 | 0 | C | T | 0.456 | 6.29E-02 | 1.17E-01 | 2.66E+00 | 2.69E+00 | 5.90E-01 | no | Gm22740 | upstream_gene_variant | MODIFIER |
| 5 | 113042319 | AX-170267255 | 0 | A | T | 0.456 | 6.29E-02 | 1.17E-01 | 2.66E+00 | 2.69E+00 | 5.90E-01 | no | Gm22740 | upstream_gene_variant | MODIFIER |
| 5 | 113042434 | AX-170278505 | 0 | A | G | 0.456 | 6.29E-02 | 1.17E-01 | 2.66E+00 | 2.69E+00 | 5.90E-01 | no | Gm22740 | upstream_gene_variant | MODIFIER |
| 5 | 113048425 | AX-170450077 | 0 | C | G | 0.456 | 6.29E-02 | 1.17E-01 | 2.66E+00 | 2.69E+00 | 5.90E-01 | no | Gm22740 | downstream_gene_variant | MODIFIER |
| 5 | 113049837 | AX-168412870 | 0 | G | A | 0.456 | 6.29E-02 | 1.17E-01 | 2.66E+00 | 2.69E+00 | 5.90E-01 | no | | regulatory_region_variant | MODIFIER |
| 5 | 113063790 | AX-170527247 | 0 | A | G | 0.456 | 6.29E-02 | 1.17E-01 | 2.66E+00 | 2.69E+00 | 5.90E-01 | no | Crybb2 | intron_variant | MODIFIER |
| 5 | 113068226 | AX-170369394 | 0 | A | G | 0.456 | 6.29E-02 | 1.17E-01 | 2.66E+00 | 2.69E+00 | 5.90E-01 | no | Crybb2 | intron_variant | MODIFIER |
| 5 | 113074131 | AX-170437290 | 0 | A | T | 0.456 | 6.29E-02 | 1.17E-01 | 2.66E+00 | 2.69E+00 | 5.90E-01 | no | Crybb3 | downstream_gene_variant | MODIFIER |
| 5 | 113089851 | AX-170176209 | 0 | G | T | 0.456 | 6.29E-02 | 1.17E-01 | 2.66E+00 | 2.69E+00 | 5.90E-01 | no | 2900026A02Rik | 3_prime_UTR_variant | MODIFIER |
| 5 | 113140808 | AX-97814145 | 0 | C | T | 0.456 | 6.29E-02 | 1.17E-01 | 2.66E+00 | 2.69E+00 | 5.90E-01 | no | 2900026A02Rik | intron_variant | MODIFIER |
| 5 | 113156032 | AX-170357467 | 0 | C | T | 0.456 | 6.29E-02 | 1.17E-01 | 2.66E+00 | 2.69E+00 | 5.90E-01 | no | 2900026A02Rik | intron_variant | MODIFIER |
| 5 | 113191266 | AX-170150909 | 0 | G | C | 0.456 | 6.29E-02 | 1.17E-01 | 2.66E+00 | 2.69E+00 | 5.90E-01 | no | 2900026A02Rik | synonymous_variant | LOW |
| 5 | 113194870 | AX-170365513 | 0 | C | T | 0.456 | 6.29E-02 | 1.17E-01 | 2.66E+00 | 2.69E+00 | 5.90E-01 | no | 2900026A02Rik | intron_variant | MODIFIER |
| 5 | 113250403 | AX-170321828 | 0 | A | G | 0.456 | 6.29E-02 | 1.17E-01 | 2.66E+00 | 2.69E+00 | 5.90E-01 | no | Sgsm1 | intron_variant | MODIFIER |
| 5 | 113253014 | AX-170376298 | 0 | T | C | 0.456 | 6.29E-02 | 1.17E-01 | 2.66E+00 | 2.69E+00 | 5.90E-01 | no | Sgsm1 | synonymous_variant | LOW |
| 5 | 113256076 | AX-170188287 | 0 | C | T | 0.456 | 6.29E-02 | 1.17E-01 | 2.66E+00 | 2.69E+00 | 5.90E-01 | no | Sgsm1 | intron_variant | MODIFIER |
| 5 | 113256769 | AX-97797128 | 0 | G | A | 0.456 | 6.29E-02 | 1.17E-01 | 2.66E+00 | 2.69E+00 | 5.90E-01 | no | Sgsm1 | intron_variant | MODIFIER |
| 5 | 113263067 | AX-170160586 | 0 | C | T | 0.456 | 6.29E-02 | 1.17E-01 | 2.66E+00 | 2.69E+00 | 5.90E-01 | no | Sgsm1 | intron_variant | MODIFIER |
| 5 | 113300048 | AX-170025914 | 0 | A | C | 0.456 | 6.29E-02 | 1.17E-01 | 2.66E+00 | 2.69E+00 | 5.90E-01 | no | Sgsm1 | intron_variant | MODIFIER |
| 5 | 113360410 | AX-170073152 | 0 | T | C | 0.456 | 6.29E-02 | 1.17E-01 | 2.66E+00 | 2.69E+00 | 5.90E-01 | no | Aym1 | downstream_gene_variant | MODIFIER |
| 5 | 113360478 | AX-170073153 | 0 | C | A | 0.456 | 6.29E-02 | 1.17E-01 | 2.66E+00 | 2.69E+00 | 5.90E-01 | no | Aym1 | downstream_gene_variant | MODIFIER |
| 5 | 113364516 | AX-97832800 | 0 | G | A | 0.456 | 6.29E-02 | 1.17E-01 | 2.66E+00 | 2.69E+00 | 5.90E-01 | no | | intergenic_variant | MODIFIER |
| 5 | 113374097 | AX-170319415 | 0 | C | T | 0.456 | 6.29E-02 | 1.17E-01 | 2.66E+00 | 2.69E+00 | 5.90E-01 | no | | regulatory_region_variant | MODIFIER |
| 5 | 113380563 | AX-170180836 | 0 | T | C | 0.456 | 6.29E-02 | 1.17E-01 | 2.66E+00 | 2.69E+00 | 5.90E-01 | no | | regulatory_region_variant | MODIFIER |
| 5 | 113393196 | AX-170352493 | 0 | C | T | 0.456 | 6.29E-02 | 1.17E-01 | 2.66E+00 | 2.69E+00 | 5.90E-01 | no | | intergenic_variant | MODIFIER |
| 5 | 113393737 | AX-170448887 | 0 | T | C | 0.456 | 6.29E-02 | 1.17E-01 | 2.66E+00 | 2.69E+00 | 5.90E-01 | no | | intergenic_variant | MODIFIER |
| 5 | 113396243 | AX-170334773 | 0 | G | T | 0.456 | 6.29E-02 | 1.17E-01 | 2.66E+00 | 2.69E+00 | 5.90E-01 | no | | regulatory_region_variant | MODIFIER |
| 5 | 113401704 | AX-170339887 | 0 | C | T | 0.456 | 6.29E-02 | 1.17E-01 | 2.66E+00 | 2.69E+00 | 5.90E-01 | no | | intergenic_variant | MODIFIER |
| 5 | 113407588 | AX-170328720 | 0 | T | C | 0.456 | 6.29E-02 | 1.17E-01 | 2.66E+00 | 2.69E+00 | 5.90E-01 | no | | regulatory_region_variant | MODIFIER |
| 5 | 113411054 | AX-170338320 | 0 | A | G | 0.456 | 6.29E-02 | 1.17E-01 | 2.66E+00 | 2.69E+00 | 5.90E-01 | no | | regulatory_region_variant | MODIFIER |
| 5 | 113411704 | AX-170484377 | 0 | T | C | 0.456 | 6.29E-02 | 1.17E-01 | 2.66E+00 | 2.69E+00 | 5.90E-01 | no | | regulatory_region_variant | MODIFIER |
| 5 | 113428855 | AX-170120744 | 0 | G | C | 0.456 | 6.29E-02 | 1.17E-01 | 2.66E+00 | 2.69E+00 | 5.90E-01 | no | | intergenic_variant | MODIFIER |
| 5 | 113434356 | AX-97832229 | 0 | C | T | 0.456 | 6.29E-02 | 1.17E-01 | 2.66E+00 | 2.69E+00 | 5.90E-01 | no | | intergenic_variant | MODIFIER |
| 5 | 113635586 | AX-170073160 | 0 | T | C | 0.456 | 6.29E-02 | 1.17E-01 | 2.66E+00 | 2.69E+00 | 5.90E-01 | no | Cmklr1 | intron_variant | MODIFIER |
| 5 | 113748337 | AX-97802769 | 0 | T | C | 0.456 | 6.29E-02 | 1.17E-01 | 2.66E+00 | 2.69E+00 | 5.90E-01 | no | Sart3 | intron_variant | MODIFIER |
| 5 | 113818928 | AX-170025928 | 0 | C | T | 0.456 | 6.29E-02 | 1.17E-01 | 2.66E+00 | 2.69E+00 | 5.90E-01 | no | Selplg | 3_prime_UTR_variant | MODIFIER |
| 5 | 113831783 | AX-97805374 | 0 | G | A | 0.456 | 6.29E-02 | 1.17E-01 | 2.66E+00 | 2.69E+00 | 5.90E-01 | no | Selplg | upstream_gene_variant | MODIFIER |
| 5 | 113838527 | AX-170025924 | 0 | C | T | 0.456 | 6.29E-02 | 1.17E-01 | 2.66E+00 | 2.69E+00 | 5.90E-01 | no | Coro1c | downstream_gene_variant | MODIFIER |

| | | | | | | | | | | | | | | | |
|---|---|---|---|---|---|---|---|---|---|---|---|---|---|---|---|
| 5 | 113856477 | AX-97828144 | 0 | A | G | 0.456 | 6.29E-02 | 1.17E-01 | 2.66E+00 | 2.69E+00 | 5.90E-01 | no | Coro1c | intron_variant | MODIFIER |
| 5 | 113858504 | AX-170197518 | 0 | T | C | 0.456 | 6.29E-02 | 1.17E-01 | 2.66E+00 | 2.69E+00 | 5.90E-01 | no | Coro1c | intron_variant | MODIFIER |
| 5 | 113859413 | AX-170486233 | 0 | A | C | 0.456 | 6.29E-02 | 1.17E-01 | 2.66E+00 | 2.69E+00 | 5.90E-01 | no | Coro1c | intron_variant | MODIFIER |
| 5 | 113876202 | AX-97819314 | 0 | T | G | 0.456 | 6.29E-02 | 1.17E-01 | 2.66E+00 | 2.69E+00 | 5.90E-01 | no | Coro1c | intron_variant | MODIFIER |
| 5 | 113883565 | AX-170025925 | 0 | T | C | 0.456 | 6.29E-02 | 1.17E-01 | 2.66E+00 | 2.69E+00 | 5.90E-01 | no | Coro1c | intron_variant | MODIFIER |
| 5 | 113895961 | AX-170293444 | 0 | A | C | 0.456 | 6.29E-02 | 1.17E-01 | 2.66E+00 | 2.69E+00 | 5.90E-01 | no | Coro1c | intron_variant | MODIFIER |
| 5 | 113915822 | AX-170173374 | 0 | C | A | 0.456 | 6.29E-02 | 1.17E-01 | 2.66E+00 | 2.69E+00 | 5.90E-01 | no | | intergenic_variant | MODIFIER |
| 5 | 113919336 | AX-170532307 | 0 | T | C | 0.456 | 6.29E-02 | 1.17E-01 | 2.66E+00 | 2.69E+00 | 5.90E-01 | no | | regulatory_region_variant | MODIFIER |
| 5 | 113933138 | AX-170145959 | 0 | G | T | 0.456 | 6.29E-02 | 1.17E-01 | 2.66E+00 | 2.69E+00 | 5.90E-01 | no | Gm22056 | upstream_gene_variant | MODIFIER |
| 5 | 113941783 | AX-170370545 | 0 | A | G | 0.456 | 6.29E-02 | 1.17E-01 | 2.66E+00 | 2.69E+00 | 5.90E-01 | no | Ssh1 | downstream_gene_variant | MODIFIER |
| 5 | 113942299 | AX-170316577 | 0 | C | T | 0.456 | 6.29E-02 | 1.17E-01 | 2.66E+00 | 2.69E+00 | 5.90E-01 | no | Ssh1 | synonymous_variant | LOW |
| 5 | 113968071 | AX-170329749 | 0 | T | C | 0.456 | 6.29E-02 | 1.17E-01 | 2.66E+00 | 2.69E+00 | 5.90E-01 | no | Ssh1 | intron_variant | MODIFIER |
| 5 | 113974912 | AX-170373175 | 0 | C | T | 0.456 | 6.29E-02 | 1.17E-01 | 2.66E+00 | 2.69E+00 | 5.90E-01 | no | Ssh1 | intron_variant | MODIFIER |
| 5 | 113978561 | AX-170458210 | 0 | T | C | 0.456 | 6.29E-02 | 1.17E-01 | 2.66E+00 | 2.69E+00 | 5.90E-01 | no | Ssh1 | intron_variant | MODIFIER |
| 5 | 113989663 | AX-170115058 | 0 | C | T | 0.456 | 6.29E-02 | 1.17E-01 | 2.66E+00 | 2.69E+00 | 5.90E-01 | no | Ssh1 | 5_prime_UTR_variant | MODIFIER |
| 5 | 114019413 | AX-170229476 | 0 | C | T | 0.456 | 6.29E-02 | 1.17E-01 | 2.66E+00 | 2.69E+00 | 5.90E-01 | no | Dao | intron_variant | MODIFIER |
| 5 | 114034692 | AX-170479575 | 0 | G | A | 0.456 | 6.29E-02 | 1.17E-01 | 2.66E+00 | 2.69E+00 | 5.90E-01 | no | Svop | intron_variant | MODIFIER |
| 5 | 114047965 | AX-170393864 | 0 | A | G | 0.456 | 6.29E-02 | 1.17E-01 | 2.66E+00 | 2.69E+00 | 5.90E-01 | no | Svop | intron_variant | MODIFIER |
| 5 | 114070257 | AX-170304452 | 0 | A | G | 0.456 | 6.29E-02 | 1.17E-01 | 2.66E+00 | 2.69E+00 | 5.90E-01 | no | Svop | intron_variant | MODIFIER |
| 5 | 114074574 | AX-170225497 | 0 | A | G | 0.456 | 6.29E-02 | 1.17E-01 | 2.66E+00 | 2.69E+00 | 5.90E-01 | no | Svop | intron_variant | MODIFIER |
| 5 | 114086894 | AX-170247864 | 0 | T | C | 0.456 | 6.29E-02 | 1.17E-01 | 2.66E+00 | 2.69E+00 | 5.90E-01 | no | Svop | intron_variant | MODIFIER |
| 5 | 114087003 | AX-170390538 | 0 | A | G | 0.456 | 6.29E-02 | 1.17E-01 | 2.66E+00 | 2.69E+00 | 5.90E-01 | no | Svop | intron_variant | MODIFIER |
| 5 | 114155214 | AX-170261732 | 0 | G | A | 0.456 | 6.29E-02 | 1.17E-01 | 2.66E+00 | 2.69E+00 | 5.90E-01 | no | Acacb | intron_variant | MODIFIER |
| 5 | 114156286 | AX-170470084 | 0 | A | G | 0.456 | 6.29E-02 | 1.17E-01 | 2.66E+00 | 2.69E+00 | 5.90E-01 | no | Acacb | intron_variant | MODIFIER |
| 5 | 114169922 | AX-97831275 | 0 | T | C | 0.456 | 6.29E-02 | 1.17E-01 | 2.66E+00 | 2.69E+00 | 5.90E-01 | no | Acacb | intron_variant | MODIFIER |
| 5 | 114220473 | AX-170467836 | 0 | G | A | 0.456 | 6.29E-02 | 1.17E-01 | 2.66E+00 | 2.69E+00 | 5.90E-01 | no | Acacb | intron_variant | MODIFIER |
| 5 | 114222053 | AX-97853579 | 0 | C | T | 0.456 | 6.29E-02 | 1.17E-01 | 2.66E+00 | 2.69E+00 | 5.90E-01 | no | Acacb | intron_variant | MODIFIER |
| 5 | 114226763 | AX-170303185 | 0 | T | C | 0.456 | 6.29E-02 | 1.17E-01 | 2.66E+00 | 2.69E+00 | 5.90E-01 | no | Acacb | synonymous_variant | LOW |
| 5 | 114227021 | AX-170485271 | 0 | T | C | 0.456 | 6.29E-02 | 1.17E-01 | 2.66E+00 | 2.69E+00 | 5.90E-01 | no | Acacb | intron_variant | MODIFIER |
| 5 | 114244270 | AX-170353601 | 0 | T | C | 0.456 | 6.29E-02 | 1.17E-01 | 2.66E+00 | 2.69E+00 | 5.90E-01 | no | Acacb | intron_variant | MODIFIER |
| 5 | 114244709 | AX-170229751 | 0 | T | C | 0.456 | 6.29E-02 | 1.17E-01 | 2.66E+00 | 2.69E+00 | 5.90E-01 | no | Acacb | intron_variant | MODIFIER |
| 5 | 114252812 | AX-170262122 | 0 | T | C | 0.456 | 6.29E-02 | 1.17E-01 | 2.66E+00 | 2.69E+00 | 5.90E-01 | no | Acacb | downstream_gene_variant | MODIFIER |
| 5 | 114267565 | AX-170025933 | 0 | A | G | 0.456 | 6.29E-02 | 1.17E-01 | 2.66E+00 | 2.69E+00 | 5.90E-01 | no | Foxn4 | intron_variant | MODIFIER |
| 5 | 114277514 | AX-170145134 | 0 | T | G | 0.456 | 6.29E-02 | 1.17E-01 | 2.66E+00 | 2.69E+00 | 5.90E-01 | no | Foxn4 | upstream_gene_variant | MODIFIER |
| 5 | 114282442 | AX-170481632 | 0 | C | T | 0.456 | 6.29E-02 | 1.17E-01 | 2.66E+00 | 2.69E+00 | 5.90E-01 | no | | intergenic_variant | MODIFIER |
| 5 | 114323570 | AX-170025931 | 0 | C | T | 0.456 | 6.29E-02 | 1.17E-01 | 2.66E+00 | 2.69E+00 | 5.90E-01 | no | Myo1h | intron_variant | MODIFIER |
| 5 | 114323621 | AX-170025930 | 0 | C | T | 0.456 | 6.29E-02 | 1.17E-01 | 2.66E+00 | 2.69E+00 | 5.90E-01 | no | Myo1h | intron_variant | MODIFIER |
| 5 | 114352019 | AX-170339744 | 0 | G | T | 0.456 | 6.29E-02 | 1.17E-01 | 2.66E+00 | 2.69E+00 | 5.90E-01 | no | Myo1h | intron_variant | MODIFIER |
| 5 | 114353508 | AX-170244926 | 0 | T | C | 0.456 | 6.29E-02 | 1.17E-01 | 2.66E+00 | 2.69E+00 | 5.90E-01 | no | Myo1h | intron_variant | MODIFIER |
| 5 | 114354243 | AX-170278919 | 0 | G | T | 0.456 | 6.29E-02 | 1.17E-01 | 2.66E+00 | 2.69E+00 | 5.90E-01 | no | Myo1h | intron_variant | MODIFIER |
| 5 | 114355114 | AX-170132675 | 0 | G | C | 0.456 | 6.29E-02 | 1.17E-01 | 2.66E+00 | 2.69E+00 | 5.90E-01 | no | Myo1h | intron_variant | MODIFIER |
| 5 | 114355277 | AX-170206615 | 0 | A | G | 0.456 | 6.29E-02 | 1.17E-01 | 2.66E+00 | 2.69E+00 | 5.90E-01 | no | Myo1h | intron_variant | MODIFIER |
| 5 | 114367081 | AX-170273431 | 0 | T | C | 0.456 | 6.29E-02 | 1.17E-01 | 2.66E+00 | 2.69E+00 | 5.90E-01 | no | Kctd10 | intron_variant | MODIFIER |
| 5 | 114398209 | AX-170520122 | 0 | G | A | 0.456 | 6.29E-02 | 1.17E-01 | 2.66E+00 | 2.69E+00 | 5.90E-01 | no | Ube3b | intron_variant | MODIFIER |
| 5 | 114400688 | AX-170470636 | 0 | A | G | 0.456 | 6.29E-02 | 1.17E-01 | 2.66E+00 | 2.69E+00 | 5.90E-01 | no | Ube3b | intron_variant | MODIFIER |
| 5 | 114403450 | AX-170095438 | 0 | T | C | 0.456 | 6.29E-02 | 1.17E-01 | 2.66E+00 | 2.69E+00 | 5.90E-01 | no | Ube3b | intron_variant | MODIFIER |
| 5 | 114406572 | AX-170284115 | 0 | G | A | 0.456 | 6.29E-02 | 1.17E-01 | 2.66E+00 | 2.69E+00 | 5.90E-01 | no | Ube3b | intron_variant | MODIFIER |
| 5 | 114409752 | AX-170358338 | 0 | T | C | 0.456 | 6.29E-02 | 1.17E-01 | 2.66E+00 | 2.69E+00 | 5.90E-01 | no | Ube3b | intron_variant | MODIFIER |
| 5 | 114435441 | AX-170475552 | 0 | A | G | 0.456 | 6.29E-02 | 1.17E-01 | 2.66E+00 | 2.69E+00 | 5.90E-01 | no | Mmab | intron_variant | MODIFIER |
| 5 | 114445042 | AX-170432498 | 0 | G | A | 0.456 | 6.29E-02 | 1.17E-01 | 2.66E+00 | 2.69E+00 | 5.90E-01 | no | Mmab | upstream_gene_variant | MODIFIER |
| 5 | 114458627 | AX-97788127 | 0 | A | G | 0.456 | 6.29E-02 | 1.17E-01 | 2.66E+00 | 2.69E+00 | 5.90E-01 | no | Mvk | intron_variant | MODIFIER |
| 5 | 114474111 | AX-97836395 | 0 | G | A | 0.456 | 6.29E-02 | 1.17E-01 | 2.66E+00 | 2.69E+00 | 5.90E-01 | no | | intergenic_variant | MODIFIER |
| 5 | 114566464 | AX-170184539 | 0 | G | A | 0.456 | 6.29E-02 | 1.17E-01 | 2.66E+00 | 2.69E+00 | 5.90E-01 | no | Fam222a | upstream_gene_variant | MODIFIER |
| 5 | 114568909 | AX-170220280 | 0 | G | A | 0.456 | 6.29E-02 | 1.17E-01 | 2.66E+00 | 2.69E+00 | 5.90E-01 | no | Fam222a | intron_variant | MODIFIER |

| | | | | | | | | | | | | | | | |
|---|---|---|---|---|---|---|---|---|---|---|---|---|---|---|---|
| 5 | 114583127 | AX-170153650 | 0 | A | G | 0.456 | 6.29E-02 | 1.17E-01 | 2.66E+00 | 2.69E+00 | 5.90E-01 | no | Fam222a | intron_variant | MODIFIER |
| 5 | 114592180 | AX-170358438 | 0 | A | G | 0.456 | 6.29E-02 | 1.17E-01 | 2.66E+00 | 2.69E+00 | 5.90E-01 | no | Fam222a | intron_variant | MODIFIER |
| 5 | 114592708 | AX-170447908 | 0 | C | T | 0.456 | 6.29E-02 | 1.17E-01 | 2.66E+00 | 2.69E+00 | 5.90E-01 | no | Fam222a | intron_variant | MODIFIER |
| 5 | 114593718 | AX-170271312 | 0 | C | A | 0.456 | 6.29E-02 | 1.17E-01 | 2.66E+00 | 2.69E+00 | 5.90E-01 | no | Fam222a | intron_variant | MODIFIER |
| 5 | 114606362 | AX-170358203 | 0 | A | G | 0.456 | 6.29E-02 | 1.17E-01 | 2.66E+00 | 2.69E+00 | 5.90E-01 | no | Fam222a | intron_variant | MODIFIER |
| 5 | 114645512 | AX-170393793 | 0 | T | C | 0.456 | 6.29E-02 | 1.17E-01 | 2.66E+00 | 2.69E+00 | 5.90E-01 | no | Trpv4 | intron_variant | MODIFIER |
| 5 | 114719606 | AX-170453593 | 0 | A | C | 0.456 | 6.29E-02 | 1.17E-01 | 2.66E+00 | 2.69E+00 | 5.90E-01 | no | Tchp | missense_variant | MODERATE |
| 5 | 114721209 | AX-170282291 | 0 | T | G | 0.456 | 6.29E-02 | 1.17E-01 | 2.66E+00 | 2.69E+00 | 5.90E-01 | no | Tchp | 3_prime_UTR_variant | MODIFIER |
| 5 | 114721341 | AX-170271374 | 0 | G | A | 0.456 | 6.29E-02 | 1.17E-01 | 2.66E+00 | 2.69E+00 | 5.90E-01 | no | Tchp | 3_prime_UTR_variant | MODIFIER |
| 5 | 114770257 | AX-170183701 | 0 | A | C | 0.456 | 6.29E-02 | 1.17E-01 | 2.66E+00 | 2.69E+00 | 5.90E-01 | no | Git2 | intron_variant | MODIFIER |
| 5 | 114795152 | AX-170317802 | 0 | A | G | 0.456 | 6.29E-02 | 1.17E-01 | 2.66E+00 | 2.69E+00 | 5.90E-01 | no | Ankrd13a | intron_variant | MODIFIER |
| 5 | 114795552 | AX-97823742 | 0 | T | C | 0.456 | 6.29E-02 | 1.17E-01 | 2.66E+00 | 2.69E+00 | 5.90E-01 | no | Ankrd13a | intron_variant | MODIFIER |
| 5 | 114809634 | AX-170170638 | 0 | A | G | 0.456 | 6.29E-02 | 1.17E-01 | 2.66E+00 | 2.69E+00 | 5.90E-01 | no | 1500011B03Rik | downstream_gene_variant | MODIFIER |
| 5 | 114875002 | AX-170268880 | 0 | G | A | 0.456 | 6.29E-02 | 1.17E-01 | 2.66E+00 | 2.69E+00 | 5.90E-01 | no | 4930519G04Rik | intron_variant | MODIFIER |
| 5 | 114877586 | AX-170114438 | 0 | A | G | 0.456 | 6.29E-02 | 1.17E-01 | 2.66E+00 | 2.69E+00 | 5.90E-01 | no | 4930519G04Rik | intron_variant | MODIFIER |
| 5 | 114879199 | AX-170073191 | 0 | A | C | 0.456 | 6.29E-02 | 1.17E-01 | 2.66E+00 | 2.69E+00 | 5.90E-01 | no | 4930519G04Rik | intron_variant | MODIFIER |
| 5 | 114879334 | AX-97787392 | 0 | T | C | 0.456 | 6.29E-02 | 1.17E-01 | 2.66E+00 | 2.69E+00 | 5.90E-01 | no | 4930519G04Rik | intron_variant | MODIFIER |
| 5 | 114899026 | AX-170073189 | 0 | C | T | 0.456 | 6.29E-02 | 1.17E-01 | 2.66E+00 | 2.69E+00 | 5.90E-01 | no | Oasl2 | intron_variant | MODIFIER |
| 5 | 114904989 | AX-170405967 | 0 | A | G | 0.456 | 6.29E-02 | 1.17E-01 | 2.66E+00 | 2.69E+00 | 5.90E-01 | no | Oasl2 | missense_variant | MODERATE |
| 5 | 114920937 | AX-170282936 | 0 | A | G | 0.456 | 6.29E-02 | 1.17E-01 | 2.66E+00 | 2.69E+00 | 5.90E-01 | no | Oasl1 | upstream_gene_variant | MODIFIER |
| 5 | 114925104 | AX-170303410 | 0 | T | C | 0.456 | 6.29E-02 | 1.17E-01 | 2.66E+00 | 2.69E+00 | 5.90E-01 | no | Oasl1 | intron_variant | MODIFIER |
| 5 | 114926421 | AX-170222676 | 0 | G | A | 0.456 | 6.29E-02 | 1.17E-01 | 2.66E+00 | 2.69E+00 | 5.90E-01 | no | Oasl1 | intron_variant | MODIFIER |
| 5 | 114942754 | AX-170025935 | 0 | A | T | 0.456 | 6.29E-02 | 1.17E-01 | 2.66E+00 | 2.69E+00 | 5.90E-01 | no | 2210016L21Rik | intron_variant | MODIFIER |
| 5 | 114984316 | AX-170025934 | 0 | A | C | 0.456 | 6.29E-02 | 1.17E-01 | 2.66E+00 | 2.69E+00 | 5.90E-01 | no | Hnf1aos1 | intron_variant,non_coding_transcript_variant | MODIFIER |
| 5 | 114993113 | AX-170025936 | 0 | C | T | 0.456 | 6.29E-02 | 1.17E-01 | 2.66E+00 | 2.69E+00 | 5.90E-01 | no | Hnf1aos1 | intron_variant,non_coding_transcript_variant | MODIFIER |
| 5 | 114993230 | AX-170025938 | 0 | C | T | 0.456 | 6.29E-02 | 1.17E-01 | 2.66E+00 | 2.69E+00 | 5.90E-01 | no | Hnf1aos1 | intron_variant,non_coding_transcript_variant | MODIFIER |
| 5 | 115031787 | AX-170025937 | 0 | A | C | 0.456 | 6.29E-02 | 1.17E-01 | 2.66E+00 | 2.69E+00 | 5.90E-01 | no | Sppl3 | intron_variant | MODIFIER |
| 5 | 115033376 | AX-170025939 | 0 | A | C | 0.456 | 6.29E-02 | 1.17E-01 | 2.66E+00 | 2.69E+00 | 5.90E-01 | no | Sppl3 | intron_variant | MODIFIER |
| 5 | 115036153 | AX-170448951 | 0 | T | C | 0.456 | 6.29E-02 | 1.17E-01 | 2.66E+00 | 2.69E+00 | 5.90E-01 | no | Sppl3 | intron_variant | MODIFIER |
| 5 | 115095804 | AX-170241019 | 0 | A | G | 0.456 | 6.29E-02 | 1.17E-01 | 2.66E+00 | 2.69E+00 | 5.90E-01 | no | Sppl3 | intron_variant | MODIFIER |
| 5 | 115096329 | AX-170222552 | 0 | A | G | 0.456 | 6.29E-02 | 1.17E-01 | 2.66E+00 | 2.69E+00 | 5.90E-01 | no | Sppl3 | intron_variant | MODIFIER |
| 5 | 115156408 | AX-170447170 | 0 | G | A | 0.456 | 6.29E-02 | 1.17E-01 | 2.66E+00 | 2.69E+00 | 5.90E-01 | no | Mlec | intron_variant | MODIFIER |
| 5 | 115196264 | AX-170164480 | 0 | G | A | 0.456 | 6.29E-02 | 1.17E-01 | 2.66E+00 | 2.69E+00 | 5.90E-01 | no | Cabp1 | upstream_gene_variant | MODIFIER |
| 5 | 115212768 | AX-170494851 | 0 | T | C | 0.456 | 6.29E-02 | 1.17E-01 | 2.66E+00 | 2.69E+00 | 5.90E-01 | no | | intergenic_variant | MODIFIER |
| 5 | 115218669 | AX-170392822 | 0 | C | T | 0.456 | 6.29E-02 | 1.17E-01 | 2.66E+00 | 2.69E+00 | 5.90E-01 | no | | intergenic_variant | MODIFIER |
| 5 | 115248571 | AX-170333312 | 0 | G | A | 0.456 | 6.29E-02 | 1.17E-01 | 2.66E+00 | 2.69E+00 | 5.90E-01 | no | Rnf10 | intron_variant | MODIFIER |
| 5 | 115250375 | AX-170109414 | 0 | C | A | 0.456 | 6.29E-02 | 1.17E-01 | 2.66E+00 | 2.69E+00 | 5.90E-01 | no | Rnf10 | intron_variant | MODIFIER |
| 5 | 115270859 | AX-170107721 | 0 | A | C | 0.456 | 6.29E-02 | 1.17E-01 | 2.66E+00 | 2.69E+00 | 5.90E-01 | no | Rnf10 | intron_variant | MODIFIER |
| 5 | 115299203 | AX-170409807 | 0 | T | C | 0.456 | 6.29E-02 | 1.17E-01 | 2.66E+00 | 2.69E+00 | 5.90E-01 | no | Dynll1 | intron_variant | MODIFIER |
| 5 | 115313501 | AX-97837649 | 0 | G | A | 0.456 | 6.29E-02 | 1.17E-01 | 2.66E+00 | 2.69E+00 | 5.90E-01 | no | Gm13830 | intron_variant,non_coding_transcript_variant | MODIFIER |
| 5 | 115384061 | AX-170196078 | 0 | G | C | 0.456 | 6.29E-02 | 1.17E-01 | 2.66E+00 | 2.69E+00 | 5.90E-01 | no | Gm13832 | intron_variant,non_coding_transcript_variant | MODIFIER |
| 5 | 115413371 | AX-170174040 | 0 | G | C | 0.456 | 6.29E-02 | 1.17E-01 | 2.66E+00 | 2.69E+00 | 5.90E-01 | no | Gm13832 | intron_variant,non_coding_transcript_variant | MODIFIER |
| 5 | 115457525 | AX-170331836 | 0 | G | A | 0.456 | 6.29E-02 | 1.17E-01 | 2.66E+00 | 2.69E+00 | 5.90E-01 | no | Msi1 | downstream_gene_variant | MODIFIER |
| 5 | 115474756 | AX-170488144 | 0 | C | G | 0.456 | 6.29E-02 | 1.17E-01 | 2.66E+00 | 2.69E+00 | 5.90E-01 | no | Pla2g1b | downstream_gene_variant | MODIFIER |
| 5 | 115525293 | AX-170356759 | 0 | C | T | 0.456 | 6.29E-02 | 1.17E-01 | 2.66E+00 | 2.69E+00 | 5.90E-01 | no | Pxn | intron_variant | MODIFIER |
| 5 | 115554285 | AX-169999728 | 0 | T | C | 0.456 | 6.29E-02 | 1.17E-01 | 2.66E+00 | 2.69E+00 | 5.90E-01 | no | Pxn | 3_prime_UTR_variant | MODIFIER |
| 5 | 115557558 | AX-97846030 | 0 | A | G | 0.456 | 6.29E-02 | 1.17E-01 | 2.66E+00 | 2.69E+00 | 5.90E-01 | no | Pxn | downstream_gene_variant | MODIFIER |
| 5 | 115578361 | AX-170159035 | 0 | T | C | 0.456 | 6.29E-02 | 1.17E-01 | 2.66E+00 | 2.69E+00 | 5.90E-01 | no | Gcn1l1 | intron_variant | MODIFIER |
| 5 | 115581298 | AX-170073209 | 0 | A | C | 0.456 | 6.29E-02 | 1.17E-01 | 2.66E+00 | 2.69E+00 | 5.90E-01 | no | Gcn1l1 | intron_variant | MODIFIER |
| 5 | 115614917 | AX-170303688 | 0 | T | C | 0.456 | 6.29E-02 | 1.17E-01 | 2.66E+00 | 2.69E+00 | 5.90E-01 | no | Gcn1l1 | synonymous_variant | LOW |
| 5 | 115615069 | AX-170189322 | 0 | C | T | 0.456 | 6.29E-02 | 1.17E-01 | 2.66E+00 | 2.69E+00 | 5.90E-01 | no | Gcn1l1 | intron_variant | MODIFIER |
| 5 | 115620518 | AX-170156565 | 0 | G | A | 0.456 | 6.29E-02 | 1.17E-01 | 2.66E+00 | 2.69E+00 | 5.90E-01 | no | Gcn1l1 | intron_variant | MODIFIER |
| 5 | 115623991 | AX-170025943 | 0 | T | C | 0.456 | 6.29E-02 | 1.17E-01 | 2.66E+00 | 2.69E+00 | 5.90E-01 | no | Gcn1l1 | downstream_gene_variant | MODIFIER |
| 5 | 115653958 | AX-97809496 | 0 | A | C | 0.456 | 6.29E-02 | 1.17E-01 | 2.66E+00 | 2.69E+00 | 5.90E-01 | no | Bicdl1 | intron_variant | MODIFIER |

| | | | | | | | | | | | | | | | |
|---|---|---|---|---|---|---|---|---|---|---|---|---|---|---|---|
| 5 | 115688321 | AX-97812809 | 0 | C | A | 0.456 | 6.29E-02 | 1.17E-01 | 2.66E+00 | 2.69E+00 | 5.90E-01 | no | Bicdl1 | intron_variant | MODIFIER |
| 5 | 115716398 | AX-170025946 | 0 | C | T | 0.456 | 6.29E-02 | 1.17E-01 | 2.66E+00 | 2.69E+00 | 5.90E-01 | no | Bicdl1 | intron_variant | MODIFIER |
| 5 | 115725905 | AX-97800156 | 0 | G | A | 0.456 | 6.29E-02 | 1.17E-01 | 2.66E+00 | 2.69E+00 | 5.90E-01 | no | Bicdl1 | intron_variant | MODIFIER |
| 5 | 115727453 | AX-170025944 | 0 | A | C | 0.456 | 6.29E-02 | 1.17E-01 | 2.66E+00 | 2.69E+00 | 5.90E-01 | no | Bicdl1 | intron_variant | MODIFIER |
| 5 | 115891586 | AX-170449949 | 0 | G | A | 0.456 | 6.29E-02 | 1.17E-01 | 2.66E+00 | 2.69E+00 | 5.90E-01 | no | Cit | intron_variant | MODIFIER |
| 5 | 115922039 | AX-170130029 | 0 | C | T | 0.456 | 6.29E-02 | 1.17E-01 | 2.66E+00 | 2.69E+00 | 5.90E-01 | no | Cit | intron_variant | MODIFIER |
| 5 | 115923667 | AX-170268827 | 0 | A | G | 0.456 | 6.29E-02 | 1.17E-01 | 2.66E+00 | 2.69E+00 | 5.90E-01 | no | Cit | intron_variant | MODIFIER |
| 5 | 115925040 | AX-170462439 | 0 | T | C | 0.456 | 6.29E-02 | 1.17E-01 | 2.66E+00 | 2.69E+00 | 5.90E-01 | no | Cit | intron_variant | MODIFIER |
| 5 | 115925110 | AX-170073216 | 0 | G | C | 0.456 | 6.29E-02 | 1.17E-01 | 2.66E+00 | 2.69E+00 | 5.90E-01 | no | Cit | intron_variant | MODIFIER |
| 5 | 115925150 | AX-170073214 | 0 | T | C | 0.456 | 6.29E-02 | 1.17E-01 | 2.66E+00 | 2.69E+00 | 5.90E-01 | no | Cit | intron_variant | MODIFIER |
| 5 | 115931115 | AX-170073215 | 0 | G | A | 0.456 | 6.29E-02 | 1.17E-01 | 2.66E+00 | 2.69E+00 | 5.90E-01 | no | Cit | intron_variant | MODIFIER |
| 5 | 115931983 | AX-170144601 | 0 | G | A | 0.456 | 6.29E-02 | 1.17E-01 | 2.66E+00 | 2.69E+00 | 5.90E-01 | no | Cit | intron_variant | MODIFIER |
| 5 | 115934340 | AX-170138745 | 0 | C | T | 0.456 | 6.29E-02 | 1.17E-01 | 2.66E+00 | 2.69E+00 | 5.90E-01 | no | Cit | intron_variant | MODIFIER |
| 5 | 115946539 | AX-170450511 | 0 | T | G | 0.456 | 6.29E-02 | 1.17E-01 | 2.66E+00 | 2.69E+00 | 5.90E-01 | no | Cit | intron_variant | MODIFIER |
| 5 | 115954959 | AX-170219055 | 0 | C | T | 0.456 | 6.29E-02 | 1.17E-01 | 2.66E+00 | 2.69E+00 | 5.90E-01 | no | Cit | intron_variant | MODIFIER |
| 5 | 115955042 | AX-97821183 | 0 | T | C | 0.456 | 6.29E-02 | 1.17E-01 | 2.66E+00 | 2.69E+00 | 5.90E-01 | no | Cit | intron_variant | MODIFIER |
| 5 | 115955128 | AX-170171732 | 0 | T | C | 0.456 | 6.29E-02 | 1.17E-01 | 2.66E+00 | 2.69E+00 | 5.90E-01 | no | Cit | intron_variant | MODIFIER |
| 5 | 115957804 | AX-170450389 | 0 | G | A | 0.456 | 6.29E-02 | 1.17E-01 | 2.66E+00 | 2.69E+00 | 5.90E-01 | no | Cit | intron_variant | MODIFIER |
| 5 | 115958277 | AX-170482535 | 0 | A | G | 0.456 | 6.29E-02 | 1.17E-01 | 2.66E+00 | 2.69E+00 | 5.90E-01 | no | Cit | intron_variant | MODIFIER |
| 5 | 115960423 | AX-170268739 | 0 | G | A | 0.456 | 6.29E-02 | 1.17E-01 | 2.66E+00 | 2.69E+00 | 5.90E-01 | no | Cit | intron_variant | MODIFIER |
| 5 | 115991372 | AX-170148818 | 0 | T | C | 0.456 | 6.29E-02 | 1.17E-01 | 2.66E+00 | 2.69E+00 | 5.90E-01 | no | Cit | intron_variant | MODIFIER |
| 5 | 115993795 | AX-170475936 | 0 | T | C | 0.456 | 6.29E-02 | 1.17E-01 | 2.66E+00 | 2.69E+00 | 5.90E-01 | no | Cit | intron_variant | MODIFIER |
| 5 | 115996212 | AX-170440363 | 0 | C | T | 0.456 | 6.29E-02 | 1.17E-01 | 2.66E+00 | 2.69E+00 | 5.90E-01 | no | Cit | intron_variant | MODIFIER |
| 5 | 116003338 | AX-170372299 | 0 | G | A | 0.456 | 6.29E-02 | 1.17E-01 | 2.66E+00 | 2.69E+00 | 5.90E-01 | no | Cit | intron_variant | MODIFIER |
| 5 | 116014356 | AX-170180941 | 0 | C | G | 0.456 | 6.29E-02 | 1.17E-01 | 2.66E+00 | 2.69E+00 | 5.90E-01 | no | Prkab1 | 3_prime_UTR_variant | MODIFIER |
| 5 | 116014379 | AX-170282883 | 0 | C | G | 0.456 | 6.29E-02 | 1.17E-01 | 2.66E+00 | 2.69E+00 | 5.90E-01 | no | Prkab1 | 3_prime_UTR_variant | MODIFIER |
| 5 | 116015427 | AX-170151730 | 0 | C | T | 0.456 | 6.29E-02 | 1.17E-01 | 2.66E+00 | 2.69E+00 | 5.90E-01 | no | Prkab1 | intron_variant | MODIFIER |
| 5 | 116015489 | AX-170419618 | 0 | A | C | 0.456 | 6.29E-02 | 1.17E-01 | 2.66E+00 | 2.69E+00 | 5.90E-01 | no | Prkab1 | intron_variant | MODIFIER |
| 5 | 116015687 | AX-97803713 | 0 | G | T | 0.456 | 6.29E-02 | 1.17E-01 | 2.66E+00 | 2.69E+00 | 5.90E-01 | no | Prkab1 | intron_variant | MODIFIER |
| 5 | 116059343 | AX-170218980 | 0 | A | C | 0.456 | 6.29E-02 | 1.17E-01 | 2.66E+00 | 2.69E+00 | 5.90E-01 | no | Tmem233 | intron_variant | MODIFIER |
| 5 | 116079186 | AX-170532626 | 0 | G | A | 0.456 | 6.29E-02 | 1.17E-01 | 2.66E+00 | 2.69E+00 | 5.90E-01 | no | Tmem233 | intron_variant | MODIFIER |
| 5 | 116086210 | AX-170392688 | 0 | T | C | 0.456 | 6.29E-02 | 1.17E-01 | 2.66E+00 | 2.69E+00 | 5.90E-01 | no | Tmem233 | upstream_gene_variant | MODIFIER |
| 5 | 116098593 | AX-170491272 | 0 | G | T | 0.456 | 6.29E-02 | 1.17E-01 | 2.66E+00 | 2.69E+00 | 5.90E-01 | no | | intergenic_variant | MODIFIER |
| 5 | 116099196 | AX-170197435 | 0 | A | G | 0.456 | 6.29E-02 | 1.17E-01 | 2.66E+00 | 2.69E+00 | 5.90E-01 | no | | intergenic_variant | MODIFIER |
| 5 | 116099699 | AX-170241730 | 0 | T | C | 0.456 | 6.29E-02 | 1.17E-01 | 2.66E+00 | 2.69E+00 | 5.90E-01 | no | | intergenic_variant | MODIFIER |
| 5 | 116101137 | AX-97836606 | 0 | T | C | 0.456 | 6.29E-02 | 1.17E-01 | 2.66E+00 | 2.69E+00 | 5.90E-01 | no | | intergenic_variant | MODIFIER |
| 5 | 116106561 | AX-170522689 | 0 | T | C | 0.456 | 6.29E-02 | 1.17E-01 | 2.66E+00 | 2.69E+00 | 5.90E-01 | no | | intergenic_variant | MODIFIER |
| 5 | 116637202 | AX-170243461 | 0 | T | C | 0.456 | 6.29E-02 | 1.17E-01 | 2.66E+00 | 2.69E+00 | 5.90E-01 | no | Gstm2-ps1 | upstream_gene_variant | MODIFIER |
| 5 | 116679379 | AX-170157745 | 0 | G | T | 0.456 | 6.29E-02 | 1.17E-01 | 2.66E+00 | 2.69E+00 | 5.90E-01 | no | Gm43122 | intron_variant,non_coding_transcript_variant | MODIFIER |
| 5 | 116729403 | AX-170359061 | 0 | G | A | 0.456 | 6.29E-02 | 1.17E-01 | 2.66E+00 | 2.69E+00 | 5.90E-01 | no | Gm43122 | intron_variant,non_coding_transcript_variant | MODIFIER |
| 5 | 116730074 | AX-170293183 | 0 | G | A | 0.456 | 6.29E-02 | 1.17E-01 | 2.66E+00 | 2.69E+00 | 5.90E-01 | no | Gm43122 | intron_variant,non_coding_transcript_variant | MODIFIER |
| 5 | 116730119 | AX-170398061 | 0 | G | A | 0.456 | 6.29E-02 | 1.17E-01 | 2.66E+00 | 2.69E+00 | 5.90E-01 | no | Gm43122 | intron_variant,non_coding_transcript_variant | MODIFIER |
| 5 | 116730211 | AX-170520625 | 0 | G | A | 0.456 | 6.29E-02 | 1.17E-01 | 2.66E+00 | 2.69E+00 | 5.90E-01 | no | Gm43122 | intron_variant,non_coding_transcript_variant | MODIFIER |
| 5 | 116742848 | AX-170180710 | 0 | C | T | 0.456 | 6.29E-02 | 1.17E-01 | 2.66E+00 | 2.69E+00 | 5.90E-01 | no | Gm43122 | intron_variant,non_coding_transcript_variant | MODIFIER |
| 5 | 116750836 | AX-170154542 | 0 | G | A | 0.456 | 6.29E-02 | 1.17E-01 | 2.66E+00 | 2.69E+00 | 5.90E-01 | no | Gm43122 | intron_variant,non_coding_transcript_variant | MODIFIER |
| 5 | 116752767 | AX-170025954 | 0 | G | A | 0.456 | 6.29E-02 | 1.17E-01 | 2.66E+00 | 2.69E+00 | 5.90E-01 | no | Gm43122 | intron_variant,non_coding_transcript_variant | MODIFIER |
| 5 | 116756430 | AX-170347491 | 0 | G | T | 0.456 | 6.29E-02 | 1.17E-01 | 2.66E+00 | 2.69E+00 | 5.90E-01 | no | Gm43122 | intron_variant,non_coding_transcript_variant | MODIFIER |
| 5 | 116756709 | AX-97844178 | 0 | G | T | 0.456 | 6.29E-02 | 1.17E-01 | 2.66E+00 | 2.69E+00 | 5.90E-01 | no | Gm43122 | intron_variant,non_coding_transcript_variant | MODIFIER |
| 5 | 116760117 | AX-170395478 | 0 | G | A | 0.456 | 6.29E-02 | 1.17E-01 | 2.66E+00 | 2.69E+00 | 5.90E-01 | no | Gm43122 | intron_variant,non_coding_transcript_variant | MODIFIER |
| 5 | 116762369 | AX-170167309 | 0 | T | C | 0.456 | 6.29E-02 | 1.17E-01 | 2.66E+00 | 2.69E+00 | 5.90E-01 | no | Gm43122 | intron_variant,non_coding_transcript_variant | MODIFIER |
| 5 | 116762669 | AX-170182738 | 0 | T | C | 0.456 | 6.29E-02 | 1.17E-01 | 2.66E+00 | 2.69E+00 | 5.90E-01 | no | Gm43122 | intron_variant,non_coding_transcript_variant | MODIFIER |
| 5 | 116764928 | AX-170073232 | 0 | T | C | 0.456 | 6.29E-02 | 1.17E-01 | 2.66E+00 | 2.69E+00 | 5.90E-01 | no | Gm43122 | intron_variant,non_coding_transcript_variant | MODIFIER |
| 5 | 116770434 | AX-97789463 | 0 | G | A | 0.456 | 6.29E-02 | 1.17E-01 | 2.66E+00 | 2.69E+00 | 5.90E-01 | no | Gm43122 | intron_variant,non_coding_transcript_variant | MODIFIER |
| 5 | 116770836 | AX-170073235 | 0 | T | G | 0.456 | 6.29E-02 | 1.17E-01 | 2.66E+00 | 2.69E+00 | 5.90E-01 | no | Gm43122 | intron_variant,non_coding_transcript_variant | MODIFIER |

| | | | | | | | | | | | | | | | |
|---|---|---|---|---|---|---|---|---|---|---|---|---|---|---|---|
| 5 | 116791191 | AX-170281945 | 0 | T | C | 0.456 | 6.29E-02 | 1.17E-01 | 2.66E+00 | 2.69E+00 | 5.90E-01 | no | Gm43122 | intron_variant,non_coding_transcript_variant | MODIFIER |
| 5 | 116794780 | AX-170476607 | 0 | C | G | 0.456 | 6.29E-02 | 1.17E-01 | 2.66E+00 | 2.69E+00 | 5.90E-01 | no | Gm43122 | intron_variant,non_coding_transcript_variant | MODIFIER |
| 5 | 116801482 | AX-170175785 | 0 | C | T | 0.456 | 6.29E-02 | 1.17E-01 | 2.66E+00 | 2.69E+00 | 5.90E-01 | no | Gm43122 | intron_variant,non_coding_transcript_variant | MODIFIER |
| 5 | 116895539 | AX-170025955 | 0 | T | C | 0.456 | 6.29E-02 | 1.17E-01 | 2.66E+00 | 2.69E+00 | 5.90E-01 | no | | regulatory_region_variant | MODIFIER |
| 5 | 116895707 | AX-170025953 | 0 | A | G | 0.456 | 6.29E-02 | 1.17E-01 | 2.66E+00 | 2.69E+00 | 5.90E-01 | no | | regulatory_region_variant | MODIFIER |
| 5 | 116904980 | AX-170508587 | 0 | C | A | 0.456 | 6.29E-02 | 1.17E-01 | 2.66E+00 | 2.69E+00 | 5.90E-01 | no | | intergenic_variant | MODIFIER |
| 5 | 116923306 | AX-170139847 | 0 | C | A | 0.456 | 6.29E-02 | 1.17E-01 | 2.66E+00 | 2.69E+00 | 5.90E-01 | no | | intergenic_variant | MODIFIER |
| 5 | 116924452 | AX-170025959 | 0 | C | T | 0.456 | 6.29E-02 | 1.17E-01 | 2.66E+00 | 2.69E+00 | 5.90E-01 | no | | intergenic_variant | MODIFIER |
| 5 | 116928925 | AX-170349710 | 0 | G | T | 0.456 | 6.29E-02 | 1.17E-01 | 2.66E+00 | 2.69E+00 | 5.90E-01 | no | | intergenic_variant | MODIFIER |
| 5 | 116929504 | AX-97823429 | 0 | C | A | 0.456 | 6.29E-02 | 1.17E-01 | 2.66E+00 | 2.69E+00 | 5.90E-01 | no | | intergenic_variant | MODIFIER |
| 5 | 116945955 | AX-170261696 | 0 | G | C | 0.456 | 6.29E-02 | 1.17E-01 | 2.66E+00 | 2.69E+00 | 5.90E-01 | no | | regulatory_region_variant | MODIFIER |
| 5 | 116968846 | AX-97821165 | 0 | A | G | 0.456 | 6.29E-02 | 1.17E-01 | 2.66E+00 | 2.69E+00 | 5.90E-01 | no | | intergenic_variant | MODIFIER |
| 5 | 116989788 | AX-170416898 | 0 | A | T | 0.456 | 6.29E-02 | 1.17E-01 | 2.66E+00 | 2.69E+00 | 5.90E-01 | no | | intergenic_variant | MODIFIER |
| 5 | 116995489 | AX-170217866 | 0 | G | A | 0.456 | 6.29E-02 | 1.17E-01 | 2.66E+00 | 2.69E+00 | 5.90E-01 | no | | intergenic_variant | MODIFIER |
| 5 | 116999961 | AX-97793830 | 0 | G | A | 0.456 | 6.29E-02 | 1.17E-01 | 2.66E+00 | 2.69E+00 | 5.90E-01 | no | | intergenic_variant | MODIFIER |
| 5 | 117023175 | AX-170535269 | 0 | C | T | 0.456 | 6.29E-02 | 1.17E-01 | 2.66E+00 | 2.69E+00 | 5.90E-01 | no | | intergenic_variant | MODIFIER |
| 5 | 117024376 | AX-170073239 | 0 | C | T | 0.456 | 6.29E-02 | 1.17E-01 | 2.66E+00 | 2.69E+00 | 5.90E-01 | no | | intergenic_variant | MODIFIER |
| 5 | 117025799 | AX-170073241 | 0 | G | A | 0.456 | 6.29E-02 | 1.17E-01 | 2.66E+00 | 2.69E+00 | 5.90E-01 | no | | intergenic_variant | MODIFIER |
| 5 | 117041080 | AX-170331893 | 0 | T | C | 0.456 | 6.29E-02 | 1.17E-01 | 2.66E+00 | 2.69E+00 | 5.90E-01 | no | | regulatory_region_variant | MODIFIER |
| 5 | 117047842 | AX-170156478 | 0 | G | A | 0.456 | 6.29E-02 | 1.17E-01 | 2.66E+00 | 2.69E+00 | 5.90E-01 | no | | intergenic_variant | MODIFIER |
| 5 | 117048527 | AX-170440710 | 0 | A | G | 0.456 | 6.29E-02 | 1.17E-01 | 2.66E+00 | 2.69E+00 | 5.90E-01 | no | | intergenic_variant | MODIFIER |
| 5 | 117048719 | AX-170349638 | 0 | A | G | 0.456 | 6.29E-02 | 1.17E-01 | 2.66E+00 | 2.69E+00 | 5.90E-01 | no | | intergenic_variant | MODIFIER |
| 5 | 117054889 | AX-170205339 | 0 | C | T | 0.456 | 6.29E-02 | 1.17E-01 | 2.66E+00 | 2.69E+00 | 5.90E-01 | no | Gm7478 | upstream_gene_variant | MODIFIER |
| 5 | 117086315 | AX-97823590 | 0 | C | T | 0.456 | 6.29E-02 | 1.17E-01 | 2.66E+00 | 2.69E+00 | 5.90E-01 | no | | intergenic_variant | MODIFIER |
| 5 | 117105895 | AX-170304667 | 0 | T | A | 0.456 | 6.29E-02 | 1.17E-01 | 2.66E+00 | 2.69E+00 | 5.90E-01 | no | Suds3 | intron_variant | MODIFIER |
| 5 | 117131631 | AX-170497695 | 0 | T | C | 0.456 | 6.29E-02 | 1.17E-01 | 2.66E+00 | 2.69E+00 | 5.90E-01 | no | Taok3 | upstream_gene_variant | MODIFIER |
| 5 | 117136799 | AX-170461025 | 0 | G | A | 0.456 | 6.29E-02 | 1.17E-01 | 2.66E+00 | 2.69E+00 | 5.90E-01 | no | Taok3 | intron_variant | MODIFIER |
| 5 | 117144816 | AX-170359031 | 0 | G | A | 0.456 | 6.29E-02 | 1.17E-01 | 2.66E+00 | 2.69E+00 | 5.90E-01 | no | Taok3 | intron_variant | MODIFIER |
| 5 | 117153736 | AX-170370875 | 0 | C | T | 0.456 | 6.29E-02 | 1.17E-01 | 2.66E+00 | 2.69E+00 | 5.90E-01 | no | Taok3 | intron_variant | MODIFIER |
| 5 | 117155576 | AX-97856675 | 0 | T | C | 0.456 | 6.29E-02 | 1.17E-01 | 2.66E+00 | 2.69E+00 | 5.90E-01 | no | Taok3 | intron_variant | MODIFIER |
| 5 | 117169183 | AX-170281169 | 0 | C | T | 0.456 | 6.29E-02 | 1.17E-01 | 2.66E+00 | 2.69E+00 | 5.90E-01 | no | Taok3 | intron_variant | MODIFIER |
| 5 | 117174737 | AX-170462804 | 0 | G | A | 0.456 | 6.29E-02 | 1.17E-01 | 2.66E+00 | 2.69E+00 | 5.90E-01 | no | Taok3 | intron_variant | MODIFIER |
| 5 | 117178365 | AX-97822727 | 0 | C | T | 0.456 | 6.29E-02 | 1.17E-01 | 2.66E+00 | 2.69E+00 | 5.90E-01 | no | Taok3 | intron_variant | MODIFIER |
| 5 | 117183423 | AX-170374480 | 0 | C | T | 0.456 | 6.29E-02 | 1.17E-01 | 2.66E+00 | 2.69E+00 | 5.90E-01 | no | Taok3 | intron_variant | MODIFIER |
| 5 | 117201206 | AX-97806297 | 0 | G | A | 0.456 | 6.29E-02 | 1.17E-01 | 2.66E+00 | 2.69E+00 | 5.90E-01 | no | Taok3 | intron_variant | MODIFIER |
| 5 | 117206627 | AX-170479240 | 0 | C | G | 0.456 | 6.29E-02 | 1.17E-01 | 2.66E+00 | 2.69E+00 | 5.90E-01 | no | Taok3 | intron_variant | MODIFIER |
| 5 | 117217347 | AX-170299658 | 0 | G | A | 0.456 | 6.29E-02 | 1.17E-01 | 2.66E+00 | 2.69E+00 | 5.90E-01 | no | Taok3 | intron_variant | MODIFIER |
| 5 | 117220339 | AX-170475618 | 0 | T | C | 0.456 | 6.29E-02 | 1.17E-01 | 2.66E+00 | 2.69E+00 | 5.90E-01 | no | Taok3 | intron_variant | MODIFIER |
| 5 | 117225044 | AX-170308120 | 0 | C | G | 0.456 | 6.29E-02 | 1.17E-01 | 2.66E+00 | 2.69E+00 | 5.90E-01 | no | Taok3 | intron_variant | MODIFIER |
| 5 | 117235939 | AX-97839061 | 0 | T | C | 0.456 | 6.29E-02 | 1.17E-01 | 2.66E+00 | 2.69E+00 | 5.90E-01 | no | Taok3 | intron_variant | MODIFIER |
| 5 | 117284097 | AX-170148762 | 0 | A | C | 0.456 | 6.29E-02 | 1.17E-01 | 2.66E+00 | 2.69E+00 | 5.90E-01 | no | Pebp1 | intron_variant | MODIFIER |
| 5 | 117287267 | AX-97799816 | 0 | C | T | 0.456 | 6.29E-02 | 1.17E-01 | 2.66E+00 | 2.69E+00 | 5.90E-01 | no | Pebp1 | intron_variant | MODIFIER |
| 5 | 117295118 | AX-170490888 | 0 | A | G | 0.456 | 6.29E-02 | 1.17E-01 | 2.66E+00 | 2.69E+00 | 5.90E-01 | no | Gm42546 | upstream_gene_variant | MODIFIER |
| 5 | 117295205 | AX-170480228 | 0 | C | T | 0.456 | 6.29E-02 | 1.17E-01 | 2.66E+00 | 2.69E+00 | 5.90E-01 | no | Gm42546 | upstream_gene_variant | MODIFIER |
| 5 | 117302687 | AX-170149944 | 0 | C | T | 0.456 | 6.29E-02 | 1.17E-01 | 2.66E+00 | 2.69E+00 | 5.90E-01 | no | Gm42546 | downstream_gene_variant | MODIFIER |
| 5 | 117303697 | AX-170089590 | 0 | G | A | 0.456 | 6.29E-02 | 1.17E-01 | 2.66E+00 | 2.69E+00 | 5.90E-01 | no | Gm42546 | downstream_gene_variant | MODIFIER |
| 5 | 117349405 | AX-170073248 | 0 | C | G | 0.456 | 6.29E-02 | 1.17E-01 | 2.66E+00 | 2.69E+00 | 5.90E-01 | no | Vsig10 | intron_variant | MODIFIER |
| 5 | 117362331 | AX-170025966 | 0 | A | C | 0.456 | 6.29E-02 | 1.17E-01 | 2.66E+00 | 2.69E+00 | 5.90E-01 | no | Wsb2 | intron_variant | MODIFIER |
| 5 | 117374045 | AX-170451754 | 0 | G | C | 0.456 | 6.29E-02 | 1.17E-01 | 2.66E+00 | 2.69E+00 | 5.90E-01 | no | Wsb2 | intron_variant | MODIFIER |
| 5 | 117383187 | AX-170395048 | 0 | T | C | 0.456 | 6.29E-02 | 1.17E-01 | 2.66E+00 | 2.69E+00 | 5.90E-01 | no | Wsb2 | downstream_gene_variant | MODIFIER |
| 5 | 117406782 | AX-170025970 | 0 | C | A | 0.456 | 6.29E-02 | 1.17E-01 | 2.66E+00 | 2.69E+00 | 5.90E-01 | no | | intergenic_variant | MODIFIER |
| 5 | 117409484 | AX-170467870 | 0 | T | C | 0.456 | 6.29E-02 | 1.17E-01 | 2.66E+00 | 2.69E+00 | 5.90E-01 | no | Ksr2 | upstream_gene_variant | MODIFIER |
| 5 | 117409688 | AX-97794232 | 0 | T | C | 0.456 | 6.29E-02 | 1.17E-01 | 2.66E+00 | 2.69E+00 | 5.90E-01 | no | Ksr2 | upstream_gene_variant | MODIFIER |
| 5 | 117434755 | AX-170095443 | 0 | A | G | 0.456 | 6.29E-02 | 1.17E-01 | 2.66E+00 | 2.69E+00 | 5.90E-01 | no | Ksr2 | intron_variant,non_coding_transcript_variant | MODIFIER |

| | | | | | | | | | | | | | | | |
|---|---|---|---|---|---|---|---|---|---|---|---|---|---|---|---|
| 5 | 117434869 | AX-170297259 | 0 | G | A | 0.456 | 6.29E-02 | 1.17E-01 | 2.66E+00 | 2.69E+00 | 5.90E-01 | no | Ksr2 | intron_variant,non_coding_transcript_variant | MODIFIER |
| 5 | 117436363 | AX-170467024 | 0 | T | G | 0.456 | 6.29E-02 | 1.17E-01 | 2.66E+00 | 2.69E+00 | 5.90E-01 | no | Ksr2 | intron_variant,non_coding_transcript_variant | MODIFIER |
| 5 | 117436847 | AX-170025964 | 0 | A | G | 0.456 | 6.29E-02 | 1.17E-01 | 2.66E+00 | 2.69E+00 | 5.90E-01 | no | Ksr2 | intron_variant,non_coding_transcript_variant | MODIFIER |
| 5 | 117441228 | AX-170025962 | 0 | A | G | 0.456 | 6.29E-02 | 1.17E-01 | 2.66E+00 | 2.69E+00 | 5.90E-01 | no | Ksr2 | intron_variant,non_coding_transcript_variant | MODIFIER |
| 5 | 117460477 | AX-170297632 | 0 | C | T | 0.456 | 6.29E-02 | 1.17E-01 | 2.66E+00 | 2.69E+00 | 5.90E-01 | no | Ksr2 | intron_variant,non_coding_transcript_variant | MODIFIER |
| 5 | 117465148 | AX-168382794 | 0 | T | G | 0.456 | 6.29E-02 | 1.17E-01 | 2.66E+00 | 2.69E+00 | 5.90E-01 | no | Ksr2 | intron_variant,non_coding_transcript_variant | MODIFIER |
| 5 | 117471334 | AX-170095444 | 0 | T | C | 0.456 | 6.29E-02 | 1.17E-01 | 2.66E+00 | 2.69E+00 | 5.90E-01 | no | Ksr2 | intron_variant,non_coding_transcript_variant | MODIFIER |
| 5 | 117480068 | AX-170025961 | 0 | G | C | 0.456 | 6.29E-02 | 1.17E-01 | 2.66E+00 | 2.69E+00 | 5.90E-01 | no | Ksr2 | intron_variant,non_coding_transcript_variant | MODIFIER |
| 5 | 117521165 | AX-170233901 | 0 | T | C | 0.456 | 6.29E-02 | 1.17E-01 | 2.66E+00 | 2.69E+00 | 5.90E-01 | no | Ksr2 | intron_variant,non_coding_transcript_variant | MODIFIER |
| 5 | 117522816 | AX-170039088 | 0 | G | A | 0.456 | 6.29E-02 | 1.17E-01 | 2.66E+00 | 2.69E+00 | 5.90E-01 | no | Ksr2 | intron_variant,non_coding_transcript_variant | MODIFIER |
| 5 | 117527967 | AX-170497657 | 0 | G | T | 0.456 | 6.29E-02 | 1.17E-01 | 2.66E+00 | 2.69E+00 | 5.90E-01 | no | Ksr2 | intron_variant,non_coding_transcript_variant | MODIFIER |
| 5 | 117528101 | AX-97842322 | 0 | A | G | 0.456 | 6.29E-02 | 1.17E-01 | 2.66E+00 | 2.69E+00 | 5.90E-01 | no | Ksr2 | intron_variant,non_coding_transcript_variant | MODIFIER |
| 7 | 60961241 | AX-170029115 | 2 | C | T | 0.436 | -5.04E-02 | 9.54E-02 | 2.76E+00 | 2.86E+00 | 5.90E-01 | no | Gm44643 | non_coding_transcript_exon_variant | MODIFIER |
| 1 | 54725965 | AX-170393623 | 0 | C | T | 0.123 | 8.74E-02 | 1.63E-01 | 2.60E+00 | 2.72E+00 | 5.90E-01 | no | Ankrd44 | intron_variant | MODIFIER |
| 13 | 39342122 | AX-170432314 | 1 | C | T | 0.17 | -6.30E-02 | 1.11E-01 | 2.48E+00 | 2.56E+00 | 5.90E-01 | no | | intergenic_variant | MODIFIER |
| 13 | 46654692 | AX-170122597 | 0 | G | C | 0.491 | -3.28E-02 | 6.24E-02 | 2.75E+00 | 2.86E+00 | 5.90E-01 | no | Cap2 | downstream_gene_variant | MODIFIER |
| 18 | 69817162 | AX-170223246 | 2 | C | T | 0.236 | -5.41E-02 | 1.02E-01 | 2.67E+00 | 2.73E+00 | 5.90E-01 | no | | intergenic_variant | MODIFIER |
| 16 | 97754038 | AX-170199408 | 0 | C | A | 0.465 | 4.05E-02 | 7.68E-02 | 2.79E+00 | 2.90E+00 | 5.90E-01 | no | Ripk4 | intron_variant | MODIFIER |
| 8 | 81228580 | AX-170364446 | 1 | C | T | 0.205 | -6.50E-02 | 1.23E-01 | 2.73E+00 | 2.82E+00 | 5.90E-01 | no | | intergenic_variant | MODIFIER |
| 8 | 68996548 | AX-170138275 | 1 | T | A | 0.188 | -7.13E-02 | 1.34E-01 | 2.67E+00 | 2.74E+00 | 5.90E-01 | no | Gm33103 | intron_variant,non_coding_transcript_variant | MODIFIER |
| 7 | 116639374 | AX-170114098 | 1 | T | C | 0.42 | 4.94E-02 | 9.22E-02 | 2.65E+00 | 2.70E+00 | 5.90E-01 | no | | intergenic_variant | MODIFIER |
| 5 | 97823439 | AX-170025675 | 1 | T | A | 0.438 | 5.85E-02 | 1.10E-01 | 2.70E+00 | 2.75E+00 | 5.91E-01 | no | | intergenic_variant | MODIFIER |
| 6 | 139433384 | AX-97853441 | 1 | A | G | 0.054 | -1.12E-01 | 2.12E-01 | 2.64E+00 | 2.77E+00 | 5.91E-01 | no | | intergenic_variant | MODIFIER |
| 13 | 40472934 | AX-97798151 | 0 | C | T | 0.184 | -6.40E-02 | 1.12E-01 | 2.47E+00 | 2.54E+00 | 5.91E-01 | no | | regulatory_region_variant | MODIFIER |
| 7 | 91653789 | AX-170480161 | 0 | T | C | 0.132 | -9.58E-02 | 1.80E-01 | 2.63E+00 | 2.73E+00 | 5.91E-01 | no | Dlg2 | intron_variant | MODIFIER |
| 9 | 96423171 | AX-97827915 | 1 | G | T | 0.143 | -6.18E-02 | 1.14E-01 | 2.57E+00 | 2.65E+00 | 5.91E-01 | no | | intergenic_variant | MODIFIER |
| 15 | 74963989 | AX-170012003 | 0 | T | C | 0.553 | 4.48E-02 | 8.24E-02 | 2.58E+00 | 2.65E+00 | 5.91E-01 | no | Ly6e | downstream_gene_variant | MODIFIER |
| 1 | 140828402 | AX-97809406 | 0 | T | C | 0.211 | 7.01E-02 | 1.32E-01 | 2.68E+00 | 2.74E+00 | 5.91E-01 | no | Gm37371 | downstream_gene_variant | MODIFIER |
| 1 | 140874537 | AX-170457306 | 0 | C | T | 0.211 | 7.01E-02 | 1.32E-01 | 2.68E+00 | 2.74E+00 | 5.91E-01 | no | | intergenic_variant | MODIFIER |
| 1 | 141078327 | AX-97836033 | 0 | A | G | 0.211 | 7.01E-02 | 1.32E-01 | 2.68E+00 | 2.74E+00 | 5.91E-01 | no | | intergenic_variant | MODIFIER |
| 1 | 141419284 | AX-170091715 | 0 | G | A | 0.211 | 7.01E-02 | 1.32E-01 | 2.68E+00 | 2.74E+00 | 5.91E-01 | no | Gm37142 | downstream_gene_variant | MODIFIER |
| 1 | 141560316 | AX-97835054 | 0 | A | G | 0.211 | 7.01E-02 | 1.32E-01 | 2.68E+00 | 2.74E+00 | 5.91E-01 | no | | intergenic_variant | MODIFIER |
| 1 | 141633885 | AX-168419615 | 0 | T | C | 0.211 | 7.01E-02 | 1.32E-01 | 2.68E+00 | 2.74E+00 | 5.91E-01 | no | | intergenic_variant | MODIFIER |
| 1 | 141976458 | AX-170445867 | 0 | A | C | 0.211 | 7.01E-02 | 1.32E-01 | 2.68E+00 | 2.74E+00 | 5.91E-01 | no | | intergenic_variant | MODIFIER |
| 1 | 141977432 | AX-97852737 | 0 | T | G | 0.211 | 7.01E-02 | 1.32E-01 | 2.68E+00 | 2.74E+00 | 5.91E-01 | no | | intergenic_variant | MODIFIER |
| 1 | 141978062 | AX-170519589 | 0 | T | C | 0.211 | 7.01E-02 | 1.32E-01 | 2.68E+00 | 2.74E+00 | 5.91E-01 | no | | intergenic_variant | MODIFIER |
| 1 | 141991484 | AX-170327727 | 0 | A | G | 0.211 | 7.01E-02 | 1.32E-01 | 2.68E+00 | 2.74E+00 | 5.91E-01 | no | | intergenic_variant | MODIFIER |
| 1 | 142003698 | AX-170435245 | 0 | T | C | 0.211 | 7.01E-02 | 1.32E-01 | 2.68E+00 | 2.74E+00 | 5.91E-01 | no | | intergenic_variant | MODIFIER |
| 1 | 142042148 | AX-170217710 | 0 | T | C | 0.211 | 7.01E-02 | 1.32E-01 | 2.68E+00 | 2.74E+00 | 5.91E-01 | no | | intergenic_variant | MODIFIER |
| 1 | 142167123 | AX-97802434 | 0 | T | C | 0.211 | 7.01E-02 | 1.32E-01 | 2.68E+00 | 2.74E+00 | 5.91E-01 | no | | intergenic_variant | MODIFIER |
| 1 | 142167187 | AX-170188926 | 0 | A | C | 0.211 | 7.01E-02 | 1.32E-01 | 2.68E+00 | 2.74E+00 | 5.91E-01 | no | | intergenic_variant | MODIFIER |
| 1 | 142580510 | AX-97797647 | 0 | G | T | 0.211 | 7.01E-02 | 1.32E-01 | 2.68E+00 | 2.74E+00 | 5.91E-01 | no | | intergenic_variant | MODIFIER |
| 1 | 143081461 | AX-97821231 | 0 | T | C | 0.211 | 7.01E-02 | 1.32E-01 | 2.68E+00 | 2.74E+00 | 5.91E-01 | no | | intergenic_variant | MODIFIER |
| 5 | 107333856 | AX-170519312 | 0 | A | G | 0.447 | 6.17E-02 | 1.15E-01 | 2.66E+00 | 2.69E+00 | 5.91E-01 | no | Brdt | intron_variant | MODIFIER |
| 5 | 107373107 | AX-170262214 | 0 | A | C | 0.447 | 6.17E-02 | 1.15E-01 | 2.66E+00 | 2.69E+00 | 5.91E-01 | no | Brdt | intron_variant | MODIFIER |
| 5 | 107376963 | AX-170215220 | 0 | G | A | 0.447 | 6.17E-02 | 1.15E-01 | 2.66E+00 | 2.69E+00 | 5.91E-01 | no | Brdt | intron_variant | MODIFIER |
| 5 | 107378653 | AX-170372319 | 0 | C | T | 0.447 | 6.17E-02 | 1.15E-01 | 2.66E+00 | 2.69E+00 | 5.91E-01 | no | Brdt | intron_variant | MODIFIER |
| 5 | 107379465 | AX-170139719 | 0 | C | T | 0.447 | 6.17E-02 | 1.15E-01 | 2.66E+00 | 2.69E+00 | 5.91E-01 | no | Brdt | intron_variant | MODIFIER |
| 5 | 107380045 | AX-97798675 | 0 | A | C | 0.447 | 6.17E-02 | 1.15E-01 | 2.66E+00 | 2.69E+00 | 5.91E-01 | no | Brdt | intron_variant | MODIFIER |
| 5 | 107397197 | AX-170274663 | 0 | T | C | 0.447 | 6.17E-02 | 1.15E-01 | 2.66E+00 | 2.69E+00 | 5.91E-01 | no | n-R5s173 | upstream_gene_variant | MODIFIER |
| 5 | 107417494 | AX-170186296 | 0 | G | A | 0.447 | 6.17E-02 | 1.15E-01 | 2.66E+00 | 2.69E+00 | 5.91E-01 | no | Ephx4 | intron_variant | MODIFIER |
| 5 | 107467046 | AX-170469368 | 0 | G | A | 0.447 | 6.17E-02 | 1.15E-01 | 2.66E+00 | 2.69E+00 | 5.91E-01 | no | Btbd8 | intron_variant | MODIFIER |
| 5 | 107552108 | AX-170115226 | 0 | A | C | 0.447 | 6.17E-02 | 1.15E-01 | 2.66E+00 | 2.69E+00 | 5.91E-01 | no | 1700028K03Rik | downstream_gene_variant | MODIFIER |
| 5 | 108016265 | AX-97817973 | 0 | G | A | 0.447 | 6.17E-02 | 1.15E-01 | 2.66E+00 | 2.69E+00 | 5.91E-01 | no | | intergenic_variant | MODIFIER |

| | | | | | | | | | | | | | | | |
|---|---|---|---|---|---|---|---|---|---|---|---|---|---|---|---|
| 5 | 108317299 | AX-170025845 | 0 | A | G | 0.447 | 6.17E-02 | 1.15E-01 | 2.66E+00 | 2.69E+00 | 5.91E-01 | no | Pigg | missense_variant | MODERATE |
| 5 | 108347167 | AX-170089579 | 0 | G | T | 0.447 | 6.17E-02 | 1.15E-01 | 2.66E+00 | 2.69E+00 | 5.91E-01 | no | Pigg | intron_variant | MODIFIER |
| 5 | 108347377 | AX-170025841 | 0 | C | G | 0.447 | 6.17E-02 | 1.15E-01 | 2.66E+00 | 2.69E+00 | 5.91E-01 | no | Pigg | intron_variant | MODIFIER |
| 5 | 108703635 | AX-170519816 | 0 | A | G | 0.447 | 6.17E-02 | 1.15E-01 | 2.66E+00 | 2.69E+00 | 5.91E-01 | no | Fgfrl1 | intron_variant | MODIFIER |
| 5 | 108727515 | AX-170127741 | 0 | A | G | 0.447 | 6.17E-02 | 1.15E-01 | 2.66E+00 | 2.69E+00 | 5.91E-01 | no | Rnf212 | downstream_gene_variant | MODIFIER |
| 5 | 108731296 | AX-170520365 | 0 | C | G | 0.447 | 6.17E-02 | 1.15E-01 | 2.66E+00 | 2.69E+00 | 5.91E-01 | no | Rnf212 | downstream_gene_variant | MODIFIER |
| 5 | 108772727 | AX-170269788 | 0 | G | A | 0.447 | 6.17E-02 | 1.15E-01 | 2.66E+00 | 2.69E+00 | 5.91E-01 | no | Tmed11 | downstream_gene_variant | MODIFIER |
| 5 | 108773243 | AX-170073050 | 0 | T | G | 0.447 | 6.17E-02 | 1.15E-01 | 2.66E+00 | 2.69E+00 | 5.91E-01 | no | Tmed11 | downstream_gene_variant | MODIFIER |
| 5 | 108784850 | AX-170379495 | 0 | T | A | 0.447 | 6.17E-02 | 1.15E-01 | 2.66E+00 | 2.69E+00 | 5.91E-01 | no | Tmed11 | intron_variant | MODIFIER |
| 5 | 108812896 | AX-97800683 | 0 | T | C | 0.447 | 6.17E-02 | 1.15E-01 | 2.66E+00 | 2.69E+00 | 5.91E-01 | no | Vmn2r8 | upstream_gene_variant | MODIFIER |
| 5 | 108854413 | AX-170025849 | 0 | T | C | 0.447 | 6.17E-02 | 1.15E-01 | 2.66E+00 | 2.69E+00 | 5.91E-01 | no | Vmn2r9 | upstream_gene_variant | MODIFIER |
| 5 | 108854736 | AX-170025859 | 0 | A | G | 0.447 | 6.17E-02 | 1.15E-01 | 2.66E+00 | 2.69E+00 | 5.91E-01 | no | Vmn2r9 | upstream_gene_variant | MODIFIER |
| 5 | 109308643 | AX-97780757 | 0 | T | C | 0.447 | 6.17E-02 | 1.15E-01 | 2.66E+00 | 2.69E+00 | 5.91E-01 | no | | intergenic_variant | MODIFIER |
| 5 | 109320084 | AX-170160425 | 0 | C | G | 0.447 | 6.17E-02 | 1.15E-01 | 2.66E+00 | 2.69E+00 | 5.91E-01 | no | | intergenic_variant | MODIFIER |
| 5 | 109395944 | AX-168388269 | 0 | A | G | 0.447 | 6.17E-02 | 1.15E-01 | 2.66E+00 | 2.69E+00 | 5.91E-01 | no | Gm7171 | upstream_gene_variant | MODIFIER |
| 5 | 109482932 | AX-97855435 | 0 | A | G | 0.447 | 6.17E-02 | 1.15E-01 | 2.66E+00 | 2.69E+00 | 5.91E-01 | no | | intergenic_variant | MODIFIER |
| 5 | 109671186 | AX-170025865 | 0 | T | C | 0.447 | 6.17E-02 | 1.15E-01 | 2.66E+00 | 2.69E+00 | 5.91E-01 | no | 5430403G16Rik | downstream_gene_variant | MODIFIER |
| 5 | 109692504 | AX-170425609 | 0 | T | G | 0.447 | 6.17E-02 | 1.15E-01 | 2.66E+00 | 2.69E+00 | 5.91E-01 | no | 5430403G16Rik | upstream_gene_variant | MODIFIER |
| 5 | 110182120 | AX-97857032 | 0 | T | C | 0.447 | 6.17E-02 | 1.15E-01 | 2.66E+00 | 2.69E+00 | 5.91E-01 | no | Golga3 | intron_variant | MODIFIER |
| 5 | 110201053 | AX-170158521 | 0 | C | T | 0.447 | 6.17E-02 | 1.15E-01 | 2.66E+00 | 2.69E+00 | 5.91E-01 | no | Golga3 | intron_variant | MODIFIER |
| 5 | 110206922 | AX-170463042 | 0 | G | A | 0.447 | 6.17E-02 | 1.15E-01 | 2.66E+00 | 2.69E+00 | 5.91E-01 | no | Golga3 | intron_variant | MODIFIER |
| 5 | 110222791 | AX-170141462 | 0 | A | G | 0.447 | 6.17E-02 | 1.15E-01 | 2.66E+00 | 2.69E+00 | 5.91E-01 | no | Golga3 | intron_variant | MODIFIER |
| 5 | 110224140 | AX-170282168 | 0 | T | C | 0.447 | 6.17E-02 | 1.15E-01 | 2.66E+00 | 2.69E+00 | 5.91E-01 | no | Golga3 | downstream_gene_variant | MODIFIER |
| 5 | 110268605 | AX-170025876 | 0 | C | G | 0.447 | 6.17E-02 | 1.15E-01 | 2.66E+00 | 2.69E+00 | 5.91E-01 | no | Pgam5 | intron_variant | MODIFIER |
| 5 | 110268708 | AX-170025877 | 0 | G | A | 0.447 | 6.17E-02 | 1.15E-01 | 2.66E+00 | 2.69E+00 | 5.91E-01 | no | Pgam5 | intron_variant | MODIFIER |
| 5 | 110290338 | AX-170340923 | 0 | T | C | 0.447 | 6.17E-02 | 1.15E-01 | 2.66E+00 | 2.69E+00 | 5.91E-01 | no | Pole | intron_variant | MODIFIER |
| 5 | 110291232 | AX-97811920 | 0 | T | C | 0.447 | 6.17E-02 | 1.15E-01 | 2.66E+00 | 2.69E+00 | 5.91E-01 | no | Pole | intron_variant | MODIFIER |
| 5 | 110294577 | AX-168382927 | 0 | T | C | 0.447 | 6.17E-02 | 1.15E-01 | 2.66E+00 | 2.69E+00 | 5.91E-01 | no | Pole | synonymous_variant | LOW |
| 5 | 110330894 | AX-170287082 | 0 | T | C | 0.447 | 6.17E-02 | 1.15E-01 | 2.66E+00 | 2.69E+00 | 5.91E-01 | no | Pole | intron_variant | MODIFIER |
| 5 | 110332605 | AX-170025875 | 0 | A | G | 0.447 | 6.17E-02 | 1.15E-01 | 2.66E+00 | 2.69E+00 | 5.91E-01 | no | Pole | intron_variant | MODIFIER |
| 5 | 110336862 | AX-170025872 | 0 | A | G | 0.447 | 6.17E-02 | 1.15E-01 | 2.66E+00 | 2.69E+00 | 5.91E-01 | no | Pole | intron_variant | MODIFIER |
| 5 | 110374178 | AX-97803764 | 0 | A | G | 0.447 | 6.17E-02 | 1.15E-01 | 2.66E+00 | 2.69E+00 | 5.91E-01 | no | Fbrsl1 | intron_variant | MODIFIER |
| 5 | 110467922 | AX-170329840 | 0 | C | T | 0.447 | 6.17E-02 | 1.15E-01 | 2.66E+00 | 2.69E+00 | 5.91E-01 | no | | intergenic_variant | MODIFIER |
| 5 | 110478979 | AX-170401464 | 0 | C | T | 0.447 | 6.17E-02 | 1.15E-01 | 2.66E+00 | 2.69E+00 | 5.91E-01 | no | Gm32996 | downstream_gene_variant | MODIFIER |
| 5 | 110489000 | AX-170089581 | 0 | T | C | 0.447 | 6.17E-02 | 1.15E-01 | 2.66E+00 | 2.69E+00 | 5.91E-01 | no | | intergenic_variant | MODIFIER |
| 5 | 110583792 | AX-170211881 | 0 | A | G | 0.447 | 6.17E-02 | 1.15E-01 | 2.66E+00 | 2.69E+00 | 5.91E-01 | no | Galnt9 | intron_variant | MODIFIER |
| 5 | 110590239 | AX-170194182 | 0 | A | G | 0.447 | 6.17E-02 | 1.15E-01 | 2.66E+00 | 2.69E+00 | 5.91E-01 | no | Galnt9 | intron_variant | MODIFIER |
| 5 | 110590311 | AX-170494257 | 0 | G | A | 0.447 | 6.17E-02 | 1.15E-01 | 2.66E+00 | 2.69E+00 | 5.91E-01 | no | Galnt9 | intron_variant | MODIFIER |
| 5 | 110590583 | AX-170226936 | 0 | G | A | 0.447 | 6.17E-02 | 1.15E-01 | 2.66E+00 | 2.69E+00 | 5.91E-01 | no | Galnt9 | intron_variant | MODIFIER |
| 5 | 110638237 | AX-170073088 | 0 | T | C | 0.447 | 6.17E-02 | 1.15E-01 | 2.66E+00 | 2.69E+00 | 5.91E-01 | no | | intergenic_variant | MODIFIER |
| 5 | 110680772 | AX-170306478 | 0 | C | T | 0.447 | 6.17E-02 | 1.15E-01 | 2.66E+00 | 2.69E+00 | 5.91E-01 | no | Ep400 | intron_variant | MODIFIER |
| 5 | 110744293 | AX-170204026 | 0 | T | C | 0.447 | 6.17E-02 | 1.15E-01 | 2.66E+00 | 2.69E+00 | 5.91E-01 | no | Ep400 | intron_variant | MODIFIER |
| 5 | 110757786 | AX-170177408 | 0 | T | G | 0.447 | 6.17E-02 | 1.15E-01 | 2.66E+00 | 2.69E+00 | 5.91E-01 | no | Ep400 | intron_variant | MODIFIER |
| 5 | 110839787 | AX-170073092 | 0 | T | C | 0.447 | 6.17E-02 | 1.15E-01 | 2.66E+00 | 2.69E+00 | 5.91E-01 | no | Hscb | upstream_gene_variant | MODIFIER |
| 5 | 110848441 | AX-170422846 | 0 | T | C | 0.447 | 6.17E-02 | 1.15E-01 | 2.66E+00 | 2.69E+00 | 5.91E-01 | no | Chek2 | synonymous_variant | LOW |
| 5 | 110854649 | AX-170483327 | 0 | T | C | 0.447 | 6.17E-02 | 1.15E-01 | 2.66E+00 | 2.69E+00 | 5.91E-01 | no | Chek2 | intron_variant | MODIFIER |
| 5 | 110855029 | AX-170452679 | 0 | A | G | 0.447 | 6.17E-02 | 1.15E-01 | 2.66E+00 | 2.69E+00 | 5.91E-01 | no | Chek2 | intron_variant | MODIFIER |
| 5 | 110855670 | AX-170464353 | 0 | G | A | 0.447 | 6.17E-02 | 1.15E-01 | 2.66E+00 | 2.69E+00 | 5.91E-01 | no | Chek2 | intron_variant | MODIFIER |
| 5 | 110855906 | AX-170511199 | 0 | A | G | 0.447 | 6.17E-02 | 1.15E-01 | 2.66E+00 | 2.69E+00 | 5.91E-01 | no | Chek2 | intron_variant | MODIFIER |
| 5 | 110868373 | AX-170403544 | 0 | T | C | 0.447 | 6.17E-02 | 1.15E-01 | 2.66E+00 | 2.69E+00 | 5.91E-01 | no | Chek2 | intron_variant | MODIFIER |
| 5 | 110869338 | AX-170241784 | 0 | G | C | 0.447 | 6.17E-02 | 1.15E-01 | 2.66E+00 | 2.69E+00 | 5.91E-01 | no | Chek2 | intron_variant | MODIFIER |
| 5 | 110893173 | AX-170025882 | 0 | C | T | 0.447 | 6.17E-02 | 1.15E-01 | 2.66E+00 | 2.69E+00 | 5.91E-01 | no | Ttc28 | intron_variant | MODIFIER |
| 5 | 110925126 | AX-170025884 | 0 | A | G | 0.447 | 6.17E-02 | 1.15E-01 | 2.66E+00 | 2.69E+00 | 5.91E-01 | no | Ttc28 | intron_variant | MODIFIER |
| 5 | 110999185 | AX-170490022 | 0 | C | A | 0.447 | 6.17E-02 | 1.15E-01 | 2.66E+00 | 2.69E+00 | 5.91E-01 | no | Ttc28 | intron_variant | MODIFIER |

| | | | | | | | | | | | | | | | |
|---|---|---|---|---|---|---|---|---|---|---|---|---|---|---|---|
| 5 | 110999261 | AX-97839084 | 0 | C | T | 0.447 | 6.17E-02 | 1.15E-01 | 2.66E+00 | 2.69E+00 | 5.91E-01 | no | Ttc28 | intron_variant | MODIFIER |
| 5 | 111003122 | AX-170190402 | 0 | A | G | 0.447 | 6.17E-02 | 1.15E-01 | 2.66E+00 | 2.69E+00 | 5.91E-01 | no | Ttc28 | intron_variant | MODIFIER |
| 5 | 111014678 | AX-170153806 | 0 | G | T | 0.447 | 6.17E-02 | 1.15E-01 | 2.66E+00 | 2.69E+00 | 5.91E-01 | no | Ttc28 | intron_variant | MODIFIER |
| 5 | 111060239 | AX-170267950 | 0 | C | G | 0.447 | 6.17E-02 | 1.15E-01 | 2.66E+00 | 2.69E+00 | 5.91E-01 | no | Ttc28 | intron_variant | MODIFIER |
| 5 | 111060355 | AX-170525332 | 0 | A | G | 0.447 | 6.17E-02 | 1.15E-01 | 2.66E+00 | 2.69E+00 | 5.91E-01 | no | Ttc28 | intron_variant | MODIFIER |
| 5 | 111113695 | AX-170099055 | 0 | T | C | 0.447 | 6.17E-02 | 1.15E-01 | 2.66E+00 | 2.69E+00 | 5.91E-01 | no | Ttc28 | intron_variant | MODIFIER |
| 5 | 111225999 | AX-170299684 | 0 | T | C | 0.447 | 6.17E-02 | 1.15E-01 | 2.66E+00 | 2.69E+00 | 5.91E-01 | no | Ttc28 | synonymous_variant | LOW |
| 5 | 111226497 | AX-170073102 | 0 | A | G | 0.447 | 6.17E-02 | 1.15E-01 | 2.66E+00 | 2.69E+00 | 5.91E-01 | no | Ttc28 | intron_variant | MODIFIER |
| 5 | 111377202 | AX-170320865 | 0 | A | G | 0.447 | 6.17E-02 | 1.15E-01 | 2.66E+00 | 2.69E+00 | 5.91E-01 | no | Pitpnb | intron_variant | MODIFIER |
| 5 | 111582609 | AX-170248041 | 0 | T | G | 0.447 | 6.17E-02 | 1.15E-01 | 2.66E+00 | 2.69E+00 | 5.91E-01 | no | C130026L21Rik | intron_variant,non_coding_transcript_variant | MODIFIER |
| 5 | 111691895 | AX-170179991 | 0 | G | A | 0.447 | 6.17E-02 | 1.15E-01 | 2.66E+00 | 2.69E+00 | 5.91E-01 | no | | regulatory_region_variant | MODIFIER |
| 5 | 111858175 | AX-170373376 | 0 | T | C | 0.447 | 6.17E-02 | 1.15E-01 | 2.66E+00 | 2.69E+00 | 5.91E-01 | no | | regulatory_region_variant | MODIFIER |
| 5 | 111869508 | AX-170534304 | 0 | T | C | 0.447 | 6.17E-02 | 1.15E-01 | 2.66E+00 | 2.69E+00 | 5.91E-01 | no | | intergenic_variant | MODIFIER |
| 5 | 111877675 | AX-170396854 | 0 | T | C | 0.447 | 6.17E-02 | 1.15E-01 | 2.66E+00 | 2.69E+00 | 5.91E-01 | no | | regulatory_region_variant | MODIFIER |
| 5 | 111884232 | AX-97783317 | 0 | C | T | 0.447 | 6.17E-02 | 1.15E-01 | 2.66E+00 | 2.69E+00 | 5.91E-01 | no | | intergenic_variant | MODIFIER |
| 5 | 111884633 | AX-170303032 | 0 | A | G | 0.447 | 6.17E-02 | 1.15E-01 | 2.66E+00 | 2.69E+00 | 5.91E-01 | no | | intergenic_variant | MODIFIER |
| 5 | 112011349 | AX-170226787 | 0 | G | A | 0.447 | 6.17E-02 | 1.15E-01 | 2.66E+00 | 2.69E+00 | 5.91E-01 | no | | regulatory_region_variant | MODIFIER |
| 5 | 112012824 | AX-170467531 | 0 | G | A | 0.447 | 6.17E-02 | 1.15E-01 | 2.66E+00 | 2.69E+00 | 5.91E-01 | no | | regulatory_region_variant | MODIFIER |
| 5 | 112054270 | AX-170025900 | 0 | G | A | 0.447 | 6.17E-02 | 1.15E-01 | 2.66E+00 | 2.69E+00 | 5.91E-01 | no | | intergenic_variant | MODIFIER |
| 5 | 112069849 | AX-170273290 | 0 | C | T | 0.447 | 6.17E-02 | 1.15E-01 | 2.66E+00 | 2.69E+00 | 5.91E-01 | no | | intergenic_variant | MODIFIER |
| 5 | 112070122 | AX-170426187 | 0 | T | C | 0.447 | 6.17E-02 | 1.15E-01 | 2.66E+00 | 2.69E+00 | 5.91E-01 | no | | intergenic_variant | MODIFIER |
| 5 | 112072398 | AX-170364604 | 0 | A | T | 0.447 | 6.17E-02 | 1.15E-01 | 2.66E+00 | 2.69E+00 | 5.91E-01 | no | | intergenic_variant | MODIFIER |
| 5 | 112124765 | AX-170025904 | 0 | A | C | 0.447 | 6.17E-02 | 1.15E-01 | 2.66E+00 | 2.69E+00 | 5.91E-01 | no | | intergenic_variant | MODIFIER |
| 5 | 112154819 | AX-170482854 | 0 | T | C | 0.447 | 6.17E-02 | 1.15E-01 | 2.66E+00 | 2.69E+00 | 5.91E-01 | no | 1700016B01Rik | upstream_gene_variant | MODIFIER |
| 5 | 112207285 | AX-97828170 | 0 | C | T | 0.447 | 6.17E-02 | 1.15E-01 | 2.66E+00 | 2.69E+00 | 5.91E-01 | no | 1700028D13Rik | intron_variant,non_coding_transcript_variant | MODIFIER |
| 5 | 112208191 | AX-170025902 | 0 | G | T | 0.447 | 6.17E-02 | 1.15E-01 | 2.66E+00 | 2.69E+00 | 5.91E-01 | no | 1700028D13Rik | intron_variant,non_coding_transcript_variant | MODIFIER |
| 5 | 112210657 | AX-170073124 | 0 | T | C | 0.447 | 6.17E-02 | 1.15E-01 | 2.66E+00 | 2.69E+00 | 5.91E-01 | no | Miat | downstream_gene_variant | MODIFIER |
| 5 | 112259173 | AX-170507685 | 0 | C | T | 0.447 | 6.17E-02 | 1.15E-01 | 2.66E+00 | 2.69E+00 | 5.91E-01 | no | Crybb1 | intron_variant | MODIFIER |
| 5 | 112332682 | AX-170296289 | 0 | C | T | 0.447 | 6.17E-02 | 1.15E-01 | 2.66E+00 | 2.69E+00 | 5.91E-01 | no | Tfip11 | intron_variant | MODIFIER |
| 5 | 113079032 | AX-170157692 | 0 | C | T | 0.447 | 6.17E-02 | 1.15E-01 | 2.66E+00 | 2.69E+00 | 5.91E-01 | no | Crybb3 | intron_variant | MODIFIER |
| 1 | 116130483 | AX-170001480 | 1 | C | T | 0.17 | 1.05E-01 | 1.94E-01 | 2.58E+00 | 2.67E+00 | 5.91E-01 | no | Cntnap5a | intron_variant | MODIFIER |
| 18 | 57646152 | AX-170230432 | 0 | G | A | 0.395 | 5.17E-02 | 9.10E-02 | 2.47E+00 | 2.55E+00 | 5.91E-01 | no | Ccdc192 | intron_variant | MODIFIER |
| 16 | 91987690 | AX-170313088 | 1 | T | C | 0.464 | -4.16E-02 | 7.86E-02 | 2.69E+00 | 2.77E+00 | 5.91E-01 | no | | intergenic_variant | MODIFIER |
| 7 | 19930416 | AX-170125929 | 0 | T | C | 0.412 | 4.82E-02 | 8.87E-02 | 2.55E+00 | 2.67E+00 | 5.91E-01 | no | Gm16251 | downstream_gene_variant | MODIFIER |
| 1 | 148227471 | AX-170197024 | 0 | T | A | 0.211 | 6.66E-02 | 1.24E-01 | 2.62E+00 | 2.68E+00 | 5.92E-01 | no | | regulatory_region_variant | MODIFIER |
| 1 | 150823625 | AX-170318808 | 1 | C | T | 0.214 | 7.17E-02 | 1.34E-01 | 2.64E+00 | 2.69E+00 | 5.92E-01 | no | Hmcn1 | intron_variant | MODIFIER |
| 9 | 91920583 | AX-97826673 | 0 | A | T | 0.14 | -5.88E-02 | 1.09E-01 | 2.58E+00 | 2.67E+00 | 5.92E-01 | no | Gm31409 | downstream_gene_variant | MODIFIER |
| 8 | 77984287 | AX-168410286 | 0 | A | G | 0.193 | 6.77E-02 | 1.28E-01 | 2.72E+00 | 2.79E+00 | 5.92E-01 | no | | regulatory_region_variant | MODIFIER |
| 5 | 86423301 | AX-170170020 | 0 | C | T | 0.43 | 5.94E-02 | 1.12E-01 | 2.76E+00 | 2.80E+00 | 5.92E-01 | no | Tmprss11a | intron_variant | MODIFIER |
| 4 | 139059184 | AX-97795593 | 1 | T | C | 0.152 | 6.27E-02 | 1.16E-01 | 2.58E+00 | 2.64E+00 | 5.93E-01 | no | Tmco4 | downstream_gene_variant | MODIFIER |
| 18 | 80840769 | AX-170517512 | 0 | T | C | 0.289 | -4.56E-02 | 8.45E-02 | 2.57E+00 | 2.67E+00 | 5.93E-01 | no | Atp9b | intron_variant | MODIFIER |
| 5 | 78890987 | AX-170192164 | 0 | A | G | 0.412 | 5.42E-02 | 1.02E-01 | 2.66E+00 | 2.73E+00 | 5.93E-01 | no | | intergenic_variant | MODIFIER |
| 5 | 81289911 | AX-170199777 | 0 | T | A | 0.412 | 5.42E-02 | 1.02E-01 | 2.66E+00 | 2.73E+00 | 5.93E-01 | no | Adgrl3 | intron_variant | MODIFIER |
| 8 | 90138983 | AX-170303809 | 0 | A | T | 0.184 | -6.09E-02 | 1.13E-01 | 2.58E+00 | 2.66E+00 | 5.93E-01 | no | | intergenic_variant | MODIFIER |
| 3 | 155502784 | AX-170068521 | 0 | C | A | 0.333 | 3.86E-02 | 7.35E-02 | 2.71E+00 | 2.80E+00 | 5.93E-01 | no | | intergenic_variant | MODIFIER |
| 7 | 103789234 | AX-97790762 | 0 | C | T | 0.456 | -5.13E-02 | 9.60E-02 | 2.87E+00 | 2.95E+00 | 5.93E-01 | no | Olfr67 | intron_variant | MODIFIER |
| 9 | 111368583 | AX-97785397 | 1 | A | C | 0.089 | -6.40E-02 | 1.17E-01 | 2.53E+00 | 2.61E+00 | 5.93E-01 | no | Trank1 | intron_variant | MODIFIER |
| 5 | 97483407 | AX-97831632 | 0 | G | T | 0.447 | 5.56E-02 | 1.05E-01 | 2.67E+00 | 2.73E+00 | 5.93E-01 | no | 4930467D21Rik | intron_variant,non_coding_transcript_variant | MODIFIER |
| 4 | 68809678 | AX-170416878 | 0 | C | T | 0.342 | 5.70E-02 | 1.09E-01 | 2.72E+00 | 2.79E+00 | 5.93E-01 | no | Brinp1 | intron_variant | MODIFIER |
| 11 | 74262314 | AX-170459535 | 2 | A | T | 0.264 | 3.71E-02 | 7.13E-02 | 2.67E+00 | 2.80E+00 | 5.94E-01 | no | Olfr405-ps1 | intron_variant,non_coding_transcript_variant | MODIFIER |
| 12 | 106686987 | AX-170377059 | 2 | T | G | 0.127 | -9.65E-02 | 1.85E-01 | 2.71E+00 | 2.81E+00 | 5.94E-01 | no | 4933406K04Rik | intron_variant,non_coding_transcript_variant | MODIFIER |
| 8 | 83043825 | AX-97781667 | 0 | G | A | 0.202 | -6.49E-02 | 1.23E-01 | 2.68E+00 | 2.76E+00 | 5.94E-01 | no | Rnf150 | intron_variant | MODIFIER |
| 8 | 83666202 | AX-170316470 | 0 | G | A | 0.202 | -6.49E-02 | 1.23E-01 | 2.68E+00 | 2.76E+00 | 5.94E-01 | no | Pkn1 | downstream_gene_variant | MODIFIER |
| 18 | 58227255 | AX-170178009 | 0 | A | T | 0.184 | -6.47E-02 | 1.22E-01 | 2.66E+00 | 2.71E+00 | 5.94E-01 | no | Gm25660 | downstream_gene_variant | MODIFIER |

| | | | | | | | | | | | | | | | |
|---|---|---|---|---|---|---|---|---|---|---|---|---|---|---|---|
| 5 | 107655624 | AX-170345577 | 1 | C | T | 0.446 | 6.05E-02 | 1.14E-01 | 2.65E+00 | 2.69E+00 | 5.94E-01 | no | Rpap2 | intron_variant | MODIFIER |
| 7 | 65549266 | AX-170175685 | 0 | C | A | 0.421 | -5.30E-02 | 1.01E-01 | 2.76E+00 | 2.83E+00 | 5.94E-01 | no | | intergenic_variant | MODIFIER |
| 15 | 69278865 | AX-170011919 | 0 | T | A | 0.553 | 4.45E-02 | 8.26E-02 | 2.58E+00 | 2.65E+00 | 5.94E-01 | no | 4930504C09Rik | downstream_gene_variant | MODIFIER |
| 18 | 57179897 | AX-170218560 | 1 | G | A | 0.196 | -6.02E-02 | 1.11E-01 | 2.57E+00 | 2.63E+00 | 5.94E-01 | no | Megf10 | intron_variant | MODIFIER |
| 2 | 130144002 | AX-170097804 | 1 | C | A | 0.143 | 1.14E-01 | 2.17E-01 | 2.71E+00 | 2.75E+00 | 5.94E-01 | no | Tgm6 | intron_variant | MODIFIER |
| 5 | 92977937 | AX-170025625 | 0 | T | C | 0.421 | 5.70E-02 | 1.08E-01 | 2.68E+00 | 2.74E+00 | 5.94E-01 | no | | intergenic_variant | MODIFIER |
| 13 | 32309166 | AX-170092709 | 1 | G | A | 0.179 | -6.41E-02 | 1.20E-01 | 2.60E+00 | 2.66E+00 | 5.95E-01 | no | Gmds | intron_variant | MODIFIER |
| 5 | 109564604 | AX-170025861 | 1 | G | C | 0.464 | 6.14E-02 | 1.15E-01 | 2.63E+00 | 2.67E+00 | 5.95E-01 | no | | intergenic_variant | MODIFIER |
| 5 | 114286416 | AX-170209159 | 1 | C | T | 0.464 | 6.14E-02 | 1.15E-01 | 2.63E+00 | 2.67E+00 | 5.95E-01 | no | Myo1h | upstream_gene_variant | MODIFIER |
| 6 | 44399162 | AX-170255253 | 0 | G | T | 0.044 | 8.41E-02 | 1.62E-01 | 2.63E+00 | 2.80E+00 | 5.95E-01 | no | | intergenic_variant | MODIFIER |
| 20 | 92741782 | AX-168397585 | 1 | A | G | 0.259 | 3.13E-02 | 5.90E-02 | 2.59E+00 | 2.71E+00 | 5.95E-01 | no | | intergenic_variant | MODIFIER |
| 6 | 133867480 | AX-170125331 | 1 | G | C | 0.562 | 4.54E-02 | 8.07E-02 | 2.47E+00 | 2.56E+00 | 5.95E-01 | no | Gm43968 | downstream_gene_variant | MODIFIER |
| 18 | 87421300 | AX-170093990 | 0 | G | A | 0.167 | -5.64E-02 | 1.04E-01 | 2.54E+00 | 2.64E+00 | 5.95E-01 | no | | intergenic_variant | MODIFIER |
| 5 | 103402879 | AX-170491552 | 1 | G | A | 0.446 | 5.52E-02 | 1.04E-01 | 2.63E+00 | 2.69E+00 | 5.95E-01 | no | | intergenic_variant | MODIFIER |
| 2 | 160124286 | AX-170507252 | 0 | C | A | 0.149 | -9.19E-02 | 1.76E-01 | 2.69E+00 | 2.78E+00 | 5.95E-01 | no | | intergenic_variant | MODIFIER |
| 9 | 38592739 | AX-97820419 | 0 | A | G | 0.491 | -4.16E-02 | 7.95E-02 | 2.70E+00 | 2.78E+00 | 5.95E-01 | no | Olfr913 | upstream_gene_variant | MODIFIER |
| 13 | 34991216 | AX-97820457 | 1 | C | T | 0.188 | -6.40E-02 | 1.19E-01 | 2.59E+00 | 2.65E+00 | 5.95E-01 | no | Eci2 | intron_variant | MODIFIER |
| 8 | 94088097 | AX-170148364 | 1 | C | T | 0.205 | -5.81E-02 | 1.11E-01 | 2.66E+00 | 2.74E+00 | 5.95E-01 | no | Bbs2 | intron_variant | MODIFIER |
| 6 | 122830200 | AX-170095789 | 0 | G | A | 0.184 | -4.27E-02 | 8.22E-02 | 2.67E+00 | 2.80E+00 | 5.95E-01 | no | Foxj2 | intron_variant | MODIFIER |
| 6 | 122862380 | AX-170530648 | 0 | G | A | 0.184 | -4.27E-02 | 8.22E-02 | 2.67E+00 | 2.80E+00 | 5.95E-01 | no | | intergenic_variant | MODIFIER |
| 6 | 122869901 | AX-97792494 | 0 | G | A | 0.184 | -4.27E-02 | 8.22E-02 | 2.67E+00 | 2.80E+00 | 5.95E-01 | no | Necap1 | upstream_gene_variant | MODIFIER |
| 6 | 122925723 | AX-170307169 | 0 | A | G | 0.184 | -4.27E-02 | 8.22E-02 | 2.67E+00 | 2.80E+00 | 5.95E-01 | no | Clec4a1 | intron_variant | MODIFIER |
| 6 | 122933010 | AX-97786369 | 0 | C | T | 0.184 | -4.27E-02 | 8.22E-02 | 2.67E+00 | 2.80E+00 | 5.95E-01 | no | Clec4a1 | intron_variant | MODIFIER |
| 6 | 122933919 | AX-170340296 | 0 | T | C | 0.184 | -4.27E-02 | 8.22E-02 | 2.67E+00 | 2.80E+00 | 5.95E-01 | no | Clec4a1 | synonymous_variant | LOW |
| 6 | 122941298 | AX-170076099 | 0 | G | A | 0.184 | -4.27E-02 | 8.22E-02 | 2.67E+00 | 2.80E+00 | 5.95E-01 | no | | intergenic_variant | MODIFIER |
| 6 | 122941871 | AX-170076097 | 0 | C | T | 0.184 | -4.27E-02 | 8.22E-02 | 2.67E+00 | 2.80E+00 | 5.95E-01 | no | | intergenic_variant | MODIFIER |
| 6 | 123256002 | AX-97842571 | 0 | C | T | 0.184 | -4.27E-02 | 8.22E-02 | 2.67E+00 | 2.80E+00 | 5.95E-01 | no | | intergenic_variant | MODIFIER |
| 6 | 123686867 | AX-97817260 | 0 | G | C | 0.184 | -4.27E-02 | 8.22E-02 | 2.67E+00 | 2.80E+00 | 5.95E-01 | no | | intergenic_variant | MODIFIER |
| 6 | 123976304 | AX-97840946 | 0 | T | C | 0.184 | -4.27E-02 | 8.22E-02 | 2.67E+00 | 2.80E+00 | 5.95E-01 | no | A330044P14Rik | intron_variant,non_coding_transcript_variant | MODIFIER |
| 6 | 124082448 | AX-97838926 | 0 | T | C | 0.184 | -4.27E-02 | 8.22E-02 | 2.67E+00 | 2.80E+00 | 5.95E-01 | no | Vmn2r-ps33 | non_coding_transcript_exon_variant | MODIFIER |
| 6 | 124145149 | AX-97792306 | 0 | C | T | 0.184 | -4.27E-02 | 8.22E-02 | 2.67E+00 | 2.80E+00 | 5.95E-01 | no | Gm23554 | downstream_gene_variant | MODIFIER |
| 6 | 124297797 | AX-170292757 | 0 | G | T | 0.184 | -4.27E-02 | 8.22E-02 | 2.67E+00 | 2.80E+00 | 5.95E-01 | no | Gm6308 | upstream_gene_variant | MODIFIER |
| 6 | 124299053 | AX-170382956 | 0 | C | T | 0.184 | -4.27E-02 | 8.22E-02 | 2.67E+00 | 2.80E+00 | 5.95E-01 | no | Gm6308 | downstream_gene_variant | MODIFIER |
| 6 | 124328502 | AX-97849903 | 0 | T | G | 0.184 | -4.27E-02 | 8.22E-02 | 2.67E+00 | 2.80E+00 | 5.95E-01 | no | Cd163 | intron_variant | MODIFIER |
| 6 | 124376386 | AX-170155538 | 0 | C | T | 0.184 | -4.27E-02 | 8.22E-02 | 2.67E+00 | 2.80E+00 | 5.95E-01 | no | Gm44006 | downstream_gene_variant | MODIFIER |
| 6 | 124627721 | AX-170095796 | 0 | T | C | 0.184 | -4.27E-02 | 8.22E-02 | 2.67E+00 | 2.80E+00 | 5.95E-01 | no | C1s2 | intron_variant | MODIFIER |
| 6 | 124920953 | AX-97814103 | 0 | G | A | 0.184 | -4.27E-02 | 8.22E-02 | 2.67E+00 | 2.80E+00 | 5.95E-01 | no | Ptms | upstream_gene_variant | MODIFIER |
| 6 | 124952622 | AX-170028170 | 0 | A | G | 0.184 | -4.27E-02 | 8.22E-02 | 2.67E+00 | 2.80E+00 | 5.95E-01 | no | Gm32536 | upstream_gene_variant | MODIFIER |
| 6 | 124953312 | AX-170089833 | 0 | G | T | 0.184 | -4.27E-02 | 8.22E-02 | 2.67E+00 | 2.80E+00 | 5.95E-01 | no | Gm32536 | upstream_gene_variant | MODIFIER |
| 6 | 124965844 | AX-170415780 | 0 | T | C | 0.184 | -4.27E-02 | 8.22E-02 | 2.67E+00 | 2.80E+00 | 5.95E-01 | no | Cops7a | upstream_gene_variant | MODIFIER |
| 6 | 124988604 | AX-97789817 | 0 | C | T | 0.184 | -4.27E-02 | 8.22E-02 | 2.67E+00 | 2.80E+00 | 5.95E-01 | no | | intergenic_variant | MODIFIER |
| 6 | 124991374 | AX-170284635 | 0 | G | A | 0.184 | -4.27E-02 | 8.22E-02 | 2.67E+00 | 2.80E+00 | 5.95E-01 | no | | intergenic_variant | MODIFIER |
| 6 | 125019403 | AX-170187730 | 0 | C | G | 0.184 | -4.27E-02 | 8.22E-02 | 2.67E+00 | 2.80E+00 | 5.95E-01 | no | Zfp384 | intron_variant | MODIFIER |
| 6 | 125027744 | AX-170293072 | 0 | C | A | 0.184 | -4.27E-02 | 8.22E-02 | 2.67E+00 | 2.80E+00 | 5.95E-01 | no | Zfp384 | intron_variant | MODIFIER |
| 6 | 125028066 | AX-170465674 | 0 | T | C | 0.184 | -4.27E-02 | 8.22E-02 | 2.67E+00 | 2.80E+00 | 5.95E-01 | no | Zfp384 | intron_variant | MODIFIER |
| 6 | 125041263 | AX-170190891 | 0 | G | A | 0.184 | -4.27E-02 | 8.22E-02 | 2.67E+00 | 2.80E+00 | 5.95E-01 | no | Ing4 | intron_variant | MODIFIER |
| 6 | 125048885 | AX-97814957 | 0 | T | C | 0.184 | -4.27E-02 | 8.22E-02 | 2.67E+00 | 2.80E+00 | 5.95E-01 | no | Ing4 | 3_prime_UTR_variant | MODIFIER |
| 6 | 125109454 | AX-170076132 | 0 | T | C | 0.184 | -4.27E-02 | 8.22E-02 | 2.67E+00 | 2.80E+00 | 5.95E-01 | no | Chd4 | synonymous_variant | LOW |
| 6 | 125112456 | AX-170528634 | 0 | T | C | 0.184 | -4.27E-02 | 8.22E-02 | 2.67E+00 | 2.80E+00 | 5.95E-01 | no | Chd4 | intron_variant | MODIFIER |
| 6 | 125121076 | AX-170265335 | 0 | T | G | 0.184 | -4.27E-02 | 8.22E-02 | 2.67E+00 | 2.80E+00 | 5.95E-01 | no | Chd4 | intron_variant | MODIFIER |
| 6 | 125121089 | AX-170364249 | 0 | C | T | 0.184 | -4.27E-02 | 8.22E-02 | 2.67E+00 | 2.80E+00 | 5.95E-01 | no | Chd4 | intron_variant | MODIFIER |
| 6 | 125133368 | AX-170504359 | 0 | T | C | 0.184 | -4.27E-02 | 8.22E-02 | 2.67E+00 | 2.80E+00 | 5.95E-01 | no | Nop2 | intron_variant | MODIFIER |
| 6 | 125139462 | AX-170271652 | 0 | C | T | 0.184 | -4.27E-02 | 8.22E-02 | 2.67E+00 | 2.80E+00 | 5.95E-01 | no | Nop2 | intron_variant | MODIFIER |
| 6 | 125139738 | AX-170390494 | 0 | C | T | 0.184 | -4.27E-02 | 8.22E-02 | 2.67E+00 | 2.80E+00 | 5.95E-01 | no | Nop2 | intron_variant | MODIFIER |

| | | | | | | | | | | | | | | | |
|---|---|---|---|---|---|---|---|---|---|---|---|---|---|---|---|
| 6 | 125144372 | AX-170225461 | 0 | A | G | 0.184 | -4.27E-02 | 8.22E-02 | 2.67E+00 | 2.80E+00 | 5.95E-01 | no | Nop2 | missense_variant | MODERATE |
| 6 | 125169856 | AX-170292092 | 0 | C | T | 0.184 | -4.27E-02 | 8.22E-02 | 2.67E+00 | 2.80E+00 | 5.95E-01 | no | Ncapd2 | missense_variant | MODERATE |
| 6 | 125175078 | AX-170287284 | 0 | A | G | 0.184 | -4.27E-02 | 8.22E-02 | 2.67E+00 | 2.80E+00 | 5.95E-01 | no | Ncapd2 | intron_variant | MODIFIER |
| 6 | 125176741 | AX-170362311 | 0 | T | A | 0.184 | -4.27E-02 | 8.22E-02 | 2.67E+00 | 2.80E+00 | 5.95E-01 | no | Ncapd2 | synonymous_variant | LOW |
| 6 | 125176912 | AX-170264006 | 0 | C | A | 0.184 | -4.27E-02 | 8.22E-02 | 2.67E+00 | 2.80E+00 | 5.95E-01 | no | Ncapd2 | intron_variant | MODIFIER |
| 6 | 125177266 | AX-97796270 | 0 | C | A | 0.184 | -4.27E-02 | 8.22E-02 | 2.67E+00 | 2.80E+00 | 5.95E-01 | no | Ncapd2 | intron_variant | MODIFIER |
| 6 | 125179902 | AX-170251294 | 0 | A | G | 0.184 | -4.27E-02 | 8.22E-02 | 2.67E+00 | 2.80E+00 | 5.95E-01 | no | Ncapd2 | intron_variant | MODIFIER |
| 6 | 125185953 | AX-170372010 | 0 | G | A | 0.184 | -4.27E-02 | 8.22E-02 | 2.67E+00 | 2.80E+00 | 5.95E-01 | no | Ncapd2 | intron_variant | MODIFIER |
| 6 | 125204964 | AX-97835503 | 0 | G | A | 0.184 | -4.27E-02 | 8.22E-02 | 2.67E+00 | 2.80E+00 | 5.95E-01 | no | Gm44487 | downstream_gene_variant | MODIFIER |
| 6 | 125221321 | AX-170407193 | 0 | C | T | 0.184 | -4.27E-02 | 8.22E-02 | 2.67E+00 | 2.80E+00 | 5.95E-01 | no | Vamp1 | 3_prime_UTR_variant | MODIFIER |
| 6 | 125222647 | AX-170462742 | 0 | T | C | 0.184 | -4.27E-02 | 8.22E-02 | 2.67E+00 | 2.80E+00 | 5.95E-01 | no | Vamp1 | downstream_gene_variant | MODIFIER |
| 6 | 125226010 | AX-170111764 | 0 | C | G | 0.184 | -4.27E-02 | 8.22E-02 | 2.67E+00 | 2.80E+00 | 5.95E-01 | no | Vamp1 | downstream_gene_variant | MODIFIER |
| 6 | 125233180 | AX-97856253 | 0 | T | C | 0.184 | -4.27E-02 | 8.22E-02 | 2.67E+00 | 2.80E+00 | 5.95E-01 | no | Cd27 | 3_prime_UTR_variant | MODIFIER |
| 6 | 125284767 | AX-97814985 | 0 | G | A | 0.184 | -4.27E-02 | 8.22E-02 | 2.67E+00 | 2.80E+00 | 5.95E-01 | no | Tuba3a | intron_variant | MODIFIER |
| 6 | 125287814 | AX-170150760 | 0 | G | A | 0.184 | -4.27E-02 | 8.22E-02 | 2.67E+00 | 2.80E+00 | 5.95E-01 | no | Tuba3a | upstream_gene_variant | MODIFIER |
| 6 | 125331628 | AX-170028179 | 0 | T | A | 0.184 | -4.27E-02 | 8.22E-02 | 2.67E+00 | 2.80E+00 | 5.95E-01 | no | Scnn1a | intron_variant | MODIFIER |
| 6 | 125332158 | AX-97804624 | 0 | A | G | 0.184 | -4.27E-02 | 8.22E-02 | 2.67E+00 | 2.80E+00 | 5.95E-01 | no | Scnn1a | synonymous_variant | LOW |
| 6 | 125333665 | AX-170028177 | 0 | T | C | 0.184 | -4.27E-02 | 8.22E-02 | 2.67E+00 | 2.80E+00 | 5.95E-01 | no | Scnn1a | intron_variant | MODIFIER |
| 6 | 125370697 | AX-170028178 | 0 | A | G | 0.184 | -4.27E-02 | 8.22E-02 | 2.67E+00 | 2.80E+00 | 5.95E-01 | no | Plekhg6 | synonymous_variant | LOW |
| 6 | 125370844 | AX-170028172 | 0 | G | T | 0.184 | -4.27E-02 | 8.22E-02 | 2.67E+00 | 2.80E+00 | 5.95E-01 | no | Plekhg6 | intron_variant | MODIFIER |
| 6 | 125375388 | AX-170361216 | 0 | A | G | 0.184 | -4.27E-02 | 8.22E-02 | 2.67E+00 | 2.80E+00 | 5.95E-01 | no | Plekhg6 | intron_variant | MODIFIER |
| 6 | 125431709 | AX-97799108 | 0 | C | T | 0.184 | -4.27E-02 | 8.22E-02 | 2.67E+00 | 2.80E+00 | 5.95E-01 | no | | intergenic_variant | MODIFIER |
| 6 | 125433104 | AX-170185148 | 0 | A | G | 0.184 | -4.27E-02 | 8.22E-02 | 2.67E+00 | 2.80E+00 | 5.95E-01 | no | | regulatory_region_variant | MODIFIER |
| 6 | 125452357 | AX-97782697 | 0 | T | C | 0.184 | -4.27E-02 | 8.22E-02 | 2.67E+00 | 2.80E+00 | 5.95E-01 | no | | intergenic_variant | MODIFIER |
| 6 | 125809043 | AX-97811513 | 0 | A | G | 0.184 | -4.27E-02 | 8.22E-02 | 2.67E+00 | 2.80E+00 | 5.95E-01 | no | Ano2 | intron_variant,non_coding_transcript_variant | MODIFIER |
| 6 | 125812494 | AX-170469122 | 0 | G | A | 0.184 | -4.27E-02 | 8.22E-02 | 2.67E+00 | 2.80E+00 | 5.95E-01 | no | Ano2 | intron_variant,non_coding_transcript_variant | MODIFIER |
| 6 | 125812519 | AX-170238244 | 0 | A | G | 0.184 | -4.27E-02 | 8.22E-02 | 2.67E+00 | 2.80E+00 | 5.95E-01 | no | Ano2 | intron_variant,non_coding_transcript_variant | MODIFIER |
| 6 | 125969800 | AX-170520496 | 0 | T | C | 0.184 | -4.27E-02 | 8.22E-02 | 2.67E+00 | 2.80E+00 | 5.95E-01 | no | Ano2 | intron_variant,NMD_transcript_variant | MODIFIER |
| 6 | 125969957 | AX-170223055 | 0 | C | T | 0.184 | -4.27E-02 | 8.22E-02 | 2.67E+00 | 2.80E+00 | 5.95E-01 | no | Ano2 | intron_variant,NMD_transcript_variant | MODIFIER |
| 6 | 125971855 | AX-170420552 | 0 | T | C | 0.184 | -4.27E-02 | 8.22E-02 | 2.67E+00 | 2.80E+00 | 5.95E-01 | no | Ano2 | intron_variant,NMD_transcript_variant | MODIFIER |
| 6 | 125979395 | AX-97813802 | 0 | T | C | 0.184 | -4.27E-02 | 8.22E-02 | 2.67E+00 | 2.80E+00 | 5.95E-01 | no | Ano2 | intron_variant,NMD_transcript_variant | MODIFIER |
| 6 | 126143103 | AX-170487818 | 0 | T | C | 0.184 | -4.27E-02 | 8.22E-02 | 2.67E+00 | 2.80E+00 | 5.95E-01 | no | Ntf3 | intron_variant | MODIFIER |
| 6 | 126143262 | AX-170170717 | 0 | G | A | 0.184 | -4.27E-02 | 8.22E-02 | 2.67E+00 | 2.80E+00 | 5.95E-01 | no | Ntf3 | intron_variant | MODIFIER |
| 6 | 126144620 | AX-170300178 | 0 | T | G | 0.184 | -4.27E-02 | 8.22E-02 | 2.67E+00 | 2.80E+00 | 5.95E-01 | no | Ntf3 | intron_variant | MODIFIER |
| 6 | 126144845 | AX-170276725 | 0 | A | G | 0.184 | -4.27E-02 | 8.22E-02 | 2.67E+00 | 2.80E+00 | 5.95E-01 | no | Ntf3 | intron_variant | MODIFIER |
| 6 | 126246551 | AX-170286526 | 0 | T | C | 0.184 | -4.27E-02 | 8.22E-02 | 2.67E+00 | 2.80E+00 | 5.95E-01 | no | | intergenic_variant | MODIFIER |
| 6 | 126276440 | AX-170185971 | 0 | C | G | 0.184 | -4.27E-02 | 8.22E-02 | 2.67E+00 | 2.80E+00 | 5.95E-01 | no | | intergenic_variant | MODIFIER |
| 6 | 126277311 | AX-170238632 | 0 | C | T | 0.184 | -4.27E-02 | 8.22E-02 | 2.67E+00 | 2.80E+00 | 5.95E-01 | no | | intergenic_variant | MODIFIER |
| 6 | 126277722 | AX-170382828 | 0 | A | G | 0.184 | -4.27E-02 | 8.22E-02 | 2.67E+00 | 2.80E+00 | 5.95E-01 | no | | intergenic_variant | MODIFIER |
| 6 | 126282882 | AX-170473896 | 0 | A | C | 0.184 | -4.27E-02 | 8.22E-02 | 2.67E+00 | 2.80E+00 | 5.95E-01 | no | Gm10415 | downstream_gene_variant | MODIFIER |
| 6 | 126284351 | AX-170268745 | 0 | C | T | 0.184 | -4.27E-02 | 8.22E-02 | 2.67E+00 | 2.80E+00 | 5.95E-01 | no | Gm10415 | downstream_gene_variant | MODIFIER |
| 6 | 126287539 | AX-170527624 | 0 | G | C | 0.184 | -4.27E-02 | 8.22E-02 | 2.67E+00 | 2.80E+00 | 5.95E-01 | no | Gm10415 | non_coding_transcript_exon_variant | MODIFIER |
| 6 | 126288237 | AX-170076147 | 0 | T | C | 0.184 | -4.27E-02 | 8.22E-02 | 2.67E+00 | 2.80E+00 | 5.95E-01 | no | Gm10415 | non_coding_transcript_exon_variant | MODIFIER |
| 6 | 126288328 | AX-170141400 | 0 | C | T | 0.184 | -4.27E-02 | 8.22E-02 | 2.67E+00 | 2.80E+00 | 5.95E-01 | no | Gm10415 | non_coding_transcript_exon_variant | MODIFIER |
| 6 | 126290232 | AX-170450372 | 0 | T | C | 0.184 | -4.27E-02 | 8.22E-02 | 2.67E+00 | 2.80E+00 | 5.95E-01 | no | Gm10415 | intron_variant,non_coding_transcript_variant | MODIFIER |
| 6 | 126304373 | AX-97786148 | 0 | C | T | 0.184 | -4.27E-02 | 8.22E-02 | 2.67E+00 | 2.80E+00 | 5.95E-01 | no | Gm10415 | intron_variant,non_coding_transcript_variant | MODIFIER |
| 6 | 126307081 | AX-170396350 | 0 | C | T | 0.184 | -4.27E-02 | 8.22E-02 | 2.67E+00 | 2.80E+00 | 5.95E-01 | no | Gm10415 | intron_variant,non_coding_transcript_variant | MODIFIER |
| 6 | 126311455 | AX-170215956 | 0 | T | G | 0.184 | -4.27E-02 | 8.22E-02 | 2.67E+00 | 2.80E+00 | 5.95E-01 | no | Gm10415 | intron_variant,non_coding_transcript_variant | MODIFIER |
| 6 | 126319300 | AX-170281072 | 0 | A | G | 0.184 | -4.27E-02 | 8.22E-02 | 2.67E+00 | 2.80E+00 | 5.95E-01 | no | Gm10415 | intron_variant,non_coding_transcript_variant | MODIFIER |
| 8 | 39445435 | AX-170151434 | 1 | A | G | 0.152 | 6.83E-02 | 1.31E-01 | 2.72E+00 | 2.78E+00 | 5.95E-01 | no | | intergenic_variant | MODIFIER |
| 10 | 120246161 | AX-170004033 | 0 | C | G | 0.342 | 3.70E-02 | 7.12E-02 | 2.72E+00 | 2.82E+00 | 5.95E-01 | no | | regulatory_region_variant | MODIFIER |
| 6 | 93647816 | AX-170224308 | 0 | C | T | 0.439 | -3.23E-02 | 6.00E-02 | 2.54E+00 | 2.66E+00 | 5.95E-01 | no | Gm7812 | downstream_gene_variant | MODIFIER |
| 16 | 32773974 | AX-170280034 | 0 | C | T | 0.491 | 4.24E-02 | 7.85E-02 | 2.57E+00 | 2.62E+00 | 5.95E-01 | no | Muc4 | intron_variant | MODIFIER |
| 16 | 77101643 | AX-170334593 | 0 | G | C | 0.456 | 4.14E-02 | 7.68E-02 | 2.56E+00 | 2.63E+00 | 5.96E-01 | no | Usp25 | intron_variant | MODIFIER |

| | | | | | | | | | | | | | | | |
|---|---|---|---|---|---|---|---|---|---|---|---|---|---|---|---|
| 2 | 124582302 | AX-170064806 | 0 | T | C | 0.149 | 9.43E-02 | 1.80E-01 | 2.67E+00 | 2.76E+00 | 5.96E-01 | no | Sema6d | intron_variant | MODIFIER |
| 18 | 46705027 | AX-170529456 | 0 | C | T | 0.061 | 1.01E-01 | 1.94E-01 | 2.62E+00 | 2.74E+00 | 5.96E-01 | no | | intergenic_variant | MODIFIER |
| 11 | 11128557 | AX-170527591 | 0 | G | A | 0.07 | 8.56E-02 | 1.66E-01 | 2.68E+00 | 2.83E+00 | 5.96E-01 | no | Vwc2 | intron_variant | MODIFIER |
| 5 | 90315155 | AX-97834638 | 1 | G | C | 0.42 | 5.65E-02 | 1.08E-01 | 2.72E+00 | 2.77E+00 | 5.96E-01 | no | Ankrd17 | intron_variant | MODIFIER |
| 13 | 39306905 | AX-97822028 | 0 | G | A | 0.184 | -6.16E-02 | 1.09E-01 | 2.47E+00 | 2.55E+00 | 5.96E-01 | no | | regulatory_region_variant | MODIFIER |
| 13 | 39307276 | AX-170133917 | 0 | T | C | 0.184 | -6.16E-02 | 1.09E-01 | 2.47E+00 | 2.55E+00 | 5.96E-01 | no | | regulatory_region_variant | MODIFIER |
| 13 | 39307305 | AX-170305939 | 0 | C | G | 0.184 | -6.16E-02 | 1.09E-01 | 2.47E+00 | 2.55E+00 | 5.96E-01 | no | | regulatory_region_variant | MODIFIER |
| 13 | 39307349 | AX-170311898 | 0 | T | C | 0.184 | -6.16E-02 | 1.09E-01 | 2.47E+00 | 2.55E+00 | 5.96E-01 | no | | regulatory_region_variant | MODIFIER |
| 13 | 39354463 | AX-170204396 | 0 | G | A | 0.184 | -6.16E-02 | 1.09E-01 | 2.47E+00 | 2.55E+00 | 5.96E-01 | no | | intergenic_variant | MODIFIER |
| 13 | 39379110 | AX-170124917 | 0 | A | G | 0.184 | -6.16E-02 | 1.09E-01 | 2.47E+00 | 2.55E+00 | 5.96E-01 | no | | intergenic_variant | MODIFIER |
| 13 | 39379482 | AX-170270081 | 0 | C | T | 0.184 | -6.16E-02 | 1.09E-01 | 2.47E+00 | 2.55E+00 | 5.96E-01 | no | | intergenic_variant | MODIFIER |
| 13 | 39382861 | AX-170232033 | 0 | C | T | 0.184 | -6.16E-02 | 1.09E-01 | 2.47E+00 | 2.55E+00 | 5.96E-01 | no | | intergenic_variant | MODIFIER |
| 13 | 39387086 | AX-170375490 | 0 | C | A | 0.184 | -6.16E-02 | 1.09E-01 | 2.47E+00 | 2.55E+00 | 5.96E-01 | no | | intergenic_variant | MODIFIER |
| 13 | 39453168 | AX-170259390 | 0 | G | A | 0.184 | -6.16E-02 | 1.09E-01 | 2.47E+00 | 2.55E+00 | 5.96E-01 | no | Gm47348 | downstream_gene_variant | MODIFIER |
| 13 | 39586749 | AX-97849131 | 0 | G | A | 0.184 | -6.16E-02 | 1.09E-01 | 2.47E+00 | 2.55E+00 | 5.96E-01 | no | Gm47352 | intron_variant,non_coding_transcript_variant | MODIFIER |
| 13 | 39589809 | AX-170328104 | 0 | A | G | 0.184 | -6.16E-02 | 1.09E-01 | 2.47E+00 | 2.55E+00 | 5.96E-01 | no | Gm47352 | intron_variant,non_coding_transcript_variant | MODIFIER |
| 13 | 39612780 | AX-97823303 | 0 | C | A | 0.184 | -6.16E-02 | 1.09E-01 | 2.47E+00 | 2.55E+00 | 5.96E-01 | no | Gm32243 | downstream_gene_variant | MODIFIER |
| 13 | 39642452 | AX-170295924 | 0 | T | G | 0.184 | -6.16E-02 | 1.09E-01 | 2.47E+00 | 2.55E+00 | 5.96E-01 | no | Gm32243 | intron_variant,non_coding_transcript_variant | MODIFIER |
| 13 | 39642481 | AX-170370906 | 0 | T | C | 0.184 | -6.16E-02 | 1.09E-01 | 2.47E+00 | 2.55E+00 | 5.96E-01 | no | Gm32243 | intron_variant,non_coding_transcript_variant | MODIFIER |
| 13 | 39643885 | AX-97828596 | 0 | A | G | 0.184 | -6.16E-02 | 1.09E-01 | 2.47E+00 | 2.55E+00 | 5.96E-01 | no | Gm32243 | intron_variant,non_coding_transcript_variant | MODIFIER |
| 13 | 39683251 | AX-170127240 | 0 | A | G | 0.184 | -6.16E-02 | 1.09E-01 | 2.47E+00 | 2.55E+00 | 5.96E-01 | no | Gm47352 | intron_variant,non_coding_transcript_variant | MODIFIER |
| 13 | 39683267 | AX-97849277 | 0 | A | G | 0.184 | -6.16E-02 | 1.09E-01 | 2.47E+00 | 2.55E+00 | 5.96E-01 | no | Gm47352 | intron_variant,non_coding_transcript_variant | MODIFIER |
| 13 | 39707710 | AX-97782312 | 0 | A | G | 0.184 | -6.16E-02 | 1.09E-01 | 2.47E+00 | 2.55E+00 | 5.96E-01 | no | | intergenic_variant | MODIFIER |
| 13 | 39781492 | AX-170051045 | 0 | T | C | 0.184 | -6.16E-02 | 1.09E-01 | 2.47E+00 | 2.55E+00 | 5.96E-01 | no | Gm40923 | intron_variant,non_coding_transcript_variant | MODIFIER |
| 13 | 39781978 | AX-170051044 | 0 | G | A | 0.184 | -6.16E-02 | 1.09E-01 | 2.47E+00 | 2.55E+00 | 5.96E-01 | no | Gm40923 | intron_variant,non_coding_transcript_variant | MODIFIER |
| 13 | 39783888 | AX-170289794 | 0 | A | G | 0.184 | -6.16E-02 | 1.09E-01 | 2.47E+00 | 2.55E+00 | 5.96E-01 | no | Gm40923 | intron_variant,non_coding_transcript_variant | MODIFIER |
| 13 | 39789027 | AX-170292675 | 0 | G | A | 0.184 | -6.16E-02 | 1.09E-01 | 2.47E+00 | 2.55E+00 | 5.96E-01 | no | A230103O09Rik | upstream_gene_variant | MODIFIER |
| 13 | 39794000 | AX-170479301 | 0 | A | G | 0.184 | -6.16E-02 | 1.09E-01 | 2.47E+00 | 2.55E+00 | 5.96E-01 | no | A230103O09Rik | intron_variant,non_coding_transcript_variant | MODIFIER |
| 13 | 39794037 | AX-170421392 | 0 | G | A | 0.184 | -6.16E-02 | 1.09E-01 | 2.47E+00 | 2.55E+00 | 5.96E-01 | no | A230103O09Rik | intron_variant,non_coding_transcript_variant | MODIFIER |
| 13 | 40388482 | AX-170179456 | 0 | A | T | 0.184 | -6.16E-02 | 1.09E-01 | 2.47E+00 | 2.55E+00 | 5.96E-01 | no | | intergenic_variant | MODIFIER |
| 13 | 40391664 | AX-170182956 | 0 | C | T | 0.184 | -6.16E-02 | 1.09E-01 | 2.47E+00 | 2.55E+00 | 5.96E-01 | no | | intergenic_variant | MODIFIER |
| 13 | 40396365 | AX-170126856 | 0 | G | A | 0.184 | -6.16E-02 | 1.09E-01 | 2.47E+00 | 2.55E+00 | 5.96E-01 | no | | intergenic_variant | MODIFIER |
| 13 | 40475458 | AX-170455298 | 0 | T | C | 0.184 | -6.16E-02 | 1.09E-01 | 2.47E+00 | 2.55E+00 | 5.96E-01 | no | | intergenic_variant | MODIFIER |
| 13 | 40476047 | AX-170380575 | 0 | G | T | 0.184 | -6.16E-02 | 1.09E-01 | 2.47E+00 | 2.55E+00 | 5.96E-01 | no | | intergenic_variant | MODIFIER |
| 13 | 40524870 | AX-170154642 | 0 | A | G | 0.184 | -6.16E-02 | 1.09E-01 | 2.47E+00 | 2.55E+00 | 5.96E-01 | no | | intergenic_variant | MODIFIER |
| 13 | 40553240 | AX-97823302 | 0 | A | G | 0.184 | -6.16E-02 | 1.09E-01 | 2.47E+00 | 2.55E+00 | 5.96E-01 | no | | intergenic_variant | MODIFIER |
| 15 | 82764439 | AX-170225530 | 1 | G | A | 0.562 | 4.49E-02 | 8.17E-02 | 2.52E+00 | 2.59E+00 | 5.96E-01 | no | Cyp2d40 | upstream_gene_variant | MODIFIER |
| 1 | 174243530 | AX-170528136 | 1 | G | A | 0.232 | 6.60E-02 | 1.22E-01 | 2.55E+00 | 2.61E+00 | 5.96E-01 | no | Spta1 | intron_variant | MODIFIER |
| 20 | 64673311 | AX-170034009 | 0 | G | A | 0.061 | 5.16E-02 | 1.00E-01 | 2.60E+00 | 2.79E+00 | 5.96E-01 | no | | intergenic_variant | MODIFIER |
| 9 | 122692268 | AX-97792309 | 0 | T | C | 0.14 | -4.64E-02 | 8.95E-02 | 2.67E+00 | 2.80E+00 | 5.96E-01 | no | Gm35501 | downstream_gene_variant | MODIFIER |
| 9 | 122719246 | AX-170336455 | 0 | G | A | 0.14 | -4.64E-02 | 8.95E-02 | 2.67E+00 | 2.80E+00 | 5.96E-01 | no | Gm47136 | upstream_gene_variant | MODIFIER |
| 9 | 122723657 | AX-97820163 | 0 | T | C | 0.14 | -4.64E-02 | 8.95E-02 | 2.67E+00 | 2.80E+00 | 5.96E-01 | no | | intergenic_variant | MODIFIER |
| 9 | 122730769 | AX-170162142 | 0 | C | T | 0.14 | -4.64E-02 | 8.95E-02 | 2.67E+00 | 2.80E+00 | 5.96E-01 | no | | intergenic_variant | MODIFIER |
| 9 | 122737033 | AX-170408975 | 0 | T | C | 0.14 | -4.64E-02 | 8.95E-02 | 2.67E+00 | 2.80E+00 | 5.96E-01 | no | | regulatory_region_variant | MODIFIER |
| 9 | 122877718 | AX-170449573 | 0 | C | T | 0.14 | -4.64E-02 | 8.95E-02 | 2.67E+00 | 2.80E+00 | 5.96E-01 | no | | intergenic_variant | MODIFIER |
| 9 | 123148774 | AX-97784655 | 0 | T | C | 0.14 | -4.64E-02 | 8.95E-02 | 2.67E+00 | 2.80E+00 | 5.96E-01 | no | Clec3b | upstream_gene_variant | MODIFIER |
| 9 | 123148889 | AX-170517607 | 0 | G | A | 0.14 | -4.64E-02 | 8.95E-02 | 2.67E+00 | 2.80E+00 | 5.96E-01 | no | Clec3b | upstream_gene_variant | MODIFIER |
| 9 | 123150736 | AX-170121235 | 0 | G | A | 0.14 | -4.64E-02 | 8.95E-02 | 2.67E+00 | 2.80E+00 | 5.96E-01 | no | Clec3b | upstream_gene_variant | MODIFIER |
| 9 | 123153990 | AX-170451030 | 0 | C | G | 0.14 | -4.64E-02 | 8.95E-02 | 2.67E+00 | 2.80E+00 | 5.96E-01 | no | Clec3b | intron_variant | MODIFIER |
| 9 | 123206721 | AX-97831712 | 0 | C | T | 0.14 | -4.64E-02 | 8.95E-02 | 2.67E+00 | 2.80E+00 | 5.96E-01 | no | Cdcp1 | intron_variant | MODIFIER |
| 9 | 123231477 | AX-97780221 | 0 | T | C | 0.14 | -4.64E-02 | 8.95E-02 | 2.67E+00 | 2.80E+00 | 5.96E-01 | no | | intergenic_variant | MODIFIER |
| 9 | 123276880 | AX-97823962 | 0 | T | C | 0.14 | -4.64E-02 | 8.95E-02 | 2.67E+00 | 2.80E+00 | 5.96E-01 | no | | intergenic_variant | MODIFIER |
| 9 | 123277349 | AX-170467443 | 0 | A | T | 0.14 | -4.64E-02 | 8.95E-02 | 2.67E+00 | 2.80E+00 | 5.96E-01 | no | | intergenic_variant | MODIFIER |
| 9 | 123278252 | AX-170425121 | 0 | T | C | 0.14 | -4.64E-02 | 8.95E-02 | 2.67E+00 | 2.80E+00 | 5.96E-01 | no | | intergenic_variant | MODIFIER |

| | | | | | | | | | | | | | | | |
|---|---|---|---|---|---|---|---|---|---|---|---|---|---|---|---|
| 1 | 43174392 | AX-168397702 | 0 | G | C | 0.096 | -1.10E-01 | 2.12E-01 | 2.71E+00 | 2.80E+00 | 5.96E-01 | no | Fhl2 | intron_variant | MODIFIER |
| 1 | 43190813 | AX-170000375 | 0 | A | G | 0.096 | -1.10E-01 | 2.12E-01 | 2.71E+00 | 2.80E+00 | 5.96E-01 | no | Gm8210 | downstream_gene_variant | MODIFIER |
| 1 | 43197026 | AX-97833416 | 0 | G | C | 0.096 | -1.10E-01 | 2.12E-01 | 2.71E+00 | 2.80E+00 | 5.96E-01 | no | Fhl2 | upstream_gene_variant | MODIFIER |
| 1 | 43202543 | AX-170202719 | 0 | C | T | 0.096 | -1.10E-01 | 2.12E-01 | 2.71E+00 | 2.80E+00 | 5.96E-01 | no | | regulatory_region_variant | MODIFIER |
| 1 | 43202835 | AX-170293582 | 0 | A | G | 0.096 | -1.10E-01 | 2.12E-01 | 2.71E+00 | 2.80E+00 | 5.96E-01 | no | | regulatory_region_variant | MODIFIER |
| 1 | 43225309 | AX-170367943 | 0 | A | G | 0.096 | -1.10E-01 | 2.12E-01 | 2.71E+00 | 2.80E+00 | 5.96E-01 | no | | regulatory_region_variant | MODIFIER |
| 1 | 43233482 | AX-97857553 | 0 | T | C | 0.096 | -1.10E-01 | 2.12E-01 | 2.71E+00 | 2.80E+00 | 5.96E-01 | no | | intergenic_variant | MODIFIER |
| 1 | 43235907 | AX-170475417 | 0 | A | C | 0.096 | -1.10E-01 | 2.12E-01 | 2.71E+00 | 2.80E+00 | 5.96E-01 | no | | intergenic_variant | MODIFIER |
| 1 | 43258610 | AX-170121129 | 0 | C | T | 0.096 | -1.10E-01 | 2.12E-01 | 2.71E+00 | 2.80E+00 | 5.96E-01 | no | Gm29040 | intron_variant,non_coding_transcript_variant | MODIFIER |
| 1 | 43262432 | AX-170000378 | 0 | C | T | 0.096 | -1.10E-01 | 2.12E-01 | 2.71E+00 | 2.80E+00 | 5.96E-01 | no | Gm29040 | intron_variant,non_coding_transcript_variant | MODIFIER |
| 1 | 43266850 | AX-170041048 | 0 | A | G | 0.096 | -1.10E-01 | 2.12E-01 | 2.71E+00 | 2.80E+00 | 5.96E-01 | no | Gm29610 | upstream_gene_variant | MODIFIER |
| 1 | 43268122 | AX-170041050 | 0 | G | A | 0.096 | -1.10E-01 | 2.12E-01 | 2.71E+00 | 2.80E+00 | 5.96E-01 | no | Gm29610 | upstream_gene_variant | MODIFIER |
| 1 | 43279346 | AX-97811377 | 0 | T | G | 0.096 | -1.10E-01 | 2.12E-01 | 2.71E+00 | 2.80E+00 | 5.96E-01 | no | Gm29040 | intron_variant,non_coding_transcript_variant | MODIFIER |
| 1 | 43280601 | AX-170000379 | 0 | T | G | 0.096 | -1.10E-01 | 2.12E-01 | 2.71E+00 | 2.80E+00 | 5.96E-01 | no | Gm29040 | intron_variant,non_coding_transcript_variant | MODIFIER |
| 1 | 43292361 | AX-170367083 | 0 | T | C | 0.096 | -1.10E-01 | 2.12E-01 | 2.71E+00 | 2.80E+00 | 5.96E-01 | no | Gm29040 | intron_variant,non_coding_transcript_variant | MODIFIER |
| 1 | 43297017 | AX-170459526 | 0 | T | G | 0.096 | -1.10E-01 | 2.12E-01 | 2.71E+00 | 2.80E+00 | 5.96E-01 | no | Gm29040 | intron_variant,non_coding_transcript_variant | MODIFIER |
| 1 | 43297193 | AX-170483161 | 0 | T | C | 0.096 | -1.10E-01 | 2.12E-01 | 2.71E+00 | 2.80E+00 | 5.96E-01 | no | Gm29040 | intron_variant,non_coding_transcript_variant | MODIFIER |
| 1 | 43297261 | AX-170518888 | 0 | A | G | 0.096 | -1.10E-01 | 2.12E-01 | 2.71E+00 | 2.80E+00 | 5.96E-01 | no | Gm29040 | intron_variant,non_coding_transcript_variant | MODIFIER |
| 1 | 43299018 | AX-170345215 | 0 | G | C | 0.096 | -1.10E-01 | 2.12E-01 | 2.71E+00 | 2.80E+00 | 5.96E-01 | no | Gm29040 | intron_variant,non_coding_transcript_variant | MODIFIER |
| 1 | 43314672 | AX-97811400 | 0 | A | G | 0.096 | -1.10E-01 | 2.12E-01 | 2.71E+00 | 2.80E+00 | 5.96E-01 | no | | intergenic_variant | MODIFIER |
| 1 | 43314918 | AX-170232059 | 0 | A | G | 0.096 | -1.10E-01 | 2.12E-01 | 2.71E+00 | 2.80E+00 | 5.96E-01 | no | | intergenic_variant | MODIFIER |
| 1 | 43318292 | AX-170299888 | 0 | T | C | 0.096 | -1.10E-01 | 2.12E-01 | 2.71E+00 | 2.80E+00 | 5.96E-01 | no | | intergenic_variant | MODIFIER |
| 1 | 43321934 | AX-170473904 | 0 | A | G | 0.096 | -1.10E-01 | 2.12E-01 | 2.71E+00 | 2.80E+00 | 5.96E-01 | no | | intergenic_variant | MODIFIER |
| 1 | 43341969 | AX-170091483 | 0 | G | A | 0.096 | -1.10E-01 | 2.12E-01 | 2.71E+00 | 2.80E+00 | 5.96E-01 | no | | intergenic_variant | MODIFIER |
| 1 | 43342649 | AX-170381131 | 0 | G | A | 0.096 | -1.10E-01 | 2.12E-01 | 2.71E+00 | 2.80E+00 | 5.96E-01 | no | | intergenic_variant | MODIFIER |
| 1 | 43342903 | AX-170159140 | 0 | A | G | 0.096 | -1.10E-01 | 2.12E-01 | 2.71E+00 | 2.80E+00 | 5.96E-01 | no | | intergenic_variant | MODIFIER |
| 1 | 43342984 | AX-170461231 | 0 | A | G | 0.096 | -1.10E-01 | 2.12E-01 | 2.71E+00 | 2.80E+00 | 5.96E-01 | no | | intergenic_variant | MODIFIER |
| 1 | 43393242 | AX-170421980 | 0 | T | C | 0.096 | -1.10E-01 | 2.12E-01 | 2.71E+00 | 2.80E+00 | 5.96E-01 | no | | intergenic_variant | MODIFIER |
| 1 | 43394963 | AX-97786775 | 0 | G | C | 0.096 | -1.10E-01 | 2.12E-01 | 2.71E+00 | 2.80E+00 | 5.96E-01 | no | | intergenic_variant | MODIFIER |
| 1 | 43436646 | AX-97811380 | 0 | C | A | 0.096 | -1.10E-01 | 2.12E-01 | 2.71E+00 | 2.80E+00 | 5.96E-01 | no | | intergenic_variant | MODIFIER |
| 1 | 43458888 | AX-170443215 | 0 | G | T | 0.096 | -1.10E-01 | 2.12E-01 | 2.71E+00 | 2.80E+00 | 5.96E-01 | no | Nck2 | intron_variant | MODIFIER |
| 1 | 43465055 | AX-170532537 | 0 | T | C | 0.096 | -1.10E-01 | 2.12E-01 | 2.71E+00 | 2.80E+00 | 5.96E-01 | no | Nck2 | intron_variant | MODIFIER |
| 1 | 43475442 | AX-170441177 | 0 | A | G | 0.096 | -1.10E-01 | 2.12E-01 | 2.71E+00 | 2.80E+00 | 5.96E-01 | no | Nck2 | intron_variant | MODIFIER |
| 1 | 43478158 | AX-170272156 | 0 | A | G | 0.096 | -1.10E-01 | 2.12E-01 | 2.71E+00 | 2.80E+00 | 5.96E-01 | no | Nck2 | intron_variant | MODIFIER |
| 1 | 43483683 | AX-170355841 | 0 | T | G | 0.096 | -1.10E-01 | 2.12E-01 | 2.71E+00 | 2.80E+00 | 5.96E-01 | no | Nck2 | intron_variant | MODIFIER |
| 1 | 43487376 | AX-170182163 | 0 | C | T | 0.096 | -1.10E-01 | 2.12E-01 | 2.71E+00 | 2.80E+00 | 5.96E-01 | no | Nck2 | intron_variant | MODIFIER |
| 1 | 43489826 | AX-170278267 | 0 | A | G | 0.096 | -1.10E-01 | 2.12E-01 | 2.71E+00 | 2.80E+00 | 5.96E-01 | no | Nck2 | intron_variant | MODIFIER |
| 1 | 43503115 | AX-170482318 | 0 | C | T | 0.096 | -1.10E-01 | 2.12E-01 | 2.71E+00 | 2.80E+00 | 5.96E-01 | no | Nck2 | intron_variant | MODIFIER |
| 1 | 43505561 | AX-170320728 | 0 | T | A | 0.096 | -1.10E-01 | 2.12E-01 | 2.71E+00 | 2.80E+00 | 5.96E-01 | no | Nck2 | intron_variant | MODIFIER |
| 1 | 43506408 | AX-170510593 | 0 | A | G | 0.096 | -1.10E-01 | 2.12E-01 | 2.71E+00 | 2.80E+00 | 5.96E-01 | no | Nck2 | intron_variant | MODIFIER |
| 1 | 43509070 | AX-97816000 | 0 | G | T | 0.096 | -1.10E-01 | 2.12E-01 | 2.71E+00 | 2.80E+00 | 5.96E-01 | no | Nck2 | intron_variant | MODIFIER |
| 1 | 43609717 | AX-170216636 | 0 | C | T | 0.096 | -1.10E-01 | 2.12E-01 | 2.71E+00 | 2.80E+00 | 5.96E-01 | no | | intergenic_variant | MODIFIER |
| 1 | 43685749 | AX-97855839 | 0 | A | G | 0.096 | -1.10E-01 | 2.12E-01 | 2.71E+00 | 2.80E+00 | 5.96E-01 | no | | intergenic_variant | MODIFIER |
| 1 | 43686349 | AX-170313599 | 0 | A | G | 0.096 | -1.10E-01 | 2.12E-01 | 2.71E+00 | 2.80E+00 | 5.96E-01 | no | | intergenic_variant | MODIFIER |
| 1 | 43686478 | AX-170000386 | 0 | A | G | 0.096 | -1.10E-01 | 2.12E-01 | 2.71E+00 | 2.80E+00 | 5.96E-01 | no | | intergenic_variant | MODIFIER |
| 1 | 43686631 | AX-170529941 | 0 | T | C | 0.096 | -1.10E-01 | 2.12E-01 | 2.71E+00 | 2.80E+00 | 5.96E-01 | no | | intergenic_variant | MODIFIER |
| 1 | 43713483 | AX-170000381 | 0 | C | G | 0.096 | -1.10E-01 | 2.12E-01 | 2.71E+00 | 2.80E+00 | 5.96E-01 | no | | intergenic_variant | MODIFIER |
| 1 | 43713757 | AX-170000385 | 0 | A | G | 0.096 | -1.10E-01 | 2.12E-01 | 2.71E+00 | 2.80E+00 | 5.96E-01 | no | | intergenic_variant | MODIFIER |
| 1 | 43713783 | AX-170456189 | 0 | T | C | 0.096 | -1.10E-01 | 2.12E-01 | 2.71E+00 | 2.80E+00 | 5.96E-01 | no | | intergenic_variant | MODIFIER |
| 1 | 43717630 | AX-170268636 | 0 | T | G | 0.096 | -1.10E-01 | 2.12E-01 | 2.71E+00 | 2.80E+00 | 5.96E-01 | no | | intergenic_variant | MODIFIER |
| 1 | 43741897 | AX-170263617 | 0 | A | G | 0.096 | -1.10E-01 | 2.12E-01 | 2.71E+00 | 2.80E+00 | 5.96E-01 | no | 1500015O10Rik | intron_variant | MODIFIER |
| 1 | 43742395 | AX-97833117 | 0 | G | A | 0.096 | -1.10E-01 | 2.12E-01 | 2.71E+00 | 2.80E+00 | 5.96E-01 | no | 1500015O10Rik | missense_variant | MODERATE |
| 1 | 43765809 | AX-97811329 | 0 | A | G | 0.096 | -1.10E-01 | 2.12E-01 | 2.71E+00 | 2.80E+00 | 5.96E-01 | no | Uxs1 | intron_variant,NMD_transcript_variant | MODIFIER |
| 1 | 43766030 | AX-170296160 | 0 | G | A | 0.096 | -1.10E-01 | 2.12E-01 | 2.71E+00 | 2.80E+00 | 5.96E-01 | no | Uxs1 | intron_variant,NMD_transcript_variant | MODIFIER |

| | | | | | | | | | | | | | | | |
|---|---|---|---|---|---|---|---|---|---|---|---|---|---|---|---|
| 1 | 43770376 | AX-170181941 | 0 | G | T | 0.096 | -1.10E-01 | 2.12E-01 | 2.71E+00 | 2.80E+00 | 5.96E-01 | no | Uxs1 | intron_variant,NMD_transcript_variant | MODIFIER |
| 1 | 43777191 | AX-170351367 | 0 | A | G | 0.096 | -1.10E-01 | 2.12E-01 | 2.71E+00 | 2.80E+00 | 5.96E-01 | no | Uxs1 | intron_variant,NMD_transcript_variant | MODIFIER |
| 1 | 43777355 | AX-170514127 | 0 | A | C | 0.096 | -1.10E-01 | 2.12E-01 | 2.71E+00 | 2.80E+00 | 5.96E-01 | no | Uxs1 | intron_variant,NMD_transcript_variant | MODIFIER |
| 1 | 43780300 | AX-97805962 | 0 | C | T | 0.096 | -1.10E-01 | 2.12E-01 | 2.71E+00 | 2.80E+00 | 5.96E-01 | no | Uxs1 | intron_variant,NMD_transcript_variant | MODIFIER |
| 1 | 43782183 | AX-170297311 | 0 | C | G | 0.096 | -1.10E-01 | 2.12E-01 | 2.71E+00 | 2.80E+00 | 5.96E-01 | no | Uxs1 | intron_variant,NMD_transcript_variant | MODIFIER |
| 1 | 43837088 | AX-170459237 | 0 | C | T | 0.096 | -1.10E-01 | 2.12E-01 | 2.71E+00 | 2.80E+00 | 5.96E-01 | no | | intergenic_variant | MODIFIER |
| 1 | 43837668 | AX-170165343 | 0 | T | C | 0.096 | -1.10E-01 | 2.12E-01 | 2.71E+00 | 2.80E+00 | 5.96E-01 | no | | intergenic_variant | MODIFIER |
| 1 | 44061378 | AX-170430404 | 0 | A | G | 0.096 | -1.10E-01 | 2.12E-01 | 2.71E+00 | 2.80E+00 | 5.96E-01 | no | Gm8251 | missense_variant | MODERATE |
| 1 | 44067676 | AX-97841976 | 0 | T | G | 0.096 | -1.10E-01 | 2.12E-01 | 2.71E+00 | 2.80E+00 | 5.96E-01 | no | | intergenic_variant | MODIFIER |
| 1 | 44067814 | AX-170508841 | 0 | A | C | 0.096 | -1.10E-01 | 2.12E-01 | 2.71E+00 | 2.80E+00 | 5.96E-01 | no | | intergenic_variant | MODIFIER |
| 1 | 44068327 | AX-170193379 | 0 | T | G | 0.096 | -1.10E-01 | 2.12E-01 | 2.71E+00 | 2.80E+00 | 5.96E-01 | no | | intergenic_variant | MODIFIER |
| 1 | 44076267 | AX-170517925 | 0 | A | G | 0.096 | -1.10E-01 | 2.12E-01 | 2.71E+00 | 2.80E+00 | 5.96E-01 | no | | intergenic_variant | MODIFIER |
| 1 | 44080397 | AX-170428135 | 0 | G | A | 0.096 | -1.10E-01 | 2.12E-01 | 2.71E+00 | 2.80E+00 | 5.96E-01 | no | | intergenic_variant | MODIFIER |
| 1 | 44080632 | AX-170344313 | 0 | T | A | 0.096 | -1.10E-01 | 2.12E-01 | 2.71E+00 | 2.80E+00 | 5.96E-01 | no | | intergenic_variant | MODIFIER |
| 1 | 44081183 | AX-170041069 | 0 | A | T | 0.096 | -1.10E-01 | 2.12E-01 | 2.71E+00 | 2.80E+00 | 5.96E-01 | no | | intergenic_variant | MODIFIER |
| 1 | 44106513 | AX-170484912 | 0 | A | G | 0.096 | -1.10E-01 | 2.12E-01 | 2.71E+00 | 2.80E+00 | 5.96E-01 | no | Kdelc1 | downstream_gene_variant | MODIFIER |
| 1 | 44110759 | AX-170360141 | 0 | T | C | 0.096 | -1.10E-01 | 2.12E-01 | 2.71E+00 | 2.80E+00 | 5.96E-01 | no | Kdelc1 | intron_variant | MODIFIER |
| 1 | 44112255 | AX-97811478 | 0 | T | C | 0.096 | -1.10E-01 | 2.12E-01 | 2.71E+00 | 2.80E+00 | 5.96E-01 | no | Kdelc1 | intron_variant | MODIFIER |
| 1 | 44112600 | AX-170468654 | 0 | C | G | 0.096 | -1.10E-01 | 2.12E-01 | 2.71E+00 | 2.80E+00 | 5.96E-01 | no | Kdelc1 | intron_variant | MODIFIER |
| 1 | 44126939 | AX-170239233 | 0 | A | T | 0.096 | -1.10E-01 | 2.12E-01 | 2.71E+00 | 2.80E+00 | 5.96E-01 | no | Bivm | intron_variant | MODIFIER |
| 1 | 44149341 | AX-170127800 | 0 | C | T | 0.096 | -1.10E-01 | 2.12E-01 | 2.71E+00 | 2.80E+00 | 5.96E-01 | no | Ercc5 | intron_variant | MODIFIER |
| 1 | 44155784 | AX-170520468 | 0 | C | T | 0.096 | -1.10E-01 | 2.12E-01 | 2.71E+00 | 2.80E+00 | 5.96E-01 | no | Ercc5 | intron_variant | MODIFIER |
| 1 | 44156861 | AX-170410022 | 0 | C | T | 0.096 | -1.10E-01 | 2.12E-01 | 2.71E+00 | 2.80E+00 | 5.96E-01 | no | Ercc5 | intron_variant | MODIFIER |
| 1 | 44159389 | AX-170294860 | 0 | C | T | 0.096 | -1.10E-01 | 2.12E-01 | 2.71E+00 | 2.80E+00 | 5.96E-01 | no | Ercc5 | intron_variant | MODIFIER |
| 1 | 44159470 | AX-170467411 | 0 | A | G | 0.096 | -1.10E-01 | 2.12E-01 | 2.71E+00 | 2.80E+00 | 5.96E-01 | no | Ercc5 | intron_variant | MODIFIER |
| 1 | 44162120 | AX-170517453 | 0 | G | A | 0.096 | -1.10E-01 | 2.12E-01 | 2.71E+00 | 2.80E+00 | 5.96E-01 | no | Ercc5 | intron_variant | MODIFIER |
| 1 | 44165708 | AX-170041074 | 0 | G | A | 0.096 | -1.10E-01 | 2.12E-01 | 2.71E+00 | 2.80E+00 | 5.96E-01 | no | Ercc5 | intron_variant | MODIFIER |
| 1 | 44167341 | AX-170041073 | 0 | T | A | 0.096 | -1.10E-01 | 2.12E-01 | 2.71E+00 | 2.80E+00 | 5.96E-01 | no | Ercc5 | synonymous_variant | LOW |
| 1 | 44169641 | AX-170370595 | 0 | T | C | 0.096 | -1.10E-01 | 2.12E-01 | 2.71E+00 | 2.80E+00 | 5.96E-01 | no | Ercc5 | intron_variant | MODIFIER |
| 1 | 44169958 | AX-170041071 | 0 | C | A | 0.096 | -1.10E-01 | 2.12E-01 | 2.71E+00 | 2.80E+00 | 5.96E-01 | no | Ercc5 | intron_variant | MODIFIER |
| 1 | 44188755 | AX-170398387 | 0 | G | A | 0.096 | -1.10E-01 | 2.12E-01 | 2.71E+00 | 2.80E+00 | 5.96E-01 | no | | intergenic_variant | MODIFIER |
| 1 | 44189119 | AX-170000391 | 0 | A | G | 0.096 | -1.10E-01 | 2.12E-01 | 2.71E+00 | 2.80E+00 | 5.96E-01 | no | | intergenic_variant | MODIFIER |
| 1 | 44201909 | AX-170144047 | 0 | T | A | 0.096 | -1.10E-01 | 2.12E-01 | 2.71E+00 | 2.80E+00 | 5.96E-01 | no | Mettl21e | downstream_gene_variant | MODIFIER |
| 1 | 44204723 | AX-170418453 | 0 | T | C | 0.096 | -1.10E-01 | 2.12E-01 | 2.71E+00 | 2.80E+00 | 5.96E-01 | no | Mettl21e | 3_prime_UTR_variant | MODIFIER |
| 1 | 44205192 | AX-170232356 | 0 | T | C | 0.096 | -1.10E-01 | 2.12E-01 | 2.71E+00 | 2.80E+00 | 5.96E-01 | no | Mettl21e | 3_prime_UTR_variant | MODIFIER |
| 1 | 44213894 | AX-170000388 | 0 | T | C | 0.096 | -1.10E-01 | 2.12E-01 | 2.71E+00 | 2.80E+00 | 5.96E-01 | no | Mettl21e | intron_variant | MODIFIER |
| 1 | 44215866 | AX-170000387 | 0 | G | C | 0.096 | -1.10E-01 | 2.12E-01 | 2.71E+00 | 2.80E+00 | 5.96E-01 | no | Mettl21e | intron_variant | MODIFIER |
| 1 | 44382417 | AX-97791448 | 0 | T | C | 0.096 | -1.10E-01 | 2.12E-01 | 2.71E+00 | 2.80E+00 | 5.96E-01 | no | | intergenic_variant | MODIFIER |
| 1 | 44461688 | AX-170508893 | 0 | A | G | 0.096 | -1.10E-01 | 2.12E-01 | 2.71E+00 | 2.80E+00 | 5.96E-01 | no | Txn-ps1 | downstream_gene_variant | MODIFIER |
| 1 | 44464395 | AX-170103674 | 0 | A | T | 0.096 | -1.10E-01 | 2.12E-01 | 2.71E+00 | 2.80E+00 | 5.96E-01 | no | Txn-ps1 | upstream_gene_variant | MODIFIER |
| 1 | 44543431 | AX-170437960 | 0 | G | A | 0.096 | -1.10E-01 | 2.12E-01 | 2.71E+00 | 2.80E+00 | 5.96E-01 | no | | intergenic_variant | MODIFIER |
| 1 | 44551989 | AX-170190933 | 0 | G | A | 0.096 | -1.10E-01 | 2.12E-01 | 2.71E+00 | 2.80E+00 | 5.96E-01 | no | Gulp1 | intron_variant | MODIFIER |
| 1 | 44567477 | AX-170225536 | 0 | A | G | 0.096 | -1.10E-01 | 2.12E-01 | 2.71E+00 | 2.80E+00 | 5.96E-01 | no | Gulp1 | intron_variant | MODIFIER |
| 1 | 44581300 | AX-97841716 | 0 | A | G | 0.096 | -1.10E-01 | 2.12E-01 | 2.71E+00 | 2.80E+00 | 5.96E-01 | no | Gulp1 | intron_variant | MODIFIER |
| 1 | 44583918 | AX-170216142 | 0 | C | T | 0.096 | -1.10E-01 | 2.12E-01 | 2.71E+00 | 2.80E+00 | 5.96E-01 | no | Gulp1 | intron_variant | MODIFIER |
| 1 | 44598694 | AX-170416739 | 0 | A | G | 0.096 | -1.10E-01 | 2.12E-01 | 2.71E+00 | 2.80E+00 | 5.96E-01 | no | Gulp1 | intron_variant | MODIFIER |
| 1 | 44610668 | AX-170283660 | 0 | C | T | 0.096 | -1.10E-01 | 2.12E-01 | 2.71E+00 | 2.80E+00 | 5.96E-01 | no | Gulp1 | intron_variant | MODIFIER |
| 1 | 44611104 | AX-170041078 | 0 | T | C | 0.096 | -1.10E-01 | 2.12E-01 | 2.71E+00 | 2.80E+00 | 5.96E-01 | no | Gulp1 | intron_variant | MODIFIER |
| 1 | 44611381 | AX-170431604 | 0 | A | G | 0.096 | -1.10E-01 | 2.12E-01 | 2.71E+00 | 2.80E+00 | 5.96E-01 | no | Gulp1 | intron_variant | MODIFIER |
| 1 | 44623437 | AX-170091488 | 0 | T | G | 0.096 | -1.10E-01 | 2.12E-01 | 2.71E+00 | 2.80E+00 | 5.96E-01 | no | Gulp1 | intron_variant | MODIFIER |
| 1 | 44634792 | AX-170389663 | 0 | A | G | 0.096 | -1.10E-01 | 2.12E-01 | 2.71E+00 | 2.80E+00 | 5.96E-01 | no | Gulp1 | intron_variant | MODIFIER |
| 1 | 44636767 | AX-170503506 | 0 | C | G | 0.096 | -1.10E-01 | 2.12E-01 | 2.71E+00 | 2.80E+00 | 5.96E-01 | no | Gulp1 | intron_variant | MODIFIER |
| 1 | 44637667 | AX-170041079 | 0 | C | T | 0.096 | -1.10E-01 | 2.12E-01 | 2.71E+00 | 2.80E+00 | 5.96E-01 | no | Gulp1 | intron_variant | MODIFIER |
| 1 | 44646451 | AX-170429663 | 0 | G | A | 0.096 | -1.10E-01 | 2.12E-01 | 2.71E+00 | 2.80E+00 | 5.96E-01 | no | Gulp1 | intron_variant | MODIFIER |

| | | | | | | | | | | | | | | | |
|---|---|---|---|---|---|---|---|---|---|---|---|---|---|---|---|
| 1 | 44666555 | AX-97795984 | 0 | T | C | 0.096 | -1.10E-01 | 2.12E-01 | 2.71E+00 | 2.80E+00 | 5.96E-01 | no | Gulp1 | intron_variant | MODIFIER |
| 1 | 44679264 | AX-97795382 | 0 | A | G | 0.096 | -1.10E-01 | 2.12E-01 | 2.71E+00 | 2.80E+00 | 5.96E-01 | no | Gulp1 | intron_variant | MODIFIER |
| 1 | 44711084 | AX-97785237 | 0 | T | C | 0.096 | -1.10E-01 | 2.12E-01 | 2.71E+00 | 2.80E+00 | 5.96E-01 | no | Gulp1 | intron_variant | MODIFIER |
| 1 | 44760263 | AX-170000399 | 0 | A | T | 0.096 | -1.10E-01 | 2.12E-01 | 2.71E+00 | 2.80E+00 | 5.96E-01 | no | Gulp1 | intron_variant | MODIFIER |
| 1 | 44760433 | AX-170000397 | 0 | A | G | 0.096 | -1.10E-01 | 2.12E-01 | 2.71E+00 | 2.80E+00 | 5.96E-01 | no | Gulp1 | intron_variant | MODIFIER |
| 1 | 44762099 | AX-170000395 | 0 | T | C | 0.096 | -1.10E-01 | 2.12E-01 | 2.71E+00 | 2.80E+00 | 5.96E-01 | no | Gulp1 | intron_variant | MODIFIER |
| 1 | 44782122 | AX-170086458 | 0 | A | G | 0.096 | -1.10E-01 | 2.12E-01 | 2.71E+00 | 2.80E+00 | 5.96E-01 | no | Gulp1 | intron_variant | MODIFIER |
| 1 | 45099422 | AX-170535579 | 0 | G | A | 0.096 | -1.10E-01 | 2.12E-01 | 2.71E+00 | 2.80E+00 | 5.96E-01 | no | | intergenic_variant | MODIFIER |
| 1 | 45132763 | AX-170423774 | 0 | T | C | 0.096 | -1.10E-01 | 2.12E-01 | 2.71E+00 | 2.80E+00 | 5.96E-01 | no | | intergenic_variant | MODIFIER |
| 1 | 45134066 | AX-97812947 | 0 | A | G | 0.096 | -1.10E-01 | 2.12E-01 | 2.71E+00 | 2.80E+00 | 5.96E-01 | no | | intergenic_variant | MODIFIER |
| 1 | 45137124 | AX-170274049 | 0 | A | T | 0.096 | -1.10E-01 | 2.12E-01 | 2.71E+00 | 2.80E+00 | 5.96E-01 | no | | intergenic_variant | MODIFIER |
| 1 | 45137160 | AX-170324753 | 0 | A | G | 0.096 | -1.10E-01 | 2.12E-01 | 2.71E+00 | 2.80E+00 | 5.96E-01 | no | | intergenic_variant | MODIFIER |
| 1 | 45160824 | AX-97846523 | 0 | C | A | 0.096 | -1.10E-01 | 2.12E-01 | 2.71E+00 | 2.80E+00 | 5.96E-01 | no | | intergenic_variant | MODIFIER |
| 1 | 45198113 | AX-170315482 | 0 | A | G | 0.096 | -1.10E-01 | 2.12E-01 | 2.71E+00 | 2.80E+00 | 5.96E-01 | no | | intergenic_variant | MODIFIER |
| 1 | 45245917 | AX-170217864 | 0 | G | A | 0.096 | -1.10E-01 | 2.12E-01 | 2.71E+00 | 2.80E+00 | 5.96E-01 | no | | regulatory_region_variant | MODIFIER |
| 1 | 45286282 | AX-170513381 | 0 | T | C | 0.096 | -1.10E-01 | 2.12E-01 | 2.71E+00 | 2.80E+00 | 5.96E-01 | no | | intergenic_variant | MODIFIER |
| 1 | 45310196 | AX-170354990 | 0 | T | G | 0.096 | -1.10E-01 | 2.12E-01 | 2.71E+00 | 2.80E+00 | 5.96E-01 | no | Col3a1 | upstream_gene_variant | MODIFIER |
| 1 | 45310433 | AX-97811406 | 0 | A | T | 0.096 | -1.10E-01 | 2.12E-01 | 2.71E+00 | 2.80E+00 | 5.96E-01 | no | Col3a1 | upstream_gene_variant | MODIFIER |
| 1 | 45353175 | AX-170376225 | 0 | T | C | 0.096 | -1.10E-01 | 2.12E-01 | 2.71E+00 | 2.80E+00 | 5.96E-01 | no | Col3a1 | downstream_gene_variant | MODIFIER |
| 1 | 45421014 | AX-170401838 | 0 | C | A | 0.096 | -1.10E-01 | 2.12E-01 | 2.71E+00 | 2.80E+00 | 5.96E-01 | no | Col5a2 | intron_variant | MODIFIER |
| 1 | 45427258 | AX-170526529 | 0 | T | C | 0.096 | -1.10E-01 | 2.12E-01 | 2.71E+00 | 2.80E+00 | 5.96E-01 | no | Col5a2 | intron_variant | MODIFIER |
| 1 | 45440116 | AX-170172096 | 0 | A | T | 0.096 | -1.10E-01 | 2.12E-01 | 2.71E+00 | 2.80E+00 | 5.96E-01 | no | Col5a2 | intron_variant | MODIFIER |
| 1 | 45454954 | AX-170422343 | 0 | T | C | 0.096 | -1.10E-01 | 2.12E-01 | 2.71E+00 | 2.80E+00 | 5.96E-01 | no | Col5a2 | intron_variant | MODIFIER |
| 1 | 45489869 | AX-170451612 | 0 | A | G | 0.096 | -1.10E-01 | 2.12E-01 | 2.71E+00 | 2.80E+00 | 5.96E-01 | no | Col5a2 | intron_variant | MODIFIER |
| 1 | 45528416 | AX-170254002 | 0 | A | G | 0.096 | -1.10E-01 | 2.12E-01 | 2.71E+00 | 2.80E+00 | 5.96E-01 | no | | intergenic_variant | MODIFIER |
| 1 | 45573493 | AX-170145329 | 0 | T | G | 0.096 | -1.10E-01 | 2.12E-01 | 2.71E+00 | 2.80E+00 | 5.96E-01 | no | | intergenic_variant | MODIFIER |
| 1 | 45577883 | AX-170041098 | 0 | T | C | 0.096 | -1.10E-01 | 2.12E-01 | 2.71E+00 | 2.80E+00 | 5.96E-01 | no | | intergenic_variant | MODIFIER |
| 1 | 45588750 | AX-170135422 | 0 | T | C | 0.096 | -1.10E-01 | 2.12E-01 | 2.71E+00 | 2.80E+00 | 5.96E-01 | no | | intergenic_variant | MODIFIER |
| 1 | 45628497 | AX-170521947 | 0 | A | C | 0.096 | -1.10E-01 | 2.12E-01 | 2.71E+00 | 2.80E+00 | 5.96E-01 | no | | intergenic_variant | MODIFIER |
| 1 | 45908699 | AX-170176856 | 0 | C | T | 0.096 | -1.10E-01 | 2.12E-01 | 2.71E+00 | 2.80E+00 | 5.96E-01 | no | Slc40a1 | 3_prime_UTR_variant | MODIFIER |
| 1 | 45924401 | AX-170240420 | 0 | C | T | 0.096 | -1.10E-01 | 2.12E-01 | 2.71E+00 | 2.80E+00 | 5.96E-01 | no | Slc40a1 | intron_variant | MODIFIER |
| 1 | 45940156 | AX-170000405 | 0 | G | A | 0.096 | -1.10E-01 | 2.12E-01 | 2.71E+00 | 2.80E+00 | 5.96E-01 | no | | intergenic_variant | MODIFIER |
| 1 | 45940535 | AX-170000409 | 0 | C | T | 0.096 | -1.10E-01 | 2.12E-01 | 2.71E+00 | 2.80E+00 | 5.96E-01 | no | | intergenic_variant | MODIFIER |
| 1 | 45940977 | AX-170492360 | 0 | A | G | 0.096 | -1.10E-01 | 2.12E-01 | 2.71E+00 | 2.80E+00 | 5.96E-01 | no | | intergenic_variant | MODIFIER |
| 1 | 45967918 | AX-170000408 | 0 | G | C | 0.096 | -1.10E-01 | 2.12E-01 | 2.71E+00 | 2.80E+00 | 5.96E-01 | no | Gm9496 | upstream_gene_variant | MODIFIER |
| 1 | 46023844 | AX-170041112 | 0 | C | G | 0.096 | -1.10E-01 | 2.12E-01 | 2.71E+00 | 2.80E+00 | 5.96E-01 | no | Stk-ps2 | intron_variant,non_coding_transcript_variant | MODIFIER |
| 1 | 46025842 | AX-170109043 | 0 | G | A | 0.096 | -1.10E-01 | 2.12E-01 | 2.71E+00 | 2.80E+00 | 5.96E-01 | no | Stk-ps2 | intron_variant,non_coding_transcript_variant | MODIFIER |
| 1 | 46034865 | AX-170000407 | 0 | T | C | 0.096 | -1.10E-01 | 2.12E-01 | 2.71E+00 | 2.80E+00 | 5.96E-01 | no | Stk-ps2 | upstream_gene_variant | MODIFIER |
| 1 | 46035728 | AX-170359765 | 0 | G | A | 0.096 | -1.10E-01 | 2.12E-01 | 2.71E+00 | 2.80E+00 | 5.96E-01 | no | | intergenic_variant | MODIFIER |
| 1 | 46036395 | AX-170118192 | 0 | A | G | 0.096 | -1.10E-01 | 2.12E-01 | 2.71E+00 | 2.80E+00 | 5.96E-01 | no | | intergenic_variant | MODIFIER |
| 1 | 46218179 | AX-170530649 | 0 | A | G | 0.096 | -1.10E-01 | 2.12E-01 | 2.71E+00 | 2.80E+00 | 5.96E-01 | no | Dnah7b | intron_variant | MODIFIER |
| 1 | 46360940 | AX-97820764 | 0 | C | T | 0.096 | -1.10E-01 | 2.12E-01 | 2.71E+00 | 2.80E+00 | 5.96E-01 | no | Dnah7b | intron_variant | MODIFIER |
| 1 | 46362045 | AX-170134322 | 0 | C | T | 0.096 | -1.10E-01 | 2.12E-01 | 2.71E+00 | 2.80E+00 | 5.96E-01 | no | Dnah7b | intron_variant | MODIFIER |
| 1 | 46382723 | AX-170498985 | 0 | C | T | 0.096 | -1.10E-01 | 2.12E-01 | 2.71E+00 | 2.80E+00 | 5.96E-01 | no | | intergenic_variant | MODIFIER |
| 1 | 46398595 | AX-170133907 | 0 | C | G | 0.096 | -1.10E-01 | 2.12E-01 | 2.71E+00 | 2.80E+00 | 5.96E-01 | no | | intergenic_variant | MODIFIER |
| 1 | 46423176 | AX-170270647 | 0 | G | C | 0.096 | -1.10E-01 | 2.12E-01 | 2.71E+00 | 2.80E+00 | 5.96E-01 | no | Dnah7c | upstream_gene_variant | MODIFIER |
| 1 | 46424714 | AX-170453850 | 0 | T | C | 0.096 | -1.10E-01 | 2.12E-01 | 2.71E+00 | 2.80E+00 | 5.96E-01 | no | Dnah7c | upstream_gene_variant | MODIFIER |
| 1 | 46457697 | AX-170384027 | 0 | T | C | 0.096 | -1.10E-01 | 2.12E-01 | 2.71E+00 | 2.80E+00 | 5.96E-01 | no | Dnah7c | intron_variant | MODIFIER |
| 1 | 46461467 | AX-170440204 | 0 | T | G | 0.096 | -1.10E-01 | 2.12E-01 | 2.71E+00 | 2.80E+00 | 5.96E-01 | no | Dnah7c | intron_variant | MODIFIER |
| 1 | 46518526 | AX-170399289 | 0 | T | C | 0.096 | -1.10E-01 | 2.12E-01 | 2.71E+00 | 2.80E+00 | 5.96E-01 | no | Dnah7c | intron_variant | MODIFIER |
| 1 | 46518597 | AX-170482272 | 0 | C | G | 0.096 | -1.10E-01 | 2.12E-01 | 2.71E+00 | 2.80E+00 | 5.96E-01 | no | Dnah7c | intron_variant | MODIFIER |
| 1 | 46526853 | AX-170329527 | 0 | T | C | 0.096 | -1.10E-01 | 2.12E-01 | 2.71E+00 | 2.80E+00 | 5.96E-01 | no | Dnah7c | intron_variant | MODIFIER |
| 1 | 46604030 | AX-170334341 | 0 | A | G | 0.096 | -1.10E-01 | 2.12E-01 | 2.71E+00 | 2.80E+00 | 5.96E-01 | no | Dnah7c | upstream_gene_variant | MODIFIER |
| 1 | 46613206 | AX-170168250 | 0 | T | G | 0.096 | -1.10E-01 | 2.12E-01 | 2.71E+00 | 2.80E+00 | 5.96E-01 | no | Dnah7c | intron_variant | MODIFIER |

| | | | | | | | | | | | | | | | |
|---|---|---|---|---|---|---|---|---|---|---|---|---|---|---|---|
| 1 | 46629933 | AX-170000410 | 0 | T | C | 0.096 | -1.10E-01 | 2.12E-01 | 2.71E+00 | 2.80E+00 | 5.96E-01 | no | Dnah7c | downstream_gene_variant | MODIFIER |
| 1 | 46781903 | AX-170207863 | 0 | G | A | 0.096 | -1.10E-01 | 2.12E-01 | 2.71E+00 | 2.80E+00 | 5.96E-01 | no | Dnah7c | intron_variant | MODIFIER |
| 1 | 46804039 | AX-170240274 | 0 | G | A | 0.096 | -1.10E-01 | 2.12E-01 | 2.71E+00 | 2.80E+00 | 5.96E-01 | no | Slc39a10 | downstream_gene_variant | MODIFIER |
| 1 | 46813362 | AX-170438006 | 0 | T | C | 0.096 | -1.10E-01 | 2.12E-01 | 2.71E+00 | 2.80E+00 | 5.96E-01 | no | Slc39a10 | intron_variant | MODIFIER |
| 1 | 46822041 | AX-170281929 | 0 | C | T | 0.096 | -1.10E-01 | 2.12E-01 | 2.71E+00 | 2.80E+00 | 5.96E-01 | no | Slc39a10 | intron_variant | MODIFIER |
| 1 | 46840650 | AX-170407007 | 0 | T | C | 0.096 | -1.10E-01 | 2.12E-01 | 2.71E+00 | 2.80E+00 | 5.96E-01 | no | Slc39a10 | intron_variant | MODIFIER |
| 1 | 46844835 | AX-170499816 | 0 | A | C | 0.096 | -1.10E-01 | 2.12E-01 | 2.71E+00 | 2.80E+00 | 5.96E-01 | no | Slc39a10 | intron_variant | MODIFIER |
| 1 | 46847534 | AX-170407910 | 0 | G | A | 0.096 | -1.10E-01 | 2.12E-01 | 2.71E+00 | 2.80E+00 | 5.96E-01 | no | Slc39a10 | intron_variant | MODIFIER |
| 1 | 46868827 | AX-170306012 | 0 | G | A | 0.096 | -1.10E-01 | 2.12E-01 | 2.71E+00 | 2.80E+00 | 5.96E-01 | no | Slc39a10 | intron_variant | MODIFIER |
| 1 | 46874670 | AX-170041128 | 0 | G | A | 0.096 | -1.10E-01 | 2.12E-01 | 2.71E+00 | 2.80E+00 | 5.96E-01 | no | Slc39a10 | intron_variant | MODIFIER |
| 1 | 46880875 | AX-170480976 | 0 | T | C | 0.096 | -1.10E-01 | 2.12E-01 | 2.71E+00 | 2.80E+00 | 5.96E-01 | no | Slc39a10 | intron_variant | MODIFIER |
| 1 | 46895009 | AX-170478694 | 0 | T | C | 0.096 | -1.10E-01 | 2.12E-01 | 2.71E+00 | 2.80E+00 | 5.96E-01 | no | Gm28527 | downstream_gene_variant | MODIFIER |
| 1 | 46895697 | AX-170394748 | 0 | T | C | 0.096 | -1.10E-01 | 2.12E-01 | 2.71E+00 | 2.80E+00 | 5.96E-01 | no | Gm28527 | downstream_gene_variant | MODIFIER |
| 5 | 5338892 | AX-170395344 | 0 | A | T | 0.439 | -3.45E-02 | 6.35E-02 | 2.52E+00 | 2.64E+00 | 5.96E-01 | no | Cdk14 | intron_variant | MODIFIER |
| 8 | 68343478 | AX-170395980 | 1 | G | A | 0.188 | -7.06E-02 | 1.35E-01 | 2.68E+00 | 2.74E+00 | 5.97E-01 | no | Sh2d4a | intron_variant | MODIFIER |
| 16 | 94483598 | AX-170235127 | 0 | C | T | 0.412 | -4.32E-02 | 8.04E-02 | 2.56E+00 | 2.64E+00 | 5.97E-01 | no | | intergenic_variant | MODIFIER |
| 16 | 24705750 | AX-170350342 | 0 | G | A | 0.254 | 4.43E-02 | 8.56E-02 | 2.64E+00 | 2.78E+00 | 5.97E-01 | no | Lpp | intron_variant | MODIFIER |
| 9 | 69069888 | AX-170366890 | 1 | G | T | 0.188 | -6.65E-02 | 1.27E-01 | 2.69E+00 | 2.75E+00 | 5.97E-01 | no | Rora | intron_variant | MODIFIER |
| 13 | 49293834 | AX-170138035 | 0 | C | A | 0.491 | -3.44E-02 | 6.67E-02 | 2.72E+00 | 2.84E+00 | 5.97E-01 | no | Fgd3 | intron_variant | MODIFIER |
| 8 | 68070107 | AX-97854063 | 1 | C | A | 0.152 | -7.64E-02 | 1.41E-01 | 2.53E+00 | 2.61E+00 | 5.97E-01 | no | Psd3 | intron_variant | MODIFIER |
| 4 | 94738581 | AX-170301838 | 0 | G | A | 0.342 | 5.48E-02 | 1.04E-01 | 2.64E+00 | 2.72E+00 | 5.97E-01 | no | Tek | upstream_gene_variant | MODIFIER |
| 5 | 105526952 | AX-170491174 | 0 | A | C | 0.447 | 5.69E-02 | 1.08E-01 | 2.63E+00 | 2.68E+00 | 5.97E-01 | no | Lrrc8c | intron_variant | MODIFIER |
| 6 | 146932620 | AX-170320182 | 0 | C | T | 0.07 | -1.10E-01 | 2.10E-01 | 2.62E+00 | 2.71E+00 | 5.97E-01 | no | Ppfibp1 | intron_variant | MODIFIER |
| 1 | 136887753 | AX-170174832 | 1 | C | A | 0.205 | 6.14E-02 | 1.18E-01 | 2.69E+00 | 2.78E+00 | 5.97E-01 | no | Nr5a2 | intron_variant | MODIFIER |
| 9 | 56491600 | AX-97804929 | 0 | C | A | 0.211 | 5.94E-02 | 1.14E-01 | 2.69E+00 | 2.76E+00 | 5.98E-01 | no | Hmg20a | intron_variant | MODIFIER |
| 4 | 127337725 | AX-170476692 | 2 | C | G | 0.145 | -6.45E-02 | 1.22E-01 | 2.90E+00 | 3.02E+00 | 5.98E-01 | no | | intergenic_variant | MODIFIER |
| 7 | 92043887 | AX-170078070 | 2 | G | A | 0.145 | 9.19E-02 | 1.78E-01 | 2.67E+00 | 2.78E+00 | 5.98E-01 | no | Dlg2 | intron_variant | MODIFIER |
| 10 | 29433189 | AX-170205389 | 0 | A | T | 0.193 | 5.03E-02 | 9.76E-02 | 2.73E+00 | 2.84E+00 | 5.98E-01 | no | | intergenic_variant | MODIFIER |
| 8 | 30138643 | AX-170315577 | 0 | T | G | 0.167 | 5.78E-02 | 1.11E-01 | 2.70E+00 | 2.78E+00 | 5.98E-01 | no | | intergenic_variant | MODIFIER |
| 5 | 88982524 | AX-170512410 | 1 | T | C | 0.42 | 5.54E-02 | 1.06E-01 | 2.74E+00 | 2.79E+00 | 5.98E-01 | no | Slc4a4 | intron_variant | MODIFIER |
| 5 | 90741903 | AX-170439130 | 1 | T | C | 0.42 | 5.54E-02 | 1.06E-01 | 2.74E+00 | 2.79E+00 | 5.98E-01 | no | | regulatory_region_variant | MODIFIER |
| 16 | 18508027 | AX-97805874 | 1 | C | T | 0.134 | 5.00E-02 | 9.75E-02 | 2.70E+00 | 2.83E+00 | 5.98E-01 | no | Gnb1l | intron_variant,NMD_transcript_variant | MODIFIER |
| 8 | 40466504 | AX-170271768 | 1 | T | C | 0.152 | 7.67E-02 | 1.47E-01 | 2.82E+00 | 2.87E+00 | 5.98E-01 | no | Zdhhc2 | intron_variant | MODIFIER |
| 8 | 40997386 | AX-170451525 | 1 | G | A | 0.152 | 7.67E-02 | 1.47E-01 | 2.82E+00 | 2.87E+00 | 5.98E-01 | no | Mtus1 | intron_variant | MODIFIER |
| 9 | 80464145 | AX-170032797 | 1 | T | C | 0.205 | 6.01E-02 | 1.15E-01 | 2.66E+00 | 2.74E+00 | 5.98E-01 | no | Impg1 | intron_variant | MODIFIER |
| 12 | 93335988 | AX-97785900 | 1 | A | C | 0.062 | 1.23E-01 | 2.36E-01 | 2.73E+00 | 2.79E+00 | 5.99E-01 | no | | intergenic_variant | MODIFIER |
| 12 | 93435649 | AX-170336988 | 1 | G | A | 0.062 | 1.23E-01 | 2.36E-01 | 2.73E+00 | 2.79E+00 | 5.99E-01 | no | 4930559C10Rik | intron_variant,non_coding_transcript_variant | MODIFIER |
| 8 | 32392377 | AX-170345479 | 1 | A | C | 0.134 | -6.75E-02 | 1.29E-01 | 2.65E+00 | 2.72E+00 | 5.99E-01 | no | Nrg1 | intron_variant | MODIFIER |
| 1 | 90599083 | AX-170295808 | 1 | C | T | 0.152 | 1.17E-01 | 2.25E-01 | 2.70E+00 | 2.76E+00 | 5.99E-01 | no | Cops8 | upstream_gene_variant | MODIFIER |
| 5 | 107213189 | AX-170025824 | 1 | C | A | 0.464 | 5.97E-02 | 1.13E-01 | 2.62E+00 | 2.67E+00 | 5.99E-01 | no | Tgfbr3 | intron_variant | MODIFIER |
| 5 | 107623301 | AX-170160541 | 1 | C | T | 0.464 | 5.97E-02 | 1.13E-01 | 2.62E+00 | 2.67E+00 | 5.99E-01 | no | Rpap2 | intron_variant | MODIFIER |
| 5 | 110173232 | AX-170130165 | 1 | C | T | 0.464 | 5.97E-02 | 1.13E-01 | 2.62E+00 | 2.67E+00 | 5.99E-01 | no | Chfr | downstream_gene_variant | MODIFIER |
| 5 | 110672037 | AX-170340160 | 1 | G | A | 0.464 | 5.97E-02 | 1.13E-01 | 2.62E+00 | 2.67E+00 | 5.99E-01 | no | Ep400 | intron_variant | MODIFIER |
| 5 | 114184155 | AX-170507296 | 1 | C | G | 0.464 | 5.97E-02 | 1.13E-01 | 2.62E+00 | 2.67E+00 | 5.99E-01 | no | Acacb | synonymous_variant | LOW |
| 5 | 114415061 | AX-170235144 | 1 | C | T | 0.464 | 5.97E-02 | 1.13E-01 | 2.62E+00 | 2.67E+00 | 5.99E-01 | no | Ube3b | intron_variant | MODIFIER |
| 16 | 97557651 | AX-170120540 | 0 | C | T | 0.465 | 3.95E-02 | 7.67E-02 | 2.78E+00 | 2.90E+00 | 5.99E-01 | no | Mx2 | intron_variant | MODIFIER |
| 5 | 109722805 | AX-97806765 | 1 | T | C | 0.455 | 5.89E-02 | 1.12E-01 | 2.62E+00 | 2.66E+00 | 5.99E-01 | no | | intergenic_variant | MODIFIER |
| 20 | 50716258 | AX-170306266 | 0 | A | G | 0.237 | 3.09E-02 | 5.91E-02 | 2.60E+00 | 2.72E+00 | 5.99E-01 | no | | intergenic_variant | MODIFIER |
| 8 | 3160572 | AX-97803476 | 0 | A | C | 0.272 | 3.88E-02 | 7.55E-02 | 2.68E+00 | 2.79E+00 | 5.99E-01 | no | Insr | intron_variant | MODIFIER |
| 8 | 3184126 | AX-97795205 | 0 | A | G | 0.272 | 3.88E-02 | 7.55E-02 | 2.68E+00 | 2.79E+00 | 5.99E-01 | no | Insr | intron_variant | MODIFIER |
| 8 | 3213522 | AX-97851427 | 0 | G | A | 0.272 | 3.88E-02 | 7.55E-02 | 2.68E+00 | 2.79E+00 | 5.99E-01 | no | Insr | intron_variant | MODIFIER |
| 8 | 3235539 | AX-170079142 | 0 | C | T | 0.272 | 3.88E-02 | 7.55E-02 | 2.68E+00 | 2.79E+00 | 5.99E-01 | no | Insr | intron_variant | MODIFIER |
| 8 | 3254285 | AX-97803274 | 0 | G | A | 0.272 | 3.88E-02 | 7.55E-02 | 2.68E+00 | 2.79E+00 | 5.99E-01 | no | Insr | intron_variant | MODIFIER |
| 8 | 3299881 | AX-97850229 | 0 | C | T | 0.272 | 3.88E-02 | 7.55E-02 | 2.68E+00 | 2.79E+00 | 5.99E-01 | no | | intergenic_variant | MODIFIER |

| | | | | | | | | | | | | | | | |
|---|---|---|---|---|---|---|---|---|---|---|---|---|---|---|---|
| 8 | 3318053 | AX-170030185 | 0 | G | A | 0.272 | 3.88E-02 | 7.55E-02 | 2.68E+00 | 2.79E+00 | 5.99E-01 | no | | intergenic_variant | MODIFIER |
| 9 | 120897482 | AX-97850835 | 0 | G | A | 0.202 | -4.29E-02 | 8.30E-02 | 2.64E+00 | 2.76E+00 | 5.99E-01 | no | | intergenic_variant | MODIFIER |
| 1 | 25782099 | AX-170040667 | 0 | C | A | 0.412 | 4.36E-02 | 8.36E-02 | 2.65E+00 | 2.72E+00 | 5.99E-01 | no | Adgrb3 | intron_variant | MODIFIER |
| 1 | 26409352 | AX-170319613 | 0 | A | T | 0.412 | 4.36E-02 | 8.36E-02 | 2.65E+00 | 2.72E+00 | 5.99E-01 | no | | intergenic_variant | MODIFIER |
| 10 | 17117580 | AX-97814875 | 0 | A | G | 0.211 | -5.09E-02 | 9.64E-02 | 2.58E+00 | 2.68E+00 | 5.99E-01 | no | Gm25382 | downstream_gene_variant | MODIFIER |
| 10 | 17282118 | AX-170002837 | 0 | A | G | 0.211 | -5.09E-02 | 9.64E-02 | 2.58E+00 | 2.68E+00 | 5.99E-01 | no | | intergenic_variant | MODIFIER |
| 1 | 14677022 | AX-170190688 | 0 | C | T | 0.377 | 4.18E-02 | 7.93E-02 | 2.61E+00 | 2.67E+00 | 5.99E-01 | no | | intergenic_variant | MODIFIER |
| 1 | 15750848 | AX-97826083 | 0 | C | T | 0.377 | 4.18E-02 | 7.93E-02 | 2.61E+00 | 2.67E+00 | 5.99E-01 | no | | intergenic_variant | MODIFIER |
| 1 | 89486280 | AX-170375069 | 1 | T | G | 0.143 | 9.07E-02 | 1.73E-01 | 2.63E+00 | 2.70E+00 | 5.99E-01 | no | Agap1 | intron_variant | MODIFIER |
| 1 | 96871146 | AX-170001143 | 1 | G | A | 0.143 | 9.07E-02 | 1.73E-01 | 2.63E+00 | 2.70E+00 | 5.99E-01 | no | Slco4c1 | intron_variant | MODIFIER |
| 5 | 93137699 | AX-170419090 | 0 | G | A | 0.412 | 5.42E-02 | 1.05E-01 | 2.70E+00 | 2.76E+00 | 5.99E-01 | no | SEP11 | intron_variant | MODIFIER |
| 9 | 87336404 | AX-170215247 | 0 | T | G | 0.14 | 6.45E-02 | 1.25E-01 | 2.81E+00 | 2.91E+00 | 5.99E-01 | no | | intergenic_variant | MODIFIER |
| 9 | 26963418 | AX-97780531 | 2 | G | A | 0.455 | 4.87E-02 | 9.24E-02 | 2.59E+00 | 2.68E+00 | 5.99E-01 | no | Gm5120 | upstream_gene_variant | MODIFIER |
| 1 | 45577969 | AX-170041099 | 1 | T | G | 0.098 | -1.08E-01 | 2.10E-01 | 2.71E+00 | 2.80E+00 | 6.00E-01 | no | | intergenic_variant | MODIFIER |
| 20 | 74256803 | AX-170084530 | 0 | A | G | 0.263 | 3.19E-02 | 6.14E-02 | 2.64E+00 | 2.73E+00 | 6.00E-01 | no | Emd | intron_variant | MODIFIER |
| 13 | 23760025 | AX-170050755 | 1 | C | T | 0.161 | -6.88E-02 | 1.25E-01 | 2.49E+00 | 2.55E+00 | 6.00E-01 | no | Hist1h1a | upstream_gene_variant | MODIFIER |
| 15 | 71433312 | AX-97856325 | 1 | T | G | 0.518 | 4.07E-02 | 7.70E-02 | 2.58E+00 | 2.68E+00 | 6.00E-01 | no | Fam135b | 3_prime_UTR_variant | MODIFIER |
| 4 | 70282721 | AX-97794140 | 1 | T | C | 0.232 | 5.58E-02 | 1.04E-01 | 2.54E+00 | 2.62E+00 | 6.00E-01 | no | Cdk5rap2 | intron_variant,NMD_transcript_variant | MODIFIER |
| 1 | 36193380 | AX-170309602 | 0 | C | G | 0.456 | 3.98E-02 | 7.73E-02 | 2.72E+00 | 2.81E+00 | 6.00E-01 | no | Uggt1 | intron_variant | MODIFIER |
| 6 | 104299458 | AX-170263348 | 0 | C | T | 0.061 | 1.15E-01 | 2.18E-01 | 2.57E+00 | 2.68E+00 | 6.00E-01 | no | Gm18100 | upstream_gene_variant | MODIFIER |
| 8 | 51854796 | AX-170090104 | 0 | G | A | 0.167 | 6.56E-02 | 1.27E-01 | 2.81E+00 | 2.91E+00 | 6.00E-01 | no | | intergenic_variant | MODIFIER |
| 5 | 33921407 | AX-170024656 | 1 | C | T | 0.366 | 5.25E-02 | 1.02E-01 | 2.69E+00 | 2.76E+00 | 6.00E-01 | no | Nelfa | intron_variant | MODIFIER |
| 10 | 58346196 | AX-170259258 | 0 | C | A | 0.298 | 4.04E-02 | 7.82E-02 | 2.85E+00 | 2.98E+00 | 6.00E-01 | no | Lims1 | intron_variant | MODIFIER |
| 10 | 127240007 | AX-97783251 | 1 | G | A | 0.321 | -3.83E-02 | 7.35E-02 | 2.61E+00 | 2.71E+00 | 6.00E-01 | no | Kif5a | intron_variant | MODIFIER |
| 1 | 91161335 | AX-170425734 | 0 | T | C | 0.158 | 1.08E-01 | 2.07E-01 | 2.66E+00 | 2.72E+00 | 6.00E-01 | no | Rbm44 | intron_variant | MODIFIER |
| 8 | 108634002 | AX-170490345 | 1 | A | T | 0.232 | 5.52E-02 | 1.07E-01 | 2.68E+00 | 2.76E+00 | 6.00E-01 | no | Zfhx3 | intron_variant | MODIFIER |
| 1 | 193338683 | AX-170399725 | 0 | A | G | 0.272 | 5.05E-02 | 8.74E-02 | 2.40E+00 | 2.51E+00 | 6.01E-01 | no | Lamb3 | intron_variant | MODIFIER |
| 1 | 193681239 | AX-97811096 | 0 | A | G | 0.272 | 5.05E-02 | 8.74E-02 | 2.40E+00 | 2.51E+00 | 6.01E-01 | no | | intergenic_variant | MODIFIER |
| 1 | 193941090 | AX-170474472 | 0 | A | G | 0.272 | 5.05E-02 | 8.74E-02 | 2.40E+00 | 2.51E+00 | 6.01E-01 | no | | intergenic_variant | MODIFIER |
| 7 | 16138524 | AX-170180805 | 1 | A | G | 0.384 | 4.14E-02 | 7.84E-02 | 2.56E+00 | 2.68E+00 | 6.01E-01 | no | Slc8a2 | intron_variant | MODIFIER |
| 2 | 124191557 | AX-170326950 | 0 | C | A | 0.149 | 9.80E-02 | 1.86E-01 | 2.60E+00 | 2.67E+00 | 6.01E-01 | no | Sema6d | intron_variant | MODIFIER |
| 2 | 127141628 | AX-170469887 | 0 | T | A | 0.149 | 9.80E-02 | 1.86E-01 | 2.60E+00 | 2.67E+00 | 6.01E-01 | no | Itpripl1 | synonymous_variant | LOW |
| 2 | 157954489 | AX-170384689 | 0 | G | A | 0.149 | 9.80E-02 | 1.86E-01 | 2.60E+00 | 2.67E+00 | 6.01E-01 | no | | intergenic_variant | MODIFIER |
| 12 | 93419315 | AX-170227778 | 1 | G | A | 0.054 | 1.11E-01 | 2.16E-01 | 2.67E+00 | 2.77E+00 | 6.01E-01 | no | 4930559C10Rik | intron_variant,non_coding_transcript_variant | MODIFIER |
| 10 | 69515294 | AX-97815786 | 0 | G | A | 0.421 | -4.10E-02 | 7.82E-02 | 2.60E+00 | 2.69E+00 | 6.01E-01 | no | Ank3 | intron_variant,non_coding_transcript_variant | MODIFIER |
| 15 | 70865001 | AX-97820988 | 2 | C | T | 0.373 | 4.03E-02 | 7.44E-02 | 2.52E+00 | 2.61E+00 | 6.01E-01 | no | | intergenic_variant | MODIFIER |
| 1 | 141061132 | AX-170531460 | 1 | C | A | 0.205 | 6.81E-02 | 1.32E-01 | 2.67E+00 | 2.74E+00 | 6.01E-01 | no | | intergenic_variant | MODIFIER |
| 8 | 18512783 | AX-170313790 | 1 | G | A | 0.179 | -4.91E-02 | 9.64E-02 | 2.68E+00 | 2.81E+00 | 6.01E-01 | no | Gm24469 | upstream_gene_variant | MODIFIER |
| 2 | 52082247 | AX-170388241 | 1 | C | T | 0.054 | 8.89E-02 | 1.72E-01 | 2.62E+00 | 2.73E+00 | 6.01E-01 | no | Rif1 | intron_variant | MODIFIER |
| 5 | 114650738 | AX-170285654 | 1 | C | T | 0.455 | 6.11E-02 | 1.18E-01 | 2.67E+00 | 2.70E+00 | 6.01E-01 | no | Trpv4 | intron_variant | MODIFIER |
| 5 | 116962880 | AX-170417760 | 1 | C | T | 0.455 | 6.11E-02 | 1.18E-01 | 2.67E+00 | 2.70E+00 | 6.01E-01 | no | | intergenic_variant | MODIFIER |
| 20 | 67850477 | AX-97845314 | 0 | G | A | 0.246 | 3.26E-02 | 6.32E-02 | 2.67E+00 | 2.75E+00 | 6.01E-01 | no | | intergenic_variant | MODIFIER |
| 15 | 78798100 | AX-170318505 | 0 | G | A | 0.561 | 4.35E-02 | 8.18E-02 | 2.56E+00 | 2.63E+00 | 6.01E-01 | no | Card10 | intron_variant | MODIFIER |
| 15 | 78885445 | AX-170292670 | 0 | T | C | 0.561 | 4.35E-02 | 8.18E-02 | 2.56E+00 | 2.63E+00 | 6.01E-01 | no | Gga1 | intron_variant | MODIFIER |
| 15 | 78889456 | AX-170184269 | 0 | A | G | 0.561 | 4.35E-02 | 8.18E-02 | 2.56E+00 | 2.63E+00 | 6.01E-01 | no | Gga1 | intron_variant | MODIFIER |
| 15 | 78889564 | AX-97843316 | 0 | G | T | 0.561 | 4.35E-02 | 8.18E-02 | 2.56E+00 | 2.63E+00 | 6.01E-01 | no | Gga1 | intron_variant | MODIFIER |
| 15 | 78894586 | AX-170186962 | 0 | C | T | 0.561 | 4.35E-02 | 8.18E-02 | 2.56E+00 | 2.63E+00 | 6.01E-01 | no | Gga1 | downstream_gene_variant | MODIFIER |
| 15 | 78895181 | AX-170146962 | 0 | G | A | 0.561 | 4.35E-02 | 8.18E-02 | 2.56E+00 | 2.63E+00 | 6.01E-01 | no | Sh3bp1 | upstream_gene_variant | MODIFIER |
| 15 | 78936817 | AX-170378523 | 0 | A | G | 0.561 | 4.35E-02 | 8.18E-02 | 2.56E+00 | 2.63E+00 | 6.01E-01 | no | Nol12 | intron_variant | MODIFIER |
| 15 | 78937415 | AX-170171785 | 0 | A | G | 0.561 | 4.35E-02 | 8.18E-02 | 2.56E+00 | 2.63E+00 | 6.01E-01 | no | Nol12 | intron_variant | MODIFIER |
| 15 | 78939145 | AX-170442065 | 0 | G | A | 0.561 | 4.35E-02 | 8.18E-02 | 2.56E+00 | 2.63E+00 | 6.01E-01 | no | Nol12 | intron_variant | MODIFIER |
| 15 | 78951145 | AX-97829847 | 0 | A | G | 0.561 | 4.35E-02 | 8.18E-02 | 2.56E+00 | 2.63E+00 | 6.01E-01 | no | Triobp | intron_variant | MODIFIER |
| 15 | 78957922 | AX-170056069 | 0 | G | T | 0.561 | 4.35E-02 | 8.18E-02 | 2.56E+00 | 2.63E+00 | 6.01E-01 | no | Triobp | intron_variant | MODIFIER |
| 15 | 78961679 | AX-97829214 | 0 | T | C | 0.561 | 4.35E-02 | 8.18E-02 | 2.56E+00 | 2.63E+00 | 6.01E-01 | no | Triobp | intron_variant | MODIFIER |

| | | | | | | | | | | | | | | | |
|---|---|---|---|---|---|---|---|---|---|---|---|---|---|---|---|
| 15 | 79076074 | AX-170299129 | 0 | C | T | 0.561 | 4.35E-02 | 8.18E-02 | 2.56E+00 | 2.63E+00 | 6.01E-01 | no | Eif3l | intron_variant | MODIFIER |
| 15 | 79076934 | AX-170398073 | 0 | C | T | 0.561 | 4.35E-02 | 8.18E-02 | 2.56E+00 | 2.63E+00 | 6.01E-01 | no | Eif3l | synonymous_variant | LOW |
| 15 | 79081838 | AX-170447331 | 0 | G | C | 0.561 | 4.35E-02 | 8.18E-02 | 2.56E+00 | 2.63E+00 | 6.01E-01 | no | Eif3l | intron_variant | MODIFIER |
| 15 | 79084641 | AX-170397764 | 0 | C | T | 0.561 | 4.35E-02 | 8.18E-02 | 2.56E+00 | 2.63E+00 | 6.01E-01 | no | Eif3l | intron_variant | MODIFIER |
| 15 | 79092453 | AX-170311820 | 0 | C | T | 0.561 | 4.35E-02 | 8.18E-02 | 2.56E+00 | 2.63E+00 | 6.01E-01 | no | Eif3l | intron_variant | MODIFIER |
| 15 | 79111216 | AX-170484517 | 0 | G | A | 0.561 | 4.35E-02 | 8.18E-02 | 2.56E+00 | 2.63E+00 | 6.01E-01 | no | Micall1 | intron_variant | MODIFIER |
| 15 | 79115005 | AX-170460989 | 0 | G | A | 0.561 | 4.35E-02 | 8.18E-02 | 2.56E+00 | 2.63E+00 | 6.01E-01 | no | Micall1 | intron_variant | MODIFIER |
| 15 | 79117980 | AX-97806801 | 0 | T | C | 0.561 | 4.35E-02 | 8.18E-02 | 2.56E+00 | 2.63E+00 | 6.01E-01 | no | Micall1 | intron_variant | MODIFIER |
| 15 | 79124674 | AX-170178362 | 0 | A | G | 0.561 | 4.35E-02 | 8.18E-02 | 2.56E+00 | 2.63E+00 | 6.01E-01 | no | Micall1 | intron_variant | MODIFIER |
| 15 | 79140838 | AX-170510153 | 0 | G | A | 0.561 | 4.35E-02 | 8.18E-02 | 2.56E+00 | 2.63E+00 | 6.01E-01 | no | Polr2f | upstream_gene_variant | MODIFIER |
| 15 | 79147289 | AX-97829846 | 0 | G | T | 0.561 | 4.35E-02 | 8.18E-02 | 2.56E+00 | 2.63E+00 | 6.01E-01 | no | Polr2f | intron_variant | MODIFIER |
| 15 | 79153929 | AX-170012062 | 0 | C | T | 0.561 | 4.35E-02 | 8.18E-02 | 2.56E+00 | 2.63E+00 | 6.01E-01 | no | Sox10 | downstream_gene_variant | MODIFIER |
| 15 | 79153958 | AX-170012067 | 0 | C | G | 0.561 | 4.35E-02 | 8.18E-02 | 2.56E+00 | 2.63E+00 | 6.01E-01 | no | Sox10 | downstream_gene_variant | MODIFIER |
| 15 | 79171637 | AX-170334634 | 0 | G | C | 0.561 | 4.35E-02 | 8.18E-02 | 2.56E+00 | 2.63E+00 | 6.01E-01 | no | Gm10863 | intron_variant,non_coding_transcript_variant | MODIFIER |
| 15 | 79175098 | AX-170354122 | 0 | T | C | 0.561 | 4.35E-02 | 8.18E-02 | 2.56E+00 | 2.63E+00 | 6.01E-01 | no | Gm10863 | intron_variant,non_coding_transcript_variant | MODIFIER |
| 15 | 79190661 | AX-97830481 | 0 | C | T | 0.561 | 4.35E-02 | 8.18E-02 | 2.56E+00 | 2.63E+00 | 6.01E-01 | no | Gm10863 | intron_variant,non_coding_transcript_variant | MODIFIER |
| 15 | 79227246 | AX-170485212 | 0 | G | A | 0.561 | 4.35E-02 | 8.18E-02 | 2.56E+00 | 2.63E+00 | 6.01E-01 | no | Pick1 | upstream_gene_variant | MODIFIER |
| 15 | 79231084 | AX-97855273 | 0 | T | C | 0.561 | 4.35E-02 | 8.18E-02 | 2.56E+00 | 2.63E+00 | 6.01E-01 | no | Pick1 | intron_variant | MODIFIER |
| 15 | 79232662 | AX-170272391 | 0 | A | C | 0.561 | 4.35E-02 | 8.18E-02 | 2.56E+00 | 2.63E+00 | 6.01E-01 | no | Pick1 | intron_variant | MODIFIER |
| 15 | 79232929 | AX-170269694 | 0 | T | C | 0.561 | 4.35E-02 | 8.18E-02 | 2.56E+00 | 2.63E+00 | 6.01E-01 | no | Pick1 | intron_variant | MODIFIER |
| 15 | 79314544 | AX-170056074 | 0 | A | G | 0.561 | 4.35E-02 | 8.18E-02 | 2.56E+00 | 2.63E+00 | 6.01E-01 | no | Pla2g6 | intron_variant | MODIFIER |
| 15 | 79316517 | AX-170056075 | 0 | A | G | 0.561 | 4.35E-02 | 8.18E-02 | 2.56E+00 | 2.63E+00 | 6.01E-01 | no | Pla2g6 | intron_variant | MODIFIER |
| 15 | 79360994 | AX-170012060 | 0 | A | G | 0.561 | 4.35E-02 | 8.18E-02 | 2.56E+00 | 2.63E+00 | 6.01E-01 | no | Tmem184b | 3_prime_UTR_variant | MODIFIER |
| 15 | 79383134 | AX-170152710 | 0 | A | G | 0.561 | 4.35E-02 | 8.18E-02 | 2.56E+00 | 2.63E+00 | 6.01E-01 | no | Tmem184b | intron_variant | MODIFIER |
| 15 | 79386745 | AX-97788261 | 0 | T | C | 0.561 | 4.35E-02 | 8.18E-02 | 2.56E+00 | 2.63E+00 | 6.01E-01 | no | Tmem184b | intron_variant | MODIFIER |
| 15 | 79423826 | AX-170012059 | 0 | A | G | 0.561 | 4.35E-02 | 8.18E-02 | 2.56E+00 | 2.63E+00 | 6.01E-01 | no | Csnk1e | intron_variant | MODIFIER |
| 15 | 79426018 | AX-170474545 | 0 | C | T | 0.561 | 4.35E-02 | 8.18E-02 | 2.56E+00 | 2.63E+00 | 6.01E-01 | no | Csnk1e | intron_variant | MODIFIER |
| 15 | 79443193 | AX-97830506 | 0 | T | C | 0.561 | 4.35E-02 | 8.18E-02 | 2.56E+00 | 2.63E+00 | 6.01E-01 | no | Csnk1e | upstream_gene_variant | MODIFIER |
| 15 | 79526717 | AX-170056080 | 0 | C | G | 0.561 | 4.35E-02 | 8.18E-02 | 2.56E+00 | 2.63E+00 | 6.01E-01 | no | Kdelr3 | intron_variant | MODIFIER |
| 15 | 79593381 | AX-170243054 | 0 | G | C | 0.561 | 4.35E-02 | 8.18E-02 | 2.56E+00 | 2.63E+00 | 6.01E-01 | no | Dmc1 | intron_variant | MODIFIER |
| 15 | 79593683 | AX-97788114 | 0 | C | T | 0.561 | 4.35E-02 | 8.18E-02 | 2.56E+00 | 2.63E+00 | 6.01E-01 | no | Dmc1 | intron_variant | MODIFIER |
| 15 | 79596763 | AX-170461744 | 0 | C | T | 0.561 | 4.35E-02 | 8.18E-02 | 2.56E+00 | 2.63E+00 | 6.01E-01 | no | Dmc1 | intron_variant | MODIFIER |
| 15 | 79621736 | AX-97824623 | 0 | T | C | 0.561 | 4.35E-02 | 8.18E-02 | 2.56E+00 | 2.63E+00 | 6.01E-01 | no | Fam227a | intron_variant | MODIFIER |
| 15 | 79634292 | AX-97829872 | 0 | G | A | 0.561 | 4.35E-02 | 8.18E-02 | 2.56E+00 | 2.63E+00 | 6.01E-01 | no | Fam227a | downstream_gene_variant | MODIFIER |
| 15 | 79664823 | AX-97805453 | 0 | C | T | 0.561 | 4.35E-02 | 8.18E-02 | 2.56E+00 | 2.63E+00 | 6.01E-01 | no | Cby1 | intron_variant | MODIFIER |
| 15 | 79688466 | AX-170087920 | 0 | T | C | 0.561 | 4.35E-02 | 8.18E-02 | 2.56E+00 | 2.63E+00 | 6.01E-01 | no | Josd1 | upstream_gene_variant | MODIFIER |
| 15 | 79697014 | AX-170526687 | 0 | G | A | 0.561 | 4.35E-02 | 8.18E-02 | 2.56E+00 | 2.63E+00 | 6.01E-01 | no | Gtpbp1 | intron_variant | MODIFIER |
| 15 | 79713786 | AX-170277185 | 0 | C | T | 0.561 | 4.35E-02 | 8.18E-02 | 2.56E+00 | 2.63E+00 | 6.01E-01 | no | Gtpbp1 | intron_variant | MODIFIER |
| 15 | 79717222 | AX-170228257 | 0 | T | C | 0.561 | 4.35E-02 | 8.18E-02 | 2.56E+00 | 2.63E+00 | 6.01E-01 | no | Gtpbp1 | intron_variant | MODIFIER |
| 15 | 79717261 | AX-97818921 | 0 | A | C | 0.561 | 4.35E-02 | 8.18E-02 | 2.56E+00 | 2.63E+00 | 6.01E-01 | no | Gtpbp1 | intron_variant | MODIFIER |
| 15 | 79719660 | AX-170208947 | 0 | T | C | 0.561 | 4.35E-02 | 8.18E-02 | 2.56E+00 | 2.63E+00 | 6.01E-01 | no | Sun2 | downstream_gene_variant | MODIFIER |
| 15 | 79723117 | AX-170424473 | 0 | A | G | 0.561 | 4.35E-02 | 8.18E-02 | 2.56E+00 | 2.63E+00 | 6.01E-01 | no | Sun2 | downstream_gene_variant | MODIFIER |
| 15 | 79740633 | AX-170012071 | 0 | T | C | 0.561 | 4.35E-02 | 8.18E-02 | 2.56E+00 | 2.63E+00 | 6.01E-01 | no | Sun2 | intron_variant | MODIFIER |
| 15 | 79756948 | AX-97832117 | 0 | T | C | 0.561 | 4.35E-02 | 8.18E-02 | 2.56E+00 | 2.63E+00 | 6.01E-01 | no | Dnal4 | downstream_gene_variant | MODIFIER |
| 15 | 79765713 | AX-97806556 | 0 | C | T | 0.561 | 4.35E-02 | 8.18E-02 | 2.56E+00 | 2.63E+00 | 6.01E-01 | no | Dnal4 | intron_variant | MODIFIER |
| 15 | 79788348 | AX-170012076 | 0 | T | C | 0.561 | 4.35E-02 | 8.18E-02 | 2.56E+00 | 2.63E+00 | 6.01E-01 | no | Nptxr | 3_prime_UTR_variant | MODIFIER |
| 15 | 79790190 | AX-170012078 | 0 | A | G | 0.561 | 4.35E-02 | 8.18E-02 | 2.56E+00 | 2.63E+00 | 6.01E-01 | no | Nptxr | intron_variant | MODIFIER |
| 15 | 79794051 | AX-97829840 | 0 | A | C | 0.561 | 4.35E-02 | 8.18E-02 | 2.56E+00 | 2.63E+00 | 6.01E-01 | no | Nptxr | intron_variant | MODIFIER |
| 15 | 79795159 | AX-170180426 | 0 | A | G | 0.561 | 4.35E-02 | 8.18E-02 | 2.56E+00 | 2.63E+00 | 6.01E-01 | no | Nptxr | intron_variant | MODIFIER |
| 15 | 79797283 | AX-170318928 | 0 | C | A | 0.561 | 4.35E-02 | 8.18E-02 | 2.56E+00 | 2.63E+00 | 6.01E-01 | no | Nptxr | intron_variant | MODIFIER |
| 15 | 79809915 | AX-170449006 | 0 | C | T | 0.561 | 4.35E-02 | 8.18E-02 | 2.56E+00 | 2.63E+00 | 6.01E-01 | no | Npcd | intron_variant | MODIFIER |
| 15 | 79816635 | AX-170252098 | 0 | A | G | 0.561 | 4.35E-02 | 8.18E-02 | 2.56E+00 | 2.63E+00 | 6.01E-01 | no | Npcd | intron_variant | MODIFIER |
| 15 | 79829951 | AX-97854595 | 0 | T | C | 0.561 | 4.35E-02 | 8.18E-02 | 2.56E+00 | 2.63E+00 | 6.01E-01 | no | Npcd | intron_variant | MODIFIER |
| 15 | 79830165 | AX-170012074 | 0 | A | G | 0.561 | 4.35E-02 | 8.18E-02 | 2.56E+00 | 2.63E+00 | 6.01E-01 | no | Npcd | intron_variant | MODIFIER |

| | | | | | | | | | | | | | | | |
|---|---|---|---|---|---|---|---|---|---|---|---|---|---|---|---|
| 15 | 79857585 | AX-170012079 | 0 | G | T | 0.561 | 4.35E-02 | 8.18E-02 | 2.56E+00 | 2.63E+00 | 6.01E-01 | no | | intergenic_variant | MODIFIER |
| 15 | 79863771 | AX-97839482 | 0 | C | T | 0.561 | 4.35E-02 | 8.18E-02 | 2.56E+00 | 2.63E+00 | 6.01E-01 | no | | intergenic_variant | MODIFIER |
| 15 | 79875161 | AX-170012075 | 0 | T | C | 0.561 | 4.35E-02 | 8.18E-02 | 2.56E+00 | 2.63E+00 | 6.01E-01 | no | Gm49412 | non_coding_transcript_exon_variant | MODIFIER |
| 15 | 79879356 | AX-170056087 | 0 | T | C | 0.561 | 4.35E-02 | 8.18E-02 | 2.56E+00 | 2.63E+00 | 6.01E-01 | no | Gm49412 | downstream_gene_variant | MODIFIER |
| 15 | 79893743 | AX-97855527 | 0 | T | C | 0.561 | 4.35E-02 | 8.18E-02 | 2.56E+00 | 2.63E+00 | 6.01E-01 | no | Apobec3 | upstream_gene_variant | MODIFIER |
| 15 | 79893868 | AX-170056086 | 0 | C | T | 0.561 | 4.35E-02 | 8.18E-02 | 2.56E+00 | 2.63E+00 | 6.01E-01 | no | Apobec3 | upstream_gene_variant | MODIFIER |
| 15 | 79901016 | AX-170056088 | 0 | A | G | 0.561 | 4.35E-02 | 8.18E-02 | 2.56E+00 | 2.63E+00 | 6.01E-01 | no | Apobec3 | intron_variant | MODIFIER |
| 15 | 79904366 | AX-170244883 | 0 | G | A | 0.561 | 4.35E-02 | 8.18E-02 | 2.56E+00 | 2.63E+00 | 6.01E-01 | no | Apobec3 | intron_variant | MODIFIER |
| 15 | 79905925 | AX-170396166 | 0 | T | A | 0.561 | 4.35E-02 | 8.18E-02 | 2.56E+00 | 2.63E+00 | 6.01E-01 | no | Apobec3 | intron_variant | MODIFIER |
| 15 | 79906950 | AX-170367872 | 0 | A | G | 0.561 | 4.35E-02 | 8.18E-02 | 2.56E+00 | 2.63E+00 | 6.01E-01 | no | Apobec3 | missense_variant | MODERATE |
| 15 | 79914686 | AX-170403033 | 0 | A | G | 0.561 | 4.35E-02 | 8.18E-02 | 2.56E+00 | 2.63E+00 | 6.01E-01 | no | Cbx7 | downstream_gene_variant | MODIFIER |
| 15 | 79917133 | AX-170139147 | 0 | C | T | 0.561 | 4.35E-02 | 8.18E-02 | 2.56E+00 | 2.63E+00 | 6.01E-01 | no | Cbx7 | 3_prime_UTR_variant | MODIFIER |
| 15 | 79918893 | AX-170456916 | 0 | T | C | 0.561 | 4.35E-02 | 8.18E-02 | 2.56E+00 | 2.63E+00 | 6.01E-01 | no | Cbx7 | missense_variant | MODERATE |
| 15 | 79921417 | AX-170174183 | 0 | T | C | 0.561 | 4.35E-02 | 8.18E-02 | 2.56E+00 | 2.63E+00 | 6.01E-01 | no | Cbx7 | synonymous_variant | LOW |
| 15 | 79923462 | AX-170093313 | 0 | A | G | 0.561 | 4.35E-02 | 8.18E-02 | 2.56E+00 | 2.63E+00 | 6.01E-01 | no | Cbx7 | intron_variant | MODIFIER |
| 15 | 79939110 | AX-170012077 | 0 | A | G | 0.561 | 4.35E-02 | 8.18E-02 | 2.56E+00 | 2.63E+00 | 6.01E-01 | no | Cbx7 | intron_variant | MODIFIER |
| 15 | 79949498 | AX-97854506 | 0 | G | T | 0.561 | 4.35E-02 | 8.18E-02 | 2.56E+00 | 2.63E+00 | 6.01E-01 | no | Cbx7 | intron_variant | MODIFIER |
| 15 | 79969950 | AX-170348466 | 0 | G | A | 0.561 | 4.35E-02 | 8.18E-02 | 2.56E+00 | 2.63E+00 | 6.01E-01 | no | Cbx7 | intron_variant | MODIFIER |
| 15 | 79970985 | AX-97821112 | 0 | T | C | 0.561 | 4.35E-02 | 8.18E-02 | 2.56E+00 | 2.63E+00 | 6.01E-01 | no | Cbx7 | 5_prime_UTR_variant | MODIFIER |
| 15 | 79984750 | AX-170466125 | 0 | G | A | 0.561 | 4.35E-02 | 8.18E-02 | 2.56E+00 | 2.63E+00 | 6.01E-01 | no | | regulatory_region_variant | MODIFIER |
| 15 | 79996691 | AX-170188632 | 0 | T | C | 0.561 | 4.35E-02 | 8.18E-02 | 2.56E+00 | 2.63E+00 | 6.01E-01 | no | Pdgfb | 3_prime_UTR_variant | MODIFIER |
| 15 | 79997372 | AX-170451797 | 0 | G | T | 0.561 | 4.35E-02 | 8.18E-02 | 2.56E+00 | 2.63E+00 | 6.01E-01 | no | Pdgfb | intron_variant | MODIFIER |
| 15 | 79997747 | AX-170161831 | 0 | G | A | 0.561 | 4.35E-02 | 8.18E-02 | 2.56E+00 | 2.63E+00 | 6.01E-01 | no | Pdgfb | intron_variant | MODIFIER |
| 15 | 79999038 | AX-170442723 | 0 | A | G | 0.561 | 4.35E-02 | 8.18E-02 | 2.56E+00 | 2.63E+00 | 6.01E-01 | no | Pdgfb | intron_variant | MODIFIER |
| 15 | 80000501 | AX-170528163 | 0 | C | T | 0.561 | 4.35E-02 | 8.18E-02 | 2.56E+00 | 2.63E+00 | 6.01E-01 | no | Pdgfb | intron_variant | MODIFIER |
| 15 | 80008624 | AX-170253422 | 0 | A | G | 0.561 | 4.35E-02 | 8.18E-02 | 2.56E+00 | 2.63E+00 | 6.01E-01 | no | Pdgfb | intron_variant | MODIFIER |
| 15 | 80025438 | AX-170266660 | 0 | T | G | 0.561 | 4.35E-02 | 8.18E-02 | 2.56E+00 | 2.63E+00 | 6.01E-01 | no | Gm41361 | intron_variant,non_coding_transcript_variant | MODIFIER |
| 15 | 80038648 | AX-97846450 | 0 | T | C | 0.561 | 4.35E-02 | 8.18E-02 | 2.56E+00 | 2.63E+00 | 6.01E-01 | no | Gm41361 | intron_variant,non_coding_transcript_variant | MODIFIER |
| 15 | 80073066 | AX-170203702 | 0 | G | A | 0.561 | 4.35E-02 | 8.18E-02 | 2.56E+00 | 2.63E+00 | 6.01E-01 | no | Rpl3 | downstream_gene_variant | MODIFIER |
| 15 | 80098240 | AX-170380752 | 0 | G | A | 0.561 | 4.35E-02 | 8.18E-02 | 2.56E+00 | 2.63E+00 | 6.01E-01 | no | Syngr1 | intron_variant | MODIFIER |
| 15 | 80193271 | AX-170277830 | 0 | C | T | 0.561 | 4.35E-02 | 8.18E-02 | 2.56E+00 | 2.63E+00 | 6.01E-01 | no | Mgat3 | intron_variant | MODIFIER |
| 15 | 80208712 | AX-170171596 | 0 | T | C | 0.561 | 4.35E-02 | 8.18E-02 | 2.56E+00 | 2.63E+00 | 6.01E-01 | no | Mgat3 | intron_variant | MODIFIER |
| 15 | 80211657 | AX-170112953 | 0 | C | T | 0.561 | 4.35E-02 | 8.18E-02 | 2.56E+00 | 2.63E+00 | 6.01E-01 | no | Mgat3 | synonymous_variant | LOW |
| 15 | 80213791 | AX-97812067 | 0 | T | C | 0.561 | 4.35E-02 | 8.18E-02 | 2.56E+00 | 2.63E+00 | 6.01E-01 | no | Mgat3 | 3_prime_UTR_variant | MODIFIER |
| 15 | 80238358 | AX-170152685 | 0 | T | C | 0.561 | 4.35E-02 | 8.18E-02 | 2.56E+00 | 2.63E+00 | 6.01E-01 | no | Mief1 | intron_variant | MODIFIER |
| 15 | 80250826 | AX-170329377 | 0 | T | C | 0.561 | 4.35E-02 | 8.18E-02 | 2.56E+00 | 2.63E+00 | 6.01E-01 | no | Mief1 | 3_prime_UTR_variant | MODIFIER |
| 15 | 80252710 | AX-170257532 | 0 | C | A | 0.561 | 4.35E-02 | 8.18E-02 | 2.56E+00 | 2.63E+00 | 6.01E-01 | no | Mief1 | 3_prime_UTR_variant | MODIFIER |
| 15 | 80252858 | AX-97811803 | 0 | C | T | 0.561 | 4.35E-02 | 8.18E-02 | 2.56E+00 | 2.63E+00 | 6.01E-01 | no | Mief1 | 3_prime_UTR_variant | MODIFIER |
| 15 | 80259647 | AX-170012083 | 0 | C | T | 0.561 | 4.35E-02 | 8.18E-02 | 2.56E+00 | 2.63E+00 | 6.01E-01 | no | Rps19bp1 | downstream_gene_variant | MODIFIER |
| 15 | 80275882 | AX-170012089 | 0 | T | C | 0.561 | 4.35E-02 | 8.18E-02 | 2.56E+00 | 2.63E+00 | 6.01E-01 | no | | intergenic_variant | MODIFIER |
| 15 | 80276087 | AX-170157525 | 0 | G | A | 0.561 | 4.35E-02 | 8.18E-02 | 2.56E+00 | 2.63E+00 | 6.01E-01 | no | | intergenic_variant | MODIFIER |
| 15 | 80279451 | AX-170492814 | 0 | T | C | 0.561 | 4.35E-02 | 8.18E-02 | 2.56E+00 | 2.63E+00 | 6.01E-01 | no | | intergenic_variant | MODIFIER |
| 15 | 80326131 | AX-170012086 | 0 | C | T | 0.561 | 4.35E-02 | 8.18E-02 | 2.56E+00 | 2.63E+00 | 6.01E-01 | no | Cacna1i | intron_variant | MODIFIER |
| 15 | 80336969 | AX-170456564 | 0 | A | C | 0.561 | 4.35E-02 | 8.18E-02 | 2.56E+00 | 2.63E+00 | 6.01E-01 | no | Cacna1i | intron_variant | MODIFIER |
| 15 | 80388528 | AX-97812064 | 0 | T | C | 0.561 | 4.35E-02 | 8.18E-02 | 2.56E+00 | 2.63E+00 | 6.01E-01 | no | Cacna1i | upstream_gene_variant | MODIFIER |
| 15 | 80388691 | AX-170012088 | 0 | A | G | 0.561 | 4.35E-02 | 8.18E-02 | 2.56E+00 | 2.63E+00 | 6.01E-01 | no | Cacna1i | upstream_gene_variant | MODIFIER |
| 15 | 80388724 | AX-170012090 | 0 | A | G | 0.561 | 4.35E-02 | 8.18E-02 | 2.56E+00 | 2.63E+00 | 6.01E-01 | no | Cacna1i | upstream_gene_variant | MODIFIER |
| 15 | 80414538 | AX-97830539 | 0 | C | A | 0.561 | 4.35E-02 | 8.18E-02 | 2.56E+00 | 2.63E+00 | 6.01E-01 | no | | regulatory_region_variant | MODIFIER |
| 15 | 80432857 | AX-170012084 | 0 | A | G | 0.561 | 4.35E-02 | 8.18E-02 | 2.56E+00 | 2.63E+00 | 6.01E-01 | no | Gm31369 | intron_variant,non_coding_transcript_variant | MODIFIER |
| 15 | 80440030 | AX-170332170 | 0 | A | G | 0.561 | 4.35E-02 | 8.18E-02 | 2.56E+00 | 2.63E+00 | 6.01E-01 | no | Gm31369 | upstream_gene_variant | MODIFIER |
| 15 | 80461665 | AX-170172126 | 0 | T | C | 0.561 | 4.35E-02 | 8.18E-02 | 2.56E+00 | 2.63E+00 | 6.01E-01 | no | Enthd1 | intron_variant | MODIFIER |
| 15 | 80481058 | AX-170195118 | 0 | G | A | 0.561 | 4.35E-02 | 8.18E-02 | 2.56E+00 | 2.63E+00 | 6.01E-01 | no | Enthd1 | intron_variant | MODIFIER |
| 15 | 80481086 | AX-170178740 | 0 | T | C | 0.561 | 4.35E-02 | 8.18E-02 | 2.56E+00 | 2.63E+00 | 6.01E-01 | no | Enthd1 | intron_variant | MODIFIER |
| 15 | 80482211 | AX-170290244 | 0 | A | G | 0.561 | 4.35E-02 | 8.18E-02 | 2.56E+00 | 2.63E+00 | 6.01E-01 | no | Enthd1 | intron_variant | MODIFIER |

| | | | | | | | | | | | | | | | |
|---|---|---|---|---|---|---|---|---|---|---|---|---|---|---|---|
| 15 | 80482589 | AX-170182519 | 0 | A | G | 0.561 | 4.35E-02 | 8.18E-02 | 2.56E+00 | 2.63E+00 | 6.01E-01 | no | Enthd1 | intron_variant | MODIFIER |
| 15 | 80483394 | AX-170512350 | 0 | T | C | 0.561 | 4.35E-02 | 8.18E-02 | 2.56E+00 | 2.63E+00 | 6.01E-01 | no | Enthd1 | intron_variant | MODIFIER |
| 15 | 80490010 | AX-170486198 | 0 | C | T | 0.561 | 4.35E-02 | 8.18E-02 | 2.56E+00 | 2.63E+00 | 6.01E-01 | no | Enthd1 | intron_variant | MODIFIER |
| 15 | 80505589 | AX-97840839 | 0 | A | C | 0.561 | 4.35E-02 | 8.18E-02 | 2.56E+00 | 2.63E+00 | 6.01E-01 | no | Enthd1 | intron_variant | MODIFIER |
| 15 | 80545682 | AX-168414276 | 0 | A | G | 0.561 | 4.35E-02 | 8.18E-02 | 2.56E+00 | 2.63E+00 | 6.01E-01 | no | Enthd1 | intron_variant | MODIFIER |
| 15 | 80604122 | AX-170443022 | 0 | G | T | 0.561 | 4.35E-02 | 8.18E-02 | 2.56E+00 | 2.63E+00 | 6.01E-01 | no | Grap2 | intron_variant | MODIFIER |
| 15 | 80806489 | AX-97844021 | 0 | A | G | 0.561 | 4.35E-02 | 8.18E-02 | 2.56E+00 | 2.63E+00 | 6.01E-01 | no | Tnrc6b | intron_variant | MODIFIER |
| 15 | 81090807 | AX-170503275 | 0 | T | C | 0.561 | 4.35E-02 | 8.18E-02 | 2.56E+00 | 2.63E+00 | 6.01E-01 | no | Mrtfa | intron_variant | MODIFIER |
| 15 | 81090961 | AX-170179182 | 0 | G | A | 0.561 | 4.35E-02 | 8.18E-02 | 2.56E+00 | 2.63E+00 | 6.01E-01 | no | Mrtfa | intron_variant | MODIFIER |
| 15 | 81102643 | AX-170190309 | 0 | C | T | 0.561 | 4.35E-02 | 8.18E-02 | 2.56E+00 | 2.63E+00 | 6.01E-01 | no | Mrtfa | intron_variant | MODIFIER |
| 15 | 81109359 | AX-170408745 | 0 | C | A | 0.561 | 4.35E-02 | 8.18E-02 | 2.56E+00 | 2.63E+00 | 6.01E-01 | no | Mrtfa | upstream_gene_variant | MODIFIER |
| 15 | 81112992 | AX-170118591 | 0 | A | G | 0.561 | 4.35E-02 | 8.18E-02 | 2.56E+00 | 2.63E+00 | 6.01E-01 | no | Mrtfa | intron_variant,non_coding_transcript_variant | MODIFIER |
| 15 | 81113107 | AX-170259717 | 0 | G | A | 0.561 | 4.35E-02 | 8.18E-02 | 2.56E+00 | 2.63E+00 | 6.01E-01 | no | Mrtfa | intron_variant,non_coding_transcript_variant | MODIFIER |
| 15 | 81117812 | AX-170468214 | 0 | G | T | 0.561 | 4.35E-02 | 8.18E-02 | 2.56E+00 | 2.63E+00 | 6.01E-01 | no | Mrtfa | intron_variant,non_coding_transcript_variant | MODIFIER |
| 15 | 81118146 | AX-168373248 | 0 | A | G | 0.561 | 4.35E-02 | 8.18E-02 | 2.56E+00 | 2.63E+00 | 6.01E-01 | no | Mrtfa | intron_variant,non_coding_transcript_variant | MODIFIER |
| 15 | 81124507 | AX-168378761 | 0 | G | A | 0.561 | 4.35E-02 | 8.18E-02 | 2.56E+00 | 2.63E+00 | 6.01E-01 | no | Mrtfa | intron_variant,non_coding_transcript_variant | MODIFIER |
| 15 | 81126969 | AX-97804477 | 0 | A | G | 0.561 | 4.35E-02 | 8.18E-02 | 2.56E+00 | 2.63E+00 | 6.01E-01 | no | Mrtfa | intron_variant,non_coding_transcript_variant | MODIFIER |
| 15 | 81140182 | AX-97787929 | 0 | T | G | 0.561 | 4.35E-02 | 8.18E-02 | 2.56E+00 | 2.63E+00 | 6.01E-01 | no | Mrtfa | intron_variant,non_coding_transcript_variant | MODIFIER |
| 15 | 81180633 | AX-170414898 | 0 | T | C | 0.561 | 4.35E-02 | 8.18E-02 | 2.56E+00 | 2.63E+00 | 6.01E-01 | no | Mrtfa | intron_variant,non_coding_transcript_variant | MODIFIER |
| 15 | 81191247 | AX-170507222 | 0 | T | G | 0.561 | 4.35E-02 | 8.18E-02 | 2.56E+00 | 2.63E+00 | 6.01E-01 | no | Mrtfa | upstream_gene_variant | MODIFIER |
| 15 | 81191614 | AX-170116345 | 0 | C | A | 0.561 | 4.35E-02 | 8.18E-02 | 2.56E+00 | 2.63E+00 | 6.01E-01 | no | Mrtfa | upstream_gene_variant | MODIFIER |
| 15 | 81204085 | AX-170338821 | 0 | G | A | 0.561 | 4.35E-02 | 8.18E-02 | 2.56E+00 | 2.63E+00 | 6.01E-01 | no | Gm49537 | downstream_gene_variant | MODIFIER |
| 15 | 81226875 | AX-97829902 | 0 | G | A | 0.561 | 4.35E-02 | 8.18E-02 | 2.56E+00 | 2.63E+00 | 6.01E-01 | no | | intergenic_variant | MODIFIER |
| 15 | 81231186 | AX-170365864 | 0 | T | G | 0.561 | 4.35E-02 | 8.18E-02 | 2.56E+00 | 2.63E+00 | 6.01E-01 | no | Mchr1 | upstream_gene_variant | MODIFIER |
| 15 | 81291801 | AX-170114150 | 0 | A | G | 0.561 | 4.35E-02 | 8.18E-02 | 2.56E+00 | 2.63E+00 | 6.01E-01 | no | 8430426J06Rik | intron_variant,non_coding_transcript_variant | MODIFIER |
| 15 | 81299795 | AX-170248218 | 0 | A | G | 0.561 | 4.35E-02 | 8.18E-02 | 2.56E+00 | 2.63E+00 | 6.01E-01 | no | 8430426J06Rik | intron_variant,non_coding_transcript_variant | MODIFIER |
| 15 | 81305049 | AX-97829905 | 0 | A | G | 0.561 | 4.35E-02 | 8.18E-02 | 2.56E+00 | 2.63E+00 | 6.01E-01 | no | 8430426J06Rik | intron_variant,non_coding_transcript_variant | MODIFIER |
| 15 | 81533128 | AX-170280288 | 0 | C | T | 0.561 | 4.35E-02 | 8.18E-02 | 2.56E+00 | 2.63E+00 | 6.01E-01 | no | | intergenic_variant | MODIFIER |
| 15 | 81704210 | AX-170175589 | 0 | C | G | 0.561 | 4.35E-02 | 8.18E-02 | 2.56E+00 | 2.63E+00 | 6.01E-01 | no | Rangap1 | downstream_gene_variant | MODIFIER |
| 15 | 81738937 | AX-170056117 | 0 | T | A | 0.561 | 4.35E-02 | 8.18E-02 | 2.56E+00 | 2.63E+00 | 6.01E-01 | no | Rangap1 | intron_variant | MODIFIER |
| 15 | 81826674 | AX-170301374 | 0 | G | T | 0.561 | 4.35E-02 | 8.18E-02 | 2.56E+00 | 2.63E+00 | 6.01E-01 | no | Tef | 3_prime_UTR_variant | MODIFIER |
| 15 | 81896422 | AX-170103775 | 0 | C | T | 0.561 | 4.35E-02 | 8.18E-02 | 2.56E+00 | 2.63E+00 | 6.01E-01 | no | Aco2 | intron_variant | MODIFIER |
| 15 | 81959184 | AX-97829925 | 0 | A | G | 0.561 | 4.35E-02 | 8.18E-02 | 2.56E+00 | 2.63E+00 | 6.01E-01 | no | Pmm1 | intron_variant | MODIFIER |
| 15 | 81990075 | AX-170236929 | 0 | A | G | 0.561 | 4.35E-02 | 8.18E-02 | 2.56E+00 | 2.63E+00 | 6.01E-01 | no | Desi1 | downstream_gene_variant | MODIFIER |
| 15 | 81992105 | AX-170012110 | 0 | G | A | 0.561 | 4.35E-02 | 8.18E-02 | 2.56E+00 | 2.63E+00 | 6.01E-01 | no | Desi1 | downstream_gene_variant | MODIFIER |
| 15 | 81995830 | AX-170012117 | 0 | T | C | 0.561 | 4.35E-02 | 8.18E-02 | 2.56E+00 | 2.63E+00 | 6.01E-01 | no | Desi1 | intron_variant,NMD_transcript_variant | MODIFIER |
| 15 | 82048355 | AX-170012109 | 0 | C | T | 0.561 | 4.35E-02 | 8.18E-02 | 2.56E+00 | 2.63E+00 | 6.01E-01 | no | Snu13 | upstream_gene_variant | MODIFIER |
| 15 | 82058850 | AX-170012116 | 0 | G | T | 0.561 | 4.35E-02 | 8.18E-02 | 2.56E+00 | 2.63E+00 | 6.01E-01 | no | 4930407I10Rik | upstream_gene_variant | MODIFIER |
| 15 | 82072118 | AX-97823488 | 0 | A | G | 0.561 | 4.35E-02 | 8.18E-02 | 2.56E+00 | 2.63E+00 | 6.01E-01 | no | Mei1 | intron_variant | MODIFIER |
| 15 | 82106218 | AX-170173756 | 0 | T | C | 0.561 | 4.35E-02 | 8.18E-02 | 2.56E+00 | 2.63E+00 | 6.01E-01 | no | Mei1 | intron_variant | MODIFIER |
| 15 | 82377300 | AX-170513278 | 0 | A | G | 0.561 | 4.35E-02 | 8.18E-02 | 2.56E+00 | 2.63E+00 | 6.01E-01 | no | Cyp2d22 | intron_variant | MODIFIER |
| 15 | 82539698 | AX-170436402 | 0 | A | G | 0.561 | 4.35E-02 | 8.18E-02 | 2.56E+00 | 2.63E+00 | 6.01E-01 | no | Gm46524 | downstream_gene_variant | MODIFIER |
| 15 | 82917159 | AX-170274740 | 0 | G | A | 0.561 | 4.35E-02 | 8.18E-02 | 2.56E+00 | 2.63E+00 | 6.01E-01 | no | Tcf20 | intron_variant | MODIFIER |
| 1 | 155866631 | AX-97786280 | 0 | C | T | 0.228 | 6.92E-02 | 1.31E-01 | 2.59E+00 | 2.65E+00 | 6.01E-01 | no | Cep350 | intron_variant | MODIFIER |
| 12 | 92544424 | AX-170431583 | 0 | T | G | 0.053 | 1.08E-01 | 2.11E-01 | 2.66E+00 | 2.77E+00 | 6.02E-01 | no | | intergenic_variant | MODIFIER |
| 11 | 32289156 | AX-170308915 | 1 | A | G | 0.071 | 9.82E-02 | 1.90E-01 | 2.64E+00 | 2.72E+00 | 6.02E-01 | no | Hba-a1 | downstream_gene_variant | MODIFIER |
| 1 | 24049930 | AX-170235446 | 0 | T | G | 0.439 | 4.40E-02 | 8.33E-02 | 2.58E+00 | 2.65E+00 | 6.02E-01 | no | Fam135a | intron_variant | MODIFIER |
| 6 | 129876217 | AX-170500940 | 0 | G | C | 0.412 | 5.50E-02 | 1.01E-01 | 2.49E+00 | 2.62E+00 | 6.02E-01 | no | Klra17 | intron_variant | MODIFIER |
| 6 | 130813903 | AX-170445103 | 0 | C | T | 0.412 | 5.50E-02 | 1.01E-01 | 2.49E+00 | 2.62E+00 | 6.02E-01 | no | | intergenic_variant | MODIFIER |
| 16 | 75698236 | AX-170416980 | 2 | T | G | 0.482 | 4.03E-02 | 7.47E-02 | 2.52E+00 | 2.61E+00 | 6.02E-01 | no | Gm30162 | intron_variant,non_coding_transcript_variant | MODIFIER |
| 13 | 31217639 | AX-170356087 | 1 | T | C | 0.17 | -6.39E-02 | 1.21E-01 | 2.59E+00 | 2.65E+00 | 6.02E-01 | no | 4930401O12Rik | intron_variant,non_coding_transcript_variant | MODIFIER |
| 18 | 58498737 | AX-97831664 | 0 | G | A | 0.395 | 5.35E-02 | 9.71E-02 | 2.49E+00 | 2.54E+00 | 6.02E-01 | no | | intergenic_variant | MODIFIER |
| 18 | 58514749 | AX-170247339 | 0 | G | T | 0.395 | 5.35E-02 | 9.71E-02 | 2.49E+00 | 2.54E+00 | 6.02E-01 | no | | intergenic_variant | MODIFIER |
| 18 | 58528888 | AX-170201124 | 0 | A | G | 0.395 | 5.35E-02 | 9.71E-02 | 2.49E+00 | 2.54E+00 | 6.02E-01 | no | | intergenic_variant | MODIFIER |

| | | | | | | | | | | | | | | | |
|---|---|---|---|---|---|---|---|---|---|---|---|---|---|---|---|
| 18 | 58535749 | AX-170131115 | 0 | C | G | 0.395 | 5.35E-02 | 9.71E-02 | 2.49E+00 | 2.54E+00 | 6.02E-01 | no | | intergenic_variant | MODIFIER |
| 18 | 58539608 | AX-97824283 | 0 | G | T | 0.395 | 5.35E-02 | 9.71E-02 | 2.49E+00 | 2.54E+00 | 6.02E-01 | no | | intergenic_variant | MODIFIER |
| 18 | 58540212 | AX-170167664 | 0 | A | G | 0.395 | 5.35E-02 | 9.71E-02 | 2.49E+00 | 2.54E+00 | 6.02E-01 | no | | intergenic_variant | MODIFIER |
| 18 | 58547624 | AX-170112213 | 0 | A | G | 0.395 | 5.35E-02 | 9.71E-02 | 2.49E+00 | 2.54E+00 | 6.02E-01 | no | | intergenic_variant | MODIFIER |
| 18 | 58553594 | AX-170471521 | 0 | C | T | 0.395 | 5.35E-02 | 9.71E-02 | 2.49E+00 | 2.54E+00 | 6.02E-01 | no | Slc27a6 | upstream_gene_variant | MODIFIER |
| 18 | 58553865 | AX-170351837 | 0 | G | A | 0.395 | 5.35E-02 | 9.71E-02 | 2.49E+00 | 2.54E+00 | 6.02E-01 | no | Slc27a6 | upstream_gene_variant | MODIFIER |
| 18 | 58554198 | AX-170454759 | 0 | G | A | 0.395 | 5.35E-02 | 9.71E-02 | 2.49E+00 | 2.54E+00 | 6.02E-01 | no | Slc27a6 | upstream_gene_variant | MODIFIER |
| 18 | 58558959 | AX-170124498 | 0 | T | C | 0.395 | 5.35E-02 | 9.71E-02 | 2.49E+00 | 2.54E+00 | 6.02E-01 | no | Slc27a6 | intron_variant | MODIFIER |
| 18 | 58559794 | AX-170461868 | 0 | T | C | 0.395 | 5.35E-02 | 9.71E-02 | 2.49E+00 | 2.54E+00 | 6.02E-01 | no | Slc27a6 | intron_variant | MODIFIER |
| 18 | 58577869 | AX-170532089 | 0 | G | A | 0.395 | 5.35E-02 | 9.71E-02 | 2.49E+00 | 2.54E+00 | 6.02E-01 | no | Slc27a6 | intron_variant | MODIFIER |
| 18 | 58581445 | AX-97835581 | 0 | A | G | 0.395 | 5.35E-02 | 9.71E-02 | 2.49E+00 | 2.54E+00 | 6.02E-01 | no | Slc27a6 | intron_variant | MODIFIER |
| 18 | 58593505 | AX-170413440 | 0 | G | T | 0.395 | 5.35E-02 | 9.71E-02 | 2.49E+00 | 2.54E+00 | 6.02E-01 | no | Slc27a6 | intron_variant | MODIFIER |
| 18 | 58593543 | AX-170286188 | 0 | G | A | 0.395 | 5.35E-02 | 9.71E-02 | 2.49E+00 | 2.54E+00 | 6.02E-01 | no | Slc27a6 | intron_variant | MODIFIER |
| 18 | 58602632 | AX-170287503 | 0 | T | C | 0.395 | 5.35E-02 | 9.71E-02 | 2.49E+00 | 2.54E+00 | 6.02E-01 | no | Slc27a6 | intron_variant | MODIFIER |
| 18 | 58602973 | AX-97826995 | 0 | G | T | 0.395 | 5.35E-02 | 9.71E-02 | 2.49E+00 | 2.54E+00 | 6.02E-01 | no | Slc27a6 | intron_variant | MODIFIER |
| 18 | 58609023 | AX-170105758 | 0 | C | T | 0.395 | 5.35E-02 | 9.71E-02 | 2.49E+00 | 2.54E+00 | 6.02E-01 | no | Slc27a6 | intron_variant | MODIFIER |
| 18 | 58613935 | AX-170412242 | 0 | T | C | 0.395 | 5.35E-02 | 9.71E-02 | 2.49E+00 | 2.54E+00 | 6.02E-01 | no | Slc27a6 | downstream_gene_variant | MODIFIER |
| 18 | 58655690 | AX-170387327 | 0 | C | T | 0.395 | 5.35E-02 | 9.71E-02 | 2.49E+00 | 2.54E+00 | 6.02E-01 | no | Isoc1 | upstream_gene_variant | MODIFIER |
| 18 | 58668701 | AX-170060971 | 0 | A | G | 0.395 | 5.35E-02 | 9.71E-02 | 2.49E+00 | 2.54E+00 | 6.02E-01 | no | Isoc1 | intron_variant | MODIFIER |
| 18 | 58708878 | AX-170304869 | 0 | C | G | 0.395 | 5.35E-02 | 9.71E-02 | 2.49E+00 | 2.54E+00 | 6.02E-01 | no | | intergenic_variant | MODIFIER |
| 18 | 58709168 | AX-170513054 | 0 | T | C | 0.395 | 5.35E-02 | 9.71E-02 | 2.49E+00 | 2.54E+00 | 6.02E-01 | no | | intergenic_variant | MODIFIER |
| 18 | 58742079 | AX-170333814 | 0 | A | G | 0.395 | 5.35E-02 | 9.71E-02 | 2.49E+00 | 2.54E+00 | 6.02E-01 | no | | regulatory_region_variant | MODIFIER |
| 18 | 58748436 | AX-170529623 | 0 | G | A | 0.395 | 5.35E-02 | 9.71E-02 | 2.49E+00 | 2.54E+00 | 6.02E-01 | no | | intergenic_variant | MODIFIER |
| 18 | 58764913 | AX-170224431 | 0 | T | C | 0.395 | 5.35E-02 | 9.71E-02 | 2.49E+00 | 2.54E+00 | 6.02E-01 | no | | intergenic_variant | MODIFIER |
| 18 | 58764962 | AX-170493759 | 0 | T | C | 0.395 | 5.35E-02 | 9.71E-02 | 2.49E+00 | 2.54E+00 | 6.02E-01 | no | | intergenic_variant | MODIFIER |
| 18 | 58778103 | AX-170368708 | 0 | T | G | 0.395 | 5.35E-02 | 9.71E-02 | 2.49E+00 | 2.54E+00 | 6.02E-01 | no | AC158924.1 | upstream_gene_variant | MODIFIER |
| 18 | 58818597 | AX-170224913 | 0 | T | C | 0.395 | 5.35E-02 | 9.71E-02 | 2.49E+00 | 2.54E+00 | 6.02E-01 | no | | intergenic_variant | MODIFIER |
| 18 | 58821012 | AX-170490109 | 0 | A | G | 0.395 | 5.35E-02 | 9.71E-02 | 2.49E+00 | 2.54E+00 | 6.02E-01 | no | | intergenic_variant | MODIFIER |
| 18 | 58871888 | AX-170478650 | 0 | G | A | 0.395 | 5.35E-02 | 9.71E-02 | 2.49E+00 | 2.54E+00 | 6.02E-01 | no | Adamts19 | intron_variant | MODIFIER |
| 18 | 58871963 | AX-170421023 | 0 | T | C | 0.395 | 5.35E-02 | 9.71E-02 | 2.49E+00 | 2.54E+00 | 6.02E-01 | no | Adamts19 | intron_variant | MODIFIER |
| 18 | 58880294 | AX-170371318 | 0 | G | A | 0.395 | 5.35E-02 | 9.71E-02 | 2.49E+00 | 2.54E+00 | 6.02E-01 | no | Adamts19 | intron_variant | MODIFIER |
| 18 | 58880578 | AX-170306156 | 0 | C | T | 0.395 | 5.35E-02 | 9.71E-02 | 2.49E+00 | 2.54E+00 | 6.02E-01 | no | Adamts19 | intron_variant | MODIFIER |
| 18 | 58882374 | AX-170060977 | 0 | A | G | 0.395 | 5.35E-02 | 9.71E-02 | 2.49E+00 | 2.54E+00 | 6.02E-01 | no | Adamts19 | intron_variant | MODIFIER |
| 18 | 58933545 | AX-97812768 | 0 | C | T | 0.395 | 5.35E-02 | 9.71E-02 | 2.49E+00 | 2.54E+00 | 6.02E-01 | no | Adamts19 | intron_variant | MODIFIER |
| 18 | 58950979 | AX-170461324 | 0 | A | T | 0.395 | 5.35E-02 | 9.71E-02 | 2.49E+00 | 2.54E+00 | 6.02E-01 | no | Adamts19 | intron_variant | MODIFIER |
| 18 | 58970760 | AX-170414565 | 0 | C | G | 0.395 | 5.35E-02 | 9.71E-02 | 2.49E+00 | 2.54E+00 | 6.02E-01 | no | Adamts19 | intron_variant | MODIFIER |
| 18 | 58976425 | AX-170435709 | 0 | A | G | 0.395 | 5.35E-02 | 9.71E-02 | 2.49E+00 | 2.54E+00 | 6.02E-01 | no | Adamts19 | intron_variant | MODIFIER |
| 18 | 59004076 | AX-170451176 | 0 | C | G | 0.395 | 5.35E-02 | 9.71E-02 | 2.49E+00 | 2.54E+00 | 6.02E-01 | no | Adamts19 | intron_variant | MODIFIER |
| 18 | 59031760 | AX-97792337 | 0 | C | G | 0.395 | 5.35E-02 | 9.71E-02 | 2.49E+00 | 2.54E+00 | 6.02E-01 | no | Adamts19 | intron_variant | MODIFIER |
| 18 | 59035377 | AX-170511818 | 0 | T | C | 0.395 | 5.35E-02 | 9.71E-02 | 2.49E+00 | 2.54E+00 | 6.02E-01 | no | Adamts19 | intron_variant | MODIFIER |
| 18 | 59039473 | AX-170288860 | 0 | A | G | 0.395 | 5.35E-02 | 9.71E-02 | 2.49E+00 | 2.54E+00 | 6.02E-01 | no | Adamts19 | intron_variant | MODIFIER |
| 18 | 59043501 | AX-170240035 | 0 | C | T | 0.395 | 5.35E-02 | 9.71E-02 | 2.49E+00 | 2.54E+00 | 6.02E-01 | no | Adamts19 | intron_variant | MODIFIER |
| 18 | 59044086 | AX-170419022 | 0 | A | G | 0.395 | 5.35E-02 | 9.71E-02 | 2.49E+00 | 2.54E+00 | 6.02E-01 | no | Adamts19 | intron_variant | MODIFIER |
| 18 | 59053527 | AX-170449350 | 0 | G | A | 0.395 | 5.35E-02 | 9.71E-02 | 2.49E+00 | 2.54E+00 | 6.02E-01 | no | Adamts19 | 3_prime_UTR_variant | MODIFIER |
| 18 | 59055092 | AX-170472406 | 0 | A | G | 0.395 | 5.35E-02 | 9.71E-02 | 2.49E+00 | 2.54E+00 | 6.02E-01 | no | Adamts19 | downstream_gene_variant | MODIFIER |
| 18 | 59061208 | AX-170522032 | 0 | T | C | 0.395 | 5.35E-02 | 9.71E-02 | 2.49E+00 | 2.54E+00 | 6.02E-01 | no | Minar2 | upstream_gene_variant | MODIFIER |
| 18 | 59065864 | AX-170526590 | 0 | A | G | 0.395 | 5.35E-02 | 9.71E-02 | 2.49E+00 | 2.54E+00 | 6.02E-01 | no | Minar2 | 5_prime_UTR_variant | MODIFIER |
| 18 | 59073381 | AX-170530481 | 0 | C | A | 0.395 | 5.35E-02 | 9.71E-02 | 2.49E+00 | 2.54E+00 | 6.02E-01 | no | Minar2 | intron_variant | MODIFIER |
| 18 | 59078272 | AX-170502735 | 0 | A | G | 0.395 | 5.35E-02 | 9.71E-02 | 2.49E+00 | 2.54E+00 | 6.02E-01 | no | Minar2 | downstream_gene_variant | MODIFIER |
| 18 | 59079893 | AX-170514866 | 0 | C | T | 0.395 | 5.35E-02 | 9.71E-02 | 2.49E+00 | 2.54E+00 | 6.02E-01 | no | Minar2 | downstream_gene_variant | MODIFIER |
| 18 | 59104656 | AX-170489806 | 0 | G | T | 0.395 | 5.35E-02 | 9.71E-02 | 2.49E+00 | 2.54E+00 | 6.02E-01 | no | | intergenic_variant | MODIFIER |
| 18 | 59136086 | AX-170471456 | 0 | T | C | 0.395 | 5.35E-02 | 9.71E-02 | 2.49E+00 | 2.54E+00 | 6.02E-01 | no | | intergenic_variant | MODIFIER |
| 18 | 59136105 | AX-170173465 | 0 | C | T | 0.395 | 5.35E-02 | 9.71E-02 | 2.49E+00 | 2.54E+00 | 6.02E-01 | no | | intergenic_variant | MODIFIER |
| 18 | 59146298 | AX-170413964 | 0 | G | A | 0.395 | 5.35E-02 | 9.71E-02 | 2.49E+00 | 2.54E+00 | 6.02E-01 | no | | intergenic_variant | MODIFIER |

| | | | | | | | | | | | | | | | |
|---|---|---|---|---|---|---|---|---|---|---|---|---|---|---|---|
| 18 | 59248131 | AX-170258392 | 0 | T | C | 0.395 | 5.35E-02 | 9.71E-02 | 2.49E+00 | 2.54E+00 | 6.02E-01 | no | Chsy3 | intron_variant | MODIFIER |
| 18 | 61846074 | AX-170160259 | 0 | T | C | 0.395 | 5.35E-02 | 9.71E-02 | 2.49E+00 | 2.54E+00 | 6.02E-01 | no | Ablim3 | intron_variant | MODIFIER |
| 1 | 174202545 | AX-170002313 | 0 | G | A | 0.228 | -5.50E-02 | 1.07E-01 | 2.87E+00 | 3.01E+00 | 6.02E-01 | no | Spta1 | intron_variant | MODIFIER |
| 1 | 175296563 | AX-170400357 | 0 | T | C | 0.228 | -5.50E-02 | 1.07E-01 | 2.87E+00 | 3.01E+00 | 6.02E-01 | no | Rgs7 | intron_variant | MODIFIER |
| 4 | 82387631 | AX-170227715 | 0 | A | T | 0.237 | 5.69E-02 | 1.07E-01 | 2.55E+00 | 2.63E+00 | 6.02E-01 | no | Nfib | intron_variant | MODIFIER |
| 20 | 69748622 | AX-170116172 | 0 | A | T | 0.254 | 3.24E-02 | 6.29E-02 | 2.66E+00 | 2.74E+00 | 6.02E-01 | no | Aff2 | intron_variant | MODIFIER |
| 20 | 75804753 | AX-170090545 | 0 | A | G | 0.254 | 3.24E-02 | 6.29E-02 | 2.66E+00 | 2.74E+00 | 6.02E-01 | no | Pls3 | intron_variant | MODIFIER |
| 8 | 39682070 | AX-170030538 | 0 | T | A | 0.149 | 7.63E-02 | 1.48E-01 | 2.79E+00 | 2.84E+00 | 6.02E-01 | no | | regulatory_region_variant | MODIFIER |
| 8 | 42289840 | AX-170115854 | 0 | A | T | 0.149 | 7.63E-02 | 1.48E-01 | 2.79E+00 | 2.84E+00 | 6.02E-01 | no | | intergenic_variant | MODIFIER |
| 4 | 83432168 | AX-170089295 | 1 | C | G | 0.241 | 5.61E-02 | 1.05E-01 | 2.54E+00 | 2.61E+00 | 6.03E-01 | no | Snapc3 | intron_variant | MODIFIER |
| 4 | 83603440 | AX-97846411 | 1 | G | A | 0.241 | 5.61E-02 | 1.05E-01 | 2.54E+00 | 2.61E+00 | 6.03E-01 | no | Ccdc171 | intron_variant | MODIFIER |
| 15 | 79393388 | AX-170012061 | 1 | C | T | 0.562 | 4.33E-02 | 8.17E-02 | 2.56E+00 | 2.63E+00 | 6.03E-01 | no | Tmem184b | intron_variant | MODIFIER |
| 18 | 58990069 | AX-170296692 | 1 | C | T | 0.402 | 5.38E-02 | 9.79E-02 | 2.49E+00 | 2.54E+00 | 6.03E-01 | no | Adamts19 | intron_variant | MODIFIER |
| 2 | 123938657 | AX-170122151 | 0 | C | A | 0.149 | 1.00E-01 | 1.95E-01 | 2.69E+00 | 2.75E+00 | 6.03E-01 | no | | intergenic_variant | MODIFIER |
| 2 | 147357231 | AX-97793854 | 0 | C | A | 0.149 | 1.00E-01 | 1.95E-01 | 2.69E+00 | 2.75E+00 | 6.03E-01 | no | Pax1 | upstream_gene_variant | MODIFIER |
| 2 | 149393601 | AX-170343582 | 0 | G | A | 0.149 | 1.00E-01 | 1.95E-01 | 2.69E+00 | 2.75E+00 | 6.03E-01 | no | | intergenic_variant | MODIFIER |
| 12 | 89313302 | AX-168406515 | 0 | T | C | 0.079 | 9.54E-02 | 1.87E-01 | 2.72E+00 | 2.80E+00 | 6.03E-01 | no | Nrxn3 | intron_variant | MODIFIER |
| 18 | 58673563 | AX-170365928 | 1 | T | A | 0.384 | 4.99E-02 | 9.04E-02 | 2.47E+00 | 2.55E+00 | 6.03E-01 | no | Isoc1 | intron_variant | MODIFIER |
| 1 | 174241778 | AX-170192790 | 0 | G | A | 0.228 | 6.55E-02 | 1.23E-01 | 2.56E+00 | 2.62E+00 | 6.03E-01 | no | Spta1 | intron_variant | MODIFIER |
| 1 | 99325013 | AX-170522507 | 2 | C | T | 0.155 | 1.14E-01 | 2.20E-01 | 2.63E+00 | 2.69E+00 | 6.03E-01 | no | | intergenic_variant | MODIFIER |
| 16 | 75520948 | AX-97818613 | 0 | C | T | 0.5 | 4.11E-02 | 7.52E-02 | 2.49E+00 | 2.57E+00 | 6.03E-01 | no | | intergenic_variant | MODIFIER |
| 12 | 90232354 | AX-170165346 | 0 | G | A | 0.088 | 8.50E-02 | 1.66E-01 | 2.67E+00 | 2.76E+00 | 6.03E-01 | no | Nrxn3 | intron_variant | MODIFIER |
| 8 | 73499548 | AX-170170356 | 1 | A | C | 0.188 | -6.87E-02 | 1.34E-01 | 2.68E+00 | 2.75E+00 | 6.03E-01 | no | | intergenic_variant | MODIFIER |
| 11 | 116683210 | AX-170006006 | 0 | A | G | 0.044 | -1.26E-01 | 2.46E-01 | 2.68E+00 | 2.75E+00 | 6.03E-01 | no | St6galnac2 | intron_variant | MODIFIER |
| 6 | 104921918 | AX-170340999 | 1 | G | A | 0.071 | 9.43E-02 | 1.85E-01 | 2.67E+00 | 2.79E+00 | 6.03E-01 | no | | intergenic_variant | MODIFIER |
| 12 | 108815887 | AX-170518527 | 1 | C | T | 0.125 | -8.31E-02 | 1.63E-01 | 2.69E+00 | 2.80E+00 | 6.03E-01 | no | Yy1 | 3_prime_UTR_variant | MODIFIER |
| 18 | 53726822 | AX-170139208 | 0 | A | G | 0.246 | -5.30E-02 | 1.01E-01 | 2.60E+00 | 2.66E+00 | 6.03E-01 | no | Cep120 | intron_variant | MODIFIER |
| 1 | 20894882 | AX-170127654 | 0 | G | A | 0.404 | 4.36E-02 | 8.32E-02 | 2.59E+00 | 2.65E+00 | 6.03E-01 | no | 6720483E21Rik | upstream_gene_variant | MODIFIER |
| 18 | 55837873 | AX-170509795 | 1 | A | G | 0.196 | -5.94E-02 | 1.13E-01 | 2.59E+00 | 2.65E+00 | 6.03E-01 | no | Gm8614 | downstream_gene_variant | MODIFIER |
| 5 | 88916960 | AX-170421293 | 0 | T | C | 0.404 | 5.44E-02 | 1.06E-01 | 2.75E+00 | 2.82E+00 | 6.03E-01 | no | Slc4a4 | intron_variant | MODIFIER |
| 2 | 87776302 | AX-170064240 | 0 | C | G | 0.053 | -1.54E-01 | 2.99E-01 | 2.77E+00 | 2.80E+00 | 6.03E-01 | no | Olfr152 | upstream_gene_variant | MODIFIER |
| 2 | 112728048 | AX-170160776 | 0 | C | T | 0.053 | -1.54E-01 | 2.99E-01 | 2.77E+00 | 2.80E+00 | 6.03E-01 | no | Ryr3 | intron_variant | MODIFIER |
| 4 | 81237051 | AX-170237980 | 0 | A | T | 0.237 | 5.65E-02 | 1.06E-01 | 2.55E+00 | 2.62E+00 | 6.03E-01 | no | | intergenic_variant | MODIFIER |
| 4 | 81252874 | AX-170393301 | 0 | C | A | 0.237 | 5.65E-02 | 1.06E-01 | 2.55E+00 | 2.62E+00 | 6.03E-01 | no | | regulatory_region_variant | MODIFIER |
| 4 | 81254533 | AX-170069806 | 0 | A | G | 0.237 | 5.65E-02 | 1.06E-01 | 2.55E+00 | 2.62E+00 | 6.03E-01 | no | | intergenic_variant | MODIFIER |
| 4 | 81256734 | AX-170189030 | 0 | C | T | 0.237 | 5.65E-02 | 1.06E-01 | 2.55E+00 | 2.62E+00 | 6.03E-01 | no | | intergenic_variant | MODIFIER |
| 4 | 81263966 | AX-97835919 | 0 | G | A | 0.237 | 5.65E-02 | 1.06E-01 | 2.55E+00 | 2.62E+00 | 6.03E-01 | no | | intergenic_variant | MODIFIER |
| 4 | 81278946 | AX-170152088 | 0 | C | T | 0.237 | 5.65E-02 | 1.06E-01 | 2.55E+00 | 2.62E+00 | 6.03E-01 | no | Mpdz | 3_prime_UTR_variant | MODIFIER |
| 4 | 81281142 | AX-170435405 | 0 | T | C | 0.237 | 5.65E-02 | 1.06E-01 | 2.55E+00 | 2.62E+00 | 6.03E-01 | no | Mpdz | intron_variant | MODIFIER |
| 4 | 81285127 | AX-170189506 | 0 | C | T | 0.237 | 5.65E-02 | 1.06E-01 | 2.55E+00 | 2.62E+00 | 6.03E-01 | no | Mpdz | intron_variant | MODIFIER |
| 4 | 81287625 | AX-170297045 | 0 | A | G | 0.237 | 5.65E-02 | 1.06E-01 | 2.55E+00 | 2.62E+00 | 6.03E-01 | no | Mpdz | intron_variant | MODIFIER |
| 4 | 81288165 | AX-97800244 | 0 | C | T | 0.237 | 5.65E-02 | 1.06E-01 | 2.55E+00 | 2.62E+00 | 6.03E-01 | no | Mpdz | intron_variant | MODIFIER |
| 4 | 81356528 | AX-97851587 | 0 | G | A | 0.237 | 5.65E-02 | 1.06E-01 | 2.55E+00 | 2.62E+00 | 6.03E-01 | no | Mpdz | intron_variant | MODIFIER |
| 4 | 81432084 | AX-170023230 | 0 | A | G | 0.237 | 5.65E-02 | 1.06E-01 | 2.55E+00 | 2.62E+00 | 6.03E-01 | no | Mpdz | intron_variant | MODIFIER |
| 4 | 81455696 | AX-97837884 | 0 | A | G | 0.237 | 5.65E-02 | 1.06E-01 | 2.55E+00 | 2.62E+00 | 6.03E-01 | no | Gm11765 | downstream_gene_variant | MODIFIER |
| 4 | 81463082 | AX-170023238 | 0 | C | G | 0.237 | 5.65E-02 | 1.06E-01 | 2.55E+00 | 2.62E+00 | 6.03E-01 | no | Gm11765 | upstream_gene_variant | MODIFIER |
| 4 | 81466543 | AX-170428256 | 0 | A | G | 0.237 | 5.65E-02 | 1.06E-01 | 2.55E+00 | 2.62E+00 | 6.03E-01 | no | Gm11765 | upstream_gene_variant | MODIFIER |
| 4 | 81466708 | AX-170202149 | 0 | C | T | 0.237 | 5.65E-02 | 1.06E-01 | 2.55E+00 | 2.62E+00 | 6.03E-01 | no | Gm11765 | upstream_gene_variant | MODIFIER |
| 4 | 81467107 | AX-170513638 | 0 | C | T | 0.237 | 5.65E-02 | 1.06E-01 | 2.55E+00 | 2.62E+00 | 6.03E-01 | no | | intergenic_variant | MODIFIER |
| 4 | 81658082 | AX-170299221 | 0 | T | C | 0.237 | 5.65E-02 | 1.06E-01 | 2.55E+00 | 2.62E+00 | 6.03E-01 | no | | intergenic_variant | MODIFIER |
| 4 | 81660832 | AX-97796745 | 0 | C | T | 0.237 | 5.65E-02 | 1.06E-01 | 2.55E+00 | 2.62E+00 | 6.03E-01 | no | | intergenic_variant | MODIFIER |
| 4 | 81746942 | AX-97840832 | 0 | A | G | 0.237 | 5.65E-02 | 1.06E-01 | 2.55E+00 | 2.62E+00 | 6.03E-01 | no | Gm11412 | downstream_gene_variant | MODIFIER |
| 4 | 81970308 | AX-170023246 | 0 | T | A | 0.237 | 5.65E-02 | 1.06E-01 | 2.55E+00 | 2.62E+00 | 6.03E-01 | no | | regulatory_region_variant | MODIFIER |
| 4 | 81971995 | AX-170023244 | 0 | C | T | 0.237 | 5.65E-02 | 1.06E-01 | 2.55E+00 | 2.62E+00 | 6.03E-01 | no | | intergenic_variant | MODIFIER |

| | | | | | | | | | | | | | | | |
|---|---|---|---|---|---|---|---|---|---|---|---|---|---|---|---|
| 4 | 82221122 | AX-170491713 | 0 | G | C | 0.237 | 5.65E-02 | 1.06E-01 | 2.55E+00 | 2.62E+00 | 6.03E-01 | no | | regulatory_region_variant | MODIFIER |
| 4 | 82306314 | AX-170462808 | 0 | T | C | 0.237 | 5.65E-02 | 1.06E-01 | 2.55E+00 | 2.62E+00 | 6.03E-01 | no | Nfib | intron_variant | MODIFIER |
| 4 | 82312068 | AX-170447602 | 0 | A | G | 0.237 | 5.65E-02 | 1.06E-01 | 2.55E+00 | 2.62E+00 | 6.03E-01 | no | Nfib | intron_variant | MODIFIER |
| 4 | 82322720 | AX-170399120 | 0 | G | A | 0.237 | 5.65E-02 | 1.06E-01 | 2.55E+00 | 2.62E+00 | 6.03E-01 | no | Nfib | intron_variant | MODIFIER |
| 4 | 82324225 | AX-170069820 | 0 | A | G | 0.237 | 5.65E-02 | 1.06E-01 | 2.55E+00 | 2.62E+00 | 6.03E-01 | no | Nfib | intron_variant | MODIFIER |
| 4 | 82377867 | AX-170175709 | 0 | A | G | 0.237 | 5.65E-02 | 1.06E-01 | 2.55E+00 | 2.62E+00 | 6.03E-01 | no | Nfib | intron_variant | MODIFIER |
| 4 | 82377898 | AX-170226682 | 0 | T | C | 0.237 | 5.65E-02 | 1.06E-01 | 2.55E+00 | 2.62E+00 | 6.03E-01 | no | Nfib | intron_variant | MODIFIER |
| 4 | 82437402 | AX-170418493 | 0 | G | T | 0.237 | 5.65E-02 | 1.06E-01 | 2.55E+00 | 2.62E+00 | 6.03E-01 | no | Nfib | intron_variant | MODIFIER |
| 4 | 82494163 | AX-170023253 | 0 | T | C | 0.237 | 5.65E-02 | 1.06E-01 | 2.55E+00 | 2.62E+00 | 6.03E-01 | no | Nfib | intron_variant | MODIFIER |
| 4 | 83469669 | AX-97782041 | 0 | G | A | 0.237 | 5.65E-02 | 1.06E-01 | 2.55E+00 | 2.62E+00 | 6.03E-01 | no | Psip1 | intron_variant | MODIFIER |
| 4 | 83589402 | AX-170178696 | 0 | C | T | 0.237 | 5.65E-02 | 1.06E-01 | 2.55E+00 | 2.62E+00 | 6.03E-01 | no | Ccdc171 | intron_variant | MODIFIER |
| 4 | 83600034 | AX-170023268 | 0 | T | C | 0.237 | 5.65E-02 | 1.06E-01 | 2.55E+00 | 2.62E+00 | 6.03E-01 | no | Ccdc171 | intron_variant | MODIFIER |
| 4 | 83708966 | AX-170023267 | 0 | C | G | 0.237 | 5.65E-02 | 1.06E-01 | 2.55E+00 | 2.62E+00 | 6.03E-01 | no | Ccdc171 | intron_variant | MODIFIER |
| 4 | 83709059 | AX-170023265 | 0 | T | A | 0.237 | 5.65E-02 | 1.06E-01 | 2.55E+00 | 2.62E+00 | 6.03E-01 | no | Ccdc171 | intron_variant | MODIFIER |
| 4 | 83717843 | AX-170533585 | 0 | T | C | 0.237 | 5.65E-02 | 1.06E-01 | 2.55E+00 | 2.62E+00 | 6.03E-01 | no | Ccdc171 | intron_variant | MODIFIER |
| 4 | 83731676 | AX-170023266 | 0 | A | G | 0.237 | 5.65E-02 | 1.06E-01 | 2.55E+00 | 2.62E+00 | 6.03E-01 | no | Ccdc171 | intron_variant | MODIFIER |
| 4 | 83731737 | AX-170023269 | 0 | G | C | 0.237 | 5.65E-02 | 1.06E-01 | 2.55E+00 | 2.62E+00 | 6.03E-01 | no | Ccdc171 | intron_variant | MODIFIER |
| 4 | 83739040 | AX-97839202 | 0 | T | C | 0.237 | 5.65E-02 | 1.06E-01 | 2.55E+00 | 2.62E+00 | 6.03E-01 | no | Ccdc171 | intron_variant | MODIFIER |
| 4 | 83769032 | AX-170089296 | 0 | T | G | 0.237 | 5.65E-02 | 1.06E-01 | 2.55E+00 | 2.62E+00 | 6.03E-01 | no | Ccdc171 | intron_variant | MODIFIER |
| 4 | 83769073 | AX-170023270 | 0 | A | T | 0.237 | 5.65E-02 | 1.06E-01 | 2.55E+00 | 2.62E+00 | 6.03E-01 | no | Ccdc171 | intron_variant | MODIFIER |
| 4 | 83770227 | AX-170126009 | 0 | G | A | 0.237 | 5.65E-02 | 1.06E-01 | 2.55E+00 | 2.62E+00 | 6.03E-01 | no | Ccdc171 | intron_variant | MODIFIER |
| 4 | 83834138 | AX-97825093 | 0 | G | A | 0.237 | 5.65E-02 | 1.06E-01 | 2.55E+00 | 2.62E+00 | 6.03E-01 | no | Ccdc171 | intron_variant | MODIFIER |
| 4 | 83876016 | AX-170315559 | 0 | T | C | 0.237 | 5.65E-02 | 1.06E-01 | 2.55E+00 | 2.62E+00 | 6.03E-01 | no | | intergenic_variant | MODIFIER |
| 4 | 83918074 | AX-170456094 | 0 | A | G | 0.237 | 5.65E-02 | 1.06E-01 | 2.55E+00 | 2.62E+00 | 6.03E-01 | no | | intergenic_variant | MODIFIER |
| 4 | 83945953 | AX-97833398 | 0 | C | T | 0.237 | 5.65E-02 | 1.06E-01 | 2.55E+00 | 2.62E+00 | 6.03E-01 | no | | intergenic_variant | MODIFIER |
| 4 | 83946544 | AX-170403217 | 0 | G | A | 0.237 | 5.65E-02 | 1.06E-01 | 2.55E+00 | 2.62E+00 | 6.03E-01 | no | | intergenic_variant | MODIFIER |
| 4 | 84006994 | AX-170525798 | 0 | T | C | 0.237 | 5.65E-02 | 1.06E-01 | 2.55E+00 | 2.62E+00 | 6.03E-01 | no | | intergenic_variant | MODIFIER |
| 4 | 84022943 | AX-170157156 | 0 | C | A | 0.237 | 5.65E-02 | 1.06E-01 | 2.55E+00 | 2.62E+00 | 6.03E-01 | no | | intergenic_variant | MODIFIER |
| 4 | 84047150 | AX-170294229 | 0 | T | G | 0.237 | 5.65E-02 | 1.06E-01 | 2.55E+00 | 2.62E+00 | 6.03E-01 | no | 6030471H07Rik | upstream_gene_variant | MODIFIER |
| 4 | 84047870 | AX-170265937 | 0 | A | C | 0.237 | 5.65E-02 | 1.06E-01 | 2.55E+00 | 2.62E+00 | 6.03E-01 | no | 6030471H07Rik | upstream_gene_variant | MODIFIER |
| 4 | 84049911 | AX-170229853 | 0 | A | G | 0.237 | 5.65E-02 | 1.06E-01 | 2.55E+00 | 2.62E+00 | 6.03E-01 | no | | intergenic_variant | MODIFIER |
| 4 | 84056121 | AX-170113432 | 0 | A | C | 0.237 | 5.65E-02 | 1.06E-01 | 2.55E+00 | 2.62E+00 | 6.03E-01 | no | | intergenic_variant | MODIFIER |
| 4 | 84059419 | AX-170089297 | 0 | A | C | 0.237 | 5.65E-02 | 1.06E-01 | 2.55E+00 | 2.62E+00 | 6.03E-01 | no | | intergenic_variant | MODIFIER |
| 5 | 123909518 | AX-170260471 | 0 | T | A | 0.447 | 5.65E-02 | 1.08E-01 | 2.60E+00 | 2.66E+00 | 6.04E-01 | no | Denr | intron_variant | MODIFIER |
| 4 | 149123428 | AX-170242611 | 0 | C | T | 0.105 | 8.70E-02 | 1.68E-01 | 2.60E+00 | 2.69E+00 | 6.04E-01 | no | Cenps | downstream_gene_variant | MODIFIER |
| 4 | 97153375 | AX-97838921 | 0 | A | T | 0.439 | -4.16E-02 | 7.85E-02 | 2.55E+00 | 2.63E+00 | 6.04E-01 | no | | intergenic_variant | MODIFIER |
| 18 | 58714251 | AX-170238496 | 1 | A | C | 0.393 | 5.31E-02 | 9.72E-02 | 2.50E+00 | 2.55E+00 | 6.04E-01 | no | | regulatory_region_variant | MODIFIER |
| 13 | 38123192 | AX-170051010 | 0 | A | T | 0.175 | -6.18E-02 | 1.18E-01 | 2.57E+00 | 2.64E+00 | 6.04E-01 | no | | intergenic_variant | MODIFIER |
| 18 | 53548059 | AX-170110834 | 0 | T | A | 0.237 | -5.76E-02 | 1.08E-01 | 2.55E+00 | 2.61E+00 | 6.04E-01 | no | Prdm6 | intron_variant | MODIFIER |
| 4 | 151664283 | AX-170165555 | 0 | A | G | 0.114 | 9.37E-02 | 1.84E-01 | 2.69E+00 | 2.77E+00 | 6.04E-01 | no | Camta1 | intron_variant | MODIFIER |
| 15 | 86068412 | AX-170121430 | 0 | G | A | 0.518 | 3.78E-02 | 7.02E-02 | 2.50E+00 | 2.61E+00 | 6.04E-01 | no | Gramd4 | intron_variant | MODIFIER |
| 10 | 92272988 | AX-170395770 | 0 | C | T | 0.316 | -4.31E-02 | 8.22E-02 | 2.58E+00 | 2.65E+00 | 6.04E-01 | no | Gm20757 | intron_variant,non_coding_transcript_variant | MODIFIER |
| 15 | 3466656 | AX-170498439 | 0 | C | T | 0.246 | -5.36E-02 | 1.03E-01 | 2.61E+00 | 2.67E+00 | 6.05E-01 | no | Ghr | intron_variant | MODIFIER |
| 18 | 57291163 | AX-170298940 | 0 | G | A | 0.404 | 5.24E-02 | 9.60E-02 | 2.49E+00 | 2.55E+00 | 6.05E-01 | no | Megf10 | intron_variant | MODIFIER |
| 6 | 135020633 | AX-170501750 | 0 | A | T | 0.57 | 4.51E-02 | 8.50E-02 | 2.55E+00 | 2.62E+00 | 6.05E-01 | no | Ddx47 | synonymous_variant | LOW |
| 5 | 42926591 | AX-170188195 | 0 | G | A | 0.211 | -5.47E-02 | 1.05E-01 | 2.58E+00 | 2.67E+00 | 6.05E-01 | no | | intergenic_variant | MODIFIER |
| 1 | 24391791 | AX-170503696 | 1 | C | T | 0.438 | 4.33E-02 | 8.39E-02 | 2.63E+00 | 2.69E+00 | 6.05E-01 | no | Col19a1 | intron_variant | MODIFIER |
| 15 | 25391200 | AX-170500874 | 0 | C | T | 0.254 | -5.30E-02 | 1.03E-01 | 2.62E+00 | 2.69E+00 | 6.05E-01 | no | Basp1 | intron_variant | MODIFIER |
| 15 | 25550239 | AX-170320067 | 0 | T | C | 0.254 | -5.30E-02 | 1.03E-01 | 2.62E+00 | 2.69E+00 | 6.05E-01 | no | Gm36642 | intron_variant,non_coding_transcript_variant | MODIFIER |
| 1 | 101658752 | AX-170430758 | 0 | G | C | 0.149 | 1.03E-01 | 2.01E-01 | 2.65E+00 | 2.71E+00 | 6.05E-01 | no | | intergenic_variant | MODIFIER |
| 1 | 101808196 | AX-170511073 | 0 | T | A | 0.149 | 1.03E-01 | 2.01E-01 | 2.65E+00 | 2.71E+00 | 6.05E-01 | no | | intergenic_variant | MODIFIER |
| 1 | 174149042 | AX-170311400 | 1 | G | A | 0.232 | 6.43E-02 | 1.21E-01 | 2.54E+00 | 2.60E+00 | 6.05E-01 | no | Olfr421-ps1 | upstream_gene_variant | MODIFIER |
| 16 | 95234019 | AX-170533681 | 0 | T | G | 0.465 | 3.53E-02 | 7.00E-02 | 2.76E+00 | 2.88E+00 | 6.05E-01 | no | Gm31641 | intron_variant,non_coding_transcript_variant | MODIFIER |
| 8 | 22124896 | AX-170353202 | 1 | T | C | 0.188 | 5.42E-02 | 1.07E-01 | 2.69E+00 | 2.78E+00 | 6.05E-01 | no | Nek3 | downstream_gene_variant | MODIFIER |

| | | | | | | | | | | | | | | | |
|---|---|---|---|---|---|---|---|---|---|---|---|---|---|---|---|
| 11 | 46354828 | AX-170463969 | 1 | G | A | 0.089 | 8.64E-02 | 1.71E-01 | 2.68E+00 | 2.80E+00 | 6.05E-01 | no | Itk | intron_variant | MODIFIER |
| 6 | 93590145 | AX-170191613 | 0 | G | A | 0.421 | -3.04E-02 | 5.83E-02 | 2.56E+00 | 2.68E+00 | 6.05E-01 | no | | intergenic_variant | MODIFIER |
| 10 | 99656612 | AX-170045720 | 0 | T | C | 0.272 | -4.57E-02 | 8.80E-02 | 2.60E+00 | 2.67E+00 | 6.05E-01 | no | Gm20110 | non_coding_transcript_exon_variant | MODIFIER |
| 1 | 151311523 | AX-170282888 | 1 | T | G | 0.205 | 7.57E-02 | 1.47E-01 | 2.65E+00 | 2.68E+00 | 6.05E-01 | no | | regulatory_region_variant | MODIFIER |
| 15 | 82759635 | AX-170460634 | 2 | A | C | 0.536 | 4.12E-02 | 7.84E-02 | 2.56E+00 | 2.64E+00 | 6.05E-01 | no | Cyp2d40 | downstream_gene_variant | MODIFIER |
| 5 | 39425793 | AX-170221276 | 0 | A | G | 0.219 | 5.38E-02 | 1.06E-01 | 2.76E+00 | 2.86E+00 | 6.05E-01 | no | | intergenic_variant | MODIFIER |
| 9 | 38254216 | AX-170433709 | 1 | A | T | 0.5 | -4.05E-02 | 8.00E-02 | 2.74E+00 | 2.83E+00 | 6.05E-01 | no | Olfr143 | synonymous_variant | LOW |
| 1 | 173289497 | AX-170256706 | 0 | G | T | 0.219 | -6.83E-02 | 1.30E-01 | 2.92E+00 | 3.04E+00 | 6.05E-01 | no | | intergenic_variant | MODIFIER |
| 5 | 104173883 | AX-170025782 | 0 | C | T | 0.465 | 5.75E-02 | 1.11E-01 | 2.61E+00 | 2.66E+00 | 6.06E-01 | no | Dspp | intron_variant | MODIFIER |
| 5 | 110757599 | AX-170487155 | 0 | G | A | 0.465 | 5.75E-02 | 1.11E-01 | 2.61E+00 | 2.66E+00 | 6.06E-01 | no | Ep400 | intron_variant | MODIFIER |
| 5 | 117525070 | AX-170299385 | 0 | C | T | 0.465 | 5.75E-02 | 1.11E-01 | 2.61E+00 | 2.66E+00 | 6.06E-01 | no | Ksr2 | intron_variant,non_coding_transcript_variant | MODIFIER |
| 18 | 57259476 | AX-170154869 | 0 | A | G | 0.404 | 5.32E-02 | 9.76E-02 | 2.49E+00 | 2.54E+00 | 6.06E-01 | no | Megf10 | intron_variant | MODIFIER |
| 18 | 57263246 | AX-170495918 | 0 | A | G | 0.404 | 5.32E-02 | 9.76E-02 | 2.49E+00 | 2.54E+00 | 6.06E-01 | no | Megf10 | intron_variant | MODIFIER |
| 18 | 57282789 | AX-170352585 | 0 | C | T | 0.404 | 5.32E-02 | 9.76E-02 | 2.49E+00 | 2.54E+00 | 6.06E-01 | no | Megf10 | intron_variant | MODIFIER |
| 18 | 57285593 | AX-97844526 | 0 | G | A | 0.404 | 5.32E-02 | 9.76E-02 | 2.49E+00 | 2.54E+00 | 6.06E-01 | no | Megf10 | intron_variant | MODIFIER |
| 18 | 57298040 | AX-97836430 | 0 | T | G | 0.404 | 5.32E-02 | 9.76E-02 | 2.49E+00 | 2.54E+00 | 6.06E-01 | no | Megf10 | downstream_gene_variant | MODIFIER |
| 18 | 57320564 | AX-170295448 | 0 | T | C | 0.404 | 5.32E-02 | 9.76E-02 | 2.49E+00 | 2.54E+00 | 6.06E-01 | no | AC157904.2 | downstream_gene_variant | MODIFIER |
| 18 | 57339980 | AX-170105133 | 0 | A | G | 0.404 | 5.32E-02 | 9.76E-02 | 2.49E+00 | 2.54E+00 | 6.06E-01 | no | AC122117.1 | downstream_gene_variant | MODIFIER |
| 18 | 57340930 | AX-97793206 | 0 | C | T | 0.404 | 5.32E-02 | 9.76E-02 | 2.49E+00 | 2.54E+00 | 6.06E-01 | no | AC122117.1 | downstream_gene_variant | MODIFIER |
| 18 | 57352439 | AX-170421556 | 0 | T | C | 0.404 | 5.32E-02 | 9.76E-02 | 2.49E+00 | 2.54E+00 | 6.06E-01 | no | Prrc1 | upstream_gene_variant | MODIFIER |
| 18 | 57365180 | AX-170222468 | 0 | A | G | 0.404 | 5.32E-02 | 9.76E-02 | 2.49E+00 | 2.54E+00 | 6.06E-01 | no | Prrc1 | intron_variant | MODIFIER |
| 18 | 57365574 | AX-170297463 | 0 | C | T | 0.404 | 5.32E-02 | 9.76E-02 | 2.49E+00 | 2.54E+00 | 6.06E-01 | no | Prrc1 | intron_variant | MODIFIER |
| 18 | 57400129 | AX-170060939 | 0 | C | A | 0.404 | 5.32E-02 | 9.76E-02 | 2.49E+00 | 2.54E+00 | 6.06E-01 | no | 4930511M06Rik | intron_variant,non_coding_transcript_variant | MODIFIER |
| 18 | 57402593 | AX-170205827 | 0 | C | T | 0.404 | 5.32E-02 | 9.76E-02 | 2.49E+00 | 2.54E+00 | 6.06E-01 | no | 4930511M06Rik | intron_variant,non_coding_transcript_variant | MODIFIER |
| 18 | 57402643 | AX-170060938 | 0 | G | A | 0.404 | 5.32E-02 | 9.76E-02 | 2.49E+00 | 2.54E+00 | 6.06E-01 | no | 4930511M06Rik | intron_variant,non_coding_transcript_variant | MODIFIER |
| 18 | 57403393 | AX-170345543 | 0 | C | T | 0.404 | 5.32E-02 | 9.76E-02 | 2.49E+00 | 2.54E+00 | 6.06E-01 | no | 4930511M06Rik | intron_variant,non_coding_transcript_variant | MODIFIER |
| 18 | 57403464 | AX-170503437 | 0 | G | T | 0.404 | 5.32E-02 | 9.76E-02 | 2.49E+00 | 2.54E+00 | 6.06E-01 | no | 4930511M06Rik | intron_variant,non_coding_transcript_variant | MODIFIER |
| 18 | 57406291 | AX-170446912 | 0 | A | G | 0.404 | 5.32E-02 | 9.76E-02 | 2.49E+00 | 2.54E+00 | 6.06E-01 | no | 4930511M06Rik | intron_variant,non_coding_transcript_variant | MODIFIER |
| 18 | 57434314 | AX-170357214 | 0 | T | C | 0.404 | 5.32E-02 | 9.76E-02 | 2.49E+00 | 2.54E+00 | 6.06E-01 | no | 4930511M06Rik | intron_variant,non_coding_transcript_variant | MODIFIER |
| 18 | 57439481 | AX-170278366 | 0 | T | C | 0.404 | 5.32E-02 | 9.76E-02 | 2.49E+00 | 2.54E+00 | 6.06E-01 | no | 4930511M06Rik | intron_variant,non_coding_transcript_variant | MODIFIER |
| 18 | 57451090 | AX-170126293 | 0 | C | T | 0.404 | 5.32E-02 | 9.76E-02 | 2.49E+00 | 2.54E+00 | 6.06E-01 | no | 4930511M06Rik | intron_variant,non_coding_transcript_variant | MODIFIER |
| 18 | 57455091 | AX-170380179 | 0 | C | A | 0.404 | 5.32E-02 | 9.76E-02 | 2.49E+00 | 2.54E+00 | 6.06E-01 | no | 4930511M06Rik | intron_variant,non_coding_transcript_variant | MODIFIER |
| 18 | 57492722 | AX-170401370 | 0 | G | A | 0.404 | 5.32E-02 | 9.76E-02 | 2.49E+00 | 2.54E+00 | 6.06E-01 | no | 4930511M06Rik | intron_variant,non_coding_transcript_variant | MODIFIER |
| 18 | 57493190 | AX-170434000 | 0 | C | T | 0.404 | 5.32E-02 | 9.76E-02 | 2.49E+00 | 2.54E+00 | 6.06E-01 | no | 4930511M06Rik | intron_variant,non_coding_transcript_variant | MODIFIER |
| 18 | 57502519 | AX-170410815 | 0 | T | G | 0.404 | 5.32E-02 | 9.76E-02 | 2.49E+00 | 2.54E+00 | 6.06E-01 | no | 4930511M06Rik | intron_variant,non_coding_transcript_variant | MODIFIER |
| 18 | 57503852 | AX-170147612 | 0 | G | C | 0.404 | 5.32E-02 | 9.76E-02 | 2.49E+00 | 2.54E+00 | 6.06E-01 | no | 4930511M06Rik | intron_variant,non_coding_transcript_variant | MODIFIER |
| 18 | 57533080 | AX-170165172 | 0 | A | T | 0.404 | 5.32E-02 | 9.76E-02 | 2.49E+00 | 2.54E+00 | 6.06E-01 | no | Ccdc192 | upstream_gene_variant | MODIFIER |
| 18 | 57543439 | AX-170152830 | 0 | C | T | 0.404 | 5.32E-02 | 9.76E-02 | 2.49E+00 | 2.54E+00 | 6.06E-01 | no | Ccdc192 | intron_variant | MODIFIER |
| 18 | 57543577 | AX-170491916 | 0 | G | T | 0.404 | 5.32E-02 | 9.76E-02 | 2.49E+00 | 2.54E+00 | 6.06E-01 | no | Ccdc192 | intron_variant | MODIFIER |
| 18 | 57560291 | AX-97816368 | 0 | T | C | 0.404 | 5.32E-02 | 9.76E-02 | 2.49E+00 | 2.54E+00 | 6.06E-01 | no | Ccdc192 | intron_variant | MODIFIER |
| 18 | 57579112 | AX-170493041 | 0 | C | T | 0.404 | 5.32E-02 | 9.76E-02 | 2.49E+00 | 2.54E+00 | 6.06E-01 | no | Ccdc192 | intron_variant | MODIFIER |
| 18 | 57579683 | AX-97797192 | 0 | T | C | 0.404 | 5.32E-02 | 9.76E-02 | 2.49E+00 | 2.54E+00 | 6.06E-01 | no | Ccdc192 | intron_variant | MODIFIER |
| 18 | 57593955 | AX-170178182 | 0 | C | T | 0.404 | 5.32E-02 | 9.76E-02 | 2.49E+00 | 2.54E+00 | 6.06E-01 | no | Ccdc192 | intron_variant | MODIFIER |
| 18 | 57594188 | AX-170270460 | 0 | T | C | 0.404 | 5.32E-02 | 9.76E-02 | 2.49E+00 | 2.54E+00 | 6.06E-01 | no | Ccdc192 | intron_variant | MODIFIER |
| 18 | 57597850 | AX-170392866 | 0 | C | T | 0.404 | 5.32E-02 | 9.76E-02 | 2.49E+00 | 2.54E+00 | 6.06E-01 | no | Ccdc192 | intron_variant | MODIFIER |
| 18 | 57607057 | AX-170060946 | 0 | A | T | 0.404 | 5.32E-02 | 9.76E-02 | 2.49E+00 | 2.54E+00 | 6.06E-01 | no | Ccdc192 | intron_variant | MODIFIER |
| 18 | 57704639 | AX-170015777 | 0 | G | A | 0.404 | 5.32E-02 | 9.76E-02 | 2.49E+00 | 2.54E+00 | 6.06E-01 | no | Ccdc192 | intron_variant | MODIFIER |
| 18 | 57755409 | AX-170060949 | 0 | A | G | 0.404 | 5.32E-02 | 9.76E-02 | 2.49E+00 | 2.54E+00 | 6.06E-01 | no | AC102232.1 | intron_variant,non_coding_transcript_variant | MODIFIER |
| 18 | 57770495 | AX-170221808 | 0 | C | G | 0.404 | 5.32E-02 | 9.76E-02 | 2.49E+00 | 2.54E+00 | 6.06E-01 | no | AC102232.1 | downstream_gene_variant | MODIFIER |
| 18 | 57775667 | AX-170312336 | 0 | T | G | 0.404 | 5.32E-02 | 9.76E-02 | 2.49E+00 | 2.54E+00 | 6.06E-01 | no | AC102232.1 | non_coding_transcript_exon_variant | MODIFIER |
| 18 | 57777987 | AX-170263935 | 0 | T | C | 0.404 | 5.32E-02 | 9.76E-02 | 2.49E+00 | 2.54E+00 | 6.06E-01 | no | AC102232.1 | intron_variant,non_coding_transcript_variant | MODIFIER |
| 18 | 57788401 | AX-170297211 | 0 | G | T | 0.404 | 5.32E-02 | 9.76E-02 | 2.49E+00 | 2.54E+00 | 6.06E-01 | no | AC102232.1 | intron_variant,non_coding_transcript_variant | MODIFIER |
| 18 | 57788569 | AX-170515192 | 0 | C | G | 0.404 | 5.32E-02 | 9.76E-02 | 2.49E+00 | 2.54E+00 | 6.06E-01 | no | AC102232.1 | intron_variant,non_coding_transcript_variant | MODIFIER |
| 18 | 57806345 | AX-170167001 | 0 | C | A | 0.404 | 5.32E-02 | 9.76E-02 | 2.49E+00 | 2.54E+00 | 6.06E-01 | no | | intergenic_variant | MODIFIER |

| | | | | | | | | | | | | | | | |
|---|---|---|---|---|---|---|---|---|---|---|---|---|---|---|---|
| 18 | 57855431 | AX-170250817 | 0 | C | T | 0.404 | 5.32E-02 | 9.76E-02 | 2.49E+00 | 2.54E+00 | 6.06E-01 | no | | intergenic_variant | MODIFIER |
| 18 | 57856250 | AX-170442601 | 0 | G | A | 0.404 | 5.32E-02 | 9.76E-02 | 2.49E+00 | 2.54E+00 | 6.06E-01 | no | | regulatory_region_variant | MODIFIER |
| 18 | 57872311 | AX-170060952 | 0 | T | C | 0.404 | 5.32E-02 | 9.76E-02 | 2.49E+00 | 2.54E+00 | 6.06E-01 | no | | intergenic_variant | MODIFIER |
| 18 | 57875233 | AX-170157556 | 0 | T | C | 0.404 | 5.32E-02 | 9.76E-02 | 2.49E+00 | 2.54E+00 | 6.06E-01 | no | Slc12a2 | upstream_gene_variant | MODIFIER |
| 18 | 57899252 | AX-170290311 | 0 | T | C | 0.404 | 5.32E-02 | 9.76E-02 | 2.49E+00 | 2.54E+00 | 6.06E-01 | no | Slc12a2 | splice_region_variant,intron_variant | LOW |
| 18 | 57909273 | AX-170191865 | 0 | T | C | 0.404 | 5.32E-02 | 9.76E-02 | 2.49E+00 | 2.54E+00 | 6.06E-01 | no | Slc12a2 | intron_variant | MODIFIER |
| 18 | 57929860 | AX-170365486 | 0 | A | G | 0.404 | 5.32E-02 | 9.76E-02 | 2.49E+00 | 2.54E+00 | 6.06E-01 | no | Slc12a2 | intron_variant | MODIFIER |
| 18 | 57965161 | AX-170315896 | 0 | A | G | 0.404 | 5.32E-02 | 9.76E-02 | 2.49E+00 | 2.54E+00 | 6.06E-01 | no | | intergenic_variant | MODIFIER |
| 18 | 58001386 | AX-170060955 | 0 | G | T | 0.404 | 5.32E-02 | 9.76E-02 | 2.49E+00 | 2.54E+00 | 6.06E-01 | no | | intergenic_variant | MODIFIER |
| 18 | 58007339 | AX-97791606 | 0 | C | T | 0.404 | 5.32E-02 | 9.76E-02 | 2.49E+00 | 2.54E+00 | 6.06E-01 | no | Fbn2 | downstream_gene_variant | MODIFIER |
| 18 | 58010495 | AX-170060953 | 0 | G | A | 0.404 | 5.32E-02 | 9.76E-02 | 2.49E+00 | 2.54E+00 | 6.06E-01 | no | Fbn2 | synonymous_variant | LOW |
| 18 | 58018509 | AX-170091040 | 0 | C | T | 0.404 | 5.32E-02 | 9.76E-02 | 2.49E+00 | 2.54E+00 | 6.06E-01 | no | Fbn2 | intron_variant | MODIFIER |
| 18 | 58047770 | AX-170101242 | 0 | C | G | 0.404 | 5.32E-02 | 9.76E-02 | 2.49E+00 | 2.54E+00 | 6.06E-01 | no | Fbn2 | intron_variant | MODIFIER |
| 18 | 58050951 | AX-170371485 | 0 | G | A | 0.404 | 5.32E-02 | 9.76E-02 | 2.49E+00 | 2.54E+00 | 6.06E-01 | no | Fbn2 | intron_variant | MODIFIER |
| 18 | 58066301 | AX-170317204 | 0 | C | T | 0.404 | 5.32E-02 | 9.76E-02 | 2.49E+00 | 2.54E+00 | 6.06E-01 | no | Fbn2 | intron_variant | MODIFIER |
| 18 | 58073760 | AX-97795272 | 0 | G | T | 0.404 | 5.32E-02 | 9.76E-02 | 2.49E+00 | 2.54E+00 | 6.06E-01 | no | Fbn2 | intron_variant | MODIFIER |
| 18 | 58117207 | AX-170316801 | 0 | T | C | 0.404 | 5.32E-02 | 9.76E-02 | 2.49E+00 | 2.54E+00 | 6.06E-01 | no | Fbn2 | intron_variant | MODIFIER |
| 18 | 58122079 | AX-170398939 | 0 | A | G | 0.404 | 5.32E-02 | 9.76E-02 | 2.49E+00 | 2.54E+00 | 6.06E-01 | no | Fbn2 | intron_variant | MODIFIER |
| 18 | 58126634 | AX-170254624 | 0 | C | T | 0.404 | 5.32E-02 | 9.76E-02 | 2.49E+00 | 2.54E+00 | 6.06E-01 | no | Fbn2 | intron_variant | MODIFIER |
| 18 | 58127861 | AX-170496997 | 0 | C | A | 0.404 | 5.32E-02 | 9.76E-02 | 2.49E+00 | 2.54E+00 | 6.06E-01 | no | Fbn2 | intron_variant | MODIFIER |
| 18 | 58171572 | AX-170380789 | 0 | T | G | 0.404 | 5.32E-02 | 9.76E-02 | 2.49E+00 | 2.54E+00 | 6.06E-01 | no | Fbn2 | intron_variant | MODIFIER |
| 18 | 58181815 | AX-170221078 | 0 | A | G | 0.404 | 5.32E-02 | 9.76E-02 | 2.49E+00 | 2.54E+00 | 6.06E-01 | no | Fbn2 | intron_variant | MODIFIER |
| 18 | 58196512 | AX-170145854 | 0 | T | C | 0.404 | 5.32E-02 | 9.76E-02 | 2.49E+00 | 2.54E+00 | 6.06E-01 | no | Fbn2 | intron_variant | MODIFIER |
| 18 | 58234036 | AX-170387623 | 0 | A | G | 0.404 | 5.32E-02 | 9.76E-02 | 2.49E+00 | 2.54E+00 | 6.06E-01 | no | Gm25660 | upstream_gene_variant | MODIFIER |
| 18 | 58235174 | AX-170334059 | 0 | T | C | 0.404 | 5.32E-02 | 9.76E-02 | 2.49E+00 | 2.54E+00 | 6.06E-01 | no | Gm25660 | upstream_gene_variant | MODIFIER |
| 18 | 58248436 | AX-170530745 | 0 | A | G | 0.404 | 5.32E-02 | 9.76E-02 | 2.49E+00 | 2.54E+00 | 6.06E-01 | no | | intergenic_variant | MODIFIER |
| 18 | 58248869 | AX-170424334 | 0 | T | C | 0.404 | 5.32E-02 | 9.76E-02 | 2.49E+00 | 2.54E+00 | 6.06E-01 | no | | intergenic_variant | MODIFIER |
| 18 | 58292663 | AX-170037827 | 0 | T | C | 0.404 | 5.32E-02 | 9.76E-02 | 2.49E+00 | 2.54E+00 | 6.06E-01 | no | | intergenic_variant | MODIFIER |
| 18 | 58299718 | AX-170245760 | 0 | G | A | 0.404 | 5.32E-02 | 9.76E-02 | 2.49E+00 | 2.54E+00 | 6.06E-01 | no | | regulatory_region_variant | MODIFIER |
| 18 | 58341684 | AX-170309046 | 0 | C | T | 0.404 | 5.32E-02 | 9.76E-02 | 2.49E+00 | 2.54E+00 | 6.06E-01 | no | | intergenic_variant | MODIFIER |
| 18 | 58348598 | AX-170213419 | 0 | G | A | 0.404 | 5.32E-02 | 9.76E-02 | 2.49E+00 | 2.54E+00 | 6.06E-01 | no | | intergenic_variant | MODIFIER |
| 18 | 58384202 | AX-170341437 | 0 | C | T | 0.404 | 5.32E-02 | 9.76E-02 | 2.49E+00 | 2.54E+00 | 6.06E-01 | no | | intergenic_variant | MODIFIER |
| 18 | 58391873 | AX-170211562 | 0 | T | C | 0.404 | 5.32E-02 | 9.76E-02 | 2.49E+00 | 2.54E+00 | 6.06E-01 | no | | intergenic_variant | MODIFIER |
| 18 | 58396973 | AX-170510029 | 0 | C | T | 0.404 | 5.32E-02 | 9.76E-02 | 2.49E+00 | 2.54E+00 | 6.06E-01 | no | | intergenic_variant | MODIFIER |
| 18 | 58402441 | AX-97827462 | 0 | G | A | 0.404 | 5.32E-02 | 9.76E-02 | 2.49E+00 | 2.54E+00 | 6.06E-01 | no | | intergenic_variant | MODIFIER |
| 18 | 58417464 | AX-170530925 | 0 | C | T | 0.404 | 5.32E-02 | 9.76E-02 | 2.49E+00 | 2.54E+00 | 6.06E-01 | no | | intergenic_variant | MODIFIER |
| 18 | 58417592 | AX-97824281 | 0 | G | T | 0.404 | 5.32E-02 | 9.76E-02 | 2.49E+00 | 2.54E+00 | 6.06E-01 | no | | intergenic_variant | MODIFIER |
| 18 | 58419025 | AX-170506717 | 0 | G | A | 0.404 | 5.32E-02 | 9.76E-02 | 2.49E+00 | 2.54E+00 | 6.06E-01 | no | | intergenic_variant | MODIFIER |
| 18 | 58436856 | AX-170505547 | 0 | G | C | 0.404 | 5.32E-02 | 9.76E-02 | 2.49E+00 | 2.54E+00 | 6.06E-01 | no | | intergenic_variant | MODIFIER |
| 18 | 58440870 | AX-170407190 | 0 | G | C | 0.404 | 5.32E-02 | 9.76E-02 | 2.49E+00 | 2.54E+00 | 6.06E-01 | no | | intergenic_variant | MODIFIER |
| 18 | 58442466 | AX-170166232 | 0 | C | T | 0.404 | 5.32E-02 | 9.76E-02 | 2.49E+00 | 2.54E+00 | 6.06E-01 | no | | regulatory_region_variant | MODIFIER |
| 18 | 58450398 | AX-170361665 | 0 | A | C | 0.404 | 5.32E-02 | 9.76E-02 | 2.49E+00 | 2.54E+00 | 6.06E-01 | no | | intergenic_variant | MODIFIER |
| 18 | 58464964 | AX-170037825 | 0 | T | A | 0.404 | 5.32E-02 | 9.76E-02 | 2.49E+00 | 2.54E+00 | 6.06E-01 | no | | intergenic_variant | MODIFIER |
| 18 | 58464987 | AX-97788692 | 0 | A | G | 0.404 | 5.32E-02 | 9.76E-02 | 2.49E+00 | 2.54E+00 | 6.06E-01 | no | | intergenic_variant | MODIFIER |
| 18 | 58465132 | AX-170037826 | 0 | C | T | 0.404 | 5.32E-02 | 9.76E-02 | 2.49E+00 | 2.54E+00 | 6.06E-01 | no | | intergenic_variant | MODIFIER |
| 18 | 58468622 | AX-170210691 | 0 | C | T | 0.404 | 5.32E-02 | 9.76E-02 | 2.49E+00 | 2.54E+00 | 6.06E-01 | no | | intergenic_variant | MODIFIER |
| 18 | 58489362 | AX-170060969 | 0 | G | C | 0.404 | 5.32E-02 | 9.76E-02 | 2.49E+00 | 2.54E+00 | 6.06E-01 | no | | intergenic_variant | MODIFIER |
| 15 | 71499862 | AX-97840245 | 1 | C | T | 0.366 | 3.95E-02 | 7.44E-02 | 2.54E+00 | 2.62E+00 | 6.06E-01 | no | Fam135b | intron_variant | MODIFIER |
| 15 | 82940265 | AX-97808894 | 1 | T | C | 0.545 | 4.30E-02 | 8.17E-02 | 2.56E+00 | 2.63E+00 | 6.06E-01 | no | Tcf20 | intron_variant | MODIFIER |
| 8 | 36966157 | AX-170481696 | 0 | C | T | 0.158 | 6.96E-02 | 1.37E-01 | 2.75E+00 | 2.82E+00 | 6.06E-01 | no | | intergenic_variant | MODIFIER |
| 12 | 93628121 | AX-170172456 | 1 | T | A | 0.062 | 1.21E-01 | 2.37E-01 | 2.73E+00 | 2.78E+00 | 6.06E-01 | no | | intergenic_variant | MODIFIER |
| 11 | 29661064 | AX-170046674 | 0 | C | T | 0.07 | 9.58E-02 | 1.87E-01 | 2.64E+00 | 2.72E+00 | 6.06E-01 | no | | regulatory_region_variant | MODIFIER |
| 1 | 169731404 | AX-170478491 | 0 | T | A | 0.219 | -6.14E-02 | 1.21E-01 | 2.77E+00 | 2.87E+00 | 6.06E-01 | no | | intergenic_variant | MODIFIER |
| 9 | 83886335 | AX-170032847 | 1 | C | A | 0.205 | -5.98E-02 | 1.17E-01 | 2.65E+00 | 2.74E+00 | 6.06E-01 | no | | regulatory_region_variant | MODIFIER |

| | | | | | | | | | | | | | | | |
|---|---|---|---|---|---|---|---|---|---|---|---|---|---|---|---|
| 8 | 100402637 | AX-170376812 | 0 | C | A | 0.228 | -5.20E-02 | 1.03E-01 | 2.71E+00 | 2.82E+00 | 6.06E-01 | no | | intergenic_variant | MODIFIER |
| 8 | 101683604 | AX-170460033 | 0 | A | G | 0.228 | -5.20E-02 | 1.03E-01 | 2.71E+00 | 2.82E+00 | 6.06E-01 | no | | intergenic_variant | MODIFIER |
| 8 | 95808259 | AX-170316372 | 0 | A | T | 0.228 | -5.16E-02 | 1.03E-01 | 2.72E+00 | 2.83E+00 | 6.06E-01 | no | Cnot1 | upstream_gene_variant | MODIFIER |
| 1 | 176530209 | AX-170195436 | 0 | T | G | 0.228 | 5.87E-02 | 1.12E-01 | 2.56E+00 | 2.63E+00 | 6.06E-01 | no | Gm36536 | upstream_gene_variant | MODIFIER |
| 9 | 21210862 | AX-170274827 | 0 | G | A | 0.465 | 4.79E-02 | 9.13E-02 | 2.57E+00 | 2.63E+00 | 6.06E-01 | no | Pde4a | missense_variant | MODERATE |
| 4 | 83599860 | AX-170023272 | 1 | T | A | 0.241 | 5.61E-02 | 1.06E-01 | 2.55E+00 | 2.62E+00 | 6.06E-01 | no | Ccdc171 | intron_variant | MODIFIER |
| 16 | 97716111 | AX-170258698 | 0 | G | A | 0.465 | 3.87E-02 | 7.68E-02 | 2.78E+00 | 2.89E+00 | 6.06E-01 | no | | intergenic_variant | MODIFIER |
| 1 | 99657570 | AX-170001191 | 1 | T | C | 0.152 | 1.08E-01 | 2.10E-01 | 2.65E+00 | 2.71E+00 | 6.06E-01 | no | | intergenic_variant | MODIFIER |
| 2 | 164499575 | AX-170379230 | 0 | A | G | 0.175 | 7.18E-02 | 1.41E-01 | 2.65E+00 | 2.75E+00 | 6.06E-01 | no | Pigt | intron_variant | MODIFIER |
| 9 | 92340266 | AX-170321391 | 0 | G | A | 0.132 | -6.39E-02 | 1.23E-01 | 2.59E+00 | 2.67E+00 | 6.06E-01 | no | 1700057G04Rik | intron_variant | MODIFIER |
| 1 | 126729134 | AX-170301409 | 0 | T | C | 0.544 | 4.24E-02 | 8.21E-02 | 2.61E+00 | 2.67E+00 | 6.06E-01 | no | Nckap5 | intron_variant | MODIFIER |
| 6 | 104318907 | AX-170143621 | 1 | G | A | 0.062 | 1.14E-01 | 2.25E-01 | 2.64E+00 | 2.75E+00 | 6.07E-01 | no | | intergenic_variant | MODIFIER |
| 3 | 37791344 | AX-170369941 | 0 | C | T | 0.193 | 4.77E-02 | 9.45E-02 | 2.73E+00 | 2.81E+00 | 6.07E-01 | no | | intergenic_variant | MODIFIER |
| 5 | 43808117 | AX-170449317 | 1 | C | A | 0.205 | -5.23E-02 | 1.02E-01 | 2.60E+00 | 2.70E+00 | 6.07E-01 | no | Fbxl5 | intron_variant | MODIFIER |
| 1 | 43394067 | AX-170388675 | 1 | C | T | 0.098 | -1.08E-01 | 2.14E-01 | 2.72E+00 | 2.81E+00 | 6.07E-01 | no | | intergenic_variant | MODIFIER |
| 14 | 65433317 | AX-170275201 | 0 | A | T | 0.061 | -7.40E-02 | 1.46E-01 | 2.60E+00 | 2.75E+00 | 6.07E-01 | no | | regulatory_region_variant | MODIFIER |
| 18 | 46411162 | AX-170362034 | 1 | A | C | 0.054 | 1.13E-01 | 2.22E-01 | 2.61E+00 | 2.74E+00 | 6.07E-01 | no | AC148019.2 | intron_variant,non_coding_transcript_variant | MODIFIER |
| 8 | 85726847 | AX-170031172 | 1 | C | A | 0.205 | -6.58E-02 | 1.30E-01 | 2.72E+00 | 2.79E+00 | 6.07E-01 | no | Itfg1 | intron_variant | MODIFIER |
| 8 | 86811223 | AX-97805526 | 1 | G | A | 0.205 | -6.58E-02 | 1.30E-01 | 2.72E+00 | 2.79E+00 | 6.07E-01 | no | N4bp1 | non_coding_transcript_exon_variant | MODIFIER |
| 7 | 35793192 | AX-170402504 | 0 | G | A | 0.351 | 4.59E-02 | 8.83E-02 | 2.57E+00 | 2.66E+00 | 6.07E-01 | no | Zfp507 | intron_variant | MODIFIER |
| 18 | 58047842 | AX-170355535 | 1 | G | T | 0.402 | 5.28E-02 | 9.77E-02 | 2.50E+00 | 2.55E+00 | 6.07E-01 | no | Fbn2 | intron_variant | MODIFIER |
| 8 | 88756563 | AX-170497756 | 0 | A | G | 0.175 | -6.46E-02 | 1.27E-01 | 2.66E+00 | 2.74E+00 | 6.07E-01 | no | Cyld | downstream_gene_variant | MODIFIER |
| 1 | 169987853 | AX-170043533 | 0 | C | A | 0.211 | -5.88E-02 | 1.17E-01 | 2.80E+00 | 2.93E+00 | 6.07E-01 | no | Ddr2 | intron_variant | MODIFIER |
| 2 | 164642841 | AX-170276458 | 0 | C | T | 0.158 | 9.89E-02 | 1.95E-01 | 2.68E+00 | 2.76E+00 | 6.08E-01 | no | Wfdc16 | upstream_gene_variant | MODIFIER |
| 13 | 31210353 | AX-170152975 | 1 | G | A | 0.179 | -6.28E-02 | 1.21E-01 | 2.59E+00 | 2.65E+00 | 6.08E-01 | no | 4930401O12Rik | downstream_gene_variant | MODIFIER |
| 13 | 33901164 | AX-97829749 | 1 | G | T | 0.179 | -6.28E-02 | 1.21E-01 | 2.59E+00 | 2.65E+00 | 6.08E-01 | no | Serpinb6c | intron_variant | MODIFIER |
| 5 | 88203906 | AX-170497907 | 1 | A | C | 0.446 | 5.50E-02 | 1.09E-01 | 2.72E+00 | 2.78E+00 | 6.08E-01 | no | Gm42621 | downstream_gene_variant | MODIFIER |
| 15 | 80250775 | AX-170269947 | 0 | A | G | 0.553 | 4.22E-02 | 8.13E-02 | 2.57E+00 | 2.65E+00 | 6.08E-01 | no | Mief1 | 3_prime_UTR_variant | MODIFIER |
| 10 | 115075421 | AX-170333783 | 0 | G | A | 0.368 | 3.77E-02 | 7.51E-02 | 2.71E+00 | 2.81E+00 | 6.08E-01 | no | Tph2 | downstream_gene_variant | MODIFIER |
| 8 | 39021224 | AX-170411872 | 1 | G | A | 0.152 | 7.51E-02 | 1.47E-01 | 2.81E+00 | 2.86E+00 | 6.08E-01 | no | Tusc3 | intron_variant | MODIFIER |
| 4 | 89361573 | AX-170196388 | 0 | A | G | 0.333 | 5.66E-02 | 1.11E-01 | 2.64E+00 | 2.71E+00 | 6.08E-01 | no | | intergenic_variant | MODIFIER |
| 4 | 89929488 | AX-170534333 | 0 | G | T | 0.333 | 5.66E-02 | 1.11E-01 | 2.64E+00 | 2.71E+00 | 6.08E-01 | no | | intergenic_variant | MODIFIER |
| 4 | 92664894 | AX-170261129 | 0 | C | A | 0.333 | 5.66E-02 | 1.11E-01 | 2.64E+00 | 2.71E+00 | 6.08E-01 | no | | intergenic_variant | MODIFIER |
| 1 | 180892626 | AX-170043772 | 0 | T | G | 0.237 | 5.40E-02 | 1.00E-01 | 2.48E+00 | 2.59E+00 | 6.08E-01 | no | Lefty2 | upstream_gene_variant | MODIFIER |
| 1 | 180916675 | AX-170306287 | 0 | G | A | 0.237 | 5.40E-02 | 1.00E-01 | 2.48E+00 | 2.59E+00 | 6.08E-01 | no | | regulatory_region_variant | MODIFIER |
| 18 | 57742684 | AX-170526491 | 0 | G | A | 0.412 | 5.27E-02 | 9.59E-02 | 2.47E+00 | 2.53E+00 | 6.08E-01 | no | AC102232.1 | intron_variant,non_coding_transcript_variant | MODIFIER |
| 1 | 68027818 | AX-170501978 | 0 | A | G | 0.123 | 1.01E-01 | 2.01E-01 | 2.68E+00 | 2.77E+00 | 6.08E-01 | no | Erbb4 | downstream_gene_variant | MODIFIER |
| 1 | 71649201 | AX-170041610 | 0 | C | T | 0.123 | 1.01E-01 | 2.01E-01 | 2.68E+00 | 2.77E+00 | 6.08E-01 | no | Fn1 | synonymous_variant | LOW |
| 18 | 57454961 | AX-170486960 | 1 | G | A | 0.411 | 5.30E-02 | 9.70E-02 | 2.47E+00 | 2.53E+00 | 6.08E-01 | no | 4930511M06Rik | intron_variant,non_coding_transcript_variant | MODIFIER |
| 18 | 39311536 | AX-170523127 | 1 | C | A | 0.071 | 1.11E-01 | 2.19E-01 | 2.65E+00 | 2.74E+00 | 6.09E-01 | no | Arhgap26 | intron_variant | MODIFIER |
| 1 | 24248370 | AX-97846028 | 1 | G | A | 0.446 | 4.31E-02 | 8.41E-02 | 2.62E+00 | 2.68E+00 | 6.09E-01 | no | Col9a1 | intron_variant | MODIFIER |
| 3 | 56058241 | AX-170505833 | 1 | C | T | 0.045 | -1.13E-01 | 2.24E-01 | 2.68E+00 | 2.76E+00 | 6.09E-01 | no | Nbea | intron_variant | MODIFIER |
| 14 | 112419850 | AX-170101036 | 0 | T | G | 0.316 | 3.73E-02 | 7.21E-02 | 2.56E+00 | 2.67E+00 | 6.09E-01 | no | | intergenic_variant | MODIFIER |
| 8 | 21619607 | AX-170106998 | 1 | G | A | 0.179 | 5.38E-02 | 1.06E-01 | 2.65E+00 | 2.74E+00 | 6.09E-01 | no | Defa26 | downstream_gene_variant | MODIFIER |
| 2 | 148945348 | AX-97786809 | 0 | C | T | 0.14 | 9.66E-02 | 1.92E-01 | 2.70E+00 | 2.78E+00 | 6.09E-01 | no | | intergenic_variant | MODIFIER |
| 16 | 76895249 | AX-170471747 | 1 | C | T | 0.455 | 3.99E-02 | 7.68E-02 | 2.57E+00 | 2.64E+00 | 6.09E-01 | no | 1700041M19Rik | intron_variant,non_coding_transcript_variant | MODIFIER |
| 13 | 40243016 | AX-170051054 | 1 | T | C | 0.161 | -6.15E-02 | 1.12E-01 | 2.45E+00 | 2.54E+00 | 6.09E-01 | no | Ofcc1 | intron_variant | MODIFIER |
| 3 | 28618295 | AX-97853043 | 0 | A | G | 0.175 | -5.03E-02 | 9.94E-02 | 2.66E+00 | 2.73E+00 | 6.09E-01 | no | Tnik | intron_variant | MODIFIER |
| 18 | 63164175 | AX-97848532 | 0 | A | C | 0.395 | 4.88E-02 | 9.08E-02 | 2.50E+00 | 2.56E+00 | 6.09E-01 | no | Piezo2 | intron_variant | MODIFIER |
| 7 | 7291840 | AX-170191218 | 0 | A | G | 0.175 | 3.93E-02 | 7.93E-02 | 2.74E+00 | 2.88E+00 | 6.09E-01 | no | Clcn4 | synonymous_variant | LOW |
| 3 | 156097289 | AX-170068533 | 1 | G | C | 0.33 | 3.76E-02 | 7.49E-02 | 2.70E+00 | 2.79E+00 | 6.09E-01 | no | | intergenic_variant | MODIFIER |
| 18 | 89425392 | AX-170198155 | 0 | G | A | 0.518 | -3.12E-02 | 6.29E-02 | 2.69E+00 | 2.82E+00 | 6.09E-01 | no | Dok6 | intron_variant | MODIFIER |
| 5 | 75049908 | AX-170035712 | 1 | G | A | 0.473 | 4.88E-02 | 9.62E-02 | 2.63E+00 | 2.73E+00 | 6.09E-01 | no | Gm18593 | non_coding_transcript_exon_variant | MODIFIER |
| 1 | 35337810 | AX-170000266 | 0 | C | T | 0.456 | 3.96E-02 | 7.89E-02 | 2.75E+00 | 2.84E+00 | 6.09E-01 | no | | intergenic_variant | MODIFIER |

| | | | | | | | | | | | | | | | |
|---|---|---|---|---|---|---|---|---|---|---|---|---|---|---|---|
| 1 | 35521397 | AX-170221653 | 0 | C | T | 0.456 | 3.96E-02 | 7.89E-02 | 2.75E+00 | 2.84E+00 | 6.09E-01 | no | | intergenic_variant | MODIFIER |
| 1 | 35549493 | AX-170329348 | 0 | A | G | 0.456 | 3.96E-02 | 7.89E-02 | 2.75E+00 | 2.84E+00 | 6.09E-01 | no | | intergenic_variant | MODIFIER |
| 1 | 35550725 | AX-170254470 | 0 | T | A | 0.456 | 3.96E-02 | 7.89E-02 | 2.75E+00 | 2.84E+00 | 6.09E-01 | no | | intergenic_variant | MODIFIER |
| 1 | 35559505 | AX-170421251 | 0 | T | C | 0.456 | 3.96E-02 | 7.89E-02 | 2.75E+00 | 2.84E+00 | 6.09E-01 | no | | intergenic_variant | MODIFIER |
| 1 | 35562284 | AX-170114516 | 0 | C | T | 0.456 | 3.96E-02 | 7.89E-02 | 2.75E+00 | 2.84E+00 | 6.09E-01 | no | | intergenic_variant | MODIFIER |
| 1 | 35563401 | AX-170105408 | 0 | C | A | 0.456 | 3.96E-02 | 7.89E-02 | 2.75E+00 | 2.84E+00 | 6.09E-01 | no | | intergenic_variant | MODIFIER |
| 1 | 35587885 | AX-97821789 | 0 | A | G | 0.456 | 3.96E-02 | 7.89E-02 | 2.75E+00 | 2.84E+00 | 6.09E-01 | no | | intergenic_variant | MODIFIER |
| 1 | 35589356 | AX-170273014 | 0 | C | T | 0.456 | 3.96E-02 | 7.89E-02 | 2.75E+00 | 2.84E+00 | 6.09E-01 | no | | intergenic_variant | MODIFIER |
| 1 | 35589368 | AX-170099893 | 0 | G | A | 0.456 | 3.96E-02 | 7.89E-02 | 2.75E+00 | 2.84E+00 | 6.09E-01 | no | | intergenic_variant | MODIFIER |
| 1 | 35589582 | AX-170484646 | 0 | C | T | 0.456 | 3.96E-02 | 7.89E-02 | 2.75E+00 | 2.84E+00 | 6.09E-01 | no | | intergenic_variant | MODIFIER |
| 1 | 35604160 | AX-170117760 | 0 | T | G | 0.456 | 3.96E-02 | 7.89E-02 | 2.75E+00 | 2.84E+00 | 6.09E-01 | no | | intergenic_variant | MODIFIER |
| 1 | 35604620 | AX-97825637 | 0 | A | C | 0.456 | 3.96E-02 | 7.89E-02 | 2.75E+00 | 2.84E+00 | 6.09E-01 | no | | intergenic_variant | MODIFIER |
| 1 | 35652429 | AX-97826256 | 0 | C | T | 0.456 | 3.96E-02 | 7.89E-02 | 2.75E+00 | 2.84E+00 | 6.09E-01 | no | Gm37068 | upstream_gene_variant | MODIFIER |
| 1 | 35683685 | AX-170254724 | 0 | G | A | 0.456 | 3.96E-02 | 7.89E-02 | 2.75E+00 | 2.84E+00 | 6.09E-01 | no | | intergenic_variant | MODIFIER |
| 8 | 78106890 | AX-170296281 | 0 | A | G | 0.202 | -6.60E-02 | 1.30E-01 | 2.65E+00 | 2.72E+00 | 6.09E-01 | no | Gm45288 | upstream_gene_variant | MODIFIER |
| 1 | 148484606 | AX-170468004 | 2 | T | A | 0.2 | 6.75E-02 | 1.33E-01 | 2.65E+00 | 2.71E+00 | 6.09E-01 | no | | intergenic_variant | MODIFIER |
| 6 | 124139849 | AX-97802078 | 1 | A | G | 0.188 | -4.09E-02 | 8.16E-02 | 2.65E+00 | 2.78E+00 | 6.09E-01 | no | Gm23554 | upstream_gene_variant | MODIFIER |
| 6 | 134401089 | AX-170118217 | 2 | C | G | 0.555 | 4.60E-02 | 8.65E-02 | 2.52E+00 | 2.58E+00 | 6.09E-01 | no | Bcl2l14 | intron_variant | MODIFIER |
| 15 | 79983846 | AX-170264544 | 2 | A | G | 0.555 | 4.19E-02 | 8.10E-02 | 2.57E+00 | 2.65E+00 | 6.10E-01 | no | | regulatory_region_variant | MODIFIER |
| 15 | 78858245 | AX-170346061 | 1 | A | G | 0.562 | 4.25E-02 | 8.18E-02 | 2.56E+00 | 2.63E+00 | 6.10E-01 | no | Lgals2 | upstream_gene_variant | MODIFIER |
| 1 | 140765628 | AX-170001841 | 1 | T | G | 0.196 | 6.71E-02 | 1.33E-01 | 2.67E+00 | 2.73E+00 | 6.10E-01 | no | | intergenic_variant | MODIFIER |
| 8 | 103504349 | AX-170146807 | 2 | C | T | 0.218 | 5.54E-02 | 1.10E-01 | 2.68E+00 | 2.76E+00 | 6.10E-01 | no | | intergenic_variant | MODIFIER |
| 9 | 78751757 | AX-170177025 | 1 | A | G | 0.196 | -6.38E-02 | 1.26E-01 | 2.65E+00 | 2.70E+00 | 6.10E-01 | no | | intergenic_variant | MODIFIER |
| 1 | 76280536 | AX-170000838 | 0 | G | A | 0.14 | 9.16E-02 | 1.81E-01 | 2.62E+00 | 2.73E+00 | 6.10E-01 | no | | intergenic_variant | MODIFIER |
| 6 | 130314759 | AX-170161750 | 0 | C | T | 0.553 | 4.18E-02 | 8.13E-02 | 2.59E+00 | 2.66E+00 | 6.10E-01 | no | | intergenic_variant | MODIFIER |
| 5 | 126131854 | AX-170127259 | 1 | G | A | 0.455 | 5.06E-02 | 1.01E-01 | 2.67E+00 | 2.75E+00 | 6.10E-01 | no | | intergenic_variant | MODIFIER |
| 2 | 130122786 | AX-170111300 | 0 | C | G | 0.14 | -9.19E-02 | 1.83E-01 | 2.72E+00 | 2.80E+00 | 6.10E-01 | no | Tgm6 | upstream_gene_variant | MODIFIER |
| 2 | 145724156 | AX-170065170 | 0 | G | A | 0.14 | -9.19E-02 | 1.83E-01 | 2.72E+00 | 2.80E+00 | 6.10E-01 | no | Rin2 | intron_variant,non_coding_transcript_variant | MODIFIER |
| 10 | 67829483 | AX-170376335 | 0 | C | T | 0.421 | -3.94E-02 | 7.71E-02 | 2.61E+00 | 2.69E+00 | 6.10E-01 | no | Gm49766 | intron_variant,non_coding_transcript_variant | MODIFIER |
| 2 | 37731455 | AX-170242962 | 1 | G | T | 0.062 | 7.71E-02 | 1.55E-01 | 2.68E+00 | 2.81E+00 | 6.10E-01 | no | | regulatory_region_variant | MODIFIER |
| 3 | 144138917 | AX-170307826 | 0 | G | A | 0.377 | 3.44E-02 | 6.80E-02 | 2.63E+00 | 2.73E+00 | 6.10E-01 | no | Gm34078 | upstream_gene_variant | MODIFIER |
| 1 | 88536509 | AX-97823054 | 0 | G | A | 0.158 | 8.68E-02 | 1.68E-01 | 2.56E+00 | 2.66E+00 | 6.10E-01 | no | | regulatory_region_variant | MODIFIER |
| 8 | 100772856 | AX-170389890 | 0 | C | T | 0.237 | -5.07E-02 | 1.02E-01 | 2.71E+00 | 2.82E+00 | 6.10E-01 | no | | regulatory_region_variant | MODIFIER |
| 1 | 152327677 | AX-170245936 | 0 | T | C | 0.202 | 6.34E-02 | 1.26E-01 | 2.70E+00 | 2.77E+00 | 6.11E-01 | no | | intergenic_variant | MODIFIER |
| 4 | 95745370 | AX-170259014 | 1 | G | A | 0.268 | 5.22E-02 | 9.86E-02 | 2.51E+00 | 2.60E+00 | 6.11E-01 | no | Fggy | intron_variant | MODIFIER |
| 6 | 131332725 | AX-170028256 | 2 | C | T | 0.555 | 4.37E-02 | 8.39E-02 | 2.55E+00 | 2.63E+00 | 6.11E-01 | no | Styk1 | intron_variant,non_coding_transcript_variant | MODIFIER |
| 11 | 14024057 | AX-170410660 | 1 | C | T | 0.071 | 9.00E-02 | 1.82E-01 | 2.71E+00 | 2.83E+00 | 6.11E-01 | no | | intergenic_variant | MODIFIER |
| 13 | 25451107 | AX-97827011 | 1 | G | A | 0.17 | -6.66E-02 | 1.23E-01 | 2.48E+00 | 2.54E+00 | 6.11E-01 | no | Tpi-rs8 | upstream_gene_variant | MODIFIER |
| 13 | 38740082 | AX-170008150 | 0 | G | C | 0.184 | -5.67E-02 | 1.09E-01 | 2.55E+00 | 2.64E+00 | 6.11E-01 | no | Gm46392 | intron_variant,non_coding_transcript_variant | MODIFIER |
| 18 | 80757183 | AX-170061461 | 0 | T | C | 0.298 | -4.44E-02 | 8.70E-02 | 2.59E+00 | 2.69E+00 | 6.11E-01 | no | Atp9b | intron_variant | MODIFIER |
| 7 | 36432900 | AX-170077064 | 0 | C | A | 0.351 | 4.30E-02 | 8.39E-02 | 2.58E+00 | 2.68E+00 | 6.11E-01 | no | Gm25247 | downstream_gene_variant | MODIFIER |
| 1 | 151838660 | AX-97813711 | 0 | C | T | 0.202 | -5.79E-02 | 1.17E-01 | 2.78E+00 | 2.91E+00 | 6.11E-01 | no | | intergenic_variant | MODIFIER |
| 13 | 116531262 | AX-170166301 | 1 | C | A | 0.045 | -8.19E-02 | 1.63E-01 | 2.61E+00 | 2.74E+00 | 6.11E-01 | no | | intergenic_variant | MODIFIER |
| 6 | 125127357 | AX-97788967 | 0 | C | T | 0.193 | -3.91E-02 | 7.90E-02 | 2.67E+00 | 2.81E+00 | 6.11E-01 | no | Nop2 | upstream_gene_variant | MODIFIER |
| 6 | 125281207 | AX-170477962 | 0 | G | A | 0.193 | -3.91E-02 | 7.90E-02 | 2.67E+00 | 2.81E+00 | 6.11E-01 | no | Tuba3a | synonymous_variant | LOW |
| 6 | 125507369 | AX-170028176 | 0 | C | T | 0.193 | -3.91E-02 | 7.90E-02 | 2.67E+00 | 2.81E+00 | 6.11E-01 | no | Gm26728 | non_coding_transcript_exon_variant | MODIFIER |
| 10 | 92146572 | AX-170104609 | 1 | C | A | 0.304 | -4.45E-02 | 8.49E-02 | 2.54E+00 | 2.60E+00 | 6.11E-01 | no | Rmst | intron_variant,non_coding_transcript_variant | MODIFIER |
| 10 | 93122894 | AX-97795559 | 1 | C | A | 0.304 | -4.45E-02 | 8.49E-02 | 2.54E+00 | 2.60E+00 | 6.11E-01 | no | | regulatory_region_variant | MODIFIER |
| 1 | 25227659 | AX-170000123 | 0 | T | C | 0.43 | 4.17E-02 | 8.16E-02 | 2.60E+00 | 2.67E+00 | 6.11E-01 | no | Adgrb3 | intron_variant | MODIFIER |
| 10 | 23619450 | AX-170002920 | 0 | G | A | 0.579 | -3.46E-02 | 7.00E-02 | 2.82E+00 | 2.97E+00 | 6.11E-01 | no | 4930520K02Rik | intron_variant,non_coding_transcript_variant | MODIFIER |
| 10 | 23627973 | AX-170347737 | 0 | A | G | 0.579 | -3.46E-02 | 7.00E-02 | 2.82E+00 | 2.97E+00 | 6.11E-01 | no | | intergenic_variant | MODIFIER |
| 10 | 23643006 | AX-97794853 | 0 | G | A | 0.579 | -3.46E-02 | 7.00E-02 | 2.82E+00 | 2.97E+00 | 6.11E-01 | no | | intergenic_variant | MODIFIER |
| 10 | 23643160 | AX-170263674 | 0 | T | G | 0.579 | -3.46E-02 | 7.00E-02 | 2.82E+00 | 2.97E+00 | 6.11E-01 | no | | intergenic_variant | MODIFIER |
| 1 | 99446814 | AX-170444952 | 1 | T | C | 0.134 | 7.69E-02 | 1.51E-01 | 2.60E+00 | 2.70E+00 | 6.11E-01 | no | | intergenic_variant | MODIFIER |

| | | | | | | | | | | | | | | | |
|---|---|---|---|---|---|---|---|---|---|---|---|---|---|---|---|
| 17 | 22209370 | AX-170220750 | 0 | C | G | 0.044 | 8.44E-02 | 1.71E-01 | 2.78E+00 | 2.93E+00 | 6.12E-01 | no | Zfp994 | intron_variant | MODIFIER |
| 13 | 22857819 | AX-170007917 | 0 | T | C | 0.167 | -6.58E-02 | 1.22E-01 | 2.48E+00 | 2.54E+00 | 6.12E-01 | no | Vmn1r211 | upstream_gene_variant | MODIFIER |
| 13 | 22858531 | AX-97817611 | 0 | C | T | 0.167 | -6.58E-02 | 1.22E-01 | 2.48E+00 | 2.54E+00 | 6.12E-01 | no | Vmn1r211 | upstream_gene_variant | MODIFIER |
| 13 | 23008852 | AX-170007913 | 0 | G | A | 0.167 | -6.58E-02 | 1.22E-01 | 2.48E+00 | 2.54E+00 | 6.12E-01 | no | Vmn1r213 | upstream_gene_variant | MODIFIER |
| 13 | 23012959 | AX-170007911 | 0 | T | C | 0.167 | -6.58E-02 | 1.22E-01 | 2.48E+00 | 2.54E+00 | 6.12E-01 | no | Vmn1r213 | downstream_gene_variant | MODIFIER |
| 13 | 23014021 | AX-170007918 | 0 | G | T | 0.167 | -6.58E-02 | 1.22E-01 | 2.48E+00 | 2.54E+00 | 6.12E-01 | no | Vmn1r213 | downstream_gene_variant | MODIFIER |
| 13 | 23042558 | AX-170007914 | 0 | A | G | 0.167 | -6.58E-02 | 1.22E-01 | 2.48E+00 | 2.54E+00 | 6.12E-01 | no | Gm19153 | non_coding_transcript_exon_variant | MODIFIER |
| 13 | 23042663 | AX-170007912 | 0 | C | G | 0.167 | -6.58E-02 | 1.22E-01 | 2.48E+00 | 2.54E+00 | 6.12E-01 | no | Gm19153 | non_coding_transcript_exon_variant | MODIFIER |
| 13 | 23043236 | AX-97827274 | 0 | G | T | 0.167 | -6.58E-02 | 1.22E-01 | 2.48E+00 | 2.54E+00 | 6.12E-01 | no | Gm19153 | upstream_gene_variant | MODIFIER |
| 13 | 23068844 | AX-170135142 | 0 | A | G | 0.167 | -6.58E-02 | 1.22E-01 | 2.48E+00 | 2.54E+00 | 6.12E-01 | no | Vmn1r215 | intron_variant | MODIFIER |
| 13 | 23075914 | AX-170050741 | 0 | A | G | 0.167 | -6.58E-02 | 1.22E-01 | 2.48E+00 | 2.54E+00 | 6.12E-01 | no | Vmn1r215 | synonymous_variant | LOW |
| 13 | 23123599 | AX-97835970 | 0 | C | T | 0.167 | -6.58E-02 | 1.22E-01 | 2.48E+00 | 2.54E+00 | 6.12E-01 | no | Vmn1r-ps128 | upstream_gene_variant | MODIFIER |
| 13 | 23123765 | AX-170050745 | 0 | T | G | 0.167 | -6.58E-02 | 1.22E-01 | 2.48E+00 | 2.54E+00 | 6.12E-01 | no | Vmn1r-ps128 | upstream_gene_variant | MODIFIER |
| 13 | 23137971 | AX-170440847 | 0 | C | T | 0.167 | -6.58E-02 | 1.22E-01 | 2.48E+00 | 2.54E+00 | 6.12E-01 | no | Vmn1r218 | downstream_gene_variant | MODIFIER |
| 13 | 23137997 | AX-170193906 | 0 | A | G | 0.167 | -6.58E-02 | 1.22E-01 | 2.48E+00 | 2.54E+00 | 6.12E-01 | no | Vmn1r218 | downstream_gene_variant | MODIFIER |
| 13 | 23139942 | AX-97822922 | 0 | C | A | 0.167 | -6.58E-02 | 1.22E-01 | 2.48E+00 | 2.54E+00 | 6.12E-01 | no | Vmn1r218 | downstream_gene_variant | MODIFIER |
| 13 | 23164023 | AX-170050744 | 0 | T | C | 0.167 | -6.58E-02 | 1.22E-01 | 2.48E+00 | 2.54E+00 | 6.12E-01 | no | Vmn1r219 | downstream_gene_variant | MODIFIER |
| 13 | 23165363 | AX-170327040 | 0 | G | C | 0.167 | -6.58E-02 | 1.22E-01 | 2.48E+00 | 2.54E+00 | 6.12E-01 | no | Vmn1r219 | downstream_gene_variant | MODIFIER |
| 13 | 23178658 | AX-170412390 | 0 | C | T | 0.167 | -6.58E-02 | 1.22E-01 | 2.48E+00 | 2.54E+00 | 6.12E-01 | no | Vmn1r220 | downstream_gene_variant | MODIFIER |
| 13 | 23227641 | AX-97787018 | 0 | A | G | 0.167 | -6.58E-02 | 1.22E-01 | 2.48E+00 | 2.54E+00 | 6.12E-01 | no | Vmn1r222 | downstream_gene_variant | MODIFIER |
| 13 | 23276483 | AX-170264534 | 0 | C | T | 0.167 | -6.58E-02 | 1.22E-01 | 2.48E+00 | 2.54E+00 | 6.12E-01 | no | 4930557F10Rik | intron_variant,non_coding_transcript_variant | MODIFIER |
| 13 | 23285904 | AX-170087386 | 0 | T | C | 0.167 | -6.58E-02 | 1.22E-01 | 2.48E+00 | 2.54E+00 | 6.12E-01 | no | 4930557F10Rik | intron_variant,non_coding_transcript_variant | MODIFIER |
| 13 | 23286345 | AX-170462448 | 0 | G | A | 0.167 | -6.58E-02 | 1.22E-01 | 2.48E+00 | 2.54E+00 | 6.12E-01 | no | 4930557F10Rik | intron_variant,non_coding_transcript_variant | MODIFIER |
| 13 | 23340434 | AX-170454595 | 0 | T | G | 0.167 | -6.58E-02 | 1.22E-01 | 2.48E+00 | 2.54E+00 | 6.12E-01 | no | 4930586N03Rik | intron_variant,non_coding_transcript_variant | MODIFIER |
| 13 | 23353117 | AX-97796508 | 0 | A | C | 0.167 | -6.58E-02 | 1.22E-01 | 2.48E+00 | 2.54E+00 | 6.12E-01 | no | Zfp322a | 3_prime_UTR_variant | MODIFIER |
| 13 | 23353322 | AX-170115909 | 0 | G | A | 0.167 | -6.58E-02 | 1.22E-01 | 2.48E+00 | 2.54E+00 | 6.12E-01 | no | Zfp322a | 3_prime_UTR_variant | MODIFIER |
| 13 | 23353454 | AX-170203451 | 0 | T | C | 0.167 | -6.58E-02 | 1.22E-01 | 2.48E+00 | 2.54E+00 | 6.12E-01 | no | Zfp322a | 3_prime_UTR_variant | MODIFIER |
| 13 | 23370523 | AX-170036874 | 0 | G | A | 0.167 | -6.58E-02 | 1.22E-01 | 2.48E+00 | 2.54E+00 | 6.12E-01 | no | Zfp322a | upstream_gene_variant | MODIFIER |
| 13 | 23398604 | AX-170532110 | 0 | A | C | 0.167 | -6.58E-02 | 1.22E-01 | 2.48E+00 | 2.54E+00 | 6.12E-01 | no | | intergenic_variant | MODIFIER |
| 13 | 23400255 | AX-170105821 | 0 | A | G | 0.167 | -6.58E-02 | 1.22E-01 | 2.48E+00 | 2.54E+00 | 6.12E-01 | no | | regulatory_region_variant | MODIFIER |
| 13 | 23403982 | AX-170248457 | 0 | C | T | 0.167 | -6.58E-02 | 1.22E-01 | 2.48E+00 | 2.54E+00 | 6.12E-01 | no | | intergenic_variant | MODIFIER |
| 13 | 23406951 | AX-97788954 | 0 | A | G | 0.167 | -6.58E-02 | 1.22E-01 | 2.48E+00 | 2.54E+00 | 6.12E-01 | no | | intergenic_variant | MODIFIER |
| 13 | 23413388 | AX-97787364 | 0 | T | G | 0.167 | -6.58E-02 | 1.22E-01 | 2.48E+00 | 2.54E+00 | 6.12E-01 | no | Abt1 | downstream_gene_variant | MODIFIER |
| 13 | 23433090 | AX-170454787 | 0 | T | C | 0.167 | -6.58E-02 | 1.22E-01 | 2.48E+00 | 2.54E+00 | 6.12E-01 | no | C230035I16Rik | upstream_gene_variant | MODIFIER |
| 13 | 23439409 | AX-97783490 | 0 | A | G | 0.167 | -6.58E-02 | 1.22E-01 | 2.48E+00 | 2.54E+00 | 6.12E-01 | no | Gm46404 | downstream_gene_variant | MODIFIER |
| 13 | 23442091 | AX-170431816 | 0 | A | G | 0.167 | -6.58E-02 | 1.22E-01 | 2.48E+00 | 2.54E+00 | 6.12E-01 | no | Gm46404 | non_coding_transcript_exon_variant | MODIFIER |
| 13 | 23461656 | AX-170480903 | 0 | C | G | 0.167 | -6.58E-02 | 1.22E-01 | 2.48E+00 | 2.54E+00 | 6.12E-01 | no | Btn1a1 | missense_variant | MODERATE |
| 13 | 23468471 | AX-97844363 | 0 | A | G | 0.167 | -6.58E-02 | 1.22E-01 | 2.48E+00 | 2.54E+00 | 6.12E-01 | no | Btn1a1 | upstream_gene_variant | MODIFIER |
| 13 | 23506069 | AX-170216568 | 0 | C | T | 0.167 | -6.58E-02 | 1.22E-01 | 2.48E+00 | 2.54E+00 | 6.12E-01 | no | | regulatory_region_variant | MODIFIER |
| 13 | 23528560 | AX-97846199 | 0 | T | C | 0.167 | -6.58E-02 | 1.22E-01 | 2.48E+00 | 2.54E+00 | 6.12E-01 | no | Hist1h4h | upstream_gene_variant | MODIFIER |
| 13 | 23531916 | AX-170133364 | 0 | T | A | 0.167 | -6.58E-02 | 1.22E-01 | 2.48E+00 | 2.54E+00 | 6.12E-01 | no | Hist1h2af | upstream_gene_variant | MODIFIER |
| 13 | 23536000 | AX-97807474 | 0 | C | T | 0.167 | -6.58E-02 | 1.22E-01 | 2.48E+00 | 2.54E+00 | 6.12E-01 | no | Hist1h2af | downstream_gene_variant | MODIFIER |
| 13 | 23560159 | AX-170050752 | 0 | T | C | 0.167 | -6.58E-02 | 1.22E-01 | 2.48E+00 | 2.54E+00 | 6.12E-01 | no | Hist1h1d | 3_prime_UTR_variant | MODIFIER |
| 13 | 23564784 | AX-170050747 | 0 | C | A | 0.167 | -6.58E-02 | 1.22E-01 | 2.48E+00 | 2.54E+00 | 6.12E-01 | no | Hist1h1d | downstream_gene_variant | MODIFIER |
| 13 | 23570366 | AX-170007927 | 0 | G | A | 0.167 | -6.58E-02 | 1.22E-01 | 2.48E+00 | 2.54E+00 | 6.12E-01 | no | Hist1h2bg | upstream_gene_variant | MODIFIER |
| 13 | 23593509 | AX-97835821 | 0 | C | T | 0.167 | -6.58E-02 | 1.22E-01 | 2.48E+00 | 2.54E+00 | 6.12E-01 | no | Hist1h2be | intron_variant | MODIFIER |
| 13 | 23594425 | AX-170254573 | 0 | C | T | 0.167 | -6.58E-02 | 1.22E-01 | 2.48E+00 | 2.54E+00 | 6.12E-01 | no | Hist1h2be | intron_variant | MODIFIER |
| 13 | 23605382 | AX-170007923 | 0 | A | G | 0.167 | -6.58E-02 | 1.22E-01 | 2.48E+00 | 2.54E+00 | 6.12E-01 | no | Hist1h2be | intron_variant | MODIFIER |
| 13 | 23606569 | AX-170007922 | 0 | C | T | 0.167 | -6.58E-02 | 1.22E-01 | 2.48E+00 | 2.54E+00 | 6.12E-01 | no | Hist1h2be | intron_variant | MODIFIER |
| 13 | 23606592 | AX-170007926 | 0 | C | G | 0.167 | -6.58E-02 | 1.22E-01 | 2.48E+00 | 2.54E+00 | 6.12E-01 | no | Hist1h2be | intron_variant | MODIFIER |
| 13 | 23606791 | AX-170036879 | 0 | G | A | 0.167 | -6.58E-02 | 1.22E-01 | 2.48E+00 | 2.54E+00 | 6.12E-01 | no | Hist1h2be | intron_variant | MODIFIER |
| 13 | 23620278 | AX-170395287 | 0 | A | G | 0.167 | -6.58E-02 | 1.22E-01 | 2.48E+00 | 2.54E+00 | 6.12E-01 | no | Hist1h2be | intron_variant | MODIFIER |
| 13 | 23621646 | AX-170249453 | 0 | A | G | 0.167 | -6.58E-02 | 1.22E-01 | 2.48E+00 | 2.54E+00 | 6.12E-01 | no | Hist1h2be | upstream_gene_variant | MODIFIER |
| 13 | 23637943 | AX-170323755 | 0 | A | G | 0.167 | -6.58E-02 | 1.22E-01 | 2.48E+00 | 2.54E+00 | 6.12E-01 | no | | intergenic_variant | MODIFIER |

| | | | | | | | | | | | | | | | |
|---|---|---|---|---|---|---|---|---|---|---|---|---|---|---|---|
| 13 | 23655655 | AX-97836625 | 0 | C | A | 0.167 | -6.58E-02 | 1.22E-01 | 2.48E+00 | 2.54E+00 | 6.12E-01 | no | | intergenic_variant | MODIFIER |
| 13 | 23658249 | AX-170412016 | 0 | A | C | 0.167 | -6.58E-02 | 1.22E-01 | 2.48E+00 | 2.54E+00 | 6.12E-01 | no | Gm22452 | upstream_gene_variant | MODIFIER |
| 13 | 23669941 | AX-97841912 | 0 | G | A | 0.167 | -6.58E-02 | 1.22E-01 | 2.48E+00 | 2.54E+00 | 6.12E-01 | no | | intergenic_variant | MODIFIER |
| 13 | 23682811 | AX-170301877 | 0 | C | G | 0.167 | -6.58E-02 | 1.22E-01 | 2.48E+00 | 2.54E+00 | 6.12E-01 | no | Hist1h2bc | upstream_gene_variant | MODIFIER |
| 13 | 23683207 | AX-170453532 | 0 | C | T | 0.167 | -6.58E-02 | 1.22E-01 | 2.48E+00 | 2.54E+00 | 6.12E-01 | no | Hist1h2bc | upstream_gene_variant | MODIFIER |
| 13 | 23693959 | AX-170254330 | 0 | G | T | 0.167 | -6.58E-02 | 1.22E-01 | 2.48E+00 | 2.54E+00 | 6.12E-01 | no | Hist1h2bc | downstream_gene_variant | MODIFIER |
| 13 | 23695160 | AX-97810630 | 0 | G | T | 0.167 | -6.58E-02 | 1.22E-01 | 2.48E+00 | 2.54E+00 | 6.12E-01 | no | Hist1h2bc | downstream_gene_variant | MODIFIER |
| 13 | 23695889 | AX-170243889 | 0 | C | T | 0.167 | -6.58E-02 | 1.22E-01 | 2.48E+00 | 2.54E+00 | 6.12E-01 | no | Hist1h2bc | downstream_gene_variant | MODIFIER |
| 13 | 23699131 | AX-170509489 | 0 | C | G | 0.167 | -6.58E-02 | 1.22E-01 | 2.48E+00 | 2.54E+00 | 6.12E-01 | no | Hist1h1t | downstream_gene_variant | MODIFIER |
| 13 | 23730033 | AX-170210852 | 0 | A | G | 0.167 | -6.58E-02 | 1.22E-01 | 2.48E+00 | 2.54E+00 | 6.12E-01 | no | Gm11338 | non_coding_transcript_exon_variant | MODIFIER |
| 1 | 95244166 | AX-170515932 | 0 | T | A | 0.158 | 1.04E-01 | 2.07E-01 | 2.65E+00 | 2.71E+00 | 6.12E-01 | no | | intergenic_variant | MODIFIER |
| 8 | 41631595 | AX-170161287 | 0 | G | T | 0.14 | -7.18E-02 | 1.42E-01 | 2.65E+00 | 2.70E+00 | 6.12E-01 | no | | intergenic_variant | MODIFIER |
| 7 | 37091110 | AX-170490370 | 0 | T | C | 0.342 | 4.40E-02 | 8.71E-02 | 2.63E+00 | 2.71E+00 | 6.12E-01 | no | | intergenic_variant | MODIFIER |
| 6 | 76952354 | AX-170214967 | 1 | C | A | 0.045 | 7.02E-02 | 1.44E-01 | 2.63E+00 | 2.82E+00 | 6.12E-01 | no | Ctnna2 | intron_variant | MODIFIER |
| 16 | 97879970 | AX-170357101 | 1 | A | G | 0.464 | 3.81E-02 | 7.68E-02 | 2.78E+00 | 2.89E+00 | 6.12E-01 | no | C2cd2 | intron_variant | MODIFIER |
| 8 | 42652884 | AX-170079897 | 1 | G | C | 0.152 | 7.43E-02 | 1.48E-01 | 2.80E+00 | 2.85E+00 | 6.12E-01 | no | | intergenic_variant | MODIFIER |
| 18 | 70892084 | AX-170015945 | 0 | A | G | 0.193 | -5.67E-02 | 1.12E-01 | 2.62E+00 | 2.68E+00 | 6.12E-01 | no | | intergenic_variant | MODIFIER |
| 18 | 59839214 | AX-170015803 | 0 | A | G | 0.421 | -4.27E-02 | 8.38E-02 | 2.61E+00 | 2.67E+00 | 6.12E-01 | no | AC138134.1 | intron_variant,non_coding_transcript_variant | MODIFIER |
| 18 | 57088108 | AX-170481200 | 1 | C | T | 0.402 | 5.41E-02 | 9.72E-02 | 2.43E+00 | 2.50E+00 | 6.12E-01 | no | | intergenic_variant | MODIFIER |
| 3 | 124427119 | AX-170412822 | 0 | C | T | 0.053 | 8.71E-02 | 1.75E-01 | 2.67E+00 | 2.77E+00 | 6.12E-01 | no | 1700006A11Rik | upstream_gene_variant | MODIFIER |
| 7 | 107183409 | AX-170142740 | 0 | T | A | 0.447 | 4.52E-02 | 8.91E-02 | 2.62E+00 | 2.68E+00 | 6.12E-01 | no | Nlrp14 | intron_variant | MODIFIER |
| 5 | 110613392 | AX-97781235 | 0 | T | G | 0.439 | 5.41E-02 | 1.07E-01 | 2.65E+00 | 2.69E+00 | 6.13E-01 | no | Galnt9 | intron_variant | MODIFIER |
| 1 | 30832628 | AX-170216939 | 1 | A | G | 0.393 | 3.97E-02 | 7.97E-02 | 2.66E+00 | 2.76E+00 | 6.13E-01 | no | Phf3 | intron_variant | MODIFIER |
| 7 | 93025398 | AX-170346877 | 0 | G | A | 0.14 | 8.63E-02 | 1.73E-01 | 2.65E+00 | 2.76E+00 | 6.13E-01 | no | Gm26862 | intron_variant,non_coding_transcript_variant | MODIFIER |
| 6 | 130892139 | AX-170465292 | 1 | A | G | 0.554 | 4.24E-02 | 8.20E-02 | 2.56E+00 | 2.63E+00 | 6.13E-01 | no | Gm33962 | upstream_gene_variant | MODIFIER |
| 5 | 47116190 | AX-170360155 | 0 | C | T | 0.5 | 5.43E-02 | 1.02E-01 | 2.50E+00 | 2.56E+00 | 6.13E-01 | no | | intergenic_variant | MODIFIER |
| 9 | 62669812 | AX-170307739 | 0 | T | C | 0.202 | -6.63E-02 | 1.33E-01 | 2.70E+00 | 2.74E+00 | 6.13E-01 | no | | intergenic_variant | MODIFIER |
| 16 | 76303998 | AX-170278842 | 0 | A | T | 0.456 | 3.96E-02 | 7.69E-02 | 2.57E+00 | 2.64E+00 | 6.13E-01 | no | Nrip1 | intron_variant | MODIFIER |
| 16 | 76327828 | AX-170340785 | 0 | T | C | 0.456 | 3.96E-02 | 7.69E-02 | 2.57E+00 | 2.64E+00 | 6.13E-01 | no | Nrip1 | intron_variant | MODIFIER |
| 16 | 76332144 | AX-170302370 | 0 | C | T | 0.456 | 3.96E-02 | 7.69E-02 | 2.57E+00 | 2.64E+00 | 6.13E-01 | no | Nrip1 | intron_variant | MODIFIER |
| 16 | 76342515 | AX-170459847 | 0 | C | T | 0.456 | 3.96E-02 | 7.69E-02 | 2.57E+00 | 2.64E+00 | 6.13E-01 | no | Nrip1 | intron_variant | MODIFIER |
| 16 | 76349249 | AX-170122325 | 0 | G | T | 0.456 | 3.96E-02 | 7.69E-02 | 2.57E+00 | 2.64E+00 | 6.13E-01 | no | Nrip1 | intron_variant | MODIFIER |
| 16 | 76360338 | AX-97822259 | 0 | A | G | 0.456 | 3.96E-02 | 7.69E-02 | 2.57E+00 | 2.64E+00 | 6.13E-01 | no | Nrip1 | intron_variant | MODIFIER |
| 16 | 76366105 | AX-170271845 | 0 | C | A | 0.456 | 3.96E-02 | 7.69E-02 | 2.57E+00 | 2.64E+00 | 6.13E-01 | no | Nrip1 | intron_variant | MODIFIER |
| 16 | 76366695 | AX-170534483 | 0 | A | G | 0.456 | 3.96E-02 | 7.69E-02 | 2.57E+00 | 2.64E+00 | 6.13E-01 | no | Nrip1 | intron_variant | MODIFIER |
| 16 | 76370340 | AX-170507117 | 0 | T | C | 0.456 | 3.96E-02 | 7.69E-02 | 2.57E+00 | 2.64E+00 | 6.13E-01 | no | Nrip1 | intron_variant | MODIFIER |
| 16 | 76383221 | AX-97843246 | 0 | A | G | 0.456 | 3.96E-02 | 7.69E-02 | 2.57E+00 | 2.64E+00 | 6.13E-01 | no | Gm49587 | intron_variant,non_coding_transcript_variant | MODIFIER |
| 16 | 76427240 | AX-170387349 | 0 | T | C | 0.456 | 3.96E-02 | 7.69E-02 | 2.57E+00 | 2.64E+00 | 6.13E-01 | no | Gm45030 | downstream_gene_variant | MODIFIER |
| 16 | 76451724 | AX-170057845 | 0 | G | A | 0.456 | 3.96E-02 | 7.69E-02 | 2.57E+00 | 2.64E+00 | 6.13E-01 | no | Gm45030 | intron_variant,non_coding_transcript_variant | MODIFIER |
| 16 | 76453444 | AX-170468678 | 0 | C | T | 0.456 | 3.96E-02 | 7.69E-02 | 2.57E+00 | 2.64E+00 | 6.13E-01 | no | Gm45030 | intron_variant,non_coding_transcript_variant | MODIFIER |
| 16 | 76462352 | AX-170316738 | 0 | A | T | 0.456 | 3.96E-02 | 7.69E-02 | 2.57E+00 | 2.64E+00 | 6.13E-01 | no | Gm45030 | intron_variant,non_coding_transcript_variant | MODIFIER |
| 16 | 76467275 | AX-170210187 | 0 | C | A | 0.456 | 3.96E-02 | 7.69E-02 | 2.57E+00 | 2.64E+00 | 6.13E-01 | no | Gm45030 | intron_variant,non_coding_transcript_variant | MODIFIER |
| 16 | 76520290 | AX-170475954 | 0 | A | G | 0.456 | 3.96E-02 | 7.69E-02 | 2.57E+00 | 2.64E+00 | 6.13E-01 | no | | intergenic_variant | MODIFIER |
| 16 | 76520596 | AX-170216989 | 0 | G | A | 0.456 | 3.96E-02 | 7.69E-02 | 2.57E+00 | 2.64E+00 | 6.13E-01 | no | | intergenic_variant | MODIFIER |
| 16 | 76526847 | AX-170239697 | 0 | G | A | 0.456 | 3.96E-02 | 7.69E-02 | 2.57E+00 | 2.64E+00 | 6.13E-01 | no | | intergenic_variant | MODIFIER |
| 16 | 76579527 | AX-170105890 | 0 | A | T | 0.456 | 3.96E-02 | 7.69E-02 | 2.57E+00 | 2.64E+00 | 6.13E-01 | no | | intergenic_variant | MODIFIER |
| 16 | 76585506 | AX-170211402 | 0 | A | G | 0.456 | 3.96E-02 | 7.69E-02 | 2.57E+00 | 2.64E+00 | 6.13E-01 | no | | intergenic_variant | MODIFIER |
| 16 | 76587927 | AX-170279478 | 0 | A | C | 0.456 | 3.96E-02 | 7.69E-02 | 2.57E+00 | 2.64E+00 | 6.13E-01 | no | | intergenic_variant | MODIFIER |
| 16 | 76588028 | AX-170286432 | 0 | G | A | 0.456 | 3.96E-02 | 7.69E-02 | 2.57E+00 | 2.64E+00 | 6.13E-01 | no | | intergenic_variant | MODIFIER |
| 16 | 76591731 | AX-170409861 | 0 | A | G | 0.456 | 3.96E-02 | 7.69E-02 | 2.57E+00 | 2.64E+00 | 6.13E-01 | no | | intergenic_variant | MODIFIER |
| 16 | 76598983 | AX-170238985 | 0 | G | A | 0.456 | 3.96E-02 | 7.69E-02 | 2.57E+00 | 2.64E+00 | 6.13E-01 | no | | intergenic_variant | MODIFIER |
| 16 | 76600939 | AX-170292822 | 0 | C | G | 0.456 | 3.96E-02 | 7.69E-02 | 2.57E+00 | 2.64E+00 | 6.13E-01 | no | | intergenic_variant | MODIFIER |
| 16 | 76603150 | AX-170057850 | 0 | G | C | 0.456 | 3.96E-02 | 7.69E-02 | 2.57E+00 | 2.64E+00 | 6.13E-01 | no | | regulatory_region_variant | MODIFIER |
| 16 | 76608654 | AX-170057851 | 0 | T | C | 0.456 | 3.96E-02 | 7.69E-02 | 2.57E+00 | 2.64E+00 | 6.13E-01 | no | | intergenic_variant | MODIFIER |

| | | | | | | | | | | | | | | | |
|---|---|---|---|---|---|---|---|---|---|---|---|---|---|---|---|
| 16 | 76621628 | AX-170170486 | 0 | A | G | 0.456 | 3.96E-02 | 7.69E-02 | 2.57E+00 | 2.64E+00 | 6.13E-01 | no | | regulatory_region_variant | MODIFIER |
| 16 | 76626918 | AX-170340718 | 0 | C | T | 0.456 | 3.96E-02 | 7.69E-02 | 2.57E+00 | 2.64E+00 | 6.13E-01 | no | | intergenic_variant | MODIFIER |
| 16 | 76655517 | AX-170196716 | 0 | C | T | 0.456 | 3.96E-02 | 7.69E-02 | 2.57E+00 | 2.64E+00 | 6.13E-01 | no | | intergenic_variant | MODIFIER |
| 16 | 76749084 | AX-170463016 | 0 | C | T | 0.456 | 3.96E-02 | 7.69E-02 | 2.57E+00 | 2.64E+00 | 6.13E-01 | no | Gm30790 | downstream_gene_variant | MODIFIER |
| 16 | 76749740 | AX-170518145 | 0 | C | T | 0.456 | 3.96E-02 | 7.69E-02 | 2.57E+00 | 2.64E+00 | 6.13E-01 | no | Gm30790 | downstream_gene_variant | MODIFIER |
| 16 | 76754649 | AX-170057853 | 0 | C | T | 0.456 | 3.96E-02 | 7.69E-02 | 2.57E+00 | 2.64E+00 | 6.13E-01 | no | Gm30790 | intron_variant,non_coding_transcript_variant | MODIFIER |
| 16 | 76754693 | AX-170057855 | 0 | T | C | 0.456 | 3.96E-02 | 7.69E-02 | 2.57E+00 | 2.64E+00 | 6.13E-01 | no | Gm30790 | intron_variant,non_coding_transcript_variant | MODIFIER |
| 16 | 76762797 | AX-170405774 | 0 | C | T | 0.456 | 3.96E-02 | 7.69E-02 | 2.57E+00 | 2.64E+00 | 6.13E-01 | no | Gm30790 | intron_variant,non_coding_transcript_variant | MODIFIER |
| 16 | 76806256 | AX-97831803 | 0 | T | C | 0.456 | 3.96E-02 | 7.69E-02 | 2.57E+00 | 2.64E+00 | 6.13E-01 | no | | intergenic_variant | MODIFIER |
| 16 | 76835194 | AX-97845875 | 0 | T | C | 0.456 | 3.96E-02 | 7.69E-02 | 2.57E+00 | 2.64E+00 | 6.13E-01 | no | 1700063K16Rik | intron_variant,non_coding_transcript_variant | MODIFIER |
| 16 | 76835380 | AX-170164198 | 0 | C | T | 0.456 | 3.96E-02 | 7.69E-02 | 2.57E+00 | 2.64E+00 | 6.13E-01 | no | 1700063K16Rik | intron_variant,non_coding_transcript_variant | MODIFIER |
| 16 | 76859310 | AX-170493855 | 0 | T | G | 0.456 | 3.96E-02 | 7.69E-02 | 2.57E+00 | 2.64E+00 | 6.13E-01 | no | 1700041M19Rik | intron_variant,non_coding_transcript_variant | MODIFIER |
| 16 | 76870448 | AX-170378864 | 0 | A | G | 0.456 | 3.96E-02 | 7.69E-02 | 2.57E+00 | 2.64E+00 | 6.13E-01 | no | 1700041M19Rik | intron_variant,non_coding_transcript_variant | MODIFIER |
| 16 | 76875789 | AX-170505617 | 0 | C | A | 0.456 | 3.96E-02 | 7.69E-02 | 2.57E+00 | 2.64E+00 | 6.13E-01 | no | 1700041M19Rik | intron_variant,non_coding_transcript_variant | MODIFIER |
| 16 | 76880730 | AX-170277107 | 0 | T | G | 0.456 | 3.96E-02 | 7.69E-02 | 2.57E+00 | 2.64E+00 | 6.13E-01 | no | 1700041M19Rik | intron_variant,non_coding_transcript_variant | MODIFIER |
| 16 | 76887660 | AX-170407608 | 0 | A | G | 0.456 | 3.96E-02 | 7.69E-02 | 2.57E+00 | 2.64E+00 | 6.13E-01 | no | 1700041M19Rik | intron_variant,non_coding_transcript_variant | MODIFIER |
| 16 | 76888075 | AX-170344836 | 0 | C | T | 0.456 | 3.96E-02 | 7.69E-02 | 2.57E+00 | 2.64E+00 | 6.13E-01 | no | 1700041M19Rik | intron_variant,non_coding_transcript_variant | MODIFIER |
| 16 | 76888198 | AX-170093530 | 0 | A | C | 0.456 | 3.96E-02 | 7.69E-02 | 2.57E+00 | 2.64E+00 | 6.13E-01 | no | 1700041M19Rik | intron_variant,non_coding_transcript_variant | MODIFIER |
| 16 | 76889420 | AX-170254758 | 0 | T | G | 0.456 | 3.96E-02 | 7.69E-02 | 2.57E+00 | 2.64E+00 | 6.13E-01 | no | 1700041M19Rik | upstream_gene_variant | MODIFIER |
| 16 | 76891047 | AX-170344088 | 0 | C | T | 0.456 | 3.96E-02 | 7.69E-02 | 2.57E+00 | 2.64E+00 | 6.13E-01 | no | 1700041M19Rik | upstream_gene_variant | MODIFIER |
| 16 | 76900159 | AX-170404131 | 0 | A | G | 0.456 | 3.96E-02 | 7.69E-02 | 2.57E+00 | 2.64E+00 | 6.13E-01 | no | 1700041M19Rik | intron_variant,non_coding_transcript_variant | MODIFIER |
| 16 | 76900380 | AX-170442133 | 0 | C | T | 0.456 | 3.96E-02 | 7.69E-02 | 2.57E+00 | 2.64E+00 | 6.13E-01 | no | 1700041M19Rik | intron_variant,non_coding_transcript_variant | MODIFIER |
| 16 | 76910650 | AX-97781695 | 0 | C | T | 0.456 | 3.96E-02 | 7.69E-02 | 2.57E+00 | 2.64E+00 | 6.13E-01 | no | 1700041M19Rik | intron_variant,non_coding_transcript_variant | MODIFIER |
| 16 | 76924830 | AX-170057859 | 0 | T | C | 0.456 | 3.96E-02 | 7.69E-02 | 2.57E+00 | 2.64E+00 | 6.13E-01 | no | 1700041M19Rik | intron_variant,non_coding_transcript_variant | MODIFIER |
| 16 | 76924878 | AX-170102232 | 0 | T | C | 0.456 | 3.96E-02 | 7.69E-02 | 2.57E+00 | 2.64E+00 | 6.13E-01 | no | 1700041M19Rik | intron_variant,non_coding_transcript_variant | MODIFIER |
| 16 | 77056163 | AX-170147896 | 0 | G | A | 0.456 | 3.96E-02 | 7.69E-02 | 2.57E+00 | 2.64E+00 | 6.13E-01 | no | Usp25 | intron_variant | MODIFIER |
| 16 | 77069403 | AX-97850143 | 0 | T | C | 0.456 | 3.96E-02 | 7.69E-02 | 2.57E+00 | 2.64E+00 | 6.13E-01 | no | Usp25 | intron_variant | MODIFIER |
| 16 | 77072069 | AX-170013426 | 0 | T | C | 0.456 | 3.96E-02 | 7.69E-02 | 2.57E+00 | 2.64E+00 | 6.13E-01 | no | Usp25 | intron_variant | MODIFIER |
| 16 | 77092286 | AX-170325925 | 0 | A | T | 0.456 | 3.96E-02 | 7.69E-02 | 2.57E+00 | 2.64E+00 | 6.13E-01 | no | Usp25 | intron_variant | MODIFIER |
| 16 | 77098850 | AX-170507979 | 0 | A | G | 0.456 | 3.96E-02 | 7.69E-02 | 2.57E+00 | 2.64E+00 | 6.13E-01 | no | Usp25 | intron_variant | MODIFIER |
| 16 | 77099091 | AX-170013429 | 0 | C | T | 0.456 | 3.96E-02 | 7.69E-02 | 2.57E+00 | 2.64E+00 | 6.13E-01 | no | Usp25 | intron_variant | MODIFIER |
| 16 | 77101352 | AX-170057861 | 0 | T | C | 0.456 | 3.96E-02 | 7.69E-02 | 2.57E+00 | 2.64E+00 | 6.13E-01 | no | Usp25 | intron_variant | MODIFIER |
| 16 | 77101398 | AX-170388526 | 0 | T | A | 0.456 | 3.96E-02 | 7.69E-02 | 2.57E+00 | 2.64E+00 | 6.13E-01 | no | Usp25 | intron_variant | MODIFIER |
| 16 | 77103794 | AX-97835825 | 0 | G | A | 0.456 | 3.96E-02 | 7.69E-02 | 2.57E+00 | 2.64E+00 | 6.13E-01 | no | Usp25 | intron_variant | MODIFIER |
| 16 | 77104136 | AX-170146486 | 0 | C | T | 0.456 | 3.96E-02 | 7.69E-02 | 2.57E+00 | 2.64E+00 | 6.13E-01 | no | Usp25 | intron_variant | MODIFIER |
| 16 | 77121978 | AX-170165435 | 0 | T | G | 0.456 | 3.96E-02 | 7.69E-02 | 2.57E+00 | 2.64E+00 | 6.13E-01 | no | | intergenic_variant | MODIFIER |
| 16 | 77122057 | AX-97833302 | 0 | T | C | 0.456 | 3.96E-02 | 7.69E-02 | 2.57E+00 | 2.64E+00 | 6.13E-01 | no | | intergenic_variant | MODIFIER |
| 16 | 77127333 | AX-170013421 | 0 | A | G | 0.456 | 3.96E-02 | 7.69E-02 | 2.57E+00 | 2.64E+00 | 6.13E-01 | no | | intergenic_variant | MODIFIER |
| 16 | 77127563 | AX-170415284 | 0 | T | G | 0.456 | 3.96E-02 | 7.69E-02 | 2.57E+00 | 2.64E+00 | 6.13E-01 | no | | intergenic_variant | MODIFIER |
| 16 | 77136920 | AX-97796570 | 0 | C | T | 0.456 | 3.96E-02 | 7.69E-02 | 2.57E+00 | 2.64E+00 | 6.13E-01 | no | | intergenic_variant | MODIFIER |
| 16 | 77174653 | AX-170350031 | 0 | C | G | 0.456 | 3.96E-02 | 7.69E-02 | 2.57E+00 | 2.64E+00 | 6.13E-01 | no | | intergenic_variant | MODIFIER |
| 16 | 77175147 | AX-170500188 | 0 | A | G | 0.456 | 3.96E-02 | 7.69E-02 | 2.57E+00 | 2.64E+00 | 6.13E-01 | no | | intergenic_variant | MODIFIER |
| 16 | 77178312 | AX-170013425 | 0 | C | A | 0.456 | 3.96E-02 | 7.69E-02 | 2.57E+00 | 2.64E+00 | 6.13E-01 | no | | intergenic_variant | MODIFIER |
| 16 | 77186745 | AX-170408107 | 0 | C | T | 0.456 | 3.96E-02 | 7.69E-02 | 2.57E+00 | 2.64E+00 | 6.13E-01 | no | | intergenic_variant | MODIFIER |
| 16 | 77198240 | AX-97830768 | 0 | A | G | 0.456 | 3.96E-02 | 7.69E-02 | 2.57E+00 | 2.64E+00 | 6.13E-01 | no | | intergenic_variant | MODIFIER |
| 16 | 77205894 | AX-170533998 | 0 | T | G | 0.456 | 3.96E-02 | 7.69E-02 | 2.57E+00 | 2.64E+00 | 6.13E-01 | no | | intergenic_variant | MODIFIER |
| 16 | 77327192 | AX-170201405 | 0 | G | A | 0.456 | 3.96E-02 | 7.69E-02 | 2.57E+00 | 2.64E+00 | 6.13E-01 | no | Mir99ahg | upstream_gene_variant | MODIFIER |
| 16 | 77344524 | AX-170190040 | 0 | C | A | 0.456 | 3.96E-02 | 7.69E-02 | 2.57E+00 | 2.64E+00 | 6.13E-01 | no | Mir99ahg | intron_variant,non_coding_transcript_variant | MODIFIER |
| 16 | 77352013 | AX-170471399 | 0 | C | A | 0.456 | 3.96E-02 | 7.69E-02 | 2.57E+00 | 2.64E+00 | 6.13E-01 | no | Mir99ahg | intron_variant,non_coding_transcript_variant | MODIFIER |
| 16 | 77384860 | AX-170174410 | 0 | G | A | 0.456 | 3.96E-02 | 7.69E-02 | 2.57E+00 | 2.64E+00 | 6.13E-01 | no | Mir99ahg | intron_variant,non_coding_transcript_variant | MODIFIER |
| 16 | 77488655 | AX-97834595 | 0 | T | C | 0.456 | 3.96E-02 | 7.69E-02 | 2.57E+00 | 2.64E+00 | 6.13E-01 | no | Mir99ahg | intron_variant,non_coding_transcript_variant | MODIFIER |
| 16 | 77525154 | AX-168376404 | 0 | G | T | 0.456 | 3.96E-02 | 7.69E-02 | 2.57E+00 | 2.64E+00 | 6.13E-01 | no | Mir99ahg | intron_variant,non_coding_transcript_variant | MODIFIER |
| 16 | 77577202 | AX-170013430 | 0 | C | T | 0.456 | 3.96E-02 | 7.69E-02 | 2.57E+00 | 2.64E+00 | 6.13E-01 | no | Mir99ahg | intron_variant,non_coding_transcript_variant | MODIFIER |
| 6 | 102653001 | AX-170249781 | 1 | G | A | 0.071 | 9.48E-02 | 1.89E-01 | 2.59E+00 | 2.73E+00 | 6.13E-01 | no | | intergenic_variant | MODIFIER |

| | | | | | | | | | | | | | | | |
|---|---|---|---|---|---|---|---|---|---|---|---|---|---|---|---|
| 9 | 19409316 | AX-97812958 | 0 | G | A | 0.465 | 4.55E-02 | 8.85E-02 | 2.57E+00 | 2.64E+00 | 6.13E-01 | no | Olfr848-ps1 | upstream_gene_variant | MODIFIER |
| 9 | 22790071 | AX-97843417 | 0 | C | T | 0.465 | 4.55E-02 | 8.85E-02 | 2.57E+00 | 2.64E+00 | 6.13E-01 | no | Bbs9 | intron_variant | MODIFIER |
| 4 | 81157901 | AX-170265778 | 0 | T | G | 0.246 | 5.43E-02 | 1.05E-01 | 2.55E+00 | 2.62E+00 | 6.13E-01 | no | | intergenic_variant | MODIFIER |
| 4 | 81187356 | AX-97811418 | 0 | C | T | 0.246 | 5.43E-02 | 1.05E-01 | 2.55E+00 | 2.62E+00 | 6.13E-01 | no | | intergenic_variant | MODIFIER |
| 4 | 81277359 | AX-170415909 | 0 | C | A | 0.246 | 5.43E-02 | 1.05E-01 | 2.55E+00 | 2.62E+00 | 6.13E-01 | no | Mpdz | downstream_gene_variant | MODIFIER |
| 4 | 81366294 | AX-170473586 | 0 | T | C | 0.246 | 5.43E-02 | 1.05E-01 | 2.55E+00 | 2.62E+00 | 6.13E-01 | no | Mpdz | intron_variant | MODIFIER |
| 4 | 81759048 | AX-170352548 | 0 | T | G | 0.246 | 5.43E-02 | 1.05E-01 | 2.55E+00 | 2.62E+00 | 6.13E-01 | no | | intergenic_variant | MODIFIER |
| 4 | 81783687 | AX-170341899 | 0 | C | G | 0.246 | 5.43E-02 | 1.05E-01 | 2.55E+00 | 2.62E+00 | 6.13E-01 | no | | intergenic_variant | MODIFIER |
| 4 | 82124172 | AX-97807367 | 0 | A | T | 0.246 | 5.43E-02 | 1.05E-01 | 2.55E+00 | 2.62E+00 | 6.13E-01 | no | | intergenic_variant | MODIFIER |
| 4 | 82169573 | AX-170221097 | 0 | C | G | 0.246 | 5.43E-02 | 1.05E-01 | 2.55E+00 | 2.62E+00 | 6.13E-01 | no | | intergenic_variant | MODIFIER |
| 4 | 82180428 | AX-170466170 | 0 | T | G | 0.246 | 5.43E-02 | 1.05E-01 | 2.55E+00 | 2.62E+00 | 6.13E-01 | no | | intergenic_variant | MODIFIER |
| 4 | 82180603 | AX-97841032 | 0 | T | C | 0.246 | 5.43E-02 | 1.05E-01 | 2.55E+00 | 2.62E+00 | 6.13E-01 | no | | intergenic_variant | MODIFIER |
| 4 | 82224272 | AX-170113261 | 0 | A | G | 0.246 | 5.43E-02 | 1.05E-01 | 2.55E+00 | 2.62E+00 | 6.13E-01 | no | | intergenic_variant | MODIFIER |
| 4 | 82246741 | AX-170107895 | 0 | A | G | 0.246 | 5.43E-02 | 1.05E-01 | 2.55E+00 | 2.62E+00 | 6.13E-01 | no | | intergenic_variant | MODIFIER |
| 4 | 82254751 | AX-170315644 | 0 | G | A | 0.246 | 5.43E-02 | 1.05E-01 | 2.55E+00 | 2.62E+00 | 6.13E-01 | no | | intergenic_variant | MODIFIER |
| 4 | 82254944 | AX-170429551 | 0 | G | A | 0.246 | 5.43E-02 | 1.05E-01 | 2.55E+00 | 2.62E+00 | 6.13E-01 | no | | intergenic_variant | MODIFIER |
| 4 | 82272673 | AX-97804394 | 0 | T | C | 0.246 | 5.43E-02 | 1.05E-01 | 2.55E+00 | 2.62E+00 | 6.13E-01 | no | | regulatory_region_variant | MODIFIER |
| 4 | 82286614 | AX-170343484 | 0 | C | T | 0.246 | 5.43E-02 | 1.05E-01 | 2.55E+00 | 2.62E+00 | 6.13E-01 | no | Nfib | downstream_gene_variant | MODIFIER |
| 4 | 82287523 | AX-97819745 | 0 | A | G | 0.246 | 5.43E-02 | 1.05E-01 | 2.55E+00 | 2.62E+00 | 6.13E-01 | no | Nfib | downstream_gene_variant | MODIFIER |
| 4 | 82288476 | AX-170262613 | 0 | C | T | 0.246 | 5.43E-02 | 1.05E-01 | 2.55E+00 | 2.62E+00 | 6.13E-01 | no | Nfib | downstream_gene_variant | MODIFIER |
| 4 | 82402887 | AX-170151249 | 0 | G | C | 0.246 | 5.43E-02 | 1.05E-01 | 2.55E+00 | 2.62E+00 | 6.13E-01 | no | Nfib | intron_variant | MODIFIER |
| 4 | 82473041 | AX-170023250 | 0 | G | A | 0.246 | 5.43E-02 | 1.05E-01 | 2.55E+00 | 2.62E+00 | 6.13E-01 | no | Nfib | intron_variant | MODIFIER |
| 4 | 82475191 | AX-170023249 | 0 | G | A | 0.246 | 5.43E-02 | 1.05E-01 | 2.55E+00 | 2.62E+00 | 6.13E-01 | no | Nfib | intron_variant | MODIFIER |
| 4 | 82476203 | AX-170069823 | 0 | G | A | 0.246 | 5.43E-02 | 1.05E-01 | 2.55E+00 | 2.62E+00 | 6.13E-01 | no | Nfib | intron_variant | MODIFIER |
| 4 | 82476503 | AX-170492587 | 0 | C | T | 0.246 | 5.43E-02 | 1.05E-01 | 2.55E+00 | 2.62E+00 | 6.13E-01 | no | Nfib | intron_variant | MODIFIER |
| 4 | 82476589 | AX-170460004 | 0 | A | G | 0.246 | 5.43E-02 | 1.05E-01 | 2.55E+00 | 2.62E+00 | 6.13E-01 | no | Nfib | intron_variant | MODIFIER |
| 4 | 83176438 | AX-170311300 | 0 | G | C | 0.246 | 5.43E-02 | 1.05E-01 | 2.55E+00 | 2.62E+00 | 6.13E-01 | no | | intergenic_variant | MODIFIER |
| 4 | 83188461 | AX-170023260 | 0 | T | C | 0.246 | 5.43E-02 | 1.05E-01 | 2.55E+00 | 2.62E+00 | 6.13E-01 | no | | intergenic_variant | MODIFIER |
| 4 | 83192297 | AX-170403024 | 0 | A | G | 0.246 | 5.43E-02 | 1.05E-01 | 2.55E+00 | 2.62E+00 | 6.13E-01 | no | Gm11185 | upstream_gene_variant | MODIFIER |
| 4 | 83200552 | AX-170247269 | 0 | A | G | 0.246 | 5.43E-02 | 1.05E-01 | 2.55E+00 | 2.62E+00 | 6.13E-01 | no | | intergenic_variant | MODIFIER |
| 4 | 83201963 | AX-170363532 | 0 | A | G | 0.246 | 5.43E-02 | 1.05E-01 | 2.55E+00 | 2.62E+00 | 6.13E-01 | no | | intergenic_variant | MODIFIER |
| 4 | 83215649 | AX-97830324 | 0 | C | T | 0.246 | 5.43E-02 | 1.05E-01 | 2.55E+00 | 2.62E+00 | 6.13E-01 | no | Ttc39b | downstream_gene_variant | MODIFIER |
| 4 | 83217687 | AX-170023264 | 0 | A | G | 0.246 | 5.43E-02 | 1.05E-01 | 2.55E+00 | 2.62E+00 | 6.13E-01 | no | Ttc39b | downstream_gene_variant | MODIFIER |
| 1 | 74577358 | AX-170000809 | 0 | A | G | 0.149 | 9.50E-02 | 1.89E-01 | 2.62E+00 | 2.72E+00 | 6.13E-01 | no | Zfp142 | intron_variant | MODIFIER |
| 16 | 77557127 | AX-170383912 | 1 | A | G | 0.455 | 3.96E-02 | 7.70E-02 | 2.57E+00 | 2.64E+00 | 6.13E-01 | no | Mir99ahg | intron_variant,non_coding_transcript_variant | MODIFIER |
| 1 | 33740146 | AX-170521124 | 1 | A | C | 0.411 | 4.18E-02 | 8.35E-02 | 2.66E+00 | 2.73E+00 | 6.13E-01 | no | Rab23 | 3_prime_UTR_variant | MODIFIER |
| 4 | 88062125 | AX-170140753 | 0 | T | A | 0.254 | 5.33E-02 | 1.03E-01 | 2.55E+00 | 2.62E+00 | 6.13E-01 | no | | intergenic_variant | MODIFIER |
| 4 | 95583627 | AX-170516316 | 0 | G | A | 0.333 | 5.01E-02 | 1.01E-01 | 2.72E+00 | 2.80E+00 | 6.14E-01 | no | Fggy | intron_variant | MODIFIER |
| 8 | 39693934 | AX-170138099 | 2 | G | A | 0.155 | 7.23E-02 | 1.45E-01 | 2.79E+00 | 2.84E+00 | 6.14E-01 | no | | intergenic_variant | MODIFIER |
| 7 | 103905178 | AX-170345959 | 0 | G | A | 0.456 | -4.65E-02 | 9.35E-02 | 2.79E+00 | 2.87E+00 | 6.14E-01 | no | Olfr65 | intron_variant | MODIFIER |
| 7 | 107182959 | AX-97791683 | 0 | C | A | 0.456 | -4.65E-02 | 9.35E-02 | 2.79E+00 | 2.87E+00 | 6.14E-01 | no | Nlrp14 | synonymous_variant | LOW |
| 13 | 35754506 | AX-97827129 | 0 | A | G | 0.175 | -5.82E-02 | 1.11E-01 | 2.52E+00 | 2.60E+00 | 6.14E-01 | no | Cdyl | intron_variant | MODIFIER |
| 3 | 29156158 | AX-170319685 | 1 | C | A | 0.161 | -4.95E-02 | 9.87E-02 | 2.63E+00 | 2.71E+00 | 6.14E-01 | no | Egfem1 | intron_variant | MODIFIER |
| 8 | 77953760 | AX-170031074 | 0 | A | G | 0.193 | -6.62E-02 | 1.32E-01 | 2.65E+00 | 2.72E+00 | 6.14E-01 | no | | intergenic_variant | MODIFIER |
| 5 | 114878934 | AX-170475702 | 1 | C | T | 0.455 | 5.84E-02 | 1.17E-01 | 2.67E+00 | 2.70E+00 | 6.14E-01 | no | 4930519G04Rik | intron_variant | MODIFIER |
| 7 | 96251250 | AX-170212112 | 0 | G | A | 0.132 | -8.61E-02 | 1.73E-01 | 2.63E+00 | 2.74E+00 | 6.14E-01 | no | Mir708 | downstream_gene_variant | MODIFIER |
| 2 | 159645109 | AX-170467732 | 2 | T | A | 0.145 | 9.58E-02 | 1.93E-01 | 2.67E+00 | 2.75E+00 | 6.14E-01 | no | | intergenic_variant | MODIFIER |
| 6 | 125014488 | AX-97814955 | 1 | C | T | 0.196 | -3.94E-02 | 8.02E-02 | 2.66E+00 | 2.80E+00 | 6.14E-01 | no | Zfp384 | intron_variant | MODIFIER |
| 4 | 144306772 | AX-170089393 | 0 | A | C | 0.114 | 8.58E-02 | 1.73E-01 | 2.68E+00 | 2.77E+00 | 6.14E-01 | no | Gm13130 | upstream_gene_variant | MODIFIER |
| 4 | 144528995 | AX-170253341 | 0 | C | A | 0.114 | 8.58E-02 | 1.73E-01 | 2.68E+00 | 2.77E+00 | 6.14E-01 | no | 9430007A20Rik | synonymous_variant | LOW |
| 5 | 125030345 | AX-170253421 | 0 | C | T | 0.456 | -5.51E-02 | 1.11E-01 | 2.80E+00 | 2.86E+00 | 6.15E-01 | no | Ncor2 | intron_variant | MODIFIER |
| 9 | 113502486 | AX-170234352 | 0 | C | T | 0.132 | -5.12E-02 | 1.03E-01 | 2.66E+00 | 2.75E+00 | 6.15E-01 | no | AU023762 | intron_variant,non_coding_transcript_variant | MODIFIER |
| 2 | 101488948 | AX-97848154 | 0 | G | A | 0.053 | -1.50E-01 | 3.01E-01 | 2.77E+00 | 2.80E+00 | 6.15E-01 | no | | intergenic_variant | MODIFIER |
| 8 | 3262487 | AX-97856203 | 1 | G | A | 0.277 | 3.77E-02 | 7.65E-02 | 2.67E+00 | 2.79E+00 | 6.15E-01 | no | Insr | intron_variant | MODIFIER |

| | | | | | | | | | | | | | | | |
|---|---|---|---|---|---|---|---|---|---|---|---|---|---|---|---|
| 1 | 43443194 | AX-170385009 | 0 | T | C | 0.184 | -5.32E-02 | 1.08E-01 | 2.71E+00 | 2.80E+00 | 6.15E-01 | no | Nck2 | upstream_gene_variant | MODIFIER |
| 9 | 70281641 | AX-170167164 | 1 | A | T | 0.188 | -6.17E-02 | 1.24E-01 | 2.70E+00 | 2.77E+00 | 6.15E-01 | no | Myo1e | intron_variant | MODIFIER |
| 5 | 76783431 | AX-170025327 | 0 | A | G | 0.447 | 5.20E-02 | 1.04E-01 | 2.66E+00 | 2.72E+00 | 6.15E-01 | no | C530008M17Rik | intron_variant | MODIFIER |
| 5 | 76823425 | AX-170025326 | 0 | T | C | 0.447 | 5.20E-02 | 1.04E-01 | 2.66E+00 | 2.72E+00 | 6.15E-01 | no | C530008M17Rik | intron_variant | MODIFIER |
| 5 | 76823522 | AX-170025329 | 0 | G | A | 0.447 | 5.20E-02 | 1.04E-01 | 2.66E+00 | 2.72E+00 | 6.15E-01 | no | C530008M17Rik | intron_variant | MODIFIER |
| 5 | 76823601 | AX-170181681 | 0 | G | A | 0.447 | 5.20E-02 | 1.04E-01 | 2.66E+00 | 2.72E+00 | 6.15E-01 | no | C530008M17Rik | intron_variant | MODIFIER |
| 5 | 76826450 | AX-97852763 | 0 | A | G | 0.447 | 5.20E-02 | 1.04E-01 | 2.66E+00 | 2.72E+00 | 6.15E-01 | no | C530008M17Rik | intron_variant | MODIFIER |
| 5 | 76865494 | AX-170025333 | 0 | A | G | 0.447 | 5.20E-02 | 1.04E-01 | 2.66E+00 | 2.72E+00 | 6.15E-01 | no | C530008M17Rik | intron_variant | MODIFIER |
| 5 | 76866100 | AX-170358352 | 0 | G | A | 0.447 | 5.20E-02 | 1.04E-01 | 2.66E+00 | 2.72E+00 | 6.15E-01 | no | C530008M17Rik | synonymous_variant | LOW |
| 5 | 76866319 | AX-97801971 | 0 | C | T | 0.447 | 5.20E-02 | 1.04E-01 | 2.66E+00 | 2.72E+00 | 6.15E-01 | no | C530008M17Rik | intron_variant | MODIFIER |
| 5 | 76889373 | AX-97786409 | 0 | T | G | 0.447 | 5.20E-02 | 1.04E-01 | 2.66E+00 | 2.72E+00 | 6.15E-01 | no | Aasdh | intron_variant | MODIFIER |
| 5 | 76950261 | AX-170025336 | 0 | C | T | 0.447 | 5.20E-02 | 1.04E-01 | 2.66E+00 | 2.72E+00 | 6.15E-01 | no | Ppat | intron_variant,non_coding_transcript_variant | MODIFIER |
| 14 | 117688066 | AX-170115511 | 0 | C | T | 0.316 | 3.69E-02 | 7.24E-02 | 2.56E+00 | 2.66E+00 | 6.15E-01 | no | Gpc6 | intron_variant | MODIFIER |
| 16 | 92560025 | AX-170442019 | 0 | C | T | 0.474 | -3.83E-02 | 7.73E-02 | 2.67E+00 | 2.76E+00 | 6.15E-01 | no | | intergenic_variant | MODIFIER |
| 16 | 92893160 | AX-170187465 | 0 | A | G | 0.474 | -3.83E-02 | 7.73E-02 | 2.67E+00 | 2.76E+00 | 6.15E-01 | no | | intergenic_variant | MODIFIER |
| 8 | 65913191 | AX-170402291 | 1 | G | C | 0.143 | -7.19E-02 | 1.42E-01 | 2.59E+00 | 2.66E+00 | 6.15E-01 | no | MAR1 | intron_variant | MODIFIER |
| 18 | 57259400 | AX-170462102 | 0 | T | C | 0.395 | 5.07E-02 | 9.64E-02 | 2.51E+00 | 2.57E+00 | 6.16E-01 | no | Megf10 | intron_variant | MODIFIER |
| 10 | 23254730 | AX-170300176 | 1 | G | A | 0.571 | -3.40E-02 | 6.96E-02 | 2.82E+00 | 2.97E+00 | 6.16E-01 | no | Eya4 | intron_variant | MODIFIER |
| 5 | 77122980 | AX-170420657 | 0 | G | T | 0.43 | 5.24E-02 | 1.06E-01 | 2.69E+00 | 2.75E+00 | 6.16E-01 | no | Gm15831 | upstream_gene_variant | MODIFIER |
| 1 | 141050576 | AX-170435292 | 0 | G | T | 0.219 | 6.09E-02 | 1.22E-01 | 2.63E+00 | 2.71E+00 | 6.16E-01 | no | | intergenic_variant | MODIFIER |
| 12 | 98978832 | AX-170007364 | 0 | G | C | 0.088 | 8.13E-02 | 1.65E-01 | 2.66E+00 | 2.76E+00 | 6.16E-01 | no | Ttc8 | intron_variant | MODIFIER |
| 16 | 87567393 | AX-97852258 | 0 | C | T | 0.43 | -3.97E-02 | 8.03E-02 | 2.71E+00 | 2.78E+00 | 6.16E-01 | no | Map3k7cl | intron_variant | MODIFIER |
| 9 | 40137370 | AX-170418109 | 0 | G | A | 0.43 | -3.80E-02 | 7.67E-02 | 2.66E+00 | 2.74E+00 | 6.16E-01 | no | | intergenic_variant | MODIFIER |
| 20 | 75799299 | AX-170413799 | 0 | C | G | 0.254 | 3.10E-02 | 6.26E-02 | 2.65E+00 | 2.74E+00 | 6.16E-01 | no | Pls3 | intron_variant | MODIFIER |
| 13 | 31969040 | AX-97805337 | 1 | G | T | 0.179 | -5.94E-02 | 1.17E-01 | 2.59E+00 | 2.65E+00 | 6.16E-01 | no | Gmds | intron_variant | MODIFIER |
| 7 | 37902740 | AX-170150438 | 0 | C | G | 0.351 | 4.88E-02 | 9.57E-02 | 2.57E+00 | 2.64E+00 | 6.16E-01 | no | | intergenic_variant | MODIFIER |
| 5 | 34356979 | AX-97789082 | 1 | T | A | 0.339 | 5.26E-02 | 1.05E-01 | 2.64E+00 | 2.70E+00 | 6.16E-01 | no | Rnf4 | downstream_gene_variant | MODIFIER |
| 7 | 7174204 | AX-170437108 | 0 | A | G | 0.167 | 3.88E-02 | 8.01E-02 | 2.74E+00 | 2.88E+00 | 6.16E-01 | no | Zfp418 | intron_variant | MODIFIER |
| 5 | 88437973 | AX-170209028 | 2 | A | T | 0.436 | 4.98E-02 | 1.00E-01 | 2.67E+00 | 2.74E+00 | 6.16E-01 | no | | intergenic_variant | MODIFIER |
| 5 | 37321774 | AX-97784560 | 1 | T | G | 0.33 | 5.13E-02 | 1.03E-01 | 2.66E+00 | 2.72E+00 | 6.16E-01 | no | Evc | intron_variant | MODIFIER |
| 1 | 120066854 | AX-97816831 | 0 | G | T | 0.579 | 4.19E-02 | 8.09E-02 | 2.53E+00 | 2.60E+00 | 6.16E-01 | no | Tmem37 | downstream_gene_variant | MODIFIER |
| 10 | 22893855 | AX-170350276 | 0 | C | T | 0.57 | -3.39E-02 | 6.96E-02 | 2.82E+00 | 2.97E+00 | 6.16E-01 | no | | regulatory_region_variant | MODIFIER |
| 10 | 22894060 | AX-170299687 | 0 | A | C | 0.57 | -3.39E-02 | 6.96E-02 | 2.82E+00 | 2.97E+00 | 6.16E-01 | no | | regulatory_region_variant | MODIFIER |
| 10 | 22894616 | AX-170247431 | 0 | G | A | 0.57 | -3.39E-02 | 6.96E-02 | 2.82E+00 | 2.97E+00 | 6.16E-01 | no | | intergenic_variant | MODIFIER |
| 10 | 22896005 | AX-170378958 | 0 | G | A | 0.57 | -3.39E-02 | 6.96E-02 | 2.82E+00 | 2.97E+00 | 6.16E-01 | no | | intergenic_variant | MODIFIER |
| 10 | 22906870 | AX-170487822 | 0 | T | C | 0.57 | -3.39E-02 | 6.96E-02 | 2.82E+00 | 2.97E+00 | 6.16E-01 | no | | intergenic_variant | MODIFIER |
| 10 | 22918984 | AX-170480452 | 0 | G | T | 0.57 | -3.39E-02 | 6.96E-02 | 2.82E+00 | 2.97E+00 | 6.16E-01 | no | | intergenic_variant | MODIFIER |
| 10 | 22924201 | AX-170002913 | 0 | T | C | 0.57 | -3.39E-02 | 6.96E-02 | 2.82E+00 | 2.97E+00 | 6.16E-01 | no | | intergenic_variant | MODIFIER |
| 10 | 22956726 | AX-170002912 | 0 | T | C | 0.57 | -3.39E-02 | 6.96E-02 | 2.82E+00 | 2.97E+00 | 6.16E-01 | no | Gm47787 | upstream_gene_variant | MODIFIER |
| 10 | 22960189 | AX-97852568 | 0 | C | T | 0.57 | -3.39E-02 | 6.96E-02 | 2.82E+00 | 2.97E+00 | 6.16E-01 | no | Gm47787 | upstream_gene_variant | MODIFIER |
| 10 | 22961452 | AX-170002911 | 0 | A | G | 0.57 | -3.39E-02 | 6.96E-02 | 2.82E+00 | 2.97E+00 | 6.16E-01 | no | Gm47787 | upstream_gene_variant | MODIFIER |
| 10 | 23069444 | AX-170002915 | 0 | G | T | 0.57 | -3.39E-02 | 6.96E-02 | 2.82E+00 | 2.97E+00 | 6.16E-01 | no | | intergenic_variant | MODIFIER |
| 10 | 23106028 | AX-170104940 | 0 | T | C | 0.57 | -3.39E-02 | 6.96E-02 | 2.82E+00 | 2.97E+00 | 6.16E-01 | no | Eya4 | 3_prime_UTR_variant | MODIFIER |
| 10 | 23113807 | AX-170285934 | 0 | T | C | 0.57 | -3.39E-02 | 6.96E-02 | 2.82E+00 | 2.97E+00 | 6.16E-01 | no | Eya4 | intron_variant | MODIFIER |
| 10 | 23125302 | AX-170113938 | 0 | C | T | 0.57 | -3.39E-02 | 6.96E-02 | 2.82E+00 | 2.97E+00 | 6.16E-01 | no | Eya4 | intron_variant | MODIFIER |
| 10 | 23167724 | AX-170452425 | 0 | A | T | 0.57 | -3.39E-02 | 6.96E-02 | 2.82E+00 | 2.97E+00 | 6.16E-01 | no | Eya4 | intron_variant | MODIFIER |
| 10 | 23176904 | AX-170385490 | 0 | C | T | 0.57 | -3.39E-02 | 6.96E-02 | 2.82E+00 | 2.97E+00 | 6.16E-01 | no | Eya4 | intron_variant | MODIFIER |
| 10 | 23225650 | AX-170479830 | 0 | T | C | 0.57 | -3.39E-02 | 6.96E-02 | 2.82E+00 | 2.97E+00 | 6.16E-01 | no | Eya4 | intron_variant | MODIFIER |
| 10 | 23227847 | AX-170498806 | 0 | T | C | 0.57 | -3.39E-02 | 6.96E-02 | 2.82E+00 | 2.97E+00 | 6.16E-01 | no | Eya4 | intron_variant | MODIFIER |
| 10 | 23232800 | AX-97782292 | 0 | T | C | 0.57 | -3.39E-02 | 6.96E-02 | 2.82E+00 | 2.97E+00 | 6.16E-01 | no | Eya4 | intron_variant | MODIFIER |
| 10 | 23252375 | AX-170483626 | 0 | T | C | 0.57 | -3.39E-02 | 6.96E-02 | 2.82E+00 | 2.97E+00 | 6.16E-01 | no | Eya4 | intron_variant | MODIFIER |
| 10 | 23264002 | AX-170500718 | 0 | G | T | 0.57 | -3.39E-02 | 6.96E-02 | 2.82E+00 | 2.97E+00 | 6.16E-01 | no | Eya4 | intron_variant | MODIFIER |
| 10 | 23264999 | AX-97854102 | 0 | C | A | 0.57 | -3.39E-02 | 6.96E-02 | 2.82E+00 | 2.97E+00 | 6.16E-01 | no | Eya4 | intron_variant | MODIFIER |
| 10 | 23341890 | AX-170308209 | 0 | C | G | 0.57 | -3.39E-02 | 6.96E-02 | 2.82E+00 | 2.97E+00 | 6.16E-01 | no | Eya4 | intron_variant | MODIFIER |

| | | | | | | | | | | | | | | | |
|---|---|---|---|---|---|---|---|---|---|---|---|---|---|---|---|
| 10 | 23346352 | AX-97841699 | 0 | G | A | 0.57 | -3.39E-02 | 6.96E-02 | 2.82E+00 | 2.97E+00 | 6.16E-01 | no | Eya4 | intron_variant | MODIFIER |
| 10 | 23357530 | AX-170277084 | 0 | A | C | 0.57 | -3.39E-02 | 6.96E-02 | 2.82E+00 | 2.97E+00 | 6.16E-01 | no | | intergenic_variant | MODIFIER |
| 10 | 23377902 | AX-170111427 | 0 | G | A | 0.57 | -3.39E-02 | 6.96E-02 | 2.82E+00 | 2.97E+00 | 6.16E-01 | no | | intergenic_variant | MODIFIER |
| 10 | 23436366 | AX-97802748 | 0 | C | T | 0.57 | -3.39E-02 | 6.96E-02 | 2.82E+00 | 2.97E+00 | 6.16E-01 | no | | intergenic_variant | MODIFIER |
| 10 | 23436622 | AX-170381889 | 0 | A | G | 0.57 | -3.39E-02 | 6.96E-02 | 2.82E+00 | 2.97E+00 | 6.16E-01 | no | | intergenic_variant | MODIFIER |
| 10 | 23493934 | AX-170002921 | 0 | G | C | 0.57 | -3.39E-02 | 6.96E-02 | 2.82E+00 | 2.97E+00 | 6.16E-01 | no | | intergenic_variant | MODIFIER |
| 5 | 83051834 | AX-170344278 | 0 | A | C | 0.439 | 5.22E-02 | 1.05E-01 | 2.69E+00 | 2.75E+00 | 6.16E-01 | no | | intergenic_variant | MODIFIER |
| 18 | 19992278 | AX-170264757 | 1 | G | A | 0.045 | -7.44E-02 | 1.48E-01 | 2.55E+00 | 2.70E+00 | 6.17E-01 | no | Dsc3 | intron_variant | MODIFIER |
| 13 | 47605776 | AX-170301469 | 0 | C | T | 0.526 | 3.18E-02 | 6.38E-02 | 2.61E+00 | 2.72E+00 | 6.17E-01 | no | | regulatory_region_variant | MODIFIER |
| 13 | 36314308 | AX-170225176 | 0 | G | A | 0.193 | -6.07E-02 | 1.19E-01 | 2.58E+00 | 2.64E+00 | 6.17E-01 | no | Fars2 | intron_variant | MODIFIER |
| 13 | 36434556 | AX-170356351 | 0 | A | G | 0.193 | -6.07E-02 | 1.19E-01 | 2.58E+00 | 2.64E+00 | 6.17E-01 | no | Fars2 | intron_variant | MODIFIER |
| 13 | 36435978 | AX-97798844 | 0 | T | C | 0.193 | -6.07E-02 | 1.19E-01 | 2.58E+00 | 2.64E+00 | 6.17E-01 | no | Fars2 | intron_variant | MODIFIER |
| 13 | 36439484 | AX-170475000 | 0 | T | C | 0.193 | -6.07E-02 | 1.19E-01 | 2.58E+00 | 2.64E+00 | 6.17E-01 | no | Fars2 | intron_variant | MODIFIER |
| 13 | 36439978 | AX-170499436 | 0 | T | C | 0.193 | -6.07E-02 | 1.19E-01 | 2.58E+00 | 2.64E+00 | 6.17E-01 | no | Fars2 | intron_variant | MODIFIER |
| 13 | 36443066 | AX-170336408 | 0 | C | G | 0.193 | -6.07E-02 | 1.19E-01 | 2.58E+00 | 2.64E+00 | 6.17E-01 | no | Fars2 | intron_variant | MODIFIER |
| 13 | 36443328 | AX-170008117 | 0 | T | C | 0.193 | -6.07E-02 | 1.19E-01 | 2.58E+00 | 2.64E+00 | 6.17E-01 | no | Fars2 | intron_variant | MODIFIER |
| 13 | 36443510 | AX-170148948 | 0 | G | T | 0.193 | -6.07E-02 | 1.19E-01 | 2.58E+00 | 2.64E+00 | 6.17E-01 | no | Fars2 | intron_variant | MODIFIER |
| 13 | 36448938 | AX-170315042 | 0 | T | G | 0.193 | -6.07E-02 | 1.19E-01 | 2.58E+00 | 2.64E+00 | 6.17E-01 | no | Fars2 | intron_variant | MODIFIER |
| 13 | 36454273 | AX-170157872 | 0 | T | G | 0.193 | -6.07E-02 | 1.19E-01 | 2.58E+00 | 2.64E+00 | 6.17E-01 | no | Fars2 | intron_variant | MODIFIER |
| 13 | 36456793 | AX-170008115 | 0 | T | C | 0.193 | -6.07E-02 | 1.19E-01 | 2.58E+00 | 2.64E+00 | 6.17E-01 | no | Fars2 | intron_variant | MODIFIER |
| 13 | 36484174 | AX-170259439 | 0 | G | A | 0.193 | -6.07E-02 | 1.19E-01 | 2.58E+00 | 2.64E+00 | 6.17E-01 | no | Fars2 | intron_variant | MODIFIER |
| 13 | 36489773 | AX-170008114 | 0 | T | C | 0.193 | -6.07E-02 | 1.19E-01 | 2.58E+00 | 2.64E+00 | 6.17E-01 | no | Fars2 | intron_variant | MODIFIER |
| 13 | 36497892 | AX-170419175 | 0 | T | C | 0.193 | -6.07E-02 | 1.19E-01 | 2.58E+00 | 2.64E+00 | 6.17E-01 | no | Fars2 | intron_variant | MODIFIER |
| 13 | 36504251 | AX-170008120 | 0 | C | T | 0.193 | -6.07E-02 | 1.19E-01 | 2.58E+00 | 2.64E+00 | 6.17E-01 | no | Fars2 | intron_variant | MODIFIER |
| 13 | 36530021 | AX-97846071 | 0 | A | C | 0.193 | -6.07E-02 | 1.19E-01 | 2.58E+00 | 2.64E+00 | 6.17E-01 | no | Fars2 | intron_variant | MODIFIER |
| 13 | 36562714 | AX-170367404 | 0 | T | C | 0.193 | -6.07E-02 | 1.19E-01 | 2.58E+00 | 2.64E+00 | 6.17E-01 | no | Gm8277 | upstream_gene_variant | MODIFIER |
| 13 | 36571998 | AX-97848090 | 0 | G | A | 0.193 | -6.07E-02 | 1.19E-01 | 2.58E+00 | 2.64E+00 | 6.17E-01 | no | Gm48767 | intron_variant,non_coding_transcript_variant | MODIFIER |
| 13 | 36575273 | AX-170255813 | 0 | T | A | 0.193 | -6.07E-02 | 1.19E-01 | 2.58E+00 | 2.64E+00 | 6.17E-01 | no | Gm48767 | non_coding_transcript_exon_variant | MODIFIER |
| 13 | 36607459 | AX-170129453 | 0 | A | G | 0.193 | -6.07E-02 | 1.19E-01 | 2.58E+00 | 2.64E+00 | 6.17E-01 | no | Fars2 | intron_variant,NMD_transcript_variant | MODIFIER |
| 13 | 36611272 | AX-170423657 | 0 | A | G | 0.193 | -6.07E-02 | 1.19E-01 | 2.58E+00 | 2.64E+00 | 6.17E-01 | no | Gm46409 | upstream_gene_variant | MODIFIER |
| 13 | 36615435 | AX-170188819 | 0 | A | T | 0.193 | -6.07E-02 | 1.19E-01 | 2.58E+00 | 2.64E+00 | 6.17E-01 | no | Gm46409 | intron_variant,non_coding_transcript_variant | MODIFIER |
| 13 | 36642539 | AX-170183652 | 0 | A | G | 0.193 | -6.07E-02 | 1.19E-01 | 2.58E+00 | 2.64E+00 | 6.17E-01 | no | Fars2 | intron_variant,NMD_transcript_variant | MODIFIER |
| 13 | 36643356 | AX-97830012 | 0 | C | A | 0.193 | -6.07E-02 | 1.19E-01 | 2.58E+00 | 2.64E+00 | 6.17E-01 | no | Fars2 | intron_variant,NMD_transcript_variant | MODIFIER |
| 13 | 36684010 | AX-170298614 | 0 | A | G | 0.193 | -6.07E-02 | 1.19E-01 | 2.58E+00 | 2.64E+00 | 6.17E-01 | no | Fars2 | intron_variant,NMD_transcript_variant | MODIFIER |
| 13 | 36685760 | AX-170087407 | 0 | G | A | 0.193 | -6.07E-02 | 1.19E-01 | 2.58E+00 | 2.64E+00 | 6.17E-01 | no | Fars2 | intron_variant,NMD_transcript_variant | MODIFIER |
| 13 | 36721721 | AX-170101304 | 0 | C | T | 0.193 | -6.07E-02 | 1.19E-01 | 2.58E+00 | 2.64E+00 | 6.17E-01 | no | Nrn1 | downstream_gene_variant | MODIFIER |
| 13 | 37598296 | AX-170483485 | 0 | C | T | 0.193 | -6.07E-02 | 1.19E-01 | 2.58E+00 | 2.64E+00 | 6.17E-01 | no | | regulatory_region_variant | MODIFIER |
| 7 | 118374623 | AX-170480298 | 1 | T | C | 0.393 | 4.24E-02 | 8.57E-02 | 2.67E+00 | 2.74E+00 | 6.17E-01 | no | | intergenic_variant | MODIFIER |
| 8 | 62726785 | AX-170216613 | 0 | G | A | 0.158 | -5.19E-02 | 1.05E-01 | 2.60E+00 | 2.73E+00 | 6.17E-01 | no | Gm45459 | intron_variant,non_coding_transcript_variant | MODIFIER |
| 16 | 78091369 | AX-170013436 | 2 | T | A | 0.455 | 3.59E-02 | 7.16E-02 | 2.60E+00 | 2.69E+00 | 6.17E-01 | no | | intergenic_variant | MODIFIER |
| 8 | 95867694 | AX-170292351 | 1 | G | A | 0.232 | 5.62E-02 | 1.14E-01 | 2.68E+00 | 2.75E+00 | 6.17E-01 | no | Got2 | intron_variant | MODIFIER |
| 4 | 144440445 | AX-97850381 | 0 | C | T | 0.114 | 8.52E-02 | 1.72E-01 | 2.64E+00 | 2.74E+00 | 6.17E-01 | no | 1700012P22Rik | upstream_gene_variant | MODIFIER |
| 4 | 145411040 | AX-170325040 | 0 | A | G | 0.114 | 8.52E-02 | 1.72E-01 | 2.64E+00 | 2.74E+00 | 6.17E-01 | no | Platr16 | intron_variant,non_coding_transcript_variant | MODIFIER |
| 2 | 90870917 | AX-170336850 | 0 | G | A | 0.053 | -1.48E-01 | 2.99E-01 | 2.76E+00 | 2.79E+00 | 6.17E-01 | no | Mtch2 | downstream_gene_variant | MODIFIER |
| 13 | 6801109 | AX-170092647 | 0 | A | G | 0.211 | -5.73E-02 | 1.13E-01 | 2.58E+00 | 2.66E+00 | 6.17E-01 | no | | intergenic_variant | MODIFIER |
| 7 | 18749900 | AX-170446415 | 1 | G | C | 0.357 | 4.10E-02 | 8.16E-02 | 2.57E+00 | 2.70E+00 | 6.17E-01 | no | Gm24464 | upstream_gene_variant | MODIFIER |
| 16 | 46417129 | AX-170134206 | 0 | T | C | 0.509 | 4.20E-02 | 8.19E-02 | 2.56E+00 | 2.60E+00 | 6.17E-01 | no | Nectin3 | intron_variant | MODIFIER |
| 16 | 46417935 | AX-170385789 | 0 | A | G | 0.509 | 4.20E-02 | 8.19E-02 | 2.56E+00 | 2.60E+00 | 6.17E-01 | no | Nectin3 | intron_variant | MODIFIER |
| 16 | 46418020 | AX-170469573 | 0 | T | C | 0.509 | 4.20E-02 | 8.19E-02 | 2.56E+00 | 2.60E+00 | 6.17E-01 | no | Nectin3 | intron_variant | MODIFIER |
| 16 | 46421290 | AX-170126662 | 0 | A | G | 0.509 | 4.20E-02 | 8.19E-02 | 2.56E+00 | 2.60E+00 | 6.17E-01 | no | Nectin3 | intron_variant | MODIFIER |
| 16 | 46421897 | AX-170419680 | 0 | C | T | 0.509 | 4.20E-02 | 8.19E-02 | 2.56E+00 | 2.60E+00 | 6.17E-01 | no | Nectin3 | intron_variant | MODIFIER |
| 16 | 46425003 | AX-170291547 | 0 | C | T | 0.509 | 4.20E-02 | 8.19E-02 | 2.56E+00 | 2.60E+00 | 6.17E-01 | no | Nectin3 | intron_variant | MODIFIER |
| 16 | 46433122 | AX-170274129 | 0 | T | C | 0.509 | 4.20E-02 | 8.19E-02 | 2.56E+00 | 2.60E+00 | 6.17E-01 | no | Nectin3 | intron_variant | MODIFIER |
| 16 | 46434618 | AX-170176503 | 0 | C | T | 0.509 | 4.20E-02 | 8.19E-02 | 2.56E+00 | 2.60E+00 | 6.17E-01 | no | Nectin3 | intron_variant | MODIFIER |

| | | | | | | | | | | | | | | | |
|---|---|---|---|---|---|---|---|---|---|---|---|---|---|---|---|
| 16 | 46436319 | AX-97855618 | 0 | T | C | 0.509 | 4.20E-02 | 8.19E-02 | 2.56E+00 | 2.60E+00 | 6.17E-01 | no | Nectin3 | intron_variant | MODIFIER |
| 16 | 49089467 | AX-97853472 | 0 | C | T | 0.509 | 4.20E-02 | 8.19E-02 | 2.56E+00 | 2.60E+00 | 6.17E-01 | no | Myh15 | intron_variant | MODIFIER |
| 16 | 49190856 | AX-97823517 | 0 | A | G | 0.509 | 4.20E-02 | 8.19E-02 | 2.56E+00 | 2.60E+00 | 6.17E-01 | no | Myh15 | intron_variant | MODIFIER |
| 9 | 38062290 | AX-170325492 | 0 | G | A | 0.491 | -3.94E-02 | 8.04E-02 | 2.71E+00 | 2.80E+00 | 6.17E-01 | no | Olfr885 | downstream_gene_variant | MODIFIER |
| 7 | 37498494 | AX-170386081 | 1 | A | G | 0.339 | 4.86E-02 | 9.53E-02 | 2.57E+00 | 2.64E+00 | 6.17E-01 | no | Zfp536 | intron_variant | MODIFIER |
| 7 | 70870390 | AX-170343964 | 2 | A | C | 0.391 | -4.99E-02 | 1.02E-01 | 2.72E+00 | 2.82E+00 | 6.17E-01 | no | Gm42397 | upstream_gene_variant | MODIFIER |
| 10 | 107898426 | AX-170310710 | 0 | G | A | 0.333 | -4.10E-02 | 8.24E-02 | 2.63E+00 | 2.70E+00 | 6.17E-01 | no | Otogl | intron_variant | MODIFIER |
| 7 | 36546865 | AX-170458646 | 0 | G | A | 0.333 | 4.48E-02 | 8.90E-02 | 2.59E+00 | 2.67E+00 | 6.17E-01 | no | | intergenic_variant | MODIFIER |
| 2 | 115867851 | AX-170144565 | 0 | C | A | 0.14 | -8.80E-02 | 1.80E-01 | 2.71E+00 | 2.80E+00 | 6.17E-01 | no | Meis2 | intron_variant | MODIFIER |
| 2 | 155227564 | AX-170472983 | 0 | G | A | 0.14 | -8.80E-02 | 1.80E-01 | 2.71E+00 | 2.80E+00 | 6.17E-01 | no | Itch | downstream_gene_variant | MODIFIER |
| 4 | 149128774 | AX-170400441 | 0 | C | T | 0.105 | 8.32E-02 | 1.68E-01 | 2.64E+00 | 2.73E+00 | 6.17E-01 | no | Cenps | stop_retained_variant | LOW |
| 5 | 118005975 | AX-170025972 | 1 | T | C | 0.455 | 5.52E-02 | 1.10E-01 | 2.62E+00 | 2.67E+00 | 6.17E-01 | no | Fbxo21 | intron_variant | MODIFIER |
| 1 | 22195681 | AX-170138066 | 0 | C | T | 0.412 | 4.31E-02 | 8.60E-02 | 2.62E+00 | 2.67E+00 | 6.17E-01 | no | | intergenic_variant | MODIFIER |
| 1 | 22271553 | AX-97821498 | 0 | G | A | 0.412 | 4.31E-02 | 8.60E-02 | 2.62E+00 | 2.67E+00 | 6.17E-01 | no | | intergenic_variant | MODIFIER |
| 1 | 22292269 | AX-170199867 | 0 | G | A | 0.412 | 4.31E-02 | 8.60E-02 | 2.62E+00 | 2.67E+00 | 6.17E-01 | no | Rims1 | intron_variant | MODIFIER |
| 1 | 22294472 | AX-170227407 | 0 | C | T | 0.412 | 4.31E-02 | 8.60E-02 | 2.62E+00 | 2.67E+00 | 6.17E-01 | no | Rims1 | intron_variant | MODIFIER |
| 1 | 22400165 | AX-97785514 | 0 | T | G | 0.412 | 4.31E-02 | 8.60E-02 | 2.62E+00 | 2.67E+00 | 6.17E-01 | no | Rims1 | intron_variant | MODIFIER |
| 1 | 22635402 | AX-97803809 | 0 | A | G | 0.412 | 4.31E-02 | 8.60E-02 | 2.62E+00 | 2.67E+00 | 6.17E-01 | no | Rims1 | intron_variant | MODIFIER |
| 1 | 22635675 | AX-170420383 | 0 | G | A | 0.412 | 4.31E-02 | 8.60E-02 | 2.62E+00 | 2.67E+00 | 6.17E-01 | no | Rims1 | intron_variant | MODIFIER |
| 1 | 22640887 | AX-170347105 | 0 | T | A | 0.412 | 4.31E-02 | 8.60E-02 | 2.62E+00 | 2.67E+00 | 6.17E-01 | no | Rims1 | intron_variant | MODIFIER |
| 1 | 22664189 | AX-97836693 | 0 | A | C | 0.412 | 4.31E-02 | 8.60E-02 | 2.62E+00 | 2.67E+00 | 6.17E-01 | no | Rims1 | intron_variant | MODIFIER |
| 1 | 22711632 | AX-170450884 | 0 | A | G | 0.412 | 4.31E-02 | 8.60E-02 | 2.62E+00 | 2.67E+00 | 6.17E-01 | no | Rims1 | intron_variant | MODIFIER |
| 1 | 22765308 | AX-170219972 | 0 | A | G | 0.412 | 4.31E-02 | 8.60E-02 | 2.62E+00 | 2.67E+00 | 6.17E-01 | no | Rims1 | intron_variant | MODIFIER |
| 1 | 22780997 | AX-168419909 | 0 | C | T | 0.412 | 4.31E-02 | 8.60E-02 | 2.62E+00 | 2.67E+00 | 6.17E-01 | no | Rims1 | intron_variant | MODIFIER |
| 1 | 22781204 | AX-170180438 | 0 | G | C | 0.412 | 4.31E-02 | 8.60E-02 | 2.62E+00 | 2.67E+00 | 6.17E-01 | no | Rims1 | intron_variant | MODIFIER |
| 1 | 22812750 | AX-170142004 | 0 | A | G | 0.412 | 4.31E-02 | 8.60E-02 | 2.62E+00 | 2.67E+00 | 6.17E-01 | no | | intergenic_variant | MODIFIER |
| 16 | 76454104 | AX-170483653 | 0 | T | C | 0.447 | 3.83E-02 | 7.56E-02 | 2.57E+00 | 2.65E+00 | 6.18E-01 | no | Gm45030 | intron_variant,non_coding_transcript_variant | MODIFIER |
| 1 | 170583860 | AX-170121994 | 1 | A | G | 0.205 | 6.93E-02 | 1.40E-01 | 2.66E+00 | 2.70E+00 | 6.18E-01 | no | Nos1ap | intron_variant,non_coding_transcript_variant | MODIFIER |
| 5 | 86830925 | AX-170072694 | 1 | T | C | 0.429 | 5.54E-02 | 1.12E-01 | 2.76E+00 | 2.80E+00 | 6.18E-01 | no | Ythdc1 | intron_variant | MODIFIER |
| 18 | 69695408 | AX-170132844 | 1 | C | A | 0.268 | -5.10E-02 | 1.03E-01 | 2.68E+00 | 2.73E+00 | 6.18E-01 | no | | intergenic_variant | MODIFIER |
| 1 | 46112450 | AX-97849735 | 2 | A | C | 0.336 | 3.55E-02 | 7.30E-02 | 2.71E+00 | 2.83E+00 | 6.18E-01 | no | Dnah7b | intron_variant | MODIFIER |
| 8 | 106789205 | AX-170442292 | 0 | C | T | 0.237 | 5.19E-02 | 1.06E-01 | 2.70E+00 | 2.78E+00 | 6.18E-01 | no | Tango6 | intron_variant | MODIFIER |
| 15 | 69260333 | AX-170479264 | 0 | T | C | 0.553 | 4.17E-02 | 8.25E-02 | 2.58E+00 | 2.65E+00 | 6.18E-01 | no | | intergenic_variant | MODIFIER |
| 15 | 69261054 | AX-170196608 | 0 | C | T | 0.553 | 4.17E-02 | 8.25E-02 | 2.58E+00 | 2.65E+00 | 6.18E-01 | no | | intergenic_variant | MODIFIER |
| 15 | 69261897 | AX-170443735 | 0 | G | A | 0.553 | 4.17E-02 | 8.25E-02 | 2.58E+00 | 2.65E+00 | 6.18E-01 | no | | intergenic_variant | MODIFIER |
| 15 | 69274288 | AX-170011921 | 0 | C | T | 0.553 | 4.17E-02 | 8.25E-02 | 2.58E+00 | 2.65E+00 | 6.18E-01 | no | 4930504C09Rik | non_coding_transcript_exon_variant | MODIFIER |
| 15 | 69277954 | AX-97820959 | 0 | T | C | 0.553 | 4.17E-02 | 8.25E-02 | 2.58E+00 | 2.65E+00 | 6.18E-01 | no | 4930504C09Rik | downstream_gene_variant | MODIFIER |
| 15 | 69284071 | AX-170329270 | 0 | A | G | 0.553 | 4.17E-02 | 8.25E-02 | 2.58E+00 | 2.65E+00 | 6.18E-01 | no | | intergenic_variant | MODIFIER |
| 15 | 69305477 | AX-97855378 | 0 | T | C | 0.553 | 4.17E-02 | 8.25E-02 | 2.58E+00 | 2.65E+00 | 6.18E-01 | no | | intergenic_variant | MODIFIER |
| 15 | 70229885 | AX-170011939 | 0 | G | A | 0.553 | 4.17E-02 | 8.25E-02 | 2.58E+00 | 2.65E+00 | 6.18E-01 | no | | intergenic_variant | MODIFIER |
| 15 | 70237404 | AX-170087899 | 0 | T | C | 0.553 | 4.17E-02 | 8.25E-02 | 2.58E+00 | 2.65E+00 | 6.18E-01 | no | | intergenic_variant | MODIFIER |
| 15 | 70346181 | AX-170413714 | 0 | A | G | 0.553 | 4.17E-02 | 8.25E-02 | 2.58E+00 | 2.65E+00 | 6.18E-01 | no | | intergenic_variant | MODIFIER |
| 15 | 70366184 | AX-170202850 | 0 | C | A | 0.553 | 4.17E-02 | 8.25E-02 | 2.58E+00 | 2.65E+00 | 6.18E-01 | no | | intergenic_variant | MODIFIER |
| 15 | 70370677 | AX-170055893 | 0 | G | A | 0.553 | 4.17E-02 | 8.25E-02 | 2.58E+00 | 2.65E+00 | 6.18E-01 | no | | intergenic_variant | MODIFIER |
| 15 | 70371551 | AX-170213878 | 0 | G | A | 0.553 | 4.17E-02 | 8.25E-02 | 2.58E+00 | 2.65E+00 | 6.18E-01 | no | | intergenic_variant | MODIFIER |
| 15 | 70374618 | AX-97847144 | 0 | A | G | 0.553 | 4.17E-02 | 8.25E-02 | 2.58E+00 | 2.65E+00 | 6.18E-01 | no | | intergenic_variant | MODIFIER |
| 15 | 70374647 | AX-170284453 | 0 | G | C | 0.553 | 4.17E-02 | 8.25E-02 | 2.58E+00 | 2.65E+00 | 6.18E-01 | no | | intergenic_variant | MODIFIER |
| 15 | 70745400 | AX-168410925 | 0 | G | A | 0.553 | 4.17E-02 | 8.25E-02 | 2.58E+00 | 2.65E+00 | 6.18E-01 | no | | intergenic_variant | MODIFIER |
| 15 | 70797663 | AX-168407503 | 0 | T | C | 0.553 | 4.17E-02 | 8.25E-02 | 2.58E+00 | 2.65E+00 | 6.18E-01 | no | | intergenic_variant | MODIFIER |
| 15 | 70805073 | AX-170011946 | 0 | G | C | 0.553 | 4.17E-02 | 8.25E-02 | 2.58E+00 | 2.65E+00 | 6.18E-01 | no | | intergenic_variant | MODIFIER |
| 15 | 71315826 | AX-170235981 | 0 | T | C | 0.553 | 4.17E-02 | 8.25E-02 | 2.58E+00 | 2.65E+00 | 6.18E-01 | no | Gm5217 | downstream_gene_variant | MODIFIER |
| 15 | 71400451 | AX-170415943 | 0 | G | A | 0.553 | 4.17E-02 | 8.25E-02 | 2.58E+00 | 2.65E+00 | 6.18E-01 | no | | intergenic_variant | MODIFIER |
| 15 | 71438880 | AX-170472978 | 0 | T | G | 0.553 | 4.17E-02 | 8.25E-02 | 2.58E+00 | 2.65E+00 | 6.18E-01 | no | Fam135b | 3_prime_UTR_variant | MODIFIER |
| 15 | 71468583 | AX-97856872 | 0 | A | G | 0.553 | 4.17E-02 | 8.25E-02 | 2.58E+00 | 2.65E+00 | 6.18E-01 | no | Fam135b | intron_variant | MODIFIER |

| | | | | | | | | | | | | | | | |
|---|---|---|---|---|---|---|---|---|---|---|---|---|---|---|---|
| 15 | 74090857 | AX-97793733 | 0 | T | C | 0.553 | 4.17E-02 | 8.25E-02 | 2.58E+00 | 2.65E+00 | 6.18E-01 | no | Gm15387 | downstream_gene_variant | MODIFIER |
| 15 | 74154426 | AX-97855682 | 0 | T | G | 0.553 | 4.17E-02 | 8.25E-02 | 2.58E+00 | 2.65E+00 | 6.18E-01 | no | | intergenic_variant | MODIFIER |
| 15 | 74168933 | AX-170488759 | 0 | G | A | 0.553 | 4.17E-02 | 8.25E-02 | 2.58E+00 | 2.65E+00 | 6.18E-01 | no | | intergenic_variant | MODIFIER |
| 15 | 74169227 | AX-97811548 | 0 | C | T | 0.553 | 4.17E-02 | 8.25E-02 | 2.58E+00 | 2.65E+00 | 6.18E-01 | no | | intergenic_variant | MODIFIER |
| 15 | 74188642 | AX-170212742 | 0 | G | A | 0.553 | 4.17E-02 | 8.25E-02 | 2.58E+00 | 2.65E+00 | 6.18E-01 | no | | regulatory_region_variant | MODIFIER |
| 15 | 74190770 | AX-170293454 | 0 | C | T | 0.553 | 4.17E-02 | 8.25E-02 | 2.58E+00 | 2.65E+00 | 6.18E-01 | no | | intergenic_variant | MODIFIER |
| 15 | 74191899 | AX-97848655 | 0 | T | C | 0.553 | 4.17E-02 | 8.25E-02 | 2.58E+00 | 2.65E+00 | 6.18E-01 | no | | intergenic_variant | MODIFIER |
| 15 | 74216919 | AX-170055978 | 0 | C | T | 0.553 | 4.17E-02 | 8.25E-02 | 2.58E+00 | 2.65E+00 | 6.18E-01 | no | | intergenic_variant | MODIFIER |
| 15 | 74217960 | AX-170391669 | 0 | A | G | 0.553 | 4.17E-02 | 8.25E-02 | 2.58E+00 | 2.65E+00 | 6.18E-01 | no | | regulatory_region_variant | MODIFIER |
| 15 | 74219269 | AX-170337155 | 0 | G | A | 0.553 | 4.17E-02 | 8.25E-02 | 2.58E+00 | 2.65E+00 | 6.18E-01 | no | | intergenic_variant | MODIFIER |
| 15 | 74221159 | AX-97832896 | 0 | T | C | 0.553 | 4.17E-02 | 8.25E-02 | 2.58E+00 | 2.65E+00 | 6.18E-01 | no | | intergenic_variant | MODIFIER |
| 15 | 74224923 | AX-170450223 | 0 | C | T | 0.553 | 4.17E-02 | 8.25E-02 | 2.58E+00 | 2.65E+00 | 6.18E-01 | no | | intergenic_variant | MODIFIER |
| 15 | 74229618 | AX-170181503 | 0 | T | C | 0.553 | 4.17E-02 | 8.25E-02 | 2.58E+00 | 2.65E+00 | 6.18E-01 | no | | intergenic_variant | MODIFIER |
| 15 | 74244824 | AX-170486230 | 0 | T | C | 0.553 | 4.17E-02 | 8.25E-02 | 2.58E+00 | 2.65E+00 | 6.18E-01 | no | | intergenic_variant | MODIFIER |
| 15 | 74244851 | AX-170486461 | 0 | T | C | 0.553 | 4.17E-02 | 8.25E-02 | 2.58E+00 | 2.65E+00 | 6.18E-01 | no | | intergenic_variant | MODIFIER |
| 15 | 74244898 | AX-170135126 | 0 | A | T | 0.553 | 4.17E-02 | 8.25E-02 | 2.58E+00 | 2.65E+00 | 6.18E-01 | no | | intergenic_variant | MODIFIER |
| 15 | 74246297 | AX-170384064 | 0 | A | G | 0.553 | 4.17E-02 | 8.25E-02 | 2.58E+00 | 2.65E+00 | 6.18E-01 | no | | intergenic_variant | MODIFIER |
| 15 | 74248116 | AX-170480209 | 0 | T | A | 0.553 | 4.17E-02 | 8.25E-02 | 2.58E+00 | 2.65E+00 | 6.18E-01 | no | | intergenic_variant | MODIFIER |
| 15 | 74248242 | AX-170396345 | 0 | A | C | 0.553 | 4.17E-02 | 8.25E-02 | 2.58E+00 | 2.65E+00 | 6.18E-01 | no | | intergenic_variant | MODIFIER |
| 15 | 74261286 | AX-170495294 | 0 | G | A | 0.553 | 4.17E-02 | 8.25E-02 | 2.58E+00 | 2.65E+00 | 6.18E-01 | no | | intergenic_variant | MODIFIER |
| 15 | 74261610 | AX-170360081 | 0 | C | T | 0.553 | 4.17E-02 | 8.25E-02 | 2.58E+00 | 2.65E+00 | 6.18E-01 | no | | intergenic_variant | MODIFIER |
| 15 | 74262019 | AX-170251568 | 0 | G | C | 0.553 | 4.17E-02 | 8.25E-02 | 2.58E+00 | 2.65E+00 | 6.18E-01 | no | | intergenic_variant | MODIFIER |
| 15 | 74276366 | AX-170151342 | 0 | G | A | 0.553 | 4.17E-02 | 8.25E-02 | 2.58E+00 | 2.65E+00 | 6.18E-01 | no | | intergenic_variant | MODIFIER |
| 15 | 74281328 | AX-170513948 | 0 | A | C | 0.553 | 4.17E-02 | 8.25E-02 | 2.58E+00 | 2.65E+00 | 6.18E-01 | no | | intergenic_variant | MODIFIER |
| 15 | 74281452 | AX-170415535 | 0 | C | A | 0.553 | 4.17E-02 | 8.25E-02 | 2.58E+00 | 2.65E+00 | 6.18E-01 | no | | intergenic_variant | MODIFIER |
| 15 | 74292117 | AX-97798729 | 0 | A | G | 0.553 | 4.17E-02 | 8.25E-02 | 2.58E+00 | 2.65E+00 | 6.18E-01 | no | | intergenic_variant | MODIFIER |
| 15 | 74311380 | AX-170408580 | 0 | A | G | 0.553 | 4.17E-02 | 8.25E-02 | 2.58E+00 | 2.65E+00 | 6.18E-01 | no | | intergenic_variant | MODIFIER |
| 15 | 74320674 | AX-170359797 | 0 | C | T | 0.553 | 4.17E-02 | 8.25E-02 | 2.58E+00 | 2.65E+00 | 6.18E-01 | no | | regulatory_region_variant | MODIFIER |
| 15 | 74320926 | AX-97822713 | 0 | T | C | 0.553 | 4.17E-02 | 8.25E-02 | 2.58E+00 | 2.65E+00 | 6.18E-01 | no | | intergenic_variant | MODIFIER |
| 15 | 74324823 | AX-170274161 | 0 | A | G | 0.553 | 4.17E-02 | 8.25E-02 | 2.58E+00 | 2.65E+00 | 6.18E-01 | no | | intergenic_variant | MODIFIER |
| 15 | 74329453 | AX-170106068 | 0 | C | T | 0.553 | 4.17E-02 | 8.25E-02 | 2.58E+00 | 2.65E+00 | 6.18E-01 | no | | regulatory_region_variant | MODIFIER |
| 15 | 74331481 | AX-170275260 | 0 | C | A | 0.553 | 4.17E-02 | 8.25E-02 | 2.58E+00 | 2.65E+00 | 6.18E-01 | no | | intergenic_variant | MODIFIER |
| 15 | 74335205 | AX-97848954 | 0 | C | A | 0.553 | 4.17E-02 | 8.25E-02 | 2.58E+00 | 2.65E+00 | 6.18E-01 | no | | intergenic_variant | MODIFIER |
| 15 | 74360502 | AX-97818867 | 0 | A | C | 0.553 | 4.17E-02 | 8.25E-02 | 2.58E+00 | 2.65E+00 | 6.18E-01 | no | | intergenic_variant | MODIFIER |
| 15 | 74368380 | AX-170165818 | 0 | C | T | 0.553 | 4.17E-02 | 8.25E-02 | 2.58E+00 | 2.65E+00 | 6.18E-01 | no | | intergenic_variant | MODIFIER |
| 15 | 74479341 | AX-170475520 | 0 | A | G | 0.553 | 4.17E-02 | 8.25E-02 | 2.58E+00 | 2.65E+00 | 6.18E-01 | no | | intergenic_variant | MODIFIER |
| 15 | 74479411 | AX-170011994 | 0 | A | G | 0.553 | 4.17E-02 | 8.25E-02 | 2.58E+00 | 2.65E+00 | 6.18E-01 | no | | intergenic_variant | MODIFIER |
| 15 | 74479439 | AX-97830375 | 0 | C | T | 0.553 | 4.17E-02 | 8.25E-02 | 2.58E+00 | 2.65E+00 | 6.18E-01 | no | | intergenic_variant | MODIFIER |
| 15 | 74491307 | AX-170269852 | 0 | A | G | 0.553 | 4.17E-02 | 8.25E-02 | 2.58E+00 | 2.65E+00 | 6.18E-01 | no | | intergenic_variant | MODIFIER |
| 15 | 74500910 | AX-97856586 | 0 | C | T | 0.553 | 4.17E-02 | 8.25E-02 | 2.58E+00 | 2.65E+00 | 6.18E-01 | no | | intergenic_variant | MODIFIER |
| 15 | 74508813 | AX-170323423 | 0 | C | T | 0.553 | 4.17E-02 | 8.25E-02 | 2.58E+00 | 2.65E+00 | 6.18E-01 | no | | intergenic_variant | MODIFIER |
| 15 | 74511318 | AX-170395425 | 0 | A | G | 0.553 | 4.17E-02 | 8.25E-02 | 2.58E+00 | 2.65E+00 | 6.18E-01 | no | Adgrb1 | upstream_gene_variant | MODIFIER |
| 15 | 74547260 | AX-170365121 | 0 | C | T | 0.553 | 4.17E-02 | 8.25E-02 | 2.58E+00 | 2.65E+00 | 6.18E-01 | no | Adgrb1 | intron_variant | MODIFIER |
| 15 | 74550214 | AX-97844331 | 0 | C | A | 0.553 | 4.17E-02 | 8.25E-02 | 2.58E+00 | 2.65E+00 | 6.18E-01 | no | Adgrb1 | intron_variant | MODIFIER |
| 15 | 74582319 | AX-170352268 | 0 | G | A | 0.553 | 4.17E-02 | 8.25E-02 | 2.58E+00 | 2.65E+00 | 6.18E-01 | no | Adgrb1 | intron_variant | MODIFIER |
| 15 | 74655134 | AX-170087907 | 0 | C | T | 0.553 | 4.17E-02 | 8.25E-02 | 2.58E+00 | 2.65E+00 | 6.18E-01 | no | | intergenic_variant | MODIFIER |
| 15 | 74693698 | AX-170363120 | 0 | A | G | 0.553 | 4.17E-02 | 8.25E-02 | 2.58E+00 | 2.65E+00 | 6.18E-01 | no | | intergenic_variant | MODIFIER |
| 15 | 74800257 | AX-170269100 | 0 | T | C | 0.553 | 4.17E-02 | 8.25E-02 | 2.58E+00 | 2.65E+00 | 6.18E-01 | no | Ly6k | upstream_gene_variant | MODIFIER |
| 15 | 74801830 | AX-97827504 | 0 | C | T | 0.553 | 4.17E-02 | 8.25E-02 | 2.58E+00 | 2.65E+00 | 6.18E-01 | no | Ly6k | upstream_gene_variant | MODIFIER |
| 15 | 74835611 | AX-170376276 | 0 | A | G | 0.553 | 4.17E-02 | 8.25E-02 | 2.58E+00 | 2.65E+00 | 6.18E-01 | no | Cyp11b1 | 3_prime_UTR_variant | MODIFIER |
| 15 | 74838664 | AX-170312517 | 0 | C | T | 0.553 | 4.17E-02 | 8.25E-02 | 2.58E+00 | 2.65E+00 | 6.18E-01 | no | Cyp11b1 | intron_variant | MODIFIER |
| 15 | 74840346 | AX-170398504 | 0 | C | T | 0.553 | 4.17E-02 | 8.25E-02 | 2.58E+00 | 2.65E+00 | 6.18E-01 | no | Cyp11b1 | intron_variant | MODIFIER |
| 15 | 74848301 | AX-170087908 | 0 | C | T | 0.553 | 4.17E-02 | 8.25E-02 | 2.58E+00 | 2.65E+00 | 6.18E-01 | no | Cyp11b2 | downstream_gene_variant | MODIFIER |
| 15 | 74854131 | AX-170055988 | 0 | C | T | 0.553 | 4.17E-02 | 8.25E-02 | 2.58E+00 | 2.65E+00 | 6.18E-01 | no | Cyp11b2 | intron_variant | MODIFIER |

| | | | | | | | | | | | | | | | |
|---|---|---|---|---|---|---|---|---|---|---|---|---|---|---|---|
| 15 | 74856890 | AX-170204787 | 0 | T | C | 0.553 | 4.17E-02 | 8.25E-02 | 2.58E+00 | 2.65E+00 | 6.18E-01 | no | Cyp11b2 | upstream_gene_variant | MODIFIER |
| 15 | 74857603 | AX-170055991 | 0 | A | G | 0.553 | 4.17E-02 | 8.25E-02 | 2.58E+00 | 2.65E+00 | 6.18E-01 | no | Cyp11b2 | upstream_gene_variant | MODIFIER |
| 15 | 74876672 | AX-170140823 | 0 | A | G | 0.553 | 4.17E-02 | 8.25E-02 | 2.58E+00 | 2.65E+00 | 6.18E-01 | no | 2010109I03Rik | downstream_gene_variant | MODIFIER |
| 15 | 74879273 | AX-170447963 | 0 | C | T | 0.553 | 4.17E-02 | 8.25E-02 | 2.58E+00 | 2.65E+00 | 6.18E-01 | no | 2010109I03Rik | 3_prime_UTR_variant | MODIFIER |
| 15 | 74879777 | AX-170338415 | 0 | T | C | 0.553 | 4.17E-02 | 8.25E-02 | 2.58E+00 | 2.65E+00 | 6.18E-01 | no | 2010109I03Rik | 3_prime_UTR_variant | MODIFIER |
| 15 | 74880766 | AX-170247356 | 0 | T | C | 0.553 | 4.17E-02 | 8.25E-02 | 2.58E+00 | 2.65E+00 | 6.18E-01 | no | 2010109I03Rik | synonymous_variant | LOW |
| 15 | 74882223 | AX-97845007 | 0 | G | A | 0.553 | 4.17E-02 | 8.25E-02 | 2.58E+00 | 2.65E+00 | 6.18E-01 | no | 2010109I03Rik | upstream_gene_variant | MODIFIER |
| 15 | 74887348 | AX-97798353 | 0 | G | A | 0.553 | 4.17E-02 | 8.25E-02 | 2.58E+00 | 2.65E+00 | 6.18E-01 | no | 2010109I03Rik | intron_variant,non_coding_transcript_variant | MODIFIER |
| 15 | 74887376 | AX-170526412 | 0 | T | C | 0.553 | 4.17E-02 | 8.25E-02 | 2.58E+00 | 2.65E+00 | 6.18E-01 | no | 2010109I03Rik | intron_variant,non_coding_transcript_variant | MODIFIER |
| 15 | 74911720 | AX-97830405 | 0 | C | T | 0.553 | 4.17E-02 | 8.25E-02 | 2.58E+00 | 2.65E+00 | 6.18E-01 | no | | intergenic_variant | MODIFIER |
| 15 | 74923278 | AX-170410139 | 0 | T | C | 0.553 | 4.17E-02 | 8.25E-02 | 2.58E+00 | 2.65E+00 | 6.18E-01 | no | Gm6610 | upstream_gene_variant | MODIFIER |
| 15 | 74926089 | AX-170152491 | 0 | G | C | 0.553 | 4.17E-02 | 8.25E-02 | 2.58E+00 | 2.65E+00 | 6.18E-01 | no | Gm6610 | downstream_gene_variant | MODIFIER |
| 15 | 74948698 | AX-170534884 | 0 | C | T | 0.553 | 4.17E-02 | 8.25E-02 | 2.58E+00 | 2.65E+00 | 6.18E-01 | no | Gm29126 | downstream_gene_variant | MODIFIER |
| 15 | 74952995 | AX-170193925 | 0 | T | C | 0.553 | 4.17E-02 | 8.25E-02 | 2.58E+00 | 2.65E+00 | 6.18E-01 | no | Ly6e | upstream_gene_variant | MODIFIER |
| 15 | 74954837 | AX-97843596 | 0 | A | G | 0.553 | 4.17E-02 | 8.25E-02 | 2.58E+00 | 2.65E+00 | 6.18E-01 | no | Ly6e | upstream_gene_variant | MODIFIER |
| 15 | 74970223 | AX-170203414 | 0 | C | T | 0.553 | 4.17E-02 | 8.25E-02 | 2.58E+00 | 2.65E+00 | 6.18E-01 | no | | intergenic_variant | MODIFIER |
| 15 | 74985808 | AX-170012005 | 0 | G | C | 0.553 | 4.17E-02 | 8.25E-02 | 2.58E+00 | 2.65E+00 | 6.18E-01 | no | Ly6i | upstream_gene_variant | MODIFIER |
| 15 | 75114844 | AX-97790260 | 0 | G | A | 0.553 | 4.17E-02 | 8.25E-02 | 2.58E+00 | 2.65E+00 | 6.18E-01 | no | Ly6c2 | upstream_gene_variant | MODIFIER |
| 15 | 75154721 | AX-97829732 | 0 | C | T | 0.553 | 4.17E-02 | 8.25E-02 | 2.58E+00 | 2.65E+00 | 6.18E-01 | no | Ly6g | upstream_gene_variant | MODIFIER |
| 15 | 75159510 | AX-97841150 | 0 | A | G | 0.553 | 4.17E-02 | 8.25E-02 | 2.58E+00 | 2.65E+00 | 6.18E-01 | no | Ly6g | downstream_gene_variant | MODIFIER |
| 15 | 75181640 | AX-170455094 | 0 | C | T | 0.553 | 4.17E-02 | 8.25E-02 | 2.58E+00 | 2.65E+00 | 6.18E-01 | no | | intergenic_variant | MODIFIER |
| 15 | 75193043 | AX-170428562 | 0 | A | G | 0.553 | 4.17E-02 | 8.25E-02 | 2.58E+00 | 2.65E+00 | 6.18E-01 | no | Gm28117 | downstream_gene_variant | MODIFIER |
| 15 | 75216939 | AX-170478866 | 0 | T | C | 0.553 | 4.17E-02 | 8.25E-02 | 2.58E+00 | 2.65E+00 | 6.18E-01 | no | BC025446 | intron_variant | MODIFIER |
| 15 | 75228182 | AX-170137903 | 0 | T | A | 0.553 | 4.17E-02 | 8.25E-02 | 2.58E+00 | 2.65E+00 | 6.18E-01 | no | | intergenic_variant | MODIFIER |
| 15 | 75287822 | AX-97800318 | 0 | T | C | 0.553 | 4.17E-02 | 8.25E-02 | 2.58E+00 | 2.65E+00 | 6.18E-01 | no | | intergenic_variant | MODIFIER |
| 15 | 75330227 | AX-170055999 | 0 | A | G | 0.553 | 4.17E-02 | 8.25E-02 | 2.58E+00 | 2.65E+00 | 6.18E-01 | no | Gm28113 | upstream_gene_variant | MODIFIER |
| 15 | 75332235 | AX-170055998 | 0 | T | G | 0.553 | 4.17E-02 | 8.25E-02 | 2.58E+00 | 2.65E+00 | 6.18E-01 | no | Gm29517 | downstream_gene_variant | MODIFIER |
| 15 | 75342815 | AX-170179006 | 0 | A | G | 0.553 | 4.17E-02 | 8.25E-02 | 2.58E+00 | 2.65E+00 | 6.18E-01 | no | | intergenic_variant | MODIFIER |
| 15 | 75352224 | AX-170203576 | 0 | G | T | 0.553 | 4.17E-02 | 8.25E-02 | 2.58E+00 | 2.65E+00 | 6.18E-01 | no | Gm28114 | upstream_gene_variant | MODIFIER |
| 15 | 75353449 | AX-170466784 | 0 | T | C | 0.553 | 4.17E-02 | 8.25E-02 | 2.58E+00 | 2.65E+00 | 6.18E-01 | no | Gm28114 | upstream_gene_variant | MODIFIER |
| 15 | 75406128 | AX-97819385 | 0 | G | A | 0.553 | 4.17E-02 | 8.25E-02 | 2.58E+00 | 2.65E+00 | 6.18E-01 | no | | intergenic_variant | MODIFIER |
| 15 | 75501720 | AX-170093305 | 0 | A | G | 0.553 | 4.17E-02 | 8.25E-02 | 2.58E+00 | 2.65E+00 | 6.18E-01 | no | Gm34593 | downstream_gene_variant | MODIFIER |
| 15 | 75517574 | AX-170417296 | 0 | G | A | 0.553 | 4.17E-02 | 8.25E-02 | 2.58E+00 | 2.65E+00 | 6.18E-01 | no | Gm34646 | non_coding_transcript_exon_variant | MODIFIER |
| 15 | 75550779 | AX-170270896 | 0 | C | T | 0.553 | 4.17E-02 | 8.25E-02 | 2.58E+00 | 2.65E+00 | 6.18E-01 | no | Ly6h | downstream_gene_variant | MODIFIER |
| 15 | 75551117 | AX-97836083 | 0 | G | A | 0.553 | 4.17E-02 | 8.25E-02 | 2.58E+00 | 2.65E+00 | 6.18E-01 | no | Ly6h | downstream_gene_variant | MODIFIER |
| 15 | 75571074 | AX-97808050 | 0 | G | A | 0.553 | 4.17E-02 | 8.25E-02 | 2.58E+00 | 2.65E+00 | 6.18E-01 | no | Ly6h | upstream_gene_variant | MODIFIER |
| 15 | 75576594 | AX-170286153 | 0 | T | A | 0.553 | 4.17E-02 | 8.25E-02 | 2.58E+00 | 2.65E+00 | 6.18E-01 | no | | regulatory_region_variant | MODIFIER |
| 15 | 75593128 | AX-97837650 | 0 | A | G | 0.553 | 4.17E-02 | 8.25E-02 | 2.58E+00 | 2.65E+00 | 6.18E-01 | no | Gpihbp1 | upstream_gene_variant | MODIFIER |
| 15 | 75659332 | AX-97804715 | 0 | A | G | 0.553 | 4.17E-02 | 8.25E-02 | 2.58E+00 | 2.65E+00 | 6.18E-01 | no | Top1mt | intron_variant | MODIFIER |
| 15 | 75716537 | AX-170533007 | 0 | A | G | 0.553 | 4.17E-02 | 8.25E-02 | 2.58E+00 | 2.65E+00 | 6.18E-01 | no | Rhpn1 | downstream_gene_variant | MODIFIER |
| 15 | 75722658 | AX-97824632 | 0 | T | C | 0.553 | 4.17E-02 | 8.25E-02 | 2.58E+00 | 2.65E+00 | 6.18E-01 | no | | intergenic_variant | MODIFIER |
| 15 | 75757223 | AX-97829728 | 0 | A | G | 0.553 | 4.17E-02 | 8.25E-02 | 2.58E+00 | 2.65E+00 | 6.18E-01 | no | Zc3h3 | intron_variant | MODIFIER |
| 15 | 75765251 | AX-97828573 | 0 | T | C | 0.553 | 4.17E-02 | 8.25E-02 | 2.58E+00 | 2.65E+00 | 6.18E-01 | no | Zc3h3 | intron_variant | MODIFIER |
| 15 | 75778854 | AX-170120897 | 0 | G | A | 0.553 | 4.17E-02 | 8.25E-02 | 2.58E+00 | 2.65E+00 | 6.18E-01 | no | Zc3h3 | intron_variant | MODIFIER |
| 15 | 75781580 | AX-170354313 | 0 | T | C | 0.553 | 4.17E-02 | 8.25E-02 | 2.58E+00 | 2.65E+00 | 6.18E-01 | no | Zc3h3 | intron_variant | MODIFIER |
| 15 | 75786304 | AX-170323452 | 0 | G | C | 0.553 | 4.17E-02 | 8.25E-02 | 2.58E+00 | 2.65E+00 | 6.18E-01 | no | Zc3h3 | intron_variant | MODIFIER |
| 15 | 75792280 | AX-97847046 | 0 | A | G | 0.553 | 4.17E-02 | 8.25E-02 | 2.58E+00 | 2.65E+00 | 6.18E-01 | no | Zc3h3 | intron_variant | MODIFIER |
| 15 | 75804217 | AX-170056011 | 0 | A | G | 0.553 | 4.17E-02 | 8.25E-02 | 2.58E+00 | 2.65E+00 | 6.18E-01 | no | Zc3h3 | intron_variant | MODIFIER |
| 15 | 75811267 | AX-168404186 | 0 | A | G | 0.553 | 4.17E-02 | 8.25E-02 | 2.58E+00 | 2.65E+00 | 6.18E-01 | no | Zc3h3 | intron_variant | MODIFIER |
| 15 | 75816817 | AX-97814287 | 0 | C | T | 0.553 | 4.17E-02 | 8.25E-02 | 2.58E+00 | 2.65E+00 | 6.18E-01 | no | Zc3h3 | intron_variant | MODIFIER |
| 15 | 75832695 | AX-170105543 | 0 | G | T | 0.553 | 4.17E-02 | 8.25E-02 | 2.58E+00 | 2.65E+00 | 6.18E-01 | no | Zc3h3 | intron_variant | MODIFIER |
| 15 | 75834320 | AX-170523669 | 0 | A | G | 0.553 | 4.17E-02 | 8.25E-02 | 2.58E+00 | 2.65E+00 | 6.18E-01 | no | Zc3h3 | intron_variant | MODIFIER |
| 15 | 75834488 | AX-170302492 | 0 | A | G | 0.553 | 4.17E-02 | 8.25E-02 | 2.58E+00 | 2.65E+00 | 6.18E-01 | no | Zc3h3 | intron_variant | MODIFIER |
| 15 | 75834653 | AX-97850271 | 0 | A | G | 0.553 | 4.17E-02 | 8.25E-02 | 2.58E+00 | 2.65E+00 | 6.18E-01 | no | Zc3h3 | intron_variant | MODIFIER |

| | | | | | | | | | | | | | | | |
|---|---|---|---|---|---|---|---|---|---|---|---|---|---|---|---|
| 15 | 75858194 | AX-170056009 | 0 | C | T | 0.553 | 4.17E-02 | 8.25E-02 | 2.58E+00 | 2.65E+00 | 6.18E-01 | no | Gsdmd | upstream_gene_variant | MODIFIER |
| 15 | 75862886 | AX-170148319 | 0 | C | G | 0.553 | 4.17E-02 | 8.25E-02 | 2.58E+00 | 2.65E+00 | 6.18E-01 | no | Gsdmd | intron_variant | MODIFIER |
| 15 | 75918363 | AX-97829893 | 0 | C | G | 0.553 | 4.17E-02 | 8.25E-02 | 2.58E+00 | 2.65E+00 | 6.18E-01 | no | Pycrl | missense_variant | MODERATE |
| 15 | 75987767 | AX-97803190 | 0 | A | G | 0.553 | 4.17E-02 | 8.25E-02 | 2.58E+00 | 2.65E+00 | 6.18E-01 | no | | intergenic_variant | MODIFIER |
| 11 | 24507496 | AX-97845012 | 1 | G | A | 0.071 | 8.67E-02 | 1.79E-01 | 2.70E+00 | 2.82E+00 | 6.18E-01 | no | 4933430M04Rik | intron_variant,non_coding_transcript_variant | MODIFIER |
| 12 | 93428239 | AX-97787959 | 1 | C | T | 0.062 | 1.16E-01 | 2.37E-01 | 2.72E+00 | 2.78E+00 | 6.18E-01 | no | 4930559C10Rik | intron_variant,non_coding_transcript_variant | MODIFIER |
| 1 | 22379976 | AX-170312731 | 1 | A | T | 0.411 | 4.28E-02 | 8.56E-02 | 2.62E+00 | 2.67E+00 | 6.18E-01 | no | Rims1 | intron_variant | MODIFIER |
| 1 | 41938943 | AX-170533601 | 1 | C | G | 0.446 | 3.82E-02 | 7.79E-02 | 2.69E+00 | 2.78E+00 | 6.18E-01 | no | Gm29260 | non_coding_transcript_exon_variant | MODIFIER |
| 15 | 91497100 | AX-170163941 | 0 | A | T | 0.123 | -6.58E-02 | 1.35E-01 | 2.72E+00 | 2.82E+00 | 6.18E-01 | no | Slc2a13 | intron_variant | MODIFIER |
| 16 | 95610249 | AX-97782856 | 0 | C | T | 0.465 | 3.73E-02 | 7.68E-02 | 2.77E+00 | 2.89E+00 | 6.19E-01 | no | | intergenic_variant | MODIFIER |
| 16 | 96118274 | AX-170305353 | 0 | T | A | 0.465 | 3.73E-02 | 7.68E-02 | 2.77E+00 | 2.89E+00 | 6.19E-01 | no | Hmgn1 | downstream_gene_variant | MODIFIER |
| 16 | 96119869 | AX-170124524 | 0 | C | T | 0.465 | 3.73E-02 | 7.68E-02 | 2.77E+00 | 2.89E+00 | 6.19E-01 | no | Hmgn1 | downstream_gene_variant | MODIFIER |
| 16 | 96122784 | AX-97839416 | 0 | A | G | 0.465 | 3.73E-02 | 7.68E-02 | 2.77E+00 | 2.89E+00 | 6.19E-01 | no | Hmgn1 | intron_variant | MODIFIER |
| 16 | 96148724 | AX-170370613 | 0 | T | C | 0.465 | 3.73E-02 | 7.68E-02 | 2.77E+00 | 2.89E+00 | 6.19E-01 | no | Wrb | intron_variant | MODIFIER |
| 16 | 96149117 | AX-170252255 | 0 | C | T | 0.465 | 3.73E-02 | 7.68E-02 | 2.77E+00 | 2.89E+00 | 6.19E-01 | no | Wrb | intron_variant | MODIFIER |
| 16 | 96157341 | AX-170483740 | 0 | C | T | 0.465 | 3.73E-02 | 7.68E-02 | 2.77E+00 | 2.89E+00 | 6.19E-01 | no | Wrb | 3_prime_UTR_variant | MODIFIER |
| 16 | 96158643 | AX-170210277 | 0 | T | C | 0.465 | 3.73E-02 | 7.68E-02 | 2.77E+00 | 2.89E+00 | 6.19E-01 | no | Wrb | downstream_gene_variant | MODIFIER |
| 16 | 96158730 | AX-170279209 | 0 | A | G | 0.465 | 3.73E-02 | 7.68E-02 | 2.77E+00 | 2.89E+00 | 6.19E-01 | no | Wrb | downstream_gene_variant | MODIFIER |
| 16 | 96205554 | AX-170480103 | 0 | A | G | 0.465 | 3.73E-02 | 7.68E-02 | 2.77E+00 | 2.89E+00 | 6.19E-01 | no | Sh3bgr | intron_variant | MODIFIER |
| 16 | 96275731 | AX-97845524 | 0 | T | C | 0.465 | 3.73E-02 | 7.68E-02 | 2.77E+00 | 2.89E+00 | 6.19E-01 | no | B3galt5 | intron_variant | MODIFIER |
| 16 | 96276142 | AX-170177136 | 0 | C | A | 0.465 | 3.73E-02 | 7.68E-02 | 2.77E+00 | 2.89E+00 | 6.19E-01 | no | B3galt5 | upstream_gene_variant | MODIFIER |
| 16 | 96281515 | AX-170141774 | 0 | G | T | 0.465 | 3.73E-02 | 7.68E-02 | 2.77E+00 | 2.89E+00 | 6.19E-01 | no | B3galt5 | intron_variant | MODIFIER |
| 16 | 96282311 | AX-170296543 | 0 | C | T | 0.465 | 3.73E-02 | 7.68E-02 | 2.77E+00 | 2.89E+00 | 6.19E-01 | no | B3galt5 | intron_variant | MODIFIER |
| 16 | 96353711 | AX-170174374 | 0 | G | A | 0.465 | 3.73E-02 | 7.68E-02 | 2.77E+00 | 2.89E+00 | 6.19E-01 | no | | regulatory_region_variant | MODIFIER |
| 16 | 96466120 | AX-170148334 | 0 | G | T | 0.465 | 3.73E-02 | 7.68E-02 | 2.77E+00 | 2.89E+00 | 6.19E-01 | no | Pcp4 | upstream_gene_variant | MODIFIER |
| 16 | 96475434 | AX-170013712 | 0 | T | C | 0.465 | 3.73E-02 | 7.68E-02 | 2.77E+00 | 2.89E+00 | 6.19E-01 | no | Pcp4 | intron_variant | MODIFIER |
| 16 | 96479096 | AX-170204911 | 0 | T | C | 0.465 | 3.73E-02 | 7.68E-02 | 2.77E+00 | 2.89E+00 | 6.19E-01 | no | Pcp4 | intron_variant | MODIFIER |
| 16 | 96482858 | AX-170489678 | 0 | G | C | 0.465 | 3.73E-02 | 7.68E-02 | 2.77E+00 | 2.89E+00 | 6.19E-01 | no | Pcp4 | intron_variant | MODIFIER |
| 16 | 96487009 | AX-170431630 | 0 | A | G | 0.465 | 3.73E-02 | 7.68E-02 | 2.77E+00 | 2.89E+00 | 6.19E-01 | no | Pcp4 | intron_variant | MODIFIER |
| 16 | 96487706 | AX-170400935 | 0 | G | A | 0.465 | 3.73E-02 | 7.68E-02 | 2.77E+00 | 2.89E+00 | 6.19E-01 | no | Pcp4 | intron_variant | MODIFIER |
| 16 | 96495816 | AX-170229229 | 0 | T | A | 0.465 | 3.73E-02 | 7.68E-02 | 2.77E+00 | 2.89E+00 | 6.19E-01 | no | Pcp4 | intron_variant | MODIFIER |
| 16 | 96498449 | AX-170459290 | 0 | T | G | 0.465 | 3.73E-02 | 7.68E-02 | 2.77E+00 | 2.89E+00 | 6.19E-01 | no | Pcp4 | intron_variant | MODIFIER |
| 16 | 96521014 | AX-170296009 | 0 | G | A | 0.465 | 3.73E-02 | 7.68E-02 | 2.77E+00 | 2.89E+00 | 6.19E-01 | no | Pcp4 | intron_variant | MODIFIER |
| 16 | 96534878 | AX-170311934 | 0 | T | C | 0.465 | 3.73E-02 | 7.68E-02 | 2.77E+00 | 2.89E+00 | 6.19E-01 | no | | intergenic_variant | MODIFIER |
| 16 | 96636205 | AX-170301705 | 0 | C | T | 0.465 | 3.73E-02 | 7.68E-02 | 2.77E+00 | 2.89E+00 | 6.19E-01 | no | Dscam | intron_variant | MODIFIER |
| 16 | 96715932 | AX-170141840 | 0 | C | T | 0.465 | 3.73E-02 | 7.68E-02 | 2.77E+00 | 2.89E+00 | 6.19E-01 | no | Dscam | intron_variant | MODIFIER |
| 16 | 96716316 | AX-170256300 | 0 | G | A | 0.465 | 3.73E-02 | 7.68E-02 | 2.77E+00 | 2.89E+00 | 6.19E-01 | no | Dscam | intron_variant | MODIFIER |
| 16 | 96716771 | AX-170140384 | 0 | C | A | 0.465 | 3.73E-02 | 7.68E-02 | 2.77E+00 | 2.89E+00 | 6.19E-01 | no | Dscam | intron_variant | MODIFIER |
| 16 | 96717008 | AX-170448584 | 0 | G | A | 0.465 | 3.73E-02 | 7.68E-02 | 2.77E+00 | 2.89E+00 | 6.19E-01 | no | Dscam | intron_variant | MODIFIER |
| 16 | 96721109 | AX-170505493 | 0 | C | T | 0.465 | 3.73E-02 | 7.68E-02 | 2.77E+00 | 2.89E+00 | 6.19E-01 | no | Dscam | intron_variant | MODIFIER |
| 16 | 96730943 | AX-170291713 | 0 | C | T | 0.465 | 3.73E-02 | 7.68E-02 | 2.77E+00 | 2.89E+00 | 6.19E-01 | no | Dscam | intron_variant | MODIFIER |
| 16 | 96737818 | AX-97790139 | 0 | T | C | 0.465 | 3.73E-02 | 7.68E-02 | 2.77E+00 | 2.89E+00 | 6.19E-01 | no | Dscam | intron_variant | MODIFIER |
| 16 | 96737997 | AX-170317351 | 0 | G | T | 0.465 | 3.73E-02 | 7.68E-02 | 2.77E+00 | 2.89E+00 | 6.19E-01 | no | Dscam | intron_variant | MODIFIER |
| 16 | 96738715 | AX-170452984 | 0 | A | G | 0.465 | 3.73E-02 | 7.68E-02 | 2.77E+00 | 2.89E+00 | 6.19E-01 | no | Dscam | intron_variant | MODIFIER |
| 16 | 96955226 | AX-170366060 | 0 | G | A | 0.465 | 3.73E-02 | 7.68E-02 | 2.77E+00 | 2.89E+00 | 6.19E-01 | no | Dscam | intron_variant | MODIFIER |
| 16 | 96996851 | AX-170013715 | 0 | C | T | 0.465 | 3.73E-02 | 7.68E-02 | 2.77E+00 | 2.89E+00 | 6.19E-01 | no | Dscam | intron_variant | MODIFIER |
| 16 | 97028356 | AX-170013717 | 0 | T | C | 0.465 | 3.73E-02 | 7.68E-02 | 2.77E+00 | 2.89E+00 | 6.19E-01 | no | Dscam | intron_variant | MODIFIER |
| 16 | 97090341 | AX-170322302 | 0 | T | G | 0.465 | 3.73E-02 | 7.68E-02 | 2.77E+00 | 2.89E+00 | 6.19E-01 | no | Dscam | intron_variant | MODIFIER |
| 16 | 97175681 | AX-170484606 | 0 | A | G | 0.465 | 3.73E-02 | 7.68E-02 | 2.77E+00 | 2.89E+00 | 6.19E-01 | no | Dscam | upstream_gene_variant | MODIFIER |
| 16 | 97216181 | AX-170238887 | 0 | C | T | 0.465 | 3.73E-02 | 7.68E-02 | 2.77E+00 | 2.89E+00 | 6.19E-01 | no | | intergenic_variant | MODIFIER |
| 16 | 97216372 | AX-170441479 | 0 | G | A | 0.465 | 3.73E-02 | 7.68E-02 | 2.77E+00 | 2.89E+00 | 6.19E-01 | no | | intergenic_variant | MODIFIER |
| 16 | 97226203 | AX-170109377 | 0 | G | A | 0.465 | 3.73E-02 | 7.68E-02 | 2.77E+00 | 2.89E+00 | 6.19E-01 | no | | intergenic_variant | MODIFIER |
| 16 | 97229253 | AX-170210047 | 0 | A | G | 0.465 | 3.73E-02 | 7.68E-02 | 2.77E+00 | 2.89E+00 | 6.19E-01 | no | | intergenic_variant | MODIFIER |
| 16 | 97247353 | AX-170162730 | 0 | T | G | 0.465 | 3.73E-02 | 7.68E-02 | 2.77E+00 | 2.89E+00 | 6.19E-01 | no | | intergenic_variant | MODIFIER |

| | | | | | | | | | | | | | | | |
|---|---|---|---|---|---|---|---|---|---|---|---|---|---|---|---|
| 16 | 97248205 | AX-170247141 | 0 | T | C | 0.465 | 3.73E-02 | 7.68E-02 | 2.77E+00 | 2.89E+00 | 6.19E-01 | no | | intergenic_variant | MODIFIER |
| 16 | 97261526 | AX-97832236 | 0 | T | G | 0.465 | 3.73E-02 | 7.68E-02 | 2.77E+00 | 2.89E+00 | 6.19E-01 | no | | intergenic_variant | MODIFIER |
| 16 | 97368149 | AX-170148489 | 0 | T | C | 0.465 | 3.73E-02 | 7.68E-02 | 2.77E+00 | 2.89E+00 | 6.19E-01 | no | Bace2 | intron_variant | MODIFIER |
| 16 | 97369962 | AX-170149240 | 0 | T | C | 0.465 | 3.73E-02 | 7.68E-02 | 2.77E+00 | 2.89E+00 | 6.19E-01 | no | Bace2 | intron_variant | MODIFIER |
| 16 | 97373270 | AX-97803724 | 0 | G | A | 0.465 | 3.73E-02 | 7.68E-02 | 2.77E+00 | 2.89E+00 | 6.19E-01 | no | Bace2 | intron_variant | MODIFIER |
| 16 | 97388616 | AX-170307395 | 0 | T | C | 0.465 | 3.73E-02 | 7.68E-02 | 2.77E+00 | 2.89E+00 | 6.19E-01 | no | Bace2 | intron_variant | MODIFIER |
| 16 | 97517402 | AX-170245049 | 0 | T | C | 0.465 | 3.73E-02 | 7.68E-02 | 2.77E+00 | 2.89E+00 | 6.19E-01 | no | Fam3b | upstream_gene_variant | MODIFIER |
| 16 | 97523741 | AX-170343072 | 0 | C | T | 0.465 | 3.73E-02 | 7.68E-02 | 2.77E+00 | 2.89E+00 | 6.19E-01 | no | | intergenic_variant | MODIFIER |
| 16 | 97523942 | AX-170317597 | 0 | A | G | 0.465 | 3.73E-02 | 7.68E-02 | 2.77E+00 | 2.89E+00 | 6.19E-01 | no | | intergenic_variant | MODIFIER |
| 16 | 97532611 | AX-170013718 | 0 | G | A | 0.465 | 3.73E-02 | 7.68E-02 | 2.77E+00 | 2.89E+00 | 6.19E-01 | no | Mx2 | upstream_gene_variant | MODIFIER |
| 16 | 97537819 | AX-170148624 | 0 | C | A | 0.465 | 3.73E-02 | 7.68E-02 | 2.77E+00 | 2.89E+00 | 6.19E-01 | no | Mx2 | intron_variant | MODIFIER |
| 16 | 97538641 | AX-97807410 | 0 | G | A | 0.465 | 3.73E-02 | 7.68E-02 | 2.77E+00 | 2.89E+00 | 6.19E-01 | no | Mx2 | 5_prime_UTR_variant | MODIFIER |
| 16 | 97575427 | AX-170530743 | 0 | A | C | 0.465 | 3.73E-02 | 7.68E-02 | 2.77E+00 | 2.89E+00 | 6.19E-01 | no | Tmprss2 | intron_variant | MODIFIER |
| 16 | 97577095 | AX-170372500 | 0 | C | T | 0.465 | 3.73E-02 | 7.68E-02 | 2.77E+00 | 2.89E+00 | 6.19E-01 | no | Tmprss2 | intron_variant | MODIFIER |
| 16 | 97582153 | AX-170351685 | 0 | G | C | 0.465 | 3.73E-02 | 7.68E-02 | 2.77E+00 | 2.89E+00 | 6.19E-01 | no | Tmprss2 | intron_variant | MODIFIER |
| 16 | 97584238 | AX-170429948 | 0 | G | A | 0.465 | 3.73E-02 | 7.68E-02 | 2.77E+00 | 2.89E+00 | 6.19E-01 | no | Tmprss2 | intron_variant | MODIFIER |
| 16 | 97596751 | AX-170471407 | 0 | G | A | 0.465 | 3.73E-02 | 7.68E-02 | 2.77E+00 | 2.89E+00 | 6.19E-01 | no | Tmprss2 | missense_variant | MODERATE |
| 16 | 97603623 | AX-170262563 | 0 | C | T | 0.465 | 3.73E-02 | 7.68E-02 | 2.77E+00 | 2.89E+00 | 6.19E-01 | no | Tmprss2 | intron_variant | MODIFIER |
| 16 | 97610432 | AX-170269601 | 0 | T | C | 0.465 | 3.73E-02 | 7.68E-02 | 2.77E+00 | 2.89E+00 | 6.19E-01 | no | Tmprss2 | intron_variant | MODIFIER |
| 16 | 97614542 | AX-170448872 | 0 | T | C | 0.465 | 3.73E-02 | 7.68E-02 | 2.77E+00 | 2.89E+00 | 6.19E-01 | no | Tmprss2 | upstream_gene_variant | MODIFIER |
| 16 | 97615416 | AX-170520307 | 0 | C | A | 0.465 | 3.73E-02 | 7.68E-02 | 2.77E+00 | 2.89E+00 | 6.19E-01 | no | Tmprss2 | upstream_gene_variant | MODIFIER |
| 16 | 97620933 | AX-170170013 | 0 | G | A | 0.465 | 3.73E-02 | 7.68E-02 | 2.77E+00 | 2.89E+00 | 6.19E-01 | no | | intergenic_variant | MODIFIER |
| 16 | 97621141 | AX-170418845 | 0 | C | T | 0.465 | 3.73E-02 | 7.68E-02 | 2.77E+00 | 2.89E+00 | 6.19E-01 | no | | intergenic_variant | MODIFIER |
| 16 | 97624286 | AX-170159766 | 0 | A | G | 0.465 | 3.73E-02 | 7.68E-02 | 2.77E+00 | 2.89E+00 | 6.19E-01 | no | | intergenic_variant | MODIFIER |
| 16 | 97625559 | AX-170223845 | 0 | C | T | 0.465 | 3.73E-02 | 7.68E-02 | 2.77E+00 | 2.89E+00 | 6.19E-01 | no | | intergenic_variant | MODIFIER |
| 16 | 97656887 | AX-170058227 | 0 | T | C | 0.465 | 3.73E-02 | 7.68E-02 | 2.77E+00 | 2.89E+00 | 6.19E-01 | no | | intergenic_variant | MODIFIER |
| 16 | 97663021 | AX-170058230 | 0 | T | C | 0.465 | 3.73E-02 | 7.68E-02 | 2.77E+00 | 2.89E+00 | 6.19E-01 | no | | intergenic_variant | MODIFIER |
| 16 | 97671710 | AX-170058229 | 0 | C | T | 0.465 | 3.73E-02 | 7.68E-02 | 2.77E+00 | 2.89E+00 | 6.19E-01 | no | | intergenic_variant | MODIFIER |
| 16 | 97673417 | AX-97850809 | 0 | C | T | 0.465 | 3.73E-02 | 7.68E-02 | 2.77E+00 | 2.89E+00 | 6.19E-01 | no | | intergenic_variant | MODIFIER |
| 16 | 97684431 | AX-170372141 | 0 | G | A | 0.465 | 3.73E-02 | 7.68E-02 | 2.77E+00 | 2.89E+00 | 6.19E-01 | no | | intergenic_variant | MODIFIER |
| 16 | 97692122 | AX-170173050 | 0 | G | T | 0.465 | 3.73E-02 | 7.68E-02 | 2.77E+00 | 2.89E+00 | 6.19E-01 | no | | intergenic_variant | MODIFIER |
| 16 | 97717466 | AX-170300353 | 0 | A | C | 0.465 | 3.73E-02 | 7.68E-02 | 2.77E+00 | 2.89E+00 | 6.19E-01 | no | | regulatory_region_variant | MODIFIER |
| 16 | 97732883 | AX-170167227 | 0 | A | G | 0.465 | 3.73E-02 | 7.68E-02 | 2.77E+00 | 2.89E+00 | 6.19E-01 | no | | intergenic_variant | MODIFIER |
| 16 | 97735550 | AX-97781190 | 0 | G | A | 0.465 | 3.73E-02 | 7.68E-02 | 2.77E+00 | 2.89E+00 | 6.19E-01 | no | | intergenic_variant | MODIFIER |
| 16 | 97735778 | AX-170494123 | 0 | G | A | 0.465 | 3.73E-02 | 7.68E-02 | 2.77E+00 | 2.89E+00 | 6.19E-01 | no | | intergenic_variant | MODIFIER |
| 16 | 97744340 | AX-170357055 | 0 | A | C | 0.465 | 3.73E-02 | 7.68E-02 | 2.77E+00 | 2.89E+00 | 6.19E-01 | no | Ripk4 | intron_variant | MODIFIER |
| 16 | 97753437 | AX-170255705 | 0 | G | A | 0.465 | 3.73E-02 | 7.68E-02 | 2.77E+00 | 2.89E+00 | 6.19E-01 | no | Ripk4 | intron_variant | MODIFIER |
| 16 | 97754725 | AX-170161331 | 0 | A | G | 0.465 | 3.73E-02 | 7.68E-02 | 2.77E+00 | 2.89E+00 | 6.19E-01 | no | Ripk4 | intron_variant | MODIFIER |
| 16 | 97755571 | AX-170119522 | 0 | T | A | 0.465 | 3.73E-02 | 7.68E-02 | 2.77E+00 | 2.89E+00 | 6.19E-01 | no | Ripk4 | intron_variant | MODIFIER |
| 16 | 97770165 | AX-170463897 | 0 | G | A | 0.465 | 3.73E-02 | 7.68E-02 | 2.77E+00 | 2.89E+00 | 6.19E-01 | no | | intergenic_variant | MODIFIER |
| 16 | 97789502 | AX-170485457 | 0 | G | A | 0.465 | 3.73E-02 | 7.68E-02 | 2.77E+00 | 2.89E+00 | 6.19E-01 | no | Prdm15 | downstream_gene_variant | MODIFIER |
| 16 | 97792490 | AX-170364799 | 0 | A | G | 0.465 | 3.73E-02 | 7.68E-02 | 2.77E+00 | 2.89E+00 | 6.19E-01 | no | Prdm15 | 3_prime_UTR_variant | MODIFIER |
| 16 | 97798105 | AX-170511127 | 0 | A | G | 0.465 | 3.73E-02 | 7.68E-02 | 2.77E+00 | 2.89E+00 | 6.19E-01 | no | Prdm15 | intron_variant | MODIFIER |
| 16 | 97812912 | AX-170058233 | 0 | T | C | 0.465 | 3.73E-02 | 7.68E-02 | 2.77E+00 | 2.89E+00 | 6.19E-01 | no | Prdm15 | intron_variant | MODIFIER |
| 16 | 97820612 | AX-170058231 | 0 | C | T | 0.465 | 3.73E-02 | 7.68E-02 | 2.77E+00 | 2.89E+00 | 6.19E-01 | no | Prdm15 | intron_variant | MODIFIER |
| 16 | 97823463 | AX-170118914 | 0 | C | T | 0.465 | 3.73E-02 | 7.68E-02 | 2.77E+00 | 2.89E+00 | 6.19E-01 | no | Prdm15 | intron_variant | MODIFIER |
| 16 | 97827530 | AX-170159216 | 0 | G | A | 0.465 | 3.73E-02 | 7.68E-02 | 2.77E+00 | 2.89E+00 | 6.19E-01 | no | Prdm15 | intron_variant | MODIFIER |
| 16 | 97833682 | AX-170156236 | 0 | A | G | 0.465 | 3.73E-02 | 7.68E-02 | 2.77E+00 | 2.89E+00 | 6.19E-01 | no | Prdm15 | intron_variant | MODIFIER |
| 16 | 97847585 | AX-170271325 | 0 | C | T | 0.465 | 3.73E-02 | 7.68E-02 | 2.77E+00 | 2.89E+00 | 6.19E-01 | no | Prdm15 | intron_variant | MODIFIER |
| 16 | 97847646 | AX-170494168 | 0 | C | A | 0.465 | 3.73E-02 | 7.68E-02 | 2.77E+00 | 2.89E+00 | 6.19E-01 | no | Prdm15 | intron_variant | MODIFIER |
| 16 | 97879892 | AX-170197207 | 0 | C | T | 0.465 | 3.73E-02 | 7.68E-02 | 2.77E+00 | 2.89E+00 | 6.19E-01 | no | C2cd2 | intron_variant | MODIFIER |
| 16 | 97884468 | AX-170427080 | 0 | T | C | 0.465 | 3.73E-02 | 7.68E-02 | 2.77E+00 | 2.89E+00 | 6.19E-01 | no | C2cd2 | intron_variant | MODIFIER |
| 16 | 97887058 | AX-170383401 | 0 | C | T | 0.465 | 3.73E-02 | 7.68E-02 | 2.77E+00 | 2.89E+00 | 6.19E-01 | no | C2cd2 | intron_variant | MODIFIER |
| 16 | 97888989 | AX-170218176 | 0 | T | C | 0.465 | 3.73E-02 | 7.68E-02 | 2.77E+00 | 2.89E+00 | 6.19E-01 | no | C2cd2 | intron_variant | MODIFIER |

| | | | | | | | | | | | | | | | |
|---|---|---|---|---|---|---|---|---|---|---|---|---|---|---|---|
| 16 | 97890131 | AX-170513167 | 0 | G | A | 0.465 | 3.73E-02 | 7.68E-02 | 2.77E+00 | 2.89E+00 | 6.19E-01 | no | C2cd2 | intron_variant | MODIFIER |
| 16 | 97891336 | AX-170440344 | 0 | C | T | 0.465 | 3.73E-02 | 7.68E-02 | 2.77E+00 | 2.89E+00 | 6.19E-01 | no | C2cd2 | intron_variant | MODIFIER |
| 16 | 97897807 | AX-170291290 | 0 | A | G | 0.465 | 3.73E-02 | 7.68E-02 | 2.77E+00 | 2.89E+00 | 6.19E-01 | no | C2cd2 | intron_variant | MODIFIER |
| 16 | 97928261 | AX-170310884 | 0 | A | C | 0.465 | 3.73E-02 | 7.68E-02 | 2.77E+00 | 2.89E+00 | 6.19E-01 | no | C2cd2 | intron_variant | MODIFIER |
| 16 | 97938122 | AX-170158530 | 0 | G | A | 0.465 | 3.73E-02 | 7.68E-02 | 2.77E+00 | 2.89E+00 | 6.19E-01 | no | C2cd2 | intron_variant | MODIFIER |
| 16 | 97945680 | AX-170528654 | 0 | T | C | 0.465 | 3.73E-02 | 7.68E-02 | 2.77E+00 | 2.89E+00 | 6.19E-01 | no | Zbtb21 | downstream_gene_variant | MODIFIER |
| 16 | 97968214 | AX-97803304 | 0 | A | T | 0.465 | 3.73E-02 | 7.68E-02 | 2.77E+00 | 2.89E+00 | 6.19E-01 | no | | intergenic_variant | MODIFIER |
| 16 | 97985710 | AX-170483652 | 0 | G | A | 0.465 | 3.73E-02 | 7.68E-02 | 2.77E+00 | 2.89E+00 | 6.19E-01 | no | | intergenic_variant | MODIFIER |
| 16 | 97997969 | AX-170343041 | 0 | A | G | 0.465 | 3.73E-02 | 7.68E-02 | 2.77E+00 | 2.89E+00 | 6.19E-01 | no | B230307C23Rik | intron_variant | MODIFIER |
| 16 | 98022563 | AX-170490174 | 0 | A | C | 0.465 | 3.73E-02 | 7.68E-02 | 2.77E+00 | 2.89E+00 | 6.19E-01 | no | B230307C23Rik | intron_variant | MODIFIER |
| 16 | 98030190 | AX-170436200 | 0 | A | G | 0.465 | 3.73E-02 | 7.68E-02 | 2.77E+00 | 2.89E+00 | 6.19E-01 | no | B230307C23Rik | intron_variant | MODIFIER |
| 16 | 98069653 | AX-97843185 | 0 | C | T | 0.465 | 3.73E-02 | 7.68E-02 | 2.77E+00 | 2.89E+00 | 6.19E-01 | no | A630089N07Rik | intron_variant | MODIFIER |
| 7 | 98895722 | AX-170353019 | 1 | A | T | 0.125 | 9.36E-02 | 1.91E-01 | 2.65E+00 | 2.76E+00 | 6.19E-01 | no | Uvrag | intron_variant | MODIFIER |
| 3 | 31602368 | AX-170094568 | 1 | G | A | 0.143 | -4.99E-02 | 1.01E-01 | 2.65E+00 | 2.73E+00 | 6.19E-01 | no | | intergenic_variant | MODIFIER |
| 5 | 38604928 | AX-170255247 | 1 | T | A | 0.304 | 4.99E-02 | 1.02E-01 | 2.70E+00 | 2.79E+00 | 6.19E-01 | no | | intergenic_variant | MODIFIER |
| 9 | 85437201 | AX-170238637 | 0 | T | C | 0.211 | -5.29E-02 | 1.08E-01 | 2.67E+00 | 2.78E+00 | 6.19E-01 | no | | intergenic_variant | MODIFIER |
| 5 | 125961923 | AX-170287851 | 0 | C | A | 0.447 | -5.34E-02 | 1.09E-01 | 2.76E+00 | 2.82E+00 | 6.19E-01 | no | Gm42952 | upstream_gene_variant | MODIFIER |
| 8 | 39896957 | AX-170293173 | 2 | G | C | 0.155 | 7.11E-02 | 1.45E-01 | 2.78E+00 | 2.83E+00 | 6.19E-01 | no | | intergenic_variant | MODIFIER |
| 1 | 189487102 | AX-170119700 | 0 | A | G | 0.509 | -2.28E-02 | 4.65E-02 | 2.87E+00 | 3.04E+00 | 6.19E-01 | no | | intergenic_variant | MODIFIER |
| 1 | 16650794 | AX-97814054 | 0 | T | A | 0.386 | 3.89E-02 | 7.62E-02 | 2.55E+00 | 2.62E+00 | 6.19E-01 | no | Eloc | intron_variant | MODIFIER |
| 1 | 23633610 | AX-170376993 | 1 | T | G | 0.429 | 4.05E-02 | 8.18E-02 | 2.63E+00 | 2.71E+00 | 6.19E-01 | no | | intergenic_variant | MODIFIER |
| 10 | 6362322 | AX-170534740 | 0 | A | G | 0.211 | -4.76E-02 | 9.61E-02 | 2.61E+00 | 2.71E+00 | 6.19E-01 | no | | intergenic_variant | MODIFIER |
| 8 | 128419945 | AX-97782660 | 0 | G | A | 0.053 | -8.58E-02 | 1.76E-01 | 2.65E+00 | 2.77E+00 | 6.19E-01 | no | Nrp1 | intron_variant | MODIFIER |
| 1 | 100402483 | AX-170154129 | 1 | A | G | 0.152 | 9.30E-02 | 1.90E-01 | 2.67E+00 | 2.75E+00 | 6.19E-01 | no | Cntnap5b | intron_variant | MODIFIER |
| 14 | 112389778 | AX-170321571 | 0 | G | A | 0.316 | 3.66E-02 | 7.37E-02 | 2.60E+00 | 2.71E+00 | 6.19E-01 | no | | intergenic_variant | MODIFIER |
| 20 | 49231624 | AX-170492552 | 0 | T | A | 0.237 | 2.88E-02 | 5.83E-02 | 2.59E+00 | 2.72E+00 | 6.19E-01 | no | Enox2 | intron_variant | MODIFIER |
| 2 | 161279054 | AX-170301311 | 1 | G | C | 0.152 | 1.05E-01 | 2.14E-01 | 2.72E+00 | 2.77E+00 | 6.19E-01 | no | | intergenic_variant | MODIFIER |
| 8 | 83909204 | AX-170031146 | 0 | T | C | 0.202 | -6.28E-02 | 1.29E-01 | 2.73E+00 | 2.80E+00 | 6.20E-01 | no | Adgrl1 | intron_variant | MODIFIER |
| 8 | 85302041 | AX-170404024 | 0 | C | T | 0.202 | -6.28E-02 | 1.29E-01 | 2.73E+00 | 2.80E+00 | 6.20E-01 | no | Vps35 | upstream_gene_variant | MODIFIER |
| 10 | 12504147 | AX-170296265 | 0 | A | G | 0.211 | -5.16E-02 | 1.04E-01 | 2.62E+00 | 2.70E+00 | 6.20E-01 | no | Utrn | intron_variant | MODIFIER |
| 15 | 79152023 | AX-170327197 | 0 | C | T | 0.553 | 3.93E-02 | 7.82E-02 | 2.58E+00 | 2.66E+00 | 6.20E-01 | no | Sox10 | downstream_gene_variant | MODIFIER |
| 1 | 72059842 | AX-170370910 | 0 | T | C | 0.123 | -8.16E-02 | 1.69E-01 | 2.67E+00 | 2.81E+00 | 6.20E-01 | no | | regulatory_region_variant | MODIFIER |
| 12 | 60696702 | AX-170416325 | 0 | A | G | 0.07 | 1.00E-01 | 2.08E-01 | 2.72E+00 | 2.84E+00 | 6.20E-01 | no | | intergenic_variant | MODIFIER |
| 9 | 61218458 | AX-97821681 | 0 | G | A | 0.193 | -6.66E-02 | 1.36E-01 | 2.71E+00 | 2.74E+00 | 6.20E-01 | no | | intergenic_variant | MODIFIER |
| 9 | 77593567 | AX-170194024 | 0 | G | A | 0.193 | -6.66E-02 | 1.36E-01 | 2.71E+00 | 2.74E+00 | 6.20E-01 | no | | intergenic_variant | MODIFIER |
| 15 | 79133304 | AX-170056072 | 1 | G | A | 0.562 | 4.13E-02 | 8.18E-02 | 2.57E+00 | 2.64E+00 | 6.20E-01 | no | Micall1 | 3_prime_UTR_variant | MODIFIER |
| 1 | 145544042 | AX-170001918 | 0 | A | T | 0.211 | 6.79E-02 | 1.37E-01 | 2.64E+00 | 2.69E+00 | 6.20E-01 | no | | intergenic_variant | MODIFIER |
| 1 | 146681904 | AX-170001933 | 0 | C | A | 0.211 | 6.79E-02 | 1.37E-01 | 2.64E+00 | 2.69E+00 | 6.20E-01 | no | Brinp3 | intron_variant | MODIFIER |
| 1 | 152327819 | AX-170139512 | 0 | G | A | 0.211 | 6.79E-02 | 1.37E-01 | 2.64E+00 | 2.69E+00 | 6.20E-01 | no | | intergenic_variant | MODIFIER |
| 1 | 152694787 | AX-170383863 | 0 | A | T | 0.211 | 6.79E-02 | 1.37E-01 | 2.64E+00 | 2.69E+00 | 6.20E-01 | no | Rgl1 | intron_variant | MODIFIER |
| 1 | 105617440 | AX-170411092 | 0 | C | A | 0.158 | 1.02E-01 | 2.07E-01 | 2.64E+00 | 2.70E+00 | 6.20E-01 | no | Pign | intron_variant | MODIFIER |
| 8 | 98384838 | AX-170205783 | 0 | G | T | 0.237 | 5.35E-02 | 1.10E-01 | 2.71E+00 | 2.78E+00 | 6.20E-01 | no | | intergenic_variant | MODIFIER |
| 1 | 189357154 | AX-170462867 | 0 | C | T | 0.491 | 2.36E-02 | 4.84E-02 | 2.87E+00 | 3.03E+00 | 6.20E-01 | no | Kcnk2 | intron_variant,NMD_transcript_variant | MODIFIER |
| 5 | 117491015 | AX-170025969 | 0 | G | A | 0.447 | 5.47E-02 | 1.11E-01 | 2.67E+00 | 2.71E+00 | 6.20E-01 | no | Ksr2 | intron_variant,non_coding_transcript_variant | MODIFIER |
| 8 | 102123943 | AX-97813328 | 1 | A | G | 0.214 | 5.21E-02 | 1.07E-01 | 2.67E+00 | 2.76E+00 | 6.20E-01 | no | | intergenic_variant | MODIFIER |
| 20 | 87246422 | AX-97806596 | 0 | G | A | 0.254 | -2.97E-02 | 6.12E-02 | 2.69E+00 | 2.80E+00 | 6.21E-01 | no | Il1rapl1 | intron_variant | MODIFIER |
| 8 | 108663834 | AX-170367335 | 0 | C | T | 0.228 | 5.28E-02 | 1.08E-01 | 2.69E+00 | 2.77E+00 | 6.21E-01 | no | Zfhx3 | intron_variant | MODIFIER |
| 8 | 108671585 | AX-170374120 | 0 | C | G | 0.228 | 5.28E-02 | 1.08E-01 | 2.69E+00 | 2.77E+00 | 6.21E-01 | no | Zfhx3 | intron_variant | MODIFIER |
| 8 | 108672344 | AX-170212325 | 0 | G | C | 0.228 | 5.28E-02 | 1.08E-01 | 2.69E+00 | 2.77E+00 | 6.21E-01 | no | Zfhx3 | intron_variant | MODIFIER |
| 8 | 108674300 | AX-170081124 | 0 | T | G | 0.228 | 5.28E-02 | 1.08E-01 | 2.69E+00 | 2.77E+00 | 6.21E-01 | no | Zfhx3 | 5_prime_UTR_variant | MODIFIER |
| 8 | 109054851 | AX-170244270 | 0 | A | G | 0.228 | 5.28E-02 | 1.08E-01 | 2.69E+00 | 2.77E+00 | 6.21E-01 | no | | intergenic_variant | MODIFIER |
| 8 | 109068603 | AX-170114704 | 0 | T | C | 0.228 | 5.28E-02 | 1.08E-01 | 2.69E+00 | 2.77E+00 | 6.21E-01 | no | | intergenic_variant | MODIFIER |
| 8 | 109117788 | AX-170327014 | 0 | C | T | 0.228 | 5.28E-02 | 1.08E-01 | 2.69E+00 | 2.77E+00 | 6.21E-01 | no | D030068K23Rik | intron_variant,non_coding_transcript_variant | MODIFIER |
| 8 | 109371523 | AX-170523430 | 0 | T | C | 0.228 | 5.28E-02 | 1.08E-01 | 2.69E+00 | 2.77E+00 | 6.21E-01 | no | | intergenic_variant | MODIFIER |

| | | | | | | | | | | | | | | | |
|---|---|---|---|---|---|---|---|---|---|---|---|---|---|---|---|
| 8 | 109402386 | AX-170511838 | 0 | A | G | 0.228 | 5.28E-02 | 1.08E-01 | 2.69E+00 | 2.77E+00 | 6.21E-01 | no | | intergenic_variant | MODIFIER |
| 8 | 109405018 | AX-170428025 | 0 | A | G | 0.228 | 5.28E-02 | 1.08E-01 | 2.69E+00 | 2.77E+00 | 6.21E-01 | no | | intergenic_variant | MODIFIER |
| 8 | 109406313 | AX-170031470 | 0 | T | G | 0.228 | 5.28E-02 | 1.08E-01 | 2.69E+00 | 2.77E+00 | 6.21E-01 | no | | intergenic_variant | MODIFIER |
| 5 | 48686434 | AX-170403059 | 2 | T | C | 0.482 | 5.04E-02 | 9.43E-02 | 2.45E+00 | 2.54E+00 | 6.21E-01 | no | Kcnip4 | intron_variant | MODIFIER |
| 7 | 92113899 | AX-170127268 | 1 | C | T | 0.143 | 8.02E-02 | 1.66E-01 | 2.65E+00 | 2.78E+00 | 6.21E-01 | no | Dlg2 | intron_variant | MODIFIER |
| 15 | 92190329 | AX-170125563 | 1 | C | T | 0.134 | 7.64E-02 | 1.55E-01 | 2.65E+00 | 2.71E+00 | 6.21E-01 | no | Cntn1 | intron_variant | MODIFIER |
| 1 | 149121356 | AX-170452327 | 1 | T | C | 0.196 | 6.79E-02 | 1.38E-01 | 2.67E+00 | 2.72E+00 | 6.21E-01 | no | Gm29020 | intron_variant,non_coding_transcript_variant | MODIFIER |
| 7 | 82100577 | AX-170463289 | 1 | T | A | 0.125 | 7.19E-02 | 1.48E-01 | 2.63E+00 | 2.76E+00 | 6.21E-01 | no | | intergenic_variant | MODIFIER |
| 13 | 23569588 | AX-170007925 | 1 | C | T | 0.17 | -6.31E-02 | 1.21E-01 | 2.50E+00 | 2.56E+00 | 6.21E-01 | no | Hist1h2bg | upstream_gene_variant | MODIFIER |
| 10 | 41140217 | AX-170375984 | 0 | T | A | 0.395 | 3.54E-02 | 7.35E-02 | 2.72E+00 | 2.84E+00 | 6.21E-01 | no | Gm32338 | upstream_gene_variant | MODIFIER |
| 8 | 49333827 | AX-170127691 | 0 | C | T | 0.149 | -7.55E-02 | 1.50E-01 | 2.58E+00 | 2.63E+00 | 6.21E-01 | no | | intergenic_variant | MODIFIER |
| 2 | 132866633 | AX-97854069 | 1 | C | T | 0.161 | -8.26E-02 | 1.71E-01 | 2.74E+00 | 2.83E+00 | 6.21E-01 | no | Crls1 | 3_prime_UTR_variant | MODIFIER |
| 18 | 40958212 | AX-97812536 | 1 | C | T | 0.08 | 8.35E-02 | 1.70E-01 | 2.59E+00 | 2.72E+00 | 6.21E-01 | no | | intergenic_variant | MODIFIER |
| 1 | 189666443 | AX-170188603 | 0 | C | T | 0.254 | 5.24E-02 | 1.00E-01 | 2.48E+00 | 2.58E+00 | 6.21E-01 | no | Cenpf | intron_variant | MODIFIER |
| 5 | 100627296 | AX-170112906 | 1 | A | C | 0.42 | -4.99E-02 | 1.03E-01 | 2.76E+00 | 2.84E+00 | 6.21E-01 | no | | intergenic_variant | MODIFIER |
| 16 | 80362631 | AX-170296657 | 0 | C | G | 0.456 | 3.70E-02 | 7.43E-02 | 2.59E+00 | 2.67E+00 | 6.21E-01 | no | | intergenic_variant | MODIFIER |
| 1 | 116691447 | AX-170001492 | 1 | G | T | 0.17 | 9.79E-02 | 1.97E-01 | 2.59E+00 | 2.67E+00 | 6.21E-01 | no | | intergenic_variant | MODIFIER |
| 12 | 95409490 | AX-97817058 | 1 | C | T | 0.08 | 9.21E-02 | 1.90E-01 | 2.69E+00 | 2.78E+00 | 6.21E-01 | no | | intergenic_variant | MODIFIER |
| 9 | 22044418 | AX-170374868 | 1 | A | G | 0.455 | 4.47E-02 | 8.81E-02 | 2.55E+00 | 2.62E+00 | 6.21E-01 | no | Elavl3 | intron_variant | MODIFIER |
| 2 | 161325000 | AX-170086193 | 0 | T | A | 0.149 | 8.23E-02 | 1.68E-01 | 2.63E+00 | 2.74E+00 | 6.21E-01 | no | | intergenic_variant | MODIFIER |
| 3 | 150400683 | AX-170021911 | 1 | T | A | 0.33 | 3.61E-02 | 7.42E-02 | 2.67E+00 | 2.76E+00 | 6.21E-01 | no | | intergenic_variant | MODIFIER |
| 18 | 60154286 | AX-97796535 | 0 | A | G | 0.386 | 4.87E-02 | 9.40E-02 | 2.51E+00 | 2.57E+00 | 6.22E-01 | no | | regulatory_region_variant | MODIFIER |
| 4 | 57064790 | AX-97780572 | 0 | C | A | 0.368 | 5.33E-02 | 1.10E-01 | 2.73E+00 | 2.81E+00 | 6.22E-01 | no | Epb41l4b | intron_variant | MODIFIER |
| 20 | 72319780 | AX-97853716 | 1 | A | T | 0.259 | 3.06E-02 | 6.28E-02 | 2.66E+00 | 2.75E+00 | 6.22E-01 | no | | intergenic_variant | MODIFIER |
| 13 | 37104352 | AX-97814600 | 1 | C | T | 0.17 | -6.29E-02 | 1.25E-01 | 2.57E+00 | 2.63E+00 | 6.22E-01 | no | | regulatory_region_variant | MODIFIER |
| 15 | 85954353 | AX-170359910 | 1 | C | T | 0.509 | 3.50E-02 | 6.75E-02 | 2.50E+00 | 2.60E+00 | 6.22E-01 | no | Celsr1 | intron_variant | MODIFIER |
| 9 | 101007226 | AX-170233952 | 1 | T | C | 0.143 | -5.43E-02 | 1.11E-01 | 2.64E+00 | 2.73E+00 | 6.22E-01 | no | Pccb | intron_variant | MODIFIER |
| 10 | 92294493 | AX-170045579 | 0 | G | A | 0.298 | -4.26E-02 | 8.38E-02 | 2.54E+00 | 2.61E+00 | 6.22E-01 | no | Gm20757 | intron_variant,non_coding_transcript_variant | MODIFIER |
| 7 | 122666210 | AX-170204384 | 0 | C | T | 0.404 | 3.98E-02 | 8.24E-02 | 2.71E+00 | 2.80E+00 | 6.22E-01 | no | Cacng3 | upstream_gene_variant | MODIFIER |
| 13 | 51362383 | AX-170092747 | 0 | T | C | 0.509 | -3.14E-02 | 6.53E-02 | 2.72E+00 | 2.82E+00 | 6.22E-01 | no | | intergenic_variant | MODIFIER |
| 10 | 23225636 | AX-97815063 | 1 | T | G | 0.562 | -3.27E-02 | 6.87E-02 | 2.79E+00 | 2.94E+00 | 6.22E-01 | no | Eya4 | intron_variant | MODIFIER |
| 15 | 81995608 | AX-170146613 | 1 | G | A | 0.562 | 4.12E-02 | 8.21E-02 | 2.57E+00 | 2.64E+00 | 6.22E-01 | no | Desi1 | intron_variant,NMD_transcript_variant | MODIFIER |
| 5 | 89270944 | AX-170443447 | 1 | A | G | 0.42 | 5.14E-02 | 1.05E-01 | 2.65E+00 | 2.73E+00 | 6.22E-01 | no | | regulatory_region_variant | MODIFIER |
| 1 | 140917784 | AX-170298437 | 2 | G | T | 0.209 | 6.38E-02 | 1.30E-01 | 2.65E+00 | 2.71E+00 | 6.22E-01 | no | | intergenic_variant | MODIFIER |
| 8 | 88608225 | AX-170511906 | 0 | A | G | 0.175 | -6.07E-02 | 1.25E-01 | 2.65E+00 | 2.74E+00 | 6.22E-01 | no | | intergenic_variant | MODIFIER |
| 9 | 21251451 | AX-170272024 | 0 | A | G | 0.465 | 4.42E-02 | 8.85E-02 | 2.58E+00 | 2.65E+00 | 6.22E-01 | no | S1pr5 | upstream_gene_variant | MODIFIER |
| 1 | 103289402 | AX-170326772 | 0 | C | G | 0.149 | 9.73E-02 | 1.99E-01 | 2.64E+00 | 2.71E+00 | 6.23E-01 | no | | intergenic_variant | MODIFIER |
| 3 | 54576304 | AX-170443876 | 1 | C | T | 0.036 | -6.53E-02 | 1.34E-01 | 2.57E+00 | 2.73E+00 | 6.23E-01 | no | | intergenic_variant | MODIFIER |
| 8 | 36243008 | AX-97825636 | 1 | A | G | 0.152 | 7.32E-02 | 1.50E-01 | 2.81E+00 | 2.86E+00 | 6.23E-01 | no | Lonrf1 | intron_variant | MODIFIER |
| 8 | 41280236 | AX-170030560 | 1 | A | G | 0.152 | 7.32E-02 | 1.50E-01 | 2.81E+00 | 2.86E+00 | 6.23E-01 | no | Pcm1 | intron_variant | MODIFIER |
| 15 | 81552029 | AX-170412422 | 1 | A | T | 0.545 | 4.07E-02 | 8.08E-02 | 2.56E+00 | 2.63E+00 | 6.23E-01 | no | | intergenic_variant | MODIFIER |
| 6 | 138606368 | AX-97784082 | 0 | A | G | 0.553 | 4.30E-02 | 8.24E-02 | 2.49E+00 | 2.57E+00 | 6.23E-01 | no | | intergenic_variant | MODIFIER |
| 9 | 15778412 | AX-170128554 | 0 | G | A | 0.474 | 4.33E-02 | 8.60E-02 | 2.56E+00 | 2.63E+00 | 6.23E-01 | no | | intergenic_variant | MODIFIER |
| 15 | 81440460 | AX-170056115 | 0 | C | T | 0.553 | 4.03E-02 | 8.13E-02 | 2.60E+00 | 2.67E+00 | 6.23E-01 | no | Xpnpep3 | intron_variant | MODIFIER |
| 5 | 35349670 | AX-170151578 | 0 | A | G | 0.342 | 5.10E-02 | 1.05E-01 | 2.66E+00 | 2.72E+00 | 6.23E-01 | no | | intergenic_variant | MODIFIER |
| 3 | 28469150 | AX-170339938 | 1 | T | A | 0.179 | -4.83E-02 | 9.97E-02 | 2.69E+00 | 2.76E+00 | 6.23E-01 | no | Tnik | intron_variant | MODIFIER |
| 1 | 105963543 | AX-97782296 | 0 | G | A | 0.158 | 1.01E-01 | 2.06E-01 | 2.65E+00 | 2.71E+00 | 6.23E-01 | no | | intergenic_variant | MODIFIER |
| 2 | 74214667 | AX-97790743 | 0 | G | T | 0.053 | -9.17E-02 | 1.92E-01 | 2.69E+00 | 2.82E+00 | 6.23E-01 | no | | intergenic_variant | MODIFIER |
| 8 | 39099540 | AX-170369780 | 0 | C | T | 0.149 | 7.29E-02 | 1.50E-01 | 2.79E+00 | 2.84E+00 | 6.23E-01 | no | Tusc3 | intron_variant | MODIFIER |
| 16 | 97655672 | AX-170392931 | 1 | T | G | 0.464 | 3.68E-02 | 7.68E-02 | 2.77E+00 | 2.89E+00 | 6.23E-01 | no | | regulatory_region_variant | MODIFIER |
| 18 | 59089945 | AX-170274784 | 1 | C | T | 0.384 | 4.80E-02 | 9.35E-02 | 2.51E+00 | 2.59E+00 | 6.23E-01 | no | | intergenic_variant | MODIFIER |
| 8 | 65229608 | AX-170528041 | 1 | C | T | 0.143 | -7.22E-02 | 1.45E-01 | 2.57E+00 | 2.65E+00 | 6.23E-01 | no | | intergenic_variant | MODIFIER |
| 13 | 33498359 | AX-97823152 | 0 | T | C | 0.184 | -5.69E-02 | 1.15E-01 | 2.60E+00 | 2.67E+00 | 6.24E-01 | no | Serpinb9g | downstream_gene_variant | MODIFIER |
| 11 | 31718059 | AX-170233625 | 0 | A | G | 0.079 | 9.82E-02 | 2.02E-01 | 2.65E+00 | 2.73E+00 | 6.24E-01 | no | | regulatory_region_variant | MODIFIER |

| | | | | | | | | | | | | | | | |
|---|---|---|---|---|---|---|---|---|---|---|---|---|---|---|---|
| 1 | 89261915 | AX-97781675 | 0 | C | T | 0.158 | 1.01E-01 | 2.08E-01 | 2.65E+00 | 2.71E+00 | 6.24E-01 | no | Gm5259 | upstream_gene_variant | MODIFIER |
| 3 | 28336716 | AX-170329809 | 0 | G | A | 0.175 | -4.81E-02 | 9.93E-02 | 2.68E+00 | 2.75E+00 | 6.24E-01 | no | Tnik | intron_variant | MODIFIER |
| 3 | 28336761 | AX-170109629 | 0 | T | G | 0.175 | -4.81E-02 | 9.93E-02 | 2.68E+00 | 2.75E+00 | 6.24E-01 | no | Tnik | intron_variant | MODIFIER |
| 3 | 28336817 | AX-170211394 | 0 | A | G | 0.175 | -4.81E-02 | 9.93E-02 | 2.68E+00 | 2.75E+00 | 6.24E-01 | no | Tnik | intron_variant | MODIFIER |
| 3 | 28340245 | AX-97835158 | 0 | G | A | 0.175 | -4.81E-02 | 9.93E-02 | 2.68E+00 | 2.75E+00 | 6.24E-01 | no | Tnik | intron_variant | MODIFIER |
| 3 | 28382083 | AX-170020198 | 0 | A | G | 0.175 | -4.81E-02 | 9.93E-02 | 2.68E+00 | 2.75E+00 | 6.24E-01 | no | Tnik | intron_variant | MODIFIER |
| 3 | 28384923 | AX-170020200 | 0 | A | G | 0.175 | -4.81E-02 | 9.93E-02 | 2.68E+00 | 2.75E+00 | 6.24E-01 | no | Tnik | intron_variant | MODIFIER |
| 3 | 28385225 | AX-170170749 | 0 | G | C | 0.175 | -4.81E-02 | 9.93E-02 | 2.68E+00 | 2.75E+00 | 6.24E-01 | no | Tnik | intron_variant | MODIFIER |
| 3 | 28453475 | AX-97837244 | 0 | T | C | 0.175 | -4.81E-02 | 9.93E-02 | 2.68E+00 | 2.75E+00 | 6.24E-01 | no | Tnik | intron_variant | MODIFIER |
| 3 | 28458385 | AX-170280207 | 0 | T | C | 0.175 | -4.81E-02 | 9.93E-02 | 2.68E+00 | 2.75E+00 | 6.24E-01 | no | Tnik | intron_variant | MODIFIER |
| 3 | 28461004 | AX-170468677 | 0 | A | G | 0.175 | -4.81E-02 | 9.93E-02 | 2.68E+00 | 2.75E+00 | 6.24E-01 | no | Tnik | intron_variant | MODIFIER |
| 3 | 28461442 | AX-170274439 | 0 | G | C | 0.175 | -4.81E-02 | 9.93E-02 | 2.68E+00 | 2.75E+00 | 6.24E-01 | no | Tnik | intron_variant | MODIFIER |
| 3 | 28461999 | AX-170439110 | 0 | T | C | 0.175 | -4.81E-02 | 9.93E-02 | 2.68E+00 | 2.75E+00 | 6.24E-01 | no | Tnik | intron_variant | MODIFIER |
| 3 | 28462115 | AX-170314795 | 0 | A | G | 0.175 | -4.81E-02 | 9.93E-02 | 2.68E+00 | 2.75E+00 | 6.24E-01 | no | Tnik | intron_variant | MODIFIER |
| 3 | 28462154 | AX-170534360 | 0 | A | G | 0.175 | -4.81E-02 | 9.93E-02 | 2.68E+00 | 2.75E+00 | 6.24E-01 | no | Tnik | intron_variant | MODIFIER |
| 3 | 28463611 | AX-97836850 | 0 | T | C | 0.175 | -4.81E-02 | 9.93E-02 | 2.68E+00 | 2.75E+00 | 6.24E-01 | no | Tnik | intron_variant | MODIFIER |
| 3 | 28468867 | AX-170265726 | 0 | G | A | 0.175 | -4.81E-02 | 9.93E-02 | 2.68E+00 | 2.75E+00 | 6.24E-01 | no | Tnik | intron_variant | MODIFIER |
| 3 | 28498481 | AX-170439525 | 0 | T | C | 0.175 | -4.81E-02 | 9.93E-02 | 2.68E+00 | 2.75E+00 | 6.24E-01 | no | Tnik | intron_variant | MODIFIER |
| 3 | 28507859 | AX-97824969 | 0 | A | G | 0.175 | -4.81E-02 | 9.93E-02 | 2.68E+00 | 2.75E+00 | 6.24E-01 | no | Tnik | intron_variant | MODIFIER |
| 3 | 28543341 | AX-97795941 | 0 | T | C | 0.175 | -4.81E-02 | 9.93E-02 | 2.68E+00 | 2.75E+00 | 6.24E-01 | no | Tnik | intron_variant | MODIFIER |
| 3 | 28579745 | AX-97828033 | 0 | C | T | 0.175 | -4.81E-02 | 9.93E-02 | 2.68E+00 | 2.75E+00 | 6.24E-01 | no | Tnik | intron_variant | MODIFIER |
| 3 | 28581586 | AX-170529007 | 0 | A | C | 0.175 | -4.81E-02 | 9.93E-02 | 2.68E+00 | 2.75E+00 | 6.24E-01 | no | Tnik | intron_variant | MODIFIER |
| 3 | 28619571 | AX-170342089 | 0 | G | T | 0.175 | -4.81E-02 | 9.93E-02 | 2.68E+00 | 2.75E+00 | 6.24E-01 | no | Tnik | intron_variant | MODIFIER |
| 3 | 28620989 | AX-170141178 | 0 | A | G | 0.175 | -4.81E-02 | 9.93E-02 | 2.68E+00 | 2.75E+00 | 6.24E-01 | no | Tnik | synonymous_variant | LOW |
| 3 | 28629132 | AX-170144696 | 0 | C | G | 0.175 | -4.81E-02 | 9.93E-02 | 2.68E+00 | 2.75E+00 | 6.24E-01 | no | Tnik | intron_variant | MODIFIER |
| 3 | 28629998 | AX-170307698 | 0 | T | C | 0.175 | -4.81E-02 | 9.93E-02 | 2.68E+00 | 2.75E+00 | 6.24E-01 | no | Tnik | intron_variant | MODIFIER |
| 3 | 28638445 | AX-170131466 | 0 | T | C | 0.175 | -4.81E-02 | 9.93E-02 | 2.68E+00 | 2.75E+00 | 6.24E-01 | no | Tnik | synonymous_variant | LOW |
| 3 | 28641669 | AX-97826389 | 0 | A | G | 0.175 | -4.81E-02 | 9.93E-02 | 2.68E+00 | 2.75E+00 | 6.24E-01 | no | Tnik | intron_variant | MODIFIER |
| 3 | 28727955 | AX-170299376 | 0 | T | A | 0.175 | -4.81E-02 | 9.93E-02 | 2.68E+00 | 2.75E+00 | 6.24E-01 | no | Slc2a2 | 3_prime_UTR_variant | MODIFIER |
| 3 | 28734643 | AX-170518389 | 0 | T | G | 0.175 | -4.81E-02 | 9.93E-02 | 2.68E+00 | 2.75E+00 | 6.24E-01 | no | Slc2a2 | downstream_gene_variant | MODIFIER |
| 3 | 28736726 | AX-170133236 | 0 | C | T | 0.175 | -4.81E-02 | 9.93E-02 | 2.68E+00 | 2.75E+00 | 6.24E-01 | no | 1700112D23Rik | intron_variant,non_coding_transcript_variant | MODIFIER |
| 3 | 28737226 | AX-170533873 | 0 | C | T | 0.175 | -4.81E-02 | 9.93E-02 | 2.68E+00 | 2.75E+00 | 6.24E-01 | no | 1700112D23Rik | intron_variant,non_coding_transcript_variant | MODIFIER |
| 3 | 28796779 | AX-170385821 | 0 | T | G | 0.175 | -4.81E-02 | 9.93E-02 | 2.68E+00 | 2.75E+00 | 6.24E-01 | no | Eif5a2 | 3_prime_UTR_variant | MODIFIER |
| 3 | 28846518 | AX-97857268 | 0 | A | G | 0.175 | -4.81E-02 | 9.93E-02 | 2.68E+00 | 2.75E+00 | 6.24E-01 | no | | intergenic_variant | MODIFIER |
| 3 | 28989197 | AX-170204968 | 0 | T | C | 0.175 | -4.81E-02 | 9.93E-02 | 2.68E+00 | 2.75E+00 | 6.24E-01 | no | | intergenic_variant | MODIFIER |
| 3 | 29047646 | AX-170320059 | 0 | C | T | 0.175 | -4.81E-02 | 9.93E-02 | 2.68E+00 | 2.75E+00 | 6.24E-01 | no | | regulatory_region_variant | MODIFIER |
| 3 | 29070163 | AX-170281527 | 0 | C | T | 0.175 | -4.81E-02 | 9.93E-02 | 2.68E+00 | 2.75E+00 | 6.24E-01 | no | | intergenic_variant | MODIFIER |
| 3 | 29075012 | AX-170112871 | 0 | A | T | 0.175 | -4.81E-02 | 9.93E-02 | 2.68E+00 | 2.75E+00 | 6.24E-01 | no | | intergenic_variant | MODIFIER |
| 3 | 29082594 | AX-170293470 | 0 | T | G | 0.175 | -4.81E-02 | 9.93E-02 | 2.68E+00 | 2.75E+00 | 6.24E-01 | no | Egfem1 | upstream_gene_variant | MODIFIER |
| 3 | 29083152 | AX-170520114 | 0 | A | G | 0.175 | -4.81E-02 | 9.93E-02 | 2.68E+00 | 2.75E+00 | 6.24E-01 | no | Egfem1 | intron_variant | MODIFIER |
| 3 | 29104400 | AX-170528345 | 0 | A | G | 0.175 | -4.81E-02 | 9.93E-02 | 2.68E+00 | 2.75E+00 | 6.24E-01 | no | Egfem1 | intron_variant | MODIFIER |
| 3 | 29105772 | AX-170470964 | 0 | G | A | 0.175 | -4.81E-02 | 9.93E-02 | 2.68E+00 | 2.75E+00 | 6.24E-01 | no | Egfem1 | intron_variant | MODIFIER |
| 3 | 29105811 | AX-170501559 | 0 | T | G | 0.175 | -4.81E-02 | 9.93E-02 | 2.68E+00 | 2.75E+00 | 6.24E-01 | no | Egfem1 | intron_variant | MODIFIER |
| 3 | 29105957 | AX-170201296 | 0 | G | A | 0.175 | -4.81E-02 | 9.93E-02 | 2.68E+00 | 2.75E+00 | 6.24E-01 | no | Egfem1 | intron_variant | MODIFIER |
| 3 | 29110757 | AX-170495784 | 0 | T | A | 0.175 | -4.81E-02 | 9.93E-02 | 2.68E+00 | 2.75E+00 | 6.24E-01 | no | Egfem1 | intron_variant | MODIFIER |
| 3 | 29110918 | AX-170256542 | 0 | C | A | 0.175 | -4.81E-02 | 9.93E-02 | 2.68E+00 | 2.75E+00 | 6.24E-01 | no | Egfem1 | intron_variant | MODIFIER |
| 3 | 29430991 | AX-97854139 | 0 | A | G | 0.175 | -4.81E-02 | 9.93E-02 | 2.68E+00 | 2.75E+00 | 6.24E-01 | no | Egfem1 | intron_variant | MODIFIER |
| 3 | 29431035 | AX-170020210 | 0 | A | G | 0.175 | -4.81E-02 | 9.93E-02 | 2.68E+00 | 2.75E+00 | 6.24E-01 | no | Egfem1 | intron_variant | MODIFIER |
| 3 | 29434256 | AX-97795942 | 0 | G | A | 0.175 | -4.81E-02 | 9.93E-02 | 2.68E+00 | 2.75E+00 | 6.24E-01 | no | Egfem1 | intron_variant | MODIFIER |
| 3 | 29460861 | AX-97802041 | 0 | C | T | 0.175 | -4.81E-02 | 9.93E-02 | 2.68E+00 | 2.75E+00 | 6.24E-01 | no | Egfem1 | intron_variant | MODIFIER |
| 3 | 29486812 | AX-170020213 | 0 | T | C | 0.175 | -4.81E-02 | 9.93E-02 | 2.68E+00 | 2.75E+00 | 6.24E-01 | no | Egfem1 | intron_variant | MODIFIER |
| 3 | 29500111 | AX-170020212 | 0 | C | T | 0.175 | -4.81E-02 | 9.93E-02 | 2.68E+00 | 2.75E+00 | 6.24E-01 | no | Egfem1 | intron_variant | MODIFIER |
| 3 | 29533841 | AX-170020206 | 0 | G | A | 0.175 | -4.81E-02 | 9.93E-02 | 2.68E+00 | 2.75E+00 | 6.24E-01 | no | Egfem1 | intron_variant | MODIFIER |
| 3 | 29541814 | AX-97801963 | 0 | T | C | 0.175 | -4.81E-02 | 9.93E-02 | 2.68E+00 | 2.75E+00 | 6.24E-01 | no | Egfem1 | intron_variant | MODIFIER |

| | | | | | | | | | | | | | | | |
|---|---|---|---|---|---|---|---|---|---|---|---|---|---|---|---|
| 3 | 29542035 | AX-170264200 | 0 | A | G | 0.175 | -4.81E-02 | 9.93E-02 | 2.68E+00 | 2.75E+00 | 6.24E-01 | no | Egfem1 | intron_variant | MODIFIER |
| 3 | 29579377 | AX-170020205 | 0 | A | G | 0.175 | -4.81E-02 | 9.93E-02 | 2.68E+00 | 2.75E+00 | 6.24E-01 | no | Egfem1 | intron_variant | MODIFIER |
| 3 | 29586549 | AX-170493044 | 0 | T | C | 0.175 | -4.81E-02 | 9.93E-02 | 2.68E+00 | 2.75E+00 | 6.24E-01 | no | Egfem1 | intron_variant | MODIFIER |
| 3 | 29588831 | AX-170281715 | 0 | C | T | 0.175 | -4.81E-02 | 9.93E-02 | 2.68E+00 | 2.75E+00 | 6.24E-01 | no | Egfem1 | intron_variant | MODIFIER |
| 3 | 29594742 | AX-170410750 | 0 | C | A | 0.175 | -4.81E-02 | 9.93E-02 | 2.68E+00 | 2.75E+00 | 6.24E-01 | no | Egfem1 | intron_variant | MODIFIER |
| 3 | 29595464 | AX-170168463 | 0 | A | C | 0.175 | -4.81E-02 | 9.93E-02 | 2.68E+00 | 2.75E+00 | 6.24E-01 | no | Egfem1 | intron_variant | MODIFIER |
| 3 | 29759236 | AX-170277450 | 0 | G | A | 0.175 | -4.81E-02 | 9.93E-02 | 2.68E+00 | 2.75E+00 | 6.24E-01 | no | | intergenic_variant | MODIFIER |
| 3 | 29765977 | AX-170220152 | 0 | C | A | 0.175 | -4.81E-02 | 9.93E-02 | 2.68E+00 | 2.75E+00 | 6.24E-01 | no | | intergenic_variant | MODIFIER |
| 3 | 29766405 | AX-170196296 | 0 | G | A | 0.175 | -4.81E-02 | 9.93E-02 | 2.68E+00 | 2.75E+00 | 6.24E-01 | no | Gm37557 | upstream_gene_variant | MODIFIER |
| 3 | 29766760 | AX-170236491 | 0 | G | A | 0.175 | -4.81E-02 | 9.93E-02 | 2.68E+00 | 2.75E+00 | 6.24E-01 | no | Gm37557 | upstream_gene_variant | MODIFIER |
| 3 | 29787029 | AX-170430945 | 0 | A | T | 0.175 | -4.81E-02 | 9.93E-02 | 2.68E+00 | 2.75E+00 | 6.24E-01 | no | Gm33206 | intron_variant,non_coding_transcript_variant | MODIFIER |
| 3 | 29829308 | AX-170066223 | 0 | G | A | 0.175 | -4.81E-02 | 9.93E-02 | 2.68E+00 | 2.75E+00 | 6.24E-01 | no | Gm33206 | intron_variant,non_coding_transcript_variant | MODIFIER |
| 3 | 29830725 | AX-170328061 | 0 | G | A | 0.175 | -4.81E-02 | 9.93E-02 | 2.68E+00 | 2.75E+00 | 6.24E-01 | no | Gm33206 | intron_variant,non_coding_transcript_variant | MODIFIER |
| 3 | 29902445 | AX-170314348 | 0 | T | G | 0.175 | -4.81E-02 | 9.93E-02 | 2.68E+00 | 2.75E+00 | 6.24E-01 | no | Mannr | intron_variant,non_coding_transcript_variant | MODIFIER |
| 3 | 30060808 | AX-170202786 | 0 | C | A | 0.175 | -4.81E-02 | 9.93E-02 | 2.68E+00 | 2.75E+00 | 6.24E-01 | no | Mecom | intron_variant | MODIFIER |
| 3 | 30060975 | AX-170461512 | 0 | C | T | 0.175 | -4.81E-02 | 9.93E-02 | 2.68E+00 | 2.75E+00 | 6.24E-01 | no | Mecom | intron_variant | MODIFIER |
| 3 | 30216139 | AX-97855144 | 0 | C | A | 0.175 | -4.81E-02 | 9.93E-02 | 2.68E+00 | 2.75E+00 | 6.24E-01 | no | Mecom | intron_variant | MODIFIER |
| 3 | 30216562 | AX-170378923 | 0 | C | T | 0.175 | -4.81E-02 | 9.93E-02 | 2.68E+00 | 2.75E+00 | 6.24E-01 | no | Mecom | intron_variant | MODIFIER |
| 3 | 30333842 | AX-170262095 | 0 | G | A | 0.175 | -4.81E-02 | 9.93E-02 | 2.68E+00 | 2.75E+00 | 6.24E-01 | no | Mecom | intron_variant | MODIFIER |
| 3 | 30335391 | AX-170526490 | 0 | A | G | 0.175 | -4.81E-02 | 9.93E-02 | 2.68E+00 | 2.75E+00 | 6.24E-01 | no | Mecom | intron_variant | MODIFIER |
| 3 | 30336223 | AX-170428703 | 0 | C | T | 0.175 | -4.81E-02 | 9.93E-02 | 2.68E+00 | 2.75E+00 | 6.24E-01 | no | Mecom | intron_variant | MODIFIER |
| 3 | 30339938 | AX-170099826 | 0 | C | G | 0.175 | -4.81E-02 | 9.93E-02 | 2.68E+00 | 2.75E+00 | 6.24E-01 | no | Mecom | intron_variant | MODIFIER |
| 3 | 30342868 | AX-170419363 | 0 | A | T | 0.175 | -4.81E-02 | 9.93E-02 | 2.68E+00 | 2.75E+00 | 6.24E-01 | no | Mecom | intron_variant | MODIFIER |
| 3 | 30351302 | AX-170371243 | 0 | A | G | 0.175 | -4.81E-02 | 9.93E-02 | 2.68E+00 | 2.75E+00 | 6.24E-01 | no | Mecom | intron_variant | MODIFIER |
| 3 | 30353561 | AX-170400011 | 0 | C | T | 0.175 | -4.81E-02 | 9.93E-02 | 2.68E+00 | 2.75E+00 | 6.24E-01 | no | Mecom | intron_variant | MODIFIER |
| 3 | 30359967 | AX-170169580 | 0 | G | T | 0.175 | -4.81E-02 | 9.93E-02 | 2.68E+00 | 2.75E+00 | 6.24E-01 | no | Mecom | intron_variant | MODIFIER |
| 3 | 30651845 | AX-170336465 | 0 | G | A | 0.175 | -4.81E-02 | 9.93E-02 | 2.68E+00 | 2.75E+00 | 6.24E-01 | no | Lrrc34 | upstream_gene_variant | MODIFIER |
| 3 | 30658187 | AX-170336189 | 0 | T | C | 0.175 | -4.81E-02 | 9.93E-02 | 2.68E+00 | 2.75E+00 | 6.24E-01 | no | Lrriq4 | intron_variant | MODIFIER |
| 3 | 30674462 | AX-170473999 | 0 | C | T | 0.175 | -4.81E-02 | 9.93E-02 | 2.68E+00 | 2.75E+00 | 6.24E-01 | no | Lrriq4 | downstream_gene_variant | MODIFIER |
| 3 | 30681194 | AX-170328485 | 0 | G | T | 0.175 | -4.81E-02 | 9.93E-02 | 2.68E+00 | 2.75E+00 | 6.24E-01 | no | Lrrc31 | downstream_gene_variant | MODIFIER |
| 3 | 30719772 | AX-170088902 | 0 | A | C | 0.175 | -4.81E-02 | 9.93E-02 | 2.68E+00 | 2.75E+00 | 6.24E-01 | no | | intergenic_variant | MODIFIER |
| 3 | 30719810 | AX-170511867 | 0 | A | G | 0.175 | -4.81E-02 | 9.93E-02 | 2.68E+00 | 2.75E+00 | 6.24E-01 | no | | intergenic_variant | MODIFIER |
| 3 | 30756519 | AX-97808995 | 0 | G | A | 0.175 | -4.81E-02 | 9.93E-02 | 2.68E+00 | 2.75E+00 | 6.24E-01 | no | Samd7 | synonymous_variant | LOW |
| 3 | 30756582 | AX-170239690 | 0 | A | G | 0.175 | -4.81E-02 | 9.93E-02 | 2.68E+00 | 2.75E+00 | 6.24E-01 | no | Samd7 | synonymous_variant | LOW |
| 3 | 30944972 | AX-170429845 | 0 | T | C | 0.175 | -4.81E-02 | 9.93E-02 | 2.68E+00 | 2.75E+00 | 6.24E-01 | no | Phc3 | intron_variant,NMD_transcript_variant | MODIFIER |
| 3 | 31223622 | AX-170066250 | 0 | C | T | 0.175 | -4.81E-02 | 9.93E-02 | 2.68E+00 | 2.75E+00 | 6.24E-01 | no | Slc7a14 | synonymous_variant | LOW |
| 3 | 31277954 | AX-170512474 | 0 | T | C | 0.175 | -4.81E-02 | 9.93E-02 | 2.68E+00 | 2.75E+00 | 6.24E-01 | no | Slc7a14 | intron_variant | MODIFIER |
| 3 | 31368143 | AX-170436853 | 0 | T | C | 0.175 | -4.81E-02 | 9.93E-02 | 2.68E+00 | 2.75E+00 | 6.24E-01 | no | | intergenic_variant | MODIFIER |
| 3 | 31682676 | AX-170399866 | 0 | G | T | 0.175 | -4.81E-02 | 9.93E-02 | 2.68E+00 | 2.75E+00 | 6.24E-01 | no | Gm6558 | intron_variant,non_coding_transcript_variant | MODIFIER |
| 5 | 121363336 | AX-97847179 | 0 | C | T | 0.447 | -5.30E-02 | 1.09E-01 | 2.79E+00 | 2.85E+00 | 6.24E-01 | no | Hectd4 | intron_variant | MODIFIER |
| 16 | 78209170 | AX-170135374 | 1 | T | C | 0.446 | 3.74E-02 | 7.60E-02 | 2.60E+00 | 2.68E+00 | 6.24E-01 | no | | intergenic_variant | MODIFIER |
| 1 | 44733533 | AX-170000400 | 1 | T | C | 0.098 | -1.02E-01 | 2.11E-01 | 2.71E+00 | 2.80E+00 | 6.24E-01 | no | Gulp1 | intron_variant | MODIFIER |
| 16 | 44870928 | AX-97813948 | 0 | C | T | 0.351 | -5.21E-02 | 1.04E-01 | 2.53E+00 | 2.66E+00 | 6.24E-01 | no | Cd200r2 | intron_variant | MODIFIER |
| 5 | 117602348 | AX-170284714 | 0 | T | C | 0.456 | -5.27E-02 | 1.09E-01 | 2.76E+00 | 2.83E+00 | 6.24E-01 | no | Ksr2 | intron_variant | MODIFIER |
| 1 | 101391117 | AX-170146080 | 0 | C | A | 0.149 | 9.76E-02 | 2.00E-01 | 2.64E+00 | 2.71E+00 | 6.24E-01 | no | Psmb6-ps2 | downstream_gene_variant | MODIFIER |
| 15 | 70244699 | AX-170522918 | 0 | A | T | 0.544 | 3.82E-02 | 7.69E-02 | 2.57E+00 | 2.66E+00 | 6.24E-01 | no | | intergenic_variant | MODIFIER |
| 13 | 23546011 | AX-170007924 | 1 | C | G | 0.17 | -6.22E-02 | 1.20E-01 | 2.50E+00 | 2.56E+00 | 6.25E-01 | no | Hist1h3f | downstream_gene_variant | MODIFIER |
| 5 | 127379036 | AX-170102828 | 0 | C | T | 0.421 | -3.91E-02 | 8.21E-02 | 2.71E+00 | 2.83E+00 | 6.25E-01 | no | Tmem132c | intron_variant | MODIFIER |
| 5 | 86415453 | AX-170390493 | 1 | G | T | 0.438 | 5.12E-02 | 1.06E-01 | 2.71E+00 | 2.77E+00 | 6.25E-01 | no | Tmprss11a | intron_variant | MODIFIER |
| 3 | 31769389 | AX-170066259 | 0 | C | A | 0.167 | -4.78E-02 | 9.97E-02 | 2.71E+00 | 2.79E+00 | 6.25E-01 | no | | regulatory_region_variant | MODIFIER |
| 13 | 34149951 | AX-97833690 | 0 | C | T | 0.184 | -5.87E-02 | 1.18E-01 | 2.58E+00 | 2.64E+00 | 6.25E-01 | no | | regulatory_region_variant | MODIFIER |
| 13 | 34156833 | AX-170389652 | 0 | C | T | 0.184 | -5.87E-02 | 1.18E-01 | 2.58E+00 | 2.64E+00 | 6.25E-01 | no | | regulatory_region_variant | MODIFIER |
| 13 | 34157473 | AX-170302886 | 0 | A | C | 0.184 | -5.87E-02 | 1.18E-01 | 2.58E+00 | 2.64E+00 | 6.25E-01 | no | | regulatory_region_variant | MODIFIER |
| 13 | 34157591 | AX-170478063 | 0 | G | A | 0.184 | -5.87E-02 | 1.18E-01 | 2.58E+00 | 2.64E+00 | 6.25E-01 | no | | regulatory_region_variant | MODIFIER |

| | | | | | | | | | | | | | | | |
|---|---|---|---|---|---|---|---|---|---|---|---|---|---|---|---|
| 13 | 34162404 | AX-170295782 | 0 | T | C | 0.184 | -5.87E-02 | 1.18E-01 | 2.58E+00 | 2.64E+00 | 6.25E-01 | no | Psmg4 | upstream_gene_variant | MODIFIER |
| 13 | 34166619 | AX-97789159 | 0 | A | G | 0.184 | -5.87E-02 | 1.18E-01 | 2.58E+00 | 2.64E+00 | 6.25E-01 | no | Psmg4 | intron_variant | MODIFIER |
| 13 | 34186681 | AX-170528387 | 0 | A | G | 0.184 | -5.87E-02 | 1.18E-01 | 2.58E+00 | 2.64E+00 | 6.25E-01 | no | Slc22a23 | intron_variant | MODIFIER |
| 13 | 34187059 | AX-170384340 | 0 | T | C | 0.184 | -5.87E-02 | 1.18E-01 | 2.58E+00 | 2.64E+00 | 6.25E-01 | no | Slc22a23 | intron_variant | MODIFIER |
| 13 | 34187313 | AX-170050923 | 0 | C | T | 0.184 | -5.87E-02 | 1.18E-01 | 2.58E+00 | 2.64E+00 | 6.25E-01 | no | Slc22a23 | intron_variant | MODIFIER |
| 13 | 34191462 | AX-168400104 | 0 | A | G | 0.184 | -5.87E-02 | 1.18E-01 | 2.58E+00 | 2.64E+00 | 6.25E-01 | no | Slc22a23 | intron_variant | MODIFIER |
| 13 | 34192770 | AX-170254096 | 0 | C | A | 0.184 | -5.87E-02 | 1.18E-01 | 2.58E+00 | 2.64E+00 | 6.25E-01 | no | Slc22a23 | intron_variant | MODIFIER |
| 13 | 34193947 | AX-97809677 | 0 | G | A | 0.184 | -5.87E-02 | 1.18E-01 | 2.58E+00 | 2.64E+00 | 6.25E-01 | no | Slc22a23 | intron_variant | MODIFIER |
| 13 | 34197822 | AX-170527372 | 0 | A | G | 0.184 | -5.87E-02 | 1.18E-01 | 2.58E+00 | 2.64E+00 | 6.25E-01 | no | Slc22a23 | synonymous_variant | LOW |
| 13 | 34212565 | AX-170371501 | 0 | C | A | 0.184 | -5.87E-02 | 1.18E-01 | 2.58E+00 | 2.64E+00 | 6.25E-01 | no | Slc22a23 | intron_variant | MODIFIER |
| 13 | 34213421 | AX-97813357 | 0 | A | G | 0.184 | -5.87E-02 | 1.18E-01 | 2.58E+00 | 2.64E+00 | 6.25E-01 | no | Slc22a23 | intron_variant | MODIFIER |
| 13 | 34221044 | AX-170008077 | 0 | A | T | 0.184 | -5.87E-02 | 1.18E-01 | 2.58E+00 | 2.64E+00 | 6.25E-01 | no | Slc22a23 | intron_variant | MODIFIER |
| 13 | 34253476 | AX-97827102 | 0 | C | T | 0.184 | -5.87E-02 | 1.18E-01 | 2.58E+00 | 2.64E+00 | 6.25E-01 | no | Slc22a23 | intron_variant | MODIFIER |
| 13 | 34267658 | AX-97826018 | 0 | A | G | 0.184 | -5.87E-02 | 1.18E-01 | 2.58E+00 | 2.64E+00 | 6.25E-01 | no | Slc22a23 | intron_variant | MODIFIER |
| 13 | 34296985 | AX-97827101 | 0 | T | A | 0.184 | -5.87E-02 | 1.18E-01 | 2.58E+00 | 2.64E+00 | 6.25E-01 | no | Slc22a23 | intron_variant | MODIFIER |
| 13 | 34313477 | AX-97823178 | 0 | T | G | 0.184 | -5.87E-02 | 1.18E-01 | 2.58E+00 | 2.64E+00 | 6.25E-01 | no | Slc22a23 | intron_variant | MODIFIER |
| 13 | 34338262 | AX-97827107 | 0 | G | A | 0.184 | -5.87E-02 | 1.18E-01 | 2.58E+00 | 2.64E+00 | 6.25E-01 | no | Slc22a23 | intron_variant | MODIFIER |
| 13 | 34371344 | AX-97818318 | 0 | T | C | 0.184 | -5.87E-02 | 1.18E-01 | 2.58E+00 | 2.64E+00 | 6.25E-01 | no | | intergenic_variant | MODIFIER |
| 13 | 34405788 | AX-97846845 | 0 | A | G | 0.184 | -5.87E-02 | 1.18E-01 | 2.58E+00 | 2.64E+00 | 6.25E-01 | no | | intergenic_variant | MODIFIER |
| 13 | 34490885 | AX-97813585 | 0 | G | A | 0.184 | -5.87E-02 | 1.18E-01 | 2.58E+00 | 2.64E+00 | 6.25E-01 | no | | intergenic_variant | MODIFIER |
| 13 | 34662733 | AX-97848780 | 0 | C | T | 0.184 | -5.87E-02 | 1.18E-01 | 2.58E+00 | 2.64E+00 | 6.25E-01 | no | Gm15908 | intron_variant,non_coding_transcript_variant | MODIFIER |
| 13 | 35385671 | AX-97783074 | 0 | C | A | 0.184 | -5.87E-02 | 1.18E-01 | 2.58E+00 | 2.64E+00 | 6.25E-01 | no | | intergenic_variant | MODIFIER |
| 13 | 35417935 | AX-170008097 | 0 | A | G | 0.184 | -5.87E-02 | 1.18E-01 | 2.58E+00 | 2.64E+00 | 6.25E-01 | no | Gm48703 | intron_variant,non_coding_transcript_variant | MODIFIER |
| 13 | 35423657 | AX-97823176 | 0 | C | T | 0.184 | -5.87E-02 | 1.18E-01 | 2.58E+00 | 2.64E+00 | 6.25E-01 | no | Gm48703 | intron_variant,non_coding_transcript_variant | MODIFIER |
| 13 | 35424244 | AX-170008091 | 0 | T | C | 0.184 | -5.87E-02 | 1.18E-01 | 2.58E+00 | 2.64E+00 | 6.25E-01 | no | Gm48703 | intron_variant,non_coding_transcript_variant | MODIFIER |
| 13 | 35429470 | AX-170326323 | 0 | G | A | 0.184 | -5.87E-02 | 1.18E-01 | 2.58E+00 | 2.64E+00 | 6.25E-01 | no | Gm48703 | intron_variant,non_coding_transcript_variant | MODIFIER |
| 13 | 35435759 | AX-97836366 | 0 | A | G | 0.184 | -5.87E-02 | 1.18E-01 | 2.58E+00 | 2.64E+00 | 6.25E-01 | no | Gm48703 | intron_variant,non_coding_transcript_variant | MODIFIER |
| 13 | 36004269 | AX-170008112 | 0 | C | T | 0.184 | -5.87E-02 | 1.18E-01 | 2.58E+00 | 2.64E+00 | 6.25E-01 | no | Lyrm4 | intron_variant | MODIFIER |
| 1 | 127457636 | AX-97824760 | 0 | G | A | 0.553 | 4.14E-02 | 8.48E-02 | 2.62E+00 | 2.69E+00 | 6.25E-01 | no | Mgat5 | intron_variant | MODIFIER |
| 8 | 39321869 | AX-170113565 | 1 | A | C | 0.152 | 7.22E-02 | 1.49E-01 | 2.79E+00 | 2.85E+00 | 6.25E-01 | no | Gm6213 | intron_variant,non_coding_transcript_variant | MODIFIER |
| 5 | 6503984 | AX-170306779 | 1 | G | A | 0.446 | 3.10E-02 | 6.58E-02 | 2.70E+00 | 2.85E+00 | 6.25E-01 | no | | intergenic_variant | MODIFIER |
| 20 | 116231726 | AX-170318175 | 0 | A | G | 0.307 | 2.46E-02 | 5.07E-02 | 2.59E+00 | 2.72E+00 | 6.25E-01 | no | | intergenic_variant | MODIFIER |
| 14 | 67283822 | AX-170509341 | 1 | T | C | 0.045 | -9.53E-02 | 1.91E-01 | 2.55E+00 | 2.65E+00 | 6.25E-01 | no | Ebf2 | intron_variant | MODIFIER |
| 6 | 125110244 | AX-170404813 | 0 | A | G | 0.167 | -3.98E-02 | 8.35E-02 | 2.67E+00 | 2.80E+00 | 6.25E-01 | no | Chd4 | intron_variant | MODIFIER |
| 6 | 125512069 | AX-170028171 | 0 | T | C | 0.167 | -3.98E-02 | 8.35E-02 | 2.67E+00 | 2.80E+00 | 6.25E-01 | no | Gm26728 | downstream_gene_variant | MODIFIER |
| 6 | 125558257 | AX-170137595 | 0 | T | C | 0.167 | -3.98E-02 | 8.35E-02 | 2.67E+00 | 2.80E+00 | 6.25E-01 | no | Vwf | intron_variant | MODIFIER |
| 6 | 125565870 | AX-170533911 | 0 | C | T | 0.167 | -3.98E-02 | 8.35E-02 | 2.67E+00 | 2.80E+00 | 6.25E-01 | no | Vwf | missense_variant | MODERATE |
| 6 | 125687763 | AX-170316587 | 0 | G | A | 0.167 | -3.98E-02 | 8.35E-02 | 2.67E+00 | 2.80E+00 | 6.25E-01 | no | Vwf | downstream_gene_variant | MODIFIER |
| 6 | 125772261 | AX-170491142 | 0 | G | A | 0.167 | -3.98E-02 | 8.35E-02 | 2.67E+00 | 2.80E+00 | 6.25E-01 | no | Ano2 | intron_variant,non_coding_transcript_variant | MODIFIER |
| 6 | 125774111 | AX-97842233 | 0 | T | C | 0.167 | -3.98E-02 | 8.35E-02 | 2.67E+00 | 2.80E+00 | 6.25E-01 | no | Ano2 | intron_variant,non_coding_transcript_variant | MODIFIER |
| 6 | 125854758 | AX-170028181 | 0 | G | T | 0.167 | -3.98E-02 | 8.35E-02 | 2.67E+00 | 2.80E+00 | 6.25E-01 | no | Ano2 | intron_variant,non_coding_transcript_variant | MODIFIER |
| 6 | 125864154 | AX-170219520 | 0 | G | C | 0.167 | -3.98E-02 | 8.35E-02 | 2.67E+00 | 2.80E+00 | 6.25E-01 | no | Ano2 | intron_variant,non_coding_transcript_variant | MODIFIER |
| 6 | 126089711 | AX-170363083 | 0 | T | C | 0.167 | -3.98E-02 | 8.35E-02 | 2.67E+00 | 2.80E+00 | 6.25E-01 | no | | intergenic_variant | MODIFIER |
| 6 | 126113800 | AX-97790289 | 0 | A | G | 0.167 | -3.98E-02 | 8.35E-02 | 2.67E+00 | 2.80E+00 | 6.25E-01 | no | Ntf3 | intron_variant | MODIFIER |
| 6 | 126137280 | AX-170497068 | 0 | G | A | 0.167 | -3.98E-02 | 8.35E-02 | 2.67E+00 | 2.80E+00 | 6.25E-01 | no | Ntf3 | intron_variant | MODIFIER |
| 6 | 126138148 | AX-170373808 | 0 | G | A | 0.167 | -3.98E-02 | 8.35E-02 | 2.67E+00 | 2.80E+00 | 6.25E-01 | no | Ntf3 | intron_variant | MODIFIER |
| 6 | 126147746 | AX-170185287 | 0 | T | C | 0.167 | -3.98E-02 | 8.35E-02 | 2.67E+00 | 2.80E+00 | 6.25E-01 | no | Ntf3 | intron_variant | MODIFIER |
| 6 | 126231067 | AX-97852959 | 0 | T | C | 0.167 | -3.98E-02 | 8.35E-02 | 2.67E+00 | 2.80E+00 | 6.25E-01 | no | | intergenic_variant | MODIFIER |
| 6 | 126236750 | AX-170424003 | 0 | T | C | 0.167 | -3.98E-02 | 8.35E-02 | 2.67E+00 | 2.80E+00 | 6.25E-01 | no | | intergenic_variant | MODIFIER |
| 6 | 126245208 | AX-170225062 | 0 | G | A | 0.167 | -3.98E-02 | 8.35E-02 | 2.67E+00 | 2.80E+00 | 6.25E-01 | no | | intergenic_variant | MODIFIER |
| 6 | 126246029 | AX-170169678 | 0 | T | C | 0.167 | -3.98E-02 | 8.35E-02 | 2.67E+00 | 2.80E+00 | 6.25E-01 | no | | intergenic_variant | MODIFIER |
| 6 | 126325674 | AX-97791207 | 0 | C | T | 0.167 | -3.98E-02 | 8.35E-02 | 2.67E+00 | 2.80E+00 | 6.25E-01 | no | Gm10415 | intron_variant,non_coding_transcript_variant | MODIFIER |
| 5 | 122678112 | AX-170026049 | 0 | G | A | 0.456 | -5.37E-02 | 1.11E-01 | 2.79E+00 | 2.85E+00 | 6.26E-01 | no | P2rx7 | intron_variant | MODIFIER |
| 9 | 21397605 | AX-170253543 | 1 | T | C | 0.455 | 4.34E-02 | 8.82E-02 | 2.59E+00 | 2.67E+00 | 6.26E-01 | no | Ilf3 | synonymous_variant | LOW |

| | | | | | | | | | | | | | | | |
|---|---|---|---|---|---|---|---|---|---|---|---|---|---|---|---|
| 1 | 66704896 | AX-170414306 | 1 | C | T | 0.116 | -7.33E-02 | 1.46E-01 | 2.53E+00 | 2.65E+00 | 6.26E-01 | no | Rpe | intron_variant | MODIFIER |
| 5 | 125502504 | AX-170400878 | 2 | A | G | 0.445 | -4.63E-02 | 9.67E-02 | 2.76E+00 | 2.84E+00 | 6.26E-01 | no | Aacs | intron_variant | MODIFIER |
| 1 | 22403770 | AX-170158307 | 0 | G | C | 0.421 | 4.16E-02 | 8.50E-02 | 2.62E+00 | 2.67E+00 | 6.26E-01 | no | Rims1 | intron_variant | MODIFIER |
| 1 | 22405883 | AX-170311638 | 0 | T | C | 0.421 | 4.16E-02 | 8.50E-02 | 2.62E+00 | 2.67E+00 | 6.26E-01 | no | Rims1 | intron_variant | MODIFIER |
| 1 | 22406343 | AX-170123726 | 0 | T | G | 0.421 | 4.16E-02 | 8.50E-02 | 2.62E+00 | 2.67E+00 | 6.26E-01 | no | Rims1 | intron_variant | MODIFIER |
| 1 | 22575478 | AX-97792462 | 0 | C | T | 0.421 | 4.16E-02 | 8.50E-02 | 2.62E+00 | 2.67E+00 | 6.26E-01 | no | Rims1 | intron_variant | MODIFIER |
| 1 | 22671850 | AX-170486065 | 0 | C | T | 0.421 | 4.16E-02 | 8.50E-02 | 2.62E+00 | 2.67E+00 | 6.26E-01 | no | Rims1 | intron_variant | MODIFIER |
| 1 | 22674465 | AX-170125340 | 0 | C | T | 0.421 | 4.16E-02 | 8.50E-02 | 2.62E+00 | 2.67E+00 | 6.26E-01 | no | Rims1 | intron_variant | MODIFIER |
| 1 | 22739878 | AX-170353575 | 0 | C | T | 0.421 | 4.16E-02 | 8.50E-02 | 2.62E+00 | 2.67E+00 | 6.26E-01 | no | Rims1 | intron_variant | MODIFIER |
| 1 | 22799064 | AX-170258241 | 0 | G | A | 0.421 | 4.16E-02 | 8.50E-02 | 2.62E+00 | 2.67E+00 | 6.26E-01 | no | Rims1 | intron_variant | MODIFIER |
| 1 | 22799715 | AX-170240766 | 0 | A | G | 0.421 | 4.16E-02 | 8.50E-02 | 2.62E+00 | 2.67E+00 | 6.26E-01 | no | Rims1 | intron_variant | MODIFIER |
| 1 | 22799976 | AX-170163116 | 0 | T | A | 0.421 | 4.16E-02 | 8.50E-02 | 2.62E+00 | 2.67E+00 | 6.26E-01 | no | Rims1 | intron_variant | MODIFIER |
| 1 | 22800419 | AX-170332870 | 0 | G | C | 0.421 | 4.16E-02 | 8.50E-02 | 2.62E+00 | 2.67E+00 | 6.26E-01 | no | Rims1 | intron_variant | MODIFIER |
| 1 | 22800467 | AX-97816970 | 0 | G | A | 0.421 | 4.16E-02 | 8.50E-02 | 2.62E+00 | 2.67E+00 | 6.26E-01 | no | Rims1 | intron_variant | MODIFIER |
| 1 | 22801625 | AX-170102875 | 0 | C | T | 0.421 | 4.16E-02 | 8.50E-02 | 2.62E+00 | 2.67E+00 | 6.26E-01 | no | Rims1 | intron_variant | MODIFIER |
| 1 | 22809394 | AX-170308166 | 0 | T | C | 0.421 | 4.16E-02 | 8.50E-02 | 2.62E+00 | 2.67E+00 | 6.26E-01 | no | Rims1 | upstream_gene_variant | MODIFIER |
| 16 | 96159061 | AX-97842960 | 1 | A | G | 0.464 | 3.66E-02 | 7.69E-02 | 2.77E+00 | 2.88E+00 | 6.26E-01 | no | Wrb | downstream_gene_variant | MODIFIER |
| 1 | 152630040 | AX-170358446 | 0 | G | A | 0.211 | 6.24E-02 | 1.28E-01 | 2.64E+00 | 2.69E+00 | 6.26E-01 | no | Rgl1 | upstream_gene_variant | MODIFIER |
| 8 | 96787458 | AX-170031323 | 0 | C | T | 0.228 | 5.35E-02 | 1.11E-01 | 2.65E+00 | 2.72E+00 | 6.26E-01 | no | | intergenic_variant | MODIFIER |
| 9 | 75063503 | AX-170343437 | 2 | T | A | 0.209 | 5.92E-02 | 1.23E-01 | 2.71E+00 | 2.78E+00 | 6.26E-01 | no | Arpp19 | downstream_gene_variant | MODIFIER |
| 20 | 72132608 | AX-97848109 | 0 | A | C | 0.254 | 3.00E-02 | 6.24E-02 | 2.65E+00 | 2.74E+00 | 6.26E-01 | no | | intergenic_variant | MODIFIER |
| 12 | 9520062 | AX-170420547 | 0 | A | T | 0.035 | -9.03E-02 | 1.86E-01 | 2.59E+00 | 2.72E+00 | 6.26E-01 | no | | intergenic_variant | MODIFIER |
| 4 | 83978930 | AX-97852437 | 0 | C | T | 0.246 | 4.96E-02 | 9.83E-02 | 2.53E+00 | 2.61E+00 | 6.26E-01 | no | | intergenic_variant | MODIFIER |
| 4 | 56716077 | AX-97819213 | 1 | T | C | 0.366 | -5.14E-02 | 1.07E-01 | 2.71E+00 | 2.79E+00 | 6.26E-01 | no | | intergenic_variant | MODIFIER |
| 20 | 77613580 | AX-170134360 | 1 | T | A | 0.259 | 3.00E-02 | 6.24E-02 | 2.66E+00 | 2.74E+00 | 6.27E-01 | no | Tbl1x | intron_variant | MODIFIER |
| 7 | 11117101 | AX-97784468 | 1 | G | A | 0.161 | -3.81E-02 | 7.66E-02 | 2.52E+00 | 2.67E+00 | 6.27E-01 | no | Gm3774 | upstream_gene_variant | MODIFIER |
| 18 | 66384688 | AX-170061151 | 1 | T | C | 0.286 | -4.81E-02 | 1.00E-01 | 2.69E+00 | 2.76E+00 | 6.27E-01 | no | | intergenic_variant | MODIFIER |
| 11 | 32976643 | AX-170344405 | 0 | A | T | 0.105 | -7.03E-02 | 1.47E-01 | 2.64E+00 | 2.76E+00 | 6.27E-01 | no | | intergenic_variant | MODIFIER |
| 1 | 43454739 | AX-170208255 | 1 | G | A | 0.089 | -9.50E-02 | 1.99E-01 | 2.71E+00 | 2.80E+00 | 6.27E-01 | no | Nck2 | intron_variant | MODIFIER |
| 11 | 31499465 | AX-97837982 | 0 | G | A | 0.07 | 8.43E-02 | 1.74E-01 | 2.61E+00 | 2.71E+00 | 6.27E-01 | no | | intergenic_variant | MODIFIER |
| 8 | 103490185 | AX-170292838 | 1 | A | G | 0.214 | 5.37E-02 | 1.12E-01 | 2.70E+00 | 2.77E+00 | 6.27E-01 | no | | intergenic_variant | MODIFIER |
| 1 | 152601240 | AX-170259642 | 1 | G | A | 0.214 | 6.52E-02 | 1.34E-01 | 2.62E+00 | 2.68E+00 | 6.27E-01 | no | Rgl1 | intron_variant | MODIFIER |
| 9 | 87861023 | AX-97842508 | 1 | G | A | 0.134 | -6.38E-02 | 1.31E-01 | 2.60E+00 | 2.68E+00 | 6.27E-01 | no | | intergenic_variant | MODIFIER |
| 10 | 28957718 | AX-97821309 | 0 | C | G | 0.175 | 4.63E-02 | 9.76E-02 | 2.72E+00 | 2.83E+00 | 6.27E-01 | no | Gm22811 | upstream_gene_variant | MODIFIER |
| 10 | 30706002 | AX-170002993 | 0 | A | G | 0.175 | 4.63E-02 | 9.76E-02 | 2.72E+00 | 2.83E+00 | 6.27E-01 | no | Ncoa7 | intron_variant | MODIFIER |
| 10 | 30710844 | AX-170002994 | 0 | G | A | 0.175 | 4.63E-02 | 9.76E-02 | 2.72E+00 | 2.83E+00 | 6.27E-01 | no | Ncoa7 | intron_variant | MODIFIER |
| 10 | 30724778 | AX-170002995 | 0 | G | A | 0.175 | 4.63E-02 | 9.76E-02 | 2.72E+00 | 2.83E+00 | 6.27E-01 | no | Ncoa7 | intron_variant | MODIFIER |
| 10 | 30780383 | AX-170002990 | 0 | A | G | 0.175 | 4.63E-02 | 9.76E-02 | 2.72E+00 | 2.83E+00 | 6.27E-01 | no | Ncoa7 | intron_variant | MODIFIER |
| 10 | 30807727 | AX-170160826 | 0 | A | G | 0.175 | 4.63E-02 | 9.76E-02 | 2.72E+00 | 2.83E+00 | 6.27E-01 | no | Ncoa7 | upstream_gene_variant | MODIFIER |
| 10 | 30849137 | AX-170086774 | 0 | A | G | 0.175 | 4.63E-02 | 9.76E-02 | 2.72E+00 | 2.83E+00 | 6.27E-01 | no | | intergenic_variant | MODIFIER |
| 10 | 30866226 | AX-170478385 | 0 | T | C | 0.175 | 4.63E-02 | 9.76E-02 | 2.72E+00 | 2.83E+00 | 6.27E-01 | no | | intergenic_variant | MODIFIER |
| 10 | 30895204 | AX-170400841 | 0 | A | G | 0.175 | 4.63E-02 | 9.76E-02 | 2.72E+00 | 2.83E+00 | 6.27E-01 | no | | intergenic_variant | MODIFIER |
| 10 | 30938975 | AX-170325979 | 0 | C | G | 0.175 | 4.63E-02 | 9.76E-02 | 2.72E+00 | 2.83E+00 | 6.27E-01 | no | Gm24168 | downstream_gene_variant | MODIFIER |
| 1 | 172248434 | AX-170492373 | 0 | C | A | 0.211 | 6.63E-02 | 1.35E-01 | 2.60E+00 | 2.65E+00 | 6.27E-01 | no | Atp1a4 | intron_variant | MODIFIER |
| 6 | 134145275 | AX-170439861 | 0 | A | T | 0.553 | 4.20E-02 | 8.36E-02 | 2.54E+00 | 2.60E+00 | 6.27E-01 | no | Etv6 | intron_variant | MODIFIER |
| 3 | 29863656 | AX-170102526 | 1 | C | T | 0.161 | -4.78E-02 | 9.97E-02 | 2.69E+00 | 2.76E+00 | 6.27E-01 | no | Gm33206 | intron_variant,non_coding_transcript_variant | MODIFIER |
| 3 | 32009515 | AX-97856057 | 1 | C | T | 0.161 | -4.78E-02 | 9.97E-02 | 2.69E+00 | 2.76E+00 | 6.27E-01 | no | Kcnmb2 | intron_variant | MODIFIER |
| 8 | 38327947 | AX-170314700 | 0 | A | T | 0.149 | -6.45E-02 | 1.35E-01 | 2.68E+00 | 2.76E+00 | 6.27E-01 | no | Sgcz | intron_variant | MODIFIER |
| 8 | 39377753 | AX-170238610 | 0 | C | T | 0.149 | -6.45E-02 | 1.35E-01 | 2.68E+00 | 2.76E+00 | 6.27E-01 | no | | intergenic_variant | MODIFIER |
| 8 | 42165649 | AX-97807156 | 0 | A | T | 0.149 | -6.45E-02 | 1.35E-01 | 2.68E+00 | 2.76E+00 | 6.27E-01 | no | | intergenic_variant | MODIFIER |
| 11 | 7172636 | AX-170391650 | 0 | T | C | 0.044 | -9.06E-02 | 1.87E-01 | 2.58E+00 | 2.72E+00 | 6.27E-01 | no | Adcy1 | 3_prime_UTR_variant | MODIFIER |
| 11 | 32665480 | AX-170134794 | 1 | C | T | 0.089 | 8.57E-02 | 1.79E-01 | 2.65E+00 | 2.75E+00 | 6.27E-01 | no | Fbxw11 | intron_variant | MODIFIER |
| 10 | 108195127 | AX-170271292 | 0 | C | T | 0.325 | -3.92E-02 | 8.11E-02 | 2.63E+00 | 2.71E+00 | 6.27E-01 | no | Ppp1r12a | intron_variant | MODIFIER |
| 1 | 95801321 | AX-170471395 | 0 | A | T | 0.175 | 6.85E-02 | 1.34E-01 | 2.49E+00 | 2.61E+00 | 6.27E-01 | no | | intergenic_variant | MODIFIER |

| | | | | | | | | | | | | | | | |
|---|---|---|---|---|---|---|---|---|---|---|---|---|---|---|---|
| 2 | 141037933 | AX-97807016 | 0 | A | G | 0.149 | 8.79E-02 | 1.82E-01 | 2.63E+00 | 2.71E+00 | 6.27E-01 | no | Macrod2 | intron_variant | MODIFIER |
| 2 | 149470396 | AX-170250863 | 0 | G | A | 0.149 | 8.79E-02 | 1.82E-01 | 2.63E+00 | 2.71E+00 | 6.27E-01 | no | | intergenic_variant | MODIFIER |
| 11 | 52939857 | AX-170047064 | 0 | T | G | 0.053 | 6.77E-02 | 1.39E-01 | 2.57E+00 | 2.70E+00 | 6.27E-01 | no | Fstl4 | intron_variant | MODIFIER |
| 1 | 14434030 | AX-170226165 | 0 | A | G | 0.377 | 3.87E-02 | 7.82E-02 | 2.57E+00 | 2.63E+00 | 6.27E-01 | no | | intergenic_variant | MODIFIER |
| 5 | 32809269 | AX-170432521 | 1 | G | A | 0.366 | 4.99E-02 | 1.04E-01 | 2.70E+00 | 2.77E+00 | 6.27E-01 | no | Prr14l | intron_variant | MODIFIER |
| 18 | 81029992 | AX-170395525 | 0 | T | G | 0.298 | -4.23E-02 | 8.63E-02 | 2.57E+00 | 2.68E+00 | 6.27E-01 | no | Gm25718 | downstream_gene_variant | MODIFIER |
| 6 | 134613939 | AX-170028305 | 0 | A | G | 0.553 | 4.19E-02 | 8.35E-02 | 2.54E+00 | 2.60E+00 | 6.27E-01 | no | Mansc1 | intron_variant | MODIFIER |
| 15 | 101244788 | AX-170110061 | 2 | C | T | 0.082 | 7.35E-02 | 1.55E-01 | 2.69E+00 | 2.81E+00 | 6.28E-01 | no | | regulatory_region_variant | MODIFIER |
| 9 | 23336504 | AX-170187591 | 1 | C | T | 0.473 | 4.38E-02 | 8.79E-02 | 2.55E+00 | 2.62E+00 | 6.28E-01 | no | Bmper | intron_variant | MODIFIER |
| 1 | 148507324 | AX-97798427 | 0 | T | C | 0.219 | 6.01E-02 | 1.24E-01 | 2.63E+00 | 2.69E+00 | 6.28E-01 | no | | intergenic_variant | MODIFIER |
| 11 | 104465880 | AX-168419940 | 0 | C | T | 0.044 | -1.18E-01 | 2.46E-01 | 2.67E+00 | 2.75E+00 | 6.28E-01 | no | Kansl1 | intron_variant | MODIFIER |
| 11 | 104706878 | AX-170152408 | 0 | A | T | 0.044 | -1.18E-01 | 2.46E-01 | 2.67E+00 | 2.75E+00 | 6.28E-01 | no | Gm11639 | intron_variant | MODIFIER |
| 11 | 110165151 | AX-170423709 | 0 | A | G | 0.044 | -1.18E-01 | 2.46E-01 | 2.67E+00 | 2.75E+00 | 6.28E-01 | no | Abca9 | intron_variant | MODIFIER |
| 11 | 115824026 | AX-170378338 | 0 | C | A | 0.044 | -1.18E-01 | 2.46E-01 | 2.67E+00 | 2.75E+00 | 6.28E-01 | no | Tsen54 | downstream_gene_variant | MODIFIER |
| 7 | 118577299 | AX-170351765 | 0 | T | A | 0.377 | 4.10E-02 | 8.50E-02 | 2.63E+00 | 2.71E+00 | 6.28E-01 | no | Tmc7 | intron_variant | MODIFIER |
| 15 | 78871860 | AX-97814410 | 1 | G | T | 0.571 | 4.13E-02 | 8.22E-02 | 2.53E+00 | 2.60E+00 | 6.28E-01 | no | Gm26634 | upstream_gene_variant | MODIFIER |
| 20 | 116503786 | AX-170090648 | 0 | A | G | 0.035 | -6.42E-02 | 1.37E-01 | 2.64E+00 | 2.80E+00 | 6.28E-01 | no | | intergenic_variant | MODIFIER |
| 3 | 31942439 | AX-97804202 | 0 | T | G | 0.149 | -4.61E-02 | 9.61E-02 | 2.64E+00 | 2.73E+00 | 6.28E-01 | no | Kcnmb2 | intron_variant | MODIFIER |
| 3 | 29750072 | AX-170130716 | 1 | G | A | 0.17 | -4.72E-02 | 9.88E-02 | 2.68E+00 | 2.75E+00 | 6.28E-01 | no | | intergenic_variant | MODIFIER |
| 9 | 77992730 | AX-170406543 | 0 | C | G | 0.202 | -6.43E-02 | 1.34E-01 | 2.70E+00 | 2.73E+00 | 6.28E-01 | no | Gm34829 | intron_variant,non_coding_transcript_variant | MODIFIER |
| 4 | 137541305 | AX-97814202 | 0 | G | A | 0.175 | 4.82E-02 | 9.95E-02 | 2.61E+00 | 2.69E+00 | 6.28E-01 | no | Hspg2 | intron_variant | MODIFIER |
| 1 | 65680453 | AX-97810940 | 1 | A | G | 0.125 | -7.53E-02 | 1.61E-01 | 2.70E+00 | 2.85E+00 | 6.28E-01 | no | | intergenic_variant | MODIFIER |
| 10 | 16119398 | AX-170044210 | 0 | A | G | 0.193 | -4.89E-02 | 1.01E-01 | 2.59E+00 | 2.69E+00 | 6.29E-01 | no | | intergenic_variant | MODIFIER |
| 1 | 170308733 | AX-170086683 | 1 | C | T | 0.205 | 6.77E-02 | 1.40E-01 | 2.64E+00 | 2.68E+00 | 6.29E-01 | no | Spata46 | upstream_gene_variant | MODIFIER |
| 18 | 48705339 | AX-170138000 | 0 | C | A | 0.044 | -1.16E-01 | 2.39E-01 | 2.59E+00 | 2.69E+00 | 6.29E-01 | no | | intergenic_variant | MODIFIER |
| 5 | 50905900 | AX-170353703 | 0 | C | G | 0.5 | 5.39E-02 | 1.05E-01 | 2.49E+00 | 2.55E+00 | 6.29E-01 | no | | intergenic_variant | MODIFIER |
| 6 | 131662672 | AX-170439685 | 0 | C | T | 0.57 | 4.38E-02 | 8.77E-02 | 2.54E+00 | 2.60E+00 | 6.29E-01 | no | Tas2r107 | upstream_gene_variant | MODIFIER |
| 8 | 109188988 | AX-170101815 | 0 | A | G | 0.237 | 5.09E-02 | 1.07E-01 | 2.69E+00 | 2.77E+00 | 6.29E-01 | no | D030068K23Rik | intron_variant,non_coding_transcript_variant | MODIFIER |
| 8 | 109206006 | AX-170150762 | 0 | T | G | 0.237 | 5.09E-02 | 1.07E-01 | 2.69E+00 | 2.77E+00 | 6.29E-01 | no | D030068K23Rik | intron_variant,non_coding_transcript_variant | MODIFIER |
| 8 | 109206481 | AX-170494877 | 0 | A | G | 0.237 | 5.09E-02 | 1.07E-01 | 2.69E+00 | 2.77E+00 | 6.29E-01 | no | D030068K23Rik | intron_variant,non_coding_transcript_variant | MODIFIER |
| 8 | 109209341 | AX-170172134 | 0 | C | T | 0.237 | 5.09E-02 | 1.07E-01 | 2.69E+00 | 2.77E+00 | 6.29E-01 | no | D030068K23Rik | intron_variant,non_coding_transcript_variant | MODIFIER |
| 8 | 109209917 | AX-170081135 | 0 | C | T | 0.237 | 5.09E-02 | 1.07E-01 | 2.69E+00 | 2.77E+00 | 6.29E-01 | no | D030068K23Rik | intron_variant,non_coding_transcript_variant | MODIFIER |
| 8 | 109216178 | AX-170164508 | 0 | A | G | 0.237 | 5.09E-02 | 1.07E-01 | 2.69E+00 | 2.77E+00 | 6.29E-01 | no | D030068K23Rik | intron_variant,non_coding_transcript_variant | MODIFIER |
| 8 | 109243245 | AX-170157884 | 0 | A | G | 0.237 | 5.09E-02 | 1.07E-01 | 2.69E+00 | 2.77E+00 | 6.29E-01 | no | D030068K23Rik | intron_variant,non_coding_transcript_variant | MODIFIER |
| 8 | 109257244 | AX-97838929 | 0 | A | G | 0.237 | 5.09E-02 | 1.07E-01 | 2.69E+00 | 2.77E+00 | 6.29E-01 | no | D030068K23Rik | downstream_gene_variant | MODIFIER |
| 7 | 61026172 | AX-170029108 | 1 | A | T | 0.438 | 4.50E-02 | 9.42E-02 | 2.65E+00 | 2.74E+00 | 6.29E-01 | no | | intergenic_variant | MODIFIER |
| 15 | 24914774 | AX-170105879 | 0 | C | T | 0.254 | -4.97E-02 | 1.03E-01 | 2.64E+00 | 2.71E+00 | 6.29E-01 | no | | intergenic_variant | MODIFIER |
| 18 | 51292979 | AX-97835449 | 0 | G | T | 0.237 | -5.34E-02 | 1.09E-01 | 2.59E+00 | 2.64E+00 | 6.29E-01 | no | Prr16 | intron_variant | MODIFIER |
| 18 | 51309021 | AX-170015690 | 0 | C | A | 0.237 | -5.34E-02 | 1.09E-01 | 2.59E+00 | 2.64E+00 | 6.29E-01 | no | Prr16 | downstream_gene_variant | MODIFIER |
| 18 | 51433670 | AX-97852539 | 0 | G | A | 0.237 | -5.34E-02 | 1.09E-01 | 2.59E+00 | 2.64E+00 | 6.29E-01 | no | | intergenic_variant | MODIFIER |
| 18 | 51455968 | AX-170375478 | 0 | G | A | 0.237 | -5.34E-02 | 1.09E-01 | 2.59E+00 | 2.64E+00 | 6.29E-01 | no | | intergenic_variant | MODIFIER |
| 18 | 51456157 | AX-170149805 | 0 | A | G | 0.237 | -5.34E-02 | 1.09E-01 | 2.59E+00 | 2.64E+00 | 6.29E-01 | no | | intergenic_variant | MODIFIER |
| 18 | 51471465 | AX-97837936 | 0 | A | C | 0.237 | -5.34E-02 | 1.09E-01 | 2.59E+00 | 2.64E+00 | 6.29E-01 | no | | regulatory_region_variant | MODIFIER |
| 18 | 51482215 | AX-170452412 | 0 | G | A | 0.237 | -5.34E-02 | 1.09E-01 | 2.59E+00 | 2.64E+00 | 6.29E-01 | no | | intergenic_variant | MODIFIER |
| 18 | 52180315 | AX-170144000 | 0 | G | C | 0.237 | -5.34E-02 | 1.09E-01 | 2.59E+00 | 2.64E+00 | 6.29E-01 | no | | intergenic_variant | MODIFIER |
| 18 | 52210347 | AX-170168606 | 0 | T | G | 0.237 | -5.34E-02 | 1.09E-01 | 2.59E+00 | 2.64E+00 | 6.29E-01 | no | | intergenic_variant | MODIFIER |
| 18 | 52210530 | AX-170228420 | 0 | A | G | 0.237 | -5.34E-02 | 1.09E-01 | 2.59E+00 | 2.64E+00 | 6.29E-01 | no | | intergenic_variant | MODIFIER |
| 18 | 52226903 | AX-170093884 | 0 | C | T | 0.237 | -5.34E-02 | 1.09E-01 | 2.59E+00 | 2.64E+00 | 6.29E-01 | no | | intergenic_variant | MODIFIER |
| 18 | 52233628 | AX-170278655 | 0 | C | T | 0.237 | -5.34E-02 | 1.09E-01 | 2.59E+00 | 2.64E+00 | 6.29E-01 | no | | intergenic_variant | MODIFIER |
| 18 | 52271206 | AX-170060850 | 0 | G | A | 0.237 | -5.34E-02 | 1.09E-01 | 2.59E+00 | 2.64E+00 | 6.29E-01 | no | | intergenic_variant | MODIFIER |
| 18 | 52283591 | AX-97816012 | 0 | C | T | 0.237 | -5.34E-02 | 1.09E-01 | 2.59E+00 | 2.64E+00 | 6.29E-01 | no | | intergenic_variant | MODIFIER |
| 18 | 52395273 | AX-170305403 | 0 | G | A | 0.237 | -5.34E-02 | 1.09E-01 | 2.59E+00 | 2.64E+00 | 6.29E-01 | no | | intergenic_variant | MODIFIER |
| 18 | 52397759 | AX-170457859 | 0 | A | C | 0.237 | -5.34E-02 | 1.09E-01 | 2.59E+00 | 2.64E+00 | 6.29E-01 | no | | intergenic_variant | MODIFIER |
| 18 | 52397886 | AX-170524595 | 0 | A | G | 0.237 | -5.34E-02 | 1.09E-01 | 2.59E+00 | 2.64E+00 | 6.29E-01 | no | | intergenic_variant | MODIFIER |

| | | | | | | | | | | | | | | | |
|---|---|---|---|---|---|---|---|---|---|---|---|---|---|---|---|
| 18 | 52894324 | AX-97802690 | 0 | A | G | 0.237 | -5.34E-02 | 1.09E-01 | 2.59E+00 | 2.64E+00 | 6.29E-01 | no | Sncaip | intron_variant | MODIFIER |
| 18 | 52907653 | AX-170176798 | 0 | C | T | 0.237 | -5.34E-02 | 1.09E-01 | 2.59E+00 | 2.64E+00 | 6.29E-01 | no | Sncaip | intron_variant | MODIFIER |
| 18 | 52908346 | AX-170324681 | 0 | G | A | 0.237 | -5.34E-02 | 1.09E-01 | 2.59E+00 | 2.64E+00 | 6.29E-01 | no | Sncaip | intron_variant | MODIFIER |
| 18 | 52912230 | AX-170037814 | 0 | G | A | 0.237 | -5.34E-02 | 1.09E-01 | 2.59E+00 | 2.64E+00 | 6.29E-01 | no | Sncaip | intron_variant | MODIFIER |
| 18 | 52913042 | AX-170060862 | 0 | G | T | 0.237 | -5.34E-02 | 1.09E-01 | 2.59E+00 | 2.64E+00 | 6.29E-01 | no | Sncaip | intron_variant | MODIFIER |
| 18 | 52914964 | AX-170060861 | 0 | A | G | 0.237 | -5.34E-02 | 1.09E-01 | 2.59E+00 | 2.64E+00 | 6.29E-01 | no | Sncaip | intron_variant | MODIFIER |
| 18 | 52935568 | AX-170037817 | 0 | A | G | 0.237 | -5.34E-02 | 1.09E-01 | 2.59E+00 | 2.64E+00 | 6.29E-01 | no | | regulatory_region_variant | MODIFIER |
| 18 | 52935581 | AX-170037816 | 0 | G | A | 0.237 | -5.34E-02 | 1.09E-01 | 2.59E+00 | 2.64E+00 | 6.29E-01 | no | | regulatory_region_variant | MODIFIER |
| 18 | 52936134 | AX-170015716 | 0 | A | C | 0.237 | -5.34E-02 | 1.09E-01 | 2.59E+00 | 2.64E+00 | 6.29E-01 | no | | intergenic_variant | MODIFIER |
| 18 | 52936333 | AX-170015714 | 0 | T | C | 0.237 | -5.34E-02 | 1.09E-01 | 2.59E+00 | 2.64E+00 | 6.29E-01 | no | | intergenic_variant | MODIFIER |
| 18 | 52936485 | AX-170015713 | 0 | A | G | 0.237 | -5.34E-02 | 1.09E-01 | 2.59E+00 | 2.64E+00 | 6.29E-01 | no | | intergenic_variant | MODIFIER |
| 18 | 52938543 | AX-170294567 | 0 | A | C | 0.237 | -5.34E-02 | 1.09E-01 | 2.59E+00 | 2.64E+00 | 6.29E-01 | no | | intergenic_variant | MODIFIER |
| 18 | 52955213 | AX-170015718 | 0 | G | A | 0.237 | -5.34E-02 | 1.09E-01 | 2.59E+00 | 2.64E+00 | 6.29E-01 | no | AC124126.1 | downstream_gene_variant | MODIFIER |
| 18 | 52960992 | AX-170422734 | 0 | T | G | 0.237 | -5.34E-02 | 1.09E-01 | 2.59E+00 | 2.64E+00 | 6.29E-01 | no | | intergenic_variant | MODIFIER |
| 18 | 52975233 | AX-170289347 | 0 | T | C | 0.237 | -5.34E-02 | 1.09E-01 | 2.59E+00 | 2.64E+00 | 6.29E-01 | no | | intergenic_variant | MODIFIER |
| 18 | 52999424 | AX-170378650 | 0 | C | T | 0.237 | -5.34E-02 | 1.09E-01 | 2.59E+00 | 2.64E+00 | 6.29E-01 | no | | intergenic_variant | MODIFIER |
| 18 | 53002339 | AX-97789055 | 0 | A | T | 0.237 | -5.34E-02 | 1.09E-01 | 2.59E+00 | 2.64E+00 | 6.29E-01 | no | | intergenic_variant | MODIFIER |
| 18 | 53002648 | AX-170417858 | 0 | G | A | 0.237 | -5.34E-02 | 1.09E-01 | 2.59E+00 | 2.64E+00 | 6.29E-01 | no | | intergenic_variant | MODIFIER |
| 18 | 53006516 | AX-170350986 | 0 | G | A | 0.237 | -5.34E-02 | 1.09E-01 | 2.59E+00 | 2.64E+00 | 6.29E-01 | no | | intergenic_variant | MODIFIER |
| 18 | 53014505 | AX-170530116 | 0 | T | C | 0.237 | -5.34E-02 | 1.09E-01 | 2.59E+00 | 2.64E+00 | 6.29E-01 | no | | intergenic_variant | MODIFIER |
| 18 | 53032416 | AX-170269954 | 0 | A | G | 0.237 | -5.34E-02 | 1.09E-01 | 2.59E+00 | 2.64E+00 | 6.29E-01 | no | | intergenic_variant | MODIFIER |
| 18 | 53037390 | AX-170383292 | 0 | C | T | 0.237 | -5.34E-02 | 1.09E-01 | 2.59E+00 | 2.64E+00 | 6.29E-01 | no | | regulatory_region_variant | MODIFIER |
| 18 | 53039569 | AX-170507188 | 0 | A | G | 0.237 | -5.34E-02 | 1.09E-01 | 2.59E+00 | 2.64E+00 | 6.29E-01 | no | | intergenic_variant | MODIFIER |
| 18 | 53079414 | AX-168409515 | 0 | T | C | 0.237 | -5.34E-02 | 1.09E-01 | 2.59E+00 | 2.64E+00 | 6.29E-01 | no | | intergenic_variant | MODIFIER |
| 18 | 53079445 | AX-170429033 | 0 | T | C | 0.237 | -5.34E-02 | 1.09E-01 | 2.59E+00 | 2.64E+00 | 6.29E-01 | no | | intergenic_variant | MODIFIER |
| 18 | 53082119 | AX-170478047 | 0 | G | T | 0.237 | -5.34E-02 | 1.09E-01 | 2.59E+00 | 2.64E+00 | 6.29E-01 | no | | regulatory_region_variant | MODIFIER |
| 18 | 53107864 | AX-170398370 | 0 | A | G | 0.237 | -5.34E-02 | 1.09E-01 | 2.59E+00 | 2.64E+00 | 6.29E-01 | no | | intergenic_variant | MODIFIER |
| 18 | 53110422 | AX-170060864 | 0 | C | T | 0.237 | -5.34E-02 | 1.09E-01 | 2.59E+00 | 2.64E+00 | 6.29E-01 | no | | intergenic_variant | MODIFIER |
| 18 | 53137377 | AX-170464306 | 0 | G | C | 0.237 | -5.34E-02 | 1.09E-01 | 2.59E+00 | 2.64E+00 | 6.29E-01 | no | | regulatory_region_variant | MODIFIER |
| 18 | 53194808 | AX-170183922 | 0 | G | A | 0.237 | -5.34E-02 | 1.09E-01 | 2.59E+00 | 2.64E+00 | 6.29E-01 | no | Snx2 | intron_variant | MODIFIER |
| 18 | 53195764 | AX-170185181 | 0 | C | T | 0.237 | -5.34E-02 | 1.09E-01 | 2.59E+00 | 2.64E+00 | 6.29E-01 | no | Snx2 | intron_variant | MODIFIER |
| 18 | 53195917 | AX-170356264 | 0 | T | A | 0.237 | -5.34E-02 | 1.09E-01 | 2.59E+00 | 2.64E+00 | 6.29E-01 | no | Snx2 | intron_variant | MODIFIER |
| 18 | 53199928 | AX-170428246 | 0 | G | A | 0.237 | -5.34E-02 | 1.09E-01 | 2.59E+00 | 2.64E+00 | 6.29E-01 | no | Snx2 | intron_variant | MODIFIER |
| 18 | 53203330 | AX-170119563 | 0 | C | T | 0.237 | -5.34E-02 | 1.09E-01 | 2.59E+00 | 2.64E+00 | 6.29E-01 | no | Snx2 | intron_variant | MODIFIER |
| 18 | 53210143 | AX-97786110 | 0 | A | G | 0.237 | -5.34E-02 | 1.09E-01 | 2.59E+00 | 2.64E+00 | 6.29E-01 | no | Snx2 | intron_variant | MODIFIER |
| 18 | 53259928 | AX-170215783 | 0 | C | T | 0.237 | -5.34E-02 | 1.09E-01 | 2.59E+00 | 2.64E+00 | 6.29E-01 | no | Snx24 | intron_variant | MODIFIER |
| 18 | 53261164 | AX-170130736 | 0 | T | C | 0.237 | -5.34E-02 | 1.09E-01 | 2.59E+00 | 2.64E+00 | 6.29E-01 | no | Snx24 | intron_variant | MODIFIER |
| 18 | 53261374 | AX-170518701 | 0 | C | T | 0.237 | -5.34E-02 | 1.09E-01 | 2.59E+00 | 2.64E+00 | 6.29E-01 | no | Snx24 | intron_variant | MODIFIER |
| 18 | 53268813 | AX-170184573 | 0 | C | A | 0.237 | -5.34E-02 | 1.09E-01 | 2.59E+00 | 2.64E+00 | 6.29E-01 | no | Snx24 | intron_variant | MODIFIER |
| 18 | 53273836 | AX-170481510 | 0 | A | G | 0.237 | -5.34E-02 | 1.09E-01 | 2.59E+00 | 2.64E+00 | 6.29E-01 | no | Snx24 | intron_variant | MODIFIER |
| 18 | 53309959 | AX-170289946 | 0 | A | G | 0.237 | -5.34E-02 | 1.09E-01 | 2.59E+00 | 2.64E+00 | 6.29E-01 | no | Snx24 | intron_variant | MODIFIER |
| 18 | 53313893 | AX-170212548 | 0 | T | C | 0.237 | -5.34E-02 | 1.09E-01 | 2.59E+00 | 2.64E+00 | 6.29E-01 | no | Snx24 | intron_variant | MODIFIER |
| 18 | 53333922 | AX-170428838 | 0 | A | G | 0.237 | -5.34E-02 | 1.09E-01 | 2.59E+00 | 2.64E+00 | 6.29E-01 | no | Snx24 | intron_variant | MODIFIER |
| 18 | 53335282 | AX-97792991 | 0 | G | A | 0.237 | -5.34E-02 | 1.09E-01 | 2.59E+00 | 2.64E+00 | 6.29E-01 | no | Snx24 | intron_variant | MODIFIER |
| 18 | 53351086 | AX-170503740 | 0 | C | T | 0.237 | -5.34E-02 | 1.09E-01 | 2.59E+00 | 2.64E+00 | 6.29E-01 | no | Snx24 | intron_variant | MODIFIER |
| 18 | 53384316 | AX-170060867 | 0 | A | G | 0.237 | -5.34E-02 | 1.09E-01 | 2.59E+00 | 2.64E+00 | 6.29E-01 | no | Snx24 | intron_variant | MODIFIER |
| 18 | 53392220 | AX-97781635 | 0 | A | T | 0.237 | -5.34E-02 | 1.09E-01 | 2.59E+00 | 2.64E+00 | 6.29E-01 | no | Snx24 | downstream_gene_variant | MODIFIER |
| 18 | 53392286 | AX-170486391 | 0 | G | T | 0.237 | -5.34E-02 | 1.09E-01 | 2.59E+00 | 2.64E+00 | 6.29E-01 | no | Snx24 | downstream_gene_variant | MODIFIER |
| 18 | 53415103 | AX-170297263 | 0 | A | G | 0.237 | -5.34E-02 | 1.09E-01 | 2.59E+00 | 2.64E+00 | 6.29E-01 | no | Ppic | intron_variant | MODIFIER |
| 18 | 53415239 | AX-170015720 | 0 | A | C | 0.237 | -5.34E-02 | 1.09E-01 | 2.59E+00 | 2.64E+00 | 6.29E-01 | no | Ppic | intron_variant | MODIFIER |
| 18 | 53439644 | AX-170165403 | 0 | A | G | 0.237 | -5.34E-02 | 1.09E-01 | 2.59E+00 | 2.64E+00 | 6.29E-01 | no | | intergenic_variant | MODIFIER |
| 18 | 53454984 | AX-170015721 | 0 | A | G | 0.237 | -5.34E-02 | 1.09E-01 | 2.59E+00 | 2.64E+00 | 6.29E-01 | no | | intergenic_variant | MODIFIER |
| 18 | 53458798 | AX-170261607 | 0 | T | C | 0.237 | -5.34E-02 | 1.09E-01 | 2.59E+00 | 2.64E+00 | 6.29E-01 | no | Prdm6 | upstream_gene_variant | MODIFIER |
| 18 | 53458810 | AX-170183869 | 0 | G | T | 0.237 | -5.34E-02 | 1.09E-01 | 2.59E+00 | 2.64E+00 | 6.29E-01 | no | Prdm6 | upstream_gene_variant | MODIFIER |

| | | | | | | | | | | | | | | | |
|---|---|---|---|---|---|---|---|---|---|---|---|---|---|---|---|
| 18 | 53487784 | AX-97809438 | 0 | T | C | 0.237 | -5.34E-02 | 1.09E-01 | 2.59E+00 | 2.64E+00 | 6.29E-01 | no | Prdm6 | intron_variant | MODIFIER |
| 18 | 53494077 | AX-170329596 | 0 | C | T | 0.237 | -5.34E-02 | 1.09E-01 | 2.59E+00 | 2.64E+00 | 6.29E-01 | no | Prdm6 | intron_variant | MODIFIER |
| 18 | 53495849 | AX-170393640 | 0 | G | C | 0.237 | -5.34E-02 | 1.09E-01 | 2.59E+00 | 2.64E+00 | 6.29E-01 | no | Prdm6 | intron_variant | MODIFIER |
| 18 | 53535039 | AX-97829567 | 0 | G | A | 0.237 | -5.34E-02 | 1.09E-01 | 2.59E+00 | 2.64E+00 | 6.29E-01 | no | Prdm6 | intron_variant | MODIFIER |
| 18 | 53540994 | AX-170401069 | 0 | T | C | 0.237 | -5.34E-02 | 1.09E-01 | 2.59E+00 | 2.64E+00 | 6.29E-01 | no | Prdm6 | intron_variant | MODIFIER |
| 18 | 53542981 | AX-170338486 | 0 | G | A | 0.237 | -5.34E-02 | 1.09E-01 | 2.59E+00 | 2.64E+00 | 6.29E-01 | no | Prdm6 | intron_variant | MODIFIER |
| 18 | 53556515 | AX-170060871 | 0 | A | C | 0.237 | -5.34E-02 | 1.09E-01 | 2.59E+00 | 2.64E+00 | 6.29E-01 | no | Prdm6 | intron_variant | MODIFIER |
| 18 | 53570156 | AX-170529946 | 0 | C | A | 0.237 | -5.34E-02 | 1.09E-01 | 2.59E+00 | 2.64E+00 | 6.29E-01 | no | Prdm6 | intron_variant | MODIFIER |
| 18 | 53598121 | AX-170309411 | 0 | G | C | 0.237 | -5.34E-02 | 1.09E-01 | 2.59E+00 | 2.64E+00 | 6.29E-01 | no | | regulatory_region_variant | MODIFIER |
| 18 | 53600688 | AX-170113934 | 0 | A | T | 0.237 | -5.34E-02 | 1.09E-01 | 2.59E+00 | 2.64E+00 | 6.29E-01 | no | | intergenic_variant | MODIFIER |
| 18 | 53650949 | AX-170282916 | 0 | T | C | 0.237 | -5.34E-02 | 1.09E-01 | 2.59E+00 | 2.64E+00 | 6.29E-01 | no | AC116597.1 | intron_variant,non_coding_transcript_variant | MODIFIER |
| 18 | 53682870 | AX-97820446 | 0 | G | A | 0.237 | -5.34E-02 | 1.09E-01 | 2.59E+00 | 2.64E+00 | 6.29E-01 | no | Cep120 | 3_prime_UTR_variant | MODIFIER |
| 18 | 53699875 | AX-170318629 | 0 | G | A | 0.237 | -5.34E-02 | 1.09E-01 | 2.59E+00 | 2.64E+00 | 6.29E-01 | no | Cep120 | intron_variant | MODIFIER |
| 18 | 53710561 | AX-170239665 | 0 | A | G | 0.237 | -5.34E-02 | 1.09E-01 | 2.59E+00 | 2.64E+00 | 6.29E-01 | no | Cep120 | intron_variant | MODIFIER |
| 18 | 53730845 | AX-170325747 | 0 | T | C | 0.237 | -5.34E-02 | 1.09E-01 | 2.59E+00 | 2.64E+00 | 6.29E-01 | no | Cep120 | intron_variant | MODIFIER |
| 18 | 53736298 | AX-170479248 | 0 | C | T | 0.237 | -5.34E-02 | 1.09E-01 | 2.59E+00 | 2.64E+00 | 6.29E-01 | no | Cep120 | intron_variant | MODIFIER |
| 18 | 53740195 | AX-170060874 | 0 | G | A | 0.237 | -5.34E-02 | 1.09E-01 | 2.59E+00 | 2.64E+00 | 6.29E-01 | no | Cep120 | synonymous_variant | LOW |
| 18 | 53755102 | AX-170207254 | 0 | G | A | 0.237 | -5.34E-02 | 1.09E-01 | 2.59E+00 | 2.64E+00 | 6.29E-01 | no | | intergenic_variant | MODIFIER |
| 18 | 53757690 | AX-170416860 | 0 | G | A | 0.237 | -5.34E-02 | 1.09E-01 | 2.59E+00 | 2.64E+00 | 6.29E-01 | no | | intergenic_variant | MODIFIER |
| 18 | 53757712 | AX-170342006 | 0 | A | C | 0.237 | -5.34E-02 | 1.09E-01 | 2.59E+00 | 2.64E+00 | 6.29E-01 | no | | intergenic_variant | MODIFIER |
| 11 | 22618085 | AX-170239604 | 1 | G | A | 0.054 | 9.16E-02 | 1.95E-01 | 2.65E+00 | 2.79E+00 | 6.29E-01 | no | Gm12052 | non_coding_transcript_exon_variant | MODIFIER |
| 1 | 149739799 | AX-170505621 | 1 | T | A | 0.196 | 6.72E-02 | 1.40E-01 | 2.65E+00 | 2.70E+00 | 6.29E-01 | no | n-R5s218 | upstream_gene_variant | MODIFIER |
| 1 | 152709671 | AX-97838595 | 1 | A | G | 0.196 | 6.72E-02 | 1.40E-01 | 2.65E+00 | 2.70E+00 | 6.29E-01 | no | Rgl1 | intron_variant | MODIFIER |
| 1 | 103203317 | AX-170210379 | 0 | C | T | 0.149 | 1.04E-01 | 2.19E-01 | 2.70E+00 | 2.77E+00 | 6.29E-01 | no | Gm18728 | intron_variant,non_coding_transcript_variant | MODIFIER |
| 3 | 32392808 | AX-170299630 | 0 | T | G | 0.175 | 4.67E-02 | 9.84E-02 | 2.71E+00 | 2.78E+00 | 6.29E-01 | no | Pik3ca | upstream_gene_variant | MODIFIER |
| 3 | 32405771 | AX-170142218 | 0 | C | T | 0.175 | 4.67E-02 | 9.84E-02 | 2.71E+00 | 2.78E+00 | 6.29E-01 | no | Pik3ca | intron_variant | MODIFIER |
| 3 | 32405858 | AX-170420263 | 0 | T | C | 0.175 | 4.67E-02 | 9.84E-02 | 2.71E+00 | 2.78E+00 | 6.29E-01 | no | Pik3ca | intron_variant | MODIFIER |
| 3 | 32412325 | AX-170236860 | 0 | T | C | 0.175 | 4.67E-02 | 9.84E-02 | 2.71E+00 | 2.78E+00 | 6.29E-01 | no | Pik3ca | intron_variant | MODIFIER |
| 3 | 32430342 | AX-170525986 | 0 | C | T | 0.175 | 4.67E-02 | 9.84E-02 | 2.71E+00 | 2.78E+00 | 6.29E-01 | no | Pik3ca | intron_variant | MODIFIER |
| 3 | 32531937 | AX-170194809 | 0 | T | C | 0.175 | 4.67E-02 | 9.84E-02 | 2.71E+00 | 2.78E+00 | 6.29E-01 | no | Mfn1 | synonymous_variant | LOW |
| 3 | 32578185 | AX-170237996 | 0 | G | A | 0.175 | 4.67E-02 | 9.84E-02 | 2.71E+00 | 2.78E+00 | 6.29E-01 | no | Mfn1 | 3_prime_UTR_variant | MODIFIER |
| 3 | 32578516 | AX-170164889 | 0 | A | G | 0.175 | 4.67E-02 | 9.84E-02 | 2.71E+00 | 2.78E+00 | 6.29E-01 | no | Mfn1 | 3_prime_UTR_variant | MODIFIER |
| 3 | 32579218 | AX-170186780 | 0 | C | T | 0.175 | 4.67E-02 | 9.84E-02 | 2.71E+00 | 2.78E+00 | 6.29E-01 | no | Mfn1 | 3_prime_UTR_variant | MODIFIER |
| 3 | 32592434 | AX-170066281 | 0 | C | G | 0.175 | 4.67E-02 | 9.84E-02 | 2.71E+00 | 2.78E+00 | 6.29E-01 | no | Gnb4 | intron_variant | MODIFIER |
| 3 | 32597517 | AX-97808687 | 0 | A | G | 0.175 | 4.67E-02 | 9.84E-02 | 2.71E+00 | 2.78E+00 | 6.29E-01 | no | Gnb4 | intron_variant | MODIFIER |
| 3 | 32612213 | AX-170393252 | 0 | C | G | 0.175 | 4.67E-02 | 9.84E-02 | 2.71E+00 | 2.78E+00 | 6.29E-01 | no | Gnb4 | intron_variant | MODIFIER |
| 3 | 32645240 | AX-97856978 | 0 | C | A | 0.175 | 4.67E-02 | 9.84E-02 | 2.71E+00 | 2.78E+00 | 6.29E-01 | no | | intergenic_variant | MODIFIER |
| 3 | 32648212 | AX-170486924 | 0 | G | A | 0.175 | 4.67E-02 | 9.84E-02 | 2.71E+00 | 2.78E+00 | 6.29E-01 | no | | intergenic_variant | MODIFIER |
| 3 | 32668158 | AX-170515469 | 0 | C | T | 0.175 | 4.67E-02 | 9.84E-02 | 2.71E+00 | 2.78E+00 | 6.29E-01 | no | | regulatory_region_variant | MODIFIER |
| 18 | 57249442 | AX-170481264 | 1 | G | A | 0.196 | -5.62E-02 | 1.15E-01 | 2.59E+00 | 2.65E+00 | 6.30E-01 | no | Megf10 | intron_variant | MODIFIER |
| 1 | 168820109 | AX-97853394 | 0 | C | G | 0.219 | -6.51E-02 | 1.36E-01 | 2.84E+00 | 2.92E+00 | 6.30E-01 | no | | intergenic_variant | MODIFIER |
| 1 | 137483475 | AX-170288482 | 1 | C | T | 0.205 | 5.66E-02 | 1.18E-01 | 2.62E+00 | 2.71E+00 | 6.30E-01 | no | | intergenic_variant | MODIFIER |
| 9 | 38331946 | AX-168373027 | 2 | A | G | 0.464 | -3.49E-02 | 7.43E-02 | 2.71E+00 | 2.82E+00 | 6.30E-01 | no | Olfr25 | downstream_gene_variant | MODIFIER |
| 2 | 122920407 | AX-170168364 | 0 | A | G | 0.149 | 8.50E-02 | 1.76E-01 | 2.61E+00 | 2.70E+00 | 6.30E-01 | no | | intergenic_variant | MODIFIER |
| 7 | 76537259 | AX-170095989 | 0 | C | T | 0.36 | -4.63E-02 | 9.82E-02 | 2.72E+00 | 2.82E+00 | 6.30E-01 | no | Agbl1 | intron_variant | MODIFIER |
| 5 | 106165668 | AX-170394472 | 0 | T | C | 0.439 | 5.23E-02 | 1.09E-01 | 2.65E+00 | 2.70E+00 | 6.30E-01 | no | | intergenic_variant | MODIFIER |
| 2 | 168521306 | AX-170019637 | 0 | T | A | 0.158 | 9.29E-02 | 1.94E-01 | 2.63E+00 | 2.70E+00 | 6.31E-01 | no | Nfatc2 | intron_variant | MODIFIER |
| 8 | 97201006 | AX-170170100 | 0 | C | G | 0.228 | 5.29E-02 | 1.12E-01 | 2.68E+00 | 2.76E+00 | 6.31E-01 | no | | intergenic_variant | MODIFIER |
| 5 | 85234573 | AX-170479088 | 0 | A | C | 0.43 | 5.36E-02 | 1.13E-01 | 2.75E+00 | 2.80E+00 | 6.31E-01 | no | | intergenic_variant | MODIFIER |
| 5 | 85235692 | AX-170072677 | 0 | A | C | 0.43 | 5.36E-02 | 1.13E-01 | 2.75E+00 | 2.80E+00 | 6.31E-01 | no | | intergenic_variant | MODIFIER |
| 5 | 85263801 | AX-97850017 | 0 | A | G | 0.43 | 5.36E-02 | 1.13E-01 | 2.75E+00 | 2.80E+00 | 6.31E-01 | no | | intergenic_variant | MODIFIER |
| 5 | 85287712 | AX-97841839 | 0 | G | C | 0.43 | 5.36E-02 | 1.13E-01 | 2.75E+00 | 2.80E+00 | 6.31E-01 | no | | intergenic_variant | MODIFIER |
| 5 | 85346260 | AX-170233680 | 0 | A | G | 0.43 | 5.36E-02 | 1.13E-01 | 2.75E+00 | 2.80E+00 | 6.31E-01 | no | | regulatory_region_variant | MODIFIER |
| 5 | 85346373 | AX-170462351 | 0 | A | G | 0.43 | 5.36E-02 | 1.13E-01 | 2.75E+00 | 2.80E+00 | 6.31E-01 | no | | regulatory_region_variant | MODIFIER |

| | | | | | | | | | | | | | | | |
|---|---|---|---|---|---|---|---|---|---|---|---|---|---|---|---|
| 5 | 85351226 | AX-97831926 | 0 | T | C | 0.43 | 5.36E-02 | 1.13E-01 | 2.75E+00 | 2.80E+00 | 6.31E-01 | no | | intergenic_variant | MODIFIER |
| 5 | 85391466 | AX-97810166 | 0 | A | G | 0.43 | 5.36E-02 | 1.13E-01 | 2.75E+00 | 2.80E+00 | 6.31E-01 | no | | intergenic_variant | MODIFIER |
| 5 | 85398896 | AX-170507859 | 0 | C | T | 0.43 | 5.36E-02 | 1.13E-01 | 2.75E+00 | 2.80E+00 | 6.31E-01 | no | | intergenic_variant | MODIFIER |
| 5 | 85446196 | AX-170072671 | 0 | A | G | 0.43 | 5.36E-02 | 1.13E-01 | 2.75E+00 | 2.80E+00 | 6.31E-01 | no | | intergenic_variant | MODIFIER |
| 5 | 85446716 | AX-170072678 | 0 | A | G | 0.43 | 5.36E-02 | 1.13E-01 | 2.75E+00 | 2.80E+00 | 6.31E-01 | no | | intergenic_variant | MODIFIER |
| 5 | 85455109 | AX-170440524 | 0 | C | A | 0.43 | 5.36E-02 | 1.13E-01 | 2.75E+00 | 2.80E+00 | 6.31E-01 | no | | intergenic_variant | MODIFIER |
| 5 | 85455838 | AX-170404981 | 0 | T | C | 0.43 | 5.36E-02 | 1.13E-01 | 2.75E+00 | 2.80E+00 | 6.31E-01 | no | | intergenic_variant | MODIFIER |
| 5 | 85456286 | AX-170153839 | 0 | C | G | 0.43 | 5.36E-02 | 1.13E-01 | 2.75E+00 | 2.80E+00 | 6.31E-01 | no | | intergenic_variant | MODIFIER |
| 5 | 85470185 | AX-170025481 | 0 | A | G | 0.43 | 5.36E-02 | 1.13E-01 | 2.75E+00 | 2.80E+00 | 6.31E-01 | no | | intergenic_variant | MODIFIER |
| 5 | 85470274 | AX-170025482 | 0 | A | T | 0.43 | 5.36E-02 | 1.13E-01 | 2.75E+00 | 2.80E+00 | 6.31E-01 | no | | intergenic_variant | MODIFIER |
| 5 | 85503634 | AX-170025488 | 0 | T | G | 0.43 | 5.36E-02 | 1.13E-01 | 2.75E+00 | 2.80E+00 | 6.31E-01 | no | | intergenic_variant | MODIFIER |
| 5 | 85537706 | AX-170025483 | 0 | G | A | 0.43 | 5.36E-02 | 1.13E-01 | 2.75E+00 | 2.80E+00 | 6.31E-01 | no | | intergenic_variant | MODIFIER |
| 5 | 85573138 | AX-170442949 | 0 | C | T | 0.43 | 5.36E-02 | 1.13E-01 | 2.75E+00 | 2.80E+00 | 6.31E-01 | no | | intergenic_variant | MODIFIER |
| 5 | 85605100 | AX-170286027 | 0 | A | G | 0.43 | 5.36E-02 | 1.13E-01 | 2.75E+00 | 2.80E+00 | 6.31E-01 | no | | intergenic_variant | MODIFIER |
| 5 | 85622353 | AX-170408411 | 0 | T | C | 0.43 | 5.36E-02 | 1.13E-01 | 2.75E+00 | 2.80E+00 | 6.31E-01 | no | | intergenic_variant | MODIFIER |
| 5 | 85625640 | AX-97800525 | 0 | C | A | 0.43 | 5.36E-02 | 1.13E-01 | 2.75E+00 | 2.80E+00 | 6.31E-01 | no | | intergenic_variant | MODIFIER |
| 5 | 85625899 | AX-170377687 | 0 | A | C | 0.43 | 5.36E-02 | 1.13E-01 | 2.75E+00 | 2.80E+00 | 6.31E-01 | no | | intergenic_variant | MODIFIER |
| 5 | 85695417 | AX-170123053 | 0 | A | G | 0.43 | 5.36E-02 | 1.13E-01 | 2.75E+00 | 2.80E+00 | 6.31E-01 | no | | intergenic_variant | MODIFIER |
| 5 | 85705083 | AX-170436974 | 0 | T | G | 0.43 | 5.36E-02 | 1.13E-01 | 2.75E+00 | 2.80E+00 | 6.31E-01 | no | | intergenic_variant | MODIFIER |
| 5 | 85728214 | AX-170361654 | 0 | C | T | 0.43 | 5.36E-02 | 1.13E-01 | 2.75E+00 | 2.80E+00 | 6.31E-01 | no | | intergenic_variant | MODIFIER |
| 5 | 85743837 | AX-170101049 | 0 | C | G | 0.43 | 5.36E-02 | 1.13E-01 | 2.75E+00 | 2.80E+00 | 6.31E-01 | no | | intergenic_variant | MODIFIER |
| 5 | 85791428 | AX-97808264 | 0 | G | A | 0.43 | 5.36E-02 | 1.13E-01 | 2.75E+00 | 2.80E+00 | 6.31E-01 | no | | intergenic_variant | MODIFIER |
| 5 | 85979806 | AX-170025501 | 0 | A | T | 0.43 | 5.36E-02 | 1.13E-01 | 2.75E+00 | 2.80E+00 | 6.31E-01 | no | | intergenic_variant | MODIFIER |
| 5 | 85980112 | AX-97857361 | 0 | C | T | 0.43 | 5.36E-02 | 1.13E-01 | 2.75E+00 | 2.80E+00 | 6.31E-01 | no | | intergenic_variant | MODIFIER |
| 5 | 86028624 | AX-170197811 | 0 | T | C | 0.43 | 5.36E-02 | 1.13E-01 | 2.75E+00 | 2.80E+00 | 6.31E-01 | no | Cenpc1 | intron_variant | MODIFIER |
| 5 | 86037398 | AX-170025491 | 0 | G | T | 0.43 | 5.36E-02 | 1.13E-01 | 2.75E+00 | 2.80E+00 | 6.31E-01 | no | Cenpc1 | missense_variant | MODERATE |
| 5 | 86344588 | AX-170238741 | 0 | C | T | 0.43 | 5.36E-02 | 1.13E-01 | 2.75E+00 | 2.80E+00 | 6.31E-01 | no | Tmprss11d | intron_variant | MODIFIER |
| 5 | 86345866 | AX-170072685 | 0 | G | A | 0.43 | 5.36E-02 | 1.13E-01 | 2.75E+00 | 2.80E+00 | 6.31E-01 | no | Tmprss11d | intron_variant | MODIFIER |
| 5 | 86351848 | AX-170387279 | 0 | T | G | 0.43 | 5.36E-02 | 1.13E-01 | 2.75E+00 | 2.80E+00 | 6.31E-01 | no | Tmprss11d | intron_variant | MODIFIER |
| 5 | 86371285 | AX-170233499 | 0 | T | C | 0.43 | 5.36E-02 | 1.13E-01 | 2.75E+00 | 2.80E+00 | 6.31E-01 | no | Tmprss11d | intron_variant | MODIFIER |
| 5 | 86371675 | AX-168398113 | 0 | C | T | 0.43 | 5.36E-02 | 1.13E-01 | 2.75E+00 | 2.80E+00 | 6.31E-01 | no | Tmprss11d | intron_variant | MODIFIER |
| 5 | 86389799 | AX-170231543 | 0 | G | A | 0.43 | 5.36E-02 | 1.13E-01 | 2.75E+00 | 2.80E+00 | 6.31E-01 | no | | intergenic_variant | MODIFIER |
| 5 | 86408518 | AX-170334049 | 0 | T | C | 0.43 | 5.36E-02 | 1.13E-01 | 2.75E+00 | 2.80E+00 | 6.31E-01 | no | Tmprss11a | downstream_gene_variant | MODIFIER |
| 5 | 86427363 | AX-170133661 | 0 | G | A | 0.43 | 5.36E-02 | 1.13E-01 | 2.75E+00 | 2.80E+00 | 6.31E-01 | no | Tmprss11a | intron_variant | MODIFIER |
| 5 | 86431437 | AX-170283213 | 0 | C | T | 0.43 | 5.36E-02 | 1.13E-01 | 2.75E+00 | 2.80E+00 | 6.31E-01 | no | Tmprss11a | intron_variant | MODIFIER |
| 5 | 86487690 | AX-170139868 | 0 | A | G | 0.43 | 5.36E-02 | 1.13E-01 | 2.75E+00 | 2.80E+00 | 6.31E-01 | no | Tmprss11g | intron_variant | MODIFIER |
| 5 | 86515365 | AX-170525495 | 0 | T | C | 0.43 | 5.36E-02 | 1.13E-01 | 2.75E+00 | 2.80E+00 | 6.31E-01 | no | Tmprss11g | intron_variant | MODIFIER |
| 5 | 86519359 | AX-170170369 | 0 | A | G | 0.43 | 5.36E-02 | 1.13E-01 | 2.75E+00 | 2.80E+00 | 6.31E-01 | no | Tmprss11f | downstream_gene_variant | MODIFIER |
| 5 | 86529080 | AX-97798865 | 0 | T | C | 0.43 | 5.36E-02 | 1.13E-01 | 2.75E+00 | 2.80E+00 | 6.31E-01 | no | Tmprss11f | intron_variant | MODIFIER |
| 5 | 86529399 | AX-170072688 | 0 | T | C | 0.43 | 5.36E-02 | 1.13E-01 | 2.75E+00 | 2.80E+00 | 6.31E-01 | no | Tmprss11f | intron_variant | MODIFIER |
| 5 | 86590518 | AX-97822148 | 0 | T | C | 0.43 | 5.36E-02 | 1.13E-01 | 2.75E+00 | 2.80E+00 | 6.31E-01 | no | Tmprss11f | intron_variant | MODIFIER |
| 5 | 86707226 | AX-97802649 | 0 | T | C | 0.43 | 5.36E-02 | 1.13E-01 | 2.75E+00 | 2.80E+00 | 6.31E-01 | no | Tmprss11e | 3_prime_UTR_variant | MODIFIER |
| 5 | 86708607 | AX-170473659 | 0 | T | C | 0.43 | 5.36E-02 | 1.13E-01 | 2.75E+00 | 2.80E+00 | 6.31E-01 | no | Tmprss11e | intron_variant | MODIFIER |
| 5 | 86727342 | AX-170238007 | 0 | G | T | 0.43 | 5.36E-02 | 1.13E-01 | 2.75E+00 | 2.80E+00 | 6.31E-01 | no | Tmprss11e | missense_variant | MODERATE |
| 5 | 86748696 | AX-170212985 | 0 | A | G | 0.43 | 5.36E-02 | 1.13E-01 | 2.75E+00 | 2.80E+00 | 6.31E-01 | no | Tmprss11e | upstream_gene_variant | MODIFIER |
| 5 | 86749303 | AX-170527534 | 0 | G | A | 0.43 | 5.36E-02 | 1.13E-01 | 2.75E+00 | 2.80E+00 | 6.31E-01 | no | Tmprss11e | upstream_gene_variant | MODIFIER |
| 5 | 86762386 | AX-170420885 | 0 | G | T | 0.43 | 5.36E-02 | 1.13E-01 | 2.75E+00 | 2.80E+00 | 6.31E-01 | no | Gm16579 | downstream_gene_variant | MODIFIER |
| 5 | 86820906 | AX-170526410 | 0 | G | T | 0.43 | 5.36E-02 | 1.13E-01 | 2.75E+00 | 2.80E+00 | 6.31E-01 | no | Ythdc1 | intron_variant | MODIFIER |
| 5 | 86876111 | AX-170205578 | 0 | C | A | 0.43 | 5.36E-02 | 1.13E-01 | 2.75E+00 | 2.80E+00 | 6.31E-01 | no | | intergenic_variant | MODIFIER |
| 5 | 86966458 | AX-97850601 | 0 | G | A | 0.43 | 5.36E-02 | 1.13E-01 | 2.75E+00 | 2.80E+00 | 6.31E-01 | no | Gm20121 | upstream_gene_variant | MODIFIER |
| 5 | 87085179 | AX-97784430 | 0 | A | G | 0.43 | 5.36E-02 | 1.13E-01 | 2.75E+00 | 2.80E+00 | 6.31E-01 | no | Ugt2b36 | intron_variant | MODIFIER |
| 5 | 87478740 | AX-97795524 | 0 | A | T | 0.43 | 5.36E-02 | 1.13E-01 | 2.75E+00 | 2.80E+00 | 6.31E-01 | no | Ugt2a2 | intron_variant | MODIFIER |
| 5 | 87779364 | AX-170359678 | 0 | A | C | 0.43 | 5.36E-02 | 1.13E-01 | 2.75E+00 | 2.80E+00 | 6.31E-01 | no | Csn1s2a | intron_variant | MODIFIER |
| 5 | 87820054 | AX-170025524 | 0 | G | A | 0.43 | 5.36E-02 | 1.13E-01 | 2.75E+00 | 2.80E+00 | 6.31E-01 | no | Csn1s2b | intron_variant | MODIFIER |

| | | | | | | | | | | | | | | | |
|---|---|---|---|---|---|---|---|---|---|---|---|---|---|---|---|
| 5 | 87850439 | AX-170201537 | 0 | T | C | 0.43 | 5.36E-02 | 1.13E-01 | 2.75E+00 | 2.80E+00 | 6.31E-01 | no | Prr27 | downstream_gene_variant | MODIFIER |
| 5 | 87851076 | AX-170472417 | 0 | T | C | 0.43 | 5.36E-02 | 1.13E-01 | 2.75E+00 | 2.80E+00 | 6.31E-01 | no | Prr27 | downstream_gene_variant | MODIFIER |
| 5 | 87851157 | AX-170149989 | 0 | T | G | 0.43 | 5.36E-02 | 1.13E-01 | 2.75E+00 | 2.80E+00 | 6.31E-01 | no | Prr27 | downstream_gene_variant | MODIFIER |
| 5 | 87863845 | AX-170025531 | 0 | C | T | 0.43 | 5.36E-02 | 1.13E-01 | 2.75E+00 | 2.80E+00 | 6.31E-01 | no | | intergenic_variant | MODIFIER |
| 5 | 87866971 | AX-170025532 | 0 | A | T | 0.43 | 5.36E-02 | 1.13E-01 | 2.75E+00 | 2.80E+00 | 6.31E-01 | no | | intergenic_variant | MODIFIER |
| 5 | 88052271 | AX-97783670 | 0 | A | G | 0.43 | 5.36E-02 | 1.13E-01 | 2.75E+00 | 2.80E+00 | 6.31E-01 | no | | intergenic_variant | MODIFIER |
| 5 | 88052386 | AX-170171130 | 0 | T | C | 0.43 | 5.36E-02 | 1.13E-01 | 2.75E+00 | 2.80E+00 | 6.31E-01 | no | | intergenic_variant | MODIFIER |
| 5 | 88054013 | AX-170153393 | 0 | T | C | 0.43 | 5.36E-02 | 1.13E-01 | 2.75E+00 | 2.80E+00 | 6.31E-01 | no | | intergenic_variant | MODIFIER |
| 5 | 88139335 | AX-168395181 | 0 | T | C | 0.43 | 5.36E-02 | 1.13E-01 | 2.75E+00 | 2.80E+00 | 6.31E-01 | no | | intergenic_variant | MODIFIER |
| 5 | 88140263 | AX-97824790 | 0 | T | C | 0.43 | 5.36E-02 | 1.13E-01 | 2.75E+00 | 2.80E+00 | 6.31E-01 | no | Gm42622 | upstream_gene_variant | MODIFIER |
| 5 | 88212466 | AX-170164055 | 0 | G | C | 0.43 | 5.36E-02 | 1.13E-01 | 2.75E+00 | 2.80E+00 | 6.31E-01 | no | Gm42164 | downstream_gene_variant | MODIFIER |
| 5 | 88293555 | AX-97781247 | 0 | G | A | 0.43 | 5.36E-02 | 1.13E-01 | 2.75E+00 | 2.80E+00 | 6.31E-01 | no | | intergenic_variant | MODIFIER |
| 5 | 88300956 | AX-97784759 | 0 | T | C | 0.43 | 5.36E-02 | 1.13E-01 | 2.75E+00 | 2.80E+00 | 6.31E-01 | no | | intergenic_variant | MODIFIER |
| 5 | 88348061 | AX-97854861 | 0 | A | G | 0.43 | 5.36E-02 | 1.13E-01 | 2.75E+00 | 2.80E+00 | 6.31E-01 | no | | intergenic_variant | MODIFIER |
| 5 | 88379256 | AX-170089548 | 0 | G | A | 0.43 | 5.36E-02 | 1.13E-01 | 2.75E+00 | 2.80E+00 | 6.31E-01 | no | Amtn | intron_variant | MODIFIER |
| 5 | 88422241 | AX-170025538 | 0 | A | C | 0.43 | 5.36E-02 | 1.13E-01 | 2.75E+00 | 2.80E+00 | 6.31E-01 | no | Gm43431 | upstream_gene_variant | MODIFIER |
| 5 | 88422557 | AX-170025536 | 0 | T | C | 0.43 | 5.36E-02 | 1.13E-01 | 2.75E+00 | 2.80E+00 | 6.31E-01 | no | Gm43431 | upstream_gene_variant | MODIFIER |
| 5 | 88438506 | AX-170095388 | 0 | T | C | 0.43 | 5.36E-02 | 1.13E-01 | 2.75E+00 | 2.80E+00 | 6.31E-01 | no | | intergenic_variant | MODIFIER |
| 5 | 88500258 | AX-170168690 | 0 | G | A | 0.43 | 5.36E-02 | 1.13E-01 | 2.75E+00 | 2.80E+00 | 6.31E-01 | no | Enam | intron_variant | MODIFIER |
| 5 | 88559426 | AX-170234628 | 0 | A | G | 0.43 | 5.36E-02 | 1.13E-01 | 2.75E+00 | 2.80E+00 | 6.31E-01 | no | Utp3 | downstream_gene_variant | MODIFIER |
| 5 | 88843674 | AX-97802691 | 0 | A | G | 0.43 | 5.36E-02 | 1.13E-01 | 2.75E+00 | 2.80E+00 | 6.31E-01 | no | Mkrn1-ps1 | upstream_gene_variant | MODIFIER |
| 5 | 88843819 | AX-170412368 | 0 | C | T | 0.43 | 5.36E-02 | 1.13E-01 | 2.75E+00 | 2.80E+00 | 6.31E-01 | no | Mkrn1-ps1 | upstream_gene_variant | MODIFIER |
| 5 | 88853351 | AX-170464557 | 0 | C | A | 0.43 | 5.36E-02 | 1.13E-01 | 2.75E+00 | 2.80E+00 | 6.31E-01 | no | | intergenic_variant | MODIFIER |
| 5 | 88854879 | AX-170025549 | 0 | A | C | 0.43 | 5.36E-02 | 1.13E-01 | 2.75E+00 | 2.80E+00 | 6.31E-01 | no | | intergenic_variant | MODIFIER |
| 5 | 88855990 | AX-170407671 | 0 | C | G | 0.43 | 5.36E-02 | 1.13E-01 | 2.75E+00 | 2.80E+00 | 6.31E-01 | no | | intergenic_variant | MODIFIER |
| 5 | 88865153 | AX-97831956 | 0 | T | C | 0.43 | 5.36E-02 | 1.13E-01 | 2.75E+00 | 2.80E+00 | 6.31E-01 | no | | intergenic_variant | MODIFIER |
| 5 | 88903619 | AX-170025547 | 0 | T | C | 0.43 | 5.36E-02 | 1.13E-01 | 2.75E+00 | 2.80E+00 | 6.31E-01 | no | Slc4a4 | intron_variant | MODIFIER |
| 5 | 88930284 | AX-97809587 | 0 | C | T | 0.43 | 5.36E-02 | 1.13E-01 | 2.75E+00 | 2.80E+00 | 6.31E-01 | no | Slc4a4 | intron_variant | MODIFIER |
| 5 | 88931572 | AX-170506827 | 0 | G | C | 0.43 | 5.36E-02 | 1.13E-01 | 2.75E+00 | 2.80E+00 | 6.31E-01 | no | Slc4a4 | intron_variant | MODIFIER |
| 5 | 88931663 | AX-170378618 | 0 | A | G | 0.43 | 5.36E-02 | 1.13E-01 | 2.75E+00 | 2.80E+00 | 6.31E-01 | no | Slc4a4 | intron_variant | MODIFIER |
| 5 | 88935584 | AX-170180529 | 0 | C | T | 0.43 | 5.36E-02 | 1.13E-01 | 2.75E+00 | 2.80E+00 | 6.31E-01 | no | Slc4a4 | intron_variant | MODIFIER |
| 5 | 88937131 | AX-170221421 | 0 | T | C | 0.43 | 5.36E-02 | 1.13E-01 | 2.75E+00 | 2.80E+00 | 6.31E-01 | no | Slc4a4 | intron_variant | MODIFIER |
| 5 | 88950189 | AX-170409064 | 0 | A | C | 0.43 | 5.36E-02 | 1.13E-01 | 2.75E+00 | 2.80E+00 | 6.31E-01 | no | Slc4a4 | intron_variant | MODIFIER |
| 5 | 88958314 | AX-170089551 | 0 | A | G | 0.43 | 5.36E-02 | 1.13E-01 | 2.75E+00 | 2.80E+00 | 6.31E-01 | no | Slc4a4 | intron_variant | MODIFIER |
| 5 | 88963933 | AX-170289783 | 0 | T | G | 0.43 | 5.36E-02 | 1.13E-01 | 2.75E+00 | 2.80E+00 | 6.31E-01 | no | Slc4a4 | intron_variant | MODIFIER |
| 5 | 88981919 | AX-170116503 | 0 | G | T | 0.43 | 5.36E-02 | 1.13E-01 | 2.75E+00 | 2.80E+00 | 6.31E-01 | no | Slc4a4 | intron_variant | MODIFIER |
| 5 | 88994046 | AX-170025550 | 0 | C | T | 0.43 | 5.36E-02 | 1.13E-01 | 2.75E+00 | 2.80E+00 | 6.31E-01 | no | Slc4a4 | intron_variant | MODIFIER |
| 5 | 89011703 | AX-97791315 | 0 | A | G | 0.43 | 5.36E-02 | 1.13E-01 | 2.75E+00 | 2.80E+00 | 6.31E-01 | no | Slc4a4 | intron_variant | MODIFIER |
| 5 | 89011807 | AX-170282920 | 0 | G | A | 0.43 | 5.36E-02 | 1.13E-01 | 2.75E+00 | 2.80E+00 | 6.31E-01 | no | Slc4a4 | intron_variant | MODIFIER |
| 5 | 89027292 | AX-170492516 | 0 | T | C | 0.43 | 5.36E-02 | 1.13E-01 | 2.75E+00 | 2.80E+00 | 6.31E-01 | no | Slc4a4 | upstream_gene_variant | MODIFIER |
| 5 | 89036372 | AX-170300414 | 0 | A | C | 0.43 | 5.36E-02 | 1.13E-01 | 2.75E+00 | 2.80E+00 | 6.31E-01 | no | Slc4a4 | intron_variant | MODIFIER |
| 5 | 89054016 | AX-97800509 | 0 | A | C | 0.43 | 5.36E-02 | 1.13E-01 | 2.75E+00 | 2.80E+00 | 6.31E-01 | no | Slc4a4 | intron_variant | MODIFIER |
| 5 | 89078585 | AX-97856886 | 0 | A | G | 0.43 | 5.36E-02 | 1.13E-01 | 2.75E+00 | 2.80E+00 | 6.31E-01 | no | Slc4a4 | intron_variant | MODIFIER |
| 5 | 89101857 | AX-97800633 | 0 | G | A | 0.43 | 5.36E-02 | 1.13E-01 | 2.75E+00 | 2.80E+00 | 6.31E-01 | no | Slc4a4 | intron_variant | MODIFIER |
| 5 | 89269364 | AX-168409749 | 0 | T | C | 0.43 | 5.36E-02 | 1.13E-01 | 2.75E+00 | 2.80E+00 | 6.31E-01 | no | | regulatory_region_variant | MODIFIER |
| 5 | 89271411 | AX-170140352 | 0 | T | G | 0.43 | 5.36E-02 | 1.13E-01 | 2.75E+00 | 2.80E+00 | 6.31E-01 | no | | intergenic_variant | MODIFIER |
| 5 | 89271745 | AX-170217935 | 0 | C | A | 0.43 | 5.36E-02 | 1.13E-01 | 2.75E+00 | 2.80E+00 | 6.31E-01 | no | | intergenic_variant | MODIFIER |
| 5 | 89272000 | AX-170247161 | 0 | G | A | 0.43 | 5.36E-02 | 1.13E-01 | 2.75E+00 | 2.80E+00 | 6.31E-01 | no | | intergenic_variant | MODIFIER |
| 5 | 89272295 | AX-170161609 | 0 | C | G | 0.43 | 5.36E-02 | 1.13E-01 | 2.75E+00 | 2.80E+00 | 6.31E-01 | no | | intergenic_variant | MODIFIER |
| 5 | 89273269 | AX-170443942 | 0 | A | C | 0.43 | 5.36E-02 | 1.13E-01 | 2.75E+00 | 2.80E+00 | 6.31E-01 | no | | intergenic_variant | MODIFIER |
| 5 | 89280480 | AX-170434638 | 0 | C | T | 0.43 | 5.36E-02 | 1.13E-01 | 2.75E+00 | 2.80E+00 | 6.31E-01 | no | | intergenic_variant | MODIFIER |
| 5 | 89280647 | AX-170455496 | 0 | T | A | 0.43 | 5.36E-02 | 1.13E-01 | 2.75E+00 | 2.80E+00 | 6.31E-01 | no | | intergenic_variant | MODIFIER |
| 5 | 89281075 | AX-170072730 | 0 | A | G | 0.43 | 5.36E-02 | 1.13E-01 | 2.75E+00 | 2.80E+00 | 6.31E-01 | no | | intergenic_variant | MODIFIER |
| 5 | 89369167 | AX-170250396 | 0 | G | A | 0.43 | 5.36E-02 | 1.13E-01 | 2.75E+00 | 2.80E+00 | 6.31E-01 | no | | intergenic_variant | MODIFIER |

| | | | | | | | | | | | | | | | |
|---|---|---|---|---|---|---|---|---|---|---|---|---|---|---|---|
| 5 | 89371113 | AX-170144855 | 0 | T | C | 0.43 | 5.36E-02 | 1.13E-01 | 2.75E+00 | 2.80E+00 | 6.31E-01 | no | | intergenic_variant | MODIFIER |
| 5 | 89373314 | AX-170288810 | 0 | A | T | 0.43 | 5.36E-02 | 1.13E-01 | 2.75E+00 | 2.80E+00 | 6.31E-01 | no | | intergenic_variant | MODIFIER |
| 5 | 89384391 | AX-170025556 | 0 | C | T | 0.43 | 5.36E-02 | 1.13E-01 | 2.75E+00 | 2.80E+00 | 6.31E-01 | no | | regulatory_region_variant | MODIFIER |
| 5 | 89390911 | AX-170490218 | 0 | T | C | 0.43 | 5.36E-02 | 1.13E-01 | 2.75E+00 | 2.80E+00 | 6.31E-01 | no | | intergenic_variant | MODIFIER |
| 5 | 89448898 | AX-97823434 | 0 | C | T | 0.43 | 5.36E-02 | 1.13E-01 | 2.75E+00 | 2.80E+00 | 6.31E-01 | no | Gc | intron_variant | MODIFIER |
| 5 | 89520851 | AX-170025553 | 0 | G | C | 0.43 | 5.36E-02 | 1.13E-01 | 2.75E+00 | 2.80E+00 | 6.31E-01 | no | Gm43515 | upstream_gene_variant | MODIFIER |
| 5 | 89536923 | AX-97791851 | 0 | T | C | 0.43 | 5.36E-02 | 1.13E-01 | 2.75E+00 | 2.80E+00 | 6.31E-01 | no | Npffr2 | intron_variant | MODIFIER |
| 5 | 89543065 | AX-170025558 | 0 | C | T | 0.43 | 5.36E-02 | 1.13E-01 | 2.75E+00 | 2.80E+00 | 6.31E-01 | no | Npffr2 | intron_variant | MODIFIER |
| 5 | 89579848 | AX-97844783 | 0 | A | G | 0.43 | 5.36E-02 | 1.13E-01 | 2.75E+00 | 2.80E+00 | 6.31E-01 | no | Npffr2 | intron_variant | MODIFIER |
| 5 | 89590929 | AX-170187009 | 0 | C | T | 0.43 | 5.36E-02 | 1.13E-01 | 2.75E+00 | 2.80E+00 | 6.31E-01 | no | | intergenic_variant | MODIFIER |
| 5 | 89622150 | AX-170108131 | 0 | C | A | 0.43 | 5.36E-02 | 1.13E-01 | 2.75E+00 | 2.80E+00 | 6.31E-01 | no | | intergenic_variant | MODIFIER |
| 5 | 89683123 | AX-170089552 | 0 | C | T | 0.43 | 5.36E-02 | 1.13E-01 | 2.75E+00 | 2.80E+00 | 6.31E-01 | no | Adamts3 | intron_variant | MODIFIER |
| 5 | 89685274 | AX-170377025 | 0 | G | A | 0.43 | 5.36E-02 | 1.13E-01 | 2.75E+00 | 2.80E+00 | 6.31E-01 | no | Adamts3 | intron_variant | MODIFIER |
| 5 | 89705458 | AX-97833895 | 0 | T | G | 0.43 | 5.36E-02 | 1.13E-01 | 2.75E+00 | 2.80E+00 | 6.31E-01 | no | Adamts3 | intron_variant | MODIFIER |
| 5 | 89738260 | AX-97813262 | 0 | T | C | 0.43 | 5.36E-02 | 1.13E-01 | 2.75E+00 | 2.80E+00 | 6.31E-01 | no | Adamts3 | intron_variant | MODIFIER |
| 5 | 89741253 | AX-170335443 | 0 | C | T | 0.43 | 5.36E-02 | 1.13E-01 | 2.75E+00 | 2.80E+00 | 6.31E-01 | no | Adamts3 | intron_variant | MODIFIER |
| 5 | 89741634 | AX-170379166 | 0 | T | A | 0.43 | 5.36E-02 | 1.13E-01 | 2.75E+00 | 2.80E+00 | 6.31E-01 | no | Adamts3 | intron_variant | MODIFIER |
| 5 | 89754030 | AX-170404857 | 0 | G | A | 0.43 | 5.36E-02 | 1.13E-01 | 2.75E+00 | 2.80E+00 | 6.31E-01 | no | Adamts3 | intron_variant | MODIFIER |
| 5 | 89764247 | AX-170311645 | 0 | C | T | 0.43 | 5.36E-02 | 1.13E-01 | 2.75E+00 | 2.80E+00 | 6.31E-01 | no | Adamts3 | intron_variant | MODIFIER |
| 5 | 89778611 | AX-170256110 | 0 | G | A | 0.43 | 5.36E-02 | 1.13E-01 | 2.75E+00 | 2.80E+00 | 6.31E-01 | no | Adamts3 | intron_variant | MODIFIER |
| 5 | 89780359 | AX-97804224 | 0 | C | T | 0.43 | 5.36E-02 | 1.13E-01 | 2.75E+00 | 2.80E+00 | 6.31E-01 | no | Adamts3 | intron_variant | MODIFIER |
| 5 | 89784781 | AX-170372150 | 0 | A | G | 0.43 | 5.36E-02 | 1.13E-01 | 2.75E+00 | 2.80E+00 | 6.31E-01 | no | Adamts3 | intron_variant | MODIFIER |
| 5 | 89799501 | AX-170339735 | 0 | C | T | 0.43 | 5.36E-02 | 1.13E-01 | 2.75E+00 | 2.80E+00 | 6.31E-01 | no | Adamts3 | intron_variant | MODIFIER |
| 5 | 89804584 | AX-170391500 | 0 | A | G | 0.43 | 5.36E-02 | 1.13E-01 | 2.75E+00 | 2.80E+00 | 6.31E-01 | no | Adamts3 | intron_variant | MODIFIER |
| 5 | 89804717 | AX-170317409 | 0 | A | T | 0.43 | 5.36E-02 | 1.13E-01 | 2.75E+00 | 2.80E+00 | 6.31E-01 | no | Adamts3 | intron_variant | MODIFIER |
| 5 | 89818182 | AX-170165786 | 0 | A | T | 0.43 | 5.36E-02 | 1.13E-01 | 2.75E+00 | 2.80E+00 | 6.31E-01 | no | Adamts3 | intron_variant | MODIFIER |
| 5 | 89821554 | AX-170534068 | 0 | C | G | 0.43 | 5.36E-02 | 1.13E-01 | 2.75E+00 | 2.80E+00 | 6.31E-01 | no | Adamts3 | intron_variant | MODIFIER |
| 5 | 89841199 | AX-170153536 | 0 | G | C | 0.43 | 5.36E-02 | 1.13E-01 | 2.75E+00 | 2.80E+00 | 6.31E-01 | no | Adamts3 | intron_variant | MODIFIER |
| 5 | 89841796 | AX-97815104 | 0 | C | T | 0.43 | 5.36E-02 | 1.13E-01 | 2.75E+00 | 2.80E+00 | 6.31E-01 | no | Adamts3 | intron_variant | MODIFIER |
| 5 | 89842011 | AX-170500109 | 0 | A | G | 0.43 | 5.36E-02 | 1.13E-01 | 2.75E+00 | 2.80E+00 | 6.31E-01 | no | Adamts3 | intron_variant | MODIFIER |
| 5 | 89843445 | AX-170299472 | 0 | A | G | 0.43 | 5.36E-02 | 1.13E-01 | 2.75E+00 | 2.80E+00 | 6.31E-01 | no | Adamts3 | intron_variant | MODIFIER |
| 5 | 89862117 | AX-170153181 | 0 | C | A | 0.43 | 5.36E-02 | 1.13E-01 | 2.75E+00 | 2.80E+00 | 6.31E-01 | no | Adamts3 | intron_variant | MODIFIER |
| 5 | 89911481 | AX-170397733 | 0 | A | C | 0.43 | 5.36E-02 | 1.13E-01 | 2.75E+00 | 2.80E+00 | 6.31E-01 | no | | intergenic_variant | MODIFIER |
| 5 | 89918672 | AX-170025563 | 0 | G | A | 0.43 | 5.36E-02 | 1.13E-01 | 2.75E+00 | 2.80E+00 | 6.31E-01 | no | | intergenic_variant | MODIFIER |
| 5 | 89931062 | AX-97835397 | 0 | C | A | 0.43 | 5.36E-02 | 1.13E-01 | 2.75E+00 | 2.80E+00 | 6.31E-01 | no | | intergenic_variant | MODIFIER |
| 5 | 89945487 | AX-170025562 | 0 | T | G | 0.43 | 5.36E-02 | 1.13E-01 | 2.75E+00 | 2.80E+00 | 6.31E-01 | no | | intergenic_variant | MODIFIER |
| 5 | 89945739 | AX-170089554 | 0 | G | T | 0.43 | 5.36E-02 | 1.13E-01 | 2.75E+00 | 2.80E+00 | 6.31E-01 | no | | regulatory_region_variant | MODIFIER |
| 5 | 89952518 | AX-170025568 | 0 | T | C | 0.43 | 5.36E-02 | 1.13E-01 | 2.75E+00 | 2.80E+00 | 6.31E-01 | no | | intergenic_variant | MODIFIER |
| 5 | 89979048 | AX-170025566 | 0 | G | T | 0.43 | 5.36E-02 | 1.13E-01 | 2.75E+00 | 2.80E+00 | 6.31E-01 | no | | intergenic_variant | MODIFIER |
| 5 | 90016215 | AX-170170921 | 0 | T | G | 0.43 | 5.36E-02 | 1.13E-01 | 2.75E+00 | 2.80E+00 | 6.31E-01 | no | | intergenic_variant | MODIFIER |
| 5 | 90016773 | AX-97850813 | 0 | A | G | 0.43 | 5.36E-02 | 1.13E-01 | 2.75E+00 | 2.80E+00 | 6.31E-01 | no | | intergenic_variant | MODIFIER |
| 5 | 90021013 | AX-170287018 | 0 | A | G | 0.43 | 5.36E-02 | 1.13E-01 | 2.75E+00 | 2.80E+00 | 6.31E-01 | no | | intergenic_variant | MODIFIER |
| 5 | 90023141 | AX-170417386 | 0 | T | C | 0.43 | 5.36E-02 | 1.13E-01 | 2.75E+00 | 2.80E+00 | 6.31E-01 | no | | intergenic_variant | MODIFIER |
| 5 | 90025977 | AX-97784816 | 0 | A | G | 0.43 | 5.36E-02 | 1.13E-01 | 2.75E+00 | 2.80E+00 | 6.31E-01 | no | | intergenic_variant | MODIFIER |
| 5 | 90044220 | AX-170025567 | 0 | A | G | 0.43 | 5.36E-02 | 1.13E-01 | 2.75E+00 | 2.80E+00 | 6.31E-01 | no | | intergenic_variant | MODIFIER |
| 5 | 90044271 | AX-170025561 | 0 | T | C | 0.43 | 5.36E-02 | 1.13E-01 | 2.75E+00 | 2.80E+00 | 6.31E-01 | no | | intergenic_variant | MODIFIER |
| 5 | 90049981 | AX-170228712 | 0 | C | G | 0.43 | 5.36E-02 | 1.13E-01 | 2.75E+00 | 2.80E+00 | 6.31E-01 | no | | intergenic_variant | MODIFIER |
| 5 | 90056593 | AX-170453126 | 0 | G | A | 0.43 | 5.36E-02 | 1.13E-01 | 2.75E+00 | 2.80E+00 | 6.31E-01 | no | | intergenic_variant | MODIFIER |
| 5 | 90060875 | AX-170025565 | 0 | C | T | 0.43 | 5.36E-02 | 1.13E-01 | 2.75E+00 | 2.80E+00 | 6.31E-01 | no | | intergenic_variant | MODIFIER |
| 5 | 90071241 | AX-170025560 | 0 | T | C | 0.43 | 5.36E-02 | 1.13E-01 | 2.75E+00 | 2.80E+00 | 6.31E-01 | no | | intergenic_variant | MODIFIER |
| 5 | 90073125 | AX-97792086 | 0 | G | A | 0.43 | 5.36E-02 | 1.13E-01 | 2.75E+00 | 2.80E+00 | 6.31E-01 | no | | intergenic_variant | MODIFIER |
| 5 | 90112929 | AX-170025570 | 0 | G | A | 0.43 | 5.36E-02 | 1.13E-01 | 2.75E+00 | 2.80E+00 | 6.31E-01 | no | | intergenic_variant | MODIFIER |
| 5 | 90127416 | AX-170025564 | 0 | A | T | 0.43 | 5.36E-02 | 1.13E-01 | 2.75E+00 | 2.80E+00 | 6.31E-01 | no | Eif5al3-ps | upstream_gene_variant | MODIFIER |
| 5 | 90145416 | AX-170321318 | 0 | T | C | 0.43 | 5.36E-02 | 1.13E-01 | 2.75E+00 | 2.80E+00 | 6.31E-01 | no | | intergenic_variant | MODIFIER |

| | | | | | | | | | | | | | | | |
|---|---|---|---|---|---|---|---|---|---|---|---|---|---|---|---|
| 5 | 90151460 | AX-170506567 | 0 | G | A | 0.43 | 5.36E-02 | 1.13E-01 | 2.75E+00 | 2.80E+00 | 6.31E-01 | no | | intergenic_variant | MODIFIER |
| 5 | 90168073 | AX-170517423 | 0 | T | C | 0.43 | 5.36E-02 | 1.13E-01 | 2.75E+00 | 2.80E+00 | 6.31E-01 | no | | intergenic_variant | MODIFIER |
| 5 | 90169220 | AX-170506753 | 0 | G | A | 0.43 | 5.36E-02 | 1.13E-01 | 2.75E+00 | 2.80E+00 | 6.31E-01 | no | | intergenic_variant | MODIFIER |
| 5 | 90177904 | AX-170137033 | 0 | C | T | 0.43 | 5.36E-02 | 1.13E-01 | 2.75E+00 | 2.80E+00 | 6.31E-01 | no | | regulatory_region_variant | MODIFIER |
| 5 | 90261664 | AX-170240198 | 0 | C | T | 0.43 | 5.36E-02 | 1.13E-01 | 2.75E+00 | 2.80E+00 | 6.31E-01 | no | Ankrd17 | intron_variant | MODIFIER |
| 5 | 90269497 | AX-170371972 | 0 | G | A | 0.43 | 5.36E-02 | 1.13E-01 | 2.75E+00 | 2.80E+00 | 6.31E-01 | no | Ankrd17 | intron_variant | MODIFIER |
| 5 | 90307164 | AX-170310391 | 0 | A | T | 0.43 | 5.36E-02 | 1.13E-01 | 2.75E+00 | 2.80E+00 | 6.31E-01 | no | Ankrd17 | intron_variant | MODIFIER |
| 5 | 90467710 | AX-170146229 | 0 | T | C | 0.43 | 5.36E-02 | 1.13E-01 | 2.75E+00 | 2.80E+00 | 6.31E-01 | no | Alb | intron_variant | MODIFIER |
| 5 | 90575386 | AX-97821216 | 0 | A | G | 0.43 | 5.36E-02 | 1.13E-01 | 2.75E+00 | 2.80E+00 | 6.31E-01 | no | 5830473C10Rik | intron_variant | MODIFIER |
| 5 | 90599978 | AX-97785665 | 0 | A | G | 0.43 | 5.36E-02 | 1.13E-01 | 2.75E+00 | 2.80E+00 | 6.31E-01 | no | Rassf6 | downstream_gene_variant | MODIFIER |
| 5 | 90619708 | AX-170025571 | 0 | G | A | 0.43 | 5.36E-02 | 1.13E-01 | 2.75E+00 | 2.80E+00 | 6.31E-01 | no | Rassf6 | intron_variant | MODIFIER |
| 5 | 90621319 | AX-170025579 | 0 | T | C | 0.43 | 5.36E-02 | 1.13E-01 | 2.75E+00 | 2.80E+00 | 6.31E-01 | no | Rassf6 | intron_variant | MODIFIER |
| 5 | 90622478 | AX-170025572 | 0 | C | T | 0.43 | 5.36E-02 | 1.13E-01 | 2.75E+00 | 2.80E+00 | 6.31E-01 | no | Rassf6 | intron_variant | MODIFIER |
| 5 | 90640890 | AX-170157408 | 0 | A | T | 0.43 | 5.36E-02 | 1.13E-01 | 2.75E+00 | 2.80E+00 | 6.31E-01 | no | Rassf6 | upstream_gene_variant | MODIFIER |
| 5 | 90641584 | AX-170086121 | 0 | G | A | 0.43 | 5.36E-02 | 1.13E-01 | 2.75E+00 | 2.80E+00 | 6.31E-01 | no | Rassf6 | upstream_gene_variant | MODIFIER |
| 5 | 90644465 | AX-170195817 | 0 | C | T | 0.43 | 5.36E-02 | 1.13E-01 | 2.75E+00 | 2.80E+00 | 6.31E-01 | no | Rassf6 | upstream_gene_variant | MODIFIER |
| 5 | 90644923 | AX-170025580 | 0 | C | T | 0.43 | 5.36E-02 | 1.13E-01 | 2.75E+00 | 2.80E+00 | 6.31E-01 | no | Rassf6 | upstream_gene_variant | MODIFIER |
| 5 | 90645112 | AX-170025573 | 0 | C | A | 0.43 | 5.36E-02 | 1.13E-01 | 2.75E+00 | 2.80E+00 | 6.31E-01 | no | Rassf6 | upstream_gene_variant | MODIFIER |
| 5 | 90652487 | AX-170341208 | 0 | A | G | 0.43 | 5.36E-02 | 1.13E-01 | 2.75E+00 | 2.80E+00 | 6.31E-01 | no | | regulatory_region_variant | MODIFIER |
| 5 | 90655619 | AX-170072752 | 0 | G | C | 0.43 | 5.36E-02 | 1.13E-01 | 2.75E+00 | 2.80E+00 | 6.31E-01 | no | | intergenic_variant | MODIFIER |
| 5 | 90656626 | AX-170072753 | 0 | C | G | 0.43 | 5.36E-02 | 1.13E-01 | 2.75E+00 | 2.80E+00 | 6.31E-01 | no | | regulatory_region_variant | MODIFIER |
| 5 | 90656666 | AX-170072751 | 0 | C | T | 0.43 | 5.36E-02 | 1.13E-01 | 2.75E+00 | 2.80E+00 | 6.31E-01 | no | | regulatory_region_variant | MODIFIER |
| 5 | 90656710 | AX-97850743 | 0 | G | A | 0.43 | 5.36E-02 | 1.13E-01 | 2.75E+00 | 2.80E+00 | 6.31E-01 | no | | regulatory_region_variant | MODIFIER |
| 5 | 90680330 | AX-97818348 | 0 | T | A | 0.43 | 5.36E-02 | 1.13E-01 | 2.75E+00 | 2.80E+00 | 6.31E-01 | no | | intergenic_variant | MODIFIER |
| 5 | 90689646 | AX-170487401 | 0 | G | C | 0.43 | 5.36E-02 | 1.13E-01 | 2.75E+00 | 2.80E+00 | 6.31E-01 | no | | regulatory_region_variant | MODIFIER |
| 5 | 90741598 | AX-97810993 | 0 | T | C | 0.43 | 5.36E-02 | 1.13E-01 | 2.75E+00 | 2.80E+00 | 6.31E-01 | no | | intergenic_variant | MODIFIER |
| 5 | 90741728 | AX-170402012 | 0 | A | C | 0.43 | 5.36E-02 | 1.13E-01 | 2.75E+00 | 2.80E+00 | 6.31E-01 | no | | intergenic_variant | MODIFIER |
| 5 | 90742276 | AX-170123408 | 0 | G | A | 0.43 | 5.36E-02 | 1.13E-01 | 2.75E+00 | 2.80E+00 | 6.31E-01 | no | | regulatory_region_variant | MODIFIER |
| 5 | 90763645 | AX-97827655 | 0 | T | C | 0.43 | 5.36E-02 | 1.13E-01 | 2.75E+00 | 2.80E+00 | 6.31E-01 | no | Cxcl5 | downstream_gene_variant | MODIFIER |
| 5 | 90774388 | AX-97832155 | 0 | G | T | 0.43 | 5.36E-02 | 1.13E-01 | 2.75E+00 | 2.80E+00 | 6.31E-01 | no | Ppbp | downstream_gene_variant | MODIFIER |
| 5 | 92081078 | AX-170535362 | 0 | G | A | 0.43 | 5.36E-02 | 1.13E-01 | 2.75E+00 | 2.80E+00 | 6.31E-01 | no | G3bp2 | intron_variant | MODIFIER |
| 6 | 146649483 | AX-170458051 | 0 | C | T | 0.07 | -9.86E-02 | 2.08E-01 | 2.67E+00 | 2.76E+00 | 6.31E-01 | no | Med21 | intron_variant | MODIFIER |
| 1 | 149236996 | AX-170223231 | 1 | A | G | 0.196 | 6.27E-02 | 1.30E-01 | 2.62E+00 | 2.69E+00 | 6.31E-01 | no | | intergenic_variant | MODIFIER |
| 8 | 34463139 | AX-170357990 | 1 | C | T | 0.143 | -6.44E-02 | 1.34E-01 | 2.64E+00 | 2.70E+00 | 6.31E-01 | no | Gm45349 | non_coding_transcript_exon_variant | MODIFIER |
| 4 | 151352444 | AX-170450417 | 0 | C | T | 0.114 | 8.67E-02 | 1.83E-01 | 2.69E+00 | 2.77E+00 | 6.31E-01 | no | Camta1 | intron_variant | MODIFIER |
| 9 | 33582181 | AX-170032093 | 0 | C | T | 0.447 | 4.02E-02 | 8.39E-02 | 2.62E+00 | 2.70E+00 | 6.31E-01 | no | | intergenic_variant | MODIFIER |
| 15 | 81297734 | AX-170429899 | 0 | C | G | 0.561 | 4.00E-02 | 8.26E-02 | 2.60E+00 | 2.67E+00 | 6.31E-01 | no | 8430426J06Rik | intron_variant,non_coding_transcript_variant | MODIFIER |
| 4 | 82240270 | AX-170262293 | 1 | G | A | 0.25 | 5.09E-02 | 1.03E-01 | 2.56E+00 | 2.63E+00 | 6.31E-01 | no | | intergenic_variant | MODIFIER |
| 1 | 128149002 | AX-170001669 | 1 | A | G | 0.536 | 4.08E-02 | 8.41E-02 | 2.59E+00 | 2.66E+00 | 6.31E-01 | no | R3hdm1 | intron_variant | MODIFIER |
| 1 | 19455084 | AX-170194976 | 0 | T | C | 0.395 | 3.95E-02 | 8.22E-02 | 2.63E+00 | 2.68E+00 | 6.32E-01 | no | | intergenic_variant | MODIFIER |
| 7 | 112893053 | AX-170107993 | 1 | G | A | 0.446 | 4.38E-02 | 9.26E-02 | 2.70E+00 | 2.76E+00 | 6.32E-01 | no | Tead1 | intron_variant | MODIFIER |
| 1 | 60213495 | AX-170041385 | 0 | A | G | 0.228 | 4.66E-02 | 9.91E-02 | 2.71E+00 | 2.80E+00 | 6.32E-01 | no | Nbeal1 | intron_variant | MODIFIER |
| 5 | 39638873 | AX-170446479 | 0 | A | G | 0.211 | -5.16E-02 | 1.07E-01 | 2.60E+00 | 2.68E+00 | 6.32E-01 | no | Hs3st1 | intron_variant | MODIFIER |
| 5 | 39640182 | AX-170481987 | 0 | G | T | 0.211 | -5.16E-02 | 1.07E-01 | 2.60E+00 | 2.68E+00 | 6.32E-01 | no | Hs3st1 | intron_variant | MODIFIER |
| 5 | 39655604 | AX-97833108 | 0 | G | A | 0.211 | -5.16E-02 | 1.07E-01 | 2.60E+00 | 2.68E+00 | 6.32E-01 | no | Hs3st1 | intron_variant | MODIFIER |
| 5 | 39666314 | AX-170429603 | 0 | T | G | 0.211 | -5.16E-02 | 1.07E-01 | 2.60E+00 | 2.68E+00 | 6.32E-01 | no | Hs3st1 | intron_variant | MODIFIER |
| 5 | 39666391 | AX-170386846 | 0 | C | T | 0.211 | -5.16E-02 | 1.07E-01 | 2.60E+00 | 2.68E+00 | 6.32E-01 | no | Hs3st1 | intron_variant | MODIFIER |
| 5 | 39680485 | AX-170110647 | 0 | T | C | 0.211 | -5.16E-02 | 1.07E-01 | 2.60E+00 | 2.68E+00 | 6.32E-01 | no | Hs3st1 | intron_variant | MODIFIER |
| 5 | 39681816 | AX-170095292 | 0 | A | C | 0.211 | -5.16E-02 | 1.07E-01 | 2.60E+00 | 2.68E+00 | 6.32E-01 | no | Hs3st1 | intron_variant | MODIFIER |
| 5 | 39954104 | AX-170277913 | 0 | A | G | 0.211 | -5.16E-02 | 1.07E-01 | 2.60E+00 | 2.68E+00 | 6.32E-01 | no | | intergenic_variant | MODIFIER |
| 5 | 40033093 | AX-170520540 | 0 | A | T | 0.211 | -5.16E-02 | 1.07E-01 | 2.60E+00 | 2.68E+00 | 6.32E-01 | no | | intergenic_variant | MODIFIER |
| 5 | 40035576 | AX-170397042 | 0 | T | G | 0.211 | -5.16E-02 | 1.07E-01 | 2.60E+00 | 2.68E+00 | 6.32E-01 | no | | intergenic_variant | MODIFIER |
| 5 | 40042356 | AX-170329152 | 0 | T | C | 0.211 | -5.16E-02 | 1.07E-01 | 2.60E+00 | 2.68E+00 | 6.32E-01 | no | | intergenic_variant | MODIFIER |
| 5 | 40044227 | AX-97803132 | 0 | A | G | 0.211 | -5.16E-02 | 1.07E-01 | 2.60E+00 | 2.68E+00 | 6.32E-01 | no | | intergenic_variant | MODIFIER |

| | | | | | | | | | | | | | | | |
|---|---|---|---|---|---|---|---|---|---|---|---|---|---|---|---|
| 5 | 40095218 | AX-170300679 | 0 | C | T | 0.211 | -5.16E-02 | 1.07E-01 | 2.60E+00 | 2.68E+00 | 6.32E-01 | no | | intergenic_variant | MODIFIER |
| 5 | 40101526 | AX-170466384 | 0 | C | T | 0.211 | -5.16E-02 | 1.07E-01 | 2.60E+00 | 2.68E+00 | 6.32E-01 | no | | intergenic_variant | MODIFIER |
| 5 | 40115847 | AX-170525833 | 0 | A | C | 0.211 | -5.16E-02 | 1.07E-01 | 2.60E+00 | 2.68E+00 | 6.32E-01 | no | | intergenic_variant | MODIFIER |
| 5 | 40119767 | AX-170071867 | 0 | A | G | 0.211 | -5.16E-02 | 1.07E-01 | 2.60E+00 | 2.68E+00 | 6.32E-01 | no | | intergenic_variant | MODIFIER |
| 5 | 40123298 | AX-170362599 | 0 | T | C | 0.211 | -5.16E-02 | 1.07E-01 | 2.60E+00 | 2.68E+00 | 6.32E-01 | no | | intergenic_variant | MODIFIER |
| 5 | 40154156 | AX-97794371 | 0 | C | T | 0.211 | -5.16E-02 | 1.07E-01 | 2.60E+00 | 2.68E+00 | 6.32E-01 | no | | intergenic_variant | MODIFIER |
| 5 | 40178016 | AX-170317308 | 0 | T | A | 0.211 | -5.16E-02 | 1.07E-01 | 2.60E+00 | 2.68E+00 | 6.32E-01 | no | | intergenic_variant | MODIFIER |
| 5 | 40202993 | AX-170351194 | 0 | A | G | 0.211 | -5.16E-02 | 1.07E-01 | 2.60E+00 | 2.68E+00 | 6.32E-01 | no | | intergenic_variant | MODIFIER |
| 5 | 40248073 | AX-170114138 | 0 | C | T | 0.211 | -5.16E-02 | 1.07E-01 | 2.60E+00 | 2.68E+00 | 6.32E-01 | no | | intergenic_variant | MODIFIER |
| 5 | 40269429 | AX-170132058 | 0 | T | C | 0.211 | -5.16E-02 | 1.07E-01 | 2.60E+00 | 2.68E+00 | 6.32E-01 | no | | intergenic_variant | MODIFIER |
| 5 | 40290521 | AX-170436726 | 0 | T | G | 0.211 | -5.16E-02 | 1.07E-01 | 2.60E+00 | 2.68E+00 | 6.32E-01 | no | | intergenic_variant | MODIFIER |
| 5 | 40295031 | AX-170156505 | 0 | C | G | 0.211 | -5.16E-02 | 1.07E-01 | 2.60E+00 | 2.68E+00 | 6.32E-01 | no | | intergenic_variant | MODIFIER |
| 5 | 40345332 | AX-170024763 | 0 | A | G | 0.211 | -5.16E-02 | 1.07E-01 | 2.60E+00 | 2.68E+00 | 6.32E-01 | no | | intergenic_variant | MODIFIER |
| 5 | 40345400 | AX-170024758 | 0 | T | C | 0.211 | -5.16E-02 | 1.07E-01 | 2.60E+00 | 2.68E+00 | 6.32E-01 | no | | intergenic_variant | MODIFIER |
| 5 | 40367822 | AX-170071871 | 0 | A | G | 0.211 | -5.16E-02 | 1.07E-01 | 2.60E+00 | 2.68E+00 | 6.32E-01 | no | | intergenic_variant | MODIFIER |
| 5 | 40374076 | AX-170024762 | 0 | A | G | 0.211 | -5.16E-02 | 1.07E-01 | 2.60E+00 | 2.68E+00 | 6.32E-01 | no | | intergenic_variant | MODIFIER |
| 5 | 40377184 | AX-170024764 | 0 | T | G | 0.211 | -5.16E-02 | 1.07E-01 | 2.60E+00 | 2.68E+00 | 6.32E-01 | no | | intergenic_variant | MODIFIER |
| 5 | 40377556 | AX-170071873 | 0 | A | T | 0.211 | -5.16E-02 | 1.07E-01 | 2.60E+00 | 2.68E+00 | 6.32E-01 | no | | intergenic_variant | MODIFIER |
| 5 | 40377723 | AX-170071870 | 0 | C | T | 0.211 | -5.16E-02 | 1.07E-01 | 2.60E+00 | 2.68E+00 | 6.32E-01 | no | | intergenic_variant | MODIFIER |
| 5 | 40615329 | AX-97796110 | 0 | G | A | 0.211 | -5.16E-02 | 1.07E-01 | 2.60E+00 | 2.68E+00 | 6.32E-01 | no | | intergenic_variant | MODIFIER |
| 5 | 40666282 | AX-97847579 | 0 | A | G | 0.211 | -5.16E-02 | 1.07E-01 | 2.60E+00 | 2.68E+00 | 6.32E-01 | no | | intergenic_variant | MODIFIER |
| 5 | 40676054 | AX-170118570 | 0 | A | G | 0.211 | -5.16E-02 | 1.07E-01 | 2.60E+00 | 2.68E+00 | 6.32E-01 | no | | intergenic_variant | MODIFIER |
| 5 | 40710488 | AX-170287912 | 0 | T | G | 0.211 | -5.16E-02 | 1.07E-01 | 2.60E+00 | 2.68E+00 | 6.32E-01 | no | | intergenic_variant | MODIFIER |
| 5 | 40714555 | AX-170192033 | 0 | C | T | 0.211 | -5.16E-02 | 1.07E-01 | 2.60E+00 | 2.68E+00 | 6.32E-01 | no | | intergenic_variant | MODIFIER |
| 5 | 40714738 | AX-170351236 | 0 | C | T | 0.211 | -5.16E-02 | 1.07E-01 | 2.60E+00 | 2.68E+00 | 6.32E-01 | no | | intergenic_variant | MODIFIER |
| 5 | 40724067 | AX-170118290 | 0 | A | T | 0.211 | -5.16E-02 | 1.07E-01 | 2.60E+00 | 2.68E+00 | 6.32E-01 | no | | intergenic_variant | MODIFIER |
| 5 | 40732289 | AX-97842701 | 0 | G | A | 0.211 | -5.16E-02 | 1.07E-01 | 2.60E+00 | 2.68E+00 | 6.32E-01 | no | | intergenic_variant | MODIFIER |
| 5 | 40740016 | AX-170366539 | 0 | C | G | 0.211 | -5.16E-02 | 1.07E-01 | 2.60E+00 | 2.68E+00 | 6.32E-01 | no | | intergenic_variant | MODIFIER |
| 15 | 80190063 | AX-97840186 | 2 | C | T | 0.555 | 3.93E-02 | 8.09E-02 | 2.58E+00 | 2.66E+00 | 6.32E-01 | no | Mgat3 | intron_variant | MODIFIER |
| 5 | 76759415 | AX-97790084 | 1 | C | T | 0.42 | 4.74E-02 | 9.88E-02 | 2.61E+00 | 2.69E+00 | 6.32E-01 | no | C530008M17Rik | intron_variant | MODIFIER |
| 3 | 122608030 | AX-170485488 | 0 | G | C | 0.035 | -8.31E-02 | 1.74E-01 | 2.59E+00 | 2.72E+00 | 6.32E-01 | no | Fnbp1l | intron_variant | MODIFIER |
| 9 | 35373801 | AX-97825334 | 0 | T | C | 0.43 | -4.14E-02 | 8.78E-02 | 2.78E+00 | 2.86E+00 | 6.32E-01 | no | | intergenic_variant | MODIFIER |
| 5 | 97086641 | AX-170491513 | 1 | A | G | 0.438 | 5.21E-02 | 1.10E-01 | 2.71E+00 | 2.76E+00 | 6.32E-01 | no | Bmp2k | intron_variant | MODIFIER |
| 5 | 82047113 | AX-97800352 | 0 | C | T | 0.412 | 4.73E-02 | 1.01E-01 | 2.70E+00 | 2.77E+00 | 6.32E-01 | no | | intergenic_variant | MODIFIER |
| 15 | 78293341 | AX-97810157 | 0 | C | A | 0.553 | 3.88E-02 | 8.04E-02 | 2.59E+00 | 2.67E+00 | 6.32E-01 | no | Csf2rb2 | intron_variant | MODIFIER |
| 12 | 94414327 | AX-170178986 | 1 | G | A | 0.089 | 9.56E-02 | 2.03E-01 | 2.72E+00 | 2.77E+00 | 6.32E-01 | no | | intergenic_variant | MODIFIER |
| 1 | 173240632 | AX-170530119 | 0 | C | T | 0.237 | 5.77E-02 | 1.18E-01 | 2.57E+00 | 2.63E+00 | 6.32E-01 | no | | intergenic_variant | MODIFIER |
| 8 | 37672441 | AX-97847011 | 2 | C | T | 0.155 | 6.89E-02 | 1.46E-01 | 2.77E+00 | 2.83E+00 | 6.32E-01 | no | Sgcz | intron_variant | MODIFIER |
| 5 | 30985501 | AX-170024617 | 0 | A | G | 0.368 | 5.00E-02 | 1.06E-01 | 2.69E+00 | 2.75E+00 | 6.33E-01 | no | | intergenic_variant | MODIFIER |
| 11 | 15369550 | AX-170046463 | 0 | A | C | 0.044 | -8.73E-02 | 1.84E-01 | 2.58E+00 | 2.73E+00 | 6.33E-01 | no | | intergenic_variant | MODIFIER |
| 11 | 18745915 | AX-170396459 | 0 | T | C | 0.044 | -8.73E-02 | 1.84E-01 | 2.58E+00 | 2.73E+00 | 6.33E-01 | no | | regulatory_region_variant | MODIFIER |
| 8 | 65934274 | AX-170528032 | 0 | G | A | 0.158 | -6.89E-02 | 1.44E-01 | 2.63E+00 | 2.70E+00 | 6.33E-01 | no | MAR1 | intron_variant | MODIFIER |
| 20 | 67123629 | AX-170034059 | 0 | T | A | 0.246 | 2.98E-02 | 6.32E-02 | 2.65E+00 | 2.74E+00 | 6.33E-01 | no | | intergenic_variant | MODIFIER |
| 8 | 116565846 | AX-170235265 | 0 | A | C | 0.254 | 4.49E-02 | 9.66E-02 | 2.69E+00 | 2.80E+00 | 6.33E-01 | no | | intergenic_variant | MODIFIER |
| 4 | 151589252 | AX-170141749 | 2 | A | G | 0.109 | 8.57E-02 | 1.84E-01 | 2.70E+00 | 2.80E+00 | 6.34E-01 | no | Camta1 | intron_variant | MODIFIER |
| 13 | 25026715 | AX-170163365 | 1 | T | G | 0.161 | -6.18E-02 | 1.20E-01 | 2.47E+00 | 2.54E+00 | 6.34E-01 | no | | intergenic_variant | MODIFIER |
| 8 | 9757872 | AX-97820458 | 0 | C | G | 0.263 | 3.49E-02 | 7.52E-02 | 2.70E+00 | 2.82E+00 | 6.34E-01 | no | Fam155a | intron_variant | MODIFIER |
| 4 | 118939566 | AX-170373302 | 0 | C | T | 0.211 | 5.54E-02 | 1.15E-01 | 2.59E+00 | 2.67E+00 | 6.34E-01 | no | Olfr1328 | upstream_gene_variant | MODIFIER |
| 5 | 86626749 | AX-170025511 | 1 | G | C | 0.429 | 5.29E-02 | 1.13E-01 | 2.76E+00 | 2.80E+00 | 6.34E-01 | no | Tmprss11f | intron_variant | MODIFIER |
| 2 | 94524862 | AX-170064332 | 0 | A | G | 0.053 | 9.81E-02 | 2.09E-01 | 2.64E+00 | 2.75E+00 | 6.34E-01 | no | | regulatory_region_variant | MODIFIER |
| 4 | 92791113 | AX-170195755 | 0 | C | T | 0.333 | 5.16E-02 | 1.09E-01 | 2.63E+00 | 2.70E+00 | 6.34E-01 | no | | intergenic_variant | MODIFIER |
| 5 | 104107716 | AX-97819055 | 1 | T | C | 0.438 | 5.52E-02 | 1.16E-01 | 2.65E+00 | 2.67E+00 | 6.34E-01 | no | Sparcl1 | intron_variant | MODIFIER |
| 11 | 34790284 | AX-97805684 | 0 | C | A | 0.096 | -7.47E-02 | 1.61E-01 | 2.67E+00 | 2.79E+00 | 6.34E-01 | no | | intergenic_variant | MODIFIER |
| 7 | 46157305 | AX-97838599 | 0 | T | C | 0.333 | 4.43E-02 | 9.33E-02 | 2.62E+00 | 2.70E+00 | 6.34E-01 | no | Abcc8 | intron_variant | MODIFIER |

| | | | | | | | | | | | | | | | |
|---|---|---|---|---|---|---|---|---|---|---|---|---|---|---|---|
| 2 | 155945797 | AX-170183066 | 1 | C | T | 0.143 | 1.03E-01 | 2.18E-01 | 2.71E+00 | 2.74E+00 | 6.34E-01 | no | Gdf5 | upstream_gene_variant | MODIFIER |
| 7 | 10308515 | AX-97856061 | 1 | C | T | 0.17 | 3.78E-02 | 8.24E-02 | 2.77E+00 | 2.91E+00 | 6.35E-01 | no | | intergenic_variant | MODIFIER |
| 1 | 69977804 | AX-97791588 | 0 | A | G | 0.14 | -8.80E-02 | 1.89E-01 | 2.67E+00 | 2.77E+00 | 6.35E-01 | no | Spag16 | intron_variant | MODIFIER |
| 1 | 74433450 | AX-170104946 | 0 | T | C | 0.14 | -8.80E-02 | 1.89E-01 | 2.67E+00 | 2.77E+00 | 6.35E-01 | no | Vil1 | intron_variant | MODIFIER |
| 20 | 81925945 | AX-97780312 | 0 | C | T | 0.053 | 4.87E-02 | 1.03E-01 | 2.58E+00 | 2.73E+00 | 6.35E-01 | no | | intergenic_variant | MODIFIER |
| 1 | 35563447 | AX-170222641 | 2 | A | G | 0.455 | 3.70E-02 | 7.93E-02 | 2.75E+00 | 2.83E+00 | 6.35E-01 | no | | intergenic_variant | MODIFIER |
| 5 | 40711011 | AX-97846914 | 1 | G | A | 0.214 | -5.09E-02 | 1.07E-01 | 2.60E+00 | 2.69E+00 | 6.35E-01 | no | | intergenic_variant | MODIFIER |
| 1 | 143350272 | AX-170091724 | 0 | C | T | 0.219 | 6.35E-02 | 1.34E-01 | 2.65E+00 | 2.70E+00 | 6.35E-01 | no | | intergenic_variant | MODIFIER |
| 5 | 93006915 | AX-97806379 | 1 | A | G | 0.42 | 5.09E-02 | 1.08E-01 | 2.69E+00 | 2.74E+00 | 6.35E-01 | no | Gm6450 | downstream_gene_variant | MODIFIER |
| 9 | 38331855 | AX-170286481 | 1 | G | A | 0.491 | -3.66E-02 | 7.87E-02 | 2.72E+00 | 2.81E+00 | 6.35E-01 | no | Olfr25 | downstream_gene_variant | MODIFIER |
| 20 | 49256367 | AX-170420561 | 0 | C | G | 0.237 | 2.76E-02 | 5.83E-02 | 2.59E+00 | 2.72E+00 | 6.35E-01 | no | Enox2 | intron_variant | MODIFIER |
| 6 | 134447983 | AX-170076300 | 0 | G | C | 0.561 | 4.31E-02 | 8.71E-02 | 2.53E+00 | 2.59E+00 | 6.35E-01 | no | Lrp6 | 3_prime_UTR_variant | MODIFIER |
| 8 | 11279405 | AX-170035771 | 0 | G | T | 0.254 | 3.45E-02 | 7.47E-02 | 2.72E+00 | 2.84E+00 | 6.35E-01 | no | Col4a1 | intron_variant | MODIFIER |
| 8 | 12958088 | AX-170475873 | 0 | G | A | 0.254 | 3.45E-02 | 7.47E-02 | 2.72E+00 | 2.84E+00 | 6.35E-01 | no | Mcf2l | intron_variant | MODIFIER |
| 5 | 34536152 | AX-170371771 | 1 | C | T | 0.339 | 5.01E-02 | 1.07E-01 | 2.67E+00 | 2.72E+00 | 6.35E-01 | no | Sh3bp2 | intron_variant | MODIFIER |
| 5 | 86644859 | AX-170095386 | 1 | A | C | 0.42 | 5.10E-02 | 1.09E-01 | 2.74E+00 | 2.79E+00 | 6.36E-01 | no | | intergenic_variant | MODIFIER |
| 5 | 110762591 | AX-97821957 | 2 | A | G | 0.455 | 5.50E-02 | 1.17E-01 | 2.67E+00 | 2.70E+00 | 6.36E-01 | no | Ep400 | intron_variant | MODIFIER |
| 5 | 82794346 | AX-170251197 | 0 | C | A | 0.43 | 4.71E-02 | 1.01E-01 | 2.68E+00 | 2.76E+00 | 6.36E-01 | no | Mir1187 | downstream_gene_variant | MODIFIER |
| 8 | 101677788 | AX-170080992 | 1 | A | G | 0.232 | 5.34E-02 | 1.14E-01 | 2.69E+00 | 2.76E+00 | 6.36E-01 | no | | intergenic_variant | MODIFIER |
| 16 | 97518985 | AX-97829677 | 1 | T | C | 0.464 | 3.53E-02 | 7.66E-02 | 2.76E+00 | 2.88E+00 | 6.36E-01 | no | Fam3b | upstream_gene_variant | MODIFIER |
| 1 | 45355981 | AX-170143917 | 1 | G | T | 0.098 | -9.82E-02 | 2.12E-01 | 2.70E+00 | 2.79E+00 | 6.36E-01 | no | | regulatory_region_variant | MODIFIER |
| 8 | 67732528 | AX-170335641 | 0 | G | A | 0.167 | -5.23E-02 | 1.10E-01 | 2.57E+00 | 2.69E+00 | 6.36E-01 | no | Psd3 | intron_variant | MODIFIER |
| 18 | 68696297 | AX-170508799 | 1 | G | T | 0.277 | -4.66E-02 | 9.86E-02 | 2.63E+00 | 2.69E+00 | 6.36E-01 | no | | intergenic_variant | MODIFIER |
| 11 | 32164909 | AX-170197311 | 0 | A | G | 0.105 | 7.96E-02 | 1.69E-01 | 2.62E+00 | 2.71E+00 | 6.36E-01 | no | | regulatory_region_variant | MODIFIER |
| 8 | 107135825 | AX-170081092 | 1 | T | C | 0.223 | 5.11E-02 | 1.10E-01 | 2.69E+00 | 2.76E+00 | 6.37E-01 | no | | regulatory_region_variant | MODIFIER |
| 4 | 71092912 | AX-170517124 | 0 | C | T | 0.237 | 5.08E-02 | 1.02E-01 | 2.51E+00 | 2.59E+00 | 6.37E-01 | no | Gm11228 | intron_variant,non_coding_transcript_variant | MODIFIER |
| 1 | 101164870 | AX-170105054 | 0 | T | A | 0.14 | 9.71E-02 | 2.09E-01 | 2.69E+00 | 2.76E+00 | 6.37E-01 | no | Gm20268 | intron_variant,non_coding_transcript_variant | MODIFIER |
| 5 | 97198512 | AX-170480500 | 0 | T | C | 0.412 | 4.93E-02 | 1.05E-01 | 2.67E+00 | 2.73E+00 | 6.37E-01 | no | Gm2861 | intron_variant,non_coding_transcript_variant | MODIFIER |
| 6 | 133332345 | AX-170155908 | 0 | A | T | 0.561 | 3.92E-02 | 8.03E-02 | 2.54E+00 | 2.62E+00 | 6.37E-01 | no | | intergenic_variant | MODIFIER |
| 16 | 25063883 | AX-170241823 | 0 | C | T | 0.246 | -3.79E-02 | 8.29E-02 | 2.65E+00 | 2.80E+00 | 6.37E-01 | no | A230028O05Rik | intron_variant,non_coding_transcript_variant | MODIFIER |
| 2 | 104120379 | AX-170035643 | 1 | G | A | 0.054 | -1.40E-01 | 3.01E-01 | 2.76E+00 | 2.79E+00 | 6.37E-01 | no | A930018P22Rik | upstream_gene_variant | MODIFIER |
| 8 | 96983281 | AX-168394401 | 1 | C | T | 0.223 | 5.40E-02 | 1.16E-01 | 2.70E+00 | 2.76E+00 | 6.37E-01 | no | | regulatory_region_variant | MODIFIER |
| 8 | 97611893 | AX-170402187 | 1 | G | A | 0.223 | 5.40E-02 | 1.16E-01 | 2.70E+00 | 2.76E+00 | 6.37E-01 | no | | intergenic_variant | MODIFIER |
| 1 | 101232685 | AX-170350080 | 0 | G | C | 0.14 | 9.21E-02 | 1.96E-01 | 2.64E+00 | 2.71E+00 | 6.37E-01 | no | | intergenic_variant | MODIFIER |
| 1 | 105464667 | AX-170001268 | 0 | C | T | 0.14 | 9.21E-02 | 1.96E-01 | 2.64E+00 | 2.71E+00 | 6.37E-01 | no | AC124553.1 | intron_variant,non_coding_transcript_variant | MODIFIER |
| 1 | 21729648 | AX-170162256 | 1 | T | G | 0.393 | 3.95E-02 | 8.30E-02 | 2.60E+00 | 2.66E+00 | 6.37E-01 | no | Kcnq5 | intron_variant | MODIFIER |
| 1 | 149647834 | AX-170111468 | 0 | A | C | 0.202 | 6.59E-02 | 1.41E-01 | 2.66E+00 | 2.71E+00 | 6.37E-01 | no | | intergenic_variant | MODIFIER |
| 1 | 20539887 | AX-170287686 | 2 | C | G | 0.391 | 3.93E-02 | 8.28E-02 | 2.60E+00 | 2.66E+00 | 6.38E-01 | no | Pkhd1 | intron_variant | MODIFIER |
| 15 | 6245656 | AX-170354228 | 0 | G | C | 0.237 | -5.03E-02 | 1.06E-01 | 2.60E+00 | 2.66E+00 | 6.38E-01 | no | | intergenic_variant | MODIFIER |
| 7 | 40488210 | AX-170391844 | 1 | A | G | 0.321 | 4.47E-02 | 9.52E-02 | 2.62E+00 | 2.71E+00 | 6.38E-01 | no | | intergenic_variant | MODIFIER |
| 7 | 44484904 | AX-170317205 | 1 | T | C | 0.321 | 4.47E-02 | 9.52E-02 | 2.62E+00 | 2.71E+00 | 6.38E-01 | no | 5430431A17Rik | non_coding_transcript_exon_variant | MODIFIER |
| 15 | 75353693 | AX-97829698 | 2 | A | C | 0.555 | 3.94E-02 | 8.25E-02 | 2.58E+00 | 2.66E+00 | 6.38E-01 | no | Gm28114 | upstream_gene_variant | MODIFIER |
| 8 | 91125298 | AX-170220489 | 0 | A | G | 0.184 | -5.53E-02 | 1.15E-01 | 2.56E+00 | 2.65E+00 | 6.38E-01 | no | Rbl2 | downstream_gene_variant | MODIFIER |
| 12 | 89660173 | AX-170325927 | 0 | G | T | 0.088 | 8.75E-02 | 1.89E-01 | 2.74E+00 | 2.81E+00 | 6.38E-01 | no | Nrxn3 | intron_variant | MODIFIER |
| 7 | 16089702 | AX-170076769 | 0 | C | T | 0.368 | 3.88E-02 | 8.32E-02 | 2.62E+00 | 2.74E+00 | 6.38E-01 | no | Zfp541 | intron_variant | MODIFIER |
| 2 | 91472165 | AX-170515597 | 1 | C | T | 0.062 | 7.89E-02 | 1.72E-01 | 2.65E+00 | 2.79E+00 | 6.38E-01 | no | Lrp4 | intron_variant | MODIFIER |
| 18 | 50097844 | AX-170449637 | 1 | T | A | 0.196 | -4.99E-02 | 1.04E-01 | 2.57E+00 | 2.66E+00 | 6.38E-01 | no | Tnfaip8 | intron_variant,non_coding_transcript_variant | MODIFIER |
| 3 | 60130255 | AX-97781738 | 1 | G | A | 0.071 | 6.53E-02 | 1.42E-01 | 2.62E+00 | 2.76E+00 | 6.38E-01 | no | Gm24382 | upstream_gene_variant | MODIFIER |
| 1 | 187240789 | AX-170190901 | 0 | G | A | 0.228 | -4.87E-02 | 1.05E-01 | 2.85E+00 | 2.97E+00 | 6.38E-01 | no | Gpatch2 | intron_variant | MODIFIER |
| 13 | 30888370 | AX-170008008 | 1 | G | A | 0.179 | -5.71E-02 | 1.20E-01 | 2.59E+00 | 2.65E+00 | 6.38E-01 | no | Exoc2 | intron_variant | MODIFIER |
| 2 | 142828484 | AX-170356095 | 0 | T | G | 0.14 | -8.28E-02 | 1.80E-01 | 2.72E+00 | 2.80E+00 | 6.38E-01 | no | Kif16b | intron_variant | MODIFIER |
| 2 | 147241990 | AX-170184324 | 0 | A | G | 0.14 | -8.28E-02 | 1.80E-01 | 2.72E+00 | 2.80E+00 | 6.38E-01 | no | Nkx2-2os | intron_variant,non_coding_transcript_variant | MODIFIER |
| 11 | 50949471 | AX-97780534 | 0 | C | T | 0.07 | -9.19E-02 | 1.97E-01 | 2.64E+00 | 2.72E+00 | 6.38E-01 | no | Prop1 | downstream_gene_variant | MODIFIER |
| 18 | 57433691 | AX-170363803 | 0 | A | G | 0.193 | -5.47E-02 | 1.17E-01 | 2.64E+00 | 2.70E+00 | 6.38E-01 | no | 4930511M06Rik | intron_variant,non_coding_transcript_variant | MODIFIER |

| | | | | | | | | | | | | | | | |
|---|---|---|---|---|---|---|---|---|---|---|---|---|---|---|---|
| 9 | 35058981 | AX-170032107 | 0 | G | A | 0.439 | -4.03E-02 | 8.71E-02 | 2.78E+00 | 2.85E+00 | 6.38E-01 | no | St3gal4 | intron_variant | MODIFIER |
| 6 | 145276221 | AX-170444653 | 0 | G | A | 0.079 | 9.83E-02 | 2.10E-01 | 2.62E+00 | 2.71E+00 | 6.39E-01 | no | | regulatory_region_variant | MODIFIER |
| 10 | 115078995 | AX-170509158 | 0 | A | G | 0.377 | 3.51E-02 | 7.62E-02 | 2.70E+00 | 2.79E+00 | 6.39E-01 | no | Tph2 | 3_prime_UTR_variant | MODIFIER |
| 10 | 115079925 | AX-170151921 | 0 | C | G | 0.377 | 3.51E-02 | 7.62E-02 | 2.70E+00 | 2.79E+00 | 6.39E-01 | no | Tph2 | intron_variant | MODIFIER |
| 10 | 115080245 | AX-97840795 | 0 | A | C | 0.377 | 3.51E-02 | 7.62E-02 | 2.70E+00 | 2.79E+00 | 6.39E-01 | no | Tph2 | intron_variant | MODIFIER |
| 10 | 115080681 | AX-170136562 | 0 | G | A | 0.377 | 3.51E-02 | 7.62E-02 | 2.70E+00 | 2.79E+00 | 6.39E-01 | no | Tph2 | intron_variant | MODIFIER |
| 10 | 115104399 | AX-170476485 | 0 | G | A | 0.377 | 3.51E-02 | 7.62E-02 | 2.70E+00 | 2.79E+00 | 6.39E-01 | no | Tph2 | intron_variant | MODIFIER |
| 10 | 115173395 | AX-170233326 | 0 | T | C | 0.377 | 3.51E-02 | 7.62E-02 | 2.70E+00 | 2.79E+00 | 6.39E-01 | no | Tph2 | intron_variant | MODIFIER |
| 10 | 115393940 | AX-170414637 | 0 | C | T | 0.377 | 3.51E-02 | 7.62E-02 | 2.70E+00 | 2.79E+00 | 6.39E-01 | no | Zfc3h1 | intron_variant | MODIFIER |
| 10 | 115399322 | AX-170416821 | 0 | T | C | 0.377 | 3.51E-02 | 7.62E-02 | 2.70E+00 | 2.79E+00 | 6.39E-01 | no | Zfc3h1 | intron_variant | MODIFIER |
| 10 | 115430596 | AX-170316151 | 0 | T | G | 0.377 | 3.51E-02 | 7.62E-02 | 2.70E+00 | 2.79E+00 | 6.39E-01 | no | Zfc3h1 | intron_variant | MODIFIER |
| 10 | 115433344 | AX-170278822 | 0 | C | G | 0.377 | 3.51E-02 | 7.62E-02 | 2.70E+00 | 2.79E+00 | 6.39E-01 | no | Zfc3h1 | downstream_gene_variant | MODIFIER |
| 10 | 115449408 | AX-170106535 | 0 | T | C | 0.377 | 3.51E-02 | 7.62E-02 | 2.70E+00 | 2.79E+00 | 6.39E-01 | no | Lgr5 | downstream_gene_variant | MODIFIER |
| 5 | 73724562 | AX-170280522 | 0 | T | G | 0.439 | 4.79E-02 | 1.02E-01 | 2.64E+00 | 2.71E+00 | 6.39E-01 | no | | intergenic_variant | MODIFIER |
| 5 | 73726338 | AX-170447117 | 0 | T | A | 0.439 | 4.79E-02 | 1.02E-01 | 2.64E+00 | 2.71E+00 | 6.39E-01 | no | | intergenic_variant | MODIFIER |
| 5 | 75162055 | AX-170246796 | 0 | T | C | 0.439 | 4.79E-02 | 1.02E-01 | 2.64E+00 | 2.71E+00 | 6.39E-01 | no | Pdgfra | intron_variant | MODIFIER |
| 12 | 109653104 | AX-170370022 | 0 | C | T | 0.123 | -7.74E-02 | 1.68E-01 | 2.70E+00 | 2.79E+00 | 6.39E-01 | no | DQ267102 | upstream_gene_variant | MODIFIER |
| 13 | 39627016 | AX-170051039 | 2 | C | T | 0.191 | -5.32E-02 | 1.08E-01 | 2.50E+00 | 2.59E+00 | 6.39E-01 | no | Gm32243 | intron_variant,non_coding_transcript_variant | MODIFIER |
| 8 | 54330957 | AX-170099579 | 0 | C | T | 0.158 | -5.70E-02 | 1.21E-01 | 2.59E+00 | 2.69E+00 | 6.39E-01 | no | | intergenic_variant | MODIFIER |
| 8 | 18556678 | AX-170467010 | 0 | C | T | 0.193 | 4.36E-02 | 9.38E-02 | 2.62E+00 | 2.73E+00 | 6.39E-01 | no | | intergenic_variant | MODIFIER |
| 1 | 65919243 | AX-170041486 | 1 | G | A | 0.134 | 9.69E-02 | 2.11E-01 | 2.69E+00 | 2.79E+00 | 6.39E-01 | no | | intergenic_variant | MODIFIER |
| 15 | 100718510 | AX-170237087 | 1 | T | A | 0.116 | -6.30E-02 | 1.37E-01 | 2.69E+00 | 2.78E+00 | 6.39E-01 | no | Galnt6 | intron_variant | MODIFIER |
| 7 | 82507065 | AX-97798923 | 1 | C | A | 0.116 | 7.81E-02 | 1.68E-01 | 2.62E+00 | 2.74E+00 | 6.39E-01 | no | Adamtsl3 | intron_variant | MODIFIER |
| 8 | 109209171 | AX-170276162 | 0 | C | T | 0.246 | 4.77E-02 | 1.03E-01 | 2.65E+00 | 2.74E+00 | 6.39E-01 | no | D030068K23Rik | intron_variant,non_coding_transcript_variant | MODIFIER |
| 11 | 74664301 | AX-170140382 | 0 | A | G | 0.263 | 3.30E-02 | 7.21E-02 | 2.65E+00 | 2.78E+00 | 6.40E-01 | no | Cluh | intron_variant | MODIFIER |
| 15 | 6994875 | AX-170330215 | 2 | G | A | 0.255 | -4.82E-02 | 1.02E-01 | 2.60E+00 | 2.67E+00 | 6.40E-01 | no | Gm38282 | upstream_gene_variant | MODIFIER |
| 4 | 48804398 | AX-170117693 | 0 | A | T | 0.36 | -4.21E-02 | 9.21E-02 | 2.70E+00 | 2.80E+00 | 6.40E-01 | no | | regulatory_region_variant | MODIFIER |
| 12 | 72849842 | AX-170146418 | 0 | C | T | 0.07 | 9.45E-02 | 2.08E-01 | 2.71E+00 | 2.83E+00 | 6.40E-01 | no | | intergenic_variant | MODIFIER |
| 5 | 51358757 | AX-170072076 | 1 | C | T | 0.491 | 4.91E-02 | 1.01E-01 | 2.54E+00 | 2.60E+00 | 6.40E-01 | no | | intergenic_variant | MODIFIER |
| 2 | 137147942 | AX-97780580 | 0 | C | T | 0.149 | -9.31E-02 | 2.03E-01 | 2.77E+00 | 2.85E+00 | 6.40E-01 | no | | intergenic_variant | MODIFIER |
| 13 | 38782765 | AX-170276403 | 0 | G | A | 0.167 | -5.58E-02 | 1.12E-01 | 2.49E+00 | 2.57E+00 | 6.40E-01 | no | Gm47509 | non_coding_transcript_exon_variant | MODIFIER |
| 13 | 38793536 | AX-97787332 | 0 | A | G | 0.167 | -5.58E-02 | 1.12E-01 | 2.49E+00 | 2.57E+00 | 6.40E-01 | no | Gm47509 | intron_variant,non_coding_transcript_variant | MODIFIER |
| 13 | 38795155 | AX-170117925 | 0 | C | A | 0.167 | -5.58E-02 | 1.12E-01 | 2.49E+00 | 2.57E+00 | 6.40E-01 | no | Gm47509 | upstream_gene_variant | MODIFIER |
| 13 | 39373030 | AX-170451979 | 0 | T | C | 0.167 | -5.58E-02 | 1.12E-01 | 2.49E+00 | 2.57E+00 | 6.40E-01 | no | | intergenic_variant | MODIFIER |
| 13 | 39406980 | AX-170119732 | 0 | C | T | 0.167 | -5.58E-02 | 1.12E-01 | 2.49E+00 | 2.57E+00 | 6.40E-01 | no | | intergenic_variant | MODIFIER |
| 13 | 39441839 | AX-170518691 | 0 | T | C | 0.167 | -5.58E-02 | 1.12E-01 | 2.49E+00 | 2.57E+00 | 6.40E-01 | no | Gm47348 | intron_variant,non_coding_transcript_variant | MODIFIER |
| 13 | 39469558 | AX-170249010 | 0 | A | G | 0.167 | -5.58E-02 | 1.12E-01 | 2.49E+00 | 2.57E+00 | 6.40E-01 | no | Gm3509 | upstream_gene_variant | MODIFIER |
| 13 | 39501486 | AX-170327779 | 0 | C | G | 0.167 | -5.58E-02 | 1.12E-01 | 2.49E+00 | 2.57E+00 | 6.40E-01 | no | | intergenic_variant | MODIFIER |
| 13 | 39503518 | AX-170470664 | 0 | A | C | 0.167 | -5.58E-02 | 1.12E-01 | 2.49E+00 | 2.57E+00 | 6.40E-01 | no | | intergenic_variant | MODIFIER |
| 13 | 39517131 | AX-97811739 | 0 | C | T | 0.167 | -5.58E-02 | 1.12E-01 | 2.49E+00 | 2.57E+00 | 6.40E-01 | no | | intergenic_variant | MODIFIER |
| 13 | 39532239 | AX-170333170 | 0 | T | G | 0.167 | -5.58E-02 | 1.12E-01 | 2.49E+00 | 2.57E+00 | 6.40E-01 | no | Gm47351 | intron_variant,non_coding_transcript_variant | MODIFIER |
| 13 | 39562158 | AX-170008158 | 0 | A | G | 0.167 | -5.58E-02 | 1.12E-01 | 2.49E+00 | 2.57E+00 | 6.40E-01 | no | | intergenic_variant | MODIFIER |
| 13 | 39756442 | AX-170369786 | 0 | C | T | 0.167 | -5.58E-02 | 1.12E-01 | 2.49E+00 | 2.57E+00 | 6.40E-01 | no | Gm5195 | upstream_gene_variant | MODIFIER |
| 13 | 39804064 | AX-170154023 | 0 | T | A | 0.167 | -5.58E-02 | 1.12E-01 | 2.49E+00 | 2.57E+00 | 6.40E-01 | no | A230103O09Rik | intron_variant,non_coding_transcript_variant | MODIFIER |
| 13 | 39891947 | AX-170217625 | 0 | T | C | 0.167 | -5.58E-02 | 1.12E-01 | 2.49E+00 | 2.57E+00 | 6.40E-01 | no | | intergenic_variant | MODIFIER |
| 13 | 39998868 | AX-170432319 | 0 | C | G | 0.167 | -5.58E-02 | 1.12E-01 | 2.49E+00 | 2.57E+00 | 6.40E-01 | no | Ofcc1 | downstream_gene_variant | MODIFIER |
| 13 | 40032396 | AX-170111154 | 0 | T | C | 0.167 | -5.58E-02 | 1.12E-01 | 2.49E+00 | 2.57E+00 | 6.40E-01 | no | Ofcc1 | intron_variant | MODIFIER |
| 13 | 40037120 | AX-170310980 | 0 | T | C | 0.167 | -5.58E-02 | 1.12E-01 | 2.49E+00 | 2.57E+00 | 6.40E-01 | no | Ofcc1 | intron_variant | MODIFIER |
| 13 | 40049160 | AX-170257285 | 0 | T | C | 0.167 | -5.58E-02 | 1.12E-01 | 2.49E+00 | 2.57E+00 | 6.40E-01 | no | Ofcc1 | intron_variant | MODIFIER |
| 13 | 40074342 | AX-97808560 | 0 | T | C | 0.167 | -5.58E-02 | 1.12E-01 | 2.49E+00 | 2.57E+00 | 6.40E-01 | no | Ofcc1 | intron_variant | MODIFIER |
| 13 | 40078982 | AX-170051051 | 0 | A | G | 0.167 | -5.58E-02 | 1.12E-01 | 2.49E+00 | 2.57E+00 | 6.40E-01 | no | Ofcc1 | intron_variant | MODIFIER |
| 13 | 40126259 | AX-170120509 | 0 | C | T | 0.167 | -5.58E-02 | 1.12E-01 | 2.49E+00 | 2.57E+00 | 6.40E-01 | no | Ofcc1 | intron_variant | MODIFIER |
| 13 | 40130085 | AX-170008167 | 0 | C | T | 0.167 | -5.58E-02 | 1.12E-01 | 2.49E+00 | 2.57E+00 | 6.40E-01 | no | Ofcc1 | intron_variant | MODIFIER |
| 13 | 40132336 | AX-170136133 | 0 | A | G | 0.167 | -5.58E-02 | 1.12E-01 | 2.49E+00 | 2.57E+00 | 6.40E-01 | no | Ofcc1 | intron_variant | MODIFIER |

| | | | | | | | | | | | | | | | |
|---|---|---|---|---|---|---|---|---|---|---|---|---|---|---|---|
| 13 | 40166324 | AX-170008166 | 0 | T | C | 0.167 | -5.58E-02 | 1.12E-01 | 2.49E+00 | 2.57E+00 | 6.40E-01 | no | Ofcc1 | intron_variant | MODIFIER |
| 13 | 40167507 | AX-170497276 | 0 | A | T | 0.167 | -5.58E-02 | 1.12E-01 | 2.49E+00 | 2.57E+00 | 6.40E-01 | no | Ofcc1 | intron_variant | MODIFIER |
| 13 | 40172757 | AX-97793786 | 0 | A | C | 0.167 | -5.58E-02 | 1.12E-01 | 2.49E+00 | 2.57E+00 | 6.40E-01 | no | Ofcc1 | intron_variant | MODIFIER |
| 13 | 40190816 | AX-170008168 | 0 | G | A | 0.167 | -5.58E-02 | 1.12E-01 | 2.49E+00 | 2.57E+00 | 6.40E-01 | no | Ofcc1 | intron_variant | MODIFIER |
| 13 | 40206992 | AX-170388111 | 0 | A | G | 0.167 | -5.58E-02 | 1.12E-01 | 2.49E+00 | 2.57E+00 | 6.40E-01 | no | Ofcc1 | intron_variant | MODIFIER |
| 13 | 40237926 | AX-170051055 | 0 | T | C | 0.167 | -5.58E-02 | 1.12E-01 | 2.49E+00 | 2.57E+00 | 6.40E-01 | no | Ofcc1 | intron_variant | MODIFIER |
| 13 | 40243143 | AX-170358812 | 0 | T | C | 0.167 | -5.58E-02 | 1.12E-01 | 2.49E+00 | 2.57E+00 | 6.40E-01 | no | Ofcc1 | intron_variant | MODIFIER |
| 13 | 40249149 | AX-170008163 | 0 | C | G | 0.167 | -5.58E-02 | 1.12E-01 | 2.49E+00 | 2.57E+00 | 6.40E-01 | no | Ofcc1 | intron_variant | MODIFIER |
| 13 | 40256044 | AX-170276513 | 0 | C | T | 0.167 | -5.58E-02 | 1.12E-01 | 2.49E+00 | 2.57E+00 | 6.40E-01 | no | Ofcc1 | intron_variant | MODIFIER |
| 13 | 40521031 | AX-170433682 | 0 | A | C | 0.167 | -5.58E-02 | 1.12E-01 | 2.49E+00 | 2.57E+00 | 6.40E-01 | no | | intergenic_variant | MODIFIER |
| 13 | 40522499 | AX-170415580 | 0 | A | G | 0.167 | -5.58E-02 | 1.12E-01 | 2.49E+00 | 2.57E+00 | 6.40E-01 | no | | intergenic_variant | MODIFIER |
| 13 | 40522730 | AX-170326497 | 0 | G | T | 0.167 | -5.58E-02 | 1.12E-01 | 2.49E+00 | 2.57E+00 | 6.40E-01 | no | | intergenic_variant | MODIFIER |
| 13 | 40524656 | AX-170419616 | 0 | A | G | 0.167 | -5.58E-02 | 1.12E-01 | 2.49E+00 | 2.57E+00 | 6.40E-01 | no | | intergenic_variant | MODIFIER |
| 13 | 40525297 | AX-170260871 | 0 | C | G | 0.167 | -5.58E-02 | 1.12E-01 | 2.49E+00 | 2.57E+00 | 6.40E-01 | no | | intergenic_variant | MODIFIER |
| 13 | 40526141 | AX-170117478 | 0 | T | C | 0.167 | -5.58E-02 | 1.12E-01 | 2.49E+00 | 2.57E+00 | 6.40E-01 | no | | intergenic_variant | MODIFIER |
| 13 | 40566167 | AX-170290242 | 0 | A | G | 0.167 | -5.58E-02 | 1.12E-01 | 2.49E+00 | 2.57E+00 | 6.40E-01 | no | Gm47039 | intron_variant,non_coding_transcript_variant | MODIFIER |
| 13 | 40668568 | AX-170497395 | 0 | G | A | 0.167 | -5.58E-02 | 1.12E-01 | 2.49E+00 | 2.57E+00 | 6.40E-01 | no | | intergenic_variant | MODIFIER |
| 13 | 40668789 | AX-170278333 | 0 | T | C | 0.167 | -5.58E-02 | 1.12E-01 | 2.49E+00 | 2.57E+00 | 6.40E-01 | no | | intergenic_variant | MODIFIER |
| 13 | 40670130 | AX-170111083 | 0 | A | G | 0.167 | -5.58E-02 | 1.12E-01 | 2.49E+00 | 2.57E+00 | 6.40E-01 | no | | intergenic_variant | MODIFIER |
| 13 | 40672822 | AX-170205136 | 0 | G | A | 0.167 | -5.58E-02 | 1.12E-01 | 2.49E+00 | 2.57E+00 | 6.40E-01 | no | | intergenic_variant | MODIFIER |
| 13 | 40679607 | AX-170315150 | 0 | C | T | 0.167 | -5.58E-02 | 1.12E-01 | 2.49E+00 | 2.57E+00 | 6.40E-01 | no | | intergenic_variant | MODIFIER |
| 13 | 40701372 | AX-170008174 | 0 | A | G | 0.167 | -5.58E-02 | 1.12E-01 | 2.49E+00 | 2.57E+00 | 6.40E-01 | no | Gm9979 | upstream_gene_variant | MODIFIER |
| 13 | 40823419 | AX-170286289 | 0 | T | G | 0.167 | -5.58E-02 | 1.12E-01 | 2.49E+00 | 2.57E+00 | 6.40E-01 | no | | intergenic_variant | MODIFIER |
| 13 | 40852172 | AX-170206743 | 0 | G | A | 0.167 | -5.58E-02 | 1.12E-01 | 2.49E+00 | 2.57E+00 | 6.40E-01 | no | Gm35160 | downstream_gene_variant | MODIFIER |
| 13 | 40887347 | AX-170331717 | 0 | T | A | 0.167 | -5.58E-02 | 1.12E-01 | 2.49E+00 | 2.57E+00 | 6.40E-01 | no | Gcnt2 | 5_prime_UTR_variant | MODIFIER |
| 13 | 40889000 | AX-170346057 | 0 | G | A | 0.167 | -5.58E-02 | 1.12E-01 | 2.49E+00 | 2.57E+00 | 6.40E-01 | no | Gcnt2 | intron_variant | MODIFIER |
| 13 | 40894626 | AX-97818197 | 0 | T | C | 0.167 | -5.58E-02 | 1.12E-01 | 2.49E+00 | 2.57E+00 | 6.40E-01 | no | Gcnt2 | intron_variant | MODIFIER |
| 13 | 40899904 | AX-170413299 | 0 | G | C | 0.167 | -5.58E-02 | 1.12E-01 | 2.49E+00 | 2.57E+00 | 6.40E-01 | no | Gcnt2 | intron_variant | MODIFIER |
| 13 | 40911892 | AX-170139694 | 0 | G | T | 0.167 | -5.58E-02 | 1.12E-01 | 2.49E+00 | 2.57E+00 | 6.40E-01 | no | Gcnt2 | intron_variant | MODIFIER |
| 13 | 40921419 | AX-170142409 | 0 | A | C | 0.167 | -5.58E-02 | 1.12E-01 | 2.49E+00 | 2.57E+00 | 6.40E-01 | no | Gcnt2 | intron_variant | MODIFIER |
| 1 | 90237015 | AX-170267877 | 1 | G | C | 0.152 | 8.73E-02 | 1.85E-01 | 2.60E+00 | 2.68E+00 | 6.40E-01 | no | | intergenic_variant | MODIFIER |
| 5 | 86544952 | AX-170072689 | 1 | C | A | 0.438 | 5.03E-02 | 1.09E-01 | 2.72E+00 | 2.77E+00 | 6.40E-01 | no | Tmprss11f | intron_variant | MODIFIER |
| 5 | 90052713 | AX-170259150 | 1 | G | A | 0.438 | 5.03E-02 | 1.09E-01 | 2.72E+00 | 2.77E+00 | 6.40E-01 | no | | intergenic_variant | MODIFIER |
| 5 | 39989625 | AX-170253507 | 1 | T | C | 0.214 | -4.99E-02 | 1.06E-01 | 2.60E+00 | 2.69E+00 | 6.40E-01 | no | Gm35191 | upstream_gene_variant | MODIFIER |
| 3 | 30191760 | AX-170035656 | 1 | A | G | 0.179 | -4.58E-02 | 9.91E-02 | 2.67E+00 | 2.73E+00 | 6.40E-01 | no | Mecom | intron_variant | MODIFIER |
| 2 | 101919221 | AX-170298216 | 0 | T | C | 0.061 | 9.97E-02 | 2.17E-01 | 2.67E+00 | 2.76E+00 | 6.40E-01 | no | | intergenic_variant | MODIFIER |
| 2 | 109684194 | AX-97832385 | 0 | C | T | 0.061 | 9.97E-02 | 2.17E-01 | 2.67E+00 | 2.76E+00 | 6.40E-01 | no | Bdnf | intron_variant | MODIFIER |
| 1 | 144989305 | AX-170001908 | 0 | G | A | 0.202 | 6.12E-02 | 1.31E-01 | 2.64E+00 | 2.70E+00 | 6.40E-01 | no | | intergenic_variant | MODIFIER |
| 5 | 92933922 | AX-97854698 | 0 | C | T | 0.421 | 5.02E-02 | 1.09E-01 | 2.69E+00 | 2.74E+00 | 6.40E-01 | no | Shroom3 | intron_variant | MODIFIER |
| 5 | 92950208 | AX-97800770 | 0 | T | C | 0.421 | 5.02E-02 | 1.09E-01 | 2.69E+00 | 2.74E+00 | 6.40E-01 | no | Shroom3 | intron_variant | MODIFIER |
| 5 | 92965961 | AX-170025618 | 0 | G | A | 0.421 | 5.02E-02 | 1.09E-01 | 2.69E+00 | 2.74E+00 | 6.40E-01 | no | Shroom3 | downstream_gene_variant | MODIFIER |
| 5 | 92981658 | AX-170382077 | 0 | C | T | 0.421 | 5.02E-02 | 1.09E-01 | 2.69E+00 | 2.74E+00 | 6.40E-01 | no | | intergenic_variant | MODIFIER |
| 5 | 92997626 | AX-170025619 | 0 | A | G | 0.421 | 5.02E-02 | 1.09E-01 | 2.69E+00 | 2.74E+00 | 6.40E-01 | no | | regulatory_region_variant | MODIFIER |
| 5 | 93022557 | AX-170250031 | 0 | A | T | 0.421 | 5.02E-02 | 1.09E-01 | 2.69E+00 | 2.74E+00 | 6.40E-01 | no | | regulatory_region_variant | MODIFIER |
| 5 | 93023070 | AX-170307129 | 0 | G | A | 0.421 | 5.02E-02 | 1.09E-01 | 2.69E+00 | 2.74E+00 | 6.40E-01 | no | | regulatory_region_variant | MODIFIER |
| 5 | 93023967 | AX-170352085 | 0 | C | T | 0.421 | 5.02E-02 | 1.09E-01 | 2.69E+00 | 2.74E+00 | 6.40E-01 | no | | intergenic_variant | MODIFIER |
| 5 | 93025501 | AX-170274782 | 0 | A | G | 0.421 | 5.02E-02 | 1.09E-01 | 2.69E+00 | 2.74E+00 | 6.40E-01 | no | | regulatory_region_variant | MODIFIER |
| 5 | 93029944 | AX-170412477 | 0 | T | C | 0.421 | 5.02E-02 | 1.09E-01 | 2.69E+00 | 2.74E+00 | 6.40E-01 | no | | intergenic_variant | MODIFIER |
| 5 | 93031496 | AX-170314878 | 0 | T | C | 0.421 | 5.02E-02 | 1.09E-01 | 2.69E+00 | 2.74E+00 | 6.40E-01 | no | | regulatory_region_variant | MODIFIER |
| 5 | 93047093 | AX-170518681 | 0 | A | C | 0.421 | 5.02E-02 | 1.09E-01 | 2.69E+00 | 2.74E+00 | 6.40E-01 | no | Sowahb | upstream_gene_variant | MODIFIER |
| 5 | 93052141 | AX-97799943 | 0 | C | T | 0.421 | 5.02E-02 | 1.09E-01 | 2.69E+00 | 2.74E+00 | 6.40E-01 | no | | intergenic_variant | MODIFIER |
| 5 | 93052274 | AX-170198434 | 0 | A | G | 0.421 | 5.02E-02 | 1.09E-01 | 2.69E+00 | 2.74E+00 | 6.40E-01 | no | | intergenic_variant | MODIFIER |
| 5 | 93052827 | AX-170072797 | 0 | G | A | 0.421 | 5.02E-02 | 1.09E-01 | 2.69E+00 | 2.74E+00 | 6.40E-01 | no | | intergenic_variant | MODIFIER |
| 5 | 93055405 | AX-170072795 | 0 | T | C | 0.421 | 5.02E-02 | 1.09E-01 | 2.69E+00 | 2.74E+00 | 6.40E-01 | no | | intergenic_variant | MODIFIER |

| | | | | | | | | | | | | | | | |
|---|---|---|---|---|---|---|---|---|---|---|---|---|---|---|---|
| 5 | 93062977 | AX-170072798 | 0 | C | T | 0.421 | 5.02E-02 | 1.09E-01 | 2.69E+00 | 2.74E+00 | 6.40E-01 | no | | intergenic_variant | MODIFIER |
| 5 | 93071246 | AX-97834691 | 0 | T | C | 0.421 | 5.02E-02 | 1.09E-01 | 2.69E+00 | 2.74E+00 | 6.40E-01 | no | Gm22915 | upstream_gene_variant | MODIFIER |
| 5 | 93099334 | AX-170345434 | 0 | T | C | 0.421 | 5.02E-02 | 1.09E-01 | 2.69E+00 | 2.74E+00 | 6.40E-01 | no | SEP11 | intron_variant | MODIFIER |
| 5 | 93099917 | AX-170469014 | 0 | C | T | 0.421 | 5.02E-02 | 1.09E-01 | 2.69E+00 | 2.74E+00 | 6.40E-01 | no | SEP11 | intron_variant | MODIFIER |
| 5 | 93124804 | AX-170399313 | 0 | C | G | 0.421 | 5.02E-02 | 1.09E-01 | 2.69E+00 | 2.74E+00 | 6.40E-01 | no | SEP11 | intron_variant | MODIFIER |
| 5 | 93144019 | AX-170443570 | 0 | G | T | 0.421 | 5.02E-02 | 1.09E-01 | 2.69E+00 | 2.74E+00 | 6.40E-01 | no | SEP11 | intron_variant | MODIFIER |
| 5 | 93165787 | AX-170255798 | 0 | C | T | 0.421 | 5.02E-02 | 1.09E-01 | 2.69E+00 | 2.74E+00 | 6.40E-01 | no | SEP11 | intron_variant | MODIFIER |
| 5 | 93175725 | AX-170283080 | 0 | T | C | 0.421 | 5.02E-02 | 1.09E-01 | 2.69E+00 | 2.74E+00 | 6.40E-01 | no | SEP11 | downstream_gene_variant | MODIFIER |
| 5 | 93180416 | AX-97808651 | 0 | C | T | 0.421 | 5.02E-02 | 1.09E-01 | 2.69E+00 | 2.74E+00 | 6.40E-01 | no | Ccni | downstream_gene_variant | MODIFIER |
| 5 | 93187797 | AX-170231788 | 0 | T | C | 0.421 | 5.02E-02 | 1.09E-01 | 2.69E+00 | 2.74E+00 | 6.40E-01 | no | Ccni | missense_variant | MODERATE |
| 5 | 93228944 | AX-170072799 | 0 | C | T | 0.421 | 5.02E-02 | 1.09E-01 | 2.69E+00 | 2.74E+00 | 6.40E-01 | no | | intergenic_variant | MODIFIER |
| 5 | 93240680 | AX-170360793 | 0 | A | G | 0.421 | 5.02E-02 | 1.09E-01 | 2.69E+00 | 2.74E+00 | 6.40E-01 | no | | regulatory_region_variant | MODIFIER |
| 5 | 93245424 | AX-170072800 | 0 | T | A | 0.421 | 5.02E-02 | 1.09E-01 | 2.69E+00 | 2.74E+00 | 6.40E-01 | no | | intergenic_variant | MODIFIER |
| 5 | 93249856 | AX-170453092 | 0 | T | C | 0.421 | 5.02E-02 | 1.09E-01 | 2.69E+00 | 2.74E+00 | 6.40E-01 | no | | intergenic_variant | MODIFIER |
| 5 | 93261259 | AX-170296788 | 0 | T | G | 0.421 | 5.02E-02 | 1.09E-01 | 2.69E+00 | 2.74E+00 | 6.40E-01 | no | | intergenic_variant | MODIFIER |
| 5 | 93266348 | AX-170025628 | 0 | C | A | 0.421 | 5.02E-02 | 1.09E-01 | 2.69E+00 | 2.74E+00 | 6.40E-01 | no | Ccng2 | upstream_gene_variant | MODIFIER |
| 5 | 93278730 | AX-170269087 | 0 | A | C | 0.421 | 5.02E-02 | 1.09E-01 | 2.69E+00 | 2.74E+00 | 6.40E-01 | no | Ccng2 | downstream_gene_variant | MODIFIER |
| 5 | 93285373 | AX-170162004 | 0 | C | A | 0.421 | 5.02E-02 | 1.09E-01 | 2.69E+00 | 2.74E+00 | 6.40E-01 | no | | intergenic_variant | MODIFIER |
| 5 | 93328283 | AX-170246477 | 0 | C | T | 0.421 | 5.02E-02 | 1.09E-01 | 2.69E+00 | 2.74E+00 | 6.40E-01 | no | | regulatory_region_variant | MODIFIER |
| 5 | 93328536 | AX-170193848 | 0 | T | C | 0.421 | 5.02E-02 | 1.09E-01 | 2.69E+00 | 2.74E+00 | 6.40E-01 | no | | regulatory_region_variant | MODIFIER |
| 5 | 93371462 | AX-170234002 | 0 | G | C | 0.421 | 5.02E-02 | 1.09E-01 | 2.69E+00 | 2.74E+00 | 6.40E-01 | no | | intergenic_variant | MODIFIER |
| 5 | 93389221 | AX-170025629 | 0 | G | A | 0.421 | 5.02E-02 | 1.09E-01 | 2.69E+00 | 2.74E+00 | 6.40E-01 | no | Kat2b-ps | downstream_gene_variant | MODIFIER |
| 5 | 93429136 | AX-97836162 | 0 | C | T | 0.421 | 5.02E-02 | 1.09E-01 | 2.69E+00 | 2.74E+00 | 6.40E-01 | no | | intergenic_variant | MODIFIER |
| 5 | 96343334 | AX-170353951 | 0 | C | T | 0.421 | 5.02E-02 | 1.09E-01 | 2.69E+00 | 2.74E+00 | 6.40E-01 | no | | intergenic_variant | MODIFIER |
| 5 | 96348134 | AX-97855215 | 0 | A | G | 0.421 | 5.02E-02 | 1.09E-01 | 2.69E+00 | 2.74E+00 | 6.40E-01 | no | | intergenic_variant | MODIFIER |
| 5 | 96403745 | AX-170470471 | 0 | C | T | 0.421 | 5.02E-02 | 1.09E-01 | 2.69E+00 | 2.74E+00 | 6.40E-01 | no | Fras1 | intron_variant | MODIFIER |
| 5 | 96409186 | AX-170121740 | 0 | C | T | 0.421 | 5.02E-02 | 1.09E-01 | 2.69E+00 | 2.74E+00 | 6.40E-01 | no | Fras1 | intron_variant | MODIFIER |
| 5 | 96413054 | AX-170072814 | 0 | T | G | 0.421 | 5.02E-02 | 1.09E-01 | 2.69E+00 | 2.74E+00 | 6.40E-01 | no | Fras1 | intron_variant | MODIFIER |
| 5 | 96413880 | AX-170512495 | 0 | A | G | 0.421 | 5.02E-02 | 1.09E-01 | 2.69E+00 | 2.74E+00 | 6.40E-01 | no | Fras1 | intron_variant | MODIFIER |
| 5 | 96441148 | AX-170279557 | 0 | A | G | 0.421 | 5.02E-02 | 1.09E-01 | 2.69E+00 | 2.74E+00 | 6.40E-01 | no | Fras1 | intron_variant | MODIFIER |
| 5 | 96457565 | AX-170176719 | 0 | G | A | 0.421 | 5.02E-02 | 1.09E-01 | 2.69E+00 | 2.74E+00 | 6.40E-01 | no | Fras1 | intron_variant | MODIFIER |
| 5 | 96459997 | AX-97820774 | 0 | C | T | 0.421 | 5.02E-02 | 1.09E-01 | 2.69E+00 | 2.74E+00 | 6.40E-01 | no | Fras1 | intron_variant | MODIFIER |
| 5 | 96460948 | AX-170162773 | 0 | C | T | 0.421 | 5.02E-02 | 1.09E-01 | 2.69E+00 | 2.74E+00 | 6.40E-01 | no | Fras1 | intron_variant | MODIFIER |
| 5 | 96521569 | AX-97836579 | 0 | C | T | 0.421 | 5.02E-02 | 1.09E-01 | 2.69E+00 | 2.74E+00 | 6.40E-01 | no | Fras1 | intron_variant | MODIFIER |
| 5 | 96576251 | AX-170308752 | 0 | A | G | 0.421 | 5.02E-02 | 1.09E-01 | 2.69E+00 | 2.74E+00 | 6.40E-01 | no | Fras1 | intron_variant | MODIFIER |
| 5 | 96581471 | AX-170151884 | 0 | G | A | 0.421 | 5.02E-02 | 1.09E-01 | 2.69E+00 | 2.74E+00 | 6.40E-01 | no | Fras1 | intron_variant | MODIFIER |
| 5 | 96662688 | AX-170025651 | 0 | A | G | 0.421 | 5.02E-02 | 1.09E-01 | 2.69E+00 | 2.74E+00 | 6.40E-01 | no | Fras1 | intron_variant | MODIFIER |
| 5 | 97068040 | AX-97835506 | 0 | T | C | 0.421 | 5.02E-02 | 1.09E-01 | 2.69E+00 | 2.74E+00 | 6.40E-01 | no | Bmp2k | intron_variant | MODIFIER |
| 5 | 97152851 | AX-170072828 | 0 | A | T | 0.421 | 5.02E-02 | 1.09E-01 | 2.69E+00 | 2.74E+00 | 6.40E-01 | no | Gm33609 | downstream_gene_variant | MODIFIER |
| 5 | 97198500 | AX-170497546 | 0 | T | C | 0.421 | 5.02E-02 | 1.09E-01 | 2.69E+00 | 2.74E+00 | 6.40E-01 | no | Gm2861 | intron_variant,non_coding_transcript_variant | MODIFIER |
| 5 | 97299871 | AX-170025661 | 0 | A | C | 0.421 | 5.02E-02 | 1.09E-01 | 2.69E+00 | 2.74E+00 | 6.40E-01 | no | | intergenic_variant | MODIFIER |
| 5 | 97304136 | AX-170305933 | 0 | T | C | 0.421 | 5.02E-02 | 1.09E-01 | 2.69E+00 | 2.74E+00 | 6.40E-01 | no | | intergenic_variant | MODIFIER |
| 5 | 97304650 | AX-170324868 | 0 | G | A | 0.421 | 5.02E-02 | 1.09E-01 | 2.69E+00 | 2.74E+00 | 6.40E-01 | no | | intergenic_variant | MODIFIER |
| 5 | 97690049 | AX-170464165 | 0 | C | A | 0.421 | 5.02E-02 | 1.09E-01 | 2.69E+00 | 2.74E+00 | 6.40E-01 | no | | intergenic_variant | MODIFIER |
| 5 | 97786840 | AX-170162923 | 0 | G | A | 0.421 | 5.02E-02 | 1.09E-01 | 2.69E+00 | 2.74E+00 | 6.40E-01 | no | | intergenic_variant | MODIFIER |
| 5 | 97877562 | AX-170305930 | 0 | A | G | 0.421 | 5.02E-02 | 1.09E-01 | 2.69E+00 | 2.74E+00 | 6.40E-01 | no | | intergenic_variant | MODIFIER |
| 5 | 97879520 | AX-170291772 | 0 | C | T | 0.421 | 5.02E-02 | 1.09E-01 | 2.69E+00 | 2.74E+00 | 6.40E-01 | no | Antxr2 | downstream_gene_variant | MODIFIER |
| 5 | 97880073 | AX-97836193 | 0 | T | C | 0.421 | 5.02E-02 | 1.09E-01 | 2.69E+00 | 2.74E+00 | 6.40E-01 | no | Antxr2 | downstream_gene_variant | MODIFIER |
| 5 | 97906100 | AX-170375544 | 0 | G | A | 0.421 | 5.02E-02 | 1.09E-01 | 2.69E+00 | 2.74E+00 | 6.40E-01 | no | Antxr2 | intron_variant | MODIFIER |
| 7 | 82246188 | AX-170077881 | 2 | A | G | 0.2 | 4.34E-02 | 9.51E-02 | 2.65E+00 | 2.78E+00 | 6.41E-01 | no | Sh3gl3 | intron_variant | MODIFIER |
| 18 | 59066820 | AX-170170565 | 1 | C | T | 0.42 | 4.72E-02 | 9.25E-02 | 2.45E+00 | 2.54E+00 | 6.41E-01 | no | Minar2 | intron_variant | MODIFIER |
| 15 | 3833008 | AX-170432089 | 2 | G | C | 0.264 | -4.60E-02 | 9.84E-02 | 2.61E+00 | 2.69E+00 | 6.41E-01 | no | | intergenic_variant | MODIFIER |
| 8 | 32178238 | AX-170455772 | 0 | A | T | 0.149 | -5.53E-02 | 1.20E-01 | 2.64E+00 | 2.73E+00 | 6.41E-01 | no | Nrg1 | intron_variant | MODIFIER |
| 9 | 81264555 | AX-97813317 | 0 | C | T | 0.219 | 5.39E-02 | 1.18E-01 | 2.73E+00 | 2.81E+00 | 6.41E-01 | no | | intergenic_variant | MODIFIER |

| | | | | | | | | | | | | | | | |
|---|---|---|---|---|---|---|---|---|---|---|---|---|---|---|---|
| 1 | 36235074 | AX-170299493 | 1 | T | A | 0.455 | 3.49E-02 | 7.64E-02 | 2.71E+00 | 2.80E+00 | 6.41E-01 | no | Uggt1 | intron_variant | MODIFIER |
| 8 | 36048816 | AX-170286004 | 0 | A | C | 0.149 | -6.80E-02 | 1.44E-01 | 2.60E+00 | 2.65E+00 | 6.41E-01 | no | | intergenic_variant | MODIFIER |
| 1 | 176043880 | AX-97849203 | 0 | C | T | 0.219 | 5.68E-02 | 1.20E-01 | 2.59E+00 | 2.65E+00 | 6.41E-01 | no | Pld5 | intron_variant | MODIFIER |
| 1 | 176044015 | AX-170161911 | 0 | G | A | 0.219 | 5.68E-02 | 1.20E-01 | 2.59E+00 | 2.65E+00 | 6.41E-01 | no | Pld5 | intron_variant | MODIFIER |
| 1 | 176044093 | AX-170221665 | 0 | A | G | 0.219 | 5.68E-02 | 1.20E-01 | 2.59E+00 | 2.65E+00 | 6.41E-01 | no | Pld5 | intron_variant | MODIFIER |
| 1 | 176045027 | AX-170446923 | 0 | T | C | 0.219 | 5.68E-02 | 1.20E-01 | 2.59E+00 | 2.65E+00 | 6.41E-01 | no | Pld5 | intron_variant | MODIFIER |
| 1 | 176045166 | AX-170256548 | 0 | A | G | 0.219 | 5.68E-02 | 1.20E-01 | 2.59E+00 | 2.65E+00 | 6.41E-01 | no | Pld5 | intron_variant | MODIFIER |
| 1 | 176067985 | AX-97854917 | 0 | G | A | 0.219 | 5.68E-02 | 1.20E-01 | 2.59E+00 | 2.65E+00 | 6.41E-01 | no | Pld5 | intron_variant | MODIFIER |
| 1 | 176069949 | AX-170043675 | 0 | T | C | 0.219 | 5.68E-02 | 1.20E-01 | 2.59E+00 | 2.65E+00 | 6.41E-01 | no | Pld5 | intron_variant | MODIFIER |
| 1 | 176070502 | AX-170043678 | 0 | T | C | 0.219 | 5.68E-02 | 1.20E-01 | 2.59E+00 | 2.65E+00 | 6.41E-01 | no | Pld5 | intron_variant | MODIFIER |
| 1 | 176070638 | AX-170043677 | 0 | A | G | 0.219 | 5.68E-02 | 1.20E-01 | 2.59E+00 | 2.65E+00 | 6.41E-01 | no | Pld5 | intron_variant | MODIFIER |
| 1 | 176092628 | AX-170254509 | 0 | T | C | 0.219 | 5.68E-02 | 1.20E-01 | 2.59E+00 | 2.65E+00 | 6.41E-01 | no | Pld5 | intron_variant | MODIFIER |
| 1 | 176095672 | AX-170115559 | 0 | T | G | 0.219 | 5.68E-02 | 1.20E-01 | 2.59E+00 | 2.65E+00 | 6.41E-01 | no | Pld5 | intron_variant | MODIFIER |
| 1 | 176096092 | AX-170387342 | 0 | A | C | 0.219 | 5.68E-02 | 1.20E-01 | 2.59E+00 | 2.65E+00 | 6.41E-01 | no | Pld5 | intron_variant | MODIFIER |
| 1 | 176098531 | AX-170251499 | 0 | T | A | 0.219 | 5.68E-02 | 1.20E-01 | 2.59E+00 | 2.65E+00 | 6.41E-01 | no | Pld5 | intron_variant | MODIFIER |
| 1 | 176112387 | AX-170523597 | 0 | A | G | 0.219 | 5.68E-02 | 1.20E-01 | 2.59E+00 | 2.65E+00 | 6.41E-01 | no | Pld5 | intron_variant | MODIFIER |
| 1 | 176112564 | AX-170199017 | 0 | G | C | 0.219 | 5.68E-02 | 1.20E-01 | 2.59E+00 | 2.65E+00 | 6.41E-01 | no | Pld5 | intron_variant | MODIFIER |
| 1 | 176113213 | AX-170477948 | 0 | T | C | 0.219 | 5.68E-02 | 1.20E-01 | 2.59E+00 | 2.65E+00 | 6.41E-01 | no | Pld5 | intron_variant | MODIFIER |
| 1 | 176117187 | AX-170002344 | 0 | C | T | 0.219 | 5.68E-02 | 1.20E-01 | 2.59E+00 | 2.65E+00 | 6.41E-01 | no | Pld5 | intron_variant | MODIFIER |
| 1 | 176117343 | AX-170381848 | 0 | T | C | 0.219 | 5.68E-02 | 1.20E-01 | 2.59E+00 | 2.65E+00 | 6.41E-01 | no | Pld5 | intron_variant | MODIFIER |
| 1 | 176122341 | AX-97853890 | 0 | G | A | 0.219 | 5.68E-02 | 1.20E-01 | 2.59E+00 | 2.65E+00 | 6.41E-01 | no | Pld5 | intron_variant | MODIFIER |
| 1 | 176124118 | AX-170002347 | 0 | G | C | 0.219 | 5.68E-02 | 1.20E-01 | 2.59E+00 | 2.65E+00 | 6.41E-01 | no | Pld5 | intron_variant | MODIFIER |
| 1 | 176143402 | AX-170391576 | 0 | C | T | 0.219 | 5.68E-02 | 1.20E-01 | 2.59E+00 | 2.65E+00 | 6.41E-01 | no | Pld5 | intron_variant | MODIFIER |
| 1 | 176171541 | AX-170341849 | 0 | G | T | 0.219 | 5.68E-02 | 1.20E-01 | 2.59E+00 | 2.65E+00 | 6.41E-01 | no | Pld5 | intron_variant | MODIFIER |
| 1 | 176171691 | AX-170228672 | 0 | A | G | 0.219 | 5.68E-02 | 1.20E-01 | 2.59E+00 | 2.65E+00 | 6.41E-01 | no | Pld5 | intron_variant | MODIFIER |
| 1 | 176173708 | AX-170435318 | 0 | C | T | 0.219 | 5.68E-02 | 1.20E-01 | 2.59E+00 | 2.65E+00 | 6.41E-01 | no | Pld5 | intron_variant | MODIFIER |
| 1 | 176185175 | AX-170002343 | 0 | C | T | 0.219 | 5.68E-02 | 1.20E-01 | 2.59E+00 | 2.65E+00 | 6.41E-01 | no | Pld5 | intron_variant | MODIFIER |
| 1 | 176186289 | AX-170300641 | 0 | G | A | 0.219 | 5.68E-02 | 1.20E-01 | 2.59E+00 | 2.65E+00 | 6.41E-01 | no | Pld5 | intron_variant | MODIFIER |
| 1 | 176192397 | AX-170197582 | 0 | A | T | 0.219 | 5.68E-02 | 1.20E-01 | 2.59E+00 | 2.65E+00 | 6.41E-01 | no | Pld5 | intron_variant | MODIFIER |
| 1 | 176193200 | AX-170472337 | 0 | C | T | 0.219 | 5.68E-02 | 1.20E-01 | 2.59E+00 | 2.65E+00 | 6.41E-01 | no | Pld5 | intron_variant | MODIFIER |
| 1 | 176193242 | AX-170309146 | 0 | C | T | 0.219 | 5.68E-02 | 1.20E-01 | 2.59E+00 | 2.65E+00 | 6.41E-01 | no | Pld5 | intron_variant | MODIFIER |
| 1 | 176289208 | AX-170339437 | 0 | A | G | 0.219 | 5.68E-02 | 1.20E-01 | 2.59E+00 | 2.65E+00 | 6.41E-01 | no | | intergenic_variant | MODIFIER |
| 1 | 176506593 | AX-170268177 | 0 | T | A | 0.219 | 5.68E-02 | 1.20E-01 | 2.59E+00 | 2.65E+00 | 6.41E-01 | no | Gm36536 | intron_variant,non_coding_transcript_variant | MODIFIER |
| 1 | 176530441 | AX-170395095 | 0 | G | A | 0.219 | 5.68E-02 | 1.20E-01 | 2.59E+00 | 2.65E+00 | 6.41E-01 | no | Gm36536 | upstream_gene_variant | MODIFIER |
| 1 | 176557307 | AX-170528494 | 0 | A | G | 0.219 | 5.68E-02 | 1.20E-01 | 2.59E+00 | 2.65E+00 | 6.41E-01 | no | | intergenic_variant | MODIFIER |
| 1 | 176563915 | AX-170120826 | 0 | G | A | 0.219 | 5.68E-02 | 1.20E-01 | 2.59E+00 | 2.65E+00 | 6.41E-01 | no | | intergenic_variant | MODIFIER |
| 1 | 176565909 | AX-170375671 | 0 | T | C | 0.219 | 5.68E-02 | 1.20E-01 | 2.59E+00 | 2.65E+00 | 6.41E-01 | no | | intergenic_variant | MODIFIER |
| 1 | 176612871 | AX-170002351 | 0 | T | G | 0.219 | 5.68E-02 | 1.20E-01 | 2.59E+00 | 2.65E+00 | 6.41E-01 | no | | intergenic_variant | MODIFIER |
| 1 | 176613011 | AX-97830939 | 0 | G | A | 0.219 | 5.68E-02 | 1.20E-01 | 2.59E+00 | 2.65E+00 | 6.41E-01 | no | | intergenic_variant | MODIFIER |
| 1 | 176664830 | AX-170002348 | 0 | T | C | 0.219 | 5.68E-02 | 1.20E-01 | 2.59E+00 | 2.65E+00 | 6.41E-01 | no | | intergenic_variant | MODIFIER |
| 1 | 176665237 | AX-170318910 | 0 | T | C | 0.219 | 5.68E-02 | 1.20E-01 | 2.59E+00 | 2.65E+00 | 6.41E-01 | no | | intergenic_variant | MODIFIER |
| 1 | 176698760 | AX-170458219 | 0 | T | G | 0.219 | 5.68E-02 | 1.20E-01 | 2.59E+00 | 2.65E+00 | 6.41E-01 | no | Gm26104 | upstream_gene_variant | MODIFIER |
| 1 | 176704717 | AX-170463987 | 0 | T | C | 0.219 | 5.68E-02 | 1.20E-01 | 2.59E+00 | 2.65E+00 | 6.41E-01 | no | Gm26104 | downstream_gene_variant | MODIFIER |
| 1 | 176707119 | AX-170376287 | 0 | T | C | 0.219 | 5.68E-02 | 1.20E-01 | 2.59E+00 | 2.65E+00 | 6.41E-01 | no | | intergenic_variant | MODIFIER |
| 1 | 176720277 | AX-170043690 | 0 | T | G | 0.219 | 5.68E-02 | 1.20E-01 | 2.59E+00 | 2.65E+00 | 6.41E-01 | no | | intergenic_variant | MODIFIER |
| 1 | 176720878 | AX-170043691 | 0 | A | C | 0.219 | 5.68E-02 | 1.20E-01 | 2.59E+00 | 2.65E+00 | 6.41E-01 | no | | intergenic_variant | MODIFIER |
| 1 | 176724068 | AX-170391243 | 0 | G | A | 0.219 | 5.68E-02 | 1.20E-01 | 2.59E+00 | 2.65E+00 | 6.41E-01 | no | | regulatory_region_variant | MODIFIER |
| 1 | 176740778 | AX-170376144 | 0 | A | T | 0.219 | 5.68E-02 | 1.20E-01 | 2.59E+00 | 2.65E+00 | 6.41E-01 | no | Cep170 | intron_variant | MODIFIER |
| 1 | 176779108 | AX-170480551 | 0 | T | G | 0.219 | 5.68E-02 | 1.20E-01 | 2.59E+00 | 2.65E+00 | 6.41E-01 | no | Cep170 | intron_variant | MODIFIER |
| 1 | 176779378 | AX-170107092 | 0 | G | A | 0.219 | 5.68E-02 | 1.20E-01 | 2.59E+00 | 2.65E+00 | 6.41E-01 | no | Cep170 | intron_variant | MODIFIER |
| 1 | 176782026 | AX-170123747 | 0 | C | T | 0.219 | 5.68E-02 | 1.20E-01 | 2.59E+00 | 2.65E+00 | 6.41E-01 | no | Cep170 | intron_variant | MODIFIER |
| 1 | 176790795 | AX-170242211 | 0 | C | T | 0.219 | 5.68E-02 | 1.20E-01 | 2.59E+00 | 2.65E+00 | 6.41E-01 | no | Cep170 | intron_variant | MODIFIER |
| 1 | 176826202 | AX-170180903 | 0 | A | G | 0.219 | 5.68E-02 | 1.20E-01 | 2.59E+00 | 2.65E+00 | 6.41E-01 | no | Sdccag8 | missense_variant | MODERATE |
| 1 | 176826901 | AX-170490127 | 0 | T | C | 0.219 | 5.68E-02 | 1.20E-01 | 2.59E+00 | 2.65E+00 | 6.41E-01 | no | Sdccag8 | intron_variant | MODIFIER |

| | | | | | | | | | | | | | | | |
|---|---|---|---|---|---|---|---|---|---|---|---|---|---|---|---|
| 1 | 176839855 | AX-97819110 | 0 | C | T | 0.219 | 5.68E-02 | 1.20E-01 | 2.59E+00 | 2.65E+00 | 6.41E-01 | no | Sdccag8 | intron_variant | MODIFIER |
| 1 | 176840252 | AX-170265701 | 0 | A | G | 0.219 | 5.68E-02 | 1.20E-01 | 2.59E+00 | 2.65E+00 | 6.41E-01 | no | Sdccag8 | intron_variant | MODIFIER |
| 1 | 176853645 | AX-170351818 | 0 | G | A | 0.219 | 5.68E-02 | 1.20E-01 | 2.59E+00 | 2.65E+00 | 6.41E-01 | no | Sdccag8 | intron_variant | MODIFIER |
| 1 | 176854010 | AX-170137200 | 0 | T | C | 0.219 | 5.68E-02 | 1.20E-01 | 2.59E+00 | 2.65E+00 | 6.41E-01 | no | Sdccag8 | intron_variant | MODIFIER |
| 1 | 176906035 | AX-170219886 | 0 | T | C | 0.219 | 5.68E-02 | 1.20E-01 | 2.59E+00 | 2.65E+00 | 6.41E-01 | no | Sdccag8 | intron_variant | MODIFIER |
| 1 | 176906468 | AX-97826893 | 0 | C | T | 0.219 | 5.68E-02 | 1.20E-01 | 2.59E+00 | 2.65E+00 | 6.41E-01 | no | Sdccag8 | intron_variant | MODIFIER |
| 1 | 176949024 | AX-170515772 | 0 | G | A | 0.219 | 5.68E-02 | 1.20E-01 | 2.59E+00 | 2.65E+00 | 6.41E-01 | no | Sdccag8 | intron_variant | MODIFIER |
| 1 | 176970165 | AX-170209935 | 0 | A | G | 0.219 | 5.68E-02 | 1.20E-01 | 2.59E+00 | 2.65E+00 | 6.41E-01 | no | Sdccag8 | intron_variant | MODIFIER |
| 1 | 176971089 | AX-170210740 | 0 | G | T | 0.219 | 5.68E-02 | 1.20E-01 | 2.59E+00 | 2.65E+00 | 6.41E-01 | no | Sdccag8 | intron_variant | MODIFIER |
| 1 | 176976794 | AX-170427812 | 0 | G | A | 0.219 | 5.68E-02 | 1.20E-01 | 2.59E+00 | 2.65E+00 | 6.41E-01 | no | Sdccag8 | intron_variant | MODIFIER |
| 19 | 6211819 | AX-170150839 | 0 | G | A | 0.053 | -8.64E-02 | 1.89E-01 | 2.66E+00 | 2.77E+00 | 6.41E-01 | no | Majin | intron_variant | MODIFIER |
| 5 | 111578878 | AX-170454161 | 1 | T | C | 0.446 | 4.76E-02 | 1.01E-01 | 2.60E+00 | 2.67E+00 | 6.41E-01 | no | C130026L21Rik | upstream_gene_variant | MODIFIER |
| 11 | 74270606 | AX-170208373 | 0 | C | A | 0.263 | 3.28E-02 | 7.20E-02 | 2.64E+00 | 2.77E+00 | 6.42E-01 | no | Olfr405-ps1 | upstream_gene_variant | MODIFIER |
| 2 | 77837584 | AX-170400195 | 0 | A | C | 0.053 | -1.39E-01 | 3.01E-01 | 2.76E+00 | 2.79E+00 | 6.42E-01 | no | Gm13661 | upstream_gene_variant | MODIFIER |
| 18 | 58747285 | AX-170097121 | 0 | T | C | 0.395 | 4.64E-02 | 9.53E-02 | 2.53E+00 | 2.59E+00 | 6.42E-01 | no | | intergenic_variant | MODIFIER |
| 9 | 40213083 | AX-168387651 | 1 | C | A | 0.42 | -3.47E-02 | 7.59E-02 | 2.68E+00 | 2.77E+00 | 6.42E-01 | no | Zfp202 | 3_prime_UTR_variant | MODIFIER |
| 16 | 45800377 | AX-170185107 | 1 | T | C | 0.473 | -3.74E-02 | 7.86E-02 | 2.57E+00 | 2.64E+00 | 6.42E-01 | no | Phldb2 | intron_variant | MODIFIER |
| 5 | 112050384 | AX-170166618 | 1 | A | G | 0.446 | 5.34E-02 | 1.15E-01 | 2.65E+00 | 2.69E+00 | 6.42E-01 | no | | regulatory_region_variant | MODIFIER |
| 5 | 117023705 | AX-170073242 | 1 | G | A | 0.455 | 5.50E-02 | 1.19E-01 | 2.66E+00 | 2.69E+00 | 6.42E-01 | no | | intergenic_variant | MODIFIER |
| 8 | 74391506 | AX-170428900 | 2 | T | A | 0.2 | -5.57E-02 | 1.21E-01 | 2.65E+00 | 2.74E+00 | 6.42E-01 | no | | intergenic_variant | MODIFIER |
| 12 | 111943922 | AX-97830383 | 0 | C | T | 0.14 | -8.76E-02 | 1.92E-01 | 2.73E+00 | 2.80E+00 | 6.42E-01 | no | 5033406O09Rik | non_coding_transcript_exon_variant | MODIFIER |
| 4 | 128259896 | AX-170174177 | 1 | G | A | 0.143 | 5.67E-02 | 1.21E-01 | 2.61E+00 | 2.67E+00 | 6.42E-01 | no | Hmgb4 | downstream_gene_variant | MODIFIER |
| 16 | 92379079 | AX-170159400 | 1 | C | T | 0.482 | 3.54E-02 | 7.78E-02 | 2.70E+00 | 2.79E+00 | 6.42E-01 | no | Gm29805 | intron_variant,non_coding_transcript_variant | MODIFIER |
| 7 | 39981505 | AX-170238703 | 1 | G | A | 0.348 | 4.47E-02 | 9.57E-02 | 2.61E+00 | 2.68E+00 | 6.42E-01 | no | Gm44992 | upstream_gene_variant | MODIFIER |
| 7 | 40101393 | AX-97857057 | 1 | G | A | 0.348 | 4.47E-02 | 9.57E-02 | 2.61E+00 | 2.68E+00 | 6.42E-01 | no | | intergenic_variant | MODIFIER |
| 5 | 81415851 | AX-170155561 | 0 | G | A | 0.412 | 4.72E-02 | 1.02E-01 | 2.65E+00 | 2.72E+00 | 6.42E-01 | no | Adgrl3 | intron_variant | MODIFIER |
| 1 | 190320332 | AX-97811141 | 0 | G | A | 0.254 | 4.82E-02 | 9.70E-02 | 2.48E+00 | 2.57E+00 | 6.43E-01 | no | | intergenic_variant | MODIFIER |
| 5 | 77130756 | AX-170346215 | 0 | C | T | 0.43 | 4.57E-02 | 9.93E-02 | 2.64E+00 | 2.72E+00 | 6.43E-01 | no | | regulatory_region_variant | MODIFIER |
| 10 | 67841206 | AX-97854091 | 0 | C | T | 0.412 | -3.60E-02 | 7.67E-02 | 2.59E+00 | 2.67E+00 | 6.43E-01 | no | Gm49766 | downstream_gene_variant | MODIFIER |
| 11 | 11087678 | AX-170004281 | 1 | C | T | 0.062 | 7.64E-02 | 1.69E-01 | 2.68E+00 | 2.80E+00 | 6.43E-01 | no | | intergenic_variant | MODIFIER |
| 7 | 6786201 | AX-170416196 | 0 | A | C | 0.167 | 3.59E-02 | 8.02E-02 | 2.75E+00 | 2.89E+00 | 6.43E-01 | no | Usp29 | intron_variant | MODIFIER |
| 7 | 6786254 | AX-170249985 | 0 | A | G | 0.167 | 3.59E-02 | 8.02E-02 | 2.75E+00 | 2.89E+00 | 6.43E-01 | no | Usp29 | intron_variant | MODIFIER |
| 7 | 6801303 | AX-170076684 | 0 | A | T | 0.167 | 3.59E-02 | 8.02E-02 | 2.75E+00 | 2.89E+00 | 6.43E-01 | no | Mir3099 | upstream_gene_variant | MODIFIER |
| 7 | 6812499 | AX-170446490 | 0 | C | T | 0.167 | 3.59E-02 | 8.02E-02 | 2.75E+00 | 2.89E+00 | 6.43E-01 | no | Usp29 | intron_variant | MODIFIER |
| 7 | 6812576 | AX-170474326 | 0 | C | T | 0.167 | 3.59E-02 | 8.02E-02 | 2.75E+00 | 2.89E+00 | 6.43E-01 | no | Usp29 | intron_variant | MODIFIER |
| 7 | 6842366 | AX-170279642 | 0 | C | A | 0.167 | 3.59E-02 | 8.02E-02 | 2.75E+00 | 2.89E+00 | 6.43E-01 | no | Gm22649 | upstream_gene_variant | MODIFIER |
| 7 | 6857620 | AX-170148724 | 0 | G | A | 0.167 | 3.59E-02 | 8.02E-02 | 2.75E+00 | 2.89E+00 | 6.43E-01 | no | Usp29 | intron_variant | MODIFIER |
| 7 | 6878316 | AX-170530087 | 0 | A | G | 0.167 | 3.59E-02 | 8.02E-02 | 2.75E+00 | 2.89E+00 | 6.43E-01 | no | Usp29 | intron_variant | MODIFIER |
| 7 | 6900969 | AX-170110965 | 0 | C | T | 0.167 | 3.59E-02 | 8.02E-02 | 2.75E+00 | 2.89E+00 | 6.43E-01 | no | Usp29 | intron_variant | MODIFIER |
| 7 | 6916885 | AX-170452109 | 0 | T | C | 0.167 | 3.59E-02 | 8.02E-02 | 2.75E+00 | 2.89E+00 | 6.43E-01 | no | Usp29 | intron_variant | MODIFIER |
| 7 | 6957605 | AX-97815509 | 0 | C | A | 0.167 | 3.59E-02 | 8.02E-02 | 2.75E+00 | 2.89E+00 | 6.43E-01 | no | Usp29 | upstream_gene_variant | MODIFIER |
| 7 | 6962862 | AX-170182305 | 0 | T | C | 0.167 | 3.59E-02 | 8.02E-02 | 2.75E+00 | 2.89E+00 | 6.43E-01 | no | Usp29 | missense_variant | MODERATE |
| 7 | 6967169 | AX-170028563 | 0 | A | G | 0.167 | 3.59E-02 | 8.02E-02 | 2.75E+00 | 2.89E+00 | 6.43E-01 | no | Usp29 | downstream_gene_variant | MODIFIER |
| 7 | 7042380 | AX-170303647 | 0 | G | A | 0.167 | 3.59E-02 | 8.02E-02 | 2.75E+00 | 2.89E+00 | 6.43E-01 | no | | intergenic_variant | MODIFIER |
| 7 | 7042708 | AX-170431858 | 0 | C | T | 0.167 | 3.59E-02 | 8.02E-02 | 2.75E+00 | 2.89E+00 | 6.43E-01 | no | | intergenic_variant | MODIFIER |
| 7 | 7092157 | AX-170202217 | 0 | T | A | 0.167 | 3.59E-02 | 8.02E-02 | 2.75E+00 | 2.89E+00 | 6.43E-01 | no | Gm6828 | downstream_gene_variant | MODIFIER |
| 7 | 7111746 | AX-170125741 | 0 | G | A | 0.167 | 3.59E-02 | 8.02E-02 | 2.75E+00 | 2.89E+00 | 6.43E-01 | no | Zfp954 | downstream_gene_variant | MODIFIER |
| 7 | 7124512 | AX-170329127 | 0 | T | C | 0.167 | 3.59E-02 | 8.02E-02 | 2.75E+00 | 2.89E+00 | 6.43E-01 | no | Zfp954 | upstream_gene_variant | MODIFIER |
| 7 | 7173133 | AX-170354790 | 0 | C | T | 0.167 | 3.59E-02 | 8.02E-02 | 2.75E+00 | 2.89E+00 | 6.43E-01 | no | Zfp418 | intron_variant | MODIFIER |
| 7 | 7179023 | AX-170151192 | 0 | G | A | 0.167 | 3.59E-02 | 8.02E-02 | 2.75E+00 | 2.89E+00 | 6.43E-01 | no | Zfp418 | intron_variant | MODIFIER |
| 7 | 7281035 | AX-170076697 | 0 | T | C | 0.167 | 3.59E-02 | 8.02E-02 | 2.75E+00 | 2.89E+00 | 6.43E-01 | no | Clcn4 | downstream_gene_variant | MODIFIER |
| 7 | 7286111 | AX-170448582 | 0 | C | T | 0.167 | 3.59E-02 | 8.02E-02 | 2.75E+00 | 2.89E+00 | 6.43E-01 | no | Clcn4 | intron_variant | MODIFIER |
| 7 | 7288409 | AX-170290059 | 0 | C | G | 0.167 | 3.59E-02 | 8.02E-02 | 2.75E+00 | 2.89E+00 | 6.43E-01 | no | Clcn4 | intron_variant | MODIFIER |
| 7 | 7295703 | AX-170457412 | 0 | C | T | 0.167 | 3.59E-02 | 8.02E-02 | 2.75E+00 | 2.89E+00 | 6.43E-01 | no | Clcn4 | intron_variant | MODIFIER |

| | | | | | | | | | | | | | | | |
|---|---|---|---|---|---|---|---|---|---|---|---|---|---|---|---|
| 7 | 10133209 | AX-97853984 | 0 | T | C | 0.167 | 3.59E-02 | 8.02E-02 | 2.75E+00 | 2.89E+00 | 6.43E-01 | no | | intergenic_variant | MODIFIER |
| 7 | 10198193 | AX-170342922 | 0 | A | G | 0.167 | 3.59E-02 | 8.02E-02 | 2.75E+00 | 2.89E+00 | 6.43E-01 | no | Gm20679 | intron_variant,non_coding_transcript_variant | MODIFIER |
| 7 | 10202889 | AX-170140986 | 0 | T | G | 0.167 | 3.59E-02 | 8.02E-02 | 2.75E+00 | 2.89E+00 | 6.43E-01 | no | Gm20679 | intron_variant,non_coding_transcript_variant | MODIFIER |
| 7 | 10207913 | AX-170125069 | 0 | T | C | 0.167 | 3.59E-02 | 8.02E-02 | 2.75E+00 | 2.89E+00 | 6.43E-01 | no | Gm20679 | upstream_gene_variant | MODIFIER |
| 7 | 10213844 | AX-170076708 | 0 | T | C | 0.167 | 3.59E-02 | 8.02E-02 | 2.75E+00 | 2.89E+00 | 6.43E-01 | no | | intergenic_variant | MODIFIER |
| 7 | 10233943 | AX-170076704 | 0 | A | G | 0.167 | 3.59E-02 | 8.02E-02 | 2.75E+00 | 2.89E+00 | 6.43E-01 | no | | intergenic_variant | MODIFIER |
| 7 | 10235739 | AX-97856000 | 0 | G | A | 0.167 | 3.59E-02 | 8.02E-02 | 2.75E+00 | 2.89E+00 | 6.43E-01 | no | | intergenic_variant | MODIFIER |
| 7 | 10238152 | AX-170508147 | 0 | C | T | 0.167 | 3.59E-02 | 8.02E-02 | 2.75E+00 | 2.89E+00 | 6.43E-01 | no | | intergenic_variant | MODIFIER |
| 7 | 10278509 | AX-97854731 | 0 | C | A | 0.167 | 3.59E-02 | 8.02E-02 | 2.75E+00 | 2.89E+00 | 6.43E-01 | no | Vmn1r66 | upstream_gene_variant | MODIFIER |
| 7 | 10285346 | AX-170168452 | 0 | A | G | 0.167 | 3.59E-02 | 8.02E-02 | 2.75E+00 | 2.89E+00 | 6.43E-01 | no | Vmn1r-ps47 | upstream_gene_variant | MODIFIER |
| 7 | 10294689 | AX-170461474 | 0 | C | G | 0.167 | 3.59E-02 | 8.02E-02 | 2.75E+00 | 2.89E+00 | 6.43E-01 | no | | intergenic_variant | MODIFIER |
| 7 | 10345008 | AX-170226217 | 0 | A | T | 0.167 | 3.59E-02 | 8.02E-02 | 2.75E+00 | 2.89E+00 | 6.43E-01 | no | Gm45400 | intron_variant,non_coding_transcript_variant | MODIFIER |
| 7 | 10355946 | AX-170154686 | 0 | G | C | 0.167 | 3.59E-02 | 8.02E-02 | 2.75E+00 | 2.89E+00 | 6.43E-01 | no | Nlrp4d | downstream_gene_variant | MODIFIER |
| 7 | 10377250 | AX-170532170 | 0 | T | A | 0.167 | 3.59E-02 | 8.02E-02 | 2.75E+00 | 2.89E+00 | 6.43E-01 | no | Nlrp4d | intron_variant,non_coding_transcript_variant | MODIFIER |
| 7 | 10410879 | AX-170234414 | 0 | T | C | 0.167 | 3.59E-02 | 8.02E-02 | 2.75E+00 | 2.89E+00 | 6.43E-01 | no | Gm3726 | intron_variant,non_coding_transcript_variant | MODIFIER |
| 7 | 10411420 | AX-170219091 | 0 | A | G | 0.167 | 3.59E-02 | 8.02E-02 | 2.75E+00 | 2.89E+00 | 6.43E-01 | no | Gm3726 | intron_variant,non_coding_transcript_variant | MODIFIER |
| 7 | 10412077 | AX-170514448 | 0 | A | G | 0.167 | 3.59E-02 | 8.02E-02 | 2.75E+00 | 2.89E+00 | 6.43E-01 | no | Gm3726 | intron_variant,non_coding_transcript_variant | MODIFIER |
| 7 | 10414874 | AX-170076712 | 0 | C | T | 0.167 | 3.59E-02 | 8.02E-02 | 2.75E+00 | 2.89E+00 | 6.43E-01 | no | Gm3726 | upstream_gene_variant | MODIFIER |
| 7 | 10456430 | AX-170028572 | 0 | A | T | 0.167 | 3.59E-02 | 8.02E-02 | 2.75E+00 | 2.89E+00 | 6.43E-01 | no | | intergenic_variant | MODIFIER |
| 7 | 10468039 | AX-170028575 | 0 | C | T | 0.167 | 3.59E-02 | 8.02E-02 | 2.75E+00 | 2.89E+00 | 6.43E-01 | no | | intergenic_variant | MODIFIER |
| 7 | 10491658 | AX-170095862 | 0 | A | G | 0.167 | 3.59E-02 | 8.02E-02 | 2.75E+00 | 2.89E+00 | 6.43E-01 | no | Zik1 | intron_variant | MODIFIER |
| 7 | 10494630 | AX-170028578 | 0 | G | A | 0.167 | 3.59E-02 | 8.02E-02 | 2.75E+00 | 2.89E+00 | 6.43E-01 | no | Zik1 | intron_variant | MODIFIER |
| 7 | 10501457 | AX-170028577 | 0 | G | A | 0.167 | 3.59E-02 | 8.02E-02 | 2.75E+00 | 2.89E+00 | 6.43E-01 | no | Gm18193 | upstream_gene_variant | MODIFIER |
| 7 | 10519813 | AX-170028569 | 0 | T | C | 0.167 | 3.59E-02 | 8.02E-02 | 2.75E+00 | 2.89E+00 | 6.43E-01 | no | Vmn1r68 | downstream_gene_variant | MODIFIER |
| 7 | 10522943 | AX-170334538 | 0 | C | A | 0.167 | 3.59E-02 | 8.02E-02 | 2.75E+00 | 2.89E+00 | 6.43E-01 | no | Vmn1r68 | downstream_gene_variant | MODIFIER |
| 7 | 10527765 | AX-170181783 | 0 | G | T | 0.167 | 3.59E-02 | 8.02E-02 | 2.75E+00 | 2.89E+00 | 6.43E-01 | no | Vmn1r68 | synonymous_variant | LOW |
| 7 | 10532171 | AX-170436785 | 0 | T | C | 0.167 | 3.59E-02 | 8.02E-02 | 2.75E+00 | 2.89E+00 | 6.43E-01 | no | Vmn1r68 | upstream_gene_variant | MODIFIER |
| 7 | 10536716 | AX-170028570 | 0 | G | A | 0.167 | 3.59E-02 | 8.02E-02 | 2.75E+00 | 2.89E+00 | 6.43E-01 | no | Vmn1r68 | intron_variant,NMD_transcript_variant | MODIFIER |
| 7 | 10543734 | AX-97805756 | 0 | A | C | 0.167 | 3.59E-02 | 8.02E-02 | 2.75E+00 | 2.89E+00 | 6.43E-01 | no | Vmn1r68 | intron_variant,NMD_transcript_variant | MODIFIER |
| 7 | 10585544 | AX-170028579 | 0 | T | C | 0.167 | 3.59E-02 | 8.02E-02 | 2.75E+00 | 2.89E+00 | 6.43E-01 | no | Vmn1r69 | upstream_gene_variant | MODIFIER |
| 7 | 10592119 | AX-170028573 | 0 | G | A | 0.167 | 3.59E-02 | 8.02E-02 | 2.75E+00 | 2.89E+00 | 6.43E-01 | no | Vmn1r69 | intron_variant | MODIFIER |
| 7 | 10604911 | AX-97842963 | 0 | A | G | 0.167 | 3.59E-02 | 8.02E-02 | 2.75E+00 | 2.89E+00 | 6.43E-01 | no | Vmn1r69 | intron_variant | MODIFIER |
| 7 | 10628599 | AX-170028574 | 0 | A | G | 0.167 | 3.59E-02 | 8.02E-02 | 2.75E+00 | 2.89E+00 | 6.43E-01 | no | Vmn1r70 | upstream_gene_variant | MODIFIER |
| 7 | 10630990 | AX-97805433 | 0 | A | G | 0.167 | 3.59E-02 | 8.02E-02 | 2.75E+00 | 2.89E+00 | 6.43E-01 | no | Vmn1r70 | upstream_gene_variant | MODIFIER |
| 7 | 10656199 | AX-170028580 | 0 | G | C | 0.167 | 3.59E-02 | 8.02E-02 | 2.75E+00 | 2.89E+00 | 6.43E-01 | no | Gm6900 | downstream_gene_variant | MODIFIER |
| 7 | 10661877 | AX-170199399 | 0 | C | T | 0.167 | 3.59E-02 | 8.02E-02 | 2.75E+00 | 2.89E+00 | 6.43E-01 | no | Vmn1r-ps52 | downstream_gene_variant | MODIFIER |
| 7 | 10668165 | AX-170254730 | 0 | A | G | 0.167 | 3.59E-02 | 8.02E-02 | 2.75E+00 | 2.89E+00 | 6.43E-01 | no | Vmn1r-ps52 | upstream_gene_variant | MODIFIER |
| 7 | 10690142 | AX-170349664 | 0 | C | T | 0.167 | 3.59E-02 | 8.02E-02 | 2.75E+00 | 2.89E+00 | 6.43E-01 | no | Nlrp4b | intron_variant | MODIFIER |
| 7 | 10727941 | AX-170445915 | 0 | T | C | 0.167 | 3.59E-02 | 8.02E-02 | 2.75E+00 | 2.89E+00 | 6.43E-01 | no | Nlrp4b | intron_variant | MODIFIER |
| 7 | 10734237 | AX-170483977 | 0 | G | C | 0.167 | 3.59E-02 | 8.02E-02 | 2.75E+00 | 2.89E+00 | 6.43E-01 | no | Nlrp4b | downstream_gene_variant | MODIFIER |
| 7 | 10734448 | AX-170149691 | 0 | T | C | 0.167 | 3.59E-02 | 8.02E-02 | 2.75E+00 | 2.89E+00 | 6.43E-01 | no | Nlrp4b | downstream_gene_variant | MODIFIER |
| 7 | 10742638 | AX-170403101 | 0 | C | A | 0.167 | 3.59E-02 | 8.02E-02 | 2.75E+00 | 2.89E+00 | 6.43E-01 | no | Vmn1r71 | downstream_gene_variant | MODIFIER |
| 7 | 10747346 | AX-170398199 | 0 | G | A | 0.167 | 3.59E-02 | 8.02E-02 | 2.75E+00 | 2.89E+00 | 6.43E-01 | no | Vmn1r71 | 3_prime_UTR_variant | MODIFIER |
| 7 | 11183700 | AX-97850974 | 0 | C | A | 0.167 | 3.59E-02 | 8.02E-02 | 2.75E+00 | 2.89E+00 | 6.43E-01 | no | Gm6909 | non_coding_transcript_exon_variant | MODIFIER |
| 7 | 11231737 | AX-170234390 | 0 | G | C | 0.167 | 3.59E-02 | 8.02E-02 | 2.75E+00 | 2.89E+00 | 6.43E-01 | no | Gm39481 | upstream_gene_variant | MODIFIER |
| 7 | 11456079 | AX-97817043 | 0 | A | G | 0.167 | 3.59E-02 | 8.02E-02 | 2.75E+00 | 2.89E+00 | 6.43E-01 | no | Gm45382 | upstream_gene_variant | MODIFIER |
| 7 | 11652215 | AX-97785621 | 0 | A | G | 0.167 | 3.59E-02 | 8.02E-02 | 2.75E+00 | 2.89E+00 | 6.43E-01 | no | Vmn1r-ps53 | downstream_gene_variant | MODIFIER |
| 7 | 11712749 | AX-170409981 | 0 | T | C | 0.167 | 3.59E-02 | 8.02E-02 | 2.75E+00 | 2.89E+00 | 6.43E-01 | no | Gm19093 | upstream_gene_variant | MODIFIER |
| 7 | 11720219 | AX-170142276 | 0 | A | C | 0.167 | 3.59E-02 | 8.02E-02 | 2.75E+00 | 2.89E+00 | 6.43E-01 | no | Gm19093 | non_coding_transcript_exon_variant | MODIFIER |
| 1 | 186926232 | AX-170115240 | 0 | T | C | 0.228 | 5.30E-02 | 1.08E-01 | 2.51E+00 | 2.58E+00 | 6.43E-01 | no | 1700023G09Rik | upstream_gene_variant | MODIFIER |
| 1 | 186935313 | AX-170091823 | 0 | A | G | 0.228 | 5.30E-02 | 1.08E-01 | 2.51E+00 | 2.58E+00 | 6.43E-01 | no | | intergenic_variant | MODIFIER |
| 1 | 186968925 | AX-170002503 | 0 | A | G | 0.228 | 5.30E-02 | 1.08E-01 | 2.51E+00 | 2.58E+00 | 6.43E-01 | no | D1Pas1 | synonymous_variant | LOW |
| 1 | 186973220 | AX-97786639 | 0 | G | A | 0.228 | 5.30E-02 | 1.08E-01 | 2.51E+00 | 2.58E+00 | 6.43E-01 | no | D1Pas1 | downstream_gene_variant | MODIFIER |
| 1 | 186991064 | AX-170381418 | 0 | A | G | 0.228 | 5.30E-02 | 1.08E-01 | 2.51E+00 | 2.58E+00 | 6.43E-01 | no | | intergenic_variant | MODIFIER |

| | | | | | | | | | | | | | | | |
|---|---|---|---|---|---|---|---|---|---|---|---|---|---|---|---|
| 1 | 186991432 | AX-170305192 | 0 | C | T | 0.228 | 5.30E-02 | 1.08E-01 | 2.51E+00 | 2.58E+00 | 6.43E-01 | no | | intergenic_variant | MODIFIER |
| 1 | 186997353 | AX-170002504 | 0 | G | A | 0.228 | 5.30E-02 | 1.08E-01 | 2.51E+00 | 2.58E+00 | 6.43E-01 | no | | intergenic_variant | MODIFIER |
| 1 | 187046066 | AX-170086718 | 0 | G | A | 0.228 | 5.30E-02 | 1.08E-01 | 2.51E+00 | 2.58E+00 | 6.43E-01 | no | Spata17 | downstream_gene_variant | MODIFIER |
| 1 | 187049375 | AX-170323581 | 0 | C | G | 0.228 | 5.30E-02 | 1.08E-01 | 2.51E+00 | 2.58E+00 | 6.43E-01 | no | Spata17 | intron_variant | MODIFIER |
| 1 | 187052607 | AX-170379735 | 0 | G | C | 0.228 | 5.30E-02 | 1.08E-01 | 2.51E+00 | 2.58E+00 | 6.43E-01 | no | Spata17 | intron_variant | MODIFIER |
| 1 | 187053680 | AX-170002498 | 0 | T | C | 0.228 | 5.30E-02 | 1.08E-01 | 2.51E+00 | 2.58E+00 | 6.43E-01 | no | Spata17 | intron_variant | MODIFIER |
| 1 | 187053749 | AX-170043889 | 0 | C | T | 0.228 | 5.30E-02 | 1.08E-01 | 2.51E+00 | 2.58E+00 | 6.43E-01 | no | Spata17 | intron_variant | MODIFIER |
| 1 | 187057623 | AX-97836585 | 0 | G | A | 0.228 | 5.30E-02 | 1.08E-01 | 2.51E+00 | 2.58E+00 | 6.43E-01 | no | Spata17 | intron_variant | MODIFIER |
| 1 | 187074429 | AX-170086719 | 0 | G | C | 0.228 | 5.30E-02 | 1.08E-01 | 2.51E+00 | 2.58E+00 | 6.43E-01 | no | Spata17 | intron_variant | MODIFIER |
| 1 | 187083711 | AX-170002505 | 0 | G | A | 0.228 | 5.30E-02 | 1.08E-01 | 2.51E+00 | 2.58E+00 | 6.43E-01 | no | Spata17 | intron_variant | MODIFIER |
| 1 | 187102128 | AX-170043887 | 0 | A | G | 0.228 | 5.30E-02 | 1.08E-01 | 2.51E+00 | 2.58E+00 | 6.43E-01 | no | Spata17 | intron_variant | MODIFIER |
| 1 | 187111637 | AX-170264101 | 0 | G | C | 0.228 | 5.30E-02 | 1.08E-01 | 2.51E+00 | 2.58E+00 | 6.43E-01 | no | Spata17 | intron_variant | MODIFIER |
| 1 | 187118570 | AX-170299197 | 0 | T | C | 0.228 | 5.30E-02 | 1.08E-01 | 2.51E+00 | 2.58E+00 | 6.43E-01 | no | Spata17 | intron_variant | MODIFIER |
| 1 | 187139192 | AX-170244601 | 0 | C | G | 0.228 | 5.30E-02 | 1.08E-01 | 2.51E+00 | 2.58E+00 | 6.43E-01 | no | Spata17 | intron_variant | MODIFIER |
| 1 | 187162722 | AX-170449722 | 0 | C | G | 0.228 | 5.30E-02 | 1.08E-01 | 2.51E+00 | 2.58E+00 | 6.43E-01 | no | Spata17 | intron_variant | MODIFIER |
| 1 | 187206687 | AX-170104458 | 0 | C | G | 0.228 | 5.30E-02 | 1.08E-01 | 2.51E+00 | 2.58E+00 | 6.43E-01 | no | Spata17 | intron_variant | MODIFIER |
| 1 | 187223281 | AX-170515796 | 0 | A | G | 0.228 | 5.30E-02 | 1.08E-01 | 2.51E+00 | 2.58E+00 | 6.43E-01 | no | Gpatch2 | intron_variant | MODIFIER |
| 1 | 187228589 | AX-170186250 | 0 | C | T | 0.228 | 5.30E-02 | 1.08E-01 | 2.51E+00 | 2.58E+00 | 6.43E-01 | no | Gpatch2 | intron_variant | MODIFIER |
| 1 | 187232610 | AX-97812453 | 0 | C | G | 0.228 | 5.30E-02 | 1.08E-01 | 2.51E+00 | 2.58E+00 | 6.43E-01 | no | Gpatch2 | intron_variant | MODIFIER |
| 1 | 187261549 | AX-170401168 | 0 | T | C | 0.228 | 5.30E-02 | 1.08E-01 | 2.51E+00 | 2.58E+00 | 6.43E-01 | no | Gpatch2 | intron_variant | MODIFIER |
| 1 | 187261833 | AX-170142043 | 0 | T | A | 0.228 | 5.30E-02 | 1.08E-01 | 2.51E+00 | 2.58E+00 | 6.43E-01 | no | Gpatch2 | intron_variant | MODIFIER |
| 1 | 187275633 | AX-170208402 | 0 | C | T | 0.228 | 5.30E-02 | 1.08E-01 | 2.51E+00 | 2.58E+00 | 6.43E-01 | no | Gpatch2 | intron_variant | MODIFIER |
| 1 | 187341078 | AX-97848744 | 0 | C | T | 0.228 | 5.30E-02 | 1.08E-01 | 2.51E+00 | 2.58E+00 | 6.43E-01 | no | Gpatch2 | intron_variant | MODIFIER |
| 1 | 187391358 | AX-170043890 | 0 | A | G | 0.228 | 5.30E-02 | 1.08E-01 | 2.51E+00 | 2.58E+00 | 6.43E-01 | no | | intergenic_variant | MODIFIER |
| 1 | 187403014 | AX-170256305 | 0 | C | T | 0.228 | 5.30E-02 | 1.08E-01 | 2.51E+00 | 2.58E+00 | 6.43E-01 | no | | intergenic_variant | MODIFIER |
| 1 | 187403041 | AX-170313014 | 0 | A | G | 0.228 | 5.30E-02 | 1.08E-01 | 2.51E+00 | 2.58E+00 | 6.43E-01 | no | | intergenic_variant | MODIFIER |
| 1 | 187408195 | AX-170274726 | 0 | C | T | 0.228 | 5.30E-02 | 1.08E-01 | 2.51E+00 | 2.58E+00 | 6.43E-01 | no | | intergenic_variant | MODIFIER |
| 7 | 96408668 | AX-170314585 | 0 | G | A | 0.123 | 7.53E-02 | 1.66E-01 | 2.63E+00 | 2.77E+00 | 6.43E-01 | no | Tenm4 | intron_variant | MODIFIER |
| 15 | 75328403 | AX-170055996 | 1 | G | A | 0.554 | 3.89E-02 | 8.27E-02 | 2.58E+00 | 2.65E+00 | 6.43E-01 | no | Gm28113 | upstream_gene_variant | MODIFIER |
| 6 | 139502134 | AX-170095829 | 0 | C | T | 0.061 | 1.03E-01 | 2.25E-01 | 2.64E+00 | 2.75E+00 | 6.43E-01 | no | Rergl | upstream_gene_variant | MODIFIER |
| 5 | 86351928 | AX-170392732 | 1 | G | A | 0.429 | 5.16E-02 | 1.13E-01 | 2.76E+00 | 2.80E+00 | 6.43E-01 | no | Tmprss11d | intron_variant | MODIFIER |
| 1 | 99337108 | AX-170418486 | 2 | C | T | 0.127 | 7.85E-02 | 1.72E-01 | 2.63E+00 | 2.75E+00 | 6.44E-01 | no | | intergenic_variant | MODIFIER |
| 1 | 45595354 | AX-170138994 | 1 | G | A | 0.098 | -9.58E-02 | 2.11E-01 | 2.71E+00 | 2.80E+00 | 6.44E-01 | no | | intergenic_variant | MODIFIER |
| 13 | 6269859 | AX-97849134 | 2 | C | T | 0.218 | -5.38E-02 | 1.14E-01 | 2.56E+00 | 2.65E+00 | 6.44E-01 | no | Gm47549 | intron_variant,non_coding_transcript_variant | MODIFIER |
| 10 | 127452133 | AX-170004130 | 0 | T | A | 0.316 | -3.36E-02 | 7.30E-02 | 2.62E+00 | 2.72E+00 | 6.44E-01 | no | R3hdm2 | intron_variant | MODIFIER |
| 5 | 114353477 | AX-170419568 | 1 | G | A | 0.455 | 5.40E-02 | 1.17E-01 | 2.66E+00 | 2.69E+00 | 6.44E-01 | no | Myo1h | intron_variant | MODIFIER |
| 1 | 114785551 | AX-170164232 | 2 | G | A | 0.173 | 7.75E-02 | 1.66E-01 | 2.57E+00 | 2.68E+00 | 6.44E-01 | no | | intergenic_variant | MODIFIER |
| 4 | 80247920 | AX-97842204 | 1 | A | G | 0.241 | 4.91E-02 | 1.04E-01 | 2.56E+00 | 2.63E+00 | 6.44E-01 | no | | intergenic_variant | MODIFIER |
| 12 | 89014378 | AX-170049726 | 1 | G | C | 0.071 | 8.00E-02 | 1.77E-01 | 2.68E+00 | 2.78E+00 | 6.44E-01 | no | Nrxn3 | intron_variant | MODIFIER |
| 1 | 153002006 | AX-170101044 | 0 | T | C | 0.202 | 6.48E-02 | 1.41E-01 | 2.66E+00 | 2.71E+00 | 6.44E-01 | no | Nmnat2 | intron_variant | MODIFIER |
| 8 | 19295389 | AX-170248820 | 1 | T | G | 0.179 | 4.74E-02 | 1.03E-01 | 2.60E+00 | 2.70E+00 | 6.44E-01 | no | Defb3 | downstream_gene_variant | MODIFIER |
| 1 | 20440701 | AX-170279251 | 0 | C | T | 0.395 | 3.82E-02 | 8.25E-02 | 2.63E+00 | 2.68E+00 | 6.44E-01 | no | Pkhd1 | intron_variant | MODIFIER |
| 9 | 80536015 | AX-170082924 | 1 | C | T | 0.205 | 5.78E-02 | 1.27E-01 | 2.79E+00 | 2.86E+00 | 6.44E-01 | no | | intergenic_variant | MODIFIER |
| 13 | 23413359 | AX-170233718 | 0 | A | T | 0.175 | -5.58E-02 | 1.14E-01 | 2.51E+00 | 2.59E+00 | 6.44E-01 | no | | regulatory_region_variant | MODIFIER |
| 5 | 73332538 | AX-170327378 | 1 | T | C | 0.429 | -4.26E-02 | 9.45E-02 | 2.73E+00 | 2.83E+00 | 6.44E-01 | no | Ociad2 | intron_variant | MODIFIER |
| 9 | 19457675 | AX-170031902 | 0 | G | T | 0.474 | 4.04E-02 | 8.49E-02 | 2.55E+00 | 2.63E+00 | 6.44E-01 | no | | intergenic_variant | MODIFIER |
| 8 | 41558837 | AX-170417314 | 1 | A | G | 0.143 | 6.89E-02 | 1.51E-01 | 2.77E+00 | 2.83E+00 | 6.44E-01 | no | | intergenic_variant | MODIFIER |
| 8 | 96132822 | AX-170275121 | 1 | C | T | 0.223 | 5.22E-02 | 1.15E-01 | 2.71E+00 | 2.78E+00 | 6.44E-01 | no | | intergenic_variant | MODIFIER |
| 7 | 124187300 | AX-170420210 | 0 | C | G | 0.377 | 3.79E-02 | 8.36E-02 | 2.70E+00 | 2.77E+00 | 6.45E-01 | no | Hs3st4 | intron_variant | MODIFIER |
| 18 | 81089228 | AX-170381531 | 1 | C | T | 0.304 | -3.95E-02 | 8.47E-02 | 2.57E+00 | 2.68E+00 | 6.45E-01 | no | AC105065.1 | intron_variant,non_coding_transcript_variant | MODIFIER |
| 2 | 123458688 | AX-170443567 | 2 | A | G | 0.145 | -8.04E-02 | 1.78E-01 | 2.71E+00 | 2.80E+00 | 6.45E-01 | no | | regulatory_region_variant | MODIFIER |
| 1 | 68431202 | AX-170041537 | 1 | T | A | 0.107 | 8.18E-02 | 1.81E-01 | 2.65E+00 | 2.77E+00 | 6.45E-01 | no | Erbb4 | intron_variant | MODIFIER |
| 8 | 31917034 | AX-170200503 | 0 | C | A | 0.14 | -5.89E-02 | 1.29E-01 | 2.66E+00 | 2.73E+00 | 6.45E-01 | no | Nrg1 | intron_variant | MODIFIER |
| 8 | 31917751 | AX-170356079 | 0 | G | A | 0.14 | -5.89E-02 | 1.29E-01 | 2.66E+00 | 2.73E+00 | 6.45E-01 | no | Nrg1 | synonymous_variant | LOW |

| | | | | | | | | | | | | | | | |
|---|---|---|---|---|---|---|---|---|---|---|---|---|---|---|---|
| 8 | 31921172 | AX-97840541 | 0 | T | C | 0.14 | -5.89E-02 | 1.29E-01 | 2.66E+00 | 2.73E+00 | 6.45E-01 | no | Nrg1 | upstream_gene_variant | MODIFIER |
| 8 | 31937226 | AX-170414986 | 0 | G | A | 0.14 | -5.89E-02 | 1.29E-01 | 2.66E+00 | 2.73E+00 | 6.45E-01 | no | Nrg1 | intron_variant | MODIFIER |
| 8 | 31945004 | AX-170410740 | 0 | T | C | 0.14 | -5.89E-02 | 1.29E-01 | 2.66E+00 | 2.73E+00 | 6.45E-01 | no | Nrg1 | intron_variant | MODIFIER |
| 8 | 31956726 | AX-170404139 | 0 | C | T | 0.14 | -5.89E-02 | 1.29E-01 | 2.66E+00 | 2.73E+00 | 6.45E-01 | no | Nrg1 | intron_variant | MODIFIER |
| 8 | 31973612 | AX-97840594 | 0 | A | G | 0.14 | -5.89E-02 | 1.29E-01 | 2.66E+00 | 2.73E+00 | 6.45E-01 | no | Nrg1 | intron_variant | MODIFIER |
| 8 | 31974562 | AX-170258426 | 0 | T | C | 0.14 | -5.89E-02 | 1.29E-01 | 2.66E+00 | 2.73E+00 | 6.45E-01 | no | Nrg1 | intron_variant | MODIFIER |
| 8 | 31979121 | AX-170181337 | 0 | A | G | 0.14 | -5.89E-02 | 1.29E-01 | 2.66E+00 | 2.73E+00 | 6.45E-01 | no | Nrg1 | intron_variant | MODIFIER |
| 8 | 31979182 | AX-170326378 | 0 | G | A | 0.14 | -5.89E-02 | 1.29E-01 | 2.66E+00 | 2.73E+00 | 6.45E-01 | no | Nrg1 | intron_variant | MODIFIER |
| 8 | 31979686 | AX-170201421 | 0 | T | G | 0.14 | -5.89E-02 | 1.29E-01 | 2.66E+00 | 2.73E+00 | 6.45E-01 | no | Nrg1 | intron_variant | MODIFIER |
| 8 | 31994482 | AX-170420824 | 0 | A | G | 0.14 | -5.89E-02 | 1.29E-01 | 2.66E+00 | 2.73E+00 | 6.45E-01 | no | Nrg1 | intron_variant | MODIFIER |
| 8 | 32000924 | AX-170145130 | 0 | T | C | 0.14 | -5.89E-02 | 1.29E-01 | 2.66E+00 | 2.73E+00 | 6.45E-01 | no | Nrg1 | intron_variant | MODIFIER |
| 8 | 32001653 | AX-170506990 | 0 | T | C | 0.14 | -5.89E-02 | 1.29E-01 | 2.66E+00 | 2.73E+00 | 6.45E-01 | no | Nrg1 | intron_variant | MODIFIER |
| 8 | 32031485 | AX-170473884 | 0 | T | C | 0.14 | -5.89E-02 | 1.29E-01 | 2.66E+00 | 2.73E+00 | 6.45E-01 | no | Nrg1 | intron_variant | MODIFIER |
| 8 | 32033013 | AX-97789230 | 0 | G | C | 0.14 | -5.89E-02 | 1.29E-01 | 2.66E+00 | 2.73E+00 | 6.45E-01 | no | Nrg1 | intron_variant | MODIFIER |
| 8 | 32035661 | AX-170258109 | 0 | A | G | 0.14 | -5.89E-02 | 1.29E-01 | 2.66E+00 | 2.73E+00 | 6.45E-01 | no | Nrg1 | intron_variant | MODIFIER |
| 8 | 32046071 | AX-170030450 | 0 | T | G | 0.14 | -5.89E-02 | 1.29E-01 | 2.66E+00 | 2.73E+00 | 6.45E-01 | no | Nrg1 | intron_variant | MODIFIER |
| 8 | 32058866 | AX-170030452 | 0 | C | T | 0.14 | -5.89E-02 | 1.29E-01 | 2.66E+00 | 2.73E+00 | 6.45E-01 | no | Nrg1 | intron_variant | MODIFIER |
| 8 | 32058883 | AX-170030454 | 0 | A | G | 0.14 | -5.89E-02 | 1.29E-01 | 2.66E+00 | 2.73E+00 | 6.45E-01 | no | Nrg1 | intron_variant | MODIFIER |
| 8 | 32059794 | AX-170190847 | 0 | C | A | 0.14 | -5.89E-02 | 1.29E-01 | 2.66E+00 | 2.73E+00 | 6.45E-01 | no | Nrg1 | intron_variant | MODIFIER |
| 8 | 32063851 | AX-170482680 | 0 | C | G | 0.14 | -5.89E-02 | 1.29E-01 | 2.66E+00 | 2.73E+00 | 6.45E-01 | no | Nrg1 | intron_variant | MODIFIER |
| 8 | 32087611 | AX-170147628 | 0 | T | C | 0.14 | -5.89E-02 | 1.29E-01 | 2.66E+00 | 2.73E+00 | 6.45E-01 | no | Nrg1 | intron_variant | MODIFIER |
| 8 | 32120597 | AX-170030455 | 0 | T | C | 0.14 | -5.89E-02 | 1.29E-01 | 2.66E+00 | 2.73E+00 | 6.45E-01 | no | Nrg1 | intron_variant | MODIFIER |
| 8 | 32137849 | AX-170198802 | 0 | C | T | 0.14 | -5.89E-02 | 1.29E-01 | 2.66E+00 | 2.73E+00 | 6.45E-01 | no | Nrg1 | intron_variant | MODIFIER |
| 8 | 32152745 | AX-170137244 | 0 | G | A | 0.14 | -5.89E-02 | 1.29E-01 | 2.66E+00 | 2.73E+00 | 6.45E-01 | no | Nrg1 | intron_variant | MODIFIER |
| 8 | 32170672 | AX-170450250 | 0 | G | A | 0.14 | -5.89E-02 | 1.29E-01 | 2.66E+00 | 2.73E+00 | 6.45E-01 | no | Nrg1 | intron_variant | MODIFIER |
| 8 | 32178577 | AX-97805595 | 0 | G | A | 0.14 | -5.89E-02 | 1.29E-01 | 2.66E+00 | 2.73E+00 | 6.45E-01 | no | Nrg1 | intron_variant | MODIFIER |
| 8 | 32198562 | AX-170146360 | 0 | T | G | 0.14 | -5.89E-02 | 1.29E-01 | 2.66E+00 | 2.73E+00 | 6.45E-01 | no | Nrg1 | intron_variant | MODIFIER |
| 8 | 32213028 | AX-170194770 | 0 | A | G | 0.14 | -5.89E-02 | 1.29E-01 | 2.66E+00 | 2.73E+00 | 6.45E-01 | no | Nrg1 | intron_variant | MODIFIER |
| 8 | 32213060 | AX-170516526 | 0 | T | C | 0.14 | -5.89E-02 | 1.29E-01 | 2.66E+00 | 2.73E+00 | 6.45E-01 | no | Nrg1 | intron_variant | MODIFIER |
| 8 | 32219646 | AX-170136273 | 0 | T | A | 0.14 | -5.89E-02 | 1.29E-01 | 2.66E+00 | 2.73E+00 | 6.45E-01 | no | Nrg1 | intron_variant | MODIFIER |
| 8 | 32220137 | AX-170096216 | 0 | C | A | 0.14 | -5.89E-02 | 1.29E-01 | 2.66E+00 | 2.73E+00 | 6.45E-01 | no | Nrg1 | intron_variant | MODIFIER |
| 8 | 32245850 | AX-170247679 | 0 | T | C | 0.14 | -5.89E-02 | 1.29E-01 | 2.66E+00 | 2.73E+00 | 6.45E-01 | no | Nrg1 | intron_variant | MODIFIER |
| 8 | 32250310 | AX-170236468 | 0 | T | C | 0.14 | -5.89E-02 | 1.29E-01 | 2.66E+00 | 2.73E+00 | 6.45E-01 | no | Nrg1 | intron_variant | MODIFIER |
| 8 | 32271638 | AX-170117766 | 0 | T | C | 0.14 | -5.89E-02 | 1.29E-01 | 2.66E+00 | 2.73E+00 | 6.45E-01 | no | Nrg1 | intron_variant | MODIFIER |
| 8 | 32284171 | AX-170147094 | 0 | T | C | 0.14 | -5.89E-02 | 1.29E-01 | 2.66E+00 | 2.73E+00 | 6.45E-01 | no | Nrg1 | intron_variant | MODIFIER |
| 8 | 32287534 | AX-170186457 | 0 | G | C | 0.14 | -5.89E-02 | 1.29E-01 | 2.66E+00 | 2.73E+00 | 6.45E-01 | no | Nrg1 | intron_variant | MODIFIER |
| 8 | 32293848 | AX-170387554 | 0 | T | G | 0.14 | -5.89E-02 | 1.29E-01 | 2.66E+00 | 2.73E+00 | 6.45E-01 | no | Nrg1 | intron_variant | MODIFIER |
| 8 | 32295068 | AX-170118249 | 0 | T | C | 0.14 | -5.89E-02 | 1.29E-01 | 2.66E+00 | 2.73E+00 | 6.45E-01 | no | Nrg1 | intron_variant | MODIFIER |
| 8 | 32300023 | AX-170348570 | 0 | A | C | 0.14 | -5.89E-02 | 1.29E-01 | 2.66E+00 | 2.73E+00 | 6.45E-01 | no | Nrg1 | intron_variant | MODIFIER |
| 8 | 32315017 | AX-170482155 | 0 | G | A | 0.14 | -5.89E-02 | 1.29E-01 | 2.66E+00 | 2.73E+00 | 6.45E-01 | no | Nrg1 | intron_variant | MODIFIER |
| 8 | 32315123 | AX-170416814 | 0 | A | G | 0.14 | -5.89E-02 | 1.29E-01 | 2.66E+00 | 2.73E+00 | 6.45E-01 | no | Nrg1 | intron_variant | MODIFIER |
| 8 | 32316809 | AX-170208080 | 0 | T | C | 0.14 | -5.89E-02 | 1.29E-01 | 2.66E+00 | 2.73E+00 | 6.45E-01 | no | Nrg1 | intron_variant | MODIFIER |
| 8 | 32326593 | AX-170079672 | 0 | G | A | 0.14 | -5.89E-02 | 1.29E-01 | 2.66E+00 | 2.73E+00 | 6.45E-01 | no | Nrg1 | intron_variant | MODIFIER |
| 8 | 32346336 | AX-170111494 | 0 | G | A | 0.14 | -5.89E-02 | 1.29E-01 | 2.66E+00 | 2.73E+00 | 6.45E-01 | no | Nrg1 | intron_variant | MODIFIER |
| 8 | 32363284 | AX-170079671 | 0 | T | C | 0.14 | -5.89E-02 | 1.29E-01 | 2.66E+00 | 2.73E+00 | 6.45E-01 | no | Nrg1 | intron_variant | MODIFIER |
| 8 | 32363612 | AX-170458910 | 0 | C | T | 0.14 | -5.89E-02 | 1.29E-01 | 2.66E+00 | 2.73E+00 | 6.45E-01 | no | Nrg1 | intron_variant | MODIFIER |
| 8 | 32365450 | AX-170466087 | 0 | T | C | 0.14 | -5.89E-02 | 1.29E-01 | 2.66E+00 | 2.73E+00 | 6.45E-01 | no | Nrg1 | intron_variant | MODIFIER |
| 8 | 32368563 | AX-170173640 | 0 | G | A | 0.14 | -5.89E-02 | 1.29E-01 | 2.66E+00 | 2.73E+00 | 6.45E-01 | no | Nrg1 | intron_variant | MODIFIER |
| 8 | 32376259 | AX-170435040 | 0 | A | C | 0.14 | -5.89E-02 | 1.29E-01 | 2.66E+00 | 2.73E+00 | 6.45E-01 | no | Nrg1 | intron_variant | MODIFIER |
| 8 | 32377221 | AX-170201812 | 0 | C | T | 0.14 | -5.89E-02 | 1.29E-01 | 2.66E+00 | 2.73E+00 | 6.45E-01 | no | Nrg1 | intron_variant | MODIFIER |
| 8 | 32391194 | AX-170370499 | 0 | T | C | 0.14 | -5.89E-02 | 1.29E-01 | 2.66E+00 | 2.73E+00 | 6.45E-01 | no | Nrg1 | intron_variant | MODIFIER |
| 8 | 32392416 | AX-97820899 | 0 | C | T | 0.14 | -5.89E-02 | 1.29E-01 | 2.66E+00 | 2.73E+00 | 6.45E-01 | no | Nrg1 | intron_variant | MODIFIER |
| 8 | 32429372 | AX-170509886 | 0 | A | G | 0.14 | -5.89E-02 | 1.29E-01 | 2.66E+00 | 2.73E+00 | 6.45E-01 | no | Nrg1 | intron_variant | MODIFIER |
| 8 | 32431835 | AX-170454156 | 0 | G | T | 0.14 | -5.89E-02 | 1.29E-01 | 2.66E+00 | 2.73E+00 | 6.45E-01 | no | Nrg1 | intron_variant | MODIFIER |

| | | | | | | | | | | | | | | | |
|---|---|---|---|---|---|---|---|---|---|---|---|---|---|---|---|
| 8 | 32469357 | AX-170129126 | 0 | C | T | 0.14 | -5.89E-02 | 1.29E-01 | 2.66E+00 | 2.73E+00 | 6.45E-01 | no | Nrg1 | intron_variant | MODIFIER |
| 8 | 32469897 | AX-97800586 | 0 | G | A | 0.14 | -5.89E-02 | 1.29E-01 | 2.66E+00 | 2.73E+00 | 6.45E-01 | no | Nrg1 | intron_variant | MODIFIER |
| 8 | 32472993 | AX-170512017 | 0 | T | C | 0.14 | -5.89E-02 | 1.29E-01 | 2.66E+00 | 2.73E+00 | 6.45E-01 | no | Nrg1 | intron_variant | MODIFIER |
| 8 | 32503202 | AX-170337538 | 0 | A | G | 0.14 | -5.89E-02 | 1.29E-01 | 2.66E+00 | 2.73E+00 | 6.45E-01 | no | Nrg1 | intron_variant | MODIFIER |
| 8 | 32510570 | AX-170108297 | 0 | G | A | 0.14 | -5.89E-02 | 1.29E-01 | 2.66E+00 | 2.73E+00 | 6.45E-01 | no | Nrg1 | intron_variant | MODIFIER |
| 8 | 32513418 | AX-170101914 | 0 | A | C | 0.14 | -5.89E-02 | 1.29E-01 | 2.66E+00 | 2.73E+00 | 6.45E-01 | no | Nrg1 | intron_variant | MODIFIER |
| 8 | 32537910 | AX-170175264 | 0 | T | C | 0.14 | -5.89E-02 | 1.29E-01 | 2.66E+00 | 2.73E+00 | 6.45E-01 | no | Nrg1 | intron_variant | MODIFIER |
| 8 | 32550844 | AX-170079675 | 0 | T | A | 0.14 | -5.89E-02 | 1.29E-01 | 2.66E+00 | 2.73E+00 | 6.45E-01 | no | Nrg1 | intron_variant | MODIFIER |
| 8 | 32559778 | AX-97783667 | 0 | A | C | 0.14 | -5.89E-02 | 1.29E-01 | 2.66E+00 | 2.73E+00 | 6.45E-01 | no | Nrg1 | intron_variant | MODIFIER |
| 8 | 32564584 | AX-170030456 | 0 | G | A | 0.14 | -5.89E-02 | 1.29E-01 | 2.66E+00 | 2.73E+00 | 6.45E-01 | no | Nrg1 | intron_variant | MODIFIER |
| 8 | 32568202 | AX-170030459 | 0 | T | C | 0.14 | -5.89E-02 | 1.29E-01 | 2.66E+00 | 2.73E+00 | 6.45E-01 | no | Nrg1 | intron_variant | MODIFIER |
| 8 | 32568394 | AX-170454074 | 0 | C | T | 0.14 | -5.89E-02 | 1.29E-01 | 2.66E+00 | 2.73E+00 | 6.45E-01 | no | Nrg1 | intron_variant | MODIFIER |
| 8 | 32597490 | AX-170456049 | 0 | C | A | 0.14 | -5.89E-02 | 1.29E-01 | 2.66E+00 | 2.73E+00 | 6.45E-01 | no | Nrg1 | intron_variant | MODIFIER |
| 8 | 32610869 | AX-170464347 | 0 | C | T | 0.14 | -5.89E-02 | 1.29E-01 | 2.66E+00 | 2.73E+00 | 6.45E-01 | no | Nrg1 | intron_variant | MODIFIER |
| 8 | 32648282 | AX-170343367 | 0 | A | T | 0.14 | -5.89E-02 | 1.29E-01 | 2.66E+00 | 2.73E+00 | 6.45E-01 | no | Nrg1 | intron_variant | MODIFIER |
| 8 | 32657970 | AX-170417404 | 0 | A | G | 0.14 | -5.89E-02 | 1.29E-01 | 2.66E+00 | 2.73E+00 | 6.45E-01 | no | Nrg1 | intron_variant | MODIFIER |
| 8 | 32665922 | AX-170364768 | 0 | A | G | 0.14 | -5.89E-02 | 1.29E-01 | 2.66E+00 | 2.73E+00 | 6.45E-01 | no | Nrg1 | intron_variant | MODIFIER |
| 8 | 32675442 | AX-170170393 | 0 | A | G | 0.14 | -5.89E-02 | 1.29E-01 | 2.66E+00 | 2.73E+00 | 6.45E-01 | no | Nrg1 | intron_variant | MODIFIER |
| 8 | 32675983 | AX-170125146 | 0 | T | C | 0.14 | -5.89E-02 | 1.29E-01 | 2.66E+00 | 2.73E+00 | 6.45E-01 | no | Nrg1 | intron_variant | MODIFIER |
| 8 | 32721441 | AX-97849838 | 0 | T | C | 0.14 | -5.89E-02 | 1.29E-01 | 2.66E+00 | 2.73E+00 | 6.45E-01 | no | Nrg1 | intron_variant | MODIFIER |
| 8 | 32721466 | AX-170351227 | 0 | G | A | 0.14 | -5.89E-02 | 1.29E-01 | 2.66E+00 | 2.73E+00 | 6.45E-01 | no | Nrg1 | intron_variant | MODIFIER |
| 8 | 32748119 | AX-170468540 | 0 | G | A | 0.14 | -5.89E-02 | 1.29E-01 | 2.66E+00 | 2.73E+00 | 6.45E-01 | no | Nrg1 | intron_variant | MODIFIER |
| 8 | 32748328 | AX-170347541 | 0 | A | G | 0.14 | -5.89E-02 | 1.29E-01 | 2.66E+00 | 2.73E+00 | 6.45E-01 | no | Nrg1 | intron_variant | MODIFIER |
| 8 | 32749066 | AX-170292026 | 0 | G | C | 0.14 | -5.89E-02 | 1.29E-01 | 2.66E+00 | 2.73E+00 | 6.45E-01 | no | Nrg1 | intron_variant | MODIFIER |
| 8 | 32749637 | AX-170377196 | 0 | C | T | 0.14 | -5.89E-02 | 1.29E-01 | 2.66E+00 | 2.73E+00 | 6.45E-01 | no | Nrg1 | intron_variant | MODIFIER |
| 8 | 32749832 | AX-170307384 | 0 | A | G | 0.14 | -5.89E-02 | 1.29E-01 | 2.66E+00 | 2.73E+00 | 6.45E-01 | no | Nrg1 | intron_variant | MODIFIER |
| 8 | 32750147 | AX-170420537 | 0 | C | G | 0.14 | -5.89E-02 | 1.29E-01 | 2.66E+00 | 2.73E+00 | 6.45E-01 | no | Nrg1 | intron_variant | MODIFIER |
| 8 | 32754387 | AX-170090078 | 0 | A | G | 0.14 | -5.89E-02 | 1.29E-01 | 2.66E+00 | 2.73E+00 | 6.45E-01 | no | Nrg1 | intron_variant | MODIFIER |
| 8 | 32757452 | AX-170365522 | 0 | C | T | 0.14 | -5.89E-02 | 1.29E-01 | 2.66E+00 | 2.73E+00 | 6.45E-01 | no | Nrg1 | intron_variant | MODIFIER |
| 8 | 32767915 | AX-170079677 | 0 | G | T | 0.14 | -5.89E-02 | 1.29E-01 | 2.66E+00 | 2.73E+00 | 6.45E-01 | no | Nrg1 | intron_variant | MODIFIER |
| 8 | 32769292 | AX-97802243 | 0 | A | G | 0.14 | -5.89E-02 | 1.29E-01 | 2.66E+00 | 2.73E+00 | 6.45E-01 | no | Nrg1 | intron_variant | MODIFIER |
| 8 | 32792391 | AX-170169984 | 0 | T | A | 0.14 | -5.89E-02 | 1.29E-01 | 2.66E+00 | 2.73E+00 | 6.45E-01 | no | Nrg1 | intron_variant | MODIFIER |
| 8 | 32792445 | AX-170399545 | 0 | A | G | 0.14 | -5.89E-02 | 1.29E-01 | 2.66E+00 | 2.73E+00 | 6.45E-01 | no | Nrg1 | intron_variant | MODIFIER |
| 8 | 32823578 | AX-170343892 | 0 | A | G | 0.14 | -5.89E-02 | 1.29E-01 | 2.66E+00 | 2.73E+00 | 6.45E-01 | no | Nrg1 | intron_variant | MODIFIER |
| 8 | 32829722 | AX-170194252 | 0 | A | G | 0.14 | -5.89E-02 | 1.29E-01 | 2.66E+00 | 2.73E+00 | 6.45E-01 | no | Nrg1 | intron_variant | MODIFIER |
| 8 | 32833773 | AX-170504012 | 0 | T | C | 0.14 | -5.89E-02 | 1.29E-01 | 2.66E+00 | 2.73E+00 | 6.45E-01 | no | Nrg1 | intron_variant | MODIFIER |
| 8 | 32834432 | AX-170313426 | 0 | T | C | 0.14 | -5.89E-02 | 1.29E-01 | 2.66E+00 | 2.73E+00 | 6.45E-01 | no | Nrg1 | intron_variant | MODIFIER |
| 8 | 32845020 | AX-170132328 | 0 | A | G | 0.14 | -5.89E-02 | 1.29E-01 | 2.66E+00 | 2.73E+00 | 6.45E-01 | no | Nrg1 | intron_variant | MODIFIER |
| 8 | 32852780 | AX-170185982 | 0 | G | A | 0.14 | -5.89E-02 | 1.29E-01 | 2.66E+00 | 2.73E+00 | 6.45E-01 | no | Nrg1 | intron_variant | MODIFIER |
| 8 | 32852840 | AX-170112401 | 0 | T | C | 0.14 | -5.89E-02 | 1.29E-01 | 2.66E+00 | 2.73E+00 | 6.45E-01 | no | Nrg1 | intron_variant | MODIFIER |
| 8 | 32866444 | AX-170455652 | 0 | G | A | 0.14 | -5.89E-02 | 1.29E-01 | 2.66E+00 | 2.73E+00 | 6.45E-01 | no | Nrg1 | intron_variant | MODIFIER |
| 8 | 32869948 | AX-170375502 | 0 | T | C | 0.14 | -5.89E-02 | 1.29E-01 | 2.66E+00 | 2.73E+00 | 6.45E-01 | no | Nrg1 | intron_variant | MODIFIER |
| 8 | 32875652 | AX-170455933 | 0 | T | C | 0.14 | -5.89E-02 | 1.29E-01 | 2.66E+00 | 2.73E+00 | 6.45E-01 | no | Nrg1 | intron_variant | MODIFIER |
| 8 | 32876535 | AX-170449262 | 0 | T | C | 0.14 | -5.89E-02 | 1.29E-01 | 2.66E+00 | 2.73E+00 | 6.45E-01 | no | Nrg1 | intron_variant | MODIFIER |
| 8 | 32882556 | AX-170265351 | 0 | G | A | 0.14 | -5.89E-02 | 1.29E-01 | 2.66E+00 | 2.73E+00 | 6.45E-01 | no | Nrg1 | intron_variant | MODIFIER |
| 16 | 77129805 | AX-170013423 | 1 | T | C | 0.446 | 3.71E-02 | 7.99E-02 | 2.60E+00 | 2.67E+00 | 6.45E-01 | no | | regulatory_region_variant | MODIFIER |
| 7 | 40124489 | AX-170445827 | 0 | T | A | 0.351 | 4.08E-02 | 8.77E-02 | 2.59E+00 | 2.68E+00 | 6.45E-01 | no | | intergenic_variant | MODIFIER |
| 8 | 21823677 | AX-170030336 | 1 | C | T | 0.179 | 4.75E-02 | 1.05E-01 | 2.66E+00 | 2.75E+00 | 6.45E-01 | no | Defb50 | intron_variant | MODIFIER |
| 6 | 134589595 | AX-170028304 | 0 | T | C | 0.561 | 4.15E-02 | 8.66E-02 | 2.54E+00 | 2.60E+00 | 6.45E-01 | no | | intergenic_variant | MODIFIER |
| 5 | 96338734 | AX-97836091 | 1 | C | T | 0.429 | 4.88E-02 | 1.07E-01 | 2.67E+00 | 2.72E+00 | 6.45E-01 | no | | intergenic_variant | MODIFIER |
| 3 | 62088401 | AX-170443901 | 1 | G | A | 0.054 | -6.60E-02 | 1.46E-01 | 2.62E+00 | 2.76E+00 | 6.45E-01 | no | | intergenic_variant | MODIFIER |
| 3 | 38329517 | AX-97844120 | 0 | C | T | 0.202 | 3.91E-02 | 8.65E-02 | 2.65E+00 | 2.76E+00 | 6.45E-01 | no | | intergenic_variant | MODIFIER |
| 10 | 128015241 | AX-170259438 | 1 | C | T | 0.321 | -3.38E-02 | 7.37E-02 | 2.61E+00 | 2.71E+00 | 6.45E-01 | no | Prim1 | 5_prime_UTR_variant | MODIFIER |
| 18 | 56963314 | AX-170450843 | 1 | G | A | 0.196 | -5.30E-02 | 1.11E-01 | 2.55E+00 | 2.62E+00 | 6.45E-01 | no | C330018D20Rik | intron_variant | MODIFIER |

| | | | | | | | | | | | | | | | |
|---|---|---|---|---|---|---|---|---|---|---|---|---|---|---|---|
| 2 | 104830888 | AX-170459150 | 0 | G | A | 0.053 | -9.74E-02 | 2.14E-01 | 2.64E+00 | 2.73E+00 | 6.46E-01 | no | Prrg4 | 3_prime_UTR_variant | MODIFIER |
| 1 | 12918473 | AX-97800844 | 1 | A | T | 0.393 | 3.53E-02 | 7.62E-02 | 2.61E+00 | 2.67E+00 | 6.46E-01 | no | Slco5a1 | intron_variant | MODIFIER |
| 1 | 43879656 | AX-170472974 | 1 | G | T | 0.098 | -7.84E-02 | 1.73E-01 | 2.63E+00 | 2.74E+00 | 6.46E-01 | no | | regulatory_region_variant | MODIFIER |
| 2 | 124322605 | AX-97839852 | 0 | A | C | 0.14 | 6.79E-02 | 1.52E-01 | 2.68E+00 | 2.80E+00 | 6.46E-01 | no | Sema6d | intron_variant | MODIFIER |
| 5 | 28362508 | AX-170491723 | 0 | G | T | 0.36 | -4.47E-02 | 9.95E-02 | 2.75E+00 | 2.84E+00 | 6.46E-01 | no | Rbm33 | intron_variant | MODIFIER |
| 2 | 164338852 | AX-170172305 | 1 | G | A | 0.143 | 8.76E-02 | 1.94E-01 | 2.68E+00 | 2.76E+00 | 6.46E-01 | no | Svs5 | downstream_gene_variant | MODIFIER |
| 17 | 3894953 | AX-170285430 | 0 | G | C | 0.044 | 5.90E-02 | 1.33E-01 | 2.66E+00 | 2.80E+00 | 6.46E-01 | no | | intergenic_variant | MODIFIER |
| 20 | 49482722 | AX-170033817 | 0 | T | C | 0.228 | 2.70E-02 | 5.87E-02 | 2.57E+00 | 2.71E+00 | 6.46E-01 | no | Arhgap36 | intron_variant | MODIFIER |
| 15 | 7765756 | AX-97832479 | 1 | G | A | 0.25 | -4.76E-02 | 1.03E-01 | 2.60E+00 | 2.66E+00 | 6.46E-01 | no | Gm49207 | downstream_gene_variant | MODIFIER |
| 15 | 3676932 | AX-170141059 | 0 | C | G | 0.246 | -4.84E-02 | 1.04E-01 | 2.59E+00 | 2.65E+00 | 6.46E-01 | no | | intergenic_variant | MODIFIER |
| 7 | 101743401 | AX-170029646 | 0 | G | A | 0.456 | 4.20E-02 | 9.16E-02 | 2.63E+00 | 2.70E+00 | 6.46E-01 | no | Clpb | intron_variant | MODIFIER |
| 5 | 90635069 | AX-170532441 | 0 | T | C | 0.439 | 4.53E-02 | 1.01E-01 | 2.71E+00 | 2.80E+00 | 6.46E-01 | no | Rassf6 | intron_variant | MODIFIER |
| 2 | 144187532 | AX-97840454 | 1 | G | A | 0.152 | 8.16E-02 | 1.79E-01 | 2.64E+00 | 2.72E+00 | 6.46E-01 | no | Gm5535 | intron_variant,non_coding_transcript_variant | MODIFIER |
| 1 | 99067091 | AX-170277824 | 0 | C | G | 0.158 | 9.48E-02 | 2.07E-01 | 2.64E+00 | 2.71E+00 | 6.46E-01 | no | | intergenic_variant | MODIFIER |
| 1 | 101721460 | AX-170001209 | 0 | C | T | 0.158 | 9.48E-02 | 2.07E-01 | 2.64E+00 | 2.71E+00 | 6.46E-01 | no | | intergenic_variant | MODIFIER |
| 1 | 102663722 | AX-170308500 | 0 | G | A | 0.158 | 9.48E-02 | 2.07E-01 | 2.64E+00 | 2.71E+00 | 6.46E-01 | no | | intergenic_variant | MODIFIER |
| 1 | 105245229 | AX-170447235 | 0 | G | A | 0.158 | 9.48E-02 | 2.07E-01 | 2.64E+00 | 2.71E+00 | 6.46E-01 | no | Gm29013 | downstream_gene_variant | MODIFIER |
| 10 | 12146550 | AX-170044154 | 0 | A | T | 0.211 | -4.52E-02 | 1.00E-01 | 2.65E+00 | 2.75E+00 | 6.46E-01 | no | | intergenic_variant | MODIFIER |
| 16 | 77592969 | AX-170013433 | 0 | G | A | 0.465 | 3.53E-02 | 7.56E-02 | 2.58E+00 | 2.65E+00 | 6.46E-01 | no | Mir99ahg | intron_variant,non_coding_transcript_variant | MODIFIER |
| 15 | 100326562 | AX-170286919 | 0 | C | A | 0.114 | 7.41E-02 | 1.63E-01 | 2.64E+00 | 2.72E+00 | 6.47E-01 | no | Mettl7a1 | intron_variant | MODIFIER |
| 18 | 52863777 | AX-170447678 | 0 | G | A | 0.237 | -5.05E-02 | 1.09E-01 | 2.60E+00 | 2.66E+00 | 6.47E-01 | no | Sncaip | intron_variant | MODIFIER |
| 18 | 52381753 | AX-97835496 | 1 | G | A | 0.196 | -5.32E-02 | 1.13E-01 | 2.56E+00 | 2.63E+00 | 6.47E-01 | no | | intergenic_variant | MODIFIER |
| 11 | 5979110 | AX-97823475 | 0 | G | A | 0.044 | -8.56E-02 | 1.88E-01 | 2.59E+00 | 2.73E+00 | 6.47E-01 | no | Camk2b | intron_variant | MODIFIER |
| 5 | 34516193 | AX-170369733 | 1 | G | A | 0.339 | 4.82E-02 | 1.06E-01 | 2.64E+00 | 2.70E+00 | 6.47E-01 | no | Tnip2 | upstream_gene_variant | MODIFIER |
| 5 | 35538420 | AX-170159276 | 1 | C | T | 0.339 | 4.82E-02 | 1.06E-01 | 2.64E+00 | 2.70E+00 | 6.47E-01 | no | | intergenic_variant | MODIFIER |
| 12 | 111593002 | AX-168397495 | 0 | T | C | 0.132 | 7.33E-02 | 1.62E-01 | 2.64E+00 | 2.75E+00 | 6.47E-01 | no | Mark3 | intron_variant | MODIFIER |
| 4 | 105987589 | AX-168396171 | 0 | C | T | 0.272 | 4.64E-02 | 9.82E-02 | 2.53E+00 | 2.64E+00 | 6.47E-01 | no | | intergenic_variant | MODIFIER |
| 5 | 39886253 | AX-170024753 | 0 | C | A | 0.219 | -4.74E-02 | 1.03E-01 | 2.60E+00 | 2.70E+00 | 6.47E-01 | no | | intergenic_variant | MODIFIER |
| 5 | 40394121 | AX-97799759 | 0 | A | T | 0.219 | -4.74E-02 | 1.03E-01 | 2.60E+00 | 2.70E+00 | 6.47E-01 | no | | intergenic_variant | MODIFIER |
| 5 | 88939409 | AX-170025545 | 0 | C | A | 0.439 | 4.80E-02 | 1.06E-01 | 2.70E+00 | 2.76E+00 | 6.47E-01 | no | Slc4a4 | intron_variant | MODIFIER |
| 5 | 89903244 | AX-170257908 | 0 | G | A | 0.439 | 4.80E-02 | 1.06E-01 | 2.70E+00 | 2.76E+00 | 6.47E-01 | no | | intergenic_variant | MODIFIER |
| 1 | 78262146 | AX-97813777 | 0 | C | A | 0.132 | 9.86E-02 | 2.19E-01 | 2.67E+00 | 2.76E+00 | 6.47E-01 | no | | intergenic_variant | MODIFIER |
| 2 | 87779912 | AX-170527727 | 0 | T | A | 0.053 | -1.35E-01 | 2.99E-01 | 2.75E+00 | 2.78E+00 | 6.47E-01 | no | Olfr152 | upstream_gene_variant | MODIFIER |
| 2 | 149814008 | AX-170206184 | 0 | C | T | 0.14 | 7.83E-02 | 1.71E-01 | 2.62E+00 | 2.70E+00 | 6.48E-01 | no | | intergenic_variant | MODIFIER |
| 4 | 73110843 | AX-170211369 | 1 | T | C | 0.446 | -3.86E-02 | 8.23E-02 | 2.56E+00 | 2.64E+00 | 6.48E-01 | no | Gm25769 | upstream_gene_variant | MODIFIER |
| 6 | 130562976 | AX-170492967 | 0 | A | G | 0.43 | 4.80E-02 | 1.01E-01 | 2.52E+00 | 2.64E+00 | 6.48E-01 | no | Gm8816 | upstream_gene_variant | MODIFIER |
| 3 | 122904579 | AX-170264700 | 0 | T | C | 0.044 | 9.15E-02 | 2.04E-01 | 2.65E+00 | 2.76E+00 | 6.48E-01 | no | 4933405D12Rik | intron_variant,non_coding_transcript_variant | MODIFIER |
| 5 | 104566177 | AX-170428554 | 1 | C | A | 0.464 | 4.99E-02 | 1.09E-01 | 2.61E+00 | 2.66E+00 | 6.48E-01 | no | | intergenic_variant | MODIFIER |
| 15 | 81743512 | AX-170381992 | 0 | A | C | 0.553 | 3.74E-02 | 8.11E-02 | 2.59E+00 | 2.67E+00 | 6.48E-01 | no | Zc3h7b | upstream_gene_variant | MODIFIER |
| 6 | 94439735 | AX-170269981 | 0 | G | A | 0.43 | -2.65E-02 | 5.69E-02 | 2.55E+00 | 2.67E+00 | 6.48E-01 | no | | intergenic_variant | MODIFIER |
| 2 | 120685242 | AX-170129586 | 0 | A | G | 0.149 | 8.03E-02 | 1.76E-01 | 2.61E+00 | 2.70E+00 | 6.48E-01 | no | Stard9 | intron_variant,NMD_transcript_variant | MODIFIER |
| 7 | 52776796 | AX-170077355 | 0 | C | T | 0.439 | 4.39E-02 | 9.62E-02 | 2.61E+00 | 2.70E+00 | 6.48E-01 | no | | intergenic_variant | MODIFIER |
| 1 | 22735360 | AX-170040602 | 1 | T | A | 0.429 | 3.87E-02 | 8.42E-02 | 2.61E+00 | 2.67E+00 | 6.48E-01 | no | Rims1 | intron_variant | MODIFIER |
| 10 | 70407962 | AX-97819425 | 0 | A | C | 0.412 | -3.59E-02 | 7.86E-02 | 2.61E+00 | 2.69E+00 | 6.49E-01 | no | Gm47946 | downstream_gene_variant | MODIFIER |
| 9 | 36867297 | AX-170032138 | 0 | A | G | 0.5 | -3.47E-02 | 7.78E-02 | 2.71E+00 | 2.80E+00 | 6.49E-01 | no | Fez1 | intron_variant | MODIFIER |
| 9 | 36909010 | AX-97830814 | 0 | C | T | 0.5 | -3.47E-02 | 7.78E-02 | 2.71E+00 | 2.80E+00 | 6.49E-01 | no | Pknox2 | intron_variant | MODIFIER |
| 9 | 36909053 | AX-170032136 | 0 | G | C | 0.5 | -3.47E-02 | 7.78E-02 | 2.71E+00 | 2.80E+00 | 6.49E-01 | no | Pknox2 | intron_variant | MODIFIER |
| 9 | 36909192 | AX-170090276 | 0 | G | C | 0.5 | -3.47E-02 | 7.78E-02 | 2.71E+00 | 2.80E+00 | 6.49E-01 | no | Pknox2 | intron_variant | MODIFIER |
| 9 | 36941972 | AX-97841104 | 0 | G | A | 0.5 | -3.47E-02 | 7.78E-02 | 2.71E+00 | 2.80E+00 | 6.49E-01 | no | Pknox2 | intron_variant | MODIFIER |
| 9 | 36952319 | AX-170032135 | 0 | C | T | 0.5 | -3.47E-02 | 7.78E-02 | 2.71E+00 | 2.80E+00 | 6.49E-01 | no | Pknox2 | intron_variant | MODIFIER |
| 9 | 36957902 | AX-170032139 | 0 | G | A | 0.5 | -3.47E-02 | 7.78E-02 | 2.71E+00 | 2.80E+00 | 6.49E-01 | no | Pknox2 | intron_variant | MODIFIER |
| 9 | 36987495 | AX-170273336 | 0 | T | C | 0.5 | -3.47E-02 | 7.78E-02 | 2.71E+00 | 2.80E+00 | 6.49E-01 | no | Pknox2 | intron_variant | MODIFIER |
| 9 | 36990511 | AX-170270340 | 0 | G | C | 0.5 | -3.47E-02 | 7.78E-02 | 2.71E+00 | 2.80E+00 | 6.49E-01 | no | Pknox2 | intron_variant | MODIFIER |
| 9 | 36991219 | AX-170456858 | 0 | G | A | 0.5 | -3.47E-02 | 7.78E-02 | 2.71E+00 | 2.80E+00 | 6.49E-01 | no | Pknox2 | intron_variant | MODIFIER |

| | | | | | | | | | | | | | | | |
|---|---|---|---|---|---|---|---|---|---|---|---|---|---|---|---|
| 9 | 36994707 | AX-170312937 | 0 | A | G | 0.5 | -3.47E-02 | 7.78E-02 | 2.71E+00 | 2.80E+00 | 6.49E-01 | no | Pknox2 | intron_variant | MODIFIER |
| 9 | 36996892 | AX-170126338 | 0 | C | G | 0.5 | -3.47E-02 | 7.78E-02 | 2.71E+00 | 2.80E+00 | 6.49E-01 | no | Pknox2 | intron_variant | MODIFIER |
| 9 | 37002384 | AX-170336439 | 0 | C | T | 0.5 | -3.47E-02 | 7.78E-02 | 2.71E+00 | 2.80E+00 | 6.49E-01 | no | Pknox2 | intron_variant | MODIFIER |
| 9 | 37045807 | AX-170369811 | 0 | G | A | 0.5 | -3.47E-02 | 7.78E-02 | 2.71E+00 | 2.80E+00 | 6.49E-01 | no | Pknox2 | intron_variant | MODIFIER |
| 9 | 37199608 | AX-170158077 | 0 | A | G | 0.5 | -3.47E-02 | 7.78E-02 | 2.71E+00 | 2.80E+00 | 6.49E-01 | no | | intergenic_variant | MODIFIER |
| 9 | 37199846 | AX-170120458 | 0 | C | A | 0.5 | -3.47E-02 | 7.78E-02 | 2.71E+00 | 2.80E+00 | 6.49E-01 | no | | intergenic_variant | MODIFIER |
| 9 | 37200415 | AX-170082127 | 0 | C | A | 0.5 | -3.47E-02 | 7.78E-02 | 2.71E+00 | 2.80E+00 | 6.49E-01 | no | | intergenic_variant | MODIFIER |
| 9 | 37262620 | AX-170090278 | 0 | A | G | 0.5 | -3.47E-02 | 7.78E-02 | 2.71E+00 | 2.80E+00 | 6.49E-01 | no | Gm47633 | downstream_gene_variant | MODIFIER |
| 9 | 37291651 | AX-170363980 | 0 | C | G | 0.5 | -3.47E-02 | 7.78E-02 | 2.71E+00 | 2.80E+00 | 6.49E-01 | no | Ccdc15 | intron_variant | MODIFIER |
| 9 | 37298696 | AX-97832787 | 0 | A | C | 0.5 | -3.47E-02 | 7.78E-02 | 2.71E+00 | 2.80E+00 | 6.49E-01 | no | Ccdc15 | intron_variant | MODIFIER |
| 9 | 37308558 | AX-170438335 | 0 | G | A | 0.5 | -3.47E-02 | 7.78E-02 | 2.71E+00 | 2.80E+00 | 6.49E-01 | no | Ccdc15 | intron_variant | MODIFIER |
| 9 | 37311258 | AX-170359038 | 0 | A | T | 0.5 | -3.47E-02 | 7.78E-02 | 2.71E+00 | 2.80E+00 | 6.49E-01 | no | Ccdc15 | intron_variant | MODIFIER |
| 9 | 37342479 | AX-170314195 | 0 | G | A | 0.5 | -3.47E-02 | 7.78E-02 | 2.71E+00 | 2.80E+00 | 6.49E-01 | no | Ccdc15 | missense_variant | MODERATE |
| 9 | 37363661 | AX-170032143 | 0 | C | T | 0.5 | -3.47E-02 | 7.78E-02 | 2.71E+00 | 2.80E+00 | 6.49E-01 | no | Hepacam | upstream_gene_variant | MODIFIER |
| 9 | 37363758 | AX-170032145 | 0 | G | A | 0.5 | -3.47E-02 | 7.78E-02 | 2.71E+00 | 2.80E+00 | 6.49E-01 | no | Hepacam | upstream_gene_variant | MODIFIER |
| 9 | 37372669 | AX-170096519 | 0 | G | T | 0.5 | -3.47E-02 | 7.78E-02 | 2.71E+00 | 2.80E+00 | 6.49E-01 | no | Hepacam | intron_variant | MODIFIER |
| 9 | 37401595 | AX-97821916 | 0 | A | G | 0.5 | -3.47E-02 | 7.78E-02 | 2.71E+00 | 2.80E+00 | 6.49E-01 | no | Robo4 | upstream_gene_variant | MODIFIER |
| 9 | 37403362 | AX-168372958 | 0 | A | G | 0.5 | -3.47E-02 | 7.78E-02 | 2.71E+00 | 2.80E+00 | 6.49E-01 | no | Robo4 | intron_variant | MODIFIER |
| 9 | 37417257 | AX-97804699 | 0 | A | C | 0.5 | -3.47E-02 | 7.78E-02 | 2.71E+00 | 2.80E+00 | 6.49E-01 | no | Robo3 | intron_variant | MODIFIER |
| 9 | 37569550 | AX-170207287 | 0 | C | A | 0.5 | -3.47E-02 | 7.78E-02 | 2.71E+00 | 2.80E+00 | 6.49E-01 | no | Siae | intron_variant | MODIFIER |
| 9 | 37575217 | AX-170393649 | 0 | C | T | 0.5 | -3.47E-02 | 7.78E-02 | 2.71E+00 | 2.80E+00 | 6.49E-01 | no | Siae | intron_variant | MODIFIER |
| 9 | 37594055 | AX-168411782 | 0 | G | A | 0.5 | -3.47E-02 | 7.78E-02 | 2.71E+00 | 2.80E+00 | 6.49E-01 | no | Siae | intron_variant | MODIFIER |
| 9 | 37626824 | AX-170160454 | 0 | T | C | 0.5 | -3.47E-02 | 7.78E-02 | 2.71E+00 | 2.80E+00 | 6.49E-01 | no | Siae | intron_variant | MODIFIER |
| 9 | 37644085 | AX-170035804 | 0 | G | A | 0.5 | -3.47E-02 | 7.78E-02 | 2.71E+00 | 2.80E+00 | 6.49E-01 | no | Siae | intron_variant | MODIFIER |
| 9 | 37824037 | AX-170463185 | 0 | T | A | 0.5 | -3.47E-02 | 7.78E-02 | 2.71E+00 | 2.80E+00 | 6.49E-01 | no | Gm24764 | upstream_gene_variant | MODIFIER |
| 9 | 37872147 | AX-97822862 | 0 | G | A | 0.5 | -3.47E-02 | 7.78E-02 | 2.71E+00 | 2.80E+00 | 6.49E-01 | no | | intergenic_variant | MODIFIER |
| 9 | 37999824 | AX-170032157 | 0 | T | C | 0.5 | -3.47E-02 | 7.78E-02 | 2.71E+00 | 2.80E+00 | 6.49E-01 | no | Olfr882-ps1 | upstream_gene_variant | MODIFIER |
| 9 | 37999859 | AX-97812240 | 0 | C | T | 0.5 | -3.47E-02 | 7.78E-02 | 2.71E+00 | 2.80E+00 | 6.49E-01 | no | Olfr882-ps1 | upstream_gene_variant | MODIFIER |
| 9 | 37999944 | AX-170032153 | 0 | T | C | 0.5 | -3.47E-02 | 7.78E-02 | 2.71E+00 | 2.80E+00 | 6.49E-01 | no | Olfr882-ps1 | upstream_gene_variant | MODIFIER |
| 9 | 38003852 | AX-170082153 | 0 | G | C | 0.5 | -3.47E-02 | 7.78E-02 | 2.71E+00 | 2.80E+00 | 6.49E-01 | no | Olfr882-ps1 | intron_variant,non_coding_transcript_variant | MODIFIER |
| 9 | 38112088 | AX-170171200 | 0 | G | A | 0.5 | -3.47E-02 | 7.78E-02 | 2.71E+00 | 2.80E+00 | 6.49E-01 | no | Olfr889 | upstream_gene_variant | MODIFIER |
| 9 | 38185476 | AX-170474914 | 0 | G | A | 0.5 | -3.47E-02 | 7.78E-02 | 2.71E+00 | 2.80E+00 | 6.49E-01 | no | Olfr891 | upstream_gene_variant | MODIFIER |
| 9 | 38185527 | AX-170244451 | 0 | C | T | 0.5 | -3.47E-02 | 7.78E-02 | 2.71E+00 | 2.80E+00 | 6.49E-01 | no | Olfr891 | upstream_gene_variant | MODIFIER |
| 9 | 38587061 | AX-170035807 | 0 | C | T | 0.5 | -3.47E-02 | 7.78E-02 | 2.71E+00 | 2.80E+00 | 6.49E-01 | no | Olfr912 | downstream_gene_variant | MODIFIER |
| 9 | 38603616 | AX-170032163 | 0 | T | A | 0.5 | -3.47E-02 | 7.78E-02 | 2.71E+00 | 2.80E+00 | 6.49E-01 | no | Olfr914 | upstream_gene_variant | MODIFIER |
| 1 | 54840146 | AX-170379390 | 0 | G | T | 0.105 | -8.30E-02 | 1.86E-01 | 2.68E+00 | 2.77E+00 | 6.49E-01 | no | Ankrd44 | intron_variant | MODIFIER |
| 11 | 110395782 | AX-97788655 | 0 | A | G | 0.053 | 8.31E-02 | 1.86E-01 | 2.63E+00 | 2.76E+00 | 6.49E-01 | no | Map2k6 | upstream_gene_variant | MODIFIER |
| 1 | 89118887 | AX-170001023 | 0 | A | C | 0.149 | 8.04E-02 | 1.77E-01 | 2.61E+00 | 2.70E+00 | 6.49E-01 | no | Sh3bp4 | intron_variant | MODIFIER |
| 2 | 117437394 | AX-170353007 | 0 | T | C | 0.132 | 8.86E-02 | 1.97E-01 | 2.71E+00 | 2.76E+00 | 6.49E-01 | no | | intergenic_variant | MODIFIER |
| 10 | 128482382 | AX-97810782 | 1 | G | C | 0.33 | -3.33E-02 | 7.31E-02 | 2.59E+00 | 2.70E+00 | 6.50E-01 | no | Smarcc2 | intron_variant | MODIFIER |
| 7 | 25027671 | AX-170300890 | 1 | G | A | 0.402 | 4.37E-02 | 9.48E-02 | 2.57E+00 | 2.67E+00 | 6.50E-01 | no | Grik5 | intron_variant | MODIFIER |
| 10 | 92949020 | AX-170471513 | 0 | A | G | 0.298 | -3.94E-02 | 8.49E-02 | 2.58E+00 | 2.65E+00 | 6.50E-01 | no | Cfap54 | intron_variant | MODIFIER |
| 10 | 92954139 | AX-170496824 | 0 | T | C | 0.298 | -3.94E-02 | 8.49E-02 | 2.58E+00 | 2.65E+00 | 6.50E-01 | no | Cfap54 | intron_variant | MODIFIER |
| 9 | 35875069 | AX-170252175 | 0 | C | T | 0.491 | -3.49E-02 | 7.84E-02 | 2.71E+00 | 2.80E+00 | 6.50E-01 | no | | intergenic_variant | MODIFIER |
| 9 | 36951738 | AX-170396558 | 0 | T | C | 0.491 | -3.49E-02 | 7.84E-02 | 2.71E+00 | 2.80E+00 | 6.50E-01 | no | Pknox2 | intron_variant | MODIFIER |
| 9 | 36987773 | AX-170378801 | 0 | T | C | 0.491 | -3.49E-02 | 7.84E-02 | 2.71E+00 | 2.80E+00 | 6.50E-01 | no | Pknox2 | intron_variant | MODIFIER |
| 9 | 37035931 | AX-170516642 | 0 | G | C | 0.491 | -3.49E-02 | 7.84E-02 | 2.71E+00 | 2.80E+00 | 6.50E-01 | no | Pknox2 | intron_variant | MODIFIER |
| 9 | 37049145 | AX-170353310 | 0 | A | G | 0.491 | -3.49E-02 | 7.84E-02 | 2.71E+00 | 2.80E+00 | 6.50E-01 | no | Pknox2 | intron_variant | MODIFIER |
| 9 | 37049475 | AX-170155976 | 0 | C | T | 0.491 | -3.49E-02 | 7.84E-02 | 2.71E+00 | 2.80E+00 | 6.50E-01 | no | Pknox2 | intron_variant | MODIFIER |
| 9 | 37053054 | AX-170314499 | 0 | A | G | 0.491 | -3.49E-02 | 7.84E-02 | 2.71E+00 | 2.80E+00 | 6.50E-01 | no | Pknox2 | intron_variant | MODIFIER |
| 9 | 37053786 | AX-170284950 | 0 | A | T | 0.491 | -3.49E-02 | 7.84E-02 | 2.71E+00 | 2.80E+00 | 6.50E-01 | no | Pknox2 | intron_variant | MODIFIER |
| 9 | 37053918 | AX-170266402 | 0 | C | A | 0.491 | -3.49E-02 | 7.84E-02 | 2.71E+00 | 2.80E+00 | 6.50E-01 | no | Pknox2 | intron_variant | MODIFIER |
| 9 | 37055313 | AX-170243217 | 0 | C | T | 0.491 | -3.49E-02 | 7.84E-02 | 2.71E+00 | 2.80E+00 | 6.50E-01 | no | Pknox2 | intron_variant | MODIFIER |
| 9 | 37100981 | AX-170489856 | 0 | A | G | 0.491 | -3.49E-02 | 7.84E-02 | 2.71E+00 | 2.80E+00 | 6.50E-01 | no | Pknox2 | intron_variant | MODIFIER |

| | | | | | | | | | | | | | | |
|---|---|---|---|---|---|---|---|---|---|---|---|---|---|---|
| 9 | 37106518 | AX-97802006 | 0 | C | T | 0.491 | -3.49E-02 | 7.84E-02 | 2.71E+00 | 2.80E+00 | 6.50E-01 | no | Pknox2 | intron_variant | MODIFIER |
| 9 | 37107769 | AX-170482732 | 0 | A | C | 0.491 | -3.49E-02 | 7.84E-02 | 2.71E+00 | 2.80E+00 | 6.50E-01 | no | Pknox2 | intron_variant | MODIFIER |
| 9 | 37114669 | AX-170082129 | 0 | A | G | 0.491 | -3.49E-02 | 7.84E-02 | 2.71E+00 | 2.80E+00 | 6.50E-01 | no | Pknox2 | intron_variant | MODIFIER |
| 9 | 37217967 | AX-170096518 | 0 | T | C | 0.491 | -3.49E-02 | 7.84E-02 | 2.71E+00 | 2.80E+00 | 6.50E-01 | no | Tmem218 | intron_variant | MODIFIER |
| 9 | 37304920 | AX-170184536 | 0 | T | C | 0.491 | -3.49E-02 | 7.84E-02 | 2.71E+00 | 2.80E+00 | 6.50E-01 | no | Ccdc15 | intron_variant | MODIFIER |
| 9 | 37363685 | AX-170459915 | 0 | G | A | 0.491 | -3.49E-02 | 7.84E-02 | 2.71E+00 | 2.80E+00 | 6.50E-01 | no | Hepacam | upstream_gene_variant | MODIFIER |
| 9 | 37371179 | AX-170250131 | 0 | G | A | 0.491 | -3.49E-02 | 7.84E-02 | 2.71E+00 | 2.80E+00 | 6.50E-01 | no | Hepacam | intron_variant | MODIFIER |
| 9 | 37374774 | AX-170279752 | 0 | T | G | 0.491 | -3.49E-02 | 7.84E-02 | 2.71E+00 | 2.80E+00 | 6.50E-01 | no | Hepacam | intron_variant | MODIFIER |
| 9 | 37413671 | AX-170280561 | 0 | G | A | 0.491 | -3.49E-02 | 7.84E-02 | 2.71E+00 | 2.80E+00 | 6.50E-01 | no | Robo3 | downstream_gene_variant | MODIFIER |
| 9 | 37441636 | AX-168404159 | 0 | G | A | 0.491 | -3.49E-02 | 7.84E-02 | 2.71E+00 | 2.80E+00 | 6.50E-01 | no | | intergenic_variant | MODIFIER |
| 9 | 37441692 | AX-170096520 | 0 | G | A | 0.491 | -3.49E-02 | 7.84E-02 | 2.71E+00 | 2.80E+00 | 6.50E-01 | no | | intergenic_variant | MODIFIER |
| 9 | 37441718 | AX-170211575 | 0 | A | G | 0.491 | -3.49E-02 | 7.84E-02 | 2.71E+00 | 2.80E+00 | 6.50E-01 | no | | intergenic_variant | MODIFIER |
| 9 | 37467850 | AX-170082137 | 0 | C | G | 0.491 | -3.49E-02 | 7.84E-02 | 2.71E+00 | 2.80E+00 | 6.50E-01 | no | | intergenic_variant | MODIFIER |
| 9 | 37645242 | AX-170082144 | 0 | C | T | 0.491 | -3.49E-02 | 7.84E-02 | 2.71E+00 | 2.80E+00 | 6.50E-01 | no | Siae | intron_variant | MODIFIER |
| 9 | 37648162 | AX-170090280 | 0 | A | G | 0.491 | -3.49E-02 | 7.84E-02 | 2.71E+00 | 2.80E+00 | 6.50E-01 | no | Siae | 3_prime_UTR_variant | MODIFIER |
| 9 | 37728862 | AX-168390407 | 0 | A | G | 0.491 | -3.49E-02 | 7.84E-02 | 2.71E+00 | 2.80E+00 | 6.50E-01 | no | Olfr151 | downstream_gene_variant | MODIFIER |
| 9 | 37729002 | AX-170297093 | 0 | A | T | 0.491 | -3.49E-02 | 7.84E-02 | 2.71E+00 | 2.80E+00 | 6.50E-01 | no | Olfr151 | downstream_gene_variant | MODIFIER |
| 9 | 37745481 | AX-170362319 | 0 | T | C | 0.491 | -3.49E-02 | 7.84E-02 | 2.71E+00 | 2.80E+00 | 6.50E-01 | no | Olfr874 | upstream_gene_variant | MODIFIER |
| 9 | 37746696 | AX-170090281 | 0 | C | T | 0.491 | -3.49E-02 | 7.84E-02 | 2.71E+00 | 2.80E+00 | 6.50E-01 | no | Olfr874 | synonymous_variant | LOW |
| 9 | 37761892 | AX-170205469 | 0 | A | G | 0.491 | -3.49E-02 | 7.84E-02 | 2.71E+00 | 2.80E+00 | 6.50E-01 | no | | intergenic_variant | MODIFIER |
| 9 | 37769941 | AX-170534015 | 0 | A | T | 0.491 | -3.49E-02 | 7.84E-02 | 2.71E+00 | 2.80E+00 | 6.50E-01 | no | Olfr875 | upstream_gene_variant | MODIFIER |
| 9 | 37786545 | AX-170534920 | 0 | T | C | 0.491 | -3.49E-02 | 7.84E-02 | 2.71E+00 | 2.80E+00 | 6.50E-01 | no | | intergenic_variant | MODIFIER |
| 9 | 37800169 | AX-97835100 | 0 | T | C | 0.491 | -3.49E-02 | 7.84E-02 | 2.71E+00 | 2.80E+00 | 6.50E-01 | no | Olfr876 | upstream_gene_variant | MODIFIER |
| 9 | 37800280 | AX-170310576 | 0 | C | T | 0.491 | -3.49E-02 | 7.84E-02 | 2.71E+00 | 2.80E+00 | 6.50E-01 | no | Olfr876 | upstream_gene_variant | MODIFIER |
| 9 | 37903320 | AX-170517059 | 0 | T | G | 0.491 | -3.49E-02 | 7.84E-02 | 2.71E+00 | 2.80E+00 | 6.50E-01 | no | Olfr145 | 3_prime_UTR_variant | MODIFIER |
| 9 | 37904591 | AX-170519927 | 0 | G | A | 0.491 | -3.49E-02 | 7.84E-02 | 2.71E+00 | 2.80E+00 | 6.50E-01 | no | Olfr145 | downstream_gene_variant | MODIFIER |
| 9 | 37904625 | AX-170327540 | 0 | A | G | 0.491 | -3.49E-02 | 7.84E-02 | 2.71E+00 | 2.80E+00 | 6.50E-01 | no | Olfr145 | downstream_gene_variant | MODIFIER |
| 9 | 37921465 | AX-170209041 | 0 | A | G | 0.491 | -3.49E-02 | 7.84E-02 | 2.71E+00 | 2.80E+00 | 6.50E-01 | no | Olfr878 | downstream_gene_variant | MODIFIER |
| 9 | 37921498 | AX-170484534 | 0 | C | T | 0.491 | -3.49E-02 | 7.84E-02 | 2.71E+00 | 2.80E+00 | 6.50E-01 | no | Olfr878 | downstream_gene_variant | MODIFIER |
| 9 | 37923872 | AX-170248870 | 0 | A | C | 0.491 | -3.49E-02 | 7.84E-02 | 2.71E+00 | 2.80E+00 | 6.50E-01 | no | Olfr878 | downstream_gene_variant | MODIFIER |
| 9 | 37933286 | AX-170475429 | 0 | C | T | 0.491 | -3.49E-02 | 7.84E-02 | 2.71E+00 | 2.80E+00 | 6.50E-01 | no | | intergenic_variant | MODIFIER |
| 9 | 37934890 | AX-97797478 | 0 | C | T | 0.491 | -3.49E-02 | 7.84E-02 | 2.71E+00 | 2.80E+00 | 6.50E-01 | no | | intergenic_variant | MODIFIER |
| 9 | 38209978 | AX-170082155 | 0 | G | T | 0.491 | -3.49E-02 | 7.84E-02 | 2.71E+00 | 2.80E+00 | 6.50E-01 | no | Olfr893 | missense_variant | MODERATE |
| 9 | 38221081 | AX-170316060 | 0 | C | T | 0.491 | -3.49E-02 | 7.84E-02 | 2.71E+00 | 2.80E+00 | 6.50E-01 | no | Olfr894 | downstream_gene_variant | MODIFIER |
| 9 | 38221490 | AX-170363066 | 0 | T | C | 0.491 | -3.49E-02 | 7.84E-02 | 2.71E+00 | 2.80E+00 | 6.50E-01 | no | Olfr894 | downstream_gene_variant | MODIFIER |
| 9 | 38255681 | AX-170335766 | 0 | A | G | 0.491 | -3.49E-02 | 7.84E-02 | 2.71E+00 | 2.80E+00 | 6.50E-01 | no | Olfr143 | downstream_gene_variant | MODIFIER |
| 9 | 38292218 | AX-170357117 | 0 | A | C | 0.491 | -3.49E-02 | 7.84E-02 | 2.71E+00 | 2.80E+00 | 6.50E-01 | no | Olfr896-ps1 | non_coding_transcript_exon_variant | MODIFIER |
| 9 | 38310589 | AX-170193565 | 0 | T | C | 0.491 | -3.49E-02 | 7.84E-02 | 2.71E+00 | 2.80E+00 | 6.50E-01 | no | Olfr897-ps1 | downstream_gene_variant | MODIFIER |
| 9 | 38329446 | AX-170368507 | 0 | G | C | 0.491 | -3.49E-02 | 7.84E-02 | 2.71E+00 | 2.80E+00 | 6.50E-01 | no | Olfr25 | upstream_gene_variant | MODIFIER |
| 9 | 38333867 | AX-170406977 | 0 | G | A | 0.491 | -3.49E-02 | 7.84E-02 | 2.71E+00 | 2.80E+00 | 6.50E-01 | no | Olfr25 | downstream_gene_variant | MODIFIER |
| 9 | 38376964 | AX-170521833 | 0 | C | T | 0.491 | -3.49E-02 | 7.84E-02 | 2.71E+00 | 2.80E+00 | 6.50E-01 | no | Olfr251 | upstream_gene_variant | MODIFIER |
| 9 | 38458358 | AX-170082161 | 0 | T | C | 0.491 | -3.49E-02 | 7.84E-02 | 2.71E+00 | 2.80E+00 | 6.50E-01 | no | Olfr904 | upstream_gene_variant | MODIFIER |
| 9 | 38463898 | AX-170458218 | 0 | G | C | 0.491 | -3.49E-02 | 7.84E-02 | 2.71E+00 | 2.80E+00 | 6.50E-01 | no | Olfr904 | upstream_gene_variant | MODIFIER |
| 9 | 38464063 | AX-170308950 | 0 | C | T | 0.491 | -3.49E-02 | 7.84E-02 | 2.71E+00 | 2.80E+00 | 6.50E-01 | no | Olfr904 | synonymous_variant | LOW |
| 9 | 38465640 | AX-170116031 | 0 | C | T | 0.491 | -3.49E-02 | 7.84E-02 | 2.71E+00 | 2.80E+00 | 6.50E-01 | no | Olfr904 | downstream_gene_variant | MODIFIER |
| 9 | 38529368 | AX-170109958 | 0 | T | A | 0.491 | -3.49E-02 | 7.84E-02 | 2.71E+00 | 2.80E+00 | 6.50E-01 | no | Olfr911-ps1 | downstream_gene_variant | MODIFIER |
| 9 | 38552184 | AX-97838151 | 0 | T | C | 0.491 | -3.49E-02 | 7.84E-02 | 2.71E+00 | 2.80E+00 | 6.50E-01 | no | | intergenic_variant | MODIFIER |
| 12 | 106617530 | AX-170313551 | 0 | A | G | 0.132 | -8.57E-02 | 1.92E-01 | 2.72E+00 | 2.79E+00 | 6.50E-01 | no | | intergenic_variant | MODIFIER |
| 1 | 126891246 | AX-170139564 | 1 | C | T | 0.545 | 3.83E-02 | 8.42E-02 | 2.62E+00 | 2.69E+00 | 6.50E-01 | no | | intergenic_variant | MODIFIER |
| 7 | 37333905 | AX-170315631 | 0 | A | C | 0.342 | 4.21E-02 | 9.16E-02 | 2.59E+00 | 2.67E+00 | 6.50E-01 | no | | intergenic_variant | MODIFIER |
| 8 | 82652182 | AX-170161855 | 0 | A | G | 0.202 | 5.50E-02 | 1.23E-01 | 2.67E+00 | 2.75E+00 | 6.50E-01 | no | | intergenic_variant | MODIFIER |
| 3 | 32250661 | AX-97798213 | 1 | G | A | 0.161 | -4.47E-02 | 9.98E-02 | 2.68E+00 | 2.75E+00 | 6.50E-01 | no | | intergenic_variant | MODIFIER |
| 6 | 126281041 | AX-97850026 | 1 | C | T | 0.188 | -3.64E-02 | 8.25E-02 | 2.66E+00 | 2.79E+00 | 6.50E-01 | no | | intergenic_variant | MODIFIER |
| 13 | 47736542 | AX-170051198 | 1 | A | G | 0.536 | 2.93E-02 | 6.47E-02 | 2.61E+00 | 2.71E+00 | 6.50E-01 | no | | intergenic_variant | MODIFIER |

| | | | | | | | | | | | | | | | |
|---|---|---|---|---|---|---|---|---|---|---|---|---|---|---|---|
| 10 | 28670582 | AX-97805341 | 1 | C | T | 0.196 | 4.25E-02 | 9.63E-02 | 2.72E+00 | 2.82E+00 | 6.50E-01 | no | Themis | intron_variant | MODIFIER |
| 8 | 49915447 | AX-97807074 | 1 | A | G | 0.143 | -6.80E-02 | 1.47E-01 | 2.59E+00 | 2.64E+00 | 6.50E-01 | no | Gm45321 | intron_variant,non_coding_transcript_variant | MODIFIER |
| 9 | 58936273 | AX-170457718 | 0 | G | A | 0.211 | -5.46E-02 | 1.22E-01 | 2.72E+00 | 2.78E+00 | 6.50E-01 | no | Neo1 | intron_variant | MODIFIER |
| 7 | 57582458 | AX-170449663 | 0 | G | A | 0.439 | 4.42E-02 | 9.77E-02 | 2.62E+00 | 2.71E+00 | 6.50E-01 | no | Gabrb3 | intron_variant | MODIFIER |
| 15 | 100694921 | AX-170250957 | 1 | C | T | 0.098 | 7.68E-02 | 1.70E-01 | 2.63E+00 | 2.72E+00 | 6.50E-01 | no | Galnt6 | intron_variant | MODIFIER |
| 7 | 118574577 | AX-170115553 | 0 | G | A | 0.404 | 3.87E-02 | 8.54E-02 | 2.63E+00 | 2.70E+00 | 6.51E-01 | no | Tmc7 | intron_variant | MODIFIER |
| 8 | 95524000 | AX-97788887 | 0 | A | C | 0.237 | 4.83E-02 | 1.09E-01 | 2.71E+00 | 2.79E+00 | 6.51E-01 | no | | intergenic_variant | MODIFIER |
| 16 | 78801756 | AX-170181150 | 0 | A | T | 0.456 | 3.53E-02 | 7.71E-02 | 2.60E+00 | 2.66E+00 | 6.51E-01 | no | | intergenic_variant | MODIFIER |
| 2 | 159718391 | AX-170512195 | 2 | C | A | 0.127 | 8.46E-02 | 1.89E-01 | 2.65E+00 | 2.74E+00 | 6.51E-01 | no | | intergenic_variant | MODIFIER |
| 10 | 27939573 | AX-97828890 | 1 | G | A | 0.179 | 4.42E-02 | 1.00E-01 | 2.73E+00 | 2.83E+00 | 6.51E-01 | no | Gm10145 | downstream_gene_variant | MODIFIER |
| 3 | 156017391 | AX-170129430 | 1 | C | A | 0.321 | 3.31E-02 | 7.47E-02 | 2.70E+00 | 2.78E+00 | 6.51E-01 | no | | intergenic_variant | MODIFIER |
| 1 | 110791442 | AX-170119683 | 1 | C | T | 0.161 | 8.56E-02 | 1.88E-01 | 2.60E+00 | 2.68E+00 | 6.51E-01 | no | | intergenic_variant | MODIFIER |
| 9 | 89954562 | AX-97847803 | 0 | T | C | 0.132 | -6.07E-02 | 1.32E-01 | 2.59E+00 | 2.66E+00 | 6.51E-01 | no | Rasgrf1 | intron_variant | MODIFIER |
| 13 | 24977872 | AX-170516887 | 0 | G | A | 0.167 | -5.83E-02 | 1.22E-01 | 2.52E+00 | 2.58E+00 | 6.51E-01 | no | Gpld1 | intron_variant | MODIFIER |
| 18 | 64292757 | AX-170015854 | 1 | C | T | 0.393 | 4.44E-02 | 9.45E-02 | 2.55E+00 | 2.60E+00 | 6.51E-01 | no | St8sia3os | intron_variant,non_coding_transcript_variant | MODIFIER |
| 7 | 117830389 | AX-170078568 | 1 | T | A | 0.384 | 3.79E-02 | 8.46E-02 | 2.65E+00 | 2.73E+00 | 6.51E-01 | no | | intergenic_variant | MODIFIER |
| 5 | 90118009 | AX-97799918 | 0 | A | G | 0.404 | 4.97E-02 | 1.12E-01 | 2.74E+00 | 2.79E+00 | 6.51E-01 | no | | intergenic_variant | MODIFIER |
| 13 | 53853879 | AX-170008335 | 0 | C | A | 0.509 | -2.96E-02 | 6.77E-02 | 2.77E+00 | 2.90E+00 | 6.51E-01 | no | | intergenic_variant | MODIFIER |
| 3 | 79236132 | AX-170120164 | 0 | T | C | 0.053 | -7.73E-02 | 1.75E-01 | 2.65E+00 | 2.77E+00 | 6.52E-01 | no | Rapgef2 | intron_variant | MODIFIER |
| 6 | 130341767 | AX-97841954 | 2 | G | A | 0.091 | -4.75E-02 | 1.01E-01 | 2.51E+00 | 2.64E+00 | 6.52E-01 | no | Klra3 | upstream_gene_variant | MODIFIER |
| 16 | 19100682 | AX-170162714 | 1 | A | G | 0.125 | 3.99E-02 | 9.12E-02 | 2.69E+00 | 2.82E+00 | 6.52E-01 | no | 2010309G21Rik | intron_variant,non_coding_transcript_variant | MODIFIER |
| 11 | 74780485 | AX-97798508 | 1 | C | T | 0.241 | 3.20E-02 | 7.25E-02 | 2.64E+00 | 2.77E+00 | 6.52E-01 | no | Mettl16 | intron_variant | MODIFIER |
| 1 | 172592457 | AX-170403909 | 0 | A | C | 0.237 | 5.39E-02 | 1.15E-01 | 2.54E+00 | 2.61E+00 | 6.52E-01 | no | Slamf8 | upstream_gene_variant | MODIFIER |
| 7 | 113840371 | AX-97800601 | 1 | G | T | 0.429 | 4.17E-02 | 9.35E-02 | 2.68E+00 | 2.73E+00 | 6.52E-01 | no | Spon1 | intron_variant | MODIFIER |
| 1 | 118700749 | AX-170147401 | 2 | C | T | 0.164 | 9.27E-02 | 2.07E-01 | 2.64E+00 | 2.73E+00 | 6.52E-01 | no | | intergenic_variant | MODIFIER |
| 18 | 59342120 | AX-170234391 | 0 | G | T | 0.412 | -3.90E-02 | 8.45E-02 | 2.57E+00 | 2.63E+00 | 6.52E-01 | no | Chsy3 | intron_variant | MODIFIER |
| 8 | 108633943 | AX-170312793 | 0 | G | A | 0.228 | 4.83E-02 | 1.09E-01 | 2.69E+00 | 2.77E+00 | 6.52E-01 | no | Zfhx3 | intron_variant | MODIFIER |
| 2 | 94465503 | AX-170018378 | 0 | C | A | 0.053 | -1.00E-01 | 2.19E-01 | 2.57E+00 | 2.67E+00 | 6.53E-01 | no | | regulatory_region_variant | MODIFIER |
| 1 | 176615703 | AX-170359376 | 0 | A | G | 0.219 | -4.60E-02 | 1.05E-01 | 2.77E+00 | 2.89E+00 | 6.53E-01 | no | | intergenic_variant | MODIFIER |
| 1 | 176779550 | AX-170306134 | 0 | T | C | 0.219 | -4.60E-02 | 1.05E-01 | 2.77E+00 | 2.89E+00 | 6.53E-01 | no | Cep170 | intron_variant | MODIFIER |
| 4 | 72218953 | AX-97819047 | 2 | T | G | 0.345 | 2.96E-02 | 6.24E-02 | 2.51E+00 | 2.60E+00 | 6.53E-01 | no | C630043F03Rik | downstream_gene_variant | MODIFIER |
| 16 | 97535160 | AX-170013719 | 1 | C | A | 0.473 | 3.36E-02 | 7.70E-02 | 2.74E+00 | 2.86E+00 | 6.53E-01 | no | Mx2 | upstream_gene_variant | MODIFIER |
| 9 | 40947184 | AX-170090286 | 0 | A | T | 0.421 | -3.20E-02 | 7.23E-02 | 2.67E+00 | 2.77E+00 | 6.53E-01 | no | Jhy | intron_variant | MODIFIER |
| 1 | 74970829 | AX-170163951 | 0 | T | C | 0.132 | 7.77E-02 | 1.73E-01 | 2.61E+00 | 2.70E+00 | 6.53E-01 | no | Nhej1 | intron_variant | MODIFIER |
| 11 | 8263594 | AX-170117985 | 0 | A | T | 0.053 | 1.01E-01 | 2.31E-01 | 2.70E+00 | 2.81E+00 | 6.53E-01 | no | | intergenic_variant | MODIFIER |
| 1 | 22708065 | AX-97811000 | 1 | A | G | 0.429 | 3.60E-02 | 7.84E-02 | 2.57E+00 | 2.66E+00 | 6.53E-01 | no | Rims1 | intron_variant | MODIFIER |
| 15 | 79145396 | AX-170413935 | 1 | A | G | 0.554 | 3.68E-02 | 8.08E-02 | 2.59E+00 | 2.67E+00 | 6.53E-01 | no | Polr2f | intron_variant | MODIFIER |
| 4 | 127756757 | AX-170394107 | 0 | C | T | 0.14 | 5.15E-02 | 1.14E-01 | 2.61E+00 | 2.69E+00 | 6.53E-01 | no | | regulatory_region_variant | MODIFIER |
| 10 | 9424091 | AX-170117752 | 0 | T | A | 0.193 | -4.58E-02 | 1.02E-01 | 2.61E+00 | 2.70E+00 | 6.53E-01 | no | | intergenic_variant | MODIFIER |
| 16 | 51575570 | AX-170013034 | 0 | C | G | 0.482 | -3.90E-02 | 8.38E-02 | 2.56E+00 | 2.60E+00 | 6.53E-01 | no | | intergenic_variant | MODIFIER |
| 1 | 25748713 | AX-97811014 | 2 | A | G | 0.427 | 3.65E-02 | 8.17E-02 | 2.65E+00 | 2.72E+00 | 6.53E-01 | no | Adgrb3 | intron_variant | MODIFIER |
| 5 | 96643759 | AX-97788976 | 2 | A | G | 0.418 | -4.35E-02 | 9.93E-02 | 2.73E+00 | 2.82E+00 | 6.53E-01 | no | Fras1 | intron_variant | MODIFIER |
| 7 | 118741325 | AX-170120905 | 0 | T | C | 0.404 | 3.99E-02 | 8.93E-02 | 2.65E+00 | 2.71E+00 | 6.53E-01 | no | Ccp110 | downstream_gene_variant | MODIFIER |
| 1 | 89228344 | AX-97812377 | 0 | C | T | 0.158 | 9.24E-02 | 2.07E-01 | 2.64E+00 | 2.71E+00 | 6.53E-01 | no | | intergenic_variant | MODIFIER |
| 1 | 90220508 | AX-170041980 | 0 | G | A | 0.158 | 9.24E-02 | 2.07E-01 | 2.64E+00 | 2.71E+00 | 6.53E-01 | no | Ackr3 | downstream_gene_variant | MODIFIER |
| 1 | 99892771 | AX-168380334 | 0 | T | C | 0.158 | 9.24E-02 | 2.07E-01 | 2.64E+00 | 2.71E+00 | 6.53E-01 | no | Cntnap5b | intron_variant | MODIFIER |
| 12 | 98450056 | AX-170007354 | 0 | G | T | 0.088 | 7.29E-02 | 1.65E-01 | 2.64E+00 | 2.75E+00 | 6.53E-01 | no | Kcnk10 | intron_variant | MODIFIER |
| 7 | 40123903 | AX-170226486 | 0 | G | A | 0.351 | 4.08E-02 | 8.99E-02 | 2.59E+00 | 2.68E+00 | 6.53E-01 | no | | intergenic_variant | MODIFIER |
| 7 | 40155283 | AX-170097998 | 0 | G | C | 0.351 | 4.08E-02 | 8.99E-02 | 2.59E+00 | 2.68E+00 | 6.53E-01 | no | | intergenic_variant | MODIFIER |
| 7 | 40164780 | AX-170461641 | 0 | T | C | 0.351 | 4.08E-02 | 8.99E-02 | 2.59E+00 | 2.68E+00 | 6.53E-01 | no | | intergenic_variant | MODIFIER |
| 7 | 40183021 | AX-170330481 | 0 | G | A | 0.351 | 4.08E-02 | 8.99E-02 | 2.59E+00 | 2.68E+00 | 6.53E-01 | no | | intergenic_variant | MODIFIER |
| 7 | 40183134 | AX-97807468 | 0 | C | T | 0.351 | 4.08E-02 | 8.99E-02 | 2.59E+00 | 2.68E+00 | 6.53E-01 | no | | intergenic_variant | MODIFIER |
| 7 | 135901206 | AX-170304326 | 1 | C | T | 0.259 | -3.67E-02 | 8.11E-02 | 2.59E+00 | 2.69E+00 | 6.53E-01 | no | | regulatory_region_variant | MODIFIER |
| 5 | 34850929 | AX-170168247 | 0 | G | A | 0.333 | 4.83E-02 | 1.09E-01 | 2.67E+00 | 2.72E+00 | 6.54E-01 | no | Htt | intron_variant | MODIFIER |

| | | | | | | | | | | | | | | | |
|---|---|---|---|---|---|---|---|---|---|---|---|---|---|---|---|
| 8 | 68321536 | AX-170494913 | 0 | T | C | 0.184 | -5.99E-02 | 1.35E-01 | 2.66E+00 | 2.72E+00 | 6.54E-01 | no | Sh2d4a | intron_variant | MODIFIER |
| 8 | 68328557 | AX-170275968 | 0 | T | C | 0.184 | -5.99E-02 | 1.35E-01 | 2.66E+00 | 2.72E+00 | 6.54E-01 | no | Sh2d4a | intron_variant | MODIFIER |
| 8 | 68335255 | AX-170347494 | 0 | C | T | 0.184 | -5.99E-02 | 1.35E-01 | 2.66E+00 | 2.72E+00 | 6.54E-01 | no | Sh2d4a | intron_variant | MODIFIER |
| 8 | 68335285 | AX-170136340 | 0 | C | T | 0.184 | -5.99E-02 | 1.35E-01 | 2.66E+00 | 2.72E+00 | 6.54E-01 | no | Sh2d4a | intron_variant | MODIFIER |
| 8 | 68348368 | AX-170096305 | 0 | T | C | 0.184 | -5.99E-02 | 1.35E-01 | 2.66E+00 | 2.72E+00 | 6.54E-01 | no | Sh2d4a | downstream_gene_variant | MODIFIER |
| 8 | 68349918 | AX-170454126 | 0 | A | G | 0.184 | -5.99E-02 | 1.35E-01 | 2.66E+00 | 2.72E+00 | 6.54E-01 | no | Sh2d4a | downstream_gene_variant | MODIFIER |
| 8 | 68356153 | AX-170484941 | 0 | T | A | 0.184 | -5.99E-02 | 1.35E-01 | 2.66E+00 | 2.72E+00 | 6.54E-01 | no | Csgalnact1 | downstream_gene_variant | MODIFIER |
| 8 | 68361881 | AX-170030971 | 0 | A | G | 0.184 | -5.99E-02 | 1.35E-01 | 2.66E+00 | 2.72E+00 | 6.54E-01 | no | Csgalnact1 | intron_variant | MODIFIER |
| 8 | 68457279 | AX-170030972 | 0 | G | T | 0.184 | -5.99E-02 | 1.35E-01 | 2.66E+00 | 2.72E+00 | 6.54E-01 | no | Csgalnact1 | intron_variant | MODIFIER |
| 8 | 68484091 | AX-168403607 | 0 | T | C | 0.184 | -5.99E-02 | 1.35E-01 | 2.66E+00 | 2.72E+00 | 6.54E-01 | no | Csgalnact1 | intron_variant | MODIFIER |
| 8 | 68486587 | AX-168417759 | 0 | G | A | 0.184 | -5.99E-02 | 1.35E-01 | 2.66E+00 | 2.72E+00 | 6.54E-01 | no | Csgalnact1 | intron_variant | MODIFIER |
| 8 | 68492569 | AX-170312453 | 0 | C | A | 0.184 | -5.99E-02 | 1.35E-01 | 2.66E+00 | 2.72E+00 | 6.54E-01 | no | Csgalnact1 | intron_variant | MODIFIER |
| 8 | 68496458 | AX-170430274 | 0 | A | G | 0.184 | -5.99E-02 | 1.35E-01 | 2.66E+00 | 2.72E+00 | 6.54E-01 | no | Csgalnact1 | intron_variant | MODIFIER |
| 8 | 68509727 | AX-97856272 | 0 | T | C | 0.184 | -5.99E-02 | 1.35E-01 | 2.66E+00 | 2.72E+00 | 6.54E-01 | no | Csgalnact1 | intron_variant | MODIFIER |
| 8 | 68540931 | AX-170251637 | 0 | T | C | 0.184 | -5.99E-02 | 1.35E-01 | 2.66E+00 | 2.72E+00 | 6.54E-01 | no | Csgalnact1 | intron_variant | MODIFIER |
| 8 | 68553471 | AX-170518301 | 0 | G | A | 0.184 | -5.99E-02 | 1.35E-01 | 2.66E+00 | 2.72E+00 | 6.54E-01 | no | Csgalnact1 | intron_variant | MODIFIER |
| 8 | 68561333 | AX-170227751 | 0 | A | G | 0.184 | -5.99E-02 | 1.35E-01 | 2.66E+00 | 2.72E+00 | 6.54E-01 | no | Csgalnact1 | intron_variant | MODIFIER |
| 8 | 68562686 | AX-170349054 | 0 | T | C | 0.184 | -5.99E-02 | 1.35E-01 | 2.66E+00 | 2.72E+00 | 6.54E-01 | no | Csgalnact1 | intron_variant | MODIFIER |
| 8 | 68605105 | AX-170080330 | 0 | C | T | 0.184 | -5.99E-02 | 1.35E-01 | 2.66E+00 | 2.72E+00 | 6.54E-01 | no | Csgalnact1 | intron_variant | MODIFIER |
| 8 | 68632798 | AX-170364626 | 0 | G | A | 0.184 | -5.99E-02 | 1.35E-01 | 2.66E+00 | 2.72E+00 | 6.54E-01 | no | Csgalnact1 | intron_variant | MODIFIER |
| 8 | 68701959 | AX-170411470 | 0 | G | C | 0.184 | -5.99E-02 | 1.35E-01 | 2.66E+00 | 2.72E+00 | 6.54E-01 | no | Csgalnact1 | intron_variant | MODIFIER |
| 8 | 68740057 | AX-170339915 | 0 | G | A | 0.184 | -5.99E-02 | 1.35E-01 | 2.66E+00 | 2.72E+00 | 6.54E-01 | no | Csgalnact1 | upstream_gene_variant | MODIFIER |
| 8 | 68741581 | AX-170331137 | 0 | T | G | 0.184 | -5.99E-02 | 1.35E-01 | 2.66E+00 | 2.72E+00 | 6.54E-01 | no | | intergenic_variant | MODIFIER |
| 8 | 68812919 | AX-170502081 | 0 | G | A | 0.184 | -5.99E-02 | 1.35E-01 | 2.66E+00 | 2.72E+00 | 6.54E-01 | no | Ints10 | intron_variant | MODIFIER |
| 8 | 68818493 | AX-170364451 | 0 | A | G | 0.184 | -5.99E-02 | 1.35E-01 | 2.66E+00 | 2.72E+00 | 6.54E-01 | no | Ints10 | intron_variant | MODIFIER |
| 8 | 68819023 | AX-170346249 | 0 | T | G | 0.184 | -5.99E-02 | 1.35E-01 | 2.66E+00 | 2.72E+00 | 6.54E-01 | no | Ints10 | intron_variant | MODIFIER |
| 8 | 68823990 | AX-170519853 | 0 | C | T | 0.184 | -5.99E-02 | 1.35E-01 | 2.66E+00 | 2.72E+00 | 6.54E-01 | no | Ints10 | intron_variant | MODIFIER |
| 8 | 68849184 | AX-170080334 | 0 | A | G | 0.184 | -5.99E-02 | 1.35E-01 | 2.66E+00 | 2.72E+00 | 6.54E-01 | no | | intergenic_variant | MODIFIER |
| 8 | 68853038 | AX-170111852 | 0 | C | T | 0.184 | -5.99E-02 | 1.35E-01 | 2.66E+00 | 2.72E+00 | 6.54E-01 | no | | intergenic_variant | MODIFIER |
| 8 | 68862394 | AX-170389887 | 0 | A | G | 0.184 | -5.99E-02 | 1.35E-01 | 2.66E+00 | 2.72E+00 | 6.54E-01 | no | | intergenic_variant | MODIFIER |
| 8 | 68887740 | AX-170519411 | 0 | A | G | 0.184 | -5.99E-02 | 1.35E-01 | 2.66E+00 | 2.72E+00 | 6.54E-01 | no | Lpl | intron_variant | MODIFIER |
| 8 | 68894449 | AX-170409172 | 0 | T | C | 0.184 | -5.99E-02 | 1.35E-01 | 2.66E+00 | 2.72E+00 | 6.54E-01 | no | Lpl | intron_variant | MODIFIER |
| 8 | 68895640 | AX-97831833 | 0 | G | A | 0.184 | -5.99E-02 | 1.35E-01 | 2.66E+00 | 2.72E+00 | 6.54E-01 | no | Lpl | intron_variant | MODIFIER |
| 8 | 68939479 | AX-97800566 | 0 | A | G | 0.184 | -5.99E-02 | 1.35E-01 | 2.66E+00 | 2.72E+00 | 6.54E-01 | no | | intergenic_variant | MODIFIER |
| 8 | 68955756 | AX-170182308 | 0 | G | A | 0.184 | -5.99E-02 | 1.35E-01 | 2.66E+00 | 2.72E+00 | 6.54E-01 | no | | regulatory_region_variant | MODIFIER |
| 8 | 68957484 | AX-170255459 | 0 | C | A | 0.184 | -5.99E-02 | 1.35E-01 | 2.66E+00 | 2.72E+00 | 6.54E-01 | no | | intergenic_variant | MODIFIER |
| 8 | 68961251 | AX-170155851 | 0 | C | T | 0.184 | -5.99E-02 | 1.35E-01 | 2.66E+00 | 2.72E+00 | 6.54E-01 | no | | intergenic_variant | MODIFIER |
| 8 | 68962316 | AX-170302712 | 0 | C | T | 0.184 | -5.99E-02 | 1.35E-01 | 2.66E+00 | 2.72E+00 | 6.54E-01 | no | | intergenic_variant | MODIFIER |
| 8 | 68976408 | AX-170155973 | 0 | C | T | 0.184 | -5.99E-02 | 1.35E-01 | 2.66E+00 | 2.72E+00 | 6.54E-01 | no | Gm33103 | intron_variant,non_coding_transcript_variant | MODIFIER |
| 8 | 68988973 | AX-170148854 | 0 | T | C | 0.184 | -5.99E-02 | 1.35E-01 | 2.66E+00 | 2.72E+00 | 6.54E-01 | no | Gm33103 | intron_variant,non_coding_transcript_variant | MODIFIER |
| 8 | 69005121 | AX-170422431 | 0 | C | T | 0.184 | -5.99E-02 | 1.35E-01 | 2.66E+00 | 2.72E+00 | 6.54E-01 | no | Gm33103 | upstream_gene_variant | MODIFIER |
| 8 | 69005316 | AX-170167046 | 0 | G | A | 0.184 | -5.99E-02 | 1.35E-01 | 2.66E+00 | 2.72E+00 | 6.54E-01 | no | Gm33103 | upstream_gene_variant | MODIFIER |
| 8 | 69011385 | AX-170137713 | 0 | T | G | 0.184 | -5.99E-02 | 1.35E-01 | 2.66E+00 | 2.72E+00 | 6.54E-01 | no | | intergenic_variant | MODIFIER |
| 8 | 69016799 | AX-170096308 | 0 | T | A | 0.184 | -5.99E-02 | 1.35E-01 | 2.66E+00 | 2.72E+00 | 6.54E-01 | no | | regulatory_region_variant | MODIFIER |
| 8 | 69039770 | AX-170080340 | 0 | C | T | 0.184 | -5.99E-02 | 1.35E-01 | 2.66E+00 | 2.72E+00 | 6.54E-01 | no | Slc18a1 | intron_variant | MODIFIER |
| 8 | 69040622 | AX-170030978 | 0 | A | G | 0.184 | -5.99E-02 | 1.35E-01 | 2.66E+00 | 2.72E+00 | 6.54E-01 | no | Slc18a1 | synonymous_variant | LOW |
| 8 | 69066948 | AX-170030975 | 0 | T | C | 0.184 | -5.99E-02 | 1.35E-01 | 2.66E+00 | 2.72E+00 | 6.54E-01 | no | Slc18a1 | intron_variant | MODIFIER |
| 8 | 69068503 | AX-170304693 | 0 | G | T | 0.184 | -5.99E-02 | 1.35E-01 | 2.66E+00 | 2.72E+00 | 6.54E-01 | no | Slc18a1 | intron_variant | MODIFIER |
| 1 | 59792885 | AX-97825392 | 0 | T | G | 0.132 | 6.12E-02 | 1.38E-01 | 2.65E+00 | 2.75E+00 | 6.54E-01 | no | Bmpr2 | intron_variant | MODIFIER |
| 11 | 74083754 | AX-97837879 | 1 | G | A | 0.268 | 3.13E-02 | 7.17E-02 | 2.65E+00 | 2.79E+00 | 6.54E-01 | no | Zfp616 | 3_prime_UTR_variant | MODIFIER |
| 12 | 93428424 | AX-170162346 | 1 | T | C | 0.062 | 1.04E-01 | 2.36E-01 | 2.71E+00 | 2.77E+00 | 6.54E-01 | no | 4930559C10Rik | intron_variant,non_coding_transcript_variant | MODIFIER |
| 3 | 148562987 | AX-170021886 | 0 | G | A | 0.316 | 3.38E-02 | 7.61E-02 | 2.65E+00 | 2.73E+00 | 6.54E-01 | no | | intergenic_variant | MODIFIER |
| 5 | 92927172 | AX-170025616 | 1 | T | C | 0.42 | 4.81E-02 | 1.09E-01 | 2.69E+00 | 2.74E+00 | 6.54E-01 | no | Shroom3 | intron_variant | MODIFIER |
| 12 | 116949528 | AX-170268969 | 1 | G | T | 0.143 | -7.16E-02 | 1.62E-01 | 2.64E+00 | 2.74E+00 | 6.54E-01 | no | Ptprn2 | intron_variant | MODIFIER |

| | | | | | | | | | | | | | | | |
|---|---|---|---|---|---|---|---|---|---|---|---|---|---|---|---|
| 8 | 51458903 | AX-170296866 | 0 | C | T | 0.14 | -6.34E-02 | 1.36E-01 | 2.54E+00 | 2.62E+00 | 6.54E-01 | no | Gm17911 | downstream_gene_variant | MODIFIER |
| 1 | 107019009 | AX-170534310 | 0 | C | T | 0.149 | 8.51E-02 | 1.87E-01 | 2.57E+00 | 2.66E+00 | 6.54E-01 | no | Gm8080 | downstream_gene_variant | MODIFIER |
| 1 | 110441552 | AX-170001356 | 0 | A | G | 0.149 | 8.51E-02 | 1.87E-01 | 2.57E+00 | 2.66E+00 | 6.54E-01 | no | | intergenic_variant | MODIFIER |
| 1 | 114756557 | AX-170278101 | 0 | T | A | 0.149 | 8.51E-02 | 1.87E-01 | 2.57E+00 | 2.66E+00 | 6.54E-01 | no | | intergenic_variant | MODIFIER |
| 1 | 116074605 | AX-170352705 | 0 | A | G | 0.149 | 8.51E-02 | 1.87E-01 | 2.57E+00 | 2.66E+00 | 6.54E-01 | no | Cntnap5a | intron_variant | MODIFIER |
| 1 | 119237190 | AX-97782254 | 0 | T | C | 0.149 | 8.51E-02 | 1.87E-01 | 2.57E+00 | 2.66E+00 | 6.54E-01 | no | Gm3508 | downstream_gene_variant | MODIFIER |
| 5 | 109498160 | AX-170025867 | 2 | C | T | 0.464 | -4.70E-02 | 1.07E-01 | 2.79E+00 | 2.87E+00 | 6.54E-01 | no | | intergenic_variant | MODIFIER |
| 7 | 40123201 | AX-170186864 | 0 | C | T | 0.36 | 4.10E-02 | 9.08E-02 | 2.60E+00 | 2.68E+00 | 6.54E-01 | no | | intergenic_variant | MODIFIER |
| 7 | 40124166 | AX-170528532 | 0 | T | C | 0.36 | 4.10E-02 | 9.08E-02 | 2.60E+00 | 2.68E+00 | 6.54E-01 | no | | intergenic_variant | MODIFIER |
| 7 | 40146848 | AX-170507637 | 0 | G | A | 0.36 | 4.10E-02 | 9.08E-02 | 2.60E+00 | 2.68E+00 | 6.54E-01 | no | | intergenic_variant | MODIFIER |
| 7 | 40158929 | AX-170319877 | 0 | C | T | 0.36 | 4.10E-02 | 9.08E-02 | 2.60E+00 | 2.68E+00 | 6.54E-01 | no | | intergenic_variant | MODIFIER |
| 7 | 40159229 | AX-170162740 | 0 | T | C | 0.36 | 4.10E-02 | 9.08E-02 | 2.60E+00 | 2.68E+00 | 6.54E-01 | no | | intergenic_variant | MODIFIER |
| 7 | 40163840 | AX-170472570 | 0 | G | C | 0.36 | 4.10E-02 | 9.08E-02 | 2.60E+00 | 2.68E+00 | 6.54E-01 | no | | intergenic_variant | MODIFIER |
| 7 | 40195683 | AX-170486424 | 0 | G | A | 0.36 | 4.10E-02 | 9.08E-02 | 2.60E+00 | 2.68E+00 | 6.54E-01 | no | Gm6851 | downstream_gene_variant | MODIFIER |
| 7 | 40195947 | AX-170172512 | 0 | A | T | 0.36 | 4.10E-02 | 9.08E-02 | 2.60E+00 | 2.68E+00 | 6.54E-01 | no | Gm6851 | downstream_gene_variant | MODIFIER |
| 7 | 40196387 | AX-170077122 | 0 | C | T | 0.36 | 4.10E-02 | 9.08E-02 | 2.60E+00 | 2.68E+00 | 6.54E-01 | no | | intergenic_variant | MODIFIER |
| 7 | 40209630 | AX-170028877 | 0 | T | C | 0.36 | 4.10E-02 | 9.08E-02 | 2.60E+00 | 2.68E+00 | 6.54E-01 | no | | intergenic_variant | MODIFIER |
| 7 | 40209690 | AX-170077123 | 0 | T | A | 0.36 | 4.10E-02 | 9.08E-02 | 2.60E+00 | 2.68E+00 | 6.54E-01 | no | | intergenic_variant | MODIFIER |
| 7 | 40211289 | AX-97841908 | 0 | C | T | 0.36 | 4.10E-02 | 9.08E-02 | 2.60E+00 | 2.68E+00 | 6.54E-01 | no | | intergenic_variant | MODIFIER |
| 18 | 64838587 | AX-170482137 | 1 | A | T | 0.429 | 3.69E-02 | 8.35E-02 | 2.79E+00 | 2.86E+00 | 6.54E-01 | no | | intergenic_variant | MODIFIER |
| 9 | 36995437 | AX-170177698 | 0 | C | A | 0.491 | -3.44E-02 | 7.86E-02 | 2.71E+00 | 2.80E+00 | 6.54E-01 | no | Pknox2 | intron_variant | MODIFIER |
| 8 | 73376649 | AX-97837671 | 0 | T | C | 0.184 | -5.94E-02 | 1.34E-01 | 2.68E+00 | 2.75E+00 | 6.55E-01 | no | | intergenic_variant | MODIFIER |
| 15 | 79232301 | AX-170392196 | 0 | G | C | 0.561 | 3.67E-02 | 8.07E-02 | 2.58E+00 | 2.66E+00 | 6.55E-01 | no | Pick1 | intron_variant | MODIFIER |
| 5 | 100063159 | AX-170319386 | 0 | T | G | 0.412 | 4.45E-02 | 1.00E-01 | 2.66E+00 | 2.73E+00 | 6.55E-01 | no | Enoph1 | intron_variant | MODIFIER |
| 16 | 80374763 | AX-170057923 | 0 | C | T | 0.465 | 3.42E-02 | 7.53E-02 | 2.59E+00 | 2.66E+00 | 6.55E-01 | no | | intergenic_variant | MODIFIER |
| 5 | 39079012 | AX-97818848 | 0 | G | A | 0.211 | -4.53E-02 | 1.02E-01 | 2.61E+00 | 2.72E+00 | 6.55E-01 | no | | intergenic_variant | MODIFIER |
| 5 | 39382065 | AX-170104282 | 0 | G | C | 0.211 | -4.53E-02 | 1.02E-01 | 2.61E+00 | 2.72E+00 | 6.55E-01 | no | | intergenic_variant | MODIFIER |
| 15 | 81952968 | AX-97828014 | 0 | C | T | 0.561 | 3.68E-02 | 8.17E-02 | 2.60E+00 | 2.68E+00 | 6.55E-01 | no | Pmm1 | intron_variant | MODIFIER |
| 11 | 34705629 | AX-170376553 | 0 | G | C | 0.105 | 7.15E-02 | 1.61E-01 | 2.62E+00 | 2.73E+00 | 6.55E-01 | no | Dock2 | intron_variant | MODIFIER |
| 4 | 79927209 | AX-170454999 | 0 | A | G | 0.447 | -3.97E-02 | 8.55E-02 | 2.55E+00 | 2.61E+00 | 6.55E-01 | no | | intergenic_variant | MODIFIER |
| 5 | 124850307 | AX-97834072 | 0 | G | A | 0.456 | -5.02E-02 | 1.14E-01 | 2.78E+00 | 2.84E+00 | 6.55E-01 | no | Ccdc92 | intron_variant | MODIFIER |
| 10 | 17569800 | AX-170286465 | 1 | C | T | 0.196 | -4.50E-02 | 1.00E-01 | 2.61E+00 | 2.69E+00 | 6.55E-01 | no | Gm47771 | intron_variant,non_coding_transcript_variant | MODIFIER |
| 12 | 115396410 | AX-170050299 | 1 | C | G | 0.152 | 7.27E-02 | 1.65E-01 | 2.64E+00 | 2.75E+00 | 6.55E-01 | no | Gm19331 | intron_variant,non_coding_transcript_variant | MODIFIER |
| 8 | 37167450 | AX-170097663 | 1 | G | A | 0.152 | 6.52E-02 | 1.47E-01 | 2.79E+00 | 2.84E+00 | 6.55E-01 | no | Gm45524 | intron_variant,non_coding_transcript_variant | MODIFIER |
| 12 | 99674334 | AX-170087322 | 1 | C | T | 0.054 | 9.75E-02 | 2.22E-01 | 2.70E+00 | 2.76E+00 | 6.55E-01 | no | | intergenic_variant | MODIFIER |
| 6 | 147335009 | AX-170120101 | 0 | A | G | 0.079 | 9.20E-02 | 2.10E-01 | 2.69E+00 | 2.79E+00 | 6.55E-01 | no | | intergenic_variant | MODIFIER |
| 16 | 92013214 | AX-170192109 | 1 | T | G | 0.455 | -3.55E-02 | 8.07E-02 | 2.67E+00 | 2.75E+00 | 6.55E-01 | no | Gm18020 | upstream_gene_variant | MODIFIER |
| 8 | 102090448 | AX-170510929 | 1 | G | T | 0.214 | 4.76E-02 | 1.09E-01 | 2.69E+00 | 2.79E+00 | 6.55E-01 | no | | intergenic_variant | MODIFIER |
| 4 | 94524925 | AX-170122449 | 1 | C | G | 0.33 | 4.83E-02 | 1.08E-01 | 2.61E+00 | 2.68E+00 | 6.55E-01 | no | Caap1 | intron_variant | MODIFIER |
| 6 | 131408940 | AX-170076253 | 1 | T | A | 0.545 | 3.79E-02 | 8.17E-02 | 2.54E+00 | 2.63E+00 | 6.55E-01 | no | | intergenic_variant | MODIFIER |
| 12 | 109829746 | AX-170366298 | 1 | A | G | 0.125 | -7.92E-02 | 1.81E-01 | 2.68E+00 | 2.78E+00 | 6.55E-01 | no | Gm9661 | downstream_gene_variant | MODIFIER |
| 6 | 124316051 | AX-97806654 | 1 | G | C | 0.188 | -3.60E-02 | 8.28E-02 | 2.66E+00 | 2.79E+00 | 6.56E-01 | no | Cd163 | intron_variant | MODIFIER |
| 3 | 32320599 | AX-97846655 | 0 | C | T | 0.167 | -4.23E-02 | 9.61E-02 | 2.67E+00 | 2.75E+00 | 6.56E-01 | no | | intergenic_variant | MODIFIER |
| 13 | 47212960 | AX-170181464 | 1 | C | G | 0.482 | -2.78E-02 | 6.41E-02 | 2.72E+00 | 2.83E+00 | 6.56E-01 | no | Rnf144b | intron_variant | MODIFIER |
| 12 | 92948065 | AX-170389460 | 0 | A | G | 0.07 | 8.50E-02 | 1.93E-01 | 2.65E+00 | 2.75E+00 | 6.56E-01 | no | | intergenic_variant | MODIFIER |
| 1 | 151446833 | AX-97818000 | 0 | C | T | 0.211 | -5.51E-02 | 1.26E-01 | 2.76E+00 | 2.84E+00 | 6.56E-01 | no | Trmt1l | intron_variant | MODIFIER |
| 13 | 22220923 | AX-97800579 | 1 | G | T | 0.161 | -5.95E-02 | 1.26E-01 | 2.52E+00 | 2.58E+00 | 6.56E-01 | no | Vmn1r193 | upstream_gene_variant | MODIFIER |
| 6 | 146864029 | AX-170445321 | 0 | G | A | 0.079 | 9.25E-02 | 2.11E-01 | 2.66E+00 | 2.75E+00 | 6.56E-01 | no | Smco2 | intron_variant | MODIFIER |
| 8 | 28360955 | AX-170452547 | 1 | T | C | 0.17 | 5.16E-02 | 1.17E-01 | 2.70E+00 | 2.76E+00 | 6.56E-01 | no | | intergenic_variant | MODIFIER |
| 1 | 127640320 | AX-170170107 | 0 | G | A | 0.544 | 3.67E-02 | 8.30E-02 | 2.66E+00 | 2.73E+00 | 6.56E-01 | no | Tmem163 | intron_variant,NMD_transcript_variant | MODIFIER |
| 13 | 21936866 | AX-97820590 | 0 | T | C | 0.158 | -5.95E-02 | 1.25E-01 | 2.51E+00 | 2.56E+00 | 6.56E-01 | no | Gm11280 | downstream_gene_variant | MODIFIER |
| 13 | 21940517 | AX-170475886 | 0 | C | G | 0.158 | -5.95E-02 | 1.25E-01 | 2.51E+00 | 2.56E+00 | 6.56E-01 | no | Zfp184 | upstream_gene_variant | MODIFIER |
| 13 | 21957867 | AX-97806762 | 0 | C | T | 0.158 | -5.95E-02 | 1.25E-01 | 2.51E+00 | 2.56E+00 | 6.56E-01 | no | Zfp184 | intron_variant | MODIFIER |
| 13 | 22019787 | AX-97850545 | 0 | C | T | 0.158 | -5.95E-02 | 1.25E-01 | 2.51E+00 | 2.56E+00 | 6.56E-01 | no | Gm11290 | downstream_gene_variant | MODIFIER |

| | | | | | | | | | | | | | | | |
|---|---|---|---|---|---|---|---|---|---|---|---|---|---|---|---|
| 13 | 22052474 | AX-97794393 | 0 | T | C | 0.158 | -5.95E-02 | 1.25E-01 | 2.51E+00 | 2.56E+00 | 6.56E-01 | no | Vmn1r-ps92 | downstream_gene_variant | MODIFIER |
| 13 | 22064243 | AX-97856532 | 0 | C | T | 0.158 | -5.95E-02 | 1.25E-01 | 2.51E+00 | 2.56E+00 | 6.56E-01 | no | | intergenic_variant | MODIFIER |
| 13 | 22096626 | AX-170524109 | 0 | G | T | 0.158 | -5.95E-02 | 1.25E-01 | 2.51E+00 | 2.56E+00 | 6.56E-01 | no | Vmn1r-ps94 | downstream_gene_variant | MODIFIER |
| 13 | 22128833 | AX-170372124 | 0 | C | T | 0.158 | -5.95E-02 | 1.25E-01 | 2.51E+00 | 2.56E+00 | 6.56E-01 | no | Vmn1r-ps95 | downstream_gene_variant | MODIFIER |
| 13 | 22140410 | AX-97815907 | 0 | C | T | 0.158 | -5.95E-02 | 1.25E-01 | 2.51E+00 | 2.56E+00 | 6.56E-01 | no | Vmn1r-ps97 | upstream_gene_variant | MODIFIER |
| 13 | 22224441 | AX-170007907 | 0 | T | C | 0.158 | -5.95E-02 | 1.25E-01 | 2.51E+00 | 2.56E+00 | 6.56E-01 | no | Vmn1r193 | upstream_gene_variant | MODIFIER |
| 13 | 22236108 | AX-170007908 | 0 | G | A | 0.158 | -5.95E-02 | 1.25E-01 | 2.51E+00 | 2.56E+00 | 6.56E-01 | no | | intergenic_variant | MODIFIER |
| 13 | 22245808 | AX-170050727 | 0 | C | G | 0.158 | -5.95E-02 | 1.25E-01 | 2.51E+00 | 2.56E+00 | 6.56E-01 | no | Vmn1r194 | downstream_gene_variant | MODIFIER |
| 13 | 22253995 | AX-170050730 | 0 | T | C | 0.158 | -5.95E-02 | 1.25E-01 | 2.51E+00 | 2.56E+00 | 6.56E-01 | no | | intergenic_variant | MODIFIER |
| 13 | 22256155 | AX-170007910 | 0 | C | T | 0.158 | -5.95E-02 | 1.25E-01 | 2.51E+00 | 2.56E+00 | 6.56E-01 | no | | intergenic_variant | MODIFIER |
| 13 | 22276815 | AX-170007902 | 0 | G | T | 0.158 | -5.95E-02 | 1.25E-01 | 2.51E+00 | 2.56E+00 | 6.56E-01 | no | Vmn1r195 | upstream_gene_variant | MODIFIER |
| 13 | 22283412 | AX-170007906 | 0 | A | G | 0.158 | -5.95E-02 | 1.25E-01 | 2.51E+00 | 2.56E+00 | 6.56E-01 | no | Vmn1r195 | downstream_gene_variant | MODIFIER |
| 13 | 22293617 | AX-97829944 | 0 | T | C | 0.158 | -5.95E-02 | 1.25E-01 | 2.51E+00 | 2.56E+00 | 6.56E-01 | no | Vmn1r196 | missense_variant | MODERATE |
| 13 | 23760204 | AX-170050753 | 0 | T | C | 0.158 | -5.95E-02 | 1.25E-01 | 2.51E+00 | 2.56E+00 | 6.56E-01 | no | Hist1h1a | upstream_gene_variant | MODIFIER |
| 13 | 23782838 | AX-170147483 | 0 | G | A | 0.158 | -5.95E-02 | 1.25E-01 | 2.51E+00 | 2.56E+00 | 6.56E-01 | no | Trim38 | synonymous_variant | LOW |
| 13 | 23783643 | AX-170464715 | 0 | A | G | 0.158 | -5.95E-02 | 1.25E-01 | 2.51E+00 | 2.56E+00 | 6.56E-01 | no | Trim38 | intron_variant | MODIFIER |
| 13 | 23783784 | AX-97817610 | 0 | C | T | 0.158 | -5.95E-02 | 1.25E-01 | 2.51E+00 | 2.56E+00 | 6.56E-01 | no | Trim38 | intron_variant | MODIFIER |
| 13 | 23784404 | AX-170530300 | 0 | A | G | 0.158 | -5.95E-02 | 1.25E-01 | 2.51E+00 | 2.56E+00 | 6.56E-01 | no | Trim38 | intron_variant | MODIFIER |
| 13 | 23798230 | AX-170198694 | 0 | T | C | 0.158 | -5.95E-02 | 1.25E-01 | 2.51E+00 | 2.56E+00 | 6.56E-01 | no | Gm11337 | intron_variant,non_coding_transcript_variant | MODIFIER |
| 13 | 23811285 | AX-170221192 | 0 | G | A | 0.158 | -5.95E-02 | 1.25E-01 | 2.51E+00 | 2.56E+00 | 6.56E-01 | no | Slc17a2 | intron_variant | MODIFIER |
| 13 | 23819633 | AX-97844808 | 0 | C | T | 0.158 | -5.95E-02 | 1.25E-01 | 2.51E+00 | 2.56E+00 | 6.56E-01 | no | Slc17a2 | intron_variant | MODIFIER |
| 13 | 23827833 | AX-170166681 | 0 | A | G | 0.158 | -5.95E-02 | 1.25E-01 | 2.51E+00 | 2.56E+00 | 6.56E-01 | no | Slc17a2 | downstream_gene_variant | MODIFIER |
| 13 | 23828006 | AX-170202663 | 0 | A | G | 0.158 | -5.95E-02 | 1.25E-01 | 2.51E+00 | 2.56E+00 | 6.56E-01 | no | Slc17a2 | downstream_gene_variant | MODIFIER |
| 13 | 23858856 | AX-97830074 | 0 | A | G | 0.158 | -5.95E-02 | 1.25E-01 | 2.51E+00 | 2.56E+00 | 6.56E-01 | no | Slc17a3 | 3_prime_UTR_variant | MODIFIER |
| 13 | 23860129 | AX-170393870 | 0 | A | T | 0.158 | -5.95E-02 | 1.25E-01 | 2.51E+00 | 2.56E+00 | 6.56E-01 | no | Slc17a3 | 3_prime_UTR_variant | MODIFIER |
| 13 | 23873142 | AX-170194484 | 0 | C | T | 0.158 | -5.95E-02 | 1.25E-01 | 2.51E+00 | 2.56E+00 | 6.56E-01 | no | Slc17a1 | intron_variant | MODIFIER |
| 13 | 23888574 | AX-170491580 | 0 | C | G | 0.158 | -5.95E-02 | 1.25E-01 | 2.51E+00 | 2.56E+00 | 6.56E-01 | no | Slc17a1 | intron_variant | MODIFIER |
| 13 | 23891844 | AX-97783833 | 0 | T | C | 0.158 | -5.95E-02 | 1.25E-01 | 2.51E+00 | 2.56E+00 | 6.56E-01 | no | Slc17a1 | intron_variant | MODIFIER |
| 13 | 23905200 | AX-170330955 | 0 | T | C | 0.158 | -5.95E-02 | 1.25E-01 | 2.51E+00 | 2.56E+00 | 6.56E-01 | no | Slc17a4 | intron_variant | MODIFIER |
| 13 | 23913369 | AX-170391238 | 0 | A | G | 0.158 | -5.95E-02 | 1.25E-01 | 2.51E+00 | 2.56E+00 | 6.56E-01 | no | Slc17a4 | intron_variant | MODIFIER |
| 13 | 23913405 | AX-170188266 | 0 | C | G | 0.158 | -5.95E-02 | 1.25E-01 | 2.51E+00 | 2.56E+00 | 6.56E-01 | no | Slc17a4 | intron_variant | MODIFIER |
| 13 | 23924273 | AX-97852183 | 0 | G | A | 0.158 | -5.95E-02 | 1.25E-01 | 2.51E+00 | 2.56E+00 | 6.56E-01 | no | Snrp1c-ps2 | upstream_gene_variant | MODIFIER |
| 13 | 23928198 | AX-170467525 | 0 | A | C | 0.158 | -5.95E-02 | 1.25E-01 | 2.51E+00 | 2.56E+00 | 6.56E-01 | no | Snrp1c-ps2 | upstream_gene_variant | MODIFIER |
| 13 | 23931535 | AX-170251899 | 0 | T | C | 0.158 | -5.95E-02 | 1.25E-01 | 2.51E+00 | 2.56E+00 | 6.56E-01 | no | Hist1h2ba | downstream_gene_variant | MODIFIER |
| 13 | 23932021 | AX-170263258 | 0 | C | T | 0.158 | -5.95E-02 | 1.25E-01 | 2.51E+00 | 2.56E+00 | 6.56E-01 | no | Hist1h2ba | downstream_gene_variant | MODIFIER |
| 13 | 23932133 | AX-170147152 | 0 | T | C | 0.158 | -5.95E-02 | 1.25E-01 | 2.51E+00 | 2.56E+00 | 6.56E-01 | no | Hist1h2ba | downstream_gene_variant | MODIFIER |
| 13 | 23945390 | AX-97787495 | 0 | G | A | 0.158 | -5.95E-02 | 1.25E-01 | 2.51E+00 | 2.56E+00 | 6.56E-01 | no | | intergenic_variant | MODIFIER |
| 13 | 23969135 | AX-97856711 | 0 | T | G | 0.158 | -5.95E-02 | 1.25E-01 | 2.51E+00 | 2.56E+00 | 6.56E-01 | no | Scgn | intron_variant | MODIFIER |
| 13 | 23971599 | AX-170288346 | 0 | C | T | 0.158 | -5.95E-02 | 1.25E-01 | 2.51E+00 | 2.56E+00 | 6.56E-01 | no | Scgn | intron_variant | MODIFIER |
| 13 | 23976217 | AX-170495708 | 0 | T | C | 0.158 | -5.95E-02 | 1.25E-01 | 2.51E+00 | 2.56E+00 | 6.56E-01 | no | Scgn | intron_variant | MODIFIER |
| 13 | 23981610 | AX-170318196 | 0 | C | T | 0.158 | -5.95E-02 | 1.25E-01 | 2.51E+00 | 2.56E+00 | 6.56E-01 | no | Scgn | intron_variant | MODIFIER |
| 13 | 24009429 | AX-170433449 | 0 | G | A | 0.158 | -5.95E-02 | 1.25E-01 | 2.51E+00 | 2.56E+00 | 6.56E-01 | no | Carmil1 | downstream_gene_variant | MODIFIER |
| 13 | 24011086 | AX-170335907 | 0 | T | C | 0.158 | -5.95E-02 | 1.25E-01 | 2.51E+00 | 2.56E+00 | 6.56E-01 | no | Carmil1 | downstream_gene_variant | MODIFIER |
| 13 | 24013126 | AX-170438680 | 0 | G | C | 0.158 | -5.95E-02 | 1.25E-01 | 2.51E+00 | 2.56E+00 | 6.56E-01 | no | Carmil1 | 3_prime_UTR_variant | MODIFIER |
| 13 | 24018392 | AX-170050759 | 0 | G | C | 0.158 | -5.95E-02 | 1.25E-01 | 2.51E+00 | 2.56E+00 | 6.56E-01 | no | Carmil1 | intron_variant | MODIFIER |
| 13 | 24039021 | AX-170050761 | 0 | G | T | 0.158 | -5.95E-02 | 1.25E-01 | 2.51E+00 | 2.56E+00 | 6.56E-01 | no | Carmil1 | intron_variant | MODIFIER |
| 13 | 24048935 | AX-170474399 | 0 | A | G | 0.158 | -5.95E-02 | 1.25E-01 | 2.51E+00 | 2.56E+00 | 6.56E-01 | no | Carmil1 | intron_variant | MODIFIER |
| 13 | 24094160 | AX-97823105 | 0 | T | C | 0.158 | -5.95E-02 | 1.25E-01 | 2.51E+00 | 2.56E+00 | 6.56E-01 | no | Carmil1 | intron_variant | MODIFIER |
| 13 | 24102680 | AX-170120408 | 0 | A | G | 0.158 | -5.95E-02 | 1.25E-01 | 2.51E+00 | 2.56E+00 | 6.56E-01 | no | Carmil1 | intron_variant | MODIFIER |
| 13 | 24110457 | AX-170347150 | 0 | A | G | 0.158 | -5.95E-02 | 1.25E-01 | 2.51E+00 | 2.56E+00 | 6.56E-01 | no | Carmil1 | intron_variant | MODIFIER |
| 13 | 24120277 | AX-97817637 | 0 | A | G | 0.158 | -5.95E-02 | 1.25E-01 | 2.51E+00 | 2.56E+00 | 6.56E-01 | no | Carmil1 | intron_variant | MODIFIER |
| 13 | 24133154 | AX-97780632 | 0 | T | G | 0.158 | -5.95E-02 | 1.25E-01 | 2.51E+00 | 2.56E+00 | 6.56E-01 | no | Carmil1 | intron_variant | MODIFIER |
| 13 | 24134410 | AX-170280836 | 0 | A | G | 0.158 | -5.95E-02 | 1.25E-01 | 2.51E+00 | 2.56E+00 | 6.56E-01 | no | Carmil1 | intron_variant | MODIFIER |
| 4 | 137353683 | AX-170143541 | 0 | C | T | 0.158 | -4.96E-02 | 1.14E-01 | 2.81E+00 | 2.93E+00 | 6.56E-01 | no | Cdc42 | intron_variant | MODIFIER |

| | | | | | | | | | | | | | | | |
|---|---|---|---|---|---|---|---|---|---|---|---|---|---|---|---|
| 7 | 6886612 | AX-170456133 | 0 | T | A | 0.167 | -3.66E-02 | 8.02E-02 | 2.53E+00 | 2.67E+00 | 6.56E-01 | no | Usp29 | intron_variant | MODIFIER |
| 7 | 7857490 | AX-170223921 | 0 | T | C | 0.167 | -3.66E-02 | 8.02E-02 | 2.53E+00 | 2.67E+00 | 6.56E-01 | no | Vmn2r-ps40 | intron_variant,non_coding_transcript_variant | MODIFIER |
| 7 | 11079313 | AX-170099378 | 0 | C | G | 0.167 | -3.66E-02 | 8.02E-02 | 2.53E+00 | 2.67E+00 | 6.56E-01 | no | Gm33601 | downstream_gene_variant | MODIFIER |
| 7 | 11083232 | AX-97791283 | 0 | G | T | 0.167 | -3.66E-02 | 8.02E-02 | 2.53E+00 | 2.67E+00 | 6.56E-01 | no | Gm7151 | intron_variant,non_coding_transcript_variant | MODIFIER |
| 7 | 11460727 | AX-170028584 | 0 | T | C | 0.167 | -3.66E-02 | 8.02E-02 | 2.53E+00 | 2.67E+00 | 6.56E-01 | no | Gm7207 | downstream_gene_variant | MODIFIER |
| 7 | 70581099 | AX-170509802 | 1 | A | C | 0.366 | -4.41E-02 | 1.01E-01 | 2.75E+00 | 2.84E+00 | 6.56E-01 | no | Gm44812 | intron_variant,non_coding_transcript_variant | MODIFIER |
| 5 | 119233291 | AX-170025992 | 0 | C | T | 0.447 | 4.69E-02 | 1.06E-01 | 2.66E+00 | 2.72E+00 | 6.56E-01 | no | | intergenic_variant | MODIFIER |
| 9 | 70980125 | AX-170124117 | 0 | T | A | 0.211 | -5.44E-02 | 1.23E-01 | 2.65E+00 | 2.71E+00 | 6.56E-01 | no | | intergenic_variant | MODIFIER |
| 2 | 87955684 | AX-170195379 | 1 | C | G | 0.054 | -1.33E-01 | 3.02E-01 | 2.75E+00 | 2.78E+00 | 6.56E-01 | no | Olfr1157 | downstream_gene_variant | MODIFIER |
| 5 | 125140064 | AX-170175198 | 0 | C | T | 0.456 | 4.76E-02 | 1.08E-01 | 2.66E+00 | 2.72E+00 | 6.56E-01 | no | Ncor2 | intron_variant | MODIFIER |
| 1 | 116788282 | AX-170001487 | 1 | G | A | 0.17 | 8.98E-02 | 2.00E-01 | 2.60E+00 | 2.68E+00 | 6.56E-01 | no | | intergenic_variant | MODIFIER |
| 12 | 99668393 | AX-170253990 | 1 | A | G | 0.054 | 9.71E-02 | 2.22E-01 | 2.70E+00 | 2.76E+00 | 6.57E-01 | no | | intergenic_variant | MODIFIER |
| 9 | 37235136 | AX-170201599 | 0 | A | G | 0.482 | -3.35E-02 | 7.69E-02 | 2.69E+00 | 2.78E+00 | 6.57E-01 | no | Slc37a2 | synonymous_variant | LOW |
| 9 | 38428394 | AX-170190952 | 0 | G | A | 0.482 | -3.35E-02 | 7.69E-02 | 2.69E+00 | 2.78E+00 | 6.57E-01 | no | Olfr901 | upstream_gene_variant | MODIFIER |
| 5 | 107071340 | AX-97799038 | 0 | G | A | 0.456 | 5.00E-02 | 1.13E-01 | 2.66E+00 | 2.71E+00 | 6.57E-01 | no | | intergenic_variant | MODIFIER |
| 5 | 112102788 | AX-97817831 | 0 | G | A | 0.456 | 5.00E-02 | 1.13E-01 | 2.66E+00 | 2.71E+00 | 6.57E-01 | no | | regulatory_region_variant | MODIFIER |
| 5 | 113378446 | AX-170251136 | 0 | C | T | 0.456 | 5.00E-02 | 1.13E-01 | 2.66E+00 | 2.71E+00 | 6.57E-01 | no | | intergenic_variant | MODIFIER |
| 13 | 32381079 | AX-170129931 | 1 | A | G | 0.179 | -5.39E-02 | 1.20E-01 | 2.59E+00 | 2.65E+00 | 6.57E-01 | no | Gm11381 | intron_variant,non_coding_transcript_variant | MODIFIER |
| 13 | 35965049 | AX-97794523 | 1 | A | G | 0.179 | -5.39E-02 | 1.20E-01 | 2.59E+00 | 2.65E+00 | 6.57E-01 | no | Ppp1r3g | upstream_gene_variant | MODIFIER |
| 9 | 77305551 | AX-170429924 | 1 | C | G | 0.196 | -5.70E-02 | 1.29E-01 | 2.66E+00 | 2.72E+00 | 6.57E-01 | no | Mlip | intron_variant | MODIFIER |
| 20 | 50713729 | AX-170210347 | 0 | T | C | 0.237 | 2.54E-02 | 5.73E-02 | 2.59E+00 | 2.72E+00 | 6.57E-01 | no | | intergenic_variant | MODIFIER |
| 12 | 90492705 | AX-170219317 | 0 | G | A | 0.07 | 7.55E-02 | 1.74E-01 | 2.67E+00 | 2.78E+00 | 6.57E-01 | no | Gm47686 | downstream_gene_variant | MODIFIER |
| 17 | 54397337 | AX-170441303 | 1 | T | C | 0.036 | -9.30E-02 | 2.01E-01 | 2.52E+00 | 2.63E+00 | 6.57E-01 | no | | intergenic_variant | MODIFIER |
| 3 | 30429620 | AX-170275579 | 1 | C | T | 0.161 | -4.37E-02 | 9.98E-02 | 2.68E+00 | 2.75E+00 | 6.57E-01 | no | Mecom | intron_variant | MODIFIER |
| 2 | 159649936 | AX-170065427 | 0 | A | G | 0.14 | 8.34E-02 | 1.90E-01 | 2.68E+00 | 2.74E+00 | 6.57E-01 | no | | intergenic_variant | MODIFIER |
| 3 | 155496109 | AX-170392701 | 0 | G | T | 0.325 | 3.25E-02 | 7.47E-02 | 2.70E+00 | 2.78E+00 | 6.57E-01 | no | | intergenic_variant | MODIFIER |
| 3 | 155497074 | AX-170412517 | 0 | A | G | 0.325 | 3.25E-02 | 7.47E-02 | 2.70E+00 | 2.78E+00 | 6.57E-01 | no | | intergenic_variant | MODIFIER |
| 3 | 155523393 | AX-170021989 | 0 | G | A | 0.325 | 3.25E-02 | 7.47E-02 | 2.70E+00 | 2.78E+00 | 6.57E-01 | no | | intergenic_variant | MODIFIER |
| 3 | 155547485 | AX-170353716 | 0 | G | C | 0.325 | 3.25E-02 | 7.47E-02 | 2.70E+00 | 2.78E+00 | 6.57E-01 | no | | intergenic_variant | MODIFIER |
| 3 | 155578054 | AX-170127008 | 0 | C | T | 0.325 | 3.25E-02 | 7.47E-02 | 2.70E+00 | 2.78E+00 | 6.57E-01 | no | | intergenic_variant | MODIFIER |
| 3 | 155605687 | AX-170346231 | 0 | C | T | 0.325 | 3.25E-02 | 7.47E-02 | 2.70E+00 | 2.78E+00 | 6.57E-01 | no | | intergenic_variant | MODIFIER |
| 3 | 155659154 | AX-170401586 | 0 | T | C | 0.325 | 3.25E-02 | 7.47E-02 | 2.70E+00 | 2.78E+00 | 6.57E-01 | no | | intergenic_variant | MODIFIER |
| 3 | 155725727 | AX-170068529 | 0 | C | G | 0.325 | 3.25E-02 | 7.47E-02 | 2.70E+00 | 2.78E+00 | 6.57E-01 | no | | intergenic_variant | MODIFIER |
| 3 | 155864665 | AX-170471969 | 0 | C | T | 0.325 | 3.25E-02 | 7.47E-02 | 2.70E+00 | 2.78E+00 | 6.57E-01 | no | | intergenic_variant | MODIFIER |
| 3 | 155944154 | AX-170089124 | 0 | C | G | 0.325 | 3.25E-02 | 7.47E-02 | 2.70E+00 | 2.78E+00 | 6.57E-01 | no | | intergenic_variant | MODIFIER |
| 3 | 155966019 | AX-97791583 | 0 | T | C | 0.325 | 3.25E-02 | 7.47E-02 | 2.70E+00 | 2.78E+00 | 6.57E-01 | no | | intergenic_variant | MODIFIER |
| 3 | 155984335 | AX-170021999 | 0 | C | T | 0.325 | 3.25E-02 | 7.47E-02 | 2.70E+00 | 2.78E+00 | 6.57E-01 | no | 9330178D15Rik | intron_variant,non_coding_transcript_variant | MODIFIER |
| 3 | 155988967 | AX-170022002 | 0 | A | G | 0.325 | 3.25E-02 | 7.47E-02 | 2.70E+00 | 2.78E+00 | 6.57E-01 | no | 9330178D15Rik | intron_variant,non_coding_transcript_variant | MODIFIER |
| 3 | 155995361 | AX-97832507 | 0 | C | T | 0.325 | 3.25E-02 | 7.47E-02 | 2.70E+00 | 2.78E+00 | 6.57E-01 | no | 9330178D15Rik | intron_variant,non_coding_transcript_variant | MODIFIER |
| 3 | 156003138 | AX-170021995 | 0 | T | C | 0.325 | 3.25E-02 | 7.47E-02 | 2.70E+00 | 2.78E+00 | 6.57E-01 | no | 9330178D15Rik | intron_variant,non_coding_transcript_variant | MODIFIER |
| 3 | 156026863 | AX-170117103 | 0 | C | T | 0.325 | 3.25E-02 | 7.47E-02 | 2.70E+00 | 2.78E+00 | 6.57E-01 | no | | intergenic_variant | MODIFIER |
| 3 | 156027478 | AX-97857508 | 0 | C | T | 0.325 | 3.25E-02 | 7.47E-02 | 2.70E+00 | 2.78E+00 | 6.57E-01 | no | | intergenic_variant | MODIFIER |
| 3 | 156028449 | AX-170378327 | 0 | G | A | 0.325 | 3.25E-02 | 7.47E-02 | 2.70E+00 | 2.78E+00 | 6.57E-01 | no | | regulatory_region_variant | MODIFIER |
| 3 | 156028589 | AX-170320805 | 0 | T | C | 0.325 | 3.25E-02 | 7.47E-02 | 2.70E+00 | 2.78E+00 | 6.57E-01 | no | | regulatory_region_variant | MODIFIER |
| 3 | 156030714 | AX-170345937 | 0 | T | C | 0.325 | 3.25E-02 | 7.47E-02 | 2.70E+00 | 2.78E+00 | 6.57E-01 | no | | intergenic_variant | MODIFIER |
| 3 | 156042571 | AX-170022003 | 0 | A | G | 0.325 | 3.25E-02 | 7.47E-02 | 2.70E+00 | 2.78E+00 | 6.57E-01 | no | Gm42947 | intron_variant,non_coding_transcript_variant | MODIFIER |
| 3 | 156072682 | AX-170182366 | 0 | C | G | 0.325 | 3.25E-02 | 7.47E-02 | 2.70E+00 | 2.78E+00 | 6.57E-01 | no | | intergenic_variant | MODIFIER |
| 3 | 156080252 | AX-170132193 | 0 | C | G | 0.325 | 3.25E-02 | 7.47E-02 | 2.70E+00 | 2.78E+00 | 6.57E-01 | no | | intergenic_variant | MODIFIER |
| 3 | 156080375 | AX-170506399 | 0 | T | A | 0.325 | 3.25E-02 | 7.47E-02 | 2.70E+00 | 2.78E+00 | 6.57E-01 | no | | intergenic_variant | MODIFIER |
| 3 | 156085923 | AX-170022001 | 0 | G | A | 0.325 | 3.25E-02 | 7.47E-02 | 2.70E+00 | 2.78E+00 | 6.57E-01 | no | | intergenic_variant | MODIFIER |
| 3 | 156087968 | AX-170089125 | 0 | G | A | 0.325 | 3.25E-02 | 7.47E-02 | 2.70E+00 | 2.78E+00 | 6.57E-01 | no | | intergenic_variant | MODIFIER |
| 3 | 156105858 | AX-170474754 | 0 | C | T | 0.325 | 3.25E-02 | 7.47E-02 | 2.70E+00 | 2.78E+00 | 6.57E-01 | no | | intergenic_variant | MODIFIER |
| 8 | 68738803 | AX-97782492 | 1 | C | G | 0.188 | -5.85E-02 | 1.33E-01 | 2.67E+00 | 2.74E+00 | 6.57E-01 | no | Csgalnact1 | upstream_gene_variant | MODIFIER |
| 8 | 68809759 | AX-170472017 | 1 | T | C | 0.188 | -5.85E-02 | 1.33E-01 | 2.67E+00 | 2.74E+00 | 6.57E-01 | no | Ints10 | intron_variant | MODIFIER |

| | | | | | | | | | | | | | | | |
|---|---|---|---|---|---|---|---|---|---|---|---|---|---|---|---|
| 8 | 69016488 | AX-170519114 | 1 | G | C | 0.188 | -5.85E-02 | 1.33E-01 | 2.67E+00 | 2.74E+00 | 6.57E-01 | no | | intergenic_variant | MODIFIER |
| 1 | 43393589 | AX-170145409 | 1 | A | C | 0.098 | -9.15E-02 | 2.10E-01 | 2.69E+00 | 2.78E+00 | 6.57E-01 | no | | intergenic_variant | MODIFIER |
| 20 | 75711517 | AX-170459064 | 1 | G | T | 0.259 | 2.67E-02 | 6.11E-02 | 2.65E+00 | 2.75E+00 | 6.58E-01 | no | 4933407K13Rik | intron_variant,non_coding_transcript_variant | MODIFIER |
| 13 | 38793870 | AX-170331970 | 0 | A | G | 0.175 | -5.07E-02 | 1.09E-01 | 2.52E+00 | 2.61E+00 | 6.58E-01 | no | Gm47509 | upstream_gene_variant | MODIFIER |
| 8 | 42447627 | AX-170525178 | 1 | C | T | 0.152 | 6.49E-02 | 1.48E-01 | 2.78E+00 | 2.82E+00 | 6.58E-01 | no | | intergenic_variant | MODIFIER |
| 5 | 104505142 | AX-170519682 | 0 | T | A | 0.474 | 4.27E-02 | 9.42E-02 | 2.56E+00 | 2.66E+00 | 6.58E-01 | no | Pkd2 | 3_prime_UTR_variant | MODIFIER |
| 1 | 30828910 | AX-97787326 | 0 | G | A | 0.404 | 3.57E-02 | 8.12E-02 | 2.65E+00 | 2.73E+00 | 6.58E-01 | no | Phf3 | intron_variant | MODIFIER |
| 4 | 101511133 | AX-170428152 | 1 | C | T | 0.357 | 4.33E-02 | 9.82E-02 | 2.63E+00 | 2.72E+00 | 6.58E-01 | no | Dnajc6 | intron_variant | MODIFIER |
| 8 | 14564779 | AX-170325439 | 1 | G | C | 0.25 | -3.35E-02 | 7.45E-02 | 2.57E+00 | 2.67E+00 | 6.58E-01 | no | Dlgap2 | intron_variant | MODIFIER |
| 9 | 34939799 | AX-170330401 | 0 | G | A | 0.439 | 3.80E-02 | 8.47E-02 | 2.60E+00 | 2.67E+00 | 6.58E-01 | no | Kirrel3 | intron_variant | MODIFIER |
| 1 | 161363497 | AX-170509422 | 0 | T | C | 0.211 | -5.82E-02 | 1.34E-01 | 2.81E+00 | 2.90E+00 | 6.58E-01 | no | | intergenic_variant | MODIFIER |
| 8 | 94968136 | AX-170517633 | 0 | T | C | 0.219 | 5.16E-02 | 1.18E-01 | 2.70E+00 | 2.76E+00 | 6.58E-01 | no | Gm10286 | non_coding_transcript_exon_variant | MODIFIER |
| 8 | 96014039 | AX-170103680 | 0 | G | A | 0.219 | 5.16E-02 | 1.18E-01 | 2.70E+00 | 2.76E+00 | 6.58E-01 | no | | intergenic_variant | MODIFIER |
| 8 | 96034379 | AX-170124853 | 0 | T | C | 0.219 | 5.16E-02 | 1.18E-01 | 2.70E+00 | 2.76E+00 | 6.58E-01 | no | | intergenic_variant | MODIFIER |
| 8 | 96039380 | AX-170218143 | 0 | A | G | 0.219 | 5.16E-02 | 1.18E-01 | 2.70E+00 | 2.76E+00 | 6.58E-01 | no | | intergenic_variant | MODIFIER |
| 8 | 96049161 | AX-97811260 | 0 | C | G | 0.219 | 5.16E-02 | 1.18E-01 | 2.70E+00 | 2.76E+00 | 6.58E-01 | no | | intergenic_variant | MODIFIER |
| 8 | 96074557 | AX-170460668 | 0 | G | A | 0.219 | 5.16E-02 | 1.18E-01 | 2.70E+00 | 2.76E+00 | 6.58E-01 | no | | intergenic_variant | MODIFIER |
| 8 | 96103711 | AX-97805929 | 0 | C | T | 0.219 | 5.16E-02 | 1.18E-01 | 2.70E+00 | 2.76E+00 | 6.58E-01 | no | | intergenic_variant | MODIFIER |
| 8 | 96123175 | AX-170126842 | 0 | T | C | 0.219 | 5.16E-02 | 1.18E-01 | 2.70E+00 | 2.76E+00 | 6.58E-01 | no | | intergenic_variant | MODIFIER |
| 8 | 96129407 | AX-170308982 | 0 | C | G | 0.219 | 5.16E-02 | 1.18E-01 | 2.70E+00 | 2.76E+00 | 6.58E-01 | no | | intergenic_variant | MODIFIER |
| 8 | 96130172 | AX-170201867 | 0 | C | G | 0.219 | 5.16E-02 | 1.18E-01 | 2.70E+00 | 2.76E+00 | 6.58E-01 | no | | intergenic_variant | MODIFIER |
| 8 | 96131373 | AX-170418757 | 0 | G | A | 0.219 | 5.16E-02 | 1.18E-01 | 2.70E+00 | 2.76E+00 | 6.58E-01 | no | | intergenic_variant | MODIFIER |
| 8 | 96131813 | AX-170429062 | 0 | T | C | 0.219 | 5.16E-02 | 1.18E-01 | 2.70E+00 | 2.76E+00 | 6.58E-01 | no | | intergenic_variant | MODIFIER |
| 8 | 96132212 | AX-170124412 | 0 | G | A | 0.219 | 5.16E-02 | 1.18E-01 | 2.70E+00 | 2.76E+00 | 6.58E-01 | no | | intergenic_variant | MODIFIER |
| 8 | 96132467 | AX-170200998 | 0 | G | A | 0.219 | 5.16E-02 | 1.18E-01 | 2.70E+00 | 2.76E+00 | 6.58E-01 | no | | regulatory_region_variant | MODIFIER |
| 8 | 96141593 | AX-170290132 | 0 | A | G | 0.219 | 5.16E-02 | 1.18E-01 | 2.70E+00 | 2.76E+00 | 6.58E-01 | no | | intergenic_variant | MODIFIER |
| 8 | 96144288 | AX-97804109 | 0 | T | A | 0.219 | 5.16E-02 | 1.18E-01 | 2.70E+00 | 2.76E+00 | 6.58E-01 | no | | intergenic_variant | MODIFIER |
| 8 | 96155930 | AX-170516786 | 0 | A | G | 0.219 | 5.16E-02 | 1.18E-01 | 2.70E+00 | 2.76E+00 | 6.58E-01 | no | | intergenic_variant | MODIFIER |
| 8 | 96163363 | AX-170098861 | 0 | C | T | 0.219 | 5.16E-02 | 1.18E-01 | 2.70E+00 | 2.76E+00 | 6.58E-01 | no | | intergenic_variant | MODIFIER |
| 8 | 96164788 | AX-170347348 | 0 | G | T | 0.219 | 5.16E-02 | 1.18E-01 | 2.70E+00 | 2.76E+00 | 6.58E-01 | no | | intergenic_variant | MODIFIER |
| 8 | 96170833 | AX-170215448 | 0 | T | A | 0.219 | 5.16E-02 | 1.18E-01 | 2.70E+00 | 2.76E+00 | 6.58E-01 | no | | intergenic_variant | MODIFIER |
| 8 | 96179974 | AX-170525532 | 0 | C | A | 0.219 | 5.16E-02 | 1.18E-01 | 2.70E+00 | 2.76E+00 | 6.58E-01 | no | | intergenic_variant | MODIFIER |
| 8 | 96186804 | AX-170200483 | 0 | C | T | 0.219 | 5.16E-02 | 1.18E-01 | 2.70E+00 | 2.76E+00 | 6.58E-01 | no | | intergenic_variant | MODIFIER |
| 8 | 96191908 | AX-97839124 | 0 | T | G | 0.219 | 5.16E-02 | 1.18E-01 | 2.70E+00 | 2.76E+00 | 6.58E-01 | no | | intergenic_variant | MODIFIER |
| 8 | 96209788 | AX-170171783 | 0 | A | G | 0.219 | 5.16E-02 | 1.18E-01 | 2.70E+00 | 2.76E+00 | 6.58E-01 | no | | intergenic_variant | MODIFIER |
| 8 | 96245984 | AX-170031312 | 0 | A | G | 0.219 | 5.16E-02 | 1.18E-01 | 2.70E+00 | 2.76E+00 | 6.58E-01 | no | | regulatory_region_variant | MODIFIER |
| 8 | 96294530 | AX-170090184 | 0 | G | A | 0.219 | 5.16E-02 | 1.18E-01 | 2.70E+00 | 2.76E+00 | 6.58E-01 | no | | intergenic_variant | MODIFIER |
| 8 | 96328819 | AX-170031320 | 0 | T | C | 0.219 | 5.16E-02 | 1.18E-01 | 2.70E+00 | 2.76E+00 | 6.58E-01 | no | Gm32005 | intron_variant,non_coding_transcript_variant | MODIFIER |
| 8 | 96367005 | AX-97852355 | 0 | C | T | 0.219 | 5.16E-02 | 1.18E-01 | 2.70E+00 | 2.76E+00 | 6.58E-01 | no | Gm32122 | downstream_gene_variant | MODIFIER |
| 8 | 96640326 | AX-170161289 | 0 | G | A | 0.219 | 5.16E-02 | 1.18E-01 | 2.70E+00 | 2.76E+00 | 6.58E-01 | no | | intergenic_variant | MODIFIER |
| 8 | 96657514 | AX-97788687 | 0 | T | C | 0.219 | 5.16E-02 | 1.18E-01 | 2.70E+00 | 2.76E+00 | 6.58E-01 | no | Gm24132 | downstream_gene_variant | MODIFIER |
| 8 | 96686755 | AX-170198534 | 0 | C | A | 0.219 | 5.16E-02 | 1.18E-01 | 2.70E+00 | 2.76E+00 | 6.58E-01 | no | | regulatory_region_variant | MODIFIER |
| 8 | 96686793 | AX-170177716 | 0 | C | T | 0.219 | 5.16E-02 | 1.18E-01 | 2.70E+00 | 2.76E+00 | 6.58E-01 | no | | regulatory_region_variant | MODIFIER |
| 8 | 96698746 | AX-170343816 | 0 | G | A | 0.219 | 5.16E-02 | 1.18E-01 | 2.70E+00 | 2.76E+00 | 6.58E-01 | no | | intergenic_variant | MODIFIER |
| 8 | 96716349 | AX-170200389 | 0 | T | C | 0.219 | 5.16E-02 | 1.18E-01 | 2.70E+00 | 2.76E+00 | 6.58E-01 | no | | intergenic_variant | MODIFIER |
| 8 | 96722988 | AX-170433241 | 0 | G | A | 0.219 | 5.16E-02 | 1.18E-01 | 2.70E+00 | 2.76E+00 | 6.58E-01 | no | | intergenic_variant | MODIFIER |
| 8 | 96724004 | AX-170393470 | 0 | C | T | 0.219 | 5.16E-02 | 1.18E-01 | 2.70E+00 | 2.76E+00 | 6.58E-01 | no | | intergenic_variant | MODIFIER |
| 8 | 96724337 | AX-170115983 | 0 | G | C | 0.219 | 5.16E-02 | 1.18E-01 | 2.70E+00 | 2.76E+00 | 6.58E-01 | no | | intergenic_variant | MODIFIER |
| 8 | 96724707 | AX-170308590 | 0 | A | C | 0.219 | 5.16E-02 | 1.18E-01 | 2.70E+00 | 2.76E+00 | 6.58E-01 | no | | intergenic_variant | MODIFIER |
| 8 | 96726357 | AX-170218382 | 0 | A | G | 0.219 | 5.16E-02 | 1.18E-01 | 2.70E+00 | 2.76E+00 | 6.58E-01 | no | | intergenic_variant | MODIFIER |
| 8 | 96739843 | AX-170031321 | 0 | G | A | 0.219 | 5.16E-02 | 1.18E-01 | 2.70E+00 | 2.76E+00 | 6.58E-01 | no | | intergenic_variant | MODIFIER |
| 8 | 96739873 | AX-170416769 | 0 | A | G | 0.219 | 5.16E-02 | 1.18E-01 | 2.70E+00 | 2.76E+00 | 6.58E-01 | no | | intergenic_variant | MODIFIER |
| 8 | 96751042 | AX-170031325 | 0 | A | G | 0.219 | 5.16E-02 | 1.18E-01 | 2.70E+00 | 2.76E+00 | 6.58E-01 | no | | intergenic_variant | MODIFIER |
| 8 | 96782050 | AX-170121147 | 0 | T | G | 0.219 | 5.16E-02 | 1.18E-01 | 2.70E+00 | 2.76E+00 | 6.58E-01 | no | | intergenic_variant | MODIFIER |

| | | | | | | | | | | | | | | | |
|---|---|---|---|---|---|---|---|---|---|---|---|---|---|---|---|
| 8 | 96847268 | AX-170031322 | 0 | A | G | 0.219 | 5.16E-02 | 1.18E-01 | 2.70E+00 | 2.76E+00 | 6.58E-01 | no | | intergenic_variant | MODIFIER |
| 8 | 96886065 | AX-97790730 | 0 | G | A | 0.219 | 5.16E-02 | 1.18E-01 | 2.70E+00 | 2.76E+00 | 6.58E-01 | no | | intergenic_variant | MODIFIER |
| 8 | 96890936 | AX-170411265 | 0 | T | C | 0.219 | 5.16E-02 | 1.18E-01 | 2.70E+00 | 2.76E+00 | 6.58E-01 | no | | intergenic_variant | MODIFIER |
| 8 | 96899189 | AX-170183221 | 0 | T | C | 0.219 | 5.16E-02 | 1.18E-01 | 2.70E+00 | 2.76E+00 | 6.58E-01 | no | | intergenic_variant | MODIFIER |
| 8 | 96914600 | AX-170419735 | 0 | T | A | 0.219 | 5.16E-02 | 1.18E-01 | 2.70E+00 | 2.76E+00 | 6.58E-01 | no | | intergenic_variant | MODIFIER |
| 8 | 96918863 | AX-170126336 | 0 | C | T | 0.219 | 5.16E-02 | 1.18E-01 | 2.70E+00 | 2.76E+00 | 6.58E-01 | no | | intergenic_variant | MODIFIER |
| 8 | 96919149 | AX-170146178 | 0 | T | C | 0.219 | 5.16E-02 | 1.18E-01 | 2.70E+00 | 2.76E+00 | 6.58E-01 | no | | intergenic_variant | MODIFIER |
| 8 | 96955780 | AX-170256654 | 0 | T | C | 0.219 | 5.16E-02 | 1.18E-01 | 2.70E+00 | 2.76E+00 | 6.58E-01 | no | | intergenic_variant | MODIFIER |
| 8 | 96982132 | AX-170147448 | 0 | C | T | 0.219 | 5.16E-02 | 1.18E-01 | 2.70E+00 | 2.76E+00 | 6.58E-01 | no | | intergenic_variant | MODIFIER |
| 8 | 97001186 | AX-97804112 | 0 | G | A | 0.219 | 5.16E-02 | 1.18E-01 | 2.70E+00 | 2.76E+00 | 6.58E-01 | no | | intergenic_variant | MODIFIER |
| 8 | 97055502 | AX-170448954 | 0 | T | C | 0.219 | 5.16E-02 | 1.18E-01 | 2.70E+00 | 2.76E+00 | 6.58E-01 | no | | intergenic_variant | MODIFIER |
| 8 | 97091608 | AX-170490779 | 0 | C | T | 0.219 | 5.16E-02 | 1.18E-01 | 2.70E+00 | 2.76E+00 | 6.58E-01 | no | | intergenic_variant | MODIFIER |
| 8 | 97091668 | AX-170436858 | 0 | C | A | 0.219 | 5.16E-02 | 1.18E-01 | 2.70E+00 | 2.76E+00 | 6.58E-01 | no | | intergenic_variant | MODIFIER |
| 8 | 97094207 | AX-97837901 | 0 | C | T | 0.219 | 5.16E-02 | 1.18E-01 | 2.70E+00 | 2.76E+00 | 6.58E-01 | no | | intergenic_variant | MODIFIER |
| 8 | 97101047 | AX-170365713 | 0 | A | G | 0.219 | 5.16E-02 | 1.18E-01 | 2.70E+00 | 2.76E+00 | 6.58E-01 | no | | intergenic_variant | MODIFIER |
| 8 | 97101665 | AX-170080909 | 0 | C | G | 0.219 | 5.16E-02 | 1.18E-01 | 2.70E+00 | 2.76E+00 | 6.58E-01 | no | | intergenic_variant | MODIFIER |
| 8 | 97133228 | AX-170080910 | 0 | A | G | 0.219 | 5.16E-02 | 1.18E-01 | 2.70E+00 | 2.76E+00 | 6.58E-01 | no | | intergenic_variant | MODIFIER |
| 8 | 97142298 | AX-170503329 | 0 | T | C | 0.219 | 5.16E-02 | 1.18E-01 | 2.70E+00 | 2.76E+00 | 6.58E-01 | no | | intergenic_variant | MODIFIER |
| 8 | 97165789 | AX-170298341 | 0 | T | G | 0.219 | 5.16E-02 | 1.18E-01 | 2.70E+00 | 2.76E+00 | 6.58E-01 | no | Gm45703 | upstream_gene_variant | MODIFIER |
| 8 | 97206372 | AX-170486846 | 0 | C | T | 0.219 | 5.16E-02 | 1.18E-01 | 2.70E+00 | 2.76E+00 | 6.58E-01 | no | | intergenic_variant | MODIFIER |
| 8 | 97225459 | AX-97842880 | 0 | T | C | 0.219 | 5.16E-02 | 1.18E-01 | 2.70E+00 | 2.76E+00 | 6.58E-01 | no | Gm25160 | downstream_gene_variant | MODIFIER |
| 8 | 97235413 | AX-170192802 | 0 | A | G | 0.219 | 5.16E-02 | 1.18E-01 | 2.70E+00 | 2.76E+00 | 6.58E-01 | no | | intergenic_variant | MODIFIER |
| 8 | 97284692 | AX-170031340 | 0 | T | C | 0.219 | 5.16E-02 | 1.18E-01 | 2.70E+00 | 2.76E+00 | 6.58E-01 | no | | intergenic_variant | MODIFIER |
| 8 | 97390832 | AX-97816679 | 0 | C | T | 0.219 | 5.16E-02 | 1.18E-01 | 2.70E+00 | 2.76E+00 | 6.58E-01 | no | | intergenic_variant | MODIFIER |
| 8 | 97392825 | AX-170031336 | 0 | A | G | 0.219 | 5.16E-02 | 1.18E-01 | 2.70E+00 | 2.76E+00 | 6.58E-01 | no | | intergenic_variant | MODIFIER |
| 8 | 97418829 | AX-170031338 | 0 | A | G | 0.219 | 5.16E-02 | 1.18E-01 | 2.70E+00 | 2.76E+00 | 6.58E-01 | no | | intergenic_variant | MODIFIER |
| 8 | 97433805 | AX-170031332 | 0 | G | A | 0.219 | 5.16E-02 | 1.18E-01 | 2.70E+00 | 2.76E+00 | 6.58E-01 | no | Gm5913 | upstream_gene_variant | MODIFIER |
| 8 | 97468506 | AX-97823603 | 0 | G | A | 0.219 | 5.16E-02 | 1.18E-01 | 2.70E+00 | 2.76E+00 | 6.58E-01 | no | | intergenic_variant | MODIFIER |
| 8 | 97528971 | AX-170168767 | 0 | G | A | 0.219 | 5.16E-02 | 1.18E-01 | 2.70E+00 | 2.76E+00 | 6.58E-01 | no | | intergenic_variant | MODIFIER |
| 8 | 97531493 | AX-170385106 | 0 | A | T | 0.219 | 5.16E-02 | 1.18E-01 | 2.70E+00 | 2.76E+00 | 6.58E-01 | no | | intergenic_variant | MODIFIER |
| 8 | 97533816 | AX-170361109 | 0 | T | C | 0.219 | 5.16E-02 | 1.18E-01 | 2.70E+00 | 2.76E+00 | 6.58E-01 | no | | intergenic_variant | MODIFIER |
| 8 | 97534101 | AX-170494568 | 0 | C | T | 0.219 | 5.16E-02 | 1.18E-01 | 2.70E+00 | 2.76E+00 | 6.58E-01 | no | | intergenic_variant | MODIFIER |
| 8 | 97553808 | AX-170402292 | 0 | T | G | 0.219 | 5.16E-02 | 1.18E-01 | 2.70E+00 | 2.76E+00 | 6.58E-01 | no | | intergenic_variant | MODIFIER |
| 8 | 97556826 | AX-170090186 | 0 | G | T | 0.219 | 5.16E-02 | 1.18E-01 | 2.70E+00 | 2.76E+00 | 6.58E-01 | no | | intergenic_variant | MODIFIER |
| 8 | 97564948 | AX-170338588 | 0 | A | G | 0.219 | 5.16E-02 | 1.18E-01 | 2.70E+00 | 2.76E+00 | 6.58E-01 | no | Gm7191 | downstream_gene_variant | MODIFIER |
| 8 | 97597448 | AX-170374237 | 0 | G | A | 0.219 | 5.16E-02 | 1.18E-01 | 2.70E+00 | 2.76E+00 | 6.58E-01 | no | | intergenic_variant | MODIFIER |
| 8 | 97598387 | AX-170373940 | 0 | T | C | 0.219 | 5.16E-02 | 1.18E-01 | 2.70E+00 | 2.76E+00 | 6.58E-01 | no | | intergenic_variant | MODIFIER |
| 8 | 97600311 | AX-97808689 | 0 | T | C | 0.219 | 5.16E-02 | 1.18E-01 | 2.70E+00 | 2.76E+00 | 6.58E-01 | no | | intergenic_variant | MODIFIER |
| 8 | 97611960 | AX-170104089 | 0 | T | C | 0.219 | 5.16E-02 | 1.18E-01 | 2.70E+00 | 2.76E+00 | 6.58E-01 | no | | intergenic_variant | MODIFIER |
| 8 | 97616601 | AX-170453039 | 0 | G | A | 0.219 | 5.16E-02 | 1.18E-01 | 2.70E+00 | 2.76E+00 | 6.58E-01 | no | | intergenic_variant | MODIFIER |
| 8 | 97616878 | AX-170245080 | 0 | A | G | 0.219 | 5.16E-02 | 1.18E-01 | 2.70E+00 | 2.76E+00 | 6.58E-01 | no | | intergenic_variant | MODIFIER |
| 8 | 97646764 | AX-170234925 | 0 | A | G | 0.219 | 5.16E-02 | 1.18E-01 | 2.70E+00 | 2.76E+00 | 6.58E-01 | no | | intergenic_variant | MODIFIER |
| 8 | 97650252 | AX-170090187 | 0 | A | G | 0.219 | 5.16E-02 | 1.18E-01 | 2.70E+00 | 2.76E+00 | 6.58E-01 | no | | intergenic_variant | MODIFIER |
| 8 | 97661035 | AX-170516918 | 0 | G | A | 0.219 | 5.16E-02 | 1.18E-01 | 2.70E+00 | 2.76E+00 | 6.58E-01 | no | | intergenic_variant | MODIFIER |
| 8 | 97662388 | AX-170196948 | 0 | T | A | 0.219 | 5.16E-02 | 1.18E-01 | 2.70E+00 | 2.76E+00 | 6.58E-01 | no | | intergenic_variant | MODIFIER |
| 8 | 97683920 | AX-170450916 | 0 | A | G | 0.219 | 5.16E-02 | 1.18E-01 | 2.70E+00 | 2.76E+00 | 6.58E-01 | no | | intergenic_variant | MODIFIER |
| 8 | 97753547 | AX-170478739 | 0 | G | A | 0.219 | 5.16E-02 | 1.18E-01 | 2.70E+00 | 2.76E+00 | 6.58E-01 | no | | intergenic_variant | MODIFIER |
| 8 | 97781357 | AX-170090188 | 0 | G | C | 0.219 | 5.16E-02 | 1.18E-01 | 2.70E+00 | 2.76E+00 | 6.58E-01 | no | | intergenic_variant | MODIFIER |
| 8 | 97886958 | AX-170031343 | 0 | G | A | 0.219 | 5.16E-02 | 1.18E-01 | 2.70E+00 | 2.76E+00 | 6.58E-01 | no | | intergenic_variant | MODIFIER |
| 8 | 97938040 | AX-170031346 | 0 | G | A | 0.219 | 5.16E-02 | 1.18E-01 | 2.70E+00 | 2.76E+00 | 6.58E-01 | no | | intergenic_variant | MODIFIER |
| 8 | 97941597 | AX-170286360 | 0 | T | C | 0.219 | 5.16E-02 | 1.18E-01 | 2.70E+00 | 2.76E+00 | 6.58E-01 | no | | intergenic_variant | MODIFIER |
| 8 | 97966987 | AX-170339024 | 0 | A | G | 0.219 | 5.16E-02 | 1.18E-01 | 2.70E+00 | 2.76E+00 | 6.58E-01 | no | | intergenic_variant | MODIFIER |
| 8 | 97969610 | AX-170507891 | 0 | C | G | 0.219 | 5.16E-02 | 1.18E-01 | 2.70E+00 | 2.76E+00 | 6.58E-01 | no | | intergenic_variant | MODIFIER |
| 8 | 97975867 | AX-170140161 | 0 | T | G | 0.219 | 5.16E-02 | 1.18E-01 | 2.70E+00 | 2.76E+00 | 6.58E-01 | no | | intergenic_variant | MODIFIER |

| | | | | | | | | | | | | | | | |
|---|---|---|---|---|---|---|---|---|---|---|---|---|---|---|---|
| 8 | 98041069 | AX-170230333 | 0 | C | A | 0.219 | 5.16E-02 | 1.18E-01 | 2.70E+00 | 2.76E+00 | 6.58E-01 | no | | intergenic_variant | MODIFIER |
| 8 | 98085600 | AX-170410432 | 0 | A | G | 0.219 | 5.16E-02 | 1.18E-01 | 2.70E+00 | 2.76E+00 | 6.58E-01 | no | | intergenic_variant | MODIFIER |
| 8 | 98092664 | AX-170414599 | 0 | G | C | 0.219 | 5.16E-02 | 1.18E-01 | 2.70E+00 | 2.76E+00 | 6.58E-01 | no | | intergenic_variant | MODIFIER |
| 8 | 98260506 | AX-170393174 | 0 | A | C | 0.219 | 5.16E-02 | 1.18E-01 | 2.70E+00 | 2.76E+00 | 6.58E-01 | no | | intergenic_variant | MODIFIER |
| 8 | 98265215 | AX-170031352 | 0 | C | T | 0.219 | 5.16E-02 | 1.18E-01 | 2.70E+00 | 2.76E+00 | 6.58E-01 | no | | intergenic_variant | MODIFIER |
| 8 | 98281473 | AX-170526155 | 0 | A | G | 0.219 | 5.16E-02 | 1.18E-01 | 2.70E+00 | 2.76E+00 | 6.58E-01 | no | | intergenic_variant | MODIFIER |
| 8 | 98291916 | AX-97786187 | 0 | G | A | 0.219 | 5.16E-02 | 1.18E-01 | 2.70E+00 | 2.76E+00 | 6.58E-01 | no | | intergenic_variant | MODIFIER |
| 8 | 98318956 | AX-170031351 | 0 | T | C | 0.219 | 5.16E-02 | 1.18E-01 | 2.70E+00 | 2.76E+00 | 6.58E-01 | no | | intergenic_variant | MODIFIER |
| 8 | 98319293 | AX-170192445 | 0 | G | A | 0.219 | 5.16E-02 | 1.18E-01 | 2.70E+00 | 2.76E+00 | 6.58E-01 | no | | intergenic_variant | MODIFIER |
| 8 | 98320147 | AX-97824799 | 0 | A | G | 0.219 | 5.16E-02 | 1.18E-01 | 2.70E+00 | 2.76E+00 | 6.58E-01 | no | | intergenic_variant | MODIFIER |
| 8 | 98321869 | AX-170363621 | 0 | T | C | 0.219 | 5.16E-02 | 1.18E-01 | 2.70E+00 | 2.76E+00 | 6.58E-01 | no | | intergenic_variant | MODIFIER |
| 8 | 98350826 | AX-170182290 | 0 | T | C | 0.219 | 5.16E-02 | 1.18E-01 | 2.70E+00 | 2.76E+00 | 6.58E-01 | no | | intergenic_variant | MODIFIER |
| 8 | 98351864 | AX-97821950 | 0 | A | G | 0.219 | 5.16E-02 | 1.18E-01 | 2.70E+00 | 2.76E+00 | 6.58E-01 | no | | intergenic_variant | MODIFIER |
| 8 | 98365602 | AX-170186700 | 0 | T | C | 0.219 | 5.16E-02 | 1.18E-01 | 2.70E+00 | 2.76E+00 | 6.58E-01 | no | | intergenic_variant | MODIFIER |
| 8 | 98405264 | AX-170377357 | 0 | T | C | 0.219 | 5.16E-02 | 1.18E-01 | 2.70E+00 | 2.76E+00 | 6.58E-01 | no | | intergenic_variant | MODIFIER |
| 8 | 98405434 | AX-170458774 | 0 | T | A | 0.219 | 5.16E-02 | 1.18E-01 | 2.70E+00 | 2.76E+00 | 6.58E-01 | no | | intergenic_variant | MODIFIER |
| 8 | 98488594 | AX-97800133 | 0 | A | G | 0.219 | 5.16E-02 | 1.18E-01 | 2.70E+00 | 2.76E+00 | 6.58E-01 | no | | intergenic_variant | MODIFIER |
| 8 | 98604694 | AX-97783998 | 0 | C | T | 0.219 | 5.16E-02 | 1.18E-01 | 2.70E+00 | 2.76E+00 | 6.58E-01 | no | | intergenic_variant | MODIFIER |
| 8 | 98630705 | AX-170338836 | 0 | C | T | 0.219 | 5.16E-02 | 1.18E-01 | 2.70E+00 | 2.76E+00 | 6.58E-01 | no | | intergenic_variant | MODIFIER |
| 8 | 98667610 | AX-97847192 | 0 | A | G | 0.219 | 5.16E-02 | 1.18E-01 | 2.70E+00 | 2.76E+00 | 6.58E-01 | no | | intergenic_variant | MODIFIER |
| 8 | 98678808 | AX-170080930 | 0 | A | C | 0.219 | 5.16E-02 | 1.18E-01 | 2.70E+00 | 2.76E+00 | 6.58E-01 | no | Gm23494 | upstream_gene_variant | MODIFIER |
| 8 | 98681078 | AX-170080936 | 0 | A | G | 0.219 | 5.16E-02 | 1.18E-01 | 2.70E+00 | 2.76E+00 | 6.58E-01 | no | | intergenic_variant | MODIFIER |
| 8 | 98832548 | AX-170399611 | 0 | A | G | 0.219 | 5.16E-02 | 1.18E-01 | 2.70E+00 | 2.76E+00 | 6.58E-01 | no | | intergenic_variant | MODIFIER |
| 8 | 98860256 | AX-170220445 | 0 | T | A | 0.219 | 5.16E-02 | 1.18E-01 | 2.70E+00 | 2.76E+00 | 6.58E-01 | no | | intergenic_variant | MODIFIER |
| 8 | 98955223 | AX-170507681 | 0 | T | C | 0.219 | 5.16E-02 | 1.18E-01 | 2.70E+00 | 2.76E+00 | 6.58E-01 | no | | intergenic_variant | MODIFIER |
| 8 | 98956475 | AX-170080938 | 0 | C | T | 0.219 | 5.16E-02 | 1.18E-01 | 2.70E+00 | 2.76E+00 | 6.58E-01 | no | | intergenic_variant | MODIFIER |
| 8 | 98957212 | AX-170521092 | 0 | C | A | 0.219 | 5.16E-02 | 1.18E-01 | 2.70E+00 | 2.76E+00 | 6.58E-01 | no | | intergenic_variant | MODIFIER |
| 8 | 98957305 | AX-170207108 | 0 | G | A | 0.219 | 5.16E-02 | 1.18E-01 | 2.70E+00 | 2.76E+00 | 6.58E-01 | no | | intergenic_variant | MODIFIER |
| 8 | 98967400 | AX-170125524 | 0 | C | T | 0.219 | 5.16E-02 | 1.18E-01 | 2.70E+00 | 2.76E+00 | 6.58E-01 | no | | intergenic_variant | MODIFIER |
| 8 | 98997506 | AX-170090191 | 0 | T | C | 0.219 | 5.16E-02 | 1.18E-01 | 2.70E+00 | 2.76E+00 | 6.58E-01 | no | | intergenic_variant | MODIFIER |
| 8 | 99027651 | AX-170412706 | 0 | T | G | 0.219 | 5.16E-02 | 1.18E-01 | 2.70E+00 | 2.76E+00 | 6.58E-01 | no | Cdh8 | downstream_gene_variant | MODIFIER |
| 8 | 99027865 | AX-170200475 | 0 | C | T | 0.219 | 5.16E-02 | 1.18E-01 | 2.70E+00 | 2.76E+00 | 6.58E-01 | no | Cdh8 | downstream_gene_variant | MODIFIER |
| 8 | 99041993 | AX-170291945 | 0 | A | G | 0.219 | 5.16E-02 | 1.18E-01 | 2.70E+00 | 2.76E+00 | 6.58E-01 | no | Cdh8 | intron_variant | MODIFIER |
| 8 | 99053984 | AX-170333619 | 0 | T | A | 0.219 | 5.16E-02 | 1.18E-01 | 2.70E+00 | 2.76E+00 | 6.58E-01 | no | Cdh8 | intron_variant | MODIFIER |
| 8 | 99057968 | AX-170217698 | 0 | G | A | 0.219 | 5.16E-02 | 1.18E-01 | 2.70E+00 | 2.76E+00 | 6.58E-01 | no | Cdh8 | intron_variant | MODIFIER |
| 8 | 99099489 | AX-170403442 | 0 | T | C | 0.219 | 5.16E-02 | 1.18E-01 | 2.70E+00 | 2.76E+00 | 6.58E-01 | no | Cdh8 | intron_variant | MODIFIER |
| 8 | 99100547 | AX-170444312 | 0 | T | G | 0.219 | 5.16E-02 | 1.18E-01 | 2.70E+00 | 2.76E+00 | 6.58E-01 | no | Cdh8 | intron_variant | MODIFIER |
| 8 | 99105548 | AX-170200853 | 0 | G | A | 0.219 | 5.16E-02 | 1.18E-01 | 2.70E+00 | 2.76E+00 | 6.58E-01 | no | Cdh8 | intron_variant | MODIFIER |
| 8 | 99118842 | AX-170134450 | 0 | G | A | 0.219 | 5.16E-02 | 1.18E-01 | 2.70E+00 | 2.76E+00 | 6.58E-01 | no | Cdh8 | intron_variant | MODIFIER |
| 8 | 99174364 | AX-97804138 | 0 | T | G | 0.219 | 5.16E-02 | 1.18E-01 | 2.70E+00 | 2.76E+00 | 6.58E-01 | no | Cdh8 | intron_variant | MODIFIER |
| 8 | 102742732 | AX-170153438 | 0 | C | T | 0.219 | 4.83E-02 | 1.11E-01 | 2.69E+00 | 2.76E+00 | 6.58E-01 | no | Cdh11 | intron_variant | MODIFIER |
| 8 | 102744322 | AX-97851576 | 0 | T | C | 0.219 | 4.83E-02 | 1.11E-01 | 2.69E+00 | 2.76E+00 | 6.58E-01 | no | Cdh11 | intron_variant | MODIFIER |
| 8 | 102836723 | AX-170433413 | 0 | G | A | 0.219 | 4.83E-02 | 1.11E-01 | 2.69E+00 | 2.76E+00 | 6.58E-01 | no | | intergenic_variant | MODIFIER |
| 8 | 103078212 | AX-170166188 | 0 | T | C | 0.219 | 4.83E-02 | 1.11E-01 | 2.69E+00 | 2.76E+00 | 6.58E-01 | no | | regulatory_region_variant | MODIFIER |
| 8 | 103085026 | AX-170475799 | 0 | A | C | 0.219 | 4.83E-02 | 1.11E-01 | 2.69E+00 | 2.76E+00 | 6.58E-01 | no | | intergenic_variant | MODIFIER |
| 8 | 103088305 | AX-170215945 | 0 | A | G | 0.219 | 4.83E-02 | 1.11E-01 | 2.69E+00 | 2.76E+00 | 6.58E-01 | no | | regulatory_region_variant | MODIFIER |
| 8 | 103233690 | AX-170421926 | 0 | G | A | 0.219 | 4.83E-02 | 1.11E-01 | 2.69E+00 | 2.76E+00 | 6.58E-01 | no | | intergenic_variant | MODIFIER |
| 8 | 103244266 | AX-170301885 | 0 | C | T | 0.219 | 4.83E-02 | 1.11E-01 | 2.69E+00 | 2.76E+00 | 6.58E-01 | no | | intergenic_variant | MODIFIER |
| 8 | 103280430 | AX-170455645 | 0 | T | C | 0.219 | 4.83E-02 | 1.11E-01 | 2.69E+00 | 2.76E+00 | 6.58E-01 | no | Gm32531 | intron_variant,non_coding_transcript_variant | MODIFIER |
| 8 | 103290754 | AX-170480982 | 0 | T | C | 0.219 | 4.83E-02 | 1.11E-01 | 2.69E+00 | 2.76E+00 | 6.58E-01 | no | Gm32531 | intron_variant,non_coding_transcript_variant | MODIFIER |
| 8 | 103290820 | AX-170367244 | 0 | T | C | 0.219 | 4.83E-02 | 1.11E-01 | 2.69E+00 | 2.76E+00 | 6.58E-01 | no | Gm32531 | intron_variant,non_coding_transcript_variant | MODIFIER |
| 8 | 103292594 | AX-170311996 | 0 | C | T | 0.219 | 4.83E-02 | 1.11E-01 | 2.69E+00 | 2.76E+00 | 6.58E-01 | no | Gm32531 | intron_variant,non_coding_transcript_variant | MODIFIER |
| 8 | 103305441 | AX-170413496 | 0 | A | G | 0.219 | 4.83E-02 | 1.11E-01 | 2.69E+00 | 2.76E+00 | 6.58E-01 | no | Gm32531 | intron_variant,non_coding_transcript_variant | MODIFIER |
| 8 | 103308931 | AX-170204508 | 0 | A | G | 0.219 | 4.83E-02 | 1.11E-01 | 2.69E+00 | 2.76E+00 | 6.58E-01 | no | Gm32531 | intron_variant,non_coding_transcript_variant | MODIFIER |

| | | | | | | | | | | | | | | | |
|---|---|---|---|---|---|---|---|---|---|---|---|---|---|---|---|
| 8 | 103312168 | AX-170081033 | 0 | T | C | 0.219 | 4.83E-02 | 1.11E-01 | 2.69E+00 | 2.76E+00 | 6.58E-01 | no | Gm32531 | intron_variant,non_coding_transcript_variant | MODIFIER |
| 8 | 103312619 | AX-170081030 | 0 | A | G | 0.219 | 4.83E-02 | 1.11E-01 | 2.69E+00 | 2.76E+00 | 6.58E-01 | no | Gm32531 | intron_variant,non_coding_transcript_variant | MODIFIER |
| 8 | 103312679 | AX-170081032 | 0 | T | C | 0.219 | 4.83E-02 | 1.11E-01 | 2.69E+00 | 2.76E+00 | 6.58E-01 | no | Gm32531 | intron_variant,non_coding_transcript_variant | MODIFIER |
| 8 | 103321937 | AX-170440845 | 0 | A | G | 0.219 | 4.83E-02 | 1.11E-01 | 2.69E+00 | 2.76E+00 | 6.58E-01 | no | Gm32531 | intron_variant,non_coding_transcript_variant | MODIFIER |
| 8 | 103338702 | AX-170181084 | 0 | C | T | 0.219 | 4.83E-02 | 1.11E-01 | 2.69E+00 | 2.76E+00 | 6.58E-01 | no | Gm32531 | intron_variant,non_coding_transcript_variant | MODIFIER |
| 8 | 103344591 | AX-170318912 | 0 | C | A | 0.219 | 4.83E-02 | 1.11E-01 | 2.69E+00 | 2.76E+00 | 6.58E-01 | no | Gm32531 | downstream_gene_variant | MODIFIER |
| 8 | 103347285 | AX-97850421 | 0 | A | G | 0.219 | 4.83E-02 | 1.11E-01 | 2.69E+00 | 2.76E+00 | 6.58E-01 | no | 1600027J07Rik | non_coding_transcript_exon_variant | MODIFIER |
| 8 | 103353359 | AX-170154980 | 0 | C | T | 0.219 | 4.83E-02 | 1.11E-01 | 2.69E+00 | 2.76E+00 | 6.58E-01 | no | | regulatory_region_variant | MODIFIER |
| 8 | 103364447 | AX-170393313 | 0 | C | T | 0.219 | 4.83E-02 | 1.11E-01 | 2.69E+00 | 2.76E+00 | 6.58E-01 | no | | intergenic_variant | MODIFIER |
| 8 | 103371479 | AX-170412615 | 0 | T | C | 0.219 | 4.83E-02 | 1.11E-01 | 2.69E+00 | 2.76E+00 | 6.58E-01 | no | | intergenic_variant | MODIFIER |
| 8 | 103372271 | AX-170195832 | 0 | T | C | 0.219 | 4.83E-02 | 1.11E-01 | 2.69E+00 | 2.76E+00 | 6.58E-01 | no | | intergenic_variant | MODIFIER |
| 8 | 103376654 | AX-170382886 | 0 | C | A | 0.219 | 4.83E-02 | 1.11E-01 | 2.69E+00 | 2.76E+00 | 6.58E-01 | no | | intergenic_variant | MODIFIER |
| 8 | 103382596 | AX-170334810 | 0 | C | G | 0.219 | 4.83E-02 | 1.11E-01 | 2.69E+00 | 2.76E+00 | 6.58E-01 | no | | intergenic_variant | MODIFIER |
| 8 | 103382828 | AX-170397578 | 0 | C | A | 0.219 | 4.83E-02 | 1.11E-01 | 2.69E+00 | 2.76E+00 | 6.58E-01 | no | | intergenic_variant | MODIFIER |
| 8 | 103415212 | AX-170403805 | 0 | C | T | 0.219 | 4.83E-02 | 1.11E-01 | 2.69E+00 | 2.76E+00 | 6.58E-01 | no | | intergenic_variant | MODIFIER |
| 8 | 103420481 | AX-170274715 | 0 | G | A | 0.219 | 4.83E-02 | 1.11E-01 | 2.69E+00 | 2.76E+00 | 6.58E-01 | no | | intergenic_variant | MODIFIER |
| 8 | 103420634 | AX-170340760 | 0 | G | A | 0.219 | 4.83E-02 | 1.11E-01 | 2.69E+00 | 2.76E+00 | 6.58E-01 | no | | intergenic_variant | MODIFIER |
| 8 | 103429527 | AX-170081034 | 0 | A | T | 0.219 | 4.83E-02 | 1.11E-01 | 2.69E+00 | 2.76E+00 | 6.58E-01 | no | | intergenic_variant | MODIFIER |
| 8 | 103429864 | AX-170081036 | 0 | A | G | 0.219 | 4.83E-02 | 1.11E-01 | 2.69E+00 | 2.76E+00 | 6.58E-01 | no | | intergenic_variant | MODIFIER |
| 8 | 103430252 | AX-170081035 | 0 | A | G | 0.219 | 4.83E-02 | 1.11E-01 | 2.69E+00 | 2.76E+00 | 6.58E-01 | no | | regulatory_region_variant | MODIFIER |
| 8 | 103430477 | AX-170221973 | 0 | G | C | 0.219 | 4.83E-02 | 1.11E-01 | 2.69E+00 | 2.76E+00 | 6.58E-01 | no | | intergenic_variant | MODIFIER |
| 8 | 103436510 | AX-170116984 | 0 | A | T | 0.219 | 4.83E-02 | 1.11E-01 | 2.69E+00 | 2.76E+00 | 6.58E-01 | no | | intergenic_variant | MODIFIER |
| 8 | 103449399 | AX-170115093 | 0 | A | G | 0.219 | 4.83E-02 | 1.11E-01 | 2.69E+00 | 2.76E+00 | 6.58E-01 | no | | intergenic_variant | MODIFIER |
| 8 | 103456297 | AX-170343336 | 0 | T | C | 0.219 | 4.83E-02 | 1.11E-01 | 2.69E+00 | 2.76E+00 | 6.58E-01 | no | | intergenic_variant | MODIFIER |
| 8 | 103463348 | AX-170458397 | 0 | C | T | 0.219 | 4.83E-02 | 1.11E-01 | 2.69E+00 | 2.76E+00 | 6.58E-01 | no | | intergenic_variant | MODIFIER |
| 8 | 103477214 | AX-170201054 | 0 | T | C | 0.219 | 4.83E-02 | 1.11E-01 | 2.69E+00 | 2.76E+00 | 6.58E-01 | no | | regulatory_region_variant | MODIFIER |
| 8 | 103491361 | AX-170269165 | 0 | A | G | 0.219 | 4.83E-02 | 1.11E-01 | 2.69E+00 | 2.76E+00 | 6.58E-01 | no | | intergenic_variant | MODIFIER |
| 8 | 103492425 | AX-170193145 | 0 | G | A | 0.219 | 4.83E-02 | 1.11E-01 | 2.69E+00 | 2.76E+00 | 6.58E-01 | no | | regulatory_region_variant | MODIFIER |
| 8 | 103505054 | AX-170398295 | 0 | T | C | 0.219 | 4.83E-02 | 1.11E-01 | 2.69E+00 | 2.76E+00 | 6.58E-01 | no | | intergenic_variant | MODIFIER |
| 8 | 103524792 | AX-170350983 | 0 | A | C | 0.219 | 4.83E-02 | 1.11E-01 | 2.69E+00 | 2.76E+00 | 6.58E-01 | no | | intergenic_variant | MODIFIER |
| 8 | 103524830 | AX-170208801 | 0 | G | A | 0.219 | 4.83E-02 | 1.11E-01 | 2.69E+00 | 2.76E+00 | 6.58E-01 | no | | intergenic_variant | MODIFIER |
| 8 | 103533939 | AX-170153963 | 0 | T | G | 0.219 | 4.83E-02 | 1.11E-01 | 2.69E+00 | 2.76E+00 | 6.58E-01 | no | Gm45277 | intron_variant,non_coding_transcript_variant | MODIFIER |
| 8 | 103554233 | AX-170353778 | 0 | A | G | 0.219 | 4.83E-02 | 1.11E-01 | 2.69E+00 | 2.76E+00 | 6.58E-01 | no | Gm45277 | intron_variant,non_coding_transcript_variant | MODIFIER |
| 8 | 103557495 | AX-170140777 | 0 | T | C | 0.219 | 4.83E-02 | 1.11E-01 | 2.69E+00 | 2.76E+00 | 6.58E-01 | no | Gm45277 | intron_variant,non_coding_transcript_variant | MODIFIER |
| 8 | 103564210 | AX-170081040 | 0 | C | T | 0.219 | 4.83E-02 | 1.11E-01 | 2.69E+00 | 2.76E+00 | 6.58E-01 | no | Gm45277 | upstream_gene_variant | MODIFIER |
| 8 | 103829713 | AX-170219412 | 0 | C | T | 0.219 | 4.83E-02 | 1.11E-01 | 2.69E+00 | 2.76E+00 | 6.58E-01 | no | | intergenic_variant | MODIFIER |
| 8 | 103832656 | AX-170222096 | 0 | T | C | 0.219 | 4.83E-02 | 1.11E-01 | 2.69E+00 | 2.76E+00 | 6.58E-01 | no | | intergenic_variant | MODIFIER |
| 8 | 103848332 | AX-97837374 | 0 | T | C | 0.219 | 4.83E-02 | 1.11E-01 | 2.69E+00 | 2.76E+00 | 6.58E-01 | no | | intergenic_variant | MODIFIER |
| 8 | 103849810 | AX-170254342 | 0 | C | A | 0.219 | 4.83E-02 | 1.11E-01 | 2.69E+00 | 2.76E+00 | 6.58E-01 | no | | regulatory_region_variant | MODIFIER |
| 8 | 103850162 | AX-170525512 | 0 | C | G | 0.219 | 4.83E-02 | 1.11E-01 | 2.69E+00 | 2.76E+00 | 6.58E-01 | no | | regulatory_region_variant | MODIFIER |
| 8 | 103850209 | AX-170533176 | 0 | T | C | 0.219 | 4.83E-02 | 1.11E-01 | 2.69E+00 | 2.76E+00 | 6.58E-01 | no | | regulatory_region_variant | MODIFIER |
| 8 | 103865902 | AX-170449889 | 0 | C | A | 0.219 | 4.83E-02 | 1.11E-01 | 2.69E+00 | 2.76E+00 | 6.58E-01 | no | | intergenic_variant | MODIFIER |
| 8 | 103885305 | AX-170475453 | 0 | G | A | 0.219 | 4.83E-02 | 1.11E-01 | 2.69E+00 | 2.76E+00 | 6.58E-01 | no | | regulatory_region_variant | MODIFIER |
| 8 | 103892326 | AX-97802463 | 0 | C | T | 0.219 | 4.83E-02 | 1.11E-01 | 2.69E+00 | 2.76E+00 | 6.58E-01 | no | | intergenic_variant | MODIFIER |
| 8 | 103940768 | AX-170300686 | 0 | G | A | 0.219 | 4.83E-02 | 1.11E-01 | 2.69E+00 | 2.76E+00 | 6.58E-01 | no | | regulatory_region_variant | MODIFIER |
| 8 | 106659727 | AX-170149391 | 0 | T | C | 0.219 | 4.83E-02 | 1.11E-01 | 2.69E+00 | 2.76E+00 | 6.58E-01 | no | Cdh1 | synonymous_variant | LOW |
| 8 | 106684162 | AX-170477338 | 0 | T | C | 0.219 | 4.83E-02 | 1.11E-01 | 2.69E+00 | 2.76E+00 | 6.58E-01 | no | Tango6 | intron_variant | MODIFIER |
| 8 | 106699908 | AX-97851483 | 0 | T | C | 0.219 | 4.83E-02 | 1.11E-01 | 2.69E+00 | 2.76E+00 | 6.58E-01 | no | Tango6 | intron_variant | MODIFIER |
| 8 | 106816584 | AX-168409331 | 0 | C | A | 0.219 | 4.83E-02 | 1.11E-01 | 2.69E+00 | 2.76E+00 | 6.58E-01 | no | Tango6 | intron_variant | MODIFIER |
| 8 | 106819183 | AX-97837421 | 0 | C | A | 0.219 | 4.83E-02 | 1.11E-01 | 2.69E+00 | 2.76E+00 | 6.58E-01 | no | Tango6 | intron_variant | MODIFIER |
| 8 | 106819978 | AX-170218229 | 0 | T | C | 0.219 | 4.83E-02 | 1.11E-01 | 2.69E+00 | 2.76E+00 | 6.58E-01 | no | Tango6 | intron_variant | MODIFIER |
| 8 | 106980296 | AX-170224536 | 0 | G | A | 0.219 | 4.83E-02 | 1.11E-01 | 2.69E+00 | 2.76E+00 | 6.58E-01 | no | Sntb2 | intron_variant | MODIFIER |
| 8 | 106988749 | AX-170313322 | 0 | A | G | 0.219 | 4.83E-02 | 1.11E-01 | 2.69E+00 | 2.76E+00 | 6.58E-01 | no | Sntb2 | intron_variant | MODIFIER |
| 8 | 107035749 | AX-170148179 | 0 | A | G | 0.219 | 4.83E-02 | 1.11E-01 | 2.69E+00 | 2.76E+00 | 6.58E-01 | no | Vps4a | intron_variant | MODIFIER |

| | | | | | | | | | | | | | | | |
|---|---|---|---|---|---|---|---|---|---|---|---|---|---|---|---|
| 8 | 107125227 | AX-170081093 | 0 | C | T | 0.219 | 4.83E-02 | 1.11E-01 | 2.69E+00 | 2.76E+00 | 6.58E-01 | no | | intergenic_variant | MODIFIER |
| 8 | 107133683 | AX-97791969 | 0 | C | T | 0.219 | 4.83E-02 | 1.11E-01 | 2.69E+00 | 2.76E+00 | 6.58E-01 | no | | regulatory_region_variant | MODIFIER |
| 8 | 107165328 | AX-170357133 | 0 | C | T | 0.219 | 4.83E-02 | 1.11E-01 | 2.69E+00 | 2.76E+00 | 6.58E-01 | no | Cyb5b | intron_variant | MODIFIER |
| 8 | 107173825 | AX-170386332 | 0 | A | G | 0.219 | 4.83E-02 | 1.11E-01 | 2.69E+00 | 2.76E+00 | 6.58E-01 | no | Cyb5b | intron_variant | MODIFIER |
| 8 | 107683417 | AX-170119839 | 0 | G | T | 0.219 | 4.83E-02 | 1.11E-01 | 2.69E+00 | 2.76E+00 | 6.58E-01 | no | | intergenic_variant | MODIFIER |
| 8 | 107687531 | AX-170515104 | 0 | T | C | 0.219 | 4.83E-02 | 1.11E-01 | 2.69E+00 | 2.76E+00 | 6.58E-01 | no | Gm8940 | downstream_gene_variant | MODIFIER |
| 8 | 108489287 | AX-170282679 | 0 | A | G | 0.219 | 4.83E-02 | 1.11E-01 | 2.69E+00 | 2.76E+00 | 6.58E-01 | no | Zfhx3 | intron_variant | MODIFIER |
| 8 | 108490657 | AX-170429740 | 0 | A | G | 0.219 | 4.83E-02 | 1.11E-01 | 2.69E+00 | 2.76E+00 | 6.58E-01 | no | Zfhx3 | intron_variant | MODIFIER |
| 8 | 108545317 | AX-170342318 | 0 | A | G | 0.219 | 4.83E-02 | 1.11E-01 | 2.69E+00 | 2.76E+00 | 6.58E-01 | no | Zfhx3 | intron_variant | MODIFIER |
| 8 | 108555254 | AX-170166786 | 0 | G | A | 0.219 | 4.83E-02 | 1.11E-01 | 2.69E+00 | 2.76E+00 | 6.58E-01 | no | Lncbate1 | intron_variant,non_coding_transcript_variant | MODIFIER |
| 1 | 76271347 | AX-97840893 | 1 | T | G | 0.134 | 8.27E-02 | 1.89E-01 | 2.64E+00 | 2.73E+00 | 6.58E-01 | no | | intergenic_variant | MODIFIER |
| 1 | 45544189 | AX-97815949 | 0 | C | G | 0.43 | -3.34E-02 | 7.60E-02 | 2.64E+00 | 2.73E+00 | 6.58E-01 | no | | intergenic_variant | MODIFIER |
| 1 | 45598872 | AX-170404919 | 0 | G | T | 0.43 | -3.34E-02 | 7.60E-02 | 2.64E+00 | 2.73E+00 | 6.58E-01 | no | Gm8304 | upstream_gene_variant | MODIFIER |
| 1 | 45598947 | AX-170130802 | 0 | G | A | 0.43 | -3.34E-02 | 7.60E-02 | 2.64E+00 | 2.73E+00 | 6.58E-01 | no | Gm8304 | upstream_gene_variant | MODIFIER |
| 1 | 45629648 | AX-170230891 | 0 | G | A | 0.43 | -3.34E-02 | 7.60E-02 | 2.64E+00 | 2.73E+00 | 6.58E-01 | no | | intergenic_variant | MODIFIER |
| 1 | 45685714 | AX-170314894 | 0 | C | T | 0.43 | -3.34E-02 | 7.60E-02 | 2.64E+00 | 2.73E+00 | 6.58E-01 | no | Gm23216 | upstream_gene_variant | MODIFIER |
| 1 | 45686043 | AX-170136034 | 0 | G | A | 0.43 | -3.34E-02 | 7.60E-02 | 2.64E+00 | 2.73E+00 | 6.58E-01 | no | Gm23216 | upstream_gene_variant | MODIFIER |
| 1 | 45689961 | AX-170334613 | 0 | C | T | 0.43 | -3.34E-02 | 7.60E-02 | 2.64E+00 | 2.73E+00 | 6.58E-01 | no | Gm23216 | upstream_gene_variant | MODIFIER |
| 1 | 45695467 | AX-170221214 | 0 | T | C | 0.43 | -3.34E-02 | 7.60E-02 | 2.64E+00 | 2.73E+00 | 6.58E-01 | no | | intergenic_variant | MODIFIER |
| 1 | 45729220 | AX-170265449 | 0 | A | T | 0.43 | -3.34E-02 | 7.60E-02 | 2.64E+00 | 2.73E+00 | 6.58E-01 | no | | intergenic_variant | MODIFIER |
| 1 | 45739897 | AX-170484824 | 0 | G | A | 0.43 | -3.34E-02 | 7.60E-02 | 2.64E+00 | 2.73E+00 | 6.58E-01 | no | | intergenic_variant | MODIFIER |
| 1 | 45764986 | AX-170522821 | 0 | T | C | 0.43 | -3.34E-02 | 7.60E-02 | 2.64E+00 | 2.73E+00 | 6.58E-01 | no | Gm8307 | upstream_gene_variant | MODIFIER |
| 1 | 45767266 | AX-170188059 | 0 | G | A | 0.43 | -3.34E-02 | 7.60E-02 | 2.64E+00 | 2.73E+00 | 6.58E-01 | no | Gm8307 | upstream_gene_variant | MODIFIER |
| 1 | 45780584 | AX-170227779 | 0 | G | T | 0.43 | -3.34E-02 | 7.60E-02 | 2.64E+00 | 2.73E+00 | 6.58E-01 | no | | intergenic_variant | MODIFIER |
| 1 | 45805990 | AX-170243589 | 0 | A | G | 0.43 | -3.34E-02 | 7.60E-02 | 2.64E+00 | 2.73E+00 | 6.58E-01 | no | Wdr75 | intron_variant | MODIFIER |
| 1 | 46629442 | AX-170086459 | 0 | G | T | 0.43 | -3.34E-02 | 7.60E-02 | 2.64E+00 | 2.73E+00 | 6.58E-01 | no | Dnah7c | downstream_gene_variant | MODIFIER |
| 1 | 46676267 | AX-97824402 | 0 | C | T | 0.43 | -3.34E-02 | 7.60E-02 | 2.64E+00 | 2.73E+00 | 6.58E-01 | no | Dnah7c | intron_variant | MODIFIER |
| 1 | 46827110 | AX-170308459 | 0 | C | T | 0.43 | -3.34E-02 | 7.60E-02 | 2.64E+00 | 2.73E+00 | 6.58E-01 | no | Slc39a10 | intron_variant | MODIFIER |
| 16 | 77791894 | AX-170361431 | 1 | C | T | 0.473 | 3.36E-02 | 7.41E-02 | 2.56E+00 | 2.65E+00 | 6.58E-01 | no | | intergenic_variant | MODIFIER |
| 11 | 74207416 | AX-170205494 | 1 | A | C | 0.268 | 3.10E-02 | 7.17E-02 | 2.64E+00 | 2.77E+00 | 6.58E-01 | no | Olfr43 | upstream_gene_variant | MODIFIER |
| 5 | 76865129 | AX-170427918 | 0 | T | G | 0.447 | 4.69E-02 | 1.08E-01 | 2.70E+00 | 2.76E+00 | 6.58E-01 | no | C530008M17Rik | intron_variant | MODIFIER |
| 1 | 3072668 | AX-169999826 | 1 | C | A | 0.429 | 3.05E-02 | 6.68E-02 | 2.55E+00 | 2.63E+00 | 6.58E-01 | no | 4933401J01Rik | upstream_gene_variant | MODIFIER |
| 2 | 124166447 | AX-97818686 | 1 | C | T | 0.152 | 7.84E-02 | 1.75E-01 | 2.59E+00 | 2.67E+00 | 6.59E-01 | no | Sema6d | intron_variant | MODIFIER |
| 8 | 68217622 | AX-170169488 | 0 | A | G | 0.158 | -6.19E-02 | 1.38E-01 | 2.59E+00 | 2.67E+00 | 6.59E-01 | no | Gm26659 | downstream_gene_variant | MODIFIER |
| 9 | 38507536 | AX-170517246 | 1 | A | C | 0.482 | -3.30E-02 | 7.64E-02 | 2.69E+00 | 2.78E+00 | 6.59E-01 | no | Gm23271 | upstream_gene_variant | MODIFIER |
| 18 | 69011158 | AX-170163382 | 0 | C | A | 0.272 | -4.32E-02 | 9.86E-02 | 2.66E+00 | 2.73E+00 | 6.59E-01 | no | | intergenic_variant | MODIFIER |
| 9 | 33058063 | AX-97830308 | 1 | C | T | 0.446 | 3.65E-02 | 8.23E-02 | 2.61E+00 | 2.69E+00 | 6.59E-01 | no | Gm27166 | intron_variant,non_coding_transcript_variant | MODIFIER |
| 15 | 81191905 | AX-170102937 | 0 | C | T | 0.561 | 3.63E-02 | 8.15E-02 | 2.61E+00 | 2.68E+00 | 6.59E-01 | no | Mrtfa | upstream_gene_variant | MODIFIER |
| 5 | 127051727 | AX-170253973 | 0 | T | C | 0.421 | 3.69E-02 | 8.45E-02 | 2.62E+00 | 2.73E+00 | 6.59E-01 | no | 4933438B17Rik | intron_variant,non_coding_transcript_variant | MODIFIER |
| 8 | 42710204 | AX-170468501 | 0 | A | T | 0.149 | -5.79E-02 | 1.32E-01 | 2.67E+00 | 2.73E+00 | 6.59E-01 | no | | intergenic_variant | MODIFIER |
| 16 | 7951026 | AX-170056609 | 0 | A | G | 0.035 | -9.26E-02 | 2.02E-01 | 2.53E+00 | 2.63E+00 | 6.59E-01 | no | | intergenic_variant | MODIFIER |
| 5 | 113367508 | AX-170025916 | 1 | A | T | 0.446 | 4.89E-02 | 1.10E-01 | 2.62E+00 | 2.67E+00 | 6.59E-01 | no | | intergenic_variant | MODIFIER |
| 8 | 96366897 | AX-170031319 | 1 | C | T | 0.223 | 5.10E-02 | 1.17E-01 | 2.68E+00 | 2.75E+00 | 6.59E-01 | no | Gm32122 | downstream_gene_variant | MODIFIER |
| 1 | 25777286 | AX-170209310 | 1 | C | A | 0.42 | 3.59E-02 | 8.22E-02 | 2.65E+00 | 2.73E+00 | 6.59E-01 | no | Adgrb3 | intron_variant | MODIFIER |
| 17 | 72566316 | AX-170447494 | 1 | C | T | 0.062 | -5.68E-02 | 1.30E-01 | 2.59E+00 | 2.73E+00 | 6.59E-01 | no | Alk | intron_variant | MODIFIER |
| 6 | 22674842 | AX-170392316 | 0 | A | G | 0.026 | 6.22E-02 | 1.42E-01 | 2.55E+00 | 2.73E+00 | 6.60E-01 | no | | intergenic_variant | MODIFIER |
| 5 | 30938493 | AX-170336049 | 0 | T | C | 0.36 | 4.60E-02 | 1.06E-01 | 2.69E+00 | 2.74E+00 | 6.60E-01 | no | Cgref1 | intron_variant | MODIFIER |
| 8 | 39999965 | AX-170462173 | 0 | A | G | 0.149 | 5.99E-02 | 1.39E-01 | 2.73E+00 | 2.80E+00 | 6.60E-01 | no | | intergenic_variant | MODIFIER |
| 8 | 45961422 | AX-170289204 | 0 | A | G | 0.149 | 5.99E-02 | 1.39E-01 | 2.73E+00 | 2.80E+00 | 6.60E-01 | no | 1700029J07Rik | intron_variant | MODIFIER |
| 8 | 97383621 | AX-170031334 | 1 | G | A | 0.223 | 5.09E-02 | 1.17E-01 | 2.69E+00 | 2.75E+00 | 6.60E-01 | no | | intergenic_variant | MODIFIER |
| 1 | 19975972 | AX-97819915 | 0 | G | A | 0.404 | 3.70E-02 | 8.33E-02 | 2.60E+00 | 2.66E+00 | 6.60E-01 | no | | intergenic_variant | MODIFIER |
| 1 | 20613452 | AX-170496318 | 0 | A | T | 0.404 | 3.70E-02 | 8.33E-02 | 2.60E+00 | 2.66E+00 | 6.60E-01 | no | Pkhd1 | missense_variant | MODERATE |
| 1 | 59990134 | AX-170314024 | 0 | G | A | 0.14 | 8.66E-02 | 2.01E-01 | 2.68E+00 | 2.78E+00 | 6.61E-01 | no | Ica1l | 3_prime_UTR_variant | MODIFIER |
| 5 | 81415351 | AX-170172133 | 2 | A | T | 0.455 | 4.25E-02 | 9.79E-02 | 2.65E+00 | 2.74E+00 | 6.61E-01 | no | Adgrl3 | intron_variant | MODIFIER |

| | | | | | | | | | | | | | | | |
|---|---|---|---|---|---|---|---|---|---|---|---|---|---|---|---|
| 15 | 10761285 | AX-170215365 | 2 | A | G | 0.236 | -4.57E-02 | 1.01E-01 | 2.56E+00 | 2.64E+00 | 6.61E-01 | no | 4930556M19Rik | intron_variant,non_coding_transcript_variant | MODIFIER |
| 13 | 33246573 | AX-97785940 | 1 | C | G | 0.179 | -5.44E-02 | 1.21E-01 | 2.58E+00 | 2.64E+00 | 6.61E-01 | no | Serpinb9e | upstream_gene_variant | MODIFIER |
| 11 | 60347172 | AX-97842328 | 1 | G | A | 0.045 | 6.86E-02 | 1.63E-01 | 2.70E+00 | 2.86E+00 | 6.61E-01 | no | Tom1l2 | intron_variant | MODIFIER |
| 7 | 10251063 | AX-97839364 | 0 | A | C | 0.167 | -3.43E-02 | 7.58E-02 | 2.52E+00 | 2.67E+00 | 6.61E-01 | no | | intergenic_variant | MODIFIER |
| 9 | 83611581 | AX-170491286 | 0 | T | C | 0.219 | 5.15E-02 | 1.20E-01 | 2.73E+00 | 2.80E+00 | 6.61E-01 | no | Sh3bgrl2 | intron_variant | MODIFIER |
| 9 | 54178296 | AX-170522928 | 1 | C | T | 0.196 | -4.86E-02 | 1.13E-01 | 2.70E+00 | 2.79E+00 | 6.61E-01 | no | Cyp19a1 | intron_variant | MODIFIER |
| 1 | 22309279 | AX-170227685 | 0 | C | T | 0.43 | 3.73E-02 | 8.48E-02 | 2.63E+00 | 2.68E+00 | 6.61E-01 | no | Rims1 | intron_variant | MODIFIER |
| 1 | 22314619 | AX-170526783 | 0 | C | T | 0.43 | 3.73E-02 | 8.48E-02 | 2.63E+00 | 2.68E+00 | 6.61E-01 | no | Rims1 | intron_variant | MODIFIER |
| 7 | 116738972 | AX-170494356 | 0 | G | A | 0.404 | 3.91E-02 | 9.00E-02 | 2.68E+00 | 2.73E+00 | 6.61E-01 | no | | intergenic_variant | MODIFIER |
| 8 | 30542180 | AX-170120362 | 0 | A | C | 0.167 | -4.84E-02 | 1.13E-01 | 2.69E+00 | 2.78E+00 | 6.61E-01 | no | | intergenic_variant | MODIFIER |
| 1 | 22185385 | AX-170260065 | 0 | G | A | 0.412 | 3.76E-02 | 8.58E-02 | 2.63E+00 | 2.69E+00 | 6.62E-01 | no | | intergenic_variant | MODIFIER |
| 5 | 46862951 | AX-170024859 | 0 | T | A | 0.5 | -4.14E-02 | 9.61E-02 | 2.83E+00 | 2.94E+00 | 6.62E-01 | no | | intergenic_variant | MODIFIER |
| 18 | 60484421 | AX-170524713 | 0 | C | T | 0.404 | -3.71E-02 | 8.27E-02 | 2.58E+00 | 2.63E+00 | 6.62E-01 | no | Smim3 | intron_variant | MODIFIER |
| 1 | 9588735 | AX-170384306 | 0 | T | C | 0.079 | 7.98E-02 | 1.86E-01 | 2.63E+00 | 2.75E+00 | 6.62E-01 | no | 2610203C22Rik | intron_variant,non_coding_transcript_variant | MODIFIER |
| 16 | 85933613 | AX-170119333 | 0 | C | G | 0.447 | 3.46E-02 | 7.97E-02 | 2.65E+00 | 2.72E+00 | 6.62E-01 | no | | intergenic_variant | MODIFIER |
| 16 | 85933745 | AX-170262004 | 0 | A | G | 0.447 | 3.46E-02 | 7.97E-02 | 2.65E+00 | 2.72E+00 | 6.62E-01 | no | | intergenic_variant | MODIFIER |
| 16 | 85936118 | AX-97792825 | 0 | T | C | 0.447 | 3.46E-02 | 7.97E-02 | 2.65E+00 | 2.72E+00 | 6.62E-01 | no | | intergenic_variant | MODIFIER |
| 16 | 85948111 | AX-170178410 | 0 | C | T | 0.447 | 3.46E-02 | 7.97E-02 | 2.65E+00 | 2.72E+00 | 6.62E-01 | no | | intergenic_variant | MODIFIER |
| 16 | 85952064 | AX-170362203 | 0 | A | G | 0.447 | 3.46E-02 | 7.97E-02 | 2.65E+00 | 2.72E+00 | 6.62E-01 | no | | intergenic_variant | MODIFIER |
| 16 | 85959023 | AX-170343503 | 0 | A | C | 0.447 | 3.46E-02 | 7.97E-02 | 2.65E+00 | 2.72E+00 | 6.62E-01 | no | | intergenic_variant | MODIFIER |
| 16 | 86143513 | AX-97791412 | 0 | C | T | 0.447 | 3.46E-02 | 7.97E-02 | 2.65E+00 | 2.72E+00 | 6.62E-01 | no | 1700007H22Rik | intron_variant,non_coding_transcript_variant | MODIFIER |
| 16 | 86148433 | AX-170200182 | 0 | C | T | 0.447 | 3.46E-02 | 7.97E-02 | 2.65E+00 | 2.72E+00 | 6.62E-01 | no | 1700007H22Rik | intron_variant,non_coding_transcript_variant | MODIFIER |
| 16 | 86171158 | AX-97808801 | 0 | A | G | 0.447 | 3.46E-02 | 7.97E-02 | 2.65E+00 | 2.72E+00 | 6.62E-01 | no | | intergenic_variant | MODIFIER |
| 16 | 86207929 | AX-170058015 | 0 | A | G | 0.447 | 3.46E-02 | 7.97E-02 | 2.65E+00 | 2.72E+00 | 6.62E-01 | no | | intergenic_variant | MODIFIER |
| 16 | 86229298 | AX-170230302 | 0 | G | A | 0.447 | 3.46E-02 | 7.97E-02 | 2.65E+00 | 2.72E+00 | 6.62E-01 | no | | intergenic_variant | MODIFIER |
| 10 | 128471605 | AX-97808908 | 0 | G | A | 0.325 | -3.29E-02 | 7.46E-02 | 2.59E+00 | 2.69E+00 | 6.62E-01 | no | Smarcc2 | intron_variant | MODIFIER |
| 8 | 19283037 | AX-170418554 | 1 | G | A | 0.179 | -4.39E-02 | 1.04E-01 | 2.68E+00 | 2.81E+00 | 6.62E-01 | no | Gm15357 | upstream_gene_variant | MODIFIER |
| 8 | 69040340 | AX-170030976 | 1 | G | A | 0.188 | -5.71E-02 | 1.32E-01 | 2.66E+00 | 2.74E+00 | 6.62E-01 | no | Slc18a1 | synonymous_variant | LOW |
| 3 | 120936263 | AX-170162899 | 0 | T | C | 0.044 | -7.16E-02 | 1.65E-01 | 2.59E+00 | 2.72E+00 | 6.62E-01 | no | | intergenic_variant | MODIFIER |
| 8 | 19140002 | AX-170113541 | 0 | A | G | 0.184 | 4.26E-02 | 9.86E-02 | 2.63E+00 | 2.73E+00 | 6.62E-01 | no | Spag11b | upstream_gene_variant | MODIFIER |
| 18 | 81102828 | AX-170230578 | 2 | C | T | 0.291 | -3.68E-02 | 8.21E-02 | 2.54E+00 | 2.66E+00 | 6.62E-01 | no | AC105065.1 | downstream_gene_variant | MODIFIER |
| 9 | 35362435 | AX-170232093 | 0 | G | T | 0.43 | -3.76E-02 | 8.76E-02 | 2.77E+00 | 2.84E+00 | 6.63E-01 | no | 2610105M22Rik | non_coding_transcript_exon_variant | MODIFIER |
| 4 | 97806403 | AX-170172863 | 0 | T | A | 0.237 | -4.97E-02 | 1.15E-01 | 2.84E+00 | 2.95E+00 | 6.63E-01 | no | Nfia | intron_variant | MODIFIER |
| 4 | 127737645 | AX-97854480 | 1 | T | C | 0.134 | 5.31E-02 | 1.20E-01 | 2.59E+00 | 2.66E+00 | 6.63E-01 | no | | intergenic_variant | MODIFIER |
| 4 | 127840520 | AX-97824287 | 1 | C | T | 0.134 | 5.31E-02 | 1.20E-01 | 2.59E+00 | 2.66E+00 | 6.63E-01 | no | | intergenic_variant | MODIFIER |
| 10 | 115742938 | AX-97821585 | 1 | G | A | 0.393 | 3.31E-02 | 7.75E-02 | 2.69E+00 | 2.78E+00 | 6.63E-01 | no | | intergenic_variant | MODIFIER |
| 10 | 127820954 | AX-170307278 | 1 | C | T | 0.33 | -3.15E-02 | 7.25E-02 | 2.61E+00 | 2.72E+00 | 6.63E-01 | no | Rdh16 | intron_variant,NMD_transcript_variant | MODIFIER |
| 9 | 34773381 | AX-170473685 | 0 | A | T | 0.439 | 3.78E-02 | 8.64E-02 | 2.63E+00 | 2.69E+00 | 6.63E-01 | no | Kirrel3 | intron_variant | MODIFIER |
| 5 | 127971388 | AX-170229101 | 2 | C | T | 0.436 | -3.24E-02 | 7.68E-02 | 2.68E+00 | 2.81E+00 | 6.63E-01 | no | Tmem132d | intron_variant | MODIFIER |
| 20 | 165526737 | AX-170513692 | 1 | C | T | 0.134 | 5.47E-02 | 1.28E-01 | 2.65E+00 | 2.77E+00 | 6.63E-01 | no | | intergenic_variant | MODIFIER |
| 10 | 12494451 | AX-97817820 | 0 | A | G | 0.211 | -4.37E-02 | 1.01E-01 | 2.64E+00 | 2.73E+00 | 6.63E-01 | no | Utrn | intron_variant | MODIFIER |
| 2 | 117560577 | AX-170223828 | 0 | A | T | 0.14 | 6.37E-02 | 1.49E-01 | 2.67E+00 | 2.78E+00 | 6.63E-01 | no | | intergenic_variant | MODIFIER |
| 8 | 37880081 | AX-170107988 | 1 | G | A | 0.152 | 6.32E-02 | 1.47E-01 | 2.79E+00 | 2.84E+00 | 6.63E-01 | no | Sgcz | intron_variant | MODIFIER |
| 8 | 40621028 | AX-170331859 | 1 | A | G | 0.152 | 6.32E-02 | 1.47E-01 | 2.79E+00 | 2.84E+00 | 6.63E-01 | no | Mtmr7 | intron_variant,NMD_transcript_variant | MODIFIER |
| 18 | 89110577 | AX-170061648 | 0 | T | A | 0.518 | -2.60E-02 | 6.18E-02 | 2.68E+00 | 2.82E+00 | 6.63E-01 | no | Rttn | intron_variant | MODIFIER |
| 14 | 111132759 | AX-170128987 | 0 | T | C | 0.316 | 3.22E-02 | 7.21E-02 | 2.55E+00 | 2.66E+00 | 6.63E-01 | no | | intergenic_variant | MODIFIER |
| 4 | 143499168 | AX-170137946 | 0 | A | C | 0.105 | -7.35E-02 | 1.72E-01 | 2.66E+00 | 2.76E+00 | 6.63E-01 | no | Gm13041 | upstream_gene_variant | MODIFIER |
| 15 | 4052703 | AX-170493831 | 0 | G | A | 0.246 | -4.58E-02 | 1.04E-01 | 2.60E+00 | 2.66E+00 | 6.63E-01 | no | Oxct1 | intron_variant | MODIFIER |
| 1 | 66642721 | AX-170041504 | 1 | G | T | 0.125 | 7.90E-02 | 1.82E-01 | 2.60E+00 | 2.71E+00 | 6.63E-01 | no | Unc80 | intron_variant | MODIFIER |
| 20 | 70843285 | AX-97789754 | 1 | G | A | 0.25 | 2.70E-02 | 6.29E-02 | 2.66E+00 | 2.74E+00 | 6.63E-01 | no | | intergenic_variant | MODIFIER |
| 9 | 35280173 | AX-170144681 | 0 | A | G | 0.421 | -3.76E-02 | 8.80E-02 | 2.78E+00 | 2.86E+00 | 6.64E-01 | no | Rpusd4 | downstream_gene_variant | MODIFIER |
| 20 | 67888451 | AX-170034067 | 1 | C | T | 0.232 | 2.72E-02 | 6.31E-02 | 2.64E+00 | 2.73E+00 | 6.64E-01 | no | | intergenic_variant | MODIFIER |
| 15 | 4106870 | AX-170010927 | 0 | G | A | 0.237 | -4.52E-02 | 1.02E-01 | 2.59E+00 | 2.66E+00 | 6.64E-01 | no | Oxct1 | intron_variant | MODIFIER |
| 15 | 6432390 | AX-170530474 | 0 | T | C | 0.237 | -4.52E-02 | 1.02E-01 | 2.59E+00 | 2.66E+00 | 6.64E-01 | no | Dab2 | intron_variant | MODIFIER |

| | | | | | | | | | | | | | | | |
|---|---|---|---|---|---|---|---|---|---|---|---|---|---|---|---|
| 1 | 39319081 | AX-170228781 | 0 | C | T | 0.404 | -3.28E-02 | 7.55E-02 | 2.62E+00 | 2.70E+00 | 6.64E-01 | no | Npas2 | intron_variant | MODIFIER |
| 1 | 101124472 | AX-170001205 | 0 | A | C | 0.158 | 6.65E-02 | 1.53E-01 | 2.61E+00 | 2.70E+00 | 6.64E-01 | no | | intergenic_variant | MODIFIER |
| 10 | 23321082 | AX-97794122 | 0 | G | T | 0.561 | -2.84E-02 | 6.81E-02 | 2.78E+00 | 2.93E+00 | 6.64E-01 | no | Eya4 | intron_variant | MODIFIER |
| 1 | 131505395 | AX-170129088 | 0 | G | A | 0.544 | 3.70E-02 | 8.45E-02 | 2.61E+00 | 2.68E+00 | 6.64E-01 | no | Srgap2 | intron_variant | MODIFIER |
| 15 | 79875848 | AX-170012080 | 0 | G | A | 0.561 | 3.60E-02 | 8.14E-02 | 2.58E+00 | 2.66E+00 | 6.64E-01 | no | Gm49412 | non_coding_transcript_exon_variant | MODIFIER |
| 5 | 113268967 | AX-170025921 | 0 | T | A | 0.456 | 4.84E-02 | 1.11E-01 | 2.63E+00 | 2.68E+00 | 6.64E-01 | no | Sgsm1 | intron_variant | MODIFIER |
| 1 | 110441310 | AX-170086573 | 1 | G | A | 0.161 | 8.42E-02 | 1.92E-01 | 2.60E+00 | 2.68E+00 | 6.64E-01 | no | | intergenic_variant | MODIFIER |
| 1 | 95995440 | AX-170428426 | 1 | C | T | 0.152 | 7.47E-02 | 1.69E-01 | 2.57E+00 | 2.68E+00 | 6.64E-01 | no | 1700063A18Rik | non_coding_transcript_exon_variant | MODIFIER |
| 1 | 191515909 | AX-170486234 | 0 | A | T | 0.263 | 4.52E-02 | 9.28E-02 | 2.43E+00 | 2.54E+00 | 6.64E-01 | no | | intergenic_variant | MODIFIER |
| 9 | 86361885 | AX-170083025 | 0 | C | A | 0.149 | 5.07E-02 | 1.20E-01 | 2.78E+00 | 2.88E+00 | 6.64E-01 | no | Ube2cbp | intron_variant | MODIFIER |
| 5 | 98085798 | AX-170072838 | 0 | G | A | 0.421 | -4.49E-02 | 1.05E-01 | 2.75E+00 | 2.83E+00 | 6.64E-01 | no | | intergenic_variant | MODIFIER |
| 18 | 63487966 | AX-97825218 | 0 | A | G | 0.395 | 4.28E-02 | 9.41E-02 | 2.54E+00 | 2.59E+00 | 6.64E-01 | no | | intergenic_variant | MODIFIER |
| 18 | 63488643 | AX-170418337 | 0 | C | T | 0.395 | 4.28E-02 | 9.41E-02 | 2.54E+00 | 2.59E+00 | 6.64E-01 | no | | intergenic_variant | MODIFIER |
| 18 | 63526459 | AX-170127209 | 0 | G | A | 0.395 | 4.28E-02 | 9.41E-02 | 2.54E+00 | 2.59E+00 | 6.64E-01 | no | | intergenic_variant | MODIFIER |
| 18 | 63528795 | AX-170488499 | 0 | A | T | 0.395 | 4.28E-02 | 9.41E-02 | 2.54E+00 | 2.59E+00 | 6.64E-01 | no | Gm22210 | upstream_gene_variant | MODIFIER |
| 18 | 63530968 | AX-170306894 | 0 | T | C | 0.395 | 4.28E-02 | 9.41E-02 | 2.54E+00 | 2.59E+00 | 6.64E-01 | no | Gm22210 | upstream_gene_variant | MODIFIER |
| 18 | 63530996 | AX-97785031 | 0 | C | T | 0.395 | 4.28E-02 | 9.41E-02 | 2.54E+00 | 2.59E+00 | 6.64E-01 | no | Gm22210 | upstream_gene_variant | MODIFIER |
| 18 | 63613948 | AX-170454233 | 0 | C | A | 0.395 | 4.28E-02 | 9.41E-02 | 2.54E+00 | 2.59E+00 | 6.64E-01 | no | | intergenic_variant | MODIFIER |
| 18 | 63921098 | AX-170146066 | 0 | C | T | 0.395 | 4.28E-02 | 9.41E-02 | 2.54E+00 | 2.59E+00 | 6.64E-01 | no | Wdr7 | intron_variant | MODIFIER |
| 18 | 63921417 | AX-170345750 | 0 | T | C | 0.395 | 4.28E-02 | 9.41E-02 | 2.54E+00 | 2.59E+00 | 6.64E-01 | no | Wdr7 | intron_variant | MODIFIER |
| 18 | 63992852 | AX-170201481 | 0 | C | T | 0.395 | 4.28E-02 | 9.41E-02 | 2.54E+00 | 2.59E+00 | 6.64E-01 | no | Wdr7 | downstream_gene_variant | MODIFIER |
| 18 | 64111996 | AX-170109706 | 0 | G | A | 0.395 | 4.28E-02 | 9.41E-02 | 2.54E+00 | 2.59E+00 | 6.64E-01 | no | | intergenic_variant | MODIFIER |
| 18 | 64277606 | AX-170476529 | 0 | A | G | 0.395 | 4.28E-02 | 9.41E-02 | 2.54E+00 | 2.59E+00 | 6.64E-01 | no | St8sia3 | 3_prime_UTR_variant | MODIFIER |
| 18 | 64278703 | AX-170299189 | 0 | C | A | 0.395 | 4.28E-02 | 9.41E-02 | 2.54E+00 | 2.59E+00 | 6.64E-01 | no | St8sia3 | 3_prime_UTR_variant | MODIFIER |
| 18 | 64280503 | AX-170363951 | 0 | T | C | 0.395 | 4.28E-02 | 9.41E-02 | 2.54E+00 | 2.59E+00 | 6.64E-01 | no | St8sia3 | downstream_gene_variant | MODIFIER |
| 18 | 64292817 | AX-170287999 | 0 | T | C | 0.395 | 4.28E-02 | 9.41E-02 | 2.54E+00 | 2.59E+00 | 6.64E-01 | no | St8sia3os | intron_variant,non_coding_transcript_variant | MODIFIER |
| 18 | 64294664 | AX-170230683 | 0 | C | G | 0.395 | 4.28E-02 | 9.41E-02 | 2.54E+00 | 2.59E+00 | 6.64E-01 | no | St8sia3os | intron_variant,non_coding_transcript_variant | MODIFIER |
| 18 | 64299849 | AX-170177339 | 0 | G | A | 0.395 | 4.28E-02 | 9.41E-02 | 2.54E+00 | 2.59E+00 | 6.64E-01 | no | St8sia3os | intron_variant,non_coding_transcript_variant | MODIFIER |
| 18 | 64387788 | AX-170215610 | 0 | A | C | 0.395 | 4.28E-02 | 9.41E-02 | 2.54E+00 | 2.59E+00 | 6.64E-01 | no | Onecut2 | 3_prime_UTR_variant | MODIFIER |
| 18 | 64420929 | AX-170118356 | 0 | C | A | 0.395 | 4.28E-02 | 9.41E-02 | 2.54E+00 | 2.59E+00 | 6.64E-01 | no | | intergenic_variant | MODIFIER |
| 18 | 64421041 | AX-170461002 | 0 | G | A | 0.395 | 4.28E-02 | 9.41E-02 | 2.54E+00 | 2.59E+00 | 6.64E-01 | no | | intergenic_variant | MODIFIER |
| 18 | 64431934 | AX-97835363 | 0 | C | T | 0.395 | 4.28E-02 | 9.41E-02 | 2.54E+00 | 2.59E+00 | 6.64E-01 | no | AC102106.2 | upstream_gene_variant | MODIFIER |
| 18 | 64695669 | AX-170450149 | 0 | G | A | 0.395 | 4.28E-02 | 9.41E-02 | 2.54E+00 | 2.59E+00 | 6.64E-01 | no | AC137585.1 | upstream_gene_variant | MODIFIER |
| 18 | 64699968 | AX-170245079 | 0 | C | G | 0.395 | 4.28E-02 | 9.41E-02 | 2.54E+00 | 2.59E+00 | 6.64E-01 | no | AC137585.1 | upstream_gene_variant | MODIFIER |
| 18 | 64702194 | AX-170428877 | 0 | C | G | 0.395 | 4.28E-02 | 9.41E-02 | 2.54E+00 | 2.59E+00 | 6.64E-01 | no | AC137585.1 | downstream_gene_variant | MODIFIER |
| 18 | 64705353 | AX-170439949 | 0 | T | C | 0.395 | 4.28E-02 | 9.41E-02 | 2.54E+00 | 2.59E+00 | 6.64E-01 | no | | intergenic_variant | MODIFIER |
| 18 | 64707412 | AX-170315925 | 0 | T | A | 0.395 | 4.28E-02 | 9.41E-02 | 2.54E+00 | 2.59E+00 | 6.64E-01 | no | | intergenic_variant | MODIFIER |
| 18 | 64711456 | AX-170318757 | 0 | C | T | 0.395 | 4.28E-02 | 9.41E-02 | 2.54E+00 | 2.59E+00 | 6.64E-01 | no | | intergenic_variant | MODIFIER |
| 18 | 64721108 | AX-170037836 | 0 | G | A | 0.395 | 4.28E-02 | 9.41E-02 | 2.54E+00 | 2.59E+00 | 6.64E-01 | no | | intergenic_variant | MODIFIER |
| 18 | 64732657 | AX-97844937 | 0 | G | A | 0.395 | 4.28E-02 | 9.41E-02 | 2.54E+00 | 2.59E+00 | 6.64E-01 | no | Gm23016 | downstream_gene_variant | MODIFIER |
| 18 | 64746645 | AX-170213112 | 0 | G | A | 0.395 | 4.28E-02 | 9.41E-02 | 2.54E+00 | 2.59E+00 | 6.64E-01 | no | | intergenic_variant | MODIFIER |
| 18 | 64749181 | AX-170289746 | 0 | A | G | 0.395 | 4.28E-02 | 9.41E-02 | 2.54E+00 | 2.59E+00 | 6.64E-01 | no | | intergenic_variant | MODIFIER |
| 18 | 64751342 | AX-170281750 | 0 | T | A | 0.395 | 4.28E-02 | 9.41E-02 | 2.54E+00 | 2.59E+00 | 6.64E-01 | no | | regulatory_region_variant | MODIFIER |
| 18 | 64756946 | AX-170228442 | 0 | A | G | 0.395 | 4.28E-02 | 9.41E-02 | 2.54E+00 | 2.59E+00 | 6.64E-01 | no | | intergenic_variant | MODIFIER |
| 18 | 64757848 | AX-170527341 | 0 | T | C | 0.395 | 4.28E-02 | 9.41E-02 | 2.54E+00 | 2.59E+00 | 6.64E-01 | no | | intergenic_variant | MODIFIER |
| 18 | 64759242 | AX-170162188 | 0 | G | C | 0.395 | 4.28E-02 | 9.41E-02 | 2.54E+00 | 2.59E+00 | 6.64E-01 | no | | intergenic_variant | MODIFIER |
| 18 | 64760768 | AX-170061112 | 0 | C | G | 0.395 | 4.28E-02 | 9.41E-02 | 2.54E+00 | 2.59E+00 | 6.64E-01 | no | | intergenic_variant | MODIFIER |
| 18 | 64760854 | AX-170061114 | 0 | T | C | 0.395 | 4.28E-02 | 9.41E-02 | 2.54E+00 | 2.59E+00 | 6.64E-01 | no | | intergenic_variant | MODIFIER |
| 18 | 64893763 | AX-170270666 | 0 | T | G | 0.395 | 4.28E-02 | 9.41E-02 | 2.54E+00 | 2.59E+00 | 6.64E-01 | no | Nedd4l | intron_variant | MODIFIER |
| 18 | 64898146 | AX-170117927 | 0 | A | G | 0.395 | 4.28E-02 | 9.41E-02 | 2.54E+00 | 2.59E+00 | 6.64E-01 | no | Nedd4l | intron_variant | MODIFIER |
| 1 | 140606100 | AX-97825663 | 0 | C | T | 0.211 | 5.22E-02 | 1.20E-01 | 2.61E+00 | 2.69E+00 | 6.64E-01 | no | Kcnt2 | intron_variant | MODIFIER |
| 8 | 94640790 | AX-97844511 | 1 | T | G | 0.214 | 4.92E-02 | 1.15E-01 | 2.72E+00 | 2.79E+00 | 6.64E-01 | no | Rspry1 | intron_variant | MODIFIER |
| 13 | 27037150 | AX-170007969 | 0 | A | G | 0.175 | -5.38E-02 | 1.22E-01 | 2.60E+00 | 2.66E+00 | 6.64E-01 | no | Gm47888 | intron_variant,non_coding_transcript_variant | MODIFIER |
| 13 | 30459929 | AX-97815901 | 0 | A | G | 0.175 | -5.38E-02 | 1.22E-01 | 2.60E+00 | 2.66E+00 | 6.64E-01 | no | | intergenic_variant | MODIFIER |

| | | | | | | | | | | | | | | | |
|---|---|---|---|---|---|---|---|---|---|---|---|---|---|---|---|
| 13 | 30485115 | AX-97790620 | 0 | C | T | 0.175 | -5.38E-02 | 1.22E-01 | 2.60E+00 | 2.66E+00 | 6.64E-01 | no | | regulatory_region_variant | MODIFIER |
| 13 | 30720147 | AX-170216122 | 0 | T | C | 0.175 | -5.38E-02 | 1.22E-01 | 2.60E+00 | 2.66E+00 | 6.64E-01 | no | | intergenic_variant | MODIFIER |
| 13 | 30816969 | AX-170087397 | 0 | C | T | 0.175 | -5.38E-02 | 1.22E-01 | 2.60E+00 | 2.66E+00 | 6.64E-01 | no | Exoc2 | intron_variant | MODIFIER |
| 13 | 30842238 | AX-97786086 | 0 | T | C | 0.175 | -5.38E-02 | 1.22E-01 | 2.60E+00 | 2.66E+00 | 6.64E-01 | no | Exoc2 | intron_variant | MODIFIER |
| 13 | 30992336 | AX-97823154 | 0 | G | A | 0.175 | -5.38E-02 | 1.22E-01 | 2.60E+00 | 2.66E+00 | 6.64E-01 | no | | intergenic_variant | MODIFIER |
| 13 | 31073033 | AX-97835942 | 0 | A | C | 0.175 | -5.38E-02 | 1.22E-01 | 2.60E+00 | 2.66E+00 | 6.64E-01 | no | Gm48662 | non_coding_transcript_exon_variant | MODIFIER |
| 13 | 31107921 | AX-170275770 | 0 | C | T | 0.175 | -5.38E-02 | 1.22E-01 | 2.60E+00 | 2.66E+00 | 6.64E-01 | no | | regulatory_region_variant | MODIFIER |
| 13 | 31108597 | AX-170162535 | 0 | T | C | 0.175 | -5.38E-02 | 1.22E-01 | 2.60E+00 | 2.66E+00 | 6.64E-01 | no | | intergenic_variant | MODIFIER |
| 13 | 31110202 | AX-170458322 | 0 | A | G | 0.175 | -5.38E-02 | 1.22E-01 | 2.60E+00 | 2.66E+00 | 6.64E-01 | no | | intergenic_variant | MODIFIER |
| 13 | 31118629 | AX-170103950 | 0 | C | A | 0.175 | -5.38E-02 | 1.22E-01 | 2.60E+00 | 2.66E+00 | 6.64E-01 | no | | intergenic_variant | MODIFIER |
| 13 | 31118908 | AX-170389715 | 0 | G | T | 0.175 | -5.38E-02 | 1.22E-01 | 2.60E+00 | 2.66E+00 | 6.64E-01 | no | | intergenic_variant | MODIFIER |
| 13 | 31120261 | AX-97798400 | 0 | C | A | 0.175 | -5.38E-02 | 1.22E-01 | 2.60E+00 | 2.66E+00 | 6.64E-01 | no | | intergenic_variant | MODIFIER |
| 13 | 31121595 | AX-170287030 | 0 | A | G | 0.175 | -5.38E-02 | 1.22E-01 | 2.60E+00 | 2.66E+00 | 6.64E-01 | no | | intergenic_variant | MODIFIER |
| 13 | 31122651 | AX-170303723 | 0 | G | C | 0.175 | -5.38E-02 | 1.22E-01 | 2.60E+00 | 2.66E+00 | 6.64E-01 | no | | intergenic_variant | MODIFIER |
| 13 | 31124264 | AX-170381630 | 0 | C | G | 0.175 | -5.38E-02 | 1.22E-01 | 2.60E+00 | 2.66E+00 | 6.64E-01 | no | | intergenic_variant | MODIFIER |
| 13 | 31124922 | AX-170102830 | 0 | T | G | 0.175 | -5.38E-02 | 1.22E-01 | 2.60E+00 | 2.66E+00 | 6.64E-01 | no | | intergenic_variant | MODIFIER |
| 13 | 31153421 | AX-170271188 | 0 | T | G | 0.175 | -5.38E-02 | 1.22E-01 | 2.60E+00 | 2.66E+00 | 6.64E-01 | no | | intergenic_variant | MODIFIER |
| 13 | 31187013 | AX-97783557 | 0 | A | G | 0.175 | -5.38E-02 | 1.22E-01 | 2.60E+00 | 2.66E+00 | 6.64E-01 | no | | regulatory_region_variant | MODIFIER |
| 13 | 31189235 | AX-170513218 | 0 | C | G | 0.175 | -5.38E-02 | 1.22E-01 | 2.60E+00 | 2.66E+00 | 6.64E-01 | no | | regulatory_region_variant | MODIFIER |
| 13 | 31209224 | AX-170176826 | 0 | G | A | 0.175 | -5.38E-02 | 1.22E-01 | 2.60E+00 | 2.66E+00 | 6.64E-01 | no | 4930401O12Rik | downstream_gene_variant | MODIFIER |
| 13 | 31210084 | AX-170254133 | 0 | C | A | 0.175 | -5.38E-02 | 1.22E-01 | 2.60E+00 | 2.66E+00 | 6.64E-01 | no | 4930401O12Rik | downstream_gene_variant | MODIFIER |
| 13 | 31210156 | AX-170316044 | 0 | G | A | 0.175 | -5.38E-02 | 1.22E-01 | 2.60E+00 | 2.66E+00 | 6.64E-01 | no | 4930401O12Rik | downstream_gene_variant | MODIFIER |
| 13 | 31217704 | AX-170385519 | 0 | G | A | 0.175 | -5.38E-02 | 1.22E-01 | 2.60E+00 | 2.66E+00 | 6.64E-01 | no | 4930401O12Rik | intron_variant,non_coding_transcript_variant | MODIFIER |
| 13 | 31218046 | AX-97820410 | 0 | A | G | 0.175 | -5.38E-02 | 1.22E-01 | 2.60E+00 | 2.66E+00 | 6.64E-01 | no | 4930401O12Rik | intron_variant,non_coding_transcript_variant | MODIFIER |
| 13 | 31232543 | AX-170386499 | 0 | C | A | 0.175 | -5.38E-02 | 1.22E-01 | 2.60E+00 | 2.66E+00 | 6.64E-01 | no | 4930401O12Rik | intron_variant,non_coding_transcript_variant | MODIFIER |
| 13 | 31241584 | AX-97794056 | 0 | G | A | 0.175 | -5.38E-02 | 1.22E-01 | 2.60E+00 | 2.66E+00 | 6.64E-01 | no | 4930401O12Rik | upstream_gene_variant | MODIFIER |
| 13 | 31258940 | AX-170092704 | 0 | C | T | 0.175 | -5.38E-02 | 1.22E-01 | 2.60E+00 | 2.66E+00 | 6.64E-01 | no | | intergenic_variant | MODIFIER |
| 13 | 31260992 | AX-170426492 | 0 | T | G | 0.175 | -5.38E-02 | 1.22E-01 | 2.60E+00 | 2.66E+00 | 6.64E-01 | no | | intergenic_variant | MODIFIER |
| 13 | 31261514 | AX-170100110 | 0 | C | G | 0.175 | -5.38E-02 | 1.22E-01 | 2.60E+00 | 2.66E+00 | 6.64E-01 | no | | intergenic_variant | MODIFIER |
| 13 | 31279210 | AX-170128753 | 0 | G | T | 0.175 | -5.38E-02 | 1.22E-01 | 2.60E+00 | 2.66E+00 | 6.64E-01 | no | | intergenic_variant | MODIFIER |
| 13 | 31297731 | AX-97823153 | 0 | A | G | 0.175 | -5.38E-02 | 1.22E-01 | 2.60E+00 | 2.66E+00 | 6.64E-01 | no | | intergenic_variant | MODIFIER |
| 13 | 31297962 | AX-170350628 | 0 | A | T | 0.175 | -5.38E-02 | 1.22E-01 | 2.60E+00 | 2.66E+00 | 6.64E-01 | no | | intergenic_variant | MODIFIER |
| 13 | 31299260 | AX-170227938 | 0 | G | C | 0.175 | -5.38E-02 | 1.22E-01 | 2.60E+00 | 2.66E+00 | 6.64E-01 | no | | intergenic_variant | MODIFIER |
| 13 | 31307994 | AX-170371514 | 0 | T | C | 0.175 | -5.38E-02 | 1.22E-01 | 2.60E+00 | 2.66E+00 | 6.64E-01 | no | | intergenic_variant | MODIFIER |
| 13 | 31309042 | AX-170360448 | 0 | C | T | 0.175 | -5.38E-02 | 1.22E-01 | 2.60E+00 | 2.66E+00 | 6.64E-01 | no | | regulatory_region_variant | MODIFIER |
| 13 | 31311287 | AX-170092705 | 0 | C | T | 0.175 | -5.38E-02 | 1.22E-01 | 2.60E+00 | 2.66E+00 | 6.64E-01 | no | | intergenic_variant | MODIFIER |
| 13 | 31317171 | AX-170351036 | 0 | G | C | 0.175 | -5.38E-02 | 1.22E-01 | 2.60E+00 | 2.66E+00 | 6.64E-01 | no | | regulatory_region_variant | MODIFIER |
| 13 | 31365981 | AX-170008016 | 0 | G | T | 0.175 | -5.38E-02 | 1.22E-01 | 2.60E+00 | 2.66E+00 | 6.64E-01 | no | G630018N14Rik | intron_variant,non_coding_transcript_variant | MODIFIER |
| 13 | 31380706 | AX-170008015 | 0 | G | A | 0.175 | -5.38E-02 | 1.22E-01 | 2.60E+00 | 2.66E+00 | 6.64E-01 | no | G630018N14Rik | intron_variant,non_coding_transcript_variant | MODIFIER |
| 13 | 31428451 | AX-170008019 | 0 | C | G | 0.175 | -5.38E-02 | 1.22E-01 | 2.60E+00 | 2.66E+00 | 6.64E-01 | no | Gm23351 | downstream_gene_variant | MODIFIER |
| 13 | 31429000 | AX-170008021 | 0 | T | C | 0.175 | -5.38E-02 | 1.22E-01 | 2.60E+00 | 2.66E+00 | 6.64E-01 | no | Gm23351 | downstream_gene_variant | MODIFIER |
| 13 | 31429210 | AX-97823203 | 0 | C | A | 0.175 | -5.38E-02 | 1.22E-01 | 2.60E+00 | 2.66E+00 | 6.64E-01 | no | Gm23351 | downstream_gene_variant | MODIFIER |
| 13 | 31439919 | AX-170362110 | 0 | T | C | 0.175 | -5.38E-02 | 1.22E-01 | 2.60E+00 | 2.66E+00 | 6.64E-01 | no | | regulatory_region_variant | MODIFIER |
| 13 | 31440702 | AX-170125004 | 0 | G | A | 0.175 | -5.38E-02 | 1.22E-01 | 2.60E+00 | 2.66E+00 | 6.64E-01 | no | | intergenic_variant | MODIFIER |
| 13 | 31440951 | AX-170358647 | 0 | G | A | 0.175 | -5.38E-02 | 1.22E-01 | 2.60E+00 | 2.66E+00 | 6.64E-01 | no | | intergenic_variant | MODIFIER |
| 13 | 31448663 | AX-170008017 | 0 | T | A | 0.175 | -5.38E-02 | 1.22E-01 | 2.60E+00 | 2.66E+00 | 6.64E-01 | no | | intergenic_variant | MODIFIER |
| 13 | 31462293 | AX-170050894 | 0 | G | A | 0.175 | -5.38E-02 | 1.22E-01 | 2.60E+00 | 2.66E+00 | 6.64E-01 | no | | intergenic_variant | MODIFIER |
| 13 | 31462396 | AX-97844492 | 0 | G | A | 0.175 | -5.38E-02 | 1.22E-01 | 2.60E+00 | 2.66E+00 | 6.64E-01 | no | | intergenic_variant | MODIFIER |
| 13 | 31488247 | AX-170277955 | 0 | T | C | 0.175 | -5.38E-02 | 1.22E-01 | 2.60E+00 | 2.66E+00 | 6.64E-01 | no | | intergenic_variant | MODIFIER |
| 13 | 31494057 | AX-97856165 | 0 | T | C | 0.175 | -5.38E-02 | 1.22E-01 | 2.60E+00 | 2.66E+00 | 6.64E-01 | no | | intergenic_variant | MODIFIER |
| 13 | 31495667 | AX-170008022 | 0 | A | C | 0.175 | -5.38E-02 | 1.22E-01 | 2.60E+00 | 2.66E+00 | 6.64E-01 | no | | intergenic_variant | MODIFIER |
| 13 | 31495688 | AX-170322996 | 0 | A | G | 0.175 | -5.38E-02 | 1.22E-01 | 2.60E+00 | 2.66E+00 | 6.64E-01 | no | | intergenic_variant | MODIFIER |
| 13 | 31497192 | AX-170328510 | 0 | A | G | 0.175 | -5.38E-02 | 1.22E-01 | 2.60E+00 | 2.66E+00 | 6.64E-01 | no | | intergenic_variant | MODIFIER |
| 13 | 31580292 | AX-170140576 | 0 | A | G | 0.175 | -5.38E-02 | 1.22E-01 | 2.60E+00 | 2.66E+00 | 6.64E-01 | no | 1700018A04Rik | intron_variant,non_coding_transcript_variant | MODIFIER |

| | | | | | | | | | | | | | | | |
|---|---|---|---|---|---|---|---|---|---|---|---|---|---|---|---|
| 13 | 31641977 | AX-170453157 | 0 | G | A | 0.175 | -5.38E-02 | 1.22E-01 | 2.60E+00 | 2.66E+00 | 6.64E-01 | no | | intergenic_variant | MODIFIER |
| 13 | 31644586 | AX-97835436 | 0 | G | A | 0.175 | -5.38E-02 | 1.22E-01 | 2.60E+00 | 2.66E+00 | 6.64E-01 | no | | intergenic_variant | MODIFIER |
| 13 | 31660239 | AX-170523155 | 0 | G | T | 0.175 | -5.38E-02 | 1.22E-01 | 2.60E+00 | 2.66E+00 | 6.64E-01 | no | | intergenic_variant | MODIFIER |
| 13 | 31667700 | AX-97793742 | 0 | T | C | 0.175 | -5.38E-02 | 1.22E-01 | 2.60E+00 | 2.66E+00 | 6.64E-01 | no | | intergenic_variant | MODIFIER |
| 13 | 31683278 | AX-97804010 | 0 | T | A | 0.175 | -5.38E-02 | 1.22E-01 | 2.60E+00 | 2.66E+00 | 6.64E-01 | no | | intergenic_variant | MODIFIER |
| 13 | 31689527 | AX-170160033 | 0 | T | A | 0.175 | -5.38E-02 | 1.22E-01 | 2.60E+00 | 2.66E+00 | 6.64E-01 | no | | regulatory_region_variant | MODIFIER |
| 13 | 31884165 | AX-170137685 | 0 | A | C | 0.175 | -5.38E-02 | 1.22E-01 | 2.60E+00 | 2.66E+00 | 6.64E-01 | no | Gmds | intron_variant | MODIFIER |
| 13 | 31886152 | AX-97823155 | 0 | T | G | 0.175 | -5.38E-02 | 1.22E-01 | 2.60E+00 | 2.66E+00 | 6.64E-01 | no | Gmds | intron_variant | MODIFIER |
| 13 | 31904484 | AX-170008027 | 0 | T | C | 0.175 | -5.38E-02 | 1.22E-01 | 2.60E+00 | 2.66E+00 | 6.64E-01 | no | Gmds | intron_variant | MODIFIER |
| 13 | 31911266 | AX-97856472 | 0 | T | C | 0.175 | -5.38E-02 | 1.22E-01 | 2.60E+00 | 2.66E+00 | 6.64E-01 | no | Gmds | intron_variant | MODIFIER |
| 13 | 31941920 | AX-170008024 | 0 | T | C | 0.175 | -5.38E-02 | 1.22E-01 | 2.60E+00 | 2.66E+00 | 6.64E-01 | no | Gmds | intron_variant | MODIFIER |
| 13 | 31941983 | AX-97827075 | 0 | T | C | 0.175 | -5.38E-02 | 1.22E-01 | 2.60E+00 | 2.66E+00 | 6.64E-01 | no | Gmds | intron_variant | MODIFIER |
| 13 | 31965233 | AX-170496458 | 0 | C | T | 0.175 | -5.38E-02 | 1.22E-01 | 2.60E+00 | 2.66E+00 | 6.64E-01 | no | Gmds | intron_variant | MODIFIER |
| 13 | 31970394 | AX-170008026 | 0 | A | T | 0.175 | -5.38E-02 | 1.22E-01 | 2.60E+00 | 2.66E+00 | 6.64E-01 | no | Gmds | intron_variant | MODIFIER |
| 13 | 31970908 | AX-170008033 | 0 | T | C | 0.175 | -5.38E-02 | 1.22E-01 | 2.60E+00 | 2.66E+00 | 6.64E-01 | no | Gmds | intron_variant | MODIFIER |
| 13 | 31971592 | AX-170418398 | 0 | C | T | 0.175 | -5.38E-02 | 1.22E-01 | 2.60E+00 | 2.66E+00 | 6.64E-01 | no | Gmds | intron_variant | MODIFIER |
| 13 | 31995283 | AX-170138820 | 0 | G | T | 0.175 | -5.38E-02 | 1.22E-01 | 2.60E+00 | 2.66E+00 | 6.64E-01 | no | Gmds | intron_variant | MODIFIER |
| 13 | 32011733 | AX-170008035 | 0 | A | G | 0.175 | -5.38E-02 | 1.22E-01 | 2.60E+00 | 2.66E+00 | 6.64E-01 | no | Gmds | intron_variant | MODIFIER |
| 13 | 32016891 | AX-97823222 | 0 | G | T | 0.175 | -5.38E-02 | 1.22E-01 | 2.60E+00 | 2.66E+00 | 6.64E-01 | no | Gmds | intron_variant | MODIFIER |
| 13 | 32038081 | AX-170150826 | 0 | G | A | 0.175 | -5.38E-02 | 1.22E-01 | 2.60E+00 | 2.66E+00 | 6.64E-01 | no | Gmds | intron_variant | MODIFIER |
| 13 | 32055964 | AX-170306617 | 0 | C | T | 0.175 | -5.38E-02 | 1.22E-01 | 2.60E+00 | 2.66E+00 | 6.64E-01 | no | Gmds | intron_variant | MODIFIER |
| 13 | 32065095 | AX-170087399 | 0 | A | G | 0.175 | -5.38E-02 | 1.22E-01 | 2.60E+00 | 2.66E+00 | 6.64E-01 | no | Gmds | intron_variant | MODIFIER |
| 13 | 32073265 | AX-170212760 | 0 | T | C | 0.175 | -5.38E-02 | 1.22E-01 | 2.60E+00 | 2.66E+00 | 6.64E-01 | no | Gmds | intron_variant | MODIFIER |
| 13 | 32097586 | AX-170087400 | 0 | A | C | 0.175 | -5.38E-02 | 1.22E-01 | 2.60E+00 | 2.66E+00 | 6.64E-01 | no | Gmds | intron_variant | MODIFIER |
| 13 | 32103531 | AX-170008034 | 0 | A | G | 0.175 | -5.38E-02 | 1.22E-01 | 2.60E+00 | 2.66E+00 | 6.64E-01 | no | Gmds | intron_variant | MODIFIER |
| 13 | 32121190 | AX-97789326 | 0 | A | G | 0.175 | -5.38E-02 | 1.22E-01 | 2.60E+00 | 2.66E+00 | 6.64E-01 | no | Gmds | intron_variant | MODIFIER |
| 13 | 32144755 | AX-97815186 | 0 | A | C | 0.175 | -5.38E-02 | 1.22E-01 | 2.60E+00 | 2.66E+00 | 6.64E-01 | no | Gmds | intron_variant | MODIFIER |
| 13 | 32147600 | AX-170050903 | 0 | G | C | 0.175 | -5.38E-02 | 1.22E-01 | 2.60E+00 | 2.66E+00 | 6.64E-01 | no | Gmds | intron_variant | MODIFIER |
| 13 | 32153985 | AX-170221781 | 0 | T | C | 0.175 | -5.38E-02 | 1.22E-01 | 2.60E+00 | 2.66E+00 | 6.64E-01 | no | Gmds | intron_variant | MODIFIER |
| 13 | 32168156 | AX-170008023 | 0 | A | G | 0.175 | -5.38E-02 | 1.22E-01 | 2.60E+00 | 2.66E+00 | 6.64E-01 | no | Gmds | intron_variant | MODIFIER |
| 13 | 32222647 | AX-170437887 | 0 | T | C | 0.175 | -5.38E-02 | 1.22E-01 | 2.60E+00 | 2.66E+00 | 6.64E-01 | no | Gmds | intron_variant | MODIFIER |
| 13 | 32222974 | AX-170249377 | 0 | T | C | 0.175 | -5.38E-02 | 1.22E-01 | 2.60E+00 | 2.66E+00 | 6.64E-01 | no | Gmds | intron_variant | MODIFIER |
| 13 | 32232460 | AX-170243424 | 0 | C | T | 0.175 | -5.38E-02 | 1.22E-01 | 2.60E+00 | 2.66E+00 | 6.64E-01 | no | Gmds | intron_variant | MODIFIER |
| 13 | 32232853 | AX-170450935 | 0 | G | C | 0.175 | -5.38E-02 | 1.22E-01 | 2.60E+00 | 2.66E+00 | 6.64E-01 | no | Gmds | intron_variant | MODIFIER |
| 13 | 32234815 | AX-170365586 | 0 | G | A | 0.175 | -5.38E-02 | 1.22E-01 | 2.60E+00 | 2.66E+00 | 6.64E-01 | no | Gmds | intron_variant | MODIFIER |
| 13 | 32244527 | AX-97857162 | 0 | C | A | 0.175 | -5.38E-02 | 1.22E-01 | 2.60E+00 | 2.66E+00 | 6.64E-01 | no | Gmds | intron_variant | MODIFIER |
| 13 | 32301655 | AX-97792143 | 0 | A | C | 0.175 | -5.38E-02 | 1.22E-01 | 2.60E+00 | 2.66E+00 | 6.64E-01 | no | Gmds | intron_variant | MODIFIER |
| 13 | 32301737 | AX-170293964 | 0 | A | C | 0.175 | -5.38E-02 | 1.22E-01 | 2.60E+00 | 2.66E+00 | 6.64E-01 | no | Gmds | intron_variant | MODIFIER |
| 13 | 32308594 | AX-168416985 | 0 | A | C | 0.175 | -5.38E-02 | 1.22E-01 | 2.60E+00 | 2.66E+00 | 6.64E-01 | no | Gmds | intron_variant | MODIFIER |
| 13 | 32314107 | AX-170492662 | 0 | G | A | 0.175 | -5.38E-02 | 1.22E-01 | 2.60E+00 | 2.66E+00 | 6.64E-01 | no | Gmds | intron_variant | MODIFIER |
| 13 | 32335790 | AX-97841632 | 0 | A | G | 0.175 | -5.38E-02 | 1.22E-01 | 2.60E+00 | 2.66E+00 | 6.64E-01 | no | Gmds | intron_variant | MODIFIER |
| 13 | 32380619 | AX-170414895 | 0 | A | G | 0.175 | -5.38E-02 | 1.22E-01 | 2.60E+00 | 2.66E+00 | 6.64E-01 | no | Gm11381 | intron_variant,non_coding_transcript_variant | MODIFIER |
| 13 | 32381397 | AX-170008039 | 0 | T | C | 0.175 | -5.38E-02 | 1.22E-01 | 2.60E+00 | 2.66E+00 | 6.64E-01 | no | Gm11381 | intron_variant,non_coding_transcript_variant | MODIFIER |
| 13 | 32411687 | AX-170008042 | 0 | A | T | 0.175 | -5.38E-02 | 1.22E-01 | 2.60E+00 | 2.66E+00 | 6.64E-01 | no | | regulatory_region_variant | MODIFIER |
| 13 | 32411721 | AX-97823220 | 0 | T | C | 0.175 | -5.38E-02 | 1.22E-01 | 2.60E+00 | 2.66E+00 | 6.64E-01 | no | | regulatory_region_variant | MODIFIER |
| 13 | 32478667 | AX-97823221 | 0 | T | C | 0.175 | -5.38E-02 | 1.22E-01 | 2.60E+00 | 2.66E+00 | 6.64E-01 | no | | intergenic_variant | MODIFIER |
| 13 | 32532662 | AX-170008041 | 0 | T | C | 0.175 | -5.38E-02 | 1.22E-01 | 2.60E+00 | 2.66E+00 | 6.64E-01 | no | Gm48073 | downstream_gene_variant | MODIFIER |
| 13 | 32534646 | AX-97850843 | 0 | G | A | 0.175 | -5.38E-02 | 1.22E-01 | 2.60E+00 | 2.66E+00 | 6.64E-01 | no | Gm48073 | non_coding_transcript_exon_variant | MODIFIER |
| 13 | 32643832 | AX-170273489 | 0 | C | G | 0.175 | -5.38E-02 | 1.22E-01 | 2.60E+00 | 2.66E+00 | 6.64E-01 | no | | intergenic_variant | MODIFIER |
| 13 | 32724588 | AX-97822431 | 0 | C | T | 0.175 | -5.38E-02 | 1.22E-01 | 2.60E+00 | 2.66E+00 | 6.64E-01 | no | Mylk4 | intron_variant | MODIFIER |
| 13 | 32766908 | AX-97823173 | 0 | A | G | 0.175 | -5.38E-02 | 1.22E-01 | 2.60E+00 | 2.66E+00 | 6.64E-01 | no | Mylk4 | intron_variant | MODIFIER |
| 13 | 32808928 | AX-97816338 | 0 | T | C | 0.175 | -5.38E-02 | 1.22E-01 | 2.60E+00 | 2.66E+00 | 6.64E-01 | no | Wrnip1 | intron_variant | MODIFIER |
| 13 | 32943624 | AX-170008054 | 0 | A | C | 0.175 | -5.38E-02 | 1.22E-01 | 2.60E+00 | 2.66E+00 | 6.64E-01 | no | Gm11382 | upstream_gene_variant | MODIFIER |
| 13 | 32945297 | AX-170087401 | 0 | T | C | 0.175 | -5.38E-02 | 1.22E-01 | 2.60E+00 | 2.66E+00 | 6.64E-01 | no | | intergenic_variant | MODIFIER |

| | | | | | | | | | | | | | | | |
|---|---|---|---|---|---|---|---|---|---|---|---|---|---|---|---|
| 13 | 32955028 | AX-170453515 | 0 | G | A | 0.175 | -5.38E-02 | 1.22E-01 | 2.60E+00 | 2.66E+00 | 6.64E-01 | no | | intergenic_variant | MODIFIER |
| 13 | 32955214 | AX-170467939 | 0 | C | G | 0.175 | -5.38E-02 | 1.22E-01 | 2.60E+00 | 2.66E+00 | 6.64E-01 | no | | intergenic_variant | MODIFIER |
| 13 | 33004990 | AX-97832649 | 0 | A | C | 0.175 | -5.38E-02 | 1.22E-01 | 2.60E+00 | 2.66E+00 | 6.64E-01 | no | Serpinb9 | intron_variant | MODIFIER |
| 13 | 33005135 | AX-170008052 | 0 | T | G | 0.175 | -5.38E-02 | 1.22E-01 | 2.60E+00 | 2.66E+00 | 6.64E-01 | no | Serpinb9 | intron_variant | MODIFIER |
| 13 | 33005556 | AX-170008057 | 0 | C | G | 0.175 | -5.38E-02 | 1.22E-01 | 2.60E+00 | 2.66E+00 | 6.64E-01 | no | Serpinb9 | intron_variant | MODIFIER |
| 13 | 33035686 | AX-170008048 | 0 | C | G | 0.175 | -5.38E-02 | 1.22E-01 | 2.60E+00 | 2.66E+00 | 6.64E-01 | no | Serpinb9b | intron_variant | MODIFIER |
| 13 | 33070806 | AX-170092712 | 0 | T | C | 0.175 | -5.38E-02 | 1.22E-01 | 2.60E+00 | 2.66E+00 | 6.64E-01 | no | | intergenic_variant | MODIFIER |
| 13 | 33071258 | AX-170391310 | 0 | G | A | 0.175 | -5.38E-02 | 1.22E-01 | 2.60E+00 | 2.66E+00 | 6.64E-01 | no | | intergenic_variant | MODIFIER |
| 13 | 33071339 | AX-170465261 | 0 | A | G | 0.175 | -5.38E-02 | 1.22E-01 | 2.60E+00 | 2.66E+00 | 6.64E-01 | no | | intergenic_variant | MODIFIER |
| 13 | 33133503 | AX-170517186 | 0 | T | G | 0.175 | -5.38E-02 | 1.22E-01 | 2.60E+00 | 2.66E+00 | 6.64E-01 | no | Gm11384 | upstream_gene_variant | MODIFIER |
| 13 | 33136590 | AX-170391466 | 0 | A | C | 0.175 | -5.38E-02 | 1.22E-01 | 2.60E+00 | 2.66E+00 | 6.64E-01 | no | Gm11384 | upstream_gene_variant | MODIFIER |
| 13 | 33151252 | AX-170092713 | 0 | T | C | 0.175 | -5.38E-02 | 1.22E-01 | 2.60E+00 | 2.66E+00 | 6.64E-01 | no | Serpinb9c | intron_variant | MODIFIER |
| 13 | 33214364 | AX-97794306 | 0 | C | G | 0.175 | -5.38E-02 | 1.22E-01 | 2.60E+00 | 2.66E+00 | 6.64E-01 | no | Gm11385 | downstream_gene_variant | MODIFIER |
| 13 | 33407888 | AX-170179827 | 0 | G | T | 0.175 | -5.38E-02 | 1.22E-01 | 2.60E+00 | 2.66E+00 | 6.64E-01 | no | Gm11397 | downstream_gene_variant | MODIFIER |
| 13 | 33514038 | AX-168416041 | 0 | C | T | 0.175 | -5.38E-02 | 1.22E-01 | 2.60E+00 | 2.66E+00 | 6.64E-01 | no | Gm11387 | intron_variant,non_coding_transcript_variant | MODIFIER |
| 13 | 34017349 | AX-97856966 | 0 | T | C | 0.175 | -5.38E-02 | 1.22E-01 | 2.60E+00 | 2.66E+00 | 6.64E-01 | no | Ripk1 | intron_variant | MODIFIER |
| 13 | 34042272 | AX-170427921 | 0 | G | C | 0.175 | -5.38E-02 | 1.22E-01 | 2.60E+00 | 2.66E+00 | 6.64E-01 | no | Bphl | intron_variant | MODIFIER |
| 13 | 34042303 | AX-170211572 | 0 | G | T | 0.175 | -5.38E-02 | 1.22E-01 | 2.60E+00 | 2.66E+00 | 6.64E-01 | no | Bphl | intron_variant | MODIFIER |
| 13 | 34044216 | AX-97845807 | 0 | G | A | 0.175 | -5.38E-02 | 1.22E-01 | 2.60E+00 | 2.66E+00 | 6.64E-01 | no | Bphl | intron_variant | MODIFIER |
| 13 | 34058666 | AX-168385718 | 0 | C | T | 0.175 | -5.38E-02 | 1.22E-01 | 2.60E+00 | 2.66E+00 | 6.64E-01 | no | Bphl | intron_variant | MODIFIER |
| 13 | 34061677 | AX-170132216 | 0 | G | C | 0.175 | -5.38E-02 | 1.22E-01 | 2.60E+00 | 2.66E+00 | 6.64E-01 | no | Bphl | intron_variant | MODIFIER |
| 13 | 34086106 | AX-97827100 | 0 | A | G | 0.175 | -5.38E-02 | 1.22E-01 | 2.60E+00 | 2.66E+00 | 6.64E-01 | no | Gm47065 | downstream_gene_variant | MODIFIER |
| 13 | 34930939 | AX-170449775 | 0 | G | C | 0.175 | -5.38E-02 | 1.22E-01 | 2.60E+00 | 2.66E+00 | 6.64E-01 | no | 4933417A18Rik | intron_variant | MODIFIER |
| 13 | 34940673 | AX-97825913 | 0 | A | C | 0.175 | -5.38E-02 | 1.22E-01 | 2.60E+00 | 2.66E+00 | 6.64E-01 | no | 4933417A18Rik | intron_variant | MODIFIER |
| 13 | 34956880 | AX-170008088 | 0 | T | C | 0.175 | -5.38E-02 | 1.22E-01 | 2.60E+00 | 2.66E+00 | 6.64E-01 | no | Eci3 | synonymous_variant | LOW |
| 13 | 34960545 | AX-97803478 | 0 | T | G | 0.175 | -5.38E-02 | 1.22E-01 | 2.60E+00 | 2.66E+00 | 6.64E-01 | no | Eci3 | intron_variant | MODIFIER |
| 13 | 35070745 | AX-170406251 | 0 | T | C | 0.175 | -5.38E-02 | 1.22E-01 | 2.60E+00 | 2.66E+00 | 6.64E-01 | no | Gm40910 | intron_variant,non_coding_transcript_variant | MODIFIER |
| 13 | 35088922 | AX-170220767 | 0 | A | G | 0.175 | -5.38E-02 | 1.22E-01 | 2.60E+00 | 2.66E+00 | 6.64E-01 | no | | intergenic_variant | MODIFIER |
| 13 | 35089099 | AX-97839393 | 0 | A | G | 0.175 | -5.38E-02 | 1.22E-01 | 2.60E+00 | 2.66E+00 | 6.64E-01 | no | | intergenic_variant | MODIFIER |
| 13 | 35101185 | AX-170175986 | 0 | C | G | 0.175 | -5.38E-02 | 1.22E-01 | 2.60E+00 | 2.66E+00 | 6.64E-01 | no | | intergenic_variant | MODIFIER |
| 13 | 35122486 | AX-97823201 | 0 | C | T | 0.175 | -5.38E-02 | 1.22E-01 | 2.60E+00 | 2.66E+00 | 6.64E-01 | no | | intergenic_variant | MODIFIER |
| 13 | 35131309 | AX-170179760 | 0 | C | T | 0.175 | -5.38E-02 | 1.22E-01 | 2.60E+00 | 2.66E+00 | 6.64E-01 | no | | regulatory_region_variant | MODIFIER |
| 13 | 35159255 | AX-170209031 | 0 | T | G | 0.175 | -5.38E-02 | 1.22E-01 | 2.60E+00 | 2.66E+00 | 6.64E-01 | no | | intergenic_variant | MODIFIER |
| 13 | 35167305 | AX-170300797 | 0 | G | C | 0.175 | -5.38E-02 | 1.22E-01 | 2.60E+00 | 2.66E+00 | 6.64E-01 | no | 4930529N20Rik | non_coding_transcript_exon_variant | MODIFIER |
| 13 | 35167382 | AX-97823279 | 0 | C | A | 0.175 | -5.38E-02 | 1.22E-01 | 2.60E+00 | 2.66E+00 | 6.64E-01 | no | 4930529N20Rik | non_coding_transcript_exon_variant | MODIFIER |
| 13 | 35241873 | AX-170087403 | 0 | C | T | 0.175 | -5.38E-02 | 1.22E-01 | 2.60E+00 | 2.66E+00 | 6.64E-01 | no | Gm48694 | downstream_gene_variant | MODIFIER |
| 13 | 35251702 | AX-170375545 | 0 | T | C | 0.175 | -5.38E-02 | 1.22E-01 | 2.60E+00 | 2.66E+00 | 6.64E-01 | no | Gm48694 | intron_variant,non_coding_transcript_variant | MODIFIER |
| 13 | 35259786 | AX-170494172 | 0 | T | C | 0.175 | -5.38E-02 | 1.22E-01 | 2.60E+00 | 2.66E+00 | 6.64E-01 | no | Gm48694 | upstream_gene_variant | MODIFIER |
| 13 | 35268894 | AX-170092720 | 0 | C | T | 0.175 | -5.38E-02 | 1.22E-01 | 2.60E+00 | 2.66E+00 | 6.64E-01 | no | | intergenic_variant | MODIFIER |
| 13 | 35270030 | AX-170326214 | 0 | A | C | 0.175 | -5.38E-02 | 1.22E-01 | 2.60E+00 | 2.66E+00 | 6.64E-01 | no | | intergenic_variant | MODIFIER |
| 13 | 35285470 | AX-170211008 | 0 | T | C | 0.175 | -5.38E-02 | 1.22E-01 | 2.60E+00 | 2.66E+00 | 6.64E-01 | no | | intergenic_variant | MODIFIER |
| 13 | 35286243 | AX-170370610 | 0 | C | A | 0.175 | -5.38E-02 | 1.22E-01 | 2.60E+00 | 2.66E+00 | 6.64E-01 | no | | intergenic_variant | MODIFIER |
| 13 | 35286758 | AX-97829316 | 0 | G | A | 0.175 | -5.38E-02 | 1.22E-01 | 2.60E+00 | 2.66E+00 | 6.64E-01 | no | | intergenic_variant | MODIFIER |
| 13 | 35315462 | AX-170168350 | 0 | A | G | 0.175 | -5.38E-02 | 1.22E-01 | 2.60E+00 | 2.66E+00 | 6.64E-01 | no | | intergenic_variant | MODIFIER |
| 13 | 35315712 | AX-170163809 | 0 | C | T | 0.175 | -5.38E-02 | 1.22E-01 | 2.60E+00 | 2.66E+00 | 6.64E-01 | no | | intergenic_variant | MODIFIER |
| 13 | 35316492 | AX-170008095 | 0 | A | G | 0.175 | -5.38E-02 | 1.22E-01 | 2.60E+00 | 2.66E+00 | 6.64E-01 | no | | intergenic_variant | MODIFIER |
| 13 | 35319410 | AX-97823196 | 0 | A | G | 0.175 | -5.38E-02 | 1.22E-01 | 2.60E+00 | 2.66E+00 | 6.64E-01 | no | | intergenic_variant | MODIFIER |
| 13 | 35345310 | AX-170376242 | 0 | A | G | 0.175 | -5.38E-02 | 1.22E-01 | 2.60E+00 | 2.66E+00 | 6.64E-01 | no | | intergenic_variant | MODIFIER |
| 13 | 35382961 | AX-170008099 | 0 | C | T | 0.175 | -5.38E-02 | 1.22E-01 | 2.60E+00 | 2.66E+00 | 6.64E-01 | no | | regulatory_region_variant | MODIFIER |
| 13 | 35466716 | AX-170008100 | 0 | T | C | 0.175 | -5.38E-02 | 1.22E-01 | 2.60E+00 | 2.66E+00 | 6.64E-01 | no | | intergenic_variant | MODIFIER |
| 13 | 35474843 | AX-97823199 | 0 | G | A | 0.175 | -5.38E-02 | 1.22E-01 | 2.60E+00 | 2.66E+00 | 6.64E-01 | no | | intergenic_variant | MODIFIER |
| 13 | 35479105 | AX-170008096 | 0 | T | A | 0.175 | -5.38E-02 | 1.22E-01 | 2.60E+00 | 2.66E+00 | 6.64E-01 | no | | intergenic_variant | MODIFIER |
| 13 | 35483446 | AX-97799022 | 0 | A | G | 0.175 | -5.38E-02 | 1.22E-01 | 2.60E+00 | 2.66E+00 | 6.64E-01 | no | | intergenic_variant | MODIFIER |
| 13 | 35541324 | AX-97823200 | 0 | C | T | 0.175 | -5.38E-02 | 1.22E-01 | 2.60E+00 | 2.66E+00 | 6.64E-01 | no | Gm48704 | upstream_gene_variant | MODIFIER |

| | | | | | | | | | | | | | | | |
|---|---|---|---|---|---|---|---|---|---|---|---|---|---|---|---|
| 13 | 35609627 | AX-97827106 | 0 | T | C | 0.175 | -5.38E-02 | 1.22E-01 | 2.60E+00 | 2.66E+00 | 6.64E-01 | no | Gm48707 | intron_variant,non_coding_transcript_variant | MODIFIER |
| 13 | 35648629 | AX-97847857 | 0 | A | C | 0.175 | -5.38E-02 | 1.22E-01 | 2.60E+00 | 2.66E+00 | 6.64E-01 | no | Gm48707 | intron_variant,non_coding_transcript_variant | MODIFIER |
| 13 | 35667295 | AX-97827103 | 0 | T | A | 0.175 | -5.38E-02 | 1.22E-01 | 2.60E+00 | 2.66E+00 | 6.64E-01 | no | Cdyl | intron_variant | MODIFIER |
| 13 | 35786932 | AX-170440678 | 0 | G | C | 0.175 | -5.38E-02 | 1.22E-01 | 2.60E+00 | 2.66E+00 | 6.64E-01 | no | Cdyl | intron_variant | MODIFIER |
| 13 | 35803458 | AX-97831890 | 0 | G | A | 0.175 | -5.38E-02 | 1.22E-01 | 2.60E+00 | 2.66E+00 | 6.64E-01 | no | Cdyl | intron_variant | MODIFIER |
| 13 | 35904413 | AX-170008104 | 0 | T | C | 0.175 | -5.38E-02 | 1.22E-01 | 2.60E+00 | 2.66E+00 | 6.64E-01 | no | Rpp40 | splice_region_variant,intron_variant | LOW |
| 13 | 35914560 | AX-170283184 | 0 | A | G | 0.175 | -5.38E-02 | 1.22E-01 | 2.60E+00 | 2.66E+00 | 6.64E-01 | no | Gm48746 | intron_variant,non_coding_transcript_variant | MODIFIER |
| 13 | 35921100 | AX-97820994 | 0 | G | A | 0.175 | -5.38E-02 | 1.22E-01 | 2.60E+00 | 2.66E+00 | 6.64E-01 | no | Gm48746 | intron_variant,non_coding_transcript_variant | MODIFIER |
| 13 | 35930261 | AX-170415694 | 0 | G | T | 0.175 | -5.38E-02 | 1.22E-01 | 2.60E+00 | 2.66E+00 | 6.64E-01 | no | | intergenic_variant | MODIFIER |
| 13 | 35943569 | AX-170305984 | 0 | C | T | 0.175 | -5.38E-02 | 1.22E-01 | 2.60E+00 | 2.66E+00 | 6.64E-01 | no | | intergenic_variant | MODIFIER |
| 13 | 35953746 | AX-170249428 | 0 | C | T | 0.175 | -5.38E-02 | 1.22E-01 | 2.60E+00 | 2.66E+00 | 6.64E-01 | no | Gm30127 | downstream_gene_variant | MODIFIER |
| 13 | 35954256 | AX-170268243 | 0 | G | A | 0.175 | -5.38E-02 | 1.22E-01 | 2.60E+00 | 2.66E+00 | 6.64E-01 | no | Gm30127 | downstream_gene_variant | MODIFIER |
| 13 | 35954325 | AX-170403311 | 0 | T | C | 0.175 | -5.38E-02 | 1.22E-01 | 2.60E+00 | 2.66E+00 | 6.64E-01 | no | Gm30127 | downstream_gene_variant | MODIFIER |
| 13 | 35979464 | AX-170511005 | 0 | G | T | 0.175 | -5.38E-02 | 1.22E-01 | 2.60E+00 | 2.66E+00 | 6.64E-01 | no | Lyrm4 | 3_prime_UTR_variant | MODIFIER |
| 13 | 35982031 | AX-170008103 | 0 | A | G | 0.175 | -5.38E-02 | 1.22E-01 | 2.60E+00 | 2.66E+00 | 6.64E-01 | no | Lyrm4 | intron_variant | MODIFIER |
| 13 | 35996229 | AX-170144053 | 0 | T | C | 0.175 | -5.38E-02 | 1.22E-01 | 2.60E+00 | 2.66E+00 | 6.64E-01 | no | Lyrm4 | intron_variant | MODIFIER |
| 13 | 36004210 | AX-97805761 | 0 | G | A | 0.175 | -5.38E-02 | 1.22E-01 | 2.60E+00 | 2.66E+00 | 6.64E-01 | no | Lyrm4 | intron_variant | MODIFIER |
| 13 | 36015897 | AX-97819434 | 0 | A | G | 0.175 | -5.38E-02 | 1.22E-01 | 2.60E+00 | 2.66E+00 | 6.64E-01 | no | Lyrm4 | intron_variant | MODIFIER |
| 13 | 36028186 | AX-170525021 | 0 | C | A | 0.175 | -5.38E-02 | 1.22E-01 | 2.60E+00 | 2.66E+00 | 6.64E-01 | no | Lyrm4 | intron_variant | MODIFIER |
| 13 | 36050583 | AX-97780862 | 0 | T | C | 0.175 | -5.38E-02 | 1.22E-01 | 2.60E+00 | 2.66E+00 | 6.64E-01 | no | Lyrm4 | intron_variant | MODIFIER |
| 13 | 36061907 | AX-170008106 | 0 | T | C | 0.175 | -5.38E-02 | 1.22E-01 | 2.60E+00 | 2.66E+00 | 6.64E-01 | no | Lyrm4 | intron_variant | MODIFIER |
| 13 | 36062253 | AX-170008107 | 0 | G | A | 0.175 | -5.38E-02 | 1.22E-01 | 2.60E+00 | 2.66E+00 | 6.64E-01 | no | Lyrm4 | intron_variant | MODIFIER |
| 13 | 36070207 | AX-170107440 | 0 | C | G | 0.175 | -5.38E-02 | 1.22E-01 | 2.60E+00 | 2.66E+00 | 6.64E-01 | no | Lyrm4 | intron_variant | MODIFIER |
| 13 | 36072330 | AX-97806104 | 0 | A | C | 0.175 | -5.38E-02 | 1.22E-01 | 2.60E+00 | 2.66E+00 | 6.64E-01 | no | Lyrm4 | intron_variant | MODIFIER |
| 13 | 36088015 | AX-170361331 | 0 | C | T | 0.175 | -5.38E-02 | 1.22E-01 | 2.60E+00 | 2.66E+00 | 6.64E-01 | no | Lyrm4 | intron_variant | MODIFIER |
| 13 | 36090606 | AX-97803011 | 0 | G | A | 0.175 | -5.38E-02 | 1.22E-01 | 2.60E+00 | 2.66E+00 | 6.64E-01 | no | Lyrm4 | intron_variant | MODIFIER |
| 13 | 36125562 | AX-170463266 | 0 | G | A | 0.175 | -5.38E-02 | 1.22E-01 | 2.60E+00 | 2.66E+00 | 6.64E-01 | no | Fars2 | intron_variant | MODIFIER |
| 13 | 36128527 | AX-170050953 | 0 | A | G | 0.175 | -5.38E-02 | 1.22E-01 | 2.60E+00 | 2.66E+00 | 6.64E-01 | no | Fars2 | intron_variant | MODIFIER |
| 13 | 36129106 | AX-170008109 | 0 | G | A | 0.175 | -5.38E-02 | 1.22E-01 | 2.60E+00 | 2.66E+00 | 6.64E-01 | no | Fars2 | intron_variant | MODIFIER |
| 13 | 36129806 | AX-170388956 | 0 | C | T | 0.175 | -5.38E-02 | 1.22E-01 | 2.60E+00 | 2.66E+00 | 6.64E-01 | no | Fars2 | intron_variant | MODIFIER |
| 13 | 36137059 | AX-170169821 | 0 | A | G | 0.175 | -5.38E-02 | 1.22E-01 | 2.60E+00 | 2.66E+00 | 6.64E-01 | no | Fars2 | intron_variant | MODIFIER |
| 13 | 36151002 | AX-170092721 | 0 | T | C | 0.175 | -5.38E-02 | 1.22E-01 | 2.60E+00 | 2.66E+00 | 6.64E-01 | no | Fars2 | intron_variant | MODIFIER |
| 13 | 36153999 | AX-170302150 | 0 | A | C | 0.175 | -5.38E-02 | 1.22E-01 | 2.60E+00 | 2.66E+00 | 6.64E-01 | no | Fars2 | intron_variant | MODIFIER |
| 13 | 36174287 | AX-97823301 | 0 | T | C | 0.175 | -5.38E-02 | 1.22E-01 | 2.60E+00 | 2.66E+00 | 6.64E-01 | no | Fars2 | intron_variant | MODIFIER |
| 13 | 36174369 | AX-170395523 | 0 | G | A | 0.175 | -5.38E-02 | 1.22E-01 | 2.60E+00 | 2.66E+00 | 6.64E-01 | no | Fars2 | intron_variant | MODIFIER |
| 13 | 36176533 | AX-170306195 | 0 | T | C | 0.175 | -5.38E-02 | 1.22E-01 | 2.60E+00 | 2.66E+00 | 6.64E-01 | no | Fars2 | intron_variant | MODIFIER |
| 13 | 36176698 | AX-170491095 | 0 | C | G | 0.175 | -5.38E-02 | 1.22E-01 | 2.60E+00 | 2.66E+00 | 6.64E-01 | no | Fars2 | intron_variant | MODIFIER |
| 13 | 36179486 | AX-170185633 | 0 | G | A | 0.175 | -5.38E-02 | 1.22E-01 | 2.60E+00 | 2.66E+00 | 6.64E-01 | no | Fars2 | intron_variant | MODIFIER |
| 13 | 36201090 | AX-97824243 | 0 | A | C | 0.175 | -5.38E-02 | 1.22E-01 | 2.60E+00 | 2.66E+00 | 6.64E-01 | no | Fars2 | intron_variant | MODIFIER |
| 13 | 36205437 | AX-170479259 | 0 | A | G | 0.175 | -5.38E-02 | 1.22E-01 | 2.60E+00 | 2.66E+00 | 6.64E-01 | no | Fars2 | intron_variant | MODIFIER |
| 13 | 36218461 | AX-170114523 | 0 | C | T | 0.175 | -5.38E-02 | 1.22E-01 | 2.60E+00 | 2.66E+00 | 6.64E-01 | no | Fars2 | intron_variant | MODIFIER |
| 13 | 36237614 | AX-170087406 | 0 | T | G | 0.175 | -5.38E-02 | 1.22E-01 | 2.60E+00 | 2.66E+00 | 6.64E-01 | no | Fars2 | intron_variant | MODIFIER |
| 13 | 36264825 | AX-170430264 | 0 | G | A | 0.175 | -5.38E-02 | 1.22E-01 | 2.60E+00 | 2.66E+00 | 6.64E-01 | no | Fars2 | intron_variant | MODIFIER |
| 13 | 36276849 | AX-170275541 | 0 | C | A | 0.175 | -5.38E-02 | 1.22E-01 | 2.60E+00 | 2.66E+00 | 6.64E-01 | no | Fars2 | intron_variant | MODIFIER |
| 13 | 36278368 | AX-170107082 | 0 | T | C | 0.175 | -5.38E-02 | 1.22E-01 | 2.60E+00 | 2.66E+00 | 6.64E-01 | no | Fars2 | intron_variant | MODIFIER |
| 13 | 36302589 | AX-97812164 | 0 | C | T | 0.175 | -5.38E-02 | 1.22E-01 | 2.60E+00 | 2.66E+00 | 6.64E-01 | no | Fars2 | intron_variant | MODIFIER |
| 13 | 36317726 | AX-97849123 | 0 | A | C | 0.175 | -5.38E-02 | 1.22E-01 | 2.60E+00 | 2.66E+00 | 6.64E-01 | no | Fars2 | intron_variant | MODIFIER |
| 13 | 36324402 | AX-170528656 | 0 | C | G | 0.175 | -5.38E-02 | 1.22E-01 | 2.60E+00 | 2.66E+00 | 6.64E-01 | no | Fars2 | intron_variant | MODIFIER |
| 13 | 36324739 | AX-170424595 | 0 | G | A | 0.175 | -5.38E-02 | 1.22E-01 | 2.60E+00 | 2.66E+00 | 6.64E-01 | no | Fars2 | intron_variant | MODIFIER |
| 13 | 36328339 | AX-170377077 | 0 | G | A | 0.175 | -5.38E-02 | 1.22E-01 | 2.60E+00 | 2.66E+00 | 6.64E-01 | no | Fars2 | intron_variant | MODIFIER |
| 13 | 36357902 | AX-170200472 | 0 | C | T | 0.175 | -5.38E-02 | 1.22E-01 | 2.60E+00 | 2.66E+00 | 6.64E-01 | no | Fars2 | intron_variant | MODIFIER |
| 13 | 36382511 | AX-170129843 | 0 | G | T | 0.175 | -5.38E-02 | 1.22E-01 | 2.60E+00 | 2.66E+00 | 6.64E-01 | no | Fars2 | intron_variant | MODIFIER |
| 13 | 36416324 | AX-170050958 | 0 | G | A | 0.175 | -5.38E-02 | 1.22E-01 | 2.60E+00 | 2.66E+00 | 6.64E-01 | no | Fars2 | intron_variant | MODIFIER |
| 13 | 36421107 | AX-97852492 | 0 | G | A | 0.175 | -5.38E-02 | 1.22E-01 | 2.60E+00 | 2.66E+00 | 6.64E-01 | no | Fars2 | intron_variant | MODIFIER |

| | | | | | | | | | | | | | | | |
|---|---|---|---|---|---|---|---|---|---|---|---|---|---|---|---|
| 13 | 36421623 | AX-170050961 | 0 | G | C | 0.175 | -5.38E-02 | 1.22E-01 | 2.60E+00 | 2.66E+00 | 6.64E-01 | no | Fars2 | intron_variant | MODIFIER |
| 13 | 36431426 | AX-168395948 | 0 | G | A | 0.175 | -5.38E-02 | 1.22E-01 | 2.60E+00 | 2.66E+00 | 6.64E-01 | no | Fars2 | intron_variant | MODIFIER |
| 13 | 36519569 | AX-170008118 | 0 | T | C | 0.175 | -5.38E-02 | 1.22E-01 | 2.60E+00 | 2.66E+00 | 6.64E-01 | no | Fars2 | intron_variant | MODIFIER |
| 13 | 36542556 | AX-97823323 | 0 | T | C | 0.175 | -5.38E-02 | 1.22E-01 | 2.60E+00 | 2.66E+00 | 6.64E-01 | no | Fars2 | downstream_gene_variant | MODIFIER |
| 13 | 36685498 | AX-170104606 | 0 | G | C | 0.175 | -5.38E-02 | 1.22E-01 | 2.60E+00 | 2.66E+00 | 6.64E-01 | no | Fars2 | intron_variant,NMD_transcript_variant | MODIFIER |
| 13 | 36943692 | AX-170087408 | 0 | T | C | 0.175 | -5.38E-02 | 1.22E-01 | 2.60E+00 | 2.66E+00 | 6.64E-01 | no | F13a1 | intron_variant | MODIFIER |
| 13 | 36972100 | AX-97814793 | 0 | T | C | 0.175 | -5.38E-02 | 1.22E-01 | 2.60E+00 | 2.66E+00 | 6.64E-01 | no | F13a1 | intron_variant | MODIFIER |
| 13 | 37104661 | AX-170331256 | 0 | T | C | 0.175 | -5.38E-02 | 1.22E-01 | 2.60E+00 | 2.66E+00 | 6.64E-01 | no | | regulatory_region_variant | MODIFIER |
| 13 | 37205941 | AX-170207558 | 0 | A | G | 0.175 | -5.38E-02 | 1.22E-01 | 2.60E+00 | 2.66E+00 | 6.64E-01 | no | | intergenic_variant | MODIFIER |
| 13 | 37216533 | AX-170403684 | 0 | T | A | 0.175 | -5.38E-02 | 1.22E-01 | 2.60E+00 | 2.66E+00 | 6.64E-01 | no | | intergenic_variant | MODIFIER |
| 13 | 37232176 | AX-170393626 | 0 | G | A | 0.175 | -5.38E-02 | 1.22E-01 | 2.60E+00 | 2.66E+00 | 6.64E-01 | no | | intergenic_variant | MODIFIER |
| 13 | 37234188 | AX-170430125 | 0 | C | T | 0.175 | -5.38E-02 | 1.22E-01 | 2.60E+00 | 2.66E+00 | 6.64E-01 | no | Gm30600 | downstream_gene_variant | MODIFIER |
| 13 | 37247118 | AX-97823251 | 0 | A | G | 0.175 | -5.38E-02 | 1.22E-01 | 2.60E+00 | 2.66E+00 | 6.64E-01 | no | Gm30600 | upstream_gene_variant | MODIFIER |
| 13 | 37247445 | AX-170050981 | 0 | G | C | 0.175 | -5.38E-02 | 1.22E-01 | 2.60E+00 | 2.66E+00 | 6.64E-01 | no | Gm30600 | upstream_gene_variant | MODIFIER |
| 13 | 37248364 | AX-170279955 | 0 | G | T | 0.175 | -5.38E-02 | 1.22E-01 | 2.60E+00 | 2.66E+00 | 6.64E-01 | no | Gm30600 | upstream_gene_variant | MODIFIER |
| 13 | 37251843 | AX-170468414 | 0 | T | C | 0.175 | -5.38E-02 | 1.22E-01 | 2.60E+00 | 2.66E+00 | 6.64E-01 | no | Gm47709 | non_coding_transcript_exon_variant | MODIFIER |
| 13 | 37253438 | AX-170493669 | 0 | T | C | 0.175 | -5.38E-02 | 1.22E-01 | 2.60E+00 | 2.66E+00 | 6.64E-01 | no | Gm47709 | downstream_gene_variant | MODIFIER |
| 13 | 37253828 | AX-170264238 | 0 | G | A | 0.175 | -5.38E-02 | 1.22E-01 | 2.60E+00 | 2.66E+00 | 6.64E-01 | no | Gm47709 | downstream_gene_variant | MODIFIER |
| 13 | 37253941 | AX-170200875 | 0 | A | G | 0.175 | -5.38E-02 | 1.22E-01 | 2.60E+00 | 2.66E+00 | 6.64E-01 | no | Gm47709 | downstream_gene_variant | MODIFIER |
| 13 | 37253965 | AX-170483723 | 0 | A | G | 0.175 | -5.38E-02 | 1.22E-01 | 2.60E+00 | 2.66E+00 | 6.64E-01 | no | Gm47709 | downstream_gene_variant | MODIFIER |
| 13 | 37272076 | AX-170306913 | 0 | C | A | 0.175 | -5.38E-02 | 1.22E-01 | 2.60E+00 | 2.66E+00 | 6.64E-01 | no | | intergenic_variant | MODIFIER |
| 13 | 37277216 | AX-97804986 | 0 | T | G | 0.175 | -5.38E-02 | 1.22E-01 | 2.60E+00 | 2.66E+00 | 6.64E-01 | no | | intergenic_variant | MODIFIER |
| 13 | 37277682 | AX-170515325 | 0 | A | G | 0.175 | -5.38E-02 | 1.22E-01 | 2.60E+00 | 2.66E+00 | 6.64E-01 | no | | intergenic_variant | MODIFIER |
| 13 | 37277731 | AX-170508132 | 0 | C | T | 0.175 | -5.38E-02 | 1.22E-01 | 2.60E+00 | 2.66E+00 | 6.64E-01 | no | | intergenic_variant | MODIFIER |
| 13 | 37282420 | AX-170176921 | 0 | T | C | 0.175 | -5.38E-02 | 1.22E-01 | 2.60E+00 | 2.66E+00 | 6.64E-01 | no | | intergenic_variant | MODIFIER |
| 13 | 37285082 | AX-170185840 | 0 | T | C | 0.175 | -5.38E-02 | 1.22E-01 | 2.60E+00 | 2.66E+00 | 6.64E-01 | no | | intergenic_variant | MODIFIER |
| 13 | 37297316 | AX-170175044 | 0 | T | C | 0.175 | -5.38E-02 | 1.22E-01 | 2.60E+00 | 2.66E+00 | 6.64E-01 | no | | intergenic_variant | MODIFIER |
| 13 | 37300089 | AX-170527786 | 0 | T | C | 0.175 | -5.38E-02 | 1.22E-01 | 2.60E+00 | 2.66E+00 | 6.64E-01 | no | | intergenic_variant | MODIFIER |
| 13 | 37300148 | AX-170318588 | 0 | A | G | 0.175 | -5.38E-02 | 1.22E-01 | 2.60E+00 | 2.66E+00 | 6.64E-01 | no | | intergenic_variant | MODIFIER |
| 9 | 84146988 | AX-170246454 | 1 | T | G | 0.214 | 5.33E-02 | 1.25E-01 | 2.75E+00 | 2.82E+00 | 6.64E-01 | no | | intergenic_variant | MODIFIER |
| 15 | 92626219 | AX-97830233 | 1 | A | G | 0.161 | -5.68E-02 | 1.34E-01 | 2.76E+00 | 2.86E+00 | 6.64E-01 | no | Pdzrn4 | intron_variant | MODIFIER |
| 20 | 69734800 | AX-170084450 | 0 | A | G | 0.254 | 2.69E-02 | 6.27E-02 | 2.65E+00 | 2.74E+00 | 6.64E-01 | no | Aff2 | intron_variant | MODIFIER |
| 20 | 69736416 | AX-170034100 | 0 | A | G | 0.254 | 2.69E-02 | 6.27E-02 | 2.65E+00 | 2.74E+00 | 6.64E-01 | no | Aff2 | intron_variant | MODIFIER |
| 20 | 69738019 | AX-97800432 | 0 | A | G | 0.254 | 2.69E-02 | 6.27E-02 | 2.65E+00 | 2.74E+00 | 6.64E-01 | no | Aff2 | intron_variant | MODIFIER |
| 20 | 69768170 | AX-97806082 | 0 | G | A | 0.254 | 2.69E-02 | 6.27E-02 | 2.65E+00 | 2.74E+00 | 6.64E-01 | no | Aff2 | intron_variant | MODIFIER |
| 20 | 69774326 | AX-170084451 | 0 | G | A | 0.254 | 2.69E-02 | 6.27E-02 | 2.65E+00 | 2.74E+00 | 6.64E-01 | no | Aff2 | intron_variant | MODIFIER |
| 20 | 69785121 | AX-97785296 | 0 | T | C | 0.254 | 2.69E-02 | 6.27E-02 | 2.65E+00 | 2.74E+00 | 6.64E-01 | no | Aff2 | intron_variant | MODIFIER |
| 20 | 69801710 | AX-170357842 | 0 | C | T | 0.254 | 2.69E-02 | 6.27E-02 | 2.65E+00 | 2.74E+00 | 6.64E-01 | no | Aff2 | intron_variant | MODIFIER |
| 20 | 69826919 | AX-97784698 | 0 | C | T | 0.254 | 2.69E-02 | 6.27E-02 | 2.65E+00 | 2.74E+00 | 6.64E-01 | no | Aff2 | intron_variant | MODIFIER |
| 20 | 69833941 | AX-170374303 | 0 | C | A | 0.254 | 2.69E-02 | 6.27E-02 | 2.65E+00 | 2.74E+00 | 6.64E-01 | no | Aff2 | intron_variant | MODIFIER |
| 20 | 69834798 | AX-170312303 | 0 | G | A | 0.254 | 2.69E-02 | 6.27E-02 | 2.65E+00 | 2.74E+00 | 6.64E-01 | no | Aff2 | missense_variant | MODERATE |
| 20 | 69837652 | AX-97834537 | 0 | C | T | 0.254 | 2.69E-02 | 6.27E-02 | 2.65E+00 | 2.74E+00 | 6.64E-01 | no | Aff2 | intron_variant | MODIFIER |
| 20 | 69859326 | AX-170286204 | 0 | T | C | 0.254 | 2.69E-02 | 6.27E-02 | 2.65E+00 | 2.74E+00 | 6.64E-01 | no | Aff2 | intron_variant | MODIFIER |
| 20 | 69859660 | AX-97802455 | 0 | A | C | 0.254 | 2.69E-02 | 6.27E-02 | 2.65E+00 | 2.74E+00 | 6.64E-01 | no | Aff2 | intron_variant | MODIFIER |
| 20 | 69865006 | AX-170279921 | 0 | A | C | 0.254 | 2.69E-02 | 6.27E-02 | 2.65E+00 | 2.74E+00 | 6.64E-01 | no | Aff2 | intron_variant | MODIFIER |
| 20 | 70167850 | AX-170346736 | 0 | A | G | 0.254 | 2.69E-02 | 6.27E-02 | 2.65E+00 | 2.74E+00 | 6.64E-01 | no | | intergenic_variant | MODIFIER |
| 20 | 70379297 | AX-170481296 | 0 | A | C | 0.254 | 2.69E-02 | 6.27E-02 | 2.65E+00 | 2.74E+00 | 6.64E-01 | no | | intergenic_variant | MODIFIER |
| 20 | 70773407 | AX-170151009 | 0 | C | T | 0.254 | 2.69E-02 | 6.27E-02 | 2.65E+00 | 2.74E+00 | 6.64E-01 | no | Gm8410 | downstream_gene_variant | MODIFIER |
| 20 | 70811983 | AX-170263205 | 0 | T | C | 0.254 | 2.69E-02 | 6.27E-02 | 2.65E+00 | 2.74E+00 | 6.64E-01 | no | | intergenic_variant | MODIFIER |
| 20 | 70812863 | AX-170267087 | 0 | C | T | 0.254 | 2.69E-02 | 6.27E-02 | 2.65E+00 | 2.74E+00 | 6.64E-01 | no | | intergenic_variant | MODIFIER |
| 20 | 70873112 | AX-170034108 | 0 | C | T | 0.254 | 2.69E-02 | 6.27E-02 | 2.65E+00 | 2.74E+00 | 6.64E-01 | no | | intergenic_variant | MODIFIER |
| 20 | 70886730 | AX-97804834 | 0 | A | G | 0.254 | 2.69E-02 | 6.27E-02 | 2.65E+00 | 2.74E+00 | 6.64E-01 | no | | intergenic_variant | MODIFIER |
| 20 | 70915957 | AX-170176623 | 0 | G | A | 0.254 | 2.69E-02 | 6.27E-02 | 2.65E+00 | 2.74E+00 | 6.64E-01 | no | | intergenic_variant | MODIFIER |
| 20 | 70935148 | AX-170213865 | 0 | G | C | 0.254 | 2.69E-02 | 6.27E-02 | 2.65E+00 | 2.74E+00 | 6.64E-01 | no | | intergenic_variant | MODIFIER |

| | | | | | | | | | | | | | | | |
|---|---|---|---|---|---|---|---|---|---|---|---|---|---|---|---|
| 20 | 70948498 | AX-170327721 | 0 | G | C | 0.254 | 2.69E-02 | 6.27E-02 | 2.65E+00 | 2.74E+00 | 6.64E-01 | no | | intergenic_variant | MODIFIER |
| 20 | 70952255 | AX-97816162 | 0 | G | A | 0.254 | 2.69E-02 | 6.27E-02 | 2.65E+00 | 2.74E+00 | 6.64E-01 | no | | intergenic_variant | MODIFIER |
| 20 | 70972409 | AX-170388645 | 0 | T | C | 0.254 | 2.69E-02 | 6.27E-02 | 2.65E+00 | 2.74E+00 | 6.64E-01 | no | | intergenic_variant | MODIFIER |
| 20 | 71000224 | AX-170136465 | 0 | G | A | 0.254 | 2.69E-02 | 6.27E-02 | 2.65E+00 | 2.74E+00 | 6.64E-01 | no | | intergenic_variant | MODIFIER |
| 20 | 71007175 | AX-170213508 | 0 | G | A | 0.254 | 2.69E-02 | 6.27E-02 | 2.65E+00 | 2.74E+00 | 6.64E-01 | no | | intergenic_variant | MODIFIER |
| 20 | 71028755 | AX-170133855 | 0 | G | A | 0.254 | 2.69E-02 | 6.27E-02 | 2.65E+00 | 2.74E+00 | 6.64E-01 | no | | intergenic_variant | MODIFIER |
| 20 | 71032112 | AX-170437770 | 0 | T | C | 0.254 | 2.69E-02 | 6.27E-02 | 2.65E+00 | 2.74E+00 | 6.64E-01 | no | | intergenic_variant | MODIFIER |
| 20 | 71032181 | AX-97850387 | 0 | C | T | 0.254 | 2.69E-02 | 6.27E-02 | 2.65E+00 | 2.74E+00 | 6.64E-01 | no | | intergenic_variant | MODIFIER |
| 20 | 71185860 | AX-170480003 | 0 | T | G | 0.254 | 2.69E-02 | 6.27E-02 | 2.65E+00 | 2.74E+00 | 6.64E-01 | no | | intergenic_variant | MODIFIER |
| 20 | 71188514 | AX-170084470 | 0 | G | T | 0.254 | 2.69E-02 | 6.27E-02 | 2.65E+00 | 2.74E+00 | 6.64E-01 | no | | intergenic_variant | MODIFIER |
| 20 | 71202868 | AX-170084469 | 0 | G | A | 0.254 | 2.69E-02 | 6.27E-02 | 2.65E+00 | 2.74E+00 | 6.64E-01 | no | Gm14730 | downstream_gene_variant | MODIFIER |
| 20 | 71380278 | AX-170273612 | 0 | A | G | 0.254 | 2.69E-02 | 6.27E-02 | 2.65E+00 | 2.74E+00 | 6.64E-01 | no | Mtmr1 | intron_variant | MODIFIER |
| 20 | 71398311 | AX-170034116 | 0 | C | A | 0.254 | 2.69E-02 | 6.27E-02 | 2.65E+00 | 2.74E+00 | 6.64E-01 | no | Mtmr1 | intron_variant | MODIFIER |
| 20 | 71413430 | AX-170406344 | 0 | G | A | 0.254 | 2.69E-02 | 6.27E-02 | 2.65E+00 | 2.74E+00 | 6.64E-01 | no | Mtmr1 | intron_variant | MODIFIER |
| 20 | 71414219 | AX-170378803 | 0 | T | G | 0.254 | 2.69E-02 | 6.27E-02 | 2.65E+00 | 2.74E+00 | 6.64E-01 | no | Mtmr1 | intron_variant | MODIFIER |
| 20 | 71423175 | AX-97807452 | 0 | T | C | 0.254 | 2.69E-02 | 6.27E-02 | 2.65E+00 | 2.74E+00 | 6.64E-01 | no | Mtmr1 | downstream_gene_variant | MODIFIER |
| 20 | 71427731 | AX-170262748 | 0 | T | C | 0.254 | 2.69E-02 | 6.27E-02 | 2.65E+00 | 2.74E+00 | 6.64E-01 | no | Cd99l2 | intron_variant | MODIFIER |
| 20 | 71476173 | AX-170084481 | 0 | A | C | 0.254 | 2.69E-02 | 6.27E-02 | 2.65E+00 | 2.74E+00 | 6.64E-01 | no | Cd99l2 | intron_variant | MODIFIER |
| 20 | 71479378 | AX-170179753 | 0 | C | T | 0.254 | 2.69E-02 | 6.27E-02 | 2.65E+00 | 2.74E+00 | 6.64E-01 | no | Cd99l2 | intron_variant | MODIFIER |
| 20 | 71479421 | AX-170443463 | 0 | C | T | 0.254 | 2.69E-02 | 6.27E-02 | 2.65E+00 | 2.74E+00 | 6.64E-01 | no | Cd99l2 | intron_variant | MODIFIER |
| 20 | 71479660 | AX-97832908 | 0 | G | A | 0.254 | 2.69E-02 | 6.27E-02 | 2.65E+00 | 2.74E+00 | 6.64E-01 | no | Cd99l2 | intron_variant | MODIFIER |
| 20 | 71479708 | AX-170084479 | 0 | C | T | 0.254 | 2.69E-02 | 6.27E-02 | 2.65E+00 | 2.74E+00 | 6.64E-01 | no | Cd99l2 | intron_variant | MODIFIER |
| 20 | 71761646 | AX-170435102 | 0 | A | C | 0.254 | 2.69E-02 | 6.27E-02 | 2.65E+00 | 2.74E+00 | 6.64E-01 | no | | intergenic_variant | MODIFIER |
| 20 | 71946109 | AX-170111815 | 0 | A | G | 0.254 | 2.69E-02 | 6.27E-02 | 2.65E+00 | 2.74E+00 | 6.64E-01 | no | | intergenic_variant | MODIFIER |
| 20 | 71972907 | AX-170164143 | 0 | C | T | 0.254 | 2.69E-02 | 6.27E-02 | 2.65E+00 | 2.74E+00 | 6.64E-01 | no | Prrg3 | downstream_gene_variant | MODIFIER |
| 20 | 71991427 | AX-170187178 | 0 | C | A | 0.254 | 2.69E-02 | 6.27E-02 | 2.65E+00 | 2.74E+00 | 6.64E-01 | no | Cnga2 | upstream_gene_variant | MODIFIER |
| 20 | 72007017 | AX-170034124 | 0 | G | A | 0.254 | 2.69E-02 | 6.27E-02 | 2.65E+00 | 2.74E+00 | 6.64E-01 | no | Cnga2 | intron_variant | MODIFIER |
| 20 | 72092990 | AX-170277985 | 0 | G | T | 0.254 | 2.69E-02 | 6.27E-02 | 2.65E+00 | 2.74E+00 | 6.64E-01 | no | | intergenic_variant | MODIFIER |
| 20 | 72133460 | AX-170084494 | 0 | G | A | 0.254 | 2.69E-02 | 6.27E-02 | 2.65E+00 | 2.74E+00 | 6.64E-01 | no | | intergenic_variant | MODIFIER |
| 20 | 72171210 | AX-170176051 | 0 | G | A | 0.254 | 2.69E-02 | 6.27E-02 | 2.65E+00 | 2.74E+00 | 6.64E-01 | no | | intergenic_variant | MODIFIER |
| 20 | 72248351 | AX-97840488 | 0 | T | C | 0.254 | 2.69E-02 | 6.27E-02 | 2.65E+00 | 2.74E+00 | 6.64E-01 | no | | intergenic_variant | MODIFIER |
| 20 | 72248991 | AX-170324306 | 0 | A | C | 0.254 | 2.69E-02 | 6.27E-02 | 2.65E+00 | 2.74E+00 | 6.64E-01 | no | | intergenic_variant | MODIFIER |
| 20 | 72250592 | AX-170201928 | 0 | C | T | 0.254 | 2.69E-02 | 6.27E-02 | 2.65E+00 | 2.74E+00 | 6.64E-01 | no | | intergenic_variant | MODIFIER |
| 20 | 72320050 | AX-170444663 | 0 | C | T | 0.254 | 2.69E-02 | 6.27E-02 | 2.65E+00 | 2.74E+00 | 6.64E-01 | no | | intergenic_variant | MODIFIER |
| 20 | 72326032 | AX-170404386 | 0 | C | T | 0.254 | 2.69E-02 | 6.27E-02 | 2.65E+00 | 2.74E+00 | 6.64E-01 | no | | intergenic_variant | MODIFIER |
| 20 | 72342817 | AX-170471795 | 0 | C | T | 0.254 | 2.69E-02 | 6.27E-02 | 2.65E+00 | 2.74E+00 | 6.64E-01 | no | | intergenic_variant | MODIFIER |
| 20 | 72351755 | AX-170204034 | 0 | G | A | 0.254 | 2.69E-02 | 6.27E-02 | 2.65E+00 | 2.74E+00 | 6.64E-01 | no | | intergenic_variant | MODIFIER |
| 20 | 72360450 | AX-97852231 | 0 | C | A | 0.254 | 2.69E-02 | 6.27E-02 | 2.65E+00 | 2.74E+00 | 6.64E-01 | no | | intergenic_variant | MODIFIER |
| 20 | 72383857 | AX-170196809 | 0 | T | G | 0.254 | 2.69E-02 | 6.27E-02 | 2.65E+00 | 2.74E+00 | 6.64E-01 | no | Magea10 | intron_variant | MODIFIER |
| 20 | 72384270 | AX-170333066 | 0 | C | T | 0.254 | 2.69E-02 | 6.27E-02 | 2.65E+00 | 2.74E+00 | 6.64E-01 | no | Magea10 | intron_variant | MODIFIER |
| 20 | 72447635 | AX-170431018 | 0 | T | C | 0.254 | 2.69E-02 | 6.27E-02 | 2.65E+00 | 2.74E+00 | 6.64E-01 | no | Gabra3 | intron_variant | MODIFIER |
| 20 | 72455868 | AX-170258126 | 0 | A | T | 0.254 | 2.69E-02 | 6.27E-02 | 2.65E+00 | 2.74E+00 | 6.64E-01 | no | Gabra3 | intron_variant | MODIFIER |
| 20 | 72466579 | AX-170406729 | 0 | C | T | 0.254 | 2.69E-02 | 6.27E-02 | 2.65E+00 | 2.74E+00 | 6.64E-01 | no | Gabra3 | intron_variant | MODIFIER |
| 20 | 72495002 | AX-170268142 | 0 | G | T | 0.254 | 2.69E-02 | 6.27E-02 | 2.65E+00 | 2.74E+00 | 6.64E-01 | no | Gabra3 | intron_variant | MODIFIER |
| 20 | 72832260 | AX-170216411 | 0 | A | C | 0.254 | 2.69E-02 | 6.27E-02 | 2.65E+00 | 2.74E+00 | 6.64E-01 | no | Gabrq | intron_variant | MODIFIER |
| 20 | 72897861 | AX-170176672 | 0 | G | A | 0.254 | 2.69E-02 | 6.27E-02 | 2.65E+00 | 2.74E+00 | 6.64E-01 | no | | intergenic_variant | MODIFIER |
| 20 | 72925679 | AX-170307064 | 0 | G | A | 0.254 | 2.69E-02 | 6.27E-02 | 2.65E+00 | 2.74E+00 | 6.64E-01 | no | Nsdhl | intron_variant | MODIFIER |
| 20 | 72931361 | AX-170435468 | 0 | A | G | 0.254 | 2.69E-02 | 6.27E-02 | 2.65E+00 | 2.74E+00 | 6.64E-01 | no | Nsdhl | intron_variant | MODIFIER |
| 20 | 72958730 | AX-170366350 | 0 | A | T | 0.254 | 2.69E-02 | 6.27E-02 | 2.65E+00 | 2.74E+00 | 6.64E-01 | no | Nsdhl | downstream_gene_variant | MODIFIER |
| 20 | 72959244 | AX-170247335 | 0 | G | A | 0.254 | 2.69E-02 | 6.27E-02 | 2.65E+00 | 2.74E+00 | 6.64E-01 | no | Nsdhl | downstream_gene_variant | MODIFIER |
| 20 | 72966785 | AX-170501613 | 0 | A | C | 0.254 | 2.69E-02 | 6.27E-02 | 2.65E+00 | 2.74E+00 | 6.64E-01 | no | | intergenic_variant | MODIFIER |
| 20 | 72969593 | AX-170147234 | 0 | A | G | 0.254 | 2.69E-02 | 6.27E-02 | 2.65E+00 | 2.74E+00 | 6.64E-01 | no | | intergenic_variant | MODIFIER |
| 20 | 72988543 | AX-170383983 | 0 | A | G | 0.254 | 2.69E-02 | 6.27E-02 | 2.65E+00 | 2.74E+00 | 6.64E-01 | no | Zfp185 | intron_variant | MODIFIER |
| 20 | 72996078 | AX-170279163 | 0 | A | G | 0.254 | 2.69E-02 | 6.27E-02 | 2.65E+00 | 2.74E+00 | 6.64E-01 | no | Zfp185 | intron_variant | MODIFIER |

| | | | | | | | | | | | | | | | |
|---|---|---|---|---|---|---|---|---|---|---|---|---|---|---|---|
| 20 | 73000553 | AX-170349515 | 0 | G | T | 0.254 | 2.69E-02 | 6.27E-02 | 2.65E+00 | 2.74E+00 | 6.64E-01 | no | Zfp185 | intron_variant | MODIFIER |
| 20 | 73004587 | AX-170115146 | 0 | A | G | 0.254 | 2.69E-02 | 6.27E-02 | 2.65E+00 | 2.74E+00 | 6.64E-01 | no | Zfp185 | intron_variant | MODIFIER |
| 20 | 73015403 | AX-170406141 | 0 | T | C | 0.254 | 2.69E-02 | 6.27E-02 | 2.65E+00 | 2.74E+00 | 6.64E-01 | no | Zfp185 | intron_variant | MODIFIER |
| 20 | 73015776 | AX-170472127 | 0 | A | G | 0.254 | 2.69E-02 | 6.27E-02 | 2.65E+00 | 2.74E+00 | 6.64E-01 | no | Zfp185 | missense_variant | MODERATE |
| 20 | 73021004 | AX-170183719 | 0 | G | A | 0.254 | 2.69E-02 | 6.27E-02 | 2.65E+00 | 2.74E+00 | 6.64E-01 | no | Zfp185 | intron_variant | MODIFIER |
| 20 | 73345736 | AX-170034138 | 0 | C | T | 0.254 | 2.69E-02 | 6.27E-02 | 2.65E+00 | 2.74E+00 | 6.64E-01 | no | Zfp275 | intron_variant | MODIFIER |
| 20 | 73346438 | AX-170034134 | 0 | C | T | 0.254 | 2.69E-02 | 6.27E-02 | 2.65E+00 | 2.74E+00 | 6.64E-01 | no | Zfp275 | intron_variant | MODIFIER |
| 20 | 73353417 | AX-170034133 | 0 | C | T | 0.254 | 2.69E-02 | 6.27E-02 | 2.65E+00 | 2.74E+00 | 6.64E-01 | no | Zfp275 | synonymous_variant | LOW |
| 20 | 73481911 | AX-97806249 | 0 | A | G | 0.254 | 2.69E-02 | 6.27E-02 | 2.65E+00 | 2.74E+00 | 6.64E-01 | no | Bgn | upstream_gene_variant | MODIFIER |
| 20 | 73510995 | AX-170266351 | 0 | G | A | 0.254 | 2.69E-02 | 6.27E-02 | 2.65E+00 | 2.74E+00 | 6.64E-01 | no | Atp2b3 | intron_variant | MODIFIER |
| 20 | 73524792 | AX-170276066 | 0 | C | T | 0.254 | 2.69E-02 | 6.27E-02 | 2.65E+00 | 2.74E+00 | 6.64E-01 | no | Atp2b3 | intron_variant | MODIFIER |
| 20 | 73525198 | AX-170382352 | 0 | C | T | 0.254 | 2.69E-02 | 6.27E-02 | 2.65E+00 | 2.74E+00 | 6.64E-01 | no | Atp2b3 | intron_variant | MODIFIER |
| 20 | 73646920 | AX-97819664 | 0 | C | T | 0.254 | 2.69E-02 | 6.27E-02 | 2.65E+00 | 2.74E+00 | 6.64E-01 | no | Dusp9 | downstream_gene_variant | MODIFIER |
| 20 | 73700780 | AX-170090541 | 0 | C | T | 0.254 | 2.69E-02 | 6.27E-02 | 2.65E+00 | 2.74E+00 | 6.64E-01 | no | Bcap31 | intron_variant | MODIFIER |
| 20 | 73734837 | AX-170158324 | 0 | A | G | 0.254 | 2.69E-02 | 6.27E-02 | 2.65E+00 | 2.74E+00 | 6.64E-01 | no | Abcd1 | intron_variant | MODIFIER |
| 20 | 73757122 | AX-170269762 | 0 | A | G | 0.254 | 2.69E-02 | 6.27E-02 | 2.65E+00 | 2.74E+00 | 6.64E-01 | no | Plxnb3 | 5_prime_UTR_variant | MODIFIER |
| 20 | 73869491 | AX-170040139 | 0 | A | G | 0.254 | 2.69E-02 | 6.27E-02 | 2.65E+00 | 2.74E+00 | 6.64E-01 | no | L1cam | intron_variant | MODIFIER |
| 20 | 73876580 | AX-170034140 | 0 | G | A | 0.254 | 2.69E-02 | 6.27E-02 | 2.65E+00 | 2.74E+00 | 6.64E-01 | no | L1cam | intron_variant | MODIFIER |
| 20 | 73887894 | AX-170393710 | 0 | A | C | 0.254 | 2.69E-02 | 6.27E-02 | 2.65E+00 | 2.74E+00 | 6.64E-01 | no | Avpr2 | upstream_gene_variant | MODIFIER |
| 20 | 73888740 | AX-170436812 | 0 | T | C | 0.254 | 2.69E-02 | 6.27E-02 | 2.65E+00 | 2.74E+00 | 6.64E-01 | no | Avpr2 | upstream_gene_variant | MODIFIER |
| 20 | 74086701 | AX-170290140 | 0 | G | A | 0.254 | 2.69E-02 | 6.27E-02 | 2.65E+00 | 2.74E+00 | 6.64E-01 | no | Mecp2 | upstream_gene_variant | MODIFIER |
| 20 | 74113117 | AX-170233086 | 0 | A | T | 0.254 | 2.69E-02 | 6.27E-02 | 2.65E+00 | 2.74E+00 | 6.64E-01 | no | Mecp2 | intron_variant | MODIFIER |
| 20 | 74123235 | AX-170505838 | 0 | G | A | 0.254 | 2.69E-02 | 6.27E-02 | 2.65E+00 | 2.74E+00 | 6.64E-01 | no | Opn1mw | upstream_gene_variant | MODIFIER |
| 20 | 74126306 | AX-170315505 | 0 | A | G | 0.254 | 2.69E-02 | 6.27E-02 | 2.65E+00 | 2.74E+00 | 6.64E-01 | no | Opn1mw | upstream_gene_variant | MODIFIER |
| 20 | 74142125 | AX-170402248 | 0 | C | A | 0.254 | 2.69E-02 | 6.27E-02 | 2.65E+00 | 2.74E+00 | 6.64E-01 | no | Opn1mw | intron_variant | MODIFIER |
| 20 | 74172005 | AX-170333923 | 0 | C | T | 0.254 | 2.69E-02 | 6.27E-02 | 2.65E+00 | 2.74E+00 | 6.64E-01 | no | Tex28 | upstream_gene_variant | MODIFIER |
| 20 | 74174287 | AX-97806251 | 0 | A | G | 0.254 | 2.69E-02 | 6.27E-02 | 2.65E+00 | 2.74E+00 | 6.64E-01 | no | Tktl1 | upstream_gene_variant | MODIFIER |
| 20 | 74177399 | AX-170366569 | 0 | A | G | 0.254 | 2.69E-02 | 6.27E-02 | 2.65E+00 | 2.74E+00 | 6.64E-01 | no | Tktl1 | missense_variant | MODERATE |
| 20 | 74207346 | AX-170430210 | 0 | G | A | 0.254 | 2.69E-02 | 6.27E-02 | 2.65E+00 | 2.74E+00 | 6.64E-01 | no | Tktl1 | intron_variant | MODIFIER |
| 20 | 74207542 | AX-170133749 | 0 | G | A | 0.254 | 2.69E-02 | 6.27E-02 | 2.65E+00 | 2.74E+00 | 6.64E-01 | no | Tktl1 | intron_variant | MODIFIER |
| 20 | 74228582 | AX-170111329 | 0 | C | T | 0.254 | 2.69E-02 | 6.27E-02 | 2.65E+00 | 2.74E+00 | 6.64E-01 | no | Flna | intron_variant | MODIFIER |
| 20 | 74242415 | AX-97850155 | 0 | A | G | 0.254 | 2.69E-02 | 6.27E-02 | 2.65E+00 | 2.74E+00 | 6.64E-01 | no | Flna | intron_variant | MODIFIER |
| 20 | 74247872 | AX-170163208 | 0 | C | A | 0.254 | 2.69E-02 | 6.27E-02 | 2.65E+00 | 2.74E+00 | 6.64E-01 | no | Flna | upstream_gene_variant | MODIFIER |
| 20 | 74281313 | AX-170156906 | 0 | G | A | 0.254 | 2.69E-02 | 6.27E-02 | 2.65E+00 | 2.74E+00 | 6.64E-01 | no | Dnase1l1 | intron_variant | MODIFIER |
| 20 | 74287380 | AX-97840577 | 0 | G | T | 0.254 | 2.69E-02 | 6.27E-02 | 2.65E+00 | 2.74E+00 | 6.64E-01 | no | Taz | intron_variant | MODIFIER |
| 20 | 74289448 | AX-170247533 | 0 | C | T | 0.254 | 2.69E-02 | 6.27E-02 | 2.65E+00 | 2.74E+00 | 6.64E-01 | no | Taz | 3_prime_UTR_variant | MODIFIER |
| 20 | 74314216 | AX-170090542 | 0 | T | C | 0.254 | 2.69E-02 | 6.27E-02 | 2.65E+00 | 2.74E+00 | 6.64E-01 | no | Gdi1 | downstream_gene_variant | MODIFIER |
| 20 | 74317666 | AX-170461912 | 0 | G | A | 0.254 | 2.69E-02 | 6.27E-02 | 2.65E+00 | 2.74E+00 | 6.64E-01 | no | Fam50a | synonymous_variant | LOW |
| 20 | 74333186 | AX-170407451 | 0 | A | G | 0.254 | 2.69E-02 | 6.27E-02 | 2.65E+00 | 2.74E+00 | 6.64E-01 | no | Plxna3 | intron_variant | MODIFIER |
| 20 | 74448039 | AX-170338518 | 0 | T | C | 0.254 | 2.69E-02 | 6.27E-02 | 2.65E+00 | 2.74E+00 | 6.64E-01 | no | Ikbkg | intron_variant | MODIFIER |
| 20 | 74456942 | AX-170232411 | 0 | T | C | 0.254 | 2.69E-02 | 6.27E-02 | 2.65E+00 | 2.74E+00 | 6.64E-01 | no | Ikbkg | downstream_gene_variant | MODIFIER |
| 20 | 74570294 | AX-170177118 | 0 | G | A | 0.254 | 2.69E-02 | 6.27E-02 | 2.65E+00 | 2.74E+00 | 6.64E-01 | no | Olfr1326-ps1 | 3_prime_UTR_variant | MODIFIER |
| 20 | 74791501 | AX-170336111 | 0 | A | C | 0.254 | 2.69E-02 | 6.27E-02 | 2.65E+00 | 2.74E+00 | 6.64E-01 | no | Gm8617 | non_coding_transcript_exon_variant | MODIFIER |
| 20 | 75701700 | AX-97785988 | 0 | C | T | 0.254 | 2.69E-02 | 6.27E-02 | 2.65E+00 | 2.74E+00 | 6.64E-01 | no | Gm15065 | upstream_gene_variant | MODIFIER |
| 20 | 75783420 | AX-170398425 | 0 | G | A | 0.254 | 2.69E-02 | 6.27E-02 | 2.65E+00 | 2.74E+00 | 6.64E-01 | no | Pls3 | downstream_gene_variant | MODIFIER |
| 20 | 75793894 | AX-170439049 | 0 | G | A | 0.254 | 2.69E-02 | 6.27E-02 | 2.65E+00 | 2.74E+00 | 6.64E-01 | no | Pls3 | intron_variant | MODIFIER |
| 20 | 75798983 | AX-170483038 | 0 | A | G | 0.254 | 2.69E-02 | 6.27E-02 | 2.65E+00 | 2.74E+00 | 6.64E-01 | no | Pls3 | intron_variant | MODIFIER |
| 20 | 75852356 | AX-170318669 | 0 | A | G | 0.254 | 2.69E-02 | 6.27E-02 | 2.65E+00 | 2.74E+00 | 6.64E-01 | no | Pls3 | intron_variant | MODIFIER |
| 20 | 75859411 | AX-170035825 | 0 | T | C | 0.254 | 2.69E-02 | 6.27E-02 | 2.65E+00 | 2.74E+00 | 6.64E-01 | no | Pls3 | intron_variant | MODIFIER |
| 20 | 75867752 | AX-97806250 | 0 | T | C | 0.254 | 2.69E-02 | 6.27E-02 | 2.65E+00 | 2.74E+00 | 6.64E-01 | no | Pls3 | intron_variant | MODIFIER |
| 20 | 75927738 | AX-170034159 | 0 | T | C | 0.254 | 2.69E-02 | 6.27E-02 | 2.65E+00 | 2.74E+00 | 6.64E-01 | no | Gm14715 | intron_variant,non_coding_transcript_variant | MODIFIER |
| 20 | 76015920 | AX-170084559 | 0 | G | A | 0.254 | 2.69E-02 | 6.27E-02 | 2.65E+00 | 2.74E+00 | 6.64E-01 | no | | intergenic_variant | MODIFIER |
| 20 | 76083257 | AX-170034162 | 0 | G | T | 0.254 | 2.69E-02 | 6.27E-02 | 2.65E+00 | 2.74E+00 | 6.64E-01 | no | | intergenic_variant | MODIFIER |
| 20 | 76126787 | AX-170443467 | 0 | G | A | 0.254 | 2.69E-02 | 6.27E-02 | 2.65E+00 | 2.74E+00 | 6.64E-01 | no | | intergenic_variant | MODIFIER |

| | | | | | | | | | | | | | | | |
|---|---|---|---|---|---|---|---|---|---|---|---|---|---|---|---|
| 20 | 76128156 | AX-168400028 | 0 | C | A | 0.254 | 2.69E-02 | 6.27E-02 | 2.65E+00 | 2.74E+00 | 6.64E-01 | no | | intergenic_variant | MODIFIER |
| 20 | 76179020 | AX-170518568 | 0 | C | A | 0.254 | 2.69E-02 | 6.27E-02 | 2.65E+00 | 2.74E+00 | 6.64E-01 | no | | intergenic_variant | MODIFIER |
| 20 | 76182115 | AX-170292547 | 0 | C | A | 0.254 | 2.69E-02 | 6.27E-02 | 2.65E+00 | 2.74E+00 | 6.64E-01 | no | | intergenic_variant | MODIFIER |
| 20 | 76196094 | AX-170422088 | 0 | T | C | 0.254 | 2.69E-02 | 6.27E-02 | 2.65E+00 | 2.74E+00 | 6.64E-01 | no | Gm5397 | upstream_gene_variant | MODIFIER |
| 20 | 76211277 | AX-170084563 | 0 | C | T | 0.254 | 2.69E-02 | 6.27E-02 | 2.65E+00 | 2.74E+00 | 6.64E-01 | no | | intergenic_variant | MODIFIER |
| 20 | 76290540 | AX-170204693 | 0 | T | G | 0.254 | 2.69E-02 | 6.27E-02 | 2.65E+00 | 2.74E+00 | 6.64E-01 | no | | intergenic_variant | MODIFIER |
| 20 | 76330186 | AX-170118063 | 0 | C | G | 0.254 | 2.69E-02 | 6.27E-02 | 2.65E+00 | 2.74E+00 | 6.64E-01 | no | | intergenic_variant | MODIFIER |
| 20 | 76423999 | AX-170247272 | 0 | A | C | 0.254 | 2.69E-02 | 6.27E-02 | 2.65E+00 | 2.74E+00 | 6.64E-01 | no | | intergenic_variant | MODIFIER |
| 20 | 76539647 | AX-170436186 | 0 | G | A | 0.254 | 2.69E-02 | 6.27E-02 | 2.65E+00 | 2.74E+00 | 6.64E-01 | no | Gm5793 | upstream_gene_variant | MODIFIER |
| 20 | 76539784 | AX-170226254 | 0 | T | C | 0.254 | 2.69E-02 | 6.27E-02 | 2.65E+00 | 2.74E+00 | 6.64E-01 | no | Gm5793 | upstream_gene_variant | MODIFIER |
| 20 | 76621473 | AX-170084568 | 0 | C | A | 0.254 | 2.69E-02 | 6.27E-02 | 2.65E+00 | 2.74E+00 | 6.64E-01 | no | | intergenic_variant | MODIFIER |
| 20 | 76621797 | AX-170034171 | 0 | C | T | 0.254 | 2.69E-02 | 6.27E-02 | 2.65E+00 | 2.74E+00 | 6.64E-01 | no | | intergenic_variant | MODIFIER |
| 20 | 76645315 | AX-170034167 | 0 | T | C | 0.254 | 2.69E-02 | 6.27E-02 | 2.65E+00 | 2.74E+00 | 6.64E-01 | no | | intergenic_variant | MODIFIER |
| 20 | 76646044 | AX-97830837 | 0 | G | T | 0.254 | 2.69E-02 | 6.27E-02 | 2.65E+00 | 2.74E+00 | 6.64E-01 | no | | intergenic_variant | MODIFIER |
| 20 | 76714815 | AX-97813755 | 0 | C | T | 0.254 | 2.69E-02 | 6.27E-02 | 2.65E+00 | 2.74E+00 | 6.64E-01 | no | | intergenic_variant | MODIFIER |
| 20 | 76747905 | AX-170463349 | 0 | C | T | 0.254 | 2.69E-02 | 6.27E-02 | 2.65E+00 | 2.74E+00 | 6.64E-01 | no | | intergenic_variant | MODIFIER |
| 20 | 76754071 | AX-170268639 | 0 | T | G | 0.254 | 2.69E-02 | 6.27E-02 | 2.65E+00 | 2.74E+00 | 6.64E-01 | no | | intergenic_variant | MODIFIER |
| 20 | 76806252 | AX-170380426 | 0 | A | G | 0.254 | 2.69E-02 | 6.27E-02 | 2.65E+00 | 2.74E+00 | 6.64E-01 | no | | intergenic_variant | MODIFIER |
| 20 | 76830139 | AX-170336185 | 0 | C | A | 0.254 | 2.69E-02 | 6.27E-02 | 2.65E+00 | 2.74E+00 | 6.64E-01 | no | Gm23280 | non_coding_transcript_exon_variant | MODIFIER |
| 20 | 76871382 | AX-170259458 | 0 | G | C | 0.254 | 2.69E-02 | 6.27E-02 | 2.65E+00 | 2.74E+00 | 6.64E-01 | no | | intergenic_variant | MODIFIER |
| 20 | 76918283 | AX-170265572 | 0 | A | G | 0.254 | 2.69E-02 | 6.27E-02 | 2.65E+00 | 2.74E+00 | 6.64E-01 | no | Gm14706 | downstream_gene_variant | MODIFIER |
| 20 | 76942812 | AX-170090547 | 0 | G | T | 0.254 | 2.69E-02 | 6.27E-02 | 2.65E+00 | 2.74E+00 | 6.64E-01 | no | | intergenic_variant | MODIFIER |
| 20 | 77091952 | AX-170397974 | 0 | G | C | 0.254 | 2.69E-02 | 6.27E-02 | 2.65E+00 | 2.74E+00 | 6.64E-01 | no | | intergenic_variant | MODIFIER |
| 20 | 77411059 | AX-170291946 | 0 | T | C | 0.254 | 2.69E-02 | 6.27E-02 | 2.65E+00 | 2.74E+00 | 6.64E-01 | no | | intergenic_variant | MODIFIER |
| 20 | 77509126 | AX-170431437 | 0 | C | T | 0.254 | 2.69E-02 | 6.27E-02 | 2.65E+00 | 2.74E+00 | 6.64E-01 | no | Tbl1x | upstream_gene_variant | MODIFIER |
| 20 | 77509268 | AX-170176405 | 0 | T | C | 0.254 | 2.69E-02 | 6.27E-02 | 2.65E+00 | 2.74E+00 | 6.64E-01 | no | Tbl1x | upstream_gene_variant | MODIFIER |
| 20 | 77531500 | AX-97811411 | 0 | T | A | 0.254 | 2.69E-02 | 6.27E-02 | 2.65E+00 | 2.74E+00 | 6.64E-01 | no | Tbl1x | intron_variant | MODIFIER |
| 20 | 77577776 | AX-97838597 | 0 | T | C | 0.254 | 2.69E-02 | 6.27E-02 | 2.65E+00 | 2.74E+00 | 6.64E-01 | no | Tbl1x | intron_variant | MODIFIER |
| 20 | 77606689 | AX-170132603 | 0 | T | G | 0.254 | 2.69E-02 | 6.27E-02 | 2.65E+00 | 2.74E+00 | 6.64E-01 | no | Tbl1x | intron_variant | MODIFIER |
| 20 | 77609261 | AX-170260262 | 0 | C | T | 0.254 | 2.69E-02 | 6.27E-02 | 2.65E+00 | 2.74E+00 | 6.64E-01 | no | Tbl1x | intron_variant | MODIFIER |
| 20 | 77635154 | AX-170271076 | 0 | G | A | 0.254 | 2.69E-02 | 6.27E-02 | 2.65E+00 | 2.74E+00 | 6.64E-01 | no | Tbl1x | intron_variant | MODIFIER |
| 2 | 142259450 | AX-170065103 | 0 | G | A | 0.14 | -7.76E-02 | 1.82E-01 | 2.69E+00 | 2.76E+00 | 6.64E-01 | no | Macrod2 | intron_variant | MODIFIER |
| 12 | 66724336 | AX-170006948 | 0 | C | G | 0.07 | 9.14E-02 | 2.16E-01 | 2.72E+00 | 2.82E+00 | 6.64E-01 | no | Mdga2 | intron_variant | MODIFIER |
| 12 | 72884769 | AX-97806115 | 0 | C | A | 0.07 | 9.14E-02 | 2.16E-01 | 2.72E+00 | 2.82E+00 | 6.64E-01 | no | 4930447C04Rik | intron_variant | MODIFIER |
| 18 | 33673901 | AX-170203956 | 0 | C | A | 0.07 | 9.14E-02 | 2.16E-01 | 2.72E+00 | 2.82E+00 | 6.64E-01 | no | | intergenic_variant | MODIFIER |
| 16 | 93321779 | AX-170264450 | 2 | C | T | 0.391 | -3.53E-02 | 8.04E-02 | 2.59E+00 | 2.68E+00 | 6.65E-01 | no | | intergenic_variant | MODIFIER |
| 3 | 38878164 | AX-170128244 | 1 | T | A | 0.196 | 4.21E-02 | 9.91E-02 | 2.72E+00 | 2.80E+00 | 6.65E-01 | no | C230034O21Rik | intron_variant,non_coding_transcript_variant | MODIFIER |
| 7 | 6882962 | AX-170280256 | 1 | G | A | 0.17 | 3.36E-02 | 8.07E-02 | 2.72E+00 | 2.87E+00 | 6.65E-01 | no | Usp29 | intron_variant | MODIFIER |
| 4 | 120792657 | AX-170122641 | 0 | A | T | 0.211 | 5.00E-02 | 1.11E-01 | 2.55E+00 | 2.63E+00 | 6.65E-01 | no | Nfyc | intron_variant | MODIFIER |
| 15 | 4763013 | AX-170463496 | 1 | A | C | 0.241 | -4.55E-02 | 1.04E-01 | 2.60E+00 | 2.67E+00 | 6.65E-01 | no | C6 | intron_variant | MODIFIER |
| 3 | 28726732 | AX-170066194 | 0 | A | G | 0.158 | -4.29E-02 | 1.00E-01 | 2.68E+00 | 2.75E+00 | 6.65E-01 | no | Slc2a2 | intron_variant | MODIFIER |
| 3 | 28728279 | AX-170066198 | 0 | C | G | 0.158 | -4.29E-02 | 1.00E-01 | 2.68E+00 | 2.75E+00 | 6.65E-01 | no | Slc2a2 | 3_prime_UTR_variant | MODIFIER |
| 3 | 28800914 | AX-170020203 | 0 | T | A | 0.158 | -4.29E-02 | 1.00E-01 | 2.68E+00 | 2.75E+00 | 6.65E-01 | no | Rpl22l1 | upstream_gene_variant | MODIFIER |
| 3 | 28816457 | AX-170020202 | 0 | T | G | 0.158 | -4.29E-02 | 1.00E-01 | 2.68E+00 | 2.75E+00 | 6.65E-01 | no | Gm33051 | upstream_gene_variant | MODIFIER |
| 3 | 28825123 | AX-170341356 | 0 | A | G | 0.158 | -4.29E-02 | 1.00E-01 | 2.68E+00 | 2.75E+00 | 6.65E-01 | no | | intergenic_variant | MODIFIER |
| 3 | 28839209 | AX-170517143 | 0 | G | A | 0.158 | -4.29E-02 | 1.00E-01 | 2.68E+00 | 2.75E+00 | 6.65E-01 | no | | intergenic_variant | MODIFIER |
| 3 | 28844145 | AX-170307780 | 0 | G | A | 0.158 | -4.29E-02 | 1.00E-01 | 2.68E+00 | 2.75E+00 | 6.65E-01 | no | | intergenic_variant | MODIFIER |
| 3 | 28844175 | AX-170362812 | 0 | G | A | 0.158 | -4.29E-02 | 1.00E-01 | 2.68E+00 | 2.75E+00 | 6.65E-01 | no | | intergenic_variant | MODIFIER |
| 3 | 28846845 | AX-170066201 | 0 | C | G | 0.158 | -4.29E-02 | 1.00E-01 | 2.68E+00 | 2.75E+00 | 6.65E-01 | no | | intergenic_variant | MODIFIER |
| 3 | 29039939 | AX-170486637 | 0 | T | C | 0.158 | -4.29E-02 | 1.00E-01 | 2.68E+00 | 2.75E+00 | 6.65E-01 | no | | intergenic_variant | MODIFIER |
| 3 | 29040175 | AX-170227513 | 0 | C | A | 0.158 | -4.29E-02 | 1.00E-01 | 2.68E+00 | 2.75E+00 | 6.65E-01 | no | | intergenic_variant | MODIFIER |
| 3 | 29040493 | AX-170113130 | 0 | A | G | 0.158 | -4.29E-02 | 1.00E-01 | 2.68E+00 | 2.75E+00 | 6.65E-01 | no | | intergenic_variant | MODIFIER |
| 3 | 29042732 | AX-170460781 | 0 | C | T | 0.158 | -4.29E-02 | 1.00E-01 | 2.68E+00 | 2.75E+00 | 6.65E-01 | no | | intergenic_variant | MODIFIER |
| 3 | 29043069 | AX-170301337 | 0 | C | A | 0.158 | -4.29E-02 | 1.00E-01 | 2.68E+00 | 2.75E+00 | 6.65E-01 | no | | intergenic_variant | MODIFIER |

| | | | | | | | | | | | | | | | |
|---|---|---|---|---|---|---|---|---|---|---|---|---|---|---|---|
| 3 | 29043297 | AX-170480902 | 0 | A | G | 0.158 | -4.29E-02 | 1.00E-01 | 2.68E+00 | 2.75E+00 | 6.65E-01 | no | | intergenic_variant | MODIFIER |
| 3 | 29046325 | AX-170478456 | 0 | G | A | 0.158 | -4.29E-02 | 1.00E-01 | 2.68E+00 | 2.75E+00 | 6.65E-01 | no | | intergenic_variant | MODIFIER |
| 3 | 29047478 | AX-170066203 | 0 | T | G | 0.158 | -4.29E-02 | 1.00E-01 | 2.68E+00 | 2.75E+00 | 6.65E-01 | no | | regulatory_region_variant | MODIFIER |
| 3 | 29047610 | AX-170066204 | 0 | A | C | 0.158 | -4.29E-02 | 1.00E-01 | 2.68E+00 | 2.75E+00 | 6.65E-01 | no | | regulatory_region_variant | MODIFIER |
| 3 | 29047855 | AX-170370828 | 0 | G | A | 0.158 | -4.29E-02 | 1.00E-01 | 2.68E+00 | 2.75E+00 | 6.65E-01 | no | | regulatory_region_variant | MODIFIER |
| 3 | 29083323 | AX-170507482 | 0 | C | T | 0.158 | -4.29E-02 | 1.00E-01 | 2.68E+00 | 2.75E+00 | 6.65E-01 | no | Egfem1 | intron_variant | MODIFIER |
| 3 | 29103165 | AX-170321407 | 0 | T | C | 0.158 | -4.29E-02 | 1.00E-01 | 2.68E+00 | 2.75E+00 | 6.65E-01 | no | Egfem1 | intron_variant | MODIFIER |
| 3 | 29104155 | AX-170204864 | 0 | A | G | 0.158 | -4.29E-02 | 1.00E-01 | 2.68E+00 | 2.75E+00 | 6.65E-01 | no | Egfem1 | intron_variant | MODIFIER |
| 3 | 29110942 | AX-170517230 | 0 | A | G | 0.158 | -4.29E-02 | 1.00E-01 | 2.68E+00 | 2.75E+00 | 6.65E-01 | no | Egfem1 | intron_variant | MODIFIER |
| 3 | 29112723 | AX-170534703 | 0 | A | G | 0.158 | -4.29E-02 | 1.00E-01 | 2.68E+00 | 2.75E+00 | 6.65E-01 | no | Egfem1 | intron_variant | MODIFIER |
| 3 | 29115048 | AX-170403418 | 0 | C | G | 0.158 | -4.29E-02 | 1.00E-01 | 2.68E+00 | 2.75E+00 | 6.65E-01 | no | Egfem1 | intron_variant | MODIFIER |
| 3 | 29115944 | AX-170492488 | 0 | C | T | 0.158 | -4.29E-02 | 1.00E-01 | 2.68E+00 | 2.75E+00 | 6.65E-01 | no | Egfem1 | intron_variant | MODIFIER |
| 3 | 29179621 | AX-170506568 | 0 | A | C | 0.158 | -4.29E-02 | 1.00E-01 | 2.68E+00 | 2.75E+00 | 6.65E-01 | no | Egfem1 | intron_variant | MODIFIER |
| 3 | 29181350 | AX-170398056 | 0 | C | G | 0.158 | -4.29E-02 | 1.00E-01 | 2.68E+00 | 2.75E+00 | 6.65E-01 | no | Egfem1 | intron_variant | MODIFIER |
| 3 | 29181420 | AX-170155379 | 0 | G | A | 0.158 | -4.29E-02 | 1.00E-01 | 2.68E+00 | 2.75E+00 | 6.65E-01 | no | Egfem1 | intron_variant | MODIFIER |
| 3 | 29181660 | AX-170194998 | 0 | A | G | 0.158 | -4.29E-02 | 1.00E-01 | 2.68E+00 | 2.75E+00 | 6.65E-01 | no | Egfem1 | intron_variant | MODIFIER |
| 3 | 29192452 | AX-170532586 | 0 | T | C | 0.158 | -4.29E-02 | 1.00E-01 | 2.68E+00 | 2.75E+00 | 6.65E-01 | no | Egfem1 | intron_variant | MODIFIER |
| 3 | 29746874 | AX-170419180 | 0 | A | C | 0.158 | -4.29E-02 | 1.00E-01 | 2.68E+00 | 2.75E+00 | 6.65E-01 | no | | intergenic_variant | MODIFIER |
| 3 | 29762902 | AX-170482299 | 0 | T | A | 0.158 | -4.29E-02 | 1.00E-01 | 2.68E+00 | 2.75E+00 | 6.65E-01 | no | | intergenic_variant | MODIFIER |
| 3 | 29763078 | AX-170163111 | 0 | T | C | 0.158 | -4.29E-02 | 1.00E-01 | 2.68E+00 | 2.75E+00 | 6.65E-01 | no | | intergenic_variant | MODIFIER |
| 3 | 29765027 | AX-170516870 | 0 | G | A | 0.158 | -4.29E-02 | 1.00E-01 | 2.68E+00 | 2.75E+00 | 6.65E-01 | no | | intergenic_variant | MODIFIER |
| 3 | 29828094 | AX-170106035 | 0 | T | C | 0.158 | -4.29E-02 | 1.00E-01 | 2.68E+00 | 2.75E+00 | 6.65E-01 | no | Gm33206 | intron_variant,non_coding_transcript_variant | MODIFIER |
| 3 | 30335477 | AX-170147506 | 0 | T | C | 0.158 | -4.29E-02 | 1.00E-01 | 2.68E+00 | 2.75E+00 | 6.65E-01 | no | Mecom | intron_variant | MODIFIER |
| 3 | 30394907 | AX-168392553 | 0 | C | T | 0.158 | -4.29E-02 | 1.00E-01 | 2.68E+00 | 2.75E+00 | 6.65E-01 | no | Mecom | intron_variant | MODIFIER |
| 3 | 30395450 | AX-170088901 | 0 | G | A | 0.158 | -4.29E-02 | 1.00E-01 | 2.68E+00 | 2.75E+00 | 6.65E-01 | no | Mecom | intron_variant | MODIFIER |
| 3 | 30409191 | AX-170354519 | 0 | G | C | 0.158 | -4.29E-02 | 1.00E-01 | 2.68E+00 | 2.75E+00 | 6.65E-01 | no | Mecom | intron_variant | MODIFIER |
| 3 | 30411056 | AX-170269012 | 0 | A | G | 0.158 | -4.29E-02 | 1.00E-01 | 2.68E+00 | 2.75E+00 | 6.65E-01 | no | Mecom | intron_variant | MODIFIER |
| 3 | 30429688 | AX-170154746 | 0 | T | G | 0.158 | -4.29E-02 | 1.00E-01 | 2.68E+00 | 2.75E+00 | 6.65E-01 | no | Mecom | intron_variant | MODIFIER |
| 3 | 30429752 | AX-170302416 | 0 | G | A | 0.158 | -4.29E-02 | 1.00E-01 | 2.68E+00 | 2.75E+00 | 6.65E-01 | no | Mecom | intron_variant | MODIFIER |
| 3 | 30429819 | AX-170434774 | 0 | G | T | 0.158 | -4.29E-02 | 1.00E-01 | 2.68E+00 | 2.75E+00 | 6.65E-01 | no | Mecom | intron_variant | MODIFIER |
| 3 | 30429905 | AX-170113273 | 0 | C | G | 0.158 | -4.29E-02 | 1.00E-01 | 2.68E+00 | 2.75E+00 | 6.65E-01 | no | Mecom | intron_variant | MODIFIER |
| 3 | 30432425 | AX-170109424 | 0 | G | A | 0.158 | -4.29E-02 | 1.00E-01 | 2.68E+00 | 2.75E+00 | 6.65E-01 | no | Mecom | intron_variant | MODIFIER |
| 3 | 30453306 | AX-170020235 | 0 | T | C | 0.158 | -4.29E-02 | 1.00E-01 | 2.68E+00 | 2.75E+00 | 6.65E-01 | no | Mecom | intron_variant | MODIFIER |
| 3 | 30654990 | AX-97839595 | 0 | T | C | 0.158 | -4.29E-02 | 1.00E-01 | 2.68E+00 | 2.75E+00 | 6.65E-01 | no | Lrriq4 | intron_variant | MODIFIER |
| 3 | 30776194 | AX-97806720 | 0 | A | G | 0.158 | -4.29E-02 | 1.00E-01 | 2.68E+00 | 2.75E+00 | 6.65E-01 | no | | intergenic_variant | MODIFIER |
| 3 | 30777027 | AX-170066237 | 0 | T | C | 0.158 | -4.29E-02 | 1.00E-01 | 2.68E+00 | 2.75E+00 | 6.65E-01 | no | | intergenic_variant | MODIFIER |
| 3 | 30778308 | AX-170115439 | 0 | A | G | 0.158 | -4.29E-02 | 1.00E-01 | 2.68E+00 | 2.75E+00 | 6.65E-01 | no | | intergenic_variant | MODIFIER |
| 3 | 30778613 | AX-170341726 | 0 | G | C | 0.158 | -4.29E-02 | 1.00E-01 | 2.68E+00 | 2.75E+00 | 6.65E-01 | no | | intergenic_variant | MODIFIER |
| 3 | 30778694 | AX-170302236 | 0 | G | A | 0.158 | -4.29E-02 | 1.00E-01 | 2.68E+00 | 2.75E+00 | 6.65E-01 | no | | intergenic_variant | MODIFIER |
| 3 | 30809002 | AX-170273469 | 0 | A | G | 0.158 | -4.29E-02 | 1.00E-01 | 2.68E+00 | 2.75E+00 | 6.65E-01 | no | Sec62 | intron_variant | MODIFIER |
| 3 | 30836609 | AX-97798434 | 0 | T | C | 0.158 | -4.29E-02 | 1.00E-01 | 2.68E+00 | 2.75E+00 | 6.65E-01 | no | | intergenic_variant | MODIFIER |
| 3 | 30870788 | AX-170182012 | 0 | C | T | 0.158 | -4.29E-02 | 1.00E-01 | 2.68E+00 | 2.75E+00 | 6.65E-01 | no | Gpr160 | intron_variant | MODIFIER |
| 3 | 30871528 | AX-170229767 | 0 | G | A | 0.158 | -4.29E-02 | 1.00E-01 | 2.68E+00 | 2.75E+00 | 6.65E-01 | no | Gpr160 | intron_variant | MODIFIER |
| 3 | 31269948 | AX-170498083 | 0 | T | C | 0.158 | -4.29E-02 | 1.00E-01 | 2.68E+00 | 2.75E+00 | 6.65E-01 | no | Slc7a14 | intron_variant | MODIFIER |
| 3 | 31463841 | AX-97803203 | 0 | T | C | 0.158 | -4.29E-02 | 1.00E-01 | 2.68E+00 | 2.75E+00 | 6.65E-01 | no | | intergenic_variant | MODIFIER |
| 3 | 31477773 | AX-170236403 | 0 | A | C | 0.158 | -4.29E-02 | 1.00E-01 | 2.68E+00 | 2.75E+00 | 6.65E-01 | no | | intergenic_variant | MODIFIER |
| 3 | 31480578 | AX-170487615 | 0 | G | A | 0.158 | -4.29E-02 | 1.00E-01 | 2.68E+00 | 2.75E+00 | 6.65E-01 | no | | intergenic_variant | MODIFIER |
| 3 | 31501521 | AX-97788284 | 0 | A | G | 0.158 | -4.29E-02 | 1.00E-01 | 2.68E+00 | 2.75E+00 | 6.65E-01 | no | | intergenic_variant | MODIFIER |
| 3 | 31518533 | AX-170199203 | 0 | T | C | 0.158 | -4.29E-02 | 1.00E-01 | 2.68E+00 | 2.75E+00 | 6.65E-01 | no | | intergenic_variant | MODIFIER |
| 3 | 31518548 | AX-97786055 | 0 | G | A | 0.158 | -4.29E-02 | 1.00E-01 | 2.68E+00 | 2.75E+00 | 6.65E-01 | no | | intergenic_variant | MODIFIER |
| 3 | 31588607 | AX-170345867 | 0 | C | A | 0.158 | -4.29E-02 | 1.00E-01 | 2.68E+00 | 2.75E+00 | 6.65E-01 | no | | intergenic_variant | MODIFIER |
| 3 | 31588632 | AX-170326720 | 0 | C | G | 0.158 | -4.29E-02 | 1.00E-01 | 2.68E+00 | 2.75E+00 | 6.65E-01 | no | | intergenic_variant | MODIFIER |
| 3 | 31589554 | AX-170301256 | 0 | C | T | 0.158 | -4.29E-02 | 1.00E-01 | 2.68E+00 | 2.75E+00 | 6.65E-01 | no | | intergenic_variant | MODIFIER |
| 3 | 31631463 | AX-97852264 | 0 | T | C | 0.158 | -4.29E-02 | 1.00E-01 | 2.68E+00 | 2.75E+00 | 6.65E-01 | no | | intergenic_variant | MODIFIER |

| | | | | | | | | | | | | | | | |
|---|---|---|---|---|---|---|---|---|---|---|---|---|---|---|---|
| 3 | 31724223 | AX-170316953 | 0 | G | A | 0.158 | -4.29E-02 | 1.00E-01 | 2.68E+00 | 2.75E+00 | 6.65E-01 | no | | intergenic_variant | MODIFIER |
| 3 | 31753420 | AX-170020246 | 0 | T | C | 0.158 | -4.29E-02 | 1.00E-01 | 2.68E+00 | 2.75E+00 | 6.65E-01 | no | | intergenic_variant | MODIFIER |
| 3 | 31753958 | AX-97845450 | 0 | T | C | 0.158 | -4.29E-02 | 1.00E-01 | 2.68E+00 | 2.75E+00 | 6.65E-01 | no | | intergenic_variant | MODIFIER |
| 3 | 31756221 | AX-170303115 | 0 | A | G | 0.158 | -4.29E-02 | 1.00E-01 | 2.68E+00 | 2.75E+00 | 6.65E-01 | no | | intergenic_variant | MODIFIER |
| 3 | 31781481 | AX-170020247 | 0 | C | A | 0.158 | -4.29E-02 | 1.00E-01 | 2.68E+00 | 2.75E+00 | 6.65E-01 | no | | intergenic_variant | MODIFIER |
| 3 | 31781570 | AX-170381306 | 0 | T | C | 0.158 | -4.29E-02 | 1.00E-01 | 2.68E+00 | 2.75E+00 | 6.65E-01 | no | | intergenic_variant | MODIFIER |
| 3 | 31792314 | AX-170486181 | 0 | A | G | 0.158 | -4.29E-02 | 1.00E-01 | 2.68E+00 | 2.75E+00 | 6.65E-01 | no | | intergenic_variant | MODIFIER |
| 3 | 31796541 | AX-170177441 | 0 | A | G | 0.158 | -4.29E-02 | 1.00E-01 | 2.68E+00 | 2.75E+00 | 6.65E-01 | no | | intergenic_variant | MODIFIER |
| 3 | 31807747 | AX-97787881 | 0 | A | C | 0.158 | -4.29E-02 | 1.00E-01 | 2.68E+00 | 2.75E+00 | 6.65E-01 | no | | intergenic_variant | MODIFIER |
| 3 | 31829493 | AX-170334164 | 0 | G | A | 0.158 | -4.29E-02 | 1.00E-01 | 2.68E+00 | 2.75E+00 | 6.65E-01 | no | | intergenic_variant | MODIFIER |
| 3 | 31853490 | AX-170374651 | 0 | A | T | 0.158 | -4.29E-02 | 1.00E-01 | 2.68E+00 | 2.75E+00 | 6.65E-01 | no | | intergenic_variant | MODIFIER |
| 3 | 31867179 | AX-170232325 | 0 | G | A | 0.158 | -4.29E-02 | 1.00E-01 | 2.68E+00 | 2.75E+00 | 6.65E-01 | no | | intergenic_variant | MODIFIER |
| 3 | 31868553 | AX-97785009 | 0 | T | G | 0.158 | -4.29E-02 | 1.00E-01 | 2.68E+00 | 2.75E+00 | 6.65E-01 | no | | intergenic_variant | MODIFIER |
| 3 | 31895959 | AX-170088903 | 0 | C | T | 0.158 | -4.29E-02 | 1.00E-01 | 2.68E+00 | 2.75E+00 | 6.65E-01 | no | | intergenic_variant | MODIFIER |
| 3 | 31917007 | AX-170456843 | 0 | C | T | 0.158 | -4.29E-02 | 1.00E-01 | 2.68E+00 | 2.75E+00 | 6.65E-01 | no | Kcnmb2 | intron_variant | MODIFIER |
| 3 | 31917115 | AX-170285560 | 0 | A | G | 0.158 | -4.29E-02 | 1.00E-01 | 2.68E+00 | 2.75E+00 | 6.65E-01 | no | Kcnmb2 | intron_variant | MODIFIER |
| 3 | 31932068 | AX-170506476 | 0 | G | A | 0.158 | -4.29E-02 | 1.00E-01 | 2.68E+00 | 2.75E+00 | 6.65E-01 | no | Kcnmb2 | intron_variant | MODIFIER |
| 3 | 31933956 | AX-170303087 | 0 | G | A | 0.158 | -4.29E-02 | 1.00E-01 | 2.68E+00 | 2.75E+00 | 6.65E-01 | no | Kcnmb2 | intron_variant | MODIFIER |
| 3 | 31933990 | AX-170283695 | 0 | C | T | 0.158 | -4.29E-02 | 1.00E-01 | 2.68E+00 | 2.75E+00 | 6.65E-01 | no | Kcnmb2 | intron_variant | MODIFIER |
| 3 | 31951781 | AX-170302790 | 0 | C | T | 0.158 | -4.29E-02 | 1.00E-01 | 2.68E+00 | 2.75E+00 | 6.65E-01 | no | Kcnmb2 | intron_variant | MODIFIER |
| 3 | 31970712 | AX-97801075 | 0 | C | T | 0.158 | -4.29E-02 | 1.00E-01 | 2.68E+00 | 2.75E+00 | 6.65E-01 | no | Kcnmb2 | intron_variant | MODIFIER |
| 3 | 31971646 | AX-170301568 | 0 | A | G | 0.158 | -4.29E-02 | 1.00E-01 | 2.68E+00 | 2.75E+00 | 6.65E-01 | no | Kcnmb2 | intron_variant | MODIFIER |
| 3 | 31971666 | AX-170186320 | 0 | A | C | 0.158 | -4.29E-02 | 1.00E-01 | 2.68E+00 | 2.75E+00 | 6.65E-01 | no | Kcnmb2 | intron_variant | MODIFIER |
| 3 | 31987602 | AX-170138295 | 0 | C | T | 0.158 | -4.29E-02 | 1.00E-01 | 2.68E+00 | 2.75E+00 | 6.65E-01 | no | Kcnmb2 | intron_variant | MODIFIER |
| 3 | 31990915 | AX-170154539 | 0 | T | G | 0.158 | -4.29E-02 | 1.00E-01 | 2.68E+00 | 2.75E+00 | 6.65E-01 | no | Kcnmb2 | intron_variant | MODIFIER |
| 3 | 31990988 | AX-170235601 | 0 | A | T | 0.158 | -4.29E-02 | 1.00E-01 | 2.68E+00 | 2.75E+00 | 6.65E-01 | no | Kcnmb2 | intron_variant | MODIFIER |
| 3 | 31991858 | AX-97803367 | 0 | C | T | 0.158 | -4.29E-02 | 1.00E-01 | 2.68E+00 | 2.75E+00 | 6.65E-01 | no | Kcnmb2 | intron_variant | MODIFIER |
| 3 | 31997858 | AX-170126629 | 0 | T | C | 0.158 | -4.29E-02 | 1.00E-01 | 2.68E+00 | 2.75E+00 | 6.65E-01 | no | Kcnmb2 | intron_variant | MODIFIER |
| 3 | 32008351 | AX-170506373 | 0 | C | T | 0.158 | -4.29E-02 | 1.00E-01 | 2.68E+00 | 2.75E+00 | 6.65E-01 | no | Kcnmb2 | intron_variant | MODIFIER |
| 3 | 32009498 | AX-170443410 | 0 | C | T | 0.158 | -4.29E-02 | 1.00E-01 | 2.68E+00 | 2.75E+00 | 6.65E-01 | no | Kcnmb2 | intron_variant | MODIFIER |
| 3 | 32018393 | AX-170231140 | 0 | A | G | 0.158 | -4.29E-02 | 1.00E-01 | 2.68E+00 | 2.75E+00 | 6.65E-01 | no | Kcnmb2 | intron_variant | MODIFIER |
| 3 | 32105572 | AX-97784331 | 0 | C | T | 0.158 | -4.29E-02 | 1.00E-01 | 2.68E+00 | 2.75E+00 | 6.65E-01 | no | Kcnmb2 | intron_variant | MODIFIER |
| 3 | 32105714 | AX-170066269 | 0 | T | C | 0.158 | -4.29E-02 | 1.00E-01 | 2.68E+00 | 2.75E+00 | 6.65E-01 | no | Kcnmb2 | intron_variant | MODIFIER |
| 3 | 32105840 | AX-170066268 | 0 | G | C | 0.158 | -4.29E-02 | 1.00E-01 | 2.68E+00 | 2.75E+00 | 6.65E-01 | no | Kcnmb2 | intron_variant | MODIFIER |
| 3 | 32105931 | AX-170285025 | 0 | C | T | 0.158 | -4.29E-02 | 1.00E-01 | 2.68E+00 | 2.75E+00 | 6.65E-01 | no | Kcnmb2 | intron_variant | MODIFIER |
| 3 | 32107531 | AX-170158322 | 0 | G | A | 0.158 | -4.29E-02 | 1.00E-01 | 2.68E+00 | 2.75E+00 | 6.65E-01 | no | Kcnmb2 | intron_variant | MODIFIER |
| 3 | 32107805 | AX-170514589 | 0 | T | C | 0.158 | -4.29E-02 | 1.00E-01 | 2.68E+00 | 2.75E+00 | 6.65E-01 | no | Kcnmb2 | intron_variant | MODIFIER |
| 3 | 32112501 | AX-170127750 | 0 | A | T | 0.158 | -4.29E-02 | 1.00E-01 | 2.68E+00 | 2.75E+00 | 6.65E-01 | no | Kcnmb2 | intron_variant | MODIFIER |
| 3 | 32181686 | AX-170225795 | 0 | T | C | 0.158 | -4.29E-02 | 1.00E-01 | 2.68E+00 | 2.75E+00 | 6.65E-01 | no | Kcnmb2 | intron_variant | MODIFIER |
| 3 | 32181951 | AX-170496887 | 0 | G | C | 0.158 | -4.29E-02 | 1.00E-01 | 2.68E+00 | 2.75E+00 | 6.65E-01 | no | Kcnmb2 | synonymous_variant | LOW |
| 3 | 32182173 | AX-170478024 | 0 | A | G | 0.158 | -4.29E-02 | 1.00E-01 | 2.68E+00 | 2.75E+00 | 6.65E-01 | no | Kcnmb2 | intron_variant | MODIFIER |
| 3 | 32183680 | AX-170145664 | 0 | A | G | 0.158 | -4.29E-02 | 1.00E-01 | 2.68E+00 | 2.75E+00 | 6.65E-01 | no | Kcnmb2 | intron_variant | MODIFIER |
| 3 | 32184194 | AX-170451125 | 0 | G | A | 0.158 | -4.29E-02 | 1.00E-01 | 2.68E+00 | 2.75E+00 | 6.65E-01 | no | Kcnmb2 | intron_variant | MODIFIER |
| 3 | 32190113 | AX-170020251 | 0 | T | C | 0.158 | -4.29E-02 | 1.00E-01 | 2.68E+00 | 2.75E+00 | 6.65E-01 | no | Kcnmb2 | intron_variant | MODIFIER |
| 3 | 32195019 | AX-170066272 | 0 | T | A | 0.158 | -4.29E-02 | 1.00E-01 | 2.68E+00 | 2.75E+00 | 6.65E-01 | no | Kcnmb2 | intron_variant | MODIFIER |
| 3 | 32195503 | AX-97845321 | 0 | C | T | 0.158 | -4.29E-02 | 1.00E-01 | 2.68E+00 | 2.75E+00 | 6.65E-01 | no | Kcnmb2 | intron_variant | MODIFIER |
| 3 | 32199276 | AX-170516390 | 0 | G | T | 0.158 | -4.29E-02 | 1.00E-01 | 2.68E+00 | 2.75E+00 | 6.65E-01 | no | Kcnmb2 | 3_prime_UTR_variant | MODIFIER |
| 3 | 32212164 | AX-170470885 | 0 | C | T | 0.158 | -4.29E-02 | 1.00E-01 | 2.68E+00 | 2.75E+00 | 6.65E-01 | no | | intergenic_variant | MODIFIER |
| 3 | 32225255 | AX-97820401 | 0 | G | A | 0.158 | -4.29E-02 | 1.00E-01 | 2.68E+00 | 2.75E+00 | 6.65E-01 | no | | intergenic_variant | MODIFIER |
| 3 | 32225786 | AX-170020250 | 0 | T | C | 0.158 | -4.29E-02 | 1.00E-01 | 2.68E+00 | 2.75E+00 | 6.65E-01 | no | | intergenic_variant | MODIFIER |
| 3 | 32226499 | AX-170020252 | 0 | C | G | 0.158 | -4.29E-02 | 1.00E-01 | 2.68E+00 | 2.75E+00 | 6.65E-01 | no | | intergenic_variant | MODIFIER |
| 3 | 32252580 | AX-170020248 | 0 | T | C | 0.158 | -4.29E-02 | 1.00E-01 | 2.68E+00 | 2.75E+00 | 6.65E-01 | no | | intergenic_variant | MODIFIER |
| 3 | 32285016 | AX-170066274 | 0 | G | A | 0.158 | -4.29E-02 | 1.00E-01 | 2.68E+00 | 2.75E+00 | 6.65E-01 | no | | intergenic_variant | MODIFIER |
| 3 | 32285996 | AX-170066277 | 0 | C | G | 0.158 | -4.29E-02 | 1.00E-01 | 2.68E+00 | 2.75E+00 | 6.65E-01 | no | | intergenic_variant | MODIFIER |

| | | | | | | | | | | | | | | | |
|---|---|---|---|---|---|---|---|---|---|---|---|---|---|---|---|
| 3 | 32301386 | AX-97782327 | 0 | T | C | 0.158 | -4.29E-02 | 1.00E-01 | 2.68E+00 | 2.75E+00 | 6.65E-01 | no | | intergenic_variant | MODIFIER |
| 3 | 32318069 | AX-170020254 | 0 | A | T | 0.158 | -4.29E-02 | 1.00E-01 | 2.68E+00 | 2.75E+00 | 6.65E-01 | no | | regulatory_region_variant | MODIFIER |
| 3 | 32345340 | AX-170020255 | 0 | T | C | 0.158 | -4.29E-02 | 1.00E-01 | 2.68E+00 | 2.75E+00 | 6.65E-01 | no | Zmat3 | intron_variant | MODIFIER |
| 3 | 32345732 | AX-170020256 | 0 | T | C | 0.158 | -4.29E-02 | 1.00E-01 | 2.68E+00 | 2.75E+00 | 6.65E-01 | no | Zmat3 | intron_variant | MODIFIER |
| 3 | 32356621 | AX-170130428 | 0 | C | G | 0.158 | -4.29E-02 | 1.00E-01 | 2.68E+00 | 2.75E+00 | 6.65E-01 | no | Zmat3 | intron_variant | MODIFIER |
| 3 | 32356682 | AX-97834807 | 0 | C | T | 0.158 | -4.29E-02 | 1.00E-01 | 2.68E+00 | 2.75E+00 | 6.65E-01 | no | Zmat3 | intron_variant | MODIFIER |
| 8 | 37200469 | AX-170221513 | 0 | G | A | 0.158 | 6.02E-02 | 1.41E-01 | 2.73E+00 | 2.79E+00 | 6.65E-01 | no | | intergenic_variant | MODIFIER |
| 8 | 38742200 | AX-170135633 | 0 | C | T | 0.158 | 6.02E-02 | 1.41E-01 | 2.73E+00 | 2.79E+00 | 6.65E-01 | no | | intergenic_variant | MODIFIER |
| 7 | 80359190 | AX-97817129 | 0 | C | A | 0.123 | 8.16E-02 | 1.91E-01 | 2.65E+00 | 2.75E+00 | 6.65E-01 | no | Man2a2 | intron_variant | MODIFIER |
| 7 | 94937739 | AX-170029552 | 0 | T | G | 0.123 | 8.16E-02 | 1.91E-01 | 2.65E+00 | 2.75E+00 | 6.65E-01 | no | | intergenic_variant | MODIFIER |
| 7 | 98875196 | AX-170372347 | 0 | T | C | 0.123 | 8.16E-02 | 1.91E-01 | 2.65E+00 | 2.75E+00 | 6.65E-01 | no | | regulatory_region_variant | MODIFIER |
| 7 | 98879267 | AX-170360314 | 0 | T | C | 0.123 | 8.16E-02 | 1.91E-01 | 2.65E+00 | 2.75E+00 | 6.65E-01 | no | | intergenic_variant | MODIFIER |
| 7 | 98882236 | AX-170475657 | 0 | C | T | 0.123 | 8.16E-02 | 1.91E-01 | 2.65E+00 | 2.75E+00 | 6.65E-01 | no | Uvrag | downstream_gene_variant | MODIFIER |
| 7 | 98900181 | AX-170229601 | 0 | T | C | 0.123 | 8.16E-02 | 1.91E-01 | 2.65E+00 | 2.75E+00 | 6.65E-01 | no | Uvrag | intron_variant | MODIFIER |
| 7 | 98902692 | AX-170322116 | 0 | C | T | 0.123 | 8.16E-02 | 1.91E-01 | 2.65E+00 | 2.75E+00 | 6.65E-01 | no | Uvrag | intron_variant | MODIFIER |
| 7 | 98903031 | AX-97785633 | 0 | C | T | 0.123 | 8.16E-02 | 1.91E-01 | 2.65E+00 | 2.75E+00 | 6.65E-01 | no | Uvrag | intron_variant | MODIFIER |
| 7 | 98913607 | AX-170272333 | 0 | G | A | 0.123 | 8.16E-02 | 1.91E-01 | 2.65E+00 | 2.75E+00 | 6.65E-01 | no | Uvrag | intron_variant | MODIFIER |
| 7 | 98922914 | AX-170244011 | 0 | T | C | 0.123 | 8.16E-02 | 1.91E-01 | 2.65E+00 | 2.75E+00 | 6.65E-01 | no | Uvrag | intron_variant | MODIFIER |
| 7 | 98940035 | AX-170439012 | 0 | C | A | 0.123 | 8.16E-02 | 1.91E-01 | 2.65E+00 | 2.75E+00 | 6.65E-01 | no | Uvrag | intron_variant | MODIFIER |
| 7 | 98947699 | AX-97847903 | 0 | T | C | 0.123 | 8.16E-02 | 1.91E-01 | 2.65E+00 | 2.75E+00 | 6.65E-01 | no | Uvrag | intron_variant | MODIFIER |
| 7 | 98988052 | AX-97809940 | 0 | G | A | 0.123 | 8.16E-02 | 1.91E-01 | 2.65E+00 | 2.75E+00 | 6.65E-01 | no | Uvrag | intron_variant | MODIFIER |
| 7 | 98992032 | AX-170101199 | 0 | G | A | 0.123 | 8.16E-02 | 1.91E-01 | 2.65E+00 | 2.75E+00 | 6.65E-01 | no | Uvrag | intron_variant | MODIFIER |
| 7 | 99001686 | AX-170457434 | 0 | A | G | 0.123 | 8.16E-02 | 1.91E-01 | 2.65E+00 | 2.75E+00 | 6.65E-01 | no | Uvrag | intron_variant | MODIFIER |
| 7 | 99002233 | AX-170507682 | 0 | C | T | 0.123 | 8.16E-02 | 1.91E-01 | 2.65E+00 | 2.75E+00 | 6.65E-01 | no | Uvrag | intron_variant | MODIFIER |
| 7 | 99007410 | AX-170463666 | 0 | G | A | 0.123 | 8.16E-02 | 1.91E-01 | 2.65E+00 | 2.75E+00 | 6.65E-01 | no | Uvrag | intron_variant | MODIFIER |
| 7 | 99009023 | AX-97846763 | 0 | C | A | 0.123 | 8.16E-02 | 1.91E-01 | 2.65E+00 | 2.75E+00 | 6.65E-01 | no | Uvrag | intron_variant | MODIFIER |
| 7 | 99023260 | AX-170109912 | 0 | G | A | 0.123 | 8.16E-02 | 1.91E-01 | 2.65E+00 | 2.75E+00 | 6.65E-01 | no | Uvrag | intron_variant | MODIFIER |
| 7 | 99046819 | AX-170286632 | 0 | T | C | 0.123 | 8.16E-02 | 1.91E-01 | 2.65E+00 | 2.75E+00 | 6.65E-01 | no | Uvrag | intron_variant | MODIFIER |
| 7 | 99056293 | AX-170187433 | 0 | T | G | 0.123 | 8.16E-02 | 1.91E-01 | 2.65E+00 | 2.75E+00 | 6.65E-01 | no | Uvrag | intron_variant | MODIFIER |
| 3 | 30353139 | AX-170377905 | 2 | A | G | 0.173 | -4.34E-02 | 1.01E-01 | 2.67E+00 | 2.74E+00 | 6.65E-01 | no | Mecom | intron_variant | MODIFIER |
| 15 | 74693106 | AX-97818895 | 0 | G | T | 0.553 | 3.59E-02 | 8.21E-02 | 2.60E+00 | 2.68E+00 | 6.65E-01 | no | | intergenic_variant | MODIFIER |
| 15 | 75709751 | AX-170359224 | 0 | G | A | 0.553 | 3.59E-02 | 8.21E-02 | 2.60E+00 | 2.68E+00 | 6.65E-01 | no | Rhpn1 | intron_variant | MODIFIER |
| 5 | 81117335 | AX-97781620 | 0 | A | C | 0.439 | 4.66E-02 | 1.09E-01 | 2.70E+00 | 2.75E+00 | 6.65E-01 | no | Adgrl3 | intron_variant | MODIFIER |
| 1 | 151171846 | AX-170001983 | 0 | T | C | 0.211 | 5.75E-02 | 1.32E-01 | 2.62E+00 | 2.69E+00 | 6.65E-01 | no | | regulatory_region_variant | MODIFIER |
| 7 | 45941278 | AX-170262245 | 1 | A | C | 0.321 | 4.06E-02 | 9.27E-02 | 2.59E+00 | 2.67E+00 | 6.66E-01 | no | Ccdc114 | intron_variant | MODIFIER |
| 7 | 89749947 | AX-170244111 | 0 | T | C | 0.14 | 7.00E-02 | 1.66E-01 | 2.64E+00 | 2.78E+00 | 6.66E-01 | no | Me3 | intron_variant | MODIFIER |
| 10 | 32433678 | AX-170528863 | 0 | C | A | 0.395 | 3.12E-02 | 7.42E-02 | 2.67E+00 | 2.79E+00 | 6.66E-01 | no | Nkain2 | intron_variant | MODIFIER |
| 12 | 99487034 | AX-97850790 | 1 | G | A | 0.054 | 9.41E-02 | 2.21E-01 | 2.69E+00 | 2.76E+00 | 6.66E-01 | no | Foxn3 | intron_variant | MODIFIER |
| 20 | 74332564 | AX-170491234 | 0 | C | T | 0.263 | 2.67E-02 | 6.23E-02 | 2.64E+00 | 2.73E+00 | 6.66E-01 | no | Plxna3 | intron_variant | MODIFIER |
| 9 | 37832294 | AX-170157594 | 0 | A | C | 0.491 | -3.32E-02 | 7.86E-02 | 2.71E+00 | 2.80E+00 | 6.66E-01 | no | | intergenic_variant | MODIFIER |
| 5 | 112986892 | AX-170473845 | 0 | G | A | 0.456 | 4.93E-02 | 1.15E-01 | 2.66E+00 | 2.70E+00 | 6.66E-01 | no | Grk3 | intron_variant | MODIFIER |
| 5 | 114281114 | AX-170025932 | 0 | T | C | 0.456 | 4.93E-02 | 1.15E-01 | 2.66E+00 | 2.70E+00 | 6.66E-01 | no | | intergenic_variant | MODIFIER |
| 8 | 106855553 | AX-170031440 | 1 | G | C | 0.223 | 4.63E-02 | 1.09E-01 | 2.67E+00 | 2.75E+00 | 6.66E-01 | no | Tango6 | downstream_gene_variant | MODIFIER |
| 9 | 39723463 | AX-170035803 | 1 | G | A | 0.438 | -3.21E-02 | 7.57E-02 | 2.68E+00 | 2.77E+00 | 6.66E-01 | no | Olfr965 | downstream_gene_variant | MODIFIER |
| 7 | 118060632 | AX-170265298 | 1 | C | T | 0.402 | 3.92E-02 | 9.15E-02 | 2.66E+00 | 2.71E+00 | 6.66E-01 | no | | intergenic_variant | MODIFIER |
| 18 | 66289713 | AX-170061148 | 0 | G | A | 0.088 | 6.63E-02 | 1.55E-01 | 2.61E+00 | 2.73E+00 | 6.66E-01 | no | Ccbe1 | intron_variant,NMD_transcript_variant | MODIFIER |
| 5 | 114396427 | AX-170189404 | 1 | C | T | 0.473 | 4.15E-02 | 9.44E-02 | 2.57E+00 | 2.66E+00 | 6.66E-01 | no | Ube3b | intron_variant | MODIFIER |
| 12 | 99566949 | AX-97800241 | 1 | T | C | 0.054 | 9.42E-02 | 2.22E-01 | 2.70E+00 | 2.76E+00 | 6.66E-01 | no | Foxn3 | upstream_gene_variant | MODIFIER |
| 1 | 168990062 | AX-170209375 | 0 | G | C | 0.439 | -2.91E-02 | 6.80E-02 | 2.83E+00 | 2.91E+00 | 6.67E-01 | no | | intergenic_variant | MODIFIER |
| 10 | 111159959 | AX-170527486 | 0 | C | T | 0.368 | -3.35E-02 | 7.84E-02 | 2.64E+00 | 2.72E+00 | 6.67E-01 | no | Osbpl8 | upstream_gene_variant | MODIFIER |
| 1 | 153165344 | AX-170466886 | 1 | G | A | 0.214 | 5.23E-02 | 1.22E-01 | 2.63E+00 | 2.72E+00 | 6.67E-01 | no | Lamc2 | intron_variant | MODIFIER |
| 14 | 124271760 | AX-97803946 | 0 | C | A | 0.289 | 3.69E-02 | 8.45E-02 | 2.57E+00 | 2.68E+00 | 6.67E-01 | no | Fgf14 | intron_variant | MODIFIER |
| 5 | 123642244 | AX-170132464 | 0 | A | G | 0.456 | -4.81E-02 | 1.13E-01 | 2.79E+00 | 2.84E+00 | 6.67E-01 | no | Clip1 | intron_variant | MODIFIER |
| 5 | 123650999 | AX-97827596 | 0 | C | T | 0.456 | -4.81E-02 | 1.13E-01 | 2.79E+00 | 2.84E+00 | 6.67E-01 | no | Clip1 | intron_variant | MODIFIER |

| | | | | | | | | | | | | | | | |
|---|---|---|---|---|---|---|---|---|---|---|---|---|---|---|---|
| 15 | 74231791 | AX-170104872 | 0 | T | G | 0.544 | 3.58E-02 | 8.18E-02 | 2.58E+00 | 2.66E+00 | 6.68E-01 | no | | intergenic_variant | MODIFIER |
| 8 | 118612835 | AX-170296090 | 0 | G | A | 0.246 | 4.08E-02 | 9.79E-02 | 2.69E+00 | 2.81E+00 | 6.68E-01 | no | Cdh13 | intron_variant | MODIFIER |
| 8 | 118613623 | AX-170139408 | 0 | T | G | 0.246 | 4.08E-02 | 9.79E-02 | 2.69E+00 | 2.81E+00 | 6.68E-01 | no | Cdh13 | intron_variant | MODIFIER |
| 8 | 118645044 | AX-170325827 | 0 | A | C | 0.246 | 4.08E-02 | 9.79E-02 | 2.69E+00 | 2.81E+00 | 6.68E-01 | no | Cdh13 | intron_variant | MODIFIER |
| 8 | 118651112 | AX-170469275 | 0 | G | A | 0.246 | 4.08E-02 | 9.79E-02 | 2.69E+00 | 2.81E+00 | 6.68E-01 | no | Cdh13 | intron_variant | MODIFIER |
| 8 | 118652382 | AX-168407412 | 0 | C | T | 0.246 | 4.08E-02 | 9.79E-02 | 2.69E+00 | 2.81E+00 | 6.68E-01 | no | Cdh13 | intron_variant | MODIFIER |
| 8 | 118655512 | AX-170389589 | 0 | A | G | 0.246 | 4.08E-02 | 9.79E-02 | 2.69E+00 | 2.81E+00 | 6.68E-01 | no | Cdh13 | intron_variant | MODIFIER |
| 8 | 118663498 | AX-170462757 | 0 | G | T | 0.246 | 4.08E-02 | 9.79E-02 | 2.69E+00 | 2.81E+00 | 6.68E-01 | no | Cdh13 | intron_variant | MODIFIER |
| 8 | 118675070 | AX-97837575 | 0 | C | T | 0.246 | 4.08E-02 | 9.79E-02 | 2.69E+00 | 2.81E+00 | 6.68E-01 | no | Cdh13 | synonymous_variant | LOW |
| 8 | 118676496 | AX-170354013 | 0 | T | C | 0.246 | 4.08E-02 | 9.79E-02 | 2.69E+00 | 2.81E+00 | 6.68E-01 | no | Cdh13 | intron_variant | MODIFIER |
| 8 | 118676622 | AX-170512874 | 0 | G | A | 0.246 | 4.08E-02 | 9.79E-02 | 2.69E+00 | 2.81E+00 | 6.68E-01 | no | Cdh13 | intron_variant | MODIFIER |
| 8 | 118684034 | AX-170109115 | 0 | A | G | 0.246 | 4.08E-02 | 9.79E-02 | 2.69E+00 | 2.81E+00 | 6.68E-01 | no | Cdh13 | intron_variant | MODIFIER |
| 3 | 31754848 | AX-170020242 | 1 | C | G | 0.161 | -4.22E-02 | 9.99E-02 | 2.68E+00 | 2.76E+00 | 6.68E-01 | no | | intergenic_variant | MODIFIER |
| 13 | 31368937 | AX-170087398 | 1 | G | A | 0.179 | -5.29E-02 | 1.22E-01 | 2.61E+00 | 2.67E+00 | 6.68E-01 | no | G630018N14Rik | intron_variant,non_coding_transcript_variant | MODIFIER |
| 13 | 31456972 | AX-170212933 | 1 | G | A | 0.179 | -5.29E-02 | 1.22E-01 | 2.61E+00 | 2.67E+00 | 6.68E-01 | no | | intergenic_variant | MODIFIER |
| 14 | 101284024 | AX-170430701 | 0 | C | T | 0.053 | -6.43E-02 | 1.49E-01 | 2.57E+00 | 2.71E+00 | 6.68E-01 | no | Gm26173 | downstream_gene_variant | MODIFIER |
| 1 | 108358879 | AX-170400234 | 1 | C | A | 0.17 | 8.46E-02 | 1.94E-01 | 2.59E+00 | 2.67E+00 | 6.68E-01 | no | | intergenic_variant | MODIFIER |
| 8 | 103988825 | AX-170371757 | 0 | A | G | 0.228 | 4.61E-02 | 1.09E-01 | 2.69E+00 | 2.76E+00 | 6.68E-01 | no | | intergenic_variant | MODIFIER |
| 8 | 103990273 | AX-170358354 | 0 | G | C | 0.228 | 4.61E-02 | 1.09E-01 | 2.69E+00 | 2.76E+00 | 6.68E-01 | no | | intergenic_variant | MODIFIER |
| 8 | 104003790 | AX-170188989 | 0 | A | C | 0.228 | 4.61E-02 | 1.09E-01 | 2.69E+00 | 2.76E+00 | 6.68E-01 | no | | intergenic_variant | MODIFIER |
| 8 | 104005225 | AX-170130743 | 0 | T | G | 0.228 | 4.61E-02 | 1.09E-01 | 2.69E+00 | 2.76E+00 | 6.68E-01 | no | | intergenic_variant | MODIFIER |
| 8 | 104008923 | AX-170031407 | 0 | T | C | 0.228 | 4.61E-02 | 1.09E-01 | 2.69E+00 | 2.76E+00 | 6.68E-01 | no | | intergenic_variant | MODIFIER |
| 8 | 104014779 | AX-170381572 | 0 | A | G | 0.228 | 4.61E-02 | 1.09E-01 | 2.69E+00 | 2.76E+00 | 6.68E-01 | no | | intergenic_variant | MODIFIER |
| 8 | 104018304 | AX-170184701 | 0 | C | A | 0.228 | 4.61E-02 | 1.09E-01 | 2.69E+00 | 2.76E+00 | 6.68E-01 | no | | intergenic_variant | MODIFIER |
| 8 | 106636687 | AX-170352975 | 0 | A | G | 0.228 | 4.61E-02 | 1.09E-01 | 2.69E+00 | 2.76E+00 | 6.68E-01 | no | Cdh1 | intron_variant | MODIFIER |
| 8 | 106653394 | AX-170523742 | 0 | T | C | 0.228 | 4.61E-02 | 1.09E-01 | 2.69E+00 | 2.76E+00 | 6.68E-01 | no | Cdh1 | intron_variant | MODIFIER |
| 8 | 106653460 | AX-170399969 | 0 | G | A | 0.228 | 4.61E-02 | 1.09E-01 | 2.69E+00 | 2.76E+00 | 6.68E-01 | no | Cdh1 | intron_variant | MODIFIER |
| 8 | 106659570 | AX-170455815 | 0 | A | T | 0.228 | 4.61E-02 | 1.09E-01 | 2.69E+00 | 2.76E+00 | 6.68E-01 | no | Cdh1 | intron_variant | MODIFIER |
| 8 | 106700264 | AX-170496639 | 0 | G | A | 0.228 | 4.61E-02 | 1.09E-01 | 2.69E+00 | 2.76E+00 | 6.68E-01 | no | Tango6 | intron_variant | MODIFIER |
| 8 | 106712791 | AX-97795097 | 0 | G | A | 0.228 | 4.61E-02 | 1.09E-01 | 2.69E+00 | 2.76E+00 | 6.68E-01 | no | Tango6 | intron_variant | MODIFIER |
| 8 | 106735368 | AX-170165978 | 0 | T | C | 0.228 | 4.61E-02 | 1.09E-01 | 2.69E+00 | 2.76E+00 | 6.68E-01 | no | Tango6 | intron_variant | MODIFIER |
| 8 | 106756202 | AX-170313250 | 0 | G | A | 0.228 | 4.61E-02 | 1.09E-01 | 2.69E+00 | 2.76E+00 | 6.68E-01 | no | Tango6 | intron_variant | MODIFIER |
| 8 | 106761655 | AX-170031439 | 0 | G | A | 0.228 | 4.61E-02 | 1.09E-01 | 2.69E+00 | 2.76E+00 | 6.68E-01 | no | Tango6 | intron_variant | MODIFIER |
| 8 | 106788338 | AX-170340894 | 0 | G | A | 0.228 | 4.61E-02 | 1.09E-01 | 2.69E+00 | 2.76E+00 | 6.68E-01 | no | Tango6 | intron_variant | MODIFIER |
| 8 | 106790086 | AX-97811394 | 0 | A | G | 0.228 | 4.61E-02 | 1.09E-01 | 2.69E+00 | 2.76E+00 | 6.68E-01 | no | Tango6 | intron_variant | MODIFIER |
| 8 | 106806405 | AX-170031437 | 0 | T | C | 0.228 | 4.61E-02 | 1.09E-01 | 2.69E+00 | 2.76E+00 | 6.68E-01 | no | Tango6 | intron_variant | MODIFIER |
| 8 | 106844660 | AX-97815265 | 0 | G | A | 0.228 | 4.61E-02 | 1.09E-01 | 2.69E+00 | 2.76E+00 | 6.68E-01 | no | Tango6 | intron_variant | MODIFIER |
| 8 | 107042704 | AX-97835374 | 0 | T | C | 0.228 | 4.61E-02 | 1.09E-01 | 2.69E+00 | 2.76E+00 | 6.68E-01 | no | Vps4a | intron_variant | MODIFIER |
| 8 | 107208935 | AX-170457610 | 0 | A | G | 0.228 | 4.61E-02 | 1.09E-01 | 2.69E+00 | 2.76E+00 | 6.68E-01 | no | | regulatory_region_variant | MODIFIER |
| 8 | 107214220 | AX-170178957 | 0 | C | T | 0.228 | 4.61E-02 | 1.09E-01 | 2.69E+00 | 2.76E+00 | 6.68E-01 | no | | intergenic_variant | MODIFIER |
| 8 | 107247085 | AX-170157122 | 0 | A | G | 0.228 | 4.61E-02 | 1.09E-01 | 2.69E+00 | 2.76E+00 | 6.68E-01 | no | | intergenic_variant | MODIFIER |
| 8 | 107251048 | AX-170031447 | 0 | T | C | 0.228 | 4.61E-02 | 1.09E-01 | 2.69E+00 | 2.76E+00 | 6.68E-01 | no | | intergenic_variant | MODIFIER |
| 8 | 107658353 | AX-170081101 | 0 | T | C | 0.228 | 4.61E-02 | 1.09E-01 | 2.69E+00 | 2.76E+00 | 6.68E-01 | no | | intergenic_variant | MODIFIER |
| 8 | 107658465 | AX-170482530 | 0 | C | G | 0.228 | 4.61E-02 | 1.09E-01 | 2.69E+00 | 2.76E+00 | 6.68E-01 | no | | intergenic_variant | MODIFIER |
| 8 | 107675178 | AX-170169128 | 0 | A | G | 0.228 | 4.61E-02 | 1.09E-01 | 2.69E+00 | 2.76E+00 | 6.68E-01 | no | | intergenic_variant | MODIFIER |
| 7 | 13238299 | AX-97806873 | 1 | A | C | 0.339 | 3.43E-02 | 7.89E-02 | 2.57E+00 | 2.69E+00 | 6.68E-01 | no | | intergenic_variant | MODIFIER |
| 12 | 104174232 | AX-170175864 | 1 | A | G | 0.107 | -8.42E-02 | 2.01E-01 | 2.75E+00 | 2.84E+00 | 6.68E-01 | no | Serpina3d-ps | intron_variant,non_coding_transcript_variant | MODIFIER |
| 18 | 69298427 | AX-170278611 | 1 | C | T | 0.268 | -4.01E-02 | 9.45E-02 | 2.65E+00 | 2.73E+00 | 6.68E-01 | no | | intergenic_variant | MODIFIER |
| 12 | 116831193 | AX-170050329 | 0 | C | T | 0.149 | -6.57E-02 | 1.58E-01 | 2.71E+00 | 2.81E+00 | 6.68E-01 | no | Ptprn2 | intron_variant | MODIFIER |
| 5 | 121180677 | AX-170140397 | 0 | G | A | 0.447 | -4.74E-02 | 1.12E-01 | 2.78E+00 | 2.84E+00 | 6.68E-01 | no | Ptpn11 | intron_variant | MODIFIER |
| 5 | 121568471 | AX-97810451 | 0 | C | T | 0.447 | -4.74E-02 | 1.12E-01 | 2.78E+00 | 2.84E+00 | 6.68E-01 | no | Aldh2 | intron_variant | MODIFIER |
| 5 | 122042972 | AX-170229445 | 0 | G | A | 0.447 | -4.74E-02 | 1.12E-01 | 2.78E+00 | 2.84E+00 | 6.68E-01 | no | Cux2 | intron_variant | MODIFIER |
| 5 | 124041084 | AX-170507369 | 0 | T | C | 0.447 | -4.74E-02 | 1.12E-01 | 2.78E+00 | 2.84E+00 | 6.68E-01 | no | Gm34086 | intron_variant,non_coding_transcript_variant | MODIFIER |
| 16 | 96480861 | AX-170356650 | 0 | C | T | 0.456 | 3.09E-02 | 7.45E-02 | 2.75E+00 | 2.87E+00 | 6.68E-01 | no | Pcp4 | intron_variant | MODIFIER |

| | | | | | | | | | | | | | | | |
|---|---|---|---|---|---|---|---|---|---|---|---|---|---|---|---|
| 1 | 68783289 | AX-170111422 | 0 | C | G | 0.114 | -7.77E-02 | 1.85E-01 | 2.68E+00 | 2.77E+00 | 6.69E-01 | no | Erbb4 | intron_variant | MODIFIER |
| 1 | 4624584 | AX-170162149 | 0 | T | C | 0.526 | 3.01E-02 | 6.73E-02 | 2.54E+00 | 2.62E+00 | 6.69E-01 | no | | intergenic_variant | MODIFIER |
| 8 | 106649805 | AX-170360455 | 0 | C | T | 0.228 | 4.49E-02 | 1.06E-01 | 2.66E+00 | 2.74E+00 | 6.69E-01 | no | Cdh1 | intron_variant | MODIFIER |
| 8 | 107132852 | AX-170248807 | 0 | G | A | 0.228 | 4.49E-02 | 1.06E-01 | 2.66E+00 | 2.74E+00 | 6.69E-01 | no | | regulatory_region_variant | MODIFIER |
| 15 | 74294955 | AX-170312164 | 2 | C | T | 0.564 | 3.54E-02 | 8.07E-02 | 2.57E+00 | 2.65E+00 | 6.69E-01 | no | | regulatory_region_variant | MODIFIER |
| 18 | 66202165 | AX-170061144 | 1 | C | T | 0.241 | -3.98E-02 | 9.42E-02 | 2.67E+00 | 2.74E+00 | 6.69E-01 | no | Ccbe1 | intron_variant,NMD_transcript_variant | MODIFIER |
| 9 | 34153078 | AX-170377489 | 1 | C | T | 0.438 | 3.68E-02 | 8.59E-02 | 2.62E+00 | 2.69E+00 | 6.69E-01 | no | | intergenic_variant | MODIFIER |
| 15 | 101256639 | AX-170012395 | 2 | T | C | 0.127 | 5.48E-02 | 1.31E-01 | 2.67E+00 | 2.77E+00 | 6.69E-01 | no | Nr4a1 | intron_variant | MODIFIER |
| 5 | 88854679 | AX-170181164 | 0 | C | T | 0.43 | 4.67E-02 | 1.11E-01 | 2.77E+00 | 2.82E+00 | 6.69E-01 | no | | intergenic_variant | MODIFIER |
| 8 | 94947118 | AX-97830223 | 0 | T | C | 0.228 | 4.89E-02 | 1.16E-01 | 2.70E+00 | 2.77E+00 | 6.69E-01 | no | Adgrg5 | downstream_gene_variant | MODIFIER |
| 8 | 94947585 | AX-170309540 | 0 | A | G | 0.228 | 4.89E-02 | 1.16E-01 | 2.70E+00 | 2.77E+00 | 6.69E-01 | no | Adgrg5 | downstream_gene_variant | MODIFIER |
| 8 | 94959676 | AX-170200280 | 0 | A | T | 0.228 | 4.89E-02 | 1.16E-01 | 2.70E+00 | 2.77E+00 | 6.69E-01 | no | | intergenic_variant | MODIFIER |
| 8 | 94959814 | AX-170386316 | 0 | A | C | 0.228 | 4.89E-02 | 1.16E-01 | 2.70E+00 | 2.77E+00 | 6.69E-01 | no | Gm10286 | downstream_gene_variant | MODIFIER |
| 8 | 94985943 | AX-170251438 | 0 | C | G | 0.228 | 4.89E-02 | 1.16E-01 | 2.70E+00 | 2.77E+00 | 6.69E-01 | no | Adgrg1 | intron_variant | MODIFIER |
| 8 | 94989977 | AX-170333034 | 0 | G | A | 0.228 | 4.89E-02 | 1.16E-01 | 2.70E+00 | 2.77E+00 | 6.69E-01 | no | Adgrg1 | intron_variant | MODIFIER |
| 8 | 94990633 | AX-170106083 | 0 | T | C | 0.228 | 4.89E-02 | 1.16E-01 | 2.70E+00 | 2.77E+00 | 6.69E-01 | no | Adgrg1 | intron_variant | MODIFIER |
| 8 | 94994103 | AX-170190343 | 0 | T | C | 0.228 | 4.89E-02 | 1.16E-01 | 2.70E+00 | 2.77E+00 | 6.69E-01 | no | Adgrg1 | intron_variant | MODIFIER |
| 8 | 95016024 | AX-97856731 | 0 | A | G | 0.228 | 4.89E-02 | 1.16E-01 | 2.70E+00 | 2.77E+00 | 6.69E-01 | no | Adgrg3 | upstream_gene_variant | MODIFIER |
| 8 | 95050713 | AX-170145988 | 0 | C | T | 0.228 | 4.89E-02 | 1.16E-01 | 2.70E+00 | 2.77E+00 | 6.69E-01 | no | Drc7 | upstream_gene_variant | MODIFIER |
| 8 | 95061203 | AX-170314930 | 0 | T | G | 0.228 | 4.89E-02 | 1.16E-01 | 2.70E+00 | 2.77E+00 | 6.69E-01 | no | Drc7 | intron_variant | MODIFIER |
| 8 | 95084109 | AX-170122129 | 0 | A | G | 0.228 | 4.89E-02 | 1.16E-01 | 2.70E+00 | 2.77E+00 | 6.69E-01 | no | Katnb1 | intron_variant | MODIFIER |
| 8 | 95111001 | AX-170080869 | 0 | C | A | 0.228 | 4.89E-02 | 1.16E-01 | 2.70E+00 | 2.77E+00 | 6.69E-01 | no | Kifc3 | intron_variant | MODIFIER |
| 8 | 95151389 | AX-170080870 | 0 | T | C | 0.228 | 4.89E-02 | 1.16E-01 | 2.70E+00 | 2.77E+00 | 6.69E-01 | no | Kifc3 | intron_variant | MODIFIER |
| 8 | 95153805 | AX-170080866 | 0 | A | G | 0.228 | 4.89E-02 | 1.16E-01 | 2.70E+00 | 2.77E+00 | 6.69E-01 | no | Kifc3 | intron_variant | MODIFIER |
| 8 | 95175266 | AX-97816568 | 0 | G | A | 0.228 | 4.89E-02 | 1.16E-01 | 2.70E+00 | 2.77E+00 | 6.69E-01 | no | Kifc3 | intron_variant | MODIFIER |
| 8 | 95176011 | AX-170166865 | 0 | C | T | 0.228 | 4.89E-02 | 1.16E-01 | 2.70E+00 | 2.77E+00 | 6.69E-01 | no | Kifc3 | intron_variant | MODIFIER |
| 8 | 95178126 | AX-170288191 | 0 | C | T | 0.228 | 4.89E-02 | 1.16E-01 | 2.70E+00 | 2.77E+00 | 6.69E-01 | no | Kifc3 | intron_variant | MODIFIER |
| 8 | 95209797 | AX-170306895 | 0 | A | C | 0.228 | 4.89E-02 | 1.16E-01 | 2.70E+00 | 2.77E+00 | 6.69E-01 | no | | regulatory_region_variant | MODIFIER |
| 8 | 95218596 | AX-97789910 | 0 | C | T | 0.228 | 4.89E-02 | 1.16E-01 | 2.70E+00 | 2.77E+00 | 6.69E-01 | no | | intergenic_variant | MODIFIER |
| 8 | 95219125 | AX-170252406 | 0 | A | G | 0.228 | 4.89E-02 | 1.16E-01 | 2.70E+00 | 2.77E+00 | 6.69E-01 | no | | intergenic_variant | MODIFIER |
| 8 | 95219359 | AX-170524743 | 0 | C | T | 0.228 | 4.89E-02 | 1.16E-01 | 2.70E+00 | 2.77E+00 | 6.69E-01 | no | | intergenic_variant | MODIFIER |
| 8 | 95228435 | AX-170031301 | 0 | G | A | 0.228 | 4.89E-02 | 1.16E-01 | 2.70E+00 | 2.77E+00 | 6.69E-01 | no | | intergenic_variant | MODIFIER |
| 8 | 95248264 | AX-170113168 | 0 | T | C | 0.228 | 4.89E-02 | 1.16E-01 | 2.70E+00 | 2.77E+00 | 6.69E-01 | no | Cngb1 | intron_variant | MODIFIER |
| 8 | 95256264 | AX-170465121 | 0 | G | C | 0.228 | 4.89E-02 | 1.16E-01 | 2.70E+00 | 2.77E+00 | 6.69E-01 | no | Cngb1 | intron_variant | MODIFIER |
| 8 | 95260823 | AX-97833983 | 0 | G | A | 0.228 | 4.89E-02 | 1.16E-01 | 2.70E+00 | 2.77E+00 | 6.69E-01 | no | Cngb1 | intron_variant | MODIFIER |
| 8 | 95264890 | AX-170031299 | 0 | A | G | 0.228 | 4.89E-02 | 1.16E-01 | 2.70E+00 | 2.77E+00 | 6.69E-01 | no | Cngb1 | intron_variant | MODIFIER |
| 8 | 95267977 | AX-170080871 | 0 | C | T | 0.228 | 4.89E-02 | 1.16E-01 | 2.70E+00 | 2.77E+00 | 6.69E-01 | no | Cngb1 | intron_variant | MODIFIER |
| 8 | 95268068 | AX-170247285 | 0 | C | T | 0.228 | 4.89E-02 | 1.16E-01 | 2.70E+00 | 2.77E+00 | 6.69E-01 | no | Cngb1 | intron_variant | MODIFIER |
| 8 | 95268090 | AX-170452670 | 0 | T | C | 0.228 | 4.89E-02 | 1.16E-01 | 2.70E+00 | 2.77E+00 | 6.69E-01 | no | Cngb1 | intron_variant | MODIFIER |
| 8 | 95269407 | AX-170193779 | 0 | G | T | 0.228 | 4.89E-02 | 1.16E-01 | 2.70E+00 | 2.77E+00 | 6.69E-01 | no | Cngb1 | intron_variant | MODIFIER |
| 8 | 95270133 | AX-170508233 | 0 | A | T | 0.228 | 4.89E-02 | 1.16E-01 | 2.70E+00 | 2.77E+00 | 6.69E-01 | no | Cngb1 | intron_variant | MODIFIER |
| 8 | 95270562 | AX-170236697 | 0 | G | A | 0.228 | 4.89E-02 | 1.16E-01 | 2.70E+00 | 2.77E+00 | 6.69E-01 | no | Cngb1 | intron_variant | MODIFIER |
| 8 | 95274768 | AX-170496664 | 0 | C | T | 0.228 | 4.89E-02 | 1.16E-01 | 2.70E+00 | 2.77E+00 | 6.69E-01 | no | Cngb1 | intron_variant | MODIFIER |
| 8 | 95295170 | AX-170031298 | 0 | G | A | 0.228 | 4.89E-02 | 1.16E-01 | 2.70E+00 | 2.77E+00 | 6.69E-01 | no | Cngb1 | synonymous_variant | LOW |
| 8 | 95295634 | AX-170273583 | 0 | A | G | 0.228 | 4.89E-02 | 1.16E-01 | 2.70E+00 | 2.77E+00 | 6.69E-01 | no | Cngb1 | intron_variant | MODIFIER |
| 8 | 95305847 | AX-170331410 | 0 | G | T | 0.228 | 4.89E-02 | 1.16E-01 | 2.70E+00 | 2.77E+00 | 6.69E-01 | no | Cngb1 | intron_variant | MODIFIER |
| 8 | 95313188 | AX-170352360 | 0 | C | G | 0.228 | 4.89E-02 | 1.16E-01 | 2.70E+00 | 2.77E+00 | 6.69E-01 | no | Tepp | intron_variant | MODIFIER |
| 8 | 95318315 | AX-170035790 | 0 | C | G | 0.228 | 4.89E-02 | 1.16E-01 | 2.70E+00 | 2.77E+00 | 6.69E-01 | no | Zfp319 | downstream_gene_variant | MODIFIER |
| 8 | 95321417 | AX-170119087 | 0 | A | G | 0.228 | 4.89E-02 | 1.16E-01 | 2.70E+00 | 2.77E+00 | 6.69E-01 | no | Zfp319 | downstream_gene_variant | MODIFIER |
| 8 | 95322273 | AX-97851752 | 0 | G | A | 0.228 | 4.89E-02 | 1.16E-01 | 2.70E+00 | 2.77E+00 | 6.69E-01 | no | Zfp319 | downstream_gene_variant | MODIFIER |
| 8 | 95388370 | AX-170483868 | 0 | G | T | 0.228 | 4.89E-02 | 1.16E-01 | 2.70E+00 | 2.77E+00 | 6.69E-01 | no | | intergenic_variant | MODIFIER |
| 8 | 95388397 | AX-170382765 | 0 | G | A | 0.228 | 4.89E-02 | 1.16E-01 | 2.70E+00 | 2.77E+00 | 6.69E-01 | no | | intergenic_variant | MODIFIER |
| 8 | 95408494 | AX-170106167 | 0 | C | T | 0.228 | 4.89E-02 | 1.16E-01 | 2.70E+00 | 2.77E+00 | 6.69E-01 | no | | intergenic_variant | MODIFIER |
| 8 | 95411858 | AX-170378089 | 0 | C | T | 0.228 | 4.89E-02 | 1.16E-01 | 2.70E+00 | 2.77E+00 | 6.69E-01 | no | | regulatory_region_variant | MODIFIER |

| | | | | | | | | | | | | | | | |
|---|---|---|---|---|---|---|---|---|---|---|---|---|---|---|---|
| 8 | 95419070 | AX-97799874 | 0 | T | C | 0.228 | 4.89E-02 | 1.16E-01 | 2.70E+00 | 2.77E+00 | 6.69E-01 | no | Cfap20 | downstream_gene_variant | MODIFIER |
| 8 | 95428150 | AX-170498097 | 0 | A | C | 0.228 | 4.89E-02 | 1.16E-01 | 2.70E+00 | 2.77E+00 | 6.69E-01 | no | Cfap20 | intron_variant | MODIFIER |
| 8 | 95455889 | AX-170439566 | 0 | G | A | 0.228 | 4.89E-02 | 1.16E-01 | 2.70E+00 | 2.77E+00 | 6.69E-01 | no | Csnk2a2 | synonymous_variant | LOW |
| 8 | 95460311 | AX-170454427 | 0 | T | C | 0.228 | 4.89E-02 | 1.16E-01 | 2.70E+00 | 2.77E+00 | 6.69E-01 | no | Csnk2a2 | intron_variant | MODIFIER |
| 8 | 95461089 | AX-170420035 | 0 | G | T | 0.228 | 4.89E-02 | 1.16E-01 | 2.70E+00 | 2.77E+00 | 6.69E-01 | no | Csnk2a2 | intron_variant | MODIFIER |
| 8 | 95463207 | AX-170090183 | 0 | C | T | 0.228 | 4.89E-02 | 1.16E-01 | 2.70E+00 | 2.77E+00 | 6.69E-01 | no | Csnk2a2 | intron_variant | MODIFIER |
| 8 | 95470463 | AX-170103423 | 0 | A | C | 0.228 | 4.89E-02 | 1.16E-01 | 2.70E+00 | 2.77E+00 | 6.69E-01 | no | Csnk2a2 | intron_variant | MODIFIER |
| 8 | 95470963 | AX-170200188 | 0 | C | T | 0.228 | 4.89E-02 | 1.16E-01 | 2.70E+00 | 2.77E+00 | 6.69E-01 | no | Csnk2a2 | intron_variant | MODIFIER |
| 8 | 95471297 | AX-170319763 | 0 | A | G | 0.228 | 4.89E-02 | 1.16E-01 | 2.70E+00 | 2.77E+00 | 6.69E-01 | no | Csnk2a2 | intron_variant | MODIFIER |
| 8 | 95472984 | AX-170155073 | 0 | A | G | 0.228 | 4.89E-02 | 1.16E-01 | 2.70E+00 | 2.77E+00 | 6.69E-01 | no | Csnk2a2 | intron_variant | MODIFIER |
| 8 | 95480318 | AX-170107909 | 0 | T | C | 0.228 | 4.89E-02 | 1.16E-01 | 2.70E+00 | 2.77E+00 | 6.69E-01 | no | Csnk2a2 | intron_variant | MODIFIER |
| 8 | 95480788 | AX-170157101 | 0 | G | C | 0.228 | 4.89E-02 | 1.16E-01 | 2.70E+00 | 2.77E+00 | 6.69E-01 | no | Csnk2a2 | intron_variant | MODIFIER |
| 8 | 95481054 | AX-170446483 | 0 | G | A | 0.228 | 4.89E-02 | 1.16E-01 | 2.70E+00 | 2.77E+00 | 6.69E-01 | no | Csnk2a2 | intron_variant | MODIFIER |
| 8 | 95481142 | AX-170210736 | 0 | A | G | 0.228 | 4.89E-02 | 1.16E-01 | 2.70E+00 | 2.77E+00 | 6.69E-01 | no | Csnk2a2 | intron_variant | MODIFIER |
| 8 | 95499318 | AX-170335249 | 0 | T | C | 0.228 | 4.89E-02 | 1.16E-01 | 2.70E+00 | 2.77E+00 | 6.69E-01 | no | 4933406B17Rik | intron_variant,non_coding_transcript_variant | MODIFIER |
| 8 | 95541386 | AX-170479230 | 0 | A | G | 0.228 | 4.89E-02 | 1.16E-01 | 2.70E+00 | 2.77E+00 | 6.69E-01 | no | Ccdc113 | intron_variant | MODIFIER |
| 8 | 95547607 | AX-170366843 | 0 | C | T | 0.228 | 4.89E-02 | 1.16E-01 | 2.70E+00 | 2.77E+00 | 6.69E-01 | no | Ccdc113 | intron_variant | MODIFIER |
| 8 | 95553804 | AX-170210345 | 0 | C | T | 0.228 | 4.89E-02 | 1.16E-01 | 2.70E+00 | 2.77E+00 | 6.69E-01 | no | Ccdc113 | intron_variant | MODIFIER |
| 8 | 95567739 | AX-170346212 | 0 | G | A | 0.228 | 4.89E-02 | 1.16E-01 | 2.70E+00 | 2.77E+00 | 6.69E-01 | no | Prss54 | intron_variant | MODIFIER |
| 8 | 95570840 | AX-170232333 | 0 | C | T | 0.228 | 4.89E-02 | 1.16E-01 | 2.70E+00 | 2.77E+00 | 6.69E-01 | no | Prss54 | intron_variant | MODIFIER |
| 8 | 95572114 | AX-170357018 | 0 | C | T | 0.228 | 4.89E-02 | 1.16E-01 | 2.70E+00 | 2.77E+00 | 6.69E-01 | no | Prss54 | intron_variant | MODIFIER |
| 8 | 95573778 | AX-170423965 | 0 | A | G | 0.228 | 4.89E-02 | 1.16E-01 | 2.70E+00 | 2.77E+00 | 6.69E-01 | no | Prss54 | intron_variant | MODIFIER |
| 8 | 95599931 | AX-170096368 | 0 | A | G | 0.228 | 4.89E-02 | 1.16E-01 | 2.70E+00 | 2.77E+00 | 6.69E-01 | no | Gm31518 | intron_variant,non_coding_transcript_variant | MODIFIER |
| 8 | 95609532 | AX-170175980 | 0 | C | T | 0.228 | 4.89E-02 | 1.16E-01 | 2.70E+00 | 2.77E+00 | 6.69E-01 | no | Gm31518 | intron_variant,non_coding_transcript_variant | MODIFIER |
| 8 | 95613319 | AX-170165628 | 0 | C | G | 0.228 | 4.89E-02 | 1.16E-01 | 2.70E+00 | 2.77E+00 | 6.69E-01 | no | Gm31518 | non_coding_transcript_exon_variant | MODIFIER |
| 8 | 95624428 | AX-170188556 | 0 | A | G | 0.228 | 4.89E-02 | 1.16E-01 | 2.70E+00 | 2.77E+00 | 6.69E-01 | no | Gm45702 | downstream_gene_variant | MODIFIER |
| 8 | 95669591 | AX-170080878 | 0 | T | C | 0.228 | 4.89E-02 | 1.16E-01 | 2.70E+00 | 2.77E+00 | 6.69E-01 | no | | intergenic_variant | MODIFIER |
| 8 | 95671928 | AX-170121439 | 0 | G | A | 0.228 | 4.89E-02 | 1.16E-01 | 2.70E+00 | 2.77E+00 | 6.69E-01 | no | Gm6013 | downstream_gene_variant | MODIFIER |
| 8 | 95674311 | AX-170234888 | 0 | T | C | 0.228 | 4.89E-02 | 1.16E-01 | 2.70E+00 | 2.77E+00 | 6.69E-01 | no | Ndrg4 | upstream_gene_variant | MODIFIER |
| 8 | 95674894 | AX-170295653 | 0 | C | G | 0.228 | 4.89E-02 | 1.16E-01 | 2.70E+00 | 2.77E+00 | 6.69E-01 | no | Ndrg4 | upstream_gene_variant | MODIFIER |
| 8 | 95688950 | AX-170242021 | 0 | A | G | 0.228 | 4.89E-02 | 1.16E-01 | 2.70E+00 | 2.77E+00 | 6.69E-01 | no | Ndrg4 | intron_variant | MODIFIER |
| 8 | 95713685 | AX-97825245 | 0 | A | G | 0.228 | 4.89E-02 | 1.16E-01 | 2.70E+00 | 2.77E+00 | 6.69E-01 | no | Setd6 | upstream_gene_variant | MODIFIER |
| 8 | 95725407 | AX-170100137 | 0 | C | T | 0.228 | 4.89E-02 | 1.16E-01 | 2.70E+00 | 2.77E+00 | 6.69E-01 | no | Cnot1 | intron_variant | MODIFIER |
| 8 | 95729367 | AX-170031309 | 0 | T | C | 0.228 | 4.89E-02 | 1.16E-01 | 2.70E+00 | 2.77E+00 | 6.69E-01 | no | Cnot1 | intron_variant | MODIFIER |
| 8 | 95729476 | AX-170031307 | 0 | T | C | 0.228 | 4.89E-02 | 1.16E-01 | 2.70E+00 | 2.77E+00 | 6.69E-01 | no | Cnot1 | intron_variant | MODIFIER |
| 8 | 95729969 | AX-170472767 | 0 | C | T | 0.228 | 4.89E-02 | 1.16E-01 | 2.70E+00 | 2.77E+00 | 6.69E-01 | no | Cnot1 | intron_variant | MODIFIER |
| 8 | 95735738 | AX-170346639 | 0 | G | A | 0.228 | 4.89E-02 | 1.16E-01 | 2.70E+00 | 2.77E+00 | 6.69E-01 | no | Cnot1 | synonymous_variant | LOW |
| 8 | 95771158 | AX-170031310 | 0 | A | C | 0.228 | 4.89E-02 | 1.16E-01 | 2.70E+00 | 2.77E+00 | 6.69E-01 | no | Cnot1 | intron_variant | MODIFIER |
| 8 | 95808511 | AX-97789060 | 0 | A | G | 0.228 | 4.89E-02 | 1.16E-01 | 2.70E+00 | 2.77E+00 | 6.69E-01 | no | Cnot1 | upstream_gene_variant | MODIFIER |
| 8 | 95829704 | AX-170463842 | 0 | T | C | 0.228 | 4.89E-02 | 1.16E-01 | 2.70E+00 | 2.77E+00 | 6.69E-01 | no | Sap18b | downstream_gene_variant | MODIFIER |
| 8 | 95837879 | AX-170328218 | 0 | C | T | 0.228 | 4.89E-02 | 1.16E-01 | 2.70E+00 | 2.77E+00 | 6.69E-01 | no | Slc38a7 | intron_variant | MODIFIER |
| 8 | 95852775 | AX-170031311 | 0 | C | T | 0.228 | 4.89E-02 | 1.16E-01 | 2.70E+00 | 2.77E+00 | 6.69E-01 | no | Slc38a7 | intron_variant | MODIFIER |
| 8 | 95869407 | AX-170196144 | 0 | T | C | 0.228 | 4.89E-02 | 1.16E-01 | 2.70E+00 | 2.77E+00 | 6.69E-01 | no | Got2 | intron_variant | MODIFIER |
| 8 | 95880062 | AX-170223938 | 0 | A | G | 0.228 | 4.89E-02 | 1.16E-01 | 2.70E+00 | 2.77E+00 | 6.69E-01 | no | Got2 | intron_variant | MODIFIER |
| 8 | 95886714 | AX-97805480 | 0 | C | T | 0.228 | 4.89E-02 | 1.16E-01 | 2.70E+00 | 2.77E+00 | 6.69E-01 | no | Got2 | intron_variant | MODIFIER |
| 8 | 95937779 | AX-170356015 | 0 | T | C | 0.228 | 4.89E-02 | 1.16E-01 | 2.70E+00 | 2.77E+00 | 6.69E-01 | no | Gm45895 | downstream_gene_variant | MODIFIER |
| 8 | 95939675 | AX-170436560 | 0 | T | C | 0.228 | 4.89E-02 | 1.16E-01 | 2.70E+00 | 2.77E+00 | 6.69E-01 | no | | intergenic_variant | MODIFIER |
| 8 | 95964224 | AX-170293393 | 0 | T | G | 0.228 | 4.89E-02 | 1.16E-01 | 2.70E+00 | 2.77E+00 | 6.69E-01 | no | | intergenic_variant | MODIFIER |
| 8 | 95972126 | AX-170272877 | 0 | T | C | 0.228 | 4.89E-02 | 1.16E-01 | 2.70E+00 | 2.77E+00 | 6.69E-01 | no | | intergenic_variant | MODIFIER |
| 8 | 95976873 | AX-170080888 | 0 | T | C | 0.228 | 4.89E-02 | 1.16E-01 | 2.70E+00 | 2.77E+00 | 6.69E-01 | no | | intergenic_variant | MODIFIER |
| 8 | 95978613 | AX-170316515 | 0 | G | A | 0.228 | 4.89E-02 | 1.16E-01 | 2.70E+00 | 2.77E+00 | 6.69E-01 | no | | intergenic_variant | MODIFIER |
| 8 | 95983177 | AX-170322841 | 0 | G | A | 0.228 | 4.89E-02 | 1.16E-01 | 2.70E+00 | 2.77E+00 | 6.69E-01 | no | | intergenic_variant | MODIFIER |
| 8 | 95993708 | AX-170428247 | 0 | G | A | 0.228 | 4.89E-02 | 1.16E-01 | 2.70E+00 | 2.77E+00 | 6.69E-01 | no | | intergenic_variant | MODIFIER |
| 8 | 95995450 | AX-170262209 | 0 | A | G | 0.228 | 4.89E-02 | 1.16E-01 | 2.70E+00 | 2.77E+00 | 6.69E-01 | no | | intergenic_variant | MODIFIER |

| | | | | | | | | | | | | | | | |
|---|---|---|---|---|---|---|---|---|---|---|---|---|---|---|---|
| 8 | 96006470 | AX-170477755 | 0 | T | C | 0.228 | 4.89E-02 | 1.16E-01 | 2.70E+00 | 2.77E+00 | 6.69E-01 | no | | intergenic_variant | MODIFIER |
| 6 | 110488321 | AX-170027967 | 0 | C | A | 0.079 | -7.06E-02 | 1.70E-01 | 2.67E+00 | 2.80E+00 | 6.69E-01 | no | | intergenic_variant | MODIFIER |
| 1 | 37160623 | AX-170254451 | 2 | G | T | 0.464 | 3.21E-02 | 7.66E-02 | 2.68E+00 | 2.77E+00 | 6.69E-01 | no | Vwa3b | intron_variant,NMD_transcript_variant | MODIFIER |
| 20 | 67831060 | AX-170034074 | 0 | T | C | 0.246 | 2.65E-02 | 6.29E-02 | 2.66E+00 | 2.75E+00 | 6.69E-01 | no | | intergenic_variant | MODIFIER |
| 20 | 71187122 | AX-97843542 | 2 | C | T | 0.255 | 2.66E-02 | 6.29E-02 | 2.65E+00 | 2.73E+00 | 6.69E-01 | no | | intergenic_variant | MODIFIER |
| 1 | 167877279 | AX-170114819 | 0 | A | T | 0.237 | -5.04E-02 | 1.21E-01 | 2.78E+00 | 2.88E+00 | 6.69E-01 | no | | intergenic_variant | MODIFIER |
| 7 | 106873456 | AX-170029718 | 0 | C | T | 0.474 | -3.96E-02 | 9.44E-02 | 2.77E+00 | 2.85E+00 | 6.70E-01 | no | Olfr705 | missense_variant | MODERATE |
| 15 | 70227133 | AX-97857546 | 0 | C | A | 0.553 | 3.53E-02 | 8.19E-02 | 2.60E+00 | 2.68E+00 | 6.70E-01 | no | | intergenic_variant | MODIFIER |
| 10 | 115231669 | AX-170003957 | 0 | G | A | 0.395 | 3.21E-02 | 7.70E-02 | 2.70E+00 | 2.79E+00 | 6.70E-01 | no | Tbc1d15 | intron_variant | MODIFIER |
| 10 | 118847315 | AX-170317220 | 1 | T | C | 0.348 | 3.12E-02 | 7.47E-02 | 2.68E+00 | 2.78E+00 | 6.70E-01 | no | | regulatory_region_variant | MODIFIER |
| 10 | 115462749 | AX-170534256 | 0 | C | T | 0.386 | 3.18E-02 | 7.61E-02 | 2.70E+00 | 2.79E+00 | 6.70E-01 | no | Lgr5 | synonymous_variant | LOW |
| 10 | 115501728 | AX-170208687 | 0 | A | G | 0.386 | 3.18E-02 | 7.61E-02 | 2.70E+00 | 2.79E+00 | 6.70E-01 | no | Lgr5 | intron_variant | MODIFIER |
| 10 | 115503508 | AX-170238818 | 0 | A | C | 0.386 | 3.18E-02 | 7.61E-02 | 2.70E+00 | 2.79E+00 | 6.70E-01 | no | Lgr5 | intron_variant | MODIFIER |
| 10 | 115534749 | AX-170152900 | 0 | C | T | 0.386 | 3.18E-02 | 7.61E-02 | 2.70E+00 | 2.79E+00 | 6.70E-01 | no | Lgr5 | intron_variant | MODIFIER |
| 10 | 115535337 | AX-170397018 | 0 | G | T | 0.386 | 3.18E-02 | 7.61E-02 | 2.70E+00 | 2.79E+00 | 6.70E-01 | no | Lgr5 | intron_variant | MODIFIER |
| 10 | 115563954 | AX-97793208 | 0 | T | C | 0.386 | 3.18E-02 | 7.61E-02 | 2.70E+00 | 2.79E+00 | 6.70E-01 | no | Lgr5 | intron_variant | MODIFIER |
| 10 | 115589382 | AX-170045977 | 0 | T | G | 0.386 | 3.18E-02 | 7.61E-02 | 2.70E+00 | 2.79E+00 | 6.70E-01 | no | Lgr5 | upstream_gene_variant | MODIFIER |
| 10 | 115624098 | AX-170238758 | 0 | G | C | 0.386 | 3.18E-02 | 7.61E-02 | 2.70E+00 | 2.79E+00 | 6.70E-01 | no | | regulatory_region_variant | MODIFIER |
| 10 | 115636838 | AX-170325490 | 0 | T | C | 0.386 | 3.18E-02 | 7.61E-02 | 2.70E+00 | 2.79E+00 | 6.70E-01 | no | | intergenic_variant | MODIFIER |
| 10 | 115637671 | AX-170156179 | 0 | T | G | 0.386 | 3.18E-02 | 7.61E-02 | 2.70E+00 | 2.79E+00 | 6.70E-01 | no | | intergenic_variant | MODIFIER |
| 10 | 115662859 | AX-170003961 | 0 | G | C | 0.386 | 3.18E-02 | 7.61E-02 | 2.70E+00 | 2.79E+00 | 6.70E-01 | no | | intergenic_variant | MODIFIER |
| 10 | 115725916 | AX-170279299 | 0 | T | C | 0.386 | 3.18E-02 | 7.61E-02 | 2.70E+00 | 2.79E+00 | 6.70E-01 | no | | intergenic_variant | MODIFIER |
| 10 | 115743481 | AX-170003966 | 0 | C | T | 0.386 | 3.18E-02 | 7.61E-02 | 2.70E+00 | 2.79E+00 | 6.70E-01 | no | | intergenic_variant | MODIFIER |
| 10 | 115800759 | AX-170003969 | 0 | C | T | 0.386 | 3.18E-02 | 7.61E-02 | 2.70E+00 | 2.79E+00 | 6.70E-01 | no | 4930422I22Rik | downstream_gene_variant | MODIFIER |
| 10 | 115832263 | AX-170331570 | 0 | A | G | 0.386 | 3.18E-02 | 7.61E-02 | 2.70E+00 | 2.79E+00 | 6.70E-01 | no | Tspan8 | intron_variant | MODIFIER |
| 10 | 115852251 | AX-170373813 | 0 | G | A | 0.386 | 3.18E-02 | 7.61E-02 | 2.70E+00 | 2.79E+00 | 6.70E-01 | no | Tspan8 | downstream_gene_variant | MODIFIER |
| 10 | 115952455 | AX-170045983 | 0 | G | A | 0.386 | 3.18E-02 | 7.61E-02 | 2.70E+00 | 2.79E+00 | 6.70E-01 | no | | intergenic_variant | MODIFIER |
| 10 | 115962383 | AX-170372553 | 0 | A | G | 0.386 | 3.18E-02 | 7.61E-02 | 2.70E+00 | 2.79E+00 | 6.70E-01 | no | | intergenic_variant | MODIFIER |
| 10 | 115964010 | AX-170497740 | 0 | G | A | 0.386 | 3.18E-02 | 7.61E-02 | 2.70E+00 | 2.79E+00 | 6.70E-01 | no | | intergenic_variant | MODIFIER |
| 10 | 115964427 | AX-170346102 | 0 | T | G | 0.386 | 3.18E-02 | 7.61E-02 | 2.70E+00 | 2.79E+00 | 6.70E-01 | no | | intergenic_variant | MODIFIER |
| 10 | 115968896 | AX-170428361 | 0 | C | T | 0.386 | 3.18E-02 | 7.61E-02 | 2.70E+00 | 2.79E+00 | 6.70E-01 | no | | intergenic_variant | MODIFIER |
| 10 | 115991321 | AX-170396150 | 0 | A | C | 0.386 | 3.18E-02 | 7.61E-02 | 2.70E+00 | 2.79E+00 | 6.70E-01 | no | | intergenic_variant | MODIFIER |
| 10 | 116033496 | AX-170512384 | 0 | C | T | 0.386 | 3.18E-02 | 7.61E-02 | 2.70E+00 | 2.79E+00 | 6.70E-01 | no | Ptprr | intron_variant | MODIFIER |
| 10 | 118703934 | AX-170004001 | 1 | T | A | 0.348 | 3.13E-02 | 7.47E-02 | 2.68E+00 | 2.77E+00 | 6.70E-01 | no | | intergenic_variant | MODIFIER |
| 10 | 28423116 | AX-170433595 | 0 | G | T | 0.193 | -3.72E-02 | 8.81E-02 | 2.61E+00 | 2.74E+00 | 6.70E-01 | no | Ptprk | intron_variant | MODIFIER |
| 10 | 5494582 | AX-97807561 | 0 | C | G | 0.211 | 3.82E-02 | 9.30E-02 | 2.71E+00 | 2.84E+00 | 6.70E-01 | no | Syne1 | intron_variant | MODIFIER |
| 15 | 78240756 | AX-170374343 | 0 | C | T | 0.553 | -3.35E-02 | 8.07E-02 | 2.76E+00 | 2.86E+00 | 6.70E-01 | no | Ncf4 | upstream_gene_variant | MODIFIER |
| 7 | 92699804 | AX-170388351 | 0 | C | A | 0.114 | -7.57E-02 | 1.80E-01 | 2.64E+00 | 2.74E+00 | 6.70E-01 | no | | intergenic_variant | MODIFIER |
| 8 | 95982730 | AX-170329448 | 1 | C | T | 0.232 | 4.82E-02 | 1.15E-01 | 2.69E+00 | 2.75E+00 | 6.71E-01 | no | | intergenic_variant | MODIFIER |
| 8 | 96598724 | AX-97826600 | 0 | C | T | 0.228 | 4.73E-02 | 1.13E-01 | 2.66E+00 | 2.74E+00 | 6.71E-01 | no | | intergenic_variant | MODIFIER |
| 8 | 96842858 | AX-170031324 | 0 | C | T | 0.228 | 4.73E-02 | 1.13E-01 | 2.66E+00 | 2.74E+00 | 6.71E-01 | no | | intergenic_variant | MODIFIER |
| 9 | 50398113 | AX-170207041 | 2 | A | T | 0.209 | -4.62E-02 | 1.09E-01 | 2.64E+00 | 2.72E+00 | 6.71E-01 | no | | intergenic_variant | MODIFIER |
| 15 | 79232215 | AX-170012064 | 1 | A | G | 0.545 | 3.37E-02 | 7.72E-02 | 2.56E+00 | 2.65E+00 | 6.71E-01 | no | Pick1 | intron_variant | MODIFIER |
| 13 | 24387040 | AX-170050767 | 0 | T | G | 0.184 | 4.10E-02 | 9.88E-02 | 2.84E+00 | 2.99E+00 | 6.71E-01 | no | Cmah | intron_variant | MODIFIER |
| 5 | 101130354 | AX-170345170 | 1 | A | T | 0.402 | -4.13E-02 | 9.92E-02 | 2.71E+00 | 2.79E+00 | 6.71E-01 | no | | intergenic_variant | MODIFIER |
| 18 | 31770935 | AX-170276818 | 0 | C | G | 0.07 | 8.86E-02 | 2.13E-01 | 2.67E+00 | 2.78E+00 | 6.71E-01 | no | Ammecr1l | intron_variant | MODIFIER |
| 18 | 38937168 | AX-170246372 | 0 | T | G | 0.07 | 8.86E-02 | 2.13E-01 | 2.67E+00 | 2.78E+00 | 6.71E-01 | no | AC110730.1 | intron_variant,non_coding_transcript_variant | MODIFIER |
| 9 | 38507494 | AX-97782792 | 0 | C | T | 0.491 | -3.27E-02 | 7.86E-02 | 2.71E+00 | 2.80E+00 | 6.71E-01 | no | Gm23271 | upstream_gene_variant | MODIFIER |
| 7 | 115580498 | AX-97800144 | 1 | A | C | 0.411 | 3.92E-02 | 9.24E-02 | 2.64E+00 | 2.70E+00 | 6.71E-01 | no | Sox6 | intron_variant | MODIFIER |
| 13 | 31447174 | AX-97823148 | 1 | G | A | 0.179 | -5.15E-02 | 1.20E-01 | 2.60E+00 | 2.67E+00 | 6.71E-01 | no | | intergenic_variant | MODIFIER |
| 7 | 123166685 | AX-170339103 | 2 | A | G | 0.382 | 3.42E-02 | 8.31E-02 | 2.74E+00 | 2.86E+00 | 6.71E-01 | no | Tnrc6a | intron_variant | MODIFIER |
| 8 | 35665173 | AX-170446242 | 1 | C | T | 0.143 | -6.31E-02 | 1.47E-01 | 2.62E+00 | 2.67E+00 | 6.71E-01 | no | Mfhas1 | intron_variant | MODIFIER |
| 4 | 92500032 | AX-170108390 | 0 | C | A | 0.333 | -4.43E-02 | 1.07E-01 | 2.74E+00 | 2.83E+00 | 6.71E-01 | no | | intergenic_variant | MODIFIER |
| 13 | 31998798 | AX-170136342 | 0 | T | C | 0.184 | -5.06E-02 | 1.17E-01 | 2.59E+00 | 2.66E+00 | 6.71E-01 | no | Gmds | intron_variant | MODIFIER |

| | | | | | | | | | | | | | | | |
|---|---|---|---|---|---|---|---|---|---|---|---|---|---|---|---|
| 1 | 174639596 | AX-170221743 | 1 | A | G | 0.223 | -4.43E-02 | 1.08E-01 | 2.78E+00 | 2.90E+00 | 6.71E-01 | no | Fmn2 | intron_variant | MODIFIER |
| 20 | 75882182 | AX-170034158 | 0 | G | A | 0.254 | 2.57E-02 | 6.14E-02 | 2.65E+00 | 2.75E+00 | 6.71E-01 | no | Gm23320 | upstream_gene_variant | MODIFIER |
| 11 | 46489859 | AX-170231738 | 0 | G | A | 0.088 | -6.82E-02 | 1.59E-01 | 2.56E+00 | 2.69E+00 | 6.71E-01 | no | | intergenic_variant | MODIFIER |
| 1 | 45565886 | AX-170381148 | 0 | G | A | 0.105 | -7.07E-02 | 1.72E-01 | 2.67E+00 | 2.80E+00 | 6.72E-01 | no | | intergenic_variant | MODIFIER |
| 18 | 80149993 | AX-170172248 | 0 | G | A | 0.386 | -3.61E-02 | 8.46E-02 | 2.60E+00 | 2.69E+00 | 6.72E-01 | no | Adnp2 | intron_variant | MODIFIER |
| 11 | 22811073 | AX-170086177 | 0 | G | A | 0.053 | 8.17E-02 | 1.97E-01 | 2.63E+00 | 2.77E+00 | 6.72E-01 | no | B3gnt2 | intron_variant,non_coding_transcript_variant | MODIFIER |
| 11 | 22873395 | AX-170162312 | 0 | C | T | 0.053 | 8.17E-02 | 1.97E-01 | 2.63E+00 | 2.77E+00 | 6.72E-01 | no | Gm24917 | downstream_gene_variant | MODIFIER |
| 1 | 126620573 | AX-170195341 | 0 | G | A | 0.57 | 3.62E-02 | 8.30E-02 | 2.57E+00 | 2.64E+00 | 6.72E-01 | no | Nckap5 | intron_variant | MODIFIER |
| 1 | 76450997 | AX-170150872 | 1 | T | G | 0.134 | 9.56E-02 | 2.30E-01 | 2.68E+00 | 2.76E+00 | 6.72E-01 | no | Gm29069 | intron_variant,non_coding_transcript_variant | MODIFIER |
| 15 | 86074413 | AX-170316141 | 1 | A | G | 0.491 | 3.07E-02 | 6.92E-02 | 2.53E+00 | 2.62E+00 | 6.72E-01 | no | Gramd4 | intron_variant | MODIFIER |
| 5 | 111580987 | AX-170172716 | 0 | G | C | 0.439 | 4.67E-02 | 1.11E-01 | 2.65E+00 | 2.70E+00 | 6.72E-01 | no | C130026L21Rik | upstream_gene_variant | MODIFIER |
| 5 | 125661621 | AX-170026103 | 1 | C | T | 0.455 | 4.28E-02 | 1.02E-01 | 2.67E+00 | 2.74E+00 | 6.72E-01 | no | Tmem132b | intron_variant | MODIFIER |
| 5 | 48311390 | AX-170280101 | 1 | C | T | 0.027 | -5.50E-02 | 1.33E-01 | 2.59E+00 | 2.76E+00 | 6.72E-01 | no | Slit2 | downstream_gene_variant | MODIFIER |
| 3 | 147788042 | AX-170472563 | 0 | G | A | 0.342 | 3.06E-02 | 7.32E-02 | 2.66E+00 | 2.73E+00 | 6.73E-01 | no | | intergenic_variant | MODIFIER |
| 7 | 102791322 | AX-170029661 | 0 | C | A | 0.465 | -3.90E-02 | 9.35E-02 | 2.81E+00 | 2.91E+00 | 6.73E-01 | no | Olfr563-ps1 | non_coding_transcript_exon_variant | MODIFIER |
| 5 | 111941686 | AX-97814413 | 1 | T | C | 0.455 | 4.96E-02 | 1.18E-01 | 2.66E+00 | 2.69E+00 | 6.73E-01 | no | Gm42488 | non_coding_transcript_exon_variant | MODIFIER |
| 1 | 33477908 | AX-170086439 | 0 | A | G | 0.421 | 3.51E-02 | 8.33E-02 | 2.63E+00 | 2.71E+00 | 6.73E-01 | no | Prim2 | intron_variant | MODIFIER |
| 5 | 118718101 | AX-170532316 | 1 | T | C | 0.455 | -4.74E-02 | 1.14E-01 | 2.77E+00 | 2.83E+00 | 6.73E-01 | no | Med13l | intron_variant | MODIFIER |
| 5 | 110605121 | AX-170185439 | 1 | A | T | 0.438 | -4.46E-02 | 1.08E-01 | 2.77E+00 | 2.85E+00 | 6.73E-01 | no | Galnt9 | intron_variant | MODIFIER |
| 1 | 149720337 | AX-170179999 | 0 | G | A | 0.202 | 5.55E-02 | 1.33E-01 | 2.68E+00 | 2.75E+00 | 6.73E-01 | no | | intergenic_variant | MODIFIER |
| 1 | 151229567 | AX-97834617 | 0 | T | C | 0.202 | 5.55E-02 | 1.33E-01 | 2.68E+00 | 2.75E+00 | 6.73E-01 | no | | intergenic_variant | MODIFIER |
| 4 | 95584679 | AX-170214674 | 1 | T | C | 0.339 | 4.58E-02 | 1.09E-01 | 2.64E+00 | 2.71E+00 | 6.73E-01 | no | Fggy | intron_variant | MODIFIER |
| 10 | 27680482 | AX-170044441 | 0 | G | A | 0.175 | 4.12E-02 | 1.00E-01 | 2.74E+00 | 2.84E+00 | 6.73E-01 | no | Gm30165 | intron_variant,non_coding_transcript_variant | MODIFIER |
| 10 | 27681632 | AX-170044440 | 0 | G | A | 0.175 | 4.12E-02 | 1.00E-01 | 2.74E+00 | 2.84E+00 | 6.73E-01 | no | Gm30165 | intron_variant,non_coding_transcript_variant | MODIFIER |
| 10 | 27701266 | AX-97783486 | 0 | C | T | 0.175 | 4.12E-02 | 1.00E-01 | 2.74E+00 | 2.84E+00 | 6.73E-01 | no | Gm30165 | upstream_gene_variant | MODIFIER |
| 10 | 27726940 | AX-170481341 | 0 | A | G | 0.175 | 4.12E-02 | 1.00E-01 | 2.74E+00 | 2.84E+00 | 6.73E-01 | no | | intergenic_variant | MODIFIER |
| 10 | 27777167 | AX-170455031 | 0 | A | G | 0.175 | 4.12E-02 | 1.00E-01 | 2.74E+00 | 2.84E+00 | 6.73E-01 | no | Gm18041 | upstream_gene_variant | MODIFIER |
| 10 | 27777268 | AX-170140679 | 0 | C | A | 0.175 | 4.12E-02 | 1.00E-01 | 2.74E+00 | 2.84E+00 | 6.73E-01 | no | Gm18041 | upstream_gene_variant | MODIFIER |
| 10 | 27832688 | AX-170351439 | 0 | T | C | 0.175 | 4.12E-02 | 1.00E-01 | 2.74E+00 | 2.84E+00 | 6.73E-01 | no | | intergenic_variant | MODIFIER |
| 10 | 27837292 | AX-170472180 | 0 | A | G | 0.175 | 4.12E-02 | 1.00E-01 | 2.74E+00 | 2.84E+00 | 6.73E-01 | no | | intergenic_variant | MODIFIER |
| 10 | 27843764 | AX-170290816 | 0 | A | G | 0.175 | 4.12E-02 | 1.00E-01 | 2.74E+00 | 2.84E+00 | 6.73E-01 | no | | intergenic_variant | MODIFIER |
| 10 | 27844166 | AX-170279711 | 0 | T | C | 0.175 | 4.12E-02 | 1.00E-01 | 2.74E+00 | 2.84E+00 | 6.73E-01 | no | | intergenic_variant | MODIFIER |
| 10 | 27862406 | AX-170165514 | 0 | A | G | 0.175 | 4.12E-02 | 1.00E-01 | 2.74E+00 | 2.84E+00 | 6.73E-01 | no | | intergenic_variant | MODIFIER |
| 10 | 27975901 | AX-170525280 | 0 | A | G | 0.175 | 4.12E-02 | 1.00E-01 | 2.74E+00 | 2.84E+00 | 6.73E-01 | no | | intergenic_variant | MODIFIER |
| 10 | 27976195 | AX-97840193 | 0 | T | C | 0.175 | 4.12E-02 | 1.00E-01 | 2.74E+00 | 2.84E+00 | 6.73E-01 | no | | intergenic_variant | MODIFIER |
| 10 | 27983601 | AX-170493060 | 0 | A | G | 0.175 | 4.12E-02 | 1.00E-01 | 2.74E+00 | 2.84E+00 | 6.73E-01 | no | | intergenic_variant | MODIFIER |
| 10 | 27989210 | AX-170281307 | 0 | G | A | 0.175 | 4.12E-02 | 1.00E-01 | 2.74E+00 | 2.84E+00 | 6.73E-01 | no | | regulatory_region_variant | MODIFIER |
| 10 | 28003721 | AX-170131356 | 0 | A | G | 0.175 | 4.12E-02 | 1.00E-01 | 2.74E+00 | 2.84E+00 | 6.73E-01 | no | | intergenic_variant | MODIFIER |
| 10 | 28004312 | AX-170356134 | 0 | C | T | 0.175 | 4.12E-02 | 1.00E-01 | 2.74E+00 | 2.84E+00 | 6.73E-01 | no | | intergenic_variant | MODIFIER |
| 10 | 28004385 | AX-170372915 | 0 | A | G | 0.175 | 4.12E-02 | 1.00E-01 | 2.74E+00 | 2.84E+00 | 6.73E-01 | no | | intergenic_variant | MODIFIER |
| 10 | 28022950 | AX-170417425 | 0 | T | C | 0.175 | 4.12E-02 | 1.00E-01 | 2.74E+00 | 2.84E+00 | 6.73E-01 | no | | intergenic_variant | MODIFIER |
| 10 | 28024052 | AX-97815222 | 0 | A | G | 0.175 | 4.12E-02 | 1.00E-01 | 2.74E+00 | 2.84E+00 | 6.73E-01 | no | | intergenic_variant | MODIFIER |
| 10 | 28034720 | AX-170389770 | 0 | C | T | 0.175 | 4.12E-02 | 1.00E-01 | 2.74E+00 | 2.84E+00 | 6.73E-01 | no | | intergenic_variant | MODIFIER |
| 10 | 28081458 | AX-170002955 | 0 | C | G | 0.175 | 4.12E-02 | 1.00E-01 | 2.74E+00 | 2.84E+00 | 6.73E-01 | no | Ptprk | intron_variant | MODIFIER |
| 10 | 28081640 | AX-170002958 | 0 | C | T | 0.175 | 4.12E-02 | 1.00E-01 | 2.74E+00 | 2.84E+00 | 6.73E-01 | no | Ptprk | intron_variant | MODIFIER |
| 10 | 28103619 | AX-170522634 | 0 | G | T | 0.175 | 4.12E-02 | 1.00E-01 | 2.74E+00 | 2.84E+00 | 6.73E-01 | no | Ptprk | intron_variant | MODIFIER |
| 5 | 39470702 | AX-170286177 | 0 | A | G | 0.193 | -4.64E-02 | 1.09E-01 | 2.60E+00 | 2.69E+00 | 6.74E-01 | no | 4930513D17Rik | intron_variant,non_coding_transcript_variant | MODIFIER |
| 5 | 39486850 | AX-170329219 | 0 | C | T | 0.193 | -4.64E-02 | 1.09E-01 | 2.60E+00 | 2.69E+00 | 6.74E-01 | no | 4930513D17Rik | intron_variant,non_coding_transcript_variant | MODIFIER |
| 5 | 39508197 | AX-97780535 | 0 | C | T | 0.193 | -4.64E-02 | 1.09E-01 | 2.60E+00 | 2.69E+00 | 6.74E-01 | no | 4930513D17Rik | intron_variant,non_coding_transcript_variant | MODIFIER |
| 5 | 39518399 | AX-170509778 | 0 | T | C | 0.193 | -4.64E-02 | 1.09E-01 | 2.60E+00 | 2.69E+00 | 6.74E-01 | no | Gm25377 | downstream_gene_variant | MODIFIER |
| 5 | 39559557 | AX-97855179 | 0 | A | G | 0.193 | -4.64E-02 | 1.09E-01 | 2.60E+00 | 2.69E+00 | 6.74E-01 | no | 4930513D17Rik | intron_variant,non_coding_transcript_variant | MODIFIER |
| 5 | 39563037 | AX-170506430 | 0 | A | G | 0.193 | -4.64E-02 | 1.09E-01 | 2.60E+00 | 2.69E+00 | 6.74E-01 | no | 4930513D17Rik | intron_variant,non_coding_transcript_variant | MODIFIER |
| 4 | 147137174 | AX-170110637 | 0 | C | G | 0.114 | 7.10E-02 | 1.72E-01 | 2.66E+00 | 2.76E+00 | 6.74E-01 | no | Zfp991 | intron_variant | MODIFIER |
| 1 | 20184798 | AX-170413497 | 1 | T | C | 0.402 | 3.56E-02 | 8.37E-02 | 2.61E+00 | 2.66E+00 | 6.74E-01 | no | Pkhd1 | intron_variant | MODIFIER |

| | | | | | | | | | | | | | | | |
|---|---|---|---|---|---|---|---|---|---|---|---|---|---|---|---|
| 5 | 117528313 | AX-170247745 | 0 | G | C | 0.447 | 4.70E-02 | 1.12E-01 | 2.66E+00 | 2.70E+00 | 6.74E-01 | no | Ksr2 | intron_variant,non_coding_transcript_variant | MODIFIER |
| 5 | 122837506 | AX-170026050 | 0 | T | A | 0.439 | -4.63E-02 | 1.11E-01 | 2.80E+00 | 2.86E+00 | 6.74E-01 | no | | intergenic_variant | MODIFIER |
| 8 | 26690449 | AX-170362563 | 0 | G | A | 0.167 | -4.67E-02 | 1.12E-01 | 2.65E+00 | 2.73E+00 | 6.74E-01 | no | | regulatory_region_variant | MODIFIER |
| 1 | 38720283 | AX-170348557 | 0 | C | T | 0.447 | -3.14E-02 | 7.55E-02 | 2.66E+00 | 2.74E+00 | 6.74E-01 | no | Gm38135 | upstream_gene_variant | MODIFIER |
| 8 | 22985756 | AX-170348694 | 0 | G | A | 0.167 | -4.75E-02 | 1.14E-01 | 2.64E+00 | 2.72E+00 | 6.75E-01 | no | Ank1 | intron_variant | MODIFIER |
| 8 | 41053404 | AX-170266257 | 2 | A | T | 0.155 | 5.98E-02 | 1.45E-01 | 2.77E+00 | 2.82E+00 | 6.75E-01 | no | Mtus1 | intron_variant | MODIFIER |
| 5 | 124943443 | AX-97798718 | 0 | C | T | 0.456 | 4.56E-02 | 1.10E-01 | 2.67E+00 | 2.73E+00 | 6.75E-01 | no | | regulatory_region_variant | MODIFIER |
| 20 | 161976412 | AX-170237713 | 0 | T | G | 0.132 | 5.28E-02 | 1.29E-01 | 2.65E+00 | 2.77E+00 | 6.75E-01 | no | Nhs | intron_variant | MODIFIER |
| 20 | 164095620 | AX-97784465 | 0 | T | G | 0.132 | 5.28E-02 | 1.29E-01 | 2.65E+00 | 2.77E+00 | 6.75E-01 | no | Cltrn | intron_variant | MODIFIER |
| 20 | 164095967 | AX-170531009 | 0 | G | T | 0.132 | 5.28E-02 | 1.29E-01 | 2.65E+00 | 2.77E+00 | 6.75E-01 | no | Cltrn | intron_variant | MODIFIER |
| 20 | 164103566 | AX-170180474 | 0 | C | T | 0.132 | 5.28E-02 | 1.29E-01 | 2.65E+00 | 2.77E+00 | 6.75E-01 | no | Cltrn | intron_variant | MODIFIER |
| 20 | 164307053 | AX-97830643 | 0 | G | A | 0.132 | 5.28E-02 | 1.29E-01 | 2.65E+00 | 2.77E+00 | 6.75E-01 | no | Pir | intron_variant | MODIFIER |
| 20 | 164314597 | AX-170035402 | 0 | T | C | 0.132 | 5.28E-02 | 1.29E-01 | 2.65E+00 | 2.77E+00 | 6.75E-01 | no | Pir | intron_variant | MODIFIER |
| 20 | 164316461 | AX-170035398 | 0 | T | C | 0.132 | 5.28E-02 | 1.29E-01 | 2.65E+00 | 2.77E+00 | 6.75E-01 | no | Pir | intron_variant | MODIFIER |
| 20 | 164418212 | AX-170408554 | 0 | A | G | 0.132 | 5.28E-02 | 1.29E-01 | 2.65E+00 | 2.77E+00 | 6.75E-01 | no | Piga | upstream_gene_variant | MODIFIER |
| 20 | 164437917 | AX-97804613 | 0 | A | G | 0.132 | 5.28E-02 | 1.29E-01 | 2.65E+00 | 2.77E+00 | 6.75E-01 | no | Piga | downstream_gene_variant | MODIFIER |
| 20 | 164512818 | AX-170421629 | 0 | G | A | 0.132 | 5.28E-02 | 1.29E-01 | 2.65E+00 | 2.77E+00 | 6.75E-01 | no | Asb9 | intron_variant | MODIFIER |
| 20 | 164571441 | AX-170090742 | 0 | A | G | 0.132 | 5.28E-02 | 1.29E-01 | 2.65E+00 | 2.77E+00 | 6.75E-01 | no | | intergenic_variant | MODIFIER |
| 20 | 164722435 | AX-170474516 | 0 | T | C | 0.132 | 5.28E-02 | 1.29E-01 | 2.65E+00 | 2.77E+00 | 6.75E-01 | no | | intergenic_variant | MODIFIER |
| 20 | 164722721 | AX-170164724 | 0 | A | C | 0.132 | 5.28E-02 | 1.29E-01 | 2.65E+00 | 2.77E+00 | 6.75E-01 | no | | intergenic_variant | MODIFIER |
| 20 | 164758618 | AX-170527070 | 0 | T | C | 0.132 | 5.28E-02 | 1.29E-01 | 2.65E+00 | 2.77E+00 | 6.75E-01 | no | | intergenic_variant | MODIFIER |
| 20 | 164918920 | AX-170035407 | 0 | C | G | 0.132 | 5.28E-02 | 1.29E-01 | 2.65E+00 | 2.77E+00 | 6.75E-01 | no | | intergenic_variant | MODIFIER |
| 20 | 165007670 | AX-170477876 | 0 | A | T | 0.132 | 5.28E-02 | 1.29E-01 | 2.65E+00 | 2.77E+00 | 6.75E-01 | no | Gm17604 | upstream_gene_variant | MODIFIER |
| 20 | 165064629 | AX-170314310 | 0 | T | C | 0.132 | 5.28E-02 | 1.29E-01 | 2.65E+00 | 2.77E+00 | 6.75E-01 | no | | intergenic_variant | MODIFIER |
| 20 | 165074702 | AX-170183473 | 0 | G | A | 0.132 | 5.28E-02 | 1.29E-01 | 2.65E+00 | 2.77E+00 | 6.75E-01 | no | | intergenic_variant | MODIFIER |
| 20 | 165076211 | AX-170430643 | 0 | G | A | 0.132 | 5.28E-02 | 1.29E-01 | 2.65E+00 | 2.77E+00 | 6.75E-01 | no | | intergenic_variant | MODIFIER |
| 20 | 165081504 | AX-97828839 | 0 | C | T | 0.132 | 5.28E-02 | 1.29E-01 | 2.65E+00 | 2.77E+00 | 6.75E-01 | no | | intergenic_variant | MODIFIER |
| 20 | 165139107 | AX-97784664 | 0 | G | T | 0.132 | 5.28E-02 | 1.29E-01 | 2.65E+00 | 2.77E+00 | 6.75E-01 | no | Glra2 | intron_variant | MODIFIER |
| 20 | 165163405 | AX-170471170 | 0 | T | C | 0.132 | 5.28E-02 | 1.29E-01 | 2.65E+00 | 2.77E+00 | 6.75E-01 | no | Glra2 | intron_variant | MODIFIER |
| 20 | 165164050 | AX-170282444 | 0 | T | G | 0.132 | 5.28E-02 | 1.29E-01 | 2.65E+00 | 2.77E+00 | 6.75E-01 | no | Glra2 | intron_variant | MODIFIER |
| 20 | 165169128 | AX-170517825 | 0 | A | C | 0.132 | 5.28E-02 | 1.29E-01 | 2.65E+00 | 2.77E+00 | 6.75E-01 | no | Glra2 | intron_variant | MODIFIER |
| 20 | 165195097 | AX-170345910 | 0 | A | T | 0.132 | 5.28E-02 | 1.29E-01 | 2.65E+00 | 2.77E+00 | 6.75E-01 | no | Glra2 | intron_variant | MODIFIER |
| 20 | 165197733 | AX-170341964 | 0 | A | G | 0.132 | 5.28E-02 | 1.29E-01 | 2.65E+00 | 2.77E+00 | 6.75E-01 | no | Glra2 | intron_variant | MODIFIER |
| 20 | 165229215 | AX-97791273 | 0 | G | A | 0.132 | 5.28E-02 | 1.29E-01 | 2.65E+00 | 2.77E+00 | 6.75E-01 | no | Glra2 | intron_variant | MODIFIER |
| 20 | 165233511 | AX-170504629 | 0 | A | G | 0.132 | 5.28E-02 | 1.29E-01 | 2.65E+00 | 2.77E+00 | 6.75E-01 | no | Glra2 | intron_variant | MODIFIER |
| 20 | 165236151 | AX-170217561 | 0 | C | A | 0.132 | 5.28E-02 | 1.29E-01 | 2.65E+00 | 2.77E+00 | 6.75E-01 | no | Glra2 | intron_variant | MODIFIER |
| 20 | 165253989 | AX-170085743 | 0 | A | G | 0.132 | 5.28E-02 | 1.29E-01 | 2.65E+00 | 2.77E+00 | 6.75E-01 | no | Glra2 | intron_variant | MODIFIER |
| 20 | 165271808 | AX-97819287 | 0 | A | G | 0.132 | 5.28E-02 | 1.29E-01 | 2.65E+00 | 2.77E+00 | 6.75E-01 | no | Glra2 | intron_variant | MODIFIER |
| 20 | 165276499 | AX-170329856 | 0 | T | C | 0.132 | 5.28E-02 | 1.29E-01 | 2.65E+00 | 2.77E+00 | 6.75E-01 | no | Glra2 | intron_variant | MODIFIER |
| 20 | 165321089 | AX-170085746 | 0 | G | A | 0.132 | 5.28E-02 | 1.29E-01 | 2.65E+00 | 2.77E+00 | 6.75E-01 | no | Glra2 | intron_variant | MODIFIER |
| 20 | 165359943 | AX-97850925 | 0 | C | T | 0.132 | 5.28E-02 | 1.29E-01 | 2.65E+00 | 2.77E+00 | 6.75E-01 | no | | intergenic_variant | MODIFIER |
| 20 | 165366989 | AX-170303327 | 0 | T | C | 0.132 | 5.28E-02 | 1.29E-01 | 2.65E+00 | 2.77E+00 | 6.75E-01 | no | | intergenic_variant | MODIFIER |
| 20 | 165368446 | AX-170428369 | 0 | G | A | 0.132 | 5.28E-02 | 1.29E-01 | 2.65E+00 | 2.77E+00 | 6.75E-01 | no | | intergenic_variant | MODIFIER |
| 20 | 165430226 | AX-170035412 | 0 | A | G | 0.132 | 5.28E-02 | 1.29E-01 | 2.65E+00 | 2.77E+00 | 6.75E-01 | no | | intergenic_variant | MODIFIER |
| 20 | 165441635 | AX-170499371 | 0 | G | T | 0.132 | 5.28E-02 | 1.29E-01 | 2.65E+00 | 2.77E+00 | 6.75E-01 | no | | intergenic_variant | MODIFIER |
| 20 | 165499641 | AX-170506713 | 0 | C | T | 0.132 | 5.28E-02 | 1.29E-01 | 2.65E+00 | 2.77E+00 | 6.75E-01 | no | | intergenic_variant | MODIFIER |
| 20 | 165573051 | AX-170504594 | 0 | C | G | 0.132 | 5.28E-02 | 1.29E-01 | 2.65E+00 | 2.77E+00 | 6.75E-01 | no | | intergenic_variant | MODIFIER |
| 20 | 165600037 | AX-170321413 | 0 | G | A | 0.132 | 5.28E-02 | 1.29E-01 | 2.65E+00 | 2.77E+00 | 6.75E-01 | no | | intergenic_variant | MODIFIER |
| 20 | 165600304 | AX-170219238 | 0 | A | G | 0.132 | 5.28E-02 | 1.29E-01 | 2.65E+00 | 2.77E+00 | 6.75E-01 | no | | intergenic_variant | MODIFIER |
| 20 | 165660852 | AX-170085749 | 0 | C | T | 0.132 | 5.28E-02 | 1.29E-01 | 2.65E+00 | 2.77E+00 | 6.75E-01 | no | | intergenic_variant | MODIFIER |
| 20 | 165678667 | AX-170361465 | 0 | A | G | 0.132 | 5.28E-02 | 1.29E-01 | 2.65E+00 | 2.77E+00 | 6.75E-01 | no | Gm15236 | upstream_gene_variant | MODIFIER |
| 20 | 165681010 | AX-170266814 | 0 | T | C | 0.132 | 5.28E-02 | 1.29E-01 | 2.65E+00 | 2.77E+00 | 6.75E-01 | no | Gm15236 | downstream_gene_variant | MODIFIER |
| 20 | 165708295 | AX-97780345 | 0 | C | G | 0.132 | 5.28E-02 | 1.29E-01 | 2.65E+00 | 2.77E+00 | 6.75E-01 | no | | intergenic_variant | MODIFIER |
| 20 | 165765887 | AX-170444869 | 0 | C | T | 0.132 | 5.28E-02 | 1.29E-01 | 2.65E+00 | 2.77E+00 | 6.75E-01 | no | | intergenic_variant | MODIFIER |

| | | | | | | | | | | | | | | | |
|---|---|---|---|---|---|---|---|---|---|---|---|---|---|---|---|
| 20 | 165782310 | AX-170340279 | 0 | A | G | 0.132 | 5.28E-02 | 1.29E-01 | 2.65E+00 | 2.77E+00 | 6.75E-01 | no | | intergenic_variant | MODIFIER |
| 20 | 165782474 | AX-170097000 | 0 | C | G | 0.132 | 5.28E-02 | 1.29E-01 | 2.65E+00 | 2.77E+00 | 6.75E-01 | no | | intergenic_variant | MODIFIER |
| 20 | 165823933 | AX-97825702 | 0 | C | T | 0.132 | 5.28E-02 | 1.29E-01 | 2.65E+00 | 2.77E+00 | 6.75E-01 | no | | intergenic_variant | MODIFIER |
| 20 | 165859509 | AX-170301978 | 0 | G | A | 0.132 | 5.28E-02 | 1.29E-01 | 2.65E+00 | 2.77E+00 | 6.75E-01 | no | | intergenic_variant | MODIFIER |
| 20 | 165876089 | AX-170441176 | 0 | G | A | 0.132 | 5.28E-02 | 1.29E-01 | 2.65E+00 | 2.77E+00 | 6.75E-01 | no | | intergenic_variant | MODIFIER |
| 20 | 165997693 | AX-170106637 | 0 | A | G | 0.132 | 5.28E-02 | 1.29E-01 | 2.65E+00 | 2.77E+00 | 6.75E-01 | no | | intergenic_variant | MODIFIER |
| 20 | 166034842 | AX-170035422 | 0 | T | C | 0.132 | 5.28E-02 | 1.29E-01 | 2.65E+00 | 2.77E+00 | 6.75E-01 | no | | intergenic_variant | MODIFIER |
| 20 | 166061674 | AX-170461748 | 0 | A | G | 0.132 | 5.28E-02 | 1.29E-01 | 2.65E+00 | 2.77E+00 | 6.75E-01 | no | Gm15225 | upstream_gene_variant | MODIFIER |
| 20 | 166198345 | AX-170118338 | 0 | T | C | 0.132 | 5.28E-02 | 1.29E-01 | 2.65E+00 | 2.77E+00 | 6.75E-01 | no | | regulatory_region_variant | MODIFIER |
| 20 | 166315319 | AX-97807280 | 0 | C | A | 0.132 | 5.28E-02 | 1.29E-01 | 2.65E+00 | 2.77E+00 | 6.75E-01 | no | Gpm6b | intron_variant | MODIFIER |
| 20 | 166319912 | AX-170333556 | 0 | C | G | 0.132 | 5.28E-02 | 1.29E-01 | 2.65E+00 | 2.77E+00 | 6.75E-01 | no | Gpm6b | intron_variant | MODIFIER |
| 20 | 166332776 | AX-170531646 | 0 | A | G | 0.132 | 5.28E-02 | 1.29E-01 | 2.65E+00 | 2.77E+00 | 6.75E-01 | no | Gpm6b | intron_variant | MODIFIER |
| 20 | 166334324 | AX-170519984 | 0 | T | G | 0.132 | 5.28E-02 | 1.29E-01 | 2.65E+00 | 2.77E+00 | 6.75E-01 | no | Gpm6b | intron_variant | MODIFIER |
| 20 | 166348315 | AX-170097002 | 0 | A | G | 0.132 | 5.28E-02 | 1.29E-01 | 2.65E+00 | 2.77E+00 | 6.75E-01 | no | Gpm6b | intron_variant | MODIFIER |
| 20 | 166358620 | AX-170164936 | 0 | C | T | 0.132 | 5.28E-02 | 1.29E-01 | 2.65E+00 | 2.77E+00 | 6.75E-01 | no | Gpm6b | intron_variant | MODIFIER |
| 20 | 166387243 | AX-97807799 | 0 | A | G | 0.132 | 5.28E-02 | 1.29E-01 | 2.65E+00 | 2.77E+00 | 6.75E-01 | no | Ofd1 | downstream_gene_variant | MODIFIER |
| 20 | 166466714 | AX-97791236 | 0 | A | G | 0.132 | 5.28E-02 | 1.29E-01 | 2.65E+00 | 2.77E+00 | 6.75E-01 | no | Rab9 | intron_variant | MODIFIER |
| 20 | 166686631 | AX-170035429 | 0 | C | T | 0.132 | 5.28E-02 | 1.29E-01 | 2.65E+00 | 2.77E+00 | 6.75E-01 | no | Gm1720 | intron_variant,non_coding_transcript_variant | MODIFIER |
| 5 | 27856118 | AX-97797293 | 0 | T | C | 0.333 | 4.50E-02 | 1.08E-01 | 2.68E+00 | 2.73E+00 | 6.75E-01 | no | Htr5a | downstream_gene_variant | MODIFIER |
| 5 | 27881466 | AX-97804518 | 0 | C | T | 0.333 | 4.50E-02 | 1.08E-01 | 2.68E+00 | 2.73E+00 | 6.75E-01 | no | | intergenic_variant | MODIFIER |
| 5 | 27906502 | AX-170071626 | 0 | A | G | 0.333 | 4.50E-02 | 1.08E-01 | 2.68E+00 | 2.73E+00 | 6.75E-01 | no | | intergenic_variant | MODIFIER |
| 5 | 27906913 | AX-170071625 | 0 | A | G | 0.333 | 4.50E-02 | 1.08E-01 | 2.68E+00 | 2.73E+00 | 6.75E-01 | no | | intergenic_variant | MODIFIER |
| 5 | 27995926 | AX-170398222 | 0 | C | T | 0.333 | 4.50E-02 | 1.08E-01 | 2.68E+00 | 2.73E+00 | 6.75E-01 | no | Gm4865 | downstream_gene_variant | MODIFIER |
| 5 | 28003060 | AX-170253909 | 0 | A | G | 0.333 | 4.50E-02 | 1.08E-01 | 2.68E+00 | 2.73E+00 | 6.75E-01 | no | Gm4865 | upstream_gene_variant | MODIFIER |
| 5 | 28004513 | AX-170328254 | 0 | C | G | 0.333 | 4.50E-02 | 1.08E-01 | 2.68E+00 | 2.73E+00 | 6.75E-01 | no | Gm4865 | upstream_gene_variant | MODIFIER |
| 5 | 28007545 | AX-170403330 | 0 | A | G | 0.333 | 4.50E-02 | 1.08E-01 | 2.68E+00 | 2.73E+00 | 6.75E-01 | no | | intergenic_variant | MODIFIER |
| 5 | 28008137 | AX-97848649 | 0 | C | T | 0.333 | 4.50E-02 | 1.08E-01 | 2.68E+00 | 2.73E+00 | 6.75E-01 | no | | intergenic_variant | MODIFIER |
| 5 | 28009018 | AX-170421179 | 0 | T | C | 0.333 | 4.50E-02 | 1.08E-01 | 2.68E+00 | 2.73E+00 | 6.75E-01 | no | | intergenic_variant | MODIFIER |
| 5 | 28044072 | AX-97843997 | 0 | A | G | 0.333 | 4.50E-02 | 1.08E-01 | 2.68E+00 | 2.73E+00 | 6.75E-01 | no | Gm26608 | intron_variant,non_coding_transcript_variant | MODIFIER |
| 5 | 28064273 | AX-97802211 | 0 | T | C | 0.333 | 4.50E-02 | 1.08E-01 | 2.68E+00 | 2.73E+00 | 6.75E-01 | no | Gm26608 | intron_variant,non_coding_transcript_variant | MODIFIER |
| 5 | 32326797 | AX-170365206 | 0 | T | C | 0.333 | 4.50E-02 | 1.08E-01 | 2.68E+00 | 2.73E+00 | 6.75E-01 | no | Plb1 | intron_variant | MODIFIER |
| 5 | 33556550 | AX-170346305 | 0 | T | A | 0.333 | 4.50E-02 | 1.08E-01 | 2.68E+00 | 2.73E+00 | 6.75E-01 | no | | intergenic_variant | MODIFIER |
| 5 | 33560286 | AX-170188351 | 0 | T | C | 0.333 | 4.50E-02 | 1.08E-01 | 2.68E+00 | 2.73E+00 | 6.75E-01 | no | | intergenic_variant | MODIFIER |
| 5 | 33602148 | AX-97812227 | 0 | T | C | 0.333 | 4.50E-02 | 1.08E-01 | 2.68E+00 | 2.73E+00 | 6.75E-01 | no | Fam53a | intron_variant | MODIFIER |
| 5 | 33802049 | AX-170024652 | 0 | A | C | 0.333 | 4.50E-02 | 1.08E-01 | 2.68E+00 | 2.73E+00 | 6.75E-01 | no | | intergenic_variant | MODIFIER |
| 5 | 33826699 | AX-170024651 | 0 | T | C | 0.333 | 4.50E-02 | 1.08E-01 | 2.68E+00 | 2.73E+00 | 6.75E-01 | no | Nsd2 | intron_variant | MODIFIER |
| 5 | 33921209 | AX-170235022 | 0 | C | T | 0.333 | 4.50E-02 | 1.08E-01 | 2.68E+00 | 2.73E+00 | 6.75E-01 | no | Nelfa | intron_variant | MODIFIER |
| 5 | 33984632 | AX-170024658 | 0 | C | G | 0.333 | 4.50E-02 | 1.08E-01 | 2.68E+00 | 2.73E+00 | 6.75E-01 | no | Gm1673 | intron_variant | MODIFIER |
| 5 | 34021820 | AX-170535268 | 0 | C | G | 0.333 | 4.50E-02 | 1.08E-01 | 2.68E+00 | 2.73E+00 | 6.75E-01 | no | Poln | intron_variant | MODIFIER |
| 5 | 34029032 | AX-97811044 | 0 | T | G | 0.333 | 4.50E-02 | 1.08E-01 | 2.68E+00 | 2.73E+00 | 6.75E-01 | no | Poln | intron_variant | MODIFIER |
| 5 | 34077432 | AX-170308522 | 0 | A | G | 0.333 | 4.50E-02 | 1.08E-01 | 2.68E+00 | 2.73E+00 | 6.75E-01 | no | Poln | intron_variant | MODIFIER |
| 5 | 34110314 | AX-97811166 | 0 | T | G | 0.333 | 4.50E-02 | 1.08E-01 | 2.68E+00 | 2.73E+00 | 6.75E-01 | no | Poln | intron_variant | MODIFIER |
| 5 | 34115946 | AX-170300618 | 0 | G | A | 0.333 | 4.50E-02 | 1.08E-01 | 2.68E+00 | 2.73E+00 | 6.75E-01 | no | Poln | intron_variant | MODIFIER |
| 5 | 34162620 | AX-170499927 | 0 | G | A | 0.333 | 4.50E-02 | 1.08E-01 | 2.68E+00 | 2.73E+00 | 6.75E-01 | no | Poln | intron_variant | MODIFIER |
| 5 | 34174955 | AX-170385023 | 0 | T | C | 0.333 | 4.50E-02 | 1.08E-01 | 2.68E+00 | 2.73E+00 | 6.75E-01 | no | Mxd4 | 3_prime_UTR_variant | MODIFIER |
| 5 | 34185209 | AX-97835735 | 0 | T | G | 0.333 | 4.50E-02 | 1.08E-01 | 2.68E+00 | 2.73E+00 | 6.75E-01 | no | Mxd4 | intron_variant | MODIFIER |
| 5 | 34332495 | AX-170411060 | 0 | A | G | 0.333 | 4.50E-02 | 1.08E-01 | 2.68E+00 | 2.73E+00 | 6.75E-01 | no | Rnf4 | upstream_gene_variant | MODIFIER |
| 5 | 34355125 | AX-170519754 | 0 | T | G | 0.333 | 4.50E-02 | 1.08E-01 | 2.68E+00 | 2.73E+00 | 6.75E-01 | no | Rnf4 | downstream_gene_variant | MODIFIER |
| 5 | 34360094 | AX-170202897 | 0 | T | A | 0.333 | 4.50E-02 | 1.08E-01 | 2.68E+00 | 2.73E+00 | 6.75E-01 | no | Rnf4 | downstream_gene_variant | MODIFIER |
| 5 | 34376257 | AX-170455967 | 0 | C | T | 0.333 | 4.50E-02 | 1.08E-01 | 2.68E+00 | 2.73E+00 | 6.75E-01 | no | Fam193a | intron_variant | MODIFIER |
| 5 | 34380395 | AX-170089446 | 0 | A | G | 0.333 | 4.50E-02 | 1.08E-01 | 2.68E+00 | 2.73E+00 | 6.75E-01 | no | Fam193a | intron_variant | MODIFIER |
| 5 | 34385103 | AX-170024663 | 0 | G | A | 0.333 | 4.50E-02 | 1.08E-01 | 2.68E+00 | 2.73E+00 | 6.75E-01 | no | Fam193a | intron_variant | MODIFIER |
| 5 | 34385208 | AX-170024661 | 0 | T | G | 0.333 | 4.50E-02 | 1.08E-01 | 2.68E+00 | 2.73E+00 | 6.75E-01 | no | Fam193a | intron_variant | MODIFIER |
| 5 | 34436629 | AX-170024660 | 0 | T | C | 0.333 | 4.50E-02 | 1.08E-01 | 2.68E+00 | 2.73E+00 | 6.75E-01 | no | Fam193a | intron_variant | MODIFIER |

| | | | | | | | | | | | | | | | |
|---|---|---|---|---|---|---|---|---|---|---|---|---|---|---|---|
| 5 | 34461698 | AX-170024664 | 0 | A | T | 0.333 | 4.50E-02 | 1.08E-01 | 2.68E+00 | 2.73E+00 | 6.75E-01 | no | Fam193a | intron_variant | MODIFIER |
| 5 | 34462771 | AX-170288871 | 0 | G | A | 0.333 | 4.50E-02 | 1.08E-01 | 2.68E+00 | 2.73E+00 | 6.75E-01 | no | Fam193a | intron_variant | MODIFIER |
| 5 | 34527139 | AX-170273038 | 0 | G | A | 0.333 | 4.50E-02 | 1.08E-01 | 2.68E+00 | 2.73E+00 | 6.75E-01 | no | Sh3bp2 | intron_variant | MODIFIER |
| 5 | 34531316 | AX-170299866 | 0 | C | A | 0.333 | 4.50E-02 | 1.08E-01 | 2.68E+00 | 2.73E+00 | 6.75E-01 | no | Sh3bp2 | intron_variant | MODIFIER |
| 5 | 34531850 | AX-97847930 | 0 | T | G | 0.333 | 4.50E-02 | 1.08E-01 | 2.68E+00 | 2.73E+00 | 6.75E-01 | no | Sh3bp2 | intron_variant | MODIFIER |
| 5 | 34535387 | AX-170123546 | 0 | A | G | 0.333 | 4.50E-02 | 1.08E-01 | 2.68E+00 | 2.73E+00 | 6.75E-01 | no | Sh3bp2 | intron_variant | MODIFIER |
| 5 | 34546999 | AX-170211184 | 0 | A | G | 0.333 | 4.50E-02 | 1.08E-01 | 2.68E+00 | 2.73E+00 | 6.75E-01 | no | Sh3bp2 | intron_variant | MODIFIER |
| 5 | 34556456 | AX-170207891 | 0 | A | G | 0.333 | 4.50E-02 | 1.08E-01 | 2.68E+00 | 2.73E+00 | 6.75E-01 | no | Sh3bp2 | intron_variant | MODIFIER |
| 5 | 34556478 | AX-170189827 | 0 | G | C | 0.333 | 4.50E-02 | 1.08E-01 | 2.68E+00 | 2.73E+00 | 6.75E-01 | no | Sh3bp2 | intron_variant | MODIFIER |
| 5 | 34563137 | AX-170260652 | 0 | T | C | 0.333 | 4.50E-02 | 1.08E-01 | 2.68E+00 | 2.73E+00 | 6.75E-01 | no | Sh3bp2 | 3_prime_UTR_variant | MODIFIER |
| 5 | 34563161 | AX-170481285 | 0 | T | C | 0.333 | 4.50E-02 | 1.08E-01 | 2.68E+00 | 2.73E+00 | 6.75E-01 | no | Sh3bp2 | 3_prime_UTR_variant | MODIFIER |
| 5 | 34563923 | AX-97815974 | 0 | T | C | 0.333 | 4.50E-02 | 1.08E-01 | 2.68E+00 | 2.73E+00 | 6.75E-01 | no | Sh3bp2 | downstream_gene_variant | MODIFIER |
| 5 | 34581808 | AX-170348417 | 0 | C | A | 0.333 | 4.50E-02 | 1.08E-01 | 2.68E+00 | 2.73E+00 | 6.75E-01 | no | Add1 | intron_variant | MODIFIER |
| 5 | 34582279 | AX-170089447 | 0 | G | A | 0.333 | 4.50E-02 | 1.08E-01 | 2.68E+00 | 2.73E+00 | 6.75E-01 | no | Add1 | intron_variant | MODIFIER |
| 5 | 34602970 | AX-170095285 | 0 | A | G | 0.333 | 4.50E-02 | 1.08E-01 | 2.68E+00 | 2.73E+00 | 6.75E-01 | no | Add1 | intron_variant | MODIFIER |
| 5 | 34613326 | AX-170459684 | 0 | G | A | 0.333 | 4.50E-02 | 1.08E-01 | 2.68E+00 | 2.73E+00 | 6.75E-01 | no | Add1 | synonymous_variant | LOW |
| 5 | 34623590 | AX-170111362 | 0 | T | C | 0.333 | 4.50E-02 | 1.08E-01 | 2.68E+00 | 2.73E+00 | 6.75E-01 | no | Add1 | intron_variant | MODIFIER |
| 5 | 34629965 | AX-170443504 | 0 | T | C | 0.333 | 4.50E-02 | 1.08E-01 | 2.68E+00 | 2.73E+00 | 6.75E-01 | no | Add1 | intron_variant | MODIFIER |
| 5 | 34630916 | AX-170495307 | 0 | A | G | 0.333 | 4.50E-02 | 1.08E-01 | 2.68E+00 | 2.73E+00 | 6.75E-01 | no | Add1 | 3_prime_UTR_variant | MODIFIER |
| 5 | 34633637 | AX-170140482 | 0 | G | T | 0.333 | 4.50E-02 | 1.08E-01 | 2.68E+00 | 2.73E+00 | 6.75E-01 | no | Add1 | downstream_gene_variant | MODIFIER |
| 5 | 34637341 | AX-170120530 | 0 | G | C | 0.333 | 4.50E-02 | 1.08E-01 | 2.68E+00 | 2.73E+00 | 6.75E-01 | no | Mfsd10 | upstream_gene_variant | MODIFIER |
| 5 | 34647694 | AX-170511570 | 0 | T | C | 0.333 | 4.50E-02 | 1.08E-01 | 2.68E+00 | 2.73E+00 | 6.75E-01 | no | Nop14 | intron_variant | MODIFIER |
| 5 | 34649273 | AX-170258844 | 0 | C | G | 0.333 | 4.50E-02 | 1.08E-01 | 2.68E+00 | 2.73E+00 | 6.75E-01 | no | Nop14 | intron_variant | MODIFIER |
| 5 | 34651146 | AX-170071764 | 0 | T | C | 0.333 | 4.50E-02 | 1.08E-01 | 2.68E+00 | 2.73E+00 | 6.75E-01 | no | Nop14 | intron_variant | MODIFIER |
| 5 | 34651661 | AX-170071766 | 0 | G | T | 0.333 | 4.50E-02 | 1.08E-01 | 2.68E+00 | 2.73E+00 | 6.75E-01 | no | Nop14 | intron_variant | MODIFIER |
| 5 | 34661173 | AX-170219630 | 0 | T | A | 0.333 | 4.50E-02 | 1.08E-01 | 2.68E+00 | 2.73E+00 | 6.75E-01 | no | Grk4 | intron_variant | MODIFIER |
| 5 | 34661302 | AX-170298281 | 0 | A | G | 0.333 | 4.50E-02 | 1.08E-01 | 2.68E+00 | 2.73E+00 | 6.75E-01 | no | Grk4 | intron_variant | MODIFIER |
| 5 | 34678369 | AX-170197652 | 0 | A | G | 0.333 | 4.50E-02 | 1.08E-01 | 2.68E+00 | 2.73E+00 | 6.75E-01 | no | Grk4 | intron_variant | MODIFIER |
| 5 | 34703577 | AX-170178931 | 0 | A | G | 0.333 | 4.50E-02 | 1.08E-01 | 2.68E+00 | 2.73E+00 | 6.75E-01 | no | Grk4 | intron_variant | MODIFIER |
| 5 | 34719112 | AX-170136026 | 0 | A | G | 0.333 | 4.50E-02 | 1.08E-01 | 2.68E+00 | 2.73E+00 | 6.75E-01 | no | Grk4 | intron_variant | MODIFIER |
| 5 | 34772018 | AX-170333708 | 0 | A | G | 0.333 | 4.50E-02 | 1.08E-01 | 2.68E+00 | 2.73E+00 | 6.75E-01 | no | Htt | intron_variant | MODIFIER |
| 5 | 34794054 | AX-170095286 | 0 | T | A | 0.333 | 4.50E-02 | 1.08E-01 | 2.68E+00 | 2.73E+00 | 6.75E-01 | no | Htt | intron_variant | MODIFIER |
| 5 | 34794927 | AX-170423163 | 0 | T | C | 0.333 | 4.50E-02 | 1.08E-01 | 2.68E+00 | 2.73E+00 | 6.75E-01 | no | Htt | intron_variant | MODIFIER |
| 5 | 34804677 | AX-170319905 | 0 | C | T | 0.333 | 4.50E-02 | 1.08E-01 | 2.68E+00 | 2.73E+00 | 6.75E-01 | no | Htt | synonymous_variant | LOW |
| 5 | 34835750 | AX-97849616 | 0 | T | C | 0.333 | 4.50E-02 | 1.08E-01 | 2.68E+00 | 2.73E+00 | 6.75E-01 | no | Htt | intron_variant | MODIFIER |
| 5 | 34875139 | AX-170384043 | 0 | A | G | 0.333 | 4.50E-02 | 1.08E-01 | 2.68E+00 | 2.73E+00 | 6.75E-01 | no | Htt | intron_variant | MODIFIER |
| 5 | 34896475 | AX-97837950 | 0 | G | T | 0.333 | 4.50E-02 | 1.08E-01 | 2.68E+00 | 2.73E+00 | 6.75E-01 | no | Htt | intron_variant | MODIFIER |
| 5 | 34905251 | AX-170197028 | 0 | C | G | 0.333 | 4.50E-02 | 1.08E-01 | 2.68E+00 | 2.73E+00 | 6.75E-01 | no | Htt | intron_variant | MODIFIER |
| 5 | 34917781 | AX-170209622 | 0 | T | C | 0.333 | 4.50E-02 | 1.08E-01 | 2.68E+00 | 2.73E+00 | 6.75E-01 | no | Msantd1 | synonymous_variant | LOW |
| 5 | 34922560 | AX-170395843 | 0 | T | C | 0.333 | 4.50E-02 | 1.08E-01 | 2.68E+00 | 2.73E+00 | 6.75E-01 | no | Msantd1 | intron_variant | MODIFIER |
| 5 | 34927204 | AX-97795165 | 0 | T | C | 0.333 | 4.50E-02 | 1.08E-01 | 2.68E+00 | 2.73E+00 | 6.75E-01 | no | Msantd1 | downstream_gene_variant | MODIFIER |
| 5 | 34940259 | AX-170174764 | 0 | A | C | 0.333 | 4.50E-02 | 1.08E-01 | 2.68E+00 | 2.73E+00 | 6.75E-01 | no | | intergenic_variant | MODIFIER |
| 5 | 34940409 | AX-170278808 | 0 | T | C | 0.333 | 4.50E-02 | 1.08E-01 | 2.68E+00 | 2.73E+00 | 6.75E-01 | no | | intergenic_variant | MODIFIER |
| 5 | 34940531 | AX-170247954 | 0 | A | C | 0.333 | 4.50E-02 | 1.08E-01 | 2.68E+00 | 2.73E+00 | 6.75E-01 | no | | intergenic_variant | MODIFIER |
| 5 | 34947153 | AX-170024665 | 0 | T | C | 0.333 | 4.50E-02 | 1.08E-01 | 2.68E+00 | 2.73E+00 | 6.75E-01 | no | Rgs12 | upstream_gene_variant | MODIFIER |
| 5 | 34951554 | AX-170328269 | 0 | T | C | 0.333 | 4.50E-02 | 1.08E-01 | 2.68E+00 | 2.73E+00 | 6.75E-01 | no | Rgs12 | intron_variant | MODIFIER |
| 5 | 34954305 | AX-170465822 | 0 | A | T | 0.333 | 4.50E-02 | 1.08E-01 | 2.68E+00 | 2.73E+00 | 6.75E-01 | no | Rgs12 | intron_variant | MODIFIER |
| 5 | 34960203 | AX-170379282 | 0 | T | C | 0.333 | 4.50E-02 | 1.08E-01 | 2.68E+00 | 2.73E+00 | 6.75E-01 | no | Rgs12 | intron_variant | MODIFIER |
| 5 | 34965813 | AX-170349172 | 0 | A | G | 0.333 | 4.50E-02 | 1.08E-01 | 2.68E+00 | 2.73E+00 | 6.75E-01 | no | Rgs12 | synonymous_variant | LOW |
| 5 | 34983177 | AX-170262588 | 0 | C | T | 0.333 | 4.50E-02 | 1.08E-01 | 2.68E+00 | 2.73E+00 | 6.75E-01 | no | Rgs12 | intron_variant | MODIFIER |
| 5 | 35008303 | AX-170401555 | 0 | G | A | 0.333 | 4.50E-02 | 1.08E-01 | 2.68E+00 | 2.73E+00 | 6.75E-01 | no | Rgs12 | intron_variant | MODIFIER |
| 5 | 35011842 | AX-97810336 | 0 | A | G | 0.333 | 4.50E-02 | 1.08E-01 | 2.68E+00 | 2.73E+00 | 6.75E-01 | no | Rgs12 | intron_variant | MODIFIER |
| 5 | 35013503 | AX-170153346 | 0 | A | T | 0.333 | 4.50E-02 | 1.08E-01 | 2.68E+00 | 2.73E+00 | 6.75E-01 | no | Rgs12 | intron_variant | MODIFIER |
| 5 | 35022045 | AX-170024668 | 0 | T | C | 0.333 | 4.50E-02 | 1.08E-01 | 2.68E+00 | 2.73E+00 | 6.75E-01 | no | Rgs12 | intron_variant | MODIFIER |

| | | | | | | | | | | | | | | | |
|---|---|---|---|---|---|---|---|---|---|---|---|---|---|---|---|
| 5 | 35030083 | AX-97781390 | 0 | T | C | 0.333 | 4.50E-02 | 1.08E-01 | 2.68E+00 | 2.73E+00 | 6.75E-01 | no | Rgs12 | missense_variant | MODERATE |
| 5 | 35039212 | AX-170392550 | 0 | T | C | 0.333 | 4.50E-02 | 1.08E-01 | 2.68E+00 | 2.73E+00 | 6.75E-01 | no | Hgfac | upstream_gene_variant | MODIFIER |
| 5 | 35082561 | AX-170071777 | 0 | G | A | 0.333 | 4.50E-02 | 1.08E-01 | 2.68E+00 | 2.73E+00 | 6.75E-01 | no | Dok7 | intron_variant | MODIFIER |
| 5 | 35105132 | AX-170165238 | 0 | C | T | 0.333 | 4.50E-02 | 1.08E-01 | 2.68E+00 | 2.73E+00 | 6.75E-01 | no | Lrpap1 | intron_variant | MODIFIER |
| 5 | 35110134 | AX-97818034 | 0 | T | C | 0.333 | 4.50E-02 | 1.08E-01 | 2.68E+00 | 2.73E+00 | 6.75E-01 | no | Lrpap1 | upstream_gene_variant | MODIFIER |
| 5 | 35121991 | AX-170314765 | 0 | A | G | 0.333 | 4.50E-02 | 1.08E-01 | 2.68E+00 | 2.73E+00 | 6.75E-01 | no | | intergenic_variant | MODIFIER |
| 5 | 35137417 | AX-97787165 | 0 | T | C | 0.333 | 4.50E-02 | 1.08E-01 | 2.68E+00 | 2.73E+00 | 6.75E-01 | no | | regulatory_region_variant | MODIFIER |
| 5 | 35145920 | AX-170381068 | 0 | T | C | 0.333 | 4.50E-02 | 1.08E-01 | 2.68E+00 | 2.73E+00 | 6.75E-01 | no | | intergenic_variant | MODIFIER |
| 5 | 35149935 | AX-97805359 | 0 | T | G | 0.333 | 4.50E-02 | 1.08E-01 | 2.68E+00 | 2.73E+00 | 6.75E-01 | no | | intergenic_variant | MODIFIER |
| 5 | 35163855 | AX-170038970 | 0 | A | G | 0.333 | 4.50E-02 | 1.08E-01 | 2.68E+00 | 2.73E+00 | 6.75E-01 | no | | intergenic_variant | MODIFIER |
| 5 | 35195638 | AX-170400500 | 0 | A | G | 0.333 | 4.50E-02 | 1.08E-01 | 2.68E+00 | 2.73E+00 | 6.75E-01 | no | | intergenic_variant | MODIFIER |
| 5 | 35197749 | AX-170246221 | 0 | C | G | 0.333 | 4.50E-02 | 1.08E-01 | 2.68E+00 | 2.73E+00 | 6.75E-01 | no | | intergenic_variant | MODIFIER |
| 5 | 35198203 | AX-170333095 | 0 | T | C | 0.333 | 4.50E-02 | 1.08E-01 | 2.68E+00 | 2.73E+00 | 6.75E-01 | no | | intergenic_variant | MODIFIER |
| 5 | 35199443 | AX-170493192 | 0 | T | C | 0.333 | 4.50E-02 | 1.08E-01 | 2.68E+00 | 2.73E+00 | 6.75E-01 | no | | intergenic_variant | MODIFIER |
| 5 | 35199574 | AX-97839022 | 0 | T | C | 0.333 | 4.50E-02 | 1.08E-01 | 2.68E+00 | 2.73E+00 | 6.75E-01 | no | | intergenic_variant | MODIFIER |
| 5 | 35208659 | AX-170427632 | 0 | T | C | 0.333 | 4.50E-02 | 1.08E-01 | 2.68E+00 | 2.73E+00 | 6.75E-01 | no | | intergenic_variant | MODIFIER |
| 5 | 35215824 | AX-170475590 | 0 | C | T | 0.333 | 4.50E-02 | 1.08E-01 | 2.68E+00 | 2.73E+00 | 6.75E-01 | no | | regulatory_region_variant | MODIFIER |
| 5 | 35217729 | AX-170417087 | 0 | A | C | 0.333 | 4.50E-02 | 1.08E-01 | 2.68E+00 | 2.73E+00 | 6.75E-01 | no | | intergenic_variant | MODIFIER |
| 5 | 35227736 | AX-170384582 | 0 | C | T | 0.333 | 4.50E-02 | 1.08E-01 | 2.68E+00 | 2.73E+00 | 6.75E-01 | no | | intergenic_variant | MODIFIER |
| 5 | 35239313 | AX-170274778 | 0 | T | C | 0.333 | 4.50E-02 | 1.08E-01 | 2.68E+00 | 2.73E+00 | 6.75E-01 | no | | intergenic_variant | MODIFIER |
| 5 | 35247331 | AX-170407818 | 0 | C | T | 0.333 | 4.50E-02 | 1.08E-01 | 2.68E+00 | 2.73E+00 | 6.75E-01 | no | | intergenic_variant | MODIFIER |
| 5 | 35264358 | AX-170071780 | 0 | G | A | 0.333 | 4.50E-02 | 1.08E-01 | 2.68E+00 | 2.73E+00 | 6.75E-01 | no | | intergenic_variant | MODIFIER |
| 5 | 35268401 | AX-170071779 | 0 | T | A | 0.333 | 4.50E-02 | 1.08E-01 | 2.68E+00 | 2.73E+00 | 6.75E-01 | no | | intergenic_variant | MODIFIER |
| 5 | 35275289 | AX-170333721 | 0 | C | G | 0.333 | 4.50E-02 | 1.08E-01 | 2.68E+00 | 2.73E+00 | 6.75E-01 | no | Adra2c | upstream_gene_variant | MODIFIER |
| 5 | 35286139 | AX-170150113 | 0 | T | A | 0.333 | 4.50E-02 | 1.08E-01 | 2.68E+00 | 2.73E+00 | 6.75E-01 | no | Adra2c | downstream_gene_variant | MODIFIER |
| 5 | 35305175 | AX-170368200 | 0 | G | A | 0.333 | 4.50E-02 | 1.08E-01 | 2.68E+00 | 2.73E+00 | 6.75E-01 | no | | regulatory_region_variant | MODIFIER |
| 5 | 35307027 | AX-170267547 | 0 | T | C | 0.333 | 4.50E-02 | 1.08E-01 | 2.68E+00 | 2.73E+00 | 6.75E-01 | no | | intergenic_variant | MODIFIER |
| 5 | 35339393 | AX-170164644 | 0 | T | A | 0.333 | 4.50E-02 | 1.08E-01 | 2.68E+00 | 2.73E+00 | 6.75E-01 | no | | intergenic_variant | MODIFIER |
| 5 | 35345059 | AX-170504053 | 0 | G | A | 0.333 | 4.50E-02 | 1.08E-01 | 2.68E+00 | 2.73E+00 | 6.75E-01 | no | | intergenic_variant | MODIFIER |
| 5 | 35364299 | AX-170175922 | 0 | G | A | 0.333 | 4.50E-02 | 1.08E-01 | 2.68E+00 | 2.73E+00 | 6.75E-01 | no | 4930478P22Rik | intron_variant,non_coding_transcript_variant | MODIFIER |
| 5 | 35365899 | AX-170264641 | 0 | G | A | 0.333 | 4.50E-02 | 1.08E-01 | 2.68E+00 | 2.73E+00 | 6.75E-01 | no | 4930478P22Rik | intron_variant,non_coding_transcript_variant | MODIFIER |
| 5 | 35413881 | AX-170239702 | 0 | G | A | 0.333 | 4.50E-02 | 1.08E-01 | 2.68E+00 | 2.73E+00 | 6.75E-01 | no | | intergenic_variant | MODIFIER |
| 5 | 35414027 | AX-170302115 | 0 | G | T | 0.333 | 4.50E-02 | 1.08E-01 | 2.68E+00 | 2.73E+00 | 6.75E-01 | no | | intergenic_variant | MODIFIER |
| 5 | 35422350 | AX-170422965 | 0 | G | A | 0.333 | 4.50E-02 | 1.08E-01 | 2.68E+00 | 2.73E+00 | 6.75E-01 | no | | intergenic_variant | MODIFIER |
| 5 | 35423092 | AX-170259931 | 0 | A | T | 0.333 | 4.50E-02 | 1.08E-01 | 2.68E+00 | 2.73E+00 | 6.75E-01 | no | | intergenic_variant | MODIFIER |
| 5 | 35429084 | AX-170225808 | 0 | G | A | 0.333 | 4.50E-02 | 1.08E-01 | 2.68E+00 | 2.73E+00 | 6.75E-01 | no | | intergenic_variant | MODIFIER |
| 5 | 35433182 | AX-170071784 | 0 | G | C | 0.333 | 4.50E-02 | 1.08E-01 | 2.68E+00 | 2.73E+00 | 6.75E-01 | no | | regulatory_region_variant | MODIFIER |
| 5 | 35438599 | AX-170071785 | 0 | A | G | 0.333 | 4.50E-02 | 1.08E-01 | 2.68E+00 | 2.73E+00 | 6.75E-01 | no | | intergenic_variant | MODIFIER |
| 5 | 35440994 | AX-170448057 | 0 | C | A | 0.333 | 4.50E-02 | 1.08E-01 | 2.68E+00 | 2.73E+00 | 6.75E-01 | no | | intergenic_variant | MODIFIER |
| 5 | 35494016 | AX-170497369 | 0 | A | G | 0.333 | 4.50E-02 | 1.08E-01 | 2.68E+00 | 2.73E+00 | 6.75E-01 | no | | intergenic_variant | MODIFIER |
| 5 | 35496449 | AX-97857424 | 0 | A | G | 0.333 | 4.50E-02 | 1.08E-01 | 2.68E+00 | 2.73E+00 | 6.75E-01 | no | | intergenic_variant | MODIFIER |
| 5 | 35502539 | AX-170024675 | 0 | A | C | 0.333 | 4.50E-02 | 1.08E-01 | 2.68E+00 | 2.73E+00 | 6.75E-01 | no | Cpz | missense_variant | MODERATE |
| 5 | 35506142 | AX-170207729 | 0 | A | G | 0.333 | 4.50E-02 | 1.08E-01 | 2.68E+00 | 2.73E+00 | 6.75E-01 | no | Cpz | intron_variant | MODIFIER |
| 5 | 35507716 | AX-170398745 | 0 | T | C | 0.333 | 4.50E-02 | 1.08E-01 | 2.68E+00 | 2.73E+00 | 6.75E-01 | no | Cpz | intron_variant | MODIFIER |
| 5 | 35513697 | AX-170372977 | 0 | G | A | 0.333 | 4.50E-02 | 1.08E-01 | 2.68E+00 | 2.73E+00 | 6.75E-01 | no | Cpz | intron_variant | MODIFIER |
| 5 | 35515340 | AX-170496269 | 0 | T | A | 0.333 | 4.50E-02 | 1.08E-01 | 2.68E+00 | 2.73E+00 | 6.75E-01 | no | Cpz | intron_variant | MODIFIER |
| 5 | 35536587 | AX-170473128 | 0 | C | T | 0.333 | 4.50E-02 | 1.08E-01 | 2.68E+00 | 2.73E+00 | 6.75E-01 | no | | regulatory_region_variant | MODIFIER |
| 5 | 35538594 | AX-170475532 | 0 | C | T | 0.333 | 4.50E-02 | 1.08E-01 | 2.68E+00 | 2.73E+00 | 6.75E-01 | no | | intergenic_variant | MODIFIER |
| 5 | 35541138 | AX-170148218 | 0 | T | C | 0.333 | 4.50E-02 | 1.08E-01 | 2.68E+00 | 2.73E+00 | 6.75E-01 | no | | regulatory_region_variant | MODIFIER |
| 5 | 35542953 | AX-170089448 | 0 | T | C | 0.333 | 4.50E-02 | 1.08E-01 | 2.68E+00 | 2.73E+00 | 6.75E-01 | no | | regulatory_region_variant | MODIFIER |
| 5 | 35544555 | AX-97827676 | 0 | T | C | 0.333 | 4.50E-02 | 1.08E-01 | 2.68E+00 | 2.73E+00 | 6.75E-01 | no | | intergenic_variant | MODIFIER |
| 5 | 35563618 | AX-170071786 | 0 | G | A | 0.333 | 4.50E-02 | 1.08E-01 | 2.68E+00 | 2.73E+00 | 6.75E-01 | no | Trmt44 | intron_variant | MODIFIER |
| 5 | 35576855 | AX-97811852 | 0 | T | C | 0.333 | 4.50E-02 | 1.08E-01 | 2.68E+00 | 2.73E+00 | 6.75E-01 | no | Trmt44 | upstream_gene_variant | MODIFIER |
| 5 | 35579036 | AX-170153126 | 0 | A | C | 0.333 | 4.50E-02 | 1.08E-01 | 2.68E+00 | 2.73E+00 | 6.75E-01 | no | Trmt44 | upstream_gene_variant | MODIFIER |

| | | | | | | | | | | | | | | | |
|---|---|---|---|---|---|---|---|---|---|---|---|---|---|---|---|
| 5 | 35581506 | AX-170363790 | 0 | A | G | 0.333 | 4.50E-02 | 1.08E-01 | 2.68E+00 | 2.73E+00 | 6.75E-01 | no | Acox3 | upstream_gene_variant | MODIFIER |
| 5 | 35612451 | AX-170478376 | 0 | T | C | 0.333 | 4.50E-02 | 1.08E-01 | 2.68E+00 | 2.73E+00 | 6.75E-01 | no | Acox3 | downstream_gene_variant | MODIFIER |
| 5 | 35619958 | AX-170483276 | 0 | A | G | 0.333 | 4.50E-02 | 1.08E-01 | 2.68E+00 | 2.73E+00 | 6.75E-01 | no | Acox3 | downstream_gene_variant | MODIFIER |
| 5 | 35633303 | AX-170459994 | 0 | G | A | 0.333 | 4.50E-02 | 1.08E-01 | 2.68E+00 | 2.73E+00 | 6.75E-01 | no | | intergenic_variant | MODIFIER |
| 5 | 35648687 | AX-170024673 | 0 | G | C | 0.333 | 4.50E-02 | 1.08E-01 | 2.68E+00 | 2.73E+00 | 6.75E-01 | no | Htra3 | downstream_gene_variant | MODIFIER |
| 5 | 35661484 | AX-170024681 | 0 | A | G | 0.333 | 4.50E-02 | 1.08E-01 | 2.68E+00 | 2.73E+00 | 6.75E-01 | no | Htra3 | intron_variant | MODIFIER |
| 9 | 117254377 | AX-170204851 | 0 | C | T | 0.132 | -3.87E-02 | 9.22E-02 | 2.60E+00 | 2.71E+00 | 6.75E-01 | no | Rbms3 | upstream_gene_variant | MODIFIER |
| 1 | 91108691 | AX-170118444 | 0 | T | C | 0.167 | 8.03E-02 | 1.91E-01 | 2.62E+00 | 2.68E+00 | 6.75E-01 | no | Lrrfip1 | intron_variant | MODIFIER |
| 10 | 28409266 | AX-170472590 | 0 | T | A | 0.193 | 3.91E-02 | 9.61E-02 | 2.72E+00 | 2.82E+00 | 6.75E-01 | no | Ptprk | intron_variant | MODIFIER |
| 10 | 28421448 | AX-97857306 | 0 | G | A | 0.193 | 3.91E-02 | 9.61E-02 | 2.72E+00 | 2.82E+00 | 6.75E-01 | no | Ptprk | intron_variant | MODIFIER |
| 10 | 28444448 | AX-170227932 | 0 | T | C | 0.193 | 3.91E-02 | 9.61E-02 | 2.72E+00 | 2.82E+00 | 6.75E-01 | no | Ptprk | intron_variant | MODIFIER |
| 10 | 28452810 | AX-170466127 | 0 | C | T | 0.193 | 3.91E-02 | 9.61E-02 | 2.72E+00 | 2.82E+00 | 6.75E-01 | no | Ptprk | intron_variant | MODIFIER |
| 10 | 28453459 | AX-170451703 | 0 | G | A | 0.193 | 3.91E-02 | 9.61E-02 | 2.72E+00 | 2.82E+00 | 6.75E-01 | no | Ptprk | intron_variant | MODIFIER |
| 10 | 28499960 | AX-170086771 | 0 | A | T | 0.193 | 3.91E-02 | 9.61E-02 | 2.72E+00 | 2.82E+00 | 6.75E-01 | no | Ptprk | intron_variant | MODIFIER |
| 10 | 28505018 | AX-170275499 | 0 | C | T | 0.193 | 3.91E-02 | 9.61E-02 | 2.72E+00 | 2.82E+00 | 6.75E-01 | no | Ptprk | intron_variant | MODIFIER |
| 10 | 28505329 | AX-170376503 | 0 | G | A | 0.193 | 3.91E-02 | 9.61E-02 | 2.72E+00 | 2.82E+00 | 6.75E-01 | no | Ptprk | intron_variant | MODIFIER |
| 10 | 28519712 | AX-97851695 | 0 | C | T | 0.193 | 3.91E-02 | 9.61E-02 | 2.72E+00 | 2.82E+00 | 6.75E-01 | no | Ptprk | intron_variant | MODIFIER |
| 10 | 28546830 | AX-170414165 | 0 | T | C | 0.193 | 3.91E-02 | 9.61E-02 | 2.72E+00 | 2.82E+00 | 6.75E-01 | no | Ptprk | intron_variant | MODIFIER |
| 10 | 28554740 | AX-97811552 | 0 | T | C | 0.193 | 3.91E-02 | 9.61E-02 | 2.72E+00 | 2.82E+00 | 6.75E-01 | no | Ptprk | intron_variant | MODIFIER |
| 10 | 28574429 | AX-170127712 | 0 | T | C | 0.193 | 3.91E-02 | 9.61E-02 | 2.72E+00 | 2.82E+00 | 6.75E-01 | no | Ptprk | intron_variant | MODIFIER |
| 10 | 28579374 | AX-170002970 | 0 | T | C | 0.193 | 3.91E-02 | 9.61E-02 | 2.72E+00 | 2.82E+00 | 6.75E-01 | no | Ptprk | intron_variant | MODIFIER |
| 10 | 28579882 | AX-170366921 | 0 | G | C | 0.193 | 3.91E-02 | 9.61E-02 | 2.72E+00 | 2.82E+00 | 6.75E-01 | no | Ptprk | intron_variant | MODIFIER |
| 10 | 28591834 | AX-170002966 | 0 | A | T | 0.193 | 3.91E-02 | 9.61E-02 | 2.72E+00 | 2.82E+00 | 6.75E-01 | no | Ptprk | intron_variant | MODIFIER |
| 10 | 28596047 | AX-97852752 | 0 | G | A | 0.193 | 3.91E-02 | 9.61E-02 | 2.72E+00 | 2.82E+00 | 6.75E-01 | no | Ptprk | 3_prime_UTR_variant | MODIFIER |
| 10 | 28600009 | AX-170250058 | 0 | A | G | 0.193 | 3.91E-02 | 9.61E-02 | 2.72E+00 | 2.82E+00 | 6.75E-01 | no | Ptprk | downstream_gene_variant | MODIFIER |
| 10 | 28602158 | AX-170412698 | 0 | G | C | 0.193 | 3.91E-02 | 9.61E-02 | 2.72E+00 | 2.82E+00 | 6.75E-01 | no | Ptprk | downstream_gene_variant | MODIFIER |
| 10 | 28617530 | AX-97828013 | 0 | G | T | 0.193 | 3.91E-02 | 9.61E-02 | 2.72E+00 | 2.82E+00 | 6.75E-01 | no | | intergenic_variant | MODIFIER |
| 10 | 28643282 | AX-97853059 | 0 | G | A | 0.193 | 3.91E-02 | 9.61E-02 | 2.72E+00 | 2.82E+00 | 6.75E-01 | no | | intergenic_variant | MODIFIER |
| 10 | 28645184 | AX-170508278 | 0 | C | T | 0.193 | 3.91E-02 | 9.61E-02 | 2.72E+00 | 2.82E+00 | 6.75E-01 | no | | intergenic_variant | MODIFIER |
| 10 | 28645784 | AX-170002969 | 0 | G | A | 0.193 | 3.91E-02 | 9.61E-02 | 2.72E+00 | 2.82E+00 | 6.75E-01 | no | | intergenic_variant | MODIFIER |
| 10 | 28649009 | AX-170002968 | 0 | A | C | 0.193 | 3.91E-02 | 9.61E-02 | 2.72E+00 | 2.82E+00 | 6.75E-01 | no | | intergenic_variant | MODIFIER |
| 10 | 28650589 | AX-170002965 | 0 | T | G | 0.193 | 3.91E-02 | 9.61E-02 | 2.72E+00 | 2.82E+00 | 6.75E-01 | no | | intergenic_variant | MODIFIER |
| 10 | 28677686 | AX-170221915 | 0 | A | G | 0.193 | 3.91E-02 | 9.61E-02 | 2.72E+00 | 2.82E+00 | 6.75E-01 | no | Themis | intron_variant | MODIFIER |
| 10 | 28717110 | AX-170195015 | 0 | G | A | 0.193 | 3.91E-02 | 9.61E-02 | 2.72E+00 | 2.82E+00 | 6.75E-01 | no | Themis | intron_variant | MODIFIER |
| 10 | 28754928 | AX-170394965 | 0 | A | T | 0.193 | 3.91E-02 | 9.61E-02 | 2.72E+00 | 2.82E+00 | 6.75E-01 | no | Themis | intron_variant | MODIFIER |
| 10 | 28761722 | AX-170206625 | 0 | T | G | 0.193 | 3.91E-02 | 9.61E-02 | 2.72E+00 | 2.82E+00 | 6.75E-01 | no | Themis | intron_variant | MODIFIER |
| 10 | 28793561 | AX-97815037 | 0 | G | T | 0.193 | 3.91E-02 | 9.61E-02 | 2.72E+00 | 2.82E+00 | 6.75E-01 | no | Themis | intron_variant | MODIFIER |
| 10 | 28793604 | AX-170198779 | 0 | A | T | 0.193 | 3.91E-02 | 9.61E-02 | 2.72E+00 | 2.82E+00 | 6.75E-01 | no | Themis | intron_variant | MODIFIER |
| 10 | 28793714 | AX-170044456 | 0 | T | G | 0.193 | 3.91E-02 | 9.61E-02 | 2.72E+00 | 2.82E+00 | 6.75E-01 | no | Themis | intron_variant | MODIFIER |
| 10 | 28843661 | AX-170479587 | 0 | C | T | 0.193 | 3.91E-02 | 9.61E-02 | 2.72E+00 | 2.82E+00 | 6.75E-01 | no | Themis | intron_variant | MODIFIER |
| 10 | 28861595 | AX-170508629 | 0 | G | A | 0.193 | 3.91E-02 | 9.61E-02 | 2.72E+00 | 2.82E+00 | 6.75E-01 | no | Themis | intron_variant | MODIFIER |
| 10 | 29124947 | AX-170002973 | 0 | G | A | 0.193 | 3.91E-02 | 9.61E-02 | 2.72E+00 | 2.82E+00 | 6.75E-01 | no | | intergenic_variant | MODIFIER |
| 10 | 29325732 | AX-170145192 | 0 | A | C | 0.193 | 3.91E-02 | 9.61E-02 | 2.72E+00 | 2.82E+00 | 6.75E-01 | no | Echdc1 | intron_variant | MODIFIER |
| 10 | 29325893 | AX-97852722 | 0 | G | T | 0.193 | 3.91E-02 | 9.61E-02 | 2.72E+00 | 2.82E+00 | 6.75E-01 | no | Echdc1 | intron_variant | MODIFIER |
| 10 | 29344545 | AX-170044464 | 0 | G | A | 0.193 | 3.91E-02 | 9.61E-02 | 2.72E+00 | 2.82E+00 | 6.75E-01 | no | Echdc1 | synonymous_variant | LOW |
| 10 | 29348135 | AX-170044468 | 0 | G | T | 0.193 | 3.91E-02 | 9.61E-02 | 2.72E+00 | 2.82E+00 | 6.75E-01 | no | Echdc1 | downstream_gene_variant | MODIFIER |
| 10 | 29348638 | AX-170044467 | 0 | A | T | 0.193 | 3.91E-02 | 9.61E-02 | 2.72E+00 | 2.82E+00 | 6.75E-01 | no | Echdc1 | downstream_gene_variant | MODIFIER |
| 10 | 29348681 | AX-170044465 | 0 | G | A | 0.193 | 3.91E-02 | 9.61E-02 | 2.72E+00 | 2.82E+00 | 6.75E-01 | no | Echdc1 | downstream_gene_variant | MODIFIER |
| 10 | 29349269 | AX-170308016 | 0 | C | T | 0.193 | 3.91E-02 | 9.61E-02 | 2.72E+00 | 2.82E+00 | 6.75E-01 | no | Echdc1 | downstream_gene_variant | MODIFIER |
| 10 | 29408061 | AX-170276101 | 0 | C | T | 0.193 | 3.91E-02 | 9.61E-02 | 2.72E+00 | 2.82E+00 | 6.75E-01 | no | | intergenic_variant | MODIFIER |
| 10 | 29409136 | AX-170348681 | 0 | A | G | 0.193 | 3.91E-02 | 9.61E-02 | 2.72E+00 | 2.82E+00 | 6.75E-01 | no | | intergenic_variant | MODIFIER |
| 10 | 29409457 | AX-97815109 | 0 | A | G | 0.193 | 3.91E-02 | 9.61E-02 | 2.72E+00 | 2.82E+00 | 6.75E-01 | no | | intergenic_variant | MODIFIER |
| 10 | 29452992 | AX-170219736 | 0 | A | C | 0.193 | 3.91E-02 | 9.61E-02 | 2.72E+00 | 2.82E+00 | 6.75E-01 | no | Rspo3 | 3_prime_UTR_variant | MODIFIER |
| 10 | 29458004 | AX-97796460 | 0 | G | A | 0.193 | 3.91E-02 | 9.61E-02 | 2.72E+00 | 2.82E+00 | 6.75E-01 | no | Rspo3 | intron_variant | MODIFIER |

| | | | | | | | | | | | | | | | |
|---|---|---|---|---|---|---|---|---|---|---|---|---|---|---|---|
| 10 | 29458413 | AX-170261577 | 0 | A | G | 0.193 | 3.91E-02 | 9.61E-02 | 2.72E+00 | 2.82E+00 | 6.75E-01 | no | Rspo3 | intron_variant | MODIFIER |
| 10 | 29469765 | AX-170328391 | 0 | G | C | 0.193 | 3.91E-02 | 9.61E-02 | 2.72E+00 | 2.82E+00 | 6.75E-01 | no | Rspo3 | intron_variant | MODIFIER |
| 10 | 29476707 | AX-170310483 | 0 | T | C | 0.193 | 3.91E-02 | 9.61E-02 | 2.72E+00 | 2.82E+00 | 6.75E-01 | no | Rspo3 | intron_variant | MODIFIER |
| 10 | 29481179 | AX-170276070 | 0 | T | C | 0.193 | 3.91E-02 | 9.61E-02 | 2.72E+00 | 2.82E+00 | 6.75E-01 | no | Rspo3 | intron_variant | MODIFIER |
| 10 | 29499441 | AX-170036156 | 0 | T | G | 0.193 | 3.91E-02 | 9.61E-02 | 2.72E+00 | 2.82E+00 | 6.75E-01 | no | Rspo3 | intron_variant | MODIFIER |
| 10 | 29507982 | AX-170339148 | 0 | G | C | 0.193 | 3.91E-02 | 9.61E-02 | 2.72E+00 | 2.82E+00 | 6.75E-01 | no | Rspo3 | intron_variant | MODIFIER |
| 10 | 29510614 | AX-170502959 | 0 | A | G | 0.193 | 3.91E-02 | 9.61E-02 | 2.72E+00 | 2.82E+00 | 6.75E-01 | no | Rspo3 | intron_variant | MODIFIER |
| 10 | 29510753 | AX-170199505 | 0 | G | A | 0.193 | 3.91E-02 | 9.61E-02 | 2.72E+00 | 2.82E+00 | 6.75E-01 | no | Rspo3 | intron_variant | MODIFIER |
| 10 | 29518761 | AX-170091905 | 0 | A | G | 0.193 | 3.91E-02 | 9.61E-02 | 2.72E+00 | 2.82E+00 | 6.75E-01 | no | Rspo3 | intron_variant | MODIFIER |
| 10 | 30167950 | AX-170002985 | 0 | A | G | 0.193 | 3.91E-02 | 9.61E-02 | 2.72E+00 | 2.82E+00 | 6.75E-01 | no | | intergenic_variant | MODIFIER |
| 10 | 30631067 | AX-170308427 | 0 | C | T | 0.193 | 3.91E-02 | 9.61E-02 | 2.72E+00 | 2.82E+00 | 6.75E-01 | no | Ncoa7 | intron_variant,NMD_transcript_variant | MODIFIER |
| 10 | 30662703 | AX-170044485 | 0 | G | A | 0.193 | 3.91E-02 | 9.61E-02 | 2.72E+00 | 2.82E+00 | 6.75E-01 | no | Ncoa7 | intron_variant | MODIFIER |
| 4 | 148903871 | AX-97852714 | 0 | C | G | 0.088 | -5.99E-02 | 1.46E-01 | 2.67E+00 | 2.77E+00 | 6.75E-01 | no | Casz1 | intron_variant | MODIFIER |
| 3 | 28736568 | AX-170126044 | 1 | G | C | 0.179 | -4.09E-02 | 9.91E-02 | 2.68E+00 | 2.75E+00 | 6.76E-01 | no | 1700112D23Rik | intron_variant,non_coding_transcript_variant | MODIFIER |
| 18 | 69589310 | AX-170061226 | 0 | A | G | 0.263 | -4.22E-02 | 1.01E-01 | 2.66E+00 | 2.72E+00 | 6.76E-01 | no | Tcf4 | intron_variant | MODIFIER |
| 4 | 148640637 | AX-170479950 | 0 | T | C | 0.096 | -6.97E-02 | 1.70E-01 | 2.66E+00 | 2.76E+00 | 6.76E-01 | no | Gm572 | upstream_gene_variant | MODIFIER |
| 10 | 27986727 | AX-170257676 | 1 | T | C | 0.179 | 4.04E-02 | 9.95E-02 | 2.72E+00 | 2.83E+00 | 6.76E-01 | no | | intergenic_variant | MODIFIER |
| 10 | 128504103 | AX-170290275 | 1 | C | T | 0.33 | -3.09E-02 | 7.45E-02 | 2.63E+00 | 2.73E+00 | 6.76E-01 | no | A430046D13Rik | non_coding_transcript_exon_variant | MODIFIER |
| 8 | 46260182 | AX-170252953 | 0 | T | C | 0.158 | 5.38E-02 | 1.31E-01 | 2.76E+00 | 2.81E+00 | 6.76E-01 | no | Gm10313 | upstream_gene_variant | MODIFIER |
| 8 | 96954177 | AX-97837304 | 1 | T | A | 0.223 | 4.88E-02 | 1.19E-01 | 2.70E+00 | 2.77E+00 | 6.76E-01 | no | | intergenic_variant | MODIFIER |
| 14 | 49342476 | AX-170108674 | 0 | T | C | 0.035 | -8.61E-02 | 2.03E-01 | 2.58E+00 | 2.68E+00 | 6.76E-01 | no | Slc35f4 | intron_variant | MODIFIER |
| 9 | 50362834 | AX-170032322 | 0 | G | A | 0.219 | -4.57E-02 | 1.10E-01 | 2.64E+00 | 2.73E+00 | 6.76E-01 | no | | regulatory_region_variant | MODIFIER |
| 2 | 114379883 | AX-170462979 | 0 | C | T | 0.14 | -6.46E-02 | 1.57E-01 | 2.65E+00 | 2.75E+00 | 6.76E-01 | no | | intergenic_variant | MODIFIER |
| 2 | 129413705 | AX-170166108 | 2 | C | G | 0.109 | 6.77E-02 | 1.64E-01 | 2.64E+00 | 2.75E+00 | 6.76E-01 | no | | regulatory_region_variant | MODIFIER |
| 2 | 76104452 | AX-170357809 | 0 | T | C | 0.061 | -6.87E-02 | 1.70E-01 | 2.65E+00 | 2.79E+00 | 6.76E-01 | no | Pde11a | intron_variant | MODIFIER |
| 2 | 162448252 | AX-170384116 | 0 | G | A | 0.158 | 7.94E-02 | 1.94E-01 | 2.68E+00 | 2.76E+00 | 6.77E-01 | no | Ptprt | intron_variant | MODIFIER |
| 8 | 80952542 | AX-170285564 | 0 | C | T | 0.211 | 5.02E-02 | 1.23E-01 | 2.68E+00 | 2.77E+00 | 6.77E-01 | no | | regulatory_region_variant | MODIFIER |
| 8 | 37464878 | AX-170111464 | 0 | G | A | 0.149 | 6.05E-02 | 1.47E-01 | 2.76E+00 | 2.81E+00 | 6.77E-01 | no | | intergenic_variant | MODIFIER |
| 9 | 98087738 | AX-170201077 | 0 | T | C | 0.14 | -4.57E-02 | 1.09E-01 | 2.61E+00 | 2.69E+00 | 6.77E-01 | no | | intergenic_variant | MODIFIER |
| 8 | 64644324 | AX-170030905 | 0 | G | A | 0.158 | -5.36E-02 | 1.26E-01 | 2.58E+00 | 2.67E+00 | 6.77E-01 | no | Cpe | intron_variant | MODIFIER |
| 16 | 31496639 | AX-170240178 | 0 | G | A | 0.5 | 3.45E-02 | 8.09E-02 | 2.59E+00 | 2.64E+00 | 6.77E-01 | no | | intergenic_variant | MODIFIER |
| 5 | 104702097 | AX-170520949 | 1 | A | C | 0.455 | 4.77E-02 | 1.15E-01 | 2.67E+00 | 2.71E+00 | 6.77E-01 | no | | intergenic_variant | MODIFIER |
| 9 | 73930460 | AX-170324847 | 0 | G | A | 0.219 | 5.49E-02 | 1.33E-01 | 2.77E+00 | 2.82E+00 | 6.77E-01 | no | Unc13c | intron_variant | MODIFIER |
| 13 | 41114492 | AX-170511265 | 0 | T | C | 0.518 | 2.67E-02 | 6.19E-02 | 2.54E+00 | 2.65E+00 | 6.77E-01 | no | Gcm2 | upstream_gene_variant | MODIFIER |
| 7 | 39762986 | AX-170028867 | 0 | G | A | 0.342 | 3.97E-02 | 9.43E-02 | 2.60E+00 | 2.68E+00 | 6.77E-01 | no | | intergenic_variant | MODIFIER |
| 7 | 39770804 | AX-170297057 | 0 | G | A | 0.342 | 3.97E-02 | 9.43E-02 | 2.60E+00 | 2.68E+00 | 6.77E-01 | no | | intergenic_variant | MODIFIER |
| 7 | 39774067 | AX-97794460 | 0 | A | G | 0.342 | 3.97E-02 | 9.43E-02 | 2.60E+00 | 2.68E+00 | 6.77E-01 | no | | intergenic_variant | MODIFIER |
| 7 | 39795388 | AX-170462640 | 0 | A | T | 0.342 | 3.97E-02 | 9.43E-02 | 2.60E+00 | 2.68E+00 | 6.77E-01 | no | 4930558N11Rik | downstream_gene_variant | MODIFIER |
| 7 | 39806772 | AX-97839958 | 0 | C | T | 0.342 | 3.97E-02 | 9.43E-02 | 2.60E+00 | 2.68E+00 | 6.77E-01 | no | 4930435C17Rik | intron_variant,non_coding_transcript_variant | MODIFIER |
| 7 | 39847566 | AX-170532797 | 0 | C | T | 0.342 | 3.97E-02 | 9.43E-02 | 2.60E+00 | 2.68E+00 | 6.77E-01 | no | 4930435C17Rik | intron_variant,non_coding_transcript_variant | MODIFIER |
| 7 | 39866874 | AX-170171728 | 0 | T | C | 0.342 | 3.97E-02 | 9.43E-02 | 2.60E+00 | 2.68E+00 | 6.77E-01 | no | 4930435C17Rik | intron_variant,non_coding_transcript_variant | MODIFIER |
| 7 | 39870207 | AX-170077115 | 0 | T | C | 0.342 | 3.97E-02 | 9.43E-02 | 2.60E+00 | 2.68E+00 | 6.77E-01 | no | 4930435C17Rik | intron_variant,non_coding_transcript_variant | MODIFIER |
| 7 | 39887412 | AX-97795530 | 0 | A | G | 0.342 | 3.97E-02 | 9.43E-02 | 2.60E+00 | 2.68E+00 | 6.77E-01 | no | 4930435C17Rik | intron_variant,non_coding_transcript_variant | MODIFIER |
| 7 | 39940688 | AX-97789576 | 0 | T | C | 0.342 | 3.97E-02 | 9.43E-02 | 2.60E+00 | 2.68E+00 | 6.77E-01 | no | Gm44992 | intron_variant,non_coding_transcript_variant | MODIFIER |
| 7 | 39959641 | AX-170103056 | 0 | A | C | 0.342 | 3.97E-02 | 9.43E-02 | 2.60E+00 | 2.68E+00 | 6.77E-01 | no | Gm44992 | intron_variant,non_coding_transcript_variant | MODIFIER |
| 7 | 39982508 | AX-170110542 | 0 | T | C | 0.342 | 3.97E-02 | 9.43E-02 | 2.60E+00 | 2.68E+00 | 6.77E-01 | no | Gm44992 | upstream_gene_variant | MODIFIER |
| 7 | 40013188 | AX-170515368 | 0 | C | G | 0.342 | 3.97E-02 | 9.43E-02 | 2.60E+00 | 2.68E+00 | 6.77E-01 | no | | intergenic_variant | MODIFIER |
| 7 | 40051180 | AX-170411339 | 0 | A | T | 0.342 | 3.97E-02 | 9.43E-02 | 2.60E+00 | 2.68E+00 | 6.77E-01 | no | | intergenic_variant | MODIFIER |
| 5 | 81653832 | AX-170025408 | 1 | C | T | 0.429 | 4.19E-02 | 1.02E-01 | 2.66E+00 | 2.73E+00 | 6.77E-01 | no | Adgrl3 | intron_variant | MODIFIER |
| 5 | 81507276 | AX-170177234 | 0 | G | A | 0.439 | 4.32E-02 | 1.05E-01 | 2.68E+00 | 2.75E+00 | 6.77E-01 | no | Adgrl3 | intron_variant | MODIFIER |
| 3 | 19501962 | AX-170020065 | 0 | A | C | 0.14 | -3.99E-02 | 9.47E-02 | 2.59E+00 | 2.69E+00 | 6.77E-01 | no | | intergenic_variant | MODIFIER |
| 7 | 70160525 | AX-97832314 | 0 | T | C | 0.368 | -4.04E-02 | 9.95E-02 | 2.71E+00 | 2.81E+00 | 6.77E-01 | no | | intergenic_variant | MODIFIER |
| 20 | 169021976 | AX-170085806 | 1 | C | T | 0.188 | 4.89E-02 | 1.17E-01 | 2.60E+00 | 2.69E+00 | 6.77E-01 | no | Arhgap6 | intron_variant | MODIFIER |
| 4 | 88051978 | AX-170357077 | 1 | A | G | 0.232 | 4.25E-02 | 9.71E-02 | 2.52E+00 | 2.63E+00 | 6.77E-01 | no | | intergenic_variant | MODIFIER |

| | | | | | | | | | | | | | | | |
|---|---|---|---|---|---|---|---|---|---|---|---|---|---|---|---|
| 5 | 127826731 | AX-170488218 | 0 | G | A | 0.421 | -3.38E-02 | 8.38E-02 | 2.70E+00 | 2.82E+00 | 6.77E-01 | no | Tmem132d | intron_variant | MODIFIER |
| 10 | 117700767 | AX-170168014 | 0 | C | T | 0.342 | -3.13E-02 | 7.52E-02 | 2.62E+00 | 2.71E+00 | 6.77E-01 | no | Mdm2 | intron_variant | MODIFIER |
| 7 | 78869551 | AX-97832096 | 1 | A | C | 0.098 | 7.25E-02 | 1.78E-01 | 2.65E+00 | 2.77E+00 | 6.78E-01 | no | | intergenic_variant | MODIFIER |
| 3 | 32671741 | AX-170483446 | 0 | G | A | 0.184 | 3.90E-02 | 9.57E-02 | 2.70E+00 | 2.78E+00 | 6.78E-01 | no | | intergenic_variant | MODIFIER |
| 11 | 45153594 | AX-170431403 | 1 | T | C | 0.098 | 6.53E-02 | 1.60E-01 | 2.65E+00 | 2.76E+00 | 6.78E-01 | no | | intergenic_variant | MODIFIER |
| 3 | 16469915 | AX-170271414 | 0 | A | G | 0.412 | -2.62E-02 | 6.51E-02 | 2.66E+00 | 2.80E+00 | 6.78E-01 | no | | intergenic_variant | MODIFIER |
| 10 | 108554020 | AX-97781600 | 0 | A | G | 0.342 | -3.32E-02 | 8.04E-02 | 2.63E+00 | 2.72E+00 | 6.78E-01 | no | Syt1 | intron_variant | MODIFIER |
| 8 | 25350170 | AX-170468350 | 0 | A | G | 0.351 | -2.78E-02 | 6.55E-02 | 2.57E+00 | 2.67E+00 | 6.78E-01 | no | | intergenic_variant | MODIFIER |
| 4 | 144312649 | AX-170141908 | 0 | G | A | 0.079 | 7.60E-02 | 1.84E-01 | 2.61E+00 | 2.73E+00 | 6.78E-01 | no | | intergenic_variant | MODIFIER |
| 1 | 190660884 | AX-168410636 | 2 | T | C | 0.255 | 4.26E-02 | 9.42E-02 | 2.48E+00 | 2.58E+00 | 6.78E-01 | no | | intergenic_variant | MODIFIER |
| 1 | 174466191 | AX-97825926 | 0 | C | T | 0.228 | 5.36E-02 | 1.26E-01 | 2.58E+00 | 2.64E+00 | 6.78E-01 | no | | regulatory_region_variant | MODIFIER |
| 10 | 126944557 | AX-170283853 | 1 | C | T | 0.33 | -3.00E-02 | 7.24E-02 | 2.61E+00 | 2.71E+00 | 6.78E-01 | no | | intergenic_variant | MODIFIER |
| 5 | 83781635 | AX-170025453 | 0 | G | A | 0.439 | 4.42E-02 | 1.08E-01 | 2.73E+00 | 2.79E+00 | 6.78E-01 | no | | intergenic_variant | MODIFIER |
| 4 | 92605196 | AX-170157098 | 1 | G | T | 0.321 | 4.37E-02 | 1.06E-01 | 2.65E+00 | 2.72E+00 | 6.79E-01 | no | | intergenic_variant | MODIFIER |
| 8 | 32258416 | AX-170325381 | 0 | A | G | 0.14 | 4.92E-02 | 1.22E-01 | 2.71E+00 | 2.80E+00 | 6.79E-01 | no | Nrg1 | intron_variant | MODIFIER |
| 5 | 33437744 | AX-170387196 | 1 | T | C | 0.339 | 4.44E-02 | 1.08E-01 | 2.68E+00 | 2.73E+00 | 6.79E-01 | no | Nkx1-1 | upstream_gene_variant | MODIFIER |
| 5 | 33545560 | AX-170262763 | 1 | A | C | 0.339 | 4.44E-02 | 1.08E-01 | 2.68E+00 | 2.73E+00 | 6.79E-01 | no | | intergenic_variant | MODIFIER |
| 2 | 93767591 | AX-97813866 | 1 | C | T | 0.054 | -1.22E-01 | 2.99E-01 | 2.75E+00 | 2.78E+00 | 6.79E-01 | no | Ext2 | intron_variant | MODIFIER |
| 2 | 104548213 | AX-170441751 | 1 | A | G | 0.054 | -1.22E-01 | 2.99E-01 | 2.75E+00 | 2.78E+00 | 6.79E-01 | no | | intergenic_variant | MODIFIER |
| 2 | 97213456 | AX-170423180 | 0 | T | C | 0.053 | -1.23E-01 | 3.00E-01 | 2.74E+00 | 2.77E+00 | 6.79E-01 | no | Lrrc4c | intron_variant | MODIFIER |
| 11 | 21877705 | AX-170369325 | 0 | A | G | 0.053 | 9.23E-02 | 2.30E-01 | 2.69E+00 | 2.81E+00 | 6.79E-01 | no | Wdpcp | intron_variant | MODIFIER |
| 6 | 44738194 | AX-170089710 | 0 | C | T | 0.044 | -6.85E-02 | 1.66E-01 | 2.60E+00 | 2.73E+00 | 6.79E-01 | no | | intergenic_variant | MODIFIER |
| 10 | 16791902 | AX-170002823 | 1 | G | A | 0.214 | 4.27E-02 | 1.05E-01 | 2.78E+00 | 2.87E+00 | 6.79E-01 | no | | intergenic_variant | MODIFIER |
| 14 | 97242062 | AX-170125417 | 0 | A | T | 0.053 | 7.15E-02 | 1.69E-01 | 2.53E+00 | 2.68E+00 | 6.79E-01 | no | | intergenic_variant | MODIFIER |
| 12 | 93592383 | AX-170049816 | 0 | T | C | 0.07 | 6.97E-02 | 1.69E-01 | 2.64E+00 | 2.71E+00 | 6.79E-01 | no | | intergenic_variant | MODIFIER |
| 15 | 96830839 | AX-97812300 | 1 | T | C | 0.161 | 6.91E-02 | 1.67E-01 | 2.61E+00 | 2.70E+00 | 6.79E-01 | no | | regulatory_region_variant | MODIFIER |
| 2 | 92486621 | AX-170064293 | 0 | T | C | 0.053 | -1.22E-01 | 2.99E-01 | 2.74E+00 | 2.77E+00 | 6.79E-01 | no | Gm13814 | intron_variant,non_coding_transcript_variant | MODIFIER |
| 7 | 39747171 | AX-170028868 | 0 | G | A | 0.351 | 3.97E-02 | 9.51E-02 | 2.60E+00 | 2.68E+00 | 6.79E-01 | no | | intergenic_variant | MODIFIER |
| 7 | 39751941 | AX-170387582 | 0 | G | A | 0.351 | 3.97E-02 | 9.51E-02 | 2.60E+00 | 2.68E+00 | 6.79E-01 | no | | intergenic_variant | MODIFIER |
| 7 | 39754341 | AX-170372404 | 0 | G | C | 0.351 | 3.97E-02 | 9.51E-02 | 2.60E+00 | 2.68E+00 | 6.79E-01 | no | | intergenic_variant | MODIFIER |
| 7 | 39776185 | AX-170115485 | 0 | G | T | 0.351 | 3.97E-02 | 9.51E-02 | 2.60E+00 | 2.68E+00 | 6.79E-01 | no | | intergenic_variant | MODIFIER |
| 7 | 39789618 | AX-170477869 | 0 | C | T | 0.351 | 3.97E-02 | 9.51E-02 | 2.60E+00 | 2.68E+00 | 6.79E-01 | no | Gm37302 | upstream_gene_variant | MODIFIER |
| 7 | 39792309 | AX-170028872 | 0 | C | T | 0.351 | 3.97E-02 | 9.51E-02 | 2.60E+00 | 2.68E+00 | 6.79E-01 | no | 4930558N11Rik | downstream_gene_variant | MODIFIER |
| 7 | 39794868 | AX-170402734 | 0 | C | T | 0.351 | 3.97E-02 | 9.51E-02 | 2.60E+00 | 2.68E+00 | 6.79E-01 | no | 4930558N11Rik | downstream_gene_variant | MODIFIER |
| 7 | 39817362 | AX-170238667 | 0 | A | G | 0.351 | 3.97E-02 | 9.51E-02 | 2.60E+00 | 2.68E+00 | 6.79E-01 | no | 4930435C17Rik | intron_variant,non_coding_transcript_variant | MODIFIER |
| 7 | 39827078 | AX-170454821 | 0 | G | A | 0.351 | 3.97E-02 | 9.51E-02 | 2.60E+00 | 2.68E+00 | 6.79E-01 | no | 4930435C17Rik | intron_variant,non_coding_transcript_variant | MODIFIER |
| 7 | 39841287 | AX-170089906 | 0 | G | T | 0.351 | 3.97E-02 | 9.51E-02 | 2.60E+00 | 2.68E+00 | 6.79E-01 | no | 4930435C17Rik | intron_variant,non_coding_transcript_variant | MODIFIER |
| 7 | 39870010 | AX-170077118 | 0 | G | C | 0.351 | 3.97E-02 | 9.51E-02 | 2.60E+00 | 2.68E+00 | 6.79E-01 | no | 4930435C17Rik | intron_variant,non_coding_transcript_variant | MODIFIER |
| 7 | 39887336 | AX-170226063 | 0 | C | A | 0.351 | 3.97E-02 | 9.51E-02 | 2.60E+00 | 2.68E+00 | 6.79E-01 | no | 4930435C17Rik | intron_variant,non_coding_transcript_variant | MODIFIER |
| 7 | 39887364 | AX-170469340 | 0 | A | C | 0.351 | 3.97E-02 | 9.51E-02 | 2.60E+00 | 2.68E+00 | 6.79E-01 | no | 4930435C17Rik | intron_variant,non_coding_transcript_variant | MODIFIER |
| 7 | 39894396 | AX-170361965 | 0 | A | T | 0.351 | 3.97E-02 | 9.51E-02 | 2.60E+00 | 2.68E+00 | 6.79E-01 | no | 4930435C17Rik | intron_variant,non_coding_transcript_variant | MODIFIER |
| 7 | 39894584 | AX-170384149 | 0 | T | A | 0.351 | 3.97E-02 | 9.51E-02 | 2.60E+00 | 2.68E+00 | 6.79E-01 | no | 4930435C17Rik | intron_variant,non_coding_transcript_variant | MODIFIER |
| 7 | 39904930 | AX-170322954 | 0 | G | A | 0.351 | 3.97E-02 | 9.51E-02 | 2.60E+00 | 2.68E+00 | 6.79E-01 | no | Gm44992 | intron_variant,non_coding_transcript_variant | MODIFIER |
| 7 | 39917795 | AX-170473394 | 0 | T | C | 0.351 | 3.97E-02 | 9.51E-02 | 2.60E+00 | 2.68E+00 | 6.79E-01 | no | Gm44992 | intron_variant,non_coding_transcript_variant | MODIFIER |
| 7 | 39919408 | AX-97853556 | 0 | T | C | 0.351 | 3.97E-02 | 9.51E-02 | 2.60E+00 | 2.68E+00 | 6.79E-01 | no | Gm44992 | intron_variant,non_coding_transcript_variant | MODIFIER |
| 7 | 40060993 | AX-170077119 | 0 | G | A | 0.351 | 3.97E-02 | 9.51E-02 | 2.60E+00 | 2.68E+00 | 6.79E-01 | no | | intergenic_variant | MODIFIER |
| 7 | 40088584 | AX-170077120 | 0 | C | T | 0.351 | 3.97E-02 | 9.51E-02 | 2.60E+00 | 2.68E+00 | 6.79E-01 | no | | intergenic_variant | MODIFIER |
| 7 | 40238584 | AX-170095901 | 0 | T | C | 0.351 | 3.97E-02 | 9.51E-02 | 2.60E+00 | 2.68E+00 | 6.79E-01 | no | | intergenic_variant | MODIFIER |
| 7 | 40242740 | AX-170117019 | 0 | C | G | 0.351 | 3.97E-02 | 9.51E-02 | 2.60E+00 | 2.68E+00 | 6.79E-01 | no | | intergenic_variant | MODIFIER |
| 7 | 40246558 | AX-170340219 | 0 | G | A | 0.351 | 3.97E-02 | 9.51E-02 | 2.60E+00 | 2.68E+00 | 6.79E-01 | no | | intergenic_variant | MODIFIER |
| 7 | 40246646 | AX-170378812 | 0 | C | T | 0.351 | 3.97E-02 | 9.51E-02 | 2.60E+00 | 2.68E+00 | 6.79E-01 | no | | intergenic_variant | MODIFIER |
| 7 | 40253721 | AX-170525182 | 0 | C | G | 0.351 | 3.97E-02 | 9.51E-02 | 2.60E+00 | 2.68E+00 | 6.79E-01 | no | | intergenic_variant | MODIFIER |
| 7 | 40253978 | AX-97795888 | 0 | A | G | 0.351 | 3.97E-02 | 9.51E-02 | 2.60E+00 | 2.68E+00 | 6.79E-01 | no | | intergenic_variant | MODIFIER |
| 7 | 40256880 | AX-170346273 | 0 | A | G | 0.351 | 3.97E-02 | 9.51E-02 | 2.60E+00 | 2.68E+00 | 6.79E-01 | no | | intergenic_variant | MODIFIER |

| | | | | | | | | | | | | | | | |
|---|---|---|---|---|---|---|---|---|---|---|---|---|---|---|---|
| 7 | 40258631 | AX-170336069 | 0 | G | A | 0.351 | 3.97E-02 | 9.51E-02 | 2.60E+00 | 2.68E+00 | 6.79E-01 | no | 4933404I11Rik | upstream_gene_variant | MODIFIER |
| 7 | 40258886 | AX-170211788 | 0 | T | C | 0.351 | 3.97E-02 | 9.51E-02 | 2.60E+00 | 2.68E+00 | 6.79E-01 | no | 4933404I11Rik | upstream_gene_variant | MODIFIER |
| 7 | 40261964 | AX-170481998 | 0 | G | A | 0.351 | 3.97E-02 | 9.51E-02 | 2.60E+00 | 2.68E+00 | 6.79E-01 | no | 4933404I11Rik | upstream_gene_variant | MODIFIER |
| 7 | 40268308 | AX-170028880 | 0 | C | A | 0.351 | 3.97E-02 | 9.51E-02 | 2.60E+00 | 2.68E+00 | 6.79E-01 | no | 4933404I11Rik | intron_variant,non_coding_transcript_variant | MODIFIER |
| 3 | 29473777 | AX-170396149 | 1 | A | T | 0.179 | -3.78E-02 | 9.21E-02 | 2.64E+00 | 2.73E+00 | 6.79E-01 | no | Egfem1 | intron_variant | MODIFIER |
| 5 | 73249698 | AX-170190135 | 2 | C | T | 0.436 | 4.15E-02 | 1.00E-01 | 2.62E+00 | 2.70E+00 | 6.79E-01 | no | Fryl | intron_variant | MODIFIER |
| 10 | 13836130 | AX-170375823 | 0 | T | A | 0.202 | -4.13E-02 | 9.98E-02 | 2.61E+00 | 2.71E+00 | 6.79E-01 | no | Aig1 | intron_variant | MODIFIER |
| 5 | 108151398 | AX-170449464 | 1 | C | T | 0.464 | 4.51E-02 | 1.09E-01 | 2.64E+00 | 2.70E+00 | 6.80E-01 | no | Ccdc18 | intron_variant | MODIFIER |
| 18 | 45654889 | AX-170060741 | 0 | T | C | 0.07 | -9.17E-02 | 2.27E-01 | 2.68E+00 | 2.77E+00 | 6.80E-01 | no | Kcnn2 | intron_variant | MODIFIER |
| 20 | 130693227 | AX-170417297 | 1 | A | G | 0.134 | 5.34E-02 | 1.32E-01 | 2.65E+00 | 2.76E+00 | 6.80E-01 | no | | intergenic_variant | MODIFIER |
| 1 | 123000666 | AX-97783310 | 0 | C | T | 0.561 | 3.52E-02 | 8.27E-02 | 2.57E+00 | 2.64E+00 | 6.80E-01 | no | | intergenic_variant | MODIFIER |
| 7 | 39795276 | AX-170259422 | 1 | G | A | 0.357 | 3.95E-02 | 9.41E-02 | 2.59E+00 | 2.67E+00 | 6.80E-01 | no | 4930558N11Rik | downstream_gene_variant | MODIFIER |
| 7 | 117245761 | AX-170327831 | 0 | G | A | 0.412 | -3.75E-02 | 9.20E-02 | 2.79E+00 | 2.86E+00 | 6.80E-01 | no | Gm39075 | intron_variant,non_coding_transcript_variant | MODIFIER |
| 8 | 93577114 | AX-170282907 | 2 | A | C | 0.2 | -4.86E-02 | 1.20E-01 | 2.69E+00 | 2.76E+00 | 6.80E-01 | no | Gm45708 | upstream_gene_variant | MODIFIER |
| 5 | 83705411 | AX-170201140 | 0 | A | G | 0.447 | 4.52E-02 | 1.11E-01 | 2.72E+00 | 2.78E+00 | 6.80E-01 | no | | intergenic_variant | MODIFIER |
| 1 | 76990243 | AX-170206724 | 1 | A | T | 0.143 | 9.64E-02 | 2.37E-01 | 2.70E+00 | 2.75E+00 | 6.80E-01 | no | Gm816 | intron_variant,non_coding_transcript_variant | MODIFIER |
| 4 | 74233531 | AX-170302937 | 0 | A | C | 0.456 | -3.60E-02 | 8.37E-02 | 2.55E+00 | 2.62E+00 | 6.80E-01 | no | Gm11405 | downstream_gene_variant | MODIFIER |
| 12 | 106577083 | AX-170500933 | 2 | T | C | 0.127 | -6.75E-02 | 1.65E-01 | 2.64E+00 | 2.74E+00 | 6.80E-01 | no | Gm17033 | downstream_gene_variant | MODIFIER |
| 12 | 94970540 | AX-170189340 | 1 | G | C | 0.062 | 9.64E-02 | 2.37E-01 | 2.71E+00 | 2.76E+00 | 6.80E-01 | no | | intergenic_variant | MODIFIER |
| 9 | 19438099 | AX-170031900 | 0 | C | G | 0.465 | 3.71E-02 | 8.81E-02 | 2.58E+00 | 2.66E+00 | 6.80E-01 | no | Olfr849 | upstream_gene_variant | MODIFIER |
| 10 | 17752122 | AX-170272969 | 0 | T | C | 0.202 | 3.93E-02 | 9.83E-02 | 2.74E+00 | 2.84E+00 | 6.80E-01 | no | | intergenic_variant | MODIFIER |
| 3 | 148083724 | AX-168409009 | 1 | C | A | 0.348 | 3.05E-02 | 7.44E-02 | 2.64E+00 | 2.71E+00 | 6.80E-01 | no | | intergenic_variant | MODIFIER |
| 2 | 88475250 | AX-170205486 | 0 | C | G | 0.053 | -1.22E-01 | 3.00E-01 | 2.75E+00 | 2.77E+00 | 6.81E-01 | no | | intergenic_variant | MODIFIER |
| 2 | 90250434 | AX-170349652 | 0 | C | A | 0.053 | -1.22E-01 | 3.00E-01 | 2.75E+00 | 2.77E+00 | 6.81E-01 | no | Olfr142 | downstream_gene_variant | MODIFIER |
| 12 | 95120281 | AX-170049849 | 1 | G | T | 0.062 | 9.60E-02 | 2.37E-01 | 2.71E+00 | 2.76E+00 | 6.81E-01 | no | | intergenic_variant | MODIFIER |
| 12 | 98165198 | AX-170511608 | 1 | C | A | 0.062 | 9.60E-02 | 2.37E-01 | 2.71E+00 | 2.76E+00 | 6.81E-01 | no | | intergenic_variant | MODIFIER |
| 4 | 153371015 | AX-97790327 | 0 | G | A | 0.105 | 6.33E-02 | 1.57E-01 | 2.67E+00 | 2.78E+00 | 6.81E-01 | no | Ajap1 | downstream_gene_variant | MODIFIER |
| 12 | 92537386 | AX-97851589 | 0 | A | G | 0.061 | 9.61E-02 | 2.37E-01 | 2.71E+00 | 2.76E+00 | 6.81E-01 | no | | intergenic_variant | MODIFIER |
| 12 | 92554864 | AX-170460113 | 0 | C | T | 0.061 | 9.61E-02 | 2.37E-01 | 2.71E+00 | 2.76E+00 | 6.81E-01 | no | | intergenic_variant | MODIFIER |
| 12 | 92572900 | AX-97847878 | 0 | A | G | 0.061 | 9.61E-02 | 2.37E-01 | 2.71E+00 | 2.76E+00 | 6.81E-01 | no | | intergenic_variant | MODIFIER |
| 12 | 92578179 | AX-170225959 | 0 | A | T | 0.061 | 9.61E-02 | 2.37E-01 | 2.71E+00 | 2.76E+00 | 6.81E-01 | no | | intergenic_variant | MODIFIER |
| 12 | 92751165 | AX-170250792 | 0 | T | C | 0.061 | 9.61E-02 | 2.37E-01 | 2.71E+00 | 2.76E+00 | 6.81E-01 | no | | intergenic_variant | MODIFIER |
| 12 | 92763139 | AX-170007276 | 0 | G | A | 0.061 | 9.61E-02 | 2.37E-01 | 2.71E+00 | 2.76E+00 | 6.81E-01 | no | | intergenic_variant | MODIFIER |
| 12 | 93199666 | AX-97844325 | 0 | T | C | 0.061 | 9.61E-02 | 2.37E-01 | 2.71E+00 | 2.76E+00 | 6.81E-01 | no | | intergenic_variant | MODIFIER |
| 12 | 93203215 | AX-170224359 | 0 | C | T | 0.061 | 9.61E-02 | 2.37E-01 | 2.71E+00 | 2.76E+00 | 6.81E-01 | no | | intergenic_variant | MODIFIER |
| 12 | 93266825 | AX-97801005 | 0 | C | T | 0.061 | 9.61E-02 | 2.37E-01 | 2.71E+00 | 2.76E+00 | 6.81E-01 | no | | intergenic_variant | MODIFIER |
| 12 | 93281527 | AX-170134332 | 0 | C | T | 0.061 | 9.61E-02 | 2.37E-01 | 2.71E+00 | 2.76E+00 | 6.81E-01 | no | | intergenic_variant | MODIFIER |
| 12 | 93324319 | AX-170007294 | 0 | T | C | 0.061 | 9.61E-02 | 2.37E-01 | 2.71E+00 | 2.76E+00 | 6.81E-01 | no | | intergenic_variant | MODIFIER |
| 12 | 93354050 | AX-170253461 | 0 | C | T | 0.061 | 9.61E-02 | 2.37E-01 | 2.71E+00 | 2.76E+00 | 6.81E-01 | no | | intergenic_variant | MODIFIER |
| 12 | 93434694 | AX-170343445 | 0 | C | G | 0.061 | 9.61E-02 | 2.37E-01 | 2.71E+00 | 2.76E+00 | 6.81E-01 | no | 4930559C10Rik | intron_variant,non_coding_transcript_variant | MODIFIER |
| 12 | 93435103 | AX-170495668 | 0 | C | A | 0.061 | 9.61E-02 | 2.37E-01 | 2.71E+00 | 2.76E+00 | 6.81E-01 | no | 4930559C10Rik | intron_variant,non_coding_transcript_variant | MODIFIER |
| 12 | 93450308 | AX-168386045 | 0 | A | G | 0.061 | 9.61E-02 | 2.37E-01 | 2.71E+00 | 2.76E+00 | 6.81E-01 | no | | intergenic_variant | MODIFIER |
| 12 | 93466568 | AX-170288963 | 0 | C | G | 0.061 | 9.61E-02 | 2.37E-01 | 2.71E+00 | 2.76E+00 | 6.81E-01 | no | | intergenic_variant | MODIFIER |
| 12 | 93473745 | AX-97854934 | 0 | A | G | 0.061 | 9.61E-02 | 2.37E-01 | 2.71E+00 | 2.76E+00 | 6.81E-01 | no | | intergenic_variant | MODIFIER |
| 12 | 93534663 | AX-170007287 | 0 | T | C | 0.061 | 9.61E-02 | 2.37E-01 | 2.71E+00 | 2.76E+00 | 6.81E-01 | no | | intergenic_variant | MODIFIER |
| 12 | 93544034 | AX-97794391 | 0 | A | C | 0.061 | 9.61E-02 | 2.37E-01 | 2.71E+00 | 2.76E+00 | 6.81E-01 | no | | intergenic_variant | MODIFIER |
| 12 | 93621203 | AX-170476546 | 0 | G | A | 0.061 | 9.61E-02 | 2.37E-01 | 2.71E+00 | 2.76E+00 | 6.81E-01 | no | | intergenic_variant | MODIFIER |
| 12 | 93678326 | AX-97853138 | 0 | T | C | 0.061 | 9.61E-02 | 2.37E-01 | 2.71E+00 | 2.76E+00 | 6.81E-01 | no | | intergenic_variant | MODIFIER |
| 12 | 94108659 | AX-170362452 | 0 | G | A | 0.061 | 9.61E-02 | 2.37E-01 | 2.71E+00 | 2.76E+00 | 6.81E-01 | no | | intergenic_variant | MODIFIER |
| 12 | 94109666 | AX-170149331 | 0 | A | G | 0.061 | 9.61E-02 | 2.37E-01 | 2.71E+00 | 2.76E+00 | 6.81E-01 | no | | intergenic_variant | MODIFIER |
| 12 | 94479433 | AX-97800614 | 0 | G | T | 0.061 | 9.61E-02 | 2.37E-01 | 2.71E+00 | 2.76E+00 | 6.81E-01 | no | | intergenic_variant | MODIFIER |
| 12 | 94584412 | AX-170049838 | 0 | G | A | 0.061 | 9.61E-02 | 2.37E-01 | 2.71E+00 | 2.76E+00 | 6.81E-01 | no | | intergenic_variant | MODIFIER |
| 12 | 94585152 | AX-97853411 | 0 | C | T | 0.061 | 9.61E-02 | 2.37E-01 | 2.71E+00 | 2.76E+00 | 6.81E-01 | no | | regulatory_region_variant | MODIFIER |
| 12 | 94585256 | AX-170049839 | 0 | T | A | 0.061 | 9.61E-02 | 2.37E-01 | 2.71E+00 | 2.76E+00 | 6.81E-01 | no | | regulatory_region_variant | MODIFIER |

| | | | | | | | | | | | | | | | |
|---|---|---|---|---|---|---|---|---|---|---|---|---|---|---|---|
| 12 | 94614371 | AX-97831650 | 0 | C | T | 0.061 | 9.61E-02 | 2.37E-01 | 2.71E+00 | 2.76E+00 | 6.81E-01 | no | | intergenic_variant | MODIFIER |
| 12 | 94970642 | AX-170470115 | 0 | G | A | 0.061 | 9.61E-02 | 2.37E-01 | 2.71E+00 | 2.76E+00 | 6.81E-01 | no | | intergenic_variant | MODIFIER |
| 12 | 95120339 | AX-170049848 | 0 | A | G | 0.061 | 9.61E-02 | 2.37E-01 | 2.71E+00 | 2.76E+00 | 6.81E-01 | no | | intergenic_variant | MODIFIER |
| 12 | 95153769 | AX-170382561 | 0 | A | G | 0.061 | 9.61E-02 | 2.37E-01 | 2.71E+00 | 2.76E+00 | 6.81E-01 | no | | intergenic_variant | MODIFIER |
| 12 | 95171236 | AX-170092580 | 0 | T | C | 0.061 | 9.61E-02 | 2.37E-01 | 2.71E+00 | 2.76E+00 | 6.81E-01 | no | | intergenic_variant | MODIFIER |
| 12 | 95229658 | AX-170108836 | 0 | C | G | 0.061 | 9.61E-02 | 2.37E-01 | 2.71E+00 | 2.76E+00 | 6.81E-01 | no | | intergenic_variant | MODIFIER |
| 12 | 95445210 | AX-170087312 | 0 | G | A | 0.061 | 9.61E-02 | 2.37E-01 | 2.71E+00 | 2.76E+00 | 6.81E-01 | no | | intergenic_variant | MODIFIER |
| 12 | 95458619 | AX-97804107 | 0 | A | G | 0.061 | 9.61E-02 | 2.37E-01 | 2.71E+00 | 2.76E+00 | 6.81E-01 | no | | intergenic_variant | MODIFIER |
| 12 | 95506625 | AX-97852406 | 0 | G | A | 0.061 | 9.61E-02 | 2.37E-01 | 2.71E+00 | 2.76E+00 | 6.81E-01 | no | | intergenic_variant | MODIFIER |
| 12 | 95506711 | AX-97856930 | 0 | A | G | 0.061 | 9.61E-02 | 2.37E-01 | 2.71E+00 | 2.76E+00 | 6.81E-01 | no | | intergenic_variant | MODIFIER |
| 12 | 95509290 | AX-170417375 | 0 | A | C | 0.061 | 9.61E-02 | 2.37E-01 | 2.71E+00 | 2.76E+00 | 6.81E-01 | no | | intergenic_variant | MODIFIER |
| 12 | 95519890 | AX-170049859 | 0 | A | C | 0.061 | 9.61E-02 | 2.37E-01 | 2.71E+00 | 2.76E+00 | 6.81E-01 | no | | intergenic_variant | MODIFIER |
| 12 | 98207632 | AX-170261968 | 0 | A | G | 0.061 | 9.61E-02 | 2.37E-01 | 2.71E+00 | 2.76E+00 | 6.81E-01 | no | Galc | intron_variant | MODIFIER |
| 12 | 98295858 | AX-170476389 | 0 | G | T | 0.061 | 9.61E-02 | 2.37E-01 | 2.71E+00 | 2.76E+00 | 6.81E-01 | no | | intergenic_variant | MODIFIER |
| 12 | 98302145 | AX-170049912 | 0 | G | A | 0.061 | 9.61E-02 | 2.37E-01 | 2.71E+00 | 2.76E+00 | 6.81E-01 | no | | intergenic_variant | MODIFIER |
| 12 | 98324515 | AX-170007348 | 0 | T | G | 0.061 | 9.61E-02 | 2.37E-01 | 2.71E+00 | 2.76E+00 | 6.81E-01 | no | | regulatory_region_variant | MODIFIER |
| 12 | 98382080 | AX-170007351 | 0 | G | A | 0.061 | 9.61E-02 | 2.37E-01 | 2.71E+00 | 2.76E+00 | 6.81E-01 | no | | intergenic_variant | MODIFIER |
| 12 | 98395000 | AX-170350365 | 0 | T | C | 0.061 | 9.61E-02 | 2.37E-01 | 2.71E+00 | 2.76E+00 | 6.81E-01 | no | | intergenic_variant | MODIFIER |
| 12 | 98418986 | AX-170007349 | 0 | G | A | 0.061 | 9.61E-02 | 2.37E-01 | 2.71E+00 | 2.76E+00 | 6.81E-01 | no | | intergenic_variant | MODIFIER |
| 12 | 98441290 | AX-170310328 | 0 | T | A | 0.061 | 9.61E-02 | 2.37E-01 | 2.71E+00 | 2.76E+00 | 6.81E-01 | no | Kcnk10 | intron_variant | MODIFIER |
| 12 | 98895213 | AX-170007363 | 0 | G | T | 0.061 | 9.61E-02 | 2.37E-01 | 2.71E+00 | 2.76E+00 | 6.81E-01 | no | Eml5 | intron_variant | MODIFIER |
| 12 | 98902343 | AX-170007366 | 0 | C | T | 0.061 | 9.61E-02 | 2.37E-01 | 2.71E+00 | 2.76E+00 | 6.81E-01 | no | Eml5 | upstream_gene_variant | MODIFIER |
| 12 | 99220600 | AX-97828360 | 0 | C | T | 0.061 | 9.61E-02 | 2.37E-01 | 2.71E+00 | 2.76E+00 | 6.81E-01 | no | Foxn3 | intron_variant | MODIFIER |
| 12 | 99249056 | AX-170417461 | 0 | G | A | 0.061 | 9.61E-02 | 2.37E-01 | 2.71E+00 | 2.76E+00 | 6.81E-01 | no | Foxn3 | intron_variant | MODIFIER |
| 20 | 166061734 | AX-170348679 | 1 | C | T | 0.134 | 5.16E-02 | 1.28E-01 | 2.66E+00 | 2.77E+00 | 6.81E-01 | no | Gm15225 | upstream_gene_variant | MODIFIER |
| 16 | 45221713 | AX-97836831 | 0 | T | G | 0.482 | -3.41E-02 | 8.10E-02 | 2.59E+00 | 2.64E+00 | 6.81E-01 | no | Btla | upstream_gene_variant | MODIFIER |
| 5 | 122845826 | AX-170026055 | 0 | A | G | 0.447 | -4.53E-02 | 1.11E-01 | 2.78E+00 | 2.83E+00 | 6.81E-01 | no | Rnf34 | upstream_gene_variant | MODIFIER |
| 2 | 168802135 | AX-170360924 | 0 | T | C | 0.167 | 6.69E-02 | 1.63E-01 | 2.62E+00 | 2.72E+00 | 6.81E-01 | no | | intergenic_variant | MODIFIER |
| 9 | 85135880 | AX-170314480 | 0 | C | T | 0.211 | 5.07E-02 | 1.25E-01 | 2.73E+00 | 2.78E+00 | 6.81E-01 | no | | intergenic_variant | MODIFIER |
| 9 | 85139682 | AX-97783852 | 0 | G | A | 0.211 | 5.07E-02 | 1.25E-01 | 2.73E+00 | 2.78E+00 | 6.81E-01 | no | | regulatory_region_variant | MODIFIER |
| 9 | 85180160 | AX-170470013 | 0 | T | C | 0.211 | 5.07E-02 | 1.25E-01 | 2.73E+00 | 2.78E+00 | 6.81E-01 | no | | intergenic_variant | MODIFIER |
| 9 | 85318229 | AX-170476259 | 0 | C | A | 0.211 | 5.07E-02 | 1.25E-01 | 2.73E+00 | 2.78E+00 | 6.81E-01 | no | Tent5a | downstream_gene_variant | MODIFIER |
| 9 | 85402323 | AX-170388941 | 0 | C | T | 0.211 | 5.07E-02 | 1.25E-01 | 2.73E+00 | 2.78E+00 | 6.81E-01 | no | | intergenic_variant | MODIFIER |
| 9 | 85436675 | AX-170527896 | 0 | A | T | 0.211 | 5.07E-02 | 1.25E-01 | 2.73E+00 | 2.78E+00 | 6.81E-01 | no | | intergenic_variant | MODIFIER |
| 9 | 85437178 | AX-170175460 | 0 | T | G | 0.211 | 5.07E-02 | 1.25E-01 | 2.73E+00 | 2.78E+00 | 6.81E-01 | no | | intergenic_variant | MODIFIER |
| 9 | 85464293 | AX-170032876 | 0 | G | A | 0.211 | 5.07E-02 | 1.25E-01 | 2.73E+00 | 2.78E+00 | 6.81E-01 | no | | intergenic_variant | MODIFIER |
| 9 | 85474025 | AX-170192054 | 0 | C | T | 0.211 | 5.07E-02 | 1.25E-01 | 2.73E+00 | 2.78E+00 | 6.81E-01 | no | Gm25125 | downstream_gene_variant | MODIFIER |
| 9 | 85482789 | AX-170032873 | 0 | T | C | 0.211 | 5.07E-02 | 1.25E-01 | 2.73E+00 | 2.78E+00 | 6.81E-01 | no | | intergenic_variant | MODIFIER |
| 11 | 33785228 | AX-170036457 | 0 | T | C | 0.105 | -6.53E-02 | 1.63E-01 | 2.68E+00 | 2.79E+00 | 6.81E-01 | no | Kcnip1 | intron_variant | MODIFIER |
| 7 | 135315860 | AX-170204927 | 0 | A | G | 0.246 | -3.44E-02 | 8.21E-02 | 2.58E+00 | 2.68E+00 | 6.81E-01 | no | | intergenic_variant | MODIFIER |
| 18 | 72384322 | AX-97851306 | 0 | G | A | 0.184 | -4.82E-02 | 1.18E-01 | 2.65E+00 | 2.71E+00 | 6.81E-01 | no | | intergenic_variant | MODIFIER |
| 8 | 95982697 | AX-170127667 | 0 | C | T | 0.237 | 4.49E-02 | 1.11E-01 | 2.66E+00 | 2.74E+00 | 6.81E-01 | no | | intergenic_variant | MODIFIER |
| 9 | 117904564 | AX-170189792 | 1 | C | T | 0.134 | -3.90E-02 | 9.41E-02 | 2.58E+00 | 2.70E+00 | 6.81E-01 | no | Zcwpw2 | intron_variant,non_coding_transcript_variant | MODIFIER |
| 1 | 156042139 | AX-170002051 | 1 | G | A | 0.223 | 5.79E-02 | 1.40E-01 | 2.62E+00 | 2.66E+00 | 6.81E-01 | no | Tor1aip2 | intron_variant | MODIFIER |
| 16 | 19742484 | AX-170362736 | 0 | C | A | 0.123 | 3.91E-02 | 9.80E-02 | 2.67E+00 | 2.79E+00 | 6.81E-01 | no | | intergenic_variant | MODIFIER |
| 4 | 72953971 | AX-170224933 | 0 | G | A | 0.316 | 4.56E-02 | 1.12E-01 | 2.66E+00 | 2.72E+00 | 6.81E-01 | no | | intergenic_variant | MODIFIER |
| 7 | 40047527 | AX-170095900 | 0 | G | A | 0.351 | 3.92E-02 | 9.47E-02 | 2.61E+00 | 2.69E+00 | 6.81E-01 | no | | intergenic_variant | MODIFIER |
| 8 | 95963794 | AX-170345881 | 1 | G | A | 0.223 | 4.67E-02 | 1.16E-01 | 2.69E+00 | 2.76E+00 | 6.81E-01 | no | | intergenic_variant | MODIFIER |
| 1 | 77673736 | AX-170516427 | 1 | T | C | 0.134 | 8.97E-02 | 2.21E-01 | 2.68E+00 | 2.75E+00 | 6.81E-01 | no | | intergenic_variant | MODIFIER |
| 15 | 85279227 | AX-170265368 | 0 | C | T | 0.518 | 3.14E-02 | 7.20E-02 | 2.52E+00 | 2.61E+00 | 6.81E-01 | no | Fbln1 | intron_variant | MODIFIER |
| 18 | 52226657 | AX-170525852 | 1 | C | A | 0.196 | -4.75E-02 | 1.13E-01 | 2.59E+00 | 2.65E+00 | 6.81E-01 | no | | intergenic_variant | MODIFIER |
| 1 | 76119261 | AX-97837414 | 1 | A | C | 0.125 | 8.62E-02 | 2.13E-01 | 2.66E+00 | 2.75E+00 | 6.82E-01 | no | | intergenic_variant | MODIFIER |
| 1 | 76146028 | AX-170189707 | 1 | G | A | 0.125 | 8.62E-02 | 2.13E-01 | 2.66E+00 | 2.75E+00 | 6.82E-01 | no | | intergenic_variant | MODIFIER |
| 9 | 77554719 | AX-97844823 | 0 | C | T | 0.211 | 5.21E-02 | 1.29E-01 | 2.76E+00 | 2.81E+00 | 6.82E-01 | no | Gm21013 | upstream_gene_variant | MODIFIER |

| | | | | | | | | | | | | | | | |
|---|---|---|---|---|---|---|---|---|---|---|---|---|---|---|---|
| 1 | 106618606 | AX-170001283 | 0 | C | T | 0.167 | 7.01E-02 | 1.63E-01 | 2.53E+00 | 2.63E+00 | 6.82E-01 | no | Bcl2 | intron_variant | MODIFIER |
| 14 | 112647242 | AX-170010717 | 0 | T | C | 0.307 | 3.01E-02 | 7.19E-02 | 2.56E+00 | 2.68E+00 | 6.82E-01 | no | | intergenic_variant | MODIFIER |
| 1 | 9721434 | AX-170520403 | 0 | T | C | 0.096 | -5.93E-02 | 1.48E-01 | 2.64E+00 | 2.77E+00 | 6.82E-01 | no | Vcpip1 | 3_prime_UTR_variant | MODIFIER |
| 1 | 186997989 | AX-170002499 | 0 | A | G | 0.237 | 4.27E-02 | 9.72E-02 | 2.50E+00 | 2.60E+00 | 6.82E-01 | no | | regulatory_region_variant | MODIFIER |
| 5 | 93032037 | AX-97793095 | 1 | G | A | 0.42 | 4.41E-02 | 1.09E-01 | 2.69E+00 | 2.74E+00 | 6.82E-01 | no | | regulatory_region_variant | MODIFIER |
| 7 | 117349123 | AX-170268080 | 1 | G | A | 0.42 | 3.58E-02 | 8.84E-02 | 2.68E+00 | 2.74E+00 | 6.82E-01 | no | | intergenic_variant | MODIFIER |
| 8 | 68568138 | AX-97845795 | 0 | T | G | 0.184 | -5.42E-02 | 1.34E-01 | 2.68E+00 | 2.75E+00 | 6.82E-01 | no | Csgalnact1 | intron_variant | MODIFIER |
| 11 | 50930436 | AX-170269354 | 0 | C | A | 0.07 | -8.05E-02 | 1.98E-01 | 2.64E+00 | 2.72E+00 | 6.82E-01 | no | Zfp354b | intron_variant | MODIFIER |
| 11 | 50936816 | AX-170387993 | 0 | T | A | 0.07 | -8.05E-02 | 1.98E-01 | 2.64E+00 | 2.72E+00 | 6.82E-01 | no | 4933414I15Rik | downstream_gene_variant | MODIFIER |
| 11 | 50962622 | AX-170372287 | 0 | G | A | 0.07 | -8.05E-02 | 1.98E-01 | 2.64E+00 | 2.72E+00 | 6.82E-01 | no | Olfr1379-ps1 | downstream_gene_variant | MODIFIER |
| 11 | 50984697 | AX-170103314 | 0 | G | A | 0.07 | -8.05E-02 | 1.98E-01 | 2.64E+00 | 2.72E+00 | 6.82E-01 | no | Olfr1377 | 5_prime_UTR_variant | MODIFIER |
| 11 | 50986911 | AX-170004918 | 0 | C | T | 0.07 | -8.05E-02 | 1.98E-01 | 2.64E+00 | 2.72E+00 | 6.82E-01 | no | Olfr1377 | downstream_gene_variant | MODIFIER |
| 11 | 50991613 | AX-170215063 | 0 | A | G | 0.07 | -8.05E-02 | 1.98E-01 | 2.64E+00 | 2.72E+00 | 6.82E-01 | no | Olfr1376-ps1 | upstream_gene_variant | MODIFIER |
| 11 | 51208401 | AX-170311547 | 0 | A | G | 0.07 | -8.05E-02 | 1.98E-01 | 2.64E+00 | 2.72E+00 | 6.82E-01 | no | Platr20 | intron_variant,non_coding_transcript_variant | MODIFIER |
| 11 | 51211550 | AX-170150192 | 0 | G | A | 0.07 | -8.05E-02 | 1.98E-01 | 2.64E+00 | 2.72E+00 | 6.82E-01 | no | Platr20 | intron_variant,non_coding_transcript_variant | MODIFIER |
| 11 | 51213075 | AX-170255023 | 0 | A | G | 0.07 | -8.05E-02 | 1.98E-01 | 2.64E+00 | 2.72E+00 | 6.82E-01 | no | Platr20 | intron_variant,non_coding_transcript_variant | MODIFIER |
| 11 | 51216630 | AX-170389917 | 0 | G | A | 0.07 | -8.05E-02 | 1.98E-01 | 2.64E+00 | 2.72E+00 | 6.82E-01 | no | Gm12569 | upstream_gene_variant | MODIFIER |
| 11 | 51221831 | AX-170296404 | 0 | G | A | 0.07 | -8.05E-02 | 1.98E-01 | 2.64E+00 | 2.72E+00 | 6.82E-01 | no | Gm12569 | intron_variant | MODIFIER |
| 11 | 51231266 | AX-170342035 | 0 | C | G | 0.07 | -8.05E-02 | 1.98E-01 | 2.64E+00 | 2.72E+00 | 6.82E-01 | no | Gm12569 | upstream_gene_variant | MODIFIER |
| 11 | 51238422 | AX-170536182 | 0 | C | G | 0.07 | -8.05E-02 | 1.98E-01 | 2.64E+00 | 2.72E+00 | 6.82E-01 | no | Gm12569 | downstream_gene_variant | MODIFIER |
| 10 | 17255872 | AX-170255825 | 1 | G | A | 0.214 | 4.27E-02 | 1.06E-01 | 2.79E+00 | 2.87E+00 | 6.82E-01 | no | | intergenic_variant | MODIFIER |
| 1 | 147231627 | AX-97813617 | 1 | A | G | 0.196 | -4.60E-02 | 1.16E-01 | 2.71E+00 | 2.83E+00 | 6.82E-01 | no | | intergenic_variant | MODIFIER |
| 2 | 113548344 | AX-97852579 | 0 | C | T | 0.07 | -7.12E-02 | 1.80E-01 | 2.68E+00 | 2.81E+00 | 6.83E-01 | no | Fmn1 | intron_variant | MODIFIER |
| 2 | 90042427 | AX-170205873 | 1 | T | C | 0.054 | -8.67E-02 | 2.14E-01 | 2.64E+00 | 2.73E+00 | 6.83E-01 | no | Olfr1265 | downstream_gene_variant | MODIFIER |
| 8 | 116148105 | AX-97805996 | 1 | G | A | 0.259 | 4.00E-02 | 1.00E-01 | 2.67E+00 | 2.78E+00 | 6.83E-01 | no | | intergenic_variant | MODIFIER |
| 1 | 78484053 | AX-170451824 | 0 | C | T | 0.14 | 9.52E-02 | 2.36E-01 | 2.70E+00 | 2.75E+00 | 6.83E-01 | no | Farsb | intron_variant | MODIFIER |
| 5 | 35462819 | AX-170454308 | 0 | C | T | 0.342 | 4.35E-02 | 1.07E-01 | 2.68E+00 | 2.73E+00 | 6.83E-01 | no | | intergenic_variant | MODIFIER |
| 9 | 52804251 | AX-170201672 | 0 | T | C | 0.219 | -4.70E-02 | 1.17E-01 | 2.69E+00 | 2.75E+00 | 6.83E-01 | no | | intergenic_variant | MODIFIER |
| 1 | 46803950 | AX-170293847 | 2 | G | A | 0.1 | -8.37E-02 | 2.09E-01 | 2.68E+00 | 2.77E+00 | 6.83E-01 | no | Slc39a10 | downstream_gene_variant | MODIFIER |
| 8 | 69029101 | AX-170080339 | 1 | A | G | 0.179 | -5.59E-02 | 1.38E-01 | 2.66E+00 | 2.72E+00 | 6.83E-01 | no | | intergenic_variant | MODIFIER |
| 9 | 20468028 | AX-170419946 | 0 | T | C | 0.465 | 3.78E-02 | 9.10E-02 | 2.59E+00 | 2.66E+00 | 6.83E-01 | no | Zfp426 | downstream_gene_variant | MODIFIER |
| 5 | 111856595 | AX-170203125 | 0 | T | C | 0.447 | 4.67E-02 | 1.14E-01 | 2.66E+00 | 2.69E+00 | 6.83E-01 | no | | intergenic_variant | MODIFIER |
| 6 | 17503748 | AX-97785909 | 0 | T | C | 0.053 | 5.82E-02 | 1.47E-01 | 2.66E+00 | 2.79E+00 | 6.83E-01 | no | Met | intron_variant | MODIFIER |
| 1 | 124677933 | AX-170193725 | 0 | G | A | 0.57 | 3.49E-02 | 8.29E-02 | 2.57E+00 | 2.64E+00 | 6.83E-01 | no | Gm28928 | intron_variant,non_coding_transcript_variant | MODIFIER |
| 10 | 57788315 | AX-170003292 | 1 | C | T | 0.295 | 3.17E-02 | 7.99E-02 | 2.79E+00 | 2.91E+00 | 6.83E-01 | no | Fabp7 | 3_prime_UTR_variant | MODIFIER |
| 1 | 168205484 | AX-170508151 | 0 | C | T | 0.219 | 5.53E-02 | 1.37E-01 | 2.66E+00 | 2.72E+00 | 6.83E-01 | no | Pbx1 | intron_variant | MODIFIER |
| 7 | 117586016 | AX-170202869 | 2 | A | G | 0.391 | 3.46E-02 | 8.49E-02 | 2.63E+00 | 2.71E+00 | 6.84E-01 | no | Xylt1 | intron_variant | MODIFIER |
| 8 | 68894228 | AX-170260136 | 0 | G | T | 0.193 | -4.99E-02 | 1.24E-01 | 2.66E+00 | 2.74E+00 | 6.84E-01 | no | Lpl | intron_variant | MODIFIER |
| 3 | 65373626 | AX-170020708 | 0 | A | G | 0.053 | 5.54E-02 | 1.35E-01 | 2.58E+00 | 2.71E+00 | 6.84E-01 | no | Kcnab1 | intron_variant | MODIFIER |
| 1 | 47891251 | AX-170198382 | 0 | G | C | 0.114 | -7.93E-02 | 1.99E-01 | 2.71E+00 | 2.79E+00 | 6.84E-01 | no | | intergenic_variant | MODIFIER |
| 13 | 37230107 | AX-170357476 | 0 | A | T | 0.184 | -4.81E-02 | 1.17E-01 | 2.61E+00 | 2.68E+00 | 6.84E-01 | no | | intergenic_variant | MODIFIER |
| 2 | 96889883 | AX-170252738 | 0 | A | G | 0.07 | 5.59E-02 | 1.41E-01 | 2.65E+00 | 2.78E+00 | 6.84E-01 | no | Lrrc4c | intron_variant | MODIFIER |
| 13 | 45372632 | AX-170376668 | 0 | G | A | 0.474 | -2.45E-02 | 6.21E-02 | 2.71E+00 | 2.82E+00 | 6.84E-01 | no | | intergenic_variant | MODIFIER |
| 9 | 113242279 | AX-170033301 | 1 | T | G | 0.125 | -4.39E-02 | 1.09E-01 | 2.65E+00 | 2.73E+00 | 6.84E-01 | no | | intergenic_variant | MODIFIER |
| 9 | 36994842 | AX-170317652 | 1 | G | A | 0.509 | -3.09E-02 | 7.78E-02 | 2.71E+00 | 2.81E+00 | 6.84E-01 | no | Pknox2 | intron_variant | MODIFIER |
| 5 | 85455765 | AX-97833410 | 0 | A | C | 0.43 | 4.48E-02 | 1.12E-01 | 2.75E+00 | 2.80E+00 | 6.84E-01 | no | | intergenic_variant | MODIFIER |
| 20 | 87312537 | AX-97813391 | 0 | C | T | 0.246 | -2.47E-02 | 6.21E-02 | 2.67E+00 | 2.77E+00 | 6.84E-01 | no | Il1rapl1 | intron_variant | MODIFIER |
| 20 | 87451703 | AX-97854938 | 0 | A | T | 0.246 | -2.47E-02 | 6.21E-02 | 2.67E+00 | 2.77E+00 | 6.84E-01 | no | Il1rapl1 | intron_variant | MODIFIER |
| 8 | 45629039 | AX-170206779 | 0 | A | G | 0.158 | -4.39E-02 | 1.10E-01 | 2.66E+00 | 2.77E+00 | 6.84E-01 | no | Sorbs2 | intron_variant | MODIFIER |
| 5 | 100226749 | AX-170227284 | 0 | C | T | 0.43 | -4.14E-02 | 1.04E-01 | 2.73E+00 | 2.80E+00 | 6.84E-01 | no | Gm8091 | intron_variant,non_coding_transcript_variant | MODIFIER |
| 5 | 100258382 | AX-170298913 | 0 | G | A | 0.43 | -4.14E-02 | 1.04E-01 | 2.73E+00 | 2.80E+00 | 6.84E-01 | no | Gm8091 | intron_variant,non_coding_transcript_variant | MODIFIER |
| 4 | 151847958 | AX-170131663 | 0 | T | A | 0.105 | 5.71E-02 | 1.40E-01 | 2.60E+00 | 2.70E+00 | 6.84E-01 | no | Camta1 | intron_variant | MODIFIER |
| 10 | 27671366 | AX-170263537 | 0 | A | C | 0.202 | 3.52E-02 | 8.93E-02 | 2.74E+00 | 2.85E+00 | 6.84E-01 | no | Gm30165 | intron_variant,non_coding_transcript_variant | MODIFIER |
| 9 | 70451673 | AX-97844588 | 0 | A | T | 0.193 | -5.51E-02 | 1.37E-01 | 2.71E+00 | 2.73E+00 | 6.85E-01 | no | Rnf111 | intron_variant | MODIFIER |

| | | | | | | | | | | | | | | | |
|---|---|---|---|---|---|---|---|---|---|---|---|---|---|---|---|
| 7 | 73890781 | AX-170364970 | 1 | A | G | 0.366 | -3.57E-02 | 9.02E-02 | 2.69E+00 | 2.79E+00 | 6.85E-01 | no | | intergenic_variant | MODIFIER |
| 7 | 93734475 | AX-170029538 | 2 | G | A | 0.145 | 7.10E-02 | 1.77E-01 | 2.63E+00 | 2.74E+00 | 6.85E-01 | no | | intergenic_variant | MODIFIER |
| 18 | 40074395 | AX-170438360 | 0 | G | A | 0.061 | -9.29E-02 | 2.23E-01 | 2.57E+00 | 2.65E+00 | 6.85E-01 | no | AC102519.1 | intron_variant,non_coding_transcript_variant | MODIFIER |
| 8 | 32792514 | AX-170507017 | 0 | C | A | 0.158 | -4.58E-02 | 1.15E-01 | 2.67E+00 | 2.77E+00 | 6.85E-01 | no | Nrg1 | intron_variant | MODIFIER |
| 13 | 53341901 | AX-170092753 | 0 | C | T | 0.5 | -2.58E-02 | 6.60E-02 | 2.75E+00 | 2.88E+00 | 6.85E-01 | no | Sptlc1 | intron_variant | MODIFIER |
| 5 | 96597644 | AX-170256497 | 0 | A | G | 0.439 | 4.42E-02 | 1.10E-01 | 2.69E+00 | 2.74E+00 | 6.85E-01 | no | Fras1 | intron_variant | MODIFIER |
| 5 | 96614995 | AX-170025656 | 0 | G | A | 0.439 | 4.42E-02 | 1.10E-01 | 2.69E+00 | 2.74E+00 | 6.85E-01 | no | Fras1 | intron_variant | MODIFIER |
| 5 | 96615071 | AX-170025649 | 0 | C | T | 0.439 | 4.42E-02 | 1.10E-01 | 2.69E+00 | 2.74E+00 | 6.85E-01 | no | Fras1 | intron_variant | MODIFIER |
| 5 | 96620361 | AX-170025652 | 0 | A | G | 0.439 | 4.42E-02 | 1.10E-01 | 2.69E+00 | 2.74E+00 | 6.85E-01 | no | Fras1 | intron_variant | MODIFIER |
| 5 | 96639967 | AX-170494880 | 0 | C | G | 0.439 | 4.42E-02 | 1.10E-01 | 2.69E+00 | 2.74E+00 | 6.85E-01 | no | Fras1 | intron_variant | MODIFIER |
| 5 | 96644976 | AX-170501160 | 0 | T | C | 0.439 | 4.42E-02 | 1.10E-01 | 2.69E+00 | 2.74E+00 | 6.85E-01 | no | Fras1 | intron_variant | MODIFIER |
| 5 | 96647446 | AX-170513880 | 0 | C | A | 0.439 | 4.42E-02 | 1.10E-01 | 2.69E+00 | 2.74E+00 | 6.85E-01 | no | Fras1 | intron_variant | MODIFIER |
| 5 | 96650662 | AX-97807484 | 0 | A | G | 0.439 | 4.42E-02 | 1.10E-01 | 2.69E+00 | 2.74E+00 | 6.85E-01 | no | Fras1 | synonymous_variant | LOW |
| 5 | 96658590 | AX-170235853 | 0 | A | G | 0.439 | 4.42E-02 | 1.10E-01 | 2.69E+00 | 2.74E+00 | 6.85E-01 | no | Fras1 | intron_variant | MODIFIER |
| 5 | 96678491 | AX-170470568 | 0 | G | A | 0.439 | 4.42E-02 | 1.10E-01 | 2.69E+00 | 2.74E+00 | 6.85E-01 | no | Fras1 | intron_variant | MODIFIER |
| 5 | 96699568 | AX-170342900 | 0 | G | T | 0.439 | 4.42E-02 | 1.10E-01 | 2.69E+00 | 2.74E+00 | 6.85E-01 | no | Fras1 | intron_variant | MODIFIER |
| 5 | 96702288 | AX-170360309 | 0 | C | G | 0.439 | 4.42E-02 | 1.10E-01 | 2.69E+00 | 2.74E+00 | 6.85E-01 | no | Fras1 | intron_variant | MODIFIER |
| 5 | 96704485 | AX-170316790 | 0 | A | G | 0.439 | 4.42E-02 | 1.10E-01 | 2.69E+00 | 2.74E+00 | 6.85E-01 | no | Fras1 | intron_variant | MODIFIER |
| 5 | 96707369 | AX-170321496 | 0 | C | A | 0.439 | 4.42E-02 | 1.10E-01 | 2.69E+00 | 2.74E+00 | 6.85E-01 | no | Fras1 | synonymous_variant | LOW |
| 5 | 96707456 | AX-170025653 | 0 | C | T | 0.439 | 4.42E-02 | 1.10E-01 | 2.69E+00 | 2.74E+00 | 6.85E-01 | no | Fras1 | synonymous_variant | LOW |
| 5 | 96731528 | AX-170458806 | 0 | C | T | 0.439 | 4.42E-02 | 1.10E-01 | 2.69E+00 | 2.74E+00 | 6.85E-01 | no | Fras1 | intron_variant | MODIFIER |
| 5 | 96731564 | AX-97845797 | 0 | C | T | 0.439 | 4.42E-02 | 1.10E-01 | 2.69E+00 | 2.74E+00 | 6.85E-01 | no | Fras1 | intron_variant | MODIFIER |
| 5 | 96731767 | AX-170477934 | 0 | A | G | 0.439 | 4.42E-02 | 1.10E-01 | 2.69E+00 | 2.74E+00 | 6.85E-01 | no | Fras1 | intron_variant | MODIFIER |
| 5 | 96732146 | AX-170431997 | 0 | A | G | 0.439 | 4.42E-02 | 1.10E-01 | 2.69E+00 | 2.74E+00 | 6.85E-01 | no | Fras1 | intron_variant | MODIFIER |
| 5 | 96739497 | AX-170025650 | 0 | C | T | 0.439 | 4.42E-02 | 1.10E-01 | 2.69E+00 | 2.74E+00 | 6.85E-01 | no | Fras1 | intron_variant | MODIFIER |
| 5 | 96741008 | AX-170025654 | 0 | A | G | 0.439 | 4.42E-02 | 1.10E-01 | 2.69E+00 | 2.74E+00 | 6.85E-01 | no | Fras1 | synonymous_variant | LOW |
| 5 | 96752356 | AX-170455601 | 0 | A | G | 0.439 | 4.42E-02 | 1.10E-01 | 2.69E+00 | 2.74E+00 | 6.85E-01 | no | Fras1 | intron_variant | MODIFIER |
| 5 | 96754703 | AX-170140062 | 0 | G | A | 0.439 | 4.42E-02 | 1.10E-01 | 2.69E+00 | 2.74E+00 | 6.85E-01 | no | Fras1 | intron_variant | MODIFIER |
| 5 | 96761007 | AX-170473988 | 0 | C | A | 0.439 | 4.42E-02 | 1.10E-01 | 2.69E+00 | 2.74E+00 | 6.85E-01 | no | Fras1 | intron_variant | MODIFIER |
| 5 | 96761463 | AX-170025655 | 0 | A | G | 0.439 | 4.42E-02 | 1.10E-01 | 2.69E+00 | 2.74E+00 | 6.85E-01 | no | Fras1 | intron_variant | MODIFIER |
| 5 | 96778485 | AX-170025658 | 0 | C | T | 0.439 | 4.42E-02 | 1.10E-01 | 2.69E+00 | 2.74E+00 | 6.85E-01 | no | Fras1 | intron_variant | MODIFIER |
| 5 | 96780060 | AX-170281843 | 0 | T | C | 0.439 | 4.42E-02 | 1.10E-01 | 2.69E+00 | 2.74E+00 | 6.85E-01 | no | Fras1 | synonymous_variant | LOW |
| 5 | 96780099 | AX-170025657 | 0 | T | C | 0.439 | 4.42E-02 | 1.10E-01 | 2.69E+00 | 2.74E+00 | 6.85E-01 | no | Fras1 | synonymous_variant | LOW |
| 5 | 96804634 | AX-170141186 | 0 | A | G | 0.439 | 4.42E-02 | 1.10E-01 | 2.69E+00 | 2.74E+00 | 6.85E-01 | no | Anxa3 | intron_variant | MODIFIER |
| 5 | 96876857 | AX-97838323 | 0 | A | G | 0.439 | 4.42E-02 | 1.10E-01 | 2.69E+00 | 2.74E+00 | 6.85E-01 | no | Gm42601 | upstream_gene_variant | MODIFIER |
| 5 | 96877955 | AX-170130829 | 0 | C | A | 0.439 | 4.42E-02 | 1.10E-01 | 2.69E+00 | 2.74E+00 | 6.85E-01 | no | Gm42601 | upstream_gene_variant | MODIFIER |
| 5 | 96897986 | AX-97807052 | 0 | A | C | 0.439 | 4.42E-02 | 1.10E-01 | 2.69E+00 | 2.74E+00 | 6.85E-01 | no | | intergenic_variant | MODIFIER |
| 5 | 96901061 | AX-170178456 | 0 | G | A | 0.439 | 4.42E-02 | 1.10E-01 | 2.69E+00 | 2.74E+00 | 6.85E-01 | no | | regulatory_region_variant | MODIFIER |
| 5 | 96942013 | AX-170102548 | 0 | G | A | 0.439 | 4.42E-02 | 1.10E-01 | 2.69E+00 | 2.74E+00 | 6.85E-01 | no | Gm43148 | downstream_gene_variant | MODIFIER |
| 5 | 96943375 | AX-170496779 | 0 | C | T | 0.439 | 4.42E-02 | 1.10E-01 | 2.69E+00 | 2.74E+00 | 6.85E-01 | no | Gm43144 | downstream_gene_variant | MODIFIER |
| 5 | 96990288 | AX-170332680 | 0 | A | G | 0.439 | 4.42E-02 | 1.10E-01 | 2.69E+00 | 2.74E+00 | 6.85E-01 | no | | intergenic_variant | MODIFIER |
| 5 | 96993339 | AX-170112039 | 0 | G | A | 0.439 | 4.42E-02 | 1.10E-01 | 2.69E+00 | 2.74E+00 | 6.85E-01 | no | Bmp2k | upstream_gene_variant | MODIFIER |
| 5 | 97039496 | AX-170244123 | 0 | A | G | 0.439 | 4.42E-02 | 1.10E-01 | 2.69E+00 | 2.74E+00 | 6.85E-01 | no | Bmp2k | intron_variant | MODIFIER |
| 5 | 97066542 | AX-170191383 | 0 | A | G | 0.439 | 4.42E-02 | 1.10E-01 | 2.69E+00 | 2.74E+00 | 6.85E-01 | no | Bmp2k | intron_variant | MODIFIER |
| 5 | 97082043 | AX-97840772 | 0 | C | T | 0.439 | 4.42E-02 | 1.10E-01 | 2.69E+00 | 2.74E+00 | 6.85E-01 | no | Bmp2k | intron_variant | MODIFIER |
| 5 | 97082258 | AX-170496283 | 0 | G | C | 0.439 | 4.42E-02 | 1.10E-01 | 2.69E+00 | 2.74E+00 | 6.85E-01 | no | Bmp2k | intron_variant | MODIFIER |
| 5 | 97083082 | AX-170168035 | 0 | A | G | 0.439 | 4.42E-02 | 1.10E-01 | 2.69E+00 | 2.74E+00 | 6.85E-01 | no | Bmp2k | intron_variant | MODIFIER |
| 5 | 97096377 | AX-97782029 | 0 | C | T | 0.439 | 4.42E-02 | 1.10E-01 | 2.69E+00 | 2.74E+00 | 6.85E-01 | no | Bmp2k | downstream_gene_variant | MODIFIER |
| 5 | 97097303 | AX-170108585 | 0 | A | G | 0.439 | 4.42E-02 | 1.10E-01 | 2.69E+00 | 2.74E+00 | 6.85E-01 | no | Paqr3 | intron_variant | MODIFIER |
| 5 | 97125989 | AX-97836138 | 0 | T | A | 0.439 | 4.42E-02 | 1.10E-01 | 2.69E+00 | 2.74E+00 | 6.85E-01 | no | | regulatory_region_variant | MODIFIER |
| 5 | 97157757 | AX-97806267 | 0 | T | C | 0.439 | 4.42E-02 | 1.10E-01 | 2.69E+00 | 2.74E+00 | 6.85E-01 | no | | intergenic_variant | MODIFIER |
| 5 | 97170643 | AX-170025671 | 0 | C | T | 0.439 | 4.42E-02 | 1.10E-01 | 2.69E+00 | 2.74E+00 | 6.85E-01 | no | | intergenic_variant | MODIFIER |
| 5 | 97174429 | AX-170072824 | 0 | A | G | 0.439 | 4.42E-02 | 1.10E-01 | 2.69E+00 | 2.74E+00 | 6.85E-01 | no | | intergenic_variant | MODIFIER |
| 5 | 97221267 | AX-170025664 | 0 | G | A | 0.439 | 4.42E-02 | 1.10E-01 | 2.69E+00 | 2.74E+00 | 6.85E-01 | no | | intergenic_variant | MODIFIER |

| | | | | | | | | | | | | | | | |
|---|---|---|---|---|---|---|---|---|---|---|---|---|---|---|---|
| 5 | 97221956 | AX-97802087 | 0 | G | A | 0.439 | 4.42E-02 | 1.10E-01 | 2.69E+00 | 2.74E+00 | 6.85E-01 | no | | intergenic_variant | MODIFIER |
| 5 | 97264972 | AX-170025668 | 0 | C | A | 0.439 | 4.42E-02 | 1.10E-01 | 2.69E+00 | 2.74E+00 | 6.85E-01 | no | Gm5560 | non_coding_transcript_exon_variant | MODIFIER |
| 5 | 97277971 | AX-97857482 | 0 | T | C | 0.439 | 4.42E-02 | 1.10E-01 | 2.69E+00 | 2.74E+00 | 6.85E-01 | no | | regulatory_region_variant | MODIFIER |
| 5 | 97322024 | AX-170136601 | 0 | G | A | 0.439 | 4.42E-02 | 1.10E-01 | 2.69E+00 | 2.74E+00 | 6.85E-01 | no | Gm33969 | upstream_gene_variant | MODIFIER |
| 5 | 97355060 | AX-170317346 | 0 | A | G | 0.439 | 4.42E-02 | 1.10E-01 | 2.69E+00 | 2.74E+00 | 6.85E-01 | no | | intergenic_variant | MODIFIER |
| 5 | 97371496 | AX-170204806 | 0 | G | A | 0.439 | 4.42E-02 | 1.10E-01 | 2.69E+00 | 2.74E+00 | 6.85E-01 | no | | intergenic_variant | MODIFIER |
| 5 | 97384496 | AX-97851301 | 0 | A | G | 0.439 | 4.42E-02 | 1.10E-01 | 2.69E+00 | 2.74E+00 | 6.85E-01 | no | Naa11 | 3_prime_UTR_variant | MODIFIER |
| 5 | 97388417 | AX-170274504 | 0 | T | C | 0.439 | 4.42E-02 | 1.10E-01 | 2.69E+00 | 2.74E+00 | 6.85E-01 | no | Naa11 | intron_variant | MODIFIER |
| 5 | 97422470 | AX-170517659 | 0 | A | G | 0.439 | 4.42E-02 | 1.10E-01 | 2.69E+00 | 2.74E+00 | 6.85E-01 | no | 4930467D21Rik | intron_variant,non_coding_transcript_variant | MODIFIER |
| 5 | 97429803 | AX-170101454 | 0 | T | C | 0.439 | 4.42E-02 | 1.10E-01 | 2.69E+00 | 2.74E+00 | 6.85E-01 | no | 4930467D21Rik | intron_variant,non_coding_transcript_variant | MODIFIER |
| 5 | 97429839 | AX-170507579 | 0 | G | A | 0.439 | 4.42E-02 | 1.10E-01 | 2.69E+00 | 2.74E+00 | 6.85E-01 | no | 4930467D21Rik | intron_variant,non_coding_transcript_variant | MODIFIER |
| 5 | 97476552 | AX-170268947 | 0 | C | G | 0.439 | 4.42E-02 | 1.10E-01 | 2.69E+00 | 2.74E+00 | 6.85E-01 | no | 4930467D21Rik | intron_variant,non_coding_transcript_variant | MODIFIER |
| 5 | 97478526 | AX-170457251 | 0 | T | G | 0.439 | 4.42E-02 | 1.10E-01 | 2.69E+00 | 2.74E+00 | 6.85E-01 | no | 4930467D21Rik | intron_variant,non_coding_transcript_variant | MODIFIER |
| 5 | 97498703 | AX-170183909 | 0 | T | C | 0.439 | 4.42E-02 | 1.10E-01 | 2.69E+00 | 2.74E+00 | 6.85E-01 | no | 4930467D21Rik | intron_variant,non_coding_transcript_variant | MODIFIER |
| 5 | 97500450 | AX-170198843 | 0 | C | G | 0.439 | 4.42E-02 | 1.10E-01 | 2.69E+00 | 2.74E+00 | 6.85E-01 | no | 4930467D21Rik | intron_variant,non_coding_transcript_variant | MODIFIER |
| 5 | 97502814 | AX-170127178 | 0 | G | C | 0.439 | 4.42E-02 | 1.10E-01 | 2.69E+00 | 2.74E+00 | 6.85E-01 | no | 4930467D21Rik | intron_variant,non_coding_transcript_variant | MODIFIER |
| 5 | 97513127 | AX-170450096 | 0 | A | T | 0.439 | 4.42E-02 | 1.10E-01 | 2.69E+00 | 2.74E+00 | 6.85E-01 | no | 4930467D21Rik | intron_variant,non_coding_transcript_variant | MODIFIER |
| 5 | 97518282 | AX-170146935 | 0 | G | A | 0.439 | 4.42E-02 | 1.10E-01 | 2.69E+00 | 2.74E+00 | 6.85E-01 | no | 4930467D21Rik | intron_variant,non_coding_transcript_variant | MODIFIER |
| 5 | 97533482 | AX-170478028 | 0 | G | A | 0.439 | 4.42E-02 | 1.10E-01 | 2.69E+00 | 2.74E+00 | 6.85E-01 | no | 4930467D21Rik | intron_variant,non_coding_transcript_variant | MODIFIER |
| 5 | 97599930 | AX-170454825 | 0 | A | C | 0.439 | 4.42E-02 | 1.10E-01 | 2.69E+00 | 2.74E+00 | 6.85E-01 | no | 4930467D21Rik | intron_variant,non_coding_transcript_variant | MODIFIER |
| 5 | 97814893 | AX-97781779 | 0 | A | G | 0.439 | 4.42E-02 | 1.10E-01 | 2.69E+00 | 2.74E+00 | 6.85E-01 | no | | regulatory_region_variant | MODIFIER |
| 1 | 115905782 | AX-170191558 | 1 | A | T | 0.161 | 7.11E-02 | 1.75E-01 | 2.61E+00 | 2.70E+00 | 6.85E-01 | no | Cntnap5a | intron_variant | MODIFIER |
| 1 | 184384939 | AX-170480406 | 0 | A | T | 0.219 | 5.04E-02 | 1.16E-01 | 2.51E+00 | 2.60E+00 | 6.85E-01 | no | Gm37223 | intron_variant,non_coding_transcript_variant | MODIFIER |
| 8 | 69011623 | AX-170292610 | 0 | G | T | 0.184 | -4.88E-02 | 1.22E-01 | 2.65E+00 | 2.74E+00 | 6.85E-01 | no | | intergenic_variant | MODIFIER |
| 1 | 118955201 | AX-170371293 | 2 | T | C | 0.155 | 7.49E-02 | 1.83E-01 | 2.59E+00 | 2.69E+00 | 6.85E-01 | no | Gli2 | intron_variant | MODIFIER |
| 15 | 74522566 | AX-97845715 | 1 | C | T | 0.545 | 3.29E-02 | 8.06E-02 | 2.60E+00 | 2.69E+00 | 6.86E-01 | no | Adgrb1 | intron_variant | MODIFIER |
| 1 | 108828371 | AX-170379874 | 0 | G | A | 0.158 | 7.17E-02 | 1.74E-01 | 2.58E+00 | 2.67E+00 | 6.86E-01 | no | | intergenic_variant | MODIFIER |
| 1 | 118750536 | AX-170001529 | 0 | A | G | 0.158 | 7.17E-02 | 1.74E-01 | 2.58E+00 | 2.67E+00 | 6.86E-01 | no | | intergenic_variant | MODIFIER |
| 7 | 118045872 | AX-170078578 | 1 | A | G | 0.402 | 3.65E-02 | 9.08E-02 | 2.66E+00 | 2.71E+00 | 6.86E-01 | no | | intergenic_variant | MODIFIER |
| 7 | 118458187 | AX-170149882 | 1 | C | T | 0.402 | 3.65E-02 | 9.08E-02 | 2.66E+00 | 2.71E+00 | 6.86E-01 | no | | intergenic_variant | MODIFIER |
| 7 | 118969057 | AX-170078600 | 1 | C | T | 0.402 | 3.65E-02 | 9.08E-02 | 2.66E+00 | 2.71E+00 | 6.86E-01 | no | Gprc5b | downstream_gene_variant | MODIFIER |
| 7 | 119168586 | AX-170300658 | 1 | C | G | 0.402 | 3.65E-02 | 9.08E-02 | 2.66E+00 | 2.71E+00 | 6.86E-01 | no | Gpr139 | intron_variant | MODIFIER |
| 1 | 107486168 | AX-170001301 | 1 | C | G | 0.027 | -6.08E-02 | 1.54E-01 | 2.58E+00 | 2.76E+00 | 6.86E-01 | no | | intergenic_variant | MODIFIER |
| 20 | 72443175 | AX-97856867 | 0 | G | A | 0.263 | 2.46E-02 | 6.20E-02 | 2.68E+00 | 2.77E+00 | 6.86E-01 | no | Gabra3 | intron_variant | MODIFIER |
| 5 | 117504840 | AX-97800657 | 0 | G | T | 0.456 | 4.68E-02 | 1.16E-01 | 2.66E+00 | 2.69E+00 | 6.86E-01 | no | Ksr2 | intron_variant,non_coding_transcript_variant | MODIFIER |
| 2 | 113677379 | AX-97831367 | 0 | C | T | 0.149 | 7.21E-02 | 1.77E-01 | 2.61E+00 | 2.70E+00 | 6.86E-01 | no | Fmn1 | intron_variant | MODIFIER |
| 2 | 158531321 | AX-170287704 | 0 | G | A | 0.149 | 7.21E-02 | 1.77E-01 | 2.61E+00 | 2.70E+00 | 6.86E-01 | no | Arhgap40 | synonymous_variant | LOW |
| 1 | 108671667 | AX-170243928 | 1 | G | C | 0.17 | 6.59E-02 | 1.59E-01 | 2.57E+00 | 2.66E+00 | 6.86E-01 | no | | intergenic_variant | MODIFIER |
| 8 | 47381845 | AX-170185386 | 1 | C | T | 0.17 | 5.00E-02 | 1.27E-01 | 2.71E+00 | 2.80E+00 | 6.86E-01 | no | Stox2 | intron_variant,non_coding_transcript_variant | MODIFIER |
| 11 | 30143941 | AX-97853970 | 0 | G | A | 0.096 | 7.80E-02 | 1.95E-01 | 2.68E+00 | 2.74E+00 | 6.86E-01 | no | Sptbn1 | intron_variant | MODIFIER |
| 11 | 30144193 | AX-170505749 | 0 | C | G | 0.096 | 7.80E-02 | 1.95E-01 | 2.68E+00 | 2.74E+00 | 6.86E-01 | no | Sptbn1 | synonymous_variant | LOW |
| 11 | 32165578 | AX-170530469 | 0 | G | A | 0.096 | 7.80E-02 | 1.95E-01 | 2.68E+00 | 2.74E+00 | 6.86E-01 | no | | intergenic_variant | MODIFIER |
| 11 | 32196037 | AX-97817961 | 0 | C | T | 0.096 | 7.80E-02 | 1.95E-01 | 2.68E+00 | 2.74E+00 | 6.86E-01 | no | Il9r | intron_variant,NMD_transcript_variant | MODIFIER |
| 11 | 32196074 | AX-170495569 | 0 | A | G | 0.096 | 7.80E-02 | 1.95E-01 | 2.68E+00 | 2.74E+00 | 6.86E-01 | no | Il9r | intron_variant,NMD_transcript_variant | MODIFIER |
| 11 | 32324980 | AX-170525218 | 0 | T | C | 0.096 | 7.80E-02 | 1.95E-01 | 2.68E+00 | 2.74E+00 | 6.86E-01 | no | | regulatory_region_variant | MODIFIER |
| 11 | 32325158 | AX-170402446 | 0 | A | C | 0.096 | 7.80E-02 | 1.95E-01 | 2.68E+00 | 2.74E+00 | 6.86E-01 | no | | regulatory_region_variant | MODIFIER |
| 11 | 32325288 | AX-170509314 | 0 | T | A | 0.096 | 7.80E-02 | 1.95E-01 | 2.68E+00 | 2.74E+00 | 6.86E-01 | no | | intergenic_variant | MODIFIER |
| 11 | 32325509 | AX-170420006 | 0 | G | A | 0.096 | 7.80E-02 | 1.95E-01 | 2.68E+00 | 2.74E+00 | 6.86E-01 | no | | intergenic_variant | MODIFIER |
| 11 | 32329001 | AX-170503776 | 0 | A | C | 0.096 | 7.80E-02 | 1.95E-01 | 2.68E+00 | 2.74E+00 | 6.86E-01 | no | | intergenic_variant | MODIFIER |
| 11 | 32359676 | AX-170340413 | 0 | C | A | 0.096 | 7.80E-02 | 1.95E-01 | 2.68E+00 | 2.74E+00 | 6.86E-01 | no | Sh3pxd2b | intron_variant | MODIFIER |
| 11 | 32396076 | AX-170467172 | 0 | G | A | 0.096 | 7.80E-02 | 1.95E-01 | 2.68E+00 | 2.74E+00 | 6.86E-01 | no | Sh3pxd2b | intron_variant | MODIFIER |
| 11 | 32548049 | AX-170259819 | 0 | G | A | 0.096 | 7.80E-02 | 1.95E-01 | 2.68E+00 | 2.74E+00 | 6.86E-01 | no | Stk10 | intron_variant | MODIFIER |
| 11 | 32552734 | AX-170046723 | 0 | G | A | 0.096 | 7.80E-02 | 1.95E-01 | 2.68E+00 | 2.74E+00 | 6.86E-01 | no | Stk10 | intron_variant | MODIFIER |
| 11 | 32553134 | AX-170181816 | 0 | G | A | 0.096 | 7.80E-02 | 1.95E-01 | 2.68E+00 | 2.74E+00 | 6.86E-01 | no | Stk10 | intron_variant | MODIFIER |

| | | | | | | | | | | | | | | | |
|---|---|---|---|---|---|---|---|---|---|---|---|---|---|---|---|
| 11 | 32564318 | AX-170004650 | 0 | G | A | 0.096 | 7.80E-02 | 1.95E-01 | 2.68E+00 | 2.74E+00 | 6.86E-01 | no | Stk10 | intron_variant | MODIFIER |
| 11 | 32566722 | AX-170452118 | 0 | C | T | 0.096 | 7.80E-02 | 1.95E-01 | 2.68E+00 | 2.74E+00 | 6.86E-01 | no | Stk10 | intron_variant | MODIFIER |
| 11 | 32577333 | AX-97801755 | 0 | C | T | 0.096 | 7.80E-02 | 1.95E-01 | 2.68E+00 | 2.74E+00 | 6.86E-01 | no | Stk10 | intron_variant | MODIFIER |
| 11 | 32605021 | AX-170287969 | 0 | T | G | 0.096 | 7.80E-02 | 1.95E-01 | 2.68E+00 | 2.74E+00 | 6.86E-01 | no | Stk10 | intron_variant | MODIFIER |
| 11 | 32649598 | AX-97820535 | 0 | A | T | 0.096 | 7.80E-02 | 1.95E-01 | 2.68E+00 | 2.74E+00 | 6.86E-01 | no | Fbxw11 | intron_variant | MODIFIER |
| 11 | 32665446 | AX-170363556 | 0 | T | A | 0.096 | 7.80E-02 | 1.95E-01 | 2.68E+00 | 2.74E+00 | 6.86E-01 | no | Fbxw11 | intron_variant | MODIFIER |
| 11 | 32674623 | AX-170142414 | 0 | C | T | 0.096 | 7.80E-02 | 1.95E-01 | 2.68E+00 | 2.74E+00 | 6.86E-01 | no | Fbxw11 | intron_variant | MODIFIER |
| 11 | 32674737 | AX-170524340 | 0 | T | A | 0.096 | 7.80E-02 | 1.95E-01 | 2.68E+00 | 2.74E+00 | 6.86E-01 | no | Fbxw11 | intron_variant | MODIFIER |
| 11 | 32724371 | AX-170165290 | 0 | A | G | 0.096 | 7.80E-02 | 1.95E-01 | 2.68E+00 | 2.74E+00 | 6.86E-01 | no | Fbxw11 | intron_variant | MODIFIER |
| 11 | 32727215 | AX-170362295 | 0 | T | C | 0.096 | 7.80E-02 | 1.95E-01 | 2.68E+00 | 2.74E+00 | 6.86E-01 | no | Fbxw11 | intron_variant | MODIFIER |
| 11 | 32765937 | AX-170259394 | 0 | C | G | 0.096 | 7.80E-02 | 1.95E-01 | 2.68E+00 | 2.74E+00 | 6.86E-01 | no | | intergenic_variant | MODIFIER |
| 11 | 32787811 | AX-170514074 | 0 | G | T | 0.096 | 7.80E-02 | 1.95E-01 | 2.68E+00 | 2.74E+00 | 6.86E-01 | no | 4930555O08Rik | downstream_gene_variant | MODIFIER |
| 11 | 32790322 | AX-170454704 | 0 | C | T | 0.096 | 7.80E-02 | 1.95E-01 | 2.68E+00 | 2.74E+00 | 6.86E-01 | no | 4930555O08Rik | intron_variant,non_coding_transcript_variant | MODIFIER |
| 11 | 32795593 | AX-170385867 | 0 | C | T | 0.096 | 7.80E-02 | 1.95E-01 | 2.68E+00 | 2.74E+00 | 6.86E-01 | no | 4930555O08Rik | upstream_gene_variant | MODIFIER |
| 11 | 32795630 | AX-97795412 | 0 | G | T | 0.096 | 7.80E-02 | 1.95E-01 | 2.68E+00 | 2.74E+00 | 6.86E-01 | no | 4930555O08Rik | upstream_gene_variant | MODIFIER |
| 11 | 32796632 | AX-170118329 | 0 | G | A | 0.096 | 7.80E-02 | 1.95E-01 | 2.68E+00 | 2.74E+00 | 6.86E-01 | no | | intergenic_variant | MODIFIER |
| 11 | 32796652 | AX-170391794 | 0 | T | C | 0.096 | 7.80E-02 | 1.95E-01 | 2.68E+00 | 2.74E+00 | 6.86E-01 | no | | intergenic_variant | MODIFIER |
| 11 | 32797965 | AX-170335690 | 0 | C | A | 0.096 | 7.80E-02 | 1.95E-01 | 2.68E+00 | 2.74E+00 | 6.86E-01 | no | | intergenic_variant | MODIFIER |
| 11 | 32801315 | AX-170046730 | 0 | G | A | 0.096 | 7.80E-02 | 1.95E-01 | 2.68E+00 | 2.74E+00 | 6.86E-01 | no | | regulatory_region_variant | MODIFIER |
| 11 | 32804597 | AX-170425842 | 0 | C | A | 0.096 | 7.80E-02 | 1.95E-01 | 2.68E+00 | 2.74E+00 | 6.86E-01 | no | | intergenic_variant | MODIFIER |
| 11 | 32822734 | AX-170090881 | 0 | T | C | 0.096 | 7.80E-02 | 1.95E-01 | 2.68E+00 | 2.74E+00 | 6.86E-01 | no | Smim23 | intron_variant | MODIFIER |
| 11 | 32830801 | AX-170316486 | 0 | G | A | 0.096 | 7.80E-02 | 1.95E-01 | 2.68E+00 | 2.74E+00 | 6.86E-01 | no | | intergenic_variant | MODIFIER |
| 11 | 32843816 | AX-97839694 | 0 | C | A | 0.096 | 7.80E-02 | 1.95E-01 | 2.68E+00 | 2.74E+00 | 6.86E-01 | no | | intergenic_variant | MODIFIER |
| 11 | 32859343 | AX-170463117 | 0 | A | G | 0.096 | 7.80E-02 | 1.95E-01 | 2.68E+00 | 2.74E+00 | 6.86E-01 | no | | intergenic_variant | MODIFIER |
| 11 | 32859551 | AX-170530096 | 0 | A | G | 0.096 | 7.80E-02 | 1.95E-01 | 2.68E+00 | 2.74E+00 | 6.86E-01 | no | | intergenic_variant | MODIFIER |
| 11 | 32867096 | AX-170173296 | 0 | C | T | 0.096 | 7.80E-02 | 1.95E-01 | 2.68E+00 | 2.74E+00 | 6.86E-01 | no | | intergenic_variant | MODIFIER |
| 11 | 32892114 | AX-97834202 | 0 | T | C | 0.096 | 7.80E-02 | 1.95E-01 | 2.68E+00 | 2.74E+00 | 6.86E-01 | no | | intergenic_variant | MODIFIER |
| 11 | 32892154 | AX-170263509 | 0 | G | C | 0.096 | 7.80E-02 | 1.95E-01 | 2.68E+00 | 2.74E+00 | 6.86E-01 | no | | intergenic_variant | MODIFIER |
| 11 | 32893328 | AX-170522828 | 0 | G | A | 0.096 | 7.80E-02 | 1.95E-01 | 2.68E+00 | 2.74E+00 | 6.86E-01 | no | | intergenic_variant | MODIFIER |
| 11 | 32898358 | AX-170326380 | 0 | C | G | 0.096 | 7.80E-02 | 1.95E-01 | 2.68E+00 | 2.74E+00 | 6.86E-01 | no | Gm12111 | downstream_gene_variant | MODIFIER |
| 11 | 32908603 | AX-170452062 | 0 | C | T | 0.096 | 7.80E-02 | 1.95E-01 | 2.68E+00 | 2.74E+00 | 6.86E-01 | no | Gm12112 | upstream_gene_variant | MODIFIER |
| 11 | 32912933 | AX-170395613 | 0 | C | T | 0.096 | 7.80E-02 | 1.95E-01 | 2.68E+00 | 2.74E+00 | 6.86E-01 | no | Gm12112 | downstream_gene_variant | MODIFIER |
| 12 | 104978447 | AX-170107519 | 0 | G | T | 0.123 | -7.39E-02 | 1.87E-01 | 2.72E+00 | 2.79E+00 | 6.86E-01 | no | Syne3 | intron_variant | MODIFIER |
| 20 | 75728354 | AX-170110111 | 0 | A | G | 0.254 | 2.52E-02 | 6.33E-02 | 2.66E+00 | 2.75E+00 | 6.86E-01 | no | Gm7153 | downstream_gene_variant | MODIFIER |
| 20 | 75834848 | AX-170278379 | 0 | G | C | 0.254 | 2.52E-02 | 6.33E-02 | 2.66E+00 | 2.75E+00 | 6.86E-01 | no | Pls3 | intron_variant | MODIFIER |
| 20 | 77604499 | AX-170418576 | 0 | A | G | 0.254 | 2.52E-02 | 6.33E-02 | 2.66E+00 | 2.75E+00 | 6.86E-01 | no | Tbl1x | intron_variant | MODIFIER |
| 1 | 169148037 | AX-170130636 | 0 | G | C | 0.237 | -4.39E-02 | 1.12E-01 | 2.73E+00 | 2.83E+00 | 6.86E-01 | no | | intergenic_variant | MODIFIER |
| 1 | 184916067 | AX-170511576 | 0 | T | G | 0.237 | -4.09E-02 | 1.03E-01 | 2.84E+00 | 2.97E+00 | 6.86E-01 | no | Mark1 | intron_variant | MODIFIER |
| 9 | 35518115 | AX-170291383 | 1 | A | G | 0.429 | -3.54E-02 | 8.91E-02 | 2.77E+00 | 2.84E+00 | 6.87E-01 | no | Gm1113 | missense_variant | MODERATE |
| 13 | 53857768 | AX-170051328 | 1 | C | A | 0.509 | 2.82E-02 | 6.79E-02 | 2.56E+00 | 2.67E+00 | 6.87E-01 | no | | intergenic_variant | MODIFIER |
| 6 | 114717597 | AX-170028032 | 0 | G | A | 0.079 | -6.71E-02 | 1.68E-01 | 2.61E+00 | 2.74E+00 | 6.87E-01 | no | Atg7 | intron_variant | MODIFIER |
| 1 | 26046456 | AX-170334878 | 0 | T | G | 0.404 | 3.43E-02 | 8.53E-02 | 2.65E+00 | 2.72E+00 | 6.87E-01 | no | | intergenic_variant | MODIFIER |
| 11 | 32675858 | AX-170046725 | 1 | G | T | 0.089 | 8.18E-02 | 2.06E-01 | 2.71E+00 | 2.76E+00 | 6.87E-01 | no | Fbxw11 | intron_variant | MODIFIER |
| 1 | 25891881 | AX-97824297 | 0 | A | C | 0.412 | 3.31E-02 | 8.23E-02 | 2.64E+00 | 2.71E+00 | 6.87E-01 | no | | intergenic_variant | MODIFIER |
| 8 | 19271523 | AX-97819317 | 0 | T | C | 0.175 | 3.95E-02 | 1.00E-01 | 2.65E+00 | 2.76E+00 | 6.87E-01 | no | | intergenic_variant | MODIFIER |
| 12 | 100805830 | AX-170363156 | 1 | G | A | 0.08 | -6.75E-02 | 1.69E-01 | 2.66E+00 | 2.74E+00 | 6.87E-01 | no | Dglucy | intron_variant | MODIFIER |
| 1 | 175268090 | AX-170496185 | 0 | A | G | 0.237 | 4.77E-02 | 1.12E-01 | 2.53E+00 | 2.60E+00 | 6.87E-01 | no | Rgs7 | intron_variant | MODIFIER |
| 7 | 102441389 | AX-170029653 | 0 | T | G | 0.456 | 3.71E-02 | 9.17E-02 | 2.63E+00 | 2.69E+00 | 6.87E-01 | no | Rrm1 | upstream_gene_variant | MODIFIER |
| 5 | 78791486 | AX-170341111 | 0 | C | T | 0.421 | 4.04E-02 | 1.01E-01 | 2.67E+00 | 2.74E+00 | 6.87E-01 | no | | intergenic_variant | MODIFIER |
| 1 | 189716989 | AX-170391437 | 0 | G | A | 0.254 | 4.19E-02 | 9.57E-02 | 2.48E+00 | 2.59E+00 | 6.87E-01 | no | | intergenic_variant | MODIFIER |
| 5 | 83479707 | AX-97835103 | 0 | G | A | 0.43 | 4.27E-02 | 1.07E-01 | 2.69E+00 | 2.74E+00 | 6.87E-01 | no | | intergenic_variant | MODIFIER |
| 7 | 52321354 | AX-170270967 | 0 | T | C | 0.43 | 4.02E-02 | 9.97E-02 | 2.63E+00 | 2.70E+00 | 6.87E-01 | no | | intergenic_variant | MODIFIER |
| 1 | 106399412 | AX-170166812 | 1 | G | A | 0.161 | 7.72E-02 | 1.89E-01 | 2.59E+00 | 2.67E+00 | 6.87E-01 | no | A530053M12Rik | downstream_gene_variant | MODIFIER |
| 7 | 10285438 | AX-170286743 | 0 | T | A | 0.175 | 3.09E-02 | 8.02E-02 | 2.72E+00 | 2.86E+00 | 6.88E-01 | no | Vmn1r-ps47 | upstream_gene_variant | MODIFIER |

| | | | | | | | | | | | | | | | |
|---|---|---|---|---|---|---|---|---|---|---|---|---|---|---|---|
| 13 | 36350974 | AX-170106468 | 1 | C | T | 0.188 | -4.64E-02 | 1.15E-01 | 2.60E+00 | 2.69E+00 | 6.88E-01 | no | Fars2 | intron_variant | MODIFIER |
| 18 | 72241098 | AX-170379788 | 0 | A | G | 0.184 | -4.53E-02 | 1.13E-01 | 2.63E+00 | 2.70E+00 | 6.88E-01 | no | Dcc | intron_variant | MODIFIER |
| 13 | 46233564 | AX-170498210 | 0 | T | A | 0.491 | -2.42E-02 | 6.22E-02 | 2.70E+00 | 2.81E+00 | 6.88E-01 | no | | intergenic_variant | MODIFIER |
| 5 | 73660280 | AX-170244893 | 0 | T | C | 0.456 | 4.06E-02 | 1.02E-01 | 2.65E+00 | 2.72E+00 | 6.88E-01 | no | Spata18 | intron_variant | MODIFIER |
| 5 | 73661989 | AX-170089521 | 0 | C | T | 0.456 | 4.06E-02 | 1.02E-01 | 2.65E+00 | 2.72E+00 | 6.88E-01 | no | Spata18 | intron_variant | MODIFIER |
| 5 | 73671384 | AX-97797511 | 0 | A | G | 0.456 | 4.06E-02 | 1.02E-01 | 2.65E+00 | 2.72E+00 | 6.88E-01 | no | Spata18 | intron_variant | MODIFIER |
| 5 | 73766161 | AX-170204727 | 0 | C | T | 0.456 | 4.06E-02 | 1.02E-01 | 2.65E+00 | 2.72E+00 | 6.88E-01 | no | | intergenic_variant | MODIFIER |
| 5 | 73964668 | AX-170025293 | 0 | G | A | 0.456 | 4.06E-02 | 1.02E-01 | 2.65E+00 | 2.72E+00 | 6.88E-01 | no | | intergenic_variant | MODIFIER |
| 5 | 73975051 | AX-170025289 | 0 | C | T | 0.456 | 4.06E-02 | 1.02E-01 | 2.65E+00 | 2.72E+00 | 6.88E-01 | no | | intergenic_variant | MODIFIER |
| 5 | 74985752 | AX-170382083 | 0 | T | C | 0.456 | 4.06E-02 | 1.02E-01 | 2.65E+00 | 2.72E+00 | 6.88E-01 | no | Gm42577 | downstream_gene_variant | MODIFIER |
| 4 | 72014259 | AX-170511645 | 1 | C | T | 0.339 | 2.67E-02 | 6.23E-02 | 2.51E+00 | 2.61E+00 | 6.88E-01 | no | | regulatory_region_variant | MODIFIER |
| 2 | 130120547 | AX-170246696 | 1 | G | A | 0.152 | -6.02E-02 | 1.55E-01 | 2.69E+00 | 2.81E+00 | 6.88E-01 | no | Tgm6 | upstream_gene_variant | MODIFIER |
| 9 | 17493453 | AX-170466957 | 0 | G | A | 0.482 | 3.64E-02 | 8.74E-02 | 2.56E+00 | 2.64E+00 | 6.88E-01 | no | | intergenic_variant | MODIFIER |
| 7 | 77136668 | AX-170029351 | 0 | C | T | 0.377 | -3.81E-02 | 9.71E-02 | 2.67E+00 | 2.77E+00 | 6.88E-01 | no | | intergenic_variant | MODIFIER |
| 8 | 116322009 | AX-170169456 | 1 | C | T | 0.259 | 3.72E-02 | 9.53E-02 | 2.67E+00 | 2.79E+00 | 6.88E-01 | no | | intergenic_variant | MODIFIER |
| 7 | 38018856 | AX-170266017 | 0 | C | T | 0.351 | 3.81E-02 | 9.28E-02 | 2.58E+00 | 2.66E+00 | 6.88E-01 | no | Uri1 | intron_variant | MODIFIER |
| 2 | 122822865 | AX-170225726 | 0 | C | T | 0.114 | 7.97E-02 | 2.01E-01 | 2.67E+00 | 2.73E+00 | 6.88E-01 | no | | intergenic_variant | MODIFIER |
| 9 | 60479468 | AX-97825679 | 0 | C | T | 0.228 | 4.43E-02 | 1.13E-01 | 2.72E+00 | 2.81E+00 | 6.88E-01 | no | Thsd4 | intron_variant | MODIFIER |
| 8 | 41915760 | AX-170460729 | 1 | A | C | 0.152 | 5.92E-02 | 1.50E-01 | 2.76E+00 | 2.81E+00 | 6.88E-01 | no | 2810404M03Rik | intron_variant,non_coding_transcript_variant | MODIFIER |
| 8 | 42815456 | AX-97798397 | 1 | G | C | 0.152 | 5.92E-02 | 1.50E-01 | 2.76E+00 | 2.81E+00 | 6.88E-01 | no | | intergenic_variant | MODIFIER |
| 15 | 74261540 | AX-170150296 | 0 | A | G | 0.535 | 3.33E-02 | 8.16E-02 | 2.59E+00 | 2.67E+00 | 6.89E-01 | no | | intergenic_variant | MODIFIER |
| 1 | 60821317 | AX-97807655 | 0 | A | G | 0.105 | -7.38E-02 | 1.86E-01 | 2.65E+00 | 2.74E+00 | 6.89E-01 | no | | regulatory_region_variant | MODIFIER |
| 9 | 19205060 | AX-97854704 | 0 | T | C | 0.456 | 3.71E-02 | 9.01E-02 | 2.57E+00 | 2.65E+00 | 6.89E-01 | no | Olfr840-ps1 | non_coding_transcript_exon_variant | MODIFIER |
| 5 | 126123697 | AX-170412600 | 0 | C | A | 0.447 | -4.21E-02 | 1.07E-01 | 2.74E+00 | 2.80E+00 | 6.89E-01 | no | | intergenic_variant | MODIFIER |
| 5 | 126204508 | AX-170026111 | 0 | T | C | 0.447 | -4.21E-02 | 1.07E-01 | 2.74E+00 | 2.80E+00 | 6.89E-01 | no | | intergenic_variant | MODIFIER |
| 13 | 53116651 | AX-170272286 | 0 | C | A | 0.518 | 2.67E-02 | 6.53E-02 | 2.57E+00 | 2.68E+00 | 6.89E-01 | no | Ror2 | intron_variant | MODIFIER |
| 7 | 46190983 | AX-170481885 | 1 | C | T | 0.33 | 3.80E-02 | 9.42E-02 | 2.61E+00 | 2.69E+00 | 6.89E-01 | no | Ush1c | downstream_gene_variant | MODIFIER |
| 16 | 18360588 | AX-170344918 | 0 | T | G | 0.132 | 3.66E-02 | 9.42E-02 | 2.65E+00 | 2.78E+00 | 6.89E-01 | no | Arvcf | intron_variant | MODIFIER |
| 16 | 18871812 | AX-170364478 | 0 | T | C | 0.132 | 3.66E-02 | 9.42E-02 | 2.65E+00 | 2.78E+00 | 6.89E-01 | no | Mrpl40 | downstream_gene_variant | MODIFIER |
| 18 | 72244090 | AX-168412495 | 0 | T | C | 0.193 | -4.13E-02 | 1.04E-01 | 2.65E+00 | 2.73E+00 | 6.89E-01 | no | Dcc | intron_variant | MODIFIER |
| 18 | 43321805 | AX-170361453 | 0 | C | T | 0.07 | 8.20E-02 | 2.09E-01 | 2.65E+00 | 2.76E+00 | 6.89E-01 | no | Dpysl3 | downstream_gene_variant | MODIFIER |
| 15 | 94143138 | AX-170352350 | 1 | A | G | 0.134 | 6.62E-02 | 1.65E-01 | 2.61E+00 | 2.70E+00 | 6.89E-01 | no | | intergenic_variant | MODIFIER |
| 8 | 68498423 | AX-170397290 | 1 | A | G | 0.188 | -5.27E-02 | 1.33E-01 | 2.68E+00 | 2.74E+00 | 6.89E-01 | no | Csgalnact1 | intron_variant | MODIFIER |
| 3 | 48911447 | AX-170020490 | 0 | C | G | 0.044 | -6.82E-02 | 1.74E-01 | 2.64E+00 | 2.75E+00 | 6.90E-01 | no | | intergenic_variant | MODIFIER |
| 6 | 122633765 | AX-170243510 | 0 | C | T | 0.053 | -8.70E-02 | 2.20E-01 | 2.63E+00 | 2.74E+00 | 6.90E-01 | no | Dppa3 | downstream_gene_variant | MODIFIER |
| 7 | 40144591 | AX-170417234 | 0 | T | C | 0.36 | 3.61E-02 | 8.91E-02 | 2.59E+00 | 2.68E+00 | 6.90E-01 | no | | intergenic_variant | MODIFIER |
| 12 | 99243472 | AX-170357275 | 0 | A | T | 0.053 | 8.72E-02 | 2.22E-01 | 2.69E+00 | 2.76E+00 | 6.90E-01 | no | Foxn3 | intron_variant | MODIFIER |
| 12 | 99250651 | AX-170296567 | 0 | T | C | 0.053 | 8.72E-02 | 2.22E-01 | 2.69E+00 | 2.76E+00 | 6.90E-01 | no | Foxn3 | intron_variant | MODIFIER |
| 12 | 99251016 | AX-170168946 | 0 | G | A | 0.053 | 8.72E-02 | 2.22E-01 | 2.69E+00 | 2.76E+00 | 6.90E-01 | no | Foxn3 | intron_variant | MODIFIER |
| 12 | 99264516 | AX-97828801 | 0 | G | A | 0.053 | 8.72E-02 | 2.22E-01 | 2.69E+00 | 2.76E+00 | 6.90E-01 | no | Foxn3 | intron_variant | MODIFIER |
| 12 | 99279903 | AX-170303753 | 0 | G | A | 0.053 | 8.72E-02 | 2.22E-01 | 2.69E+00 | 2.76E+00 | 6.90E-01 | no | Foxn3 | intron_variant | MODIFIER |
| 12 | 99282248 | AX-170284898 | 0 | C | T | 0.053 | 8.72E-02 | 2.22E-01 | 2.69E+00 | 2.76E+00 | 6.90E-01 | no | Foxn3 | intron_variant | MODIFIER |
| 12 | 99312047 | AX-170155456 | 0 | T | C | 0.053 | 8.72E-02 | 2.22E-01 | 2.69E+00 | 2.76E+00 | 6.90E-01 | no | Foxn3 | intron_variant | MODIFIER |
| 12 | 99314870 | AX-170120723 | 0 | G | A | 0.053 | 8.72E-02 | 2.22E-01 | 2.69E+00 | 2.76E+00 | 6.90E-01 | no | Foxn3 | intron_variant | MODIFIER |
| 12 | 99322135 | AX-97801589 | 0 | C | T | 0.053 | 8.72E-02 | 2.22E-01 | 2.69E+00 | 2.76E+00 | 6.90E-01 | no | Foxn3 | intron_variant | MODIFIER |
| 12 | 99328912 | AX-170386034 | 0 | A | G | 0.053 | 8.72E-02 | 2.22E-01 | 2.69E+00 | 2.76E+00 | 6.90E-01 | no | Foxn3 | intron_variant | MODIFIER |
| 12 | 99436379 | AX-170007376 | 0 | A | G | 0.053 | 8.72E-02 | 2.22E-01 | 2.69E+00 | 2.76E+00 | 6.90E-01 | no | Foxn3 | intron_variant | MODIFIER |
| 12 | 99437422 | AX-170049928 | 0 | C | T | 0.053 | 8.72E-02 | 2.22E-01 | 2.69E+00 | 2.76E+00 | 6.90E-01 | no | Foxn3 | intron_variant | MODIFIER |
| 12 | 99443147 | AX-170514434 | 0 | A | G | 0.053 | 8.72E-02 | 2.22E-01 | 2.69E+00 | 2.76E+00 | 6.90E-01 | no | Foxn3 | intron_variant | MODIFIER |
| 12 | 99445089 | AX-170332688 | 0 | G | A | 0.053 | 8.72E-02 | 2.22E-01 | 2.69E+00 | 2.76E+00 | 6.90E-01 | no | Foxn3 | intron_variant | MODIFIER |
| 12 | 99471061 | AX-170294054 | 0 | C | A | 0.053 | 8.72E-02 | 2.22E-01 | 2.69E+00 | 2.76E+00 | 6.90E-01 | no | Foxn3 | intron_variant | MODIFIER |
| 12 | 99473623 | AX-170123595 | 0 | T | G | 0.053 | 8.72E-02 | 2.22E-01 | 2.69E+00 | 2.76E+00 | 6.90E-01 | no | Foxn3 | intron_variant | MODIFIER |
| 12 | 99474731 | AX-170432121 | 0 | G | T | 0.053 | 8.72E-02 | 2.22E-01 | 2.69E+00 | 2.76E+00 | 6.90E-01 | no | Foxn3 | intron_variant | MODIFIER |
| 12 | 99474797 | AX-97826715 | 0 | C | T | 0.053 | 8.72E-02 | 2.22E-01 | 2.69E+00 | 2.76E+00 | 6.90E-01 | no | Foxn3 | intron_variant | MODIFIER |

| | | | | | | | | | | | | | | | |
|---|---|---|---|---|---|---|---|---|---|---|---|---|---|---|---|
| 12 | 99474856 | AX-170007369 | 0 | G | A | 0.053 | 8.72E-02 | 2.22E-01 | 2.69E+00 | 2.76E+00 | 6.90E-01 | no | Foxn3 | intron_variant | MODIFIER |
| 12 | 99500461 | AX-170007372 | 0 | A | G | 0.053 | 8.72E-02 | 2.22E-01 | 2.69E+00 | 2.76E+00 | 6.90E-01 | no | Foxn3 | intron_variant | MODIFIER |
| 12 | 99514841 | AX-170240352 | 0 | G | C | 0.053 | 8.72E-02 | 2.22E-01 | 2.69E+00 | 2.76E+00 | 6.90E-01 | no | Foxn3 | intron_variant | MODIFIER |
| 12 | 99517689 | AX-97786163 | 0 | T | C | 0.053 | 8.72E-02 | 2.22E-01 | 2.69E+00 | 2.76E+00 | 6.90E-01 | no | Foxn3 | intron_variant | MODIFIER |
| 12 | 99529618 | AX-170007375 | 0 | C | T | 0.053 | 8.72E-02 | 2.22E-01 | 2.69E+00 | 2.76E+00 | 6.90E-01 | no | Foxn3 | intron_variant | MODIFIER |
| 12 | 99532941 | AX-170049932 | 0 | T | G | 0.053 | 8.72E-02 | 2.22E-01 | 2.69E+00 | 2.76E+00 | 6.90E-01 | no | Foxn3 | intron_variant | MODIFIER |
| 12 | 99533071 | AX-170049933 | 0 | G | A | 0.053 | 8.72E-02 | 2.22E-01 | 2.69E+00 | 2.76E+00 | 6.90E-01 | no | Foxn3 | intron_variant | MODIFIER |
| 12 | 99545283 | AX-170276873 | 0 | C | T | 0.053 | 8.72E-02 | 2.22E-01 | 2.69E+00 | 2.76E+00 | 6.90E-01 | no | Foxn3 | intron_variant | MODIFIER |
| 12 | 99573134 | AX-170182525 | 0 | A | G | 0.053 | 8.72E-02 | 2.22E-01 | 2.69E+00 | 2.76E+00 | 6.90E-01 | no | Gm26839 | intron_variant,non_coding_transcript_variant | MODIFIER |
| 12 | 99589109 | AX-170099068 | 0 | G | A | 0.053 | 8.72E-02 | 2.22E-01 | 2.69E+00 | 2.76E+00 | 6.90E-01 | no | Gm26839 | intron_variant,non_coding_transcript_variant | MODIFIER |
| 12 | 99596477 | AX-170240495 | 0 | C | T | 0.053 | 8.72E-02 | 2.22E-01 | 2.69E+00 | 2.76E+00 | 6.90E-01 | no | Gm26839 | intron_variant,non_coding_transcript_variant | MODIFIER |
| 12 | 99606610 | AX-170190417 | 0 | A | G | 0.053 | 8.72E-02 | 2.22E-01 | 2.69E+00 | 2.76E+00 | 6.90E-01 | no | Gm26839 | intron_variant,non_coding_transcript_variant | MODIFIER |
| 12 | 99607344 | AX-170400362 | 0 | T | C | 0.053 | 8.72E-02 | 2.22E-01 | 2.69E+00 | 2.76E+00 | 6.90E-01 | no | Gm26839 | intron_variant,non_coding_transcript_variant | MODIFIER |
| 12 | 99607419 | AX-170451892 | 0 | G | A | 0.053 | 8.72E-02 | 2.22E-01 | 2.69E+00 | 2.76E+00 | 6.90E-01 | no | Gm26839 | intron_variant,non_coding_transcript_variant | MODIFIER |
| 12 | 99609017 | AX-170184264 | 0 | G | A | 0.053 | 8.72E-02 | 2.22E-01 | 2.69E+00 | 2.76E+00 | 6.90E-01 | no | Gm26839 | intron_variant,non_coding_transcript_variant | MODIFIER |
| 12 | 99609735 | AX-170230158 | 0 | T | C | 0.053 | 8.72E-02 | 2.22E-01 | 2.69E+00 | 2.76E+00 | 6.90E-01 | no | Gm26839 | intron_variant,non_coding_transcript_variant | MODIFIER |
| 12 | 99610486 | AX-170171880 | 0 | T | C | 0.053 | 8.72E-02 | 2.22E-01 | 2.69E+00 | 2.76E+00 | 6.90E-01 | no | Gm26839 | intron_variant,non_coding_transcript_variant | MODIFIER |
| 12 | 99617597 | AX-170524168 | 0 | T | C | 0.053 | 8.72E-02 | 2.22E-01 | 2.69E+00 | 2.76E+00 | 6.90E-01 | no | Gm26839 | intron_variant,non_coding_transcript_variant | MODIFIER |
| 12 | 99618420 | AX-170467628 | 0 | T | C | 0.053 | 8.72E-02 | 2.22E-01 | 2.69E+00 | 2.76E+00 | 6.90E-01 | no | Gm26839 | intron_variant,non_coding_transcript_variant | MODIFIER |
| 12 | 99626115 | AX-170286265 | 0 | G | A | 0.053 | 8.72E-02 | 2.22E-01 | 2.69E+00 | 2.76E+00 | 6.90E-01 | no | 1700064M15Rik | non_coding_transcript_exon_variant | MODIFIER |
| 12 | 99680767 | AX-170340397 | 0 | T | G | 0.053 | 8.72E-02 | 2.22E-01 | 2.69E+00 | 2.76E+00 | 6.90E-01 | no | | intergenic_variant | MODIFIER |
| 12 | 99697156 | AX-170382786 | 0 | C | T | 0.053 | 8.72E-02 | 2.22E-01 | 2.69E+00 | 2.76E+00 | 6.90E-01 | no | Efcab11 | intron_variant | MODIFIER |
| 12 | 99700137 | AX-97789010 | 0 | T | C | 0.053 | 8.72E-02 | 2.22E-01 | 2.69E+00 | 2.76E+00 | 6.90E-01 | no | Efcab11 | intron_variant | MODIFIER |
| 12 | 99703092 | AX-170394154 | 0 | C | T | 0.053 | 8.72E-02 | 2.22E-01 | 2.69E+00 | 2.76E+00 | 6.90E-01 | no | Efcab11 | intron_variant | MODIFIER |
| 12 | 99704416 | AX-170430525 | 0 | T | G | 0.053 | 8.72E-02 | 2.22E-01 | 2.69E+00 | 2.76E+00 | 6.90E-01 | no | Efcab11 | intron_variant | MODIFIER |
| 12 | 99715387 | AX-170384660 | 0 | G | C | 0.053 | 8.72E-02 | 2.22E-01 | 2.69E+00 | 2.76E+00 | 6.90E-01 | no | Efcab11 | downstream_gene_variant | MODIFIER |
| 12 | 99733362 | AX-170294313 | 0 | G | T | 0.053 | 8.72E-02 | 2.22E-01 | 2.69E+00 | 2.76E+00 | 6.90E-01 | no | Efcab11 | intron_variant | MODIFIER |
| 12 | 99740720 | AX-97846097 | 0 | A | G | 0.053 | 8.72E-02 | 2.22E-01 | 2.69E+00 | 2.76E+00 | 6.90E-01 | no | Efcab11 | intron_variant | MODIFIER |
| 12 | 99755117 | AX-170049936 | 0 | G | C | 0.053 | 8.72E-02 | 2.22E-01 | 2.69E+00 | 2.76E+00 | 6.90E-01 | no | Efcab11 | intron_variant | MODIFIER |
| 12 | 99763578 | AX-97822216 | 0 | G | A | 0.053 | 8.72E-02 | 2.22E-01 | 2.69E+00 | 2.76E+00 | 6.90E-01 | no | Efcab11 | intron_variant | MODIFIER |
| 12 | 99764253 | AX-170481912 | 0 | G | A | 0.053 | 8.72E-02 | 2.22E-01 | 2.69E+00 | 2.76E+00 | 6.90E-01 | no | Efcab11 | intron_variant | MODIFIER |
| 12 | 99764433 | AX-170049934 | 0 | T | C | 0.053 | 8.72E-02 | 2.22E-01 | 2.69E+00 | 2.76E+00 | 6.90E-01 | no | Efcab11 | intron_variant | MODIFIER |
| 12 | 99775814 | AX-170197791 | 0 | C | T | 0.053 | 8.72E-02 | 2.22E-01 | 2.69E+00 | 2.76E+00 | 6.90E-01 | no | Efcab11 | intron_variant | MODIFIER |
| 12 | 99792016 | AX-170188472 | 0 | T | C | 0.053 | 8.72E-02 | 2.22E-01 | 2.69E+00 | 2.76E+00 | 6.90E-01 | no | Efcab11 | intron_variant | MODIFIER |
| 1 | 172194217 | AX-170043591 | 1 | C | A | 0.214 | 5.24E-02 | 1.26E-01 | 2.55E+00 | 2.63E+00 | 6.90E-01 | no | Pea15a | downstream_gene_variant | MODIFIER |
| 13 | 37390439 | AX-170228460 | 2 | G | A | 0.173 | -4.35E-02 | 1.06E-01 | 2.56E+00 | 2.66E+00 | 6.90E-01 | no | Ly86 | intron_variant | MODIFIER |
| 1 | 45811205 | AX-170528072 | 1 | C | G | 0.438 | -3.01E-02 | 7.65E-02 | 2.66E+00 | 2.74E+00 | 6.90E-01 | no | Wdr75 | intron_variant | MODIFIER |
| 1 | 9981148 | AX-170391124 | 0 | T | A | 0.088 | 6.81E-02 | 1.74E-01 | 2.64E+00 | 2.76E+00 | 6.90E-01 | no | Ppp1r42 | intron_variant | MODIFIER |
| 10 | 26564806 | AX-170091895 | 0 | G | A | 0.184 | 3.91E-02 | 1.01E-01 | 2.72E+00 | 2.83E+00 | 6.90E-01 | no | | regulatory_region_variant | MODIFIER |
| 5 | 32918436 | AX-170183195 | 0 | A | T | 0.351 | -3.81E-02 | 9.80E-02 | 2.69E+00 | 2.79E+00 | 6.90E-01 | no | Depdc5 | intron_variant | MODIFIER |
| 12 | 99294970 | AX-97822137 | 1 | C | T | 0.054 | 8.81E-02 | 2.24E-01 | 2.70E+00 | 2.76E+00 | 6.90E-01 | no | Foxn3 | intron_variant | MODIFIER |
| 11 | 19002542 | AX-170216398 | 0 | G | A | 0.07 | 6.94E-02 | 1.79E-01 | 2.68E+00 | 2.80E+00 | 6.90E-01 | no | Meis1 | intron_variant | MODIFIER |
| 7 | 89961354 | AX-170448453 | 0 | A | G | 0.132 | -8.35E-02 | 2.13E-01 | 2.69E+00 | 2.77E+00 | 6.90E-01 | no | Eed | intron_variant | MODIFIER |
| 7 | 94934716 | AX-97847384 | 0 | C | T | 0.132 | -8.35E-02 | 2.13E-01 | 2.69E+00 | 2.77E+00 | 6.90E-01 | no | | intergenic_variant | MODIFIER |
| 7 | 112733690 | AX-170418381 | 1 | G | A | 0.446 | 3.69E-02 | 9.37E-02 | 2.68E+00 | 2.74E+00 | 6.91E-01 | no | Tead1 | intron_variant | MODIFIER |
| 13 | 37295444 | AX-97853570 | 1 | T | C | 0.179 | -5.06E-02 | 1.23E-01 | 2.58E+00 | 2.64E+00 | 6.91E-01 | no | | intergenic_variant | MODIFIER |
| 8 | 63437893 | AX-97823144 | 0 | G | A | 0.035 | 4.84E-02 | 1.28E-01 | 2.61E+00 | 2.80E+00 | 6.91E-01 | no | | intergenic_variant | MODIFIER |
| 8 | 97663983 | AX-170368417 | 1 | G | A | 0.223 | 4.55E-02 | 1.16E-01 | 2.70E+00 | 2.76E+00 | 6.91E-01 | no | | intergenic_variant | MODIFIER |
| 7 | 38043732 | AX-170274757 | 1 | C | A | 0.348 | 3.75E-02 | 9.23E-02 | 2.59E+00 | 2.67E+00 | 6.91E-01 | no | | intergenic_variant | MODIFIER |
| 10 | 43049376 | AX-97791639 | 0 | C | A | 0.368 | -2.95E-02 | 7.37E-02 | 2.59E+00 | 2.70E+00 | 6.91E-01 | no | Sobp | intron_variant | MODIFIER |
| 19 | 57847721 | AX-170116487 | 0 | C | A | 0.044 | 5.68E-02 | 1.45E-01 | 2.59E+00 | 2.75E+00 | 6.91E-01 | no | Atrnl1 | intron_variant | MODIFIER |
| 5 | 113373954 | AX-170134975 | 1 | A | T | 0.455 | 4.66E-02 | 1.17E-01 | 2.66E+00 | 2.69E+00 | 6.91E-01 | no | | regulatory_region_variant | MODIFIER |
| 9 | 110968852 | AX-170307420 | 0 | T | G | 0.096 | -4.87E-02 | 1.19E-01 | 2.57E+00 | 2.65E+00 | 6.91E-01 | no | Lrrc2 | intron_variant | MODIFIER |
| 9 | 21604532 | AX-170156092 | 0 | T | C | 0.465 | 3.64E-02 | 8.90E-02 | 2.58E+00 | 2.65E+00 | 6.91E-01 | no | Gm7904 | downstream_gene_variant | MODIFIER |

| | | | | | | | | | | | | | | | |
|---|---|---|---|---|---|---|---|---|---|---|---|---|---|---|---|
| 9 | 22870458 | AX-97807337 | 0 | A | G | 0.465 | 3.64E-02 | 8.90E-02 | 2.58E+00 | 2.65E+00 | 6.91E-01 | no | Bbs9 | intron_variant | MODIFIER |
| 7 | 95485893 | AX-170456093 | 1 | T | G | 0.116 | -5.59E-02 | 1.43E-01 | 2.65E+00 | 2.76E+00 | 6.91E-01 | no | | intergenic_variant | MODIFIER |
| 5 | 88386830 | AX-170072715 | 1 | C | A | 0.438 | 4.28E-02 | 1.09E-01 | 2.73E+00 | 2.78E+00 | 6.91E-01 | no | Amtn | downstream_gene_variant | MODIFIER |
| 5 | 101570365 | AX-170025737 | 0 | T | C | 0.439 | 4.35E-02 | 1.11E-01 | 2.68E+00 | 2.73E+00 | 6.91E-01 | no | | intergenic_variant | MODIFIER |
| 5 | 101570501 | AX-170025739 | 0 | C | T | 0.439 | 4.35E-02 | 1.11E-01 | 2.68E+00 | 2.73E+00 | 6.91E-01 | no | | intergenic_variant | MODIFIER |
| 5 | 101570689 | AX-170123077 | 0 | G | A | 0.439 | 4.35E-02 | 1.11E-01 | 2.68E+00 | 2.73E+00 | 6.91E-01 | no | | intergenic_variant | MODIFIER |
| 5 | 101570843 | AX-170380792 | 0 | C | T | 0.439 | 4.35E-02 | 1.11E-01 | 2.68E+00 | 2.73E+00 | 6.91E-01 | no | | intergenic_variant | MODIFIER |
| 5 | 101570881 | AX-170348024 | 0 | C | T | 0.439 | 4.35E-02 | 1.11E-01 | 2.68E+00 | 2.73E+00 | 6.91E-01 | no | | intergenic_variant | MODIFIER |
| 5 | 101571417 | AX-170476021 | 0 | G | A | 0.439 | 4.35E-02 | 1.11E-01 | 2.68E+00 | 2.73E+00 | 6.91E-01 | no | | intergenic_variant | MODIFIER |
| 5 | 101589543 | AX-170109874 | 0 | A | G | 0.439 | 4.35E-02 | 1.11E-01 | 2.68E+00 | 2.73E+00 | 6.91E-01 | no | | intergenic_variant | MODIFIER |
| 5 | 101601273 | AX-97797321 | 0 | C | A | 0.439 | 4.35E-02 | 1.11E-01 | 2.68E+00 | 2.73E+00 | 6.91E-01 | no | Gm20484 | intron_variant,non_coding_transcript_variant | MODIFIER |
| 5 | 101615177 | AX-170515454 | 0 | A | G | 0.439 | 4.35E-02 | 1.11E-01 | 2.68E+00 | 2.73E+00 | 6.91E-01 | no | | intergenic_variant | MODIFIER |
| 7 | 78000552 | AX-170029369 | 0 | C | T | 0.377 | -3.74E-02 | 9.52E-02 | 2.64E+00 | 2.74E+00 | 6.91E-01 | no | | intergenic_variant | MODIFIER |
| 2 | 107085355 | AX-168381205 | 0 | G | A | 0.053 | -1.18E-01 | 3.00E-01 | 2.74E+00 | 2.77E+00 | 6.91E-01 | no | | intergenic_variant | MODIFIER |
| 9 | 61279218 | AX-170315858 | 1 | C | G | 0.196 | 5.35E-02 | 1.36E-01 | 2.79E+00 | 2.82E+00 | 6.91E-01 | no | B930092H01Rik | intron_variant,non_coding_transcript_variant | MODIFIER |
| 9 | 67508227 | AX-170468352 | 1 | C | T | 0.196 | 5.35E-02 | 1.36E-01 | 2.79E+00 | 2.82E+00 | 6.91E-01 | no | Tln2 | intron_variant | MODIFIER |
| 9 | 69790466 | AX-170159692 | 1 | T | C | 0.196 | 5.35E-02 | 1.36E-01 | 2.79E+00 | 2.82E+00 | 6.91E-01 | no | B230323A14Rik | intron_variant,non_coding_transcript_variant | MODIFIER |
| 8 | 72813100 | AX-170253622 | 0 | C | T | 0.193 | -5.10E-02 | 1.28E-01 | 2.61E+00 | 2.69E+00 | 6.91E-01 | no | Large1 | downstream_gene_variant | MODIFIER |
| 9 | 79584619 | AX-170137783 | 0 | G | A | 0.193 | 4.96E-02 | 1.27E-01 | 2.79E+00 | 2.85E+00 | 6.91E-01 | no | | intergenic_variant | MODIFIER |
| 2 | 142257088 | AX-170019228 | 2 | C | T | 0.145 | 8.56E-02 | 2.18E-01 | 2.69E+00 | 2.72E+00 | 6.92E-01 | no | Macrod2 | intron_variant | MODIFIER |
| 5 | 73737381 | AX-170463617 | 0 | G | C | 0.439 | 3.96E-02 | 1.00E-01 | 2.65E+00 | 2.73E+00 | 6.92E-01 | no | | intergenic_variant | MODIFIER |
| 1 | 74487664 | AX-170000808 | 1 | T | A | 0.116 | 7.87E-02 | 2.01E-01 | 2.65E+00 | 2.74E+00 | 6.92E-01 | no | Usp37 | intron_variant | MODIFIER |
| 18 | 53773979 | AX-170526521 | 1 | G | A | 0.438 | 4.14E-02 | 9.29E-02 | 2.46E+00 | 2.54E+00 | 6.92E-01 | no | | intergenic_variant | MODIFIER |
| 7 | 117099646 | AX-170486928 | 1 | G | A | 0.42 | 3.66E-02 | 9.25E-02 | 2.65E+00 | 2.70E+00 | 6.92E-01 | no | | intergenic_variant | MODIFIER |
| 20 | 165114496 | AX-170035408 | 1 | G | A | 0.134 | 5.04E-02 | 1.30E-01 | 2.65E+00 | 2.76E+00 | 6.92E-01 | no | | intergenic_variant | MODIFIER |
| 2 | 96346995 | AX-170245046 | 0 | T | G | 0.053 | -1.17E-01 | 3.00E-01 | 2.74E+00 | 2.77E+00 | 6.92E-01 | no | Lrrc4c | intron_variant | MODIFIER |
| 12 | 89695958 | AX-170264140 | 2 | G | C | 0.055 | 7.62E-02 | 1.96E-01 | 2.65E+00 | 2.76E+00 | 6.92E-01 | no | Nrxn3 | intron_variant | MODIFIER |
| 18 | 47823561 | AX-170357939 | 2 | C | A | 0.055 | 9.78E-02 | 2.47E-01 | 2.61E+00 | 2.71E+00 | 6.92E-01 | no | | intergenic_variant | MODIFIER |
| 14 | 85367143 | AX-97819747 | 2 | C | T | 0.055 | 3.94E-02 | 1.04E-01 | 2.62E+00 | 2.81E+00 | 6.92E-01 | no | | intergenic_variant | MODIFIER |
| 5 | 125374706 | AX-170089603 | 0 | T | C | 0.456 | -4.27E-02 | 1.10E-01 | 2.76E+00 | 2.82E+00 | 6.92E-01 | no | Gm40323 | downstream_gene_variant | MODIFIER |
| 9 | 87026881 | AX-170032902 | 0 | T | G | 0.132 | -4.84E-02 | 1.20E-01 | 2.58E+00 | 2.66E+00 | 6.92E-01 | no | Cyb5r4 | intron_variant | MODIFIER |
| 1 | 96837363 | AX-170001149 | 0 | A | C | 0.149 | 7.81E-02 | 1.97E-01 | 2.63E+00 | 2.70E+00 | 6.92E-01 | no | Slco4c1 | intron_variant | MODIFIER |
| 1 | 104797518 | AX-170478897 | 0 | T | G | 0.149 | 7.81E-02 | 1.97E-01 | 2.63E+00 | 2.70E+00 | 6.92E-01 | no | Cdh20 | intron_variant | MODIFIER |
| 5 | 43929005 | AX-97854372 | 0 | A | G | 0.193 | -3.97E-02 | 1.01E-01 | 2.63E+00 | 2.72E+00 | 6.92E-01 | no | | intergenic_variant | MODIFIER |
| 4 | 153325611 | AX-170274274 | 0 | T | A | 0.105 | 5.99E-02 | 1.56E-01 | 2.69E+00 | 2.81E+00 | 6.93E-01 | no | | regulatory_region_variant | MODIFIER |
| 1 | 67307069 | AX-97844264 | 1 | T | G | 0.098 | 7.14E-02 | 1.83E-01 | 2.61E+00 | 2.74E+00 | 6.93E-01 | no | Gm15668 | intron_variant,non_coding_transcript_variant | MODIFIER |
| 18 | 69521824 | AX-170439585 | 0 | A | G | 0.272 | -3.73E-02 | 9.49E-02 | 2.65E+00 | 2.72E+00 | 6.93E-01 | no | Tcf4 | intron_variant | MODIFIER |
| 12 | 94390803 | AX-170049835 | 0 | G | T | 0.088 | 7.92E-02 | 2.04E-01 | 2.71E+00 | 2.77E+00 | 6.93E-01 | no | | intergenic_variant | MODIFIER |
| 12 | 94392049 | AX-170049833 | 0 | T | C | 0.088 | 7.92E-02 | 2.04E-01 | 2.71E+00 | 2.77E+00 | 6.93E-01 | no | | intergenic_variant | MODIFIER |
| 12 | 94392089 | AX-97800486 | 0 | G | A | 0.088 | 7.92E-02 | 2.04E-01 | 2.71E+00 | 2.77E+00 | 6.93E-01 | no | | intergenic_variant | MODIFIER |
| 12 | 94414373 | AX-170414187 | 0 | C | T | 0.088 | 7.92E-02 | 2.04E-01 | 2.71E+00 | 2.77E+00 | 6.93E-01 | no | | intergenic_variant | MODIFIER |
| 12 | 94422104 | AX-170049834 | 0 | G | T | 0.088 | 7.92E-02 | 2.04E-01 | 2.71E+00 | 2.77E+00 | 6.93E-01 | no | | regulatory_region_variant | MODIFIER |
| 12 | 94422156 | AX-170160252 | 0 | G | A | 0.088 | 7.92E-02 | 2.04E-01 | 2.71E+00 | 2.77E+00 | 6.93E-01 | no | | regulatory_region_variant | MODIFIER |
| 12 | 94424788 | AX-170505975 | 0 | T | G | 0.088 | 7.92E-02 | 2.04E-01 | 2.71E+00 | 2.77E+00 | 6.93E-01 | no | | regulatory_region_variant | MODIFIER |
| 12 | 94427535 | AX-170464773 | 0 | T | A | 0.088 | 7.92E-02 | 2.04E-01 | 2.71E+00 | 2.77E+00 | 6.93E-01 | no | | intergenic_variant | MODIFIER |
| 12 | 94456792 | AX-170007300 | 0 | G | A | 0.088 | 7.92E-02 | 2.04E-01 | 2.71E+00 | 2.77E+00 | 6.93E-01 | no | | intergenic_variant | MODIFIER |
| 12 | 94478730 | AX-170218894 | 0 | T | G | 0.088 | 7.92E-02 | 2.04E-01 | 2.71E+00 | 2.77E+00 | 6.93E-01 | no | | intergenic_variant | MODIFIER |
| 12 | 94686866 | AX-97817720 | 0 | T | A | 0.088 | 7.92E-02 | 2.04E-01 | 2.71E+00 | 2.77E+00 | 6.93E-01 | no | | intergenic_variant | MODIFIER |
| 12 | 95127313 | AX-170279649 | 0 | C | T | 0.088 | 7.92E-02 | 2.04E-01 | 2.71E+00 | 2.77E+00 | 6.93E-01 | no | | intergenic_variant | MODIFIER |
| 12 | 95155658 | AX-170391433 | 0 | G | A | 0.088 | 7.92E-02 | 2.04E-01 | 2.71E+00 | 2.77E+00 | 6.93E-01 | no | | intergenic_variant | MODIFIER |
| 12 | 95177421 | AX-170500322 | 0 | C | G | 0.088 | 7.92E-02 | 2.04E-01 | 2.71E+00 | 2.77E+00 | 6.93E-01 | no | | intergenic_variant | MODIFIER |
| 12 | 95385102 | AX-170409050 | 0 | A | G | 0.088 | 7.92E-02 | 2.04E-01 | 2.71E+00 | 2.77E+00 | 6.93E-01 | no | | regulatory_region_variant | MODIFIER |
| 12 | 95387116 | AX-170147491 | 0 | A | T | 0.088 | 7.92E-02 | 2.04E-01 | 2.71E+00 | 2.77E+00 | 6.93E-01 | no | | intergenic_variant | MODIFIER |
| 12 | 95398854 | AX-97802425 | 0 | A | G | 0.088 | 7.92E-02 | 2.04E-01 | 2.71E+00 | 2.77E+00 | 6.93E-01 | no | | intergenic_variant | MODIFIER |

| | | | | | | | | | | | | | | | |
|---|---|---|---|---|---|---|---|---|---|---|---|---|---|---|---|
| 12 | 95399001 | AX-170222742 | 0 | C | T | 0.088 | 7.92E-02 | 2.04E-01 | 2.71E+00 | 2.77E+00 | 6.93E-01 | no | | intergenic_variant | MODIFIER |
| 12 | 95412752 | AX-170148678 | 0 | T | C | 0.088 | 7.92E-02 | 2.04E-01 | 2.71E+00 | 2.77E+00 | 6.93E-01 | no | | intergenic_variant | MODIFIER |
| 17 | 72582174 | AX-97783352 | 1 | C | T | 0.054 | -5.73E-02 | 1.49E-01 | 2.62E+00 | 2.76E+00 | 6.93E-01 | no | Alk | intron_variant | MODIFIER |
| 12 | 93294638 | AX-97826657 | 0 | T | C | 0.07 | 8.30E-02 | 2.12E-01 | 2.66E+00 | 2.73E+00 | 6.93E-01 | no | | intergenic_variant | MODIFIER |
| 8 | 96248399 | AX-97843896 | 0 | C | A | 0.219 | 4.66E-02 | 1.20E-01 | 2.70E+00 | 2.77E+00 | 6.93E-01 | no | | intergenic_variant | MODIFIER |
| 8 | 96606635 | AX-170177238 | 0 | A | G | 0.219 | 4.66E-02 | 1.20E-01 | 2.70E+00 | 2.77E+00 | 6.93E-01 | no | | intergenic_variant | MODIFIER |
| 1 | 8083615 | AX-170169772 | 1 | T | G | 0.411 | -2.72E-02 | 7.03E-02 | 2.79E+00 | 2.88E+00 | 6.93E-01 | no | | intergenic_variant | MODIFIER |
| 16 | 23467605 | AX-170137425 | 0 | T | C | 0.158 | 3.53E-02 | 9.22E-02 | 2.66E+00 | 2.79E+00 | 6.93E-01 | no | Masp1 | intron_variant | MODIFIER |
| 15 | 94702772 | AX-97813382 | 0 | G | T | 0.149 | -5.98E-02 | 1.56E-01 | 2.71E+00 | 2.81E+00 | 6.93E-01 | no | Tmem117 | intron_variant | MODIFIER |
| 20 | 155593969 | AX-170223722 | 1 | A | T | 0.116 | 4.67E-02 | 1.20E-01 | 2.66E+00 | 2.76E+00 | 6.93E-01 | no | Ptchd1 | intron_variant | MODIFIER |
| 8 | 37005186 | AX-170079778 | 1 | T | C | 0.152 | -5.18E-02 | 1.32E-01 | 2.65E+00 | 2.73E+00 | 6.93E-01 | no | G630064G18Rik | intron_variant,non_coding_transcript_variant | MODIFIER |
| 20 | 164796437 | AX-170133445 | 0 | T | G | 0.132 | 4.66E-02 | 1.21E-01 | 2.66E+00 | 2.78E+00 | 6.93E-01 | no | | intergenic_variant | MODIFIER |
| 8 | 95329472 | AX-170267386 | 1 | G | A | 0.232 | 4.39E-02 | 1.13E-01 | 2.69E+00 | 2.76E+00 | 6.93E-01 | no | Usb1 | upstream_gene_variant | MODIFIER |
| 8 | 95777576 | AX-170080886 | 1 | A | C | 0.232 | 4.39E-02 | 1.13E-01 | 2.69E+00 | 2.76E+00 | 6.93E-01 | no | Cnot1 | intron_variant | MODIFIER |
| 8 | 38471171 | AX-170495280 | 2 | A | T | 0.155 | 5.73E-02 | 1.47E-01 | 2.77E+00 | 2.81E+00 | 6.93E-01 | no | Sgcz | intron_variant | MODIFIER |
| 8 | 91160078 | AX-170453240 | 1 | G | A | 0.179 | -4.74E-02 | 1.20E-01 | 2.63E+00 | 2.71E+00 | 6.93E-01 | no | Aktip | intron_variant | MODIFIER |
| 1 | 23834559 | AX-170458650 | 0 | T | A | 0.421 | 3.32E-02 | 8.43E-02 | 2.64E+00 | 2.70E+00 | 6.93E-01 | no | B3gat2 | intron_variant | MODIFIER |
| 1 | 68643871 | AX-170394217 | 0 | G | T | 0.114 | -7.39E-02 | 1.90E-01 | 2.65E+00 | 2.74E+00 | 6.93E-01 | no | Erbb4 | intron_variant | MODIFIER |
| 7 | 116914596 | AX-170418536 | 0 | G | A | 0.421 | 3.47E-02 | 8.79E-02 | 2.64E+00 | 2.70E+00 | 6.93E-01 | no | | intergenic_variant | MODIFIER |
| 5 | 33648325 | AX-170089444 | 1 | G | A | 0.071 | 4.69E-02 | 1.22E-01 | 2.61E+00 | 2.77E+00 | 6.94E-01 | no | Tmem129 | downstream_gene_variant | MODIFIER |
| 1 | 118850367 | AX-170042454 | 0 | A | T | 0.158 | 6.32E-02 | 1.58E-01 | 2.57E+00 | 2.69E+00 | 6.94E-01 | no | Gli2 | intron_variant | MODIFIER |
| 11 | 74212580 | AX-170417735 | 0 | G | A | 0.263 | 2.75E-02 | 7.14E-02 | 2.63E+00 | 2.77E+00 | 6.94E-01 | no | Olfr43 | upstream_gene_variant | MODIFIER |
| 15 | 88129869 | AX-170201760 | 0 | G | A | 0.132 | -5.35E-02 | 1.40E-01 | 2.72E+00 | 2.82E+00 | 6.94E-01 | no | | intergenic_variant | MODIFIER |
| 5 | 127818423 | AX-170139159 | 2 | T | C | 0.4 | -3.10E-02 | 8.09E-02 | 2.67E+00 | 2.78E+00 | 6.94E-01 | no | Tmem132d | intron_variant | MODIFIER |
| 18 | 66094169 | AX-170100531 | 2 | A | G | 0.4 | 3.52E-02 | 8.23E-02 | 2.50E+00 | 2.58E+00 | 6.94E-01 | no | Ccbe1 | intron_variant,NMD_transcript_variant | MODIFIER |
| 3 | 37381448 | AX-170422923 | 1 | T | C | 0.188 | 3.48E-02 | 9.00E-02 | 2.65E+00 | 2.76E+00 | 6.94E-01 | no | Fgf2 | intron_variant | MODIFIER |
| 1 | 108742845 | AX-170535290 | 1 | G | A | 0.161 | 6.73E-02 | 1.68E-01 | 2.57E+00 | 2.69E+00 | 6.94E-01 | no | | intergenic_variant | MODIFIER |
| 8 | 75224901 | AX-170460287 | 0 | A | G | 0.202 | 4.71E-02 | 1.23E-01 | 2.73E+00 | 2.81E+00 | 6.94E-01 | no | Rasd2 | downstream_gene_variant | MODIFIER |
| 12 | 93268860 | AX-170380713 | 0 | G | C | 0.061 | 9.17E-02 | 2.37E-01 | 2.71E+00 | 2.77E+00 | 6.94E-01 | no | | intergenic_variant | MODIFIER |
| 5 | 86371875 | AX-97791769 | 1 | T | C | 0.429 | 4.39E-02 | 1.13E-01 | 2.74E+00 | 2.78E+00 | 6.94E-01 | no | Tmprss11d | intron_variant | MODIFIER |
| 5 | 87681001 | AX-170192170 | 1 | A | G | 0.429 | 4.39E-02 | 1.13E-01 | 2.74E+00 | 2.78E+00 | 6.94E-01 | no | Csn1s1 | intron_variant | MODIFIER |
| 5 | 97456759 | AX-97799628 | 0 | C | T | 0.447 | 4.17E-02 | 1.07E-01 | 2.67E+00 | 2.73E+00 | 6.94E-01 | no | Gk2 | synonymous_variant | LOW |
| 2 | 142672630 | AX-97798458 | 0 | C | T | 0.158 | 6.08E-02 | 1.51E-01 | 2.56E+00 | 2.68E+00 | 6.94E-01 | no | Kif16b | intron_variant | MODIFIER |
| 7 | 55484912 | AX-170444645 | 2 | C | T | 0.436 | -3.49E-02 | 9.13E-02 | 2.69E+00 | 2.80E+00 | 6.94E-01 | no | | intergenic_variant | MODIFIER |
| 2 | 118939999 | AX-97833911 | 0 | G | T | 0.14 | 8.54E-02 | 2.20E-01 | 2.71E+00 | 2.74E+00 | 6.94E-01 | no | | intergenic_variant | MODIFIER |
| 2 | 119172660 | AX-170301870 | 0 | T | A | 0.14 | 8.54E-02 | 2.20E-01 | 2.71E+00 | 2.74E+00 | 6.94E-01 | no | Dnajc17 | missense_variant | MODERATE |
| 2 | 119194522 | AX-170373196 | 0 | C | T | 0.14 | 8.54E-02 | 2.20E-01 | 2.71E+00 | 2.74E+00 | 6.94E-01 | no | Dnajc17 | intron_variant | MODIFIER |
| 2 | 121547262 | AX-170064758 | 1 | C | T | 0.143 | 8.62E-02 | 2.22E-01 | 2.70E+00 | 2.73E+00 | 6.94E-01 | no | Wdr76 | downstream_gene_variant | MODIFIER |
| 2 | 129497042 | AX-170320229 | 1 | C | T | 0.143 | 8.62E-02 | 2.22E-01 | 2.70E+00 | 2.73E+00 | 6.94E-01 | no | F830045P16Rik | intron_variant | MODIFIER |
| 2 | 136844971 | AX-170065008 | 1 | C | T | 0.143 | 8.62E-02 | 2.22E-01 | 2.70E+00 | 2.73E+00 | 6.94E-01 | no | | intergenic_variant | MODIFIER |
| 2 | 139493090 | AX-97853485 | 1 | C | T | 0.143 | 8.62E-02 | 2.22E-01 | 2.70E+00 | 2.73E+00 | 6.94E-01 | no | Sptlc3 | upstream_gene_variant | MODIFIER |
| 2 | 143655742 | AX-97840453 | 1 | G | T | 0.143 | 8.62E-02 | 2.22E-01 | 2.70E+00 | 2.73E+00 | 6.94E-01 | no | Pcsk2 | intron_variant | MODIFIER |
| 2 | 145649209 | AX-170437594 | 1 | T | A | 0.143 | 8.62E-02 | 2.22E-01 | 2.70E+00 | 2.73E+00 | 6.94E-01 | no | | intergenic_variant | MODIFIER |
| 2 | 145840587 | AX-170245548 | 1 | C | T | 0.143 | 8.62E-02 | 2.22E-01 | 2.70E+00 | 2.73E+00 | 6.94E-01 | no | Rin2 | intron_variant | MODIFIER |
| 11 | 74513947 | AX-170005316 | 0 | T | A | 0.263 | 2.73E-02 | 7.13E-02 | 2.63E+00 | 2.76E+00 | 6.94E-01 | no | Rap1gap2 | intron_variant,NMD_transcript_variant | MODIFIER |
| 18 | 69078798 | AX-170339886 | 0 | C | T | 0.272 | -3.77E-02 | 9.60E-02 | 2.64E+00 | 2.70E+00 | 6.94E-01 | no | | intergenic_variant | MODIFIER |
| 1 | 51074008 | AX-170000484 | 0 | G | A | 0.114 | 7.66E-02 | 1.99E-01 | 2.68E+00 | 2.77E+00 | 6.94E-01 | no | Tmeff2 | intron_variant | MODIFIER |
| 1 | 53977226 | AX-170000512 | 0 | C | G | 0.114 | 7.66E-02 | 1.99E-01 | 2.68E+00 | 2.77E+00 | 6.94E-01 | no | Hecw2 | intron_variant | MODIFIER |
| 1 | 55665372 | AX-170403673 | 0 | G | A | 0.114 | 7.66E-02 | 1.99E-01 | 2.68E+00 | 2.77E+00 | 6.94E-01 | no | Plcl1 | intron_variant | MODIFIER |
| 8 | 95427497 | AX-170413392 | 1 | A | G | 0.232 | 4.52E-02 | 1.17E-01 | 2.70E+00 | 2.77E+00 | 6.95E-01 | no | Cfap20 | intron_variant | MODIFIER |
| 12 | 94931172 | AX-97843042 | 1 | C | T | 0.062 | 9.27E-02 | 2.39E-01 | 2.71E+00 | 2.76E+00 | 6.95E-01 | no | | intergenic_variant | MODIFIER |
| 2 | 91472926 | AX-97838944 | 0 | G | T | 0.053 | -1.16E-01 | 2.99E-01 | 2.74E+00 | 2.77E+00 | 6.95E-01 | no | Lrp4 | intron_variant | MODIFIER |
| 7 | 82237295 | AX-170274568 | 1 | G | A | 0.098 | 7.04E-02 | 1.84E-01 | 2.64E+00 | 2.77E+00 | 6.95E-01 | no | Sh3gl3 | intron_variant | MODIFIER |
| 1 | 100798279 | AX-170472477 | 0 | G | A | 0.158 | 8.12E-02 | 2.08E-01 | 2.65E+00 | 2.71E+00 | 6.95E-01 | no | | intergenic_variant | MODIFIER |

| | | | | | | | | | | | | | | | |
|---|---|---|---|---|---|---|---|---|---|---|---|---|---|---|---|
| 16 | 45696030 | AX-170097787 | 1 | C | T | 0.518 | 3.21E-02 | 7.89E-02 | 2.57E+00 | 2.63E+00 | 6.95E-01 | no | Tmprss7 | upstream_gene_variant | MODIFIER |
| 18 | 51180613 | AX-170183430 | 1 | G | A | 0.241 | -4.31E-02 | 1.07E-01 | 2.59E+00 | 2.65E+00 | 6.95E-01 | no | Prr16 | intron_variant | MODIFIER |
| 9 | 62460098 | AX-170162109 | 0 | T | G | 0.202 | -4.11E-02 | 1.06E-01 | 2.65E+00 | 2.75E+00 | 6.95E-01 | no | Coro2b | intron_variant | MODIFIER |
| 5 | 104862877 | AX-170362076 | 1 | G | A | 0.446 | 4.29E-02 | 1.10E-01 | 2.65E+00 | 2.70E+00 | 6.95E-01 | no | Zfp951 | upstream_gene_variant | MODIFIER |
| 2 | 97371944 | AX-170216164 | 0 | A | G | 0.053 | -1.16E-01 | 2.99E-01 | 2.74E+00 | 2.77E+00 | 6.95E-01 | no | Lrrc4c | intron_variant | MODIFIER |
| 2 | 106454292 | AX-97831567 | 0 | G | A | 0.053 | -1.16E-01 | 2.99E-01 | 2.74E+00 | 2.77E+00 | 6.95E-01 | no | | regulatory_region_variant | MODIFIER |
| 5 | 27155589 | AX-170450307 | 0 | A | G | 0.053 | -1.16E-01 | 2.99E-01 | 2.74E+00 | 2.77E+00 | 6.95E-01 | no | Dpp6 | intron_variant | MODIFIER |
| 4 | 93961760 | AX-170023428 | 1 | T | C | 0.321 | 4.06E-02 | 1.04E-01 | 2.65E+00 | 2.72E+00 | 6.95E-01 | no | | intergenic_variant | MODIFIER |
| 8 | 87200142 | AX-170031204 | 1 | C | T | 0.205 | -4.91E-02 | 1.28E-01 | 2.69E+00 | 2.77E+00 | 6.95E-01 | no | Gm27169 | upstream_gene_variant | MODIFIER |
| 9 | 17217744 | AX-97826240 | 0 | C | T | 0.482 | 3.48E-02 | 8.73E-02 | 2.59E+00 | 2.67E+00 | 6.95E-01 | no | | intergenic_variant | MODIFIER |
| 1 | 63727433 | AX-170000663 | 1 | C | T | 0.134 | -7.56E-02 | 1.95E-01 | 2.64E+00 | 2.73E+00 | 6.95E-01 | no | Fastkd2 | upstream_gene_variant | MODIFIER |
| 10 | 4963931 | AX-170204825 | 2 | C | A | 0.191 | 3.78E-02 | 9.94E-02 | 2.68E+00 | 2.79E+00 | 6.96E-01 | no | Esr1 | intron_variant | MODIFIER |
| 17 | 55439786 | AX-170014434 | 0 | T | C | 0.053 | 5.42E-02 | 1.45E-01 | 2.66E+00 | 2.82E+00 | 6.96E-01 | no | | intergenic_variant | MODIFIER |
| 1 | 4164403 | AX-170515304 | 0 | A | G | 0.105 | -4.83E-02 | 1.27E-01 | 2.62E+00 | 2.76E+00 | 6.96E-01 | no | Rp1 | intron_variant | MODIFIER |
| 8 | 47480930 | AX-170447496 | 0 | A | G | 0.14 | -4.83E-02 | 1.20E-01 | 2.56E+00 | 2.66E+00 | 6.96E-01 | no | | intergenic_variant | MODIFIER |
| 1 | 25696173 | AX-170275949 | 1 | G | C | 0.411 | 3.24E-02 | 8.30E-02 | 2.64E+00 | 2.71E+00 | 6.96E-01 | no | Adgrb3 | intron_variant | MODIFIER |
| 1 | 30795968 | AX-170475211 | 0 | G | T | 0.404 | 3.20E-02 | 8.29E-02 | 2.67E+00 | 2.74E+00 | 6.96E-01 | no | Gm2914 | upstream_gene_variant | MODIFIER |
| 1 | 105153160 | AX-170375503 | 0 | G | A | 0.167 | 6.85E-02 | 1.72E-01 | 2.58E+00 | 2.68E+00 | 6.96E-01 | no | | intergenic_variant | MODIFIER |
| 5 | 110862219 | AX-170166507 | 1 | C | T | 0.455 | 4.20E-02 | 1.07E-01 | 2.63E+00 | 2.69E+00 | 6.96E-01 | no | Chek2 | intron_variant | MODIFIER |
| 15 | 86599761 | AX-170141713 | 0 | A | G | 0.491 | 2.87E-02 | 6.92E-02 | 2.52E+00 | 2.62E+00 | 6.96E-01 | no | | intergenic_variant | MODIFIER |
| 1 | 45625409 | AX-170100640 | 0 | G | A | 0.439 | -2.94E-02 | 7.63E-02 | 2.66E+00 | 2.75E+00 | 6.96E-01 | no | | intergenic_variant | MODIFIER |
| 9 | 111857993 | AX-170506126 | 0 | C | T | 0.105 | 4.45E-02 | 1.17E-01 | 2.74E+00 | 2.82E+00 | 6.96E-01 | no | | intergenic_variant | MODIFIER |
| 10 | 118896580 | AX-170263748 | 1 | C | A | 0.339 | 2.82E-02 | 7.38E-02 | 2.68E+00 | 2.77E+00 | 6.97E-01 | no | Gm26976 | downstream_gene_variant | MODIFIER |
| 12 | 111370214 | AX-170149793 | 0 | A | C | 0.132 | 6.40E-02 | 1.66E-01 | 2.64E+00 | 2.75E+00 | 6.97E-01 | no | Cdc42bpb | intron_variant | MODIFIER |
| 7 | 10456369 | AX-170487728 | 1 | C | T | 0.17 | 3.01E-02 | 8.09E-02 | 2.73E+00 | 2.87E+00 | 6.97E-01 | no | | intergenic_variant | MODIFIER |
| 1 | 169077719 | AX-170491209 | 2 | C | G | 0.218 | 4.86E-02 | 1.24E-01 | 2.61E+00 | 2.69E+00 | 6.97E-01 | no | | intergenic_variant | MODIFIER |
| 2 | 71044297 | AX-170374064 | 0 | C | T | 0.053 | -7.55E-02 | 1.98E-01 | 2.66E+00 | 2.77E+00 | 6.97E-01 | no | Mettl8 | intron_variant | MODIFIER |
| 1 | 100523459 | AX-170507403 | 1 | G | A | 0.161 | 8.02E-02 | 2.06E-01 | 2.64E+00 | 2.70E+00 | 6.97E-01 | no | | intergenic_variant | MODIFIER |
| 15 | 79927044 | AX-170485779 | 1 | G | C | 0.554 | 3.18E-02 | 7.92E-02 | 2.57E+00 | 2.65E+00 | 6.97E-01 | no | Cbx7 | intron_variant | MODIFIER |
| 12 | 105940478 | AX-170300064 | 0 | G | A | 0.123 | -6.94E-02 | 1.82E-01 | 2.69E+00 | 2.78E+00 | 6.97E-01 | no | | intergenic_variant | MODIFIER |
| 5 | 26202199 | AX-170243951 | 0 | C | T | 0.061 | 8.03E-02 | 2.12E-01 | 2.67E+00 | 2.78E+00 | 6.97E-01 | no | | intergenic_variant | MODIFIER |
| 10 | 8896485 | AX-170002682 | 0 | A | G | 0.202 | 3.81E-02 | 9.97E-02 | 2.66E+00 | 2.76E+00 | 6.97E-01 | no | | intergenic_variant | MODIFIER |
| 3 | 154942003 | AX-170021980 | 0 | T | C | 0.342 | 2.79E-02 | 7.32E-02 | 2.70E+00 | 2.78E+00 | 6.97E-01 | no | Tnni3k | intron_variant | MODIFIER |
| 3 | 155126410 | AX-170409879 | 0 | G | A | 0.342 | 2.79E-02 | 7.32E-02 | 2.70E+00 | 2.78E+00 | 6.97E-01 | no | Lrriq3 | intron_variant | MODIFIER |
| 3 | 155186484 | AX-170089122 | 0 | T | C | 0.342 | 2.79E-02 | 7.32E-02 | 2.70E+00 | 2.78E+00 | 6.97E-01 | no | Lrriq3 | intron_variant | MODIFIER |
| 3 | 155291992 | AX-97809202 | 0 | T | C | 0.342 | 2.79E-02 | 7.32E-02 | 2.70E+00 | 2.78E+00 | 6.97E-01 | no | | intergenic_variant | MODIFIER |
| 3 | 155367490 | AX-97798830 | 0 | G | T | 0.342 | 2.79E-02 | 7.32E-02 | 2.70E+00 | 2.78E+00 | 6.97E-01 | no | | intergenic_variant | MODIFIER |
| 3 | 155552913 | AX-97834679 | 0 | G | A | 0.342 | 2.79E-02 | 7.32E-02 | 2.70E+00 | 2.78E+00 | 6.97E-01 | no | | intergenic_variant | MODIFIER |
| 3 | 155782709 | AX-97797751 | 0 | G | A | 0.342 | 2.79E-02 | 7.32E-02 | 2.70E+00 | 2.78E+00 | 6.97E-01 | no | | intergenic_variant | MODIFIER |
| 8 | 47538695 | AX-170271909 | 0 | T | C | 0.149 | -5.94E-02 | 1.53E-01 | 2.65E+00 | 2.69E+00 | 6.97E-01 | no | Rwdd4a | intron_variant | MODIFIER |
| 9 | 33926100 | AX-170163207 | 0 | A | T | 0.43 | 3.33E-02 | 8.57E-02 | 2.64E+00 | 2.71E+00 | 6.97E-01 | no | Gm47785 | downstream_gene_variant | MODIFIER |
| 9 | 105077218 | AX-170083349 | 0 | G | T | 0.114 | 4.05E-02 | 1.08E-01 | 2.77E+00 | 2.88E+00 | 6.97E-01 | no | Mrpl3 | 3_prime_UTR_variant | MODIFIER |
| 2 | 147684219 | AX-170279097 | 1 | C | T | 0.134 | 7.49E-02 | 1.96E-01 | 2.68E+00 | 2.76E+00 | 6.97E-01 | no | | intergenic_variant | MODIFIER |
| 9 | 81220513 | AX-170090349 | 1 | A | G | 0.214 | -4.56E-02 | 1.17E-01 | 2.63E+00 | 2.71E+00 | 6.98E-01 | no | | intergenic_variant | MODIFIER |
| 13 | 53168147 | AX-170319291 | 0 | C | T | 0.518 | -2.43E-02 | 6.50E-02 | 2.73E+00 | 2.85E+00 | 6.98E-01 | no | Ror2 | intron_variant | MODIFIER |
| 5 | 97260215 | AX-97857143 | 0 | T | C | 0.447 | 4.03E-02 | 1.04E-01 | 2.66E+00 | 2.72E+00 | 6.98E-01 | no | Gm5560 | upstream_gene_variant | MODIFIER |
| 1 | 3475736 | AX-170286062 | 1 | A | G | 0.527 | 2.80E-02 | 6.86E-02 | 2.55E+00 | 2.62E+00 | 6.98E-01 | no | Xkr4 | intron_variant | MODIFIER |
| 2 | 72345481 | AX-170427349 | 0 | G | A | 0.053 | -7.29E-02 | 1.92E-01 | 2.64E+00 | 2.76E+00 | 6.98E-01 | no | Map3k20 | intron_variant | MODIFIER |
| 5 | 127843873 | AX-170111027 | 1 | C | T | 0.429 | -3.19E-02 | 8.49E-02 | 2.71E+00 | 2.82E+00 | 6.98E-01 | no | Tmem132d | intron_variant | MODIFIER |
| 12 | 109627267 | AX-170410337 | 0 | C | T | 0.132 | 6.36E-02 | 1.66E-01 | 2.63E+00 | 2.74E+00 | 6.98E-01 | no | Gm24564 | upstream_gene_variant | MODIFIER |
| 7 | 27298127 | AX-170028729 | 1 | C | T | 0.402 | 3.46E-02 | 8.74E-02 | 2.57E+00 | 2.68E+00 | 6.98E-01 | no | | regulatory_region_variant | MODIFIER |
| 7 | 40165427 | AX-170220863 | 1 | T | C | 0.348 | 3.72E-02 | 9.47E-02 | 2.61E+00 | 2.68E+00 | 6.98E-01 | no | | intergenic_variant | MODIFIER |
| 7 | 40209922 | AX-170028875 | 1 | A | C | 0.348 | 3.72E-02 | 9.47E-02 | 2.61E+00 | 2.68E+00 | 6.98E-01 | no | | intergenic_variant | MODIFIER |
| 1 | 30553867 | AX-97823599 | 1 | G | A | 0.429 | 3.07E-02 | 8.06E-02 | 2.68E+00 | 2.76E+00 | 6.98E-01 | no | | intergenic_variant | MODIFIER |

| | | | | | | | | | | | | | | | |
|---|---|---|---|---|---|---|---|---|---|---|---|---|---|---|---|
| 4 | 80552826 | AX-170023215 | 0 | C | A | 0.228 | 3.97E-02 | 9.94E-02 | 2.57E+00 | 2.66E+00 | 6.98E-01 | no | | regulatory_region_variant | MODIFIER |
| 16 | 88986072 | AX-170098744 | 0 | T | C | 0.465 | -3.07E-02 | 8.07E-02 | 2.69E+00 | 2.77E+00 | 6.98E-01 | no | | intergenic_variant | MODIFIER |
| 16 | 88992144 | AX-170513972 | 0 | T | A | 0.465 | -3.07E-02 | 8.07E-02 | 2.69E+00 | 2.77E+00 | 6.98E-01 | no | | intergenic_variant | MODIFIER |
| 19 | 39436017 | AX-170467162 | 1 | T | G | 0.036 | -4.61E-02 | 1.11E-01 | 2.46E+00 | 2.64E+00 | 6.98E-01 | no | Cyp2c38 | intron_variant | MODIFIER |
| 12 | 64258145 | AX-168391426 | 1 | C | T | 0.071 | -5.76E-02 | 1.52E-01 | 2.62E+00 | 2.75E+00 | 6.99E-01 | no | | intergenic_variant | MODIFIER |
| 5 | 78102856 | AX-170318574 | 1 | C | T | 0.429 | 4.10E-02 | 1.07E-01 | 2.68E+00 | 2.74E+00 | 6.99E-01 | no | | intergenic_variant | MODIFIER |
| 9 | 37200661 | AX-170412667 | 1 | C | A | 0.509 | -3.03E-02 | 8.01E-02 | 2.68E+00 | 2.78E+00 | 6.99E-01 | no | | intergenic_variant | MODIFIER |
| 20 | 165367385 | AX-170101799 | 0 | C | T | 0.132 | 4.86E-02 | 1.28E-01 | 2.65E+00 | 2.76E+00 | 6.99E-01 | no | | intergenic_variant | MODIFIER |
| 10 | 31396438 | AX-97845100 | 0 | C | T | 0.386 | -2.89E-02 | 7.46E-02 | 2.60E+00 | 2.71E+00 | 6.99E-01 | no | Tpd52l1 | intron_variant | MODIFIER |
| 10 | 32571411 | AX-170513312 | 0 | C | G | 0.386 | -2.89E-02 | 7.46E-02 | 2.60E+00 | 2.71E+00 | 6.99E-01 | no | Nkain2 | intron_variant | MODIFIER |
| 1 | 168211409 | AX-170242181 | 0 | T | C | 0.219 | 5.15E-02 | 1.32E-01 | 2.61E+00 | 2.68E+00 | 6.99E-01 | no | Pbx1 | intron_variant | MODIFIER |
| 1 | 168933860 | AX-97846721 | 0 | G | A | 0.219 | 5.15E-02 | 1.32E-01 | 2.61E+00 | 2.68E+00 | 6.99E-01 | no | | intergenic_variant | MODIFIER |
| 18 | 80235825 | AX-170534985 | 0 | C | G | 0.386 | -3.23E-02 | 8.34E-02 | 2.61E+00 | 2.71E+00 | 6.99E-01 | no | Hsbp1l1 | intron_variant | MODIFIER |
| 4 | 141181309 | AX-170070891 | 1 | A | T | 0.071 | 7.11E-02 | 1.85E-01 | 2.64E+00 | 2.71E+00 | 6.99E-01 | no | Fbxo42 | intron_variant | MODIFIER |
| 4 | 142080098 | AX-97833835 | 2 | T | C | 0.073 | 7.13E-02 | 1.85E-01 | 2.64E+00 | 2.71E+00 | 6.99E-01 | no | Tmem51 | intron_variant | MODIFIER |
| 4 | 144343831 | AX-97813813 | 2 | T | C | 0.073 | 7.13E-02 | 1.85E-01 | 2.64E+00 | 2.71E+00 | 6.99E-01 | no | Gm13122 | downstream_gene_variant | MODIFIER |
| 5 | 45056672 | AX-170159822 | 0 | T | C | 0.184 | -4.27E-02 | 1.11E-01 | 2.62E+00 | 2.70E+00 | 7.00E-01 | no | D5Ertd615e | intron_variant,non_coding_transcript_variant | MODIFIER |
| 5 | 28562038 | AX-97799160 | 0 | A | T | 0.368 | 3.86E-02 | 1.00E-01 | 2.64E+00 | 2.71E+00 | 7.00E-01 | no | 9530036O11Rik | intron_variant,non_coding_transcript_variant | MODIFIER |
| 7 | 77530095 | AX-170029352 | 0 | G | C | 0.377 | -3.62E-02 | 9.65E-02 | 2.68E+00 | 2.78E+00 | 7.00E-01 | no | | intergenic_variant | MODIFIER |
| 8 | 39034058 | AX-170216344 | 0 | T | C | 0.149 | -5.19E-02 | 1.36E-01 | 2.65E+00 | 2.73E+00 | 7.00E-01 | no | Tusc3 | intron_variant | MODIFIER |
| 8 | 39931390 | AX-170121039 | 0 | C | G | 0.149 | -5.19E-02 | 1.36E-01 | 2.65E+00 | 2.73E+00 | 7.00E-01 | no | | intergenic_variant | MODIFIER |
| 8 | 40462896 | AX-170264745 | 0 | T | C | 0.149 | -5.19E-02 | 1.36E-01 | 2.65E+00 | 2.73E+00 | 7.00E-01 | no | Zdhhc2 | intron_variant | MODIFIER |
| 8 | 41132028 | AX-170378918 | 0 | T | C | 0.149 | -5.19E-02 | 1.36E-01 | 2.65E+00 | 2.73E+00 | 7.00E-01 | no | Mtus1 | intron_variant | MODIFIER |
| 8 | 42263843 | AX-170079894 | 0 | T | G | 0.149 | -5.19E-02 | 1.36E-01 | 2.65E+00 | 2.73E+00 | 7.00E-01 | no | | intergenic_variant | MODIFIER |
| 8 | 43033239 | AX-170198835 | 0 | A | G | 0.149 | -5.19E-02 | 1.36E-01 | 2.65E+00 | 2.73E+00 | 7.00E-01 | no | | intergenic_variant | MODIFIER |
| 8 | 45574679 | AX-170079957 | 0 | A | G | 0.149 | -5.19E-02 | 1.36E-01 | 2.65E+00 | 2.73E+00 | 7.00E-01 | no | Sorbs2 | intron_variant | MODIFIER |
| 9 | 39630915 | AX-97783178 | 0 | A | G | 0.43 | -2.86E-02 | 7.55E-02 | 2.67E+00 | 2.76E+00 | 7.00E-01 | no | | intergenic_variant | MODIFIER |
| 9 | 39643758 | AX-170168593 | 0 | C | T | 0.43 | -2.86E-02 | 7.55E-02 | 2.67E+00 | 2.76E+00 | 7.00E-01 | no | Olfr961 | upstream_gene_variant | MODIFIER |
| 9 | 39675897 | AX-170109143 | 0 | G | A | 0.43 | -2.86E-02 | 7.55E-02 | 2.67E+00 | 2.76E+00 | 7.00E-01 | no | | intergenic_variant | MODIFIER |
| 9 | 39713287 | AX-170134203 | 0 | C | A | 0.43 | -2.86E-02 | 7.55E-02 | 2.67E+00 | 2.76E+00 | 7.00E-01 | no | Olfr965 | intron_variant | MODIFIER |
| 9 | 39746528 | AX-97783312 | 0 | A | C | 0.43 | -2.86E-02 | 7.55E-02 | 2.67E+00 | 2.76E+00 | 7.00E-01 | no | Olfr967 | upstream_gene_variant | MODIFIER |
| 9 | 39750160 | AX-168383045 | 0 | A | G | 0.43 | -2.86E-02 | 7.55E-02 | 2.67E+00 | 2.76E+00 | 7.00E-01 | no | Olfr967 | upstream_gene_variant | MODIFIER |
| 9 | 40137298 | AX-170353402 | 0 | G | C | 0.43 | -2.86E-02 | 7.55E-02 | 2.67E+00 | 2.76E+00 | 7.00E-01 | no | | intergenic_variant | MODIFIER |
| 9 | 40140621 | AX-170315088 | 0 | C | T | 0.43 | -2.86E-02 | 7.55E-02 | 2.67E+00 | 2.76E+00 | 7.00E-01 | no | | intergenic_variant | MODIFIER |
| 9 | 40148186 | AX-170368393 | 0 | T | C | 0.43 | -2.86E-02 | 7.55E-02 | 2.67E+00 | 2.76E+00 | 7.00E-01 | no | Tmem225 | 5_prime_UTR_variant | MODIFIER |
| 9 | 40404073 | AX-170082202 | 0 | G | C | 0.43 | -2.86E-02 | 7.55E-02 | 2.67E+00 | 2.76E+00 | 7.00E-01 | no | Gramd1b | intron_variant | MODIFIER |
| 9 | 40408637 | AX-170364462 | 0 | C | T | 0.43 | -2.86E-02 | 7.55E-02 | 2.67E+00 | 2.76E+00 | 7.00E-01 | no | Gramd1b | intron_variant | MODIFIER |
| 9 | 40408693 | AX-170112664 | 0 | A | G | 0.43 | -2.86E-02 | 7.55E-02 | 2.67E+00 | 2.76E+00 | 7.00E-01 | no | Gramd1b | intron_variant | MODIFIER |
| 9 | 40430128 | AX-170316303 | 0 | A | G | 0.43 | -2.86E-02 | 7.55E-02 | 2.67E+00 | 2.76E+00 | 7.00E-01 | no | Gramd1b | intron_variant | MODIFIER |
| 9 | 40438203 | AX-170353609 | 0 | G | A | 0.43 | -2.86E-02 | 7.55E-02 | 2.67E+00 | 2.76E+00 | 7.00E-01 | no | Gramd1b | intron_variant | MODIFIER |
| 9 | 40473610 | AX-170436487 | 0 | A | G | 0.43 | -2.86E-02 | 7.55E-02 | 2.67E+00 | 2.76E+00 | 7.00E-01 | no | Gramd1b | intron_variant | MODIFIER |
| 9 | 40484898 | AX-170468159 | 0 | C | T | 0.43 | -2.86E-02 | 7.55E-02 | 2.67E+00 | 2.76E+00 | 7.00E-01 | no | Gramd1b | intron_variant | MODIFIER |
| 9 | 40503368 | AX-170498911 | 0 | A | C | 0.43 | -2.86E-02 | 7.55E-02 | 2.67E+00 | 2.76E+00 | 7.00E-01 | no | Gramd1b | intron_variant | MODIFIER |
| 9 | 40508137 | AX-97784431 | 0 | A | G | 0.43 | -2.86E-02 | 7.55E-02 | 2.67E+00 | 2.76E+00 | 7.00E-01 | no | Gramd1b | intron_variant | MODIFIER |
| 9 | 40510871 | AX-170082206 | 0 | A | C | 0.43 | -2.86E-02 | 7.55E-02 | 2.67E+00 | 2.76E+00 | 7.00E-01 | no | Gramd1b | intron_variant | MODIFIER |
| 9 | 40539059 | AX-170082207 | 0 | C | G | 0.43 | -2.86E-02 | 7.55E-02 | 2.67E+00 | 2.76E+00 | 7.00E-01 | no | | regulatory_region_variant | MODIFIER |
| 9 | 40540057 | AX-170514993 | 0 | A | G | 0.43 | -2.86E-02 | 7.55E-02 | 2.67E+00 | 2.76E+00 | 7.00E-01 | no | | regulatory_region_variant | MODIFIER |
| 9 | 40548026 | AX-170269716 | 0 | G | T | 0.43 | -2.86E-02 | 7.55E-02 | 2.67E+00 | 2.76E+00 | 7.00E-01 | no | Gm35177 | intron_variant,non_coding_transcript_variant | MODIFIER |
| 9 | 40548276 | AX-170238821 | 0 | A | C | 0.43 | -2.86E-02 | 7.55E-02 | 2.67E+00 | 2.76E+00 | 7.00E-01 | no | Gm35177 | intron_variant,non_coding_transcript_variant | MODIFIER |
| 9 | 40549051 | AX-170452487 | 0 | A | C | 0.43 | -2.86E-02 | 7.55E-02 | 2.67E+00 | 2.76E+00 | 7.00E-01 | no | Gm35177 | intron_variant,non_coding_transcript_variant | MODIFIER |
| 9 | 40561368 | AX-170174728 | 0 | G | C | 0.43 | -2.86E-02 | 7.55E-02 | 2.67E+00 | 2.76E+00 | 7.00E-01 | no | Gm48277 | upstream_gene_variant | MODIFIER |
| 9 | 40566478 | AX-170321130 | 0 | G | A | 0.43 | -2.86E-02 | 7.55E-02 | 2.67E+00 | 2.76E+00 | 7.00E-01 | no | | intergenic_variant | MODIFIER |
| 9 | 40580629 | AX-170301457 | 0 | C | A | 0.43 | -2.86E-02 | 7.55E-02 | 2.67E+00 | 2.76E+00 | 7.00E-01 | no | | intergenic_variant | MODIFIER |
| 9 | 40585392 | AX-170455669 | 0 | T | C | 0.43 | -2.86E-02 | 7.55E-02 | 2.67E+00 | 2.76E+00 | 7.00E-01 | no | | intergenic_variant | MODIFIER |

| | | | | | | | | | | | | | | | |
|---|---|---|---|---|---|---|---|---|---|---|---|---|---|---|---|
| 9 | 40586413 | AX-170510790 | 0 | G | A | 0.43 | -2.86E-02 | 7.55E-02 | 2.67E+00 | 2.76E+00 | 7.00E-01 | no | | intergenic_variant | MODIFIER |
| 9 | 40646961 | AX-170097389 | 0 | G | C | 0.43 | -2.86E-02 | 7.55E-02 | 2.67E+00 | 2.76E+00 | 7.00E-01 | no | | intergenic_variant | MODIFIER |
| 9 | 40651333 | AX-97834905 | 0 | G | A | 0.43 | -2.86E-02 | 7.55E-02 | 2.67E+00 | 2.76E+00 | 7.00E-01 | no | | intergenic_variant | MODIFIER |
| 9 | 40673316 | AX-170082209 | 0 | A | G | 0.43 | -2.86E-02 | 7.55E-02 | 2.67E+00 | 2.76E+00 | 7.00E-01 | no | Gm21915 | downstream_gene_variant | MODIFIER |
| 9 | 40710538 | AX-170272709 | 0 | G | A | 0.43 | -2.86E-02 | 7.55E-02 | 2.67E+00 | 2.76E+00 | 7.00E-01 | no | Clmp | intron_variant | MODIFIER |
| 9 | 40718452 | AX-170420518 | 0 | G | C | 0.43 | -2.86E-02 | 7.55E-02 | 2.67E+00 | 2.76E+00 | 7.00E-01 | no | Clmp | intron_variant | MODIFIER |
| 9 | 40722043 | AX-97839296 | 0 | G | A | 0.43 | -2.86E-02 | 7.55E-02 | 2.67E+00 | 2.76E+00 | 7.00E-01 | no | Clmp | intron_variant | MODIFIER |
| 9 | 40725437 | AX-170315049 | 0 | G | T | 0.43 | -2.86E-02 | 7.55E-02 | 2.67E+00 | 2.76E+00 | 7.00E-01 | no | Clmp | intron_variant | MODIFIER |
| 9 | 40744780 | AX-170192915 | 0 | G | A | 0.43 | -2.86E-02 | 7.55E-02 | 2.67E+00 | 2.76E+00 | 7.00E-01 | no | Clmp | intron_variant | MODIFIER |
| 9 | 40747600 | AX-170148151 | 0 | T | G | 0.43 | -2.86E-02 | 7.55E-02 | 2.67E+00 | 2.76E+00 | 7.00E-01 | no | Clmp | intron_variant | MODIFIER |
| 9 | 40774081 | AX-170330050 | 0 | C | T | 0.43 | -2.86E-02 | 7.55E-02 | 2.67E+00 | 2.76E+00 | 7.00E-01 | no | Clmp | intron_variant | MODIFIER |
| 9 | 40810066 | AX-170158549 | 0 | A | G | 0.43 | -2.86E-02 | 7.55E-02 | 2.67E+00 | 2.76E+00 | 7.00E-01 | no | Hspa8 | downstream_gene_variant | MODIFIER |
| 9 | 40857095 | AX-170255434 | 0 | G | A | 0.43 | -2.86E-02 | 7.55E-02 | 2.67E+00 | 2.76E+00 | 7.00E-01 | no | | intergenic_variant | MODIFIER |
| 9 | 40861965 | AX-170533960 | 0 | T | C | 0.43 | -2.86E-02 | 7.55E-02 | 2.67E+00 | 2.76E+00 | 7.00E-01 | no | | intergenic_variant | MODIFIER |
| 9 | 40862963 | AX-97840526 | 0 | G | A | 0.43 | -2.86E-02 | 7.55E-02 | 2.67E+00 | 2.76E+00 | 7.00E-01 | no | | intergenic_variant | MODIFIER |
| 9 | 40870023 | AX-170391743 | 0 | G | A | 0.43 | -2.86E-02 | 7.55E-02 | 2.67E+00 | 2.76E+00 | 7.00E-01 | no | Bsx | upstream_gene_variant | MODIFIER |
| 9 | 40897081 | AX-170500528 | 0 | T | C | 0.43 | -2.86E-02 | 7.55E-02 | 2.67E+00 | 2.76E+00 | 7.00E-01 | no | Jhy | intron_variant | MODIFIER |
| 9 | 40908748 | AX-170032191 | 0 | A | C | 0.43 | -2.86E-02 | 7.55E-02 | 2.67E+00 | 2.76E+00 | 7.00E-01 | no | Jhy | intron_variant | MODIFIER |
| 9 | 40910406 | AX-170032192 | 0 | G | A | 0.43 | -2.86E-02 | 7.55E-02 | 2.67E+00 | 2.76E+00 | 7.00E-01 | no | Jhy | intron_variant | MODIFIER |
| 9 | 40910484 | AX-170032187 | 0 | T | A | 0.43 | -2.86E-02 | 7.55E-02 | 2.67E+00 | 2.76E+00 | 7.00E-01 | no | Jhy | intron_variant | MODIFIER |
| 9 | 40933933 | AX-97832189 | 0 | G | A | 0.43 | -2.86E-02 | 7.55E-02 | 2.67E+00 | 2.76E+00 | 7.00E-01 | no | Jhy | intron_variant | MODIFIER |
| 9 | 40984136 | AX-170032189 | 0 | C | T | 0.43 | -2.86E-02 | 7.55E-02 | 2.67E+00 | 2.76E+00 | 7.00E-01 | no | Crtam | intron_variant | MODIFIER |
| 9 | 41002587 | AX-170374653 | 0 | A | G | 0.43 | -2.86E-02 | 7.55E-02 | 2.67E+00 | 2.76E+00 | 7.00E-01 | no | Crtam | intron_variant | MODIFIER |
| 9 | 41031978 | AX-97848416 | 0 | A | C | 0.43 | -2.86E-02 | 7.55E-02 | 2.67E+00 | 2.76E+00 | 7.00E-01 | no | Ubash3b | intron_variant | MODIFIER |
| 9 | 41040054 | AX-170126443 | 0 | A | G | 0.43 | -2.86E-02 | 7.55E-02 | 2.67E+00 | 2.76E+00 | 7.00E-01 | no | Ubash3b | intron_variant | MODIFIER |
| 9 | 41050740 | AX-170178457 | 0 | C | A | 0.43 | -2.86E-02 | 7.55E-02 | 2.67E+00 | 2.76E+00 | 7.00E-01 | no | Ubash3b | intron_variant | MODIFIER |
| 9 | 41051378 | AX-170451215 | 0 | A | G | 0.43 | -2.86E-02 | 7.55E-02 | 2.67E+00 | 2.76E+00 | 7.00E-01 | no | Ubash3b | splice_region_variant,intron_variant | LOW |
| 9 | 41062708 | AX-170379498 | 0 | G | A | 0.43 | -2.86E-02 | 7.55E-02 | 2.67E+00 | 2.76E+00 | 7.00E-01 | no | Ubash3b | intron_variant | MODIFIER |
| 9 | 41085406 | AX-170359888 | 0 | A | G | 0.43 | -2.86E-02 | 7.55E-02 | 2.67E+00 | 2.76E+00 | 7.00E-01 | no | Ubash3b | intron_variant | MODIFIER |
| 15 | 90072911 | AX-170377991 | 0 | T | C | 0.14 | 6.60E-02 | 1.71E-01 | 2.62E+00 | 2.71E+00 | 7.00E-01 | no | Gm36330 | upstream_gene_variant | MODIFIER |
| 4 | 150433481 | AX-170403975 | 0 | C | G | 0.096 | -6.32E-02 | 1.67E-01 | 2.67E+00 | 2.77E+00 | 7.00E-01 | no | Rere | intron_variant | MODIFIER |
| 9 | 40539388 | AX-170405340 | 1 | T | C | 0.429 | -2.87E-02 | 7.56E-02 | 2.67E+00 | 2.76E+00 | 7.00E-01 | no | | regulatory_region_variant | MODIFIER |
| 3 | 148025347 | AX-97828117 | 0 | T | C | 0.342 | 2.80E-02 | 7.32E-02 | 2.65E+00 | 2.72E+00 | 7.00E-01 | no | | intergenic_variant | MODIFIER |
| 12 | 105910420 | AX-170092604 | 0 | A | G | 0.132 | -5.92E-02 | 1.58E-01 | 2.70E+00 | 2.79E+00 | 7.00E-01 | no | | intergenic_variant | MODIFIER |
| 1 | 43717497 | AX-170000383 | 1 | A | G | 0.089 | -7.23E-02 | 1.92E-01 | 2.70E+00 | 2.78E+00 | 7.00E-01 | no | | intergenic_variant | MODIFIER |
| 20 | 114270615 | AX-170529890 | 1 | C | A | 0.321 | 1.95E-02 | 5.10E-02 | 2.61E+00 | 2.73E+00 | 7.00E-01 | no | | intergenic_variant | MODIFIER |
| 4 | 57010280 | AX-170022850 | 1 | G | T | 0.366 | -4.05E-02 | 1.07E-01 | 2.69E+00 | 2.76E+00 | 7.00E-01 | no | Epb41l4b | intron_variant | MODIFIER |
| 1 | 114007297 | AX-97807249 | 1 | T | C | 0.188 | 5.25E-02 | 1.36E-01 | 2.60E+00 | 2.72E+00 | 7.01E-01 | no | | intergenic_variant | MODIFIER |
| 7 | 45972360 | AX-170229320 | 0 | G | C | 0.325 | 3.49E-02 | 9.00E-02 | 2.60E+00 | 2.69E+00 | 7.01E-01 | no | Abcc6 | downstream_gene_variant | MODIFIER |
| 1 | 76998234 | AX-97825703 | 0 | G | A | 0.132 | 8.42E-02 | 2.22E-01 | 2.68E+00 | 2.74E+00 | 7.01E-01 | no | Gm816 | intron_variant,non_coding_transcript_variant | MODIFIER |
| 8 | 78901206 | AX-170080524 | 0 | C | T | 0.202 | -5.06E-02 | 1.33E-01 | 2.69E+00 | 2.75E+00 | 7.01E-01 | no | | intergenic_variant | MODIFIER |
| 4 | 88610839 | AX-170198443 | 0 | A | G | 0.333 | 4.18E-02 | 1.09E-01 | 2.65E+00 | 2.71E+00 | 7.01E-01 | no | Gm12601 | downstream_gene_variant | MODIFIER |
| 4 | 88686089 | AX-170130583 | 0 | A | G | 0.333 | 4.18E-02 | 1.09E-01 | 2.65E+00 | 2.71E+00 | 7.01E-01 | no | Ifnab | downstream_gene_variant | MODIFIER |
| 4 | 89051267 | AX-170127153 | 0 | A | G | 0.333 | 4.18E-02 | 1.09E-01 | 2.65E+00 | 2.71E+00 | 7.01E-01 | no | Wincr1 | intron_variant,non_coding_transcript_variant | MODIFIER |
| 4 | 89130810 | AX-170198762 | 0 | T | C | 0.333 | 4.18E-02 | 1.09E-01 | 2.65E+00 | 2.71E+00 | 7.01E-01 | no | Gm12602 | upstream_gene_variant | MODIFIER |
| 4 | 89141979 | AX-170089310 | 0 | T | A | 0.333 | 4.18E-02 | 1.09E-01 | 2.65E+00 | 2.71E+00 | 7.01E-01 | no | Mtap | intron_variant | MODIFIER |
| 4 | 89701055 | AX-170206167 | 0 | A | G | 0.333 | 4.18E-02 | 1.09E-01 | 2.65E+00 | 2.71E+00 | 7.01E-01 | no | | intergenic_variant | MODIFIER |
| 4 | 89805275 | AX-170410090 | 0 | G | A | 0.333 | 4.18E-02 | 1.09E-01 | 2.65E+00 | 2.71E+00 | 7.01E-01 | no | | intergenic_variant | MODIFIER |
| 4 | 89812788 | AX-170332519 | 0 | C | T | 0.333 | 4.18E-02 | 1.09E-01 | 2.65E+00 | 2.71E+00 | 7.01E-01 | no | | intergenic_variant | MODIFIER |
| 4 | 89939632 | AX-97823229 | 0 | C | G | 0.333 | 4.18E-02 | 1.09E-01 | 2.65E+00 | 2.71E+00 | 7.01E-01 | no | | intergenic_variant | MODIFIER |
| 4 | 89948684 | AX-170351435 | 0 | C | T | 0.333 | 4.18E-02 | 1.09E-01 | 2.65E+00 | 2.71E+00 | 7.01E-01 | no | | intergenic_variant | MODIFIER |
| 4 | 89959549 | AX-170446568 | 0 | C | T | 0.333 | 4.18E-02 | 1.09E-01 | 2.65E+00 | 2.71E+00 | 7.01E-01 | no | | intergenic_variant | MODIFIER |
| 4 | 90051624 | AX-170485159 | 0 | A | G | 0.333 | 4.18E-02 | 1.09E-01 | 2.65E+00 | 2.71E+00 | 7.01E-01 | no | | intergenic_variant | MODIFIER |
| 4 | 90052502 | AX-97821407 | 0 | T | G | 0.333 | 4.18E-02 | 1.09E-01 | 2.65E+00 | 2.71E+00 | 7.01E-01 | no | | intergenic_variant | MODIFIER |

| | | | | | | | | | | | | | | | |
|---|---|---|---|---|---|---|---|---|---|---|---|---|---|---|---|
| 4 | 90069174 | AX-170023373 | 0 | A | C | 0.333 | 4.18E-02 | 1.09E-01 | 2.65E+00 | 2.71E+00 | 7.01E-01 | no | | intergenic_variant | MODIFIER |
| 4 | 90069222 | AX-170069961 | 0 | G | C | 0.333 | 4.18E-02 | 1.09E-01 | 2.65E+00 | 2.71E+00 | 7.01E-01 | no | | intergenic_variant | MODIFIER |
| 4 | 90076936 | AX-170023378 | 0 | A | G | 0.333 | 4.18E-02 | 1.09E-01 | 2.65E+00 | 2.71E+00 | 7.01E-01 | no | | intergenic_variant | MODIFIER |
| 4 | 90188153 | AX-170471719 | 0 | A | G | 0.333 | 4.18E-02 | 1.09E-01 | 2.65E+00 | 2.71E+00 | 7.01E-01 | no | Gm12629 | intron_variant,non_coding_transcript_variant | MODIFIER |
| 4 | 90190624 | AX-97785220 | 0 | G | A | 0.333 | 4.18E-02 | 1.09E-01 | 2.65E+00 | 2.71E+00 | 7.01E-01 | no | Gm12629 | downstream_gene_variant | MODIFIER |
| 4 | 90192284 | AX-170023374 | 0 | A | G | 0.333 | 4.18E-02 | 1.09E-01 | 2.65E+00 | 2.71E+00 | 7.01E-01 | no | Gm12629 | downstream_gene_variant | MODIFIER |
| 4 | 91311085 | AX-97797059 | 0 | C | T | 0.333 | 4.18E-02 | 1.09E-01 | 2.65E+00 | 2.71E+00 | 7.01E-01 | no | Elavl2 | intron_variant | MODIFIER |
| 4 | 91503070 | AX-170158642 | 0 | C | T | 0.333 | 4.18E-02 | 1.09E-01 | 2.65E+00 | 2.71E+00 | 7.01E-01 | no | | intergenic_variant | MODIFIER |
| 4 | 91549258 | AX-170216003 | 0 | G | A | 0.333 | 4.18E-02 | 1.09E-01 | 2.65E+00 | 2.71E+00 | 7.01E-01 | no | | intergenic_variant | MODIFIER |
| 4 | 91940232 | AX-170319773 | 0 | G | C | 0.333 | 4.18E-02 | 1.09E-01 | 2.65E+00 | 2.71E+00 | 7.01E-01 | no | 4930577H14Rik | upstream_gene_variant | MODIFIER |
| 4 | 92340970 | AX-97782717 | 0 | A | G | 0.333 | 4.18E-02 | 1.09E-01 | 2.65E+00 | 2.71E+00 | 7.01E-01 | no | | intergenic_variant | MODIFIER |
| 4 | 92585716 | AX-170123937 | 0 | C | T | 0.333 | 4.18E-02 | 1.09E-01 | 2.65E+00 | 2.71E+00 | 7.01E-01 | no | | intergenic_variant | MODIFIER |
| 4 | 92599815 | AX-170427141 | 0 | G | T | 0.333 | 4.18E-02 | 1.09E-01 | 2.65E+00 | 2.71E+00 | 7.01E-01 | no | | intergenic_variant | MODIFIER |
| 4 | 92605128 | AX-170309877 | 0 | G | A | 0.333 | 4.18E-02 | 1.09E-01 | 2.65E+00 | 2.71E+00 | 7.01E-01 | no | | intergenic_variant | MODIFIER |
| 4 | 92714231 | AX-170023410 | 0 | T | C | 0.333 | 4.18E-02 | 1.09E-01 | 2.65E+00 | 2.71E+00 | 7.01E-01 | no | | intergenic_variant | MODIFIER |
| 4 | 92750049 | AX-170338878 | 0 | T | C | 0.333 | 4.18E-02 | 1.09E-01 | 2.65E+00 | 2.71E+00 | 7.01E-01 | no | | intergenic_variant | MODIFIER |
| 4 | 92750422 | AX-170345498 | 0 | G | A | 0.333 | 4.18E-02 | 1.09E-01 | 2.65E+00 | 2.71E+00 | 7.01E-01 | no | | intergenic_variant | MODIFIER |
| 4 | 92947230 | AX-170165458 | 0 | G | A | 0.333 | 4.18E-02 | 1.09E-01 | 2.65E+00 | 2.71E+00 | 7.01E-01 | no | | intergenic_variant | MODIFIER |
| 4 | 93050582 | AX-170184227 | 0 | T | C | 0.333 | 4.18E-02 | 1.09E-01 | 2.65E+00 | 2.71E+00 | 7.01E-01 | no | | intergenic_variant | MODIFIER |
| 4 | 93051420 | AX-170193682 | 0 | G | T | 0.333 | 4.18E-02 | 1.09E-01 | 2.65E+00 | 2.71E+00 | 7.01E-01 | no | | intergenic_variant | MODIFIER |
| 4 | 93058589 | AX-170460565 | 0 | A | G | 0.333 | 4.18E-02 | 1.09E-01 | 2.65E+00 | 2.71E+00 | 7.01E-01 | no | | intergenic_variant | MODIFIER |
| 4 | 93079962 | AX-170162552 | 0 | T | G | 0.333 | 4.18E-02 | 1.09E-01 | 2.65E+00 | 2.71E+00 | 7.01E-01 | no | | intergenic_variant | MODIFIER |
| 4 | 93083802 | AX-170262252 | 0 | A | G | 0.333 | 4.18E-02 | 1.09E-01 | 2.65E+00 | 2.71E+00 | 7.01E-01 | no | | intergenic_variant | MODIFIER |
| 4 | 93090291 | AX-97781006 | 0 | T | C | 0.333 | 4.18E-02 | 1.09E-01 | 2.65E+00 | 2.71E+00 | 7.01E-01 | no | | intergenic_variant | MODIFIER |
| 4 | 93130614 | AX-170355850 | 0 | C | T | 0.333 | 4.18E-02 | 1.09E-01 | 2.65E+00 | 2.71E+00 | 7.01E-01 | no | | intergenic_variant | MODIFIER |
| 4 | 93319402 | AX-97784826 | 0 | A | G | 0.333 | 4.18E-02 | 1.09E-01 | 2.65E+00 | 2.71E+00 | 7.01E-01 | no | | intergenic_variant | MODIFIER |
| 4 | 93327101 | AX-170103009 | 0 | T | C | 0.333 | 4.18E-02 | 1.09E-01 | 2.65E+00 | 2.71E+00 | 7.01E-01 | no | | intergenic_variant | MODIFIER |
| 4 | 93612746 | AX-170414480 | 0 | G | A | 0.333 | 4.18E-02 | 1.09E-01 | 2.65E+00 | 2.71E+00 | 7.01E-01 | no | Gm12649 | intron_variant,non_coding_transcript_variant | MODIFIER |
| 4 | 93617343 | AX-170327073 | 0 | T | G | 0.333 | 4.18E-02 | 1.09E-01 | 2.65E+00 | 2.71E+00 | 7.01E-01 | no | Gm12649 | intron_variant,non_coding_transcript_variant | MODIFIER |
| 4 | 93632650 | AX-170441830 | 0 | T | C | 0.333 | 4.18E-02 | 1.09E-01 | 2.65E+00 | 2.71E+00 | 7.01E-01 | no | Gm12649 | intron_variant,non_coding_transcript_variant | MODIFIER |
| 4 | 93667245 | AX-97855133 | 0 | G | C | 0.333 | 4.18E-02 | 1.09E-01 | 2.65E+00 | 2.71E+00 | 7.01E-01 | no | Gm12649 | intron_variant,non_coding_transcript_variant | MODIFIER |
| 4 | 93717603 | AX-170335095 | 0 | A | T | 0.333 | 4.18E-02 | 1.09E-01 | 2.65E+00 | 2.71E+00 | 7.01E-01 | no | | intergenic_variant | MODIFIER |
| 4 | 93721078 | AX-170322356 | 0 | G | T | 0.333 | 4.18E-02 | 1.09E-01 | 2.65E+00 | 2.71E+00 | 7.01E-01 | no | | intergenic_variant | MODIFIER |
| 4 | 93728931 | AX-170320251 | 0 | G | A | 0.333 | 4.18E-02 | 1.09E-01 | 2.65E+00 | 2.71E+00 | 7.01E-01 | no | | intergenic_variant | MODIFIER |
| 4 | 93756185 | AX-170172862 | 0 | G | A | 0.333 | 4.18E-02 | 1.09E-01 | 2.65E+00 | 2.71E+00 | 7.01E-01 | no | | intergenic_variant | MODIFIER |
| 4 | 93882938 | AX-170023424 | 0 | G | A | 0.333 | 4.18E-02 | 1.09E-01 | 2.65E+00 | 2.71E+00 | 7.01E-01 | no | | intergenic_variant | MODIFIER |
| 4 | 93899166 | AX-170070024 | 0 | C | T | 0.333 | 4.18E-02 | 1.09E-01 | 2.65E+00 | 2.71E+00 | 7.01E-01 | no | | intergenic_variant | MODIFIER |
| 4 | 93929116 | AX-170023423 | 0 | G | A | 0.333 | 4.18E-02 | 1.09E-01 | 2.65E+00 | 2.71E+00 | 7.01E-01 | no | | intergenic_variant | MODIFIER |
| 4 | 93972322 | AX-170340220 | 0 | G | A | 0.333 | 4.18E-02 | 1.09E-01 | 2.65E+00 | 2.71E+00 | 7.01E-01 | no | | intergenic_variant | MODIFIER |
| 4 | 94014079 | AX-168392605 | 0 | C | T | 0.333 | 4.18E-02 | 1.09E-01 | 2.65E+00 | 2.71E+00 | 7.01E-01 | no | | intergenic_variant | MODIFIER |
| 4 | 94152723 | AX-170233229 | 0 | C | T | 0.333 | 4.18E-02 | 1.09E-01 | 2.65E+00 | 2.71E+00 | 7.01E-01 | no | Gm12648 | intron_variant,non_coding_transcript_variant | MODIFIER |
| 4 | 94311045 | AX-170023435 | 0 | G | A | 0.333 | 4.18E-02 | 1.09E-01 | 2.65E+00 | 2.71E+00 | 7.01E-01 | no | Gm12648 | intron_variant,non_coding_transcript_variant | MODIFIER |
| 4 | 94432389 | AX-170402246 | 0 | G | A | 0.333 | 4.18E-02 | 1.09E-01 | 2.65E+00 | 2.71E+00 | 7.01E-01 | no | Gm26410 | downstream_gene_variant | MODIFIER |
| 4 | 94445003 | AX-170090782 | 0 | A | G | 0.333 | 4.18E-02 | 1.09E-01 | 2.65E+00 | 2.71E+00 | 7.01E-01 | no | | intergenic_variant | MODIFIER |
| 4 | 94445069 | AX-170135226 | 0 | A | T | 0.333 | 4.18E-02 | 1.09E-01 | 2.65E+00 | 2.71E+00 | 7.01E-01 | no | | intergenic_variant | MODIFIER |
| 4 | 94447779 | AX-170409072 | 0 | T | C | 0.333 | 4.18E-02 | 1.09E-01 | 2.65E+00 | 2.71E+00 | 7.01E-01 | no | | intergenic_variant | MODIFIER |
| 4 | 94451348 | AX-170500932 | 0 | G | A | 0.333 | 4.18E-02 | 1.09E-01 | 2.65E+00 | 2.71E+00 | 7.01E-01 | no | | intergenic_variant | MODIFIER |
| 4 | 94453616 | AX-170363927 | 0 | T | A | 0.333 | 4.18E-02 | 1.09E-01 | 2.65E+00 | 2.71E+00 | 7.01E-01 | no | | intergenic_variant | MODIFIER |
| 4 | 94465145 | AX-170288435 | 0 | A | G | 0.333 | 4.18E-02 | 1.09E-01 | 2.65E+00 | 2.71E+00 | 7.01E-01 | no | Gm12655 | intron_variant,non_coding_transcript_variant | MODIFIER |
| 4 | 94472195 | AX-170442678 | 0 | C | T | 0.333 | 4.18E-02 | 1.09E-01 | 2.65E+00 | 2.71E+00 | 7.01E-01 | no | Gm12655 | intron_variant,non_coding_transcript_variant | MODIFIER |
| 4 | 94487316 | AX-170218871 | 0 | C | T | 0.333 | 4.18E-02 | 1.09E-01 | 2.65E+00 | 2.71E+00 | 7.01E-01 | no | Gm22804 | upstream_gene_variant | MODIFIER |
| 4 | 94492733 | AX-170159016 | 0 | T | C | 0.333 | 4.18E-02 | 1.09E-01 | 2.65E+00 | 2.71E+00 | 7.01E-01 | no | Gm12656 | upstream_gene_variant | MODIFIER |
| 4 | 94516525 | AX-97804423 | 0 | A | T | 0.333 | 4.18E-02 | 1.09E-01 | 2.65E+00 | 2.71E+00 | 7.01E-01 | no | Caap1 | intron_variant | MODIFIER |
| 4 | 94545876 | AX-170189418 | 0 | C | G | 0.333 | 4.18E-02 | 1.09E-01 | 2.65E+00 | 2.71E+00 | 7.01E-01 | no | Caap1 | intron_variant | MODIFIER |

| | | | | | | | | | | | | | | | |
|---|---|---|---|---|---|---|---|---|---|---|---|---|---|---|---|
| 4 | 94546079 | AX-170140088 | 0 | T | C | 0.333 | 4.18E-02 | 1.09E-01 | 2.65E+00 | 2.71E+00 | 7.01E-01 | no | Caap1 | intron_variant | MODIFIER |
| 4 | 94557675 | AX-170132483 | 0 | A | C | 0.333 | 4.18E-02 | 1.09E-01 | 2.65E+00 | 2.71E+00 | 7.01E-01 | no | Caap1 | upstream_gene_variant | MODIFIER |
| 4 | 94582904 | AX-170402530 | 0 | A | G | 0.333 | 4.18E-02 | 1.09E-01 | 2.65E+00 | 2.71E+00 | 7.01E-01 | no | Plaa | intron_variant | MODIFIER |
| 4 | 94588702 | AX-170368536 | 0 | A | C | 0.333 | 4.18E-02 | 1.09E-01 | 2.65E+00 | 2.71E+00 | 7.01E-01 | no | Plaa | intron_variant | MODIFIER |
| 4 | 94639240 | AX-97788956 | 0 | C | T | 0.333 | 4.18E-02 | 1.09E-01 | 2.65E+00 | 2.71E+00 | 7.01E-01 | no | Ift74 | intron_variant | MODIFIER |
| 4 | 94661389 | AX-170495238 | 0 | A | C | 0.333 | 4.18E-02 | 1.09E-01 | 2.65E+00 | 2.71E+00 | 7.01E-01 | no | Ift74 | intron_variant | MODIFIER |
| 4 | 94676592 | AX-97857265 | 0 | T | C | 0.333 | 4.18E-02 | 1.09E-01 | 2.65E+00 | 2.71E+00 | 7.01E-01 | no | Ift74 | intron_variant | MODIFIER |
| 4 | 94677526 | AX-170344324 | 0 | C | T | 0.333 | 4.18E-02 | 1.09E-01 | 2.65E+00 | 2.71E+00 | 7.01E-01 | no | Ift74 | intron_variant | MODIFIER |
| 4 | 94686991 | AX-170158675 | 0 | C | A | 0.333 | 4.18E-02 | 1.09E-01 | 2.65E+00 | 2.71E+00 | 7.01E-01 | no | Ift74 | intron_variant | MODIFIER |
| 4 | 94748052 | AX-170158631 | 0 | C | G | 0.333 | 4.18E-02 | 1.09E-01 | 2.65E+00 | 2.71E+00 | 7.01E-01 | no | Tek | intron_variant | MODIFIER |
| 4 | 94758443 | AX-170164363 | 0 | A | G | 0.333 | 4.18E-02 | 1.09E-01 | 2.65E+00 | 2.71E+00 | 7.01E-01 | no | Tek | intron_variant | MODIFIER |
| 4 | 94780717 | AX-170353228 | 0 | G | T | 0.333 | 4.18E-02 | 1.09E-01 | 2.65E+00 | 2.71E+00 | 7.01E-01 | no | Tek | intron_variant | MODIFIER |
| 4 | 94781206 | AX-170232757 | 0 | C | A | 0.333 | 4.18E-02 | 1.09E-01 | 2.65E+00 | 2.71E+00 | 7.01E-01 | no | Tek | intron_variant | MODIFIER |
| 4 | 94830182 | AX-97829711 | 0 | G | T | 0.333 | 4.18E-02 | 1.09E-01 | 2.65E+00 | 2.71E+00 | 7.01E-01 | no | Tek | intron_variant | MODIFIER |
| 4 | 94834952 | AX-170023443 | 0 | A | G | 0.333 | 4.18E-02 | 1.09E-01 | 2.65E+00 | 2.71E+00 | 7.01E-01 | no | Tek | intron_variant | MODIFIER |
| 4 | 94852928 | AX-170070038 | 0 | C | T | 0.333 | 4.18E-02 | 1.09E-01 | 2.65E+00 | 2.71E+00 | 7.01E-01 | no | Tek | intron_variant | MODIFIER |
| 4 | 94854023 | AX-170023441 | 0 | C | T | 0.333 | 4.18E-02 | 1.09E-01 | 2.65E+00 | 2.71E+00 | 7.01E-01 | no | Tek | intron_variant | MODIFIER |
| 4 | 94902209 | AX-170023437 | 0 | G | A | 0.333 | 4.18E-02 | 1.09E-01 | 2.65E+00 | 2.71E+00 | 7.01E-01 | no | | intergenic_variant | MODIFIER |
| 4 | 94904813 | AX-170228954 | 0 | G | A | 0.333 | 4.18E-02 | 1.09E-01 | 2.65E+00 | 2.71E+00 | 7.01E-01 | no | Eqtn | downstream_gene_variant | MODIFIER |
| 4 | 94943593 | AX-170168039 | 0 | C | T | 0.333 | 4.18E-02 | 1.09E-01 | 2.65E+00 | 2.71E+00 | 7.01E-01 | no | Mysm1 | 3_prime_UTR_variant | MODIFIER |
| 4 | 94951120 | AX-170221225 | 0 | A | G | 0.333 | 4.18E-02 | 1.09E-01 | 2.65E+00 | 2.71E+00 | 7.01E-01 | no | Mysm1 | intron_variant | MODIFIER |
| 4 | 94959578 | AX-170494293 | 0 | A | T | 0.333 | 4.18E-02 | 1.09E-01 | 2.65E+00 | 2.71E+00 | 7.01E-01 | no | Mysm1 | intron_variant | MODIFIER |
| 4 | 94960553 | AX-170268748 | 0 | C | T | 0.333 | 4.18E-02 | 1.09E-01 | 2.65E+00 | 2.71E+00 | 7.01E-01 | no | Mysm1 | intron_variant | MODIFIER |
| 4 | 94970099 | AX-97848506 | 0 | C | T | 0.333 | 4.18E-02 | 1.09E-01 | 2.65E+00 | 2.71E+00 | 7.01E-01 | no | Mysm1 | intron_variant | MODIFIER |
| 4 | 94972445 | AX-170070041 | 0 | A | G | 0.333 | 4.18E-02 | 1.09E-01 | 2.65E+00 | 2.71E+00 | 7.01E-01 | no | Mysm1 | intron_variant | MODIFIER |
| 4 | 95002707 | AX-170202922 | 0 | G | A | 0.333 | 4.18E-02 | 1.09E-01 | 2.65E+00 | 2.71E+00 | 7.01E-01 | no | Gm12694 | intron_variant,non_coding_transcript_variant | MODIFIER |
| 4 | 95020773 | AX-170466436 | 0 | C | T | 0.333 | 4.18E-02 | 1.09E-01 | 2.65E+00 | 2.71E+00 | 7.01E-01 | no | Gm12694 | intron_variant,non_coding_transcript_variant | MODIFIER |
| 4 | 95030493 | AX-170288434 | 0 | C | T | 0.333 | 4.18E-02 | 1.09E-01 | 2.65E+00 | 2.71E+00 | 7.01E-01 | no | Gm12703 | non_coding_transcript_exon_variant | MODIFIER |
| 4 | 95287593 | AX-170239680 | 0 | A | G | 0.333 | 4.18E-02 | 1.09E-01 | 2.65E+00 | 2.71E+00 | 7.01E-01 | no | | intergenic_variant | MODIFIER |
| 4 | 95288748 | AX-97801116 | 0 | T | C | 0.333 | 4.18E-02 | 1.09E-01 | 2.65E+00 | 2.71E+00 | 7.01E-01 | no | | intergenic_variant | MODIFIER |
| 4 | 95289151 | AX-170189417 | 0 | T | C | 0.333 | 4.18E-02 | 1.09E-01 | 2.65E+00 | 2.71E+00 | 7.01E-01 | no | | intergenic_variant | MODIFIER |
| 4 | 95290592 | AX-170468773 | 0 | T | A | 0.333 | 4.18E-02 | 1.09E-01 | 2.65E+00 | 2.71E+00 | 7.01E-01 | no | | intergenic_variant | MODIFIER |
| 4 | 95291148 | AX-170242922 | 0 | G | A | 0.333 | 4.18E-02 | 1.09E-01 | 2.65E+00 | 2.71E+00 | 7.01E-01 | no | | intergenic_variant | MODIFIER |
| 4 | 95296437 | AX-170163135 | 0 | G | T | 0.333 | 4.18E-02 | 1.09E-01 | 2.65E+00 | 2.71E+00 | 7.01E-01 | no | | intergenic_variant | MODIFIER |
| 4 | 95309255 | AX-170023448 | 0 | C | T | 0.333 | 4.18E-02 | 1.09E-01 | 2.65E+00 | 2.71E+00 | 7.01E-01 | no | | intergenic_variant | MODIFIER |
| 4 | 95456802 | AX-170023444 | 0 | C | T | 0.333 | 4.18E-02 | 1.09E-01 | 2.65E+00 | 2.71E+00 | 7.01E-01 | no | | intergenic_variant | MODIFIER |
| 4 | 95462161 | AX-170376479 | 0 | C | G | 0.333 | 4.18E-02 | 1.09E-01 | 2.65E+00 | 2.71E+00 | 7.01E-01 | no | Gm830 | downstream_gene_variant | MODIFIER |
| 4 | 95462396 | AX-168404895 | 0 | G | A | 0.333 | 4.18E-02 | 1.09E-01 | 2.65E+00 | 2.71E+00 | 7.01E-01 | no | Gm830 | downstream_gene_variant | MODIFIER |
| 4 | 95494291 | AX-170289786 | 0 | G | C | 0.333 | 4.18E-02 | 1.09E-01 | 2.65E+00 | 2.71E+00 | 7.01E-01 | no | | intergenic_variant | MODIFIER |
| 4 | 95494309 | AX-97835463 | 0 | C | G | 0.333 | 4.18E-02 | 1.09E-01 | 2.65E+00 | 2.71E+00 | 7.01E-01 | no | | intergenic_variant | MODIFIER |
| 4 | 95494727 | AX-170273766 | 0 | G | A | 0.333 | 4.18E-02 | 1.09E-01 | 2.65E+00 | 2.71E+00 | 7.01E-01 | no | | intergenic_variant | MODIFIER |
| 4 | 95504458 | AX-170473641 | 0 | T | C | 0.333 | 4.18E-02 | 1.09E-01 | 2.65E+00 | 2.71E+00 | 7.01E-01 | no | | intergenic_variant | MODIFIER |
| 4 | 95504912 | AX-170208266 | 0 | T | C | 0.333 | 4.18E-02 | 1.09E-01 | 2.65E+00 | 2.71E+00 | 7.01E-01 | no | | intergenic_variant | MODIFIER |
| 4 | 95519559 | AX-170346190 | 0 | A | G | 0.333 | 4.18E-02 | 1.09E-01 | 2.65E+00 | 2.71E+00 | 7.01E-01 | no | Gm25004 | upstream_gene_variant | MODIFIER |
| 4 | 95543156 | AX-168386779 | 0 | G | A | 0.333 | 4.18E-02 | 1.09E-01 | 2.65E+00 | 2.71E+00 | 7.01E-01 | no | | intergenic_variant | MODIFIER |
| 4 | 95543318 | AX-170495414 | 0 | G | A | 0.333 | 4.18E-02 | 1.09E-01 | 2.65E+00 | 2.71E+00 | 7.01E-01 | no | | intergenic_variant | MODIFIER |
| 4 | 95543740 | AX-170115774 | 0 | T | C | 0.333 | 4.18E-02 | 1.09E-01 | 2.65E+00 | 2.71E+00 | 7.01E-01 | no | | regulatory_region_variant | MODIFIER |
| 4 | 95571154 | AX-170089319 | 0 | T | G | 0.333 | 4.18E-02 | 1.09E-01 | 2.65E+00 | 2.71E+00 | 7.01E-01 | no | Fggy | intron_variant | MODIFIER |
| 4 | 95571566 | AX-170109176 | 0 | A | G | 0.333 | 4.18E-02 | 1.09E-01 | 2.65E+00 | 2.71E+00 | 7.01E-01 | no | Fggy | intron_variant | MODIFIER |
| 4 | 95578036 | AX-170403186 | 0 | A | G | 0.333 | 4.18E-02 | 1.09E-01 | 2.65E+00 | 2.71E+00 | 7.01E-01 | no | Fggy | upstream_gene_variant | MODIFIER |
| 4 | 95581204 | AX-170461923 | 0 | G | A | 0.333 | 4.18E-02 | 1.09E-01 | 2.65E+00 | 2.71E+00 | 7.01E-01 | no | Fggy | intron_variant | MODIFIER |
| 4 | 95584137 | AX-170387356 | 0 | C | T | 0.333 | 4.18E-02 | 1.09E-01 | 2.65E+00 | 2.71E+00 | 7.01E-01 | no | Fggy | intron_variant | MODIFIER |
| 4 | 95584753 | AX-170502589 | 0 | G | C | 0.333 | 4.18E-02 | 1.09E-01 | 2.65E+00 | 2.71E+00 | 7.01E-01 | no | Fggy | intron_variant | MODIFIER |
| 4 | 95601041 | AX-170361364 | 0 | A | G | 0.333 | 4.18E-02 | 1.09E-01 | 2.65E+00 | 2.71E+00 | 7.01E-01 | no | Fggy | synonymous_variant | LOW |

| | | | | | | | | | | | | | | | |
|---|---|---|---|---|---|---|---|---|---|---|---|---|---|---|---|
| 4 | 88892236 | AX-170218739 | 0 | C | T | 0.325 | 4.10E-02 | 1.07E-01 | 2.65E+00 | 2.71E+00 | 7.01E-01 | no | | intergenic_variant | MODIFIER |
| 4 | 88892571 | AX-170355581 | 0 | T | G | 0.325 | 4.10E-02 | 1.07E-01 | 2.65E+00 | 2.71E+00 | 7.01E-01 | no | | intergenic_variant | MODIFIER |
| 4 | 88894454 | AX-170388839 | 0 | C | A | 0.325 | 4.10E-02 | 1.07E-01 | 2.65E+00 | 2.71E+00 | 7.01E-01 | no | | intergenic_variant | MODIFIER |
| 4 | 88895556 | AX-170267922 | 0 | G | C | 0.325 | 4.10E-02 | 1.07E-01 | 2.65E+00 | 2.71E+00 | 7.01E-01 | no | | intergenic_variant | MODIFIER |
| 4 | 88906713 | AX-170242438 | 0 | A | G | 0.325 | 4.10E-02 | 1.07E-01 | 2.65E+00 | 2.71E+00 | 7.01E-01 | no | Mir31 | downstream_gene_variant | MODIFIER |
| 4 | 88910112 | AX-170307615 | 0 | G | A | 0.325 | 4.10E-02 | 1.07E-01 | 2.65E+00 | 2.71E+00 | 7.01E-01 | no | Mir31 | downstream_gene_variant | MODIFIER |
| 4 | 88964008 | AX-170524907 | 0 | A | G | 0.325 | 4.10E-02 | 1.07E-01 | 2.65E+00 | 2.71E+00 | 7.01E-01 | no | | intergenic_variant | MODIFIER |
| 4 | 88967749 | AX-170210729 | 0 | C | A | 0.325 | 4.10E-02 | 1.07E-01 | 2.65E+00 | 2.71E+00 | 7.01E-01 | no | | intergenic_variant | MODIFIER |
| 4 | 88970938 | AX-170453586 | 0 | C | T | 0.325 | 4.10E-02 | 1.07E-01 | 2.65E+00 | 2.71E+00 | 7.01E-01 | no | | intergenic_variant | MODIFIER |
| 4 | 88984741 | AX-170106051 | 0 | A | T | 0.325 | 4.10E-02 | 1.07E-01 | 2.65E+00 | 2.71E+00 | 7.01E-01 | no | | intergenic_variant | MODIFIER |
| 4 | 88985046 | AX-170023358 | 0 | G | A | 0.325 | 4.10E-02 | 1.07E-01 | 2.65E+00 | 2.71E+00 | 7.01E-01 | no | | intergenic_variant | MODIFIER |
| 4 | 88992553 | AX-170419655 | 0 | A | G | 0.325 | 4.10E-02 | 1.07E-01 | 2.65E+00 | 2.71E+00 | 7.01E-01 | no | AL805899.2 | downstream_gene_variant | MODIFIER |
| 4 | 89012990 | AX-170069938 | 0 | A | T | 0.325 | 4.10E-02 | 1.07E-01 | 2.65E+00 | 2.71E+00 | 7.01E-01 | no | AL805899.2 | intron_variant,non_coding_transcript_variant | MODIFIER |
| 4 | 89022820 | AX-170211691 | 0 | G | A | 0.325 | 4.10E-02 | 1.07E-01 | 2.65E+00 | 2.71E+00 | 7.01E-01 | no | AL805899.2 | intron_variant,non_coding_transcript_variant | MODIFIER |
| 4 | 89052516 | AX-97802559 | 0 | C | A | 0.325 | 4.10E-02 | 1.07E-01 | 2.65E+00 | 2.71E+00 | 7.01E-01 | no | Wincr1 | intron_variant,non_coding_transcript_variant | MODIFIER |
| 4 | 89113006 | AX-170023361 | 0 | T | G | 0.325 | 4.10E-02 | 1.07E-01 | 2.65E+00 | 2.71E+00 | 7.01E-01 | no | Gm12602 | intron_variant,non_coding_transcript_variant | MODIFIER |
| 4 | 89120338 | AX-170306873 | 0 | A | G | 0.325 | 4.10E-02 | 1.07E-01 | 2.65E+00 | 2.71E+00 | 7.01E-01 | no | Gm12602 | downstream_gene_variant | MODIFIER |
| 4 | 89403312 | AX-170097946 | 0 | G | C | 0.325 | 4.10E-02 | 1.07E-01 | 2.65E+00 | 2.71E+00 | 7.01E-01 | no | Gm12609 | downstream_gene_variant | MODIFIER |
| 4 | 89404761 | AX-170493779 | 0 | T | G | 0.325 | 4.10E-02 | 1.07E-01 | 2.65E+00 | 2.71E+00 | 7.01E-01 | no | Gm12609 | downstream_gene_variant | MODIFIER |
| 4 | 89405580 | AX-170418050 | 0 | A | C | 0.325 | 4.10E-02 | 1.07E-01 | 2.65E+00 | 2.71E+00 | 7.01E-01 | no | | regulatory_region_variant | MODIFIER |
| 4 | 89405849 | AX-170356881 | 0 | T | C | 0.325 | 4.10E-02 | 1.07E-01 | 2.65E+00 | 2.71E+00 | 7.01E-01 | no | | regulatory_region_variant | MODIFIER |
| 4 | 89436104 | AX-170530605 | 0 | C | T | 0.325 | 4.10E-02 | 1.07E-01 | 2.65E+00 | 2.71E+00 | 7.01E-01 | no | Gm12610 | intron_variant,non_coding_transcript_variant | MODIFIER |
| 4 | 89444960 | AX-170493954 | 0 | A | G | 0.325 | 4.10E-02 | 1.07E-01 | 2.65E+00 | 2.71E+00 | 7.01E-01 | no | Gm12608 | upstream_gene_variant | MODIFIER |
| 4 | 89447874 | AX-170431339 | 0 | A | G | 0.325 | 4.10E-02 | 1.07E-01 | 2.65E+00 | 2.71E+00 | 7.01E-01 | no | Gm12608 | upstream_gene_variant | MODIFIER |
| 4 | 89468113 | AX-170069946 | 0 | C | A | 0.325 | 4.10E-02 | 1.07E-01 | 2.65E+00 | 2.71E+00 | 7.01E-01 | no | Gm12610 | downstream_gene_variant | MODIFIER |
| 4 | 89470846 | AX-170069948 | 0 | A | T | 0.325 | 4.10E-02 | 1.07E-01 | 2.65E+00 | 2.71E+00 | 7.01E-01 | no | | regulatory_region_variant | MODIFIER |
| 4 | 89476740 | AX-170254882 | 0 | T | A | 0.325 | 4.10E-02 | 1.07E-01 | 2.65E+00 | 2.71E+00 | 7.01E-01 | no | | intergenic_variant | MODIFIER |
| 4 | 89774404 | AX-170069954 | 0 | T | C | 0.325 | 4.10E-02 | 1.07E-01 | 2.65E+00 | 2.71E+00 | 7.01E-01 | no | | intergenic_variant | MODIFIER |
| 4 | 89774456 | AX-170370702 | 0 | G | T | 0.325 | 4.10E-02 | 1.07E-01 | 2.65E+00 | 2.71E+00 | 7.01E-01 | no | | intergenic_variant | MODIFIER |
| 4 | 89997582 | AX-97796928 | 0 | G | A | 0.325 | 4.10E-02 | 1.07E-01 | 2.65E+00 | 2.71E+00 | 7.01E-01 | no | | intergenic_variant | MODIFIER |
| 4 | 89997832 | AX-170131865 | 0 | C | T | 0.325 | 4.10E-02 | 1.07E-01 | 2.65E+00 | 2.71E+00 | 7.01E-01 | no | | intergenic_variant | MODIFIER |
| 4 | 90946115 | AX-170404835 | 0 | A | G | 0.325 | 4.10E-02 | 1.07E-01 | 2.65E+00 | 2.71E+00 | 7.01E-01 | no | | intergenic_variant | MODIFIER |
| 4 | 90997628 | AX-170425592 | 0 | A | C | 0.325 | 4.10E-02 | 1.07E-01 | 2.65E+00 | 2.71E+00 | 7.01E-01 | no | Gm25244 | downstream_gene_variant | MODIFIER |
| 4 | 90998468 | AX-170209237 | 0 | T | G | 0.325 | 4.10E-02 | 1.07E-01 | 2.65E+00 | 2.71E+00 | 7.01E-01 | no | Gm25244 | upstream_gene_variant | MODIFIER |
| 4 | 91041678 | AX-170432063 | 0 | A | G | 0.325 | 4.10E-02 | 1.07E-01 | 2.65E+00 | 2.71E+00 | 7.01E-01 | no | Gm12643 | downstream_gene_variant | MODIFIER |
| 4 | 91093858 | AX-170146256 | 0 | C | T | 0.325 | 4.10E-02 | 1.07E-01 | 2.65E+00 | 2.71E+00 | 7.01E-01 | no | | intergenic_variant | MODIFIER |
| 4 | 91158735 | AX-170150804 | 0 | C | T | 0.325 | 4.10E-02 | 1.07E-01 | 2.65E+00 | 2.71E+00 | 7.01E-01 | no | Gm12644 | downstream_gene_variant | MODIFIER |
| 4 | 91205189 | AX-97798911 | 0 | G | A | 0.325 | 4.10E-02 | 1.07E-01 | 2.65E+00 | 2.71E+00 | 7.01E-01 | no | | intergenic_variant | MODIFIER |
| 4 | 92468992 | AX-170459280 | 0 | A | T | 0.325 | 4.10E-02 | 1.07E-01 | 2.65E+00 | 2.71E+00 | 7.01E-01 | no | | intergenic_variant | MODIFIER |
| 4 | 92483757 | AX-170070000 | 0 | A | T | 0.325 | 4.10E-02 | 1.07E-01 | 2.65E+00 | 2.71E+00 | 7.01E-01 | no | | intergenic_variant | MODIFIER |
| 4 | 92547694 | AX-170199472 | 0 | C | G | 0.325 | 4.10E-02 | 1.07E-01 | 2.65E+00 | 2.71E+00 | 7.01E-01 | no | Gm12637 | intron_variant,non_coding_transcript_variant | MODIFIER |
| 4 | 92557352 | AX-170089316 | 0 | A | C | 0.325 | 4.10E-02 | 1.07E-01 | 2.65E+00 | 2.71E+00 | 7.01E-01 | no | | intergenic_variant | MODIFIER |
| 4 | 92578271 | AX-97795229 | 0 | A | G | 0.325 | 4.10E-02 | 1.07E-01 | 2.65E+00 | 2.71E+00 | 7.01E-01 | no | | intergenic_variant | MODIFIER |
| 4 | 92605427 | AX-97843978 | 0 | G | A | 0.325 | 4.10E-02 | 1.07E-01 | 2.65E+00 | 2.71E+00 | 7.01E-01 | no | | intergenic_variant | MODIFIER |
| 4 | 92752551 | AX-170070005 | 0 | A | T | 0.325 | 4.10E-02 | 1.07E-01 | 2.65E+00 | 2.71E+00 | 7.01E-01 | no | | intergenic_variant | MODIFIER |
| 4 | 92946109 | AX-170147101 | 0 | T | C | 0.325 | 4.10E-02 | 1.07E-01 | 2.65E+00 | 2.71E+00 | 7.01E-01 | no | | intergenic_variant | MODIFIER |
| 4 | 93016526 | AX-97842903 | 0 | A | C | 0.325 | 4.10E-02 | 1.07E-01 | 2.65E+00 | 2.71E+00 | 7.01E-01 | no | | intergenic_variant | MODIFIER |
| 4 | 93217190 | AX-97801447 | 0 | G | A | 0.325 | 4.10E-02 | 1.07E-01 | 2.65E+00 | 2.71E+00 | 7.01E-01 | no | | intergenic_variant | MODIFIER |
| 4 | 93218809 | AX-168406321 | 0 | G | T | 0.325 | 4.10E-02 | 1.07E-01 | 2.65E+00 | 2.71E+00 | 7.01E-01 | no | | intergenic_variant | MODIFIER |
| 4 | 93717845 | AX-97810013 | 0 | G | C | 0.325 | 4.10E-02 | 1.07E-01 | 2.65E+00 | 2.71E+00 | 7.01E-01 | no | | intergenic_variant | MODIFIER |
| 4 | 93755104 | AX-170401564 | 0 | A | G | 0.325 | 4.10E-02 | 1.07E-01 | 2.65E+00 | 2.71E+00 | 7.01E-01 | no | | intergenic_variant | MODIFIER |
| 4 | 93809791 | AX-170023426 | 0 | C | A | 0.325 | 4.10E-02 | 1.07E-01 | 2.65E+00 | 2.71E+00 | 7.01E-01 | no | | intergenic_variant | MODIFIER |
| 4 | 93900636 | AX-170227569 | 0 | T | C | 0.325 | 4.10E-02 | 1.07E-01 | 2.65E+00 | 2.71E+00 | 7.01E-01 | no | | intergenic_variant | MODIFIER |
| 4 | 93961915 | AX-97849755 | 0 | C | T | 0.325 | 4.10E-02 | 1.07E-01 | 2.65E+00 | 2.71E+00 | 7.01E-01 | no | | intergenic_variant | MODIFIER |

| | | | | | | | | | | | | | | | |
|---|---|---|---|---|---|---|---|---|---|---|---|---|---|---|---|
| 4 | 93990983 | AX-97807179 | 0 | T | C | 0.325 | 4.10E-02 | 1.07E-01 | 2.65E+00 | 2.71E+00 | 7.01E-01 | no | | intergenic_variant | MODIFIER |
| 4 | 93992034 | AX-170431283 | 0 | A | G | 0.325 | 4.10E-02 | 1.07E-01 | 2.65E+00 | 2.71E+00 | 7.01E-01 | no | | intergenic_variant | MODIFIER |
| 4 | 94011485 | AX-170239534 | 0 | T | C | 0.325 | 4.10E-02 | 1.07E-01 | 2.65E+00 | 2.71E+00 | 7.01E-01 | no | | intergenic_variant | MODIFIER |
| 4 | 94011658 | AX-97822293 | 0 | T | C | 0.325 | 4.10E-02 | 1.07E-01 | 2.65E+00 | 2.71E+00 | 7.01E-01 | no | | intergenic_variant | MODIFIER |
| 4 | 94853437 | AX-170070036 | 0 | A | G | 0.325 | 4.10E-02 | 1.07E-01 | 2.65E+00 | 2.71E+00 | 7.01E-01 | no | Tek | synonymous_variant | LOW |
| 4 | 95002311 | AX-170349377 | 0 | T | C | 0.325 | 4.10E-02 | 1.07E-01 | 2.65E+00 | 2.71E+00 | 7.01E-01 | no | Gm12694 | intron_variant,non_coding_transcript_variant | MODIFIER |
| 4 | 95004630 | AX-170163566 | 0 | A | G | 0.325 | 4.10E-02 | 1.07E-01 | 2.65E+00 | 2.71E+00 | 7.01E-01 | no | Gm12694 | intron_variant,non_coding_transcript_variant | MODIFIER |
| 4 | 95046422 | AX-97848153 | 0 | G | A | 0.325 | 4.10E-02 | 1.07E-01 | 2.65E+00 | 2.71E+00 | 7.01E-01 | no | Jun | downstream_gene_variant | MODIFIER |
| 4 | 95136670 | AX-170370859 | 0 | T | A | 0.325 | 4.10E-02 | 1.07E-01 | 2.65E+00 | 2.71E+00 | 7.01E-01 | no | Junos | intron_variant,non_coding_transcript_variant | MODIFIER |
| 4 | 95188678 | AX-97850264 | 0 | A | G | 0.325 | 4.10E-02 | 1.07E-01 | 2.65E+00 | 2.71E+00 | 7.01E-01 | no | Gm12708 | upstream_gene_variant | MODIFIER |
| 4 | 95194603 | AX-170070047 | 0 | T | C | 0.325 | 4.10E-02 | 1.07E-01 | 2.65E+00 | 2.71E+00 | 7.01E-01 | no | Gm12708 | intron_variant,non_coding_transcript_variant | MODIFIER |
| 4 | 95199438 | AX-170070048 | 0 | C | G | 0.325 | 4.10E-02 | 1.07E-01 | 2.65E+00 | 2.71E+00 | 7.01E-01 | no | Gm12708 | intron_variant,non_coding_transcript_variant | MODIFIER |
| 4 | 95214278 | AX-97795067 | 0 | A | G | 0.325 | 4.10E-02 | 1.07E-01 | 2.65E+00 | 2.71E+00 | 7.01E-01 | no | Gm12708 | downstream_gene_variant | MODIFIER |
| 4 | 95350335 | AX-170392863 | 0 | T | A | 0.325 | 4.10E-02 | 1.07E-01 | 2.65E+00 | 2.71E+00 | 7.01E-01 | no | | intergenic_variant | MODIFIER |
| 4 | 95352556 | AX-170023445 | 0 | T | C | 0.325 | 4.10E-02 | 1.07E-01 | 2.65E+00 | 2.71E+00 | 7.01E-01 | no | | intergenic_variant | MODIFIER |
| 4 | 95353020 | AX-97850444 | 0 | C | T | 0.325 | 4.10E-02 | 1.07E-01 | 2.65E+00 | 2.71E+00 | 7.01E-01 | no | | intergenic_variant | MODIFIER |
| 4 | 95379481 | AX-170454376 | 0 | G | A | 0.325 | 4.10E-02 | 1.07E-01 | 2.65E+00 | 2.71E+00 | 7.01E-01 | no | | intergenic_variant | MODIFIER |
| 4 | 95380587 | AX-170317648 | 0 | T | C | 0.325 | 4.10E-02 | 1.07E-01 | 2.65E+00 | 2.71E+00 | 7.01E-01 | no | | intergenic_variant | MODIFIER |
| 4 | 95550516 | AX-170095060 | 0 | G | A | 0.325 | 4.10E-02 | 1.07E-01 | 2.65E+00 | 2.71E+00 | 7.01E-01 | no | | intergenic_variant | MODIFIER |
| 2 | 180452139 | AX-170178636 | 0 | C | T | 0.053 | -5.50E-02 | 1.45E-01 | 2.60E+00 | 2.74E+00 | 7.01E-01 | no | Slco4a1 | upstream_gene_variant | MODIFIER |
| 12 | 99444640 | AX-170407429 | 1 | C | T | 0.054 | 8.45E-02 | 2.23E-01 | 2.69E+00 | 2.75E+00 | 7.01E-01 | no | Foxn3 | intron_variant | MODIFIER |
| 12 | 102571964 | AX-170293763 | 1 | A | G | 0.054 | 8.45E-02 | 2.23E-01 | 2.69E+00 | 2.75E+00 | 7.01E-01 | no | Itpk1 | intron_variant | MODIFIER |
| 3 | 144043819 | AX-97793306 | 0 | C | T | 0.377 | 2.64E-02 | 6.90E-02 | 2.63E+00 | 2.72E+00 | 7.01E-01 | no | | intergenic_variant | MODIFIER |
| 10 | 27777125 | AX-97834733 | 1 | A | C | 0.179 | 3.72E-02 | 1.00E-01 | 2.72E+00 | 2.82E+00 | 7.01E-01 | no | Gm18041 | upstream_gene_variant | MODIFIER |
| 18 | 69718806 | AX-170145940 | 0 | C | T | 0.263 | -3.91E-02 | 1.02E-01 | 2.66E+00 | 2.71E+00 | 7.01E-01 | no | | intergenic_variant | MODIFIER |
| 7 | 114517232 | AX-97800976 | 1 | C | T | 0.446 | 3.57E-02 | 9.36E-02 | 2.65E+00 | 2.72E+00 | 7.01E-01 | no | Pde3b | intron_variant | MODIFIER |
| 15 | 86127486 | AX-97831347 | 0 | G | T | 0.5 | 2.85E-02 | 6.99E-02 | 2.53E+00 | 2.62E+00 | 7.01E-01 | no | Gramd4 | intron_variant | MODIFIER |
| 16 | 19154604 | AX-170142822 | 0 | T | C | 0.132 | 3.47E-02 | 9.45E-02 | 2.68E+00 | 2.82E+00 | 7.01E-01 | no | | intergenic_variant | MODIFIER |
| 2 | 161453098 | AX-170088822 | 0 | C | A | 0.149 | 5.65E-02 | 1.50E-01 | 2.63E+00 | 2.75E+00 | 7.01E-01 | no | | intergenic_variant | MODIFIER |
| 8 | 16495025 | AX-170120271 | 0 | A | G | 0.228 | -3.04E-02 | 7.62E-02 | 2.55E+00 | 2.65E+00 | 7.02E-01 | no | Csmd1 | intron_variant | MODIFIER |
| 9 | 38080594 | AX-170275718 | 1 | T | C | 0.473 | -2.84E-02 | 7.56E-02 | 2.66E+00 | 2.76E+00 | 7.02E-01 | no | Olfr887 | upstream_gene_variant | MODIFIER |
| 8 | 95659354 | AX-170080879 | 0 | C | A | 0.219 | 4.44E-02 | 1.18E-01 | 2.71E+00 | 2.78E+00 | 7.02E-01 | no | | intergenic_variant | MODIFIER |
| 1 | 101167383 | AX-170439829 | 1 | T | C | 0.134 | 6.98E-02 | 1.80E-01 | 2.59E+00 | 2.69E+00 | 7.02E-01 | no | Gm20268 | intron_variant,non_coding_transcript_variant | MODIFIER |
| 7 | 53913913 | AX-170336993 | 1 | A | C | 0.429 | 3.79E-02 | 9.93E-02 | 2.64E+00 | 2.71E+00 | 7.02E-01 | no | | intergenic_variant | MODIFIER |
| 4 | 75008988 | AX-170295833 | 0 | T | G | 0.246 | 3.99E-02 | 1.01E-01 | 2.57E+00 | 2.65E+00 | 7.02E-01 | no | | intergenic_variant | MODIFIER |
| 12 | 98301901 | AX-170419591 | 0 | A | T | 0.088 | -6.36E-02 | 1.70E-01 | 2.69E+00 | 2.78E+00 | 7.02E-01 | no | | intergenic_variant | MODIFIER |
| 9 | 39716044 | AX-170197686 | 0 | A | G | 0.421 | -2.85E-02 | 7.59E-02 | 2.67E+00 | 2.76E+00 | 7.02E-01 | no | Olfr965 | upstream_gene_variant | MODIFIER |
| 9 | 39811514 | AX-97805229 | 0 | G | T | 0.421 | -2.85E-02 | 7.59E-02 | 2.67E+00 | 2.76E+00 | 7.02E-01 | no | Olfr970 | intron_variant | MODIFIER |
| 9 | 39838410 | AX-168377679 | 0 | T | C | 0.421 | -2.85E-02 | 7.59E-02 | 2.67E+00 | 2.76E+00 | 7.02E-01 | no | Olfr971 | upstream_gene_variant | MODIFIER |
| 9 | 39908958 | AX-170497609 | 0 | G | A | 0.421 | -2.85E-02 | 7.59E-02 | 2.67E+00 | 2.76E+00 | 7.02E-01 | no | Olfr229 | upstream_gene_variant | MODIFIER |
| 9 | 40140551 | AX-170415721 | 0 | T | C | 0.421 | -2.85E-02 | 7.59E-02 | 2.67E+00 | 2.76E+00 | 7.02E-01 | no | | intergenic_variant | MODIFIER |
| 9 | 40140782 | AX-170250501 | 0 | C | G | 0.421 | -2.85E-02 | 7.59E-02 | 2.67E+00 | 2.76E+00 | 7.02E-01 | no | | intergenic_variant | MODIFIER |
| 9 | 40426782 | AX-170449888 | 0 | G | A | 0.421 | -2.85E-02 | 7.59E-02 | 2.67E+00 | 2.76E+00 | 7.02E-01 | no | Gramd1b | intron_variant | MODIFIER |
| 9 | 40433270 | AX-97815776 | 0 | A | G | 0.421 | -2.85E-02 | 7.59E-02 | 2.67E+00 | 2.76E+00 | 7.02E-01 | no | Gramd1b | intron_variant | MODIFIER |
| 9 | 40436073 | AX-97813289 | 0 | A | G | 0.421 | -2.85E-02 | 7.59E-02 | 2.67E+00 | 2.76E+00 | 7.02E-01 | no | Gramd1b | intron_variant | MODIFIER |
| 9 | 40438526 | AX-170225986 | 0 | T | G | 0.421 | -2.85E-02 | 7.59E-02 | 2.67E+00 | 2.76E+00 | 7.02E-01 | no | Gramd1b | intron_variant | MODIFIER |
| 9 | 40438677 | AX-168387313 | 0 | G | A | 0.421 | -2.85E-02 | 7.59E-02 | 2.67E+00 | 2.76E+00 | 7.02E-01 | no | Gramd1b | intron_variant | MODIFIER |
| 9 | 40456607 | AX-170032185 | 0 | A | C | 0.421 | -2.85E-02 | 7.59E-02 | 2.67E+00 | 2.76E+00 | 7.02E-01 | no | Gramd1b | upstream_gene_variant | MODIFIER |
| 9 | 40550296 | AX-168373416 | 0 | A | G | 0.421 | -2.85E-02 | 7.59E-02 | 2.67E+00 | 2.76E+00 | 7.02E-01 | no | Gm48277 | downstream_gene_variant | MODIFIER |
| 9 | 40585106 | AX-97842682 | 0 | G | A | 0.421 | -2.85E-02 | 7.59E-02 | 2.67E+00 | 2.76E+00 | 7.02E-01 | no | | intergenic_variant | MODIFIER |
| 9 | 40585231 | AX-170134554 | 0 | T | C | 0.421 | -2.85E-02 | 7.59E-02 | 2.67E+00 | 2.76E+00 | 7.02E-01 | no | | intergenic_variant | MODIFIER |
| 9 | 40993284 | AX-170457651 | 0 | T | C | 0.421 | -2.85E-02 | 7.59E-02 | 2.67E+00 | 2.76E+00 | 7.02E-01 | no | Crtam | intron_variant | MODIFIER |
| 9 | 40993578 | AX-170311467 | 0 | T | C | 0.421 | -2.85E-02 | 7.59E-02 | 2.67E+00 | 2.76E+00 | 7.02E-01 | no | Crtam | intron_variant | MODIFIER |
| 9 | 41004548 | AX-170100787 | 0 | T | C | 0.421 | -2.85E-02 | 7.59E-02 | 2.67E+00 | 2.76E+00 | 7.02E-01 | no | Crtam | synonymous_variant | LOW |

| | | | | | | | | | | | | | | | |
|---|---|---|---|---|---|---|---|---|---|---|---|---|---|---|---|
| 9 | 41088680 | AX-170362328 | 0 | C | G | 0.421 | -2.85E-02 | 7.59E-02 | 2.67E+00 | 2.76E+00 | 7.02E-01 | no | Ubash3b | intron_variant | MODIFIER |
| 9 | 41100403 | AX-170327897 | 0 | C | T | 0.421 | -2.85E-02 | 7.59E-02 | 2.67E+00 | 2.76E+00 | 7.02E-01 | no | Ubash3b | intron_variant | MODIFIER |
| 9 | 41110693 | AX-170082219 | 0 | T | G | 0.421 | -2.85E-02 | 7.59E-02 | 2.67E+00 | 2.76E+00 | 7.02E-01 | no | Ubash3b | intron_variant | MODIFIER |
| 1 | 46868702 | AX-170525439 | 1 | G | A | 0.098 | -7.93E-02 | 2.12E-01 | 2.69E+00 | 2.77E+00 | 7.02E-01 | no | Slc39a10 | intron_variant | MODIFIER |
| 4 | 124628026 | AX-170492410 | 0 | T | C | 0.219 | 4.13E-02 | 1.02E-01 | 2.53E+00 | 2.64E+00 | 7.02E-01 | no | Gm2164 | intron_variant,non_coding_transcript_variant | MODIFIER |
| 3 | 148065034 | AX-97857726 | 0 | T | C | 0.351 | 2.76E-02 | 7.22E-02 | 2.63E+00 | 2.70E+00 | 7.02E-01 | no | | intergenic_variant | MODIFIER |
| 10 | 126768198 | AX-170004122 | 0 | G | T | 0.325 | -2.80E-02 | 7.33E-02 | 2.61E+00 | 2.71E+00 | 7.02E-01 | no | | intergenic_variant | MODIFIER |
| 10 | 126773653 | AX-97817084 | 0 | A | G | 0.325 | -2.80E-02 | 7.33E-02 | 2.61E+00 | 2.71E+00 | 7.02E-01 | no | | intergenic_variant | MODIFIER |
| 10 | 126800033 | AX-170132055 | 0 | C | T | 0.325 | -2.80E-02 | 7.33E-02 | 2.61E+00 | 2.71E+00 | 7.02E-01 | no | | intergenic_variant | MODIFIER |
| 10 | 126803042 | AX-97836576 | 0 | G | A | 0.325 | -2.80E-02 | 7.33E-02 | 2.61E+00 | 2.71E+00 | 7.02E-01 | no | | intergenic_variant | MODIFIER |
| 10 | 126807507 | AX-170004123 | 0 | G | A | 0.325 | -2.80E-02 | 7.33E-02 | 2.61E+00 | 2.71E+00 | 7.02E-01 | no | Gm9111 | downstream_gene_variant | MODIFIER |
| 10 | 126821435 | AX-170004124 | 0 | T | C | 0.325 | -2.80E-02 | 7.33E-02 | 2.61E+00 | 2.71E+00 | 7.02E-01 | no | | intergenic_variant | MODIFIER |
| 10 | 126821481 | AX-170294009 | 0 | A | G | 0.325 | -2.80E-02 | 7.33E-02 | 2.61E+00 | 2.71E+00 | 7.02E-01 | no | | intergenic_variant | MODIFIER |
| 10 | 126850399 | AX-170389401 | 0 | A | G | 0.325 | -2.80E-02 | 7.33E-02 | 2.61E+00 | 2.71E+00 | 7.02E-01 | no | | intergenic_variant | MODIFIER |
| 10 | 126851658 | AX-170160926 | 0 | T | C | 0.325 | -2.80E-02 | 7.33E-02 | 2.61E+00 | 2.71E+00 | 7.02E-01 | no | | intergenic_variant | MODIFIER |
| 10 | 126852217 | AX-97810617 | 0 | G | A | 0.325 | -2.80E-02 | 7.33E-02 | 2.61E+00 | 2.71E+00 | 7.02E-01 | no | | intergenic_variant | MODIFIER |
| 10 | 126864771 | AX-97801026 | 0 | C | A | 0.325 | -2.80E-02 | 7.33E-02 | 2.61E+00 | 2.71E+00 | 7.02E-01 | no | Atp23 | downstream_gene_variant | MODIFIER |
| 10 | 126864808 | AX-170004121 | 0 | G | A | 0.325 | -2.80E-02 | 7.33E-02 | 2.61E+00 | 2.71E+00 | 7.02E-01 | no | Atp23 | downstream_gene_variant | MODIFIER |
| 10 | 126865089 | AX-170362424 | 0 | G | A | 0.325 | -2.80E-02 | 7.33E-02 | 2.61E+00 | 2.71E+00 | 7.02E-01 | no | Atp23 | downstream_gene_variant | MODIFIER |
| 10 | 126867311 | AX-170144348 | 0 | A | G | 0.325 | -2.80E-02 | 7.33E-02 | 2.61E+00 | 2.71E+00 | 7.02E-01 | no | Atp23 | downstream_gene_variant | MODIFIER |
| 10 | 126869782 | AX-170046204 | 0 | G | A | 0.325 | -2.80E-02 | 7.33E-02 | 2.61E+00 | 2.71E+00 | 7.02E-01 | no | Atp23 | intron_variant | MODIFIER |
| 10 | 126887677 | AX-170299223 | 0 | T | C | 0.325 | -2.80E-02 | 7.33E-02 | 2.61E+00 | 2.71E+00 | 7.02E-01 | no | Atp23 | intron_variant | MODIFIER |
| 10 | 126888547 | AX-170092148 | 0 | T | C | 0.325 | -2.80E-02 | 7.33E-02 | 2.61E+00 | 2.71E+00 | 7.02E-01 | no | Atp23 | intron_variant | MODIFIER |
| 10 | 126891715 | AX-170454016 | 0 | A | G | 0.325 | -2.80E-02 | 7.33E-02 | 2.61E+00 | 2.71E+00 | 7.02E-01 | no | Atp23 | synonymous_variant | LOW |
| 10 | 126910230 | AX-97803242 | 0 | C | T | 0.325 | -2.80E-02 | 7.33E-02 | 2.61E+00 | 2.71E+00 | 7.02E-01 | no | | intergenic_variant | MODIFIER |
| 10 | 126931266 | AX-170465644 | 0 | C | T | 0.325 | -2.80E-02 | 7.33E-02 | 2.61E+00 | 2.71E+00 | 7.02E-01 | no | | regulatory_region_variant | MODIFIER |
| 10 | 126931734 | AX-170386479 | 0 | G | C | 0.325 | -2.80E-02 | 7.33E-02 | 2.61E+00 | 2.71E+00 | 7.02E-01 | no | | regulatory_region_variant | MODIFIER |
| 10 | 126938380 | AX-170442860 | 0 | C | T | 0.325 | -2.80E-02 | 7.33E-02 | 2.61E+00 | 2.71E+00 | 7.02E-01 | no | | intergenic_variant | MODIFIER |
| 10 | 126938795 | AX-170520578 | 0 | T | G | 0.325 | -2.80E-02 | 7.33E-02 | 2.61E+00 | 2.71E+00 | 7.02E-01 | no | | intergenic_variant | MODIFIER |
| 10 | 126939286 | AX-170428649 | 0 | C | T | 0.325 | -2.80E-02 | 7.33E-02 | 2.61E+00 | 2.71E+00 | 7.02E-01 | no | | regulatory_region_variant | MODIFIER |
| 10 | 126943892 | AX-170301571 | 0 | C | T | 0.325 | -2.80E-02 | 7.33E-02 | 2.61E+00 | 2.71E+00 | 7.02E-01 | no | | intergenic_variant | MODIFIER |
| 10 | 126948319 | AX-170412747 | 0 | G | T | 0.325 | -2.80E-02 | 7.33E-02 | 2.61E+00 | 2.71E+00 | 7.02E-01 | no | | intergenic_variant | MODIFIER |
| 10 | 127003732 | AX-170405784 | 0 | G | A | 0.325 | -2.80E-02 | 7.33E-02 | 2.61E+00 | 2.71E+00 | 7.02E-01 | no | Avil | intron_variant | MODIFIER |
| 10 | 127007086 | AX-170320201 | 0 | T | C | 0.325 | -2.80E-02 | 7.33E-02 | 2.61E+00 | 2.71E+00 | 7.02E-01 | no | Avil | intron_variant | MODIFIER |
| 10 | 127007523 | AX-170114538 | 0 | G | A | 0.325 | -2.80E-02 | 7.33E-02 | 2.61E+00 | 2.71E+00 | 7.02E-01 | no | Avil | intron_variant | MODIFIER |
| 10 | 127009303 | AX-170204150 | 0 | G | A | 0.325 | -2.80E-02 | 7.33E-02 | 2.61E+00 | 2.71E+00 | 7.02E-01 | no | Avil | intron_variant | MODIFIER |
| 10 | 127011575 | AX-170358809 | 0 | G | A | 0.325 | -2.80E-02 | 7.33E-02 | 2.61E+00 | 2.71E+00 | 7.02E-01 | no | Avil | intron_variant | MODIFIER |
| 10 | 127013573 | AX-170122862 | 0 | A | G | 0.325 | -2.80E-02 | 7.33E-02 | 2.61E+00 | 2.71E+00 | 7.02E-01 | no | Avil | synonymous_variant | LOW |
| 10 | 127014168 | AX-170298052 | 0 | C | T | 0.325 | -2.80E-02 | 7.33E-02 | 2.61E+00 | 2.71E+00 | 7.02E-01 | no | Avil | synonymous_variant | LOW |
| 10 | 127014312 | AX-170106466 | 0 | G | A | 0.325 | -2.80E-02 | 7.33E-02 | 2.61E+00 | 2.71E+00 | 7.02E-01 | no | Avil | intron_variant | MODIFIER |
| 10 | 127173742 | AX-170046211 | 0 | T | C | 0.325 | -2.80E-02 | 7.33E-02 | 2.61E+00 | 2.71E+00 | 7.02E-01 | no | B4galnt1 | downstream_gene_variant | MODIFIER |
| 10 | 127174043 | AX-170150447 | 0 | C | T | 0.325 | -2.80E-02 | 7.33E-02 | 2.61E+00 | 2.71E+00 | 7.02E-01 | no | B4galnt1 | downstream_gene_variant | MODIFIER |
| 10 | 127174816 | AX-170114756 | 0 | C | T | 0.325 | -2.80E-02 | 7.33E-02 | 2.61E+00 | 2.71E+00 | 7.02E-01 | no | B4galnt1 | downstream_gene_variant | MODIFIER |
| 10 | 127188117 | AX-170442038 | 0 | C | T | 0.325 | -2.80E-02 | 7.33E-02 | 2.61E+00 | 2.71E+00 | 7.02E-01 | no | Arhgef25 | intron_variant | MODIFIER |
| 10 | 127209665 | AX-97817085 | 0 | A | C | 0.325 | -2.80E-02 | 7.33E-02 | 2.61E+00 | 2.71E+00 | 7.02E-01 | no | Pip4k2c | intron_variant | MODIFIER |
| 10 | 127230474 | AX-170233905 | 0 | T | G | 0.325 | -2.80E-02 | 7.33E-02 | 2.61E+00 | 2.71E+00 | 7.02E-01 | no | Kif5a | intron_variant | MODIFIER |
| 10 | 127246953 | AX-170097149 | 0 | C | G | 0.325 | -2.80E-02 | 7.33E-02 | 2.61E+00 | 2.71E+00 | 7.02E-01 | no | Kif5a | intron_variant | MODIFIER |
| 10 | 127247365 | AX-170352608 | 0 | T | C | 0.325 | -2.80E-02 | 7.33E-02 | 2.61E+00 | 2.71E+00 | 7.02E-01 | no | Kif5a | intron_variant | MODIFIER |
| 10 | 127247648 | AX-170394626 | 0 | A | G | 0.325 | -2.80E-02 | 7.33E-02 | 2.61E+00 | 2.71E+00 | 7.02E-01 | no | Kif5a | intron_variant | MODIFIER |
| 10 | 127247895 | AX-170299953 | 0 | A | C | 0.325 | -2.80E-02 | 7.33E-02 | 2.61E+00 | 2.71E+00 | 7.02E-01 | no | Kif5a | intron_variant | MODIFIER |
| 10 | 127248161 | AX-170301879 | 0 | G | A | 0.325 | -2.80E-02 | 7.33E-02 | 2.61E+00 | 2.71E+00 | 7.02E-01 | no | Kif5a | intron_variant | MODIFIER |
| 10 | 127283027 | AX-170359049 | 0 | T | G | 0.325 | -2.80E-02 | 7.33E-02 | 2.61E+00 | 2.71E+00 | 7.02E-01 | no | Mbd6 | splice_region_variant,synonymous_variant | LOW |
| 10 | 127305306 | AX-170270059 | 0 | A | G | 0.325 | -2.80E-02 | 7.33E-02 | 2.61E+00 | 2.71E+00 | 7.02E-01 | no | Mars | intron_variant | MODIFIER |
| 10 | 127307356 | AX-170107774 | 0 | C | G | 0.325 | -2.80E-02 | 7.33E-02 | 2.61E+00 | 2.71E+00 | 7.02E-01 | no | Mars | intron_variant | MODIFIER |

| | | | | | | | | | | | | | | | |
|---|---|---|---|---|---|---|---|---|---|---|---|---|---|---|---|
| 10 | 127307598 | AX-170262247 | 0 | C | T | 0.325 | -2.80E-02 | 7.33E-02 | 2.61E+00 | 2.71E+00 | 7.02E-01 | no | Mars | intron_variant | MODIFIER |
| 10 | 127325823 | AX-97857096 | 0 | A | G | 0.325 | -2.80E-02 | 7.33E-02 | 2.61E+00 | 2.71E+00 | 7.02E-01 | no | Gli1 | downstream_gene_variant | MODIFIER |
| 10 | 127335843 | AX-170113439 | 0 | C | G | 0.325 | -2.80E-02 | 7.33E-02 | 2.61E+00 | 2.71E+00 | 7.02E-01 | no | Gli1 | intron_variant | MODIFIER |
| 10 | 127352182 | AX-170004127 | 0 | A | G | 0.325 | -2.80E-02 | 7.33E-02 | 2.61E+00 | 2.71E+00 | 7.02E-01 | no | Inhbc | downstream_gene_variant | MODIFIER |
| 10 | 127354223 | AX-170389117 | 0 | A | G | 0.325 | -2.80E-02 | 7.33E-02 | 2.61E+00 | 2.71E+00 | 7.02E-01 | no | Inhbc | downstream_gene_variant | MODIFIER |
| 10 | 127357771 | AX-170117848 | 0 | A | G | 0.325 | -2.80E-02 | 7.33E-02 | 2.61E+00 | 2.71E+00 | 7.02E-01 | no | Inhbc | synonymous_variant | LOW |
| 10 | 127363719 | AX-170092150 | 0 | T | C | 0.325 | -2.80E-02 | 7.33E-02 | 2.61E+00 | 2.71E+00 | 7.02E-01 | no | Inhbc | intron_variant | MODIFIER |
| 10 | 127408411 | AX-170158111 | 0 | A | G | 0.325 | -2.80E-02 | 7.33E-02 | 2.61E+00 | 2.71E+00 | 7.02E-01 | no | R3hdm2 | intron_variant | MODIFIER |
| 10 | 127411506 | AX-97830161 | 0 | G | T | 0.325 | -2.80E-02 | 7.33E-02 | 2.61E+00 | 2.71E+00 | 7.02E-01 | no | R3hdm2 | intron_variant | MODIFIER |
| 10 | 127447538 | AX-170046217 | 0 | G | T | 0.325 | -2.80E-02 | 7.33E-02 | 2.61E+00 | 2.71E+00 | 7.02E-01 | no | R3hdm2 | intron_variant | MODIFIER |
| 10 | 127451886 | AX-170004129 | 0 | T | G | 0.325 | -2.80E-02 | 7.33E-02 | 2.61E+00 | 2.71E+00 | 7.02E-01 | no | R3hdm2 | intron_variant | MODIFIER |
| 10 | 127468822 | AX-170046215 | 0 | C | T | 0.325 | -2.80E-02 | 7.33E-02 | 2.61E+00 | 2.71E+00 | 7.02E-01 | no | R3hdm2 | intron_variant | MODIFIER |
| 10 | 127471474 | AX-97810729 | 0 | C | T | 0.325 | -2.80E-02 | 7.33E-02 | 2.61E+00 | 2.71E+00 | 7.02E-01 | no | R3hdm2 | intron_variant | MODIFIER |
| 10 | 127471792 | AX-170386334 | 0 | C | T | 0.325 | -2.80E-02 | 7.33E-02 | 2.61E+00 | 2.71E+00 | 7.02E-01 | no | R3hdm2 | synonymous_variant | LOW |
| 10 | 127472994 | AX-170361602 | 0 | G | C | 0.325 | -2.80E-02 | 7.33E-02 | 2.61E+00 | 2.71E+00 | 7.02E-01 | no | R3hdm2 | intron_variant | MODIFIER |
| 10 | 127477862 | AX-97805026 | 0 | G | A | 0.325 | -2.80E-02 | 7.33E-02 | 2.61E+00 | 2.71E+00 | 7.02E-01 | no | R3hdm2 | intron_variant | MODIFIER |
| 10 | 127498092 | AX-170246595 | 0 | C | T | 0.325 | -2.80E-02 | 7.33E-02 | 2.61E+00 | 2.71E+00 | 7.02E-01 | no | Stac3 | upstream_gene_variant | MODIFIER |
| 10 | 127511005 | AX-170126724 | 0 | G | A | 0.325 | -2.80E-02 | 7.33E-02 | 2.61E+00 | 2.71E+00 | 7.02E-01 | no | Ndufa4l2 | upstream_gene_variant | MODIFIER |
| 10 | 127606250 | AX-170361851 | 0 | G | A | 0.325 | -2.80E-02 | 7.33E-02 | 2.61E+00 | 2.71E+00 | 7.02E-01 | no | Lrp1 | intron_variant | MODIFIER |
| 10 | 127608814 | AX-170243783 | 0 | C | T | 0.325 | -2.80E-02 | 7.33E-02 | 2.61E+00 | 2.71E+00 | 7.02E-01 | no | Lrp1 | intron_variant | MODIFIER |
| 10 | 127624497 | AX-170194802 | 0 | C | A | 0.325 | -2.80E-02 | 7.33E-02 | 2.61E+00 | 2.71E+00 | 7.02E-01 | no | Lrp1 | upstream_gene_variant | MODIFIER |
| 10 | 127676327 | AX-170317203 | 0 | A | G | 0.325 | -2.80E-02 | 7.33E-02 | 2.61E+00 | 2.71E+00 | 7.02E-01 | no | Nemp1 | upstream_gene_variant | MODIFIER |
| 10 | 127703587 | AX-170046220 | 0 | A | G | 0.325 | -2.80E-02 | 7.33E-02 | 2.61E+00 | 2.71E+00 | 7.02E-01 | no | Nemp1 | downstream_gene_variant | MODIFIER |
| 10 | 127704764 | AX-168385387 | 0 | T | C | 0.325 | -2.80E-02 | 7.33E-02 | 2.61E+00 | 2.71E+00 | 7.02E-01 | no | Nemp1 | downstream_gene_variant | MODIFIER |
| 10 | 127708480 | AX-170246452 | 0 | T | C | 0.325 | -2.80E-02 | 7.33E-02 | 2.61E+00 | 2.71E+00 | 7.02E-01 | no | Myo1a | intron_variant | MODIFIER |
| 10 | 127709375 | AX-168407848 | 0 | A | G | 0.325 | -2.80E-02 | 7.33E-02 | 2.61E+00 | 2.71E+00 | 7.02E-01 | no | Myo1a | intron_variant | MODIFIER |
| 10 | 127709539 | AX-170173525 | 0 | G | A | 0.325 | -2.80E-02 | 7.33E-02 | 2.61E+00 | 2.71E+00 | 7.02E-01 | no | Myo1a | intron_variant | MODIFIER |
| 10 | 127711922 | AX-170046226 | 0 | A | G | 0.325 | -2.80E-02 | 7.33E-02 | 2.61E+00 | 2.71E+00 | 7.02E-01 | no | Myo1a | intron_variant | MODIFIER |
| 10 | 127712387 | AX-170046223 | 0 | A | C | 0.325 | -2.80E-02 | 7.33E-02 | 2.61E+00 | 2.71E+00 | 7.02E-01 | no | Myo1a | intron_variant | MODIFIER |
| 10 | 127814481 | AX-97802734 | 0 | C | T | 0.325 | -2.80E-02 | 7.33E-02 | 2.61E+00 | 2.71E+00 | 7.02E-01 | no | Rdh16 | 3_prime_UTR_variant | MODIFIER |
| 10 | 127841979 | AX-170424797 | 0 | G | C | 0.325 | -2.80E-02 | 7.33E-02 | 2.61E+00 | 2.71E+00 | 7.02E-01 | no | Rdh16 | intron_variant,NMD_transcript_variant | MODIFIER |
| 10 | 127857383 | AX-170534785 | 0 | G | A | 0.325 | -2.80E-02 | 7.33E-02 | 2.61E+00 | 2.71E+00 | 7.02E-01 | no | Rdh19 | intron_variant | MODIFIER |
| 10 | 127859269 | AX-170419027 | 0 | C | A | 0.325 | -2.80E-02 | 7.33E-02 | 2.61E+00 | 2.71E+00 | 7.02E-01 | no | Rdh19 | intron_variant | MODIFIER |
| 10 | 127862278 | AX-170456303 | 0 | A | G | 0.325 | -2.80E-02 | 7.33E-02 | 2.61E+00 | 2.71E+00 | 7.02E-01 | no | Rdh19 | downstream_gene_variant | MODIFIER |
| 10 | 127874743 | AX-170463475 | 0 | T | C | 0.325 | -2.80E-02 | 7.33E-02 | 2.61E+00 | 2.71E+00 | 7.02E-01 | no | Rdh16f2 | intron_variant | MODIFIER |
| 10 | 127877211 | AX-170046229 | 0 | C | T | 0.325 | -2.80E-02 | 7.33E-02 | 2.61E+00 | 2.71E+00 | 7.02E-01 | no | Rdh16f2 | 3_prime_UTR_variant | MODIFIER |
| 10 | 127894809 | AX-170046230 | 0 | G | A | 0.325 | -2.80E-02 | 7.33E-02 | 2.61E+00 | 2.71E+00 | 7.02E-01 | no | Sdr9c7 | upstream_gene_variant | MODIFIER |
| 10 | 127898352 | AX-170448669 | 0 | C | A | 0.325 | -2.80E-02 | 7.33E-02 | 2.61E+00 | 2.71E+00 | 7.02E-01 | no | Sdr9c7 | upstream_gene_variant | MODIFIER |
| 10 | 127901786 | AX-170004134 | 0 | A | G | 0.325 | -2.80E-02 | 7.33E-02 | 2.61E+00 | 2.71E+00 | 7.02E-01 | no | Sdr9c7 | intron_variant | MODIFIER |
| 10 | 127904910 | AX-170046232 | 0 | A | T | 0.325 | -2.80E-02 | 7.33E-02 | 2.61E+00 | 2.71E+00 | 7.02E-01 | no | Sdr9c7 | intron_variant | MODIFIER |
| 10 | 127910378 | AX-170433648 | 0 | G | A | 0.325 | -2.80E-02 | 7.33E-02 | 2.61E+00 | 2.71E+00 | 7.02E-01 | no | Sdr9c7 | 3_prime_UTR_variant | MODIFIER |
| 10 | 127910449 | AX-170315851 | 0 | C | T | 0.325 | -2.80E-02 | 7.33E-02 | 2.61E+00 | 2.71E+00 | 7.02E-01 | no | Sdr9c7 | 3_prime_UTR_variant | MODIFIER |
| 10 | 127916401 | AX-170106762 | 0 | T | C | 0.325 | -2.80E-02 | 7.33E-02 | 2.61E+00 | 2.71E+00 | 7.02E-01 | no | Sdr9c7 | downstream_gene_variant | MODIFIER |
| 10 | 127919111 | AX-170372764 | 0 | A | G | 0.325 | -2.80E-02 | 7.33E-02 | 2.61E+00 | 2.71E+00 | 7.02E-01 | no | Gm15905 | upstream_gene_variant | MODIFIER |
| 10 | 127925814 | AX-170004136 | 0 | C | T | 0.325 | -2.80E-02 | 7.33E-02 | 2.61E+00 | 2.71E+00 | 7.02E-01 | no | Gm15905 | downstream_gene_variant | MODIFIER |
| 10 | 127931423 | AX-97825857 | 0 | T | C | 0.325 | -2.80E-02 | 7.33E-02 | 2.61E+00 | 2.71E+00 | 7.02E-01 | no | | intergenic_variant | MODIFIER |
| 10 | 127956061 | AX-170247722 | 0 | A | G | 0.325 | -2.80E-02 | 7.33E-02 | 2.61E+00 | 2.71E+00 | 7.02E-01 | no | Gm23996 | upstream_gene_variant | MODIFIER |
| 10 | 127967679 | AX-170092153 | 0 | G | A | 0.325 | -2.80E-02 | 7.33E-02 | 2.61E+00 | 2.71E+00 | 7.02E-01 | no | | intergenic_variant | MODIFIER |
| 10 | 127976443 | AX-170004132 | 0 | G | A | 0.325 | -2.80E-02 | 7.33E-02 | 2.61E+00 | 2.71E+00 | 7.02E-01 | no | | intergenic_variant | MODIFIER |
| 10 | 127977696 | AX-170314626 | 0 | T | C | 0.325 | -2.80E-02 | 7.33E-02 | 2.61E+00 | 2.71E+00 | 7.02E-01 | no | | intergenic_variant | MODIFIER |
| 10 | 128015123 | AX-170100083 | 0 | C | T | 0.325 | -2.80E-02 | 7.33E-02 | 2.61E+00 | 2.71E+00 | 7.02E-01 | no | Prim1 | upstream_gene_variant | MODIFIER |
| 10 | 128017589 | AX-170092154 | 0 | T | C | 0.325 | -2.80E-02 | 7.33E-02 | 2.61E+00 | 2.71E+00 | 7.02E-01 | no | Prim1 | intron_variant | MODIFIER |
| 10 | 128069892 | AX-170046237 | 0 | T | C | 0.325 | -2.80E-02 | 7.33E-02 | 2.61E+00 | 2.71E+00 | 7.02E-01 | no | Ptges3 | intron_variant | MODIFIER |
| 10 | 128074164 | AX-97825578 | 0 | G | A | 0.325 | -2.80E-02 | 7.33E-02 | 2.61E+00 | 2.71E+00 | 7.02E-01 | no | Ptges3 | intron_variant | MODIFIER |

| | | | | | | | | | | | | | | | |
|---|---|---|---|---|---|---|---|---|---|---|---|---|---|---|---|
| 10 | 128104832 | AX-170364336 | 0 | A | G | 0.325 | -2.80E-02 | 7.33E-02 | 2.61E+00 | 2.71E+00 | 7.02E-01 | no | Baz2a | intron_variant | MODIFIER |
| 10 | 128114704 | AX-170262171 | 0 | T | G | 0.325 | -2.80E-02 | 7.33E-02 | 2.61E+00 | 2.71E+00 | 7.02E-01 | no | Baz2a | missense_variant | MODERATE |
| 10 | 128124024 | AX-170397597 | 0 | T | G | 0.325 | -2.80E-02 | 7.33E-02 | 2.61E+00 | 2.71E+00 | 7.02E-01 | no | Baz2a | intron_variant | MODIFIER |
| 10 | 128135365 | AX-170278297 | 0 | A | T | 0.325 | -2.80E-02 | 7.33E-02 | 2.61E+00 | 2.71E+00 | 7.02E-01 | no | Rbms2 | intron_variant | MODIFIER |
| 10 | 128152972 | AX-170128656 | 0 | A | G | 0.325 | -2.80E-02 | 7.33E-02 | 2.61E+00 | 2.71E+00 | 7.02E-01 | no | Rbms2 | intron_variant | MODIFIER |
| 10 | 128153162 | AX-170483840 | 0 | G | A | 0.325 | -2.80E-02 | 7.33E-02 | 2.61E+00 | 2.71E+00 | 7.02E-01 | no | Rbms2 | intron_variant | MODIFIER |
| 10 | 128162877 | AX-97783128 | 0 | G | A | 0.325 | -2.80E-02 | 7.33E-02 | 2.61E+00 | 2.71E+00 | 7.02E-01 | no | Rbms2 | intron_variant | MODIFIER |
| 10 | 128179106 | AX-170330694 | 0 | C | T | 0.325 | -2.80E-02 | 7.33E-02 | 2.61E+00 | 2.71E+00 | 7.02E-01 | no | Rbms2 | upstream_gene_variant | MODIFIER |
| 10 | 128179283 | AX-170158298 | 0 | T | C | 0.325 | -2.80E-02 | 7.33E-02 | 2.61E+00 | 2.71E+00 | 7.02E-01 | no | Rbms2 | upstream_gene_variant | MODIFIER |
| 10 | 128190729 | AX-170405978 | 0 | T | G | 0.325 | -2.80E-02 | 7.33E-02 | 2.61E+00 | 2.71E+00 | 7.02E-01 | no | Gls2 | upstream_gene_variant | MODIFIER |
| 10 | 128210048 | AX-170256438 | 0 | T | A | 0.325 | -2.80E-02 | 7.33E-02 | 2.61E+00 | 2.71E+00 | 7.02E-01 | no | Gls2 | downstream_gene_variant | MODIFIER |
| 10 | 128210787 | AX-170240031 | 0 | T | G | 0.325 | -2.80E-02 | 7.33E-02 | 2.61E+00 | 2.71E+00 | 7.02E-01 | no | Gls2 | downstream_gene_variant | MODIFIER |
| 10 | 128221248 | AX-170499958 | 0 | C | A | 0.325 | -2.80E-02 | 7.33E-02 | 2.61E+00 | 2.71E+00 | 7.02E-01 | no | Mip | upstream_gene_variant | MODIFIER |
| 10 | 128224413 | AX-170419274 | 0 | A | G | 0.325 | -2.80E-02 | 7.33E-02 | 2.61E+00 | 2.71E+00 | 7.02E-01 | no | Mip | upstream_gene_variant | MODIFIER |
| 10 | 128225861 | AX-170262857 | 0 | G | A | 0.325 | -2.80E-02 | 7.33E-02 | 2.61E+00 | 2.71E+00 | 7.02E-01 | no | Mip | 5_prime_UTR_variant | MODIFIER |
| 10 | 128226510 | AX-170371918 | 0 | C | T | 0.325 | -2.80E-02 | 7.33E-02 | 2.61E+00 | 2.71E+00 | 7.02E-01 | no | Mip | intron_variant | MODIFIER |
| 10 | 128239367 | AX-170334765 | 0 | A | C | 0.325 | -2.80E-02 | 7.33E-02 | 2.61E+00 | 2.71E+00 | 7.02E-01 | no | Timeless | intron_variant | MODIFIER |
| 10 | 128243181 | AX-97829991 | 0 | A | G | 0.325 | -2.80E-02 | 7.33E-02 | 2.61E+00 | 2.71E+00 | 7.02E-01 | no | Timeless | intron_variant | MODIFIER |
| 10 | 128282696 | AX-170046240 | 0 | C | T | 0.325 | -2.80E-02 | 7.33E-02 | 2.61E+00 | 2.71E+00 | 7.02E-01 | no | Stat2 | intron_variant | MODIFIER |
| 10 | 128283501 | AX-170046241 | 0 | C | T | 0.325 | -2.80E-02 | 7.33E-02 | 2.61E+00 | 2.71E+00 | 7.02E-01 | no | Stat2 | synonymous_variant | LOW |
| 10 | 128289788 | AX-97809946 | 0 | C | A | 0.325 | -2.80E-02 | 7.33E-02 | 2.61E+00 | 2.71E+00 | 7.02E-01 | no | Stat2 | intron_variant | MODIFIER |
| 10 | 128292645 | AX-170368143 | 0 | T | C | 0.325 | -2.80E-02 | 7.33E-02 | 2.61E+00 | 2.71E+00 | 7.02E-01 | no | Il23a | downstream_gene_variant | MODIFIER |
| 10 | 128293069 | AX-170306142 | 0 | G | A | 0.325 | -2.80E-02 | 7.33E-02 | 2.61E+00 | 2.71E+00 | 7.02E-01 | no | Il23a | downstream_gene_variant | MODIFIER |
| 10 | 128342094 | AX-170205379 | 0 | A | G | 0.325 | -2.80E-02 | 7.33E-02 | 2.61E+00 | 2.71E+00 | 7.02E-01 | no | Cs | intron_variant | MODIFIER |
| 10 | 128342188 | AX-170365812 | 0 | G | C | 0.325 | -2.80E-02 | 7.33E-02 | 2.61E+00 | 2.71E+00 | 7.02E-01 | no | Cs | intron_variant | MODIFIER |
| 10 | 128348227 | AX-170326151 | 0 | G | A | 0.325 | -2.80E-02 | 7.33E-02 | 2.61E+00 | 2.71E+00 | 7.02E-01 | no | Cs | intron_variant | MODIFIER |
| 10 | 128351246 | AX-170207578 | 0 | A | G | 0.325 | -2.80E-02 | 7.33E-02 | 2.61E+00 | 2.71E+00 | 7.02E-01 | no | Cs | synonymous_variant | LOW |
| 10 | 128356905 | AX-170481326 | 0 | C | T | 0.325 | -2.80E-02 | 7.33E-02 | 2.61E+00 | 2.71E+00 | 7.02E-01 | no | Cs | intron_variant | MODIFIER |
| 10 | 128358883 | AX-170102632 | 0 | G | A | 0.325 | -2.80E-02 | 7.33E-02 | 2.61E+00 | 2.71E+00 | 7.02E-01 | no | Cs | intron_variant | MODIFIER |
| 10 | 128359068 | AX-170389923 | 0 | T | C | 0.325 | -2.80E-02 | 7.33E-02 | 2.61E+00 | 2.71E+00 | 7.02E-01 | no | Cs | intron_variant | MODIFIER |
| 10 | 128379143 | AX-170263146 | 0 | G | A | 0.325 | -2.80E-02 | 7.33E-02 | 2.61E+00 | 2.71E+00 | 7.02E-01 | no | Ankrd52 | intron_variant | MODIFIER |
| 10 | 128398995 | AX-170450349 | 0 | G | A | 0.325 | -2.80E-02 | 7.33E-02 | 2.61E+00 | 2.71E+00 | 7.02E-01 | no | Ankrd52 | downstream_gene_variant | MODIFIER |
| 10 | 128435607 | AX-170172202 | 0 | G | A | 0.325 | -2.80E-02 | 7.33E-02 | 2.61E+00 | 2.71E+00 | 7.02E-01 | no | Rnf41 | intron_variant | MODIFIER |
| 10 | 128435869 | AX-170295277 | 0 | A | T | 0.325 | -2.80E-02 | 7.33E-02 | 2.61E+00 | 2.71E+00 | 7.02E-01 | no | Rnf41 | intron_variant | MODIFIER |
| 10 | 128437400 | AX-97783568 | 0 | C | T | 0.325 | -2.80E-02 | 7.33E-02 | 2.61E+00 | 2.71E+00 | 7.02E-01 | no | Rnf41 | synonymous_variant | LOW |
| 10 | 128475290 | AX-170276624 | 0 | A | C | 0.325 | -2.80E-02 | 7.33E-02 | 2.61E+00 | 2.71E+00 | 7.02E-01 | no | Smarcc2 | intron_variant | MODIFIER |
| 10 | 128475347 | AX-170418145 | 0 | A | G | 0.325 | -2.80E-02 | 7.33E-02 | 2.61E+00 | 2.71E+00 | 7.02E-01 | no | Smarcc2 | intron_variant | MODIFIER |
| 10 | 128510115 | AX-170046243 | 0 | G | T | 0.325 | -2.80E-02 | 7.33E-02 | 2.61E+00 | 2.71E+00 | 7.02E-01 | no | Esyt1 | 3_prime_UTR_variant | MODIFIER |
| 2 | 159292325 | AX-170525055 | 0 | C | T | 0.149 | 8.15E-02 | 2.16E-01 | 2.71E+00 | 2.75E+00 | 7.02E-01 | no | | intergenic_variant | MODIFIER |
| 2 | 159602158 | AX-170331310 | 0 | G | T | 0.149 | 8.15E-02 | 2.16E-01 | 2.71E+00 | 2.75E+00 | 7.02E-01 | no | | intergenic_variant | MODIFIER |
| 2 | 159606275 | AX-170270869 | 0 | A | G | 0.149 | 8.15E-02 | 2.16E-01 | 2.71E+00 | 2.75E+00 | 7.02E-01 | no | | intergenic_variant | MODIFIER |
| 2 | 159606348 | AX-170143475 | 0 | C | T | 0.149 | 8.15E-02 | 2.16E-01 | 2.71E+00 | 2.75E+00 | 7.02E-01 | no | | intergenic_variant | MODIFIER |
| 2 | 159645437 | AX-170199781 | 0 | T | C | 0.149 | 8.15E-02 | 2.16E-01 | 2.71E+00 | 2.75E+00 | 7.02E-01 | no | | intergenic_variant | MODIFIER |
| 2 | 159687846 | AX-170476244 | 0 | A | G | 0.149 | 8.15E-02 | 2.16E-01 | 2.71E+00 | 2.75E+00 | 7.02E-01 | no | | intergenic_variant | MODIFIER |
| 2 | 159688190 | AX-170362795 | 0 | T | C | 0.149 | 8.15E-02 | 2.16E-01 | 2.71E+00 | 2.75E+00 | 7.02E-01 | no | | intergenic_variant | MODIFIER |
| 2 | 159698008 | AX-170279367 | 0 | T | C | 0.149 | 8.15E-02 | 2.16E-01 | 2.71E+00 | 2.75E+00 | 7.02E-01 | no | | intergenic_variant | MODIFIER |
| 2 | 159709236 | AX-170336898 | 0 | A | C | 0.149 | 8.15E-02 | 2.16E-01 | 2.71E+00 | 2.75E+00 | 7.02E-01 | no | | intergenic_variant | MODIFIER |
| 2 | 159709338 | AX-170347344 | 0 | A | T | 0.149 | 8.15E-02 | 2.16E-01 | 2.71E+00 | 2.75E+00 | 7.02E-01 | no | | intergenic_variant | MODIFIER |
| 2 | 159711935 | AX-170228802 | 0 | C | G | 0.149 | 8.15E-02 | 2.16E-01 | 2.71E+00 | 2.75E+00 | 7.02E-01 | no | | intergenic_variant | MODIFIER |
| 2 | 159712552 | AX-97794924 | 0 | T | C | 0.149 | 8.15E-02 | 2.16E-01 | 2.71E+00 | 2.75E+00 | 7.02E-01 | no | | intergenic_variant | MODIFIER |
| 2 | 159713226 | AX-170373796 | 0 | C | T | 0.149 | 8.15E-02 | 2.16E-01 | 2.71E+00 | 2.75E+00 | 7.02E-01 | no | | intergenic_variant | MODIFIER |
| 2 | 159713390 | AX-170132568 | 0 | A | G | 0.149 | 8.15E-02 | 2.16E-01 | 2.71E+00 | 2.75E+00 | 7.02E-01 | no | | intergenic_variant | MODIFIER |
| 2 | 159750201 | AX-170187691 | 0 | A | G | 0.149 | 8.15E-02 | 2.16E-01 | 2.71E+00 | 2.75E+00 | 7.02E-01 | no | | intergenic_variant | MODIFIER |
| 2 | 159750336 | AX-170471077 | 0 | T | C | 0.149 | 8.15E-02 | 2.16E-01 | 2.71E+00 | 2.75E+00 | 7.02E-01 | no | | intergenic_variant | MODIFIER |

| | | | | | | | | | | | | | | | |
|---|---|---|---|---|---|---|---|---|---|---|---|---|---|---|---|
| 2 | 159750595 | AX-170387217 | 0 | T | C | 0.149 | 8.15E-02 | 2.16E-01 | 2.71E+00 | 2.75E+00 | 7.02E-01 | no | | intergenic_variant | MODIFIER |
| 2 | 159761412 | AX-97811745 | 0 | C | T | 0.149 | 8.15E-02 | 2.16E-01 | 2.71E+00 | 2.75E+00 | 7.02E-01 | no | | intergenic_variant | MODIFIER |
| 2 | 159761579 | AX-170121852 | 0 | T | C | 0.149 | 8.15E-02 | 2.16E-01 | 2.71E+00 | 2.75E+00 | 7.02E-01 | no | | intergenic_variant | MODIFIER |
| 2 | 159761964 | AX-170462407 | 0 | C | T | 0.149 | 8.15E-02 | 2.16E-01 | 2.71E+00 | 2.75E+00 | 7.02E-01 | no | | intergenic_variant | MODIFIER |
| 2 | 159762533 | AX-170498220 | 0 | G | A | 0.149 | 8.15E-02 | 2.16E-01 | 2.71E+00 | 2.75E+00 | 7.02E-01 | no | | regulatory_region_variant | MODIFIER |
| 2 | 159764020 | AX-170393719 | 0 | A | C | 0.149 | 8.15E-02 | 2.16E-01 | 2.71E+00 | 2.75E+00 | 7.02E-01 | no | | intergenic_variant | MODIFIER |
| 2 | 159764510 | AX-170234154 | 0 | T | G | 0.149 | 8.15E-02 | 2.16E-01 | 2.71E+00 | 2.75E+00 | 7.02E-01 | no | | intergenic_variant | MODIFIER |
| 2 | 159785049 | AX-97831488 | 0 | A | G | 0.149 | 8.15E-02 | 2.16E-01 | 2.71E+00 | 2.75E+00 | 7.02E-01 | no | | intergenic_variant | MODIFIER |
| 2 | 159849851 | AX-170363905 | 0 | G | A | 0.149 | 8.15E-02 | 2.16E-01 | 2.71E+00 | 2.75E+00 | 7.02E-01 | no | | intergenic_variant | MODIFIER |
| 2 | 159855067 | AX-170427966 | 0 | C | T | 0.149 | 8.15E-02 | 2.16E-01 | 2.71E+00 | 2.75E+00 | 7.02E-01 | no | Gm11446 | downstream_gene_variant | MODIFIER |
| 2 | 159860338 | AX-97847025 | 0 | T | C | 0.149 | 8.15E-02 | 2.16E-01 | 2.71E+00 | 2.75E+00 | 7.02E-01 | no | Gm11446 | upstream_gene_variant | MODIFIER |
| 2 | 159869124 | AX-170461905 | 0 | C | T | 0.149 | 8.15E-02 | 2.16E-01 | 2.71E+00 | 2.75E+00 | 7.02E-01 | no | | intergenic_variant | MODIFIER |
| 2 | 159871747 | AX-170480617 | 0 | C | T | 0.149 | 8.15E-02 | 2.16E-01 | 2.71E+00 | 2.75E+00 | 7.02E-01 | no | | intergenic_variant | MODIFIER |
| 2 | 159884294 | AX-97839735 | 0 | T | C | 0.149 | 8.15E-02 | 2.16E-01 | 2.71E+00 | 2.75E+00 | 7.02E-01 | no | | intergenic_variant | MODIFIER |
| 2 | 159891111 | AX-170156038 | 0 | A | G | 0.149 | 8.15E-02 | 2.16E-01 | 2.71E+00 | 2.75E+00 | 7.02E-01 | no | | intergenic_variant | MODIFIER |
| 2 | 159974973 | AX-170201029 | 0 | A | G | 0.149 | 8.15E-02 | 2.16E-01 | 2.71E+00 | 2.75E+00 | 7.02E-01 | no | Gm14220 | upstream_gene_variant | MODIFIER |
| 2 | 159976270 | AX-170473491 | 0 | A | G | 0.149 | 8.15E-02 | 2.16E-01 | 2.71E+00 | 2.75E+00 | 7.02E-01 | no | Gm14220 | upstream_gene_variant | MODIFIER |
| 2 | 159982190 | AX-170455811 | 0 | A | G | 0.149 | 8.15E-02 | 2.16E-01 | 2.71E+00 | 2.75E+00 | 7.02E-01 | no | Gm14220 | downstream_gene_variant | MODIFIER |
| 2 | 159982929 | AX-170354623 | 0 | G | A | 0.149 | 8.15E-02 | 2.16E-01 | 2.71E+00 | 2.75E+00 | 7.02E-01 | no | Gm14220 | downstream_gene_variant | MODIFIER |
| 2 | 160001104 | AX-170088814 | 0 | A | C | 0.149 | 8.15E-02 | 2.16E-01 | 2.71E+00 | 2.75E+00 | 7.02E-01 | no | | intergenic_variant | MODIFIER |
| 2 | 160058754 | AX-170019516 | 0 | G | C | 0.149 | 8.15E-02 | 2.16E-01 | 2.71E+00 | 2.75E+00 | 7.02E-01 | no | | intergenic_variant | MODIFIER |
| 2 | 160069100 | AX-170401704 | 0 | T | C | 0.149 | 8.15E-02 | 2.16E-01 | 2.71E+00 | 2.75E+00 | 7.02E-01 | no | | intergenic_variant | MODIFIER |
| 2 | 160110857 | AX-170305146 | 0 | T | C | 0.149 | 8.15E-02 | 2.16E-01 | 2.71E+00 | 2.75E+00 | 7.02E-01 | no | | intergenic_variant | MODIFIER |
| 2 | 160110940 | AX-170209625 | 0 | T | C | 0.149 | 8.15E-02 | 2.16E-01 | 2.71E+00 | 2.75E+00 | 7.02E-01 | no | | intergenic_variant | MODIFIER |
| 2 | 160124516 | AX-170310099 | 0 | T | C | 0.149 | 8.15E-02 | 2.16E-01 | 2.71E+00 | 2.75E+00 | 7.02E-01 | no | | intergenic_variant | MODIFIER |
| 2 | 160285278 | AX-97839733 | 0 | G | A | 0.149 | 8.15E-02 | 2.16E-01 | 2.71E+00 | 2.75E+00 | 7.02E-01 | no | | intergenic_variant | MODIFIER |
| 2 | 160342408 | AX-97850584 | 0 | A | T | 0.149 | 8.15E-02 | 2.16E-01 | 2.71E+00 | 2.75E+00 | 7.02E-01 | no | | intergenic_variant | MODIFIER |
| 2 | 160351025 | AX-170120779 | 0 | A | G | 0.149 | 8.15E-02 | 2.16E-01 | 2.71E+00 | 2.75E+00 | 7.02E-01 | no | | regulatory_region_variant | MODIFIER |
| 2 | 160354969 | AX-170259622 | 0 | G | C | 0.149 | 8.15E-02 | 2.16E-01 | 2.71E+00 | 2.75E+00 | 7.02E-01 | no | | intergenic_variant | MODIFIER |
| 2 | 160355016 | AX-170532140 | 0 | T | G | 0.149 | 8.15E-02 | 2.16E-01 | 2.71E+00 | 2.75E+00 | 7.02E-01 | no | | intergenic_variant | MODIFIER |
| 2 | 160355322 | AX-170065439 | 0 | T | C | 0.149 | 8.15E-02 | 2.16E-01 | 2.71E+00 | 2.75E+00 | 7.02E-01 | no | | intergenic_variant | MODIFIER |
| 2 | 160515865 | AX-170019525 | 0 | C | G | 0.149 | 8.15E-02 | 2.16E-01 | 2.71E+00 | 2.75E+00 | 7.02E-01 | no | | intergenic_variant | MODIFIER |
| 2 | 160811177 | AX-170251139 | 0 | G | A | 0.149 | 8.15E-02 | 2.16E-01 | 2.71E+00 | 2.75E+00 | 7.02E-01 | no | Zhx3 | intron_variant | MODIFIER |
| 2 | 161042742 | AX-170088818 | 0 | A | G | 0.149 | 8.15E-02 | 2.16E-01 | 2.71E+00 | 2.75E+00 | 7.02E-01 | no | Chd6 | intron_variant | MODIFIER |
| 2 | 161056792 | AX-97828516 | 0 | G | A | 0.149 | 8.15E-02 | 2.16E-01 | 2.71E+00 | 2.75E+00 | 7.02E-01 | no | Chd6 | intron_variant | MODIFIER |
| 2 | 161096263 | AX-170019530 | 0 | C | G | 0.149 | 8.15E-02 | 2.16E-01 | 2.71E+00 | 2.75E+00 | 7.02E-01 | no | Chd6 | intron_variant | MODIFIER |
| 2 | 161117794 | AX-170088819 | 0 | C | T | 0.149 | 8.15E-02 | 2.16E-01 | 2.71E+00 | 2.75E+00 | 7.02E-01 | no | Gm45447 | downstream_gene_variant | MODIFIER |
| 2 | 161215435 | AX-170245884 | 0 | G | A | 0.149 | 8.15E-02 | 2.16E-01 | 2.71E+00 | 2.75E+00 | 7.02E-01 | no | Gm14243 | non_coding_transcript_exon_variant | MODIFIER |
| 2 | 161215659 | AX-170271122 | 0 | G | A | 0.149 | 8.15E-02 | 2.16E-01 | 2.71E+00 | 2.75E+00 | 7.02E-01 | no | Gm14243 | non_coding_transcript_exon_variant | MODIFIER |
| 2 | 161219896 | AX-97805730 | 0 | T | C | 0.149 | 8.15E-02 | 2.16E-01 | 2.71E+00 | 2.75E+00 | 7.02E-01 | no | Gm14243 | downstream_gene_variant | MODIFIER |
| 2 | 161289622 | AX-170500489 | 0 | C | T | 0.149 | 8.15E-02 | 2.16E-01 | 2.71E+00 | 2.75E+00 | 7.02E-01 | no | Gm11448 | downstream_gene_variant | MODIFIER |
| 2 | 161302503 | AX-170136359 | 0 | C | T | 0.149 | 8.15E-02 | 2.16E-01 | 2.71E+00 | 2.75E+00 | 7.02E-01 | no | | intergenic_variant | MODIFIER |
| 2 | 161303721 | AX-170347567 | 0 | A | G | 0.149 | 8.15E-02 | 2.16E-01 | 2.71E+00 | 2.75E+00 | 7.02E-01 | no | | intergenic_variant | MODIFIER |
| 2 | 161305909 | AX-170065455 | 0 | G | C | 0.149 | 8.15E-02 | 2.16E-01 | 2.71E+00 | 2.75E+00 | 7.02E-01 | no | | intergenic_variant | MODIFIER |
| 2 | 161305950 | AX-170065454 | 0 | C | T | 0.149 | 8.15E-02 | 2.16E-01 | 2.71E+00 | 2.75E+00 | 7.02E-01 | no | | intergenic_variant | MODIFIER |
| 2 | 161322752 | AX-170237730 | 0 | G | C | 0.149 | 8.15E-02 | 2.16E-01 | 2.71E+00 | 2.75E+00 | 7.02E-01 | no | | intergenic_variant | MODIFIER |
| 2 | 161327042 | AX-170502881 | 0 | T | C | 0.149 | 8.15E-02 | 2.16E-01 | 2.71E+00 | 2.75E+00 | 7.02E-01 | no | Gm14241 | upstream_gene_variant | MODIFIER |
| 2 | 161341322 | AX-97800609 | 0 | G | A | 0.149 | 8.15E-02 | 2.16E-01 | 2.71E+00 | 2.75E+00 | 7.02E-01 | no | | regulatory_region_variant | MODIFIER |
| 8 | 14706968 | AX-170030299 | 0 | A | T | 0.246 | -2.91E-02 | 7.44E-02 | 2.57E+00 | 2.67E+00 | 7.02E-01 | no | Dlgap2 | intron_variant | MODIFIER |
| 8 | 15267335 | AX-170368085 | 1 | C | T | 0.241 | -3.03E-02 | 7.67E-02 | 2.57E+00 | 2.66E+00 | 7.02E-01 | no | Gm7706 | intron_variant,non_coding_transcript_variant | MODIFIER |
| 1 | 172680923 | AX-170362413 | 0 | T | C | 0.228 | -3.96E-02 | 1.08E-01 | 2.77E+00 | 2.90E+00 | 7.02E-01 | no | Gm22383 | downstream_gene_variant | MODIFIER |
| 1 | 172989821 | AX-170216325 | 0 | A | T | 0.228 | -3.96E-02 | 1.08E-01 | 2.77E+00 | 2.90E+00 | 7.02E-01 | no | | regulatory_region_variant | MODIFIER |
| 1 | 174580918 | AX-170346065 | 0 | G | A | 0.228 | -3.96E-02 | 1.08E-01 | 2.77E+00 | 2.90E+00 | 7.02E-01 | no | Fmn2 | intron_variant | MODIFIER |
| 1 | 174637540 | AX-170106707 | 0 | A | G | 0.228 | -3.96E-02 | 1.08E-01 | 2.77E+00 | 2.90E+00 | 7.02E-01 | no | Fmn2 | intron_variant | MODIFIER |

| | | | | | | | | | | | | | | | |
|---|---|---|---|---|---|---|---|---|---|---|---|---|---|---|---|
| 15 | 86144427 | AX-170056189 | 1 | G | A | 0.518 | 2.93E-02 | 7.21E-02 | 2.52E+00 | 2.62E+00 | 7.02E-01 | no | Cerk | intron_variant | MODIFIER |
| 4 | 94927735 | AX-170418352 | 1 | T | G | 0.321 | 4.07E-02 | 1.07E-01 | 2.65E+00 | 2.72E+00 | 7.02E-01 | no | Eqtn | intron_variant | MODIFIER |
| 10 | 121222367 | AX-170517273 | 0 | G | A | 0.351 | 2.75E-02 | 7.35E-02 | 2.68E+00 | 2.77E+00 | 7.03E-01 | no | | intergenic_variant | MODIFIER |
| 1 | 174650604 | AX-170391349 | 1 | A | C | 0.223 | 4.81E-02 | 1.21E-01 | 2.57E+00 | 2.63E+00 | 7.03E-01 | no | Fmn2 | intron_variant | MODIFIER |
| 6 | 125775115 | AX-170508801 | 0 | A | C | 0.167 | -3.06E-02 | 8.28E-02 | 2.65E+00 | 2.79E+00 | 7.03E-01 | no | Ano2 | intron_variant,non_coding_transcript_variant | MODIFIER |
| 4 | 148415678 | AX-170070993 | 1 | G | C | 0.089 | 7.83E-02 | 2.07E-01 | 2.66E+00 | 2.73E+00 | 7.03E-01 | no | | regulatory_region_variant | MODIFIER |
| 5 | 112987372 | AX-170073144 | 1 | C | T | 0.446 | -4.01E-02 | 1.07E-01 | 2.77E+00 | 2.84E+00 | 7.03E-01 | no | Grk3 | intron_variant | MODIFIER |
| 8 | 41014038 | AX-170146021 | 0 | G | A | 0.167 | 4.73E-02 | 1.26E-01 | 2.67E+00 | 2.75E+00 | 7.03E-01 | no | Mtus1 | intron_variant | MODIFIER |
| 1 | 75353925 | AX-170335973 | 2 | A | G | 0.118 | 7.47E-02 | 1.99E-01 | 2.67E+00 | 2.75E+00 | 7.04E-01 | no | | intergenic_variant | MODIFIER |
| 12 | 107134115 | AX-97789901 | 2 | T | C | 0.109 | -7.43E-02 | 2.00E-01 | 2.74E+00 | 2.83E+00 | 7.04E-01 | no | Gm16085 | downstream_gene_variant | MODIFIER |
| 1 | 176075248 | AX-170461559 | 0 | C | T | 0.228 | 4.45E-02 | 1.15E-01 | 2.61E+00 | 2.68E+00 | 7.04E-01 | no | Pld5 | intron_variant | MODIFIER |
| 10 | 17575431 | AX-170402922 | 1 | C | G | 0.205 | 3.83E-02 | 1.03E-01 | 2.76E+00 | 2.85E+00 | 7.04E-01 | no | Gm47771 | intron_variant,non_coding_transcript_variant | MODIFIER |
| 8 | 86400742 | AX-97785140 | 1 | T | C | 0.196 | -4.88E-02 | 1.30E-01 | 2.69E+00 | 2.75E+00 | 7.04E-01 | no | | intergenic_variant | MODIFIER |
| 3 | 37957241 | AX-170398858 | 0 | T | C | 0.193 | 3.43E-02 | 9.25E-02 | 2.68E+00 | 2.77E+00 | 7.04E-01 | no | Gm7799 | downstream_gene_variant | MODIFIER |
| 7 | 113952841 | AX-170078482 | 1 | G | A | 0.42 | 3.64E-02 | 9.67E-02 | 2.69E+00 | 2.73E+00 | 7.04E-01 | no | Spon1 | intron_variant | MODIFIER |
| 5 | 88888631 | AX-170465962 | 0 | G | A | 0.43 | 4.10E-02 | 1.10E-01 | 2.74E+00 | 2.80E+00 | 7.04E-01 | no | Slc4a4 | intron_variant | MODIFIER |
| 5 | 88942627 | AX-170025544 | 0 | C | T | 0.43 | 4.10E-02 | 1.10E-01 | 2.74E+00 | 2.80E+00 | 7.04E-01 | no | Slc4a4 | intron_variant | MODIFIER |
| 7 | 93897408 | AX-170127659 | 0 | A | T | 0.132 | -6.76E-02 | 1.76E-01 | 2.60E+00 | 2.70E+00 | 7.04E-01 | no | | intergenic_variant | MODIFIER |
| 9 | 39723793 | AX-97829559 | 1 | A | G | 0.42 | -2.79E-02 | 7.43E-02 | 2.64E+00 | 2.74E+00 | 7.04E-01 | no | Olfr965 | downstream_gene_variant | MODIFIER |
| 20 | 139848768 | AX-170085390 | 1 | A | T | 0.152 | 5.05E-02 | 1.35E-01 | 2.64E+00 | 2.74E+00 | 7.04E-01 | no | Morc4 | intron_variant | MODIFIER |
| 5 | 38720342 | AX-170343498 | 0 | G | A | 0.211 | 4.16E-02 | 1.12E-01 | 2.73E+00 | 2.82E+00 | 7.04E-01 | no | Clnk | intron_variant | MODIFIER |
| 5 | 38734306 | AX-97843737 | 0 | A | G | 0.211 | 4.16E-02 | 1.12E-01 | 2.73E+00 | 2.82E+00 | 7.04E-01 | no | Clnkos | upstream_gene_variant | MODIFIER |
| 5 | 38776074 | AX-97792187 | 0 | A | G | 0.211 | 4.16E-02 | 1.12E-01 | 2.73E+00 | 2.82E+00 | 7.04E-01 | no | Clnk | intron_variant | MODIFIER |
| 5 | 38790442 | AX-170024726 | 0 | A | C | 0.211 | 4.16E-02 | 1.12E-01 | 2.73E+00 | 2.82E+00 | 7.04E-01 | no | Clnk | intron_variant | MODIFIER |
| 5 | 39076482 | AX-170329795 | 0 | C | T | 0.211 | 4.16E-02 | 1.12E-01 | 2.73E+00 | 2.82E+00 | 7.04E-01 | no | | regulatory_region_variant | MODIFIER |
| 5 | 39076596 | AX-170150569 | 0 | T | C | 0.211 | 4.16E-02 | 1.12E-01 | 2.73E+00 | 2.82E+00 | 7.04E-01 | no | | regulatory_region_variant | MODIFIER |
| 5 | 39082197 | AX-170110526 | 0 | T | C | 0.211 | 4.16E-02 | 1.12E-01 | 2.73E+00 | 2.82E+00 | 7.04E-01 | no | | intergenic_variant | MODIFIER |
| 5 | 39082277 | AX-170318826 | 0 | T | G | 0.211 | 4.16E-02 | 1.12E-01 | 2.73E+00 | 2.82E+00 | 7.04E-01 | no | | intergenic_variant | MODIFIER |
| 5 | 39148437 | AX-170089455 | 0 | T | C | 0.211 | 4.16E-02 | 1.12E-01 | 2.73E+00 | 2.82E+00 | 7.04E-01 | no | | intergenic_variant | MODIFIER |
| 5 | 39218494 | AX-97794797 | 0 | T | C | 0.211 | 4.16E-02 | 1.12E-01 | 2.73E+00 | 2.82E+00 | 7.04E-01 | no | | intergenic_variant | MODIFIER |
| 5 | 39321463 | AX-170024738 | 0 | A | G | 0.211 | 4.16E-02 | 1.12E-01 | 2.73E+00 | 2.82E+00 | 7.04E-01 | no | | intergenic_variant | MODIFIER |
| 5 | 39322836 | AX-97814271 | 0 | G | A | 0.211 | 4.16E-02 | 1.12E-01 | 2.73E+00 | 2.82E+00 | 7.04E-01 | no | | intergenic_variant | MODIFIER |
| 5 | 39343600 | AX-170024742 | 0 | T | C | 0.211 | 4.16E-02 | 1.12E-01 | 2.73E+00 | 2.82E+00 | 7.04E-01 | no | Gm25505 | upstream_gene_variant | MODIFIER |
| 5 | 39352938 | AX-97784878 | 0 | C | T | 0.211 | 4.16E-02 | 1.12E-01 | 2.73E+00 | 2.82E+00 | 7.04E-01 | no | | intergenic_variant | MODIFIER |
| 5 | 39362197 | AX-170463350 | 0 | T | G | 0.211 | 4.16E-02 | 1.12E-01 | 2.73E+00 | 2.82E+00 | 7.04E-01 | no | | intergenic_variant | MODIFIER |
| 5 | 39365009 | AX-170210822 | 0 | C | T | 0.211 | 4.16E-02 | 1.12E-01 | 2.73E+00 | 2.82E+00 | 7.04E-01 | no | | intergenic_variant | MODIFIER |
| 5 | 39366789 | AX-170024736 | 0 | A | G | 0.211 | 4.16E-02 | 1.12E-01 | 2.73E+00 | 2.82E+00 | 7.04E-01 | no | | intergenic_variant | MODIFIER |
| 5 | 39383971 | AX-97811490 | 0 | C | T | 0.211 | 4.16E-02 | 1.12E-01 | 2.73E+00 | 2.82E+00 | 7.04E-01 | no | | intergenic_variant | MODIFIER |
| 5 | 39416939 | AX-170363563 | 0 | A | G | 0.211 | 4.16E-02 | 1.12E-01 | 2.73E+00 | 2.82E+00 | 7.04E-01 | no | | regulatory_region_variant | MODIFIER |
| 5 | 39417140 | AX-170398602 | 0 | T | C | 0.211 | 4.16E-02 | 1.12E-01 | 2.73E+00 | 2.82E+00 | 7.04E-01 | no | | regulatory_region_variant | MODIFIER |
| 5 | 39427694 | AX-97831401 | 0 | A | C | 0.211 | 4.16E-02 | 1.12E-01 | 2.73E+00 | 2.82E+00 | 7.04E-01 | no | | intergenic_variant | MODIFIER |
| 20 | 71479858 | AX-97823856 | 0 | A | C | 0.254 | 2.32E-02 | 6.20E-02 | 2.65E+00 | 2.75E+00 | 7.04E-01 | no | Cd99l2 | intron_variant | MODIFIER |
| 20 | 75917865 | AX-170034161 | 0 | C | T | 0.254 | 2.32E-02 | 6.20E-02 | 2.65E+00 | 2.75E+00 | 7.04E-01 | no | Oat-rs1 | downstream_gene_variant | MODIFIER |
| 15 | 3236252 | AX-97837455 | 0 | C | T | 0.088 | -4.56E-02 | 9.51E-02 | 2.35E+00 | 2.49E+00 | 7.05E-01 | no | | intergenic_variant | MODIFIER |
| 9 | 32971790 | AX-97791594 | 0 | C | T | 0.439 | 3.35E-02 | 8.80E-02 | 2.63E+00 | 2.69E+00 | 7.05E-01 | no | Gm37192 | non_coding_transcript_exon_variant | MODIFIER |
| 15 | 92102289 | AX-170012266 | 0 | T | C | 0.149 | 6.77E-02 | 1.79E-01 | 2.65E+00 | 2.71E+00 | 7.05E-01 | no | Cntn1 | intron_variant,non_coding_transcript_variant | MODIFIER |
| 9 | 106842567 | AX-170489683 | 0 | A | T | 0.132 | -3.56E-02 | 8.77E-02 | 2.51E+00 | 2.62E+00 | 7.05E-01 | no | Dcaf1 | intron_variant | MODIFIER |
| 1 | 173831291 | AX-170273641 | 1 | C | T | 0.223 | 4.64E-02 | 1.19E-01 | 2.58E+00 | 2.66E+00 | 7.05E-01 | no | Gm7897 | intron_variant,non_coding_transcript_variant | MODIFIER |
| 11 | 5313298 | AX-170244304 | 0 | G | T | 0.061 | -6.47E-02 | 1.71E-01 | 2.60E+00 | 2.72E+00 | 7.05E-01 | no | Znrf3 | intron_variant | MODIFIER |
| 13 | 36554405 | AX-170466096 | 1 | G | C | 0.179 | -4.64E-02 | 1.20E-01 | 2.60E+00 | 2.66E+00 | 7.05E-01 | no | Gm8277 | downstream_gene_variant | MODIFIER |
| 10 | 118800346 | AX-170454559 | 2 | A | G | 0.327 | 2.72E-02 | 7.36E-02 | 2.67E+00 | 2.77E+00 | 7.05E-01 | no | Gm4065 | downstream_gene_variant | MODIFIER |
| 9 | 35022155 | AX-97827299 | 0 | T | C | 0.439 | -3.28E-02 | 8.86E-02 | 2.74E+00 | 2.81E+00 | 7.05E-01 | no | Kirrel3 | intron_variant | MODIFIER |
| 2 | 124335907 | AX-170311211 | 1 | A | G | 0.125 | 6.80E-02 | 1.82E-01 | 2.64E+00 | 2.74E+00 | 7.05E-01 | no | Sema6d | intron_variant | MODIFIER |
| 8 | 84428916 | AX-97785866 | 1 | C | T | 0.205 | 4.74E-02 | 1.27E-01 | 2.70E+00 | 2.76E+00 | 7.05E-01 | no | Cacna1a | intron_variant | MODIFIER |

| | | | | | | | | | | | | | | | |
|---|---|---|---|---|---|---|---|---|---|---|---|---|---|---|---|
| 5 | 34871744 | AX-170451810 | 0 | C | G | 0.333 | -3.88E-02 | 1.05E-01 | 2.73E+00 | 2.81E+00 | 7.05E-01 | no | Htt | intron_variant | MODIFIER |
| 11 | 51244013 | AX-170047034 | 1 | G | A | 0.071 | -7.40E-02 | 1.97E-01 | 2.64E+00 | 2.72E+00 | 7.05E-01 | no | | intergenic_variant | MODIFIER |
| 2 | 100111811 | AX-97783324 | 0 | A | G | 0.061 | -7.50E-02 | 2.03E-01 | 2.70E+00 | 2.78E+00 | 7.05E-01 | no | | intergenic_variant | MODIFIER |
| 4 | 107557947 | AX-170338430 | 0 | A | T | 0.254 | 4.14E-02 | 1.02E-01 | 2.51E+00 | 2.62E+00 | 7.05E-01 | no | Glis1 | intron_variant | MODIFIER |
| 8 | 45155191 | AX-170383383 | 0 | C | T | 0.167 | 4.47E-02 | 1.20E-01 | 2.65E+00 | 2.75E+00 | 7.05E-01 | no | Gm30504 | intron_variant,non_coding_transcript_variant | MODIFIER |
| 10 | 114836175 | AX-97798155 | 0 | G | A | 0.368 | 2.76E-02 | 7.51E-02 | 2.70E+00 | 2.79E+00 | 7.06E-01 | no | | intergenic_variant | MODIFIER |
| 8 | 30697938 | AX-170187902 | 0 | G | A | 0.175 | 4.20E-02 | 1.13E-01 | 2.70E+00 | 2.77E+00 | 7.06E-01 | no | | intergenic_variant | MODIFIER |
| 6 | 139834129 | AX-170323648 | 1 | C | A | 0.08 | -8.45E-02 | 2.30E-01 | 2.71E+00 | 2.80E+00 | 7.06E-01 | no | Pik3c2g | intron_variant | MODIFIER |
| 7 | 137450046 | AX-170275329 | 0 | C | T | 0.289 | -3.06E-02 | 8.04E-02 | 2.60E+00 | 2.70E+00 | 7.06E-01 | no | Glrx3 | intron_variant | MODIFIER |
| 15 | 78281404 | AX-170056058 | 0 | G | A | 0.553 | -2.85E-02 | 7.75E-02 | 2.79E+00 | 2.90E+00 | 7.06E-01 | no | Csf2rb2 | downstream_gene_variant | MODIFIER |
| 5 | 32972222 | AX-170122760 | 0 | A | T | 0.368 | 3.75E-02 | 1.01E-01 | 2.67E+00 | 2.73E+00 | 7.06E-01 | no | Depdc5 | intron_variant | MODIFIER |
| 18 | 65309156 | AX-170509689 | 1 | T | C | 0.402 | 3.40E-02 | 8.28E-02 | 2.51E+00 | 2.59E+00 | 7.06E-01 | no | Alpk2 | intron_variant | MODIFIER |
| 5 | 77866914 | AX-170490254 | 1 | G | A | 0.411 | 3.74E-02 | 9.95E-02 | 2.64E+00 | 2.72E+00 | 7.06E-01 | no | | intergenic_variant | MODIFIER |
| 15 | 93682989 | AX-170056335 | 1 | G | A | 0.143 | 6.04E-02 | 1.59E-01 | 2.61E+00 | 2.70E+00 | 7.06E-01 | no | Zfp42-ps1 | upstream_gene_variant | MODIFIER |
| 7 | 40146182 | AX-170520070 | 1 | A | T | 0.357 | 3.26E-02 | 8.62E-02 | 2.61E+00 | 2.71E+00 | 7.06E-01 | no | | intergenic_variant | MODIFIER |
| 14 | 105293080 | AX-170330681 | 0 | C | A | 0.053 | 5.66E-02 | 1.49E-01 | 2.53E+00 | 2.71E+00 | 7.06E-01 | no | Ndfip2 | non_coding_transcript_exon_variant | MODIFIER |
| 18 | 67864609 | AX-170061180 | 0 | A | G | 0.272 | -3.93E-02 | 1.05E-01 | 2.66E+00 | 2.72E+00 | 7.07E-01 | no | Cep192 | intron_variant | MODIFIER |
| 5 | 86529802 | AX-170072687 | 0 | G | T | 0.439 | 3.63E-02 | 9.87E-02 | 2.69E+00 | 2.79E+00 | 7.07E-01 | no | Tmprss11f | intron_variant | MODIFIER |
| 8 | 49248578 | AX-97786176 | 0 | G | A | 0.149 | 5.11E-02 | 1.38E-01 | 2.78E+00 | 2.86E+00 | 7.07E-01 | no | | intergenic_variant | MODIFIER |
| 18 | 67598798 | AX-170128699 | 2 | G | A | 0.291 | -3.72E-02 | 1.00E-01 | 2.67E+00 | 2.74E+00 | 7.07E-01 | no | Spire1 | intron_variant | MODIFIER |
| 15 | 88234352 | AX-170423839 | 1 | G | T | 0.152 | 6.72E-02 | 1.79E-01 | 2.64E+00 | 2.70E+00 | 7.07E-01 | no | | intergenic_variant | MODIFIER |
| 12 | 93127337 | AX-170192772 | 0 | T | A | 0.061 | 8.76E-02 | 2.37E-01 | 2.71E+00 | 2.77E+00 | 7.07E-01 | no | | intergenic_variant | MODIFIER |
| 10 | 126893288 | AX-170224152 | 0 | G | A | 0.333 | -2.70E-02 | 7.22E-02 | 2.62E+00 | 2.72E+00 | 7.07E-01 | no | Atp23 | intron_variant | MODIFIER |
| 2 | 93784672 | AX-170179915 | 0 | A | G | 0.061 | 8.06E-02 | 2.19E-01 | 2.67E+00 | 2.76E+00 | 7.07E-01 | no | Ext2 | intron_variant | MODIFIER |
| 7 | 123857426 | AX-170263670 | 1 | A | G | 0.393 | 3.07E-02 | 8.31E-02 | 2.67E+00 | 2.76E+00 | 7.07E-01 | no | | intergenic_variant | MODIFIER |
| 15 | 85863648 | AX-170012174 | 1 | T | G | 0.509 | 2.86E-02 | 7.16E-02 | 2.53E+00 | 2.62E+00 | 7.07E-01 | no | Ttc38 | downstream_gene_variant | MODIFIER |
| 1 | 106960831 | AX-97846915 | 1 | G | A | 0.17 | 7.51E-02 | 1.96E-01 | 2.59E+00 | 2.67E+00 | 7.07E-01 | no | Serpinb12 | downstream_gene_variant | MODIFIER |
| 18 | 80138608 | AX-97792930 | 0 | G | A | 0.158 | -4.49E-02 | 1.14E-01 | 2.55E+00 | 2.64E+00 | 7.07E-01 | no | Adnp2 | intron_variant | MODIFIER |
| 8 | 95931620 | AX-170414172 | 1 | C | T | 0.232 | 4.27E-02 | 1.16E-01 | 2.69E+00 | 2.76E+00 | 7.07E-01 | no | Gm45895 | intron_variant,non_coding_transcript_variant | MODIFIER |
| 7 | 92208455 | AX-170029515 | 0 | G | A | 0.123 | -5.80E-02 | 1.60E-01 | 2.63E+00 | 2.78E+00 | 7.08E-01 | no | Dlg2 | intron_variant | MODIFIER |
| 7 | 94879544 | AX-170230764 | 1 | C | A | 0.107 | -7.36E-02 | 2.02E-01 | 2.68E+00 | 2.78E+00 | 7.08E-01 | no | | intergenic_variant | MODIFIER |
| 7 | 135303360 | AX-170030040 | 1 | C | T | 0.25 | -3.14E-02 | 8.20E-02 | 2.58E+00 | 2.68E+00 | 7.08E-01 | no | | intergenic_variant | MODIFIER |
| 3 | 31970598 | AX-170234898 | 0 | C | G | 0.158 | -3.65E-02 | 9.94E-02 | 2.70E+00 | 2.78E+00 | 7.08E-01 | no | Kcnmb2 | intron_variant | MODIFIER |
| 1 | 5276181 | AX-169999859 | 1 | C | T | 0.098 | -5.38E-02 | 1.48E-01 | 2.64E+00 | 2.77E+00 | 7.08E-01 | no | Gm7182 | non_coding_transcript_exon_variant | MODIFIER |
| 9 | 27772028 | AX-97843563 | 0 | T | C | 0.456 | 3.50E-02 | 9.24E-02 | 2.62E+00 | 2.68E+00 | 7.08E-01 | no | | intergenic_variant | MODIFIER |
| 9 | 27793953 | AX-97809632 | 0 | T | G | 0.456 | 3.50E-02 | 9.24E-02 | 2.62E+00 | 2.68E+00 | 7.08E-01 | no | Opcml | intron_variant | MODIFIER |
| 9 | 27805896 | AX-170205958 | 0 | T | G | 0.456 | 3.50E-02 | 9.24E-02 | 2.62E+00 | 2.68E+00 | 7.08E-01 | no | Opcml | intron_variant | MODIFIER |
| 9 | 27822342 | AX-170493096 | 0 | T | G | 0.456 | 3.50E-02 | 9.24E-02 | 2.62E+00 | 2.68E+00 | 7.08E-01 | no | Opcml | intron_variant | MODIFIER |
| 9 | 27826158 | AX-170285569 | 0 | T | C | 0.456 | 3.50E-02 | 9.24E-02 | 2.62E+00 | 2.68E+00 | 7.08E-01 | no | Opcml | intron_variant | MODIFIER |
| 9 | 27831778 | AX-97790193 | 0 | T | C | 0.456 | 3.50E-02 | 9.24E-02 | 2.62E+00 | 2.68E+00 | 7.08E-01 | no | Opcml | intron_variant | MODIFIER |
| 9 | 27843201 | AX-170374224 | 0 | T | G | 0.456 | 3.50E-02 | 9.24E-02 | 2.62E+00 | 2.68E+00 | 7.08E-01 | no | Opcml | intron_variant | MODIFIER |
| 9 | 27888856 | AX-170145541 | 0 | C | A | 0.456 | 3.50E-02 | 9.24E-02 | 2.62E+00 | 2.68E+00 | 7.08E-01 | no | Opcml | intron_variant | MODIFIER |
| 9 | 27899268 | AX-170213930 | 0 | G | A | 0.456 | 3.50E-02 | 9.24E-02 | 2.62E+00 | 2.68E+00 | 7.08E-01 | no | Opcml | intron_variant | MODIFIER |
| 9 | 27918426 | AX-170081937 | 0 | A | G | 0.456 | 3.50E-02 | 9.24E-02 | 2.62E+00 | 2.68E+00 | 7.08E-01 | no | Opcml | intron_variant | MODIFIER |
| 9 | 27918886 | AX-170212027 | 0 | G | C | 0.456 | 3.50E-02 | 9.24E-02 | 2.62E+00 | 2.68E+00 | 7.08E-01 | no | Opcml | intron_variant | MODIFIER |
| 9 | 27920120 | AX-97846713 | 0 | G | A | 0.456 | 3.50E-02 | 9.24E-02 | 2.62E+00 | 2.68E+00 | 7.08E-01 | no | Opcml | intron_variant | MODIFIER |
| 9 | 27929041 | AX-170350267 | 0 | T | G | 0.456 | 3.50E-02 | 9.24E-02 | 2.62E+00 | 2.68E+00 | 7.08E-01 | no | Opcml | intron_variant | MODIFIER |
| 9 | 27935323 | AX-170474833 | 0 | A | G | 0.456 | 3.50E-02 | 9.24E-02 | 2.62E+00 | 2.68E+00 | 7.08E-01 | no | Opcml | intron_variant | MODIFIER |
| 9 | 27937311 | AX-170352900 | 0 | A | G | 0.456 | 3.50E-02 | 9.24E-02 | 2.62E+00 | 2.68E+00 | 7.08E-01 | no | Opcml | intron_variant | MODIFIER |
| 9 | 27938022 | AX-170494814 | 0 | C | T | 0.456 | 3.50E-02 | 9.24E-02 | 2.62E+00 | 2.68E+00 | 7.08E-01 | no | Opcml | intron_variant | MODIFIER |
| 9 | 27938387 | AX-170096497 | 0 | G | T | 0.456 | 3.50E-02 | 9.24E-02 | 2.62E+00 | 2.68E+00 | 7.08E-01 | no | Opcml | intron_variant | MODIFIER |
| 9 | 27941470 | AX-170428468 | 0 | C | G | 0.456 | 3.50E-02 | 9.24E-02 | 2.62E+00 | 2.68E+00 | 7.08E-01 | no | Opcml | intron_variant | MODIFIER |
| 15 | 68064855 | AX-170266678 | 1 | G | A | 0.348 | -3.17E-02 | 7.64E-02 | 2.49E+00 | 2.58E+00 | 7.08E-01 | no | | intergenic_variant | MODIFIER |
| 5 | 122569121 | AX-170160389 | 1 | G | T | 0.455 | 4.10E-02 | 1.10E-01 | 2.67E+00 | 2.72E+00 | 7.08E-01 | no | Ift81 | intron_variant | MODIFIER |

| | | | | | | | | | | | | | | | |
|---|---|---|---|---|---|---|---|---|---|---|---|---|---|---|---|
| 4 | 85731098 | AX-170023303 | 0 | C | T | 0.333 | 4.10E-02 | 1.10E-01 | 2.65E+00 | 2.72E+00 | 7.08E-01 | no | Adamtsl1 | intron_variant,non_coding_transcript_variant | MODIFIER |
| 4 | 58519745 | AX-170069466 | 1 | C | T | 0.375 | 3.74E-02 | 1.01E-01 | 2.64E+00 | 2.74E+00 | 7.08E-01 | no | Lpar1 | intron_variant | MODIFIER |
| 5 | 27453090 | AX-170261422 | 0 | A | G | 0.061 | -7.77E-02 | 2.15E-01 | 2.71E+00 | 2.82E+00 | 7.08E-01 | no | Dpp6 | intron_variant | MODIFIER |
| 16 | 89224669 | AX-97832703 | 0 | C | T | 0.465 | -2.90E-02 | 7.88E-02 | 2.68E+00 | 2.75E+00 | 7.08E-01 | no | | intergenic_variant | MODIFIER |
| 8 | 90542301 | AX-170181216 | 0 | C | A | 0.193 | -4.41E-02 | 1.14E-01 | 2.57E+00 | 2.66E+00 | 7.09E-01 | no | | intergenic_variant | MODIFIER |
| 4 | 140693843 | AX-170287001 | 0 | T | C | 0.158 | 4.12E-02 | 1.09E-01 | 2.61E+00 | 2.67E+00 | 7.09E-01 | no | Gm23045 | downstream_gene_variant | MODIFIER |
| 16 | 27570303 | AX-170148561 | 0 | T | C | 0.246 | 3.27E-02 | 8.90E-02 | 2.63E+00 | 2.75E+00 | 7.09E-01 | no | | intergenic_variant | MODIFIER |
| 4 | 101445322 | AX-170201628 | 2 | C | T | 0.364 | 3.11E-02 | 8.50E-02 | 2.63E+00 | 2.76E+00 | 7.09E-01 | no | Ak4 | intron_variant | MODIFIER |
| 1 | 11464498 | AX-97832935 | 0 | T | G | 0.386 | -2.62E-02 | 7.17E-02 | 2.77E+00 | 2.85E+00 | 7.09E-01 | no | A830018L16Rik | intron_variant | MODIFIER |
| 20 | 52940727 | AX-170090504 | 1 | G | T | 0.25 | 2.27E-02 | 6.16E-02 | 2.65E+00 | 2.75E+00 | 7.09E-01 | no | Phf6 | intron_variant | MODIFIER |
| 12 | 101741906 | AX-170007402 | 0 | C | T | 0.096 | -5.30E-02 | 1.45E-01 | 2.64E+00 | 2.76E+00 | 7.09E-01 | no | Fbln5 | downstream_gene_variant | MODIFIER |
| 8 | 101282436 | AX-170096385 | 0 | C | A | 0.228 | -3.97E-02 | 1.09E-01 | 2.71E+00 | 2.81E+00 | 7.09E-01 | no | | intergenic_variant | MODIFIER |
| 1 | 190557422 | AX-170002559 | 0 | C | T | 0.254 | 3.88E-02 | 9.45E-02 | 2.49E+00 | 2.59E+00 | 7.09E-01 | no | | intergenic_variant | MODIFIER |
| 4 | 94758785 | AX-170023436 | 0 | C | T | 0.333 | -3.78E-02 | 1.04E-01 | 2.70E+00 | 2.79E+00 | 7.09E-01 | no | Tek | intron_variant | MODIFIER |
| 13 | 47231187 | AX-170051187 | 0 | C | T | 0.465 | -2.34E-02 | 6.46E-02 | 2.69E+00 | 2.80E+00 | 7.09E-01 | no | Rnf144b | intron_variant | MODIFIER |
| 1 | 51065313 | AX-97784134 | 0 | C | G | 0.105 | -6.83E-02 | 1.84E-01 | 2.62E+00 | 2.72E+00 | 7.09E-01 | no | Tmeff2 | intron_variant | MODIFIER |
| 1 | 60205710 | AX-170292199 | 0 | T | C | 0.105 | -6.83E-02 | 1.84E-01 | 2.62E+00 | 2.72E+00 | 7.09E-01 | no | Nbeal1 | intron_variant | MODIFIER |
| 5 | 73720043 | AX-170459634 | 1 | T | C | 0.438 | 3.81E-02 | 1.02E-01 | 2.63E+00 | 2.70E+00 | 7.09E-01 | no | | intergenic_variant | MODIFIER |
| 1 | 182433895 | AX-170131384 | 0 | G | A | 0.228 | 4.04E-02 | 1.01E-01 | 2.51E+00 | 2.62E+00 | 7.09E-01 | no | Trp53bp2 | intron_variant | MODIFIER |
| 1 | 184161293 | AX-170138277 | 0 | A | G | 0.228 | 4.04E-02 | 1.01E-01 | 2.51E+00 | 2.62E+00 | 7.09E-01 | no | | intergenic_variant | MODIFIER |
| 2 | 137196038 | AX-170463268 | 1 | T | G | 0.143 | 5.54E-02 | 1.52E-01 | 2.65E+00 | 2.76E+00 | 7.09E-01 | no | Gm28214 | upstream_gene_variant | MODIFIER |
| 12 | 89172674 | AX-170502733 | 0 | G | C | 0.079 | 5.76E-02 | 1.59E-01 | 2.68E+00 | 2.79E+00 | 7.09E-01 | no | Nrxn3 | intron_variant | MODIFIER |
| 1 | 47824266 | AX-170468805 | 0 | T | C | 0.114 | 6.80E-02 | 1.84E-01 | 2.62E+00 | 2.73E+00 | 7.09E-01 | no | | intergenic_variant | MODIFIER |
| 1 | 161245853 | AX-170184746 | 1 | A | G | 0.214 | 5.36E-02 | 1.42E-01 | 2.62E+00 | 2.68E+00 | 7.09E-01 | no | Prdx6 | intron_variant | MODIFIER |
| 11 | 74298056 | AX-170160829 | 1 | G | C | 0.268 | 2.59E-02 | 7.14E-02 | 2.63E+00 | 2.77E+00 | 7.09E-01 | no | Olfr408-ps1 | downstream_gene_variant | MODIFIER |
| 15 | 84806335 | AX-170056169 | 0 | G | C | 0.518 | 2.87E-02 | 7.20E-02 | 2.53E+00 | 2.62E+00 | 7.09E-01 | no | Phf21b | intron_variant | MODIFIER |
| 15 | 84827584 | AX-170441525 | 0 | C | T | 0.518 | 2.87E-02 | 7.20E-02 | 2.53E+00 | 2.62E+00 | 7.09E-01 | no | Phf21b | intron_variant | MODIFIER |
| 15 | 84829333 | AX-170012154 | 0 | T | C | 0.518 | 2.87E-02 | 7.20E-02 | 2.53E+00 | 2.62E+00 | 7.09E-01 | no | Phf21b | intron_variant | MODIFIER |
| 15 | 84836199 | AX-170313480 | 0 | C | T | 0.518 | 2.87E-02 | 7.20E-02 | 2.53E+00 | 2.62E+00 | 7.09E-01 | no | Phf21b | intron_variant | MODIFIER |
| 15 | 85279253 | AX-170283420 | 0 | G | C | 0.518 | 2.87E-02 | 7.20E-02 | 2.53E+00 | 2.62E+00 | 7.09E-01 | no | Fbln1 | intron_variant | MODIFIER |
| 15 | 85313568 | AX-97850300 | 0 | T | C | 0.518 | 2.87E-02 | 7.20E-02 | 2.53E+00 | 2.62E+00 | 7.09E-01 | no | | intergenic_variant | MODIFIER |
| 15 | 85357153 | AX-170308194 | 0 | C | G | 0.518 | 2.87E-02 | 7.20E-02 | 2.53E+00 | 2.62E+00 | 7.09E-01 | no | Atxn10 | intron_variant | MODIFIER |
| 15 | 85360778 | AX-170491658 | 0 | C | A | 0.518 | 2.87E-02 | 7.20E-02 | 2.53E+00 | 2.62E+00 | 7.09E-01 | no | Atxn10 | intron_variant | MODIFIER |
| 15 | 85361788 | AX-170186891 | 0 | A | T | 0.518 | 2.87E-02 | 7.20E-02 | 2.53E+00 | 2.62E+00 | 7.09E-01 | no | Atxn10 | intron_variant | MODIFIER |
| 15 | 85362773 | AX-170056175 | 0 | T | C | 0.518 | 2.87E-02 | 7.20E-02 | 2.53E+00 | 2.62E+00 | 7.09E-01 | no | Atxn10 | intron_variant | MODIFIER |
| 15 | 85372634 | AX-170405341 | 0 | G | T | 0.518 | 2.87E-02 | 7.20E-02 | 2.53E+00 | 2.62E+00 | 7.09E-01 | no | Atxn10 | intron_variant | MODIFIER |
| 15 | 85657968 | AX-170093323 | 0 | C | A | 0.518 | 2.87E-02 | 7.20E-02 | 2.53E+00 | 2.62E+00 | 7.09E-01 | no | Gm49539 | intron_variant,non_coding_transcript_variant | MODIFIER |
| 15 | 85662335 | AX-170363376 | 0 | G | A | 0.518 | 2.87E-02 | 7.20E-02 | 2.53E+00 | 2.62E+00 | 7.09E-01 | no | Gm49539 | intron_variant,non_coding_transcript_variant | MODIFIER |
| 15 | 85691144 | AX-170407824 | 0 | A | T | 0.518 | 2.87E-02 | 7.20E-02 | 2.53E+00 | 2.62E+00 | 7.09E-01 | no | Lncppara | intron_variant,non_coding_transcript_variant | MODIFIER |
| 15 | 85691533 | AX-170134634 | 0 | C | A | 0.518 | 2.87E-02 | 7.20E-02 | 2.53E+00 | 2.62E+00 | 7.09E-01 | no | Lncppara | intron_variant,non_coding_transcript_variant | MODIFIER |
| 15 | 85733754 | AX-170374210 | 0 | A | G | 0.518 | 2.87E-02 | 7.20E-02 | 2.53E+00 | 2.62E+00 | 7.09E-01 | no | Ppara | upstream_gene_variant | MODIFIER |
| 15 | 85791470 | AX-170035557 | 0 | G | A | 0.518 | 2.87E-02 | 7.20E-02 | 2.53E+00 | 2.62E+00 | 7.09E-01 | no | Ppara | intron_variant | MODIFIER |
| 15 | 85818574 | AX-170012176 | 0 | T | C | 0.518 | 2.87E-02 | 7.20E-02 | 2.53E+00 | 2.62E+00 | 7.09E-01 | no | Pkdrej | missense_variant | MODERATE |
| 15 | 85819106 | AX-170531261 | 0 | T | C | 0.518 | 2.87E-02 | 7.20E-02 | 2.53E+00 | 2.62E+00 | 7.09E-01 | no | Pkdrej | synonymous_variant | LOW |
| 15 | 85905510 | AX-170035558 | 0 | C | T | 0.518 | 2.87E-02 | 7.20E-02 | 2.53E+00 | 2.62E+00 | 7.09E-01 | no | Celsr1 | synonymous_variant | LOW |
| 15 | 85910120 | AX-170484393 | 0 | T | G | 0.518 | 2.87E-02 | 7.20E-02 | 2.53E+00 | 2.62E+00 | 7.09E-01 | no | Celsr1 | intron_variant | MODIFIER |
| 15 | 85958381 | AX-170349097 | 0 | A | G | 0.518 | 2.87E-02 | 7.20E-02 | 2.53E+00 | 2.62E+00 | 7.09E-01 | no | Celsr1 | intron_variant | MODIFIER |
| 15 | 85976221 | AX-170213802 | 0 | C | T | 0.518 | 2.87E-02 | 7.20E-02 | 2.53E+00 | 2.62E+00 | 7.09E-01 | no | Celsr1 | intron_variant | MODIFIER |
| 15 | 86003757 | AX-170103928 | 0 | A | G | 0.518 | 2.87E-02 | 7.20E-02 | 2.53E+00 | 2.62E+00 | 7.09E-01 | no | Celsr1 | intron_variant | MODIFIER |
| 15 | 86005311 | AX-170103190 | 0 | T | C | 0.518 | 2.87E-02 | 7.20E-02 | 2.53E+00 | 2.62E+00 | 7.09E-01 | no | Celsr1 | intron_variant | MODIFIER |
| 15 | 86069268 | AX-170142832 | 0 | A | G | 0.518 | 2.87E-02 | 7.20E-02 | 2.53E+00 | 2.62E+00 | 7.09E-01 | no | Gramd4 | intron_variant | MODIFIER |
| 15 | 86074939 | AX-170367318 | 0 | A | G | 0.518 | 2.87E-02 | 7.20E-02 | 2.53E+00 | 2.62E+00 | 7.09E-01 | no | Gramd4 | intron_variant | MODIFIER |
| 15 | 86078890 | AX-170296464 | 0 | G | A | 0.518 | 2.87E-02 | 7.20E-02 | 2.53E+00 | 2.62E+00 | 7.09E-01 | no | Gramd4 | intron_variant | MODIFIER |
| 15 | 86143319 | AX-170097949 | 0 | C | T | 0.518 | 2.87E-02 | 7.20E-02 | 2.53E+00 | 2.62E+00 | 7.09E-01 | no | Cerk | intron_variant | MODIFIER |

| | | | | | | | | | | | | | | | |
|---|---|---|---|---|---|---|---|---|---|---|---|---|---|---|---|
| 15 | 86145247 | AX-170056190 | 0 | C | T | 0.518 | 2.87E-02 | 7.20E-02 | 2.53E+00 | 2.62E+00 | 7.09E-01 | no | Cerk | intron_variant | MODIFIER |
| 15 | 86148278 | AX-170056192 | 0 | C | T | 0.518 | 2.87E-02 | 7.20E-02 | 2.53E+00 | 2.62E+00 | 7.09E-01 | no | Cerk | intron_variant | MODIFIER |
| 15 | 86550440 | AX-170270774 | 0 | A | G | 0.518 | 2.87E-02 | 7.20E-02 | 2.53E+00 | 2.62E+00 | 7.09E-01 | no | | intergenic_variant | MODIFIER |
| 15 | 86559167 | AX-170406908 | 0 | C | T | 0.518 | 2.87E-02 | 7.20E-02 | 2.53E+00 | 2.62E+00 | 7.09E-01 | no | | intergenic_variant | MODIFIER |
| 15 | 86599464 | AX-170372180 | 0 | T | C | 0.518 | 2.87E-02 | 7.20E-02 | 2.53E+00 | 2.62E+00 | 7.09E-01 | no | | intergenic_variant | MODIFIER |
| 15 | 86606809 | AX-170035556 | 0 | C | T | 0.518 | 2.87E-02 | 7.20E-02 | 2.53E+00 | 2.62E+00 | 7.09E-01 | no | | intergenic_variant | MODIFIER |
| 15 | 86644765 | AX-97821472 | 0 | C | T | 0.518 | 2.87E-02 | 7.20E-02 | 2.53E+00 | 2.62E+00 | 7.09E-01 | no | | intergenic_variant | MODIFIER |
| 15 | 86647405 | AX-170306296 | 0 | T | C | 0.518 | 2.87E-02 | 7.20E-02 | 2.53E+00 | 2.62E+00 | 7.09E-01 | no | | intergenic_variant | MODIFIER |
| 15 | 86657216 | AX-170490904 | 0 | T | C | 0.518 | 2.87E-02 | 7.20E-02 | 2.53E+00 | 2.62E+00 | 7.09E-01 | no | | intergenic_variant | MODIFIER |
| 15 | 86659463 | AX-170527820 | 0 | C | T | 0.518 | 2.87E-02 | 7.20E-02 | 2.53E+00 | 2.62E+00 | 7.09E-01 | no | | intergenic_variant | MODIFIER |
| 15 | 86663073 | AX-170532452 | 0 | T | C | 0.518 | 2.87E-02 | 7.20E-02 | 2.53E+00 | 2.62E+00 | 7.09E-01 | no | | intergenic_variant | MODIFIER |
| 5 | 101072968 | AX-97795565 | 1 | G | T | 0.402 | -3.53E-02 | 9.75E-02 | 2.73E+00 | 2.82E+00 | 7.09E-01 | no | | intergenic_variant | MODIFIER |
| 7 | 112388388 | AX-170263099 | 0 | G | A | 0.456 | 3.43E-02 | 9.29E-02 | 2.66E+00 | 2.73E+00 | 7.10E-01 | no | Mical2 | intron_variant | MODIFIER |
| 7 | 135887466 | AX-170218698 | 1 | A | C | 0.25 | -3.06E-02 | 8.08E-02 | 2.59E+00 | 2.69E+00 | 7.10E-01 | no | | intergenic_variant | MODIFIER |
| 18 | 50130790 | AX-170394526 | 2 | G | A | 0.2 | -3.94E-02 | 1.03E-01 | 2.57E+00 | 2.66E+00 | 7.10E-01 | no | Hsd17b4 | intron_variant | MODIFIER |
| 12 | 109712143 | AX-170399714 | 1 | C | T | 0.134 | -7.08E-02 | 1.94E-01 | 2.72E+00 | 2.79E+00 | 7.10E-01 | no | Mir411 | downstream_gene_variant | MODIFIER |
| 1 | 60225473 | AX-170157989 | 1 | T | G | 0.098 | 7.72E-02 | 2.11E-01 | 2.66E+00 | 2.76E+00 | 7.10E-01 | no | Nbeal1 | intron_variant | MODIFIER |
| 7 | 116830241 | AX-170165546 | 1 | A | G | 0.438 | 3.46E-02 | 9.33E-02 | 2.66E+00 | 2.71E+00 | 7.10E-01 | no | Gm4366 | upstream_gene_variant | MODIFIER |
| 5 | 102834494 | AX-170512257 | 0 | T | G | 0.43 | 3.90E-02 | 1.05E-01 | 2.65E+00 | 2.72E+00 | 7.10E-01 | no | Arhgap24 | intron_variant | MODIFIER |
| 5 | 28009123 | AX-170265018 | 1 | G | A | 0.339 | 3.94E-02 | 1.07E-01 | 2.68E+00 | 2.73E+00 | 7.10E-01 | no | | intergenic_variant | MODIFIER |
| 1 | 70681722 | AX-170097643 | 2 | T | G | 0.109 | 8.10E-02 | 2.21E-01 | 2.66E+00 | 2.75E+00 | 7.10E-01 | no | Spag16 | intron_variant | MODIFIER |
| 15 | 98446101 | AX-170458199 | 0 | C | T | 0.14 | 5.26E-02 | 1.39E-01 | 2.59E+00 | 2.68E+00 | 7.10E-01 | no | Gm6979 | upstream_gene_variant | MODIFIER |
| 1 | 176613157 | AX-170002350 | 0 | G | A | 0.219 | -4.19E-02 | 1.16E-01 | 2.77E+00 | 2.86E+00 | 7.10E-01 | no | | intergenic_variant | MODIFIER |
| 14 | 92973595 | AX-170102972 | 0 | G | A | 0.044 | -6.26E-02 | 1.68E-01 | 2.59E+00 | 2.71E+00 | 7.10E-01 | no | | intergenic_variant | MODIFIER |
| 17 | 25072321 | AX-170371812 | 0 | G | A | 0.044 | -6.26E-02 | 1.68E-01 | 2.59E+00 | 2.71E+00 | 7.10E-01 | no | Ift140 | intron_variant | MODIFIER |
| 12 | 108834225 | AX-170369440 | 0 | G | A | 0.132 | -7.04E-02 | 1.93E-01 | 2.70E+00 | 2.77E+00 | 7.10E-01 | no | Slc25a29 | intron_variant | MODIFIER |
| 8 | 48599882 | AX-170510837 | 0 | C | G | 0.149 | -5.15E-02 | 1.33E-01 | 2.57E+00 | 2.64E+00 | 7.10E-01 | no | Tenm3 | intron_variant | MODIFIER |
| 8 | 49296379 | AX-170119792 | 0 | A | G | 0.149 | -5.15E-02 | 1.33E-01 | 2.57E+00 | 2.64E+00 | 7.10E-01 | no | | intergenic_variant | MODIFIER |
| 1 | 170774911 | AX-170356626 | 0 | C | T | 0.211 | -5.19E-02 | 1.41E-01 | 2.82E+00 | 2.88E+00 | 7.10E-01 | no | Atf6 | intron_variant | MODIFIER |
| 3 | 148210431 | AX-97847174 | 1 | G | A | 0.33 | 2.76E-02 | 7.45E-02 | 2.64E+00 | 2.71E+00 | 7.11E-01 | no | | intergenic_variant | MODIFIER |
| 3 | 148642508 | AX-170021890 | 1 | A | G | 0.33 | 2.76E-02 | 7.45E-02 | 2.64E+00 | 2.71E+00 | 7.11E-01 | no | | regulatory_region_variant | MODIFIER |
| 9 | 85427027 | AX-170289501 | 0 | C | T | 0.211 | 4.49E-02 | 1.23E-01 | 2.73E+00 | 2.80E+00 | 7.11E-01 | no | | regulatory_region_variant | MODIFIER |
| 1 | 67474751 | AX-170000726 | 1 | C | G | 0.107 | 8.04E-02 | 2.20E-01 | 2.67E+00 | 2.75E+00 | 7.11E-01 | no | | intergenic_variant | MODIFIER |
| 1 | 67488306 | AX-97811906 | 1 | G | T | 0.107 | 8.04E-02 | 2.20E-01 | 2.67E+00 | 2.75E+00 | 7.11E-01 | no | | intergenic_variant | MODIFIER |
| 1 | 70192200 | AX-170484012 | 1 | A | G | 0.107 | 8.04E-02 | 2.20E-01 | 2.67E+00 | 2.75E+00 | 7.11E-01 | no | Spag16 | intron_variant | MODIFIER |
| 10 | 92703150 | AX-170296702 | 0 | G | A | 0.298 | -3.13E-02 | 8.14E-02 | 2.56E+00 | 2.65E+00 | 7.11E-01 | no | Nedd1 | intron_variant | MODIFIER |
| 6 | 130379017 | AX-170231490 | 1 | T | C | 0.429 | 3.92E-02 | 1.01E-01 | 2.53E+00 | 2.66E+00 | 7.11E-01 | no | Klra1 | missense_variant | MODERATE |
| 7 | 13132562 | AX-97853971 | 0 | A | G | 0.351 | 3.01E-02 | 8.01E-02 | 2.58E+00 | 2.70E+00 | 7.11E-01 | no | Vmn1r87 | upstream_gene_variant | MODIFIER |
| 7 | 13215934 | AX-97785779 | 0 | T | C | 0.351 | 3.01E-02 | 8.01E-02 | 2.58E+00 | 2.70E+00 | 7.11E-01 | no | Vmn1r89 | upstream_gene_variant | MODIFIER |
| 7 | 13250293 | AX-170028611 | 0 | G | A | 0.351 | 3.01E-02 | 8.01E-02 | 2.58E+00 | 2.70E+00 | 7.11E-01 | no | | intergenic_variant | MODIFIER |
| 7 | 13277052 | AX-170028606 | 0 | T | C | 0.351 | 3.01E-02 | 8.01E-02 | 2.58E+00 | 2.70E+00 | 7.11E-01 | no | Lig1 | upstream_gene_variant | MODIFIER |
| 7 | 13290569 | AX-97792349 | 0 | T | C | 0.351 | 3.01E-02 | 8.01E-02 | 2.58E+00 | 2.70E+00 | 7.11E-01 | no | Lig1 | intron_variant | MODIFIER |
| 7 | 13292421 | AX-170492439 | 0 | C | T | 0.351 | 3.01E-02 | 8.01E-02 | 2.58E+00 | 2.70E+00 | 7.11E-01 | no | Lig1 | intron_variant | MODIFIER |
| 7 | 13305746 | AX-97856041 | 0 | A | C | 0.351 | 3.01E-02 | 8.01E-02 | 2.58E+00 | 2.70E+00 | 7.11E-01 | no | Lig1 | intron_variant | MODIFIER |
| 7 | 13343707 | AX-97786257 | 0 | A | G | 0.351 | 3.01E-02 | 8.01E-02 | 2.58E+00 | 2.70E+00 | 7.11E-01 | no | Pla2g4c | intron_variant | MODIFIER |
| 7 | 13433926 | AX-97849921 | 0 | T | G | 0.351 | 3.01E-02 | 8.01E-02 | 2.58E+00 | 2.70E+00 | 7.11E-01 | no | | intergenic_variant | MODIFIER |
| 7 | 13480197 | AX-170414382 | 0 | A | G | 0.351 | 3.01E-02 | 8.01E-02 | 2.58E+00 | 2.70E+00 | 7.11E-01 | no | Gm44254 | upstream_gene_variant | MODIFIER |
| 7 | 13546628 | AX-170451721 | 0 | G | T | 0.351 | 3.01E-02 | 8.01E-02 | 2.58E+00 | 2.70E+00 | 7.11E-01 | no | | intergenic_variant | MODIFIER |
| 7 | 14515300 | AX-97790421 | 0 | T | C | 0.351 | 3.01E-02 | 8.01E-02 | 2.58E+00 | 2.70E+00 | 7.11E-01 | no | Obox4-ps1 | downstream_gene_variant | MODIFIER |
| 7 | 14544160 | AX-170295185 | 0 | G | T | 0.351 | 3.01E-02 | 8.01E-02 | 2.58E+00 | 2.70E+00 | 7.11E-01 | no | Nlrp5-ps | intron_variant,non_coding_transcript_variant | MODIFIER |
| 7 | 14548495 | AX-170464328 | 0 | G | A | 0.351 | 3.01E-02 | 8.01E-02 | 2.58E+00 | 2.70E+00 | 7.11E-01 | no | Nlrp5-ps | intron_variant,non_coding_transcript_variant | MODIFIER |
| 7 | 14569506 | AX-170098155 | 0 | G | C | 0.351 | 3.01E-02 | 8.01E-02 | 2.58E+00 | 2.70E+00 | 7.11E-01 | no | Nlrp5-ps | intron_variant,non_coding_transcript_variant | MODIFIER |
| 6 | 112924514 | AX-170216759 | 2 | G | T | 0.055 | 1.08E-01 | 2.96E-01 | 2.68E+00 | 2.74E+00 | 7.11E-01 | no | Srgap3 | intron_variant,non_coding_transcript_variant | MODIFIER |
| 5 | 110567839 | AX-170073081 | 2 | T | G | 0.427 | -3.78E-02 | 1.04E-01 | 2.77E+00 | 2.86E+00 | 7.11E-01 | no | Galnt9 | intron_variant | MODIFIER |

| | | | | | | | | | | | | | | | |
|---|---|---|---|---|---|---|---|---|---|---|---|---|---|---|---|
| 20 | 58709777 | AX-170101035 | 2 | G | A | 0.036 | 4.43E-02 | 1.26E-01 | 2.65E+00 | 2.82E+00 | 7.11E-01 | no | Gm7071 | upstream_gene_variant | MODIFIER |
| 12 | 99322166 | AX-170007368 | 1 | G | A | 0.054 | 8.09E-02 | 2.22E-01 | 2.69E+00 | 2.75E+00 | 7.11E-01 | no | Foxn3 | intron_variant | MODIFIER |
| 2 | 119271435 | AX-170388198 | 0 | C | T | 0.149 | 6.72E-02 | 1.80E-01 | 2.62E+00 | 2.70E+00 | 7.11E-01 | no | Rhov | upstream_gene_variant | MODIFIER |
| 7 | 40620591 | AX-170099481 | 1 | C | T | 0.33 | 3.56E-02 | 9.49E-02 | 2.61E+00 | 2.69E+00 | 7.11E-01 | no | | intergenic_variant | MODIFIER |
| 9 | 88117246 | AX-170122157 | 1 | T | C | 0.143 | -4.68E-02 | 1.25E-01 | 2.61E+00 | 2.68E+00 | 7.11E-01 | no | Gm2324 | downstream_gene_variant | MODIFIER |
| 2 | 101978363 | AX-170534889 | 1 | G | A | 0.054 | -1.10E-01 | 3.00E-01 | 2.74E+00 | 2.77E+00 | 7.11E-01 | no | Ldlrad3 | intron_variant | MODIFIER |
| 2 | 110793167 | AX-170018666 | 1 | A | G | 0.054 | -1.10E-01 | 3.00E-01 | 2.74E+00 | 2.77E+00 | 7.11E-01 | no | Ano3 | intron_variant | MODIFIER |
| 5 | 30987751 | AX-170024615 | 1 | A | G | 0.357 | 3.85E-02 | 1.05E-01 | 2.69E+00 | 2.74E+00 | 7.11E-01 | no | | intergenic_variant | MODIFIER |
| 5 | 32014932 | AX-170109677 | 1 | T | A | 0.357 | 3.85E-02 | 1.05E-01 | 2.69E+00 | 2.74E+00 | 7.11E-01 | no | Babam2 | intron_variant | MODIFIER |
| 5 | 33916133 | AX-170024655 | 1 | T | A | 0.357 | 3.85E-02 | 1.05E-01 | 2.69E+00 | 2.74E+00 | 7.11E-01 | no | Nelfa | intron_variant | MODIFIER |
| 1 | 38477112 | AX-170331479 | 0 | A | G | 0.456 | 2.75E-02 | 7.64E-02 | 2.72E+00 | 2.81E+00 | 7.11E-01 | no | Aff3 | intron_variant,non_coding_transcript_variant | MODIFIER |
| 15 | 84719381 | AX-170129125 | 0 | A | G | 0.5 | 2.74E-02 | 6.98E-02 | 2.53E+00 | 2.62E+00 | 7.11E-01 | no | Arhgap8 | upstream_gene_variant | MODIFIER |
| 15 | 84722244 | AX-97783363 | 0 | G | A | 0.5 | 2.74E-02 | 6.98E-02 | 2.53E+00 | 2.62E+00 | 7.11E-01 | no | Arhgap8 | intron_variant | MODIFIER |
| 15 | 84776435 | AX-97830022 | 0 | A | G | 0.5 | 2.74E-02 | 6.98E-02 | 2.53E+00 | 2.62E+00 | 7.11E-01 | no | Arhgap8 | downstream_gene_variant | MODIFIER |
| 15 | 84785110 | AX-170246248 | 0 | G | C | 0.5 | 2.74E-02 | 6.98E-02 | 2.53E+00 | 2.62E+00 | 7.11E-01 | no | Phf21b | downstream_gene_variant | MODIFIER |
| 15 | 84794858 | AX-170012159 | 0 | A | C | 0.5 | 2.74E-02 | 6.98E-02 | 2.53E+00 | 2.62E+00 | 7.11E-01 | no | Phf21b | intron_variant | MODIFIER |
| 15 | 84795611 | AX-97788885 | 0 | T | C | 0.5 | 2.74E-02 | 6.98E-02 | 2.53E+00 | 2.62E+00 | 7.11E-01 | no | Phf21b | intron_variant | MODIFIER |
| 15 | 84795679 | AX-170363270 | 0 | G | A | 0.5 | 2.74E-02 | 6.98E-02 | 2.53E+00 | 2.62E+00 | 7.11E-01 | no | Phf21b | intron_variant | MODIFIER |
| 15 | 84798859 | AX-170214694 | 0 | T | C | 0.5 | 2.74E-02 | 6.98E-02 | 2.53E+00 | 2.62E+00 | 7.11E-01 | no | Phf21b | intron_variant | MODIFIER |
| 15 | 84801852 | AX-170012157 | 0 | A | G | 0.5 | 2.74E-02 | 6.98E-02 | 2.53E+00 | 2.62E+00 | 7.11E-01 | no | Phf21b | intron_variant | MODIFIER |
| 15 | 84820466 | AX-170087927 | 0 | T | C | 0.5 | 2.74E-02 | 6.98E-02 | 2.53E+00 | 2.62E+00 | 7.11E-01 | no | Phf21b | intron_variant | MODIFIER |
| 15 | 84827459 | AX-170481019 | 0 | A | G | 0.5 | 2.74E-02 | 6.98E-02 | 2.53E+00 | 2.62E+00 | 7.11E-01 | no | Phf21b | intron_variant | MODIFIER |
| 15 | 85312450 | AX-170280812 | 0 | A | C | 0.5 | 2.74E-02 | 6.98E-02 | 2.53E+00 | 2.62E+00 | 7.11E-01 | no | | regulatory_region_variant | MODIFIER |
| 15 | 85357227 | AX-170305646 | 0 | C | T | 0.5 | 2.74E-02 | 6.98E-02 | 2.53E+00 | 2.62E+00 | 7.11E-01 | no | Atxn10 | intron_variant | MODIFIER |
| 15 | 85361356 | AX-170123533 | 0 | T | C | 0.5 | 2.74E-02 | 6.98E-02 | 2.53E+00 | 2.62E+00 | 7.11E-01 | no | Atxn10 | intron_variant | MODIFIER |
| 15 | 85512941 | AX-170379656 | 0 | A | G | 0.5 | 2.74E-02 | 6.98E-02 | 2.53E+00 | 2.62E+00 | 7.11E-01 | no | Gm4825 | downstream_gene_variant | MODIFIER |
| 15 | 85661428 | AX-170240153 | 0 | T | C | 0.5 | 2.74E-02 | 6.98E-02 | 2.53E+00 | 2.62E+00 | 7.11E-01 | no | Gm49539 | intron_variant,non_coding_transcript_variant | MODIFIER |
| 15 | 85680084 | AX-170012169 | 0 | A | C | 0.5 | 2.74E-02 | 6.98E-02 | 2.53E+00 | 2.62E+00 | 7.11E-01 | no | Lncppara | intron_variant,non_coding_transcript_variant | MODIFIER |
| 15 | 85686339 | AX-170292146 | 0 | A | G | 0.5 | 2.74E-02 | 6.98E-02 | 2.53E+00 | 2.62E+00 | 7.11E-01 | no | Lncppara | intron_variant,non_coding_transcript_variant | MODIFIER |
| 15 | 85687419 | AX-170288528 | 0 | C | T | 0.5 | 2.74E-02 | 6.98E-02 | 2.53E+00 | 2.62E+00 | 7.11E-01 | no | Lncppara | intron_variant,non_coding_transcript_variant | MODIFIER |
| 15 | 85688559 | AX-170056181 | 0 | C | T | 0.5 | 2.74E-02 | 6.98E-02 | 2.53E+00 | 2.62E+00 | 7.11E-01 | no | Lncppara | intron_variant,non_coding_transcript_variant | MODIFIER |
| 15 | 85693778 | AX-170012172 | 0 | T | C | 0.5 | 2.74E-02 | 6.98E-02 | 2.53E+00 | 2.62E+00 | 7.11E-01 | no | Lncppara | intron_variant,non_coding_transcript_variant | MODIFIER |
| 15 | 85710999 | AX-97816746 | 0 | T | C | 0.5 | 2.74E-02 | 6.98E-02 | 2.53E+00 | 2.62E+00 | 7.11E-01 | no | Mirlet7b | downstream_gene_variant | MODIFIER |
| 15 | 85755577 | AX-97809435 | 0 | A | G | 0.5 | 2.74E-02 | 6.98E-02 | 2.53E+00 | 2.62E+00 | 7.11E-01 | no | Ppara | intron_variant | MODIFIER |
| 15 | 85756747 | AX-170414771 | 0 | A | G | 0.5 | 2.74E-02 | 6.98E-02 | 2.53E+00 | 2.62E+00 | 7.11E-01 | no | Ppara | intron_variant | MODIFIER |
| 15 | 85795233 | AX-170434416 | 0 | C | T | 0.5 | 2.74E-02 | 6.98E-02 | 2.53E+00 | 2.62E+00 | 7.11E-01 | no | Ppara | intron_variant | MODIFIER |
| 15 | 85854194 | AX-170170848 | 0 | T | C | 0.5 | 2.74E-02 | 6.98E-02 | 2.53E+00 | 2.62E+00 | 7.11E-01 | no | Ttc38 | intron_variant,non_coding_transcript_variant | MODIFIER |
| 15 | 85912263 | AX-170379248 | 0 | C | T | 0.5 | 2.74E-02 | 6.98E-02 | 2.53E+00 | 2.62E+00 | 7.11E-01 | no | Celsr1 | intron_variant | MODIFIER |
| 15 | 85920552 | AX-170146209 | 0 | T | C | 0.5 | 2.74E-02 | 6.98E-02 | 2.53E+00 | 2.62E+00 | 7.11E-01 | no | Celsr1 | intron_variant | MODIFIER |
| 15 | 85929858 | AX-170395413 | 0 | T | G | 0.5 | 2.74E-02 | 6.98E-02 | 2.53E+00 | 2.62E+00 | 7.11E-01 | no | Celsr1 | synonymous_variant | LOW |
| 15 | 85935518 | AX-97788478 | 0 | A | G | 0.5 | 2.74E-02 | 6.98E-02 | 2.53E+00 | 2.62E+00 | 7.11E-01 | no | Celsr1 | intron_variant | MODIFIER |
| 15 | 85976201 | AX-170056185 | 0 | T | C | 0.5 | 2.74E-02 | 6.98E-02 | 2.53E+00 | 2.62E+00 | 7.11E-01 | no | Celsr1 | intron_variant | MODIFIER |
| 15 | 85987020 | AX-170243774 | 0 | T | C | 0.5 | 2.74E-02 | 6.98E-02 | 2.53E+00 | 2.62E+00 | 7.11E-01 | no | Celsr1 | intron_variant | MODIFIER |
| 15 | 85994878 | AX-97813294 | 0 | T | G | 0.5 | 2.74E-02 | 6.98E-02 | 2.53E+00 | 2.62E+00 | 7.11E-01 | no | Celsr1 | intron_variant | MODIFIER |
| 15 | 86004743 | AX-170499464 | 0 | T | G | 0.5 | 2.74E-02 | 6.98E-02 | 2.53E+00 | 2.62E+00 | 7.11E-01 | no | Celsr1 | intron_variant | MODIFIER |
| 15 | 86018849 | AX-170242737 | 0 | G | C | 0.5 | 2.74E-02 | 6.98E-02 | 2.53E+00 | 2.62E+00 | 7.11E-01 | no | Celsr1 | intron_variant | MODIFIER |
| 15 | 86018924 | AX-170353920 | 0 | C | A | 0.5 | 2.74E-02 | 6.98E-02 | 2.53E+00 | 2.62E+00 | 7.11E-01 | no | Celsr1 | intron_variant | MODIFIER |
| 15 | 86065546 | AX-170436900 | 0 | T | C | 0.5 | 2.74E-02 | 6.98E-02 | 2.53E+00 | 2.62E+00 | 7.11E-01 | no | Gramd4 | intron_variant | MODIFIER |
| 15 | 86066022 | AX-170056187 | 0 | T | C | 0.5 | 2.74E-02 | 6.98E-02 | 2.53E+00 | 2.62E+00 | 7.11E-01 | no | Gramd4 | intron_variant | MODIFIER |
| 15 | 86076792 | AX-170484191 | 0 | A | G | 0.5 | 2.74E-02 | 6.98E-02 | 2.53E+00 | 2.62E+00 | 7.11E-01 | no | Gramd4 | intron_variant | MODIFIER |
| 15 | 86092523 | AX-97805092 | 0 | G | A | 0.5 | 2.74E-02 | 6.98E-02 | 2.53E+00 | 2.62E+00 | 7.11E-01 | no | Gramd4 | intron_variant | MODIFIER |
| 15 | 86095537 | AX-170313584 | 0 | A | G | 0.5 | 2.74E-02 | 6.98E-02 | 2.53E+00 | 2.62E+00 | 7.11E-01 | no | Gramd4 | intron_variant | MODIFIER |
| 15 | 86099593 | AX-170245821 | 0 | T | G | 0.5 | 2.74E-02 | 6.98E-02 | 2.53E+00 | 2.62E+00 | 7.11E-01 | no | Gramd4 | intron_variant | MODIFIER |
| 15 | 86120274 | AX-170451293 | 0 | A | G | 0.5 | 2.74E-02 | 6.98E-02 | 2.53E+00 | 2.62E+00 | 7.11E-01 | no | Gramd4 | intron_variant | MODIFIER |

| | | | | | | | | | | | | | | | |
|---|---|---|---|---|---|---|---|---|---|---|---|---|---|---|---|
| 15 | 86133008 | AX-170448674 | 0 | T | C | 0.5 | 2.74E-02 | 6.98E-02 | 2.53E+00 | 2.62E+00 | 7.11E-01 | no | Gramd4 | intron_variant | MODIFIER |
| 15 | 86145348 | AX-170093324 | 0 | A | G | 0.5 | 2.74E-02 | 6.98E-02 | 2.53E+00 | 2.62E+00 | 7.11E-01 | no | Cerk | intron_variant | MODIFIER |
| 15 | 86591362 | AX-170358173 | 0 | T | C | 0.5 | 2.74E-02 | 6.98E-02 | 2.53E+00 | 2.62E+00 | 7.11E-01 | no | | regulatory_region_variant | MODIFIER |
| 15 | 86597100 | AX-97830640 | 0 | C | T | 0.5 | 2.74E-02 | 6.98E-02 | 2.53E+00 | 2.62E+00 | 7.11E-01 | no | | intergenic_variant | MODIFIER |
| 15 | 86602896 | AX-170356139 | 0 | G | A | 0.5 | 2.74E-02 | 6.98E-02 | 2.53E+00 | 2.62E+00 | 7.11E-01 | no | | intergenic_variant | MODIFIER |
| 15 | 86608373 | AX-170445853 | 0 | A | G | 0.5 | 2.74E-02 | 6.98E-02 | 2.53E+00 | 2.62E+00 | 7.11E-01 | no | | intergenic_variant | MODIFIER |
| 15 | 86644639 | AX-170202785 | 0 | A | G | 0.5 | 2.74E-02 | 6.98E-02 | 2.53E+00 | 2.62E+00 | 7.11E-01 | no | | intergenic_variant | MODIFIER |
| 15 | 86663437 | AX-170056201 | 0 | G | A | 0.5 | 2.74E-02 | 6.98E-02 | 2.53E+00 | 2.62E+00 | 7.11E-01 | no | | intergenic_variant | MODIFIER |
| 15 | 86664705 | AX-170056202 | 0 | A | G | 0.5 | 2.74E-02 | 6.98E-02 | 2.53E+00 | 2.62E+00 | 7.11E-01 | no | | intergenic_variant | MODIFIER |
| 5 | 109502489 | AX-170073061 | 0 | A | G | 0.447 | 4.02E-02 | 1.07E-01 | 2.62E+00 | 2.68E+00 | 7.11E-01 | no | | intergenic_variant | MODIFIER |
| 7 | 117950308 | AX-170029838 | 0 | C | G | 0.395 | 3.37E-02 | 9.16E-02 | 2.67E+00 | 2.72E+00 | 7.11E-01 | no | | intergenic_variant | MODIFIER |
| 1 | 156015349 | AX-170002044 | 0 | A | T | 0.219 | 5.25E-02 | 1.41E-01 | 2.63E+00 | 2.67E+00 | 7.12E-01 | no | Tor1aip1 | intron_variant | MODIFIER |
| 8 | 29794974 | AX-97845242 | 0 | G | A | 0.175 | 4.05E-02 | 1.11E-01 | 2.67E+00 | 2.75E+00 | 7.12E-01 | no | | intergenic_variant | MODIFIER |
| 7 | 51042153 | AX-170028993 | 1 | C | T | 0.438 | 3.65E-02 | 9.83E-02 | 2.62E+00 | 2.70E+00 | 7.12E-01 | no | | intergenic_variant | MODIFIER |
| 2 | 100201266 | AX-170461545 | 0 | C | T | 0.044 | -5.83E-02 | 1.63E-01 | 2.62E+00 | 2.77E+00 | 7.12E-01 | no | | intergenic_variant | MODIFIER |
| 1 | 19704291 | AX-170000033 | 0 | C | A | 0.404 | 3.09E-02 | 8.33E-02 | 2.63E+00 | 2.69E+00 | 7.12E-01 | no | | intergenic_variant | MODIFIER |
| 3 | 148548940 | AX-170068400 | 0 | T | G | 0.325 | 2.81E-02 | 7.62E-02 | 2.63E+00 | 2.71E+00 | 7.12E-01 | no | | intergenic_variant | MODIFIER |
| 3 | 148564434 | AX-97839724 | 0 | C | T | 0.325 | 2.81E-02 | 7.62E-02 | 2.63E+00 | 2.71E+00 | 7.12E-01 | no | Gm38068 | upstream_gene_variant | MODIFIER |
| 3 | 148604727 | AX-97825227 | 0 | T | C | 0.325 | 2.81E-02 | 7.62E-02 | 2.63E+00 | 2.71E+00 | 7.12E-01 | no | | intergenic_variant | MODIFIER |
| 3 | 148611352 | AX-170109375 | 0 | A | G | 0.325 | 2.81E-02 | 7.62E-02 | 2.63E+00 | 2.71E+00 | 7.12E-01 | no | | intergenic_variant | MODIFIER |
| 3 | 148611554 | AX-170364090 | 0 | G | A | 0.325 | 2.81E-02 | 7.62E-02 | 2.63E+00 | 2.71E+00 | 7.12E-01 | no | | intergenic_variant | MODIFIER |
| 3 | 148611833 | AX-170468314 | 0 | T | C | 0.325 | 2.81E-02 | 7.62E-02 | 2.63E+00 | 2.71E+00 | 7.12E-01 | no | | intergenic_variant | MODIFIER |
| 3 | 148613780 | AX-170187791 | 0 | G | T | 0.325 | 2.81E-02 | 7.62E-02 | 2.63E+00 | 2.71E+00 | 7.12E-01 | no | | intergenic_variant | MODIFIER |
| 3 | 148614758 | AX-170424686 | 0 | A | G | 0.325 | 2.81E-02 | 7.62E-02 | 2.63E+00 | 2.71E+00 | 7.12E-01 | no | | regulatory_region_variant | MODIFIER |
| 3 | 148616112 | AX-170107979 | 0 | A | G | 0.325 | 2.81E-02 | 7.62E-02 | 2.63E+00 | 2.71E+00 | 7.12E-01 | no | | intergenic_variant | MODIFIER |
| 3 | 148678226 | AX-170511613 | 0 | A | C | 0.325 | 2.81E-02 | 7.62E-02 | 2.63E+00 | 2.71E+00 | 7.12E-01 | no | | intergenic_variant | MODIFIER |
| 3 | 148678268 | AX-170292703 | 0 | A | T | 0.325 | 2.81E-02 | 7.62E-02 | 2.63E+00 | 2.71E+00 | 7.12E-01 | no | | intergenic_variant | MODIFIER |
| 3 | 148726955 | AX-170172875 | 0 | A | G | 0.325 | 2.81E-02 | 7.62E-02 | 2.63E+00 | 2.71E+00 | 7.12E-01 | no | | regulatory_region_variant | MODIFIER |
| 3 | 148731470 | AX-170225587 | 0 | A | T | 0.325 | 2.81E-02 | 7.62E-02 | 2.63E+00 | 2.71E+00 | 7.12E-01 | no | | intergenic_variant | MODIFIER |
| 3 | 148731701 | AX-170306997 | 0 | C | T | 0.325 | 2.81E-02 | 7.62E-02 | 2.63E+00 | 2.71E+00 | 7.12E-01 | no | | intergenic_variant | MODIFIER |
| 3 | 148733260 | AX-170527975 | 0 | T | C | 0.325 | 2.81E-02 | 7.62E-02 | 2.63E+00 | 2.71E+00 | 7.12E-01 | no | | regulatory_region_variant | MODIFIER |
| 3 | 148736355 | AX-97842164 | 0 | A | G | 0.325 | 2.81E-02 | 7.62E-02 | 2.63E+00 | 2.71E+00 | 7.12E-01 | no | | intergenic_variant | MODIFIER |
| 3 | 148740070 | AX-170397645 | 0 | G | A | 0.325 | 2.81E-02 | 7.62E-02 | 2.63E+00 | 2.71E+00 | 7.12E-01 | no | | intergenic_variant | MODIFIER |
| 3 | 148757355 | AX-170216990 | 0 | T | C | 0.325 | 2.81E-02 | 7.62E-02 | 2.63E+00 | 2.71E+00 | 7.12E-01 | no | | intergenic_variant | MODIFIER |
| 3 | 148758514 | AX-170068407 | 0 | G | A | 0.325 | 2.81E-02 | 7.62E-02 | 2.63E+00 | 2.71E+00 | 7.12E-01 | no | | intergenic_variant | MODIFIER |
| 3 | 148759300 | AX-170089105 | 0 | T | G | 0.325 | 2.81E-02 | 7.62E-02 | 2.63E+00 | 2.71E+00 | 7.12E-01 | no | | intergenic_variant | MODIFIER |
| 3 | 148786956 | AX-97818142 | 0 | C | G | 0.325 | 2.81E-02 | 7.62E-02 | 2.63E+00 | 2.71E+00 | 7.12E-01 | no | | intergenic_variant | MODIFIER |
| 3 | 148821735 | AX-170168812 | 0 | G | T | 0.325 | 2.81E-02 | 7.62E-02 | 2.63E+00 | 2.71E+00 | 7.12E-01 | no | Adgrl2 | intron_variant | MODIFIER |
| 3 | 148831818 | AX-170499091 | 0 | T | A | 0.325 | 2.81E-02 | 7.62E-02 | 2.63E+00 | 2.71E+00 | 7.12E-01 | no | Adgrl2 | intron_variant | MODIFIER |
| 3 | 148839085 | AX-170355694 | 0 | T | C | 0.325 | 2.81E-02 | 7.62E-02 | 2.63E+00 | 2.71E+00 | 7.12E-01 | no | Adgrl2 | intron_variant | MODIFIER |
| 3 | 148850195 | AX-170476175 | 0 | C | T | 0.325 | 2.81E-02 | 7.62E-02 | 2.63E+00 | 2.71E+00 | 7.12E-01 | no | Adgrl2 | intron_variant | MODIFIER |
| 3 | 148853634 | AX-170168385 | 0 | G | T | 0.325 | 2.81E-02 | 7.62E-02 | 2.63E+00 | 2.71E+00 | 7.12E-01 | no | Adgrl2 | intron_variant | MODIFIER |
| 3 | 148855398 | AX-170315711 | 0 | G | A | 0.325 | 2.81E-02 | 7.62E-02 | 2.63E+00 | 2.71E+00 | 7.12E-01 | no | Adgrl2 | intron_variant | MODIFIER |
| 3 | 148888812 | AX-170251032 | 0 | G | A | 0.325 | 2.81E-02 | 7.62E-02 | 2.63E+00 | 2.71E+00 | 7.12E-01 | no | Adgrl2 | downstream_gene_variant | MODIFIER |
| 3 | 148895346 | AX-97794714 | 0 | C | T | 0.325 | 2.81E-02 | 7.62E-02 | 2.63E+00 | 2.71E+00 | 7.12E-01 | no | Adgrl2 | intron_variant,non_coding_transcript_variant | MODIFIER |
| 3 | 148896330 | AX-170203932 | 0 | C | T | 0.325 | 2.81E-02 | 7.62E-02 | 2.63E+00 | 2.71E+00 | 7.12E-01 | no | Adgrl2 | intron_variant,non_coding_transcript_variant | MODIFIER |
| 3 | 148912431 | AX-170255887 | 0 | C | T | 0.325 | 2.81E-02 | 7.62E-02 | 2.63E+00 | 2.71E+00 | 7.12E-01 | no | Adgrl2 | intron_variant,non_coding_transcript_variant | MODIFIER |
| 3 | 148913928 | AX-170207499 | 0 | T | C | 0.325 | 2.81E-02 | 7.62E-02 | 2.63E+00 | 2.71E+00 | 7.12E-01 | no | Adgrl2 | intron_variant,non_coding_transcript_variant | MODIFIER |
| 3 | 148914311 | AX-170377768 | 0 | G | T | 0.325 | 2.81E-02 | 7.62E-02 | 2.63E+00 | 2.71E+00 | 7.12E-01 | no | Adgrl2 | intron_variant,non_coding_transcript_variant | MODIFIER |
| 3 | 148917383 | AX-170193828 | 0 | G | A | 0.325 | 2.81E-02 | 7.62E-02 | 2.63E+00 | 2.71E+00 | 7.12E-01 | no | Adgrl2 | intron_variant,non_coding_transcript_variant | MODIFIER |
| 3 | 148930546 | AX-170358292 | 0 | T | C | 0.325 | 2.81E-02 | 7.62E-02 | 2.63E+00 | 2.71E+00 | 7.12E-01 | no | Adgrl2 | intron_variant,non_coding_transcript_variant | MODIFIER |
| 3 | 148935119 | AX-97783801 | 0 | G | A | 0.325 | 2.81E-02 | 7.62E-02 | 2.63E+00 | 2.71E+00 | 7.12E-01 | no | Adgrl2 | intron_variant,non_coding_transcript_variant | MODIFIER |
| 3 | 148960715 | AX-170417150 | 0 | A | G | 0.325 | 2.81E-02 | 7.62E-02 | 2.63E+00 | 2.71E+00 | 7.12E-01 | no | Adgrl2 | intron_variant,non_coding_transcript_variant | MODIFIER |
| 3 | 148968689 | AX-170388599 | 0 | A | G | 0.325 | 2.81E-02 | 7.62E-02 | 2.63E+00 | 2.71E+00 | 7.12E-01 | no | Adgrl2 | intron_variant,non_coding_transcript_variant | MODIFIER |

| | | | | | | | | | | | | | | | |
|---|---|---|---|---|---|---|---|---|---|---|---|---|---|---|---|
| 3 | 148970795 | AX-170329758 | 0 | A | G | 0.325 | 2.81E-02 | 7.62E-02 | 2.63E+00 | 2.71E+00 | 7.12E-01 | no | Adgrl2 | intron_variant,non_coding_transcript_variant | MODIFIER |
| 12 | 93473406 | AX-170357886 | 1 | T | A | 0.071 | 7.03E-02 | 1.92E-01 | 2.64E+00 | 2.73E+00 | 7.12E-01 | no | | intergenic_variant | MODIFIER |
| 8 | 4959138 | AX-170124717 | 1 | A | G | 0.268 | 2.76E-02 | 7.68E-02 | 2.66E+00 | 2.77E+00 | 7.12E-01 | no | | intergenic_variant | MODIFIER |
| 4 | 148149701 | AX-170277506 | 1 | G | A | 0.098 | 5.58E-02 | 1.50E-01 | 2.57E+00 | 2.70E+00 | 7.12E-01 | no | Fbxo6 | intron_variant | MODIFIER |
| 7 | 8419130 | AX-170028567 | 0 | C | T | 0.175 | -2.86E-02 | 7.39E-02 | 2.51E+00 | 2.67E+00 | 7.12E-01 | no | Vmn2r-ps46 | intron_variant,non_coding_transcript_variant | MODIFIER |
| 5 | 32265482 | AX-97800085 | 0 | G | A | 0.36 | 3.67E-02 | 9.95E-02 | 2.63E+00 | 2.70E+00 | 7.13E-01 | no | Plb1 | intron_variant | MODIFIER |
| 5 | 106785201 | AX-170368456 | 0 | T | A | 0.439 | 4.10E-02 | 1.11E-01 | 2.64E+00 | 2.69E+00 | 7.13E-01 | no | | regulatory_region_variant | MODIFIER |
| 16 | 27156081 | AX-97809726 | 1 | C | T | 0.25 | 3.15E-02 | 8.73E-02 | 2.63E+00 | 2.76E+00 | 7.13E-01 | no | | intergenic_variant | MODIFIER |
| 18 | 58171284 | AX-170329536 | 0 | T | C | 0.395 | 3.90E-02 | 9.75E-02 | 2.52E+00 | 2.58E+00 | 7.13E-01 | no | Fbn2 | intron_variant | MODIFIER |
| 13 | 31676000 | AX-170248214 | 1 | A | T | 0.179 | -4.21E-02 | 1.11E-01 | 2.57E+00 | 2.66E+00 | 7.13E-01 | no | | intergenic_variant | MODIFIER |
| 13 | 23734285 | AX-170050757 | 1 | C | A | 0.17 | -4.77E-02 | 1.21E-01 | 2.54E+00 | 2.60E+00 | 7.13E-01 | no | Hist1h1c | upstream_gene_variant | MODIFIER |
| 11 | 73785130 | AX-170005295 | 1 | T | A | 0.241 | 2.57E-02 | 7.13E-02 | 2.62E+00 | 2.76E+00 | 7.13E-01 | no | Olfr390 | upstream_gene_variant | MODIFIER |
| 10 | 28551934 | AX-170290451 | 1 | C | T | 0.196 | 3.47E-02 | 9.72E-02 | 2.70E+00 | 2.81E+00 | 7.13E-01 | no | Ptprk | intron_variant | MODIFIER |
| 10 | 27877642 | AX-170501951 | 1 | A | G | 0.179 | 3.61E-02 | 1.01E-01 | 2.72E+00 | 2.82E+00 | 7.13E-01 | no | | intergenic_variant | MODIFIER |
| 10 | 31773336 | AX-170467853 | 0 | T | C | 0.386 | -2.73E-02 | 7.44E-02 | 2.61E+00 | 2.72E+00 | 7.13E-01 | no | Nkain2 | intron_variant | MODIFIER |
| 1 | 19982791 | AX-170465719 | 0 | G | A | 0.395 | 3.05E-02 | 8.21E-02 | 2.62E+00 | 2.68E+00 | 7.13E-01 | no | | intergenic_variant | MODIFIER |
| 8 | 94981941 | AX-170035786 | 1 | C | A | 0.223 | 4.31E-02 | 1.19E-01 | 2.69E+00 | 2.75E+00 | 7.13E-01 | no | Adgrg1 | upstream_gene_variant | MODIFIER |
| 2 | 105088816 | AX-170494612 | 1 | C | G | 0.054 | -1.09E-01 | 3.01E-01 | 2.74E+00 | 2.77E+00 | 7.13E-01 | no | Gm11060 | upstream_gene_variant | MODIFIER |
| 2 | 106228690 | AX-170352516 | 1 | C | T | 0.054 | -1.09E-01 | 3.01E-01 | 2.74E+00 | 2.77E+00 | 7.13E-01 | no | Dcdc5 | intron_variant,non_coding_transcript_variant | MODIFIER |
| 2 | 107856347 | AX-97800367 | 1 | A | C | 0.054 | -1.09E-01 | 3.01E-01 | 2.74E+00 | 2.77E+00 | 7.13E-01 | no | | intergenic_variant | MODIFIER |
| 2 | 110520332 | AX-170398682 | 1 | A | C | 0.054 | -1.09E-01 | 3.01E-01 | 2.74E+00 | 2.77E+00 | 7.13E-01 | no | | intergenic_variant | MODIFIER |
| 9 | 34957003 | AX-170211323 | 1 | G | T | 0.446 | 2.99E-02 | 8.18E-02 | 2.64E+00 | 2.72E+00 | 7.14E-01 | no | Kirrel3 | intron_variant | MODIFIER |
| 9 | 72057597 | AX-170187371 | 0 | C | G | 0.193 | -4.72E-02 | 1.30E-01 | 2.67E+00 | 2.72E+00 | 7.14E-01 | no | Tcf12 | intron_variant | MODIFIER |
| 9 | 74799697 | AX-170032718 | 0 | T | C | 0.193 | -4.72E-02 | 1.30E-01 | 2.67E+00 | 2.72E+00 | 7.14E-01 | no | | intergenic_variant | MODIFIER |
| 1 | 164894145 | AX-97814061 | 0 | T | C | 0.228 | 4.74E-02 | 1.28E-01 | 2.62E+00 | 2.69E+00 | 7.14E-01 | no | Gm32569 | intron_variant,non_coding_transcript_variant | MODIFIER |
| 1 | 167821238 | AX-170375208 | 0 | T | A | 0.228 | 4.74E-02 | 1.28E-01 | 2.62E+00 | 2.69E+00 | 7.14E-01 | no | Lmx1a | intron_variant | MODIFIER |
| 1 | 13444281 | AX-170035608 | 1 | T | G | 0.366 | 2.66E-02 | 7.22E-02 | 2.62E+00 | 2.70E+00 | 7.14E-01 | no | | intergenic_variant | MODIFIER |
| 9 | 35516112 | AX-170422933 | 0 | G | A | 0.439 | -3.23E-02 | 9.00E-02 | 2.77E+00 | 2.84E+00 | 7.14E-01 | no | Gm1113 | intron_variant | MODIFIER |
| 8 | 34066020 | AX-97819584 | 0 | A | G | 0.149 | -4.53E-02 | 1.23E-01 | 2.61E+00 | 2.70E+00 | 7.14E-01 | no | | regulatory_region_variant | MODIFIER |
| 4 | 149158118 | AX-170274502 | 0 | C | G | 0.105 | 5.80E-02 | 1.60E-01 | 2.67E+00 | 2.75E+00 | 7.14E-01 | no | Pgd | intron_variant | MODIFIER |
| 8 | 65935695 | AX-170226794 | 1 | T | C | 0.161 | -4.46E-02 | 1.21E-01 | 2.61E+00 | 2.71E+00 | 7.14E-01 | no | MAR1 | intron_variant | MODIFIER |
| 3 | 21111042 | AX-170329006 | 1 | C | T | 0.152 | -3.86E-02 | 1.02E-01 | 2.58E+00 | 2.66E+00 | 7.14E-01 | no | Gm43160 | downstream_gene_variant | MODIFIER |
| 5 | 100683038 | AX-170409908 | 1 | A | G | 0.438 | -3.67E-02 | 1.03E-01 | 2.72E+00 | 2.80E+00 | 7.14E-01 | no | Hpse | intron_variant | MODIFIER |
| 7 | 77854354 | AX-170525811 | 1 | G | C | 0.384 | -3.54E-02 | 9.92E-02 | 2.68E+00 | 2.78E+00 | 7.14E-01 | no | | intergenic_variant | MODIFIER |
| 3 | 148676814 | AX-170285366 | 1 | T | A | 0.339 | 2.71E-02 | 7.35E-02 | 2.61E+00 | 2.69E+00 | 7.14E-01 | no | | intergenic_variant | MODIFIER |
| 1 | 13040242 | AX-170380999 | 1 | T | C | 0.393 | 2.73E-02 | 7.28E-02 | 2.60E+00 | 2.66E+00 | 7.14E-01 | no | | intergenic_variant | MODIFIER |
| 4 | 82325152 | AX-170069818 | 0 | G | T | 0.246 | 3.76E-02 | 9.83E-02 | 2.56E+00 | 2.64E+00 | 7.15E-01 | no | Nfib | intron_variant | MODIFIER |
| 9 | 35479324 | AX-170360383 | 2 | G | A | 0.427 | -3.13E-02 | 8.76E-02 | 2.73E+00 | 2.80E+00 | 7.15E-01 | no | Cdon | intron_variant | MODIFIER |
| 5 | 122852469 | AX-170311450 | 0 | G | A | 0.456 | 3.92E-02 | 1.07E-01 | 2.64E+00 | 2.70E+00 | 7.15E-01 | no | Rnf34 | intron_variant | MODIFIER |
| 5 | 123481275 | AX-170026064 | 0 | T | C | 0.456 | 3.92E-02 | 1.07E-01 | 2.64E+00 | 2.70E+00 | 7.15E-01 | no | Il31 | intron_variant | MODIFIER |
| 16 | 40192383 | AX-97825472 | 0 | G | A | 0.491 | 3.11E-02 | 8.18E-02 | 2.58E+00 | 2.63E+00 | 7.15E-01 | no | Lsamp | intron_variant | MODIFIER |
| 8 | 29613389 | AX-97844895 | 1 | C | T | 0.17 | 4.12E-02 | 1.15E-01 | 2.71E+00 | 2.79E+00 | 7.15E-01 | no | | intergenic_variant | MODIFIER |
| 8 | 106665160 | AX-170379531 | 1 | T | C | 0.223 | 3.97E-02 | 1.11E-01 | 2.69E+00 | 2.76E+00 | 7.15E-01 | no | Cdh1 | intron_variant | MODIFIER |
| 6 | 133204164 | AX-170339715 | 2 | A | C | 0.555 | 3.30E-02 | 8.51E-02 | 2.54E+00 | 2.62E+00 | 7.15E-01 | no | | intergenic_variant | MODIFIER |
| 9 | 69078070 | AX-170523484 | 1 | G | A | 0.196 | -4.88E-02 | 1.35E-01 | 2.70E+00 | 2.73E+00 | 7.15E-01 | no | Rora | intron_variant | MODIFIER |
| 5 | 108648762 | AX-170073046 | 1 | A | T | 0.464 | 4.27E-02 | 1.16E-01 | 2.64E+00 | 2.67E+00 | 7.15E-01 | no | Dgkq | missense_variant | MODERATE |
| 1 | 89409635 | AX-170146173 | 0 | C | T | 0.149 | 7.28E-02 | 1.99E-01 | 2.64E+00 | 2.70E+00 | 7.15E-01 | no | C030007H22Rik | upstream_gene_variant | MODIFIER |
| 14 | 120038324 | AX-170010828 | 0 | G | A | 0.333 | 2.77E-02 | 7.64E-02 | 2.62E+00 | 2.73E+00 | 7.15E-01 | no | Oxgr1 | intron_variant | MODIFIER |
| 2 | 75513781 | AX-170099907 | 1 | C | T | 0.054 | -6.57E-02 | 1.85E-01 | 2.65E+00 | 2.77E+00 | 7.15E-01 | no | | intergenic_variant | MODIFIER |
| 5 | 40415074 | AX-170024757 | 1 | A | G | 0.214 | -3.97E-02 | 1.07E-01 | 2.60E+00 | 2.68E+00 | 7.15E-01 | no | | intergenic_variant | MODIFIER |
| 1 | 148914216 | AX-170001963 | 0 | A | G | 0.202 | -4.13E-02 | 1.18E-01 | 2.72E+00 | 2.83E+00 | 7.15E-01 | no | | intergenic_variant | MODIFIER |
| 1 | 149739444 | AX-170455444 | 0 | A | G | 0.202 | -4.13E-02 | 1.18E-01 | 2.72E+00 | 2.83E+00 | 7.15E-01 | no | n-R5s218 | upstream_gene_variant | MODIFIER |
| 1 | 151431834 | AX-170344192 | 0 | T | C | 0.202 | -4.13E-02 | 1.18E-01 | 2.72E+00 | 2.83E+00 | 7.15E-01 | no | Swt1 | upstream_gene_variant | MODIFIER |
| 8 | 75620658 | AX-97784213 | 1 | G | A | 0.188 | -4.85E-02 | 1.34E-01 | 2.67E+00 | 2.73E+00 | 7.15E-01 | no | Iqcm | intron_variant | MODIFIER |

| | | | | | | | | | | | | | | | |
|---|---|---|---|---|---|---|---|---|---|---|---|---|---|---|---|
| 5 | 27902582 | AX-170071623 | 1 | G | C | 0.348 | 3.90E-02 | 1.07E-01 | 2.63E+00 | 2.70E+00 | 7.15E-01 | no | | intergenic_variant | MODIFIER |
| 5 | 98286500 | AX-97815695 | 0 | C | T | 0.421 | -3.72E-02 | 1.04E-01 | 2.71E+00 | 2.77E+00 | 7.16E-01 | no | | intergenic_variant | MODIFIER |
| 2 | 114721999 | AX-97841952 | 1 | C | T | 0.143 | 7.94E-02 | 2.20E-01 | 2.70E+00 | 2.73E+00 | 7.16E-01 | no | | intergenic_variant | MODIFIER |
| 2 | 121382676 | AX-170510565 | 1 | G | C | 0.143 | 7.94E-02 | 2.20E-01 | 2.70E+00 | 2.73E+00 | 7.16E-01 | no | Strc | upstream_gene_variant | MODIFIER |
| 2 | 123023963 | AX-170390084 | 1 | G | T | 0.143 | 7.94E-02 | 2.20E-01 | 2.70E+00 | 2.73E+00 | 7.16E-01 | no | | intergenic_variant | MODIFIER |
| 2 | 149952001 | AX-97840585 | 1 | G | T | 0.143 | 7.94E-02 | 2.20E-01 | 2.70E+00 | 2.73E+00 | 7.16E-01 | no | Syndig1 | intron_variant | MODIFIER |
| 13 | 35322290 | AX-170008092 | 0 | A | T | 0.175 | -4.46E-02 | 1.21E-01 | 2.61E+00 | 2.67E+00 | 7.16E-01 | no | | intergenic_variant | MODIFIER |
| 8 | 107209033 | AX-170188047 | 0 | A | G | 0.228 | -3.50E-02 | 9.86E-02 | 2.67E+00 | 2.78E+00 | 7.16E-01 | no | | regulatory_region_variant | MODIFIER |
| 15 | 74979530 | AX-170154163 | 0 | G | A | 0.553 | 2.94E-02 | 7.73E-02 | 2.56E+00 | 2.64E+00 | 7.16E-01 | no | Ly6i | downstream_gene_variant | MODIFIER |
| 4 | 142523672 | AX-170484727 | 1 | C | T | 0.08 | 7.23E-02 | 2.02E-01 | 2.66E+00 | 2.74E+00 | 7.16E-01 | no | | intergenic_variant | MODIFIER |
| 18 | 62209376 | AX-170430842 | 1 | C | T | 0.188 | -3.94E-02 | 1.06E-01 | 2.59E+00 | 2.67E+00 | 7.16E-01 | no | | intergenic_variant | MODIFIER |
| 6 | 105081107 | AX-170128156 | 0 | C | T | 0.053 | -7.67E-02 | 2.16E-01 | 2.66E+00 | 2.76E+00 | 7.16E-01 | no | | intergenic_variant | MODIFIER |
| 6 | 139894921 | AX-170110007 | 0 | C | G | 0.053 | -7.67E-02 | 2.16E-01 | 2.66E+00 | 2.76E+00 | 7.16E-01 | no | Pik3c2g | intron_variant | MODIFIER |
| 7 | 10100346 | AX-170207144 | 0 | G | T | 0.175 | 2.75E-02 | 7.96E-02 | 2.71E+00 | 2.84E+00 | 7.16E-01 | no | Vmn2r51 | missense_variant | MODERATE |
| 17 | 22029844 | AX-170341493 | 0 | A | G | 0.053 | 6.04E-02 | 1.72E-01 | 2.68E+00 | 2.80E+00 | 7.17E-01 | no | Zfp943 | intron_variant,NMD_transcript_variant | MODIFIER |
| 1 | 15760547 | AX-170513988 | 1 | T | C | 0.366 | 3.03E-02 | 8.18E-02 | 2.61E+00 | 2.66E+00 | 7.17E-01 | no | Gm25168 | downstream_gene_variant | MODIFIER |
| 6 | 134501736 | AX-170328990 | 0 | T | C | 0.57 | 3.26E-02 | 8.64E-02 | 2.58E+00 | 2.64E+00 | 7.17E-01 | no | Lrp6 | intron_variant | MODIFIER |
| 15 | 87315125 | AX-170012199 | 2 | G | A | 0.127 | 5.67E-02 | 1.56E-01 | 2.63E+00 | 2.71E+00 | 7.17E-01 | no | | intergenic_variant | MODIFIER |
| 18 | 34503829 | AX-97845357 | 1 | A | G | 0.071 | 7.62E-02 | 2.16E-01 | 2.71E+00 | 2.80E+00 | 7.17E-01 | no | Fam13b | intron_variant | MODIFIER |
| 13 | 36455255 | AX-170008119 | 1 | G | C | 0.17 | -4.56E-02 | 1.22E-01 | 2.60E+00 | 2.66E+00 | 7.17E-01 | no | Fars2 | intron_variant | MODIFIER |
| 5 | 107134873 | AX-170224714 | 0 | G | A | 0.439 | 4.17E-02 | 1.15E-01 | 2.67E+00 | 2.70E+00 | 7.17E-01 | no | Tgfbr3 | intron_variant | MODIFIER |
| 1 | 101808032 | AX-170230058 | 0 | T | C | 0.149 | 6.01E-02 | 1.68E-01 | 2.64E+00 | 2.74E+00 | 7.17E-01 | no | | intergenic_variant | MODIFIER |
| 1 | 20160322 | AX-170301896 | 2 | G | T | 0.409 | 3.14E-02 | 8.57E-02 | 2.63E+00 | 2.68E+00 | 7.17E-01 | no | Pkhd1 | intron_variant | MODIFIER |
| 1 | 46560899 | AX-170511936 | 1 | T | C | 0.089 | -7.56E-02 | 2.13E-01 | 2.66E+00 | 2.76E+00 | 7.17E-01 | no | Dnah7c | intron_variant | MODIFIER |
| 1 | 173480286 | AX-170407947 | 0 | C | G | 0.228 | 4.59E-02 | 1.22E-01 | 2.58E+00 | 2.64E+00 | 7.17E-01 | no | Ifi206 | intron_variant | MODIFIER |
| 10 | 17553983 | AX-170356038 | 1 | C | T | 0.214 | 3.76E-02 | 1.06E-01 | 2.76E+00 | 2.84E+00 | 7.17E-01 | no | | intergenic_variant | MODIFIER |
| 13 | 35385399 | AX-170008098 | 1 | C | T | 0.188 | -4.36E-02 | 1.18E-01 | 2.60E+00 | 2.67E+00 | 7.17E-01 | no | | intergenic_variant | MODIFIER |
| 8 | 39208994 | AX-170320353 | 1 | A | T | 0.152 | 5.33E-02 | 1.49E-01 | 2.76E+00 | 2.81E+00 | 7.17E-01 | no | Gm17766 | downstream_gene_variant | MODIFIER |
| 8 | 43124997 | AX-170411107 | 1 | G | C | 0.152 | 5.33E-02 | 1.49E-01 | 2.76E+00 | 2.81E+00 | 7.17E-01 | no | Triml1 | downstream_gene_variant | MODIFIER |
| 4 | 128281019 | AX-170227548 | 0 | G | A | 0.14 | 4.24E-02 | 1.15E-01 | 2.60E+00 | 2.68E+00 | 7.18E-01 | no | Csmd2 | intron_variant,non_coding_transcript_variant | MODIFIER |
| 6 | 136963997 | AX-170028333 | 0 | G | A | 0.579 | 3.48E-02 | 8.86E-02 | 2.53E+00 | 2.58E+00 | 7.18E-01 | no | Pde6h | downstream_gene_variant | MODIFIER |
| 2 | 104180039 | AX-170238930 | 0 | C | T | 0.053 | -1.07E-01 | 2.99E-01 | 2.74E+00 | 2.76E+00 | 7.18E-01 | no | D430041D05Rik | intron_variant | MODIFIER |
| 5 | 108719273 | AX-97831996 | 1 | T | C | 0.438 | 4.01E-02 | 1.10E-01 | 2.64E+00 | 2.67E+00 | 7.18E-01 | no | | regulatory_region_variant | MODIFIER |
| 1 | 25646552 | AX-97785282 | 1 | C | A | 0.411 | 2.94E-02 | 8.06E-02 | 2.62E+00 | 2.69E+00 | 7.18E-01 | no | Adgrb3 | intron_variant | MODIFIER |
| 1 | 23068978 | AX-170396008 | 0 | G | C | 0.439 | 3.00E-02 | 8.32E-02 | 2.64E+00 | 2.71E+00 | 7.18E-01 | no | Gm29506 | intron_variant,non_coding_transcript_variant | MODIFIER |
| 9 | 91297993 | AX-170249262 | 0 | T | C | 0.123 | -4.45E-02 | 1.23E-01 | 2.63E+00 | 2.72E+00 | 7.18E-01 | no | | intergenic_variant | MODIFIER |
| 2 | 119687519 | AX-170088730 | 0 | G | A | 0.14 | 7.87E-02 | 2.20E-01 | 2.70E+00 | 2.73E+00 | 7.18E-01 | no | Rtf1 | intron_variant | MODIFIER |
| 2 | 120767086 | AX-170358416 | 0 | T | G | 0.14 | 7.87E-02 | 2.20E-01 | 2.70E+00 | 2.73E+00 | 7.18E-01 | no | Ttbk2 | intron_variant | MODIFIER |
| 2 | 124339284 | AX-170535042 | 0 | T | A | 0.14 | 7.87E-02 | 2.20E-01 | 2.70E+00 | 2.73E+00 | 7.18E-01 | no | Sema6d | intron_variant | MODIFIER |
| 2 | 138572723 | AX-97852229 | 0 | C | T | 0.14 | 7.87E-02 | 2.20E-01 | 2.70E+00 | 2.73E+00 | 7.18E-01 | no | Btbd3 | intron_variant,non_coding_transcript_variant | MODIFIER |
| 12 | 111798984 | AX-97856533 | 0 | A | G | 0.14 | -6.76E-02 | 1.91E-01 | 2.69E+00 | 2.76E+00 | 7.18E-01 | no | Xrcc3 | downstream_gene_variant | MODIFIER |
| 1 | 147248357 | AX-170218564 | 1 | T | C | 0.196 | 4.95E-02 | 1.37E-01 | 2.65E+00 | 2.71E+00 | 7.18E-01 | no | | intergenic_variant | MODIFIER |
| 1 | 149695246 | AX-170208171 | 1 | T | C | 0.196 | 4.95E-02 | 1.37E-01 | 2.65E+00 | 2.71E+00 | 7.18E-01 | no | | intergenic_variant | MODIFIER |
| 1 | 41920635 | AX-170359457 | 1 | C | A | 0.446 | -2.68E-02 | 7.51E-02 | 2.65E+00 | 2.74E+00 | 7.18E-01 | no | Gm29260 | intron_variant,non_coding_transcript_variant | MODIFIER |
| 5 | 37648447 | AX-170173152 | 0 | T | G | 0.333 | 3.66E-02 | 1.03E-01 | 2.68E+00 | 2.75E+00 | 7.18E-01 | no | Stk32b | intron_variant | MODIFIER |
| 9 | 106489939 | AX-97805238 | 2 | G | A | 0.127 | 4.14E-02 | 1.17E-01 | 2.79E+00 | 2.87E+00 | 7.18E-01 | no | Rrp9 | downstream_gene_variant | MODIFIER |
| 9 | 70297336 | AX-170255607 | 0 | G | A | 0.202 | 4.43E-02 | 1.25E-01 | 2.71E+00 | 2.76E+00 | 7.18E-01 | no | Myo1e | missense_variant | MODERATE |
| 18 | 89749866 | AX-170275768 | 1 | A | G | 0.5 | -2.24E-02 | 6.45E-02 | 2.67E+00 | 2.79E+00 | 7.18E-01 | no | Dok6 | intron_variant | MODIFIER |
| 18 | 88516355 | AX-97796427 | 1 | A | G | 0.17 | -3.60E-02 | 9.83E-02 | 2.58E+00 | 2.70E+00 | 7.18E-01 | no | | intergenic_variant | MODIFIER |
| 7 | 79161002 | AX-170420621 | 0 | G | T | 0.132 | -7.44E-02 | 2.11E-01 | 2.69E+00 | 2.77E+00 | 7.19E-01 | no | | intergenic_variant | MODIFIER |
| 1 | 91033676 | AX-170183883 | 0 | G | A | 0.158 | 7.46E-02 | 2.08E-01 | 2.65E+00 | 2.71E+00 | 7.19E-01 | no | Lrrfip1 | intron_variant | MODIFIER |
| 1 | 100789027 | AX-170215240 | 0 | C | T | 0.158 | 7.46E-02 | 2.08E-01 | 2.65E+00 | 2.71E+00 | 7.19E-01 | no | | intergenic_variant | MODIFIER |
| 5 | 35633341 | AX-170376882 | 1 | A | G | 0.339 | 3.82E-02 | 1.07E-01 | 2.66E+00 | 2.72E+00 | 7.19E-01 | no | | intergenic_variant | MODIFIER |
| 9 | 87568494 | AX-170100215 | 0 | A | C | 0.14 | 4.50E-02 | 1.27E-01 | 2.79E+00 | 2.87E+00 | 7.19E-01 | no | | intergenic_variant | MODIFIER |

| | | | | | | | | | | | | | | | |
|---|---|---|---|---|---|---|---|---|---|---|---|---|---|---|---|
| 7 | 94641521 | AX-170524154 | 0 | G | T | 0.132 | -7.45E-02 | 2.11E-01 | 2.68E+00 | 2.76E+00 | 7.19E-01 | no | Gm32647 | intron_variant,non_coding_transcript_variant | MODIFIER |
| 4 | 148581395 | AX-170024206 | 1 | A | G | 0.08 | 6.39E-02 | 1.78E-01 | 2.63E+00 | 2.72E+00 | 7.19E-01 | no | Exosc10 | intron_variant | MODIFIER |
| 4 | 149336845 | AX-170191481 | 1 | A | T | 0.08 | 6.39E-02 | 1.78E-01 | 2.63E+00 | 2.72E+00 | 7.19E-01 | no | Ube4b | intron_variant | MODIFIER |
| 8 | 35257976 | AX-170103905 | 0 | A | T | 0.158 | -4.78E-02 | 1.31E-01 | 2.61E+00 | 2.68E+00 | 7.19E-01 | no | | intergenic_variant | MODIFIER |
| 15 | 92516221 | AX-170526131 | 1 | T | C | 0.134 | -4.44E-02 | 1.29E-01 | 2.70E+00 | 2.82E+00 | 7.19E-01 | no | Pdzrn4 | intron_variant | MODIFIER |
| 8 | 102123830 | AX-170295954 | 0 | A | G | 0.219 | -3.82E-02 | 1.08E-01 | 2.67E+00 | 2.76E+00 | 7.19E-01 | no | | intergenic_variant | MODIFIER |
| 9 | 38219961 | AX-170248935 | 0 | A | G | 0.482 | -2.69E-02 | 7.67E-02 | 2.68E+00 | 2.77E+00 | 7.19E-01 | no | Olfr894 | downstream_gene_variant | MODIFIER |
| 2 | 94519963 | AX-170377637 | 1 | C | T | 0.054 | -1.07E-01 | 3.02E-01 | 2.74E+00 | 2.77E+00 | 7.19E-01 | no | | intergenic_variant | MODIFIER |
| 10 | 9099568 | AX-170448544 | 0 | G | A | 0.219 | -3.58E-02 | 9.94E-02 | 2.62E+00 | 2.71E+00 | 7.20E-01 | no | | intergenic_variant | MODIFIER |
| 7 | 102939613 | AX-97842528 | 1 | T | C | 0.464 | 3.39E-02 | 9.38E-02 | 2.63E+00 | 2.69E+00 | 7.20E-01 | no | Olfr573-ps1 | downstream_gene_variant | MODIFIER |
| 1 | 25944975 | AX-170532079 | 1 | A | G | 0.393 | 3.01E-02 | 8.48E-02 | 2.66E+00 | 2.74E+00 | 7.20E-01 | no | | intergenic_variant | MODIFIER |
| 15 | 92596884 | AX-97790441 | 1 | C | T | 0.161 | 6.03E-02 | 1.63E-01 | 2.58E+00 | 2.66E+00 | 7.20E-01 | no | Pdzrn4 | upstream_gene_variant | MODIFIER |
| 2 | 89123780 | AX-97782321 | 0 | C | T | 0.053 | -1.06E-01 | 3.00E-01 | 2.73E+00 | 2.76E+00 | 7.20E-01 | no | Olfr1222 | downstream_gene_variant | MODIFIER |
| 2 | 92727308 | AX-170187251 | 0 | T | C | 0.053 | -1.06E-01 | 3.00E-01 | 2.73E+00 | 2.76E+00 | 7.20E-01 | no | | regulatory_region_variant | MODIFIER |
| 1 | 144983719 | AX-170001905 | 2 | T | C | 0.2 | 4.82E-02 | 1.34E-01 | 2.64E+00 | 2.70E+00 | 7.20E-01 | no | Gm25708 | upstream_gene_variant | MODIFIER |
| 1 | 69886652 | AX-170459152 | 1 | T | C | 0.143 | 8.33E-02 | 2.36E-01 | 2.70E+00 | 2.76E+00 | 7.20E-01 | no | Spag16 | intron_variant | MODIFIER |
| 8 | 96251394 | AX-170031316 | 0 | C | T | 0.228 | 3.97E-02 | 1.13E-01 | 2.69E+00 | 2.76E+00 | 7.20E-01 | no | | intergenic_variant | MODIFIER |
| 1 | 9051580 | AX-170190604 | 0 | T | G | 0.518 | -2.34E-02 | 6.73E-02 | 2.77E+00 | 2.87E+00 | 7.20E-01 | no | Sntg1 | intron_variant,non_coding_transcript_variant | MODIFIER |
| 10 | 119058168 | AX-170356057 | 0 | T | G | 0.342 | 2.55E-02 | 7.28E-02 | 2.69E+00 | 2.78E+00 | 7.20E-01 | no | | regulatory_region_variant | MODIFIER |
| 10 | 120285597 | AX-97835836 | 0 | A | G | 0.342 | 2.55E-02 | 7.28E-02 | 2.69E+00 | 2.78E+00 | 7.20E-01 | no | Gm48118 | downstream_gene_variant | MODIFIER |
| 9 | 65280507 | AX-97843532 | 1 | T | A | 0.188 | -4.77E-02 | 1.34E-01 | 2.69E+00 | 2.73E+00 | 7.20E-01 | no | Cilp | 3_prime_UTR_variant | MODIFIER |
| 1 | 82276100 | AX-97852540 | 2 | A | C | 0.082 | 5.81E-02 | 1.70E-01 | 2.65E+00 | 2.81E+00 | 7.20E-01 | no | Irs1 | intron_variant | MODIFIER |
| 8 | 40676684 | AX-170493982 | 1 | A | G | 0.143 | 5.01E-02 | 1.43E-01 | 2.73E+00 | 2.79E+00 | 7.20E-01 | no | Adam24 | intron_variant | MODIFIER |
| 1 | 68658569 | AX-170237773 | 0 | G | T | 0.123 | 6.65E-02 | 1.86E-01 | 2.61E+00 | 2.71E+00 | 7.21E-01 | no | Erbb4 | intron_variant | MODIFIER |
| 2 | 119923124 | AX-97835666 | 0 | T | C | 0.132 | 6.98E-02 | 1.98E-01 | 2.67E+00 | 2.74E+00 | 7.21E-01 | no | Mga | intron_variant | MODIFIER |
| 1 | 137141160 | AX-170295526 | 1 | G | A | 0.205 | 4.46E-02 | 1.25E-01 | 2.64E+00 | 2.71E+00 | 7.21E-01 | no | | intergenic_variant | MODIFIER |
| 5 | 46857320 | AX-97803716 | 0 | C | T | 0.491 | 3.73E-02 | 9.64E-02 | 2.52E+00 | 2.60E+00 | 7.21E-01 | no | | intergenic_variant | MODIFIER |
| 12 | 91998518 | AX-170528757 | 0 | A | T | 0.079 | 6.80E-02 | 1.93E-01 | 2.66E+00 | 2.75E+00 | 7.21E-01 | no | | intergenic_variant | MODIFIER |
| 3 | 148409824 | AX-170498414 | 0 | A | G | 0.333 | 2.67E-02 | 7.48E-02 | 2.64E+00 | 2.71E+00 | 7.21E-01 | no | | intergenic_variant | MODIFIER |
| 8 | 91098245 | AX-97837225 | 0 | G | A | 0.175 | -4.35E-02 | 1.19E-01 | 2.60E+00 | 2.68E+00 | 7.21E-01 | no | Rbl2 | intron_variant | MODIFIER |
| 20 | 133734147 | AX-97788362 | 1 | G | T | 0.152 | 4.77E-02 | 1.35E-01 | 2.64E+00 | 2.74E+00 | 7.21E-01 | no | | intergenic_variant | MODIFIER |
| 5 | 107015887 | AX-97797900 | 0 | A | G | 0.43 | 4.02E-02 | 1.11E-01 | 2.63E+00 | 2.68E+00 | 7.21E-01 | no | | regulatory_region_variant | MODIFIER |
| 7 | 60194531 | AX-170099272 | 1 | G | C | 0.42 | -3.38E-02 | 9.74E-02 | 2.70E+00 | 2.80E+00 | 7.21E-01 | no | | intergenic_variant | MODIFIER |
| 8 | 36107599 | AX-170442568 | 0 | G | A | 0.149 | 5.31E-02 | 1.50E-01 | 2.77E+00 | 2.81E+00 | 7.21E-01 | no | Prag1 | intron_variant | MODIFIER |
| 8 | 36110201 | AX-170229627 | 0 | T | C | 0.149 | 5.31E-02 | 1.50E-01 | 2.77E+00 | 2.81E+00 | 7.21E-01 | no | Prag1 | intron_variant | MODIFIER |
| 8 | 36113809 | AX-170341495 | 0 | T | G | 0.149 | 5.31E-02 | 1.50E-01 | 2.77E+00 | 2.81E+00 | 7.21E-01 | no | Prag1 | intron_variant | MODIFIER |
| 8 | 36120092 | AX-170168736 | 0 | G | A | 0.149 | 5.31E-02 | 1.50E-01 | 2.77E+00 | 2.81E+00 | 7.21E-01 | no | Prag1 | intron_variant | MODIFIER |
| 8 | 36131984 | AX-170266368 | 0 | A | C | 0.149 | 5.31E-02 | 1.50E-01 | 2.77E+00 | 2.81E+00 | 7.21E-01 | no | Prag1 | intron_variant | MODIFIER |
| 8 | 36132144 | AX-170299965 | 0 | A | G | 0.149 | 5.31E-02 | 1.50E-01 | 2.77E+00 | 2.81E+00 | 7.21E-01 | no | Prag1 | intron_variant | MODIFIER |
| 8 | 36153558 | AX-170145983 | 0 | G | A | 0.149 | 5.31E-02 | 1.50E-01 | 2.77E+00 | 2.81E+00 | 7.21E-01 | no | Gm38414 | upstream_gene_variant | MODIFIER |
| 8 | 36159795 | AX-170210812 | 0 | A | G | 0.149 | 5.31E-02 | 1.50E-01 | 2.77E+00 | 2.81E+00 | 7.21E-01 | no | Gm38414 | non_coding_transcript_exon_variant | MODIFIER |
| 8 | 36166740 | AX-170389491 | 0 | A | G | 0.149 | 5.31E-02 | 1.50E-01 | 2.77E+00 | 2.81E+00 | 7.21E-01 | no | | intergenic_variant | MODIFIER |
| 8 | 36172784 | AX-170531179 | 0 | A | G | 0.149 | 5.31E-02 | 1.50E-01 | 2.77E+00 | 2.81E+00 | 7.21E-01 | no | Gm39164 | downstream_gene_variant | MODIFIER |
| 8 | 36173072 | AX-170364273 | 0 | A | G | 0.149 | 5.31E-02 | 1.50E-01 | 2.77E+00 | 2.81E+00 | 7.21E-01 | no | Gm39164 | downstream_gene_variant | MODIFIER |
| 8 | 36183338 | AX-170079762 | 0 | T | C | 0.149 | 5.31E-02 | 1.50E-01 | 2.77E+00 | 2.81E+00 | 7.21E-01 | no | 4933430A20Rik | downstream_gene_variant | MODIFIER |
| 8 | 36184192 | AX-170079758 | 0 | C | A | 0.149 | 5.31E-02 | 1.50E-01 | 2.77E+00 | 2.81E+00 | 7.21E-01 | no | Gm35520 | upstream_gene_variant | MODIFIER |
| 8 | 36194317 | AX-170438794 | 0 | T | C | 0.149 | 5.31E-02 | 1.50E-01 | 2.77E+00 | 2.81E+00 | 7.21E-01 | no | Gm35520 | intron_variant,non_coding_transcript_variant | MODIFIER |
| 8 | 36195593 | AX-170079763 | 0 | T | C | 0.149 | 5.31E-02 | 1.50E-01 | 2.77E+00 | 2.81E+00 | 7.21E-01 | no | Gm35520 | intron_variant,non_coding_transcript_variant | MODIFIER |
| 8 | 36197215 | AX-170079757 | 0 | G | A | 0.149 | 5.31E-02 | 1.50E-01 | 2.77E+00 | 2.81E+00 | 7.21E-01 | no | Gm35520 | intron_variant,non_coding_transcript_variant | MODIFIER |
| 8 | 36205963 | AX-97823181 | 0 | G | A | 0.149 | 5.31E-02 | 1.50E-01 | 2.77E+00 | 2.81E+00 | 7.21E-01 | no | Gm35520 | downstream_gene_variant | MODIFIER |
| 8 | 36211494 | AX-97797068 | 0 | A | G | 0.149 | 5.31E-02 | 1.50E-01 | 2.77E+00 | 2.81E+00 | 7.21E-01 | no | Lonrf1 | downstream_gene_variant | MODIFIER |
| 8 | 36219305 | AX-170486171 | 0 | C | T | 0.149 | 5.31E-02 | 1.50E-01 | 2.77E+00 | 2.81E+00 | 7.21E-01 | no | Lonrf1 | intron_variant | MODIFIER |
| 8 | 36219607 | AX-170348608 | 0 | G | A | 0.149 | 5.31E-02 | 1.50E-01 | 2.77E+00 | 2.81E+00 | 7.21E-01 | no | Lonrf1 | intron_variant | MODIFIER |
| 8 | 36266888 | AX-170221040 | 0 | T | C | 0.149 | 5.31E-02 | 1.50E-01 | 2.77E+00 | 2.81E+00 | 7.21E-01 | no | | intergenic_variant | MODIFIER |

| | | | | | | | | | | | | | | | |
|---|---|---|---|---|---|---|---|---|---|---|---|---|---|---|---|
| 8 | 36287334 | AX-170104319 | 0 | G | A | 0.149 | 5.31E-02 | 1.50E-01 | 2.77E+00 | 2.81E+00 | 7.21E-01 | no | | intergenic_variant | MODIFIER |
| 8 | 36289644 | AX-97794425 | 0 | C | T | 0.149 | 5.31E-02 | 1.50E-01 | 2.77E+00 | 2.81E+00 | 7.21E-01 | no | | intergenic_variant | MODIFIER |
| 8 | 36290433 | AX-170376838 | 0 | C | T | 0.149 | 5.31E-02 | 1.50E-01 | 2.77E+00 | 2.81E+00 | 7.21E-01 | no | | regulatory_region_variant | MODIFIER |
| 8 | 36294660 | AX-170446530 | 0 | G | A | 0.149 | 5.31E-02 | 1.50E-01 | 2.77E+00 | 2.81E+00 | 7.21E-01 | no | | intergenic_variant | MODIFIER |
| 8 | 36317644 | AX-97839977 | 0 | C | T | 0.149 | 5.31E-02 | 1.50E-01 | 2.77E+00 | 2.81E+00 | 7.21E-01 | no | | intergenic_variant | MODIFIER |
| 8 | 36325972 | AX-170447244 | 0 | C | A | 0.149 | 5.31E-02 | 1.50E-01 | 2.77E+00 | 2.81E+00 | 7.21E-01 | no | | regulatory_region_variant | MODIFIER |
| 8 | 36347040 | AX-170483355 | 0 | T | A | 0.149 | 5.31E-02 | 1.50E-01 | 2.77E+00 | 2.81E+00 | 7.21E-01 | no | | regulatory_region_variant | MODIFIER |
| 8 | 36347302 | AX-170126233 | 0 | G | C | 0.149 | 5.31E-02 | 1.50E-01 | 2.77E+00 | 2.81E+00 | 7.21E-01 | no | | regulatory_region_variant | MODIFIER |
| 8 | 36347940 | AX-170333614 | 0 | A | G | 0.149 | 5.31E-02 | 1.50E-01 | 2.77E+00 | 2.81E+00 | 7.21E-01 | no | | intergenic_variant | MODIFIER |
| 8 | 36362946 | AX-170273125 | 0 | G | A | 0.149 | 5.31E-02 | 1.50E-01 | 2.77E+00 | 2.81E+00 | 7.21E-01 | no | | intergenic_variant | MODIFIER |
| 8 | 36363673 | AX-170531010 | 0 | A | G | 0.149 | 5.31E-02 | 1.50E-01 | 2.77E+00 | 2.81E+00 | 7.21E-01 | no | | regulatory_region_variant | MODIFIER |
| 8 | 36363811 | AX-170196403 | 0 | T | G | 0.149 | 5.31E-02 | 1.50E-01 | 2.77E+00 | 2.81E+00 | 7.21E-01 | no | | intergenic_variant | MODIFIER |
| 8 | 36364287 | AX-170477540 | 0 | C | T | 0.149 | 5.31E-02 | 1.50E-01 | 2.77E+00 | 2.81E+00 | 7.21E-01 | no | | intergenic_variant | MODIFIER |
| 8 | 36376239 | AX-170327310 | 0 | C | T | 0.149 | 5.31E-02 | 1.50E-01 | 2.77E+00 | 2.81E+00 | 7.21E-01 | no | Gm5787 | upstream_gene_variant | MODIFIER |
| 8 | 36376364 | AX-170333098 | 0 | C | G | 0.149 | 5.31E-02 | 1.50E-01 | 2.77E+00 | 2.81E+00 | 7.21E-01 | no | Gm5787 | upstream_gene_variant | MODIFIER |
| 8 | 36397513 | AX-97811880 | 0 | A | G | 0.149 | 5.31E-02 | 1.50E-01 | 2.77E+00 | 2.81E+00 | 7.21E-01 | no | | intergenic_variant | MODIFIER |
| 8 | 36397792 | AX-170373842 | 0 | T | C | 0.149 | 5.31E-02 | 1.50E-01 | 2.77E+00 | 2.81E+00 | 7.21E-01 | no | | intergenic_variant | MODIFIER |
| 8 | 36418415 | AX-97852802 | 0 | G | C | 0.149 | 5.31E-02 | 1.50E-01 | 2.77E+00 | 2.81E+00 | 7.21E-01 | no | | intergenic_variant | MODIFIER |
| 8 | 36426856 | AX-170469958 | 0 | A | G | 0.149 | 5.31E-02 | 1.50E-01 | 2.77E+00 | 2.81E+00 | 7.21E-01 | no | | regulatory_region_variant | MODIFIER |
| 8 | 36428436 | AX-170268103 | 0 | A | C | 0.149 | 5.31E-02 | 1.50E-01 | 2.77E+00 | 2.81E+00 | 7.21E-01 | no | | intergenic_variant | MODIFIER |
| 8 | 36435077 | AX-170283744 | 0 | T | C | 0.149 | 5.31E-02 | 1.50E-01 | 2.77E+00 | 2.81E+00 | 7.21E-01 | no | | intergenic_variant | MODIFIER |
| 8 | 36435676 | AX-97848486 | 0 | A | G | 0.149 | 5.31E-02 | 1.50E-01 | 2.77E+00 | 2.81E+00 | 7.21E-01 | no | | regulatory_region_variant | MODIFIER |
| 8 | 36438275 | AX-170326517 | 0 | C | G | 0.149 | 5.31E-02 | 1.50E-01 | 2.77E+00 | 2.81E+00 | 7.21E-01 | no | | intergenic_variant | MODIFIER |
| 8 | 36444109 | AX-170167942 | 0 | T | G | 0.149 | 5.31E-02 | 1.50E-01 | 2.77E+00 | 2.81E+00 | 7.21E-01 | no | | intergenic_variant | MODIFIER |
| 8 | 36444325 | AX-170346042 | 0 | G | A | 0.149 | 5.31E-02 | 1.50E-01 | 2.77E+00 | 2.81E+00 | 7.21E-01 | no | | intergenic_variant | MODIFIER |
| 8 | 36474981 | AX-170153193 | 0 | A | G | 0.149 | 5.31E-02 | 1.50E-01 | 2.77E+00 | 2.81E+00 | 7.21E-01 | no | Trmt9b | intron_variant | MODIFIER |
| 8 | 36501243 | AX-170488200 | 0 | C | G | 0.149 | 5.31E-02 | 1.50E-01 | 2.77E+00 | 2.81E+00 | 7.21E-01 | no | Trmt9b | upstream_gene_variant | MODIFIER |
| 8 | 36501410 | AX-97850898 | 0 | T | A | 0.149 | 5.31E-02 | 1.50E-01 | 2.77E+00 | 2.81E+00 | 7.21E-01 | no | Trmt9b | upstream_gene_variant | MODIFIER |
| 8 | 36504861 | AX-170247426 | 0 | A | G | 0.149 | 5.31E-02 | 1.50E-01 | 2.77E+00 | 2.81E+00 | 7.21E-01 | no | Trmt9b | upstream_gene_variant | MODIFIER |
| 8 | 36507674 | AX-170452473 | 0 | A | G | 0.149 | 5.31E-02 | 1.50E-01 | 2.77E+00 | 2.81E+00 | 7.21E-01 | no | Trmt9b | intron_variant,non_coding_transcript_variant | MODIFIER |
| 8 | 36510743 | AX-170234128 | 0 | A | G | 0.149 | 5.31E-02 | 1.50E-01 | 2.77E+00 | 2.81E+00 | 7.21E-01 | no | Trmt9b | intron_variant,non_coding_transcript_variant | MODIFIER |
| 8 | 36510831 | AX-170379763 | 0 | C | T | 0.149 | 5.31E-02 | 1.50E-01 | 2.77E+00 | 2.81E+00 | 7.21E-01 | no | Trmt9b | intron_variant,non_coding_transcript_variant | MODIFIER |
| 8 | 36548326 | AX-170304201 | 0 | C | T | 0.149 | 5.31E-02 | 1.50E-01 | 2.77E+00 | 2.81E+00 | 7.21E-01 | no | | regulatory_region_variant | MODIFIER |
| 8 | 36548404 | AX-170030512 | 0 | A | G | 0.149 | 5.31E-02 | 1.50E-01 | 2.77E+00 | 2.81E+00 | 7.21E-01 | no | | regulatory_region_variant | MODIFIER |
| 8 | 36548468 | AX-170030513 | 0 | T | C | 0.149 | 5.31E-02 | 1.50E-01 | 2.77E+00 | 2.81E+00 | 7.21E-01 | no | | regulatory_region_variant | MODIFIER |
| 8 | 36558714 | AX-170199442 | 0 | A | G | 0.149 | 5.31E-02 | 1.50E-01 | 2.77E+00 | 2.81E+00 | 7.21E-01 | no | | intergenic_variant | MODIFIER |
| 8 | 36559606 | AX-170174027 | 0 | C | T | 0.149 | 5.31E-02 | 1.50E-01 | 2.77E+00 | 2.81E+00 | 7.21E-01 | no | | regulatory_region_variant | MODIFIER |
| 8 | 36567931 | AX-170142809 | 0 | T | C | 0.149 | 5.31E-02 | 1.50E-01 | 2.77E+00 | 2.81E+00 | 7.21E-01 | no | Dlc1 | 3_prime_UTR_variant | MODIFIER |
| 8 | 36573307 | AX-170079767 | 0 | C | G | 0.149 | 5.31E-02 | 1.50E-01 | 2.77E+00 | 2.81E+00 | 7.21E-01 | no | Dlc1 | intron_variant | MODIFIER |
| 8 | 36580749 | AX-170079770 | 0 | A | C | 0.149 | 5.31E-02 | 1.50E-01 | 2.77E+00 | 2.81E+00 | 7.21E-01 | no | Dlc1 | intron_variant | MODIFIER |
| 8 | 36584410 | AX-170030514 | 0 | T | C | 0.149 | 5.31E-02 | 1.50E-01 | 2.77E+00 | 2.81E+00 | 7.21E-01 | no | Dlc1 | synonymous_variant | LOW |
| 8 | 36613355 | AX-170230373 | 0 | C | A | 0.149 | 5.31E-02 | 1.50E-01 | 2.77E+00 | 2.81E+00 | 7.21E-01 | no | Dlc1 | intron_variant | MODIFIER |
| 8 | 36614633 | AX-170244132 | 0 | C | T | 0.149 | 5.31E-02 | 1.50E-01 | 2.77E+00 | 2.81E+00 | 7.21E-01 | no | Dlc1 | upstream_gene_variant | MODIFIER |
| 8 | 36616465 | AX-170389107 | 0 | G | C | 0.149 | 5.31E-02 | 1.50E-01 | 2.77E+00 | 2.81E+00 | 7.21E-01 | no | Dlc1 | upstream_gene_variant | MODIFIER |
| 8 | 36645806 | AX-170366252 | 0 | A | C | 0.149 | 5.31E-02 | 1.50E-01 | 2.77E+00 | 2.81E+00 | 7.21E-01 | no | Dlc1 | intron_variant | MODIFIER |
| 8 | 36661363 | AX-170283803 | 0 | T | C | 0.149 | 5.31E-02 | 1.50E-01 | 2.77E+00 | 2.81E+00 | 7.21E-01 | no | Dlc1 | intron_variant | MODIFIER |
| 8 | 36675087 | AX-97799970 | 0 | G | A | 0.149 | 5.31E-02 | 1.50E-01 | 2.77E+00 | 2.81E+00 | 7.21E-01 | no | Dlc1 | intron_variant | MODIFIER |
| 8 | 36677122 | AX-170216817 | 0 | G | A | 0.149 | 5.31E-02 | 1.50E-01 | 2.77E+00 | 2.81E+00 | 7.21E-01 | no | Dlc1 | intron_variant | MODIFIER |
| 8 | 36698572 | AX-97847326 | 0 | A | G | 0.149 | 5.31E-02 | 1.50E-01 | 2.77E+00 | 2.81E+00 | 7.21E-01 | no | Dlc1 | intron_variant | MODIFIER |
| 8 | 36704831 | AX-170377296 | 0 | T | C | 0.149 | 5.31E-02 | 1.50E-01 | 2.77E+00 | 2.81E+00 | 7.21E-01 | no | Dlc1 | intron_variant | MODIFIER |
| 8 | 36706963 | AX-170030510 | 0 | A | G | 0.149 | 5.31E-02 | 1.50E-01 | 2.77E+00 | 2.81E+00 | 7.21E-01 | no | Dlc1 | intron_variant | MODIFIER |
| 8 | 36715341 | AX-97839574 | 0 | C | A | 0.149 | 5.31E-02 | 1.50E-01 | 2.77E+00 | 2.81E+00 | 7.21E-01 | no | Dlc1 | intron_variant | MODIFIER |
| 8 | 36751690 | AX-170365421 | 0 | A | G | 0.149 | 5.31E-02 | 1.50E-01 | 2.77E+00 | 2.81E+00 | 7.21E-01 | no | Dlc1 | intron_variant | MODIFIER |
| 8 | 36753937 | AX-170030508 | 0 | G | C | 0.149 | 5.31E-02 | 1.50E-01 | 2.77E+00 | 2.81E+00 | 7.21E-01 | no | Dlc1 | intron_variant | MODIFIER |

| | | | | | | | | | | | | | | | |
|---|---|---|---|---|---|---|---|---|---|---|---|---|---|---|---|
| 8 | 36767384 | AX-170219214 | 0 | C | T | 0.149 | 5.31E-02 | 1.50E-01 | 2.77E+00 | 2.81E+00 | 7.21E-01 | no | Dlc1 | intron_variant | MODIFIER |
| 8 | 36782646 | AX-170383545 | 0 | T | A | 0.149 | 5.31E-02 | 1.50E-01 | 2.77E+00 | 2.81E+00 | 7.21E-01 | no | Dlc1 | intron_variant | MODIFIER |
| 8 | 36787985 | AX-170365934 | 0 | A | G | 0.149 | 5.31E-02 | 1.50E-01 | 2.77E+00 | 2.81E+00 | 7.21E-01 | no | Dlc1 | intron_variant | MODIFIER |
| 8 | 36788253 | AX-170449035 | 0 | A | C | 0.149 | 5.31E-02 | 1.50E-01 | 2.77E+00 | 2.81E+00 | 7.21E-01 | no | Dlc1 | intron_variant | MODIFIER |
| 8 | 36791124 | AX-170524460 | 0 | C | T | 0.149 | 5.31E-02 | 1.50E-01 | 2.77E+00 | 2.81E+00 | 7.21E-01 | no | Dlc1 | intron_variant | MODIFIER |
| 8 | 36801776 | AX-97807022 | 0 | T | C | 0.149 | 5.31E-02 | 1.50E-01 | 2.77E+00 | 2.81E+00 | 7.21E-01 | no | Dlc1 | intron_variant | MODIFIER |
| 8 | 36816037 | AX-170079775 | 0 | T | C | 0.149 | 5.31E-02 | 1.50E-01 | 2.77E+00 | 2.81E+00 | 7.21E-01 | no | Dlc1 | intron_variant | MODIFIER |
| 8 | 36838774 | AX-170030509 | 0 | A | G | 0.149 | 5.31E-02 | 1.50E-01 | 2.77E+00 | 2.81E+00 | 7.21E-01 | no | Dlc1 | intron_variant | MODIFIER |
| 8 | 36838905 | AX-170030516 | 0 | C | G | 0.149 | 5.31E-02 | 1.50E-01 | 2.77E+00 | 2.81E+00 | 7.21E-01 | no | Dlc1 | intron_variant | MODIFIER |
| 8 | 36841350 | AX-97794116 | 0 | C | T | 0.149 | 5.31E-02 | 1.50E-01 | 2.77E+00 | 2.81E+00 | 7.21E-01 | no | Dlc1 | intron_variant | MODIFIER |
| 8 | 36841374 | AX-170338399 | 0 | G | A | 0.149 | 5.31E-02 | 1.50E-01 | 2.77E+00 | 2.81E+00 | 7.21E-01 | no | Dlc1 | intron_variant | MODIFIER |
| 8 | 36863021 | AX-170307830 | 0 | T | C | 0.149 | 5.31E-02 | 1.50E-01 | 2.77E+00 | 2.81E+00 | 7.21E-01 | no | Dlc1 | intron_variant,non_coding_transcript_variant | MODIFIER |
| 8 | 36867688 | AX-170098867 | 0 | T | C | 0.149 | 5.31E-02 | 1.50E-01 | 2.77E+00 | 2.81E+00 | 7.21E-01 | no | Dlc1 | intron_variant,non_coding_transcript_variant | MODIFIER |
| 8 | 36868805 | AX-97838845 | 0 | T | G | 0.149 | 5.31E-02 | 1.50E-01 | 2.77E+00 | 2.81E+00 | 7.21E-01 | no | Dlc1 | intron_variant,non_coding_transcript_variant | MODIFIER |
| 8 | 36891971 | AX-170361163 | 0 | G | T | 0.149 | 5.31E-02 | 1.50E-01 | 2.77E+00 | 2.81E+00 | 7.21E-01 | no | Dlc1 | intron_variant,non_coding_transcript_variant | MODIFIER |
| 8 | 36892939 | AX-170231894 | 0 | A | G | 0.149 | 5.31E-02 | 1.50E-01 | 2.77E+00 | 2.81E+00 | 7.21E-01 | no | Dlc1 | intron_variant,non_coding_transcript_variant | MODIFIER |
| 8 | 36909761 | AX-170133073 | 0 | A | G | 0.149 | 5.31E-02 | 1.50E-01 | 2.77E+00 | 2.81E+00 | 7.21E-01 | no | Dlc1 | intron_variant,non_coding_transcript_variant | MODIFIER |
| 8 | 36912719 | AX-170436605 | 0 | C | T | 0.149 | 5.31E-02 | 1.50E-01 | 2.77E+00 | 2.81E+00 | 7.21E-01 | no | Dlc1 | intron_variant,non_coding_transcript_variant | MODIFIER |
| 8 | 36946330 | AX-170443829 | 0 | T | G | 0.149 | 5.31E-02 | 1.50E-01 | 2.77E+00 | 2.81E+00 | 7.21E-01 | no | Dlc1 | intron_variant,non_coding_transcript_variant | MODIFIER |
| 8 | 36953665 | AX-97845821 | 0 | G | T | 0.149 | 5.31E-02 | 1.50E-01 | 2.77E+00 | 2.81E+00 | 7.21E-01 | no | Dlc1 | upstream_gene_variant | MODIFIER |
| 8 | 36953943 | AX-170310556 | 0 | T | C | 0.149 | 5.31E-02 | 1.50E-01 | 2.77E+00 | 2.81E+00 | 7.21E-01 | no | Dlc1 | upstream_gene_variant | MODIFIER |
| 8 | 36980411 | AX-170406555 | 0 | T | C | 0.149 | 5.31E-02 | 1.50E-01 | 2.77E+00 | 2.81E+00 | 7.21E-01 | no | | intergenic_variant | MODIFIER |
| 8 | 36986274 | AX-170508432 | 0 | A | G | 0.149 | 5.31E-02 | 1.50E-01 | 2.77E+00 | 2.81E+00 | 7.21E-01 | no | G630064G18Rik | non_coding_transcript_exon_variant | MODIFIER |
| 8 | 37022421 | AX-170090083 | 0 | A | G | 0.149 | 5.31E-02 | 1.50E-01 | 2.77E+00 | 2.81E+00 | 7.21E-01 | no | G630064G18Rik | intron_variant,non_coding_transcript_variant | MODIFIER |
| 8 | 37024633 | AX-170409718 | 0 | C | G | 0.149 | 5.31E-02 | 1.50E-01 | 2.77E+00 | 2.81E+00 | 7.21E-01 | no | G630064G18Rik | upstream_gene_variant | MODIFIER |
| 8 | 37050804 | AX-170232264 | 0 | C | T | 0.149 | 5.31E-02 | 1.50E-01 | 2.77E+00 | 2.81E+00 | 7.21E-01 | no | | intergenic_variant | MODIFIER |
| 8 | 37051057 | AX-97818938 | 0 | T | C | 0.149 | 5.31E-02 | 1.50E-01 | 2.77E+00 | 2.81E+00 | 7.21E-01 | no | | intergenic_variant | MODIFIER |
| 8 | 37079891 | AX-97798414 | 0 | T | C | 0.149 | 5.31E-02 | 1.50E-01 | 2.77E+00 | 2.81E+00 | 7.21E-01 | no | | intergenic_variant | MODIFIER |
| 8 | 37094843 | AX-170202176 | 0 | A | C | 0.149 | 5.31E-02 | 1.50E-01 | 2.77E+00 | 2.81E+00 | 7.21E-01 | no | Gm8342 | non_coding_transcript_exon_variant | MODIFIER |
| 8 | 37100559 | AX-170530408 | 0 | T | C | 0.149 | 5.31E-02 | 1.50E-01 | 2.77E+00 | 2.81E+00 | 7.21E-01 | no | Gm3433 | upstream_gene_variant | MODIFIER |
| 8 | 37107176 | AX-170531376 | 0 | T | C | 0.149 | 5.31E-02 | 1.50E-01 | 2.77E+00 | 2.81E+00 | 7.21E-01 | no | Gm3433 | downstream_gene_variant | MODIFIER |
| 8 | 37109672 | AX-170490148 | 0 | G | C | 0.149 | 5.31E-02 | 1.50E-01 | 2.77E+00 | 2.81E+00 | 7.21E-01 | no | Gm3433 | downstream_gene_variant | MODIFIER |
| 8 | 37136397 | AX-170506991 | 0 | G | C | 0.149 | 5.31E-02 | 1.50E-01 | 2.77E+00 | 2.81E+00 | 7.21E-01 | no | Gm45524 | intron_variant,non_coding_transcript_variant | MODIFIER |
| 8 | 37161835 | AX-170434738 | 0 | A | G | 0.149 | 5.31E-02 | 1.50E-01 | 2.77E+00 | 2.81E+00 | 7.21E-01 | no | Gm45524 | intron_variant,non_coding_transcript_variant | MODIFIER |
| 8 | 37167855 | AX-170264312 | 0 | G | A | 0.149 | 5.31E-02 | 1.50E-01 | 2.77E+00 | 2.81E+00 | 7.21E-01 | no | Gm45524 | intron_variant,non_coding_transcript_variant | MODIFIER |
| 8 | 37178169 | AX-170399298 | 0 | G | A | 0.149 | 5.31E-02 | 1.50E-01 | 2.77E+00 | 2.81E+00 | 7.21E-01 | no | Gm45524 | intron_variant,non_coding_transcript_variant | MODIFIER |
| 8 | 37195450 | AX-170096228 | 0 | A | C | 0.149 | 5.31E-02 | 1.50E-01 | 2.77E+00 | 2.81E+00 | 7.21E-01 | no | | intergenic_variant | MODIFIER |
| 8 | 37200170 | AX-170224955 | 0 | A | T | 0.149 | 5.31E-02 | 1.50E-01 | 2.77E+00 | 2.81E+00 | 7.21E-01 | no | | intergenic_variant | MODIFIER |
| 8 | 37206861 | AX-170468160 | 0 | G | T | 0.149 | 5.31E-02 | 1.50E-01 | 2.77E+00 | 2.81E+00 | 7.21E-01 | no | | intergenic_variant | MODIFIER |
| 8 | 37213467 | AX-170392066 | 0 | A | G | 0.149 | 5.31E-02 | 1.50E-01 | 2.77E+00 | 2.81E+00 | 7.21E-01 | no | | intergenic_variant | MODIFIER |
| 8 | 37232603 | AX-97803353 | 0 | T | G | 0.149 | 5.31E-02 | 1.50E-01 | 2.77E+00 | 2.81E+00 | 7.21E-01 | no | | intergenic_variant | MODIFIER |
| 8 | 37264674 | AX-170477723 | 0 | T | G | 0.149 | 5.31E-02 | 1.50E-01 | 2.77E+00 | 2.81E+00 | 7.21E-01 | no | | intergenic_variant | MODIFIER |
| 8 | 37267197 | AX-170519590 | 0 | A | G | 0.149 | 5.31E-02 | 1.50E-01 | 2.77E+00 | 2.81E+00 | 7.21E-01 | no | | intergenic_variant | MODIFIER |
| 8 | 37274041 | AX-170170233 | 0 | G | C | 0.149 | 5.31E-02 | 1.50E-01 | 2.77E+00 | 2.81E+00 | 7.21E-01 | no | | intergenic_variant | MODIFIER |
| 8 | 37277202 | AX-170410666 | 0 | T | C | 0.149 | 5.31E-02 | 1.50E-01 | 2.77E+00 | 2.81E+00 | 7.21E-01 | no | | intergenic_variant | MODIFIER |
| 8 | 37277836 | AX-170296133 | 0 | G | T | 0.149 | 5.31E-02 | 1.50E-01 | 2.77E+00 | 2.81E+00 | 7.21E-01 | no | | intergenic_variant | MODIFIER |
| 8 | 37279845 | AX-170198964 | 0 | C | T | 0.149 | 5.31E-02 | 1.50E-01 | 2.77E+00 | 2.81E+00 | 7.21E-01 | no | | intergenic_variant | MODIFIER |
| 8 | 37298238 | AX-170198687 | 0 | T | C | 0.149 | 5.31E-02 | 1.50E-01 | 2.77E+00 | 2.81E+00 | 7.21E-01 | no | | intergenic_variant | MODIFIER |
| 8 | 37301806 | AX-170090084 | 0 | C | T | 0.149 | 5.31E-02 | 1.50E-01 | 2.77E+00 | 2.81E+00 | 7.21E-01 | no | | intergenic_variant | MODIFIER |
| 8 | 37341345 | AX-170177644 | 0 | A | G | 0.149 | 5.31E-02 | 1.50E-01 | 2.77E+00 | 2.81E+00 | 7.21E-01 | no | Gm45368 | intron_variant,non_coding_transcript_variant | MODIFIER |
| 8 | 37347675 | AX-97803534 | 0 | A | C | 0.149 | 5.31E-02 | 1.50E-01 | 2.77E+00 | 2.81E+00 | 7.21E-01 | no | Gm45368 | downstream_gene_variant | MODIFIER |
| 8 | 37347699 | AX-170030519 | 0 | A | G | 0.149 | 5.31E-02 | 1.50E-01 | 2.77E+00 | 2.81E+00 | 7.21E-01 | no | Gm45368 | downstream_gene_variant | MODIFIER |
| 8 | 37350218 | AX-170030520 | 0 | A | G | 0.149 | 5.31E-02 | 1.50E-01 | 2.77E+00 | 2.81E+00 | 7.21E-01 | no | Gm45368 | intron_variant,non_coding_transcript_variant | MODIFIER |
| 8 | 37370966 | AX-170158894 | 0 | G | C | 0.149 | 5.31E-02 | 1.50E-01 | 2.77E+00 | 2.81E+00 | 7.21E-01 | no | Gm6203 | downstream_gene_variant | MODIFIER |

| | | | | | | | | | | | | | | | |
|---|---|---|---|---|---|---|---|---|---|---|---|---|---|---|---|
| 8 | 37370988 | AX-170115672 | 0 | C | T | 0.149 | 5.31E-02 | 1.50E-01 | 2.77E+00 | 2.81E+00 | 7.21E-01 | no | Gm6203 | downstream_gene_variant | MODIFIER |
| 8 | 37392690 | AX-170030521 | 0 | A | G | 0.149 | 5.31E-02 | 1.50E-01 | 2.77E+00 | 2.81E+00 | 7.21E-01 | no | | intergenic_variant | MODIFIER |
| 8 | 37442587 | AX-170030517 | 0 | G | A | 0.149 | 5.31E-02 | 1.50E-01 | 2.77E+00 | 2.81E+00 | 7.21E-01 | no | | intergenic_variant | MODIFIER |
| 8 | 37468392 | AX-170079786 | 0 | C | A | 0.149 | 5.31E-02 | 1.50E-01 | 2.77E+00 | 2.81E+00 | 7.21E-01 | no | | intergenic_variant | MODIFIER |
| 8 | 37468673 | AX-170079789 | 0 | A | T | 0.149 | 5.31E-02 | 1.50E-01 | 2.77E+00 | 2.81E+00 | 7.21E-01 | no | | intergenic_variant | MODIFIER |
| 8 | 37469511 | AX-170522492 | 0 | T | C | 0.149 | 5.31E-02 | 1.50E-01 | 2.77E+00 | 2.81E+00 | 7.21E-01 | no | | intergenic_variant | MODIFIER |
| 8 | 37471814 | AX-170398393 | 0 | A | G | 0.149 | 5.31E-02 | 1.50E-01 | 2.77E+00 | 2.81E+00 | 7.21E-01 | no | | intergenic_variant | MODIFIER |
| 8 | 37473543 | AX-97847396 | 0 | G | A | 0.149 | 5.31E-02 | 1.50E-01 | 2.77E+00 | 2.81E+00 | 7.21E-01 | no | | intergenic_variant | MODIFIER |
| 8 | 37505366 | AX-170079784 | 0 | A | G | 0.149 | 5.31E-02 | 1.50E-01 | 2.77E+00 | 2.81E+00 | 7.21E-01 | no | | intergenic_variant | MODIFIER |
| 8 | 37507923 | AX-170420095 | 0 | A | G | 0.149 | 5.31E-02 | 1.50E-01 | 2.77E+00 | 2.81E+00 | 7.21E-01 | no | | intergenic_variant | MODIFIER |
| 8 | 37508314 | AX-170464791 | 0 | A | G | 0.149 | 5.31E-02 | 1.50E-01 | 2.77E+00 | 2.81E+00 | 7.21E-01 | no | | intergenic_variant | MODIFIER |
| 8 | 37510246 | AX-170223081 | 0 | G | T | 0.149 | 5.31E-02 | 1.50E-01 | 2.77E+00 | 2.81E+00 | 7.21E-01 | no | | intergenic_variant | MODIFIER |
| 8 | 37514177 | AX-170165606 | 0 | A | G | 0.149 | 5.31E-02 | 1.50E-01 | 2.77E+00 | 2.81E+00 | 7.21E-01 | no | | intergenic_variant | MODIFIER |
| 8 | 37541538 | AX-97803536 | 0 | A | G | 0.149 | 5.31E-02 | 1.50E-01 | 2.77E+00 | 2.81E+00 | 7.21E-01 | no | Sgcz | intron_variant | MODIFIER |
| 8 | 37552848 | AX-170477820 | 0 | C | T | 0.149 | 5.31E-02 | 1.50E-01 | 2.77E+00 | 2.81E+00 | 7.21E-01 | no | Sgcz | intron_variant | MODIFIER |
| 8 | 37566013 | AX-97852609 | 0 | A | G | 0.149 | 5.31E-02 | 1.50E-01 | 2.77E+00 | 2.81E+00 | 7.21E-01 | no | Sgcz | intron_variant | MODIFIER |
| 8 | 37568142 | AX-170141759 | 0 | C | T | 0.149 | 5.31E-02 | 1.50E-01 | 2.77E+00 | 2.81E+00 | 7.21E-01 | no | Sgcz | intron_variant | MODIFIER |
| 8 | 37587768 | AX-170413502 | 0 | A | G | 0.149 | 5.31E-02 | 1.50E-01 | 2.77E+00 | 2.81E+00 | 7.21E-01 | no | Sgcz | intron_variant | MODIFIER |
| 8 | 37589184 | AX-170298529 | 0 | A | C | 0.149 | 5.31E-02 | 1.50E-01 | 2.77E+00 | 2.81E+00 | 7.21E-01 | no | Sgcz | intron_variant | MODIFIER |
| 8 | 37591920 | AX-170256695 | 0 | T | C | 0.149 | 5.31E-02 | 1.50E-01 | 2.77E+00 | 2.81E+00 | 7.21E-01 | no | Sgcz | intron_variant | MODIFIER |
| 8 | 37593298 | AX-170490107 | 0 | G | A | 0.149 | 5.31E-02 | 1.50E-01 | 2.77E+00 | 2.81E+00 | 7.21E-01 | no | Sgcz | intron_variant | MODIFIER |
| 8 | 37707157 | AX-170244868 | 0 | C | T | 0.149 | 5.31E-02 | 1.50E-01 | 2.77E+00 | 2.81E+00 | 7.21E-01 | no | Sgcz | intron_variant | MODIFIER |
| 8 | 37726857 | AX-170176226 | 0 | A | T | 0.149 | 5.31E-02 | 1.50E-01 | 2.77E+00 | 2.81E+00 | 7.21E-01 | no | Sgcz | intron_variant | MODIFIER |
| 8 | 37768314 | AX-170387863 | 0 | G | C | 0.149 | 5.31E-02 | 1.50E-01 | 2.77E+00 | 2.81E+00 | 7.21E-01 | no | Sgcz | intron_variant | MODIFIER |
| 8 | 37787619 | AX-170079793 | 0 | A | T | 0.149 | 5.31E-02 | 1.50E-01 | 2.77E+00 | 2.81E+00 | 7.21E-01 | no | Gm16205 | downstream_gene_variant | MODIFIER |
| 8 | 37795344 | AX-97844304 | 0 | C | T | 0.149 | 5.31E-02 | 1.50E-01 | 2.77E+00 | 2.81E+00 | 7.21E-01 | no | Sgcz | intron_variant | MODIFIER |
| 8 | 37832618 | AX-170239151 | 0 | A | G | 0.149 | 5.31E-02 | 1.50E-01 | 2.77E+00 | 2.81E+00 | 7.21E-01 | no | Sgcz | intron_variant | MODIFIER |
| 8 | 37877238 | AX-170294026 | 0 | T | A | 0.149 | 5.31E-02 | 1.50E-01 | 2.77E+00 | 2.81E+00 | 7.21E-01 | no | Sgcz | intron_variant | MODIFIER |
| 8 | 37881892 | AX-170264753 | 0 | G | A | 0.149 | 5.31E-02 | 1.50E-01 | 2.77E+00 | 2.81E+00 | 7.21E-01 | no | Sgcz | intron_variant | MODIFIER |
| 8 | 37891214 | AX-170451257 | 0 | G | A | 0.149 | 5.31E-02 | 1.50E-01 | 2.77E+00 | 2.81E+00 | 7.21E-01 | no | Sgcz | intron_variant | MODIFIER |
| 8 | 37905413 | AX-170122591 | 0 | G | A | 0.149 | 5.31E-02 | 1.50E-01 | 2.77E+00 | 2.81E+00 | 7.21E-01 | no | Sgcz | intron_variant | MODIFIER |
| 8 | 37917413 | AX-97822121 | 0 | A | G | 0.149 | 5.31E-02 | 1.50E-01 | 2.77E+00 | 2.81E+00 | 7.21E-01 | no | Sgcz | intron_variant | MODIFIER |
| 8 | 37949788 | AX-170030523 | 0 | A | G | 0.149 | 5.31E-02 | 1.50E-01 | 2.77E+00 | 2.81E+00 | 7.21E-01 | no | Sgcz | intron_variant | MODIFIER |
| 8 | 37963452 | AX-170126027 | 0 | T | C | 0.149 | 5.31E-02 | 1.50E-01 | 2.77E+00 | 2.81E+00 | 7.21E-01 | no | Sgcz | intron_variant | MODIFIER |
| 8 | 37965483 | AX-170330910 | 0 | A | G | 0.149 | 5.31E-02 | 1.50E-01 | 2.77E+00 | 2.81E+00 | 7.21E-01 | no | Sgcz | intron_variant | MODIFIER |
| 8 | 37982438 | AX-170079803 | 0 | T | C | 0.149 | 5.31E-02 | 1.50E-01 | 2.77E+00 | 2.81E+00 | 7.21E-01 | no | Sgcz | intron_variant | MODIFIER |
| 8 | 37983454 | AX-170079802 | 0 | T | A | 0.149 | 5.31E-02 | 1.50E-01 | 2.77E+00 | 2.81E+00 | 7.21E-01 | no | Sgcz | intron_variant | MODIFIER |
| 8 | 37992178 | AX-170522756 | 0 | T | C | 0.149 | 5.31E-02 | 1.50E-01 | 2.77E+00 | 2.81E+00 | 7.21E-01 | no | Sgcz | intron_variant | MODIFIER |
| 8 | 38012849 | AX-170138204 | 0 | A | G | 0.149 | 5.31E-02 | 1.50E-01 | 2.77E+00 | 2.81E+00 | 7.21E-01 | no | Sgcz | intron_variant | MODIFIER |
| 8 | 38027839 | AX-170326865 | 0 | T | G | 0.149 | 5.31E-02 | 1.50E-01 | 2.77E+00 | 2.81E+00 | 7.21E-01 | no | Sgcz | intron_variant | MODIFIER |
| 8 | 38029027 | AX-170524100 | 0 | T | A | 0.149 | 5.31E-02 | 1.50E-01 | 2.77E+00 | 2.81E+00 | 7.21E-01 | no | Sgcz | intron_variant | MODIFIER |
| 8 | 38057252 | AX-97836597 | 0 | C | T | 0.149 | 5.31E-02 | 1.50E-01 | 2.77E+00 | 2.81E+00 | 7.21E-01 | no | Sgcz | intron_variant | MODIFIER |
| 8 | 38080716 | AX-170328515 | 0 | T | C | 0.149 | 5.31E-02 | 1.50E-01 | 2.77E+00 | 2.81E+00 | 7.21E-01 | no | Gm16204 | downstream_gene_variant | MODIFIER |
| 8 | 38089337 | AX-170164159 | 0 | C | T | 0.149 | 5.31E-02 | 1.50E-01 | 2.77E+00 | 2.81E+00 | 7.21E-01 | no | Sgcz | intron_variant | MODIFIER |
| 8 | 38116379 | AX-170154015 | 0 | A | G | 0.149 | 5.31E-02 | 1.50E-01 | 2.77E+00 | 2.81E+00 | 7.21E-01 | no | Sgcz | intron_variant | MODIFIER |
| 8 | 38128337 | AX-170321831 | 0 | T | C | 0.149 | 5.31E-02 | 1.50E-01 | 2.77E+00 | 2.81E+00 | 7.21E-01 | no | Sgcz | intron_variant | MODIFIER |
| 8 | 38150201 | AX-170169318 | 0 | C | G | 0.149 | 5.31E-02 | 1.50E-01 | 2.77E+00 | 2.81E+00 | 7.21E-01 | no | Sgcz | intron_variant | MODIFIER |
| 8 | 38150260 | AX-170488794 | 0 | A | G | 0.149 | 5.31E-02 | 1.50E-01 | 2.77E+00 | 2.81E+00 | 7.21E-01 | no | Sgcz | intron_variant | MODIFIER |
| 8 | 38191910 | AX-170299439 | 0 | A | G | 0.149 | 5.31E-02 | 1.50E-01 | 2.77E+00 | 2.81E+00 | 7.21E-01 | no | Sgcz | intron_variant | MODIFIER |
| 8 | 38229846 | AX-170152533 | 0 | T | C | 0.149 | 5.31E-02 | 1.50E-01 | 2.77E+00 | 2.81E+00 | 7.21E-01 | no | Sgcz | intron_variant | MODIFIER |
| 8 | 38232187 | AX-170328554 | 0 | T | C | 0.149 | 5.31E-02 | 1.50E-01 | 2.77E+00 | 2.81E+00 | 7.21E-01 | no | Sgcz | intron_variant | MODIFIER |
| 8 | 38259912 | AX-170079809 | 0 | T | C | 0.149 | 5.31E-02 | 1.50E-01 | 2.77E+00 | 2.81E+00 | 7.21E-01 | no | Sgcz | intron_variant | MODIFIER |
| 8 | 38263262 | AX-170442370 | 0 | G | A | 0.149 | 5.31E-02 | 1.50E-01 | 2.77E+00 | 2.81E+00 | 7.21E-01 | no | Sgcz | intron_variant | MODIFIER |
| 8 | 38321288 | AX-170526988 | 0 | G | T | 0.149 | 5.31E-02 | 1.50E-01 | 2.77E+00 | 2.81E+00 | 7.21E-01 | no | Sgcz | intron_variant | MODIFIER |

| | | | | | | | | | | | | | | | |
|---|---|---|---|---|---|---|---|---|---|---|---|---|---|---|---|
| 8 | 38336626 | AX-170370580 | 0 | T | A | 0.149 | 5.31E-02 | 1.50E-01 | 2.77E+00 | 2.81E+00 | 7.21E-01 | no | Sgcz | intron_variant | MODIFIER |
| 8 | 38336687 | AX-97837574 | 0 | A | C | 0.149 | 5.31E-02 | 1.50E-01 | 2.77E+00 | 2.81E+00 | 7.21E-01 | no | Sgcz | intron_variant | MODIFIER |
| 8 | 38338738 | AX-170144078 | 0 | A | G | 0.149 | 5.31E-02 | 1.50E-01 | 2.77E+00 | 2.81E+00 | 7.21E-01 | no | Sgcz | intron_variant | MODIFIER |
| 8 | 38350059 | AX-170202464 | 0 | A | C | 0.149 | 5.31E-02 | 1.50E-01 | 2.77E+00 | 2.81E+00 | 7.21E-01 | no | Sgcz | intron_variant | MODIFIER |
| 8 | 38371404 | AX-170506975 | 0 | C | T | 0.149 | 5.31E-02 | 1.50E-01 | 2.77E+00 | 2.81E+00 | 7.21E-01 | no | Sgcz | intron_variant | MODIFIER |
| 8 | 38371551 | AX-170454453 | 0 | C | T | 0.149 | 5.31E-02 | 1.50E-01 | 2.77E+00 | 2.81E+00 | 7.21E-01 | no | Sgcz | intron_variant | MODIFIER |
| 8 | 38371636 | AX-170109755 | 0 | G | A | 0.149 | 5.31E-02 | 1.50E-01 | 2.77E+00 | 2.81E+00 | 7.21E-01 | no | Sgcz | intron_variant | MODIFIER |
| 8 | 38384350 | AX-170102551 | 0 | G | A | 0.149 | 5.31E-02 | 1.50E-01 | 2.77E+00 | 2.81E+00 | 7.21E-01 | no | Sgcz | intron_variant | MODIFIER |
| 8 | 38398959 | AX-170448695 | 0 | G | A | 0.149 | 5.31E-02 | 1.50E-01 | 2.77E+00 | 2.81E+00 | 7.21E-01 | no | Sgcz | intron_variant | MODIFIER |
| 8 | 38416933 | AX-170200012 | 0 | T | A | 0.149 | 5.31E-02 | 1.50E-01 | 2.77E+00 | 2.81E+00 | 7.21E-01 | no | Sgcz | intron_variant | MODIFIER |
| 8 | 38427075 | AX-97852894 | 0 | C | T | 0.149 | 5.31E-02 | 1.50E-01 | 2.77E+00 | 2.81E+00 | 7.21E-01 | no | Sgcz | intron_variant | MODIFIER |
| 8 | 38457124 | AX-97836595 | 0 | A | G | 0.149 | 5.31E-02 | 1.50E-01 | 2.77E+00 | 2.81E+00 | 7.21E-01 | no | Sgcz | intron_variant | MODIFIER |
| 8 | 38457143 | AX-170153039 | 0 | C | T | 0.149 | 5.31E-02 | 1.50E-01 | 2.77E+00 | 2.81E+00 | 7.21E-01 | no | Sgcz | intron_variant | MODIFIER |
| 8 | 38468948 | AX-170172800 | 0 | T | C | 0.149 | 5.31E-02 | 1.50E-01 | 2.77E+00 | 2.81E+00 | 7.21E-01 | no | Sgcz | intron_variant | MODIFIER |
| 8 | 38485826 | AX-170413623 | 0 | A | G | 0.149 | 5.31E-02 | 1.50E-01 | 2.77E+00 | 2.81E+00 | 7.21E-01 | no | Sgcz | intron_variant | MODIFIER |
| 8 | 38492654 | AX-170412864 | 0 | G | A | 0.149 | 5.31E-02 | 1.50E-01 | 2.77E+00 | 2.81E+00 | 7.21E-01 | no | Sgcz | intron_variant | MODIFIER |
| 8 | 38511065 | AX-170504297 | 0 | C | T | 0.149 | 5.31E-02 | 1.50E-01 | 2.77E+00 | 2.81E+00 | 7.21E-01 | no | Sgcz | intron_variant | MODIFIER |
| 8 | 38512702 | AX-170315647 | 0 | T | A | 0.149 | 5.31E-02 | 1.50E-01 | 2.77E+00 | 2.81E+00 | 7.21E-01 | no | Sgcz | intron_variant | MODIFIER |
| 8 | 38545771 | AX-170294659 | 0 | T | C | 0.149 | 5.31E-02 | 1.50E-01 | 2.77E+00 | 2.81E+00 | 7.21E-01 | no | Sgcz | intron_variant | MODIFIER |
| 8 | 38545825 | AX-170030529 | 0 | A | G | 0.149 | 5.31E-02 | 1.50E-01 | 2.77E+00 | 2.81E+00 | 7.21E-01 | no | Sgcz | intron_variant | MODIFIER |
| 8 | 38555376 | AX-170506475 | 0 | T | C | 0.149 | 5.31E-02 | 1.50E-01 | 2.77E+00 | 2.81E+00 | 7.21E-01 | no | Sgcz | intron_variant | MODIFIER |
| 8 | 38600762 | AX-170090085 | 0 | C | T | 0.149 | 5.31E-02 | 1.50E-01 | 2.77E+00 | 2.81E+00 | 7.21E-01 | no | Sgcz | intron_variant | MODIFIER |
| 8 | 38601123 | AX-170183911 | 0 | A | G | 0.149 | 5.31E-02 | 1.50E-01 | 2.77E+00 | 2.81E+00 | 7.21E-01 | no | Sgcz | intron_variant | MODIFIER |
| 8 | 38601429 | AX-170397489 | 0 | T | C | 0.149 | 5.31E-02 | 1.50E-01 | 2.77E+00 | 2.81E+00 | 7.21E-01 | no | Sgcz | intron_variant | MODIFIER |
| 8 | 38605162 | AX-170030526 | 0 | G | T | 0.149 | 5.31E-02 | 1.50E-01 | 2.77E+00 | 2.81E+00 | 7.21E-01 | no | Sgcz | intron_variant | MODIFIER |
| 8 | 38611549 | AX-170340133 | 0 | C | T | 0.149 | 5.31E-02 | 1.50E-01 | 2.77E+00 | 2.81E+00 | 7.21E-01 | no | Sgcz | intron_variant | MODIFIER |
| 8 | 38624141 | AX-170474607 | 0 | A | G | 0.149 | 5.31E-02 | 1.50E-01 | 2.77E+00 | 2.81E+00 | 7.21E-01 | no | Sgcz | intron_variant | MODIFIER |
| 8 | 38630627 | AX-97836602 | 0 | A | G | 0.149 | 5.31E-02 | 1.50E-01 | 2.77E+00 | 2.81E+00 | 7.21E-01 | no | Sgcz | intron_variant | MODIFIER |
| 8 | 38630685 | AX-170413670 | 0 | T | C | 0.149 | 5.31E-02 | 1.50E-01 | 2.77E+00 | 2.81E+00 | 7.21E-01 | no | Sgcz | intron_variant | MODIFIER |
| 8 | 38657511 | AX-170139350 | 0 | G | C | 0.149 | 5.31E-02 | 1.50E-01 | 2.77E+00 | 2.81E+00 | 7.21E-01 | no | Sgcz | intron_variant | MODIFIER |
| 8 | 38669925 | AX-170524035 | 0 | A | G | 0.149 | 5.31E-02 | 1.50E-01 | 2.77E+00 | 2.81E+00 | 7.21E-01 | no | Gm40493 | intron_variant,non_coding_transcript_variant | MODIFIER |
| 8 | 38685025 | AX-170390313 | 0 | A | G | 0.149 | 5.31E-02 | 1.50E-01 | 2.77E+00 | 2.81E+00 | 7.21E-01 | no | | intergenic_variant | MODIFIER |
| 8 | 38696751 | AX-97849114 | 0 | G | T | 0.149 | 5.31E-02 | 1.50E-01 | 2.77E+00 | 2.81E+00 | 7.21E-01 | no | | intergenic_variant | MODIFIER |
| 8 | 38707629 | AX-170504552 | 0 | T | C | 0.149 | 5.31E-02 | 1.50E-01 | 2.77E+00 | 2.81E+00 | 7.21E-01 | no | | intergenic_variant | MODIFIER |
| 8 | 38739785 | AX-170079815 | 0 | A | C | 0.149 | 5.31E-02 | 1.50E-01 | 2.77E+00 | 2.81E+00 | 7.21E-01 | no | | intergenic_variant | MODIFIER |
| 8 | 38759182 | AX-170079816 | 0 | A | G | 0.149 | 5.31E-02 | 1.50E-01 | 2.77E+00 | 2.81E+00 | 7.21E-01 | no | | intergenic_variant | MODIFIER |
| 8 | 38759450 | AX-170532329 | 0 | T | C | 0.149 | 5.31E-02 | 1.50E-01 | 2.77E+00 | 2.81E+00 | 7.21E-01 | no | | intergenic_variant | MODIFIER |
| 8 | 38763418 | AX-170239037 | 0 | C | T | 0.149 | 5.31E-02 | 1.50E-01 | 2.77E+00 | 2.81E+00 | 7.21E-01 | no | | intergenic_variant | MODIFIER |
| 8 | 38790369 | AX-170273753 | 0 | G | A | 0.149 | 5.31E-02 | 1.50E-01 | 2.77E+00 | 2.81E+00 | 7.21E-01 | no | | intergenic_variant | MODIFIER |
| 8 | 38813968 | AX-170238160 | 0 | A | G | 0.149 | 5.31E-02 | 1.50E-01 | 2.77E+00 | 2.81E+00 | 7.21E-01 | no | | intergenic_variant | MODIFIER |
| 8 | 38813996 | AX-170156134 | 0 | C | T | 0.149 | 5.31E-02 | 1.50E-01 | 2.77E+00 | 2.81E+00 | 7.21E-01 | no | | intergenic_variant | MODIFIER |
| 8 | 38837474 | AX-170391115 | 0 | A | C | 0.149 | 5.31E-02 | 1.50E-01 | 2.77E+00 | 2.81E+00 | 7.21E-01 | no | | intergenic_variant | MODIFIER |
| 8 | 38840209 | AX-170405344 | 0 | G | C | 0.149 | 5.31E-02 | 1.50E-01 | 2.77E+00 | 2.81E+00 | 7.21E-01 | no | | intergenic_variant | MODIFIER |
| 8 | 38861327 | AX-170130551 | 0 | A | G | 0.149 | 5.31E-02 | 1.50E-01 | 2.77E+00 | 2.81E+00 | 7.21E-01 | no | | intergenic_variant | MODIFIER |
| 8 | 38867330 | AX-170276026 | 0 | T | G | 0.149 | 5.31E-02 | 1.50E-01 | 2.77E+00 | 2.81E+00 | 7.21E-01 | no | | intergenic_variant | MODIFIER |
| 8 | 38892171 | AX-170220799 | 0 | C | T | 0.149 | 5.31E-02 | 1.50E-01 | 2.77E+00 | 2.81E+00 | 7.21E-01 | no | | intergenic_variant | MODIFIER |
| 8 | 38895585 | AX-170474095 | 0 | G | C | 0.149 | 5.31E-02 | 1.50E-01 | 2.77E+00 | 2.81E+00 | 7.21E-01 | no | | intergenic_variant | MODIFIER |
| 8 | 38909640 | AX-170318222 | 0 | A | G | 0.149 | 5.31E-02 | 1.50E-01 | 2.77E+00 | 2.81E+00 | 7.21E-01 | no | | intergenic_variant | MODIFIER |
| 8 | 38909878 | AX-170390145 | 0 | A | T | 0.149 | 5.31E-02 | 1.50E-01 | 2.77E+00 | 2.81E+00 | 7.21E-01 | no | | intergenic_variant | MODIFIER |
| 8 | 38910099 | AX-97787448 | 0 | T | C | 0.149 | 5.31E-02 | 1.50E-01 | 2.77E+00 | 2.81E+00 | 7.21E-01 | no | | intergenic_variant | MODIFIER |
| 8 | 38945017 | AX-170109042 | 0 | G | A | 0.149 | 5.31E-02 | 1.50E-01 | 2.77E+00 | 2.81E+00 | 7.21E-01 | no | | intergenic_variant | MODIFIER |
| 8 | 38945343 | AX-170391165 | 0 | C | T | 0.149 | 5.31E-02 | 1.50E-01 | 2.77E+00 | 2.81E+00 | 7.21E-01 | no | | intergenic_variant | MODIFIER |
| 8 | 38979291 | AX-170256102 | 0 | C | T | 0.149 | 5.31E-02 | 1.50E-01 | 2.77E+00 | 2.81E+00 | 7.21E-01 | no | Gm23128 | downstream_gene_variant | MODIFIER |
| 8 | 38980713 | AX-170188942 | 0 | C | T | 0.149 | 5.31E-02 | 1.50E-01 | 2.77E+00 | 2.81E+00 | 7.21E-01 | no | Gm23128 | downstream_gene_variant | MODIFIER |

| | | | | | | | | | | | | | | | |
|---|---|---|---|---|---|---|---|---|---|---|---|---|---|---|---|
| 8 | 38987763 | AX-170219735 | 0 | A | C | 0.149 | 5.31E-02 | 1.50E-01 | 2.77E+00 | 2.81E+00 | 7.21E-01 | no | Gm23128 | upstream_gene_variant | MODIFIER |
| 8 | 39006918 | AX-170530423 | 0 | T | C | 0.149 | 5.31E-02 | 1.50E-01 | 2.77E+00 | 2.81E+00 | 7.21E-01 | no | Tusc3 | intron_variant | MODIFIER |
| 8 | 39008286 | AX-170283557 | 0 | A | C | 0.149 | 5.31E-02 | 1.50E-01 | 2.77E+00 | 2.81E+00 | 7.21E-01 | no | Tusc3 | intron_variant | MODIFIER |
| 8 | 39031328 | AX-170115471 | 0 | C | T | 0.149 | 5.31E-02 | 1.50E-01 | 2.77E+00 | 2.81E+00 | 7.21E-01 | no | Tusc3 | intron_variant | MODIFIER |
| 8 | 39034077 | AX-170099015 | 0 | T | C | 0.149 | 5.31E-02 | 1.50E-01 | 2.77E+00 | 2.81E+00 | 7.21E-01 | no | Tusc3 | intron_variant | MODIFIER |
| 8 | 39037117 | AX-170419683 | 0 | T | G | 0.149 | 5.31E-02 | 1.50E-01 | 2.77E+00 | 2.81E+00 | 7.21E-01 | no | Tusc3 | intron_variant | MODIFIER |
| 8 | 39046583 | AX-170325313 | 0 | C | T | 0.149 | 5.31E-02 | 1.50E-01 | 2.77E+00 | 2.81E+00 | 7.21E-01 | no | Tusc3 | synonymous_variant | LOW |
| 8 | 39053398 | AX-170264971 | 0 | A | T | 0.149 | 5.31E-02 | 1.50E-01 | 2.77E+00 | 2.81E+00 | 7.21E-01 | no | Tusc3 | intron_variant | MODIFIER |
| 8 | 39060595 | AX-170030534 | 0 | G | A | 0.149 | 5.31E-02 | 1.50E-01 | 2.77E+00 | 2.81E+00 | 7.21E-01 | no | Tusc3 | intron_variant | MODIFIER |
| 8 | 39064157 | AX-170209318 | 0 | A | G | 0.149 | 5.31E-02 | 1.50E-01 | 2.77E+00 | 2.81E+00 | 7.21E-01 | no | Tusc3 | intron_variant | MODIFIER |
| 8 | 39092453 | AX-170079830 | 0 | A | G | 0.149 | 5.31E-02 | 1.50E-01 | 2.77E+00 | 2.81E+00 | 7.21E-01 | no | Tusc3 | intron_variant | MODIFIER |
| 8 | 39098302 | AX-170079829 | 0 | A | G | 0.149 | 5.31E-02 | 1.50E-01 | 2.77E+00 | 2.81E+00 | 7.21E-01 | no | Tusc3 | intron_variant | MODIFIER |
| 8 | 39098892 | AX-170269051 | 0 | A | G | 0.149 | 5.31E-02 | 1.50E-01 | 2.77E+00 | 2.81E+00 | 7.21E-01 | no | Tusc3 | intron_variant | MODIFIER |
| 8 | 39099981 | AX-170481791 | 0 | G | A | 0.149 | 5.31E-02 | 1.50E-01 | 2.77E+00 | 2.81E+00 | 7.21E-01 | no | Tusc3 | intron_variant | MODIFIER |
| 8 | 39100447 | AX-170357741 | 0 | C | A | 0.149 | 5.31E-02 | 1.50E-01 | 2.77E+00 | 2.81E+00 | 7.21E-01 | no | Tusc3 | intron_variant | MODIFIER |
| 8 | 39112863 | AX-170114559 | 0 | C | G | 0.149 | 5.31E-02 | 1.50E-01 | 2.77E+00 | 2.81E+00 | 7.21E-01 | no | Tusc3 | intron_variant | MODIFIER |
| 8 | 39113221 | AX-170030532 | 0 | C | G | 0.149 | 5.31E-02 | 1.50E-01 | 2.77E+00 | 2.81E+00 | 7.21E-01 | no | Tusc3 | intron_variant | MODIFIER |
| 8 | 39113542 | AX-170208057 | 0 | C | A | 0.149 | 5.31E-02 | 1.50E-01 | 2.77E+00 | 2.81E+00 | 7.21E-01 | no | Tusc3 | intron_variant | MODIFIER |
| 8 | 39121961 | AX-170030533 | 0 | C | T | 0.149 | 5.31E-02 | 1.50E-01 | 2.77E+00 | 2.81E+00 | 7.21E-01 | no | Tusc3 | intron_variant | MODIFIER |
| 8 | 39123421 | AX-170322489 | 0 | G | C | 0.149 | 5.31E-02 | 1.50E-01 | 2.77E+00 | 2.81E+00 | 7.21E-01 | no | Tusc3 | intron_variant | MODIFIER |
| 8 | 39163663 | AX-97843443 | 0 | T | C | 0.149 | 5.31E-02 | 1.50E-01 | 2.77E+00 | 2.81E+00 | 7.21E-01 | no | Tusc3 | intron_variant | MODIFIER |
| 8 | 39165199 | AX-170386902 | 0 | C | T | 0.149 | 5.31E-02 | 1.50E-01 | 2.77E+00 | 2.81E+00 | 7.21E-01 | no | Tusc3 | downstream_gene_variant | MODIFIER |
| 8 | 39169862 | AX-170152646 | 0 | C | A | 0.149 | 5.31E-02 | 1.50E-01 | 2.77E+00 | 2.81E+00 | 7.21E-01 | no | Tusc3 | downstream_gene_variant | MODIFIER |
| 8 | 39170560 | AX-170208091 | 0 | A | G | 0.149 | 5.31E-02 | 1.50E-01 | 2.77E+00 | 2.81E+00 | 7.21E-01 | no | | intergenic_variant | MODIFIER |
| 8 | 39193081 | AX-170482493 | 0 | A | T | 0.149 | 5.31E-02 | 1.50E-01 | 2.77E+00 | 2.81E+00 | 7.21E-01 | no | | intergenic_variant | MODIFIER |
| 8 | 39211267 | AX-97844253 | 0 | A | G | 0.149 | 5.31E-02 | 1.50E-01 | 2.77E+00 | 2.81E+00 | 7.21E-01 | no | Gm45398 | downstream_gene_variant | MODIFIER |
| 8 | 39211473 | AX-170246847 | 0 | G | T | 0.149 | 5.31E-02 | 1.50E-01 | 2.77E+00 | 2.81E+00 | 7.21E-01 | no | Gm45398 | downstream_gene_variant | MODIFIER |
| 8 | 39211586 | AX-170079834 | 0 | T | C | 0.149 | 5.31E-02 | 1.50E-01 | 2.77E+00 | 2.81E+00 | 7.21E-01 | no | Gm45398 | downstream_gene_variant | MODIFIER |
| 8 | 39211606 | AX-170079833 | 0 | C | T | 0.149 | 5.31E-02 | 1.50E-01 | 2.77E+00 | 2.81E+00 | 7.21E-01 | no | Gm45398 | downstream_gene_variant | MODIFIER |
| 8 | 39211642 | AX-170079831 | 0 | T | C | 0.149 | 5.31E-02 | 1.50E-01 | 2.77E+00 | 2.81E+00 | 7.21E-01 | no | | intergenic_variant | MODIFIER |
| 8 | 39285944 | AX-97816942 | 0 | G | A | 0.149 | 5.31E-02 | 1.50E-01 | 2.77E+00 | 2.81E+00 | 7.21E-01 | no | Zfp148-ps1 | downstream_gene_variant | MODIFIER |
| 8 | 39310256 | AX-170116087 | 0 | C | A | 0.149 | 5.31E-02 | 1.50E-01 | 2.77E+00 | 2.81E+00 | 7.21E-01 | no | Gm6213 | intron_variant,non_coding_transcript_variant | MODIFIER |
| 8 | 39310423 | AX-170352927 | 0 | C | A | 0.149 | 5.31E-02 | 1.50E-01 | 2.77E+00 | 2.81E+00 | 7.21E-01 | no | Gm6213 | intron_variant,non_coding_transcript_variant | MODIFIER |
| 8 | 39310768 | AX-170524269 | 0 | G | C | 0.149 | 5.31E-02 | 1.50E-01 | 2.77E+00 | 2.81E+00 | 7.21E-01 | no | Gm6213 | intron_variant,non_coding_transcript_variant | MODIFIER |
| 8 | 39313746 | AX-170300286 | 0 | G | A | 0.149 | 5.31E-02 | 1.50E-01 | 2.77E+00 | 2.81E+00 | 7.21E-01 | no | Gm6213 | intron_variant,non_coding_transcript_variant | MODIFIER |
| 8 | 39316126 | AX-170442094 | 0 | T | C | 0.149 | 5.31E-02 | 1.50E-01 | 2.77E+00 | 2.81E+00 | 7.21E-01 | no | Gm6213 | intron_variant,non_coding_transcript_variant | MODIFIER |
| 8 | 39316446 | AX-170403474 | 0 | C | T | 0.149 | 5.31E-02 | 1.50E-01 | 2.77E+00 | 2.81E+00 | 7.21E-01 | no | Gm6213 | intron_variant,non_coding_transcript_variant | MODIFIER |
| 8 | 39340058 | AX-170116883 | 0 | G | T | 0.149 | 5.31E-02 | 1.50E-01 | 2.77E+00 | 2.81E+00 | 7.21E-01 | no | Gm6213 | intron_variant,non_coding_transcript_variant | MODIFIER |
| 8 | 39340648 | AX-97834051 | 0 | T | G | 0.149 | 5.31E-02 | 1.50E-01 | 2.77E+00 | 2.81E+00 | 7.21E-01 | no | Gm6213 | intron_variant,non_coding_transcript_variant | MODIFIER |
| 8 | 39404742 | AX-170188175 | 0 | C | G | 0.149 | 5.31E-02 | 1.50E-01 | 2.77E+00 | 2.81E+00 | 7.21E-01 | no | | intergenic_variant | MODIFIER |
| 8 | 39408566 | AX-170166717 | 0 | C | G | 0.149 | 5.31E-02 | 1.50E-01 | 2.77E+00 | 2.81E+00 | 7.21E-01 | no | | intergenic_variant | MODIFIER |
| 8 | 39419093 | AX-170516295 | 0 | G | A | 0.149 | 5.31E-02 | 1.50E-01 | 2.77E+00 | 2.81E+00 | 7.21E-01 | no | | intergenic_variant | MODIFIER |
| 8 | 39419173 | AX-170185928 | 0 | A | C | 0.149 | 5.31E-02 | 1.50E-01 | 2.77E+00 | 2.81E+00 | 7.21E-01 | no | | intergenic_variant | MODIFIER |
| 8 | 39427390 | AX-170514176 | 0 | C | A | 0.149 | 5.31E-02 | 1.50E-01 | 2.77E+00 | 2.81E+00 | 7.21E-01 | no | | intergenic_variant | MODIFIER |
| 8 | 39445413 | AX-170397994 | 0 | A | C | 0.149 | 5.31E-02 | 1.50E-01 | 2.77E+00 | 2.81E+00 | 7.21E-01 | no | | intergenic_variant | MODIFIER |
| 8 | 39445456 | AX-97853120 | 0 | T | C | 0.149 | 5.31E-02 | 1.50E-01 | 2.77E+00 | 2.81E+00 | 7.21E-01 | no | | intergenic_variant | MODIFIER |
| 8 | 39475374 | AX-170257272 | 0 | C | T | 0.149 | 5.31E-02 | 1.50E-01 | 2.77E+00 | 2.81E+00 | 7.21E-01 | no | | intergenic_variant | MODIFIER |
| 8 | 39483752 | AX-97824169 | 0 | A | C | 0.149 | 5.31E-02 | 1.50E-01 | 2.77E+00 | 2.81E+00 | 7.21E-01 | no | | intergenic_variant | MODIFIER |
| 8 | 39499258 | AX-170439365 | 0 | A | G | 0.149 | 5.31E-02 | 1.50E-01 | 2.77E+00 | 2.81E+00 | 7.21E-01 | no | | intergenic_variant | MODIFIER |
| 8 | 39511303 | AX-170158625 | 0 | T | A | 0.149 | 5.31E-02 | 1.50E-01 | 2.77E+00 | 2.81E+00 | 7.21E-01 | no | | intergenic_variant | MODIFIER |
| 8 | 39537900 | AX-170408347 | 0 | C | T | 0.149 | 5.31E-02 | 1.50E-01 | 2.77E+00 | 2.81E+00 | 7.21E-01 | no | | intergenic_variant | MODIFIER |
| 8 | 39558951 | AX-97837335 | 0 | T | A | 0.149 | 5.31E-02 | 1.50E-01 | 2.77E+00 | 2.81E+00 | 7.21E-01 | no | | intergenic_variant | MODIFIER |
| 8 | 39560891 | AX-170496809 | 0 | T | C | 0.149 | 5.31E-02 | 1.50E-01 | 2.77E+00 | 2.81E+00 | 7.21E-01 | no | | intergenic_variant | MODIFIER |
| 8 | 39592079 | AX-170249265 | 0 | A | G | 0.149 | 5.31E-02 | 1.50E-01 | 2.77E+00 | 2.81E+00 | 7.21E-01 | no | Msr1 | intron_variant | MODIFIER |

| | | | | | | | | | | | | | | | |
|---|---|---|---|---|---|---|---|---|---|---|---|---|---|---|---|
| 8 | 39592176 | AX-170489585 | 0 | A | G | 0.149 | 5.31E-02 | 1.50E-01 | 2.77E+00 | 2.81E+00 | 7.21E-01 | no | Msr1 | intron_variant | MODIFIER |
| 8 | 39628676 | AX-170079840 | 0 | T | C | 0.149 | 5.31E-02 | 1.50E-01 | 2.77E+00 | 2.81E+00 | 7.21E-01 | no | Msr1 | intron_variant | MODIFIER |
| 8 | 39646998 | AX-170030539 | 0 | T | C | 0.149 | 5.31E-02 | 1.50E-01 | 2.77E+00 | 2.81E+00 | 7.21E-01 | no | Msr1 | upstream_gene_variant | MODIFIER |
| 8 | 39668016 | AX-170030540 | 0 | T | C | 0.149 | 5.31E-02 | 1.50E-01 | 2.77E+00 | 2.81E+00 | 7.21E-01 | no | | intergenic_variant | MODIFIER |
| 8 | 39693970 | AX-97837606 | 0 | A | G | 0.149 | 5.31E-02 | 1.50E-01 | 2.77E+00 | 2.81E+00 | 7.21E-01 | no | | intergenic_variant | MODIFIER |
| 8 | 39694727 | AX-170386048 | 0 | T | C | 0.149 | 5.31E-02 | 1.50E-01 | 2.77E+00 | 2.81E+00 | 7.21E-01 | no | | intergenic_variant | MODIFIER |
| 8 | 39694787 | AX-170246783 | 0 | C | T | 0.149 | 5.31E-02 | 1.50E-01 | 2.77E+00 | 2.81E+00 | 7.21E-01 | no | | intergenic_variant | MODIFIER |
| 8 | 39717657 | AX-170030541 | 0 | C | T | 0.149 | 5.31E-02 | 1.50E-01 | 2.77E+00 | 2.81E+00 | 7.21E-01 | no | Gm2033 | downstream_gene_variant | MODIFIER |
| 8 | 39752614 | AX-97849761 | 0 | G | T | 0.149 | 5.31E-02 | 1.50E-01 | 2.77E+00 | 2.81E+00 | 7.21E-01 | no | | intergenic_variant | MODIFIER |
| 8 | 39762399 | AX-170030542 | 0 | G | C | 0.149 | 5.31E-02 | 1.50E-01 | 2.77E+00 | 2.81E+00 | 7.21E-01 | no | | intergenic_variant | MODIFIER |
| 8 | 39771067 | AX-170218717 | 0 | C | G | 0.149 | 5.31E-02 | 1.50E-01 | 2.77E+00 | 2.81E+00 | 7.21E-01 | no | | intergenic_variant | MODIFIER |
| 8 | 39841990 | AX-170287678 | 0 | G | A | 0.149 | 5.31E-02 | 1.50E-01 | 2.77E+00 | 2.81E+00 | 7.21E-01 | no | | intergenic_variant | MODIFIER |
| 8 | 39842637 | AX-170438915 | 0 | G | A | 0.149 | 5.31E-02 | 1.50E-01 | 2.77E+00 | 2.81E+00 | 7.21E-01 | no | | intergenic_variant | MODIFIER |
| 8 | 39853399 | AX-170079841 | 0 | T | C | 0.149 | 5.31E-02 | 1.50E-01 | 2.77E+00 | 2.81E+00 | 7.21E-01 | no | | intergenic_variant | MODIFIER |
| 8 | 39853604 | AX-170079842 | 0 | C | T | 0.149 | 5.31E-02 | 1.50E-01 | 2.77E+00 | 2.81E+00 | 7.21E-01 | no | | intergenic_variant | MODIFIER |
| 8 | 39879679 | AX-170156516 | 0 | G | A | 0.149 | 5.31E-02 | 1.50E-01 | 2.77E+00 | 2.81E+00 | 7.21E-01 | no | | intergenic_variant | MODIFIER |
| 8 | 39929400 | AX-97841926 | 0 | T | C | 0.149 | 5.31E-02 | 1.50E-01 | 2.77E+00 | 2.81E+00 | 7.21E-01 | no | | intergenic_variant | MODIFIER |
| 8 | 39939749 | AX-170476636 | 0 | C | A | 0.149 | 5.31E-02 | 1.50E-01 | 2.77E+00 | 2.81E+00 | 7.21E-01 | no | | intergenic_variant | MODIFIER |
| 8 | 39939781 | AX-170114056 | 0 | G | A | 0.149 | 5.31E-02 | 1.50E-01 | 2.77E+00 | 2.81E+00 | 7.21E-01 | no | | intergenic_variant | MODIFIER |
| 8 | 39941236 | AX-170398652 | 0 | T | C | 0.149 | 5.31E-02 | 1.50E-01 | 2.77E+00 | 2.81E+00 | 7.21E-01 | no | | intergenic_variant | MODIFIER |
| 8 | 39949601 | AX-97788036 | 0 | G | A | 0.149 | 5.31E-02 | 1.50E-01 | 2.77E+00 | 2.81E+00 | 7.21E-01 | no | | intergenic_variant | MODIFIER |
| 8 | 39957537 | AX-170226072 | 0 | T | C | 0.149 | 5.31E-02 | 1.50E-01 | 2.77E+00 | 2.81E+00 | 7.21E-01 | no | | intergenic_variant | MODIFIER |
| 8 | 39960386 | AX-170450840 | 0 | C | T | 0.149 | 5.31E-02 | 1.50E-01 | 2.77E+00 | 2.81E+00 | 7.21E-01 | no | | intergenic_variant | MODIFIER |
| 8 | 39992349 | AX-170192158 | 0 | G | A | 0.149 | 5.31E-02 | 1.50E-01 | 2.77E+00 | 2.81E+00 | 7.21E-01 | no | | intergenic_variant | MODIFIER |
| 8 | 40004001 | AX-170217344 | 0 | G | A | 0.149 | 5.31E-02 | 1.50E-01 | 2.77E+00 | 2.81E+00 | 7.21E-01 | no | | intergenic_variant | MODIFIER |
| 8 | 40063645 | AX-170143830 | 0 | C | T | 0.149 | 5.31E-02 | 1.50E-01 | 2.77E+00 | 2.81E+00 | 7.21E-01 | no | | intergenic_variant | MODIFIER |
| 8 | 40084268 | AX-170165262 | 0 | A | G | 0.149 | 5.31E-02 | 1.50E-01 | 2.77E+00 | 2.81E+00 | 7.21E-01 | no | | intergenic_variant | MODIFIER |
| 8 | 40089164 | AX-170493995 | 0 | G | A | 0.149 | 5.31E-02 | 1.50E-01 | 2.77E+00 | 2.81E+00 | 7.21E-01 | no | | intergenic_variant | MODIFIER |
| 8 | 40089192 | AX-170433687 | 0 | T | G | 0.149 | 5.31E-02 | 1.50E-01 | 2.77E+00 | 2.81E+00 | 7.21E-01 | no | | intergenic_variant | MODIFIER |
| 8 | 40111155 | AX-170079845 | 0 | C | A | 0.149 | 5.31E-02 | 1.50E-01 | 2.77E+00 | 2.81E+00 | 7.21E-01 | no | | intergenic_variant | MODIFIER |
| 8 | 40111639 | AX-170079847 | 0 | A | G | 0.149 | 5.31E-02 | 1.50E-01 | 2.77E+00 | 2.81E+00 | 7.21E-01 | no | | intergenic_variant | MODIFIER |
| 8 | 40115062 | AX-170409780 | 0 | G | C | 0.149 | 5.31E-02 | 1.50E-01 | 2.77E+00 | 2.81E+00 | 7.21E-01 | no | | intergenic_variant | MODIFIER |
| 8 | 40123164 | AX-170206173 | 0 | G | A | 0.149 | 5.31E-02 | 1.50E-01 | 2.77E+00 | 2.81E+00 | 7.21E-01 | no | | intergenic_variant | MODIFIER |
| 8 | 40160049 | AX-170534347 | 0 | T | C | 0.149 | 5.31E-02 | 1.50E-01 | 2.77E+00 | 2.81E+00 | 7.21E-01 | no | | intergenic_variant | MODIFIER |
| 8 | 40185269 | AX-170387436 | 0 | C | T | 0.149 | 5.31E-02 | 1.50E-01 | 2.77E+00 | 2.81E+00 | 7.21E-01 | no | | intergenic_variant | MODIFIER |
| 8 | 40205411 | AX-170504550 | 0 | A | G | 0.149 | 5.31E-02 | 1.50E-01 | 2.77E+00 | 2.81E+00 | 7.21E-01 | no | | intergenic_variant | MODIFIER |
| 8 | 40205500 | AX-170392511 | 0 | A | G | 0.149 | 5.31E-02 | 1.50E-01 | 2.77E+00 | 2.81E+00 | 7.21E-01 | no | | intergenic_variant | MODIFIER |
| 8 | 40205610 | AX-170203753 | 0 | G | A | 0.149 | 5.31E-02 | 1.50E-01 | 2.77E+00 | 2.81E+00 | 7.21E-01 | no | | intergenic_variant | MODIFIER |
| 8 | 40228413 | AX-170224475 | 0 | A | C | 0.149 | 5.31E-02 | 1.50E-01 | 2.77E+00 | 2.81E+00 | 7.21E-01 | no | | intergenic_variant | MODIFIER |
| 8 | 40233013 | AX-170030544 | 0 | A | C | 0.149 | 5.31E-02 | 1.50E-01 | 2.77E+00 | 2.81E+00 | 7.21E-01 | no | | intergenic_variant | MODIFIER |
| 8 | 40239409 | AX-170030549 | 0 | G | A | 0.149 | 5.31E-02 | 1.50E-01 | 2.77E+00 | 2.81E+00 | 7.21E-01 | no | | intergenic_variant | MODIFIER |
| 8 | 40242912 | AX-97793330 | 0 | G | A | 0.149 | 5.31E-02 | 1.50E-01 | 2.77E+00 | 2.81E+00 | 7.21E-01 | no | | intergenic_variant | MODIFIER |
| 8 | 40271445 | AX-170090086 | 0 | T | C | 0.149 | 5.31E-02 | 1.50E-01 | 2.77E+00 | 2.81E+00 | 7.21E-01 | no | | regulatory_region_variant | MODIFIER |
| 8 | 40296555 | AX-170222240 | 0 | G | C | 0.149 | 5.31E-02 | 1.50E-01 | 2.77E+00 | 2.81E+00 | 7.21E-01 | no | Fgf20 | intron_variant | MODIFIER |
| 8 | 40314490 | AX-170168645 | 0 | G | A | 0.149 | 5.31E-02 | 1.50E-01 | 2.77E+00 | 2.81E+00 | 7.21E-01 | no | Micu3 | intron_variant | MODIFIER |
| 8 | 40316998 | AX-170030547 | 0 | T | C | 0.149 | 5.31E-02 | 1.50E-01 | 2.77E+00 | 2.81E+00 | 7.21E-01 | no | Micu3 | intron_variant | MODIFIER |
| 8 | 40317858 | AX-170030551 | 0 | T | A | 0.149 | 5.31E-02 | 1.50E-01 | 2.77E+00 | 2.81E+00 | 7.21E-01 | no | Micu3 | intron_variant | MODIFIER |
| 8 | 40323888 | AX-97788187 | 0 | G | A | 0.149 | 5.31E-02 | 1.50E-01 | 2.77E+00 | 2.81E+00 | 7.21E-01 | no | Micu3 | intron_variant | MODIFIER |
| 8 | 40345851 | AX-97837608 | 0 | G | A | 0.149 | 5.31E-02 | 1.50E-01 | 2.77E+00 | 2.81E+00 | 7.21E-01 | no | Micu3 | intron_variant | MODIFIER |
| 8 | 40378840 | AX-170030552 | 0 | A | G | 0.149 | 5.31E-02 | 1.50E-01 | 2.77E+00 | 2.81E+00 | 7.21E-01 | no | Micu3 | synonymous_variant | LOW |
| 8 | 40393729 | AX-170079848 | 0 | G | A | 0.149 | 5.31E-02 | 1.50E-01 | 2.77E+00 | 2.81E+00 | 7.21E-01 | no | | intergenic_variant | MODIFIER |
| 8 | 40402098 | AX-170030545 | 0 | A | T | 0.149 | 5.31E-02 | 1.50E-01 | 2.77E+00 | 2.81E+00 | 7.21E-01 | no | | intergenic_variant | MODIFIER |
| 8 | 40402133 | AX-170030546 | 0 | C | T | 0.149 | 5.31E-02 | 1.50E-01 | 2.77E+00 | 2.81E+00 | 7.21E-01 | no | | intergenic_variant | MODIFIER |
| 8 | 40406261 | AX-97842485 | 0 | C | T | 0.149 | 5.31E-02 | 1.50E-01 | 2.77E+00 | 2.81E+00 | 7.21E-01 | no | | intergenic_variant | MODIFIER |

| | | | | | | | | | | | | | | | |
|---|---|---|---|---|---|---|---|---|---|---|---|---|---|---|---|
| 8 | 40421907 | AX-170030548 | 0 | G | A | 0.149 | 5.31E-02 | 1.50E-01 | 2.77E+00 | 2.81E+00 | 7.21E-01 | no | Zdhhc2 | upstream_gene_variant | MODIFIER |
| 8 | 40428539 | AX-170079849 | 0 | C | T | 0.149 | 5.31E-02 | 1.50E-01 | 2.77E+00 | 2.81E+00 | 7.21E-01 | no | Zdhhc2 | intron_variant | MODIFIER |
| 8 | 40435247 | AX-170357817 | 0 | G | A | 0.149 | 5.31E-02 | 1.50E-01 | 2.77E+00 | 2.81E+00 | 7.21E-01 | no | Zdhhc2 | intron_variant | MODIFIER |
| 8 | 40439426 | AX-170484583 | 0 | G | T | 0.149 | 5.31E-02 | 1.50E-01 | 2.77E+00 | 2.81E+00 | 7.21E-01 | no | Zdhhc2 | intron_variant | MODIFIER |
| 8 | 40461216 | AX-170334376 | 0 | T | C | 0.149 | 5.31E-02 | 1.50E-01 | 2.77E+00 | 2.81E+00 | 7.21E-01 | no | Zdhhc2 | intron_variant | MODIFIER |
| 8 | 40486242 | AX-170130581 | 0 | C | T | 0.149 | 5.31E-02 | 1.50E-01 | 2.77E+00 | 2.81E+00 | 7.21E-01 | no | Zdhhc2 | downstream_gene_variant | MODIFIER |
| 8 | 40529051 | AX-170487638 | 0 | A | G | 0.149 | 5.31E-02 | 1.50E-01 | 2.77E+00 | 2.81E+00 | 7.21E-01 | no | Vps37a | intron_variant | MODIFIER |
| 8 | 40529305 | AX-170192069 | 0 | C | T | 0.149 | 5.31E-02 | 1.50E-01 | 2.77E+00 | 2.81E+00 | 7.21E-01 | no | Vps37a | intron_variant | MODIFIER |
| 8 | 40546122 | AX-97803563 | 0 | G | T | 0.149 | 5.31E-02 | 1.50E-01 | 2.77E+00 | 2.81E+00 | 7.21E-01 | no | Mtmr7 | downstream_gene_variant | MODIFIER |
| 8 | 40548461 | AX-170288594 | 0 | C | T | 0.149 | 5.31E-02 | 1.50E-01 | 2.77E+00 | 2.81E+00 | 7.21E-01 | no | Mtmr7 | downstream_gene_variant | MODIFIER |
| 8 | 40550797 | AX-170279976 | 0 | A | G | 0.149 | 5.31E-02 | 1.50E-01 | 2.77E+00 | 2.81E+00 | 7.21E-01 | no | Mtmr7 | downstream_gene_variant | MODIFIER |
| 8 | 40551270 | AX-170179271 | 0 | A | G | 0.149 | 5.31E-02 | 1.50E-01 | 2.77E+00 | 2.81E+00 | 7.21E-01 | no | Mtmr7 | 3_prime_UTR_variant,NMD_transcript_variant | MODIFIER |
| 8 | 40597989 | AX-97837605 | 0 | C | A | 0.149 | 5.31E-02 | 1.50E-01 | 2.77E+00 | 2.81E+00 | 7.21E-01 | no | Mtmr7 | intron_variant,NMD_transcript_variant | MODIFIER |
| 8 | 40603601 | AX-170079855 | 0 | A | G | 0.149 | 5.31E-02 | 1.50E-01 | 2.77E+00 | 2.81E+00 | 7.21E-01 | no | Mtmr7 | intron_variant,NMD_transcript_variant | MODIFIER |
| 8 | 40630127 | AX-97840721 | 0 | T | C | 0.149 | 5.31E-02 | 1.50E-01 | 2.77E+00 | 2.81E+00 | 7.21E-01 | no | Mtmr7 | intron_variant,NMD_transcript_variant | MODIFIER |
| 8 | 40637771 | AX-170516532 | 0 | C | G | 0.149 | 5.31E-02 | 1.50E-01 | 2.77E+00 | 2.81E+00 | 7.21E-01 | no | Mtmr7 | upstream_gene_variant | MODIFIER |
| 8 | 40639510 | AX-97788864 | 0 | T | C | 0.149 | 5.31E-02 | 1.50E-01 | 2.77E+00 | 2.81E+00 | 7.21E-01 | no | Mtmr7 | upstream_gene_variant | MODIFIER |
| 8 | 40674951 | AX-170487340 | 0 | T | C | 0.149 | 5.31E-02 | 1.50E-01 | 2.77E+00 | 2.81E+00 | 7.21E-01 | no | Adam24 | upstream_gene_variant | MODIFIER |
| 8 | 40680556 | AX-170333252 | 0 | G | A | 0.149 | 5.31E-02 | 1.50E-01 | 2.77E+00 | 2.81E+00 | 7.21E-01 | no | Adam24 | synonymous_variant | LOW |
| 8 | 40680855 | AX-170412122 | 0 | G | A | 0.149 | 5.31E-02 | 1.50E-01 | 2.77E+00 | 2.81E+00 | 7.21E-01 | no | Adam24 | missense_variant | MODERATE |
| 8 | 40710397 | AX-170128204 | 0 | A | G | 0.149 | 5.31E-02 | 1.50E-01 | 2.77E+00 | 2.81E+00 | 7.21E-01 | no | Gm8423 | downstream_gene_variant | MODIFIER |
| 8 | 40723065 | AX-170190783 | 0 | C | T | 0.149 | 5.31E-02 | 1.50E-01 | 2.77E+00 | 2.81E+00 | 7.21E-01 | no | Gm2080 | upstream_gene_variant | MODIFIER |
| 8 | 40756617 | AX-170351880 | 0 | A | T | 0.149 | 5.31E-02 | 1.50E-01 | 2.77E+00 | 2.81E+00 | 7.21E-01 | no | Adam25 | downstream_gene_variant | MODIFIER |
| 8 | 40797188 | AX-170030553 | 0 | A | G | 0.149 | 5.31E-02 | 1.50E-01 | 2.77E+00 | 2.81E+00 | 7.21E-01 | no | Adam20 | 3_prime_UTR_variant | MODIFIER |
| 8 | 40801161 | AX-170030556 | 0 | G | C | 0.149 | 5.31E-02 | 1.50E-01 | 2.77E+00 | 2.81E+00 | 7.21E-01 | no | Adam20 | downstream_gene_variant | MODIFIER |
| 8 | 40836457 | AX-170090087 | 0 | T | C | 0.149 | 5.31E-02 | 1.50E-01 | 2.77E+00 | 2.81E+00 | 7.21E-01 | no | | intergenic_variant | MODIFIER |
| 8 | 40836537 | AX-170030554 | 0 | G | T | 0.149 | 5.31E-02 | 1.50E-01 | 2.77E+00 | 2.81E+00 | 7.21E-01 | no | | intergenic_variant | MODIFIER |
| 8 | 40876069 | AX-170142789 | 0 | C | T | 0.149 | 5.31E-02 | 1.50E-01 | 2.77E+00 | 2.81E+00 | 7.21E-01 | no | Slc7a2 | intron_variant | MODIFIER |
| 8 | 40879416 | AX-170490510 | 0 | A | G | 0.149 | 5.31E-02 | 1.50E-01 | 2.77E+00 | 2.81E+00 | 7.21E-01 | no | Slc7a2 | intron_variant | MODIFIER |
| 8 | 40881491 | AX-170526285 | 0 | A | G | 0.149 | 5.31E-02 | 1.50E-01 | 2.77E+00 | 2.81E+00 | 7.21E-01 | no | Slc7a2 | intron_variant | MODIFIER |
| 8 | 40881684 | AX-170096238 | 0 | G | A | 0.149 | 5.31E-02 | 1.50E-01 | 2.77E+00 | 2.81E+00 | 7.21E-01 | no | Slc7a2 | intron_variant | MODIFIER |
| 8 | 40887134 | AX-170356236 | 0 | G | A | 0.149 | 5.31E-02 | 1.50E-01 | 2.77E+00 | 2.81E+00 | 7.21E-01 | no | Slc7a2 | intron_variant | MODIFIER |
| 8 | 40889202 | AX-170525812 | 0 | A | G | 0.149 | 5.31E-02 | 1.50E-01 | 2.77E+00 | 2.81E+00 | 7.21E-01 | no | Slc7a2 | intron_variant | MODIFIER |
| 8 | 40910746 | AX-170149800 | 0 | T | C | 0.149 | 5.31E-02 | 1.50E-01 | 2.77E+00 | 2.81E+00 | 7.21E-01 | no | Slc7a2 | intron_variant | MODIFIER |
| 8 | 40914215 | AX-170528251 | 0 | T | C | 0.149 | 5.31E-02 | 1.50E-01 | 2.77E+00 | 2.81E+00 | 7.21E-01 | no | Slc7a2 | intron_variant | MODIFIER |
| 8 | 40915371 | AX-170210107 | 0 | G | C | 0.149 | 5.31E-02 | 1.50E-01 | 2.77E+00 | 2.81E+00 | 7.21E-01 | no | Slc7a2 | intron_variant | MODIFIER |
| 8 | 40915460 | AX-170274009 | 0 | T | C | 0.149 | 5.31E-02 | 1.50E-01 | 2.77E+00 | 2.81E+00 | 7.21E-01 | no | Slc7a2 | intron_variant | MODIFIER |
| 8 | 40917038 | AX-170384653 | 0 | C | A | 0.149 | 5.31E-02 | 1.50E-01 | 2.77E+00 | 2.81E+00 | 7.21E-01 | no | Slc7a2 | 3_prime_UTR_variant | MODIFIER |
| 8 | 40921672 | AX-170274393 | 0 | T | G | 0.149 | 5.31E-02 | 1.50E-01 | 2.77E+00 | 2.81E+00 | 7.21E-01 | no | Pdgfrl | upstream_gene_variant | MODIFIER |
| 8 | 40943392 | AX-97821705 | 0 | A | G | 0.149 | 5.31E-02 | 1.50E-01 | 2.77E+00 | 2.81E+00 | 7.21E-01 | no | Pdgfrl | intron_variant | MODIFIER |
| 8 | 40949397 | AX-170487466 | 0 | T | G | 0.149 | 5.31E-02 | 1.50E-01 | 2.77E+00 | 2.81E+00 | 7.21E-01 | no | Pdgfrl | intron_variant | MODIFIER |
| 8 | 40950109 | AX-170161631 | 0 | T | C | 0.149 | 5.31E-02 | 1.50E-01 | 2.77E+00 | 2.81E+00 | 7.21E-01 | no | Pdgfrl | intron_variant | MODIFIER |
| 8 | 40959337 | AX-170451280 | 0 | A | G | 0.149 | 5.31E-02 | 1.50E-01 | 2.77E+00 | 2.81E+00 | 7.21E-01 | no | Pdgfrl | intron_variant | MODIFIER |
| 8 | 40968685 | AX-170212457 | 0 | A | T | 0.149 | 5.31E-02 | 1.50E-01 | 2.77E+00 | 2.81E+00 | 7.21E-01 | no | Pdgfrl | intron_variant | MODIFIER |
| 8 | 40977235 | AX-170380956 | 0 | T | C | 0.149 | 5.31E-02 | 1.50E-01 | 2.77E+00 | 2.81E+00 | 7.21E-01 | no | Pdgfrl | intron_variant | MODIFIER |
| 8 | 40977260 | AX-170507966 | 0 | T | C | 0.149 | 5.31E-02 | 1.50E-01 | 2.77E+00 | 2.81E+00 | 7.21E-01 | no | Pdgfrl | intron_variant | MODIFIER |
| 8 | 40986575 | AX-170280130 | 0 | T | C | 0.149 | 5.31E-02 | 1.50E-01 | 2.77E+00 | 2.81E+00 | 7.21E-01 | no | Pdgfrl | intron_variant | MODIFIER |
| 8 | 40986680 | AX-170243183 | 0 | T | C | 0.149 | 5.31E-02 | 1.50E-01 | 2.77E+00 | 2.81E+00 | 7.21E-01 | no | Pdgfrl | intron_variant | MODIFIER |
| 8 | 41006193 | AX-170079861 | 0 | G | A | 0.149 | 5.31E-02 | 1.50E-01 | 2.77E+00 | 2.81E+00 | 7.21E-01 | no | Mtus1 | intron_variant | MODIFIER |
| 8 | 41006391 | AX-170425561 | 0 | T | C | 0.149 | 5.31E-02 | 1.50E-01 | 2.77E+00 | 2.81E+00 | 7.21E-01 | no | Mtus1 | intron_variant | MODIFIER |
| 8 | 41013658 | AX-170159543 | 0 | G | A | 0.149 | 5.31E-02 | 1.50E-01 | 2.77E+00 | 2.81E+00 | 7.21E-01 | no | Mtus1 | intron_variant | MODIFIER |
| 8 | 41044672 | AX-170446602 | 0 | C | T | 0.149 | 5.31E-02 | 1.50E-01 | 2.77E+00 | 2.81E+00 | 7.21E-01 | no | Mtus1 | upstream_gene_variant | MODIFIER |
| 8 | 41044732 | AX-170271309 | 0 | C | T | 0.149 | 5.31E-02 | 1.50E-01 | 2.77E+00 | 2.81E+00 | 7.21E-01 | no | Mtus1 | upstream_gene_variant | MODIFIER |
| 8 | 41044822 | AX-170351579 | 0 | A | G | 0.149 | 5.31E-02 | 1.50E-01 | 2.77E+00 | 2.81E+00 | 7.21E-01 | no | Mtus1 | upstream_gene_variant | MODIFIER |

| | | | | | | | | | | | | | | | |
|---|---|---|---|---|---|---|---|---|---|---|---|---|---|---|---|
| 8 | 41045657 | AX-170474685 | 0 | A | G | 0.149 | 5.31E-02 | 1.50E-01 | 2.77E+00 | 2.81E+00 | 7.21E-01 | no | Mtus1 | upstream_gene_variant | MODIFIER |
| 8 | 41045827 | AX-170389903 | 0 | G | A | 0.149 | 5.31E-02 | 1.50E-01 | 2.77E+00 | 2.81E+00 | 7.21E-01 | no | Mtus1 | upstream_gene_variant | MODIFIER |
| 8 | 41065263 | AX-170227385 | 0 | T | C | 0.149 | 5.31E-02 | 1.50E-01 | 2.77E+00 | 2.81E+00 | 7.21E-01 | no | Mtus1 | intron_variant | MODIFIER |
| 8 | 41065778 | AX-170215034 | 0 | C | A | 0.149 | 5.31E-02 | 1.50E-01 | 2.77E+00 | 2.81E+00 | 7.21E-01 | no | Mtus1 | intron_variant | MODIFIER |
| 8 | 41079409 | AX-97842130 | 0 | C | T | 0.149 | 5.31E-02 | 1.50E-01 | 2.77E+00 | 2.81E+00 | 7.21E-01 | no | Mtus1 | intron_variant | MODIFIER |
| 8 | 41079523 | AX-170303865 | 0 | C | T | 0.149 | 5.31E-02 | 1.50E-01 | 2.77E+00 | 2.81E+00 | 7.21E-01 | no | Mtus1 | intron_variant | MODIFIER |
| 8 | 41084750 | AX-170297549 | 0 | A | G | 0.149 | 5.31E-02 | 1.50E-01 | 2.77E+00 | 2.81E+00 | 7.21E-01 | no | Mtus1 | 5_prime_UTR_variant | MODIFIER |
| 8 | 41101754 | AX-170119721 | 0 | T | G | 0.149 | 5.31E-02 | 1.50E-01 | 2.77E+00 | 2.81E+00 | 7.21E-01 | no | Mtus1 | intron_variant | MODIFIER |
| 8 | 41115565 | AX-97799552 | 0 | A | G | 0.149 | 5.31E-02 | 1.50E-01 | 2.77E+00 | 2.81E+00 | 7.21E-01 | no | Mtus1 | intron_variant | MODIFIER |
| 8 | 41131900 | AX-170318052 | 0 | G | A | 0.149 | 5.31E-02 | 1.50E-01 | 2.77E+00 | 2.81E+00 | 7.21E-01 | no | Mtus1 | intron_variant | MODIFIER |
| 8 | 41138779 | AX-170079870 | 0 | A | G | 0.149 | 5.31E-02 | 1.50E-01 | 2.77E+00 | 2.81E+00 | 7.21E-01 | no | | intergenic_variant | MODIFIER |
| 8 | 41162694 | AX-170224805 | 0 | A | G | 0.149 | 5.31E-02 | 1.50E-01 | 2.77E+00 | 2.81E+00 | 7.21E-01 | no | | intergenic_variant | MODIFIER |
| 8 | 41162887 | AX-170134497 | 0 | T | C | 0.149 | 5.31E-02 | 1.50E-01 | 2.77E+00 | 2.81E+00 | 7.21E-01 | no | | intergenic_variant | MODIFIER |
| 8 | 41162912 | AX-170326941 | 0 | A | G | 0.149 | 5.31E-02 | 1.50E-01 | 2.77E+00 | 2.81E+00 | 7.21E-01 | no | | intergenic_variant | MODIFIER |
| 8 | 41171021 | AX-170079866 | 0 | G | A | 0.149 | 5.31E-02 | 1.50E-01 | 2.77E+00 | 2.81E+00 | 7.21E-01 | no | | intergenic_variant | MODIFIER |
| 8 | 41173318 | AX-170079864 | 0 | G | T | 0.149 | 5.31E-02 | 1.50E-01 | 2.77E+00 | 2.81E+00 | 7.21E-01 | no | | intergenic_variant | MODIFIER |
| 8 | 41190613 | AX-170338405 | 0 | C | T | 0.149 | 5.31E-02 | 1.50E-01 | 2.77E+00 | 2.81E+00 | 7.21E-01 | no | Fgl1 | downstream_gene_variant | MODIFIER |
| 8 | 41196090 | AX-170209097 | 0 | T | C | 0.149 | 5.31E-02 | 1.50E-01 | 2.77E+00 | 2.81E+00 | 7.21E-01 | no | Fgl1 | intron_variant | MODIFIER |
| 8 | 41197008 | AX-170261706 | 0 | A | G | 0.149 | 5.31E-02 | 1.50E-01 | 2.77E+00 | 2.81E+00 | 7.21E-01 | no | Fgl1 | synonymous_variant | LOW |
| 8 | 41224474 | AX-170430092 | 0 | T | C | 0.149 | 5.31E-02 | 1.50E-01 | 2.77E+00 | 2.81E+00 | 7.21E-01 | no | Gm16348 | downstream_gene_variant | MODIFIER |
| 8 | 41224832 | AX-170480404 | 0 | T | G | 0.149 | 5.31E-02 | 1.50E-01 | 2.77E+00 | 2.81E+00 | 7.21E-01 | no | Gm16348 | downstream_gene_variant | MODIFIER |
| 8 | 41236292 | AX-170137706 | 0 | G | A | 0.149 | 5.31E-02 | 1.50E-01 | 2.77E+00 | 2.81E+00 | 7.21E-01 | no | Pcm1 | upstream_gene_variant | MODIFIER |
| 8 | 41252594 | AX-170098462 | 0 | C | T | 0.149 | 5.31E-02 | 1.50E-01 | 2.77E+00 | 2.81E+00 | 7.21E-01 | no | Pcm1 | intron_variant | MODIFIER |
| 8 | 41291250 | AX-97843629 | 0 | A | G | 0.149 | 5.31E-02 | 1.50E-01 | 2.77E+00 | 2.81E+00 | 7.21E-01 | no | Pcm1 | intron_variant | MODIFIER |
| 8 | 41322924 | AX-170030559 | 0 | C | T | 0.149 | 5.31E-02 | 1.50E-01 | 2.77E+00 | 2.81E+00 | 7.21E-01 | no | Pcm1 | intron_variant | MODIFIER |
| 8 | 41324502 | AX-97826423 | 0 | A | G | 0.149 | 5.31E-02 | 1.50E-01 | 2.77E+00 | 2.81E+00 | 7.21E-01 | no | Pcm1 | intron_variant | MODIFIER |
| 8 | 41379081 | AX-170505442 | 0 | A | T | 0.149 | 5.31E-02 | 1.50E-01 | 2.77E+00 | 2.81E+00 | 7.21E-01 | no | Asah1 | upstream_gene_variant | MODIFIER |
| 8 | 41387794 | AX-170262185 | 0 | A | C | 0.149 | 5.31E-02 | 1.50E-01 | 2.77E+00 | 2.81E+00 | 7.21E-01 | no | | intergenic_variant | MODIFIER |
| 8 | 41410560 | AX-170301295 | 0 | C | T | 0.149 | 5.31E-02 | 1.50E-01 | 2.77E+00 | 2.81E+00 | 7.21E-01 | no | Frg1 | intron_variant | MODIFIER |
| 8 | 41414978 | AX-170154213 | 0 | G | A | 0.149 | 5.31E-02 | 1.50E-01 | 2.77E+00 | 2.81E+00 | 7.21E-01 | no | Frg1 | intron_variant | MODIFIER |
| 8 | 41442857 | AX-97781912 | 0 | G | A | 0.149 | 5.31E-02 | 1.50E-01 | 2.77E+00 | 2.81E+00 | 7.21E-01 | no | | intergenic_variant | MODIFIER |
| 8 | 41455438 | AX-170174796 | 0 | A | G | 0.149 | 5.31E-02 | 1.50E-01 | 2.77E+00 | 2.81E+00 | 7.21E-01 | no | | intergenic_variant | MODIFIER |
| 8 | 41457436 | AX-170291333 | 0 | C | A | 0.149 | 5.31E-02 | 1.50E-01 | 2.77E+00 | 2.81E+00 | 7.21E-01 | no | | intergenic_variant | MODIFIER |
| 8 | 41473801 | AX-170363132 | 0 | G | C | 0.149 | 5.31E-02 | 1.50E-01 | 2.77E+00 | 2.81E+00 | 7.21E-01 | no | | regulatory_region_variant | MODIFIER |
| 8 | 41485216 | AX-170362966 | 0 | T | G | 0.149 | 5.31E-02 | 1.50E-01 | 2.77E+00 | 2.81E+00 | 7.21E-01 | no | | intergenic_variant | MODIFIER |
| 8 | 41486093 | AX-170329010 | 0 | A | C | 0.149 | 5.31E-02 | 1.50E-01 | 2.77E+00 | 2.81E+00 | 7.21E-01 | no | | intergenic_variant | MODIFIER |
| 8 | 41492844 | AX-170495524 | 0 | C | T | 0.149 | 5.31E-02 | 1.50E-01 | 2.77E+00 | 2.81E+00 | 7.21E-01 | no | | intergenic_variant | MODIFIER |
| 8 | 41493137 | AX-170526165 | 0 | T | C | 0.149 | 5.31E-02 | 1.50E-01 | 2.77E+00 | 2.81E+00 | 7.21E-01 | no | | intergenic_variant | MODIFIER |
| 8 | 41526825 | AX-97803562 | 0 | A | G | 0.149 | 5.31E-02 | 1.50E-01 | 2.77E+00 | 2.81E+00 | 7.21E-01 | no | | intergenic_variant | MODIFIER |
| 8 | 41528825 | AX-170146904 | 0 | T | C | 0.149 | 5.31E-02 | 1.50E-01 | 2.77E+00 | 2.81E+00 | 7.21E-01 | no | | intergenic_variant | MODIFIER |
| 8 | 41530482 | AX-170176755 | 0 | A | C | 0.149 | 5.31E-02 | 1.50E-01 | 2.77E+00 | 2.81E+00 | 7.21E-01 | no | | intergenic_variant | MODIFIER |
| 8 | 41530536 | AX-170079877 | 0 | T | C | 0.149 | 5.31E-02 | 1.50E-01 | 2.77E+00 | 2.81E+00 | 7.21E-01 | no | | intergenic_variant | MODIFIER |
| 8 | 41554284 | AX-170329675 | 0 | C | T | 0.149 | 5.31E-02 | 1.50E-01 | 2.77E+00 | 2.81E+00 | 7.21E-01 | no | | intergenic_variant | MODIFIER |
| 8 | 41558632 | AX-170447535 | 0 | A | C | 0.149 | 5.31E-02 | 1.50E-01 | 2.77E+00 | 2.81E+00 | 7.21E-01 | no | | intergenic_variant | MODIFIER |
| 8 | 41586861 | AX-170403606 | 0 | T | G | 0.149 | 5.31E-02 | 1.50E-01 | 2.77E+00 | 2.81E+00 | 7.21E-01 | no | | intergenic_variant | MODIFIER |
| 8 | 41587589 | AX-170181507 | 0 | A | C | 0.149 | 5.31E-02 | 1.50E-01 | 2.77E+00 | 2.81E+00 | 7.21E-01 | no | | intergenic_variant | MODIFIER |
| 8 | 41599352 | AX-170334854 | 0 | G | A | 0.149 | 5.31E-02 | 1.50E-01 | 2.77E+00 | 2.81E+00 | 7.21E-01 | no | | intergenic_variant | MODIFIER |
| 8 | 41612157 | AX-170317230 | 0 | C | T | 0.149 | 5.31E-02 | 1.50E-01 | 2.77E+00 | 2.81E+00 | 7.21E-01 | no | Gapdh-ps14 | upstream_gene_variant | MODIFIER |
| 8 | 41612456 | AX-170176563 | 0 | G | A | 0.149 | 5.31E-02 | 1.50E-01 | 2.77E+00 | 2.81E+00 | 7.21E-01 | no | Gapdh-ps14 | upstream_gene_variant | MODIFIER |
| 8 | 41631302 | AX-170494105 | 0 | C | T | 0.149 | 5.31E-02 | 1.50E-01 | 2.77E+00 | 2.81E+00 | 7.21E-01 | no | | intergenic_variant | MODIFIER |
| 8 | 41631836 | AX-170372620 | 0 | A | G | 0.149 | 5.31E-02 | 1.50E-01 | 2.77E+00 | 2.81E+00 | 7.21E-01 | no | | intergenic_variant | MODIFIER |
| 8 | 41645061 | AX-97830417 | 0 | G | C | 0.149 | 5.31E-02 | 1.50E-01 | 2.77E+00 | 2.81E+00 | 7.21E-01 | no | | intergenic_variant | MODIFIER |
| 8 | 41645232 | AX-170288005 | 0 | T | C | 0.149 | 5.31E-02 | 1.50E-01 | 2.77E+00 | 2.81E+00 | 7.21E-01 | no | | intergenic_variant | MODIFIER |
| 8 | 41691468 | AX-97837056 | 0 | G | A | 0.149 | 5.31E-02 | 1.50E-01 | 2.77E+00 | 2.81E+00 | 7.21E-01 | no | | intergenic_variant | MODIFIER |

| | | | | | | | | | | | | | | | |
|---|---|---|---|---|---|---|---|---|---|---|---|---|---|---|---|
| 8 | 41699793 | AX-170185231 | 0 | G | A | 0.149 | 5.31E-02 | 1.50E-01 | 2.77E+00 | 2.81E+00 | 7.21E-01 | no | | intergenic_variant | MODIFIER |
| 8 | 41707138 | AX-170475182 | 0 | T | G | 0.149 | 5.31E-02 | 1.50E-01 | 2.77E+00 | 2.81E+00 | 7.21E-01 | no | | intergenic_variant | MODIFIER |
| 8 | 41709863 | AX-170383248 | 0 | G | A | 0.149 | 5.31E-02 | 1.50E-01 | 2.77E+00 | 2.81E+00 | 7.21E-01 | no | | intergenic_variant | MODIFIER |
| 8 | 41720544 | AX-170265271 | 0 | T | C | 0.149 | 5.31E-02 | 1.50E-01 | 2.77E+00 | 2.81E+00 | 7.21E-01 | no | | intergenic_variant | MODIFIER |
| 8 | 41720567 | AX-170423127 | 0 | A | G | 0.149 | 5.31E-02 | 1.50E-01 | 2.77E+00 | 2.81E+00 | 7.21E-01 | no | | intergenic_variant | MODIFIER |
| 8 | 41740333 | AX-170528034 | 0 | T | G | 0.149 | 5.31E-02 | 1.50E-01 | 2.77E+00 | 2.81E+00 | 7.21E-01 | no | | intergenic_variant | MODIFIER |
| 8 | 41749676 | AX-170188438 | 0 | T | C | 0.149 | 5.31E-02 | 1.50E-01 | 2.77E+00 | 2.81E+00 | 7.21E-01 | no | | intergenic_variant | MODIFIER |
| 8 | 41749896 | AX-97839652 | 0 | G | A | 0.149 | 5.31E-02 | 1.50E-01 | 2.77E+00 | 2.81E+00 | 7.21E-01 | no | | intergenic_variant | MODIFIER |
| 8 | 41774608 | AX-170169953 | 0 | A | G | 0.149 | 5.31E-02 | 1.50E-01 | 2.77E+00 | 2.81E+00 | 7.21E-01 | no | | intergenic_variant | MODIFIER |
| 8 | 41777747 | AX-170079881 | 0 | T | C | 0.149 | 5.31E-02 | 1.50E-01 | 2.77E+00 | 2.81E+00 | 7.21E-01 | no | | intergenic_variant | MODIFIER |
| 8 | 41778496 | AX-170079883 | 0 | G | A | 0.149 | 5.31E-02 | 1.50E-01 | 2.77E+00 | 2.81E+00 | 7.21E-01 | no | | intergenic_variant | MODIFIER |
| 8 | 41789102 | AX-170225605 | 0 | C | T | 0.149 | 5.31E-02 | 1.50E-01 | 2.77E+00 | 2.81E+00 | 7.21E-01 | no | | intergenic_variant | MODIFIER |
| 8 | 41794749 | AX-170079880 | 0 | A | C | 0.149 | 5.31E-02 | 1.50E-01 | 2.77E+00 | 2.81E+00 | 7.21E-01 | no | | intergenic_variant | MODIFIER |
| 8 | 41796186 | AX-170372114 | 0 | C | T | 0.149 | 5.31E-02 | 1.50E-01 | 2.77E+00 | 2.81E+00 | 7.21E-01 | no | | intergenic_variant | MODIFIER |
| 8 | 41801443 | AX-170399710 | 0 | A | G | 0.149 | 5.31E-02 | 1.50E-01 | 2.77E+00 | 2.81E+00 | 7.21E-01 | no | | intergenic_variant | MODIFIER |
| 8 | 41830858 | AX-170239484 | 0 | G | C | 0.149 | 5.31E-02 | 1.50E-01 | 2.77E+00 | 2.81E+00 | 7.21E-01 | no | 2810404M03Rik | intron_variant,non_coding_transcript_variant | MODIFIER |
| 8 | 41832537 | AX-170306861 | 0 | G | A | 0.149 | 5.31E-02 | 1.50E-01 | 2.77E+00 | 2.81E+00 | 7.21E-01 | no | 2810404M03Rik | intron_variant,non_coding_transcript_variant | MODIFIER |
| 8 | 41854092 | AX-170195720 | 0 | T | C | 0.149 | 5.31E-02 | 1.50E-01 | 2.77E+00 | 2.81E+00 | 7.21E-01 | no | 2810404M03Rik | intron_variant,non_coding_transcript_variant | MODIFIER |
| 8 | 41859136 | AX-170263038 | 0 | T | C | 0.149 | 5.31E-02 | 1.50E-01 | 2.77E+00 | 2.81E+00 | 7.21E-01 | no | 2810404M03Rik | intron_variant,non_coding_transcript_variant | MODIFIER |
| 8 | 41863010 | AX-170307823 | 0 | A | C | 0.149 | 5.31E-02 | 1.50E-01 | 2.77E+00 | 2.81E+00 | 7.21E-01 | no | 2810404M03Rik | intron_variant,non_coding_transcript_variant | MODIFIER |
| 8 | 41874113 | AX-170105289 | 0 | T | G | 0.149 | 5.31E-02 | 1.50E-01 | 2.77E+00 | 2.81E+00 | 7.21E-01 | no | 2810404M03Rik | intron_variant,non_coding_transcript_variant | MODIFIER |
| 8 | 41879407 | AX-170030568 | 0 | C | T | 0.149 | 5.31E-02 | 1.50E-01 | 2.77E+00 | 2.81E+00 | 7.21E-01 | no | 2810404M03Rik | intron_variant,non_coding_transcript_variant | MODIFIER |
| 8 | 41884997 | AX-170030563 | 0 | C | T | 0.149 | 5.31E-02 | 1.50E-01 | 2.77E+00 | 2.81E+00 | 7.21E-01 | no | 2810404M03Rik | intron_variant,non_coding_transcript_variant | MODIFIER |
| 8 | 41893065 | AX-170430819 | 0 | C | T | 0.149 | 5.31E-02 | 1.50E-01 | 2.77E+00 | 2.81E+00 | 7.21E-01 | no | 2810404M03Rik | intron_variant,non_coding_transcript_variant | MODIFIER |
| 8 | 41916239 | AX-170423016 | 0 | G | C | 0.149 | 5.31E-02 | 1.50E-01 | 2.77E+00 | 2.81E+00 | 7.21E-01 | no | 2810404M03Rik | intron_variant,non_coding_transcript_variant | MODIFIER |
| 8 | 41919039 | AX-97844657 | 0 | T | C | 0.149 | 5.31E-02 | 1.50E-01 | 2.77E+00 | 2.81E+00 | 7.21E-01 | no | 2810404M03Rik | intron_variant,non_coding_transcript_variant | MODIFIER |
| 8 | 41919076 | AX-170030566 | 0 | T | C | 0.149 | 5.31E-02 | 1.50E-01 | 2.77E+00 | 2.81E+00 | 7.21E-01 | no | 2810404M03Rik | intron_variant,non_coding_transcript_variant | MODIFIER |
| 8 | 41919718 | AX-170472546 | 0 | C | T | 0.149 | 5.31E-02 | 1.50E-01 | 2.77E+00 | 2.81E+00 | 7.21E-01 | no | 2810404M03Rik | intron_variant,non_coding_transcript_variant | MODIFIER |
| 8 | 41919806 | AX-170030562 | 0 | T | C | 0.149 | 5.31E-02 | 1.50E-01 | 2.77E+00 | 2.81E+00 | 7.21E-01 | no | 2810404M03Rik | intron_variant,non_coding_transcript_variant | MODIFIER |
| 8 | 41941141 | AX-97825886 | 0 | C | A | 0.149 | 5.31E-02 | 1.50E-01 | 2.77E+00 | 2.81E+00 | 7.21E-01 | no | 2810404M03Rik | intron_variant,non_coding_transcript_variant | MODIFIER |
| 8 | 41946112 | AX-170227082 | 0 | C | G | 0.149 | 5.31E-02 | 1.50E-01 | 2.77E+00 | 2.81E+00 | 7.21E-01 | no | 2810404M03Rik | intron_variant,non_coding_transcript_variant | MODIFIER |
| 8 | 41964645 | AX-97850621 | 0 | G | A | 0.149 | 5.31E-02 | 1.50E-01 | 2.77E+00 | 2.81E+00 | 7.21E-01 | no | 2810404M03Rik | intron_variant,non_coding_transcript_variant | MODIFIER |
| 8 | 41964760 | AX-170483280 | 0 | A | G | 0.149 | 5.31E-02 | 1.50E-01 | 2.77E+00 | 2.81E+00 | 7.21E-01 | no | 2810404M03Rik | intron_variant,non_coding_transcript_variant | MODIFIER |
| 8 | 42030017 | AX-170099778 | 0 | A | C | 0.149 | 5.31E-02 | 1.50E-01 | 2.77E+00 | 2.81E+00 | 7.21E-01 | no | 2810404M03Rik | intron_variant,non_coding_transcript_variant | MODIFIER |
| 8 | 42030097 | AX-170079885 | 0 | G | A | 0.149 | 5.31E-02 | 1.50E-01 | 2.77E+00 | 2.81E+00 | 7.21E-01 | no | 2810404M03Rik | intron_variant,non_coding_transcript_variant | MODIFIER |
| 8 | 42039795 | AX-170168264 | 0 | C | G | 0.149 | 5.31E-02 | 1.50E-01 | 2.77E+00 | 2.81E+00 | 7.21E-01 | no | 2810404M03Rik | intron_variant,non_coding_transcript_variant | MODIFIER |
| 8 | 42041620 | AX-170176397 | 0 | A | C | 0.149 | 5.31E-02 | 1.50E-01 | 2.77E+00 | 2.81E+00 | 7.21E-01 | no | 2810404M03Rik | intron_variant,non_coding_transcript_variant | MODIFIER |
| 8 | 42041649 | AX-97847915 | 0 | C | T | 0.149 | 5.31E-02 | 1.50E-01 | 2.77E+00 | 2.81E+00 | 7.21E-01 | no | 2810404M03Rik | intron_variant,non_coding_transcript_variant | MODIFIER |
| 8 | 42044871 | AX-170105880 | 0 | G | C | 0.149 | 5.31E-02 | 1.50E-01 | 2.77E+00 | 2.81E+00 | 7.21E-01 | no | 2810404M03Rik | splice_donor_variant,non_coding_transcript_variant | HIGH |
| 8 | 42067238 | AX-170492498 | 0 | T | C | 0.149 | 5.31E-02 | 1.50E-01 | 2.77E+00 | 2.81E+00 | 7.21E-01 | no | | intergenic_variant | MODIFIER |
| 8 | 42068223 | AX-170387934 | 0 | G | T | 0.149 | 5.31E-02 | 1.50E-01 | 2.77E+00 | 2.81E+00 | 7.21E-01 | no | | intergenic_variant | MODIFIER |
| 8 | 42069848 | AX-97839947 | 0 | C | A | 0.149 | 5.31E-02 | 1.50E-01 | 2.77E+00 | 2.81E+00 | 7.21E-01 | no | | intergenic_variant | MODIFIER |
| 8 | 42070847 | AX-170454209 | 0 | C | T | 0.149 | 5.31E-02 | 1.50E-01 | 2.77E+00 | 2.81E+00 | 7.21E-01 | no | | intergenic_variant | MODIFIER |
| 8 | 42100282 | AX-170534379 | 0 | G | C | 0.149 | 5.31E-02 | 1.50E-01 | 2.77E+00 | 2.81E+00 | 7.21E-01 | no | | intergenic_variant | MODIFIER |
| 8 | 42109858 | AX-170111676 | 0 | C | A | 0.149 | 5.31E-02 | 1.50E-01 | 2.77E+00 | 2.81E+00 | 7.21E-01 | no | | intergenic_variant | MODIFIER |
| 8 | 42110099 | AX-170483826 | 0 | A | G | 0.149 | 5.31E-02 | 1.50E-01 | 2.77E+00 | 2.81E+00 | 7.21E-01 | no | | intergenic_variant | MODIFIER |
| 8 | 42115292 | AX-170108938 | 0 | G | A | 0.149 | 5.31E-02 | 1.50E-01 | 2.77E+00 | 2.81E+00 | 7.21E-01 | no | | intergenic_variant | MODIFIER |
| 8 | 42115318 | AX-170446038 | 0 | C | T | 0.149 | 5.31E-02 | 1.50E-01 | 2.77E+00 | 2.81E+00 | 7.21E-01 | no | | intergenic_variant | MODIFIER |
| 8 | 42154511 | AX-170242491 | 0 | A | G | 0.149 | 5.31E-02 | 1.50E-01 | 2.77E+00 | 2.81E+00 | 7.21E-01 | no | | intergenic_variant | MODIFIER |
| 8 | 42165575 | AX-170365064 | 0 | A | C | 0.149 | 5.31E-02 | 1.50E-01 | 2.77E+00 | 2.81E+00 | 7.21E-01 | no | | intergenic_variant | MODIFIER |
| 8 | 42165744 | AX-170292841 | 0 | G | A | 0.149 | 5.31E-02 | 1.50E-01 | 2.77E+00 | 2.81E+00 | 7.21E-01 | no | | intergenic_variant | MODIFIER |
| 8 | 42168805 | AX-170347964 | 0 | A | C | 0.149 | 5.31E-02 | 1.50E-01 | 2.77E+00 | 2.81E+00 | 7.21E-01 | no | | intergenic_variant | MODIFIER |
| 8 | 42207846 | AX-170516415 | 0 | G | C | 0.149 | 5.31E-02 | 1.50E-01 | 2.77E+00 | 2.81E+00 | 7.21E-01 | no | | intergenic_variant | MODIFIER |
| 8 | 42207874 | AX-97850536 | 0 | A | C | 0.149 | 5.31E-02 | 1.50E-01 | 2.77E+00 | 2.81E+00 | 7.21E-01 | no | | intergenic_variant | MODIFIER |

| | | | | | | | | | | | | | | | |
|---|---|---|---|---|---|---|---|---|---|---|---|---|---|---|---|
| 8 | 42258284 | AX-170498835 | 0 | T | C | 0.149 | 5.31E-02 | 1.50E-01 | 2.77E+00 | 2.81E+00 | 7.21E-01 | no | | intergenic_variant | MODIFIER |
| 8 | 42263985 | AX-170079892 | 0 | C | T | 0.149 | 5.31E-02 | 1.50E-01 | 2.77E+00 | 2.81E+00 | 7.21E-01 | no | | intergenic_variant | MODIFIER |
| 8 | 42265735 | AX-170079891 | 0 | A | G | 0.149 | 5.31E-02 | 1.50E-01 | 2.77E+00 | 2.81E+00 | 7.21E-01 | no | | intergenic_variant | MODIFIER |
| 8 | 42281315 | AX-97840020 | 0 | C | T | 0.149 | 5.31E-02 | 1.50E-01 | 2.77E+00 | 2.81E+00 | 7.21E-01 | no | | intergenic_variant | MODIFIER |
| 8 | 42292519 | AX-170526578 | 0 | T | G | 0.149 | 5.31E-02 | 1.50E-01 | 2.77E+00 | 2.81E+00 | 7.21E-01 | no | | intergenic_variant | MODIFIER |
| 8 | 42317944 | AX-170170080 | 0 | G | A | 0.149 | 5.31E-02 | 1.50E-01 | 2.77E+00 | 2.81E+00 | 7.21E-01 | no | | intergenic_variant | MODIFIER |
| 8 | 42349575 | AX-97810056 | 0 | G | C | 0.149 | 5.31E-02 | 1.50E-01 | 2.77E+00 | 2.81E+00 | 7.21E-01 | no | | intergenic_variant | MODIFIER |
| 8 | 42351758 | AX-170231088 | 0 | A | G | 0.149 | 5.31E-02 | 1.50E-01 | 2.77E+00 | 2.81E+00 | 7.21E-01 | no | | intergenic_variant | MODIFIER |
| 8 | 42372807 | AX-170121954 | 0 | T | A | 0.149 | 5.31E-02 | 1.50E-01 | 2.77E+00 | 2.81E+00 | 7.21E-01 | no | | intergenic_variant | MODIFIER |
| 8 | 42402557 | AX-170196844 | 0 | C | G | 0.149 | 5.31E-02 | 1.50E-01 | 2.77E+00 | 2.81E+00 | 7.21E-01 | no | | intergenic_variant | MODIFIER |
| 8 | 42415066 | AX-170357854 | 0 | C | T | 0.149 | 5.31E-02 | 1.50E-01 | 2.77E+00 | 2.81E+00 | 7.21E-01 | no | | intergenic_variant | MODIFIER |
| 8 | 42415135 | AX-170133105 | 0 | G | A | 0.149 | 5.31E-02 | 1.50E-01 | 2.77E+00 | 2.81E+00 | 7.21E-01 | no | | intergenic_variant | MODIFIER |
| 8 | 42424190 | AX-170030571 | 0 | G | A | 0.149 | 5.31E-02 | 1.50E-01 | 2.77E+00 | 2.81E+00 | 7.21E-01 | no | | intergenic_variant | MODIFIER |
| 8 | 42443402 | AX-170030569 | 0 | G | T | 0.149 | 5.31E-02 | 1.50E-01 | 2.77E+00 | 2.81E+00 | 7.21E-01 | no | | intergenic_variant | MODIFIER |
| 8 | 42476246 | AX-97804137 | 0 | T | C | 0.149 | 5.31E-02 | 1.50E-01 | 2.77E+00 | 2.81E+00 | 7.21E-01 | no | | intergenic_variant | MODIFIER |
| 8 | 42481290 | AX-170030575 | 0 | T | A | 0.149 | 5.31E-02 | 1.50E-01 | 2.77E+00 | 2.81E+00 | 7.21E-01 | no | | intergenic_variant | MODIFIER |
| 8 | 42524645 | AX-170148400 | 0 | G | A | 0.149 | 5.31E-02 | 1.50E-01 | 2.77E+00 | 2.81E+00 | 7.21E-01 | no | Gm23531 | upstream_gene_variant | MODIFIER |
| 8 | 42537968 | AX-170226549 | 0 | C | A | 0.149 | 5.31E-02 | 1.50E-01 | 2.77E+00 | 2.81E+00 | 7.21E-01 | no | | intergenic_variant | MODIFIER |
| 8 | 42558882 | AX-170287654 | 0 | G | A | 0.149 | 5.31E-02 | 1.50E-01 | 2.77E+00 | 2.81E+00 | 7.21E-01 | no | | intergenic_variant | MODIFIER |
| 8 | 42559387 | AX-170096241 | 0 | A | G | 0.149 | 5.31E-02 | 1.50E-01 | 2.77E+00 | 2.81E+00 | 7.21E-01 | no | | intergenic_variant | MODIFIER |
| 8 | 42577052 | AX-170253098 | 0 | A | G | 0.149 | 5.31E-02 | 1.50E-01 | 2.77E+00 | 2.81E+00 | 7.21E-01 | no | | intergenic_variant | MODIFIER |
| 8 | 42591468 | AX-97853116 | 0 | G | A | 0.149 | 5.31E-02 | 1.50E-01 | 2.77E+00 | 2.81E+00 | 7.21E-01 | no | | intergenic_variant | MODIFIER |
| 8 | 42598053 | AX-170250242 | 0 | G | A | 0.149 | 5.31E-02 | 1.50E-01 | 2.77E+00 | 2.81E+00 | 7.21E-01 | no | | intergenic_variant | MODIFIER |
| 8 | 42614626 | AX-170079898 | 0 | C | T | 0.149 | 5.31E-02 | 1.50E-01 | 2.77E+00 | 2.81E+00 | 7.21E-01 | no | | intergenic_variant | MODIFIER |
| 8 | 42640541 | AX-170294231 | 0 | C | T | 0.149 | 5.31E-02 | 1.50E-01 | 2.77E+00 | 2.81E+00 | 7.21E-01 | no | | intergenic_variant | MODIFIER |
| 8 | 42670217 | AX-97804136 | 0 | G | C | 0.149 | 5.31E-02 | 1.50E-01 | 2.77E+00 | 2.81E+00 | 7.21E-01 | no | | intergenic_variant | MODIFIER |
| 8 | 42680797 | AX-170455700 | 0 | A | G | 0.149 | 5.31E-02 | 1.50E-01 | 2.77E+00 | 2.81E+00 | 7.21E-01 | no | | intergenic_variant | MODIFIER |
| 8 | 42699305 | AX-170362496 | 0 | G | C | 0.149 | 5.31E-02 | 1.50E-01 | 2.77E+00 | 2.81E+00 | 7.21E-01 | no | | intergenic_variant | MODIFIER |
| 8 | 42764142 | AX-170151709 | 0 | A | C | 0.149 | 5.31E-02 | 1.50E-01 | 2.77E+00 | 2.81E+00 | 7.21E-01 | no | | intergenic_variant | MODIFIER |
| 8 | 42764171 | AX-170111156 | 0 | A | G | 0.149 | 5.31E-02 | 1.50E-01 | 2.77E+00 | 2.81E+00 | 7.21E-01 | no | | intergenic_variant | MODIFIER |
| 8 | 42764979 | AX-170315208 | 0 | T | C | 0.149 | 5.31E-02 | 1.50E-01 | 2.77E+00 | 2.81E+00 | 7.21E-01 | no | Gm18414 | upstream_gene_variant | MODIFIER |
| 8 | 42765556 | AX-170185252 | 0 | C | T | 0.149 | 5.31E-02 | 1.50E-01 | 2.77E+00 | 2.81E+00 | 7.21E-01 | no | Gm18414 | upstream_gene_variant | MODIFIER |
| 8 | 42769958 | AX-170460118 | 0 | C | T | 0.149 | 5.31E-02 | 1.50E-01 | 2.77E+00 | 2.81E+00 | 7.21E-01 | no | Gm18414 | non_coding_transcript_exon_variant | MODIFIER |
| 8 | 42771346 | AX-170118379 | 0 | C | T | 0.149 | 5.31E-02 | 1.50E-01 | 2.77E+00 | 2.81E+00 | 7.21E-01 | no | Gm18414 | downstream_gene_variant | MODIFIER |
| 8 | 42774237 | AX-170195560 | 0 | G | A | 0.149 | 5.31E-02 | 1.50E-01 | 2.77E+00 | 2.81E+00 | 7.21E-01 | no | Gm18414 | downstream_gene_variant | MODIFIER |
| 8 | 42795500 | AX-97838211 | 0 | T | C | 0.149 | 5.31E-02 | 1.50E-01 | 2.77E+00 | 2.81E+00 | 7.21E-01 | no | | intergenic_variant | MODIFIER |
| 8 | 42810258 | AX-170207325 | 0 | G | C | 0.149 | 5.31E-02 | 1.50E-01 | 2.77E+00 | 2.81E+00 | 7.21E-01 | no | | intergenic_variant | MODIFIER |
| 8 | 42837633 | AX-170140635 | 0 | A | C | 0.149 | 5.31E-02 | 1.50E-01 | 2.77E+00 | 2.81E+00 | 7.21E-01 | no | | intergenic_variant | MODIFIER |
| 8 | 42838036 | AX-170333920 | 0 | G | A | 0.149 | 5.31E-02 | 1.50E-01 | 2.77E+00 | 2.81E+00 | 7.21E-01 | no | | intergenic_variant | MODIFIER |
| 8 | 42856654 | AX-170424869 | 0 | C | G | 0.149 | 5.31E-02 | 1.50E-01 | 2.77E+00 | 2.81E+00 | 7.21E-01 | no | | intergenic_variant | MODIFIER |
| 8 | 42857064 | AX-97837854 | 0 | C | T | 0.149 | 5.31E-02 | 1.50E-01 | 2.77E+00 | 2.81E+00 | 7.21E-01 | no | | intergenic_variant | MODIFIER |
| 8 | 42870001 | AX-170322779 | 0 | C | T | 0.149 | 5.31E-02 | 1.50E-01 | 2.77E+00 | 2.81E+00 | 7.21E-01 | no | | intergenic_variant | MODIFIER |
| 8 | 42874942 | AX-97822366 | 0 | T | C | 0.149 | 5.31E-02 | 1.50E-01 | 2.77E+00 | 2.81E+00 | 7.21E-01 | no | | intergenic_variant | MODIFIER |
| 8 | 42888122 | AX-170079903 | 0 | G | A | 0.149 | 5.31E-02 | 1.50E-01 | 2.77E+00 | 2.81E+00 | 7.21E-01 | no | | intergenic_variant | MODIFIER |
| 8 | 42921729 | AX-170401665 | 0 | A | G | 0.149 | 5.31E-02 | 1.50E-01 | 2.77E+00 | 2.81E+00 | 7.21E-01 | no | | intergenic_variant | MODIFIER |
| 8 | 42921994 | AX-170079906 | 0 | G | T | 0.149 | 5.31E-02 | 1.50E-01 | 2.77E+00 | 2.81E+00 | 7.21E-01 | no | | intergenic_variant | MODIFIER |
| 8 | 42932989 | AX-97804135 | 0 | A | C | 0.149 | 5.31E-02 | 1.50E-01 | 2.77E+00 | 2.81E+00 | 7.21E-01 | no | | intergenic_variant | MODIFIER |
| 8 | 42935787 | AX-170434462 | 0 | T | C | 0.149 | 5.31E-02 | 1.50E-01 | 2.77E+00 | 2.81E+00 | 7.21E-01 | no | | intergenic_variant | MODIFIER |
| 8 | 42945611 | AX-170030577 | 0 | C | T | 0.149 | 5.31E-02 | 1.50E-01 | 2.77E+00 | 2.81E+00 | 7.21E-01 | no | | intergenic_variant | MODIFIER |
| 8 | 42946339 | AX-170288540 | 0 | C | T | 0.149 | 5.31E-02 | 1.50E-01 | 2.77E+00 | 2.81E+00 | 7.21E-01 | no | | intergenic_variant | MODIFIER |
| 8 | 43021021 | AX-170496742 | 0 | C | T | 0.149 | 5.31E-02 | 1.50E-01 | 2.77E+00 | 2.81E+00 | 7.21E-01 | no | | intergenic_variant | MODIFIER |
| 8 | 43021124 | AX-170278987 | 0 | G | A | 0.149 | 5.31E-02 | 1.50E-01 | 2.77E+00 | 2.81E+00 | 7.21E-01 | no | | intergenic_variant | MODIFIER |
| 8 | 43022964 | AX-170338439 | 0 | C | T | 0.149 | 5.31E-02 | 1.50E-01 | 2.77E+00 | 2.81E+00 | 7.21E-01 | no | | intergenic_variant | MODIFIER |
| 8 | 43059128 | AX-170390712 | 0 | G | A | 0.149 | 5.31E-02 | 1.50E-01 | 2.77E+00 | 2.81E+00 | 7.21E-01 | no | | intergenic_variant | MODIFIER |

| | | | | | | | | | | | | | | | |
|---|---|---|---|---|---|---|---|---|---|---|---|---|---|---|---|
| 8 | 43064111 | AX-170162948 | 0 | G | A | 0.149 | 5.31E-02 | 1.50E-01 | 2.77E+00 | 2.81E+00 | 7.21E-01 | no | | intergenic_variant | MODIFIER |
| 8 | 43071246 | AX-170235487 | 0 | C | G | 0.149 | 5.31E-02 | 1.50E-01 | 2.77E+00 | 2.81E+00 | 7.21E-01 | no | | intergenic_variant | MODIFIER |
| 8 | 43076648 | AX-170357136 | 0 | T | G | 0.149 | 5.31E-02 | 1.50E-01 | 2.77E+00 | 2.81E+00 | 7.21E-01 | no | | intergenic_variant | MODIFIER |
| 8 | 43102127 | AX-170298420 | 0 | G | A | 0.149 | 5.31E-02 | 1.50E-01 | 2.77E+00 | 2.81E+00 | 7.21E-01 | no | | intergenic_variant | MODIFIER |
| 8 | 43117589 | AX-170491615 | 0 | A | G | 0.149 | 5.31E-02 | 1.50E-01 | 2.77E+00 | 2.81E+00 | 7.21E-01 | no | | intergenic_variant | MODIFIER |
| 8 | 43130285 | AX-170122133 | 0 | G | A | 0.149 | 5.31E-02 | 1.50E-01 | 2.77E+00 | 2.81E+00 | 7.21E-01 | no | Triml1 | synonymous_variant | LOW |
| 8 | 43136741 | AX-170426975 | 0 | T | C | 0.149 | 5.31E-02 | 1.50E-01 | 2.77E+00 | 2.81E+00 | 7.21E-01 | no | Triml1 | intron_variant | MODIFIER |
| 8 | 43137952 | AX-97836677 | 0 | A | G | 0.149 | 5.31E-02 | 1.50E-01 | 2.77E+00 | 2.81E+00 | 7.21E-01 | no | Triml1 | intron_variant | MODIFIER |
| 8 | 43159977 | AX-170173311 | 0 | T | C | 0.149 | 5.31E-02 | 1.50E-01 | 2.77E+00 | 2.81E+00 | 7.21E-01 | no | Gm6284 | non_coding_transcript_exon_variant | MODIFIER |
| 8 | 43167268 | AX-170490621 | 0 | A | G | 0.149 | 5.31E-02 | 1.50E-01 | 2.77E+00 | 2.81E+00 | 7.21E-01 | no | Gm6284 | downstream_gene_variant | MODIFIER |
| 8 | 43187053 | AX-170276069 | 0 | A | G | 0.149 | 5.31E-02 | 1.50E-01 | 2.77E+00 | 2.81E+00 | 7.21E-01 | no | Triml2 | intron_variant | MODIFIER |
| 8 | 43188814 | AX-170090088 | 0 | T | C | 0.149 | 5.31E-02 | 1.50E-01 | 2.77E+00 | 2.81E+00 | 7.21E-01 | no | Triml2 | intron_variant | MODIFIER |
| 8 | 43215947 | AX-170363540 | 0 | T | C | 0.149 | 5.31E-02 | 1.50E-01 | 2.77E+00 | 2.81E+00 | 7.21E-01 | no | | intergenic_variant | MODIFIER |
| 8 | 43217608 | AX-170372849 | 0 | T | C | 0.149 | 5.31E-02 | 1.50E-01 | 2.77E+00 | 2.81E+00 | 7.21E-01 | no | | intergenic_variant | MODIFIER |
| 8 | 43231546 | AX-170497196 | 0 | G | T | 0.149 | 5.31E-02 | 1.50E-01 | 2.77E+00 | 2.81E+00 | 7.21E-01 | no | | intergenic_variant | MODIFIER |
| 8 | 43243951 | AX-170487443 | 0 | G | A | 0.149 | 5.31E-02 | 1.50E-01 | 2.77E+00 | 2.81E+00 | 7.21E-01 | no | | intergenic_variant | MODIFIER |
| 8 | 43274527 | AX-170519483 | 0 | T | C | 0.149 | 5.31E-02 | 1.50E-01 | 2.77E+00 | 2.81E+00 | 7.21E-01 | no | | intergenic_variant | MODIFIER |
| 8 | 43274699 | AX-170308599 | 0 | C | G | 0.149 | 5.31E-02 | 1.50E-01 | 2.77E+00 | 2.81E+00 | 7.21E-01 | no | | intergenic_variant | MODIFIER |
| 8 | 43277791 | AX-170167510 | 0 | A | G | 0.149 | 5.31E-02 | 1.50E-01 | 2.77E+00 | 2.81E+00 | 7.21E-01 | no | | intergenic_variant | MODIFIER |
| 8 | 43286405 | AX-97845272 | 0 | G | A | 0.149 | 5.31E-02 | 1.50E-01 | 2.77E+00 | 2.81E+00 | 7.21E-01 | no | | intergenic_variant | MODIFIER |
| 8 | 43300389 | AX-170317715 | 0 | C | G | 0.149 | 5.31E-02 | 1.50E-01 | 2.77E+00 | 2.81E+00 | 7.21E-01 | no | Zfp42 | intron_variant | MODIFIER |
| 8 | 43309272 | AX-170230411 | 0 | C | T | 0.149 | 5.31E-02 | 1.50E-01 | 2.77E+00 | 2.81E+00 | 7.21E-01 | no | Zfp42 | upstream_gene_variant | MODIFIER |
| 8 | 43309586 | AX-170079911 | 0 | C | T | 0.149 | 5.31E-02 | 1.50E-01 | 2.77E+00 | 2.81E+00 | 7.21E-01 | no | Zfp42 | upstream_gene_variant | MODIFIER |
| 8 | 43332204 | AX-170420180 | 0 | A | G | 0.149 | 5.31E-02 | 1.50E-01 | 2.77E+00 | 2.81E+00 | 7.21E-01 | no | | intergenic_variant | MODIFIER |
| 8 | 43336297 | AX-170435787 | 0 | A | G | 0.149 | 5.31E-02 | 1.50E-01 | 2.77E+00 | 2.81E+00 | 7.21E-01 | no | | intergenic_variant | MODIFIER |
| 8 | 43340877 | AX-170228628 | 0 | T | C | 0.149 | 5.31E-02 | 1.50E-01 | 2.77E+00 | 2.81E+00 | 7.21E-01 | no | | intergenic_variant | MODIFIER |
| 8 | 43354948 | AX-170327036 | 0 | C | T | 0.149 | 5.31E-02 | 1.50E-01 | 2.77E+00 | 2.81E+00 | 7.21E-01 | no | | intergenic_variant | MODIFIER |
| 8 | 44812025 | AX-97807955 | 0 | T | C | 0.149 | 5.31E-02 | 1.50E-01 | 2.77E+00 | 2.81E+00 | 7.21E-01 | no | | intergenic_variant | MODIFIER |
| 8 | 45043434 | AX-170209639 | 0 | C | A | 0.149 | 5.31E-02 | 1.50E-01 | 2.77E+00 | 2.81E+00 | 7.21E-01 | no | Fat1 | intron_variant | MODIFIER |
| 8 | 45072486 | AX-170430516 | 0 | T | G | 0.149 | 5.31E-02 | 1.50E-01 | 2.77E+00 | 2.81E+00 | 7.21E-01 | no | Mtnr1a | intron_variant | MODIFIER |
| 8 | 45157116 | AX-170339247 | 0 | G | C | 0.149 | 5.31E-02 | 1.50E-01 | 2.77E+00 | 2.81E+00 | 7.21E-01 | no | Gm6329 | downstream_gene_variant | MODIFIER |
| 8 | 45159459 | AX-170316728 | 0 | A | G | 0.149 | 5.31E-02 | 1.50E-01 | 2.77E+00 | 2.81E+00 | 7.21E-01 | no | Gm6329 | downstream_gene_variant | MODIFIER |
| 8 | 45159667 | AX-170210829 | 0 | A | T | 0.149 | 5.31E-02 | 1.50E-01 | 2.77E+00 | 2.81E+00 | 7.21E-01 | no | Gm6329 | downstream_gene_variant | MODIFIER |
| 8 | 45405903 | AX-170463413 | 0 | A | G | 0.149 | 5.31E-02 | 1.50E-01 | 2.77E+00 | 2.81E+00 | 7.21E-01 | no | Tlr3 | intron_variant | MODIFIER |
| 8 | 45473732 | AX-170153601 | 0 | A | G | 0.149 | 5.31E-02 | 1.50E-01 | 2.77E+00 | 2.81E+00 | 7.21E-01 | no | | intergenic_variant | MODIFIER |
| 8 | 45475764 | AX-170310244 | 0 | C | T | 0.149 | 5.31E-02 | 1.50E-01 | 2.77E+00 | 2.81E+00 | 7.21E-01 | no | | intergenic_variant | MODIFIER |
| 8 | 45526964 | AX-170465418 | 0 | T | C | 0.149 | 5.31E-02 | 1.50E-01 | 2.77E+00 | 2.81E+00 | 7.21E-01 | no | Sorbs2 | intron_variant | MODIFIER |
| 8 | 45545649 | AX-170397955 | 0 | A | G | 0.149 | 5.31E-02 | 1.50E-01 | 2.77E+00 | 2.81E+00 | 7.21E-01 | no | Sorbs2 | intron_variant | MODIFIER |
| 8 | 45573078 | AX-170079956 | 0 | T | C | 0.149 | 5.31E-02 | 1.50E-01 | 2.77E+00 | 2.81E+00 | 7.21E-01 | no | Sorbs2 | intron_variant | MODIFIER |
| 8 | 45574304 | AX-97835014 | 0 | T | C | 0.149 | 5.31E-02 | 1.50E-01 | 2.77E+00 | 2.81E+00 | 7.21E-01 | no | Sorbs2 | intron_variant | MODIFIER |
| 8 | 45584783 | AX-170243000 | 0 | C | A | 0.149 | 5.31E-02 | 1.50E-01 | 2.77E+00 | 2.81E+00 | 7.21E-01 | no | Sorbs2 | intron_variant | MODIFIER |
| 8 | 45586358 | AX-170030608 | 0 | T | G | 0.149 | 5.31E-02 | 1.50E-01 | 2.77E+00 | 2.81E+00 | 7.21E-01 | no | Sorbs2 | intron_variant | MODIFIER |
| 8 | 45610559 | AX-170530560 | 0 | A | G | 0.149 | 5.31E-02 | 1.50E-01 | 2.77E+00 | 2.81E+00 | 7.21E-01 | no | Sorbs2 | intron_variant | MODIFIER |
| 8 | 45709584 | AX-170533012 | 0 | C | T | 0.149 | 5.31E-02 | 1.50E-01 | 2.77E+00 | 2.81E+00 | 7.21E-01 | no | Sorbs2 | intron_variant | MODIFIER |
| 8 | 45825912 | AX-170180850 | 0 | A | G | 0.149 | 5.31E-02 | 1.50E-01 | 2.77E+00 | 2.81E+00 | 7.21E-01 | no | Sorbs2 | 3_prime_UTR_variant | MODIFIER |
| 8 | 45842054 | AX-170117710 | 0 | G | A | 0.149 | 5.31E-02 | 1.50E-01 | 2.77E+00 | 2.81E+00 | 7.21E-01 | no | | intergenic_variant | MODIFIER |
| 8 | 45846123 | AX-97830527 | 0 | G | A | 0.149 | 5.31E-02 | 1.50E-01 | 2.77E+00 | 2.81E+00 | 7.21E-01 | no | | regulatory_region_variant | MODIFIER |
| 8 | 45878008 | AX-170470504 | 0 | T | G | 0.149 | 5.31E-02 | 1.50E-01 | 2.77E+00 | 2.81E+00 | 7.21E-01 | no | | intergenic_variant | MODIFIER |
| 8 | 45889240 | AX-170124093 | 0 | T | C | 0.149 | 5.31E-02 | 1.50E-01 | 2.77E+00 | 2.81E+00 | 7.21E-01 | no | Pdlim3 | intron_variant | MODIFIER |
| 8 | 45902163 | AX-170480912 | 0 | A | T | 0.149 | 5.31E-02 | 1.50E-01 | 2.77E+00 | 2.81E+00 | 7.21E-01 | no | Pdlim3 | intron_variant | MODIFIER |
| 8 | 45905358 | AX-170183746 | 0 | C | T | 0.149 | 5.31E-02 | 1.50E-01 | 2.77E+00 | 2.81E+00 | 7.21E-01 | no | Pdlim3 | intron_variant | MODIFIER |
| 8 | 45907197 | AX-170442839 | 0 | T | G | 0.149 | 5.31E-02 | 1.50E-01 | 2.77E+00 | 2.81E+00 | 7.21E-01 | no | Pdlim3 | intron_variant | MODIFIER |
| 8 | 45919078 | AX-170234622 | 0 | T | C | 0.149 | 5.31E-02 | 1.50E-01 | 2.77E+00 | 2.81E+00 | 7.21E-01 | no | Pdlim3 | 3_prime_UTR_variant | MODIFIER |
| 8 | 45919375 | AX-97833442 | 0 | A | G | 0.149 | 5.31E-02 | 1.50E-01 | 2.77E+00 | 2.81E+00 | 7.21E-01 | no | Pdlim3 | 3_prime_UTR_variant | MODIFIER |

| | | | | | | | | | | | | | | | |
|---|---|---|---|---|---|---|---|---|---|---|---|---|---|---|---|
| 8 | 45919573 | AX-170498246 | 0 | G | T | 0.149 | 5.31E-02 | 1.50E-01 | 2.77E+00 | 2.81E+00 | 7.21E-01 | no | Pdlim3 | downstream_gene_variant | MODIFIER |
| 8 | 45920934 | AX-170214283 | 0 | T | C | 0.149 | 5.31E-02 | 1.50E-01 | 2.77E+00 | 2.81E+00 | 7.21E-01 | no | Pdlim3 | downstream_gene_variant | MODIFIER |
| 8 | 45953605 | AX-170412756 | 0 | T | C | 0.149 | 5.31E-02 | 1.50E-01 | 2.77E+00 | 2.81E+00 | 7.21E-01 | no | 1700029J07Rik | downstream_gene_variant | MODIFIER |
| 8 | 45965040 | AX-170181229 | 0 | G | T | 0.149 | 5.31E-02 | 1.50E-01 | 2.77E+00 | 2.81E+00 | 7.21E-01 | no | 1700029J07Rik | intron_variant | MODIFIER |
| 8 | 45967756 | AX-97816221 | 0 | T | C | 0.149 | 5.31E-02 | 1.50E-01 | 2.77E+00 | 2.81E+00 | 7.21E-01 | no | 1700029J07Rik | intron_variant | MODIFIER |
| 8 | 45976446 | AX-170407358 | 0 | A | G | 0.149 | 5.31E-02 | 1.50E-01 | 2.77E+00 | 2.81E+00 | 7.21E-01 | no | Ufsp2 | intron_variant | MODIFIER |
| 8 | 45982238 | AX-170407802 | 0 | G | A | 0.149 | 5.31E-02 | 1.50E-01 | 2.77E+00 | 2.81E+00 | 7.21E-01 | no | Ufsp2 | intron_variant | MODIFIER |
| 8 | 45989730 | AX-97787139 | 0 | C | T | 0.149 | 5.31E-02 | 1.50E-01 | 2.77E+00 | 2.81E+00 | 7.21E-01 | no | Ufsp2 | intron_variant | MODIFIER |
| 8 | 45995888 | AX-170315819 | 0 | C | T | 0.149 | 5.31E-02 | 1.50E-01 | 2.77E+00 | 2.81E+00 | 7.21E-01 | no | Ufsp2 | intron_variant | MODIFIER |
| 8 | 46065547 | AX-170228875 | 0 | T | C | 0.149 | 5.31E-02 | 1.50E-01 | 2.77E+00 | 2.81E+00 | 7.21E-01 | no | Snx25 | intron_variant | MODIFIER |
| 8 | 46276876 | AX-170281238 | 0 | G | A | 0.149 | 5.31E-02 | 1.50E-01 | 2.77E+00 | 2.81E+00 | 7.21E-01 | no | Gm24669 | downstream_gene_variant | MODIFIER |
| 8 | 46277201 | AX-170174841 | 0 | G | A | 0.149 | 5.31E-02 | 1.50E-01 | 2.77E+00 | 2.81E+00 | 7.21E-01 | no | Gm8575 | upstream_gene_variant | MODIFIER |
| 8 | 46277732 | AX-170251057 | 0 | A | C | 0.149 | 5.31E-02 | 1.50E-01 | 2.77E+00 | 2.81E+00 | 7.21E-01 | no | Gm8575 | upstream_gene_variant | MODIFIER |
| 8 | 46281362 | AX-97842533 | 0 | T | C | 0.149 | 5.31E-02 | 1.50E-01 | 2.77E+00 | 2.81E+00 | 7.21E-01 | no | Gm8575 | downstream_gene_variant | MODIFIER |
| 8 | 46293984 | AX-170428432 | 0 | T | C | 0.149 | 5.31E-02 | 1.50E-01 | 2.77E+00 | 2.81E+00 | 7.21E-01 | no | Helt | intron_variant | MODIFIER |
| 8 | 46316173 | AX-170232376 | 0 | C | A | 0.149 | 5.31E-02 | 1.50E-01 | 2.77E+00 | 2.81E+00 | 7.21E-01 | no | Gm45255 | downstream_gene_variant | MODIFIER |
| 8 | 46318636 | AX-170369215 | 0 | G | A | 0.149 | 5.31E-02 | 1.50E-01 | 2.77E+00 | 2.81E+00 | 7.21E-01 | no | Gm45255 | downstream_gene_variant | MODIFIER |
| 8 | 46329396 | AX-170484896 | 0 | A | G | 0.149 | 5.31E-02 | 1.50E-01 | 2.77E+00 | 2.81E+00 | 7.21E-01 | no | Gm39168 | downstream_gene_variant | MODIFIER |
| 8 | 46352031 | AX-170096252 | 0 | T | C | 0.149 | 5.31E-02 | 1.50E-01 | 2.77E+00 | 2.81E+00 | 7.21E-01 | no | | intergenic_variant | MODIFIER |
| 8 | 46363339 | AX-170419328 | 0 | T | C | 0.149 | 5.31E-02 | 1.50E-01 | 2.77E+00 | 2.81E+00 | 7.21E-01 | no | Gm30931 | non_coding_transcript_exon_variant | MODIFIER |
| 8 | 46364023 | AX-170487074 | 0 | T | C | 0.149 | 5.31E-02 | 1.50E-01 | 2.77E+00 | 2.81E+00 | 7.21E-01 | no | Gm30931 | intron_variant,non_coding_transcript_variant | MODIFIER |
| 8 | 46394247 | AX-170324332 | 0 | T | C | 0.149 | 5.31E-02 | 1.50E-01 | 2.77E+00 | 2.81E+00 | 7.21E-01 | no | | intergenic_variant | MODIFIER |
| 8 | 46399801 | AX-170518488 | 0 | G | A | 0.149 | 5.31E-02 | 1.50E-01 | 2.77E+00 | 2.81E+00 | 7.21E-01 | no | | intergenic_variant | MODIFIER |
| 8 | 46414812 | AX-170534751 | 0 | A | G | 0.149 | 5.31E-02 | 1.50E-01 | 2.77E+00 | 2.81E+00 | 7.21E-01 | no | | regulatory_region_variant | MODIFIER |
| 8 | 46431280 | AX-170157372 | 0 | T | A | 0.149 | 5.31E-02 | 1.50E-01 | 2.77E+00 | 2.81E+00 | 7.21E-01 | no | | intergenic_variant | MODIFIER |
| 8 | 46438869 | AX-170320892 | 0 | G | T | 0.149 | 5.31E-02 | 1.50E-01 | 2.77E+00 | 2.81E+00 | 7.21E-01 | no | | intergenic_variant | MODIFIER |
| 8 | 46446472 | AX-170079974 | 0 | T | A | 0.149 | 5.31E-02 | 1.50E-01 | 2.77E+00 | 2.81E+00 | 7.21E-01 | no | | regulatory_region_variant | MODIFIER |
| 8 | 46448383 | AX-170079973 | 0 | C | T | 0.149 | 5.31E-02 | 1.50E-01 | 2.77E+00 | 2.81E+00 | 7.21E-01 | no | Gm45245 | downstream_gene_variant | MODIFIER |
| 8 | 46449660 | AX-170079971 | 0 | G | A | 0.149 | 5.31E-02 | 1.50E-01 | 2.77E+00 | 2.81E+00 | 7.21E-01 | no | Gm45245 | downstream_gene_variant | MODIFIER |
| 8 | 46474399 | AX-170478278 | 0 | G | T | 0.149 | 5.31E-02 | 1.50E-01 | 2.77E+00 | 2.81E+00 | 7.21E-01 | no | Acsl1 | intron_variant | MODIFIER |
| 8 | 46492038 | AX-170492159 | 0 | A | G | 0.149 | 5.31E-02 | 1.50E-01 | 2.77E+00 | 2.81E+00 | 7.21E-01 | no | Acsl1 | intron_variant | MODIFIER |
| 8 | 46495797 | AX-170424811 | 0 | T | C | 0.149 | 5.31E-02 | 1.50E-01 | 2.77E+00 | 2.81E+00 | 7.21E-01 | no | Acsl1 | intron_variant | MODIFIER |
| 8 | 46517678 | AX-170509182 | 0 | A | G | 0.149 | 5.31E-02 | 1.50E-01 | 2.77E+00 | 2.81E+00 | 7.21E-01 | no | Acsl1 | intron_variant | MODIFIER |
| 8 | 46522683 | AX-170291508 | 0 | A | G | 0.149 | 5.31E-02 | 1.50E-01 | 2.77E+00 | 2.81E+00 | 7.21E-01 | no | Acsl1 | intron_variant | MODIFIER |
| 8 | 46530071 | AX-170517585 | 0 | G | A | 0.149 | 5.31E-02 | 1.50E-01 | 2.77E+00 | 2.81E+00 | 7.21E-01 | no | Acsl1 | intron_variant | MODIFIER |
| 18 | 65094152 | AX-170178935 | 0 | T | A | 0.421 | 2.86E-02 | 8.17E-02 | 2.78E+00 | 2.86E+00 | 7.21E-01 | no | Nedd4l | intron_variant | MODIFIER |
| 1 | 37906176 | AX-97831768 | 0 | T | C | 0.456 | 2.64E-02 | 7.60E-02 | 2.70E+00 | 2.79E+00 | 7.21E-01 | no | Lyg2 | 3_prime_UTR_variant | MODIFIER |
| 8 | 83794292 | AX-170080617 | 2 | A | T | 0.209 | 4.55E-02 | 1.30E-01 | 2.70E+00 | 2.76E+00 | 7.21E-01 | no | | intergenic_variant | MODIFIER |
| 2 | 144040026 | AX-170266926 | 1 | C | T | 0.143 | 7.83E-02 | 2.21E-01 | 2.70E+00 | 2.73E+00 | 7.21E-01 | no | Banf2 | intron_variant | MODIFIER |
| 2 | 145731262 | AX-170423091 | 1 | T | A | 0.143 | 7.83E-02 | 2.21E-01 | 2.70E+00 | 2.73E+00 | 7.21E-01 | no | Rin2 | intron_variant,non_coding_transcript_variant | MODIFIER |
| 7 | 35019568 | AX-170128851 | 0 | A | C | 0.333 | 3.31E-02 | 8.94E-02 | 2.57E+00 | 2.66E+00 | 7.21E-01 | no | Pepd | intron_variant | MODIFIER |
| 18 | 53559938 | AX-170060869 | 1 | G | A | 0.241 | -4.00E-02 | 1.10E-01 | 2.60E+00 | 2.66E+00 | 7.21E-01 | no | Prdm6 | intron_variant | MODIFIER |
| 18 | 53763011 | AX-97823824 | 1 | C | T | 0.241 | -4.00E-02 | 1.10E-01 | 2.60E+00 | 2.66E+00 | 7.21E-01 | no | | regulatory_region_variant | MODIFIER |
| 4 | 83714246 | AX-97790305 | 1 | G | T | 0.232 | 3.94E-02 | 1.05E-01 | 2.56E+00 | 2.64E+00 | 7.21E-01 | no | Ccdc171 | intron_variant | MODIFIER |
| 6 | 126507960 | AX-170076149 | 0 | G | A | 0.553 | 3.21E-02 | 8.61E-02 | 2.57E+00 | 2.63E+00 | 7.21E-01 | no | | intergenic_variant | MODIFIER |
| 11 | 108465774 | AX-170228865 | 0 | C | T | 0.053 | -6.44E-02 | 1.85E-01 | 2.65E+00 | 2.77E+00 | 7.21E-01 | no | Cep112 | intron_variant | MODIFIER |
| 11 | 108646182 | AX-170202351 | 0 | G | A | 0.053 | -6.44E-02 | 1.85E-01 | 2.65E+00 | 2.77E+00 | 7.21E-01 | no | Cep112 | intron_variant | MODIFIER |
| 11 | 109463233 | AX-170180433 | 0 | G | A | 0.053 | -6.44E-02 | 1.85E-01 | 2.65E+00 | 2.77E+00 | 7.21E-01 | no | Slc16a6 | intron_variant | MODIFIER |
| 11 | 116995616 | AX-170392069 | 0 | C | T | 0.053 | -6.44E-02 | 1.85E-01 | 2.65E+00 | 2.77E+00 | 7.21E-01 | no | | intergenic_variant | MODIFIER |
| 2 | 95246196 | AX-170416262 | 0 | A | G | 0.053 | -7.93E-02 | 2.21E-01 | 2.62E+00 | 2.71E+00 | 7.21E-01 | no | | intergenic_variant | MODIFIER |
| 8 | 62987523 | AX-97842061 | 0 | C | T | 0.14 | -4.52E-02 | 1.24E-01 | 2.57E+00 | 2.69E+00 | 7.21E-01 | no | Spock3 | intron_variant | MODIFIER |
| 5 | 32808415 | AX-170413028 | 1 | C | T | 0.366 | -3.54E-02 | 1.02E-01 | 2.71E+00 | 2.79E+00 | 7.21E-01 | no | Prr14l | intron_variant | MODIFIER |
| 11 | 32560095 | AX-170351312 | 0 | C | T | 0.096 | 6.83E-02 | 1.94E-01 | 2.68E+00 | 2.75E+00 | 7.21E-01 | no | Stk10 | intron_variant | MODIFIER |
| 1 | 96730402 | AX-170001146 | 1 | C | T | 0.143 | 6.75E-02 | 1.88E-01 | 2.61E+00 | 2.70E+00 | 7.22E-01 | no | | intergenic_variant | MODIFIER |

| | | | | | | | | | | | | | | | |
|---|---|---|---|---|---|---|---|---|---|---|---|---|---|---|---|
| 16 | 97813298 | AX-97809573 | 1 | C | T | 0.482 | 2.67E-02 | 7.77E-02 | 2.69E+00 | 2.81E+00 | 7.22E-01 | no | Prdm15 | intron_variant | MODIFIER |
| 8 | 115617501 | AX-97821596 | 0 | T | C | 0.254 | 3.38E-02 | 9.76E-02 | 2.67E+00 | 2.78E+00 | 7.22E-01 | no | | intergenic_variant | MODIFIER |
| 8 | 115899860 | AX-170031543 | 0 | T | C | 0.254 | 3.38E-02 | 9.76E-02 | 2.67E+00 | 2.78E+00 | 7.22E-01 | no | | intergenic_variant | MODIFIER |
| 8 | 115911823 | AX-170502309 | 0 | T | C | 0.254 | 3.38E-02 | 9.76E-02 | 2.67E+00 | 2.78E+00 | 7.22E-01 | no | | intergenic_variant | MODIFIER |
| 8 | 115916313 | AX-170219684 | 0 | T | C | 0.254 | 3.38E-02 | 9.76E-02 | 2.67E+00 | 2.78E+00 | 7.22E-01 | no | | intergenic_variant | MODIFIER |
| 8 | 115918099 | AX-170339884 | 0 | G | A | 0.254 | 3.38E-02 | 9.76E-02 | 2.67E+00 | 2.78E+00 | 7.22E-01 | no | | intergenic_variant | MODIFIER |
| 8 | 115918932 | AX-170237111 | 0 | G | A | 0.254 | 3.38E-02 | 9.76E-02 | 2.67E+00 | 2.78E+00 | 7.22E-01 | no | | intergenic_variant | MODIFIER |
| 8 | 115941781 | AX-170081266 | 0 | T | G | 0.254 | 3.38E-02 | 9.76E-02 | 2.67E+00 | 2.78E+00 | 7.22E-01 | no | | intergenic_variant | MODIFIER |
| 8 | 115945559 | AX-170096420 | 0 | T | C | 0.254 | 3.38E-02 | 9.76E-02 | 2.67E+00 | 2.78E+00 | 7.22E-01 | no | | intergenic_variant | MODIFIER |
| 8 | 115956862 | AX-170249980 | 0 | G | C | 0.254 | 3.38E-02 | 9.76E-02 | 2.67E+00 | 2.78E+00 | 7.22E-01 | no | | intergenic_variant | MODIFIER |
| 8 | 115974119 | AX-170528898 | 0 | A | G | 0.254 | 3.38E-02 | 9.76E-02 | 2.67E+00 | 2.78E+00 | 7.22E-01 | no | Gm45733 | upstream_gene_variant | MODIFIER |
| 8 | 115975610 | AX-170296948 | 0 | C | T | 0.254 | 3.38E-02 | 9.76E-02 | 2.67E+00 | 2.78E+00 | 7.22E-01 | no | Gm45733 | upstream_gene_variant | MODIFIER |
| 8 | 115981340 | AX-170169311 | 0 | A | C | 0.254 | 3.38E-02 | 9.76E-02 | 2.67E+00 | 2.78E+00 | 7.22E-01 | no | | intergenic_variant | MODIFIER |
| 8 | 116005815 | AX-97785221 | 0 | A | G | 0.254 | 3.38E-02 | 9.76E-02 | 2.67E+00 | 2.78E+00 | 7.22E-01 | no | | intergenic_variant | MODIFIER |
| 8 | 116014633 | AX-170283824 | 0 | T | C | 0.254 | 3.38E-02 | 9.76E-02 | 2.67E+00 | 2.78E+00 | 7.22E-01 | no | | intergenic_variant | MODIFIER |
| 8 | 116028016 | AX-170193680 | 0 | A | C | 0.254 | 3.38E-02 | 9.76E-02 | 2.67E+00 | 2.78E+00 | 7.22E-01 | no | | intergenic_variant | MODIFIER |
| 8 | 116063150 | AX-170446777 | 0 | G | A | 0.254 | 3.38E-02 | 9.76E-02 | 2.67E+00 | 2.78E+00 | 7.22E-01 | no | | intergenic_variant | MODIFIER |
| 8 | 116064898 | AX-97782455 | 0 | T | C | 0.254 | 3.38E-02 | 9.76E-02 | 2.67E+00 | 2.78E+00 | 7.22E-01 | no | | intergenic_variant | MODIFIER |
| 8 | 116075756 | AX-170243761 | 0 | G | C | 0.254 | 3.38E-02 | 9.76E-02 | 2.67E+00 | 2.78E+00 | 7.22E-01 | no | | intergenic_variant | MODIFIER |
| 8 | 116084212 | AX-170140273 | 0 | T | C | 0.254 | 3.38E-02 | 9.76E-02 | 2.67E+00 | 2.78E+00 | 7.22E-01 | no | | intergenic_variant | MODIFIER |
| 8 | 116119187 | AX-170484328 | 0 | T | C | 0.254 | 3.38E-02 | 9.76E-02 | 2.67E+00 | 2.78E+00 | 7.22E-01 | no | | intergenic_variant | MODIFIER |
| 8 | 116120224 | AX-170235014 | 0 | T | C | 0.254 | 3.38E-02 | 9.76E-02 | 2.67E+00 | 2.78E+00 | 7.22E-01 | no | | regulatory_region_variant | MODIFIER |
| 8 | 116120555 | AX-170529731 | 0 | T | C | 0.254 | 3.38E-02 | 9.76E-02 | 2.67E+00 | 2.78E+00 | 7.22E-01 | no | | regulatory_region_variant | MODIFIER |
| 8 | 116121848 | AX-170081267 | 0 | T | G | 0.254 | 3.38E-02 | 9.76E-02 | 2.67E+00 | 2.78E+00 | 7.22E-01 | no | | intergenic_variant | MODIFIER |
| 8 | 116148240 | AX-170245021 | 0 | G | A | 0.254 | 3.38E-02 | 9.76E-02 | 2.67E+00 | 2.78E+00 | 7.22E-01 | no | | intergenic_variant | MODIFIER |
| 8 | 116256261 | AX-170031553 | 0 | T | C | 0.254 | 3.38E-02 | 9.76E-02 | 2.67E+00 | 2.78E+00 | 7.22E-01 | no | | intergenic_variant | MODIFIER |
| 8 | 116288356 | AX-170031556 | 0 | T | C | 0.254 | 3.38E-02 | 9.76E-02 | 2.67E+00 | 2.78E+00 | 7.22E-01 | no | | intergenic_variant | MODIFIER |
| 8 | 116288502 | AX-170031554 | 0 | G | T | 0.254 | 3.38E-02 | 9.76E-02 | 2.67E+00 | 2.78E+00 | 7.22E-01 | no | | intergenic_variant | MODIFIER |
| 8 | 116288740 | AX-97850865 | 0 | A | G | 0.254 | 3.38E-02 | 9.76E-02 | 2.67E+00 | 2.78E+00 | 7.22E-01 | no | | intergenic_variant | MODIFIER |
| 8 | 116311068 | AX-170221835 | 0 | A | G | 0.254 | 3.38E-02 | 9.76E-02 | 2.67E+00 | 2.78E+00 | 7.22E-01 | no | | intergenic_variant | MODIFIER |
| 8 | 116313504 | AX-170476657 | 0 | C | A | 0.254 | 3.38E-02 | 9.76E-02 | 2.67E+00 | 2.78E+00 | 7.22E-01 | no | | intergenic_variant | MODIFIER |
| 8 | 116315832 | AX-170081272 | 0 | A | G | 0.254 | 3.38E-02 | 9.76E-02 | 2.67E+00 | 2.78E+00 | 7.22E-01 | no | | regulatory_region_variant | MODIFIER |
| 8 | 116317673 | AX-170081270 | 0 | T | C | 0.254 | 3.38E-02 | 9.76E-02 | 2.67E+00 | 2.78E+00 | 7.22E-01 | no | | intergenic_variant | MODIFIER |
| 8 | 116318401 | AX-170081271 | 0 | C | G | 0.254 | 3.38E-02 | 9.76E-02 | 2.67E+00 | 2.78E+00 | 7.22E-01 | no | | intergenic_variant | MODIFIER |
| 8 | 116322113 | AX-170401253 | 0 | C | T | 0.254 | 3.38E-02 | 9.76E-02 | 2.67E+00 | 2.78E+00 | 7.22E-01 | no | | intergenic_variant | MODIFIER |
| 8 | 116322789 | AX-170031550 | 0 | C | T | 0.254 | 3.38E-02 | 9.76E-02 | 2.67E+00 | 2.78E+00 | 7.22E-01 | no | | intergenic_variant | MODIFIER |
| 8 | 116322882 | AX-170031552 | 0 | T | C | 0.254 | 3.38E-02 | 9.76E-02 | 2.67E+00 | 2.78E+00 | 7.22E-01 | no | | intergenic_variant | MODIFIER |
| 8 | 116331752 | AX-170031551 | 0 | A | C | 0.254 | 3.38E-02 | 9.76E-02 | 2.67E+00 | 2.78E+00 | 7.22E-01 | no | | intergenic_variant | MODIFIER |
| 8 | 116346520 | AX-170372341 | 0 | C | T | 0.254 | 3.38E-02 | 9.76E-02 | 2.67E+00 | 2.78E+00 | 7.22E-01 | no | | intergenic_variant | MODIFIER |
| 8 | 116353280 | AX-170181000 | 0 | C | A | 0.254 | 3.38E-02 | 9.76E-02 | 2.67E+00 | 2.78E+00 | 7.22E-01 | no | 1700018P08Rik | upstream_gene_variant | MODIFIER |
| 8 | 116353459 | AX-170193207 | 0 | C | T | 0.254 | 3.38E-02 | 9.76E-02 | 2.67E+00 | 2.78E+00 | 7.22E-01 | no | 1700018P08Rik | upstream_gene_variant | MODIFIER |
| 8 | 116353577 | AX-170388793 | 0 | T | C | 0.254 | 3.38E-02 | 9.76E-02 | 2.67E+00 | 2.78E+00 | 7.22E-01 | no | 1700018P08Rik | upstream_gene_variant | MODIFIER |
| 8 | 116360061 | AX-170190522 | 0 | C | T | 0.254 | 3.38E-02 | 9.76E-02 | 2.67E+00 | 2.78E+00 | 7.22E-01 | no | 1700018P08Rik | downstream_gene_variant | MODIFIER |
| 8 | 116361198 | AX-170229594 | 0 | A | G | 0.254 | 3.38E-02 | 9.76E-02 | 2.67E+00 | 2.78E+00 | 7.22E-01 | no | 1700018P08Rik | downstream_gene_variant | MODIFIER |
| 8 | 116403351 | AX-170090218 | 0 | C | A | 0.254 | 3.38E-02 | 9.76E-02 | 2.67E+00 | 2.78E+00 | 7.22E-01 | no | | intergenic_variant | MODIFIER |
| 8 | 116408741 | AX-170431076 | 0 | T | C | 0.254 | 3.38E-02 | 9.76E-02 | 2.67E+00 | 2.78E+00 | 7.22E-01 | no | | intergenic_variant | MODIFIER |
| 8 | 116408894 | AX-170173571 | 0 | A | G | 0.254 | 3.38E-02 | 9.76E-02 | 2.67E+00 | 2.78E+00 | 7.22E-01 | no | | intergenic_variant | MODIFIER |
| 8 | 116529221 | AX-170368823 | 0 | T | C | 0.254 | 3.38E-02 | 9.76E-02 | 2.67E+00 | 2.78E+00 | 7.22E-01 | no | | intergenic_variant | MODIFIER |
| 8 | 116563971 | AX-170128520 | 0 | A | T | 0.254 | 3.38E-02 | 9.76E-02 | 2.67E+00 | 2.78E+00 | 7.22E-01 | no | | intergenic_variant | MODIFIER |
| 8 | 116566791 | AX-170457085 | 0 | A | G | 0.254 | 3.38E-02 | 9.76E-02 | 2.67E+00 | 2.78E+00 | 7.22E-01 | no | | intergenic_variant | MODIFIER |
| 8 | 116567949 | AX-170327200 | 0 | G | A | 0.254 | 3.38E-02 | 9.76E-02 | 2.67E+00 | 2.78E+00 | 7.22E-01 | no | | intergenic_variant | MODIFIER |
| 8 | 116569216 | AX-97785260 | 0 | A | G | 0.254 | 3.38E-02 | 9.76E-02 | 2.67E+00 | 2.78E+00 | 7.22E-01 | no | | intergenic_variant | MODIFIER |
| 8 | 116667076 | AX-97781254 | 0 | A | G | 0.254 | 3.38E-02 | 9.76E-02 | 2.67E+00 | 2.78E+00 | 7.22E-01 | no | Cdyl2 | intron_variant | MODIFIER |
| 8 | 116763767 | AX-170218428 | 0 | C | T | 0.254 | 3.38E-02 | 9.76E-02 | 2.67E+00 | 2.78E+00 | 7.22E-01 | no | Gm10620 | downstream_gene_variant | MODIFIER |

| | | | | | | | | | | | | | | | |
|---|---|---|---|---|---|---|---|---|---|---|---|---|---|---|---|
| 8 | 116767274 | AX-170355097 | 0 | A | G | 0.254 | 3.38E-02 | 9.76E-02 | 2.67E+00 | 2.78E+00 | 7.22E-01 | no | Gm10620 | upstream_gene_variant | MODIFIER |
| 8 | 116767365 | AX-97809328 | 0 | A | G | 0.254 | 3.38E-02 | 9.76E-02 | 2.67E+00 | 2.78E+00 | 7.22E-01 | no | Gm10620 | upstream_gene_variant | MODIFIER |
| 8 | 116767384 | AX-170293557 | 0 | T | C | 0.254 | 3.38E-02 | 9.76E-02 | 2.67E+00 | 2.78E+00 | 7.22E-01 | no | Gm10620 | upstream_gene_variant | MODIFIER |
| 8 | 116813730 | AX-170310627 | 0 | A | C | 0.254 | 3.38E-02 | 9.76E-02 | 2.67E+00 | 2.78E+00 | 7.22E-01 | no | | intergenic_variant | MODIFIER |
| 8 | 116814045 | AX-170459727 | 0 | G | A | 0.254 | 3.38E-02 | 9.76E-02 | 2.67E+00 | 2.78E+00 | 7.22E-01 | no | | regulatory_region_variant | MODIFIER |
| 8 | 116814222 | AX-170207936 | 0 | C | T | 0.254 | 3.38E-02 | 9.76E-02 | 2.67E+00 | 2.78E+00 | 7.22E-01 | no | | regulatory_region_variant | MODIFIER |
| 8 | 116814315 | AX-170377570 | 0 | C | T | 0.254 | 3.38E-02 | 9.76E-02 | 2.67E+00 | 2.78E+00 | 7.22E-01 | no | | regulatory_region_variant | MODIFIER |
| 8 | 116820187 | AX-170163106 | 0 | G | T | 0.254 | 3.38E-02 | 9.76E-02 | 2.67E+00 | 2.78E+00 | 7.22E-01 | no | | intergenic_variant | MODIFIER |
| 8 | 116820859 | AX-170165798 | 0 | A | G | 0.254 | 3.38E-02 | 9.76E-02 | 2.67E+00 | 2.78E+00 | 7.22E-01 | no | | intergenic_variant | MODIFIER |
| 8 | 116838652 | AX-97839165 | 0 | A | G | 0.254 | 3.38E-02 | 9.76E-02 | 2.67E+00 | 2.78E+00 | 7.22E-01 | no | | intergenic_variant | MODIFIER |
| 8 | 116858179 | AX-170242221 | 0 | G | A | 0.254 | 3.38E-02 | 9.76E-02 | 2.67E+00 | 2.78E+00 | 7.22E-01 | no | | intergenic_variant | MODIFIER |
| 8 | 116933268 | AX-170269786 | 0 | C | A | 0.254 | 3.38E-02 | 9.76E-02 | 2.67E+00 | 2.78E+00 | 7.22E-01 | no | Cenpn | intron_variant | MODIFIER |
| 8 | 116945986 | AX-170143731 | 0 | A | G | 0.254 | 3.38E-02 | 9.76E-02 | 2.67E+00 | 2.78E+00 | 7.22E-01 | no | Cenpn | downstream_gene_variant | MODIFIER |
| 8 | 116946371 | AX-170438437 | 0 | G | A | 0.254 | 3.38E-02 | 9.76E-02 | 2.67E+00 | 2.78E+00 | 7.22E-01 | no | Cenpn | downstream_gene_variant | MODIFIER |
| 8 | 116946401 | AX-170081285 | 0 | C | G | 0.254 | 3.38E-02 | 9.76E-02 | 2.67E+00 | 2.78E+00 | 7.22E-01 | no | Cenpn | downstream_gene_variant | MODIFIER |
| 8 | 116946699 | AX-170081286 | 0 | C | T | 0.254 | 3.38E-02 | 9.76E-02 | 2.67E+00 | 2.78E+00 | 7.22E-01 | no | Atmin | intron_variant | MODIFIER |
| 8 | 116953705 | AX-170081284 | 0 | C | T | 0.254 | 3.38E-02 | 9.76E-02 | 2.67E+00 | 2.78E+00 | 7.22E-01 | no | Atmin | intron_variant | MODIFIER |
| 8 | 116972836 | AX-97804273 | 0 | T | C | 0.254 | 3.38E-02 | 9.76E-02 | 2.67E+00 | 2.78E+00 | 7.22E-01 | no | 1700030J22Rik | intron_variant | MODIFIER |
| 8 | 116988274 | AX-170090220 | 0 | G | C | 0.254 | 3.38E-02 | 9.76E-02 | 2.67E+00 | 2.78E+00 | 7.22E-01 | no | Gcsh | intron_variant | MODIFIER |
| 8 | 117032929 | AX-97823777 | 0 | A | G | 0.254 | 3.38E-02 | 9.76E-02 | 2.67E+00 | 2.78E+00 | 7.22E-01 | no | Pkd1l2 | intron_variant | MODIFIER |
| 8 | 117034447 | AX-170357882 | 0 | G | A | 0.254 | 3.38E-02 | 9.76E-02 | 2.67E+00 | 2.78E+00 | 7.22E-01 | no | Pkd1l2 | intron_variant | MODIFIER |
| 8 | 117057275 | AX-97854903 | 0 | C | T | 0.254 | 3.38E-02 | 9.76E-02 | 2.67E+00 | 2.78E+00 | 7.22E-01 | no | Pkd1l2 | intron_variant | MODIFIER |
| 8 | 117058635 | AX-170172793 | 0 | A | G | 0.254 | 3.38E-02 | 9.76E-02 | 2.67E+00 | 2.78E+00 | 7.22E-01 | no | Pkd1l2 | intron_variant | MODIFIER |
| 8 | 117078875 | AX-170346250 | 0 | G | A | 0.254 | 3.38E-02 | 9.76E-02 | 2.67E+00 | 2.78E+00 | 7.22E-01 | no | Pkd1l2 | intron_variant | MODIFIER |
| 8 | 117081571 | AX-170276875 | 0 | C | T | 0.254 | 3.38E-02 | 9.76E-02 | 2.67E+00 | 2.78E+00 | 7.22E-01 | no | Pkd1l2 | intron_variant | MODIFIER |
| 8 | 117082345 | AX-170124562 | 0 | A | G | 0.254 | 3.38E-02 | 9.76E-02 | 2.67E+00 | 2.78E+00 | 7.22E-01 | no | Pkd1l2 | synonymous_variant | LOW |
| 8 | 117272751 | AX-170486606 | 0 | A | G | 0.254 | 3.38E-02 | 9.76E-02 | 2.67E+00 | 2.78E+00 | 7.22E-01 | no | Cmip | intron_variant | MODIFIER |
| 8 | 117292215 | AX-170031567 | 0 | A | G | 0.254 | 3.38E-02 | 9.76E-02 | 2.67E+00 | 2.78E+00 | 7.22E-01 | no | Cmip | intron_variant | MODIFIER |
| 8 | 117384268 | AX-170081293 | 0 | T | C | 0.254 | 3.38E-02 | 9.76E-02 | 2.67E+00 | 2.78E+00 | 7.22E-01 | no | Cmip | intron_variant | MODIFIER |
| 8 | 117413225 | AX-170371698 | 0 | T | C | 0.254 | 3.38E-02 | 9.76E-02 | 2.67E+00 | 2.78E+00 | 7.22E-01 | no | Cmip | intron_variant | MODIFIER |
| 8 | 117637991 | AX-170205814 | 0 | C | T | 0.254 | 3.38E-02 | 9.76E-02 | 2.67E+00 | 2.78E+00 | 7.22E-01 | no | Plcg2 | downstream_gene_variant | MODIFIER |
| 8 | 117818911 | AX-170081303 | 0 | T | A | 0.254 | 3.38E-02 | 9.76E-02 | 2.67E+00 | 2.78E+00 | 7.22E-01 | no | | regulatory_region_variant | MODIFIER |
| 8 | 117819355 | AX-97830138 | 0 | G | A | 0.254 | 3.38E-02 | 9.76E-02 | 2.67E+00 | 2.78E+00 | 7.22E-01 | no | | intergenic_variant | MODIFIER |
| 8 | 117824243 | AX-170081308 | 0 | G | A | 0.254 | 3.38E-02 | 9.76E-02 | 2.67E+00 | 2.78E+00 | 7.22E-01 | no | | intergenic_variant | MODIFIER |
| 8 | 117838676 | AX-170081307 | 0 | G | A | 0.254 | 3.38E-02 | 9.76E-02 | 2.67E+00 | 2.78E+00 | 7.22E-01 | no | | intergenic_variant | MODIFIER |
| 8 | 117840421 | AX-170323440 | 0 | A | T | 0.254 | 3.38E-02 | 9.76E-02 | 2.67E+00 | 2.78E+00 | 7.22E-01 | no | | intergenic_variant | MODIFIER |
| 8 | 117844491 | AX-97805950 | 0 | A | G | 0.254 | 3.38E-02 | 9.76E-02 | 2.67E+00 | 2.78E+00 | 7.22E-01 | no | | intergenic_variant | MODIFIER |
| 8 | 117852080 | AX-170031570 | 0 | A | G | 0.254 | 3.38E-02 | 9.76E-02 | 2.67E+00 | 2.78E+00 | 7.22E-01 | no | | intergenic_variant | MODIFIER |
| 8 | 117854875 | AX-170031573 | 0 | C | T | 0.254 | 3.38E-02 | 9.76E-02 | 2.67E+00 | 2.78E+00 | 7.22E-01 | no | | regulatory_region_variant | MODIFIER |
| 8 | 117855494 | AX-170477970 | 0 | C | T | 0.254 | 3.38E-02 | 9.76E-02 | 2.67E+00 | 2.78E+00 | 7.22E-01 | no | | regulatory_region_variant | MODIFIER |
| 2 | 118467463 | AX-170148235 | 0 | A | G | 0.123 | 6.49E-02 | 1.84E-01 | 2.67E+00 | 2.73E+00 | 7.22E-01 | no | Eif2ak4 | intron_variant | MODIFIER |
| 5 | 44993848 | AX-170336094 | 1 | A | G | 0.188 | -3.99E-02 | 1.11E-01 | 2.62E+00 | 2.70E+00 | 7.22E-01 | no | | intergenic_variant | MODIFIER |
| 9 | 38069217 | AX-97784894 | 1 | C | A | 0.5 | -2.71E-02 | 7.82E-02 | 2.70E+00 | 2.79E+00 | 7.22E-01 | no | Olfr886-ps1 | upstream_gene_variant | MODIFIER |
| 8 | 74142027 | AX-170080428 | 0 | C | T | 0.202 | 4.38E-02 | 1.26E-01 | 2.71E+00 | 2.79E+00 | 7.22E-01 | no | | intergenic_variant | MODIFIER |
| 1 | 19742816 | AX-170337781 | 0 | C | T | 0.404 | 3.08E-02 | 8.48E-02 | 2.61E+00 | 2.66E+00 | 7.22E-01 | no | | intergenic_variant | MODIFIER |
| 16 | 92362157 | AX-97857106 | 0 | C | T | 0.474 | 2.73E-02 | 7.86E-02 | 2.68E+00 | 2.77E+00 | 7.22E-01 | no | Kcne1 | upstream_gene_variant | MODIFIER |
| 9 | 55942510 | AX-170223429 | 0 | G | A | 0.211 | -4.08E-02 | 1.17E-01 | 2.71E+00 | 2.78E+00 | 7.22E-01 | no | Gm7435 | intron_variant,non_coding_transcript_variant | MODIFIER |
| 6 | 94298049 | AX-170404908 | 1 | C | T | 0.438 | -2.19E-02 | 5.90E-02 | 2.54E+00 | 2.67E+00 | 7.22E-01 | no | | intergenic_variant | MODIFIER |
| 4 | 145229399 | AX-170535814 | 0 | A | G | 0.105 | 7.14E-02 | 2.04E-01 | 2.69E+00 | 2.75E+00 | 7.22E-01 | no | Tnfrsf1b | intron_variant | MODIFIER |
| 10 | 3444708 | AX-170002656 | 0 | G | A | 0.202 | -3.32E-02 | 9.44E-02 | 2.62E+00 | 2.74E+00 | 7.22E-01 | no | Ppp1r14c | intron_variant | MODIFIER |
| 18 | 89776478 | AX-97835195 | 0 | A | T | 0.518 | 2.23E-02 | 6.17E-02 | 2.57E+00 | 2.70E+00 | 7.22E-01 | no | | intergenic_variant | MODIFIER |
| 5 | 146576598 | AX-170484822 | 0 | G | A | 0.053 | 5.71E-02 | 1.64E-01 | 2.63E+00 | 2.75E+00 | 7.22E-01 | no | Gm8494 | downstream_gene_variant | MODIFIER |
| 3 | 148620777 | AX-170436778 | 0 | T | G | 0.333 | 2.66E-02 | 7.42E-02 | 2.61E+00 | 2.70E+00 | 7.22E-01 | no | | intergenic_variant | MODIFIER |
| 8 | 90010035 | AX-170480707 | 1 | C | T | 0.188 | -4.28E-02 | 1.19E-01 | 2.61E+00 | 2.68E+00 | 7.22E-01 | no | | intergenic_variant | MODIFIER |

| | | | | | | | | | | | | | | | |
|---|---|---|---|---|---|---|---|---|---|---|---|---|---|---|---|
| 18 | 90133781 | AX-170016173 | 0 | T | C | 0.491 | -2.28E-02 | 6.68E-02 | 2.67E+00 | 2.80E+00 | 7.22E-01 | no | | intergenic_variant | MODIFIER |
| 2 | 110603086 | AX-97838509 | 1 | G | A | 0.054 | -1.05E-01 | 3.00E-01 | 2.74E+00 | 2.77E+00 | 7.22E-01 | no | Slc5a12 | intron_variant | MODIFIER |
| 7 | 112733976 | AX-170369791 | 2 | G | T | 0.445 | 3.21E-02 | 9.16E-02 | 2.67E+00 | 2.74E+00 | 7.23E-01 | no | Tead1 | intron_variant | MODIFIER |
| 15 | 88469701 | AX-170357478 | 0 | A | G | 0.123 | 5.30E-02 | 1.49E-01 | 2.62E+00 | 2.71E+00 | 7.23E-01 | no | | intergenic_variant | MODIFIER |
| 18 | 72274448 | AX-170504348 | 0 | T | C | 0.184 | -4.22E-02 | 1.19E-01 | 2.65E+00 | 2.71E+00 | 7.23E-01 | no | Dcc | intron_variant | MODIFIER |
| 5 | 92892721 | AX-170025621 | 1 | T | A | 0.42 | 3.76E-02 | 1.07E-01 | 2.69E+00 | 2.75E+00 | 7.23E-01 | no | Shroom3 | intron_variant | MODIFIER |
| 3 | 32743919 | AX-170020264 | 1 | G | A | 0.179 | 3.43E-02 | 9.86E-02 | 2.69E+00 | 2.77E+00 | 7.23E-01 | no | Ndufb5 | intron_variant,NMD_transcript_variant | MODIFIER |
| 9 | 55817617 | AX-170223685 | 0 | G | C | 0.202 | 4.01E-02 | 1.14E-01 | 2.67E+00 | 2.74E+00 | 7.23E-01 | no | Scaper | intron_variant | MODIFIER |
| 2 | 108561205 | AX-97839346 | 0 | C | T | 0.07 | 5.96E-02 | 1.70E-01 | 2.61E+00 | 2.73E+00 | 7.23E-01 | no | | intergenic_variant | MODIFIER |
| 11 | 34508842 | AX-170046760 | 0 | A | T | 0.096 | -5.67E-02 | 1.62E-01 | 2.63E+00 | 2.74E+00 | 7.23E-01 | no | Dock2 | intron_variant | MODIFIER |
| 1 | 3474289 | AX-170160383 | 0 | G | A | 0.518 | 2.65E-02 | 6.95E-02 | 2.53E+00 | 2.61E+00 | 7.23E-01 | no | Xkr4 | intron_variant | MODIFIER |
| 2 | 95586673 | AX-168379873 | 0 | T | G | 0.061 | -6.22E-02 | 1.78E-01 | 2.61E+00 | 2.74E+00 | 7.23E-01 | no | | intergenic_variant | MODIFIER |
| 15 | 82141519 | AX-170244406 | 1 | T | A | 0.545 | 2.93E-02 | 7.96E-02 | 2.57E+00 | 2.65E+00 | 7.23E-01 | no | Ccdc134 | 3_prime_UTR_variant | MODIFIER |
| 18 | 58947982 | AX-97822897 | 2 | A | G | 0.418 | 2.87E-02 | 8.27E-02 | 2.79E+00 | 2.87E+00 | 7.23E-01 | no | Adamts19 | intron_variant | MODIFIER |
| 10 | 5349572 | AX-170204514 | 0 | C | T | 0.219 | 3.31E-02 | 9.62E-02 | 2.67E+00 | 2.78E+00 | 7.24E-01 | no | Syne1 | intron_variant | MODIFIER |
| 2 | 130121224 | AX-170358671 | 2 | A | T | 0.145 | 7.78E-02 | 2.22E-01 | 2.70E+00 | 2.73E+00 | 7.24E-01 | no | Tgm6 | upstream_gene_variant | MODIFIER |
| 16 | 95360715 | AX-170244171 | 0 | T | C | 0.456 | 2.57E-02 | 7.57E-02 | 2.73E+00 | 2.85E+00 | 7.24E-01 | no | Erg | 3_prime_UTR_variant | MODIFIER |
| 10 | 28854042 | AX-170227041 | 1 | T | C | 0.188 | 3.29E-02 | 9.57E-02 | 2.68E+00 | 2.78E+00 | 7.24E-01 | no | Themis | intron_variant | MODIFIER |
| 9 | 66169158 | AX-170161506 | 0 | T | A | 0.202 | 4.37E-02 | 1.26E-01 | 2.73E+00 | 2.78E+00 | 7.24E-01 | no | Dapk2 | intron_variant | MODIFIER |
| 8 | 39315249 | AX-97804078 | 1 | G | T | 0.152 | 5.14E-02 | 1.48E-01 | 2.75E+00 | 2.80E+00 | 7.24E-01 | no | Gm6213 | intron_variant,non_coding_transcript_variant | MODIFIER |
| 8 | 39718810 | AX-170210783 | 1 | G | T | 0.152 | 5.14E-02 | 1.48E-01 | 2.75E+00 | 2.80E+00 | 7.24E-01 | no | Gm2033 | non_coding_transcript_exon_variant | MODIFIER |
| 8 | 41393640 | AX-170079874 | 1 | T | C | 0.152 | 5.14E-02 | 1.48E-01 | 2.75E+00 | 2.80E+00 | 7.24E-01 | no | Frg1 | downstream_gene_variant | MODIFIER |
| 8 | 45956506 | AX-170151103 | 1 | C | T | 0.152 | 5.14E-02 | 1.48E-01 | 2.75E+00 | 2.80E+00 | 7.24E-01 | no | 1700029J07Rik | missense_variant | MODERATE |
| 9 | 50413376 | AX-170324077 | 1 | A | T | 0.214 | -3.78E-02 | 1.08E-01 | 2.66E+00 | 2.74E+00 | 7.24E-01 | no | | intergenic_variant | MODIFIER |
| 5 | 97190707 | AX-170274226 | 1 | C | A | 0.438 | 3.85E-02 | 1.11E-01 | 2.70E+00 | 2.75E+00 | 7.24E-01 | no | Gm2861 | upstream_gene_variant | MODIFIER |
| 2 | 114748751 | AX-97841200 | 1 | G | A | 0.143 | 7.72E-02 | 2.21E-01 | 2.70E+00 | 2.73E+00 | 7.24E-01 | no | | intergenic_variant | MODIFIER |
| 10 | 27608314 | AX-170416863 | 0 | T | C | 0.193 | 3.38E-02 | 9.89E-02 | 2.73E+00 | 2.83E+00 | 7.24E-01 | no | Lama2 | intron_variant | MODIFIER |
| 10 | 27608637 | AX-170136673 | 0 | A | T | 0.193 | 3.38E-02 | 9.89E-02 | 2.73E+00 | 2.83E+00 | 7.24E-01 | no | Lama2 | intron_variant | MODIFIER |
| 10 | 27608688 | AX-170150406 | 0 | G | A | 0.193 | 3.38E-02 | 9.89E-02 | 2.73E+00 | 2.83E+00 | 7.24E-01 | no | Lama2 | intron_variant | MODIFIER |
| 10 | 27609274 | AX-170242782 | 0 | C | T | 0.193 | 3.38E-02 | 9.89E-02 | 2.73E+00 | 2.83E+00 | 7.24E-01 | no | Lama2 | intron_variant | MODIFIER |
| 10 | 27618262 | AX-170327524 | 0 | G | C | 0.193 | 3.38E-02 | 9.89E-02 | 2.73E+00 | 2.83E+00 | 7.24E-01 | no | Lama2 | upstream_gene_variant | MODIFIER |
| 10 | 27620992 | AX-170355849 | 0 | G | T | 0.193 | 3.38E-02 | 9.89E-02 | 2.73E+00 | 2.83E+00 | 7.24E-01 | no | Lama2 | upstream_gene_variant | MODIFIER |
| 10 | 27634603 | AX-97843228 | 0 | C | G | 0.193 | 3.38E-02 | 9.89E-02 | 2.73E+00 | 2.83E+00 | 7.24E-01 | no | | intergenic_variant | MODIFIER |
| 10 | 27664165 | AX-97820305 | 0 | A | C | 0.193 | 3.38E-02 | 9.89E-02 | 2.73E+00 | 2.83E+00 | 7.24E-01 | no | Gm30165 | intron_variant,non_coding_transcript_variant | MODIFIER |
| 10 | 27669599 | AX-170195654 | 0 | A | G | 0.193 | 3.38E-02 | 9.89E-02 | 2.73E+00 | 2.83E+00 | 7.24E-01 | no | Gm30165 | intron_variant,non_coding_transcript_variant | MODIFIER |
| 10 | 27670258 | AX-170416112 | 0 | A | T | 0.193 | 3.38E-02 | 9.89E-02 | 2.73E+00 | 2.83E+00 | 7.24E-01 | no | Gm30165 | intron_variant,non_coding_transcript_variant | MODIFIER |
| 10 | 27670874 | AX-170520883 | 0 | A | G | 0.193 | 3.38E-02 | 9.89E-02 | 2.73E+00 | 2.83E+00 | 7.24E-01 | no | Gm30165 | intron_variant,non_coding_transcript_variant | MODIFIER |
| 10 | 27673998 | AX-170471623 | 0 | C | G | 0.193 | 3.38E-02 | 9.89E-02 | 2.73E+00 | 2.83E+00 | 7.24E-01 | no | Gm30165 | intron_variant,non_coding_transcript_variant | MODIFIER |
| 10 | 27674194 | AX-170044439 | 0 | A | G | 0.193 | 3.38E-02 | 9.89E-02 | 2.73E+00 | 2.83E+00 | 7.24E-01 | no | Gm30165 | intron_variant,non_coding_transcript_variant | MODIFIER |
| 10 | 27709157 | AX-97812559 | 0 | G | A | 0.193 | 3.38E-02 | 9.89E-02 | 2.73E+00 | 2.83E+00 | 7.24E-01 | no | | intergenic_variant | MODIFIER |
| 10 | 27712820 | AX-170193311 | 0 | T | C | 0.193 | 3.38E-02 | 9.89E-02 | 2.73E+00 | 2.83E+00 | 7.24E-01 | no | | intergenic_variant | MODIFIER |
| 10 | 27729486 | AX-170445189 | 0 | A | G | 0.193 | 3.38E-02 | 9.89E-02 | 2.73E+00 | 2.83E+00 | 7.24E-01 | no | | regulatory_region_variant | MODIFIER |
| 10 | 27830809 | AX-170515957 | 0 | T | C | 0.193 | 3.38E-02 | 9.89E-02 | 2.73E+00 | 2.83E+00 | 7.24E-01 | no | | intergenic_variant | MODIFIER |
| 10 | 27877955 | AX-170147375 | 0 | A | G | 0.193 | 3.38E-02 | 9.89E-02 | 2.73E+00 | 2.83E+00 | 7.24E-01 | no | | intergenic_variant | MODIFIER |
| 10 | 27878708 | AX-170379758 | 0 | A | C | 0.193 | 3.38E-02 | 9.89E-02 | 2.73E+00 | 2.83E+00 | 7.24E-01 | no | | intergenic_variant | MODIFIER |
| 10 | 27926043 | AX-170238280 | 0 | C | A | 0.193 | 3.38E-02 | 9.89E-02 | 2.73E+00 | 2.83E+00 | 7.24E-01 | no | | intergenic_variant | MODIFIER |
| 10 | 28250367 | AX-170086770 | 0 | G | T | 0.193 | 3.38E-02 | 9.89E-02 | 2.73E+00 | 2.83E+00 | 7.24E-01 | no | Ptprk | intron_variant | MODIFIER |
| 1 | 53846645 | AX-170322267 | 0 | C | T | 0.096 | 6.85E-02 | 2.00E-01 | 2.68E+00 | 2.78E+00 | 7.24E-01 | no | Hecw2 | intron_variant | MODIFIER |
| 1 | 54386634 | AX-170454299 | 0 | T | C | 0.096 | 6.85E-02 | 2.00E-01 | 2.68E+00 | 2.78E+00 | 7.24E-01 | no | | intergenic_variant | MODIFIER |
| 10 | 13013785 | AX-97814773 | 1 | C | T | 0.196 | -3.84E-02 | 1.08E-01 | 2.62E+00 | 2.69E+00 | 7.24E-01 | no | Sf3b5 | downstream_gene_variant | MODIFIER |
| 3 | 145431690 | AX-170068352 | 0 | G | A | 0.36 | -2.38E-02 | 6.94E-02 | 2.69E+00 | 2.78E+00 | 7.24E-01 | no | Col24a1 | intron_variant | MODIFIER |
| 7 | 54097358 | AX-170249270 | 1 | T | C | 0.42 | 3.57E-02 | 1.02E-01 | 2.65E+00 | 2.73E+00 | 7.24E-01 | no | | intergenic_variant | MODIFIER |
| 1 | 187679641 | AX-170114918 | 2 | A | C | 0.236 | 3.78E-02 | 9.77E-02 | 2.50E+00 | 2.61E+00 | 7.24E-01 | no | Esrrg | intron_variant | MODIFIER |
| 5 | 118658052 | AX-170487951 | 0 | C | T | 0.456 | 3.86E-02 | 1.10E-01 | 2.66E+00 | 2.71E+00 | 7.25E-01 | no | Med13l | intron_variant | MODIFIER |

| | | | | | | | | | | | | | | | |
|---|---|---|---|---|---|---|---|---|---|---|---|---|---|---|---|
| 5 | 122624104 | AX-170100565 | 0 | C | T | 0.456 | 3.86E-02 | 1.10E-01 | 2.66E+00 | 2.71E+00 | 7.25E-01 | no | | regulatory_region_variant | MODIFIER |
| 5 | 124787889 | AX-97849401 | 0 | A | G | 0.456 | 3.86E-02 | 1.10E-01 | 2.66E+00 | 2.71E+00 | 7.25E-01 | no | Dnah10 | intron_variant | MODIFIER |
| 5 | 114355170 | AX-170365760 | 0 | G | A | 0.447 | 3.73E-02 | 1.07E-01 | 2.66E+00 | 2.73E+00 | 7.25E-01 | no | Myo1h | intron_variant | MODIFIER |
| 1 | 99571940 | AX-97855053 | 0 | A | C | 0.158 | 7.30E-02 | 2.07E-01 | 2.65E+00 | 2.71E+00 | 7.25E-01 | no | | intergenic_variant | MODIFIER |
| 1 | 170988751 | AX-170255176 | 0 | C | T | 0.219 | 4.55E-02 | 1.27E-01 | 2.60E+00 | 2.68E+00 | 7.25E-01 | no | Gm10522 | downstream_gene_variant | MODIFIER |
| 1 | 145032074 | AX-170119917 | 1 | G | A | 0.205 | 4.73E-02 | 1.35E-01 | 2.66E+00 | 2.71E+00 | 7.25E-01 | no | | intergenic_variant | MODIFIER |
| 8 | 34825914 | AX-97843037 | 0 | A | G | 0.149 | -5.18E-02 | 1.46E-01 | 2.64E+00 | 2.68E+00 | 7.25E-01 | no | Tnks | downstream_gene_variant | MODIFIER |
| 8 | 34833494 | AX-170030494 | 0 | C | T | 0.149 | -5.18E-02 | 1.46E-01 | 2.64E+00 | 2.68E+00 | 7.25E-01 | no | Tnks | intron_variant | MODIFIER |
| 8 | 34836726 | AX-170079727 | 0 | A | T | 0.149 | -5.18E-02 | 1.46E-01 | 2.64E+00 | 2.68E+00 | 7.25E-01 | no | Tnks | intron_variant | MODIFIER |
| 8 | 34837928 | AX-170079726 | 0 | A | G | 0.149 | -5.18E-02 | 1.46E-01 | 2.64E+00 | 2.68E+00 | 7.25E-01 | no | Tnks | intron_variant | MODIFIER |
| 8 | 34838254 | AX-170079725 | 0 | C | T | 0.149 | -5.18E-02 | 1.46E-01 | 2.64E+00 | 2.68E+00 | 7.25E-01 | no | Tnks | intron_variant | MODIFIER |
| 8 | 34838345 | AX-97821838 | 0 | T | C | 0.149 | -5.18E-02 | 1.46E-01 | 2.64E+00 | 2.68E+00 | 7.25E-01 | no | Tnks | intron_variant | MODIFIER |
| 8 | 34854897 | AX-170079723 | 0 | G | A | 0.149 | -5.18E-02 | 1.46E-01 | 2.64E+00 | 2.68E+00 | 7.25E-01 | no | Tnks | intron_variant | MODIFIER |
| 8 | 34865874 | AX-97820608 | 0 | T | C | 0.149 | -5.18E-02 | 1.46E-01 | 2.64E+00 | 2.68E+00 | 7.25E-01 | no | Tnks | intron_variant | MODIFIER |
| 8 | 34898004 | AX-97815853 | 0 | A | G | 0.149 | -5.18E-02 | 1.46E-01 | 2.64E+00 | 2.68E+00 | 7.25E-01 | no | Tnks | intron_variant | MODIFIER |
| 8 | 34913560 | AX-170346922 | 0 | A | G | 0.149 | -5.18E-02 | 1.46E-01 | 2.64E+00 | 2.68E+00 | 7.25E-01 | no | Tnks | intron_variant | MODIFIER |
| 8 | 34914248 | AX-97849490 | 0 | T | C | 0.149 | -5.18E-02 | 1.46E-01 | 2.64E+00 | 2.68E+00 | 7.25E-01 | no | Tnks | intron_variant | MODIFIER |
| 8 | 34935345 | AX-170516731 | 0 | A | T | 0.149 | -5.18E-02 | 1.46E-01 | 2.64E+00 | 2.68E+00 | 7.25E-01 | no | Tnks | intron_variant | MODIFIER |
| 8 | 34941298 | AX-170079724 | 0 | T | A | 0.149 | -5.18E-02 | 1.46E-01 | 2.64E+00 | 2.68E+00 | 7.25E-01 | no | Tnks | intron_variant | MODIFIER |
| 8 | 34950697 | AX-97788969 | 0 | T | C | 0.149 | -5.18E-02 | 1.46E-01 | 2.64E+00 | 2.68E+00 | 7.25E-01 | no | Tnks | intron_variant | MODIFIER |
| 8 | 34962524 | AX-170388175 | 0 | G | A | 0.149 | -5.18E-02 | 1.46E-01 | 2.64E+00 | 2.68E+00 | 7.25E-01 | no | Tnks | intron_variant | MODIFIER |
| 8 | 34974989 | AX-170455582 | 0 | A | G | 0.149 | -5.18E-02 | 1.46E-01 | 2.64E+00 | 2.68E+00 | 7.25E-01 | no | Gm34368 | intron_variant,non_coding_transcript_variant | MODIFIER |
| 8 | 34977741 | AX-97811992 | 0 | T | C | 0.149 | -5.18E-02 | 1.46E-01 | 2.64E+00 | 2.68E+00 | 7.25E-01 | no | Gm34368 | intron_variant,non_coding_transcript_variant | MODIFIER |
| 8 | 34996349 | AX-97844245 | 0 | T | G | 0.149 | -5.18E-02 | 1.46E-01 | 2.64E+00 | 2.68E+00 | 7.25E-01 | no | | regulatory_region_variant | MODIFIER |
| 8 | 35010740 | AX-170306452 | 0 | A | G | 0.149 | -5.18E-02 | 1.46E-01 | 2.64E+00 | 2.68E+00 | 7.25E-01 | no | | intergenic_variant | MODIFIER |
| 8 | 35014594 | AX-97785339 | 0 | T | C | 0.149 | -5.18E-02 | 1.46E-01 | 2.64E+00 | 2.68E+00 | 7.25E-01 | no | | intergenic_variant | MODIFIER |
| 8 | 35016026 | AX-170109718 | 0 | G | A | 0.149 | -5.18E-02 | 1.46E-01 | 2.64E+00 | 2.68E+00 | 7.25E-01 | no | | intergenic_variant | MODIFIER |
| 8 | 35016956 | AX-170402896 | 0 | C | T | 0.149 | -5.18E-02 | 1.46E-01 | 2.64E+00 | 2.68E+00 | 7.25E-01 | no | | intergenic_variant | MODIFIER |
| 8 | 35022593 | AX-170035779 | 0 | G | T | 0.149 | -5.18E-02 | 1.46E-01 | 2.64E+00 | 2.68E+00 | 7.25E-01 | no | | regulatory_region_variant | MODIFIER |
| 8 | 35077356 | AX-97847943 | 0 | A | C | 0.149 | -5.18E-02 | 1.46E-01 | 2.64E+00 | 2.68E+00 | 7.25E-01 | no | | intergenic_variant | MODIFIER |
| 8 | 35087133 | AX-170128795 | 0 | C | T | 0.149 | -5.18E-02 | 1.46E-01 | 2.64E+00 | 2.68E+00 | 7.25E-01 | no | | regulatory_region_variant | MODIFIER |
| 8 | 35089001 | AX-170079734 | 0 | C | T | 0.149 | -5.18E-02 | 1.46E-01 | 2.64E+00 | 2.68E+00 | 7.25E-01 | no | | regulatory_region_variant | MODIFIER |
| 8 | 35105521 | AX-170079733 | 0 | A | T | 0.149 | -5.18E-02 | 1.46E-01 | 2.64E+00 | 2.68E+00 | 7.25E-01 | no | Gm34419 | downstream_gene_variant | MODIFIER |
| 8 | 35106178 | AX-170079729 | 0 | C | T | 0.149 | -5.18E-02 | 1.46E-01 | 2.64E+00 | 2.68E+00 | 7.25E-01 | no | Gm34419 | downstream_gene_variant | MODIFIER |
| 8 | 35114804 | AX-170079731 | 0 | G | C | 0.149 | -5.18E-02 | 1.46E-01 | 2.64E+00 | 2.68E+00 | 7.25E-01 | no | | intergenic_variant | MODIFIER |
| 8 | 35122551 | AX-170178416 | 0 | A | T | 0.149 | -5.18E-02 | 1.46E-01 | 2.64E+00 | 2.68E+00 | 7.25E-01 | no | | intergenic_variant | MODIFIER |
| 8 | 35122696 | AX-97804106 | 0 | A | G | 0.149 | -5.18E-02 | 1.46E-01 | 2.64E+00 | 2.68E+00 | 7.25E-01 | no | | regulatory_region_variant | MODIFIER |
| 8 | 35126526 | AX-170151497 | 0 | A | G | 0.149 | -5.18E-02 | 1.46E-01 | 2.64E+00 | 2.68E+00 | 7.25E-01 | no | | regulatory_region_variant | MODIFIER |
| 8 | 35133877 | AX-170456810 | 0 | T | C | 0.149 | -5.18E-02 | 1.46E-01 | 2.64E+00 | 2.68E+00 | 7.25E-01 | no | Rpl31-ps23 | upstream_gene_variant | MODIFIER |
| 8 | 35158874 | AX-170457364 | 0 | C | T | 0.149 | -5.18E-02 | 1.46E-01 | 2.64E+00 | 2.68E+00 | 7.25E-01 | no | | regulatory_region_variant | MODIFIER |
| 8 | 35170273 | AX-170155257 | 0 | C | G | 0.149 | -5.18E-02 | 1.46E-01 | 2.64E+00 | 2.68E+00 | 7.25E-01 | no | | intergenic_variant | MODIFIER |
| 8 | 35197108 | AX-170311765 | 0 | A | T | 0.149 | -5.18E-02 | 1.46E-01 | 2.64E+00 | 2.68E+00 | 7.25E-01 | no | | intergenic_variant | MODIFIER |
| 8 | 35198792 | AX-170228088 | 0 | C | T | 0.149 | -5.18E-02 | 1.46E-01 | 2.64E+00 | 2.68E+00 | 7.25E-01 | no | | regulatory_region_variant | MODIFIER |
| 8 | 35201379 | AX-170529409 | 0 | G | A | 0.149 | -5.18E-02 | 1.46E-01 | 2.64E+00 | 2.68E+00 | 7.25E-01 | no | | regulatory_region_variant | MODIFIER |
| 8 | 35205726 | AX-170354954 | 0 | C | T | 0.149 | -5.18E-02 | 1.46E-01 | 2.64E+00 | 2.68E+00 | 7.25E-01 | no | | regulatory_region_variant | MODIFIER |
| 8 | 35205796 | AX-170286227 | 0 | T | C | 0.149 | -5.18E-02 | 1.46E-01 | 2.64E+00 | 2.68E+00 | 7.25E-01 | no | | regulatory_region_variant | MODIFIER |
| 8 | 35209087 | AX-170122709 | 0 | G | A | 0.149 | -5.18E-02 | 1.46E-01 | 2.64E+00 | 2.68E+00 | 7.25E-01 | no | | regulatory_region_variant | MODIFIER |
| 8 | 35213322 | AX-170478291 | 0 | C | T | 0.149 | -5.18E-02 | 1.46E-01 | 2.64E+00 | 2.68E+00 | 7.25E-01 | no | | intergenic_variant | MODIFIER |
| 8 | 35214045 | AX-170412432 | 0 | G | A | 0.149 | -5.18E-02 | 1.46E-01 | 2.64E+00 | 2.68E+00 | 7.25E-01 | no | Gm34474 | upstream_gene_variant | MODIFIER |
| 8 | 35226192 | AX-170417618 | 0 | A | G | 0.149 | -5.18E-02 | 1.46E-01 | 2.64E+00 | 2.68E+00 | 7.25E-01 | no | Gm34533 | downstream_gene_variant | MODIFIER |
| 8 | 35244192 | AX-170345722 | 0 | A | C | 0.149 | -5.18E-02 | 1.46E-01 | 2.64E+00 | 2.68E+00 | 7.25E-01 | no | Gm34597 | intron_variant,non_coding_transcript_variant | MODIFIER |
| 8 | 35258267 | AX-170344344 | 0 | T | C | 0.149 | -5.18E-02 | 1.46E-01 | 2.64E+00 | 2.68E+00 | 7.25E-01 | no | | intergenic_variant | MODIFIER |
| 8 | 35268635 | AX-97804053 | 0 | G | A | 0.149 | -5.18E-02 | 1.46E-01 | 2.64E+00 | 2.68E+00 | 7.25E-01 | no | | regulatory_region_variant | MODIFIER |
| 8 | 35295455 | AX-170126574 | 0 | T | C | 0.149 | -5.18E-02 | 1.46E-01 | 2.64E+00 | 2.68E+00 | 7.25E-01 | no | | intergenic_variant | MODIFIER |

| | | | | | | | | | | | | | | | |
|---|---|---|---|---|---|---|---|---|---|---|---|---|---|---|---|
| 8 | 35301007 | AX-97852304 | 0 | A | G | 0.149 | -5.18E-02 | 1.46E-01 | 2.64E+00 | 2.68E+00 | 7.25E-01 | no | | regulatory_region_variant | MODIFIER |
| 8 | 35301641 | AX-170237475 | 0 | T | C | 0.149 | -5.18E-02 | 1.46E-01 | 2.64E+00 | 2.68E+00 | 7.25E-01 | no | | intergenic_variant | MODIFIER |
| 8 | 35301736 | AX-170282645 | 0 | C | T | 0.149 | -5.18E-02 | 1.46E-01 | 2.64E+00 | 2.68E+00 | 7.25E-01 | no | | intergenic_variant | MODIFIER |
| 8 | 35343110 | AX-97786706 | 0 | G | T | 0.149 | -5.18E-02 | 1.46E-01 | 2.64E+00 | 2.68E+00 | 7.25E-01 | no | | intergenic_variant | MODIFIER |
| 8 | 35349906 | AX-170136203 | 0 | C | G | 0.149 | -5.18E-02 | 1.46E-01 | 2.64E+00 | 2.68E+00 | 7.25E-01 | no | | regulatory_region_variant | MODIFIER |
| 8 | 35357538 | AX-170079736 | 0 | C | T | 0.149 | -5.18E-02 | 1.46E-01 | 2.64E+00 | 2.68E+00 | 7.25E-01 | no | | intergenic_variant | MODIFIER |
| 8 | 35359643 | AX-170079737 | 0 | C | G | 0.149 | -5.18E-02 | 1.46E-01 | 2.64E+00 | 2.68E+00 | 7.25E-01 | no | | regulatory_region_variant | MODIFIER |
| 8 | 35361080 | AX-170477394 | 0 | T | C | 0.149 | -5.18E-02 | 1.46E-01 | 2.64E+00 | 2.68E+00 | 7.25E-01 | no | | intergenic_variant | MODIFIER |
| 8 | 35366524 | AX-170499741 | 0 | A | G | 0.149 | -5.18E-02 | 1.46E-01 | 2.64E+00 | 2.68E+00 | 7.25E-01 | no | Gm45469 | downstream_gene_variant | MODIFIER |
| 8 | 35372853 | AX-170030502 | 0 | T | C | 0.149 | -5.18E-02 | 1.46E-01 | 2.64E+00 | 2.68E+00 | 7.25E-01 | no | Ppp1r3b | upstream_gene_variant | MODIFIER |
| 8 | 35387370 | AX-170030501 | 0 | C | T | 0.149 | -5.18E-02 | 1.46E-01 | 2.64E+00 | 2.68E+00 | 7.25E-01 | no | Ppp1r3b | 3_prime_UTR_variant | MODIFIER |
| 8 | 35391555 | AX-170283348 | 0 | A | G | 0.149 | -5.18E-02 | 1.46E-01 | 2.64E+00 | 2.68E+00 | 7.25E-01 | no | Ppp1r3b | downstream_gene_variant | MODIFIER |
| 8 | 35392384 | AX-170181265 | 0 | T | C | 0.149 | -5.18E-02 | 1.46E-01 | 2.64E+00 | 2.68E+00 | 7.25E-01 | no | Ppp1r3b | downstream_gene_variant | MODIFIER |
| 8 | 35392832 | AX-97854238 | 0 | A | G | 0.149 | -5.18E-02 | 1.46E-01 | 2.64E+00 | 2.68E+00 | 7.25E-01 | no | Ppp1r3b | downstream_gene_variant | MODIFIER |
| 8 | 35405269 | AX-170118848 | 0 | T | C | 0.149 | -5.18E-02 | 1.46E-01 | 2.64E+00 | 2.68E+00 | 7.25E-01 | no | Gm45301 | downstream_gene_variant | MODIFIER |
| 8 | 35414710 | AX-170493357 | 0 | A | G | 0.149 | -5.18E-02 | 1.46E-01 | 2.64E+00 | 2.68E+00 | 7.25E-01 | no | | intergenic_variant | MODIFIER |
| 8 | 35415176 | AX-170030500 | 0 | A | G | 0.149 | -5.18E-02 | 1.46E-01 | 2.64E+00 | 2.68E+00 | 7.25E-01 | no | | intergenic_variant | MODIFIER |
| 8 | 35440725 | AX-170341223 | 0 | T | G | 0.149 | -5.18E-02 | 1.46E-01 | 2.64E+00 | 2.68E+00 | 7.25E-01 | no | Gm34853 | intron_variant,non_coding_transcript_variant | MODIFIER |
| 8 | 35448261 | AX-170234063 | 0 | C | T | 0.149 | -5.18E-02 | 1.46E-01 | 2.64E+00 | 2.68E+00 | 7.25E-01 | no | Gm34853 | downstream_gene_variant | MODIFIER |
| 8 | 35451387 | AX-170488816 | 0 | C | T | 0.149 | -5.18E-02 | 1.46E-01 | 2.64E+00 | 2.68E+00 | 7.25E-01 | no | | intergenic_variant | MODIFIER |
| 8 | 35456742 | AX-170190625 | 0 | T | C | 0.149 | -5.18E-02 | 1.46E-01 | 2.64E+00 | 2.68E+00 | 7.25E-01 | no | | intergenic_variant | MODIFIER |
| 8 | 35461909 | AX-170208269 | 0 | A | G | 0.149 | -5.18E-02 | 1.46E-01 | 2.64E+00 | 2.68E+00 | 7.25E-01 | no | Eri1 | downstream_gene_variant | MODIFIER |
| 8 | 35473508 | AX-170356503 | 0 | G | A | 0.149 | -5.18E-02 | 1.46E-01 | 2.64E+00 | 2.68E+00 | 7.25E-01 | no | Eri1 | intron_variant | MODIFIER |
| 8 | 35490905 | AX-97788200 | 0 | G | C | 0.149 | -5.18E-02 | 1.46E-01 | 2.64E+00 | 2.68E+00 | 7.25E-01 | no | Eri1 | intron_variant | MODIFIER |
| 8 | 35514572 | AX-170079739 | 0 | G | A | 0.149 | -5.18E-02 | 1.46E-01 | 2.64E+00 | 2.68E+00 | 7.25E-01 | no | | intergenic_variant | MODIFIER |
| 8 | 35514767 | AX-170079740 | 0 | G | A | 0.149 | -5.18E-02 | 1.46E-01 | 2.64E+00 | 2.68E+00 | 7.25E-01 | no | | intergenic_variant | MODIFIER |
| 8 | 35527448 | AX-170469113 | 0 | A | G | 0.149 | -5.18E-02 | 1.46E-01 | 2.64E+00 | 2.68E+00 | 7.25E-01 | no | | intergenic_variant | MODIFIER |
| 8 | 35530190 | AX-97809609 | 0 | C | T | 0.149 | -5.18E-02 | 1.46E-01 | 2.64E+00 | 2.68E+00 | 7.25E-01 | no | | intergenic_variant | MODIFIER |
| 8 | 35533470 | AX-170190023 | 0 | G | A | 0.149 | -5.18E-02 | 1.46E-01 | 2.64E+00 | 2.68E+00 | 7.25E-01 | no | | intergenic_variant | MODIFIER |
| 8 | 35533700 | AX-170485069 | 0 | C | T | 0.149 | -5.18E-02 | 1.46E-01 | 2.64E+00 | 2.68E+00 | 7.25E-01 | no | | intergenic_variant | MODIFIER |
| 8 | 35536094 | AX-170344716 | 0 | T | A | 0.149 | -5.18E-02 | 1.46E-01 | 2.64E+00 | 2.68E+00 | 7.25E-01 | no | | intergenic_variant | MODIFIER |
| 8 | 35549966 | AX-170490960 | 0 | T | A | 0.149 | -5.18E-02 | 1.46E-01 | 2.64E+00 | 2.68E+00 | 7.25E-01 | no | | intergenic_variant | MODIFIER |
| 8 | 35550576 | AX-170463940 | 0 | A | G | 0.149 | -5.18E-02 | 1.46E-01 | 2.64E+00 | 2.68E+00 | 7.25E-01 | no | | intergenic_variant | MODIFIER |
| 8 | 35551729 | AX-170257603 | 0 | G | C | 0.149 | -5.18E-02 | 1.46E-01 | 2.64E+00 | 2.68E+00 | 7.25E-01 | no | | regulatory_region_variant | MODIFIER |
| 8 | 35554554 | AX-170125448 | 0 | C | T | 0.149 | -5.18E-02 | 1.46E-01 | 2.64E+00 | 2.68E+00 | 7.25E-01 | no | | regulatory_region_variant | MODIFIER |
| 8 | 35570374 | AX-170503076 | 0 | C | T | 0.149 | -5.18E-02 | 1.46E-01 | 2.64E+00 | 2.68E+00 | 7.25E-01 | no | | regulatory_region_variant | MODIFIER |
| 8 | 35600053 | AX-170204654 | 0 | C | T | 0.149 | -5.18E-02 | 1.46E-01 | 2.64E+00 | 2.68E+00 | 7.25E-01 | no | Mfhas1 | intron_variant | MODIFIER |
| 8 | 35603013 | AX-170278993 | 0 | C | T | 0.149 | -5.18E-02 | 1.46E-01 | 2.64E+00 | 2.68E+00 | 7.25E-01 | no | Mfhas1 | intron_variant | MODIFIER |
| 8 | 35603469 | AX-170337951 | 0 | T | C | 0.149 | -5.18E-02 | 1.46E-01 | 2.64E+00 | 2.68E+00 | 7.25E-01 | no | Mfhas1 | intron_variant | MODIFIER |
| 8 | 35605108 | AX-170511498 | 0 | G | A | 0.149 | -5.18E-02 | 1.46E-01 | 2.64E+00 | 2.68E+00 | 7.25E-01 | no | Mfhas1 | intron_variant | MODIFIER |
| 8 | 35605217 | AX-170474271 | 0 | G | A | 0.149 | -5.18E-02 | 1.46E-01 | 2.64E+00 | 2.68E+00 | 7.25E-01 | no | Mfhas1 | intron_variant | MODIFIER |
| 8 | 35608178 | AX-170503007 | 0 | C | A | 0.149 | -5.18E-02 | 1.46E-01 | 2.64E+00 | 2.68E+00 | 7.25E-01 | no | Mfhas1 | intron_variant | MODIFIER |
| 8 | 35609546 | AX-170316789 | 0 | G | T | 0.149 | -5.18E-02 | 1.46E-01 | 2.64E+00 | 2.68E+00 | 7.25E-01 | no | Mfhas1 | intron_variant | MODIFIER |
| 8 | 35616875 | AX-170489065 | 0 | T | C | 0.149 | -5.18E-02 | 1.46E-01 | 2.64E+00 | 2.68E+00 | 7.25E-01 | no | Mfhas1 | intron_variant | MODIFIER |
| 8 | 35618636 | AX-170372568 | 0 | T | C | 0.149 | -5.18E-02 | 1.46E-01 | 2.64E+00 | 2.68E+00 | 7.25E-01 | no | Mfhas1 | intron_variant | MODIFIER |
| 8 | 35621467 | AX-97854894 | 0 | T | C | 0.149 | -5.18E-02 | 1.46E-01 | 2.64E+00 | 2.68E+00 | 7.25E-01 | no | Mfhas1 | intron_variant | MODIFIER |
| 8 | 35623122 | AX-170141643 | 0 | A | G | 0.149 | -5.18E-02 | 1.46E-01 | 2.64E+00 | 2.68E+00 | 7.25E-01 | no | Mfhas1 | intron_variant | MODIFIER |
| 8 | 35623754 | AX-170501234 | 0 | T | C | 0.149 | -5.18E-02 | 1.46E-01 | 2.64E+00 | 2.68E+00 | 7.25E-01 | no | Mfhas1 | intron_variant | MODIFIER |
| 8 | 35641710 | AX-170443287 | 0 | T | C | 0.149 | -5.18E-02 | 1.46E-01 | 2.64E+00 | 2.68E+00 | 7.25E-01 | no | Mfhas1 | intron_variant | MODIFIER |
| 8 | 35641832 | AX-170443242 | 0 | C | T | 0.149 | -5.18E-02 | 1.46E-01 | 2.64E+00 | 2.68E+00 | 7.25E-01 | no | Mfhas1 | intron_variant | MODIFIER |
| 8 | 35649086 | AX-170251839 | 0 | C | T | 0.149 | -5.18E-02 | 1.46E-01 | 2.64E+00 | 2.68E+00 | 7.25E-01 | no | Mfhas1 | intron_variant | MODIFIER |
| 8 | 35651461 | AX-170401471 | 0 | A | G | 0.149 | -5.18E-02 | 1.46E-01 | 2.64E+00 | 2.68E+00 | 7.25E-01 | no | Mfhas1 | intron_variant | MODIFIER |
| 8 | 35652499 | AX-170347672 | 0 | C | T | 0.149 | -5.18E-02 | 1.46E-01 | 2.64E+00 | 2.68E+00 | 7.25E-01 | no | Mfhas1 | intron_variant | MODIFIER |
| 8 | 35652548 | AX-170208082 | 0 | A | T | 0.149 | -5.18E-02 | 1.46E-01 | 2.64E+00 | 2.68E+00 | 7.25E-01 | no | Mfhas1 | intron_variant | MODIFIER |

| | | | | | | | | | | | | | | | |
|---|---|---|---|---|---|---|---|---|---|---|---|---|---|---|---|
| 8 | 35655667 | AX-170158411 | 0 | G | A | 0.149 | -5.18E-02 | 1.46E-01 | 2.64E+00 | 2.68E+00 | 7.25E-01 | no | Mfhas1 | intron_variant | MODIFIER |
| 8 | 35672585 | AX-97826741 | 0 | A | C | 0.149 | -5.18E-02 | 1.46E-01 | 2.64E+00 | 2.68E+00 | 7.25E-01 | no | Mfhas1 | intron_variant | MODIFIER |
| 8 | 35676016 | AX-170378380 | 0 | C | T | 0.149 | -5.18E-02 | 1.46E-01 | 2.64E+00 | 2.68E+00 | 7.25E-01 | no | Mfhas1 | intron_variant | MODIFIER |
| 8 | 35678111 | AX-170330590 | 0 | T | C | 0.149 | -5.18E-02 | 1.46E-01 | 2.64E+00 | 2.68E+00 | 7.25E-01 | no | Mfhas1 | 3_prime_UTR_variant | MODIFIER |
| 8 | 35681499 | AX-170331482 | 0 | C | T | 0.149 | -5.18E-02 | 1.46E-01 | 2.64E+00 | 2.68E+00 | 7.25E-01 | no | Mfhas1 | downstream_gene_variant | MODIFIER |
| 8 | 35700979 | AX-170481405 | 0 | T | G | 0.149 | -5.18E-02 | 1.46E-01 | 2.64E+00 | 2.68E+00 | 7.25E-01 | no | | regulatory_region_variant | MODIFIER |
| 8 | 35713900 | AX-170258294 | 0 | T | C | 0.149 | -5.18E-02 | 1.46E-01 | 2.64E+00 | 2.68E+00 | 7.25E-01 | no | 1700015I17Rik | intron_variant,non_coding_transcript_variant | MODIFIER |
| 8 | 35716094 | AX-170103852 | 0 | A | G | 0.149 | -5.18E-02 | 1.46E-01 | 2.64E+00 | 2.68E+00 | 7.25E-01 | no | 1700015I17Rik | intron_variant,non_coding_transcript_variant | MODIFIER |
| 8 | 35718863 | AX-170338660 | 0 | A | T | 0.149 | -5.18E-02 | 1.46E-01 | 2.64E+00 | 2.68E+00 | 7.25E-01 | no | 1700015I17Rik | intron_variant,non_coding_transcript_variant | MODIFIER |
| 8 | 35730449 | AX-170426720 | 0 | G | C | 0.149 | -5.18E-02 | 1.46E-01 | 2.64E+00 | 2.68E+00 | 7.25E-01 | no | Gm35021 | intron_variant,non_coding_transcript_variant | MODIFIER |
| 8 | 35735763 | AX-170295751 | 0 | T | G | 0.149 | -5.18E-02 | 1.46E-01 | 2.64E+00 | 2.68E+00 | 7.25E-01 | no | Gm35021 | intron_variant,non_coding_transcript_variant | MODIFIER |
| 8 | 35743540 | AX-170079744 | 0 | A | G | 0.149 | -5.18E-02 | 1.46E-01 | 2.64E+00 | 2.68E+00 | 7.25E-01 | no | Gm22216 | upstream_gene_variant | MODIFIER |
| 8 | 35755040 | AX-170079745 | 0 | G | A | 0.149 | -5.18E-02 | 1.46E-01 | 2.64E+00 | 2.68E+00 | 7.25E-01 | no | Gm35021 | intron_variant,non_coding_transcript_variant | MODIFIER |
| 8 | 35759224 | AX-170261346 | 0 | T | G | 0.149 | -5.18E-02 | 1.46E-01 | 2.64E+00 | 2.68E+00 | 7.25E-01 | no | Gm44474 | downstream_gene_variant | MODIFIER |
| 8 | 35759266 | AX-170462988 | 0 | G | C | 0.149 | -5.18E-02 | 1.46E-01 | 2.64E+00 | 2.68E+00 | 7.25E-01 | no | Gm44474 | downstream_gene_variant | MODIFIER |
| 8 | 35762930 | AX-170394318 | 0 | C | G | 0.149 | -5.18E-02 | 1.46E-01 | 2.64E+00 | 2.68E+00 | 7.25E-01 | no | Gm44474 | downstream_gene_variant | MODIFIER |
| 8 | 35772891 | AX-170347665 | 0 | T | C | 0.149 | -5.18E-02 | 1.46E-01 | 2.64E+00 | 2.68E+00 | 7.25E-01 | no | Gm19410 | intron_variant | MODIFIER |
| 8 | 35773291 | AX-170532144 | 0 | G | T | 0.149 | -5.18E-02 | 1.46E-01 | 2.64E+00 | 2.68E+00 | 7.25E-01 | no | Gm19410 | intron_variant | MODIFIER |
| 8 | 35776082 | AX-170468248 | 0 | G | T | 0.149 | -5.18E-02 | 1.46E-01 | 2.64E+00 | 2.68E+00 | 7.25E-01 | no | Gm19410 | intron_variant | MODIFIER |
| 8 | 35779294 | AX-170532687 | 0 | C | T | 0.149 | -5.18E-02 | 1.46E-01 | 2.64E+00 | 2.68E+00 | 7.25E-01 | no | Gm19410 | intron_variant | MODIFIER |
| 8 | 35788464 | AX-97821065 | 0 | A | G | 0.149 | -5.18E-02 | 1.46E-01 | 2.64E+00 | 2.68E+00 | 7.25E-01 | no | Gm19410 | intron_variant | MODIFIER |
| 8 | 35788870 | AX-170325053 | 0 | A | G | 0.149 | -5.18E-02 | 1.46E-01 | 2.64E+00 | 2.68E+00 | 7.25E-01 | no | Gm19410 | intron_variant | MODIFIER |
| 8 | 35796075 | AX-170455105 | 0 | T | C | 0.149 | -5.18E-02 | 1.46E-01 | 2.64E+00 | 2.68E+00 | 7.25E-01 | no | Gm19410 | intron_variant | MODIFIER |
| 8 | 35807095 | AX-170486956 | 0 | G | A | 0.149 | -5.18E-02 | 1.46E-01 | 2.64E+00 | 2.68E+00 | 7.25E-01 | no | Gm19410 | synonymous_variant | LOW |
| 8 | 35832996 | AX-170319986 | 0 | C | T | 0.149 | -5.18E-02 | 1.46E-01 | 2.64E+00 | 2.68E+00 | 7.25E-01 | no | | intergenic_variant | MODIFIER |
| 8 | 35836494 | AX-170348676 | 0 | G | A | 0.149 | -5.18E-02 | 1.46E-01 | 2.64E+00 | 2.68E+00 | 7.25E-01 | no | | intergenic_variant | MODIFIER |
| 8 | 35836766 | AX-170363230 | 0 | C | T | 0.149 | -5.18E-02 | 1.46E-01 | 2.64E+00 | 2.68E+00 | 7.25E-01 | no | | intergenic_variant | MODIFIER |
| 8 | 35848873 | AX-97824752 | 0 | C | A | 0.149 | -5.18E-02 | 1.46E-01 | 2.64E+00 | 2.68E+00 | 7.25E-01 | no | | intergenic_variant | MODIFIER |
| 8 | 35861473 | AX-170315843 | 0 | C | T | 0.149 | -5.18E-02 | 1.46E-01 | 2.64E+00 | 2.68E+00 | 7.25E-01 | no | 5430403N17Rik | downstream_gene_variant | MODIFIER |
| 8 | 35861741 | AX-170351841 | 0 | T | C | 0.149 | -5.18E-02 | 1.46E-01 | 2.64E+00 | 2.68E+00 | 7.25E-01 | no | 5430403N17Rik | downstream_gene_variant | MODIFIER |
| 8 | 35864620 | AX-170449571 | 0 | G | A | 0.149 | -5.18E-02 | 1.46E-01 | 2.64E+00 | 2.68E+00 | 7.25E-01 | no | 5430403N17Rik | non_coding_transcript_exon_variant | MODIFIER |
| 8 | 35869696 | AX-170451121 | 0 | C | T | 0.149 | -5.18E-02 | 1.46E-01 | 2.64E+00 | 2.68E+00 | 7.25E-01 | no | 5430403N17Rik | intron_variant,non_coding_transcript_variant | MODIFIER |
| 8 | 35889762 | AX-170482417 | 0 | A | C | 0.149 | -5.18E-02 | 1.46E-01 | 2.64E+00 | 2.68E+00 | 7.25E-01 | no | | intergenic_variant | MODIFIER |
| 8 | 35906932 | AX-170488189 | 0 | C | T | 0.149 | -5.18E-02 | 1.46E-01 | 2.64E+00 | 2.68E+00 | 7.25E-01 | no | | intergenic_variant | MODIFIER |
| 8 | 35935448 | AX-170328405 | 0 | T | C | 0.149 | -5.18E-02 | 1.46E-01 | 2.64E+00 | 2.68E+00 | 7.25E-01 | no | | intergenic_variant | MODIFIER |
| 8 | 35943662 | AX-170205466 | 0 | A | G | 0.149 | -5.18E-02 | 1.46E-01 | 2.64E+00 | 2.68E+00 | 7.25E-01 | no | | regulatory_region_variant | MODIFIER |
| 8 | 35951777 | AX-170030507 | 0 | T | C | 0.149 | -5.18E-02 | 1.46E-01 | 2.64E+00 | 2.68E+00 | 7.25E-01 | no | | intergenic_variant | MODIFIER |
| 8 | 35952356 | AX-170491208 | 0 | A | G | 0.149 | -5.18E-02 | 1.46E-01 | 2.64E+00 | 2.68E+00 | 7.25E-01 | no | | intergenic_variant | MODIFIER |
| 8 | 35953950 | AX-170445763 | 0 | G | A | 0.149 | -5.18E-02 | 1.46E-01 | 2.64E+00 | 2.68E+00 | 7.25E-01 | no | | intergenic_variant | MODIFIER |
| 8 | 35961614 | AX-170147287 | 0 | C | A | 0.149 | -5.18E-02 | 1.46E-01 | 2.64E+00 | 2.68E+00 | 7.25E-01 | no | | regulatory_region_variant | MODIFIER |
| 8 | 35964206 | AX-97795721 | 0 | C | T | 0.149 | -5.18E-02 | 1.46E-01 | 2.64E+00 | 2.68E+00 | 7.25E-01 | no | | intergenic_variant | MODIFIER |
| 8 | 35968377 | AX-170290620 | 0 | T | C | 0.149 | -5.18E-02 | 1.46E-01 | 2.64E+00 | 2.68E+00 | 7.25E-01 | no | | regulatory_region_variant | MODIFIER |
| 8 | 35968941 | AX-170108400 | 0 | T | C | 0.149 | -5.18E-02 | 1.46E-01 | 2.64E+00 | 2.68E+00 | 7.25E-01 | no | | regulatory_region_variant | MODIFIER |
| 8 | 35991273 | AX-170306529 | 0 | A | G | 0.149 | -5.18E-02 | 1.46E-01 | 2.64E+00 | 2.68E+00 | 7.25E-01 | no | | regulatory_region_variant | MODIFIER |
| 8 | 35991805 | AX-170163864 | 0 | C | T | 0.149 | -5.18E-02 | 1.46E-01 | 2.64E+00 | 2.68E+00 | 7.25E-01 | no | | regulatory_region_variant | MODIFIER |
| 8 | 35997641 | AX-170371331 | 0 | T | C | 0.149 | -5.18E-02 | 1.46E-01 | 2.64E+00 | 2.68E+00 | 7.25E-01 | no | | intergenic_variant | MODIFIER |
| 8 | 35998305 | AX-97782060 | 0 | C | T | 0.149 | -5.18E-02 | 1.46E-01 | 2.64E+00 | 2.68E+00 | 7.25E-01 | no | | intergenic_variant | MODIFIER |
| 8 | 36001987 | AX-170079756 | 0 | T | G | 0.149 | -5.18E-02 | 1.46E-01 | 2.64E+00 | 2.68E+00 | 7.25E-01 | no | | intergenic_variant | MODIFIER |
| 8 | 36011111 | AX-170079755 | 0 | C | A | 0.149 | -5.18E-02 | 1.46E-01 | 2.64E+00 | 2.68E+00 | 7.25E-01 | no | | regulatory_region_variant | MODIFIER |
| 8 | 36026290 | AX-170035777 | 0 | C | T | 0.149 | -5.18E-02 | 1.46E-01 | 2.64E+00 | 2.68E+00 | 7.25E-01 | no | Rps12-ps24 | downstream_gene_variant | MODIFIER |
| 8 | 36032321 | AX-170030506 | 0 | T | C | 0.149 | -5.18E-02 | 1.46E-01 | 2.64E+00 | 2.68E+00 | 7.25E-01 | no | | intergenic_variant | MODIFIER |
| 8 | 36033223 | AX-170469937 | 0 | T | C | 0.149 | -5.18E-02 | 1.46E-01 | 2.64E+00 | 2.68E+00 | 7.25E-01 | no | | intergenic_variant | MODIFIER |
| 8 | 36040185 | AX-170205158 | 0 | A | G | 0.149 | -5.18E-02 | 1.46E-01 | 2.64E+00 | 2.68E+00 | 7.25E-01 | no | | intergenic_variant | MODIFIER |
| 8 | 36058034 | AX-170296489 | 0 | A | G | 0.149 | -5.18E-02 | 1.46E-01 | 2.64E+00 | 2.68E+00 | 7.25E-01 | no | | intergenic_variant | MODIFIER |

| | | | | | | | | | | | | | | | |
|---|---|---|---|---|---|---|---|---|---|---|---|---|---|---|---|
| 8 | 36069255 | AX-170391291 | 0 | A | G | 0.149 | -5.18E-02 | 1.46E-01 | 2.64E+00 | 2.68E+00 | 7.25E-01 | no | | intergenic_variant | MODIFIER |
| 8 | 36086887 | AX-170280360 | 0 | T | C | 0.149 | -5.18E-02 | 1.46E-01 | 2.64E+00 | 2.68E+00 | 7.25E-01 | no | | intergenic_variant | MODIFIER |
| 15 | 93687716 | AX-170012290 | 1 | C | T | 0.143 | 6.21E-02 | 1.76E-01 | 2.63E+00 | 2.71E+00 | 7.25E-01 | no | | intergenic_variant | MODIFIER |
| 5 | 39402304 | AX-170071853 | 2 | G | A | 0.218 | 3.88E-02 | 1.13E-01 | 2.70E+00 | 2.79E+00 | 7.25E-01 | no | Gm5291 | upstream_gene_variant | MODIFIER |
| 15 | 92187393 | AX-97839486 | 0 | C | A | 0.149 | 6.30E-02 | 1.79E-01 | 2.64E+00 | 2.70E+00 | 7.25E-01 | no | Cntn1 | intron_variant | MODIFIER |
| 15 | 93239137 | AX-170012278 | 0 | G | A | 0.149 | 6.30E-02 | 1.79E-01 | 2.64E+00 | 2.70E+00 | 7.25E-01 | no | Gxylt1 | downstream_gene_variant | MODIFIER |
| 15 | 96634814 | AX-170514418 | 0 | G | A | 0.149 | 6.30E-02 | 1.79E-01 | 2.64E+00 | 2.70E+00 | 7.25E-01 | no | Slc38a1 | intron_variant | MODIFIER |
| 8 | 15130261 | AX-97787123 | 0 | C | T | 0.237 | -2.77E-02 | 7.60E-02 | 2.57E+00 | 2.66E+00 | 7.25E-01 | no | Myom2 | intron_variant | MODIFIER |
| 7 | 118682436 | AX-170327161 | 1 | A | C | 0.384 | 3.10E-02 | 8.89E-02 | 2.67E+00 | 2.73E+00 | 7.25E-01 | no | | intergenic_variant | MODIFIER |
| 10 | 19371780 | AX-170470562 | 1 | T | G | 0.58 | 2.50E-02 | 6.59E-02 | 2.50E+00 | 2.64E+00 | 7.25E-01 | no | | intergenic_variant | MODIFIER |
| 8 | 101591274 | AX-170309177 | 0 | G | A | 0.237 | 3.88E-02 | 1.11E-01 | 2.66E+00 | 2.73E+00 | 7.25E-01 | no | | intergenic_variant | MODIFIER |
| 2 | 93159695 | AX-170375731 | 0 | G | A | 0.053 | -1.04E-01 | 3.00E-01 | 2.73E+00 | 2.76E+00 | 7.25E-01 | no | | intergenic_variant | MODIFIER |
| 8 | 41126022 | AX-170079868 | 0 | T | A | 0.158 | 4.87E-02 | 1.41E-01 | 2.74E+00 | 2.79E+00 | 7.25E-01 | no | Mtus1 | intron_variant | MODIFIER |
| 11 | 49859822 | AX-170383688 | 0 | C | T | 0.088 | -5.59E-02 | 1.56E-01 | 2.57E+00 | 2.69E+00 | 7.26E-01 | no | Mapk9 | intron_variant | MODIFIER |
| 1 | 3404187 | AX-170486258 | 1 | A | T | 0.527 | 2.60E-02 | 6.91E-02 | 2.54E+00 | 2.61E+00 | 7.26E-01 | no | Xkr4 | intron_variant | MODIFIER |
| 6 | 133204377 | AX-170506103 | 0 | G | A | 0.421 | 4.14E-02 | 1.12E-01 | 2.54E+00 | 2.65E+00 | 7.26E-01 | no | | intergenic_variant | MODIFIER |
| 8 | 39898918 | AX-97807762 | 1 | G | A | 0.152 | 5.10E-02 | 1.48E-01 | 2.75E+00 | 2.80E+00 | 7.26E-01 | no | | intergenic_variant | MODIFIER |
| 9 | 50306646 | AX-170032321 | 0 | C | G | 0.211 | -3.77E-02 | 1.08E-01 | 2.65E+00 | 2.73E+00 | 7.26E-01 | no | Gm39336 | upstream_gene_variant | MODIFIER |
| 9 | 50362112 | AX-170032320 | 0 | G | C | 0.211 | -3.77E-02 | 1.08E-01 | 2.65E+00 | 2.73E+00 | 7.26E-01 | no | | intergenic_variant | MODIFIER |
| 9 | 50378621 | AX-97781941 | 0 | T | C | 0.211 | -3.77E-02 | 1.08E-01 | 2.65E+00 | 2.73E+00 | 7.26E-01 | no | | intergenic_variant | MODIFIER |
| 9 | 50387534 | AX-170264885 | 0 | C | T | 0.211 | -3.77E-02 | 1.08E-01 | 2.65E+00 | 2.73E+00 | 7.26E-01 | no | | intergenic_variant | MODIFIER |
| 9 | 50387633 | AX-170120577 | 0 | C | A | 0.211 | -3.77E-02 | 1.08E-01 | 2.65E+00 | 2.73E+00 | 7.26E-01 | no | | intergenic_variant | MODIFIER |
| 9 | 50388129 | AX-170032318 | 0 | A | G | 0.211 | -3.77E-02 | 1.08E-01 | 2.65E+00 | 2.73E+00 | 7.26E-01 | no | | intergenic_variant | MODIFIER |
| 9 | 50388333 | AX-170354110 | 0 | A | C | 0.211 | -3.77E-02 | 1.08E-01 | 2.65E+00 | 2.73E+00 | 7.26E-01 | no | | intergenic_variant | MODIFIER |
| 9 | 50390680 | AX-170459878 | 0 | T | C | 0.211 | -3.77E-02 | 1.08E-01 | 2.65E+00 | 2.73E+00 | 7.26E-01 | no | | intergenic_variant | MODIFIER |
| 9 | 50390868 | AX-170112902 | 0 | T | C | 0.211 | -3.77E-02 | 1.08E-01 | 2.65E+00 | 2.73E+00 | 7.26E-01 | no | | intergenic_variant | MODIFIER |
| 9 | 50396362 | AX-170082422 | 0 | A | G | 0.211 | -3.77E-02 | 1.08E-01 | 2.65E+00 | 2.73E+00 | 7.26E-01 | no | | regulatory_region_variant | MODIFIER |
| 9 | 50398096 | AX-170472942 | 0 | G | A | 0.211 | -3.77E-02 | 1.08E-01 | 2.65E+00 | 2.73E+00 | 7.26E-01 | no | | intergenic_variant | MODIFIER |
| 9 | 50423272 | AX-97837954 | 0 | G | T | 0.211 | -3.77E-02 | 1.08E-01 | 2.65E+00 | 2.73E+00 | 7.26E-01 | no | | intergenic_variant | MODIFIER |
| 9 | 50449391 | AX-170282419 | 0 | G | A | 0.211 | -3.77E-02 | 1.08E-01 | 2.65E+00 | 2.73E+00 | 7.26E-01 | no | | regulatory_region_variant | MODIFIER |
| 8 | 39099603 | AX-97836620 | 0 | C | A | 0.149 | 5.09E-02 | 1.48E-01 | 2.75E+00 | 2.80E+00 | 7.26E-01 | no | Tusc3 | intron_variant | MODIFIER |
| 18 | 58655865 | AX-170351305 | 0 | C | T | 0.395 | 3.66E-02 | 9.70E-02 | 2.54E+00 | 2.59E+00 | 7.26E-01 | no | Isoc1 | upstream_gene_variant | MODIFIER |
| 11 | 74283604 | AX-170005298 | 1 | G | A | 0.268 | 2.46E-02 | 7.20E-02 | 2.63E+00 | 2.76E+00 | 7.26E-01 | no | Olfr407-ps1 | upstream_gene_variant | MODIFIER |
| 4 | 106799888 | AX-170023611 | 0 | T | A | 0.246 | 4.29E-02 | 1.11E-01 | 2.50E+00 | 2.60E+00 | 7.26E-01 | no | Acot11 | intron_variant | MODIFIER |
| 1 | 150226549 | AX-170388733 | 0 | G | T | 0.202 | -4.29E-02 | 1.26E-01 | 2.73E+00 | 2.82E+00 | 7.27E-01 | no | | intergenic_variant | MODIFIER |
| 18 | 31856396 | AX-170015470 | 0 | C | T | 0.053 | 7.84E-02 | 2.22E-01 | 2.58E+00 | 2.70E+00 | 7.27E-01 | no | Wdr33 | intron_variant | MODIFIER |
| 2 | 90490797 | AX-170412913 | 1 | C | T | 0.054 | -1.04E-01 | 3.01E-01 | 2.74E+00 | 2.76E+00 | 7.27E-01 | no | 4933423P22Rik | downstream_gene_variant | MODIFIER |
| 2 | 94815409 | AX-168382292 | 1 | G | A | 0.054 | -1.04E-01 | 3.01E-01 | 2.74E+00 | 2.76E+00 | 7.27E-01 | no | | intergenic_variant | MODIFIER |
| 2 | 105113057 | AX-170484351 | 1 | G | A | 0.054 | -1.04E-01 | 3.01E-01 | 2.74E+00 | 2.76E+00 | 7.27E-01 | no | Wt1os | intron_variant,non_coding_transcript_variant | MODIFIER |
| 5 | 27473957 | AX-170241910 | 1 | G | A | 0.054 | -1.04E-01 | 3.01E-01 | 2.74E+00 | 2.76E+00 | 7.27E-01 | no | Dpp6 | intron_variant | MODIFIER |
| 2 | 127214715 | AX-170314496 | 0 | G | T | 0.149 | 5.46E-02 | 1.60E-01 | 2.67E+00 | 2.77E+00 | 7.27E-01 | no | Snrnp200 | intron_variant | MODIFIER |
| 7 | 50421752 | AX-170406190 | 0 | A | G | 0.412 | -3.38E-02 | 9.94E-02 | 2.74E+00 | 2.82E+00 | 7.27E-01 | no | Nell1 | intron_variant | MODIFIER |
| 12 | 99354339 | AX-97795486 | 1 | G | A | 0.062 | 6.23E-02 | 1.78E-01 | 2.61E+00 | 2.72E+00 | 7.27E-01 | no | Foxn3 | intron_variant | MODIFIER |
| 8 | 40912031 | AX-170357644 | 1 | A | G | 0.152 | 5.05E-02 | 1.47E-01 | 2.75E+00 | 2.80E+00 | 7.27E-01 | no | Slc7a2 | intron_variant | MODIFIER |
| 9 | 110983428 | AX-170352201 | 1 | G | A | 0.089 | -4.20E-02 | 1.17E-01 | 2.58E+00 | 2.67E+00 | 7.27E-01 | no | Lrrc2 | 3_prime_UTR_variant | MODIFIER |
| 4 | 133626638 | AX-170097499 | 0 | A | G | 0.158 | 4.12E-02 | 1.13E-01 | 2.57E+00 | 2.64E+00 | 7.27E-01 | no | Zdhhc18 | intron_variant | MODIFIER |
| 4 | 136889926 | AX-170147824 | 0 | T | C | 0.158 | 4.12E-02 | 1.13E-01 | 2.57E+00 | 2.64E+00 | 7.27E-01 | no | C1qc | 3_prime_UTR_variant | MODIFIER |
| 8 | 117824826 | AX-170081305 | 0 | A | T | 0.263 | -3.09E-02 | 9.09E-02 | 2.63E+00 | 2.76E+00 | 7.27E-01 | no | | intergenic_variant | MODIFIER |
| 15 | 82153368 | AX-170535720 | 0 | A | G | 0.544 | 2.68E-02 | 7.64E-02 | 2.62E+00 | 2.70E+00 | 7.27E-01 | no | Srebf2 | intron_variant | MODIFIER |
| 1 | 114780171 | AX-170091647 | 2 | T | G | 0.164 | 7.40E-02 | 2.13E-01 | 2.64E+00 | 2.72E+00 | 7.27E-01 | no | | intergenic_variant | MODIFIER |
| 8 | 85560840 | AX-170122316 | 0 | C | T | 0.211 | 4.14E-02 | 1.21E-01 | 2.67E+00 | 2.75E+00 | 7.28E-01 | no | | regulatory_region_variant | MODIFIER |
| 1 | 16697050 | AX-170424295 | 0 | T | G | 0.377 | 2.77E-02 | 7.72E-02 | 2.59E+00 | 2.66E+00 | 7.28E-01 | no | Ly96 | intron_variant | MODIFIER |
| 4 | 117457022 | AX-168381149 | 0 | C | T | 0.211 | 3.95E-02 | 1.12E-01 | 2.60E+00 | 2.69E+00 | 7.28E-01 | no | Rnf220 | intron_variant | MODIFIER |
| 4 | 118372010 | AX-170375847 | 0 | A | G | 0.211 | 3.95E-02 | 1.12E-01 | 2.60E+00 | 2.69E+00 | 7.28E-01 | no | Szt2 | missense_variant | MODERATE |

| | | | | | | | | | | | | | | | |
|---|---|---|---|---|---|---|---|---|---|---|---|---|---|---|---|
| 5 | 119918329 | AX-170161900 | 0 | C | T | 0.447 | 3.76E-02 | 1.09E-01 | 2.66E+00 | 2.71E+00 | 7.28E-01 | no | Gm43269 | downstream_gene_variant | MODIFIER |
| 1 | 13780160 | AX-170091403 | 1 | T | C | 0.384 | 2.72E-02 | 7.75E-02 | 2.63E+00 | 2.69E+00 | 7.28E-01 | no | | intergenic_variant | MODIFIER |
| 1 | 4918544 | AX-170165954 | 0 | G | A | 0.43 | 2.46E-02 | 6.78E-02 | 2.57E+00 | 2.64E+00 | 7.28E-01 | no | Rgs20 | intron_variant | MODIFIER |
| 8 | 36045391 | AX-170246583 | 1 | T | C | 0.152 | -5.05E-02 | 1.45E-01 | 2.65E+00 | 2.70E+00 | 7.28E-01 | no | | intergenic_variant | MODIFIER |
| 18 | 58250699 | AX-170427899 | 1 | G | T | 0.393 | 3.51E-02 | 9.42E-02 | 2.54E+00 | 2.61E+00 | 7.28E-01 | no | | intergenic_variant | MODIFIER |
| 15 | 87860828 | AX-170442985 | 1 | G | A | 0.116 | 5.89E-02 | 1.66E-01 | 2.61E+00 | 2.68E+00 | 7.28E-01 | no | | intergenic_variant | MODIFIER |
| 10 | 8449696 | AX-170086736 | 1 | C | T | 0.205 | -3.63E-02 | 1.06E-01 | 2.65E+00 | 2.74E+00 | 7.28E-01 | no | Ust | intron_variant | MODIFIER |
| 2 | 93027852 | AX-170393118 | 2 | C | T | 0.055 | -1.03E-01 | 3.00E-01 | 2.74E+00 | 2.77E+00 | 7.28E-01 | no | 4631405J19Rik | upstream_gene_variant | MODIFIER |
| 5 | 122638179 | AX-170026051 | 0 | T | C | 0.456 | -3.69E-02 | 1.08E-01 | 2.74E+00 | 2.80E+00 | 7.28E-01 | no | | regulatory_region_variant | MODIFIER |
| 1 | 108498863 | AX-170480910 | 1 | G | A | 0.17 | 6.97E-02 | 1.96E-01 | 2.60E+00 | 2.68E+00 | 7.29E-01 | no | | intergenic_variant | MODIFIER |
| 2 | 121225591 | AX-170279573 | 2 | G | A | 0.145 | 7.50E-02 | 2.17E-01 | 2.68E+00 | 2.71E+00 | 7.29E-01 | no | Trp53bp1 | intron_variant | MODIFIER |
| 3 | 148544805 | AX-97785712 | 1 | C | T | 0.321 | 2.62E-02 | 7.57E-02 | 2.64E+00 | 2.72E+00 | 7.29E-01 | no | | intergenic_variant | MODIFIER |
| 5 | 37661221 | AX-170089451 | 0 | T | C | 0.333 | 3.55E-02 | 1.03E-01 | 2.65E+00 | 2.73E+00 | 7.29E-01 | no | Stk32b | intron_variant | MODIFIER |
| 1 | 106357062 | AX-170249550 | 0 | C | T | 0.167 | 6.94E-02 | 1.97E-01 | 2.60E+00 | 2.68E+00 | 7.29E-01 | no | Phlpp1 | intron_variant | MODIFIER |
| 1 | 106452282 | AX-170384604 | 0 | A | G | 0.167 | 6.94E-02 | 1.97E-01 | 2.60E+00 | 2.68E+00 | 7.29E-01 | no | | intergenic_variant | MODIFIER |
| 1 | 106609946 | AX-170192202 | 0 | G | C | 0.167 | 6.94E-02 | 1.97E-01 | 2.60E+00 | 2.68E+00 | 7.29E-01 | no | Bcl2 | intron_variant | MODIFIER |
| 1 | 106758507 | AX-170001282 | 0 | A | G | 0.167 | 6.94E-02 | 1.97E-01 | 2.60E+00 | 2.68E+00 | 7.29E-01 | no | Kdsr | intron_variant | MODIFIER |
| 1 | 106761956 | AX-97812612 | 0 | G | T | 0.167 | 6.94E-02 | 1.97E-01 | 2.60E+00 | 2.68E+00 | 7.29E-01 | no | Kdsr | upstream_gene_variant | MODIFIER |
| 1 | 106767813 | AX-170042296 | 0 | A | G | 0.167 | 6.94E-02 | 1.97E-01 | 2.60E+00 | 2.68E+00 | 7.29E-01 | no | Vps4b | downstream_gene_variant | MODIFIER |
| 1 | 106768328 | AX-170042293 | 0 | T | C | 0.167 | 6.94E-02 | 1.97E-01 | 2.60E+00 | 2.68E+00 | 7.29E-01 | no | Vps4b | downstream_gene_variant | MODIFIER |
| 1 | 106774142 | AX-170524248 | 0 | A | T | 0.167 | 6.94E-02 | 1.97E-01 | 2.60E+00 | 2.68E+00 | 7.29E-01 | no | Vps4b | synonymous_variant | LOW |
| 1 | 106820542 | AX-170494883 | 0 | C | T | 0.167 | 6.94E-02 | 1.97E-01 | 2.60E+00 | 2.68E+00 | 7.29E-01 | no | | intergenic_variant | MODIFIER |
| 1 | 106902616 | AX-170136621 | 0 | T | C | 0.167 | 6.94E-02 | 1.97E-01 | 2.60E+00 | 2.68E+00 | 7.29E-01 | no | | intergenic_variant | MODIFIER |
| 1 | 106999550 | AX-97844899 | 0 | G | T | 0.167 | 6.94E-02 | 1.97E-01 | 2.60E+00 | 2.68E+00 | 7.29E-01 | no | Serpinb13 | intron_variant | MODIFIER |
| 1 | 107024838 | AX-97812680 | 0 | G | A | 0.167 | 6.94E-02 | 1.97E-01 | 2.60E+00 | 2.68E+00 | 7.29E-01 | no | Gm8080 | intron_variant,non_coding_transcript_variant | MODIFIER |
| 1 | 107094538 | AX-170001299 | 0 | G | A | 0.167 | 6.94E-02 | 1.97E-01 | 2.60E+00 | 2.68E+00 | 7.29E-01 | no | | intergenic_variant | MODIFIER |
| 1 | 107192452 | AX-170086564 | 0 | A | T | 0.167 | 6.94E-02 | 1.97E-01 | 2.60E+00 | 2.68E+00 | 7.29E-01 | no | Gm8084 | intron_variant,non_coding_transcript_variant | MODIFIER |
| 1 | 107321627 | AX-170163650 | 0 | T | G | 0.167 | 6.94E-02 | 1.97E-01 | 2.60E+00 | 2.68E+00 | 7.29E-01 | no | | intergenic_variant | MODIFIER |
| 1 | 107336666 | AX-97830270 | 0 | T | C | 0.167 | 6.94E-02 | 1.97E-01 | 2.60E+00 | 2.68E+00 | 7.29E-01 | no | | intergenic_variant | MODIFIER |
| 1 | 107351514 | AX-170362912 | 0 | T | C | 0.167 | 6.94E-02 | 1.97E-01 | 2.60E+00 | 2.68E+00 | 7.29E-01 | no | | intergenic_variant | MODIFIER |
| 1 | 107358099 | AX-97791274 | 0 | T | C | 0.167 | 6.94E-02 | 1.97E-01 | 2.60E+00 | 2.68E+00 | 7.29E-01 | no | Serpinb11 | upstream_gene_variant | MODIFIER |
| 1 | 107360988 | AX-170001293 | 0 | G | A | 0.167 | 6.94E-02 | 1.97E-01 | 2.60E+00 | 2.68E+00 | 7.29E-01 | no | Serpinb11 | upstream_gene_variant | MODIFIER |
| 1 | 107373389 | AX-97783851 | 0 | C | T | 0.167 | 6.94E-02 | 1.97E-01 | 2.60E+00 | 2.68E+00 | 7.29E-01 | no | Serpinb11 | intron_variant | MODIFIER |
| 1 | 107411390 | AX-97796726 | 0 | T | G | 0.167 | 6.94E-02 | 1.97E-01 | 2.60E+00 | 2.68E+00 | 7.29E-01 | no | Serpinb7 | intron_variant | MODIFIER |
| 1 | 107420860 | AX-170001297 | 0 | G | A | 0.167 | 6.94E-02 | 1.97E-01 | 2.60E+00 | 2.68E+00 | 7.29E-01 | no | Serpinb7 | upstream_gene_variant | MODIFIER |
| 1 | 107482112 | AX-170035625 | 0 | C | T | 0.167 | 6.94E-02 | 1.97E-01 | 2.60E+00 | 2.68E+00 | 7.29E-01 | no | | intergenic_variant | MODIFIER |
| 1 | 107674021 | AX-170035626 | 0 | A | T | 0.167 | 6.94E-02 | 1.97E-01 | 2.60E+00 | 2.68E+00 | 7.29E-01 | no | | intergenic_variant | MODIFIER |
| 1 | 107775649 | AX-170001307 | 0 | A | C | 0.167 | 6.94E-02 | 1.97E-01 | 2.60E+00 | 2.68E+00 | 7.29E-01 | no | | intergenic_variant | MODIFIER |
| 1 | 108319896 | AX-170001316 | 0 | C | G | 0.167 | 6.94E-02 | 1.97E-01 | 2.60E+00 | 2.68E+00 | 7.29E-01 | no | | intergenic_variant | MODIFIER |
| 1 | 108329244 | AX-97812681 | 0 | A | G | 0.167 | 6.94E-02 | 1.97E-01 | 2.60E+00 | 2.68E+00 | 7.29E-01 | no | | intergenic_variant | MODIFIER |
| 1 | 108339823 | AX-170035628 | 0 | C | T | 0.167 | 6.94E-02 | 1.97E-01 | 2.60E+00 | 2.68E+00 | 7.29E-01 | no | | intergenic_variant | MODIFIER |
| 1 | 108358764 | AX-170042320 | 0 | T | C | 0.167 | 6.94E-02 | 1.97E-01 | 2.60E+00 | 2.68E+00 | 7.29E-01 | no | | intergenic_variant | MODIFIER |
| 1 | 108358983 | AX-170500385 | 0 | G | A | 0.167 | 6.94E-02 | 1.97E-01 | 2.60E+00 | 2.68E+00 | 7.29E-01 | no | | intergenic_variant | MODIFIER |
| 1 | 108369043 | AX-170001318 | 0 | T | C | 0.167 | 6.94E-02 | 1.97E-01 | 2.60E+00 | 2.68E+00 | 7.29E-01 | no | | intergenic_variant | MODIFIER |
| 1 | 108369694 | AX-170001321 | 0 | T | C | 0.167 | 6.94E-02 | 1.97E-01 | 2.60E+00 | 2.68E+00 | 7.29E-01 | no | | intergenic_variant | MODIFIER |
| 1 | 108369835 | AX-170001314 | 0 | C | G | 0.167 | 6.94E-02 | 1.97E-01 | 2.60E+00 | 2.68E+00 | 7.29E-01 | no | | intergenic_variant | MODIFIER |
| 1 | 108370035 | AX-170524482 | 0 | T | G | 0.167 | 6.94E-02 | 1.97E-01 | 2.60E+00 | 2.68E+00 | 7.29E-01 | no | | intergenic_variant | MODIFIER |
| 1 | 108378999 | AX-170381859 | 0 | T | C | 0.167 | 6.94E-02 | 1.97E-01 | 2.60E+00 | 2.68E+00 | 7.29E-01 | no | Gm5702 | non_coding_transcript_exon_variant | MODIFIER |
| 1 | 108379083 | AX-170142932 | 0 | G | T | 0.167 | 6.94E-02 | 1.97E-01 | 2.60E+00 | 2.68E+00 | 7.29E-01 | no | Gm5702 | non_coding_transcript_exon_variant | MODIFIER |
| 1 | 108379266 | AX-170312616 | 0 | G | A | 0.167 | 6.94E-02 | 1.97E-01 | 2.60E+00 | 2.68E+00 | 7.29E-01 | no | Gm5702 | non_coding_transcript_exon_variant | MODIFIER |
| 1 | 108379632 | AX-170519790 | 0 | G | A | 0.167 | 6.94E-02 | 1.97E-01 | 2.60E+00 | 2.68E+00 | 7.29E-01 | no | Gm5702 | upstream_gene_variant | MODIFIER |
| 1 | 108379780 | AX-170197646 | 0 | A | G | 0.167 | 6.94E-02 | 1.97E-01 | 2.60E+00 | 2.68E+00 | 7.29E-01 | no | Gm5702 | upstream_gene_variant | MODIFIER |
| 1 | 108579036 | AX-97794956 | 0 | C | T | 0.167 | 6.94E-02 | 1.97E-01 | 2.60E+00 | 2.68E+00 | 7.29E-01 | no | | intergenic_variant | MODIFIER |
| 1 | 108587995 | AX-170211749 | 0 | A | G | 0.167 | 6.94E-02 | 1.97E-01 | 2.60E+00 | 2.68E+00 | 7.29E-01 | no | | intergenic_variant | MODIFIER |

| | | | | | | | | | | | | | | | |
|---|---|---|---|---|---|---|---|---|---|---|---|---|---|---|---|
| 1 | 108656196 | AX-170228778 | 0 | C | T | 0.167 | 6.94E-02 | 1.97E-01 | 2.60E+00 | 2.68E+00 | 7.29E-01 | no | | intergenic_variant | MODIFIER |
| 1 | 108658141 | AX-97793681 | 0 | C | T | 0.167 | 6.94E-02 | 1.97E-01 | 2.60E+00 | 2.68E+00 | 7.29E-01 | no | | intergenic_variant | MODIFIER |
| 1 | 108828955 | AX-170303920 | 0 | A | T | 0.167 | 6.94E-02 | 1.97E-01 | 2.60E+00 | 2.68E+00 | 7.29E-01 | no | | intergenic_variant | MODIFIER |
| 1 | 109121290 | AX-97812704 | 0 | C | A | 0.167 | 6.94E-02 | 1.97E-01 | 2.60E+00 | 2.68E+00 | 7.29E-01 | no | | intergenic_variant | MODIFIER |
| 1 | 110328501 | AX-170281516 | 0 | C | T | 0.167 | 6.94E-02 | 1.97E-01 | 2.60E+00 | 2.68E+00 | 7.29E-01 | no | | intergenic_variant | MODIFIER |
| 1 | 113479268 | AX-170001422 | 0 | A | G | 0.167 | 6.94E-02 | 1.97E-01 | 2.60E+00 | 2.68E+00 | 7.29E-01 | no | | intergenic_variant | MODIFIER |
| 1 | 113479571 | AX-97801131 | 0 | T | G | 0.167 | 6.94E-02 | 1.97E-01 | 2.60E+00 | 2.68E+00 | 7.29E-01 | no | | intergenic_variant | MODIFIER |
| 1 | 113482458 | AX-170388780 | 0 | T | G | 0.167 | 6.94E-02 | 1.97E-01 | 2.60E+00 | 2.68E+00 | 7.29E-01 | no | | intergenic_variant | MODIFIER |
| 1 | 113508476 | AX-170042389 | 0 | C | T | 0.167 | 6.94E-02 | 1.97E-01 | 2.60E+00 | 2.68E+00 | 7.29E-01 | no | | intergenic_variant | MODIFIER |
| 1 | 113519328 | AX-170001420 | 0 | G | A | 0.167 | 6.94E-02 | 1.97E-01 | 2.60E+00 | 2.68E+00 | 7.29E-01 | no | | intergenic_variant | MODIFIER |
| 1 | 113529295 | AX-170042388 | 0 | T | C | 0.167 | 6.94E-02 | 1.97E-01 | 2.60E+00 | 2.68E+00 | 7.29E-01 | no | | intergenic_variant | MODIFIER |
| 1 | 113544405 | AX-170001418 | 0 | T | A | 0.167 | 6.94E-02 | 1.97E-01 | 2.60E+00 | 2.68E+00 | 7.29E-01 | no | | intergenic_variant | MODIFIER |
| 1 | 113544448 | AX-170001425 | 0 | T | A | 0.167 | 6.94E-02 | 1.97E-01 | 2.60E+00 | 2.68E+00 | 7.29E-01 | no | | intergenic_variant | MODIFIER |
| 1 | 113545453 | AX-170336029 | 0 | C | G | 0.167 | 6.94E-02 | 1.97E-01 | 2.60E+00 | 2.68E+00 | 7.29E-01 | no | | intergenic_variant | MODIFIER |
| 1 | 113545832 | AX-170480431 | 0 | G | A | 0.167 | 6.94E-02 | 1.97E-01 | 2.60E+00 | 2.68E+00 | 7.29E-01 | no | | intergenic_variant | MODIFIER |
| 1 | 113719172 | AX-170280741 | 0 | C | T | 0.167 | 6.94E-02 | 1.97E-01 | 2.60E+00 | 2.68E+00 | 7.29E-01 | no | Gm24937 | upstream_gene_variant | MODIFIER |
| 1 | 114385434 | AX-97824678 | 0 | A | T | 0.167 | 6.94E-02 | 1.97E-01 | 2.60E+00 | 2.68E+00 | 7.29E-01 | no | | intergenic_variant | MODIFIER |
| 1 | 114419047 | AX-170505575 | 0 | T | C | 0.167 | 6.94E-02 | 1.97E-01 | 2.60E+00 | 2.68E+00 | 7.29E-01 | no | | intergenic_variant | MODIFIER |
| 1 | 114656464 | AX-170001447 | 0 | T | C | 0.167 | 6.94E-02 | 1.97E-01 | 2.60E+00 | 2.68E+00 | 7.29E-01 | no | | intergenic_variant | MODIFIER |
| 1 | 114755913 | AX-170515016 | 0 | G | A | 0.167 | 6.94E-02 | 1.97E-01 | 2.60E+00 | 2.68E+00 | 7.29E-01 | no | | intergenic_variant | MODIFIER |
| 1 | 114756058 | AX-170340675 | 0 | A | G | 0.167 | 6.94E-02 | 1.97E-01 | 2.60E+00 | 2.68E+00 | 7.29E-01 | no | | intergenic_variant | MODIFIER |
| 1 | 114758075 | AX-170091646 | 0 | G | A | 0.167 | 6.94E-02 | 1.97E-01 | 2.60E+00 | 2.68E+00 | 7.29E-01 | no | | intergenic_variant | MODIFIER |
| 1 | 114758617 | AX-170355125 | 0 | C | G | 0.167 | 6.94E-02 | 1.97E-01 | 2.60E+00 | 2.68E+00 | 7.29E-01 | no | | intergenic_variant | MODIFIER |
| 1 | 114774699 | AX-170494593 | 0 | C | T | 0.167 | 6.94E-02 | 1.97E-01 | 2.60E+00 | 2.68E+00 | 7.29E-01 | no | | intergenic_variant | MODIFIER |
| 1 | 114784460 | AX-170188669 | 0 | C | T | 0.167 | 6.94E-02 | 1.97E-01 | 2.60E+00 | 2.68E+00 | 7.29E-01 | no | | intergenic_variant | MODIFIER |
| 1 | 114811847 | AX-170441284 | 0 | T | C | 0.167 | 6.94E-02 | 1.97E-01 | 2.60E+00 | 2.68E+00 | 7.29E-01 | no | | intergenic_variant | MODIFIER |
| 1 | 114826603 | AX-170178390 | 0 | G | A | 0.167 | 6.94E-02 | 1.97E-01 | 2.60E+00 | 2.68E+00 | 7.29E-01 | no | | intergenic_variant | MODIFIER |
| 1 | 114882167 | AX-97786175 | 0 | T | G | 0.167 | 6.94E-02 | 1.97E-01 | 2.60E+00 | 2.68E+00 | 7.29E-01 | no | | intergenic_variant | MODIFIER |
| 1 | 114905549 | AX-170477020 | 0 | A | G | 0.167 | 6.94E-02 | 1.97E-01 | 2.60E+00 | 2.68E+00 | 7.29E-01 | no | | intergenic_variant | MODIFIER |
| 1 | 114909459 | AX-97853775 | 0 | T | C | 0.167 | 6.94E-02 | 1.97E-01 | 2.60E+00 | 2.68E+00 | 7.29E-01 | no | | intergenic_variant | MODIFIER |
| 1 | 114921664 | AX-170153711 | 0 | G | A | 0.167 | 6.94E-02 | 1.97E-01 | 2.60E+00 | 2.68E+00 | 7.29E-01 | no | | intergenic_variant | MODIFIER |
| 1 | 114967428 | AX-170120430 | 0 | C | T | 0.167 | 6.94E-02 | 1.97E-01 | 2.60E+00 | 2.68E+00 | 7.29E-01 | no | | intergenic_variant | MODIFIER |
| 1 | 114980364 | AX-170325089 | 0 | A | C | 0.167 | 6.94E-02 | 1.97E-01 | 2.60E+00 | 2.68E+00 | 7.29E-01 | no | | intergenic_variant | MODIFIER |
| 1 | 115001262 | AX-170001463 | 0 | C | T | 0.167 | 6.94E-02 | 1.97E-01 | 2.60E+00 | 2.68E+00 | 7.29E-01 | no | | intergenic_variant | MODIFIER |
| 1 | 115052265 | AX-97826861 | 0 | G | A | 0.167 | 6.94E-02 | 1.97E-01 | 2.60E+00 | 2.68E+00 | 7.29E-01 | no | | intergenic_variant | MODIFIER |
| 1 | 115075751 | AX-97813015 | 0 | G | A | 0.167 | 6.94E-02 | 1.97E-01 | 2.60E+00 | 2.68E+00 | 7.29E-01 | no | | intergenic_variant | MODIFIER |
| 1 | 115136598 | AX-170001456 | 0 | A | T | 0.167 | 6.94E-02 | 1.97E-01 | 2.60E+00 | 2.68E+00 | 7.29E-01 | no | | intergenic_variant | MODIFIER |
| 1 | 115190666 | AX-170118585 | 0 | C | T | 0.167 | 6.94E-02 | 1.97E-01 | 2.60E+00 | 2.68E+00 | 7.29E-01 | no | | intergenic_variant | MODIFIER |
| 1 | 115230898 | AX-170001458 | 0 | C | T | 0.167 | 6.94E-02 | 1.97E-01 | 2.60E+00 | 2.68E+00 | 7.29E-01 | no | | intergenic_variant | MODIFIER |
| 1 | 115253080 | AX-97835642 | 0 | G | A | 0.167 | 6.94E-02 | 1.97E-01 | 2.60E+00 | 2.68E+00 | 7.29E-01 | no | | intergenic_variant | MODIFIER |
| 1 | 115268302 | AX-170042409 | 0 | A | C | 0.167 | 6.94E-02 | 1.97E-01 | 2.60E+00 | 2.68E+00 | 7.29E-01 | no | | intergenic_variant | MODIFIER |
| 1 | 115271376 | AX-170042408 | 0 | T | G | 0.167 | 6.94E-02 | 1.97E-01 | 2.60E+00 | 2.68E+00 | 7.29E-01 | no | | intergenic_variant | MODIFIER |
| 1 | 115272956 | AX-170001452 | 0 | G | C | 0.167 | 6.94E-02 | 1.97E-01 | 2.60E+00 | 2.68E+00 | 7.29E-01 | no | | intergenic_variant | MODIFIER |
| 1 | 115275769 | AX-170001455 | 0 | A | G | 0.167 | 6.94E-02 | 1.97E-01 | 2.60E+00 | 2.68E+00 | 7.29E-01 | no | | intergenic_variant | MODIFIER |
| 1 | 115276584 | AX-170001460 | 0 | G | C | 0.167 | 6.94E-02 | 1.97E-01 | 2.60E+00 | 2.68E+00 | 7.29E-01 | no | | intergenic_variant | MODIFIER |
| 1 | 115285524 | AX-97811958 | 0 | A | G | 0.167 | 6.94E-02 | 1.97E-01 | 2.60E+00 | 2.68E+00 | 7.29E-01 | no | | intergenic_variant | MODIFIER |
| 1 | 115285695 | AX-170257018 | 0 | A | C | 0.167 | 6.94E-02 | 1.97E-01 | 2.60E+00 | 2.68E+00 | 7.29E-01 | no | | intergenic_variant | MODIFIER |
| 1 | 115589173 | AX-97848972 | 0 | A | C | 0.167 | 6.94E-02 | 1.97E-01 | 2.60E+00 | 2.68E+00 | 7.29E-01 | no | | intergenic_variant | MODIFIER |
| 1 | 115659440 | AX-97813018 | 0 | A | T | 0.167 | 6.94E-02 | 1.97E-01 | 2.60E+00 | 2.68E+00 | 7.29E-01 | no | | intergenic_variant | MODIFIER |
| 1 | 115665798 | AX-170427647 | 0 | C | G | 0.167 | 6.94E-02 | 1.97E-01 | 2.60E+00 | 2.68E+00 | 7.29E-01 | no | Gm17916 | downstream_gene_variant | MODIFIER |
| 1 | 115689473 | AX-170001472 | 0 | T | C | 0.167 | 6.94E-02 | 1.97E-01 | 2.60E+00 | 2.68E+00 | 7.29E-01 | no | Cntnap5a | intron_variant | MODIFIER |
| 1 | 115692155 | AX-170495865 | 0 | C | T | 0.167 | 6.94E-02 | 1.97E-01 | 2.60E+00 | 2.68E+00 | 7.29E-01 | no | Cntnap5a | intron_variant | MODIFIER |
| 1 | 115693280 | AX-170362724 | 0 | T | C | 0.167 | 6.94E-02 | 1.97E-01 | 2.60E+00 | 2.68E+00 | 7.29E-01 | no | Cntnap5a | intron_variant | MODIFIER |
| 1 | 115707249 | AX-170410035 | 0 | A | C | 0.167 | 6.94E-02 | 1.97E-01 | 2.60E+00 | 2.68E+00 | 7.29E-01 | no | Cntnap5a | intron_variant | MODIFIER |

| | | | | | | | | | | | | | | | |
|---|---|---|---|---|---|---|---|---|---|---|---|---|---|---|---|
| 1 | 115742139 | AX-170001476 | 0 | C | T | 0.167 | 6.94E-02 | 1.97E-01 | 2.60E+00 | 2.68E+00 | 7.29E-01 | no | Cntnap5a | intron_variant | MODIFIER |
| 1 | 115743268 | AX-170001474 | 0 | G | A | 0.167 | 6.94E-02 | 1.97E-01 | 2.60E+00 | 2.68E+00 | 7.29E-01 | no | Cntnap5a | intron_variant | MODIFIER |
| 1 | 115776232 | AX-170001470 | 0 | C | A | 0.167 | 6.94E-02 | 1.97E-01 | 2.60E+00 | 2.68E+00 | 7.29E-01 | no | Cntnap5a | intron_variant | MODIFIER |
| 1 | 115927019 | AX-170344470 | 0 | G | C | 0.167 | 6.94E-02 | 1.97E-01 | 2.60E+00 | 2.68E+00 | 7.29E-01 | no | Cntnap5a | intron_variant | MODIFIER |
| 1 | 115951738 | AX-170325274 | 0 | G | T | 0.167 | 6.94E-02 | 1.97E-01 | 2.60E+00 | 2.68E+00 | 7.29E-01 | no | Cntnap5a | intron_variant | MODIFIER |
| 1 | 115952059 | AX-170343155 | 0 | T | G | 0.167 | 6.94E-02 | 1.97E-01 | 2.60E+00 | 2.68E+00 | 7.29E-01 | no | Cntnap5a | intron_variant | MODIFIER |
| 1 | 115975643 | AX-97786777 | 0 | G | A | 0.167 | 6.94E-02 | 1.97E-01 | 2.60E+00 | 2.68E+00 | 7.29E-01 | no | Cntnap5a | intron_variant | MODIFIER |
| 1 | 115996849 | AX-170517751 | 0 | T | C | 0.167 | 6.94E-02 | 1.97E-01 | 2.60E+00 | 2.68E+00 | 7.29E-01 | no | Cntnap5a | intron_variant | MODIFIER |
| 1 | 116034230 | AX-97813017 | 0 | C | T | 0.167 | 6.94E-02 | 1.97E-01 | 2.60E+00 | 2.68E+00 | 7.29E-01 | no | Cntnap5a | intron_variant | MODIFIER |
| 1 | 116035434 | AX-170245164 | 0 | A | C | 0.167 | 6.94E-02 | 1.97E-01 | 2.60E+00 | 2.68E+00 | 7.29E-01 | no | Cntnap5a | intron_variant | MODIFIER |
| 1 | 116057033 | AX-170152505 | 0 | A | C | 0.167 | 6.94E-02 | 1.97E-01 | 2.60E+00 | 2.68E+00 | 7.29E-01 | no | Cntnap5a | intron_variant | MODIFIER |
| 1 | 116130637 | AX-170001483 | 0 | A | G | 0.167 | 6.94E-02 | 1.97E-01 | 2.60E+00 | 2.68E+00 | 7.29E-01 | no | Cntnap5a | intron_variant | MODIFIER |
| 1 | 116131165 | AX-97814220 | 0 | A | G | 0.167 | 6.94E-02 | 1.97E-01 | 2.60E+00 | 2.68E+00 | 7.29E-01 | no | Cntnap5a | intron_variant | MODIFIER |
| 1 | 116232807 | AX-170001479 | 0 | G | C | 0.167 | 6.94E-02 | 1.97E-01 | 2.60E+00 | 2.68E+00 | 7.29E-01 | no | Cntnap5a | intron_variant | MODIFIER |
| 1 | 116312380 | AX-170236988 | 0 | G | A | 0.167 | 6.94E-02 | 1.97E-01 | 2.60E+00 | 2.68E+00 | 7.29E-01 | no | Cntnap5a | intron_variant | MODIFIER |
| 1 | 116312466 | AX-170261884 | 0 | C | T | 0.167 | 6.94E-02 | 1.97E-01 | 2.60E+00 | 2.68E+00 | 7.29E-01 | no | Cntnap5a | intron_variant | MODIFIER |
| 1 | 116352601 | AX-170365585 | 0 | G | T | 0.167 | 6.94E-02 | 1.97E-01 | 2.60E+00 | 2.68E+00 | 7.29E-01 | no | Cntnap5a | intron_variant | MODIFIER |
| 1 | 116366810 | AX-170313780 | 0 | T | G | 0.167 | 6.94E-02 | 1.97E-01 | 2.60E+00 | 2.68E+00 | 7.29E-01 | no | Cntnap5a | intron_variant | MODIFIER |
| 1 | 116380970 | AX-97789394 | 0 | A | C | 0.167 | 6.94E-02 | 1.97E-01 | 2.60E+00 | 2.68E+00 | 7.29E-01 | no | Cntnap5a | intron_variant | MODIFIER |
| 1 | 116381759 | AX-170153166 | 0 | A | G | 0.167 | 6.94E-02 | 1.97E-01 | 2.60E+00 | 2.68E+00 | 7.29E-01 | no | Cntnap5a | intron_variant | MODIFIER |
| 1 | 116403820 | AX-170220744 | 0 | G | A | 0.167 | 6.94E-02 | 1.97E-01 | 2.60E+00 | 2.68E+00 | 7.29E-01 | no | Cntnap5a | intron_variant | MODIFIER |
| 1 | 116438886 | AX-170471091 | 0 | A | C | 0.167 | 6.94E-02 | 1.97E-01 | 2.60E+00 | 2.68E+00 | 7.29E-01 | no | Cntnap5a | intron_variant | MODIFIER |
| 1 | 116446190 | AX-170393326 | 0 | G | T | 0.167 | 6.94E-02 | 1.97E-01 | 2.60E+00 | 2.68E+00 | 7.29E-01 | no | Cntnap5a | intron_variant | MODIFIER |
| 1 | 116468252 | AX-170042427 | 0 | G | T | 0.167 | 6.94E-02 | 1.97E-01 | 2.60E+00 | 2.68E+00 | 7.29E-01 | no | Cntnap5a | intron_variant | MODIFIER |
| 1 | 116504378 | AX-97792068 | 0 | A | T | 0.167 | 6.94E-02 | 1.97E-01 | 2.60E+00 | 2.68E+00 | 7.29E-01 | no | Cntnap5a | intron_variant | MODIFIER |
| 1 | 116512157 | AX-170385102 | 0 | C | A | 0.167 | 6.94E-02 | 1.97E-01 | 2.60E+00 | 2.68E+00 | 7.29E-01 | no | Cntnap5a | intron_variant | MODIFIER |
| 1 | 116673034 | AX-170118031 | 0 | T | C | 0.167 | 6.94E-02 | 1.97E-01 | 2.60E+00 | 2.68E+00 | 7.29E-01 | no | | intergenic_variant | MODIFIER |
| 1 | 116673557 | AX-170134848 | 0 | C | T | 0.167 | 6.94E-02 | 1.97E-01 | 2.60E+00 | 2.68E+00 | 7.29E-01 | no | | intergenic_variant | MODIFIER |
| 1 | 116674352 | AX-170001496 | 0 | C | T | 0.167 | 6.94E-02 | 1.97E-01 | 2.60E+00 | 2.68E+00 | 7.29E-01 | no | | intergenic_variant | MODIFIER |
| 1 | 116715728 | AX-170001493 | 0 | G | A | 0.167 | 6.94E-02 | 1.97E-01 | 2.60E+00 | 2.68E+00 | 7.29E-01 | no | | intergenic_variant | MODIFIER |
| 1 | 116719644 | AX-170141934 | 0 | T | C | 0.167 | 6.94E-02 | 1.97E-01 | 2.60E+00 | 2.68E+00 | 7.29E-01 | no | | intergenic_variant | MODIFIER |
| 1 | 116719974 | AX-170001485 | 0 | T | C | 0.167 | 6.94E-02 | 1.97E-01 | 2.60E+00 | 2.68E+00 | 7.29E-01 | no | | intergenic_variant | MODIFIER |
| 1 | 116787987 | AX-170001494 | 0 | C | T | 0.167 | 6.94E-02 | 1.97E-01 | 2.60E+00 | 2.68E+00 | 7.29E-01 | no | | intergenic_variant | MODIFIER |
| 1 | 116844203 | AX-97781147 | 0 | A | G | 0.167 | 6.94E-02 | 1.97E-01 | 2.60E+00 | 2.68E+00 | 7.29E-01 | no | | intergenic_variant | MODIFIER |
| 1 | 116853451 | AX-170393461 | 0 | G | T | 0.167 | 6.94E-02 | 1.97E-01 | 2.60E+00 | 2.68E+00 | 7.29E-01 | no | | intergenic_variant | MODIFIER |
| 1 | 116864495 | AX-170509731 | 0 | G | C | 0.167 | 6.94E-02 | 1.97E-01 | 2.60E+00 | 2.68E+00 | 7.29E-01 | no | | intergenic_variant | MODIFIER |
| 1 | 116865281 | AX-170232856 | 0 | T | C | 0.167 | 6.94E-02 | 1.97E-01 | 2.60E+00 | 2.68E+00 | 7.29E-01 | no | | intergenic_variant | MODIFIER |
| 1 | 116865348 | AX-170266297 | 0 | T | C | 0.167 | 6.94E-02 | 1.97E-01 | 2.60E+00 | 2.68E+00 | 7.29E-01 | no | | intergenic_variant | MODIFIER |
| 1 | 116871480 | AX-170436234 | 0 | C | T | 0.167 | 6.94E-02 | 1.97E-01 | 2.60E+00 | 2.68E+00 | 7.29E-01 | no | | intergenic_variant | MODIFIER |
| 1 | 116917078 | AX-97822905 | 0 | C | T | 0.167 | 6.94E-02 | 1.97E-01 | 2.60E+00 | 2.68E+00 | 7.29E-01 | no | | intergenic_variant | MODIFIER |
| 1 | 116962958 | AX-170412868 | 0 | T | C | 0.167 | 6.94E-02 | 1.97E-01 | 2.60E+00 | 2.68E+00 | 7.29E-01 | no | | intergenic_variant | MODIFIER |
| 1 | 116964579 | AX-97812790 | 0 | C | T | 0.167 | 6.94E-02 | 1.97E-01 | 2.60E+00 | 2.68E+00 | 7.29E-01 | no | | intergenic_variant | MODIFIER |
| 1 | 116975781 | AX-170042432 | 0 | G | A | 0.167 | 6.94E-02 | 1.97E-01 | 2.60E+00 | 2.68E+00 | 7.29E-01 | no | | intergenic_variant | MODIFIER |
| 1 | 116975860 | AX-97825940 | 0 | G | A | 0.167 | 6.94E-02 | 1.97E-01 | 2.60E+00 | 2.68E+00 | 7.29E-01 | no | | intergenic_variant | MODIFIER |
| 1 | 116985245 | AX-170111479 | 0 | A | C | 0.167 | 6.94E-02 | 1.97E-01 | 2.60E+00 | 2.68E+00 | 7.29E-01 | no | | intergenic_variant | MODIFIER |
| 1 | 117000518 | AX-170177622 | 0 | A | G | 0.167 | 6.94E-02 | 1.97E-01 | 2.60E+00 | 2.68E+00 | 7.29E-01 | no | | intergenic_variant | MODIFIER |
| 1 | 117000606 | AX-97857012 | 0 | A | G | 0.167 | 6.94E-02 | 1.97E-01 | 2.60E+00 | 2.68E+00 | 7.29E-01 | no | | intergenic_variant | MODIFIER |
| 1 | 117001976 | AX-170042434 | 0 | C | G | 0.167 | 6.94E-02 | 1.97E-01 | 2.60E+00 | 2.68E+00 | 7.29E-01 | no | | intergenic_variant | MODIFIER |
| 1 | 117750868 | AX-170149136 | 0 | A | G | 0.167 | 6.94E-02 | 1.97E-01 | 2.60E+00 | 2.68E+00 | 7.29E-01 | no | Gm28359 | intron_variant,non_coding_transcript_variant | MODIFIER |
| 1 | 117773022 | AX-170168640 | 0 | C | T | 0.167 | 6.94E-02 | 1.97E-01 | 2.60E+00 | 2.68E+00 | 7.29E-01 | no | Gm28359 | downstream_gene_variant | MODIFIER |
| 1 | 117900925 | AX-170376935 | 0 | G | A | 0.167 | 6.94E-02 | 1.97E-01 | 2.60E+00 | 2.68E+00 | 7.29E-01 | no | Gm7195 | downstream_gene_variant | MODIFIER |
| 1 | 118449012 | AX-170102748 | 0 | T | C | 0.167 | 6.94E-02 | 1.97E-01 | 2.60E+00 | 2.68E+00 | 7.29E-01 | no | Clasp1 | intron_variant | MODIFIER |
| 1 | 118471313 | AX-170536085 | 0 | G | A | 0.167 | 6.94E-02 | 1.97E-01 | 2.60E+00 | 2.68E+00 | 7.29E-01 | no | Clasp1 | intron_variant | MODIFIER |
| 1 | 118510943 | AX-168400294 | 0 | T | C | 0.167 | 6.94E-02 | 1.97E-01 | 2.60E+00 | 2.68E+00 | 7.29E-01 | no | Clasp1 | intron_variant | MODIFIER |

| | | | | | | | | | | | | | | | |
|---|---|---|---|---|---|---|---|---|---|---|---|---|---|---|---|
| 1 | 118511775 | AX-170421490 | 0 | A | G | 0.167 | 6.94E-02 | 1.97E-01 | 2.60E+00 | 2.68E+00 | 7.29E-01 | no | Clasp1 | intron_variant | MODIFIER |
| 1 | 118626277 | AX-170301456 | 0 | C | T | 0.167 | 6.94E-02 | 1.97E-01 | 2.60E+00 | 2.68E+00 | 7.29E-01 | no | Tfcp2l1 | upstream_gene_variant | MODIFIER |
| 1 | 118628333 | AX-170268303 | 0 | C | T | 0.167 | 6.94E-02 | 1.97E-01 | 2.60E+00 | 2.68E+00 | 7.29E-01 | no | Tfcp2l1 | intron_variant | MODIFIER |
| 1 | 118702671 | AX-170277804 | 0 | G | A | 0.167 | 6.94E-02 | 1.97E-01 | 2.60E+00 | 2.68E+00 | 7.29E-01 | no | | intergenic_variant | MODIFIER |
| 1 | 118713129 | AX-170001528 | 0 | C | T | 0.167 | 6.94E-02 | 1.97E-01 | 2.60E+00 | 2.68E+00 | 7.29E-01 | no | | intergenic_variant | MODIFIER |
| 1 | 118716234 | AX-170504260 | 0 | C | T | 0.167 | 6.94E-02 | 1.97E-01 | 2.60E+00 | 2.68E+00 | 7.29E-01 | no | | intergenic_variant | MODIFIER |
| 1 | 118723442 | AX-170461439 | 0 | G | C | 0.167 | 6.94E-02 | 1.97E-01 | 2.60E+00 | 2.68E+00 | 7.29E-01 | no | | intergenic_variant | MODIFIER |
| 1 | 118750497 | AX-170001522 | 0 | A | T | 0.167 | 6.94E-02 | 1.97E-01 | 2.60E+00 | 2.68E+00 | 7.29E-01 | no | | intergenic_variant | MODIFIER |
| 1 | 118751216 | AX-170335049 | 0 | T | C | 0.167 | 6.94E-02 | 1.97E-01 | 2.60E+00 | 2.68E+00 | 7.29E-01 | no | | intergenic_variant | MODIFIER |
| 1 | 118754464 | AX-97838761 | 0 | A | C | 0.167 | 6.94E-02 | 1.97E-01 | 2.60E+00 | 2.68E+00 | 7.29E-01 | no | | intergenic_variant | MODIFIER |
| 1 | 118780699 | AX-170263656 | 0 | T | C | 0.167 | 6.94E-02 | 1.97E-01 | 2.60E+00 | 2.68E+00 | 7.29E-01 | no | | intergenic_variant | MODIFIER |
| 1 | 118797064 | AX-170378579 | 0 | G | T | 0.167 | 6.94E-02 | 1.97E-01 | 2.60E+00 | 2.68E+00 | 7.29E-01 | no | | intergenic_variant | MODIFIER |
| 1 | 118797114 | AX-170265639 | 0 | A | C | 0.167 | 6.94E-02 | 1.97E-01 | 2.60E+00 | 2.68E+00 | 7.29E-01 | no | | intergenic_variant | MODIFIER |
| 1 | 118813287 | AX-170289258 | 0 | G | A | 0.167 | 6.94E-02 | 1.97E-01 | 2.60E+00 | 2.68E+00 | 7.29E-01 | no | Gm28467 | downstream_gene_variant | MODIFIER |
| 1 | 118815524 | AX-170478996 | 0 | T | G | 0.167 | 6.94E-02 | 1.97E-01 | 2.60E+00 | 2.68E+00 | 7.29E-01 | no | | intergenic_variant | MODIFIER |
| 1 | 118850330 | AX-170042455 | 0 | T | C | 0.167 | 6.94E-02 | 1.97E-01 | 2.60E+00 | 2.68E+00 | 7.29E-01 | no | Gli2 | intron_variant | MODIFIER |
| 1 | 118852804 | AX-170042453 | 0 | G | A | 0.167 | 6.94E-02 | 1.97E-01 | 2.60E+00 | 2.68E+00 | 7.29E-01 | no | Gli2 | intron_variant | MODIFIER |
| 1 | 118863577 | AX-170173235 | 0 | G | A | 0.167 | 6.94E-02 | 1.97E-01 | 2.60E+00 | 2.68E+00 | 7.29E-01 | no | Gli2 | intron_variant | MODIFIER |
| 1 | 118872655 | AX-170473972 | 0 | G | A | 0.167 | 6.94E-02 | 1.97E-01 | 2.60E+00 | 2.68E+00 | 7.29E-01 | no | Gli2 | intron_variant | MODIFIER |
| 1 | 119016101 | AX-170042457 | 0 | T | C | 0.167 | 6.94E-02 | 1.97E-01 | 2.60E+00 | 2.68E+00 | 7.29E-01 | no | Gli2 | intron_variant | MODIFIER |
| 1 | 119031229 | AX-170120914 | 0 | C | A | 0.167 | 6.94E-02 | 1.97E-01 | 2.60E+00 | 2.68E+00 | 7.29E-01 | no | Gli2 | intron_variant | MODIFIER |
| 1 | 119054852 | AX-170288938 | 0 | C | A | 0.167 | 6.94E-02 | 1.97E-01 | 2.60E+00 | 2.68E+00 | 7.29E-01 | no | Gli2 | upstream_gene_variant | MODIFIER |
| 1 | 119153798 | AX-170173133 | 0 | A | G | 0.167 | 6.94E-02 | 1.97E-01 | 2.60E+00 | 2.68E+00 | 7.29E-01 | no | Gm8321 | upstream_gene_variant | MODIFIER |
| 6 | 148069784 | AX-170225300 | 0 | A | G | 0.044 | -6.29E-02 | 1.80E-01 | 2.60E+00 | 2.71E+00 | 7.29E-01 | no | Far2 | intron_variant | MODIFIER |
| 7 | 118795109 | AX-170474168 | 0 | T | C | 0.395 | 3.17E-02 | 9.18E-02 | 2.66E+00 | 2.71E+00 | 7.29E-01 | no | Vps35l | intron_variant | MODIFIER |
| 5 | 87790767 | AX-170039061 | 1 | A | G | 0.429 | 3.88E-02 | 1.14E-01 | 2.73E+00 | 2.77E+00 | 7.29E-01 | no | Csn1s2a | downstream_gene_variant | MODIFIER |
| 3 | 30897988 | AX-170230258 | 0 | C | G | 0.167 | -3.26E-02 | 9.59E-02 | 2.68E+00 | 2.76E+00 | 7.29E-01 | no | Phc3 | downstream_gene_variant | MODIFIER |
| 1 | 108900588 | AX-170001332 | 1 | G | A | 0.17 | 6.91E-02 | 1.95E-01 | 2.59E+00 | 2.68E+00 | 7.29E-01 | no | | intergenic_variant | MODIFIER |
| 1 | 115793989 | AX-170001467 | 1 | T | A | 0.17 | 6.91E-02 | 1.95E-01 | 2.59E+00 | 2.68E+00 | 7.29E-01 | no | Cntnap5a | intron_variant | MODIFIER |
| 1 | 116690999 | AX-170086587 | 1 | G | T | 0.17 | 6.91E-02 | 1.95E-01 | 2.59E+00 | 2.68E+00 | 7.29E-01 | no | | intergenic_variant | MODIFIER |
| 8 | 43356948 | AX-170436500 | 1 | A | G | 0.152 | 5.02E-02 | 1.48E-01 | 2.75E+00 | 2.80E+00 | 7.29E-01 | no | | intergenic_variant | MODIFIER |
| 12 | 103668052 | AX-170162419 | 0 | C | T | 0.132 | -6.60E-02 | 1.95E-01 | 2.70E+00 | 2.77E+00 | 7.29E-01 | no | Serpina16 | downstream_gene_variant | MODIFIER |
| 12 | 103674783 | AX-170318867 | 0 | T | C | 0.132 | -6.60E-02 | 1.95E-01 | 2.70E+00 | 2.77E+00 | 7.29E-01 | no | Serpina16 | intron_variant | MODIFIER |
| 12 | 103675193 | AX-170306499 | 0 | T | C | 0.132 | -6.60E-02 | 1.95E-01 | 2.70E+00 | 2.77E+00 | 7.29E-01 | no | Serpina16 | missense_variant | MODERATE |
| 12 | 103675653 | AX-170344482 | 0 | G | A | 0.132 | -6.60E-02 | 1.95E-01 | 2.70E+00 | 2.77E+00 | 7.29E-01 | no | Serpina16 | upstream_gene_variant | MODIFIER |
| 12 | 103681854 | AX-170098023 | 0 | A | C | 0.132 | -6.60E-02 | 1.95E-01 | 2.70E+00 | 2.77E+00 | 7.29E-01 | no | | intergenic_variant | MODIFIER |
| 12 | 103685173 | AX-170157668 | 0 | C | T | 0.132 | -6.60E-02 | 1.95E-01 | 2.70E+00 | 2.77E+00 | 7.29E-01 | no | Serpina1f | downstream_gene_variant | MODIFIER |
| 12 | 103692440 | AX-170087326 | 0 | C | A | 0.132 | -6.60E-02 | 1.95E-01 | 2.70E+00 | 2.77E+00 | 7.29E-01 | no | Serpina1f | intron_variant | MODIFIER |
| 12 | 103693428 | AX-170265487 | 0 | C | T | 0.132 | -6.60E-02 | 1.95E-01 | 2.70E+00 | 2.77E+00 | 7.29E-01 | no | Serpina1f | synonymous_variant | LOW |
| 12 | 103696203 | AX-170373437 | 0 | T | C | 0.132 | -6.60E-02 | 1.95E-01 | 2.70E+00 | 2.77E+00 | 7.29E-01 | no | Serpina1f | upstream_gene_variant | MODIFIER |
| 12 | 103705600 | AX-170343333 | 0 | G | T | 0.132 | -6.60E-02 | 1.95E-01 | 2.70E+00 | 2.77E+00 | 7.29E-01 | no | Gm5656 | upstream_gene_variant | MODIFIER |
| 12 | 103710055 | AX-170116471 | 0 | A | G | 0.132 | -6.60E-02 | 1.95E-01 | 2.70E+00 | 2.77E+00 | 7.29E-01 | no | Gm17142 | downstream_gene_variant | MODIFIER |
| 12 | 103738324 | AX-170007435 | 0 | C | G | 0.132 | -6.60E-02 | 1.95E-01 | 2.70E+00 | 2.77E+00 | 7.29E-01 | no | Serpina1b | upstream_gene_variant | MODIFIER |
| 12 | 103738432 | AX-170007437 | 0 | T | C | 0.132 | -6.60E-02 | 1.95E-01 | 2.70E+00 | 2.77E+00 | 7.29E-01 | no | Serpina1b | upstream_gene_variant | MODIFIER |
| 12 | 103940737 | AX-170263519 | 0 | C | T | 0.132 | -6.60E-02 | 1.95E-01 | 2.70E+00 | 2.77E+00 | 7.29E-01 | no | | intergenic_variant | MODIFIER |
| 12 | 104015042 | AX-170438634 | 0 | C | G | 0.132 | -6.60E-02 | 1.95E-01 | 2.70E+00 | 2.77E+00 | 7.29E-01 | no | Serpina9 | upstream_gene_variant | MODIFIER |
| 12 | 104037462 | AX-170314955 | 0 | C | T | 0.132 | -6.60E-02 | 1.95E-01 | 2.70E+00 | 2.77E+00 | 7.29E-01 | no | Serpina12 | intron_variant | MODIFIER |
| 12 | 104063073 | AX-97826731 | 0 | A | G | 0.132 | -6.60E-02 | 1.95E-01 | 2.70E+00 | 2.77E+00 | 7.29E-01 | no | | intergenic_variant | MODIFIER |
| 12 | 104110937 | AX-170442993 | 0 | C | G | 0.132 | -6.60E-02 | 1.95E-01 | 2.70E+00 | 2.77E+00 | 7.29E-01 | no | Serpina5 | downstream_gene_variant | MODIFIER |
| 12 | 104112011 | AX-170136329 | 0 | C | G | 0.132 | -6.60E-02 | 1.95E-01 | 2.70E+00 | 2.77E+00 | 7.29E-01 | no | Serpina3a | upstream_gene_variant | MODIFIER |
| 12 | 104114686 | AX-170437174 | 0 | T | G | 0.132 | -6.60E-02 | 1.95E-01 | 2.70E+00 | 2.77E+00 | 7.29E-01 | no | Serpina3a | upstream_gene_variant | MODIFIER |
| 12 | 104130560 | AX-170441865 | 0 | A | G | 0.132 | -6.60E-02 | 1.95E-01 | 2.70E+00 | 2.77E+00 | 7.29E-01 | no | Serpina3b | synonymous_variant | LOW |
| 12 | 104139910 | AX-170518894 | 0 | A | G | 0.132 | -6.60E-02 | 1.95E-01 | 2.70E+00 | 2.77E+00 | 7.29E-01 | no | Serpina3b | downstream_gene_variant | MODIFIER |
| 12 | 104141723 | AX-170335272 | 0 | A | G | 0.132 | -6.60E-02 | 1.95E-01 | 2.70E+00 | 2.77E+00 | 7.29E-01 | no | Serpina3c | downstream_gene_variant | MODIFIER |

| | | | | | | | | | | | | | | | |
|---|---|---|---|---|---|---|---|---|---|---|---|---|---|---|---|
| 12 | 104154183 | AX-170146950 | 0 | T | C | 0.132 | -6.60E-02 | 1.95E-01 | 2.70E+00 | 2.77E+00 | 7.29E-01 | no | Serpina3c | upstream_gene_variant | MODIFIER |
| 12 | 104174830 | AX-170276299 | 0 | T | G | 0.132 | -6.60E-02 | 1.95E-01 | 2.70E+00 | 2.77E+00 | 7.29E-01 | no | Serpina3d-ps | intron_variant,non_coding_transcript_variant | MODIFIER |
| 12 | 104197342 | AX-170440494 | 0 | T | C | 0.132 | -6.60E-02 | 1.95E-01 | 2.70E+00 | 2.77E+00 | 7.29E-01 | no | Serpina3e-ps | upstream_gene_variant | MODIFIER |
| 12 | 104197433 | AX-170332191 | 0 | T | C | 0.132 | -6.60E-02 | 1.95E-01 | 2.70E+00 | 2.77E+00 | 7.29E-01 | no | Serpina3e-ps | upstream_gene_variant | MODIFIER |
| 12 | 104209134 | AX-170050036 | 0 | T | C | 0.132 | -6.60E-02 | 1.95E-01 | 2.70E+00 | 2.77E+00 | 7.29E-01 | no | | intergenic_variant | MODIFIER |
| 12 | 104209899 | AX-170050034 | 0 | C | T | 0.132 | -6.60E-02 | 1.95E-01 | 2.70E+00 | 2.77E+00 | 7.29E-01 | no | Serpina3f | upstream_gene_variant | MODIFIER |
| 12 | 104211235 | AX-170279375 | 0 | G | A | 0.132 | -6.60E-02 | 1.95E-01 | 2.70E+00 | 2.77E+00 | 7.29E-01 | no | Serpina3f | upstream_gene_variant | MODIFIER |
| 12 | 104211312 | AX-170170844 | 0 | G | A | 0.132 | -6.60E-02 | 1.95E-01 | 2.70E+00 | 2.77E+00 | 7.29E-01 | no | Serpina3f | upstream_gene_variant | MODIFIER |
| 12 | 104211430 | AX-170299961 | 0 | C | G | 0.132 | -6.60E-02 | 1.95E-01 | 2.70E+00 | 2.77E+00 | 7.29E-01 | no | Serpina3f | upstream_gene_variant | MODIFIER |
| 12 | 104216685 | AX-170303297 | 0 | G | T | 0.132 | -6.60E-02 | 1.95E-01 | 2.70E+00 | 2.77E+00 | 7.29E-01 | no | Serpina3f | intron_variant | MODIFIER |
| 12 | 104218550 | AX-170269751 | 0 | G | T | 0.132 | -6.60E-02 | 1.95E-01 | 2.70E+00 | 2.77E+00 | 7.29E-01 | no | Serpina3f | intron_variant | MODIFIER |
| 12 | 104219426 | AX-170321423 | 0 | G | A | 0.132 | -6.60E-02 | 1.95E-01 | 2.70E+00 | 2.77E+00 | 7.29E-01 | no | Serpina3f | missense_variant | MODERATE |
| 12 | 104222928 | AX-170227032 | 0 | C | T | 0.132 | -6.60E-02 | 1.95E-01 | 2.70E+00 | 2.77E+00 | 7.29E-01 | no | Serpina3f | downstream_gene_variant | MODIFIER |
| 12 | 104224193 | AX-170195643 | 0 | A | G | 0.132 | -6.60E-02 | 1.95E-01 | 2.70E+00 | 2.77E+00 | 7.29E-01 | no | Serpina3f | downstream_gene_variant | MODIFIER |
| 12 | 104224362 | AX-170521410 | 0 | G | A | 0.132 | -6.60E-02 | 1.95E-01 | 2.70E+00 | 2.77E+00 | 7.29E-01 | no | Serpina3f | downstream_gene_variant | MODIFIER |
| 12 | 104244778 | AX-170199590 | 0 | T | C | 0.132 | -6.60E-02 | 1.95E-01 | 2.70E+00 | 2.77E+00 | 7.29E-01 | no | Serpina3g | downstream_gene_variant | MODIFIER |
| 12 | 104260095 | AX-170169762 | 0 | G | A | 0.132 | -6.60E-02 | 1.95E-01 | 2.70E+00 | 2.77E+00 | 7.29E-01 | no | Serpina3i | upstream_gene_variant | MODIFIER |
| 12 | 104294481 | AX-170495217 | 0 | G | A | 0.132 | -6.60E-02 | 1.95E-01 | 2.70E+00 | 2.77E+00 | 7.29E-01 | no | | intergenic_variant | MODIFIER |
| 12 | 104294652 | AX-170442619 | 0 | T | C | 0.132 | -6.60E-02 | 1.95E-01 | 2.70E+00 | 2.77E+00 | 7.29E-01 | no | | intergenic_variant | MODIFIER |
| 12 | 104296200 | AX-170292455 | 0 | A | G | 0.132 | -6.60E-02 | 1.95E-01 | 2.70E+00 | 2.77E+00 | 7.29E-01 | no | | intergenic_variant | MODIFIER |
| 12 | 104297556 | AX-170397566 | 0 | G | T | 0.132 | -6.60E-02 | 1.95E-01 | 2.70E+00 | 2.77E+00 | 7.29E-01 | no | | intergenic_variant | MODIFIER |
| 12 | 104297975 | AX-170490412 | 0 | C | T | 0.132 | -6.60E-02 | 1.95E-01 | 2.70E+00 | 2.77E+00 | 7.29E-01 | no | | intergenic_variant | MODIFIER |
| 12 | 104305908 | AX-170050040 | 0 | G | A | 0.132 | -6.60E-02 | 1.95E-01 | 2.70E+00 | 2.77E+00 | 7.29E-01 | no | | regulatory_region_variant | MODIFIER |
| 12 | 104314127 | AX-170050039 | 0 | T | G | 0.132 | -6.60E-02 | 1.95E-01 | 2.70E+00 | 2.77E+00 | 7.29E-01 | no | Serpina3j | upstream_gene_variant | MODIFIER |
| 12 | 104318643 | AX-170179835 | 0 | G | A | 0.132 | -6.60E-02 | 1.95E-01 | 2.70E+00 | 2.77E+00 | 7.29E-01 | no | Serpina3j | intron_variant | MODIFIER |
| 12 | 104320197 | AX-170531204 | 0 | T | C | 0.132 | -6.60E-02 | 1.95E-01 | 2.70E+00 | 2.77E+00 | 7.29E-01 | no | Serpina3j | 3_prime_UTR_variant | MODIFIER |
| 12 | 104321995 | AX-170375869 | 0 | A | C | 0.132 | -6.60E-02 | 1.95E-01 | 2.70E+00 | 2.77E+00 | 7.29E-01 | no | Serpina3j | downstream_gene_variant | MODIFIER |
| 12 | 104322260 | AX-170007439 | 0 | G | A | 0.132 | -6.60E-02 | 1.95E-01 | 2.70E+00 | 2.77E+00 | 7.29E-01 | no | Serpina3j | downstream_gene_variant | MODIFIER |
| 12 | 104323614 | AX-170007442 | 0 | T | C | 0.132 | -6.60E-02 | 1.95E-01 | 2.70E+00 | 2.77E+00 | 7.29E-01 | no | Serpina3j | downstream_gene_variant | MODIFIER |
| 12 | 104357723 | AX-170177062 | 0 | A | T | 0.132 | -6.60E-02 | 1.95E-01 | 2.70E+00 | 2.77E+00 | 7.29E-01 | no | Serpina3m | intron_variant | MODIFIER |
| 12 | 104357754 | AX-170525585 | 0 | T | G | 0.132 | -6.60E-02 | 1.95E-01 | 2.70E+00 | 2.77E+00 | 7.29E-01 | no | Serpina3m | intron_variant | MODIFIER |
| 12 | 104357780 | AX-170273955 | 0 | G | C | 0.132 | -6.60E-02 | 1.95E-01 | 2.70E+00 | 2.77E+00 | 7.29E-01 | no | Serpina3m | intron_variant | MODIFIER |
| 12 | 104357877 | AX-170300167 | 0 | G | A | 0.132 | -6.60E-02 | 1.95E-01 | 2.70E+00 | 2.77E+00 | 7.29E-01 | no | Serpina3m | intron_variant | MODIFIER |
| 12 | 104363519 | AX-170007444 | 0 | C | T | 0.132 | -6.60E-02 | 1.95E-01 | 2.70E+00 | 2.77E+00 | 7.29E-01 | no | Serpina3m | intron_variant | MODIFIER |
| 12 | 104363684 | AX-170007440 | 0 | A | G | 0.132 | -6.60E-02 | 1.95E-01 | 2.70E+00 | 2.77E+00 | 7.29E-01 | no | Serpina3m | intron_variant | MODIFIER |
| 12 | 104417583 | AX-170007443 | 0 | A | G | 0.132 | -6.60E-02 | 1.95E-01 | 2.70E+00 | 2.77E+00 | 7.29E-01 | no | Serpina3n | downstream_gene_variant | MODIFIER |
| 12 | 104429611 | AX-170165404 | 0 | T | C | 0.132 | -6.60E-02 | 1.95E-01 | 2.70E+00 | 2.77E+00 | 7.29E-01 | no | | intergenic_variant | MODIFIER |
| 12 | 104429664 | AX-170266893 | 0 | A | G | 0.132 | -6.60E-02 | 1.95E-01 | 2.70E+00 | 2.77E+00 | 7.29E-01 | no | | intergenic_variant | MODIFIER |
| 12 | 104485733 | AX-170470843 | 0 | A | G | 0.132 | -6.60E-02 | 1.95E-01 | 2.70E+00 | 2.77E+00 | 7.29E-01 | no | | intergenic_variant | MODIFIER |
| 12 | 104944424 | AX-97822284 | 0 | T | C | 0.132 | -6.60E-02 | 1.95E-01 | 2.70E+00 | 2.77E+00 | 7.29E-01 | no | Syne3 | intron_variant | MODIFIER |
| 12 | 104970671 | AX-170296760 | 0 | C | T | 0.132 | -6.60E-02 | 1.95E-01 | 2.70E+00 | 2.77E+00 | 7.29E-01 | no | Syne3 | intron_variant | MODIFIER |
| 12 | 105065151 | AX-170352009 | 0 | G | A | 0.132 | -6.60E-02 | 1.95E-01 | 2.70E+00 | 2.77E+00 | 7.29E-01 | no | | regulatory_region_variant | MODIFIER |
| 12 | 106475298 | AX-170364668 | 0 | C | T | 0.132 | -6.60E-02 | 1.95E-01 | 2.70E+00 | 2.77E+00 | 7.29E-01 | no | | regulatory_region_variant | MODIFIER |
| 12 | 106493008 | AX-170239330 | 0 | A | G | 0.132 | -6.60E-02 | 1.95E-01 | 2.70E+00 | 2.77E+00 | 7.29E-01 | no | | intergenic_variant | MODIFIER |
| 12 | 106493900 | AX-170309585 | 0 | A | G | 0.132 | -6.60E-02 | 1.95E-01 | 2.70E+00 | 2.77E+00 | 7.29E-01 | no | | intergenic_variant | MODIFIER |
| 12 | 106530917 | AX-170191649 | 0 | T | C | 0.132 | -6.60E-02 | 1.95E-01 | 2.70E+00 | 2.77E+00 | 7.29E-01 | no | | intergenic_variant | MODIFIER |
| 12 | 106577461 | AX-170509865 | 0 | A | T | 0.132 | -6.60E-02 | 1.95E-01 | 2.70E+00 | 2.77E+00 | 7.29E-01 | no | Gm17033 | downstream_gene_variant | MODIFIER |
| 12 | 106580396 | AX-170149788 | 0 | A | T | 0.132 | -6.60E-02 | 1.95E-01 | 2.70E+00 | 2.77E+00 | 7.29E-01 | no | | intergenic_variant | MODIFIER |
| 12 | 106582692 | AX-170196601 | 0 | C | G | 0.132 | -6.60E-02 | 1.95E-01 | 2.70E+00 | 2.77E+00 | 7.29E-01 | no | | intergenic_variant | MODIFIER |
| 12 | 106590343 | AX-170496661 | 0 | A | T | 0.132 | -6.60E-02 | 1.95E-01 | 2.70E+00 | 2.77E+00 | 7.29E-01 | no | | intergenic_variant | MODIFIER |
| 12 | 106593120 | AX-170092606 | 0 | C | T | 0.132 | -6.60E-02 | 1.95E-01 | 2.70E+00 | 2.77E+00 | 7.29E-01 | no | | intergenic_variant | MODIFIER |
| 12 | 106601543 | AX-170146403 | 0 | C | T | 0.132 | -6.60E-02 | 1.95E-01 | 2.70E+00 | 2.77E+00 | 7.29E-01 | no | | intergenic_variant | MODIFIER |
| 12 | 106616150 | AX-170493741 | 0 | G | A | 0.132 | -6.60E-02 | 1.95E-01 | 2.70E+00 | 2.77E+00 | 7.29E-01 | no | | intergenic_variant | MODIFIER |
| 12 | 106616300 | AX-97814129 | 0 | A | G | 0.132 | -6.60E-02 | 1.95E-01 | 2.70E+00 | 2.77E+00 | 7.29E-01 | no | | intergenic_variant | MODIFIER |

| | | | | | | | | | | | | | | | |
|---|---|---|---|---|---|---|---|---|---|---|---|---|---|---|---|
| 12 | 106617634 | AX-170299188 | 0 | C | T | 0.132 | -6.60E-02 | 1.95E-01 | 2.70E+00 | 2.77E+00 | 7.29E-01 | no | | intergenic_variant | MODIFIER |
| 12 | 106618351 | AX-170390083 | 0 | G | A | 0.132 | -6.60E-02 | 1.95E-01 | 2.70E+00 | 2.77E+00 | 7.29E-01 | no | | intergenic_variant | MODIFIER |
| 12 | 106638667 | AX-97802264 | 0 | T | C | 0.132 | -6.60E-02 | 1.95E-01 | 2.70E+00 | 2.77E+00 | 7.29E-01 | no | | intergenic_variant | MODIFIER |
| 12 | 106657877 | AX-170007469 | 0 | C | T | 0.132 | -6.60E-02 | 1.95E-01 | 2.70E+00 | 2.77E+00 | 7.29E-01 | no | | intergenic_variant | MODIFIER |
| 12 | 106661450 | AX-170446834 | 0 | G | A | 0.132 | -6.60E-02 | 1.95E-01 | 2.70E+00 | 2.77E+00 | 7.29E-01 | no | | intergenic_variant | MODIFIER |
| 12 | 106666008 | AX-97834789 | 0 | C | T | 0.132 | -6.60E-02 | 1.95E-01 | 2.70E+00 | 2.77E+00 | 7.29E-01 | no | | intergenic_variant | MODIFIER |
| 12 | 106687045 | AX-170519280 | 0 | A | C | 0.132 | -6.60E-02 | 1.95E-01 | 2.70E+00 | 2.77E+00 | 7.29E-01 | no | 4933406K04Rik | intron_variant,non_coding_transcript_variant | MODIFIER |
| 12 | 106689383 | AX-170201893 | 0 | A | C | 0.132 | -6.60E-02 | 1.95E-01 | 2.70E+00 | 2.77E+00 | 7.29E-01 | no | 4933406K04Rik | intron_variant,non_coding_transcript_variant | MODIFIER |
| 12 | 106691880 | AX-170007471 | 0 | G | A | 0.132 | -6.60E-02 | 1.95E-01 | 2.70E+00 | 2.77E+00 | 7.29E-01 | no | 4933406K04Rik | downstream_gene_variant | MODIFIER |
| 12 | 106693009 | AX-97792102 | 0 | C | T | 0.132 | -6.60E-02 | 1.95E-01 | 2.70E+00 | 2.77E+00 | 7.29E-01 | no | 4933406K04Rik | downstream_gene_variant | MODIFIER |
| 12 | 106706456 | AX-170147334 | 0 | G | A | 0.132 | -6.60E-02 | 1.95E-01 | 2.70E+00 | 2.77E+00 | 7.29E-01 | no | 4933406K04Rik | intron_variant,non_coding_transcript_variant | MODIFIER |
| 12 | 106714867 | AX-97794208 | 0 | G | A | 0.132 | -6.60E-02 | 1.95E-01 | 2.70E+00 | 2.77E+00 | 7.29E-01 | no | 4933406K04Rik | intron_variant,non_coding_transcript_variant | MODIFIER |
| 12 | 106719387 | AX-170314010 | 0 | G | A | 0.132 | -6.60E-02 | 1.95E-01 | 2.70E+00 | 2.77E+00 | 7.29E-01 | no | 4933406K04Rik | upstream_gene_variant | MODIFIER |
| 12 | 106867060 | AX-170357572 | 0 | C | T | 0.132 | -6.60E-02 | 1.95E-01 | 2.70E+00 | 2.77E+00 | 7.29E-01 | no | 1700013N06Rik | intron_variant,non_coding_transcript_variant | MODIFIER |
| 12 | 106873087 | AX-170503424 | 0 | C | A | 0.132 | -6.60E-02 | 1.95E-01 | 2.70E+00 | 2.77E+00 | 7.29E-01 | no | 1700013N06Rik | intron_variant,non_coding_transcript_variant | MODIFIER |
| 12 | 106876135 | AX-170278476 | 0 | A | G | 0.132 | -6.60E-02 | 1.95E-01 | 2.70E+00 | 2.77E+00 | 7.29E-01 | no | 1700013N06Rik | intron_variant,non_coding_transcript_variant | MODIFIER |
| 12 | 106897986 | AX-170152808 | 0 | G | A | 0.132 | -6.60E-02 | 1.95E-01 | 2.70E+00 | 2.77E+00 | 7.29E-01 | no | | intergenic_variant | MODIFIER |
| 12 | 107018055 | AX-170214790 | 0 | C | T | 0.132 | -6.60E-02 | 1.95E-01 | 2.70E+00 | 2.77E+00 | 7.29E-01 | no | | intergenic_variant | MODIFIER |
| 12 | 108490372 | AX-170277645 | 0 | C | G | 0.132 | -6.60E-02 | 1.95E-01 | 2.70E+00 | 2.77E+00 | 7.29E-01 | no | Eml1 | intron_variant | MODIFIER |
| 12 | 108532038 | AX-97826831 | 0 | T | C | 0.132 | -6.60E-02 | 1.95E-01 | 2.70E+00 | 2.77E+00 | 7.29E-01 | no | Eml1 | intron_variant | MODIFIER |
| 12 | 108567105 | AX-97856624 | 0 | T | C | 0.132 | -6.60E-02 | 1.95E-01 | 2.70E+00 | 2.77E+00 | 7.29E-01 | no | Evl | intron_variant | MODIFIER |
| 12 | 108585327 | AX-170491177 | 0 | C | A | 0.132 | -6.60E-02 | 1.95E-01 | 2.70E+00 | 2.77E+00 | 7.29E-01 | no | Evl | intron_variant | MODIFIER |
| 12 | 108585481 | AX-170361707 | 0 | A | T | 0.132 | -6.60E-02 | 1.95E-01 | 2.70E+00 | 2.77E+00 | 7.29E-01 | no | Evl | intron_variant | MODIFIER |
| 12 | 108587278 | AX-170315990 | 0 | A | G | 0.132 | -6.60E-02 | 1.95E-01 | 2.70E+00 | 2.77E+00 | 7.29E-01 | no | Evl | intron_variant | MODIFIER |
| 12 | 108606500 | AX-170243667 | 0 | G | A | 0.132 | -6.60E-02 | 1.95E-01 | 2.70E+00 | 2.77E+00 | 7.29E-01 | no | Evl | intron_variant | MODIFIER |
| 12 | 108623642 | AX-170382084 | 0 | T | C | 0.132 | -6.60E-02 | 1.95E-01 | 2.70E+00 | 2.77E+00 | 7.29E-01 | no | Evl | intron_variant | MODIFIER |
| 12 | 108641050 | AX-170105129 | 0 | A | G | 0.132 | -6.60E-02 | 1.95E-01 | 2.70E+00 | 2.77E+00 | 7.29E-01 | no | Evl | intron_variant | MODIFIER |
| 12 | 108641086 | AX-170469905 | 0 | C | T | 0.132 | -6.60E-02 | 1.95E-01 | 2.70E+00 | 2.77E+00 | 7.29E-01 | no | Evl | intron_variant | MODIFIER |
| 12 | 108641730 | AX-170358139 | 0 | A | G | 0.132 | -6.60E-02 | 1.95E-01 | 2.70E+00 | 2.77E+00 | 7.29E-01 | no | Evl | intron_variant | MODIFIER |
| 12 | 108645557 | AX-97790284 | 0 | A | G | 0.132 | -6.60E-02 | 1.95E-01 | 2.70E+00 | 2.77E+00 | 7.29E-01 | no | Evl | intron_variant | MODIFIER |
| 12 | 108651497 | AX-170346759 | 0 | C | T | 0.132 | -6.60E-02 | 1.95E-01 | 2.70E+00 | 2.77E+00 | 7.29E-01 | no | Evl | intron_variant | MODIFIER |
| 12 | 108656178 | AX-170164109 | 0 | G | C | 0.132 | -6.60E-02 | 1.95E-01 | 2.70E+00 | 2.77E+00 | 7.29E-01 | no | Evl | intron_variant | MODIFIER |
| 12 | 108660937 | AX-170322215 | 0 | T | C | 0.132 | -6.60E-02 | 1.95E-01 | 2.70E+00 | 2.77E+00 | 7.29E-01 | no | Evl | intron_variant | MODIFIER |
| 12 | 108665357 | AX-170275191 | 0 | C | T | 0.132 | -6.60E-02 | 1.95E-01 | 2.70E+00 | 2.77E+00 | 7.29E-01 | no | Evl | intron_variant | MODIFIER |
| 12 | 108682168 | AX-170466083 | 0 | G | A | 0.132 | -6.60E-02 | 1.95E-01 | 2.70E+00 | 2.77E+00 | 7.29E-01 | no | Evl | intron_variant | MODIFIER |
| 12 | 108766559 | AX-170007497 | 0 | T | C | 0.132 | -6.60E-02 | 1.95E-01 | 2.70E+00 | 2.77E+00 | 7.29E-01 | no | Gm33682 | upstream_gene_variant | MODIFIER |
| 12 | 108828391 | AX-170404421 | 0 | C | T | 0.132 | -6.60E-02 | 1.95E-01 | 2.70E+00 | 2.77E+00 | 7.29E-01 | no | Slc25a29 | intron_variant | MODIFIER |
| 12 | 108829963 | AX-170233274 | 0 | C | T | 0.132 | -6.60E-02 | 1.95E-01 | 2.70E+00 | 2.77E+00 | 7.29E-01 | no | Slc25a29 | intron_variant | MODIFIER |
| 12 | 108837594 | AX-97839860 | 0 | A | G | 0.132 | -6.60E-02 | 1.95E-01 | 2.70E+00 | 2.77E+00 | 7.29E-01 | no | Slc25a29 | upstream_gene_variant | MODIFIER |
| 2 | 71271651 | AX-170117193 | 0 | T | C | 0.053 | -5.47E-02 | 1.58E-01 | 2.58E+00 | 2.72E+00 | 7.29E-01 | no | Slc25a12 | downstream_gene_variant | MODIFIER |
| 6 | 129514528 | AX-170195552 | 0 | C | T | 0.553 | 2.86E-02 | 8.11E-02 | 2.60E+00 | 2.68E+00 | 7.29E-01 | no | Tmem52b | intron_variant | MODIFIER |
| 1 | 118754380 | AX-170459145 | 2 | G | A | 0.164 | 6.55E-02 | 1.86E-01 | 2.59E+00 | 2.69E+00 | 7.30E-01 | no | | intergenic_variant | MODIFIER |
| 9 | 62466952 | AX-170032517 | 1 | C | T | 0.205 | 4.55E-02 | 1.34E-01 | 2.75E+00 | 2.79E+00 | 7.30E-01 | no | Coro2b | intron_variant | MODIFIER |
| 7 | 14588143 | AX-97809797 | 0 | C | A | 0.368 | 2.87E-02 | 8.21E-02 | 2.58E+00 | 2.71E+00 | 7.30E-01 | no | Nlrp5-ps | intron_variant,non_coding_transcript_variant | MODIFIER |
| 2 | 124208622 | AX-170344234 | 0 | T | C | 0.149 | 6.32E-02 | 1.78E-01 | 2.58E+00 | 2.67E+00 | 7.30E-01 | no | Sema6d | intron_variant | MODIFIER |
| 1 | 106406374 | AX-170394839 | 0 | C | A | 0.158 | 6.63E-02 | 1.89E-01 | 2.60E+00 | 2.68E+00 | 7.30E-01 | no | A530053M12Rik | upstream_gene_variant | MODIFIER |
| 1 | 106422210 | AX-170510425 | 0 | T | C | 0.158 | 6.63E-02 | 1.89E-01 | 2.60E+00 | 2.68E+00 | 7.30E-01 | no | | intergenic_variant | MODIFIER |
| 1 | 106459750 | AX-97840629 | 0 | A | G | 0.158 | 6.63E-02 | 1.89E-01 | 2.60E+00 | 2.68E+00 | 7.30E-01 | no | | intergenic_variant | MODIFIER |
| 1 | 106480347 | AX-170086563 | 0 | A | T | 0.158 | 6.63E-02 | 1.89E-01 | 2.60E+00 | 2.68E+00 | 7.30E-01 | no | | intergenic_variant | MODIFIER |
| 1 | 106505071 | AX-170121469 | 0 | C | G | 0.158 | 6.63E-02 | 1.89E-01 | 2.60E+00 | 2.68E+00 | 7.30E-01 | no | | intergenic_variant | MODIFIER |
| 1 | 106508160 | AX-97839587 | 0 | T | C | 0.158 | 6.63E-02 | 1.89E-01 | 2.60E+00 | 2.68E+00 | 7.30E-01 | no | | intergenic_variant | MODIFIER |
| 1 | 106554591 | AX-170312437 | 0 | G | A | 0.158 | 6.63E-02 | 1.89E-01 | 2.60E+00 | 2.68E+00 | 7.30E-01 | no | Bcl2 | intron_variant | MODIFIER |
| 1 | 106555394 | AX-170177896 | 0 | G | A | 0.158 | 6.63E-02 | 1.89E-01 | 2.60E+00 | 2.68E+00 | 7.30E-01 | no | Bcl2 | intron_variant | MODIFIER |
| 1 | 106571444 | AX-170001285 | 0 | T | C | 0.158 | 6.63E-02 | 1.89E-01 | 2.60E+00 | 2.68E+00 | 7.30E-01 | no | Bcl2 | intron_variant | MODIFIER |

| | | | | | | | | | | | | | | | |
|---|---|---|---|---|---|---|---|---|---|---|---|---|---|---|---|
| 1 | 106574774 | AX-97854157 | 0 | T | C | 0.158 | 6.63E-02 | 1.89E-01 | 2.60E+00 | 2.68E+00 | 7.30E-01 | no | Bcl2 | intron_variant | MODIFIER |
| 1 | 106591534 | AX-170496591 | 0 | G | A | 0.158 | 6.63E-02 | 1.89E-01 | 2.60E+00 | 2.68E+00 | 7.30E-01 | no | Bcl2 | intron_variant | MODIFIER |
| 1 | 106591714 | AX-170406192 | 0 | G | A | 0.158 | 6.63E-02 | 1.89E-01 | 2.60E+00 | 2.68E+00 | 7.30E-01 | no | Bcl2 | intron_variant | MODIFIER |
| 1 | 106592746 | AX-170042291 | 0 | C | T | 0.158 | 6.63E-02 | 1.89E-01 | 2.60E+00 | 2.68E+00 | 7.30E-01 | no | Bcl2 | intron_variant | MODIFIER |
| 1 | 106595403 | AX-170001286 | 0 | T | C | 0.158 | 6.63E-02 | 1.89E-01 | 2.60E+00 | 2.68E+00 | 7.30E-01 | no | Bcl2 | intron_variant | MODIFIER |
| 1 | 106603714 | AX-170254012 | 0 | G | T | 0.158 | 6.63E-02 | 1.89E-01 | 2.60E+00 | 2.68E+00 | 7.30E-01 | no | Bcl2 | intron_variant | MODIFIER |
| 1 | 106605734 | AX-97787297 | 0 | T | C | 0.158 | 6.63E-02 | 1.89E-01 | 2.60E+00 | 2.68E+00 | 7.30E-01 | no | Bcl2 | intron_variant | MODIFIER |
| 1 | 106607311 | AX-170190889 | 0 | C | G | 0.158 | 6.63E-02 | 1.89E-01 | 2.60E+00 | 2.68E+00 | 7.30E-01 | no | Bcl2 | intron_variant | MODIFIER |
| 1 | 106610454 | AX-170001284 | 0 | T | C | 0.158 | 6.63E-02 | 1.89E-01 | 2.60E+00 | 2.68E+00 | 7.30E-01 | no | Bcl2 | intron_variant | MODIFIER |
| 1 | 106614577 | AX-170001288 | 0 | T | C | 0.158 | 6.63E-02 | 1.89E-01 | 2.60E+00 | 2.68E+00 | 7.30E-01 | no | Bcl2 | intron_variant | MODIFIER |
| 1 | 106648709 | AX-170433539 | 0 | C | T | 0.158 | 6.63E-02 | 1.89E-01 | 2.60E+00 | 2.68E+00 | 7.30E-01 | no | Bcl2 | intron_variant | MODIFIER |
| 1 | 106648942 | AX-170001287 | 0 | G | C | 0.158 | 6.63E-02 | 1.89E-01 | 2.60E+00 | 2.68E+00 | 7.30E-01 | no | Bcl2 | intron_variant | MODIFIER |
| 1 | 106649546 | AX-170091628 | 0 | A | C | 0.158 | 6.63E-02 | 1.89E-01 | 2.60E+00 | 2.68E+00 | 7.30E-01 | no | Bcl2 | intron_variant | MODIFIER |
| 1 | 108658217 | AX-170270659 | 0 | T | C | 0.158 | 6.63E-02 | 1.89E-01 | 2.60E+00 | 2.68E+00 | 7.30E-01 | no | | intergenic_variant | MODIFIER |
| 1 | 108719095 | AX-170201801 | 0 | A | C | 0.158 | 6.63E-02 | 1.89E-01 | 2.60E+00 | 2.68E+00 | 7.30E-01 | no | | intergenic_variant | MODIFIER |
| 1 | 108719701 | AX-97791118 | 0 | A | G | 0.158 | 6.63E-02 | 1.89E-01 | 2.60E+00 | 2.68E+00 | 7.30E-01 | no | | intergenic_variant | MODIFIER |
| 1 | 108721978 | AX-170459205 | 0 | T | C | 0.158 | 6.63E-02 | 1.89E-01 | 2.60E+00 | 2.68E+00 | 7.30E-01 | no | | intergenic_variant | MODIFIER |
| 1 | 108744645 | AX-170115160 | 0 | A | G | 0.158 | 6.63E-02 | 1.89E-01 | 2.60E+00 | 2.68E+00 | 7.30E-01 | no | | intergenic_variant | MODIFIER |
| 1 | 108937963 | AX-97824810 | 0 | A | G | 0.158 | 6.63E-02 | 1.89E-01 | 2.60E+00 | 2.68E+00 | 7.30E-01 | no | | intergenic_variant | MODIFIER |
| 1 | 109114411 | AX-97827670 | 0 | A | G | 0.158 | 6.63E-02 | 1.89E-01 | 2.60E+00 | 2.68E+00 | 7.30E-01 | no | | intergenic_variant | MODIFIER |
| 1 | 109211670 | AX-170209077 | 0 | T | A | 0.158 | 6.63E-02 | 1.89E-01 | 2.60E+00 | 2.68E+00 | 7.30E-01 | no | | intergenic_variant | MODIFIER |
| 1 | 109225220 | AX-170172696 | 0 | G | C | 0.158 | 6.63E-02 | 1.89E-01 | 2.60E+00 | 2.68E+00 | 7.30E-01 | no | | intergenic_variant | MODIFIER |
| 1 | 110384380 | AX-170457128 | 0 | G | A | 0.158 | 6.63E-02 | 1.89E-01 | 2.60E+00 | 2.68E+00 | 7.30E-01 | no | | intergenic_variant | MODIFIER |
| 1 | 110407349 | AX-170001365 | 0 | G | C | 0.158 | 6.63E-02 | 1.89E-01 | 2.60E+00 | 2.68E+00 | 7.30E-01 | no | Gm18552 | downstream_gene_variant | MODIFIER |
| 1 | 110415224 | AX-97812220 | 0 | G | A | 0.158 | 6.63E-02 | 1.89E-01 | 2.60E+00 | 2.68E+00 | 7.30E-01 | no | Gm18552 | upstream_gene_variant | MODIFIER |
| 1 | 110437109 | AX-170001358 | 0 | A | T | 0.158 | 6.63E-02 | 1.89E-01 | 2.60E+00 | 2.68E+00 | 7.30E-01 | no | | intergenic_variant | MODIFIER |
| 1 | 110446752 | AX-170411128 | 0 | T | C | 0.158 | 6.63E-02 | 1.89E-01 | 2.60E+00 | 2.68E+00 | 7.30E-01 | no | | intergenic_variant | MODIFIER |
| 1 | 110452749 | AX-170392957 | 0 | C | T | 0.158 | 6.63E-02 | 1.89E-01 | 2.60E+00 | 2.68E+00 | 7.30E-01 | no | | intergenic_variant | MODIFIER |
| 1 | 110478763 | AX-170001364 | 0 | G | A | 0.158 | 6.63E-02 | 1.89E-01 | 2.60E+00 | 2.68E+00 | 7.30E-01 | no | Gm3332 | downstream_gene_variant | MODIFIER |
| 1 | 110529598 | AX-97788585 | 0 | A | C | 0.158 | 6.63E-02 | 1.89E-01 | 2.60E+00 | 2.68E+00 | 7.30E-01 | no | | intergenic_variant | MODIFIER |
| 1 | 110551626 | AX-170001362 | 0 | T | C | 0.158 | 6.63E-02 | 1.89E-01 | 2.60E+00 | 2.68E+00 | 7.30E-01 | no | | intergenic_variant | MODIFIER |
| 1 | 110563707 | AX-170086574 | 0 | A | G | 0.158 | 6.63E-02 | 1.89E-01 | 2.60E+00 | 2.68E+00 | 7.30E-01 | no | | intergenic_variant | MODIFIER |
| 1 | 110684678 | AX-170416132 | 0 | T | C | 0.158 | 6.63E-02 | 1.89E-01 | 2.60E+00 | 2.68E+00 | 7.30E-01 | no | | intergenic_variant | MODIFIER |
| 1 | 113569088 | AX-170192769 | 0 | T | G | 0.158 | 6.63E-02 | 1.89E-01 | 2.60E+00 | 2.68E+00 | 7.30E-01 | no | Gm17946 | upstream_gene_variant | MODIFIER |
| 1 | 113705178 | AX-170382939 | 0 | G | A | 0.158 | 6.63E-02 | 1.89E-01 | 2.60E+00 | 2.68E+00 | 7.30E-01 | no | | intergenic_variant | MODIFIER |
| 1 | 113732939 | AX-170180702 | 0 | A | T | 0.158 | 6.63E-02 | 1.89E-01 | 2.60E+00 | 2.68E+00 | 7.30E-01 | no | | intergenic_variant | MODIFIER |
| 1 | 113733010 | AX-170117281 | 0 | C | T | 0.158 | 6.63E-02 | 1.89E-01 | 2.60E+00 | 2.68E+00 | 7.30E-01 | no | | intergenic_variant | MODIFIER |
| 1 | 113736514 | AX-170107941 | 0 | A | C | 0.158 | 6.63E-02 | 1.89E-01 | 2.60E+00 | 2.68E+00 | 7.30E-01 | no | | intergenic_variant | MODIFIER |
| 1 | 113802961 | AX-170194953 | 0 | A | G | 0.158 | 6.63E-02 | 1.89E-01 | 2.60E+00 | 2.68E+00 | 7.30E-01 | no | | intergenic_variant | MODIFIER |
| 1 | 113809767 | AX-97824638 | 0 | C | T | 0.158 | 6.63E-02 | 1.89E-01 | 2.60E+00 | 2.68E+00 | 7.30E-01 | no | | intergenic_variant | MODIFIER |
| 1 | 113834083 | AX-97787762 | 0 | C | T | 0.158 | 6.63E-02 | 1.89E-01 | 2.60E+00 | 2.68E+00 | 7.30E-01 | no | | intergenic_variant | MODIFIER |
| 1 | 113846158 | AX-170469524 | 0 | G | T | 0.158 | 6.63E-02 | 1.89E-01 | 2.60E+00 | 2.68E+00 | 7.30E-01 | no | | intergenic_variant | MODIFIER |
| 1 | 113847564 | AX-170124482 | 0 | A | G | 0.158 | 6.63E-02 | 1.89E-01 | 2.60E+00 | 2.68E+00 | 7.30E-01 | no | | intergenic_variant | MODIFIER |
| 1 | 113861902 | AX-97794080 | 0 | G | A | 0.158 | 6.63E-02 | 1.89E-01 | 2.60E+00 | 2.68E+00 | 7.30E-01 | no | | intergenic_variant | MODIFIER |
| 1 | 113865139 | AX-170001432 | 0 | A | G | 0.158 | 6.63E-02 | 1.89E-01 | 2.60E+00 | 2.68E+00 | 7.30E-01 | no | | intergenic_variant | MODIFIER |
| 1 | 113887315 | AX-97824682 | 0 | T | C | 0.158 | 6.63E-02 | 1.89E-01 | 2.60E+00 | 2.68E+00 | 7.30E-01 | no | 9330185C12Rik | downstream_gene_variant | MODIFIER |
| 1 | 113927718 | AX-170001427 | 0 | A | G | 0.158 | 6.63E-02 | 1.89E-01 | 2.60E+00 | 2.68E+00 | 7.30E-01 | no | 9330185C12Rik | intron_variant,non_coding_transcript_variant | MODIFIER |
| 1 | 113928481 | AX-170001434 | 0 | A | G | 0.158 | 6.63E-02 | 1.89E-01 | 2.60E+00 | 2.68E+00 | 7.30E-01 | no | 9330185C12Rik | intron_variant,non_coding_transcript_variant | MODIFIER |
| 1 | 113928963 | AX-170001430 | 0 | C | T | 0.158 | 6.63E-02 | 1.89E-01 | 2.60E+00 | 2.68E+00 | 7.30E-01 | no | 9330185C12Rik | intron_variant,non_coding_transcript_variant | MODIFIER |
| 1 | 113932685 | AX-170244041 | 0 | T | A | 0.158 | 6.63E-02 | 1.89E-01 | 2.60E+00 | 2.68E+00 | 7.30E-01 | no | 9330185C12Rik | intron_variant,non_coding_transcript_variant | MODIFIER |
| 1 | 113934768 | AX-170106817 | 0 | C | T | 0.158 | 6.63E-02 | 1.89E-01 | 2.60E+00 | 2.68E+00 | 7.30E-01 | no | 9330185C12Rik | intron_variant,non_coding_transcript_variant | MODIFIER |
| 1 | 113934998 | AX-97836349 | 0 | T | G | 0.158 | 6.63E-02 | 1.89E-01 | 2.60E+00 | 2.68E+00 | 7.30E-01 | no | 9330185C12Rik | intron_variant,non_coding_transcript_variant | MODIFIER |
| 1 | 113938328 | AX-170442407 | 0 | C | T | 0.158 | 6.63E-02 | 1.89E-01 | 2.60E+00 | 2.68E+00 | 7.30E-01 | no | 9330185C12Rik | intron_variant,non_coding_transcript_variant | MODIFIER |
| 1 | 113938867 | AX-168395466 | 0 | T | C | 0.158 | 6.63E-02 | 1.89E-01 | 2.60E+00 | 2.68E+00 | 7.30E-01 | no | 9330185C12Rik | intron_variant,non_coding_transcript_variant | MODIFIER |

| | | | | | | | | | | | | | | | |
|---|---|---|---|---|---|---|---|---|---|---|---|---|---|---|---|
| 1 | 113947758 | AX-170001437 | 0 | T | C | 0.158 | 6.63E-02 | 1.89E-01 | 2.60E+00 | 2.68E+00 | 7.30E-01 | no | 9330185C12Rik | intron_variant,non_coding_transcript_variant | MODIFIER |
| 1 | 113948032 | AX-170001431 | 0 | C | A | 0.158 | 6.63E-02 | 1.89E-01 | 2.60E+00 | 2.68E+00 | 7.30E-01 | no | 9330185C12Rik | intron_variant,non_coding_transcript_variant | MODIFIER |
| 1 | 113948087 | AX-97831992 | 0 | G | A | 0.158 | 6.63E-02 | 1.89E-01 | 2.60E+00 | 2.68E+00 | 7.30E-01 | no | 9330185C12Rik | intron_variant,non_coding_transcript_variant | MODIFIER |
| 1 | 115837652 | AX-170191716 | 0 | T | A | 0.158 | 6.63E-02 | 1.89E-01 | 2.60E+00 | 2.68E+00 | 7.30E-01 | no | Cntnap5a | intron_variant | MODIFIER |
| 1 | 115932151 | AX-97813019 | 0 | A | G | 0.158 | 6.63E-02 | 1.89E-01 | 2.60E+00 | 2.68E+00 | 7.30E-01 | no | Cntnap5a | intron_variant | MODIFIER |
| 1 | 116074385 | AX-97792097 | 0 | G | T | 0.158 | 6.63E-02 | 1.89E-01 | 2.60E+00 | 2.68E+00 | 7.30E-01 | no | Cntnap5a | intron_variant | MODIFIER |
| 1 | 116472620 | AX-170345622 | 0 | T | A | 0.158 | 6.63E-02 | 1.89E-01 | 2.60E+00 | 2.68E+00 | 7.30E-01 | no | Cntnap5a | intron_variant | MODIFIER |
| 1 | 116473147 | AX-170189530 | 0 | G | C | 0.158 | 6.63E-02 | 1.89E-01 | 2.60E+00 | 2.68E+00 | 7.30E-01 | no | Cntnap5a | intron_variant | MODIFIER |
| 1 | 116609473 | AX-170091652 | 0 | G | T | 0.158 | 6.63E-02 | 1.89E-01 | 2.60E+00 | 2.68E+00 | 7.30E-01 | no | | intergenic_variant | MODIFIER |
| 1 | 116613534 | AX-170487866 | 0 | G | A | 0.158 | 6.63E-02 | 1.89E-01 | 2.60E+00 | 2.68E+00 | 7.30E-01 | no | | intergenic_variant | MODIFIER |
| 1 | 116613905 | AX-170303900 | 0 | A | C | 0.158 | 6.63E-02 | 1.89E-01 | 2.60E+00 | 2.68E+00 | 7.30E-01 | no | | intergenic_variant | MODIFIER |
| 1 | 117026516 | AX-97804951 | 0 | G | T | 0.158 | 6.63E-02 | 1.89E-01 | 2.60E+00 | 2.68E+00 | 7.30E-01 | no | | intergenic_variant | MODIFIER |
| 1 | 118360109 | AX-170120638 | 0 | A | G | 0.158 | 6.63E-02 | 1.89E-01 | 2.60E+00 | 2.68E+00 | 7.30E-01 | no | Gm28867 | intron_variant,non_coding_transcript_variant | MODIFIER |
| 1 | 118483713 | AX-170466662 | 0 | C | G | 0.158 | 6.63E-02 | 1.89E-01 | 2.60E+00 | 2.68E+00 | 7.30E-01 | no | Clasp1 | intron_variant | MODIFIER |
| 1 | 118564485 | AX-170346422 | 0 | C | A | 0.158 | 6.63E-02 | 1.89E-01 | 2.60E+00 | 2.68E+00 | 7.30E-01 | no | Clasp1 | intron_variant | MODIFIER |
| 1 | 118581368 | AX-170175054 | 0 | C | T | 0.158 | 6.63E-02 | 1.89E-01 | 2.60E+00 | 2.68E+00 | 7.30E-01 | no | Clasp1 | synonymous_variant | LOW |
| 1 | 118602685 | AX-170138931 | 0 | G | A | 0.158 | 6.63E-02 | 1.89E-01 | 2.60E+00 | 2.68E+00 | 7.30E-01 | no | Clasp1 | intron_variant | MODIFIER |
| 1 | 118602971 | AX-97851134 | 0 | C | T | 0.158 | 6.63E-02 | 1.89E-01 | 2.60E+00 | 2.68E+00 | 7.30E-01 | no | Clasp1 | intron_variant | MODIFIER |
| 1 | 118603625 | AX-170199447 | 0 | A | G | 0.158 | 6.63E-02 | 1.89E-01 | 2.60E+00 | 2.68E+00 | 7.30E-01 | no | Clasp1 | intron_variant | MODIFIER |
| 1 | 118839199 | AX-170138502 | 0 | A | G | 0.158 | 6.63E-02 | 1.89E-01 | 2.60E+00 | 2.68E+00 | 7.30E-01 | no | Gli2 | intron_variant | MODIFIER |
| 1 | 118892085 | AX-170495001 | 0 | C | T | 0.158 | 6.63E-02 | 1.89E-01 | 2.60E+00 | 2.68E+00 | 7.30E-01 | no | Gli2 | intron_variant | MODIFIER |
| 1 | 118909046 | AX-170350266 | 0 | T | C | 0.158 | 6.63E-02 | 1.89E-01 | 2.60E+00 | 2.68E+00 | 7.30E-01 | no | Gli2 | intron_variant | MODIFIER |
| 1 | 118909445 | AX-170279354 | 0 | C | T | 0.158 | 6.63E-02 | 1.89E-01 | 2.60E+00 | 2.68E+00 | 7.30E-01 | no | Gli2 | intron_variant | MODIFIER |
| 1 | 118934353 | AX-170463401 | 0 | T | C | 0.158 | 6.63E-02 | 1.89E-01 | 2.60E+00 | 2.68E+00 | 7.30E-01 | no | Gli2 | intron_variant | MODIFIER |
| 1 | 118962327 | AX-170143298 | 0 | C | T | 0.158 | 6.63E-02 | 1.89E-01 | 2.60E+00 | 2.68E+00 | 7.30E-01 | no | Gli2 | intron_variant | MODIFIER |
| 1 | 118965789 | AX-170246211 | 0 | G | A | 0.158 | 6.63E-02 | 1.89E-01 | 2.60E+00 | 2.68E+00 | 7.30E-01 | no | Gli2 | intron_variant | MODIFIER |
| 1 | 118975178 | AX-170470090 | 0 | C | T | 0.158 | 6.63E-02 | 1.89E-01 | 2.60E+00 | 2.68E+00 | 7.30E-01 | no | Gli2 | intron_variant | MODIFIER |
| 1 | 118976407 | AX-170424635 | 0 | A | C | 0.158 | 6.63E-02 | 1.89E-01 | 2.60E+00 | 2.68E+00 | 7.30E-01 | no | Gli2 | intron_variant | MODIFIER |
| 1 | 118978473 | AX-170131723 | 0 | G | A | 0.158 | 6.63E-02 | 1.89E-01 | 2.60E+00 | 2.68E+00 | 7.30E-01 | no | Gli2 | intron_variant | MODIFIER |
| 1 | 118978536 | AX-170288158 | 0 | G | C | 0.158 | 6.63E-02 | 1.89E-01 | 2.60E+00 | 2.68E+00 | 7.30E-01 | no | Gli2 | intron_variant | MODIFIER |
| 1 | 118981019 | AX-170259820 | 0 | A | G | 0.158 | 6.63E-02 | 1.89E-01 | 2.60E+00 | 2.68E+00 | 7.30E-01 | no | Gli2 | intron_variant | MODIFIER |
| 1 | 118986108 | AX-97813089 | 0 | C | T | 0.158 | 6.63E-02 | 1.89E-01 | 2.60E+00 | 2.68E+00 | 7.30E-01 | no | Gli2 | intron_variant | MODIFIER |
| 1 | 119068785 | AX-170261329 | 0 | T | A | 0.158 | 6.63E-02 | 1.89E-01 | 2.60E+00 | 2.68E+00 | 7.30E-01 | no | | intergenic_variant | MODIFIER |
| 1 | 119153871 | AX-170001534 | 0 | G | A | 0.158 | 6.63E-02 | 1.89E-01 | 2.60E+00 | 2.68E+00 | 7.30E-01 | no | Gm8321 | upstream_gene_variant | MODIFIER |
| 1 | 119170468 | AX-170001531 | 0 | T | C | 0.158 | 6.63E-02 | 1.89E-01 | 2.60E+00 | 2.68E+00 | 7.30E-01 | no | | intergenic_variant | MODIFIER |
| 1 | 119187411 | AX-170253627 | 0 | C | T | 0.158 | 6.63E-02 | 1.89E-01 | 2.60E+00 | 2.68E+00 | 7.30E-01 | no | | intergenic_variant | MODIFIER |
| 1 | 119199867 | AX-170294578 | 0 | A | C | 0.158 | 6.63E-02 | 1.89E-01 | 2.60E+00 | 2.68E+00 | 7.30E-01 | no | | intergenic_variant | MODIFIER |
| 1 | 119202244 | AX-170001536 | 0 | C | T | 0.158 | 6.63E-02 | 1.89E-01 | 2.60E+00 | 2.68E+00 | 7.30E-01 | no | | intergenic_variant | MODIFIER |
| 1 | 119207042 | AX-170001532 | 0 | G | A | 0.158 | 6.63E-02 | 1.89E-01 | 2.60E+00 | 2.68E+00 | 7.30E-01 | no | | intergenic_variant | MODIFIER |
| 1 | 119223017 | AX-170001538 | 0 | G | A | 0.158 | 6.63E-02 | 1.89E-01 | 2.60E+00 | 2.68E+00 | 7.30E-01 | no | | intergenic_variant | MODIFIER |
| 1 | 119224364 | AX-170001537 | 0 | A | G | 0.158 | 6.63E-02 | 1.89E-01 | 2.60E+00 | 2.68E+00 | 7.30E-01 | no | | intergenic_variant | MODIFIER |
| 1 | 119234357 | AX-170001533 | 0 | G | A | 0.158 | 6.63E-02 | 1.89E-01 | 2.60E+00 | 2.68E+00 | 7.30E-01 | no | | intergenic_variant | MODIFIER |
| 1 | 119257335 | AX-97808185 | 0 | G | A | 0.158 | 6.63E-02 | 1.89E-01 | 2.60E+00 | 2.68E+00 | 7.30E-01 | no | | regulatory_region_variant | MODIFIER |
| 1 | 119260297 | AX-170384426 | 0 | A | G | 0.158 | 6.63E-02 | 1.89E-01 | 2.60E+00 | 2.68E+00 | 7.30E-01 | no | | intergenic_variant | MODIFIER |
| 1 | 119267219 | AX-170001535 | 0 | A | G | 0.158 | 6.63E-02 | 1.89E-01 | 2.60E+00 | 2.68E+00 | 7.30E-01 | no | | intergenic_variant | MODIFIER |
| 1 | 119272980 | AX-97813062 | 0 | G | A | 0.158 | 6.63E-02 | 1.89E-01 | 2.60E+00 | 2.68E+00 | 7.30E-01 | no | | intergenic_variant | MODIFIER |
| 8 | 35747384 | AX-170263156 | 1 | G | A | 0.152 | -4.98E-02 | 1.44E-01 | 2.64E+00 | 2.69E+00 | 7.30E-01 | no | Gm22216 | upstream_gene_variant | MODIFIER |
| 4 | 70224694 | AX-170023054 | 0 | T | A | 0.456 | -2.88E-02 | 7.97E-02 | 2.57E+00 | 2.64E+00 | 7.30E-01 | no | Cdk5rap2 | intron_variant,NMD_transcript_variant | MODIFIER |
| 11 | 118403672 | AX-170323338 | 0 | G | A | 0.053 | -6.51E-02 | 1.95E-01 | 2.67E+00 | 2.79E+00 | 7.30E-01 | no | Cant1 | downstream_gene_variant | MODIFIER |
| 4 | 136880502 | AX-170153111 | 1 | A | C | 0.152 | -3.37E-02 | 1.02E-01 | 2.75E+00 | 2.87E+00 | 7.30E-01 | no | C1qb | synonymous_variant | LOW |
| 5 | 35449610 | AX-170326884 | 1 | A | G | 0.33 | 3.66E-02 | 1.07E-01 | 2.67E+00 | 2.73E+00 | 7.30E-01 | no | | intergenic_variant | MODIFIER |
| 5 | 84963415 | AX-170134540 | 0 | A | G | 0.439 | 3.69E-02 | 1.09E-01 | 2.72E+00 | 2.78E+00 | 7.30E-01 | no | | intergenic_variant | MODIFIER |
| 10 | 128206955 | AX-170388562 | 1 | A | G | 0.33 | -2.50E-02 | 7.29E-02 | 2.62E+00 | 2.72E+00 | 7.30E-01 | no | Gls2 | synonymous_variant | LOW |
| 5 | 114048375 | AX-170484349 | 1 | C | T | 0.464 | 3.74E-02 | 1.09E-01 | 2.65E+00 | 2.70E+00 | 7.31E-01 | no | Svop | intron_variant | MODIFIER |

| | | | | | | | | | | | | | | | |
|---|---|---|---|---|---|---|---|---|---|---|---|---|---|---|---|
| 8 | 31917000 | AX-170367226 | 0 | T | C | 0.149 | -4.06E-02 | 1.20E-01 | 2.66E+00 | 2.75E+00 | 7.31E-01 | no | Nrg1 | intron_variant | MODIFIER |
| 7 | 54838125 | AX-170384843 | 1 | T | C | 0.438 | 3.44E-02 | 9.97E-02 | 2.63E+00 | 2.71E+00 | 7.31E-01 | no | Luzp2 | intron_variant | MODIFIER |
| 1 | 95201679 | AX-97797747 | 1 | C | T | 0.152 | 4.66E-02 | 1.34E-01 | 2.60E+00 | 2.70E+00 | 7.31E-01 | no | | intergenic_variant | MODIFIER |
| 9 | 113356625 | AX-170130610 | 1 | G | A | 0.125 | -3.62E-02 | 1.06E-01 | 2.64E+00 | 2.73E+00 | 7.31E-01 | no | Gm39436 | upstream_gene_variant | MODIFIER |
| 10 | 17185775 | AX-97814879 | 2 | C | T | 0.227 | 3.57E-02 | 1.07E-01 | 2.74E+00 | 2.83E+00 | 7.31E-01 | no | | intergenic_variant | MODIFIER |
| 1 | 101839635 | AX-170086555 | 0 | C | A | 0.149 | 5.60E-02 | 1.67E-01 | 2.66E+00 | 2.76E+00 | 7.31E-01 | no | | intergenic_variant | MODIFIER |
| 4 | 85452057 | AX-170450553 | 1 | C | T | 0.071 | 3.21E-02 | 9.50E-02 | 2.56E+00 | 2.75E+00 | 7.31E-01 | no | Sh3gl2 | intron_variant | MODIFIER |
| 2 | 75936665 | AX-170185708 | 2 | A | G | 0.482 | 1.83E-01 | 5.36E-01 | 2.67E+00 | 2.72E+00 | 7.31E-01 | Yes | Ttc30b | synonymous_variant | LOW |
| 8 | 109935693 | AX-170031477 | 0 | T | C | 0.237 | -3.06E-02 | 9.09E-02 | 2.64E+00 | 2.76E+00 | 7.31E-01 | no | Phlpp2 | intron_variant | MODIFIER |
| 13 | 38230175 | AX-170403597 | 0 | G | A | 0.175 | -4.26E-02 | 1.17E-01 | 2.56E+00 | 2.63E+00 | 7.31E-01 | no | Snrnp48 | downstream_gene_variant | MODIFIER |
| 8 | 21898456 | AX-170134157 | 0 | C | T | 0.175 | -3.76E-02 | 1.11E-01 | 2.67E+00 | 2.75E+00 | 7.31E-01 | no | | intergenic_variant | MODIFIER |
| 1 | 82066542 | AX-170447021 | 1 | C | G | 0.152 | 7.53E-02 | 2.20E-01 | 2.65E+00 | 2.71E+00 | 7.31E-01 | no | | intergenic_variant | MODIFIER |
| 5 | 28520709 | AX-170444452 | 1 | C | T | 0.366 | 3.64E-02 | 1.07E-01 | 2.67E+00 | 2.72E+00 | 7.32E-01 | no | 9530036O11Rik | intron_variant,non_coding_transcript_variant | MODIFIER |
| 5 | 32321895 | AX-170296820 | 1 | T | C | 0.366 | 3.64E-02 | 1.07E-01 | 2.67E+00 | 2.72E+00 | 7.32E-01 | no | Plb1 | intron_variant | MODIFIER |
| 1 | 82197278 | AX-170452930 | 0 | C | T | 0.158 | 5.74E-02 | 1.71E-01 | 2.66E+00 | 2.77E+00 | 7.32E-01 | no | | intergenic_variant | MODIFIER |
| 1 | 95245398 | AX-170207282 | 0 | G | A | 0.158 | 5.74E-02 | 1.71E-01 | 2.66E+00 | 2.77E+00 | 7.32E-01 | no | | intergenic_variant | MODIFIER |
| 1 | 23338636 | AX-170228801 | 0 | A | C | 0.43 | 2.86E-02 | 8.11E-02 | 2.59E+00 | 2.67E+00 | 7.32E-01 | no | | intergenic_variant | MODIFIER |
| 8 | 78654122 | AX-170495121 | 0 | T | C | 0.211 | 4.06E-02 | 1.21E-01 | 2.70E+00 | 2.78E+00 | 7.32E-01 | no | Slc10a7 | intron_variant | MODIFIER |
| 9 | 40641914 | AX-170518113 | 1 | G | A | 0.438 | -2.54E-02 | 7.57E-02 | 2.68E+00 | 2.76E+00 | 7.32E-01 | no | | regulatory_region_variant | MODIFIER |
| 12 | 104005212 | AX-170106915 | 1 | A | G | 0.134 | -6.47E-02 | 1.93E-01 | 2.71E+00 | 2.78E+00 | 7.32E-01 | no | Serpina9 | intron_variant | MODIFIER |
| 12 | 106453375 | AX-97786221 | 1 | G | A | 0.134 | -6.47E-02 | 1.93E-01 | 2.71E+00 | 2.78E+00 | 7.32E-01 | no | | intergenic_variant | MODIFIER |
| 7 | 50870737 | AX-170303515 | 1 | A | G | 0.429 | 3.41E-02 | 9.93E-02 | 2.63E+00 | 2.71E+00 | 7.32E-01 | no | Nell1 | downstream_gene_variant | MODIFIER |
| 1 | 138667651 | AX-170458988 | 0 | A | G | 0.202 | 4.03E-02 | 1.18E-01 | 2.63E+00 | 2.71E+00 | 7.32E-01 | no | Gm5833 | downstream_gene_variant | MODIFIER |
| 11 | 73242433 | AX-170260028 | 1 | G | A | 0.241 | 2.42E-02 | 7.20E-02 | 2.62E+00 | 2.75E+00 | 7.32E-01 | no | Trpv1 | intron_variant | MODIFIER |
| 1 | 53123376 | AX-170267110 | 1 | T | C | 0.116 | 6.47E-02 | 1.91E-01 | 2.64E+00 | 2.73E+00 | 7.32E-01 | no | | intergenic_variant | MODIFIER |
| 7 | 97659342 | AX-97803425 | 0 | C | T | 0.132 | -7.10E-02 | 2.12E-01 | 2.68E+00 | 2.76E+00 | 7.32E-01 | no | Rsf1 | intron_variant | MODIFIER |
| 6 | 137459282 | AX-170028344 | 0 | A | G | 0.579 | 3.21E-02 | 8.71E-02 | 2.54E+00 | 2.61E+00 | 7.32E-01 | no | Ptpro | intron_variant | MODIFIER |
| 7 | 117637179 | AX-170323923 | 1 | A | G | 0.384 | 3.00E-02 | 8.86E-02 | 2.66E+00 | 2.73E+00 | 7.32E-01 | no | Xylt1 | intron_variant | MODIFIER |
| 2 | 149087392 | AX-97780669 | 0 | T | C | 0.114 | -5.88E-02 | 1.78E-01 | 2.71E+00 | 2.81E+00 | 7.32E-01 | no | | intergenic_variant | MODIFIER |
| 19 | 40090806 | AX-170062444 | 0 | T | C | 0.482 | 1.81E-01 | 5.31E-01 | 2.67E+00 | 2.71E+00 | 7.32E-01 | Yes | Cyp2c50 | intron_variant | MODIFIER |
| 2 | 77194760 | AX-170210481 | 0 | G | A | 0.053 | -1.01E-01 | 3.00E-01 | 2.74E+00 | 2.76E+00 | 7.33E-01 | no | Sestd1 | intron_variant | MODIFIER |
| 2 | 77203285 | AX-170132077 | 0 | T | G | 0.053 | -1.01E-01 | 3.00E-01 | 2.74E+00 | 2.76E+00 | 7.33E-01 | no | Sestd1 | intron_variant | MODIFIER |
| 2 | 77365487 | AX-170094293 | 0 | T | C | 0.053 | -1.01E-01 | 3.00E-01 | 2.74E+00 | 2.76E+00 | 7.33E-01 | no | | intergenic_variant | MODIFIER |
| 2 | 77370479 | AX-170365899 | 0 | C | A | 0.053 | -1.01E-01 | 3.00E-01 | 2.74E+00 | 2.76E+00 | 7.33E-01 | no | | intergenic_variant | MODIFIER |
| 2 | 77389538 | AX-170248612 | 0 | A | C | 0.053 | -1.01E-01 | 3.00E-01 | 2.74E+00 | 2.76E+00 | 7.33E-01 | no | | intergenic_variant | MODIFIER |
| 2 | 77389594 | AX-170505170 | 0 | C | T | 0.053 | -1.01E-01 | 3.00E-01 | 2.74E+00 | 2.76E+00 | 7.33E-01 | no | | intergenic_variant | MODIFIER |
| 2 | 77395296 | AX-170308434 | 0 | G | A | 0.053 | -1.01E-01 | 3.00E-01 | 2.74E+00 | 2.76E+00 | 7.33E-01 | no | | intergenic_variant | MODIFIER |
| 2 | 77414628 | AX-97838609 | 0 | G | A | 0.053 | -1.01E-01 | 3.00E-01 | 2.74E+00 | 2.76E+00 | 7.33E-01 | no | Zfp385b | missense_variant | MODERATE |
| 2 | 77455148 | AX-170137679 | 0 | G | C | 0.053 | -1.01E-01 | 3.00E-01 | 2.74E+00 | 2.76E+00 | 7.33E-01 | no | Zfp385b | intron_variant | MODIFIER |
| 2 | 77483550 | AX-170094294 | 0 | A | G | 0.053 | -1.01E-01 | 3.00E-01 | 2.74E+00 | 2.76E+00 | 7.33E-01 | no | Zfp385b | intron_variant | MODIFIER |
| 2 | 77484921 | AX-170064060 | 0 | C | T | 0.053 | -1.01E-01 | 3.00E-01 | 2.74E+00 | 2.76E+00 | 7.33E-01 | no | Zfp385b | intron_variant | MODIFIER |
| 2 | 77485353 | AX-170064063 | 0 | A | G | 0.053 | -1.01E-01 | 3.00E-01 | 2.74E+00 | 2.76E+00 | 7.33E-01 | no | Zfp385b | intron_variant | MODIFIER |
| 2 | 77486737 | AX-170064061 | 0 | A | T | 0.053 | -1.01E-01 | 3.00E-01 | 2.74E+00 | 2.76E+00 | 7.33E-01 | no | Zfp385b | intron_variant | MODIFIER |
| 2 | 77487168 | AX-170362764 | 0 | C | G | 0.053 | -1.01E-01 | 3.00E-01 | 2.74E+00 | 2.76E+00 | 7.33E-01 | no | Zfp385b | intron_variant | MODIFIER |
| 2 | 77487185 | AX-170140061 | 0 | T | A | 0.053 | -1.01E-01 | 3.00E-01 | 2.74E+00 | 2.76E+00 | 7.33E-01 | no | Zfp385b | intron_variant | MODIFIER |
| 2 | 77514457 | AX-97833098 | 0 | A | G | 0.053 | -1.01E-01 | 3.00E-01 | 2.74E+00 | 2.76E+00 | 7.33E-01 | no | Zfp385b | intron_variant | MODIFIER |
| 2 | 77516097 | AX-170145215 | 0 | C | T | 0.053 | -1.01E-01 | 3.00E-01 | 2.74E+00 | 2.76E+00 | 7.33E-01 | no | Zfp385b | intron_variant | MODIFIER |
| 2 | 77530236 | AX-170088637 | 0 | T | C | 0.053 | -1.01E-01 | 3.00E-01 | 2.74E+00 | 2.76E+00 | 7.33E-01 | no | Zfp385b | intron_variant | MODIFIER |
| 2 | 77543037 | AX-170265661 | 0 | C | T | 0.053 | -1.01E-01 | 3.00E-01 | 2.74E+00 | 2.76E+00 | 7.33E-01 | no | Zfp385b | intron_variant | MODIFIER |
| 2 | 77566410 | AX-170451177 | 0 | G | A | 0.053 | -1.01E-01 | 3.00E-01 | 2.74E+00 | 2.76E+00 | 7.33E-01 | no | Zfp385b | intron_variant | MODIFIER |
| 2 | 77573877 | AX-170115053 | 0 | T | G | 0.053 | -1.01E-01 | 3.00E-01 | 2.74E+00 | 2.76E+00 | 7.33E-01 | no | Zfp385b | intron_variant | MODIFIER |
| 2 | 77606940 | AX-170360572 | 0 | T | C | 0.053 | -1.01E-01 | 3.00E-01 | 2.74E+00 | 2.76E+00 | 7.33E-01 | no | Zfp385b | intron_variant | MODIFIER |
| 2 | 77610678 | AX-170210505 | 0 | T | C | 0.053 | -1.01E-01 | 3.00E-01 | 2.74E+00 | 2.76E+00 | 7.33E-01 | no | Zfp385b | intron_variant | MODIFIER |
| 2 | 77610844 | AX-170361185 | 0 | A | G | 0.053 | -1.01E-01 | 3.00E-01 | 2.74E+00 | 2.76E+00 | 7.33E-01 | no | Zfp385b | intron_variant | MODIFIER |

| | | | | | | | | | | | | | | | |
|---|---|---|---|---|---|---|---|---|---|---|---|---|---|---|---|
| 2 | 77615489 | AX-170164493 | 0 | A | T | 0.053 | -1.01E-01 | 3.00E-01 | 2.74E+00 | 2.76E+00 | 7.33E-01 | no | Zfp385b | intron_variant | MODIFIER |
| 2 | 77623117 | AX-170147113 | 0 | T | G | 0.053 | -1.01E-01 | 3.00E-01 | 2.74E+00 | 2.76E+00 | 7.33E-01 | no | Zfp385b | intron_variant | MODIFIER |
| 2 | 77675987 | AX-170400928 | 0 | T | C | 0.053 | -1.01E-01 | 3.00E-01 | 2.74E+00 | 2.76E+00 | 7.33E-01 | no | Zfp385b | intron_variant | MODIFIER |
| 2 | 77680607 | AX-170168214 | 0 | G | A | 0.053 | -1.01E-01 | 3.00E-01 | 2.74E+00 | 2.76E+00 | 7.33E-01 | no | Zfp385b | intron_variant | MODIFIER |
| 2 | 77689018 | AX-170439147 | 0 | G | A | 0.053 | -1.01E-01 | 3.00E-01 | 2.74E+00 | 2.76E+00 | 7.33E-01 | no | Zfp385b | intron_variant | MODIFIER |
| 2 | 77727562 | AX-170289362 | 0 | A | G | 0.053 | -1.01E-01 | 3.00E-01 | 2.74E+00 | 2.76E+00 | 7.33E-01 | no | Zfp385b | intron_variant | MODIFIER |
| 2 | 77837814 | AX-170383720 | 0 | C | T | 0.053 | -1.01E-01 | 3.00E-01 | 2.74E+00 | 2.76E+00 | 7.33E-01 | no | Gm13661 | upstream_gene_variant | MODIFIER |
| 2 | 77837892 | AX-170233814 | 0 | G | A | 0.053 | -1.01E-01 | 3.00E-01 | 2.74E+00 | 2.76E+00 | 7.33E-01 | no | Gm13661 | upstream_gene_variant | MODIFIER |
| 2 | 77841522 | AX-170237748 | 0 | G | A | 0.053 | -1.01E-01 | 3.00E-01 | 2.74E+00 | 2.76E+00 | 7.33E-01 | no | Gm13661 | intron_variant,non_coding_transcript_variant | MODIFIER |
| 2 | 77842410 | AX-170323548 | 0 | G | T | 0.053 | -1.01E-01 | 3.00E-01 | 2.74E+00 | 2.76E+00 | 7.33E-01 | no | Gm13661 | intron_variant,non_coding_transcript_variant | MODIFIER |
| 2 | 77889425 | AX-97837711 | 0 | T | C | 0.053 | -1.01E-01 | 3.00E-01 | 2.74E+00 | 2.76E+00 | 7.33E-01 | no | Cwc22 | intron_variant | MODIFIER |
| 2 | 77932003 | AX-170101943 | 0 | G | A | 0.053 | -1.01E-01 | 3.00E-01 | 2.74E+00 | 2.76E+00 | 7.33E-01 | no | Cwc22 | intron_variant | MODIFIER |
| 2 | 77974829 | AX-170209784 | 0 | A | G | 0.053 | -1.01E-01 | 3.00E-01 | 2.74E+00 | 2.76E+00 | 7.33E-01 | no | | intergenic_variant | MODIFIER |
| 2 | 78222243 | AX-170127480 | 0 | A | G | 0.053 | -1.01E-01 | 3.00E-01 | 2.74E+00 | 2.76E+00 | 7.33E-01 | no | | intergenic_variant | MODIFIER |
| 2 | 78492065 | AX-170272717 | 0 | A | G | 0.053 | -1.01E-01 | 3.00E-01 | 2.74E+00 | 2.76E+00 | 7.33E-01 | no | | intergenic_variant | MODIFIER |
| 2 | 78497450 | AX-97805541 | 0 | G | A | 0.053 | -1.01E-01 | 3.00E-01 | 2.74E+00 | 2.76E+00 | 7.33E-01 | no | | intergenic_variant | MODIFIER |
| 2 | 78529937 | AX-170471955 | 0 | A | C | 0.053 | -1.01E-01 | 3.00E-01 | 2.74E+00 | 2.76E+00 | 7.33E-01 | no | | intergenic_variant | MODIFIER |
| 2 | 78547235 | AX-97813657 | 0 | T | C | 0.053 | -1.01E-01 | 3.00E-01 | 2.74E+00 | 2.76E+00 | 7.33E-01 | no | Gm23149 | downstream_gene_variant | MODIFIER |
| 2 | 78567258 | AX-170523842 | 0 | G | T | 0.053 | -1.01E-01 | 3.00E-01 | 2.74E+00 | 2.76E+00 | 7.33E-01 | no | | intergenic_variant | MODIFIER |
| 2 | 78567571 | AX-170064079 | 0 | T | C | 0.053 | -1.01E-01 | 3.00E-01 | 2.74E+00 | 2.76E+00 | 7.33E-01 | no | | intergenic_variant | MODIFIER |
| 2 | 78567745 | AX-170064078 | 0 | G | T | 0.053 | -1.01E-01 | 3.00E-01 | 2.74E+00 | 2.76E+00 | 7.33E-01 | no | | intergenic_variant | MODIFIER |
| 2 | 78568303 | AX-170220500 | 0 | A | G | 0.053 | -1.01E-01 | 3.00E-01 | 2.74E+00 | 2.76E+00 | 7.33E-01 | no | | intergenic_variant | MODIFIER |
| 2 | 78579233 | AX-170168346 | 0 | A | G | 0.053 | -1.01E-01 | 3.00E-01 | 2.74E+00 | 2.76E+00 | 7.33E-01 | no | | intergenic_variant | MODIFIER |
| 2 | 78591136 | AX-170168101 | 0 | T | C | 0.053 | -1.01E-01 | 3.00E-01 | 2.74E+00 | 2.76E+00 | 7.33E-01 | no | | intergenic_variant | MODIFIER |
| 2 | 78604270 | AX-97847286 | 0 | G | A | 0.053 | -1.01E-01 | 3.00E-01 | 2.74E+00 | 2.76E+00 | 7.33E-01 | no | | intergenic_variant | MODIFIER |
| 2 | 78604307 | AX-170187144 | 0 | T | A | 0.053 | -1.01E-01 | 3.00E-01 | 2.74E+00 | 2.76E+00 | 7.33E-01 | no | | intergenic_variant | MODIFIER |
| 2 | 78606115 | AX-170273700 | 0 | T | C | 0.053 | -1.01E-01 | 3.00E-01 | 2.74E+00 | 2.76E+00 | 7.33E-01 | no | | regulatory_region_variant | MODIFIER |
| 2 | 78612667 | AX-170134498 | 0 | C | G | 0.053 | -1.01E-01 | 3.00E-01 | 2.74E+00 | 2.76E+00 | 7.33E-01 | no | | intergenic_variant | MODIFIER |
| 2 | 78612876 | AX-170509558 | 0 | T | C | 0.053 | -1.01E-01 | 3.00E-01 | 2.74E+00 | 2.76E+00 | 7.33E-01 | no | | intergenic_variant | MODIFIER |
| 2 | 78613494 | AX-170314553 | 0 | T | C | 0.053 | -1.01E-01 | 3.00E-01 | 2.74E+00 | 2.76E+00 | 7.33E-01 | no | | intergenic_variant | MODIFIER |
| 2 | 78630013 | AX-97806623 | 0 | C | A | 0.053 | -1.01E-01 | 3.00E-01 | 2.74E+00 | 2.76E+00 | 7.33E-01 | no | | intergenic_variant | MODIFIER |
| 2 | 78649195 | AX-170453392 | 0 | T | C | 0.053 | -1.01E-01 | 3.00E-01 | 2.74E+00 | 2.76E+00 | 7.33E-01 | no | | intergenic_variant | MODIFIER |
| 2 | 78649735 | AX-97830903 | 0 | A | G | 0.053 | -1.01E-01 | 3.00E-01 | 2.74E+00 | 2.76E+00 | 7.33E-01 | no | | intergenic_variant | MODIFIER |
| 2 | 78652887 | AX-170018129 | 0 | C | T | 0.053 | -1.01E-01 | 3.00E-01 | 2.74E+00 | 2.76E+00 | 7.33E-01 | no | Gm14463 | upstream_gene_variant | MODIFIER |
| 2 | 78682984 | AX-97821339 | 0 | A | G | 0.053 | -1.01E-01 | 3.00E-01 | 2.74E+00 | 2.76E+00 | 7.33E-01 | no | | intergenic_variant | MODIFIER |
| 2 | 78684506 | AX-170018130 | 0 | A | C | 0.053 | -1.01E-01 | 3.00E-01 | 2.74E+00 | 2.76E+00 | 7.33E-01 | no | | intergenic_variant | MODIFIER |
| 2 | 78686046 | AX-170018123 | 0 | G | A | 0.053 | -1.01E-01 | 3.00E-01 | 2.74E+00 | 2.76E+00 | 7.33E-01 | no | | intergenic_variant | MODIFIER |
| 2 | 78726401 | AX-170276444 | 0 | T | A | 0.053 | -1.01E-01 | 3.00E-01 | 2.74E+00 | 2.76E+00 | 7.33E-01 | no | | intergenic_variant | MODIFIER |
| 2 | 78726467 | AX-170103142 | 0 | T | C | 0.053 | -1.01E-01 | 3.00E-01 | 2.74E+00 | 2.76E+00 | 7.33E-01 | no | | intergenic_variant | MODIFIER |
| 2 | 78735648 | AX-97827204 | 0 | C | T | 0.053 | -1.01E-01 | 3.00E-01 | 2.74E+00 | 2.76E+00 | 7.33E-01 | no | | intergenic_variant | MODIFIER |
| 2 | 78738020 | AX-170018124 | 0 | A | G | 0.053 | -1.01E-01 | 3.00E-01 | 2.74E+00 | 2.76E+00 | 7.33E-01 | no | | intergenic_variant | MODIFIER |
| 2 | 78738437 | AX-170018125 | 0 | A | G | 0.053 | -1.01E-01 | 3.00E-01 | 2.74E+00 | 2.76E+00 | 7.33E-01 | no | | intergenic_variant | MODIFIER |
| 2 | 78738605 | AX-170018121 | 0 | G | A | 0.053 | -1.01E-01 | 3.00E-01 | 2.74E+00 | 2.76E+00 | 7.33E-01 | no | | intergenic_variant | MODIFIER |
| 2 | 78919426 | AX-97856294 | 0 | C | T | 0.053 | -1.01E-01 | 3.00E-01 | 2.74E+00 | 2.76E+00 | 7.33E-01 | no | Ube2e3 | intron_variant | MODIFIER |
| 2 | 80328322 | AX-170264970 | 0 | T | G | 0.053 | -1.01E-01 | 3.00E-01 | 2.74E+00 | 2.76E+00 | 7.33E-01 | no | Dnajc10 | intron_variant | MODIFIER |
| 2 | 80338469 | AX-170351931 | 0 | A | G | 0.053 | -1.01E-01 | 3.00E-01 | 2.74E+00 | 2.76E+00 | 7.33E-01 | no | Dnajc10 | intron_variant | MODIFIER |
| 2 | 80339916 | AX-170094300 | 0 | G | A | 0.053 | -1.01E-01 | 3.00E-01 | 2.74E+00 | 2.76E+00 | 7.33E-01 | no | Dnajc10 | intron_variant | MODIFIER |
| 2 | 80384442 | AX-170018153 | 0 | C | T | 0.053 | -1.01E-01 | 3.00E-01 | 2.74E+00 | 2.76E+00 | 7.33E-01 | no | | intergenic_variant | MODIFIER |
| 2 | 80386831 | AX-170064105 | 0 | G | A | 0.053 | -1.01E-01 | 3.00E-01 | 2.74E+00 | 2.76E+00 | 7.33E-01 | no | | intergenic_variant | MODIFIER |
| 2 | 80387743 | AX-97839811 | 0 | A | C | 0.053 | -1.01E-01 | 3.00E-01 | 2.74E+00 | 2.76E+00 | 7.33E-01 | no | | regulatory_region_variant | MODIFIER |
| 2 | 80388977 | AX-170320879 | 0 | A | G | 0.053 | -1.01E-01 | 3.00E-01 | 2.74E+00 | 2.76E+00 | 7.33E-01 | no | | regulatory_region_variant | MODIFIER |
| 2 | 80397126 | AX-170345527 | 0 | G | A | 0.053 | -1.01E-01 | 3.00E-01 | 2.74E+00 | 2.76E+00 | 7.33E-01 | no | Gm13752 | intron_variant,non_coding_transcript_variant | MODIFIER |
| 2 | 80397207 | AX-170094301 | 0 | A | G | 0.053 | -1.01E-01 | 3.00E-01 | 2.74E+00 | 2.76E+00 | 7.33E-01 | no | Gm13752 | intron_variant,non_coding_transcript_variant | MODIFIER |
| 2 | 80397787 | AX-170320237 | 0 | C | G | 0.053 | -1.01E-01 | 3.00E-01 | 2.74E+00 | 2.76E+00 | 7.33E-01 | no | Gm13752 | intron_variant,non_coding_transcript_variant | MODIFIER |

| | | | | | | | | | | | | | | | |
|---|---|---|---|---|---|---|---|---|---|---|---|---|---|---|---|
| 2 | 80408815 | AX-170018159 | 0 | T | C | 0.053 | -1.01E-01 | 3.00E-01 | 2.74E+00 | 2.76E+00 | 7.33E-01 | no | Frzb | downstream_gene_variant | MODIFIER |
| 2 | 80408843 | AX-170018157 | 0 | T | C | 0.053 | -1.01E-01 | 3.00E-01 | 2.74E+00 | 2.76E+00 | 7.33E-01 | no | Frzb | downstream_gene_variant | MODIFIER |
| 2 | 87593775 | AX-170018256 | 0 | G | T | 0.053 | -1.01E-01 | 3.00E-01 | 2.74E+00 | 2.76E+00 | 7.33E-01 | no | | intergenic_variant | MODIFIER |
| 2 | 87600626 | AX-170110325 | 0 | C | T | 0.053 | -1.01E-01 | 3.00E-01 | 2.74E+00 | 2.76E+00 | 7.33E-01 | no | | intergenic_variant | MODIFIER |
| 2 | 87601194 | AX-170242416 | 0 | G | C | 0.053 | -1.01E-01 | 3.00E-01 | 2.74E+00 | 2.76E+00 | 7.33E-01 | no | | intergenic_variant | MODIFIER |
| 2 | 87630427 | AX-170180124 | 0 | A | G | 0.053 | -1.01E-01 | 3.00E-01 | 2.74E+00 | 2.76E+00 | 7.33E-01 | no | Olfr1132 | downstream_gene_variant | MODIFIER |
| 2 | 87640501 | AX-170202551 | 0 | C | T | 0.053 | -1.01E-01 | 3.00E-01 | 2.74E+00 | 2.76E+00 | 7.33E-01 | no | Olfr1132 | upstream_gene_variant | MODIFIER |
| 2 | 87663107 | AX-170155606 | 0 | T | C | 0.053 | -1.01E-01 | 3.00E-01 | 2.74E+00 | 2.76E+00 | 7.33E-01 | no | Olfr1134 | upstream_gene_variant | MODIFIER |
| 2 | 87691661 | AX-170137845 | 0 | G | T | 0.053 | -1.01E-01 | 3.00E-01 | 2.74E+00 | 2.76E+00 | 7.33E-01 | no | Olfr1136 | downstream_gene_variant | MODIFIER |
| 2 | 87693774 | AX-170213715 | 0 | A | G | 0.053 | -1.01E-01 | 3.00E-01 | 2.74E+00 | 2.76E+00 | 7.33E-01 | no | Olfr1136 | missense_variant | MODERATE |
| 2 | 87734514 | AX-170134656 | 0 | T | C | 0.053 | -1.01E-01 | 3.00E-01 | 2.74E+00 | 2.76E+00 | 7.33E-01 | no | Olfr1138 | downstream_gene_variant | MODIFIER |
| 2 | 87782370 | AX-170231345 | 0 | A | G | 0.053 | -1.01E-01 | 3.00E-01 | 2.74E+00 | 2.76E+00 | 7.33E-01 | no | Olfr152 | upstream_gene_variant | MODIFIER |
| 2 | 87782384 | AX-170327877 | 0 | C | T | 0.053 | -1.01E-01 | 3.00E-01 | 2.74E+00 | 2.76E+00 | 7.33E-01 | no | Olfr152 | upstream_gene_variant | MODIFIER |
| 2 | 87790145 | AX-97854000 | 0 | T | G | 0.053 | -1.01E-01 | 3.00E-01 | 2.74E+00 | 2.76E+00 | 7.33E-01 | no | Olfr1142-ps1 | downstream_gene_variant | MODIFIER |
| 2 | 87799751 | AX-170197453 | 0 | G | C | 0.053 | -1.01E-01 | 3.00E-01 | 2.74E+00 | 2.76E+00 | 7.33E-01 | no | Olfr1143 | upstream_gene_variant | MODIFIER |
| 2 | 87817728 | AX-170134082 | 0 | G | C | 0.053 | -1.01E-01 | 3.00E-01 | 2.74E+00 | 2.76E+00 | 7.33E-01 | no | Olfr1146-ps1 | downstream_gene_variant | MODIFIER |
| 2 | 87838333 | AX-170212253 | 0 | C | T | 0.053 | -1.01E-01 | 3.00E-01 | 2.74E+00 | 2.76E+00 | 7.33E-01 | no | Olfr1148 | downstream_gene_variant | MODIFIER |
| 2 | 87867188 | AX-170252729 | 0 | T | C | 0.053 | -1.01E-01 | 3.00E-01 | 2.74E+00 | 2.76E+00 | 7.33E-01 | no | Olfr1152 | upstream_gene_variant | MODIFIER |
| 2 | 87895494 | AX-170430845 | 0 | G | A | 0.053 | -1.01E-01 | 3.00E-01 | 2.74E+00 | 2.76E+00 | 7.33E-01 | no | Olfr1153 | upstream_gene_variant | MODIFIER |
| 2 | 87950514 | AX-170159838 | 0 | A | T | 0.053 | -1.01E-01 | 3.00E-01 | 2.74E+00 | 2.76E+00 | 7.33E-01 | no | Olfr1156 | upstream_gene_variant | MODIFIER |
| 2 | 87952554 | AX-170490014 | 0 | G | A | 0.053 | -1.01E-01 | 3.00E-01 | 2.74E+00 | 2.76E+00 | 7.33E-01 | no | Olfr1156 | upstream_gene_variant | MODIFIER |
| 2 | 87956482 | AX-170218935 | 0 | C | A | 0.053 | -1.01E-01 | 3.00E-01 | 2.74E+00 | 2.76E+00 | 7.33E-01 | no | Olfr1157 | downstream_gene_variant | MODIFIER |
| 2 | 88033253 | AX-170064245 | 0 | G | C | 0.053 | -1.01E-01 | 3.00E-01 | 2.74E+00 | 2.76E+00 | 7.33E-01 | no | Olfr73 | downstream_gene_variant | MODIFIER |
| 2 | 88068231 | AX-170266869 | 0 | T | C | 0.053 | -1.01E-01 | 3.00E-01 | 2.74E+00 | 2.76E+00 | 7.33E-01 | no | Olfr1163 | downstream_gene_variant | MODIFIER |
| 2 | 88073164 | AX-170157848 | 0 | T | C | 0.053 | -1.01E-01 | 3.00E-01 | 2.74E+00 | 2.76E+00 | 7.33E-01 | no | Olfr1163 | upstream_gene_variant | MODIFIER |
| 2 | 88237372 | AX-170018260 | 0 | T | C | 0.053 | -1.01E-01 | 3.00E-01 | 2.74E+00 | 2.76E+00 | 7.33E-01 | no | | intergenic_variant | MODIFIER |
| 2 | 88248570 | AX-170187543 | 0 | A | C | 0.053 | -1.01E-01 | 3.00E-01 | 2.74E+00 | 2.76E+00 | 7.33E-01 | no | Olfr1171-ps1 | downstream_gene_variant | MODIFIER |
| 2 | 88248736 | AX-97835893 | 0 | G | A | 0.053 | -1.01E-01 | 3.00E-01 | 2.74E+00 | 2.76E+00 | 7.33E-01 | no | Olfr1171-ps1 | downstream_gene_variant | MODIFIER |
| 2 | 88309360 | AX-170018267 | 0 | A | C | 0.053 | -1.01E-01 | 3.00E-01 | 2.74E+00 | 2.76E+00 | 7.33E-01 | no | Olfr1174-ps | downstream_gene_variant | MODIFIER |
| 2 | 88355914 | AX-97846665 | 0 | G | T | 0.053 | -1.01E-01 | 3.00E-01 | 2.74E+00 | 2.76E+00 | 7.33E-01 | no | | intergenic_variant | MODIFIER |
| 2 | 88456088 | AX-170064249 | 0 | G | C | 0.053 | -1.01E-01 | 3.00E-01 | 2.74E+00 | 2.76E+00 | 7.33E-01 | no | Olfr1183 | intron_variant | MODIFIER |
| 2 | 88523746 | AX-97794528 | 0 | C | A | 0.053 | -1.01E-01 | 3.00E-01 | 2.74E+00 | 2.76E+00 | 7.33E-01 | no | Olfr1186 | upstream_gene_variant | MODIFIER |
| 2 | 88523857 | AX-170232945 | 0 | A | T | 0.053 | -1.01E-01 | 3.00E-01 | 2.74E+00 | 2.76E+00 | 7.33E-01 | no | Olfr1186 | upstream_gene_variant | MODIFIER |
| 2 | 88580050 | AX-97832552 | 0 | A | C | 0.053 | -1.01E-01 | 3.00E-01 | 2.74E+00 | 2.76E+00 | 7.33E-01 | no | Gm13758 | downstream_gene_variant | MODIFIER |
| 2 | 88580647 | AX-170339675 | 0 | T | G | 0.053 | -1.01E-01 | 3.00E-01 | 2.74E+00 | 2.76E+00 | 7.33E-01 | no | Gm13758 | downstream_gene_variant | MODIFIER |
| 2 | 88586325 | AX-170426399 | 0 | G | A | 0.053 | -1.01E-01 | 3.00E-01 | 2.74E+00 | 2.76E+00 | 7.33E-01 | no | Olfr1189 | upstream_gene_variant | MODIFIER |
| 2 | 88678276 | AX-97839789 | 0 | T | G | 0.053 | -1.01E-01 | 3.00E-01 | 2.74E+00 | 2.76E+00 | 7.33E-01 | no | Olfr1193 | missense_variant | MODERATE |
| 2 | 88700217 | AX-170203101 | 0 | A | C | 0.053 | -1.01E-01 | 3.00E-01 | 2.74E+00 | 2.76E+00 | 7.33E-01 | no | Olfr1196 | downstream_gene_variant | MODIFIER |
| 2 | 88864141 | AX-170018277 | 0 | A | C | 0.053 | -1.01E-01 | 3.00E-01 | 2.74E+00 | 2.76E+00 | 7.33E-01 | no | Olfr1206 | upstream_gene_variant | MODIFIER |
| 2 | 88864630 | AX-170480916 | 0 | C | G | 0.053 | -1.01E-01 | 3.00E-01 | 2.74E+00 | 2.76E+00 | 7.33E-01 | no | Olfr1206 | synonymous_variant | LOW |
| 2 | 88865329 | AX-170376541 | 0 | T | C | 0.053 | -1.01E-01 | 3.00E-01 | 2.74E+00 | 2.76E+00 | 7.33E-01 | no | Olfr1206 | synonymous_variant | LOW |
| 2 | 88911308 | AX-170018273 | 0 | T | A | 0.053 | -1.01E-01 | 3.00E-01 | 2.74E+00 | 2.76E+00 | 7.33E-01 | no | Olfr1209 | upstream_gene_variant | MODIFIER |
| 2 | 88992477 | AX-170458570 | 0 | C | A | 0.053 | -1.01E-01 | 3.00E-01 | 2.74E+00 | 2.76E+00 | 7.33E-01 | no | Olfr1214 | upstream_gene_variant | MODIFIER |
| 2 | 89002257 | AX-170438223 | 0 | A | G | 0.053 | -1.01E-01 | 3.00E-01 | 2.74E+00 | 2.76E+00 | 7.33E-01 | no | Olfr1215 | synonymous_variant | LOW |
| 2 | 89002412 | AX-170309805 | 0 | T | C | 0.053 | -1.01E-01 | 3.00E-01 | 2.74E+00 | 2.76E+00 | 7.33E-01 | no | Olfr1215 | upstream_gene_variant | MODIFIER |
| 2 | 89143151 | AX-170463227 | 0 | C | A | 0.053 | -1.01E-01 | 3.00E-01 | 2.74E+00 | 2.76E+00 | 7.33E-01 | no | Olfr1223 | downstream_gene_variant | MODIFIER |
| 2 | 89143897 | AX-170380201 | 0 | A | T | 0.053 | -1.01E-01 | 3.00E-01 | 2.74E+00 | 2.76E+00 | 7.33E-01 | no | Olfr1223 | 3_prime_UTR_variant | MODIFIER |
| 2 | 89171929 | AX-97838886 | 0 | G | A | 0.053 | -1.01E-01 | 3.00E-01 | 2.74E+00 | 2.76E+00 | 7.33E-01 | no | Olfr1225 | upstream_gene_variant | MODIFIER |
| 2 | 89257890 | AX-170128113 | 0 | C | T | 0.053 | -1.01E-01 | 3.00E-01 | 2.74E+00 | 2.76E+00 | 7.33E-01 | no | Olfr1228 | upstream_gene_variant | MODIFIER |
| 2 | 89610770 | AX-97848783 | 0 | C | T | 0.053 | -1.01E-01 | 3.00E-01 | 2.74E+00 | 2.76E+00 | 7.33E-01 | no | Olfr1247 | upstream_gene_variant | MODIFIER |
| 2 | 89751782 | AX-170118216 | 0 | C | A | 0.053 | -1.01E-01 | 3.00E-01 | 2.74E+00 | 2.76E+00 | 7.33E-01 | no | Olfr1253 | downstream_gene_variant | MODIFIER |
| 2 | 89978637 | AX-97839177 | 0 | T | C | 0.053 | -1.01E-01 | 3.00E-01 | 2.74E+00 | 2.76E+00 | 7.33E-01 | no | Olfr1260 | synonymous_variant | LOW |
| 2 | 90000878 | AX-170206193 | 0 | T | A | 0.053 | -1.01E-01 | 3.00E-01 | 2.74E+00 | 2.76E+00 | 7.33E-01 | no | Olfr1262 | upstream_gene_variant | MODIFIER |
| 2 | 90015329 | AX-170018291 | 0 | G | A | 0.053 | -1.01E-01 | 3.00E-01 | 2.74E+00 | 2.76E+00 | 7.33E-01 | no | Olfr1263 | missense_variant | MODERATE |

| | | | | | | | | | | | | | | | |
|---|---|---|---|---|---|---|---|---|---|---|---|---|---|---|---|
| 2 | 90034279 | AX-97840950 | 0 | G | A | 0.053 | -1.01E-01 | 3.00E-01 | 2.74E+00 | 2.76E+00 | 7.33E-01 | no | Olfr1265 | upstream_gene_variant | MODIFIER |
| 2 | 90057527 | AX-170100714 | 0 | A | G | 0.053 | -1.01E-01 | 3.00E-01 | 2.74E+00 | 2.76E+00 | 7.33E-01 | no | Olfr140 | intron_variant | MODIFIER |
| 2 | 90063466 | AX-170018295 | 0 | C | T | 0.053 | -1.01E-01 | 3.00E-01 | 2.74E+00 | 2.76E+00 | 7.33E-01 | no | Olfr140 | 5_prime_UTR_variant | MODIFIER |
| 2 | 90083440 | AX-97838002 | 0 | A | G | 0.053 | -1.01E-01 | 3.00E-01 | 2.74E+00 | 2.76E+00 | 7.33E-01 | no | Olfr1267-ps1 | downstream_gene_variant | MODIFIER |
| 2 | 90111491 | AX-170319167 | 0 | G | T | 0.053 | -1.01E-01 | 3.00E-01 | 2.74E+00 | 2.76E+00 | 7.33E-01 | no | Olfr1268-ps1 | intron_variant | MODIFIER |
| 2 | 90266411 | AX-170064271 | 0 | A | G | 0.053 | -1.01E-01 | 3.00E-01 | 2.74E+00 | 2.76E+00 | 7.33E-01 | no | Olfr1271 | synonymous_variant | LOW |
| 2 | 90267377 | AX-97790348 | 0 | T | C | 0.053 | -1.01E-01 | 3.00E-01 | 2.74E+00 | 2.76E+00 | 7.33E-01 | no | Olfr1271 | upstream_gene_variant | MODIFIER |
| 2 | 90296836 | AX-97837965 | 0 | T | C | 0.053 | -1.01E-01 | 3.00E-01 | 2.74E+00 | 2.76E+00 | 7.33E-01 | no | Olfr1272 | upstream_gene_variant | MODIFIER |
| 2 | 90442646 | AX-170352227 | 0 | A | G | 0.053 | -1.01E-01 | 3.00E-01 | 2.74E+00 | 2.76E+00 | 7.33E-01 | no | Ptprj | intron_variant | MODIFIER |
| 2 | 90443965 | AX-170169252 | 0 | C | T | 0.053 | -1.01E-01 | 3.00E-01 | 2.74E+00 | 2.76E+00 | 7.33E-01 | no | Ptprj | intron_variant | MODIFIER |
| 2 | 90444049 | AX-170468363 | 0 | A | G | 0.053 | -1.01E-01 | 3.00E-01 | 2.74E+00 | 2.76E+00 | 7.33E-01 | no | Ptprj | intron_variant | MODIFIER |
| 2 | 90467880 | AX-170259027 | 0 | G | A | 0.053 | -1.01E-01 | 3.00E-01 | 2.74E+00 | 2.76E+00 | 7.33E-01 | no | Ptprj | intron_variant | MODIFIER |
| 2 | 90468050 | AX-170272465 | 0 | C | T | 0.053 | -1.01E-01 | 3.00E-01 | 2.74E+00 | 2.76E+00 | 7.33E-01 | no | Ptprj | intron_variant | MODIFIER |
| 2 | 90481670 | AX-97793757 | 0 | G | A | 0.053 | -1.01E-01 | 3.00E-01 | 2.74E+00 | 2.76E+00 | 7.33E-01 | no | Ptprj | upstream_gene_variant | MODIFIER |
| 2 | 90483516 | AX-170222206 | 0 | T | C | 0.053 | -1.01E-01 | 3.00E-01 | 2.74E+00 | 2.76E+00 | 7.33E-01 | no | Ptprj | upstream_gene_variant | MODIFIER |
| 2 | 90484351 | AX-170435200 | 0 | G | A | 0.053 | -1.01E-01 | 3.00E-01 | 2.74E+00 | 2.76E+00 | 7.33E-01 | no | 4933423P22Rik | non_coding_transcript_exon_variant | MODIFIER |
| 2 | 90515752 | AX-170064274 | 0 | C | T | 0.053 | -1.01E-01 | 3.00E-01 | 2.74E+00 | 2.76E+00 | 7.33E-01 | no | Ptprj | intron_variant | MODIFIER |
| 2 | 90518806 | AX-170191635 | 0 | A | G | 0.053 | -1.01E-01 | 3.00E-01 | 2.74E+00 | 2.76E+00 | 7.33E-01 | no | Ptprj | intron_variant | MODIFIER |
| 2 | 90523224 | AX-170240919 | 0 | T | G | 0.053 | -1.01E-01 | 3.00E-01 | 2.74E+00 | 2.76E+00 | 7.33E-01 | no | Ptprj | intron_variant | MODIFIER |
| 2 | 90523672 | AX-170411596 | 0 | T | C | 0.053 | -1.01E-01 | 3.00E-01 | 2.74E+00 | 2.76E+00 | 7.33E-01 | no | Ptprj | intron_variant | MODIFIER |
| 2 | 90546543 | AX-170204610 | 0 | G | C | 0.053 | -1.01E-01 | 3.00E-01 | 2.74E+00 | 2.76E+00 | 7.33E-01 | no | Ptprj | intron_variant | MODIFIER |
| 2 | 90548406 | AX-170146349 | 0 | G | A | 0.053 | -1.01E-01 | 3.00E-01 | 2.74E+00 | 2.76E+00 | 7.33E-01 | no | Ptprj | intron_variant | MODIFIER |
| 2 | 90548689 | AX-97843979 | 0 | C | T | 0.053 | -1.01E-01 | 3.00E-01 | 2.74E+00 | 2.76E+00 | 7.33E-01 | no | Ptprj | intron_variant | MODIFIER |
| 2 | 90550180 | AX-170268784 | 0 | T | A | 0.053 | -1.01E-01 | 3.00E-01 | 2.74E+00 | 2.76E+00 | 7.33E-01 | no | Ptprj | intron_variant | MODIFIER |
| 2 | 90552817 | AX-170257968 | 0 | G | A | 0.053 | -1.01E-01 | 3.00E-01 | 2.74E+00 | 2.76E+00 | 7.33E-01 | no | Ptprj | intron_variant | MODIFIER |
| 2 | 90553651 | AX-170331204 | 0 | A | G | 0.053 | -1.01E-01 | 3.00E-01 | 2.74E+00 | 2.76E+00 | 7.33E-01 | no | Ptprj | intron_variant | MODIFIER |
| 2 | 90554558 | AX-170341985 | 0 | A | G | 0.053 | -1.01E-01 | 3.00E-01 | 2.74E+00 | 2.76E+00 | 7.33E-01 | no | Ptprj | intron_variant | MODIFIER |
| 2 | 90559338 | AX-170258536 | 0 | C | G | 0.053 | -1.01E-01 | 3.00E-01 | 2.74E+00 | 2.76E+00 | 7.33E-01 | no | Ptprj | intron_variant | MODIFIER |
| 2 | 90562035 | AX-170393364 | 0 | T | C | 0.053 | -1.01E-01 | 3.00E-01 | 2.74E+00 | 2.76E+00 | 7.33E-01 | no | Ptprj | intron_variant | MODIFIER |
| 2 | 90613680 | AX-97838918 | 0 | G | T | 0.053 | -1.01E-01 | 3.00E-01 | 2.74E+00 | 2.76E+00 | 7.33E-01 | no | | intergenic_variant | MODIFIER |
| 2 | 90632050 | AX-170335343 | 0 | C | A | 0.053 | -1.01E-01 | 3.00E-01 | 2.74E+00 | 2.76E+00 | 7.33E-01 | no | | intergenic_variant | MODIFIER |
| 2 | 90632148 | AX-170295087 | 0 | T | C | 0.053 | -1.01E-01 | 3.00E-01 | 2.74E+00 | 2.76E+00 | 7.33E-01 | no | | intergenic_variant | MODIFIER |
| 2 | 90637672 | AX-170506422 | 0 | A | G | 0.053 | -1.01E-01 | 3.00E-01 | 2.74E+00 | 2.76E+00 | 7.33E-01 | no | | intergenic_variant | MODIFIER |
| 2 | 90768607 | AX-170064277 | 0 | C | A | 0.053 | -1.01E-01 | 3.00E-01 | 2.74E+00 | 2.76E+00 | 7.33E-01 | no | Fnbp4 | missense_variant | MODERATE |
| 2 | 90777959 | AX-97794795 | 0 | A | G | 0.053 | -1.01E-01 | 3.00E-01 | 2.74E+00 | 2.76E+00 | 7.33E-01 | no | Fnbp4 | intron_variant | MODIFIER |
| 2 | 90802800 | AX-170108401 | 0 | G | A | 0.053 | -1.01E-01 | 3.00E-01 | 2.74E+00 | 2.76E+00 | 7.33E-01 | no | Agbl2 | intron_variant | MODIFIER |
| 2 | 90820505 | AX-170088660 | 0 | G | A | 0.053 | -1.01E-01 | 3.00E-01 | 2.74E+00 | 2.76E+00 | 7.33E-01 | no | Agbl2 | downstream_gene_variant | MODIFIER |
| 2 | 90839710 | AX-170359733 | 0 | A | G | 0.053 | -1.01E-01 | 3.00E-01 | 2.74E+00 | 2.76E+00 | 7.33E-01 | no | | intergenic_variant | MODIFIER |
| 2 | 90860960 | AX-170414286 | 0 | T | C | 0.053 | -1.01E-01 | 3.00E-01 | 2.74E+00 | 2.76E+00 | 7.33E-01 | no | Mtch2 | intron_variant,non_coding_transcript_variant | MODIFIER |
| 2 | 90861014 | AX-170333466 | 0 | G | T | 0.053 | -1.01E-01 | 3.00E-01 | 2.74E+00 | 2.76E+00 | 7.33E-01 | no | Mtch2 | intron_variant,non_coding_transcript_variant | MODIFIER |
| 2 | 90863538 | AX-170491725 | 0 | T | G | 0.053 | -1.01E-01 | 3.00E-01 | 2.74E+00 | 2.76E+00 | 7.33E-01 | no | Mtch2 | intron_variant,non_coding_transcript_variant | MODIFIER |
| 2 | 90866746 | AX-170278019 | 0 | T | G | 0.053 | -1.01E-01 | 3.00E-01 | 2.74E+00 | 2.76E+00 | 7.33E-01 | no | Mtch2 | downstream_gene_variant | MODIFIER |
| 2 | 90870425 | AX-170121422 | 0 | G | A | 0.053 | -1.01E-01 | 3.00E-01 | 2.74E+00 | 2.76E+00 | 7.33E-01 | no | Mtch2 | downstream_gene_variant | MODIFIER |
| 2 | 90870976 | AX-170469102 | 0 | T | C | 0.053 | -1.01E-01 | 3.00E-01 | 2.74E+00 | 2.76E+00 | 7.33E-01 | no | Mtch2 | downstream_gene_variant | MODIFIER |
| 2 | 90928899 | AX-170064279 | 0 | G | A | 0.053 | -1.01E-01 | 3.00E-01 | 2.74E+00 | 2.76E+00 | 7.33E-01 | no | | intergenic_variant | MODIFIER |
| 2 | 90932632 | AX-170064280 | 0 | C | T | 0.053 | -1.01E-01 | 3.00E-01 | 2.74E+00 | 2.76E+00 | 7.33E-01 | no | | intergenic_variant | MODIFIER |
| 2 | 90944271 | AX-170340695 | 0 | G | T | 0.053 | -1.01E-01 | 3.00E-01 | 2.74E+00 | 2.76E+00 | 7.33E-01 | no | Celf1 | intron_variant | MODIFIER |
| 2 | 91021224 | AX-170109836 | 0 | G | A | 0.053 | -1.01E-01 | 3.00E-01 | 2.74E+00 | 2.76E+00 | 7.33E-01 | no | Celf1 | downstream_gene_variant | MODIFIER |
| 2 | 91059677 | AX-97818636 | 0 | A | G | 0.053 | -1.01E-01 | 3.00E-01 | 2.74E+00 | 2.76E+00 | 7.33E-01 | no | Psmc3 | intron_variant | MODIFIER |
| 2 | 91060132 | AX-170383580 | 0 | G | A | 0.053 | -1.01E-01 | 3.00E-01 | 2.74E+00 | 2.76E+00 | 7.33E-01 | no | Psmc3 | intron_variant | MODIFIER |
| 2 | 91085930 | AX-170124487 | 0 | C | G | 0.053 | -1.01E-01 | 3.00E-01 | 2.74E+00 | 2.76E+00 | 7.33E-01 | no | Spi1 | intron_variant,non_coding_transcript_variant | MODIFIER |
| 2 | 91086352 | AX-170270203 | 0 | G | C | 0.053 | -1.01E-01 | 3.00E-01 | 2.74E+00 | 2.76E+00 | 7.33E-01 | no | Spi1 | intron_variant,non_coding_transcript_variant | MODIFIER |
| 2 | 91087223 | AX-170094327 | 0 | G | A | 0.053 | -1.01E-01 | 3.00E-01 | 2.74E+00 | 2.76E+00 | 7.33E-01 | no | Spi1 | intron_variant,non_coding_transcript_variant | MODIFIER |
| 2 | 91113202 | AX-170353137 | 0 | C | A | 0.053 | -1.01E-01 | 3.00E-01 | 2.74E+00 | 2.76E+00 | 7.33E-01 | no | Spi1 | intron_variant | MODIFIER |

| | | | | | | | | | | | | | | | |
|---|---|---|---|---|---|---|---|---|---|---|---|---|---|---|---|
| 2 | 91114231 | AX-170174111 | 0 | C | T | 0.053 | -1.01E-01 | 3.00E-01 | 2.74E+00 | 2.76E+00 | 7.33E-01 | no | Spi1 | intron_variant | MODIFIER |
| 2 | 91122150 | AX-170452454 | 0 | C | T | 0.053 | -1.01E-01 | 3.00E-01 | 2.74E+00 | 2.76E+00 | 7.33E-01 | no | Mybpc3 | intron_variant | MODIFIER |
| 2 | 91136790 | AX-97836662 | 0 | G | A | 0.053 | -1.01E-01 | 3.00E-01 | 2.74E+00 | 2.76E+00 | 7.33E-01 | no | Madd | downstream_gene_variant | MODIFIER |
| 2 | 91176115 | AX-97791399 | 0 | A | G | 0.053 | -1.01E-01 | 3.00E-01 | 2.74E+00 | 2.76E+00 | 7.33E-01 | no | Madd | intron_variant | MODIFIER |
| 2 | 91181295 | AX-170018306 | 0 | T | C | 0.053 | -1.01E-01 | 3.00E-01 | 2.74E+00 | 2.76E+00 | 7.33E-01 | no | Nr1h3 | downstream_gene_variant | MODIFIER |
| 2 | 91190465 | AX-170018313 | 0 | A | G | 0.053 | -1.01E-01 | 3.00E-01 | 2.74E+00 | 2.76E+00 | 7.33E-01 | no | Nr1h3 | synonymous_variant | LOW |
| 2 | 91212157 | AX-97838915 | 0 | G | A | 0.053 | -1.01E-01 | 3.00E-01 | 2.74E+00 | 2.76E+00 | 7.33E-01 | no | Acp2 | 3_prime_UTR_variant | MODIFIER |
| 2 | 91271304 | AX-170018308 | 0 | T | C | 0.053 | -1.01E-01 | 3.00E-01 | 2.74E+00 | 2.76E+00 | 7.33E-01 | no | Arfgap2 | intron_variant | MODIFIER |
| 2 | 91275142 | AX-97826460 | 0 | G | A | 0.053 | -1.01E-01 | 3.00E-01 | 2.74E+00 | 2.76E+00 | 7.33E-01 | no | Arfgap2 | missense_variant | MODERATE |
| 2 | 91299661 | AX-170018315 | 0 | A | G | 0.053 | -1.01E-01 | 3.00E-01 | 2.74E+00 | 2.76E+00 | 7.33E-01 | no | 1110051M20Rik | intron_variant | MODIFIER |
| 2 | 91317933 | AX-97814157 | 0 | C | T | 0.053 | -1.01E-01 | 3.00E-01 | 2.74E+00 | 2.76E+00 | 7.33E-01 | no | 1110051M20Rik | intron_variant | MODIFIER |
| 2 | 91322235 | AX-170018305 | 0 | C | T | 0.053 | -1.01E-01 | 3.00E-01 | 2.74E+00 | 2.76E+00 | 7.33E-01 | no | 1110051M20Rik | intron_variant | MODIFIER |
| 2 | 91327897 | AX-170018312 | 0 | G | C | 0.053 | -1.01E-01 | 3.00E-01 | 2.74E+00 | 2.76E+00 | 7.33E-01 | no | 1110051M20Rik | intron_variant | MODIFIER |
| 2 | 91370509 | AX-170018309 | 0 | C | T | 0.053 | -1.01E-01 | 3.00E-01 | 2.74E+00 | 2.76E+00 | 7.33E-01 | no | 1110051M20Rik | intron_variant | MODIFIER |
| 2 | 91373013 | AX-97849249 | 0 | C | T | 0.053 | -1.01E-01 | 3.00E-01 | 2.74E+00 | 2.76E+00 | 7.33E-01 | no | 1110051M20Rik | intron_variant | MODIFIER |
| 2 | 91389920 | AX-170414355 | 0 | T | C | 0.053 | -1.01E-01 | 3.00E-01 | 2.74E+00 | 2.76E+00 | 7.33E-01 | no | 1110051M20Rik | intron_variant | MODIFIER |
| 2 | 91399200 | AX-170212918 | 0 | G | C | 0.053 | -1.01E-01 | 3.00E-01 | 2.74E+00 | 2.76E+00 | 7.33E-01 | no | 1110051M20Rik | intron_variant | MODIFIER |
| 2 | 91420000 | AX-170294369 | 0 | T | C | 0.053 | -1.01E-01 | 3.00E-01 | 2.74E+00 | 2.76E+00 | 7.33E-01 | no | 1110051M20Rik | intron_variant | MODIFIER |
| 2 | 91475050 | AX-170181100 | 0 | T | G | 0.053 | -1.01E-01 | 3.00E-01 | 2.74E+00 | 2.76E+00 | 7.33E-01 | no | Lrp4 | intron_variant | MODIFIER |
| 2 | 91495040 | AX-170432987 | 0 | C | T | 0.053 | -1.01E-01 | 3.00E-01 | 2.74E+00 | 2.76E+00 | 7.33E-01 | no | Lrp4 | synonymous_variant | LOW |
| 2 | 91498910 | AX-97804390 | 0 | G | A | 0.053 | -1.01E-01 | 3.00E-01 | 2.74E+00 | 2.76E+00 | 7.33E-01 | no | Lrp4 | intron_variant | MODIFIER |
| 2 | 91630472 | AX-170528612 | 0 | G | A | 0.053 | -1.01E-01 | 3.00E-01 | 2.74E+00 | 2.76E+00 | 7.33E-01 | no | F2 | intron_variant | MODIFIER |
| 2 | 91709815 | AX-170088663 | 0 | G | A | 0.053 | -1.01E-01 | 3.00E-01 | 2.74E+00 | 2.76E+00 | 7.33E-01 | no | Atg13 | intron_variant | MODIFIER |
| 2 | 91794796 | AX-170018327 | 0 | G | A | 0.053 | -1.01E-01 | 3.00E-01 | 2.74E+00 | 2.76E+00 | 7.33E-01 | no | Ambra1 | intron_variant | MODIFIER |
| 2 | 91849874 | AX-170018329 | 0 | C | T | 0.053 | -1.01E-01 | 3.00E-01 | 2.74E+00 | 2.76E+00 | 7.33E-01 | no | Ambra1 | intron_variant | MODIFIER |
| 2 | 91853666 | AX-170350062 | 0 | A | G | 0.053 | -1.01E-01 | 3.00E-01 | 2.74E+00 | 2.76E+00 | 7.33E-01 | no | Ambra1 | intron_variant | MODIFIER |
| 2 | 91857652 | AX-97838064 | 0 | G | A | 0.053 | -1.01E-01 | 3.00E-01 | 2.74E+00 | 2.76E+00 | 7.33E-01 | no | Ambra1 | intron_variant | MODIFIER |
| 2 | 91899787 | AX-97814760 | 0 | C | T | 0.053 | -1.01E-01 | 3.00E-01 | 2.74E+00 | 2.76E+00 | 7.33E-01 | no | Ambra1 | intron_variant | MODIFIER |
| 2 | 91903052 | AX-170400422 | 0 | C | G | 0.053 | -1.01E-01 | 3.00E-01 | 2.74E+00 | 2.76E+00 | 7.33E-01 | no | Ambra1 | intron_variant | MODIFIER |
| 2 | 91919965 | AX-97786599 | 0 | T | C | 0.053 | -1.01E-01 | 3.00E-01 | 2.74E+00 | 2.76E+00 | 7.33E-01 | no | Ambra1 | downstream_gene_variant | MODIFIER |
| 2 | 91976911 | AX-97838016 | 0 | G | C | 0.053 | -1.01E-01 | 3.00E-01 | 2.74E+00 | 2.76E+00 | 7.33E-01 | no | Dgkz | upstream_gene_variant | MODIFIER |
| 2 | 92015853 | AX-170352088 | 0 | A | T | 0.053 | -1.01E-01 | 3.00E-01 | 2.74E+00 | 2.76E+00 | 7.33E-01 | no | Creb3l1 | intron_variant | MODIFIER |
| 2 | 92018052 | AX-170018330 | 0 | C | T | 0.053 | -1.01E-01 | 3.00E-01 | 2.74E+00 | 2.76E+00 | 7.33E-01 | no | Creb3l1 | intron_variant | MODIFIER |
| 2 | 92020480 | AX-170088665 | 0 | T | C | 0.053 | -1.01E-01 | 3.00E-01 | 2.74E+00 | 2.76E+00 | 7.33E-01 | no | Creb3l1 | intron_variant | MODIFIER |
| 2 | 92030041 | AX-170103944 | 0 | G | A | 0.053 | -1.01E-01 | 3.00E-01 | 2.74E+00 | 2.76E+00 | 7.33E-01 | no | | regulatory_region_variant | MODIFIER |
| 2 | 92036275 | AX-170155989 | 0 | C | T | 0.053 | -1.01E-01 | 3.00E-01 | 2.74E+00 | 2.76E+00 | 7.33E-01 | no | | intergenic_variant | MODIFIER |
| 2 | 92037097 | AX-170374899 | 0 | T | A | 0.053 | -1.01E-01 | 3.00E-01 | 2.74E+00 | 2.76E+00 | 7.33E-01 | no | | intergenic_variant | MODIFIER |
| 2 | 92037168 | AX-170329226 | 0 | C | T | 0.053 | -1.01E-01 | 3.00E-01 | 2.74E+00 | 2.76E+00 | 7.33E-01 | no | | intergenic_variant | MODIFIER |
| 2 | 92051218 | AX-170347912 | 0 | A | G | 0.053 | -1.01E-01 | 3.00E-01 | 2.74E+00 | 2.76E+00 | 7.33E-01 | no | 2900072N19Rik | downstream_gene_variant | MODIFIER |
| 2 | 92054877 | AX-170240638 | 0 | A | G | 0.053 | -1.01E-01 | 3.00E-01 | 2.74E+00 | 2.76E+00 | 7.33E-01 | no | 2900072N19Rik | downstream_gene_variant | MODIFIER |
| 2 | 92058733 | AX-170018323 | 0 | G | C | 0.053 | -1.01E-01 | 3.00E-01 | 2.74E+00 | 2.76E+00 | 7.33E-01 | no | 2900072N19Rik | intron_variant,non_coding_transcript_variant | MODIFIER |
| 2 | 92064145 | AX-170214979 | 0 | T | C | 0.053 | -1.01E-01 | 3.00E-01 | 2.74E+00 | 2.76E+00 | 7.33E-01 | no | 2900072N19Rik | upstream_gene_variant | MODIFIER |
| 2 | 92073134 | AX-170207487 | 0 | A | T | 0.053 | -1.01E-01 | 3.00E-01 | 2.74E+00 | 2.76E+00 | 7.33E-01 | no | Gm13780 | intron_variant,non_coding_transcript_variant | MODIFIER |
| 2 | 92078981 | AX-170114298 | 0 | A | G | 0.053 | -1.01E-01 | 3.00E-01 | 2.74E+00 | 2.76E+00 | 7.33E-01 | no | Gm13780 | intron_variant,non_coding_transcript_variant | MODIFIER |
| 2 | 92087067 | AX-170495518 | 0 | T | C | 0.053 | -1.01E-01 | 3.00E-01 | 2.74E+00 | 2.76E+00 | 7.33E-01 | no | Gm13780 | intron_variant,non_coding_transcript_variant | MODIFIER |
| 2 | 92090332 | AX-170088666 | 0 | C | A | 0.053 | -1.01E-01 | 3.00E-01 | 2.74E+00 | 2.76E+00 | 7.33E-01 | no | Phf21a | upstream_gene_variant | MODIFIER |
| 2 | 92092379 | AX-170018331 | 0 | A | G | 0.053 | -1.01E-01 | 3.00E-01 | 2.74E+00 | 2.76E+00 | 7.33E-01 | no | Phf21a | upstream_gene_variant | MODIFIER |
| 2 | 92138725 | AX-170406678 | 0 | C | T | 0.053 | -1.01E-01 | 3.00E-01 | 2.74E+00 | 2.76E+00 | 7.33E-01 | no | Phf21a | intron_variant | MODIFIER |
| 2 | 92147975 | AX-97838035 | 0 | A | G | 0.053 | -1.01E-01 | 3.00E-01 | 2.74E+00 | 2.76E+00 | 7.33E-01 | no | Phf21a | intron_variant | MODIFIER |
| 2 | 92165964 | AX-170256359 | 0 | T | C | 0.053 | -1.01E-01 | 3.00E-01 | 2.74E+00 | 2.76E+00 | 7.33E-01 | no | Phf21a | intron_variant | MODIFIER |
| 2 | 92171010 | AX-170115083 | 0 | G | C | 0.053 | -1.01E-01 | 3.00E-01 | 2.74E+00 | 2.76E+00 | 7.33E-01 | no | Phf21a | intron_variant | MODIFIER |
| 2 | 92174119 | AX-97852969 | 0 | G | A | 0.053 | -1.01E-01 | 3.00E-01 | 2.74E+00 | 2.76E+00 | 7.33E-01 | no | Phf21a | intron_variant | MODIFIER |
| 2 | 92176288 | AX-170018321 | 0 | T | C | 0.053 | -1.01E-01 | 3.00E-01 | 2.74E+00 | 2.76E+00 | 7.33E-01 | no | Phf21a | intron_variant | MODIFIER |
| 2 | 92221674 | AX-170473783 | 0 | T | C | 0.053 | -1.01E-01 | 3.00E-01 | 2.74E+00 | 2.76E+00 | 7.33E-01 | no | Phf21a | intron_variant | MODIFIER |

| | | | | | | | | | | | | | | | |
|---|---|---|---|---|---|---|---|---|---|---|---|---|---|---|---|
| 2 | 92239433 | AX-170064289 | 0 | C | T | 0.053 | -1.01E-01 | 3.00E-01 | 2.74E+00 | 2.76E+00 | 7.33E-01 | no | Phf21a | intron_variant | MODIFIER |
| 2 | 92253544 | AX-170018340 | 0 | C | T | 0.053 | -1.01E-01 | 3.00E-01 | 2.74E+00 | 2.76E+00 | 7.33E-01 | no | Phf21a | intron_variant | MODIFIER |
| 2 | 92288292 | AX-170094330 | 0 | C | T | 0.053 | -1.01E-01 | 3.00E-01 | 2.74E+00 | 2.76E+00 | 7.33E-01 | no | Phf21a | intron_variant | MODIFIER |
| 2 | 92288338 | AX-170496705 | 0 | C | G | 0.053 | -1.01E-01 | 3.00E-01 | 2.74E+00 | 2.76E+00 | 7.33E-01 | no | Phf21a | intron_variant | MODIFIER |
| 2 | 92316730 | AX-170018335 | 0 | G | A | 0.053 | -1.01E-01 | 3.00E-01 | 2.74E+00 | 2.76E+00 | 7.33E-01 | no | Phf21a | intron_variant | MODIFIER |
| 2 | 92331271 | AX-97821655 | 0 | A | G | 0.053 | -1.01E-01 | 3.00E-01 | 2.74E+00 | 2.76E+00 | 7.33E-01 | no | Phf21a | intron_variant | MODIFIER |
| 2 | 92351941 | AX-97857235 | 0 | G | A | 0.053 | -1.01E-01 | 3.00E-01 | 2.74E+00 | 2.76E+00 | 7.33E-01 | no | Phf21a | intron_variant | MODIFIER |
| 2 | 92388497 | AX-170277858 | 0 | T | C | 0.053 | -1.01E-01 | 3.00E-01 | 2.74E+00 | 2.76E+00 | 7.33E-01 | no | Mapk8ip1 | intron_variant | MODIFIER |
| 2 | 92399529 | AX-97806563 | 0 | A | G | 0.053 | -1.01E-01 | 3.00E-01 | 2.74E+00 | 2.76E+00 | 7.33E-01 | no | Mapk8ip1 | intron_variant | MODIFIER |
| 2 | 92454070 | AX-97833644 | 0 | C | T | 0.053 | -1.01E-01 | 3.00E-01 | 2.74E+00 | 2.76E+00 | 7.33E-01 | no | Slc35c1 | 3_prime_UTR_variant | MODIFIER |
| 2 | 92487524 | AX-97831301 | 0 | T | G | 0.053 | -1.01E-01 | 3.00E-01 | 2.74E+00 | 2.76E+00 | 7.33E-01 | no | Gm13814 | intron_variant,non_coding_transcript_variant | MODIFIER |
| 2 | 92493356 | AX-170402739 | 0 | A | G | 0.053 | -1.01E-01 | 3.00E-01 | 2.74E+00 | 2.76E+00 | 7.33E-01 | no | Gm13814 | intron_variant,non_coding_transcript_variant | MODIFIER |
| 2 | 92494869 | AX-170328696 | 0 | T | C | 0.053 | -1.01E-01 | 3.00E-01 | 2.74E+00 | 2.76E+00 | 7.33E-01 | no | Gm13814 | intron_variant,non_coding_transcript_variant | MODIFIER |
| 2 | 92497923 | AX-170453615 | 0 | T | C | 0.053 | -1.01E-01 | 3.00E-01 | 2.74E+00 | 2.76E+00 | 7.33E-01 | no | Gm13814 | intron_variant,non_coding_transcript_variant | MODIFIER |
| 2 | 92500889 | AX-170358501 | 0 | T | C | 0.053 | -1.01E-01 | 3.00E-01 | 2.74E+00 | 2.76E+00 | 7.33E-01 | no | Gm13814 | intron_variant,non_coding_transcript_variant | MODIFIER |
| 2 | 92566139 | AX-97840395 | 0 | G | A | 0.053 | -1.01E-01 | 3.00E-01 | 2.74E+00 | 2.76E+00 | 7.33E-01 | no | Gm13815 | downstream_gene_variant | MODIFIER |
| 2 | 92584955 | AX-170470991 | 0 | T | C | 0.053 | -1.01E-01 | 3.00E-01 | 2.74E+00 | 2.76E+00 | 7.33E-01 | no | | intergenic_variant | MODIFIER |
| 2 | 92589058 | AX-170258379 | 0 | G | A | 0.053 | -1.01E-01 | 3.00E-01 | 2.74E+00 | 2.76E+00 | 7.33E-01 | no | Mir7221 | upstream_gene_variant | MODIFIER |
| 2 | 92618112 | AX-170495373 | 0 | T | C | 0.053 | -1.01E-01 | 3.00E-01 | 2.74E+00 | 2.76E+00 | 7.33E-01 | no | Chst1 | downstream_gene_variant | MODIFIER |
| 2 | 92618158 | AX-97850886 | 0 | T | C | 0.053 | -1.01E-01 | 3.00E-01 | 2.74E+00 | 2.76E+00 | 7.33E-01 | no | Chst1 | downstream_gene_variant | MODIFIER |
| 2 | 92653803 | AX-170064300 | 0 | T | C | 0.053 | -1.01E-01 | 3.00E-01 | 2.74E+00 | 2.76E+00 | 7.33E-01 | no | | intergenic_variant | MODIFIER |
| 2 | 92758858 | AX-170365955 | 0 | G | A | 0.053 | -1.01E-01 | 3.00E-01 | 2.74E+00 | 2.76E+00 | 7.33E-01 | no | Gm13816 | intron_variant,non_coding_transcript_variant | MODIFIER |
| 2 | 92759584 | AX-170175136 | 0 | A | G | 0.053 | -1.01E-01 | 3.00E-01 | 2.74E+00 | 2.76E+00 | 7.33E-01 | no | Gm13816 | downstream_gene_variant | MODIFIER |
| 2 | 92761881 | AX-97851009 | 0 | T | C | 0.053 | -1.01E-01 | 3.00E-01 | 2.74E+00 | 2.76E+00 | 7.33E-01 | no | Gm13816 | downstream_gene_variant | MODIFIER |
| 2 | 92761904 | AX-170194474 | 0 | G | A | 0.053 | -1.01E-01 | 3.00E-01 | 2.74E+00 | 2.76E+00 | 7.33E-01 | no | Gm13816 | downstream_gene_variant | MODIFIER |
| 2 | 92764526 | AX-170172077 | 0 | T | A | 0.053 | -1.01E-01 | 3.00E-01 | 2.74E+00 | 2.76E+00 | 7.33E-01 | no | | intergenic_variant | MODIFIER |
| 2 | 92764641 | AX-170064304 | 0 | T | C | 0.053 | -1.01E-01 | 3.00E-01 | 2.74E+00 | 2.76E+00 | 7.33E-01 | no | | intergenic_variant | MODIFIER |
| 2 | 92764897 | AX-170064302 | 0 | G | A | 0.053 | -1.01E-01 | 3.00E-01 | 2.74E+00 | 2.76E+00 | 7.33E-01 | no | | intergenic_variant | MODIFIER |
| 2 | 92795876 | AX-170425710 | 0 | A | G | 0.053 | -1.01E-01 | 3.00E-01 | 2.74E+00 | 2.76E+00 | 7.33E-01 | no | | intergenic_variant | MODIFIER |
| 2 | 92795999 | AX-170258058 | 0 | C | T | 0.053 | -1.01E-01 | 3.00E-01 | 2.74E+00 | 2.76E+00 | 7.33E-01 | no | | intergenic_variant | MODIFIER |
| 2 | 92796245 | AX-97807287 | 0 | A | G | 0.053 | -1.01E-01 | 3.00E-01 | 2.74E+00 | 2.76E+00 | 7.33E-01 | no | | intergenic_variant | MODIFIER |
| 2 | 92799135 | AX-170442711 | 0 | T | A | 0.053 | -1.01E-01 | 3.00E-01 | 2.74E+00 | 2.76E+00 | 7.33E-01 | no | | intergenic_variant | MODIFIER |
| 2 | 92799205 | AX-170154693 | 0 | G | A | 0.053 | -1.01E-01 | 3.00E-01 | 2.74E+00 | 2.76E+00 | 7.33E-01 | no | | intergenic_variant | MODIFIER |
| 2 | 92812224 | AX-170018347 | 0 | T | C | 0.053 | -1.01E-01 | 3.00E-01 | 2.74E+00 | 2.76E+00 | 7.33E-01 | no | | intergenic_variant | MODIFIER |
| 2 | 92813655 | AX-170018341 | 0 | C | T | 0.053 | -1.01E-01 | 3.00E-01 | 2.74E+00 | 2.76E+00 | 7.33E-01 | no | | regulatory_region_variant | MODIFIER |
| 2 | 92824934 | AX-170184408 | 0 | A | G | 0.053 | -1.01E-01 | 3.00E-01 | 2.74E+00 | 2.76E+00 | 7.33E-01 | no | | regulatory_region_variant | MODIFIER |
| 2 | 92826811 | AX-170018344 | 0 | G | A | 0.053 | -1.01E-01 | 3.00E-01 | 2.74E+00 | 2.76E+00 | 7.33E-01 | no | | intergenic_variant | MODIFIER |
| 2 | 92832090 | AX-170467311 | 0 | C | T | 0.053 | -1.01E-01 | 3.00E-01 | 2.74E+00 | 2.76E+00 | 7.33E-01 | no | | intergenic_variant | MODIFIER |
| 2 | 92869022 | AX-170018345 | 0 | A | G | 0.053 | -1.01E-01 | 3.00E-01 | 2.74E+00 | 2.76E+00 | 7.33E-01 | no | Gm13792 | downstream_gene_variant | MODIFIER |
| 2 | 92869138 | AX-170018346 | 0 | A | G | 0.053 | -1.01E-01 | 3.00E-01 | 2.74E+00 | 2.76E+00 | 7.33E-01 | no | Gm13792 | downstream_gene_variant | MODIFIER |
| 2 | 92933563 | AX-170466347 | 0 | C | T | 0.053 | -1.01E-01 | 3.00E-01 | 2.74E+00 | 2.76E+00 | 7.33E-01 | no | Syt13 | intron_variant | MODIFIER |
| 2 | 92954889 | AX-97838039 | 0 | C | T | 0.053 | -1.01E-01 | 3.00E-01 | 2.74E+00 | 2.76E+00 | 7.33E-01 | no | Syt13 | 3_prime_UTR_variant | MODIFIER |
| 2 | 92968264 | AX-170111476 | 0 | A | G | 0.053 | -1.01E-01 | 3.00E-01 | 2.74E+00 | 2.76E+00 | 7.33E-01 | no | Prdm11 | 3_prime_UTR_variant | MODIFIER |
| 2 | 92968850 | AX-170506651 | 0 | A | G | 0.053 | -1.01E-01 | 3.00E-01 | 2.74E+00 | 2.76E+00 | 7.33E-01 | no | Prdm11 | 3_prime_UTR_variant | MODIFIER |
| 2 | 92969058 | AX-170229139 | 0 | A | C | 0.053 | -1.01E-01 | 3.00E-01 | 2.74E+00 | 2.76E+00 | 7.33E-01 | no | Prdm11 | 3_prime_UTR_variant | MODIFIER |
| 2 | 92974142 | AX-170141460 | 0 | A | G | 0.053 | -1.01E-01 | 3.00E-01 | 2.74E+00 | 2.76E+00 | 7.33E-01 | no | Prdm11 | 3_prime_UTR_variant | MODIFIER |
| 2 | 92974665 | AX-170112394 | 0 | G | A | 0.053 | -1.01E-01 | 3.00E-01 | 2.74E+00 | 2.76E+00 | 7.33E-01 | no | Prdm11 | 3_prime_UTR_variant | MODIFIER |
| 2 | 92978912 | AX-170364955 | 0 | C | A | 0.053 | -1.01E-01 | 3.00E-01 | 2.74E+00 | 2.76E+00 | 7.33E-01 | no | Prdm11 | intron_variant | MODIFIER |
| 2 | 92979049 | AX-170302760 | 0 | T | C | 0.053 | -1.01E-01 | 3.00E-01 | 2.74E+00 | 2.76E+00 | 7.33E-01 | no | Prdm11 | intron_variant | MODIFIER |
| 2 | 92979370 | AX-170292601 | 0 | C | T | 0.053 | -1.01E-01 | 3.00E-01 | 2.74E+00 | 2.76E+00 | 7.33E-01 | no | Prdm11 | intron_variant | MODIFIER |
| 2 | 92979667 | AX-170469842 | 0 | A | G | 0.053 | -1.01E-01 | 3.00E-01 | 2.74E+00 | 2.76E+00 | 7.33E-01 | no | Prdm11 | intron_variant | MODIFIER |
| 2 | 92983380 | AX-170466780 | 0 | T | C | 0.053 | -1.01E-01 | 3.00E-01 | 2.74E+00 | 2.76E+00 | 7.33E-01 | no | Prdm11 | downstream_gene_variant | MODIFIER |
| 2 | 92984948 | AX-97795061 | 0 | A | G | 0.053 | -1.01E-01 | 3.00E-01 | 2.74E+00 | 2.76E+00 | 7.33E-01 | no | Prdm11 | 3_prime_UTR_variant | MODIFIER |
| 2 | 92985288 | AX-170360970 | 0 | A | C | 0.053 | -1.01E-01 | 3.00E-01 | 2.74E+00 | 2.76E+00 | 7.33E-01 | no | Prdm11 | 3_prime_UTR_variant | MODIFIER |

| | | | | | | | | | | | | | | | |
|---|---|---|---|---|---|---|---|---|---|---|---|---|---|---|---|
| 2 | 92998929 | AX-170139595 | 0 | G | A | 0.053 | -1.01E-01 | 3.00E-01 | 2.74E+00 | 2.76E+00 | 7.33E-01 | no | Prdm11 | intron_variant | MODIFIER |
| 2 | 93021231 | AX-170346743 | 0 | A | G | 0.053 | -1.01E-01 | 3.00E-01 | 2.74E+00 | 2.76E+00 | 7.33E-01 | no | Prdm11 | intron_variant | MODIFIER |
| 2 | 93021335 | AX-170463953 | 0 | G | A | 0.053 | -1.01E-01 | 3.00E-01 | 2.74E+00 | 2.76E+00 | 7.33E-01 | no | Prdm11 | intron_variant | MODIFIER |
| 2 | 93023954 | AX-170534496 | 0 | C | T | 0.053 | -1.01E-01 | 3.00E-01 | 2.74E+00 | 2.76E+00 | 7.33E-01 | no | 4631405J19Rik | upstream_gene_variant | MODIFIER |
| 2 | 93026268 | AX-170261920 | 0 | C | G | 0.053 | -1.01E-01 | 3.00E-01 | 2.74E+00 | 2.76E+00 | 7.33E-01 | no | 4631405J19Rik | upstream_gene_variant | MODIFIER |
| 2 | 93028557 | AX-170247184 | 0 | A | G | 0.053 | -1.01E-01 | 3.00E-01 | 2.74E+00 | 2.76E+00 | 7.33E-01 | no | 4631405J19Rik | upstream_gene_variant | MODIFIER |
| 2 | 93053147 | AX-170523586 | 0 | T | G | 0.053 | -1.01E-01 | 3.00E-01 | 2.74E+00 | 2.76E+00 | 7.33E-01 | no | | intergenic_variant | MODIFIER |
| 2 | 93054875 | AX-97849071 | 0 | G | A | 0.053 | -1.01E-01 | 3.00E-01 | 2.74E+00 | 2.76E+00 | 7.33E-01 | no | | intergenic_variant | MODIFIER |
| 2 | 93065335 | AX-170201956 | 0 | T | C | 0.053 | -1.01E-01 | 3.00E-01 | 2.74E+00 | 2.76E+00 | 7.33E-01 | no | | regulatory_region_variant | MODIFIER |
| 2 | 93066233 | AX-170353382 | 0 | T | G | 0.053 | -1.01E-01 | 3.00E-01 | 2.74E+00 | 2.76E+00 | 7.33E-01 | no | | intergenic_variant | MODIFIER |
| 2 | 93100354 | AX-170319931 | 0 | T | C | 0.053 | -1.01E-01 | 3.00E-01 | 2.74E+00 | 2.76E+00 | 7.33E-01 | no | Gm13802 | upstream_gene_variant | MODIFIER |
| 2 | 93114093 | AX-170354450 | 0 | G | C | 0.053 | -1.01E-01 | 3.00E-01 | 2.74E+00 | 2.76E+00 | 7.33E-01 | no | | regulatory_region_variant | MODIFIER |
| 2 | 93141060 | AX-97838038 | 0 | A | G | 0.053 | -1.01E-01 | 3.00E-01 | 2.74E+00 | 2.76E+00 | 7.33E-01 | no | | regulatory_region_variant | MODIFIER |
| 2 | 93157093 | AX-170460585 | 0 | A | G | 0.053 | -1.01E-01 | 3.00E-01 | 2.74E+00 | 2.76E+00 | 7.33E-01 | no | | intergenic_variant | MODIFIER |
| 2 | 93168142 | AX-170348431 | 0 | C | T | 0.053 | -1.01E-01 | 3.00E-01 | 2.74E+00 | 2.76E+00 | 7.33E-01 | no | | intergenic_variant | MODIFIER |
| 2 | 93175494 | AX-97838082 | 0 | A | G | 0.053 | -1.01E-01 | 3.00E-01 | 2.74E+00 | 2.76E+00 | 7.33E-01 | no | | regulatory_region_variant | MODIFIER |
| 2 | 93192570 | AX-170233075 | 0 | C | T | 0.053 | -1.01E-01 | 3.00E-01 | 2.74E+00 | 2.76E+00 | 7.33E-01 | no | Trp53i11 | intron_variant | MODIFIER |
| 2 | 93245707 | AX-170427396 | 0 | T | G | 0.053 | -1.01E-01 | 3.00E-01 | 2.74E+00 | 2.76E+00 | 7.33E-01 | no | Tspan18 | intron_variant | MODIFIER |
| 2 | 93253865 | AX-170351782 | 0 | C | T | 0.053 | -1.01E-01 | 3.00E-01 | 2.74E+00 | 2.76E+00 | 7.33E-01 | no | Tspan18 | intron_variant | MODIFIER |
| 2 | 93255039 | AX-170156698 | 0 | T | C | 0.053 | -1.01E-01 | 3.00E-01 | 2.74E+00 | 2.76E+00 | 7.33E-01 | no | Tspan18 | intron_variant | MODIFIER |
| 2 | 93255390 | AX-170323810 | 0 | T | C | 0.053 | -1.01E-01 | 3.00E-01 | 2.74E+00 | 2.76E+00 | 7.33E-01 | no | Tspan18 | intron_variant | MODIFIER |
| 2 | 93255423 | AX-170064314 | 0 | T | C | 0.053 | -1.01E-01 | 3.00E-01 | 2.74E+00 | 2.76E+00 | 7.33E-01 | no | Tspan18 | intron_variant | MODIFIER |
| 2 | 93255698 | AX-170064315 | 0 | C | T | 0.053 | -1.01E-01 | 3.00E-01 | 2.74E+00 | 2.76E+00 | 7.33E-01 | no | Tspan18 | intron_variant | MODIFIER |
| 2 | 93257619 | AX-97853334 | 0 | G | A | 0.053 | -1.01E-01 | 3.00E-01 | 2.74E+00 | 2.76E+00 | 7.33E-01 | no | Tspan18 | intron_variant | MODIFIER |
| 2 | 93258157 | AX-170447539 | 0 | T | C | 0.053 | -1.01E-01 | 3.00E-01 | 2.74E+00 | 2.76E+00 | 7.33E-01 | no | Tspan18 | intron_variant | MODIFIER |
| 2 | 93259672 | AX-170242365 | 0 | C | T | 0.053 | -1.01E-01 | 3.00E-01 | 2.74E+00 | 2.76E+00 | 7.33E-01 | no | Tspan18 | intron_variant | MODIFIER |
| 2 | 93286156 | AX-97787367 | 0 | T | G | 0.053 | -1.01E-01 | 3.00E-01 | 2.74E+00 | 2.76E+00 | 7.33E-01 | no | Tspan18 | intron_variant | MODIFIER |
| 2 | 93303076 | AX-97838948 | 0 | A | C | 0.053 | -1.01E-01 | 3.00E-01 | 2.74E+00 | 2.76E+00 | 7.33E-01 | no | Tspan18 | intron_variant | MODIFIER |
| 2 | 93522967 | AX-170511307 | 0 | T | C | 0.053 | -1.01E-01 | 3.00E-01 | 2.74E+00 | 2.76E+00 | 7.33E-01 | no | | intergenic_variant | MODIFIER |
| 2 | 93778768 | AX-97854195 | 0 | C | T | 0.053 | -1.01E-01 | 3.00E-01 | 2.74E+00 | 2.76E+00 | 7.33E-01 | no | Ext2 | intron_variant | MODIFIER |
| 2 | 93780995 | AX-170153992 | 0 | T | A | 0.053 | -1.01E-01 | 3.00E-01 | 2.74E+00 | 2.76E+00 | 7.33E-01 | no | Ext2 | intron_variant | MODIFIER |
| 2 | 93786519 | AX-170418067 | 0 | A | G | 0.053 | -1.01E-01 | 3.00E-01 | 2.74E+00 | 2.76E+00 | 7.33E-01 | no | Ext2 | intron_variant | MODIFIER |
| 2 | 93789311 | AX-170111822 | 0 | G | C | 0.053 | -1.01E-01 | 3.00E-01 | 2.74E+00 | 2.76E+00 | 7.33E-01 | no | Ext2 | intron_variant | MODIFIER |
| 2 | 93791859 | AX-170348388 | 0 | A | G | 0.053 | -1.01E-01 | 3.00E-01 | 2.74E+00 | 2.76E+00 | 7.33E-01 | no | Ext2 | intron_variant | MODIFIER |
| 2 | 93805291 | AX-170018370 | 0 | T | C | 0.053 | -1.01E-01 | 3.00E-01 | 2.74E+00 | 2.76E+00 | 7.33E-01 | no | Ext2 | intron_variant | MODIFIER |
| 2 | 93807490 | AX-97813721 | 0 | C | A | 0.053 | -1.01E-01 | 3.00E-01 | 2.74E+00 | 2.76E+00 | 7.33E-01 | no | Ext2 | intron_variant | MODIFIER |
| 2 | 93807900 | AX-170172266 | 0 | T | C | 0.053 | -1.01E-01 | 3.00E-01 | 2.74E+00 | 2.76E+00 | 7.33E-01 | no | Ext2 | intron_variant | MODIFIER |
| 2 | 93808244 | AX-170374084 | 0 | C | T | 0.053 | -1.01E-01 | 3.00E-01 | 2.74E+00 | 2.76E+00 | 7.33E-01 | no | Ext2 | intron_variant | MODIFIER |
| 2 | 93813302 | AX-170347506 | 0 | A | G | 0.053 | -1.01E-01 | 3.00E-01 | 2.74E+00 | 2.76E+00 | 7.33E-01 | no | Ext2 | intron_variant | MODIFIER |
| 2 | 93818788 | AX-170401534 | 0 | C | G | 0.053 | -1.01E-01 | 3.00E-01 | 2.74E+00 | 2.76E+00 | 7.33E-01 | no | Ext2 | intron_variant | MODIFIER |
| 2 | 93818881 | AX-170144181 | 0 | A | G | 0.053 | -1.01E-01 | 3.00E-01 | 2.74E+00 | 2.76E+00 | 7.33E-01 | no | Ext2 | intron_variant | MODIFIER |
| 2 | 93835084 | AX-170310742 | 0 | T | C | 0.053 | -1.01E-01 | 3.00E-01 | 2.74E+00 | 2.76E+00 | 7.33E-01 | no | Accs | downstream_gene_variant | MODIFIER |
| 2 | 94098587 | AX-97822670 | 0 | A | G | 0.053 | -1.01E-01 | 3.00E-01 | 2.74E+00 | 2.76E+00 | 7.33E-01 | no | Hsd17b12 | intron_variant | MODIFIER |
| 2 | 94295092 | AX-170283834 | 0 | T | C | 0.053 | -1.01E-01 | 3.00E-01 | 2.74E+00 | 2.76E+00 | 7.33E-01 | no | | regulatory_region_variant | MODIFIER |
| 2 | 94297820 | AX-170373130 | 0 | T | C | 0.053 | -1.01E-01 | 3.00E-01 | 2.74E+00 | 2.76E+00 | 7.33E-01 | no | Ttc17 | downstream_gene_variant | MODIFIER |
| 2 | 94319215 | AX-170178616 | 0 | T | C | 0.053 | -1.01E-01 | 3.00E-01 | 2.74E+00 | 2.76E+00 | 7.33E-01 | no | Ttc17 | intron_variant | MODIFIER |
| 2 | 94324673 | AX-170370598 | 0 | C | T | 0.053 | -1.01E-01 | 3.00E-01 | 2.74E+00 | 2.76E+00 | 7.33E-01 | no | Ttc17 | intron_variant | MODIFIER |
| 2 | 94329041 | AX-168410960 | 0 | A | G | 0.053 | -1.01E-01 | 3.00E-01 | 2.74E+00 | 2.76E+00 | 7.33E-01 | no | Ttc17 | intron_variant | MODIFIER |
| 2 | 94333147 | AX-170216950 | 0 | T | C | 0.053 | -1.01E-01 | 3.00E-01 | 2.74E+00 | 2.76E+00 | 7.33E-01 | no | Ttc17 | intron_variant | MODIFIER |
| 2 | 94337103 | AX-170173259 | 0 | C | T | 0.053 | -1.01E-01 | 3.00E-01 | 2.74E+00 | 2.76E+00 | 7.33E-01 | no | Ttc17 | intron_variant | MODIFIER |
| 2 | 94361454 | AX-170338181 | 0 | G | C | 0.053 | -1.01E-01 | 3.00E-01 | 2.74E+00 | 2.76E+00 | 7.33E-01 | no | Ttc17 | intron_variant | MODIFIER |
| 2 | 94390718 | AX-170258310 | 0 | C | T | 0.053 | -1.01E-01 | 3.00E-01 | 2.74E+00 | 2.76E+00 | 7.33E-01 | no | Ttc17 | intron_variant | MODIFIER |
| 2 | 94395298 | AX-170455063 | 0 | C | T | 0.053 | -1.01E-01 | 3.00E-01 | 2.74E+00 | 2.76E+00 | 7.33E-01 | no | Ttc17 | intron_variant | MODIFIER |
| 2 | 94412440 | AX-97804934 | 0 | T | C | 0.053 | -1.01E-01 | 3.00E-01 | 2.74E+00 | 2.76E+00 | 7.33E-01 | no | Api5 | 3_prime_UTR_variant | MODIFIER |

| | | | | | | | | | | | | | | | |
|---|---|---|---|---|---|---|---|---|---|---|---|---|---|---|---|
| 2 | 94498593 | AX-170512987 | 0 | G | T | 0.053 | -1.01E-01 | 3.00E-01 | 2.74E+00 | 2.76E+00 | 7.33E-01 | no | | regulatory_region_variant | MODIFIER |
| 2 | 94503564 | AX-170511439 | 0 | T | C | 0.053 | -1.01E-01 | 3.00E-01 | 2.74E+00 | 2.76E+00 | 7.33E-01 | no | | intergenic_variant | MODIFIER |
| 2 | 94509208 | AX-170261294 | 0 | A | C | 0.053 | -1.01E-01 | 3.00E-01 | 2.74E+00 | 2.76E+00 | 7.33E-01 | no | | intergenic_variant | MODIFIER |
| 2 | 94518932 | AX-170436721 | 0 | T | G | 0.053 | -1.01E-01 | 3.00E-01 | 2.74E+00 | 2.76E+00 | 7.33E-01 | no | | intergenic_variant | MODIFIER |
| 2 | 94524604 | AX-170318731 | 0 | T | C | 0.053 | -1.01E-01 | 3.00E-01 | 2.74E+00 | 2.76E+00 | 7.33E-01 | no | | regulatory_region_variant | MODIFIER |
| 2 | 94527258 | AX-170064330 | 0 | T | C | 0.053 | -1.01E-01 | 3.00E-01 | 2.74E+00 | 2.76E+00 | 7.33E-01 | no | | intergenic_variant | MODIFIER |
| 2 | 94551363 | AX-170153016 | 0 | C | T | 0.053 | -1.01E-01 | 3.00E-01 | 2.74E+00 | 2.76E+00 | 7.33E-01 | no | | intergenic_variant | MODIFIER |
| 2 | 94582057 | AX-97852384 | 0 | C | T | 0.053 | -1.01E-01 | 3.00E-01 | 2.74E+00 | 2.76E+00 | 7.33E-01 | no | | intergenic_variant | MODIFIER |
| 2 | 94585115 | AX-170496622 | 0 | G | T | 0.053 | -1.01E-01 | 3.00E-01 | 2.74E+00 | 2.76E+00 | 7.33E-01 | no | | intergenic_variant | MODIFIER |
| 2 | 94700247 | AX-170522591 | 0 | T | G | 0.053 | -1.01E-01 | 3.00E-01 | 2.74E+00 | 2.76E+00 | 7.33E-01 | no | | intergenic_variant | MODIFIER |
| 2 | 94731864 | AX-170129732 | 0 | C | G | 0.053 | -1.01E-01 | 3.00E-01 | 2.74E+00 | 2.76E+00 | 7.33E-01 | no | Gm13793 | upstream_gene_variant | MODIFIER |
| 2 | 94779167 | AX-170480588 | 0 | A | G | 0.053 | -1.01E-01 | 3.00E-01 | 2.74E+00 | 2.76E+00 | 7.33E-01 | no | | intergenic_variant | MODIFIER |
| 2 | 94783736 | AX-170088672 | 0 | C | A | 0.053 | -1.01E-01 | 3.00E-01 | 2.74E+00 | 2.76E+00 | 7.33E-01 | no | | intergenic_variant | MODIFIER |
| 2 | 94799237 | AX-170064333 | 0 | T | C | 0.053 | -1.01E-01 | 3.00E-01 | 2.74E+00 | 2.76E+00 | 7.33E-01 | no | | intergenic_variant | MODIFIER |
| 2 | 94799514 | AX-170064335 | 0 | G | A | 0.053 | -1.01E-01 | 3.00E-01 | 2.74E+00 | 2.76E+00 | 7.33E-01 | no | | intergenic_variant | MODIFIER |
| 2 | 94800799 | AX-170064334 | 0 | A | G | 0.053 | -1.01E-01 | 3.00E-01 | 2.74E+00 | 2.76E+00 | 7.33E-01 | no | | intergenic_variant | MODIFIER |
| 2 | 94814341 | AX-97828688 | 0 | T | C | 0.053 | -1.01E-01 | 3.00E-01 | 2.74E+00 | 2.76E+00 | 7.33E-01 | no | | intergenic_variant | MODIFIER |
| 2 | 94815816 | AX-170231734 | 0 | C | T | 0.053 | -1.01E-01 | 3.00E-01 | 2.74E+00 | 2.76E+00 | 7.33E-01 | no | | intergenic_variant | MODIFIER |
| 2 | 94875050 | AX-97838085 | 0 | G | T | 0.053 | -1.01E-01 | 3.00E-01 | 2.74E+00 | 2.76E+00 | 7.33E-01 | no | | intergenic_variant | MODIFIER |
| 2 | 94915845 | AX-170018383 | 0 | C | T | 0.053 | -1.01E-01 | 3.00E-01 | 2.74E+00 | 2.76E+00 | 7.33E-01 | no | | intergenic_variant | MODIFIER |
| 2 | 94917960 | AX-170109904 | 0 | C | T | 0.053 | -1.01E-01 | 3.00E-01 | 2.74E+00 | 2.76E+00 | 7.33E-01 | no | | intergenic_variant | MODIFIER |
| 2 | 94920183 | AX-170174344 | 0 | G | A | 0.053 | -1.01E-01 | 3.00E-01 | 2.74E+00 | 2.76E+00 | 7.33E-01 | no | | intergenic_variant | MODIFIER |
| 2 | 94939817 | AX-170018385 | 0 | C | T | 0.053 | -1.01E-01 | 3.00E-01 | 2.74E+00 | 2.76E+00 | 7.33E-01 | no | | intergenic_variant | MODIFIER |
| 2 | 94967834 | AX-170018382 | 0 | G | A | 0.053 | -1.01E-01 | 3.00E-01 | 2.74E+00 | 2.76E+00 | 7.33E-01 | no | | intergenic_variant | MODIFIER |
| 2 | 94982299 | AX-170018388 | 0 | C | T | 0.053 | -1.01E-01 | 3.00E-01 | 2.74E+00 | 2.76E+00 | 7.33E-01 | no | | intergenic_variant | MODIFIER |
| 2 | 94982538 | AX-170018387 | 0 | C | T | 0.053 | -1.01E-01 | 3.00E-01 | 2.74E+00 | 2.76E+00 | 7.33E-01 | no | | intergenic_variant | MODIFIER |
| 2 | 95023226 | AX-170241459 | 0 | G | A | 0.053 | -1.01E-01 | 3.00E-01 | 2.74E+00 | 2.76E+00 | 7.33E-01 | no | | intergenic_variant | MODIFIER |
| 2 | 95026917 | AX-170203921 | 0 | T | G | 0.053 | -1.01E-01 | 3.00E-01 | 2.74E+00 | 2.76E+00 | 7.33E-01 | no | | intergenic_variant | MODIFIER |
| 2 | 95027154 | AX-168382501 | 0 | A | C | 0.053 | -1.01E-01 | 3.00E-01 | 2.74E+00 | 2.76E+00 | 7.33E-01 | no | | intergenic_variant | MODIFIER |
| 2 | 95059671 | AX-170474831 | 0 | G | A | 0.053 | -1.01E-01 | 3.00E-01 | 2.74E+00 | 2.76E+00 | 7.33E-01 | no | | intergenic_variant | MODIFIER |
| 2 | 95059911 | AX-170444749 | 0 | C | G | 0.053 | -1.01E-01 | 3.00E-01 | 2.74E+00 | 2.76E+00 | 7.33E-01 | no | | intergenic_variant | MODIFIER |
| 2 | 95077535 | AX-170064341 | 0 | C | T | 0.053 | -1.01E-01 | 3.00E-01 | 2.74E+00 | 2.76E+00 | 7.33E-01 | no | | intergenic_variant | MODIFIER |
| 2 | 95131905 | AX-170440395 | 0 | G | A | 0.053 | -1.01E-01 | 3.00E-01 | 2.74E+00 | 2.76E+00 | 7.33E-01 | no | | intergenic_variant | MODIFIER |
| 2 | 95134624 | AX-170337436 | 0 | T | A | 0.053 | -1.01E-01 | 3.00E-01 | 2.74E+00 | 2.76E+00 | 7.33E-01 | no | | intergenic_variant | MODIFIER |
| 2 | 95142762 | AX-170494622 | 0 | C | G | 0.053 | -1.01E-01 | 3.00E-01 | 2.74E+00 | 2.76E+00 | 7.33E-01 | no | | intergenic_variant | MODIFIER |
| 2 | 95165019 | AX-170088674 | 0 | A | G | 0.053 | -1.01E-01 | 3.00E-01 | 2.74E+00 | 2.76E+00 | 7.33E-01 | no | | intergenic_variant | MODIFIER |
| 2 | 95169808 | AX-170191208 | 0 | G | A | 0.053 | -1.01E-01 | 3.00E-01 | 2.74E+00 | 2.76E+00 | 7.33E-01 | no | | intergenic_variant | MODIFIER |
| 2 | 95230093 | AX-97788903 | 0 | T | C | 0.053 | -1.01E-01 | 3.00E-01 | 2.74E+00 | 2.76E+00 | 7.33E-01 | no | | intergenic_variant | MODIFIER |
| 2 | 95230811 | AX-170294752 | 0 | C | T | 0.053 | -1.01E-01 | 3.00E-01 | 2.74E+00 | 2.76E+00 | 7.33E-01 | no | | intergenic_variant | MODIFIER |
| 2 | 95264715 | AX-97794345 | 0 | A | G | 0.053 | -1.01E-01 | 3.00E-01 | 2.74E+00 | 2.76E+00 | 7.33E-01 | no | | intergenic_variant | MODIFIER |
| 2 | 95272572 | AX-170209308 | 0 | G | C | 0.053 | -1.01E-01 | 3.00E-01 | 2.74E+00 | 2.76E+00 | 7.33E-01 | no | | intergenic_variant | MODIFIER |
| 2 | 95362161 | AX-97794203 | 0 | T | C | 0.053 | -1.01E-01 | 3.00E-01 | 2.74E+00 | 2.76E+00 | 7.33E-01 | no | | intergenic_variant | MODIFIER |
| 2 | 95511227 | AX-170018391 | 0 | C | G | 0.053 | -1.01E-01 | 3.00E-01 | 2.74E+00 | 2.76E+00 | 7.33E-01 | no | | intergenic_variant | MODIFIER |
| 2 | 95522753 | AX-170018393 | 0 | A | T | 0.053 | -1.01E-01 | 3.00E-01 | 2.74E+00 | 2.76E+00 | 7.33E-01 | no | | intergenic_variant | MODIFIER |
| 2 | 95586134 | AX-97825234 | 0 | A | G | 0.053 | -1.01E-01 | 3.00E-01 | 2.74E+00 | 2.76E+00 | 7.33E-01 | no | | intergenic_variant | MODIFIER |
| 2 | 95607142 | AX-170286766 | 0 | G | A | 0.053 | -1.01E-01 | 3.00E-01 | 2.74E+00 | 2.76E+00 | 7.33E-01 | no | | intergenic_variant | MODIFIER |
| 2 | 95645504 | AX-170373495 | 0 | G | A | 0.053 | -1.01E-01 | 3.00E-01 | 2.74E+00 | 2.76E+00 | 7.33E-01 | no | | intergenic_variant | MODIFIER |
| 2 | 95742447 | AX-170088675 | 0 | A | G | 0.053 | -1.01E-01 | 3.00E-01 | 2.74E+00 | 2.76E+00 | 7.33E-01 | no | | intergenic_variant | MODIFIER |
| 2 | 96313826 | AX-97836664 | 0 | A | G | 0.053 | -1.01E-01 | 3.00E-01 | 2.74E+00 | 2.76E+00 | 7.33E-01 | no | Lrrc4c | upstream_gene_variant | MODIFIER |
| 2 | 96316587 | AX-170443059 | 0 | A | G | 0.053 | -1.01E-01 | 3.00E-01 | 2.74E+00 | 2.76E+00 | 7.33E-01 | no | Lrrc4c | upstream_gene_variant | MODIFIER |
| 2 | 96340085 | AX-170259396 | 0 | C | A | 0.053 | -1.01E-01 | 3.00E-01 | 2.74E+00 | 2.76E+00 | 7.33E-01 | no | Lrrc4c | intron_variant | MODIFIER |
| 2 | 96459831 | AX-170433946 | 0 | T | C | 0.053 | -1.01E-01 | 3.00E-01 | 2.74E+00 | 2.76E+00 | 7.33E-01 | no | Lrrc4c | intron_variant | MODIFIER |
| 2 | 96472029 | AX-170097670 | 0 | A | G | 0.053 | -1.01E-01 | 3.00E-01 | 2.74E+00 | 2.76E+00 | 7.33E-01 | no | Lrrc4c | intron_variant | MODIFIER |
| 2 | 96474185 | AX-170288180 | 0 | T | C | 0.053 | -1.01E-01 | 3.00E-01 | 2.74E+00 | 2.76E+00 | 7.33E-01 | no | Lrrc4c | intron_variant | MODIFIER |

| | | | | | | | | | | | | | | | |
|---|---|---|---|---|---|---|---|---|---|---|---|---|---|---|---|
| 2 | 96481253 | AX-97791623 | 0 | A | G | 0.053 | -1.01E-01 | 3.00E-01 | 2.74E+00 | 2.76E+00 | 7.33E-01 | no | Lrrc4c | intron_variant | MODIFIER |
| 2 | 96481621 | AX-170313384 | 0 | A | T | 0.053 | -1.01E-01 | 3.00E-01 | 2.74E+00 | 2.76E+00 | 7.33E-01 | no | Lrrc4c | intron_variant | MODIFIER |
| 2 | 96490084 | AX-170018410 | 0 | G | T | 0.053 | -1.01E-01 | 3.00E-01 | 2.74E+00 | 2.76E+00 | 7.33E-01 | no | Lrrc4c | intron_variant | MODIFIER |
| 2 | 96494554 | AX-170530417 | 0 | G | T | 0.053 | -1.01E-01 | 3.00E-01 | 2.74E+00 | 2.76E+00 | 7.33E-01 | no | Lrrc4c | intron_variant | MODIFIER |
| 2 | 96496185 | AX-170018411 | 0 | G | T | 0.053 | -1.01E-01 | 3.00E-01 | 2.74E+00 | 2.76E+00 | 7.33E-01 | no | Lrrc4c | intron_variant | MODIFIER |
| 2 | 96497703 | AX-170194762 | 0 | C | G | 0.053 | -1.01E-01 | 3.00E-01 | 2.74E+00 | 2.76E+00 | 7.33E-01 | no | Lrrc4c | intron_variant | MODIFIER |
| 2 | 96514881 | AX-97844831 | 0 | A | G | 0.053 | -1.01E-01 | 3.00E-01 | 2.74E+00 | 2.76E+00 | 7.33E-01 | no | Lrrc4c | intron_variant | MODIFIER |
| 2 | 96539319 | AX-170530443 | 0 | C | T | 0.053 | -1.01E-01 | 3.00E-01 | 2.74E+00 | 2.76E+00 | 7.33E-01 | no | Lrrc4c | intron_variant | MODIFIER |
| 2 | 96539516 | AX-170466665 | 0 | T | C | 0.053 | -1.01E-01 | 3.00E-01 | 2.74E+00 | 2.76E+00 | 7.33E-01 | no | Lrrc4c | intron_variant | MODIFIER |
| 2 | 96583060 | AX-97838167 | 0 | A | G | 0.053 | -1.01E-01 | 3.00E-01 | 2.74E+00 | 2.76E+00 | 7.33E-01 | no | Lrrc4c | intron_variant | MODIFIER |
| 2 | 96623459 | AX-170172933 | 0 | T | C | 0.053 | -1.01E-01 | 3.00E-01 | 2.74E+00 | 2.76E+00 | 7.33E-01 | no | Lrrc4c | intron_variant | MODIFIER |
| 2 | 96640533 | AX-170251183 | 0 | A | T | 0.053 | -1.01E-01 | 3.00E-01 | 2.74E+00 | 2.76E+00 | 7.33E-01 | no | Lrrc4c | intron_variant | MODIFIER |
| 2 | 96666366 | AX-97826504 | 0 | C | T | 0.053 | -1.01E-01 | 3.00E-01 | 2.74E+00 | 2.76E+00 | 7.33E-01 | no | Lrrc4c | intron_variant | MODIFIER |
| 2 | 96672107 | AX-170342989 | 0 | C | T | 0.053 | -1.01E-01 | 3.00E-01 | 2.74E+00 | 2.76E+00 | 7.33E-01 | no | Lrrc4c | intron_variant | MODIFIER |
| 2 | 96716911 | AX-170321541 | 0 | T | C | 0.053 | -1.01E-01 | 3.00E-01 | 2.74E+00 | 2.76E+00 | 7.33E-01 | no | Lrrc4c | intron_variant | MODIFIER |
| 2 | 96776881 | AX-170417528 | 0 | A | G | 0.053 | -1.01E-01 | 3.00E-01 | 2.74E+00 | 2.76E+00 | 7.33E-01 | no | Lrrc4c | intron_variant | MODIFIER |
| 2 | 96796203 | AX-170225749 | 0 | T | C | 0.053 | -1.01E-01 | 3.00E-01 | 2.74E+00 | 2.76E+00 | 7.33E-01 | no | Lrrc4c | intron_variant | MODIFIER |
| 2 | 96820094 | AX-170422017 | 0 | T | C | 0.053 | -1.01E-01 | 3.00E-01 | 2.74E+00 | 2.76E+00 | 7.33E-01 | no | Lrrc4c | intron_variant | MODIFIER |
| 2 | 96820775 | AX-170465918 | 0 | G | A | 0.053 | -1.01E-01 | 3.00E-01 | 2.74E+00 | 2.76E+00 | 7.33E-01 | no | Lrrc4c | intron_variant | MODIFIER |
| 2 | 96823311 | AX-97843951 | 0 | T | C | 0.053 | -1.01E-01 | 3.00E-01 | 2.74E+00 | 2.76E+00 | 7.33E-01 | no | Lrrc4c | intron_variant | MODIFIER |
| 2 | 96823768 | AX-170507629 | 0 | A | T | 0.053 | -1.01E-01 | 3.00E-01 | 2.74E+00 | 2.76E+00 | 7.33E-01 | no | Lrrc4c | intron_variant | MODIFIER |
| 2 | 96854447 | AX-97820917 | 0 | T | C | 0.053 | -1.01E-01 | 3.00E-01 | 2.74E+00 | 2.76E+00 | 7.33E-01 | no | Lrrc4c | intron_variant | MODIFIER |
| 2 | 96857512 | AX-170361482 | 0 | A | G | 0.053 | -1.01E-01 | 3.00E-01 | 2.74E+00 | 2.76E+00 | 7.33E-01 | no | Lrrc4c | intron_variant | MODIFIER |
| 2 | 96872248 | AX-170064369 | 0 | G | A | 0.053 | -1.01E-01 | 3.00E-01 | 2.74E+00 | 2.76E+00 | 7.33E-01 | no | Lrrc4c | intron_variant | MODIFIER |
| 2 | 96881893 | AX-170281714 | 0 | C | T | 0.053 | -1.01E-01 | 3.00E-01 | 2.74E+00 | 2.76E+00 | 7.33E-01 | no | Lrrc4c | intron_variant | MODIFIER |
| 2 | 96881973 | AX-170416439 | 0 | A | G | 0.053 | -1.01E-01 | 3.00E-01 | 2.74E+00 | 2.76E+00 | 7.33E-01 | no | Lrrc4c | intron_variant | MODIFIER |
| 2 | 96882316 | AX-170338390 | 0 | C | T | 0.053 | -1.01E-01 | 3.00E-01 | 2.74E+00 | 2.76E+00 | 7.33E-01 | no | Lrrc4c | intron_variant | MODIFIER |
| 2 | 96882433 | AX-170411384 | 0 | A | G | 0.053 | -1.01E-01 | 3.00E-01 | 2.74E+00 | 2.76E+00 | 7.33E-01 | no | Lrrc4c | intron_variant | MODIFIER |
| 2 | 96903154 | AX-170250209 | 0 | G | A | 0.053 | -1.01E-01 | 3.00E-01 | 2.74E+00 | 2.76E+00 | 7.33E-01 | no | Lrrc4c | intron_variant | MODIFIER |
| 2 | 96922489 | AX-170475162 | 0 | T | G | 0.053 | -1.01E-01 | 3.00E-01 | 2.74E+00 | 2.76E+00 | 7.33E-01 | no | Lrrc4c | intron_variant | MODIFIER |
| 2 | 96923002 | AX-170303588 | 0 | T | G | 0.053 | -1.01E-01 | 3.00E-01 | 2.74E+00 | 2.76E+00 | 7.33E-01 | no | Lrrc4c | intron_variant | MODIFIER |
| 2 | 96923768 | AX-97795109 | 0 | T | C | 0.053 | -1.01E-01 | 3.00E-01 | 2.74E+00 | 2.76E+00 | 7.33E-01 | no | Lrrc4c | intron_variant | MODIFIER |
| 2 | 96946272 | AX-170129254 | 0 | A | G | 0.053 | -1.01E-01 | 3.00E-01 | 2.74E+00 | 2.76E+00 | 7.33E-01 | no | Lrrc4c | intron_variant | MODIFIER |
| 2 | 97018579 | AX-97852823 | 0 | T | C | 0.053 | -1.01E-01 | 3.00E-01 | 2.74E+00 | 2.76E+00 | 7.33E-01 | no | Lrrc4c | intron_variant | MODIFIER |
| 2 | 97018652 | AX-170064374 | 0 | T | C | 0.053 | -1.01E-01 | 3.00E-01 | 2.74E+00 | 2.76E+00 | 7.33E-01 | no | Lrrc4c | intron_variant | MODIFIER |
| 2 | 97025984 | AX-97838999 | 0 | G | T | 0.053 | -1.01E-01 | 3.00E-01 | 2.74E+00 | 2.76E+00 | 7.33E-01 | no | Lrrc4c | intron_variant | MODIFIER |
| 2 | 97047014 | AX-170018413 | 0 | G | C | 0.053 | -1.01E-01 | 3.00E-01 | 2.74E+00 | 2.76E+00 | 7.33E-01 | no | Lrrc4c | intron_variant | MODIFIER |
| 2 | 97080207 | AX-97840631 | 0 | T | G | 0.053 | -1.01E-01 | 3.00E-01 | 2.74E+00 | 2.76E+00 | 7.33E-01 | no | Lrrc4c | intron_variant | MODIFIER |
| 2 | 97128110 | AX-170270166 | 0 | C | T | 0.053 | -1.01E-01 | 3.00E-01 | 2.74E+00 | 2.76E+00 | 7.33E-01 | no | Lrrc4c | intron_variant | MODIFIER |
| 2 | 97161627 | AX-170424255 | 0 | T | C | 0.053 | -1.01E-01 | 3.00E-01 | 2.74E+00 | 2.76E+00 | 7.33E-01 | no | Lrrc4c | intron_variant | MODIFIER |
| 2 | 97166240 | AX-170411483 | 0 | C | T | 0.053 | -1.01E-01 | 3.00E-01 | 2.74E+00 | 2.76E+00 | 7.33E-01 | no | Lrrc4c | intron_variant | MODIFIER |
| 2 | 97191609 | AX-170350514 | 0 | C | G | 0.053 | -1.01E-01 | 3.00E-01 | 2.74E+00 | 2.76E+00 | 7.33E-01 | no | Lrrc4c | intron_variant | MODIFIER |
| 2 | 97223697 | AX-170261340 | 0 | T | G | 0.053 | -1.01E-01 | 3.00E-01 | 2.74E+00 | 2.76E+00 | 7.33E-01 | no | Lrrc4c | intron_variant | MODIFIER |
| 2 | 97224541 | AX-170281947 | 0 | C | T | 0.053 | -1.01E-01 | 3.00E-01 | 2.74E+00 | 2.76E+00 | 7.33E-01 | no | Lrrc4c | intron_variant | MODIFIER |
| 2 | 97232637 | AX-97845685 | 0 | T | C | 0.053 | -1.01E-01 | 3.00E-01 | 2.74E+00 | 2.76E+00 | 7.33E-01 | no | Lrrc4c | intron_variant | MODIFIER |
| 2 | 97262939 | AX-170339173 | 0 | C | A | 0.053 | -1.01E-01 | 3.00E-01 | 2.74E+00 | 2.76E+00 | 7.33E-01 | no | Lrrc4c | intron_variant | MODIFIER |
| 2 | 97292405 | AX-170489580 | 0 | C | G | 0.053 | -1.01E-01 | 3.00E-01 | 2.74E+00 | 2.76E+00 | 7.33E-01 | no | Lrrc4c | intron_variant | MODIFIER |
| 2 | 97335615 | AX-97807542 | 0 | A | G | 0.053 | -1.01E-01 | 3.00E-01 | 2.74E+00 | 2.76E+00 | 7.33E-01 | no | Lrrc4c | intron_variant | MODIFIER |
| 2 | 97335855 | AX-170345995 | 0 | C | T | 0.053 | -1.01E-01 | 3.00E-01 | 2.74E+00 | 2.76E+00 | 7.33E-01 | no | Lrrc4c | intron_variant | MODIFIER |
| 2 | 97348157 | AX-170064376 | 0 | T | G | 0.053 | -1.01E-01 | 3.00E-01 | 2.74E+00 | 2.76E+00 | 7.33E-01 | no | Lrrc4c | intron_variant | MODIFIER |
| 2 | 97395866 | AX-170516861 | 0 | T | C | 0.053 | -1.01E-01 | 3.00E-01 | 2.74E+00 | 2.76E+00 | 7.33E-01 | no | Lrrc4c | intron_variant | MODIFIER |
| 2 | 97413201 | AX-170395845 | 0 | T | G | 0.053 | -1.01E-01 | 3.00E-01 | 2.74E+00 | 2.76E+00 | 7.33E-01 | no | Lrrc4c | intron_variant | MODIFIER |
| 2 | 97413733 | AX-170320796 | 0 | A | G | 0.053 | -1.01E-01 | 3.00E-01 | 2.74E+00 | 2.76E+00 | 7.33E-01 | no | Lrrc4c | intron_variant | MODIFIER |
| 2 | 97418535 | AX-97838166 | 0 | G | A | 0.053 | -1.01E-01 | 3.00E-01 | 2.74E+00 | 2.76E+00 | 7.33E-01 | no | Lrrc4c | intron_variant | MODIFIER |

| | | | | | | | | | | | | | | |
|---|---|---|---|---|---|---|---|---|---|---|---|---|---|---|
| 2 | 97449037 | AX-170390269 | 0 | G | C | 0.053 | -1.01E-01 | 3.00E-01 | 2.74E+00 | 2.76E+00 | 7.33E-01 | no | Lrrc4c | intron_variant | MODIFIER |
| 2 | 97505681 | AX-170306180 | 0 | A | T | 0.053 | -1.01E-01 | 3.00E-01 | 2.74E+00 | 2.76E+00 | 7.33E-01 | no | Lrrc4c | intron_variant | MODIFIER |
| 2 | 97506390 | AX-170094342 | 0 | T | C | 0.053 | -1.01E-01 | 3.00E-01 | 2.74E+00 | 2.76E+00 | 7.33E-01 | no | Lrrc4c | intron_variant | MODIFIER |
| 2 | 97506744 | AX-170445169 | 0 | A | G | 0.053 | -1.01E-01 | 3.00E-01 | 2.74E+00 | 2.76E+00 | 7.33E-01 | no | Lrrc4c | intron_variant | MODIFIER |
| 2 | 97525007 | AX-170324531 | 0 | G | T | 0.053 | -1.01E-01 | 3.00E-01 | 2.74E+00 | 2.76E+00 | 7.33E-01 | no | Lrrc4c | intron_variant | MODIFIER |
| 2 | 97535898 | AX-170250730 | 0 | G | A | 0.053 | -1.01E-01 | 3.00E-01 | 2.74E+00 | 2.76E+00 | 7.33E-01 | no | Lrrc4c | intron_variant | MODIFIER |
| 2 | 97537000 | AX-170192321 | 0 | G | A | 0.053 | -1.01E-01 | 3.00E-01 | 2.74E+00 | 2.76E+00 | 7.33E-01 | no | Lrrc4c | intron_variant | MODIFIER |
| 2 | 97555081 | AX-170221014 | 0 | A | G | 0.053 | -1.01E-01 | 3.00E-01 | 2.74E+00 | 2.76E+00 | 7.33E-01 | no | Lrrc4c | intron_variant | MODIFIER |
| 2 | 97555233 | AX-170498679 | 0 | T | G | 0.053 | -1.01E-01 | 3.00E-01 | 2.74E+00 | 2.76E+00 | 7.33E-01 | no | Lrrc4c | intron_variant | MODIFIER |
| 2 | 97576822 | AX-170201509 | 0 | G | A | 0.053 | -1.01E-01 | 3.00E-01 | 2.74E+00 | 2.76E+00 | 7.33E-01 | no | Lrrc4c | intron_variant | MODIFIER |
| 2 | 97580418 | AX-97786355 | 0 | A | G | 0.053 | -1.01E-01 | 3.00E-01 | 2.74E+00 | 2.76E+00 | 7.33E-01 | no | Lrrc4c | intron_variant | MODIFIER |
| 2 | 97975740 | AX-170018432 | 0 | G | T | 0.053 | -1.01E-01 | 3.00E-01 | 2.74E+00 | 2.76E+00 | 7.33E-01 | no | | intergenic_variant | MODIFIER |
| 2 | 97981147 | AX-97819671 | 0 | T | C | 0.053 | -1.01E-01 | 3.00E-01 | 2.74E+00 | 2.76E+00 | 7.33E-01 | no | | intergenic_variant | MODIFIER |
| 2 | 97991643 | AX-170157175 | 0 | G | T | 0.053 | -1.01E-01 | 3.00E-01 | 2.74E+00 | 2.76E+00 | 7.33E-01 | no | | intergenic_variant | MODIFIER |
| 2 | 98521164 | AX-97824579 | 0 | A | G | 0.053 | -1.01E-01 | 3.00E-01 | 2.74E+00 | 2.76E+00 | 7.33E-01 | no | | intergenic_variant | MODIFIER |
| 2 | 99217002 | AX-97852096 | 0 | C | A | 0.053 | -1.01E-01 | 3.00E-01 | 2.74E+00 | 2.76E+00 | 7.33E-01 | no | | intergenic_variant | MODIFIER |
| 2 | 99239681 | AX-170321208 | 0 | G | A | 0.053 | -1.01E-01 | 3.00E-01 | 2.74E+00 | 2.76E+00 | 7.33E-01 | no | | intergenic_variant | MODIFIER |
| 2 | 99310049 | AX-170256677 | 0 | C | G | 0.053 | -1.01E-01 | 3.00E-01 | 2.74E+00 | 2.76E+00 | 7.33E-01 | no | | intergenic_variant | MODIFIER |
| 2 | 99389137 | AX-97813108 | 0 | A | G | 0.053 | -1.01E-01 | 3.00E-01 | 2.74E+00 | 2.76E+00 | 7.33E-01 | no | | intergenic_variant | MODIFIER |
| 2 | 99469825 | AX-170018466 | 0 | G | A | 0.053 | -1.01E-01 | 3.00E-01 | 2.74E+00 | 2.76E+00 | 7.33E-01 | no | | intergenic_variant | MODIFIER |
| 2 | 99471492 | AX-170018463 | 0 | G | A | 0.053 | -1.01E-01 | 3.00E-01 | 2.74E+00 | 2.76E+00 | 7.33E-01 | no | | intergenic_variant | MODIFIER |
| 2 | 99493591 | AX-97783751 | 0 | C | A | 0.053 | -1.01E-01 | 3.00E-01 | 2.74E+00 | 2.76E+00 | 7.33E-01 | no | | intergenic_variant | MODIFIER |
| 2 | 99497446 | AX-170197894 | 0 | G | A | 0.053 | -1.01E-01 | 3.00E-01 | 2.74E+00 | 2.76E+00 | 7.33E-01 | no | | intergenic_variant | MODIFIER |
| 2 | 99498042 | AX-170302898 | 0 | T | C | 0.053 | -1.01E-01 | 3.00E-01 | 2.74E+00 | 2.76E+00 | 7.33E-01 | no | | intergenic_variant | MODIFIER |
| 2 | 99501834 | AX-97847727 | 0 | A | G | 0.053 | -1.01E-01 | 3.00E-01 | 2.74E+00 | 2.76E+00 | 7.33E-01 | no | | intergenic_variant | MODIFIER |
| 2 | 99505292 | AX-170434657 | 0 | T | C | 0.053 | -1.01E-01 | 3.00E-01 | 2.74E+00 | 2.76E+00 | 7.33E-01 | no | | intergenic_variant | MODIFIER |
| 2 | 99521055 | AX-170517287 | 0 | C | T | 0.053 | -1.01E-01 | 3.00E-01 | 2.74E+00 | 2.76E+00 | 7.33E-01 | no | | intergenic_variant | MODIFIER |
| 2 | 99614851 | AX-97847778 | 0 | T | C | 0.053 | -1.01E-01 | 3.00E-01 | 2.74E+00 | 2.76E+00 | 7.33E-01 | no | | intergenic_variant | MODIFIER |
| 2 | 99624856 | AX-170018464 | 0 | T | C | 0.053 | -1.01E-01 | 3.00E-01 | 2.74E+00 | 2.76E+00 | 7.33E-01 | no | | intergenic_variant | MODIFIER |
| 2 | 99665228 | AX-97838217 | 0 | T | G | 0.053 | -1.01E-01 | 3.00E-01 | 2.74E+00 | 2.76E+00 | 7.33E-01 | no | | intergenic_variant | MODIFIER |
| 2 | 99667401 | AX-170366123 | 0 | T | C | 0.053 | -1.01E-01 | 3.00E-01 | 2.74E+00 | 2.76E+00 | 7.33E-01 | no | | intergenic_variant | MODIFIER |
| 2 | 99675799 | AX-170360121 | 0 | A | G | 0.053 | -1.01E-01 | 3.00E-01 | 2.74E+00 | 2.76E+00 | 7.33E-01 | no | | intergenic_variant | MODIFIER |
| 2 | 99680603 | AX-170381441 | 0 | T | C | 0.053 | -1.01E-01 | 3.00E-01 | 2.74E+00 | 2.76E+00 | 7.33E-01 | no | | intergenic_variant | MODIFIER |
| 2 | 99703427 | AX-97800502 | 0 | A | G | 0.053 | -1.01E-01 | 3.00E-01 | 2.74E+00 | 2.76E+00 | 7.33E-01 | no | | intergenic_variant | MODIFIER |
| 2 | 99739903 | AX-170205436 | 0 | C | T | 0.053 | -1.01E-01 | 3.00E-01 | 2.74E+00 | 2.76E+00 | 7.33E-01 | no | | intergenic_variant | MODIFIER |
| 2 | 99739978 | AX-168386465 | 0 | T | C | 0.053 | -1.01E-01 | 3.00E-01 | 2.74E+00 | 2.76E+00 | 7.33E-01 | no | | intergenic_variant | MODIFIER |
| 2 | 99772290 | AX-168410667 | 0 | A | G | 0.053 | -1.01E-01 | 3.00E-01 | 2.74E+00 | 2.76E+00 | 7.33E-01 | no | | intergenic_variant | MODIFIER |
| 2 | 99810417 | AX-170342882 | 0 | A | G | 0.053 | -1.01E-01 | 3.00E-01 | 2.74E+00 | 2.76E+00 | 7.33E-01 | no | | intergenic_variant | MODIFIER |
| 2 | 99810439 | AX-170414608 | 0 | A | T | 0.053 | -1.01E-01 | 3.00E-01 | 2.74E+00 | 2.76E+00 | 7.33E-01 | no | | intergenic_variant | MODIFIER |
| 2 | 99869763 | AX-170018474 | 0 | A | G | 0.053 | -1.01E-01 | 3.00E-01 | 2.74E+00 | 2.76E+00 | 7.33E-01 | no | | intergenic_variant | MODIFIER |
| 2 | 99979990 | AX-170127185 | 0 | A | G | 0.053 | -1.01E-01 | 3.00E-01 | 2.74E+00 | 2.76E+00 | 7.33E-01 | no | | intergenic_variant | MODIFIER |
| 2 | 100019113 | AX-170018472 | 0 | G | A | 0.053 | -1.01E-01 | 3.00E-01 | 2.74E+00 | 2.76E+00 | 7.33E-01 | no | | intergenic_variant | MODIFIER |
| 2 | 100072147 | AX-170064403 | 0 | G | A | 0.053 | -1.01E-01 | 3.00E-01 | 2.74E+00 | 2.76E+00 | 7.33E-01 | no | | intergenic_variant | MODIFIER |
| 2 | 100077444 | AX-170329817 | 0 | T | C | 0.053 | -1.01E-01 | 3.00E-01 | 2.74E+00 | 2.76E+00 | 7.33E-01 | no | | intergenic_variant | MODIFIER |
| 2 | 100093062 | AX-170198572 | 0 | G | A | 0.053 | -1.01E-01 | 3.00E-01 | 2.74E+00 | 2.76E+00 | 7.33E-01 | no | | intergenic_variant | MODIFIER |
| 2 | 100153705 | AX-170404439 | 0 | T | C | 0.053 | -1.01E-01 | 3.00E-01 | 2.74E+00 | 2.76E+00 | 7.33E-01 | no | | intergenic_variant | MODIFIER |
| 2 | 100187612 | AX-97839082 | 0 | T | C | 0.053 | -1.01E-01 | 3.00E-01 | 2.74E+00 | 2.76E+00 | 7.33E-01 | no | | intergenic_variant | MODIFIER |
| 2 | 100200016 | AX-97802100 | 0 | G | A | 0.053 | -1.01E-01 | 3.00E-01 | 2.74E+00 | 2.76E+00 | 7.33E-01 | no | | intergenic_variant | MODIFIER |
| 2 | 100917904 | AX-170064421 | 0 | T | C | 0.053 | -1.01E-01 | 3.00E-01 | 2.74E+00 | 2.76E+00 | 7.33E-01 | no | | intergenic_variant | MODIFIER |
| 2 | 100956592 | AX-170231215 | 0 | A | G | 0.053 | -1.01E-01 | 3.00E-01 | 2.74E+00 | 2.76E+00 | 7.33E-01 | no | | intergenic_variant | MODIFIER |
| 2 | 100980390 | AX-170094350 | 0 | T | C | 0.053 | -1.01E-01 | 3.00E-01 | 2.74E+00 | 2.76E+00 | 7.33E-01 | no | | intergenic_variant | MODIFIER |
| 2 | 101009312 | AX-170415413 | 0 | C | G | 0.053 | -1.01E-01 | 3.00E-01 | 2.74E+00 | 2.76E+00 | 7.33E-01 | no | | intergenic_variant | MODIFIER |
| 2 | 101010078 | AX-170408913 | 0 | A | G | 0.053 | -1.01E-01 | 3.00E-01 | 2.74E+00 | 2.76E+00 | 7.33E-01 | no | | intergenic_variant | MODIFIER |
| 2 | 101060868 | AX-170472022 | 0 | T | C | 0.053 | -1.01E-01 | 3.00E-01 | 2.74E+00 | 2.76E+00 | 7.33E-01 | no | | intergenic_variant | MODIFIER |

| | | | | | | | | | | | | | | | |
|---|---|---|---|---|---|---|---|---|---|---|---|---|---|---|---|
| 2 | 101100586 | AX-170140106 | 0 | A | G | 0.053 | -1.01E-01 | 3.00E-01 | 2.74E+00 | 2.76E+00 | 7.33E-01 | no | | intergenic_variant | MODIFIER |
| 2 | 101112714 | AX-170514517 | 0 | G | A | 0.053 | -1.01E-01 | 3.00E-01 | 2.74E+00 | 2.76E+00 | 7.33E-01 | no | | intergenic_variant | MODIFIER |
| 2 | 101135833 | AX-97848545 | 0 | A | G | 0.053 | -1.01E-01 | 3.00E-01 | 2.74E+00 | 2.76E+00 | 7.33E-01 | no | | intergenic_variant | MODIFIER |
| 2 | 101139742 | AX-168411736 | 0 | C | T | 0.053 | -1.01E-01 | 3.00E-01 | 2.74E+00 | 2.76E+00 | 7.33E-01 | no | | intergenic_variant | MODIFIER |
| 2 | 101141059 | AX-170161281 | 0 | T | C | 0.053 | -1.01E-01 | 3.00E-01 | 2.74E+00 | 2.76E+00 | 7.33E-01 | no | | intergenic_variant | MODIFIER |
| 2 | 101154751 | AX-170438910 | 0 | G | A | 0.053 | -1.01E-01 | 3.00E-01 | 2.74E+00 | 2.76E+00 | 7.33E-01 | no | | intergenic_variant | MODIFIER |
| 2 | 101165447 | AX-168381680 | 0 | C | T | 0.053 | -1.01E-01 | 3.00E-01 | 2.74E+00 | 2.76E+00 | 7.33E-01 | no | | intergenic_variant | MODIFIER |
| 2 | 101167708 | AX-97798321 | 0 | A | G | 0.053 | -1.01E-01 | 3.00E-01 | 2.74E+00 | 2.76E+00 | 7.33E-01 | no | | intergenic_variant | MODIFIER |
| 2 | 101206884 | AX-170514574 | 0 | A | G | 0.053 | -1.01E-01 | 3.00E-01 | 2.74E+00 | 2.76E+00 | 7.33E-01 | no | | intergenic_variant | MODIFIER |
| 2 | 101214945 | AX-170216248 | 0 | G | A | 0.053 | -1.01E-01 | 3.00E-01 | 2.74E+00 | 2.76E+00 | 7.33E-01 | no | | intergenic_variant | MODIFIER |
| 2 | 101255483 | AX-170195942 | 0 | C | T | 0.053 | -1.01E-01 | 3.00E-01 | 2.74E+00 | 2.76E+00 | 7.33E-01 | no | | intergenic_variant | MODIFIER |
| 2 | 101324911 | AX-170187929 | 0 | C | T | 0.053 | -1.01E-01 | 3.00E-01 | 2.74E+00 | 2.76E+00 | 7.33E-01 | no | | intergenic_variant | MODIFIER |
| 2 | 101332205 | AX-170309609 | 0 | C | T | 0.053 | -1.01E-01 | 3.00E-01 | 2.74E+00 | 2.76E+00 | 7.33E-01 | no | | intergenic_variant | MODIFIER |
| 2 | 101332303 | AX-170352871 | 0 | C | A | 0.053 | -1.01E-01 | 3.00E-01 | 2.74E+00 | 2.76E+00 | 7.33E-01 | no | | intergenic_variant | MODIFIER |
| 2 | 101354651 | AX-170244148 | 0 | G | A | 0.053 | -1.01E-01 | 3.00E-01 | 2.74E+00 | 2.76E+00 | 7.33E-01 | no | | intergenic_variant | MODIFIER |
| 2 | 101354820 | AX-170094351 | 0 | A | G | 0.053 | -1.01E-01 | 3.00E-01 | 2.74E+00 | 2.76E+00 | 7.33E-01 | no | | intergenic_variant | MODIFIER |
| 2 | 101354873 | AX-170265382 | 0 | A | C | 0.053 | -1.01E-01 | 3.00E-01 | 2.74E+00 | 2.76E+00 | 7.33E-01 | no | | intergenic_variant | MODIFIER |
| 2 | 101362176 | AX-170492350 | 0 | T | G | 0.053 | -1.01E-01 | 3.00E-01 | 2.74E+00 | 2.76E+00 | 7.33E-01 | no | | intergenic_variant | MODIFIER |
| 2 | 101369029 | AX-170064431 | 0 | A | G | 0.053 | -1.01E-01 | 3.00E-01 | 2.74E+00 | 2.76E+00 | 7.33E-01 | no | | intergenic_variant | MODIFIER |
| 2 | 101373460 | AX-170064434 | 0 | C | T | 0.053 | -1.01E-01 | 3.00E-01 | 2.74E+00 | 2.76E+00 | 7.33E-01 | no | | intergenic_variant | MODIFIER |
| 2 | 101377228 | AX-170018485 | 0 | G | A | 0.053 | -1.01E-01 | 3.00E-01 | 2.74E+00 | 2.76E+00 | 7.33E-01 | no | | intergenic_variant | MODIFIER |
| 2 | 101410893 | AX-170018489 | 0 | A | G | 0.053 | -1.01E-01 | 3.00E-01 | 2.74E+00 | 2.76E+00 | 7.33E-01 | no | | intergenic_variant | MODIFIER |
| 2 | 101438154 | AX-97852157 | 0 | T | C | 0.053 | -1.01E-01 | 3.00E-01 | 2.74E+00 | 2.76E+00 | 7.33E-01 | no | | intergenic_variant | MODIFIER |
| 2 | 101438216 | AX-170179123 | 0 | A | G | 0.053 | -1.01E-01 | 3.00E-01 | 2.74E+00 | 2.76E+00 | 7.33E-01 | no | | intergenic_variant | MODIFIER |
| 2 | 101440722 | AX-170234167 | 0 | C | T | 0.053 | -1.01E-01 | 3.00E-01 | 2.74E+00 | 2.76E+00 | 7.33E-01 | no | | intergenic_variant | MODIFIER |
| 2 | 101462193 | AX-97836054 | 0 | T | G | 0.053 | -1.01E-01 | 3.00E-01 | 2.74E+00 | 2.76E+00 | 7.33E-01 | no | | intergenic_variant | MODIFIER |
| 2 | 101462273 | AX-170018483 | 0 | C | T | 0.053 | -1.01E-01 | 3.00E-01 | 2.74E+00 | 2.76E+00 | 7.33E-01 | no | | intergenic_variant | MODIFIER |
| 2 | 101462327 | AX-170198775 | 0 | T | G | 0.053 | -1.01E-01 | 3.00E-01 | 2.74E+00 | 2.76E+00 | 7.33E-01 | no | | intergenic_variant | MODIFIER |
| 2 | 101464384 | AX-170468183 | 0 | A | G | 0.053 | -1.01E-01 | 3.00E-01 | 2.74E+00 | 2.76E+00 | 7.33E-01 | no | | intergenic_variant | MODIFIER |
| 2 | 101551232 | AX-170169162 | 0 | G | A | 0.053 | -1.01E-01 | 3.00E-01 | 2.74E+00 | 2.76E+00 | 7.33E-01 | no | | intergenic_variant | MODIFIER |
| 2 | 101564155 | AX-170239996 | 0 | G | C | 0.053 | -1.01E-01 | 3.00E-01 | 2.74E+00 | 2.76E+00 | 7.33E-01 | no | B230118H07Rik | intron_variant | MODIFIER |
| 2 | 101565513 | AX-170356842 | 0 | A | G | 0.053 | -1.01E-01 | 3.00E-01 | 2.74E+00 | 2.76E+00 | 7.33E-01 | no | B230118H07Rik | intron_variant | MODIFIER |
| 2 | 101576322 | AX-170396208 | 0 | T | C | 0.053 | -1.01E-01 | 3.00E-01 | 2.74E+00 | 2.76E+00 | 7.33E-01 | no | B230118H07Rik | intron_variant | MODIFIER |
| 2 | 101577131 | AX-170114825 | 0 | C | T | 0.053 | -1.01E-01 | 3.00E-01 | 2.74E+00 | 2.76E+00 | 7.33E-01 | no | B230118H07Rik | intron_variant | MODIFIER |
| 2 | 101578239 | AX-97830617 | 0 | G | A | 0.053 | -1.01E-01 | 3.00E-01 | 2.74E+00 | 2.76E+00 | 7.33E-01 | no | B230118H07Rik | intron_variant | MODIFIER |
| 2 | 101600765 | AX-170176630 | 0 | T | C | 0.053 | -1.01E-01 | 3.00E-01 | 2.74E+00 | 2.76E+00 | 7.33E-01 | no | B230118H07Rik | intron_variant | MODIFIER |
| 2 | 101611523 | AX-97838241 | 0 | A | G | 0.053 | -1.01E-01 | 3.00E-01 | 2.74E+00 | 2.76E+00 | 7.33E-01 | no | B230118H07Rik | intron_variant | MODIFIER |
| 2 | 101619773 | AX-97843043 | 0 | A | G | 0.053 | -1.01E-01 | 3.00E-01 | 2.74E+00 | 2.76E+00 | 7.33E-01 | no | Rag2 | upstream_gene_variant | MODIFIER |
| 2 | 101621887 | AX-170511988 | 0 | T | C | 0.053 | -1.01E-01 | 3.00E-01 | 2.74E+00 | 2.76E+00 | 7.33E-01 | no | Rag2 | upstream_gene_variant | MODIFIER |
| 2 | 101640399 | AX-170064436 | 0 | G | A | 0.053 | -1.01E-01 | 3.00E-01 | 2.74E+00 | 2.76E+00 | 7.33E-01 | no | Rag1 | 3_prime_UTR_variant | MODIFIER |
| 2 | 101641931 | AX-170064437 | 0 | T | G | 0.053 | -1.01E-01 | 3.00E-01 | 2.74E+00 | 2.76E+00 | 7.33E-01 | no | Rag1 | synonymous_variant | LOW |
| 2 | 101648294 | AX-170484037 | 0 | G | A | 0.053 | -1.01E-01 | 3.00E-01 | 2.74E+00 | 2.76E+00 | 7.33E-01 | no | Rag1 | intron_variant | MODIFIER |
| 2 | 101651197 | AX-170502323 | 0 | C | T | 0.053 | -1.01E-01 | 3.00E-01 | 2.74E+00 | 2.76E+00 | 7.33E-01 | no | Rag1 | upstream_gene_variant | MODIFIER |
| 2 | 101652876 | AX-170402174 | 0 | G | C | 0.053 | -1.01E-01 | 3.00E-01 | 2.74E+00 | 2.76E+00 | 7.33E-01 | no | Rag1 | upstream_gene_variant | MODIFIER |
| 2 | 101653858 | AX-170319020 | 0 | A | G | 0.053 | -1.01E-01 | 3.00E-01 | 2.74E+00 | 2.76E+00 | 7.33E-01 | no | Rag1 | upstream_gene_variant | MODIFIER |
| 2 | 101654644 | AX-97789233 | 0 | T | C | 0.053 | -1.01E-01 | 3.00E-01 | 2.74E+00 | 2.76E+00 | 7.33E-01 | no | | intergenic_variant | MODIFIER |
| 2 | 101672271 | AX-170471115 | 0 | G | A | 0.053 | -1.01E-01 | 3.00E-01 | 2.74E+00 | 2.76E+00 | 7.33E-01 | no | | intergenic_variant | MODIFIER |
| 2 | 101672716 | AX-170343157 | 0 | A | G | 0.053 | -1.01E-01 | 3.00E-01 | 2.74E+00 | 2.76E+00 | 7.33E-01 | no | | intergenic_variant | MODIFIER |
| 2 | 101674929 | AX-170275737 | 0 | C | T | 0.053 | -1.01E-01 | 3.00E-01 | 2.74E+00 | 2.76E+00 | 7.33E-01 | no | Traf6 | upstream_gene_variant | MODIFIER |
| 2 | 101675104 | AX-168390151 | 0 | G | T | 0.053 | -1.01E-01 | 3.00E-01 | 2.74E+00 | 2.76E+00 | 7.33E-01 | no | Traf6 | upstream_gene_variant | MODIFIER |
| 2 | 101683348 | AX-170505928 | 0 | T | G | 0.053 | -1.01E-01 | 3.00E-01 | 2.74E+00 | 2.76E+00 | 7.33E-01 | no | Traf6 | intron_variant | MODIFIER |
| 2 | 101693914 | AX-170489227 | 0 | T | C | 0.053 | -1.01E-01 | 3.00E-01 | 2.74E+00 | 2.76E+00 | 7.33E-01 | no | Traf6 | intron_variant | MODIFIER |
| 2 | 101697377 | AX-170519748 | 0 | C | A | 0.053 | -1.01E-01 | 3.00E-01 | 2.74E+00 | 2.76E+00 | 7.33E-01 | no | Traf6 | synonymous_variant | LOW |
| 2 | 101706732 | AX-170532659 | 0 | T | C | 0.053 | -1.01E-01 | 3.00E-01 | 2.74E+00 | 2.76E+00 | 7.33E-01 | no | | intergenic_variant | MODIFIER |

| | | | | | | | | | | | | | | | |
|---|---|---|---|---|---|---|---|---|---|---|---|---|---|---|---|
| 2 | 101707287 | AX-97781122 | 0 | G | A | 0.053 | -1.01E-01 | 3.00E-01 | 2.74E+00 | 2.76E+00 | 7.33E-01 | no | | intergenic_variant | MODIFIER |
| 2 | 101707393 | AX-170249210 | 0 | G | C | 0.053 | -1.01E-01 | 3.00E-01 | 2.74E+00 | 2.76E+00 | 7.33E-01 | no | | intergenic_variant | MODIFIER |
| 2 | 101708338 | AX-170433116 | 0 | A | C | 0.053 | -1.01E-01 | 3.00E-01 | 2.74E+00 | 2.76E+00 | 7.33E-01 | no | | intergenic_variant | MODIFIER |
| 2 | 101709262 | AX-170190754 | 0 | T | G | 0.053 | -1.01E-01 | 3.00E-01 | 2.74E+00 | 2.76E+00 | 7.33E-01 | no | | intergenic_variant | MODIFIER |
| 2 | 101717297 | AX-170400369 | 0 | A | G | 0.053 | -1.01E-01 | 3.00E-01 | 2.74E+00 | 2.76E+00 | 7.33E-01 | no | Prr5l | synonymous_variant | LOW |
| 2 | 101722539 | AX-170133400 | 0 | A | C | 0.053 | -1.01E-01 | 3.00E-01 | 2.74E+00 | 2.76E+00 | 7.33E-01 | no | Prr5l | intron_variant | MODIFIER |
| 2 | 101722610 | AX-170371580 | 0 | C | T | 0.053 | -1.01E-01 | 3.00E-01 | 2.74E+00 | 2.76E+00 | 7.33E-01 | no | Prr5l | intron_variant | MODIFIER |
| 2 | 101722835 | AX-170452854 | 0 | G | T | 0.053 | -1.01E-01 | 3.00E-01 | 2.74E+00 | 2.76E+00 | 7.33E-01 | no | Prr5l | intron_variant | MODIFIER |
| 2 | 101723112 | AX-170123397 | 0 | A | G | 0.053 | -1.01E-01 | 3.00E-01 | 2.74E+00 | 2.76E+00 | 7.33E-01 | no | Prr5l | intron_variant | MODIFIER |
| 2 | 101723685 | AX-170064441 | 0 | T | C | 0.053 | -1.01E-01 | 3.00E-01 | 2.74E+00 | 2.76E+00 | 7.33E-01 | no | Prr5l | intron_variant | MODIFIER |
| 2 | 101734846 | AX-170337672 | 0 | A | G | 0.053 | -1.01E-01 | 3.00E-01 | 2.74E+00 | 2.76E+00 | 7.33E-01 | no | Prr5l | intron_variant | MODIFIER |
| 2 | 101734912 | AX-97838251 | 0 | T | C | 0.053 | -1.01E-01 | 3.00E-01 | 2.74E+00 | 2.76E+00 | 7.33E-01 | no | Prr5l | intron_variant | MODIFIER |
| 2 | 101744327 | AX-170242445 | 0 | T | C | 0.053 | -1.01E-01 | 3.00E-01 | 2.74E+00 | 2.76E+00 | 7.33E-01 | no | Prr5l | intron_variant | MODIFIER |
| 2 | 101782611 | AX-97804254 | 0 | T | C | 0.053 | -1.01E-01 | 3.00E-01 | 2.74E+00 | 2.76E+00 | 7.33E-01 | no | Prr5l | intron_variant | MODIFIER |
| 2 | 101783092 | AX-170269284 | 0 | G | A | 0.053 | -1.01E-01 | 3.00E-01 | 2.74E+00 | 2.76E+00 | 7.33E-01 | no | Prr5l | intron_variant | MODIFIER |
| 2 | 101790644 | AX-170018494 | 0 | T | C | 0.053 | -1.01E-01 | 3.00E-01 | 2.74E+00 | 2.76E+00 | 7.33E-01 | no | Prr5l | intron_variant | MODIFIER |
| 2 | 101802511 | AX-170209616 | 0 | T | G | 0.053 | -1.01E-01 | 3.00E-01 | 2.74E+00 | 2.76E+00 | 7.33E-01 | no | Prr5l | upstream_gene_variant | MODIFIER |
| 2 | 101803532 | AX-97842650 | 0 | A | C | 0.053 | -1.01E-01 | 3.00E-01 | 2.74E+00 | 2.76E+00 | 7.33E-01 | no | Prr5l | intron_variant | MODIFIER |
| 2 | 101824784 | AX-97835419 | 0 | T | C | 0.053 | -1.01E-01 | 3.00E-01 | 2.74E+00 | 2.76E+00 | 7.33E-01 | no | Prr5l | intron_variant | MODIFIER |
| 2 | 101846570 | AX-170449780 | 0 | A | G | 0.053 | -1.01E-01 | 3.00E-01 | 2.74E+00 | 2.76E+00 | 7.33E-01 | no | Prr5l | intron_variant,non_coding_transcript_variant | MODIFIER |
| 2 | 101852159 | AX-170018493 | 0 | C | A | 0.053 | -1.01E-01 | 3.00E-01 | 2.74E+00 | 2.76E+00 | 7.33E-01 | no | Prr5l | intron_variant,non_coding_transcript_variant | MODIFIER |
| 2 | 101854509 | AX-170018496 | 0 | G | A | 0.053 | -1.01E-01 | 3.00E-01 | 2.74E+00 | 2.76E+00 | 7.33E-01 | no | Prr5l | intron_variant,non_coding_transcript_variant | MODIFIER |
| 2 | 101854801 | AX-170448694 | 0 | G | A | 0.053 | -1.01E-01 | 3.00E-01 | 2.74E+00 | 2.76E+00 | 7.33E-01 | no | Prr5l | intron_variant,non_coding_transcript_variant | MODIFIER |
| 2 | 101877738 | AX-97814075 | 0 | C | T | 0.053 | -1.01E-01 | 3.00E-01 | 2.74E+00 | 2.76E+00 | 7.33E-01 | no | Prr5l | intron_variant,non_coding_transcript_variant | MODIFIER |
| 2 | 101891825 | AX-170324635 | 0 | T | C | 0.053 | -1.01E-01 | 3.00E-01 | 2.74E+00 | 2.76E+00 | 7.33E-01 | no | Commd9 | intron_variant | MODIFIER |
| 2 | 101895447 | AX-170432968 | 0 | C | T | 0.053 | -1.01E-01 | 3.00E-01 | 2.74E+00 | 2.76E+00 | 7.33E-01 | no | Commd9 | intron_variant | MODIFIER |
| 2 | 101897356 | AX-97804752 | 0 | T | C | 0.053 | -1.01E-01 | 3.00E-01 | 2.74E+00 | 2.76E+00 | 7.33E-01 | no | Commd9 | intron_variant | MODIFIER |
| 2 | 101920763 | AX-170064444 | 0 | G | A | 0.053 | -1.01E-01 | 3.00E-01 | 2.74E+00 | 2.76E+00 | 7.33E-01 | no | | regulatory_region_variant | MODIFIER |
| 2 | 101925083 | AX-170064447 | 0 | A | G | 0.053 | -1.01E-01 | 3.00E-01 | 2.74E+00 | 2.76E+00 | 7.33E-01 | no | | intergenic_variant | MODIFIER |
| 2 | 101931736 | AX-170064446 | 0 | C | T | 0.053 | -1.01E-01 | 3.00E-01 | 2.74E+00 | 2.76E+00 | 7.33E-01 | no | | intergenic_variant | MODIFIER |
| 2 | 101956943 | AX-170356834 | 0 | G | A | 0.053 | -1.01E-01 | 3.00E-01 | 2.74E+00 | 2.76E+00 | 7.33E-01 | no | Ldlrad3 | intron_variant | MODIFIER |
| 2 | 101973666 | AX-170522046 | 0 | G | A | 0.053 | -1.01E-01 | 3.00E-01 | 2.74E+00 | 2.76E+00 | 7.33E-01 | no | Ldlrad3 | intron_variant | MODIFIER |
| 2 | 101980486 | AX-170269066 | 0 | G | C | 0.053 | -1.01E-01 | 3.00E-01 | 2.74E+00 | 2.76E+00 | 7.33E-01 | no | Ldlrad3 | intron_variant | MODIFIER |
| 2 | 101981076 | AX-97815456 | 0 | G | T | 0.053 | -1.01E-01 | 3.00E-01 | 2.74E+00 | 2.76E+00 | 7.33E-01 | no | Ldlrad3 | intron_variant | MODIFIER |
| 2 | 101984219 | AX-170531116 | 0 | C | T | 0.053 | -1.01E-01 | 3.00E-01 | 2.74E+00 | 2.76E+00 | 7.33E-01 | no | Ldlrad3 | intron_variant | MODIFIER |
| 2 | 101985386 | AX-170512300 | 0 | G | A | 0.053 | -1.01E-01 | 3.00E-01 | 2.74E+00 | 2.76E+00 | 7.33E-01 | no | Ldlrad3 | intron_variant | MODIFIER |
| 2 | 102017978 | AX-97799405 | 0 | C | T | 0.053 | -1.01E-01 | 3.00E-01 | 2.74E+00 | 2.76E+00 | 7.33E-01 | no | Ldlrad3 | intron_variant | MODIFIER |
| 2 | 102022352 | AX-170445788 | 0 | G | A | 0.053 | -1.01E-01 | 3.00E-01 | 2.74E+00 | 2.76E+00 | 7.33E-01 | no | Ldlrad3 | intron_variant | MODIFIER |
| 2 | 102024135 | AX-170387742 | 0 | T | C | 0.053 | -1.01E-01 | 3.00E-01 | 2.74E+00 | 2.76E+00 | 7.33E-01 | no | Ldlrad3 | intron_variant | MODIFIER |
| 2 | 102024585 | AX-170354344 | 0 | C | T | 0.053 | -1.01E-01 | 3.00E-01 | 2.74E+00 | 2.76E+00 | 7.33E-01 | no | Ldlrad3 | intron_variant | MODIFIER |
| 2 | 102030297 | AX-170526428 | 0 | G | A | 0.053 | -1.01E-01 | 3.00E-01 | 2.74E+00 | 2.76E+00 | 7.33E-01 | no | Ldlrad3 | intron_variant | MODIFIER |
| 2 | 102031619 | AX-170265059 | 0 | C | T | 0.053 | -1.01E-01 | 3.00E-01 | 2.74E+00 | 2.76E+00 | 7.33E-01 | no | Ldlrad3 | intron_variant | MODIFIER |
| 2 | 102032359 | AX-97827493 | 0 | A | G | 0.053 | -1.01E-01 | 3.00E-01 | 2.74E+00 | 2.76E+00 | 7.33E-01 | no | Ldlrad3 | intron_variant | MODIFIER |
| 2 | 102032852 | AX-170290329 | 0 | G | A | 0.053 | -1.01E-01 | 3.00E-01 | 2.74E+00 | 2.76E+00 | 7.33E-01 | no | Ldlrad3 | intron_variant | MODIFIER |
| 2 | 102045485 | AX-170106431 | 0 | A | G | 0.053 | -1.01E-01 | 3.00E-01 | 2.74E+00 | 2.76E+00 | 7.33E-01 | no | Ldlrad3 | intron_variant | MODIFIER |
| 2 | 102053501 | AX-170396892 | 0 | A | G | 0.053 | -1.01E-01 | 3.00E-01 | 2.74E+00 | 2.76E+00 | 7.33E-01 | no | Ldlrad3 | intron_variant | MODIFIER |
| 2 | 102057397 | AX-97839520 | 0 | A | G | 0.053 | -1.01E-01 | 3.00E-01 | 2.74E+00 | 2.76E+00 | 7.33E-01 | no | Ldlrad3 | intron_variant | MODIFIER |
| 2 | 102069294 | AX-170175644 | 0 | C | T | 0.053 | -1.01E-01 | 3.00E-01 | 2.74E+00 | 2.76E+00 | 7.33E-01 | no | Ldlrad3 | intron_variant | MODIFIER |
| 2 | 102069838 | AX-170268193 | 0 | A | C | 0.053 | -1.01E-01 | 3.00E-01 | 2.74E+00 | 2.76E+00 | 7.33E-01 | no | Ldlrad3 | intron_variant | MODIFIER |
| 2 | 102073421 | AX-97792007 | 0 | A | G | 0.053 | -1.01E-01 | 3.00E-01 | 2.74E+00 | 2.76E+00 | 7.33E-01 | no | Ldlrad3 | intron_variant | MODIFIER |
| 2 | 102074350 | AX-170187485 | 0 | T | C | 0.053 | -1.01E-01 | 3.00E-01 | 2.74E+00 | 2.76E+00 | 7.33E-01 | no | Ldlrad3 | intron_variant | MODIFIER |
| 2 | 102097406 | AX-170345500 | 0 | G | T | 0.053 | -1.01E-01 | 3.00E-01 | 2.74E+00 | 2.76E+00 | 7.33E-01 | no | Ldlrad3 | intron_variant | MODIFIER |
| 2 | 102102451 | AX-170142651 | 0 | C | A | 0.053 | -1.01E-01 | 3.00E-01 | 2.74E+00 | 2.76E+00 | 7.33E-01 | no | Ldlrad3 | intron_variant | MODIFIER |
| 2 | 102102972 | AX-170116522 | 0 | G | A | 0.053 | -1.01E-01 | 3.00E-01 | 2.74E+00 | 2.76E+00 | 7.33E-01 | no | Ldlrad3 | intron_variant | MODIFIER |

| | | | | | | | | | | | | | | |
|---|---|---|---|---|---|---|---|---|---|---|---|---|---|---|
| 2 | 102103259 | AX-97852862 | 0 | T | G | 0.053 | -1.01E-01 | 3.00E-01 | 2.74E+00 | 2.76E+00 | 7.33E-01 | no | Ldlrad3 | intron_variant | MODIFIER |
| 2 | 102103347 | AX-170467986 | 0 | C | T | 0.053 | -1.01E-01 | 3.00E-01 | 2.74E+00 | 2.76E+00 | 7.33E-01 | no | Ldlrad3 | intron_variant | MODIFIER |
| 2 | 102103749 | AX-170218163 | 0 | A | C | 0.053 | -1.01E-01 | 3.00E-01 | 2.74E+00 | 2.76E+00 | 7.33E-01 | no | Ldlrad3 | intron_variant | MODIFIER |
| 2 | 102105015 | AX-170235291 | 0 | T | A | 0.053 | -1.01E-01 | 3.00E-01 | 2.74E+00 | 2.76E+00 | 7.33E-01 | no | Ldlrad3 | intron_variant | MODIFIER |
| 2 | 102107192 | AX-170172240 | 0 | C | T | 0.053 | -1.01E-01 | 3.00E-01 | 2.74E+00 | 2.76E+00 | 7.33E-01 | no | Ldlrad3 | intron_variant | MODIFIER |
| 2 | 102113322 | AX-170454177 | 0 | A | G | 0.053 | -1.01E-01 | 3.00E-01 | 2.74E+00 | 2.76E+00 | 7.33E-01 | no | Ldlrad3 | intron_variant | MODIFIER |
| 2 | 102115838 | AX-170258309 | 0 | G | A | 0.053 | -1.01E-01 | 3.00E-01 | 2.74E+00 | 2.76E+00 | 7.33E-01 | no | Ldlrad3 | intron_variant | MODIFIER |
| 2 | 102122423 | AX-170188834 | 0 | A | G | 0.053 | -1.01E-01 | 3.00E-01 | 2.74E+00 | 2.76E+00 | 7.33E-01 | no | Ldlrad3 | intron_variant | MODIFIER |
| 2 | 102126451 | AX-170115482 | 0 | T | G | 0.053 | -1.01E-01 | 3.00E-01 | 2.74E+00 | 2.76E+00 | 7.33E-01 | no | Ldlrad3 | intron_variant | MODIFIER |
| 2 | 102126738 | AX-97827492 | 0 | A | C | 0.053 | -1.01E-01 | 3.00E-01 | 2.74E+00 | 2.76E+00 | 7.33E-01 | no | Ldlrad3 | intron_variant | MODIFIER |
| 2 | 102127418 | AX-170430042 | 0 | T | C | 0.053 | -1.01E-01 | 3.00E-01 | 2.74E+00 | 2.76E+00 | 7.33E-01 | no | Ldlrad3 | intron_variant | MODIFIER |
| 2 | 102131207 | AX-170243339 | 0 | A | G | 0.053 | -1.01E-01 | 3.00E-01 | 2.74E+00 | 2.76E+00 | 7.33E-01 | no | Ldlrad3 | intron_variant | MODIFIER |
| 2 | 102132503 | AX-170102092 | 0 | G | T | 0.053 | -1.01E-01 | 3.00E-01 | 2.74E+00 | 2.76E+00 | 7.33E-01 | no | Ldlrad3 | intron_variant | MODIFIER |
| 2 | 102138436 | AX-170334873 | 0 | A | G | 0.053 | -1.01E-01 | 3.00E-01 | 2.74E+00 | 2.76E+00 | 7.33E-01 | no | Ldlrad3 | intron_variant | MODIFIER |
| 2 | 102138495 | AX-170265664 | 0 | T | C | 0.053 | -1.01E-01 | 3.00E-01 | 2.74E+00 | 2.76E+00 | 7.33E-01 | no | Ldlrad3 | intron_variant | MODIFIER |
| 2 | 102139299 | AX-170172113 | 0 | C | A | 0.053 | -1.01E-01 | 3.00E-01 | 2.74E+00 | 2.76E+00 | 7.33E-01 | no | Ldlrad3 | intron_variant | MODIFIER |
| 2 | 102179687 | AX-97835877 | 0 | A | G | 0.053 | -1.01E-01 | 3.00E-01 | 2.74E+00 | 2.76E+00 | 7.33E-01 | no | Ldlrad3 | intron_variant | MODIFIER |
| 2 | 102184769 | AX-170283787 | 0 | A | G | 0.053 | -1.01E-01 | 3.00E-01 | 2.74E+00 | 2.76E+00 | 7.33E-01 | no | Ldlrad3 | intron_variant | MODIFIER |
| 2 | 102193314 | AX-170446406 | 0 | A | G | 0.053 | -1.01E-01 | 3.00E-01 | 2.74E+00 | 2.76E+00 | 7.33E-01 | no | | intergenic_variant | MODIFIER |
| 2 | 102193590 | AX-170439612 | 0 | G | A | 0.053 | -1.01E-01 | 3.00E-01 | 2.74E+00 | 2.76E+00 | 7.33E-01 | no | | regulatory_region_variant | MODIFIER |
| 2 | 102203668 | AX-170175146 | 0 | G | A | 0.053 | -1.01E-01 | 3.00E-01 | 2.74E+00 | 2.76E+00 | 7.33E-01 | no | | regulatory_region_variant | MODIFIER |
| 2 | 102210936 | AX-170338396 | 0 | C | T | 0.053 | -1.01E-01 | 3.00E-01 | 2.74E+00 | 2.76E+00 | 7.33E-01 | no | Gm13866 | upstream_gene_variant | MODIFIER |
| 2 | 102213523 | AX-170220924 | 0 | A | G | 0.053 | -1.01E-01 | 3.00E-01 | 2.74E+00 | 2.76E+00 | 7.33E-01 | no | Gm13866 | upstream_gene_variant | MODIFIER |
| 2 | 102214217 | AX-168411617 | 0 | G | A | 0.053 | -1.01E-01 | 3.00E-01 | 2.74E+00 | 2.76E+00 | 7.33E-01 | no | Gm13866 | upstream_gene_variant | MODIFIER |
| 2 | 102222858 | AX-170321847 | 0 | C | T | 0.053 | -1.01E-01 | 3.00E-01 | 2.74E+00 | 2.76E+00 | 7.33E-01 | no | | regulatory_region_variant | MODIFIER |
| 2 | 102223447 | AX-97806169 | 0 | G | T | 0.053 | -1.01E-01 | 3.00E-01 | 2.74E+00 | 2.76E+00 | 7.33E-01 | no | | regulatory_region_variant | MODIFIER |
| 2 | 102284971 | AX-97832482 | 0 | G | T | 0.053 | -1.01E-01 | 3.00E-01 | 2.74E+00 | 2.76E+00 | 7.33E-01 | no | | intergenic_variant | MODIFIER |
| 2 | 102298749 | AX-170204860 | 0 | T | C | 0.053 | -1.01E-01 | 3.00E-01 | 2.74E+00 | 2.76E+00 | 7.33E-01 | no | Trim44 | downstream_gene_variant | MODIFIER |
| 2 | 102299163 | AX-170335047 | 0 | G | A | 0.053 | -1.01E-01 | 3.00E-01 | 2.74E+00 | 2.76E+00 | 7.33E-01 | no | Trim44 | downstream_gene_variant | MODIFIER |
| 2 | 102303725 | AX-97838265 | 0 | T | G | 0.053 | -1.01E-01 | 3.00E-01 | 2.74E+00 | 2.76E+00 | 7.33E-01 | no | Trim44 | 3_prime_UTR_variant | MODIFIER |
| 2 | 102303787 | AX-170519004 | 0 | C | G | 0.053 | -1.01E-01 | 3.00E-01 | 2.74E+00 | 2.76E+00 | 7.33E-01 | no | Trim44 | 3_prime_UTR_variant | MODIFIER |
| 2 | 102313635 | AX-170446469 | 0 | G | A | 0.053 | -1.01E-01 | 3.00E-01 | 2.74E+00 | 2.76E+00 | 7.33E-01 | no | Trim44 | intron_variant | MODIFIER |
| 2 | 102327326 | AX-97800464 | 0 | A | G | 0.053 | -1.01E-01 | 3.00E-01 | 2.74E+00 | 2.76E+00 | 7.33E-01 | no | Trim44 | intron_variant | MODIFIER |
| 2 | 102338717 | AX-170018508 | 0 | A | G | 0.053 | -1.01E-01 | 3.00E-01 | 2.74E+00 | 2.76E+00 | 7.33E-01 | no | Trim44 | intron_variant | MODIFIER |
| 2 | 102338774 | AX-170018499 | 0 | A | G | 0.053 | -1.01E-01 | 3.00E-01 | 2.74E+00 | 2.76E+00 | 7.33E-01 | no | Trim44 | intron_variant | MODIFIER |
| 2 | 102341580 | AX-170287817 | 0 | A | G | 0.053 | -1.01E-01 | 3.00E-01 | 2.74E+00 | 2.76E+00 | 7.33E-01 | no | Trim44 | intron_variant | MODIFIER |
| 2 | 102342362 | AX-170018501 | 0 | G | T | 0.053 | -1.01E-01 | 3.00E-01 | 2.74E+00 | 2.76E+00 | 7.33E-01 | no | Trim44 | intron_variant | MODIFIER |
| 2 | 102348621 | AX-170387092 | 0 | C | T | 0.053 | -1.01E-01 | 3.00E-01 | 2.74E+00 | 2.76E+00 | 7.33E-01 | no | Trim44 | intron_variant | MODIFIER |
| 2 | 102361389 | AX-97845787 | 0 | C | T | 0.053 | -1.01E-01 | 3.00E-01 | 2.74E+00 | 2.76E+00 | 7.33E-01 | no | Trim44 | intron_variant | MODIFIER |
| 2 | 102365153 | AX-170315884 | 0 | A | G | 0.053 | -1.01E-01 | 3.00E-01 | 2.74E+00 | 2.76E+00 | 7.33E-01 | no | Trim44 | intron_variant | MODIFIER |
| 2 | 102374017 | AX-170018505 | 0 | T | C | 0.053 | -1.01E-01 | 3.00E-01 | 2.74E+00 | 2.76E+00 | 7.33E-01 | no | Trim44 | intron_variant | MODIFIER |
| 2 | 102374070 | AX-170018500 | 0 | T | C | 0.053 | -1.01E-01 | 3.00E-01 | 2.74E+00 | 2.76E+00 | 7.33E-01 | no | Trim44 | intron_variant | MODIFIER |
| 2 | 102387006 | AX-97795172 | 0 | C | T | 0.053 | -1.01E-01 | 3.00E-01 | 2.74E+00 | 2.76E+00 | 7.33E-01 | no | Trim44 | intron_variant | MODIFIER |
| 2 | 102414185 | AX-97838267 | 0 | T | C | 0.053 | -1.01E-01 | 3.00E-01 | 2.74E+00 | 2.76E+00 | 7.33E-01 | no | | intergenic_variant | MODIFIER |
| 2 | 102422477 | AX-170018503 | 0 | G | A | 0.053 | -1.01E-01 | 3.00E-01 | 2.74E+00 | 2.76E+00 | 7.33E-01 | no | | intergenic_variant | MODIFIER |
| 2 | 102427409 | AX-170315348 | 0 | A | C | 0.053 | -1.01E-01 | 3.00E-01 | 2.74E+00 | 2.76E+00 | 7.33E-01 | no | | intergenic_variant | MODIFIER |
| 2 | 102429894 | AX-170413013 | 0 | G | C | 0.053 | -1.01E-01 | 3.00E-01 | 2.74E+00 | 2.76E+00 | 7.33E-01 | no | | intergenic_variant | MODIFIER |
| 2 | 102432380 | AX-97782729 | 0 | C | T | 0.053 | -1.01E-01 | 3.00E-01 | 2.74E+00 | 2.76E+00 | 7.33E-01 | no | | intergenic_variant | MODIFIER |
| 2 | 102439125 | AX-170117308 | 0 | T | C | 0.053 | -1.01E-01 | 3.00E-01 | 2.74E+00 | 2.76E+00 | 7.33E-01 | no | | intergenic_variant | MODIFIER |
| 2 | 102468770 | AX-170520438 | 0 | C | G | 0.053 | -1.01E-01 | 3.00E-01 | 2.74E+00 | 2.76E+00 | 7.33E-01 | no | | intergenic_variant | MODIFIER |
| 2 | 102469093 | AX-170018502 | 0 | G | C | 0.053 | -1.01E-01 | 3.00E-01 | 2.74E+00 | 2.76E+00 | 7.33E-01 | no | | intergenic_variant | MODIFIER |
| 2 | 102480656 | AX-170221337 | 0 | G | A | 0.053 | -1.01E-01 | 3.00E-01 | 2.74E+00 | 2.76E+00 | 7.33E-01 | no | | intergenic_variant | MODIFIER |
| 2 | 102482481 | AX-97787688 | 0 | A | G | 0.053 | -1.01E-01 | 3.00E-01 | 2.74E+00 | 2.76E+00 | 7.33E-01 | no | | intergenic_variant | MODIFIER |
| 2 | 102498926 | AX-170195219 | 0 | G | A | 0.053 | -1.01E-01 | 3.00E-01 | 2.74E+00 | 2.76E+00 | 7.33E-01 | no | | intergenic_variant | MODIFIER |

| | | | | | | | | | | | | | | | |
|---|---|---|---|---|---|---|---|---|---|---|---|---|---|---|---|
| 2 | 102507690 | AX-170088685 | 0 | C | T | 0.053 | -1.01E-01 | 3.00E-01 | 2.74E+00 | 2.76E+00 | 7.33E-01 | no | | intergenic_variant | MODIFIER |
| 2 | 102529366 | AX-170356010 | 0 | G | A | 0.053 | -1.01E-01 | 3.00E-01 | 2.74E+00 | 2.76E+00 | 7.33E-01 | no | | intergenic_variant | MODIFIER |
| 2 | 102530809 | AX-97792459 | 0 | T | C | 0.053 | -1.01E-01 | 3.00E-01 | 2.74E+00 | 2.76E+00 | 7.33E-01 | no | | intergenic_variant | MODIFIER |
| 2 | 102570000 | AX-170275421 | 0 | T | C | 0.053 | -1.01E-01 | 3.00E-01 | 2.74E+00 | 2.76E+00 | 7.33E-01 | no | Pamr1 | intron_variant | MODIFIER |
| 2 | 102571035 | AX-170478689 | 0 | C | T | 0.053 | -1.01E-01 | 3.00E-01 | 2.74E+00 | 2.76E+00 | 7.33E-01 | no | Pamr1 | intron_variant | MODIFIER |
| 2 | 102580182 | AX-170386908 | 0 | C | T | 0.053 | -1.01E-01 | 3.00E-01 | 2.74E+00 | 2.76E+00 | 7.33E-01 | no | Pamr1 | intron_variant | MODIFIER |
| 2 | 102581758 | AX-170064458 | 0 | A | T | 0.053 | -1.01E-01 | 3.00E-01 | 2.74E+00 | 2.76E+00 | 7.33E-01 | no | Pamr1 | intron_variant | MODIFIER |
| 2 | 102585755 | AX-170253041 | 0 | T | C | 0.053 | -1.01E-01 | 3.00E-01 | 2.74E+00 | 2.76E+00 | 7.33E-01 | no | Pamr1 | intron_variant | MODIFIER |
| 2 | 102590470 | AX-170112666 | 0 | A | G | 0.053 | -1.01E-01 | 3.00E-01 | 2.74E+00 | 2.76E+00 | 7.33E-01 | no | Pamr1 | intron_variant | MODIFIER |
| 2 | 102600994 | AX-170430613 | 0 | T | C | 0.053 | -1.01E-01 | 3.00E-01 | 2.74E+00 | 2.76E+00 | 7.33E-01 | no | Pamr1 | intron_variant | MODIFIER |
| 2 | 102610922 | AX-97807870 | 0 | A | G | 0.053 | -1.01E-01 | 3.00E-01 | 2.74E+00 | 2.76E+00 | 7.33E-01 | no | Pamr1 | intron_variant | MODIFIER |
| 2 | 102613054 | AX-170385760 | 0 | G | A | 0.053 | -1.01E-01 | 3.00E-01 | 2.74E+00 | 2.76E+00 | 7.33E-01 | no | Pamr1 | intron_variant | MODIFIER |
| 2 | 102615857 | AX-170138162 | 0 | C | A | 0.053 | -1.01E-01 | 3.00E-01 | 2.74E+00 | 2.76E+00 | 7.33E-01 | no | Pamr1 | intron_variant | MODIFIER |
| 2 | 102617314 | AX-170344152 | 0 | C | A | 0.053 | -1.01E-01 | 3.00E-01 | 2.74E+00 | 2.76E+00 | 7.33E-01 | no | Pamr1 | intron_variant | MODIFIER |
| 2 | 102618807 | AX-170367703 | 0 | C | T | 0.053 | -1.01E-01 | 3.00E-01 | 2.74E+00 | 2.76E+00 | 7.33E-01 | no | Pamr1 | intron_variant | MODIFIER |
| 2 | 102625036 | AX-97832550 | 0 | A | G | 0.053 | -1.01E-01 | 3.00E-01 | 2.74E+00 | 2.76E+00 | 7.33E-01 | no | Pamr1 | intron_variant | MODIFIER |
| 2 | 102632371 | AX-170234218 | 0 | A | T | 0.053 | -1.01E-01 | 3.00E-01 | 2.74E+00 | 2.76E+00 | 7.33E-01 | no | Pamr1 | intron_variant | MODIFIER |
| 2 | 102641737 | AX-168398380 | 0 | A | G | 0.053 | -1.01E-01 | 3.00E-01 | 2.74E+00 | 2.76E+00 | 7.33E-01 | no | Pamr1 | intron_variant | MODIFIER |
| 2 | 102641836 | AX-170246972 | 0 | A | G | 0.053 | -1.01E-01 | 3.00E-01 | 2.74E+00 | 2.76E+00 | 7.33E-01 | no | Pamr1 | intron_variant | MODIFIER |
| 2 | 102646200 | AX-168415652 | 0 | C | A | 0.053 | -1.01E-01 | 3.00E-01 | 2.74E+00 | 2.76E+00 | 7.33E-01 | no | Pamr1 | downstream_gene_variant | MODIFIER |
| 2 | 102649620 | AX-170361213 | 0 | C | T | 0.053 | -1.01E-01 | 3.00E-01 | 2.74E+00 | 2.76E+00 | 7.33E-01 | no | | intergenic_variant | MODIFIER |
| 2 | 102650165 | AX-170335478 | 0 | A | G | 0.053 | -1.01E-01 | 3.00E-01 | 2.74E+00 | 2.76E+00 | 7.33E-01 | no | | intergenic_variant | MODIFIER |
| 2 | 102650208 | AX-170064464 | 0 | A | G | 0.053 | -1.01E-01 | 3.00E-01 | 2.74E+00 | 2.76E+00 | 7.33E-01 | no | | intergenic_variant | MODIFIER |
| 2 | 102660380 | AX-97839166 | 0 | T | C | 0.053 | -1.01E-01 | 3.00E-01 | 2.74E+00 | 2.76E+00 | 7.33E-01 | no | Slc1a2 | intron_variant | MODIFIER |
| 2 | 102660525 | AX-170234877 | 0 | A | G | 0.053 | -1.01E-01 | 3.00E-01 | 2.74E+00 | 2.76E+00 | 7.33E-01 | no | Slc1a2 | intron_variant | MODIFIER |
| 2 | 102671210 | AX-170103598 | 0 | G | A | 0.053 | -1.01E-01 | 3.00E-01 | 2.74E+00 | 2.76E+00 | 7.33E-01 | no | Slc1a2 | intron_variant | MODIFIER |
| 2 | 102672579 | AX-170450062 | 0 | G | A | 0.053 | -1.01E-01 | 3.00E-01 | 2.74E+00 | 2.76E+00 | 7.33E-01 | no | Slc1a2 | intron_variant | MODIFIER |
| 2 | 102677177 | AX-170094353 | 0 | T | C | 0.053 | -1.01E-01 | 3.00E-01 | 2.74E+00 | 2.76E+00 | 7.33E-01 | no | Slc1a2 | intron_variant | MODIFIER |
| 2 | 102680488 | AX-170244540 | 0 | G | A | 0.053 | -1.01E-01 | 3.00E-01 | 2.74E+00 | 2.76E+00 | 7.33E-01 | no | Slc1a2 | intron_variant | MODIFIER |
| 2 | 102690196 | AX-170247085 | 0 | C | T | 0.053 | -1.01E-01 | 3.00E-01 | 2.74E+00 | 2.76E+00 | 7.33E-01 | no | Slc1a2 | intron_variant | MODIFIER |
| 2 | 102694280 | AX-170402585 | 0 | T | A | 0.053 | -1.01E-01 | 3.00E-01 | 2.74E+00 | 2.76E+00 | 7.33E-01 | no | Slc1a2 | intron_variant | MODIFIER |
| 2 | 102704748 | AX-97823631 | 0 | C | A | 0.053 | -1.01E-01 | 3.00E-01 | 2.74E+00 | 2.76E+00 | 7.33E-01 | no | Slc1a2 | upstream_gene_variant | MODIFIER |
| 2 | 102707086 | AX-170377004 | 0 | A | T | 0.053 | -1.01E-01 | 3.00E-01 | 2.74E+00 | 2.76E+00 | 7.33E-01 | no | Slc1a2 | intron_variant | MODIFIER |
| 2 | 102724632 | AX-170415819 | 0 | A | G | 0.053 | -1.01E-01 | 3.00E-01 | 2.74E+00 | 2.76E+00 | 7.33E-01 | no | Slc1a2 | intron_variant | MODIFIER |
| 2 | 102726587 | AX-170179649 | 0 | C | G | 0.053 | -1.01E-01 | 3.00E-01 | 2.74E+00 | 2.76E+00 | 7.33E-01 | no | Slc1a2 | intron_variant | MODIFIER |
| 2 | 102726936 | AX-170477550 | 0 | G | A | 0.053 | -1.01E-01 | 3.00E-01 | 2.74E+00 | 2.76E+00 | 7.33E-01 | no | Slc1a2 | intron_variant | MODIFIER |
| 2 | 102729105 | AX-170304341 | 0 | C | G | 0.053 | -1.01E-01 | 3.00E-01 | 2.74E+00 | 2.76E+00 | 7.33E-01 | no | Slc1a2 | intron_variant | MODIFIER |
| 2 | 102729245 | AX-170335075 | 0 | T | A | 0.053 | -1.01E-01 | 3.00E-01 | 2.74E+00 | 2.76E+00 | 7.33E-01 | no | Slc1a2 | intron_variant | MODIFIER |
| 2 | 102731991 | AX-170528229 | 0 | C | T | 0.053 | -1.01E-01 | 3.00E-01 | 2.74E+00 | 2.76E+00 | 7.33E-01 | no | Slc1a2 | intron_variant | MODIFIER |
| 2 | 102736854 | AX-170268256 | 0 | A | G | 0.053 | -1.01E-01 | 3.00E-01 | 2.74E+00 | 2.76E+00 | 7.33E-01 | no | Slc1a2 | intron_variant | MODIFIER |
| 2 | 102742204 | AX-170361111 | 0 | A | G | 0.053 | -1.01E-01 | 3.00E-01 | 2.74E+00 | 2.76E+00 | 7.33E-01 | no | Slc1a2 | intron_variant | MODIFIER |
| 2 | 102747774 | AX-170490365 | 0 | T | G | 0.053 | -1.01E-01 | 3.00E-01 | 2.74E+00 | 2.76E+00 | 7.33E-01 | no | Slc1a2 | intron_variant | MODIFIER |
| 2 | 102757707 | AX-97784793 | 0 | A | C | 0.053 | -1.01E-01 | 3.00E-01 | 2.74E+00 | 2.76E+00 | 7.33E-01 | no | Slc1a2 | intron_variant | MODIFIER |
| 2 | 102764263 | AX-170398510 | 0 | G | A | 0.053 | -1.01E-01 | 3.00E-01 | 2.74E+00 | 2.76E+00 | 7.33E-01 | no | Slc1a2 | intron_variant | MODIFIER |
| 2 | 102765028 | AX-170064467 | 0 | C | A | 0.053 | -1.01E-01 | 3.00E-01 | 2.74E+00 | 2.76E+00 | 7.33E-01 | no | Slc1a2 | intron_variant | MODIFIER |
| 2 | 102779840 | AX-170094354 | 0 | A | G | 0.053 | -1.01E-01 | 3.00E-01 | 2.74E+00 | 2.76E+00 | 7.33E-01 | no | Slc1a2 | downstream_gene_variant | MODIFIER |
| 2 | 102790481 | AX-170247317 | 0 | T | C | 0.053 | -1.01E-01 | 3.00E-01 | 2.74E+00 | 2.76E+00 | 7.33E-01 | no | Slc1a2 | 3_prime_UTR_variant | MODIFIER |
| 2 | 102793248 | AX-170490080 | 0 | G | A | 0.053 | -1.01E-01 | 3.00E-01 | 2.74E+00 | 2.76E+00 | 7.33E-01 | no | Slc1a2 | downstream_gene_variant | MODIFIER |
| 2 | 102820392 | AX-170018511 | 0 | T | G | 0.053 | -1.01E-01 | 3.00E-01 | 2.74E+00 | 2.76E+00 | 7.33E-01 | no | Cd44 | intron_variant | MODIFIER |
| 2 | 102872332 | AX-170018512 | 0 | C | T | 0.053 | -1.01E-01 | 3.00E-01 | 2.74E+00 | 2.76E+00 | 7.33E-01 | no | Cd44 | intron_variant | MODIFIER |
| 2 | 102872523 | AX-170119424 | 0 | C | T | 0.053 | -1.01E-01 | 3.00E-01 | 2.74E+00 | 2.76E+00 | 7.33E-01 | no | Cd44 | intron_variant | MODIFIER |
| 2 | 102882839 | AX-97821028 | 0 | C | T | 0.053 | -1.01E-01 | 3.00E-01 | 2.74E+00 | 2.76E+00 | 7.33E-01 | no | Cd44 | intron_variant | MODIFIER |
| 2 | 102918310 | AX-170239406 | 0 | G | A | 0.053 | -1.01E-01 | 3.00E-01 | 2.74E+00 | 2.76E+00 | 7.33E-01 | no | | intergenic_variant | MODIFIER |
| 2 | 102932487 | AX-170018513 | 0 | A | G | 0.053 | -1.01E-01 | 3.00E-01 | 2.74E+00 | 2.76E+00 | 7.33E-01 | no | | intergenic_variant | MODIFIER |

| | | | | | | | | | | | | | | | |
|---|---|---|---|---|---|---|---|---|---|---|---|---|---|---|---|
| 2 | 102934394 | AX-97847249 | 0 | T | C | 0.053 | -1.01E-01 | 3.00E-01 | 2.74E+00 | 2.76E+00 | 7.33E-01 | no | | intergenic_variant | MODIFIER |
| 2 | 102943200 | AX-170018514 | 0 | G | A | 0.053 | -1.01E-01 | 3.00E-01 | 2.74E+00 | 2.76E+00 | 7.33E-01 | no | | regulatory_region_variant | MODIFIER |
| 2 | 102949192 | AX-170018515 | 0 | G | A | 0.053 | -1.01E-01 | 3.00E-01 | 2.74E+00 | 2.76E+00 | 7.33E-01 | no | | regulatory_region_variant | MODIFIER |
| 2 | 102956675 | AX-97780569 | 0 | T | G | 0.053 | -1.01E-01 | 3.00E-01 | 2.74E+00 | 2.76E+00 | 7.33E-01 | no | | regulatory_region_variant | MODIFIER |
| 2 | 102970334 | AX-170018509 | 0 | G | A | 0.053 | -1.01E-01 | 3.00E-01 | 2.74E+00 | 2.76E+00 | 7.33E-01 | no | | intergenic_variant | MODIFIER |
| 2 | 102970366 | AX-170147790 | 0 | T | C | 0.053 | -1.01E-01 | 3.00E-01 | 2.74E+00 | 2.76E+00 | 7.33E-01 | no | | intergenic_variant | MODIFIER |
| 2 | 102975927 | AX-170164181 | 0 | G | T | 0.053 | -1.01E-01 | 3.00E-01 | 2.74E+00 | 2.76E+00 | 7.33E-01 | no | | regulatory_region_variant | MODIFIER |
| 2 | 102976622 | AX-170116304 | 0 | T | C | 0.053 | -1.01E-01 | 3.00E-01 | 2.74E+00 | 2.76E+00 | 7.33E-01 | no | | intergenic_variant | MODIFIER |
| 2 | 103043796 | AX-170382854 | 0 | C | G | 0.053 | -1.01E-01 | 3.00E-01 | 2.74E+00 | 2.76E+00 | 7.33E-01 | no | Pdhx | intron_variant | MODIFIER |
| 2 | 103130407 | AX-170354439 | 0 | C | T | 0.053 | -1.01E-01 | 3.00E-01 | 2.74E+00 | 2.76E+00 | 7.33E-01 | no | | intergenic_variant | MODIFIER |
| 2 | 103141238 | AX-170088687 | 0 | T | C | 0.053 | -1.01E-01 | 3.00E-01 | 2.74E+00 | 2.76E+00 | 7.33E-01 | no | | intergenic_variant | MODIFIER |
| 2 | 103267259 | AX-97839164 | 0 | C | T | 0.053 | -1.01E-01 | 3.00E-01 | 2.74E+00 | 2.76E+00 | 7.33E-01 | no | Ehf | intron_variant | MODIFIER |
| 2 | 103267498 | AX-168379259 | 0 | T | C | 0.053 | -1.01E-01 | 3.00E-01 | 2.74E+00 | 2.76E+00 | 7.33E-01 | no | Ehf | intron_variant | MODIFIER |
| 2 | 103418740 | AX-170400476 | 0 | G | T | 0.053 | -1.01E-01 | 3.00E-01 | 2.74E+00 | 2.76E+00 | 7.33E-01 | no | Elf5 | intron_variant,non_coding_transcript_variant | MODIFIER |
| 2 | 103420096 | AX-170018522 | 0 | T | C | 0.053 | -1.01E-01 | 3.00E-01 | 2.74E+00 | 2.76E+00 | 7.33E-01 | no | Elf5 | upstream_gene_variant | MODIFIER |
| 2 | 103423310 | AX-170018519 | 0 | C | A | 0.053 | -1.01E-01 | 3.00E-01 | 2.74E+00 | 2.76E+00 | 7.33E-01 | no | Elf5 | upstream_gene_variant | MODIFIER |
| 2 | 103439276 | AX-170121697 | 0 | T | C | 0.053 | -1.01E-01 | 3.00E-01 | 2.74E+00 | 2.76E+00 | 7.33E-01 | no | Elf5 | synonymous_variant | LOW |
| 2 | 103442246 | AX-170018527 | 0 | T | A | 0.053 | -1.01E-01 | 3.00E-01 | 2.74E+00 | 2.76E+00 | 7.33E-01 | no | Elf5 | intron_variant | MODIFIER |
| 2 | 103444544 | AX-97816002 | 0 | T | C | 0.053 | -1.01E-01 | 3.00E-01 | 2.74E+00 | 2.76E+00 | 7.33E-01 | no | Elf5 | intron_variant | MODIFIER |
| 2 | 103450038 | AX-170526407 | 0 | T | C | 0.053 | -1.01E-01 | 3.00E-01 | 2.74E+00 | 2.76E+00 | 7.33E-01 | no | Elf5 | 3_prime_UTR_variant | MODIFIER |
| 2 | 103450501 | AX-168417714 | 0 | A | G | 0.053 | -1.01E-01 | 3.00E-01 | 2.74E+00 | 2.76E+00 | 7.33E-01 | no | Elf5 | 3_prime_UTR_variant | MODIFIER |
| 2 | 103459050 | AX-170334324 | 0 | T | C | 0.053 | -1.01E-01 | 3.00E-01 | 2.74E+00 | 2.76E+00 | 7.33E-01 | no | Cat | intron_variant | MODIFIER |
| 2 | 103461918 | AX-97783236 | 0 | A | G | 0.053 | -1.01E-01 | 3.00E-01 | 2.74E+00 | 2.76E+00 | 7.33E-01 | no | Cat | intron_variant | MODIFIER |
| 2 | 103469500 | AX-170378306 | 0 | T | C | 0.053 | -1.01E-01 | 3.00E-01 | 2.74E+00 | 2.76E+00 | 7.33E-01 | no | Cat | intron_variant | MODIFIER |
| 2 | 103568378 | AX-170365910 | 0 | T | C | 0.053 | -1.01E-01 | 3.00E-01 | 2.74E+00 | 2.76E+00 | 7.33E-01 | no | Abtb2 | intron_variant | MODIFIER |
| 2 | 103581353 | AX-170018529 | 0 | T | C | 0.053 | -1.01E-01 | 3.00E-01 | 2.74E+00 | 2.76E+00 | 7.33E-01 | no | Abtb2 | intron_variant | MODIFIER |
| 2 | 103877838 | AX-97809512 | 0 | C | T | 0.053 | -1.01E-01 | 3.00E-01 | 2.74E+00 | 2.76E+00 | 7.33E-01 | no | | intergenic_variant | MODIFIER |
| 2 | 103911067 | AX-170301585 | 0 | C | T | 0.053 | -1.01E-01 | 3.00E-01 | 2.74E+00 | 2.76E+00 | 7.33E-01 | no | Gm13881 | upstream_gene_variant | MODIFIER |
| 2 | 103924035 | AX-170118547 | 0 | T | G | 0.053 | -1.01E-01 | 3.00E-01 | 2.74E+00 | 2.76E+00 | 7.33E-01 | no | | intergenic_variant | MODIFIER |
| 2 | 103925883 | AX-170322254 | 0 | T | C | 0.053 | -1.01E-01 | 3.00E-01 | 2.74E+00 | 2.76E+00 | 7.33E-01 | no | | intergenic_variant | MODIFIER |
| 2 | 104060034 | AX-97838353 | 0 | A | G | 0.053 | -1.01E-01 | 3.00E-01 | 2.74E+00 | 2.76E+00 | 7.33E-01 | no | Fbxo3 | synonymous_variant | LOW |
| 2 | 104120567 | AX-170237218 | 0 | T | C | 0.053 | -1.01E-01 | 3.00E-01 | 2.74E+00 | 2.76E+00 | 7.33E-01 | no | A930018P22Rik | upstream_gene_variant | MODIFIER |
| 2 | 104139810 | AX-170302213 | 0 | G | T | 0.053 | -1.01E-01 | 3.00E-01 | 2.74E+00 | 2.76E+00 | 7.33E-01 | no | D430041D05Rik | downstream_gene_variant | MODIFIER |
| 2 | 104180513 | AX-170238285 | 0 | G | A | 0.053 | -1.01E-01 | 3.00E-01 | 2.74E+00 | 2.76E+00 | 7.33E-01 | no | D430041D05Rik | intron_variant | MODIFIER |
| 2 | 104182091 | AX-170401072 | 0 | G | A | 0.053 | -1.01E-01 | 3.00E-01 | 2.74E+00 | 2.76E+00 | 7.33E-01 | no | D430041D05Rik | intron_variant | MODIFIER |
| 2 | 104184891 | AX-170494648 | 0 | C | G | 0.053 | -1.01E-01 | 3.00E-01 | 2.74E+00 | 2.76E+00 | 7.33E-01 | no | D430041D05Rik | intron_variant | MODIFIER |
| 2 | 104188842 | AX-170462404 | 0 | A | G | 0.053 | -1.01E-01 | 3.00E-01 | 2.74E+00 | 2.76E+00 | 7.33E-01 | no | D430041D05Rik | intron_variant | MODIFIER |
| 2 | 104205250 | AX-170449749 | 0 | A | G | 0.053 | -1.01E-01 | 3.00E-01 | 2.74E+00 | 2.76E+00 | 7.33E-01 | no | D430041D05Rik | intron_variant | MODIFIER |
| 2 | 104205749 | AX-170064499 | 0 | G | C | 0.053 | -1.01E-01 | 3.00E-01 | 2.74E+00 | 2.76E+00 | 7.33E-01 | no | D430041D05Rik | intron_variant | MODIFIER |
| 2 | 104206358 | AX-170064500 | 0 | G | C | 0.053 | -1.01E-01 | 3.00E-01 | 2.74E+00 | 2.76E+00 | 7.33E-01 | no | D430041D05Rik | intron_variant | MODIFIER |
| 2 | 104220855 | AX-170434621 | 0 | T | C | 0.053 | -1.01E-01 | 3.00E-01 | 2.74E+00 | 2.76E+00 | 7.33E-01 | no | D430041D05Rik | intron_variant | MODIFIER |
| 2 | 104232567 | AX-170339040 | 0 | G | A | 0.053 | -1.01E-01 | 3.00E-01 | 2.74E+00 | 2.76E+00 | 7.33E-01 | no | D430041D05Rik | intron_variant | MODIFIER |
| 2 | 104247132 | AX-170194939 | 0 | T | C | 0.053 | -1.01E-01 | 3.00E-01 | 2.74E+00 | 2.76E+00 | 7.33E-01 | no | D430041D05Rik | intron_variant | MODIFIER |
| 2 | 104253552 | AX-170349139 | 0 | G | T | 0.053 | -1.01E-01 | 3.00E-01 | 2.74E+00 | 2.76E+00 | 7.33E-01 | no | D430041D05Rik | intron_variant | MODIFIER |
| 2 | 104260184 | AX-170481824 | 0 | A | G | 0.053 | -1.01E-01 | 3.00E-01 | 2.74E+00 | 2.76E+00 | 7.33E-01 | no | D430041D05Rik | intron_variant | MODIFIER |
| 2 | 104427308 | AX-170035642 | 0 | C | A | 0.053 | -1.01E-01 | 3.00E-01 | 2.74E+00 | 2.76E+00 | 7.33E-01 | no | Hipk3 | 3_prime_UTR_variant | MODIFIER |
| 2 | 104470657 | AX-168408962 | 0 | G | A | 0.053 | -1.01E-01 | 3.00E-01 | 2.74E+00 | 2.76E+00 | 7.33E-01 | no | Hipk3 | intron_variant | MODIFIER |
| 2 | 104582736 | AX-170192009 | 0 | G | T | 0.053 | -1.01E-01 | 3.00E-01 | 2.74E+00 | 2.76E+00 | 7.33E-01 | no | Gm22439 | downstream_gene_variant | MODIFIER |
| 2 | 104583791 | AX-170346570 | 0 | T | C | 0.053 | -1.01E-01 | 3.00E-01 | 2.74E+00 | 2.76E+00 | 7.33E-01 | no | Gm22439 | downstream_gene_variant | MODIFIER |
| 2 | 104624482 | AX-170428285 | 0 | A | G | 0.053 | -1.01E-01 | 3.00E-01 | 2.74E+00 | 2.76E+00 | 7.33E-01 | no | Cstf3 | intron_variant | MODIFIER |
| 2 | 104626331 | AX-97785094 | 0 | C | A | 0.053 | -1.01E-01 | 3.00E-01 | 2.74E+00 | 2.76E+00 | 7.33E-01 | no | Cstf3 | intron_variant | MODIFIER |
| 2 | 104643517 | AX-170111694 | 0 | C | T | 0.053 | -1.01E-01 | 3.00E-01 | 2.74E+00 | 2.76E+00 | 7.33E-01 | no | Cstf3 | intron_variant | MODIFIER |
| 2 | 104644077 | AX-170248674 | 0 | G | A | 0.053 | -1.01E-01 | 3.00E-01 | 2.74E+00 | 2.76E+00 | 7.33E-01 | no | Cstf3 | intron_variant | MODIFIER |
| 2 | 104698402 | AX-170035641 | 0 | G | A | 0.053 | -1.01E-01 | 3.00E-01 | 2.74E+00 | 2.76E+00 | 7.33E-01 | no | Tcp11l1 | intron_variant | MODIFIER |

| | | | | | | | | | | | | | | | |
|---|---|---|---|---|---|---|---|---|---|---|---|---|---|---|---|
| 2 | 104698701 | AX-97838374 | 0 | A | G | 0.053 | -1.01E-01 | 3.00E-01 | 2.74E+00 | 2.76E+00 | 7.33E-01 | no | Tcp11l1 | intron_variant | MODIFIER |
| 2 | 104716155 | AX-170400013 | 0 | T | C | 0.053 | -1.01E-01 | 3.00E-01 | 2.74E+00 | 2.76E+00 | 7.33E-01 | no | Tcp11l1 | upstream_gene_variant | MODIFIER |
| 2 | 104716820 | AX-170391700 | 0 | A | G | 0.053 | -1.01E-01 | 3.00E-01 | 2.74E+00 | 2.76E+00 | 7.33E-01 | no | Depdc7 | downstream_gene_variant | MODIFIER |
| 2 | 104727424 | AX-170454314 | 0 | C | G | 0.053 | -1.01E-01 | 3.00E-01 | 2.74E+00 | 2.76E+00 | 7.33E-01 | no | Depdc7 | intron_variant | MODIFIER |
| 2 | 104736844 | AX-97795198 | 0 | A | G | 0.053 | -1.01E-01 | 3.00E-01 | 2.74E+00 | 2.76E+00 | 7.33E-01 | no | Depdc7 | intron_variant | MODIFIER |
| 2 | 104736894 | AX-170147357 | 0 | C | G | 0.053 | -1.01E-01 | 3.00E-01 | 2.74E+00 | 2.76E+00 | 7.33E-01 | no | Depdc7 | intron_variant | MODIFIER |
| 2 | 104778067 | AX-97839585 | 0 | T | C | 0.053 | -1.01E-01 | 3.00E-01 | 2.74E+00 | 2.76E+00 | 7.33E-01 | no | Qser1 | intron_variant | MODIFIER |
| 2 | 104779125 | AX-170371839 | 0 | C | T | 0.053 | -1.01E-01 | 3.00E-01 | 2.74E+00 | 2.76E+00 | 7.33E-01 | no | Qser1 | intron_variant | MODIFIER |
| 2 | 104830118 | AX-170455465 | 0 | G | T | 0.053 | -1.01E-01 | 3.00E-01 | 2.74E+00 | 2.76E+00 | 7.33E-01 | no | Prrg4 | downstream_gene_variant | MODIFIER |
| 2 | 104845300 | AX-170018547 | 0 | T | C | 0.053 | -1.01E-01 | 3.00E-01 | 2.74E+00 | 2.76E+00 | 7.33E-01 | no | Prrg4 | intron_variant | MODIFIER |
| 2 | 104992164 | AX-170064513 | 0 | G | A | 0.053 | -1.01E-01 | 3.00E-01 | 2.74E+00 | 2.76E+00 | 7.33E-01 | no | Ccdc73 | synonymous_variant | LOW |
| 2 | 104992180 | AX-170311127 | 0 | C | T | 0.053 | -1.01E-01 | 3.00E-01 | 2.74E+00 | 2.76E+00 | 7.33E-01 | no | Ccdc73 | missense_variant | MODERATE |
| 2 | 104993952 | AX-170404056 | 0 | C | T | 0.053 | -1.01E-01 | 3.00E-01 | 2.74E+00 | 2.76E+00 | 7.33E-01 | no | Ccdc73 | intron_variant | MODIFIER |
| 2 | 105007258 | AX-97821632 | 0 | T | C | 0.053 | -1.01E-01 | 3.00E-01 | 2.74E+00 | 2.76E+00 | 7.33E-01 | no | Eif3m | intron_variant | MODIFIER |
| 2 | 105036040 | AX-170385908 | 0 | A | C | 0.053 | -1.01E-01 | 3.00E-01 | 2.74E+00 | 2.76E+00 | 7.33E-01 | no | Gm13888 | upstream_gene_variant | MODIFIER |
| 2 | 105038624 | AX-97799737 | 0 | G | T | 0.053 | -1.01E-01 | 3.00E-01 | 2.74E+00 | 2.76E+00 | 7.33E-01 | no | | intergenic_variant | MODIFIER |
| 2 | 105038957 | AX-170511679 | 0 | G | A | 0.053 | -1.01E-01 | 3.00E-01 | 2.74E+00 | 2.76E+00 | 7.33E-01 | no | | intergenic_variant | MODIFIER |
| 2 | 105039458 | AX-168392074 | 0 | C | T | 0.053 | -1.01E-01 | 3.00E-01 | 2.74E+00 | 2.76E+00 | 7.33E-01 | no | | regulatory_region_variant | MODIFIER |
| 2 | 105070001 | AX-170218902 | 0 | C | A | 0.053 | -1.01E-01 | 3.00E-01 | 2.74E+00 | 2.76E+00 | 7.33E-01 | no | | intergenic_variant | MODIFIER |
| 2 | 105070097 | AX-170297351 | 0 | C | T | 0.053 | -1.01E-01 | 3.00E-01 | 2.74E+00 | 2.76E+00 | 7.33E-01 | no | | intergenic_variant | MODIFIER |
| 2 | 105076924 | AX-170315903 | 0 | A | G | 0.053 | -1.01E-01 | 3.00E-01 | 2.74E+00 | 2.76E+00 | 7.33E-01 | no | Wt1os | downstream_gene_variant | MODIFIER |
| 2 | 105106351 | AX-170263414 | 0 | C | A | 0.053 | -1.01E-01 | 3.00E-01 | 2.74E+00 | 2.76E+00 | 7.33E-01 | no | Wt1os | intron_variant,non_coding_transcript_variant | MODIFIER |
| 2 | 105112032 | AX-97800334 | 0 | T | C | 0.053 | -1.01E-01 | 3.00E-01 | 2.74E+00 | 2.76E+00 | 7.33E-01 | no | Wt1os | intron_variant,non_coding_transcript_variant | MODIFIER |
| 2 | 105112149 | AX-170370090 | 0 | A | G | 0.053 | -1.01E-01 | 3.00E-01 | 2.74E+00 | 2.76E+00 | 7.33E-01 | no | Wt1os | intron_variant,non_coding_transcript_variant | MODIFIER |
| 2 | 105164649 | AX-170172510 | 0 | G | A | 0.053 | -1.01E-01 | 3.00E-01 | 2.74E+00 | 2.76E+00 | 7.33E-01 | no | Wt1 | intron_variant | MODIFIER |
| 2 | 105166668 | AX-170289454 | 0 | C | A | 0.053 | -1.01E-01 | 3.00E-01 | 2.74E+00 | 2.76E+00 | 7.33E-01 | no | Wt1 | intron_variant | MODIFIER |
| 2 | 105168085 | AX-170530513 | 0 | A | G | 0.053 | -1.01E-01 | 3.00E-01 | 2.74E+00 | 2.76E+00 | 7.33E-01 | no | Wt1 | intron_variant | MODIFIER |
| 2 | 105180964 | AX-168387544 | 0 | A | G | 0.053 | -1.01E-01 | 3.00E-01 | 2.74E+00 | 2.76E+00 | 7.33E-01 | no | | intergenic_variant | MODIFIER |
| 2 | 105209465 | AX-170253336 | 0 | C | A | 0.053 | -1.01E-01 | 3.00E-01 | 2.74E+00 | 2.76E+00 | 7.33E-01 | no | | intergenic_variant | MODIFIER |
| 2 | 105209885 | AX-170297596 | 0 | T | C | 0.053 | -1.01E-01 | 3.00E-01 | 2.74E+00 | 2.76E+00 | 7.33E-01 | no | | intergenic_variant | MODIFIER |
| 2 | 105241251 | AX-170319984 | 0 | C | T | 0.053 | -1.01E-01 | 3.00E-01 | 2.74E+00 | 2.76E+00 | 7.33E-01 | no | Them7 | intron_variant | MODIFIER |
| 2 | 105326883 | AX-170282520 | 0 | G | A | 0.053 | -1.01E-01 | 3.00E-01 | 2.74E+00 | 2.76E+00 | 7.33E-01 | no | Them7 | intron_variant | MODIFIER |
| 2 | 105633014 | AX-170279784 | 0 | G | A | 0.053 | -1.01E-01 | 3.00E-01 | 2.74E+00 | 2.76E+00 | 7.33E-01 | no | Pax6os1 | intron_variant,non_coding_transcript_variant | MODIFIER |
| 2 | 105634443 | AX-97821689 | 0 | C | T | 0.053 | -1.01E-01 | 3.00E-01 | 2.74E+00 | 2.76E+00 | 7.33E-01 | no | Pax6os1 | intron_variant,non_coding_transcript_variant | MODIFIER |
| 2 | 105654231 | AX-97796897 | 0 | C | G | 0.053 | -1.01E-01 | 3.00E-01 | 2.74E+00 | 2.76E+00 | 7.33E-01 | no | Pax6os1 | intron_variant,non_coding_transcript_variant | MODIFIER |
| 2 | 105654518 | AX-170147095 | 0 | A | C | 0.053 | -1.01E-01 | 3.00E-01 | 2.74E+00 | 2.76E+00 | 7.33E-01 | no | Pax6os1 | intron_variant,non_coding_transcript_variant | MODIFIER |
| 2 | 106006333 | AX-170445747 | 0 | C | A | 0.053 | -1.01E-01 | 3.00E-01 | 2.74E+00 | 2.76E+00 | 7.33E-01 | no | Dnajc24 | upstream_gene_variant | MODIFIER |
| 2 | 106011367 | AX-170375854 | 0 | C | G | 0.053 | -1.01E-01 | 3.00E-01 | 2.74E+00 | 2.76E+00 | 7.33E-01 | no | Gm13992 | upstream_gene_variant | MODIFIER |
| 2 | 106042154 | AX-168379504 | 0 | T | C | 0.053 | -1.01E-01 | 3.00E-01 | 2.74E+00 | 2.76E+00 | 7.33E-01 | no | Dcdc5 | intron_variant,non_coding_transcript_variant | MODIFIER |
| 2 | 106042528 | AX-170401804 | 0 | C | T | 0.053 | -1.01E-01 | 3.00E-01 | 2.74E+00 | 2.76E+00 | 7.33E-01 | no | Dcdc5 | intron_variant,non_coding_transcript_variant | MODIFIER |
| 2 | 106061983 | AX-170336020 | 0 | A | G | 0.053 | -1.01E-01 | 3.00E-01 | 2.74E+00 | 2.76E+00 | 7.33E-01 | no | Dcdc5 | intron_variant,non_coding_transcript_variant | MODIFIER |
| 2 | 106068971 | AX-97797687 | 0 | C | T | 0.053 | -1.01E-01 | 3.00E-01 | 2.74E+00 | 2.76E+00 | 7.33E-01 | no | Dcdc5 | intron_variant,non_coding_transcript_variant | MODIFIER |
| 2 | 106072958 | AX-170412554 | 0 | C | A | 0.053 | -1.01E-01 | 3.00E-01 | 2.74E+00 | 2.76E+00 | 7.33E-01 | no | Dcdc5 | intron_variant,non_coding_transcript_variant | MODIFIER |
| 2 | 106074005 | AX-170344774 | 0 | T | C | 0.053 | -1.01E-01 | 3.00E-01 | 2.74E+00 | 2.76E+00 | 7.33E-01 | no | Dcdc5 | intron_variant,non_coding_transcript_variant | MODIFIER |
| 2 | 106110496 | AX-170064533 | 0 | G | A | 0.053 | -1.01E-01 | 3.00E-01 | 2.74E+00 | 2.76E+00 | 7.33E-01 | no | Dcdc5 | intron_variant,non_coding_transcript_variant | MODIFIER |
| 2 | 106113851 | AX-97802321 | 0 | T | C | 0.053 | -1.01E-01 | 3.00E-01 | 2.74E+00 | 2.76E+00 | 7.33E-01 | no | Dcdc5 | intron_variant,non_coding_transcript_variant | MODIFIER |
| 2 | 106188791 | AX-170410712 | 0 | G | A | 0.053 | -1.01E-01 | 3.00E-01 | 2.74E+00 | 2.76E+00 | 7.33E-01 | no | Dcdc5 | intron_variant,non_coding_transcript_variant | MODIFIER |
| 2 | 106188833 | AX-97839271 | 0 | C | T | 0.053 | -1.01E-01 | 3.00E-01 | 2.74E+00 | 2.76E+00 | 7.33E-01 | no | Dcdc5 | intron_variant,non_coding_transcript_variant | MODIFIER |
| 2 | 106192421 | AX-170361327 | 0 | A | G | 0.053 | -1.01E-01 | 3.00E-01 | 2.74E+00 | 2.76E+00 | 7.33E-01 | no | Dcdc5 | intron_variant,non_coding_transcript_variant | MODIFIER |
| 2 | 106205043 | AX-170234705 | 0 | T | A | 0.053 | -1.01E-01 | 3.00E-01 | 2.74E+00 | 2.76E+00 | 7.33E-01 | no | Dcdc5 | intron_variant,non_coding_transcript_variant | MODIFIER |
| 2 | 106227740 | AX-170275020 | 0 | C | G | 0.053 | -1.01E-01 | 3.00E-01 | 2.74E+00 | 2.76E+00 | 7.33E-01 | no | Dcdc5 | intron_variant,non_coding_transcript_variant | MODIFIER |
| 2 | 106228752 | AX-170507210 | 0 | T | C | 0.053 | -1.01E-01 | 3.00E-01 | 2.74E+00 | 2.76E+00 | 7.33E-01 | no | Dcdc5 | intron_variant,non_coding_transcript_variant | MODIFIER |
| 2 | 106232534 | AX-170064537 | 0 | C | T | 0.053 | -1.01E-01 | 3.00E-01 | 2.74E+00 | 2.76E+00 | 7.33E-01 | no | Dcdc5 | intron_variant,non_coding_transcript_variant | MODIFIER |
| 2 | 106238731 | AX-170506043 | 0 | A | C | 0.053 | -1.01E-01 | 3.00E-01 | 2.74E+00 | 2.76E+00 | 7.33E-01 | no | Dcdc5 | intron_variant,non_coding_transcript_variant | MODIFIER |

| | | | | | | | | | | | | | | | |
|---|---|---|---|---|---|---|---|---|---|---|---|---|---|---|---|
| 2 | 106266592 | AX-97837648 | 0 | C | A | 0.053 | -1.01E-01 | 3.00E-01 | 2.74E+00 | 2.76E+00 | 7.33E-01 | no | Gm14013 | downstream_gene_variant | MODIFIER |
| 2 | 106282582 | AX-170140898 | 0 | T | G | 0.053 | -1.01E-01 | 3.00E-01 | 2.74E+00 | 2.76E+00 | 7.33E-01 | no | Dcdc5 | intron_variant,non_coding_transcript_variant | MODIFIER |
| 2 | 106308969 | AX-97846448 | 0 | G | A | 0.053 | -1.01E-01 | 3.00E-01 | 2.74E+00 | 2.76E+00 | 7.33E-01 | no | Dcdc5 | intron_variant,non_coding_transcript_variant | MODIFIER |
| 2 | 106363672 | AX-170410002 | 0 | T | C | 0.053 | -1.01E-01 | 3.00E-01 | 2.74E+00 | 2.76E+00 | 7.33E-01 | no | Dcdc5 | intron_variant,non_coding_transcript_variant | MODIFIER |
| 2 | 106365592 | AX-170152498 | 0 | T | C | 0.053 | -1.01E-01 | 3.00E-01 | 2.74E+00 | 2.76E+00 | 7.33E-01 | no | Dcdc5 | intron_variant,non_coding_transcript_variant | MODIFIER |
| 2 | 106370149 | AX-170482985 | 0 | A | G | 0.053 | -1.01E-01 | 3.00E-01 | 2.74E+00 | 2.76E+00 | 7.33E-01 | no | Dcdc5 | intron_variant,non_coding_transcript_variant | MODIFIER |
| 2 | 106382991 | AX-97794755 | 0 | C | T | 0.053 | -1.01E-01 | 3.00E-01 | 2.74E+00 | 2.76E+00 | 7.33E-01 | no | Dcdc5 | upstream_gene_variant | MODIFIER |
| 2 | 106383008 | AX-170442684 | 0 | A | G | 0.053 | -1.01E-01 | 3.00E-01 | 2.74E+00 | 2.76E+00 | 7.33E-01 | no | Dcdc5 | upstream_gene_variant | MODIFIER |
| 2 | 106384866 | AX-170496068 | 0 | G | A | 0.053 | -1.01E-01 | 3.00E-01 | 2.74E+00 | 2.76E+00 | 7.33E-01 | no | Dcdc5 | intron_variant,non_coding_transcript_variant | MODIFIER |
| 2 | 106385259 | AX-170391920 | 0 | A | G | 0.053 | -1.01E-01 | 3.00E-01 | 2.74E+00 | 2.76E+00 | 7.33E-01 | no | Dcdc5 | intron_variant,non_coding_transcript_variant | MODIFIER |
| 2 | 106385328 | AX-170064538 | 0 | G | A | 0.053 | -1.01E-01 | 3.00E-01 | 2.74E+00 | 2.76E+00 | 7.33E-01 | no | Dcdc5 | intron_variant,non_coding_transcript_variant | MODIFIER |
| 2 | 106429885 | AX-170324940 | 0 | A | G | 0.053 | -1.01E-01 | 3.00E-01 | 2.74E+00 | 2.76E+00 | 7.33E-01 | no | | intergenic_variant | MODIFIER |
| 2 | 106435423 | AX-170146754 | 0 | C | T | 0.053 | -1.01E-01 | 3.00E-01 | 2.74E+00 | 2.76E+00 | 7.33E-01 | no | | intergenic_variant | MODIFIER |
| 2 | 106436459 | AX-170524352 | 0 | C | G | 0.053 | -1.01E-01 | 3.00E-01 | 2.74E+00 | 2.76E+00 | 7.33E-01 | no | | intergenic_variant | MODIFIER |
| 2 | 106665335 | AX-97843680 | 0 | G | A | 0.053 | -1.01E-01 | 3.00E-01 | 2.74E+00 | 2.76E+00 | 7.33E-01 | no | | regulatory_region_variant | MODIFIER |
| 2 | 106698454 | AX-170207512 | 0 | C | T | 0.053 | -1.01E-01 | 3.00E-01 | 2.74E+00 | 2.76E+00 | 7.33E-01 | no | Mpped2 | intron_variant | MODIFIER |
| 2 | 106727537 | AX-170314161 | 0 | A | G | 0.053 | -1.01E-01 | 3.00E-01 | 2.74E+00 | 2.76E+00 | 7.33E-01 | no | Mpped2 | intron_variant | MODIFIER |
| 2 | 106727593 | AX-170088696 | 0 | G | A | 0.053 | -1.01E-01 | 3.00E-01 | 2.74E+00 | 2.76E+00 | 7.33E-01 | no | Mpped2 | intron_variant | MODIFIER |
| 2 | 106752168 | AX-168398241 | 0 | G | A | 0.053 | -1.01E-01 | 3.00E-01 | 2.74E+00 | 2.76E+00 | 7.33E-01 | no | Mpped2 | intron_variant | MODIFIER |
| 2 | 106753096 | AX-97827675 | 0 | G | A | 0.053 | -1.01E-01 | 3.00E-01 | 2.74E+00 | 2.76E+00 | 7.33E-01 | no | Mpped2 | intron_variant | MODIFIER |
| 2 | 106753256 | AX-170407119 | 0 | A | G | 0.053 | -1.01E-01 | 3.00E-01 | 2.74E+00 | 2.76E+00 | 7.33E-01 | no | Mpped2 | intron_variant | MODIFIER |
| 2 | 106764200 | AX-170340668 | 0 | C | T | 0.053 | -1.01E-01 | 3.00E-01 | 2.74E+00 | 2.76E+00 | 7.33E-01 | no | Mpped2 | intron_variant | MODIFIER |
| 2 | 106771341 | AX-170147980 | 0 | A | C | 0.053 | -1.01E-01 | 3.00E-01 | 2.74E+00 | 2.76E+00 | 7.33E-01 | no | Mpped2 | intron_variant | MODIFIER |
| 2 | 106773355 | AX-97787341 | 0 | C | T | 0.053 | -1.01E-01 | 3.00E-01 | 2.74E+00 | 2.76E+00 | 7.33E-01 | no | Mpped2 | intron_variant | MODIFIER |
| 2 | 106785577 | AX-170151769 | 0 | T | C | 0.053 | -1.01E-01 | 3.00E-01 | 2.74E+00 | 2.76E+00 | 7.33E-01 | no | Mpped2 | intron_variant | MODIFIER |
| 2 | 106809075 | AX-97839278 | 0 | C | T | 0.053 | -1.01E-01 | 3.00E-01 | 2.74E+00 | 2.76E+00 | 7.33E-01 | no | Mpped2 | intron_variant | MODIFIER |
| 2 | 106819622 | AX-170238361 | 0 | A | G | 0.053 | -1.01E-01 | 3.00E-01 | 2.74E+00 | 2.76E+00 | 7.33E-01 | no | Mpped2 | intron_variant | MODIFIER |
| 2 | 106821396 | AX-170064545 | 0 | A | T | 0.053 | -1.01E-01 | 3.00E-01 | 2.74E+00 | 2.76E+00 | 7.33E-01 | no | Mpped2 | intron_variant | MODIFIER |
| 2 | 106821999 | AX-170237008 | 0 | T | A | 0.053 | -1.01E-01 | 3.00E-01 | 2.74E+00 | 2.76E+00 | 7.33E-01 | no | Mpped2 | intron_variant | MODIFIER |
| 2 | 106822128 | AX-170394351 | 0 | A | G | 0.053 | -1.01E-01 | 3.00E-01 | 2.74E+00 | 2.76E+00 | 7.33E-01 | no | Mpped2 | intron_variant | MODIFIER |
| 2 | 106822192 | AX-170524847 | 0 | T | C | 0.053 | -1.01E-01 | 3.00E-01 | 2.74E+00 | 2.76E+00 | 7.33E-01 | no | Mpped2 | intron_variant | MODIFIER |
| 2 | 106830959 | AX-170018588 | 0 | C | T | 0.053 | -1.01E-01 | 3.00E-01 | 2.74E+00 | 2.76E+00 | 7.33E-01 | no | Mpped2 | intron_variant | MODIFIER |
| 2 | 106841364 | AX-97788929 | 0 | T | C | 0.053 | -1.01E-01 | 3.00E-01 | 2.74E+00 | 2.76E+00 | 7.33E-01 | no | Mpped2 | intron_variant | MODIFIER |
| 2 | 106850027 | AX-170018596 | 0 | C | T | 0.053 | -1.01E-01 | 3.00E-01 | 2.74E+00 | 2.76E+00 | 7.33E-01 | no | Mpped2 | intron_variant | MODIFIER |
| 2 | 106851733 | AX-97813596 | 0 | C | T | 0.053 | -1.01E-01 | 3.00E-01 | 2.74E+00 | 2.76E+00 | 7.33E-01 | no | Mpped2 | intron_variant | MODIFIER |
| 2 | 106868614 | AX-170453898 | 0 | C | T | 0.053 | -1.01E-01 | 3.00E-01 | 2.74E+00 | 2.76E+00 | 7.33E-01 | no | Mpped2 | downstream_gene_variant | MODIFIER |
| 2 | 106872125 | AX-170270102 | 0 | G | A | 0.053 | -1.01E-01 | 3.00E-01 | 2.74E+00 | 2.76E+00 | 7.33E-01 | no | Mpped2 | downstream_gene_variant | MODIFIER |
| 2 | 107048679 | AX-170018595 | 0 | C | A | 0.053 | -1.01E-01 | 3.00E-01 | 2.74E+00 | 2.76E+00 | 7.33E-01 | no | | intergenic_variant | MODIFIER |
| 2 | 107048755 | AX-170349314 | 0 | T | C | 0.053 | -1.01E-01 | 3.00E-01 | 2.74E+00 | 2.76E+00 | 7.33E-01 | no | | intergenic_variant | MODIFIER |
| 2 | 107052332 | AX-170088698 | 0 | C | T | 0.053 | -1.01E-01 | 3.00E-01 | 2.74E+00 | 2.76E+00 | 7.33E-01 | no | Fshb | downstream_gene_variant | MODIFIER |
| 2 | 107055388 | AX-170335021 | 0 | T | C | 0.053 | -1.01E-01 | 3.00E-01 | 2.74E+00 | 2.76E+00 | 7.33E-01 | no | Fshb | downstream_gene_variant | MODIFIER |
| 2 | 107085976 | AX-170018590 | 0 | A | G | 0.053 | -1.01E-01 | 3.00E-01 | 2.74E+00 | 2.76E+00 | 7.33E-01 | no | | intergenic_variant | MODIFIER |
| 2 | 107118511 | AX-97856931 | 0 | C | T | 0.053 | -1.01E-01 | 3.00E-01 | 2.74E+00 | 2.76E+00 | 7.33E-01 | no | | intergenic_variant | MODIFIER |
| 2 | 107118879 | AX-170476419 | 0 | T | G | 0.053 | -1.01E-01 | 3.00E-01 | 2.74E+00 | 2.76E+00 | 7.33E-01 | no | Gm13903 | downstream_gene_variant | MODIFIER |
| 2 | 107123153 | AX-168420022 | 0 | T | C | 0.053 | -1.01E-01 | 3.00E-01 | 2.74E+00 | 2.76E+00 | 7.33E-01 | no | Gm13903 | downstream_gene_variant | MODIFIER |
| 2 | 107150801 | AX-170200965 | 0 | G | A | 0.053 | -1.01E-01 | 3.00E-01 | 2.74E+00 | 2.76E+00 | 7.33E-01 | no | | intergenic_variant | MODIFIER |
| 2 | 107153805 | AX-170165945 | 0 | T | C | 0.053 | -1.01E-01 | 3.00E-01 | 2.74E+00 | 2.76E+00 | 7.33E-01 | no | | intergenic_variant | MODIFIER |
| 2 | 107160080 | AX-170498845 | 0 | T | C | 0.053 | -1.01E-01 | 3.00E-01 | 2.74E+00 | 2.76E+00 | 7.33E-01 | no | | intergenic_variant | MODIFIER |
| 2 | 107160114 | AX-97826860 | 0 | A | C | 0.053 | -1.01E-01 | 3.00E-01 | 2.74E+00 | 2.76E+00 | 7.33E-01 | no | | intergenic_variant | MODIFIER |
| 2 | 107160235 | AX-170189275 | 0 | T | C | 0.053 | -1.01E-01 | 3.00E-01 | 2.74E+00 | 2.76E+00 | 7.33E-01 | no | | intergenic_variant | MODIFIER |
| 2 | 107181715 | AX-97788720 | 0 | C | T | 0.053 | -1.01E-01 | 3.00E-01 | 2.74E+00 | 2.76E+00 | 7.33E-01 | no | | intergenic_variant | MODIFIER |
| 2 | 107222183 | AX-170490240 | 0 | C | T | 0.053 | -1.01E-01 | 3.00E-01 | 2.74E+00 | 2.76E+00 | 7.33E-01 | no | | intergenic_variant | MODIFIER |
| 2 | 107227039 | AX-170128733 | 0 | C | G | 0.053 | -1.01E-01 | 3.00E-01 | 2.74E+00 | 2.76E+00 | 7.33E-01 | no | | regulatory_region_variant | MODIFIER |
| 2 | 107227487 | AX-170162288 | 0 | G | A | 0.053 | -1.01E-01 | 3.00E-01 | 2.74E+00 | 2.76E+00 | 7.33E-01 | no | | regulatory_region_variant | MODIFIER |

| | | | | | | | | | | | | | | | |
|---|---|---|---|---|---|---|---|---|---|---|---|---|---|---|---|
| 2 | 107228030 | AX-170131176 | 0 | T | G | 0.053 | -1.01E-01 | 3.00E-01 | 2.74E+00 | 2.76E+00 | 7.33E-01 | no | | regulatory_region_variant | MODIFIER |
| 2 | 107233333 | AX-97846042 | 0 | C | T | 0.053 | -1.01E-01 | 3.00E-01 | 2.74E+00 | 2.76E+00 | 7.33E-01 | no | | intergenic_variant | MODIFIER |
| 2 | 107234081 | AX-170116205 | 0 | C | A | 0.053 | -1.01E-01 | 3.00E-01 | 2.74E+00 | 2.76E+00 | 7.33E-01 | no | | intergenic_variant | MODIFIER |
| 2 | 107236717 | AX-170064556 | 0 | A | G | 0.053 | -1.01E-01 | 3.00E-01 | 2.74E+00 | 2.76E+00 | 7.33E-01 | no | | intergenic_variant | MODIFIER |
| 2 | 107241372 | AX-170064555 | 0 | G | A | 0.053 | -1.01E-01 | 3.00E-01 | 2.74E+00 | 2.76E+00 | 7.33E-01 | no | | intergenic_variant | MODIFIER |
| 2 | 107465137 | AX-170018603 | 0 | A | T | 0.053 | -1.01E-01 | 3.00E-01 | 2.74E+00 | 2.76E+00 | 7.33E-01 | no | | intergenic_variant | MODIFIER |
| 2 | 107547837 | AX-170018604 | 0 | G | C | 0.053 | -1.01E-01 | 3.00E-01 | 2.74E+00 | 2.76E+00 | 7.33E-01 | no | | intergenic_variant | MODIFIER |
| 2 | 107549292 | AX-170038219 | 0 | G | A | 0.053 | -1.01E-01 | 3.00E-01 | 2.74E+00 | 2.76E+00 | 7.33E-01 | no | | intergenic_variant | MODIFIER |
| 2 | 107569680 | AX-97824343 | 0 | G | A | 0.053 | -1.01E-01 | 3.00E-01 | 2.74E+00 | 2.76E+00 | 7.33E-01 | no | | intergenic_variant | MODIFIER |
| 2 | 107612181 | AX-97798233 | 0 | T | C | 0.053 | -1.01E-01 | 3.00E-01 | 2.74E+00 | 2.76E+00 | 7.33E-01 | no | | intergenic_variant | MODIFIER |
| 2 | 107615961 | AX-170514115 | 0 | T | C | 0.053 | -1.01E-01 | 3.00E-01 | 2.74E+00 | 2.76E+00 | 7.33E-01 | no | | intergenic_variant | MODIFIER |
| 2 | 107616053 | AX-170343110 | 0 | T | G | 0.053 | -1.01E-01 | 3.00E-01 | 2.74E+00 | 2.76E+00 | 7.33E-01 | no | | intergenic_variant | MODIFIER |
| 2 | 107616644 | AX-170170999 | 0 | C | T | 0.053 | -1.01E-01 | 3.00E-01 | 2.74E+00 | 2.76E+00 | 7.33E-01 | no | | intergenic_variant | MODIFIER |
| 2 | 107617058 | AX-170100793 | 0 | T | C | 0.053 | -1.01E-01 | 3.00E-01 | 2.74E+00 | 2.76E+00 | 7.33E-01 | no | | intergenic_variant | MODIFIER |
| 2 | 107620397 | AX-170257353 | 0 | C | G | 0.053 | -1.01E-01 | 3.00E-01 | 2.74E+00 | 2.76E+00 | 7.33E-01 | no | | intergenic_variant | MODIFIER |
| 2 | 107624537 | AX-170018607 | 0 | T | G | 0.053 | -1.01E-01 | 3.00E-01 | 2.74E+00 | 2.76E+00 | 7.33E-01 | no | | intergenic_variant | MODIFIER |
| 2 | 107631043 | AX-97796656 | 0 | A | C | 0.053 | -1.01E-01 | 3.00E-01 | 2.74E+00 | 2.76E+00 | 7.33E-01 | no | | intergenic_variant | MODIFIER |
| 2 | 107656336 | AX-97855001 | 0 | A | G | 0.053 | -1.01E-01 | 3.00E-01 | 2.74E+00 | 2.76E+00 | 7.33E-01 | no | | intergenic_variant | MODIFIER |
| 2 | 107699197 | AX-170120088 | 0 | T | A | 0.053 | -1.01E-01 | 3.00E-01 | 2.74E+00 | 2.76E+00 | 7.33E-01 | no | | intergenic_variant | MODIFIER |
| 2 | 107699319 | AX-170116297 | 0 | A | C | 0.053 | -1.01E-01 | 3.00E-01 | 2.74E+00 | 2.76E+00 | 7.33E-01 | no | | intergenic_variant | MODIFIER |
| 2 | 107700949 | AX-97840965 | 0 | G | A | 0.053 | -1.01E-01 | 3.00E-01 | 2.74E+00 | 2.76E+00 | 7.33E-01 | no | | intergenic_variant | MODIFIER |
| 2 | 107728500 | AX-97839314 | 0 | C | T | 0.053 | -1.01E-01 | 3.00E-01 | 2.74E+00 | 2.76E+00 | 7.33E-01 | no | | intergenic_variant | MODIFIER |
| 2 | 107734956 | AX-170187430 | 0 | C | T | 0.053 | -1.01E-01 | 3.00E-01 | 2.74E+00 | 2.76E+00 | 7.33E-01 | no | | intergenic_variant | MODIFIER |
| 2 | 107748426 | AX-170305766 | 0 | C | T | 0.053 | -1.01E-01 | 3.00E-01 | 2.74E+00 | 2.76E+00 | 7.33E-01 | no | | intergenic_variant | MODIFIER |
| 2 | 107788114 | AX-170166562 | 0 | C | A | 0.053 | -1.01E-01 | 3.00E-01 | 2.74E+00 | 2.76E+00 | 7.33E-01 | no | | intergenic_variant | MODIFIER |
| 2 | 107809379 | AX-97791940 | 0 | C | T | 0.053 | -1.01E-01 | 3.00E-01 | 2.74E+00 | 2.76E+00 | 7.33E-01 | no | | intergenic_variant | MODIFIER |
| 2 | 107816394 | AX-170337268 | 0 | G | C | 0.053 | -1.01E-01 | 3.00E-01 | 2.74E+00 | 2.76E+00 | 7.33E-01 | no | | intergenic_variant | MODIFIER |
| 2 | 107819494 | AX-170131816 | 0 | G | A | 0.053 | -1.01E-01 | 3.00E-01 | 2.74E+00 | 2.76E+00 | 7.33E-01 | no | | intergenic_variant | MODIFIER |
| 2 | 107819966 | AX-170064562 | 0 | T | G | 0.053 | -1.01E-01 | 3.00E-01 | 2.74E+00 | 2.76E+00 | 7.33E-01 | no | | intergenic_variant | MODIFIER |
| 2 | 107827545 | AX-170359098 | 0 | T | A | 0.053 | -1.01E-01 | 3.00E-01 | 2.74E+00 | 2.76E+00 | 7.33E-01 | no | | intergenic_variant | MODIFIER |
| 2 | 107827572 | AX-170482802 | 0 | A | T | 0.053 | -1.01E-01 | 3.00E-01 | 2.74E+00 | 2.76E+00 | 7.33E-01 | no | | intergenic_variant | MODIFIER |
| 2 | 107854385 | AX-170245944 | 0 | C | T | 0.053 | -1.01E-01 | 3.00E-01 | 2.74E+00 | 2.76E+00 | 7.33E-01 | no | | intergenic_variant | MODIFIER |
| 2 | 107861512 | AX-170328425 | 0 | C | G | 0.053 | -1.01E-01 | 3.00E-01 | 2.74E+00 | 2.76E+00 | 7.33E-01 | no | | intergenic_variant | MODIFIER |
| 2 | 107903796 | AX-170458129 | 0 | A | C | 0.053 | -1.01E-01 | 3.00E-01 | 2.74E+00 | 2.76E+00 | 7.33E-01 | no | | intergenic_variant | MODIFIER |
| 2 | 107906650 | AX-170088700 | 0 | T | C | 0.053 | -1.01E-01 | 3.00E-01 | 2.74E+00 | 2.76E+00 | 7.33E-01 | no | | intergenic_variant | MODIFIER |
| 2 | 107924471 | AX-168400751 | 0 | A | G | 0.053 | -1.01E-01 | 3.00E-01 | 2.74E+00 | 2.76E+00 | 7.33E-01 | no | | intergenic_variant | MODIFIER |
| 2 | 107970047 | AX-97839316 | 0 | C | A | 0.053 | -1.01E-01 | 3.00E-01 | 2.74E+00 | 2.76E+00 | 7.33E-01 | no | | intergenic_variant | MODIFIER |
| 2 | 107972397 | AX-170018617 | 0 | C | T | 0.053 | -1.01E-01 | 3.00E-01 | 2.74E+00 | 2.76E+00 | 7.33E-01 | no | | intergenic_variant | MODIFIER |
| 2 | 107985458 | AX-170018612 | 0 | C | T | 0.053 | -1.01E-01 | 3.00E-01 | 2.74E+00 | 2.76E+00 | 7.33E-01 | no | | intergenic_variant | MODIFIER |
| 2 | 107988777 | AX-170064564 | 0 | G | A | 0.053 | -1.01E-01 | 3.00E-01 | 2.74E+00 | 2.76E+00 | 7.33E-01 | no | | intergenic_variant | MODIFIER |
| 2 | 108014173 | AX-170018618 | 0 | C | G | 0.053 | -1.01E-01 | 3.00E-01 | 2.74E+00 | 2.76E+00 | 7.33E-01 | no | | intergenic_variant | MODIFIER |
| 2 | 108050244 | AX-170416442 | 0 | A | G | 0.053 | -1.01E-01 | 3.00E-01 | 2.74E+00 | 2.76E+00 | 7.33E-01 | no | | intergenic_variant | MODIFIER |
| 2 | 108138205 | AX-170478956 | 0 | T | G | 0.053 | -1.01E-01 | 3.00E-01 | 2.74E+00 | 2.76E+00 | 7.33E-01 | no | | intergenic_variant | MODIFIER |
| 2 | 108139286 | AX-170188749 | 0 | A | G | 0.053 | -1.01E-01 | 3.00E-01 | 2.74E+00 | 2.76E+00 | 7.33E-01 | no | | intergenic_variant | MODIFIER |
| 2 | 108143386 | AX-97838398 | 0 | T | C | 0.053 | -1.01E-01 | 3.00E-01 | 2.74E+00 | 2.76E+00 | 7.33E-01 | no | | intergenic_variant | MODIFIER |
| 2 | 108168037 | AX-97851194 | 0 | T | G | 0.053 | -1.01E-01 | 3.00E-01 | 2.74E+00 | 2.76E+00 | 7.33E-01 | no | | intergenic_variant | MODIFIER |
| 2 | 108174519 | AX-170170059 | 0 | A | G | 0.053 | -1.01E-01 | 3.00E-01 | 2.74E+00 | 2.76E+00 | 7.33E-01 | no | | intergenic_variant | MODIFIER |
| 2 | 108183533 | AX-170526946 | 0 | T | G | 0.053 | -1.01E-01 | 3.00E-01 | 2.74E+00 | 2.76E+00 | 7.33E-01 | no | Gm13912 | downstream_gene_variant | MODIFIER |
| 2 | 108223073 | AX-170285032 | 0 | G | C | 0.053 | -1.01E-01 | 3.00E-01 | 2.74E+00 | 2.76E+00 | 7.33E-01 | no | Gm13912 | intron_variant,non_coding_transcript_variant | MODIFIER |
| 2 | 108223224 | AX-97800933 | 0 | T | C | 0.053 | -1.01E-01 | 3.00E-01 | 2.74E+00 | 2.76E+00 | 7.33E-01 | no | Gm13912 | intron_variant,non_coding_transcript_variant | MODIFIER |
| 2 | 108258687 | AX-170232980 | 0 | C | T | 0.053 | -1.01E-01 | 3.00E-01 | 2.74E+00 | 2.76E+00 | 7.33E-01 | no | Gm13912 | intron_variant,non_coding_transcript_variant | MODIFIER |
| 2 | 108266382 | AX-170289036 | 0 | C | A | 0.053 | -1.01E-01 | 3.00E-01 | 2.74E+00 | 2.76E+00 | 7.33E-01 | no | Gm13912 | intron_variant,non_coding_transcript_variant | MODIFIER |
| 2 | 108267839 | AX-97853315 | 0 | C | T | 0.053 | -1.01E-01 | 3.00E-01 | 2.74E+00 | 2.76E+00 | 7.33E-01 | no | Gm13912 | intron_variant,non_coding_transcript_variant | MODIFIER |
| 2 | 108275542 | AX-97852662 | 0 | A | G | 0.053 | -1.01E-01 | 3.00E-01 | 2.74E+00 | 2.76E+00 | 7.33E-01 | no | Gm13912 | intron_variant,non_coding_transcript_variant | MODIFIER |

| | | | | | | | | | | | | | | | |
|---|---|---|---|---|---|---|---|---|---|---|---|---|---|---|---|
| 2 | 108300954 | AX-97793282 | 0 | T | G | 0.053 | -1.01E-01 | 3.00E-01 | 2.74E+00 | 2.76E+00 | 7.33E-01 | no | | intergenic_variant | MODIFIER |
| 2 | 108319997 | AX-170310792 | 0 | T | G | 0.053 | -1.01E-01 | 3.00E-01 | 2.74E+00 | 2.76E+00 | 7.33E-01 | no | | intergenic_variant | MODIFIER |
| 2 | 108320253 | AX-170293652 | 0 | T | G | 0.053 | -1.01E-01 | 3.00E-01 | 2.74E+00 | 2.76E+00 | 7.33E-01 | no | | intergenic_variant | MODIFIER |
| 2 | 108322998 | AX-170345906 | 0 | T | C | 0.053 | -1.01E-01 | 3.00E-01 | 2.74E+00 | 2.76E+00 | 7.33E-01 | no | | intergenic_variant | MODIFIER |
| 2 | 108334842 | AX-170317001 | 0 | G | A | 0.053 | -1.01E-01 | 3.00E-01 | 2.74E+00 | 2.76E+00 | 7.33E-01 | no | | intergenic_variant | MODIFIER |
| 2 | 108342897 | AX-170210571 | 0 | A | G | 0.053 | -1.01E-01 | 3.00E-01 | 2.74E+00 | 2.76E+00 | 7.33E-01 | no | | intergenic_variant | MODIFIER |
| 2 | 108402223 | AX-170413881 | 0 | G | A | 0.053 | -1.01E-01 | 3.00E-01 | 2.74E+00 | 2.76E+00 | 7.33E-01 | no | | intergenic_variant | MODIFIER |
| 2 | 108405893 | AX-170401636 | 0 | T | C | 0.053 | -1.01E-01 | 3.00E-01 | 2.74E+00 | 2.76E+00 | 7.33E-01 | no | | intergenic_variant | MODIFIER |
| 2 | 108406815 | AX-170102863 | 0 | T | C | 0.053 | -1.01E-01 | 3.00E-01 | 2.74E+00 | 2.76E+00 | 7.33E-01 | no | | intergenic_variant | MODIFIER |
| 2 | 108454305 | AX-170018625 | 0 | A | G | 0.053 | -1.01E-01 | 3.00E-01 | 2.74E+00 | 2.76E+00 | 7.33E-01 | no | | intergenic_variant | MODIFIER |
| 2 | 108461549 | AX-97839318 | 0 | T | C | 0.053 | -1.01E-01 | 3.00E-01 | 2.74E+00 | 2.76E+00 | 7.33E-01 | no | | intergenic_variant | MODIFIER |
| 2 | 108519078 | AX-170018628 | 0 | A | G | 0.053 | -1.01E-01 | 3.00E-01 | 2.74E+00 | 2.76E+00 | 7.33E-01 | no | | intergenic_variant | MODIFIER |
| 2 | 108936768 | AX-170359921 | 0 | A | T | 0.053 | -1.01E-01 | 3.00E-01 | 2.74E+00 | 2.76E+00 | 7.33E-01 | no | | regulatory_region_variant | MODIFIER |
| 2 | 109398655 | AX-97839375 | 0 | A | C | 0.053 | -1.01E-01 | 3.00E-01 | 2.74E+00 | 2.76E+00 | 7.33E-01 | no | | intergenic_variant | MODIFIER |
| 2 | 109640587 | AX-170518458 | 0 | T | C | 0.053 | -1.01E-01 | 3.00E-01 | 2.74E+00 | 2.76E+00 | 7.33E-01 | no | | intergenic_variant | MODIFIER |
| 2 | 109661253 | AX-170018645 | 0 | T | A | 0.053 | -1.01E-01 | 3.00E-01 | 2.74E+00 | 2.76E+00 | 7.33E-01 | no | | intergenic_variant | MODIFIER |
| 2 | 109661351 | AX-170018649 | 0 | T | A | 0.053 | -1.01E-01 | 3.00E-01 | 2.74E+00 | 2.76E+00 | 7.33E-01 | no | | intergenic_variant | MODIFIER |
| 2 | 109702851 | AX-170018646 | 0 | A | G | 0.053 | -1.01E-01 | 3.00E-01 | 2.74E+00 | 2.76E+00 | 7.33E-01 | no | Bdnf | intron_variant | MODIFIER |
| 2 | 109713244 | AX-170377765 | 0 | A | G | 0.053 | -1.01E-01 | 3.00E-01 | 2.74E+00 | 2.76E+00 | 7.33E-01 | no | Bdnf | intron_variant | MODIFIER |
| 2 | 109720212 | AX-170437605 | 0 | A | G | 0.053 | -1.01E-01 | 3.00E-01 | 2.74E+00 | 2.76E+00 | 7.33E-01 | no | Bdnf | intron_variant | MODIFIER |
| 2 | 109720277 | AX-170209815 | 0 | G | A | 0.053 | -1.01E-01 | 3.00E-01 | 2.74E+00 | 2.76E+00 | 7.33E-01 | no | Bdnf | intron_variant | MODIFIER |
| 2 | 109732500 | AX-97850286 | 0 | A | G | 0.053 | -1.01E-01 | 3.00E-01 | 2.74E+00 | 2.76E+00 | 7.33E-01 | no | | intergenic_variant | MODIFIER |
| 2 | 109735632 | AX-170018648 | 0 | A | G | 0.053 | -1.01E-01 | 3.00E-01 | 2.74E+00 | 2.76E+00 | 7.33E-01 | no | | regulatory_region_variant | MODIFIER |
| 2 | 109755211 | AX-170369737 | 0 | A | G | 0.053 | -1.01E-01 | 3.00E-01 | 2.74E+00 | 2.76E+00 | 7.33E-01 | no | | intergenic_variant | MODIFIER |
| 2 | 109779236 | AX-97787513 | 0 | G | A | 0.053 | -1.01E-01 | 3.00E-01 | 2.74E+00 | 2.76E+00 | 7.33E-01 | no | | intergenic_variant | MODIFIER |
| 2 | 109794883 | AX-97850170 | 0 | G | A | 0.053 | -1.01E-01 | 3.00E-01 | 2.74E+00 | 2.76E+00 | 7.33E-01 | no | Gm13932 | upstream_gene_variant | MODIFIER |
| 2 | 109798018 | AX-170430111 | 0 | A | G | 0.053 | -1.01E-01 | 3.00E-01 | 2.74E+00 | 2.76E+00 | 7.33E-01 | no | | intergenic_variant | MODIFIER |
| 2 | 109823273 | AX-170398241 | 0 | G | A | 0.053 | -1.01E-01 | 3.00E-01 | 2.74E+00 | 2.76E+00 | 7.33E-01 | no | | intergenic_variant | MODIFIER |
| 2 | 109832308 | AX-97796133 | 0 | T | C | 0.053 | -1.01E-01 | 3.00E-01 | 2.74E+00 | 2.76E+00 | 7.33E-01 | no | | intergenic_variant | MODIFIER |
| 2 | 109858203 | AX-168387102 | 0 | T | G | 0.053 | -1.01E-01 | 3.00E-01 | 2.74E+00 | 2.76E+00 | 7.33E-01 | no | | intergenic_variant | MODIFIER |
| 2 | 109859194 | AX-170144491 | 0 | G | A | 0.053 | -1.01E-01 | 3.00E-01 | 2.74E+00 | 2.76E+00 | 7.33E-01 | no | | intergenic_variant | MODIFIER |
| 2 | 109866779 | AX-170208791 | 0 | T | C | 0.053 | -1.01E-01 | 3.00E-01 | 2.74E+00 | 2.76E+00 | 7.33E-01 | no | | intergenic_variant | MODIFIER |
| 2 | 109908872 | AX-170159930 | 0 | C | T | 0.053 | -1.01E-01 | 3.00E-01 | 2.74E+00 | 2.76E+00 | 7.33E-01 | no | Platr8 | intron_variant,non_coding_transcript_variant | MODIFIER |
| 2 | 109908924 | AX-170389370 | 0 | T | C | 0.053 | -1.01E-01 | 3.00E-01 | 2.74E+00 | 2.76E+00 | 7.33E-01 | no | Platr8 | intron_variant,non_coding_transcript_variant | MODIFIER |
| 2 | 109909187 | AX-170233434 | 0 | A | C | 0.053 | -1.01E-01 | 3.00E-01 | 2.74E+00 | 2.76E+00 | 7.33E-01 | no | Platr8 | intron_variant,non_coding_transcript_variant | MODIFIER |
| 2 | 109929202 | AX-170466925 | 0 | A | G | 0.053 | -1.01E-01 | 3.00E-01 | 2.74E+00 | 2.76E+00 | 7.33E-01 | no | Lgr4 | intron_variant | MODIFIER |
| 2 | 109936366 | AX-170454477 | 0 | T | G | 0.053 | -1.01E-01 | 3.00E-01 | 2.74E+00 | 2.76E+00 | 7.33E-01 | no | Lgr4 | intron_variant | MODIFIER |
| 2 | 109940028 | AX-170501345 | 0 | G | T | 0.053 | -1.01E-01 | 3.00E-01 | 2.74E+00 | 2.76E+00 | 7.33E-01 | no | Lgr4 | intron_variant | MODIFIER |
| 2 | 109943485 | AX-170533209 | 0 | A | G | 0.053 | -1.01E-01 | 3.00E-01 | 2.74E+00 | 2.76E+00 | 7.33E-01 | no | Lgr4 | intron_variant | MODIFIER |
| 2 | 110193610 | AX-170018652 | 0 | C | A | 0.053 | -1.01E-01 | 3.00E-01 | 2.74E+00 | 2.76E+00 | 7.33E-01 | no | Gm13936 | upstream_gene_variant | MODIFIER |
| 2 | 110311644 | AX-97839402 | 0 | C | A | 0.053 | -1.01E-01 | 3.00E-01 | 2.74E+00 | 2.76E+00 | 7.33E-01 | no | Bbox1 | intron_variant | MODIFIER |
| 2 | 110319816 | AX-97797403 | 0 | G | T | 0.053 | -1.01E-01 | 3.00E-01 | 2.74E+00 | 2.76E+00 | 7.33E-01 | no | | intergenic_variant | MODIFIER |
| 2 | 110360537 | AX-97795312 | 0 | T | C | 0.053 | -1.01E-01 | 3.00E-01 | 2.74E+00 | 2.76E+00 | 7.33E-01 | no | Fibin | downstream_gene_variant | MODIFIER |
| 2 | 110389085 | AX-97792122 | 0 | G | A | 0.053 | -1.01E-01 | 3.00E-01 | 2.74E+00 | 2.76E+00 | 7.33E-01 | no | | intergenic_variant | MODIFIER |
| 2 | 110389483 | AX-170018660 | 0 | G | C | 0.053 | -1.01E-01 | 3.00E-01 | 2.74E+00 | 2.76E+00 | 7.33E-01 | no | | intergenic_variant | MODIFIER |
| 2 | 110411422 | AX-97805717 | 0 | G | A | 0.053 | -1.01E-01 | 3.00E-01 | 2.74E+00 | 2.76E+00 | 7.33E-01 | no | | intergenic_variant | MODIFIER |
| 2 | 110416878 | AX-170385113 | 0 | A | C | 0.053 | -1.01E-01 | 3.00E-01 | 2.74E+00 | 2.76E+00 | 7.33E-01 | no | Gm13935 | downstream_gene_variant | MODIFIER |
| 2 | 110416910 | AX-170475810 | 0 | T | C | 0.053 | -1.01E-01 | 3.00E-01 | 2.74E+00 | 2.76E+00 | 7.33E-01 | no | Gm13935 | downstream_gene_variant | MODIFIER |
| 2 | 110440852 | AX-170088706 | 0 | C | T | 0.053 | -1.01E-01 | 3.00E-01 | 2.74E+00 | 2.76E+00 | 7.33E-01 | no | | intergenic_variant | MODIFIER |
| 2 | 110455265 | AX-97793119 | 0 | C | G | 0.053 | -1.01E-01 | 3.00E-01 | 2.74E+00 | 2.76E+00 | 7.33E-01 | no | | intergenic_variant | MODIFIER |
| 2 | 110484676 | AX-170088707 | 0 | T | C | 0.053 | -1.01E-01 | 3.00E-01 | 2.74E+00 | 2.76E+00 | 7.33E-01 | no | | intergenic_variant | MODIFIER |
| 2 | 110518160 | AX-170435244 | 0 | A | G | 0.053 | -1.01E-01 | 3.00E-01 | 2.74E+00 | 2.76E+00 | 7.33E-01 | no | | intergenic_variant | MODIFIER |
| 2 | 110593970 | AX-170453137 | 0 | T | C | 0.053 | -1.01E-01 | 3.00E-01 | 2.74E+00 | 2.76E+00 | 7.33E-01 | no | Slc5a12 | upstream_gene_variant | MODIFIER |
| 2 | 110594006 | AX-97795127 | 0 | C | T | 0.053 | -1.01E-01 | 3.00E-01 | 2.74E+00 | 2.76E+00 | 7.33E-01 | no | Slc5a12 | upstream_gene_variant | MODIFIER |

| | | | | | | | | | | | | | | |
|---|---|---|---|---|---|---|---|---|---|---|---|---|---|---|
| 2 | 110597359 | AX-170507928 | 0 | A | G | 0.053 | -1.01E-01 | 3.00E-01 | 2.74E+00 | 2.76E+00 | 7.33E-01 | no | Slc5a12 | 5_prime_UTR_variant | MODIFIER |
| 2 | 110598894 | AX-170474264 | 0 | A | T | 0.053 | -1.01E-01 | 3.00E-01 | 2.74E+00 | 2.76E+00 | 7.33E-01 | no | Slc5a12 | intron_variant | MODIFIER |
| 2 | 110602810 | AX-170018670 | 0 | G | A | 0.053 | -1.01E-01 | 3.00E-01 | 2.74E+00 | 2.76E+00 | 7.33E-01 | no | Slc5a12 | intron_variant | MODIFIER |
| 2 | 110667461 | AX-170018675 | 0 | A | G | 0.053 | -1.01E-01 | 3.00E-01 | 2.74E+00 | 2.76E+00 | 7.33E-01 | no | Ano3 | intron_variant | MODIFIER |
| 2 | 110710230 | AX-170018672 | 0 | G | C | 0.053 | -1.01E-01 | 3.00E-01 | 2.74E+00 | 2.76E+00 | 7.33E-01 | no | Ano3 | intron_variant | MODIFIER |
| 2 | 110710778 | AX-97839418 | 0 | T | C | 0.053 | -1.01E-01 | 3.00E-01 | 2.74E+00 | 2.76E+00 | 7.33E-01 | no | Ano3 | intron_variant | MODIFIER |
| 2 | 110737285 | AX-97839449 | 0 | A | G | 0.053 | -1.01E-01 | 3.00E-01 | 2.74E+00 | 2.76E+00 | 7.33E-01 | no | Muc15 | intron_variant | MODIFIER |
| 2 | 110788361 | AX-97801055 | 0 | T | C | 0.053 | -1.01E-01 | 3.00E-01 | 2.74E+00 | 2.76E+00 | 7.33E-01 | no | Ano3 | intron_variant | MODIFIER |
| 2 | 110793143 | AX-170018671 | 0 | G | T | 0.053 | -1.01E-01 | 3.00E-01 | 2.74E+00 | 2.76E+00 | 7.33E-01 | no | Ano3 | intron_variant | MODIFIER |
| 2 | 110793294 | AX-170199706 | 0 | T | C | 0.053 | -1.01E-01 | 3.00E-01 | 2.74E+00 | 2.76E+00 | 7.33E-01 | no | Ano3 | intron_variant | MODIFIER |
| 2 | 110829010 | AX-170186903 | 0 | G | C | 0.053 | -1.01E-01 | 3.00E-01 | 2.74E+00 | 2.76E+00 | 7.33E-01 | no | Ano3 | intron_variant | MODIFIER |
| 2 | 110829087 | AX-97841656 | 0 | T | A | 0.053 | -1.01E-01 | 3.00E-01 | 2.74E+00 | 2.76E+00 | 7.33E-01 | no | Ano3 | intron_variant | MODIFIER |
| 2 | 110851998 | AX-97839446 | 0 | T | C | 0.053 | -1.01E-01 | 3.00E-01 | 2.74E+00 | 2.76E+00 | 7.33E-01 | no | Ano3 | intron_variant | MODIFIER |
| 2 | 110914374 | AX-170210002 | 0 | G | C | 0.053 | -1.01E-01 | 3.00E-01 | 2.74E+00 | 2.76E+00 | 7.33E-01 | no | Ano3 | intron_variant | MODIFIER |
| 2 | 110951682 | AX-168411394 | 0 | T | C | 0.053 | -1.01E-01 | 3.00E-01 | 2.74E+00 | 2.76E+00 | 7.33E-01 | no | Ano3 | upstream_gene_variant | MODIFIER |
| 2 | 110954205 | AX-170038226 | 0 | G | C | 0.053 | -1.01E-01 | 3.00E-01 | 2.74E+00 | 2.76E+00 | 7.33E-01 | no | Ano3 | upstream_gene_variant | MODIFIER |
| 2 | 112526636 | AX-170018699 | 0 | C | T | 0.053 | -1.01E-01 | 3.00E-01 | 2.74E+00 | 2.76E+00 | 7.33E-01 | no | Aven | intron_variant | MODIFIER |
| 2 | 112565050 | AX-170018698 | 0 | T | C | 0.053 | -1.01E-01 | 3.00E-01 | 2.74E+00 | 2.76E+00 | 7.33E-01 | no | Aven | intron_variant | MODIFIER |
| 2 | 112621594 | AX-170143717 | 0 | T | C | 0.053 | -1.01E-01 | 3.00E-01 | 2.74E+00 | 2.76E+00 | 7.33E-01 | no | Aven | intron_variant | MODIFIER |
| 2 | 112629897 | AX-170395382 | 0 | C | T | 0.053 | -1.01E-01 | 3.00E-01 | 2.74E+00 | 2.76E+00 | 7.33E-01 | no | Aven | missense_variant | MODERATE |
| 2 | 112651183 | AX-170131610 | 0 | T | C | 0.053 | -1.01E-01 | 3.00E-01 | 2.74E+00 | 2.76E+00 | 7.33E-01 | no | Ryr3 | intron_variant | MODIFIER |
| 2 | 112651700 | AX-170350541 | 0 | G | A | 0.053 | -1.01E-01 | 3.00E-01 | 2.74E+00 | 2.76E+00 | 7.33E-01 | no | Ryr3 | intron_variant | MODIFIER |
| 2 | 112652154 | AX-170163978 | 0 | G | A | 0.053 | -1.01E-01 | 3.00E-01 | 2.74E+00 | 2.76E+00 | 7.33E-01 | no | Ryr3 | intron_variant | MODIFIER |
| 2 | 112720956 | AX-170311778 | 0 | T | C | 0.053 | -1.01E-01 | 3.00E-01 | 2.74E+00 | 2.76E+00 | 7.33E-01 | no | Ryr3 | intron_variant | MODIFIER |
| 2 | 112727968 | AX-170306842 | 0 | T | C | 0.053 | -1.01E-01 | 3.00E-01 | 2.74E+00 | 2.76E+00 | 7.33E-01 | no | Ryr3 | intron_variant | MODIFIER |
| 2 | 112729603 | AX-170258900 | 0 | T | G | 0.053 | -1.01E-01 | 3.00E-01 | 2.74E+00 | 2.76E+00 | 7.33E-01 | no | Ryr3 | intron_variant | MODIFIER |
| 2 | 112732933 | AX-97838606 | 0 | C | T | 0.053 | -1.01E-01 | 3.00E-01 | 2.74E+00 | 2.76E+00 | 7.33E-01 | no | Ryr3 | intron_variant | MODIFIER |
| 2 | 112734262 | AX-170064620 | 0 | G | A | 0.053 | -1.01E-01 | 3.00E-01 | 2.74E+00 | 2.76E+00 | 7.33E-01 | no | Ryr3 | intron_variant | MODIFIER |
| 2 | 112755365 | AX-97780602 | 0 | T | C | 0.053 | -1.01E-01 | 3.00E-01 | 2.74E+00 | 2.76E+00 | 7.33E-01 | no | Ryr3 | intron_variant | MODIFIER |
| 2 | 112759846 | AX-170338236 | 0 | A | G | 0.053 | -1.01E-01 | 3.00E-01 | 2.74E+00 | 2.76E+00 | 7.33E-01 | no | Ryr3 | intron_variant | MODIFIER |
| 2 | 112771983 | AX-170348860 | 0 | T | C | 0.053 | -1.01E-01 | 3.00E-01 | 2.74E+00 | 2.76E+00 | 7.33E-01 | no | Ryr3 | intron_variant | MODIFIER |
| 2 | 112778898 | AX-170099756 | 0 | A | C | 0.053 | -1.01E-01 | 3.00E-01 | 2.74E+00 | 2.76E+00 | 7.33E-01 | no | Ryr3 | intron_variant | MODIFIER |
| 2 | 113070190 | AX-170333627 | 0 | T | A | 0.053 | -1.01E-01 | 3.00E-01 | 2.74E+00 | 2.76E+00 | 7.33E-01 | no | Ryr3 | intron_variant | MODIFIER |
| 2 | 113184012 | AX-97780921 | 0 | G | A | 0.053 | -1.01E-01 | 3.00E-01 | 2.74E+00 | 2.76E+00 | 7.33E-01 | no | Ryr3 | intron_variant | MODIFIER |
| 2 | 113242182 | AX-97835569 | 0 | T | C | 0.053 | -1.01E-01 | 3.00E-01 | 2.74E+00 | 2.76E+00 | 7.33E-01 | no | | intergenic_variant | MODIFIER |
| 2 | 113251426 | AX-170365203 | 0 | T | C | 0.053 | -1.01E-01 | 3.00E-01 | 2.74E+00 | 2.76E+00 | 7.33E-01 | no | | intergenic_variant | MODIFIER |
| 2 | 113258733 | AX-170267851 | 0 | T | C | 0.053 | -1.01E-01 | 3.00E-01 | 2.74E+00 | 2.76E+00 | 7.33E-01 | no | | intergenic_variant | MODIFIER |
| 2 | 113258891 | AX-170208172 | 0 | G | C | 0.053 | -1.01E-01 | 3.00E-01 | 2.74E+00 | 2.76E+00 | 7.33E-01 | no | | intergenic_variant | MODIFIER |
| 2 | 113260069 | AX-170424229 | 0 | G | T | 0.053 | -1.01E-01 | 3.00E-01 | 2.74E+00 | 2.76E+00 | 7.33E-01 | no | | intergenic_variant | MODIFIER |
| 2 | 113343061 | AX-97802571 | 0 | G | C | 0.053 | -1.01E-01 | 3.00E-01 | 2.74E+00 | 2.76E+00 | 7.33E-01 | no | Fmn1 | intron_variant | MODIFIER |
| 2 | 113381723 | AX-170018711 | 0 | T | C | 0.053 | -1.01E-01 | 3.00E-01 | 2.74E+00 | 2.76E+00 | 7.33E-01 | no | Fmn1 | intron_variant | MODIFIER |
| 2 | 113396165 | AX-170018712 | 0 | T | C | 0.053 | -1.01E-01 | 3.00E-01 | 2.74E+00 | 2.76E+00 | 7.33E-01 | no | Fmn1 | intron_variant | MODIFIER |
| 2 | 113407440 | AX-170094375 | 0 | C | T | 0.053 | -1.01E-01 | 3.00E-01 | 2.74E+00 | 2.76E+00 | 7.33E-01 | no | Fmn1 | intron_variant | MODIFIER |
| 2 | 113416620 | AX-170185397 | 0 | C | T | 0.053 | -1.01E-01 | 3.00E-01 | 2.74E+00 | 2.76E+00 | 7.33E-01 | no | Fmn1 | intron_variant | MODIFIER |
| 2 | 113416957 | AX-170064630 | 0 | G | A | 0.053 | -1.01E-01 | 3.00E-01 | 2.74E+00 | 2.76E+00 | 7.33E-01 | no | Fmn1 | intron_variant | MODIFIER |
| 2 | 113428636 | AX-97854621 | 0 | G | T | 0.053 | -1.01E-01 | 3.00E-01 | 2.74E+00 | 2.76E+00 | 7.33E-01 | no | Fmn1 | intron_variant | MODIFIER |
| 2 | 113445953 | AX-170018717 | 0 | T | A | 0.053 | -1.01E-01 | 3.00E-01 | 2.74E+00 | 2.76E+00 | 7.33E-01 | no | Fmn1 | intron_variant | MODIFIER |
| 2 | 113470716 | AX-170205218 | 0 | A | G | 0.053 | -1.01E-01 | 3.00E-01 | 2.74E+00 | 2.76E+00 | 7.33E-01 | no | Fmn1 | intron_variant | MODIFIER |
| 2 | 113472758 | AX-170313511 | 0 | T | A | 0.053 | -1.01E-01 | 3.00E-01 | 2.74E+00 | 2.76E+00 | 7.33E-01 | no | Fmn1 | intron_variant | MODIFIER |
| 2 | 113482979 | AX-170018708 | 0 | G | A | 0.053 | -1.01E-01 | 3.00E-01 | 2.74E+00 | 2.76E+00 | 7.33E-01 | no | Fmn1 | intron_variant | MODIFIER |
| 2 | 113483011 | AX-97798550 | 0 | C | A | 0.053 | -1.01E-01 | 3.00E-01 | 2.74E+00 | 2.76E+00 | 7.33E-01 | no | Fmn1 | intron_variant | MODIFIER |
| 2 | 113483135 | AX-170018710 | 0 | C | T | 0.053 | -1.01E-01 | 3.00E-01 | 2.74E+00 | 2.76E+00 | 7.33E-01 | no | Fmn1 | intron_variant | MODIFIER |
| 2 | 113528988 | AX-170203723 | 0 | G | A | 0.053 | -1.01E-01 | 3.00E-01 | 2.74E+00 | 2.76E+00 | 7.33E-01 | no | Fmn1 | intron_variant | MODIFIER |
| 2 | 113531660 | AX-97834538 | 0 | C | T | 0.053 | -1.01E-01 | 3.00E-01 | 2.74E+00 | 2.76E+00 | 7.33E-01 | no | Fmn1 | intron_variant | MODIFIER |

| | | | | | | | | | | | | | | | |
|---|---|---|---|---|---|---|---|---|---|---|---|---|---|---|---|
| 2 | 113546050 | AX-170018709 | 0 | T | C | 0.053 | -1.01E-01 | 3.00E-01 | 2.74E+00 | 2.76E+00 | 7.33E-01 | no | Fmn1 | intron_variant | MODIFIER |
| 2 | 113573661 | AX-170018714 | 0 | T | C | 0.053 | -1.01E-01 | 3.00E-01 | 2.74E+00 | 2.76E+00 | 7.33E-01 | no | Fmn1 | intron_variant | MODIFIER |
| 2 | 113577404 | AX-170318503 | 0 | T | A | 0.053 | -1.01E-01 | 3.00E-01 | 2.74E+00 | 2.76E+00 | 7.33E-01 | no | Fmn1 | intron_variant | MODIFIER |
| 2 | 113577483 | AX-170018707 | 0 | G | A | 0.053 | -1.01E-01 | 3.00E-01 | 2.74E+00 | 2.76E+00 | 7.33E-01 | no | Fmn1 | intron_variant | MODIFIER |
| 2 | 113583497 | AX-97801901 | 0 | G | A | 0.053 | -1.01E-01 | 3.00E-01 | 2.74E+00 | 2.76E+00 | 7.33E-01 | no | Fmn1 | intron_variant | MODIFIER |
| 2 | 113608326 | AX-97839579 | 0 | C | G | 0.053 | -1.01E-01 | 3.00E-01 | 2.74E+00 | 2.76E+00 | 7.33E-01 | no | Fmn1 | intron_variant | MODIFIER |
| 2 | 113616590 | AX-170018718 | 0 | T | A | 0.053 | -1.01E-01 | 3.00E-01 | 2.74E+00 | 2.76E+00 | 7.33E-01 | no | Fmn1 | intron_variant | MODIFIER |
| 5 | 23860767 | AX-170024527 | 0 | C | T | 0.053 | -1.01E-01 | 3.00E-01 | 2.74E+00 | 2.76E+00 | 7.33E-01 | no | | intergenic_variant | MODIFIER |
| 5 | 23911125 | AX-170024520 | 0 | C | T | 0.053 | -1.01E-01 | 3.00E-01 | 2.74E+00 | 2.76E+00 | 7.33E-01 | no | Fam126a | downstream_gene_variant | MODIFIER |
| 5 | 23985831 | AX-170123237 | 0 | C | T | 0.053 | -1.01E-01 | 3.00E-01 | 2.74E+00 | 2.76E+00 | 7.33E-01 | no | Fam126a | synonymous_variant | LOW |
| 5 | 24013407 | AX-170488640 | 0 | C | T | 0.053 | -1.01E-01 | 3.00E-01 | 2.74E+00 | 2.76E+00 | 7.33E-01 | no | Fam126a | intron_variant | MODIFIER |
| 5 | 24018866 | AX-170310976 | 0 | C | T | 0.053 | -1.01E-01 | 3.00E-01 | 2.74E+00 | 2.76E+00 | 7.33E-01 | no | Fam126a | intron_variant | MODIFIER |
| 5 | 24025091 | AX-97821038 | 0 | A | G | 0.053 | -1.01E-01 | 3.00E-01 | 2.74E+00 | 2.76E+00 | 7.33E-01 | no | Fam126a | intron_variant | MODIFIER |
| 5 | 24034175 | AX-170328204 | 0 | T | C | 0.053 | -1.01E-01 | 3.00E-01 | 2.74E+00 | 2.76E+00 | 7.33E-01 | no | Fam126a | upstream_gene_variant | MODIFIER |
| 5 | 24066837 | AX-170172900 | 0 | G | T | 0.053 | -1.01E-01 | 3.00E-01 | 2.74E+00 | 2.76E+00 | 7.33E-01 | no | | intergenic_variant | MODIFIER |
| 5 | 24068293 | AX-97831040 | 0 | A | G | 0.053 | -1.01E-01 | 3.00E-01 | 2.74E+00 | 2.76E+00 | 7.33E-01 | no | | intergenic_variant | MODIFIER |
| 5 | 24083892 | AX-97793572 | 0 | A | G | 0.053 | -1.01E-01 | 3.00E-01 | 2.74E+00 | 2.76E+00 | 7.33E-01 | no | Gm18038 | upstream_gene_variant | MODIFIER |
| 5 | 24087305 | AX-170435707 | 0 | T | G | 0.053 | -1.01E-01 | 3.00E-01 | 2.74E+00 | 2.76E+00 | 7.33E-01 | no | Gm18038 | downstream_gene_variant | MODIFIER |
| 5 | 24092345 | AX-170196995 | 0 | G | A | 0.053 | -1.01E-01 | 3.00E-01 | 2.74E+00 | 2.76E+00 | 7.33E-01 | no | | intergenic_variant | MODIFIER |
| 5 | 24111387 | AX-97835650 | 0 | G | A | 0.053 | -1.01E-01 | 3.00E-01 | 2.74E+00 | 2.76E+00 | 7.33E-01 | no | Klhl7 | intron_variant | MODIFIER |
| 5 | 24144336 | AX-97844581 | 0 | C | A | 0.053 | -1.01E-01 | 3.00E-01 | 2.74E+00 | 2.76E+00 | 7.33E-01 | no | Klhl7 | intron_variant | MODIFIER |
| 5 | 24146441 | AX-170338012 | 0 | A | G | 0.053 | -1.01E-01 | 3.00E-01 | 2.74E+00 | 2.76E+00 | 7.33E-01 | no | Klhl7 | intron_variant | MODIFIER |
| 5 | 24156387 | AX-168401700 | 0 | G | A | 0.053 | -1.01E-01 | 3.00E-01 | 2.74E+00 | 2.76E+00 | 7.33E-01 | no | Klhl7 | intron_variant | MODIFIER |
| 5 | 24156727 | AX-170149833 | 0 | A | G | 0.053 | -1.01E-01 | 3.00E-01 | 2.74E+00 | 2.76E+00 | 7.33E-01 | no | Klhl7 | intron_variant | MODIFIER |
| 5 | 24157151 | AX-170217355 | 0 | T | A | 0.053 | -1.01E-01 | 3.00E-01 | 2.74E+00 | 2.76E+00 | 7.33E-01 | no | Klhl7 | intron_variant | MODIFIER |
| 5 | 24160531 | AX-170287946 | 0 | G | A | 0.053 | -1.01E-01 | 3.00E-01 | 2.74E+00 | 2.76E+00 | 7.33E-01 | no | Klhl7 | 3_prime_UTR_variant | MODIFIER |
| 5 | 24182136 | AX-170395683 | 0 | C | G | 0.053 | -1.01E-01 | 3.00E-01 | 2.74E+00 | 2.76E+00 | 7.33E-01 | no | Nupl2 | missense_variant | MODERATE |
| 5 | 24195455 | AX-170024536 | 0 | A | G | 0.053 | -1.01E-01 | 3.00E-01 | 2.74E+00 | 2.76E+00 | 7.33E-01 | no | | intergenic_variant | MODIFIER |
| 5 | 24196567 | AX-170024540 | 0 | A | T | 0.053 | -1.01E-01 | 3.00E-01 | 2.74E+00 | 2.76E+00 | 7.33E-01 | no | | intergenic_variant | MODIFIER |
| 5 | 24213704 | AX-170024533 | 0 | A | G | 0.053 | -1.01E-01 | 3.00E-01 | 2.74E+00 | 2.76E+00 | 7.33E-01 | no | Gm5575 | upstream_gene_variant | MODIFIER |
| 5 | 24223090 | AX-170071550 | 0 | T | A | 0.053 | -1.01E-01 | 3.00E-01 | 2.74E+00 | 2.76E+00 | 7.33E-01 | no | | regulatory_region_variant | MODIFIER |
| 5 | 24236731 | AX-97835654 | 0 | A | G | 0.053 | -1.01E-01 | 3.00E-01 | 2.74E+00 | 2.76E+00 | 7.33E-01 | no | Gm43828 | downstream_gene_variant | MODIFIER |
| 5 | 24264150 | AX-97830933 | 0 | C | T | 0.053 | -1.01E-01 | 3.00E-01 | 2.74E+00 | 2.76E+00 | 7.33E-01 | no | | regulatory_region_variant | MODIFIER |
| 5 | 24271180 | AX-170071551 | 0 | C | G | 0.053 | -1.01E-01 | 3.00E-01 | 2.74E+00 | 2.76E+00 | 7.33E-01 | no | | intergenic_variant | MODIFIER |
| 5 | 24282042 | AX-97835657 | 0 | A | G | 0.053 | -1.01E-01 | 3.00E-01 | 2.74E+00 | 2.76E+00 | 7.33E-01 | no | | intergenic_variant | MODIFIER |
| 5 | 24288477 | AX-170484580 | 0 | A | G | 0.053 | -1.01E-01 | 3.00E-01 | 2.74E+00 | 2.76E+00 | 7.33E-01 | no | | intergenic_variant | MODIFIER |
| 5 | 24289171 | AX-170024539 | 0 | T | C | 0.053 | -1.01E-01 | 3.00E-01 | 2.74E+00 | 2.76E+00 | 7.33E-01 | no | | intergenic_variant | MODIFIER |
| 5 | 24289686 | AX-170327947 | 0 | G | A | 0.053 | -1.01E-01 | 3.00E-01 | 2.74E+00 | 2.76E+00 | 7.33E-01 | no | | intergenic_variant | MODIFIER |
| 5 | 24291151 | AX-170157798 | 0 | A | T | 0.053 | -1.01E-01 | 3.00E-01 | 2.74E+00 | 2.76E+00 | 7.33E-01 | no | | intergenic_variant | MODIFIER |
| 5 | 24291432 | AX-170516912 | 0 | A | G | 0.053 | -1.01E-01 | 3.00E-01 | 2.74E+00 | 2.76E+00 | 7.33E-01 | no | | intergenic_variant | MODIFIER |
| 5 | 26548606 | AX-97784545 | 0 | T | G | 0.053 | -1.01E-01 | 3.00E-01 | 2.74E+00 | 2.76E+00 | 7.33E-01 | no | | intergenic_variant | MODIFIER |
| 5 | 26616122 | AX-170134959 | 0 | A | G | 0.053 | -1.01E-01 | 3.00E-01 | 2.74E+00 | 2.76E+00 | 7.33E-01 | no | | intergenic_variant | MODIFIER |
| 5 | 26616158 | AX-170445204 | 0 | T | G | 0.053 | -1.01E-01 | 3.00E-01 | 2.74E+00 | 2.76E+00 | 7.33E-01 | no | | intergenic_variant | MODIFIER |
| 5 | 26635710 | AX-97822535 | 0 | T | C | 0.053 | -1.01E-01 | 3.00E-01 | 2.74E+00 | 2.76E+00 | 7.33E-01 | no | | intergenic_variant | MODIFIER |
| 5 | 26729468 | AX-170299956 | 0 | C | T | 0.053 | -1.01E-01 | 3.00E-01 | 2.74E+00 | 2.76E+00 | 7.33E-01 | no | | intergenic_variant | MODIFIER |
| 5 | 26729906 | AX-97800195 | 0 | A | G | 0.053 | -1.01E-01 | 3.00E-01 | 2.74E+00 | 2.76E+00 | 7.33E-01 | no | | intergenic_variant | MODIFIER |
| 5 | 26729956 | AX-170119749 | 0 | G | A | 0.053 | -1.01E-01 | 3.00E-01 | 2.74E+00 | 2.76E+00 | 7.33E-01 | no | | intergenic_variant | MODIFIER |
| 5 | 26730294 | AX-170381478 | 0 | T | A | 0.053 | -1.01E-01 | 3.00E-01 | 2.74E+00 | 2.76E+00 | 7.33E-01 | no | | intergenic_variant | MODIFIER |
| 5 | 26755219 | AX-97827771 | 0 | G | A | 0.053 | -1.01E-01 | 3.00E-01 | 2.74E+00 | 2.76E+00 | 7.33E-01 | no | | intergenic_variant | MODIFIER |
| 5 | 27100253 | AX-170024565 | 0 | T | C | 0.053 | -1.01E-01 | 3.00E-01 | 2.74E+00 | 2.76E+00 | 7.33E-01 | no | Dpp6 | intron_variant | MODIFIER |
| 5 | 27114401 | AX-170024569 | 0 | C | T | 0.053 | -1.01E-01 | 3.00E-01 | 2.74E+00 | 2.76E+00 | 7.33E-01 | no | Dpp6 | intron_variant | MODIFIER |
| 5 | 27125231 | AX-170024567 | 0 | C | T | 0.053 | -1.01E-01 | 3.00E-01 | 2.74E+00 | 2.76E+00 | 7.33E-01 | no | Dpp6 | intron_variant | MODIFIER |
| 5 | 27125986 | AX-97845042 | 0 | T | C | 0.053 | -1.01E-01 | 3.00E-01 | 2.74E+00 | 2.76E+00 | 7.33E-01 | no | Dpp6 | intron_variant | MODIFIER |
| 5 | 27129337 | AX-170238183 | 0 | G | T | 0.053 | -1.01E-01 | 3.00E-01 | 2.74E+00 | 2.76E+00 | 7.33E-01 | no | Dpp6 | intron_variant | MODIFIER |

| | | | | | | | | | | | | | | | |
|---|---|---|---|---|---|---|---|---|---|---|---|---|---|---|---|
| 5 | 27147152 | AX-170183204 | 0 | C | T | 0.053 | -1.01E-01 | 3.00E-01 | 2.74E+00 | 2.76E+00 | 7.33E-01 | no | Dpp6 | intron_variant | MODIFIER |
| 5 | 27154219 | AX-170333610 | 0 | T | C | 0.053 | -1.01E-01 | 3.00E-01 | 2.74E+00 | 2.76E+00 | 7.33E-01 | no | Dpp6 | intron_variant | MODIFIER |
| 5 | 27155641 | AX-170121558 | 0 | A | G | 0.053 | -1.01E-01 | 3.00E-01 | 2.74E+00 | 2.76E+00 | 7.33E-01 | no | Dpp6 | intron_variant | MODIFIER |
| 5 | 27156380 | AX-97794499 | 0 | A | G | 0.053 | -1.01E-01 | 3.00E-01 | 2.74E+00 | 2.76E+00 | 7.33E-01 | no | Dpp6 | intron_variant | MODIFIER |
| 5 | 27157919 | AX-170388888 | 0 | C | G | 0.053 | -1.01E-01 | 3.00E-01 | 2.74E+00 | 2.76E+00 | 7.33E-01 | no | Dpp6 | intron_variant | MODIFIER |
| 5 | 27157952 | AX-170453252 | 0 | T | C | 0.053 | -1.01E-01 | 3.00E-01 | 2.74E+00 | 2.76E+00 | 7.33E-01 | no | Dpp6 | intron_variant | MODIFIER |
| 5 | 27162066 | AX-170269315 | 0 | G | A | 0.053 | -1.01E-01 | 3.00E-01 | 2.74E+00 | 2.76E+00 | 7.33E-01 | no | Dpp6 | intron_variant | MODIFIER |
| 5 | 27164427 | AX-170398651 | 0 | A | G | 0.053 | -1.01E-01 | 3.00E-01 | 2.74E+00 | 2.76E+00 | 7.33E-01 | no | Dpp6 | intron_variant | MODIFIER |
| 5 | 27174756 | AX-170306099 | 0 | A | G | 0.053 | -1.01E-01 | 3.00E-01 | 2.74E+00 | 2.76E+00 | 7.33E-01 | no | Dpp6 | intron_variant | MODIFIER |
| 5 | 27176625 | AX-170490230 | 0 | A | T | 0.053 | -1.01E-01 | 3.00E-01 | 2.74E+00 | 2.76E+00 | 7.33E-01 | no | Dpp6 | intron_variant | MODIFIER |
| 5 | 27180205 | AX-170071611 | 0 | G | T | 0.053 | -1.01E-01 | 3.00E-01 | 2.74E+00 | 2.76E+00 | 7.33E-01 | no | Dpp6 | intron_variant | MODIFIER |
| 5 | 27180840 | AX-170071608 | 0 | T | C | 0.053 | -1.01E-01 | 3.00E-01 | 2.74E+00 | 2.76E+00 | 7.33E-01 | no | Dpp6 | intron_variant | MODIFIER |
| 5 | 27181213 | AX-97799019 | 0 | A | G | 0.053 | -1.01E-01 | 3.00E-01 | 2.74E+00 | 2.76E+00 | 7.33E-01 | no | Dpp6 | intron_variant | MODIFIER |
| 5 | 27357250 | AX-170095268 | 0 | C | T | 0.053 | -1.01E-01 | 3.00E-01 | 2.74E+00 | 2.76E+00 | 7.33E-01 | no | Dpp6 | intron_variant | MODIFIER |
| 5 | 27369646 | AX-170531051 | 0 | A | G | 0.053 | -1.01E-01 | 3.00E-01 | 2.74E+00 | 2.76E+00 | 7.33E-01 | no | Dpp6 | intron_variant | MODIFIER |
| 5 | 27369682 | AX-170322311 | 0 | A | G | 0.053 | -1.01E-01 | 3.00E-01 | 2.74E+00 | 2.76E+00 | 7.33E-01 | no | Dpp6 | intron_variant | MODIFIER |
| 5 | 27382620 | AX-97805510 | 0 | T | C | 0.053 | -1.01E-01 | 3.00E-01 | 2.74E+00 | 2.76E+00 | 7.33E-01 | no | Dpp6 | intron_variant | MODIFIER |
| 5 | 27382894 | AX-170406681 | 0 | T | C | 0.053 | -1.01E-01 | 3.00E-01 | 2.74E+00 | 2.76E+00 | 7.33E-01 | no | Dpp6 | intron_variant | MODIFIER |
| 5 | 27383065 | AX-170492137 | 0 | A | G | 0.053 | -1.01E-01 | 3.00E-01 | 2.74E+00 | 2.76E+00 | 7.33E-01 | no | Dpp6 | intron_variant | MODIFIER |
| 5 | 27383520 | AX-170202071 | 0 | T | C | 0.053 | -1.01E-01 | 3.00E-01 | 2.74E+00 | 2.76E+00 | 7.33E-01 | no | Dpp6 | intron_variant | MODIFIER |
| 5 | 27386258 | AX-170330330 | 0 | T | C | 0.053 | -1.01E-01 | 3.00E-01 | 2.74E+00 | 2.76E+00 | 7.33E-01 | no | Dpp6 | intron_variant | MODIFIER |
| 5 | 27390724 | AX-170442483 | 0 | T | C | 0.053 | -1.01E-01 | 3.00E-01 | 2.74E+00 | 2.76E+00 | 7.33E-01 | no | Dpp6 | intron_variant | MODIFIER |
| 5 | 27416183 | AX-97814486 | 0 | T | C | 0.053 | -1.01E-01 | 3.00E-01 | 2.74E+00 | 2.76E+00 | 7.33E-01 | no | Dpp6 | intron_variant | MODIFIER |
| 5 | 27448069 | AX-170467144 | 0 | G | T | 0.053 | -1.01E-01 | 3.00E-01 | 2.74E+00 | 2.76E+00 | 7.33E-01 | no | Dpp6 | intron_variant | MODIFIER |
| 5 | 27457120 | AX-97841688 | 0 | A | G | 0.053 | -1.01E-01 | 3.00E-01 | 2.74E+00 | 2.76E+00 | 7.33E-01 | no | Dpp6 | intron_variant | MODIFIER |
| 5 | 27470298 | AX-170295339 | 0 | T | C | 0.053 | -1.01E-01 | 3.00E-01 | 2.74E+00 | 2.76E+00 | 7.33E-01 | no | Dpp6 | intron_variant | MODIFIER |
| 5 | 27476771 | AX-170148089 | 0 | T | C | 0.053 | -1.01E-01 | 3.00E-01 | 2.74E+00 | 2.76E+00 | 7.33E-01 | no | Dpp6 | intron_variant | MODIFIER |
| 5 | 27478618 | AX-170353510 | 0 | T | A | 0.053 | -1.01E-01 | 3.00E-01 | 2.74E+00 | 2.76E+00 | 7.33E-01 | no | Dpp6 | intron_variant | MODIFIER |
| 5 | 27480471 | AX-97809578 | 0 | G | T | 0.053 | -1.01E-01 | 3.00E-01 | 2.74E+00 | 2.76E+00 | 7.33E-01 | no | Dpp6 | intron_variant | MODIFIER |
| 5 | 27538393 | AX-97817247 | 0 | G | A | 0.053 | -1.01E-01 | 3.00E-01 | 2.74E+00 | 2.76E+00 | 7.33E-01 | no | Dpp6 | intron_variant | MODIFIER |
| 5 | 27539035 | AX-170235113 | 0 | T | C | 0.053 | -1.01E-01 | 3.00E-01 | 2.74E+00 | 2.76E+00 | 7.33E-01 | no | Dpp6 | intron_variant | MODIFIER |
| 5 | 27572111 | AX-170024580 | 0 | G | A | 0.053 | -1.01E-01 | 3.00E-01 | 2.74E+00 | 2.76E+00 | 7.33E-01 | no | Dpp6 | intron_variant | MODIFIER |
| 5 | 27572315 | AX-170024579 | 0 | C | T | 0.053 | -1.01E-01 | 3.00E-01 | 2.74E+00 | 2.76E+00 | 7.33E-01 | no | Dpp6 | intron_variant | MODIFIER |
| 5 | 27572380 | AX-170024581 | 0 | T | G | 0.053 | -1.01E-01 | 3.00E-01 | 2.74E+00 | 2.76E+00 | 7.33E-01 | no | Dpp6 | intron_variant | MODIFIER |
| 5 | 27575410 | AX-170095269 | 0 | T | C | 0.053 | -1.01E-01 | 3.00E-01 | 2.74E+00 | 2.76E+00 | 7.33E-01 | no | Dpp6 | intron_variant | MODIFIER |
| 5 | 27625317 | AX-170314792 | 0 | G | A | 0.053 | -1.01E-01 | 3.00E-01 | 2.74E+00 | 2.76E+00 | 7.33E-01 | no | Dpp6 | intron_variant | MODIFIER |
| 5 | 27629000 | AX-170024572 | 0 | C | A | 0.053 | -1.01E-01 | 3.00E-01 | 2.74E+00 | 2.76E+00 | 7.33E-01 | no | Dpp6 | intron_variant | MODIFIER |
| 5 | 27634421 | AX-170112294 | 0 | G | A | 0.053 | -1.01E-01 | 3.00E-01 | 2.74E+00 | 2.76E+00 | 7.33E-01 | no | Dpp6 | intron_variant | MODIFIER |
| 5 | 27635290 | AX-170460522 | 0 | A | T | 0.053 | -1.01E-01 | 3.00E-01 | 2.74E+00 | 2.76E+00 | 7.33E-01 | no | Dpp6 | intron_variant | MODIFIER |
| 5 | 27651857 | AX-97818092 | 0 | T | C | 0.053 | -1.01E-01 | 3.00E-01 | 2.74E+00 | 2.76E+00 | 7.33E-01 | no | Dpp6 | intron_variant | MODIFIER |
| 5 | 27652285 | AX-170133160 | 0 | A | G | 0.053 | -1.01E-01 | 3.00E-01 | 2.74E+00 | 2.76E+00 | 7.33E-01 | no | Dpp6 | synonymous_variant | LOW |
| 5 | 27710032 | AX-97854213 | 0 | C | T | 0.053 | -1.01E-01 | 3.00E-01 | 2.74E+00 | 2.76E+00 | 7.33E-01 | no | Dpp6 | intron_variant | MODIFIER |
| 5 | 27710143 | AX-170327319 | 0 | A | G | 0.053 | -1.01E-01 | 3.00E-01 | 2.74E+00 | 2.76E+00 | 7.33E-01 | no | Dpp6 | intron_variant | MODIFIER |
| 5 | 27710161 | AX-170148049 | 0 | T | G | 0.053 | -1.01E-01 | 3.00E-01 | 2.74E+00 | 2.76E+00 | 7.33E-01 | no | Dpp6 | intron_variant | MODIFIER |
| 5 | 27717954 | AX-170114152 | 0 | G | A | 0.053 | -1.01E-01 | 3.00E-01 | 2.74E+00 | 2.76E+00 | 7.33E-01 | no | Dpp6 | intron_variant | MODIFIER |
| 5 | 27718139 | AX-170387304 | 0 | T | G | 0.053 | -1.01E-01 | 3.00E-01 | 2.74E+00 | 2.76E+00 | 7.33E-01 | no | Dpp6 | intron_variant | MODIFIER |
| 5 | 27720897 | AX-170024575 | 0 | G | A | 0.053 | -1.01E-01 | 3.00E-01 | 2.74E+00 | 2.76E+00 | 7.33E-01 | no | Dpp6 | intron_variant | MODIFIER |
| 5 | 27723655 | AX-97821123 | 0 | A | C | 0.053 | -1.01E-01 | 3.00E-01 | 2.74E+00 | 2.76E+00 | 7.33E-01 | no | Dpp6 | intron_variant | MODIFIER |
| 5 | 27754526 | AX-170024577 | 0 | C | T | 0.053 | -1.01E-01 | 3.00E-01 | 2.74E+00 | 2.76E+00 | 7.33E-01 | no | Paxip1 | synonymous_variant | LOW |
| 5 | 27759076 | AX-170224610 | 0 | A | G | 0.053 | -1.01E-01 | 3.00E-01 | 2.74E+00 | 2.76E+00 | 7.33E-01 | no | Paxip1 | intron_variant | MODIFIER |
| 5 | 27769714 | AX-170302148 | 0 | G | A | 0.053 | -1.01E-01 | 3.00E-01 | 2.74E+00 | 2.76E+00 | 7.33E-01 | no | Paxip1 | intron_variant | MODIFIER |
| 5 | 27775946 | AX-97821618 | 0 | C | T | 0.053 | -1.01E-01 | 3.00E-01 | 2.74E+00 | 2.76E+00 | 7.33E-01 | no | Paxip1 | intron_variant | MODIFIER |
| 5 | 27817723 | AX-97813370 | 0 | G | A | 0.053 | -1.01E-01 | 3.00E-01 | 2.74E+00 | 2.76E+00 | 7.33E-01 | no | Gm43433 | non_coding_transcript_exon_variant | MODIFIER |
| 5 | 27836865 | AX-170095270 | 0 | T | G | 0.053 | -1.01E-01 | 3.00E-01 | 2.74E+00 | 2.76E+00 | 7.33E-01 | no | | intergenic_variant | MODIFIER |

| | | | | | | | | | | | | | | | |
|---|---|---|---|---|---|---|---|---|---|---|---|---|---|---|---|
| 5 | 27837365 | AX-170369142 | 0 | T | G | 0.053 | -1.01E-01 | 3.00E-01 | 2.74E+00 | 2.76E+00 | 7.33E-01 | no | Htr5a | upstream_gene_variant | MODIFIER |
| 12 | 64871549 | AX-170532796 | 0 | C | G | 0.053 | -1.01E-01 | 3.00E-01 | 2.74E+00 | 2.76E+00 | 7.33E-01 | no | | intergenic_variant | MODIFIER |
| 12 | 70326471 | AX-97837487 | 0 | G | A | 0.053 | -1.01E-01 | 3.00E-01 | 2.74E+00 | 2.76E+00 | 7.33E-01 | no | Trim9 | intron_variant | MODIFIER |
| 4 | 142269080 | AX-170488669 | 2 | A | T | 0.073 | 5.84E-02 | 1.74E-01 | 2.68E+00 | 2.75E+00 | 7.33E-01 | no | | intergenic_variant | MODIFIER |
| 16 | 78395760 | AX-170258011 | 0 | C | T | 0.456 | 2.63E-02 | 7.51E-02 | 2.59E+00 | 2.67E+00 | 7.33E-01 | no | | intergenic_variant | MODIFIER |
| 2 | 102936814 | AX-170064474 | 1 | G | T | 0.054 | -1.01E-01 | 3.00E-01 | 2.74E+00 | 2.77E+00 | 7.33E-01 | no | | intergenic_variant | MODIFIER |
| 8 | 38259664 | AX-170079805 | 2 | C | A | 0.155 | 4.94E-02 | 1.47E-01 | 2.75E+00 | 2.80E+00 | 7.33E-01 | no | Sgcz | intron_variant | MODIFIER |
| 11 | 32331100 | AX-170242600 | 0 | C | T | 0.096 | 6.57E-02 | 1.95E-01 | 2.67E+00 | 2.74E+00 | 7.33E-01 | no | | regulatory_region_variant | MODIFIER |
| 1 | 41720191 | AX-170258442 | 1 | A | T | 0.42 | -2.52E-02 | 7.42E-02 | 2.63E+00 | 2.73E+00 | 7.33E-01 | no | | intergenic_variant | MODIFIER |
| 10 | 6617751 | AX-170466434 | 2 | G | A | 0.209 | -3.26E-02 | 9.74E-02 | 2.65E+00 | 2.75E+00 | 7.33E-01 | no | | intergenic_variant | MODIFIER |
| 8 | 23707562 | AX-170115697 | 1 | G | A | 0.179 | 3.62E-02 | 1.09E-01 | 2.69E+00 | 2.78E+00 | 7.33E-01 | no | Zmat4 | intron_variant | MODIFIER |
| 18 | 57807052 | AX-170472797 | 0 | A | T | 0.193 | -4.01E-02 | 1.18E-01 | 2.64E+00 | 2.71E+00 | 7.33E-01 | no | | intergenic_variant | MODIFIER |
| 18 | 59328622 | AX-170189256 | 0 | A | G | 0.193 | -4.01E-02 | 1.18E-01 | 2.64E+00 | 2.71E+00 | 7.33E-01 | no | Chsy3 | intron_variant | MODIFIER |
| 15 | 74687339 | AX-168416437 | 2 | C | A | 0.545 | 2.81E-02 | 8.03E-02 | 2.59E+00 | 2.67E+00 | 7.33E-01 | no | | intergenic_variant | MODIFIER |
| 9 | 54169394 | AX-170142937 | 2 | C | T | 0.227 | 3.79E-02 | 1.12E-01 | 2.66E+00 | 2.73E+00 | 7.33E-01 | no | Cyp19a1 | intron_variant | MODIFIER |
| 9 | 40098969 | AX-170035806 | 1 | C | T | 0.438 | -2.50E-02 | 7.50E-02 | 2.68E+00 | 2.77E+00 | 7.33E-01 | no | Olfr984 | downstream_gene_variant | MODIFIER |
| 7 | 110056726 | AX-170125246 | 0 | C | A | 0.044 | 4.24E-02 | 1.22E-01 | 2.54E+00 | 2.70E+00 | 7.34E-01 | no | Zfp143 | upstream_gene_variant | MODIFIER |
| 5 | 128536132 | AX-170477108 | 0 | T | C | 0.158 | -3.61E-02 | 8.91E-02 | 2.44E+00 | 2.55E+00 | 7.34E-01 | no | | regulatory_region_variant | MODIFIER |
| 1 | 39017100 | AX-170397655 | 0 | T | C | 0.456 | 2.68E-02 | 8.05E-02 | 2.69E+00 | 2.77E+00 | 7.34E-01 | no | | intergenic_variant | MODIFIER |
| 3 | 47566335 | AX-170020471 | 0 | T | C | 0.044 | 6.92E-02 | 2.10E-01 | 2.70E+00 | 2.79E+00 | 7.34E-01 | no | | intergenic_variant | MODIFIER |
| 16 | 77164136 | AX-170504485 | 2 | T | G | 0.464 | 2.64E-02 | 7.53E-02 | 2.58E+00 | 2.67E+00 | 7.34E-01 | no | | intergenic_variant | MODIFIER |
| 18 | 66858746 | AX-170320317 | 1 | A | G | 0.286 | -3.40E-02 | 1.01E-01 | 2.68E+00 | 2.74E+00 | 7.34E-01 | no | Mc4r | 3_prime_UTR_variant | MODIFIER |
| 11 | 73093085 | AX-170211913 | 0 | G | A | 0.237 | 2.39E-02 | 7.19E-02 | 2.62E+00 | 2.75E+00 | 7.34E-01 | no | Itgae | intron_variant | MODIFIER |
| 11 | 73095454 | AX-170405180 | 0 | T | C | 0.237 | 2.39E-02 | 7.19E-02 | 2.62E+00 | 2.75E+00 | 7.34E-01 | no | Itgae | intron_variant | MODIFIER |
| 11 | 73095956 | AX-97830490 | 0 | A | G | 0.237 | 2.39E-02 | 7.19E-02 | 2.62E+00 | 2.75E+00 | 7.34E-01 | no | Itgae | intron_variant | MODIFIER |
| 11 | 73111873 | AX-170005290 | 0 | A | G | 0.237 | 2.39E-02 | 7.19E-02 | 2.62E+00 | 2.75E+00 | 7.34E-01 | no | Itgae | intron_variant | MODIFIER |
| 11 | 73115640 | AX-170175884 | 0 | A | G | 0.237 | 2.39E-02 | 7.19E-02 | 2.62E+00 | 2.75E+00 | 7.34E-01 | no | Itgae | missense_variant | MODERATE |
| 11 | 73119579 | AX-170394393 | 0 | C | T | 0.237 | 2.39E-02 | 7.19E-02 | 2.62E+00 | 2.75E+00 | 7.34E-01 | no | Itgae | intron_variant | MODIFIER |
| 11 | 73119705 | AX-169999717 | 0 | G | A | 0.237 | 2.39E-02 | 7.19E-02 | 2.62E+00 | 2.75E+00 | 7.34E-01 | no | Itgae | intron_variant | MODIFIER |
| 11 | 73121957 | AX-170005289 | 0 | G | T | 0.237 | 2.39E-02 | 7.19E-02 | 2.62E+00 | 2.75E+00 | 7.34E-01 | no | Itgae | missense_variant | MODERATE |
| 11 | 73124393 | AX-97816262 | 0 | A | G | 0.237 | 2.39E-02 | 7.19E-02 | 2.62E+00 | 2.75E+00 | 7.34E-01 | no | Itgae | intron_variant | MODIFIER |
| 11 | 73236014 | AX-170219250 | 0 | C | A | 0.237 | 2.39E-02 | 7.19E-02 | 2.62E+00 | 2.75E+00 | 7.34E-01 | no | Trpv1 | upstream_gene_variant | MODIFIER |
| 11 | 73236239 | AX-170005286 | 0 | A | C | 0.237 | 2.39E-02 | 7.19E-02 | 2.62E+00 | 2.75E+00 | 7.34E-01 | no | Trpv1 | upstream_gene_variant | MODIFIER |
| 11 | 73237631 | AX-170170582 | 0 | T | C | 0.237 | 2.39E-02 | 7.19E-02 | 2.62E+00 | 2.75E+00 | 7.34E-01 | no | Trpv1 | upstream_gene_variant | MODIFIER |
| 11 | 73244223 | AX-168397851 | 0 | C | T | 0.237 | 2.39E-02 | 7.19E-02 | 2.62E+00 | 2.75E+00 | 7.34E-01 | no | Trpv1 | synonymous_variant | LOW |
| 11 | 73250469 | AX-168408145 | 0 | C | T | 0.237 | 2.39E-02 | 7.19E-02 | 2.62E+00 | 2.75E+00 | 7.34E-01 | no | Trpv1 | intron_variant | MODIFIER |
| 11 | 73267180 | AX-170321309 | 0 | A | G | 0.237 | 2.39E-02 | 7.19E-02 | 2.62E+00 | 2.75E+00 | 7.34E-01 | no | Trpv3 | upstream_gene_variant | MODIFIER |
| 11 | 73322187 | AX-97819019 | 0 | G | T | 0.237 | 2.39E-02 | 7.19E-02 | 2.62E+00 | 2.75E+00 | 7.34E-01 | no | Aspa | missense_variant | MODERATE |
| 11 | 73950116 | AX-97846736 | 0 | G | T | 0.237 | 2.39E-02 | 7.19E-02 | 2.62E+00 | 2.75E+00 | 7.34E-01 | no | Olfr22-ps1 | upstream_gene_variant | MODIFIER |
| 11 | 74128320 | AX-170087059 | 0 | A | G | 0.237 | 2.39E-02 | 7.19E-02 | 2.62E+00 | 2.75E+00 | 7.34E-01 | no | | intergenic_variant | MODIFIER |
| 11 | 74652381 | AX-97851243 | 0 | T | C | 0.237 | 2.39E-02 | 7.19E-02 | 2.62E+00 | 2.75E+00 | 7.34E-01 | no | Cluh | intron_variant | MODIFIER |
| 11 | 74674871 | AX-170234593 | 0 | A | T | 0.237 | 2.39E-02 | 7.19E-02 | 2.62E+00 | 2.75E+00 | 7.34E-01 | no | Pafah1b1 | 3_prime_UTR_variant | MODIFIER |
| 11 | 74677995 | AX-97842283 | 0 | T | C | 0.237 | 2.39E-02 | 7.19E-02 | 2.62E+00 | 2.75E+00 | 7.34E-01 | no | Pafah1b1 | intron_variant | MODIFIER |
| 11 | 74692899 | AX-97789834 | 0 | T | C | 0.237 | 2.39E-02 | 7.19E-02 | 2.62E+00 | 2.75E+00 | 7.34E-01 | no | Pafah1b1 | intron_variant | MODIFIER |
| 11 | 74693468 | AX-170005311 | 0 | G | C | 0.237 | 2.39E-02 | 7.19E-02 | 2.62E+00 | 2.75E+00 | 7.34E-01 | no | Pafah1b1 | intron_variant | MODIFIER |
| 11 | 74709010 | AX-170278674 | 0 | G | A | 0.237 | 2.39E-02 | 7.19E-02 | 2.62E+00 | 2.75E+00 | 7.34E-01 | no | Pafah1b1 | intron_variant | MODIFIER |
| 11 | 74709216 | AX-170263836 | 0 | A | G | 0.237 | 2.39E-02 | 7.19E-02 | 2.62E+00 | 2.75E+00 | 7.34E-01 | no | Pafah1b1 | intron_variant | MODIFIER |
| 11 | 74709374 | AX-170157166 | 0 | T | C | 0.237 | 2.39E-02 | 7.19E-02 | 2.62E+00 | 2.75E+00 | 7.34E-01 | no | Pafah1b1 | intron_variant | MODIFIER |
| 11 | 74735137 | AX-170352661 | 0 | A | C | 0.237 | 2.39E-02 | 7.19E-02 | 2.62E+00 | 2.75E+00 | 7.34E-01 | no | Gm16032 | upstream_gene_variant | MODIFIER |
| 11 | 74749160 | AX-97794129 | 0 | G | A | 0.237 | 2.39E-02 | 7.19E-02 | 2.62E+00 | 2.75E+00 | 7.34E-01 | no | | regulatory_region_variant | MODIFIER |
| 11 | 74780745 | AX-170481015 | 0 | A | G | 0.237 | 2.39E-02 | 7.19E-02 | 2.62E+00 | 2.75E+00 | 7.34E-01 | no | Mettl16 | intron_variant | MODIFIER |
| 11 | 74813056 | AX-170425991 | 0 | C | T | 0.237 | 2.39E-02 | 7.19E-02 | 2.62E+00 | 2.75E+00 | 7.34E-01 | no | Mettl16 | intron_variant | MODIFIER |
| 11 | 74818973 | AX-170451899 | 0 | T | G | 0.237 | 2.39E-02 | 7.19E-02 | 2.62E+00 | 2.75E+00 | 7.34E-01 | no | Mettl16 | downstream_gene_variant | MODIFIER |
| 11 | 74827564 | AX-170390173 | 0 | G | A | 0.237 | 2.39E-02 | 7.19E-02 | 2.62E+00 | 2.75E+00 | 7.34E-01 | no | Mnt | upstream_gene_variant | MODIFIER |

| | | | | | | | | | | | | | | | |
|---|---|---|---|---|---|---|---|---|---|---|---|---|---|---|---|
| 11 | 74933050 | AX-170097061 | 0 | T | C | 0.237 | 2.39E-02 | 7.19E-02 | 2.62E+00 | 2.75E+00 | 7.34E-01 | no | Smg6 | intron_variant | MODIFIER |
| 8 | 35861691 | AX-170464293 | 1 | A | G | 0.152 | -4.95E-02 | 1.45E-01 | 2.65E+00 | 2.69E+00 | 7.34E-01 | no | 5430403N17Rik | downstream_gene_variant | MODIFIER |
| 8 | 83498974 | AX-170265805 | 1 | T | C | 0.205 | 4.28E-02 | 1.27E-01 | 2.67E+00 | 2.74E+00 | 7.34E-01 | no | | intergenic_variant | MODIFIER |
| 8 | 86922521 | AX-170128555 | 1 | G | A | 0.205 | -4.30E-02 | 1.29E-01 | 2.69E+00 | 2.76E+00 | 7.34E-01 | no | Gm27168 | upstream_gene_variant | MODIFIER |
| 5 | 98370248 | AX-170089565 | 0 | A | G | 0.43 | -3.47E-02 | 1.04E-01 | 2.72E+00 | 2.79E+00 | 7.34E-01 | no | Cfap299 | intron_variant | MODIFIER |
| 16 | 92540581 | AX-170440486 | 0 | A | G | 0.474 | 2.69E-02 | 8.05E-02 | 2.67E+00 | 2.75E+00 | 7.34E-01 | no | Clic6 | 3_prime_UTR_variant | MODIFIER |
| 2 | 94368467 | AX-170330108 | 1 | G | A | 0.054 | -1.00E-01 | 2.99E-01 | 2.74E+00 | 2.77E+00 | 7.34E-01 | no | Ttc17 | upstream_gene_variant | MODIFIER |
| 2 | 94679308 | AX-170123082 | 1 | G | T | 0.054 | -1.00E-01 | 2.99E-01 | 2.74E+00 | 2.77E+00 | 7.34E-01 | no | | intergenic_variant | MODIFIER |
| 2 | 96797036 | AX-170397611 | 1 | C | T | 0.054 | -1.00E-01 | 2.99E-01 | 2.74E+00 | 2.77E+00 | 7.34E-01 | no | Lrrc4c | intron_variant | MODIFIER |
| 2 | 102581671 | AX-170064462 | 1 | G | A | 0.054 | -1.00E-01 | 2.99E-01 | 2.74E+00 | 2.77E+00 | 7.34E-01 | no | Pamr1 | intron_variant | MODIFIER |
| 6 | 110032240 | AX-170027958 | 1 | C | T | 0.054 | 9.91E-02 | 2.96E-01 | 2.68E+00 | 2.74E+00 | 7.34E-01 | no | Gm44162 | upstream_gene_variant | MODIFIER |
| 6 | 139845305 | AX-170420619 | 1 | A | G | 0.054 | 9.91E-02 | 2.96E-01 | 2.68E+00 | 2.74E+00 | 7.34E-01 | no | Pik3c2g | intron_variant | MODIFIER |
| 4 | 56306965 | AX-170528895 | 1 | A | G | 0.357 | -3.47E-02 | 1.04E-01 | 2.68E+00 | 2.76E+00 | 7.34E-01 | no | | intergenic_variant | MODIFIER |
| 5 | 30176536 | AX-170154268 | 0 | G | A | 0.368 | 3.55E-02 | 1.05E-01 | 2.66E+00 | 2.72E+00 | 7.34E-01 | no | Hadhb | intron_variant | MODIFIER |
| 16 | 93047704 | AX-170160495 | 1 | C | T | 0.393 | -2.73E-02 | 7.95E-02 | 2.61E+00 | 2.69E+00 | 7.35E-01 | no | | intergenic_variant | MODIFIER |
| 2 | 127534337 | AX-170064854 | 0 | A | C | 0.14 | -5.96E-02 | 1.79E-01 | 2.69E+00 | 2.76E+00 | 7.35E-01 | no | Prom2 | intron_variant | MODIFIER |
| 2 | 147066591 | AX-97855552 | 0 | A | G | 0.14 | -5.96E-02 | 1.79E-01 | 2.69E+00 | 2.76E+00 | 7.35E-01 | no | Xrn2 | intron_variant | MODIFIER |
| 6 | 148520235 | AX-170223548 | 0 | G | A | 0.053 | 5.87E-02 | 1.78E-01 | 2.66E+00 | 2.77E+00 | 7.35E-01 | no | | intergenic_variant | MODIFIER |
| 1 | 95242832 | AX-170391009 | 0 | G | A | 0.158 | 7.02E-02 | 2.08E-01 | 2.65E+00 | 2.72E+00 | 7.35E-01 | no | | intergenic_variant | MODIFIER |
| 6 | 139716116 | AX-170437222 | 0 | G | A | 0.07 | -9.21E-02 | 2.76E-01 | 2.71E+00 | 2.75E+00 | 7.35E-01 | no | Pik3c2g | intron_variant | MODIFIER |
| 15 | 90812528 | AX-170463888 | 0 | T | G | 0.132 | -4.63E-02 | 1.41E-01 | 2.70E+00 | 2.79E+00 | 7.35E-01 | no | | intergenic_variant | MODIFIER |
| 1 | 21690169 | AX-170331558 | 1 | C | T | 0.402 | 2.91E-02 | 8.47E-02 | 2.62E+00 | 2.67E+00 | 7.35E-01 | no | Kcnq5 | intron_variant | MODIFIER |
| 5 | 124011984 | AX-170408174 | 0 | T | G | 0.439 | -3.67E-02 | 1.10E-01 | 2.76E+00 | 2.82E+00 | 7.35E-01 | no | Vps37b | intron_variant | MODIFIER |
| 18 | 61011530 | AX-170230936 | 0 | T | C | 0.412 | 2.79E-02 | 8.41E-02 | 2.78E+00 | 2.85E+00 | 7.35E-01 | no | Slc6a7 | intron_variant | MODIFIER |
| 2 | 121848091 | AX-97856952 | 2 | G | A | 0.145 | 7.42E-02 | 2.21E-01 | 2.69E+00 | 2.73E+00 | 7.35E-01 | no | | intergenic_variant | MODIFIER |
| 2 | 143098062 | AX-97808377 | 2 | G | A | 0.145 | 7.42E-02 | 2.21E-01 | 2.69E+00 | 2.73E+00 | 7.35E-01 | no | | intergenic_variant | MODIFIER |
| 4 | 140238230 | AX-97810398 | 0 | T | C | 0.158 | 3.89E-02 | 1.11E-01 | 2.58E+00 | 2.66E+00 | 7.35E-01 | no | Igsf21 | intron_variant | MODIFIER |
| 15 | 95909609 | AX-170056387 | 2 | G | A | 0.145 | 5.06E-02 | 1.52E-01 | 2.64E+00 | 2.75E+00 | 7.35E-01 | no | Ano6 | intron_variant | MODIFIER |
| 9 | 101283556 | AX-170083268 | 0 | T | C | 0.149 | -3.69E-02 | 1.08E-01 | 2.61E+00 | 2.70E+00 | 7.35E-01 | no | Gm3121 | upstream_gene_variant | MODIFIER |
| 10 | 12580165 | AX-97814012 | 0 | G | T | 0.228 | 3.28E-02 | 9.96E-02 | 2.69E+00 | 2.78E+00 | 7.35E-01 | no | Utrn | intron_variant | MODIFIER |
| 8 | 103292651 | AX-97805382 | 1 | T | C | 0.214 | 3.69E-02 | 1.11E-01 | 2.67E+00 | 2.75E+00 | 7.35E-01 | no | Gm32531 | intron_variant,non_coding_transcript_variant | MODIFIER |
| 10 | 18300054 | AX-170002856 | 0 | T | C | 0.202 | 3.22E-02 | 9.84E-02 | 2.69E+00 | 2.79E+00 | 7.35E-01 | no | Gm48229 | downstream_gene_variant | MODIFIER |
| 9 | 69794991 | AX-170266785 | 2 | G | A | 0.209 | 4.14E-02 | 1.25E-01 | 2.69E+00 | 2.76E+00 | 7.35E-01 | no | B230323A14Rik | intron_variant,non_coding_transcript_variant | MODIFIER |
| 5 | 28425791 | AX-170266296 | 1 | A | G | 0.357 | 3.53E-02 | 1.06E-01 | 2.68E+00 | 2.74E+00 | 7.35E-01 | no | | intergenic_variant | MODIFIER |
| 5 | 33107805 | AX-170244179 | 1 | C | A | 0.357 | 3.53E-02 | 1.06E-01 | 2.68E+00 | 2.74E+00 | 7.35E-01 | no | Slc5a1 | intron_variant | MODIFIER |
| 9 | 77959192 | AX-170230968 | 1 | G | A | 0.205 | 4.47E-02 | 1.34E-01 | 2.77E+00 | 2.81E+00 | 7.35E-01 | no | Elovl5 | intron_variant | MODIFIER |
| 6 | 105629134 | AX-170218868 | 0 | T | A | 0.079 | 5.40E-02 | 1.68E-01 | 2.67E+00 | 2.81E+00 | 7.36E-01 | no | | intergenic_variant | MODIFIER |
| 1 | 3322824 | AX-170085959 | 1 | A | G | 0.518 | 2.50E-02 | 7.02E-02 | 2.57E+00 | 2.64E+00 | 7.36E-01 | no | Xkr4 | intron_variant | MODIFIER |
| 8 | 78656225 | AX-170277086 | 0 | C | T | 0.193 | -4.15E-02 | 1.23E-01 | 2.64E+00 | 2.72E+00 | 7.36E-01 | no | Slc10a7 | intron_variant | MODIFIER |
| 15 | 97834537 | AX-170500895 | 0 | C | T | 0.123 | 5.20E-02 | 1.54E-01 | 2.64E+00 | 2.72E+00 | 7.36E-01 | no | Hdac7 | upstream_gene_variant | MODIFIER |
| 8 | 49676497 | AX-170464779 | 0 | A | C | 0.158 | 4.51E-02 | 1.38E-01 | 2.77E+00 | 2.87E+00 | 7.36E-01 | no | | intergenic_variant | MODIFIER |
| 2 | 146985650 | AX-97809359 | 1 | C | T | 0.152 | 6.63E-02 | 1.97E-01 | 2.65E+00 | 2.72E+00 | 7.36E-01 | no | | regulatory_region_variant | MODIFIER |
| 18 | 53724979 | AX-170423589 | 1 | G | T | 0.25 | -3.58E-02 | 1.02E-01 | 2.57E+00 | 2.65E+00 | 7.36E-01 | no | Cep120 | intron_variant | MODIFIER |
| 2 | 156524951 | AX-170224808 | 0 | T | C | 0.132 | 6.86E-02 | 2.05E-01 | 2.68E+00 | 2.72E+00 | 7.36E-01 | no | Epb41l1 | missense_variant | MODERATE |
| 10 | 27672233 | AX-170134025 | 0 | C | A | 0.202 | 3.06E-02 | 9.40E-02 | 2.69E+00 | 2.80E+00 | 7.36E-01 | no | Gm30165 | intron_variant,non_coding_transcript_variant | MODIFIER |
| 5 | 100701979 | AX-97839674 | 0 | T | A | 0.439 | -3.47E-02 | 1.05E-01 | 2.74E+00 | 2.81E+00 | 7.36E-01 | no | Hpse | intron_variant | MODIFIER |
| 9 | 85297340 | AX-170338126 | 0 | T | A | 0.219 | 4.10E-02 | 1.24E-01 | 2.72E+00 | 2.78E+00 | 7.36E-01 | no | | intergenic_variant | MODIFIER |
| 9 | 85332175 | AX-170445448 | 0 | C | T | 0.219 | 4.10E-02 | 1.24E-01 | 2.72E+00 | 2.78E+00 | 7.36E-01 | no | Tent5a | upstream_gene_variant | MODIFIER |
| 9 | 85359896 | AX-170405210 | 0 | A | C | 0.219 | 4.10E-02 | 1.24E-01 | 2.72E+00 | 2.78E+00 | 7.36E-01 | no | | intergenic_variant | MODIFIER |
| 9 | 85494523 | AX-97821088 | 0 | T | C | 0.219 | 4.10E-02 | 1.24E-01 | 2.72E+00 | 2.78E+00 | 7.36E-01 | no | | intergenic_variant | MODIFIER |
| 5 | 28097656 | AX-97805348 | 0 | C | T | 0.36 | 3.53E-02 | 1.06E-01 | 2.68E+00 | 2.74E+00 | 7.36E-01 | no | | intergenic_variant | MODIFIER |
| 5 | 28132247 | AX-170389243 | 0 | C | G | 0.36 | 3.53E-02 | 1.06E-01 | 2.68E+00 | 2.74E+00 | 7.36E-01 | no | | intergenic_variant | MODIFIER |
| 5 | 28133773 | AX-97830674 | 0 | G | A | 0.36 | 3.53E-02 | 1.06E-01 | 2.68E+00 | 2.74E+00 | 7.36E-01 | no | | intergenic_variant | MODIFIER |
| 5 | 28139093 | AX-170189027 | 0 | G | A | 0.36 | 3.53E-02 | 1.06E-01 | 2.68E+00 | 2.74E+00 | 7.36E-01 | no | | intergenic_variant | MODIFIER |

| | | | | | | | | | | | | | | | |
|---|---|---|---|---|---|---|---|---|---|---|---|---|---|---|---|
| 5 | 28158453 | AX-170089437 | 0 | G | T | 0.36 | 3.53E-02 | 1.06E-01 | 2.68E+00 | 2.74E+00 | 7.36E-01 | no | | regulatory_region_variant | MODIFIER |
| 5 | 28158519 | AX-170358460 | 0 | T | A | 0.36 | 3.53E-02 | 1.06E-01 | 2.68E+00 | 2.74E+00 | 7.36E-01 | no | | regulatory_region_variant | MODIFIER |
| 5 | 28184306 | AX-170381210 | 0 | A | G | 0.36 | 3.53E-02 | 1.06E-01 | 2.68E+00 | 2.74E+00 | 7.36E-01 | no | | intergenic_variant | MODIFIER |
| 5 | 28184512 | AX-170024588 | 0 | C | G | 0.36 | 3.53E-02 | 1.06E-01 | 2.68E+00 | 2.74E+00 | 7.36E-01 | no | | intergenic_variant | MODIFIER |
| 5 | 28239321 | AX-97815310 | 0 | G | T | 0.36 | 3.53E-02 | 1.06E-01 | 2.68E+00 | 2.74E+00 | 7.36E-01 | no | Cnpy1 | upstream_gene_variant | MODIFIER |
| 5 | 28313567 | AX-170071633 | 0 | T | A | 0.36 | 3.53E-02 | 1.06E-01 | 2.68E+00 | 2.74E+00 | 7.36E-01 | no | Rbm33 | upstream_gene_variant | MODIFIER |
| 5 | 28359953 | AX-97836491 | 0 | G | A | 0.36 | 3.53E-02 | 1.06E-01 | 2.68E+00 | 2.74E+00 | 7.36E-01 | no | Rbm33 | downstream_gene_variant | MODIFIER |
| 5 | 28364804 | AX-170109727 | 0 | T | C | 0.36 | 3.53E-02 | 1.06E-01 | 2.68E+00 | 2.74E+00 | 7.36E-01 | no | Rbm33 | intron_variant | MODIFIER |
| 5 | 28365037 | AX-170387138 | 0 | A | G | 0.36 | 3.53E-02 | 1.06E-01 | 2.68E+00 | 2.74E+00 | 7.36E-01 | no | Rbm33 | intron_variant | MODIFIER |
| 5 | 28365142 | AX-170480566 | 0 | G | A | 0.36 | 3.53E-02 | 1.06E-01 | 2.68E+00 | 2.74E+00 | 7.36E-01 | no | Rbm33 | intron_variant | MODIFIER |
| 5 | 28376975 | AX-170099898 | 0 | G | A | 0.36 | 3.53E-02 | 1.06E-01 | 2.68E+00 | 2.74E+00 | 7.36E-01 | no | Rbm33 | intron_variant | MODIFIER |
| 5 | 28379409 | AX-97847480 | 0 | C | T | 0.36 | 3.53E-02 | 1.06E-01 | 2.68E+00 | 2.74E+00 | 7.36E-01 | no | Rbm33 | intron_variant | MODIFIER |
| 5 | 28403413 | AX-170440345 | 0 | C | T | 0.36 | 3.53E-02 | 1.06E-01 | 2.68E+00 | 2.74E+00 | 7.36E-01 | no | Rbm33 | intron_variant | MODIFIER |
| 5 | 28420465 | AX-170278203 | 0 | G | A | 0.36 | 3.53E-02 | 1.06E-01 | 2.68E+00 | 2.74E+00 | 7.36E-01 | no | Rbm33 | downstream_gene_variant | MODIFIER |
| 5 | 28423329 | AX-170178589 | 0 | A | G | 0.36 | 3.53E-02 | 1.06E-01 | 2.68E+00 | 2.74E+00 | 7.36E-01 | no | Rbm33 | downstream_gene_variant | MODIFIER |
| 5 | 28423401 | AX-170161117 | 0 | A | C | 0.36 | 3.53E-02 | 1.06E-01 | 2.68E+00 | 2.74E+00 | 7.36E-01 | no | Rbm33 | downstream_gene_variant | MODIFIER |
| 5 | 28423510 | AX-170532345 | 0 | T | C | 0.36 | 3.53E-02 | 1.06E-01 | 2.68E+00 | 2.74E+00 | 7.36E-01 | no | Rbm33 | downstream_gene_variant | MODIFIER |
| 5 | 28447914 | AX-97786364 | 0 | C | T | 0.36 | 3.53E-02 | 1.06E-01 | 2.68E+00 | 2.74E+00 | 7.36E-01 | no | | intergenic_variant | MODIFIER |
| 5 | 28449456 | AX-170430853 | 0 | T | C | 0.36 | 3.53E-02 | 1.06E-01 | 2.68E+00 | 2.74E+00 | 7.36E-01 | no | | intergenic_variant | MODIFIER |
| 5 | 28479304 | AX-170518277 | 0 | C | T | 0.36 | 3.53E-02 | 1.06E-01 | 2.68E+00 | 2.74E+00 | 7.36E-01 | no | Gm26894 | intron_variant,non_coding_transcript_variant | MODIFIER |
| 5 | 28499217 | AX-97798828 | 0 | G | A | 0.36 | 3.53E-02 | 1.06E-01 | 2.68E+00 | 2.74E+00 | 7.36E-01 | no | 9530036O11Rik | intron_variant,non_coding_transcript_variant | MODIFIER |
| 5 | 28522623 | AX-170246694 | 0 | T | C | 0.36 | 3.53E-02 | 1.06E-01 | 2.68E+00 | 2.74E+00 | 7.36E-01 | no | 9530036O11Rik | intron_variant,non_coding_transcript_variant | MODIFIER |
| 5 | 28524797 | AX-170171289 | 0 | G | A | 0.36 | 3.53E-02 | 1.06E-01 | 2.68E+00 | 2.74E+00 | 7.36E-01 | no | 9530036O11Rik | intron_variant,non_coding_transcript_variant | MODIFIER |
| 5 | 28525891 | AX-170095271 | 0 | G | A | 0.36 | 3.53E-02 | 1.06E-01 | 2.68E+00 | 2.74E+00 | 7.36E-01 | no | 9530036O11Rik | intron_variant,non_coding_transcript_variant | MODIFIER |
| 5 | 28562491 | AX-170174109 | 0 | T | C | 0.36 | 3.53E-02 | 1.06E-01 | 2.68E+00 | 2.74E+00 | 7.36E-01 | no | 9530036O11Rik | intron_variant,non_coding_transcript_variant | MODIFIER |
| 5 | 28575479 | AX-170135564 | 0 | A | T | 0.36 | 3.53E-02 | 1.06E-01 | 2.68E+00 | 2.74E+00 | 7.36E-01 | no | 9530036O11Rik | intron_variant,non_coding_transcript_variant | MODIFIER |
| 5 | 28577215 | AX-170120427 | 0 | C | T | 0.36 | 3.53E-02 | 1.06E-01 | 2.68E+00 | 2.74E+00 | 7.36E-01 | no | 9530036O11Rik | intron_variant,non_coding_transcript_variant | MODIFIER |
| 5 | 28577383 | AX-97844661 | 0 | A | G | 0.36 | 3.53E-02 | 1.06E-01 | 2.68E+00 | 2.74E+00 | 7.36E-01 | no | 9530036O11Rik | intron_variant,non_coding_transcript_variant | MODIFIER |
| 5 | 28577798 | AX-170337002 | 0 | A | T | 0.36 | 3.53E-02 | 1.06E-01 | 2.68E+00 | 2.74E+00 | 7.36E-01 | no | 9530036O11Rik | intron_variant,non_coding_transcript_variant | MODIFIER |
| 5 | 28618229 | AX-170095272 | 0 | A | G | 0.36 | 3.53E-02 | 1.06E-01 | 2.68E+00 | 2.74E+00 | 7.36E-01 | no | 9530036O11Rik | intron_variant,non_coding_transcript_variant | MODIFIER |
| 5 | 28622078 | AX-170504923 | 0 | C | G | 0.36 | 3.53E-02 | 1.06E-01 | 2.68E+00 | 2.74E+00 | 7.36E-01 | no | 9530036O11Rik | intron_variant,non_coding_transcript_variant | MODIFIER |
| 5 | 28639693 | AX-97838626 | 0 | C | T | 0.36 | 3.53E-02 | 1.06E-01 | 2.68E+00 | 2.74E+00 | 7.36E-01 | no | 9530036O11Rik | intron_variant,non_coding_transcript_variant | MODIFIER |
| 5 | 28639892 | AX-170512584 | 0 | A | T | 0.36 | 3.53E-02 | 1.06E-01 | 2.68E+00 | 2.74E+00 | 7.36E-01 | no | 9530036O11Rik | intron_variant,non_coding_transcript_variant | MODIFIER |
| 5 | 28664593 | AX-97816965 | 0 | A | G | 0.36 | 3.53E-02 | 1.06E-01 | 2.68E+00 | 2.74E+00 | 7.36E-01 | no | 9530036O11Rik | intron_variant,non_coding_transcript_variant | MODIFIER |
| 5 | 28664605 | AX-170189013 | 0 | C | T | 0.36 | 3.53E-02 | 1.06E-01 | 2.68E+00 | 2.74E+00 | 7.36E-01 | no | 9530036O11Rik | intron_variant,non_coding_transcript_variant | MODIFIER |
| 5 | 28665224 | AX-170261343 | 0 | G | C | 0.36 | 3.53E-02 | 1.06E-01 | 2.68E+00 | 2.74E+00 | 7.36E-01 | no | 9530036O11Rik | intron_variant,non_coding_transcript_variant | MODIFIER |
| 5 | 28665893 | AX-170339349 | 0 | G | A | 0.36 | 3.53E-02 | 1.06E-01 | 2.68E+00 | 2.74E+00 | 7.36E-01 | no | 9530036O11Rik | intron_variant,non_coding_transcript_variant | MODIFIER |
| 5 | 28666064 | AX-170474468 | 0 | T | A | 0.36 | 3.53E-02 | 1.06E-01 | 2.68E+00 | 2.74E+00 | 7.36E-01 | no | 9530036O11Rik | intron_variant,non_coding_transcript_variant | MODIFIER |
| 5 | 28667868 | AX-170455260 | 0 | G | A | 0.36 | 3.53E-02 | 1.06E-01 | 2.68E+00 | 2.74E+00 | 7.36E-01 | no | 9530036O11Rik | intron_variant,non_coding_transcript_variant | MODIFIER |
| 5 | 28668831 | AX-170341317 | 0 | T | G | 0.36 | 3.53E-02 | 1.06E-01 | 2.68E+00 | 2.74E+00 | 7.36E-01 | no | 9530036O11Rik | intron_variant,non_coding_transcript_variant | MODIFIER |
| 5 | 28710506 | AX-97821368 | 0 | A | G | 0.36 | 3.53E-02 | 1.06E-01 | 2.68E+00 | 2.74E+00 | 7.36E-01 | no | 9530036O11Rik | intron_variant,non_coding_transcript_variant | MODIFIER |
| 5 | 29361892 | AX-170163537 | 0 | G | T | 0.36 | 3.53E-02 | 1.06E-01 | 2.68E+00 | 2.74E+00 | 7.36E-01 | no | Lmbr1 | intron_variant | MODIFIER |
| 5 | 30646487 | AX-168395187 | 0 | T | C | 0.36 | 3.53E-02 | 1.06E-01 | 2.68E+00 | 2.74E+00 | 7.36E-01 | no | Slc35f6 | upstream_gene_variant | MODIFIER |
| 5 | 30710476 | AX-170071682 | 0 | G | A | 0.36 | 3.53E-02 | 1.06E-01 | 2.68E+00 | 2.74E+00 | 7.36E-01 | no | Dpysl5 | upstream_gene_variant | MODIFIER |
| 5 | 30911677 | AX-170265155 | 0 | T | C | 0.36 | 3.53E-02 | 1.06E-01 | 2.68E+00 | 2.74E+00 | 7.36E-01 | no | Emilin1 | upstream_gene_variant | MODIFIER |
| 5 | 30917520 | AX-97808223 | 0 | G | A | 0.36 | 3.53E-02 | 1.06E-01 | 2.68E+00 | 2.74E+00 | 7.36E-01 | no | Khk | upstream_gene_variant | MODIFIER |
| 5 | 30923979 | AX-168408209 | 0 | T | C | 0.36 | 3.53E-02 | 1.06E-01 | 2.68E+00 | 2.74E+00 | 7.36E-01 | no | Khk | intron_variant | MODIFIER |
| 5 | 30926870 | AX-170086208 | 0 | A | G | 0.36 | 3.53E-02 | 1.06E-01 | 2.68E+00 | 2.74E+00 | 7.36E-01 | no | Khk | intron_variant | MODIFIER |
| 5 | 30938474 | AX-170371751 | 0 | C | T | 0.36 | 3.53E-02 | 1.06E-01 | 2.68E+00 | 2.74E+00 | 7.36E-01 | no | Cgref1 | intron_variant | MODIFIER |
| 5 | 30938607 | AX-170404496 | 0 | G | A | 0.36 | 3.53E-02 | 1.06E-01 | 2.68E+00 | 2.74E+00 | 7.36E-01 | no | Cgref1 | intron_variant | MODIFIER |
| 5 | 30938682 | AX-170370962 | 0 | A | G | 0.36 | 3.53E-02 | 1.06E-01 | 2.68E+00 | 2.74E+00 | 7.36E-01 | no | Cgref1 | intron_variant | MODIFIER |
| 5 | 30955785 | AX-97802651 | 0 | A | G | 0.36 | 3.53E-02 | 1.06E-01 | 2.68E+00 | 2.74E+00 | 7.36E-01 | no | Abhd1 | downstream_gene_variant | MODIFIER |
| 5 | 30958581 | AX-170183806 | 0 | G | T | 0.36 | 3.53E-02 | 1.06E-01 | 2.68E+00 | 2.74E+00 | 7.36E-01 | no | Abhd1 | downstream_gene_variant | MODIFIER |
| 5 | 30981759 | AX-97846849 | 0 | T | C | 0.36 | 3.53E-02 | 1.06E-01 | 2.68E+00 | 2.74E+00 | 7.36E-01 | no | Tcf23 | downstream_gene_variant | MODIFIER |

| | | | | | | | | | | | | | | | |
|---|---|---|---|---|---|---|---|---|---|---|---|---|---|---|---|
| 5 | 31000234 | AX-170272541 | 0 | T | C | 0.36 | 3.53E-02 | 1.06E-01 | 2.68E+00 | 2.74E+00 | 7.36E-01 | no | | intergenic_variant | MODIFIER |
| 5 | 31006191 | AX-170071687 | 0 | C | T | 0.36 | 3.53E-02 | 1.06E-01 | 2.68E+00 | 2.74E+00 | 7.36E-01 | no | | intergenic_variant | MODIFIER |
| 5 | 31030075 | AX-170024619 | 0 | C | T | 0.36 | 3.53E-02 | 1.06E-01 | 2.68E+00 | 2.74E+00 | 7.36E-01 | no | | intergenic_variant | MODIFIER |
| 5 | 31034697 | AX-170024614 | 0 | A | G | 0.36 | 3.53E-02 | 1.06E-01 | 2.68E+00 | 2.74E+00 | 7.36E-01 | no | Slc5a6 | downstream_gene_variant | MODIFIER |
| 5 | 31038032 | AX-170368573 | 0 | G | A | 0.36 | 3.53E-02 | 1.06E-01 | 2.68E+00 | 2.74E+00 | 7.36E-01 | no | Slc5a6 | synonymous_variant | LOW |
| 5 | 31058566 | AX-170496595 | 0 | G | C | 0.36 | 3.53E-02 | 1.06E-01 | 2.68E+00 | 2.74E+00 | 7.36E-01 | no | Atraid | downstream_gene_variant | MODIFIER |
| 5 | 31067852 | AX-170024616 | 0 | A | G | 0.36 | 3.53E-02 | 1.06E-01 | 2.68E+00 | 2.74E+00 | 7.36E-01 | no | Cad | intron_variant | MODIFIER |
| 5 | 31068965 | AX-97833060 | 0 | A | G | 0.36 | 3.53E-02 | 1.06E-01 | 2.68E+00 | 2.74E+00 | 7.36E-01 | no | Cad | intron_variant | MODIFIER |
| 5 | 31072074 | AX-97812025 | 0 | T | C | 0.36 | 3.53E-02 | 1.06E-01 | 2.68E+00 | 2.74E+00 | 7.36E-01 | no | Cad | synonymous_variant | LOW |
| 5 | 31074104 | AX-170272080 | 0 | C | T | 0.36 | 3.53E-02 | 1.06E-01 | 2.68E+00 | 2.74E+00 | 7.36E-01 | no | Cad | synonymous_variant | LOW |
| 5 | 31077915 | AX-170024612 | 0 | T | C | 0.36 | 3.53E-02 | 1.06E-01 | 2.68E+00 | 2.74E+00 | 7.36E-01 | no | Cad | synonymous_variant | LOW |
| 5 | 31096231 | AX-97825226 | 0 | G | A | 0.36 | 3.53E-02 | 1.06E-01 | 2.68E+00 | 2.74E+00 | 7.36E-01 | no | Slc30a3 | upstream_gene_variant | MODIFIER |
| 5 | 31161270 | AX-170249775 | 0 | T | C | 0.36 | 3.53E-02 | 1.06E-01 | 2.68E+00 | 2.74E+00 | 7.36E-01 | no | Gtf3c2 | intron_variant | MODIFIER |
| 5 | 31162431 | AX-168374642 | 0 | T | G | 0.36 | 3.53E-02 | 1.06E-01 | 2.68E+00 | 2.74E+00 | 7.36E-01 | no | Gtf3c2 | intron_variant | MODIFIER |
| 5 | 31222735 | AX-170100174 | 0 | A | G | 0.36 | 3.53E-02 | 1.06E-01 | 2.68E+00 | 2.74E+00 | 7.36E-01 | no | Ppm1g | upstream_gene_variant | MODIFIER |
| 5 | 31227098 | AX-170280238 | 0 | G | T | 0.36 | 3.53E-02 | 1.06E-01 | 2.68E+00 | 2.74E+00 | 7.36E-01 | no | | intergenic_variant | MODIFIER |
| 5 | 31228246 | AX-97838525 | 0 | T | C | 0.36 | 3.53E-02 | 1.06E-01 | 2.68E+00 | 2.74E+00 | 7.36E-01 | no | | intergenic_variant | MODIFIER |
| 5 | 31260583 | AX-170201701 | 0 | T | C | 0.36 | 3.53E-02 | 1.06E-01 | 2.68E+00 | 2.74E+00 | 7.36E-01 | no | Ift172 | intron_variant | MODIFIER |
| 5 | 31452114 | AX-97820798 | 0 | T | C | 0.36 | 3.53E-02 | 1.06E-01 | 2.68E+00 | 2.74E+00 | 7.36E-01 | no | Zfp512 | upstream_gene_variant | MODIFIER |
| 5 | 31506617 | AX-97808907 | 0 | C | T | 0.36 | 3.53E-02 | 1.06E-01 | 2.68E+00 | 2.74E+00 | 7.36E-01 | no | Gpn1 | intron_variant | MODIFIER |
| 5 | 31730628 | AX-97799701 | 0 | T | C | 0.36 | 3.53E-02 | 1.06E-01 | 2.68E+00 | 2.74E+00 | 7.36E-01 | no | Babam2 | intron_variant | MODIFIER |
| 5 | 31782466 | AX-97841841 | 0 | G | A | 0.36 | 3.53E-02 | 1.06E-01 | 2.68E+00 | 2.74E+00 | 7.36E-01 | no | Babam2 | intron_variant | MODIFIER |
| 5 | 31792769 | AX-168393690 | 0 | A | G | 0.36 | 3.53E-02 | 1.06E-01 | 2.68E+00 | 2.74E+00 | 7.36E-01 | no | Babam2 | intron_variant | MODIFIER |
| 5 | 31814430 | AX-168396278 | 0 | C | T | 0.36 | 3.53E-02 | 1.06E-01 | 2.68E+00 | 2.74E+00 | 7.36E-01 | no | Babam2 | intron_variant | MODIFIER |
| 5 | 31924821 | AX-170316398 | 0 | T | C | 0.36 | 3.53E-02 | 1.06E-01 | 2.68E+00 | 2.74E+00 | 7.36E-01 | no | Babam2 | intron_variant | MODIFIER |
| 5 | 31933761 | AX-97831089 | 0 | T | C | 0.36 | 3.53E-02 | 1.06E-01 | 2.68E+00 | 2.74E+00 | 7.36E-01 | no | Babam2 | intron_variant | MODIFIER |
| 5 | 31936656 | AX-170360582 | 0 | G | T | 0.36 | 3.53E-02 | 1.06E-01 | 2.68E+00 | 2.74E+00 | 7.36E-01 | no | Babam2 | intron_variant | MODIFIER |
| 5 | 31938116 | AX-170350358 | 0 | G | T | 0.36 | 3.53E-02 | 1.06E-01 | 2.68E+00 | 2.74E+00 | 7.36E-01 | no | Babam2 | intron_variant | MODIFIER |
| 5 | 31952690 | AX-170534829 | 0 | G | A | 0.36 | 3.53E-02 | 1.06E-01 | 2.68E+00 | 2.74E+00 | 7.36E-01 | no | Babam2 | intron_variant | MODIFIER |
| 5 | 31968949 | AX-170107018 | 0 | T | C | 0.36 | 3.53E-02 | 1.06E-01 | 2.68E+00 | 2.74E+00 | 7.36E-01 | no | Babam2 | intron_variant | MODIFIER |
| 5 | 32004334 | AX-170411516 | 0 | T | C | 0.36 | 3.53E-02 | 1.06E-01 | 2.68E+00 | 2.74E+00 | 7.36E-01 | no | Babam2 | intron_variant | MODIFIER |
| 5 | 32004656 | AX-170273240 | 0 | C | A | 0.36 | 3.53E-02 | 1.06E-01 | 2.68E+00 | 2.74E+00 | 7.36E-01 | no | Babam2 | intron_variant | MODIFIER |
| 5 | 32010477 | AX-170275510 | 0 | C | T | 0.36 | 3.53E-02 | 1.06E-01 | 2.68E+00 | 2.74E+00 | 7.36E-01 | no | Babam2 | intron_variant | MODIFIER |
| 5 | 32014184 | AX-170164206 | 0 | T | C | 0.36 | 3.53E-02 | 1.06E-01 | 2.68E+00 | 2.74E+00 | 7.36E-01 | no | Babam2 | intron_variant | MODIFIER |
| 5 | 32015867 | AX-97852370 | 0 | G | A | 0.36 | 3.53E-02 | 1.06E-01 | 2.68E+00 | 2.74E+00 | 7.36E-01 | no | Babam2 | intron_variant | MODIFIER |
| 5 | 32049267 | AX-170071708 | 0 | C | T | 0.36 | 3.53E-02 | 1.06E-01 | 2.68E+00 | 2.74E+00 | 7.36E-01 | no | Babam2 | intron_variant | MODIFIER |
| 5 | 32049398 | AX-170071715 | 0 | C | G | 0.36 | 3.53E-02 | 1.06E-01 | 2.68E+00 | 2.74E+00 | 7.36E-01 | no | Babam2 | intron_variant | MODIFIER |
| 5 | 32052932 | AX-170071714 | 0 | T | G | 0.36 | 3.53E-02 | 1.06E-01 | 2.68E+00 | 2.74E+00 | 7.36E-01 | no | Babam2 | intron_variant | MODIFIER |
| 5 | 32073723 | AX-170071712 | 0 | C | T | 0.36 | 3.53E-02 | 1.06E-01 | 2.68E+00 | 2.74E+00 | 7.36E-01 | no | Babam2 | intron_variant | MODIFIER |
| 5 | 32084054 | AX-170071713 | 0 | G | C | 0.36 | 3.53E-02 | 1.06E-01 | 2.68E+00 | 2.74E+00 | 7.36E-01 | no | Babam2 | intron_variant | MODIFIER |
| 5 | 32085462 | AX-170071709 | 0 | C | T | 0.36 | 3.53E-02 | 1.06E-01 | 2.68E+00 | 2.74E+00 | 7.36E-01 | no | Babam2 | downstream_gene_variant | MODIFIER |
| 5 | 32089970 | AX-170317509 | 0 | G | C | 0.36 | 3.53E-02 | 1.06E-01 | 2.68E+00 | 2.74E+00 | 7.36E-01 | no | | regulatory_region_variant | MODIFIER |
| 5 | 32102851 | AX-170243755 | 0 | A | G | 0.36 | 3.53E-02 | 1.06E-01 | 2.68E+00 | 2.74E+00 | 7.36E-01 | no | | regulatory_region_variant | MODIFIER |
| 5 | 32103217 | AX-170306208 | 0 | T | C | 0.36 | 3.53E-02 | 1.06E-01 | 2.68E+00 | 2.74E+00 | 7.36E-01 | no | | regulatory_region_variant | MODIFIER |
| 5 | 32113259 | AX-170256640 | 0 | C | T | 0.36 | 3.53E-02 | 1.06E-01 | 2.68E+00 | 2.74E+00 | 7.36E-01 | no | | regulatory_region_variant | MODIFIER |
| 5 | 32122814 | AX-170320538 | 0 | G | A | 0.36 | 3.53E-02 | 1.06E-01 | 2.68E+00 | 2.74E+00 | 7.36E-01 | no | | intergenic_variant | MODIFIER |
| 5 | 32126934 | AX-170393765 | 0 | T | C | 0.36 | 3.53E-02 | 1.06E-01 | 2.68E+00 | 2.74E+00 | 7.36E-01 | no | Gm10463 | downstream_gene_variant | MODIFIER |
| 5 | 32140833 | AX-170024634 | 0 | A | G | 0.36 | 3.53E-02 | 1.06E-01 | 2.68E+00 | 2.74E+00 | 7.36E-01 | no | Fosl2 | intron_variant | MODIFIER |
| 5 | 32141085 | AX-170024633 | 0 | G | A | 0.36 | 3.53E-02 | 1.06E-01 | 2.68E+00 | 2.74E+00 | 7.36E-01 | no | Fosl2 | intron_variant | MODIFIER |
| 5 | 32141132 | AX-170024632 | 0 | G | A | 0.36 | 3.53E-02 | 1.06E-01 | 2.68E+00 | 2.74E+00 | 7.36E-01 | no | Fosl2 | intron_variant | MODIFIER |
| 5 | 32150230 | AX-97849448 | 0 | C | G | 0.36 | 3.53E-02 | 1.06E-01 | 2.68E+00 | 2.74E+00 | 7.36E-01 | no | Fosl2 | intron_variant | MODIFIER |
| 5 | 32216411 | AX-170367207 | 0 | A | C | 0.36 | 3.53E-02 | 1.06E-01 | 2.68E+00 | 2.74E+00 | 7.36E-01 | no | | intergenic_variant | MODIFIER |
| 5 | 32222495 | AX-170329407 | 0 | C | T | 0.36 | 3.53E-02 | 1.06E-01 | 2.68E+00 | 2.74E+00 | 7.36E-01 | no | | intergenic_variant | MODIFIER |
| 5 | 32235460 | AX-170287510 | 0 | G | A | 0.36 | 3.53E-02 | 1.06E-01 | 2.68E+00 | 2.74E+00 | 7.36E-01 | no | Plb1 | intron_variant,non_coding_transcript_variant | MODIFIER |

| | | | | | | | | | | | | | | | |
|---|---|---|---|---|---|---|---|---|---|---|---|---|---|---|---|
| 5 | 32238852 | AX-97783897 | 0 | C | T | 0.36 | 3.53E-02 | 1.06E-01 | 2.68E+00 | 2.74E+00 | 7.36E-01 | no | Plb1 | intron_variant,non_coding_transcript_variant | MODIFIER |
| 5 | 32244095 | AX-170477792 | 0 | G | A | 0.36 | 3.53E-02 | 1.06E-01 | 2.68E+00 | 2.74E+00 | 7.36E-01 | no | Plb1 | upstream_gene_variant | MODIFIER |
| 5 | 32272989 | AX-170210746 | 0 | G | A | 0.36 | 3.53E-02 | 1.06E-01 | 2.68E+00 | 2.74E+00 | 7.36E-01 | no | Plb1 | intron_variant | MODIFIER |
| 5 | 32274427 | AX-170485523 | 0 | T | C | 0.36 | 3.53E-02 | 1.06E-01 | 2.68E+00 | 2.74E+00 | 7.36E-01 | no | Plb1 | intron_variant | MODIFIER |
| 5 | 32274870 | AX-170415018 | 0 | T | G | 0.36 | 3.53E-02 | 1.06E-01 | 2.68E+00 | 2.74E+00 | 7.36E-01 | no | Plb1 | intron_variant | MODIFIER |
| 5 | 32278625 | AX-170302172 | 0 | C | T | 0.36 | 3.53E-02 | 1.06E-01 | 2.68E+00 | 2.74E+00 | 7.36E-01 | no | Plb1 | intron_variant | MODIFIER |
| 5 | 32285856 | AX-170303733 | 0 | C | T | 0.36 | 3.53E-02 | 1.06E-01 | 2.68E+00 | 2.74E+00 | 7.36E-01 | no | Plb1 | synonymous_variant | LOW |
| 5 | 32299567 | AX-168385114 | 0 | T | G | 0.36 | 3.53E-02 | 1.06E-01 | 2.68E+00 | 2.74E+00 | 7.36E-01 | no | Plb1 | intron_variant | MODIFIER |
| 5 | 32319918 | AX-170089440 | 0 | T | C | 0.36 | 3.53E-02 | 1.06E-01 | 2.68E+00 | 2.74E+00 | 7.36E-01 | no | Plb1 | intron_variant | MODIFIER |
| 5 | 32386387 | AX-170071726 | 0 | T | C | 0.36 | 3.53E-02 | 1.06E-01 | 2.68E+00 | 2.74E+00 | 7.36E-01 | no | Gm7596 | downstream_gene_variant | MODIFIER |
| 5 | 32392349 | AX-170118357 | 0 | T | C | 0.36 | 3.53E-02 | 1.06E-01 | 2.68E+00 | 2.74E+00 | 7.36E-01 | no | Gm7596 | intron_variant,non_coding_transcript_variant | MODIFIER |
| 5 | 32393125 | AX-170353757 | 0 | T | C | 0.36 | 3.53E-02 | 1.06E-01 | 2.68E+00 | 2.74E+00 | 7.36E-01 | no | Gm7596 | intron_variant,non_coding_transcript_variant | MODIFIER |
| 5 | 32411725 | AX-170144918 | 0 | G | T | 0.36 | 3.53E-02 | 1.06E-01 | 2.68E+00 | 2.74E+00 | 7.36E-01 | no | | regulatory_region_variant | MODIFIER |
| 5 | 32416953 | AX-97826590 | 0 | T | C | 0.36 | 3.53E-02 | 1.06E-01 | 2.68E+00 | 2.74E+00 | 7.36E-01 | no | Gm5864 | upstream_gene_variant | MODIFIER |
| 5 | 32472308 | AX-170228730 | 0 | T | C | 0.36 | 3.53E-02 | 1.06E-01 | 2.68E+00 | 2.74E+00 | 7.36E-01 | no | Ppp1cb | intron_variant | MODIFIER |
| 5 | 32473592 | AX-97829860 | 0 | A | G | 0.36 | 3.53E-02 | 1.06E-01 | 2.68E+00 | 2.74E+00 | 7.36E-01 | no | Ppp1cb | intron_variant | MODIFIER |
| 5 | 32477811 | AX-170471914 | 0 | T | A | 0.36 | 3.53E-02 | 1.06E-01 | 2.68E+00 | 2.74E+00 | 7.36E-01 | no | Ppp1cb | intron_variant | MODIFIER |
| 5 | 32498805 | AX-97833969 | 0 | G | A | 0.36 | 3.53E-02 | 1.06E-01 | 2.68E+00 | 2.74E+00 | 7.36E-01 | no | Ppp1cb | downstream_gene_variant | MODIFIER |
| 5 | 32508131 | AX-170532075 | 0 | T | C | 0.36 | 3.53E-02 | 1.06E-01 | 2.68E+00 | 2.74E+00 | 7.36E-01 | no | Gm15614 | non_coding_transcript_exon_variant | MODIFIER |
| 5 | 32516630 | AX-170103276 | 0 | C | T | 0.36 | 3.53E-02 | 1.06E-01 | 2.68E+00 | 2.74E+00 | 7.36E-01 | no | Gm15614 | intron_variant,non_coding_transcript_variant | MODIFIER |
| 5 | 32564992 | AX-170089441 | 0 | G | A | 0.36 | 3.53E-02 | 1.06E-01 | 2.68E+00 | 2.74E+00 | 7.36E-01 | no | Gm15614 | intron_variant,non_coding_transcript_variant | MODIFIER |
| 5 | 32618126 | AX-170095280 | 0 | A | G | 0.36 | 3.53E-02 | 1.06E-01 | 2.68E+00 | 2.74E+00 | 7.36E-01 | no | Yes1 | intron_variant | MODIFIER |
| 5 | 32640728 | AX-170425652 | 0 | T | C | 0.36 | 3.53E-02 | 1.06E-01 | 2.68E+00 | 2.74E+00 | 7.36E-01 | no | Yes1 | intron_variant | MODIFIER |
| 5 | 32643800 | AX-170445933 | 0 | T | C | 0.36 | 3.53E-02 | 1.06E-01 | 2.68E+00 | 2.74E+00 | 7.36E-01 | no | Yes1 | intron_variant | MODIFIER |
| 5 | 32661955 | AX-97813126 | 0 | T | C | 0.36 | 3.53E-02 | 1.06E-01 | 2.68E+00 | 2.74E+00 | 7.36E-01 | no | Yes1 | intron_variant | MODIFIER |
| 5 | 32666718 | AX-170176467 | 0 | G | C | 0.36 | 3.53E-02 | 1.06E-01 | 2.68E+00 | 2.74E+00 | 7.36E-01 | no | Yes1 | intron_variant | MODIFIER |
| 5 | 32691732 | AX-170371280 | 0 | T | C | 0.36 | 3.53E-02 | 1.06E-01 | 2.68E+00 | 2.74E+00 | 7.36E-01 | no | Yes1 | downstream_gene_variant | MODIFIER |
| 5 | 32695587 | AX-170497039 | 0 | C | T | 0.36 | 3.53E-02 | 1.06E-01 | 2.68E+00 | 2.74E+00 | 7.36E-01 | no | | intergenic_variant | MODIFIER |
| 5 | 32707102 | AX-97794702 | 0 | A | G | 0.36 | 3.53E-02 | 1.06E-01 | 2.68E+00 | 2.74E+00 | 7.36E-01 | no | | intergenic_variant | MODIFIER |
| 5 | 32708495 | AX-170024641 | 0 | C | T | 0.36 | 3.53E-02 | 1.06E-01 | 2.68E+00 | 2.74E+00 | 7.36E-01 | no | Gm43852 | downstream_gene_variant | MODIFIER |
| 5 | 32748695 | AX-170024640 | 0 | A | G | 0.36 | 3.53E-02 | 1.06E-01 | 2.68E+00 | 2.74E+00 | 7.36E-01 | no | Pisd | intron_variant | MODIFIER |
| 5 | 32785392 | AX-170024639 | 0 | G | C | 0.36 | 3.53E-02 | 1.06E-01 | 2.68E+00 | 2.74E+00 | 7.36E-01 | no | Pisd | intron_variant | MODIFIER |
| 5 | 32800988 | AX-170024637 | 0 | G | T | 0.36 | 3.53E-02 | 1.06E-01 | 2.68E+00 | 2.74E+00 | 7.36E-01 | no | Prr14l | intron_variant | MODIFIER |
| 5 | 32804345 | AX-170428519 | 0 | A | G | 0.36 | 3.53E-02 | 1.06E-01 | 2.68E+00 | 2.74E+00 | 7.36E-01 | no | Prr14l | intron_variant | MODIFIER |
| 5 | 32804519 | AX-170393693 | 0 | C | T | 0.36 | 3.53E-02 | 1.06E-01 | 2.68E+00 | 2.74E+00 | 7.36E-01 | no | Prr14l | intron_variant | MODIFIER |
| 5 | 32807528 | AX-170206386 | 0 | A | G | 0.36 | 3.53E-02 | 1.06E-01 | 2.68E+00 | 2.74E+00 | 7.36E-01 | no | Prr14l | intron_variant | MODIFIER |
| 5 | 32813483 | AX-170024636 | 0 | T | C | 0.36 | 3.53E-02 | 1.06E-01 | 2.68E+00 | 2.74E+00 | 7.36E-01 | no | Prr14l | intron_variant | MODIFIER |
| 5 | 32820422 | AX-170024638 | 0 | A | G | 0.36 | 3.53E-02 | 1.06E-01 | 2.68E+00 | 2.74E+00 | 7.36E-01 | no | Prr14l | intron_variant | MODIFIER |
| 5 | 32905179 | AX-170152922 | 0 | C | T | 0.36 | 3.53E-02 | 1.06E-01 | 2.68E+00 | 2.74E+00 | 7.36E-01 | no | Depdc5 | intron_variant | MODIFIER |
| 5 | 32918466 | AX-97796182 | 0 | C | T | 0.36 | 3.53E-02 | 1.06E-01 | 2.68E+00 | 2.74E+00 | 7.36E-01 | no | Depdc5 | intron_variant | MODIFIER |
| 5 | 32921947 | AX-170310461 | 0 | G | A | 0.36 | 3.53E-02 | 1.06E-01 | 2.68E+00 | 2.74E+00 | 7.36E-01 | no | Depdc5 | intron_variant | MODIFIER |
| 5 | 32923520 | AX-170506690 | 0 | G | A | 0.36 | 3.53E-02 | 1.06E-01 | 2.68E+00 | 2.74E+00 | 7.36E-01 | no | Depdc5 | intron_variant | MODIFIER |
| 5 | 32923949 | AX-170507176 | 0 | G | C | 0.36 | 3.53E-02 | 1.06E-01 | 2.68E+00 | 2.74E+00 | 7.36E-01 | no | Depdc5 | intron_variant | MODIFIER |
| 5 | 32927606 | AX-170314995 | 0 | G | T | 0.36 | 3.53E-02 | 1.06E-01 | 2.68E+00 | 2.74E+00 | 7.36E-01 | no | Depdc5 | intron_variant | MODIFIER |
| 5 | 32932132 | AX-97809369 | 0 | A | C | 0.36 | 3.53E-02 | 1.06E-01 | 2.68E+00 | 2.74E+00 | 7.36E-01 | no | Depdc5 | intron_variant | MODIFIER |
| 5 | 32948682 | AX-170255416 | 0 | T | A | 0.36 | 3.53E-02 | 1.06E-01 | 2.68E+00 | 2.74E+00 | 7.36E-01 | no | Depdc5 | intron_variant | MODIFIER |
| 5 | 32957439 | AX-170301347 | 0 | T | C | 0.36 | 3.53E-02 | 1.06E-01 | 2.68E+00 | 2.74E+00 | 7.36E-01 | no | Depdc5 | downstream_gene_variant | MODIFIER |
| 5 | 32977029 | AX-97792967 | 0 | C | A | 0.36 | 3.53E-02 | 1.06E-01 | 2.68E+00 | 2.74E+00 | 7.36E-01 | no | Depdc5 | intron_variant | MODIFIER |
| 5 | 32983559 | AX-170175078 | 0 | T | C | 0.36 | 3.53E-02 | 1.06E-01 | 2.68E+00 | 2.74E+00 | 7.36E-01 | no | Depdc5 | intron_variant | MODIFIER |
| 5 | 32989918 | AX-170071734 | 0 | C | T | 0.36 | 3.53E-02 | 1.06E-01 | 2.68E+00 | 2.74E+00 | 7.36E-01 | no | Depdc5 | intron_variant | MODIFIER |
| 5 | 33105328 | AX-170404594 | 0 | A | G | 0.36 | 3.53E-02 | 1.06E-01 | 2.68E+00 | 2.74E+00 | 7.36E-01 | no | Slc5a1 | intron_variant | MODIFIER |
| 5 | 33105490 | AX-170282456 | 0 | C | T | 0.36 | 3.53E-02 | 1.06E-01 | 2.68E+00 | 2.74E+00 | 7.36E-01 | no | Slc5a1 | intron_variant | MODIFIER |
| 5 | 33107311 | AX-170280310 | 0 | G | A | 0.36 | 3.53E-02 | 1.06E-01 | 2.68E+00 | 2.74E+00 | 7.36E-01 | no | Slc5a1 | intron_variant | MODIFIER |
| 5 | 33107453 | AX-97813757 | 0 | T | C | 0.36 | 3.53E-02 | 1.06E-01 | 2.68E+00 | 2.74E+00 | 7.36E-01 | no | Slc5a1 | intron_variant | MODIFIER |

| | | | | | | | | | | | | | | | |
|---|---|---|---|---|---|---|---|---|---|---|---|---|---|---|---|
| 5 | 33116746 | AX-170183092 | 0 | G | C | 0.36 | 3.53E-02 | 1.06E-01 | 2.68E+00 | 2.74E+00 | 7.36E-01 | no | Slc5a1 | intron_variant | MODIFIER |
| 5 | 33116845 | AX-170407202 | 0 | G | A | 0.36 | 3.53E-02 | 1.06E-01 | 2.68E+00 | 2.74E+00 | 7.36E-01 | no | Slc5a1 | synonymous_variant | LOW |
| 5 | 33119342 | AX-97799190 | 0 | A | G | 0.36 | 3.53E-02 | 1.06E-01 | 2.68E+00 | 2.74E+00 | 7.36E-01 | no | Slc5a1 | intron_variant | MODIFIER |
| 5 | 33365126 | AX-97847600 | 0 | G | A | 0.36 | 3.53E-02 | 1.06E-01 | 2.68E+00 | 2.74E+00 | 7.36E-01 | no | Maea | intron_variant | MODIFIER |
| 5 | 33385971 | AX-170334120 | 0 | G | A | 0.36 | 3.53E-02 | 1.06E-01 | 2.68E+00 | 2.74E+00 | 7.36E-01 | no | Uvssa | intron_variant | MODIFIER |
| 5 | 33559089 | AX-170188335 | 0 | C | T | 0.36 | 3.53E-02 | 1.06E-01 | 2.68E+00 | 2.74E+00 | 7.36E-01 | no | | intergenic_variant | MODIFIER |
| 5 | 33563313 | AX-170395477 | 0 | A | C | 0.36 | 3.53E-02 | 1.06E-01 | 2.68E+00 | 2.74E+00 | 7.36E-01 | no | | intergenic_variant | MODIFIER |
| 5 | 33588652 | AX-170259669 | 0 | G | A | 0.36 | 3.53E-02 | 1.06E-01 | 2.68E+00 | 2.74E+00 | 7.36E-01 | no | | intergenic_variant | MODIFIER |
| 5 | 33737741 | AX-97835737 | 0 | C | T | 0.36 | 3.53E-02 | 1.06E-01 | 2.68E+00 | 2.74E+00 | 7.36E-01 | no | Letm1 | downstream_gene_variant | MODIFIER |
| 5 | 33801850 | AX-170103799 | 0 | C | T | 0.36 | 3.53E-02 | 1.06E-01 | 2.68E+00 | 2.74E+00 | 7.36E-01 | no | | intergenic_variant | MODIFIER |
| 5 | 33838726 | AX-97840845 | 0 | C | T | 0.36 | 3.53E-02 | 1.06E-01 | 2.68E+00 | 2.74E+00 | 7.36E-01 | no | Nsd2 | upstream_gene_variant | MODIFIER |
| 5 | 33851779 | AX-170024657 | 0 | C | A | 0.36 | 3.53E-02 | 1.06E-01 | 2.68E+00 | 2.74E+00 | 7.36E-01 | no | Nsd2 | intron_variant | MODIFIER |
| 5 | 33859977 | AX-97817070 | 0 | G | A | 0.36 | 3.53E-02 | 1.06E-01 | 2.68E+00 | 2.74E+00 | 7.36E-01 | no | Nsd2 | intron_variant | MODIFIER |
| 5 | 33901406 | AX-97796388 | 0 | T | C | 0.36 | 3.53E-02 | 1.06E-01 | 2.68E+00 | 2.74E+00 | 7.36E-01 | no | Nelfa | intron_variant | MODIFIER |
| 5 | 34007597 | AX-170024654 | 0 | C | T | 0.36 | 3.53E-02 | 1.06E-01 | 2.68E+00 | 2.74E+00 | 7.36E-01 | no | Poln | intron_variant | MODIFIER |
| 5 | 34051690 | AX-170214745 | 0 | A | G | 0.36 | 3.53E-02 | 1.06E-01 | 2.68E+00 | 2.74E+00 | 7.36E-01 | no | Poln | intron_variant | MODIFIER |
| 5 | 34160771 | AX-170499211 | 0 | C | T | 0.36 | 3.53E-02 | 1.06E-01 | 2.68E+00 | 2.74E+00 | 7.36E-01 | no | Poln | intron_variant | MODIFIER |
| 5 | 35096125 | AX-170229821 | 0 | G | T | 0.36 | 3.53E-02 | 1.06E-01 | 2.68E+00 | 2.74E+00 | 7.36E-01 | no | Lrpap1 | intron_variant | MODIFIER |
| 5 | 35622439 | AX-170202997 | 0 | G | T | 0.36 | 3.53E-02 | 1.06E-01 | 2.68E+00 | 2.74E+00 | 7.36E-01 | no | | intergenic_variant | MODIFIER |
| 5 | 35662462 | AX-97798117 | 0 | A | G | 0.36 | 3.53E-02 | 1.06E-01 | 2.68E+00 | 2.74E+00 | 7.36E-01 | no | Htra3 | intron_variant | MODIFIER |
| 5 | 35666547 | AX-170024679 | 0 | A | G | 0.36 | 3.53E-02 | 1.06E-01 | 2.68E+00 | 2.74E+00 | 7.36E-01 | no | Htra3 | intron_variant | MODIFIER |
| 5 | 35672892 | AX-170378050 | 0 | C | T | 0.36 | 3.53E-02 | 1.06E-01 | 2.68E+00 | 2.74E+00 | 7.36E-01 | no | Htra3 | intron_variant | MODIFIER |
| 5 | 35673043 | AX-170104832 | 0 | A | T | 0.36 | 3.53E-02 | 1.06E-01 | 2.68E+00 | 2.74E+00 | 7.36E-01 | no | Htra3 | intron_variant | MODIFIER |
| 5 | 35677016 | AX-170459200 | 0 | C | T | 0.36 | 3.53E-02 | 1.06E-01 | 2.68E+00 | 2.74E+00 | 7.36E-01 | no | Htra3 | intron_variant | MODIFIER |
| 5 | 28007376 | AX-170142093 | 0 | T | G | 0.342 | 3.13E-02 | 9.31E-02 | 2.62E+00 | 2.72E+00 | 7.36E-01 | no | | intergenic_variant | MODIFIER |
| 18 | 89086126 | AX-170191934 | 0 | A | G | 0.518 | -1.95E-02 | 6.03E-02 | 2.66E+00 | 2.79E+00 | 7.36E-01 | no | Rttn | intron_variant | MODIFIER |
| 1 | 90232548 | AX-170163812 | 1 | T | A | 0.152 | 6.46E-02 | 1.93E-01 | 2.65E+00 | 2.72E+00 | 7.36E-01 | no | Gm38277 | upstream_gene_variant | MODIFIER |
| 1 | 28886254 | AX-170333546 | 0 | G | C | 0.404 | 2.75E-02 | 8.26E-02 | 2.67E+00 | 2.74E+00 | 7.36E-01 | no | | intergenic_variant | MODIFIER |
| 6 | 135908369 | AX-170372226 | 0 | G | T | 0.579 | 3.25E-02 | 8.85E-02 | 2.53E+00 | 2.59E+00 | 7.36E-01 | no | Grin2b | intron_variant | MODIFIER |
| 6 | 136271398 | AX-97794880 | 0 | C | A | 0.579 | 3.25E-02 | 8.85E-02 | 2.53E+00 | 2.59E+00 | 7.36E-01 | no | | intergenic_variant | MODIFIER |
| 20 | 116665527 | AX-97813525 | 1 | T | C | 0.312 | -1.64E-02 | 5.06E-02 | 2.65E+00 | 2.78E+00 | 7.37E-01 | no | | intergenic_variant | MODIFIER |
| 3 | 147615807 | AX-170068382 | 0 | G | T | 0.351 | 2.42E-02 | 7.23E-02 | 2.64E+00 | 2.72E+00 | 7.37E-01 | no | | intergenic_variant | MODIFIER |
| 17 | 73773396 | AX-170214986 | 0 | C | A | 0.053 | -5.82E-02 | 1.78E-01 | 2.66E+00 | 2.77E+00 | 7.37E-01 | no | | intergenic_variant | MODIFIER |
| 1 | 55277012 | AX-170041282 | 1 | G | A | 0.107 | -6.14E-02 | 1.85E-01 | 2.65E+00 | 2.75E+00 | 7.37E-01 | no | Boll | intron_variant,NMD_transcript_variant | MODIFIER |
| 3 | 30334951 | AX-170138428 | 1 | C | G | 0.161 | -3.43E-02 | 1.03E-01 | 2.67E+00 | 2.74E+00 | 7.37E-01 | no | Mecom | intron_variant | MODIFIER |
| 8 | 48335773 | AX-97837732 | 0 | T | C | 0.149 | -5.08E-02 | 1.51E-01 | 2.65E+00 | 2.70E+00 | 7.37E-01 | no | Tenm3 | splice_region_variant,synonymous_variant | LOW |
| 15 | 93837396 | AX-170222936 | 0 | C | T | 0.14 | -6.17E-02 | 1.88E-01 | 2.75E+00 | 2.83E+00 | 7.37E-01 | no | Gm49445 | intron_variant,non_coding_transcript_variant | MODIFIER |
| 7 | 11688471 | AX-170123568 | 0 | T | A | 0.167 | 2.55E-02 | 8.04E-02 | 2.70E+00 | 2.85E+00 | 7.37E-01 | no | Gm18850 | upstream_gene_variant | MODIFIER |
| 8 | 32152494 | AX-170306574 | 2 | G | A | 0.145 | -4.20E-02 | 1.26E-01 | 2.66E+00 | 2.73E+00 | 7.37E-01 | no | Nrg1 | intron_variant | MODIFIER |
| 8 | 100399179 | AX-170289225 | 1 | C | A | 0.241 | 3.74E-02 | 1.13E-01 | 2.67E+00 | 2.74E+00 | 7.37E-01 | no | | intergenic_variant | MODIFIER |
| 12 | 116438124 | AX-170149633 | 1 | A | G | 0.134 | -6.06E-02 | 1.84E-01 | 2.67E+00 | 2.76E+00 | 7.37E-01 | no | Ncapg2 | intron_variant | MODIFIER |
| 9 | 62505976 | AX-170235660 | 1 | G | A | 0.214 | 4.01E-02 | 1.22E-01 | 2.73E+00 | 2.79E+00 | 7.37E-01 | no | Coro2b | intron_variant | MODIFIER |
| 7 | 91615149 | AX-170140284 | 1 | G | A | 0.152 | 4.97E-02 | 1.54E-01 | 2.65E+00 | 2.78E+00 | 7.38E-01 | no | Dlg2 | intron_variant | MODIFIER |
| 7 | 76648848 | AX-97801869 | 0 | T | C | 0.36 | -3.20E-02 | 9.81E-02 | 2.68E+00 | 2.78E+00 | 7.38E-01 | no | Agbl1 | intron_variant | MODIFIER |
| 12 | 104578257 | AX-170263082 | 0 | G | T | 0.123 | -6.12E-02 | 1.87E-01 | 2.71E+00 | 2.78E+00 | 7.38E-01 | no | | intergenic_variant | MODIFIER |
| 12 | 108378289 | AX-170139917 | 1 | G | A | 0.116 | -6.45E-02 | 1.97E-01 | 2.72E+00 | 2.79E+00 | 7.38E-01 | no | Eml1 | intron_variant,non_coding_transcript_variant | MODIFIER |
| 15 | 89574534 | AX-170514188 | 0 | G | A | 0.149 | -5.06E-02 | 1.55E-01 | 2.68E+00 | 2.77E+00 | 7.38E-01 | no | Acr | 3_prime_UTR_variant | MODIFIER |
| 14 | 64544806 | AX-97826372 | 0 | C | T | 0.053 | 5.97E-02 | 1.85E-01 | 2.66E+00 | 2.78E+00 | 7.38E-01 | no | | intergenic_variant | MODIFIER |
| 12 | 104045602 | AX-170098322 | 0 | C | T | 0.114 | -5.85E-02 | 1.78E-01 | 2.67E+00 | 2.76E+00 | 7.38E-01 | no | Serpina12 | upstream_gene_variant | MODIFIER |
| 11 | 74239162 | AX-170085836 | 0 | T | G | 0.246 | 2.34E-02 | 7.17E-02 | 2.62E+00 | 2.76E+00 | 7.38E-01 | no | Olfr404-ps1 | upstream_gene_variant | MODIFIER |
| 12 | 112300018 | AX-97826121 | 0 | A | G | 0.123 | -6.31E-02 | 1.93E-01 | 2.70E+00 | 2.78E+00 | 7.38E-01 | no | | regulatory_region_variant | MODIFIER |
| 2 | 129978415 | AX-170454000 | 1 | T | C | 0.134 | 6.49E-02 | 1.97E-01 | 2.70E+00 | 2.75E+00 | 7.38E-01 | no | | intergenic_variant | MODIFIER |
| 9 | 80254203 | AX-170082915 | 1 | G | T | 0.205 | 4.29E-02 | 1.30E-01 | 2.76E+00 | 2.81E+00 | 7.38E-01 | no | Myo6 | intron_variant | MODIFIER |
| 1 | 4740872 | AX-169999850 | 0 | T | C | 0.509 | 2.40E-02 | 6.88E-02 | 2.57E+00 | 2.65E+00 | 7.38E-01 | no | | intergenic_variant | MODIFIER |

| | | | | | | | | | | | | | | | |
|---|---|---|---|---|---|---|---|---|---|---|---|---|---|---|---|
| 20 | 134901347 | AX-170035012 | 2 | A | G | 0.136 | 3.94E-02 | 1.19E-01 | 2.65E+00 | 2.73E+00 | 7.38E-01 | no | Gm15018 | downstream_gene_variant | MODIFIER |
| 7 | 106911349 | AX-170029720 | 1 | C | A | 0.473 | -3.00E-02 | 9.25E-02 | 2.74E+00 | 2.82E+00 | 7.38E-01 | no | Olfr1532-ps1 | upstream_gene_variant | MODIFIER |
| 20 | 164733931 | AX-170441398 | 1 | A | C | 0.125 | 4.01E-02 | 1.25E-01 | 2.65E+00 | 2.78E+00 | 7.39E-01 | no | | intergenic_variant | MODIFIER |
| 20 | 165430169 | AX-170035414 | 1 | G | A | 0.125 | 4.01E-02 | 1.25E-01 | 2.65E+00 | 2.78E+00 | 7.39E-01 | no | | intergenic_variant | MODIFIER |
| 12 | 115345276 | AX-170327778 | 0 | C | A | 0.14 | -6.30E-02 | 1.92E-01 | 2.68E+00 | 2.75E+00 | 7.39E-01 | no | Ighv1-60 | upstream_gene_variant | MODIFIER |
| 7 | 13344341 | AX-170028612 | 0 | T | A | 0.351 | 2.71E-02 | 8.05E-02 | 2.59E+00 | 2.71E+00 | 7.39E-01 | no | Pla2g4c | intron_variant | MODIFIER |
| 3 | 73225561 | AX-170217447 | 0 | C | T | 0.044 | -5.56E-02 | 1.67E-01 | 2.60E+00 | 2.72E+00 | 7.39E-01 | no | Gm20754 | intron_variant,non_coding_transcript_variant | MODIFIER |
| 18 | 18115471 | AX-170435045 | 0 | C | T | 0.044 | -5.56E-02 | 1.67E-01 | 2.60E+00 | 2.72E+00 | 7.39E-01 | no | | intergenic_variant | MODIFIER |
| 6 | 136105648 | AX-97827170 | 1 | T | A | 0.562 | 3.09E-02 | 8.61E-02 | 2.54E+00 | 2.62E+00 | 7.39E-01 | no | Grin2b | intron_variant | MODIFIER |
| 8 | 97234930 | AX-97837735 | 1 | A | G | 0.214 | 3.82E-02 | 1.17E-01 | 2.69E+00 | 2.76E+00 | 7.39E-01 | no | | intergenic_variant | MODIFIER |
| 18 | 74558476 | AX-170304828 | 0 | C | A | 0.395 | -2.93E-02 | 8.72E-02 | 2.63E+00 | 2.70E+00 | 7.39E-01 | no | Myo5b | intron_variant | MODIFIER |
| 18 | 74587860 | AX-170453773 | 0 | T | C | 0.395 | -2.93E-02 | 8.72E-02 | 2.63E+00 | 2.70E+00 | 7.39E-01 | no | Myo5b | intron_variant | MODIFIER |
| 18 | 74654196 | AX-97829495 | 0 | T | C | 0.395 | -2.93E-02 | 8.72E-02 | 2.63E+00 | 2.70E+00 | 7.39E-01 | no | Myo5b | intron_variant | MODIFIER |
| 18 | 74664583 | AX-170456160 | 0 | C | T | 0.395 | -2.93E-02 | 8.72E-02 | 2.63E+00 | 2.70E+00 | 7.39E-01 | no | Myo5b | intron_variant | MODIFIER |
| 18 | 74896090 | AX-170154480 | 0 | C | T | 0.395 | -2.93E-02 | 8.72E-02 | 2.63E+00 | 2.70E+00 | 7.39E-01 | no | | regulatory_region_variant | MODIFIER |
| 18 | 74901001 | AX-170114957 | 0 | A | C | 0.395 | -2.93E-02 | 8.72E-02 | 2.63E+00 | 2.70E+00 | 7.39E-01 | no | | intergenic_variant | MODIFIER |
| 18 | 74901145 | AX-170295181 | 0 | A | G | 0.395 | -2.93E-02 | 8.72E-02 | 2.63E+00 | 2.70E+00 | 7.39E-01 | no | | intergenic_variant | MODIFIER |
| 18 | 74901387 | AX-170486359 | 0 | C | T | 0.395 | -2.93E-02 | 8.72E-02 | 2.63E+00 | 2.70E+00 | 7.39E-01 | no | Gm24559 | downstream_gene_variant | MODIFIER |
| 13 | 23385414 | AX-170035527 | 1 | A | G | 0.17 | -4.40E-02 | 1.20E-01 | 2.53E+00 | 2.59E+00 | 7.39E-01 | no | Gm11334 | downstream_gene_variant | MODIFIER |
| 4 | 143000989 | AX-170070917 | 0 | C | A | 0.105 | 6.70E-02 | 2.04E-01 | 2.68E+00 | 2.75E+00 | 7.39E-01 | no | 6330411D24Rik | intron_variant,non_coding_transcript_variant | MODIFIER |
| 4 | 145309040 | AX-170447967 | 0 | G | A | 0.105 | 6.70E-02 | 2.04E-01 | 2.68E+00 | 2.75E+00 | 7.39E-01 | no | Tnfrsf8 | intron_variant | MODIFIER |
| 15 | 88177566 | AX-170232488 | 1 | G | C | 0.161 | 5.30E-02 | 1.53E-01 | 2.58E+00 | 2.67E+00 | 7.39E-01 | no | | intergenic_variant | MODIFIER |
| 13 | 37348692 | AX-97823349 | 0 | C | T | 0.184 | 3.40E-02 | 1.07E-01 | 2.75E+00 | 2.87E+00 | 7.39E-01 | no | Ly86 | intron_variant | MODIFIER |
| 5 | 45974480 | AX-170507301 | 0 | T | G | 0.5 | 3.43E-02 | 9.72E-02 | 2.56E+00 | 2.63E+00 | 7.39E-01 | no | | intergenic_variant | MODIFIER |
| 5 | 30925742 | AX-170146435 | 0 | T | C | 0.368 | 3.32E-02 | 1.01E-01 | 2.67E+00 | 2.73E+00 | 7.39E-01 | no | Khk | intron_variant | MODIFIER |
| 9 | 37312905 | AX-170125065 | 2 | C | T | 0.5 | -2.45E-02 | 7.56E-02 | 2.67E+00 | 2.77E+00 | 7.39E-01 | no | Ccdc15 | intron_variant | MODIFIER |
| 5 | 104321754 | AX-97781207 | 0 | T | C | 0.439 | 3.59E-02 | 1.08E-01 | 2.66E+00 | 2.70E+00 | 7.39E-01 | no | Mepe | upstream_gene_variant | MODIFIER |
| 1 | 53122920 | AX-170097132 | 1 | G | A | 0.089 | 5.85E-02 | 1.81E-01 | 2.66E+00 | 2.77E+00 | 7.39E-01 | no | | intergenic_variant | MODIFIER |
| 15 | 10958591 | AX-170510599 | 0 | C | T | 0.254 | -3.31E-02 | 9.81E-02 | 2.61E+00 | 2.69E+00 | 7.40E-01 | no | C1qtnf3 | intron_variant | MODIFIER |
| 2 | 120581723 | AX-97828684 | 1 | C | T | 0.143 | 7.24E-02 | 2.20E-01 | 2.70E+00 | 2.73E+00 | 7.40E-01 | no | Snap23 | intron_variant | MODIFIER |
| 2 | 142589368 | AX-170300985 | 1 | G | T | 0.143 | 7.24E-02 | 2.20E-01 | 2.70E+00 | 2.73E+00 | 7.40E-01 | no | | regulatory_region_variant | MODIFIER |
| 7 | 135379807 | AX-170518198 | 0 | A | G | 0.246 | -2.95E-02 | 8.57E-02 | 2.57E+00 | 2.67E+00 | 7.40E-01 | no | | intergenic_variant | MODIFIER |
| 18 | 64754755 | AX-170494166 | 1 | A | C | 0.393 | 3.32E-02 | 9.44E-02 | 2.57E+00 | 2.62E+00 | 7.40E-01 | no | | intergenic_variant | MODIFIER |
| 7 | 117137301 | AX-170415817 | 1 | A | G | 0.42 | 3.07E-02 | 9.19E-02 | 2.64E+00 | 2.69E+00 | 7.40E-01 | no | | intergenic_variant | MODIFIER |
| 2 | 101377441 | AX-170018487 | 0 | G | C | 0.061 | 6.81E-02 | 2.11E-01 | 2.66E+00 | 2.77E+00 | 7.40E-01 | no | | intergenic_variant | MODIFIER |
| 9 | 39908881 | AX-97847590 | 0 | T | C | 0.421 | -2.47E-02 | 7.58E-02 | 2.67E+00 | 2.76E+00 | 7.40E-01 | no | Olfr229 | upstream_gene_variant | MODIFIER |
| 10 | 122143675 | AX-170284245 | 0 | C | T | 0.035 | -4.71E-02 | 1.45E-01 | 2.57E+00 | 2.75E+00 | 7.40E-01 | no | | intergenic_variant | MODIFIER |
| 8 | 21746015 | AX-170422149 | 1 | A | T | 0.179 | 3.40E-02 | 1.04E-01 | 2.65E+00 | 2.74E+00 | 7.40E-01 | no | Defa-ps11 | upstream_gene_variant | MODIFIER |
| 2 | 119806839 | AX-170435579 | 2 | A | G | 0.109 | 6.07E-02 | 1.82E-01 | 2.65E+00 | 2.70E+00 | 7.40E-01 | no | Tyro3 | intron_variant | MODIFIER |
| 2 | 33711196 | AX-97852866 | 0 | C | T | 0.035 | -8.78E-02 | 2.66E-01 | 2.67E+00 | 2.72E+00 | 7.40E-01 | no | | intergenic_variant | MODIFIER |
| 2 | 122492499 | AX-170328660 | 0 | G | A | 0.482 | 1.76E-01 | 5.32E-01 | 2.67E+00 | 2.72E+00 | 7.40E-01 | Yes | Gm14085 | intron_variant | MODIFIER |
| 3 | 94944863 | AX-170067399 | 0 | A | C | 0.482 | 1.76E-01 | 5.32E-01 | 2.67E+00 | 2.72E+00 | 7.40E-01 | Yes | Selenbp1 | downstream_gene_variant | MODIFIER |
| 4 | 11092067 | AX-97783372 | 0 | A | G | 0.482 | 1.76E-01 | 5.32E-01 | 2.67E+00 | 2.72E+00 | 7.40E-01 | Yes | | intergenic_variant | MODIFIER |
| 4 | 147244977 | AX-170070969 | 0 | C | A | 0.482 | 1.76E-01 | 5.32E-01 | 2.67E+00 | 2.72E+00 | 7.40E-01 | Yes | Gm13147 | intron_variant,non_coding_transcript_variant | MODIFIER |
| 5 | 143598236 | AX-170307652 | 0 | G | C | 0.482 | 1.76E-01 | 5.32E-01 | 2.67E+00 | 2.72E+00 | 7.40E-01 | Yes | | intergenic_variant | MODIFIER |
| 5 | 145758576 | AX-170026409 | 0 | A | C | 0.482 | 1.76E-01 | 5.32E-01 | 2.67E+00 | 2.72E+00 | 7.40E-01 | Yes | Gm43626 | upstream_gene_variant | MODIFIER |
| 6 | 3180317 | AX-170399821 | 0 | G | T | 0.482 | 1.76E-01 | 5.32E-01 | 2.67E+00 | 2.72E+00 | 7.40E-01 | Yes | Gm42642 | downstream_gene_variant | MODIFIER |
| 6 | 3274631 | AX-170145340 | 0 | C | T | 0.482 | 1.76E-01 | 5.32E-01 | 2.67E+00 | 2.72E+00 | 7.40E-01 | Yes | Gm42961 | upstream_gene_variant | MODIFIER |
| 6 | 32702192 | AX-170230422 | 0 | C | T | 0.482 | 1.76E-01 | 5.32E-01 | 2.67E+00 | 2.72E+00 | 7.40E-01 | Yes | | intergenic_variant | MODIFIER |
| 11 | 52201628 | AX-97799579 | 0 | C | T | 0.482 | 1.76E-01 | 5.32E-01 | 2.67E+00 | 2.72E+00 | 7.40E-01 | Yes | Gm26551 | intron_variant,non_coding_transcript_variant | MODIFIER |
| 11 | 56191985 | AX-170035498 | 0 | A | G | 0.482 | 1.76E-01 | 5.32E-01 | 2.67E+00 | 2.72E+00 | 7.40E-01 | Yes | | intergenic_variant | MODIFIER |
| 12 | 17948456 | AX-170533468 | 0 | G | T | 0.482 | 1.76E-01 | 5.32E-01 | 2.67E+00 | 2.72E+00 | 7.40E-01 | Yes | Gm49329 | intron_variant,non_coding_transcript_variant | MODIFIER |
| 12 | 23766465 | AX-97851882 | 0 | C | G | 0.482 | 1.76E-01 | 5.32E-01 | 2.67E+00 | 2.72E+00 | 7.40E-01 | Yes | | intergenic_variant | MODIFIER |
| 12 | 30877584 | AX-170510084 | 0 | C | T | 0.482 | 1.76E-01 | 5.32E-01 | 2.67E+00 | 2.72E+00 | 7.40E-01 | Yes | | intergenic_variant | MODIFIER |

| | | | | | | | | | | | | | | | |
|---|---|---|---|---|---|---|---|---|---|---|---|---|---|---|---|
| 12 | 103766875 | AX-170302698 | 0 | C | A | 0.482 | 1.76E-01 | 5.32E-01 | 2.67E+00 | 2.72E+00 | 7.40E-01 | Yes | Serpina1d | intron_variant | MODIFIER |
| 13 | 66501917 | AX-170131053 | 0 | T | C | 0.482 | 1.76E-01 | 5.32E-01 | 2.67E+00 | 2.72E+00 | 7.40E-01 | Yes | Gm26806 | downstream_gene_variant | MODIFIER |
| 13 | 83769215 | AX-170008728 | 0 | T | C | 0.482 | 1.76E-01 | 5.32E-01 | 2.67E+00 | 2.72E+00 | 7.40E-01 | Yes | C130071C03Rik | intron_variant,non_coding_transcript_variant | MODIFIER |
| 14 | 43825311 | AX-170009700 | 0 | A | G | 0.482 | 1.76E-01 | 5.32E-01 | 2.67E+00 | 2.72E+00 | 7.40E-01 | Yes | Ear1 | intron_variant | MODIFIER |
| 14 | 45101386 | AX-170172817 | 0 | G | T | 0.482 | 1.76E-01 | 5.32E-01 | 2.67E+00 | 2.72E+00 | 7.40E-01 | Yes | Gm49180 | intron_variant,non_coding_transcript_variant | MODIFIER |
| 14 | 51410376 | AX-170009778 | 0 | C | T | 0.482 | 1.76E-01 | 5.32E-01 | 2.67E+00 | 2.72E+00 | 7.40E-01 | Yes | Vmn2r88 | upstream_gene_variant | MODIFIER |
| 14 | 51642439 | AX-170053374 | 0 | T | G | 0.482 | 1.76E-01 | 5.32E-01 | 2.67E+00 | 2.72E+00 | 7.40E-01 | Yes | Gm5622 | intron_variant | MODIFIER |
| 14 | 52348956 | AX-170531206 | 0 | C | T | 0.482 | 1.76E-01 | 5.32E-01 | 2.67E+00 | 2.72E+00 | 7.40E-01 | Yes | Olfr1513 | downstream_gene_variant | MODIFIER |
| 14 | 53489077 | AX-170296012 | 0 | G | C | 0.482 | 1.76E-01 | 5.32E-01 | 2.67E+00 | 2.72E+00 | 7.40E-01 | Yes | Trav9-1 | downstream_gene_variant | MODIFIER |
| 14 | 53620697 | AX-97816533 | 0 | T | G | 0.482 | 1.76E-01 | 5.32E-01 | 2.67E+00 | 2.72E+00 | 7.40E-01 | Yes | Trav12-3 | upstream_gene_variant | MODIFIER |
| 14 | 96353181 | AX-97827133 | 0 | G | C | 0.482 | 1.76E-01 | 5.32E-01 | 2.67E+00 | 2.72E+00 | 7.40E-01 | Yes | Klhl1 | intron_variant | MODIFIER |
| 16 | 91510253 | AX-170058104 | 0 | A | G | 0.482 | 1.76E-01 | 5.32E-01 | 2.67E+00 | 2.72E+00 | 7.40E-01 | Yes | Ifnar1 | downstream_gene_variant | MODIFIER |
| 17 | 7337230 | AX-170013771 | 0 | A | G | 0.482 | 1.76E-01 | 5.32E-01 | 2.67E+00 | 2.72E+00 | 7.40E-01 | Yes | Tcp10a | intron_variant | MODIFIER |
| 17 | 28437828 | AX-170175931 | 0 | T | C | 0.482 | 1.76E-01 | 5.32E-01 | 2.67E+00 | 2.72E+00 | 7.40E-01 | Yes | Fkbp5 | intron_variant | MODIFIER |
| 17 | 34474082 | AX-170518636 | 0 | G | A | 0.482 | 1.76E-01 | 5.32E-01 | 2.67E+00 | 2.72E+00 | 7.40E-01 | Yes | Btnl4 | missense_variant | MODERATE |
| 17 | 34534591 | AX-170148360 | 0 | C | A | 0.482 | 1.76E-01 | 5.32E-01 | 2.67E+00 | 2.72E+00 | 7.40E-01 | Yes | Btnl7-ps | intron_variant,non_coding_transcript_variant | MODIFIER |
| 17 | 36031387 | AX-97793881 | 0 | A | C | 0.482 | 1.76E-01 | 5.32E-01 | 2.67E+00 | 2.72E+00 | 7.40E-01 | Yes | H2-T23 | intron_variant | MODIFIER |
| 17 | 44902367 | AX-170247021 | 0 | G | A | 0.482 | 1.76E-01 | 5.32E-01 | 2.67E+00 | 2.72E+00 | 7.40E-01 | Yes | Supt3 | intron_variant | MODIFIER |
| 17 | 82904230 | AX-170207894 | 0 | G | A | 0.482 | 1.76E-01 | 5.32E-01 | 2.67E+00 | 2.72E+00 | 7.40E-01 | Yes | | intergenic_variant | MODIFIER |
| 18 | 20270206 | AX-97831307 | 0 | A | G | 0.482 | 1.76E-01 | 5.32E-01 | 2.67E+00 | 2.72E+00 | 7.40E-01 | Yes | Dsg1c | intron_variant | MODIFIER |
| 19 | 40089945 | AX-170429823 | 0 | G | T | 0.482 | 1.76E-01 | 5.32E-01 | 2.67E+00 | 2.72E+00 | 7.40E-01 | Yes | Cyp2c50 | intron_variant | MODIFIER |
| 12 | 107546231 | AX-170092608 | 0 | A | G | 0.105 | -6.20E-02 | 1.92E-01 | 2.68E+00 | 2.77E+00 | 7.40E-01 | no | | intergenic_variant | MODIFIER |
| 7 | 70631329 | AX-170029243 | 0 | C | T | 0.386 | -3.31E-02 | 1.03E-01 | 2.71E+00 | 2.81E+00 | 7.40E-01 | no | | intergenic_variant | MODIFIER |
| 14 | 51461024 | AX-170009783 | 1 | G | C | 0.482 | 1.75E-01 | 5.32E-01 | 2.67E+00 | 2.72E+00 | 7.40E-01 | Yes | Vmn2r89 | 3_prime_UTR_variant | MODIFIER |
| 5 | 31025751 | AX-97814109 | 1 | C | T | 0.366 | 3.43E-02 | 1.04E-01 | 2.65E+00 | 2.72E+00 | 7.40E-01 | no | | intergenic_variant | MODIFIER |
| 1 | 192753445 | AX-170269722 | 0 | G | A | 0.456 | -1.78E-02 | 4.67E-02 | 2.47E+00 | 2.58E+00 | 7.40E-01 | no | Hhat | intron_variant | MODIFIER |
| 19 | 4041905 | AX-170164597 | 0 | C | G | 0.272 | 3.57E-02 | 9.34E-02 | 2.47E+00 | 2.58E+00 | 7.40E-01 | no | Gstp2 | missense_variant,splice_region_variant | MODERATE |
| 6 | 69540123 | AX-170384178 | 0 | C | A | 0.482 | 1.75E-01 | 5.31E-01 | 2.67E+00 | 2.71E+00 | 7.40E-01 | Yes | Igkv4-57-1 | downstream_gene_variant | MODIFIER |
| 1 | 4613879 | AX-170416397 | 0 | G | A | 0.509 | 2.38E-02 | 6.77E-02 | 2.56E+00 | 2.63E+00 | 7.40E-01 | no | Gm7369 | downstream_gene_variant | MODIFIER |
| 8 | 51455652 | AX-97804217 | 1 | A | G | 0.143 | -5.10E-02 | 1.50E-01 | 2.61E+00 | 2.66E+00 | 7.40E-01 | no | Gm17911 | downstream_gene_variant | MODIFIER |
| 2 | 114139277 | AX-170383264 | 1 | C | T | 0.134 | 6.77E-02 | 2.07E-01 | 2.68E+00 | 2.74E+00 | 7.40E-01 | no | Aqr | intron_variant | MODIFIER |
| 2 | 119376522 | AX-170064717 | 1 | T | C | 0.134 | 6.77E-02 | 2.07E-01 | 2.68E+00 | 2.74E+00 | 7.40E-01 | no | Ino80 | intron_variant | MODIFIER |
| 2 | 119448644 | AX-170364752 | 1 | G | A | 0.134 | 6.77E-02 | 2.07E-01 | 2.68E+00 | 2.74E+00 | 7.40E-01 | no | Ino80 | intron_variant | MODIFIER |
| 2 | 120376975 | AX-170236597 | 1 | T | G | 0.134 | 6.77E-02 | 2.07E-01 | 2.68E+00 | 2.74E+00 | 7.40E-01 | no | Tmem87a | intron_variant | MODIFIER |
| 2 | 121140002 | AX-170283146 | 1 | T | G | 0.134 | 6.77E-02 | 2.07E-01 | 2.68E+00 | 2.74E+00 | 7.40E-01 | no | Adal | upstream_gene_variant | MODIFIER |
| 2 | 122007605 | AX-170018880 | 1 | C | T | 0.134 | 6.77E-02 | 2.07E-01 | 2.68E+00 | 2.74E+00 | 7.40E-01 | no | Ctdspl2 | intron_variant | MODIFIER |
| 2 | 137085033 | AX-170520525 | 1 | T | C | 0.134 | 6.77E-02 | 2.07E-01 | 2.68E+00 | 2.74E+00 | 7.40E-01 | no | Jag1 | synonymous_variant | LOW |
| 2 | 143588407 | AX-170164212 | 1 | A | C | 0.134 | 6.77E-02 | 2.07E-01 | 2.68E+00 | 2.74E+00 | 7.40E-01 | no | Pcsk2 | intron_variant | MODIFIER |
| 2 | 144703647 | AX-97857052 | 1 | C | T | 0.134 | 6.77E-02 | 2.07E-01 | 2.68E+00 | 2.74E+00 | 7.40E-01 | no | Dtd1 | intron_variant | MODIFIER |
| 4 | 94960954 | AX-170361932 | 0 | C | T | 0.333 | 3.58E-02 | 1.09E-01 | 2.66E+00 | 2.73E+00 | 7.40E-01 | no | Mysm1 | intron_variant | MODIFIER |
| 4 | 95040284 | AX-170502129 | 0 | C | T | 0.333 | 3.58E-02 | 1.09E-01 | 2.66E+00 | 2.73E+00 | 7.40E-01 | no | Gm12704 | upstream_gene_variant | MODIFIER |
| 8 | 50750762 | AX-97844293 | 1 | G | T | 0.152 | 4.93E-02 | 1.50E-01 | 2.81E+00 | 2.87E+00 | 7.41E-01 | no | C130073E24Rik | intron_variant,non_coding_transcript_variant | MODIFIER |
| 8 | 53834521 | AX-97844160 | 1 | A | G | 0.152 | 4.93E-02 | 1.50E-01 | 2.81E+00 | 2.87E+00 | 7.41E-01 | no | | intergenic_variant | MODIFIER |
| 8 | 54677173 | AX-97804270 | 1 | C | T | 0.152 | 4.93E-02 | 1.50E-01 | 2.81E+00 | 2.87E+00 | 7.41E-01 | no | Wdr17 | intron_variant,non_coding_transcript_variant | MODIFIER |
| 10 | 68037420 | AX-97799346 | 0 | T | C | 0.421 | -2.56E-02 | 7.67E-02 | 2.62E+00 | 2.70E+00 | 7.41E-01 | no | Rtkn2 | intron_variant,NMD_transcript_variant | MODIFIER |
| 1 | 44439631 | AX-170210464 | 0 | G | A | 0.105 | 5.67E-02 | 1.73E-01 | 2.61E+00 | 2.73E+00 | 7.41E-01 | no | | intergenic_variant | MODIFIER |
| 18 | 55508731 | AX-170314387 | 0 | G | A | 0.447 | 3.45E-02 | 9.45E-02 | 2.52E+00 | 2.59E+00 | 7.41E-01 | no | | regulatory_region_variant | MODIFIER |
| 4 | 143424482 | AX-170133963 | 1 | G | T | 0.107 | 6.67E-02 | 2.05E-01 | 2.69E+00 | 2.75E+00 | 7.41E-01 | no | Pramef8 | downstream_gene_variant | MODIFIER |
| 4 | 145524529 | AX-170343785 | 1 | G | A | 0.107 | 6.67E-02 | 2.05E-01 | 2.69E+00 | 2.75E+00 | 7.41E-01 | no | Zfp990 | intron_variant | MODIFIER |
| 12 | 114352346 | AX-170471575 | 0 | T | G | 0.114 | -6.24E-02 | 1.94E-01 | 2.69E+00 | 2.78E+00 | 7.41E-01 | no | Ighv1-1 | upstream_gene_variant | MODIFIER |
| 9 | 80588359 | AX-170082923 | 1 | A | G | 0.205 | -3.97E-02 | 1.21E-01 | 2.65E+00 | 2.72E+00 | 7.41E-01 | no | Gm48387 | intron_variant,non_coding_transcript_variant | MODIFIER |
| 8 | 50837073 | AX-170101020 | 2 | T | A | 0.155 | 4.88E-02 | 1.49E-01 | 2.81E+00 | 2.87E+00 | 7.41E-01 | no | C130073E24Rik | intron_variant,non_coding_transcript_variant | MODIFIER |
| 9 | 76717386 | AX-170032742 | 1 | G | T | 0.196 | 4.30E-02 | 1.32E-01 | 2.73E+00 | 2.77E+00 | 7.41E-01 | no | Gm22938 | upstream_gene_variant | MODIFIER |
| 16 | 86224255 | AX-97789570 | 1 | G | A | 0.455 | 2.63E-02 | 7.92E-02 | 2.63E+00 | 2.70E+00 | 7.42E-01 | no | | intergenic_variant | MODIFIER |

| | | | | | | | | | | | | | | | |
|---|---|---|---|---|---|---|---|---|---|---|---|---|---|---|---|
| 15 | 95204718 | AX-170400005 | 1 | C | T | 0.143 | 5.89E-02 | 1.79E-01 | 2.64E+00 | 2.71E+00 | 7.42E-01 | no | Nell2 | intron_variant | MODIFIER |
| 10 | 18250315 | AX-170124734 | 0 | A | G | 0.202 | 3.15E-02 | 9.92E-02 | 2.72E+00 | 2.82E+00 | 7.42E-01 | no | Gm10827 | downstream_gene_variant | MODIFIER |
| 2 | 124974521 | AX-170350445 | 1 | G | A | 0.143 | 7.21E-02 | 2.21E-01 | 2.69E+00 | 2.72E+00 | 7.42E-01 | no | | intergenic_variant | MODIFIER |
| 2 | 138164768 | AX-170088765 | 1 | C | T | 0.143 | 7.21E-02 | 2.21E-01 | 2.69E+00 | 2.72E+00 | 7.42E-01 | no | | intergenic_variant | MODIFIER |
| 2 | 155882844 | AX-170188000 | 1 | A | C | 0.143 | 7.21E-02 | 2.21E-01 | 2.69E+00 | 2.72E+00 | 7.42E-01 | no | Uqcc1 | intron_variant | MODIFIER |
| 15 | 96927469 | AX-170525624 | 1 | C | T | 0.152 | -5.06E-02 | 1.58E-01 | 2.70E+00 | 2.79E+00 | 7.42E-01 | no | Rpl10a-ps3 | upstream_gene_variant | MODIFIER |
| 2 | 95072584 | AX-97813321 | 1 | G | A | 0.054 | -9.78E-02 | 3.01E-01 | 2.73E+00 | 2.76E+00 | 7.42E-01 | no | | intergenic_variant | MODIFIER |
| 2 | 102570102 | AX-170064461 | 1 | C | T | 0.054 | -9.78E-02 | 3.01E-01 | 2.73E+00 | 2.76E+00 | 7.42E-01 | no | Pamr1 | intron_variant | MODIFIER |
| 12 | 93328188 | AX-170486880 | 0 | A | T | 0.07 | 6.10E-02 | 1.85E-01 | 2.62E+00 | 2.72E+00 | 7.42E-01 | no | | intergenic_variant | MODIFIER |
| 10 | 8372619 | AX-170086735 | 0 | G | A | 0.211 | -3.48E-02 | 1.05E-01 | 2.62E+00 | 2.71E+00 | 7.42E-01 | no | Ust | intron_variant | MODIFIER |
| 1 | 144984394 | AX-97787984 | 0 | T | C | 0.202 | 4.29E-02 | 1.30E-01 | 2.66E+00 | 2.71E+00 | 7.42E-01 | no | Gm25708 | upstream_gene_variant | MODIFIER |
| 18 | 57789881 | AX-170454845 | 0 | A | T | 0.404 | -3.00E-02 | 9.21E-02 | 2.82E+00 | 2.92E+00 | 7.42E-01 | no | AC102232.1 | intron_variant,non_coding_transcript_variant | MODIFIER |
| 8 | 98862920 | AX-170305364 | 0 | T | C | 0.219 | 3.47E-02 | 1.08E-01 | 2.67E+00 | 2.76E+00 | 7.42E-01 | no | | intergenic_variant | MODIFIER |
| 9 | 107081673 | AX-97847529 | 0 | G | A | 0.132 | -3.59E-02 | 1.07E-01 | 2.59E+00 | 2.69E+00 | 7.42E-01 | no | Dock3 | intron_variant | MODIFIER |
| 13 | 54672492 | AX-97803018 | 1 | C | T | 0.536 | 2.34E-02 | 6.87E-02 | 2.57E+00 | 2.68E+00 | 7.42E-01 | no | | intergenic_variant | MODIFIER |
| 7 | 134908805 | AX-170096137 | 0 | C | T | 0.254 | -2.68E-02 | 8.04E-02 | 2.60E+00 | 2.70E+00 | 7.42E-01 | no | Dock1 | intron_variant | MODIFIER |
| 9 | 69664307 | AX-170535853 | 1 | T | C | 0.196 | 4.36E-02 | 1.34E-01 | 2.77E+00 | 2.82E+00 | 7.42E-01 | no | | intergenic_variant | MODIFIER |
| 8 | 104019284 | AX-170501046 | 1 | A | G | 0.223 | 3.53E-02 | 1.09E-01 | 2.68E+00 | 2.75E+00 | 7.42E-01 | no | | intergenic_variant | MODIFIER |
| 18 | 68753457 | AX-97827041 | 1 | C | A | 0.089 | 5.18E-02 | 1.58E-01 | 2.61E+00 | 2.73E+00 | 7.42E-01 | no | | intergenic_variant | MODIFIER |
| 10 | 56158298 | AX-170475456 | 0 | G | A | 0.289 | 2.53E-02 | 8.03E-02 | 2.77E+00 | 2.88E+00 | 7.42E-01 | no | Tbc1d32 | intron_variant | MODIFIER |
| 10 | 56172732 | AX-97792355 | 0 | C | A | 0.289 | 2.53E-02 | 8.03E-02 | 2.77E+00 | 2.88E+00 | 7.42E-01 | no | Tbc1d32 | intron_variant | MODIFIER |
| 10 | 56177765 | AX-97805343 | 0 | T | C | 0.289 | 2.53E-02 | 8.03E-02 | 2.77E+00 | 2.88E+00 | 7.42E-01 | no | Tbc1d32 | intron_variant | MODIFIER |
| 10 | 56177957 | AX-170246898 | 0 | A | G | 0.289 | 2.53E-02 | 8.03E-02 | 2.77E+00 | 2.88E+00 | 7.42E-01 | no | Tbc1d32 | intron_variant | MODIFIER |
| 10 | 58290962 | AX-170044966 | 0 | C | T | 0.289 | 2.53E-02 | 8.03E-02 | 2.77E+00 | 2.88E+00 | 7.42E-01 | no | Gcc2 | intron_variant | MODIFIER |
| 10 | 58329767 | AX-170044968 | 0 | C | T | 0.289 | 2.53E-02 | 8.03E-02 | 2.77E+00 | 2.88E+00 | 7.42E-01 | no | Lims1 | intron_variant | MODIFIER |
| 10 | 58589364 | AX-97816322 | 0 | A | G | 0.289 | 2.53E-02 | 8.03E-02 | 2.77E+00 | 2.88E+00 | 7.42E-01 | no | | intergenic_variant | MODIFIER |
| 10 | 58635763 | AX-97801045 | 0 | G | A | 0.289 | 2.53E-02 | 8.03E-02 | 2.77E+00 | 2.88E+00 | 7.42E-01 | no | Edar | intron_variant | MODIFIER |
| 10 | 58665966 | AX-170044974 | 0 | G | C | 0.289 | 2.53E-02 | 8.03E-02 | 2.77E+00 | 2.88E+00 | 7.42E-01 | no | Edar | intron_variant | MODIFIER |
| 10 | 58737565 | AX-170348968 | 0 | A | G | 0.289 | 2.53E-02 | 8.03E-02 | 2.77E+00 | 2.88E+00 | 7.42E-01 | no | | intergenic_variant | MODIFIER |
| 10 | 58738138 | AX-97818119 | 0 | G | A | 0.289 | 2.53E-02 | 8.03E-02 | 2.77E+00 | 2.88E+00 | 7.42E-01 | no | | intergenic_variant | MODIFIER |
| 10 | 58892034 | AX-170450839 | 0 | C | G | 0.289 | 2.53E-02 | 8.03E-02 | 2.77E+00 | 2.88E+00 | 7.42E-01 | no | Sh3rf3 | intron_variant,non_coding_transcript_variant | MODIFIER |
| 10 | 58893468 | AX-170338136 | 0 | T | C | 0.289 | 2.53E-02 | 8.03E-02 | 2.77E+00 | 2.88E+00 | 7.42E-01 | no | Sh3rf3 | intron_variant,non_coding_transcript_variant | MODIFIER |
| 10 | 58921451 | AX-170435953 | 0 | C | G | 0.289 | 2.53E-02 | 8.03E-02 | 2.77E+00 | 2.88E+00 | 7.42E-01 | no | Sh3rf3 | intron_variant,non_coding_transcript_variant | MODIFIER |
| 10 | 58922328 | AX-170464195 | 0 | G | A | 0.289 | 2.53E-02 | 8.03E-02 | 2.77E+00 | 2.88E+00 | 7.42E-01 | no | Sh3rf3 | intron_variant,non_coding_transcript_variant | MODIFIER |
| 10 | 59103823 | AX-170425556 | 0 | C | T | 0.289 | 2.53E-02 | 8.03E-02 | 2.77E+00 | 2.88E+00 | 7.42E-01 | no | Sh3rf3 | intron_variant,non_coding_transcript_variant | MODIFIER |
| 10 | 59104422 | AX-170348060 | 0 | A | C | 0.289 | 2.53E-02 | 8.03E-02 | 2.77E+00 | 2.88E+00 | 7.42E-01 | no | Sh3rf3 | non_coding_transcript_exon_variant | MODIFIER |
| 10 | 59104467 | AX-170305974 | 0 | A | G | 0.289 | 2.53E-02 | 8.03E-02 | 2.77E+00 | 2.88E+00 | 7.42E-01 | no | Sh3rf3 | non_coding_transcript_exon_variant | MODIFIER |
| 10 | 59104494 | AX-170375171 | 0 | A | T | 0.289 | 2.53E-02 | 8.03E-02 | 2.77E+00 | 2.88E+00 | 7.42E-01 | no | Sh3rf3 | non_coding_transcript_exon_variant | MODIFIER |
| 10 | 59104901 | AX-170136277 | 0 | T | C | 0.289 | 2.53E-02 | 8.03E-02 | 2.77E+00 | 2.88E+00 | 7.42E-01 | no | Sh3rf3 | downstream_gene_variant | MODIFIER |
| 10 | 59106008 | AX-170174778 | 0 | T | C | 0.289 | 2.53E-02 | 8.03E-02 | 2.77E+00 | 2.88E+00 | 7.42E-01 | no | Sh3rf3 | downstream_gene_variant | MODIFIER |
| 10 | 59622782 | AX-170328722 | 0 | G | A | 0.289 | 2.53E-02 | 8.03E-02 | 2.77E+00 | 2.88E+00 | 7.42E-01 | no | | regulatory_region_variant | MODIFIER |
| 1 | 90822253 | AX-170426305 | 0 | A | G | 0.158 | 5.66E-02 | 1.74E-01 | 2.64E+00 | 2.73E+00 | 7.43E-01 | no | Col6a3 | synonymous_variant | LOW |
| 2 | 159706394 | AX-170462313 | 1 | A | T | 0.125 | 6.16E-02 | 1.89E-01 | 2.65E+00 | 2.73E+00 | 7.43E-01 | no | | intergenic_variant | MODIFIER |
| 4 | 137388339 | AX-170421388 | 0 | T | C | 0.167 | 3.74E-02 | 1.10E-01 | 2.59E+00 | 2.66E+00 | 7.43E-01 | no | 2810405F17Rik | non_coding_transcript_exon_variant | MODIFIER |
| 3 | 30335793 | AX-170215758 | 0 | A | T | 0.158 | -3.25E-02 | 1.01E-01 | 2.68E+00 | 2.76E+00 | 7.43E-01 | no | Mecom | intron_variant | MODIFIER |
| 5 | 71422036 | AX-170522365 | 1 | C | T | 0.455 | 3.19E-02 | 9.71E-02 | 2.63E+00 | 2.71E+00 | 7.43E-01 | no | | intergenic_variant | MODIFIER |
| 18 | 47565660 | AX-170494842 | 0 | C | T | 0.053 | 7.54E-02 | 2.36E-01 | 2.65E+00 | 2.77E+00 | 7.43E-01 | no | AC141639.4 | intron_variant,non_coding_transcript_variant | MODIFIER |
| 7 | 117148073 | AX-170335391 | 1 | C | T | 0.42 | -2.89E-02 | 9.03E-02 | 2.73E+00 | 2.80E+00 | 7.43E-01 | no | | intergenic_variant | MODIFIER |
| 2 | 97338980 | AX-170460732 | 0 | T | A | 0.061 | -7.21E-02 | 2.24E-01 | 2.65E+00 | 2.75E+00 | 7.43E-01 | no | Lrrc4c | intron_variant | MODIFIER |
| 2 | 100199939 | AX-170185399 | 0 | C | A | 0.061 | -7.21E-02 | 2.24E-01 | 2.65E+00 | 2.75E+00 | 7.43E-01 | no | | intergenic_variant | MODIFIER |
| 2 | 106419240 | AX-97835967 | 0 | G | A | 0.061 | -7.21E-02 | 2.24E-01 | 2.65E+00 | 2.75E+00 | 7.43E-01 | no | | intergenic_variant | MODIFIER |
| 8 | 47505492 | AX-97792019 | 1 | G | A | 0.152 | 4.76E-02 | 1.48E-01 | 2.76E+00 | 2.81E+00 | 7.43E-01 | no | Trappc11 | intron_variant | MODIFIER |
| 1 | 147193808 | AX-170036047 | 1 | A | G | 0.196 | 4.67E-02 | 1.42E-01 | 2.65E+00 | 2.69E+00 | 7.43E-01 | no | | intergenic_variant | MODIFIER |
| 1 | 30801998 | AX-170040753 | 1 | T | C | 0.402 | 2.79E-02 | 8.53E-02 | 2.64E+00 | 2.71E+00 | 7.43E-01 | no | Phf3 | downstream_gene_variant | MODIFIER |

| | | | | | | | | | | | | | | | |
|---|---|---|---|---|---|---|---|---|---|---|---|---|---|---|---|
| 8 | 96886034 | AX-170031327 | 0 | C | T | 0.219 | 3.82E-02 | 1.19E-01 | 2.70E+00 | 2.77E+00 | 7.43E-01 | no | | intergenic_variant | MODIFIER |
| 1 | 25534513 | AX-168391054 | 0 | A | G | 0.412 | 2.75E-02 | 8.32E-02 | 2.63E+00 | 2.69E+00 | 7.44E-01 | no | Adgrb3 | intron_variant | MODIFIER |
| 6 | 126537268 | AX-170229246 | 2 | T | C | 0.527 | 2.82E-02 | 7.94E-02 | 2.53E+00 | 2.63E+00 | 7.44E-01 | no | Kcna5 | upstream_gene_variant | MODIFIER |
| 9 | 112822674 | AX-170472317 | 1 | A | C | 0.125 | -3.45E-02 | 1.06E-01 | 2.65E+00 | 2.73E+00 | 7.44E-01 | no | | intergenic_variant | MODIFIER |
| 9 | 113102630 | AX-170209997 | 1 | A | G | 0.125 | -3.45E-02 | 1.06E-01 | 2.65E+00 | 2.73E+00 | 7.44E-01 | no | Gm36251 | intron_variant,non_coding_transcript_variant | MODIFIER |
| 2 | 147489172 | AX-170019309 | 0 | C | T | 0.149 | 6.24E-02 | 1.93E-01 | 2.67E+00 | 2.74E+00 | 7.44E-01 | no | | intergenic_variant | MODIFIER |
| 7 | 81982277 | AX-170216884 | 0 | T | C | 0.123 | 6.71E-02 | 2.07E-01 | 2.65E+00 | 2.74E+00 | 7.44E-01 | no | Bnc1 | intron_variant | MODIFIER |
| 9 | 35109322 | AX-170118468 | 0 | A | G | 0.43 | -2.68E-02 | 8.40E-02 | 2.73E+00 | 2.80E+00 | 7.44E-01 | no | St3gal4 | intron_variant | MODIFIER |
| 8 | 39650423 | AX-170030537 | 0 | C | A | 0.158 | 4.34E-02 | 1.36E-01 | 2.71E+00 | 2.78E+00 | 7.44E-01 | no | | intergenic_variant | MODIFIER |
| 8 | 41691347 | AX-170287793 | 0 | C | A | 0.158 | 4.34E-02 | 1.36E-01 | 2.71E+00 | 2.78E+00 | 7.44E-01 | no | | intergenic_variant | MODIFIER |
| 8 | 45597198 | AX-170079958 | 0 | G | A | 0.158 | 4.34E-02 | 1.36E-01 | 2.71E+00 | 2.78E+00 | 7.44E-01 | no | Sorbs2 | intron_variant | MODIFIER |
| 8 | 57453851 | AX-170030800 | 0 | A | G | 0.158 | -4.47E-02 | 1.35E-01 | 2.62E+00 | 2.69E+00 | 7.44E-01 | no | Scrg1 | upstream_gene_variant | MODIFIER |
| 12 | 113293025 | AX-170007563 | 1 | C | T | 0.134 | -6.38E-02 | 1.99E-01 | 2.71E+00 | 2.77E+00 | 7.44E-01 | no | Ighg2c | upstream_gene_variant | MODIFIER |
| 12 | 118155337 | AX-170501509 | 1 | G | A | 0.134 | -6.38E-02 | 1.99E-01 | 2.71E+00 | 2.77E+00 | 7.44E-01 | no | Dnah11 | intron_variant | MODIFIER |
| 2 | 104786695 | AX-170305798 | 1 | C | T | 0.054 | -9.70E-02 | 3.01E-01 | 2.74E+00 | 2.76E+00 | 7.44E-01 | no | Qser1 | synonymous_variant | LOW |
| 1 | 91167136 | AX-170001062 | 0 | G | A | 0.158 | -5.66E-02 | 1.77E-01 | 2.67E+00 | 2.76E+00 | 7.44E-01 | no | Rbm44 | intron_variant | MODIFIER |
| 5 | 27628743 | AX-170138313 | 1 | C | A | 0.054 | -9.65E-02 | 2.99E-01 | 2.74E+00 | 2.77E+00 | 7.44E-01 | no | Dpp6 | intron_variant | MODIFIER |
| 16 | 18959994 | AX-170446618 | 0 | A | G | 0.132 | 2.99E-02 | 9.41E-02 | 2.63E+00 | 2.76E+00 | 7.44E-01 | no | Hira | intron_variant | MODIFIER |
| 16 | 19883079 | AX-97830688 | 0 | A | G | 0.132 | 2.99E-02 | 9.41E-02 | 2.63E+00 | 2.76E+00 | 7.44E-01 | no | A930003A15Rik | intron_variant,non_coding_transcript_variant | MODIFIER |
| 7 | 40164095 | AX-170342449 | 0 | C | A | 0.368 | 3.05E-02 | 9.11E-02 | 2.59E+00 | 2.68E+00 | 7.44E-01 | no | | intergenic_variant | MODIFIER |
| 8 | 30550381 | AX-170030433 | 1 | C | A | 0.17 | 3.72E-02 | 1.16E-01 | 2.69E+00 | 2.76E+00 | 7.45E-01 | no | | intergenic_variant | MODIFIER |
| 8 | 97508518 | AX-97851568 | 0 | C | A | 0.219 | 3.62E-02 | 1.14E-01 | 2.70E+00 | 2.79E+00 | 7.45E-01 | no | | intergenic_variant | MODIFIER |
| 8 | 97822432 | AX-97843651 | 0 | C | T | 0.219 | 3.62E-02 | 1.14E-01 | 2.70E+00 | 2.79E+00 | 7.45E-01 | no | | intergenic_variant | MODIFIER |
| 10 | 118896803 | AX-170236282 | 0 | A | G | 0.333 | 2.36E-02 | 7.40E-02 | 2.67E+00 | 2.76E+00 | 7.45E-01 | no | Gm26976 | downstream_gene_variant | MODIFIER |
| 10 | 118898321 | AX-170131542 | 0 | G | C | 0.333 | 2.36E-02 | 7.40E-02 | 2.67E+00 | 2.76E+00 | 7.45E-01 | no | Gm26976 | downstream_gene_variant | MODIFIER |
| 10 | 128258964 | AX-170300517 | 0 | C | T | 0.333 | -2.34E-02 | 7.22E-02 | 2.62E+00 | 2.73E+00 | 7.45E-01 | no | Apon | downstream_gene_variant | MODIFIER |
| 9 | 65592500 | AX-97804191 | 0 | T | C | 0.193 | 4.09E-02 | 1.28E-01 | 2.72E+00 | 2.77E+00 | 7.45E-01 | no | Pif1 | intron_variant | MODIFIER |
| 4 | 100089468 | AX-97801774 | 0 | C | T | 0.351 | 3.22E-02 | 9.90E-02 | 2.64E+00 | 2.72E+00 | 7.45E-01 | no | | intergenic_variant | MODIFIER |
| 4 | 100519065 | AX-170089324 | 0 | C | T | 0.351 | 3.22E-02 | 9.90E-02 | 2.64E+00 | 2.72E+00 | 7.45E-01 | no | Ube2u | intron_variant | MODIFIER |
| 4 | 100546888 | AX-97830957 | 0 | C | T | 0.351 | 3.22E-02 | 9.90E-02 | 2.64E+00 | 2.72E+00 | 7.45E-01 | no | Ube2u | intron_variant | MODIFIER |
| 4 | 100548003 | AX-170183813 | 0 | T | C | 0.351 | 3.22E-02 | 9.90E-02 | 2.64E+00 | 2.72E+00 | 7.45E-01 | no | Ube2u | intron_variant | MODIFIER |
| 4 | 100829781 | AX-97792237 | 0 | G | A | 0.351 | 3.22E-02 | 9.90E-02 | 2.64E+00 | 2.72E+00 | 7.45E-01 | no | Cachd1 | intron_variant | MODIFIER |
| 4 | 101438421 | AX-170336211 | 0 | C | A | 0.351 | 3.22E-02 | 9.90E-02 | 2.64E+00 | 2.72E+00 | 7.45E-01 | no | Ak4 | intron_variant | MODIFIER |
| 4 | 101450863 | AX-170506273 | 0 | T | A | 0.351 | 3.22E-02 | 9.90E-02 | 2.64E+00 | 2.72E+00 | 7.45E-01 | no | Ak4 | intron_variant | MODIFIER |
| 4 | 101456134 | AX-170311015 | 0 | A | G | 0.351 | 3.22E-02 | 9.90E-02 | 2.64E+00 | 2.72E+00 | 7.45E-01 | no | Ak4 | intron_variant | MODIFIER |
| 4 | 101457745 | AX-170459692 | 0 | A | G | 0.351 | 3.22E-02 | 9.90E-02 | 2.64E+00 | 2.72E+00 | 7.45E-01 | no | Ak4 | intron_variant | MODIFIER |
| 4 | 101458681 | AX-170070174 | 0 | A | G | 0.351 | 3.22E-02 | 9.90E-02 | 2.64E+00 | 2.72E+00 | 7.45E-01 | no | Ak4 | intron_variant | MODIFIER |
| 4 | 101500071 | AX-168401404 | 0 | T | C | 0.351 | 3.22E-02 | 9.90E-02 | 2.64E+00 | 2.72E+00 | 7.45E-01 | no | Dnajc6 | intron_variant | MODIFIER |
| 4 | 101500145 | AX-170400442 | 0 | T | C | 0.351 | 3.22E-02 | 9.90E-02 | 2.64E+00 | 2.72E+00 | 7.45E-01 | no | Dnajc6 | intron_variant | MODIFIER |
| 4 | 101510153 | AX-170213769 | 0 | T | A | 0.351 | 3.22E-02 | 9.90E-02 | 2.64E+00 | 2.72E+00 | 7.45E-01 | no | Dnajc6 | intron_variant | MODIFIER |
| 10 | 58667200 | AX-170044975 | 0 | C | T | 0.298 | 2.47E-02 | 7.94E-02 | 2.75E+00 | 2.86E+00 | 7.45E-01 | no | Edar | intron_variant | MODIFIER |
| 1 | 156184322 | AX-170248369 | 1 | C | T | 0.223 | 4.55E-02 | 1.39E-01 | 2.64E+00 | 2.68E+00 | 7.45E-01 | no | Fam163a | intron_variant | MODIFIER |
| 8 | 40437999 | AX-170079852 | 1 | C | A | 0.152 | 4.65E-02 | 1.46E-01 | 2.75E+00 | 2.80E+00 | 7.45E-01 | no | Zdhhc2 | intron_variant | MODIFIER |
| 8 | 40619551 | AX-170528084 | 1 | G | A | 0.152 | 4.65E-02 | 1.46E-01 | 2.75E+00 | 2.80E+00 | 7.45E-01 | no | Mtmr7 | intron_variant,NMD_transcript_variant | MODIFIER |
| 8 | 43290907 | AX-170247183 | 1 | A | G | 0.152 | 4.65E-02 | 1.46E-01 | 2.75E+00 | 2.80E+00 | 7.45E-01 | no | Zfp42 | downstream_gene_variant | MODIFIER |
| 10 | 56314780 | AX-170454623 | 0 | A | G | 0.298 | 2.51E-02 | 8.05E-02 | 2.77E+00 | 2.88E+00 | 7.45E-01 | no | | intergenic_variant | MODIFIER |
| 10 | 57265804 | AX-170415782 | 0 | G | A | 0.298 | 2.51E-02 | 8.05E-02 | 2.77E+00 | 2.88E+00 | 7.45E-01 | no | Gm48055 | downstream_gene_variant | MODIFIER |
| 4 | 55706467 | AX-170022832 | 1 | T | C | 0.375 | -3.39E-02 | 1.07E-01 | 2.69E+00 | 2.77E+00 | 7.45E-01 | no | | intergenic_variant | MODIFIER |
| 5 | 35638693 | AX-170124537 | 0 | C | T | 0.333 | 3.57E-02 | 1.11E-01 | 2.67E+00 | 2.73E+00 | 7.46E-01 | no | | intergenic_variant | MODIFIER |
| 3 | 32581611 | AX-97810193 | 1 | T | C | 0.17 | 3.08E-02 | 9.64E-02 | 2.66E+00 | 2.75E+00 | 7.46E-01 | no | Mfn1 | downstream_gene_variant | MODIFIER |
| 16 | 92852787 | AX-170109529 | 1 | T | C | 0.464 | -2.48E-02 | 7.81E-02 | 2.67E+00 | 2.76E+00 | 7.46E-01 | no | | intergenic_variant | MODIFIER |
| 10 | 118754296 | AX-97823617 | 0 | T | C | 0.342 | 2.37E-02 | 7.47E-02 | 2.67E+00 | 2.76E+00 | 7.46E-01 | no | | intergenic_variant | MODIFIER |
| 10 | 118761537 | AX-170004004 | 0 | G | A | 0.342 | 2.37E-02 | 7.47E-02 | 2.67E+00 | 2.76E+00 | 7.46E-01 | no | | regulatory_region_variant | MODIFIER |
| 10 | 118767046 | AX-170004002 | 0 | G | T | 0.342 | 2.37E-02 | 7.47E-02 | 2.67E+00 | 2.76E+00 | 7.46E-01 | no | | regulatory_region_variant | MODIFIER |

| | | | | | | | | | | | | | | | |
|---|---|---|---|---|---|---|---|---|---|---|---|---|---|---|---|
| 10 | 118792960 | AX-170468903 | 0 | G | C | 0.342 | 2.37E-02 | 7.47E-02 | 2.67E+00 | 2.76E+00 | 7.46E-01 | no | | regulatory_region_variant | MODIFIER |
| 10 | 118796008 | AX-170499564 | 0 | C | T | 0.342 | 2.37E-02 | 7.47E-02 | 2.67E+00 | 2.76E+00 | 7.46E-01 | no | | intergenic_variant | MODIFIER |
| 10 | 118796749 | AX-170310048 | 0 | G | T | 0.342 | 2.37E-02 | 7.47E-02 | 2.67E+00 | 2.76E+00 | 7.46E-01 | no | | regulatory_region_variant | MODIFIER |
| 10 | 118796823 | AX-170381283 | 0 | G | C | 0.342 | 2.37E-02 | 7.47E-02 | 2.67E+00 | 2.76E+00 | 7.46E-01 | no | | regulatory_region_variant | MODIFIER |
| 10 | 118797099 | AX-170004008 | 0 | A | C | 0.342 | 2.37E-02 | 7.47E-02 | 2.67E+00 | 2.76E+00 | 7.46E-01 | no | | regulatory_region_variant | MODIFIER |
| 10 | 118797135 | AX-170273158 | 0 | T | G | 0.342 | 2.37E-02 | 7.47E-02 | 2.67E+00 | 2.76E+00 | 7.46E-01 | no | | regulatory_region_variant | MODIFIER |
| 10 | 118798472 | AX-170317632 | 0 | T | C | 0.342 | 2.37E-02 | 7.47E-02 | 2.67E+00 | 2.76E+00 | 7.46E-01 | no | | regulatory_region_variant | MODIFIER |
| 10 | 118816526 | AX-170004006 | 0 | T | G | 0.342 | 2.37E-02 | 7.47E-02 | 2.67E+00 | 2.76E+00 | 7.46E-01 | no | | intergenic_variant | MODIFIER |
| 10 | 118846218 | AX-170316424 | 0 | T | C | 0.342 | 2.37E-02 | 7.47E-02 | 2.67E+00 | 2.76E+00 | 7.46E-01 | no | | intergenic_variant | MODIFIER |
| 10 | 118846411 | AX-170275047 | 0 | G | A | 0.342 | 2.37E-02 | 7.47E-02 | 2.67E+00 | 2.76E+00 | 7.46E-01 | no | | intergenic_variant | MODIFIER |
| 10 | 118847252 | AX-170257091 | 0 | G | A | 0.342 | 2.37E-02 | 7.47E-02 | 2.67E+00 | 2.76E+00 | 7.46E-01 | no | | regulatory_region_variant | MODIFIER |
| 10 | 118851376 | AX-97830318 | 0 | G | A | 0.342 | 2.37E-02 | 7.47E-02 | 2.67E+00 | 2.76E+00 | 7.46E-01 | no | Dyrk2 | downstream_gene_variant | MODIFIER |
| 10 | 118860674 | AX-97845717 | 0 | A | G | 0.342 | 2.37E-02 | 7.47E-02 | 2.67E+00 | 2.76E+00 | 7.46E-01 | no | Dyrk2 | synonymous_variant | LOW |
| 10 | 119999113 | AX-97819328 | 0 | A | G | 0.342 | 2.37E-02 | 7.47E-02 | 2.67E+00 | 2.76E+00 | 7.46E-01 | no | Grip1 | intron_variant | MODIFIER |
| 1 | 53727762 | AX-170311971 | 1 | C | G | 0.098 | -6.95E-02 | 2.19E-01 | 2.69E+00 | 2.76E+00 | 7.46E-01 | no | | intergenic_variant | MODIFIER |
| 1 | 170416495 | AX-170171125 | 0 | C | G | 0.211 | 4.18E-02 | 1.29E-01 | 2.65E+00 | 2.70E+00 | 7.46E-01 | no | Nos1ap | intron_variant,non_coding_transcript_variant | MODIFIER |
| 18 | 40897222 | AX-170437750 | 1 | A | T | 0.054 | -9.80E-02 | 3.06E-01 | 2.73E+00 | 2.76E+00 | 7.46E-01 | no | AC107756.1 | downstream_gene_variant | MODIFIER |
| 16 | 92090297 | AX-170421266 | 0 | C | A | 0.465 | 2.51E-02 | 7.92E-02 | 2.70E+00 | 2.78E+00 | 7.46E-01 | no | Mrps6 | intron_variant | MODIFIER |
| 6 | 103331748 | AX-170289271 | 0 | G | T | 0.053 | 9.43E-02 | 2.95E-01 | 2.68E+00 | 2.74E+00 | 7.46E-01 | no | | intergenic_variant | MODIFIER |
| 7 | 55577336 | AX-170029048 | 0 | C | G | 0.43 | -3.01E-02 | 9.59E-02 | 2.71E+00 | 2.80E+00 | 7.46E-01 | no | | intergenic_variant | MODIFIER |
| 1 | 9085589 | AX-97810579 | 0 | T | C | 0.509 | -2.23E-02 | 7.08E-02 | 2.77E+00 | 2.86E+00 | 7.46E-01 | no | Sntg1 | intron_variant,non_coding_transcript_variant | MODIFIER |
| 2 | 123256562 | AX-170347971 | 0 | C | G | 0.149 | -6.12E-02 | 1.94E-01 | 2.73E+00 | 2.80E+00 | 7.46E-01 | no | | intergenic_variant | MODIFIER |
| 4 | 100593030 | AX-97799606 | 0 | A | G | 0.342 | 3.12E-02 | 9.66E-02 | 2.64E+00 | 2.72E+00 | 7.46E-01 | no | | intergenic_variant | MODIFIER |
| 4 | 100619898 | AX-170264518 | 0 | A | G | 0.342 | 3.12E-02 | 9.66E-02 | 2.64E+00 | 2.72E+00 | 7.46E-01 | no | | intergenic_variant | MODIFIER |
| 4 | 100620549 | AX-97786928 | 0 | A | G | 0.342 | 3.12E-02 | 9.66E-02 | 2.64E+00 | 2.72E+00 | 7.46E-01 | no | | intergenic_variant | MODIFIER |
| 1 | 127062951 | AX-170245898 | 1 | A | G | 0.527 | 2.79E-02 | 8.56E-02 | 2.63E+00 | 2.70E+00 | 7.46E-01 | no | | intergenic_variant | MODIFIER |
| 1 | 28097012 | AX-170318904 | 0 | A | T | 0.395 | 2.52E-02 | 7.95E-02 | 2.67E+00 | 2.76E+00 | 7.46E-01 | no | | intergenic_variant | MODIFIER |
| 15 | 91202651 | AX-170211659 | 1 | C | T | 0.125 | 5.32E-02 | 1.61E-01 | 2.60E+00 | 2.68E+00 | 7.47E-01 | no | CN725425 | intron_variant | MODIFIER |
| 15 | 94876080 | AX-170487117 | 1 | T | A | 0.125 | 5.32E-02 | 1.61E-01 | 2.60E+00 | 2.68E+00 | 7.47E-01 | no | Tmem117 | intron_variant | MODIFIER |
| 10 | 127166557 | AX-97850861 | 1 | A | G | 0.312 | -2.33E-02 | 7.23E-02 | 2.62E+00 | 2.72E+00 | 7.47E-01 | no | B4galnt1 | intron_variant | MODIFIER |
| 10 | 127911183 | AX-170438810 | 1 | A | G | 0.312 | -2.33E-02 | 7.23E-02 | 2.62E+00 | 2.72E+00 | 7.47E-01 | no | Sdr9c7 | 3_prime_UTR_variant | MODIFIER |
| 3 | 37026248 | AX-170259076 | 1 | G | A | 0.196 | 2.91E-02 | 9.23E-02 | 2.67E+00 | 2.76E+00 | 7.47E-01 | no | 4932438A13Rik | synonymous_variant | LOW |
| 9 | 113245938 | AX-170497942 | 2 | C | T | 0.136 | -3.27E-02 | 1.02E-01 | 2.65E+00 | 2.73E+00 | 7.47E-01 | no | | intergenic_variant | MODIFIER |
| 8 | 102755059 | AX-170105998 | 1 | C | A | 0.214 | 3.54E-02 | 1.12E-01 | 2.68E+00 | 2.75E+00 | 7.47E-01 | no | Cdh11 | intron_variant | MODIFIER |
| 9 | 80724517 | AX-170352445 | 1 | A | G | 0.205 | -3.67E-02 | 1.13E-01 | 2.62E+00 | 2.71E+00 | 7.47E-01 | no | | intergenic_variant | MODIFIER |
| 8 | 37950474 | AX-170030522 | 0 | G | T | 0.158 | 4.30E-02 | 1.36E-01 | 2.72E+00 | 2.78E+00 | 7.47E-01 | no | Sgcz | intron_variant | MODIFIER |
| 1 | 90822948 | AX-97812328 | 1 | A | G | 0.152 | 6.85E-02 | 2.15E-01 | 2.67E+00 | 2.74E+00 | 7.47E-01 | no | Col6a3 | intron_variant | MODIFIER |
| 4 | 137833055 | AX-170089386 | 1 | T | C | 0.161 | 3.64E-02 | 1.10E-01 | 2.59E+00 | 2.68E+00 | 7.47E-01 | no | | intergenic_variant | MODIFIER |
| 18 | 70750557 | AX-170218974 | 1 | A | T | 0.179 | -3.99E-02 | 1.23E-01 | 2.64E+00 | 2.69E+00 | 7.47E-01 | no | | intergenic_variant | MODIFIER |
| 16 | 46201516 | AX-97805594 | 1 | A | G | 0.509 | 2.69E-02 | 7.96E-02 | 2.58E+00 | 2.64E+00 | 7.47E-01 | no | | intergenic_variant | MODIFIER |
| 15 | 92515926 | AX-97848859 | 1 | C | G | 0.152 | 5.77E-02 | 1.78E-01 | 2.64E+00 | 2.70E+00 | 7.47E-01 | no | Pdzrn4 | intron_variant | MODIFIER |
| 10 | 16537809 | AX-170226384 | 0 | T | C | 0.193 | 3.38E-02 | 1.08E-01 | 2.75E+00 | 2.83E+00 | 7.47E-01 | no | | intergenic_variant | MODIFIER |
| 10 | 16552391 | AX-170035480 | 0 | C | T | 0.193 | 3.38E-02 | 1.08E-01 | 2.75E+00 | 2.83E+00 | 7.47E-01 | no | | intergenic_variant | MODIFIER |
| 8 | 95533961 | AX-170522860 | 1 | A | G | 0.223 | 3.66E-02 | 1.16E-01 | 2.68E+00 | 2.75E+00 | 7.47E-01 | no | Ccdc113 | upstream_gene_variant | MODIFIER |
| 1 | 36928774 | AX-170322601 | 0 | G | A | 0.456 | -2.50E-02 | 7.87E-02 | 2.67E+00 | 2.74E+00 | 7.48E-01 | no | Tmem131 | intron_variant | MODIFIER |
| 11 | 31912575 | AX-170502398 | 0 | C | T | 0.079 | 5.39E-02 | 1.71E-01 | 2.64E+00 | 2.75E+00 | 7.48E-01 | no | Cpeb4 | intron_variant | MODIFIER |
| 15 | 95296121 | AX-170190022 | 1 | G | A | 0.161 | 5.08E-02 | 1.54E-01 | 2.59E+00 | 2.69E+00 | 7.48E-01 | no | Nell2 | intron_variant | MODIFIER |
| 9 | 110672864 | AX-97850129 | 0 | A | G | 0.105 | -3.74E-02 | 1.13E-01 | 2.60E+00 | 2.68E+00 | 7.48E-01 | no | Ccdc12 | intron_variant | MODIFIER |
| 4 | 55612958 | AX-97792152 | 0 | C | T | 0.368 | -3.38E-02 | 1.07E-01 | 2.69E+00 | 2.76E+00 | 7.48E-01 | no | Gm12506 | intron_variant,non_coding_transcript_variant | MODIFIER |
| 4 | 55618624 | AX-170307466 | 0 | G | T | 0.368 | -3.38E-02 | 1.07E-01 | 2.69E+00 | 2.76E+00 | 7.48E-01 | no | Gm12506 | intron_variant,non_coding_transcript_variant | MODIFIER |
| 4 | 55618694 | AX-170467401 | 0 | C | T | 0.368 | -3.38E-02 | 1.07E-01 | 2.69E+00 | 2.76E+00 | 7.48E-01 | no | Gm12506 | intron_variant,non_coding_transcript_variant | MODIFIER |
| 4 | 55623277 | AX-170069417 | 0 | G | A | 0.368 | -3.38E-02 | 1.07E-01 | 2.69E+00 | 2.76E+00 | 7.48E-01 | no | Gm12506 | intron_variant,non_coding_transcript_variant | MODIFIER |
| 4 | 55624940 | AX-170069416 | 0 | G | T | 0.368 | -3.38E-02 | 1.07E-01 | 2.69E+00 | 2.76E+00 | 7.48E-01 | no | Gm12506 | intron_variant,non_coding_transcript_variant | MODIFIER |
| 4 | 55626548 | AX-170022835 | 0 | G | A | 0.368 | -3.38E-02 | 1.07E-01 | 2.69E+00 | 2.76E+00 | 7.48E-01 | no | Gm12506 | intron_variant,non_coding_transcript_variant | MODIFIER |

| | | | | | | | | | | | | | | | |
|---|---|---|---|---|---|---|---|---|---|---|---|---|---|---|---|
| 4 | 55626667 | AX-170022829 | 0 | G | T | 0.368 | -3.38E-02 | 1.07E-01 | 2.69E+00 | 2.76E+00 | 7.48E-01 | no | Gm12506 | intron_variant,non_coding_transcript_variant | MODIFIER |
| 4 | 55629348 | AX-170464577 | 0 | C | T | 0.368 | -3.38E-02 | 1.07E-01 | 2.69E+00 | 2.76E+00 | 7.48E-01 | no | Gm12506 | intron_variant,non_coding_transcript_variant | MODIFIER |
| 4 | 55672338 | AX-97816829 | 0 | A | G | 0.368 | -3.38E-02 | 1.07E-01 | 2.69E+00 | 2.76E+00 | 7.48E-01 | no | | regulatory_region_variant | MODIFIER |
| 4 | 55679317 | AX-170359104 | 0 | T | C | 0.368 | -3.38E-02 | 1.07E-01 | 2.69E+00 | 2.76E+00 | 7.48E-01 | no | | intergenic_variant | MODIFIER |
| 4 | 55680924 | AX-170089241 | 0 | A | G | 0.368 | -3.38E-02 | 1.07E-01 | 2.69E+00 | 2.76E+00 | 7.48E-01 | no | | intergenic_variant | MODIFIER |
| 4 | 55697192 | AX-170527045 | 0 | C | A | 0.368 | -3.38E-02 | 1.07E-01 | 2.69E+00 | 2.76E+00 | 7.48E-01 | no | | regulatory_region_variant | MODIFIER |
| 4 | 55697595 | AX-170335287 | 0 | G | T | 0.368 | -3.38E-02 | 1.07E-01 | 2.69E+00 | 2.76E+00 | 7.48E-01 | no | | regulatory_region_variant | MODIFIER |
| 4 | 55717283 | AX-170431892 | 0 | A | G | 0.368 | -3.38E-02 | 1.07E-01 | 2.69E+00 | 2.76E+00 | 7.48E-01 | no | | intergenic_variant | MODIFIER |
| 4 | 55720602 | AX-170022830 | 0 | A | G | 0.368 | -3.38E-02 | 1.07E-01 | 2.69E+00 | 2.76E+00 | 7.48E-01 | no | | intergenic_variant | MODIFIER |
| 4 | 55734260 | AX-170360709 | 0 | T | C | 0.368 | -3.38E-02 | 1.07E-01 | 2.69E+00 | 2.76E+00 | 7.48E-01 | no | | intergenic_variant | MODIFIER |
| 4 | 55736783 | AX-170340062 | 0 | G | A | 0.368 | -3.38E-02 | 1.07E-01 | 2.69E+00 | 2.76E+00 | 7.48E-01 | no | | intergenic_variant | MODIFIER |
| 4 | 55740958 | AX-170316249 | 0 | A | G | 0.368 | -3.38E-02 | 1.07E-01 | 2.69E+00 | 2.76E+00 | 7.48E-01 | no | | intergenic_variant | MODIFIER |
| 4 | 55741387 | AX-170349340 | 0 | A | G | 0.368 | -3.38E-02 | 1.07E-01 | 2.69E+00 | 2.76E+00 | 7.48E-01 | no | | intergenic_variant | MODIFIER |
| 4 | 55752089 | AX-97852499 | 0 | G | C | 0.368 | -3.38E-02 | 1.07E-01 | 2.69E+00 | 2.76E+00 | 7.48E-01 | no | | intergenic_variant | MODIFIER |
| 4 | 55752144 | AX-170022834 | 0 | T | C | 0.368 | -3.38E-02 | 1.07E-01 | 2.69E+00 | 2.76E+00 | 7.48E-01 | no | | intergenic_variant | MODIFIER |
| 4 | 55758108 | AX-170397519 | 0 | T | C | 0.368 | -3.38E-02 | 1.07E-01 | 2.69E+00 | 2.76E+00 | 7.48E-01 | no | | intergenic_variant | MODIFIER |
| 4 | 55760243 | AX-170022833 | 0 | A | G | 0.368 | -3.38E-02 | 1.07E-01 | 2.69E+00 | 2.76E+00 | 7.48E-01 | no | | intergenic_variant | MODIFIER |
| 4 | 55760780 | AX-170225420 | 0 | C | T | 0.368 | -3.38E-02 | 1.07E-01 | 2.69E+00 | 2.76E+00 | 7.48E-01 | no | | intergenic_variant | MODIFIER |
| 4 | 55784351 | AX-170069420 | 0 | G | A | 0.368 | -3.38E-02 | 1.07E-01 | 2.69E+00 | 2.76E+00 | 7.48E-01 | no | | intergenic_variant | MODIFIER |
| 4 | 55784441 | AX-170069419 | 0 | C | T | 0.368 | -3.38E-02 | 1.07E-01 | 2.69E+00 | 2.76E+00 | 7.48E-01 | no | | intergenic_variant | MODIFIER |
| 4 | 55785330 | AX-170382027 | 0 | C | T | 0.368 | -3.38E-02 | 1.07E-01 | 2.69E+00 | 2.76E+00 | 7.48E-01 | no | | intergenic_variant | MODIFIER |
| 4 | 55859905 | AX-170357705 | 0 | C | A | 0.368 | -3.38E-02 | 1.07E-01 | 2.69E+00 | 2.76E+00 | 7.48E-01 | no | | regulatory_region_variant | MODIFIER |
| 4 | 55974038 | AX-170129718 | 0 | C | T | 0.368 | -3.38E-02 | 1.07E-01 | 2.69E+00 | 2.76E+00 | 7.48E-01 | no | Gm12519 | intron_variant,non_coding_transcript_variant | MODIFIER |
| 4 | 56018459 | AX-170393238 | 0 | G | A | 0.368 | -3.38E-02 | 1.07E-01 | 2.69E+00 | 2.76E+00 | 7.48E-01 | no | | intergenic_variant | MODIFIER |
| 4 | 56026051 | AX-170118787 | 0 | G | A | 0.368 | -3.38E-02 | 1.07E-01 | 2.69E+00 | 2.76E+00 | 7.48E-01 | no | | intergenic_variant | MODIFIER |
| 4 | 56026097 | AX-170149363 | 0 | A | G | 0.368 | -3.38E-02 | 1.07E-01 | 2.69E+00 | 2.76E+00 | 7.48E-01 | no | | intergenic_variant | MODIFIER |
| 4 | 56037031 | AX-170373481 | 0 | G | A | 0.368 | -3.38E-02 | 1.07E-01 | 2.69E+00 | 2.76E+00 | 7.48E-01 | no | | intergenic_variant | MODIFIER |
| 4 | 56037802 | AX-170224560 | 0 | C | T | 0.368 | -3.38E-02 | 1.07E-01 | 2.69E+00 | 2.76E+00 | 7.48E-01 | no | | intergenic_variant | MODIFIER |
| 4 | 56198745 | AX-97800617 | 0 | C | A | 0.368 | -3.38E-02 | 1.07E-01 | 2.69E+00 | 2.76E+00 | 7.48E-01 | no | | intergenic_variant | MODIFIER |
| 4 | 56233354 | AX-97813530 | 0 | G | T | 0.368 | -3.38E-02 | 1.07E-01 | 2.69E+00 | 2.76E+00 | 7.48E-01 | no | | intergenic_variant | MODIFIER |
| 4 | 56255564 | AX-170022839 | 0 | C | T | 0.368 | -3.38E-02 | 1.07E-01 | 2.69E+00 | 2.76E+00 | 7.48E-01 | no | | intergenic_variant | MODIFIER |
| 4 | 56255604 | AX-170022838 | 0 | T | C | 0.368 | -3.38E-02 | 1.07E-01 | 2.69E+00 | 2.76E+00 | 7.48E-01 | no | | intergenic_variant | MODIFIER |
| 4 | 56269802 | AX-170340416 | 0 | G | A | 0.368 | -3.38E-02 | 1.07E-01 | 2.69E+00 | 2.76E+00 | 7.48E-01 | no | | intergenic_variant | MODIFIER |
| 4 | 56270225 | AX-170140733 | 0 | A | G | 0.368 | -3.38E-02 | 1.07E-01 | 2.69E+00 | 2.76E+00 | 7.48E-01 | no | | intergenic_variant | MODIFIER |
| 4 | 56272228 | AX-170312399 | 0 | A | G | 0.368 | -3.38E-02 | 1.07E-01 | 2.69E+00 | 2.76E+00 | 7.48E-01 | no | | intergenic_variant | MODIFIER |
| 4 | 56305180 | AX-170022836 | 0 | T | G | 0.368 | -3.38E-02 | 1.07E-01 | 2.69E+00 | 2.76E+00 | 7.48E-01 | no | | regulatory_region_variant | MODIFIER |
| 4 | 56413683 | AX-97834738 | 0 | T | C | 0.368 | -3.38E-02 | 1.07E-01 | 2.69E+00 | 2.76E+00 | 7.48E-01 | no | | intergenic_variant | MODIFIER |
| 4 | 56478884 | AX-170420756 | 0 | G | A | 0.368 | -3.38E-02 | 1.07E-01 | 2.69E+00 | 2.76E+00 | 7.48E-01 | no | | intergenic_variant | MODIFIER |
| 4 | 56487275 | AX-170402439 | 0 | G | C | 0.368 | -3.38E-02 | 1.07E-01 | 2.69E+00 | 2.76E+00 | 7.48E-01 | no | | intergenic_variant | MODIFIER |
| 4 | 56498483 | AX-170350412 | 0 | G | A | 0.368 | -3.38E-02 | 1.07E-01 | 2.69E+00 | 2.76E+00 | 7.48E-01 | no | | intergenic_variant | MODIFIER |
| 4 | 56504926 | AX-170436365 | 0 | A | C | 0.368 | -3.38E-02 | 1.07E-01 | 2.69E+00 | 2.76E+00 | 7.48E-01 | no | | intergenic_variant | MODIFIER |
| 4 | 56539334 | AX-97820880 | 0 | C | T | 0.368 | -3.38E-02 | 1.07E-01 | 2.69E+00 | 2.76E+00 | 7.48E-01 | no | | regulatory_region_variant | MODIFIER |
| 4 | 56711272 | AX-170022855 | 0 | C | A | 0.368 | -3.38E-02 | 1.07E-01 | 2.69E+00 | 2.76E+00 | 7.48E-01 | no | | intergenic_variant | MODIFIER |
| 4 | 56745159 | AX-97841500 | 0 | T | A | 0.368 | -3.38E-02 | 1.07E-01 | 2.69E+00 | 2.76E+00 | 7.48E-01 | no | Elp1 | downstream_gene_variant | MODIFIER |
| 4 | 56775461 | AX-170022854 | 0 | G | C | 0.368 | -3.38E-02 | 1.07E-01 | 2.69E+00 | 2.76E+00 | 7.48E-01 | no | Elp1 | synonymous_variant | LOW |
| 4 | 56821352 | AX-97804082 | 0 | C | A | 0.368 | -3.38E-02 | 1.07E-01 | 2.69E+00 | 2.76E+00 | 7.48E-01 | no | Ctnnal1 | intron_variant | MODIFIER |
| 4 | 56841251 | AX-97798418 | 0 | A | G | 0.368 | -3.38E-02 | 1.07E-01 | 2.69E+00 | 2.76E+00 | 7.48E-01 | no | Ctnnal1 | intron_variant | MODIFIER |
| 4 | 56867883 | AX-97835598 | 0 | T | G | 0.368 | -3.38E-02 | 1.07E-01 | 2.69E+00 | 2.76E+00 | 7.48E-01 | no | Ctnnal1 | upstream_gene_variant | MODIFIER |
| 4 | 56886684 | AX-170022856 | 0 | A | G | 0.368 | -3.38E-02 | 1.07E-01 | 2.69E+00 | 2.76E+00 | 7.48E-01 | no | Tmem245 | intron_variant | MODIFIER |
| 4 | 56886712 | AX-170022849 | 0 | G | A | 0.368 | -3.38E-02 | 1.07E-01 | 2.69E+00 | 2.76E+00 | 7.48E-01 | no | Tmem245 | intron_variant | MODIFIER |
| 4 | 56891893 | AX-97804212 | 0 | G | A | 0.368 | -3.38E-02 | 1.07E-01 | 2.69E+00 | 2.76E+00 | 7.48E-01 | no | Tmem245 | intron_variant | MODIFIER |
| 4 | 56914427 | AX-97797874 | 0 | C | A | 0.368 | -3.38E-02 | 1.07E-01 | 2.69E+00 | 2.76E+00 | 7.48E-01 | no | Tmem245 | intron_variant | MODIFIER |
| 4 | 56922157 | AX-170022858 | 0 | G | A | 0.368 | -3.38E-02 | 1.07E-01 | 2.69E+00 | 2.76E+00 | 7.48E-01 | no | Tmem245 | intron_variant | MODIFIER |
| 4 | 56950482 | AX-97791772 | 0 | G | C | 0.368 | -3.38E-02 | 1.07E-01 | 2.69E+00 | 2.76E+00 | 7.48E-01 | no | Tmem245 | upstream_gene_variant | MODIFIER |

| | | | | | | | | | | | | | | | |
|---|---|---|---|---|---|---|---|---|---|---|---|---|---|---|---|
| 4 | 56953014 | AX-170177138 | 0 | G | A | 0.368 | -3.38E-02 | 1.07E-01 | 2.69E+00 | 2.76E+00 | 7.48E-01 | no | Frrs1l | downstream_gene_variant | MODIFIER |
| 4 | 56960261 | AX-170180033 | 0 | C | T | 0.368 | -3.38E-02 | 1.07E-01 | 2.69E+00 | 2.76E+00 | 7.48E-01 | no | Frrs1l | 3_prime_UTR_variant | MODIFIER |
| 4 | 56960301 | AX-170455974 | 0 | G | A | 0.368 | -3.38E-02 | 1.07E-01 | 2.69E+00 | 2.76E+00 | 7.48E-01 | no | Frrs1l | 3_prime_UTR_variant | MODIFIER |
| 4 | 56966059 | AX-97782988 | 0 | A | G | 0.368 | -3.38E-02 | 1.07E-01 | 2.69E+00 | 2.76E+00 | 7.48E-01 | no | Frrs1l | intron_variant | MODIFIER |
| 4 | 56966792 | AX-170022859 | 0 | A | G | 0.368 | -3.38E-02 | 1.07E-01 | 2.69E+00 | 2.76E+00 | 7.48E-01 | no | Frrs1l | intron_variant | MODIFIER |
| 4 | 56971084 | AX-170455831 | 0 | A | G | 0.368 | -3.38E-02 | 1.07E-01 | 2.69E+00 | 2.76E+00 | 7.48E-01 | no | Frrs1l | intron_variant | MODIFIER |
| 4 | 56987545 | AX-97808252 | 0 | A | G | 0.368 | -3.38E-02 | 1.07E-01 | 2.69E+00 | 2.76E+00 | 7.48E-01 | no | Frrs1l | intron_variant | MODIFIER |
| 4 | 57008984 | AX-97803873 | 0 | C | T | 0.368 | -3.38E-02 | 1.07E-01 | 2.69E+00 | 2.76E+00 | 7.48E-01 | no | Epb41l4b | intron_variant | MODIFIER |
| 4 | 57052826 | AX-170419663 | 0 | G | A | 0.368 | -3.38E-02 | 1.07E-01 | 2.69E+00 | 2.76E+00 | 7.48E-01 | no | Epb41l4b | intron_variant | MODIFIER |
| 4 | 57053830 | AX-170441202 | 0 | C | G | 0.368 | -3.38E-02 | 1.07E-01 | 2.69E+00 | 2.76E+00 | 7.48E-01 | no | Epb41l4b | intron_variant | MODIFIER |
| 4 | 57061518 | AX-170069438 | 0 | C | T | 0.368 | -3.38E-02 | 1.07E-01 | 2.69E+00 | 2.76E+00 | 7.48E-01 | no | Epb41l4b | downstream_gene_variant | MODIFIER |
| 4 | 57063147 | AX-170356565 | 0 | T | C | 0.368 | -3.38E-02 | 1.07E-01 | 2.69E+00 | 2.76E+00 | 7.48E-01 | no | Epb41l4b | synonymous_variant | LOW |
| 4 | 57068898 | AX-170286819 | 0 | C | T | 0.368 | -3.38E-02 | 1.07E-01 | 2.69E+00 | 2.76E+00 | 7.48E-01 | no | Epb41l4b | intron_variant | MODIFIER |
| 4 | 57074426 | AX-170253347 | 0 | A | T | 0.368 | -3.38E-02 | 1.07E-01 | 2.69E+00 | 2.76E+00 | 7.48E-01 | no | Epb41l4b | intron_variant | MODIFIER |
| 4 | 57078117 | AX-170196654 | 0 | T | C | 0.368 | -3.38E-02 | 1.07E-01 | 2.69E+00 | 2.76E+00 | 7.48E-01 | no | Epb41l4b | intron_variant | MODIFIER |
| 4 | 57085528 | AX-97789840 | 0 | G | A | 0.368 | -3.38E-02 | 1.07E-01 | 2.69E+00 | 2.76E+00 | 7.48E-01 | no | Epb41l4b | intron_variant | MODIFIER |
| 4 | 57101316 | AX-170419695 | 0 | G | A | 0.368 | -3.38E-02 | 1.07E-01 | 2.69E+00 | 2.76E+00 | 7.48E-01 | no | Epb41l4b | intron_variant | MODIFIER |
| 4 | 57203906 | AX-170089247 | 0 | A | G | 0.368 | -3.38E-02 | 1.07E-01 | 2.69E+00 | 2.76E+00 | 7.48E-01 | no | Ptpn3 | intron_variant | MODIFIER |
| 5 | 126769954 | AX-170184391 | 1 | G | A | 0.473 | 3.13E-02 | 9.74E-02 | 2.64E+00 | 2.72E+00 | 7.48E-01 | no | | regulatory_region_variant | MODIFIER |
| 7 | 79019130 | AX-170369833 | 0 | C | T | 0.105 | 6.40E-02 | 2.02E-01 | 2.63E+00 | 2.74E+00 | 7.48E-01 | no | Gm26633 | non_coding_transcript_exon_variant | MODIFIER |
| 2 | 118447035 | AX-170018808 | 1 | C | T | 0.134 | 6.33E-02 | 2.00E-01 | 2.69E+00 | 2.74E+00 | 7.48E-01 | no | Eif2ak4 | intron_variant | MODIFIER |
| 9 | 76920549 | AX-170366959 | 0 | G | A | 0.193 | 4.19E-02 | 1.33E-01 | 2.72E+00 | 2.76E+00 | 7.48E-01 | no | | intergenic_variant | MODIFIER |
| 9 | 78881030 | AX-170230568 | 0 | A | T | 0.202 | 4.04E-02 | 1.28E-01 | 2.75E+00 | 2.81E+00 | 7.48E-01 | no | | regulatory_region_variant | MODIFIER |
| 2 | 139331105 | AX-170165420 | 0 | G | A | 0.14 | -6.93E-02 | 2.19E-01 | 2.75E+00 | 2.79E+00 | 7.48E-01 | no | Gm14066 | intron_variant,non_coding_transcript_variant | MODIFIER |
| 18 | 65710457 | AX-170232455 | 0 | G | A | 0.412 | 2.46E-02 | 7.89E-02 | 2.77E+00 | 2.86E+00 | 7.48E-01 | no | Oacyl | intron_variant | MODIFIER |
| 5 | 109735690 | AX-170025870 | 0 | C | G | 0.465 | 3.72E-02 | 1.15E-01 | 2.64E+00 | 2.68E+00 | 7.48E-01 | no | 4930522L14Rik | downstream_gene_variant | MODIFIER |
| 13 | 51147321 | AX-97789280 | 0 | T | C | 0.518 | 2.21E-02 | 6.74E-02 | 2.59E+00 | 2.69E+00 | 7.48E-01 | no | Spin1 | intron_variant | MODIFIER |
| 3 | 148825915 | AX-170131347 | 0 | G | A | 0.351 | 2.37E-02 | 7.38E-02 | 2.63E+00 | 2.71E+00 | 7.49E-01 | no | Adgrl2 | intron_variant | MODIFIER |
| 16 | 93728200 | AX-170148604 | 1 | T | C | 0.402 | -2.67E-02 | 8.03E-02 | 2.58E+00 | 2.67E+00 | 7.49E-01 | no | Dop1b | intron_variant | MODIFIER |
| 9 | 74304563 | AX-170344018 | 0 | A | G | 0.202 | 4.11E-02 | 1.30E-01 | 2.75E+00 | 2.80E+00 | 7.49E-01 | no | | intergenic_variant | MODIFIER |
| 5 | 33008673 | AX-170089442 | 0 | C | T | 0.36 | 3.20E-02 | 1.00E-01 | 2.65E+00 | 2.72E+00 | 7.49E-01 | no | | intergenic_variant | MODIFIER |
| 6 | 121578148 | AX-170168945 | 1 | C | A | 0.071 | 8.65E-02 | 2.74E-01 | 2.73E+00 | 2.77E+00 | 7.49E-01 | no | | intergenic_variant | MODIFIER |
| 7 | 118676478 | AX-97828675 | 1 | C | T | 0.402 | 2.91E-02 | 9.07E-02 | 2.65E+00 | 2.70E+00 | 7.49E-01 | no | Tmc5 | downstream_gene_variant | MODIFIER |
| 7 | 75211777 | AX-170103175 | 0 | C | T | 0.386 | -3.03E-02 | 9.70E-02 | 2.65E+00 | 2.76E+00 | 7.49E-01 | no | Sv2b | intron_variant | MODIFIER |
| 7 | 3131034 | AX-170277457 | 0 | A | T | 0.184 | 2.33E-02 | 7.56E-02 | 2.62E+00 | 2.77E+00 | 7.49E-01 | no | Speer9-ps1 | intron_variant,non_coding_transcript_variant | MODIFIER |
| 9 | 33773963 | AX-170230514 | 1 | A | C | 0.42 | 2.81E-02 | 8.75E-02 | 2.63E+00 | 2.71E+00 | 7.50E-01 | no | | intergenic_variant | MODIFIER |
| 4 | 56025116 | AX-170207361 | 0 | G | A | 0.377 | -3.25E-02 | 1.04E-01 | 2.68E+00 | 2.77E+00 | 7.50E-01 | no | | intergenic_variant | MODIFIER |
| 5 | 128025422 | AX-170385796 | 1 | G | A | 0.429 | 2.60E-02 | 7.99E-02 | 2.58E+00 | 2.70E+00 | 7.50E-01 | no | Tmem132d | intron_variant | MODIFIER |
| 1 | 75040083 | AX-170000816 | 0 | C | G | 0.132 | 5.46E-02 | 1.71E-01 | 2.62E+00 | 2.72E+00 | 7.50E-01 | no | Nhej1 | intron_variant | MODIFIER |
| 1 | 116482828 | AX-170482668 | 1 | G | A | 0.17 | 6.39E-02 | 1.95E-01 | 2.59E+00 | 2.68E+00 | 7.50E-01 | no | Cntnap5a | intron_variant | MODIFIER |
| 20 | 76922021 | AX-170409237 | 1 | T | A | 0.25 | 1.97E-02 | 6.24E-02 | 2.63E+00 | 2.73E+00 | 7.50E-01 | no | Gm14706 | upstream_gene_variant | MODIFIER |
| 5 | 126399391 | AX-97822478 | 1 | T | C | 0.482 | 2.59E-02 | 8.15E-02 | 2.61E+00 | 2.73E+00 | 7.50E-01 | no | | intergenic_variant | MODIFIER |
| 9 | 60921675 | AX-170032490 | 1 | T | C | 0.214 | 3.47E-02 | 1.11E-01 | 2.68E+00 | 2.76E+00 | 7.50E-01 | no | Gm47923 | intron_variant,non_coding_transcript_variant | MODIFIER |
| 4 | 90072157 | AX-170325469 | 1 | T | C | 0.33 | 3.43E-02 | 1.08E-01 | 2.64E+00 | 2.71E+00 | 7.50E-01 | no | | intergenic_variant | MODIFIER |
| 1 | 60496333 | AX-170469185 | 1 | T | G | 0.098 | 6.44E-02 | 2.08E-01 | 2.68E+00 | 2.77E+00 | 7.50E-01 | no | Raph1 | intron_variant | MODIFIER |
| 5 | 28177764 | AX-97797562 | 0 | A | G | 0.368 | -3.07E-02 | 9.98E-02 | 2.70E+00 | 2.80E+00 | 7.50E-01 | no | | intergenic_variant | MODIFIER |
| 2 | 160062864 | AX-97797685 | 0 | C | T | 0.158 | 5.58E-02 | 1.75E-01 | 2.62E+00 | 2.72E+00 | 7.50E-01 | no | | regulatory_region_variant | MODIFIER |
| 15 | 81980674 | AX-170130153 | 0 | A | G | 0.553 | 2.70E-02 | 8.17E-02 | 2.59E+00 | 2.66E+00 | 7.50E-01 | no | 1700029P11Rik | missense_variant | MODERATE |
| 8 | 117843401 | AX-170493851 | 1 | G | T | 0.259 | 3.03E-02 | 9.89E-02 | 2.68E+00 | 2.79E+00 | 7.50E-01 | no | | intergenic_variant | MODIFIER |
| 8 | 75554897 | AX-170122950 | 1 | C | A | 0.188 | -4.36E-02 | 1.38E-01 | 2.67E+00 | 2.73E+00 | 7.51E-01 | no | Iqcm | missense_variant | MODERATE |
| 20 | 68920001 | AX-170471620 | 1 | T | C | 0.223 | 2.01E-02 | 6.36E-02 | 2.63E+00 | 2.73E+00 | 7.51E-01 | no | Gm8344 | downstream_gene_variant | MODIFIER |
| 12 | 113579185 | AX-168392304 | 0 | C | T | 0.123 | -6.07E-02 | 1.96E-01 | 2.71E+00 | 2.79E+00 | 7.51E-01 | no | Ighv5-2 | upstream_gene_variant | MODIFIER |
| 5 | 111548666 | AX-170160475 | 0 | C | T | 0.456 | -3.37E-02 | 1.08E-01 | 2.76E+00 | 2.83E+00 | 7.51E-01 | no | | regulatory_region_variant | MODIFIER |
| 9 | 27918636 | AX-170478571 | 1 | C | T | 0.455 | 2.96E-02 | 9.19E-02 | 2.62E+00 | 2.69E+00 | 7.51E-01 | no | Opcml | intron_variant | MODIFIER |

| | | | | | | | | | | | | | | | |
|---|---|---|---|---|---|---|---|---|---|---|---|---|---|---|---|
| 1 | 118022406 | AX-170085856 | 0 | G | A | 0.167 | 6.24E-02 | 1.94E-01 | 2.62E+00 | 2.70E+00 | 7.51E-01 | no | Gm28358 | intron_variant,non_coding_transcript_variant | MODIFIER |
| 18 | 58581475 | AX-170109473 | 1 | A | C | 0.384 | 3.34E-02 | 9.52E-02 | 2.53E+00 | 2.60E+00 | 7.51E-01 | no | Slc27a6 | intron_variant | MODIFIER |
| 12 | 17496977 | AX-170048483 | 0 | C | T | 0.044 | -5.45E-02 | 1.71E-01 | 2.59E+00 | 2.72E+00 | 7.51E-01 | no | | intergenic_variant | MODIFIER |
| 12 | 113156847 | AX-170235055 | 0 | C | T | 0.14 | -5.99E-02 | 1.93E-01 | 2.69E+00 | 2.76E+00 | 7.51E-01 | no | Crip1 | downstream_gene_variant | MODIFIER |
| 8 | 74575898 | AX-170337518 | 1 | T | C | 0.188 | -4.32E-02 | 1.38E-01 | 2.67E+00 | 2.73E+00 | 7.51E-01 | no | | intergenic_variant | MODIFIER |
| 5 | 100240072 | AX-170072875 | 0 | G | T | 0.43 | -3.37E-02 | 1.09E-01 | 2.72E+00 | 2.77E+00 | 7.51E-01 | no | Gm8091 | intron_variant,non_coding_transcript_variant | MODIFIER |
| 1 | 156466736 | AX-170110777 | 2 | C | A | 0.218 | 4.50E-02 | 1.39E-01 | 2.62E+00 | 2.67E+00 | 7.51E-01 | no | Soat1 | synonymous_variant | LOW |
| 6 | 101094931 | AX-170337277 | 0 | G | A | 0.079 | 6.85E-02 | 2.21E-01 | 2.67E+00 | 2.76E+00 | 7.52E-01 | no | Gm43948 | intron_variant,non_coding_transcript_variant | MODIFIER |
| 6 | 120613616 | AX-170105425 | 0 | C | A | 0.079 | 6.85E-02 | 2.21E-01 | 2.67E+00 | 2.76E+00 | 7.52E-01 | no | | intergenic_variant | MODIFIER |
| 4 | 80931838 | AX-97785622 | 0 | C | T | 0.237 | 3.43E-02 | 1.01E-01 | 2.55E+00 | 2.63E+00 | 7.52E-01 | no | Lurap1l | intron_variant | MODIFIER |
| 3 | 38145541 | AX-170268709 | 0 | T | C | 0.202 | 2.90E-02 | 9.32E-02 | 2.66E+00 | 2.75E+00 | 7.52E-01 | no | | regulatory_region_variant | MODIFIER |
| 3 | 38327220 | AX-170142774 | 0 | A | T | 0.202 | 2.90E-02 | 9.32E-02 | 2.66E+00 | 2.75E+00 | 7.52E-01 | no | | intergenic_variant | MODIFIER |
| 1 | 66600241 | AX-168405609 | 0 | T | G | 0.123 | 7.30E-02 | 2.32E-01 | 2.66E+00 | 2.72E+00 | 7.52E-01 | no | Unc80 | intron_variant | MODIFIER |
| 1 | 75410705 | AX-170377211 | 0 | C | T | 0.123 | 7.30E-02 | 2.32E-01 | 2.66E+00 | 2.72E+00 | 7.52E-01 | no | Speg | synonymous_variant | LOW |
| 1 | 78825646 | AX-97812145 | 0 | G | A | 0.123 | 7.30E-02 | 2.32E-01 | 2.66E+00 | 2.72E+00 | 7.52E-01 | no | Gm29187 | downstream_gene_variant | MODIFIER |
| 5 | 127412914 | AX-170283801 | 0 | C | T | 0.421 | -2.52E-02 | 8.23E-02 | 2.66E+00 | 2.77E+00 | 7.52E-01 | no | Tmem132c | intron_variant | MODIFIER |
| 12 | 93451022 | AX-97782654 | 0 | G | A | 0.061 | -5.84E-02 | 1.87E-01 | 2.63E+00 | 2.74E+00 | 7.52E-01 | no | | intergenic_variant | MODIFIER |
| 1 | 65838702 | AX-170514847 | 0 | T | C | 0.149 | 6.93E-02 | 2.17E-01 | 2.63E+00 | 2.70E+00 | 7.52E-01 | no | | intergenic_variant | MODIFIER |
| 8 | 115974053 | AX-170090217 | 0 | C | T | 0.254 | 3.05E-02 | 1.00E-01 | 2.68E+00 | 2.79E+00 | 7.52E-01 | no | Gm45733 | upstream_gene_variant | MODIFIER |
| 11 | 50936441 | AX-170087021 | 1 | G | A | 0.071 | -6.21E-02 | 1.97E-01 | 2.64E+00 | 2.72E+00 | 7.52E-01 | no | 4933414I15Rik | downstream_gene_variant | MODIFIER |
| 11 | 33996297 | AX-170451169 | 0 | C | G | 0.079 | -5.78E-02 | 1.86E-01 | 2.63E+00 | 2.75E+00 | 7.52E-01 | no | Kcnip1 | upstream_gene_variant | MODIFIER |
| 17 | 77182679 | AX-170209791 | 0 | T | C | 0.053 | -5.65E-02 | 1.85E-01 | 2.67E+00 | 2.78E+00 | 7.52E-01 | no | | intergenic_variant | MODIFIER |
| 20 | 145212520 | AX-97808250 | 2 | A | G | 0.155 | 4.25E-02 | 1.36E-01 | 2.64E+00 | 2.73E+00 | 7.53E-01 | no | | intergenic_variant | MODIFIER |
| 1 | 11699972 | AX-170387317 | 2 | T | C | 0.382 | -2.22E-02 | 7.28E-02 | 2.76E+00 | 2.85E+00 | 7.53E-01 | no | A830018L16Rik | intron_variant | MODIFIER |
| 7 | 57652324 | AX-170494893 | 1 | T | A | 0.438 | 3.21E-02 | 1.01E-01 | 2.63E+00 | 2.70E+00 | 7.53E-01 | no | Gabrb3 | intron_variant | MODIFIER |
| 1 | 48208744 | AX-170000431 | 1 | T | A | 0.098 | -6.83E-02 | 2.21E-01 | 2.67E+00 | 2.75E+00 | 7.53E-01 | no | | intergenic_variant | MODIFIER |
| 5 | 125401101 | AX-170406950 | 1 | A | C | 0.438 | -3.33E-02 | 1.08E-01 | 2.73E+00 | 2.80E+00 | 7.53E-01 | no | | regulatory_region_variant | MODIFIER |
| 8 | 94231304 | AX-170404617 | 1 | G | A | 0.214 | -3.70E-02 | 1.20E-01 | 2.69E+00 | 2.75E+00 | 7.53E-01 | no | Nup93 | intron_variant | MODIFIER |
| 13 | 51407647 | AX-170430118 | 0 | A | G | 0.509 | -1.95E-02 | 6.43E-02 | 2.68E+00 | 2.79E+00 | 7.53E-01 | no | S1pr3 | upstream_gene_variant | MODIFIER |
| 8 | 34935893 | AX-170030495 | 0 | C | T | 0.158 | -4.21E-02 | 1.34E-01 | 2.64E+00 | 2.71E+00 | 7.53E-01 | no | Tnks | intron_variant | MODIFIER |
| 8 | 35019223 | AX-170100016 | 0 | G | A | 0.158 | -4.21E-02 | 1.34E-01 | 2.64E+00 | 2.71E+00 | 7.53E-01 | no | | intergenic_variant | MODIFIER |
| 18 | 58395228 | AX-170235403 | 1 | T | C | 0.393 | 3.32E-02 | 9.58E-02 | 2.53E+00 | 2.60E+00 | 7.53E-01 | no | | regulatory_region_variant | MODIFIER |
| 7 | 134874885 | AX-170292239 | 0 | C | T | 0.254 | -2.62E-02 | 8.16E-02 | 2.59E+00 | 2.69E+00 | 7.53E-01 | no | Dock1 | intron_variant | MODIFIER |
| 7 | 134878615 | AX-170517165 | 0 | C | G | 0.254 | -2.62E-02 | 8.16E-02 | 2.59E+00 | 2.69E+00 | 7.53E-01 | no | Dock1 | intron_variant | MODIFIER |
| 7 | 134879807 | AX-170391515 | 0 | T | C | 0.254 | -2.62E-02 | 8.16E-02 | 2.59E+00 | 2.69E+00 | 7.53E-01 | no | Dock1 | intron_variant | MODIFIER |
| 7 | 134885742 | AX-170109389 | 0 | A | G | 0.254 | -2.62E-02 | 8.16E-02 | 2.59E+00 | 2.69E+00 | 7.53E-01 | no | Dock1 | intron_variant | MODIFIER |
| 7 | 134886204 | AX-170506391 | 0 | G | A | 0.254 | -2.62E-02 | 8.16E-02 | 2.59E+00 | 2.69E+00 | 7.53E-01 | no | Dock1 | intron_variant | MODIFIER |
| 7 | 134887602 | AX-170519922 | 0 | C | T | 0.254 | -2.62E-02 | 8.16E-02 | 2.59E+00 | 2.69E+00 | 7.53E-01 | no | Dock1 | intron_variant | MODIFIER |
| 7 | 134980351 | AX-170078948 | 0 | T | C | 0.254 | -2.62E-02 | 8.16E-02 | 2.59E+00 | 2.69E+00 | 7.53E-01 | no | Dock1 | intron_variant | MODIFIER |
| 7 | 135330639 | AX-170299198 | 0 | T | C | 0.254 | -2.62E-02 | 8.16E-02 | 2.59E+00 | 2.69E+00 | 7.53E-01 | no | | intergenic_variant | MODIFIER |
| 7 | 135337591 | AX-170132289 | 0 | G | A | 0.254 | -2.62E-02 | 8.16E-02 | 2.59E+00 | 2.69E+00 | 7.53E-01 | no | | intergenic_variant | MODIFIER |
| 7 | 135378659 | AX-170279593 | 0 | T | C | 0.254 | -2.62E-02 | 8.16E-02 | 2.59E+00 | 2.69E+00 | 7.53E-01 | no | | intergenic_variant | MODIFIER |
| 7 | 135379737 | AX-170455859 | 0 | C | T | 0.254 | -2.62E-02 | 8.16E-02 | 2.59E+00 | 2.69E+00 | 7.53E-01 | no | | intergenic_variant | MODIFIER |
| 7 | 135388475 | AX-170200672 | 0 | T | C | 0.254 | -2.62E-02 | 8.16E-02 | 2.59E+00 | 2.69E+00 | 7.53E-01 | no | | intergenic_variant | MODIFIER |
| 7 | 135420523 | AX-170245098 | 0 | A | C | 0.254 | -2.62E-02 | 8.16E-02 | 2.59E+00 | 2.69E+00 | 7.53E-01 | no | | intergenic_variant | MODIFIER |
| 7 | 135430898 | AX-97837015 | 0 | G | A | 0.254 | -2.62E-02 | 8.16E-02 | 2.59E+00 | 2.69E+00 | 7.53E-01 | no | | intergenic_variant | MODIFIER |
| 7 | 135490382 | AX-170355346 | 0 | A | G | 0.254 | -2.62E-02 | 8.16E-02 | 2.59E+00 | 2.69E+00 | 7.53E-01 | no | Gm36356 | non_coding_transcript_exon_variant | MODIFIER |
| 7 | 135601719 | AX-170529207 | 0 | C | T | 0.254 | -2.62E-02 | 8.16E-02 | 2.59E+00 | 2.69E+00 | 7.53E-01 | no | Ptpre | intron_variant | MODIFIER |
| 7 | 135865972 | AX-97797166 | 0 | G | A | 0.254 | -2.62E-02 | 8.16E-02 | 2.59E+00 | 2.69E+00 | 7.53E-01 | no | | intergenic_variant | MODIFIER |
| 7 | 135897653 | AX-170352028 | 0 | T | A | 0.254 | -2.62E-02 | 8.16E-02 | 2.59E+00 | 2.69E+00 | 7.53E-01 | no | Gm45440 | downstream_gene_variant | MODIFIER |
| 7 | 135926859 | AX-170271397 | 0 | G | A | 0.254 | -2.62E-02 | 8.16E-02 | 2.59E+00 | 2.69E+00 | 7.53E-01 | no | | intergenic_variant | MODIFIER |
| 7 | 135950229 | AX-170263908 | 0 | G | T | 0.254 | -2.62E-02 | 8.16E-02 | 2.59E+00 | 2.69E+00 | 7.53E-01 | no | Gm9341 | upstream_gene_variant | MODIFIER |
| 7 | 135962301 | AX-170525788 | 0 | T | G | 0.254 | -2.62E-02 | 8.16E-02 | 2.59E+00 | 2.69E+00 | 7.53E-01 | no | | intergenic_variant | MODIFIER |
| 7 | 135987701 | AX-170489680 | 0 | A | G | 0.254 | -2.62E-02 | 8.16E-02 | 2.59E+00 | 2.69E+00 | 7.53E-01 | no | | intergenic_variant | MODIFIER |

| | | | | | | | | | | | | | | | |
|---|---|---|---|---|---|---|---|---|---|---|---|---|---|---|---|
| 12 | 105489833 | AX-170121688 | 0 | T | G | 0.123 | -5.60E-02 | 1.83E-01 | 2.69E+00 | 2.77E+00 | 7.54E-01 | no | D430019H16Rik | 3_prime_UTR_variant | MODIFIER |
| 12 | 105504476 | AX-97856070 | 0 | G | A | 0.123 | -5.60E-02 | 1.83E-01 | 2.69E+00 | 2.77E+00 | 7.54E-01 | no | | intergenic_variant | MODIFIER |
| 12 | 105506679 | AX-170115706 | 0 | A | C | 0.123 | -5.60E-02 | 1.83E-01 | 2.69E+00 | 2.77E+00 | 7.54E-01 | no | | intergenic_variant | MODIFIER |
| 12 | 105514596 | AX-170007459 | 0 | G | A | 0.123 | -5.60E-02 | 1.83E-01 | 2.69E+00 | 2.77E+00 | 7.54E-01 | no | AU015791 | intron_variant,non_coding_transcript_variant | MODIFIER |
| 12 | 105537106 | AX-170007456 | 0 | C | T | 0.123 | -5.60E-02 | 1.83E-01 | 2.69E+00 | 2.77E+00 | 7.54E-01 | no | | regulatory_region_variant | MODIFIER |
| 12 | 105556697 | AX-170210477 | 0 | G | A | 0.123 | -5.60E-02 | 1.83E-01 | 2.69E+00 | 2.77E+00 | 7.54E-01 | no | Gm47796 | upstream_gene_variant | MODIFIER |
| 12 | 105556903 | AX-170120941 | 0 | C | T | 0.123 | -5.60E-02 | 1.83E-01 | 2.69E+00 | 2.77E+00 | 7.54E-01 | no | Gm47796 | upstream_gene_variant | MODIFIER |
| 12 | 105557046 | AX-170327394 | 0 | T | C | 0.123 | -5.60E-02 | 1.83E-01 | 2.69E+00 | 2.77E+00 | 7.54E-01 | no | Gm47796 | upstream_gene_variant | MODIFIER |
| 12 | 105573790 | AX-170007458 | 0 | G | A | 0.123 | -5.60E-02 | 1.83E-01 | 2.69E+00 | 2.77E+00 | 7.54E-01 | no | Bdkrb2 | intron_variant | MODIFIER |
| 12 | 105643393 | AX-170188364 | 0 | C | T | 0.123 | -5.60E-02 | 1.83E-01 | 2.69E+00 | 2.77E+00 | 7.54E-01 | no | Atg2b | intron_variant | MODIFIER |
| 12 | 105658852 | AX-170225704 | 0 | G | A | 0.123 | -5.60E-02 | 1.83E-01 | 2.69E+00 | 2.77E+00 | 7.54E-01 | no | Atg2b | missense_variant | MODERATE |
| 12 | 105675045 | AX-170326902 | 0 | A | G | 0.123 | -5.60E-02 | 1.83E-01 | 2.69E+00 | 2.77E+00 | 7.54E-01 | no | Atg2b | intron_variant | MODIFIER |
| 12 | 105698041 | AX-170210480 | 0 | C | T | 0.123 | -5.60E-02 | 1.83E-01 | 2.69E+00 | 2.77E+00 | 7.54E-01 | no | Gskip | intron_variant | MODIFIER |
| 12 | 105710040 | AX-170087327 | 0 | T | C | 0.123 | -5.60E-02 | 1.83E-01 | 2.69E+00 | 2.77E+00 | 7.54E-01 | no | Ak7 | intron_variant | MODIFIER |
| 12 | 105746096 | AX-97807770 | 0 | A | G | 0.123 | -5.60E-02 | 1.83E-01 | 2.69E+00 | 2.77E+00 | 7.54E-01 | no | Ak7 | intron_variant | MODIFIER |
| 12 | 105747811 | AX-170397120 | 0 | C | T | 0.123 | -5.60E-02 | 1.83E-01 | 2.69E+00 | 2.77E+00 | 7.54E-01 | no | Ak7 | intron_variant | MODIFIER |
| 12 | 105748080 | AX-170171584 | 0 | C | T | 0.123 | -5.60E-02 | 1.83E-01 | 2.69E+00 | 2.77E+00 | 7.54E-01 | no | Ak7 | intron_variant | MODIFIER |
| 12 | 105748599 | AX-170050072 | 0 | T | C | 0.123 | -5.60E-02 | 1.83E-01 | 2.69E+00 | 2.77E+00 | 7.54E-01 | no | Ak7 | intron_variant | MODIFIER |
| 12 | 105752310 | AX-170050068 | 0 | T | A | 0.123 | -5.60E-02 | 1.83E-01 | 2.69E+00 | 2.77E+00 | 7.54E-01 | no | Ak7 | intron_variant | MODIFIER |
| 12 | 105752579 | AX-170050071 | 0 | T | C | 0.123 | -5.60E-02 | 1.83E-01 | 2.69E+00 | 2.77E+00 | 7.54E-01 | no | Ak7 | intron_variant | MODIFIER |
| 12 | 105761148 | AX-170050069 | 0 | T | C | 0.123 | -5.60E-02 | 1.83E-01 | 2.69E+00 | 2.77E+00 | 7.54E-01 | no | Ak7 | intron_variant | MODIFIER |
| 12 | 105768694 | AX-170115070 | 0 | T | C | 0.123 | -5.60E-02 | 1.83E-01 | 2.69E+00 | 2.77E+00 | 7.54E-01 | no | Ak7 | missense_variant | MODERATE |
| 12 | 105770776 | AX-170342818 | 0 | A | T | 0.123 | -5.60E-02 | 1.83E-01 | 2.69E+00 | 2.77E+00 | 7.54E-01 | no | Ak7 | intron_variant | MODIFIER |
| 12 | 105791836 | AX-170303331 | 0 | A | G | 0.123 | -5.60E-02 | 1.83E-01 | 2.69E+00 | 2.77E+00 | 7.54E-01 | no | Papola | intron_variant,NMD_transcript_variant | MODIFIER |
| 12 | 105798769 | AX-170386769 | 0 | G | A | 0.123 | -5.60E-02 | 1.83E-01 | 2.69E+00 | 2.77E+00 | 7.54E-01 | no | Papola | intron_variant,NMD_transcript_variant | MODIFIER |
| 12 | 105821404 | AX-170180191 | 0 | T | A | 0.123 | -5.60E-02 | 1.83E-01 | 2.69E+00 | 2.77E+00 | 7.54E-01 | no | Papola | intron_variant,NMD_transcript_variant | MODIFIER |
| 12 | 105821599 | AX-170231318 | 0 | C | T | 0.123 | -5.60E-02 | 1.83E-01 | 2.69E+00 | 2.77E+00 | 7.54E-01 | no | Papola | intron_variant,NMD_transcript_variant | MODIFIER |
| 12 | 105824701 | AX-170463478 | 0 | C | A | 0.123 | -5.60E-02 | 1.83E-01 | 2.69E+00 | 2.77E+00 | 7.54E-01 | no | Papola | intron_variant,NMD_transcript_variant | MODIFIER |
| 12 | 105829765 | AX-170268529 | 0 | T | C | 0.123 | -5.60E-02 | 1.83E-01 | 2.69E+00 | 2.77E+00 | 7.54E-01 | no | Papola | intron_variant,NMD_transcript_variant | MODIFIER |
| 12 | 105847317 | AX-97840633 | 0 | G | C | 0.123 | -5.60E-02 | 1.83E-01 | 2.69E+00 | 2.77E+00 | 7.54E-01 | no | | regulatory_region_variant | MODIFIER |
| 12 | 105861800 | AX-170340447 | 0 | A | G | 0.123 | -5.60E-02 | 1.83E-01 | 2.69E+00 | 2.77E+00 | 7.54E-01 | no | Gm22120 | downstream_gene_variant | MODIFIER |
| 12 | 105861953 | AX-170184593 | 0 | T | G | 0.123 | -5.60E-02 | 1.83E-01 | 2.69E+00 | 2.77E+00 | 7.54E-01 | no | Gm22120 | downstream_gene_variant | MODIFIER |
| 12 | 105862175 | AX-170459857 | 0 | T | C | 0.123 | -5.60E-02 | 1.83E-01 | 2.69E+00 | 2.77E+00 | 7.54E-01 | no | Gm22120 | downstream_gene_variant | MODIFIER |
| 12 | 105862197 | AX-170355386 | 0 | C | T | 0.123 | -5.60E-02 | 1.83E-01 | 2.69E+00 | 2.77E+00 | 7.54E-01 | no | Gm22120 | downstream_gene_variant | MODIFIER |
| 12 | 105862810 | AX-170206319 | 0 | C | T | 0.123 | -5.60E-02 | 1.83E-01 | 2.69E+00 | 2.77E+00 | 7.54E-01 | no | Gm22120 | downstream_gene_variant | MODIFIER |
| 12 | 105895546 | AX-97852020 | 0 | G | A | 0.123 | -5.60E-02 | 1.83E-01 | 2.69E+00 | 2.77E+00 | 7.54E-01 | no | Gm19554 | upstream_gene_variant | MODIFIER |
| 12 | 105904904 | AX-170322421 | 0 | C | T | 0.123 | -5.60E-02 | 1.83E-01 | 2.69E+00 | 2.77E+00 | 7.54E-01 | no | Gm19554 | downstream_gene_variant | MODIFIER |
| 12 | 105914971 | AX-170375751 | 0 | A | G | 0.123 | -5.60E-02 | 1.83E-01 | 2.69E+00 | 2.77E+00 | 7.54E-01 | no | | intergenic_variant | MODIFIER |
| 12 | 105943277 | AX-170356809 | 0 | T | G | 0.123 | -5.60E-02 | 1.83E-01 | 2.69E+00 | 2.77E+00 | 7.54E-01 | no | | regulatory_region_variant | MODIFIER |
| 12 | 105946208 | AX-170446966 | 0 | C | T | 0.123 | -5.60E-02 | 1.83E-01 | 2.69E+00 | 2.77E+00 | 7.54E-01 | no | | intergenic_variant | MODIFIER |
| 12 | 105953548 | AX-170530612 | 0 | G | A | 0.123 | -5.60E-02 | 1.83E-01 | 2.69E+00 | 2.77E+00 | 7.54E-01 | no | | intergenic_variant | MODIFIER |
| 12 | 105954813 | AX-170234450 | 0 | A | G | 0.123 | -5.60E-02 | 1.83E-01 | 2.69E+00 | 2.77E+00 | 7.54E-01 | no | | regulatory_region_variant | MODIFIER |
| 12 | 105955472 | AX-170356671 | 0 | T | C | 0.123 | -5.60E-02 | 1.83E-01 | 2.69E+00 | 2.77E+00 | 7.54E-01 | no | | regulatory_region_variant | MODIFIER |
| 12 | 105963342 | AX-170280732 | 0 | A | G | 0.123 | -5.60E-02 | 1.83E-01 | 2.69E+00 | 2.77E+00 | 7.54E-01 | no | | regulatory_region_variant | MODIFIER |
| 12 | 105964259 | AX-170336423 | 0 | C | A | 0.123 | -5.60E-02 | 1.83E-01 | 2.69E+00 | 2.77E+00 | 7.54E-01 | no | | intergenic_variant | MODIFIER |
| 12 | 105992323 | AX-170190024 | 0 | A | T | 0.123 | -5.60E-02 | 1.83E-01 | 2.69E+00 | 2.77E+00 | 7.54E-01 | no | | intergenic_variant | MODIFIER |
| 12 | 105992374 | AX-170484304 | 0 | T | C | 0.123 | -5.60E-02 | 1.83E-01 | 2.69E+00 | 2.77E+00 | 7.54E-01 | no | | intergenic_variant | MODIFIER |
| 12 | 105992428 | AX-170474292 | 0 | G | A | 0.123 | -5.60E-02 | 1.83E-01 | 2.69E+00 | 2.77E+00 | 7.54E-01 | no | | regulatory_region_variant | MODIFIER |
| 12 | 105994480 | AX-170270909 | 0 | G | T | 0.123 | -5.60E-02 | 1.83E-01 | 2.69E+00 | 2.77E+00 | 7.54E-01 | no | | intergenic_variant | MODIFIER |
| 12 | 106358542 | AX-170099793 | 0 | T | C | 0.123 | -5.60E-02 | 1.83E-01 | 2.69E+00 | 2.77E+00 | 7.54E-01 | no | | intergenic_variant | MODIFIER |
| 12 | 106371998 | AX-97822308 | 0 | G | A | 0.123 | -5.60E-02 | 1.83E-01 | 2.69E+00 | 2.77E+00 | 7.54E-01 | no | | intergenic_variant | MODIFIER |
| 12 | 106403924 | AX-170241618 | 0 | C | T | 0.123 | -5.60E-02 | 1.83E-01 | 2.69E+00 | 2.77E+00 | 7.54E-01 | no | | intergenic_variant | MODIFIER |
| 12 | 106417307 | AX-170122095 | 0 | G | A | 0.123 | -5.60E-02 | 1.83E-01 | 2.69E+00 | 2.77E+00 | 7.54E-01 | no | | intergenic_variant | MODIFIER |
| 12 | 108543178 | AX-170361978 | 0 | T | G | 0.123 | -5.60E-02 | 1.83E-01 | 2.69E+00 | 2.77E+00 | 7.54E-01 | no | Eml1 | downstream_gene_variant | MODIFIER |

| | | | | | | | | | | | | | | | |
|---|---|---|---|---|---|---|---|---|---|---|---|---|---|---|---|
| 12 | 108601916 | AX-170221306 | 0 | C | T | 0.123 | -5.60E-02 | 1.83E-01 | 2.69E+00 | 2.77E+00 | 7.54E-01 | no | Evl | intron_variant | MODIFIER |
| 12 | 108612573 | AX-170238382 | 0 | T | C | 0.123 | -5.60E-02 | 1.83E-01 | 2.69E+00 | 2.77E+00 | 7.54E-01 | no | Evl | intron_variant | MODIFIER |
| 12 | 108613551 | AX-170152376 | 0 | T | C | 0.123 | -5.60E-02 | 1.83E-01 | 2.69E+00 | 2.77E+00 | 7.54E-01 | no | Evl | intron_variant | MODIFIER |
| 12 | 108614593 | AX-170121497 | 0 | T | G | 0.123 | -5.60E-02 | 1.83E-01 | 2.69E+00 | 2.77E+00 | 7.54E-01 | no | Evl | intron_variant | MODIFIER |
| 12 | 108624139 | AX-170050142 | 0 | C | T | 0.123 | -5.60E-02 | 1.83E-01 | 2.69E+00 | 2.77E+00 | 7.54E-01 | no | Evl | intron_variant | MODIFIER |
| 12 | 108624607 | AX-170050143 | 0 | G | A | 0.123 | -5.60E-02 | 1.83E-01 | 2.69E+00 | 2.77E+00 | 7.54E-01 | no | Evl | intron_variant | MODIFIER |
| 12 | 108627586 | AX-170512521 | 0 | A | G | 0.123 | -5.60E-02 | 1.83E-01 | 2.69E+00 | 2.77E+00 | 7.54E-01 | no | Evl | intron_variant | MODIFIER |
| 12 | 108627766 | AX-170270684 | 0 | C | T | 0.123 | -5.60E-02 | 1.83E-01 | 2.69E+00 | 2.77E+00 | 7.54E-01 | no | Evl | intron_variant | MODIFIER |
| 12 | 108627846 | AX-170451192 | 0 | T | C | 0.123 | -5.60E-02 | 1.83E-01 | 2.69E+00 | 2.77E+00 | 7.54E-01 | no | Evl | intron_variant | MODIFIER |
| 12 | 108675087 | AX-170310184 | 0 | A | T | 0.123 | -5.60E-02 | 1.83E-01 | 2.69E+00 | 2.77E+00 | 7.54E-01 | no | Evl | intron_variant | MODIFIER |
| 12 | 108675736 | AX-170496173 | 0 | T | C | 0.123 | -5.60E-02 | 1.83E-01 | 2.69E+00 | 2.77E+00 | 7.54E-01 | no | Evl | intron_variant | MODIFIER |
| 7 | 98754742 | AX-170242098 | 0 | A | G | 0.14 | 5.87E-02 | 1.91E-01 | 2.66E+00 | 2.76E+00 | 7.54E-01 | no | | intergenic_variant | MODIFIER |
| 7 | 98775517 | AX-170298300 | 0 | G | A | 0.14 | 5.87E-02 | 1.91E-01 | 2.66E+00 | 2.76E+00 | 7.54E-01 | no | | intergenic_variant | MODIFIER |
| 7 | 98801609 | AX-170376483 | 0 | T | G | 0.14 | 5.87E-02 | 1.91E-01 | 2.66E+00 | 2.76E+00 | 7.54E-01 | no | | intergenic_variant | MODIFIER |
| 7 | 98804202 | AX-170127319 | 0 | T | C | 0.14 | 5.87E-02 | 1.91E-01 | 2.66E+00 | 2.76E+00 | 7.54E-01 | no | | intergenic_variant | MODIFIER |
| 7 | 98804983 | AX-97834711 | 0 | C | T | 0.14 | 5.87E-02 | 1.91E-01 | 2.66E+00 | 2.76E+00 | 7.54E-01 | no | | regulatory_region_variant | MODIFIER |
| 7 | 98821952 | AX-170244091 | 0 | A | C | 0.14 | 5.87E-02 | 1.91E-01 | 2.66E+00 | 2.76E+00 | 7.54E-01 | no | | intergenic_variant | MODIFIER |
| 7 | 98824212 | AX-97810252 | 0 | T | C | 0.14 | 5.87E-02 | 1.91E-01 | 2.66E+00 | 2.76E+00 | 7.54E-01 | no | | regulatory_region_variant | MODIFIER |
| 7 | 99100829 | AX-170224638 | 0 | A | G | 0.14 | 5.87E-02 | 1.91E-01 | 2.66E+00 | 2.76E+00 | 7.54E-01 | no | Uvrag | intron_variant | MODIFIER |
| 1 | 41593840 | AX-170518303 | 1 | C | T | 0.429 | -2.30E-02 | 7.44E-02 | 2.65E+00 | 2.74E+00 | 7.54E-01 | no | | intergenic_variant | MODIFIER |
| 12 | 111651828 | AX-97786334 | 1 | G | A | 0.134 | -5.52E-02 | 1.79E-01 | 2.68E+00 | 2.76E+00 | 7.54E-01 | no | Mark3 | intron_variant | MODIFIER |
| 9 | 84435165 | AX-170032851 | 0 | A | T | 0.219 | 3.66E-02 | 1.20E-01 | 2.72E+00 | 2.79E+00 | 7.54E-01 | no | | intergenic_variant | MODIFIER |
| 4 | 148087744 | AX-170306928 | 0 | T | C | 0.088 | 6.45E-02 | 2.07E-01 | 2.65E+00 | 2.72E+00 | 7.54E-01 | no | Agtrap | intron_variant | MODIFIER |
| 4 | 148352695 | AX-170264557 | 0 | G | C | 0.088 | 6.45E-02 | 2.07E-01 | 2.65E+00 | 2.72E+00 | 7.54E-01 | no | Gm13200 | downstream_gene_variant | MODIFIER |
| 4 | 149534840 | AX-170071010 | 0 | G | C | 0.088 | 6.45E-02 | 2.07E-01 | 2.65E+00 | 2.72E+00 | 7.54E-01 | no | Ctnnbip1 | intron_variant | MODIFIER |
| 9 | 57424282 | AX-170227202 | 1 | T | C | 0.223 | -3.30E-02 | 1.07E-01 | 2.67E+00 | 2.75E+00 | 7.54E-01 | no | Ppcdc | intron_variant | MODIFIER |
| 12 | 59041038 | AX-170254896 | 0 | G | A | 0.07 | -7.12E-02 | 2.30E-01 | 2.66E+00 | 2.74E+00 | 7.54E-01 | no | Trappc6b | downstream_gene_variant | MODIFIER |
| 13 | 24048519 | AX-170441792 | 1 | G | A | 0.161 | -4.21E-02 | 1.23E-01 | 2.54E+00 | 2.61E+00 | 7.54E-01 | no | Carmil1 | intron_variant | MODIFIER |
| 10 | 127879833 | AX-170378471 | 0 | A | G | 0.325 | -2.36E-02 | 7.56E-02 | 2.63E+00 | 2.73E+00 | 7.54E-01 | no | Rdh7 | downstream_gene_variant | MODIFIER |
| 10 | 128514129 | AX-170450663 | 0 | A | C | 0.325 | -2.36E-02 | 7.56E-02 | 2.63E+00 | 2.73E+00 | 7.54E-01 | no | Esyt1 | intron_variant | MODIFIER |
| 14 | 67188665 | AX-170333654 | 1 | G | A | 0.054 | 6.17E-02 | 2.04E-01 | 2.72E+00 | 2.81E+00 | 7.54E-01 | no | Gm30806 | intron_variant,non_coding_transcript_variant | MODIFIER |
| 18 | 89209578 | AX-97790223 | 1 | C | A | 0.509 | -1.92E-02 | 6.43E-02 | 2.67E+00 | 2.80E+00 | 7.54E-01 | no | Cd226 | intron_variant | MODIFIER |
| 3 | 148116728 | AX-97852042 | 0 | T | G | 0.342 | 2.32E-02 | 7.43E-02 | 2.65E+00 | 2.72E+00 | 7.54E-01 | no | | intergenic_variant | MODIFIER |
| 3 | 148137627 | AX-170021880 | 0 | A | G | 0.342 | 2.32E-02 | 7.43E-02 | 2.65E+00 | 2.72E+00 | 7.54E-01 | no | Gm43574 | intron_variant,non_coding_transcript_variant | MODIFIER |
| 3 | 148139390 | AX-97846935 | 0 | A | G | 0.342 | 2.32E-02 | 7.43E-02 | 2.65E+00 | 2.72E+00 | 7.54E-01 | no | Gm43574 | intron_variant,non_coding_transcript_variant | MODIFIER |
| 3 | 148166409 | AX-170021881 | 0 | C | G | 0.342 | 2.32E-02 | 7.43E-02 | 2.65E+00 | 2.72E+00 | 7.54E-01 | no | | intergenic_variant | MODIFIER |
| 3 | 148202600 | AX-170089104 | 0 | C | T | 0.342 | 2.32E-02 | 7.43E-02 | 2.65E+00 | 2.72E+00 | 7.54E-01 | no | | regulatory_region_variant | MODIFIER |
| 3 | 148204681 | AX-170021883 | 0 | T | G | 0.342 | 2.32E-02 | 7.43E-02 | 2.65E+00 | 2.72E+00 | 7.54E-01 | no | | intergenic_variant | MODIFIER |
| 3 | 148210240 | AX-170424847 | 0 | A | G | 0.342 | 2.32E-02 | 7.43E-02 | 2.65E+00 | 2.72E+00 | 7.54E-01 | no | | intergenic_variant | MODIFIER |
| 3 | 148285258 | AX-97840886 | 0 | G | A | 0.342 | 2.32E-02 | 7.43E-02 | 2.65E+00 | 2.72E+00 | 7.54E-01 | no | | intergenic_variant | MODIFIER |
| 3 | 148325769 | AX-170021874 | 0 | G | C | 0.342 | 2.32E-02 | 7.43E-02 | 2.65E+00 | 2.72E+00 | 7.54E-01 | no | | intergenic_variant | MODIFIER |
| 3 | 148327034 | AX-97798695 | 0 | G | A | 0.342 | 2.32E-02 | 7.43E-02 | 2.65E+00 | 2.72E+00 | 7.54E-01 | no | | intergenic_variant | MODIFIER |
| 3 | 148343926 | AX-170094847 | 0 | T | C | 0.342 | 2.32E-02 | 7.43E-02 | 2.65E+00 | 2.72E+00 | 7.54E-01 | no | | regulatory_region_variant | MODIFIER |
| 3 | 148346451 | AX-170516483 | 0 | A | G | 0.342 | 2.32E-02 | 7.43E-02 | 2.65E+00 | 2.72E+00 | 7.54E-01 | no | | intergenic_variant | MODIFIER |
| 3 | 148353855 | AX-170021877 | 0 | A | G | 0.342 | 2.32E-02 | 7.43E-02 | 2.65E+00 | 2.72E+00 | 7.54E-01 | no | Gm23131 | downstream_gene_variant | MODIFIER |
| 3 | 148379854 | AX-97818358 | 0 | T | C | 0.342 | 2.32E-02 | 7.43E-02 | 2.65E+00 | 2.72E+00 | 7.54E-01 | no | | intergenic_variant | MODIFIER |
| 3 | 148384077 | AX-97813093 | 0 | A | C | 0.342 | 2.32E-02 | 7.43E-02 | 2.65E+00 | 2.72E+00 | 7.54E-01 | no | | intergenic_variant | MODIFIER |
| 3 | 148387624 | AX-170068395 | 0 | G | A | 0.342 | 2.32E-02 | 7.43E-02 | 2.65E+00 | 2.72E+00 | 7.54E-01 | no | | regulatory_region_variant | MODIFIER |
| 3 | 148394547 | AX-170021882 | 0 | T | C | 0.342 | 2.32E-02 | 7.43E-02 | 2.65E+00 | 2.72E+00 | 7.54E-01 | no | | intergenic_variant | MODIFIER |
| 3 | 148396238 | AX-170149074 | 0 | T | C | 0.342 | 2.32E-02 | 7.43E-02 | 2.65E+00 | 2.72E+00 | 7.54E-01 | no | | intergenic_variant | MODIFIER |
| 3 | 148410345 | AX-170132257 | 0 | G | T | 0.342 | 2.32E-02 | 7.43E-02 | 2.65E+00 | 2.72E+00 | 7.54E-01 | no | | intergenic_variant | MODIFIER |
| 3 | 148410655 | AX-97787237 | 0 | A | G | 0.342 | 2.32E-02 | 7.43E-02 | 2.65E+00 | 2.72E+00 | 7.54E-01 | no | | intergenic_variant | MODIFIER |
| 3 | 148410977 | AX-168375122 | 0 | T | C | 0.342 | 2.32E-02 | 7.43E-02 | 2.65E+00 | 2.72E+00 | 7.54E-01 | no | | intergenic_variant | MODIFIER |
| 3 | 148412622 | AX-170174024 | 0 | A | T | 0.342 | 2.32E-02 | 7.43E-02 | 2.65E+00 | 2.72E+00 | 7.54E-01 | no | | intergenic_variant | MODIFIER |

| | | | | | | | | | | | | | | | |
|---|---|---|---|---|---|---|---|---|---|---|---|---|---|---|---|
| 3 | 148413507 | AX-170507992 | 0 | A | G | 0.342 | 2.32E-02 | 7.43E-02 | 2.65E+00 | 2.72E+00 | 7.54E-01 | no | | intergenic_variant | MODIFIER |
| 3 | 148530755 | AX-170375223 | 0 | C | T | 0.342 | 2.32E-02 | 7.43E-02 | 2.65E+00 | 2.72E+00 | 7.54E-01 | no | | intergenic_variant | MODIFIER |
| 3 | 148536908 | AX-170387195 | 0 | G | A | 0.342 | 2.32E-02 | 7.43E-02 | 2.65E+00 | 2.72E+00 | 7.54E-01 | no | | intergenic_variant | MODIFIER |
| 3 | 148540043 | AX-170068398 | 0 | A | G | 0.342 | 2.32E-02 | 7.43E-02 | 2.65E+00 | 2.72E+00 | 7.54E-01 | no | | intergenic_variant | MODIFIER |
| 3 | 148563560 | AX-170319779 | 0 | T | G | 0.342 | 2.32E-02 | 7.43E-02 | 2.65E+00 | 2.72E+00 | 7.54E-01 | no | | intergenic_variant | MODIFIER |
| 3 | 148576751 | AX-170021885 | 0 | C | T | 0.342 | 2.32E-02 | 7.43E-02 | 2.65E+00 | 2.72E+00 | 7.54E-01 | no | | intergenic_variant | MODIFIER |
| 3 | 148579141 | AX-170378217 | 0 | G | A | 0.342 | 2.32E-02 | 7.43E-02 | 2.65E+00 | 2.72E+00 | 7.54E-01 | no | | intergenic_variant | MODIFIER |
| 3 | 148606952 | AX-170021892 | 0 | A | G | 0.342 | 2.32E-02 | 7.43E-02 | 2.65E+00 | 2.72E+00 | 7.54E-01 | no | | intergenic_variant | MODIFIER |
| 3 | 148607369 | AX-170339781 | 0 | G | C | 0.342 | 2.32E-02 | 7.43E-02 | 2.65E+00 | 2.72E+00 | 7.54E-01 | no | | intergenic_variant | MODIFIER |
| 3 | 148607496 | AX-170021889 | 0 | T | C | 0.342 | 2.32E-02 | 7.43E-02 | 2.65E+00 | 2.72E+00 | 7.54E-01 | no | | intergenic_variant | MODIFIER |
| 3 | 148643234 | AX-170499778 | 0 | C | A | 0.342 | 2.32E-02 | 7.43E-02 | 2.65E+00 | 2.72E+00 | 7.54E-01 | no | | intergenic_variant | MODIFIER |
| 3 | 148656164 | AX-170021887 | 0 | T | C | 0.342 | 2.32E-02 | 7.43E-02 | 2.65E+00 | 2.72E+00 | 7.54E-01 | no | Gm43575 | downstream_gene_variant | MODIFIER |
| 3 | 148719466 | AX-170529147 | 0 | G | A | 0.342 | 2.32E-02 | 7.43E-02 | 2.65E+00 | 2.72E+00 | 7.54E-01 | no | | intergenic_variant | MODIFIER |
| 3 | 148720354 | AX-170385770 | 0 | G | T | 0.342 | 2.32E-02 | 7.43E-02 | 2.65E+00 | 2.72E+00 | 7.54E-01 | no | | intergenic_variant | MODIFIER |
| 3 | 148727309 | AX-170238719 | 0 | G | A | 0.342 | 2.32E-02 | 7.43E-02 | 2.65E+00 | 2.72E+00 | 7.54E-01 | no | | regulatory_region_variant | MODIFIER |
| 3 | 148727690 | AX-170218851 | 0 | T | C | 0.342 | 2.32E-02 | 7.43E-02 | 2.65E+00 | 2.72E+00 | 7.54E-01 | no | | intergenic_variant | MODIFIER |
| 3 | 148825525 | AX-170219527 | 0 | C | T | 0.342 | 2.32E-02 | 7.43E-02 | 2.65E+00 | 2.72E+00 | 7.54E-01 | no | Adgrl2 | intron_variant | MODIFIER |
| 3 | 148831759 | AX-170485303 | 0 | T | C | 0.342 | 2.32E-02 | 7.43E-02 | 2.65E+00 | 2.72E+00 | 7.54E-01 | no | Adgrl2 | intron_variant | MODIFIER |
| 3 | 148978480 | AX-170157688 | 0 | C | T | 0.342 | 2.32E-02 | 7.43E-02 | 2.65E+00 | 2.72E+00 | 7.54E-01 | no | Adgrl2 | intron_variant,non_coding_transcript_variant | MODIFIER |
| 3 | 149001012 | AX-170465894 | 0 | G | A | 0.342 | 2.32E-02 | 7.43E-02 | 2.65E+00 | 2.72E+00 | 7.54E-01 | no | Gm43573 | non_coding_transcript_exon_variant | MODIFIER |
| 3 | 149006715 | AX-170278368 | 0 | A | G | 0.342 | 2.32E-02 | 7.43E-02 | 2.65E+00 | 2.72E+00 | 7.54E-01 | no | Gm43573 | downstream_gene_variant | MODIFIER |
| 3 | 149010188 | AX-170229518 | 0 | A | G | 0.342 | 2.32E-02 | 7.43E-02 | 2.65E+00 | 2.72E+00 | 7.54E-01 | no | Gm43572 | downstream_gene_variant | MODIFIER |
| 3 | 149030122 | AX-170068414 | 0 | A | G | 0.342 | 2.32E-02 | 7.43E-02 | 2.65E+00 | 2.72E+00 | 7.54E-01 | no | Gm25127 | downstream_gene_variant | MODIFIER |
| 3 | 149049159 | AX-170462378 | 0 | A | G | 0.342 | 2.32E-02 | 7.43E-02 | 2.65E+00 | 2.72E+00 | 7.54E-01 | no | | intergenic_variant | MODIFIER |
| 3 | 149053680 | AX-170513183 | 0 | T | C | 0.342 | 2.32E-02 | 7.43E-02 | 2.65E+00 | 2.72E+00 | 7.54E-01 | no | | intergenic_variant | MODIFIER |
| 3 | 149054302 | AX-170316912 | 0 | C | T | 0.342 | 2.32E-02 | 7.43E-02 | 2.65E+00 | 2.72E+00 | 7.54E-01 | no | | intergenic_variant | MODIFIER |
| 16 | 92779138 | AX-170037543 | 0 | G | T | 0.474 | -2.38E-02 | 7.74E-02 | 2.66E+00 | 2.75E+00 | 7.54E-01 | no | Runx1 | intron_variant | MODIFIER |
| 16 | 92884040 | AX-170379550 | 0 | C | T | 0.474 | -2.38E-02 | 7.74E-02 | 2.66E+00 | 2.75E+00 | 7.54E-01 | no | | regulatory_region_variant | MODIFIER |
| 18 | 55850729 | AX-97835546 | 1 | G | T | 0.196 | -3.68E-02 | 1.14E-01 | 2.60E+00 | 2.67E+00 | 7.54E-01 | no | | intergenic_variant | MODIFIER |
| 7 | 73173908 | AX-170029286 | 1 | C | T | 0.366 | -2.96E-02 | 9.72E-02 | 2.67E+00 | 2.77E+00 | 7.54E-01 | no | | intergenic_variant | MODIFIER |
| 11 | 74323926 | AX-97855409 | 1 | T | G | 0.259 | 2.18E-02 | 7.18E-02 | 2.63E+00 | 2.76E+00 | 7.54E-01 | no | n-R5s71 | downstream_gene_variant | MODIFIER |
| 9 | 35050346 | AX-170261089 | 0 | C | T | 0.439 | -2.70E-02 | 8.83E-02 | 2.73E+00 | 2.80E+00 | 7.54E-01 | no | St3gal4 | intron_variant | MODIFIER |
| 8 | 19126606 | AX-170302895 | 1 | T | C | 0.179 | 3.24E-02 | 1.05E-01 | 2.64E+00 | 2.73E+00 | 7.54E-01 | no | Defb34 | downstream_gene_variant | MODIFIER |
| 2 | 97465616 | AX-170250379 | 1 | G | C | 0.054 | -9.22E-02 | 2.99E-01 | 2.73E+00 | 2.76E+00 | 7.54E-01 | no | Lrrc4c | upstream_gene_variant | MODIFIER |
| 2 | 94389992 | AX-170515190 | 1 | C | T | 0.054 | -9.26E-02 | 3.00E-01 | 2.73E+00 | 2.76E+00 | 7.54E-01 | no | Ttc17 | intron_variant | MODIFIER |
| 2 | 95744829 | AX-170064350 | 1 | A | G | 0.054 | -9.26E-02 | 3.00E-01 | 2.73E+00 | 2.76E+00 | 7.54E-01 | no | | intergenic_variant | MODIFIER |
| 2 | 142933209 | AX-170432855 | 1 | T | A | 0.107 | 5.73E-02 | 1.85E-01 | 2.64E+00 | 2.73E+00 | 7.54E-01 | no | | intergenic_variant | MODIFIER |
| 8 | 74581531 | AX-170474017 | 1 | A | G | 0.188 | -4.14E-02 | 1.32E-01 | 2.65E+00 | 2.72E+00 | 7.55E-01 | no | | intergenic_variant | MODIFIER |
| 18 | 57602268 | AX-170060945 | 1 | A | G | 0.393 | 3.03E-02 | 9.10E-02 | 2.55E+00 | 2.65E+00 | 7.55E-01 | no | Ccdc192 | intron_variant | MODIFIER |
| 1 | 99405602 | AX-170042152 | 0 | C | A | 0.158 | 5.21E-02 | 1.72E-01 | 2.67E+00 | 2.77E+00 | 7.55E-01 | no | | intergenic_variant | MODIFIER |
| 12 | 118199447 | AX-170050363 | 0 | C | T | 0.14 | -5.31E-02 | 1.75E-01 | 2.69E+00 | 2.78E+00 | 7.55E-01 | no | Dnah11 | upstream_gene_variant | MODIFIER |
| 9 | 107423375 | AX-97788703 | 0 | G | A | 0.123 | -3.79E-02 | 1.18E-01 | 2.60E+00 | 2.67E+00 | 7.55E-01 | no | Cacna2d2 | intron_variant | MODIFIER |
| 11 | 33237787 | AX-170261034 | 1 | G | T | 0.107 | 5.84E-02 | 1.87E-01 | 2.64E+00 | 2.72E+00 | 7.55E-01 | no | Ranbp17 | intron_variant | MODIFIER |
| 9 | 54313032 | AX-170337758 | 0 | C | T | 0.211 | 3.53E-02 | 1.14E-01 | 2.66E+00 | 2.74E+00 | 7.55E-01 | no | Gldn | intron_variant | MODIFIER |
| 7 | 113825232 | AX-170469559 | 0 | A | G | 0.421 | 2.92E-02 | 9.48E-02 | 2.69E+00 | 2.73E+00 | 7.55E-01 | no | Spon1 | intron_variant | MODIFIER |
| 7 | 113829803 | AX-170275999 | 0 | C | T | 0.421 | 2.92E-02 | 9.48E-02 | 2.69E+00 | 2.73E+00 | 7.55E-01 | no | Spon1 | intron_variant | MODIFIER |
| 7 | 113849547 | AX-170235197 | 0 | T | C | 0.421 | 2.92E-02 | 9.48E-02 | 2.69E+00 | 2.73E+00 | 7.55E-01 | no | Spon1 | intron_variant | MODIFIER |
| 7 | 113859917 | AX-170143119 | 0 | G | A | 0.421 | 2.92E-02 | 9.48E-02 | 2.69E+00 | 2.73E+00 | 7.55E-01 | no | Spon1 | intron_variant | MODIFIER |
| 16 | 88831457 | AX-170093558 | 1 | T | G | 0.455 | 2.34E-02 | 7.61E-02 | 2.65E+00 | 2.74E+00 | 7.55E-01 | no | Krtap15 | downstream_gene_variant | MODIFIER |
| 5 | 27837165 | AX-170435895 | 1 | T | C | 0.054 | -9.28E-02 | 3.02E-01 | 2.73E+00 | 2.76E+00 | 7.55E-01 | no | Htr5a | upstream_gene_variant | MODIFIER |
| 1 | 136425888 | AX-170086619 | 1 | T | G | 0.205 | 3.27E-02 | 1.05E-01 | 2.61E+00 | 2.73E+00 | 7.55E-01 | no | Ddx59 | intron_variant | MODIFIER |
| 20 | 74526985 | AX-170223208 | 2 | T | A | 0.036 | -3.30E-02 | 9.53E-02 | 2.46E+00 | 2.66E+00 | 7.56E-01 | no | | intergenic_variant | MODIFIER |
| 11 | 33703091 | AX-170230478 | 0 | C | T | 0.105 | 5.83E-02 | 1.88E-01 | 2.65E+00 | 2.72E+00 | 7.56E-01 | no | Kcnip1 | intron_variant | MODIFIER |
| 4 | 11580024 | AX-170224242 | 0 | C | T | 0.053 | -5.32E-02 | 1.78E-01 | 2.66E+00 | 2.78E+00 | 7.56E-01 | no | Rad54b | intron_variant | MODIFIER |

| | | | | | | | | | | | | | | | |
|---|---|---|---|---|---|---|---|---|---|---|---|---|---|---|---|
| 18 | 66619311 | AX-170491012 | 1 | C | T | 0.098 | 5.30E-02 | 1.70E-01 | 2.61E+00 | 2.72E+00 | 7.56E-01 | no | AC100212.8 | downstream_gene_variant | MODIFIER |
| 7 | 135421029 | AX-97819658 | 1 | A | C | 0.241 | -2.63E-02 | 8.21E-02 | 2.58E+00 | 2.69E+00 | 7.56E-01 | no | | intergenic_variant | MODIFIER |
| 10 | 109641951 | AX-170411837 | 1 | C | T | 0.357 | -2.59E-02 | 8.38E-02 | 2.64E+00 | 2.73E+00 | 7.56E-01 | no | | intergenic_variant | MODIFIER |
| 16 | 88970078 | AX-97849620 | 0 | G | A | 0.474 | 2.39E-02 | 7.78E-02 | 2.65E+00 | 2.74E+00 | 7.56E-01 | no | | intergenic_variant | MODIFIER |
| 7 | 59269741 | AX-97844836 | 1 | C | T | 0.438 | 3.01E-02 | 9.75E-02 | 2.65E+00 | 2.73E+00 | 7.56E-01 | no | Ube3a | intron_variant | MODIFIER |
| 7 | 134861110 | AX-170482730 | 0 | A | G | 0.246 | -2.61E-02 | 8.22E-02 | 2.59E+00 | 2.69E+00 | 7.56E-01 | no | Dock1 | intron_variant | MODIFIER |
| 7 | 134870307 | AX-170384853 | 0 | G | A | 0.246 | -2.61E-02 | 8.22E-02 | 2.59E+00 | 2.69E+00 | 7.56E-01 | no | Dock1 | intron_variant | MODIFIER |
| 7 | 134890572 | AX-170422263 | 0 | C | T | 0.246 | -2.61E-02 | 8.22E-02 | 2.59E+00 | 2.69E+00 | 7.56E-01 | no | Dock1 | intron_variant | MODIFIER |
| 7 | 134891241 | AX-170489892 | 0 | C | T | 0.246 | -2.61E-02 | 8.22E-02 | 2.59E+00 | 2.69E+00 | 7.56E-01 | no | Dock1 | intron_variant | MODIFIER |
| 7 | 134896723 | AX-170458865 | 0 | A | T | 0.246 | -2.61E-02 | 8.22E-02 | 2.59E+00 | 2.69E+00 | 7.56E-01 | no | Dock1 | intron_variant | MODIFIER |
| 7 | 134898703 | AX-170340342 | 0 | C | T | 0.246 | -2.61E-02 | 8.22E-02 | 2.59E+00 | 2.69E+00 | 7.56E-01 | no | Dock1 | intron_variant | MODIFIER |
| 7 | 134898883 | AX-170376741 | 0 | A | C | 0.246 | -2.61E-02 | 8.22E-02 | 2.59E+00 | 2.69E+00 | 7.56E-01 | no | Dock1 | intron_variant | MODIFIER |
| 7 | 134904945 | AX-170186865 | 0 | A | G | 0.246 | -2.61E-02 | 8.22E-02 | 2.59E+00 | 2.69E+00 | 7.56E-01 | no | Dock1 | intron_variant | MODIFIER |
| 7 | 134907891 | AX-170491473 | 0 | A | G | 0.246 | -2.61E-02 | 8.22E-02 | 2.59E+00 | 2.69E+00 | 7.56E-01 | no | Dock1 | intron_variant | MODIFIER |
| 7 | 134909201 | AX-170334581 | 0 | C | T | 0.246 | -2.61E-02 | 8.22E-02 | 2.59E+00 | 2.69E+00 | 7.56E-01 | no | Dock1 | intron_variant | MODIFIER |
| 7 | 134912561 | AX-170452164 | 0 | T | C | 0.246 | -2.61E-02 | 8.22E-02 | 2.59E+00 | 2.69E+00 | 7.56E-01 | no | Dock1 | intron_variant | MODIFIER |
| 7 | 134912854 | AX-170383191 | 0 | C | T | 0.246 | -2.61E-02 | 8.22E-02 | 2.59E+00 | 2.69E+00 | 7.56E-01 | no | Dock1 | intron_variant | MODIFIER |
| 7 | 134913542 | AX-170309112 | 0 | G | A | 0.246 | -2.61E-02 | 8.22E-02 | 2.59E+00 | 2.69E+00 | 7.56E-01 | no | Dock1 | intron_variant | MODIFIER |
| 7 | 135072661 | AX-170350205 | 0 | C | T | 0.246 | -2.61E-02 | 8.22E-02 | 2.59E+00 | 2.69E+00 | 7.56E-01 | no | Dock1 | intron_variant | MODIFIER |
| 7 | 135072862 | AX-170343063 | 0 | C | T | 0.246 | -2.61E-02 | 8.22E-02 | 2.59E+00 | 2.69E+00 | 7.56E-01 | no | Dock1 | intron_variant | MODIFIER |
| 7 | 135072879 | AX-170447788 | 0 | A | C | 0.246 | -2.61E-02 | 8.22E-02 | 2.59E+00 | 2.69E+00 | 7.56E-01 | no | Dock1 | intron_variant | MODIFIER |
| 7 | 135072939 | AX-170172977 | 0 | T | C | 0.246 | -2.61E-02 | 8.22E-02 | 2.59E+00 | 2.69E+00 | 7.56E-01 | no | Dock1 | intron_variant | MODIFIER |
| 7 | 135083124 | AX-97846672 | 0 | C | T | 0.246 | -2.61E-02 | 8.22E-02 | 2.59E+00 | 2.69E+00 | 7.56E-01 | no | Dock1 | intron_variant | MODIFIER |
| 7 | 135101462 | AX-170405298 | 0 | A | C | 0.246 | -2.61E-02 | 8.22E-02 | 2.59E+00 | 2.69E+00 | 7.56E-01 | no | Dock1 | intron_variant | MODIFIER |
| 7 | 135259074 | AX-97807501 | 0 | T | C | 0.246 | -2.61E-02 | 8.22E-02 | 2.59E+00 | 2.69E+00 | 7.56E-01 | no | Nps | intron_variant | MODIFIER |
| 7 | 135281872 | AX-97837021 | 0 | G | A | 0.246 | -2.61E-02 | 8.22E-02 | 2.59E+00 | 2.69E+00 | 7.56E-01 | no | Gm45339 | upstream_gene_variant | MODIFIER |
| 7 | 135422622 | AX-170173729 | 0 | G | C | 0.246 | -2.61E-02 | 8.22E-02 | 2.59E+00 | 2.69E+00 | 7.56E-01 | no | | intergenic_variant | MODIFIER |
| 2 | 130153531 | AX-170335625 | 0 | G | A | 0.149 | 5.57E-02 | 1.77E-01 | 2.62E+00 | 2.70E+00 | 7.56E-01 | no | Tgm6 | 3_prime_UTR_variant | MODIFIER |
| 3 | 30609241 | AX-170020233 | 0 | G | A | 0.175 | -2.95E-02 | 9.63E-02 | 2.66E+00 | 2.74E+00 | 7.56E-01 | no | Mynn | intron_variant | MODIFIER |
| 13 | 115613171 | AX-170205173 | 1 | G | A | 0.036 | 4.69E-02 | 1.61E-01 | 2.61E+00 | 2.79E+00 | 7.56E-01 | no | | intergenic_variant | MODIFIER |
| 1 | 33719127 | AX-170035609 | 0 | A | G | 0.412 | 2.49E-02 | 8.09E-02 | 2.66E+00 | 2.73E+00 | 7.56E-01 | no | Rab23 | upstream_gene_variant | MODIFIER |
| 8 | 106638854 | AX-170502880 | 0 | A | G | 0.228 | 3.45E-02 | 1.13E-01 | 2.69E+00 | 2.76E+00 | 7.56E-01 | no | Cdh1 | intron_variant | MODIFIER |
| 12 | 101048345 | AX-170007393 | 2 | C | T | 0.073 | -5.34E-02 | 1.76E-01 | 2.64E+00 | 2.76E+00 | 7.56E-01 | no | Ppp4r3a | intron_variant | MODIFIER |
| 4 | 31935763 | AX-170224462 | 1 | C | T | 0.027 | 5.04E-02 | 1.72E-01 | 2.61E+00 | 2.79E+00 | 7.56E-01 | no | | intergenic_variant | MODIFIER |
| 7 | 106650689 | AX-170309329 | 1 | A | T | 0.482 | -2.73E-02 | 9.13E-02 | 2.73E+00 | 2.82E+00 | 7.57E-01 | no | | regulatory_region_variant | MODIFIER |
| 7 | 122566590 | AX-170370113 | 1 | G | A | 0.411 | 2.66E-02 | 8.71E-02 | 2.67E+00 | 2.75E+00 | 7.57E-01 | no | Prkcb | intron_variant | MODIFIER |
| 1 | 39770275 | AX-170391114 | 1 | C | T | 0.393 | 2.24E-02 | 7.47E-02 | 2.71E+00 | 2.79E+00 | 7.57E-01 | no | | regulatory_region_variant | MODIFIER |
| 4 | 77715198 | AX-170333281 | 0 | C | G | 0.325 | 3.54E-02 | 1.14E-01 | 2.64E+00 | 2.70E+00 | 7.57E-01 | no | Ptprd | intron_variant,non_coding_transcript_variant | MODIFIER |
| 4 | 78905656 | AX-97854865 | 0 | T | C | 0.237 | 3.25E-02 | 9.82E-02 | 2.55E+00 | 2.64E+00 | 7.57E-01 | no | | intergenic_variant | MODIFIER |
| 1 | 136680909 | AX-97786994 | 1 | T | C | 0.205 | -3.17E-02 | 1.07E-01 | 2.67E+00 | 2.79E+00 | 7.57E-01 | no | Gm19705 | intron_variant,non_coding_transcript_variant | MODIFIER |
| 8 | 35405283 | AX-170306444 | 1 | C | T | 0.152 | -4.52E-02 | 1.44E-01 | 2.64E+00 | 2.69E+00 | 7.57E-01 | no | Gm45301 | downstream_gene_variant | MODIFIER |
| 4 | 134462112 | AX-170334126 | 1 | C | T | 0.143 | 3.77E-02 | 1.16E-01 | 2.59E+00 | 2.65E+00 | 7.57E-01 | no | | intergenic_variant | MODIFIER |
| 15 | 80208376 | AX-170322130 | 0 | G | C | 0.561 | 2.51E-02 | 7.86E-02 | 2.59E+00 | 2.68E+00 | 7.57E-01 | no | Mgat3 | intron_variant | MODIFIER |
| 1 | 67360694 | AX-170232894 | 0 | G | A | 0.114 | 7.05E-02 | 2.31E-01 | 2.69E+00 | 2.74E+00 | 7.57E-01 | no | | intergenic_variant | MODIFIER |
| 7 | 112903464 | AX-170167092 | 1 | A | G | 0.429 | 2.82E-02 | 9.21E-02 | 2.68E+00 | 2.73E+00 | 7.57E-01 | no | Tead1 | 3_prime_UTR_variant | MODIFIER |
| 7 | 113050309 | AX-170205508 | 1 | A | G | 0.429 | 2.82E-02 | 9.21E-02 | 2.68E+00 | 2.73E+00 | 7.57E-01 | no | | intergenic_variant | MODIFIER |
| 7 | 113051488 | AX-170227615 | 1 | A | G | 0.429 | 2.82E-02 | 9.21E-02 | 2.68E+00 | 2.73E+00 | 7.57E-01 | no | | intergenic_variant | MODIFIER |
| 16 | 97370166 | AX-170310338 | 0 | G | A | 0.465 | 2.28E-02 | 7.73E-02 | 2.71E+00 | 2.82E+00 | 7.57E-01 | no | Bace2 | intron_variant | MODIFIER |
| 7 | 116805048 | AX-170388852 | 1 | C | G | 0.402 | 2.74E-02 | 8.82E-02 | 2.64E+00 | 2.71E+00 | 7.57E-01 | no | Gm6008 | upstream_gene_variant | MODIFIER |
| 8 | 32101310 | AX-170079666 | 1 | C | G | 0.143 | -3.90E-02 | 1.26E-01 | 2.65E+00 | 2.72E+00 | 7.57E-01 | no | Nrg1 | intron_variant | MODIFIER |
| 8 | 42318000 | AX-170131239 | 1 | T | C | 0.152 | 4.58E-02 | 1.51E-01 | 2.76E+00 | 2.79E+00 | 7.57E-01 | no | | intergenic_variant | MODIFIER |
| 11 | 23646269 | AX-170141062 | 0 | G | A | 0.053 | 6.29E-02 | 2.03E-01 | 2.60E+00 | 2.72E+00 | 7.58E-01 | no | Pex13 | downstream_gene_variant | MODIFIER |
| 7 | 7140369 | AX-170376176 | 1 | T | C | 0.17 | 2.34E-02 | 8.09E-02 | 2.69E+00 | 2.83E+00 | 7.58E-01 | no | Zfp773 | upstream_gene_variant | MODIFIER |
| 4 | 58582145 | AX-170409116 | 0 | C | G | 0.377 | -3.23E-02 | 1.07E-01 | 2.70E+00 | 2.78E+00 | 7.58E-01 | no | | intergenic_variant | MODIFIER |

| | | | | | | | | | | | | | | | |
|---|---|---|---|---|---|---|---|---|---|---|---|---|---|---|---|
| 9 | 80947386 | AX-170513113 | 0 | C | T | 0.193 | 3.77E-02 | 1.25E-01 | 2.75E+00 | 2.81E+00 | 7.58E-01 | no | | intergenic_variant | MODIFIER |
| 9 | 108633611 | AX-170033245 | 2 | G | A | 0.127 | 3.51E-02 | 1.17E-01 | 2.76E+00 | 2.84E+00 | 7.58E-01 | no | Arih2 | intron_variant | MODIFIER |
| 3 | 83159446 | AX-170435555 | 0 | G | A | 0.044 | -4.53E-02 | 1.50E-01 | 2.60E+00 | 2.75E+00 | 7.58E-01 | no | Dchs2 | intron_variant | MODIFIER |
| 4 | 80355842 | AX-170470440 | 0 | T | G | 0.237 | -2.98E-02 | 1.01E-01 | 2.75E+00 | 2.86E+00 | 7.58E-01 | no | | intergenic_variant | MODIFIER |
| 7 | 116339847 | AX-170533032 | 0 | A | G | 0.404 | 2.79E-02 | 9.04E-02 | 2.65E+00 | 2.71E+00 | 7.58E-01 | no | Pik3c2a | 3_prime_UTR_variant | MODIFIER |
| 4 | 142857963 | AX-97840320 | 1 | C | T | 0.098 | 6.06E-02 | 1.99E-01 | 2.68E+00 | 2.74E+00 | 7.58E-01 | no | 6330411D24Rik | intron_variant,non_coding_transcript_variant | MODIFIER |
| 8 | 69040082 | AX-97843156 | 0 | G | A | 0.184 | -4.03E-02 | 1.32E-01 | 2.66E+00 | 2.73E+00 | 7.58E-01 | no | Slc18a1 | intron_variant | MODIFIER |
| 13 | 31232646 | AX-170199046 | 1 | C | A | 0.179 | -3.84E-02 | 1.22E-01 | 2.61E+00 | 2.68E+00 | 7.58E-01 | no | 4930401O12Rik | intron_variant,non_coding_transcript_variant | MODIFIER |
| 13 | 31998914 | AX-97801793 | 1 | A | G | 0.179 | -3.84E-02 | 1.22E-01 | 2.61E+00 | 2.68E+00 | 7.58E-01 | no | Gmds | intron_variant | MODIFIER |
| 18 | 89727492 | AX-170525135 | 1 | T | C | 0.509 | -1.86E-02 | 6.34E-02 | 2.66E+00 | 2.79E+00 | 7.58E-01 | no | Dok6 | intron_variant | MODIFIER |
| 10 | 29324819 | AX-170121848 | 0 | G | C | 0.193 | 2.72E-02 | 9.21E-02 | 2.70E+00 | 2.80E+00 | 7.58E-01 | no | Echdc1 | intron_variant | MODIFIER |
| 6 | 107094366 | AX-170438002 | 1 | G | A | 0.054 | 8.99E-02 | 2.95E-01 | 2.68E+00 | 2.74E+00 | 7.58E-01 | no | Gm6064 | upstream_gene_variant | MODIFIER |
| 12 | 113194394 | AX-170524651 | 0 | C | T | 0.149 | 4.91E-02 | 1.62E-01 | 2.64E+00 | 2.75E+00 | 7.58E-01 | no | Tmem121 | downstream_gene_variant | MODIFIER |
| 10 | 126809772 | AX-170004120 | 1 | C | A | 0.33 | -2.23E-02 | 7.23E-02 | 2.61E+00 | 2.72E+00 | 7.58E-01 | no | Gm9111 | non_coding_transcript_exon_variant | MODIFIER |
| 12 | 104550099 | AX-170374231 | 0 | T | A | 0.123 | -5.58E-02 | 1.86E-01 | 2.70E+00 | 2.77E+00 | 7.58E-01 | no | | intergenic_variant | MODIFIER |
| 1 | 26050726 | AX-170383065 | 1 | A | T | 0.411 | 2.63E-02 | 8.50E-02 | 2.63E+00 | 2.70E+00 | 7.59E-01 | no | | intergenic_variant | MODIFIER |
| 15 | 85658505 | AX-170173248 | 0 | A | T | 0.509 | 2.36E-02 | 7.01E-02 | 2.53E+00 | 2.63E+00 | 7.59E-01 | no | Gm49539 | intron_variant,non_coding_transcript_variant | MODIFIER |
| 10 | 17008030 | AX-170044224 | 0 | A | C | 0.219 | 3.01E-02 | 1.01E-01 | 2.72E+00 | 2.80E+00 | 7.59E-01 | no | Gm20125 | intron_variant,non_coding_transcript_variant | MODIFIER |
| 1 | 170733168 | AX-170108882 | 0 | G | A | 0.202 | 4.05E-02 | 1.32E-01 | 2.66E+00 | 2.72E+00 | 7.59E-01 | no | Atf6 | intron_variant | MODIFIER |
| 9 | 105798316 | AX-170501419 | 1 | A | G | 0.125 | -3.69E-02 | 1.18E-01 | 2.62E+00 | 2.69E+00 | 7.59E-01 | no | Col6a6 | intron_variant | MODIFIER |
| 7 | 68669231 | AX-170133561 | 0 | G | A | 0.404 | -2.80E-02 | 9.41E-02 | 2.71E+00 | 2.79E+00 | 7.59E-01 | no | | intergenic_variant | MODIFIER |
| 7 | 118583184 | AX-170302534 | 2 | T | C | 0.382 | 2.80E-02 | 9.27E-02 | 2.68E+00 | 2.74E+00 | 7.59E-01 | no | Tmc7 | intron_variant | MODIFIER |
| 12 | 112173823 | AX-170204073 | 0 | C | T | 0.14 | -5.72E-02 | 1.90E-01 | 2.69E+00 | 2.76E+00 | 7.59E-01 | no | Kif26a | synonymous_variant | LOW |
| 4 | 95532553 | AX-170335283 | 1 | C | T | 0.33 | 3.32E-02 | 1.09E-01 | 2.65E+00 | 2.72E+00 | 7.59E-01 | no | | intergenic_variant | MODIFIER |
| 4 | 155166735 | AX-97781692 | 0 | T | C | 0.105 | 4.40E-02 | 1.48E-01 | 2.64E+00 | 2.76E+00 | 7.59E-01 | no | Ski | intron_variant | MODIFIER |
| 4 | 92457285 | AX-170136812 | 0 | T | A | 0.316 | 3.12E-02 | 1.02E-01 | 2.64E+00 | 2.72E+00 | 7.59E-01 | no | | intergenic_variant | MODIFIER |
| 1 | 75298772 | AX-170000820 | 1 | C | A | 0.143 | 7.10E-02 | 2.35E-01 | 2.69E+00 | 2.74E+00 | 7.59E-01 | no | | intergenic_variant | MODIFIER |
| 5 | 88938948 | AX-170297490 | 0 | T | C | 0.404 | 3.23E-02 | 1.07E-01 | 2.69E+00 | 2.75E+00 | 7.59E-01 | no | Slc4a4 | intron_variant | MODIFIER |
| 3 | 150899163 | AX-170244321 | 1 | A | C | 0.339 | 2.21E-02 | 7.34E-02 | 2.66E+00 | 2.75E+00 | 7.59E-01 | no | | intergenic_variant | MODIFIER |
| 9 | 33773587 | AX-170480180 | 0 | A | C | 0.421 | 2.74E-02 | 8.86E-02 | 2.63E+00 | 2.70E+00 | 7.59E-01 | no | | intergenic_variant | MODIFIER |
| 4 | 72474542 | AX-170437044 | 0 | T | C | 0.228 | -2.99E-02 | 1.02E-01 | 2.75E+00 | 2.87E+00 | 7.60E-01 | no | | intergenic_variant | MODIFIER |
| 4 | 72800598 | AX-170023097 | 0 | A | C | 0.228 | -2.99E-02 | 1.02E-01 | 2.75E+00 | 2.87E+00 | 7.60E-01 | no | | intergenic_variant | MODIFIER |
| 4 | 74903114 | AX-170023139 | 0 | A | T | 0.228 | -2.99E-02 | 1.02E-01 | 2.75E+00 | 2.87E+00 | 7.60E-01 | no | | intergenic_variant | MODIFIER |
| 20 | 139907538 | AX-170479078 | 0 | C | T | 0.088 | -3.60E-02 | 1.22E-01 | 2.62E+00 | 2.77E+00 | 7.60E-01 | no | Rbm41 | intron_variant | MODIFIER |
| 1 | 26078707 | AX-168396812 | 0 | C | T | 0.412 | 2.61E-02 | 8.45E-02 | 2.63E+00 | 2.70E+00 | 7.60E-01 | no | | intergenic_variant | MODIFIER |
| 20 | 167884463 | AX-97809424 | 1 | G | T | 0.188 | 3.67E-02 | 1.18E-01 | 2.61E+00 | 2.69E+00 | 7.60E-01 | no | Frmpd4 | intron_variant | MODIFIER |
| 8 | 53388146 | AX-170096272 | 2 | C | T | 0.145 | -4.67E-02 | 1.48E-01 | 2.61E+00 | 2.66E+00 | 7.60E-01 | no | | intergenic_variant | MODIFIER |
| 5 | 109498223 | AX-97793598 | 2 | G | A | 0.473 | 3.34E-02 | 1.07E-01 | 2.62E+00 | 2.69E+00 | 7.60E-01 | no | | intergenic_variant | MODIFIER |
| 12 | 103986082 | AX-170050032 | 0 | G | A | 0.14 | -4.77E-02 | 1.61E-01 | 2.67E+00 | 2.78E+00 | 7.60E-01 | no | Serpina11 | missense_variant | MODERATE |
| 6 | 128216583 | AX-170283356 | 0 | T | G | 0.553 | 2.77E-02 | 8.61E-02 | 2.58E+00 | 2.65E+00 | 7.60E-01 | no | Gm19065 | downstream_gene_variant | MODIFIER |
| 6 | 129093568 | AX-170028235 | 0 | C | T | 0.553 | 2.77E-02 | 8.61E-02 | 2.58E+00 | 2.65E+00 | 7.60E-01 | no | Clec2e | intron_variant | MODIFIER |
| 4 | 147738672 | AX-170371919 | 1 | C | G | 0.482 | 1.62E-01 | 5.31E-01 | 2.67E+00 | 2.72E+00 | 7.60E-01 | Yes | Gm13136 | upstream_gene_variant | MODIFIER |
| 1 | 74441563 | AX-170501684 | 1 | G | A | 0.143 | 7.19E-02 | 2.38E-01 | 2.69E+00 | 2.74E+00 | 7.60E-01 | no | Usp37 | synonymous_variant | LOW |
| 9 | 77992941 | AX-97828759 | 0 | T | A | 0.193 | 4.12E-02 | 1.37E-01 | 2.75E+00 | 2.78E+00 | 7.60E-01 | no | Gm34829 | intron_variant,non_coding_transcript_variant | MODIFIER |
| 12 | 63881458 | AX-97849103 | 0 | G | T | 0.07 | 6.90E-02 | 2.33E-01 | 2.70E+00 | 2.78E+00 | 7.60E-01 | no | | intergenic_variant | MODIFIER |
| 18 | 35137035 | AX-97781312 | 0 | C | T | 0.07 | 6.90E-02 | 2.33E-01 | 2.70E+00 | 2.78E+00 | 7.60E-01 | no | Ctnna1 | intron_variant | MODIFIER |
| 1 | 169693259 | AX-170494761 | 1 | A | G | 0.205 | 4.30E-02 | 1.40E-01 | 2.65E+00 | 2.69E+00 | 7.60E-01 | no | Rgs5 | 3_prime_UTR_variant | MODIFIER |
| 5 | 32073021 | AX-170445446 | 0 | G | T | 0.351 | -3.00E-02 | 1.00E-01 | 2.65E+00 | 2.75E+00 | 7.60E-01 | no | Babam2 | intron_variant | MODIFIER |
| 17 | 49513917 | AX-170059209 | 1 | A | T | 0.045 | -4.72E-02 | 1.54E-01 | 2.58E+00 | 2.72E+00 | 7.60E-01 | no | Daam2 | intron_variant | MODIFIER |
| 9 | 111004051 | AX-170504435 | 1 | C | T | 0.098 | -3.70E-02 | 1.17E-01 | 2.59E+00 | 2.67E+00 | 7.61E-01 | no | Gm10030 | upstream_gene_variant | MODIFIER |
| 4 | 92831953 | AX-97825873 | 1 | C | T | 0.33 | 3.30E-02 | 1.08E-01 | 2.64E+00 | 2.71E+00 | 7.61E-01 | no | | intergenic_variant | MODIFIER |
| 1 | 25612891 | AX-97830089 | 0 | T | C | 0.404 | 2.60E-02 | 8.53E-02 | 2.65E+00 | 2.71E+00 | 7.61E-01 | no | Adgrb3 | intron_variant | MODIFIER |
| 1 | 25687599 | AX-170000127 | 0 | G | A | 0.404 | 2.60E-02 | 8.53E-02 | 2.65E+00 | 2.71E+00 | 7.61E-01 | no | Adgrb3 | intron_variant | MODIFIER |
| 1 | 25881755 | AX-170426665 | 0 | G | A | 0.404 | 2.60E-02 | 8.53E-02 | 2.65E+00 | 2.71E+00 | 7.61E-01 | no | | intergenic_variant | MODIFIER |

| | | | | | | | | | | | | | | | |
|---|---|---|---|---|---|---|---|---|---|---|---|---|---|---|---|
| 1 | 25940852 | AX-170091437 | 0 | G | T | 0.404 | 2.60E-02 | 8.53E-02 | 2.65E+00 | 2.71E+00 | 7.61E-01 | no | | intergenic_variant | MODIFIER |
| 1 | 25975579 | AX-170499328 | 0 | C | A | 0.404 | 2.60E-02 | 8.53E-02 | 2.65E+00 | 2.71E+00 | 7.61E-01 | no | | intergenic_variant | MODIFIER |
| 1 | 26044918 | AX-97841183 | 0 | G | A | 0.404 | 2.60E-02 | 8.53E-02 | 2.65E+00 | 2.71E+00 | 7.61E-01 | no | | intergenic_variant | MODIFIER |
| 1 | 26049737 | AX-170116789 | 0 | A | G | 0.404 | 2.60E-02 | 8.53E-02 | 2.65E+00 | 2.71E+00 | 7.61E-01 | no | | intergenic_variant | MODIFIER |
| 1 | 26061889 | AX-170189191 | 0 | G | A | 0.404 | 2.60E-02 | 8.53E-02 | 2.65E+00 | 2.71E+00 | 7.61E-01 | no | | intergenic_variant | MODIFIER |
| 1 | 26091444 | AX-170111223 | 0 | C | A | 0.404 | 2.60E-02 | 8.53E-02 | 2.65E+00 | 2.71E+00 | 7.61E-01 | no | | intergenic_variant | MODIFIER |
| 1 | 26100750 | AX-170108830 | 0 | C | G | 0.404 | 2.60E-02 | 8.53E-02 | 2.65E+00 | 2.71E+00 | 7.61E-01 | no | | intergenic_variant | MODIFIER |
| 1 | 26109943 | AX-170141133 | 0 | G | A | 0.404 | 2.60E-02 | 8.53E-02 | 2.65E+00 | 2.71E+00 | 7.61E-01 | no | | intergenic_variant | MODIFIER |
| 1 | 26114395 | AX-170086423 | 0 | G | A | 0.404 | 2.60E-02 | 8.53E-02 | 2.65E+00 | 2.71E+00 | 7.61E-01 | no | | intergenic_variant | MODIFIER |
| 1 | 26404334 | AX-170235386 | 0 | G | C | 0.404 | 2.60E-02 | 8.53E-02 | 2.65E+00 | 2.71E+00 | 7.61E-01 | no | | intergenic_variant | MODIFIER |
| 1 | 26463601 | AX-97811015 | 0 | G | A | 0.404 | 2.60E-02 | 8.53E-02 | 2.65E+00 | 2.71E+00 | 7.61E-01 | no | Gm19123 | upstream_gene_variant | MODIFIER |
| 1 | 26656818 | AX-170394115 | 0 | A | G | 0.404 | 2.60E-02 | 8.53E-02 | 2.65E+00 | 2.71E+00 | 7.61E-01 | no | | intergenic_variant | MODIFIER |
| 1 | 26723533 | AX-170528983 | 0 | T | C | 0.404 | 2.60E-02 | 8.53E-02 | 2.65E+00 | 2.71E+00 | 7.61E-01 | no | | intergenic_variant | MODIFIER |
| 7 | 39929244 | AX-170237884 | 1 | T | C | 0.348 | 2.80E-02 | 8.75E-02 | 2.57E+00 | 2.66E+00 | 7.61E-01 | no | Gm44992 | intron_variant,non_coding_transcript_variant | MODIFIER |
| 7 | 69520838 | AX-170138547 | 1 | G | T | 0.411 | 3.03E-02 | 9.69E-02 | 2.61E+00 | 2.68E+00 | 7.61E-01 | no | 4930402F11Rik | downstream_gene_variant | MODIFIER |
| 4 | 141099855 | AX-97844280 | 0 | G | A | 0.079 | 5.82E-02 | 1.92E-01 | 2.66E+00 | 2.72E+00 | 7.61E-01 | no | Spata21 | intron_variant | MODIFIER |
| 4 | 150484865 | AX-170441586 | 0 | T | C | 0.079 | 5.82E-02 | 1.92E-01 | 2.66E+00 | 2.72E+00 | 7.61E-01 | no | Rere | upstream_gene_variant | MODIFIER |
| 1 | 70000344 | AX-170508996 | 1 | C | G | 0.116 | 6.95E-02 | 2.31E-01 | 2.69E+00 | 2.74E+00 | 7.61E-01 | no | Spag16 | intron_variant | MODIFIER |
| 15 | 92930106 | AX-170299689 | 1 | C | T | 0.143 | 4.83E-02 | 1.55E-01 | 2.59E+00 | 2.69E+00 | 7.61E-01 | no | Gm49418 | upstream_gene_variant | MODIFIER |
| 1 | 152608313 | AX-97828101 | 1 | A | C | 0.196 | 4.24E-02 | 1.39E-01 | 2.65E+00 | 2.70E+00 | 7.61E-01 | no | Rgl1 | intron_variant | MODIFIER |
| 4 | 140371804 | AX-170294750 | 0 | T | G | 0.158 | 3.42E-02 | 1.11E-01 | 2.62E+00 | 2.69E+00 | 7.61E-01 | no | | intergenic_variant | MODIFIER |
| 9 | 65322323 | AX-97830162 | 0 | C | T | 0.193 | 4.08E-02 | 1.36E-01 | 2.75E+00 | 2.78E+00 | 7.61E-01 | no | Clpx | intron_variant | MODIFIER |
| 3 | 143928949 | AX-170311371 | 0 | C | T | 0.368 | 2.10E-02 | 6.91E-02 | 2.63E+00 | 2.72E+00 | 7.61E-01 | no | Gm42707 | intron_variant,non_coding_transcript_variant | MODIFIER |
| 1 | 171225406 | AX-170002277 | 0 | A | G | 0.211 | 4.03E-02 | 1.27E-01 | 2.59E+00 | 2.66E+00 | 7.61E-01 | no | Tomm40l | upstream_gene_variant | MODIFIER |
| 1 | 171281362 | AX-170272107 | 0 | C | T | 0.211 | 4.03E-02 | 1.27E-01 | 2.59E+00 | 2.66E+00 | 7.61E-01 | no | B4galt3 | downstream_gene_variant | MODIFIER |
| 2 | 17660723 | AX-170321532 | 0 | T | A | 0.018 | 4.10E-02 | 1.46E-01 | 2.62E+00 | 2.81E+00 | 7.62E-01 | no | Nebl | intron_variant | MODIFIER |
| 9 | 35132011 | AX-170175202 | 1 | T | C | 0.429 | -2.48E-02 | 8.43E-02 | 2.74E+00 | 2.82E+00 | 7.62E-01 | no | Dcps | intron_variant | MODIFIER |
| 2 | 92924494 | AX-170121865 | 0 | G | T | 0.07 | 5.38E-02 | 1.82E-01 | 2.65E+00 | 2.76E+00 | 7.62E-01 | no | Syt13 | intron_variant | MODIFIER |
| 1 | 32430741 | AX-170193435 | 1 | C | A | 0.429 | 2.45E-02 | 8.21E-02 | 2.67E+00 | 2.75E+00 | 7.62E-01 | no | Khdrbs2 | intron_variant | MODIFIER |
| 15 | 95488565 | AX-170123701 | 1 | A | T | 0.134 | 5.62E-02 | 1.81E-01 | 2.61E+00 | 2.68E+00 | 7.62E-01 | no | Nell2 | intron_variant | MODIFIER |
| 9 | 61762840 | AX-170138936 | 1 | G | A | 0.188 | 4.01E-02 | 1.34E-01 | 2.75E+00 | 2.79E+00 | 7.62E-01 | no | Gm47241 | upstream_gene_variant | MODIFIER |
| 9 | 76918277 | AX-170375560 | 1 | G | A | 0.188 | 4.01E-02 | 1.34E-01 | 2.75E+00 | 2.79E+00 | 7.62E-01 | no | | intergenic_variant | MODIFIER |
| 1 | 5227555 | AX-169999858 | 1 | G | C | 0.429 | 2.21E-02 | 6.84E-02 | 2.57E+00 | 2.64E+00 | 7.62E-01 | no | | intergenic_variant | MODIFIER |
| 2 | 118938588 | AX-170064706 | 2 | T | C | 0.145 | 5.72E-02 | 1.83E-01 | 2.60E+00 | 2.68E+00 | 7.62E-01 | no | | intergenic_variant | MODIFIER |
| 9 | 70429929 | AX-97851513 | 2 | T | G | 0.2 | 3.77E-02 | 1.27E-01 | 2.71E+00 | 2.77E+00 | 7.62E-01 | no | Rnf111 | intron_variant | MODIFIER |
| 16 | 19469521 | AX-97843911 | 0 | G | A | 0.132 | 2.60E-02 | 9.02E-02 | 2.67E+00 | 2.80E+00 | 7.62E-01 | no | | intergenic_variant | MODIFIER |
| 8 | 75742027 | AX-170420493 | 1 | C | G | 0.196 | -4.00E-02 | 1.34E-01 | 2.68E+00 | 2.74E+00 | 7.62E-01 | no | Iqcm | intron_variant | MODIFIER |
| 3 | 32403329 | AX-170400533 | 0 | T | C | 0.175 | -2.68E-02 | 9.07E-02 | 2.65E+00 | 2.76E+00 | 7.62E-01 | no | Pik3ca | intron_variant | MODIFIER |
| 13 | 53730624 | AX-170329370 | 0 | C | T | 0.535 | 2.17E-02 | 6.81E-02 | 2.56E+00 | 2.67E+00 | 7.62E-01 | no | | regulatory_region_variant | MODIFIER |
| 5 | 97793452 | AX-170025673 | 1 | G | C | 0.438 | 3.29E-02 | 1.10E-01 | 2.68E+00 | 2.73E+00 | 7.63E-01 | no | | intergenic_variant | MODIFIER |
| 8 | 34865851 | AX-170030491 | 1 | G | A | 0.152 | -4.41E-02 | 1.45E-01 | 2.65E+00 | 2.70E+00 | 7.63E-01 | no | Tnks | intron_variant | MODIFIER |
| 8 | 35050870 | AX-97812450 | 1 | C | A | 0.152 | -4.41E-02 | 1.45E-01 | 2.65E+00 | 2.70E+00 | 7.63E-01 | no | Gm8291 | downstream_gene_variant | MODIFIER |
| 8 | 35832497 | AX-170284383 | 1 | C | T | 0.152 | -4.41E-02 | 1.45E-01 | 2.65E+00 | 2.70E+00 | 7.63E-01 | no | | intergenic_variant | MODIFIER |
| 1 | 138661871 | AX-97824159 | 1 | C | T | 0.214 | -3.58E-02 | 1.23E-01 | 2.75E+00 | 2.85E+00 | 7.63E-01 | no | Gm5833 | intron_variant,non_coding_transcript_variant | MODIFIER |
| 8 | 40159186 | AX-97849940 | 1 | G | C | 0.152 | 4.37E-02 | 1.48E-01 | 2.75E+00 | 2.80E+00 | 7.63E-01 | no | | intergenic_variant | MODIFIER |
| 1 | 33696966 | AX-97811234 | 1 | A | G | 0.411 | 2.40E-02 | 8.07E-02 | 2.67E+00 | 2.75E+00 | 7.63E-01 | no | | intergenic_variant | MODIFIER |
| 15 | 87107626 | AX-170370117 | 1 | T | A | 0.134 | 4.74E-02 | 1.56E-01 | 2.64E+00 | 2.70E+00 | 7.63E-01 | no | | intergenic_variant | MODIFIER |
| 1 | 74611440 | AX-97816370 | 0 | A | G | 0.123 | 6.11E-02 | 2.06E-01 | 2.69E+00 | 2.75E+00 | 7.63E-01 | no | Stk36 | missense_variant | MODERATE |
| 9 | 62192213 | AX-97839509 | 0 | G | T | 0.193 | -4.03E-02 | 1.34E-01 | 2.69E+00 | 2.72E+00 | 7.63E-01 | no | | regulatory_region_variant | MODIFIER |
| 9 | 65992349 | AX-170032573 | 0 | T | C | 0.193 | -4.03E-02 | 1.34E-01 | 2.69E+00 | 2.72E+00 | 7.63E-01 | no | Csnk1g1 | intron_variant | MODIFIER |
| 1 | 44732978 | AX-170287740 | 1 | G | T | 0.098 | -6.47E-02 | 2.19E-01 | 2.68E+00 | 2.76E+00 | 7.63E-01 | no | Gulp1 | intron_variant | MODIFIER |
| 11 | 3242050 | AX-97850544 | 0 | C | T | 0.061 | 7.24E-02 | 2.47E-01 | 2.66E+00 | 2.76E+00 | 7.63E-01 | no | Eif4enif1 | intron_variant | MODIFIER |
| 12 | 105651676 | AX-170414057 | 2 | A | T | 0.127 | -5.32E-02 | 1.82E-01 | 2.69E+00 | 2.77E+00 | 7.63E-01 | no | Atg2b | intron_variant | MODIFIER |
| 5 | 38720156 | AX-170024728 | 0 | T | C | 0.228 | 3.19E-02 | 1.09E-01 | 2.72E+00 | 2.81E+00 | 7.63E-01 | no | Clnk | intron_variant | MODIFIER |

| | | | | | | | | | | | | | | | |
|---|---|---|---|---|---|---|---|---|---|---|---|---|---|---|---|
| 5 | 38829263 | AX-170024729 | 0 | A | G | 0.228 | 3.19E-02 | 1.09E-01 | 2.72E+00 | 2.81E+00 | 7.63E-01 | no | Clnk | intron_variant | MODIFIER |
| 5 | 38840291 | AX-170024735 | 0 | G | A | 0.228 | 3.19E-02 | 1.09E-01 | 2.72E+00 | 2.81E+00 | 7.63E-01 | no | Clnk | intron_variant | MODIFIER |
| 5 | 38842475 | AX-97814698 | 0 | G | T | 0.228 | 3.19E-02 | 1.09E-01 | 2.72E+00 | 2.81E+00 | 7.63E-01 | no | Clnk | intron_variant | MODIFIER |
| 5 | 39078713 | AX-170267886 | 0 | A | G | 0.228 | 3.19E-02 | 1.09E-01 | 2.72E+00 | 2.81E+00 | 7.63E-01 | no | | intergenic_variant | MODIFIER |
| 5 | 39125991 | AX-170089454 | 0 | T | C | 0.228 | 3.19E-02 | 1.09E-01 | 2.72E+00 | 2.81E+00 | 7.63E-01 | no | | regulatory_region_variant | MODIFIER |
| 5 | 39151817 | AX-170191718 | 0 | G | A | 0.228 | 3.19E-02 | 1.09E-01 | 2.72E+00 | 2.81E+00 | 7.63E-01 | no | | intergenic_variant | MODIFIER |
| 5 | 34340418 | AX-97781801 | 2 | C | T | 0.336 | 3.18E-02 | 1.07E-01 | 2.68E+00 | 2.74E+00 | 7.64E-01 | no | Rnf4 | intron_variant | MODIFIER |
| 10 | 5788480 | AX-170151919 | 1 | G | A | 0.214 | 3.00E-02 | 1.03E-01 | 2.69E+00 | 2.79E+00 | 7.64E-01 | no | | intergenic_variant | MODIFIER |
| 7 | 59986065 | AX-170135316 | 0 | T | A | 0.421 | -2.88E-02 | 9.89E-02 | 2.70E+00 | 2.78E+00 | 7.64E-01 | no | Snrpn | intron_variant | MODIFIER |
| 7 | 135304208 | AX-170030042 | 0 | G | T | 0.254 | -2.52E-02 | 8.23E-02 | 2.60E+00 | 2.70E+00 | 7.64E-01 | no | | intergenic_variant | MODIFIER |
| 8 | 97898976 | AX-97787899 | 0 | A | G | 0.228 | 3.45E-02 | 1.16E-01 | 2.67E+00 | 2.74E+00 | 7.64E-01 | no | | intergenic_variant | MODIFIER |
| 8 | 98632174 | AX-170125096 | 0 | C | T | 0.228 | 3.45E-02 | 1.16E-01 | 2.67E+00 | 2.74E+00 | 7.64E-01 | no | | intergenic_variant | MODIFIER |
| 8 | 100404048 | AX-170181898 | 0 | A | T | 0.228 | 3.45E-02 | 1.16E-01 | 2.67E+00 | 2.74E+00 | 7.64E-01 | no | | intergenic_variant | MODIFIER |
| 8 | 100692758 | AX-170137737 | 0 | C | T | 0.228 | 3.45E-02 | 1.16E-01 | 2.67E+00 | 2.74E+00 | 7.64E-01 | no | Gm39232 | intron_variant,non_coding_transcript_variant | MODIFIER |
| 8 | 100769492 | AX-170256696 | 0 | G | A | 0.228 | 3.45E-02 | 1.16E-01 | 2.67E+00 | 2.74E+00 | 7.64E-01 | no | | intergenic_variant | MODIFIER |
| 8 | 100855119 | AX-170119149 | 0 | A | G | 0.228 | 3.45E-02 | 1.16E-01 | 2.67E+00 | 2.74E+00 | 7.64E-01 | no | | intergenic_variant | MODIFIER |
| 8 | 100920045 | AX-170177878 | 0 | C | T | 0.228 | 3.45E-02 | 1.16E-01 | 2.67E+00 | 2.74E+00 | 7.64E-01 | no | | intergenic_variant | MODIFIER |
| 8 | 100934431 | AX-170158824 | 0 | G | T | 0.228 | 3.45E-02 | 1.16E-01 | 2.67E+00 | 2.74E+00 | 7.64E-01 | no | | intergenic_variant | MODIFIER |
| 8 | 100938918 | AX-170189201 | 0 | C | T | 0.228 | 3.45E-02 | 1.16E-01 | 2.67E+00 | 2.74E+00 | 7.64E-01 | no | | intergenic_variant | MODIFIER |
| 8 | 100959915 | AX-170224290 | 0 | T | C | 0.228 | 3.45E-02 | 1.16E-01 | 2.67E+00 | 2.74E+00 | 7.64E-01 | no | | intergenic_variant | MODIFIER |
| 8 | 101106977 | AX-170481859 | 0 | T | C | 0.228 | 3.45E-02 | 1.16E-01 | 2.67E+00 | 2.74E+00 | 7.64E-01 | no | | intergenic_variant | MODIFIER |
| 8 | 101145087 | AX-170339433 | 0 | T | G | 0.228 | 3.45E-02 | 1.16E-01 | 2.67E+00 | 2.74E+00 | 7.64E-01 | no | | intergenic_variant | MODIFIER |
| 8 | 101179579 | AX-170031390 | 0 | T | A | 0.228 | 3.45E-02 | 1.16E-01 | 2.67E+00 | 2.74E+00 | 7.64E-01 | no | | intergenic_variant | MODIFIER |
| 8 | 101203396 | AX-170031385 | 0 | C | T | 0.228 | 3.45E-02 | 1.16E-01 | 2.67E+00 | 2.74E+00 | 7.64E-01 | no | | intergenic_variant | MODIFIER |
| 8 | 101205674 | AX-170031391 | 0 | G | T | 0.228 | 3.45E-02 | 1.16E-01 | 2.67E+00 | 2.74E+00 | 7.64E-01 | no | | intergenic_variant | MODIFIER |
| 8 | 101218563 | AX-170031386 | 0 | C | T | 0.228 | 3.45E-02 | 1.16E-01 | 2.67E+00 | 2.74E+00 | 7.64E-01 | no | | intergenic_variant | MODIFIER |
| 8 | 101220309 | AX-170328692 | 0 | T | C | 0.228 | 3.45E-02 | 1.16E-01 | 2.67E+00 | 2.74E+00 | 7.64E-01 | no | | intergenic_variant | MODIFIER |
| 8 | 101235608 | AX-97854545 | 0 | T | A | 0.228 | 3.45E-02 | 1.16E-01 | 2.67E+00 | 2.74E+00 | 7.64E-01 | no | | intergenic_variant | MODIFIER |
| 8 | 101274545 | AX-170031389 | 0 | T | C | 0.228 | 3.45E-02 | 1.16E-01 | 2.67E+00 | 2.74E+00 | 7.64E-01 | no | | intergenic_variant | MODIFIER |
| 8 | 101278747 | AX-170080988 | 0 | C | T | 0.228 | 3.45E-02 | 1.16E-01 | 2.67E+00 | 2.74E+00 | 7.64E-01 | no | | intergenic_variant | MODIFIER |
| 8 | 101288465 | AX-170031388 | 0 | T | G | 0.228 | 3.45E-02 | 1.16E-01 | 2.67E+00 | 2.74E+00 | 7.64E-01 | no | | intergenic_variant | MODIFIER |
| 8 | 101345614 | AX-170031392 | 0 | A | G | 0.228 | 3.45E-02 | 1.16E-01 | 2.67E+00 | 2.74E+00 | 7.64E-01 | no | | intergenic_variant | MODIFIER |
| 8 | 101346737 | AX-170031387 | 0 | A | C | 0.228 | 3.45E-02 | 1.16E-01 | 2.67E+00 | 2.74E+00 | 7.64E-01 | no | | intergenic_variant | MODIFIER |
| 8 | 101439267 | AX-170481961 | 0 | T | A | 0.228 | 3.45E-02 | 1.16E-01 | 2.67E+00 | 2.74E+00 | 7.64E-01 | no | | intergenic_variant | MODIFIER |
| 8 | 101460227 | AX-170535413 | 0 | C | G | 0.228 | 3.45E-02 | 1.16E-01 | 2.67E+00 | 2.74E+00 | 7.64E-01 | no | | intergenic_variant | MODIFIER |
| 8 | 101496741 | AX-170317609 | 0 | T | C | 0.228 | 3.45E-02 | 1.16E-01 | 2.67E+00 | 2.74E+00 | 7.64E-01 | no | | intergenic_variant | MODIFIER |
| 8 | 101518429 | AX-170431774 | 0 | T | C | 0.228 | 3.45E-02 | 1.16E-01 | 2.67E+00 | 2.74E+00 | 7.64E-01 | no | | intergenic_variant | MODIFIER |
| 8 | 101530314 | AX-170185253 | 0 | A | G | 0.228 | 3.45E-02 | 1.16E-01 | 2.67E+00 | 2.74E+00 | 7.64E-01 | no | | intergenic_variant | MODIFIER |
| 8 | 101540397 | AX-170249351 | 0 | T | A | 0.228 | 3.45E-02 | 1.16E-01 | 2.67E+00 | 2.74E+00 | 7.64E-01 | no | Gm22223 | upstream_gene_variant | MODIFIER |
| 8 | 101573910 | AX-170114848 | 0 | G | C | 0.228 | 3.45E-02 | 1.16E-01 | 2.67E+00 | 2.74E+00 | 7.64E-01 | no | | intergenic_variant | MODIFIER |
| 8 | 101579617 | AX-170478546 | 0 | G | C | 0.228 | 3.45E-02 | 1.16E-01 | 2.67E+00 | 2.74E+00 | 7.64E-01 | no | | intergenic_variant | MODIFIER |
| 8 | 101583732 | AX-170172607 | 0 | A | G | 0.228 | 3.45E-02 | 1.16E-01 | 2.67E+00 | 2.74E+00 | 7.64E-01 | no | | intergenic_variant | MODIFIER |
| 8 | 101583864 | AX-170271491 | 0 | C | T | 0.228 | 3.45E-02 | 1.16E-01 | 2.67E+00 | 2.74E+00 | 7.64E-01 | no | | intergenic_variant | MODIFIER |
| 8 | 101591373 | AX-170314360 | 0 | G | T | 0.228 | 3.45E-02 | 1.16E-01 | 2.67E+00 | 2.74E+00 | 7.64E-01 | no | | intergenic_variant | MODIFIER |
| 8 | 101591503 | AX-170373550 | 0 | C | T | 0.228 | 3.45E-02 | 1.16E-01 | 2.67E+00 | 2.74E+00 | 7.64E-01 | no | | intergenic_variant | MODIFIER |
| 8 | 101651815 | AX-170031393 | 0 | A | T | 0.228 | 3.45E-02 | 1.16E-01 | 2.67E+00 | 2.74E+00 | 7.64E-01 | no | | intergenic_variant | MODIFIER |
| 8 | 101668774 | AX-97790812 | 0 | C | A | 0.228 | 3.45E-02 | 1.16E-01 | 2.67E+00 | 2.74E+00 | 7.64E-01 | no | | intergenic_variant | MODIFIER |
| 8 | 101677231 | AX-170031394 | 0 | T | C | 0.228 | 3.45E-02 | 1.16E-01 | 2.67E+00 | 2.74E+00 | 7.64E-01 | no | | intergenic_variant | MODIFIER |
| 8 | 101680398 | AX-170080993 | 0 | T | C | 0.228 | 3.45E-02 | 1.16E-01 | 2.67E+00 | 2.74E+00 | 7.64E-01 | no | | intergenic_variant | MODIFIER |
| 9 | 40865140 | AX-170433829 | 1 | T | A | 0.438 | -2.20E-02 | 7.45E-02 | 2.66E+00 | 2.75E+00 | 7.64E-01 | no | | intergenic_variant | MODIFIER |
| 11 | 46980969 | AX-97810569 | 0 | T | C | 0.096 | -4.64E-02 | 1.56E-01 | 2.61E+00 | 2.74E+00 | 7.64E-01 | no | Sgcd | intron_variant | MODIFIER |
| 7 | 112806867 | AX-97838741 | 1 | G | A | 0.446 | 2.78E-02 | 9.31E-02 | 2.67E+00 | 2.72E+00 | 7.64E-01 | no | Tead1 | intron_variant | MODIFIER |
| 1 | 190062347 | AX-170438177 | 0 | A | T | 0.254 | 3.23E-02 | 9.44E-02 | 2.50E+00 | 2.60E+00 | 7.64E-01 | no | | regulatory_region_variant | MODIFIER |
| 9 | 112202535 | AX-170328148 | 0 | T | C | 0.123 | -3.19E-02 | 1.07E-01 | 2.64E+00 | 2.73E+00 | 7.64E-01 | no | Arpp21 | intron_variant | MODIFIER |

| | | | | | | | | | | | | | | | |
|---|---|---|---|---|---|---|---|---|---|---|---|---|---|---|---|
| 9 | 112247330 | AX-170090420 | 0 | G | A | 0.123 | -3.19E-02 | 1.07E-01 | 2.64E+00 | 2.73E+00 | 7.64E-01 | no | 2900079G21Rik | intron_variant,non_coding_transcript_variant | MODIFIER |
| 9 | 112262170 | AX-170491311 | 0 | T | C | 0.123 | -3.19E-02 | 1.07E-01 | 2.64E+00 | 2.73E+00 | 7.64E-01 | no | 2900079G21Rik | intron_variant,non_coding_transcript_variant | MODIFIER |
| 9 | 112267292 | AX-170156058 | 0 | C | T | 0.123 | -3.19E-02 | 1.07E-01 | 2.64E+00 | 2.73E+00 | 7.64E-01 | no | 2900079G21Rik | intron_variant,non_coding_transcript_variant | MODIFIER |
| 9 | 112303884 | AX-170387901 | 0 | A | G | 0.123 | -3.19E-02 | 1.07E-01 | 2.64E+00 | 2.73E+00 | 7.64E-01 | no | 2900079G21Rik | intron_variant,non_coding_transcript_variant | MODIFIER |
| 9 | 112316438 | AX-170165235 | 0 | G | T | 0.123 | -3.19E-02 | 1.07E-01 | 2.64E+00 | 2.73E+00 | 7.64E-01 | no | 2900079G21Rik | intron_variant,non_coding_transcript_variant | MODIFIER |
| 9 | 112388841 | AX-170108704 | 0 | G | T | 0.123 | -3.19E-02 | 1.07E-01 | 2.64E+00 | 2.73E+00 | 7.64E-01 | no | 4933411E02Rik | downstream_gene_variant | MODIFIER |
| 9 | 112395337 | AX-170394394 | 0 | A | G | 0.123 | -3.19E-02 | 1.07E-01 | 2.64E+00 | 2.73E+00 | 7.64E-01 | no | 4933411E02Rik | intron_variant,non_coding_transcript_variant | MODIFIER |
| 9 | 112544254 | AX-97786811 | 0 | G | A | 0.123 | -3.19E-02 | 1.07E-01 | 2.64E+00 | 2.73E+00 | 7.64E-01 | no | | intergenic_variant | MODIFIER |
| 9 | 112550425 | AX-170133106 | 0 | A | G | 0.123 | -3.19E-02 | 1.07E-01 | 2.64E+00 | 2.73E+00 | 7.64E-01 | no | | intergenic_variant | MODIFIER |
| 9 | 112550739 | AX-170201108 | 0 | T | C | 0.123 | -3.19E-02 | 1.07E-01 | 2.64E+00 | 2.73E+00 | 7.64E-01 | no | | intergenic_variant | MODIFIER |
| 9 | 112552406 | AX-170194199 | 0 | T | C | 0.123 | -3.19E-02 | 1.07E-01 | 2.64E+00 | 2.73E+00 | 7.64E-01 | no | | intergenic_variant | MODIFIER |
| 9 | 112568187 | AX-170415496 | 0 | C | T | 0.123 | -3.19E-02 | 1.07E-01 | 2.64E+00 | 2.73E+00 | 7.64E-01 | no | | intergenic_variant | MODIFIER |
| 9 | 112569308 | AX-170327718 | 0 | T | C | 0.123 | -3.19E-02 | 1.07E-01 | 2.64E+00 | 2.73E+00 | 7.64E-01 | no | | intergenic_variant | MODIFIER |
| 9 | 112572358 | AX-170096671 | 0 | C | T | 0.123 | -3.19E-02 | 1.07E-01 | 2.64E+00 | 2.73E+00 | 7.64E-01 | no | | intergenic_variant | MODIFIER |
| 9 | 112578177 | AX-170352786 | 0 | T | C | 0.123 | -3.19E-02 | 1.07E-01 | 2.64E+00 | 2.73E+00 | 7.64E-01 | no | | intergenic_variant | MODIFIER |
| 9 | 112583856 | AX-170338571 | 0 | T | C | 0.123 | -3.19E-02 | 1.07E-01 | 2.64E+00 | 2.73E+00 | 7.64E-01 | no | | intergenic_variant | MODIFIER |
| 9 | 112647975 | AX-97829334 | 0 | G | T | 0.123 | -3.19E-02 | 1.07E-01 | 2.64E+00 | 2.73E+00 | 7.64E-01 | no | | intergenic_variant | MODIFIER |
| 9 | 112648142 | AX-170317378 | 0 | G | A | 0.123 | -3.19E-02 | 1.07E-01 | 2.64E+00 | 2.73E+00 | 7.64E-01 | no | | intergenic_variant | MODIFIER |
| 9 | 112678651 | AX-170307457 | 0 | C | T | 0.123 | -3.19E-02 | 1.07E-01 | 2.64E+00 | 2.73E+00 | 7.64E-01 | no | | intergenic_variant | MODIFIER |
| 9 | 112736891 | AX-170263718 | 0 | C | T | 0.123 | -3.19E-02 | 1.07E-01 | 2.64E+00 | 2.73E+00 | 7.64E-01 | no | | intergenic_variant | MODIFIER |
| 9 | 112777167 | AX-170273692 | 0 | G | T | 0.123 | -3.19E-02 | 1.07E-01 | 2.64E+00 | 2.73E+00 | 7.64E-01 | no | | intergenic_variant | MODIFIER |
| 9 | 112804997 | AX-170083491 | 0 | A | G | 0.123 | -3.19E-02 | 1.07E-01 | 2.64E+00 | 2.73E+00 | 7.64E-01 | no | | intergenic_variant | MODIFIER |
| 9 | 112822374 | AX-170276811 | 0 | C | T | 0.123 | -3.19E-02 | 1.07E-01 | 2.64E+00 | 2.73E+00 | 7.64E-01 | no | | intergenic_variant | MODIFIER |
| 9 | 112822403 | AX-170424784 | 0 | C | T | 0.123 | -3.19E-02 | 1.07E-01 | 2.64E+00 | 2.73E+00 | 7.64E-01 | no | | intergenic_variant | MODIFIER |
| 9 | 112822647 | AX-170350419 | 0 | G | A | 0.123 | -3.19E-02 | 1.07E-01 | 2.64E+00 | 2.73E+00 | 7.64E-01 | no | | intergenic_variant | MODIFIER |
| 9 | 112825604 | AX-170138593 | 0 | G | A | 0.123 | -3.19E-02 | 1.07E-01 | 2.64E+00 | 2.73E+00 | 7.64E-01 | no | | intergenic_variant | MODIFIER |
| 9 | 112827297 | AX-170145266 | 0 | T | C | 0.123 | -3.19E-02 | 1.07E-01 | 2.64E+00 | 2.73E+00 | 7.64E-01 | no | | intergenic_variant | MODIFIER |
| 9 | 112831064 | AX-170490153 | 0 | A | G | 0.123 | -3.19E-02 | 1.07E-01 | 2.64E+00 | 2.73E+00 | 7.64E-01 | no | | intergenic_variant | MODIFIER |
| 9 | 112832163 | AX-170282019 | 0 | T | C | 0.123 | -3.19E-02 | 1.07E-01 | 2.64E+00 | 2.73E+00 | 7.64E-01 | no | | intergenic_variant | MODIFIER |
| 9 | 112865119 | AX-170302245 | 0 | G | T | 0.123 | -3.19E-02 | 1.07E-01 | 2.64E+00 | 2.73E+00 | 7.64E-01 | no | | regulatory_region_variant | MODIFIER |
| 9 | 112867896 | AX-170335447 | 0 | C | T | 0.123 | -3.19E-02 | 1.07E-01 | 2.64E+00 | 2.73E+00 | 7.64E-01 | no | | intergenic_variant | MODIFIER |
| 9 | 112876151 | AX-170448155 | 0 | C | T | 0.123 | -3.19E-02 | 1.07E-01 | 2.64E+00 | 2.73E+00 | 7.64E-01 | no | | intergenic_variant | MODIFIER |
| 9 | 112921572 | AX-170300122 | 0 | C | T | 0.123 | -3.19E-02 | 1.07E-01 | 2.64E+00 | 2.73E+00 | 7.64E-01 | no | | intergenic_variant | MODIFIER |
| 9 | 112975345 | AX-97827598 | 0 | G | A | 0.123 | -3.19E-02 | 1.07E-01 | 2.64E+00 | 2.73E+00 | 7.64E-01 | no | | intergenic_variant | MODIFIER |
| 9 | 112975444 | AX-170360333 | 0 | T | C | 0.123 | -3.19E-02 | 1.07E-01 | 2.64E+00 | 2.73E+00 | 7.64E-01 | no | | intergenic_variant | MODIFIER |
| 9 | 113057851 | AX-170083498 | 0 | A | G | 0.123 | -3.19E-02 | 1.07E-01 | 2.64E+00 | 2.73E+00 | 7.64E-01 | no | | intergenic_variant | MODIFIER |
| 9 | 113057967 | AX-170186756 | 0 | T | G | 0.123 | -3.19E-02 | 1.07E-01 | 2.64E+00 | 2.73E+00 | 7.64E-01 | no | | intergenic_variant | MODIFIER |
| 9 | 113061098 | AX-97809008 | 0 | G | A | 0.123 | -3.19E-02 | 1.07E-01 | 2.64E+00 | 2.73E+00 | 7.64E-01 | no | | intergenic_variant | MODIFIER |
| 9 | 113061149 | AX-170194785 | 0 | C | G | 0.123 | -3.19E-02 | 1.07E-01 | 2.64E+00 | 2.73E+00 | 7.64E-01 | no | | intergenic_variant | MODIFIER |
| 9 | 113071709 | AX-170083496 | 0 | C | T | 0.123 | -3.19E-02 | 1.07E-01 | 2.64E+00 | 2.73E+00 | 7.64E-01 | no | Gm36251 | non_coding_transcript_exon_variant | MODIFIER |
| 9 | 113101771 | AX-170279346 | 0 | T | C | 0.123 | -3.19E-02 | 1.07E-01 | 2.64E+00 | 2.73E+00 | 7.64E-01 | no | Gm36251 | intron_variant,non_coding_transcript_variant | MODIFIER |
| 9 | 113101823 | AX-170153819 | 0 | A | G | 0.123 | -3.19E-02 | 1.07E-01 | 2.64E+00 | 2.73E+00 | 7.64E-01 | no | Gm36251 | intron_variant,non_coding_transcript_variant | MODIFIER |
| 9 | 113101955 | AX-170228252 | 0 | C | T | 0.123 | -3.19E-02 | 1.07E-01 | 2.64E+00 | 2.73E+00 | 7.64E-01 | no | Gm36251 | intron_variant,non_coding_transcript_variant | MODIFIER |
| 9 | 113102299 | AX-170215650 | 0 | A | G | 0.123 | -3.19E-02 | 1.07E-01 | 2.64E+00 | 2.73E+00 | 7.64E-01 | no | Gm36251 | intron_variant,non_coding_transcript_variant | MODIFIER |
| 9 | 113102449 | AX-170096673 | 0 | A | G | 0.123 | -3.19E-02 | 1.07E-01 | 2.64E+00 | 2.73E+00 | 7.64E-01 | no | Gm36251 | intron_variant,non_coding_transcript_variant | MODIFIER |
| 9 | 113105674 | AX-170104943 | 0 | G | T | 0.123 | -3.19E-02 | 1.07E-01 | 2.64E+00 | 2.73E+00 | 7.64E-01 | no | Gm36251 | intron_variant,non_coding_transcript_variant | MODIFIER |
| 9 | 113180166 | AX-170323535 | 0 | C | G | 0.123 | -3.19E-02 | 1.07E-01 | 2.64E+00 | 2.73E+00 | 7.64E-01 | no | Gm4657 | intron_variant,non_coding_transcript_variant | MODIFIER |
| 9 | 113185052 | AX-170033302 | 0 | T | C | 0.123 | -3.19E-02 | 1.07E-01 | 2.64E+00 | 2.73E+00 | 7.64E-01 | no | Gm4657 | intron_variant,non_coding_transcript_variant | MODIFIER |
| 9 | 113185450 | AX-170033300 | 0 | C | T | 0.123 | -3.19E-02 | 1.07E-01 | 2.64E+00 | 2.73E+00 | 7.64E-01 | no | Gm4657 | intron_variant,non_coding_transcript_variant | MODIFIER |
| 9 | 113204961 | AX-170033299 | 0 | A | G | 0.123 | -3.19E-02 | 1.07E-01 | 2.64E+00 | 2.73E+00 | 7.64E-01 | no | | intergenic_variant | MODIFIER |
| 9 | 113229524 | AX-170274876 | 0 | G | A | 0.123 | -3.19E-02 | 1.07E-01 | 2.64E+00 | 2.73E+00 | 7.64E-01 | no | | intergenic_variant | MODIFIER |
| 9 | 113244851 | AX-170083504 | 0 | G | T | 0.123 | -3.19E-02 | 1.07E-01 | 2.64E+00 | 2.73E+00 | 7.64E-01 | no | | intergenic_variant | MODIFIER |
| 9 | 113255150 | AX-170100330 | 0 | T | C | 0.123 | -3.19E-02 | 1.07E-01 | 2.64E+00 | 2.73E+00 | 7.64E-01 | no | | intergenic_variant | MODIFIER |
| 9 | 113271933 | AX-170111884 | 0 | A | C | 0.123 | -3.19E-02 | 1.07E-01 | 2.64E+00 | 2.73E+00 | 7.64E-01 | no | | intergenic_variant | MODIFIER |

| | | | | | | | | | | | | | | | |
|---|---|---|---|---|---|---|---|---|---|---|---|---|---|---|---|
| 9 | 113275248 | AX-170265586 | 0 | A | G | 0.123 | -3.19E-02 | 1.07E-01 | 2.64E+00 | 2.73E+00 | 7.64E-01 | no | | intergenic_variant | MODIFIER |
| 9 | 113291117 | AX-170090421 | 0 | G | A | 0.123 | -3.19E-02 | 1.07E-01 | 2.64E+00 | 2.73E+00 | 7.64E-01 | no | Gm36367 | intron_variant,non_coding_transcript_variant | MODIFIER |
| 9 | 113305774 | AX-170225532 | 0 | C | T | 0.123 | -3.19E-02 | 1.07E-01 | 2.64E+00 | 2.73E+00 | 7.64E-01 | no | Gm36367 | intron_variant,non_coding_transcript_variant | MODIFIER |
| 9 | 113320363 | AX-97797194 | 0 | T | C | 0.123 | -3.19E-02 | 1.07E-01 | 2.64E+00 | 2.73E+00 | 7.64E-01 | no | Gm39436 | downstream_gene_variant | MODIFIER |
| 9 | 113324235 | AX-170357768 | 0 | T | C | 0.123 | -3.19E-02 | 1.07E-01 | 2.64E+00 | 2.73E+00 | 7.64E-01 | no | Gm39436 | downstream_gene_variant | MODIFIER |
| 9 | 113344358 | AX-170446826 | 0 | T | C | 0.123 | -3.19E-02 | 1.07E-01 | 2.64E+00 | 2.73E+00 | 7.64E-01 | no | Gm39436 | intron_variant,non_coding_transcript_variant | MODIFIER |
| 9 | 113348898 | AX-170497325 | 0 | A | T | 0.123 | -3.19E-02 | 1.07E-01 | 2.64E+00 | 2.73E+00 | 7.64E-01 | no | Gm39436 | intron_variant,non_coding_transcript_variant | MODIFIER |
| 9 | 113350584 | AX-170471188 | 0 | T | C | 0.123 | -3.19E-02 | 1.07E-01 | 2.64E+00 | 2.73E+00 | 7.64E-01 | no | Gm39436 | intron_variant,non_coding_transcript_variant | MODIFIER |
| 9 | 113351959 | AX-170488540 | 0 | C | T | 0.123 | -3.19E-02 | 1.07E-01 | 2.64E+00 | 2.73E+00 | 7.64E-01 | no | Gm39436 | intron_variant,non_coding_transcript_variant | MODIFIER |
| 9 | 113352178 | AX-97830678 | 0 | T | C | 0.123 | -3.19E-02 | 1.07E-01 | 2.64E+00 | 2.73E+00 | 7.64E-01 | no | Gm39436 | intron_variant,non_coding_transcript_variant | MODIFIER |
| 9 | 113356856 | AX-170083505 | 0 | C | T | 0.123 | -3.19E-02 | 1.07E-01 | 2.64E+00 | 2.73E+00 | 7.64E-01 | no | Gm39436 | upstream_gene_variant | MODIFIER |
| 9 | 113357001 | AX-170111439 | 0 | G | A | 0.123 | -3.19E-02 | 1.07E-01 | 2.64E+00 | 2.73E+00 | 7.64E-01 | no | Gm39436 | upstream_gene_variant | MODIFIER |
| 9 | 113357996 | AX-170348481 | 0 | A | G | 0.123 | -3.19E-02 | 1.07E-01 | 2.64E+00 | 2.73E+00 | 7.64E-01 | no | | intergenic_variant | MODIFIER |
| 9 | 113368983 | AX-170391886 | 0 | A | G | 0.123 | -3.19E-02 | 1.07E-01 | 2.64E+00 | 2.73E+00 | 7.64E-01 | no | | intergenic_variant | MODIFIER |
| 9 | 113387143 | AX-170334894 | 0 | A | G | 0.123 | -3.19E-02 | 1.07E-01 | 2.64E+00 | 2.73E+00 | 7.64E-01 | no | | regulatory_region_variant | MODIFIER |
| 9 | 113387987 | AX-170410916 | 0 | G | A | 0.123 | -3.19E-02 | 1.07E-01 | 2.64E+00 | 2.73E+00 | 7.64E-01 | no | | regulatory_region_variant | MODIFIER |
| 9 | 113402439 | AX-170476835 | 0 | T | C | 0.123 | -3.19E-02 | 1.07E-01 | 2.64E+00 | 2.73E+00 | 7.64E-01 | no | Gm22492 | upstream_gene_variant | MODIFIER |
| 9 | 113403278 | AX-97816385 | 0 | T | C | 0.123 | -3.19E-02 | 1.07E-01 | 2.64E+00 | 2.73E+00 | 7.64E-01 | no | Gm22492 | upstream_gene_variant | MODIFIER |
| 9 | 113403721 | AX-170457548 | 0 | A | G | 0.123 | -3.19E-02 | 1.07E-01 | 2.64E+00 | 2.73E+00 | 7.64E-01 | no | Gm22492 | non_coding_transcript_exon_variant | MODIFIER |
| 9 | 113444442 | AX-170209922 | 0 | T | C | 0.123 | -3.19E-02 | 1.07E-01 | 2.64E+00 | 2.73E+00 | 7.64E-01 | no | | intergenic_variant | MODIFIER |
| 9 | 113446322 | AX-170163555 | 0 | A | T | 0.123 | -3.19E-02 | 1.07E-01 | 2.64E+00 | 2.73E+00 | 7.64E-01 | no | | intergenic_variant | MODIFIER |
| 9 | 113458285 | AX-170353059 | 0 | T | C | 0.123 | -3.19E-02 | 1.07E-01 | 2.64E+00 | 2.73E+00 | 7.64E-01 | no | | intergenic_variant | MODIFIER |
| 9 | 113458320 | AX-170485366 | 0 | T | A | 0.123 | -3.19E-02 | 1.07E-01 | 2.64E+00 | 2.73E+00 | 7.64E-01 | no | | intergenic_variant | MODIFIER |
| 9 | 113458349 | AX-170410805 | 0 | C | T | 0.123 | -3.19E-02 | 1.07E-01 | 2.64E+00 | 2.73E+00 | 7.64E-01 | no | | intergenic_variant | MODIFIER |
| 9 | 113458557 | AX-170388630 | 0 | A | G | 0.123 | -3.19E-02 | 1.07E-01 | 2.64E+00 | 2.73E+00 | 7.64E-01 | no | | intergenic_variant | MODIFIER |
| 9 | 113484157 | AX-170461798 | 0 | C | T | 0.123 | -3.19E-02 | 1.07E-01 | 2.64E+00 | 2.73E+00 | 7.64E-01 | no | | intergenic_variant | MODIFIER |
| 9 | 113484649 | AX-170457019 | 0 | C | T | 0.123 | -3.19E-02 | 1.07E-01 | 2.64E+00 | 2.73E+00 | 7.64E-01 | no | | intergenic_variant | MODIFIER |
| 9 | 113502568 | AX-97789108 | 0 | A | G | 0.123 | -3.19E-02 | 1.07E-01 | 2.64E+00 | 2.73E+00 | 7.64E-01 | no | AU023762 | intron_variant,non_coding_transcript_variant | MODIFIER |
| 9 | 113505553 | AX-170225164 | 0 | A | T | 0.123 | -3.19E-02 | 1.07E-01 | 2.64E+00 | 2.73E+00 | 7.64E-01 | no | AU023762 | intron_variant,non_coding_transcript_variant | MODIFIER |
| 9 | 84566352 | AX-170358641 | 1 | A | G | 0.214 | 3.74E-02 | 1.28E-01 | 2.73E+00 | 2.79E+00 | 7.64E-01 | no | | intergenic_variant | MODIFIER |
| 8 | 101523300 | AX-97780758 | 1 | A | G | 0.232 | 3.42E-02 | 1.15E-01 | 2.66E+00 | 2.73E+00 | 7.64E-01 | no | | intergenic_variant | MODIFIER |
| 7 | 119137630 | AX-170389764 | 1 | A | G | 0.384 | 2.60E-02 | 8.69E-02 | 2.64E+00 | 2.72E+00 | 7.65E-01 | no | Gpr139 | downstream_gene_variant | MODIFIER |
| 5 | 106841764 | AX-170397539 | 0 | A | G | 0.439 | 3.40E-02 | 1.13E-01 | 2.65E+00 | 2.69E+00 | 7.65E-01 | no | Hfm1 | missense_variant,splice_region_variant | MODERATE |
| 2 | 93167201 | AX-170229330 | 0 | C | A | 0.053 | -8.85E-02 | 3.00E-01 | 2.73E+00 | 2.75E+00 | 7.65E-01 | no | | regulatory_region_variant | MODIFIER |
| 2 | 80379812 | AX-170018151 | 1 | A | C | 0.054 | -8.82E-02 | 2.99E-01 | 2.73E+00 | 2.76E+00 | 7.65E-01 | no | | intergenic_variant | MODIFIER |
| 2 | 109705407 | AX-97838479 | 1 | G | A | 0.054 | -8.82E-02 | 2.99E-01 | 2.73E+00 | 2.76E+00 | 7.65E-01 | no | Bdnf | intron_variant | MODIFIER |
| 5 | 24018412 | AX-170276314 | 1 | T | C | 0.054 | -8.82E-02 | 2.99E-01 | 2.73E+00 | 2.76E+00 | 7.65E-01 | no | Fam126a | intron_variant | MODIFIER |
| 13 | 53248165 | AX-97808160 | 0 | G | T | 0.526 | 2.14E-02 | 6.82E-02 | 2.56E+00 | 2.67E+00 | 7.65E-01 | no | Ror2 | intron_variant | MODIFIER |
| 5 | 34360030 | AX-170137178 | 1 | T | C | 0.33 | 3.22E-02 | 1.09E-01 | 2.68E+00 | 2.73E+00 | 7.65E-01 | no | Rnf4 | downstream_gene_variant | MODIFIER |
| 8 | 78036935 | AX-97853469 | 0 | C | T | 0.193 | -4.06E-02 | 1.33E-01 | 2.62E+00 | 2.69E+00 | 7.65E-01 | no | | intergenic_variant | MODIFIER |
| 1 | 3129299 | AX-170190667 | 0 | C | T | 0.518 | 2.21E-02 | 6.92E-02 | 2.57E+00 | 2.64E+00 | 7.65E-01 | no | | intergenic_variant | MODIFIER |
| 1 | 3135418 | AX-169999828 | 0 | A | G | 0.518 | 2.21E-02 | 6.92E-02 | 2.57E+00 | 2.64E+00 | 7.65E-01 | no | | intergenic_variant | MODIFIER |
| 1 | 3135686 | AX-97804858 | 0 | G | A | 0.518 | 2.21E-02 | 6.92E-02 | 2.57E+00 | 2.64E+00 | 7.65E-01 | no | | intergenic_variant | MODIFIER |
| 1 | 3206491 | AX-169999832 | 0 | G | C | 0.518 | 2.21E-02 | 6.92E-02 | 2.57E+00 | 2.64E+00 | 7.65E-01 | no | Xkr4 | downstream_gene_variant | MODIFIER |
| 1 | 3211082 | AX-169999830 | 0 | T | C | 0.518 | 2.21E-02 | 6.92E-02 | 2.57E+00 | 2.64E+00 | 7.65E-01 | no | Xkr4 | downstream_gene_variant | MODIFIER |
| 1 | 3219103 | AX-169999831 | 0 | T | G | 0.518 | 2.21E-02 | 6.92E-02 | 2.57E+00 | 2.64E+00 | 7.65E-01 | no | Xkr4 | intron_variant | MODIFIER |
| 1 | 3255359 | AX-170085958 | 0 | C | G | 0.518 | 2.21E-02 | 6.92E-02 | 2.57E+00 | 2.64E+00 | 7.65E-01 | no | Xkr4 | intron_variant | MODIFIER |
| 1 | 3265516 | AX-170085963 | 0 | A | G | 0.518 | 2.21E-02 | 6.92E-02 | 2.57E+00 | 2.64E+00 | 7.65E-01 | no | Xkr4 | intron_variant | MODIFIER |
| 1 | 3320361 | AX-170085960 | 0 | T | C | 0.518 | 2.21E-02 | 6.92E-02 | 2.57E+00 | 2.64E+00 | 7.65E-01 | no | Xkr4 | intron_variant | MODIFIER |
| 1 | 3323400 | AX-170513178 | 0 | T | C | 0.518 | 2.21E-02 | 6.92E-02 | 2.57E+00 | 2.64E+00 | 7.65E-01 | no | Xkr4 | intron_variant | MODIFIER |
| 1 | 3323489 | AX-170085961 | 0 | A | G | 0.518 | 2.21E-02 | 6.92E-02 | 2.57E+00 | 2.64E+00 | 7.65E-01 | no | Xkr4 | intron_variant | MODIFIER |
| 1 | 3355656 | AX-170386943 | 0 | G | A | 0.518 | 2.21E-02 | 6.92E-02 | 2.57E+00 | 2.64E+00 | 7.65E-01 | no | Xkr4 | intron_variant | MODIFIER |
| 1 | 3355917 | AX-170128133 | 0 | G | A | 0.518 | 2.21E-02 | 6.92E-02 | 2.57E+00 | 2.64E+00 | 7.65E-01 | no | Xkr4 | intron_variant | MODIFIER |
| 1 | 3359621 | AX-170124939 | 0 | G | A | 0.518 | 2.21E-02 | 6.92E-02 | 2.57E+00 | 2.64E+00 | 7.65E-01 | no | Xkr4 | intron_variant | MODIFIER |

| | | | | | | | | | | | | | | | |
|---|---|---|---|---|---|---|---|---|---|---|---|---|---|---|---|
| 1 | 3360461 | AX-170528434 | 0 | T | C | 0.518 | 2.21E-02 | 6.92E-02 | 2.57E+00 | 2.64E+00 | 7.65E-01 | no | Xkr4 | intron_variant | MODIFIER |
| 1 | 3362907 | AX-170268413 | 0 | C | T | 0.518 | 2.21E-02 | 6.92E-02 | 2.57E+00 | 2.64E+00 | 7.65E-01 | no | Xkr4 | intron_variant | MODIFIER |
| 1 | 3362949 | AX-97797866 | 0 | A | G | 0.518 | 2.21E-02 | 6.92E-02 | 2.57E+00 | 2.64E+00 | 7.65E-01 | no | Xkr4 | intron_variant | MODIFIER |
| 1 | 3366585 | AX-170383534 | 0 | A | C | 0.518 | 2.21E-02 | 6.92E-02 | 2.57E+00 | 2.64E+00 | 7.65E-01 | no | Xkr4 | intron_variant | MODIFIER |
| 1 | 3367041 | AX-170301923 | 0 | C | T | 0.518 | 2.21E-02 | 6.92E-02 | 2.57E+00 | 2.64E+00 | 7.65E-01 | no | Xkr4 | intron_variant | MODIFIER |
| 1 | 3367883 | AX-170501399 | 0 | G | A | 0.518 | 2.21E-02 | 6.92E-02 | 2.57E+00 | 2.64E+00 | 7.65E-01 | no | Xkr4 | intron_variant | MODIFIER |
| 1 | 3381737 | AX-170517778 | 0 | T | C | 0.518 | 2.21E-02 | 6.92E-02 | 2.57E+00 | 2.64E+00 | 7.65E-01 | no | Xkr4 | intron_variant | MODIFIER |
| 1 | 3438213 | AX-170401142 | 0 | A | G | 0.518 | 2.21E-02 | 6.92E-02 | 2.57E+00 | 2.64E+00 | 7.65E-01 | no | Xkr4 | intron_variant | MODIFIER |
| 1 | 3453121 | AX-97806506 | 0 | A | G | 0.518 | 2.21E-02 | 6.92E-02 | 2.57E+00 | 2.64E+00 | 7.65E-01 | no | Xkr4 | intron_variant | MODIFIER |
| 1 | 3466623 | AX-170318095 | 0 | T | G | 0.518 | 2.21E-02 | 6.92E-02 | 2.57E+00 | 2.64E+00 | 7.65E-01 | no | Xkr4 | intron_variant | MODIFIER |
| 1 | 3466799 | AX-170376866 | 0 | A | T | 0.518 | 2.21E-02 | 6.92E-02 | 2.57E+00 | 2.64E+00 | 7.65E-01 | no | Xkr4 | intron_variant | MODIFIER |
| 1 | 3479267 | AX-170450952 | 0 | A | T | 0.518 | 2.21E-02 | 6.92E-02 | 2.57E+00 | 2.64E+00 | 7.65E-01 | no | Xkr4 | intron_variant | MODIFIER |
| 1 | 3492195 | AX-170239468 | 0 | T | C | 0.518 | 2.21E-02 | 6.92E-02 | 2.57E+00 | 2.64E+00 | 7.65E-01 | no | Xkr4 | intron_variant | MODIFIER |
| 1 | 3494927 | AX-170157377 | 0 | A | G | 0.518 | 2.21E-02 | 6.92E-02 | 2.57E+00 | 2.64E+00 | 7.65E-01 | no | Xkr4 | intron_variant | MODIFIER |
| 1 | 3497294 | AX-170129614 | 0 | T | C | 0.518 | 2.21E-02 | 6.92E-02 | 2.57E+00 | 2.64E+00 | 7.65E-01 | no | Xkr4 | intron_variant | MODIFIER |
| 1 | 3877160 | AX-170109620 | 0 | G | A | 0.518 | 2.21E-02 | 6.92E-02 | 2.57E+00 | 2.64E+00 | 7.65E-01 | no | | intergenic_variant | MODIFIER |
| 1 | 3887862 | AX-170091372 | 0 | C | T | 0.518 | 2.21E-02 | 6.92E-02 | 2.57E+00 | 2.64E+00 | 7.65E-01 | no | | intergenic_variant | MODIFIER |
| 1 | 3899239 | AX-97810598 | 0 | G | A | 0.518 | 2.21E-02 | 6.92E-02 | 2.57E+00 | 2.64E+00 | 7.65E-01 | no | | intergenic_variant | MODIFIER |
| 1 | 3953455 | AX-170494289 | 0 | A | G | 0.518 | 2.21E-02 | 6.92E-02 | 2.57E+00 | 2.64E+00 | 7.65E-01 | no | Gm37381 | intron_variant,non_coding_transcript_variant | MODIFIER |
| 1 | 3971811 | AX-170254075 | 0 | T | C | 0.518 | 2.21E-02 | 6.92E-02 | 2.57E+00 | 2.64E+00 | 7.65E-01 | no | Gm37381 | intron_variant,non_coding_transcript_variant | MODIFIER |
| 1 | 4004793 | AX-170383615 | 0 | C | A | 0.518 | 2.21E-02 | 6.92E-02 | 2.57E+00 | 2.64E+00 | 7.65E-01 | no | Rp1 | intron_variant | MODIFIER |
| 1 | 4028619 | AX-170207714 | 0 | A | G | 0.518 | 2.21E-02 | 6.92E-02 | 2.57E+00 | 2.64E+00 | 7.65E-01 | no | Rp1 | intron_variant | MODIFIER |
| 1 | 4029675 | AX-170450770 | 0 | C | T | 0.518 | 2.21E-02 | 6.92E-02 | 2.57E+00 | 2.64E+00 | 7.65E-01 | no | Rp1 | intron_variant | MODIFIER |
| 1 | 4058749 | AX-170040249 | 0 | C | A | 0.518 | 2.21E-02 | 6.92E-02 | 2.57E+00 | 2.64E+00 | 7.65E-01 | no | Rp1 | intron_variant | MODIFIER |
| 1 | 4058803 | AX-170172748 | 0 | G | C | 0.518 | 2.21E-02 | 6.92E-02 | 2.57E+00 | 2.64E+00 | 7.65E-01 | no | Rp1 | intron_variant | MODIFIER |
| 1 | 4108464 | AX-170416961 | 0 | A | G | 0.518 | 2.21E-02 | 6.92E-02 | 2.57E+00 | 2.64E+00 | 7.65E-01 | no | Rp1 | intron_variant | MODIFIER |
| 1 | 4108608 | AX-170230492 | 0 | G | A | 0.518 | 2.21E-02 | 6.92E-02 | 2.57E+00 | 2.64E+00 | 7.65E-01 | no | Rp1 | intron_variant | MODIFIER |
| 1 | 4116917 | AX-170146073 | 0 | G | A | 0.518 | 2.21E-02 | 6.92E-02 | 2.57E+00 | 2.64E+00 | 7.65E-01 | no | Rp1 | intron_variant | MODIFIER |
| 1 | 4613691 | AX-170247626 | 0 | A | G | 0.518 | 2.21E-02 | 6.92E-02 | 2.57E+00 | 2.64E+00 | 7.65E-01 | no | Gm7369 | downstream_gene_variant | MODIFIER |
| 1 | 4627892 | AX-170501731 | 0 | C | A | 0.518 | 2.21E-02 | 6.92E-02 | 2.57E+00 | 2.64E+00 | 7.65E-01 | no | | intergenic_variant | MODIFIER |
| 1 | 4627984 | AX-170236023 | 0 | T | C | 0.518 | 2.21E-02 | 6.92E-02 | 2.57E+00 | 2.64E+00 | 7.65E-01 | no | | intergenic_variant | MODIFIER |
| 1 | 4643553 | AX-97853829 | 0 | C | T | 0.518 | 2.21E-02 | 6.92E-02 | 2.57E+00 | 2.64E+00 | 7.65E-01 | no | | intergenic_variant | MODIFIER |
| 1 | 4643630 | AX-170260336 | 0 | A | G | 0.518 | 2.21E-02 | 6.92E-02 | 2.57E+00 | 2.64E+00 | 7.65E-01 | no | | intergenic_variant | MODIFIER |
| 1 | 4643661 | AX-170136623 | 0 | T | C | 0.518 | 2.21E-02 | 6.92E-02 | 2.57E+00 | 2.64E+00 | 7.65E-01 | no | | intergenic_variant | MODIFIER |
| 1 | 4643840 | AX-170353337 | 0 | C | T | 0.518 | 2.21E-02 | 6.92E-02 | 2.57E+00 | 2.64E+00 | 7.65E-01 | no | | intergenic_variant | MODIFIER |
| 1 | 4661140 | AX-170517470 | 0 | G | A | 0.518 | 2.21E-02 | 6.92E-02 | 2.57E+00 | 2.64E+00 | 7.65E-01 | no | | intergenic_variant | MODIFIER |
| 1 | 4689445 | AX-97842478 | 0 | G | C | 0.518 | 2.21E-02 | 6.92E-02 | 2.57E+00 | 2.64E+00 | 7.65E-01 | no | Gm6085 | upstream_gene_variant | MODIFIER |
| 1 | 4718171 | AX-169999854 | 0 | C | T | 0.518 | 2.21E-02 | 6.92E-02 | 2.57E+00 | 2.64E+00 | 7.65E-01 | no | | intergenic_variant | MODIFIER |
| 1 | 4721145 | AX-97833012 | 0 | A | G | 0.518 | 2.21E-02 | 6.92E-02 | 2.57E+00 | 2.64E+00 | 7.65E-01 | no | Gm25493 | downstream_gene_variant | MODIFIER |
| 1 | 4759718 | AX-169999852 | 0 | A | G | 0.518 | 2.21E-02 | 6.92E-02 | 2.57E+00 | 2.64E+00 | 7.65E-01 | no | | intergenic_variant | MODIFIER |
| 1 | 4774069 | AX-170499001 | 0 | G | C | 0.518 | 2.21E-02 | 6.92E-02 | 2.57E+00 | 2.64E+00 | 7.65E-01 | no | Mrpl15 | downstream_gene_variant | MODIFIER |
| 1 | 4774160 | AX-170404815 | 0 | T | G | 0.518 | 2.21E-02 | 6.92E-02 | 2.57E+00 | 2.64E+00 | 7.65E-01 | no | Mrpl15 | downstream_gene_variant | MODIFIER |
| 1 | 4775973 | AX-170320611 | 0 | C | G | 0.518 | 2.21E-02 | 6.92E-02 | 2.57E+00 | 2.64E+00 | 7.65E-01 | no | Mrpl15 | intron_variant,NMD_transcript_variant | MODIFIER |
| 1 | 4777026 | AX-170439895 | 0 | G | A | 0.518 | 2.21E-02 | 6.92E-02 | 2.57E+00 | 2.64E+00 | 7.65E-01 | no | Mrpl15 | intron_variant,NMD_transcript_variant | MODIFIER |
| 1 | 4777943 | AX-169999851 | 0 | A | G | 0.518 | 2.21E-02 | 6.92E-02 | 2.57E+00 | 2.64E+00 | 7.65E-01 | no | Mrpl15 | intron_variant,NMD_transcript_variant | MODIFIER |
| 1 | 4780260 | AX-170528467 | 0 | T | C | 0.518 | 2.21E-02 | 6.92E-02 | 2.57E+00 | 2.64E+00 | 7.65E-01 | no | Mrpl15 | intron_variant,NMD_transcript_variant | MODIFIER |
| 1 | 4780355 | AX-170086291 | 0 | T | C | 0.518 | 2.21E-02 | 6.92E-02 | 2.57E+00 | 2.64E+00 | 7.65E-01 | no | Mrpl15 | intron_variant,NMD_transcript_variant | MODIFIER |
| 1 | 4796014 | AX-169999849 | 0 | T | A | 0.518 | 2.21E-02 | 6.92E-02 | 2.57E+00 | 2.64E+00 | 7.65E-01 | no | | intergenic_variant | MODIFIER |
| 1 | 4867555 | AX-170163352 | 0 | C | A | 0.518 | 2.21E-02 | 6.92E-02 | 2.57E+00 | 2.64E+00 | 7.65E-01 | no | Tcea1 | intron_variant | MODIFIER |
| 1 | 4891489 | AX-170306813 | 0 | G | A | 0.518 | 2.21E-02 | 6.92E-02 | 2.57E+00 | 2.64E+00 | 7.65E-01 | no | Tcea1 | intron_variant | MODIFIER |
| 1 | 4899398 | AX-170135796 | 0 | A | G | 0.518 | 2.21E-02 | 6.92E-02 | 2.57E+00 | 2.64E+00 | 7.65E-01 | no | Tcea1 | downstream_gene_variant | MODIFIER |
| 1 | 4934863 | AX-170477344 | 0 | G | A | 0.518 | 2.21E-02 | 6.92E-02 | 2.57E+00 | 2.64E+00 | 7.65E-01 | no | Rgs20 | intron_variant | MODIFIER |
| 1 | 4971193 | AX-97846679 | 0 | C | T | 0.518 | 2.21E-02 | 6.92E-02 | 2.57E+00 | 2.64E+00 | 7.65E-01 | no | Rgs20 | intron_variant | MODIFIER |
| 1 | 4974676 | AX-170392817 | 0 | G | A | 0.518 | 2.21E-02 | 6.92E-02 | 2.57E+00 | 2.64E+00 | 7.65E-01 | no | Rgs20 | intron_variant | MODIFIER |

| | | | | | | | | | | | | | | | |
|---|---|---|---|---|---|---|---|---|---|---|---|---|---|---|---|
| 1 | 5014713 | AX-170382804 | 0 | C | G | 0.518 | 2.21E-02 | 6.92E-02 | 2.57E+00 | 2.64E+00 | 7.65E-01 | no | Rgs20 | intron_variant | MODIFIER |
| 1 | 5233486 | AX-169999860 | 0 | G | T | 0.518 | 2.21E-02 | 6.92E-02 | 2.57E+00 | 2.64E+00 | 7.65E-01 | no | | regulatory_region_variant | MODIFIER |
| 1 | 5315915 | AX-170337790 | 0 | C | G | 0.518 | 2.21E-02 | 6.92E-02 | 2.57E+00 | 2.64E+00 | 7.65E-01 | no | | intergenic_variant | MODIFIER |
| 1 | 5316051 | AX-170343800 | 0 | T | A | 0.518 | 2.21E-02 | 6.92E-02 | 2.57E+00 | 2.64E+00 | 7.65E-01 | no | | intergenic_variant | MODIFIER |
| 7 | 117744412 | AX-170136176 | 1 | G | T | 0.402 | 2.76E-02 | 9.25E-02 | 2.66E+00 | 2.71E+00 | 7.65E-01 | no | | intergenic_variant | MODIFIER |
| 15 | 51907050 | AX-170526376 | 1 | G | A | 0.054 | -3.63E-02 | 1.25E-01 | 2.60E+00 | 2.76E+00 | 7.65E-01 | no | | regulatory_region_variant | MODIFIER |
| 1 | 25876559 | AX-170528876 | 0 | A | G | 0.404 | 2.50E-02 | 8.39E-02 | 2.64E+00 | 2.72E+00 | 7.65E-01 | no | | intergenic_variant | MODIFIER |
| 20 | 72530856 | AX-170034130 | 2 | G | A | 0.255 | 1.83E-02 | 6.16E-02 | 2.63E+00 | 2.73E+00 | 7.66E-01 | no | Gabra3 | intron_variant | MODIFIER |
| 15 | 98021793 | AX-170458584 | 1 | A | T | 0.116 | 4.83E-02 | 1.58E-01 | 2.60E+00 | 2.69E+00 | 7.66E-01 | no | | regulatory_region_variant | MODIFIER |
| 1 | 66048107 | AX-170086486 | 1 | C | G | 0.143 | 6.98E-02 | 2.37E-01 | 2.69E+00 | 2.74E+00 | 7.66E-01 | no | | intergenic_variant | MODIFIER |
| 2 | 104048908 | AX-170281558 | 1 | C | T | 0.054 | -8.81E-02 | 3.00E-01 | 2.73E+00 | 2.76E+00 | 7.66E-01 | no | Fbxo3 | intron_variant | MODIFIER |
| 5 | 27702294 | AX-170024571 | 1 | A | G | 0.054 | -8.81E-02 | 3.00E-01 | 2.73E+00 | 2.76E+00 | 7.66E-01 | no | Dpp6 | intron_variant | MODIFIER |
| 3 | 29186516 | AX-170118878 | 1 | T | C | 0.161 | -2.98E-02 | 1.01E-01 | 2.67E+00 | 2.74E+00 | 7.66E-01 | no | Egfem1 | intron_variant | MODIFIER |
| 13 | 24061509 | AX-97852632 | 2 | A | G | 0.164 | -3.92E-02 | 1.21E-01 | 2.55E+00 | 2.62E+00 | 7.66E-01 | no | Carmil1 | intron_variant | MODIFIER |
| 7 | 117667607 | AX-170340461 | 0 | T | C | 0.386 | 2.65E-02 | 8.91E-02 | 2.65E+00 | 2.72E+00 | 7.66E-01 | no | Xylt1 | synonymous_variant | LOW |
| 7 | 112614918 | AX-170291182 | 0 | A | G | 0.447 | 2.71E-02 | 9.20E-02 | 2.68E+00 | 2.74E+00 | 7.66E-01 | no | 2310014F06Rik | intron_variant,non_coding_transcript_variant | MODIFIER |
| 1 | 136042503 | AX-170130447 | 1 | G | A | 0.205 | 3.64E-02 | 1.23E-01 | 2.65E+00 | 2.72E+00 | 7.66E-01 | no | | intergenic_variant | MODIFIER |
| 5 | 100126097 | AX-97829138 | 1 | G | A | 0.438 | -3.11E-02 | 1.07E-01 | 2.73E+00 | 2.80E+00 | 7.66E-01 | no | Tmem150c | intron_variant | MODIFIER |
| 15 | 95246063 | AX-170123368 | 0 | C | T | 0.149 | -4.42E-02 | 1.53E-01 | 2.68E+00 | 2.77E+00 | 7.66E-01 | no | Nell2 | intron_variant | MODIFIER |
| 9 | 84437612 | AX-170242143 | 0 | C | T | 0.211 | -3.63E-02 | 1.20E-01 | 2.62E+00 | 2.70E+00 | 7.66E-01 | no | | intergenic_variant | MODIFIER |
| 10 | 28169005 | AX-170002963 | 1 | T | C | 0.196 | 2.86E-02 | 1.00E-01 | 2.71E+00 | 2.81E+00 | 7.66E-01 | no | Ptprk | intron_variant | MODIFIER |
| 6 | 105259365 | AX-170440609 | 0 | G | T | 0.053 | -6.34E-02 | 2.15E-01 | 2.62E+00 | 2.74E+00 | 7.66E-01 | no | | intergenic_variant | MODIFIER |
| 6 | 114113503 | AX-170028028 | 0 | T | C | 0.053 | -6.34E-02 | 2.15E-01 | 2.62E+00 | 2.74E+00 | 7.66E-01 | no | | intergenic_variant | MODIFIER |
| 6 | 121423143 | AX-170316605 | 0 | G | A | 0.053 | -6.34E-02 | 2.15E-01 | 2.62E+00 | 2.74E+00 | 7.66E-01 | no | Iqsec3 | intron_variant | MODIFIER |
| 16 | 94200886 | AX-170482591 | 1 | G | C | 0.411 | -2.50E-02 | 8.11E-02 | 2.59E+00 | 2.68E+00 | 7.66E-01 | no | Hlcs | intron_variant | MODIFIER |
| 8 | 109416678 | AX-97811943 | 0 | C | T | 0.228 | 3.07E-02 | 1.06E-01 | 2.68E+00 | 2.77E+00 | 7.66E-01 | no | | regulatory_region_variant | MODIFIER |
| 8 | 109430202 | AX-170460423 | 0 | A | G | 0.228 | 3.07E-02 | 1.06E-01 | 2.68E+00 | 2.77E+00 | 7.66E-01 | no | Gm23163 | upstream_gene_variant | MODIFIER |
| 8 | 109434047 | AX-170081141 | 0 | A | T | 0.228 | 3.07E-02 | 1.06E-01 | 2.68E+00 | 2.77E+00 | 7.66E-01 | no | | intergenic_variant | MODIFIER |
| 8 | 109438340 | AX-170081139 | 0 | G | A | 0.228 | 3.07E-02 | 1.06E-01 | 2.68E+00 | 2.77E+00 | 7.66E-01 | no | | intergenic_variant | MODIFIER |
| 8 | 109442494 | AX-170081140 | 0 | T | C | 0.228 | 3.07E-02 | 1.06E-01 | 2.68E+00 | 2.77E+00 | 7.66E-01 | no | | intergenic_variant | MODIFIER |
| 8 | 109451824 | AX-97804004 | 0 | C | T | 0.228 | 3.07E-02 | 1.06E-01 | 2.68E+00 | 2.77E+00 | 7.66E-01 | no | | intergenic_variant | MODIFIER |
| 8 | 109465387 | AX-170442143 | 0 | A | G | 0.228 | 3.07E-02 | 1.06E-01 | 2.68E+00 | 2.77E+00 | 7.66E-01 | no | | regulatory_region_variant | MODIFIER |
| 8 | 109466904 | AX-170241494 | 0 | A | G | 0.228 | 3.07E-02 | 1.06E-01 | 2.68E+00 | 2.77E+00 | 7.66E-01 | no | | regulatory_region_variant | MODIFIER |
| 8 | 110024333 | AX-170031476 | 0 | C | T | 0.228 | 3.07E-02 | 1.06E-01 | 2.68E+00 | 2.77E+00 | 7.66E-01 | no | Chst4 | downstream_gene_variant | MODIFIER |
| 8 | 110029074 | AX-170098833 | 0 | T | C | 0.228 | 3.07E-02 | 1.06E-01 | 2.68E+00 | 2.77E+00 | 7.66E-01 | no | Chst4 | downstream_gene_variant | MODIFIER |
| 8 | 110055787 | AX-170513398 | 0 | A | G | 0.228 | 3.07E-02 | 1.06E-01 | 2.68E+00 | 2.77E+00 | 7.66E-01 | no | 8030455M16Rik | intron_variant,non_coding_transcript_variant | MODIFIER |
| 8 | 110055957 | AX-170380892 | 0 | T | C | 0.228 | 3.07E-02 | 1.06E-01 | 2.68E+00 | 2.77E+00 | 7.66E-01 | no | 8030455M16Rik | intron_variant,non_coding_transcript_variant | MODIFIER |
| 8 | 110066912 | AX-170081152 | 0 | T | C | 0.228 | 3.07E-02 | 1.06E-01 | 2.68E+00 | 2.77E+00 | 7.66E-01 | no | | intergenic_variant | MODIFIER |
| 8 | 110072037 | AX-170387633 | 0 | T | C | 0.228 | 3.07E-02 | 1.06E-01 | 2.68E+00 | 2.77E+00 | 7.66E-01 | no | | regulatory_region_variant | MODIFIER |
| 8 | 110073408 | AX-170081156 | 0 | G | T | 0.228 | 3.07E-02 | 1.06E-01 | 2.68E+00 | 2.77E+00 | 7.66E-01 | no | | intergenic_variant | MODIFIER |
| 8 | 110074417 | AX-170250091 | 0 | T | C | 0.228 | 3.07E-02 | 1.06E-01 | 2.68E+00 | 2.77E+00 | 7.66E-01 | no | | intergenic_variant | MODIFIER |
| 8 | 110219910 | AX-170325930 | 0 | A | T | 0.228 | 3.07E-02 | 1.06E-01 | 2.68E+00 | 2.77E+00 | 7.66E-01 | no | Cmtr2 | intron_variant | MODIFIER |
| 16 | 44259475 | AX-170178503 | 1 | G | A | 0.509 | 2.50E-02 | 8.06E-02 | 2.60E+00 | 2.66E+00 | 7.66E-01 | no | Sidt1 | synonymous_variant | LOW |
| 16 | 45500034 | AX-97831628 | 1 | A | C | 0.509 | 2.50E-02 | 8.06E-02 | 2.60E+00 | 2.66E+00 | 7.66E-01 | no | | intergenic_variant | MODIFIER |
| 2 | 77842097 | AX-97852308 | 1 | G | A | 0.054 | -8.77E-02 | 2.99E-01 | 2.73E+00 | 2.76E+00 | 7.66E-01 | no | Gm13661 | intron_variant,non_coding_transcript_variant | MODIFIER |
| 2 | 101932217 | AX-170106209 | 1 | T | C | 0.054 | -8.77E-02 | 2.99E-01 | 2.73E+00 | 2.76E+00 | 7.66E-01 | no | | regulatory_region_variant | MODIFIER |
| 2 | 102469160 | AX-170018507 | 1 | G | A | 0.054 | -8.77E-02 | 2.99E-01 | 2.73E+00 | 2.76E+00 | 7.66E-01 | no | | intergenic_variant | MODIFIER |
| 7 | 10306729 | AX-170134559 | 0 | G | C | 0.167 | 2.26E-02 | 8.19E-02 | 2.70E+00 | 2.84E+00 | 7.67E-01 | no | | intergenic_variant | MODIFIER |
| 7 | 112382856 | AX-97846201 | 0 | G | A | 0.474 | -2.52E-02 | 8.82E-02 | 2.69E+00 | 2.79E+00 | 7.67E-01 | no | Mical2 | intron_variant | MODIFIER |
| 6 | 125177961 | AX-170414702 | 2 | T | A | 0.191 | -2.35E-02 | 8.14E-02 | 2.63E+00 | 2.76E+00 | 7.67E-01 | no | Ncapd2 | intron_variant | MODIFIER |
| 15 | 86019906 | AX-97808191 | 2 | T | C | 0.5 | 2.27E-02 | 7.03E-02 | 2.54E+00 | 2.64E+00 | 7.67E-01 | no | Celsr1 | intron_variant | MODIFIER |
| 16 | 24561397 | AX-170286972 | 1 | T | C | 0.241 | 2.57E-02 | 8.85E-02 | 2.62E+00 | 2.75E+00 | 7.67E-01 | no | Lpp | intron_variant | MODIFIER |
| 5 | 105493733 | AX-170025800 | 1 | C | A | 0.429 | 3.28E-02 | 1.09E-01 | 2.63E+00 | 2.68E+00 | 7.67E-01 | no | Lrrc8b | downstream_gene_variant | MODIFIER |
| 1 | 89401801 | AX-170317432 | 0 | G | T | 0.14 | 5.42E-02 | 1.80E-01 | 2.60E+00 | 2.70E+00 | 7.67E-01 | no | C030007H22Rik | intron_variant,non_coding_transcript_variant | MODIFIER |

| | | | | | | | | | | | | | | | |
|---|---|---|---|---|---|---|---|---|---|---|---|---|---|---|---|
| 12 | 92001425 | AX-170145805 | 2 | G | A | 0.073 | -4.99E-02 | 1.71E-01 | 2.63E+00 | 2.74E+00 | 7.67E-01 | no | | intergenic_variant | MODIFIER |
| 15 | 84831450 | AX-170012160 | 1 | A | G | 0.509 | 2.25E-02 | 7.04E-02 | 2.55E+00 | 2.65E+00 | 7.67E-01 | no | Phf21b | intron_variant | MODIFIER |
| 1 | 169904257 | AX-170091784 | 1 | A | G | 0.214 | 3.82E-02 | 1.27E-01 | 2.63E+00 | 2.69E+00 | 7.68E-01 | no | | intergenic_variant | MODIFIER |
| 16 | 91419909 | AX-170241633 | 1 | T | A | 0.464 | -2.03E-02 | 6.86E-02 | 2.62E+00 | 2.72E+00 | 7.68E-01 | no | Il10rb | intron_variant | MODIFIER |
| 4 | 118780740 | AX-170023797 | 0 | A | T | 0.219 | -3.30E-02 | 1.17E-01 | 2.75E+00 | 2.86E+00 | 7.68E-01 | no | Olfr1337 | downstream_gene_variant | MODIFIER |
| 9 | 32857491 | AX-170278562 | 1 | T | G | 0.429 | 2.55E-02 | 8.51E-02 | 2.63E+00 | 2.69E+00 | 7.68E-01 | no | | intergenic_variant | MODIFIER |
| 16 | 94163482 | AX-170535575 | 1 | G | A | 0.411 | -2.49E-02 | 8.13E-02 | 2.59E+00 | 2.68E+00 | 7.68E-01 | no | Hlcs | intron_variant | MODIFIER |
| 4 | 129993718 | AX-170321007 | 0 | G | C | 0.158 | 3.45E-02 | 1.13E-01 | 2.61E+00 | 2.67E+00 | 7.68E-01 | no | Adgrb2 | intron_variant | MODIFIER |
| 8 | 78211125 | AX-170454545 | 1 | C | A | 0.205 | 3.75E-02 | 1.30E-01 | 2.71E+00 | 2.77E+00 | 7.68E-01 | no | Ttc29 | upstream_gene_variant | MODIFIER |
| 8 | 78269458 | AX-170498586 | 1 | C | T | 0.205 | 3.75E-02 | 1.30E-01 | 2.71E+00 | 2.77E+00 | 7.68E-01 | no | Ttc29 | intron_variant | MODIFIER |
| 1 | 3407348 | AX-170254782 | 1 | G | C | 0.518 | 2.18E-02 | 6.92E-02 | 2.57E+00 | 2.64E+00 | 7.68E-01 | no | Xkr4 | intron_variant | MODIFIER |
| 2 | 87643811 | AX-97820905 | 2 | G | A | 0.055 | -8.70E-02 | 2.99E-01 | 2.73E+00 | 2.76E+00 | 7.68E-01 | no | Olfr1133 | downstream_gene_variant | MODIFIER |
| 5 | 36110683 | AX-170024682 | 1 | A | T | 0.321 | -3.08E-02 | 1.07E-01 | 2.69E+00 | 2.77E+00 | 7.68E-01 | no | Sorcs2 | intron_variant | MODIFIER |
| 8 | 103516989 | AX-97806974 | 0 | G | A | 0.035 | -3.73E-02 | 1.33E-01 | 2.61E+00 | 2.78E+00 | 7.68E-01 | no | | intergenic_variant | MODIFIER |
| 2 | 103503508 | AX-170018525 | 1 | C | T | 0.054 | -8.70E-02 | 2.99E-01 | 2.73E+00 | 2.76E+00 | 7.68E-01 | no | | intergenic_variant | MODIFIER |
| 5 | 111972685 | AX-170419098 | 0 | C | T | 0.456 | 3.33E-02 | 1.12E-01 | 2.66E+00 | 2.70E+00 | 7.68E-01 | no | | intergenic_variant | MODIFIER |
| 1 | 37000023 | AX-170186207 | 0 | G | T | 0.465 | 2.28E-02 | 7.89E-02 | 2.67E+00 | 2.75E+00 | 7.68E-01 | no | | regulatory_region_variant | MODIFIER |
| 1 | 13541753 | AX-97789789 | 1 | G | A | 0.384 | 2.25E-02 | 7.50E-02 | 2.63E+00 | 2.69E+00 | 7.68E-01 | no | | intergenic_variant | MODIFIER |
| 1 | 139661990 | AX-170232753 | 0 | G | A | 0.202 | -3.54E-02 | 1.25E-01 | 2.75E+00 | 2.84E+00 | 7.68E-01 | no | Cfhr3 | upstream_gene_variant | MODIFIER |
| 7 | 106425983 | AX-170422640 | 2 | T | C | 0.445 | -2.54E-02 | 9.06E-02 | 2.73E+00 | 2.85E+00 | 7.68E-01 | no | Gm22372 | upstream_gene_variant | MODIFIER |
| 1 | 68027865 | AX-170404589 | 2 | G | A | 0.118 | 6.65E-02 | 2.29E-01 | 2.69E+00 | 2.74E+00 | 7.68E-01 | no | Erbb4 | downstream_gene_variant | MODIFIER |
| 9 | 77259048 | AX-170032751 | 1 | T | C | 0.205 | 3.12E-02 | 1.10E-01 | 2.68E+00 | 2.79E+00 | 7.68E-01 | no | Mlip | intron_variant | MODIFIER |
| 20 | 164296296 | AX-170085721 | 0 | C | G | 0.132 | 3.40E-02 | 1.20E-01 | 2.65E+00 | 2.77E+00 | 7.69E-01 | no | Pir | intron_variant | MODIFIER |
| 20 | 157865838 | AX-170115335 | 0 | A | G | 0.14 | 3.28E-02 | 1.11E-01 | 2.63E+00 | 2.71E+00 | 7.69E-01 | no | Cnksr2 | synonymous_variant | LOW |
| 20 | 157878490 | AX-170215303 | 0 | T | C | 0.14 | 3.28E-02 | 1.11E-01 | 2.63E+00 | 2.71E+00 | 7.69E-01 | no | Cnksr2 | intron_variant | MODIFIER |
| 9 | 84214035 | AX-170132113 | 0 | C | T | 0.211 | 3.65E-02 | 1.27E-01 | 2.72E+00 | 2.78E+00 | 7.69E-01 | no | | intergenic_variant | MODIFIER |
| 8 | 90009760 | AX-97784280 | 1 | G | C | 0.196 | -3.48E-02 | 1.17E-01 | 2.63E+00 | 2.71E+00 | 7.69E-01 | no | | intergenic_variant | MODIFIER |
| 16 | 45519612 | AX-170447956 | 0 | G | A | 0.509 | 2.46E-02 | 8.11E-02 | 2.61E+00 | 2.67E+00 | 7.69E-01 | no | Gm17783 | intron_variant | MODIFIER |
| 5 | 85235457 | AX-170072672 | 0 | T | G | 0.412 | 3.04E-02 | 1.05E-01 | 2.69E+00 | 2.75E+00 | 7.69E-01 | no | | intergenic_variant | MODIFIER |
| 5 | 113200726 | AX-97804828 | 2 | A | G | 0.436 | 3.13E-02 | 1.05E-01 | 2.63E+00 | 2.69E+00 | 7.69E-01 | no | 2900026A02Rik | intron_variant | MODIFIER |
| 5 | 99691232 | AX-170161121 | 2 | T | G | 0.436 | -2.73E-02 | 9.60E-02 | 2.69E+00 | 2.78E+00 | 7.69E-01 | no | Rasgef1b | intron_variant,non_coding_transcript_variant | MODIFIER |
| 1 | 9622506 | AX-170178665 | 0 | T | G | 0.096 | 4.15E-02 | 1.39E-01 | 2.58E+00 | 2.71E+00 | 7.69E-01 | no | Vxn | intron_variant | MODIFIER |
| 12 | 108316363 | AX-170007488 | 0 | C | T | 0.114 | -5.66E-02 | 1.98E-01 | 2.70E+00 | 2.78E+00 | 7.69E-01 | no | Hhipl1 | intron_variant | MODIFIER |
| 20 | 72964113 | AX-170299103 | 0 | T | C | 0.246 | 1.85E-02 | 6.33E-02 | 2.63E+00 | 2.73E+00 | 7.69E-01 | no | | intergenic_variant | MODIFIER |
| 2 | 114590778 | AX-170249165 | 2 | C | A | 0.136 | 5.87E-02 | 2.01E-01 | 2.66E+00 | 2.72E+00 | 7.69E-01 | no | Dph6 | intron_variant | MODIFIER |
| 9 | 89451897 | AX-97807861 | 0 | G | A | 0.132 | -4.00E-02 | 1.33E-01 | 2.61E+00 | 2.68E+00 | 7.69E-01 | no | | intergenic_variant | MODIFIER |
| 8 | 93398687 | AX-170245585 | 0 | T | C | 0.211 | -3.03E-02 | 1.05E-01 | 2.66E+00 | 2.75E+00 | 7.69E-01 | no | Gm45765 | intron_variant,non_coding_transcript_variant | MODIFIER |
| 15 | 81397972 | AX-97794673 | 1 | T | C | 0.545 | 2.48E-02 | 8.00E-02 | 2.58E+00 | 2.66E+00 | 7.69E-01 | no | St13 | intron_variant | MODIFIER |
| 7 | 55473149 | AX-170408146 | 1 | T | C | 0.438 | 2.73E-02 | 9.28E-02 | 2.62E+00 | 2.72E+00 | 7.69E-01 | no | | intergenic_variant | MODIFIER |
| 8 | 50272518 | AX-97823317 | 1 | C | T | 0.152 | -4.64E-02 | 1.51E-01 | 2.59E+00 | 2.65E+00 | 7.69E-01 | no | Gm45455 | intron_variant,non_coding_transcript_variant | MODIFIER |
| 11 | 33613394 | AX-170004668 | 0 | T | C | 0.088 | 5.65E-02 | 1.90E-01 | 2.61E+00 | 2.70E+00 | 7.69E-01 | no | D130052B06Rik | intron_variant | MODIFIER |
| 16 | 92563188 | AX-170101477 | 0 | A | G | 0.474 | 2.29E-02 | 7.88E-02 | 2.65E+00 | 2.74E+00 | 7.69E-01 | no | | intergenic_variant | MODIFIER |
| 9 | 111671944 | AX-170497235 | 0 | G | T | 0.105 | -3.51E-02 | 1.19E-01 | 2.63E+00 | 2.70E+00 | 7.70E-01 | no | Stac | intron_variant | MODIFIER |
| 1 | 60551119 | AX-170133246 | 0 | T | C | 0.096 | -6.14E-02 | 2.15E-01 | 2.67E+00 | 2.76E+00 | 7.70E-01 | no | Raph1 | intron_variant | MODIFIER |
| 16 | 51596455 | AX-170088015 | 0 | T | G | 0.482 | -2.58E-02 | 8.20E-02 | 2.57E+00 | 2.63E+00 | 7.70E-01 | no | | intergenic_variant | MODIFIER |
| 2 | 40669795 | AX-170017482 | 0 | G | A | 0.053 | 4.89E-02 | 1.74E-01 | 2.66E+00 | 2.79E+00 | 7.70E-01 | no | Lrp1b | intron_variant | MODIFIER |
| 10 | 8333169 | AX-170002668 | 1 | T | C | 0.205 | -3.14E-02 | 1.06E-01 | 2.62E+00 | 2.71E+00 | 7.70E-01 | no | Ust | intron_variant | MODIFIER |
| 7 | 50811843 | AX-170120745 | 0 | A | G | 0.43 | 2.97E-02 | 1.00E-01 | 2.63E+00 | 2.70E+00 | 7.70E-01 | no | Nell1 | intron_variant | MODIFIER |
| 9 | 120769440 | AX-170185494 | 1 | A | T | 0.214 | 2.28E-02 | 8.04E-02 | 2.64E+00 | 2.76E+00 | 7.70E-01 | no | | intergenic_variant | MODIFIER |
| 1 | 149355798 | AX-97840755 | 0 | C | T | 0.211 | 4.32E-02 | 1.46E-01 | 2.64E+00 | 2.68E+00 | 7.70E-01 | no | | intergenic_variant | MODIFIER |
| 1 | 155701287 | AX-97857204 | 0 | T | C | 0.211 | 4.32E-02 | 1.46E-01 | 2.64E+00 | 2.68E+00 | 7.70E-01 | no | Lhx4 | 3_prime_UTR_variant | MODIFIER |
| 1 | 155720722 | AX-170281984 | 0 | A | G | 0.211 | 4.32E-02 | 1.46E-01 | 2.64E+00 | 2.68E+00 | 7.70E-01 | no | Lhx4 | intron_variant | MODIFIER |
| 1 | 155720754 | AX-170282550 | 0 | G | A | 0.211 | 4.32E-02 | 1.46E-01 | 2.64E+00 | 2.68E+00 | 7.70E-01 | no | Lhx4 | intron_variant | MODIFIER |
| 1 | 158699489 | AX-170043306 | 0 | T | C | 0.211 | 4.32E-02 | 1.46E-01 | 2.64E+00 | 2.68E+00 | 7.70E-01 | no | | regulatory_region_variant | MODIFIER |

| | | | | | | | | | | | | | | | |
|---|---|---|---|---|---|---|---|---|---|---|---|---|---|---|---|
| 1 | 161238241 | AX-170297037 | 0 | G | A | 0.211 | 4.32E-02 | 1.46E-01 | 2.64E+00 | 2.68E+00 | 7.70E-01 | no | Prdx6 | downstream_gene_variant | MODIFIER |
| 1 | 161239736 | AX-170043340 | 0 | T | C | 0.211 | 4.32E-02 | 1.46E-01 | 2.64E+00 | 2.68E+00 | 7.70E-01 | no | Prdx6 | downstream_gene_variant | MODIFIER |
| 1 | 161244598 | AX-170043342 | 0 | C | T | 0.211 | 4.32E-02 | 1.46E-01 | 2.64E+00 | 2.68E+00 | 7.70E-01 | no | Prdx6 | intron_variant | MODIFIER |
| 1 | 161244832 | AX-97807165 | 0 | A | G | 0.211 | 4.32E-02 | 1.46E-01 | 2.64E+00 | 2.68E+00 | 7.70E-01 | no | Prdx6 | intron_variant | MODIFIER |
| 1 | 161274908 | AX-170463871 | 0 | T | C | 0.211 | 4.32E-02 | 1.46E-01 | 2.64E+00 | 2.68E+00 | 7.70E-01 | no | | intergenic_variant | MODIFIER |
| 1 | 161283462 | AX-170091768 | 0 | C | T | 0.211 | 4.32E-02 | 1.46E-01 | 2.64E+00 | 2.68E+00 | 7.70E-01 | no | | intergenic_variant | MODIFIER |
| 1 | 161284191 | AX-170104933 | 0 | G | A | 0.211 | 4.32E-02 | 1.46E-01 | 2.64E+00 | 2.68E+00 | 7.70E-01 | no | | intergenic_variant | MODIFIER |
| 1 | 161365580 | AX-170497086 | 0 | A | C | 0.211 | 4.32E-02 | 1.46E-01 | 2.64E+00 | 2.68E+00 | 7.70E-01 | no | | intergenic_variant | MODIFIER |
| 1 | 161365686 | AX-170400616 | 0 | C | T | 0.211 | 4.32E-02 | 1.46E-01 | 2.64E+00 | 2.68E+00 | 7.70E-01 | no | | intergenic_variant | MODIFIER |
| 1 | 161366660 | AX-170127231 | 0 | T | C | 0.211 | 4.32E-02 | 1.46E-01 | 2.64E+00 | 2.68E+00 | 7.70E-01 | no | | intergenic_variant | MODIFIER |
| 1 | 161452261 | AX-97816286 | 0 | A | G | 0.211 | 4.32E-02 | 1.46E-01 | 2.64E+00 | 2.68E+00 | 7.70E-01 | no | | intergenic_variant | MODIFIER |
| 1 | 161466406 | AX-170035636 | 0 | A | G | 0.211 | 4.32E-02 | 1.46E-01 | 2.64E+00 | 2.68E+00 | 7.70E-01 | no | | intergenic_variant | MODIFIER |
| 1 | 161512269 | AX-170496416 | 0 | A | G | 0.211 | 4.32E-02 | 1.46E-01 | 2.64E+00 | 2.68E+00 | 7.70E-01 | no | | intergenic_variant | MODIFIER |
| 1 | 161519746 | AX-170232906 | 0 | A | C | 0.211 | 4.32E-02 | 1.46E-01 | 2.64E+00 | 2.68E+00 | 7.70E-01 | no | | intergenic_variant | MODIFIER |
| 1 | 161576327 | AX-97801965 | 0 | G | A | 0.211 | 4.32E-02 | 1.46E-01 | 2.64E+00 | 2.68E+00 | 7.70E-01 | no | | regulatory_region_variant | MODIFIER |
| 1 | 161991182 | AX-97809184 | 0 | T | G | 0.211 | 4.32E-02 | 1.46E-01 | 2.64E+00 | 2.68E+00 | 7.70E-01 | no | Dnm3 | 3_prime_UTR_variant | MODIFIER |
| 1 | 162091885 | AX-97836019 | 0 | G | A | 0.211 | 4.32E-02 | 1.46E-01 | 2.64E+00 | 2.68E+00 | 7.70E-01 | no | Dnm3 | intron_variant | MODIFIER |
| 1 | 162292887 | AX-97821717 | 0 | C | T | 0.211 | 4.32E-02 | 1.46E-01 | 2.64E+00 | 2.68E+00 | 7.70E-01 | no | Dnm3 | intron_variant | MODIFIER |
| 10 | 94111832 | AX-97792537 | 0 | C | T | 0.289 | -2.64E-02 | 8.47E-02 | 2.57E+00 | 2.64E+00 | 7.70E-01 | no | Fgd6 | intron_variant | MODIFIER |
| 9 | 85683220 | AX-170324824 | 1 | G | A | 0.143 | -3.79E-02 | 1.27E-01 | 2.62E+00 | 2.68E+00 | 7.70E-01 | no | Ibtk | downstream_gene_variant | MODIFIER |
| 3 | 144122626 | AX-170094840 | 0 | A | C | 0.368 | 2.03E-02 | 6.91E-02 | 2.62E+00 | 2.71E+00 | 7.71E-01 | no | Gm34078 | intron_variant,non_coding_transcript_variant | MODIFIER |
| 16 | 86859020 | AX-170013580 | 0 | G | A | 0.43 | -2.29E-02 | 8.00E-02 | 2.69E+00 | 2.75E+00 | 7.71E-01 | no | Gm32624 | intron_variant,non_coding_transcript_variant | MODIFIER |
| 16 | 86860080 | AX-170109282 | 0 | C | A | 0.43 | -2.29E-02 | 8.00E-02 | 2.69E+00 | 2.75E+00 | 7.71E-01 | no | Gm32624 | intron_variant,non_coding_transcript_variant | MODIFIER |
| 16 | 86860133 | AX-170235353 | 0 | C | T | 0.43 | -2.29E-02 | 8.00E-02 | 2.69E+00 | 2.75E+00 | 7.71E-01 | no | Gm32624 | intron_variant,non_coding_transcript_variant | MODIFIER |
| 16 | 86878701 | AX-170013579 | 0 | G | A | 0.43 | -2.29E-02 | 8.00E-02 | 2.69E+00 | 2.75E+00 | 7.71E-01 | no | Gm32624 | downstream_gene_variant | MODIFIER |
| 16 | 87025379 | AX-170058026 | 0 | G | A | 0.43 | -2.29E-02 | 8.00E-02 | 2.69E+00 | 2.75E+00 | 7.71E-01 | no | Gm38477 | upstream_gene_variant | MODIFIER |
| 16 | 87071893 | AX-170186723 | 0 | A | T | 0.43 | -2.29E-02 | 8.00E-02 | 2.69E+00 | 2.75E+00 | 7.71E-01 | no | | intergenic_variant | MODIFIER |
| 16 | 87078078 | AX-97785698 | 0 | G | A | 0.43 | -2.29E-02 | 8.00E-02 | 2.69E+00 | 2.75E+00 | 7.71E-01 | no | | intergenic_variant | MODIFIER |
| 16 | 87083260 | AX-170170183 | 0 | G | A | 0.43 | -2.29E-02 | 8.00E-02 | 2.69E+00 | 2.75E+00 | 7.71E-01 | no | | intergenic_variant | MODIFIER |
| 16 | 87083300 | AX-170426360 | 0 | C | T | 0.43 | -2.29E-02 | 8.00E-02 | 2.69E+00 | 2.75E+00 | 7.71E-01 | no | | intergenic_variant | MODIFIER |
| 16 | 87099382 | AX-170130810 | 0 | G | A | 0.43 | -2.29E-02 | 8.00E-02 | 2.69E+00 | 2.75E+00 | 7.71E-01 | no | | intergenic_variant | MODIFIER |
| 16 | 87105114 | AX-170462917 | 0 | A | T | 0.43 | -2.29E-02 | 8.00E-02 | 2.69E+00 | 2.75E+00 | 7.71E-01 | no | | intergenic_variant | MODIFIER |
| 16 | 87105428 | AX-170268011 | 0 | C | T | 0.43 | -2.29E-02 | 8.00E-02 | 2.69E+00 | 2.75E+00 | 7.71E-01 | no | | intergenic_variant | MODIFIER |
| 16 | 87105558 | AX-170189746 | 0 | A | G | 0.43 | -2.29E-02 | 8.00E-02 | 2.69E+00 | 2.75E+00 | 7.71E-01 | no | | intergenic_variant | MODIFIER |
| 16 | 87124805 | AX-97833328 | 0 | C | T | 0.43 | -2.29E-02 | 8.00E-02 | 2.69E+00 | 2.75E+00 | 7.71E-01 | no | Gm32865 | downstream_gene_variant | MODIFIER |
| 16 | 87124821 | AX-170347834 | 0 | G | A | 0.43 | -2.29E-02 | 8.00E-02 | 2.69E+00 | 2.75E+00 | 7.71E-01 | no | Gm32865 | downstream_gene_variant | MODIFIER |
| 16 | 87125915 | AX-170311361 | 0 | C | T | 0.43 | -2.29E-02 | 8.00E-02 | 2.69E+00 | 2.75E+00 | 7.71E-01 | no | Gm32865 | intron_variant,non_coding_transcript_variant | MODIFIER |
| 16 | 87126377 | AX-170427034 | 0 | T | C | 0.43 | -2.29E-02 | 8.00E-02 | 2.69E+00 | 2.75E+00 | 7.71E-01 | no | Gm32865 | intron_variant,non_coding_transcript_variant | MODIFIER |
| 16 | 87156554 | AX-97831570 | 0 | T | C | 0.43 | -2.29E-02 | 8.00E-02 | 2.69E+00 | 2.75E+00 | 7.71E-01 | no | | intergenic_variant | MODIFIER |
| 16 | 87169778 | AX-170116600 | 0 | C | A | 0.43 | -2.29E-02 | 8.00E-02 | 2.69E+00 | 2.75E+00 | 7.71E-01 | no | | intergenic_variant | MODIFIER |
| 16 | 87169887 | AX-170303595 | 0 | G | A | 0.43 | -2.29E-02 | 8.00E-02 | 2.69E+00 | 2.75E+00 | 7.71E-01 | no | | intergenic_variant | MODIFIER |
| 16 | 87207137 | AX-97792740 | 0 | C | A | 0.43 | -2.29E-02 | 8.00E-02 | 2.69E+00 | 2.75E+00 | 7.71E-01 | no | | intergenic_variant | MODIFIER |
| 16 | 87225197 | AX-170188278 | 0 | G | A | 0.43 | -2.29E-02 | 8.00E-02 | 2.69E+00 | 2.75E+00 | 7.71E-01 | no | | intergenic_variant | MODIFIER |
| 16 | 87263563 | AX-97845806 | 0 | G | A | 0.43 | -2.29E-02 | 8.00E-02 | 2.69E+00 | 2.75E+00 | 7.71E-01 | no | | regulatory_region_variant | MODIFIER |
| 16 | 87268047 | AX-170233416 | 0 | A | T | 0.43 | -2.29E-02 | 8.00E-02 | 2.69E+00 | 2.75E+00 | 7.71E-01 | no | | regulatory_region_variant | MODIFIER |
| 16 | 87286476 | AX-170177918 | 0 | G | T | 0.43 | -2.29E-02 | 8.00E-02 | 2.69E+00 | 2.75E+00 | 7.71E-01 | no | | intergenic_variant | MODIFIER |
| 16 | 87293417 | AX-170418903 | 0 | A | G | 0.43 | -2.29E-02 | 8.00E-02 | 2.69E+00 | 2.75E+00 | 7.71E-01 | no | | intergenic_variant | MODIFIER |
| 16 | 87293785 | AX-97791110 | 0 | C | T | 0.43 | -2.29E-02 | 8.00E-02 | 2.69E+00 | 2.75E+00 | 7.71E-01 | no | | intergenic_variant | MODIFIER |
| 16 | 87319271 | AX-170179019 | 0 | A | G | 0.43 | -2.29E-02 | 8.00E-02 | 2.69E+00 | 2.75E+00 | 7.71E-01 | no | | regulatory_region_variant | MODIFIER |
| 16 | 87319647 | AX-170195533 | 0 | A | G | 0.43 | -2.29E-02 | 8.00E-02 | 2.69E+00 | 2.75E+00 | 7.71E-01 | no | | regulatory_region_variant | MODIFIER |
| 16 | 87323478 | AX-97828276 | 0 | C | T | 0.43 | -2.29E-02 | 8.00E-02 | 2.69E+00 | 2.75E+00 | 7.71E-01 | no | | intergenic_variant | MODIFIER |
| 16 | 87323917 | AX-170532509 | 0 | C | T | 0.43 | -2.29E-02 | 8.00E-02 | 2.69E+00 | 2.75E+00 | 7.71E-01 | no | | intergenic_variant | MODIFIER |
| 16 | 87395256 | AX-97850290 | 0 | C | T | 0.43 | -2.29E-02 | 8.00E-02 | 2.69E+00 | 2.75E+00 | 7.71E-01 | no | Ltn1 | intron_variant | MODIFIER |
| 16 | 87478370 | AX-170526856 | 0 | A | G | 0.43 | -2.29E-02 | 8.00E-02 | 2.69E+00 | 2.75E+00 | 7.71E-01 | no | Cct8 | downstream_gene_variant | MODIFIER |

| | | | | | | | | | | | | | | | |
|---|---|---|---|---|---|---|---|---|---|---|---|---|---|---|---|
| 16 | 87534104 | AX-170391656 | 0 | T | C | 0.43 | -2.29E-02 | 8.00E-02 | 2.69E+00 | 2.75E+00 | 7.71E-01 | no | | regulatory_region_variant | MODIFIER |
| 16 | 87552552 | AX-170520006 | 0 | C | T | 0.43 | -2.29E-02 | 8.00E-02 | 2.69E+00 | 2.75E+00 | 7.71E-01 | no | Map3k7cl | upstream_gene_variant | MODIFIER |
| 16 | 87561864 | AX-170370554 | 0 | G | A | 0.43 | -2.29E-02 | 8.00E-02 | 2.69E+00 | 2.75E+00 | 7.71E-01 | no | Map3k7cl | intron_variant | MODIFIER |
| 16 | 87563636 | AX-170384491 | 0 | T | C | 0.43 | -2.29E-02 | 8.00E-02 | 2.69E+00 | 2.75E+00 | 7.71E-01 | no | Map3k7cl | intron_variant | MODIFIER |
| 16 | 87566392 | AX-170058033 | 0 | C | T | 0.43 | -2.29E-02 | 8.00E-02 | 2.69E+00 | 2.75E+00 | 7.71E-01 | no | Map3k7cl | intron_variant | MODIFIER |
| 16 | 87618900 | AX-97804926 | 0 | G | A | 0.43 | -2.29E-02 | 8.00E-02 | 2.69E+00 | 2.75E+00 | 7.71E-01 | no | Gm22808 | downstream_gene_variant | MODIFIER |
| 16 | 87629934 | AX-170131337 | 0 | T | C | 0.43 | -2.29E-02 | 8.00E-02 | 2.69E+00 | 2.75E+00 | 7.71E-01 | no | | intergenic_variant | MODIFIER |
| 16 | 87629963 | AX-97832078 | 0 | T | C | 0.43 | -2.29E-02 | 8.00E-02 | 2.69E+00 | 2.75E+00 | 7.71E-01 | no | | intergenic_variant | MODIFIER |
| 16 | 87655346 | AX-170119568 | 0 | C | G | 0.43 | -2.29E-02 | 8.00E-02 | 2.69E+00 | 2.75E+00 | 7.71E-01 | no | | regulatory_region_variant | MODIFIER |
| 16 | 87663397 | AX-97846758 | 0 | C | T | 0.43 | -2.29E-02 | 8.00E-02 | 2.69E+00 | 2.75E+00 | 7.71E-01 | no | | regulatory_region_variant | MODIFIER |
| 16 | 87664971 | AX-170474953 | 0 | A | C | 0.43 | -2.29E-02 | 8.00E-02 | 2.69E+00 | 2.75E+00 | 7.71E-01 | no | | regulatory_region_variant | MODIFIER |
| 16 | 87819629 | AX-170116799 | 0 | C | T | 0.43 | -2.29E-02 | 8.00E-02 | 2.69E+00 | 2.75E+00 | 7.71E-01 | no | 2810407A14Rik | intron_variant,non_coding_transcript_variant | MODIFIER |
| 3 | 144007595 | AX-170146189 | 0 | G | A | 0.377 | 2.01E-02 | 6.88E-02 | 2.63E+00 | 2.72E+00 | 7.71E-01 | no | | intergenic_variant | MODIFIER |
| 11 | 17751111 | AX-170217457 | 0 | G | A | 0.044 | -5.31E-02 | 1.84E-01 | 2.59E+00 | 2.73E+00 | 7.71E-01 | no | | intergenic_variant | MODIFIER |
| 5 | 102542417 | AX-170072920 | 0 | T | C | 0.439 | 3.10E-02 | 1.08E-01 | 2.69E+00 | 2.76E+00 | 7.71E-01 | no | Arhgap24 | intron_variant | MODIFIER |
| 1 | 49311562 | AX-170438959 | 0 | C | T | 0.105 | -6.75E-02 | 2.36E-01 | 2.72E+00 | 2.76E+00 | 7.71E-01 | no | C230029F24Rik | intron_variant,non_coding_transcript_variant | MODIFIER |
| 1 | 52595949 | AX-170409860 | 0 | C | T | 0.105 | -6.75E-02 | 2.36E-01 | 2.72E+00 | 2.76E+00 | 7.71E-01 | no | | intergenic_variant | MODIFIER |
| 5 | 107603899 | AX-170455785 | 0 | A | C | 0.447 | 3.33E-02 | 1.13E-01 | 2.64E+00 | 2.69E+00 | 7.71E-01 | no | Rpap2 | intron_variant | MODIFIER |
| 5 | 113193521 | AX-170409922 | 0 | T | G | 0.447 | 3.33E-02 | 1.13E-01 | 2.64E+00 | 2.69E+00 | 7.71E-01 | no | 2900026A02Rik | intron_variant | MODIFIER |
| 5 | 113921099 | AX-170143827 | 0 | A | G | 0.447 | 3.33E-02 | 1.13E-01 | 2.64E+00 | 2.69E+00 | 7.71E-01 | no | | regulatory_region_variant | MODIFIER |
| 5 | 114077673 | AX-170123769 | 0 | A | G | 0.447 | 3.33E-02 | 1.13E-01 | 2.64E+00 | 2.69E+00 | 7.71E-01 | no | Svop | intron_variant | MODIFIER |
| 7 | 82318220 | AX-170434376 | 1 | C | T | 0.107 | 5.96E-02 | 2.08E-01 | 2.63E+00 | 2.74E+00 | 7.71E-01 | no | | regulatory_region_variant | MODIFIER |
| 4 | 141210827 | AX-97792693 | 1 | C | T | 0.089 | 4.86E-02 | 1.69E-01 | 2.66E+00 | 2.73E+00 | 7.71E-01 | no | Rsg1 | upstream_gene_variant | MODIFIER |
| 8 | 41989111 | AX-170030567 | 1 | C | T | 0.143 | 4.35E-02 | 1.53E-01 | 2.75E+00 | 2.80E+00 | 7.72E-01 | no | 2810404M03Rik | intron_variant,non_coding_transcript_variant | MODIFIER |
| 7 | 67635826 | AX-170138356 | 1 | A | T | 0.42 | -2.84E-02 | 1.01E-01 | 2.71E+00 | 2.79E+00 | 7.72E-01 | no | Lrrc28 | intron_variant | MODIFIER |
| 20 | 52213579 | AX-97857017 | 0 | G | A | 0.254 | 1.78E-02 | 6.23E-02 | 2.65E+00 | 2.74E+00 | 7.72E-01 | no | | intergenic_variant | MODIFIER |
| 20 | 52245618 | AX-97845568 | 0 | C | A | 0.254 | 1.78E-02 | 6.23E-02 | 2.65E+00 | 2.74E+00 | 7.72E-01 | no | | intergenic_variant | MODIFIER |
| 20 | 52291269 | AX-170237805 | 0 | C | A | 0.254 | 1.78E-02 | 6.23E-02 | 2.65E+00 | 2.74E+00 | 7.72E-01 | no | Gpc3 | intron_variant | MODIFIER |
| 20 | 52905037 | AX-170033868 | 0 | G | C | 0.254 | 1.78E-02 | 6.23E-02 | 2.65E+00 | 2.74E+00 | 7.72E-01 | no | | intergenic_variant | MODIFIER |
| 20 | 52931976 | AX-170429392 | 0 | T | C | 0.254 | 1.78E-02 | 6.23E-02 | 2.65E+00 | 2.74E+00 | 7.72E-01 | no | Phf6 | splice_region_variant,intron_variant | LOW |
| 20 | 53051523 | AX-97806042 | 0 | A | G | 0.254 | 1.78E-02 | 6.23E-02 | 2.65E+00 | 2.74E+00 | 7.72E-01 | no | Mir322 | downstream_gene_variant | MODIFIER |
| 20 | 53059139 | AX-97841635 | 0 | C | T | 0.254 | 1.78E-02 | 6.23E-02 | 2.65E+00 | 2.74E+00 | 7.72E-01 | no | Mir322 | upstream_gene_variant | MODIFIER |
| 20 | 53089412 | AX-170210299 | 0 | G | A | 0.254 | 1.78E-02 | 6.23E-02 | 2.65E+00 | 2.74E+00 | 7.72E-01 | no | Plac1 | intron_variant | MODIFIER |
| 20 | 53120779 | AX-97806638 | 0 | G | A | 0.254 | 1.78E-02 | 6.23E-02 | 2.65E+00 | 2.74E+00 | 7.72E-01 | no | Plac1 | intron_variant | MODIFIER |
| 20 | 53132019 | AX-97822757 | 0 | A | G | 0.254 | 1.78E-02 | 6.23E-02 | 2.65E+00 | 2.74E+00 | 7.72E-01 | no | Plac1 | intron_variant | MODIFIER |
| 20 | 53139257 | AX-170319743 | 0 | T | G | 0.254 | 1.78E-02 | 6.23E-02 | 2.65E+00 | 2.74E+00 | 7.72E-01 | no | Plac1 | intron_variant | MODIFIER |
| 20 | 53144129 | AX-170481856 | 0 | G | C | 0.254 | 1.78E-02 | 6.23E-02 | 2.65E+00 | 2.74E+00 | 7.72E-01 | no | Plac1 | intron_variant | MODIFIER |
| 20 | 53146797 | AX-170168166 | 0 | G | A | 0.254 | 1.78E-02 | 6.23E-02 | 2.65E+00 | 2.74E+00 | 7.72E-01 | no | Plac1 | intron_variant | MODIFIER |
| 20 | 53157033 | AX-170084252 | 0 | A | G | 0.254 | 1.78E-02 | 6.23E-02 | 2.65E+00 | 2.74E+00 | 7.72E-01 | no | Plac1 | intron_variant | MODIFIER |
| 20 | 53158958 | AX-170084250 | 0 | G | A | 0.254 | 1.78E-02 | 6.23E-02 | 2.65E+00 | 2.74E+00 | 7.72E-01 | no | Plac1 | intron_variant | MODIFIER |
| 20 | 53162235 | AX-170084253 | 0 | G | A | 0.254 | 1.78E-02 | 6.23E-02 | 2.65E+00 | 2.74E+00 | 7.72E-01 | no | Plac1 | intron_variant | MODIFIER |
| 20 | 53166335 | AX-97815429 | 0 | A | G | 0.254 | 1.78E-02 | 6.23E-02 | 2.65E+00 | 2.74E+00 | 7.72E-01 | no | Plac1 | intron_variant | MODIFIER |
| 7 | 40475748 | AX-170437081 | 1 | A | G | 0.321 | 2.75E-02 | 9.27E-02 | 2.61E+00 | 2.69E+00 | 7.72E-01 | no | | intergenic_variant | MODIFIER |
| 7 | 44611702 | AX-170028921 | 1 | A | G | 0.321 | 2.75E-02 | 9.27E-02 | 2.61E+00 | 2.69E+00 | 7.72E-01 | no | Myh14 | synonymous_variant | LOW |
| 4 | 71175213 | AX-170089272 | 1 | G | T | 0.241 | 3.19E-02 | 1.03E-01 | 2.56E+00 | 2.65E+00 | 7.72E-01 | no | | intergenic_variant | MODIFIER |
| 18 | 68670787 | AX-170182062 | 1 | C | T | 0.098 | 4.97E-02 | 1.70E-01 | 2.61E+00 | 2.71E+00 | 7.72E-01 | no | | regulatory_region_variant | MODIFIER |
| 7 | 94303725 | AX-170217991 | 0 | C | T | 0.14 | 5.10E-02 | 1.77E-01 | 2.63E+00 | 2.74E+00 | 7.72E-01 | no | Gm32647 | intron_variant,non_coding_transcript_variant | MODIFIER |
| 10 | 86656805 | AX-97788540 | 1 | G | T | 0.482 | 1.54E-01 | 5.32E-01 | 2.67E+00 | 2.72E+00 | 7.72E-01 | Yes | Gm5174 | non_coding_transcript_exon_variant | MODIFIER |
| 11 | 74083386 | AX-170005301 | 0 | T | C | 0.263 | 2.02E-02 | 7.17E-02 | 2.62E+00 | 2.76E+00 | 7.72E-01 | no | Zfp616 | 3_prime_UTR_variant | MODIFIER |
| 11 | 74185779 | AX-97817268 | 0 | A | C | 0.263 | 2.02E-02 | 7.17E-02 | 2.62E+00 | 2.76E+00 | 7.72E-01 | no | Olfr403 | upstream_gene_variant | MODIFIER |
| 11 | 74195022 | AX-170326727 | 0 | G | A | 0.263 | 2.02E-02 | 7.17E-02 | 2.62E+00 | 2.76E+00 | 7.72E-01 | no | Olfr403 | upstream_gene_variant | MODIFIER |
| 11 | 74197840 | AX-97844105 | 0 | G | A | 0.263 | 2.02E-02 | 7.17E-02 | 2.62E+00 | 2.76E+00 | 7.72E-01 | no | Olfr403 | downstream_gene_variant | MODIFIER |
| 11 | 74203943 | AX-170005299 | 0 | T | C | 0.263 | 2.02E-02 | 7.17E-02 | 2.62E+00 | 2.76E+00 | 7.72E-01 | no | Olfr43 | downstream_gene_variant | MODIFIER |
| 11 | 74206109 | AX-170005296 | 0 | G | A | 0.263 | 2.02E-02 | 7.17E-02 | 2.62E+00 | 2.76E+00 | 7.72E-01 | no | Olfr43 | downstream_gene_variant | MODIFIER |

| | | | | | | | | | | | | | | | |
|---|---|---|---|---|---|---|---|---|---|---|---|---|---|---|---|
| 11 | 74207492 | AX-170256632 | 0 | T | C | 0.263 | 2.02E-02 | 7.17E-02 | 2.62E+00 | 2.76E+00 | 7.72E-01 | no | Olfr43 | upstream_gene_variant | MODIFIER |
| 11 | 74209923 | AX-170435147 | 0 | G | C | 0.263 | 2.02E-02 | 7.17E-02 | 2.62E+00 | 2.76E+00 | 7.72E-01 | no | Olfr43 | upstream_gene_variant | MODIFIER |
| 11 | 74210641 | AX-170275575 | 0 | G | A | 0.263 | 2.02E-02 | 7.17E-02 | 2.62E+00 | 2.76E+00 | 7.72E-01 | no | Olfr43 | upstream_gene_variant | MODIFIER |
| 11 | 74210995 | AX-170426728 | 0 | C | T | 0.263 | 2.02E-02 | 7.17E-02 | 2.62E+00 | 2.76E+00 | 7.72E-01 | no | Olfr43 | upstream_gene_variant | MODIFIER |
| 11 | 74211178 | AX-170150652 | 0 | G | A | 0.263 | 2.02E-02 | 7.17E-02 | 2.62E+00 | 2.76E+00 | 7.72E-01 | no | Olfr43 | upstream_gene_variant | MODIFIER |
| 11 | 74211248 | AX-170464835 | 0 | G | A | 0.263 | 2.02E-02 | 7.17E-02 | 2.62E+00 | 2.76E+00 | 7.72E-01 | no | Olfr43 | upstream_gene_variant | MODIFIER |
| 11 | 74213858 | AX-170517373 | 0 | A | G | 0.263 | 2.02E-02 | 7.17E-02 | 2.62E+00 | 2.76E+00 | 7.72E-01 | no | Olfr43 | upstream_gene_variant | MODIFIER |
| 11 | 74213969 | AX-170190180 | 0 | T | C | 0.263 | 2.02E-02 | 7.17E-02 | 2.62E+00 | 2.76E+00 | 7.72E-01 | no | Olfr43 | upstream_gene_variant | MODIFIER |
| 11 | 74217077 | AX-170267499 | 0 | G | A | 0.263 | 2.02E-02 | 7.17E-02 | 2.62E+00 | 2.76E+00 | 7.72E-01 | no | | intergenic_variant | MODIFIER |
| 11 | 74238299 | AX-170005292 | 0 | T | C | 0.263 | 2.02E-02 | 7.17E-02 | 2.62E+00 | 2.76E+00 | 7.72E-01 | no | Olfr404-ps1 | upstream_gene_variant | MODIFIER |
| 11 | 74238369 | AX-170005306 | 0 | G | A | 0.263 | 2.02E-02 | 7.17E-02 | 2.62E+00 | 2.76E+00 | 7.72E-01 | no | Olfr404-ps1 | upstream_gene_variant | MODIFIER |
| 11 | 74238672 | AX-170085833 | 0 | A | G | 0.263 | 2.02E-02 | 7.17E-02 | 2.62E+00 | 2.76E+00 | 7.72E-01 | no | Olfr404-ps1 | upstream_gene_variant | MODIFIER |
| 11 | 74239992 | AX-97789432 | 0 | T | C | 0.263 | 2.02E-02 | 7.17E-02 | 2.62E+00 | 2.76E+00 | 7.72E-01 | no | Olfr404-ps1 | missense_variant | MODERATE |
| 11 | 74241685 | AX-170174894 | 0 | A | G | 0.263 | 2.02E-02 | 7.17E-02 | 2.62E+00 | 2.76E+00 | 7.72E-01 | no | Olfr404-ps1 | downstream_gene_variant | MODIFIER |
| 11 | 74264374 | AX-170476132 | 0 | G | A | 0.263 | 2.02E-02 | 7.17E-02 | 2.62E+00 | 2.76E+00 | 7.72E-01 | no | Olfr405-ps1 | intron_variant,non_coding_transcript_variant | MODIFIER |
| 11 | 74265385 | AX-170404182 | 0 | G | A | 0.263 | 2.02E-02 | 7.17E-02 | 2.62E+00 | 2.76E+00 | 7.72E-01 | no | Olfr405-ps1 | intron_variant,non_coding_transcript_variant | MODIFIER |
| 11 | 74269786 | AX-170340650 | 0 | T | C | 0.263 | 2.02E-02 | 7.17E-02 | 2.62E+00 | 2.76E+00 | 7.72E-01 | no | Olfr405-ps1 | upstream_gene_variant | MODIFIER |
| 11 | 74270749 | AX-170162820 | 0 | T | C | 0.263 | 2.02E-02 | 7.17E-02 | 2.62E+00 | 2.76E+00 | 7.72E-01 | no | Olfr405-ps1 | upstream_gene_variant | MODIFIER |
| 11 | 74281564 | AX-97818828 | 0 | G | A | 0.263 | 2.02E-02 | 7.17E-02 | 2.62E+00 | 2.76E+00 | 7.72E-01 | no | Olfr407-ps1 | non_coding_transcript_exon_variant | MODIFIER |
| 11 | 74315819 | AX-170519348 | 0 | G | A | 0.263 | 2.02E-02 | 7.17E-02 | 2.62E+00 | 2.76E+00 | 7.72E-01 | no | n-R5s71 | upstream_gene_variant | MODIFIER |
| 11 | 74316484 | AX-170157581 | 0 | G | C | 0.263 | 2.02E-02 | 7.17E-02 | 2.62E+00 | 2.76E+00 | 7.72E-01 | no | n-R5s71 | upstream_gene_variant | MODIFIER |
| 11 | 74327461 | AX-170297078 | 0 | A | G | 0.263 | 2.02E-02 | 7.17E-02 | 2.62E+00 | 2.76E+00 | 7.72E-01 | no | Gm12332 | downstream_gene_variant | MODIFIER |
| 11 | 74327665 | AX-170099939 | 0 | C | T | 0.263 | 2.02E-02 | 7.17E-02 | 2.62E+00 | 2.76E+00 | 7.72E-01 | no | Gm12332 | downstream_gene_variant | MODIFIER |
| 11 | 74330357 | AX-170429734 | 0 | T | C | 0.263 | 2.02E-02 | 7.17E-02 | 2.62E+00 | 2.76E+00 | 7.72E-01 | no | Olfr410 | downstream_gene_variant | MODIFIER |
| 11 | 74332963 | AX-170257342 | 0 | G | A | 0.263 | 2.02E-02 | 7.17E-02 | 2.62E+00 | 2.76E+00 | 7.72E-01 | no | Olfr410 | downstream_gene_variant | MODIFIER |
| 11 | 74333434 | AX-170186583 | 0 | T | A | 0.263 | 2.02E-02 | 7.17E-02 | 2.62E+00 | 2.76E+00 | 7.72E-01 | no | Olfr410 | downstream_gene_variant | MODIFIER |
| 11 | 74364621 | AX-170407520 | 0 | T | C | 0.263 | 2.02E-02 | 7.17E-02 | 2.62E+00 | 2.76E+00 | 7.72E-01 | no | Olfr412 | upstream_gene_variant | MODIFIER |
| 11 | 74377960 | AX-97787414 | 0 | A | G | 0.263 | 2.02E-02 | 7.17E-02 | 2.62E+00 | 2.76E+00 | 7.72E-01 | no | | intergenic_variant | MODIFIER |
| 11 | 74378763 | AX-170239325 | 0 | C | T | 0.263 | 2.02E-02 | 7.17E-02 | 2.62E+00 | 2.76E+00 | 7.72E-01 | no | Rap1gap2 | downstream_gene_variant | MODIFIER |
| 11 | 74391187 | AX-170087060 | 0 | G | A | 0.263 | 2.02E-02 | 7.17E-02 | 2.62E+00 | 2.76E+00 | 7.72E-01 | no | Rap1gap2 | intron_variant,NMD_transcript_variant | MODIFIER |
| 11 | 74426148 | AX-97805320 | 0 | G | A | 0.263 | 2.02E-02 | 7.17E-02 | 2.62E+00 | 2.76E+00 | 7.72E-01 | no | Rap1gap2 | intron_variant,NMD_transcript_variant | MODIFIER |
| 11 | 74429750 | AX-170005312 | 0 | T | C | 0.263 | 2.02E-02 | 7.17E-02 | 2.62E+00 | 2.76E+00 | 7.72E-01 | no | Rap1gap2 | intron_variant,NMD_transcript_variant | MODIFIER |
| 11 | 74462426 | AX-170320560 | 0 | A | G | 0.263 | 2.02E-02 | 7.17E-02 | 2.62E+00 | 2.76E+00 | 7.72E-01 | no | Rap1gap2 | intron_variant,NMD_transcript_variant | MODIFIER |
| 11 | 74466045 | AX-97816210 | 0 | C | T | 0.263 | 2.02E-02 | 7.17E-02 | 2.62E+00 | 2.76E+00 | 7.72E-01 | no | Rap1gap2 | intron_variant,NMD_transcript_variant | MODIFIER |
| 11 | 74485614 | AX-170158378 | 0 | G | A | 0.263 | 2.02E-02 | 7.17E-02 | 2.62E+00 | 2.76E+00 | 7.72E-01 | no | Rap1gap2 | intron_variant,NMD_transcript_variant | MODIFIER |
| 11 | 74486823 | AX-170242714 | 0 | T | C | 0.263 | 2.02E-02 | 7.17E-02 | 2.62E+00 | 2.76E+00 | 7.72E-01 | no | Rap1gap2 | intron_variant,NMD_transcript_variant | MODIFIER |
| 11 | 74497915 | AX-170279217 | 0 | A | G | 0.263 | 2.02E-02 | 7.17E-02 | 2.62E+00 | 2.76E+00 | 7.72E-01 | no | Rap1gap2 | intron_variant,NMD_transcript_variant | MODIFIER |
| 11 | 74498347 | AX-97809377 | 0 | C | T | 0.263 | 2.02E-02 | 7.17E-02 | 2.62E+00 | 2.76E+00 | 7.72E-01 | no | Rap1gap2 | intron_variant,NMD_transcript_variant | MODIFIER |
| 11 | 74517133 | AX-170005309 | 0 | C | A | 0.263 | 2.02E-02 | 7.17E-02 | 2.62E+00 | 2.76E+00 | 7.72E-01 | no | Rap1gap2 | intron_variant,NMD_transcript_variant | MODIFIER |
| 11 | 74523629 | AX-97852994 | 0 | A | G | 0.263 | 2.02E-02 | 7.17E-02 | 2.62E+00 | 2.76E+00 | 7.72E-01 | no | Rap1gap2 | intron_variant,NMD_transcript_variant | MODIFIER |
| 11 | 74531384 | AX-170245314 | 0 | G | A | 0.263 | 2.02E-02 | 7.17E-02 | 2.62E+00 | 2.76E+00 | 7.72E-01 | no | Rap1gap2 | intron_variant,NMD_transcript_variant | MODIFIER |
| 11 | 74567248 | AX-170423230 | 0 | A | G | 0.263 | 2.02E-02 | 7.17E-02 | 2.62E+00 | 2.76E+00 | 7.72E-01 | no | Rap1gap2 | synonymous_variant,NMD_transcript_variant | LOW |
| 11 | 74594376 | AX-170297554 | 0 | C | T | 0.263 | 2.02E-02 | 7.17E-02 | 2.62E+00 | 2.76E+00 | 7.72E-01 | no | Rap1gap2 | intron_variant,NMD_transcript_variant | MODIFIER |
| 1 | 99802734 | AX-170199262 | 0 | C | T | 0.149 | 5.82E-02 | 1.98E-01 | 2.63E+00 | 2.70E+00 | 7.72E-01 | no | Cntnap5b | intron_variant | MODIFIER |
| 6 | 111069294 | AX-170027982 | 1 | C | T | 0.054 | 8.52E-02 | 2.97E-01 | 2.67E+00 | 2.73E+00 | 7.72E-01 | no | Grm7 | intron_variant | MODIFIER |
| 2 | 121935402 | AX-170018876 | 1 | G | A | 0.143 | 6.32E-02 | 2.19E-01 | 2.68E+00 | 2.72E+00 | 7.72E-01 | no | Casc4 | 3_prime_UTR_variant | MODIFIER |
| 2 | 147284724 | AX-170193743 | 1 | G | T | 0.143 | 6.32E-02 | 2.19E-01 | 2.68E+00 | 2.72E+00 | 7.72E-01 | no | Gm14115 | upstream_gene_variant | MODIFIER |
| 1 | 29068208 | AX-168381516 | 0 | A | G | 0.404 | 2.37E-02 | 8.27E-02 | 2.66E+00 | 2.74E+00 | 7.72E-01 | no | | intergenic_variant | MODIFIER |
| 15 | 89777907 | AX-97797032 | 2 | G | A | 0.082 | -4.60E-02 | 1.67E-01 | 2.68E+00 | 2.80E+00 | 7.72E-01 | no | Syt10 | downstream_gene_variant | MODIFIER |
| 5 | 125744426 | AX-170363008 | 0 | G | A | 0.447 | -3.17E-02 | 1.11E-01 | 2.72E+00 | 2.77E+00 | 7.72E-01 | no | Tmem132b | intron_variant | MODIFIER |
| 18 | 81487857 | AX-170520284 | 0 | T | A | 0.43 | 1.71E-02 | 6.11E-02 | 2.64E+00 | 2.76E+00 | 7.72E-01 | no | AC102255.3 | intron_variant,non_coding_transcript_variant | MODIFIER |
| 7 | 117246027 | AX-170474096 | 1 | A | G | 0.411 | 2.70E-02 | 9.34E-02 | 2.67E+00 | 2.71E+00 | 7.72E-01 | no | Gm39075 | intron_variant,non_coding_transcript_variant | MODIFIER |
| 20 | 72761757 | AX-170442832 | 0 | A | G | 0.237 | -1.73E-02 | 6.17E-02 | 2.68E+00 | 2.77E+00 | 7.72E-01 | no | | intergenic_variant | MODIFIER |
| 5 | 73322062 | AX-170460408 | 0 | A | T | 0.456 | 2.60E-02 | 8.82E-02 | 2.60E+00 | 2.70E+00 | 7.72E-01 | no | Ociad2 | downstream_gene_variant | MODIFIER |

| | | | | | | | | | | | | | | | |
|---|---|---|---|---|---|---|---|---|---|---|---|---|---|---|---|
| 12 | 104305490 | AX-170431682 | 0 | C | A | 0.14 | -4.52E-02 | 1.63E-01 | 2.68E+00 | 2.78E+00 | 7.72E-01 | no | | intergenic_variant | MODIFIER |
| 9 | 40586062 | AX-170394800 | 1 | A | T | 0.429 | 2.09E-02 | 7.36E-02 | 2.66E+00 | 2.75E+00 | 7.73E-01 | no | | intergenic_variant | MODIFIER |
| 16 | 24983248 | AX-170280026 | 1 | C | T | 0.25 | 2.56E-02 | 8.99E-02 | 2.62E+00 | 2.74E+00 | 7.73E-01 | no | Lpp | 3_prime_UTR_variant | MODIFIER |
| 16 | 26005940 | AX-97800019 | 1 | C | T | 0.25 | 2.56E-02 | 8.99E-02 | 2.62E+00 | 2.74E+00 | 7.73E-01 | no | P3h2 | intron_variant | MODIFIER |
| 8 | 93233181 | AX-170331217 | 0 | C | G | 0.211 | 3.33E-02 | 1.18E-01 | 2.71E+00 | 2.77E+00 | 7.73E-01 | no | Ces1e | upstream_gene_variant | MODIFIER |
| 8 | 93418325 | AX-170466432 | 0 | T | C | 0.211 | 3.33E-02 | 1.18E-01 | 2.71E+00 | 2.77E+00 | 7.73E-01 | no | Gm45765 | intron_variant,non_coding_transcript_variant | MODIFIER |
| 1 | 150546149 | AX-97844696 | 1 | C | T | 0.196 | -3.62E-02 | 1.29E-01 | 2.69E+00 | 2.78E+00 | 7.73E-01 | no | Ska2l-ps | upstream_gene_variant | MODIFIER |
| 1 | 33817924 | AX-170000244 | 0 | G | T | 0.412 | 2.30E-02 | 8.06E-02 | 2.66E+00 | 2.73E+00 | 7.73E-01 | no | Zfp451 | upstream_gene_variant | MODIFIER |
| 1 | 33817955 | AX-97811228 | 0 | T | A | 0.412 | 2.30E-02 | 8.06E-02 | 2.66E+00 | 2.73E+00 | 7.73E-01 | no | Zfp451 | upstream_gene_variant | MODIFIER |
| 5 | 124324722 | AX-170026083 | 1 | C | T | 0.464 | -3.08E-02 | 1.10E-01 | 2.73E+00 | 2.79E+00 | 7.73E-01 | no | Mphosph9 | intron_variant | MODIFIER |
| 2 | 155144958 | AX-170394698 | 1 | G | T | 0.134 | -5.44E-02 | 1.94E-01 | 2.69E+00 | 2.77E+00 | 7.73E-01 | no | Itch | intron_variant | MODIFIER |
| 2 | 78629976 | AX-170175403 | 1 | G | T | 0.054 | -8.53E-02 | 3.00E-01 | 2.73E+00 | 2.76E+00 | 7.73E-01 | no | | intergenic_variant | MODIFIER |
| 7 | 116263126 | AX-170141814 | 0 | G | C | 0.412 | 2.70E-02 | 9.36E-02 | 2.67E+00 | 2.71E+00 | 7.73E-01 | no | Plekha7 | intron_variant | MODIFIER |
| 7 | 116265025 | AX-170330084 | 0 | A | C | 0.412 | 2.70E-02 | 9.36E-02 | 2.67E+00 | 2.71E+00 | 7.73E-01 | no | Plekha7 | intron_variant | MODIFIER |
| 7 | 116265177 | AX-170385430 | 0 | T | C | 0.412 | 2.70E-02 | 9.36E-02 | 2.67E+00 | 2.71E+00 | 7.73E-01 | no | Plekha7 | intron_variant | MODIFIER |
| 7 | 116270894 | AX-97805324 | 0 | A | G | 0.412 | 2.70E-02 | 9.36E-02 | 2.67E+00 | 2.71E+00 | 7.73E-01 | no | Plekha7 | intron_variant | MODIFIER |
| 7 | 116271033 | AX-170125467 | 0 | G | A | 0.412 | 2.70E-02 | 9.36E-02 | 2.67E+00 | 2.71E+00 | 7.73E-01 | no | Plekha7 | intron_variant | MODIFIER |
| 7 | 116272234 | AX-170196518 | 0 | A | G | 0.412 | 2.70E-02 | 9.36E-02 | 2.67E+00 | 2.71E+00 | 7.73E-01 | no | Plekha7 | intron_variant | MODIFIER |
| 7 | 116280978 | AX-170332051 | 0 | T | C | 0.412 | 2.70E-02 | 9.36E-02 | 2.67E+00 | 2.71E+00 | 7.73E-01 | no | Plekha7 | intron_variant | MODIFIER |
| 7 | 116288951 | AX-170457567 | 0 | T | C | 0.412 | 2.70E-02 | 9.36E-02 | 2.67E+00 | 2.71E+00 | 7.73E-01 | no | Plekha7 | intron_variant | MODIFIER |
| 7 | 116289893 | AX-170209885 | 0 | A | G | 0.412 | 2.70E-02 | 9.36E-02 | 2.67E+00 | 2.71E+00 | 7.73E-01 | no | Plekha7 | intron_variant | MODIFIER |
| 7 | 116298401 | AX-170389329 | 0 | A | G | 0.412 | 2.70E-02 | 9.36E-02 | 2.67E+00 | 2.71E+00 | 7.73E-01 | no | Plekha7 | intron_variant | MODIFIER |
| 7 | 116302216 | AX-170233010 | 0 | A | G | 0.412 | 2.70E-02 | 9.36E-02 | 2.67E+00 | 2.71E+00 | 7.73E-01 | no | Plekha7 | intron_variant | MODIFIER |
| 7 | 116325104 | AX-170353381 | 0 | A | T | 0.412 | 2.70E-02 | 9.36E-02 | 2.67E+00 | 2.71E+00 | 7.73E-01 | no | | intergenic_variant | MODIFIER |
| 7 | 116330915 | AX-97854335 | 0 | C | T | 0.412 | 2.70E-02 | 9.36E-02 | 2.67E+00 | 2.71E+00 | 7.73E-01 | no | Snord14a | downstream_gene_variant | MODIFIER |
| 7 | 116334355 | AX-170395296 | 0 | A | T | 0.412 | 2.70E-02 | 9.36E-02 | 2.67E+00 | 2.71E+00 | 7.73E-01 | no | Snord14a | upstream_gene_variant | MODIFIER |
| 7 | 116348971 | AX-170179885 | 0 | A | G | 0.412 | 2.70E-02 | 9.36E-02 | 2.67E+00 | 2.71E+00 | 7.73E-01 | no | Pik3c2a | intron_variant | MODIFIER |
| 7 | 116371828 | AX-170237209 | 0 | A | G | 0.412 | 2.70E-02 | 9.36E-02 | 2.67E+00 | 2.71E+00 | 7.73E-01 | no | Pik3c2a | intron_variant | MODIFIER |
| 7 | 116380472 | AX-170484421 | 0 | C | T | 0.412 | 2.70E-02 | 9.36E-02 | 2.67E+00 | 2.71E+00 | 7.73E-01 | no | Pik3c2a | intron_variant | MODIFIER |
| 7 | 116392096 | AX-170489777 | 0 | T | C | 0.412 | 2.70E-02 | 9.36E-02 | 2.67E+00 | 2.71E+00 | 7.73E-01 | no | Pik3c2a | intron_variant | MODIFIER |
| 7 | 116426244 | AX-97798240 | 0 | C | A | 0.412 | 2.70E-02 | 9.36E-02 | 2.67E+00 | 2.71E+00 | 7.73E-01 | no | Pik3c2a | intron_variant | MODIFIER |
| 7 | 116427400 | AX-170101775 | 0 | T | C | 0.412 | 2.70E-02 | 9.36E-02 | 2.67E+00 | 2.71E+00 | 7.73E-01 | no | Pik3c2a | intron_variant | MODIFIER |
| 7 | 116427812 | AX-170131498 | 0 | T | C | 0.412 | 2.70E-02 | 9.36E-02 | 2.67E+00 | 2.71E+00 | 7.73E-01 | no | Pik3c2a | intron_variant | MODIFIER |
| 7 | 116448261 | AX-170029829 | 0 | G | C | 0.412 | 2.70E-02 | 9.36E-02 | 2.67E+00 | 2.71E+00 | 7.73E-01 | no | Pik3c2a | upstream_gene_variant | MODIFIER |
| 7 | 116452156 | AX-170261024 | 0 | C | T | 0.412 | 2.70E-02 | 9.36E-02 | 2.67E+00 | 2.71E+00 | 7.73E-01 | no | | intergenic_variant | MODIFIER |
| 7 | 116452970 | AX-170206018 | 0 | T | A | 0.412 | 2.70E-02 | 9.36E-02 | 2.67E+00 | 2.71E+00 | 7.73E-01 | no | | intergenic_variant | MODIFIER |
| 7 | 116461566 | AX-97833665 | 0 | G | A | 0.412 | 2.70E-02 | 9.36E-02 | 2.67E+00 | 2.71E+00 | 7.73E-01 | no | | intergenic_variant | MODIFIER |
| 7 | 116479985 | AX-170029827 | 0 | G | A | 0.412 | 2.70E-02 | 9.36E-02 | 2.67E+00 | 2.71E+00 | 7.73E-01 | no | | intergenic_variant | MODIFIER |
| 7 | 116482605 | AX-170078538 | 0 | G | A | 0.412 | 2.70E-02 | 9.36E-02 | 2.67E+00 | 2.71E+00 | 7.73E-01 | no | | intergenic_variant | MODIFIER |
| 7 | 116482639 | AX-170078540 | 0 | T | C | 0.412 | 2.70E-02 | 9.36E-02 | 2.67E+00 | 2.71E+00 | 7.73E-01 | no | | intergenic_variant | MODIFIER |
| 7 | 116489685 | AX-170527553 | 0 | G | T | 0.412 | 2.70E-02 | 9.36E-02 | 2.67E+00 | 2.71E+00 | 7.73E-01 | no | Gm45031 | downstream_gene_variant | MODIFIER |
| 7 | 116493270 | AX-170165354 | 0 | A | G | 0.412 | 2.70E-02 | 9.36E-02 | 2.67E+00 | 2.71E+00 | 7.73E-01 | no | Gm45031 | upstream_gene_variant | MODIFIER |
| 7 | 116508050 | AX-97784458 | 0 | T | C | 0.412 | 2.70E-02 | 9.36E-02 | 2.67E+00 | 2.71E+00 | 7.73E-01 | no | Nucb2 | intron_variant | MODIFIER |
| 7 | 116514794 | AX-170144757 | 0 | A | G | 0.412 | 2.70E-02 | 9.36E-02 | 2.67E+00 | 2.71E+00 | 7.73E-01 | no | Nucb2 | intron_variant | MODIFIER |
| 7 | 116552972 | AX-170325260 | 0 | A | C | 0.412 | 2.70E-02 | 9.36E-02 | 2.67E+00 | 2.71E+00 | 7.73E-01 | no | 4732496C06Rik | downstream_gene_variant | MODIFIER |
| 7 | 116559795 | AX-170315764 | 0 | T | C | 0.412 | 2.70E-02 | 9.36E-02 | 2.67E+00 | 2.71E+00 | 7.73E-01 | no | | regulatory_region_variant | MODIFIER |
| 7 | 116567388 | AX-170459375 | 0 | G | A | 0.412 | 2.70E-02 | 9.36E-02 | 2.67E+00 | 2.71E+00 | 7.73E-01 | no | | intergenic_variant | MODIFIER |
| 7 | 116587384 | AX-170455322 | 0 | A | G | 0.412 | 2.70E-02 | 9.36E-02 | 2.67E+00 | 2.71E+00 | 7.73E-01 | no | | intergenic_variant | MODIFIER |
| 7 | 116645344 | AX-170206496 | 0 | C | A | 0.412 | 2.70E-02 | 9.36E-02 | 2.67E+00 | 2.71E+00 | 7.73E-01 | no | | regulatory_region_variant | MODIFIER |
| 7 | 116661453 | AX-170099124 | 0 | T | C | 0.412 | 2.70E-02 | 9.36E-02 | 2.67E+00 | 2.71E+00 | 7.73E-01 | no | | intergenic_variant | MODIFIER |
| 7 | 116684675 | AX-97786730 | 0 | A | G | 0.412 | 2.70E-02 | 9.36E-02 | 2.67E+00 | 2.71E+00 | 7.73E-01 | no | | intergenic_variant | MODIFIER |
| 7 | 116685501 | AX-170385369 | 0 | T | C | 0.412 | 2.70E-02 | 9.36E-02 | 2.67E+00 | 2.71E+00 | 7.73E-01 | no | | intergenic_variant | MODIFIER |
| 7 | 116714368 | AX-170337344 | 0 | A | G | 0.412 | 2.70E-02 | 9.36E-02 | 2.67E+00 | 2.71E+00 | 7.73E-01 | no | | intergenic_variant | MODIFIER |
| 7 | 116738495 | AX-170534742 | 0 | T | C | 0.412 | 2.70E-02 | 9.36E-02 | 2.67E+00 | 2.71E+00 | 7.73E-01 | no | | intergenic_variant | MODIFIER |

| | | | | | | | | | | | | | | | |
|---|---|---|---|---|---|---|---|---|---|---|---|---|---|---|---|
| 7 | 116764033 | AX-170294387 | 0 | C | G | 0.412 | 2.70E-02 | 9.36E-02 | 2.67E+00 | 2.71E+00 | 7.73E-01 | no | | intergenic_variant | MODIFIER |
| 7 | 116764294 | AX-170182426 | 0 | T | C | 0.412 | 2.70E-02 | 9.36E-02 | 2.67E+00 | 2.71E+00 | 7.73E-01 | no | | intergenic_variant | MODIFIER |
| 7 | 116764548 | AX-170176424 | 0 | G | T | 0.412 | 2.70E-02 | 9.36E-02 | 2.67E+00 | 2.71E+00 | 7.73E-01 | no | | intergenic_variant | MODIFIER |
| 7 | 116834498 | AX-170252336 | 0 | A | G | 0.412 | 2.70E-02 | 9.36E-02 | 2.67E+00 | 2.71E+00 | 7.73E-01 | no | | intergenic_variant | MODIFIER |
| 7 | 116836457 | AX-170078547 | 0 | T | G | 0.412 | 2.70E-02 | 9.36E-02 | 2.67E+00 | 2.71E+00 | 7.73E-01 | no | | intergenic_variant | MODIFIER |
| 7 | 116874000 | AX-170392844 | 0 | T | C | 0.412 | 2.70E-02 | 9.36E-02 | 2.67E+00 | 2.71E+00 | 7.73E-01 | no | | intergenic_variant | MODIFIER |
| 7 | 116876619 | AX-170318318 | 0 | G | T | 0.412 | 2.70E-02 | 9.36E-02 | 2.67E+00 | 2.71E+00 | 7.73E-01 | no | | intergenic_variant | MODIFIER |
| 7 | 116876854 | AX-170236940 | 0 | T | C | 0.412 | 2.70E-02 | 9.36E-02 | 2.67E+00 | 2.71E+00 | 7.73E-01 | no | | intergenic_variant | MODIFIER |
| 7 | 116877368 | AX-170281632 | 0 | G | C | 0.412 | 2.70E-02 | 9.36E-02 | 2.67E+00 | 2.71E+00 | 7.73E-01 | no | | intergenic_variant | MODIFIER |
| 7 | 116881664 | AX-170176068 | 0 | A | G | 0.412 | 2.70E-02 | 9.36E-02 | 2.67E+00 | 2.71E+00 | 7.73E-01 | no | | intergenic_variant | MODIFIER |
| 7 | 116884301 | AX-170434500 | 0 | C | T | 0.412 | 2.70E-02 | 9.36E-02 | 2.67E+00 | 2.71E+00 | 7.73E-01 | no | | intergenic_variant | MODIFIER |
| 7 | 116885064 | AX-170357159 | 0 | G | A | 0.412 | 2.70E-02 | 9.36E-02 | 2.67E+00 | 2.71E+00 | 7.73E-01 | no | | intergenic_variant | MODIFIER |
| 7 | 116887430 | AX-170335125 | 0 | A | T | 0.412 | 2.70E-02 | 9.36E-02 | 2.67E+00 | 2.71E+00 | 7.73E-01 | no | | intergenic_variant | MODIFIER |
| 7 | 116890921 | AX-170475391 | 0 | A | G | 0.412 | 2.70E-02 | 9.36E-02 | 2.67E+00 | 2.71E+00 | 7.73E-01 | no | | intergenic_variant | MODIFIER |
| 7 | 116904814 | AX-170277134 | 0 | C | T | 0.412 | 2.70E-02 | 9.36E-02 | 2.67E+00 | 2.71E+00 | 7.73E-01 | no | | intergenic_variant | MODIFIER |
| 7 | 116904939 | AX-170481247 | 0 | G | A | 0.412 | 2.70E-02 | 9.36E-02 | 2.67E+00 | 2.71E+00 | 7.73E-01 | no | | intergenic_variant | MODIFIER |
| 7 | 116904981 | AX-170499952 | 0 | A | G | 0.412 | 2.70E-02 | 9.36E-02 | 2.67E+00 | 2.71E+00 | 7.73E-01 | no | | intergenic_variant | MODIFIER |
| 7 | 116921422 | AX-170235519 | 0 | C | T | 0.412 | 2.70E-02 | 9.36E-02 | 2.67E+00 | 2.71E+00 | 7.73E-01 | no | | intergenic_variant | MODIFIER |
| 7 | 116927762 | AX-170384656 | 0 | G | C | 0.412 | 2.70E-02 | 9.36E-02 | 2.67E+00 | 2.71E+00 | 7.73E-01 | no | | intergenic_variant | MODIFIER |
| 7 | 116928456 | AX-170372179 | 0 | C | A | 0.412 | 2.70E-02 | 9.36E-02 | 2.67E+00 | 2.71E+00 | 7.73E-01 | no | | intergenic_variant | MODIFIER |
| 7 | 116947852 | AX-170457317 | 0 | G | A | 0.412 | 2.70E-02 | 9.36E-02 | 2.67E+00 | 2.71E+00 | 7.73E-01 | no | Gm26054 | downstream_gene_variant | MODIFIER |
| 7 | 116947879 | AX-170471807 | 0 | T | C | 0.412 | 2.70E-02 | 9.36E-02 | 2.67E+00 | 2.71E+00 | 7.73E-01 | no | Gm26054 | downstream_gene_variant | MODIFIER |
| 7 | 116948531 | AX-170402230 | 0 | T | C | 0.412 | 2.70E-02 | 9.36E-02 | 2.67E+00 | 2.71E+00 | 7.73E-01 | no | Gm26054 | downstream_gene_variant | MODIFIER |
| 7 | 116983218 | AX-170029833 | 0 | C | T | 0.412 | 2.70E-02 | 9.36E-02 | 2.67E+00 | 2.71E+00 | 7.73E-01 | no | | intergenic_variant | MODIFIER |
| 7 | 117000896 | AX-170078550 | 0 | C | A | 0.412 | 2.70E-02 | 9.36E-02 | 2.67E+00 | 2.71E+00 | 7.73E-01 | no | | intergenic_variant | MODIFIER |
| 7 | 117001430 | AX-170198275 | 0 | T | G | 0.412 | 2.70E-02 | 9.36E-02 | 2.67E+00 | 2.71E+00 | 7.73E-01 | no | | intergenic_variant | MODIFIER |
| 7 | 117005571 | AX-170413983 | 0 | T | G | 0.412 | 2.70E-02 | 9.36E-02 | 2.67E+00 | 2.71E+00 | 7.73E-01 | no | | intergenic_variant | MODIFIER |
| 7 | 117028455 | AX-97825257 | 0 | G | A | 0.412 | 2.70E-02 | 9.36E-02 | 2.67E+00 | 2.71E+00 | 7.73E-01 | no | | intergenic_variant | MODIFIER |
| 7 | 117028546 | AX-170300012 | 0 | T | C | 0.412 | 2.70E-02 | 9.36E-02 | 2.67E+00 | 2.71E+00 | 7.73E-01 | no | | intergenic_variant | MODIFIER |
| 7 | 117029976 | AX-170029831 | 0 | T | C | 0.412 | 2.70E-02 | 9.36E-02 | 2.67E+00 | 2.71E+00 | 7.73E-01 | no | | intergenic_variant | MODIFIER |
| 7 | 117030491 | AX-170029832 | 0 | C | T | 0.412 | 2.70E-02 | 9.36E-02 | 2.67E+00 | 2.71E+00 | 7.73E-01 | no | | intergenic_variant | MODIFIER |
| 7 | 117036124 | AX-170156117 | 0 | T | C | 0.412 | 2.70E-02 | 9.36E-02 | 2.67E+00 | 2.71E+00 | 7.73E-01 | no | | intergenic_variant | MODIFIER |
| 7 | 117066522 | AX-170214261 | 0 | G | A | 0.412 | 2.70E-02 | 9.36E-02 | 2.67E+00 | 2.71E+00 | 7.73E-01 | no | | intergenic_variant | MODIFIER |
| 7 | 117066896 | AX-170427708 | 0 | T | C | 0.412 | 2.70E-02 | 9.36E-02 | 2.67E+00 | 2.71E+00 | 7.73E-01 | no | | intergenic_variant | MODIFIER |
| 7 | 117074519 | AX-170162182 | 0 | A | G | 0.412 | 2.70E-02 | 9.36E-02 | 2.67E+00 | 2.71E+00 | 7.73E-01 | no | | intergenic_variant | MODIFIER |
| 7 | 117078419 | AX-170199999 | 0 | A | C | 0.412 | 2.70E-02 | 9.36E-02 | 2.67E+00 | 2.71E+00 | 7.73E-01 | no | | intergenic_variant | MODIFIER |
| 7 | 117082683 | AX-170143565 | 0 | A | G | 0.412 | 2.70E-02 | 9.36E-02 | 2.67E+00 | 2.71E+00 | 7.73E-01 | no | | intergenic_variant | MODIFIER |
| 7 | 117084026 | AX-170276076 | 0 | C | T | 0.412 | 2.70E-02 | 9.36E-02 | 2.67E+00 | 2.71E+00 | 7.73E-01 | no | | intergenic_variant | MODIFIER |
| 7 | 117091422 | AX-170408494 | 0 | T | C | 0.412 | 2.70E-02 | 9.36E-02 | 2.67E+00 | 2.71E+00 | 7.73E-01 | no | | intergenic_variant | MODIFIER |
| 7 | 117092524 | AX-170096090 | 0 | G | A | 0.412 | 2.70E-02 | 9.36E-02 | 2.67E+00 | 2.71E+00 | 7.73E-01 | no | | intergenic_variant | MODIFIER |
| 7 | 117099687 | AX-170394090 | 0 | A | C | 0.412 | 2.70E-02 | 9.36E-02 | 2.67E+00 | 2.71E+00 | 7.73E-01 | no | | intergenic_variant | MODIFIER |
| 7 | 117100958 | AX-97829939 | 0 | A | G | 0.412 | 2.70E-02 | 9.36E-02 | 2.67E+00 | 2.71E+00 | 7.73E-01 | no | | intergenic_variant | MODIFIER |
| 7 | 117108127 | AX-170136591 | 0 | T | C | 0.412 | 2.70E-02 | 9.36E-02 | 2.67E+00 | 2.71E+00 | 7.73E-01 | no | | intergenic_variant | MODIFIER |
| 7 | 117126316 | AX-170365400 | 0 | G | C | 0.412 | 2.70E-02 | 9.36E-02 | 2.67E+00 | 2.71E+00 | 7.73E-01 | no | | intergenic_variant | MODIFIER |
| 7 | 117159363 | AX-170437363 | 0 | A | G | 0.412 | 2.70E-02 | 9.36E-02 | 2.67E+00 | 2.71E+00 | 7.73E-01 | no | | intergenic_variant | MODIFIER |
| 7 | 117165044 | AX-170078551 | 0 | T | G | 0.412 | 2.70E-02 | 9.36E-02 | 2.67E+00 | 2.71E+00 | 7.73E-01 | no | | intergenic_variant | MODIFIER |
| 7 | 117182710 | AX-170333930 | 0 | T | C | 0.412 | 2.70E-02 | 9.36E-02 | 2.67E+00 | 2.71E+00 | 7.73E-01 | no | | intergenic_variant | MODIFIER |
| 7 | 117190233 | AX-170376499 | 0 | C | T | 0.412 | 2.70E-02 | 9.36E-02 | 2.67E+00 | 2.71E+00 | 7.73E-01 | no | | intergenic_variant | MODIFIER |
| 7 | 117190260 | AX-170191516 | 0 | A | C | 0.412 | 2.70E-02 | 9.36E-02 | 2.67E+00 | 2.71E+00 | 7.73E-01 | no | | intergenic_variant | MODIFIER |
| 7 | 117194217 | AX-170366479 | 0 | A | G | 0.412 | 2.70E-02 | 9.36E-02 | 2.67E+00 | 2.71E+00 | 7.73E-01 | no | | intergenic_variant | MODIFIER |
| 7 | 117213110 | AX-170532159 | 0 | A | G | 0.412 | 2.70E-02 | 9.36E-02 | 2.67E+00 | 2.71E+00 | 7.73E-01 | no | | intergenic_variant | MODIFIER |
| 7 | 117219268 | AX-170403954 | 0 | A | G | 0.412 | 2.70E-02 | 9.36E-02 | 2.67E+00 | 2.71E+00 | 7.73E-01 | no | | intergenic_variant | MODIFIER |
| 7 | 117236190 | AX-170467874 | 0 | C | T | 0.412 | 2.70E-02 | 9.36E-02 | 2.67E+00 | 2.71E+00 | 7.73E-01 | no | Gm39075 | intron_variant,non_coding_transcript_variant | MODIFIER |
| 7 | 117238211 | AX-170298262 | 0 | T | A | 0.412 | 2.70E-02 | 9.36E-02 | 2.67E+00 | 2.71E+00 | 7.73E-01 | no | Gm39075 | intron_variant,non_coding_transcript_variant | MODIFIER |

| | | | | | | | | | | | | | | | |
|---|---|---|---|---|---|---|---|---|---|---|---|---|---|---|---|
| 7 | 117238481 | AX-170503726 | 0 | A | C | 0.412 | 2.70E-02 | 9.36E-02 | 2.67E+00 | 2.71E+00 | 7.73E-01 | no | Gm39075 | intron_variant,non_coding_transcript_variant | MODIFIER |
| 7 | 117250876 | AX-170442762 | 0 | G | A | 0.412 | 2.70E-02 | 9.36E-02 | 2.67E+00 | 2.71E+00 | 7.73E-01 | no | Gm39075 | intron_variant,non_coding_transcript_variant | MODIFIER |
| 7 | 117251583 | AX-97790673 | 0 | C | T | 0.412 | 2.70E-02 | 9.36E-02 | 2.67E+00 | 2.71E+00 | 7.73E-01 | no | Gm39075 | intron_variant,non_coding_transcript_variant | MODIFIER |
| 7 | 117259943 | AX-170215178 | 0 | C | G | 0.412 | 2.70E-02 | 9.36E-02 | 2.67E+00 | 2.71E+00 | 7.73E-01 | no | Gm39075 | intron_variant,non_coding_transcript_variant | MODIFIER |
| 7 | 117262680 | AX-170470766 | 0 | C | T | 0.412 | 2.70E-02 | 9.36E-02 | 2.67E+00 | 2.71E+00 | 7.73E-01 | no | Gm39075 | intron_variant,non_coding_transcript_variant | MODIFIER |
| 7 | 117263191 | AX-170255757 | 0 | G | A | 0.412 | 2.70E-02 | 9.36E-02 | 2.67E+00 | 2.71E+00 | 7.73E-01 | no | Gm39075 | intron_variant,non_coding_transcript_variant | MODIFIER |
| 7 | 117264782 | AX-170275500 | 0 | G | A | 0.412 | 2.70E-02 | 9.36E-02 | 2.67E+00 | 2.71E+00 | 7.73E-01 | no | Gm39075 | intron_variant,non_coding_transcript_variant | MODIFIER |
| 7 | 117272708 | AX-170375742 | 0 | G | T | 0.412 | 2.70E-02 | 9.36E-02 | 2.67E+00 | 2.71E+00 | 7.73E-01 | no | Gm39075 | intron_variant,non_coding_transcript_variant | MODIFIER |
| 7 | 117280708 | AX-170189382 | 0 | T | C | 0.412 | 2.70E-02 | 9.36E-02 | 2.67E+00 | 2.71E+00 | 7.73E-01 | no | Gm39075 | downstream_gene_variant | MODIFIER |
| 7 | 117281067 | AX-170105031 | 0 | G | A | 0.412 | 2.70E-02 | 9.36E-02 | 2.67E+00 | 2.71E+00 | 7.73E-01 | no | Gm39075 | downstream_gene_variant | MODIFIER |
| 7 | 117284434 | AX-170345650 | 0 | C | G | 0.412 | 2.70E-02 | 9.36E-02 | 2.67E+00 | 2.71E+00 | 7.73E-01 | no | | regulatory_region_variant | MODIFIER |
| 7 | 117292070 | AX-170451569 | 0 | C | T | 0.412 | 2.70E-02 | 9.36E-02 | 2.67E+00 | 2.71E+00 | 7.73E-01 | no | | intergenic_variant | MODIFIER |
| 7 | 117293217 | AX-170370134 | 0 | A | G | 0.412 | 2.70E-02 | 9.36E-02 | 2.67E+00 | 2.71E+00 | 7.73E-01 | no | | intergenic_variant | MODIFIER |
| 7 | 117298722 | AX-170258524 | 0 | T | C | 0.412 | 2.70E-02 | 9.36E-02 | 2.67E+00 | 2.71E+00 | 7.73E-01 | no | | intergenic_variant | MODIFIER |
| 7 | 117303222 | AX-170522873 | 0 | G | A | 0.412 | 2.70E-02 | 9.36E-02 | 2.67E+00 | 2.71E+00 | 7.73E-01 | no | | intergenic_variant | MODIFIER |
| 7 | 117305188 | AX-170348698 | 0 | C | T | 0.412 | 2.70E-02 | 9.36E-02 | 2.67E+00 | 2.71E+00 | 7.73E-01 | no | | intergenic_variant | MODIFIER |
| 7 | 117309002 | AX-170154970 | 0 | A | C | 0.412 | 2.70E-02 | 9.36E-02 | 2.67E+00 | 2.71E+00 | 7.73E-01 | no | | intergenic_variant | MODIFIER |
| 7 | 117309315 | AX-170299818 | 0 | G | A | 0.412 | 2.70E-02 | 9.36E-02 | 2.67E+00 | 2.71E+00 | 7.73E-01 | no | | intergenic_variant | MODIFIER |
| 7 | 117319173 | AX-170438563 | 0 | A | G | 0.412 | 2.70E-02 | 9.36E-02 | 2.67E+00 | 2.71E+00 | 7.73E-01 | no | | intergenic_variant | MODIFIER |
| 7 | 117319933 | AX-170466242 | 0 | C | T | 0.412 | 2.70E-02 | 9.36E-02 | 2.67E+00 | 2.71E+00 | 7.73E-01 | no | | intergenic_variant | MODIFIER |
| 7 | 117333920 | AX-170103028 | 0 | C | A | 0.412 | 2.70E-02 | 9.36E-02 | 2.67E+00 | 2.71E+00 | 7.73E-01 | no | | intergenic_variant | MODIFIER |
| 7 | 117334163 | AX-170180499 | 0 | A | G | 0.412 | 2.70E-02 | 9.36E-02 | 2.67E+00 | 2.71E+00 | 7.73E-01 | no | | regulatory_region_variant | MODIFIER |
| 7 | 117341221 | AX-170294163 | 0 | G | C | 0.412 | 2.70E-02 | 9.36E-02 | 2.67E+00 | 2.71E+00 | 7.73E-01 | no | | intergenic_variant | MODIFIER |
| 7 | 117342196 | AX-170175000 | 0 | G | A | 0.412 | 2.70E-02 | 9.36E-02 | 2.67E+00 | 2.71E+00 | 7.73E-01 | no | | intergenic_variant | MODIFIER |
| 7 | 117346572 | AX-170272851 | 0 | G | C | 0.412 | 2.70E-02 | 9.36E-02 | 2.67E+00 | 2.71E+00 | 7.73E-01 | no | | intergenic_variant | MODIFIER |
| 7 | 117346602 | AX-170173321 | 0 | C | T | 0.412 | 2.70E-02 | 9.36E-02 | 2.67E+00 | 2.71E+00 | 7.73E-01 | no | | intergenic_variant | MODIFIER |
| 7 | 117347202 | AX-170265285 | 0 | T | C | 0.412 | 2.70E-02 | 9.36E-02 | 2.67E+00 | 2.71E+00 | 7.73E-01 | no | | intergenic_variant | MODIFIER |
| 7 | 117347556 | AX-170261319 | 0 | A | G | 0.412 | 2.70E-02 | 9.36E-02 | 2.67E+00 | 2.71E+00 | 7.73E-01 | no | | intergenic_variant | MODIFIER |
| 7 | 117348520 | AX-170196240 | 0 | C | T | 0.412 | 2.70E-02 | 9.36E-02 | 2.67E+00 | 2.71E+00 | 7.73E-01 | no | | intergenic_variant | MODIFIER |
| 7 | 117348835 | AX-170277160 | 0 | C | A | 0.412 | 2.70E-02 | 9.36E-02 | 2.67E+00 | 2.71E+00 | 7.73E-01 | no | | intergenic_variant | MODIFIER |
| 7 | 117362738 | AX-170424008 | 0 | G | A | 0.412 | 2.70E-02 | 9.36E-02 | 2.67E+00 | 2.71E+00 | 7.73E-01 | no | | regulatory_region_variant | MODIFIER |
| 7 | 117372445 | AX-170078555 | 0 | T | C | 0.412 | 2.70E-02 | 9.36E-02 | 2.67E+00 | 2.71E+00 | 7.73E-01 | no | | intergenic_variant | MODIFIER |
| 7 | 117399765 | AX-170168948 | 0 | T | C | 0.412 | 2.70E-02 | 9.36E-02 | 2.67E+00 | 2.71E+00 | 7.73E-01 | no | Xylt1 | intron_variant | MODIFIER |
| 7 | 117399890 | AX-170472282 | 0 | C | T | 0.412 | 2.70E-02 | 9.36E-02 | 2.67E+00 | 2.71E+00 | 7.73E-01 | no | Xylt1 | intron_variant | MODIFIER |
| 15 | 75192622 | AX-170012002 | 0 | C | T | 0.553 | 2.35E-02 | 8.10E-02 | 2.63E+00 | 2.71E+00 | 7.73E-01 | no | Gm28117 | downstream_gene_variant | MODIFIER |
| 18 | 60971270 | AX-170274019 | 1 | T | A | 0.411 | -2.48E-02 | 8.37E-02 | 2.62E+00 | 2.67E+00 | 7.73E-01 | no | Camk2a | intron_variant | MODIFIER |
| 18 | 68525893 | AX-170015916 | 1 | T | A | 0.089 | 5.31E-02 | 1.81E-01 | 2.59E+00 | 2.71E+00 | 7.73E-01 | no | | intergenic_variant | MODIFIER |
| 8 | 100214992 | AX-170287735 | 0 | G | C | 0.237 | 3.26E-02 | 1.14E-01 | 2.68E+00 | 2.74E+00 | 7.73E-01 | no | | intergenic_variant | MODIFIER |
| 8 | 100215348 | AX-170317048 | 0 | G | T | 0.237 | 3.26E-02 | 1.14E-01 | 2.68E+00 | 2.74E+00 | 7.73E-01 | no | | intergenic_variant | MODIFIER |
| 8 | 100235362 | AX-170512094 | 0 | C | T | 0.237 | 3.26E-02 | 1.14E-01 | 2.68E+00 | 2.74E+00 | 7.73E-01 | no | | intergenic_variant | MODIFIER |
| 8 | 100262353 | AX-170430019 | 0 | G | A | 0.237 | 3.26E-02 | 1.14E-01 | 2.68E+00 | 2.74E+00 | 7.73E-01 | no | | intergenic_variant | MODIFIER |
| 8 | 100286204 | AX-170102909 | 0 | T | C | 0.237 | 3.26E-02 | 1.14E-01 | 2.68E+00 | 2.74E+00 | 7.73E-01 | no | | intergenic_variant | MODIFIER |
| 8 | 100286891 | AX-170363759 | 0 | T | C | 0.237 | 3.26E-02 | 1.14E-01 | 2.68E+00 | 2.74E+00 | 7.73E-01 | no | | intergenic_variant | MODIFIER |
| 8 | 100488509 | AX-170222757 | 0 | T | C | 0.237 | 3.26E-02 | 1.14E-01 | 2.68E+00 | 2.74E+00 | 7.73E-01 | no | | intergenic_variant | MODIFIER |
| 8 | 100504677 | AX-170286424 | 0 | A | T | 0.237 | 3.26E-02 | 1.14E-01 | 2.68E+00 | 2.74E+00 | 7.73E-01 | no | | intergenic_variant | MODIFIER |
| 8 | 100536416 | AX-170031380 | 0 | G | T | 0.237 | 3.26E-02 | 1.14E-01 | 2.68E+00 | 2.74E+00 | 7.73E-01 | no | | intergenic_variant | MODIFIER |
| 8 | 100587059 | AX-97836970 | 0 | T | C | 0.237 | 3.26E-02 | 1.14E-01 | 2.68E+00 | 2.74E+00 | 7.73E-01 | no | | intergenic_variant | MODIFIER |
| 8 | 100609392 | AX-170031377 | 0 | G | A | 0.237 | 3.26E-02 | 1.14E-01 | 2.68E+00 | 2.74E+00 | 7.73E-01 | no | | intergenic_variant | MODIFIER |
| 8 | 100614185 | AX-170439820 | 0 | C | G | 0.237 | 3.26E-02 | 1.14E-01 | 2.68E+00 | 2.74E+00 | 7.73E-01 | no | | intergenic_variant | MODIFIER |
| 8 | 100614781 | AX-97819219 | 0 | T | A | 0.237 | 3.26E-02 | 1.14E-01 | 2.68E+00 | 2.74E+00 | 7.73E-01 | no | | intergenic_variant | MODIFIER |
| 8 | 100681002 | AX-170031378 | 0 | A | T | 0.237 | 3.26E-02 | 1.14E-01 | 2.68E+00 | 2.74E+00 | 7.73E-01 | no | Gm39232 | intron_variant,non_coding_transcript_variant | MODIFIER |
| 8 | 100791028 | AX-170442547 | 0 | C | T | 0.237 | 3.26E-02 | 1.14E-01 | 2.68E+00 | 2.74E+00 | 7.73E-01 | no | | regulatory_region_variant | MODIFIER |
| 5 | 98174043 | AX-170238603 | 0 | T | C | 0.404 | -2.90E-02 | 1.04E-01 | 2.72E+00 | 2.79E+00 | 7.73E-01 | no | Prdm8 | intron_variant | MODIFIER |
| 15 | 101249078 | AX-170012389 | 0 | A | T | 0.105 | 4.76E-02 | 1.64E-01 | 2.63E+00 | 2.71E+00 | 7.73E-01 | no | | regulatory_region_variant | MODIFIER |

| | | | | | | | | | | | | | | | |
|---|---|---|---|---|---|---|---|---|---|---|---|---|---|---|---|
| 6 | 112482772 | AX-97854669 | 1 | G | T | 0.054 | 8.35E-02 | 2.94E-01 | 2.68E+00 | 2.74E+00 | 7.73E-01 | no | Oxtr | intron_variant | MODIFIER |
| 6 | 114486657 | AX-170386757 | 0 | T | C | 0.061 | 6.31E-02 | 2.21E-01 | 2.62E+00 | 2.73E+00 | 7.73E-01 | no | Hrh1 | downstream_gene_variant | MODIFIER |
| 5 | 83942269 | AX-170415956 | 0 | G | C | 0.447 | 2.96E-02 | 1.04E-01 | 2.69E+00 | 2.75E+00 | 7.73E-01 | no | | intergenic_variant | MODIFIER |
| 1 | 22735271 | AX-97847904 | 0 | G | A | 0.421 | 2.43E-02 | 8.37E-02 | 2.64E+00 | 2.70E+00 | 7.73E-01 | no | Rims1 | intron_variant | MODIFIER |
| 1 | 5182828 | AX-170040266 | 0 | C | T | 0.526 | 2.05E-02 | 6.59E-02 | 2.55E+00 | 2.64E+00 | 7.74E-01 | no | | intergenic_variant | MODIFIER |
| 3 | 30411327 | AX-170511291 | 0 | T | C | 0.167 | -2.74E-02 | 9.64E-02 | 2.66E+00 | 2.74E+00 | 7.74E-01 | no | Mecom | intron_variant | MODIFIER |
| 1 | 41770820 | AX-97823849 | 0 | C | T | 0.439 | -2.16E-02 | 7.56E-02 | 2.65E+00 | 2.73E+00 | 7.74E-01 | no | | intergenic_variant | MODIFIER |
| 1 | 41878488 | AX-170133720 | 0 | A | G | 0.439 | -2.16E-02 | 7.56E-02 | 2.65E+00 | 2.73E+00 | 7.74E-01 | no | Gm29260 | intron_variant,non_coding_transcript_variant | MODIFIER |
| 1 | 41894713 | AX-170144333 | 0 | G | T | 0.439 | -2.16E-02 | 7.56E-02 | 2.65E+00 | 2.73E+00 | 7.74E-01 | no | Gm29260 | intron_variant,non_coding_transcript_variant | MODIFIER |
| 1 | 41897584 | AX-170117328 | 0 | C | T | 0.439 | -2.16E-02 | 7.56E-02 | 2.65E+00 | 2.73E+00 | 7.74E-01 | no | Gm29260 | intron_variant,non_coding_transcript_variant | MODIFIER |
| 1 | 41900454 | AX-170158460 | 0 | C | T | 0.439 | -2.16E-02 | 7.56E-02 | 2.65E+00 | 2.73E+00 | 7.74E-01 | no | Gm29260 | intron_variant,non_coding_transcript_variant | MODIFIER |
| 1 | 41908556 | AX-97839568 | 0 | A | G | 0.439 | -2.16E-02 | 7.56E-02 | 2.65E+00 | 2.73E+00 | 7.74E-01 | no | Gm29260 | intron_variant,non_coding_transcript_variant | MODIFIER |
| 1 | 41908853 | AX-170292112 | 0 | A | G | 0.439 | -2.16E-02 | 7.56E-02 | 2.65E+00 | 2.73E+00 | 7.74E-01 | no | Gm29260 | intron_variant,non_coding_transcript_variant | MODIFIER |
| 1 | 41955254 | AX-170109940 | 0 | T | G | 0.439 | -2.16E-02 | 7.56E-02 | 2.65E+00 | 2.73E+00 | 7.74E-01 | no | | intergenic_variant | MODIFIER |
| 1 | 41960305 | AX-170041019 | 0 | C | T | 0.439 | -2.16E-02 | 7.56E-02 | 2.65E+00 | 2.73E+00 | 7.74E-01 | no | | regulatory_region_variant | MODIFIER |
| 1 | 41961602 | AX-97824851 | 0 | T | C | 0.439 | -2.16E-02 | 7.56E-02 | 2.65E+00 | 2.73E+00 | 7.74E-01 | no | | intergenic_variant | MODIFIER |
| 1 | 41966532 | AX-170321745 | 0 | C | T | 0.439 | -2.16E-02 | 7.56E-02 | 2.65E+00 | 2.73E+00 | 7.74E-01 | no | | intergenic_variant | MODIFIER |
| 1 | 41967252 | AX-170041021 | 0 | A | G | 0.439 | -2.16E-02 | 7.56E-02 | 2.65E+00 | 2.73E+00 | 7.74E-01 | no | | intergenic_variant | MODIFIER |
| 1 | 42000342 | AX-170474055 | 0 | A | G | 0.439 | -2.16E-02 | 7.56E-02 | 2.65E+00 | 2.73E+00 | 7.74E-01 | no | | intergenic_variant | MODIFIER |
| 1 | 42080664 | AX-170000354 | 0 | A | C | 0.439 | -2.16E-02 | 7.56E-02 | 2.65E+00 | 2.73E+00 | 7.74E-01 | no | | intergenic_variant | MODIFIER |
| 1 | 42096274 | AX-170345153 | 0 | C | T | 0.439 | -2.16E-02 | 7.56E-02 | 2.65E+00 | 2.73E+00 | 7.74E-01 | no | | intergenic_variant | MODIFIER |
| 1 | 42108512 | AX-170000360 | 0 | C | T | 0.439 | -2.16E-02 | 7.56E-02 | 2.65E+00 | 2.73E+00 | 7.74E-01 | no | Gm5267 | downstream_gene_variant | MODIFIER |
| 1 | 42108769 | AX-170262973 | 0 | A | G | 0.439 | -2.16E-02 | 7.56E-02 | 2.65E+00 | 2.73E+00 | 7.74E-01 | no | Gm5267 | downstream_gene_variant | MODIFIER |
| 1 | 42110784 | AX-97844816 | 0 | G | A | 0.439 | -2.16E-02 | 7.56E-02 | 2.65E+00 | 2.73E+00 | 7.74E-01 | no | | intergenic_variant | MODIFIER |
| 1 | 42135288 | AX-170086452 | 0 | A | C | 0.439 | -2.16E-02 | 7.56E-02 | 2.65E+00 | 2.73E+00 | 7.74E-01 | no | | intergenic_variant | MODIFIER |
| 1 | 42151100 | AX-170307654 | 0 | C | A | 0.439 | -2.16E-02 | 7.56E-02 | 2.65E+00 | 2.73E+00 | 7.74E-01 | no | | intergenic_variant | MODIFIER |
| 1 | 42158075 | AX-170132007 | 0 | T | A | 0.439 | -2.16E-02 | 7.56E-02 | 2.65E+00 | 2.73E+00 | 7.74E-01 | no | | intergenic_variant | MODIFIER |
| 1 | 42159678 | AX-170000356 | 0 | A | G | 0.439 | -2.16E-02 | 7.56E-02 | 2.65E+00 | 2.73E+00 | 7.74E-01 | no | | intergenic_variant | MODIFIER |
| 1 | 42223027 | AX-170105424 | 0 | T | C | 0.439 | -2.16E-02 | 7.56E-02 | 2.65E+00 | 2.73E+00 | 7.74E-01 | no | | intergenic_variant | MODIFIER |
| 1 | 42231363 | AX-97816009 | 0 | C | T | 0.439 | -2.16E-02 | 7.56E-02 | 2.65E+00 | 2.73E+00 | 7.74E-01 | no | Gm9915 | downstream_gene_variant | MODIFIER |
| 1 | 42247552 | AX-170000362 | 0 | A | G | 0.439 | -2.16E-02 | 7.56E-02 | 2.65E+00 | 2.73E+00 | 7.74E-01 | no | | intergenic_variant | MODIFIER |
| 4 | 72557775 | AX-170389942 | 0 | C | G | 0.237 | 2.91E-02 | 9.33E-02 | 2.55E+00 | 2.65E+00 | 7.74E-01 | no | | intergenic_variant | MODIFIER |
| 9 | 59210032 | AX-170221640 | 0 | T | C | 0.219 | 3.41E-02 | 1.20E-01 | 2.69E+00 | 2.74E+00 | 7.74E-01 | no | Gm33053 | intron_variant,non_coding_transcript_variant | MODIFIER |
| 16 | 95440599 | AX-170350293 | 1 | G | A | 0.446 | 2.09E-02 | 7.63E-02 | 2.72E+00 | 2.82E+00 | 7.74E-01 | no | Erg | intron_variant | MODIFIER |
| 6 | 127529564 | AX-97797470 | 0 | C | T | 0.535 | 2.44E-02 | 8.09E-02 | 2.58E+00 | 2.67E+00 | 7.74E-01 | no | | regulatory_region_variant | MODIFIER |
| 8 | 90250705 | AX-170147353 | 0 | G | A | 0.193 | 3.20E-02 | 1.16E-01 | 2.70E+00 | 2.79E+00 | 7.74E-01 | no | Tox3 | intron_variant | MODIFIER |
| 16 | 24383920 | AX-97790414 | 0 | A | G | 0.246 | 2.52E-02 | 8.97E-02 | 2.63E+00 | 2.75E+00 | 7.75E-01 | no | AC169509.1 | upstream_gene_variant | MODIFIER |
| 16 | 24420976 | AX-170437354 | 0 | G | T | 0.246 | 2.52E-02 | 8.97E-02 | 2.63E+00 | 2.75E+00 | 7.75E-01 | no | Lpp | intron_variant | MODIFIER |
| 16 | 24421062 | AX-97841307 | 0 | T | C | 0.246 | 2.52E-02 | 8.97E-02 | 2.63E+00 | 2.75E+00 | 7.75E-01 | no | Lpp | intron_variant | MODIFIER |
| 16 | 24423233 | AX-170309281 | 0 | G | A | 0.246 | 2.52E-02 | 8.97E-02 | 2.63E+00 | 2.75E+00 | 7.75E-01 | no | Lpp | intron_variant | MODIFIER |
| 16 | 24541226 | AX-170329472 | 0 | T | C | 0.246 | 2.52E-02 | 8.97E-02 | 2.63E+00 | 2.75E+00 | 7.75E-01 | no | Lpp | intron_variant | MODIFIER |
| 16 | 24578497 | AX-170056920 | 0 | C | T | 0.246 | 2.52E-02 | 8.97E-02 | 2.63E+00 | 2.75E+00 | 7.75E-01 | no | Lpp | intron_variant | MODIFIER |
| 16 | 24639275 | AX-170145752 | 0 | T | C | 0.246 | 2.52E-02 | 8.97E-02 | 2.63E+00 | 2.75E+00 | 7.75E-01 | no | Lpp | intron_variant | MODIFIER |
| 16 | 24642134 | AX-170290301 | 0 | T | C | 0.246 | 2.52E-02 | 8.97E-02 | 2.63E+00 | 2.75E+00 | 7.75E-01 | no | Lpp | intron_variant | MODIFIER |
| 16 | 24674711 | AX-170338496 | 0 | G | A | 0.246 | 2.52E-02 | 8.97E-02 | 2.63E+00 | 2.75E+00 | 7.75E-01 | no | Lpp | intron_variant | MODIFIER |
| 16 | 24678928 | AX-170395879 | 0 | A | G | 0.246 | 2.52E-02 | 8.97E-02 | 2.63E+00 | 2.75E+00 | 7.75E-01 | no | Lpp | intron_variant | MODIFIER |
| 16 | 24699760 | AX-170491093 | 0 | T | C | 0.246 | 2.52E-02 | 8.97E-02 | 2.63E+00 | 2.75E+00 | 7.75E-01 | no | Lpp | intron_variant | MODIFIER |
| 16 | 24699788 | AX-170288057 | 0 | G | A | 0.246 | 2.52E-02 | 8.97E-02 | 2.63E+00 | 2.75E+00 | 7.75E-01 | no | Lpp | intron_variant | MODIFIER |
| 16 | 24707019 | AX-170482117 | 0 | C | T | 0.246 | 2.52E-02 | 8.97E-02 | 2.63E+00 | 2.75E+00 | 7.75E-01 | no | Lpp | intron_variant | MODIFIER |
| 16 | 24712048 | AX-170163256 | 0 | A | G | 0.246 | 2.52E-02 | 8.97E-02 | 2.63E+00 | 2.75E+00 | 7.75E-01 | no | Lpp | intron_variant | MODIFIER |
| 16 | 24972541 | AX-170093417 | 0 | G | A | 0.246 | 2.52E-02 | 8.97E-02 | 2.63E+00 | 2.75E+00 | 7.75E-01 | no | Lpp | intron_variant | MODIFIER |
| 16 | 24986621 | AX-170012663 | 0 | G | A | 0.246 | 2.52E-02 | 8.97E-02 | 2.63E+00 | 2.75E+00 | 7.75E-01 | no | Lpp | 3_prime_UTR_variant | MODIFIER |
| 16 | 25012998 | AX-170012661 | 0 | C | T | 0.246 | 2.52E-02 | 8.97E-02 | 2.63E+00 | 2.75E+00 | 7.75E-01 | no | | intergenic_variant | MODIFIER |
| 16 | 25016963 | AX-170380053 | 0 | C | T | 0.246 | 2.52E-02 | 8.97E-02 | 2.63E+00 | 2.75E+00 | 7.75E-01 | no | | regulatory_region_variant | MODIFIER |

| | | | | | | | | | | | | | | | |
|---|---|---|---|---|---|---|---|---|---|---|---|---|---|---|---|
| 16 | 25026659 | AX-97802475 | 0 | T | C | 0.246 | 2.52E-02 | 8.97E-02 | 2.63E+00 | 2.75E+00 | 7.75E-01 | no | | regulatory_region_variant | MODIFIER |
| 16 | 25029555 | AX-170156577 | 0 | A | T | 0.246 | 2.52E-02 | 8.97E-02 | 2.63E+00 | 2.75E+00 | 7.75E-01 | no | | intergenic_variant | MODIFIER |
| 16 | 25031925 | AX-170485657 | 0 | T | C | 0.246 | 2.52E-02 | 8.97E-02 | 2.63E+00 | 2.75E+00 | 7.75E-01 | no | | intergenic_variant | MODIFIER |
| 16 | 25061352 | AX-170177345 | 0 | C | T | 0.246 | 2.52E-02 | 8.97E-02 | 2.63E+00 | 2.75E+00 | 7.75E-01 | no | A230028O05Rik | intron_variant,non_coding_transcript_variant | MODIFIER |
| 16 | 25062760 | AX-170354305 | 0 | C | T | 0.246 | 2.52E-02 | 8.97E-02 | 2.63E+00 | 2.75E+00 | 7.75E-01 | no | A230028O05Rik | intron_variant,non_coding_transcript_variant | MODIFIER |
| 16 | 25063762 | AX-170487091 | 0 | T | A | 0.246 | 2.52E-02 | 8.97E-02 | 2.63E+00 | 2.75E+00 | 7.75E-01 | no | A230028O05Rik | intron_variant,non_coding_transcript_variant | MODIFIER |
| 16 | 25102681 | AX-170256669 | 0 | T | A | 0.246 | 2.52E-02 | 8.97E-02 | 2.63E+00 | 2.75E+00 | 7.75E-01 | no | | intergenic_variant | MODIFIER |
| 16 | 25165753 | AX-170056931 | 0 | A | G | 0.246 | 2.52E-02 | 8.97E-02 | 2.63E+00 | 2.75E+00 | 7.75E-01 | no | | intergenic_variant | MODIFIER |
| 16 | 25258471 | AX-97835591 | 0 | A | G | 0.246 | 2.52E-02 | 8.97E-02 | 2.63E+00 | 2.75E+00 | 7.75E-01 | no | | intergenic_variant | MODIFIER |
| 16 | 25306940 | AX-170512476 | 0 | G | A | 0.246 | 2.52E-02 | 8.97E-02 | 2.63E+00 | 2.75E+00 | 7.75E-01 | no | Tprg | intron_variant | MODIFIER |
| 16 | 25343191 | AX-170218390 | 0 | G | T | 0.246 | 2.52E-02 | 8.97E-02 | 2.63E+00 | 2.75E+00 | 7.75E-01 | no | Tprg | intron_variant | MODIFIER |
| 16 | 25344769 | AX-170472990 | 0 | G | A | 0.246 | 2.52E-02 | 8.97E-02 | 2.63E+00 | 2.75E+00 | 7.75E-01 | no | Tprg | intron_variant | MODIFIER |
| 16 | 25344967 | AX-170131074 | 0 | A | G | 0.246 | 2.52E-02 | 8.97E-02 | 2.63E+00 | 2.75E+00 | 7.75E-01 | no | Tprg | intron_variant | MODIFIER |
| 16 | 25350063 | AX-170486063 | 0 | G | A | 0.246 | 2.52E-02 | 8.97E-02 | 2.63E+00 | 2.75E+00 | 7.75E-01 | no | Tprg | intron_variant | MODIFIER |
| 16 | 25382747 | AX-170401451 | 0 | A | G | 0.246 | 2.52E-02 | 8.97E-02 | 2.63E+00 | 2.75E+00 | 7.75E-01 | no | Tprg | intron_variant | MODIFIER |
| 16 | 25399354 | AX-170056937 | 0 | A | G | 0.246 | 2.52E-02 | 8.97E-02 | 2.63E+00 | 2.75E+00 | 7.75E-01 | no | Tprg | intron_variant | MODIFIER |
| 16 | 25739184 | AX-170499339 | 0 | T | C | 0.246 | 2.52E-02 | 8.97E-02 | 2.63E+00 | 2.75E+00 | 7.75E-01 | no | Trp63 | intron_variant | MODIFIER |
| 16 | 25886474 | AX-170263862 | 0 | C | G | 0.246 | 2.52E-02 | 8.97E-02 | 2.63E+00 | 2.75E+00 | 7.75E-01 | no | Trp63 | intron_variant | MODIFIER |
| 16 | 25887170 | AX-97846951 | 0 | T | C | 0.246 | 2.52E-02 | 8.97E-02 | 2.63E+00 | 2.75E+00 | 7.75E-01 | no | Trp63 | intron_variant | MODIFIER |
| 16 | 25887820 | AX-170056945 | 0 | A | G | 0.246 | 2.52E-02 | 8.97E-02 | 2.63E+00 | 2.75E+00 | 7.75E-01 | no | Trp63 | intron_variant | MODIFIER |
| 16 | 25901100 | AX-170457326 | 0 | T | A | 0.246 | 2.52E-02 | 8.97E-02 | 2.63E+00 | 2.75E+00 | 7.75E-01 | no | | intergenic_variant | MODIFIER |
| 16 | 25966198 | AX-97788821 | 0 | T | G | 0.246 | 2.52E-02 | 8.97E-02 | 2.63E+00 | 2.75E+00 | 7.75E-01 | no | P3h2 | intron_variant | MODIFIER |
| 16 | 25985142 | AX-97792570 | 0 | T | G | 0.246 | 2.52E-02 | 8.97E-02 | 2.63E+00 | 2.75E+00 | 7.75E-01 | no | P3h2 | intron_variant | MODIFIER |
| 16 | 26008966 | AX-170171272 | 0 | A | G | 0.246 | 2.52E-02 | 8.97E-02 | 2.63E+00 | 2.75E+00 | 7.75E-01 | no | P3h2 | intron_variant | MODIFIER |
| 16 | 26018959 | AX-170487889 | 0 | C | G | 0.246 | 2.52E-02 | 8.97E-02 | 2.63E+00 | 2.75E+00 | 7.75E-01 | no | P3h2 | intron_variant | MODIFIER |
| 16 | 26038233 | AX-170482805 | 0 | C | G | 0.246 | 2.52E-02 | 8.97E-02 | 2.63E+00 | 2.75E+00 | 7.75E-01 | no | P3h2 | intron_variant | MODIFIER |
| 16 | 26039920 | AX-97796331 | 0 | G | A | 0.246 | 2.52E-02 | 8.97E-02 | 2.63E+00 | 2.75E+00 | 7.75E-01 | no | P3h2 | intron_variant | MODIFIER |
| 16 | 26057855 | AX-170492901 | 0 | T | C | 0.246 | 2.52E-02 | 8.97E-02 | 2.63E+00 | 2.75E+00 | 7.75E-01 | no | P3h2 | intron_variant | MODIFIER |
| 16 | 26169485 | AX-97807688 | 0 | G | A | 0.246 | 2.52E-02 | 8.97E-02 | 2.63E+00 | 2.75E+00 | 7.75E-01 | no | | intergenic_variant | MODIFIER |
| 16 | 26169551 | AX-170119298 | 0 | G | T | 0.246 | 2.52E-02 | 8.97E-02 | 2.63E+00 | 2.75E+00 | 7.75E-01 | no | | intergenic_variant | MODIFIER |
| 16 | 26529102 | AX-97822358 | 0 | C | A | 0.246 | 2.52E-02 | 8.97E-02 | 2.63E+00 | 2.75E+00 | 7.75E-01 | no | Tmem207 | upstream_gene_variant | MODIFIER |
| 16 | 26612083 | AX-97857590 | 0 | A | G | 0.246 | 2.52E-02 | 8.97E-02 | 2.63E+00 | 2.75E+00 | 7.75E-01 | no | Il1rap | intron_variant | MODIFIER |
| 16 | 26631434 | AX-170056960 | 0 | C | T | 0.246 | 2.52E-02 | 8.97E-02 | 2.63E+00 | 2.75E+00 | 7.75E-01 | no | Il1rap | intron_variant | MODIFIER |
| 16 | 26665933 | AX-97818599 | 0 | T | C | 0.246 | 2.52E-02 | 8.97E-02 | 2.63E+00 | 2.75E+00 | 7.75E-01 | no | Il1rap | intron_variant | MODIFIER |
| 16 | 26703089 | AX-97854317 | 0 | A | C | 0.246 | 2.52E-02 | 8.97E-02 | 2.63E+00 | 2.75E+00 | 7.75E-01 | no | Il1rap | intron_variant | MODIFIER |
| 16 | 26707187 | AX-97850460 | 0 | G | T | 0.246 | 2.52E-02 | 8.97E-02 | 2.63E+00 | 2.75E+00 | 7.75E-01 | no | Il1rap | intron_variant | MODIFIER |
| 16 | 26779804 | AX-170056964 | 0 | G | A | 0.246 | 2.52E-02 | 8.97E-02 | 2.63E+00 | 2.75E+00 | 7.75E-01 | no | | intergenic_variant | MODIFIER |
| 16 | 27551691 | AX-168411083 | 0 | G | A | 0.246 | 2.52E-02 | 8.97E-02 | 2.63E+00 | 2.75E+00 | 7.75E-01 | no | | intergenic_variant | MODIFIER |
| 16 | 27551720 | AX-97794710 | 0 | A | G | 0.246 | 2.52E-02 | 8.97E-02 | 2.63E+00 | 2.75E+00 | 7.75E-01 | no | | intergenic_variant | MODIFIER |
| 16 | 27568573 | AX-97846546 | 0 | C | A | 0.246 | 2.52E-02 | 8.97E-02 | 2.63E+00 | 2.75E+00 | 7.75E-01 | no | | intergenic_variant | MODIFIER |
| 16 | 27568726 | AX-170414289 | 0 | A | G | 0.246 | 2.52E-02 | 8.97E-02 | 2.63E+00 | 2.75E+00 | 7.75E-01 | no | | intergenic_variant | MODIFIER |
| 16 | 27578758 | AX-170517725 | 0 | C | T | 0.246 | 2.52E-02 | 8.97E-02 | 2.63E+00 | 2.75E+00 | 7.75E-01 | no | | intergenic_variant | MODIFIER |
| 16 | 27587008 | AX-170200484 | 0 | G | A | 0.246 | 2.52E-02 | 8.97E-02 | 2.63E+00 | 2.75E+00 | 7.75E-01 | no | | intergenic_variant | MODIFIER |
| 16 | 27587582 | AX-170175900 | 0 | C | T | 0.246 | 2.52E-02 | 8.97E-02 | 2.63E+00 | 2.75E+00 | 7.75E-01 | no | | intergenic_variant | MODIFIER |
| 16 | 27587620 | AX-170492144 | 0 | C | T | 0.246 | 2.52E-02 | 8.97E-02 | 2.63E+00 | 2.75E+00 | 7.75E-01 | no | | intergenic_variant | MODIFIER |
| 16 | 27589316 | AX-170203445 | 0 | G | A | 0.246 | 2.52E-02 | 8.97E-02 | 2.63E+00 | 2.75E+00 | 7.75E-01 | no | | intergenic_variant | MODIFIER |
| 16 | 27589454 | AX-97830563 | 0 | A | G | 0.246 | 2.52E-02 | 8.97E-02 | 2.63E+00 | 2.75E+00 | 7.75E-01 | no | | intergenic_variant | MODIFIER |
| 16 | 27640440 | AX-170172363 | 0 | A | G | 0.246 | 2.52E-02 | 8.97E-02 | 2.63E+00 | 2.75E+00 | 7.75E-01 | no | | intergenic_variant | MODIFIER |
| 16 | 27642144 | AX-170087982 | 0 | A | G | 0.246 | 2.52E-02 | 8.97E-02 | 2.63E+00 | 2.75E+00 | 7.75E-01 | no | | intergenic_variant | MODIFIER |
| 12 | 105943823 | AX-170493201 | 0 | A | G | 0.132 | -4.61E-02 | 1.67E-01 | 2.68E+00 | 2.78E+00 | 7.75E-01 | no | | regulatory_region_variant | MODIFIER |
| 18 | 69128945 | AX-170353947 | 1 | T | C | 0.089 | 4.79E-02 | 1.65E-01 | 2.59E+00 | 2.71E+00 | 7.75E-01 | no | | intergenic_variant | MODIFIER |
| 7 | 94506126 | AX-97799557 | 2 | A | G | 0.109 | 4.47E-02 | 1.58E-01 | 2.63E+00 | 2.74E+00 | 7.75E-01 | no | Gm32647 | intron_variant,non_coding_transcript_variant | MODIFIER |
| 1 | 9928899 | AX-170407420 | 2 | T | C | 0.5 | 2.16E-02 | 7.35E-02 | 2.62E+00 | 2.68E+00 | 7.75E-01 | no | Mcmdc2 | intron_variant | MODIFIER |
| 19 | 5040820 | AX-97835935 | 0 | C | T | 0.044 | -5.46E-02 | 1.81E-01 | 2.56E+00 | 2.68E+00 | 7.75E-01 | no | B4gat1 | 3_prime_UTR_variant | MODIFIER |

| | | | | | | | | | | | | | | | |
|---|---|---|---|---|---|---|---|---|---|---|---|---|---|---|---|
| 9 | 69753365 | AX-170388886 | 0 | G | T | 0.193 | -3.72E-02 | 1.30E-01 | 2.67E+00 | 2.71E+00 | 7.75E-01 | no | Foxb1 | downstream_gene_variant | MODIFIER |
| 7 | 137104356 | AX-170152272 | 1 | C | T | 0.295 | -2.32E-02 | 8.02E-02 | 2.61E+00 | 2.71E+00 | 7.75E-01 | no | Mgmt | intron_variant | MODIFIER |
| 4 | 56608785 | AX-170393431 | 1 | C | T | 0.375 | -3.07E-02 | 1.10E-01 | 2.71E+00 | 2.77E+00 | 7.75E-01 | no | | regulatory_region_variant | MODIFIER |
| 4 | 147420193 | AX-170477100 | 1 | T | G | 0.098 | 5.24E-02 | 1.86E-01 | 2.64E+00 | 2.74E+00 | 7.75E-01 | no | Gm26624 | intron_variant,non_coding_transcript_variant | MODIFIER |
| 8 | 96723035 | AX-170214585 | 1 | G | A | 0.214 | 3.35E-02 | 1.19E-01 | 2.68E+00 | 2.74E+00 | 7.76E-01 | no | | intergenic_variant | MODIFIER |
| 4 | 78739472 | AX-170023183 | 0 | G | A | 0.289 | -2.85E-02 | 1.04E-01 | 2.67E+00 | 2.78E+00 | 7.76E-01 | no | | intergenic_variant | MODIFIER |
| 7 | 135380900 | AX-170404796 | 1 | C | T | 0.259 | -2.37E-02 | 8.16E-02 | 2.60E+00 | 2.70E+00 | 7.76E-01 | no | | intergenic_variant | MODIFIER |
| 15 | 94592595 | AX-170087942 | 1 | T | C | 0.134 | 4.48E-02 | 1.56E-01 | 2.64E+00 | 2.70E+00 | 7.76E-01 | no | Twf1 | upstream_gene_variant | MODIFIER |
| 9 | 95136091 | AX-170033034 | 0 | G | A | 0.149 | -3.24E-02 | 1.12E-01 | 2.62E+00 | 2.71E+00 | 7.76E-01 | no | Slc9a9 | intron_variant | MODIFIER |
| 3 | 29497524 | AX-170020207 | 0 | G | A | 0.184 | -2.75E-02 | 9.73E-02 | 2.66E+00 | 2.73E+00 | 7.76E-01 | no | Egfem1 | intron_variant | MODIFIER |
| 3 | 29587108 | AX-170503430 | 0 | T | C | 0.184 | -2.75E-02 | 9.73E-02 | 2.66E+00 | 2.73E+00 | 7.76E-01 | no | Egfem1 | intron_variant | MODIFIER |
| 7 | 98019688 | AX-170333745 | 1 | A | G | 0.116 | 4.94E-02 | 1.80E-01 | 2.65E+00 | 2.77E+00 | 7.76E-01 | no | Gdpd4 | intron_variant | MODIFIER |
| 10 | 31769559 | AX-170387590 | 0 | C | T | 0.386 | -2.15E-02 | 7.49E-02 | 2.61E+00 | 2.72E+00 | 7.76E-01 | no | Nkain2 | intron_variant | MODIFIER |
| 18 | 64790616 | AX-170522542 | 0 | G | T | 0.421 | -2.38E-02 | 8.20E-02 | 2.62E+00 | 2.69E+00 | 7.76E-01 | no | Gm24504 | upstream_gene_variant | MODIFIER |
| 12 | 108187055 | AX-170409221 | 0 | G | A | 0.123 | -5.19E-02 | 1.87E-01 | 2.70E+00 | 2.77E+00 | 7.77E-01 | no | Ccnk | intron_variant | MODIFIER |
| 12 | 108308713 | AX-170007487 | 0 | C | G | 0.123 | -5.19E-02 | 1.87E-01 | 2.70E+00 | 2.77E+00 | 7.77E-01 | no | Hhipl1 | intron_variant | MODIFIER |
| 16 | 92796485 | AX-170142082 | 0 | C | T | 0.474 | -2.18E-02 | 7.80E-02 | 2.66E+00 | 2.74E+00 | 7.77E-01 | no | Runx1 | intron_variant | MODIFIER |
| 2 | 138227604 | AX-170168738 | 1 | G | A | 0.143 | 6.22E-02 | 2.21E-01 | 2.69E+00 | 2.72E+00 | 7.77E-01 | no | | intergenic_variant | MODIFIER |
| 4 | 139866967 | AX-170024102 | 1 | A | T | 0.143 | -2.54E-02 | 9.73E-02 | 2.67E+00 | 2.83E+00 | 7.77E-01 | no | | intergenic_variant | MODIFIER |
| 10 | 67885464 | AX-170307442 | 0 | C | T | 0.412 | -2.27E-02 | 7.83E-02 | 2.61E+00 | 2.69E+00 | 7.77E-01 | no | Zfp365 | downstream_gene_variant | MODIFIER |
| 14 | 92935033 | AX-170317164 | 1 | A | C | 0.045 | -4.98E-02 | 1.75E-01 | 2.61E+00 | 2.72E+00 | 7.77E-01 | no | | intergenic_variant | MODIFIER |
| 3 | 25527319 | AX-170418768 | 0 | C | T | 0.474 | -9.71E-03 | 3.54E-02 | 2.60E+00 | 2.76E+00 | 7.77E-01 | no | Nlgn1 | intron_variant | MODIFIER |
| 1 | 47219926 | AX-170323920 | 1 | A | G | 0.116 | 5.45E-02 | 1.96E-01 | 2.66E+00 | 2.75E+00 | 7.77E-01 | no | | intergenic_variant | MODIFIER |
| 10 | 128151082 | AX-170282087 | 0 | C | T | 0.333 | -2.07E-02 | 7.27E-02 | 2.61E+00 | 2.71E+00 | 7.77E-01 | no | Rbms2 | intron_variant | MODIFIER |
| 7 | 67691836 | AX-170265266 | 0 | C | T | 0.412 | -2.64E-02 | 9.65E-02 | 2.71E+00 | 2.79E+00 | 7.78E-01 | no | Ttc23 | intron_variant | MODIFIER |
| 18 | 41534850 | AX-170468134 | 0 | A | G | 0.07 | 5.86E-02 | 2.13E-01 | 2.66E+00 | 2.76E+00 | 7.78E-01 | no | | intergenic_variant | MODIFIER |
| 1 | 151385944 | AX-170164284 | 1 | T | C | 0.196 | -3.08E-02 | 1.16E-01 | 2.69E+00 | 2.82E+00 | 7.78E-01 | no | Swt1 | intron_variant | MODIFIER |
| 2 | 142933367 | AX-97843871 | 1 | C | T | 0.152 | 4.26E-02 | 1.53E-01 | 2.63E+00 | 2.74E+00 | 7.78E-01 | no | | intergenic_variant | MODIFIER |
| 16 | 89055873 | AX-170013614 | 0 | C | T | 0.482 | 2.22E-02 | 7.97E-02 | 2.66E+00 | 2.74E+00 | 7.78E-01 | no | | intergenic_variant | MODIFIER |
| 2 | 96597009 | AX-170355117 | 1 | A | G | 0.054 | -8.33E-02 | 2.99E-01 | 2.73E+00 | 2.75E+00 | 7.78E-01 | no | Lrrc4c | intron_variant | MODIFIER |
| 1 | 40904321 | AX-170460783 | 0 | G | A | 0.412 | -2.11E-02 | 7.52E-02 | 2.65E+00 | 2.73E+00 | 7.78E-01 | no | | intergenic_variant | MODIFIER |
| 1 | 40904493 | AX-170405686 | 0 | T | C | 0.412 | -2.11E-02 | 7.52E-02 | 2.65E+00 | 2.73E+00 | 7.78E-01 | no | | intergenic_variant | MODIFIER |
| 1 | 40919111 | AX-97798831 | 0 | C | T | 0.412 | -2.11E-02 | 7.52E-02 | 2.65E+00 | 2.73E+00 | 7.78E-01 | no | | intergenic_variant | MODIFIER |
| 1 | 40926380 | AX-170404671 | 0 | C | T | 0.412 | -2.11E-02 | 7.52E-02 | 2.65E+00 | 2.73E+00 | 7.78E-01 | no | | intergenic_variant | MODIFIER |
| 1 | 40938683 | AX-170257043 | 0 | G | T | 0.412 | -2.11E-02 | 7.52E-02 | 2.65E+00 | 2.73E+00 | 7.78E-01 | no | | intergenic_variant | MODIFIER |
| 1 | 40942631 | AX-97827840 | 0 | A | G | 0.412 | -2.11E-02 | 7.52E-02 | 2.65E+00 | 2.73E+00 | 7.78E-01 | no | | intergenic_variant | MODIFIER |
| 1 | 41061580 | AX-97810238 | 0 | C | G | 0.412 | -2.11E-02 | 7.52E-02 | 2.65E+00 | 2.73E+00 | 7.78E-01 | no | | intergenic_variant | MODIFIER |
| 1 | 68084698 | AX-97783143 | 1 | G | A | 0.116 | 6.47E-02 | 2.32E-01 | 2.69E+00 | 2.73E+00 | 7.78E-01 | no | Erbb4 | intron_variant | MODIFIER |
| 13 | 54809105 | AX-97827178 | 0 | T | C | 0.518 | -1.81E-02 | 6.74E-02 | 2.69E+00 | 2.80E+00 | 7.78E-01 | no | | intergenic_variant | MODIFIER |
| 9 | 68592685 | AX-170082728 | 1 | G | A | 0.196 | 3.83E-02 | 1.38E-01 | 2.76E+00 | 2.78E+00 | 7.78E-01 | no | | intergenic_variant | MODIFIER |
| 9 | 75236895 | AX-170304891 | 1 | C | G | 0.196 | 3.83E-02 | 1.38E-01 | 2.76E+00 | 2.78E+00 | 7.78E-01 | no | Myo5c | intron_variant | MODIFIER |
| 1 | 30891492 | AX-97852690 | 0 | G | A | 0.395 | 2.20E-02 | 7.81E-02 | 2.64E+00 | 2.72E+00 | 7.78E-01 | no | | intergenic_variant | MODIFIER |
| 16 | 87561961 | AX-170503555 | 0 | T | C | 0.43 | -2.21E-02 | 7.98E-02 | 2.69E+00 | 2.75E+00 | 7.78E-01 | no | Map3k7cl | intron_variant | MODIFIER |
| 12 | 104425478 | AX-170007441 | 0 | C | T | 0.14 | -4.38E-02 | 1.62E-01 | 2.67E+00 | 2.78E+00 | 7.78E-01 | no | | intergenic_variant | MODIFIER |
| 3 | 63440895 | AX-170379837 | 0 | C | A | 0.044 | 5.13E-02 | 1.91E-01 | 2.67E+00 | 2.79E+00 | 7.78E-01 | no | Gm34240 | intron_variant,non_coding_transcript_variant | MODIFIER |
| 5 | 120262398 | AX-170026009 | 1 | C | T | 0.455 | 3.10E-02 | 1.09E-01 | 2.65E+00 | 2.70E+00 | 7.78E-01 | no | | intergenic_variant | MODIFIER |
| 20 | 156404876 | AX-170035290 | 1 | T | A | 0.134 | 3.32E-02 | 1.19E-01 | 2.66E+00 | 2.74E+00 | 7.78E-01 | no | | intergenic_variant | MODIFIER |
| 5 | 73336759 | AX-170193360 | 1 | A | G | 0.429 | -2.59E-02 | 9.59E-02 | 2.69E+00 | 2.79E+00 | 7.79E-01 | no | Ociad2 | intron_variant | MODIFIER |
| 12 | 14164789 | AX-170211752 | 0 | G | A | 0.044 | 4.00E-02 | 1.51E-01 | 2.64E+00 | 2.79E+00 | 7.79E-01 | no | | intergenic_variant | MODIFIER |
| 20 | 104614457 | AX-170145882 | 1 | C | G | 0.339 | 1.53E-02 | 5.49E-02 | 2.62E+00 | 2.73E+00 | 7.79E-01 | no | Zdhhc15 | intron_variant | MODIFIER |
| 1 | 6472311 | AX-170344409 | 0 | G | T | 0.412 | -1.85E-02 | 6.85E-02 | 2.75E+00 | 2.84E+00 | 7.79E-01 | no | | intergenic_variant | MODIFIER |
| 8 | 36233207 | AX-170309874 | 1 | T | C | 0.143 | 3.92E-02 | 1.43E-01 | 2.74E+00 | 2.80E+00 | 7.79E-01 | no | Lonrf1 | intron_variant | MODIFIER |
| 9 | 33381905 | AX-97790593 | 1 | C | T | 0.446 | 2.44E-02 | 8.53E-02 | 2.62E+00 | 2.69E+00 | 7.79E-01 | no | | intergenic_variant | MODIFIER |
| 9 | 83013804 | AX-170390998 | 0 | T | C | 0.202 | 3.55E-02 | 1.29E-01 | 2.73E+00 | 2.78E+00 | 7.79E-01 | no | Gm39383 | intron_variant,non_coding_transcript_variant | MODIFIER |

| | | | | | | | | | | | | | | | |
|---|---|---|---|---|---|---|---|---|---|---|---|---|---|---|---|
| 18 | 46991743 | AX-170146684 | 0 | C | A | 0.053 | 6.40E-02 | 2.23E-01 | 2.58E+00 | 2.71E+00 | 7.79E-01 | no | Commd10 | intron_variant | MODIFIER |
| 9 | 73173772 | AX-170427203 | 0 | T | C | 0.219 | -3.23E-02 | 1.17E-01 | 2.65E+00 | 2.74E+00 | 7.79E-01 | no | Gm5745 | downstream_gene_variant | MODIFIER |
| 13 | 36091498 | AX-170209014 | 0 | T | A | 0.175 | -3.51E-02 | 1.22E-01 | 2.62E+00 | 2.68E+00 | 7.79E-01 | no | Lyrm4 | intron_variant | MODIFIER |
| 6 | 148670427 | AX-170106614 | 0 | A | G | 0.053 | 4.75E-02 | 1.76E-01 | 2.65E+00 | 2.77E+00 | 7.79E-01 | no | 4930528G23Rik | intron_variant,non_coding_transcript_variant | MODIFIER |
| 12 | 42657119 | AX-170454534 | 0 | A | C | 0.053 | 4.75E-02 | 1.76E-01 | 2.65E+00 | 2.77E+00 | 7.79E-01 | no | | intergenic_variant | MODIFIER |
| 12 | 114035719 | AX-170285094 | 2 | C | A | 0.482 | 1.48E-01 | 5.31E-01 | 2.67E+00 | 2.72E+00 | 7.79E-01 | Yes | Ighv3-2 | upstream_gene_variant | MODIFIER |
| 2 | 88869085 | AX-170508764 | 1 | T | C | 0.054 | -8.29E-02 | 3.00E-01 | 2.73E+00 | 2.75E+00 | 7.79E-01 | no | Olfr1206 | downstream_gene_variant | MODIFIER |
| 1 | 176615115 | AX-170257005 | 1 | A | T | 0.223 | -3.00E-02 | 1.12E-01 | 2.74E+00 | 2.84E+00 | 7.79E-01 | no | | intergenic_variant | MODIFIER |
| 8 | 95767472 | AX-170080884 | 1 | G | A | 0.241 | 3.07E-02 | 1.11E-01 | 2.66E+00 | 2.73E+00 | 7.80E-01 | no | Cnot1 | intron_variant | MODIFIER |
| 8 | 18874477 | AX-170494198 | 0 | C | T | 0.184 | 2.72E-02 | 9.83E-02 | 2.63E+00 | 2.74E+00 | 7.80E-01 | no | Agpat5 | intron_variant | MODIFIER |
| 20 | 77654610 | AX-170476956 | 1 | A | T | 0.25 | 1.74E-02 | 6.26E-02 | 2.64E+00 | 2.73E+00 | 7.80E-01 | no | Tbl1x | intron_variant | MODIFIER |
| 7 | 135922343 | AX-170156207 | 1 | T | C | 0.25 | -2.31E-02 | 7.97E-02 | 2.58E+00 | 2.69E+00 | 7.80E-01 | no | | intergenic_variant | MODIFIER |
| 2 | 120466383 | AX-170511753 | 1 | G | C | 0.134 | 6.08E-02 | 2.20E-01 | 2.68E+00 | 2.74E+00 | 7.80E-01 | no | Capn3 | intron_variant | MODIFIER |
| 10 | 114593695 | AX-97842402 | 1 | T | C | 0.357 | -2.15E-02 | 7.81E-02 | 2.66E+00 | 2.74E+00 | 7.80E-01 | no | Trhde | intron_variant | MODIFIER |
| 9 | 59319840 | AX-170183903 | 0 | C | A | 0.228 | 3.09E-02 | 1.12E-01 | 2.65E+00 | 2.73E+00 | 7.80E-01 | no | Bbs4 | downstream_gene_variant | MODIFIER |
| 5 | 104269767 | AX-170408506 | 0 | T | C | 0.43 | 3.17E-02 | 1.11E-01 | 2.64E+00 | 2.68E+00 | 7.80E-01 | no | | intergenic_variant | MODIFIER |
| 8 | 100466582 | AX-97797531 | 1 | G | A | 0.241 | 3.11E-02 | 1.12E-01 | 2.66E+00 | 2.73E+00 | 7.80E-01 | no | | intergenic_variant | MODIFIER |
| 5 | 102302937 | AX-97787691 | 2 | C | T | 0.436 | 2.99E-02 | 1.09E-01 | 2.69E+00 | 2.75E+00 | 7.80E-01 | no | | intergenic_variant | MODIFIER |
| 1 | 155959415 | AX-170002049 | 1 | C | T | 0.223 | 4.12E-02 | 1.46E-01 | 2.65E+00 | 2.68E+00 | 7.80E-01 | no | Gm23744 | downstream_gene_variant | MODIFIER |
| 1 | 41901964 | AX-170487062 | 0 | G | A | 0.439 | -2.09E-02 | 7.56E-02 | 2.65E+00 | 2.73E+00 | 7.80E-01 | no | Gm29260 | intron_variant,non_coding_transcript_variant | MODIFIER |
| 15 | 101833498 | AX-97846004 | 1 | T | C | 0.125 | 4.36E-02 | 1.58E-01 | 2.66E+00 | 2.74E+00 | 7.80E-01 | no | | intergenic_variant | MODIFIER |
| 2 | 155184967 | AX-170470139 | 2 | G | A | 0.164 | -3.82E-02 | 1.35E-01 | 2.58E+00 | 2.71E+00 | 7.80E-01 | no | Itch | intron_variant | MODIFIER |
| 15 | 93156603 | AX-170012282 | 0 | C | T | 0.132 | 5.15E-02 | 1.81E-01 | 2.62E+00 | 2.69E+00 | 7.80E-01 | no | | intergenic_variant | MODIFIER |
| 15 | 93611074 | AX-170012293 | 0 | T | A | 0.132 | 5.15E-02 | 1.81E-01 | 2.62E+00 | 2.69E+00 | 7.80E-01 | no | | regulatory_region_variant | MODIFIER |
| 15 | 94801291 | AX-170056361 | 0 | G | A | 0.132 | 5.15E-02 | 1.81E-01 | 2.62E+00 | 2.69E+00 | 7.80E-01 | no | Tmem117 | intron_variant | MODIFIER |
| 12 | 114481240 | AX-170501985 | 0 | C | T | 0.14 | -5.24E-02 | 1.92E-01 | 2.69E+00 | 2.76E+00 | 7.80E-01 | no | Ighv10-1 | upstream_gene_variant | MODIFIER |
| 12 | 117408343 | AX-170240018 | 0 | G | T | 0.14 | -5.24E-02 | 1.92E-01 | 2.69E+00 | 2.76E+00 | 7.80E-01 | no | | intergenic_variant | MODIFIER |
| 10 | 127910591 | AX-170159146 | 1 | C | G | 0.33 | -2.04E-02 | 7.28E-02 | 2.61E+00 | 2.72E+00 | 7.81E-01 | no | Sdr9c7 | 3_prime_UTR_variant | MODIFIER |
| 15 | 101772991 | AX-170419112 | 2 | G | C | 0.118 | 4.17E-02 | 1.52E-01 | 2.66E+00 | 2.75E+00 | 7.81E-01 | no | Krt72 | downstream_gene_variant | MODIFIER |
| 1 | 66960539 | AX-170164395 | 2 | C | A | 0.118 | 6.46E-02 | 2.33E-01 | 2.68E+00 | 2.72E+00 | 7.81E-01 | no | | intergenic_variant | MODIFIER |
| 7 | 136487519 | AX-170118144 | 1 | T | C | 0.295 | -2.23E-02 | 7.89E-02 | 2.61E+00 | 2.71E+00 | 7.81E-01 | no | | intergenic_variant | MODIFIER |
| 15 | 94282597 | AX-170340841 | 1 | C | A | 0.152 | 5.00E-02 | 1.78E-01 | 2.64E+00 | 2.71E+00 | 7.81E-01 | no | Adamts20 | intron_variant | MODIFIER |
| 12 | 116175097 | AX-170353648 | 0 | A | G | 0.149 | 4.40E-02 | 1.62E-01 | 2.64E+00 | 2.75E+00 | 7.81E-01 | no | Gm23732 | downstream_gene_variant | MODIFIER |
| 4 | 136975292 | AX-170422806 | 0 | G | A | 0.175 | 3.04E-02 | 1.07E-01 | 2.63E+00 | 2.69E+00 | 7.81E-01 | no | Zbtb40 | downstream_gene_variant | MODIFIER |
| 2 | 115578578 | AX-170253818 | 1 | G | C | 0.143 | 6.07E-02 | 2.20E-01 | 2.69E+00 | 2.72E+00 | 7.81E-01 | no | BC052040 | upstream_gene_variant | MODIFIER |
| 2 | 115871178 | AX-170192302 | 1 | T | C | 0.143 | 6.07E-02 | 2.20E-01 | 2.69E+00 | 2.72E+00 | 7.81E-01 | no | Meis2 | intron_variant | MODIFIER |
| 2 | 149831883 | AX-170508250 | 1 | C | G | 0.143 | 6.07E-02 | 2.20E-01 | 2.69E+00 | 2.72E+00 | 7.81E-01 | no | C530025M09Rik | upstream_gene_variant | MODIFIER |
| 4 | 145309722 | AX-170503451 | 1 | G | A | 0.107 | 5.60E-02 | 2.04E-01 | 2.68E+00 | 2.74E+00 | 7.81E-01 | no | Tnfrsf8 | intron_variant | MODIFIER |
| 1 | 25970096 | AX-170040669 | 0 | A | C | 0.395 | 2.35E-02 | 8.38E-02 | 2.63E+00 | 2.71E+00 | 7.81E-01 | no | | intergenic_variant | MODIFIER |
| 5 | 7025667 | AX-170024363 | 0 | G | A | 0.447 | 1.67E-02 | 6.33E-02 | 2.65E+00 | 2.79E+00 | 7.81E-01 | no | Zfp804b | intron_variant | MODIFIER |
| 1 | 26507758 | AX-170467673 | 1 | A | C | 0.402 | 2.37E-02 | 8.53E-02 | 2.65E+00 | 2.71E+00 | 7.81E-01 | no | | intergenic_variant | MODIFIER |
| 7 | 69043030 | AX-170144235 | 0 | G | T | 0.404 | -2.53E-02 | 9.41E-02 | 2.70E+00 | 2.78E+00 | 7.81E-01 | no | | intergenic_variant | MODIFIER |
| 8 | 90030920 | AX-170333559 | 0 | A | G | 0.193 | -3.33E-02 | 1.19E-01 | 2.64E+00 | 2.71E+00 | 7.81E-01 | no | | regulatory_region_variant | MODIFIER |
| 8 | 90031182 | AX-170136887 | 0 | A | G | 0.193 | -3.33E-02 | 1.19E-01 | 2.64E+00 | 2.71E+00 | 7.81E-01 | no | | regulatory_region_variant | MODIFIER |
| 8 | 90036580 | AX-170530193 | 0 | C | A | 0.193 | -3.33E-02 | 1.19E-01 | 2.64E+00 | 2.71E+00 | 7.81E-01 | no | | regulatory_region_variant | MODIFIER |
| 8 | 90038482 | AX-170199695 | 0 | C | G | 0.193 | -3.33E-02 | 1.19E-01 | 2.64E+00 | 2.71E+00 | 7.81E-01 | no | | intergenic_variant | MODIFIER |
| 8 | 90038687 | AX-170409845 | 0 | A | G | 0.193 | -3.33E-02 | 1.19E-01 | 2.64E+00 | 2.71E+00 | 7.81E-01 | no | | intergenic_variant | MODIFIER |
| 8 | 90039713 | AX-170449472 | 0 | G | A | 0.193 | -3.33E-02 | 1.19E-01 | 2.64E+00 | 2.71E+00 | 7.81E-01 | no | | intergenic_variant | MODIFIER |
| 8 | 90079273 | AX-170325402 | 0 | C | T | 0.193 | -3.33E-02 | 1.19E-01 | 2.64E+00 | 2.71E+00 | 7.81E-01 | no | | intergenic_variant | MODIFIER |
| 8 | 90079369 | AX-170307552 | 0 | A | G | 0.193 | -3.33E-02 | 1.19E-01 | 2.64E+00 | 2.71E+00 | 7.81E-01 | no | | intergenic_variant | MODIFIER |
| 8 | 90080087 | AX-170415470 | 0 | T | G | 0.193 | -3.33E-02 | 1.19E-01 | 2.64E+00 | 2.71E+00 | 7.81E-01 | no | | intergenic_variant | MODIFIER |
| 8 | 90080566 | AX-170080744 | 0 | T | C | 0.193 | -3.33E-02 | 1.19E-01 | 2.64E+00 | 2.71E+00 | 7.81E-01 | no | | intergenic_variant | MODIFIER |
| 8 | 90086869 | AX-170257321 | 0 | C | T | 0.193 | -3.33E-02 | 1.19E-01 | 2.64E+00 | 2.71E+00 | 7.81E-01 | no | | intergenic_variant | MODIFIER |
| 8 | 90086948 | AX-170096359 | 0 | G | T | 0.193 | -3.33E-02 | 1.19E-01 | 2.64E+00 | 2.71E+00 | 7.81E-01 | no | | intergenic_variant | MODIFIER |

| | | | | | | | | | | | | | | | |
|---|---|---|---|---|---|---|---|---|---|---|---|---|---|---|---|
| 8 | 90106290 | AX-170259015 | 0 | A | G | 0.193 | -3.33E-02 | 1.19E-01 | 2.64E+00 | 2.71E+00 | 7.81E-01 | no | | intergenic_variant | MODIFIER |
| 8 | 90106445 | AX-170530902 | 0 | C | T | 0.193 | -3.33E-02 | 1.19E-01 | 2.64E+00 | 2.71E+00 | 7.81E-01 | no | | intergenic_variant | MODIFIER |
| 8 | 90112287 | AX-170463882 | 0 | A | G | 0.193 | -3.33E-02 | 1.19E-01 | 2.64E+00 | 2.71E+00 | 7.81E-01 | no | | intergenic_variant | MODIFIER |
| 8 | 90112962 | AX-170168671 | 0 | T | G | 0.193 | -3.33E-02 | 1.19E-01 | 2.64E+00 | 2.71E+00 | 7.81E-01 | no | | intergenic_variant | MODIFIER |
| 8 | 90113092 | AX-170321589 | 0 | T | C | 0.193 | -3.33E-02 | 1.19E-01 | 2.64E+00 | 2.71E+00 | 7.81E-01 | no | | intergenic_variant | MODIFIER |
| 8 | 90113553 | AX-97795642 | 0 | T | C | 0.193 | -3.33E-02 | 1.19E-01 | 2.64E+00 | 2.71E+00 | 7.81E-01 | no | | intergenic_variant | MODIFIER |
| 8 | 90125447 | AX-170255922 | 0 | G | A | 0.193 | -3.33E-02 | 1.19E-01 | 2.64E+00 | 2.71E+00 | 7.81E-01 | no | | intergenic_variant | MODIFIER |
| 8 | 90125973 | AX-170483616 | 0 | C | T | 0.193 | -3.33E-02 | 1.19E-01 | 2.64E+00 | 2.71E+00 | 7.81E-01 | no | | intergenic_variant | MODIFIER |
| 8 | 90130615 | AX-170419542 | 0 | T | C | 0.193 | -3.33E-02 | 1.19E-01 | 2.64E+00 | 2.71E+00 | 7.81E-01 | no | | intergenic_variant | MODIFIER |
| 8 | 90143414 | AX-170452175 | 0 | T | C | 0.193 | -3.33E-02 | 1.19E-01 | 2.64E+00 | 2.71E+00 | 7.81E-01 | no | | intergenic_variant | MODIFIER |
| 8 | 90151156 | AX-170479091 | 0 | G | A | 0.193 | -3.33E-02 | 1.19E-01 | 2.64E+00 | 2.71E+00 | 7.81E-01 | no | | intergenic_variant | MODIFIER |
| 8 | 90151975 | AX-170449798 | 0 | C | G | 0.193 | -3.33E-02 | 1.19E-01 | 2.64E+00 | 2.71E+00 | 7.81E-01 | no | | intergenic_variant | MODIFIER |
| 8 | 90154613 | AX-97809877 | 0 | G | A | 0.193 | -3.33E-02 | 1.19E-01 | 2.64E+00 | 2.71E+00 | 7.81E-01 | no | | intergenic_variant | MODIFIER |
| 8 | 90161809 | AX-170080749 | 0 | A | G | 0.193 | -3.33E-02 | 1.19E-01 | 2.64E+00 | 2.71E+00 | 7.81E-01 | no | | intergenic_variant | MODIFIER |
| 8 | 90168142 | AX-97801471 | 0 | T | C | 0.193 | -3.33E-02 | 1.19E-01 | 2.64E+00 | 2.71E+00 | 7.81E-01 | no | | intergenic_variant | MODIFIER |
| 8 | 90173634 | AX-170414130 | 0 | A | G | 0.193 | -3.33E-02 | 1.19E-01 | 2.64E+00 | 2.71E+00 | 7.81E-01 | no | | regulatory_region_variant | MODIFIER |
| 8 | 90179792 | AX-170080745 | 0 | T | C | 0.193 | -3.33E-02 | 1.19E-01 | 2.64E+00 | 2.71E+00 | 7.81E-01 | no | | intergenic_variant | MODIFIER |
| 8 | 90183834 | AX-97833921 | 0 | T | C | 0.193 | -3.33E-02 | 1.19E-01 | 2.64E+00 | 2.71E+00 | 7.81E-01 | no | | intergenic_variant | MODIFIER |
| 8 | 90184251 | AX-170080750 | 0 | C | T | 0.193 | -3.33E-02 | 1.19E-01 | 2.64E+00 | 2.71E+00 | 7.81E-01 | no | | intergenic_variant | MODIFIER |
| 8 | 90187286 | AX-170513371 | 0 | T | G | 0.193 | -3.33E-02 | 1.19E-01 | 2.64E+00 | 2.71E+00 | 7.81E-01 | no | | intergenic_variant | MODIFIER |
| 8 | 90188250 | AX-170363884 | 0 | T | C | 0.193 | -3.33E-02 | 1.19E-01 | 2.64E+00 | 2.71E+00 | 7.81E-01 | no | | intergenic_variant | MODIFIER |
| 8 | 90205904 | AX-170124072 | 0 | G | A | 0.193 | -3.33E-02 | 1.19E-01 | 2.64E+00 | 2.71E+00 | 7.81E-01 | no | | intergenic_variant | MODIFIER |
| 8 | 90206020 | AX-170337036 | 0 | G | A | 0.193 | -3.33E-02 | 1.19E-01 | 2.64E+00 | 2.71E+00 | 7.81E-01 | no | | intergenic_variant | MODIFIER |
| 8 | 90209331 | AX-170339590 | 0 | T | C | 0.193 | -3.33E-02 | 1.19E-01 | 2.64E+00 | 2.71E+00 | 7.81E-01 | no | | regulatory_region_variant | MODIFIER |
| 8 | 90209748 | AX-170417689 | 0 | G | T | 0.193 | -3.33E-02 | 1.19E-01 | 2.64E+00 | 2.71E+00 | 7.81E-01 | no | | regulatory_region_variant | MODIFIER |
| 8 | 90209857 | AX-170157158 | 0 | G | A | 0.193 | -3.33E-02 | 1.19E-01 | 2.64E+00 | 2.71E+00 | 7.81E-01 | no | | regulatory_region_variant | MODIFIER |
| 8 | 90214485 | AX-170437837 | 0 | A | C | 0.193 | -3.33E-02 | 1.19E-01 | 2.64E+00 | 2.71E+00 | 7.81E-01 | no | | intergenic_variant | MODIFIER |
| 8 | 90230219 | AX-97836793 | 0 | A | C | 0.193 | -3.33E-02 | 1.19E-01 | 2.64E+00 | 2.71E+00 | 7.81E-01 | no | | intergenic_variant | MODIFIER |
| 8 | 90230362 | AX-170469865 | 0 | C | G | 0.193 | -3.33E-02 | 1.19E-01 | 2.64E+00 | 2.71E+00 | 7.81E-01 | no | | intergenic_variant | MODIFIER |
| 8 | 90248316 | AX-170196290 | 0 | C | T | 0.193 | -3.33E-02 | 1.19E-01 | 2.64E+00 | 2.71E+00 | 7.81E-01 | no | Tox3 | synonymous_variant | LOW |
| 8 | 90250923 | AX-170191531 | 0 | C | T | 0.193 | -3.33E-02 | 1.19E-01 | 2.64E+00 | 2.71E+00 | 7.81E-01 | no | Tox3 | intron_variant | MODIFIER |
| 8 | 90251194 | AX-170340588 | 0 | C | A | 0.193 | -3.33E-02 | 1.19E-01 | 2.64E+00 | 2.71E+00 | 7.81E-01 | no | Tox3 | intron_variant | MODIFIER |
| 8 | 90279057 | AX-170265646 | 0 | A | G | 0.193 | -3.33E-02 | 1.19E-01 | 2.64E+00 | 2.71E+00 | 7.81E-01 | no | Tox3 | intron_variant | MODIFIER |
| 8 | 90279300 | AX-97792073 | 0 | A | G | 0.193 | -3.33E-02 | 1.19E-01 | 2.64E+00 | 2.71E+00 | 7.81E-01 | no | Tox3 | intron_variant | MODIFIER |
| 8 | 90279647 | AX-170080753 | 0 | A | G | 0.193 | -3.33E-02 | 1.19E-01 | 2.64E+00 | 2.71E+00 | 7.81E-01 | no | Tox3 | intron_variant | MODIFIER |
| 8 | 90279849 | AX-170080752 | 0 | G | A | 0.193 | -3.33E-02 | 1.19E-01 | 2.64E+00 | 2.71E+00 | 7.81E-01 | no | Tox3 | intron_variant | MODIFIER |
| 8 | 90287994 | AX-170146714 | 0 | A | C | 0.193 | -3.33E-02 | 1.19E-01 | 2.64E+00 | 2.71E+00 | 7.81E-01 | no | Tox3 | intron_variant | MODIFIER |
| 8 | 90293080 | AX-170450195 | 0 | T | G | 0.193 | -3.33E-02 | 1.19E-01 | 2.64E+00 | 2.71E+00 | 7.81E-01 | no | Tox3 | intron_variant | MODIFIER |
| 8 | 90293581 | AX-170309479 | 0 | T | C | 0.193 | -3.33E-02 | 1.19E-01 | 2.64E+00 | 2.71E+00 | 7.81E-01 | no | Tox3 | intron_variant | MODIFIER |
| 8 | 90295803 | AX-170498298 | 0 | A | C | 0.193 | -3.33E-02 | 1.19E-01 | 2.64E+00 | 2.71E+00 | 7.81E-01 | no | Tox3 | intron_variant | MODIFIER |
| 8 | 90301533 | AX-170461305 | 0 | T | C | 0.193 | -3.33E-02 | 1.19E-01 | 2.64E+00 | 2.71E+00 | 7.81E-01 | no | Tox3 | intron_variant | MODIFIER |
| 8 | 90317845 | AX-170253203 | 0 | A | C | 0.193 | -3.33E-02 | 1.19E-01 | 2.64E+00 | 2.71E+00 | 7.81E-01 | no | Tox3 | intron_variant | MODIFIER |
| 8 | 90339123 | AX-170142783 | 0 | C | T | 0.193 | -3.33E-02 | 1.19E-01 | 2.64E+00 | 2.71E+00 | 7.81E-01 | no | Tox3 | intron_variant | MODIFIER |
| 8 | 90339301 | AX-170535687 | 0 | A | G | 0.193 | -3.33E-02 | 1.19E-01 | 2.64E+00 | 2.71E+00 | 7.81E-01 | no | Tox3 | intron_variant | MODIFIER |
| 8 | 90365799 | AX-170107059 | 0 | T | C | 0.193 | -3.33E-02 | 1.19E-01 | 2.64E+00 | 2.71E+00 | 7.81E-01 | no | | intergenic_variant | MODIFIER |
| 8 | 90373324 | AX-170259495 | 0 | T | C | 0.193 | -3.33E-02 | 1.19E-01 | 2.64E+00 | 2.71E+00 | 7.81E-01 | no | | intergenic_variant | MODIFIER |
| 8 | 90374087 | AX-170322366 | 0 | G | A | 0.193 | -3.33E-02 | 1.19E-01 | 2.64E+00 | 2.71E+00 | 7.81E-01 | no | | intergenic_variant | MODIFIER |
| 8 | 90377763 | AX-170445936 | 0 | C | T | 0.193 | -3.33E-02 | 1.19E-01 | 2.64E+00 | 2.71E+00 | 7.81E-01 | no | | intergenic_variant | MODIFIER |
| 8 | 90378281 | AX-170531316 | 0 | C | T | 0.193 | -3.33E-02 | 1.19E-01 | 2.64E+00 | 2.71E+00 | 7.81E-01 | no | | regulatory_region_variant | MODIFIER |
| 8 | 90389813 | AX-170150133 | 0 | G | A | 0.193 | -3.33E-02 | 1.19E-01 | 2.64E+00 | 2.71E+00 | 7.81E-01 | no | | intergenic_variant | MODIFIER |
| 8 | 90400733 | AX-170435712 | 0 | T | C | 0.193 | -3.33E-02 | 1.19E-01 | 2.64E+00 | 2.71E+00 | 7.81E-01 | no | | intergenic_variant | MODIFIER |
| 8 | 90410168 | AX-170376345 | 0 | G | A | 0.193 | -3.33E-02 | 1.19E-01 | 2.64E+00 | 2.71E+00 | 7.81E-01 | no | | intergenic_variant | MODIFIER |
| 8 | 90415289 | AX-170333919 | 0 | C | T | 0.193 | -3.33E-02 | 1.19E-01 | 2.64E+00 | 2.71E+00 | 7.81E-01 | no | | intergenic_variant | MODIFIER |
| 8 | 90416885 | AX-170534222 | 0 | G | T | 0.193 | -3.33E-02 | 1.19E-01 | 2.64E+00 | 2.71E+00 | 7.81E-01 | no | | intergenic_variant | MODIFIER |

| | | | | | | | | | | | | | | | |
|---|---|---|---|---|---|---|---|---|---|---|---|---|---|---|---|
| 8 | 90417243 | AX-170111952 | 0 | A | G | 0.193 | -3.33E-02 | 1.19E-01 | 2.64E+00 | 2.71E+00 | 7.81E-01 | no | | intergenic_variant | MODIFIER |
| 8 | 90417645 | AX-170513706 | 0 | T | C | 0.193 | -3.33E-02 | 1.19E-01 | 2.64E+00 | 2.71E+00 | 7.81E-01 | no | | intergenic_variant | MODIFIER |
| 8 | 90418480 | AX-170261094 | 0 | G | A | 0.193 | -3.33E-02 | 1.19E-01 | 2.64E+00 | 2.71E+00 | 7.81E-01 | no | | intergenic_variant | MODIFIER |
| 8 | 90435661 | AX-170114380 | 0 | G | A | 0.193 | -3.33E-02 | 1.19E-01 | 2.64E+00 | 2.71E+00 | 7.81E-01 | no | | intergenic_variant | MODIFIER |
| 8 | 90438905 | AX-170031248 | 0 | G | A | 0.193 | -3.33E-02 | 1.19E-01 | 2.64E+00 | 2.71E+00 | 7.81E-01 | no | | intergenic_variant | MODIFIER |
| 8 | 90486281 | AX-170122838 | 0 | G | A | 0.193 | -3.33E-02 | 1.19E-01 | 2.64E+00 | 2.71E+00 | 7.81E-01 | no | | intergenic_variant | MODIFIER |
| 8 | 90486311 | AX-170159007 | 0 | C | G | 0.193 | -3.33E-02 | 1.19E-01 | 2.64E+00 | 2.71E+00 | 7.81E-01 | no | | intergenic_variant | MODIFIER |
| 8 | 90492640 | AX-97855946 | 0 | G | A | 0.193 | -3.33E-02 | 1.19E-01 | 2.64E+00 | 2.71E+00 | 7.81E-01 | no | | regulatory_region_variant | MODIFIER |
| 8 | 90493069 | AX-170031247 | 0 | C | A | 0.193 | -3.33E-02 | 1.19E-01 | 2.64E+00 | 2.71E+00 | 7.81E-01 | no | | regulatory_region_variant | MODIFIER |
| 8 | 90493495 | AX-170204977 | 0 | G | C | 0.193 | -3.33E-02 | 1.19E-01 | 2.64E+00 | 2.71E+00 | 7.81E-01 | no | | intergenic_variant | MODIFIER |
| 8 | 90511405 | AX-170403578 | 0 | C | T | 0.193 | -3.33E-02 | 1.19E-01 | 2.64E+00 | 2.71E+00 | 7.81E-01 | no | | intergenic_variant | MODIFIER |
| 8 | 90515101 | AX-170280345 | 0 | C | T | 0.193 | -3.33E-02 | 1.19E-01 | 2.64E+00 | 2.71E+00 | 7.81E-01 | no | | intergenic_variant | MODIFIER |
| 8 | 90517998 | AX-170165390 | 0 | T | G | 0.193 | -3.33E-02 | 1.19E-01 | 2.64E+00 | 2.71E+00 | 7.81E-01 | no | | intergenic_variant | MODIFIER |
| 8 | 90537896 | AX-170031245 | 0 | G | A | 0.193 | -3.33E-02 | 1.19E-01 | 2.64E+00 | 2.71E+00 | 7.81E-01 | no | | intergenic_variant | MODIFIER |
| 8 | 90538760 | AX-170228922 | 0 | T | G | 0.193 | -3.33E-02 | 1.19E-01 | 2.64E+00 | 2.71E+00 | 7.81E-01 | no | | intergenic_variant | MODIFIER |
| 8 | 90544533 | AX-97804199 | 0 | T | C | 0.193 | -3.33E-02 | 1.19E-01 | 2.64E+00 | 2.71E+00 | 7.81E-01 | no | | intergenic_variant | MODIFIER |
| 8 | 90548166 | AX-170031244 | 0 | T | C | 0.193 | -3.33E-02 | 1.19E-01 | 2.64E+00 | 2.71E+00 | 7.81E-01 | no | | intergenic_variant | MODIFIER |
| 8 | 90566640 | AX-97855183 | 0 | A | T | 0.193 | -3.33E-02 | 1.19E-01 | 2.64E+00 | 2.71E+00 | 7.81E-01 | no | | intergenic_variant | MODIFIER |
| 8 | 90568749 | AX-170080758 | 0 | A | G | 0.193 | -3.33E-02 | 1.19E-01 | 2.64E+00 | 2.71E+00 | 7.81E-01 | no | | intergenic_variant | MODIFIER |
| 8 | 90607176 | AX-170401363 | 0 | T | C | 0.193 | -3.33E-02 | 1.19E-01 | 2.64E+00 | 2.71E+00 | 7.81E-01 | no | | regulatory_region_variant | MODIFIER |
| 8 | 90608243 | AX-170155390 | 0 | T | C | 0.193 | -3.33E-02 | 1.19E-01 | 2.64E+00 | 2.71E+00 | 7.81E-01 | no | | intergenic_variant | MODIFIER |
| 8 | 90610794 | AX-170121816 | 0 | A | C | 0.193 | -3.33E-02 | 1.19E-01 | 2.64E+00 | 2.71E+00 | 7.81E-01 | no | | intergenic_variant | MODIFIER |
| 8 | 90616689 | AX-170258784 | 0 | A | G | 0.193 | -3.33E-02 | 1.19E-01 | 2.64E+00 | 2.71E+00 | 7.81E-01 | no | | intergenic_variant | MODIFIER |
| 8 | 90648094 | AX-170357928 | 0 | G | A | 0.193 | -3.33E-02 | 1.19E-01 | 2.64E+00 | 2.71E+00 | 7.81E-01 | no | | intergenic_variant | MODIFIER |
| 8 | 90651465 | AX-170430104 | 0 | A | G | 0.193 | -3.33E-02 | 1.19E-01 | 2.64E+00 | 2.71E+00 | 7.81E-01 | no | | intergenic_variant | MODIFIER |
| 8 | 90651750 | AX-170345152 | 0 | A | G | 0.193 | -3.33E-02 | 1.19E-01 | 2.64E+00 | 2.71E+00 | 7.81E-01 | no | | intergenic_variant | MODIFIER |
| 8 | 90659129 | AX-97831198 | 0 | C | T | 0.193 | -3.33E-02 | 1.19E-01 | 2.64E+00 | 2.71E+00 | 7.81E-01 | no | | regulatory_region_variant | MODIFIER |
| 8 | 90672201 | AX-170267850 | 0 | G | A | 0.193 | -3.33E-02 | 1.19E-01 | 2.64E+00 | 2.71E+00 | 7.81E-01 | no | Gm45639 | intron_variant,non_coding_transcript_variant | MODIFIER |
| 8 | 90694918 | AX-170309128 | 0 | A | G | 0.193 | -3.33E-02 | 1.19E-01 | 2.64E+00 | 2.71E+00 | 7.81E-01 | no | | intergenic_variant | MODIFIER |
| 8 | 90703965 | AX-97780764 | 0 | A | C | 0.193 | -3.33E-02 | 1.19E-01 | 2.64E+00 | 2.71E+00 | 7.81E-01 | no | | intergenic_variant | MODIFIER |
| 8 | 90707692 | AX-170197594 | 0 | G | T | 0.193 | -3.33E-02 | 1.19E-01 | 2.64E+00 | 2.71E+00 | 7.81E-01 | no | Gm35850 | downstream_gene_variant | MODIFIER |
| 8 | 90707724 | AX-170254979 | 0 | C | T | 0.193 | -3.33E-02 | 1.19E-01 | 2.64E+00 | 2.71E+00 | 7.81E-01 | no | Gm35850 | downstream_gene_variant | MODIFIER |
| 8 | 90736938 | AX-170428757 | 0 | G | A | 0.193 | -3.33E-02 | 1.19E-01 | 2.64E+00 | 2.71E+00 | 7.81E-01 | no | Gm35850 | non_coding_transcript_exon_variant | MODIFIER |
| 8 | 90737643 | AX-170397083 | 0 | G | A | 0.193 | -3.33E-02 | 1.19E-01 | 2.64E+00 | 2.71E+00 | 7.81E-01 | no | Gm35850 | intron_variant,non_coding_transcript_variant | MODIFIER |
| 8 | 90738422 | AX-170190700 | 0 | C | T | 0.193 | -3.33E-02 | 1.19E-01 | 2.64E+00 | 2.71E+00 | 7.81E-01 | no | Gm35850 | intron_variant,non_coding_transcript_variant | MODIFIER |
| 8 | 90739726 | AX-170394930 | 0 | T | G | 0.193 | -3.33E-02 | 1.19E-01 | 2.64E+00 | 2.71E+00 | 7.81E-01 | no | Gm35850 | intron_variant,non_coding_transcript_variant | MODIFIER |
| 8 | 90763574 | AX-97785350 | 0 | T | C | 0.193 | -3.33E-02 | 1.19E-01 | 2.64E+00 | 2.71E+00 | 7.81E-01 | no | Gm35963 | upstream_gene_variant | MODIFIER |
| 8 | 90784282 | AX-170277446 | 0 | G | T | 0.193 | -3.33E-02 | 1.19E-01 | 2.64E+00 | 2.71E+00 | 7.81E-01 | no | | intergenic_variant | MODIFIER |
| 8 | 90819663 | AX-170228237 | 0 | G | A | 0.193 | -3.33E-02 | 1.19E-01 | 2.64E+00 | 2.71E+00 | 7.81E-01 | no | Gm19935 | upstream_gene_variant | MODIFIER |
| 8 | 90820631 | AX-97854701 | 0 | C | T | 0.193 | -3.33E-02 | 1.19E-01 | 2.64E+00 | 2.71E+00 | 7.81E-01 | no | Gm19935 | upstream_gene_variant | MODIFIER |
| 8 | 90821140 | AX-170227809 | 0 | C | T | 0.193 | -3.33E-02 | 1.19E-01 | 2.64E+00 | 2.71E+00 | 7.81E-01 | no | Gm19935 | upstream_gene_variant | MODIFIER |
| 8 | 90821916 | AX-170158038 | 0 | C | T | 0.193 | -3.33E-02 | 1.19E-01 | 2.64E+00 | 2.71E+00 | 7.81E-01 | no | Gm19935 | upstream_gene_variant | MODIFIER |
| 8 | 90825287 | AX-170251846 | 0 | A | G | 0.193 | -3.33E-02 | 1.19E-01 | 2.64E+00 | 2.71E+00 | 7.81E-01 | no | Chd9 | upstream_gene_variant | MODIFIER |
| 8 | 90825760 | AX-170276890 | 0 | A | G | 0.193 | -3.33E-02 | 1.19E-01 | 2.64E+00 | 2.71E+00 | 7.81E-01 | no | Chd9 | upstream_gene_variant | MODIFIER |
| 8 | 90842047 | AX-170254849 | 0 | T | C | 0.193 | -3.33E-02 | 1.19E-01 | 2.64E+00 | 2.71E+00 | 7.81E-01 | no | Chd9 | intron_variant | MODIFIER |
| 8 | 90842688 | AX-170273942 | 0 | A | C | 0.193 | -3.33E-02 | 1.19E-01 | 2.64E+00 | 2.71E+00 | 7.81E-01 | no | Chd9 | intron_variant | MODIFIER |
| 8 | 90844368 | AX-170234610 | 0 | T | A | 0.193 | -3.33E-02 | 1.19E-01 | 2.64E+00 | 2.71E+00 | 7.81E-01 | no | Chd9 | intron_variant | MODIFIER |
| 8 | 90850597 | AX-170529566 | 0 | C | A | 0.193 | -3.33E-02 | 1.19E-01 | 2.64E+00 | 2.71E+00 | 7.81E-01 | no | Chd9 | intron_variant | MODIFIER |
| 8 | 90851690 | AX-170313550 | 0 | G | A | 0.193 | -3.33E-02 | 1.19E-01 | 2.64E+00 | 2.71E+00 | 7.81E-01 | no | Chd9 | intron_variant | MODIFIER |
| 8 | 90857070 | AX-170427062 | 0 | C | T | 0.193 | -3.33E-02 | 1.19E-01 | 2.64E+00 | 2.71E+00 | 7.81E-01 | no | Chd9 | intron_variant | MODIFIER |
| 8 | 90857927 | AX-170227505 | 0 | C | T | 0.193 | -3.33E-02 | 1.19E-01 | 2.64E+00 | 2.71E+00 | 7.81E-01 | no | Chd9 | intron_variant | MODIFIER |
| 8 | 90872945 | AX-97811265 | 0 | A | G | 0.193 | -3.33E-02 | 1.19E-01 | 2.64E+00 | 2.71E+00 | 7.81E-01 | no | Chd9 | intron_variant | MODIFIER |
| 8 | 90880797 | AX-170080765 | 0 | C | G | 0.193 | -3.33E-02 | 1.19E-01 | 2.64E+00 | 2.71E+00 | 7.81E-01 | no | Chd9 | intron_variant | MODIFIER |
| 8 | 90882610 | AX-170514275 | 0 | G | A | 0.193 | -3.33E-02 | 1.19E-01 | 2.64E+00 | 2.71E+00 | 7.81E-01 | no | Chd9 | intron_variant | MODIFIER |

| | | | | | | | | | | | | | | | |
|---|---|---|---|---|---|---|---|---|---|---|---|---|---|---|---|
| 8 | 90883106 | AX-170134128 | 0 | C | T | 0.193 | -3.33E-02 | 1.19E-01 | 2.64E+00 | 2.71E+00 | 7.81E-01 | no | Chd9 | intron_variant | MODIFIER |
| 8 | 90903660 | AX-170133670 | 0 | A | G | 0.193 | -3.33E-02 | 1.19E-01 | 2.64E+00 | 2.71E+00 | 7.81E-01 | no | Chd9 | intron_variant | MODIFIER |
| 8 | 90907151 | AX-170496674 | 0 | C | G | 0.193 | -3.33E-02 | 1.19E-01 | 2.64E+00 | 2.71E+00 | 7.81E-01 | no | Chd9 | intron_variant | MODIFIER |
| 8 | 90907322 | AX-97844505 | 0 | T | C | 0.193 | -3.33E-02 | 1.19E-01 | 2.64E+00 | 2.71E+00 | 7.81E-01 | no | Chd9 | intron_variant | MODIFIER |
| 8 | 90930623 | AX-97850172 | 0 | T | A | 0.193 | -3.33E-02 | 1.19E-01 | 2.64E+00 | 2.71E+00 | 7.81E-01 | no | Chd9 | intron_variant | MODIFIER |
| 8 | 90938552 | AX-97788015 | 0 | T | C | 0.193 | -3.33E-02 | 1.19E-01 | 2.64E+00 | 2.71E+00 | 7.81E-01 | no | Chd9 | intron_variant | MODIFIER |
| 8 | 91053525 | AX-170147362 | 0 | C | T | 0.193 | -3.33E-02 | 1.19E-01 | 2.64E+00 | 2.71E+00 | 7.81E-01 | no | Chd9 | 3_prime_UTR_variant | MODIFIER |
| 8 | 91073706 | AX-170295293 | 0 | G | A | 0.193 | -3.33E-02 | 1.19E-01 | 2.64E+00 | 2.71E+00 | 7.81E-01 | no | Rbl2 | intron_variant | MODIFIER |
| 8 | 91169112 | AX-170247158 | 0 | A | G | 0.193 | -3.33E-02 | 1.19E-01 | 2.64E+00 | 2.71E+00 | 7.81E-01 | no | Aktip | intron_variant | MODIFIER |
| 6 | 144134665 | AX-170350620 | 1 | C | A | 0.054 | 8.11E-02 | 2.95E-01 | 2.68E+00 | 2.74E+00 | 7.81E-01 | no | Sox5 | intron_variant | MODIFIER |
| 2 | 97194036 | AX-170018416 | 1 | G | A | 0.054 | -8.21E-02 | 3.00E-01 | 2.73E+00 | 2.75E+00 | 7.81E-01 | no | Lrrc4c | intron_variant | MODIFIER |
| 5 | 24304650 | AX-97851692 | 1 | A | T | 0.054 | -8.21E-02 | 3.00E-01 | 2.73E+00 | 2.75E+00 | 7.81E-01 | no | | intergenic_variant | MODIFIER |
| 8 | 116017637 | AX-97836530 | 1 | G | A | 0.25 | 2.60E-02 | 9.71E-02 | 2.66E+00 | 2.77E+00 | 7.81E-01 | no | | intergenic_variant | MODIFIER |
| 8 | 116953907 | AX-170081283 | 1 | G | A | 0.25 | 2.60E-02 | 9.71E-02 | 2.66E+00 | 2.77E+00 | 7.81E-01 | no | Atmin | intron_variant | MODIFIER |
| 4 | 142912374 | AX-170221289 | 1 | A | T | 0.089 | 4.62E-02 | 1.68E-01 | 2.67E+00 | 2.74E+00 | 7.82E-01 | no | 6330411D24Rik | intron_variant,non_coding_transcript_variant | MODIFIER |
| 4 | 95572102 | AX-170524171 | 1 | C | T | 0.348 | 2.83E-02 | 1.00E-01 | 2.61E+00 | 2.70E+00 | 7.82E-01 | no | Fggy | intron_variant | MODIFIER |
| 7 | 117457813 | AX-170259734 | 0 | T | G | 0.395 | 2.54E-02 | 9.19E-02 | 2.67E+00 | 2.72E+00 | 7.82E-01 | no | Xylt1 | intron_variant | MODIFIER |
| 7 | 117462200 | AX-170366304 | 0 | A | C | 0.395 | 2.54E-02 | 9.19E-02 | 2.67E+00 | 2.72E+00 | 7.82E-01 | no | Xylt1 | intron_variant | MODIFIER |
| 7 | 117465250 | AX-170179307 | 0 | A | G | 0.395 | 2.54E-02 | 9.19E-02 | 2.67E+00 | 2.72E+00 | 7.82E-01 | no | Xylt1 | intron_variant | MODIFIER |
| 7 | 117479886 | AX-170029835 | 0 | C | T | 0.395 | 2.54E-02 | 9.19E-02 | 2.67E+00 | 2.72E+00 | 7.82E-01 | no | Xylt1 | intron_variant | MODIFIER |
| 7 | 117480149 | AX-170183018 | 0 | A | C | 0.395 | 2.54E-02 | 9.19E-02 | 2.67E+00 | 2.72E+00 | 7.82E-01 | no | Xylt1 | intron_variant | MODIFIER |
| 7 | 117480929 | AX-170484803 | 0 | A | G | 0.395 | 2.54E-02 | 9.19E-02 | 2.67E+00 | 2.72E+00 | 7.82E-01 | no | Xylt1 | intron_variant | MODIFIER |
| 7 | 117494707 | AX-170078557 | 0 | G | T | 0.395 | 2.54E-02 | 9.19E-02 | 2.67E+00 | 2.72E+00 | 7.82E-01 | no | Xylt1 | intron_variant | MODIFIER |
| 7 | 117495231 | AX-170281684 | 0 | A | G | 0.395 | 2.54E-02 | 9.19E-02 | 2.67E+00 | 2.72E+00 | 7.82E-01 | no | Xylt1 | intron_variant | MODIFIER |
| 7 | 117497507 | AX-170199896 | 0 | G | A | 0.395 | 2.54E-02 | 9.19E-02 | 2.67E+00 | 2.72E+00 | 7.82E-01 | no | Xylt1 | intron_variant | MODIFIER |
| 7 | 117502673 | AX-170167809 | 0 | C | A | 0.395 | 2.54E-02 | 9.19E-02 | 2.67E+00 | 2.72E+00 | 7.82E-01 | no | Xylt1 | intron_variant | MODIFIER |
| 7 | 117503341 | AX-170142795 | 0 | T | G | 0.395 | 2.54E-02 | 9.19E-02 | 2.67E+00 | 2.72E+00 | 7.82E-01 | no | Xylt1 | intron_variant | MODIFIER |
| 7 | 117508453 | AX-170086033 | 0 | T | C | 0.395 | 2.54E-02 | 9.19E-02 | 2.67E+00 | 2.72E+00 | 7.82E-01 | no | Xylt1 | intron_variant | MODIFIER |
| 7 | 117524415 | AX-97803593 | 0 | G | A | 0.395 | 2.54E-02 | 9.19E-02 | 2.67E+00 | 2.72E+00 | 7.82E-01 | no | Xylt1 | intron_variant | MODIFIER |
| 7 | 117531598 | AX-170396285 | 0 | A | T | 0.395 | 2.54E-02 | 9.19E-02 | 2.67E+00 | 2.72E+00 | 7.82E-01 | no | Xylt1 | intron_variant | MODIFIER |
| 7 | 117538913 | AX-170401313 | 0 | T | C | 0.395 | 2.54E-02 | 9.19E-02 | 2.67E+00 | 2.72E+00 | 7.82E-01 | no | Xylt1 | intron_variant | MODIFIER |
| 7 | 117543021 | AX-170177073 | 0 | A | G | 0.395 | 2.54E-02 | 9.19E-02 | 2.67E+00 | 2.72E+00 | 7.82E-01 | no | Xylt1 | intron_variant | MODIFIER |
| 7 | 117549438 | AX-97837174 | 0 | G | A | 0.395 | 2.54E-02 | 9.19E-02 | 2.67E+00 | 2.72E+00 | 7.82E-01 | no | Xylt1 | intron_variant | MODIFIER |
| 7 | 117549726 | AX-170507099 | 0 | T | C | 0.395 | 2.54E-02 | 9.19E-02 | 2.67E+00 | 2.72E+00 | 7.82E-01 | no | Xylt1 | intron_variant | MODIFIER |
| 7 | 117567470 | AX-170350008 | 0 | T | C | 0.395 | 2.54E-02 | 9.19E-02 | 2.67E+00 | 2.72E+00 | 7.82E-01 | no | Xylt1 | intron_variant | MODIFIER |
| 7 | 117582511 | AX-170426269 | 0 | G | T | 0.395 | 2.54E-02 | 9.19E-02 | 2.67E+00 | 2.72E+00 | 7.82E-01 | no | Xylt1 | intron_variant | MODIFIER |
| 7 | 117582533 | AX-170498048 | 0 | A | G | 0.395 | 2.54E-02 | 9.19E-02 | 2.67E+00 | 2.72E+00 | 7.82E-01 | no | Xylt1 | intron_variant | MODIFIER |
| 7 | 117591553 | AX-170124913 | 0 | T | C | 0.395 | 2.54E-02 | 9.19E-02 | 2.67E+00 | 2.72E+00 | 7.82E-01 | no | Xylt1 | intron_variant | MODIFIER |
| 7 | 117592547 | AX-170267878 | 0 | G | A | 0.395 | 2.54E-02 | 9.19E-02 | 2.67E+00 | 2.72E+00 | 7.82E-01 | no | Xylt1 | intron_variant | MODIFIER |
| 7 | 117602466 | AX-170447850 | 0 | C | T | 0.395 | 2.54E-02 | 9.19E-02 | 2.67E+00 | 2.72E+00 | 7.82E-01 | no | Xylt1 | intron_variant | MODIFIER |
| 7 | 117604183 | AX-170243146 | 0 | G | A | 0.395 | 2.54E-02 | 9.19E-02 | 2.67E+00 | 2.72E+00 | 7.82E-01 | no | Xylt1 | intron_variant | MODIFIER |
| 7 | 117604302 | AX-170394259 | 0 | T | C | 0.395 | 2.54E-02 | 9.19E-02 | 2.67E+00 | 2.72E+00 | 7.82E-01 | no | Xylt1 | intron_variant | MODIFIER |
| 7 | 117608119 | AX-170507562 | 0 | T | C | 0.395 | 2.54E-02 | 9.19E-02 | 2.67E+00 | 2.72E+00 | 7.82E-01 | no | Xylt1 | intron_variant | MODIFIER |
| 7 | 117612840 | AX-170078559 | 0 | T | C | 0.395 | 2.54E-02 | 9.19E-02 | 2.67E+00 | 2.72E+00 | 7.82E-01 | no | Xylt1 | intron_variant | MODIFIER |
| 7 | 117617704 | AX-170230489 | 0 | C | T | 0.395 | 2.54E-02 | 9.19E-02 | 2.67E+00 | 2.72E+00 | 7.82E-01 | no | Xylt1 | intron_variant | MODIFIER |
| 7 | 117618445 | AX-170424308 | 0 | A | G | 0.395 | 2.54E-02 | 9.19E-02 | 2.67E+00 | 2.72E+00 | 7.82E-01 | no | Xylt1 | intron_variant | MODIFIER |
| 7 | 117627775 | AX-170328281 | 0 | G | C | 0.395 | 2.54E-02 | 9.19E-02 | 2.67E+00 | 2.72E+00 | 7.82E-01 | no | Xylt1 | intron_variant | MODIFIER |
| 7 | 117630967 | AX-170396714 | 0 | A | G | 0.395 | 2.54E-02 | 9.19E-02 | 2.67E+00 | 2.72E+00 | 7.82E-01 | no | Xylt1 | intron_variant | MODIFIER |
| 7 | 117631259 | AX-97810941 | 0 | G | T | 0.395 | 2.54E-02 | 9.19E-02 | 2.67E+00 | 2.72E+00 | 7.82E-01 | no | Xylt1 | intron_variant | MODIFIER |
| 7 | 117637371 | AX-170339836 | 0 | G | C | 0.395 | 2.54E-02 | 9.19E-02 | 2.67E+00 | 2.72E+00 | 7.82E-01 | no | Xylt1 | intron_variant | MODIFIER |
| 7 | 117638870 | AX-170353580 | 0 | C | A | 0.395 | 2.54E-02 | 9.19E-02 | 2.67E+00 | 2.72E+00 | 7.82E-01 | no | Xylt1 | intron_variant | MODIFIER |
| 7 | 117652386 | AX-170273213 | 0 | A | G | 0.395 | 2.54E-02 | 9.19E-02 | 2.67E+00 | 2.72E+00 | 7.82E-01 | no | Xylt1 | intron_variant | MODIFIER |
| 7 | 117653327 | AX-170425325 | 0 | C | T | 0.395 | 2.54E-02 | 9.19E-02 | 2.67E+00 | 2.72E+00 | 7.82E-01 | no | Xylt1 | intron_variant | MODIFIER |
| 7 | 117657083 | AX-170530374 | 0 | G | A | 0.395 | 2.54E-02 | 9.19E-02 | 2.67E+00 | 2.72E+00 | 7.82E-01 | no | Xylt1 | intron_variant | MODIFIER |

| | | | | | | | | | | | | | | | |
|---|---|---|---|---|---|---|---|---|---|---|---|---|---|---|---|
| 7 | 117659725 | AX-170478030 | 0 | G | C | 0.395 | 2.54E-02 | 9.19E-02 | 2.67E+00 | 2.72E+00 | 7.82E-01 | no | Xylt1 | intron_variant | MODIFIER |
| 7 | 117668204 | AX-170493908 | 0 | G | T | 0.395 | 2.54E-02 | 9.19E-02 | 2.67E+00 | 2.72E+00 | 7.82E-01 | no | Xylt1 | 3_prime_UTR_variant | MODIFIER |
| 7 | 117682105 | AX-170216017 | 0 | T | C | 0.395 | 2.54E-02 | 9.19E-02 | 2.67E+00 | 2.72E+00 | 7.82E-01 | no | | intergenic_variant | MODIFIER |
| 7 | 117691544 | AX-97836149 | 0 | T | C | 0.395 | 2.54E-02 | 9.19E-02 | 2.67E+00 | 2.72E+00 | 7.82E-01 | no | | intergenic_variant | MODIFIER |
| 7 | 117699755 | AX-170421047 | 0 | T | C | 0.395 | 2.54E-02 | 9.19E-02 | 2.67E+00 | 2.72E+00 | 7.82E-01 | no | | regulatory_region_variant | MODIFIER |
| 7 | 117699969 | AX-170096097 | 0 | C | G | 0.395 | 2.54E-02 | 9.19E-02 | 2.67E+00 | 2.72E+00 | 7.82E-01 | no | | regulatory_region_variant | MODIFIER |
| 7 | 117702298 | AX-170287328 | 0 | G | A | 0.395 | 2.54E-02 | 9.19E-02 | 2.67E+00 | 2.72E+00 | 7.82E-01 | no | | intergenic_variant | MODIFIER |
| 7 | 117704390 | AX-170208835 | 0 | A | G | 0.395 | 2.54E-02 | 9.19E-02 | 2.67E+00 | 2.72E+00 | 7.82E-01 | no | | intergenic_variant | MODIFIER |
| 7 | 117704416 | AX-170390109 | 0 | G | A | 0.395 | 2.54E-02 | 9.19E-02 | 2.67E+00 | 2.72E+00 | 7.82E-01 | no | | intergenic_variant | MODIFIER |
| 7 | 117708506 | AX-170078564 | 0 | G | T | 0.395 | 2.54E-02 | 9.19E-02 | 2.67E+00 | 2.72E+00 | 7.82E-01 | no | | intergenic_variant | MODIFIER |
| 7 | 117711953 | AX-170234371 | 0 | C | T | 0.395 | 2.54E-02 | 9.19E-02 | 2.67E+00 | 2.72E+00 | 7.82E-01 | no | | intergenic_variant | MODIFIER |
| 7 | 117712800 | AX-170431146 | 0 | G | A | 0.395 | 2.54E-02 | 9.19E-02 | 2.67E+00 | 2.72E+00 | 7.82E-01 | no | | intergenic_variant | MODIFIER |
| 7 | 117729932 | AX-97833075 | 0 | T | C | 0.395 | 2.54E-02 | 9.19E-02 | 2.67E+00 | 2.72E+00 | 7.82E-01 | no | | intergenic_variant | MODIFIER |
| 7 | 117744891 | AX-170411887 | 0 | G | A | 0.395 | 2.54E-02 | 9.19E-02 | 2.67E+00 | 2.72E+00 | 7.82E-01 | no | | intergenic_variant | MODIFIER |
| 7 | 117745929 | AX-170396027 | 0 | G | C | 0.395 | 2.54E-02 | 9.19E-02 | 2.67E+00 | 2.72E+00 | 7.82E-01 | no | | intergenic_variant | MODIFIER |
| 7 | 117746063 | AX-170450965 | 0 | G | A | 0.395 | 2.54E-02 | 9.19E-02 | 2.67E+00 | 2.72E+00 | 7.82E-01 | no | | intergenic_variant | MODIFIER |
| 7 | 117759845 | AX-97800786 | 0 | C | T | 0.395 | 2.54E-02 | 9.19E-02 | 2.67E+00 | 2.72E+00 | 7.82E-01 | no | | intergenic_variant | MODIFIER |
| 7 | 117766274 | AX-170132936 | 0 | C | T | 0.395 | 2.54E-02 | 9.19E-02 | 2.67E+00 | 2.72E+00 | 7.82E-01 | no | | intergenic_variant | MODIFIER |
| 7 | 117779735 | AX-170483788 | 0 | G | A | 0.395 | 2.54E-02 | 9.19E-02 | 2.67E+00 | 2.72E+00 | 7.82E-01 | no | Gm24063 | downstream_gene_variant | MODIFIER |
| 7 | 117780378 | AX-170233364 | 0 | C | T | 0.395 | 2.54E-02 | 9.19E-02 | 2.67E+00 | 2.72E+00 | 7.82E-01 | no | Gm24063 | downstream_gene_variant | MODIFIER |
| 7 | 117781285 | AX-170379532 | 0 | C | T | 0.395 | 2.54E-02 | 9.19E-02 | 2.67E+00 | 2.72E+00 | 7.82E-01 | no | Gm24063 | downstream_gene_variant | MODIFIER |
| 7 | 117784800 | AX-170417245 | 0 | G | C | 0.395 | 2.54E-02 | 9.19E-02 | 2.67E+00 | 2.72E+00 | 7.82E-01 | no | | intergenic_variant | MODIFIER |
| 7 | 117796229 | AX-170451103 | 0 | A | G | 0.395 | 2.54E-02 | 9.19E-02 | 2.67E+00 | 2.72E+00 | 7.82E-01 | no | | intergenic_variant | MODIFIER |
| 7 | 117815137 | AX-170361814 | 0 | A | G | 0.395 | 2.54E-02 | 9.19E-02 | 2.67E+00 | 2.72E+00 | 7.82E-01 | no | | intergenic_variant | MODIFIER |
| 7 | 117815203 | AX-170105921 | 0 | T | G | 0.395 | 2.54E-02 | 9.19E-02 | 2.67E+00 | 2.72E+00 | 7.82E-01 | no | | intergenic_variant | MODIFIER |
| 7 | 117818843 | AX-170096098 | 0 | C | T | 0.395 | 2.54E-02 | 9.19E-02 | 2.67E+00 | 2.72E+00 | 7.82E-01 | no | | intergenic_variant | MODIFIER |
| 7 | 117821653 | AX-170078569 | 0 | T | C | 0.395 | 2.54E-02 | 9.19E-02 | 2.67E+00 | 2.72E+00 | 7.82E-01 | no | | intergenic_variant | MODIFIER |
| 7 | 117830835 | AX-170282521 | 0 | T | C | 0.395 | 2.54E-02 | 9.19E-02 | 2.67E+00 | 2.72E+00 | 7.82E-01 | no | | intergenic_variant | MODIFIER |
| 7 | 117834082 | AX-170488210 | 0 | G | T | 0.395 | 2.54E-02 | 9.19E-02 | 2.67E+00 | 2.72E+00 | 7.82E-01 | no | | intergenic_variant | MODIFIER |
| 7 | 117850010 | AX-170414704 | 0 | T | C | 0.395 | 2.54E-02 | 9.19E-02 | 2.67E+00 | 2.72E+00 | 7.82E-01 | no | | intergenic_variant | MODIFIER |
| 7 | 117851513 | AX-170105751 | 0 | T | C | 0.395 | 2.54E-02 | 9.19E-02 | 2.67E+00 | 2.72E+00 | 7.82E-01 | no | | intergenic_variant | MODIFIER |
| 7 | 117853124 | AX-170516246 | 0 | G | A | 0.395 | 2.54E-02 | 9.19E-02 | 2.67E+00 | 2.72E+00 | 7.82E-01 | no | | regulatory_region_variant | MODIFIER |
| 7 | 117860515 | AX-170113283 | 0 | T | C | 0.395 | 2.54E-02 | 9.19E-02 | 2.67E+00 | 2.72E+00 | 7.82E-01 | no | | intergenic_variant | MODIFIER |
| 7 | 117860668 | AX-170415573 | 0 | A | G | 0.395 | 2.54E-02 | 9.19E-02 | 2.67E+00 | 2.72E+00 | 7.82E-01 | no | | intergenic_variant | MODIFIER |
| 7 | 117868776 | AX-170142236 | 0 | C | T | 0.395 | 2.54E-02 | 9.19E-02 | 2.67E+00 | 2.72E+00 | 7.82E-01 | no | | intergenic_variant | MODIFIER |
| 7 | 117869184 | AX-170276376 | 0 | G | T | 0.395 | 2.54E-02 | 9.19E-02 | 2.67E+00 | 2.72E+00 | 7.82E-01 | no | | intergenic_variant | MODIFIER |
| 7 | 117870108 | AX-170458167 | 0 | A | G | 0.395 | 2.54E-02 | 9.19E-02 | 2.67E+00 | 2.72E+00 | 7.82E-01 | no | | intergenic_variant | MODIFIER |
| 7 | 117872407 | AX-170370014 | 0 | A | C | 0.395 | 2.54E-02 | 9.19E-02 | 2.67E+00 | 2.72E+00 | 7.82E-01 | no | | intergenic_variant | MODIFIER |
| 7 | 117872498 | AX-170086229 | 0 | T | A | 0.395 | 2.54E-02 | 9.19E-02 | 2.67E+00 | 2.72E+00 | 7.82E-01 | no | | intergenic_variant | MODIFIER |
| 7 | 117873489 | AX-170201696 | 0 | T | A | 0.395 | 2.54E-02 | 9.19E-02 | 2.67E+00 | 2.72E+00 | 7.82E-01 | no | | intergenic_variant | MODIFIER |
| 7 | 117873915 | AX-170517415 | 0 | A | G | 0.395 | 2.54E-02 | 9.19E-02 | 2.67E+00 | 2.72E+00 | 7.82E-01 | no | | intergenic_variant | MODIFIER |
| 7 | 117880815 | AX-170500275 | 0 | T | C | 0.395 | 2.54E-02 | 9.19E-02 | 2.67E+00 | 2.72E+00 | 7.82E-01 | no | | intergenic_variant | MODIFIER |
| 7 | 117881673 | AX-170145022 | 0 | C | T | 0.395 | 2.54E-02 | 9.19E-02 | 2.67E+00 | 2.72E+00 | 7.82E-01 | no | | intergenic_variant | MODIFIER |
| 7 | 117896920 | AX-170246974 | 0 | C | T | 0.395 | 2.54E-02 | 9.19E-02 | 2.67E+00 | 2.72E+00 | 7.82E-01 | no | | intergenic_variant | MODIFIER |
| 7 | 117899046 | AX-170405207 | 0 | G | A | 0.395 | 2.54E-02 | 9.19E-02 | 2.67E+00 | 2.72E+00 | 7.82E-01 | no | | intergenic_variant | MODIFIER |
| 7 | 117904824 | AX-170458132 | 0 | A | G | 0.395 | 2.54E-02 | 9.19E-02 | 2.67E+00 | 2.72E+00 | 7.82E-01 | no | | intergenic_variant | MODIFIER |
| 7 | 117912767 | AX-170489023 | 0 | G | A | 0.395 | 2.54E-02 | 9.19E-02 | 2.67E+00 | 2.72E+00 | 7.82E-01 | no | | intergenic_variant | MODIFIER |
| 7 | 117912994 | AX-170078570 | 0 | A | G | 0.395 | 2.54E-02 | 9.19E-02 | 2.67E+00 | 2.72E+00 | 7.82E-01 | no | | intergenic_variant | MODIFIER |
| 7 | 117913735 | AX-170280928 | 0 | C | T | 0.395 | 2.54E-02 | 9.19E-02 | 2.67E+00 | 2.72E+00 | 7.82E-01 | no | | intergenic_variant | MODIFIER |
| 7 | 117914108 | AX-170454716 | 0 | T | G | 0.395 | 2.54E-02 | 9.19E-02 | 2.67E+00 | 2.72E+00 | 7.82E-01 | no | | regulatory_region_variant | MODIFIER |
| 7 | 117914474 | AX-170078572 | 0 | G | A | 0.395 | 2.54E-02 | 9.19E-02 | 2.67E+00 | 2.72E+00 | 7.82E-01 | no | | regulatory_region_variant | MODIFIER |
| 7 | 117915878 | AX-97831348 | 0 | T | C | 0.395 | 2.54E-02 | 9.19E-02 | 2.67E+00 | 2.72E+00 | 7.82E-01 | no | | intergenic_variant | MODIFIER |
| 7 | 117927089 | AX-170355248 | 0 | G | A | 0.395 | 2.54E-02 | 9.19E-02 | 2.67E+00 | 2.72E+00 | 7.82E-01 | no | | intergenic_variant | MODIFIER |
| 7 | 117928722 | AX-170270074 | 0 | A | G | 0.395 | 2.54E-02 | 9.19E-02 | 2.67E+00 | 2.72E+00 | 7.82E-01 | no | | intergenic_variant | MODIFIER |

| | | | | | | | | | | | | | | | |
|---|---|---|---|---|---|---|---|---|---|---|---|---|---|---|---|
| 7 | 117943046 | AX-170029837 | 0 | C | T | 0.395 | 2.54E-02 | 9.19E-02 | 2.67E+00 | 2.72E+00 | 7.82E-01 | no | | intergenic_variant | MODIFIER |
| 7 | 117959158 | AX-170416823 | 0 | A | G | 0.395 | 2.54E-02 | 9.19E-02 | 2.67E+00 | 2.72E+00 | 7.82E-01 | no | | intergenic_variant | MODIFIER |
| 7 | 117960061 | AX-170441879 | 0 | A | G | 0.395 | 2.54E-02 | 9.19E-02 | 2.67E+00 | 2.72E+00 | 7.82E-01 | no | | intergenic_variant | MODIFIER |
| 7 | 117960228 | AX-97853483 | 0 | T | G | 0.395 | 2.54E-02 | 9.19E-02 | 2.67E+00 | 2.72E+00 | 7.82E-01 | no | | intergenic_variant | MODIFIER |
| 7 | 117969323 | AX-170303234 | 0 | A | G | 0.395 | 2.54E-02 | 9.19E-02 | 2.67E+00 | 2.72E+00 | 7.82E-01 | no | | intergenic_variant | MODIFIER |
| 7 | 117969496 | AX-170519942 | 0 | C | G | 0.395 | 2.54E-02 | 9.19E-02 | 2.67E+00 | 2.72E+00 | 7.82E-01 | no | | intergenic_variant | MODIFIER |
| 7 | 117975678 | AX-170489846 | 0 | G | A | 0.395 | 2.54E-02 | 9.19E-02 | 2.67E+00 | 2.72E+00 | 7.82E-01 | no | | intergenic_variant | MODIFIER |
| 7 | 117979855 | AX-97843990 | 0 | T | C | 0.395 | 2.54E-02 | 9.19E-02 | 2.67E+00 | 2.72E+00 | 7.82E-01 | no | | intergenic_variant | MODIFIER |
| 7 | 117980341 | AX-170029840 | 0 | A | G | 0.395 | 2.54E-02 | 9.19E-02 | 2.67E+00 | 2.72E+00 | 7.82E-01 | no | | intergenic_variant | MODIFIER |
| 7 | 117992503 | AX-170029839 | 0 | G | A | 0.395 | 2.54E-02 | 9.19E-02 | 2.67E+00 | 2.72E+00 | 7.82E-01 | no | | intergenic_variant | MODIFIER |
| 7 | 118003910 | AX-170309360 | 0 | C | G | 0.395 | 2.54E-02 | 9.19E-02 | 2.67E+00 | 2.72E+00 | 7.82E-01 | no | | intergenic_variant | MODIFIER |
| 7 | 118009420 | AX-170350743 | 0 | C | T | 0.395 | 2.54E-02 | 9.19E-02 | 2.67E+00 | 2.72E+00 | 7.82E-01 | no | | intergenic_variant | MODIFIER |
| 7 | 118012446 | AX-170099962 | 0 | A | G | 0.395 | 2.54E-02 | 9.19E-02 | 2.67E+00 | 2.72E+00 | 7.82E-01 | no | | intergenic_variant | MODIFIER |
| 7 | 118013199 | AX-170096100 | 0 | A | G | 0.395 | 2.54E-02 | 9.19E-02 | 2.67E+00 | 2.72E+00 | 7.82E-01 | no | | intergenic_variant | MODIFIER |
| 7 | 118023528 | AX-97835785 | 0 | T | C | 0.395 | 2.54E-02 | 9.19E-02 | 2.67E+00 | 2.72E+00 | 7.82E-01 | no | | intergenic_variant | MODIFIER |
| 7 | 118025225 | AX-170375521 | 0 | T | G | 0.395 | 2.54E-02 | 9.19E-02 | 2.67E+00 | 2.72E+00 | 7.82E-01 | no | | intergenic_variant | MODIFIER |
| 7 | 118045890 | AX-170078576 | 0 | C | T | 0.395 | 2.54E-02 | 9.19E-02 | 2.67E+00 | 2.72E+00 | 7.82E-01 | no | | intergenic_variant | MODIFIER |
| 7 | 118060526 | AX-170433914 | 0 | A | G | 0.395 | 2.54E-02 | 9.19E-02 | 2.67E+00 | 2.72E+00 | 7.82E-01 | no | | intergenic_variant | MODIFIER |
| 7 | 118062219 | AX-170193546 | 0 | G | A | 0.395 | 2.54E-02 | 9.19E-02 | 2.67E+00 | 2.72E+00 | 7.82E-01 | no | | intergenic_variant | MODIFIER |
| 7 | 118063347 | AX-170374956 | 0 | T | C | 0.395 | 2.54E-02 | 9.19E-02 | 2.67E+00 | 2.72E+00 | 7.82E-01 | no | | intergenic_variant | MODIFIER |
| 7 | 118064910 | AX-170405130 | 0 | A | G | 0.395 | 2.54E-02 | 9.19E-02 | 2.67E+00 | 2.72E+00 | 7.82E-01 | no | | intergenic_variant | MODIFIER |
| 7 | 118065272 | AX-170312195 | 0 | G | A | 0.395 | 2.54E-02 | 9.19E-02 | 2.67E+00 | 2.72E+00 | 7.82E-01 | no | | intergenic_variant | MODIFIER |
| 7 | 118095513 | AX-170142342 | 0 | G | A | 0.395 | 2.54E-02 | 9.19E-02 | 2.67E+00 | 2.72E+00 | 7.82E-01 | no | | intergenic_variant | MODIFIER |
| 7 | 118099066 | AX-170138724 | 0 | G | C | 0.395 | 2.54E-02 | 9.19E-02 | 2.67E+00 | 2.72E+00 | 7.82E-01 | no | | intergenic_variant | MODIFIER |
| 7 | 118099253 | AX-170271392 | 0 | A | G | 0.395 | 2.54E-02 | 9.19E-02 | 2.67E+00 | 2.72E+00 | 7.82E-01 | no | | intergenic_variant | MODIFIER |
| 7 | 118108884 | AX-170193163 | 0 | G | C | 0.395 | 2.54E-02 | 9.19E-02 | 2.67E+00 | 2.72E+00 | 7.82E-01 | no | Rps15a | downstream_gene_variant | MODIFIER |
| 7 | 118114215 | AX-97832562 | 0 | A | G | 0.395 | 2.54E-02 | 9.19E-02 | 2.67E+00 | 2.72E+00 | 7.82E-01 | no | Arl6ip1 | downstream_gene_variant | MODIFIER |
| 7 | 118114430 | AX-170105452 | 0 | C | A | 0.395 | 2.54E-02 | 9.19E-02 | 2.67E+00 | 2.72E+00 | 7.82E-01 | no | Arl6ip1 | downstream_gene_variant | MODIFIER |
| 7 | 118146787 | AX-170414375 | 0 | T | C | 0.395 | 2.54E-02 | 9.19E-02 | 2.67E+00 | 2.72E+00 | 7.82E-01 | no | Smg1 | intron_variant | MODIFIER |
| 7 | 118147416 | AX-97790449 | 0 | G | A | 0.395 | 2.54E-02 | 9.19E-02 | 2.67E+00 | 2.72E+00 | 7.82E-01 | no | Smg1 | intron_variant | MODIFIER |
| 7 | 118159997 | AX-170312648 | 0 | T | C | 0.395 | 2.54E-02 | 9.19E-02 | 2.67E+00 | 2.72E+00 | 7.82E-01 | no | Smg1 | intron_variant | MODIFIER |
| 7 | 118171786 | AX-170166754 | 0 | T | A | 0.395 | 2.54E-02 | 9.19E-02 | 2.67E+00 | 2.72E+00 | 7.82E-01 | no | Smg1 | synonymous_variant | LOW |
| 7 | 118172828 | AX-170427238 | 0 | T | C | 0.395 | 2.54E-02 | 9.19E-02 | 2.67E+00 | 2.72E+00 | 7.82E-01 | no | Smg1 | intron_variant | MODIFIER |
| 7 | 118174457 | AX-170208794 | 0 | T | A | 0.395 | 2.54E-02 | 9.19E-02 | 2.67E+00 | 2.72E+00 | 7.82E-01 | no | Smg1 | intron_variant | MODIFIER |
| 7 | 118189707 | AX-170371408 | 0 | C | A | 0.395 | 2.54E-02 | 9.19E-02 | 2.67E+00 | 2.72E+00 | 7.82E-01 | no | Smg1 | intron_variant | MODIFIER |
| 7 | 118193126 | AX-170224360 | 0 | T | C | 0.395 | 2.54E-02 | 9.19E-02 | 2.67E+00 | 2.72E+00 | 7.82E-01 | no | Smg1 | intron_variant | MODIFIER |
| 7 | 118214528 | AX-170209515 | 0 | C | T | 0.395 | 2.54E-02 | 9.19E-02 | 2.67E+00 | 2.72E+00 | 7.82E-01 | no | Smg1 | intron_variant | MODIFIER |
| 7 | 118235748 | AX-97792520 | 0 | G | C | 0.395 | 2.54E-02 | 9.19E-02 | 2.67E+00 | 2.72E+00 | 7.82E-01 | no | Smg1 | intron_variant | MODIFIER |
| 7 | 118253382 | AX-170224273 | 0 | G | A | 0.395 | 2.54E-02 | 9.19E-02 | 2.67E+00 | 2.72E+00 | 7.82E-01 | no | | regulatory_region_variant | MODIFIER |
| 7 | 118255221 | AX-170181088 | 0 | T | C | 0.395 | 2.54E-02 | 9.19E-02 | 2.67E+00 | 2.72E+00 | 7.82E-01 | no | | regulatory_region_variant | MODIFIER |
| 7 | 118267486 | AX-170170900 | 0 | A | G | 0.395 | 2.54E-02 | 9.19E-02 | 2.67E+00 | 2.72E+00 | 7.82E-01 | no | | intergenic_variant | MODIFIER |
| 7 | 118267540 | AX-97786247 | 0 | G | A | 0.395 | 2.54E-02 | 9.19E-02 | 2.67E+00 | 2.72E+00 | 7.82E-01 | no | | intergenic_variant | MODIFIER |
| 7 | 118267598 | AX-170163164 | 0 | G | C | 0.395 | 2.54E-02 | 9.19E-02 | 2.67E+00 | 2.72E+00 | 7.82E-01 | no | | intergenic_variant | MODIFIER |
| 7 | 118274713 | AX-170210289 | 0 | C | A | 0.395 | 2.54E-02 | 9.19E-02 | 2.67E+00 | 2.72E+00 | 7.82E-01 | no | Gm45155 | intron_variant,non_coding_transcript_variant | MODIFIER |
| 7 | 118293639 | AX-170262651 | 0 | T | C | 0.395 | 2.54E-02 | 9.19E-02 | 2.67E+00 | 2.72E+00 | 7.82E-01 | no | | intergenic_variant | MODIFIER |
| 7 | 118293683 | AX-170371433 | 0 | C | T | 0.395 | 2.54E-02 | 9.19E-02 | 2.67E+00 | 2.72E+00 | 7.82E-01 | no | | intergenic_variant | MODIFIER |
| 7 | 118305496 | AX-170479106 | 0 | G | A | 0.395 | 2.54E-02 | 9.19E-02 | 2.67E+00 | 2.72E+00 | 7.82E-01 | no | | regulatory_region_variant | MODIFIER |
| 7 | 118314262 | AX-170174422 | 0 | A | G | 0.395 | 2.54E-02 | 9.19E-02 | 2.67E+00 | 2.72E+00 | 7.82E-01 | no | | regulatory_region_variant | MODIFIER |
| 7 | 118320155 | AX-170258448 | 0 | T | G | 0.395 | 2.54E-02 | 9.19E-02 | 2.67E+00 | 2.72E+00 | 7.82E-01 | no | | regulatory_region_variant | MODIFIER |
| 7 | 118323021 | AX-170195120 | 0 | G | T | 0.395 | 2.54E-02 | 9.19E-02 | 2.67E+00 | 2.72E+00 | 7.82E-01 | no | | intergenic_variant | MODIFIER |
| 7 | 118326350 | AX-170449794 | 0 | A | G | 0.395 | 2.54E-02 | 9.19E-02 | 2.67E+00 | 2.72E+00 | 7.82E-01 | no | Gm35309 | upstream_gene_variant | MODIFIER |
| 7 | 118331460 | AX-170404582 | 0 | A | G | 0.395 | 2.54E-02 | 9.19E-02 | 2.67E+00 | 2.72E+00 | 7.82E-01 | no | Gm35309 | downstream_gene_variant | MODIFIER |
| 7 | 118350009 | AX-97848064 | 0 | T | C | 0.395 | 2.54E-02 | 9.19E-02 | 2.67E+00 | 2.72E+00 | 7.82E-01 | no | | intergenic_variant | MODIFIER |
| 7 | 118350048 | AX-170130561 | 0 | C | A | 0.395 | 2.54E-02 | 9.19E-02 | 2.67E+00 | 2.72E+00 | 7.82E-01 | no | | intergenic_variant | MODIFIER |

| | | | | | | | | | | | | | | | |
|---|---|---|---|---|---|---|---|---|---|---|---|---|---|---|---|
| 7 | 118352655 | AX-170535178 | 0 | G | T | 0.395 | 2.54E-02 | 9.19E-02 | 2.67E+00 | 2.72E+00 | 7.82E-01 | no | | intergenic_variant | MODIFIER |
| 7 | 118352771 | AX-170306526 | 0 | C | T | 0.395 | 2.54E-02 | 9.19E-02 | 2.67E+00 | 2.72E+00 | 7.82E-01 | no | | intergenic_variant | MODIFIER |
| 7 | 118355814 | AX-170267066 | 0 | C | T | 0.395 | 2.54E-02 | 9.19E-02 | 2.67E+00 | 2.72E+00 | 7.82E-01 | no | | intergenic_variant | MODIFIER |
| 7 | 118361001 | AX-170290392 | 0 | A | T | 0.395 | 2.54E-02 | 9.19E-02 | 2.67E+00 | 2.72E+00 | 7.82E-01 | no | | intergenic_variant | MODIFIER |
| 7 | 118364449 | AX-170375230 | 0 | G | C | 0.395 | 2.54E-02 | 9.19E-02 | 2.67E+00 | 2.72E+00 | 7.82E-01 | no | | regulatory_region_variant | MODIFIER |
| 7 | 118372786 | AX-170382043 | 0 | G | A | 0.395 | 2.54E-02 | 9.19E-02 | 2.67E+00 | 2.72E+00 | 7.82E-01 | no | | intergenic_variant | MODIFIER |
| 7 | 118381312 | AX-170510928 | 0 | T | C | 0.395 | 2.54E-02 | 9.19E-02 | 2.67E+00 | 2.72E+00 | 7.82E-01 | no | Syt17 | 3_prime_UTR_variant | MODIFIER |
| 7 | 118386561 | AX-170470790 | 0 | C | A | 0.395 | 2.54E-02 | 9.19E-02 | 2.67E+00 | 2.72E+00 | 7.82E-01 | no | Syt17 | intron_variant | MODIFIER |
| 7 | 118393137 | AX-170296894 | 0 | A | C | 0.395 | 2.54E-02 | 9.19E-02 | 2.67E+00 | 2.72E+00 | 7.82E-01 | no | Syt17 | intron_variant | MODIFIER |
| 7 | 118396915 | AX-170142657 | 0 | C | T | 0.395 | 2.54E-02 | 9.19E-02 | 2.67E+00 | 2.72E+00 | 7.82E-01 | no | Syt17 | intron_variant | MODIFIER |
| 7 | 118398002 | AX-170368212 | 0 | A | G | 0.395 | 2.54E-02 | 9.19E-02 | 2.67E+00 | 2.72E+00 | 7.82E-01 | no | Syt17 | intron_variant | MODIFIER |
| 7 | 118402620 | AX-170208431 | 0 | A | G | 0.395 | 2.54E-02 | 9.19E-02 | 2.67E+00 | 2.72E+00 | 7.82E-01 | no | Syt17 | intron_variant | MODIFIER |
| 7 | 118405635 | AX-170241711 | 0 | C | T | 0.395 | 2.54E-02 | 9.19E-02 | 2.67E+00 | 2.72E+00 | 7.82E-01 | no | Syt17 | intron_variant | MODIFIER |
| 7 | 118407848 | AX-170506719 | 0 | A | G | 0.395 | 2.54E-02 | 9.19E-02 | 2.67E+00 | 2.72E+00 | 7.82E-01 | no | Syt17 | intron_variant | MODIFIER |
| 7 | 118408212 | AX-170380629 | 0 | G | A | 0.395 | 2.54E-02 | 9.19E-02 | 2.67E+00 | 2.72E+00 | 7.82E-01 | no | Syt17 | intron_variant | MODIFIER |
| 7 | 118408626 | AX-170235179 | 0 | T | A | 0.395 | 2.54E-02 | 9.19E-02 | 2.67E+00 | 2.72E+00 | 7.82E-01 | no | Syt17 | intron_variant | MODIFIER |
| 7 | 118409293 | AX-170400564 | 0 | A | G | 0.395 | 2.54E-02 | 9.19E-02 | 2.67E+00 | 2.72E+00 | 7.82E-01 | no | Syt17 | intron_variant | MODIFIER |
| 7 | 118409410 | AX-97837894 | 0 | T | C | 0.395 | 2.54E-02 | 9.19E-02 | 2.67E+00 | 2.72E+00 | 7.82E-01 | no | Syt17 | intron_variant | MODIFIER |
| 7 | 118409646 | AX-170236068 | 0 | A | G | 0.395 | 2.54E-02 | 9.19E-02 | 2.67E+00 | 2.72E+00 | 7.82E-01 | no | Syt17 | intron_variant | MODIFIER |
| 7 | 118409686 | AX-170331188 | 0 | G | C | 0.395 | 2.54E-02 | 9.19E-02 | 2.67E+00 | 2.72E+00 | 7.82E-01 | no | Syt17 | intron_variant | MODIFIER |
| 7 | 118410187 | AX-170257555 | 0 | T | G | 0.395 | 2.54E-02 | 9.19E-02 | 2.67E+00 | 2.72E+00 | 7.82E-01 | no | Syt17 | intron_variant | MODIFIER |
| 7 | 118414600 | AX-170167198 | 0 | A | C | 0.395 | 2.54E-02 | 9.19E-02 | 2.67E+00 | 2.72E+00 | 7.82E-01 | no | Syt17 | intron_variant | MODIFIER |
| 7 | 118425172 | AX-170266057 | 0 | T | C | 0.395 | 2.54E-02 | 9.19E-02 | 2.67E+00 | 2.72E+00 | 7.82E-01 | no | Syt17 | intron_variant | MODIFIER |
| 7 | 118426447 | AX-170381167 | 0 | T | A | 0.395 | 2.54E-02 | 9.19E-02 | 2.67E+00 | 2.72E+00 | 7.82E-01 | no | Syt17 | intron_variant | MODIFIER |
| 7 | 118426473 | AX-170029843 | 0 | G | A | 0.395 | 2.54E-02 | 9.19E-02 | 2.67E+00 | 2.72E+00 | 7.82E-01 | no | Syt17 | intron_variant | MODIFIER |
| 7 | 118430573 | AX-170244471 | 0 | A | G | 0.395 | 2.54E-02 | 9.19E-02 | 2.67E+00 | 2.72E+00 | 7.82E-01 | no | Syt17 | intron_variant | MODIFIER |
| 7 | 118432644 | AX-170533385 | 0 | G | A | 0.395 | 2.54E-02 | 9.19E-02 | 2.67E+00 | 2.72E+00 | 7.82E-01 | no | Syt17 | intron_variant | MODIFIER |
| 7 | 118434372 | AX-170284252 | 0 | G | A | 0.395 | 2.54E-02 | 9.19E-02 | 2.67E+00 | 2.72E+00 | 7.82E-01 | no | Syt17 | missense_variant | MODERATE |
| 7 | 118454346 | AX-170078585 | 0 | G | T | 0.395 | 2.54E-02 | 9.19E-02 | 2.67E+00 | 2.72E+00 | 7.82E-01 | no | | intergenic_variant | MODIFIER |
| 7 | 118460983 | AX-170319733 | 0 | C | T | 0.395 | 2.54E-02 | 9.19E-02 | 2.67E+00 | 2.72E+00 | 7.82E-01 | no | | regulatory_region_variant | MODIFIER |
| 7 | 118461124 | AX-170397164 | 0 | C | G | 0.395 | 2.54E-02 | 9.19E-02 | 2.67E+00 | 2.72E+00 | 7.82E-01 | no | | regulatory_region_variant | MODIFIER |
| 7 | 118467328 | AX-170029841 | 0 | C | T | 0.395 | 2.54E-02 | 9.19E-02 | 2.67E+00 | 2.72E+00 | 7.82E-01 | no | | regulatory_region_variant | MODIFIER |
| 7 | 118467495 | AX-170029842 | 0 | T | G | 0.395 | 2.54E-02 | 9.19E-02 | 2.67E+00 | 2.72E+00 | 7.82E-01 | no | | regulatory_region_variant | MODIFIER |
| 7 | 118470408 | AX-170440139 | 0 | C | T | 0.395 | 2.54E-02 | 9.19E-02 | 2.67E+00 | 2.72E+00 | 7.82E-01 | no | | regulatory_region_variant | MODIFIER |
| 7 | 118481406 | AX-170407654 | 0 | G | T | 0.395 | 2.54E-02 | 9.19E-02 | 2.67E+00 | 2.72E+00 | 7.82E-01 | no | Itpripl2 | downstream_gene_variant | MODIFIER |
| 7 | 118481684 | AX-170188931 | 0 | T | C | 0.395 | 2.54E-02 | 9.19E-02 | 2.67E+00 | 2.72E+00 | 7.82E-01 | no | Itpripl2 | downstream_gene_variant | MODIFIER |
| 7 | 118483045 | AX-170476822 | 0 | C | T | 0.395 | 2.54E-02 | 9.19E-02 | 2.67E+00 | 2.72E+00 | 7.82E-01 | no | Itpripl2 | downstream_gene_variant | MODIFIER |
| 7 | 118488425 | AX-170179564 | 0 | G | C | 0.395 | 2.54E-02 | 9.19E-02 | 2.67E+00 | 2.72E+00 | 7.82E-01 | no | Itpripl2 | 3_prime_UTR_variant | MODIFIER |
| 7 | 118488489 | AX-170239545 | 0 | T | C | 0.395 | 2.54E-02 | 9.19E-02 | 2.67E+00 | 2.72E+00 | 7.82E-01 | no | Itpripl2 | 3_prime_UTR_variant | MODIFIER |
| 7 | 118500158 | AX-170126869 | 0 | T | G | 0.395 | 2.54E-02 | 9.19E-02 | 2.67E+00 | 2.72E+00 | 7.82E-01 | no | | regulatory_region_variant | MODIFIER |
| 7 | 118501284 | AX-170399720 | 0 | A | G | 0.395 | 2.54E-02 | 9.19E-02 | 2.67E+00 | 2.72E+00 | 7.82E-01 | no | | regulatory_region_variant | MODIFIER |
| 7 | 118502628 | AX-170307701 | 0 | G | A | 0.395 | 2.54E-02 | 9.19E-02 | 2.67E+00 | 2.72E+00 | 7.82E-01 | no | | regulatory_region_variant | MODIFIER |
| 7 | 118502735 | AX-170455212 | 0 | A | G | 0.395 | 2.54E-02 | 9.19E-02 | 2.67E+00 | 2.72E+00 | 7.82E-01 | no | | regulatory_region_variant | MODIFIER |
| 7 | 118504533 | AX-170325128 | 0 | G | A | 0.395 | 2.54E-02 | 9.19E-02 | 2.67E+00 | 2.72E+00 | 7.82E-01 | no | | intergenic_variant | MODIFIER |
| 7 | 118509362 | AX-170174076 | 0 | G | A | 0.395 | 2.54E-02 | 9.19E-02 | 2.67E+00 | 2.72E+00 | 7.82E-01 | no | Coq7 | downstream_gene_variant | MODIFIER |
| 7 | 118512135 | AX-170191183 | 0 | A | G | 0.395 | 2.54E-02 | 9.19E-02 | 2.67E+00 | 2.72E+00 | 7.82E-01 | no | Coq7 | intron_variant | MODIFIER |
| 7 | 118512301 | AX-170214189 | 0 | G | A | 0.395 | 2.54E-02 | 9.19E-02 | 2.67E+00 | 2.72E+00 | 7.82E-01 | no | Coq7 | intron_variant | MODIFIER |
| 7 | 118512437 | AX-170511243 | 0 | G | A | 0.395 | 2.54E-02 | 9.19E-02 | 2.67E+00 | 2.72E+00 | 7.82E-01 | no | Coq7 | intron_variant | MODIFIER |
| 7 | 118514848 | AX-170078586 | 0 | G | A | 0.395 | 2.54E-02 | 9.19E-02 | 2.67E+00 | 2.72E+00 | 7.82E-01 | no | Coq7 | intron_variant | MODIFIER |
| 7 | 118524757 | AX-170078588 | 0 | G | A | 0.395 | 2.54E-02 | 9.19E-02 | 2.67E+00 | 2.72E+00 | 7.82E-01 | no | Coq7 | downstream_gene_variant | MODIFIER |
| 7 | 118536891 | AX-170137599 | 0 | A | G | 0.395 | 2.54E-02 | 9.19E-02 | 2.67E+00 | 2.72E+00 | 7.82E-01 | no | Coq7 | upstream_gene_variant | MODIFIER |
| 7 | 118547213 | AX-170147115 | 0 | C | T | 0.395 | 2.54E-02 | 9.19E-02 | 2.67E+00 | 2.72E+00 | 7.82E-01 | no | Tmc7 | intron_variant | MODIFIER |
| 7 | 118548422 | AX-170269179 | 0 | G | A | 0.395 | 2.54E-02 | 9.19E-02 | 2.67E+00 | 2.72E+00 | 7.82E-01 | no | Tmc7 | intron_variant | MODIFIER |
| 7 | 118550340 | AX-170208334 | 0 | T | C | 0.395 | 2.54E-02 | 9.19E-02 | 2.67E+00 | 2.72E+00 | 7.82E-01 | no | Tmc7 | intron_variant | MODIFIER |

| | | | | | | | | | | | | | | | |
|---|---|---|---|---|---|---|---|---|---|---|---|---|---|---|---|
| 7 | 118552603 | AX-170126257 | 0 | C | T | 0.395 | 2.54E-02 | 9.19E-02 | 2.67E+00 | 2.72E+00 | 7.82E-01 | no | Tmc7 | intron_variant | MODIFIER |
| 7 | 118552855 | AX-170385874 | 0 | C | A | 0.395 | 2.54E-02 | 9.19E-02 | 2.67E+00 | 2.72E+00 | 7.82E-01 | no | Tmc7 | intron_variant | MODIFIER |
| 7 | 118553989 | AX-170325176 | 0 | A | G | 0.395 | 2.54E-02 | 9.19E-02 | 2.67E+00 | 2.72E+00 | 7.82E-01 | no | Tmc7 | intron_variant | MODIFIER |
| 7 | 118566272 | AX-170496708 | 0 | G | C | 0.395 | 2.54E-02 | 9.19E-02 | 2.67E+00 | 2.72E+00 | 7.82E-01 | no | Tmc7 | missense_variant | MODERATE |
| 7 | 118566337 | AX-170398909 | 0 | C | T | 0.395 | 2.54E-02 | 9.19E-02 | 2.67E+00 | 2.72E+00 | 7.82E-01 | no | Tmc7 | synonymous_variant | LOW |
| 7 | 118571125 | AX-170362341 | 0 | G | A | 0.395 | 2.54E-02 | 9.19E-02 | 2.67E+00 | 2.72E+00 | 7.82E-01 | no | Tmc7 | intron_variant | MODIFIER |
| 7 | 118581738 | AX-97825591 | 0 | G | A | 0.395 | 2.54E-02 | 9.19E-02 | 2.67E+00 | 2.72E+00 | 7.82E-01 | no | Tmc7 | intron_variant | MODIFIER |
| 7 | 118582952 | AX-170323795 | 0 | A | G | 0.395 | 2.54E-02 | 9.19E-02 | 2.67E+00 | 2.72E+00 | 7.82E-01 | no | Tmc7 | intron_variant | MODIFIER |
| 7 | 118592094 | AX-170115101 | 0 | A | G | 0.395 | 2.54E-02 | 9.19E-02 | 2.67E+00 | 2.72E+00 | 7.82E-01 | no | B230311B06Rik | upstream_gene_variant | MODIFIER |
| 7 | 118609099 | AX-170356429 | 0 | C | T | 0.395 | 2.54E-02 | 9.19E-02 | 2.67E+00 | 2.72E+00 | 7.82E-01 | no | Tmc5 | intron_variant | MODIFIER |
| 7 | 118628702 | AX-170078592 | 0 | T | C | 0.395 | 2.54E-02 | 9.19E-02 | 2.67E+00 | 2.72E+00 | 7.82E-01 | no | Tmc5 | intron_variant | MODIFIER |
| 7 | 118633976 | AX-170078590 | 0 | T | C | 0.395 | 2.54E-02 | 9.19E-02 | 2.67E+00 | 2.72E+00 | 7.82E-01 | no | Tmc5 | synonymous_variant | LOW |
| 7 | 118644425 | AX-170275188 | 0 | T | C | 0.395 | 2.54E-02 | 9.19E-02 | 2.67E+00 | 2.72E+00 | 7.82E-01 | no | Tmc5 | intron_variant | MODIFIER |
| 7 | 118647286 | AX-170424922 | 0 | A | G | 0.395 | 2.54E-02 | 9.19E-02 | 2.67E+00 | 2.72E+00 | 7.82E-01 | no | Tmc5 | intron_variant | MODIFIER |
| 7 | 118652928 | AX-170078591 | 0 | G | A | 0.395 | 2.54E-02 | 9.19E-02 | 2.67E+00 | 2.72E+00 | 7.82E-01 | no | Tmc5 | intron_variant | MODIFIER |
| 7 | 118654432 | AX-170186901 | 0 | A | T | 0.395 | 2.54E-02 | 9.19E-02 | 2.67E+00 | 2.72E+00 | 7.82E-01 | no | Tmc5 | intron_variant | MODIFIER |
| 7 | 118654844 | AX-170175334 | 0 | G | A | 0.395 | 2.54E-02 | 9.19E-02 | 2.67E+00 | 2.72E+00 | 7.82E-01 | no | Tmc5 | intron_variant | MODIFIER |
| 7 | 118655020 | AX-170303773 | 0 | C | T | 0.395 | 2.54E-02 | 9.19E-02 | 2.67E+00 | 2.72E+00 | 7.82E-01 | no | Tmc5 | intron_variant | MODIFIER |
| 7 | 118659528 | AX-170458021 | 0 | T | A | 0.395 | 2.54E-02 | 9.19E-02 | 2.67E+00 | 2.72E+00 | 7.82E-01 | no | Tmc5 | intron_variant | MODIFIER |
| 7 | 118659613 | AX-170529411 | 0 | T | C | 0.395 | 2.54E-02 | 9.19E-02 | 2.67E+00 | 2.72E+00 | 7.82E-01 | no | Tmc5 | synonymous_variant | LOW |
| 7 | 118660834 | AX-170385214 | 0 | A | G | 0.395 | 2.54E-02 | 9.19E-02 | 2.67E+00 | 2.72E+00 | 7.82E-01 | no | Tmc5 | intron_variant | MODIFIER |
| 7 | 118667880 | AX-170414871 | 0 | C | A | 0.395 | 2.54E-02 | 9.19E-02 | 2.67E+00 | 2.72E+00 | 7.82E-01 | no | Tmc5 | intron_variant | MODIFIER |
| 7 | 118682246 | AX-170269526 | 0 | C | T | 0.395 | 2.54E-02 | 9.19E-02 | 2.67E+00 | 2.72E+00 | 7.82E-01 | no | | intergenic_variant | MODIFIER |
| 7 | 118682309 | AX-170451165 | 0 | A | G | 0.395 | 2.54E-02 | 9.19E-02 | 2.67E+00 | 2.72E+00 | 7.82E-01 | no | | intergenic_variant | MODIFIER |
| 7 | 118682464 | AX-170441668 | 0 | T | C | 0.395 | 2.54E-02 | 9.19E-02 | 2.67E+00 | 2.72E+00 | 7.82E-01 | no | | intergenic_variant | MODIFIER |
| 7 | 118706530 | AX-170397410 | 0 | T | C | 0.395 | 2.54E-02 | 9.19E-02 | 2.67E+00 | 2.72E+00 | 7.82E-01 | no | Gde1 | upstream_gene_variant | MODIFIER |
| 7 | 118721866 | AX-97812061 | 0 | G | A | 0.395 | 2.54E-02 | 9.19E-02 | 2.67E+00 | 2.72E+00 | 7.82E-01 | no | Ccp110 | missense_variant | MODERATE |
| 7 | 118722215 | AX-170100342 | 0 | C | T | 0.395 | 2.54E-02 | 9.19E-02 | 2.67E+00 | 2.72E+00 | 7.82E-01 | no | Ccp110 | synonymous_variant | LOW |
| 7 | 118726035 | AX-170427378 | 0 | A | G | 0.395 | 2.54E-02 | 9.19E-02 | 2.67E+00 | 2.72E+00 | 7.82E-01 | no | Ccp110 | missense_variant | MODERATE |
| 7 | 118728158 | AX-170106189 | 0 | G | A | 0.395 | 2.54E-02 | 9.19E-02 | 2.67E+00 | 2.72E+00 | 7.82E-01 | no | Ccp110 | intron_variant | MODIFIER |
| 7 | 118732285 | AX-170189600 | 0 | T | C | 0.395 | 2.54E-02 | 9.19E-02 | 2.67E+00 | 2.72E+00 | 7.82E-01 | no | Ccp110 | intron_variant | MODIFIER |
| 7 | 118732505 | AX-170485284 | 0 | A | G | 0.395 | 2.54E-02 | 9.19E-02 | 2.67E+00 | 2.72E+00 | 7.82E-01 | no | Ccp110 | intron_variant | MODIFIER |
| 7 | 118737382 | AX-170345398 | 0 | G | A | 0.395 | 2.54E-02 | 9.19E-02 | 2.67E+00 | 2.72E+00 | 7.82E-01 | no | Ccp110 | downstream_gene_variant | MODIFIER |
| 7 | 118741403 | AX-170490478 | 0 | C | T | 0.395 | 2.54E-02 | 9.19E-02 | 2.67E+00 | 2.72E+00 | 7.82E-01 | no | Ccp110 | downstream_gene_variant | MODIFIER |
| 7 | 118743142 | AX-170475712 | 0 | G | A | 0.395 | 2.54E-02 | 9.19E-02 | 2.67E+00 | 2.72E+00 | 7.82E-01 | no | Vps35l | intron_variant | MODIFIER |
| 7 | 118748108 | AX-170281049 | 0 | C | T | 0.395 | 2.54E-02 | 9.19E-02 | 2.67E+00 | 2.72E+00 | 7.82E-01 | no | Vps35l | intron_variant | MODIFIER |
| 7 | 118749893 | AX-170261756 | 0 | C | T | 0.395 | 2.54E-02 | 9.19E-02 | 2.67E+00 | 2.72E+00 | 7.82E-01 | no | Vps35l | intron_variant | MODIFIER |
| 7 | 118764675 | AX-170282581 | 0 | G | T | 0.395 | 2.54E-02 | 9.19E-02 | 2.67E+00 | 2.72E+00 | 7.82E-01 | no | Vps35l | 5_prime_UTR_variant | MODIFIER |
| 7 | 118775651 | AX-170405613 | 0 | G | A | 0.395 | 2.54E-02 | 9.19E-02 | 2.67E+00 | 2.72E+00 | 7.82E-01 | no | Vps35l | intron_variant | MODIFIER |
| 7 | 118777060 | AX-170232940 | 0 | C | G | 0.395 | 2.54E-02 | 9.19E-02 | 2.67E+00 | 2.72E+00 | 7.82E-01 | no | Vps35l | intron_variant | MODIFIER |
| 7 | 118778246 | AX-170219632 | 0 | G | C | 0.395 | 2.54E-02 | 9.19E-02 | 2.67E+00 | 2.72E+00 | 7.82E-01 | no | Vps35l | intron_variant | MODIFIER |
| 7 | 118778276 | AX-170177726 | 0 | C | T | 0.395 | 2.54E-02 | 9.19E-02 | 2.67E+00 | 2.72E+00 | 7.82E-01 | no | Vps35l | intron_variant | MODIFIER |
| 7 | 118780969 | AX-170330177 | 0 | G | A | 0.395 | 2.54E-02 | 9.19E-02 | 2.67E+00 | 2.72E+00 | 7.82E-01 | no | Vps35l | intron_variant | MODIFIER |
| 7 | 118788362 | AX-170272474 | 0 | A | T | 0.395 | 2.54E-02 | 9.19E-02 | 2.67E+00 | 2.72E+00 | 7.82E-01 | no | Vps35l | intron_variant | MODIFIER |
| 7 | 118790872 | AX-170252966 | 0 | C | A | 0.395 | 2.54E-02 | 9.19E-02 | 2.67E+00 | 2.72E+00 | 7.82E-01 | no | Vps35l | intron_variant | MODIFIER |
| 7 | 118794453 | AX-170167835 | 0 | A | C | 0.395 | 2.54E-02 | 9.19E-02 | 2.67E+00 | 2.72E+00 | 7.82E-01 | no | Vps35l | intron_variant | MODIFIER |
| 7 | 118794797 | AX-170180168 | 0 | T | C | 0.395 | 2.54E-02 | 9.19E-02 | 2.67E+00 | 2.72E+00 | 7.82E-01 | no | Vps35l | intron_variant | MODIFIER |
| 7 | 118795142 | AX-170295234 | 0 | A | C | 0.395 | 2.54E-02 | 9.19E-02 | 2.67E+00 | 2.72E+00 | 7.82E-01 | no | Vps35l | intron_variant | MODIFIER |
| 7 | 118814582 | AX-170442899 | 0 | G | A | 0.395 | 2.54E-02 | 9.19E-02 | 2.67E+00 | 2.72E+00 | 7.82E-01 | no | Vps35l | intron_variant | MODIFIER |
| 7 | 118824427 | AX-170360269 | 0 | T | C | 0.395 | 2.54E-02 | 9.19E-02 | 2.67E+00 | 2.72E+00 | 7.82E-01 | no | Vps35l | intron_variant | MODIFIER |
| 7 | 118824749 | AX-170222652 | 0 | T | C | 0.395 | 2.54E-02 | 9.19E-02 | 2.67E+00 | 2.72E+00 | 7.82E-01 | no | Vps35l | intron_variant | MODIFIER |
| 7 | 118834802 | AX-170301657 | 0 | G | A | 0.395 | 2.54E-02 | 9.19E-02 | 2.67E+00 | 2.72E+00 | 7.82E-01 | no | Vps35l | intron_variant | MODIFIER |
| 7 | 118837219 | AX-170171891 | 0 | T | C | 0.395 | 2.54E-02 | 9.19E-02 | 2.67E+00 | 2.72E+00 | 7.82E-01 | no | Vps35l | intron_variant | MODIFIER |
| 7 | 118838159 | AX-170327350 | 0 | C | T | 0.395 | 2.54E-02 | 9.19E-02 | 2.67E+00 | 2.72E+00 | 7.82E-01 | no | Vps35l | synonymous_variant | LOW |

| | | | | | | | | | | | | | | | |
|---|---|---|---|---|---|---|---|---|---|---|---|---|---|---|---|
| 7 | 118843310 | AX-170102431 | 0 | T | C | 0.395 | 2.54E-02 | 9.19E-02 | 2.67E+00 | 2.72E+00 | 7.82E-01 | no | Knop1 | downstream_gene_variant | MODIFIER |
| 7 | 118865717 | AX-170429157 | 0 | G | A | 0.395 | 2.54E-02 | 9.19E-02 | 2.67E+00 | 2.72E+00 | 7.82E-01 | no | Iqck | intron_variant | MODIFIER |
| 7 | 118868711 | AX-170153397 | 0 | G | T | 0.395 | 2.54E-02 | 9.19E-02 | 2.67E+00 | 2.72E+00 | 7.82E-01 | no | Iqck | intron_variant | MODIFIER |
| 7 | 118873619 | AX-170312738 | 0 | C | T | 0.395 | 2.54E-02 | 9.19E-02 | 2.67E+00 | 2.72E+00 | 7.82E-01 | no | Iqck | intron_variant | MODIFIER |
| 7 | 118879784 | AX-170504026 | 0 | T | C | 0.395 | 2.54E-02 | 9.19E-02 | 2.67E+00 | 2.72E+00 | 7.82E-01 | no | Iqck | intron_variant | MODIFIER |
| 7 | 118903268 | AX-170116529 | 0 | C | T | 0.395 | 2.54E-02 | 9.19E-02 | 2.67E+00 | 2.72E+00 | 7.82E-01 | no | Iqck | intron_variant | MODIFIER |
| 7 | 118903784 | AX-170506073 | 0 | A | G | 0.395 | 2.54E-02 | 9.19E-02 | 2.67E+00 | 2.72E+00 | 7.82E-01 | no | Iqck | intron_variant | MODIFIER |
| 7 | 118912281 | AX-170181863 | 0 | C | A | 0.395 | 2.54E-02 | 9.19E-02 | 2.67E+00 | 2.72E+00 | 7.82E-01 | no | Iqck | intron_variant | MODIFIER |
| 7 | 118933331 | AX-170380365 | 0 | A | G | 0.395 | 2.54E-02 | 9.19E-02 | 2.67E+00 | 2.72E+00 | 7.82E-01 | no | Iqck | intron_variant | MODIFIER |
| 7 | 118944847 | AX-170452242 | 0 | T | C | 0.395 | 2.54E-02 | 9.19E-02 | 2.67E+00 | 2.72E+00 | 7.82E-01 | no | Iqck | intron_variant | MODIFIER |
| 7 | 118944994 | AX-170207338 | 0 | T | C | 0.395 | 2.54E-02 | 9.19E-02 | 2.67E+00 | 2.72E+00 | 7.82E-01 | no | Iqck | intron_variant | MODIFIER |
| 7 | 118945637 | AX-170078603 | 0 | T | C | 0.395 | 2.54E-02 | 9.19E-02 | 2.67E+00 | 2.72E+00 | 7.82E-01 | no | Iqck | intron_variant | MODIFIER |
| 7 | 118945693 | AX-170078604 | 0 | A | G | 0.395 | 2.54E-02 | 9.19E-02 | 2.67E+00 | 2.72E+00 | 7.82E-01 | no | Iqck | intron_variant | MODIFIER |
| 7 | 118950219 | AX-170078605 | 0 | G | A | 0.395 | 2.54E-02 | 9.19E-02 | 2.67E+00 | 2.72E+00 | 7.82E-01 | no | Iqck | intron_variant | MODIFIER |
| 7 | 118962207 | AX-170353290 | 0 | G | T | 0.395 | 2.54E-02 | 9.19E-02 | 2.67E+00 | 2.72E+00 | 7.82E-01 | no | Iqck | intron_variant | MODIFIER |
| 7 | 118971679 | AX-170078602 | 0 | C | T | 0.395 | 2.54E-02 | 9.19E-02 | 2.67E+00 | 2.72E+00 | 7.82E-01 | no | Gprc5b | downstream_gene_variant | MODIFIER |
| 7 | 118974474 | AX-170380266 | 0 | T | C | 0.395 | 2.54E-02 | 9.19E-02 | 2.67E+00 | 2.72E+00 | 7.82E-01 | no | Gprc5b | 3_prime_UTR_variant | MODIFIER |
| 7 | 118980329 | AX-170029844 | 0 | G | A | 0.395 | 2.54E-02 | 9.19E-02 | 2.67E+00 | 2.72E+00 | 7.82E-01 | no | Gprc5b | intron_variant | MODIFIER |
| 7 | 118984683 | AX-170160282 | 0 | A | G | 0.395 | 2.54E-02 | 9.19E-02 | 2.67E+00 | 2.72E+00 | 7.82E-01 | no | Gprc5b | intron_variant | MODIFIER |
| 7 | 118984833 | AX-170296973 | 0 | G | A | 0.395 | 2.54E-02 | 9.19E-02 | 2.67E+00 | 2.72E+00 | 7.82E-01 | no | Gprc5b | intron_variant | MODIFIER |
| 7 | 119010648 | AX-170474673 | 0 | G | A | 0.395 | 2.54E-02 | 9.19E-02 | 2.67E+00 | 2.72E+00 | 7.82E-01 | no | | intergenic_variant | MODIFIER |
| 7 | 119013880 | AX-170460526 | 0 | T | C | 0.395 | 2.54E-02 | 9.19E-02 | 2.67E+00 | 2.72E+00 | 7.82E-01 | no | | intergenic_variant | MODIFIER |
| 7 | 119048632 | AX-170253529 | 0 | T | C | 0.395 | 2.54E-02 | 9.19E-02 | 2.67E+00 | 2.72E+00 | 7.82E-01 | no | | intergenic_variant | MODIFIER |
| 7 | 119064680 | AX-170231581 | 0 | A | G | 0.395 | 2.54E-02 | 9.19E-02 | 2.67E+00 | 2.72E+00 | 7.82E-01 | no | | intergenic_variant | MODIFIER |
| 7 | 119076725 | AX-97808387 | 0 | C | A | 0.395 | 2.54E-02 | 9.19E-02 | 2.67E+00 | 2.72E+00 | 7.82E-01 | no | | regulatory_region_variant | MODIFIER |
| 7 | 119086592 | AX-170104912 | 0 | A | G | 0.395 | 2.54E-02 | 9.19E-02 | 2.67E+00 | 2.72E+00 | 7.82E-01 | no | | intergenic_variant | MODIFIER |
| 7 | 119086621 | AX-170319033 | 0 | T | C | 0.395 | 2.54E-02 | 9.19E-02 | 2.67E+00 | 2.72E+00 | 7.82E-01 | no | | intergenic_variant | MODIFIER |
| 7 | 119090538 | AX-170273750 | 0 | G | T | 0.395 | 2.54E-02 | 9.19E-02 | 2.67E+00 | 2.72E+00 | 7.82E-01 | no | | intergenic_variant | MODIFIER |
| 7 | 119090593 | AX-170143387 | 0 | T | C | 0.395 | 2.54E-02 | 9.19E-02 | 2.67E+00 | 2.72E+00 | 7.82E-01 | no | | intergenic_variant | MODIFIER |
| 7 | 119095703 | AX-170378122 | 0 | G | A | 0.395 | 2.54E-02 | 9.19E-02 | 2.67E+00 | 2.72E+00 | 7.82E-01 | no | | intergenic_variant | MODIFIER |
| 7 | 119117379 | AX-170189690 | 0 | C | G | 0.395 | 2.54E-02 | 9.19E-02 | 2.67E+00 | 2.72E+00 | 7.82E-01 | no | | intergenic_variant | MODIFIER |
| 7 | 119119942 | AX-170445522 | 0 | C | T | 0.395 | 2.54E-02 | 9.19E-02 | 2.67E+00 | 2.72E+00 | 7.82E-01 | no | | intergenic_variant | MODIFIER |
| 7 | 119136130 | AX-97828320 | 0 | T | C | 0.395 | 2.54E-02 | 9.19E-02 | 2.67E+00 | 2.72E+00 | 7.82E-01 | no | Gpr139 | downstream_gene_variant | MODIFIER |
| 7 | 119136242 | AX-170513307 | 0 | C | T | 0.395 | 2.54E-02 | 9.19E-02 | 2.67E+00 | 2.72E+00 | 7.82E-01 | no | Gpr139 | downstream_gene_variant | MODIFIER |
| 7 | 119142737 | AX-168413045 | 0 | C | T | 0.395 | 2.54E-02 | 9.19E-02 | 2.67E+00 | 2.72E+00 | 7.82E-01 | no | Gpr139 | 3_prime_UTR_variant | MODIFIER |
| 7 | 119145106 | AX-170455787 | 0 | C | G | 0.395 | 2.54E-02 | 9.19E-02 | 2.67E+00 | 2.72E+00 | 7.82E-01 | no | Gpr139 | synonymous_variant | LOW |
| 7 | 119149385 | AX-170485568 | 0 | C | T | 0.395 | 2.54E-02 | 9.19E-02 | 2.67E+00 | 2.72E+00 | 7.82E-01 | no | Gpr139 | intron_variant | MODIFIER |
| 7 | 119149567 | AX-170388568 | 0 | C | T | 0.395 | 2.54E-02 | 9.19E-02 | 2.67E+00 | 2.72E+00 | 7.82E-01 | no | Gpr139 | intron_variant | MODIFIER |
| 7 | 119158142 | AX-170459245 | 0 | A | C | 0.395 | 2.54E-02 | 9.19E-02 | 2.67E+00 | 2.72E+00 | 7.82E-01 | no | Gpr139 | intron_variant | MODIFIER |
| 7 | 119160363 | AX-170490632 | 0 | G | A | 0.395 | 2.54E-02 | 9.19E-02 | 2.67E+00 | 2.72E+00 | 7.82E-01 | no | Gpr139 | intron_variant | MODIFIER |
| 7 | 119160823 | AX-170277947 | 0 | A | G | 0.395 | 2.54E-02 | 9.19E-02 | 2.67E+00 | 2.72E+00 | 7.82E-01 | no | Gpr139 | intron_variant | MODIFIER |
| 7 | 119161594 | AX-170304074 | 0 | G | A | 0.395 | 2.54E-02 | 9.19E-02 | 2.67E+00 | 2.72E+00 | 7.82E-01 | no | Gpr139 | intron_variant | MODIFIER |
| 7 | 119169137 | AX-97844888 | 0 | A | G | 0.395 | 2.54E-02 | 9.19E-02 | 2.67E+00 | 2.72E+00 | 7.82E-01 | no | Gpr139 | intron_variant | MODIFIER |
| 7 | 119172395 | AX-170437519 | 0 | C | T | 0.395 | 2.54E-02 | 9.19E-02 | 2.67E+00 | 2.72E+00 | 7.82E-01 | no | Gpr139 | intron_variant | MODIFIER |
| 7 | 119172440 | AX-170316941 | 0 | C | T | 0.395 | 2.54E-02 | 9.19E-02 | 2.67E+00 | 2.72E+00 | 7.82E-01 | no | Gpr139 | intron_variant | MODIFIER |
| 7 | 119172578 | AX-170240566 | 0 | T | C | 0.395 | 2.54E-02 | 9.19E-02 | 2.67E+00 | 2.72E+00 | 7.82E-01 | no | Gpr139 | intron_variant | MODIFIER |
| 7 | 119173323 | AX-170389488 | 0 | A | G | 0.395 | 2.54E-02 | 9.19E-02 | 2.67E+00 | 2.72E+00 | 7.82E-01 | no | Gpr139 | intron_variant | MODIFIER |
| 2 | 163763633 | AX-170065500 | 0 | G | A | 0.158 | 4.42E-02 | 1.64E-01 | 2.66E+00 | 2.76E+00 | 7.82E-01 | no | | intergenic_variant | MODIFIER |
| 15 | 92528335 | AX-170180713 | 2 | T | C | 0.1 | 4.40E-02 | 1.57E-01 | 2.61E+00 | 2.71E+00 | 7.82E-01 | no | Pdzrn4 | intron_variant | MODIFIER |
| 8 | 52111403 | AX-97850736 | 0 | A | C | 0.158 | -3.99E-02 | 1.40E-01 | 2.62E+00 | 2.68E+00 | 7.82E-01 | no | | intergenic_variant | MODIFIER |
| 1 | 62640810 | AX-170360413 | 0 | T | C | 0.096 | 5.74E-02 | 2.12E-01 | 2.66E+00 | 2.75E+00 | 7.82E-01 | no | Pard3b | downstream_gene_variant | MODIFIER |
| 1 | 155720899 | AX-170362744 | 0 | G | A | 0.219 | 4.00E-02 | 1.43E-01 | 2.65E+00 | 2.69E+00 | 7.82E-01 | no | Lhx4 | intron_variant | MODIFIER |
| 1 | 155748114 | AX-170143203 | 0 | T | C | 0.219 | 4.00E-02 | 1.43E-01 | 2.65E+00 | 2.69E+00 | 7.82E-01 | no | Lhx4 | intron_variant | MODIFIER |
| 1 | 155772284 | AX-170430313 | 0 | A | G | 0.219 | 4.00E-02 | 1.43E-01 | 2.65E+00 | 2.69E+00 | 7.82E-01 | no | Qsox1 | downstream_gene_variant | MODIFIER |

| | | | | | | | | | | | | | | | |
|---|---|---|---|---|---|---|---|---|---|---|---|---|---|---|---|
| 1 | 155774184 | AX-97806041 | 0 | T | C | 0.219 | 4.00E-02 | 1.43E-01 | 2.65E+00 | 2.69E+00 | 7.82E-01 | no | Qsox1 | downstream_gene_variant | MODIFIER |
| 1 | 155778951 | AX-170283758 | 0 | G | A | 0.219 | 4.00E-02 | 1.43E-01 | 2.65E+00 | 2.69E+00 | 7.82E-01 | no | Qsox1 | 3_prime_UTR_variant | MODIFIER |
| 1 | 155784183 | AX-170367061 | 0 | C | A | 0.219 | 4.00E-02 | 1.43E-01 | 2.65E+00 | 2.69E+00 | 7.82E-01 | no | Qsox1 | intron_variant | MODIFIER |
| 1 | 155785795 | AX-170270188 | 0 | T | C | 0.219 | 4.00E-02 | 1.43E-01 | 2.65E+00 | 2.69E+00 | 7.82E-01 | no | Qsox1 | intron_variant | MODIFIER |
| 1 | 155804562 | AX-170391392 | 0 | C | G | 0.219 | 4.00E-02 | 1.43E-01 | 2.65E+00 | 2.69E+00 | 7.82E-01 | no | Qsox1 | intron_variant | MODIFIER |
| 1 | 155805057 | AX-170043241 | 0 | G | A | 0.219 | 4.00E-02 | 1.43E-01 | 2.65E+00 | 2.69E+00 | 7.82E-01 | no | Qsox1 | intron_variant | MODIFIER |
| 1 | 155809590 | AX-170043244 | 0 | T | A | 0.219 | 4.00E-02 | 1.43E-01 | 2.65E+00 | 2.69E+00 | 7.82E-01 | no | Qsox1 | intron_variant | MODIFIER |
| 1 | 155809714 | AX-97791612 | 0 | G | A | 0.219 | 4.00E-02 | 1.43E-01 | 2.65E+00 | 2.69E+00 | 7.82E-01 | no | Qsox1 | intron_variant | MODIFIER |
| 1 | 155826467 | AX-170043242 | 0 | C | A | 0.219 | 4.00E-02 | 1.43E-01 | 2.65E+00 | 2.69E+00 | 7.82E-01 | no | Gm37571 | intron_variant,non_coding_transcript_variant | MODIFIER |
| 1 | 155834122 | AX-97815869 | 0 | G | T | 0.219 | 4.00E-02 | 1.43E-01 | 2.65E+00 | 2.69E+00 | 7.82E-01 | no | Gm37571 | intron_variant,non_coding_transcript_variant | MODIFIER |
| 1 | 155844519 | AX-170300003 | 0 | A | T | 0.219 | 4.00E-02 | 1.43E-01 | 2.65E+00 | 2.69E+00 | 7.82E-01 | no | Cep350 | downstream_gene_variant | MODIFIER |
| 1 | 155849570 | AX-170211864 | 0 | A | G | 0.219 | 4.00E-02 | 1.43E-01 | 2.65E+00 | 2.69E+00 | 7.82E-01 | no | Cep350 | intron_variant,non_coding_transcript_variant | MODIFIER |
| 1 | 155852495 | AX-170488485 | 0 | A | G | 0.219 | 4.00E-02 | 1.43E-01 | 2.65E+00 | 2.69E+00 | 7.82E-01 | no | Cep350 | intron_variant,non_coding_transcript_variant | MODIFIER |
| 1 | 155852517 | AX-170277813 | 0 | A | G | 0.219 | 4.00E-02 | 1.43E-01 | 2.65E+00 | 2.69E+00 | 7.82E-01 | no | Cep350 | intron_variant,non_coding_transcript_variant | MODIFIER |
| 1 | 155855144 | AX-170197342 | 0 | A | G | 0.219 | 4.00E-02 | 1.43E-01 | 2.65E+00 | 2.69E+00 | 7.82E-01 | no | Cep350 | intron_variant,non_coding_transcript_variant | MODIFIER |
| 1 | 155860718 | AX-170355615 | 0 | A | G | 0.219 | 4.00E-02 | 1.43E-01 | 2.65E+00 | 2.69E+00 | 7.82E-01 | no | Cep350 | downstream_gene_variant | MODIFIER |
| 1 | 155866308 | AX-170163568 | 0 | T | C | 0.219 | 4.00E-02 | 1.43E-01 | 2.65E+00 | 2.69E+00 | 7.82E-01 | no | Cep350 | intron_variant | MODIFIER |
| 1 | 155872250 | AX-170518914 | 0 | A | C | 0.219 | 4.00E-02 | 1.43E-01 | 2.65E+00 | 2.69E+00 | 7.82E-01 | no | Cep350 | intron_variant | MODIFIER |
| 1 | 155880514 | AX-97799912 | 0 | T | C | 0.219 | 4.00E-02 | 1.43E-01 | 2.65E+00 | 2.69E+00 | 7.82E-01 | no | Cep350 | intron_variant | MODIFIER |
| 1 | 155887820 | AX-170193232 | 0 | T | C | 0.219 | 4.00E-02 | 1.43E-01 | 2.65E+00 | 2.69E+00 | 7.82E-01 | no | Cep350 | intron_variant | MODIFIER |
| 1 | 155889833 | AX-170393051 | 0 | T | C | 0.219 | 4.00E-02 | 1.43E-01 | 2.65E+00 | 2.69E+00 | 7.82E-01 | no | Cep350 | intron_variant | MODIFIER |
| 1 | 155909395 | AX-170235331 | 0 | C | T | 0.219 | 4.00E-02 | 1.43E-01 | 2.65E+00 | 2.69E+00 | 7.82E-01 | no | Cep350 | intron_variant | MODIFIER |
| 1 | 155915140 | AX-170151428 | 0 | C | T | 0.219 | 4.00E-02 | 1.43E-01 | 2.65E+00 | 2.69E+00 | 7.82E-01 | no | Cep350 | intron_variant | MODIFIER |
| 1 | 155928211 | AX-170263725 | 0 | C | T | 0.219 | 4.00E-02 | 1.43E-01 | 2.65E+00 | 2.69E+00 | 7.82E-01 | no | Cep350 | missense_variant | MODERATE |
| 1 | 155928402 | AX-170332529 | 0 | G | C | 0.219 | 4.00E-02 | 1.43E-01 | 2.65E+00 | 2.69E+00 | 7.82E-01 | no | Cep350 | synonymous_variant | LOW |
| 1 | 155938001 | AX-97780606 | 0 | A | G | 0.219 | 4.00E-02 | 1.43E-01 | 2.65E+00 | 2.69E+00 | 7.82E-01 | no | Cep350 | intron_variant | MODIFIER |
| 1 | 155938082 | AX-170242441 | 0 | C | T | 0.219 | 4.00E-02 | 1.43E-01 | 2.65E+00 | 2.69E+00 | 7.82E-01 | no | Cep350 | intron_variant | MODIFIER |
| 1 | 155940096 | AX-170002046 | 0 | G | A | 0.219 | 4.00E-02 | 1.43E-01 | 2.65E+00 | 2.69E+00 | 7.82E-01 | no | Cep350 | intron_variant | MODIFIER |
| 1 | 155951274 | AX-170155707 | 0 | G | A | 0.219 | 4.00E-02 | 1.43E-01 | 2.65E+00 | 2.69E+00 | 7.82E-01 | no | Gm23744 | upstream_gene_variant | MODIFIER |
| 1 | 155959302 | AX-97813842 | 0 | C | T | 0.219 | 4.00E-02 | 1.43E-01 | 2.65E+00 | 2.69E+00 | 7.82E-01 | no | Gm23744 | downstream_gene_variant | MODIFIER |
| 1 | 155983173 | AX-170241246 | 0 | C | T | 0.219 | 4.00E-02 | 1.43E-01 | 2.65E+00 | 2.69E+00 | 7.82E-01 | no | | intergenic_variant | MODIFIER |
| 1 | 155985673 | AX-170143019 | 0 | A | G | 0.219 | 4.00E-02 | 1.43E-01 | 2.65E+00 | 2.69E+00 | 7.82E-01 | no | | intergenic_variant | MODIFIER |
| 1 | 155995327 | AX-170420496 | 0 | T | C | 0.219 | 4.00E-02 | 1.43E-01 | 2.65E+00 | 2.69E+00 | 7.82E-01 | no | | intergenic_variant | MODIFIER |
| 1 | 155996427 | AX-97830914 | 0 | T | G | 0.219 | 4.00E-02 | 1.43E-01 | 2.65E+00 | 2.69E+00 | 7.82E-01 | no | | intergenic_variant | MODIFIER |
| 1 | 155998605 | AX-170531404 | 0 | T | C | 0.219 | 4.00E-02 | 1.43E-01 | 2.65E+00 | 2.69E+00 | 7.82E-01 | no | | intergenic_variant | MODIFIER |
| 1 | 156027160 | AX-97781999 | 0 | A | C | 0.219 | 4.00E-02 | 1.43E-01 | 2.65E+00 | 2.69E+00 | 7.82E-01 | no | Tor1aip1 | intron_variant | MODIFIER |
| 1 | 156049323 | AX-170002045 | 0 | C | G | 0.219 | 4.00E-02 | 1.43E-01 | 2.65E+00 | 2.69E+00 | 7.82E-01 | no | Tor1aip2 | intron_variant | MODIFIER |
| 1 | 156050309 | AX-170152400 | 0 | T | A | 0.219 | 4.00E-02 | 1.43E-01 | 2.65E+00 | 2.69E+00 | 7.82E-01 | no | Tor1aip2 | intron_variant | MODIFIER |
| 1 | 156054179 | AX-170522538 | 0 | C | A | 0.219 | 4.00E-02 | 1.43E-01 | 2.65E+00 | 2.69E+00 | 7.82E-01 | no | Tor1aip2 | intron_variant | MODIFIER |
| 1 | 156057353 | AX-170310860 | 0 | C | T | 0.219 | 4.00E-02 | 1.43E-01 | 2.65E+00 | 2.69E+00 | 7.82E-01 | no | Tor1aip2 | intron_variant | MODIFIER |
| 1 | 156057384 | AX-97811644 | 0 | C | T | 0.219 | 4.00E-02 | 1.43E-01 | 2.65E+00 | 2.69E+00 | 7.82E-01 | no | Tor1aip2 | intron_variant | MODIFIER |
| 1 | 156061765 | AX-170175578 | 0 | A | G | 0.219 | 4.00E-02 | 1.43E-01 | 2.65E+00 | 2.69E+00 | 7.82E-01 | no | Tor1aip2 | missense_variant | MODERATE |
| 1 | 156062093 | AX-170322958 | 0 | C | T | 0.219 | 4.00E-02 | 1.43E-01 | 2.65E+00 | 2.69E+00 | 7.82E-01 | no | Tor1aip2 | synonymous_variant | LOW |
| 1 | 156101102 | AX-170355905 | 0 | C | T | 0.219 | 4.00E-02 | 1.43E-01 | 2.65E+00 | 2.69E+00 | 7.82E-01 | no | Fam163a | intron_variant | MODIFIER |
| 1 | 156106380 | AX-170002047 | 0 | T | C | 0.219 | 4.00E-02 | 1.43E-01 | 2.65E+00 | 2.69E+00 | 7.82E-01 | no | Fam163a | intron_variant | MODIFIER |
| 1 | 156106657 | AX-170304112 | 0 | C | G | 0.219 | 4.00E-02 | 1.43E-01 | 2.65E+00 | 2.69E+00 | 7.82E-01 | no | Fam163a | intron_variant | MODIFIER |
| 1 | 156106738 | AX-97843837 | 0 | A | C | 0.219 | 4.00E-02 | 1.43E-01 | 2.65E+00 | 2.69E+00 | 7.82E-01 | no | Fam163a | intron_variant | MODIFIER |
| 1 | 156185032 | AX-170165720 | 0 | G | T | 0.219 | 4.00E-02 | 1.43E-01 | 2.65E+00 | 2.69E+00 | 7.82E-01 | no | Fam163a | intron_variant | MODIFIER |
| 1 | 156185957 | AX-170439305 | 0 | G | A | 0.219 | 4.00E-02 | 1.43E-01 | 2.65E+00 | 2.69E+00 | 7.82E-01 | no | Fam163a | intron_variant | MODIFIER |
| 1 | 156186989 | AX-170174452 | 0 | A | G | 0.219 | 4.00E-02 | 1.43E-01 | 2.65E+00 | 2.69E+00 | 7.82E-01 | no | Fam163a | intron_variant | MODIFIER |
| 1 | 156187344 | AX-170443988 | 0 | T | C | 0.219 | 4.00E-02 | 1.43E-01 | 2.65E+00 | 2.69E+00 | 7.82E-01 | no | Fam163a | intron_variant | MODIFIER |
| 1 | 156191091 | AX-170494376 | 0 | T | G | 0.219 | 4.00E-02 | 1.43E-01 | 2.65E+00 | 2.69E+00 | 7.82E-01 | no | Fam163a | intron_variant | MODIFIER |
| 1 | 156212020 | AX-170322294 | 0 | C | T | 0.219 | 4.00E-02 | 1.43E-01 | 2.65E+00 | 2.69E+00 | 7.82E-01 | no | | regulatory_region_variant | MODIFIER |
| 1 | 156219116 | AX-170430455 | 0 | T | C | 0.219 | 4.00E-02 | 1.43E-01 | 2.65E+00 | 2.69E+00 | 7.82E-01 | no | | regulatory_region_variant | MODIFIER |

| | | | | | | | | | | | | | | | |
|---|---|---|---|---|---|---|---|---|---|---|---|---|---|---|---|
| 1 | 156225080 | AX-170476740 | 0 | G | A | 0.219 | 4.00E-02 | 1.43E-01 | 2.65E+00 | 2.69E+00 | 7.82E-01 | no | Gm38113 | downstream_gene_variant | MODIFIER |
| 1 | 156241089 | AX-97800963 | 0 | C | T | 0.219 | 4.00E-02 | 1.43E-01 | 2.65E+00 | 2.69E+00 | 7.82E-01 | no | | regulatory_region_variant | MODIFIER |
| 1 | 156242305 | AX-170386503 | 0 | G | A | 0.219 | 4.00E-02 | 1.43E-01 | 2.65E+00 | 2.69E+00 | 7.82E-01 | no | | regulatory_region_variant | MODIFIER |
| 1 | 156242870 | AX-170212975 | 0 | T | G | 0.219 | 4.00E-02 | 1.43E-01 | 2.65E+00 | 2.69E+00 | 7.82E-01 | no | | intergenic_variant | MODIFIER |
| 1 | 156245435 | AX-170193374 | 0 | C | T | 0.219 | 4.00E-02 | 1.43E-01 | 2.65E+00 | 2.69E+00 | 7.82E-01 | no | | intergenic_variant | MODIFIER |
| 1 | 156262597 | AX-170501451 | 0 | C | A | 0.219 | 4.00E-02 | 1.43E-01 | 2.65E+00 | 2.69E+00 | 7.82E-01 | no | Tdrd5 | intron_variant | MODIFIER |
| 1 | 156262881 | AX-170478884 | 0 | G | C | 0.219 | 4.00E-02 | 1.43E-01 | 2.65E+00 | 2.69E+00 | 7.82E-01 | no | Tdrd5 | missense_variant | MODERATE |
| 1 | 156265387 | AX-170425421 | 0 | G | A | 0.219 | 4.00E-02 | 1.43E-01 | 2.65E+00 | 2.69E+00 | 7.82E-01 | no | Tdrd5 | intron_variant | MODIFIER |
| 1 | 156266000 | AX-97817413 | 0 | A | G | 0.219 | 4.00E-02 | 1.43E-01 | 2.65E+00 | 2.69E+00 | 7.82E-01 | no | Tdrd5 | intron_variant | MODIFIER |
| 1 | 156269920 | AX-170530199 | 0 | T | C | 0.219 | 4.00E-02 | 1.43E-01 | 2.65E+00 | 2.69E+00 | 7.82E-01 | no | Tdrd5 | intron_variant | MODIFIER |
| 1 | 156269964 | AX-170459038 | 0 | T | C | 0.219 | 4.00E-02 | 1.43E-01 | 2.65E+00 | 2.69E+00 | 7.82E-01 | no | Tdrd5 | intron_variant | MODIFIER |
| 1 | 156270646 | AX-170299678 | 0 | C | T | 0.219 | 4.00E-02 | 1.43E-01 | 2.65E+00 | 2.69E+00 | 7.82E-01 | no | Tdrd5 | missense_variant | MODERATE |
| 1 | 156287617 | AX-97852551 | 0 | C | A | 0.219 | 4.00E-02 | 1.43E-01 | 2.65E+00 | 2.69E+00 | 7.82E-01 | no | Tdrd5 | intron_variant | MODIFIER |
| 1 | 156292306 | AX-170126517 | 0 | G | A | 0.219 | 4.00E-02 | 1.43E-01 | 2.65E+00 | 2.69E+00 | 7.82E-01 | no | Tdrd5 | intron_variant | MODIFIER |
| 1 | 156298114 | AX-170320356 | 0 | T | G | 0.219 | 4.00E-02 | 1.43E-01 | 2.65E+00 | 2.69E+00 | 7.82E-01 | no | Tdrd5 | intron_variant | MODIFIER |
| 1 | 156298233 | AX-170114186 | 0 | A | C | 0.219 | 4.00E-02 | 1.43E-01 | 2.65E+00 | 2.69E+00 | 7.82E-01 | no | Tdrd5 | intron_variant | MODIFIER |
| 1 | 156301208 | AX-97782391 | 0 | A | G | 0.219 | 4.00E-02 | 1.43E-01 | 2.65E+00 | 2.69E+00 | 7.82E-01 | no | Tdrd5 | intron_variant | MODIFIER |
| 1 | 156321317 | AX-170176767 | 0 | A | G | 0.219 | 4.00E-02 | 1.43E-01 | 2.65E+00 | 2.69E+00 | 7.82E-01 | no | Nphs2 | intron_variant | MODIFIER |
| 1 | 156322458 | AX-170523827 | 0 | T | C | 0.219 | 4.00E-02 | 1.43E-01 | 2.65E+00 | 2.69E+00 | 7.82E-01 | no | Nphs2 | intron_variant | MODIFIER |
| 1 | 156329965 | AX-170167961 | 0 | A | G | 0.219 | 4.00E-02 | 1.43E-01 | 2.65E+00 | 2.69E+00 | 7.82E-01 | no | Nphs2 | downstream_gene_variant | MODIFIER |
| 1 | 156331477 | AX-170384158 | 0 | C | G | 0.219 | 4.00E-02 | 1.43E-01 | 2.65E+00 | 2.69E+00 | 7.82E-01 | no | Nphs2 | downstream_gene_variant | MODIFIER |
| 1 | 156331574 | AX-170315177 | 0 | T | C | 0.219 | 4.00E-02 | 1.43E-01 | 2.65E+00 | 2.69E+00 | 7.82E-01 | no | Nphs2 | downstream_gene_variant | MODIFIER |
| 1 | 156343494 | AX-97813812 | 0 | T | C | 0.219 | 4.00E-02 | 1.43E-01 | 2.65E+00 | 2.69E+00 | 7.82E-01 | no | Axdnd1 | intron_variant | MODIFIER |
| 1 | 156347329 | AX-170339163 | 0 | G | A | 0.219 | 4.00E-02 | 1.43E-01 | 2.65E+00 | 2.69E+00 | 7.82E-01 | no | Axdnd1 | intron_variant | MODIFIER |
| 1 | 156383898 | AX-170417012 | 0 | C | A | 0.219 | 4.00E-02 | 1.43E-01 | 2.65E+00 | 2.69E+00 | 7.82E-01 | no | Axdnd1 | intron_variant,non_coding_transcript_variant | MODIFIER |
| 1 | 156400202 | AX-170251664 | 0 | T | C | 0.219 | 4.00E-02 | 1.43E-01 | 2.65E+00 | 2.69E+00 | 7.82E-01 | no | Axdnd1 | upstream_gene_variant | MODIFIER |
| 1 | 156411507 | AX-97803532 | 0 | T | C | 0.219 | 4.00E-02 | 1.43E-01 | 2.65E+00 | 2.69E+00 | 7.82E-01 | no | Axdnd1 | intron_variant | MODIFIER |
| 1 | 156425939 | AX-170273076 | 0 | G | A | 0.219 | 4.00E-02 | 1.43E-01 | 2.65E+00 | 2.69E+00 | 7.82E-01 | no | Soat1 | downstream_gene_variant | MODIFIER |
| 1 | 156442776 | AX-170258303 | 0 | G | A | 0.219 | 4.00E-02 | 1.43E-01 | 2.65E+00 | 2.69E+00 | 7.82E-01 | no | Soat1 | intron_variant | MODIFIER |
| 1 | 156443084 | AX-170255863 | 0 | C | T | 0.219 | 4.00E-02 | 1.43E-01 | 2.65E+00 | 2.69E+00 | 7.82E-01 | no | Soat1 | intron_variant | MODIFIER |
| 1 | 156448692 | AX-170382403 | 0 | A | G | 0.219 | 4.00E-02 | 1.43E-01 | 2.65E+00 | 2.69E+00 | 7.82E-01 | no | Soat1 | intron_variant | MODIFIER |
| 1 | 156450015 | AX-170043254 | 0 | G | A | 0.219 | 4.00E-02 | 1.43E-01 | 2.65E+00 | 2.69E+00 | 7.82E-01 | no | Soat1 | intron_variant | MODIFIER |
| 1 | 156457572 | AX-97813810 | 0 | A | G | 0.219 | 4.00E-02 | 1.43E-01 | 2.65E+00 | 2.69E+00 | 7.82E-01 | no | Soat1 | intron_variant | MODIFIER |
| 1 | 156495123 | AX-170002055 | 0 | G | A | 0.219 | 4.00E-02 | 1.43E-01 | 2.65E+00 | 2.69E+00 | 7.82E-01 | no | | intergenic_variant | MODIFIER |
| 1 | 156497086 | AX-170002062 | 0 | C | T | 0.219 | 4.00E-02 | 1.43E-01 | 2.65E+00 | 2.69E+00 | 7.82E-01 | no | | intergenic_variant | MODIFIER |
| 1 | 156500078 | AX-170485176 | 0 | T | G | 0.219 | 4.00E-02 | 1.43E-01 | 2.65E+00 | 2.69E+00 | 7.82E-01 | no | | intergenic_variant | MODIFIER |
| 1 | 156500590 | AX-170260549 | 0 | T | C | 0.219 | 4.00E-02 | 1.43E-01 | 2.65E+00 | 2.69E+00 | 7.82E-01 | no | | intergenic_variant | MODIFIER |
| 1 | 156517257 | AX-170102292 | 0 | G | A | 0.219 | 4.00E-02 | 1.43E-01 | 2.65E+00 | 2.69E+00 | 7.82E-01 | no | | intergenic_variant | MODIFIER |
| 1 | 159604671 | AX-97818712 | 0 | C | T | 0.219 | 4.00E-02 | 1.43E-01 | 2.65E+00 | 2.69E+00 | 7.82E-01 | no | Tnr | intron_variant | MODIFIER |
| 1 | 159617045 | AX-170366992 | 0 | T | G | 0.219 | 4.00E-02 | 1.43E-01 | 2.65E+00 | 2.69E+00 | 7.82E-01 | no | Gm10530 | intron_variant,non_coding_transcript_variant | MODIFIER |
| 1 | 159626751 | AX-170279291 | 0 | G | A | 0.219 | 4.00E-02 | 1.43E-01 | 2.65E+00 | 2.69E+00 | 7.82E-01 | no | Gm10530 | intron_variant,non_coding_transcript_variant | MODIFIER |
| 1 | 159629540 | AX-170196704 | 0 | T | C | 0.219 | 4.00E-02 | 1.43E-01 | 2.65E+00 | 2.69E+00 | 7.82E-01 | no | Gm10530 | intron_variant,non_coding_transcript_variant | MODIFIER |
| 1 | 159641943 | AX-170002114 | 0 | C | T | 0.219 | 4.00E-02 | 1.43E-01 | 2.65E+00 | 2.69E+00 | 7.82E-01 | no | Gm10530 | intron_variant,non_coding_transcript_variant | MODIFIER |
| 1 | 159644731 | AX-170002115 | 0 | C | T | 0.219 | 4.00E-02 | 1.43E-01 | 2.65E+00 | 2.69E+00 | 7.82E-01 | no | Gm10530 | intron_variant,non_coding_transcript_variant | MODIFIER |
| 1 | 160022961 | AX-170512608 | 0 | G | T | 0.219 | 4.00E-02 | 1.43E-01 | 2.65E+00 | 2.69E+00 | 7.82E-01 | no | | intergenic_variant | MODIFIER |
| 1 | 160058871 | AX-170500490 | 0 | A | G | 0.219 | 4.00E-02 | 1.43E-01 | 2.65E+00 | 2.69E+00 | 7.82E-01 | no | 4930523C07Rik | intron_variant,non_coding_transcript_variant | MODIFIER |
| 1 | 160059257 | AX-170451750 | 0 | G | A | 0.219 | 4.00E-02 | 1.43E-01 | 2.65E+00 | 2.69E+00 | 7.82E-01 | no | 4930523C07Rik | intron_variant,non_coding_transcript_variant | MODIFIER |
| 1 | 160061383 | AX-97852468 | 0 | G | A | 0.219 | 4.00E-02 | 1.43E-01 | 2.65E+00 | 2.69E+00 | 7.82E-01 | no | 4930523C07Rik | intron_variant,non_coding_transcript_variant | MODIFIER |
| 1 | 160061395 | AX-170309064 | 0 | T | G | 0.219 | 4.00E-02 | 1.43E-01 | 2.65E+00 | 2.69E+00 | 7.82E-01 | no | 4930523C07Rik | intron_variant,non_coding_transcript_variant | MODIFIER |
| 1 | 161214262 | AX-170481675 | 0 | C | T | 0.219 | 4.00E-02 | 1.43E-01 | 2.65E+00 | 2.69E+00 | 7.82E-01 | no | Gm6185 | intron_variant,non_coding_transcript_variant | MODIFIER |
| 1 | 161986294 | AX-170185141 | 0 | G | A | 0.219 | 4.00E-02 | 1.43E-01 | 2.65E+00 | 2.69E+00 | 7.82E-01 | no | Dnm3 | downstream_gene_variant | MODIFIER |
| 1 | 161991892 | AX-170249561 | 0 | A | G | 0.219 | 4.00E-02 | 1.43E-01 | 2.65E+00 | 2.69E+00 | 7.82E-01 | no | Dnm3 | 3_prime_UTR_variant | MODIFIER |
| 1 | 162023779 | AX-170493577 | 0 | T | G | 0.219 | 4.00E-02 | 1.43E-01 | 2.65E+00 | 2.69E+00 | 7.82E-01 | no | Dnm3 | intron_variant | MODIFIER |
| 1 | 162037124 | AX-170208012 | 0 | C | G | 0.219 | 4.00E-02 | 1.43E-01 | 2.65E+00 | 2.69E+00 | 7.82E-01 | no | Dnm3 | intron_variant | MODIFIER |

| | | | | | | | | | | | | | | | |
|---|---|---|---|---|---|---|---|---|---|---|---|---|---|---|---|
| 1 | 162037144 | AX-97804057 | 0 | G | C | 0.219 | 4.00E-02 | 1.43E-01 | 2.65E+00 | 2.69E+00 | 7.82E-01 | no | Dnm3 | intron_variant | MODIFIER |
| 1 | 162042000 | AX-170412539 | 0 | C | T | 0.219 | 4.00E-02 | 1.43E-01 | 2.65E+00 | 2.69E+00 | 7.82E-01 | no | Dnm3 | intron_variant | MODIFIER |
| 1 | 162049324 | AX-170196696 | 0 | T | C | 0.219 | 4.00E-02 | 1.43E-01 | 2.65E+00 | 2.69E+00 | 7.82E-01 | no | Dnm3 | intron_variant | MODIFIER |
| 1 | 162099092 | AX-170457012 | 0 | A | G | 0.219 | 4.00E-02 | 1.43E-01 | 2.65E+00 | 2.69E+00 | 7.82E-01 | no | Dnm3 | intron_variant | MODIFIER |
| 1 | 162110133 | AX-170385900 | 0 | T | A | 0.219 | 4.00E-02 | 1.43E-01 | 2.65E+00 | 2.69E+00 | 7.82E-01 | no | Dnm3 | intron_variant | MODIFIER |
| 1 | 162128965 | AX-170239877 | 0 | C | T | 0.219 | 4.00E-02 | 1.43E-01 | 2.65E+00 | 2.69E+00 | 7.82E-01 | no | Dnm3 | intron_variant | MODIFIER |
| 1 | 162156922 | AX-97849386 | 0 | T | C | 0.219 | 4.00E-02 | 1.43E-01 | 2.65E+00 | 2.69E+00 | 7.82E-01 | no | Dnm3 | intron_variant | MODIFIER |
| 1 | 162213132 | AX-170091770 | 0 | A | G | 0.219 | 4.00E-02 | 1.43E-01 | 2.65E+00 | 2.69E+00 | 7.82E-01 | no | Dnm3 | intron_variant | MODIFIER |
| 12 | 73260065 | AX-170007042 | 0 | G | T | 0.053 | 6.41E-02 | 2.36E-01 | 2.63E+00 | 2.75E+00 | 7.82E-01 | no | Mnat1 | intron_variant | MODIFIER |
| 18 | 28358039 | AX-97822622 | 0 | T | C | 0.053 | 6.41E-02 | 2.36E-01 | 2.63E+00 | 2.75E+00 | 7.82E-01 | no | | intergenic_variant | MODIFIER |
| 1 | 77560438 | AX-170474948 | 1 | A | G | 0.143 | -3.70E-02 | 1.37E-01 | 2.61E+00 | 2.75E+00 | 7.82E-01 | no | | intergenic_variant | MODIFIER |
| 6 | 146945061 | AX-170350285 | 0 | G | C | 0.079 | -6.03E-02 | 2.23E-01 | 2.67E+00 | 2.76E+00 | 7.82E-01 | no | Ppfibp1 | intron_variant | MODIFIER |
| 7 | 117402160 | AX-170167540 | 0 | G | A | 0.404 | 2.58E-02 | 9.36E-02 | 2.67E+00 | 2.72E+00 | 7.82E-01 | no | Xylt1 | intron_variant | MODIFIER |
| 7 | 117404721 | AX-170314967 | 0 | G | A | 0.404 | 2.58E-02 | 9.36E-02 | 2.67E+00 | 2.72E+00 | 7.82E-01 | no | Xylt1 | intron_variant | MODIFIER |
| 7 | 117407525 | AX-170350946 | 0 | T | C | 0.404 | 2.58E-02 | 9.36E-02 | 2.67E+00 | 2.72E+00 | 7.82E-01 | no | Xylt1 | intron_variant | MODIFIER |
| 7 | 117415159 | AX-170349441 | 0 | A | G | 0.404 | 2.58E-02 | 9.36E-02 | 2.67E+00 | 2.72E+00 | 7.82E-01 | no | Xylt1 | intron_variant | MODIFIER |
| 7 | 117415956 | AX-170332156 | 0 | C | T | 0.404 | 2.58E-02 | 9.36E-02 | 2.67E+00 | 2.72E+00 | 7.82E-01 | no | Xylt1 | intron_variant | MODIFIER |
| 7 | 117417921 | AX-170471062 | 0 | T | C | 0.404 | 2.58E-02 | 9.36E-02 | 2.67E+00 | 2.72E+00 | 7.82E-01 | no | Xylt1 | intron_variant | MODIFIER |
| 7 | 117422283 | AX-170131761 | 0 | C | G | 0.404 | 2.58E-02 | 9.36E-02 | 2.67E+00 | 2.72E+00 | 7.82E-01 | no | Xylt1 | intron_variant | MODIFIER |
| 7 | 117428949 | AX-170112216 | 0 | A | C | 0.404 | 2.58E-02 | 9.36E-02 | 2.67E+00 | 2.72E+00 | 7.82E-01 | no | Xylt1 | intron_variant | MODIFIER |
| 7 | 117431910 | AX-170212036 | 0 | A | C | 0.404 | 2.58E-02 | 9.36E-02 | 2.67E+00 | 2.72E+00 | 7.82E-01 | no | Xylt1 | intron_variant | MODIFIER |
| 1 | 33676221 | AX-97785786 | 0 | T | A | 0.421 | -2.13E-02 | 7.90E-02 | 2.68E+00 | 2.77E+00 | 7.82E-01 | no | Gm38173 | upstream_gene_variant | MODIFIER |
| 7 | 117612399 | AX-170244932 | 1 | G | A | 0.393 | 2.53E-02 | 9.20E-02 | 2.67E+00 | 2.72E+00 | 7.82E-01 | no | Xylt1 | intron_variant | MODIFIER |
| 7 | 117943606 | AX-170029836 | 1 | C | T | 0.393 | 2.53E-02 | 9.20E-02 | 2.67E+00 | 2.72E+00 | 7.82E-01 | no | | intergenic_variant | MODIFIER |
| 4 | 89436350 | AX-170310618 | 0 | C | T | 0.325 | 2.81E-02 | 1.03E-01 | 2.67E+00 | 2.74E+00 | 7.82E-01 | no | Gm12610 | intron_variant,non_coding_transcript_variant | MODIFIER |
| 4 | 93221544 | AX-170471992 | 0 | G | T | 0.325 | 2.81E-02 | 1.03E-01 | 2.67E+00 | 2.74E+00 | 7.82E-01 | no | | intergenic_variant | MODIFIER |
| 12 | 104944870 | AX-170492230 | 1 | G | A | 0.134 | -5.19E-02 | 1.93E-01 | 2.69E+00 | 2.77E+00 | 7.82E-01 | no | Syne3 | intron_variant | MODIFIER |
| 1 | 13340946 | AX-170040420 | 0 | G | A | 0.386 | 2.11E-02 | 7.49E-02 | 2.63E+00 | 2.69E+00 | 7.82E-01 | no | Ncoa2 | intron_variant | MODIFIER |
| 8 | 41064100 | AX-170251770 | 0 | T | G | 0.149 | 4.04E-02 | 1.49E-01 | 2.78E+00 | 2.83E+00 | 7.82E-01 | no | Mtus1 | intron_variant | MODIFIER |
| 8 | 50921346 | AX-97781907 | 0 | A | G | 0.149 | 4.04E-02 | 1.49E-01 | 2.78E+00 | 2.83E+00 | 7.82E-01 | no | C130073E24Rik | upstream_gene_variant | MODIFIER |
| 10 | 38009987 | AX-170473713 | 1 | T | C | 0.045 | 3.26E-02 | 1.22E-01 | 2.58E+00 | 2.76E+00 | 7.83E-01 | no | | intergenic_variant | MODIFIER |
| 7 | 91441392 | AX-170170205 | 0 | A | T | 0.123 | 5.14E-02 | 1.93E-01 | 2.67E+00 | 2.77E+00 | 7.83E-01 | no | Dlg2 | intron_variant | MODIFIER |
| 1 | 51772807 | AX-170366985 | 1 | C | A | 0.098 | -5.99E-02 | 2.20E-01 | 2.68E+00 | 2.74E+00 | 7.83E-01 | no | Myo1b | intron_variant | MODIFIER |
| 5 | 100771178 | AX-170349668 | 1 | A | C | 0.429 | 2.87E-02 | 1.02E-01 | 2.62E+00 | 2.69E+00 | 7.83E-01 | no | Helq | intron_variant | MODIFIER |
| 2 | 117386980 | AX-170534334 | 0 | A | T | 0.123 | 5.22E-02 | 1.92E-01 | 2.68E+00 | 2.74E+00 | 7.83E-01 | no | | intergenic_variant | MODIFIER |
| 18 | 69060839 | AX-170367883 | 1 | G | C | 0.089 | 4.86E-02 | 1.76E-01 | 2.61E+00 | 2.72E+00 | 7.83E-01 | no | | intergenic_variant | MODIFIER |
| 4 | 140903313 | AX-170490821 | 0 | G | C | 0.088 | 4.58E-02 | 1.68E-01 | 2.67E+00 | 2.74E+00 | 7.83E-01 | no | Padi2 | upstream_gene_variant | MODIFIER |
| 4 | 150453207 | AX-170463343 | 0 | A | C | 0.088 | 4.58E-02 | 1.68E-01 | 2.67E+00 | 2.74E+00 | 7.83E-01 | no | Rere | intron_variant | MODIFIER |
| 2 | 161960928 | AX-170019545 | 0 | C | G | 0.158 | -4.73E-02 | 1.72E-01 | 2.63E+00 | 2.73E+00 | 7.83E-01 | no | Ptprt | intron_variant | MODIFIER |
| 1 | 74113109 | AX-170335965 | 0 | G | A | 0.123 | 5.59E-02 | 2.06E-01 | 2.69E+00 | 2.74E+00 | 7.83E-01 | no | Tns1 | intron_variant | MODIFIER |
| 7 | 134890190 | AX-170286455 | 1 | T | C | 0.241 | -2.28E-02 | 8.01E-02 | 2.58E+00 | 2.69E+00 | 7.83E-01 | no | Dock1 | intron_variant | MODIFIER |
| 1 | 156482689 | AX-97822694 | 1 | C | T | 0.223 | 3.95E-02 | 1.41E-01 | 2.64E+00 | 2.68E+00 | 7.83E-01 | no | | intergenic_variant | MODIFIER |
| 1 | 175871705 | AX-170295269 | 0 | G | T | 0.228 | 3.21E-02 | 1.12E-01 | 2.60E+00 | 2.68E+00 | 7.83E-01 | no | | intergenic_variant | MODIFIER |
| 4 | 118069025 | AX-170149874 | 1 | A | T | 0.214 | -2.64E-02 | 1.04E-01 | 2.71E+00 | 2.85E+00 | 7.83E-01 | no | St3gal3 | intron_variant | MODIFIER |
| 5 | 5897715 | AX-168379250 | 1 | T | C | 0.429 | 1.74E-02 | 6.61E-02 | 2.64E+00 | 2.77E+00 | 7.83E-01 | no | | intergenic_variant | MODIFIER |
| 7 | 58790264 | AX-170346201 | 0 | G | T | 0.43 | -2.59E-02 | 9.70E-02 | 2.68E+00 | 2.77E+00 | 7.83E-01 | no | Atp10a | intron_variant | MODIFIER |
| 8 | 77972059 | AX-170139947 | 0 | T | C | 0.193 | -3.25E-02 | 1.18E-01 | 2.63E+00 | 2.72E+00 | 7.84E-01 | no | | intergenic_variant | MODIFIER |
| 16 | 94756932 | AX-170379841 | 0 | G | A | 0.404 | -2.31E-02 | 8.03E-02 | 2.59E+00 | 2.68E+00 | 7.84E-01 | no | Kcnj6 | intron_variant | MODIFIER |
| 7 | 116487302 | AX-170078536 | 0 | A | G | 0.421 | 2.43E-02 | 8.76E-02 | 2.64E+00 | 2.70E+00 | 7.84E-01 | no | Gm45031 | downstream_gene_variant | MODIFIER |
| 18 | 57993885 | AX-170418989 | 1 | C | A | 0.411 | 2.87E-02 | 9.45E-02 | 2.55E+00 | 2.62E+00 | 7.84E-01 | no | | intergenic_variant | MODIFIER |
| 8 | 81445005 | AX-170316140 | 1 | C | T | 0.205 | 3.51E-02 | 1.30E-01 | 2.68E+00 | 2.74E+00 | 7.84E-01 | no | Inpp4b | intron_variant,NMD_transcript_variant | MODIFIER |
| 8 | 82079665 | AX-170477818 | 1 | T | G | 0.205 | 3.51E-02 | 1.30E-01 | 2.68E+00 | 2.74E+00 | 7.84E-01 | no | Inpp4b | intron_variant | MODIFIER |
| 18 | 70683713 | AX-170207362 | 1 | C | G | 0.268 | -2.68E-02 | 9.82E-02 | 2.66E+00 | 2.73E+00 | 7.84E-01 | no | | regulatory_region_variant | MODIFIER |
| 6 | 139581085 | AX-170426578 | 1 | T | C | 0.054 | 7.99E-02 | 2.95E-01 | 2.67E+00 | 2.74E+00 | 7.84E-01 | no | 4930443G03Rik | upstream_gene_variant | MODIFIER |

| | | | | | | | | | | | | | | | |
|---|---|---|---|---|---|---|---|---|---|---|---|---|---|---|---|
| 8 | 66290109 | AX-170408225 | 0 | T | G | 0.14 | -4.21E-02 | 1.47E-01 | 2.60E+00 | 2.67E+00 | 7.84E-01 | no | MAR1 | intron_variant | MODIFIER |
| 8 | 37760258 | AX-170079792 | 1 | T | A | 0.152 | 3.96E-02 | 1.47E-01 | 2.74E+00 | 2.78E+00 | 7.84E-01 | no | Sgcz | intron_variant | MODIFIER |
| 8 | 40124803 | AX-170208684 | 1 | T | C | 0.152 | 3.96E-02 | 1.47E-01 | 2.74E+00 | 2.78E+00 | 7.84E-01 | no | | intergenic_variant | MODIFIER |
| 8 | 40654151 | AX-170102060 | 1 | T | C | 0.152 | 3.96E-02 | 1.47E-01 | 2.74E+00 | 2.78E+00 | 7.84E-01 | no | | regulatory_region_variant | MODIFIER |
| 8 | 43345465 | AX-97823566 | 1 | G | A | 0.152 | 3.96E-02 | 1.47E-01 | 2.74E+00 | 2.78E+00 | 7.84E-01 | no | | intergenic_variant | MODIFIER |
| 15 | 82994125 | AX-170474465 | 0 | A | G | 0.553 | 2.25E-02 | 7.89E-02 | 2.60E+00 | 2.68E+00 | 7.84E-01 | no | Nfam1 | downstream_gene_variant | MODIFIER |
| 9 | 77685737 | AX-170468437 | 0 | C | T | 0.193 | -3.54E-02 | 1.29E-01 | 2.67E+00 | 2.71E+00 | 7.84E-01 | no | | regulatory_region_variant | MODIFIER |
| 13 | 36538943 | AX-170426958 | 1 | G | A | 0.179 | -3.44E-02 | 1.23E-01 | 2.62E+00 | 2.69E+00 | 7.84E-01 | no | Fars2 | downstream_gene_variant | MODIFIER |
| 8 | 90321722 | AX-97826350 | 0 | G | C | 0.184 | -3.32E-02 | 1.20E-01 | 2.63E+00 | 2.71E+00 | 7.84E-01 | no | Tox3 | intron_variant | MODIFIER |
| 8 | 90962789 | AX-170274403 | 0 | G | A | 0.184 | -3.32E-02 | 1.20E-01 | 2.63E+00 | 2.71E+00 | 7.84E-01 | no | Chd9 | intron_variant | MODIFIER |
| 8 | 90965899 | AX-170513043 | 0 | A | G | 0.184 | -3.32E-02 | 1.20E-01 | 2.63E+00 | 2.71E+00 | 7.84E-01 | no | Chd9 | intron_variant | MODIFIER |
| 8 | 90981316 | AX-170197523 | 0 | G | A | 0.184 | -3.32E-02 | 1.20E-01 | 2.63E+00 | 2.71E+00 | 7.84E-01 | no | Chd9 | intron_variant | MODIFIER |
| 8 | 90984600 | AX-170096361 | 0 | T | C | 0.184 | -3.32E-02 | 1.20E-01 | 2.63E+00 | 2.71E+00 | 7.84E-01 | no | Chd9 | intron_variant | MODIFIER |
| 8 | 90988907 | AX-170425127 | 0 | A | C | 0.184 | -3.32E-02 | 1.20E-01 | 2.63E+00 | 2.71E+00 | 7.84E-01 | no | Chd9 | intron_variant | MODIFIER |
| 8 | 91014957 | AX-170522442 | 0 | G | C | 0.184 | -3.32E-02 | 1.20E-01 | 2.63E+00 | 2.71E+00 | 7.84E-01 | no | Chd9 | intron_variant | MODIFIER |
| 8 | 91018509 | AX-170031252 | 0 | C | T | 0.184 | -3.32E-02 | 1.20E-01 | 2.63E+00 | 2.71E+00 | 7.84E-01 | no | Chd9 | intron_variant | MODIFIER |
| 8 | 91022463 | AX-170535565 | 0 | A | T | 0.184 | -3.32E-02 | 1.20E-01 | 2.63E+00 | 2.71E+00 | 7.84E-01 | no | Chd9 | intron_variant | MODIFIER |
| 8 | 91027886 | AX-170283875 | 0 | T | C | 0.184 | -3.32E-02 | 1.20E-01 | 2.63E+00 | 2.71E+00 | 7.84E-01 | no | Chd9 | intron_variant | MODIFIER |
| 8 | 91050809 | AX-170031249 | 0 | T | C | 0.184 | -3.32E-02 | 1.20E-01 | 2.63E+00 | 2.71E+00 | 7.84E-01 | no | Chd9 | intron_variant | MODIFIER |
| 8 | 91055612 | AX-170031250 | 0 | T | C | 0.184 | -3.32E-02 | 1.20E-01 | 2.63E+00 | 2.71E+00 | 7.84E-01 | no | Chd9 | downstream_gene_variant | MODIFIER |
| 8 | 91077215 | AX-170296372 | 0 | A | G | 0.184 | -3.32E-02 | 1.20E-01 | 2.63E+00 | 2.71E+00 | 7.84E-01 | no | Rbl2 | intron_variant | MODIFIER |
| 8 | 91090398 | AX-170417759 | 0 | C | G | 0.184 | -3.32E-02 | 1.20E-01 | 2.63E+00 | 2.71E+00 | 7.84E-01 | no | Rbl2 | intron_variant | MODIFIER |
| 8 | 91092171 | AX-170175899 | 0 | A | C | 0.184 | -3.32E-02 | 1.20E-01 | 2.63E+00 | 2.71E+00 | 7.84E-01 | no | Rbl2 | intron_variant | MODIFIER |
| 8 | 91098269 | AX-170080767 | 0 | A | G | 0.184 | -3.32E-02 | 1.20E-01 | 2.63E+00 | 2.71E+00 | 7.84E-01 | no | Rbl2 | intron_variant | MODIFIER |
| 8 | 91162135 | AX-170533692 | 0 | T | C | 0.184 | -3.32E-02 | 1.20E-01 | 2.63E+00 | 2.71E+00 | 7.84E-01 | no | Aktip | intron_variant | MODIFIER |
| 8 | 91507617 | AX-170483936 | 0 | T | C | 0.184 | -3.32E-02 | 1.20E-01 | 2.63E+00 | 2.71E+00 | 7.84E-01 | no | Fto | intron_variant | MODIFIER |
| 3 | 28638816 | AX-170334562 | 1 | C | T | 0.179 | -2.70E-02 | 9.95E-02 | 2.66E+00 | 2.73E+00 | 7.84E-01 | no | Tnik | intron_variant | MODIFIER |
| 18 | 68626998 | AX-170179586 | 0 | T | C | 0.088 | 5.04E-02 | 1.81E-01 | 2.60E+00 | 2.71E+00 | 7.84E-01 | no | | intergenic_variant | MODIFIER |
| 2 | 116722033 | AX-170501530 | 0 | T | A | 0.132 | 5.38E-02 | 1.99E-01 | 2.69E+00 | 2.74E+00 | 7.84E-01 | no | | intergenic_variant | MODIFIER |
| 2 | 116788074 | AX-97807944 | 0 | C | T | 0.132 | 5.38E-02 | 1.99E-01 | 2.69E+00 | 2.74E+00 | 7.84E-01 | no | | intergenic_variant | MODIFIER |
| 2 | 116944990 | AX-170501430 | 0 | G | A | 0.132 | 5.38E-02 | 1.99E-01 | 2.69E+00 | 2.74E+00 | 7.84E-01 | no | | intergenic_variant | MODIFIER |
| 2 | 117000383 | AX-170018783 | 0 | A | G | 0.132 | 5.38E-02 | 1.99E-01 | 2.69E+00 | 2.74E+00 | 7.84E-01 | no | | intergenic_variant | MODIFIER |
| 2 | 117361031 | AX-170238429 | 0 | T | C | 0.132 | 5.38E-02 | 1.99E-01 | 2.69E+00 | 2.74E+00 | 7.84E-01 | no | | intergenic_variant | MODIFIER |
| 2 | 117361996 | AX-170119013 | 0 | A | G | 0.132 | 5.38E-02 | 1.99E-01 | 2.69E+00 | 2.74E+00 | 7.84E-01 | no | | intergenic_variant | MODIFIER |
| 2 | 117362951 | AX-97848457 | 0 | T | G | 0.132 | 5.38E-02 | 1.99E-01 | 2.69E+00 | 2.74E+00 | 7.84E-01 | no | | intergenic_variant | MODIFIER |
| 2 | 117372224 | AX-170064681 | 0 | C | T | 0.132 | 5.38E-02 | 1.99E-01 | 2.69E+00 | 2.74E+00 | 7.84E-01 | no | | intergenic_variant | MODIFIER |
| 2 | 117372346 | AX-170064682 | 0 | C | T | 0.132 | 5.38E-02 | 1.99E-01 | 2.69E+00 | 2.74E+00 | 7.84E-01 | no | | intergenic_variant | MODIFIER |
| 2 | 117372419 | AX-170064680 | 0 | G | C | 0.132 | 5.38E-02 | 1.99E-01 | 2.69E+00 | 2.74E+00 | 7.84E-01 | no | | intergenic_variant | MODIFIER |
| 2 | 117376291 | AX-170170173 | 0 | C | A | 0.132 | 5.38E-02 | 1.99E-01 | 2.69E+00 | 2.74E+00 | 7.84E-01 | no | | intergenic_variant | MODIFIER |
| 2 | 117377830 | AX-170241657 | 0 | T | A | 0.132 | 5.38E-02 | 1.99E-01 | 2.69E+00 | 2.74E+00 | 7.84E-01 | no | | intergenic_variant | MODIFIER |
| 2 | 117377867 | AX-170303531 | 0 | C | T | 0.132 | 5.38E-02 | 1.99E-01 | 2.69E+00 | 2.74E+00 | 7.84E-01 | no | | intergenic_variant | MODIFIER |
| 2 | 117408920 | AX-170018797 | 0 | A | G | 0.132 | 5.38E-02 | 1.99E-01 | 2.69E+00 | 2.74E+00 | 7.84E-01 | no | | intergenic_variant | MODIFIER |
| 2 | 117412280 | AX-170018793 | 0 | A | G | 0.132 | 5.38E-02 | 1.99E-01 | 2.69E+00 | 2.74E+00 | 7.84E-01 | no | Gm13982 | downstream_gene_variant | MODIFIER |
| 2 | 117412444 | AX-170018795 | 0 | A | G | 0.132 | 5.38E-02 | 1.99E-01 | 2.69E+00 | 2.74E+00 | 7.84E-01 | no | Gm13982 | downstream_gene_variant | MODIFIER |
| 2 | 117414012 | AX-170385284 | 0 | C | T | 0.132 | 5.38E-02 | 1.99E-01 | 2.69E+00 | 2.74E+00 | 7.84E-01 | no | Gm13982 | downstream_gene_variant | MODIFIER |
| 2 | 117417173 | AX-170282071 | 0 | G | C | 0.132 | 5.38E-02 | 1.99E-01 | 2.69E+00 | 2.74E+00 | 7.84E-01 | no | Gm13982 | intron_variant,non_coding_transcript_variant | MODIFIER |
| 2 | 117449630 | AX-170018796 | 0 | C | T | 0.132 | 5.38E-02 | 1.99E-01 | 2.69E+00 | 2.74E+00 | 7.84E-01 | no | | intergenic_variant | MODIFIER |
| 2 | 117449711 | AX-170018791 | 0 | A | G | 0.132 | 5.38E-02 | 1.99E-01 | 2.69E+00 | 2.74E+00 | 7.84E-01 | no | | intergenic_variant | MODIFIER |
| 2 | 117474163 | AX-97850694 | 0 | C | T | 0.132 | 5.38E-02 | 1.99E-01 | 2.69E+00 | 2.74E+00 | 7.84E-01 | no | | intergenic_variant | MODIFIER |
| 2 | 117503997 | AX-170018789 | 0 | C | G | 0.132 | 5.38E-02 | 1.99E-01 | 2.69E+00 | 2.74E+00 | 7.84E-01 | no | | intergenic_variant | MODIFIER |
| 2 | 117524595 | AX-170129871 | 0 | T | C | 0.132 | 5.38E-02 | 1.99E-01 | 2.69E+00 | 2.74E+00 | 7.84E-01 | no | | regulatory_region_variant | MODIFIER |
| 2 | 117524726 | AX-97839711 | 0 | G | A | 0.132 | 5.38E-02 | 1.99E-01 | 2.69E+00 | 2.74E+00 | 7.84E-01 | no | | regulatory_region_variant | MODIFIER |
| 2 | 117609186 | AX-170255377 | 0 | T | C | 0.132 | 5.38E-02 | 1.99E-01 | 2.69E+00 | 2.74E+00 | 7.84E-01 | no | | intergenic_variant | MODIFIER |
| 2 | 117610258 | AX-97795452 | 0 | C | A | 0.132 | 5.38E-02 | 1.99E-01 | 2.69E+00 | 2.74E+00 | 7.84E-01 | no | | intergenic_variant | MODIFIER |

| | | | | | | | | | | | | | | | |
|---|---|---|---|---|---|---|---|---|---|---|---|---|---|---|---|
| 2 | 117627191 | AX-170364674 | 0 | A | G | 0.132 | 5.38E-02 | 1.99E-01 | 2.69E+00 | 2.74E+00 | 7.84E-01 | no | | intergenic_variant | MODIFIER |
| 2 | 117707564 | AX-97813894 | 0 | G | T | 0.132 | 5.38E-02 | 1.99E-01 | 2.69E+00 | 2.74E+00 | 7.84E-01 | no | | intergenic_variant | MODIFIER |
| 2 | 117824321 | AX-97841349 | 0 | A | G | 0.132 | 5.38E-02 | 1.99E-01 | 2.69E+00 | 2.74E+00 | 7.84E-01 | no | 4930412B13Rik | intron_variant,non_coding_transcript_variant | MODIFIER |
| 2 | 117826012 | AX-170214394 | 0 | T | C | 0.132 | 5.38E-02 | 1.99E-01 | 2.69E+00 | 2.74E+00 | 7.84E-01 | no | 4930412B13Rik | intron_variant,non_coding_transcript_variant | MODIFIER |
| 2 | 117841670 | AX-170211068 | 0 | C | T | 0.132 | 5.38E-02 | 1.99E-01 | 2.69E+00 | 2.74E+00 | 7.84E-01 | no | 4930412B13Rik | intron_variant,non_coding_transcript_variant | MODIFIER |
| 2 | 117843803 | AX-170365863 | 0 | T | C | 0.132 | 5.38E-02 | 1.99E-01 | 2.69E+00 | 2.74E+00 | 7.84E-01 | no | 4930412B13Rik | intron_variant,non_coding_transcript_variant | MODIFIER |
| 2 | 117875620 | AX-97838755 | 0 | C | A | 0.132 | 5.38E-02 | 1.99E-01 | 2.69E+00 | 2.74E+00 | 7.84E-01 | no | Gm13986 | intron_variant,non_coding_transcript_variant | MODIFIER |
| 2 | 117903858 | AX-170134183 | 0 | A | C | 0.132 | 5.38E-02 | 1.99E-01 | 2.69E+00 | 2.74E+00 | 7.84E-01 | no | Gm13986 | intron_variant,non_coding_transcript_variant | MODIFIER |
| 2 | 117907593 | AX-97841622 | 0 | A | G | 0.132 | 5.38E-02 | 1.99E-01 | 2.69E+00 | 2.74E+00 | 7.84E-01 | no | Gm13986 | intron_variant,non_coding_transcript_variant | MODIFIER |
| 2 | 117912820 | AX-170101750 | 0 | T | C | 0.132 | 5.38E-02 | 1.99E-01 | 2.69E+00 | 2.74E+00 | 7.84E-01 | no | Gm13986 | intron_variant,non_coding_transcript_variant | MODIFIER |
| 2 | 117914091 | AX-170114379 | 0 | T | A | 0.132 | 5.38E-02 | 1.99E-01 | 2.69E+00 | 2.74E+00 | 7.84E-01 | no | Gm13986 | intron_variant,non_coding_transcript_variant | MODIFIER |
| 2 | 117918027 | AX-170018800 | 0 | C | T | 0.132 | 5.38E-02 | 1.99E-01 | 2.69E+00 | 2.74E+00 | 7.84E-01 | no | Gm13986 | intron_variant,non_coding_transcript_variant | MODIFIER |
| 2 | 117926656 | AX-170064692 | 0 | T | C | 0.132 | 5.38E-02 | 1.99E-01 | 2.69E+00 | 2.74E+00 | 7.84E-01 | no | Gm13986 | intron_variant,non_coding_transcript_variant | MODIFIER |
| 2 | 117927326 | AX-170094383 | 0 | A | G | 0.132 | 5.38E-02 | 1.99E-01 | 2.69E+00 | 2.74E+00 | 7.84E-01 | no | Gm13986 | intron_variant,non_coding_transcript_variant | MODIFIER |
| 2 | 117927434 | AX-170222113 | 0 | G | A | 0.132 | 5.38E-02 | 1.99E-01 | 2.69E+00 | 2.74E+00 | 7.84E-01 | no | Gm13986 | intron_variant,non_coding_transcript_variant | MODIFIER |
| 2 | 117927524 | AX-170336577 | 0 | A | G | 0.132 | 5.38E-02 | 1.99E-01 | 2.69E+00 | 2.74E+00 | 7.84E-01 | no | Gm13986 | intron_variant,non_coding_transcript_variant | MODIFIER |
| 2 | 117993038 | AX-170018807 | 0 | C | T | 0.132 | 5.38E-02 | 1.99E-01 | 2.69E+00 | 2.74E+00 | 7.84E-01 | no | Gm13986 | intron_variant,non_coding_transcript_variant | MODIFIER |
| 2 | 117993061 | AX-170018802 | 0 | T | C | 0.132 | 5.38E-02 | 1.99E-01 | 2.69E+00 | 2.74E+00 | 7.84E-01 | no | Gm13986 | intron_variant,non_coding_transcript_variant | MODIFIER |
| 2 | 117994755 | AX-170183478 | 0 | C | T | 0.132 | 5.38E-02 | 1.99E-01 | 2.69E+00 | 2.74E+00 | 7.84E-01 | no | Gm13986 | intron_variant,non_coding_transcript_variant | MODIFIER |
| 2 | 118011318 | AX-170508532 | 0 | T | C | 0.132 | 5.38E-02 | 1.99E-01 | 2.69E+00 | 2.74E+00 | 7.84E-01 | no | Gm13986 | intron_variant,non_coding_transcript_variant | MODIFIER |
| 2 | 118057643 | AX-170518750 | 0 | T | C | 0.132 | 5.38E-02 | 1.99E-01 | 2.69E+00 | 2.74E+00 | 7.84E-01 | no | Gm13986 | intron_variant,non_coding_transcript_variant | MODIFIER |
| 2 | 118057697 | AX-170157817 | 0 | C | T | 0.132 | 5.38E-02 | 1.99E-01 | 2.69E+00 | 2.74E+00 | 7.84E-01 | no | Gm13986 | intron_variant,non_coding_transcript_variant | MODIFIER |
| 2 | 118057710 | AX-170480251 | 0 | T | C | 0.132 | 5.38E-02 | 1.99E-01 | 2.69E+00 | 2.74E+00 | 7.84E-01 | no | Gm13986 | intron_variant,non_coding_transcript_variant | MODIFIER |
| 2 | 118058391 | AX-170261161 | 0 | A | G | 0.132 | 5.38E-02 | 1.99E-01 | 2.69E+00 | 2.74E+00 | 7.84E-01 | no | Gm13986 | intron_variant,non_coding_transcript_variant | MODIFIER |
| 2 | 118059432 | AX-170423354 | 0 | G | C | 0.132 | 5.38E-02 | 1.99E-01 | 2.69E+00 | 2.74E+00 | 7.84E-01 | no | Gm13986 | intron_variant,non_coding_transcript_variant | MODIFIER |
| 2 | 118059503 | AX-170174633 | 0 | G | A | 0.132 | 5.38E-02 | 1.99E-01 | 2.69E+00 | 2.74E+00 | 7.84E-01 | no | Gm13986 | intron_variant,non_coding_transcript_variant | MODIFIER |
| 2 | 118059621 | AX-170064693 | 0 | A | G | 0.132 | 5.38E-02 | 1.99E-01 | 2.69E+00 | 2.74E+00 | 7.84E-01 | no | Gm13986 | intron_variant,non_coding_transcript_variant | MODIFIER |
| 2 | 118059734 | AX-170064695 | 0 | G | A | 0.132 | 5.38E-02 | 1.99E-01 | 2.69E+00 | 2.74E+00 | 7.84E-01 | no | Gm13986 | intron_variant,non_coding_transcript_variant | MODIFIER |
| 2 | 118060224 | AX-170334291 | 0 | T | C | 0.132 | 5.38E-02 | 1.99E-01 | 2.69E+00 | 2.74E+00 | 7.84E-01 | no | Gm13986 | intron_variant,non_coding_transcript_variant | MODIFIER |
| 2 | 118070933 | AX-170456617 | 0 | C | T | 0.132 | 5.38E-02 | 1.99E-01 | 2.69E+00 | 2.74E+00 | 7.84E-01 | no | Gm13986 | non_coding_transcript_exon_variant | MODIFIER |
| 2 | 118089167 | AX-170480512 | 0 | A | G | 0.132 | 5.38E-02 | 1.99E-01 | 2.69E+00 | 2.74E+00 | 7.84E-01 | no | Gm13986 | intron_variant,non_coding_transcript_variant | MODIFIER |
| 2 | 118125198 | AX-170121596 | 0 | G | A | 0.132 | 5.38E-02 | 1.99E-01 | 2.69E+00 | 2.74E+00 | 7.84E-01 | no | Thbs1 | 3_prime_UTR_variant | MODIFIER |
| 2 | 118132598 | AX-97839743 | 0 | G | A | 0.132 | 5.38E-02 | 1.99E-01 | 2.69E+00 | 2.74E+00 | 7.84E-01 | no | Fsip1 | intron_variant,non_coding_transcript_variant | MODIFIER |
| 2 | 118148368 | AX-170256758 | 0 | C | A | 0.132 | 5.38E-02 | 1.99E-01 | 2.69E+00 | 2.74E+00 | 7.84E-01 | no | Fsip1 | intron_variant | MODIFIER |
| 2 | 118149871 | AX-170102492 | 0 | T | A | 0.132 | 5.38E-02 | 1.99E-01 | 2.69E+00 | 2.74E+00 | 7.84E-01 | no | Fsip1 | intron_variant | MODIFIER |
| 2 | 118150734 | AX-170442153 | 0 | C | A | 0.132 | 5.38E-02 | 1.99E-01 | 2.69E+00 | 2.74E+00 | 7.84E-01 | no | Fsip1 | intron_variant | MODIFIER |
| 2 | 118151162 | AX-170307088 | 0 | C | T | 0.132 | 5.38E-02 | 1.99E-01 | 2.69E+00 | 2.74E+00 | 7.84E-01 | no | Fsip1 | intron_variant | MODIFIER |
| 2 | 118158877 | AX-170391957 | 0 | C | T | 0.132 | 5.38E-02 | 1.99E-01 | 2.69E+00 | 2.74E+00 | 7.84E-01 | no | Fsip1 | intron_variant | MODIFIER |
| 2 | 118198510 | AX-97794931 | 0 | A | G | 0.132 | 5.38E-02 | 1.99E-01 | 2.69E+00 | 2.74E+00 | 7.84E-01 | no | Fsip1 | intron_variant | MODIFIER |
| 2 | 118236369 | AX-170246157 | 0 | G | C | 0.132 | 5.38E-02 | 1.99E-01 | 2.69E+00 | 2.74E+00 | 7.84E-01 | no | Fsip1 | intron_variant | MODIFIER |
| 2 | 118251632 | AX-170432512 | 0 | A | G | 0.132 | 5.38E-02 | 1.99E-01 | 2.69E+00 | 2.74E+00 | 7.84E-01 | no | Fsip1 | splice_region_variant,intron_variant | LOW |
| 2 | 118251701 | AX-170336025 | 0 | A | G | 0.132 | 5.38E-02 | 1.99E-01 | 2.69E+00 | 2.74E+00 | 7.84E-01 | no | Fsip1 | missense_variant | MODERATE |
| 2 | 118251724 | AX-170334870 | 0 | C | T | 0.132 | 5.38E-02 | 1.99E-01 | 2.69E+00 | 2.74E+00 | 7.84E-01 | no | Fsip1 | synonymous_variant | LOW |
| 2 | 118252469 | AX-170508894 | 0 | C | A | 0.132 | 5.38E-02 | 1.99E-01 | 2.69E+00 | 2.74E+00 | 7.84E-01 | no | Fsip1 | intron_variant | MODIFIER |
| 2 | 118254834 | AX-170144311 | 0 | G | A | 0.132 | 5.38E-02 | 1.99E-01 | 2.69E+00 | 2.74E+00 | 7.84E-01 | no | Fsip1 | intron_variant | MODIFIER |
| 2 | 118262522 | AX-170104013 | 0 | T | C | 0.132 | 5.38E-02 | 1.99E-01 | 2.69E+00 | 2.74E+00 | 7.84E-01 | no | Gm29232 | intron_variant,non_coding_transcript_variant | MODIFIER |
| 2 | 118286569 | AX-170474962 | 0 | A | C | 0.132 | 5.38E-02 | 1.99E-01 | 2.69E+00 | 2.74E+00 | 7.84E-01 | no | Gpr176 | intron_variant | MODIFIER |
| 2 | 118287154 | AX-170435979 | 0 | C | T | 0.132 | 5.38E-02 | 1.99E-01 | 2.69E+00 | 2.74E+00 | 7.84E-01 | no | Gpr176 | intron_variant | MODIFIER |
| 2 | 118294315 | AX-170064698 | 0 | T | C | 0.132 | 5.38E-02 | 1.99E-01 | 2.69E+00 | 2.74E+00 | 7.84E-01 | no | Gpr176 | intron_variant | MODIFIER |
| 2 | 118295163 | AX-170444086 | 0 | A | G | 0.132 | 5.38E-02 | 1.99E-01 | 2.69E+00 | 2.74E+00 | 7.84E-01 | no | Gpr176 | intron_variant | MODIFIER |
| 2 | 118295865 | AX-170368788 | 0 | G | T | 0.132 | 5.38E-02 | 1.99E-01 | 2.69E+00 | 2.74E+00 | 7.84E-01 | no | Gpr176 | intron_variant | MODIFIER |
| 2 | 118298630 | AX-170099947 | 0 | A | C | 0.132 | 5.38E-02 | 1.99E-01 | 2.69E+00 | 2.74E+00 | 7.84E-01 | no | Gpr176 | intron_variant | MODIFIER |
| 2 | 118299985 | AX-170230544 | 0 | A | G | 0.132 | 5.38E-02 | 1.99E-01 | 2.69E+00 | 2.74E+00 | 7.84E-01 | no | Gpr176 | intron_variant | MODIFIER |
| 2 | 118300043 | AX-170116840 | 0 | C | T | 0.132 | 5.38E-02 | 1.99E-01 | 2.69E+00 | 2.74E+00 | 7.84E-01 | no | Gpr176 | intron_variant | MODIFIER |

| | | | | | | | | | | | | | | | |
|---|---|---|---|---|---|---|---|---|---|---|---|---|---|---|---|
| 2 | 118307620 | AX-170254004 | 0 | C | T | 0.132 | 5.38E-02 | 1.99E-01 | 2.69E+00 | 2.74E+00 | 7.84E-01 | no | Gpr176 | intron_variant | MODIFIER |
| 2 | 118327288 | AX-170523619 | 0 | A | C | 0.132 | 5.38E-02 | 1.99E-01 | 2.69E+00 | 2.74E+00 | 7.84E-01 | no | Gpr176 | intron_variant | MODIFIER |
| 2 | 118329556 | AX-170372637 | 0 | A | T | 0.132 | 5.38E-02 | 1.99E-01 | 2.69E+00 | 2.74E+00 | 7.84E-01 | no | Gpr176 | intron_variant | MODIFIER |
| 2 | 118329800 | AX-170301040 | 0 | A | G | 0.132 | 5.38E-02 | 1.99E-01 | 2.69E+00 | 2.74E+00 | 7.84E-01 | no | Gpr176 | intron_variant | MODIFIER |
| 2 | 118361281 | AX-170523911 | 0 | G | C | 0.132 | 5.38E-02 | 1.99E-01 | 2.69E+00 | 2.74E+00 | 7.84E-01 | no | Gpr176 | intron_variant | MODIFIER |
| 2 | 118369597 | AX-170210613 | 0 | G | T | 0.132 | 5.38E-02 | 1.99E-01 | 2.69E+00 | 2.74E+00 | 7.84E-01 | no | Gpr176 | intron_variant | MODIFIER |
| 2 | 118369934 | AX-97794929 | 0 | C | T | 0.132 | 5.38E-02 | 1.99E-01 | 2.69E+00 | 2.74E+00 | 7.84E-01 | no | Gpr176 | intron_variant | MODIFIER |
| 2 | 118390252 | AX-170346696 | 0 | C | T | 0.132 | 5.38E-02 | 1.99E-01 | 2.69E+00 | 2.74E+00 | 7.84E-01 | no | Eif2ak4 | intron_variant | MODIFIER |
| 2 | 118396547 | AX-170213777 | 0 | C | T | 0.132 | 5.38E-02 | 1.99E-01 | 2.69E+00 | 2.74E+00 | 7.84E-01 | no | Eif2ak4 | intron_variant | MODIFIER |
| 2 | 118399336 | AX-170220034 | 0 | T | C | 0.132 | 5.38E-02 | 1.99E-01 | 2.69E+00 | 2.74E+00 | 7.84E-01 | no | Eif2ak4 | intron_variant | MODIFIER |
| 2 | 118415815 | AX-170468505 | 0 | C | A | 0.132 | 5.38E-02 | 1.99E-01 | 2.69E+00 | 2.74E+00 | 7.84E-01 | no | Eif2ak4 | intron_variant | MODIFIER |
| 2 | 118415863 | AX-170018817 | 0 | C | T | 0.132 | 5.38E-02 | 1.99E-01 | 2.69E+00 | 2.74E+00 | 7.84E-01 | no | Eif2ak4 | intron_variant | MODIFIER |
| 2 | 118415884 | AX-170175350 | 0 | G | A | 0.132 | 5.38E-02 | 1.99E-01 | 2.69E+00 | 2.74E+00 | 7.84E-01 | no | Eif2ak4 | intron_variant | MODIFIER |
| 2 | 118416082 | AX-170429352 | 0 | G | A | 0.132 | 5.38E-02 | 1.99E-01 | 2.69E+00 | 2.74E+00 | 7.84E-01 | no | Eif2ak4 | intron_variant | MODIFIER |
| 2 | 118430315 | AX-170355139 | 0 | A | C | 0.132 | 5.38E-02 | 1.99E-01 | 2.69E+00 | 2.74E+00 | 7.84E-01 | no | Eif2ak4 | intron_variant | MODIFIER |
| 2 | 118438605 | AX-170150227 | 0 | G | A | 0.132 | 5.38E-02 | 1.99E-01 | 2.69E+00 | 2.74E+00 | 7.84E-01 | no | Eif2ak4 | intron_variant | MODIFIER |
| 2 | 118442170 | AX-170064701 | 0 | C | T | 0.132 | 5.38E-02 | 1.99E-01 | 2.69E+00 | 2.74E+00 | 7.84E-01 | no | Eif2ak4 | intron_variant | MODIFIER |
| 2 | 118442812 | AX-170064699 | 0 | G | A | 0.132 | 5.38E-02 | 1.99E-01 | 2.69E+00 | 2.74E+00 | 7.84E-01 | no | Eif2ak4 | intron_variant | MODIFIER |
| 2 | 118461935 | AX-170516001 | 0 | C | A | 0.132 | 5.38E-02 | 1.99E-01 | 2.69E+00 | 2.74E+00 | 7.84E-01 | no | Eif2ak4 | intron_variant | MODIFIER |
| 2 | 118478416 | AX-170236912 | 0 | A | T | 0.132 | 5.38E-02 | 1.99E-01 | 2.69E+00 | 2.74E+00 | 7.84E-01 | no | Eif2ak4 | downstream_gene_variant | MODIFIER |
| 2 | 118499172 | AX-170341296 | 0 | A | G | 0.132 | 5.38E-02 | 1.99E-01 | 2.69E+00 | 2.74E+00 | 7.84E-01 | no | | regulatory_region_variant | MODIFIER |
| 2 | 118513267 | AX-97850617 | 0 | C | T | 0.132 | 5.38E-02 | 1.99E-01 | 2.69E+00 | 2.74E+00 | 7.84E-01 | no | | regulatory_region_variant | MODIFIER |
| 2 | 118526291 | AX-170018813 | 0 | A | G | 0.132 | 5.38E-02 | 1.99E-01 | 2.69E+00 | 2.74E+00 | 7.84E-01 | no | Bmf | downstream_gene_variant | MODIFIER |
| 2 | 118529313 | AX-170018812 | 0 | A | G | 0.132 | 5.38E-02 | 1.99E-01 | 2.69E+00 | 2.74E+00 | 7.84E-01 | no | Bmf | 3_prime_UTR_variant | MODIFIER |
| 2 | 118537709 | AX-170365411 | 0 | G | T | 0.132 | 5.38E-02 | 1.99E-01 | 2.69E+00 | 2.74E+00 | 7.84E-01 | no | Bmf | intron_variant | MODIFIER |
| 2 | 118539260 | AX-170104812 | 0 | T | C | 0.132 | 5.38E-02 | 1.99E-01 | 2.69E+00 | 2.74E+00 | 7.84E-01 | no | Bmf | intron_variant | MODIFIER |
| 2 | 118582646 | AX-170289308 | 0 | A | G | 0.132 | 5.38E-02 | 1.99E-01 | 2.69E+00 | 2.74E+00 | 7.84E-01 | no | | intergenic_variant | MODIFIER |
| 2 | 118598800 | AX-170194400 | 0 | C | T | 0.132 | 5.38E-02 | 1.99E-01 | 2.69E+00 | 2.74E+00 | 7.84E-01 | no | Bub1b | intron_variant | MODIFIER |
| 2 | 118605380 | AX-170018810 | 0 | T | C | 0.132 | 5.38E-02 | 1.99E-01 | 2.69E+00 | 2.74E+00 | 7.84E-01 | no | Bub1b | intron_variant | MODIFIER |
| 2 | 118605543 | AX-170217136 | 0 | A | G | 0.132 | 5.38E-02 | 1.99E-01 | 2.69E+00 | 2.74E+00 | 7.84E-01 | no | Bub1b | intron_variant | MODIFIER |
| 2 | 118605688 | AX-170018809 | 0 | T | A | 0.132 | 5.38E-02 | 1.99E-01 | 2.69E+00 | 2.74E+00 | 7.84E-01 | no | Bub1b | intron_variant | MODIFIER |
| 2 | 118609588 | AX-170511149 | 0 | C | T | 0.132 | 5.38E-02 | 1.99E-01 | 2.69E+00 | 2.74E+00 | 7.84E-01 | no | Bub1b | intron_variant | MODIFIER |
| 2 | 118623263 | AX-170320234 | 0 | A | G | 0.132 | 5.38E-02 | 1.99E-01 | 2.69E+00 | 2.74E+00 | 7.84E-01 | no | Bub1b | intron_variant | MODIFIER |
| 2 | 118628982 | AX-170267391 | 0 | C | A | 0.132 | 5.38E-02 | 1.99E-01 | 2.69E+00 | 2.74E+00 | 7.84E-01 | no | Bub1b | intron_variant | MODIFIER |
| 2 | 118629030 | AX-170258847 | 0 | A | T | 0.132 | 5.38E-02 | 1.99E-01 | 2.69E+00 | 2.74E+00 | 7.84E-01 | no | Bub1b | intron_variant | MODIFIER |
| 2 | 118663319 | AX-170358998 | 0 | A | T | 0.132 | 5.38E-02 | 1.99E-01 | 2.69E+00 | 2.74E+00 | 7.84E-01 | no | Pak6 | 5_prime_UTR_variant | MODIFIER |
| 2 | 118666090 | AX-97826734 | 0 | A | G | 0.132 | 5.38E-02 | 1.99E-01 | 2.69E+00 | 2.74E+00 | 7.84E-01 | no | Pak6 | intron_variant | MODIFIER |
| 2 | 118669291 | AX-170468423 | 0 | T | G | 0.132 | 5.38E-02 | 1.99E-01 | 2.69E+00 | 2.74E+00 | 7.84E-01 | no | Pak6 | intron_variant | MODIFIER |
| 2 | 118671274 | AX-170178479 | 0 | A | G | 0.132 | 5.38E-02 | 1.99E-01 | 2.69E+00 | 2.74E+00 | 7.84E-01 | no | Pak6 | intron_variant | MODIFIER |
| 2 | 118673024 | AX-170088728 | 0 | C | T | 0.132 | 5.38E-02 | 1.99E-01 | 2.69E+00 | 2.74E+00 | 7.84E-01 | no | Pak6 | intron_variant | MODIFIER |
| 2 | 118676461 | AX-170235066 | 0 | A | T | 0.132 | 5.38E-02 | 1.99E-01 | 2.69E+00 | 2.74E+00 | 7.84E-01 | no | Pak6 | intron_variant | MODIFIER |
| 2 | 118723291 | AX-170323373 | 0 | T | C | 0.132 | 5.38E-02 | 1.99E-01 | 2.69E+00 | 2.74E+00 | 7.84E-01 | no | Plcb2 | intron_variant,non_coding_transcript_variant | MODIFIER |
| 2 | 118724059 | AX-170376201 | 0 | T | A | 0.132 | 5.38E-02 | 1.99E-01 | 2.69E+00 | 2.74E+00 | 7.84E-01 | no | Plcb2 | intron_variant,non_coding_transcript_variant | MODIFIER |
| 2 | 118725385 | AX-170296254 | 0 | C | T | 0.132 | 5.38E-02 | 1.99E-01 | 2.69E+00 | 2.74E+00 | 7.84E-01 | no | Plcb2 | intron_variant,non_coding_transcript_variant | MODIFIER |
| 2 | 118730035 | AX-170301637 | 0 | T | C | 0.132 | 5.38E-02 | 1.99E-01 | 2.69E+00 | 2.74E+00 | 7.84E-01 | no | Plcb2 | upstream_gene_variant | MODIFIER |
| 2 | 118742312 | AX-170113850 | 0 | G | A | 0.132 | 5.38E-02 | 1.99E-01 | 2.69E+00 | 2.74E+00 | 7.84E-01 | no | Inafm2 | upstream_gene_variant | MODIFIER |
| 2 | 118749783 | AX-170472694 | 0 | A | G | 0.132 | 5.38E-02 | 1.99E-01 | 2.69E+00 | 2.74E+00 | 7.84E-01 | no | Ccdc9b | downstream_gene_variant | MODIFIER |
| 2 | 118750022 | AX-170529283 | 0 | T | G | 0.132 | 5.38E-02 | 1.99E-01 | 2.69E+00 | 2.74E+00 | 7.84E-01 | no | Ccdc9b | downstream_gene_variant | MODIFIER |
| 2 | 118766172 | AX-170367702 | 0 | T | C | 0.132 | 5.38E-02 | 1.99E-01 | 2.69E+00 | 2.74E+00 | 7.84E-01 | no | Ccdc9b | upstream_gene_variant | MODIFIER |
| 2 | 118766977 | AX-170456414 | 0 | T | C | 0.132 | 5.38E-02 | 1.99E-01 | 2.69E+00 | 2.74E+00 | 7.84E-01 | no | Ccdc9b | upstream_gene_variant | MODIFIER |
| 2 | 118788502 | AX-170191073 | 0 | C | T | 0.132 | 5.38E-02 | 1.99E-01 | 2.69E+00 | 2.74E+00 | 7.84E-01 | no | Disp2 | intron_variant | MODIFIER |
| 2 | 118789003 | AX-170218494 | 0 | C | T | 0.132 | 5.38E-02 | 1.99E-01 | 2.69E+00 | 2.74E+00 | 7.84E-01 | no | Disp2 | intron_variant | MODIFIER |
| 2 | 118789958 | AX-170240527 | 0 | C | T | 0.132 | 5.38E-02 | 1.99E-01 | 2.69E+00 | 2.74E+00 | 7.84E-01 | no | Disp2 | synonymous_variant | LOW |
| 2 | 118807696 | AX-97853252 | 0 | T | C | 0.132 | 5.38E-02 | 1.99E-01 | 2.69E+00 | 2.74E+00 | 7.84E-01 | no | Disp2 | intron_variant | MODIFIER |

| | | | | | | | | | | | | | | | |
|---|---|---|---|---|---|---|---|---|---|---|---|---|---|---|---|
| 2 | 118817085 | AX-170190106 | 0 | T | C | 0.132 | 5.38E-02 | 1.99E-01 | 2.69E+00 | 2.74E+00 | 7.84E-01 | no | Knstrn | intron_variant | MODIFIER |
| 2 | 118854656 | AX-97839788 | 0 | T | C | 0.132 | 5.38E-02 | 1.99E-01 | 2.69E+00 | 2.74E+00 | 7.84E-01 | no | Knstrn | downstream_gene_variant | MODIFIER |
| 2 | 128754643 | AX-170360667 | 0 | A | T | 0.132 | 5.38E-02 | 1.99E-01 | 2.69E+00 | 2.74E+00 | 7.84E-01 | no | Mertk | intron_variant | MODIFIER |
| 2 | 142266236 | AX-170065102 | 0 | G | A | 0.132 | 5.38E-02 | 1.99E-01 | 2.69E+00 | 2.74E+00 | 7.84E-01 | no | Macrod2 | intron_variant | MODIFIER |
| 2 | 149636938 | AX-170179905 | 0 | G | A | 0.132 | 5.38E-02 | 1.99E-01 | 2.69E+00 | 2.74E+00 | 7.84E-01 | no | | intergenic_variant | MODIFIER |
| 7 | 106973965 | AX-170290514 | 1 | G | A | 0.482 | 2.56E-02 | 9.22E-02 | 2.63E+00 | 2.70E+00 | 7.85E-01 | no | Olfr711 | intron_variant | MODIFIER |
| 5 | 32958115 | AX-170476305 | 1 | A | T | 0.357 | 2.84E-02 | 1.05E-01 | 2.68E+00 | 2.74E+00 | 7.85E-01 | no | Depdc5 | downstream_gene_variant | MODIFIER |
| 5 | 30931063 | AX-170386398 | 0 | A | T | 0.368 | 2.81E-02 | 1.04E-01 | 2.68E+00 | 2.74E+00 | 7.85E-01 | no | Cgref1 | downstream_gene_variant | MODIFIER |
| 7 | 103866951 | AX-170086228 | 1 | G | A | 0.473 | 2.43E-02 | 8.58E-02 | 2.60E+00 | 2.68E+00 | 7.85E-01 | no | | regulatory_region_variant | MODIFIER |
| 4 | 91664594 | AX-170023400 | 1 | C | T | 0.33 | 2.98E-02 | 1.09E-01 | 2.65E+00 | 2.72E+00 | 7.85E-01 | no | | intergenic_variant | MODIFIER |
| 6 | 36259901 | AX-97855873 | 0 | T | C | 0.044 | -4.61E-02 | 1.69E-01 | 2.60E+00 | 2.73E+00 | 7.85E-01 | no | 9330158H04Rik | intron_variant,non_coding_transcript_variant | MODIFIER |
| 1 | 41708114 | AX-170338859 | 0 | G | A | 0.421 | -2.05E-02 | 7.48E-02 | 2.63E+00 | 2.72E+00 | 7.85E-01 | no | | intergenic_variant | MODIFIER |
| 7 | 111871886 | AX-170148876 | 2 | A | C | 0.464 | 2.41E-02 | 8.86E-02 | 2.65E+00 | 2.72E+00 | 7.85E-01 | no | | intergenic_variant | MODIFIER |
| 20 | 87209861 | AX-170090573 | 1 | G | A | 0.25 | -1.62E-02 | 6.08E-02 | 2.65E+00 | 2.76E+00 | 7.85E-01 | no | Il1rapl1 | intron_variant | MODIFIER |
| 13 | 41442786 | AX-170488493 | 0 | C | T | 0.518 | -1.60E-02 | 6.21E-02 | 2.70E+00 | 2.81E+00 | 7.85E-01 | no | Nedd9 | intron_variant | MODIFIER |
| 15 | 94265300 | AX-170205463 | 1 | A | G | 0.152 | 4.94E-02 | 1.80E-01 | 2.65E+00 | 2.71E+00 | 7.85E-01 | no | Adamts20 | downstream_gene_variant | MODIFIER |
| 13 | 42449950 | AX-97850077 | 0 | T | C | 0.491 | 1.71E-02 | 6.09E-02 | 2.59E+00 | 2.70E+00 | 7.85E-01 | no | | intergenic_variant | MODIFIER |
| 15 | 93288025 | AX-170056328 | 1 | A | G | 0.134 | 3.94E-02 | 1.41E-01 | 2.60E+00 | 2.70E+00 | 7.85E-01 | no | Yaf2 | intron_variant | MODIFIER |
| 9 | 85622946 | AX-97840152 | 0 | A | G | 0.211 | -3.26E-02 | 1.21E-01 | 2.68E+00 | 2.74E+00 | 7.85E-01 | no | | intergenic_variant | MODIFIER |
| 9 | 85651245 | AX-170317253 | 0 | A | G | 0.211 | -3.26E-02 | 1.21E-01 | 2.68E+00 | 2.74E+00 | 7.85E-01 | no | | intergenic_variant | MODIFIER |
| 9 | 85651644 | AX-170284473 | 0 | A | C | 0.211 | -3.26E-02 | 1.21E-01 | 2.68E+00 | 2.74E+00 | 7.85E-01 | no | | intergenic_variant | MODIFIER |
| 8 | 39209037 | AX-170511110 | 1 | G | A | 0.152 | -3.96E-02 | 1.42E-01 | 2.63E+00 | 2.69E+00 | 7.85E-01 | no | Gm17766 | downstream_gene_variant | MODIFIER |
| 1 | 99757005 | AX-170091607 | 0 | C | A | 0.158 | -3.77E-02 | 1.42E-01 | 2.64E+00 | 2.75E+00 | 7.85E-01 | no | | intergenic_variant | MODIFIER |
| 16 | 91333310 | AX-97832136 | 0 | G | C | 0.465 | -2.15E-02 | 8.03E-02 | 2.68E+00 | 2.75E+00 | 7.85E-01 | no | Gm15966 | upstream_gene_variant | MODIFIER |
| 16 | 91374901 | AX-170168223 | 0 | T | C | 0.465 | -2.15E-02 | 8.03E-02 | 2.68E+00 | 2.75E+00 | 7.85E-01 | no | Ifnar2 | intron_variant | MODIFIER |
| 16 | 91471460 | AX-97827017 | 0 | G | A | 0.465 | -2.15E-02 | 8.03E-02 | 2.68E+00 | 2.75E+00 | 7.85E-01 | no | A930006K02Rik | downstream_gene_variant | MODIFIER |
| 16 | 91538382 | AX-170140066 | 0 | G | C | 0.465 | -2.15E-02 | 8.03E-02 | 2.68E+00 | 2.75E+00 | 7.85E-01 | no | Gm7860 | downstream_gene_variant | MODIFIER |
| 16 | 91538455 | AX-97832134 | 0 | C | T | 0.465 | -2.15E-02 | 8.03E-02 | 2.68E+00 | 2.75E+00 | 7.85E-01 | no | Gm7860 | downstream_gene_variant | MODIFIER |
| 16 | 91620044 | AX-97832760 | 0 | A | G | 0.465 | -2.15E-02 | 8.03E-02 | 2.68E+00 | 2.75E+00 | 7.85E-01 | no | Gart | downstream_gene_variant | MODIFIER |
| 16 | 91623830 | AX-170013654 | 0 | A | G | 0.465 | -2.15E-02 | 8.03E-02 | 2.68E+00 | 2.75E+00 | 7.85E-01 | no | Gart | intron_variant | MODIFIER |
| 16 | 91636884 | AX-97797283 | 0 | G | A | 0.465 | -2.15E-02 | 8.03E-02 | 2.68E+00 | 2.75E+00 | 7.85E-01 | no | Gart | intron_variant | MODIFIER |
| 16 | 91666641 | AX-97848582 | 0 | C | T | 0.465 | -2.15E-02 | 8.03E-02 | 2.68E+00 | 2.75E+00 | 7.85E-01 | no | Son | downstream_gene_variant | MODIFIER |
| 16 | 91725005 | AX-170013656 | 0 | A | G | 0.465 | -2.15E-02 | 8.03E-02 | 2.68E+00 | 2.75E+00 | 7.85E-01 | no | Itsn1 | upstream_gene_variant | MODIFIER |
| 16 | 91833328 | AX-170297963 | 0 | T | C | 0.465 | -2.15E-02 | 8.03E-02 | 2.68E+00 | 2.75E+00 | 7.85E-01 | no | Itsn1 | intron_variant | MODIFIER |
| 16 | 91881669 | AX-170412090 | 0 | A | G | 0.465 | -2.15E-02 | 8.03E-02 | 2.68E+00 | 2.75E+00 | 7.85E-01 | no | Itsn1 | intron_variant | MODIFIER |
| 16 | 91893094 | AX-170269433 | 0 | T | C | 0.465 | -2.15E-02 | 8.03E-02 | 2.68E+00 | 2.75E+00 | 7.85E-01 | no | Itsn1 | intron_variant | MODIFIER |
| 16 | 91899208 | AX-170169160 | 0 | C | A | 0.465 | -2.15E-02 | 8.03E-02 | 2.68E+00 | 2.75E+00 | 7.85E-01 | no | Itsn1 | intron_variant | MODIFIER |
| 16 | 91900102 | AX-170267449 | 0 | C | G | 0.465 | -2.15E-02 | 8.03E-02 | 2.68E+00 | 2.75E+00 | 7.85E-01 | no | Itsn1 | intron_variant | MODIFIER |
| 16 | 91900494 | AX-170293153 | 0 | C | T | 0.465 | -2.15E-02 | 8.03E-02 | 2.68E+00 | 2.75E+00 | 7.85E-01 | no | Itsn1 | intron_variant | MODIFIER |
| 16 | 91913110 | AX-170484905 | 0 | A | T | 0.465 | -2.15E-02 | 8.03E-02 | 2.68E+00 | 2.75E+00 | 7.85E-01 | no | Itsn1 | 3_prime_UTR_variant | MODIFIER |
| 16 | 91913441 | AX-170058114 | 0 | G | T | 0.465 | -2.15E-02 | 8.03E-02 | 2.68E+00 | 2.75E+00 | 7.85E-01 | no | Itsn1 | 3_prime_UTR_variant | MODIFIER |
| 16 | 91913521 | AX-170058112 | 0 | G | A | 0.465 | -2.15E-02 | 8.03E-02 | 2.68E+00 | 2.75E+00 | 7.85E-01 | no | Itsn1 | 3_prime_UTR_variant | MODIFIER |
| 16 | 91931301 | AX-170155108 | 0 | T | C | 0.465 | -2.15E-02 | 8.03E-02 | 2.68E+00 | 2.75E+00 | 7.85E-01 | no | Atp5o | intron_variant | MODIFIER |
| 16 | 91931662 | AX-97832101 | 0 | T | C | 0.465 | -2.15E-02 | 8.03E-02 | 2.68E+00 | 2.75E+00 | 7.85E-01 | no | Atp5o | 5_prime_UTR_variant | MODIFIER |
| 16 | 91948951 | AX-170148611 | 0 | A | G | 0.465 | -2.15E-02 | 8.03E-02 | 2.68E+00 | 2.75E+00 | 7.85E-01 | no | D430001F17Rik | intron_variant,non_coding_transcript_variant | MODIFIER |
| 16 | 91970668 | AX-170380532 | 0 | C | T | 0.465 | -2.15E-02 | 8.03E-02 | 2.68E+00 | 2.75E+00 | 7.85E-01 | no | | regulatory_region_variant | MODIFIER |
| 16 | 92013201 | AX-170486679 | 0 | G | A | 0.465 | -2.15E-02 | 8.03E-02 | 2.68E+00 | 2.75E+00 | 7.85E-01 | no | Gm18020 | upstream_gene_variant | MODIFIER |
| 16 | 92014693 | AX-170158228 | 0 | T | C | 0.465 | -2.15E-02 | 8.03E-02 | 2.68E+00 | 2.75E+00 | 7.85E-01 | no | Gm18020 | upstream_gene_variant | MODIFIER |
| 16 | 92026311 | AX-170130969 | 0 | C | T | 0.465 | -2.15E-02 | 8.03E-02 | 2.68E+00 | 2.75E+00 | 7.85E-01 | no | | intergenic_variant | MODIFIER |
| 16 | 92047334 | AX-170467748 | 0 | G | C | 0.465 | -2.15E-02 | 8.03E-02 | 2.68E+00 | 2.75E+00 | 7.85E-01 | no | Gm23210 | downstream_gene_variant | MODIFIER |
| 16 | 92080132 | AX-168420009 | 0 | A | T | 0.465 | -2.15E-02 | 8.03E-02 | 2.68E+00 | 2.75E+00 | 7.85E-01 | no | Mrps6 | intron_variant | MODIFIER |
| 16 | 92081622 | AX-170475209 | 0 | G | A | 0.465 | -2.15E-02 | 8.03E-02 | 2.68E+00 | 2.75E+00 | 7.85E-01 | no | Mrps6 | intron_variant | MODIFIER |
| 16 | 92081668 | AX-170458709 | 0 | C | T | 0.465 | -2.15E-02 | 8.03E-02 | 2.68E+00 | 2.75E+00 | 7.85E-01 | no | Mrps6 | intron_variant | MODIFIER |
| 16 | 92088560 | AX-170438692 | 0 | G | C | 0.465 | -2.15E-02 | 8.03E-02 | 2.68E+00 | 2.75E+00 | 7.85E-01 | no | Mrps6 | intron_variant | MODIFIER |

| | | | | | | | | | | | | | | | |
|---|---|---|---|---|---|---|---|---|---|---|---|---|---|---|---|
| 16 | 92102249 | AX-170444773 | 0 | T | A | 0.465 | -2.15E-02 | 8.03E-02 | 2.68E+00 | 2.75E+00 | 7.85E-01 | no | Mrps6 | intron_variant | MODIFIER |
| 16 | 92105213 | AX-170360978 | 0 | A | G | 0.465 | -2.15E-02 | 8.03E-02 | 2.68E+00 | 2.75E+00 | 7.85E-01 | no | Mrps6 | intron_variant | MODIFIER |
| 16 | 92111312 | AX-170316686 | 0 | T | C | 0.465 | -2.15E-02 | 8.03E-02 | 2.68E+00 | 2.75E+00 | 7.85E-01 | no | Mrps6 | intron_variant | MODIFIER |
| 16 | 92143020 | AX-170058118 | 0 | C | A | 0.465 | -2.15E-02 | 8.03E-02 | 2.68E+00 | 2.75E+00 | 7.85E-01 | no | Gm41496 | intron_variant,non_coding_transcript_variant | MODIFIER |
| 16 | 92171726 | AX-170494143 | 0 | A | G | 0.465 | -2.15E-02 | 8.03E-02 | 2.68E+00 | 2.75E+00 | 7.85E-01 | no | Gm41496 | intron_variant,non_coding_transcript_variant | MODIFIER |
| 16 | 92172399 | AX-97832458 | 0 | C | T | 0.465 | -2.15E-02 | 8.03E-02 | 2.68E+00 | 2.75E+00 | 7.85E-01 | no | Gm41496 | intron_variant,non_coding_transcript_variant | MODIFIER |
| 16 | 92185602 | AX-170451067 | 0 | C | T | 0.465 | -2.15E-02 | 8.03E-02 | 2.68E+00 | 2.75E+00 | 7.85E-01 | no | Gm41496 | intron_variant,non_coding_transcript_variant | MODIFIER |
| 16 | 92185837 | AX-170186927 | 0 | A | G | 0.465 | -2.15E-02 | 8.03E-02 | 2.68E+00 | 2.75E+00 | 7.85E-01 | no | Gm41496 | intron_variant,non_coding_transcript_variant | MODIFIER |
| 16 | 92189948 | AX-170530176 | 0 | G | T | 0.465 | -2.15E-02 | 8.03E-02 | 2.68E+00 | 2.75E+00 | 7.85E-01 | no | Gm41496 | downstream_gene_variant | MODIFIER |
| 16 | 92189969 | AX-97819242 | 0 | A | G | 0.465 | -2.15E-02 | 8.03E-02 | 2.68E+00 | 2.75E+00 | 7.85E-01 | no | Gm41496 | downstream_gene_variant | MODIFIER |
| 16 | 92197611 | AX-170496539 | 0 | A | G | 0.465 | -2.15E-02 | 8.03E-02 | 2.68E+00 | 2.75E+00 | 7.85E-01 | no | | regulatory_region_variant | MODIFIER |
| 16 | 92210536 | AX-170013669 | 0 | A | G | 0.465 | -2.15E-02 | 8.03E-02 | 2.68E+00 | 2.75E+00 | 7.85E-01 | no | Gm29880 | upstream_gene_variant | MODIFIER |
| 16 | 92216263 | AX-170151866 | 0 | A | G | 0.465 | -2.15E-02 | 8.03E-02 | 2.68E+00 | 2.75E+00 | 7.85E-01 | no | | regulatory_region_variant | MODIFIER |
| 16 | 92224880 | AX-170532498 | 0 | T | C | 0.465 | -2.15E-02 | 8.03E-02 | 2.68E+00 | 2.75E+00 | 7.85E-01 | no | | intergenic_variant | MODIFIER |
| 16 | 92224991 | AX-170384332 | 0 | G | A | 0.465 | -2.15E-02 | 8.03E-02 | 2.68E+00 | 2.75E+00 | 7.85E-01 | no | | intergenic_variant | MODIFIER |
| 16 | 92229205 | AX-170365770 | 0 | T | C | 0.465 | -2.15E-02 | 8.03E-02 | 2.68E+00 | 2.75E+00 | 7.85E-01 | no | | intergenic_variant | MODIFIER |
| 16 | 92241299 | AX-170088099 | 0 | C | T | 0.465 | -2.15E-02 | 8.03E-02 | 2.68E+00 | 2.75E+00 | 7.85E-01 | no | | intergenic_variant | MODIFIER |
| 12 | 108623739 | AX-170153076 | 1 | G | A | 0.125 | -4.91E-02 | 1.85E-01 | 2.68E+00 | 2.76E+00 | 7.85E-01 | no | Evl | intron_variant | MODIFIER |
| 9 | 105343327 | AX-170535544 | 1 | G | A | 0.134 | 2.98E-02 | 1.13E-01 | 2.73E+00 | 2.81E+00 | 7.86E-01 | no | Nek11 | intron_variant | MODIFIER |
| 16 | 91912727 | AX-170473548 | 0 | C | T | 0.474 | -2.16E-02 | 8.11E-02 | 2.68E+00 | 2.76E+00 | 7.86E-01 | no | Itsn1 | 3_prime_UTR_variant | MODIFIER |
| 1 | 66637773 | AX-170391851 | 1 | G | A | 0.116 | 6.24E-02 | 2.32E-01 | 2.69E+00 | 2.74E+00 | 7.86E-01 | no | Unc80 | intron_variant | MODIFIER |
| 5 | 73331409 | AX-170343940 | 1 | G | T | 0.438 | 2.66E-02 | 9.57E-02 | 2.61E+00 | 2.70E+00 | 7.86E-01 | no | Ociad2 | intron_variant | MODIFIER |
| 8 | 72722040 | AX-97794141 | 1 | C | T | 0.196 | -3.62E-02 | 1.34E-01 | 2.67E+00 | 2.73E+00 | 7.86E-01 | no | Sin3b | upstream_gene_variant | MODIFIER |
| 6 | 105656093 | AX-170488694 | 0 | C | A | 0.061 | 6.18E-02 | 2.22E-01 | 2.60E+00 | 2.70E+00 | 7.86E-01 | no | | intergenic_variant | MODIFIER |
| 5 | 108191010 | AX-170132471 | 0 | T | C | 0.447 | 2.79E-02 | 1.03E-01 | 2.66E+00 | 2.72E+00 | 7.86E-01 | no | Ccdc18 | intron_variant | MODIFIER |
| 1 | 60515043 | AX-170138548 | 2 | G | A | 0.1 | 5.57E-02 | 2.09E-01 | 2.65E+00 | 2.74E+00 | 7.86E-01 | no | Raph1 | intron_variant | MODIFIER |
| 1 | 13821768 | AX-170253899 | 1 | C | T | 0.375 | 2.11E-02 | 7.64E-02 | 2.64E+00 | 2.69E+00 | 7.86E-01 | no | | intergenic_variant | MODIFIER |
| 7 | 122597784 | AX-170229138 | 0 | A | G | 0.404 | 2.35E-02 | 8.79E-02 | 2.68E+00 | 2.75E+00 | 7.86E-01 | no | Prkcb | intron_variant | MODIFIER |
| 8 | 75293606 | AX-170031031 | 2 | T | C | 0.2 | -3.33E-02 | 1.24E-01 | 2.64E+00 | 2.73E+00 | 7.86E-01 | no | Gm45907 | upstream_gene_variant | MODIFIER |
| 12 | 109655614 | AX-170152850 | 0 | C | G | 0.132 | -4.22E-02 | 1.57E-01 | 2.64E+00 | 2.74E+00 | 7.86E-01 | no | DQ267102 | upstream_gene_variant | MODIFIER |
| 15 | 93615988 | AX-170012291 | 0 | G | C | 0.158 | 5.24E-02 | 1.94E-01 | 2.66E+00 | 2.72E+00 | 7.86E-01 | no | | intergenic_variant | MODIFIER |
| 8 | 83669813 | AX-170356302 | 0 | A | G | 0.202 | -3.57E-02 | 1.34E-01 | 2.70E+00 | 2.76E+00 | 7.86E-01 | no | Pkn1 | 3_prime_UTR_variant | MODIFIER |
| 7 | 17161868 | AX-170340809 | 0 | G | A | 0.36 | 2.36E-02 | 8.50E-02 | 2.59E+00 | 2.70E+00 | 7.86E-01 | no | Ceacam3 | missense_variant | MODERATE |
| 7 | 17886774 | AX-97803262 | 0 | G | A | 0.36 | 2.36E-02 | 8.50E-02 | 2.59E+00 | 2.70E+00 | 7.86E-01 | no | Gm5155 | intron_variant | MODIFIER |
| 1 | 71301837 | AX-170041605 | 1 | G | A | 0.143 | 6.36E-02 | 2.37E-01 | 2.68E+00 | 2.73E+00 | 7.86E-01 | no | Abca12 | intron_variant | MODIFIER |
| 15 | 70340465 | AX-170363206 | 0 | T | A | 0.553 | 2.19E-02 | 7.84E-02 | 2.61E+00 | 2.69E+00 | 7.86E-01 | no | | intergenic_variant | MODIFIER |
| 9 | 59095111 | AX-97836165 | 0 | G | A | 0.211 | -3.27E-02 | 1.23E-01 | 2.70E+00 | 2.75E+00 | 7.87E-01 | no | | intergenic_variant | MODIFIER |
| 1 | 136749717 | AX-170387655 | 0 | T | C | 0.211 | -2.80E-02 | 1.09E-01 | 2.67E+00 | 2.79E+00 | 7.87E-01 | no | | regulatory_region_variant | MODIFIER |
| 8 | 116005925 | AX-170257576 | 0 | G | A | 0.254 | 2.51E-02 | 9.79E-02 | 2.68E+00 | 2.80E+00 | 7.87E-01 | no | | intergenic_variant | MODIFIER |
| 2 | 100091412 | AX-170143788 | 1 | G | T | 0.054 | -8.01E-02 | 3.00E-01 | 2.73E+00 | 2.75E+00 | 7.87E-01 | no | | intergenic_variant | MODIFIER |
| 7 | 118737066 | AX-170481891 | 1 | T | C | 0.384 | 2.36E-02 | 8.68E-02 | 2.64E+00 | 2.72E+00 | 7.87E-01 | no | Ccp110 | downstream_gene_variant | MODIFIER |
| 5 | 108583687 | AX-170448817 | 0 | A | G | 0.456 | -2.95E-02 | 1.11E-01 | 2.75E+00 | 2.81E+00 | 7.87E-01 | no | Gak | intron_variant | MODIFIER |
| 5 | 111900381 | AX-170329986 | 0 | G | A | 0.456 | -2.95E-02 | 1.11E-01 | 2.75E+00 | 2.81E+00 | 7.87E-01 | no | | intergenic_variant | MODIFIER |
| 1 | 90454797 | AX-97794685 | 0 | G | A | 0.158 | 5.66E-02 | 2.06E-01 | 2.64E+00 | 2.70E+00 | 7.87E-01 | no | Gm28722 | upstream_gene_variant | MODIFIER |
| 8 | 39121553 | AX-170030530 | 1 | T | A | 0.152 | 3.83E-02 | 1.45E-01 | 2.72E+00 | 2.77E+00 | 7.87E-01 | no | Tusc3 | intron_variant | MODIFIER |
| 1 | 100425456 | AX-97851035 | 0 | C | T | 0.149 | 5.79E-02 | 2.18E-01 | 2.68E+00 | 2.75E+00 | 7.87E-01 | no | Cntnap5b | intron_variant | MODIFIER |
| 2 | 113665383 | AX-170473505 | 0 | C | T | 0.123 | 5.71E-02 | 2.11E-01 | 2.67E+00 | 2.71E+00 | 7.87E-01 | no | Fmn1 | intron_variant | MODIFIER |
| 2 | 120864121 | AX-170114791 | 0 | C | G | 0.123 | 5.71E-02 | 2.11E-01 | 2.67E+00 | 2.71E+00 | 7.87E-01 | no | Ubr1 | intron_variant | MODIFIER |
| 2 | 121509448 | AX-170064757 | 0 | T | G | 0.123 | 5.71E-02 | 2.11E-01 | 2.67E+00 | 2.71E+00 | 7.87E-01 | no | Wdr76 | intron_variant | MODIFIER |
| 2 | 123135421 | AX-170368887 | 0 | A | T | 0.123 | 5.71E-02 | 2.11E-01 | 2.67E+00 | 2.71E+00 | 7.87E-01 | no | | intergenic_variant | MODIFIER |
| 2 | 124749635 | AX-97813000 | 0 | A | T | 0.123 | 5.71E-02 | 2.11E-01 | 2.67E+00 | 2.71E+00 | 7.87E-01 | no | | intergenic_variant | MODIFIER |
| 2 | 129632356 | AX-170351338 | 0 | T | A | 0.123 | 5.71E-02 | 2.11E-01 | 2.67E+00 | 2.71E+00 | 7.87E-01 | no | Sirpa | downstream_gene_variant | MODIFIER |
| 2 | 129827739 | AX-170530765 | 0 | T | A | 0.123 | 5.71E-02 | 2.11E-01 | 2.67E+00 | 2.71E+00 | 7.87E-01 | no | Stk35 | intron_variant | MODIFIER |
| 2 | 144092011 | AX-97827951 | 0 | C | T | 0.123 | 5.71E-02 | 2.11E-01 | 2.67E+00 | 2.71E+00 | 7.87E-01 | no | | intergenic_variant | MODIFIER |

| | | | | | | | | | | | | | | | |
|---|---|---|---|---|---|---|---|---|---|---|---|---|---|---|---|
| 2 | 146861909 | AX-170319180 | 0 | T | C | 0.123 | 5.71E-02 | 2.11E-01 | 2.67E+00 | 2.71E+00 | 7.87E-01 | no | Kiz | intron_variant | MODIFIER |
| 2 | 147300240 | AX-170276211 | 0 | T | A | 0.123 | 5.71E-02 | 2.11E-01 | 2.67E+00 | 2.71E+00 | 7.87E-01 | no | Nkx2-2os | intron_variant,non_coding_transcript_variant | MODIFIER |
| 2 | 147781632 | AX-170526601 | 0 | T | A | 0.123 | 5.71E-02 | 2.11E-01 | 2.67E+00 | 2.71E+00 | 7.87E-01 | no | | intergenic_variant | MODIFIER |
| 2 | 156338540 | AX-170449852 | 0 | C | A | 0.123 | 5.71E-02 | 2.11E-01 | 2.67E+00 | 2.71E+00 | 7.87E-01 | no | Cnbd2 | intron_variant | MODIFIER |
| 2 | 156356234 | AX-170276946 | 0 | T | A | 0.123 | 5.71E-02 | 2.11E-01 | 2.67E+00 | 2.71E+00 | 7.87E-01 | no | Cnbd2 | intron_variant | MODIFIER |
| 8 | 90099515 | AX-170169205 | 1 | G | C | 0.196 | -3.24E-02 | 1.19E-01 | 2.64E+00 | 2.71E+00 | 7.87E-01 | no | | intergenic_variant | MODIFIER |
| 1 | 75587969 | AX-97811230 | 1 | C | T | 0.143 | 6.41E-02 | 2.41E-01 | 2.69E+00 | 2.74E+00 | 7.87E-01 | no | | regulatory_region_variant | MODIFIER |
| 9 | 34779771 | AX-97841669 | 1 | A | C | 0.438 | 2.37E-02 | 8.67E-02 | 2.64E+00 | 2.70E+00 | 7.87E-01 | no | Kirrel3 | intron_variant | MODIFIER |
| 4 | 143618782 | AX-97807073 | 0 | G | C | 0.096 | 4.99E-02 | 1.86E-01 | 2.65E+00 | 2.73E+00 | 7.87E-01 | no | Gm13083 | downstream_gene_variant | MODIFIER |
| 4 | 143842377 | AX-170024175 | 0 | T | C | 0.096 | 4.99E-02 | 1.86E-01 | 2.65E+00 | 2.73E+00 | 7.87E-01 | no | Gm13103 | upstream_gene_variant | MODIFIER |
| 4 | 144224124 | AX-170402344 | 0 | T | C | 0.096 | 4.99E-02 | 1.86E-01 | 2.65E+00 | 2.73E+00 | 7.87E-01 | no | | intergenic_variant | MODIFIER |
| 4 | 145083711 | AX-170501049 | 0 | T | C | 0.096 | 4.99E-02 | 1.86E-01 | 2.65E+00 | 2.73E+00 | 7.87E-01 | no | Vps13d | intron_variant | MODIFIER |
| 1 | 116615520 | AX-170242593 | 2 | A | T | 0.145 | 4.86E-02 | 1.79E-01 | 2.61E+00 | 2.72E+00 | 7.87E-01 | no | | intergenic_variant | MODIFIER |
| 12 | 61147720 | AX-170322187 | 0 | G | A | 0.07 | -6.10E-02 | 2.30E-01 | 2.67E+00 | 2.75E+00 | 7.87E-01 | no | | intergenic_variant | MODIFIER |
| 12 | 73606962 | AX-170502139 | 0 | G | A | 0.07 | -6.10E-02 | 2.30E-01 | 2.67E+00 | 2.75E+00 | 7.87E-01 | no | Prkch | intron_variant | MODIFIER |
| 18 | 36003534 | AX-170097349 | 0 | T | A | 0.07 | -6.10E-02 | 2.30E-01 | 2.67E+00 | 2.75E+00 | 7.87E-01 | no | Psd2 | intron_variant | MODIFIER |
| 18 | 42414593 | AX-170473120 | 0 | G | A | 0.07 | -6.10E-02 | 2.30E-01 | 2.67E+00 | 2.75E+00 | 7.87E-01 | no | | intergenic_variant | MODIFIER |
| 18 | 45184417 | AX-170502918 | 0 | C | T | 0.07 | -6.10E-02 | 2.30E-01 | 2.67E+00 | 2.75E+00 | 7.87E-01 | no | | intergenic_variant | MODIFIER |
| 18 | 57898834 | AX-170437048 | 1 | T | G | 0.402 | 2.88E-02 | 9.75E-02 | 2.57E+00 | 2.63E+00 | 7.87E-01 | no | Slc12a2 | intron_variant | MODIFIER |
| 2 | 123299476 | AX-97818153 | 1 | C | T | 0.143 | 5.91E-02 | 2.21E-01 | 2.69E+00 | 2.73E+00 | 7.87E-01 | no | | intergenic_variant | MODIFIER |
| 4 | 143401867 | AX-170343349 | 1 | G | C | 0.107 | 5.44E-02 | 2.05E-01 | 2.68E+00 | 2.74E+00 | 7.88E-01 | no | Pramel1 | downstream_gene_variant | MODIFIER |
| 10 | 10755792 | AX-97832894 | 1 | A | C | 0.205 | 2.81E-02 | 1.07E-01 | 2.69E+00 | 2.77E+00 | 7.88E-01 | no | Grm1 | intron_variant | MODIFIER |
| 8 | 39164048 | AX-170030531 | 2 | A | G | 0.136 | -3.57E-02 | 1.30E-01 | 2.62E+00 | 2.71E+00 | 7.88E-01 | no | Tusc3 | intron_variant | MODIFIER |
| 13 | 46318553 | AX-170051166 | 0 | A | T | 0.491 | -1.60E-02 | 6.16E-02 | 2.66E+00 | 2.77E+00 | 7.88E-01 | no | | intergenic_variant | MODIFIER |
| 8 | 96598787 | AX-170187241 | 0 | C | G | 0.211 | 3.15E-02 | 1.18E-01 | 2.67E+00 | 2.74E+00 | 7.88E-01 | no | | intergenic_variant | MODIFIER |
| 20 | 116581209 | AX-170034791 | 0 | G | A | 0.307 | -1.35E-02 | 5.27E-02 | 2.65E+00 | 2.78E+00 | 7.88E-01 | no | | intergenic_variant | MODIFIER |
| 18 | 56459033 | AX-170383841 | 0 | A | C | 0.202 | -2.96E-02 | 1.06E-01 | 2.61E+00 | 2.68E+00 | 7.88E-01 | no | Gramd3 | intron_variant | MODIFIER |
| 7 | 116261090 | AX-170240893 | 1 | G | C | 0.402 | 2.48E-02 | 9.25E-02 | 2.67E+00 | 2.72E+00 | 7.88E-01 | no | Plekha7 | intron_variant | MODIFIER |
| 7 | 69549090 | AX-170145368 | 0 | T | A | 0.421 | 2.78E-02 | 1.01E-01 | 2.62E+00 | 2.69E+00 | 7.88E-01 | no | 4930402F11Rik | intron_variant,non_coding_transcript_variant | MODIFIER |
| 10 | 12187787 | AX-97799977 | 1 | G | A | 0.205 | 2.79E-02 | 1.07E-01 | 2.70E+00 | 2.78E+00 | 7.88E-01 | no | | regulatory_region_variant | MODIFIER |
| 4 | 128910938 | AX-170130121 | 0 | G | A | 0.149 | -2.72E-02 | 1.07E-01 | 2.72E+00 | 2.83E+00 | 7.88E-01 | no | Trim62 | 3_prime_UTR_variant | MODIFIER |
| 18 | 52620057 | AX-170015711 | 0 | T | A | 0.061 | 4.18E-02 | 1.63E-01 | 2.62E+00 | 2.77E+00 | 7.88E-01 | no | Zfp474 | intron_variant | MODIFIER |
| 12 | 118981157 | AX-170007625 | 0 | C | T | 0.149 | -4.38E-02 | 1.70E-01 | 2.69E+00 | 2.79E+00 | 7.88E-01 | no | | intergenic_variant | MODIFIER |
| 1 | 180310557 | AX-170153929 | 0 | T | C | 0.237 | 2.94E-02 | 1.02E-01 | 2.56E+00 | 2.67E+00 | 7.88E-01 | no | | intergenic_variant | MODIFIER |
| 8 | 54153631 | AX-170080101 | 0 | T | C | 0.149 | -3.77E-02 | 1.34E-01 | 2.60E+00 | 2.67E+00 | 7.88E-01 | no | Vegfc | intron_variant | MODIFIER |
| 10 | 19348068 | AX-97809233 | 0 | G | T | 0.579 | 1.99E-02 | 6.64E-02 | 2.52E+00 | 2.65E+00 | 7.88E-01 | no | | intergenic_variant | MODIFIER |
| 7 | 114551154 | AX-97841028 | 0 | A | G | 0.421 | 2.42E-02 | 9.10E-02 | 2.66E+00 | 2.74E+00 | 7.88E-01 | no | Cyp2r1 | intron_variant | MODIFIER |
| 1 | 13569005 | AX-170518795 | 1 | A | T | 0.366 | 2.05E-02 | 7.60E-02 | 2.65E+00 | 2.71E+00 | 7.88E-01 | no | Tram1 | intron_variant | MODIFIER |
| 1 | 71197132 | AX-170311874 | 1 | G | A | 0.143 | 6.25E-02 | 2.36E-01 | 2.69E+00 | 2.75E+00 | 7.88E-01 | no | | intergenic_variant | MODIFIER |
| 15 | 82784277 | AX-170328400 | 1 | T | C | 0.545 | 2.29E-02 | 8.14E-02 | 2.59E+00 | 2.67E+00 | 7.89E-01 | no | Cyp2d41-ps | upstream_gene_variant | MODIFIER |
| 2 | 119383593 | AX-170327041 | 1 | A | G | 0.125 | 4.30E-02 | 1.59E-01 | 2.64E+00 | 2.71E+00 | 7.89E-01 | no | Ino80 | intron_variant | MODIFIER |
| 2 | 130055679 | AX-170432345 | 1 | T | C | 0.125 | 4.30E-02 | 1.59E-01 | 2.64E+00 | 2.71E+00 | 7.89E-01 | no | | intergenic_variant | MODIFIER |
| 7 | 102943278 | AX-170101423 | 0 | A | G | 0.474 | 2.59E-02 | 9.50E-02 | 2.63E+00 | 2.69E+00 | 7.89E-01 | no | Olfr573-ps1 | upstream_gene_variant | MODIFIER |
| 3 | 147469915 | AX-170312605 | 0 | T | C | 0.342 | -1.89E-02 | 7.26E-02 | 2.69E+00 | 2.77E+00 | 7.89E-01 | no | | intergenic_variant | MODIFIER |
| 3 | 147723509 | AX-170227670 | 0 | C | T | 0.342 | -1.89E-02 | 7.26E-02 | 2.69E+00 | 2.77E+00 | 7.89E-01 | no | | intergenic_variant | MODIFIER |
| 6 | 112737466 | AX-170387323 | 2 | C | T | 0.055 | 7.81E-02 | 2.95E-01 | 2.67E+00 | 2.73E+00 | 7.89E-01 | no | Srgap3 | intron_variant | MODIFIER |
| 15 | 96453422 | AX-170087945 | 1 | T | C | 0.152 | 4.84E-02 | 1.79E-01 | 2.64E+00 | 2.71E+00 | 7.89E-01 | no | Scaf11 | intron_variant | MODIFIER |
| 15 | 97529109 | AX-170479685 | 1 | C | T | 0.152 | 4.84E-02 | 1.79E-01 | 2.64E+00 | 2.71E+00 | 7.89E-01 | no | | intergenic_variant | MODIFIER |
| 16 | 78281228 | AX-170184119 | 0 | C | A | 0.456 | 2.05E-02 | 7.40E-02 | 2.61E+00 | 2.69E+00 | 7.89E-01 | no | E330011O21Rik | intron_variant,non_coding_transcript_variant | MODIFIER |
| 9 | 87362222 | AX-170314434 | 0 | A | T | 0.132 | -3.64E-02 | 1.26E-01 | 2.57E+00 | 2.66E+00 | 7.89E-01 | no | | intergenic_variant | MODIFIER |
| 5 | 114322725 | AX-170025929 | 0 | A | C | 0.474 | 2.96E-02 | 1.08E-01 | 2.64E+00 | 2.69E+00 | 7.89E-01 | no | Myo1h | intron_variant | MODIFIER |
| 7 | 94168562 | AX-170277627 | 0 | C | T | 0.114 | -4.72E-02 | 1.75E-01 | 2.61E+00 | 2.72E+00 | 7.89E-01 | no | Gm32647 | intron_variant,non_coding_transcript_variant | MODIFIER |
| 2 | 115850763 | AX-170018762 | 0 | T | A | 0.175 | 3.46E-02 | 1.29E-01 | 2.64E+00 | 2.72E+00 | 7.89E-01 | no | | intergenic_variant | MODIFIER |
| 7 | 60555064 | AX-170509685 | 1 | A | G | 0.411 | -4.39E-02 | 1.67E-01 | 2.60E+00 | 2.74E+00 | 7.89E-01 | no | | intergenic_variant | MODIFIER |

| | | | | | | | | | | | | | | | |
|---|---|---|---|---|---|---|---|---|---|---|---|---|---|---|---|
| 10 | 29413328 | AX-170495246 | 1 | C | T | 0.196 | 2.32E-02 | 9.19E-02 | 2.68E+00 | 2.80E+00 | 7.89E-01 | no | | intergenic_variant | MODIFIER |
| 1 | 30539786 | AX-170430363 | 0 | A | T | 0.421 | 2.15E-02 | 8.16E-02 | 2.67E+00 | 2.74E+00 | 7.89E-01 | no | | intergenic_variant | MODIFIER |
| 18 | 58161829 | AX-170157290 | 2 | C | T | 0.409 | 2.22E-02 | 8.54E-02 | 2.76E+00 | 2.83E+00 | 7.89E-01 | no | Fbn2 | intron_variant | MODIFIER |
| 9 | 62323278 | AX-170090316 | 0 | G | T | 0.202 | 3.38E-02 | 1.29E-01 | 2.74E+00 | 2.79E+00 | 7.90E-01 | no | Gm35288 | intron_variant,non_coding_transcript_variant | MODIFIER |
| 5 | 90746778 | AX-170389528 | 1 | C | T | 0.42 | -2.78E-02 | 1.06E-01 | 2.68E+00 | 2.74E+00 | 7.90E-01 | no | Gm43090 | upstream_gene_variant | MODIFIER |
| 8 | 15089162 | AX-97785670 | 1 | T | C | 0.232 | -2.10E-02 | 7.46E-02 | 2.58E+00 | 2.68E+00 | 7.90E-01 | no | Myom2 | intron_variant | MODIFIER |
| 9 | 27827784 | AX-170259500 | 0 | G | A | 0.465 | 2.54E-02 | 9.20E-02 | 2.61E+00 | 2.68E+00 | 7.90E-01 | no | Opcml | intron_variant | MODIFIER |
| 20 | 165996631 | AX-170444926 | 1 | G | T | 0.125 | 3.41E-02 | 1.31E-01 | 2.63E+00 | 2.75E+00 | 7.90E-01 | no | | intergenic_variant | MODIFIER |
| 1 | 38163261 | AX-97811308 | 1 | C | T | 0.464 | -2.04E-02 | 7.73E-02 | 2.65E+00 | 2.73E+00 | 7.90E-01 | no | Gm5099 | downstream_gene_variant | MODIFIER |
| 7 | 122642297 | AX-170108588 | 1 | C | T | 0.393 | 2.14E-02 | 8.25E-02 | 2.67E+00 | 2.76E+00 | 7.90E-01 | no | | intergenic_variant | MODIFIER |
| 12 | 116462549 | AX-170087342 | 0 | C | T | 0.14 | -5.00E-02 | 1.91E-01 | 2.68E+00 | 2.75E+00 | 7.90E-01 | no | Ncapg2 | downstream_gene_variant | MODIFIER |
| 16 | 73609061 | AX-170438234 | 1 | A | G | 0.027 | 3.62E-02 | 1.40E-01 | 2.57E+00 | 2.75E+00 | 7.90E-01 | no | | intergenic_variant | MODIFIER |
| 1 | 78820086 | AX-170200316 | 0 | G | A | 0.14 | 4.98E-02 | 1.81E-01 | 2.59E+00 | 2.70E+00 | 7.90E-01 | no | Kcne4 | downstream_gene_variant | MODIFIER |
| 17 | 80591760 | AX-97847773 | 1 | G | T | 0.054 | 4.35E-02 | 1.70E-01 | 2.65E+00 | 2.77E+00 | 7.90E-01 | no | Map4k3 | intron_variant | MODIFIER |
| 15 | 98990851 | AX-170385367 | 1 | A | T | 0.107 | 4.78E-02 | 1.78E-01 | 2.65E+00 | 2.71E+00 | 7.90E-01 | no | 4930578M01Rik | downstream_gene_variant | MODIFIER |
| 16 | 93815921 | AX-170323527 | 0 | A | G | 0.412 | -2.25E-02 | 8.11E-02 | 2.60E+00 | 2.68E+00 | 7.90E-01 | no | Gm30881 | intron_variant,non_coding_transcript_variant | MODIFIER |
| 16 | 93851108 | AX-170361368 | 0 | A | G | 0.412 | -2.25E-02 | 8.11E-02 | 2.60E+00 | 2.68E+00 | 7.90E-01 | no | Morc3 | intron_variant | MODIFIER |
| 16 | 93854943 | AX-170184499 | 0 | T | C | 0.412 | -2.25E-02 | 8.11E-02 | 2.60E+00 | 2.68E+00 | 7.90E-01 | no | Morc3 | intron_variant | MODIFIER |
| 16 | 93866178 | AX-97808596 | 0 | G | A | 0.412 | -2.25E-02 | 8.11E-02 | 2.60E+00 | 2.68E+00 | 7.90E-01 | no | Morc3 | intron_variant | MODIFIER |
| 16 | 93879439 | AX-170107283 | 0 | G | A | 0.412 | -2.25E-02 | 8.11E-02 | 2.60E+00 | 2.68E+00 | 7.90E-01 | no | Chaf1b | upstream_gene_variant | MODIFIER |
| 16 | 93888158 | AX-170431729 | 0 | T | C | 0.412 | -2.25E-02 | 8.11E-02 | 2.60E+00 | 2.68E+00 | 7.90E-01 | no | Chaf1b | intron_variant | MODIFIER |
| 16 | 93934507 | AX-170287515 | 0 | G | C | 0.412 | -2.25E-02 | 8.11E-02 | 2.60E+00 | 2.68E+00 | 7.90E-01 | no | Cldn14 | intron_variant | MODIFIER |
| 16 | 93952840 | AX-170397972 | 0 | T | C | 0.412 | -2.25E-02 | 8.11E-02 | 2.60E+00 | 2.68E+00 | 7.90E-01 | no | Cldn14 | intron_variant | MODIFIER |
| 16 | 93957971 | AX-170324510 | 0 | G | T | 0.412 | -2.25E-02 | 8.11E-02 | 2.60E+00 | 2.68E+00 | 7.90E-01 | no | Cldn14 | intron_variant | MODIFIER |
| 16 | 93969894 | AX-170251144 | 0 | A | T | 0.412 | -2.25E-02 | 8.11E-02 | 2.60E+00 | 2.68E+00 | 7.90E-01 | no | Cldn14 | intron_variant | MODIFIER |
| 16 | 93982328 | AX-170314473 | 0 | C | G | 0.412 | -2.25E-02 | 8.11E-02 | 2.60E+00 | 2.68E+00 | 7.90E-01 | no | Cldn14 | intron_variant | MODIFIER |
| 16 | 93989953 | AX-170304780 | 0 | G | A | 0.412 | -2.25E-02 | 8.11E-02 | 2.60E+00 | 2.68E+00 | 7.90E-01 | no | Cldn14 | intron_variant | MODIFIER |
| 16 | 94024143 | AX-170449611 | 0 | C | T | 0.412 | -2.25E-02 | 8.11E-02 | 2.60E+00 | 2.68E+00 | 7.90E-01 | no | | regulatory_region_variant | MODIFIER |
| 16 | 94167562 | AX-170294398 | 0 | G | A | 0.412 | -2.25E-02 | 8.11E-02 | 2.60E+00 | 2.68E+00 | 7.90E-01 | no | Hlcs | intron_variant | MODIFIER |
| 16 | 94172529 | AX-170278623 | 0 | G | A | 0.412 | -2.25E-02 | 8.11E-02 | 2.60E+00 | 2.68E+00 | 7.90E-01 | no | Hlcs | intron_variant | MODIFIER |
| 16 | 94187576 | AX-170226925 | 0 | C | T | 0.412 | -2.25E-02 | 8.11E-02 | 2.60E+00 | 2.68E+00 | 7.90E-01 | no | Hlcs | intron_variant | MODIFIER |
| 16 | 94197013 | AX-170325699 | 0 | A | C | 0.412 | -2.25E-02 | 8.11E-02 | 2.60E+00 | 2.68E+00 | 7.90E-01 | no | Hlcs | intron_variant | MODIFIER |
| 16 | 94204088 | AX-170406275 | 0 | T | A | 0.412 | -2.25E-02 | 8.11E-02 | 2.60E+00 | 2.68E+00 | 7.90E-01 | no | Hlcs | intron_variant | MODIFIER |
| 16 | 94204829 | AX-97826351 | 0 | G | A | 0.412 | -2.25E-02 | 8.11E-02 | 2.60E+00 | 2.68E+00 | 7.90E-01 | no | Hlcs | intron_variant | MODIFIER |
| 16 | 94205101 | AX-170240965 | 0 | G | A | 0.412 | -2.25E-02 | 8.11E-02 | 2.60E+00 | 2.68E+00 | 7.90E-01 | no | Hlcs | intron_variant | MODIFIER |
| 16 | 94354852 | AX-170013686 | 0 | C | G | 0.412 | -2.25E-02 | 8.11E-02 | 2.60E+00 | 2.68E+00 | 7.90E-01 | no | Pigp | downstream_gene_variant | MODIFIER |
| 16 | 94367174 | AX-170190478 | 0 | G | T | 0.412 | -2.25E-02 | 8.11E-02 | 2.60E+00 | 2.68E+00 | 7.90E-01 | no | Pigp | intron_variant | MODIFIER |
| 16 | 94367451 | AX-170013683 | 0 | G | C | 0.412 | -2.25E-02 | 8.11E-02 | 2.60E+00 | 2.68E+00 | 7.90E-01 | no | Pigp | intron_variant | MODIFIER |
| 16 | 94380364 | AX-170058158 | 0 | C | A | 0.412 | -2.25E-02 | 8.11E-02 | 2.60E+00 | 2.68E+00 | 7.90E-01 | no | Ttc3 | intron_variant | MODIFIER |
| 16 | 94387724 | AX-170187940 | 0 | T | C | 0.412 | -2.25E-02 | 8.11E-02 | 2.60E+00 | 2.68E+00 | 7.90E-01 | no | Ttc3 | intron_variant | MODIFIER |
| 16 | 94387771 | AX-170500446 | 0 | T | C | 0.412 | -2.25E-02 | 8.11E-02 | 2.60E+00 | 2.68E+00 | 7.90E-01 | no | Ttc3 | intron_variant | MODIFIER |
| 16 | 94414490 | AX-170381042 | 0 | G | T | 0.412 | -2.25E-02 | 8.11E-02 | 2.60E+00 | 2.68E+00 | 7.90E-01 | no | Ttc3 | intron_variant | MODIFIER |
| 16 | 94426055 | AX-170464162 | 0 | T | G | 0.412 | -2.25E-02 | 8.11E-02 | 2.60E+00 | 2.68E+00 | 7.90E-01 | no | Gm23692 | upstream_gene_variant | MODIFIER |
| 16 | 94427763 | AX-170117762 | 0 | G | A | 0.412 | -2.25E-02 | 8.11E-02 | 2.60E+00 | 2.68E+00 | 7.90E-01 | no | Gm23692 | upstream_gene_variant | MODIFIER |
| 16 | 94428493 | AX-170519707 | 0 | A | G | 0.412 | -2.25E-02 | 8.11E-02 | 2.60E+00 | 2.68E+00 | 7.90E-01 | no | Gm23692 | upstream_gene_variant | MODIFIER |
| 16 | 94432601 | AX-170395507 | 0 | C | T | 0.412 | -2.25E-02 | 8.11E-02 | 2.60E+00 | 2.68E+00 | 7.90E-01 | no | Gm23692 | downstream_gene_variant | MODIFIER |
| 16 | 94441884 | AX-170369507 | 0 | A | G | 0.412 | -2.25E-02 | 8.11E-02 | 2.60E+00 | 2.68E+00 | 7.90E-01 | no | Ttc3 | intron_variant | MODIFIER |
| 16 | 94452854 | AX-170408535 | 0 | A | G | 0.412 | -2.25E-02 | 8.11E-02 | 2.60E+00 | 2.68E+00 | 7.90E-01 | no | Ttc3 | intron_variant | MODIFIER |
| 16 | 94456864 | AX-170403343 | 0 | A | G | 0.412 | -2.25E-02 | 8.11E-02 | 2.60E+00 | 2.68E+00 | 7.90E-01 | no | Ttc3 | intron_variant | MODIFIER |
| 16 | 94457969 | AX-170248782 | 0 | C | T | 0.412 | -2.25E-02 | 8.11E-02 | 2.60E+00 | 2.68E+00 | 7.90E-01 | no | Ttc3 | synonymous_variant | LOW |
| 16 | 94468768 | AX-170534502 | 0 | A | G | 0.412 | -2.25E-02 | 8.11E-02 | 2.60E+00 | 2.68E+00 | 7.90E-01 | no | Ttc3 | 3_prime_UTR_variant | MODIFIER |
| 16 | 94471812 | AX-170497090 | 0 | T | C | 0.412 | -2.25E-02 | 8.11E-02 | 2.60E+00 | 2.68E+00 | 7.90E-01 | no | Ttc3 | downstream_gene_variant | MODIFIER |
| 16 | 94474578 | AX-170353005 | 0 | G | A | 0.412 | -2.25E-02 | 8.11E-02 | 2.60E+00 | 2.68E+00 | 7.90E-01 | no | | regulatory_region_variant | MODIFIER |
| 16 | 94493764 | AX-170514407 | 0 | T | G | 0.412 | -2.25E-02 | 8.11E-02 | 2.60E+00 | 2.68E+00 | 7.90E-01 | no | Vps26c | downstream_gene_variant | MODIFIER |

| | | | | | | | | | | | | | | | |
|---|---|---|---|---|---|---|---|---|---|---|---|---|---|---|---|
| 16 | 94498167 | AX-97799782 | 0 | G | A | 0.412 | -2.25E-02 | 8.11E-02 | 2.60E+00 | 2.68E+00 | 7.90E-01 | no | Vps26c | 3_prime_UTR_variant | MODIFIER |
| 16 | 94515048 | AX-170277358 | 0 | C | G | 0.412 | -2.25E-02 | 8.11E-02 | 2.60E+00 | 2.68E+00 | 7.90E-01 | no | Vps26c | intron_variant | MODIFIER |
| 16 | 94515159 | AX-97843732 | 0 | C | A | 0.412 | -2.25E-02 | 8.11E-02 | 2.60E+00 | 2.68E+00 | 7.90E-01 | no | Vps26c | intron_variant | MODIFIER |
| 16 | 94652590 | AX-170354994 | 0 | T | G | 0.412 | -2.25E-02 | 8.11E-02 | 2.60E+00 | 2.68E+00 | 7.90E-01 | no | Dyrk1a | intron_variant | MODIFIER |
| 16 | 94663024 | AX-97831276 | 0 | T | C | 0.412 | -2.25E-02 | 8.11E-02 | 2.60E+00 | 2.68E+00 | 7.90E-01 | no | Dyrk1a | intron_variant | MODIFIER |
| 16 | 94756404 | AX-170230763 | 0 | T | C | 0.412 | -2.25E-02 | 8.11E-02 | 2.60E+00 | 2.68E+00 | 7.90E-01 | no | Kcnj6 | intron_variant | MODIFIER |
| 16 | 94756419 | AX-170426906 | 0 | G | A | 0.412 | -2.25E-02 | 8.11E-02 | 2.60E+00 | 2.68E+00 | 7.90E-01 | no | Kcnj6 | intron_variant | MODIFIER |
| 16 | 94758306 | AX-97830343 | 0 | G | A | 0.412 | -2.25E-02 | 8.11E-02 | 2.60E+00 | 2.68E+00 | 7.90E-01 | no | Kcnj6 | intron_variant | MODIFIER |
| 18 | 68659900 | AX-170523990 | 1 | G | A | 0.08 | 4.60E-02 | 1.70E-01 | 2.60E+00 | 2.71E+00 | 7.90E-01 | no | | intergenic_variant | MODIFIER |
| 5 | 123668435 | AX-170125626 | 2 | C | T | 0.464 | 2.67E-02 | 1.00E-01 | 2.64E+00 | 2.72E+00 | 7.90E-01 | no | Clip1 | intron_variant | MODIFIER |
| 4 | 150445707 | AX-170371607 | 1 | G | A | 0.098 | 5.26E-02 | 2.00E-01 | 2.67E+00 | 2.74E+00 | 7.90E-01 | no | Rere | intron_variant | MODIFIER |
| 15 | 79130277 | AX-170313430 | 1 | T | C | 0.554 | 2.29E-02 | 8.19E-02 | 2.59E+00 | 2.67E+00 | 7.90E-01 | no | Micall1 | intron_variant | MODIFIER |
| 8 | 94875861 | AX-170035791 | 0 | A | G | 0.202 | 2.99E-02 | 1.15E-01 | 2.68E+00 | 2.75E+00 | 7.91E-01 | no | Dok4 | intron_variant | MODIFIER |
| 12 | 94428551 | AX-170007305 | 1 | C | T | 0.08 | 5.15E-02 | 2.01E-01 | 2.69E+00 | 2.78E+00 | 7.91E-01 | no | | intergenic_variant | MODIFIER |
| 1 | 13347293 | AX-170234965 | 0 | C | T | 0.377 | -1.91E-02 | 7.44E-02 | 2.74E+00 | 2.82E+00 | 7.91E-01 | no | Ncoa2 | intron_variant | MODIFIER |
| 9 | 62727720 | AX-170116326 | 0 | G | A | 0.211 | -3.24E-02 | 1.22E-01 | 2.66E+00 | 2.72E+00 | 7.91E-01 | no | Itga11 | intron_variant | MODIFIER |
| 8 | 93791040 | AX-170340798 | 0 | G | T | 0.219 | 3.02E-02 | 1.16E-01 | 2.70E+00 | 2.76E+00 | 7.91E-01 | no | 4930488L21Rik | intron_variant,non_coding_transcript_variant | MODIFIER |
| 10 | 17255856 | AX-170310810 | 1 | C | T | 0.205 | -2.77E-02 | 1.02E-01 | 2.61E+00 | 2.70E+00 | 7.91E-01 | no | | intergenic_variant | MODIFIER |
| 18 | 57440411 | AX-170142747 | 1 | A | G | 0.411 | 2.84E-02 | 9.61E-02 | 2.55E+00 | 2.62E+00 | 7.91E-01 | no | 4930511M06Rik | intron_variant,non_coding_transcript_variant | MODIFIER |
| 12 | 104358224 | AX-170375463 | 1 | T | C | 0.116 | -5.14E-02 | 1.99E-01 | 2.70E+00 | 2.77E+00 | 7.91E-01 | no | Serpina3m | intron_variant | MODIFIER |
| 8 | 56940169 | AX-97845493 | 1 | G | A | 0.143 | -4.05E-02 | 1.48E-01 | 2.62E+00 | 2.68E+00 | 7.91E-01 | no | | intergenic_variant | MODIFIER |
| 9 | 66640455 | AX-170106748 | 2 | A | G | 0.182 | 3.29E-02 | 1.28E-01 | 2.75E+00 | 2.81E+00 | 7.91E-01 | no | | intergenic_variant | MODIFIER |
| 18 | 56186869 | AX-170015760 | 1 | G | T | 0.205 | 2.76E-02 | 1.09E-01 | 2.74E+00 | 2.84E+00 | 7.91E-01 | no | | intergenic_variant | MODIFIER |
| 5 | 28011892 | AX-170257863 | 0 | C | T | 0.333 | 2.89E-02 | 1.10E-01 | 2.68E+00 | 2.73E+00 | 7.91E-01 | no | | intergenic_variant | MODIFIER |
| 6 | 105762213 | AX-97848191 | 1 | G | A | 0.062 | 4.83E-02 | 1.89E-01 | 2.68E+00 | 2.78E+00 | 7.91E-01 | no | Cntn4 | intron_variant | MODIFIER |
| 4 | 150263995 | AX-97856205 | 0 | G | T | 0.096 | -4.37E-02 | 1.67E-01 | 2.64E+00 | 2.74E+00 | 7.91E-01 | no | | intergenic_variant | MODIFIER |
| 10 | 12529932 | AX-97826232 | 1 | C | A | 0.205 | 2.67E-02 | 1.04E-01 | 2.69E+00 | 2.78E+00 | 7.91E-01 | no | Utrn | intron_variant | MODIFIER |
| 1 | 47079796 | AX-97818762 | 1 | C | A | 0.098 | -5.75E-02 | 2.22E-01 | 2.68E+00 | 2.76E+00 | 7.91E-01 | no | | intergenic_variant | MODIFIER |
| 16 | 94604586 | AX-170395979 | 0 | G | C | 0.412 | -2.22E-02 | 8.13E-02 | 2.61E+00 | 2.69E+00 | 7.91E-01 | no | Dyrk1a | intron_variant | MODIFIER |
| 2 | 33368363 | AX-97836746 | 1 | A | G | 0.045 | 4.07E-02 | 1.64E-01 | 2.66E+00 | 2.80E+00 | 7.91E-01 | no | Ralgps1 | intron_variant | MODIFIER |
| 8 | 48454647 | AX-170447719 | 1 | G | T | 0.152 | 3.73E-02 | 1.45E-01 | 2.74E+00 | 2.81E+00 | 7.91E-01 | no | Tenm3 | intron_variant | MODIFIER |
| 12 | 116147263 | AX-170007586 | 0 | C | T | 0.105 | -4.48E-02 | 1.74E-01 | 2.64E+00 | 2.76E+00 | 7.91E-01 | no | Vipr2 | downstream_gene_variant | MODIFIER |
| 2 | 156530440 | AX-170451662 | 1 | C | G | 0.143 | 5.79E-02 | 2.21E-01 | 2.69E+00 | 2.72E+00 | 7.92E-01 | no | Epb41l1 | intron_variant | MODIFIER |
| 5 | 97277685 | AX-170511042 | 0 | C | G | 0.447 | 2.75E-02 | 1.05E-01 | 2.67E+00 | 2.73E+00 | 7.92E-01 | no | | regulatory_region_variant | MODIFIER |
| 12 | 93543820 | AX-170364616 | 0 | A | T | 0.061 | 6.16E-02 | 2.37E-01 | 2.69E+00 | 2.75E+00 | 7.92E-01 | no | | intergenic_variant | MODIFIER |
| 5 | 28524858 | AX-170518413 | 1 | T | C | 0.357 | -2.60E-02 | 1.01E-01 | 2.69E+00 | 2.77E+00 | 7.92E-01 | no | 9530036O11Rik | intron_variant,non_coding_transcript_variant | MODIFIER |
| 8 | 48648893 | AX-97850947 | 1 | C | T | 0.152 | 3.90E-02 | 1.50E-01 | 2.78E+00 | 2.84E+00 | 7.92E-01 | no | Tenm3 | intron_variant | MODIFIER |
| 9 | 70289356 | AX-170503340 | 1 | C | T | 0.214 | 3.24E-02 | 1.26E-01 | 2.71E+00 | 2.77E+00 | 7.92E-01 | no | Myo1e | intron_variant | MODIFIER |
| 16 | 86168639 | AX-170192645 | 2 | C | A | 0.436 | 2.06E-02 | 7.83E-02 | 2.64E+00 | 2.72E+00 | 7.92E-01 | no | | intergenic_variant | MODIFIER |
| 2 | 71807159 | AX-170018022 | 0 | A | T | 0.044 | -4.55E-02 | 1.70E-01 | 2.58E+00 | 2.72E+00 | 7.92E-01 | no | Itga6 | intron_variant | MODIFIER |
| 1 | 70890065 | AX-170267022 | 0 | G | T | 0.149 | 5.09E-02 | 1.88E-01 | 2.60E+00 | 2.70E+00 | 7.92E-01 | no | Vwc2l | downstream_gene_variant | MODIFIER |
| 1 | 78699749 | AX-170499532 | 0 | C | T | 0.149 | 5.09E-02 | 1.88E-01 | 2.60E+00 | 2.70E+00 | 7.92E-01 | no | Acsl3 | splice_region_variant,intron_variant | LOW |
| 1 | 71259132 | AX-97784792 | 1 | C | T | 0.143 | 6.21E-02 | 2.38E-01 | 2.68E+00 | 2.73E+00 | 7.92E-01 | no | Abca12 | intron_variant | MODIFIER |
| 1 | 78539270 | AX-170238332 | 1 | C | T | 0.143 | 6.21E-02 | 2.38E-01 | 2.68E+00 | 2.73E+00 | 7.92E-01 | no | Mogat1 | downstream_gene_variant | MODIFIER |
| 12 | 99668374 | AX-170395592 | 0 | T | C | 0.053 | -4.68E-02 | 1.81E-01 | 2.64E+00 | 2.74E+00 | 7.92E-01 | no | | intergenic_variant | MODIFIER |
| 9 | 66630560 | AX-97815465 | 1 | A | G | 0.188 | 3.38E-02 | 1.30E-01 | 2.71E+00 | 2.75E+00 | 7.92E-01 | no | | regulatory_region_variant | MODIFIER |
| 9 | 29470593 | AX-170090268 | 0 | C | T | 0.439 | -2.19E-02 | 8.65E-02 | 2.74E+00 | 2.82E+00 | 7.92E-01 | no | Ntm | intron_variant | MODIFIER |
| 15 | 100787801 | AX-170056483 | 0 | T | A | 0.105 | 4.66E-02 | 1.77E-01 | 2.65E+00 | 2.71E+00 | 7.92E-01 | no | Slc4a8 | intron_variant | MODIFIER |
| 8 | 36317709 | AX-170154168 | 1 | A | G | 0.152 | 3.82E-02 | 1.48E-01 | 2.74E+00 | 2.79E+00 | 7.93E-01 | no | | intergenic_variant | MODIFIER |
| 8 | 42565310 | AX-170079901 | 1 | G | A | 0.152 | 3.82E-02 | 1.48E-01 | 2.74E+00 | 2.79E+00 | 7.93E-01 | no | | intergenic_variant | MODIFIER |
| 8 | 43062580 | AX-170199692 | 1 | G | T | 0.152 | 3.82E-02 | 1.48E-01 | 2.74E+00 | 2.79E+00 | 7.93E-01 | no | | intergenic_variant | MODIFIER |
| 1 | 21973798 | AX-170149955 | 1 | T | C | 0.393 | 2.33E-02 | 8.58E-02 | 2.63E+00 | 2.67E+00 | 7.93E-01 | no | | intergenic_variant | MODIFIER |
| 5 | 125762985 | AX-170482271 | 0 | C | T | 0.447 | -2.79E-02 | 1.08E-01 | 2.70E+00 | 2.76E+00 | 7.93E-01 | no | Tmem132b | intron_variant | MODIFIER |
| 2 | 115785815 | AX-97833051 | 2 | G | A | 0.145 | -3.92E-02 | 1.54E-01 | 2.64E+00 | 2.76E+00 | 7.93E-01 | no | | regulatory_region_variant | MODIFIER |

| | | | | | | | | | | | | | | | |
|---|---|---|---|---|---|---|---|---|---|---|---|---|---|---|---|
| 18 | 47098111 | AX-170149930 | 0 | A | G | 0.053 | 6.07E-02 | 2.36E-01 | 2.63E+00 | 2.75E+00 | 7.93E-01 | no | Commd10 | intron_variant,NMD_transcript_variant | MODIFIER |
| 20 | 167049740 | AX-170389608 | 0 | T | G | 0.202 | 3.63E-02 | 1.34E-01 | 2.59E+00 | 2.70E+00 | 7.93E-01 | no | | regulatory_region_variant | MODIFIER |
| 20 | 167117452 | AX-170035431 | 0 | C | T | 0.202 | 3.63E-02 | 1.34E-01 | 2.59E+00 | 2.70E+00 | 7.93E-01 | no | | intergenic_variant | MODIFIER |
| 1 | 137368238 | AX-170169669 | 0 | A | G | 0.211 | 3.17E-02 | 1.17E-01 | 2.60E+00 | 2.69E+00 | 7.93E-01 | no | Gm23534 | upstream_gene_variant | MODIFIER |
| 1 | 76166143 | AX-170324645 | 1 | C | T | 0.134 | 5.86E-02 | 2.26E-01 | 2.67E+00 | 2.74E+00 | 7.93E-01 | no | | intergenic_variant | MODIFIER |
| 10 | 19346806 | AX-170101241 | 0 | C | A | 0.588 | 1.93E-02 | 6.59E-02 | 2.52E+00 | 2.66E+00 | 7.93E-01 | no | | intergenic_variant | MODIFIER |
| 10 | 19347443 | AX-170178070 | 0 | A | G | 0.588 | 1.93E-02 | 6.59E-02 | 2.52E+00 | 2.66E+00 | 7.93E-01 | no | | intergenic_variant | MODIFIER |
| 10 | 19351373 | AX-170324092 | 0 | C | A | 0.588 | 1.93E-02 | 6.59E-02 | 2.52E+00 | 2.66E+00 | 7.93E-01 | no | | intergenic_variant | MODIFIER |
| 10 | 19359596 | AX-170158503 | 0 | C | T | 0.588 | 1.93E-02 | 6.59E-02 | 2.52E+00 | 2.66E+00 | 7.93E-01 | no | Olig3 | downstream_gene_variant | MODIFIER |
| 10 | 19409600 | AX-170002862 | 0 | A | G | 0.588 | 1.93E-02 | 6.59E-02 | 2.52E+00 | 2.66E+00 | 7.93E-01 | no | | intergenic_variant | MODIFIER |
| 10 | 19409622 | AX-170002861 | 0 | T | C | 0.588 | 1.93E-02 | 6.59E-02 | 2.52E+00 | 2.66E+00 | 7.93E-01 | no | | intergenic_variant | MODIFIER |
| 11 | 33997057 | AX-170273670 | 0 | A | G | 0.096 | -4.74E-02 | 1.88E-01 | 2.69E+00 | 2.78E+00 | 7.93E-01 | no | Kcnip1 | upstream_gene_variant | MODIFIER |
| 1 | 155721749 | AX-170379647 | 0 | G | A | 0.228 | 3.62E-02 | 1.34E-01 | 2.63E+00 | 2.68E+00 | 7.93E-01 | no | Lhx4 | intron_variant | MODIFIER |
| 18 | 65215928 | AX-170384473 | 0 | T | C | 0.193 | -3.08E-02 | 1.15E-01 | 2.63E+00 | 2.70E+00 | 7.93E-01 | no | Nedd4l | downstream_gene_variant | MODIFIER |
| 14 | 62872010 | AX-170053573 | 0 | A | G | 0.053 | 4.85E-02 | 1.94E-01 | 2.69E+00 | 2.80E+00 | 7.93E-01 | no | Wdfy2 | intron_variant | MODIFIER |
| 15 | 89564891 | AX-170338799 | 1 | C | T | 0.152 | 4.73E-02 | 1.79E-01 | 2.64E+00 | 2.71E+00 | 7.94E-01 | no | Acr | upstream_gene_variant | MODIFIER |
| 4 | 80992165 | AX-170023223 | 0 | T | C | 0.228 | 2.73E-02 | 9.28E-02 | 2.53E+00 | 2.64E+00 | 7.94E-01 | no | | intergenic_variant | MODIFIER |
| 7 | 117961092 | AX-170138583 | 1 | A | G | 0.393 | 2.40E-02 | 9.19E-02 | 2.67E+00 | 2.72E+00 | 7.94E-01 | no | | intergenic_variant | MODIFIER |
| 13 | 36132286 | AX-170242006 | 1 | C | T | 0.179 | -3.25E-02 | 1.20E-01 | 2.61E+00 | 2.68E+00 | 7.94E-01 | no | Fars2 | intron_variant | MODIFIER |
| 4 | 133587078 | AX-170070742 | 0 | A | T | 0.158 | 3.17E-02 | 1.13E-01 | 2.58E+00 | 2.65E+00 | 7.94E-01 | no | Gpn2 | intron_variant | MODIFIER |
| 9 | 108500600 | AX-170454070 | 1 | T | C | 0.116 | 2.84E-02 | 1.14E-01 | 2.74E+00 | 2.83E+00 | 7.94E-01 | no | Usp19 | intron_variant | MODIFIER |
| 6 | 147583223 | AX-170028501 | 0 | A | G | 0.053 | -4.70E-02 | 1.81E-01 | 2.61E+00 | 2.73E+00 | 7.94E-01 | no | Ccdc91 | intron_variant | MODIFIER |
| 11 | 33394135 | AX-170205306 | 1 | A | T | 0.089 | -4.50E-02 | 1.79E-01 | 2.68E+00 | 2.77E+00 | 7.94E-01 | no | Ranbp17 | intron_variant | MODIFIER |
| 7 | 93105335 | AX-170106764 | 0 | G | A | 0.132 | -5.44E-02 | 2.13E-01 | 2.67E+00 | 2.75E+00 | 7.94E-01 | no | Gm26862 | downstream_gene_variant | MODIFIER |
| 7 | 97682986 | AX-170433228 | 0 | G | A | 0.132 | -5.44E-02 | 2.13E-01 | 2.67E+00 | 2.75E+00 | 7.94E-01 | no | Rsf1 | intron_variant | MODIFIER |
| 2 | 113658312 | AX-170246737 | 0 | A | G | 0.132 | 4.87E-02 | 1.87E-01 | 2.64E+00 | 2.72E+00 | 7.94E-01 | no | Fmn1 | intron_variant | MODIFIER |
| 9 | 117899274 | AX-170300100 | 2 | T | C | 0.127 | -2.57E-02 | 9.69E-02 | 2.60E+00 | 2.71E+00 | 7.94E-01 | no | Zcwpw2 | intron_variant,non_coding_transcript_variant | MODIFIER |
| 9 | 35000099 | AX-170096511 | 0 | T | C | 0.439 | -2.24E-02 | 8.85E-02 | 2.73E+00 | 2.80E+00 | 7.94E-01 | no | Kirrel3 | intron_variant | MODIFIER |
| 9 | 35001127 | AX-170495086 | 0 | T | C | 0.439 | -2.24E-02 | 8.85E-02 | 2.73E+00 | 2.80E+00 | 7.94E-01 | no | Kirrel3 | intron_variant | MODIFIER |
| 9 | 35022177 | AX-170032109 | 0 | T | C | 0.439 | -2.24E-02 | 8.85E-02 | 2.73E+00 | 2.80E+00 | 7.94E-01 | no | Kirrel3 | intron_variant | MODIFIER |
| 9 | 35050235 | AX-170239503 | 0 | G | A | 0.439 | -2.24E-02 | 8.85E-02 | 2.73E+00 | 2.80E+00 | 7.94E-01 | no | St3gal4 | intron_variant | MODIFIER |
| 9 | 35052611 | AX-170485566 | 0 | C | G | 0.439 | -2.24E-02 | 8.85E-02 | 2.73E+00 | 2.80E+00 | 7.94E-01 | no | St3gal4 | intron_variant | MODIFIER |
| 9 | 35061764 | AX-170032108 | 0 | A | G | 0.439 | -2.24E-02 | 8.85E-02 | 2.73E+00 | 2.80E+00 | 7.94E-01 | no | St3gal4 | intron_variant | MODIFIER |
| 9 | 35129506 | AX-170096512 | 0 | G | A | 0.439 | -2.24E-02 | 8.85E-02 | 2.73E+00 | 2.80E+00 | 7.94E-01 | no | Dcps | synonymous_variant | LOW |
| 9 | 35134206 | AX-170298489 | 0 | C | T | 0.439 | -2.24E-02 | 8.85E-02 | 2.73E+00 | 2.80E+00 | 7.94E-01 | no | Dcps | intron_variant | MODIFIER |
| 9 | 35157972 | AX-170437397 | 0 | T | C | 0.439 | -2.24E-02 | 8.85E-02 | 2.73E+00 | 2.80E+00 | 7.94E-01 | no | Dcps | intron_variant | MODIFIER |
| 9 | 35184433 | AX-170194941 | 0 | G | A | 0.439 | -2.24E-02 | 8.85E-02 | 2.73E+00 | 2.80E+00 | 7.94E-01 | no | Tirap | downstream_gene_variant | MODIFIER |
| 9 | 35192961 | AX-170082096 | 0 | C | T | 0.439 | -2.24E-02 | 8.85E-02 | 2.73E+00 | 2.80E+00 | 7.94E-01 | no | Tirap | intron_variant | MODIFIER |
| 9 | 35223385 | AX-170424447 | 0 | C | A | 0.439 | -2.24E-02 | 8.85E-02 | 2.73E+00 | 2.80E+00 | 7.94E-01 | no | Fam118b | intron_variant | MODIFIER |
| 9 | 35311693 | AX-170229317 | 0 | C | A | 0.439 | -2.24E-02 | 8.85E-02 | 2.73E+00 | 2.80E+00 | 7.94E-01 | no | Gm33838 | upstream_gene_variant | MODIFIER |
| 9 | 35311726 | AX-170098507 | 0 | A | G | 0.439 | -2.24E-02 | 8.85E-02 | 2.73E+00 | 2.80E+00 | 7.94E-01 | no | Gm33838 | upstream_gene_variant | MODIFIER |
| 17 | 50041796 | AX-170456186 | 0 | A | C | 0.035 | -5.34E-02 | 1.98E-01 | 2.59E+00 | 2.70E+00 | 7.94E-01 | no | Rftn1 | intron_variant | MODIFIER |
| 16 | 92665575 | AX-97806382 | 0 | C | A | 0.474 | -2.05E-02 | 8.00E-02 | 2.67E+00 | 2.75E+00 | 7.94E-01 | no | Runx1 | intron_variant | MODIFIER |
| 12 | 101287908 | AX-170510849 | 1 | T | C | 0.054 | 6.00E-02 | 2.39E-01 | 2.70E+00 | 2.79E+00 | 7.94E-01 | no | Gm18795 | intron_variant,non_coding_transcript_variant | MODIFIER |
| 1 | 54314733 | AX-170000529 | 0 | C | T | 0.114 | -5.01E-02 | 1.98E-01 | 2.68E+00 | 2.77E+00 | 7.94E-01 | no | Ccdc150 | intron_variant | MODIFIER |
| 8 | 78653978 | AX-170220567 | 1 | C | T | 0.205 | -3.20E-02 | 1.23E-01 | 2.64E+00 | 2.72E+00 | 7.94E-01 | no | Slc10a7 | intron_variant | MODIFIER |
| 4 | 149952554 | AX-170269720 | 1 | T | C | 0.089 | 5.41E-02 | 2.09E-01 | 2.67E+00 | 2.73E+00 | 7.94E-01 | no | Spsb1 | intron_variant | MODIFIER |
| 13 | 24856255 | AX-170474848 | 0 | G | T | 0.158 | -3.46E-02 | 1.10E-01 | 2.49E+00 | 2.59E+00 | 7.94E-01 | no | D130043K22Rik | intron_variant | MODIFIER |
| 9 | 110638628 | AX-97792396 | 0 | T | G | 0.096 | -3.22E-02 | 1.18E-01 | 2.60E+00 | 2.68E+00 | 7.95E-01 | no | Nbeal2 | upstream_gene_variant | MODIFIER |
| 9 | 110642792 | AX-170414509 | 0 | G | A | 0.096 | -3.22E-02 | 1.18E-01 | 2.60E+00 | 2.68E+00 | 7.95E-01 | no | Nbeal2 | upstream_gene_variant | MODIFIER |
| 9 | 110649429 | AX-170033270 | 0 | C | G | 0.096 | -3.22E-02 | 1.18E-01 | 2.60E+00 | 2.68E+00 | 7.95E-01 | no | Nbeal2 | intron_variant | MODIFIER |
| 9 | 110669493 | AX-170332168 | 0 | C | T | 0.096 | -3.22E-02 | 1.18E-01 | 2.60E+00 | 2.68E+00 | 7.95E-01 | no | Ccdc12 | intron_variant | MODIFIER |
| 9 | 110673100 | AX-170033271 | 0 | A | G | 0.096 | -3.22E-02 | 1.18E-01 | 2.60E+00 | 2.68E+00 | 7.95E-01 | no | Ccdc12 | intron_variant | MODIFIER |
| 9 | 110675466 | AX-170033269 | 0 | T | C | 0.096 | -3.22E-02 | 1.18E-01 | 2.60E+00 | 2.68E+00 | 7.95E-01 | no | Ccdc12 | intron_variant | MODIFIER |

| | | | | | | | | | | | | | | | |
|---|---|---|---|---|---|---|---|---|---|---|---|---|---|---|---|
| 9 | 110675541 | AX-170033268 | 0 | A | G | 0.096 | -3.22E-02 | 1.18E-01 | 2.60E+00 | 2.68E+00 | 7.95E-01 | no | Ccdc12 | intron_variant | MODIFIER |
| 9 | 110847571 | AX-97848163 | 0 | A | G | 0.096 | -3.22E-02 | 1.18E-01 | 2.60E+00 | 2.68E+00 | 7.95E-01 | no | Prss46 | upstream_gene_variant | MODIFIER |
| 9 | 110860810 | AX-170180750 | 0 | C | T | 0.096 | -3.22E-02 | 1.18E-01 | 2.60E+00 | 2.68E+00 | 7.95E-01 | no | Tmie | downstream_gene_variant | MODIFIER |
| 9 | 110865384 | AX-170393581 | 0 | T | C | 0.096 | -3.22E-02 | 1.18E-01 | 2.60E+00 | 2.68E+00 | 7.95E-01 | no | Tmie | downstream_gene_variant | MODIFIER |
| 9 | 110893369 | AX-170358427 | 0 | C | T | 0.096 | -3.22E-02 | 1.18E-01 | 2.60E+00 | 2.68E+00 | 7.95E-01 | no | Als2cl | intron_variant | MODIFIER |
| 9 | 110897289 | AX-170460040 | 0 | G | A | 0.096 | -3.22E-02 | 1.18E-01 | 2.60E+00 | 2.68E+00 | 7.95E-01 | no | Als2cl | intron_variant | MODIFIER |
| 9 | 110898487 | AX-170300908 | 0 | A | T | 0.096 | -3.22E-02 | 1.18E-01 | 2.60E+00 | 2.68E+00 | 7.95E-01 | no | Als2cl | missense_variant | MODERATE |
| 9 | 110901281 | AX-170312578 | 0 | C | T | 0.096 | -3.22E-02 | 1.18E-01 | 2.60E+00 | 2.68E+00 | 7.95E-01 | no | Als2cl | downstream_gene_variant | MODIFIER |
| 9 | 110926072 | AX-170406736 | 0 | A | G | 0.096 | -3.22E-02 | 1.18E-01 | 2.60E+00 | 2.68E+00 | 7.95E-01 | no | Gm24269 | downstream_gene_variant | MODIFIER |
| 9 | 110926825 | AX-97851603 | 0 | G | C | 0.096 | -3.22E-02 | 1.18E-01 | 2.60E+00 | 2.68E+00 | 7.95E-01 | no | Gm24269 | downstream_gene_variant | MODIFIER |
| 9 | 110954312 | AX-170497829 | 0 | T | G | 0.096 | -3.22E-02 | 1.18E-01 | 2.60E+00 | 2.68E+00 | 7.95E-01 | no | Lrrc2 | intron_variant | MODIFIER |
| 9 | 110964835 | AX-170455180 | 0 | G | T | 0.096 | -3.22E-02 | 1.18E-01 | 2.60E+00 | 2.68E+00 | 7.95E-01 | no | Lrrc2 | intron_variant | MODIFIER |
| 9 | 110964851 | AX-170172734 | 0 | A | G | 0.096 | -3.22E-02 | 1.18E-01 | 2.60E+00 | 2.68E+00 | 7.95E-01 | no | Lrrc2 | intron_variant | MODIFIER |
| 9 | 110970100 | AX-170447335 | 0 | T | C | 0.096 | -3.22E-02 | 1.18E-01 | 2.60E+00 | 2.68E+00 | 7.95E-01 | no | Lrrc2 | synonymous_variant | LOW |
| 9 | 110970242 | AX-170237860 | 0 | G | T | 0.096 | -3.22E-02 | 1.18E-01 | 2.60E+00 | 2.68E+00 | 7.95E-01 | no | Lrrc2 | intron_variant | MODIFIER |
| 9 | 110989005 | AX-170383398 | 0 | T | C | 0.096 | -3.22E-02 | 1.18E-01 | 2.60E+00 | 2.68E+00 | 7.95E-01 | no | Lrrc2 | downstream_gene_variant | MODIFIER |
| 9 | 111001927 | AX-170369984 | 0 | T | C | 0.096 | -3.22E-02 | 1.18E-01 | 2.60E+00 | 2.68E+00 | 7.95E-01 | no | Gm10030 | upstream_gene_variant | MODIFIER |
| 9 | 111007777 | AX-170456332 | 0 | T | C | 0.096 | -3.22E-02 | 1.18E-01 | 2.60E+00 | 2.68E+00 | 7.95E-01 | no | Gm10030 | downstream_gene_variant | MODIFIER |
| 9 | 111041408 | AX-170371652 | 0 | G | A | 0.096 | -3.22E-02 | 1.18E-01 | 2.60E+00 | 2.68E+00 | 7.95E-01 | no | Ltf | intron_variant | MODIFIER |
| 9 | 111058006 | AX-97781603 | 0 | C | T | 0.096 | -3.22E-02 | 1.18E-01 | 2.60E+00 | 2.68E+00 | 7.95E-01 | no | Ccrl2 | upstream_gene_variant | MODIFIER |
| 9 | 111318211 | AX-170384182 | 0 | C | T | 0.096 | -3.22E-02 | 1.18E-01 | 2.60E+00 | 2.68E+00 | 7.95E-01 | no | Trank1 | intron_variant | MODIFIER |
| 9 | 111334623 | AX-170403956 | 0 | T | G | 0.096 | -3.22E-02 | 1.18E-01 | 2.60E+00 | 2.68E+00 | 7.95E-01 | no | Trank1 | intron_variant | MODIFIER |
| 9 | 111344315 | AX-170096668 | 0 | G | T | 0.096 | -3.22E-02 | 1.18E-01 | 2.60E+00 | 2.68E+00 | 7.95E-01 | no | Trank1 | intron_variant | MODIFIER |
| 9 | 111364385 | AX-170205937 | 0 | T | C | 0.096 | -3.22E-02 | 1.18E-01 | 2.60E+00 | 2.68E+00 | 7.95E-01 | no | Trank1 | intron_variant | MODIFIER |
| 9 | 111372187 | AX-170324095 | 0 | T | G | 0.096 | -3.22E-02 | 1.18E-01 | 2.60E+00 | 2.68E+00 | 7.95E-01 | no | Trank1 | intron_variant | MODIFIER |
| 9 | 111404874 | AX-97839463 | 0 | T | C | 0.096 | -3.22E-02 | 1.18E-01 | 2.60E+00 | 2.68E+00 | 7.95E-01 | no | | intergenic_variant | MODIFIER |
| 9 | 111407792 | AX-170153577 | 0 | C | A | 0.096 | -3.22E-02 | 1.18E-01 | 2.60E+00 | 2.68E+00 | 7.95E-01 | no | Gm42527 | upstream_gene_variant | MODIFIER |
| 9 | 111408000 | AX-170306270 | 0 | T | C | 0.096 | -3.22E-02 | 1.18E-01 | 2.60E+00 | 2.68E+00 | 7.95E-01 | no | Gm42527 | upstream_gene_variant | MODIFIER |
| 9 | 111408196 | AX-170179025 | 0 | G | A | 0.096 | -3.22E-02 | 1.18E-01 | 2.60E+00 | 2.68E+00 | 7.95E-01 | no | Gm42527 | upstream_gene_variant | MODIFIER |
| 20 | 157716671 | AX-97842479 | 0 | C | T | 0.149 | 2.86E-02 | 1.09E-01 | 2.62E+00 | 2.71E+00 | 7.95E-01 | no | Smpx | intron_variant,NMD_transcript_variant | MODIFIER |
| 7 | 91824628 | AX-170029498 | 1 | T | C | 0.125 | -5.03E-02 | 1.99E-01 | 2.68E+00 | 2.77E+00 | 7.95E-01 | no | Dlg2 | intron_variant | MODIFIER |
| 7 | 97092887 | AX-170029584 | 1 | C | A | 0.125 | -5.03E-02 | 1.99E-01 | 2.68E+00 | 2.77E+00 | 7.95E-01 | no | Gab2 | intron_variant | MODIFIER |
| 16 | 93338981 | AX-170013678 | 0 | T | A | 0.386 | 1.92E-02 | 7.79E-02 | 2.70E+00 | 2.81E+00 | 7.95E-01 | no | 1810053B23Rik | downstream_gene_variant | MODIFIER |
| 18 | 86952848 | AX-170155071 | 1 | A | G | 0.188 | -2.87E-02 | 1.03E-01 | 2.57E+00 | 2.67E+00 | 7.95E-01 | no | | intergenic_variant | MODIFIER |
| 1 | 50239103 | AX-170481681 | 0 | C | G | 0.105 | -6.01E-02 | 2.35E-01 | 2.72E+00 | 2.76E+00 | 7.95E-01 | no | | intergenic_variant | MODIFIER |
| 1 | 52715534 | AX-170404387 | 0 | T | G | 0.105 | -6.01E-02 | 2.35E-01 | 2.72E+00 | 2.76E+00 | 7.95E-01 | no | Mfsd6 | intron_variant | MODIFIER |
| 1 | 55061302 | AX-170091518 | 0 | G | A | 0.105 | -6.01E-02 | 2.35E-01 | 2.72E+00 | 2.76E+00 | 7.95E-01 | no | Coq10b | missense_variant | MODERATE |
| 4 | 149950392 | AX-170270783 | 0 | T | C | 0.079 | 4.97E-02 | 1.92E-01 | 2.66E+00 | 2.73E+00 | 7.95E-01 | no | Spsb1 | intron_variant | MODIFIER |
| 5 | 85573000 | AX-170477945 | 0 | G | A | 0.43 | 2.78E-02 | 1.09E-01 | 2.71E+00 | 2.76E+00 | 7.95E-01 | no | | intergenic_variant | MODIFIER |
| 8 | 66020655 | AX-170509670 | 0 | A | G | 0.149 | -3.70E-02 | 1.36E-01 | 2.59E+00 | 2.68E+00 | 7.95E-01 | no | MAR1 | intron_variant | MODIFIER |
| 16 | 94321550 | AX-170088102 | 0 | G | A | 0.421 | -2.16E-02 | 8.06E-02 | 2.61E+00 | 2.69E+00 | 7.95E-01 | no | | intergenic_variant | MODIFIER |
| 10 | 117490769 | AX-170269016 | 2 | T | C | 0.327 | 1.88E-02 | 7.54E-02 | 2.68E+00 | 2.78E+00 | 7.95E-01 | no | | intergenic_variant | MODIFIER |
| 18 | 69559385 | AX-170373169 | 1 | G | A | 0.268 | -2.62E-02 | 9.98E-02 | 2.64E+00 | 2.71E+00 | 7.95E-01 | no | Tcf4 | intron_variant | MODIFIER |
| 16 | 91898967 | AX-170270822 | 2 | T | C | 0.464 | 1.93E-02 | 7.56E-02 | 2.65E+00 | 2.75E+00 | 7.95E-01 | no | Itsn1 | intron_variant | MODIFIER |
| 4 | 147967345 | AX-170352609 | 0 | A | T | 0.096 | 5.47E-02 | 2.13E-01 | 2.68E+00 | 2.73E+00 | 7.95E-01 | no | | intergenic_variant | MODIFIER |
| 8 | 15323566 | AX-97798043 | 0 | A | G | 0.237 | -2.10E-02 | 7.65E-02 | 2.58E+00 | 2.68E+00 | 7.95E-01 | no | Gm7706 | intron_variant,non_coding_transcript_variant | MODIFIER |
| 1 | 55195652 | AX-170120348 | 1 | A | T | 0.107 | 4.64E-02 | 1.78E-01 | 2.61E+00 | 2.72E+00 | 7.96E-01 | no | Rftn2 | intron_variant | MODIFIER |
| 2 | 115703143 | AX-170088718 | 1 | G | A | 0.143 | 5.69E-02 | 2.21E-01 | 2.69E+00 | 2.73E+00 | 7.96E-01 | no | BC052040 | intron_variant | MODIFIER |
| 2 | 115843491 | AX-97783137 | 1 | C | A | 0.143 | 5.69E-02 | 2.21E-01 | 2.69E+00 | 2.73E+00 | 7.96E-01 | no | | intergenic_variant | MODIFIER |
| 2 | 122168681 | AX-170414370 | 1 | C | T | 0.143 | 5.69E-02 | 2.21E-01 | 2.69E+00 | 2.73E+00 | 7.96E-01 | no | Patl2 | intron_variant | MODIFIER |
| 2 | 125756862 | AX-170507091 | 1 | A | T | 0.143 | 5.69E-02 | 2.21E-01 | 2.69E+00 | 2.73E+00 | 7.96E-01 | no | Secisbp2l | intron_variant | MODIFIER |
| 2 | 130429916 | AX-170357241 | 1 | C | A | 0.143 | 5.69E-02 | 2.21E-01 | 2.69E+00 | 2.73E+00 | 7.96E-01 | no | Vps16 | intron_variant | MODIFIER |
| 2 | 138228324 | AX-170065032 | 1 | C | T | 0.143 | 5.69E-02 | 2.21E-01 | 2.69E+00 | 2.73E+00 | 7.96E-01 | no | | intergenic_variant | MODIFIER |
| 2 | 142910683 | AX-170409550 | 1 | G | A | 0.143 | 5.69E-02 | 2.21E-01 | 2.69E+00 | 2.73E+00 | 7.96E-01 | no | | intergenic_variant | MODIFIER |

| | | | | | | | | | | | | | | | |
|---|---|---|---|---|---|---|---|---|---|---|---|---|---|---|---|
| 2 | 155886841 | AX-170019449 | 1 | G | A | 0.143 | 5.69E-02 | 2.21E-01 | 2.69E+00 | 2.73E+00 | 7.96E-01 | no | Uqcc1 | intron_variant | MODIFIER |
| 1 | 3046184 | AX-169999825 | 0 | A | G | 0.421 | 1.89E-02 | 6.86E-02 | 2.59E+00 | 2.66E+00 | 7.96E-01 | no | | intergenic_variant | MODIFIER |
| 1 | 3060252 | AX-170091369 | 0 | G | T | 0.421 | 1.89E-02 | 6.86E-02 | 2.59E+00 | 2.66E+00 | 7.96E-01 | no | | intergenic_variant | MODIFIER |
| 1 | 3061281 | AX-169999827 | 0 | A | G | 0.421 | 1.89E-02 | 6.86E-02 | 2.59E+00 | 2.66E+00 | 7.96E-01 | no | | intergenic_variant | MODIFIER |
| 1 | 3073457 | AX-97805837 | 0 | C | G | 0.421 | 1.89E-02 | 6.86E-02 | 2.59E+00 | 2.66E+00 | 7.96E-01 | no | 4933401J01Rik | non_coding_transcript_exon_variant | MODIFIER |
| 1 | 3101325 | AX-170040233 | 0 | G | T | 0.421 | 1.89E-02 | 6.86E-02 | 2.59E+00 | 2.66E+00 | 7.96E-01 | no | Gm26206 | upstream_gene_variant | MODIFIER |
| 1 | 3103735 | AX-169999829 | 0 | C | T | 0.421 | 1.89E-02 | 6.86E-02 | 2.59E+00 | 2.66E+00 | 7.96E-01 | no | Gm26206 | downstream_gene_variant | MODIFIER |
| 1 | 3105903 | AX-97803249 | 0 | C | A | 0.421 | 1.89E-02 | 6.86E-02 | 2.59E+00 | 2.66E+00 | 7.96E-01 | no | Gm26206 | downstream_gene_variant | MODIFIER |
| 1 | 3644739 | AX-170351609 | 0 | C | G | 0.421 | 1.89E-02 | 6.86E-02 | 2.59E+00 | 2.66E+00 | 7.96E-01 | no | Xkr4 | intron_variant | MODIFIER |
| 1 | 3668324 | AX-170274044 | 0 | A | G | 0.421 | 1.89E-02 | 6.86E-02 | 2.59E+00 | 2.66E+00 | 7.96E-01 | no | Xkr4 | intron_variant | MODIFIER |
| 1 | 3668351 | AX-97810552 | 0 | G | A | 0.421 | 1.89E-02 | 6.86E-02 | 2.59E+00 | 2.66E+00 | 7.96E-01 | no | Xkr4 | intron_variant | MODIFIER |
| 1 | 4201770 | AX-169999843 | 0 | C | T | 0.421 | 1.89E-02 | 6.86E-02 | 2.59E+00 | 2.66E+00 | 7.96E-01 | no | Rp1 | intron_variant | MODIFIER |
| 1 | 4317090 | AX-170123347 | 0 | T | G | 0.421 | 1.89E-02 | 6.86E-02 | 2.59E+00 | 2.66E+00 | 7.96E-01 | no | Rp1 | intron_variant | MODIFIER |
| 1 | 4350805 | AX-97817170 | 0 | A | G | 0.421 | 1.89E-02 | 6.86E-02 | 2.59E+00 | 2.66E+00 | 7.96E-01 | no | Rp1 | intron_variant | MODIFIER |
| 1 | 4606072 | AX-170479979 | 0 | A | G | 0.421 | 1.89E-02 | 6.86E-02 | 2.59E+00 | 2.66E+00 | 7.96E-01 | no | Gm7369 | upstream_gene_variant | MODIFIER |
| 1 | 4976580 | AX-170361819 | 0 | T | C | 0.421 | 1.89E-02 | 6.86E-02 | 2.59E+00 | 2.66E+00 | 7.96E-01 | no | Rgs20 | intron_variant | MODIFIER |
| 1 | 5065231 | AX-170086398 | 0 | T | C | 0.421 | 1.89E-02 | 6.86E-02 | 2.59E+00 | 2.66E+00 | 7.96E-01 | no | Rgs20 | intron_variant | MODIFIER |
| 1 | 5088401 | AX-170477286 | 0 | C | G | 0.421 | 1.89E-02 | 6.86E-02 | 2.59E+00 | 2.66E+00 | 7.96E-01 | no | Atp6v1h | intron_variant | MODIFIER |
| 1 | 5089449 | AX-97808344 | 0 | A | G | 0.421 | 1.89E-02 | 6.86E-02 | 2.59E+00 | 2.66E+00 | 7.96E-01 | no | Atp6v1h | intron_variant | MODIFIER |
| 1 | 5095686 | AX-170356034 | 0 | C | T | 0.421 | 1.89E-02 | 6.86E-02 | 2.59E+00 | 2.66E+00 | 7.96E-01 | no | Atp6v1h | synonymous_variant | LOW |
| 1 | 5119587 | AX-170434246 | 0 | G | A | 0.421 | 1.89E-02 | 6.86E-02 | 2.59E+00 | 2.66E+00 | 7.96E-01 | no | Atp6v1h | intron_variant | MODIFIER |
| 1 | 5119671 | AX-170487665 | 0 | T | C | 0.421 | 1.89E-02 | 6.86E-02 | 2.59E+00 | 2.66E+00 | 7.96E-01 | no | Atp6v1h | intron_variant | MODIFIER |
| 1 | 5120013 | AX-170410118 | 0 | T | C | 0.421 | 1.89E-02 | 6.86E-02 | 2.59E+00 | 2.66E+00 | 7.96E-01 | no | Atp6v1h | intron_variant | MODIFIER |
| 1 | 5133848 | AX-170414223 | 0 | C | T | 0.421 | 1.89E-02 | 6.86E-02 | 2.59E+00 | 2.66E+00 | 7.96E-01 | no | Atp6v1h | intron_variant | MODIFIER |
| 1 | 5140044 | AX-170458136 | 0 | T | C | 0.421 | 1.89E-02 | 6.86E-02 | 2.59E+00 | 2.66E+00 | 7.96E-01 | no | Atp6v1h | intron_variant | MODIFIER |
| 1 | 5142376 | AX-97848170 | 0 | T | C | 0.421 | 1.89E-02 | 6.86E-02 | 2.59E+00 | 2.66E+00 | 7.96E-01 | no | Atp6v1h | intron_variant | MODIFIER |
| 1 | 5163127 | AX-170091377 | 0 | A | G | 0.421 | 1.89E-02 | 6.86E-02 | 2.59E+00 | 2.66E+00 | 7.96E-01 | no | Atp6v1h | downstream_gene_variant | MODIFIER |
| 1 | 5177384 | AX-170097500 | 0 | A | G | 0.421 | 1.89E-02 | 6.86E-02 | 2.59E+00 | 2.66E+00 | 7.96E-01 | no | | intergenic_variant | MODIFIER |
| 1 | 5181919 | AX-170467830 | 0 | A | G | 0.421 | 1.89E-02 | 6.86E-02 | 2.59E+00 | 2.66E+00 | 7.96E-01 | no | | intergenic_variant | MODIFIER |
| 1 | 5210579 | AX-97802133 | 0 | A | G | 0.421 | 1.89E-02 | 6.86E-02 | 2.59E+00 | 2.66E+00 | 7.96E-01 | no | | intergenic_variant | MODIFIER |
| 1 | 5218963 | AX-169999856 | 0 | T | C | 0.421 | 1.89E-02 | 6.86E-02 | 2.59E+00 | 2.66E+00 | 7.96E-01 | no | | intergenic_variant | MODIFIER |
| 1 | 5233400 | AX-97822719 | 0 | C | G | 0.421 | 1.89E-02 | 6.86E-02 | 2.59E+00 | 2.66E+00 | 7.96E-01 | no | | regulatory_region_variant | MODIFIER |
| 1 | 5249721 | AX-97855603 | 0 | C | T | 0.421 | 1.89E-02 | 6.86E-02 | 2.59E+00 | 2.66E+00 | 7.96E-01 | no | | intergenic_variant | MODIFIER |
| 1 | 5273634 | AX-170286242 | 0 | T | C | 0.421 | 1.89E-02 | 6.86E-02 | 2.59E+00 | 2.66E+00 | 7.96E-01 | no | Gm7182 | downstream_gene_variant | MODIFIER |
| 1 | 5301769 | AX-170295303 | 0 | T | C | 0.421 | 1.89E-02 | 6.86E-02 | 2.59E+00 | 2.66E+00 | 7.96E-01 | no | | intergenic_variant | MODIFIER |
| 1 | 5314171 | AX-97810555 | 0 | C | A | 0.421 | 1.89E-02 | 6.86E-02 | 2.59E+00 | 2.66E+00 | 7.96E-01 | no | Gm37567 | downstream_gene_variant | MODIFIER |
| 1 | 5325874 | AX-170100229 | 0 | A | G | 0.421 | 1.89E-02 | 6.86E-02 | 2.59E+00 | 2.66E+00 | 7.96E-01 | no | | intergenic_variant | MODIFIER |
| 1 | 5334728 | AX-170294333 | 0 | C | G | 0.421 | 1.89E-02 | 6.86E-02 | 2.59E+00 | 2.66E+00 | 7.96E-01 | no | | intergenic_variant | MODIFIER |
| 1 | 5340857 | AX-170224621 | 0 | C | T | 0.421 | 1.89E-02 | 6.86E-02 | 2.59E+00 | 2.66E+00 | 7.96E-01 | no | | intergenic_variant | MODIFIER |
| 18 | 59256577 | AX-170015799 | 0 | A | G | 0.412 | -2.24E-02 | 8.38E-02 | 2.62E+00 | 2.68E+00 | 7.96E-01 | no | Chsy3 | intron_variant | MODIFIER |
| 18 | 59285024 | AX-170190191 | 0 | T | C | 0.412 | -2.24E-02 | 8.38E-02 | 2.62E+00 | 2.68E+00 | 7.96E-01 | no | Chsy3 | intron_variant | MODIFIER |
| 18 | 59340925 | AX-170152141 | 0 | C | A | 0.412 | -2.24E-02 | 8.38E-02 | 2.62E+00 | 2.68E+00 | 7.96E-01 | no | Chsy3 | intron_variant | MODIFIER |
| 18 | 59349296 | AX-170299863 | 0 | C | G | 0.412 | -2.24E-02 | 8.38E-02 | 2.62E+00 | 2.68E+00 | 7.96E-01 | no | Chsy3 | intron_variant | MODIFIER |
| 18 | 59356748 | AX-170225874 | 0 | G | A | 0.412 | -2.24E-02 | 8.38E-02 | 2.62E+00 | 2.68E+00 | 7.96E-01 | no | Chsy3 | intron_variant | MODIFIER |
| 18 | 59359908 | AX-170362303 | 0 | G | A | 0.412 | -2.24E-02 | 8.38E-02 | 2.62E+00 | 2.68E+00 | 7.96E-01 | no | Chsy3 | intron_variant | MODIFIER |
| 18 | 59377550 | AX-170348522 | 0 | C | T | 0.412 | -2.24E-02 | 8.38E-02 | 2.62E+00 | 2.68E+00 | 7.96E-01 | no | Chsy3 | intron_variant | MODIFIER |
| 18 | 59379081 | AX-170144683 | 0 | C | T | 0.412 | -2.24E-02 | 8.38E-02 | 2.62E+00 | 2.68E+00 | 7.96E-01 | no | Chsy3 | intron_variant | MODIFIER |
| 18 | 59382425 | AX-170060991 | 0 | C | G | 0.412 | -2.24E-02 | 8.38E-02 | 2.62E+00 | 2.68E+00 | 7.96E-01 | no | Chsy3 | intron_variant | MODIFIER |
| 18 | 59468827 | AX-170436122 | 0 | T | C | 0.412 | -2.24E-02 | 8.38E-02 | 2.62E+00 | 2.68E+00 | 7.96E-01 | no | Chsy3 | intron_variant,NMD_transcript_variant | MODIFIER |
| 18 | 60123787 | AX-170437914 | 0 | A | G | 0.412 | -2.24E-02 | 8.38E-02 | 2.62E+00 | 2.68E+00 | 7.96E-01 | no | | intergenic_variant | MODIFIER |
| 18 | 60128554 | AX-170513492 | 0 | G | T | 0.412 | -2.24E-02 | 8.38E-02 | 2.62E+00 | 2.68E+00 | 7.96E-01 | no | | intergenic_variant | MODIFIER |
| 18 | 60486642 | AX-168384739 | 0 | A | G | 0.412 | -2.24E-02 | 8.38E-02 | 2.62E+00 | 2.68E+00 | 7.96E-01 | no | Smim3 | intron_variant | MODIFIER |
| 18 | 60609788 | AX-170166630 | 0 | T | C | 0.412 | -2.24E-02 | 8.38E-02 | 2.62E+00 | 2.68E+00 | 7.96E-01 | no | Synpo | intron_variant | MODIFIER |
| 18 | 60823311 | AX-170372602 | 0 | G | A | 0.412 | -2.24E-02 | 8.38E-02 | 2.62E+00 | 2.68E+00 | 7.96E-01 | no | Tcof1 | intron_variant,NMD_transcript_variant | MODIFIER |

| | | | | | | | | | | | | | | | |
|---|---|---|---|---|---|---|---|---|---|---|---|---|---|---|---|
| 18 | 60828006 | AX-170093905 | 0 | A | G | 0.412 | -2.24E-02 | 8.38E-02 | 2.62E+00 | 2.68E+00 | 7.96E-01 | no | Tcof1 | intron_variant,NMD_transcript_variant | MODIFIER |
| 18 | 60837794 | AX-170116372 | 0 | A | G | 0.412 | -2.24E-02 | 8.38E-02 | 2.62E+00 | 2.68E+00 | 7.96E-01 | no | Tcof1 | intron_variant,NMD_transcript_variant | MODIFIER |
| 18 | 60838298 | AX-170118777 | 0 | T | C | 0.412 | -2.24E-02 | 8.38E-02 | 2.62E+00 | 2.68E+00 | 7.96E-01 | no | Tcof1 | intron_variant,NMD_transcript_variant | MODIFIER |
| 18 | 60860760 | AX-170421695 | 0 | T | G | 0.412 | -2.24E-02 | 8.38E-02 | 2.62E+00 | 2.68E+00 | 7.96E-01 | no | | intergenic_variant | MODIFIER |
| 18 | 60862278 | AX-170113977 | 0 | T | C | 0.412 | -2.24E-02 | 8.38E-02 | 2.62E+00 | 2.68E+00 | 7.96E-01 | no | | intergenic_variant | MODIFIER |
| 18 | 60863826 | AX-170209285 | 0 | C | T | 0.412 | -2.24E-02 | 8.38E-02 | 2.62E+00 | 2.68E+00 | 7.96E-01 | no | | regulatory_region_variant | MODIFIER |
| 18 | 60865722 | AX-170384959 | 0 | C | T | 0.412 | -2.24E-02 | 8.38E-02 | 2.62E+00 | 2.68E+00 | 7.96E-01 | no | | intergenic_variant | MODIFIER |
| 18 | 60866833 | AX-170160855 | 0 | C | T | 0.412 | -2.24E-02 | 8.38E-02 | 2.62E+00 | 2.68E+00 | 7.96E-01 | no | | regulatory_region_variant | MODIFIER |
| 18 | 60867479 | AX-97792416 | 0 | A | G | 0.412 | -2.24E-02 | 8.38E-02 | 2.62E+00 | 2.68E+00 | 7.96E-01 | no | | regulatory_region_variant | MODIFIER |
| 18 | 60888437 | AX-170433271 | 0 | T | G | 0.412 | -2.24E-02 | 8.38E-02 | 2.62E+00 | 2.68E+00 | 7.96E-01 | no | | regulatory_region_variant | MODIFIER |
| 18 | 60907768 | AX-97783518 | 0 | A | G | 0.412 | -2.24E-02 | 8.38E-02 | 2.62E+00 | 2.68E+00 | 7.96E-01 | no | Arsi | upstream_gene_variant | MODIFIER |
| 18 | 60908399 | AX-170152104 | 0 | A | G | 0.412 | -2.24E-02 | 8.38E-02 | 2.62E+00 | 2.68E+00 | 7.96E-01 | no | Arsi | upstream_gene_variant | MODIFIER |
| 18 | 60921377 | AX-170348637 | 0 | T | C | 0.412 | -2.24E-02 | 8.38E-02 | 2.62E+00 | 2.68E+00 | 7.96E-01 | no | Camk2a | upstream_gene_variant | MODIFIER |
| 18 | 60944366 | AX-170325487 | 0 | T | C | 0.412 | -2.24E-02 | 8.38E-02 | 2.62E+00 | 2.68E+00 | 7.96E-01 | no | Camk2a | intron_variant | MODIFIER |
| 18 | 60948947 | AX-170061020 | 0 | G | A | 0.412 | -2.24E-02 | 8.38E-02 | 2.62E+00 | 2.68E+00 | 7.96E-01 | no | Camk2a | intron_variant | MODIFIER |
| 18 | 60949318 | AX-170061021 | 0 | A | G | 0.412 | -2.24E-02 | 8.38E-02 | 2.62E+00 | 2.68E+00 | 7.96E-01 | no | Camk2a | intron_variant | MODIFIER |
| 18 | 60953545 | AX-97808409 | 0 | A | G | 0.412 | -2.24E-02 | 8.38E-02 | 2.62E+00 | 2.68E+00 | 7.96E-01 | no | Camk2a | intron_variant | MODIFIER |
| 18 | 60958003 | AX-170469076 | 0 | A | G | 0.412 | -2.24E-02 | 8.38E-02 | 2.62E+00 | 2.68E+00 | 7.96E-01 | no | Camk2a | intron_variant | MODIFIER |
| 18 | 60958018 | AX-170352628 | 0 | C | T | 0.412 | -2.24E-02 | 8.38E-02 | 2.62E+00 | 2.68E+00 | 7.96E-01 | no | Camk2a | intron_variant | MODIFIER |
| 18 | 60976190 | AX-170509123 | 0 | C | G | 0.412 | -2.24E-02 | 8.38E-02 | 2.62E+00 | 2.68E+00 | 7.96E-01 | no | Camk2a | intron_variant | MODIFIER |
| 18 | 60976432 | AX-170502238 | 0 | G | A | 0.412 | -2.24E-02 | 8.38E-02 | 2.62E+00 | 2.68E+00 | 7.96E-01 | no | Camk2a | intron_variant | MODIFIER |
| 18 | 60978313 | AX-170180911 | 0 | C | A | 0.412 | -2.24E-02 | 8.38E-02 | 2.62E+00 | 2.68E+00 | 7.96E-01 | no | Camk2a | intron_variant | MODIFIER |
| 18 | 60993585 | AX-170119029 | 0 | A | C | 0.412 | -2.24E-02 | 8.38E-02 | 2.62E+00 | 2.68E+00 | 7.96E-01 | no | Slc6a7 | downstream_gene_variant | MODIFIER |
| 18 | 60993966 | AX-170202763 | 0 | A | G | 0.412 | -2.24E-02 | 8.38E-02 | 2.62E+00 | 2.68E+00 | 7.96E-01 | no | Slc6a7 | downstream_gene_variant | MODIFIER |
| 18 | 61002073 | AX-170158520 | 0 | T | C | 0.412 | -2.24E-02 | 8.38E-02 | 2.62E+00 | 2.68E+00 | 7.96E-01 | no | Slc6a7 | intron_variant | MODIFIER |
| 18 | 61012294 | AX-170148937 | 0 | C | T | 0.412 | -2.24E-02 | 8.38E-02 | 2.62E+00 | 2.68E+00 | 7.96E-01 | no | Slc6a7 | intron_variant | MODIFIER |
| 18 | 61012717 | AX-170374821 | 0 | A | G | 0.412 | -2.24E-02 | 8.38E-02 | 2.62E+00 | 2.68E+00 | 7.96E-01 | no | Slc6a7 | intron_variant | MODIFIER |
| 18 | 61049334 | AX-97835605 | 0 | A | G | 0.412 | -2.24E-02 | 8.38E-02 | 2.62E+00 | 2.68E+00 | 7.96E-01 | no | Pdgfrb | intron_variant | MODIFIER |
| 18 | 61185846 | AX-170061029 | 0 | C | T | 0.412 | -2.24E-02 | 8.38E-02 | 2.62E+00 | 2.68E+00 | 7.96E-01 | no | | intergenic_variant | MODIFIER |
| 18 | 61189783 | AX-170061028 | 0 | G | A | 0.412 | -2.24E-02 | 8.38E-02 | 2.62E+00 | 2.68E+00 | 7.96E-01 | no | Slc26a2 | downstream_gene_variant | MODIFIER |
| 18 | 61190709 | AX-170061027 | 0 | T | C | 0.412 | -2.24E-02 | 8.38E-02 | 2.62E+00 | 2.68E+00 | 7.96E-01 | no | Slc26a2 | downstream_gene_variant | MODIFIER |
| 18 | 61190779 | AX-170466674 | 0 | G | A | 0.412 | -2.24E-02 | 8.38E-02 | 2.62E+00 | 2.68E+00 | 7.96E-01 | no | Slc26a2 | downstream_gene_variant | MODIFIER |
| 18 | 61221327 | AX-170088354 | 0 | C | A | 0.412 | -2.24E-02 | 8.38E-02 | 2.62E+00 | 2.68E+00 | 7.96E-01 | no | Pde6a | intron_variant | MODIFIER |
| 18 | 61386553 | AX-170454137 | 0 | A | G | 0.412 | -2.24E-02 | 8.38E-02 | 2.62E+00 | 2.68E+00 | 7.96E-01 | no | Ppargc1b | intron_variant | MODIFIER |
| 18 | 61387563 | AX-170304467 | 0 | A | G | 0.412 | -2.24E-02 | 8.38E-02 | 2.62E+00 | 2.68E+00 | 7.96E-01 | no | Ppargc1b | intron_variant | MODIFIER |
| 18 | 61476887 | AX-97828478 | 0 | A | C | 0.412 | -2.24E-02 | 8.38E-02 | 2.62E+00 | 2.68E+00 | 7.96E-01 | no | AC153155.2 | intron_variant,non_coding_transcript_variant | MODIFIER |
| 18 | 61487657 | AX-170119461 | 0 | T | C | 0.412 | -2.24E-02 | 8.38E-02 | 2.62E+00 | 2.68E+00 | 7.96E-01 | no | Arhgef37 | downstream_gene_variant | MODIFIER |
| 18 | 61746123 | AX-170401462 | 0 | A | G | 0.412 | -2.24E-02 | 8.38E-02 | 2.62E+00 | 2.68E+00 | 7.96E-01 | no | Afap1l1 | intron_variant | MODIFIER |
| 18 | 61870235 | AX-170440634 | 0 | C | T | 0.412 | -2.24E-02 | 8.38E-02 | 2.62E+00 | 2.68E+00 | 7.96E-01 | no | Ablim3 | intron_variant | MODIFIER |
| 18 | 61950873 | AX-170297421 | 0 | T | C | 0.412 | -2.24E-02 | 8.38E-02 | 2.62E+00 | 2.68E+00 | 7.96E-01 | no | Sh3tc2 | upstream_gene_variant | MODIFIER |
| 18 | 61952635 | AX-170474266 | 0 | C | T | 0.412 | -2.24E-02 | 8.38E-02 | 2.62E+00 | 2.68E+00 | 7.96E-01 | no | Sh3tc2 | upstream_gene_variant | MODIFIER |
| 18 | 61953342 | AX-97851961 | 0 | A | C | 0.412 | -2.24E-02 | 8.38E-02 | 2.62E+00 | 2.68E+00 | 7.96E-01 | no | Sh3tc2 | intron_variant | MODIFIER |
| 18 | 61958618 | AX-170261415 | 0 | G | A | 0.412 | -2.24E-02 | 8.38E-02 | 2.62E+00 | 2.68E+00 | 7.96E-01 | no | Sh3tc2 | intron_variant | MODIFIER |
| 18 | 61970794 | AX-170384470 | 0 | A | G | 0.412 | -2.24E-02 | 8.38E-02 | 2.62E+00 | 2.68E+00 | 7.96E-01 | no | Sh3tc2 | intron_variant | MODIFIER |
| 18 | 61971281 | AX-170525645 | 0 | A | G | 0.412 | -2.24E-02 | 8.38E-02 | 2.62E+00 | 2.68E+00 | 7.96E-01 | no | Sh3tc2 | intron_variant | MODIFIER |
| 18 | 62384530 | AX-170231660 | 0 | C | T | 0.412 | -2.24E-02 | 8.38E-02 | 2.62E+00 | 2.68E+00 | 7.96E-01 | no | Htr4 | intron_variant | MODIFIER |
| 18 | 62954147 | AX-97794538 | 0 | A | G | 0.412 | -2.24E-02 | 8.38E-02 | 2.62E+00 | 2.68E+00 | 7.96E-01 | no | Apcdd1 | downstream_gene_variant | MODIFIER |
| 18 | 63100398 | AX-170170585 | 0 | A | G | 0.412 | -2.24E-02 | 8.38E-02 | 2.62E+00 | 2.68E+00 | 7.96E-01 | no | Piezo2 | intron_variant | MODIFIER |
| 2 | 121058751 | AX-170428433 | 2 | A | T | 0.145 | 5.74E-02 | 2.22E-01 | 2.68E+00 | 2.71E+00 | 7.96E-01 | no | Tgm5 | intron_variant | MODIFIER |
| 1 | 60605054 | AX-170211513 | 0 | T | C | 0.114 | 4.95E-02 | 1.91E-01 | 2.63E+00 | 2.72E+00 | 7.96E-01 | no | | intergenic_variant | MODIFIER |
| 16 | 77281301 | AX-170492328 | 1 | G | T | 0.455 | 1.93E-02 | 7.17E-02 | 2.59E+00 | 2.69E+00 | 7.96E-01 | no | Mir99ahg | intron_variant,non_coding_transcript_variant | MODIFIER |
| 2 | 119496201 | AX-170439164 | 1 | G | A | 0.116 | 5.02E-02 | 1.94E-01 | 2.66E+00 | 2.71E+00 | 7.96E-01 | no | | intergenic_variant | MODIFIER |
| 2 | 128513147 | AX-170252664 | 1 | T | A | 0.116 | 5.02E-02 | 1.94E-01 | 2.66E+00 | 2.71E+00 | 7.96E-01 | no | | regulatory_region_variant | MODIFIER |
| 2 | 114360016 | AX-170126317 | 1 | A | C | 0.134 | 5.31E-02 | 2.08E-01 | 2.70E+00 | 2.74E+00 | 7.96E-01 | no | | intergenic_variant | MODIFIER |

| | | | | | | | | | | | | | | | |
|---|---|---|---|---|---|---|---|---|---|---|---|---|---|---|---|
| 2 | 123653096 | AX-170290357 | 1 | C | T | 0.134 | 5.31E-02 | 2.08E-01 | 2.70E+00 | 2.74E+00 | 7.96E-01 | no | | intergenic_variant | MODIFIER |
| 2 | 143936926 | AX-170415739 | 1 | T | C | 0.134 | 5.31E-02 | 2.08E-01 | 2.70E+00 | 2.74E+00 | 7.96E-01 | no | Dstn | intron_variant | MODIFIER |
| 2 | 150055112 | AX-170117200 | 1 | C | G | 0.134 | 5.31E-02 | 2.08E-01 | 2.70E+00 | 2.74E+00 | 7.96E-01 | no | | intergenic_variant | MODIFIER |
| 2 | 155656947 | AX-97832996 | 1 | T | C | 0.134 | 5.31E-02 | 2.08E-01 | 2.70E+00 | 2.74E+00 | 7.96E-01 | no | Trpc4ap | intron_variant | MODIFIER |
| 1 | 25924609 | AX-170292680 | 1 | G | A | 0.411 | 2.17E-02 | 8.37E-02 | 2.65E+00 | 2.72E+00 | 7.96E-01 | no | | intergenic_variant | MODIFIER |
| 8 | 84242122 | AX-170254158 | 1 | G | A | 0.196 | 3.32E-02 | 1.31E-01 | 2.68E+00 | 2.75E+00 | 7.96E-01 | no | D8Ertd738e | downstream_gene_variant | MODIFIER |
| 2 | 123840980 | AX-170231110 | 1 | C | G | 0.143 | 4.96E-02 | 1.92E-01 | 2.64E+00 | 2.72E+00 | 7.96E-01 | no | | intergenic_variant | MODIFIER |
| 1 | 71509748 | AX-170409181 | 0 | C | T | 0.123 | 5.44E-02 | 2.07E-01 | 2.63E+00 | 2.70E+00 | 7.96E-01 | no | | intergenic_variant | MODIFIER |
| 6 | 102745884 | AX-97819663 | 1 | G | A | 0.054 | 7.58E-02 | 2.96E-01 | 2.68E+00 | 2.73E+00 | 7.96E-01 | no | | intergenic_variant | MODIFIER |
| 6 | 103351271 | AX-170499022 | 1 | G | A | 0.054 | 7.58E-02 | 2.96E-01 | 2.68E+00 | 2.73E+00 | 7.96E-01 | no | | intergenic_variant | MODIFIER |
| 5 | 44042052 | AX-170253026 | 1 | C | A | 0.214 | -2.64E-02 | 9.82E-02 | 2.59E+00 | 2.69E+00 | 7.96E-01 | no | Prom1 | intron_variant | MODIFIER |
| 4 | 144901269 | AX-170517843 | 1 | T | G | 0.125 | 3.62E-02 | 1.42E-01 | 2.65E+00 | 2.74E+00 | 7.96E-01 | no | Dhrs3 | intron_variant,NMD_transcript_variant | MODIFIER |
| 4 | 58345499 | AX-170520476 | 0 | C | T | 0.377 | -2.74E-02 | 1.10E-01 | 2.71E+00 | 2.78E+00 | 7.97E-01 | no | Musk | intron_variant | MODIFIER |
| 4 | 58353669 | AX-170501788 | 0 | G | A | 0.377 | -2.74E-02 | 1.10E-01 | 2.71E+00 | 2.78E+00 | 7.97E-01 | no | Musk | intron_variant | MODIFIER |
| 4 | 58363486 | AX-170022880 | 0 | T | C | 0.377 | -2.74E-02 | 1.10E-01 | 2.71E+00 | 2.78E+00 | 7.97E-01 | no | Musk | intron_variant | MODIFIER |
| 4 | 58366092 | AX-170473345 | 0 | T | G | 0.377 | -2.74E-02 | 1.10E-01 | 2.71E+00 | 2.78E+00 | 7.97E-01 | no | Musk | intron_variant | MODIFIER |
| 4 | 58369008 | AX-170356475 | 0 | C | T | 0.377 | -2.74E-02 | 1.10E-01 | 2.71E+00 | 2.78E+00 | 7.97E-01 | no | Musk | synonymous_variant | LOW |
| 4 | 58373446 | AX-170271992 | 0 | G | T | 0.377 | -2.74E-02 | 1.10E-01 | 2.71E+00 | 2.78E+00 | 7.97E-01 | no | Musk | synonymous_variant | LOW |
| 4 | 58374025 | AX-170272310 | 0 | T | C | 0.377 | -2.74E-02 | 1.10E-01 | 2.71E+00 | 2.78E+00 | 7.97E-01 | no | Musk | 3_prime_UTR_variant | MODIFIER |
| 4 | 58507941 | AX-170288566 | 0 | G | T | 0.377 | -2.74E-02 | 1.10E-01 | 2.71E+00 | 2.78E+00 | 7.97E-01 | no | Lpar1 | intron_variant | MODIFIER |
| 4 | 58542555 | AX-97791247 | 0 | A | G | 0.377 | -2.74E-02 | 1.10E-01 | 2.71E+00 | 2.78E+00 | 7.97E-01 | no | Lpar1 | intron_variant | MODIFIER |
| 4 | 58564164 | AX-170497558 | 0 | C | G | 0.377 | -2.74E-02 | 1.10E-01 | 2.71E+00 | 2.78E+00 | 7.97E-01 | no | | intergenic_variant | MODIFIER |
| 4 | 58581326 | AX-170304551 | 0 | G | A | 0.377 | -2.74E-02 | 1.10E-01 | 2.71E+00 | 2.78E+00 | 7.97E-01 | no | | regulatory_region_variant | MODIFIER |
| 4 | 58584687 | AX-170424268 | 0 | G | A | 0.377 | -2.74E-02 | 1.10E-01 | 2.71E+00 | 2.78E+00 | 7.97E-01 | no | | intergenic_variant | MODIFIER |
| 4 | 58590104 | AX-170336620 | 0 | T | C | 0.377 | -2.74E-02 | 1.10E-01 | 2.71E+00 | 2.78E+00 | 7.97E-01 | no | Gm12578 | non_coding_transcript_exon_variant | MODIFIER |
| 4 | 58591442 | AX-170478500 | 0 | G | A | 0.377 | -2.74E-02 | 1.10E-01 | 2.71E+00 | 2.78E+00 | 7.97E-01 | no | Gm12578 | downstream_gene_variant | MODIFIER |
| 4 | 58617198 | AX-170217916 | 0 | G | A | 0.377 | -2.74E-02 | 1.10E-01 | 2.71E+00 | 2.78E+00 | 7.97E-01 | no | | intergenic_variant | MODIFIER |
| 6 | 104697245 | AX-170205839 | 1 | C | T | 0.054 | 7.53E-02 | 2.96E-01 | 2.67E+00 | 2.74E+00 | 7.97E-01 | no | Cntn6 | intron_variant | MODIFIER |
| 6 | 144411549 | AX-170281796 | 1 | G | C | 0.054 | 7.53E-02 | 2.96E-01 | 2.67E+00 | 2.74E+00 | 7.97E-01 | no | Sox5 | intron_variant,non_coding_transcript_variant | MODIFIER |
| 13 | 38299004 | AX-170051011 | 0 | G | A | 0.193 | -3.07E-02 | 1.10E-01 | 2.57E+00 | 2.65E+00 | 7.97E-01 | no | | intergenic_variant | MODIFIER |
| 13 | 38451674 | AX-97823219 | 0 | C | T | 0.193 | -3.07E-02 | 1.10E-01 | 2.57E+00 | 2.65E+00 | 7.97E-01 | no | Bmp6 | intron_variant | MODIFIER |
| 13 | 39372506 | AX-170128443 | 0 | G | A | 0.193 | -3.07E-02 | 1.10E-01 | 2.57E+00 | 2.65E+00 | 7.97E-01 | no | | intergenic_variant | MODIFIER |
| 8 | 81182526 | AX-170270864 | 0 | A | G | 0.202 | -3.00E-02 | 1.18E-01 | 2.66E+00 | 2.74E+00 | 7.97E-01 | no | | intergenic_variant | MODIFIER |
| 5 | 32382132 | AX-97823690 | 0 | C | T | 0.368 | 2.68E-02 | 1.05E-01 | 2.66E+00 | 2.73E+00 | 7.97E-01 | no | | intergenic_variant | MODIFIER |
| 9 | 34970535 | AX-170273973 | 0 | T | A | 0.43 | -2.23E-02 | 8.92E-02 | 2.73E+00 | 2.80E+00 | 7.97E-01 | no | Kirrel3 | intron_variant | MODIFIER |
| 9 | 35015901 | AX-170032110 | 0 | T | C | 0.43 | -2.23E-02 | 8.92E-02 | 2.73E+00 | 2.80E+00 | 7.97E-01 | no | Kirrel3 | intron_variant | MODIFIER |
| 9 | 35018269 | AX-170362324 | 0 | T | G | 0.43 | -2.23E-02 | 8.92E-02 | 2.73E+00 | 2.80E+00 | 7.97E-01 | no | Kirrel3 | intron_variant | MODIFIER |
| 9 | 35022121 | AX-170192590 | 0 | T | C | 0.43 | -2.23E-02 | 8.92E-02 | 2.73E+00 | 2.80E+00 | 7.97E-01 | no | Kirrel3 | intron_variant | MODIFIER |
| 9 | 35034319 | AX-170466344 | 0 | G | A | 0.43 | -2.23E-02 | 8.92E-02 | 2.73E+00 | 2.80E+00 | 7.97E-01 | no | Kirrel3 | intron_variant | MODIFIER |
| 9 | 35040959 | AX-170331914 | 0 | A | G | 0.43 | -2.23E-02 | 8.92E-02 | 2.73E+00 | 2.80E+00 | 7.97E-01 | no | Kirrel3 | downstream_gene_variant | MODIFIER |
| 9 | 35056260 | AX-170391937 | 0 | C | T | 0.43 | -2.23E-02 | 8.92E-02 | 2.73E+00 | 2.80E+00 | 7.97E-01 | no | St3gal4 | intron_variant | MODIFIER |
| 9 | 35057381 | AX-170500684 | 0 | G | A | 0.43 | -2.23E-02 | 8.92E-02 | 2.73E+00 | 2.80E+00 | 7.97E-01 | no | St3gal4 | intron_variant | MODIFIER |
| 9 | 35111841 | AX-170082092 | 0 | G | A | 0.43 | -2.23E-02 | 8.92E-02 | 2.73E+00 | 2.80E+00 | 7.97E-01 | no | St3gal4 | intron_variant | MODIFIER |
| 9 | 35136476 | AX-170096513 | 0 | G | A | 0.43 | -2.23E-02 | 8.92E-02 | 2.73E+00 | 2.80E+00 | 7.97E-01 | no | Dcps | synonymous_variant | LOW |
| 9 | 35138282 | AX-170376019 | 0 | A | G | 0.43 | -2.23E-02 | 8.92E-02 | 2.73E+00 | 2.80E+00 | 7.97E-01 | no | Dcps | intron_variant | MODIFIER |
| 9 | 35155100 | AX-170120771 | 0 | A | G | 0.43 | -2.23E-02 | 8.92E-02 | 2.73E+00 | 2.80E+00 | 7.97E-01 | no | Dcps | intron_variant | MODIFIER |
| 9 | 35159521 | AX-170285779 | 0 | A | G | 0.43 | -2.23E-02 | 8.92E-02 | 2.73E+00 | 2.80E+00 | 7.97E-01 | no | Dcps | intron_variant | MODIFIER |
| 9 | 35159727 | AX-97825302 | 0 | A | G | 0.43 | -2.23E-02 | 8.92E-02 | 2.73E+00 | 2.80E+00 | 7.97E-01 | no | Dcps | intron_variant | MODIFIER |
| 9 | 35192702 | AX-170082097 | 0 | A | G | 0.43 | -2.23E-02 | 8.92E-02 | 2.73E+00 | 2.80E+00 | 7.97E-01 | no | Tirap | intron_variant | MODIFIER |
| 9 | 35213921 | AX-170421394 | 0 | C | T | 0.43 | -2.23E-02 | 8.92E-02 | 2.73E+00 | 2.80E+00 | 7.97E-01 | no | Srpr | intron_variant | MODIFIER |
| 9 | 35224053 | AX-170316827 | 0 | A | G | 0.43 | -2.23E-02 | 8.92E-02 | 2.73E+00 | 2.80E+00 | 7.97E-01 | no | Fam118b | intron_variant | MODIFIER |
| 9 | 35262076 | AX-170142029 | 0 | C | T | 0.43 | -2.23E-02 | 8.92E-02 | 2.73E+00 | 2.80E+00 | 7.97E-01 | no | Fam118b | intron_variant | MODIFIER |
| 9 | 35277780 | AX-97848595 | 0 | G | T | 0.43 | -2.23E-02 | 8.92E-02 | 2.73E+00 | 2.80E+00 | 7.97E-01 | no | Rpusd4 | downstream_gene_variant | MODIFIER |
| 9 | 35287107 | AX-170176803 | 0 | T | G | 0.43 | -2.23E-02 | 8.92E-02 | 2.73E+00 | 2.80E+00 | 7.97E-01 | no | 4933422A05Rik | downstream_gene_variant | MODIFIER |

| | | | | | | | | | | | | | | | |
|---|---|---|---|---|---|---|---|---|---|---|---|---|---|---|---|
| 9 | 35289697 | AX-170358049 | 0 | G | A | 0.43 | -2.23E-02 | 8.92E-02 | 2.73E+00 | 2.80E+00 | 7.97E-01 | no | 4933422A05Rik | downstream_gene_variant | MODIFIER |
| 9 | 35302508 | AX-97785275 | 0 | T | A | 0.43 | -2.23E-02 | 8.92E-02 | 2.73E+00 | 2.80E+00 | 7.97E-01 | no | Gm33838 | intron_variant,non_coding_transcript_variant | MODIFIER |
| 9 | 35318934 | AX-170279195 | 0 | A | G | 0.43 | -2.23E-02 | 8.92E-02 | 2.73E+00 | 2.80E+00 | 7.97E-01 | no | | regulatory_region_variant | MODIFIER |
| 9 | 35319319 | AX-170310301 | 0 | G | C | 0.43 | -2.23E-02 | 8.92E-02 | 2.73E+00 | 2.80E+00 | 7.97E-01 | no | | regulatory_region_variant | MODIFIER |
| 9 | 35330636 | AX-170082098 | 0 | T | C | 0.43 | -2.23E-02 | 8.92E-02 | 2.73E+00 | 2.80E+00 | 7.97E-01 | no | | regulatory_region_variant | MODIFIER |
| 9 | 35332488 | AX-170082100 | 0 | T | A | 0.43 | -2.23E-02 | 8.92E-02 | 2.73E+00 | 2.80E+00 | 7.97E-01 | no | | regulatory_region_variant | MODIFIER |
| 9 | 35333113 | AX-170082099 | 0 | C | A | 0.43 | -2.23E-02 | 8.92E-02 | 2.73E+00 | 2.80E+00 | 7.97E-01 | no | | regulatory_region_variant | MODIFIER |
| 9 | 35365555 | AX-170435481 | 0 | C | G | 0.43 | -2.23E-02 | 8.92E-02 | 2.73E+00 | 2.80E+00 | 7.97E-01 | no | 2610105M22Rik | upstream_gene_variant | MODIFIER |
| 9 | 35378703 | AX-170258835 | 0 | C | T | 0.43 | -2.23E-02 | 8.92E-02 | 2.73E+00 | 2.80E+00 | 7.97E-01 | no | | regulatory_region_variant | MODIFIER |
| 9 | 35378725 | AX-170247263 | 0 | A | C | 0.43 | -2.23E-02 | 8.92E-02 | 2.73E+00 | 2.80E+00 | 7.97E-01 | no | | regulatory_region_variant | MODIFIER |
| 9 | 35378825 | AX-170480521 | 0 | C | T | 0.43 | -2.23E-02 | 8.92E-02 | 2.73E+00 | 2.80E+00 | 7.97E-01 | no | | regulatory_region_variant | MODIFIER |
| 9 | 35379597 | AX-170111368 | 0 | A | G | 0.43 | -2.23E-02 | 8.92E-02 | 2.73E+00 | 2.80E+00 | 7.97E-01 | no | | regulatory_region_variant | MODIFIER |
| 9 | 35380824 | AX-170524107 | 0 | T | C | 0.43 | -2.23E-02 | 8.92E-02 | 2.73E+00 | 2.80E+00 | 7.97E-01 | no | | regulatory_region_variant | MODIFIER |
| 9 | 35380905 | AX-170172906 | 0 | A | G | 0.43 | -2.23E-02 | 8.92E-02 | 2.73E+00 | 2.80E+00 | 7.97E-01 | no | | regulatory_region_variant | MODIFIER |
| 9 | 35381022 | AX-170168906 | 0 | G | A | 0.43 | -2.23E-02 | 8.92E-02 | 2.73E+00 | 2.80E+00 | 7.97E-01 | no | | intergenic_variant | MODIFIER |
| 9 | 35399955 | AX-170193560 | 0 | C | T | 0.43 | -2.23E-02 | 8.92E-02 | 2.73E+00 | 2.80E+00 | 7.97E-01 | no | | regulatory_region_variant | MODIFIER |
| 9 | 35402440 | AX-170333166 | 0 | C | T | 0.43 | -2.23E-02 | 8.92E-02 | 2.73E+00 | 2.80E+00 | 7.97E-01 | no | | regulatory_region_variant | MODIFIER |
| 9 | 35403276 | AX-97817282 | 0 | A | G | 0.43 | -2.23E-02 | 8.92E-02 | 2.73E+00 | 2.80E+00 | 7.97E-01 | no | | regulatory_region_variant | MODIFIER |
| 9 | 35404366 | AX-170209365 | 0 | A | G | 0.43 | -2.23E-02 | 8.92E-02 | 2.73E+00 | 2.80E+00 | 7.97E-01 | no | | regulatory_region_variant | MODIFIER |
| 9 | 35405996 | AX-170465285 | 0 | A | C | 0.43 | -2.23E-02 | 8.92E-02 | 2.73E+00 | 2.80E+00 | 7.97E-01 | no | | regulatory_region_variant | MODIFIER |
| 9 | 35440935 | AX-170121937 | 0 | A | G | 0.43 | -2.23E-02 | 8.92E-02 | 2.73E+00 | 2.80E+00 | 7.97E-01 | no | Cdon | intron_variant | MODIFIER |
| 9 | 35473363 | AX-170082103 | 0 | C | T | 0.43 | -2.23E-02 | 8.92E-02 | 2.73E+00 | 2.80E+00 | 7.97E-01 | no | Cdon | intron_variant | MODIFIER |
| 9 | 35477708 | AX-170096515 | 0 | T | C | 0.43 | -2.23E-02 | 8.92E-02 | 2.73E+00 | 2.80E+00 | 7.97E-01 | no | Cdon | intron_variant | MODIFIER |
| 9 | 35479773 | AX-170135754 | 0 | G | T | 0.43 | -2.23E-02 | 8.92E-02 | 2.73E+00 | 2.80E+00 | 7.97E-01 | no | Cdon | intron_variant | MODIFIER |
| 9 | 35480365 | AX-170192584 | 0 | T | G | 0.43 | -2.23E-02 | 8.92E-02 | 2.73E+00 | 2.80E+00 | 7.97E-01 | no | Cdon | intron_variant | MODIFIER |
| 9 | 35483809 | AX-170423785 | 0 | G | C | 0.43 | -2.23E-02 | 8.92E-02 | 2.73E+00 | 2.80E+00 | 7.97E-01 | no | Cdon | intron_variant | MODIFIER |
| 9 | 35487721 | AX-170157231 | 0 | T | C | 0.43 | -2.23E-02 | 8.92E-02 | 2.73E+00 | 2.80E+00 | 7.97E-01 | no | Cdon | intron_variant | MODIFIER |
| 9 | 35487807 | AX-170500546 | 0 | T | G | 0.43 | -2.23E-02 | 8.92E-02 | 2.73E+00 | 2.80E+00 | 7.97E-01 | no | Cdon | intron_variant | MODIFIER |
| 9 | 35489695 | AX-170382151 | 0 | T | G | 0.43 | -2.23E-02 | 8.92E-02 | 2.73E+00 | 2.80E+00 | 7.97E-01 | no | Cdon | intron_variant | MODIFIER |
| 9 | 35494689 | AX-170035802 | 0 | C | T | 0.43 | -2.23E-02 | 8.92E-02 | 2.73E+00 | 2.80E+00 | 7.97E-01 | no | Cdon | intron_variant | MODIFIER |
| 9 | 35496265 | AX-170481806 | 0 | T | G | 0.43 | -2.23E-02 | 8.92E-02 | 2.73E+00 | 2.80E+00 | 7.97E-01 | no | Cdon | intron_variant | MODIFIER |
| 9 | 35508104 | AX-170248440 | 0 | T | C | 0.43 | -2.23E-02 | 8.92E-02 | 2.73E+00 | 2.80E+00 | 7.97E-01 | no | Cdon | downstream_gene_variant | MODIFIER |
| 9 | 35517157 | AX-170461244 | 0 | G | C | 0.43 | -2.23E-02 | 8.92E-02 | 2.73E+00 | 2.80E+00 | 7.97E-01 | no | Gm1113 | intron_variant | MODIFIER |
| 9 | 35517486 | AX-170323908 | 0 | T | A | 0.43 | -2.23E-02 | 8.92E-02 | 2.73E+00 | 2.80E+00 | 7.97E-01 | no | Gm1113 | intron_variant | MODIFIER |
| 9 | 35517842 | AX-170357209 | 0 | G | A | 0.43 | -2.23E-02 | 8.92E-02 | 2.73E+00 | 2.80E+00 | 7.97E-01 | no | Gm1113 | intron_variant | MODIFIER |
| 9 | 35522596 | AX-97850203 | 0 | G | A | 0.43 | -2.23E-02 | 8.92E-02 | 2.73E+00 | 2.80E+00 | 7.97E-01 | no | Gm1113 | upstream_gene_variant | MODIFIER |
| 7 | 62187547 | AX-170274719 | 0 | C | T | 0.43 | 2.43E-02 | 9.52E-02 | 2.65E+00 | 2.73E+00 | 7.97E-01 | no | Gm9801 | intron_variant,non_coding_transcript_variant | MODIFIER |
| 5 | 69418916 | AX-170025220 | 0 | C | A | 0.447 | -2.29E-02 | 9.27E-02 | 2.68E+00 | 2.78E+00 | 7.97E-01 | no | | intergenic_variant | MODIFIER |
| 6 | 102526733 | AX-170205638 | 1 | T | C | 0.054 | -5.40E-02 | 2.15E-01 | 2.64E+00 | 2.75E+00 | 7.97E-01 | no | Cntn3 | intron_variant | MODIFIER |
| 1 | 96545407 | AX-170137351 | 0 | G | T | 0.149 | 5.21E-02 | 2.01E-01 | 2.64E+00 | 2.71E+00 | 7.97E-01 | no | Gm29601 | intron_variant,non_coding_transcript_variant | MODIFIER |
| 4 | 145083916 | AX-170267128 | 1 | G | A | 0.107 | 5.24E-02 | 2.07E-01 | 2.68E+00 | 2.74E+00 | 7.97E-01 | no | Vps13d | intron_variant | MODIFIER |
| 6 | 139438057 | AX-170250337 | 2 | G | A | 0.055 | 7.51E-02 | 2.95E-01 | 2.67E+00 | 2.73E+00 | 7.97E-01 | no | | intergenic_variant | MODIFIER |
| 6 | 5142233 | AX-97783663 | 0 | G | A | 0.044 | -4.21E-02 | 1.68E-01 | 2.63E+00 | 2.75E+00 | 7.97E-01 | no | Ppp1r9a | intron_variant | MODIFIER |
| 2 | 159826842 | AX-97839739 | 0 | C | T | 0.149 | -4.34E-02 | 1.73E-01 | 2.65E+00 | 2.75E+00 | 7.98E-01 | no | | regulatory_region_variant | MODIFIER |
| 2 | 160357111 | AX-97805202 | 0 | A | G | 0.149 | -4.34E-02 | 1.73E-01 | 2.65E+00 | 2.75E+00 | 7.98E-01 | no | | intergenic_variant | MODIFIER |
| 2 | 94502163 | AX-170527019 | 0 | A | G | 0.061 | -5.60E-02 | 2.25E-01 | 2.68E+00 | 2.77E+00 | 7.98E-01 | no | | intergenic_variant | MODIFIER |
| 2 | 96698405 | AX-170285258 | 0 | G | A | 0.061 | -5.60E-02 | 2.25E-01 | 2.68E+00 | 2.77E+00 | 7.98E-01 | no | Lrrc4c | intron_variant | MODIFIER |
| 2 | 112700792 | AX-97839514 | 0 | C | T | 0.061 | -5.60E-02 | 2.25E-01 | 2.68E+00 | 2.77E+00 | 7.98E-01 | no | Ryr3 | intron_variant | MODIFIER |
| 1 | 76032892 | AX-170302343 | 1 | G | A | 0.134 | 5.68E-02 | 2.23E-01 | 2.67E+00 | 2.73E+00 | 7.98E-01 | no | | intergenic_variant | MODIFIER |
| 20 | 168949456 | AX-170085809 | 1 | A | G | 0.179 | 3.20E-02 | 1.23E-01 | 2.62E+00 | 2.71E+00 | 7.98E-01 | no | Arhgap6 | intron_variant | MODIFIER |
| 4 | 144382277 | AX-170410592 | 1 | A | G | 0.098 | 5.51E-02 | 2.18E-01 | 2.69E+00 | 2.74E+00 | 7.98E-01 | no | Pramef20 | upstream_gene_variant | MODIFIER |
| 8 | 94189605 | AX-170133010 | 0 | C | T | 0.211 | 2.91E-02 | 1.16E-01 | 2.68E+00 | 2.75E+00 | 7.98E-01 | no | Gm39228 | downstream_gene_variant | MODIFIER |
| 8 | 75578308 | AX-170090150 | 0 | T | G | 0.175 | -3.00E-02 | 1.16E-01 | 2.61E+00 | 2.72E+00 | 7.98E-01 | no | Iqcm | intron_variant | MODIFIER |
| 15 | 87112926 | AX-170056210 | 1 | G | A | 0.134 | 4.05E-02 | 1.57E-01 | 2.64E+00 | 2.71E+00 | 7.98E-01 | no | | intergenic_variant | MODIFIER |

| | | | | | | | | | | | | | | | |
|---|---|---|---|---|---|---|---|---|---|---|---|---|---|---|---|
| 9 | 106123841 | AX-170273349 | 0 | G | A | 0.123 | -3.18E-02 | 1.19E-01 | 2.61E+00 | 2.67E+00 | 7.98E-01 | no | Gm29208 | intron_variant,non_coding_transcript_variant | MODIFIER |
| 9 | 107003767 | AX-170033231 | 0 | C | T | 0.123 | -3.18E-02 | 1.19E-01 | 2.61E+00 | 2.67E+00 | 7.98E-01 | no | Dock3 | intron_variant | MODIFIER |
| 5 | 7059370 | AX-170155717 | 0 | T | C | 0.43 | 1.49E-02 | 6.16E-02 | 2.65E+00 | 2.78E+00 | 7.98E-01 | no | Zfp804b | intron_variant | MODIFIER |
| 9 | 55941590 | AX-170507981 | 1 | C | G | 0.196 | -3.03E-02 | 1.21E-01 | 2.69E+00 | 2.76E+00 | 7.98E-01 | no | Gm7435 | intron_variant,non_coding_transcript_variant | MODIFIER |
| 8 | 36572547 | AX-170366132 | 0 | C | T | 0.158 | 3.51E-02 | 1.41E-01 | 2.74E+00 | 2.79E+00 | 7.98E-01 | no | Dlc1 | intron_variant | MODIFIER |
| 5 | 39996070 | AX-97799276 | 0 | C | A | 0.219 | -2.81E-02 | 1.09E-01 | 2.64E+00 | 2.72E+00 | 7.99E-01 | no | | intergenic_variant | MODIFIER |
| 8 | 128491097 | AX-170035799 | 1 | T | A | 0.054 | 5.09E-02 | 2.07E-01 | 2.68E+00 | 2.77E+00 | 7.99E-01 | no | Nrp1 | intron_variant | MODIFIER |
| 7 | 75641893 | AX-170133359 | 0 | T | C | 0.368 | -2.35E-02 | 9.53E-02 | 2.65E+00 | 2.76E+00 | 7.99E-01 | no | Akap13 | intron_variant | MODIFIER |
| 4 | 6258826 | AX-170022094 | 2 | C | A | 0.064 | -3.55E-02 | 1.42E-01 | 2.62E+00 | 2.75E+00 | 7.99E-01 | no | Gm11798 | downstream_gene_variant | MODIFIER |
| 6 | 126217267 | AX-170271405 | 0 | A | G | 0.167 | -2.05E-02 | 8.38E-02 | 2.64E+00 | 2.76E+00 | 7.99E-01 | no | | intergenic_variant | MODIFIER |
| 1 | 68179276 | AX-170159695 | 2 | T | A | 0.1 | -4.76E-02 | 1.94E-01 | 2.66E+00 | 2.76E+00 | 7.99E-01 | no | Erbb4 | intron_variant | MODIFIER |
| 8 | 94634637 | AX-170341717 | 1 | G | A | 0.205 | -3.03E-02 | 1.20E-01 | 2.67E+00 | 2.74E+00 | 7.99E-01 | no | Rspry1 | intron_variant | MODIFIER |
| 10 | 3920230 | AX-170396406 | 2 | A | G | 0.2 | -2.38E-02 | 9.46E-02 | 2.62E+00 | 2.74E+00 | 8.00E-01 | no | Plekhg1 | intron_variant | MODIFIER |
| 16 | 92649231 | AX-170267464 | 1 | A | G | 0.464 | 1.85E-02 | 7.49E-02 | 2.65E+00 | 2.75E+00 | 8.00E-01 | no | Runx1 | intron_variant | MODIFIER |
| 1 | 62844209 | AX-170217174 | 1 | G | A | 0.134 | 6.03E-02 | 2.41E-01 | 2.70E+00 | 2.75E+00 | 8.00E-01 | no | | intergenic_variant | MODIFIER |
| 7 | 112641494 | AX-170494221 | 0 | C | T | 0.439 | 2.39E-02 | 9.52E-02 | 2.69E+00 | 2.74E+00 | 8.00E-01 | no | 2310014F06Rik | intron_variant,non_coding_transcript_variant | MODIFIER |
| 7 | 112641650 | AX-170356383 | 0 | C | T | 0.439 | 2.39E-02 | 9.52E-02 | 2.69E+00 | 2.74E+00 | 8.00E-01 | no | 2310014F06Rik | intron_variant,non_coding_transcript_variant | MODIFIER |
| 7 | 112650562 | AX-168385766 | 0 | C | T | 0.439 | 2.39E-02 | 9.52E-02 | 2.69E+00 | 2.74E+00 | 8.00E-01 | no | 2310014F06Rik | intron_variant,non_coding_transcript_variant | MODIFIER |
| 7 | 112656425 | AX-170278867 | 0 | C | T | 0.439 | 2.39E-02 | 9.52E-02 | 2.69E+00 | 2.74E+00 | 8.00E-01 | no | 2310014F06Rik | intron_variant,non_coding_transcript_variant | MODIFIER |
| 7 | 112663383 | AX-170035768 | 0 | C | T | 0.439 | 2.39E-02 | 9.52E-02 | 2.69E+00 | 2.74E+00 | 8.00E-01 | no | 2310014F06Rik | intron_variant,non_coding_transcript_variant | MODIFIER |
| 7 | 112669905 | AX-170471029 | 0 | T | C | 0.439 | 2.39E-02 | 9.52E-02 | 2.69E+00 | 2.74E+00 | 8.00E-01 | no | 2310014F06Rik | intron_variant,non_coding_transcript_variant | MODIFIER |
| 7 | 112712542 | AX-170475176 | 0 | T | G | 0.439 | 2.39E-02 | 9.52E-02 | 2.69E+00 | 2.74E+00 | 8.00E-01 | no | Tead1 | intron_variant | MODIFIER |
| 7 | 112712589 | AX-170276394 | 0 | C | T | 0.439 | 2.39E-02 | 9.52E-02 | 2.69E+00 | 2.74E+00 | 8.00E-01 | no | Tead1 | intron_variant | MODIFIER |
| 7 | 112721581 | AX-170103966 | 0 | C | T | 0.439 | 2.39E-02 | 9.52E-02 | 2.69E+00 | 2.74E+00 | 8.00E-01 | no | Tead1 | intron_variant | MODIFIER |
| 7 | 112735922 | AX-170123617 | 0 | A | G | 0.439 | 2.39E-02 | 9.52E-02 | 2.69E+00 | 2.74E+00 | 8.00E-01 | no | Tead1 | intron_variant | MODIFIER |
| 7 | 112736553 | AX-170228900 | 0 | G | A | 0.439 | 2.39E-02 | 9.52E-02 | 2.69E+00 | 2.74E+00 | 8.00E-01 | no | Tead1 | intron_variant | MODIFIER |
| 7 | 112805721 | AX-170134819 | 0 | T | C | 0.439 | 2.39E-02 | 9.52E-02 | 2.69E+00 | 2.74E+00 | 8.00E-01 | no | Tead1 | intron_variant | MODIFIER |
| 7 | 112807353 | AX-170276060 | 0 | G | A | 0.439 | 2.39E-02 | 9.52E-02 | 2.69E+00 | 2.74E+00 | 8.00E-01 | no | Tead1 | intron_variant | MODIFIER |
| 7 | 112807964 | AX-170422202 | 0 | A | T | 0.439 | 2.39E-02 | 9.52E-02 | 2.69E+00 | 2.74E+00 | 8.00E-01 | no | Tead1 | intron_variant | MODIFIER |
| 7 | 112813095 | AX-170117031 | 0 | G | A | 0.439 | 2.39E-02 | 9.52E-02 | 2.69E+00 | 2.74E+00 | 8.00E-01 | no | Tead1 | intron_variant | MODIFIER |
| 7 | 112844051 | AX-170413825 | 0 | T | C | 0.439 | 2.39E-02 | 9.52E-02 | 2.69E+00 | 2.74E+00 | 8.00E-01 | no | Tead1 | intron_variant | MODIFIER |
| 7 | 112862033 | AX-170105606 | 0 | G | C | 0.439 | 2.39E-02 | 9.52E-02 | 2.69E+00 | 2.74E+00 | 8.00E-01 | no | Tead1 | intron_variant | MODIFIER |
| 7 | 112863974 | AX-170388125 | 0 | C | T | 0.439 | 2.39E-02 | 9.52E-02 | 2.69E+00 | 2.74E+00 | 8.00E-01 | no | Tead1 | intron_variant | MODIFIER |
| 7 | 112913045 | AX-170113612 | 0 | C | T | 0.439 | 2.39E-02 | 9.52E-02 | 2.69E+00 | 2.74E+00 | 8.00E-01 | no | | intergenic_variant | MODIFIER |
| 7 | 112942910 | AX-170078460 | 0 | A | G | 0.439 | 2.39E-02 | 9.52E-02 | 2.69E+00 | 2.74E+00 | 8.00E-01 | no | | intergenic_variant | MODIFIER |
| 7 | 112946722 | AX-170262387 | 0 | G | A | 0.439 | 2.39E-02 | 9.52E-02 | 2.69E+00 | 2.74E+00 | 8.00E-01 | no | | intergenic_variant | MODIFIER |
| 7 | 112946767 | AX-170078458 | 0 | C | T | 0.439 | 2.39E-02 | 9.52E-02 | 2.69E+00 | 2.74E+00 | 8.00E-01 | no | | intergenic_variant | MODIFIER |
| 7 | 112962149 | AX-170443090 | 0 | C | A | 0.439 | 2.39E-02 | 9.52E-02 | 2.69E+00 | 2.74E+00 | 8.00E-01 | no | Rassf10 | downstream_gene_variant | MODIFIER |
| 7 | 112971130 | AX-170167194 | 0 | A | G | 0.439 | 2.39E-02 | 9.52E-02 | 2.69E+00 | 2.74E+00 | 8.00E-01 | no | Gm5599 | downstream_gene_variant | MODIFIER |
| 7 | 112989080 | AX-170096076 | 0 | T | C | 0.439 | 2.39E-02 | 9.52E-02 | 2.69E+00 | 2.74E+00 | 8.00E-01 | no | Gm33586 | intron_variant,non_coding_transcript_variant | MODIFIER |
| 7 | 113048203 | AX-170326927 | 0 | T | C | 0.439 | 2.39E-02 | 9.52E-02 | 2.69E+00 | 2.74E+00 | 8.00E-01 | no | | regulatory_region_variant | MODIFIER |
| 7 | 113051666 | AX-170196409 | 0 | A | T | 0.439 | 2.39E-02 | 9.52E-02 | 2.69E+00 | 2.74E+00 | 8.00E-01 | no | | intergenic_variant | MODIFIER |
| 7 | 113058209 | AX-97815804 | 0 | C | T | 0.439 | 2.39E-02 | 9.52E-02 | 2.69E+00 | 2.74E+00 | 8.00E-01 | no | | regulatory_region_variant | MODIFIER |
| 7 | 113069417 | AX-170485185 | 0 | G | A | 0.439 | 2.39E-02 | 9.52E-02 | 2.69E+00 | 2.74E+00 | 8.00E-01 | no | | intergenic_variant | MODIFIER |
| 7 | 113071384 | AX-170134797 | 0 | T | C | 0.439 | 2.39E-02 | 9.52E-02 | 2.69E+00 | 2.74E+00 | 8.00E-01 | no | | intergenic_variant | MODIFIER |
| 7 | 113101179 | AX-170117537 | 0 | C | G | 0.439 | 2.39E-02 | 9.52E-02 | 2.69E+00 | 2.74E+00 | 8.00E-01 | no | 4930543E12Rik | intron_variant,non_coding_transcript_variant | MODIFIER |
| 7 | 113104217 | AX-97785325 | 0 | G | A | 0.439 | 2.39E-02 | 9.52E-02 | 2.69E+00 | 2.74E+00 | 8.00E-01 | no | 4930543E12Rik | intron_variant,non_coding_transcript_variant | MODIFIER |
| 7 | 113135073 | AX-170258914 | 0 | T | C | 0.439 | 2.39E-02 | 9.52E-02 | 2.69E+00 | 2.74E+00 | 8.00E-01 | no | 4930543E12Rik | intron_variant,non_coding_transcript_variant | MODIFIER |
| 7 | 113135445 | AX-170508786 | 0 | A | G | 0.439 | 2.39E-02 | 9.52E-02 | 2.69E+00 | 2.74E+00 | 8.00E-01 | no | 4930543E12Rik | intron_variant,non_coding_transcript_variant | MODIFIER |
| 7 | 113143280 | AX-170239631 | 0 | T | G | 0.439 | 2.39E-02 | 9.52E-02 | 2.69E+00 | 2.74E+00 | 8.00E-01 | no | 4930543E12Rik | upstream_gene_variant | MODIFIER |
| 7 | 113143397 | AX-170291406 | 0 | G | A | 0.439 | 2.39E-02 | 9.52E-02 | 2.69E+00 | 2.74E+00 | 8.00E-01 | no | 4930543E12Rik | upstream_gene_variant | MODIFIER |
| 7 | 113149608 | AX-170451835 | 0 | A | G | 0.439 | 2.39E-02 | 9.52E-02 | 2.69E+00 | 2.74E+00 | 8.00E-01 | no | | intergenic_variant | MODIFIER |
| 7 | 113165006 | AX-170419358 | 0 | G | A | 0.439 | 2.39E-02 | 9.52E-02 | 2.69E+00 | 2.74E+00 | 8.00E-01 | no | | regulatory_region_variant | MODIFIER |
| 7 | 113189972 | AX-170268173 | 0 | G | A | 0.439 | 2.39E-02 | 9.52E-02 | 2.69E+00 | 2.74E+00 | 8.00E-01 | no | | intergenic_variant | MODIFIER |

| | | | | | | | | | | | | | | | |
|---|---|---|---|---|---|---|---|---|---|---|---|---|---|---|---|
| 7 | 113254058 | AX-170480425 | 0 | A | G | 0.439 | 2.39E-02 | 9.52E-02 | 2.69E+00 | 2.74E+00 | 8.00E-01 | no | Arntl | intron_variant | MODIFIER |
| 7 | 113261071 | AX-170321965 | 0 | G | A | 0.439 | 2.39E-02 | 9.52E-02 | 2.69E+00 | 2.74E+00 | 8.00E-01 | no | Arntl | intron_variant | MODIFIER |
| 7 | 113274730 | AX-170152960 | 0 | A | G | 0.439 | 2.39E-02 | 9.52E-02 | 2.69E+00 | 2.74E+00 | 8.00E-01 | no | Arntl | 5_prime_UTR_variant | MODIFIER |
| 7 | 113304233 | AX-170163855 | 0 | T | C | 0.439 | 2.39E-02 | 9.52E-02 | 2.69E+00 | 2.74E+00 | 8.00E-01 | no | Arntl | intron_variant | MODIFIER |
| 7 | 113308166 | AX-170280461 | 0 | T | C | 0.439 | 2.39E-02 | 9.52E-02 | 2.69E+00 | 2.74E+00 | 8.00E-01 | no | Arntl | intron_variant | MODIFIER |
| 7 | 113334384 | AX-170078466 | 0 | G | T | 0.439 | 2.39E-02 | 9.52E-02 | 2.69E+00 | 2.74E+00 | 8.00E-01 | no | Btbd10 | intron_variant | MODIFIER |
| 7 | 113342762 | AX-170194712 | 0 | G | A | 0.439 | 2.39E-02 | 9.52E-02 | 2.69E+00 | 2.74E+00 | 8.00E-01 | no | Btbd10 | intron_variant | MODIFIER |
| 7 | 113345255 | AX-170340129 | 0 | G | C | 0.439 | 2.39E-02 | 9.52E-02 | 2.69E+00 | 2.74E+00 | 8.00E-01 | no | Btbd10 | intron_variant | MODIFIER |
| 7 | 113367904 | AX-170402328 | 0 | G | A | 0.439 | 2.39E-02 | 9.52E-02 | 2.69E+00 | 2.74E+00 | 8.00E-01 | no | Btbd10 | intron_variant | MODIFIER |
| 7 | 113891348 | AX-170389472 | 0 | A | G | 0.439 | 2.39E-02 | 9.52E-02 | 2.69E+00 | 2.74E+00 | 8.00E-01 | no | Spon1 | intron_variant | MODIFIER |
| 7 | 113894620 | AX-170214258 | 0 | C | T | 0.439 | 2.39E-02 | 9.52E-02 | 2.69E+00 | 2.74E+00 | 8.00E-01 | no | Spon1 | intron_variant | MODIFIER |
| 7 | 113897841 | AX-170435502 | 0 | A | T | 0.439 | 2.39E-02 | 9.52E-02 | 2.69E+00 | 2.74E+00 | 8.00E-01 | no | Spon1 | intron_variant | MODIFIER |
| 7 | 113898218 | AX-170418881 | 0 | G | A | 0.439 | 2.39E-02 | 9.52E-02 | 2.69E+00 | 2.74E+00 | 8.00E-01 | no | Spon1 | intron_variant | MODIFIER |
| 7 | 113903302 | AX-170315161 | 0 | A | G | 0.439 | 2.39E-02 | 9.52E-02 | 2.69E+00 | 2.74E+00 | 8.00E-01 | no | Spon1 | intron_variant | MODIFIER |
| 10 | 16956938 | AX-97806165 | 0 | C | G | 0.202 | 2.56E-02 | 1.05E-01 | 2.73E+00 | 2.81E+00 | 8.00E-01 | no | Gm20125 | intron_variant,non_coding_transcript_variant | MODIFIER |
| 10 | 17012594 | AX-97850430 | 0 | T | C | 0.202 | 2.56E-02 | 1.05E-01 | 2.73E+00 | 2.81E+00 | 8.00E-01 | no | Gm20125 | intron_variant,non_coding_transcript_variant | MODIFIER |
| 10 | 17261954 | AX-97825106 | 0 | T | C | 0.202 | 2.56E-02 | 1.05E-01 | 2.73E+00 | 2.81E+00 | 8.00E-01 | no | | regulatory_region_variant | MODIFIER |
| 10 | 17278317 | AX-97796623 | 0 | C | G | 0.202 | 2.56E-02 | 1.05E-01 | 2.73E+00 | 2.81E+00 | 8.00E-01 | no | | intergenic_variant | MODIFIER |
| 10 | 17450957 | AX-170035478 | 0 | T | C | 0.202 | 2.56E-02 | 1.05E-01 | 2.73E+00 | 2.81E+00 | 8.00E-01 | no | | intergenic_variant | MODIFIER |
| 10 | 17497121 | AX-170179850 | 0 | T | C | 0.202 | 2.56E-02 | 1.05E-01 | 2.73E+00 | 2.81E+00 | 8.00E-01 | no | | intergenic_variant | MODIFIER |
| 10 | 17543891 | AX-97814298 | 0 | A | G | 0.202 | 2.56E-02 | 1.05E-01 | 2.73E+00 | 2.81E+00 | 8.00E-01 | no | | intergenic_variant | MODIFIER |
| 17 | 32903731 | AX-170307954 | 0 | C | T | 0.044 | 5.11E-02 | 2.09E-01 | 2.69E+00 | 2.78E+00 | 8.00E-01 | no | Cyp4f14 | downstream_gene_variant | MODIFIER |
| 15 | 102434474 | AX-170142340 | 1 | T | A | 0.116 | 3.78E-02 | 1.50E-01 | 2.65E+00 | 2.73E+00 | 8.00E-01 | no | Sp1 | 3_prime_UTR_variant | MODIFIER |
| 13 | 24371616 | AX-170007933 | 0 | C | G | 0.175 | -3.22E-02 | 1.12E-01 | 2.54E+00 | 2.62E+00 | 8.00E-01 | no | Cmah | intron_variant | MODIFIER |
| 4 | 55965845 | AX-170291157 | 0 | G | A | 0.36 | -2.57E-02 | 1.04E-01 | 2.68E+00 | 2.76E+00 | 8.00E-01 | no | Gm12519 | downstream_gene_variant | MODIFIER |
| 4 | 55975799 | AX-170367666 | 0 | G | A | 0.36 | -2.57E-02 | 1.04E-01 | 2.68E+00 | 2.76E+00 | 8.00E-01 | no | Gm12519 | intron_variant,non_coding_transcript_variant | MODIFIER |
| 4 | 55990349 | AX-170178278 | 0 | T | C | 0.36 | -2.57E-02 | 1.04E-01 | 2.68E+00 | 2.76E+00 | 8.00E-01 | no | Gm12519 | intron_variant,non_coding_transcript_variant | MODIFIER |
| 4 | 56015714 | AX-170370989 | 0 | T | C | 0.36 | -2.57E-02 | 1.04E-01 | 2.68E+00 | 2.76E+00 | 8.00E-01 | no | | intergenic_variant | MODIFIER |
| 4 | 56060863 | AX-170414764 | 0 | G | A | 0.36 | -2.57E-02 | 1.04E-01 | 2.68E+00 | 2.76E+00 | 8.00E-01 | no | | intergenic_variant | MODIFIER |
| 4 | 56085063 | AX-170133759 | 0 | A | G | 0.36 | -2.57E-02 | 1.04E-01 | 2.68E+00 | 2.76E+00 | 8.00E-01 | no | | intergenic_variant | MODIFIER |
| 4 | 56087258 | AX-170362193 | 0 | A | T | 0.36 | -2.57E-02 | 1.04E-01 | 2.68E+00 | 2.76E+00 | 8.00E-01 | no | | intergenic_variant | MODIFIER |
| 4 | 56093549 | AX-97788812 | 0 | A | G | 0.36 | -2.57E-02 | 1.04E-01 | 2.68E+00 | 2.76E+00 | 8.00E-01 | no | | intergenic_variant | MODIFIER |
| 4 | 56099837 | AX-170069423 | 0 | G | A | 0.36 | -2.57E-02 | 1.04E-01 | 2.68E+00 | 2.76E+00 | 8.00E-01 | no | | intergenic_variant | MODIFIER |
| 4 | 56100824 | AX-170069424 | 0 | G | T | 0.36 | -2.57E-02 | 1.04E-01 | 2.68E+00 | 2.76E+00 | 8.00E-01 | no | | intergenic_variant | MODIFIER |
| 4 | 56120430 | AX-170458105 | 0 | T | A | 0.36 | -2.57E-02 | 1.04E-01 | 2.68E+00 | 2.76E+00 | 8.00E-01 | no | | intergenic_variant | MODIFIER |
| 4 | 56120563 | AX-170187882 | 0 | C | G | 0.36 | -2.57E-02 | 1.04E-01 | 2.68E+00 | 2.76E+00 | 8.00E-01 | no | | intergenic_variant | MODIFIER |
| 4 | 56153988 | AX-170521412 | 0 | A | C | 0.36 | -2.57E-02 | 1.04E-01 | 2.68E+00 | 2.76E+00 | 8.00E-01 | no | | intergenic_variant | MODIFIER |
| 4 | 56233218 | AX-170199868 | 0 | C | T | 0.36 | -2.57E-02 | 1.04E-01 | 2.68E+00 | 2.76E+00 | 8.00E-01 | no | | intergenic_variant | MODIFIER |
| 4 | 56233783 | AX-170114191 | 0 | A | G | 0.36 | -2.57E-02 | 1.04E-01 | 2.68E+00 | 2.76E+00 | 8.00E-01 | no | | intergenic_variant | MODIFIER |
| 4 | 56239415 | AX-170022840 | 0 | G | A | 0.36 | -2.57E-02 | 1.04E-01 | 2.68E+00 | 2.76E+00 | 8.00E-01 | no | | regulatory_region_variant | MODIFIER |
| 4 | 56267029 | AX-170420372 | 0 | C | A | 0.36 | -2.57E-02 | 1.04E-01 | 2.68E+00 | 2.76E+00 | 8.00E-01 | no | | intergenic_variant | MODIFIER |
| 4 | 56307909 | AX-170450435 | 0 | G | T | 0.36 | -2.57E-02 | 1.04E-01 | 2.68E+00 | 2.76E+00 | 8.00E-01 | no | | intergenic_variant | MODIFIER |
| 4 | 56307940 | AX-170523073 | 0 | T | C | 0.36 | -2.57E-02 | 1.04E-01 | 2.68E+00 | 2.76E+00 | 8.00E-01 | no | | intergenic_variant | MODIFIER |
| 4 | 56317002 | AX-170445361 | 0 | C | T | 0.36 | -2.57E-02 | 1.04E-01 | 2.68E+00 | 2.76E+00 | 8.00E-01 | no | | intergenic_variant | MODIFIER |
| 4 | 56317445 | AX-170341363 | 0 | T | G | 0.36 | -2.57E-02 | 1.04E-01 | 2.68E+00 | 2.76E+00 | 8.00E-01 | no | | intergenic_variant | MODIFIER |
| 4 | 56317889 | AX-170486848 | 0 | A | C | 0.36 | -2.57E-02 | 1.04E-01 | 2.68E+00 | 2.76E+00 | 8.00E-01 | no | | intergenic_variant | MODIFIER |
| 4 | 56319085 | AX-170452515 | 0 | C | T | 0.36 | -2.57E-02 | 1.04E-01 | 2.68E+00 | 2.76E+00 | 8.00E-01 | no | | intergenic_variant | MODIFIER |
| 4 | 56319304 | AX-170412265 | 0 | C | T | 0.36 | -2.57E-02 | 1.04E-01 | 2.68E+00 | 2.76E+00 | 8.00E-01 | no | | intergenic_variant | MODIFIER |
| 4 | 56758425 | AX-170472125 | 0 | A | T | 0.36 | -2.57E-02 | 1.04E-01 | 2.68E+00 | 2.76E+00 | 8.00E-01 | no | Elp1 | intron_variant | MODIFIER |
| 4 | 57031997 | AX-170226643 | 0 | T | C | 0.36 | -2.57E-02 | 1.04E-01 | 2.68E+00 | 2.76E+00 | 8.00E-01 | no | Epb41l4b | upstream_gene_variant | MODIFIER |
| 4 | 57032650 | AX-170317400 | 0 | G | T | 0.36 | -2.57E-02 | 1.04E-01 | 2.68E+00 | 2.76E+00 | 8.00E-01 | no | Epb41l4b | upstream_gene_variant | MODIFIER |
| 4 | 57045061 | AX-170097887 | 0 | T | C | 0.36 | -2.57E-02 | 1.04E-01 | 2.68E+00 | 2.76E+00 | 8.00E-01 | no | Epb41l4b | intron_variant | MODIFIER |
| 4 | 57046762 | AX-170486015 | 0 | T | C | 0.36 | -2.57E-02 | 1.04E-01 | 2.68E+00 | 2.76E+00 | 8.00E-01 | no | Epb41l4b | intron_variant | MODIFIER |
| 4 | 57114390 | AX-170162517 | 0 | G | A | 0.36 | -2.57E-02 | 1.04E-01 | 2.68E+00 | 2.76E+00 | 8.00E-01 | no | Epb41l4b | intron_variant | MODIFIER |

| | | | | | | | | | | | | | | | |
|---|---|---|---|---|---|---|---|---|---|---|---|---|---|---|---|
| 4 | 58319395 | AX-170022879 | 0 | G | A | 0.36 | -2.57E-02 | 1.04E-01 | 2.68E+00 | 2.76E+00 | 8.00E-01 | no | Musk | intron_variant | MODIFIER |
| 4 | 58325151 | AX-168409194 | 0 | T | C | 0.36 | -2.57E-02 | 1.04E-01 | 2.68E+00 | 2.76E+00 | 8.00E-01 | no | Musk | intron_variant | MODIFIER |
| 1 | 115263402 | AX-170164008 | 0 | A | G | 0.167 | -4.01E-02 | 1.66E-01 | 2.67E+00 | 2.78E+00 | 8.00E-01 | no | | intergenic_variant | MODIFIER |
| 5 | 126638355 | AX-97822167 | 1 | C | T | 0.464 | 2.34E-02 | 9.32E-02 | 2.65E+00 | 2.74E+00 | 8.00E-01 | no | | intergenic_variant | MODIFIER |
| 7 | 61341121 | AX-97847452 | 1 | C | A | 0.429 | -2.40E-02 | 9.80E-02 | 2.69E+00 | 2.77E+00 | 8.00E-01 | no | | intergenic_variant | MODIFIER |
| 7 | 62269674 | AX-170334052 | 1 | T | C | 0.429 | -2.40E-02 | 9.80E-02 | 2.69E+00 | 2.77E+00 | 8.00E-01 | no | | regulatory_region_variant | MODIFIER |
| 8 | 51794462 | AX-170313490 | 1 | A | C | 0.143 | -3.93E-02 | 1.49E-01 | 2.62E+00 | 2.67E+00 | 8.00E-01 | no | | intergenic_variant | MODIFIER |
| 13 | 54248721 | AX-170487529 | 1 | C | T | 0.536 | 1.82E-02 | 6.84E-02 | 2.58E+00 | 2.69E+00 | 8.00E-01 | no | Gm48615 | downstream_gene_variant | MODIFIER |
| 2 | 113632747 | AX-97838662 | 1 | G | A | 0.143 | 5.68E-02 | 2.26E-01 | 2.70E+00 | 2.72E+00 | 8.00E-01 | no | Fmn1 | intron_variant | MODIFIER |
| 2 | 123922506 | AX-170225285 | 1 | C | T | 0.143 | 5.68E-02 | 2.26E-01 | 2.70E+00 | 2.72E+00 | 8.00E-01 | no | | intergenic_variant | MODIFIER |
| 2 | 138495902 | AX-170088766 | 1 | C | T | 0.143 | 5.68E-02 | 2.26E-01 | 2.70E+00 | 2.72E+00 | 8.00E-01 | no | Btbd3 | intron_variant,non_coding_transcript_variant | MODIFIER |
| 2 | 139450173 | AX-97815514 | 1 | C | T | 0.143 | 5.68E-02 | 2.26E-01 | 2.70E+00 | 2.72E+00 | 8.00E-01 | no | Gm14066 | intron_variant,non_coding_transcript_variant | MODIFIER |
| 2 | 155569220 | AX-170342775 | 1 | T | A | 0.143 | 5.68E-02 | 2.26E-01 | 2.70E+00 | 2.72E+00 | 8.00E-01 | no | Acss2 | intron_variant | MODIFIER |
| 1 | 33114617 | AX-170000241 | 1 | T | G | 0.42 | -1.96E-02 | 7.95E-02 | 2.68E+00 | 2.76E+00 | 8.00E-01 | no | Gm4785 | upstream_gene_variant | MODIFIER |
| 9 | 35146503 | AX-170466309 | 1 | C | T | 0.429 | -2.11E-02 | 8.62E-02 | 2.70E+00 | 2.78E+00 | 8.00E-01 | no | Dcps | intron_variant | MODIFIER |
| 7 | 113849365 | AX-170297388 | 1 | A | T | 0.438 | 2.37E-02 | 9.50E-02 | 2.69E+00 | 2.74E+00 | 8.00E-01 | no | Spon1 | intron_variant | MODIFIER |
| 8 | 23557080 | AX-170030353 | 1 | C | A | 0.152 | -2.87E-02 | 1.11E-01 | 2.61E+00 | 2.71E+00 | 8.00E-01 | no | | intergenic_variant | MODIFIER |
| 10 | 127901298 | AX-170004133 | 1 | G | A | 0.33 | -1.84E-02 | 7.18E-02 | 2.61E+00 | 2.72E+00 | 8.00E-01 | no | Sdr9c7 | intron_variant | MODIFIER |
| 8 | 68215549 | AX-170141704 | 0 | T | C | 0.149 | -3.46E-02 | 1.33E-01 | 2.61E+00 | 2.70E+00 | 8.01E-01 | no | Gm26659 | downstream_gene_variant | MODIFIER |
| 2 | 139103835 | AX-170485109 | 2 | A | C | 0.127 | 4.99E-02 | 2.00E-01 | 2.65E+00 | 2.74E+00 | 8.01E-01 | no | | intergenic_variant | MODIFIER |
| 3 | 148682706 | AX-170102699 | 0 | C | T | 0.325 | 1.92E-02 | 7.52E-02 | 2.63E+00 | 2.71E+00 | 8.01E-01 | no | | regulatory_region_variant | MODIFIER |
| 1 | 184175662 | AX-97845546 | 0 | A | C | 0.228 | 2.88E-02 | 1.00E-01 | 2.53E+00 | 2.64E+00 | 8.01E-01 | no | | intergenic_variant | MODIFIER |
| 8 | 102538001 | AX-170348703 | 1 | T | C | 0.223 | 2.80E-02 | 1.12E-01 | 2.67E+00 | 2.74E+00 | 8.01E-01 | no | Gm45422 | downstream_gene_variant | MODIFIER |
| 5 | 126392135 | AX-97837080 | 0 | C | T | 0.465 | 2.33E-02 | 9.32E-02 | 2.65E+00 | 2.73E+00 | 8.01E-01 | no | Gm23151 | upstream_gene_variant | MODIFIER |
| 5 | 126392928 | AX-170360655 | 0 | C | T | 0.465 | 2.33E-02 | 9.32E-02 | 2.65E+00 | 2.73E+00 | 8.01E-01 | no | Gm23151 | upstream_gene_variant | MODIFIER |
| 5 | 126411859 | AX-170345775 | 0 | T | A | 0.465 | 2.33E-02 | 9.32E-02 | 2.65E+00 | 2.73E+00 | 8.01E-01 | no | | intergenic_variant | MODIFIER |
| 5 | 126421571 | AX-170073421 | 0 | G | A | 0.465 | 2.33E-02 | 9.32E-02 | 2.65E+00 | 2.73E+00 | 8.01E-01 | no | | intergenic_variant | MODIFIER |
| 5 | 126426714 | AX-97801058 | 0 | A | G | 0.465 | 2.33E-02 | 9.32E-02 | 2.65E+00 | 2.73E+00 | 8.01E-01 | no | | intergenic_variant | MODIFIER |
| 5 | 126449391 | AX-170495141 | 0 | G | A | 0.465 | 2.33E-02 | 9.32E-02 | 2.65E+00 | 2.73E+00 | 8.01E-01 | no | Gm37700 | non_coding_transcript_exon_variant | MODIFIER |
| 5 | 126454156 | AX-97830810 | 0 | G | A | 0.465 | 2.33E-02 | 9.32E-02 | 2.65E+00 | 2.73E+00 | 8.01E-01 | no | Gm37700 | downstream_gene_variant | MODIFIER |
| 5 | 126463584 | AX-170202784 | 0 | C | T | 0.465 | 2.33E-02 | 9.32E-02 | 2.65E+00 | 2.73E+00 | 8.01E-01 | no | | intergenic_variant | MODIFIER |
| 5 | 126494892 | AX-97829783 | 0 | G | A | 0.465 | 2.33E-02 | 9.32E-02 | 2.65E+00 | 2.73E+00 | 8.01E-01 | no | | intergenic_variant | MODIFIER |
| 5 | 126522255 | AX-170251638 | 0 | A | G | 0.465 | 2.33E-02 | 9.32E-02 | 2.65E+00 | 2.73E+00 | 8.01E-01 | no | | intergenic_variant | MODIFIER |
| 5 | 126523723 | AX-97853948 | 0 | A | G | 0.465 | 2.33E-02 | 9.32E-02 | 2.65E+00 | 2.73E+00 | 8.01E-01 | no | | intergenic_variant | MODIFIER |
| 5 | 126641044 | AX-170446668 | 0 | A | G | 0.465 | 2.33E-02 | 9.32E-02 | 2.65E+00 | 2.73E+00 | 8.01E-01 | no | | intergenic_variant | MODIFIER |
| 5 | 126641074 | AX-170454913 | 0 | T | C | 0.465 | 2.33E-02 | 9.32E-02 | 2.65E+00 | 2.73E+00 | 8.01E-01 | no | | intergenic_variant | MODIFIER |
| 5 | 126641420 | AX-170116948 | 0 | T | C | 0.465 | 2.33E-02 | 9.32E-02 | 2.65E+00 | 2.73E+00 | 8.01E-01 | no | | intergenic_variant | MODIFIER |
| 5 | 126652477 | AX-97833361 | 0 | A | G | 0.465 | 2.33E-02 | 9.32E-02 | 2.65E+00 | 2.73E+00 | 8.01E-01 | no | | intergenic_variant | MODIFIER |
| 5 | 126663897 | AX-170319485 | 0 | C | T | 0.465 | 2.33E-02 | 9.32E-02 | 2.65E+00 | 2.73E+00 | 8.01E-01 | no | | intergenic_variant | MODIFIER |
| 5 | 126663916 | AX-170217232 | 0 | A | C | 0.465 | 2.33E-02 | 9.32E-02 | 2.65E+00 | 2.73E+00 | 8.01E-01 | no | | intergenic_variant | MODIFIER |
| 5 | 126673628 | AX-170518347 | 0 | G | C | 0.465 | 2.33E-02 | 9.32E-02 | 2.65E+00 | 2.73E+00 | 8.01E-01 | no | | intergenic_variant | MODIFIER |
| 5 | 126691600 | AX-170173522 | 0 | A | G | 0.465 | 2.33E-02 | 9.32E-02 | 2.65E+00 | 2.73E+00 | 8.01E-01 | no | | intergenic_variant | MODIFIER |
| 5 | 126697217 | AX-97823978 | 0 | T | C | 0.465 | 2.33E-02 | 9.32E-02 | 2.65E+00 | 2.73E+00 | 8.01E-01 | no | | intergenic_variant | MODIFIER |
| 5 | 126769053 | AX-170089606 | 0 | G | C | 0.465 | 2.33E-02 | 9.32E-02 | 2.65E+00 | 2.73E+00 | 8.01E-01 | no | | regulatory_region_variant | MODIFIER |
| 5 | 126769520 | AX-170169774 | 0 | C | T | 0.465 | 2.33E-02 | 9.32E-02 | 2.65E+00 | 2.73E+00 | 8.01E-01 | no | | regulatory_region_variant | MODIFIER |
| 5 | 126790082 | AX-170287466 | 0 | G | A | 0.465 | 2.33E-02 | 9.32E-02 | 2.65E+00 | 2.73E+00 | 8.01E-01 | no | | intergenic_variant | MODIFIER |
| 5 | 126791812 | AX-170110304 | 0 | C | T | 0.465 | 2.33E-02 | 9.32E-02 | 2.65E+00 | 2.73E+00 | 8.01E-01 | no | | intergenic_variant | MODIFIER |
| 5 | 126792283 | AX-170464410 | 0 | C | T | 0.465 | 2.33E-02 | 9.32E-02 | 2.65E+00 | 2.73E+00 | 8.01E-01 | no | Gm42953 | downstream_gene_variant | MODIFIER |
| 5 | 126792536 | AX-170392839 | 0 | T | C | 0.465 | 2.33E-02 | 9.32E-02 | 2.65E+00 | 2.73E+00 | 8.01E-01 | no | Gm42953 | downstream_gene_variant | MODIFIER |
| 5 | 126793348 | AX-170320558 | 0 | G | A | 0.465 | 2.33E-02 | 9.32E-02 | 2.65E+00 | 2.73E+00 | 8.01E-01 | no | Gm42953 | downstream_gene_variant | MODIFIER |
| 5 | 126793922 | AX-170445227 | 0 | T | C | 0.465 | 2.33E-02 | 9.32E-02 | 2.65E+00 | 2.73E+00 | 8.01E-01 | no | Gm42953 | downstream_gene_variant | MODIFIER |
| 5 | 126809025 | AX-170285816 | 0 | C | T | 0.465 | 2.33E-02 | 9.32E-02 | 2.65E+00 | 2.73E+00 | 8.01E-01 | no | Gm42953 | intron_variant,non_coding_transcript_variant | MODIFIER |
| 2 | 96722076 | AX-170228541 | 1 | A | G | 0.054 | -7.43E-02 | 2.98E-01 | 2.72E+00 | 2.75E+00 | 8.01E-01 | no | Lrrc4c | intron_variant | MODIFIER |
| 1 | 107049093 | AX-170447613 | 1 | G | T | 0.161 | 5.33E-02 | 2.11E-01 | 2.64E+00 | 2.72E+00 | 8.01E-01 | no | Serpinb3a | intron_variant | MODIFIER |

| | | | | | | | | | | | | | | | |
|---|---|---|---|---|---|---|---|---|---|---|---|---|---|---|---|
| 6 | 102302678 | AX-170500297 | 0 | C | A | 0.079 | -4.43E-02 | 1.76E-01 | 2.61E+00 | 2.73E+00 | 8.01E-01 | no | Cntn3 | intron_variant | MODIFIER |
| 5 | 38820590 | AX-170024734 | 2 | T | C | 0.236 | 2.60E-02 | 1.08E-01 | 2.69E+00 | 2.78E+00 | 8.01E-01 | no | Clnk | intron_variant | MODIFIER |
| 5 | 85226606 | AX-170227018 | 0 | C | A | 0.421 | 2.64E-02 | 1.07E-01 | 2.69E+00 | 2.75E+00 | 8.01E-01 | no | | intergenic_variant | MODIFIER |
| 9 | 78869786 | AX-170032774 | 0 | A | T | 0.211 | 3.14E-02 | 1.28E-01 | 2.73E+00 | 2.78E+00 | 8.01E-01 | no | | intergenic_variant | MODIFIER |
| 8 | 42680109 | AX-170079896 | 1 | T | C | 0.143 | 3.57E-02 | 1.45E-01 | 2.72E+00 | 2.77E+00 | 8.01E-01 | no | | intergenic_variant | MODIFIER |
| 15 | 79708079 | AX-170387759 | 1 | A | G | 0.554 | 2.16E-02 | 8.06E-02 | 2.59E+00 | 2.67E+00 | 8.01E-01 | no | Gtpbp1 | intron_variant | MODIFIER |
| 15 | 80251476 | AX-170119550 | 1 | A | T | 0.554 | 2.16E-02 | 8.06E-02 | 2.59E+00 | 2.67E+00 | 8.01E-01 | no | Mief1 | 3_prime_UTR_variant | MODIFIER |
| 4 | 136924714 | AX-170334642 | 0 | C | T | 0.167 | 3.03E-02 | 1.12E-01 | 2.58E+00 | 2.66E+00 | 8.01E-01 | no | Epha8 | downstream_gene_variant | MODIFIER |
| 1 | 25439157 | AX-170443981 | 0 | C | T | 0.395 | -1.93E-02 | 7.95E-02 | 2.71E+00 | 2.79E+00 | 8.01E-01 | no | Adgrb3 | intron_variant | MODIFIER |
| 2 | 150132367 | AX-97840586 | 1 | C | A | 0.134 | -4.38E-02 | 1.75E-01 | 2.65E+00 | 2.73E+00 | 8.02E-01 | no | Zfp120 | intron_variant | MODIFIER |
| 4 | 70913565 | AX-170159899 | 0 | C | T | 0.228 | 2.81E-02 | 1.05E-01 | 2.59E+00 | 2.67E+00 | 8.02E-01 | no | | intergenic_variant | MODIFIER |
| 4 | 71054627 | AX-97855528 | 0 | C | T | 0.228 | 2.81E-02 | 1.05E-01 | 2.59E+00 | 2.67E+00 | 8.02E-01 | no | | intergenic_variant | MODIFIER |
| 4 | 71055170 | AX-170455193 | 0 | C | T | 0.228 | 2.81E-02 | 1.05E-01 | 2.59E+00 | 2.67E+00 | 8.02E-01 | no | | intergenic_variant | MODIFIER |
| 4 | 71105271 | AX-97818604 | 0 | G | A | 0.228 | 2.81E-02 | 1.05E-01 | 2.59E+00 | 2.67E+00 | 8.02E-01 | no | Gm11228 | intron_variant,non_coding_transcript_variant | MODIFIER |
| 4 | 71128693 | AX-170204429 | 0 | C | T | 0.228 | 2.81E-02 | 1.05E-01 | 2.59E+00 | 2.67E+00 | 8.02E-01 | no | Gm11228 | intron_variant,non_coding_transcript_variant | MODIFIER |
| 4 | 71197717 | AX-170390135 | 0 | C | G | 0.228 | 2.81E-02 | 1.05E-01 | 2.59E+00 | 2.67E+00 | 8.02E-01 | no | | intergenic_variant | MODIFIER |
| 4 | 71225659 | AX-170315770 | 0 | T | G | 0.228 | 2.81E-02 | 1.05E-01 | 2.59E+00 | 2.67E+00 | 8.02E-01 | no | | intergenic_variant | MODIFIER |
| 4 | 71225781 | AX-170089273 | 0 | T | C | 0.228 | 2.81E-02 | 1.05E-01 | 2.59E+00 | 2.67E+00 | 8.02E-01 | no | | intergenic_variant | MODIFIER |
| 4 | 71229158 | AX-170069642 | 0 | C | T | 0.228 | 2.81E-02 | 1.05E-01 | 2.59E+00 | 2.67E+00 | 8.02E-01 | no | | intergenic_variant | MODIFIER |
| 4 | 71229849 | AX-170069639 | 0 | A | G | 0.228 | 2.81E-02 | 1.05E-01 | 2.59E+00 | 2.67E+00 | 8.02E-01 | no | | regulatory_region_variant | MODIFIER |
| 4 | 71267393 | AX-97806228 | 0 | T | C | 0.228 | 2.81E-02 | 1.05E-01 | 2.59E+00 | 2.67E+00 | 8.02E-01 | no | | intergenic_variant | MODIFIER |
| 4 | 71273774 | AX-170023077 | 0 | A | G | 0.228 | 2.81E-02 | 1.05E-01 | 2.59E+00 | 2.67E+00 | 8.02E-01 | no | | intergenic_variant | MODIFIER |
| 4 | 71287465 | AX-170023075 | 0 | G | C | 0.228 | 2.81E-02 | 1.05E-01 | 2.59E+00 | 2.67E+00 | 8.02E-01 | no | Gm11229 | downstream_gene_variant | MODIFIER |
| 4 | 71308380 | AX-97840284 | 0 | T | C | 0.228 | 2.81E-02 | 1.05E-01 | 2.59E+00 | 2.67E+00 | 8.02E-01 | no | | intergenic_variant | MODIFIER |
| 4 | 71325740 | AX-170023078 | 0 | C | T | 0.228 | 2.81E-02 | 1.05E-01 | 2.59E+00 | 2.67E+00 | 8.02E-01 | no | | intergenic_variant | MODIFIER |
| 4 | 71346326 | AX-97825204 | 0 | C | A | 0.228 | 2.81E-02 | 1.05E-01 | 2.59E+00 | 2.67E+00 | 8.02E-01 | no | | intergenic_variant | MODIFIER |
| 4 | 71400909 | AX-97814811 | 0 | T | C | 0.228 | 2.81E-02 | 1.05E-01 | 2.59E+00 | 2.67E+00 | 8.02E-01 | no | | intergenic_variant | MODIFIER |
| 4 | 71542987 | AX-168396107 | 0 | T | C | 0.228 | 2.81E-02 | 1.05E-01 | 2.59E+00 | 2.67E+00 | 8.02E-01 | no | | intergenic_variant | MODIFIER |
| 4 | 71583329 | AX-170217605 | 0 | T | C | 0.228 | 2.81E-02 | 1.05E-01 | 2.59E+00 | 2.67E+00 | 8.02E-01 | no | | intergenic_variant | MODIFIER |
| 4 | 71584676 | AX-97782973 | 0 | A | G | 0.228 | 2.81E-02 | 1.05E-01 | 2.59E+00 | 2.67E+00 | 8.02E-01 | no | | intergenic_variant | MODIFIER |
| 4 | 71645734 | AX-170125321 | 0 | T | C | 0.228 | 2.81E-02 | 1.05E-01 | 2.59E+00 | 2.67E+00 | 8.02E-01 | no | | intergenic_variant | MODIFIER |
| 4 | 71687438 | AX-97790281 | 0 | C | A | 0.228 | 2.81E-02 | 1.05E-01 | 2.59E+00 | 2.67E+00 | 8.02E-01 | no | | intergenic_variant | MODIFIER |
| 4 | 71773659 | AX-170023086 | 0 | T | C | 0.228 | 2.81E-02 | 1.05E-01 | 2.59E+00 | 2.67E+00 | 8.02E-01 | no | | intergenic_variant | MODIFIER |
| 4 | 71970070 | AX-170277401 | 0 | A | G | 0.228 | 2.81E-02 | 1.05E-01 | 2.59E+00 | 2.67E+00 | 8.02E-01 | no | | intergenic_variant | MODIFIER |
| 4 | 71989174 | AX-97789723 | 0 | G | T | 0.228 | 2.81E-02 | 1.05E-01 | 2.59E+00 | 2.67E+00 | 8.02E-01 | no | Gm11234 | intron_variant,non_coding_transcript_variant | MODIFIER |
| 4 | 71989472 | AX-170465583 | 0 | T | C | 0.228 | 2.81E-02 | 1.05E-01 | 2.59E+00 | 2.67E+00 | 8.02E-01 | no | Gm11234 | intron_variant,non_coding_transcript_variant | MODIFIER |
| 4 | 72013858 | AX-170328559 | 0 | G | A | 0.228 | 2.81E-02 | 1.05E-01 | 2.59E+00 | 2.67E+00 | 8.02E-01 | no | | regulatory_region_variant | MODIFIER |
| 4 | 72016291 | AX-170452455 | 0 | T | C | 0.228 | 2.81E-02 | 1.05E-01 | 2.59E+00 | 2.67E+00 | 8.02E-01 | no | | intergenic_variant | MODIFIER |
| 4 | 72038353 | AX-170035688 | 0 | T | G | 0.228 | 2.81E-02 | 1.05E-01 | 2.59E+00 | 2.67E+00 | 8.02E-01 | no | | intergenic_variant | MODIFIER |
| 4 | 72047112 | AX-170089274 | 0 | T | C | 0.228 | 2.81E-02 | 1.05E-01 | 2.59E+00 | 2.67E+00 | 8.02E-01 | no | | intergenic_variant | MODIFIER |
| 4 | 72138328 | AX-170409708 | 0 | A | G | 0.228 | 2.81E-02 | 1.05E-01 | 2.59E+00 | 2.67E+00 | 8.02E-01 | no | Tle1 | intron_variant | MODIFIER |
| 4 | 72139536 | AX-97835936 | 0 | G | A | 0.228 | 2.81E-02 | 1.05E-01 | 2.59E+00 | 2.67E+00 | 8.02E-01 | no | Tle1 | intron_variant | MODIFIER |
| 4 | 72141010 | AX-170235581 | 0 | A | G | 0.228 | 2.81E-02 | 1.05E-01 | 2.59E+00 | 2.67E+00 | 8.02E-01 | no | Tle1 | intron_variant | MODIFIER |
| 4 | 72382641 | AX-97813920 | 0 | T | C | 0.228 | 2.81E-02 | 1.05E-01 | 2.59E+00 | 2.67E+00 | 8.02E-01 | no | | regulatory_region_variant | MODIFIER |
| 4 | 72578771 | AX-170103293 | 0 | A | G | 0.228 | 2.81E-02 | 1.05E-01 | 2.59E+00 | 2.67E+00 | 8.02E-01 | no | | intergenic_variant | MODIFIER |
| 4 | 72651258 | AX-170360113 | 0 | C | T | 0.228 | 2.81E-02 | 1.05E-01 | 2.59E+00 | 2.67E+00 | 8.02E-01 | no | | intergenic_variant | MODIFIER |
| 4 | 72652002 | AX-170430495 | 0 | T | C | 0.228 | 2.81E-02 | 1.05E-01 | 2.59E+00 | 2.67E+00 | 8.02E-01 | no | | intergenic_variant | MODIFIER |
| 4 | 72652822 | AX-97830373 | 0 | C | T | 0.228 | 2.81E-02 | 1.05E-01 | 2.59E+00 | 2.67E+00 | 8.02E-01 | no | | intergenic_variant | MODIFIER |
| 4 | 72716348 | AX-97832010 | 0 | C | G | 0.228 | 2.81E-02 | 1.05E-01 | 2.59E+00 | 2.67E+00 | 8.02E-01 | no | | intergenic_variant | MODIFIER |
| 4 | 72851242 | AX-170023100 | 0 | G | C | 0.228 | 2.81E-02 | 1.05E-01 | 2.59E+00 | 2.67E+00 | 8.02E-01 | no | Aldoart1 | 3_prime_UTR_variant | MODIFIER |
| 4 | 73355551 | AX-97837752 | 0 | A | G | 0.228 | 2.81E-02 | 1.05E-01 | 2.59E+00 | 2.67E+00 | 8.02E-01 | no | | intergenic_variant | MODIFIER |
| 4 | 73541112 | AX-97807303 | 0 | A | C | 0.228 | 2.81E-02 | 1.05E-01 | 2.59E+00 | 2.67E+00 | 8.02E-01 | no | | intergenic_variant | MODIFIER |
| 4 | 73954832 | AX-170215778 | 0 | T | C | 0.228 | 2.81E-02 | 1.05E-01 | 2.59E+00 | 2.67E+00 | 8.02E-01 | no | 2310002L09Rik | upstream_gene_variant | MODIFIER |
| 4 | 73956132 | AX-170298854 | 0 | T | G | 0.228 | 2.81E-02 | 1.05E-01 | 2.59E+00 | 2.67E+00 | 8.02E-01 | no | | intergenic_variant | MODIFIER |

| | | | | | | | | | | | | | | | |
|---|---|---|---|---|---|---|---|---|---|---|---|---|---|---|---|
| 4 | 73956185 | AX-170178197 | 0 | G | C | 0.228 | 2.81E-02 | 1.05E-01 | 2.59E+00 | 2.67E+00 | 8.02E-01 | no | | intergenic_variant | MODIFIER |
| 4 | 73962288 | AX-97843549 | 0 | T | C | 0.228 | 2.81E-02 | 1.05E-01 | 2.59E+00 | 2.67E+00 | 8.02E-01 | no | | intergenic_variant | MODIFIER |
| 4 | 74007260 | AX-170455456 | 0 | T | A | 0.228 | 2.81E-02 | 1.05E-01 | 2.59E+00 | 2.67E+00 | 8.02E-01 | no | | regulatory_region_variant | MODIFIER |
| 4 | 74020661 | AX-170023118 | 0 | C | A | 0.228 | 2.81E-02 | 1.05E-01 | 2.59E+00 | 2.67E+00 | 8.02E-01 | no | Frmd3 | intron_variant | MODIFIER |
| 4 | 74037543 | AX-170023120 | 0 | A | G | 0.228 | 2.81E-02 | 1.05E-01 | 2.59E+00 | 2.67E+00 | 8.02E-01 | no | Frmd3 | intron_variant | MODIFIER |
| 4 | 74050916 | AX-170089277 | 0 | T | C | 0.228 | 2.81E-02 | 1.05E-01 | 2.59E+00 | 2.67E+00 | 8.02E-01 | no | Frmd3 | intron_variant | MODIFIER |
| 4 | 74106443 | AX-170069686 | 0 | A | T | 0.228 | 2.81E-02 | 1.05E-01 | 2.59E+00 | 2.67E+00 | 8.02E-01 | no | Frmd3 | intron_variant | MODIFIER |
| 4 | 74119477 | AX-170023123 | 0 | A | T | 0.228 | 2.81E-02 | 1.05E-01 | 2.59E+00 | 2.67E+00 | 8.02E-01 | no | Frmd3 | intron_variant | MODIFIER |
| 4 | 74309742 | AX-170318742 | 0 | A | G | 0.228 | 2.81E-02 | 1.05E-01 | 2.59E+00 | 2.67E+00 | 8.02E-01 | no | Kdm4c | intron_variant | MODIFIER |
| 4 | 74727754 | AX-170023136 | 0 | A | G | 0.228 | 2.81E-02 | 1.05E-01 | 2.59E+00 | 2.67E+00 | 8.02E-01 | no | | intergenic_variant | MODIFIER |
| 4 | 74730418 | AX-97810368 | 0 | T | C | 0.228 | 2.81E-02 | 1.05E-01 | 2.59E+00 | 2.67E+00 | 8.02E-01 | no | | intergenic_variant | MODIFIER |
| 4 | 74764472 | AX-97816305 | 0 | A | G | 0.228 | 2.81E-02 | 1.05E-01 | 2.59E+00 | 2.67E+00 | 8.02E-01 | no | | intergenic_variant | MODIFIER |
| 4 | 74788169 | AX-170069695 | 0 | A | G | 0.228 | 2.81E-02 | 1.05E-01 | 2.59E+00 | 2.67E+00 | 8.02E-01 | no | | intergenic_variant | MODIFIER |
| 4 | 74788466 | AX-97795820 | 0 | T | C | 0.228 | 2.81E-02 | 1.05E-01 | 2.59E+00 | 2.67E+00 | 8.02E-01 | no | | intergenic_variant | MODIFIER |
| 4 | 74795440 | AX-170434593 | 0 | C | T | 0.228 | 2.81E-02 | 1.05E-01 | 2.59E+00 | 2.67E+00 | 8.02E-01 | no | | intergenic_variant | MODIFIER |
| 4 | 74811205 | AX-97854607 | 0 | C | T | 0.228 | 2.81E-02 | 1.05E-01 | 2.59E+00 | 2.67E+00 | 8.02E-01 | no | | intergenic_variant | MODIFIER |
| 4 | 74839198 | AX-97784578 | 0 | A | G | 0.228 | 2.81E-02 | 1.05E-01 | 2.59E+00 | 2.67E+00 | 8.02E-01 | no | | intergenic_variant | MODIFIER |
| 4 | 74849031 | AX-170023137 | 0 | T | C | 0.228 | 2.81E-02 | 1.05E-01 | 2.59E+00 | 2.67E+00 | 8.02E-01 | no | | intergenic_variant | MODIFIER |
| 4 | 74907930 | AX-170023134 | 0 | C | G | 0.228 | 2.81E-02 | 1.05E-01 | 2.59E+00 | 2.67E+00 | 8.02E-01 | no | | intergenic_variant | MODIFIER |
| 4 | 74943359 | AX-170429868 | 0 | A | C | 0.228 | 2.81E-02 | 1.05E-01 | 2.59E+00 | 2.67E+00 | 8.02E-01 | no | | intergenic_variant | MODIFIER |
| 4 | 74978517 | AX-170424751 | 0 | A | G | 0.228 | 2.81E-02 | 1.05E-01 | 2.59E+00 | 2.67E+00 | 8.02E-01 | no | | intergenic_variant | MODIFIER |
| 4 | 74982212 | AX-97799581 | 0 | T | C | 0.228 | 2.81E-02 | 1.05E-01 | 2.59E+00 | 2.67E+00 | 8.02E-01 | no | | intergenic_variant | MODIFIER |
| 4 | 74982289 | AX-170366045 | 0 | G | C | 0.228 | 2.81E-02 | 1.05E-01 | 2.59E+00 | 2.67E+00 | 8.02E-01 | no | | intergenic_variant | MODIFIER |
| 4 | 75018113 | AX-97852369 | 0 | T | C | 0.228 | 2.81E-02 | 1.05E-01 | 2.59E+00 | 2.67E+00 | 8.02E-01 | no | | intergenic_variant | MODIFIER |
| 4 | 75063294 | AX-170023144 | 0 | G | A | 0.228 | 2.81E-02 | 1.05E-01 | 2.59E+00 | 2.67E+00 | 8.02E-01 | no | | regulatory_region_variant | MODIFIER |
| 4 | 75086420 | AX-170023148 | 0 | C | T | 0.228 | 2.81E-02 | 1.05E-01 | 2.59E+00 | 2.67E+00 | 8.02E-01 | no | Gm11253 | upstream_gene_variant | MODIFIER |
| 4 | 75108417 | AX-170023142 | 0 | G | A | 0.228 | 2.81E-02 | 1.05E-01 | 2.59E+00 | 2.67E+00 | 8.02E-01 | no | | intergenic_variant | MODIFIER |
| 4 | 80757905 | AX-170484943 | 0 | T | C | 0.228 | 2.81E-02 | 1.05E-01 | 2.59E+00 | 2.67E+00 | 8.02E-01 | no | | intergenic_variant | MODIFIER |
| 4 | 80899803 | AX-170023225 | 0 | T | A | 0.228 | 2.81E-02 | 1.05E-01 | 2.59E+00 | 2.67E+00 | 8.02E-01 | no | | intergenic_variant | MODIFIER |
| 4 | 80899834 | AX-170023226 | 0 | A | G | 0.228 | 2.81E-02 | 1.05E-01 | 2.59E+00 | 2.67E+00 | 8.02E-01 | no | | intergenic_variant | MODIFIER |
| 4 | 80900444 | AX-170023227 | 0 | A | G | 0.228 | 2.81E-02 | 1.05E-01 | 2.59E+00 | 2.67E+00 | 8.02E-01 | no | | intergenic_variant | MODIFIER |
| 4 | 80923418 | AX-97787761 | 0 | C | T | 0.228 | 2.81E-02 | 1.05E-01 | 2.59E+00 | 2.67E+00 | 8.02E-01 | no | Lurap1l | intron_variant | MODIFIER |
| 4 | 80933812 | AX-170023228 | 0 | G | C | 0.228 | 2.81E-02 | 1.05E-01 | 2.59E+00 | 2.67E+00 | 8.02E-01 | no | Lurap1l | intron_variant | MODIFIER |
| 13 | 56794023 | AX-170518652 | 0 | T | C | 0.509 | 1.81E-02 | 6.62E-02 | 2.55E+00 | 2.67E+00 | 8.02E-01 | no | Trpc7 | intron_variant | MODIFIER |
| 12 | 103769890 | AX-170007436 | 1 | T | G | 0.08 | -4.04E-02 | 1.65E-01 | 2.61E+00 | 2.75E+00 | 8.02E-01 | no | Serpina1d | intron_variant | MODIFIER |
| 3 | 143913843 | AX-170151753 | 0 | C | T | 0.377 | 1.73E-02 | 6.87E-02 | 2.63E+00 | 2.72E+00 | 8.02E-01 | no | Gm42707 | non_coding_transcript_exon_variant | MODIFIER |
| 3 | 143914918 | AX-170418346 | 0 | G | T | 0.377 | 1.73E-02 | 6.87E-02 | 2.63E+00 | 2.72E+00 | 8.02E-01 | no | Gm42707 | intron_variant,non_coding_transcript_variant | MODIFIER |
| 3 | 143917146 | AX-170331305 | 0 | C | A | 0.377 | 1.73E-02 | 6.87E-02 | 2.63E+00 | 2.72E+00 | 8.02E-01 | no | Gm42707 | intron_variant,non_coding_transcript_variant | MODIFIER |
| 3 | 143927153 | AX-170473819 | 0 | G | A | 0.377 | 1.73E-02 | 6.87E-02 | 2.63E+00 | 2.72E+00 | 8.02E-01 | no | Gm42707 | intron_variant,non_coding_transcript_variant | MODIFIER |
| 3 | 143927804 | AX-170531819 | 0 | A | G | 0.377 | 1.73E-02 | 6.87E-02 | 2.63E+00 | 2.72E+00 | 8.02E-01 | no | Gm42707 | intron_variant,non_coding_transcript_variant | MODIFIER |
| 3 | 143936452 | AX-170371283 | 0 | A | G | 0.377 | 1.73E-02 | 6.87E-02 | 2.63E+00 | 2.72E+00 | 8.02E-01 | no | Gm42707 | intron_variant,non_coding_transcript_variant | MODIFIER |
| 3 | 143960475 | AX-97787007 | 0 | G | A | 0.377 | 1.73E-02 | 6.87E-02 | 2.63E+00 | 2.72E+00 | 8.02E-01 | no | | intergenic_variant | MODIFIER |
| 3 | 143964280 | AX-170509905 | 0 | G | A | 0.377 | 1.73E-02 | 6.87E-02 | 2.63E+00 | 2.72E+00 | 8.02E-01 | no | | intergenic_variant | MODIFIER |
| 3 | 143984889 | AX-170301521 | 0 | G | A | 0.377 | 1.73E-02 | 6.87E-02 | 2.63E+00 | 2.72E+00 | 8.02E-01 | no | | intergenic_variant | MODIFIER |
| 3 | 143985369 | AX-170535381 | 0 | T | C | 0.377 | 1.73E-02 | 6.87E-02 | 2.63E+00 | 2.72E+00 | 8.02E-01 | no | | intergenic_variant | MODIFIER |
| 3 | 143991358 | AX-170436225 | 0 | G | A | 0.377 | 1.73E-02 | 6.87E-02 | 2.63E+00 | 2.72E+00 | 8.02E-01 | no | | intergenic_variant | MODIFIER |
| 3 | 144005590 | AX-170329240 | 0 | C | T | 0.377 | 1.73E-02 | 6.87E-02 | 2.63E+00 | 2.72E+00 | 8.02E-01 | no | | intergenic_variant | MODIFIER |
| 3 | 144005787 | AX-170189448 | 0 | G | A | 0.377 | 1.73E-02 | 6.87E-02 | 2.63E+00 | 2.72E+00 | 8.02E-01 | no | | intergenic_variant | MODIFIER |
| 3 | 144008931 | AX-170257214 | 0 | A | C | 0.377 | 1.73E-02 | 6.87E-02 | 2.63E+00 | 2.72E+00 | 8.02E-01 | no | | intergenic_variant | MODIFIER |
| 3 | 144045298 | AX-170381580 | 0 | T | C | 0.377 | 1.73E-02 | 6.87E-02 | 2.63E+00 | 2.72E+00 | 8.02E-01 | no | | intergenic_variant | MODIFIER |
| 3 | 144050854 | AX-170331145 | 0 | G | T | 0.377 | 1.73E-02 | 6.87E-02 | 2.63E+00 | 2.72E+00 | 8.02E-01 | no | | intergenic_variant | MODIFIER |
| 3 | 144065512 | AX-170225246 | 0 | G | A | 0.377 | 1.73E-02 | 6.87E-02 | 2.63E+00 | 2.72E+00 | 8.02E-01 | no | | intergenic_variant | MODIFIER |
| 3 | 144066042 | AX-170348807 | 0 | G | A | 0.377 | 1.73E-02 | 6.87E-02 | 2.63E+00 | 2.72E+00 | 8.02E-01 | no | | intergenic_variant | MODIFIER |
| 3 | 144066364 | AX-170451506 | 0 | A | G | 0.377 | 1.73E-02 | 6.87E-02 | 2.63E+00 | 2.72E+00 | 8.02E-01 | no | | intergenic_variant | MODIFIER |

| | | | | | | | | | | | | | | | |
|---|---|---|---|---|---|---|---|---|---|---|---|---|---|---|---|
| 3 | 144067679 | AX-170068320 | 0 | T | C | 0.377 | 1.73E-02 | 6.87E-02 | 2.63E+00 | 2.72E+00 | 8.02E-01 | no | | intergenic_variant | MODIFIER |
| 3 | 144070873 | AX-170068321 | 0 | G | C | 0.377 | 1.73E-02 | 6.87E-02 | 2.63E+00 | 2.72E+00 | 8.02E-01 | no | | intergenic_variant | MODIFIER |
| 3 | 144078534 | AX-170211653 | 0 | C | T | 0.377 | 1.73E-02 | 6.87E-02 | 2.63E+00 | 2.72E+00 | 8.02E-01 | no | | regulatory_region_variant | MODIFIER |
| 3 | 144090151 | AX-170445873 | 0 | G | A | 0.377 | 1.73E-02 | 6.87E-02 | 2.63E+00 | 2.72E+00 | 8.02E-01 | no | | regulatory_region_variant | MODIFIER |
| 3 | 144103447 | AX-170325798 | 0 | T | G | 0.377 | 1.73E-02 | 6.87E-02 | 2.63E+00 | 2.72E+00 | 8.02E-01 | no | Gm34078 | intron_variant,non_coding_transcript_variant | MODIFIER |
| 3 | 144103464 | AX-170267864 | 0 | C | G | 0.377 | 1.73E-02 | 6.87E-02 | 2.63E+00 | 2.72E+00 | 8.02E-01 | no | Gm34078 | intron_variant,non_coding_transcript_variant | MODIFIER |
| 3 | 144103939 | AX-170474045 | 0 | C | T | 0.377 | 1.73E-02 | 6.87E-02 | 2.63E+00 | 2.72E+00 | 8.02E-01 | no | Gm34078 | intron_variant,non_coding_transcript_variant | MODIFIER |
| 3 | 144107041 | AX-97814153 | 0 | T | C | 0.377 | 1.73E-02 | 6.87E-02 | 2.63E+00 | 2.72E+00 | 8.02E-01 | no | Gm34078 | intron_variant,non_coding_transcript_variant | MODIFIER |
| 3 | 144107171 | AX-170384134 | 0 | G | T | 0.377 | 1.73E-02 | 6.87E-02 | 2.63E+00 | 2.72E+00 | 8.02E-01 | no | Gm34078 | intron_variant,non_coding_transcript_variant | MODIFIER |
| 3 | 144123723 | AX-170181169 | 0 | G | A | 0.377 | 1.73E-02 | 6.87E-02 | 2.63E+00 | 2.72E+00 | 8.02E-01 | no | Gm34078 | intron_variant,non_coding_transcript_variant | MODIFIER |
| 3 | 144129034 | AX-170109474 | 0 | G | A | 0.377 | 1.73E-02 | 6.87E-02 | 2.63E+00 | 2.72E+00 | 8.02E-01 | no | Gm34078 | intron_variant,non_coding_transcript_variant | MODIFIER |
| 3 | 144129643 | AX-170195816 | 0 | C | T | 0.377 | 1.73E-02 | 6.87E-02 | 2.63E+00 | 2.72E+00 | 8.02E-01 | no | Gm34078 | intron_variant,non_coding_transcript_variant | MODIFIER |
| 3 | 144141049 | AX-170272239 | 0 | A | G | 0.377 | 1.73E-02 | 6.87E-02 | 2.63E+00 | 2.72E+00 | 8.02E-01 | no | | intergenic_variant | MODIFIER |
| 3 | 144152733 | AX-170445494 | 0 | G | A | 0.377 | 1.73E-02 | 6.87E-02 | 2.63E+00 | 2.72E+00 | 8.02E-01 | no | | intergenic_variant | MODIFIER |
| 3 | 144156204 | AX-170509031 | 0 | A | G | 0.377 | 1.73E-02 | 6.87E-02 | 2.63E+00 | 2.72E+00 | 8.02E-01 | no | | intergenic_variant | MODIFIER |
| 3 | 144164568 | AX-97800806 | 0 | C | T | 0.377 | 1.73E-02 | 6.87E-02 | 2.63E+00 | 2.72E+00 | 8.02E-01 | no | | regulatory_region_variant | MODIFIER |
| 3 | 144174109 | AX-170474605 | 0 | G | A | 0.377 | 1.73E-02 | 6.87E-02 | 2.63E+00 | 2.72E+00 | 8.02E-01 | no | | intergenic_variant | MODIFIER |
| 3 | 144201869 | AX-170021813 | 0 | C | T | 0.377 | 1.73E-02 | 6.87E-02 | 2.63E+00 | 2.72E+00 | 8.02E-01 | no | Lmo4 | missense_variant | MODERATE |
| 8 | 95162625 | AX-170422236 | 1 | A | G | 0.214 | 2.86E-02 | 1.16E-01 | 2.68E+00 | 2.75E+00 | 8.02E-01 | no | Kifc3 | intron_variant | MODIFIER |
| 2 | 127284162 | AX-170090773 | 1 | C | T | 0.152 | 4.50E-02 | 1.77E-01 | 2.62E+00 | 2.71E+00 | 8.02E-01 | no | Stard7 | missense_variant | MODERATE |
| 11 | 47058248 | AX-170287782 | 2 | G | T | 0.082 | -3.93E-02 | 1.54E-01 | 2.59E+00 | 2.72E+00 | 8.02E-01 | no | Sgcd | intron_variant | MODIFIER |
| 5 | 105136138 | AX-170196029 | 1 | G | A | 0.438 | 2.74E-02 | 1.07E-01 | 2.63E+00 | 2.69E+00 | 8.02E-01 | no | Gbp8 | intron_variant | MODIFIER |
| 4 | 89770335 | AX-170488284 | 0 | T | A | 0.325 | 2.65E-02 | 1.06E-01 | 2.65E+00 | 2.72E+00 | 8.02E-01 | no | | intergenic_variant | MODIFIER |
| 1 | 42367595 | AX-170430801 | 0 | C | T | 0.439 | -1.89E-02 | 7.57E-02 | 2.65E+00 | 2.73E+00 | 8.02E-01 | no | | intergenic_variant | MODIFIER |
| 8 | 91125317 | AX-170105281 | 0 | C | T | 0.193 | -2.84E-02 | 1.13E-01 | 2.63E+00 | 2.72E+00 | 8.02E-01 | no | Rbl2 | downstream_gene_variant | MODIFIER |
| 16 | 93057146 | AX-97846947 | 1 | C | G | 0.384 | -2.07E-02 | 7.98E-02 | 2.61E+00 | 2.69E+00 | 8.02E-01 | no | | intergenic_variant | MODIFIER |
| 9 | 56799280 | AX-170433319 | 1 | C | A | 0.214 | 2.78E-02 | 1.12E-01 | 2.66E+00 | 2.74E+00 | 8.02E-01 | no | Lingo1 | upstream_gene_variant | MODIFIER |
| 7 | 78207559 | AX-170273520 | 1 | T | A | 0.348 | -2.27E-02 | 9.27E-02 | 2.65E+00 | 2.75E+00 | 8.02E-01 | no | Ntrk3 | intron_variant | MODIFIER |
| 9 | 104360186 | AX-170083339 | 1 | G | A | 0.125 | 2.87E-02 | 1.19E-01 | 2.75E+00 | 2.83E+00 | 8.02E-01 | no | Gm28548 | upstream_gene_variant | MODIFIER |
| 9 | 104366488 | AX-97840557 | 1 | C | A | 0.125 | 2.87E-02 | 1.19E-01 | 2.75E+00 | 2.83E+00 | 8.02E-01 | no | Gm28548 | intron_variant,non_coding_transcript_variant | MODIFIER |
| 11 | 117246863 | AX-170006009 | 0 | C | T | 0.053 | -4.56E-02 | 1.90E-01 | 2.66E+00 | 2.77E+00 | 8.02E-01 | no | SEP9 | intron_variant | MODIFIER |
| 1 | 25557293 | AX-170283661 | 0 | G | A | 0.412 | 2.09E-02 | 8.33E-02 | 2.65E+00 | 2.72E+00 | 8.02E-01 | no | Adgrb3 | intron_variant | MODIFIER |
| 1 | 25631188 | AX-170474746 | 0 | G | T | 0.412 | 2.09E-02 | 8.33E-02 | 2.65E+00 | 2.72E+00 | 8.02E-01 | no | Adgrb3 | intron_variant | MODIFIER |
| 1 | 25642764 | AX-97784774 | 0 | T | G | 0.412 | 2.09E-02 | 8.33E-02 | 2.65E+00 | 2.72E+00 | 8.02E-01 | no | Adgrb3 | intron_variant | MODIFIER |
| 1 | 25669793 | AX-170170282 | 0 | A | G | 0.412 | 2.09E-02 | 8.33E-02 | 2.65E+00 | 2.72E+00 | 8.02E-01 | no | Adgrb3 | intron_variant | MODIFIER |
| 1 | 25679781 | AX-170331870 | 0 | G | A | 0.412 | 2.09E-02 | 8.33E-02 | 2.65E+00 | 2.72E+00 | 8.02E-01 | no | Adgrb3 | intron_variant | MODIFIER |
| 1 | 25697943 | AX-170000129 | 0 | A | G | 0.412 | 2.09E-02 | 8.33E-02 | 2.65E+00 | 2.72E+00 | 8.02E-01 | no | Adgrb3 | intron_variant | MODIFIER |
| 1 | 25698150 | AX-168384030 | 0 | C | T | 0.412 | 2.09E-02 | 8.33E-02 | 2.65E+00 | 2.72E+00 | 8.02E-01 | no | Adgrb3 | intron_variant | MODIFIER |
| 1 | 25699821 | AX-170256499 | 0 | C | T | 0.412 | 2.09E-02 | 8.33E-02 | 2.65E+00 | 2.72E+00 | 8.02E-01 | no | Adgrb3 | intron_variant | MODIFIER |
| 1 | 25705150 | AX-97851640 | 0 | A | G | 0.412 | 2.09E-02 | 8.33E-02 | 2.65E+00 | 2.72E+00 | 8.02E-01 | no | Adgrb3 | intron_variant | MODIFIER |
| 1 | 25713853 | AX-170314471 | 0 | G | A | 0.412 | 2.09E-02 | 8.33E-02 | 2.65E+00 | 2.72E+00 | 8.02E-01 | no | Adgrb3 | intron_variant | MODIFIER |
| 1 | 25714393 | AX-170379123 | 0 | T | A | 0.412 | 2.09E-02 | 8.33E-02 | 2.65E+00 | 2.72E+00 | 8.02E-01 | no | Adgrb3 | intron_variant | MODIFIER |
| 1 | 25765511 | AX-170000132 | 0 | G | C | 0.412 | 2.09E-02 | 8.33E-02 | 2.65E+00 | 2.72E+00 | 8.02E-01 | no | Adgrb3 | intron_variant | MODIFIER |
| 1 | 25765958 | AX-170169928 | 0 | A | G | 0.412 | 2.09E-02 | 8.33E-02 | 2.65E+00 | 2.72E+00 | 8.02E-01 | no | Adgrb3 | intron_variant | MODIFIER |
| 1 | 25766924 | AX-170497253 | 0 | T | C | 0.412 | 2.09E-02 | 8.33E-02 | 2.65E+00 | 2.72E+00 | 8.02E-01 | no | Adgrb3 | intron_variant | MODIFIER |
| 1 | 25807128 | AX-170272620 | 0 | G | A | 0.412 | 2.09E-02 | 8.33E-02 | 2.65E+00 | 2.72E+00 | 8.02E-01 | no | Adgrb3 | intron_variant | MODIFIER |
| 1 | 25842037 | AX-170380730 | 0 | C | A | 0.412 | 2.09E-02 | 8.33E-02 | 2.65E+00 | 2.72E+00 | 8.02E-01 | no | | intergenic_variant | MODIFIER |
| 1 | 26395208 | AX-170272923 | 0 | C | T | 0.412 | 2.09E-02 | 8.33E-02 | 2.65E+00 | 2.72E+00 | 8.02E-01 | no | | intergenic_variant | MODIFIER |
| 1 | 26641448 | AX-170328004 | 0 | A | G | 0.412 | 2.09E-02 | 8.33E-02 | 2.65E+00 | 2.72E+00 | 8.02E-01 | no | | intergenic_variant | MODIFIER |
| 1 | 26880778 | AX-170000148 | 0 | A | G | 0.412 | 2.09E-02 | 8.33E-02 | 2.65E+00 | 2.72E+00 | 8.02E-01 | no | | intergenic_variant | MODIFIER |
| 6 | 126287992 | AX-170076146 | 0 | A | G | 0.193 | -1.92E-02 | 7.94E-02 | 2.62E+00 | 2.76E+00 | 8.02E-01 | no | Gm10415 | non_coding_transcript_exon_variant | MODIFIER |
| 1 | 55530711 | AX-170500265 | 2 | C | T | 0.091 | 4.88E-02 | 2.00E-01 | 2.66E+00 | 2.75E+00 | 8.03E-01 | no | Plcl1 | intron_variant | MODIFIER |
| 1 | 46036932 | AX-170106248 | 2 | A | T | 0.073 | -4.80E-02 | 1.97E-01 | 2.66E+00 | 2.75E+00 | 8.03E-01 | no | | intergenic_variant | MODIFIER |
| 1 | 3061773 | AX-170040231 | 1 | A | G | 0.429 | 1.76E-02 | 6.52E-02 | 2.57E+00 | 2.66E+00 | 8.03E-01 | no | | intergenic_variant | MODIFIER |

| | | | | | | | | | | | | | | | |
|---|---|---|---|---|---|---|---|---|---|---|---|---|---|---|---|
| 1 | 67747444 | AX-170342486 | 0 | T | C | 0.105 | 4.86E-02 | 1.96E-01 | 2.64E+00 | 2.73E+00 | 8.03E-01 | no | | intergenic_variant | MODIFIER |
| 4 | 88910068 | AX-170069935 | 0 | C | T | 0.325 | 2.59E-02 | 1.06E-01 | 2.68E+00 | 2.76E+00 | 8.03E-01 | no | Mir31 | downstream_gene_variant | MODIFIER |
| 5 | 101684860 | AX-170526839 | 1 | G | A | 0.402 | 2.47E-02 | 9.88E-02 | 2.63E+00 | 2.72E+00 | 8.03E-01 | no | | intergenic_variant | MODIFIER |
| 16 | 76895220 | AX-170272500 | 1 | A | G | 0.446 | 2.01E-02 | 7.65E-02 | 2.60E+00 | 2.68E+00 | 8.03E-01 | no | 1700041M19Rik | intron_variant,non_coding_transcript_variant | MODIFIER |
| 2 | 122607174 | AX-170018895 | 1 | C | T | 0.134 | 4.93E-02 | 1.98E-01 | 2.66E+00 | 2.72E+00 | 8.03E-01 | no | Gatm | intron_variant | MODIFIER |
| 2 | 147647239 | AX-170511873 | 1 | A | T | 0.134 | 4.93E-02 | 1.98E-01 | 2.66E+00 | 2.72E+00 | 8.03E-01 | no | Gm14107 | upstream_gene_variant | MODIFIER |
| 5 | 84126050 | AX-168419969 | 2 | C | A | 0.436 | 2.55E-02 | 1.04E-01 | 2.67E+00 | 2.74E+00 | 8.03E-01 | no | Epha5 | intron_variant | MODIFIER |
| 7 | 112378984 | AX-170285282 | 0 | G | T | 0.465 | -2.26E-02 | 9.28E-02 | 2.69E+00 | 2.76E+00 | 8.03E-01 | no | Mical2 | intron_variant | MODIFIER |
| 15 | 75918193 | AX-170342240 | 0 | T | C | 0.544 | 2.15E-02 | 8.19E-02 | 2.60E+00 | 2.68E+00 | 8.03E-01 | no | Pycrl | intron_variant | MODIFIER |
| 12 | 108390975 | AX-170140085 | 1 | G | A | 0.116 | -4.82E-02 | 1.99E-01 | 2.71E+00 | 2.78E+00 | 8.03E-01 | no | Eml1 | intron_variant,non_coding_transcript_variant | MODIFIER |
| 8 | 36021453 | AX-170035776 | 1 | G | A | 0.152 | -3.63E-02 | 1.43E-01 | 2.64E+00 | 2.69E+00 | 8.03E-01 | no | Rps12-ps24 | upstream_gene_variant | MODIFIER |
| 1 | 28415233 | AX-97820482 | 0 | C | T | 0.404 | 2.04E-02 | 8.28E-02 | 2.66E+00 | 2.74E+00 | 8.03E-01 | no | | regulatory_region_variant | MODIFIER |
| 1 | 28735447 | AX-97797979 | 0 | C | T | 0.404 | 2.04E-02 | 8.28E-02 | 2.66E+00 | 2.74E+00 | 8.03E-01 | no | | intergenic_variant | MODIFIER |
| 1 | 28780469 | AX-170473460 | 0 | A | G | 0.404 | 2.04E-02 | 8.28E-02 | 2.66E+00 | 2.74E+00 | 8.03E-01 | no | Gm597 | upstream_gene_variant | MODIFIER |
| 1 | 28819869 | AX-170303066 | 0 | C | A | 0.404 | 2.04E-02 | 8.28E-02 | 2.66E+00 | 2.74E+00 | 8.03E-01 | no | | intergenic_variant | MODIFIER |
| 1 | 28901359 | AX-170000189 | 0 | A | T | 0.404 | 2.04E-02 | 8.28E-02 | 2.66E+00 | 2.74E+00 | 8.03E-01 | no | | intergenic_variant | MODIFIER |
| 1 | 28904233 | AX-170000178 | 0 | G | A | 0.404 | 2.04E-02 | 8.28E-02 | 2.66E+00 | 2.74E+00 | 8.03E-01 | no | | intergenic_variant | MODIFIER |
| 1 | 29101457 | AX-97800702 | 0 | C | A | 0.404 | 2.04E-02 | 8.28E-02 | 2.66E+00 | 2.74E+00 | 8.03E-01 | no | | intergenic_variant | MODIFIER |
| 1 | 29796251 | AX-170252300 | 0 | A | G | 0.404 | 2.04E-02 | 8.28E-02 | 2.66E+00 | 2.74E+00 | 8.03E-01 | no | | intergenic_variant | MODIFIER |
| 1 | 29823707 | AX-170117075 | 0 | T | A | 0.404 | 2.04E-02 | 8.28E-02 | 2.66E+00 | 2.74E+00 | 8.03E-01 | no | | intergenic_variant | MODIFIER |
| 1 | 29829755 | AX-170190103 | 0 | G | T | 0.404 | 2.04E-02 | 8.28E-02 | 2.66E+00 | 2.74E+00 | 8.03E-01 | no | | intergenic_variant | MODIFIER |
| 1 | 29884159 | AX-170532961 | 0 | T | A | 0.404 | 2.04E-02 | 8.28E-02 | 2.66E+00 | 2.74E+00 | 8.03E-01 | no | | intergenic_variant | MODIFIER |
| 1 | 29885102 | AX-170509779 | 0 | G | T | 0.404 | 2.04E-02 | 8.28E-02 | 2.66E+00 | 2.74E+00 | 8.03E-01 | no | | intergenic_variant | MODIFIER |
| 1 | 29904254 | AX-170401931 | 0 | T | G | 0.404 | 2.04E-02 | 8.28E-02 | 2.66E+00 | 2.74E+00 | 8.03E-01 | no | | intergenic_variant | MODIFIER |
| 1 | 30710506 | AX-170121567 | 0 | A | C | 0.404 | 2.04E-02 | 8.28E-02 | 2.66E+00 | 2.74E+00 | 8.03E-01 | no | | intergenic_variant | MODIFIER |
| 1 | 30715214 | AX-170259301 | 0 | C | T | 0.404 | 2.04E-02 | 8.28E-02 | 2.66E+00 | 2.74E+00 | 8.03E-01 | no | Gm9898 | downstream_gene_variant | MODIFIER |
| 1 | 30718768 | AX-97808258 | 0 | G | T | 0.404 | 2.04E-02 | 8.28E-02 | 2.66E+00 | 2.74E+00 | 8.03E-01 | no | Gm9898 | non_coding_transcript_exon_variant | MODIFIER |
| 1 | 30801803 | AX-170040756 | 0 | A | G | 0.404 | 2.04E-02 | 8.28E-02 | 2.66E+00 | 2.74E+00 | 8.03E-01 | no | Phf3 | downstream_gene_variant | MODIFIER |
| 1 | 30801911 | AX-97798905 | 0 | C | T | 0.404 | 2.04E-02 | 8.28E-02 | 2.66E+00 | 2.74E+00 | 8.03E-01 | no | Phf3 | downstream_gene_variant | MODIFIER |
| 1 | 30821895 | AX-170361252 | 0 | G | C | 0.404 | 2.04E-02 | 8.28E-02 | 2.66E+00 | 2.74E+00 | 8.03E-01 | no | Phf3 | intron_variant | MODIFIER |
| 1 | 30821986 | AX-170242396 | 0 | A | C | 0.404 | 2.04E-02 | 8.28E-02 | 2.66E+00 | 2.74E+00 | 8.03E-01 | no | Phf3 | intron_variant | MODIFIER |
| 1 | 30846574 | AX-170316437 | 0 | T | C | 0.404 | 2.04E-02 | 8.28E-02 | 2.66E+00 | 2.74E+00 | 8.03E-01 | no | Phf3 | intron_variant | MODIFIER |
| 1 | 30866138 | AX-170419872 | 0 | G | C | 0.404 | 2.04E-02 | 8.28E-02 | 2.66E+00 | 2.74E+00 | 8.03E-01 | no | Phf3 | upstream_gene_variant | MODIFIER |
| 1 | 30887178 | AX-170450693 | 0 | G | C | 0.404 | 2.04E-02 | 8.28E-02 | 2.66E+00 | 2.74E+00 | 8.03E-01 | no | | intergenic_variant | MODIFIER |
| 1 | 30905570 | AX-97812718 | 0 | A | G | 0.404 | 2.04E-02 | 8.28E-02 | 2.66E+00 | 2.74E+00 | 8.03E-01 | no | | intergenic_variant | MODIFIER |
| 1 | 30941352 | AX-170000219 | 0 | C | T | 0.404 | 2.04E-02 | 8.28E-02 | 2.66E+00 | 2.74E+00 | 8.03E-01 | no | Ptp4a1 | 3_prime_UTR_variant | MODIFIER |
| 1 | 30944871 | AX-97840794 | 0 | T | C | 0.404 | 2.04E-02 | 8.28E-02 | 2.66E+00 | 2.74E+00 | 8.03E-01 | no | Ptp4a1 | intron_variant | MODIFIER |
| 1 | 31047545 | AX-170000220 | 0 | C | T | 0.404 | 2.04E-02 | 8.28E-02 | 2.66E+00 | 2.74E+00 | 8.03E-01 | no | 4931428L18Rik | intron_variant | MODIFIER |
| 1 | 31048107 | AX-170403820 | 0 | G | C | 0.404 | 2.04E-02 | 8.28E-02 | 2.66E+00 | 2.74E+00 | 8.03E-01 | no | 4931428L18Rik | intron_variant | MODIFIER |
| 1 | 31048703 | AX-170520910 | 0 | T | C | 0.404 | 2.04E-02 | 8.28E-02 | 2.66E+00 | 2.74E+00 | 8.03E-01 | no | 4931428L18Rik | intron_variant | MODIFIER |
| 1 | 31048767 | AX-170406437 | 0 | T | C | 0.404 | 2.04E-02 | 8.28E-02 | 2.66E+00 | 2.74E+00 | 8.03E-01 | no | 4931428L18Rik | intron_variant | MODIFIER |
| 1 | 31050246 | AX-170233882 | 0 | C | T | 0.404 | 2.04E-02 | 8.28E-02 | 2.66E+00 | 2.74E+00 | 8.03E-01 | no | 4931428L18Rik | intron_variant | MODIFIER |
| 1 | 31051420 | AX-170000214 | 0 | G | C | 0.404 | 2.04E-02 | 8.28E-02 | 2.66E+00 | 2.74E+00 | 8.03E-01 | no | 4931428L18Rik | intron_variant | MODIFIER |
| 1 | 31057820 | AX-170223706 | 0 | C | T | 0.404 | 2.04E-02 | 8.28E-02 | 2.66E+00 | 2.74E+00 | 8.03E-01 | no | 4931428L18Rik | intron_variant | MODIFIER |
| 1 | 31058177 | AX-170279555 | 0 | C | T | 0.404 | 2.04E-02 | 8.28E-02 | 2.66E+00 | 2.74E+00 | 8.03E-01 | no | 4931428L18Rik | intron_variant | MODIFIER |
| 1 | 31058299 | AX-170166314 | 0 | A | G | 0.404 | 2.04E-02 | 8.28E-02 | 2.66E+00 | 2.74E+00 | 8.03E-01 | no | 4931428L18Rik | intron_variant | MODIFIER |
| 1 | 31058783 | AX-170529292 | 0 | T | C | 0.404 | 2.04E-02 | 8.28E-02 | 2.66E+00 | 2.74E+00 | 8.03E-01 | no | 4931428L18Rik | intron_variant | MODIFIER |
| 1 | 31058925 | AX-97845761 | 0 | A | G | 0.404 | 2.04E-02 | 8.28E-02 | 2.66E+00 | 2.74E+00 | 8.03E-01 | no | 4931428L18Rik | intron_variant | MODIFIER |
| 1 | 31889774 | AX-170233261 | 0 | A | G | 0.404 | 2.04E-02 | 8.28E-02 | 2.66E+00 | 2.74E+00 | 8.03E-01 | no | | intergenic_variant | MODIFIER |
| 1 | 32238803 | AX-170423031 | 0 | T | C | 0.404 | 2.04E-02 | 8.28E-02 | 2.66E+00 | 2.74E+00 | 8.03E-01 | no | Khdrbs2 | intron_variant | MODIFIER |
| 1 | 32400925 | AX-170312765 | 0 | C | T | 0.404 | 2.04E-02 | 8.28E-02 | 2.66E+00 | 2.74E+00 | 8.03E-01 | no | Khdrbs2 | intron_variant | MODIFIER |
| 1 | 32423782 | AX-170370988 | 0 | T | A | 0.404 | 2.04E-02 | 8.28E-02 | 2.66E+00 | 2.74E+00 | 8.03E-01 | no | Khdrbs2 | intron_variant | MODIFIER |
| 1 | 32531095 | AX-170091456 | 0 | G | A | 0.404 | 2.04E-02 | 8.28E-02 | 2.66E+00 | 2.74E+00 | 8.03E-01 | no | Khdrbs2 | intron_variant | MODIFIER |
| 1 | 32625818 | AX-170404073 | 0 | C | A | 0.404 | 2.04E-02 | 8.28E-02 | 2.66E+00 | 2.74E+00 | 8.03E-01 | no | Khdrbs2 | intron_variant | MODIFIER |

| | | | | | | | | | | | | | | | |
|---|---|---|---|---|---|---|---|---|---|---|---|---|---|---|---|
| 1 | 32635126 | AX-168375265 | 0 | C | T | 0.404 | 2.04E-02 | 8.28E-02 | 2.66E+00 | 2.74E+00 | 8.03E-01 | no | Khdrbs2 | intron_variant | MODIFIER |
| 1 | 32635494 | AX-170208628 | 0 | A | G | 0.404 | 2.04E-02 | 8.28E-02 | 2.66E+00 | 2.74E+00 | 8.03E-01 | no | Khdrbs2 | intron_variant | MODIFIER |
| 1 | 32639709 | AX-97844713 | 0 | G | A | 0.404 | 2.04E-02 | 8.28E-02 | 2.66E+00 | 2.74E+00 | 8.03E-01 | no | Khdrbs2 | intron_variant | MODIFIER |
| 1 | 32729578 | AX-168393173 | 0 | T | C | 0.404 | 2.04E-02 | 8.28E-02 | 2.66E+00 | 2.74E+00 | 8.03E-01 | no | Gm23024 | downstream_gene_variant | MODIFIER |
| 1 | 33032669 | AX-170311809 | 0 | A | T | 0.404 | 2.04E-02 | 8.28E-02 | 2.66E+00 | 2.74E+00 | 8.03E-01 | no | Gm28624 | upstream_gene_variant | MODIFIER |
| 1 | 33033152 | AX-170348345 | 0 | G | C | 0.404 | 2.04E-02 | 8.28E-02 | 2.66E+00 | 2.74E+00 | 8.03E-01 | no | Gm28624 | upstream_gene_variant | MODIFIER |
| 1 | 33057244 | AX-170503299 | 0 | C | T | 0.404 | 2.04E-02 | 8.28E-02 | 2.66E+00 | 2.74E+00 | 8.03E-01 | no | | intergenic_variant | MODIFIER |
| 1 | 33059889 | AX-170184958 | 0 | A | G | 0.404 | 2.04E-02 | 8.28E-02 | 2.66E+00 | 2.74E+00 | 8.03E-01 | no | | intergenic_variant | MODIFIER |
| 1 | 33297352 | AX-170320512 | 0 | A | G | 0.404 | 2.04E-02 | 8.28E-02 | 2.66E+00 | 2.74E+00 | 8.03E-01 | no | Gm25792 | downstream_gene_variant | MODIFIER |
| 1 | 33313934 | AX-170097860 | 0 | G | A | 0.404 | 2.04E-02 | 8.28E-02 | 2.66E+00 | 2.74E+00 | 8.03E-01 | no | Gm18799 | downstream_gene_variant | MODIFIER |
| 1 | 33331702 | AX-170086165 | 0 | T | C | 0.404 | 2.04E-02 | 8.28E-02 | 2.66E+00 | 2.74E+00 | 8.03E-01 | no | | intergenic_variant | MODIFIER |
| 1 | 33723514 | AX-170000245 | 0 | C | T | 0.404 | 2.04E-02 | 8.28E-02 | 2.66E+00 | 2.74E+00 | 8.03E-01 | no | Rab23 | intron_variant | MODIFIER |
| 1 | 33740032 | AX-170395020 | 0 | A | C | 0.404 | 2.04E-02 | 8.28E-02 | 2.66E+00 | 2.74E+00 | 8.03E-01 | no | Rab23 | 3_prime_UTR_variant | MODIFIER |
| 12 | 94975289 | AX-170418680 | 0 | C | T | 0.061 | 5.77E-02 | 2.36E-01 | 2.69E+00 | 2.75E+00 | 8.03E-01 | no | | intergenic_variant | MODIFIER |
| 16 | 80719790 | AX-170182835 | 1 | G | A | 0.482 | 1.88E-02 | 7.23E-02 | 2.60E+00 | 2.69E+00 | 8.03E-01 | no | | intergenic_variant | MODIFIER |
| 5 | 78663894 | AX-170417323 | 0 | G | A | 0.404 | 2.48E-02 | 9.89E-02 | 2.64E+00 | 2.72E+00 | 8.03E-01 | no | | intergenic_variant | MODIFIER |
| 10 | 16621339 | AX-97853903 | 0 | G | A | 0.211 | 2.53E-02 | 1.06E-01 | 2.73E+00 | 2.81E+00 | 8.03E-01 | no | Gm47729 | intron_variant,non_coding_transcript_variant | MODIFIER |
| 10 | 16622000 | AX-170253500 | 0 | G | A | 0.211 | 2.53E-02 | 1.06E-01 | 2.73E+00 | 2.81E+00 | 8.03E-01 | no | Gm47729 | intron_variant,non_coding_transcript_variant | MODIFIER |
| 10 | 16723281 | AX-170002820 | 0 | C | T | 0.211 | 2.53E-02 | 1.06E-01 | 2.73E+00 | 2.81E+00 | 8.03E-01 | no | | intergenic_variant | MODIFIER |
| 10 | 16903459 | AX-97825110 | 0 | T | C | 0.211 | 2.53E-02 | 1.06E-01 | 2.73E+00 | 2.81E+00 | 8.03E-01 | no | Gm20125 | downstream_gene_variant | MODIFIER |
| 10 | 16906932 | AX-170140836 | 0 | C | A | 0.211 | 2.53E-02 | 1.06E-01 | 2.73E+00 | 2.81E+00 | 8.03E-01 | no | Gm20125 | intron_variant,non_coding_transcript_variant | MODIFIER |
| 10 | 16987471 | AX-170316866 | 0 | T | A | 0.211 | 2.53E-02 | 1.06E-01 | 2.73E+00 | 2.81E+00 | 8.03E-01 | no | Gm20125 | intron_variant,non_coding_transcript_variant | MODIFIER |
| 10 | 16998383 | AX-170179759 | 0 | T | G | 0.211 | 2.53E-02 | 1.06E-01 | 2.73E+00 | 2.81E+00 | 8.03E-01 | no | Gm20125 | intron_variant,non_coding_transcript_variant | MODIFIER |
| 10 | 17073039 | AX-170086757 | 0 | C | G | 0.211 | 2.53E-02 | 1.06E-01 | 2.73E+00 | 2.81E+00 | 8.03E-01 | no | Gm20125 | intron_variant,non_coding_transcript_variant | MODIFIER |
| 10 | 17085986 | AX-170148977 | 0 | A | G | 0.211 | 2.53E-02 | 1.06E-01 | 2.73E+00 | 2.81E+00 | 8.03E-01 | no | Gm20125 | intron_variant,non_coding_transcript_variant | MODIFIER |
| 10 | 17117301 | AX-170278662 | 0 | G | T | 0.211 | 2.53E-02 | 1.06E-01 | 2.73E+00 | 2.81E+00 | 8.03E-01 | no | Gm25382 | downstream_gene_variant | MODIFIER |
| 10 | 17122108 | AX-170002827 | 0 | G | A | 0.211 | 2.53E-02 | 1.06E-01 | 2.73E+00 | 2.81E+00 | 8.03E-01 | no | | intergenic_variant | MODIFIER |
| 10 | 17172324 | AX-170002838 | 0 | A | G | 0.211 | 2.53E-02 | 1.06E-01 | 2.73E+00 | 2.81E+00 | 8.03E-01 | no | | intergenic_variant | MODIFIER |
| 10 | 17197937 | AX-170002834 | 0 | T | C | 0.211 | 2.53E-02 | 1.06E-01 | 2.73E+00 | 2.81E+00 | 8.03E-01 | no | | intergenic_variant | MODIFIER |
| 10 | 17262924 | AX-170002829 | 0 | C | T | 0.211 | 2.53E-02 | 1.06E-01 | 2.73E+00 | 2.81E+00 | 8.03E-01 | no | | regulatory_region_variant | MODIFIER |
| 10 | 17513438 | AX-170289164 | 0 | T | C | 0.211 | 2.53E-02 | 1.06E-01 | 2.73E+00 | 2.81E+00 | 8.03E-01 | no | Gm47768 | upstream_gene_variant | MODIFIER |
| 10 | 17519855 | AX-170287342 | 0 | A | G | 0.211 | 2.53E-02 | 1.06E-01 | 2.73E+00 | 2.81E+00 | 8.03E-01 | no | Gm47768 | downstream_gene_variant | MODIFIER |
| 1 | 33000880 | AX-170416177 | 1 | C | G | 0.411 | 1.99E-02 | 8.18E-02 | 2.68E+00 | 2.76E+00 | 8.03E-01 | no | | intergenic_variant | MODIFIER |
| 7 | 137263180 | AX-170030071 | 1 | G | T | 0.286 | -1.99E-02 | 7.85E-02 | 2.61E+00 | 2.71E+00 | 8.03E-01 | no | Ebf3 | intron_variant | MODIFIER |
| 8 | 97055555 | AX-97792511 | 0 | G | T | 0.219 | -2.68E-02 | 1.11E-01 | 2.68E+00 | 2.77E+00 | 8.03E-01 | no | | intergenic_variant | MODIFIER |
| 5 | 7258874 | AX-97832454 | 0 | C | T | 0.439 | 1.49E-02 | 6.33E-02 | 2.65E+00 | 2.78E+00 | 8.03E-01 | no | Gm42597 | downstream_gene_variant | MODIFIER |
| 15 | 92590988 | AX-170012271 | 1 | A | T | 0.143 | 4.16E-02 | 1.67E-01 | 2.64E+00 | 2.72E+00 | 8.04E-01 | no | Pdzrn4 | intron_variant | MODIFIER |
| 8 | 78024114 | AX-170236812 | 0 | C | G | 0.184 | 2.77E-02 | 1.16E-01 | 2.66E+00 | 2.77E+00 | 8.04E-01 | no | | regulatory_region_variant | MODIFIER |
| 13 | 53727365 | AX-170222594 | 0 | G | A | 0.526 | -1.58E-02 | 6.78E-02 | 2.69E+00 | 2.81E+00 | 8.04E-01 | no | | intergenic_variant | MODIFIER |
| 10 | 9328716 | AX-170002691 | 0 | T | C | 0.202 | -2.49E-02 | 9.75E-02 | 2.61E+00 | 2.70E+00 | 8.04E-01 | no | Gm48748 | downstream_gene_variant | MODIFIER |
| 1 | 25714010 | AX-170482078 | 1 | A | C | 0.402 | 2.05E-02 | 8.29E-02 | 2.65E+00 | 2.73E+00 | 8.04E-01 | no | Adgrb3 | intron_variant | MODIFIER |
| 16 | 91559068 | AX-170495982 | 1 | C | A | 0.446 | -1.92E-02 | 7.88E-02 | 2.66E+00 | 2.75E+00 | 8.04E-01 | no | Ifngr2 | intron_variant | MODIFIER |
| 7 | 117876860 | AX-170480505 | 2 | G | A | 0.364 | 2.06E-02 | 8.31E-02 | 2.64E+00 | 2.73E+00 | 8.04E-01 | no | | intergenic_variant | MODIFIER |
| 6 | 104936654 | AX-170472704 | 1 | T | C | 0.054 | 7.25E-02 | 2.96E-01 | 2.68E+00 | 2.74E+00 | 8.04E-01 | no | | intergenic_variant | MODIFIER |
| 1 | 151641334 | AX-170244697 | 1 | T | C | 0.205 | 3.35E-02 | 1.35E-01 | 2.66E+00 | 2.72E+00 | 8.04E-01 | no | Fam129a | intron_variant,NMD_transcript_variant | MODIFIER |
| 18 | 63258094 | AX-168403812 | 0 | A | C | 0.421 | 2.04E-02 | 8.45E-02 | 2.75E+00 | 2.81E+00 | 8.04E-01 | no | Piezo2 | intron_variant | MODIFIER |
| 18 | 63847451 | AX-170061092 | 0 | T | C | 0.421 | 2.04E-02 | 8.45E-02 | 2.75E+00 | 2.81E+00 | 8.04E-01 | no | Wdr7 | intron_variant | MODIFIER |
| 18 | 63850482 | AX-170190335 | 0 | C | T | 0.421 | 2.04E-02 | 8.45E-02 | 2.75E+00 | 2.81E+00 | 8.04E-01 | no | Wdr7 | intron_variant | MODIFIER |
| 18 | 63852555 | AX-170198913 | 0 | C | T | 0.421 | 2.04E-02 | 8.45E-02 | 2.75E+00 | 2.81E+00 | 8.04E-01 | no | Wdr7 | intron_variant | MODIFIER |
| 18 | 63852785 | AX-170105422 | 0 | A | C | 0.421 | 2.04E-02 | 8.45E-02 | 2.75E+00 | 2.81E+00 | 8.04E-01 | no | Wdr7 | intron_variant | MODIFIER |
| 18 | 63896115 | AX-97835653 | 0 | A | G | 0.421 | 2.04E-02 | 8.45E-02 | 2.75E+00 | 2.81E+00 | 8.04E-01 | no | Wdr7 | intron_variant | MODIFIER |
| 18 | 63896358 | AX-170172380 | 0 | G | A | 0.421 | 2.04E-02 | 8.45E-02 | 2.75E+00 | 2.81E+00 | 8.04E-01 | no | Wdr7 | intron_variant | MODIFIER |
| 18 | 63921724 | AX-97802581 | 0 | T | G | 0.421 | 2.04E-02 | 8.45E-02 | 2.75E+00 | 2.81E+00 | 8.04E-01 | no | Wdr7 | intron_variant | MODIFIER |
| 18 | 63952956 | AX-170484493 | 0 | C | T | 0.421 | 2.04E-02 | 8.45E-02 | 2.75E+00 | 2.81E+00 | 8.04E-01 | no | Wdr7 | intron_variant | MODIFIER |

| | | | | | | | | | | | | | | | |
|---|---|---|---|---|---|---|---|---|---|---|---|---|---|---|---|
| 18 | 63982440 | AX-170200300 | 0 | T | A | 0.421 | 2.04E-02 | 8.45E-02 | 2.75E+00 | 2.81E+00 | 8.04E-01 | no | Wdr7 | intron_variant | MODIFIER |
| 18 | 63983167 | AX-170189292 | 0 | A | G | 0.421 | 2.04E-02 | 8.45E-02 | 2.75E+00 | 2.81E+00 | 8.04E-01 | no | Wdr7 | intron_variant | MODIFIER |
| 18 | 64094891 | AX-170269388 | 0 | A | G | 0.421 | 2.04E-02 | 8.45E-02 | 2.75E+00 | 2.81E+00 | 8.04E-01 | no | | intergenic_variant | MODIFIER |
| 18 | 64157461 | AX-97835675 | 0 | C | T | 0.421 | 2.04E-02 | 8.45E-02 | 2.75E+00 | 2.81E+00 | 8.04E-01 | no | St8sia3os | downstream_gene_variant | MODIFIER |
| 18 | 64175329 | AX-170447348 | 0 | C | G | 0.421 | 2.04E-02 | 8.45E-02 | 2.75E+00 | 2.81E+00 | 8.04E-01 | no | St8sia3os | intron_variant,non_coding_transcript_variant | MODIFIER |
| 18 | 64179965 | AX-170093914 | 0 | T | C | 0.421 | 2.04E-02 | 8.45E-02 | 2.75E+00 | 2.81E+00 | 8.04E-01 | no | St8sia3os | intron_variant,non_coding_transcript_variant | MODIFIER |
| 18 | 64225608 | AX-170015857 | 0 | C | A | 0.421 | 2.04E-02 | 8.45E-02 | 2.75E+00 | 2.81E+00 | 8.04E-01 | no | St8sia3os | intron_variant,non_coding_transcript_variant | MODIFIER |
| 18 | 64230628 | AX-170165791 | 0 | C | T | 0.421 | 2.04E-02 | 8.45E-02 | 2.75E+00 | 2.81E+00 | 8.04E-01 | no | St8sia3os | intron_variant,non_coding_transcript_variant | MODIFIER |
| 18 | 64230855 | AX-97812105 | 0 | G | A | 0.421 | 2.04E-02 | 8.45E-02 | 2.75E+00 | 2.81E+00 | 8.04E-01 | no | St8sia3os | intron_variant,non_coding_transcript_variant | MODIFIER |
| 18 | 64231076 | AX-170228834 | 0 | G | T | 0.421 | 2.04E-02 | 8.45E-02 | 2.75E+00 | 2.81E+00 | 8.04E-01 | no | St8sia3os | intron_variant,non_coding_transcript_variant | MODIFIER |
| 18 | 64244395 | AX-170240340 | 0 | G | T | 0.421 | 2.04E-02 | 8.45E-02 | 2.75E+00 | 2.81E+00 | 8.04E-01 | no | St8sia3os | intron_variant,non_coding_transcript_variant | MODIFIER |
| 18 | 64257179 | AX-170371434 | 0 | C | T | 0.421 | 2.04E-02 | 8.45E-02 | 2.75E+00 | 2.81E+00 | 8.04E-01 | no | St8sia3 | intron_variant | MODIFIER |
| 18 | 64263115 | AX-170323650 | 0 | T | C | 0.421 | 2.04E-02 | 8.45E-02 | 2.75E+00 | 2.81E+00 | 8.04E-01 | no | St8sia3 | intron_variant | MODIFIER |
| 18 | 64300855 | AX-170373361 | 0 | T | C | 0.421 | 2.04E-02 | 8.45E-02 | 2.75E+00 | 2.81E+00 | 8.04E-01 | no | St8sia3os | intron_variant,non_coding_transcript_variant | MODIFIER |
| 18 | 64301057 | AX-170345582 | 0 | C | T | 0.421 | 2.04E-02 | 8.45E-02 | 2.75E+00 | 2.81E+00 | 8.04E-01 | no | St8sia3os | intron_variant,non_coding_transcript_variant | MODIFIER |
| 18 | 64476033 | AX-97841946 | 0 | T | C | 0.421 | 2.04E-02 | 8.45E-02 | 2.75E+00 | 2.81E+00 | 8.04E-01 | no | Fech | intron_variant | MODIFIER |
| 18 | 64503743 | AX-170480989 | 0 | T | C | 0.421 | 2.04E-02 | 8.45E-02 | 2.75E+00 | 2.81E+00 | 8.04E-01 | no | Nars | intron_variant | MODIFIER |
| 18 | 64504522 | AX-97787666 | 0 | T | C | 0.421 | 2.04E-02 | 8.45E-02 | 2.75E+00 | 2.81E+00 | 8.04E-01 | no | Nars | intron_variant | MODIFIER |
| 18 | 64512147 | AX-170303406 | 0 | A | C | 0.421 | 2.04E-02 | 8.45E-02 | 2.75E+00 | 2.81E+00 | 8.04E-01 | no | Nars | intron_variant | MODIFIER |
| 18 | 64789150 | AX-170278875 | 0 | A | G | 0.421 | 2.04E-02 | 8.45E-02 | 2.75E+00 | 2.81E+00 | 8.04E-01 | no | Gm24504 | upstream_gene_variant | MODIFIER |
| 18 | 64789373 | AX-170015861 | 0 | C | T | 0.421 | 2.04E-02 | 8.45E-02 | 2.75E+00 | 2.81E+00 | 8.04E-01 | no | Gm24504 | upstream_gene_variant | MODIFIER |
| 18 | 64801293 | AX-97827692 | 0 | T | G | 0.421 | 2.04E-02 | 8.45E-02 | 2.75E+00 | 2.81E+00 | 8.04E-01 | no | AC131709.1 | intron_variant,non_coding_transcript_variant | MODIFIER |
| 18 | 64813693 | AX-170452759 | 0 | T | C | 0.421 | 2.04E-02 | 8.45E-02 | 2.75E+00 | 2.81E+00 | 8.04E-01 | no | | intergenic_variant | MODIFIER |
| 18 | 64820517 | AX-170015862 | 0 | T | C | 0.421 | 2.04E-02 | 8.45E-02 | 2.75E+00 | 2.81E+00 | 8.04E-01 | no | | intergenic_variant | MODIFIER |
| 18 | 64820850 | AX-97820058 | 0 | C | A | 0.421 | 2.04E-02 | 8.45E-02 | 2.75E+00 | 2.81E+00 | 8.04E-01 | no | | intergenic_variant | MODIFIER |
| 18 | 64845774 | AX-170529514 | 0 | C | G | 0.421 | 2.04E-02 | 8.45E-02 | 2.75E+00 | 2.81E+00 | 8.04E-01 | no | | regulatory_region_variant | MODIFIER |
| 18 | 64906120 | AX-170293502 | 0 | T | C | 0.421 | 2.04E-02 | 8.45E-02 | 2.75E+00 | 2.81E+00 | 8.04E-01 | no | Nedd4l | intron_variant | MODIFIER |
| 1 | 59722382 | AX-170138995 | 1 | C | T | 0.098 | 5.34E-02 | 2.19E-01 | 2.66E+00 | 2.75E+00 | 8.04E-01 | no | Nop58 | downstream_gene_variant | MODIFIER |
| 1 | 59775802 | AX-170302818 | 1 | T | C | 0.098 | 5.34E-02 | 2.19E-01 | 2.66E+00 | 2.75E+00 | 8.04E-01 | no | Bmpr2 | intron_variant | MODIFIER |
| 1 | 60478124 | AX-170000613 | 1 | A | G | 0.098 | 5.34E-02 | 2.19E-01 | 2.66E+00 | 2.75E+00 | 8.04E-01 | no | Abi2 | 3_prime_UTR_variant | MODIFIER |
| 1 | 60792374 | AX-170041392 | 1 | C | T | 0.098 | 5.34E-02 | 2.19E-01 | 2.66E+00 | 2.75E+00 | 8.04E-01 | no | Gm11609 | upstream_gene_variant | MODIFIER |
| 1 | 102593671 | AX-170420911 | 0 | C | A | 0.044 | 3.88E-02 | 1.65E-01 | 2.61E+00 | 2.77E+00 | 8.04E-01 | no | | intergenic_variant | MODIFIER |
| 18 | 53699828 | AX-170170365 | 0 | T | C | 0.202 | -2.83E-02 | 1.08E-01 | 2.60E+00 | 2.67E+00 | 8.04E-01 | no | Cep120 | intron_variant | MODIFIER |
| 1 | 69295983 | AX-170000752 | 1 | G | A | 0.143 | 5.98E-02 | 2.44E-01 | 2.69E+00 | 2.74E+00 | 8.04E-01 | no | | intergenic_variant | MODIFIER |
| 1 | 46896797 | AX-170236007 | 0 | G | A | 0.43 | -1.90E-02 | 7.74E-02 | 2.65E+00 | 2.74E+00 | 8.04E-01 | no | Gm28527 | non_coding_transcript_exon_variant | MODIFIER |
| 8 | 75272380 | AX-170444568 | 0 | G | A | 0.193 | -3.43E-02 | 1.38E-01 | 2.65E+00 | 2.71E+00 | 8.05E-01 | no | Gm9644 | downstream_gene_variant | MODIFIER |
| 18 | 53548098 | AX-170101351 | 0 | A | G | 0.237 | -2.78E-02 | 1.08E-01 | 2.62E+00 | 2.68E+00 | 8.05E-01 | no | Prdm6 | intron_variant | MODIFIER |
| 4 | 98041067 | AX-170023476 | 0 | G | T | 0.228 | 2.89E-02 | 1.14E-01 | 2.62E+00 | 2.69E+00 | 8.05E-01 | no | Nfia | intron_variant | MODIFIER |
| 8 | 51559377 | AX-170109935 | 0 | G | C | 0.158 | 3.24E-02 | 1.37E-01 | 2.72E+00 | 2.81E+00 | 8.05E-01 | no | | intergenic_variant | MODIFIER |
| 20 | 63324692 | AX-170090522 | 1 | G | A | 0.25 | 1.54E-02 | 6.24E-02 | 2.64E+00 | 2.73E+00 | 8.05E-01 | no | | intergenic_variant | MODIFIER |
| 1 | 46153182 | AX-97811408 | 1 | G | A | 0.33 | -1.84E-02 | 7.40E-02 | 2.63E+00 | 2.72E+00 | 8.05E-01 | no | Dnah7b | intron_variant | MODIFIER |
| 10 | 5078733 | AX-168418141 | 1 | C | A | 0.214 | 2.31E-02 | 9.76E-02 | 2.67E+00 | 2.78E+00 | 8.05E-01 | no | Syne1 | intron_variant | MODIFIER |
| 20 | 166777429 | AX-170085771 | 0 | C | T | 0.184 | 2.99E-02 | 1.18E-01 | 2.61E+00 | 2.70E+00 | 8.05E-01 | no | | intergenic_variant | MODIFIER |
| 20 | 166797693 | AX-170035425 | 0 | G | A | 0.184 | 2.99E-02 | 1.18E-01 | 2.61E+00 | 2.70E+00 | 8.05E-01 | no | | intergenic_variant | MODIFIER |
| 20 | 166800861 | AX-170503939 | 0 | G | C | 0.184 | 2.99E-02 | 1.18E-01 | 2.61E+00 | 2.70E+00 | 8.05E-01 | no | | intergenic_variant | MODIFIER |
| 20 | 166801343 | AX-170396458 | 0 | G | C | 0.184 | 2.99E-02 | 1.18E-01 | 2.61E+00 | 2.70E+00 | 8.05E-01 | no | | intergenic_variant | MODIFIER |
| 20 | 166805814 | AX-170306304 | 0 | T | C | 0.184 | 2.99E-02 | 1.18E-01 | 2.61E+00 | 2.70E+00 | 8.05E-01 | no | | regulatory_region_variant | MODIFIER |
| 20 | 166806207 | AX-170506397 | 0 | A | G | 0.184 | 2.99E-02 | 1.18E-01 | 2.61E+00 | 2.70E+00 | 8.05E-01 | no | | intergenic_variant | MODIFIER |
| 20 | 166823677 | AX-170425669 | 0 | C | G | 0.184 | 2.99E-02 | 1.18E-01 | 2.61E+00 | 2.70E+00 | 8.05E-01 | no | | intergenic_variant | MODIFIER |
| 20 | 166833944 | AX-170035424 | 0 | T | A | 0.184 | 2.99E-02 | 1.18E-01 | 2.61E+00 | 2.70E+00 | 8.05E-01 | no | | intergenic_variant | MODIFIER |
| 20 | 167105018 | AX-170160861 | 0 | T | C | 0.184 | 2.99E-02 | 1.18E-01 | 2.61E+00 | 2.70E+00 | 8.05E-01 | no | Gm15232 | upstream_gene_variant | MODIFIER |
| 20 | 167115305 | AX-170146315 | 0 | C | G | 0.184 | 2.99E-02 | 1.18E-01 | 2.61E+00 | 2.70E+00 | 8.05E-01 | no | | intergenic_variant | MODIFIER |
| 20 | 167119698 | AX-97841806 | 0 | T | C | 0.184 | 2.99E-02 | 1.18E-01 | 2.61E+00 | 2.70E+00 | 8.05E-01 | no | | regulatory_region_variant | MODIFIER |
| 20 | 167136220 | AX-170228470 | 0 | T | C | 0.184 | 2.99E-02 | 1.18E-01 | 2.61E+00 | 2.70E+00 | 8.05E-01 | no | | intergenic_variant | MODIFIER |

| | | | | | | | | | | | | | | | |
|---|---|---|---|---|---|---|---|---|---|---|---|---|---|---|---|
| 20 | 167147882 | AX-170503161 | 0 | A | T | 0.184 | 2.99E-02 | 1.18E-01 | 2.61E+00 | 2.70E+00 | 8.05E-01 | no | | intergenic_variant | MODIFIER |
| 20 | 167175377 | AX-170428029 | 0 | A | G | 0.184 | 2.99E-02 | 1.18E-01 | 2.61E+00 | 2.70E+00 | 8.05E-01 | no | Gm15228 | downstream_gene_variant | MODIFIER |
| 20 | 167176755 | AX-97830769 | 0 | T | A | 0.184 | 2.99E-02 | 1.18E-01 | 2.61E+00 | 2.70E+00 | 8.05E-01 | no | | intergenic_variant | MODIFIER |
| 20 | 167198750 | AX-170311602 | 0 | G | C | 0.184 | 2.99E-02 | 1.18E-01 | 2.61E+00 | 2.70E+00 | 8.05E-01 | no | | intergenic_variant | MODIFIER |
| 20 | 167198928 | AX-97807282 | 0 | G | A | 0.184 | 2.99E-02 | 1.18E-01 | 2.61E+00 | 2.70E+00 | 8.05E-01 | no | | intergenic_variant | MODIFIER |
| 20 | 167221872 | AX-97849844 | 0 | G | T | 0.184 | 2.99E-02 | 1.18E-01 | 2.61E+00 | 2.70E+00 | 8.05E-01 | no | | intergenic_variant | MODIFIER |
| 20 | 167416568 | AX-97823588 | 0 | G | A | 0.184 | 2.99E-02 | 1.18E-01 | 2.61E+00 | 2.70E+00 | 8.05E-01 | no | | intergenic_variant | MODIFIER |
| 20 | 167428075 | AX-170193373 | 0 | A | G | 0.184 | 2.99E-02 | 1.18E-01 | 2.61E+00 | 2.70E+00 | 8.05E-01 | no | | intergenic_variant | MODIFIER |
| 20 | 167434783 | AX-170196727 | 0 | A | G | 0.184 | 2.99E-02 | 1.18E-01 | 2.61E+00 | 2.70E+00 | 8.05E-01 | no | | intergenic_variant | MODIFIER |
| 20 | 167462864 | AX-97809450 | 0 | A | G | 0.184 | 2.99E-02 | 1.18E-01 | 2.61E+00 | 2.70E+00 | 8.05E-01 | no | | intergenic_variant | MODIFIER |
| 20 | 167514342 | AX-170425355 | 0 | T | C | 0.184 | 2.99E-02 | 1.18E-01 | 2.61E+00 | 2.70E+00 | 8.05E-01 | no | Frmpd4 | intron_variant | MODIFIER |
| 20 | 167516158 | AX-97808168 | 0 | A | G | 0.184 | 2.99E-02 | 1.18E-01 | 2.61E+00 | 2.70E+00 | 8.05E-01 | no | Frmpd4 | intron_variant | MODIFIER |
| 20 | 167516181 | AX-170514990 | 0 | T | C | 0.184 | 2.99E-02 | 1.18E-01 | 2.61E+00 | 2.70E+00 | 8.05E-01 | no | Frmpd4 | intron_variant | MODIFIER |
| 20 | 167557208 | AX-97807822 | 0 | A | G | 0.184 | 2.99E-02 | 1.18E-01 | 2.61E+00 | 2.70E+00 | 8.05E-01 | no | Frmpd4 | intron_variant | MODIFIER |
| 20 | 167614117 | AX-97794889 | 0 | T | G | 0.184 | 2.99E-02 | 1.18E-01 | 2.61E+00 | 2.70E+00 | 8.05E-01 | no | Frmpd4 | intron_variant | MODIFIER |
| 20 | 168048295 | AX-97815493 | 0 | C | T | 0.184 | 2.99E-02 | 1.18E-01 | 2.61E+00 | 2.70E+00 | 8.05E-01 | no | Frmpd4 | intron_variant | MODIFIER |
| 20 | 168052342 | AX-170270779 | 0 | A | G | 0.184 | 2.99E-02 | 1.18E-01 | 2.61E+00 | 2.70E+00 | 8.05E-01 | no | Frmpd4 | intron_variant | MODIFIER |
| 20 | 168460680 | AX-97803114 | 0 | G | T | 0.184 | 2.99E-02 | 1.18E-01 | 2.61E+00 | 2.70E+00 | 8.05E-01 | no | Frmpd4 | intron_variant,NMD_transcript_variant | MODIFIER |
| 20 | 168463296 | AX-170271631 | 0 | T | C | 0.184 | 2.99E-02 | 1.18E-01 | 2.61E+00 | 2.70E+00 | 8.05E-01 | no | Frmpd4 | intron_variant,NMD_transcript_variant | MODIFIER |
| 20 | 168463567 | AX-170466766 | 0 | C | G | 0.184 | 2.99E-02 | 1.18E-01 | 2.61E+00 | 2.70E+00 | 8.05E-01 | no | Frmpd4 | intron_variant,NMD_transcript_variant | MODIFIER |
| 20 | 168525833 | AX-97784883 | 0 | G | A | 0.184 | 2.99E-02 | 1.18E-01 | 2.61E+00 | 2.70E+00 | 8.05E-01 | no | Frmpd4 | intron_variant,NMD_transcript_variant | MODIFIER |
| 20 | 168587468 | AX-97821118 | 0 | C | T | 0.184 | 2.99E-02 | 1.18E-01 | 2.61E+00 | 2.70E+00 | 8.05E-01 | no | | intergenic_variant | MODIFIER |
| 20 | 168634597 | AX-170371993 | 0 | C | T | 0.184 | 2.99E-02 | 1.18E-01 | 2.61E+00 | 2.70E+00 | 8.05E-01 | no | | intergenic_variant | MODIFIER |
| 20 | 168634917 | AX-170301307 | 0 | A | G | 0.184 | 2.99E-02 | 1.18E-01 | 2.61E+00 | 2.70E+00 | 8.05E-01 | no | | intergenic_variant | MODIFIER |
| 20 | 168646467 | AX-97849484 | 0 | C | T | 0.184 | 2.99E-02 | 1.18E-01 | 2.61E+00 | 2.70E+00 | 8.05E-01 | no | | intergenic_variant | MODIFIER |
| 20 | 168664861 | AX-97802553 | 0 | A | T | 0.184 | 2.99E-02 | 1.18E-01 | 2.61E+00 | 2.70E+00 | 8.05E-01 | no | Msl3 | intron_variant | MODIFIER |
| 20 | 168704178 | AX-170097005 | 0 | A | G | 0.184 | 2.99E-02 | 1.18E-01 | 2.61E+00 | 2.70E+00 | 8.05E-01 | no | | intergenic_variant | MODIFIER |
| 20 | 168714012 | AX-97782143 | 0 | T | C | 0.184 | 2.99E-02 | 1.18E-01 | 2.61E+00 | 2.70E+00 | 8.05E-01 | no | | intergenic_variant | MODIFIER |
| 20 | 168749590 | AX-97833507 | 0 | G | A | 0.184 | 2.99E-02 | 1.18E-01 | 2.61E+00 | 2.70E+00 | 8.05E-01 | no | | intergenic_variant | MODIFIER |
| 20 | 168756699 | AX-170090748 | 0 | C | T | 0.184 | 2.99E-02 | 1.18E-01 | 2.61E+00 | 2.70E+00 | 8.05E-01 | no | | intergenic_variant | MODIFIER |
| 20 | 168916960 | AX-97807309 | 0 | T | C | 0.184 | 2.99E-02 | 1.18E-01 | 2.61E+00 | 2.70E+00 | 8.05E-01 | no | Arhgap6 | intron_variant | MODIFIER |
| 20 | 168938804 | AX-97844558 | 0 | A | C | 0.184 | 2.99E-02 | 1.18E-01 | 2.61E+00 | 2.70E+00 | 8.05E-01 | no | Arhgap6 | intron_variant | MODIFIER |
| 20 | 169014065 | AX-170097007 | 0 | C | T | 0.184 | 2.99E-02 | 1.18E-01 | 2.61E+00 | 2.70E+00 | 8.05E-01 | no | Arhgap6 | intron_variant | MODIFIER |
| 20 | 169037976 | AX-97822932 | 0 | A | C | 0.184 | 2.99E-02 | 1.18E-01 | 2.61E+00 | 2.70E+00 | 8.05E-01 | no | Arhgap6 | intron_variant | MODIFIER |
| 20 | 169323841 | AX-97783087 | 0 | C | T | 0.184 | 2.99E-02 | 1.18E-01 | 2.61E+00 | 2.70E+00 | 8.05E-01 | no | Hccs | upstream_gene_variant | MODIFIER |
| 20 | 169351569 | AX-97854624 | 0 | A | G | 0.184 | 2.99E-02 | 1.18E-01 | 2.61E+00 | 2.70E+00 | 8.05E-01 | no | Gm15246 | intron_variant,non_coding_transcript_variant | MODIFIER |
| 9 | 34016546 | AX-170032097 | 1 | C | T | 0.429 | 2.19E-02 | 8.72E-02 | 2.64E+00 | 2.70E+00 | 8.05E-01 | no | | intergenic_variant | MODIFIER |
| 1 | 45582552 | AX-170041096 | 0 | A | G | 0.333 | -1.84E-02 | 7.42E-02 | 2.63E+00 | 2.72E+00 | 8.05E-01 | no | | intergenic_variant | MODIFIER |
| 1 | 45624980 | AX-170299038 | 0 | A | C | 0.333 | -1.84E-02 | 7.42E-02 | 2.63E+00 | 2.72E+00 | 8.05E-01 | no | | intergenic_variant | MODIFIER |
| 1 | 45689179 | AX-170401857 | 0 | A | G | 0.333 | -1.84E-02 | 7.42E-02 | 2.63E+00 | 2.72E+00 | 8.05E-01 | no | Gm23216 | upstream_gene_variant | MODIFIER |
| 1 | 45690940 | AX-170041102 | 0 | A | C | 0.333 | -1.84E-02 | 7.42E-02 | 2.63E+00 | 2.72E+00 | 8.05E-01 | no | | intergenic_variant | MODIFIER |
| 1 | 46081446 | AX-170462877 | 0 | C | G | 0.333 | -1.84E-02 | 7.42E-02 | 2.63E+00 | 2.72E+00 | 8.05E-01 | no | Dnah7b | synonymous_variant | LOW |
| 1 | 46169182 | AX-170525469 | 0 | C | T | 0.333 | -1.84E-02 | 7.42E-02 | 2.63E+00 | 2.72E+00 | 8.05E-01 | no | Dnah7b | intron_variant | MODIFIER |
| 1 | 46177959 | AX-170207326 | 0 | G | C | 0.333 | -1.84E-02 | 7.42E-02 | 2.63E+00 | 2.72E+00 | 8.05E-01 | no | Dnah7b | intron_variant | MODIFIER |
| 1 | 46640920 | AX-170196738 | 0 | A | G | 0.333 | -1.84E-02 | 7.42E-02 | 2.63E+00 | 2.72E+00 | 8.05E-01 | no | Dnah7c | intron_variant,non_coding_transcript_variant | MODIFIER |
| 1 | 46819293 | AX-170382756 | 0 | A | G | 0.333 | -1.84E-02 | 7.42E-02 | 2.63E+00 | 2.72E+00 | 8.05E-01 | no | Slc39a10 | synonymous_variant | LOW |
| 1 | 46884637 | AX-170215573 | 0 | C | T | 0.333 | -1.84E-02 | 7.42E-02 | 2.63E+00 | 2.72E+00 | 8.05E-01 | no | Slc39a10 | intron_variant | MODIFIER |
| 15 | 98393025 | AX-170012355 | 1 | G | A | 0.134 | 3.91E-02 | 1.57E-01 | 2.65E+00 | 2.71E+00 | 8.05E-01 | no | | intergenic_variant | MODIFIER |
| 1 | 28637697 | AX-170130820 | 0 | A | G | 0.404 | 1.96E-02 | 8.13E-02 | 2.67E+00 | 2.75E+00 | 8.05E-01 | no | | intergenic_variant | MODIFIER |
| 10 | 119058001 | AX-170288165 | 0 | C | A | 0.342 | 1.69E-02 | 7.11E-02 | 2.67E+00 | 2.77E+00 | 8.05E-01 | no | | regulatory_region_variant | MODIFIER |
| 9 | 112809206 | AX-170391644 | 0 | G | T | 0.123 | -2.63E-02 | 1.07E-01 | 2.65E+00 | 2.73E+00 | 8.05E-01 | no | | intergenic_variant | MODIFIER |
| 9 | 32706771 | AX-170309463 | 0 | T | C | 0.43 | 2.20E-02 | 8.76E-02 | 2.64E+00 | 2.70E+00 | 8.05E-01 | no | Ets1 | intron_variant | MODIFIER |
| 9 | 32707460 | AX-170238117 | 0 | A | G | 0.43 | 2.20E-02 | 8.76E-02 | 2.64E+00 | 2.70E+00 | 8.05E-01 | no | Ets1 | intron_variant | MODIFIER |
| 9 | 32715095 | AX-170367361 | 0 | C | T | 0.43 | 2.20E-02 | 8.76E-02 | 2.64E+00 | 2.70E+00 | 8.05E-01 | no | Ets1 | intron_variant | MODIFIER |

| | | | | | | | | | | | | | | | |
|---|---|---|---|---|---|---|---|---|---|---|---|---|---|---|---|
| 9 | 32721237 | AX-170524050 | 0 | G | C | 0.43 | 2.20E-02 | 8.76E-02 | 2.64E+00 | 2.70E+00 | 8.05E-01 | no | Ets1 | intron_variant | MODIFIER |
| 9 | 32721639 | AX-170313501 | 0 | A | G | 0.43 | 2.20E-02 | 8.76E-02 | 2.64E+00 | 2.70E+00 | 8.05E-01 | no | Ets1 | intron_variant | MODIFIER |
| 9 | 32745081 | AX-170144999 | 0 | T | G | 0.43 | 2.20E-02 | 8.76E-02 | 2.64E+00 | 2.70E+00 | 8.05E-01 | no | Ets1 | intron_variant | MODIFIER |
| 9 | 32745223 | AX-170382958 | 0 | C | T | 0.43 | 2.20E-02 | 8.76E-02 | 2.64E+00 | 2.70E+00 | 8.05E-01 | no | Ets1 | intron_variant | MODIFIER |
| 9 | 32759050 | AX-170391008 | 0 | C | T | 0.43 | 2.20E-02 | 8.76E-02 | 2.64E+00 | 2.70E+00 | 8.05E-01 | no | Ets1 | downstream_gene_variant | MODIFIER |
| 9 | 32759200 | AX-97787026 | 0 | G | A | 0.43 | 2.20E-02 | 8.76E-02 | 2.64E+00 | 2.70E+00 | 8.05E-01 | no | Ets1 | downstream_gene_variant | MODIFIER |
| 9 | 32764731 | AX-170484357 | 0 | T | C | 0.43 | 2.20E-02 | 8.76E-02 | 2.64E+00 | 2.70E+00 | 8.05E-01 | no | | intergenic_variant | MODIFIER |
| 9 | 32771569 | AX-170510461 | 0 | T | G | 0.43 | 2.20E-02 | 8.76E-02 | 2.64E+00 | 2.70E+00 | 8.05E-01 | no | | intergenic_variant | MODIFIER |
| 9 | 32852433 | AX-97838013 | 0 | T | C | 0.43 | 2.20E-02 | 8.76E-02 | 2.64E+00 | 2.70E+00 | 8.05E-01 | no | | intergenic_variant | MODIFIER |
| 9 | 32853930 | AX-170344379 | 0 | T | G | 0.43 | 2.20E-02 | 8.76E-02 | 2.64E+00 | 2.70E+00 | 8.05E-01 | no | | intergenic_variant | MODIFIER |
| 9 | 32864201 | AX-170133069 | 0 | C | A | 0.43 | 2.20E-02 | 8.76E-02 | 2.64E+00 | 2.70E+00 | 8.05E-01 | no | | intergenic_variant | MODIFIER |
| 9 | 32909738 | AX-170192475 | 0 | A | G | 0.43 | 2.20E-02 | 8.76E-02 | 2.64E+00 | 2.70E+00 | 8.05E-01 | no | Gm27162 | intron_variant,non_coding_transcript_variant | MODIFIER |
| 9 | 32924730 | AX-170113423 | 0 | T | C | 0.43 | 2.20E-02 | 8.76E-02 | 2.64E+00 | 2.70E+00 | 8.05E-01 | no | Gm27162 | intron_variant,non_coding_transcript_variant | MODIFIER |
| 9 | 32971356 | AX-170032078 | 0 | A | G | 0.43 | 2.20E-02 | 8.76E-02 | 2.64E+00 | 2.70E+00 | 8.05E-01 | no | Gm37192 | non_coding_transcript_exon_variant | MODIFIER |
| 9 | 33023514 | AX-170090272 | 0 | G | T | 0.43 | 2.20E-02 | 8.76E-02 | 2.64E+00 | 2.70E+00 | 8.05E-01 | no | | intergenic_variant | MODIFIER |
| 9 | 33029567 | AX-170032082 | 0 | T | G | 0.43 | 2.20E-02 | 8.76E-02 | 2.64E+00 | 2.70E+00 | 8.05E-01 | no | Gm27166 | upstream_gene_variant | MODIFIER |
| 9 | 33070998 | AX-168409538 | 0 | G | A | 0.43 | 2.20E-02 | 8.76E-02 | 2.64E+00 | 2.70E+00 | 8.05E-01 | no | Gm27166 | intron_variant,non_coding_transcript_variant | MODIFIER |
| 9 | 33087837 | AX-170418840 | 0 | C | T | 0.43 | 2.20E-02 | 8.76E-02 | 2.64E+00 | 2.70E+00 | 8.05E-01 | no | Gm27166 | intron_variant,non_coding_transcript_variant | MODIFIER |
| 9 | 33150286 | AX-170032086 | 0 | A | G | 0.43 | 2.20E-02 | 8.76E-02 | 2.64E+00 | 2.70E+00 | 8.05E-01 | no | Gm32171 | upstream_gene_variant | MODIFIER |
| 9 | 33212734 | AX-170032079 | 0 | T | C | 0.43 | 2.20E-02 | 8.76E-02 | 2.64E+00 | 2.70E+00 | 8.05E-01 | no | Gm32171 | intron_variant,non_coding_transcript_variant | MODIFIER |
| 9 | 33225340 | AX-170032083 | 0 | T | C | 0.43 | 2.20E-02 | 8.76E-02 | 2.64E+00 | 2.70E+00 | 8.05E-01 | no | Gm32171 | intron_variant,non_coding_transcript_variant | MODIFIER |
| 9 | 33267930 | AX-170424300 | 0 | C | G | 0.43 | 2.20E-02 | 8.76E-02 | 2.64E+00 | 2.70E+00 | 8.05E-01 | no | | intergenic_variant | MODIFIER |
| 9 | 33382066 | AX-170410444 | 0 | A | C | 0.43 | 2.20E-02 | 8.76E-02 | 2.64E+00 | 2.70E+00 | 8.05E-01 | no | | intergenic_variant | MODIFIER |
| 9 | 33387310 | AX-170445034 | 0 | A | G | 0.43 | 2.20E-02 | 8.76E-02 | 2.64E+00 | 2.70E+00 | 8.05E-01 | no | | intergenic_variant | MODIFIER |
| 9 | 33414095 | AX-170513466 | 0 | G | C | 0.43 | 2.20E-02 | 8.76E-02 | 2.64E+00 | 2.70E+00 | 8.05E-01 | no | | intergenic_variant | MODIFIER |
| 9 | 33415645 | AX-170245366 | 0 | T | C | 0.43 | 2.20E-02 | 8.76E-02 | 2.64E+00 | 2.70E+00 | 8.05E-01 | no | | intergenic_variant | MODIFIER |
| 9 | 33516644 | AX-170032088 | 0 | C | T | 0.43 | 2.20E-02 | 8.76E-02 | 2.64E+00 | 2.70E+00 | 8.05E-01 | no | | intergenic_variant | MODIFIER |
| 9 | 33516817 | AX-170032091 | 0 | T | A | 0.43 | 2.20E-02 | 8.76E-02 | 2.64E+00 | 2.70E+00 | 8.05E-01 | no | | intergenic_variant | MODIFIER |
| 9 | 33537673 | AX-170431436 | 0 | T | C | 0.43 | 2.20E-02 | 8.76E-02 | 2.64E+00 | 2.70E+00 | 8.05E-01 | no | | intergenic_variant | MODIFIER |
| 9 | 33538724 | AX-170032094 | 0 | A | C | 0.43 | 2.20E-02 | 8.76E-02 | 2.64E+00 | 2.70E+00 | 8.05E-01 | no | | intergenic_variant | MODIFIER |
| 9 | 33540280 | AX-97844333 | 0 | A | C | 0.43 | 2.20E-02 | 8.76E-02 | 2.64E+00 | 2.70E+00 | 8.05E-01 | no | | intergenic_variant | MODIFIER |
| 9 | 33554031 | AX-170082053 | 0 | T | C | 0.43 | 2.20E-02 | 8.76E-02 | 2.64E+00 | 2.70E+00 | 8.05E-01 | no | | intergenic_variant | MODIFIER |
| 9 | 33554547 | AX-170082054 | 0 | G | C | 0.43 | 2.20E-02 | 8.76E-02 | 2.64E+00 | 2.70E+00 | 8.05E-01 | no | | intergenic_variant | MODIFIER |
| 9 | 33649305 | AX-170032089 | 0 | G | C | 0.43 | 2.20E-02 | 8.76E-02 | 2.64E+00 | 2.70E+00 | 8.05E-01 | no | | intergenic_variant | MODIFIER |
| 9 | 33677485 | AX-97790581 | 0 | T | C | 0.43 | 2.20E-02 | 8.76E-02 | 2.64E+00 | 2.70E+00 | 8.05E-01 | no | | intergenic_variant | MODIFIER |
| 9 | 33714565 | AX-170299526 | 0 | T | C | 0.43 | 2.20E-02 | 8.76E-02 | 2.64E+00 | 2.70E+00 | 8.05E-01 | no | | intergenic_variant | MODIFIER |
| 9 | 33749093 | AX-170525370 | 0 | A | G | 0.43 | 2.20E-02 | 8.76E-02 | 2.64E+00 | 2.70E+00 | 8.05E-01 | no | | intergenic_variant | MODIFIER |
| 9 | 33778775 | AX-97828054 | 0 | C | G | 0.43 | 2.20E-02 | 8.76E-02 | 2.64E+00 | 2.70E+00 | 8.05E-01 | no | | intergenic_variant | MODIFIER |
| 9 | 33798345 | AX-170308332 | 0 | C | G | 0.43 | 2.20E-02 | 8.76E-02 | 2.64E+00 | 2.70E+00 | 8.05E-01 | no | | intergenic_variant | MODIFIER |
| 9 | 33832078 | AX-97804675 | 0 | T | C | 0.43 | 2.20E-02 | 8.76E-02 | 2.64E+00 | 2.70E+00 | 8.05E-01 | no | | intergenic_variant | MODIFIER |
| 9 | 33832298 | AX-170168575 | 0 | A | C | 0.43 | 2.20E-02 | 8.76E-02 | 2.64E+00 | 2.70E+00 | 8.05E-01 | no | | intergenic_variant | MODIFIER |
| 9 | 33868479 | AX-97821539 | 0 | G | T | 0.43 | 2.20E-02 | 8.76E-02 | 2.64E+00 | 2.70E+00 | 8.05E-01 | no | Gm47784 | downstream_gene_variant | MODIFIER |
| 9 | 33874604 | AX-170342720 | 0 | A | G | 0.43 | 2.20E-02 | 8.76E-02 | 2.64E+00 | 2.70E+00 | 8.05E-01 | no | | intergenic_variant | MODIFIER |
| 9 | 33904281 | AX-170319997 | 0 | C | G | 0.43 | 2.20E-02 | 8.76E-02 | 2.64E+00 | 2.70E+00 | 8.05E-01 | no | Gm47785 | intron_variant,non_coding_transcript_variant | MODIFIER |
| 9 | 33989294 | AX-170389672 | 0 | G | A | 0.43 | 2.20E-02 | 8.76E-02 | 2.64E+00 | 2.70E+00 | 8.05E-01 | no | Gm27164 | intron_variant,non_coding_transcript_variant | MODIFIER |
| 9 | 34033932 | AX-170032095 | 0 | T | G | 0.43 | 2.20E-02 | 8.76E-02 | 2.64E+00 | 2.70E+00 | 8.05E-01 | no | | intergenic_variant | MODIFIER |
| 9 | 34050505 | AX-97850945 | 0 | A | G | 0.43 | 2.20E-02 | 8.76E-02 | 2.64E+00 | 2.70E+00 | 8.05E-01 | no | 7630403G23Rik | intron_variant,non_coding_transcript_variant | MODIFIER |
| 9 | 34155194 | AX-170269918 | 0 | T | C | 0.43 | 2.20E-02 | 8.76E-02 | 2.64E+00 | 2.70E+00 | 8.05E-01 | no | | intergenic_variant | MODIFIER |
| 9 | 34155323 | AX-170244963 | 0 | T | C | 0.43 | 2.20E-02 | 8.76E-02 | 2.64E+00 | 2.70E+00 | 8.05E-01 | no | | regulatory_region_variant | MODIFIER |
| 9 | 34205915 | AX-170113320 | 0 | A | G | 0.43 | 2.20E-02 | 8.76E-02 | 2.64E+00 | 2.70E+00 | 8.05E-01 | no | | intergenic_variant | MODIFIER |
| 9 | 34206033 | AX-170082066 | 0 | A | C | 0.43 | 2.20E-02 | 8.76E-02 | 2.64E+00 | 2.70E+00 | 8.05E-01 | no | | intergenic_variant | MODIFIER |
| 9 | 34243111 | AX-170417874 | 0 | A | G | 0.43 | 2.20E-02 | 8.76E-02 | 2.64E+00 | 2.70E+00 | 8.05E-01 | no | | intergenic_variant | MODIFIER |
| 9 | 34263843 | AX-170363709 | 0 | T | C | 0.43 | 2.20E-02 | 8.76E-02 | 2.64E+00 | 2.70E+00 | 8.05E-01 | no | | intergenic_variant | MODIFIER |
| 9 | 34280164 | AX-170184167 | 0 | A | G | 0.43 | 2.20E-02 | 8.76E-02 | 2.64E+00 | 2.70E+00 | 8.05E-01 | no | | regulatory_region_variant | MODIFIER |

| | | | | | | | | | | | | | | | |
|---|---|---|---|---|---|---|---|---|---|---|---|---|---|---|---|
| 9 | 34537550 | AX-170316974 | 0 | G | T | 0.43 | 2.20E-02 | 8.76E-02 | 2.64E+00 | 2.70E+00 | 8.05E-01 | no | Kirrel3 | intron_variant | MODIFIER |
| 9 | 34538606 | AX-170332642 | 0 | G | A | 0.43 | 2.20E-02 | 8.76E-02 | 2.64E+00 | 2.70E+00 | 8.05E-01 | no | Kirrel3 | intron_variant | MODIFIER |
| 9 | 34539265 | AX-170260539 | 0 | C | T | 0.43 | 2.20E-02 | 8.76E-02 | 2.64E+00 | 2.70E+00 | 8.05E-01 | no | Kirrel3 | intron_variant | MODIFIER |
| 9 | 34546077 | AX-170107019 | 0 | G | A | 0.43 | 2.20E-02 | 8.76E-02 | 2.64E+00 | 2.70E+00 | 8.05E-01 | no | Kirrel3 | intron_variant | MODIFIER |
| 9 | 34548447 | AX-168409879 | 0 | T | C | 0.43 | 2.20E-02 | 8.76E-02 | 2.64E+00 | 2.70E+00 | 8.05E-01 | no | Kirrel3 | intron_variant | MODIFIER |
| 9 | 34548826 | AX-168411657 | 0 | G | A | 0.43 | 2.20E-02 | 8.76E-02 | 2.64E+00 | 2.70E+00 | 8.05E-01 | no | Kirrel3 | intron_variant | MODIFIER |
| 9 | 34585707 | AX-168417760 | 0 | G | A | 0.43 | 2.20E-02 | 8.76E-02 | 2.64E+00 | 2.70E+00 | 8.05E-01 | no | Kirrel3 | intron_variant | MODIFIER |
| 9 | 34620021 | AX-168384879 | 0 | T | C | 0.43 | 2.20E-02 | 8.76E-02 | 2.64E+00 | 2.70E+00 | 8.05E-01 | no | Kirrel3 | intron_variant | MODIFIER |
| 9 | 34673824 | AX-170096510 | 0 | G | A | 0.43 | 2.20E-02 | 8.76E-02 | 2.64E+00 | 2.70E+00 | 8.05E-01 | no | Kirrel3 | intron_variant | MODIFIER |
| 9 | 34675086 | AX-170082081 | 0 | A | G | 0.43 | 2.20E-02 | 8.76E-02 | 2.64E+00 | 2.70E+00 | 8.05E-01 | no | Kirrel3 | intron_variant | MODIFIER |
| 9 | 34675379 | AX-170486044 | 0 | G | C | 0.43 | 2.20E-02 | 8.76E-02 | 2.64E+00 | 2.70E+00 | 8.05E-01 | no | Kirrel3 | intron_variant | MODIFIER |
| 9 | 34860126 | AX-170268798 | 0 | G | A | 0.43 | 2.20E-02 | 8.76E-02 | 2.64E+00 | 2.70E+00 | 8.05E-01 | no | Kirrel3 | intron_variant | MODIFIER |
| 9 | 34861615 | AX-170472088 | 0 | C | T | 0.43 | 2.20E-02 | 8.76E-02 | 2.64E+00 | 2.70E+00 | 8.05E-01 | no | Kirrel3 | intron_variant | MODIFIER |
| 9 | 34862065 | AX-170123227 | 0 | G | A | 0.43 | 2.20E-02 | 8.76E-02 | 2.64E+00 | 2.70E+00 | 8.05E-01 | no | Kirrel3 | intron_variant | MODIFIER |
| 9 | 34863207 | AX-170309523 | 0 | T | A | 0.43 | 2.20E-02 | 8.76E-02 | 2.64E+00 | 2.70E+00 | 8.05E-01 | no | Kirrel3 | intron_variant | MODIFIER |
| 9 | 34870108 | AX-170510889 | 0 | G | T | 0.43 | 2.20E-02 | 8.76E-02 | 2.64E+00 | 2.70E+00 | 8.05E-01 | no | Kirrel3 | intron_variant | MODIFIER |
| 9 | 34875159 | AX-97804935 | 0 | A | G | 0.43 | 2.20E-02 | 8.76E-02 | 2.64E+00 | 2.70E+00 | 8.05E-01 | no | Kirrel3 | intron_variant | MODIFIER |
| 9 | 34967507 | AX-170112941 | 0 | T | C | 0.43 | 2.20E-02 | 8.76E-02 | 2.64E+00 | 2.70E+00 | 8.05E-01 | no | Kirrel3 | intron_variant | MODIFIER |
| 3 | 32712661 | AX-170529202 | 0 | T | G | 0.175 | 2.39E-02 | 9.92E-02 | 2.68E+00 | 2.76E+00 | 8.06E-01 | no | Actl6a | intron_variant | MODIFIER |
| 3 | 32718227 | AX-97821244 | 0 | C | T | 0.175 | 2.39E-02 | 9.92E-02 | 2.68E+00 | 2.76E+00 | 8.06E-01 | no | Actl6a | intron_variant | MODIFIER |
| 3 | 32732572 | AX-170398910 | 0 | G | C | 0.175 | 2.39E-02 | 9.92E-02 | 2.68E+00 | 2.76E+00 | 8.06E-01 | no | Ndufb5 | upstream_gene_variant | MODIFIER |
| 3 | 32742160 | AX-170301957 | 0 | A | G | 0.175 | 2.39E-02 | 9.92E-02 | 2.68E+00 | 2.76E+00 | 8.06E-01 | no | Ndufb5 | intron_variant,NMD_transcript_variant | MODIFIER |
| 3 | 32742184 | AX-170330690 | 0 | T | C | 0.175 | 2.39E-02 | 9.92E-02 | 2.68E+00 | 2.76E+00 | 8.06E-01 | no | Ndufb5 | intron_variant,NMD_transcript_variant | MODIFIER |
| 3 | 32742219 | AX-97848600 | 0 | G | T | 0.175 | 2.39E-02 | 9.92E-02 | 2.68E+00 | 2.76E+00 | 8.06E-01 | no | Ndufb5 | intron_variant,NMD_transcript_variant | MODIFIER |
| 18 | 66174796 | AX-170366794 | 0 | G | C | 0.096 | 4.24E-02 | 1.70E-01 | 2.61E+00 | 2.72E+00 | 8.06E-01 | no | Ccbe1 | intron_variant,NMD_transcript_variant | MODIFIER |
| 18 | 66176207 | AX-170341909 | 0 | A | G | 0.096 | 4.24E-02 | 1.70E-01 | 2.61E+00 | 2.72E+00 | 8.06E-01 | no | Ccbe1 | intron_variant,NMD_transcript_variant | MODIFIER |
| 18 | 66235474 | AX-170368482 | 0 | C | G | 0.096 | 4.24E-02 | 1.70E-01 | 2.61E+00 | 2.72E+00 | 8.06E-01 | no | Ccbe1 | intron_variant,NMD_transcript_variant | MODIFIER |
| 18 | 66262148 | AX-170449467 | 0 | G | A | 0.096 | 4.24E-02 | 1.70E-01 | 2.61E+00 | 2.72E+00 | 8.06E-01 | no | Ccbe1 | intron_variant,NMD_transcript_variant | MODIFIER |
| 18 | 66262699 | AX-170416318 | 0 | C | A | 0.096 | 4.24E-02 | 1.70E-01 | 2.61E+00 | 2.72E+00 | 8.06E-01 | no | Ccbe1 | intron_variant,NMD_transcript_variant | MODIFIER |
| 18 | 66264342 | AX-170227171 | 0 | A | C | 0.096 | 4.24E-02 | 1.70E-01 | 2.61E+00 | 2.72E+00 | 8.06E-01 | no | Ccbe1 | intron_variant,NMD_transcript_variant | MODIFIER |
| 18 | 66264588 | AX-170301021 | 0 | T | C | 0.096 | 4.24E-02 | 1.70E-01 | 2.61E+00 | 2.72E+00 | 8.06E-01 | no | Ccbe1 | intron_variant,NMD_transcript_variant | MODIFIER |
| 18 | 66266566 | AX-170274146 | 0 | A | T | 0.096 | 4.24E-02 | 1.70E-01 | 2.61E+00 | 2.72E+00 | 8.06E-01 | no | Ccbe1 | intron_variant,NMD_transcript_variant | MODIFIER |
| 18 | 66267305 | AX-170464455 | 0 | A | G | 0.096 | 4.24E-02 | 1.70E-01 | 2.61E+00 | 2.72E+00 | 8.06E-01 | no | Ccbe1 | intron_variant,NMD_transcript_variant | MODIFIER |
| 18 | 66285378 | AX-170061147 | 0 | G | C | 0.096 | 4.24E-02 | 1.70E-01 | 2.61E+00 | 2.72E+00 | 8.06E-01 | no | Ccbe1 | intron_variant,NMD_transcript_variant | MODIFIER |
| 18 | 66289976 | AX-170061149 | 0 | A | G | 0.096 | 4.24E-02 | 1.70E-01 | 2.61E+00 | 2.72E+00 | 8.06E-01 | no | Ccbe1 | intron_variant,NMD_transcript_variant | MODIFIER |
| 18 | 66292321 | AX-97835002 | 0 | A | C | 0.096 | 4.24E-02 | 1.70E-01 | 2.61E+00 | 2.72E+00 | 8.06E-01 | no | Ccbe1 | upstream_gene_variant | MODIFIER |
| 18 | 66292587 | AX-170174010 | 0 | C | T | 0.096 | 4.24E-02 | 1.70E-01 | 2.61E+00 | 2.72E+00 | 8.06E-01 | no | Ccbe1 | upstream_gene_variant | MODIFIER |
| 18 | 66293054 | AX-170236313 | 0 | G | A | 0.096 | 4.24E-02 | 1.70E-01 | 2.61E+00 | 2.72E+00 | 8.06E-01 | no | Ccbe1 | upstream_gene_variant | MODIFIER |
| 18 | 66293657 | AX-170240133 | 0 | T | C | 0.096 | 4.24E-02 | 1.70E-01 | 2.61E+00 | 2.72E+00 | 8.06E-01 | no | Ccbe1 | upstream_gene_variant | MODIFIER |
| 18 | 66455789 | AX-170265454 | 0 | C | G | 0.096 | 4.24E-02 | 1.70E-01 | 2.61E+00 | 2.72E+00 | 8.06E-01 | no | Pmaip1 | upstream_gene_variant | MODIFIER |
| 18 | 66457142 | AX-170358858 | 0 | C | A | 0.096 | 4.24E-02 | 1.70E-01 | 2.61E+00 | 2.72E+00 | 8.06E-01 | no | Pmaip1 | upstream_gene_variant | MODIFIER |
| 18 | 66476538 | AX-170329255 | 0 | G | A | 0.096 | 4.24E-02 | 1.70E-01 | 2.61E+00 | 2.72E+00 | 8.06E-01 | no | | regulatory_region_variant | MODIFIER |
| 18 | 66551001 | AX-170498783 | 0 | T | A | 0.096 | 4.24E-02 | 1.70E-01 | 2.61E+00 | 2.72E+00 | 8.06E-01 | no | Gm24679 | upstream_gene_variant | MODIFIER |
| 18 | 66551648 | AX-170432294 | 0 | T | C | 0.096 | 4.24E-02 | 1.70E-01 | 2.61E+00 | 2.72E+00 | 8.06E-01 | no | Gm24679 | upstream_gene_variant | MODIFIER |
| 18 | 66556355 | AX-170167843 | 0 | C | T | 0.096 | 4.24E-02 | 1.70E-01 | 2.61E+00 | 2.72E+00 | 8.06E-01 | no | | intergenic_variant | MODIFIER |
| 18 | 66593825 | AX-170360346 | 0 | C | G | 0.096 | 4.24E-02 | 1.70E-01 | 2.61E+00 | 2.72E+00 | 8.06E-01 | no | | intergenic_variant | MODIFIER |
| 18 | 66632827 | AX-170384694 | 0 | G | C | 0.096 | 4.24E-02 | 1.70E-01 | 2.61E+00 | 2.72E+00 | 8.06E-01 | no | AC133607.4 | downstream_gene_variant | MODIFIER |
| 18 | 66657559 | AX-170453065 | 0 | A | T | 0.096 | 4.24E-02 | 1.70E-01 | 2.61E+00 | 2.72E+00 | 8.06E-01 | no | AC133607.2 | intron_variant,non_coding_transcript_variant | MODIFIER |
| 18 | 66658134 | AX-170130898 | 0 | A | G | 0.096 | 4.24E-02 | 1.70E-01 | 2.61E+00 | 2.72E+00 | 8.06E-01 | no | AC133607.2 | intron_variant,non_coding_transcript_variant | MODIFIER |
| 18 | 66700808 | AX-97786460 | 0 | T | C | 0.096 | 4.24E-02 | 1.70E-01 | 2.61E+00 | 2.72E+00 | 8.06E-01 | no | | intergenic_variant | MODIFIER |
| 18 | 66716488 | AX-170426448 | 0 | T | A | 0.096 | 4.24E-02 | 1.70E-01 | 2.61E+00 | 2.72E+00 | 8.06E-01 | no | | intergenic_variant | MODIFIER |
| 18 | 66733195 | AX-170146812 | 0 | A | C | 0.096 | 4.24E-02 | 1.70E-01 | 2.61E+00 | 2.72E+00 | 8.06E-01 | no | | intergenic_variant | MODIFIER |
| 18 | 66738678 | AX-170310659 | 0 | T | A | 0.096 | 4.24E-02 | 1.70E-01 | 2.61E+00 | 2.72E+00 | 8.06E-01 | no | | intergenic_variant | MODIFIER |
| 18 | 66743672 | AX-97852536 | 0 | T | C | 0.096 | 4.24E-02 | 1.70E-01 | 2.61E+00 | 2.72E+00 | 8.06E-01 | no | | intergenic_variant | MODIFIER |

| | | | | | | | | | | | | | | | |
|---|---|---|---|---|---|---|---|---|---|---|---|---|---|---|---|
| 18 | 66804449 | AX-170395692 | 0 | G | C | 0.096 | 4.24E-02 | 1.70E-01 | 2.61E+00 | 2.72E+00 | 8.06E-01 | no | AC133607.3 | downstream_gene_variant | MODIFIER |
| 18 | 66805330 | AX-97841757 | 0 | C | T | 0.096 | 4.24E-02 | 1.70E-01 | 2.61E+00 | 2.72E+00 | 8.06E-01 | no | AC133607.3 | downstream_gene_variant | MODIFIER |
| 18 | 66818290 | AX-97799359 | 0 | C | A | 0.096 | 4.24E-02 | 1.70E-01 | 2.61E+00 | 2.72E+00 | 8.06E-01 | no | | intergenic_variant | MODIFIER |
| 18 | 66836240 | AX-170525260 | 0 | C | T | 0.096 | 4.24E-02 | 1.70E-01 | 2.61E+00 | 2.72E+00 | 8.06E-01 | no | | intergenic_variant | MODIFIER |
| 18 | 66836541 | AX-170425882 | 0 | G | A | 0.096 | 4.24E-02 | 1.70E-01 | 2.61E+00 | 2.72E+00 | 8.06E-01 | no | | intergenic_variant | MODIFIER |
| 18 | 66859942 | AX-170015891 | 0 | T | C | 0.096 | 4.24E-02 | 1.70E-01 | 2.61E+00 | 2.72E+00 | 8.06E-01 | no | Mc4r | synonymous_variant | LOW |
| 18 | 66872409 | AX-170108149 | 0 | C | T | 0.096 | 4.24E-02 | 1.70E-01 | 2.61E+00 | 2.72E+00 | 8.06E-01 | no | | intergenic_variant | MODIFIER |
| 18 | 66877370 | AX-170093925 | 0 | C | T | 0.096 | 4.24E-02 | 1.70E-01 | 2.61E+00 | 2.72E+00 | 8.06E-01 | no | | intergenic_variant | MODIFIER |
| 18 | 66900335 | AX-170015892 | 0 | T | A | 0.096 | 4.24E-02 | 1.70E-01 | 2.61E+00 | 2.72E+00 | 8.06E-01 | no | | intergenic_variant | MODIFIER |
| 18 | 66907659 | AX-170015895 | 0 | G | A | 0.096 | 4.24E-02 | 1.70E-01 | 2.61E+00 | 2.72E+00 | 8.06E-01 | no | | intergenic_variant | MODIFIER |
| 18 | 66910410 | AX-170015894 | 0 | T | A | 0.096 | 4.24E-02 | 1.70E-01 | 2.61E+00 | 2.72E+00 | 8.06E-01 | no | | intergenic_variant | MODIFIER |
| 18 | 66925278 | AX-170088365 | 0 | A | G | 0.096 | 4.24E-02 | 1.70E-01 | 2.61E+00 | 2.72E+00 | 8.06E-01 | no | AC127234.1 | downstream_gene_variant | MODIFIER |
| 18 | 66925385 | AX-170169240 | 0 | T | C | 0.096 | 4.24E-02 | 1.70E-01 | 2.61E+00 | 2.72E+00 | 8.06E-01 | no | AC127234.1 | downstream_gene_variant | MODIFIER |
| 18 | 67329182 | AX-97782066 | 0 | C | T | 0.096 | 4.24E-02 | 1.70E-01 | 2.61E+00 | 2.72E+00 | 8.06E-01 | no | B430212C06Rik | intron_variant,non_coding_transcript_variant | MODIFIER |
| 18 | 67440760 | AX-170225829 | 0 | A | G | 0.096 | 4.24E-02 | 1.70E-01 | 2.61E+00 | 2.72E+00 | 8.06E-01 | no | Afg3l2 | intron_variant | MODIFIER |
| 18 | 67651265 | AX-97851465 | 0 | A | G | 0.096 | 4.24E-02 | 1.70E-01 | 2.61E+00 | 2.72E+00 | 8.06E-01 | no | Psmg2 | intron_variant | MODIFIER |
| 18 | 67664020 | AX-170224746 | 0 | A | G | 0.096 | 4.24E-02 | 1.70E-01 | 2.61E+00 | 2.72E+00 | 8.06E-01 | no | Ptpn2 | downstream_gene_variant | MODIFIER |
| 18 | 67695952 | AX-170404415 | 0 | T | C | 0.096 | 4.24E-02 | 1.70E-01 | 2.61E+00 | 2.72E+00 | 8.06E-01 | no | Ptpn2 | intron_variant | MODIFIER |
| 18 | 67764820 | AX-170360530 | 0 | C | T | 0.096 | 4.24E-02 | 1.70E-01 | 2.61E+00 | 2.72E+00 | 8.06E-01 | no | 4930549G23Rik | downstream_gene_variant | MODIFIER |
| 18 | 67845651 | AX-170268266 | 0 | T | C | 0.096 | 4.24E-02 | 1.70E-01 | 2.61E+00 | 2.72E+00 | 8.06E-01 | no | Cep192 | intron_variant | MODIFIER |
| 18 | 67848222 | AX-170219613 | 0 | A | T | 0.096 | 4.24E-02 | 1.70E-01 | 2.61E+00 | 2.72E+00 | 8.06E-01 | no | Cep192 | intron_variant | MODIFIER |
| 18 | 67903710 | AX-170518241 | 0 | T | C | 0.096 | 4.24E-02 | 1.70E-01 | 2.61E+00 | 2.72E+00 | 8.06E-01 | no | AC127236.1 | upstream_gene_variant | MODIFIER |
| 18 | 67906663 | AX-170330250 | 0 | A | G | 0.096 | 4.24E-02 | 1.70E-01 | 2.61E+00 | 2.72E+00 | 8.06E-01 | no | AC127236.1 | upstream_gene_variant | MODIFIER |
| 18 | 68185097 | AX-97785240 | 0 | A | G | 0.096 | 4.24E-02 | 1.70E-01 | 2.61E+00 | 2.72E+00 | 8.06E-01 | no | Ldlrad4 | intron_variant | MODIFIER |
| 18 | 68186613 | AX-170061190 | 0 | C | T | 0.096 | 4.24E-02 | 1.70E-01 | 2.61E+00 | 2.72E+00 | 8.06E-01 | no | Ldlrad4 | intron_variant | MODIFIER |
| 18 | 68279444 | AX-170236416 | 0 | T | G | 0.096 | 4.24E-02 | 1.70E-01 | 2.61E+00 | 2.72E+00 | 8.06E-01 | no | Fam210a | intron_variant | MODIFIER |
| 18 | 68454200 | AX-97844987 | 0 | C | A | 0.096 | 4.24E-02 | 1.70E-01 | 2.61E+00 | 2.72E+00 | 8.06E-01 | no | | intergenic_variant | MODIFIER |
| 18 | 68515501 | AX-170501780 | 0 | C | T | 0.096 | 4.24E-02 | 1.70E-01 | 2.61E+00 | 2.72E+00 | 8.06E-01 | no | | intergenic_variant | MODIFIER |
| 18 | 68521145 | AX-170238545 | 0 | A | G | 0.096 | 4.24E-02 | 1.70E-01 | 2.61E+00 | 2.72E+00 | 8.06E-01 | no | | intergenic_variant | MODIFIER |
| 18 | 68521229 | AX-170472547 | 0 | T | C | 0.096 | 4.24E-02 | 1.70E-01 | 2.61E+00 | 2.72E+00 | 8.06E-01 | no | | intergenic_variant | MODIFIER |
| 18 | 68529652 | AX-170125821 | 0 | C | A | 0.096 | 4.24E-02 | 1.70E-01 | 2.61E+00 | 2.72E+00 | 8.06E-01 | no | | intergenic_variant | MODIFIER |
| 18 | 68533248 | AX-170015915 | 0 | A | C | 0.096 | 4.24E-02 | 1.70E-01 | 2.61E+00 | 2.72E+00 | 8.06E-01 | no | | intergenic_variant | MODIFIER |
| 18 | 68604730 | AX-97832491 | 0 | T | C | 0.096 | 4.24E-02 | 1.70E-01 | 2.61E+00 | 2.72E+00 | 8.06E-01 | no | | intergenic_variant | MODIFIER |
| 18 | 68660089 | AX-170206429 | 0 | A | G | 0.096 | 4.24E-02 | 1.70E-01 | 2.61E+00 | 2.72E+00 | 8.06E-01 | no | | intergenic_variant | MODIFIER |
| 18 | 68669276 | AX-170398509 | 0 | G | T | 0.096 | 4.24E-02 | 1.70E-01 | 2.61E+00 | 2.72E+00 | 8.06E-01 | no | | intergenic_variant | MODIFIER |
| 18 | 68670962 | AX-170228244 | 0 | C | T | 0.096 | 4.24E-02 | 1.70E-01 | 2.61E+00 | 2.72E+00 | 8.06E-01 | no | | intergenic_variant | MODIFIER |
| 18 | 68671837 | AX-170396448 | 0 | G | A | 0.096 | 4.24E-02 | 1.70E-01 | 2.61E+00 | 2.72E+00 | 8.06E-01 | no | | intergenic_variant | MODIFIER |
| 18 | 68672267 | AX-170490775 | 0 | G | T | 0.096 | 4.24E-02 | 1.70E-01 | 2.61E+00 | 2.72E+00 | 8.06E-01 | no | | intergenic_variant | MODIFIER |
| 18 | 68741304 | AX-170533121 | 0 | T | G | 0.096 | 4.24E-02 | 1.70E-01 | 2.61E+00 | 2.72E+00 | 8.06E-01 | no | Mir6356 | downstream_gene_variant | MODIFIER |
| 18 | 68746382 | AX-170315838 | 0 | T | C | 0.096 | 4.24E-02 | 1.70E-01 | 2.61E+00 | 2.72E+00 | 8.06E-01 | no | | intergenic_variant | MODIFIER |
| 18 | 68857033 | AX-170061205 | 0 | A | G | 0.096 | 4.24E-02 | 1.70E-01 | 2.61E+00 | 2.72E+00 | 8.06E-01 | no | | intergenic_variant | MODIFIER |
| 18 | 69030750 | AX-170487730 | 0 | A | C | 0.096 | 4.24E-02 | 1.70E-01 | 2.61E+00 | 2.72E+00 | 8.06E-01 | no | | intergenic_variant | MODIFIER |
| 18 | 69043253 | AX-170185182 | 0 | G | T | 0.096 | 4.24E-02 | 1.70E-01 | 2.61E+00 | 2.72E+00 | 8.06E-01 | no | | intergenic_variant | MODIFIER |
| 18 | 69043307 | AX-170495517 | 0 | A | G | 0.096 | 4.24E-02 | 1.70E-01 | 2.61E+00 | 2.72E+00 | 8.06E-01 | no | | intergenic_variant | MODIFIER |
| 18 | 69046300 | AX-97786563 | 0 | T | C | 0.096 | 4.24E-02 | 1.70E-01 | 2.61E+00 | 2.72E+00 | 8.06E-01 | no | | intergenic_variant | MODIFIER |
| 18 | 69076156 | AX-170167171 | 0 | A | G | 0.096 | 4.24E-02 | 1.70E-01 | 2.61E+00 | 2.72E+00 | 8.06E-01 | no | | intergenic_variant | MODIFIER |
| 18 | 69076196 | AX-170387259 | 0 | T | C | 0.096 | 4.24E-02 | 1.70E-01 | 2.61E+00 | 2.72E+00 | 8.06E-01 | no | | intergenic_variant | MODIFIER |
| 18 | 69096775 | AX-170501540 | 0 | G | A | 0.096 | 4.24E-02 | 1.70E-01 | 2.61E+00 | 2.72E+00 | 8.06E-01 | no | | intergenic_variant | MODIFIER |
| 18 | 69097244 | AX-170181721 | 0 | C | T | 0.096 | 4.24E-02 | 1.70E-01 | 2.61E+00 | 2.72E+00 | 8.06E-01 | no | | intergenic_variant | MODIFIER |
| 18 | 69099615 | AX-170015919 | 0 | G | A | 0.096 | 4.24E-02 | 1.70E-01 | 2.61E+00 | 2.72E+00 | 8.06E-01 | no | | intergenic_variant | MODIFIER |
| 18 | 69099995 | AX-170093935 | 0 | T | G | 0.096 | 4.24E-02 | 1.70E-01 | 2.61E+00 | 2.72E+00 | 8.06E-01 | no | | intergenic_variant | MODIFIER |
| 18 | 69108398 | AX-170015920 | 0 | T | C | 0.096 | 4.24E-02 | 1.70E-01 | 2.61E+00 | 2.72E+00 | 8.06E-01 | no | | intergenic_variant | MODIFIER |
| 18 | 69124354 | AX-170134427 | 0 | T | C | 0.096 | 4.24E-02 | 1.70E-01 | 2.61E+00 | 2.72E+00 | 8.06E-01 | no | | regulatory_region_variant | MODIFIER |
| 18 | 69124377 | AX-97844190 | 0 | T | C | 0.096 | 4.24E-02 | 1.70E-01 | 2.61E+00 | 2.72E+00 | 8.06E-01 | no | | regulatory_region_variant | MODIFIER |

| | | | | | | | | | | | | | | | |
|---|---|---|---|---|---|---|---|---|---|---|---|---|---|---|---|
| 18 | 69130732 | AX-170243858 | 0 | A | G | 0.096 | 4.24E-02 | 1.70E-01 | 2.61E+00 | 2.72E+00 | 8.06E-01 | no | | intergenic_variant | MODIFIER |
| 18 | 69135422 | AX-170501762 | 0 | G | T | 0.096 | 4.24E-02 | 1.70E-01 | 2.61E+00 | 2.72E+00 | 8.06E-01 | no | | intergenic_variant | MODIFIER |
| 18 | 69135759 | AX-170175893 | 0 | C | T | 0.096 | 4.24E-02 | 1.70E-01 | 2.61E+00 | 2.72E+00 | 8.06E-01 | no | | intergenic_variant | MODIFIER |
| 18 | 69152303 | AX-170340176 | 0 | A | G | 0.096 | 4.24E-02 | 1.70E-01 | 2.61E+00 | 2.72E+00 | 8.06E-01 | no | | regulatory_region_variant | MODIFIER |
| 18 | 69173996 | AX-97820739 | 0 | G | A | 0.096 | 4.24E-02 | 1.70E-01 | 2.61E+00 | 2.72E+00 | 8.06E-01 | no | Gm23536 | non_coding_transcript_exon_variant | MODIFIER |
| 18 | 69192625 | AX-97814071 | 0 | C | T | 0.096 | 4.24E-02 | 1.70E-01 | 2.61E+00 | 2.72E+00 | 8.06E-01 | no | Gm31294 | downstream_gene_variant | MODIFIER |
| 18 | 69213849 | AX-170330275 | 0 | A | G | 0.096 | 4.24E-02 | 1.70E-01 | 2.61E+00 | 2.72E+00 | 8.06E-01 | no | AC121263.2 | upstream_gene_variant | MODIFIER |
| 18 | 69213916 | AX-170358442 | 0 | T | C | 0.096 | 4.24E-02 | 1.70E-01 | 2.61E+00 | 2.72E+00 | 8.06E-01 | no | AC121263.2 | upstream_gene_variant | MODIFIER |
| 18 | 69223561 | AX-170475043 | 0 | C | T | 0.096 | 4.24E-02 | 1.70E-01 | 2.61E+00 | 2.72E+00 | 8.06E-01 | no | AC121263.2 | downstream_gene_variant | MODIFIER |
| 18 | 69231941 | AX-170292308 | 0 | T | C | 0.096 | 4.24E-02 | 1.70E-01 | 2.61E+00 | 2.72E+00 | 8.06E-01 | no | | intergenic_variant | MODIFIER |
| 18 | 69281201 | AX-170156952 | 0 | G | A | 0.096 | 4.24E-02 | 1.70E-01 | 2.61E+00 | 2.72E+00 | 8.06E-01 | no | | intergenic_variant | MODIFIER |
| 18 | 69291248 | AX-170356546 | 0 | G | T | 0.096 | 4.24E-02 | 1.70E-01 | 2.61E+00 | 2.72E+00 | 8.06E-01 | no | | intergenic_variant | MODIFIER |
| 18 | 69298860 | AX-170417501 | 0 | T | A | 0.096 | 4.24E-02 | 1.70E-01 | 2.61E+00 | 2.72E+00 | 8.06E-01 | no | | intergenic_variant | MODIFIER |
| 18 | 69299041 | AX-170188267 | 0 | C | G | 0.096 | 4.24E-02 | 1.70E-01 | 2.61E+00 | 2.72E+00 | 8.06E-01 | no | | intergenic_variant | MODIFIER |
| 18 | 69315857 | AX-170276266 | 0 | C | T | 0.096 | 4.24E-02 | 1.70E-01 | 2.61E+00 | 2.72E+00 | 8.06E-01 | no | | intergenic_variant | MODIFIER |
| 18 | 69317235 | AX-170422844 | 0 | A | T | 0.096 | 4.24E-02 | 1.70E-01 | 2.61E+00 | 2.72E+00 | 8.06E-01 | no | | intergenic_variant | MODIFIER |
| 18 | 69317522 | AX-170404518 | 0 | C | T | 0.096 | 4.24E-02 | 1.70E-01 | 2.61E+00 | 2.72E+00 | 8.06E-01 | no | | intergenic_variant | MODIFIER |
| 18 | 69319804 | AX-170061222 | 0 | A | G | 0.096 | 4.24E-02 | 1.70E-01 | 2.61E+00 | 2.72E+00 | 8.06E-01 | no | | intergenic_variant | MODIFIER |
| 18 | 69331341 | AX-170495799 | 0 | G | A | 0.096 | 4.24E-02 | 1.70E-01 | 2.61E+00 | 2.72E+00 | 8.06E-01 | no | | intergenic_variant | MODIFIER |
| 18 | 69339695 | AX-170162787 | 0 | C | G | 0.096 | 4.24E-02 | 1.70E-01 | 2.61E+00 | 2.72E+00 | 8.06E-01 | no | Tcf4 | upstream_gene_variant | MODIFIER |
| 18 | 69363789 | AX-170444378 | 0 | A | C | 0.096 | 4.24E-02 | 1.70E-01 | 2.61E+00 | 2.72E+00 | 8.06E-01 | no | Tcf4 | intron_variant | MODIFIER |
| 18 | 69373852 | AX-97835758 | 0 | G | A | 0.096 | 4.24E-02 | 1.70E-01 | 2.61E+00 | 2.72E+00 | 8.06E-01 | no | Tcf4 | intron_variant | MODIFIER |
| 18 | 69462446 | AX-170505367 | 0 | C | G | 0.096 | 4.24E-02 | 1.70E-01 | 2.61E+00 | 2.72E+00 | 8.06E-01 | no | Tcf4 | intron_variant | MODIFIER |
| 18 | 69469988 | AX-170373819 | 0 | T | C | 0.096 | 4.24E-02 | 1.70E-01 | 2.61E+00 | 2.72E+00 | 8.06E-01 | no | Tcf4 | intron_variant | MODIFIER |
| 18 | 69476780 | AX-170281302 | 0 | T | C | 0.096 | 4.24E-02 | 1.70E-01 | 2.61E+00 | 2.72E+00 | 8.06E-01 | no | Tcf4 | intron_variant | MODIFIER |
| 18 | 69491392 | AX-170442243 | 0 | T | C | 0.096 | 4.24E-02 | 1.70E-01 | 2.61E+00 | 2.72E+00 | 8.06E-01 | no | Tcf4 | intron_variant | MODIFIER |
| 18 | 69581692 | AX-170465991 | 0 | G | A | 0.096 | 4.24E-02 | 1.70E-01 | 2.61E+00 | 2.72E+00 | 8.06E-01 | no | Tcf4 | intron_variant | MODIFIER |
| 18 | 69763903 | AX-170269392 | 0 | A | T | 0.096 | 4.24E-02 | 1.70E-01 | 2.61E+00 | 2.72E+00 | 8.06E-01 | no | | intergenic_variant | MODIFIER |
| 18 | 69819005 | AX-170386376 | 0 | T | C | 0.096 | 4.24E-02 | 1.70E-01 | 2.61E+00 | 2.72E+00 | 8.06E-01 | no | | intergenic_variant | MODIFIER |
| 18 | 69832141 | AX-170156011 | 0 | C | G | 0.096 | 4.24E-02 | 1.70E-01 | 2.61E+00 | 2.72E+00 | 8.06E-01 | no | | intergenic_variant | MODIFIER |
| 1 | 156185001 | AX-170398743 | 1 | A | G | 0.223 | 3.48E-02 | 1.39E-01 | 2.65E+00 | 2.69E+00 | 8.06E-01 | no | Fam163a | intron_variant | MODIFIER |
| 12 | 103834238 | AX-170449071 | 2 | C | A | 0.136 | -4.58E-02 | 1.91E-01 | 2.69E+00 | 2.76E+00 | 8.06E-01 | no | Serpina1b | upstream_gene_variant | MODIFIER |
| 5 | 80901588 | AX-170365051 | 2 | T | C | 0.418 | 2.52E-02 | 1.02E-01 | 2.64E+00 | 2.71E+00 | 8.06E-01 | no | Gm2199 | non_coding_transcript_exon_variant | MODIFIER |
| 15 | 11089018 | AX-170307934 | 2 | C | T | 0.264 | -2.51E-02 | 1.00E-01 | 2.63E+00 | 2.71E+00 | 8.06E-01 | no | Adamts12 | intron_variant | MODIFIER |
| 14 | 27925501 | AX-170252516 | 1 | A | G | 0.045 | -3.99E-02 | 1.51E-01 | 2.55E+00 | 2.69E+00 | 8.06E-01 | no | Erc2 | intron_variant | MODIFIER |
| 20 | 130688847 | AX-97782744 | 0 | A | G | 0.149 | 3.31E-02 | 1.35E-01 | 2.64E+00 | 2.73E+00 | 8.06E-01 | no | | intergenic_variant | MODIFIER |
| 20 | 131850300 | AX-97840563 | 0 | G | A | 0.149 | 3.31E-02 | 1.35E-01 | 2.64E+00 | 2.73E+00 | 8.06E-01 | no | Gm4994 | upstream_gene_variant | MODIFIER |
| 20 | 133505754 | AX-170503514 | 0 | A | G | 0.149 | 3.31E-02 | 1.35E-01 | 2.64E+00 | 2.73E+00 | 8.06E-01 | no | | intergenic_variant | MODIFIER |
| 20 | 133670104 | AX-170034992 | 0 | G | A | 0.149 | 3.31E-02 | 1.35E-01 | 2.64E+00 | 2.73E+00 | 8.06E-01 | no | Pcdh19 | intron_variant | MODIFIER |
| 20 | 133756507 | AX-170153720 | 0 | C | T | 0.149 | 3.31E-02 | 1.35E-01 | 2.64E+00 | 2.73E+00 | 8.06E-01 | no | | intergenic_variant | MODIFIER |
| 20 | 133759846 | AX-170403791 | 0 | G | C | 0.149 | 3.31E-02 | 1.35E-01 | 2.64E+00 | 2.73E+00 | 8.06E-01 | no | | intergenic_variant | MODIFIER |
| 20 | 133774509 | AX-170231915 | 0 | A | G | 0.149 | 3.31E-02 | 1.35E-01 | 2.64E+00 | 2.73E+00 | 8.06E-01 | no | | intergenic_variant | MODIFIER |
| 20 | 133775535 | AX-97787063 | 0 | G | A | 0.149 | 3.31E-02 | 1.35E-01 | 2.64E+00 | 2.73E+00 | 8.06E-01 | no | | intergenic_variant | MODIFIER |
| 20 | 133783044 | AX-97786475 | 0 | A | G | 0.149 | 3.31E-02 | 1.35E-01 | 2.64E+00 | 2.73E+00 | 8.06E-01 | no | | intergenic_variant | MODIFIER |
| 20 | 133814502 | AX-170396885 | 0 | G | A | 0.149 | 3.31E-02 | 1.35E-01 | 2.64E+00 | 2.73E+00 | 8.06E-01 | no | | intergenic_variant | MODIFIER |
| 20 | 133827828 | AX-170499748 | 0 | A | G | 0.149 | 3.31E-02 | 1.35E-01 | 2.64E+00 | 2.73E+00 | 8.06E-01 | no | | intergenic_variant | MODIFIER |
| 20 | 133836992 | AX-170161555 | 0 | C | T | 0.149 | 3.31E-02 | 1.35E-01 | 2.64E+00 | 2.73E+00 | 8.06E-01 | no | | intergenic_variant | MODIFIER |
| 20 | 138478499 | AX-97792168 | 0 | C | T | 0.149 | 3.31E-02 | 1.35E-01 | 2.64E+00 | 2.73E+00 | 8.06E-01 | no | Il1rapl2 | intron_variant | MODIFIER |
| 20 | 139810681 | AX-170316586 | 0 | T | C | 0.149 | 3.31E-02 | 1.35E-01 | 2.64E+00 | 2.73E+00 | 8.06E-01 | no | Cldn2 | 3_prime_UTR_variant | MODIFIER |
| 20 | 139827830 | AX-170240526 | 0 | T | A | 0.149 | 3.31E-02 | 1.35E-01 | 2.64E+00 | 2.73E+00 | 8.06E-01 | no | Morc4 | intron_variant | MODIFIER |
| 20 | 139840710 | AX-97801382 | 0 | G | A | 0.149 | 3.31E-02 | 1.35E-01 | 2.64E+00 | 2.73E+00 | 8.06E-01 | no | Morc4 | missense_variant | MODERATE |
| 20 | 139903860 | AX-170422192 | 0 | T | C | 0.149 | 3.31E-02 | 1.35E-01 | 2.64E+00 | 2.73E+00 | 8.06E-01 | no | Rbm41 | intron_variant | MODIFIER |
| 20 | 139907580 | AX-170336171 | 0 | G | C | 0.149 | 3.31E-02 | 1.35E-01 | 2.64E+00 | 2.73E+00 | 8.06E-01 | no | Rbm41 | intron_variant | MODIFIER |
| 20 | 139907735 | AX-97856705 | 0 | A | G | 0.149 | 3.31E-02 | 1.35E-01 | 2.64E+00 | 2.73E+00 | 8.06E-01 | no | Rbm41 | intron_variant | MODIFIER |

| | | | | | | | | | | | | | | | |
|---|---|---|---|---|---|---|---|---|---|---|---|---|---|---|---|
| 20 | 139909103 | AX-170278780 | 0 | T | A | 0.149 | 3.31E-02 | 1.35E-01 | 2.64E+00 | 2.73E+00 | 8.06E-01 | no | Rbm41 | intron_variant | MODIFIER |
| 20 | 139940754 | AX-97782720 | 0 | T | C | 0.149 | 3.31E-02 | 1.35E-01 | 2.64E+00 | 2.73E+00 | 8.06E-01 | no | Rbm41 | downstream_gene_variant | MODIFIER |
| 20 | 139968326 | AX-170413109 | 0 | A | G | 0.149 | 3.31E-02 | 1.35E-01 | 2.64E+00 | 2.73E+00 | 8.06E-01 | no | Rbm41 | missense_variant | MODERATE |
| 20 | 139993694 | AX-170213415 | 0 | C | T | 0.149 | 3.31E-02 | 1.35E-01 | 2.64E+00 | 2.73E+00 | 8.06E-01 | no | Rbm41 | intron_variant | MODIFIER |
| 20 | 140026942 | AX-170035080 | 0 | G | T | 0.149 | 3.31E-02 | 1.35E-01 | 2.64E+00 | 2.73E+00 | 8.06E-01 | no | Nup62cl | intron_variant | MODIFIER |
| 20 | 140037001 | AX-170035077 | 0 | C | G | 0.149 | 3.31E-02 | 1.35E-01 | 2.64E+00 | 2.73E+00 | 8.06E-01 | no | Nup62cl | intron_variant | MODIFIER |
| 20 | 140092615 | AX-168404557 | 0 | T | C | 0.149 | 3.31E-02 | 1.35E-01 | 2.64E+00 | 2.73E+00 | 8.06E-01 | no | Pih1h3b | intron_variant | MODIFIER |
| 20 | 140094434 | AX-170035081 | 0 | T | C | 0.149 | 3.31E-02 | 1.35E-01 | 2.64E+00 | 2.73E+00 | 8.06E-01 | no | Pih1h3b | intron_variant | MODIFIER |
| 20 | 140393728 | AX-170499870 | 0 | T | C | 0.149 | 3.31E-02 | 1.35E-01 | 2.64E+00 | 2.73E+00 | 8.06E-01 | no | Frmpd3 | intron_variant,non_coding_transcript_variant | MODIFIER |
| 20 | 140548414 | AX-170179244 | 0 | A | G | 0.149 | 3.31E-02 | 1.35E-01 | 2.64E+00 | 2.73E+00 | 8.06E-01 | no | Tsc22d3 | intron_variant | MODIFIER |
| 20 | 140549033 | AX-97786568 | 0 | A | G | 0.149 | 3.31E-02 | 1.35E-01 | 2.64E+00 | 2.73E+00 | 8.06E-01 | no | Tsc22d3 | intron_variant | MODIFIER |
| 20 | 140556388 | AX-170344494 | 0 | G | A | 0.149 | 3.31E-02 | 1.35E-01 | 2.64E+00 | 2.73E+00 | 8.06E-01 | no | Tsc22d3 | intron_variant | MODIFIER |
| 20 | 140571798 | AX-97810848 | 0 | T | C | 0.149 | 3.31E-02 | 1.35E-01 | 2.64E+00 | 2.73E+00 | 8.06E-01 | no | Tsc22d3 | intron_variant | MODIFIER |
| 20 | 140576010 | AX-170369978 | 0 | T | C | 0.149 | 3.31E-02 | 1.35E-01 | 2.64E+00 | 2.73E+00 | 8.06E-01 | no | Tsc22d3 | intron_variant | MODIFIER |
| 20 | 140576405 | AX-170308077 | 0 | C | T | 0.149 | 3.31E-02 | 1.35E-01 | 2.64E+00 | 2.73E+00 | 8.06E-01 | no | Tsc22d3 | intron_variant | MODIFIER |
| 20 | 140667742 | AX-170035088 | 0 | T | A | 0.149 | 3.31E-02 | 1.35E-01 | 2.64E+00 | 2.73E+00 | 8.06E-01 | no | Mid2 | intron_variant | MODIFIER |
| 20 | 140678432 | AX-170035087 | 0 | A | G | 0.149 | 3.31E-02 | 1.35E-01 | 2.64E+00 | 2.73E+00 | 8.06E-01 | no | Mid2 | synonymous_variant | LOW |
| 20 | 140687733 | AX-170316888 | 0 | G | T | 0.149 | 3.31E-02 | 1.35E-01 | 2.64E+00 | 2.73E+00 | 8.06E-01 | no | Mid2 | intron_variant | MODIFIER |
| 20 | 140696912 | AX-170448986 | 0 | G | C | 0.149 | 3.31E-02 | 1.35E-01 | 2.64E+00 | 2.73E+00 | 8.06E-01 | no | Mid2 | intron_variant | MODIFIER |
| 20 | 140704588 | AX-170458735 | 0 | C | T | 0.149 | 3.31E-02 | 1.35E-01 | 2.64E+00 | 2.73E+00 | 8.06E-01 | no | Mid2 | intron_variant | MODIFIER |
| 20 | 140704679 | AX-97786168 | 0 | A | G | 0.149 | 3.31E-02 | 1.35E-01 | 2.64E+00 | 2.73E+00 | 8.06E-01 | no | Mid2 | intron_variant | MODIFIER |
| 20 | 140715469 | AX-170035090 | 0 | G | A | 0.149 | 3.31E-02 | 1.35E-01 | 2.64E+00 | 2.73E+00 | 8.06E-01 | no | Mid2 | intron_variant | MODIFIER |
| 20 | 140976454 | AX-97788373 | 0 | A | C | 0.149 | 3.31E-02 | 1.35E-01 | 2.64E+00 | 2.73E+00 | 8.06E-01 | no | Atg4a | intron_variant | MODIFIER |
| 20 | 140976724 | AX-170121307 | 0 | A | G | 0.149 | 3.31E-02 | 1.35E-01 | 2.64E+00 | 2.73E+00 | 8.06E-01 | no | Atg4a | intron_variant | MODIFIER |
| 20 | 140980339 | AX-97786668 | 0 | C | G | 0.149 | 3.31E-02 | 1.35E-01 | 2.64E+00 | 2.73E+00 | 8.06E-01 | no | Atg4a | intron_variant | MODIFIER |
| 20 | 141165407 | AX-97794813 | 0 | C | T | 0.149 | 3.31E-02 | 1.35E-01 | 2.64E+00 | 2.73E+00 | 8.06E-01 | no | Col4a6 | 3_prime_UTR_variant | MODIFIER |
| 20 | 141175663 | AX-170311248 | 0 | C | G | 0.149 | 3.31E-02 | 1.35E-01 | 2.64E+00 | 2.73E+00 | 8.06E-01 | no | Col4a6 | intron_variant | MODIFIER |
| 20 | 141175794 | AX-170302972 | 0 | G | A | 0.149 | 3.31E-02 | 1.35E-01 | 2.64E+00 | 2.73E+00 | 8.06E-01 | no | Col4a6 | intron_variant | MODIFIER |
| 20 | 143822899 | AX-97807606 | 0 | T | C | 0.149 | 3.31E-02 | 1.35E-01 | 2.64E+00 | 2.73E+00 | 8.06E-01 | no | Capn6 | intron_variant | MODIFIER |
| 20 | 143832022 | AX-170035128 | 0 | A | C | 0.149 | 3.31E-02 | 1.35E-01 | 2.64E+00 | 2.73E+00 | 8.06E-01 | no | Capn6 | upstream_gene_variant | MODIFIER |
| 20 | 143875257 | AX-170216357 | 0 | T | G | 0.149 | 3.31E-02 | 1.35E-01 | 2.64E+00 | 2.73E+00 | 8.06E-01 | no | Dcx | intron_variant | MODIFIER |
| 20 | 144420518 | AX-170035141 | 0 | G | A | 0.149 | 3.31E-02 | 1.35E-01 | 2.64E+00 | 2.73E+00 | 8.06E-01 | no | Trpc5 | intron_variant | MODIFIER |
| 20 | 144556550 | AX-170485070 | 0 | C | G | 0.149 | 3.31E-02 | 1.35E-01 | 2.64E+00 | 2.73E+00 | 8.06E-01 | no | Trpc5 | intron_variant | MODIFIER |
| 20 | 144570718 | AX-97807607 | 0 | A | C | 0.149 | 3.31E-02 | 1.35E-01 | 2.64E+00 | 2.73E+00 | 8.06E-01 | no | Trpc5 | intron_variant | MODIFIER |
| 20 | 144578790 | AX-170303473 | 0 | T | G | 0.149 | 3.31E-02 | 1.35E-01 | 2.64E+00 | 2.73E+00 | 8.06E-01 | no | Trpc5 | intron_variant | MODIFIER |
| 20 | 144599392 | AX-97785765 | 0 | C | A | 0.149 | 3.31E-02 | 1.35E-01 | 2.64E+00 | 2.73E+00 | 8.06E-01 | no | Trpc5 | intron_variant | MODIFIER |
| 20 | 144886207 | AX-170035153 | 0 | C | G | 0.149 | 3.31E-02 | 1.35E-01 | 2.64E+00 | 2.73E+00 | 8.06E-01 | no | Rtl4 | intron_variant | MODIFIER |
| 20 | 145057499 | AX-97781676 | 0 | G | C | 0.149 | 3.31E-02 | 1.35E-01 | 2.64E+00 | 2.73E+00 | 8.06E-01 | no | Rtl4 | intron_variant | MODIFIER |
| 20 | 145187648 | AX-170143959 | 0 | A | G | 0.149 | 3.31E-02 | 1.35E-01 | 2.64E+00 | 2.73E+00 | 8.06E-01 | no | | intergenic_variant | MODIFIER |
| 20 | 145208970 | AX-170258874 | 0 | T | C | 0.149 | 3.31E-02 | 1.35E-01 | 2.64E+00 | 2.73E+00 | 8.06E-01 | no | | intergenic_variant | MODIFIER |
| 20 | 145224175 | AX-170497518 | 0 | T | A | 0.149 | 3.31E-02 | 1.35E-01 | 2.64E+00 | 2.73E+00 | 8.06E-01 | no | Gm15073 | downstream_gene_variant | MODIFIER |
| 20 | 145310988 | AX-170090710 | 0 | A | G | 0.149 | 3.31E-02 | 1.35E-01 | 2.64E+00 | 2.73E+00 | 8.06E-01 | no | Lhfpl1 | intron_variant | MODIFIER |
| 20 | 145489609 | AX-170035167 | 0 | G | A | 0.149 | 3.31E-02 | 1.35E-01 | 2.64E+00 | 2.73E+00 | 8.06E-01 | no | Amot | intron_variant | MODIFIER |
| 20 | 145489698 | AX-170035165 | 0 | T | C | 0.149 | 3.31E-02 | 1.35E-01 | 2.64E+00 | 2.73E+00 | 8.06E-01 | no | Amot | intron_variant | MODIFIER |
| 20 | 145518527 | AX-97789951 | 0 | T | C | 0.149 | 3.31E-02 | 1.35E-01 | 2.64E+00 | 2.73E+00 | 8.06E-01 | no | | intergenic_variant | MODIFIER |
| 20 | 145536273 | AX-170035156 | 0 | C | G | 0.149 | 3.31E-02 | 1.35E-01 | 2.64E+00 | 2.73E+00 | 8.06E-01 | no | | intergenic_variant | MODIFIER |
| 20 | 145551456 | AX-97835834 | 0 | C | T | 0.149 | 3.31E-02 | 1.35E-01 | 2.64E+00 | 2.73E+00 | 8.06E-01 | no | | intergenic_variant | MODIFIER |
| 20 | 145610998 | AX-97796820 | 0 | T | C | 0.149 | 3.31E-02 | 1.35E-01 | 2.64E+00 | 2.73E+00 | 8.06E-01 | no | | intergenic_variant | MODIFIER |
| 20 | 145659689 | AX-170369725 | 0 | C | T | 0.149 | 3.31E-02 | 1.35E-01 | 2.64E+00 | 2.73E+00 | 8.06E-01 | no | | intergenic_variant | MODIFIER |
| 20 | 145678028 | AX-170493441 | 0 | A | G | 0.149 | 3.31E-02 | 1.35E-01 | 2.64E+00 | 2.73E+00 | 8.06E-01 | no | | intergenic_variant | MODIFIER |
| 20 | 145696212 | AX-170468633 | 0 | T | C | 0.149 | 3.31E-02 | 1.35E-01 | 2.64E+00 | 2.73E+00 | 8.06E-01 | no | | intergenic_variant | MODIFIER |
| 20 | 145719940 | AX-170458502 | 0 | A | T | 0.149 | 3.31E-02 | 1.35E-01 | 2.64E+00 | 2.73E+00 | 8.06E-01 | no | | intergenic_variant | MODIFIER |
| 20 | 145760744 | AX-97848517 | 0 | A | G | 0.149 | 3.31E-02 | 1.35E-01 | 2.64E+00 | 2.73E+00 | 8.06E-01 | no | | intergenic_variant | MODIFIER |
| 20 | 151470124 | AX-170035823 | 0 | G | A | 0.149 | 3.31E-02 | 1.35E-01 | 2.64E+00 | 2.73E+00 | 8.06E-01 | no | Fam120c | 3_prime_UTR_variant | MODIFIER |

| | | | | | | | | | | | | | | | |
|---|---|---|---|---|---|---|---|---|---|---|---|---|---|---|---|
| 8 | 29376227 | AX-170510819 | 0 | C | T | 0.167 | 2.73E-02 | 1.11E-01 | 2.64E+00 | 2.72E+00 | 8.06E-01 | no | | intergenic_variant | MODIFIER |
| 9 | 54012972 | AX-170438017 | 1 | C | T | 0.214 | -2.91E-02 | 1.21E-01 | 2.69E+00 | 2.75E+00 | 8.06E-01 | no | | intergenic_variant | MODIFIER |
| 6 | 139800098 | AX-170479228 | 1 | C | T | 0.054 | 7.19E-02 | 2.96E-01 | 2.67E+00 | 2.74E+00 | 8.06E-01 | no | Pik3c2g | intron_variant | MODIFIER |
| 12 | 95457708 | AX-170523760 | 0 | C | T | 0.061 | 5.72E-02 | 2.36E-01 | 2.69E+00 | 2.75E+00 | 8.06E-01 | no | | intergenic_variant | MODIFIER |
| 7 | 112245491 | AX-170132031 | 1 | G | T | 0.464 | -2.25E-02 | 9.33E-02 | 2.67E+00 | 2.75E+00 | 8.06E-01 | no | Mical2 | intron_variant | MODIFIER |
| 2 | 99746252 | AX-97838220 | 1 | T | C | 0.054 | -7.22E-02 | 2.99E-01 | 2.72E+00 | 2.75E+00 | 8.06E-01 | no | | intergenic_variant | MODIFIER |
| 10 | 114629460 | AX-170201935 | 0 | G | A | 0.342 | -1.89E-02 | 7.71E-02 | 2.64E+00 | 2.73E+00 | 8.06E-01 | no | Trhde | intron_variant | MODIFIER |
| 9 | 33915569 | AX-97794985 | 1 | T | C | 0.42 | 2.18E-02 | 8.79E-02 | 2.64E+00 | 2.71E+00 | 8.06E-01 | no | Gm25439 | upstream_gene_variant | MODIFIER |
| 7 | 137218240 | AX-170145181 | 0 | A | G | 0.281 | -2.11E-02 | 8.27E-02 | 2.60E+00 | 2.70E+00 | 8.06E-01 | no | Ebf3 | intron_variant | MODIFIER |
| 6 | 18142528 | AX-170489397 | 1 | C | T | 0.062 | 3.64E-02 | 1.49E-01 | 2.61E+00 | 2.73E+00 | 8.06E-01 | no | Gm15593 | upstream_gene_variant | MODIFIER |
| 3 | 29475212 | AX-170176602 | 0 | A | G | 0.184 | -2.32E-02 | 9.56E-02 | 2.66E+00 | 2.74E+00 | 8.06E-01 | no | Egfem1 | intron_variant | MODIFIER |
| 18 | 68608350 | AX-170481051 | 1 | C | T | 0.098 | 4.23E-02 | 1.70E-01 | 2.61E+00 | 2.72E+00 | 8.07E-01 | no | | intergenic_variant | MODIFIER |
| 20 | 140007223 | AX-170192919 | 1 | C | A | 0.152 | 3.29E-02 | 1.35E-01 | 2.64E+00 | 2.73E+00 | 8.07E-01 | no | Nup62cl | 3_prime_UTR_variant | MODIFIER |
| 20 | 145719818 | AX-170477345 | 1 | G | A | 0.152 | 3.29E-02 | 1.35E-01 | 2.64E+00 | 2.73E+00 | 8.07E-01 | no | | intergenic_variant | MODIFIER |
| 9 | 67581897 | AX-170149269 | 2 | C | G | 0.182 | 3.30E-02 | 1.37E-01 | 2.75E+00 | 2.79E+00 | 8.07E-01 | no | | regulatory_region_variant | MODIFIER |
| 7 | 138071550 | AX-170269305 | 0 | A | T | 0.298 | 1.82E-02 | 7.79E-02 | 2.67E+00 | 2.78E+00 | 8.07E-01 | no | | intergenic_variant | MODIFIER |
| 7 | 138823081 | AX-170030093 | 0 | A | G | 0.298 | 1.82E-02 | 7.79E-02 | 2.67E+00 | 2.78E+00 | 8.07E-01 | no | Gm45506 | downstream_gene_variant | MODIFIER |
| 2 | 162027896 | AX-170019547 | 0 | A | G | 0.158 | -4.19E-02 | 1.73E-01 | 2.65E+00 | 2.74E+00 | 8.07E-01 | no | Ptprt | intron_variant | MODIFIER |
| 13 | 36400156 | AX-97854772 | 0 | G | A | 0.193 | -2.76E-02 | 1.10E-01 | 2.62E+00 | 2.69E+00 | 8.07E-01 | no | Fars2 | intron_variant | MODIFIER |
| 1 | 113800396 | AX-170463272 | 0 | C | A | 0.158 | 4.72E-02 | 1.87E-01 | 2.61E+00 | 2.70E+00 | 8.07E-01 | no | | intergenic_variant | MODIFIER |
| 2 | 129807266 | AX-170064903 | 1 | T | C | 0.143 | 5.29E-02 | 2.18E-01 | 2.69E+00 | 2.73E+00 | 8.07E-01 | no | Stk35 | intron_variant | MODIFIER |
| 9 | 54064854 | AX-170176667 | 1 | A | T | 0.205 | -2.84E-02 | 1.19E-01 | 2.69E+00 | 2.76E+00 | 8.07E-01 | no | Tnfaip8l3 | intron_variant | MODIFIER |
| 9 | 56067192 | AX-170032415 | 1 | C | T | 0.205 | -2.84E-02 | 1.19E-01 | 2.69E+00 | 2.76E+00 | 8.07E-01 | no | | intergenic_variant | MODIFIER |
| 8 | 109453419 | AX-170031471 | 0 | A | G | 0.228 | -2.47E-02 | 1.02E-01 | 2.63E+00 | 2.74E+00 | 8.07E-01 | no | | intergenic_variant | MODIFIER |
| 2 | 156207333 | AX-170088802 | 1 | C | T | 0.143 | 5.37E-02 | 2.21E-01 | 2.69E+00 | 2.72E+00 | 8.07E-01 | no | Phf20 | intron_variant | MODIFIER |
| 12 | 115867740 | AX-170290905 | 1 | G | A | 0.143 | -4.68E-02 | 1.96E-01 | 2.69E+00 | 2.76E+00 | 8.07E-01 | no | Ighv1-78 | downstream_gene_variant | MODIFIER |
| 8 | 94943047 | AX-170145053 | 1 | T | C | 0.223 | 2.73E-02 | 1.14E-01 | 2.68E+00 | 2.75E+00 | 8.07E-01 | no | Adgrg5 | 3_prime_UTR_variant | MODIFIER |
| 15 | 91802671 | AX-97830737 | 0 | G | A | 0.158 | 3.97E-02 | 1.54E-01 | 2.59E+00 | 2.68E+00 | 8.07E-01 | no | Lrrk2 | intron_variant | MODIFIER |
| 8 | 73398344 | AX-170149906 | 0 | T | G | 0.193 | -2.84E-02 | 1.15E-01 | 2.62E+00 | 2.72E+00 | 8.07E-01 | no | | intergenic_variant | MODIFIER |
| 1 | 51209947 | AX-170518684 | 2 | A | G | 0.091 | 4.87E-02 | 2.03E-01 | 2.66E+00 | 2.75E+00 | 8.07E-01 | no | | intergenic_variant | MODIFIER |
| 1 | 68990527 | AX-170270928 | 0 | C | T | 0.123 | 4.41E-02 | 1.79E-01 | 2.61E+00 | 2.72E+00 | 8.07E-01 | no | Erbb4 | intron_variant | MODIFIER |
| 1 | 33131775 | AX-170111785 | 0 | T | C | 0.412 | 1.96E-02 | 8.22E-02 | 2.68E+00 | 2.76E+00 | 8.07E-01 | no | | intergenic_variant | MODIFIER |
| 20 | 77651857 | AX-170513684 | 0 | T | G | 0.272 | 1.42E-02 | 5.89E-02 | 2.64E+00 | 2.74E+00 | 8.07E-01 | no | Tbl1x | intron_variant | MODIFIER |
| 12 | 104417355 | AX-170007438 | 0 | C | T | 0.14 | -4.24E-02 | 1.81E-01 | 2.70E+00 | 2.79E+00 | 8.07E-01 | no | Serpina3n | downstream_gene_variant | MODIFIER |
| 1 | 142716483 | AX-170125759 | 2 | G | A | 0.209 | 2.87E-02 | 1.17E-01 | 2.62E+00 | 2.72E+00 | 8.07E-01 | no | | intergenic_variant | MODIFIER |
| 10 | 31541083 | AX-170109954 | 0 | G | A | 0.386 | -1.85E-02 | 7.52E-02 | 2.61E+00 | 2.72E+00 | 8.07E-01 | no | Rnf217 | intron_variant | MODIFIER |
| 4 | 143915706 | AX-170024176 | 1 | C | A | 0.107 | 4.92E-02 | 2.04E-01 | 2.67E+00 | 2.73E+00 | 8.07E-01 | no | Gm13100 | intron_variant,non_coding_transcript_variant | MODIFIER |
| 11 | 74266059 | AX-170162763 | 1 | G | C | 0.268 | 1.71E-02 | 7.23E-02 | 2.62E+00 | 2.75E+00 | 8.08E-01 | no | Olfr405-ps1 | intron_variant,non_coding_transcript_variant | MODIFIER |
| 11 | 30183667 | AX-170375550 | 1 | C | T | 0.089 | 4.50E-02 | 1.82E-01 | 2.62E+00 | 2.72E+00 | 8.08E-01 | no | Sptbn1 | intron_variant | MODIFIER |
| 8 | 48787675 | AX-170030643 | 0 | C | T | 0.158 | -3.44E-02 | 1.37E-01 | 2.62E+00 | 2.69E+00 | 8.08E-01 | no | Tenm3 | intron_variant | MODIFIER |
| 6 | 118541407 | AX-170028073 | 1 | A | T | 0.054 | 7.13E-02 | 2.96E-01 | 2.67E+00 | 2.74E+00 | 8.08E-01 | no | Ankrd26 | intron_variant | MODIFIER |
| 6 | 139307766 | AX-170141098 | 1 | T | C | 0.054 | 7.13E-02 | 2.96E-01 | 2.67E+00 | 2.74E+00 | 8.08E-01 | no | | intergenic_variant | MODIFIER |
| 3 | 25541349 | AX-170186086 | 0 | C | G | 0.474 | 8.54E-03 | 3.55E-02 | 2.58E+00 | 2.74E+00 | 8.08E-01 | no | Nlgn1 | intron_variant | MODIFIER |
| 5 | 126184515 | AX-170089604 | 1 | T | C | 0.446 | -2.66E-02 | 1.12E-01 | 2.72E+00 | 2.77E+00 | 8.08E-01 | no | | intergenic_variant | MODIFIER |
| 9 | 84430211 | AX-97825249 | 0 | C | T | 0.211 | 3.04E-02 | 1.28E-01 | 2.72E+00 | 2.78E+00 | 8.08E-01 | no | | intergenic_variant | MODIFIER |
| 9 | 73479427 | AX-170131025 | 0 | C | T | 0.219 | 3.15E-02 | 1.32E-01 | 2.74E+00 | 2.79E+00 | 8.08E-01 | no | Unc13c | downstream_gene_variant | MODIFIER |
| 1 | 54279715 | AX-170126590 | 0 | G | A | 0.105 | -5.62E-02 | 2.36E-01 | 2.71E+00 | 2.76E+00 | 8.08E-01 | no | Ccdc150 | intron_variant | MODIFIER |
| 5 | 79473025 | AX-170025381 | 0 | C | T | 0.439 | 2.59E-02 | 1.08E-01 | 2.68E+00 | 2.74E+00 | 8.08E-01 | no | | intergenic_variant | MODIFIER |
| 20 | 87174018 | AX-170034345 | 0 | G | A | 0.254 | -1.48E-02 | 6.20E-02 | 2.64E+00 | 2.74E+00 | 8.08E-01 | no | Il1rapl1 | intron_variant | MODIFIER |
| 7 | 92155018 | AX-170107318 | 0 | A | G | 0.132 | -4.24E-02 | 1.76E-01 | 2.63E+00 | 2.73E+00 | 8.08E-01 | no | Dlg2 | intron_variant | MODIFIER |
| 11 | 33573397 | AX-170389581 | 2 | G | A | 0.073 | 4.61E-02 | 1.87E-01 | 2.62E+00 | 2.71E+00 | 8.08E-01 | no | Gabrp | 5_prime_UTR_variant | MODIFIER |
| 15 | 90447285 | AX-97825415 | 0 | T | C | 0.14 | 4.08E-02 | 1.64E-01 | 2.62E+00 | 2.70E+00 | 8.09E-01 | no | | intergenic_variant | MODIFIER |
| 15 | 94750708 | AX-170338748 | 0 | T | C | 0.14 | 4.08E-02 | 1.64E-01 | 2.62E+00 | 2.70E+00 | 8.09E-01 | no | Tmem117 | intron_variant | MODIFIER |
| 11 | 74483995 | AX-170005313 | 0 | T | C | 0.263 | 1.57E-02 | 6.77E-02 | 2.62E+00 | 2.76E+00 | 8.09E-01 | no | Rap1gap2 | intron_variant,NMD_transcript_variant | MODIFIER |

| | | | | | | | | | | | | | | | |
|---|---|---|---|---|---|---|---|---|---|---|---|---|---|---|---|
| 2 | 130469130 | AX-170495838 | 1 | G | A | 0.143 | 5.22E-02 | 2.18E-01 | 2.70E+00 | 2.73E+00 | 8.09E-01 | no | Ptpra | intron_variant | MODIFIER |
| 8 | 63347034 | AX-97804417 | 0 | T | G | 0.149 | 2.93E-02 | 1.28E-01 | 2.68E+00 | 2.79E+00 | 8.09E-01 | no | Spock3 | intron_variant | MODIFIER |
| 16 | 94986149 | AX-170188448 | 0 | G | A | 0.447 | 1.74E-02 | 7.63E-02 | 2.70E+00 | 2.81E+00 | 8.09E-01 | no | Kcnj6 | intron_variant | MODIFIER |
| 16 | 95018177 | AX-170088104 | 0 | T | C | 0.447 | 1.74E-02 | 7.63E-02 | 2.70E+00 | 2.81E+00 | 8.09E-01 | no | Gm7976 | intron_variant,non_coding_transcript_variant | MODIFIER |
| 16 | 95027390 | AX-170414988 | 0 | G | T | 0.447 | 1.74E-02 | 7.63E-02 | 2.70E+00 | 2.81E+00 | 8.09E-01 | no | Gm7976 | intron_variant,non_coding_transcript_variant | MODIFIER |
| 16 | 95034136 | AX-170088105 | 0 | G | A | 0.447 | 1.74E-02 | 7.63E-02 | 2.70E+00 | 2.81E+00 | 8.09E-01 | no | Gm7976 | intron_variant,non_coding_transcript_variant | MODIFIER |
| 16 | 95060241 | AX-170356597 | 0 | C | A | 0.447 | 1.74E-02 | 7.63E-02 | 2.70E+00 | 2.81E+00 | 8.09E-01 | no | Gm7976 | intron_variant,non_coding_transcript_variant | MODIFIER |
| 16 | 95062498 | AX-170124890 | 0 | C | T | 0.447 | 1.74E-02 | 7.63E-02 | 2.70E+00 | 2.81E+00 | 8.09E-01 | no | Gm7976 | intron_variant,non_coding_transcript_variant | MODIFIER |
| 16 | 95063114 | AX-170137101 | 0 | A | G | 0.447 | 1.74E-02 | 7.63E-02 | 2.70E+00 | 2.81E+00 | 8.09E-01 | no | Gm7976 | intron_variant,non_coding_transcript_variant | MODIFIER |
| 16 | 95063169 | AX-170217814 | 0 | T | C | 0.447 | 1.74E-02 | 7.63E-02 | 2.70E+00 | 2.81E+00 | 8.09E-01 | no | Gm7976 | intron_variant,non_coding_transcript_variant | MODIFIER |
| 16 | 95064919 | AX-97832181 | 0 | A | G | 0.447 | 1.74E-02 | 7.63E-02 | 2.70E+00 | 2.81E+00 | 8.09E-01 | no | Gm7976 | intron_variant,non_coding_transcript_variant | MODIFIER |
| 16 | 95075929 | AX-170285840 | 0 | T | C | 0.447 | 1.74E-02 | 7.63E-02 | 2.70E+00 | 2.81E+00 | 8.09E-01 | no | | regulatory_region_variant | MODIFIER |
| 16 | 95076086 | AX-170058173 | 0 | G | A | 0.447 | 1.74E-02 | 7.63E-02 | 2.70E+00 | 2.81E+00 | 8.09E-01 | no | | regulatory_region_variant | MODIFIER |
| 16 | 95076977 | AX-170058174 | 0 | C | G | 0.447 | 1.74E-02 | 7.63E-02 | 2.70E+00 | 2.81E+00 | 8.09E-01 | no | | regulatory_region_variant | MODIFIER |
| 16 | 95089735 | AX-97847386 | 0 | T | C | 0.447 | 1.74E-02 | 7.63E-02 | 2.70E+00 | 2.81E+00 | 8.09E-01 | no | | intergenic_variant | MODIFIER |
| 16 | 95121234 | AX-170340734 | 0 | T | C | 0.447 | 1.74E-02 | 7.63E-02 | 2.70E+00 | 2.81E+00 | 8.09E-01 | no | | intergenic_variant | MODIFIER |
| 16 | 95121616 | AX-170385689 | 0 | T | C | 0.447 | 1.74E-02 | 7.63E-02 | 2.70E+00 | 2.81E+00 | 8.09E-01 | no | | intergenic_variant | MODIFIER |
| 16 | 95122367 | AX-170455749 | 0 | C | A | 0.447 | 1.74E-02 | 7.63E-02 | 2.70E+00 | 2.81E+00 | 8.09E-01 | no | | intergenic_variant | MODIFIER |
| 16 | 95124122 | AX-170371161 | 0 | T | C | 0.447 | 1.74E-02 | 7.63E-02 | 2.70E+00 | 2.81E+00 | 8.09E-01 | no | | intergenic_variant | MODIFIER |
| 16 | 95143119 | AX-97857718 | 0 | G | A | 0.447 | 1.74E-02 | 7.63E-02 | 2.70E+00 | 2.81E+00 | 8.09E-01 | no | | intergenic_variant | MODIFIER |
| 16 | 95170502 | AX-170058177 | 0 | T | C | 0.447 | 1.74E-02 | 7.63E-02 | 2.70E+00 | 2.81E+00 | 8.09E-01 | no | Gm49642 | non_coding_transcript_exon_variant | MODIFIER |
| 16 | 95195039 | AX-97832178 | 0 | C | T | 0.447 | 1.74E-02 | 7.63E-02 | 2.70E+00 | 2.81E+00 | 8.09E-01 | no | Gm31641 | intron_variant,non_coding_transcript_variant | MODIFIER |
| 16 | 95197082 | AX-170258191 | 0 | C | T | 0.447 | 1.74E-02 | 7.63E-02 | 2.70E+00 | 2.81E+00 | 8.09E-01 | no | Gm31641 | intron_variant,non_coding_transcript_variant | MODIFIER |
| 16 | 95199622 | AX-170132142 | 0 | G | A | 0.447 | 1.74E-02 | 7.63E-02 | 2.70E+00 | 2.81E+00 | 8.09E-01 | no | Gm31641 | intron_variant,non_coding_transcript_variant | MODIFIER |
| 16 | 95203492 | AX-170369094 | 0 | G | A | 0.447 | 1.74E-02 | 7.63E-02 | 2.70E+00 | 2.81E+00 | 8.09E-01 | no | Gm31641 | intron_variant,non_coding_transcript_variant | MODIFIER |
| 16 | 95231171 | AX-170318896 | 0 | A | G | 0.447 | 1.74E-02 | 7.63E-02 | 2.70E+00 | 2.81E+00 | 8.09E-01 | no | Gm31641 | intron_variant,non_coding_transcript_variant | MODIFIER |
| 16 | 95234216 | AX-170227465 | 0 | C | T | 0.447 | 1.74E-02 | 7.63E-02 | 2.70E+00 | 2.81E+00 | 8.09E-01 | no | Gm31641 | intron_variant,non_coding_transcript_variant | MODIFIER |
| 16 | 95235049 | AX-170167688 | 0 | C | T | 0.447 | 1.74E-02 | 7.63E-02 | 2.70E+00 | 2.81E+00 | 8.09E-01 | no | Gm31641 | intron_variant,non_coding_transcript_variant | MODIFIER |
| 16 | 95235103 | AX-170496154 | 0 | A | G | 0.447 | 1.74E-02 | 7.63E-02 | 2.70E+00 | 2.81E+00 | 8.09E-01 | no | Gm31641 | intron_variant,non_coding_transcript_variant | MODIFIER |
| 16 | 95235268 | AX-170088106 | 0 | C | A | 0.447 | 1.74E-02 | 7.63E-02 | 2.70E+00 | 2.81E+00 | 8.09E-01 | no | Gm31641 | intron_variant,non_coding_transcript_variant | MODIFIER |
| 16 | 95276095 | AX-170149334 | 0 | G | A | 0.447 | 1.74E-02 | 7.63E-02 | 2.70E+00 | 2.81E+00 | 8.09E-01 | no | Kcnj15 | intron_variant | MODIFIER |
| 16 | 95276107 | AX-170166186 | 0 | G | A | 0.447 | 1.74E-02 | 7.63E-02 | 2.70E+00 | 2.81E+00 | 8.09E-01 | no | Kcnj15 | intron_variant | MODIFIER |
| 16 | 95304735 | AX-170013701 | 0 | T | A | 0.447 | 1.74E-02 | 7.63E-02 | 2.70E+00 | 2.81E+00 | 8.09E-01 | no | Kcnj15 | downstream_gene_variant | MODIFIER |
| 16 | 95386387 | AX-170498033 | 0 | G | A | 0.447 | 1.74E-02 | 7.63E-02 | 2.70E+00 | 2.81E+00 | 8.09E-01 | no | Erg | intron_variant | MODIFIER |
| 16 | 95415913 | AX-170510932 | 0 | C | T | 0.447 | 1.74E-02 | 7.63E-02 | 2.70E+00 | 2.81E+00 | 8.09E-01 | no | Erg | intron_variant | MODIFIER |
| 16 | 95421119 | AX-170366703 | 0 | G | A | 0.447 | 1.74E-02 | 7.63E-02 | 2.70E+00 | 2.81E+00 | 8.09E-01 | no | Erg | intron_variant | MODIFIER |
| 11 | 5825757 | AX-170086934 | 0 | A | G | 0.044 | -4.62E-02 | 1.88E-01 | 2.58E+00 | 2.72E+00 | 8.09E-01 | no | Polm | downstream_gene_variant | MODIFIER |
| 3 | 147445555 | AX-170068381 | 1 | C | A | 0.348 | 1.77E-02 | 7.29E-02 | 2.64E+00 | 2.72E+00 | 8.09E-01 | no | | regulatory_region_variant | MODIFIER |
| 8 | 101485084 | AX-97817292 | 0 | T | C | 0.228 | 2.77E-02 | 1.17E-01 | 2.68E+00 | 2.75E+00 | 8.09E-01 | no | | intergenic_variant | MODIFIER |
| 4 | 149014871 | AX-97819961 | 0 | A | G | 0.096 | 3.41E-02 | 1.40E-01 | 2.63E+00 | 2.71E+00 | 8.09E-01 | no | Pex14 | intron_variant | MODIFIER |
| 5 | 84028095 | AX-170494100 | 0 | T | C | 0.439 | 2.49E-02 | 1.06E-01 | 2.70E+00 | 2.76E+00 | 8.09E-01 | no | | intergenic_variant | MODIFIER |
| 1 | 22698733 | AX-170475117 | 1 | T | C | 0.429 | 2.04E-02 | 7.97E-02 | 2.59E+00 | 2.68E+00 | 8.09E-01 | no | Rims1 | intron_variant | MODIFIER |
| 7 | 116983149 | AX-97855285 | 0 | A | G | 0.404 | 2.27E-02 | 9.30E-02 | 2.65E+00 | 2.70E+00 | 8.09E-01 | no | | intergenic_variant | MODIFIER |
| 2 | 143522331 | AX-97843433 | 0 | C | A | 0.149 | -4.82E-02 | 2.07E-01 | 2.73E+00 | 2.80E+00 | 8.09E-01 | no | | intergenic_variant | MODIFIER |
| 4 | 71285934 | AX-97810139 | 0 | C | T | 0.237 | 2.67E-02 | 1.04E-01 | 2.60E+00 | 2.67E+00 | 8.09E-01 | no | Gm11229 | upstream_gene_variant | MODIFIER |
| 4 | 72140939 | AX-168410045 | 0 | G | A | 0.237 | 2.67E-02 | 1.04E-01 | 2.60E+00 | 2.67E+00 | 8.09E-01 | no | Tle1 | intron_variant | MODIFIER |
| 4 | 72141103 | AX-170120855 | 0 | C | T | 0.237 | 2.67E-02 | 1.04E-01 | 2.60E+00 | 2.67E+00 | 8.09E-01 | no | Tle1 | intron_variant | MODIFIER |
| 4 | 74001181 | AX-170023117 | 0 | G | A | 0.237 | 2.67E-02 | 1.04E-01 | 2.60E+00 | 2.67E+00 | 8.09E-01 | no | | intergenic_variant | MODIFIER |
| 4 | 74001193 | AX-170023125 | 0 | G | A | 0.237 | 2.67E-02 | 1.04E-01 | 2.60E+00 | 2.67E+00 | 8.09E-01 | no | | intergenic_variant | MODIFIER |
| 4 | 74027033 | AX-170270007 | 0 | C | T | 0.237 | 2.67E-02 | 1.04E-01 | 2.60E+00 | 2.67E+00 | 8.09E-01 | no | Frmd3 | intron_variant | MODIFIER |
| 4 | 74201062 | AX-170095014 | 0 | T | C | 0.237 | 2.67E-02 | 1.04E-01 | 2.60E+00 | 2.67E+00 | 8.09E-01 | no | Frmd3 | intron_variant | MODIFIER |
| 4 | 74204576 | AX-97829974 | 0 | G | T | 0.237 | 2.67E-02 | 1.04E-01 | 2.60E+00 | 2.67E+00 | 8.09E-01 | no | Frmd3 | downstream_gene_variant | MODIFIER |
| 4 | 74212640 | AX-170146688 | 0 | C | T | 0.237 | 2.67E-02 | 1.04E-01 | 2.60E+00 | 2.67E+00 | 8.09E-01 | no | | intergenic_variant | MODIFIER |
| 4 | 74785234 | AX-170223533 | 0 | C | T | 0.237 | 2.67E-02 | 1.04E-01 | 2.60E+00 | 2.67E+00 | 8.09E-01 | no | | intergenic_variant | MODIFIER |

| | | | | | | | | | | | | | | | |
|---|---|---|---|---|---|---|---|---|---|---|---|---|---|---|---|
| 4 | 74785731 | AX-170023130 | 0 | G | C | 0.237 | 2.67E-02 | 1.04E-01 | 2.60E+00 | 2.67E+00 | 8.09E-01 | no | | intergenic_variant | MODIFIER |
| 4 | 74907182 | AX-97834798 | 0 | G | A | 0.237 | 2.67E-02 | 1.04E-01 | 2.60E+00 | 2.67E+00 | 8.09E-01 | no | | intergenic_variant | MODIFIER |
| 4 | 78905937 | AX-170420673 | 0 | T | C | 0.237 | 2.67E-02 | 1.04E-01 | 2.60E+00 | 2.67E+00 | 8.09E-01 | no | | intergenic_variant | MODIFIER |
| 4 | 79106798 | AX-170280158 | 0 | A | G | 0.237 | 2.67E-02 | 1.04E-01 | 2.60E+00 | 2.67E+00 | 8.09E-01 | no | | intergenic_variant | MODIFIER |
| 4 | 79112108 | AX-97850654 | 0 | T | G | 0.237 | 2.67E-02 | 1.04E-01 | 2.60E+00 | 2.67E+00 | 8.09E-01 | no | | intergenic_variant | MODIFIER |
| 4 | 79157202 | AX-97790536 | 0 | A | G | 0.237 | 2.67E-02 | 1.04E-01 | 2.60E+00 | 2.67E+00 | 8.09E-01 | no | | intergenic_variant | MODIFIER |
| 4 | 79190513 | AX-170280244 | 0 | G | C | 0.237 | 2.67E-02 | 1.04E-01 | 2.60E+00 | 2.67E+00 | 8.09E-01 | no | | intergenic_variant | MODIFIER |
| 4 | 79822356 | AX-170150592 | 0 | G | A | 0.237 | 2.67E-02 | 1.04E-01 | 2.60E+00 | 2.67E+00 | 8.09E-01 | no | | intergenic_variant | MODIFIER |
| 4 | 80199223 | AX-170023204 | 0 | T | C | 0.237 | 2.67E-02 | 1.04E-01 | 2.60E+00 | 2.67E+00 | 8.09E-01 | no | | intergenic_variant | MODIFIER |
| 4 | 80209292 | AX-170023202 | 0 | T | C | 0.237 | 2.67E-02 | 1.04E-01 | 2.60E+00 | 2.67E+00 | 8.09E-01 | no | | intergenic_variant | MODIFIER |
| 4 | 80286759 | AX-170023197 | 0 | G | A | 0.237 | 2.67E-02 | 1.04E-01 | 2.60E+00 | 2.67E+00 | 8.09E-01 | no | | intergenic_variant | MODIFIER |
| 4 | 80287358 | AX-97810882 | 0 | G | T | 0.237 | 2.67E-02 | 1.04E-01 | 2.60E+00 | 2.67E+00 | 8.09E-01 | no | | intergenic_variant | MODIFIER |
| 4 | 80320483 | AX-170023201 | 0 | A | T | 0.237 | 2.67E-02 | 1.04E-01 | 2.60E+00 | 2.67E+00 | 8.09E-01 | no | | intergenic_variant | MODIFIER |
| 4 | 80324261 | AX-170089290 | 0 | C | A | 0.237 | 2.67E-02 | 1.04E-01 | 2.60E+00 | 2.67E+00 | 8.09E-01 | no | | intergenic_variant | MODIFIER |
| 4 | 80347018 | AX-170351982 | 0 | G | C | 0.237 | 2.67E-02 | 1.04E-01 | 2.60E+00 | 2.67E+00 | 8.09E-01 | no | | intergenic_variant | MODIFIER |
| 4 | 80355360 | AX-170339614 | 0 | C | A | 0.237 | 2.67E-02 | 1.04E-01 | 2.60E+00 | 2.67E+00 | 8.09E-01 | no | | intergenic_variant | MODIFIER |
| 4 | 80368026 | AX-170214804 | 0 | T | C | 0.237 | 2.67E-02 | 1.04E-01 | 2.60E+00 | 2.67E+00 | 8.09E-01 | no | | intergenic_variant | MODIFIER |
| 4 | 80369005 | AX-97791309 | 0 | A | G | 0.237 | 2.67E-02 | 1.04E-01 | 2.60E+00 | 2.67E+00 | 8.09E-01 | no | | intergenic_variant | MODIFIER |
| 4 | 80388013 | AX-97849259 | 0 | T | C | 0.237 | 2.67E-02 | 1.04E-01 | 2.60E+00 | 2.67E+00 | 8.09E-01 | no | | intergenic_variant | MODIFIER |
| 4 | 80419906 | AX-97833935 | 0 | C | T | 0.237 | 2.67E-02 | 1.04E-01 | 2.60E+00 | 2.67E+00 | 8.09E-01 | no | | intergenic_variant | MODIFIER |
| 4 | 80436433 | AX-170023213 | 0 | A | T | 0.237 | 2.67E-02 | 1.04E-01 | 2.60E+00 | 2.67E+00 | 8.09E-01 | no | Gm11406 | upstream_gene_variant | MODIFIER |
| 4 | 80437557 | AX-170089291 | 0 | A | C | 0.237 | 2.67E-02 | 1.04E-01 | 2.60E+00 | 2.67E+00 | 8.09E-01 | no | Gm11406 | upstream_gene_variant | MODIFIER |
| 4 | 80466909 | AX-97849199 | 0 | T | C | 0.237 | 2.67E-02 | 1.04E-01 | 2.60E+00 | 2.67E+00 | 8.09E-01 | no | | intergenic_variant | MODIFIER |
| 4 | 80466945 | AX-170023217 | 0 | C | T | 0.237 | 2.67E-02 | 1.04E-01 | 2.60E+00 | 2.67E+00 | 8.09E-01 | no | | intergenic_variant | MODIFIER |
| 4 | 80466982 | AX-170023214 | 0 | C | T | 0.237 | 2.67E-02 | 1.04E-01 | 2.60E+00 | 2.67E+00 | 8.09E-01 | no | | intergenic_variant | MODIFIER |
| 4 | 80493655 | AX-97815906 | 0 | A | G | 0.237 | 2.67E-02 | 1.04E-01 | 2.60E+00 | 2.67E+00 | 8.09E-01 | no | | intergenic_variant | MODIFIER |
| 4 | 80524933 | AX-97798765 | 0 | G | A | 0.237 | 2.67E-02 | 1.04E-01 | 2.60E+00 | 2.67E+00 | 8.09E-01 | no | | intergenic_variant | MODIFIER |
| 4 | 80552839 | AX-170023211 | 0 | G | A | 0.237 | 2.67E-02 | 1.04E-01 | 2.60E+00 | 2.67E+00 | 8.09E-01 | no | | regulatory_region_variant | MODIFIER |
| 4 | 80557745 | AX-97781668 | 0 | G | A | 0.237 | 2.67E-02 | 1.04E-01 | 2.60E+00 | 2.67E+00 | 8.09E-01 | no | | intergenic_variant | MODIFIER |
| 4 | 80572705 | AX-170069795 | 0 | G | C | 0.237 | 2.67E-02 | 1.04E-01 | 2.60E+00 | 2.67E+00 | 8.09E-01 | no | | intergenic_variant | MODIFIER |
| 4 | 80609318 | AX-170069796 | 0 | C | T | 0.237 | 2.67E-02 | 1.04E-01 | 2.60E+00 | 2.67E+00 | 8.09E-01 | no | | intergenic_variant | MODIFIER |
| 4 | 80617911 | AX-97790109 | 0 | T | C | 0.237 | 2.67E-02 | 1.04E-01 | 2.60E+00 | 2.67E+00 | 8.09E-01 | no | | intergenic_variant | MODIFIER |
| 4 | 80628552 | AX-170531893 | 0 | A | G | 0.237 | 2.67E-02 | 1.04E-01 | 2.60E+00 | 2.67E+00 | 8.09E-01 | no | | intergenic_variant | MODIFIER |
| 4 | 80631938 | AX-97783856 | 0 | A | G | 0.237 | 2.67E-02 | 1.04E-01 | 2.60E+00 | 2.67E+00 | 8.09E-01 | no | | intergenic_variant | MODIFIER |
| 4 | 80643774 | AX-170116191 | 0 | G | A | 0.237 | 2.67E-02 | 1.04E-01 | 2.60E+00 | 2.67E+00 | 8.09E-01 | no | | intergenic_variant | MODIFIER |
| 4 | 80661685 | AX-170144463 | 0 | G | A | 0.237 | 2.67E-02 | 1.04E-01 | 2.60E+00 | 2.67E+00 | 8.09E-01 | no | | intergenic_variant | MODIFIER |
| 4 | 80661830 | AX-170415737 | 0 | G | A | 0.237 | 2.67E-02 | 1.04E-01 | 2.60E+00 | 2.67E+00 | 8.09E-01 | no | | intergenic_variant | MODIFIER |
| 4 | 80694824 | AX-170341008 | 0 | G | A | 0.237 | 2.67E-02 | 1.04E-01 | 2.60E+00 | 2.67E+00 | 8.09E-01 | no | | intergenic_variant | MODIFIER |
| 4 | 80839166 | AX-97855329 | 0 | G | A | 0.237 | 2.67E-02 | 1.04E-01 | 2.60E+00 | 2.67E+00 | 8.09E-01 | no | Tyrp1 | intron_variant | MODIFIER |
| 4 | 80950758 | AX-170023221 | 0 | G | A | 0.237 | 2.67E-02 | 1.04E-01 | 2.60E+00 | 2.67E+00 | 8.09E-01 | no | Lurap1l | intron_variant | MODIFIER |
| 10 | 16119358 | AX-97804743 | 0 | G | T | 0.202 | 2.34E-02 | 1.02E-01 | 2.70E+00 | 2.79E+00 | 8.09E-01 | no | | intergenic_variant | MODIFIER |
| 5 | 112376703 | AX-170302487 | 0 | C | T | 0.447 | 2.64E-02 | 1.08E-01 | 2.65E+00 | 2.70E+00 | 8.10E-01 | no | Hps4 | intron_variant | MODIFIER |
| 15 | 94315305 | AX-170505173 | 1 | G | C | 0.152 | 4.39E-02 | 1.79E-01 | 2.64E+00 | 2.70E+00 | 8.10E-01 | no | Adamts20 | intron_variant | MODIFIER |
| 15 | 94802910 | AX-170123371 | 1 | A | G | 0.152 | 4.42E-02 | 1.81E-01 | 2.64E+00 | 2.70E+00 | 8.10E-01 | no | Tmem117 | intron_variant | MODIFIER |
| 11 | 73267237 | AX-170386065 | 0 | C | T | 0.246 | 1.68E-02 | 7.11E-02 | 2.61E+00 | 2.74E+00 | 8.10E-01 | no | Trpv3 | upstream_gene_variant | MODIFIER |
| 9 | 83539306 | AX-170138822 | 1 | G | A | 0.214 | 3.00E-02 | 1.28E-01 | 2.73E+00 | 2.78E+00 | 8.10E-01 | no | | intergenic_variant | MODIFIER |
| 12 | 105952744 | AX-170523585 | 1 | G | A | 0.125 | -4.26E-02 | 1.82E-01 | 2.68E+00 | 2.76E+00 | 8.10E-01 | no | | intergenic_variant | MODIFIER |
| 1 | 41173511 | AX-170362247 | 0 | T | A | 0.421 | -1.81E-02 | 7.57E-02 | 2.65E+00 | 2.73E+00 | 8.10E-01 | no | 4930448I06Rik | intron_variant,non_coding_transcript_variant | MODIFIER |
| 1 | 41228127 | AX-170454978 | 0 | A | G | 0.421 | -1.81E-02 | 7.57E-02 | 2.65E+00 | 2.73E+00 | 8.10E-01 | no | | intergenic_variant | MODIFIER |
| 1 | 41292526 | AX-170350963 | 0 | T | G | 0.421 | -1.81E-02 | 7.57E-02 | 2.65E+00 | 2.73E+00 | 8.10E-01 | no | | intergenic_variant | MODIFIER |
| 1 | 41422748 | AX-170312570 | 0 | A | G | 0.421 | -1.81E-02 | 7.57E-02 | 2.65E+00 | 2.73E+00 | 8.10E-01 | no | | intergenic_variant | MODIFIER |
| 1 | 41423874 | AX-170458182 | 0 | A | G | 0.421 | -1.81E-02 | 7.57E-02 | 2.65E+00 | 2.73E+00 | 8.10E-01 | no | | intergenic_variant | MODIFIER |
| 1 | 41437565 | AX-170335793 | 0 | T | C | 0.421 | -1.81E-02 | 7.57E-02 | 2.65E+00 | 2.73E+00 | 8.10E-01 | no | | intergenic_variant | MODIFIER |
| 1 | 41439794 | AX-170265686 | 0 | G | A | 0.421 | -1.81E-02 | 7.57E-02 | 2.65E+00 | 2.73E+00 | 8.10E-01 | no | | intergenic_variant | MODIFIER |

| | | | | | | | | | | | | | | | |
|---|---|---|---|---|---|---|---|---|---|---|---|---|---|---|---|
| 1 | 41440064 | AX-97789912 | 0 | A | C | 0.421 | -1.81E-02 | 7.57E-02 | 2.65E+00 | 2.73E+00 | 8.10E-01 | no | | intergenic_variant | MODIFIER |
| 1 | 41460988 | AX-170326348 | 0 | T | A | 0.421 | -1.81E-02 | 7.57E-02 | 2.65E+00 | 2.73E+00 | 8.10E-01 | no | | intergenic_variant | MODIFIER |
| 1 | 41466706 | AX-170423397 | 0 | T | C | 0.421 | -1.81E-02 | 7.57E-02 | 2.65E+00 | 2.73E+00 | 8.10E-01 | no | | intergenic_variant | MODIFIER |
| 1 | 41490739 | AX-97815512 | 0 | T | C | 0.421 | -1.81E-02 | 7.57E-02 | 2.65E+00 | 2.73E+00 | 8.10E-01 | no | | intergenic_variant | MODIFIER |
| 1 | 41513723 | AX-170323964 | 0 | A | G | 0.421 | -1.81E-02 | 7.57E-02 | 2.65E+00 | 2.73E+00 | 8.10E-01 | no | | intergenic_variant | MODIFIER |
| 1 | 41552304 | AX-170487877 | 0 | C | T | 0.421 | -1.81E-02 | 7.57E-02 | 2.65E+00 | 2.73E+00 | 8.10E-01 | no | | intergenic_variant | MODIFIER |
| 1 | 41590612 | AX-170000350 | 0 | G | T | 0.421 | -1.81E-02 | 7.57E-02 | 2.65E+00 | 2.73E+00 | 8.10E-01 | no | | intergenic_variant | MODIFIER |
| 1 | 41595144 | AX-170000351 | 0 | A | G | 0.421 | -1.81E-02 | 7.57E-02 | 2.65E+00 | 2.73E+00 | 8.10E-01 | no | | intergenic_variant | MODIFIER |
| 1 | 41605524 | AX-170458372 | 0 | C | A | 0.421 | -1.81E-02 | 7.57E-02 | 2.65E+00 | 2.73E+00 | 8.10E-01 | no | | regulatory_region_variant | MODIFIER |
| 1 | 41629164 | AX-170041011 | 0 | C | T | 0.421 | -1.81E-02 | 7.57E-02 | 2.65E+00 | 2.73E+00 | 8.10E-01 | no | | intergenic_variant | MODIFIER |
| 1 | 41639079 | AX-170522483 | 0 | C | A | 0.421 | -1.81E-02 | 7.57E-02 | 2.65E+00 | 2.73E+00 | 8.10E-01 | no | | intergenic_variant | MODIFIER |
| 1 | 41641263 | AX-170455557 | 0 | A | G | 0.421 | -1.81E-02 | 7.57E-02 | 2.65E+00 | 2.73E+00 | 8.10E-01 | no | | intergenic_variant | MODIFIER |
| 1 | 41642572 | AX-170471006 | 0 | G | A | 0.421 | -1.81E-02 | 7.57E-02 | 2.65E+00 | 2.73E+00 | 8.10E-01 | no | | intergenic_variant | MODIFIER |
| 1 | 41648796 | AX-170118639 | 0 | T | C | 0.421 | -1.81E-02 | 7.57E-02 | 2.65E+00 | 2.73E+00 | 8.10E-01 | no | | intergenic_variant | MODIFIER |
| 1 | 41650870 | AX-170281823 | 0 | A | G | 0.421 | -1.81E-02 | 7.57E-02 | 2.65E+00 | 2.73E+00 | 8.10E-01 | no | | intergenic_variant | MODIFIER |
| 1 | 41650935 | AX-170189354 | 0 | A | G | 0.421 | -1.81E-02 | 7.57E-02 | 2.65E+00 | 2.73E+00 | 8.10E-01 | no | | intergenic_variant | MODIFIER |
| 1 | 41650973 | AX-170441798 | 0 | T | C | 0.421 | -1.81E-02 | 7.57E-02 | 2.65E+00 | 2.73E+00 | 8.10E-01 | no | | intergenic_variant | MODIFIER |
| 1 | 41669484 | AX-170227919 | 0 | G | T | 0.421 | -1.81E-02 | 7.57E-02 | 2.65E+00 | 2.73E+00 | 8.10E-01 | no | | intergenic_variant | MODIFIER |
| 1 | 41684349 | AX-170342954 | 0 | C | T | 0.421 | -1.81E-02 | 7.57E-02 | 2.65E+00 | 2.73E+00 | 8.10E-01 | no | | intergenic_variant | MODIFIER |
| 1 | 41912677 | AX-170305920 | 0 | A | G | 0.421 | -1.81E-02 | 7.57E-02 | 2.65E+00 | 2.73E+00 | 8.10E-01 | no | Gm29260 | intron_variant,non_coding_transcript_variant | MODIFIER |
| 1 | 41988729 | AX-97824267 | 0 | C | T | 0.421 | -1.81E-02 | 7.57E-02 | 2.65E+00 | 2.73E+00 | 8.10E-01 | no | | intergenic_variant | MODIFIER |
| 1 | 41994144 | AX-170125374 | 0 | T | C | 0.421 | -1.81E-02 | 7.57E-02 | 2.65E+00 | 2.73E+00 | 8.10E-01 | no | | intergenic_variant | MODIFIER |
| 1 | 42007844 | AX-97811953 | 0 | T | C | 0.421 | -1.81E-02 | 7.57E-02 | 2.65E+00 | 2.73E+00 | 8.10E-01 | no | | intergenic_variant | MODIFIER |
| 1 | 42009743 | AX-170161575 | 0 | T | C | 0.421 | -1.81E-02 | 7.57E-02 | 2.65E+00 | 2.73E+00 | 8.10E-01 | no | | intergenic_variant | MODIFIER |
| 1 | 42010160 | AX-170207206 | 0 | C | T | 0.421 | -1.81E-02 | 7.57E-02 | 2.65E+00 | 2.73E+00 | 8.10E-01 | no | | intergenic_variant | MODIFIER |
| 5 | 34093070 | AX-97797697 | 1 | G | A | 0.366 | -2.30E-02 | 9.97E-02 | 2.68E+00 | 2.78E+00 | 8.10E-01 | no | Poln | intron_variant | MODIFIER |
| 1 | 70748595 | AX-170415779 | 1 | A | G | 0.107 | 4.90E-02 | 2.06E-01 | 2.65E+00 | 2.73E+00 | 8.10E-01 | no | Vwc2l | intron_variant | MODIFIER |
| 16 | 11991403 | AX-168399677 | 0 | G | A | 0.061 | -4.45E-02 | 1.88E-01 | 2.63E+00 | 2.74E+00 | 8.10E-01 | no | Shisa9 | intron_variant | MODIFIER |
| 16 | 12865865 | AX-97799991 | 0 | C | T | 0.061 | -4.45E-02 | 1.88E-01 | 2.63E+00 | 2.74E+00 | 8.10E-01 | no | 4930414F18Rik | intron_variant,non_coding_transcript_variant | MODIFIER |
| 9 | 32792647 | AX-170469373 | 1 | T | C | 0.42 | 2.09E-02 | 8.59E-02 | 2.64E+00 | 2.71E+00 | 8.10E-01 | no | | intergenic_variant | MODIFIER |
| 8 | 73642143 | AX-170031000 | 0 | C | A | 0.202 | -2.89E-02 | 1.22E-01 | 2.66E+00 | 2.74E+00 | 8.10E-01 | no | Gm23255 | upstream_gene_variant | MODIFIER |
| 1 | 46807987 | AX-170391643 | 0 | G | A | 0.088 | -4.92E-02 | 2.09E-01 | 2.65E+00 | 2.75E+00 | 8.10E-01 | no | Slc39a10 | 3_prime_UTR_variant | MODIFIER |
| 18 | 63991901 | AX-170419298 | 1 | C | T | 0.384 | 2.44E-02 | 9.23E-02 | 2.57E+00 | 2.64E+00 | 8.10E-01 | no | Wdr7 | downstream_gene_variant | MODIFIER |
| 1 | 3252796 | AX-170461196 | 0 | C | T | 0.518 | 1.78E-02 | 6.90E-02 | 2.59E+00 | 2.67E+00 | 8.10E-01 | no | Xkr4 | intron_variant | MODIFIER |
| 3 | 148184540 | AX-170021878 | 1 | G | A | 0.339 | 1.78E-02 | 7.39E-02 | 2.65E+00 | 2.72E+00 | 8.11E-01 | no | | intergenic_variant | MODIFIER |
| 7 | 68382716 | AX-97803034 | 0 | A | G | 0.395 | -2.28E-02 | 9.89E-02 | 2.69E+00 | 2.78E+00 | 8.11E-01 | no | Gm44889 | upstream_gene_variant | MODIFIER |
| 1 | 140936459 | AX-97815327 | 1 | T | C | 0.205 | 3.07E-02 | 1.29E-01 | 2.66E+00 | 2.73E+00 | 8.11E-01 | no | | intergenic_variant | MODIFIER |
| 15 | 91437407 | AX-170087938 | 1 | G | A | 0.134 | 3.81E-02 | 1.57E-01 | 2.65E+00 | 2.71E+00 | 8.11E-01 | no | Slc2a13 | intron_variant | MODIFIER |
| 5 | 123259690 | AX-97815368 | 1 | C | A | 0.446 | -2.68E-02 | 1.14E-01 | 2.75E+00 | 2.79E+00 | 8.11E-01 | no | Wdr66 | intron_variant | MODIFIER |
| 16 | 26522150 | AX-170388182 | 0 | A | T | 0.246 | -2.00E-02 | 8.67E-02 | 2.62E+00 | 2.76E+00 | 8.11E-01 | no | Tmem207 | intron_variant | MODIFIER |
| 7 | 52689714 | AX-170029007 | 1 | C | T | 0.438 | -2.12E-02 | 9.37E-02 | 2.70E+00 | 2.80E+00 | 8.11E-01 | no | | intergenic_variant | MODIFIER |
| 1 | 75287093 | AX-97816419 | 1 | T | C | 0.125 | 4.87E-02 | 2.07E-01 | 2.69E+00 | 2.74E+00 | 8.11E-01 | no | | intergenic_variant | MODIFIER |
| 9 | 29493976 | AX-170081975 | 0 | T | C | 0.439 | 2.11E-02 | 8.70E-02 | 2.64E+00 | 2.70E+00 | 8.11E-01 | no | Ntm | intron_variant | MODIFIER |
| 9 | 29497301 | AX-97801576 | 0 | T | C | 0.439 | 2.11E-02 | 8.70E-02 | 2.64E+00 | 2.70E+00 | 8.11E-01 | no | Ntm | intron_variant | MODIFIER |
| 9 | 29497745 | AX-170113692 | 0 | C | A | 0.439 | 2.11E-02 | 8.70E-02 | 2.64E+00 | 2.70E+00 | 8.11E-01 | no | Ntm | intron_variant | MODIFIER |
| 9 | 29509348 | AX-170137040 | 0 | G | A | 0.439 | 2.11E-02 | 8.70E-02 | 2.64E+00 | 2.70E+00 | 8.11E-01 | no | Ntm | intron_variant | MODIFIER |
| 9 | 29515360 | AX-170227646 | 0 | C | G | 0.439 | 2.11E-02 | 8.70E-02 | 2.64E+00 | 2.70E+00 | 8.11E-01 | no | Ntm | intron_variant | MODIFIER |
| 9 | 29516407 | AX-170413186 | 0 | C | T | 0.439 | 2.11E-02 | 8.70E-02 | 2.64E+00 | 2.70E+00 | 8.11E-01 | no | Ntm | intron_variant | MODIFIER |
| 9 | 29519521 | AX-97785525 | 0 | G | A | 0.439 | 2.11E-02 | 8.70E-02 | 2.64E+00 | 2.70E+00 | 8.11E-01 | no | Ntm | intron_variant | MODIFIER |
| 9 | 29524335 | AX-170299590 | 0 | G | A | 0.439 | 2.11E-02 | 8.70E-02 | 2.64E+00 | 2.70E+00 | 8.11E-01 | no | Ntm | intron_variant | MODIFIER |
| 9 | 29549787 | AX-170096503 | 0 | T | G | 0.439 | 2.11E-02 | 8.70E-02 | 2.64E+00 | 2.70E+00 | 8.11E-01 | no | Ntm | intron_variant | MODIFIER |
| 9 | 29550451 | AX-170125583 | 0 | G | A | 0.439 | 2.11E-02 | 8.70E-02 | 2.64E+00 | 2.70E+00 | 8.11E-01 | no | Ntm | intron_variant | MODIFIER |
| 9 | 29562080 | AX-170234838 | 0 | A | C | 0.439 | 2.11E-02 | 8.70E-02 | 2.64E+00 | 2.70E+00 | 8.11E-01 | no | Ntm | intron_variant | MODIFIER |
| 9 | 29563178 | AX-170373948 | 0 | G | A | 0.439 | 2.11E-02 | 8.70E-02 | 2.64E+00 | 2.70E+00 | 8.11E-01 | no | Ntm | intron_variant | MODIFIER |

| | | | | | | | | | | | | | | | |
|---|---|---|---|---|---|---|---|---|---|---|---|---|---|---|---|
| 9 | 29572593 | AX-97805304 | 0 | G | A | 0.439 | 2.11E-02 | 8.70E-02 | 2.64E+00 | 2.70E+00 | 8.11E-01 | no | Ntm | intron_variant | MODIFIER |
| 9 | 29575626 | AX-170153116 | 0 | C | A | 0.439 | 2.11E-02 | 8.70E-02 | 2.64E+00 | 2.70E+00 | 8.11E-01 | no | Ntm | intron_variant | MODIFIER |
| 9 | 29607004 | AX-97833730 | 0 | C | T | 0.439 | 2.11E-02 | 8.70E-02 | 2.64E+00 | 2.70E+00 | 8.11E-01 | no | Ntm | intron_variant | MODIFIER |
| 9 | 29610291 | AX-170354975 | 0 | T | C | 0.439 | 2.11E-02 | 8.70E-02 | 2.64E+00 | 2.70E+00 | 8.11E-01 | no | Ntm | intron_variant | MODIFIER |
| 9 | 29610375 | AX-97847438 | 0 | A | G | 0.439 | 2.11E-02 | 8.70E-02 | 2.64E+00 | 2.70E+00 | 8.11E-01 | no | Ntm | intron_variant | MODIFIER |
| 9 | 29612520 | AX-170261196 | 0 | T | C | 0.439 | 2.11E-02 | 8.70E-02 | 2.64E+00 | 2.70E+00 | 8.11E-01 | no | Ntm | intron_variant | MODIFIER |
| 9 | 29621683 | AX-170379768 | 0 | G | A | 0.439 | 2.11E-02 | 8.70E-02 | 2.64E+00 | 2.70E+00 | 8.11E-01 | no | Ntm | intron_variant | MODIFIER |
| 9 | 29622238 | AX-170373378 | 0 | T | C | 0.439 | 2.11E-02 | 8.70E-02 | 2.64E+00 | 2.70E+00 | 8.11E-01 | no | Ntm | intron_variant | MODIFIER |
| 9 | 29630794 | AX-170164115 | 0 | G | A | 0.439 | 2.11E-02 | 8.70E-02 | 2.64E+00 | 2.70E+00 | 8.11E-01 | no | Ntm | intron_variant | MODIFIER |
| 9 | 29637287 | AX-170519435 | 0 | G | T | 0.439 | 2.11E-02 | 8.70E-02 | 2.64E+00 | 2.70E+00 | 8.11E-01 | no | Ntm | intron_variant | MODIFIER |
| 9 | 29641583 | AX-170291957 | 0 | A | C | 0.439 | 2.11E-02 | 8.70E-02 | 2.64E+00 | 2.70E+00 | 8.11E-01 | no | Ntm | intron_variant | MODIFIER |
| 9 | 29642260 | AX-97845612 | 0 | A | G | 0.439 | 2.11E-02 | 8.70E-02 | 2.64E+00 | 2.70E+00 | 8.11E-01 | no | Ntm | intron_variant | MODIFIER |
| 9 | 29642407 | AX-170490448 | 0 | C | T | 0.439 | 2.11E-02 | 8.70E-02 | 2.64E+00 | 2.70E+00 | 8.11E-01 | no | Ntm | intron_variant | MODIFIER |
| 9 | 29645634 | AX-170308222 | 0 | A | C | 0.439 | 2.11E-02 | 8.70E-02 | 2.64E+00 | 2.70E+00 | 8.11E-01 | no | Ntm | intron_variant | MODIFIER |
| 9 | 29656431 | AX-170419364 | 0 | A | C | 0.439 | 2.11E-02 | 8.70E-02 | 2.64E+00 | 2.70E+00 | 8.11E-01 | no | Ntm | intron_variant | MODIFIER |
| 9 | 29673670 | AX-170354237 | 0 | T | C | 0.439 | 2.11E-02 | 8.70E-02 | 2.64E+00 | 2.70E+00 | 8.11E-01 | no | Ntm | intron_variant | MODIFIER |
| 9 | 29674014 | AX-170527137 | 0 | G | A | 0.439 | 2.11E-02 | 8.70E-02 | 2.64E+00 | 2.70E+00 | 8.11E-01 | no | Ntm | intron_variant | MODIFIER |
| 9 | 29674450 | AX-97805302 | 0 | T | C | 0.439 | 2.11E-02 | 8.70E-02 | 2.64E+00 | 2.70E+00 | 8.11E-01 | no | Ntm | intron_variant | MODIFIER |
| 9 | 29694257 | AX-97839136 | 0 | T | C | 0.439 | 2.11E-02 | 8.70E-02 | 2.64E+00 | 2.70E+00 | 8.11E-01 | no | Ntm | intron_variant | MODIFIER |
| 9 | 29708083 | AX-170185361 | 0 | T | G | 0.439 | 2.11E-02 | 8.70E-02 | 2.64E+00 | 2.70E+00 | 8.11E-01 | no | Ntm | intron_variant | MODIFIER |
| 9 | 29714634 | AX-170032048 | 0 | G | T | 0.439 | 2.11E-02 | 8.70E-02 | 2.64E+00 | 2.70E+00 | 8.11E-01 | no | Ntm | intron_variant | MODIFIER |
| 9 | 29716490 | AX-170443668 | 0 | G | T | 0.439 | 2.11E-02 | 8.70E-02 | 2.64E+00 | 2.70E+00 | 8.11E-01 | no | Ntm | intron_variant | MODIFIER |
| 9 | 29726596 | AX-170392958 | 0 | A | G | 0.439 | 2.11E-02 | 8.70E-02 | 2.64E+00 | 2.70E+00 | 8.11E-01 | no | Ntm | intron_variant | MODIFIER |
| 9 | 29731986 | AX-170142807 | 0 | G | A | 0.439 | 2.11E-02 | 8.70E-02 | 2.64E+00 | 2.70E+00 | 8.11E-01 | no | Ntm | intron_variant | MODIFIER |
| 9 | 29732784 | AX-170189456 | 0 | C | T | 0.439 | 2.11E-02 | 8.70E-02 | 2.64E+00 | 2.70E+00 | 8.11E-01 | no | Ntm | intron_variant | MODIFIER |
| 9 | 29746864 | AX-170032046 | 0 | G | T | 0.439 | 2.11E-02 | 8.70E-02 | 2.64E+00 | 2.70E+00 | 8.11E-01 | no | Ntm | intron_variant | MODIFIER |
| 9 | 29747087 | AX-170090269 | 0 | T | G | 0.439 | 2.11E-02 | 8.70E-02 | 2.64E+00 | 2.70E+00 | 8.11E-01 | no | Ntm | intron_variant | MODIFIER |
| 9 | 29747635 | AX-170032045 | 0 | C | A | 0.439 | 2.11E-02 | 8.70E-02 | 2.64E+00 | 2.70E+00 | 8.11E-01 | no | Ntm | intron_variant | MODIFIER |
| 9 | 29747714 | AX-170160899 | 0 | A | T | 0.439 | 2.11E-02 | 8.70E-02 | 2.64E+00 | 2.70E+00 | 8.11E-01 | no | Ntm | intron_variant | MODIFIER |
| 9 | 29760209 | AX-170290263 | 0 | G | A | 0.439 | 2.11E-02 | 8.70E-02 | 2.64E+00 | 2.70E+00 | 8.11E-01 | no | Ntm | intron_variant | MODIFIER |
| 9 | 29760619 | AX-170337743 | 0 | T | C | 0.439 | 2.11E-02 | 8.70E-02 | 2.64E+00 | 2.70E+00 | 8.11E-01 | no | Ntm | intron_variant | MODIFIER |
| 9 | 29760712 | AX-97818611 | 0 | G | A | 0.439 | 2.11E-02 | 8.70E-02 | 2.64E+00 | 2.70E+00 | 8.11E-01 | no | Ntm | intron_variant | MODIFIER |
| 9 | 29780487 | AX-170248984 | 0 | G | A | 0.439 | 2.11E-02 | 8.70E-02 | 2.64E+00 | 2.70E+00 | 8.11E-01 | no | Ntm | intron_variant | MODIFIER |
| 9 | 29785943 | AX-170032044 | 0 | G | A | 0.439 | 2.11E-02 | 8.70E-02 | 2.64E+00 | 2.70E+00 | 8.11E-01 | no | Ntm | intron_variant | MODIFIER |
| 9 | 29787520 | AX-170032047 | 0 | A | C | 0.439 | 2.11E-02 | 8.70E-02 | 2.64E+00 | 2.70E+00 | 8.11E-01 | no | Ntm | intron_variant | MODIFIER |
| 9 | 29796018 | AX-97811596 | 0 | C | T | 0.439 | 2.11E-02 | 8.70E-02 | 2.64E+00 | 2.70E+00 | 8.11E-01 | no | Ntm | intron_variant | MODIFIER |
| 9 | 29823035 | AX-170032049 | 0 | C | A | 0.439 | 2.11E-02 | 8.70E-02 | 2.64E+00 | 2.70E+00 | 8.11E-01 | no | Ntm | intron_variant | MODIFIER |
| 9 | 29823256 | AX-97798511 | 0 | T | G | 0.439 | 2.11E-02 | 8.70E-02 | 2.64E+00 | 2.70E+00 | 8.11E-01 | no | Ntm | intron_variant | MODIFIER |
| 9 | 29827216 | AX-170310610 | 0 | G | A | 0.439 | 2.11E-02 | 8.70E-02 | 2.64E+00 | 2.70E+00 | 8.11E-01 | no | Ntm | intron_variant | MODIFIER |
| 9 | 29864716 | AX-170232671 | 0 | G | T | 0.439 | 2.11E-02 | 8.70E-02 | 2.64E+00 | 2.70E+00 | 8.11E-01 | no | Ntm | intron_variant | MODIFIER |
| 9 | 29871609 | AX-97827246 | 0 | G | A | 0.439 | 2.11E-02 | 8.70E-02 | 2.64E+00 | 2.70E+00 | 8.11E-01 | no | Ntm | intron_variant | MODIFIER |
| 9 | 29914163 | AX-170370821 | 0 | C | T | 0.439 | 2.11E-02 | 8.70E-02 | 2.64E+00 | 2.70E+00 | 8.11E-01 | no | Ntm | intron_variant | MODIFIER |
| 9 | 29916820 | AX-170133674 | 0 | C | T | 0.439 | 2.11E-02 | 8.70E-02 | 2.64E+00 | 2.70E+00 | 8.11E-01 | no | Ntm | intron_variant | MODIFIER |
| 9 | 29916851 | AX-170312078 | 0 | T | C | 0.439 | 2.11E-02 | 8.70E-02 | 2.64E+00 | 2.70E+00 | 8.11E-01 | no | Ntm | intron_variant | MODIFIER |
| 9 | 29923306 | AX-97805198 | 0 | T | G | 0.439 | 2.11E-02 | 8.70E-02 | 2.64E+00 | 2.70E+00 | 8.11E-01 | no | Ntm | intron_variant | MODIFIER |
| 9 | 29926690 | AX-170081987 | 0 | C | G | 0.439 | 2.11E-02 | 8.70E-02 | 2.64E+00 | 2.70E+00 | 8.11E-01 | no | Ntm | intron_variant | MODIFIER |
| 9 | 29933413 | AX-170422308 | 0 | G | A | 0.439 | 2.11E-02 | 8.70E-02 | 2.64E+00 | 2.70E+00 | 8.11E-01 | no | Ntm | intron_variant | MODIFIER |
| 9 | 29940961 | AX-170470871 | 0 | G | A | 0.439 | 2.11E-02 | 8.70E-02 | 2.64E+00 | 2.70E+00 | 8.11E-01 | no | Ntm | intron_variant | MODIFIER |
| 9 | 29944918 | AX-97811894 | 0 | G | T | 0.439 | 2.11E-02 | 8.70E-02 | 2.64E+00 | 2.70E+00 | 8.11E-01 | no | Ntm | intron_variant | MODIFIER |
| 9 | 29945057 | AX-170462004 | 0 | G | C | 0.439 | 2.11E-02 | 8.70E-02 | 2.64E+00 | 2.70E+00 | 8.11E-01 | no | Ntm | intron_variant | MODIFIER |
| 9 | 29981884 | AX-97814767 | 0 | G | A | 0.439 | 2.11E-02 | 8.70E-02 | 2.64E+00 | 2.70E+00 | 8.11E-01 | no | | intergenic_variant | MODIFIER |
| 9 | 30202620 | AX-170345448 | 0 | G | A | 0.439 | 2.11E-02 | 8.70E-02 | 2.64E+00 | 2.70E+00 | 8.11E-01 | no | | intergenic_variant | MODIFIER |
| 9 | 32708612 | AX-170082037 | 0 | G | A | 0.439 | 2.11E-02 | 8.70E-02 | 2.64E+00 | 2.70E+00 | 8.11E-01 | no | Ets1 | intron_variant | MODIFIER |
| 9 | 32719309 | AX-170139902 | 0 | A | G | 0.439 | 2.11E-02 | 8.70E-02 | 2.64E+00 | 2.70E+00 | 8.11E-01 | no | Ets1 | intron_variant | MODIFIER |

| | | | | | | | | | | | | | | | |
|---|---|---|---|---|---|---|---|---|---|---|---|---|---|---|---|
| 9 | 32720505 | AX-170473629 | 0 | C | T | 0.439 | 2.11E-02 | 8.70E-02 | 2.64E+00 | 2.70E+00 | 8.11E-01 | no | Ets1 | intron_variant | MODIFIER |
| 9 | 32720759 | AX-170303523 | 0 | A | G | 0.439 | 2.11E-02 | 8.70E-02 | 2.64E+00 | 2.70E+00 | 8.11E-01 | no | Ets1 | intron_variant | MODIFIER |
| 9 | 32740709 | AX-97827117 | 0 | T | A | 0.439 | 2.11E-02 | 8.70E-02 | 2.64E+00 | 2.70E+00 | 8.11E-01 | no | Ets1 | intron_variant | MODIFIER |
| 9 | 32792520 | AX-170473948 | 0 | G | A | 0.439 | 2.11E-02 | 8.70E-02 | 2.64E+00 | 2.70E+00 | 8.11E-01 | no | | intergenic_variant | MODIFIER |
| 9 | 32793409 | AX-170403822 | 0 | T | C | 0.439 | 2.11E-02 | 8.70E-02 | 2.64E+00 | 2.70E+00 | 8.11E-01 | no | | intergenic_variant | MODIFIER |
| 9 | 32827507 | AX-97818917 | 0 | C | T | 0.439 | 2.11E-02 | 8.70E-02 | 2.64E+00 | 2.70E+00 | 8.11E-01 | no | Gm3331 | intron_variant,non_coding_transcript_variant | MODIFIER |
| 9 | 32857366 | AX-170535750 | 0 | C | T | 0.439 | 2.11E-02 | 8.70E-02 | 2.64E+00 | 2.70E+00 | 8.11E-01 | no | | intergenic_variant | MODIFIER |
| 9 | 32858543 | AX-170082043 | 0 | A | G | 0.439 | 2.11E-02 | 8.70E-02 | 2.64E+00 | 2.70E+00 | 8.11E-01 | no | | intergenic_variant | MODIFIER |
| 9 | 32905086 | AX-97857214 | 0 | C | T | 0.439 | 2.11E-02 | 8.70E-02 | 2.64E+00 | 2.70E+00 | 8.11E-01 | no | Gm27162 | non_coding_transcript_exon_variant | MODIFIER |
| 9 | 32910959 | AX-170146516 | 0 | C | T | 0.439 | 2.11E-02 | 8.70E-02 | 2.64E+00 | 2.70E+00 | 8.11E-01 | no | Gm27162 | non_coding_transcript_exon_variant | MODIFIER |
| 9 | 32932073 | AX-170182516 | 0 | C | T | 0.439 | 2.11E-02 | 8.70E-02 | 2.64E+00 | 2.70E+00 | 8.11E-01 | no | Gm27162 | upstream_gene_variant | MODIFIER |
| 9 | 32971233 | AX-170262919 | 0 | C | T | 0.439 | 2.11E-02 | 8.70E-02 | 2.64E+00 | 2.70E+00 | 8.11E-01 | no | Gm37192 | non_coding_transcript_exon_variant | MODIFIER |
| 9 | 32986790 | AX-170032080 | 0 | A | G | 0.439 | 2.11E-02 | 8.70E-02 | 2.64E+00 | 2.70E+00 | 8.11E-01 | no | | regulatory_region_variant | MODIFIER |
| 9 | 33039827 | AX-170032077 | 0 | C | G | 0.439 | 2.11E-02 | 8.70E-02 | 2.64E+00 | 2.70E+00 | 8.11E-01 | no | Gm27166 | intron_variant,non_coding_transcript_variant | MODIFIER |
| 9 | 33071183 | AX-97822826 | 0 | A | C | 0.439 | 2.11E-02 | 8.70E-02 | 2.64E+00 | 2.70E+00 | 8.11E-01 | no | Gm27166 | intron_variant,non_coding_transcript_variant | MODIFIER |
| 9 | 33150554 | AX-170032084 | 0 | G | T | 0.439 | 2.11E-02 | 8.70E-02 | 2.64E+00 | 2.70E+00 | 8.11E-01 | no | Gm32171 | upstream_gene_variant | MODIFIER |
| 9 | 33206964 | AX-170032076 | 0 | A | C | 0.439 | 2.11E-02 | 8.70E-02 | 2.64E+00 | 2.70E+00 | 8.11E-01 | no | Gm32171 | intron_variant,non_coding_transcript_variant | MODIFIER |
| 9 | 33225789 | AX-170125799 | 0 | G | A | 0.439 | 2.11E-02 | 8.70E-02 | 2.64E+00 | 2.70E+00 | 8.11E-01 | no | Gm32171 | intron_variant,non_coding_transcript_variant | MODIFIER |
| 9 | 33241631 | AX-170242896 | 0 | T | G | 0.439 | 2.11E-02 | 8.70E-02 | 2.64E+00 | 2.70E+00 | 8.11E-01 | no | Gm32171 | intron_variant,non_coding_transcript_variant | MODIFIER |
| 9 | 33262082 | AX-170212872 | 0 | G | A | 0.439 | 2.11E-02 | 8.70E-02 | 2.64E+00 | 2.70E+00 | 8.11E-01 | no | Gm32171 | non_coding_transcript_exon_variant | MODIFIER |
| 9 | 33267596 | AX-170379897 | 0 | C | T | 0.439 | 2.11E-02 | 8.70E-02 | 2.64E+00 | 2.70E+00 | 8.11E-01 | no | | intergenic_variant | MODIFIER |
| 9 | 33304434 | AX-97806780 | 0 | T | C | 0.439 | 2.11E-02 | 8.70E-02 | 2.64E+00 | 2.70E+00 | 8.11E-01 | no | Gm47775 | intron_variant,non_coding_transcript_variant | MODIFIER |
| 9 | 33335123 | AX-170451265 | 0 | C | T | 0.439 | 2.11E-02 | 8.70E-02 | 2.64E+00 | 2.70E+00 | 8.11E-01 | no | | intergenic_variant | MODIFIER |
| 9 | 33335283 | AX-170315043 | 0 | T | C | 0.439 | 2.11E-02 | 8.70E-02 | 2.64E+00 | 2.70E+00 | 8.11E-01 | no | | intergenic_variant | MODIFIER |
| 9 | 33340079 | AX-170104206 | 0 | C | T | 0.439 | 2.11E-02 | 8.70E-02 | 2.64E+00 | 2.70E+00 | 8.11E-01 | no | | intergenic_variant | MODIFIER |
| 9 | 33341032 | AX-170103657 | 0 | G | A | 0.439 | 2.11E-02 | 8.70E-02 | 2.64E+00 | 2.70E+00 | 8.11E-01 | no | | intergenic_variant | MODIFIER |
| 9 | 33361714 | AX-170417534 | 0 | T | C | 0.439 | 2.11E-02 | 8.70E-02 | 2.64E+00 | 2.70E+00 | 8.11E-01 | no | | intergenic_variant | MODIFIER |
| 9 | 33383243 | AX-170347593 | 0 | G | A | 0.439 | 2.11E-02 | 8.70E-02 | 2.64E+00 | 2.70E+00 | 8.11E-01 | no | | intergenic_variant | MODIFIER |
| 9 | 33413653 | AX-170478021 | 0 | A | G | 0.439 | 2.11E-02 | 8.70E-02 | 2.64E+00 | 2.70E+00 | 8.11E-01 | no | | intergenic_variant | MODIFIER |
| 9 | 33507569 | AX-97832193 | 0 | C | G | 0.439 | 2.11E-02 | 8.70E-02 | 2.64E+00 | 2.70E+00 | 8.11E-01 | no | | intergenic_variant | MODIFIER |
| 9 | 33585587 | AX-170120445 | 0 | G | A | 0.439 | 2.11E-02 | 8.70E-02 | 2.64E+00 | 2.70E+00 | 8.11E-01 | no | | intergenic_variant | MODIFIER |
| 9 | 33631022 | AX-97804682 | 0 | G | T | 0.439 | 2.11E-02 | 8.70E-02 | 2.64E+00 | 2.70E+00 | 8.11E-01 | no | | intergenic_variant | MODIFIER |
| 9 | 33651752 | AX-170145500 | 0 | A | T | 0.439 | 2.11E-02 | 8.70E-02 | 2.64E+00 | 2.70E+00 | 8.11E-01 | no | | intergenic_variant | MODIFIER |
| 9 | 33679585 | AX-170525410 | 0 | C | T | 0.439 | 2.11E-02 | 8.70E-02 | 2.64E+00 | 2.70E+00 | 8.11E-01 | no | | intergenic_variant | MODIFIER |
| 9 | 33688170 | AX-170144832 | 0 | T | C | 0.439 | 2.11E-02 | 8.70E-02 | 2.64E+00 | 2.70E+00 | 8.11E-01 | no | | intergenic_variant | MODIFIER |
| 9 | 33830449 | AX-170082058 | 0 | C | A | 0.439 | 2.11E-02 | 8.70E-02 | 2.64E+00 | 2.70E+00 | 8.11E-01 | no | | intergenic_variant | MODIFIER |
| 9 | 33856807 | AX-170123250 | 0 | T | C | 0.439 | 2.11E-02 | 8.70E-02 | 2.64E+00 | 2.70E+00 | 8.11E-01 | no | Gm47784 | upstream_gene_variant | MODIFIER |
| 9 | 33926270 | AX-170330060 | 0 | C | G | 0.439 | 2.11E-02 | 8.70E-02 | 2.64E+00 | 2.70E+00 | 8.11E-01 | no | Gm47785 | downstream_gene_variant | MODIFIER |
| 9 | 33930048 | AX-170517179 | 0 | G | A | 0.439 | 2.11E-02 | 8.70E-02 | 2.64E+00 | 2.70E+00 | 8.11E-01 | no | Gm27165 | upstream_gene_variant | MODIFIER |
| 9 | 33930823 | AX-170510884 | 0 | G | T | 0.439 | 2.11E-02 | 8.70E-02 | 2.64E+00 | 2.70E+00 | 8.11E-01 | no | Gm27165 | upstream_gene_variant | MODIFIER |
| 9 | 33936144 | AX-170530361 | 0 | A | G | 0.439 | 2.11E-02 | 8.70E-02 | 2.64E+00 | 2.70E+00 | 8.11E-01 | no | Gm27165 | downstream_gene_variant | MODIFIER |
| 9 | 33939690 | AX-170364299 | 0 | A | G | 0.439 | 2.11E-02 | 8.70E-02 | 2.64E+00 | 2.70E+00 | 8.11E-01 | no | Gm27164 | upstream_gene_variant | MODIFIER |
| 9 | 33945420 | AX-97804681 | 0 | T | C | 0.439 | 2.11E-02 | 8.70E-02 | 2.64E+00 | 2.70E+00 | 8.11E-01 | no | Gm27164 | intron_variant,non_coding_transcript_variant | MODIFIER |
| 9 | 34002825 | AX-170032098 | 0 | G | T | 0.439 | 2.11E-02 | 8.70E-02 | 2.64E+00 | 2.70E+00 | 8.11E-01 | no | Gm27164 | intron_variant,non_coding_transcript_variant | MODIFIER |
| 9 | 34112786 | AX-170313179 | 0 | C | T | 0.439 | 2.11E-02 | 8.70E-02 | 2.64E+00 | 2.70E+00 | 8.11E-01 | no | | intergenic_variant | MODIFIER |
| 9 | 34153204 | AX-170267912 | 0 | T | C | 0.439 | 2.11E-02 | 8.70E-02 | 2.64E+00 | 2.70E+00 | 8.11E-01 | no | | intergenic_variant | MODIFIER |
| 9 | 34153961 | AX-170453902 | 0 | G | A | 0.439 | 2.11E-02 | 8.70E-02 | 2.64E+00 | 2.70E+00 | 8.11E-01 | no | | intergenic_variant | MODIFIER |
| 9 | 34155118 | AX-170391027 | 0 | C | T | 0.439 | 2.11E-02 | 8.70E-02 | 2.64E+00 | 2.70E+00 | 8.11E-01 | no | | intergenic_variant | MODIFIER |
| 9 | 34166928 | AX-170487204 | 0 | G | A | 0.439 | 2.11E-02 | 8.70E-02 | 2.64E+00 | 2.70E+00 | 8.11E-01 | no | | intergenic_variant | MODIFIER |
| 9 | 34180111 | AX-170162044 | 0 | T | C | 0.439 | 2.11E-02 | 8.70E-02 | 2.64E+00 | 2.70E+00 | 8.11E-01 | no | | intergenic_variant | MODIFIER |
| 9 | 34181167 | AX-170082065 | 0 | A | T | 0.439 | 2.11E-02 | 8.70E-02 | 2.64E+00 | 2.70E+00 | 8.11E-01 | no | | intergenic_variant | MODIFIER |
| 9 | 34205966 | AX-170454467 | 0 | G | T | 0.439 | 2.11E-02 | 8.70E-02 | 2.64E+00 | 2.70E+00 | 8.11E-01 | no | | intergenic_variant | MODIFIER |
| 9 | 34227613 | AX-170126123 | 0 | G | T | 0.439 | 2.11E-02 | 8.70E-02 | 2.64E+00 | 2.70E+00 | 8.11E-01 | no | | intergenic_variant | MODIFIER |
| 9 | 34237613 | AX-170391614 | 0 | A | T | 0.439 | 2.11E-02 | 8.70E-02 | 2.64E+00 | 2.70E+00 | 8.11E-01 | no | | intergenic_variant | MODIFIER |

| | | | | | | | | | | | | | | | |
|---|---|---|---|---|---|---|---|---|---|---|---|---|---|---|---|
| 9 | 34237917 | AX-97786325 | 0 | G | A | 0.439 | 2.11E-02 | 8.70E-02 | 2.64E+00 | 2.70E+00 | 8.11E-01 | no | | intergenic_variant | MODIFIER |
| 9 | 34262778 | AX-170267780 | 0 | T | A | 0.439 | 2.11E-02 | 8.70E-02 | 2.64E+00 | 2.70E+00 | 8.11E-01 | no | | intergenic_variant | MODIFIER |
| 9 | 34262882 | AX-170417132 | 0 | G | A | 0.439 | 2.11E-02 | 8.70E-02 | 2.64E+00 | 2.70E+00 | 8.11E-01 | no | | intergenic_variant | MODIFIER |
| 9 | 34264662 | AX-170286415 | 0 | T | C | 0.439 | 2.11E-02 | 8.70E-02 | 2.64E+00 | 2.70E+00 | 8.11E-01 | no | | intergenic_variant | MODIFIER |
| 9 | 34442834 | AX-170461576 | 0 | T | A | 0.439 | 2.11E-02 | 8.70E-02 | 2.64E+00 | 2.70E+00 | 8.11E-01 | no | | intergenic_variant | MODIFIER |
| 9 | 34462672 | AX-170162831 | 0 | G | A | 0.439 | 2.11E-02 | 8.70E-02 | 2.64E+00 | 2.70E+00 | 8.11E-01 | no | Gm47858 | intron_variant,non_coding_transcript_variant | MODIFIER |
| 9 | 34619983 | AX-170032104 | 0 | C | T | 0.439 | 2.11E-02 | 8.70E-02 | 2.64E+00 | 2.70E+00 | 8.11E-01 | no | Kirrel3 | intron_variant | MODIFIER |
| 9 | 34658510 | AX-170035808 | 0 | C | T | 0.439 | 2.11E-02 | 8.70E-02 | 2.64E+00 | 2.70E+00 | 8.11E-01 | no | Kirrel3 | intron_variant | MODIFIER |
| 9 | 34671508 | AX-170479033 | 0 | T | C | 0.439 | 2.11E-02 | 8.70E-02 | 2.64E+00 | 2.70E+00 | 8.11E-01 | no | Kirrel3 | intron_variant | MODIFIER |
| 9 | 34704374 | AX-170178921 | 0 | G | A | 0.439 | 2.11E-02 | 8.70E-02 | 2.64E+00 | 2.70E+00 | 8.11E-01 | no | Kirrel3 | intron_variant | MODIFIER |
| 9 | 34704531 | AX-170344579 | 0 | G | A | 0.439 | 2.11E-02 | 8.70E-02 | 2.64E+00 | 2.70E+00 | 8.11E-01 | no | Kirrel3 | intron_variant | MODIFIER |
| 9 | 34709670 | AX-170197385 | 0 | T | C | 0.439 | 2.11E-02 | 8.70E-02 | 2.64E+00 | 2.70E+00 | 8.11E-01 | no | Kirrel3 | intron_variant | MODIFIER |
| 9 | 34725264 | AX-97821921 | 0 | G | A | 0.439 | 2.11E-02 | 8.70E-02 | 2.64E+00 | 2.70E+00 | 8.11E-01 | no | Kirrel3 | intron_variant | MODIFIER |
| 9 | 34743586 | AX-170503079 | 0 | T | A | 0.439 | 2.11E-02 | 8.70E-02 | 2.64E+00 | 2.70E+00 | 8.11E-01 | no | Kirrel3 | intron_variant | MODIFIER |
| 9 | 34744163 | AX-170304233 | 0 | G | A | 0.439 | 2.11E-02 | 8.70E-02 | 2.64E+00 | 2.70E+00 | 8.11E-01 | no | Kirrel3 | intron_variant | MODIFIER |
| 9 | 34746336 | AX-170266593 | 0 | A | G | 0.439 | 2.11E-02 | 8.70E-02 | 2.64E+00 | 2.70E+00 | 8.11E-01 | no | Kirrel3 | intron_variant | MODIFIER |
| 9 | 34746350 | AX-170482739 | 0 | A | G | 0.439 | 2.11E-02 | 8.70E-02 | 2.64E+00 | 2.70E+00 | 8.11E-01 | no | Kirrel3 | intron_variant | MODIFIER |
| 9 | 34751385 | AX-97790266 | 0 | T | C | 0.439 | 2.11E-02 | 8.70E-02 | 2.64E+00 | 2.70E+00 | 8.11E-01 | no | Kirrel3 | intron_variant | MODIFIER |
| 9 | 34752172 | AX-170398427 | 0 | A | G | 0.439 | 2.11E-02 | 8.70E-02 | 2.64E+00 | 2.70E+00 | 8.11E-01 | no | Kirrel3 | intron_variant | MODIFIER |
| 9 | 34753151 | AX-170407394 | 0 | C | T | 0.439 | 2.11E-02 | 8.70E-02 | 2.64E+00 | 2.70E+00 | 8.11E-01 | no | Kirrel3 | intron_variant | MODIFIER |
| 9 | 34763343 | AX-170223568 | 0 | T | C | 0.439 | 2.11E-02 | 8.70E-02 | 2.64E+00 | 2.70E+00 | 8.11E-01 | no | Kirrel3 | intron_variant | MODIFIER |
| 9 | 34774282 | AX-170472166 | 0 | A | G | 0.439 | 2.11E-02 | 8.70E-02 | 2.64E+00 | 2.70E+00 | 8.11E-01 | no | Kirrel3 | intron_variant | MODIFIER |
| 9 | 34774642 | AX-170365033 | 0 | G | T | 0.439 | 2.11E-02 | 8.70E-02 | 2.64E+00 | 2.70E+00 | 8.11E-01 | no | Kirrel3 | intron_variant | MODIFIER |
| 9 | 34793283 | AX-168387747 | 0 | G | A | 0.439 | 2.11E-02 | 8.70E-02 | 2.64E+00 | 2.70E+00 | 8.11E-01 | no | Kirrel3 | intron_variant | MODIFIER |
| 9 | 34794020 | AX-170500136 | 0 | T | G | 0.439 | 2.11E-02 | 8.70E-02 | 2.64E+00 | 2.70E+00 | 8.11E-01 | no | Kirrel3 | intron_variant | MODIFIER |
| 9 | 34794281 | AX-170524161 | 0 | G | A | 0.439 | 2.11E-02 | 8.70E-02 | 2.64E+00 | 2.70E+00 | 8.11E-01 | no | Kirrel3 | intron_variant | MODIFIER |
| 9 | 34795118 | AX-170096493 | 0 | C | T | 0.439 | 2.11E-02 | 8.70E-02 | 2.64E+00 | 2.70E+00 | 8.11E-01 | no | Kirrel3 | intron_variant | MODIFIER |
| 9 | 34837880 | AX-170280056 | 0 | A | C | 0.439 | 2.11E-02 | 8.70E-02 | 2.64E+00 | 2.70E+00 | 8.11E-01 | no | Kirrel3 | intron_variant | MODIFIER |
| 9 | 34887803 | AX-170169573 | 0 | T | G | 0.439 | 2.11E-02 | 8.70E-02 | 2.64E+00 | 2.70E+00 | 8.11E-01 | no | Kirrel3 | intron_variant | MODIFIER |
| 9 | 34922995 | AX-170082088 | 0 | C | T | 0.439 | 2.11E-02 | 8.70E-02 | 2.64E+00 | 2.70E+00 | 8.11E-01 | no | Kirrel3 | intron_variant | MODIFIER |
| 9 | 34933109 | AX-170405608 | 0 | T | C | 0.439 | 2.11E-02 | 8.70E-02 | 2.64E+00 | 2.70E+00 | 8.11E-01 | no | Kirrel3 | intron_variant | MODIFIER |
| 9 | 34938479 | AX-170523956 | 0 | C | T | 0.439 | 2.11E-02 | 8.70E-02 | 2.64E+00 | 2.70E+00 | 8.11E-01 | no | Kirrel3 | intron_variant | MODIFIER |
| 9 | 34942020 | AX-170206873 | 0 | G | A | 0.439 | 2.11E-02 | 8.70E-02 | 2.64E+00 | 2.70E+00 | 8.11E-01 | no | Kirrel3 | intron_variant | MODIFIER |
| 9 | 34942417 | AX-170162200 | 0 | A | C | 0.439 | 2.11E-02 | 8.70E-02 | 2.64E+00 | 2.70E+00 | 8.11E-01 | no | Kirrel3 | intron_variant | MODIFIER |
| 9 | 34946025 | AX-170283252 | 0 | G | C | 0.439 | 2.11E-02 | 8.70E-02 | 2.64E+00 | 2.70E+00 | 8.11E-01 | no | Kirrel3 | intron_variant | MODIFIER |
| 9 | 34948500 | AX-170281592 | 0 | G | A | 0.439 | 2.11E-02 | 8.70E-02 | 2.64E+00 | 2.70E+00 | 8.11E-01 | no | Kirrel3 | intron_variant | MODIFIER |
| 9 | 34948827 | AX-170106665 | 0 | G | C | 0.439 | 2.11E-02 | 8.70E-02 | 2.64E+00 | 2.70E+00 | 8.11E-01 | no | Kirrel3 | intron_variant | MODIFIER |
| 9 | 34951932 | AX-170438577 | 0 | A | G | 0.439 | 2.11E-02 | 8.70E-02 | 2.64E+00 | 2.70E+00 | 8.11E-01 | no | Kirrel3 | intron_variant | MODIFIER |
| 9 | 34952348 | AX-170133072 | 0 | T | C | 0.439 | 2.11E-02 | 8.70E-02 | 2.64E+00 | 2.70E+00 | 8.11E-01 | no | Kirrel3 | intron_variant | MODIFIER |
| 9 | 34953009 | AX-170292654 | 0 | G | C | 0.439 | 2.11E-02 | 8.70E-02 | 2.64E+00 | 2.70E+00 | 8.11E-01 | no | Kirrel3 | intron_variant | MODIFIER |
| 9 | 34956456 | AX-170461382 | 0 | A | T | 0.439 | 2.11E-02 | 8.70E-02 | 2.64E+00 | 2.70E+00 | 8.11E-01 | no | Kirrel3 | intron_variant | MODIFIER |
| 9 | 34957097 | AX-170232861 | 0 | C | T | 0.439 | 2.11E-02 | 8.70E-02 | 2.64E+00 | 2.70E+00 | 8.11E-01 | no | Kirrel3 | intron_variant | MODIFIER |
| 9 | 34957339 | AX-170233016 | 0 | A | G | 0.439 | 2.11E-02 | 8.70E-02 | 2.64E+00 | 2.70E+00 | 8.11E-01 | no | Kirrel3 | intron_variant | MODIFIER |
| 9 | 34961378 | AX-170182950 | 0 | T | C | 0.439 | 2.11E-02 | 8.70E-02 | 2.64E+00 | 2.70E+00 | 8.11E-01 | no | Kirrel3 | intron_variant | MODIFIER |
| 9 | 34963711 | AX-170459081 | 0 | G | C | 0.439 | 2.11E-02 | 8.70E-02 | 2.64E+00 | 2.70E+00 | 8.11E-01 | no | Kirrel3 | intron_variant | MODIFIER |
| 9 | 34963760 | AX-97821014 | 0 | A | G | 0.439 | 2.11E-02 | 8.70E-02 | 2.64E+00 | 2.70E+00 | 8.11E-01 | no | Kirrel3 | intron_variant | MODIFIER |
| 9 | 34963883 | AX-170348429 | 0 | A | T | 0.439 | 2.11E-02 | 8.70E-02 | 2.64E+00 | 2.70E+00 | 8.11E-01 | no | Kirrel3 | intron_variant | MODIFIER |
| 9 | 34966431 | AX-170396342 | 0 | C | T | 0.439 | 2.11E-02 | 8.70E-02 | 2.64E+00 | 2.70E+00 | 8.11E-01 | no | Kirrel3 | intron_variant | MODIFIER |
| 3 | 152899342 | AX-170068473 | 2 | A | G | 0.318 | 1.70E-02 | 7.29E-02 | 2.65E+00 | 2.75E+00 | 8.11E-01 | no | St6galnac5 | intron_variant | MODIFIER |
| 7 | 112470226 | AX-170332819 | 1 | G | A | 0.464 | 2.25E-02 | 9.49E-02 | 2.66E+00 | 2.73E+00 | 8.11E-01 | no | Parva | intron_variant | MODIFIER |
| 5 | 28002954 | AX-170311499 | 0 | G | A | 0.333 | 2.58E-02 | 1.08E-01 | 2.67E+00 | 2.73E+00 | 8.11E-01 | no | Gm4865 | upstream_gene_variant | MODIFIER |
| 12 | 60826292 | AX-170006847 | 0 | T | A | 0.044 | 3.91E-02 | 1.75E-01 | 2.65E+00 | 2.79E+00 | 8.11E-01 | no | | intergenic_variant | MODIFIER |
| 2 | 120353484 | AX-97815634 | 0 | A | G | 0.132 | 4.69E-02 | 1.96E-01 | 2.66E+00 | 2.72E+00 | 8.12E-01 | no | Vps39 | upstream_gene_variant | MODIFIER |
| 2 | 123665180 | AX-170094398 | 0 | A | C | 0.132 | 4.69E-02 | 1.96E-01 | 2.66E+00 | 2.72E+00 | 8.12E-01 | no | | intergenic_variant | MODIFIER |

| | | | | | | | | | | | | | | | |
|---|---|---|---|---|---|---|---|---|---|---|---|---|---|---|---|
| 2 | 125867551 | AX-170248997 | 0 | A | G | 0.132 | 4.69E-02 | 1.96E-01 | 2.66E+00 | 2.72E+00 | 8.12E-01 | no | Galk2 | intron_variant | MODIFIER |
| 2 | 127085380 | AX-170117054 | 0 | A | G | 0.132 | 4.69E-02 | 1.96E-01 | 2.66E+00 | 2.72E+00 | 8.12E-01 | no | Blvra | intron_variant | MODIFIER |
| 2 | 130122530 | AX-170064909 | 0 | T | C | 0.132 | 4.69E-02 | 1.96E-01 | 2.66E+00 | 2.72E+00 | 8.12E-01 | no | Tgm6 | upstream_gene_variant | MODIFIER |
| 2 | 145560707 | AX-170280366 | 0 | T | A | 0.132 | 4.69E-02 | 1.96E-01 | 2.66E+00 | 2.72E+00 | 8.12E-01 | no | Slc24a3 | intron_variant | MODIFIER |
| 7 | 136450088 | AX-170472813 | 0 | C | A | 0.298 | -1.93E-02 | 7.93E-02 | 2.61E+00 | 2.71E+00 | 8.12E-01 | no | | intergenic_variant | MODIFIER |
| 7 | 136457889 | AX-170389255 | 0 | G | A | 0.298 | -1.93E-02 | 7.93E-02 | 2.61E+00 | 2.71E+00 | 8.12E-01 | no | | intergenic_variant | MODIFIER |
| 7 | 136462237 | AX-170030057 | 0 | C | A | 0.298 | -1.93E-02 | 7.93E-02 | 2.61E+00 | 2.71E+00 | 8.12E-01 | no | | intergenic_variant | MODIFIER |
| 7 | 136469016 | AX-170267203 | 0 | T | C | 0.298 | -1.93E-02 | 7.93E-02 | 2.61E+00 | 2.71E+00 | 8.12E-01 | no | | intergenic_variant | MODIFIER |
| 7 | 136475041 | AX-170411331 | 0 | A | T | 0.298 | -1.93E-02 | 7.93E-02 | 2.61E+00 | 2.71E+00 | 8.12E-01 | no | | intergenic_variant | MODIFIER |
| 7 | 136504234 | AX-170227048 | 0 | G | T | 0.298 | -1.93E-02 | 7.93E-02 | 2.61E+00 | 2.71E+00 | 8.12E-01 | no | Gm24444 | downstream_gene_variant | MODIFIER |
| 7 | 136511804 | AX-170334026 | 0 | G | C | 0.298 | -1.93E-02 | 7.93E-02 | 2.61E+00 | 2.71E+00 | 8.12E-01 | no | Gm24444 | upstream_gene_variant | MODIFIER |
| 7 | 136512092 | AX-170361085 | 0 | G | A | 0.298 | -1.93E-02 | 7.93E-02 | 2.61E+00 | 2.71E+00 | 8.12E-01 | no | Gm24444 | upstream_gene_variant | MODIFIER |
| 7 | 136819243 | AX-97809600 | 0 | G | A | 0.298 | -1.93E-02 | 7.93E-02 | 2.61E+00 | 2.71E+00 | 8.12E-01 | no | | intergenic_variant | MODIFIER |
| 7 | 136838497 | AX-170181484 | 0 | C | A | 0.298 | -1.93E-02 | 7.93E-02 | 2.61E+00 | 2.71E+00 | 8.12E-01 | no | | intergenic_variant | MODIFIER |
| 7 | 136842671 | AX-170030065 | 0 | C | T | 0.298 | -1.93E-02 | 7.93E-02 | 2.61E+00 | 2.71E+00 | 8.12E-01 | no | | intergenic_variant | MODIFIER |
| 7 | 137005277 | AX-170284160 | 0 | G | A | 0.298 | -1.93E-02 | 7.93E-02 | 2.61E+00 | 2.71E+00 | 8.12E-01 | no | Mgmt | intron_variant | MODIFIER |
| 7 | 137020642 | AX-97822855 | 0 | A | G | 0.298 | -1.93E-02 | 7.93E-02 | 2.61E+00 | 2.71E+00 | 8.12E-01 | no | Mgmt | intron_variant | MODIFIER |
| 7 | 137022105 | AX-170244649 | 0 | G | T | 0.298 | -1.93E-02 | 7.93E-02 | 2.61E+00 | 2.71E+00 | 8.12E-01 | no | Mgmt | intron_variant | MODIFIER |
| 7 | 137023474 | AX-97798450 | 0 | C | T | 0.298 | -1.93E-02 | 7.93E-02 | 2.61E+00 | 2.71E+00 | 8.12E-01 | no | Mgmt | intron_variant | MODIFIER |
| 7 | 137052531 | AX-170403659 | 0 | G | A | 0.298 | -1.93E-02 | 7.93E-02 | 2.61E+00 | 2.71E+00 | 8.12E-01 | no | Mgmt | intron_variant | MODIFIER |
| 7 | 137063316 | AX-97850084 | 0 | C | T | 0.298 | -1.93E-02 | 7.93E-02 | 2.61E+00 | 2.71E+00 | 8.12E-01 | no | Mgmt | intron_variant | MODIFIER |
| 7 | 137066077 | AX-170433773 | 0 | G | A | 0.298 | -1.93E-02 | 7.93E-02 | 2.61E+00 | 2.71E+00 | 8.12E-01 | no | Mgmt | intron_variant | MODIFIER |
| 7 | 137066167 | AX-170078984 | 0 | C | T | 0.298 | -1.93E-02 | 7.93E-02 | 2.61E+00 | 2.71E+00 | 8.12E-01 | no | Mgmt | intron_variant | MODIFIER |
| 7 | 137157328 | AX-170156566 | 0 | T | C | 0.298 | -1.93E-02 | 7.93E-02 | 2.61E+00 | 2.71E+00 | 8.12E-01 | no | | intergenic_variant | MODIFIER |
| 7 | 137690216 | AX-168403330 | 0 | C | T | 0.298 | -1.93E-02 | 7.93E-02 | 2.61E+00 | 2.71E+00 | 8.12E-01 | no | | intergenic_variant | MODIFIER |
| 1 | 174004332 | AX-170518234 | 0 | G | A | 0.228 | -2.33E-02 | 1.06E-01 | 2.71E+00 | 2.83E+00 | 8.12E-01 | no | Gm18445 | upstream_gene_variant | MODIFIER |
| 12 | 99114105 | AX-97826710 | 0 | A | G | 0.07 | 4.72E-02 | 2.02E-01 | 2.69E+00 | 2.75E+00 | 8.12E-01 | no | | intergenic_variant | MODIFIER |
| 12 | 99150584 | AX-97814149 | 0 | C | T | 0.07 | 4.72E-02 | 2.02E-01 | 2.69E+00 | 2.75E+00 | 8.12E-01 | no | | regulatory_region_variant | MODIFIER |
| 12 | 99185927 | AX-170144093 | 0 | G | A | 0.07 | 4.72E-02 | 2.02E-01 | 2.69E+00 | 2.75E+00 | 8.12E-01 | no | Foxn3 | downstream_gene_variant | MODIFIER |
| 12 | 99187041 | AX-170294115 | 0 | T | C | 0.07 | 4.72E-02 | 2.02E-01 | 2.69E+00 | 2.75E+00 | 8.12E-01 | no | Foxn3 | downstream_gene_variant | MODIFIER |
| 8 | 75734885 | AX-170197694 | 0 | A | G | 0.202 | -2.74E-02 | 1.16E-01 | 2.65E+00 | 2.73E+00 | 8.12E-01 | no | Iqcm | intron_variant | MODIFIER |
| 5 | 78556224 | AX-170163223 | 0 | C | A | 0.439 | 2.24E-02 | 9.55E-02 | 2.65E+00 | 2.74E+00 | 8.12E-01 | no | | intergenic_variant | MODIFIER |
| 13 | 36461340 | AX-97785730 | 1 | G | T | 0.196 | -2.97E-02 | 1.16E-01 | 2.59E+00 | 2.66E+00 | 8.12E-01 | no | Fars2 | intron_variant | MODIFIER |
| 9 | 27785621 | AX-170274608 | 1 | T | C | 0.455 | 2.27E-02 | 9.25E-02 | 2.63E+00 | 2.69E+00 | 8.12E-01 | no | | intergenic_variant | MODIFIER |
| 5 | 30252612 | AX-170513621 | 0 | C | G | 0.36 | -2.44E-02 | 1.05E-01 | 2.71E+00 | 2.78E+00 | 8.12E-01 | no | Selenoi | intron_variant,NMD_transcript_variant | MODIFIER |
| 3 | 150890579 | AX-170021923 | 2 | G | A | 0.327 | -1.76E-02 | 7.61E-02 | 2.67E+00 | 2.76E+00 | 8.12E-01 | no | | intergenic_variant | MODIFIER |
| 1 | 22008134 | AX-97801861 | 0 | G | A | 0.404 | 2.15E-02 | 8.56E-02 | 2.61E+00 | 2.67E+00 | 8.12E-01 | no | | intergenic_variant | MODIFIER |
| 9 | 57245843 | AX-170082512 | 0 | T | C | 0.219 | -2.69E-02 | 1.16E-01 | 2.71E+00 | 2.76E+00 | 8.12E-01 | no | Trcg1 | missense_variant | MODERATE |
| 9 | 58664290 | AX-168396157 | 0 | C | T | 0.219 | -2.69E-02 | 1.16E-01 | 2.71E+00 | 2.76E+00 | 8.12E-01 | no | Rec114 | intron_variant | MODIFIER |
| 3 | 29110872 | AX-170343993 | 0 | G | A | 0.184 | -2.27E-02 | 9.59E-02 | 2.66E+00 | 2.73E+00 | 8.12E-01 | no | Egfem1 | intron_variant | MODIFIER |
| 9 | 63797831 | AX-170418099 | 1 | T | C | 0.205 | 3.07E-02 | 1.32E-01 | 2.73E+00 | 2.77E+00 | 8.12E-01 | no | Gm2553 | downstream_gene_variant | MODIFIER |
| 12 | 93428344 | AX-170103760 | 2 | G | C | 0.1 | 5.18E-02 | 2.19E-01 | 2.67E+00 | 2.73E+00 | 8.12E-01 | no | 4930559C10Rik | intron_variant,non_coding_transcript_variant | MODIFIER |
| 18 | 59195581 | AX-170361370 | 0 | G | C | 0.421 | 1.87E-02 | 8.20E-02 | 2.71E+00 | 2.79E+00 | 8.12E-01 | no | Chsy3 | intron_variant | MODIFIER |
| 12 | 104319370 | AX-170385166 | 2 | T | C | 0.091 | -3.67E-02 | 1.63E-01 | 2.63E+00 | 2.77E+00 | 8.12E-01 | no | Serpina3j | intron_variant | MODIFIER |
| 12 | 64674353 | AX-170102747 | 0 | C | T | 0.053 | 4.24E-02 | 1.78E-01 | 2.60E+00 | 2.73E+00 | 8.12E-01 | no | | intergenic_variant | MODIFIER |
| 20 | 71421158 | AX-170034115 | 0 | A | T | 0.246 | 1.51E-02 | 6.36E-02 | 2.64E+00 | 2.73E+00 | 8.12E-01 | no | Mtmr1 | downstream_gene_variant | MODIFIER |
| 7 | 116904998 | AX-170144751 | 1 | G | T | 0.411 | 2.23E-02 | 9.37E-02 | 2.67E+00 | 2.71E+00 | 8.13E-01 | no | | intergenic_variant | MODIFIER |
| 1 | 155915916 | AX-168386993 | 0 | C | T | 0.219 | -3.27E-02 | 1.41E-01 | 2.75E+00 | 2.81E+00 | 8.13E-01 | no | Cep350 | intron_variant | MODIFIER |
| 13 | 34111818 | AX-170101811 | 1 | T | A | 0.179 | -3.01E-02 | 1.22E-01 | 2.62E+00 | 2.68E+00 | 8.13E-01 | no | 4930447K03Rik | intron_variant,non_coding_transcript_variant | MODIFIER |
| 6 | 104602116 | AX-170095738 | 0 | G | A | 0.079 | -5.20E-02 | 2.25E-01 | 2.69E+00 | 2.76E+00 | 8.13E-01 | no | Cntn6 | intron_variant | MODIFIER |
| 6 | 119195391 | AX-170152230 | 0 | C | G | 0.079 | -5.20E-02 | 2.25E-01 | 2.69E+00 | 2.76E+00 | 8.13E-01 | no | Dcp1b | intron_variant | MODIFIER |
| 6 | 138922059 | AX-170028368 | 0 | C | T | 0.079 | -5.20E-02 | 2.25E-01 | 2.69E+00 | 2.76E+00 | 8.13E-01 | no | | intergenic_variant | MODIFIER |
| 4 | 58321182 | AX-97829891 | 0 | T | C | 0.368 | -2.35E-02 | 1.02E-01 | 2.68E+00 | 2.76E+00 | 8.13E-01 | no | Musk | intron_variant | MODIFIER |
| 15 | 95206411 | AX-170012313 | 0 | G | T | 0.149 | -3.54E-02 | 1.56E-01 | 2.68E+00 | 2.78E+00 | 8.13E-01 | no | Nell2 | intron_variant | MODIFIER |

| | | | | | | | | | | | | | | | |
|---|---|---|---|---|---|---|---|---|---|---|---|---|---|---|---|
| 5 | 113885235 | AX-170108073 | 1 | T | G | 0.446 | 2.66E-02 | 1.10E-01 | 2.64E+00 | 2.70E+00 | 8.13E-01 | no | Coro1c | intron_variant | MODIFIER |
| 5 | 116763367 | AX-170470652 | 1 | C | T | 0.446 | 2.66E-02 | 1.10E-01 | 2.64E+00 | 2.70E+00 | 8.13E-01 | no | Gm43122 | intron_variant,non_coding_transcript_variant | MODIFIER |
| 20 | 87423440 | AX-170084715 | 0 | C | A | 0.254 | -1.42E-02 | 6.13E-02 | 2.64E+00 | 2.75E+00 | 8.13E-01 | no | Il1rapl1 | intron_variant | MODIFIER |
| 7 | 135867840 | AX-170078964 | 1 | G | C | 0.259 | -2.02E-02 | 8.25E-02 | 2.60E+00 | 2.70E+00 | 8.13E-01 | no | | intergenic_variant | MODIFIER |
| 9 | 87517866 | AX-170516487 | 1 | T | C | 0.143 | -3.08E-02 | 1.26E-01 | 2.62E+00 | 2.69E+00 | 8.13E-01 | no | | intergenic_variant | MODIFIER |
| 8 | 37300452 | AX-170311855 | 0 | T | C | 0.149 | 3.41E-02 | 1.48E-01 | 2.75E+00 | 2.80E+00 | 8.13E-01 | no | | intergenic_variant | MODIFIER |
| 8 | 40027250 | AX-170369649 | 0 | G | T | 0.149 | 3.41E-02 | 1.48E-01 | 2.75E+00 | 2.80E+00 | 8.13E-01 | no | | intergenic_variant | MODIFIER |
| 2 | 123876072 | AX-170522834 | 1 | C | T | 0.143 | 5.23E-02 | 2.22E-01 | 2.69E+00 | 2.72E+00 | 8.13E-01 | no | | intergenic_variant | MODIFIER |
| 2 | 123967251 | AX-97803508 | 1 | A | G | 0.143 | 5.23E-02 | 2.22E-01 | 2.69E+00 | 2.72E+00 | 8.13E-01 | no | | intergenic_variant | MODIFIER |
| 2 | 124528011 | AX-170135154 | 1 | A | G | 0.143 | 5.23E-02 | 2.22E-01 | 2.69E+00 | 2.72E+00 | 8.13E-01 | no | Sema6d | intron_variant | MODIFIER |
| 2 | 126986430 | AX-170154530 | 1 | A | T | 0.143 | 5.23E-02 | 2.22E-01 | 2.69E+00 | 2.72E+00 | 8.13E-01 | no | | intergenic_variant | MODIFIER |
| 1 | 103167988 | AX-170228839 | 1 | G | C | 0.152 | 3.79E-02 | 1.63E-01 | 2.64E+00 | 2.74E+00 | 8.13E-01 | no | Gm18728 | upstream_gene_variant | MODIFIER |
| 1 | 29047621 | AX-97810762 | 0 | A | C | 0.404 | -1.84E-02 | 8.01E-02 | 2.69E+00 | 2.76E+00 | 8.13E-01 | no | | intergenic_variant | MODIFIER |
| 1 | 42233008 | AX-170262627 | 1 | A | G | 0.429 | -1.75E-02 | 7.47E-02 | 2.65E+00 | 2.73E+00 | 8.14E-01 | no | Gm9915 | downstream_gene_variant | MODIFIER |
| 13 | 55304880 | AX-170254217 | 0 | T | G | 0.526 | 1.67E-02 | 6.62E-02 | 2.57E+00 | 2.68E+00 | 8.14E-01 | no | Nsd1 | intron_variant | MODIFIER |
| 1 | 151215020 | AX-170367886 | 0 | A | T | 0.211 | 3.04E-02 | 1.30E-01 | 2.66E+00 | 2.73E+00 | 8.14E-01 | no | Gm47996 | downstream_gene_variant | MODIFIER |
| 12 | 118756024 | AX-97852564 | 0 | C | T | 0.132 | -4.19E-02 | 1.83E-01 | 2.68E+00 | 2.76E+00 | 8.14E-01 | no | Gm22537 | downstream_gene_variant | MODIFIER |
| 7 | 136433882 | AX-170030058 | 0 | T | C | 0.289 | -1.93E-02 | 8.00E-02 | 2.61E+00 | 2.71E+00 | 8.14E-01 | no | | intergenic_variant | MODIFIER |
| 7 | 136437640 | AX-170078976 | 0 | A | G | 0.289 | -1.93E-02 | 8.00E-02 | 2.61E+00 | 2.71E+00 | 8.14E-01 | no | | intergenic_variant | MODIFIER |
| 7 | 136439916 | AX-170030054 | 0 | A | G | 0.289 | -1.93E-02 | 8.00E-02 | 2.61E+00 | 2.71E+00 | 8.14E-01 | no | | intergenic_variant | MODIFIER |
| 7 | 136447805 | AX-170464209 | 0 | A | G | 0.289 | -1.93E-02 | 8.00E-02 | 2.61E+00 | 2.71E+00 | 8.14E-01 | no | | intergenic_variant | MODIFIER |
| 7 | 136511154 | AX-170529876 | 0 | G | A | 0.289 | -1.93E-02 | 8.00E-02 | 2.61E+00 | 2.71E+00 | 8.14E-01 | no | Gm24444 | upstream_gene_variant | MODIFIER |
| 7 | 136516616 | AX-170408450 | 0 | T | C | 0.289 | -1.93E-02 | 8.00E-02 | 2.61E+00 | 2.71E+00 | 8.14E-01 | no | | intergenic_variant | MODIFIER |
| 7 | 136530970 | AX-97803300 | 0 | G | T | 0.289 | -1.93E-02 | 8.00E-02 | 2.61E+00 | 2.71E+00 | 8.14E-01 | no | | intergenic_variant | MODIFIER |
| 7 | 136537210 | AX-170274203 | 0 | A | G | 0.289 | -1.93E-02 | 8.00E-02 | 2.61E+00 | 2.71E+00 | 8.14E-01 | no | | intergenic_variant | MODIFIER |
| 7 | 136551341 | AX-170387320 | 0 | T | C | 0.289 | -1.93E-02 | 8.00E-02 | 2.61E+00 | 2.71E+00 | 8.14E-01 | no | | intergenic_variant | MODIFIER |
| 7 | 136557242 | AX-170449632 | 0 | T | A | 0.289 | -1.93E-02 | 8.00E-02 | 2.61E+00 | 2.71E+00 | 8.14E-01 | no | | intergenic_variant | MODIFIER |
| 7 | 136558533 | AX-170365871 | 0 | A | G | 0.289 | -1.93E-02 | 8.00E-02 | 2.61E+00 | 2.71E+00 | 8.14E-01 | no | | intergenic_variant | MODIFIER |
| 7 | 136558581 | AX-170078978 | 0 | G | A | 0.289 | -1.93E-02 | 8.00E-02 | 2.61E+00 | 2.71E+00 | 8.14E-01 | no | | intergenic_variant | MODIFIER |
| 7 | 136586379 | AX-97811613 | 0 | C | T | 0.289 | -1.93E-02 | 8.00E-02 | 2.61E+00 | 2.71E+00 | 8.14E-01 | no | | intergenic_variant | MODIFIER |
| 7 | 136587257 | AX-170458886 | 0 | C | A | 0.289 | -1.93E-02 | 8.00E-02 | 2.61E+00 | 2.71E+00 | 8.14E-01 | no | | intergenic_variant | MODIFIER |
| 7 | 136598298 | AX-170534117 | 0 | T | C | 0.289 | -1.93E-02 | 8.00E-02 | 2.61E+00 | 2.71E+00 | 8.14E-01 | no | | intergenic_variant | MODIFIER |
| 7 | 136608408 | AX-170443670 | 0 | C | T | 0.289 | -1.93E-02 | 8.00E-02 | 2.61E+00 | 2.71E+00 | 8.14E-01 | no | | intergenic_variant | MODIFIER |
| 7 | 136638762 | AX-97791582 | 0 | T | C | 0.289 | -1.93E-02 | 8.00E-02 | 2.61E+00 | 2.71E+00 | 8.14E-01 | no | | intergenic_variant | MODIFIER |
| 7 | 136640522 | AX-170134385 | 0 | T | C | 0.289 | -1.93E-02 | 8.00E-02 | 2.61E+00 | 2.71E+00 | 8.14E-01 | no | | regulatory_region_variant | MODIFIER |
| 7 | 136814149 | AX-170407781 | 0 | G | A | 0.289 | -1.93E-02 | 8.00E-02 | 2.61E+00 | 2.71E+00 | 8.14E-01 | no | Gm9347 | downstream_gene_variant | MODIFIER |
| 7 | 136827775 | AX-170479796 | 0 | A | G | 0.289 | -1.93E-02 | 8.00E-02 | 2.61E+00 | 2.71E+00 | 8.14E-01 | no | | intergenic_variant | MODIFIER |
| 7 | 136830283 | AX-170030067 | 0 | T | G | 0.289 | -1.93E-02 | 8.00E-02 | 2.61E+00 | 2.71E+00 | 8.14E-01 | no | | intergenic_variant | MODIFIER |
| 7 | 136846741 | AX-97819443 | 0 | T | G | 0.289 | -1.93E-02 | 8.00E-02 | 2.61E+00 | 2.71E+00 | 8.14E-01 | no | | intergenic_variant | MODIFIER |
| 7 | 136913413 | AX-97855657 | 0 | A | G | 0.289 | -1.93E-02 | 8.00E-02 | 2.61E+00 | 2.71E+00 | 8.14E-01 | no | Mgmt | intron_variant | MODIFIER |
| 7 | 136961017 | AX-170531697 | 0 | G | A | 0.289 | -1.93E-02 | 8.00E-02 | 2.61E+00 | 2.71E+00 | 8.14E-01 | no | Mgmt | intron_variant | MODIFIER |
| 7 | 136961436 | AX-170491348 | 0 | T | C | 0.289 | -1.93E-02 | 8.00E-02 | 2.61E+00 | 2.71E+00 | 8.14E-01 | no | Mgmt | intron_variant | MODIFIER |
| 7 | 136962283 | AX-170171882 | 0 | A | C | 0.289 | -1.93E-02 | 8.00E-02 | 2.61E+00 | 2.71E+00 | 8.14E-01 | no | Mgmt | intron_variant | MODIFIER |
| 7 | 136962512 | AX-170150315 | 0 | A | G | 0.289 | -1.93E-02 | 8.00E-02 | 2.61E+00 | 2.71E+00 | 8.14E-01 | no | Mgmt | intron_variant | MODIFIER |
| 7 | 136962650 | AX-170459028 | 0 | G | C | 0.289 | -1.93E-02 | 8.00E-02 | 2.61E+00 | 2.71E+00 | 8.14E-01 | no | Mgmt | intron_variant | MODIFIER |
| 7 | 136963318 | AX-170182989 | 0 | T | C | 0.289 | -1.93E-02 | 8.00E-02 | 2.61E+00 | 2.71E+00 | 8.14E-01 | no | Mgmt | intron_variant | MODIFIER |
| 7 | 136964015 | AX-170383236 | 0 | T | C | 0.289 | -1.93E-02 | 8.00E-02 | 2.61E+00 | 2.71E+00 | 8.14E-01 | no | Mgmt | intron_variant | MODIFIER |
| 7 | 136966524 | AX-170356743 | 0 | A | G | 0.289 | -1.93E-02 | 8.00E-02 | 2.61E+00 | 2.71E+00 | 8.14E-01 | no | Mgmt | intron_variant | MODIFIER |
| 7 | 136967977 | AX-170354081 | 0 | T | C | 0.289 | -1.93E-02 | 8.00E-02 | 2.61E+00 | 2.71E+00 | 8.14E-01 | no | Mgmt | intron_variant | MODIFIER |
| 7 | 136978212 | AX-170276141 | 0 | C | A | 0.289 | -1.93E-02 | 8.00E-02 | 2.61E+00 | 2.71E+00 | 8.14E-01 | no | Mgmt | intron_variant | MODIFIER |
| 7 | 136978778 | AX-97821414 | 0 | G | T | 0.289 | -1.93E-02 | 8.00E-02 | 2.61E+00 | 2.71E+00 | 8.14E-01 | no | Mgmt | intron_variant | MODIFIER |
| 7 | 136980860 | AX-170533166 | 0 | T | A | 0.289 | -1.93E-02 | 8.00E-02 | 2.61E+00 | 2.71E+00 | 8.14E-01 | no | Mgmt | intron_variant | MODIFIER |
| 7 | 136996208 | AX-170445153 | 0 | A | C | 0.289 | -1.93E-02 | 8.00E-02 | 2.61E+00 | 2.71E+00 | 8.14E-01 | no | Mgmt | intron_variant | MODIFIER |
| 7 | 137021697 | AX-170309496 | 0 | C | T | 0.289 | -1.93E-02 | 8.00E-02 | 2.61E+00 | 2.71E+00 | 8.14E-01 | no | Mgmt | intron_variant | MODIFIER |

| | | | | | | | | | | | | | | | |
|---|---|---|---|---|---|---|---|---|---|---|---|---|---|---|---|
| 7 | 137070419 | AX-170122897 | 0 | A | G | 0.289 | -1.93E-02 | 8.00E-02 | 2.61E+00 | 2.71E+00 | 8.14E-01 | no | Mgmt | intron_variant | MODIFIER |
| 7 | 137070545 | AX-170370571 | 0 | C | A | 0.289 | -1.93E-02 | 8.00E-02 | 2.61E+00 | 2.71E+00 | 8.14E-01 | no | Mgmt | intron_variant | MODIFIER |
| 7 | 137071141 | AX-170429258 | 0 | G | A | 0.289 | -1.93E-02 | 8.00E-02 | 2.61E+00 | 2.71E+00 | 8.14E-01 | no | Mgmt | intron_variant | MODIFIER |
| 7 | 137082360 | AX-170166097 | 0 | A | G | 0.289 | -1.93E-02 | 8.00E-02 | 2.61E+00 | 2.71E+00 | 8.14E-01 | no | Mgmt | intron_variant | MODIFIER |
| 7 | 137085239 | AX-97792965 | 0 | T | C | 0.289 | -1.93E-02 | 8.00E-02 | 2.61E+00 | 2.71E+00 | 8.14E-01 | no | Mgmt | intron_variant | MODIFIER |
| 7 | 137088031 | AX-170155936 | 0 | A | G | 0.289 | -1.93E-02 | 8.00E-02 | 2.61E+00 | 2.71E+00 | 8.14E-01 | no | Mgmt | intron_variant | MODIFIER |
| 7 | 137090648 | AX-170440369 | 0 | A | C | 0.289 | -1.93E-02 | 8.00E-02 | 2.61E+00 | 2.71E+00 | 8.14E-01 | no | Mgmt | intron_variant | MODIFIER |
| 7 | 137100089 | AX-170211468 | 0 | G | A | 0.289 | -1.93E-02 | 8.00E-02 | 2.61E+00 | 2.71E+00 | 8.14E-01 | no | Mgmt | intron_variant | MODIFIER |
| 7 | 137106488 | AX-170213622 | 0 | G | A | 0.289 | -1.93E-02 | 8.00E-02 | 2.61E+00 | 2.71E+00 | 8.14E-01 | no | Mgmt | intron_variant | MODIFIER |
| 7 | 137107501 | AX-97801238 | 0 | C | T | 0.289 | -1.93E-02 | 8.00E-02 | 2.61E+00 | 2.71E+00 | 8.14E-01 | no | Mgmt | intron_variant | MODIFIER |
| 7 | 137109048 | AX-170388640 | 0 | G | A | 0.289 | -1.93E-02 | 8.00E-02 | 2.61E+00 | 2.71E+00 | 8.14E-01 | no | Mgmt | intron_variant | MODIFIER |
| 7 | 137109583 | AX-170398537 | 0 | G | A | 0.289 | -1.93E-02 | 8.00E-02 | 2.61E+00 | 2.71E+00 | 8.14E-01 | no | Mgmt | intron_variant | MODIFIER |
| 7 | 137119788 | AX-170458455 | 0 | T | C | 0.289 | -1.93E-02 | 8.00E-02 | 2.61E+00 | 2.71E+00 | 8.14E-01 | no | Mgmt | intron_variant | MODIFIER |
| 7 | 137121638 | AX-170427008 | 0 | A | G | 0.289 | -1.93E-02 | 8.00E-02 | 2.61E+00 | 2.71E+00 | 8.14E-01 | no | Mgmt | intron_variant | MODIFIER |
| 7 | 137122012 | AX-170278223 | 0 | T | A | 0.289 | -1.93E-02 | 8.00E-02 | 2.61E+00 | 2.71E+00 | 8.14E-01 | no | Mgmt | intron_variant | MODIFIER |
| 7 | 137122324 | AX-170124126 | 0 | G | A | 0.289 | -1.93E-02 | 8.00E-02 | 2.61E+00 | 2.71E+00 | 8.14E-01 | no | Mgmt | intron_variant | MODIFIER |
| 7 | 137122712 | AX-170163490 | 0 | C | G | 0.289 | -1.93E-02 | 8.00E-02 | 2.61E+00 | 2.71E+00 | 8.14E-01 | no | Mgmt | intron_variant | MODIFIER |
| 7 | 137132519 | AX-170078988 | 0 | A | G | 0.289 | -1.93E-02 | 8.00E-02 | 2.61E+00 | 2.71E+00 | 8.14E-01 | no | Mgmt | downstream_gene_variant | MODIFIER |
| 7 | 137132929 | AX-97821457 | 0 | C | T | 0.289 | -1.93E-02 | 8.00E-02 | 2.61E+00 | 2.71E+00 | 8.14E-01 | no | Mgmt | downstream_gene_variant | MODIFIER |
| 7 | 137174147 | AX-97831414 | 0 | G | A | 0.289 | -1.93E-02 | 8.00E-02 | 2.61E+00 | 2.71E+00 | 8.14E-01 | no | | intergenic_variant | MODIFIER |
| 7 | 137177903 | AX-170152851 | 0 | C | G | 0.289 | -1.93E-02 | 8.00E-02 | 2.61E+00 | 2.71E+00 | 8.14E-01 | no | | intergenic_variant | MODIFIER |
| 7 | 137180540 | AX-170457628 | 0 | T | G | 0.289 | -1.93E-02 | 8.00E-02 | 2.61E+00 | 2.71E+00 | 8.14E-01 | no | | intergenic_variant | MODIFIER |
| 7 | 137217876 | AX-170310161 | 0 | G | A | 0.289 | -1.93E-02 | 8.00E-02 | 2.61E+00 | 2.71E+00 | 8.14E-01 | no | Ebf3 | intron_variant | MODIFIER |
| 7 | 137218928 | AX-97834802 | 0 | C | T | 0.289 | -1.93E-02 | 8.00E-02 | 2.61E+00 | 2.71E+00 | 8.14E-01 | no | Ebf3 | intron_variant | MODIFIER |
| 7 | 137224362 | AX-170137458 | 0 | A | C | 0.289 | -1.93E-02 | 8.00E-02 | 2.61E+00 | 2.71E+00 | 8.14E-01 | no | Ebf3 | intron_variant | MODIFIER |
| 7 | 137224443 | AX-170493652 | 0 | A | G | 0.289 | -1.93E-02 | 8.00E-02 | 2.61E+00 | 2.71E+00 | 8.14E-01 | no | Ebf3 | intron_variant | MODIFIER |
| 7 | 137231084 | AX-170372045 | 0 | G | C | 0.289 | -1.93E-02 | 8.00E-02 | 2.61E+00 | 2.71E+00 | 8.14E-01 | no | Ebf3 | intron_variant | MODIFIER |
| 7 | 137249585 | AX-170030069 | 0 | A | T | 0.289 | -1.93E-02 | 8.00E-02 | 2.61E+00 | 2.71E+00 | 8.14E-01 | no | Ebf3 | intron_variant | MODIFIER |
| 7 | 137250994 | AX-170346965 | 0 | G | A | 0.289 | -1.93E-02 | 8.00E-02 | 2.61E+00 | 2.71E+00 | 8.14E-01 | no | Ebf3 | intron_variant | MODIFIER |
| 7 | 137252743 | AX-170030070 | 0 | A | C | 0.289 | -1.93E-02 | 8.00E-02 | 2.61E+00 | 2.71E+00 | 8.14E-01 | no | Ebf3 | intron_variant | MODIFIER |
| 7 | 137257260 | AX-170251284 | 0 | G | A | 0.289 | -1.93E-02 | 8.00E-02 | 2.61E+00 | 2.71E+00 | 8.14E-01 | no | Ebf3 | intron_variant | MODIFIER |
| 7 | 137290266 | AX-97821984 | 0 | G | A | 0.289 | -1.93E-02 | 8.00E-02 | 2.61E+00 | 2.71E+00 | 8.14E-01 | no | Ebf3 | intron_variant | MODIFIER |
| 7 | 137292125 | AX-170030072 | 0 | T | C | 0.289 | -1.93E-02 | 8.00E-02 | 2.61E+00 | 2.71E+00 | 8.14E-01 | no | Ebf3 | intron_variant | MODIFIER |
| 7 | 137292203 | AX-170030074 | 0 | G | A | 0.289 | -1.93E-02 | 8.00E-02 | 2.61E+00 | 2.71E+00 | 8.14E-01 | no | Ebf3 | intron_variant | MODIFIER |
| 7 | 137343282 | AX-97819079 | 0 | C | T | 0.289 | -1.93E-02 | 8.00E-02 | 2.61E+00 | 2.71E+00 | 8.14E-01 | no | | intergenic_variant | MODIFIER |
| 7 | 137348243 | AX-170030073 | 0 | G | A | 0.289 | -1.93E-02 | 8.00E-02 | 2.61E+00 | 2.71E+00 | 8.14E-01 | no | | intergenic_variant | MODIFIER |
| 7 | 137348269 | AX-170030075 | 0 | A | G | 0.289 | -1.93E-02 | 8.00E-02 | 2.61E+00 | 2.71E+00 | 8.14E-01 | no | | intergenic_variant | MODIFIER |
| 7 | 137381673 | AX-170030077 | 0 | T | C | 0.289 | -1.93E-02 | 8.00E-02 | 2.61E+00 | 2.71E+00 | 8.14E-01 | no | 9430038I01Rik | intron_variant | MODIFIER |
| 7 | 137383625 | AX-170117579 | 0 | G | C | 0.289 | -1.93E-02 | 8.00E-02 | 2.61E+00 | 2.71E+00 | 8.14E-01 | no | 9430038I01Rik | intron_variant | MODIFIER |
| 7 | 137384625 | AX-170264927 | 0 | C | G | 0.289 | -1.93E-02 | 8.00E-02 | 2.61E+00 | 2.71E+00 | 8.14E-01 | no | 9430038I01Rik | intron_variant | MODIFIER |
| 7 | 137421860 | AX-97811420 | 0 | A | G | 0.289 | -1.93E-02 | 8.00E-02 | 2.61E+00 | 2.71E+00 | 8.14E-01 | no | | intergenic_variant | MODIFIER |
| 7 | 137425675 | AX-170486529 | 0 | G | A | 0.289 | -1.93E-02 | 8.00E-02 | 2.61E+00 | 2.71E+00 | 8.14E-01 | no | | intergenic_variant | MODIFIER |
| 7 | 137452846 | AX-170030076 | 0 | C | T | 0.289 | -1.93E-02 | 8.00E-02 | 2.61E+00 | 2.71E+00 | 8.14E-01 | no | Glrx3 | intron_variant | MODIFIER |
| 7 | 137502294 | AX-170090031 | 0 | T | C | 0.289 | -1.93E-02 | 8.00E-02 | 2.61E+00 | 2.71E+00 | 8.14E-01 | no | | intergenic_variant | MODIFIER |
| 7 | 137618370 | AX-170149369 | 0 | A | T | 0.289 | -1.93E-02 | 8.00E-02 | 2.61E+00 | 2.71E+00 | 8.14E-01 | no | | intergenic_variant | MODIFIER |
| 7 | 137618888 | AX-170359479 | 0 | T | C | 0.289 | -1.93E-02 | 8.00E-02 | 2.61E+00 | 2.71E+00 | 8.14E-01 | no | | intergenic_variant | MODIFIER |
| 7 | 137619001 | AX-97856277 | 0 | C | T | 0.289 | -1.93E-02 | 8.00E-02 | 2.61E+00 | 2.71E+00 | 8.14E-01 | no | | intergenic_variant | MODIFIER |
| 7 | 137620826 | AX-170219497 | 0 | C | G | 0.289 | -1.93E-02 | 8.00E-02 | 2.61E+00 | 2.71E+00 | 8.14E-01 | no | | intergenic_variant | MODIFIER |
| 7 | 137621935 | AX-170383699 | 0 | C | T | 0.289 | -1.93E-02 | 8.00E-02 | 2.61E+00 | 2.71E+00 | 8.14E-01 | no | | intergenic_variant | MODIFIER |
| 7 | 137622928 | AX-170254232 | 0 | C | G | 0.289 | -1.93E-02 | 8.00E-02 | 2.61E+00 | 2.71E+00 | 8.14E-01 | no | | intergenic_variant | MODIFIER |
| 7 | 137690005 | AX-97814374 | 0 | T | C | 0.289 | -1.93E-02 | 8.00E-02 | 2.61E+00 | 2.71E+00 | 8.14E-01 | no | | intergenic_variant | MODIFIER |
| 2 | 161203138 | AX-170088820 | 0 | C | T | 0.149 | 4.92E-02 | 2.12E-01 | 2.69E+00 | 2.74E+00 | 8.14E-01 | no | Gm14243 | intron_variant,non_coding_transcript_variant | MODIFIER |
| 1 | 41869032 | AX-97794423 | 0 | G | A | 0.447 | -1.76E-02 | 7.60E-02 | 2.65E+00 | 2.74E+00 | 8.14E-01 | no | Gm29260 | intron_variant,non_coding_transcript_variant | MODIFIER |
| 13 | 38656313 | AX-170008143 | 1 | A | T | 0.188 | 2.31E-02 | 1.04E-01 | 2.70E+00 | 2.80E+00 | 8.14E-01 | no | Eef1e1 | intron_variant | MODIFIER |

| | | | | | | | | | | | | | | | |
|---|---|---|---|---|---|---|---|---|---|---|---|---|---|---|---|
| 18 | 58189461 | AX-170426530 | 0 | T | C | 0.395 | 2.56E-02 | 9.59E-02 | 2.56E+00 | 2.63E+00 | 8.14E-01 | no | Fbn2 | intron_variant | MODIFIER |
| 10 | 120995705 | AX-97810423 | 0 | A | C | 0.342 | 1.70E-02 | 7.42E-02 | 2.66E+00 | 2.75E+00 | 8.14E-01 | no | Gm48341 | intron_variant,non_coding_transcript_variant | MODIFIER |
| 10 | 121157274 | AX-170496597 | 0 | T | C | 0.342 | 1.70E-02 | 7.42E-02 | 2.66E+00 | 2.75E+00 | 8.14E-01 | no | Gm10741 | downstream_gene_variant | MODIFIER |
| 10 | 121180286 | AX-170432206 | 0 | G | C | 0.342 | 1.70E-02 | 7.42E-02 | 2.66E+00 | 2.75E+00 | 8.14E-01 | no | Gm48426 | downstream_gene_variant | MODIFIER |
| 10 | 121198608 | AX-97819953 | 0 | A | G | 0.342 | 1.70E-02 | 7.42E-02 | 2.66E+00 | 2.75E+00 | 8.14E-01 | no | 4930432O09Rik | downstream_gene_variant | MODIFIER |
| 10 | 121207132 | AX-170298893 | 0 | G | A | 0.342 | 1.70E-02 | 7.42E-02 | 2.66E+00 | 2.75E+00 | 8.14E-01 | no | 4930432O09Rik | upstream_gene_variant | MODIFIER |
| 10 | 121220334 | AX-97837038 | 0 | G | T | 0.342 | 1.70E-02 | 7.42E-02 | 2.66E+00 | 2.75E+00 | 8.14E-01 | no | | intergenic_variant | MODIFIER |
| 10 | 121229924 | AX-170508992 | 0 | T | C | 0.342 | 1.70E-02 | 7.42E-02 | 2.66E+00 | 2.75E+00 | 8.14E-01 | no | | intergenic_variant | MODIFIER |
| 10 | 121232081 | AX-97817842 | 0 | G | A | 0.342 | 1.70E-02 | 7.42E-02 | 2.66E+00 | 2.75E+00 | 8.14E-01 | no | | intergenic_variant | MODIFIER |
| 12 | 95384252 | AX-170505320 | 0 | A | T | 0.096 | -3.62E-02 | 1.56E-01 | 2.63E+00 | 2.74E+00 | 8.14E-01 | no | | intergenic_variant | MODIFIER |
| 1 | 46663368 | AX-170329613 | 1 | T | C | 0.33 | -1.75E-02 | 7.43E-02 | 2.63E+00 | 2.72E+00 | 8.14E-01 | no | Dnah7c | downstream_gene_variant | MODIFIER |
| 9 | 55964232 | AX-170397599 | 1 | C | T | 0.205 | -2.75E-02 | 1.19E-01 | 2.69E+00 | 2.75E+00 | 8.14E-01 | no | 4930563M21Rik | intron_variant | MODIFIER |
| 2 | 102447409 | AX-97839162 | 1 | G | A | 0.054 | -6.95E-02 | 3.00E-01 | 2.72E+00 | 2.75E+00 | 8.14E-01 | no | Fjx1 | downstream_gene_variant | MODIFIER |
| 5 | 110485108 | AX-170385383 | 0 | C | T | 0.456 | 2.49E-02 | 1.03E-01 | 2.63E+00 | 2.70E+00 | 8.14E-01 | no | Gm32996 | upstream_gene_variant | MODIFIER |
| 12 | 111755740 | AX-170050217 | 1 | A | G | 0.143 | -4.51E-02 | 1.96E-01 | 2.68E+00 | 2.75E+00 | 8.14E-01 | no | Klc1 | upstream_gene_variant | MODIFIER |
| 10 | 6266084 | AX-170288081 | 0 | A | G | 0.211 | 2.34E-02 | 1.04E-01 | 2.68E+00 | 2.77E+00 | 8.14E-01 | no | | intergenic_variant | MODIFIER |
| 10 | 17790507 | AX-170217025 | 0 | T | G | 0.202 | 2.05E-02 | 9.21E-02 | 2.66E+00 | 2.78E+00 | 8.14E-01 | no | Gm20655 | downstream_gene_variant | MODIFIER |
| 7 | 82301736 | AX-170121792 | 1 | C | T | 0.125 | 3.62E-02 | 1.50E-01 | 2.57E+00 | 2.71E+00 | 8.15E-01 | no | Sh3gl3 | intron_variant | MODIFIER |
| 9 | 106158832 | AX-170138005 | 0 | G | A | 0.132 | 2.47E-02 | 1.11E-01 | 2.73E+00 | 2.82E+00 | 8.15E-01 | no | Glyctk | upstream_gene_variant | MODIFIER |
| 1 | 156191044 | AX-170091761 | 1 | G | A | 0.223 | 3.40E-02 | 1.43E-01 | 2.66E+00 | 2.70E+00 | 8.15E-01 | no | Fam163a | intron_variant | MODIFIER |
| 7 | 118025722 | AX-170465664 | 0 | A | C | 0.395 | 2.08E-02 | 8.96E-02 | 2.67E+00 | 2.73E+00 | 8.15E-01 | no | | regulatory_region_variant | MODIFIER |
| 9 | 53589031 | AX-97793987 | 1 | A | C | 0.214 | 2.67E-02 | 1.16E-01 | 2.69E+00 | 2.75E+00 | 8.15E-01 | no | Acat1 | intron_variant | MODIFIER |
| 10 | 109574038 | AX-170354016 | 0 | A | G | 0.351 | 1.83E-02 | 8.15E-02 | 2.68E+00 | 2.78E+00 | 8.15E-01 | no | | intergenic_variant | MODIFIER |
| 7 | 117517963 | AX-170300572 | 0 | G | A | 0.395 | -1.92E-02 | 8.45E-02 | 2.68E+00 | 2.76E+00 | 8.15E-01 | no | Xylt1 | intron_variant | MODIFIER |
| 15 | 103300870 | AX-170012431 | 1 | A | T | 0.143 | 3.17E-02 | 1.33E-01 | 2.62E+00 | 2.71E+00 | 8.15E-01 | no | Copz1 | downstream_gene_variant | MODIFIER |
| 1 | 21919753 | AX-97785206 | 0 | T | C | 0.395 | 2.09E-02 | 8.60E-02 | 2.63E+00 | 2.68E+00 | 8.15E-01 | no | Kcnq5 | intron_variant | MODIFIER |
| 1 | 21921918 | AX-170000062 | 0 | G | C | 0.395 | 2.09E-02 | 8.60E-02 | 2.63E+00 | 2.68E+00 | 8.15E-01 | no | Kcnq5 | intron_variant | MODIFIER |
| 1 | 21941226 | AX-97819908 | 0 | A | G | 0.395 | 2.09E-02 | 8.60E-02 | 2.63E+00 | 2.68E+00 | 8.15E-01 | no | Kcnq5 | intron_variant | MODIFIER |
| 1 | 21948919 | AX-97845388 | 0 | T | C | 0.395 | 2.09E-02 | 8.60E-02 | 2.63E+00 | 2.68E+00 | 8.15E-01 | no | Kcnq5 | intron_variant | MODIFIER |
| 1 | 21977020 | AX-170176020 | 0 | T | C | 0.395 | 2.09E-02 | 8.60E-02 | 2.63E+00 | 2.68E+00 | 8.15E-01 | no | | intergenic_variant | MODIFIER |
| 1 | 21994653 | AX-170000059 | 0 | C | G | 0.395 | 2.09E-02 | 8.60E-02 | 2.63E+00 | 2.68E+00 | 8.15E-01 | no | | intergenic_variant | MODIFIER |
| 1 | 22003563 | AX-170000061 | 0 | A | G | 0.395 | 2.09E-02 | 8.60E-02 | 2.63E+00 | 2.68E+00 | 8.15E-01 | no | | intergenic_variant | MODIFIER |
| 1 | 22062131 | AX-170000068 | 0 | A | G | 0.395 | 2.09E-02 | 8.60E-02 | 2.63E+00 | 2.68E+00 | 8.15E-01 | no | | intergenic_variant | MODIFIER |
| 4 | 149539054 | AX-170301854 | 0 | C | A | 0.105 | 4.70E-02 | 2.04E-01 | 2.68E+00 | 2.74E+00 | 8.15E-01 | no | Ctnnbip1 | intron_variant | MODIFIER |
| 2 | 129614473 | AX-170289009 | 0 | C | T | 0.132 | 4.27E-02 | 1.76E-01 | 2.61E+00 | 2.70E+00 | 8.15E-01 | no | Sirpa | intron_variant | MODIFIER |
| 2 | 117997826 | AX-170464872 | 0 | G | A | 0.14 | 3.89E-02 | 1.65E-01 | 2.63E+00 | 2.72E+00 | 8.15E-01 | no | Gm13986 | intron_variant,non_coding_transcript_variant | MODIFIER |
| 2 | 142909722 | AX-170182566 | 1 | C | T | 0.152 | 4.28E-02 | 1.79E-01 | 2.62E+00 | 2.71E+00 | 8.15E-01 | no | | intergenic_variant | MODIFIER |
| 4 | 90717364 | AX-97795682 | 0 | T | A | 0.254 | 2.77E-02 | 1.07E-01 | 2.57E+00 | 2.65E+00 | 8.15E-01 | no | | intergenic_variant | MODIFIER |
| 2 | 119789851 | AX-170232590 | 0 | G | T | 0.149 | 4.28E-02 | 1.76E-01 | 2.61E+00 | 2.69E+00 | 8.15E-01 | no | Rpap1 | upstream_gene_variant | MODIFIER |
| 2 | 123655857 | AX-170156115 | 0 | T | A | 0.149 | 4.28E-02 | 1.76E-01 | 2.61E+00 | 2.69E+00 | 8.15E-01 | no | | intergenic_variant | MODIFIER |
| 2 | 125181253 | AX-170342476 | 0 | C | T | 0.149 | 4.28E-02 | 1.76E-01 | 2.61E+00 | 2.69E+00 | 8.15E-01 | no | Slc12a1 | intron_variant | MODIFIER |
| 2 | 126938416 | AX-170105367 | 0 | G | A | 0.149 | 4.28E-02 | 1.76E-01 | 2.61E+00 | 2.69E+00 | 8.15E-01 | no | | regulatory_region_variant | MODIFIER |
| 5 | 28309132 | AX-170104486 | 0 | C | A | 0.36 | -2.38E-02 | 1.05E-01 | 2.70E+00 | 2.77E+00 | 8.15E-01 | no | | intergenic_variant | MODIFIER |
| 5 | 33589613 | AX-170184188 | 0 | G | T | 0.36 | -2.38E-02 | 1.05E-01 | 2.70E+00 | 2.77E+00 | 8.15E-01 | no | | regulatory_region_variant | MODIFIER |
| 5 | 109330290 | AX-97845114 | 2 | T | G | 0.427 | 2.49E-02 | 1.05E-01 | 2.64E+00 | 2.70E+00 | 8.16E-01 | no | Vmn2r16 | 5_prime_UTR_variant | MODIFIER |
| 10 | 32108856 | AX-170377309 | 0 | C | T | 0.395 | 1.58E-02 | 7.10E-02 | 2.65E+00 | 2.77E+00 | 8.16E-01 | no | Nkain2 | intron_variant | MODIFIER |
| 2 | 119335763 | AX-97838779 | 1 | A | G | 0.116 | 4.63E-02 | 1.99E-01 | 2.67E+00 | 2.72E+00 | 8.16E-01 | no | Dll4 | 3_prime_UTR_variant | MODIFIER |
| 1 | 88595034 | AX-170167318 | 0 | A | T | 0.158 | 4.93E-02 | 2.09E-01 | 2.65E+00 | 2.71E+00 | 8.16E-01 | no | | regulatory_region_variant | MODIFIER |
| 1 | 100243437 | AX-97851773 | 0 | T | C | 0.158 | 4.93E-02 | 2.09E-01 | 2.65E+00 | 2.71E+00 | 8.16E-01 | no | Cntnap5b | intron_variant | MODIFIER |
| 5 | 7105241 | AX-170272916 | 0 | A | G | 0.439 | 1.38E-02 | 6.29E-02 | 2.65E+00 | 2.78E+00 | 8.16E-01 | no | Zfp804b | intron_variant | MODIFIER |
| 8 | 85603551 | AX-97837147 | 1 | C | T | 0.205 | 2.94E-02 | 1.28E-01 | 2.68E+00 | 2.75E+00 | 8.16E-01 | no | Gm45424 | downstream_gene_variant | MODIFIER |
| 1 | 190791795 | AX-170477018 | 0 | A | G | 0.254 | -2.07E-02 | 9.68E-02 | 2.75E+00 | 2.88E+00 | 8.16E-01 | no | Rps6kc1 | intron_variant | MODIFIER |
| 1 | 7566494 | AX-97791009 | 0 | T | C | 0.412 | -1.63E-02 | 7.27E-02 | 2.73E+00 | 2.81E+00 | 8.16E-01 | no | | intergenic_variant | MODIFIER |
| 5 | 5896110 | AX-168399030 | 0 | A | G | 0.43 | 1.40E-02 | 6.38E-02 | 2.64E+00 | 2.78E+00 | 8.16E-01 | no | | intergenic_variant | MODIFIER |

| | | | | | | | | | | | | | | | |
|---|---|---|---|---|---|---|---|---|---|---|---|---|---|---|---|
| 5 | 6452908 | AX-97793958 | 0 | A | C | 0.43 | 1.40E-02 | 6.38E-02 | 2.64E+00 | 2.78E+00 | 8.16E-01 | no | | intergenic_variant | MODIFIER |
| 5 | 6583416 | AX-170533706 | 0 | G | C | 0.43 | 1.40E-02 | 6.38E-02 | 2.64E+00 | 2.78E+00 | 8.16E-01 | no | | intergenic_variant | MODIFIER |
| 5 | 6613643 | AX-97855122 | 0 | T | C | 0.43 | 1.40E-02 | 6.38E-02 | 2.64E+00 | 2.78E+00 | 8.16E-01 | no | | intergenic_variant | MODIFIER |
| 5 | 6675599 | AX-170450682 | 0 | T | G | 0.43 | 1.40E-02 | 6.38E-02 | 2.64E+00 | 2.78E+00 | 8.16E-01 | no | | intergenic_variant | MODIFIER |
| 5 | 7140137 | AX-170502730 | 0 | A | T | 0.43 | 1.40E-02 | 6.38E-02 | 2.64E+00 | 2.78E+00 | 8.16E-01 | no | Zfp804b | intron_variant | MODIFIER |
| 5 | 7152050 | AX-170136618 | 0 | C | A | 0.43 | 1.40E-02 | 6.38E-02 | 2.64E+00 | 2.78E+00 | 8.16E-01 | no | Zfp804b | intron_variant | MODIFIER |
| 5 | 7171095 | AX-170323479 | 0 | T | C | 0.43 | 1.40E-02 | 6.38E-02 | 2.64E+00 | 2.78E+00 | 8.16E-01 | no | Zfp804b | intron_variant | MODIFIER |
| 5 | 7179019 | AX-170071235 | 0 | G | A | 0.43 | 1.40E-02 | 6.38E-02 | 2.64E+00 | 2.78E+00 | 8.16E-01 | no | Gm24525 | upstream_gene_variant | MODIFIER |
| 5 | 7251336 | AX-170166994 | 0 | A | G | 0.43 | 1.40E-02 | 6.38E-02 | 2.64E+00 | 2.78E+00 | 8.16E-01 | no | Zfp804b | intron_variant | MODIFIER |
| 5 | 7252509 | AX-170115677 | 0 | T | C | 0.43 | 1.40E-02 | 6.38E-02 | 2.64E+00 | 2.78E+00 | 8.16E-01 | no | Zfp804b | intron_variant | MODIFIER |
| 5 | 7252962 | AX-170193193 | 0 | A | T | 0.43 | 1.40E-02 | 6.38E-02 | 2.64E+00 | 2.78E+00 | 8.16E-01 | no | Zfp804b | intron_variant | MODIFIER |
| 8 | 101639545 | AX-97842812 | 0 | G | A | 0.228 | 2.67E-02 | 1.16E-01 | 2.67E+00 | 2.74E+00 | 8.16E-01 | no | | intergenic_variant | MODIFIER |
| 2 | 125970201 | AX-170350162 | 1 | T | C | 0.143 | 5.19E-02 | 2.24E-01 | 2.70E+00 | 2.72E+00 | 8.16E-01 | no | Galk2 | intron_variant | MODIFIER |
| 2 | 130000986 | AX-170452686 | 1 | A | G | 0.143 | 5.19E-02 | 2.24E-01 | 2.70E+00 | 2.72E+00 | 8.16E-01 | no | | intergenic_variant | MODIFIER |
| 2 | 137153153 | AX-170256608 | 1 | C | G | 0.143 | 5.19E-02 | 2.24E-01 | 2.70E+00 | 2.72E+00 | 8.16E-01 | no | | regulatory_region_variant | MODIFIER |
| 2 | 143786682 | AX-170110042 | 1 | C | T | 0.143 | 5.19E-02 | 2.24E-01 | 2.70E+00 | 2.72E+00 | 8.16E-01 | no | Pcsk2 | intron_variant | MODIFIER |
| 2 | 155784489 | AX-170521735 | 1 | G | A | 0.143 | 5.19E-02 | 2.24E-01 | 2.70E+00 | 2.72E+00 | 8.16E-01 | no | Mmp24 | intron_variant | MODIFIER |
| 20 | 130776471 | AX-170142237 | 0 | T | G | 0.132 | 2.77E-02 | 1.20E-01 | 2.65E+00 | 2.73E+00 | 8.16E-01 | no | | intergenic_variant | MODIFIER |
| 20 | 130844448 | AX-170260271 | 0 | C | T | 0.132 | 2.77E-02 | 1.20E-01 | 2.65E+00 | 2.73E+00 | 8.16E-01 | no | | intergenic_variant | MODIFIER |
| 20 | 130845662 | AX-97783070 | 0 | A | T | 0.132 | 2.77E-02 | 1.20E-01 | 2.65E+00 | 2.73E+00 | 8.16E-01 | no | | intergenic_variant | MODIFIER |
| 20 | 130846063 | AX-170190200 | 0 | A | T | 0.132 | 2.77E-02 | 1.20E-01 | 2.65E+00 | 2.73E+00 | 8.16E-01 | no | | intergenic_variant | MODIFIER |
| 20 | 130847408 | AX-170354396 | 0 | C | T | 0.132 | 2.77E-02 | 1.20E-01 | 2.65E+00 | 2.73E+00 | 8.16E-01 | no | | intergenic_variant | MODIFIER |
| 20 | 130860008 | AX-170422522 | 0 | C | T | 0.132 | 2.77E-02 | 1.20E-01 | 2.65E+00 | 2.73E+00 | 8.16E-01 | no | | intergenic_variant | MODIFIER |
| 20 | 130868223 | AX-170417918 | 0 | G | A | 0.132 | 2.77E-02 | 1.20E-01 | 2.65E+00 | 2.73E+00 | 8.16E-01 | no | | intergenic_variant | MODIFIER |
| 20 | 130869933 | AX-170463319 | 0 | C | G | 0.132 | 2.77E-02 | 1.20E-01 | 2.65E+00 | 2.73E+00 | 8.16E-01 | no | | intergenic_variant | MODIFIER |
| 20 | 130896344 | AX-170219951 | 0 | A | C | 0.132 | 2.77E-02 | 1.20E-01 | 2.65E+00 | 2.73E+00 | 8.16E-01 | no | | intergenic_variant | MODIFIER |
| 20 | 133603917 | AX-170034987 | 0 | T | C | 0.132 | 2.77E-02 | 1.20E-01 | 2.65E+00 | 2.73E+00 | 8.16E-01 | no | Pcdh19 | intron_variant | MODIFIER |
| 20 | 133670156 | AX-97787133 | 0 | A | G | 0.132 | 2.77E-02 | 1.20E-01 | 2.65E+00 | 2.73E+00 | 8.16E-01 | no | Pcdh19 | intron_variant | MODIFIER |
| 20 | 133924347 | AX-97800398 | 0 | C | T | 0.132 | 2.77E-02 | 1.20E-01 | 2.65E+00 | 2.73E+00 | 8.16E-01 | no | Srpx2 | intron_variant | MODIFIER |
| 20 | 133924406 | AX-170347911 | 0 | T | C | 0.132 | 2.77E-02 | 1.20E-01 | 2.65E+00 | 2.73E+00 | 8.16E-01 | no | Srpx2 | intron_variant | MODIFIER |
| 20 | 133924419 | AX-170509690 | 0 | A | C | 0.132 | 2.77E-02 | 1.20E-01 | 2.65E+00 | 2.73E+00 | 8.16E-01 | no | Srpx2 | intron_variant | MODIFIER |
| 20 | 133969426 | AX-170350871 | 0 | T | A | 0.132 | 2.77E-02 | 1.20E-01 | 2.65E+00 | 2.73E+00 | 8.16E-01 | no | Sytl4 | intron_variant | MODIFIER |
| 20 | 133990477 | AX-170423232 | 0 | A | T | 0.132 | 2.77E-02 | 1.20E-01 | 2.65E+00 | 2.73E+00 | 8.16E-01 | no | | intergenic_variant | MODIFIER |
| 20 | 133997474 | AX-97796912 | 0 | A | C | 0.132 | 2.77E-02 | 1.20E-01 | 2.65E+00 | 2.73E+00 | 8.16E-01 | no | | intergenic_variant | MODIFIER |
| 20 | 134115975 | AX-170034998 | 0 | G | C | 0.132 | 2.77E-02 | 1.20E-01 | 2.65E+00 | 2.73E+00 | 8.16E-01 | no | Nox1 | upstream_gene_variant | MODIFIER |
| 20 | 134124353 | AX-97837571 | 0 | C | T | 0.132 | 2.77E-02 | 1.20E-01 | 2.65E+00 | 2.73E+00 | 8.16E-01 | no | Nox1 | intron_variant | MODIFIER |
| 20 | 134184808 | AX-170302426 | 0 | G | A | 0.132 | 2.77E-02 | 1.20E-01 | 2.65E+00 | 2.73E+00 | 8.16E-01 | no | Arl13a | upstream_gene_variant | MODIFIER |
| 20 | 134192021 | AX-170034997 | 0 | C | T | 0.132 | 2.77E-02 | 1.20E-01 | 2.65E+00 | 2.73E+00 | 8.16E-01 | no | Arl13a | intron_variant | MODIFIER |
| 20 | 134234699 | AX-97832974 | 0 | C | T | 0.132 | 2.77E-02 | 1.20E-01 | 2.65E+00 | 2.73E+00 | 8.16E-01 | no | Trmt2b | intron_variant | MODIFIER |
| 20 | 134240163 | AX-170035002 | 0 | G | C | 0.132 | 2.77E-02 | 1.20E-01 | 2.65E+00 | 2.73E+00 | 8.16E-01 | no | Trmt2b | missense_variant | MODERATE |
| 20 | 134245617 | AX-170085293 | 0 | C | T | 0.132 | 2.77E-02 | 1.20E-01 | 2.65E+00 | 2.73E+00 | 8.16E-01 | no | Trmt2b | intron_variant | MODIFIER |
| 20 | 134297193 | AX-170096931 | 0 | T | C | 0.132 | 2.77E-02 | 1.20E-01 | 2.65E+00 | 2.73E+00 | 8.16E-01 | no | Tmem35a | intron_variant | MODIFIER |
| 20 | 134307971 | AX-170138847 | 0 | G | A | 0.132 | 2.77E-02 | 1.20E-01 | 2.65E+00 | 2.73E+00 | 8.16E-01 | no | Tmem35a | downstream_gene_variant | MODIFIER |
| 20 | 134364586 | AX-170035004 | 0 | A | G | 0.132 | 2.77E-02 | 1.20E-01 | 2.65E+00 | 2.73E+00 | 8.16E-01 | no | Cenpi | downstream_gene_variant | MODIFIER |
| 20 | 134367666 | AX-170096932 | 0 | T | G | 0.132 | 2.77E-02 | 1.20E-01 | 2.65E+00 | 2.73E+00 | 8.16E-01 | no | | intergenic_variant | MODIFIER |
| 20 | 134378842 | AX-170175961 | 0 | T | C | 0.132 | 2.77E-02 | 1.20E-01 | 2.65E+00 | 2.73E+00 | 8.16E-01 | no | | intergenic_variant | MODIFIER |
| 20 | 134385213 | AX-97831970 | 0 | T | C | 0.132 | 2.77E-02 | 1.20E-01 | 2.65E+00 | 2.73E+00 | 8.16E-01 | no | | regulatory_region_variant | MODIFIER |
| 20 | 134386909 | AX-170455203 | 0 | G | A | 0.132 | 2.77E-02 | 1.20E-01 | 2.65E+00 | 2.73E+00 | 8.16E-01 | no | | intergenic_variant | MODIFIER |
| 20 | 134511896 | AX-170090687 | 0 | A | G | 0.132 | 2.77E-02 | 1.20E-01 | 2.65E+00 | 2.73E+00 | 8.16E-01 | no | | intergenic_variant | MODIFIER |
| 20 | 134546838 | AX-170361194 | 0 | G | C | 0.132 | 2.77E-02 | 1.20E-01 | 2.65E+00 | 2.73E+00 | 8.16E-01 | no | Btk | splice_region_variant,intron_variant | LOW |
| 20 | 134551054 | AX-170090688 | 0 | G | A | 0.132 | 2.77E-02 | 1.20E-01 | 2.65E+00 | 2.73E+00 | 8.16E-01 | no | Btk | intron_variant | MODIFIER |
| 20 | 134552435 | AX-170152268 | 0 | C | T | 0.132 | 2.77E-02 | 1.20E-01 | 2.65E+00 | 2.73E+00 | 8.16E-01 | no | Btk | intron_variant | MODIFIER |
| 20 | 134564633 | AX-97802547 | 0 | G | A | 0.132 | 2.77E-02 | 1.20E-01 | 2.65E+00 | 2.73E+00 | 8.16E-01 | no | Btk | intron_variant | MODIFIER |
| 20 | 134568006 | AX-170085294 | 0 | G | A | 0.132 | 2.77E-02 | 1.20E-01 | 2.65E+00 | 2.73E+00 | 8.16E-01 | no | Btk | intron_variant | MODIFIER |

| | | | | | | | | | | | | | | | |
|---|---|---|---|---|---|---|---|---|---|---|---|---|---|---|---|
| 20 | 134581637 | AX-170493805 | 0 | G | A | 0.132 | 2.77E-02 | 1.20E-01 | 2.65E+00 | 2.73E+00 | 8.16E-01 | no | Btk | intron_variant | MODIFIER |
| 20 | 134588153 | AX-170090689 | 0 | A | G | 0.132 | 2.77E-02 | 1.20E-01 | 2.65E+00 | 2.73E+00 | 8.16E-01 | no | Gla | 3_prime_UTR_variant | MODIFIER |
| 20 | 134591045 | AX-170466082 | 0 | G | C | 0.132 | 2.77E-02 | 1.20E-01 | 2.65E+00 | 2.73E+00 | 8.16E-01 | no | Gla | synonymous_variant | LOW |
| 20 | 134708353 | AX-170035017 | 0 | T | C | 0.132 | 2.77E-02 | 1.20E-01 | 2.65E+00 | 2.73E+00 | 8.16E-01 | no | | intergenic_variant | MODIFIER |
| 20 | 134718366 | AX-97821326 | 0 | G | T | 0.132 | 2.77E-02 | 1.20E-01 | 2.65E+00 | 2.73E+00 | 8.16E-01 | no | Armcx1 | splice_region_variant,intron_variant | LOW |
| 20 | 134719607 | AX-170035007 | 0 | A | C | 0.132 | 2.77E-02 | 1.20E-01 | 2.65E+00 | 2.73E+00 | 8.16E-01 | no | Armcx1 | intron_variant | MODIFIER |
| 20 | 134722499 | AX-170123084 | 0 | A | G | 0.132 | 2.77E-02 | 1.20E-01 | 2.65E+00 | 2.73E+00 | 8.16E-01 | no | Armcx1 | downstream_gene_variant | MODIFIER |
| 20 | 134738573 | AX-97817978 | 0 | C | T | 0.132 | 2.77E-02 | 1.20E-01 | 2.65E+00 | 2.73E+00 | 8.16E-01 | no | | intergenic_variant | MODIFIER |
| 20 | 134818549 | AX-97857218 | 0 | T | G | 0.132 | 2.77E-02 | 1.20E-01 | 2.65E+00 | 2.73E+00 | 8.16E-01 | no | | intergenic_variant | MODIFIER |
| 20 | 134876198 | AX-97807486 | 0 | C | A | 0.132 | 2.77E-02 | 1.20E-01 | 2.65E+00 | 2.73E+00 | 8.16E-01 | no | | intergenic_variant | MODIFIER |
| 20 | 134876861 | AX-170035011 | 0 | G | A | 0.132 | 2.77E-02 | 1.20E-01 | 2.65E+00 | 2.73E+00 | 8.16E-01 | no | | intergenic_variant | MODIFIER |
| 20 | 134880549 | AX-97781461 | 0 | C | T | 0.132 | 2.77E-02 | 1.20E-01 | 2.65E+00 | 2.73E+00 | 8.16E-01 | no | | intergenic_variant | MODIFIER |
| 20 | 134928259 | AX-97847753 | 0 | A | C | 0.132 | 2.77E-02 | 1.20E-01 | 2.65E+00 | 2.73E+00 | 8.16E-01 | no | | intergenic_variant | MODIFIER |
| 20 | 134929293 | AX-170035013 | 0 | T | C | 0.132 | 2.77E-02 | 1.20E-01 | 2.65E+00 | 2.73E+00 | 8.16E-01 | no | | intergenic_variant | MODIFIER |
| 20 | 134952379 | AX-97807487 | 0 | A | C | 0.132 | 2.77E-02 | 1.20E-01 | 2.65E+00 | 2.73E+00 | 8.16E-01 | no | Nxf2 | intron_variant | MODIFIER |
| 20 | 135586393 | AX-97804952 | 0 | C | T | 0.132 | 2.77E-02 | 1.20E-01 | 2.65E+00 | 2.73E+00 | 8.16E-01 | no | Nxf7 | intron_variant | MODIFIER |
| 20 | 135589513 | AX-170035020 | 0 | G | T | 0.132 | 2.77E-02 | 1.20E-01 | 2.65E+00 | 2.73E+00 | 8.16E-01 | no | Nxf7 | intron_variant | MODIFIER |
| 20 | 135589773 | AX-170035026 | 0 | G | A | 0.132 | 2.77E-02 | 1.20E-01 | 2.65E+00 | 2.73E+00 | 8.16E-01 | no | Nxf7 | intron_variant | MODIFIER |
| 20 | 141324778 | AX-97815025 | 0 | A | G | 0.132 | 2.77E-02 | 1.20E-01 | 2.65E+00 | 2.73E+00 | 8.16E-01 | no | Col4a6 | intron_variant | MODIFIER |
| 20 | 141327075 | AX-170035095 | 0 | T | C | 0.132 | 2.77E-02 | 1.20E-01 | 2.65E+00 | 2.73E+00 | 8.16E-01 | no | Col4a6 | intron_variant | MODIFIER |
| 20 | 141327785 | AX-170035098 | 0 | T | G | 0.132 | 2.77E-02 | 1.20E-01 | 2.65E+00 | 2.73E+00 | 8.16E-01 | no | Col4a6 | intron_variant | MODIFIER |
| 20 | 141331287 | AX-170126458 | 0 | T | G | 0.132 | 2.77E-02 | 1.20E-01 | 2.65E+00 | 2.73E+00 | 8.16E-01 | no | Col4a6 | intron_variant | MODIFIER |
| 20 | 141341382 | AX-97793064 | 0 | C | T | 0.132 | 2.77E-02 | 1.20E-01 | 2.65E+00 | 2.73E+00 | 8.16E-01 | no | Col4a6 | intron_variant | MODIFIER |
| 20 | 141354228 | AX-170408633 | 0 | C | T | 0.132 | 2.77E-02 | 1.20E-01 | 2.65E+00 | 2.73E+00 | 8.16E-01 | no | Col4a6 | intron_variant | MODIFIER |
| 20 | 141354344 | AX-170401024 | 0 | C | T | 0.132 | 2.77E-02 | 1.20E-01 | 2.65E+00 | 2.73E+00 | 8.16E-01 | no | Col4a6 | intron_variant | MODIFIER |
| 20 | 141424534 | AX-97836229 | 0 | C | G | 0.132 | 2.77E-02 | 1.20E-01 | 2.65E+00 | 2.73E+00 | 8.16E-01 | no | Col4a6 | intron_variant | MODIFIER |
| 20 | 141475847 | AX-170477274 | 0 | A | G | 0.132 | 2.77E-02 | 1.20E-01 | 2.65E+00 | 2.73E+00 | 8.16E-01 | no | Col4a6 | upstream_gene_variant | MODIFIER |
| 20 | 141979879 | AX-170085426 | 0 | A | C | 0.132 | 2.77E-02 | 1.20E-01 | 2.65E+00 | 2.73E+00 | 8.16E-01 | no | Gm6368 | upstream_gene_variant | MODIFIER |
| 20 | 141996124 | AX-170096949 | 0 | C | T | 0.132 | 2.77E-02 | 1.20E-01 | 2.65E+00 | 2.73E+00 | 8.16E-01 | no | | intergenic_variant | MODIFIER |
| 20 | 141996341 | AX-170085422 | 0 | A | G | 0.132 | 2.77E-02 | 1.20E-01 | 2.65E+00 | 2.73E+00 | 8.16E-01 | no | | intergenic_variant | MODIFIER |
| 20 | 143068176 | AX-170217777 | 0 | C | G | 0.132 | 2.77E-02 | 1.20E-01 | 2.65E+00 | 2.73E+00 | 8.16E-01 | no | | intergenic_variant | MODIFIER |
| 20 | 143068892 | AX-170181861 | 0 | C | T | 0.132 | 2.77E-02 | 1.20E-01 | 2.65E+00 | 2.73E+00 | 8.16E-01 | no | | intergenic_variant | MODIFIER |
| 20 | 143071820 | AX-170035117 | 0 | A | G | 0.132 | 2.77E-02 | 1.20E-01 | 2.65E+00 | 2.73E+00 | 8.16E-01 | no | | intergenic_variant | MODIFIER |
| 20 | 143099415 | AX-170035123 | 0 | C | T | 0.132 | 2.77E-02 | 1.20E-01 | 2.65E+00 | 2.73E+00 | 8.16E-01 | no | Rtl9 | upstream_gene_variant | MODIFIER |
| 20 | 143119165 | AX-170035120 | 0 | G | A | 0.132 | 2.77E-02 | 1.20E-01 | 2.65E+00 | 2.73E+00 | 8.16E-01 | no | | intergenic_variant | MODIFIER |
| 20 | 155597211 | AX-170090725 | 0 | C | T | 0.132 | 2.77E-02 | 1.20E-01 | 2.65E+00 | 2.73E+00 | 8.16E-01 | no | Ptchd1 | intron_variant | MODIFIER |
| 20 | 155597238 | AX-170035279 | 0 | G | A | 0.132 | 2.77E-02 | 1.20E-01 | 2.65E+00 | 2.73E+00 | 8.16E-01 | no | Ptchd1 | intron_variant | MODIFIER |
| 20 | 155602285 | AX-170153997 | 0 | A | G | 0.132 | 2.77E-02 | 1.20E-01 | 2.65E+00 | 2.73E+00 | 8.16E-01 | no | Ptchd1 | intron_variant | MODIFIER |
| 20 | 155606584 | AX-170096980 | 0 | A | G | 0.132 | 2.77E-02 | 1.20E-01 | 2.65E+00 | 2.73E+00 | 8.16E-01 | no | Ptchd1 | intron_variant | MODIFIER |
| 20 | 155638903 | AX-97807654 | 0 | G | A | 0.132 | 2.77E-02 | 1.20E-01 | 2.65E+00 | 2.73E+00 | 8.16E-01 | no | Gm15155 | intron_variant | MODIFIER |
| 20 | 155676163 | AX-97830322 | 0 | T | C | 0.132 | 2.77E-02 | 1.20E-01 | 2.65E+00 | 2.73E+00 | 8.16E-01 | no | Gm15155 | intron_variant | MODIFIER |
| 20 | 155681077 | AX-170035273 | 0 | C | T | 0.132 | 2.77E-02 | 1.20E-01 | 2.65E+00 | 2.73E+00 | 8.16E-01 | no | Gm15155 | intron_variant | MODIFIER |
| 20 | 155709737 | AX-170035271 | 0 | T | A | 0.132 | 2.77E-02 | 1.20E-01 | 2.65E+00 | 2.73E+00 | 8.16E-01 | no | Gm15155 | intron_variant | MODIFIER |
| 20 | 155716622 | AX-170035275 | 0 | A | T | 0.132 | 2.77E-02 | 1.20E-01 | 2.65E+00 | 2.73E+00 | 8.16E-01 | no | Gm15155 | intron_variant | MODIFIER |
| 20 | 155716806 | AX-97853045 | 0 | T | C | 0.132 | 2.77E-02 | 1.20E-01 | 2.65E+00 | 2.73E+00 | 8.16E-01 | no | Gm15155 | intron_variant | MODIFIER |
| 20 | 155948167 | AX-97825322 | 0 | A | G | 0.132 | 2.77E-02 | 1.20E-01 | 2.65E+00 | 2.73E+00 | 8.16E-01 | no | Gm15155 | intron_variant | MODIFIER |
| 20 | 155969112 | AX-170090726 | 0 | C | T | 0.132 | 2.77E-02 | 1.20E-01 | 2.65E+00 | 2.73E+00 | 8.16E-01 | no | Gm15155 | intron_variant | MODIFIER |
| 20 | 156099058 | AX-97848687 | 0 | C | T | 0.132 | 2.77E-02 | 1.20E-01 | 2.65E+00 | 2.73E+00 | 8.16E-01 | no | Gm15155 | intron_variant | MODIFIER |
| 20 | 156287508 | AX-170035291 | 0 | A | T | 0.132 | 2.77E-02 | 1.20E-01 | 2.65E+00 | 2.73E+00 | 8.16E-01 | no | Gm15155 | intron_variant | MODIFIER |
| 20 | 156324324 | AX-97826587 | 0 | A | G | 0.132 | 2.77E-02 | 1.20E-01 | 2.65E+00 | 2.73E+00 | 8.16E-01 | no | Gm15155 | intron_variant | MODIFIER |
| 20 | 156332321 | AX-97829490 | 0 | T | C | 0.132 | 2.77E-02 | 1.20E-01 | 2.65E+00 | 2.73E+00 | 8.16E-01 | no | Gm15155 | intron_variant | MODIFIER |
| 20 | 156429273 | AX-97780828 | 0 | A | G | 0.132 | 2.77E-02 | 1.20E-01 | 2.65E+00 | 2.73E+00 | 8.16E-01 | no | | intergenic_variant | MODIFIER |
| 20 | 156475955 | AX-97846676 | 0 | G | A | 0.132 | 2.77E-02 | 1.20E-01 | 2.65E+00 | 2.73E+00 | 8.16E-01 | no | | intergenic_variant | MODIFIER |
| 20 | 156490735 | AX-170096981 | 0 | A | G | 0.132 | 2.77E-02 | 1.20E-01 | 2.65E+00 | 2.73E+00 | 8.16E-01 | no | | intergenic_variant | MODIFIER |

| | | | | | | | | | | | | | | | |
|---|---|---|---|---|---|---|---|---|---|---|---|---|---|---|---|
| 20 | 156497909 | AX-170035286 | 0 | C | G | 0.132 | 2.77E-02 | 1.20E-01 | 2.65E+00 | 2.73E+00 | 8.16E-01 | no | | intergenic_variant | MODIFIER |
| 20 | 156519501 | AX-97825609 | 0 | G | T | 0.132 | 2.77E-02 | 1.20E-01 | 2.65E+00 | 2.73E+00 | 8.16E-01 | no | | intergenic_variant | MODIFIER |
| 20 | 156612133 | AX-97787653 | 0 | T | C | 0.132 | 2.77E-02 | 1.20E-01 | 2.65E+00 | 2.73E+00 | 8.16E-01 | no | | intergenic_variant | MODIFIER |
| 20 | 156641087 | AX-170521882 | 0 | A | T | 0.132 | 2.77E-02 | 1.20E-01 | 2.65E+00 | 2.73E+00 | 8.16E-01 | no | | intergenic_variant | MODIFIER |
| 20 | 156664109 | AX-97844459 | 0 | A | C | 0.132 | 2.77E-02 | 1.20E-01 | 2.65E+00 | 2.73E+00 | 8.16E-01 | no | | intergenic_variant | MODIFIER |
| 20 | 156666835 | AX-170508541 | 0 | A | G | 0.132 | 2.77E-02 | 1.20E-01 | 2.65E+00 | 2.73E+00 | 8.16E-01 | no | | intergenic_variant | MODIFIER |
| 20 | 156888762 | AX-97833061 | 0 | A | T | 0.132 | 2.77E-02 | 1.20E-01 | 2.65E+00 | 2.73E+00 | 8.16E-01 | no | | intergenic_variant | MODIFIER |
| 20 | 156927669 | AX-97849679 | 0 | C | T | 0.132 | 2.77E-02 | 1.20E-01 | 2.65E+00 | 2.73E+00 | 8.16E-01 | no | | intergenic_variant | MODIFIER |
| 20 | 156947262 | AX-97802107 | 0 | T | C | 0.132 | 2.77E-02 | 1.20E-01 | 2.65E+00 | 2.73E+00 | 8.16E-01 | no | | intergenic_variant | MODIFIER |
| 20 | 157026977 | AX-170035299 | 0 | C | A | 0.132 | 2.77E-02 | 1.20E-01 | 2.65E+00 | 2.73E+00 | 8.16E-01 | no | | intergenic_variant | MODIFIER |
| 20 | 157084291 | AX-170035298 | 0 | T | C | 0.132 | 2.77E-02 | 1.20E-01 | 2.65E+00 | 2.73E+00 | 8.16E-01 | no | | intergenic_variant | MODIFIER |
| 20 | 157087905 | AX-170090729 | 0 | C | T | 0.132 | 2.77E-02 | 1.20E-01 | 2.65E+00 | 2.73E+00 | 8.16E-01 | no | | intergenic_variant | MODIFIER |
| 20 | 157108454 | AX-170035295 | 0 | T | A | 0.132 | 2.77E-02 | 1.20E-01 | 2.65E+00 | 2.73E+00 | 8.16E-01 | no | | intergenic_variant | MODIFIER |
| 20 | 157118465 | AX-97807681 | 0 | A | G | 0.132 | 2.77E-02 | 1.20E-01 | 2.65E+00 | 2.73E+00 | 8.16E-01 | no | | intergenic_variant | MODIFIER |
| 20 | 157137219 | AX-170418442 | 0 | C | T | 0.132 | 2.77E-02 | 1.20E-01 | 2.65E+00 | 2.73E+00 | 8.16E-01 | no | | intergenic_variant | MODIFIER |
| 20 | 157138675 | AX-168418253 | 0 | C | T | 0.132 | 2.77E-02 | 1.20E-01 | 2.65E+00 | 2.73E+00 | 8.16E-01 | no | | intergenic_variant | MODIFIER |
| 20 | 157139654 | AX-97852723 | 0 | A | G | 0.132 | 2.77E-02 | 1.20E-01 | 2.65E+00 | 2.73E+00 | 8.16E-01 | no | | intergenic_variant | MODIFIER |
| 20 | 157216237 | AX-97850225 | 0 | C | G | 0.132 | 2.77E-02 | 1.20E-01 | 2.65E+00 | 2.73E+00 | 8.16E-01 | no | Phex | intron_variant | MODIFIER |
| 20 | 157232116 | AX-170035297 | 0 | G | C | 0.132 | 2.77E-02 | 1.20E-01 | 2.65E+00 | 2.73E+00 | 8.16E-01 | no | Phex | intron_variant | MODIFIER |
| 20 | 157275768 | AX-97808207 | 0 | A | C | 0.132 | 2.77E-02 | 1.20E-01 | 2.65E+00 | 2.73E+00 | 8.16E-01 | no | Phex | intron_variant | MODIFIER |
| 20 | 157293060 | AX-97845081 | 0 | A | G | 0.132 | 2.77E-02 | 1.20E-01 | 2.65E+00 | 2.73E+00 | 8.16E-01 | no | Phex | intron_variant | MODIFIER |
| 20 | 157293303 | AX-170197461 | 0 | T | C | 0.132 | 2.77E-02 | 1.20E-01 | 2.65E+00 | 2.73E+00 | 8.16E-01 | no | Phex | intron_variant | MODIFIER |
| 20 | 157297791 | AX-170286321 | 0 | A | C | 0.132 | 2.77E-02 | 1.20E-01 | 2.65E+00 | 2.73E+00 | 8.16E-01 | no | Phex | intron_variant | MODIFIER |
| 20 | 157331891 | AX-97780731 | 0 | T | C | 0.132 | 2.77E-02 | 1.20E-01 | 2.65E+00 | 2.73E+00 | 8.16E-01 | no | Phex | intron_variant | MODIFIER |
| 20 | 157372261 | AX-97781193 | 0 | G | A | 0.132 | 2.77E-02 | 1.20E-01 | 2.65E+00 | 2.73E+00 | 8.16E-01 | no | Phex | intron_variant | MODIFIER |
| 20 | 157444417 | AX-97809640 | 0 | G | A | 0.132 | 2.77E-02 | 1.20E-01 | 2.65E+00 | 2.73E+00 | 8.16E-01 | no | Sms | 3_prime_UTR_variant | MODIFIER |
| 20 | 157445557 | AX-170035317 | 0 | C | G | 0.132 | 2.77E-02 | 1.20E-01 | 2.65E+00 | 2.73E+00 | 8.16E-01 | no | Sms | 3_prime_UTR_variant | MODIFIER |
| 20 | 157464140 | AX-170035312 | 0 | G | A | 0.132 | 2.77E-02 | 1.20E-01 | 2.65E+00 | 2.73E+00 | 8.16E-01 | no | Sms | intron_variant | MODIFIER |
| 20 | 157464826 | AX-170035311 | 0 | G | A | 0.132 | 2.77E-02 | 1.20E-01 | 2.65E+00 | 2.73E+00 | 8.16E-01 | no | Sms | intron_variant | MODIFIER |
| 20 | 157466071 | AX-97807909 | 0 | C | T | 0.132 | 2.77E-02 | 1.20E-01 | 2.65E+00 | 2.73E+00 | 8.16E-01 | no | Sms | intron_variant | MODIFIER |
| 20 | 157485243 | AX-170449279 | 0 | T | A | 0.132 | 2.77E-02 | 1.20E-01 | 2.65E+00 | 2.73E+00 | 8.16E-01 | no | Sms | intron_variant | MODIFIER |
| 20 | 157496716 | AX-97783531 | 0 | G | A | 0.132 | 2.77E-02 | 1.20E-01 | 2.65E+00 | 2.73E+00 | 8.16E-01 | no | Sms | upstream_gene_variant | MODIFIER |
| 20 | 157502004 | AX-170035318 | 0 | C | T | 0.132 | 2.77E-02 | 1.20E-01 | 2.65E+00 | 2.73E+00 | 8.16E-01 | no | | intergenic_variant | MODIFIER |
| 20 | 157736548 | AX-97789112 | 0 | C | T | 0.132 | 2.77E-02 | 1.20E-01 | 2.65E+00 | 2.73E+00 | 8.16E-01 | no | Smpx | intron_variant,NMD_transcript_variant | MODIFIER |
| 20 | 157757952 | AX-170132148 | 0 | T | C | 0.132 | 2.77E-02 | 1.20E-01 | 2.65E+00 | 2.73E+00 | 8.16E-01 | no | Gm15169 | non_coding_transcript_exon_variant | MODIFIER |
| 20 | 157758877 | AX-170258541 | 0 | A | T | 0.132 | 2.77E-02 | 1.20E-01 | 2.65E+00 | 2.73E+00 | 8.16E-01 | no | Gm15169 | non_coding_transcript_exon_variant | MODIFIER |
| 20 | 157762961 | AX-97821011 | 0 | A | G | 0.132 | 2.77E-02 | 1.20E-01 | 2.65E+00 | 2.73E+00 | 8.16E-01 | no | Gm15169 | intron_variant,non_coding_transcript_variant | MODIFIER |
| 20 | 157780964 | AX-170343368 | 0 | A | T | 0.132 | 2.77E-02 | 1.20E-01 | 2.65E+00 | 2.73E+00 | 8.16E-01 | no | Gm15169 | downstream_gene_variant | MODIFIER |
| 20 | 157782120 | AX-170455534 | 0 | A | G | 0.132 | 2.77E-02 | 1.20E-01 | 2.65E+00 | 2.73E+00 | 8.16E-01 | no | Gm15169 | downstream_gene_variant | MODIFIER |
| 20 | 157786496 | AX-97848141 | 0 | C | T | 0.132 | 2.77E-02 | 1.20E-01 | 2.65E+00 | 2.73E+00 | 8.16E-01 | no | Gm15169 | non_coding_transcript_exon_variant | MODIFIER |
| 20 | 157817250 | AX-170382823 | 0 | G | A | 0.132 | 2.77E-02 | 1.20E-01 | 2.65E+00 | 2.73E+00 | 8.16E-01 | no | Cnksr2 | downstream_gene_variant | MODIFIER |
| 20 | 157819028 | AX-170329439 | 0 | T | C | 0.132 | 2.77E-02 | 1.20E-01 | 2.65E+00 | 2.73E+00 | 8.16E-01 | no | Cnksr2 | downstream_gene_variant | MODIFIER |
| 20 | 157819181 | AX-97851167 | 0 | G | A | 0.132 | 2.77E-02 | 1.20E-01 | 2.65E+00 | 2.73E+00 | 8.16E-01 | no | Cnksr2 | downstream_gene_variant | MODIFIER |
| 20 | 157834989 | AX-170306187 | 0 | A | G | 0.132 | 2.77E-02 | 1.20E-01 | 2.65E+00 | 2.73E+00 | 8.16E-01 | no | Cnksr2 | intron_variant | MODIFIER |
| 20 | 157835679 | AX-170165804 | 0 | A | T | 0.132 | 2.77E-02 | 1.20E-01 | 2.65E+00 | 2.73E+00 | 8.16E-01 | no | Cnksr2 | intron_variant | MODIFIER |
| 20 | 157835795 | AX-170492312 | 0 | C | T | 0.132 | 2.77E-02 | 1.20E-01 | 2.65E+00 | 2.73E+00 | 8.16E-01 | no | Cnksr2 | intron_variant | MODIFIER |
| 20 | 157841177 | AX-170106491 | 0 | C | T | 0.132 | 2.77E-02 | 1.20E-01 | 2.65E+00 | 2.73E+00 | 8.16E-01 | no | Cnksr2 | intron_variant | MODIFIER |
| 20 | 157842870 | AX-97796925 | 0 | C | A | 0.132 | 2.77E-02 | 1.20E-01 | 2.65E+00 | 2.73E+00 | 8.16E-01 | no | Cnksr2 | intron_variant | MODIFIER |
| 20 | 157848307 | AX-168419407 | 0 | T | C | 0.132 | 2.77E-02 | 1.20E-01 | 2.65E+00 | 2.73E+00 | 8.16E-01 | no | Cnksr2 | intron_variant | MODIFIER |
| 20 | 157848597 | AX-168383860 | 0 | C | T | 0.132 | 2.77E-02 | 1.20E-01 | 2.65E+00 | 2.73E+00 | 8.16E-01 | no | Cnksr2 | intron_variant | MODIFIER |
| 20 | 157849240 | AX-170494310 | 0 | G | A | 0.132 | 2.77E-02 | 1.20E-01 | 2.65E+00 | 2.73E+00 | 8.16E-01 | no | Cnksr2 | intron_variant | MODIFIER |
| 20 | 157856501 | AX-170085613 | 0 | T | G | 0.132 | 2.77E-02 | 1.20E-01 | 2.65E+00 | 2.73E+00 | 8.16E-01 | no | Cnksr2 | intron_variant | MODIFIER |
| 20 | 157872900 | AX-97844686 | 0 | C | T | 0.132 | 2.77E-02 | 1.20E-01 | 2.65E+00 | 2.73E+00 | 8.16E-01 | no | Cnksr2 | intron_variant | MODIFIER |
| 20 | 157873637 | AX-170508850 | 0 | T | C | 0.132 | 2.77E-02 | 1.20E-01 | 2.65E+00 | 2.73E+00 | 8.16E-01 | no | Cnksr2 | intron_variant | MODIFIER |

| | | | | | | | | | | | | | | | |
|---|---|---|---|---|---|---|---|---|---|---|---|---|---|---|---|
| 20 | 157920140 | AX-170417580 | 0 | A | G | 0.132 | 2.77E-02 | 1.20E-01 | 2.65E+00 | 2.73E+00 | 8.16E-01 | no | Cnksr2 | intron_variant | MODIFIER |
| 13 | 51350793 | AX-170051265 | 0 | A | G | 0.509 | -1.45E-02 | 6.53E-02 | 2.66E+00 | 2.77E+00 | 8.16E-01 | no | | regulatory_region_variant | MODIFIER |
| 12 | 101313098 | AX-170469022 | 1 | G | A | 0.08 | -4.30E-02 | 1.90E-01 | 2.67E+00 | 2.75E+00 | 8.16E-01 | no | Gm18795 | intron_variant,non_coding_transcript_variant | MODIFIER |
| 1 | 13256087 | AX-170349217 | 0 | T | C | 0.377 | 1.79E-02 | 7.43E-02 | 2.62E+00 | 2.69E+00 | 8.16E-01 | no | Ncoa2 | intron_variant | MODIFIER |
| 8 | 35606145 | AX-170532242 | 0 | A | G | 0.158 | -3.21E-02 | 1.38E-01 | 2.66E+00 | 2.72E+00 | 8.17E-01 | no | Mfhas1 | intron_variant | MODIFIER |
| 20 | 93817942 | AX-170123589 | 0 | C | T | 0.053 | -2.19E-02 | 1.03E-01 | 2.60E+00 | 2.78E+00 | 8.17E-01 | no | Pdk3 | intron_variant | MODIFIER |
| 9 | 114455568 | AX-170519977 | 0 | G | A | 0.14 | 2.25E-02 | 1.01E-01 | 2.65E+00 | 2.77E+00 | 8.17E-01 | no | Glb1 | intron_variant | MODIFIER |
| 2 | 90788599 | AX-170018296 | 1 | G | T | 0.054 | -6.82E-02 | 2.98E-01 | 2.72E+00 | 2.75E+00 | 8.17E-01 | no | Agbl2 | intron_variant | MODIFIER |
| 7 | 137452402 | AX-97794906 | 0 | G | T | 0.289 | -1.88E-02 | 7.83E-02 | 2.60E+00 | 2.70E+00 | 8.17E-01 | no | Glrx3 | intron_variant | MODIFIER |
| 7 | 135073250 | AX-170381483 | 1 | C | T | 0.25 | -1.97E-02 | 8.23E-02 | 2.60E+00 | 2.70E+00 | 8.17E-01 | no | Dock1 | intron_variant | MODIFIER |
| 7 | 116323725 | AX-170527781 | 2 | T | A | 0.4 | 1.99E-02 | 8.70E-02 | 2.66E+00 | 2.74E+00 | 8.17E-01 | no | | regulatory_region_variant | MODIFIER |
| 1 | 10726118 | AX-170426821 | 0 | G | A | 0.395 | -1.70E-02 | 7.64E-02 | 2.75E+00 | 2.82E+00 | 8.17E-01 | no | | intergenic_variant | MODIFIER |
| 4 | 143371557 | AX-97802751 | 1 | G | A | 0.089 | 3.86E-02 | 1.68E-01 | 2.66E+00 | 2.74E+00 | 8.17E-01 | no | Lrrc38 | downstream_gene_variant | MODIFIER |
| 10 | 128279730 | AX-170046242 | 0 | G | T | 0.333 | -1.64E-02 | 7.05E-02 | 2.62E+00 | 2.72E+00 | 8.17E-01 | no | Stat2 | intron_variant | MODIFIER |
| 18 | 69893874 | AX-170362527 | 1 | T | A | 0.179 | -2.64E-02 | 1.13E-01 | 2.64E+00 | 2.72E+00 | 8.17E-01 | no | AC108433.1 | intron_variant,non_coding_transcript_variant | MODIFIER |
| 8 | 78839669 | AX-170099830 | 1 | C | G | 0.205 | 2.92E-02 | 1.29E-01 | 2.69E+00 | 2.76E+00 | 8.17E-01 | no | | intergenic_variant | MODIFIER |
| 12 | 116496496 | AX-170223661 | 1 | G | A | 0.143 | -3.70E-02 | 1.58E-01 | 2.62E+00 | 2.72E+00 | 8.17E-01 | no | Ptprn2 | intron_variant | MODIFIER |
| 2 | 128689337 | AX-170264458 | 1 | A | T | 0.143 | 5.08E-02 | 2.20E-01 | 2.69E+00 | 2.72E+00 | 8.17E-01 | no | Anapc1 | upstream_gene_variant | MODIFIER |
| 2 | 131451291 | AX-170528909 | 1 | C | T | 0.143 | 5.08E-02 | 2.20E-01 | 2.69E+00 | 2.72E+00 | 8.17E-01 | no | Gm14304 | downstream_gene_variant | MODIFIER |
| 2 | 138603247 | AX-97795171 | 1 | T | C | 0.143 | 5.08E-02 | 2.20E-01 | 2.69E+00 | 2.72E+00 | 8.17E-01 | no | Gm14061 | intron_variant,non_coding_transcript_variant | MODIFIER |
| 2 | 141939320 | AX-97817434 | 1 | C | T | 0.143 | 5.08E-02 | 2.20E-01 | 2.69E+00 | 2.72E+00 | 8.17E-01 | no | Macrod2 | intron_variant | MODIFIER |
| 10 | 92891733 | AX-170109064 | 1 | A | T | 0.312 | -1.92E-02 | 7.74E-02 | 2.59E+00 | 2.67E+00 | 8.17E-01 | no | Cfap54 | intron_variant | MODIFIER |
| 13 | 22988324 | AX-97836433 | 0 | G | A | 0.175 | -2.96E-02 | 1.14E-01 | 2.56E+00 | 2.64E+00 | 8.17E-01 | no | | intergenic_variant | MODIFIER |
| 7 | 116560142 | AX-170459529 | 0 | G | C | 0.43 | 2.18E-02 | 9.42E-02 | 2.67E+00 | 2.72E+00 | 8.17E-01 | no | | intergenic_variant | MODIFIER |
| 7 | 116565120 | AX-170151811 | 0 | T | G | 0.43 | 2.18E-02 | 9.42E-02 | 2.67E+00 | 2.72E+00 | 8.17E-01 | no | | intergenic_variant | MODIFIER |
| 7 | 116587093 | AX-170273844 | 0 | C | G | 0.43 | 2.18E-02 | 9.42E-02 | 2.67E+00 | 2.72E+00 | 8.17E-01 | no | | intergenic_variant | MODIFIER |
| 7 | 116643927 | AX-170276574 | 0 | T | C | 0.43 | 2.18E-02 | 9.42E-02 | 2.67E+00 | 2.72E+00 | 8.17E-01 | no | | intergenic_variant | MODIFIER |
| 7 | 116645467 | AX-170280958 | 0 | T | C | 0.43 | 2.18E-02 | 9.42E-02 | 2.67E+00 | 2.72E+00 | 8.17E-01 | no | | regulatory_region_variant | MODIFIER |
| 7 | 116657956 | AX-170466937 | 0 | C | T | 0.43 | 2.18E-02 | 9.42E-02 | 2.67E+00 | 2.72E+00 | 8.17E-01 | no | | intergenic_variant | MODIFIER |
| 7 | 116678446 | AX-170078543 | 0 | G | T | 0.43 | 2.18E-02 | 9.42E-02 | 2.67E+00 | 2.72E+00 | 8.17E-01 | no | | intergenic_variant | MODIFIER |
| 7 | 116685364 | AX-170460450 | 0 | G | A | 0.43 | 2.18E-02 | 9.42E-02 | 2.67E+00 | 2.72E+00 | 8.17E-01 | no | | intergenic_variant | MODIFIER |
| 7 | 116714766 | AX-170229895 | 0 | G | A | 0.43 | 2.18E-02 | 9.42E-02 | 2.67E+00 | 2.72E+00 | 8.17E-01 | no | | intergenic_variant | MODIFIER |
| 7 | 116715731 | AX-170393651 | 0 | G | C | 0.43 | 2.18E-02 | 9.42E-02 | 2.67E+00 | 2.72E+00 | 8.17E-01 | no | | regulatory_region_variant | MODIFIER |
| 7 | 116722856 | AX-170344734 | 0 | C | G | 0.43 | 2.18E-02 | 9.42E-02 | 2.67E+00 | 2.72E+00 | 8.17E-01 | no | | intergenic_variant | MODIFIER |
| 7 | 116731661 | AX-170226705 | 0 | G | A | 0.43 | 2.18E-02 | 9.42E-02 | 2.67E+00 | 2.72E+00 | 8.17E-01 | no | | intergenic_variant | MODIFIER |
| 7 | 116738100 | AX-97845823 | 0 | C | A | 0.43 | 2.18E-02 | 9.42E-02 | 2.67E+00 | 2.72E+00 | 8.17E-01 | no | | intergenic_variant | MODIFIER |
| 7 | 116755559 | AX-97813335 | 0 | C | T | 0.43 | 2.18E-02 | 9.42E-02 | 2.67E+00 | 2.72E+00 | 8.17E-01 | no | | intergenic_variant | MODIFIER |
| 7 | 116820942 | AX-170096093 | 0 | G | A | 0.43 | 2.18E-02 | 9.42E-02 | 2.67E+00 | 2.72E+00 | 8.17E-01 | no | Gm4366 | downstream_gene_variant | MODIFIER |
| 7 | 116895482 | AX-168419975 | 0 | T | C | 0.43 | 2.18E-02 | 9.42E-02 | 2.67E+00 | 2.72E+00 | 8.17E-01 | no | | intergenic_variant | MODIFIER |
| 15 | 78556569 | AX-170493197 | 0 | G | A | 0.553 | -1.72E-02 | 7.82E-02 | 2.71E+00 | 2.80E+00 | 8.17E-01 | no | Rac2 | downstream_gene_variant | MODIFIER |
| 1 | 36002988 | AX-170342859 | 1 | T | G | 0.446 | 1.76E-02 | 7.86E-02 | 2.68E+00 | 2.76E+00 | 8.17E-01 | no | Gm33222 | non_coding_transcript_exon_variant | MODIFIER |
| 5 | 73408644 | AX-170293466 | 0 | C | T | 0.43 | -2.14E-02 | 9.71E-02 | 2.69E+00 | 2.78E+00 | 8.18E-01 | no | Cwh43 | intron_variant | MODIFIER |
| 16 | 97914215 | AX-97808206 | 0 | G | A | 0.43 | 1.76E-02 | 8.09E-02 | 2.69E+00 | 2.80E+00 | 8.18E-01 | no | C2cd2 | intron_variant | MODIFIER |
| 9 | 52539389 | AX-170488828 | 1 | T | G | 0.214 | -2.52E-02 | 1.08E-01 | 2.64E+00 | 2.71E+00 | 8.18E-01 | no | | intergenic_variant | MODIFIER |
| 1 | 37227217 | AX-170208895 | 1 | A | G | 0.446 | 1.75E-02 | 7.80E-02 | 2.67E+00 | 2.75E+00 | 8.18E-01 | no | Cnga3 | intron_variant | MODIFIER |
| 7 | 117288798 | AX-170223589 | 0 | A | G | 0.412 | -2.08E-02 | 9.31E-02 | 2.73E+00 | 2.79E+00 | 8.18E-01 | no | | intergenic_variant | MODIFIER |
| 8 | 73973512 | AX-170524275 | 0 | A | G | 0.193 | -3.01E-02 | 1.29E-01 | 2.63E+00 | 2.71E+00 | 8.18E-01 | no | | intergenic_variant | MODIFIER |
| 8 | 36986247 | AX-170433409 | 1 | T | A | 0.152 | 3.31E-02 | 1.47E-01 | 2.73E+00 | 2.77E+00 | 8.18E-01 | no | G630064G18Rik | non_coding_transcript_exon_variant | MODIFIER |
| 1 | 177311194 | AX-170345316 | 1 | C | T | 0.232 | 2.66E-02 | 1.09E-01 | 2.59E+00 | 2.69E+00 | 8.18E-01 | no | | intergenic_variant | MODIFIER |
| 1 | 147537284 | AX-170106129 | 1 | T | C | 0.205 | 3.04E-02 | 1.27E-01 | 2.61E+00 | 2.69E+00 | 8.18E-01 | no | | intergenic_variant | MODIFIER |
| 8 | 118661348 | AX-170316487 | 1 | C | T | 0.241 | 2.19E-02 | 9.91E-02 | 2.65E+00 | 2.77E+00 | 8.18E-01 | no | Cdh13 | intron_variant | MODIFIER |
| 11 | 110577886 | AX-170389040 | 0 | G | A | 0.07 | 3.44E-02 | 1.42E-01 | 2.56E+00 | 2.70E+00 | 8.18E-01 | no | | intergenic_variant | MODIFIER |
| 1 | 108499200 | AX-97801002 | 0 | T | C | 0.167 | 4.61E-02 | 1.95E-01 | 2.62E+00 | 2.70E+00 | 8.18E-01 | no | | intergenic_variant | MODIFIER |
| 6 | 103998787 | AX-170221662 | 1 | G | A | 0.054 | 6.76E-02 | 2.96E-01 | 2.67E+00 | 2.73E+00 | 8.18E-01 | no | | intergenic_variant | MODIFIER |

| | | | | | | | | | | | | | | | |
|---|---|---|---|---|---|---|---|---|---|---|---|---|---|---|---|
| 13 | 22215886 | AX-170156721 | 0 | T | A | 0.167 | -2.91E-02 | 1.14E-01 | 2.58E+00 | 2.64E+00 | 8.18E-01 | no | Vmn1r193 | downstream_gene_variant | MODIFIER |
| 5 | 113063410 | AX-170184256 | 0 | T | C | 0.456 | 2.50E-02 | 1.09E-01 | 2.67E+00 | 2.72E+00 | 8.18E-01 | no | Crybb2 | intron_variant | MODIFIER |
| 7 | 119064593 | AX-170371426 | 1 | C | T | 0.402 | 2.13E-02 | 9.26E-02 | 2.67E+00 | 2.72E+00 | 8.18E-01 | no | | intergenic_variant | MODIFIER |
| 8 | 74757821 | AX-170031024 | 0 | A | G | 0.193 | -2.89E-02 | 1.22E-01 | 2.63E+00 | 2.70E+00 | 8.18E-01 | no | Gm11033 | intron_variant,non_coding_transcript_variant | MODIFIER |
| 4 | 143468725 | AX-97786672 | 1 | G | A | 0.098 | 4.91E-02 | 2.15E-01 | 2.68E+00 | 2.73E+00 | 8.18E-01 | no | Gm13046 | upstream_gene_variant | MODIFIER |
| 7 | 107313508 | AX-170355562 | 1 | C | T | 0.464 | 2.18E-02 | 9.23E-02 | 2.63E+00 | 2.70E+00 | 8.18E-01 | no | | intergenic_variant | MODIFIER |
| 10 | 27425429 | AX-170002953 | 1 | G | C | 0.179 | -2.12E-02 | 9.33E-02 | 2.61E+00 | 2.74E+00 | 8.18E-01 | no | Lama2 | intron_variant | MODIFIER |
| 13 | 53104052 | AX-170482233 | 0 | T | C | 0.509 | 1.58E-02 | 6.44E-02 | 2.57E+00 | 2.69E+00 | 8.18E-01 | no | | intergenic_variant | MODIFIER |
| 12 | 103765811 | AX-170251758 | 0 | A | G | 0.114 | -4.43E-02 | 1.99E-01 | 2.70E+00 | 2.77E+00 | 8.18E-01 | no | Serpina1d | missense_variant | MODERATE |
| 12 | 104259448 | AX-170147298 | 0 | G | A | 0.114 | -4.43E-02 | 1.99E-01 | 2.70E+00 | 2.77E+00 | 8.18E-01 | no | Serpina3i | upstream_gene_variant | MODIFIER |
| 12 | 105094850 | AX-97828569 | 0 | A | G | 0.114 | -4.43E-02 | 1.99E-01 | 2.70E+00 | 2.77E+00 | 8.18E-01 | no | | intergenic_variant | MODIFIER |
| 12 | 105489892 | AX-170403663 | 0 | A | G | 0.114 | -4.43E-02 | 1.99E-01 | 2.70E+00 | 2.77E+00 | 8.18E-01 | no | D430019H16Rik | 3_prime_UTR_variant | MODIFIER |
| 12 | 105514378 | AX-170050063 | 0 | T | G | 0.114 | -4.43E-02 | 1.99E-01 | 2.70E+00 | 2.77E+00 | 8.18E-01 | no | AU015791 | non_coding_transcript_exon_variant | MODIFIER |
| 12 | 105517015 | AX-170050062 | 0 | G | A | 0.114 | -4.43E-02 | 1.99E-01 | 2.70E+00 | 2.77E+00 | 8.18E-01 | no | AU015791 | downstream_gene_variant | MODIFIER |
| 12 | 105522082 | AX-97857112 | 0 | A | G | 0.114 | -4.43E-02 | 1.99E-01 | 2.70E+00 | 2.77E+00 | 8.18E-01 | no | | regulatory_region_variant | MODIFIER |
| 12 | 105555850 | AX-97851922 | 0 | A | G | 0.114 | -4.43E-02 | 1.99E-01 | 2.70E+00 | 2.77E+00 | 8.18E-01 | no | | regulatory_region_variant | MODIFIER |
| 12 | 106005871 | AX-170299705 | 0 | G | T | 0.114 | -4.43E-02 | 1.99E-01 | 2.70E+00 | 2.77E+00 | 8.18E-01 | no | Vrk1 | upstream_gene_variant | MODIFIER |
| 12 | 106358555 | AX-170176451 | 0 | G | T | 0.114 | -4.43E-02 | 1.99E-01 | 2.70E+00 | 2.77E+00 | 8.18E-01 | no | | intergenic_variant | MODIFIER |
| 12 | 106373822 | AX-170476656 | 0 | A | G | 0.114 | -4.43E-02 | 1.99E-01 | 2.70E+00 | 2.77E+00 | 8.18E-01 | no | | intergenic_variant | MODIFIER |
| 12 | 106394868 | AX-170087328 | 0 | C | T | 0.114 | -4.43E-02 | 1.99E-01 | 2.70E+00 | 2.77E+00 | 8.18E-01 | no | | intergenic_variant | MODIFIER |
| 12 | 106417378 | AX-170492248 | 0 | A | G | 0.114 | -4.43E-02 | 1.99E-01 | 2.70E+00 | 2.77E+00 | 8.18E-01 | no | | intergenic_variant | MODIFIER |
| 12 | 107141257 | AX-170478051 | 0 | C | T | 0.114 | -4.43E-02 | 1.99E-01 | 2.70E+00 | 2.77E+00 | 8.18E-01 | no | | intergenic_variant | MODIFIER |
| 12 | 107144317 | AX-170448955 | 0 | C | A | 0.114 | -4.43E-02 | 1.99E-01 | 2.70E+00 | 2.77E+00 | 8.18E-01 | no | Gm16084 | downstream_gene_variant | MODIFIER |
| 12 | 107145168 | AX-170191667 | 0 | A | G | 0.114 | -4.43E-02 | 1.99E-01 | 2.70E+00 | 2.77E+00 | 8.18E-01 | no | Gm16084 | downstream_gene_variant | MODIFIER |
| 12 | 107436315 | AX-170377242 | 0 | T | A | 0.114 | -4.43E-02 | 1.99E-01 | 2.70E+00 | 2.77E+00 | 8.18E-01 | no | | intergenic_variant | MODIFIER |
| 12 | 107436451 | AX-97808841 | 0 | C | A | 0.114 | -4.43E-02 | 1.99E-01 | 2.70E+00 | 2.77E+00 | 8.18E-01 | no | | intergenic_variant | MODIFIER |
| 12 | 107474192 | AX-170253156 | 0 | A | G | 0.114 | -4.43E-02 | 1.99E-01 | 2.70E+00 | 2.77E+00 | 8.18E-01 | no | Gm3234 | intron_variant,non_coding_transcript_variant | MODIFIER |
| 12 | 107525459 | AX-170507849 | 0 | C | T | 0.114 | -4.43E-02 | 1.99E-01 | 2.70E+00 | 2.77E+00 | 8.18E-01 | no | | intergenic_variant | MODIFIER |
| 12 | 107525496 | AX-170472537 | 0 | A | G | 0.114 | -4.43E-02 | 1.99E-01 | 2.70E+00 | 2.77E+00 | 8.18E-01 | no | | intergenic_variant | MODIFIER |
| 12 | 107548390 | AX-170160893 | 0 | C | A | 0.114 | -4.43E-02 | 1.99E-01 | 2.70E+00 | 2.77E+00 | 8.18E-01 | no | | regulatory_region_variant | MODIFIER |
| 12 | 107549247 | AX-170386488 | 0 | A | T | 0.114 | -4.43E-02 | 1.99E-01 | 2.70E+00 | 2.77E+00 | 8.18E-01 | no | | intergenic_variant | MODIFIER |
| 12 | 107549319 | AX-170162962 | 0 | T | C | 0.114 | -4.43E-02 | 1.99E-01 | 2.70E+00 | 2.77E+00 | 8.18E-01 | no | | intergenic_variant | MODIFIER |
| 12 | 107610589 | AX-97791737 | 0 | T | C | 0.114 | -4.43E-02 | 1.99E-01 | 2.70E+00 | 2.77E+00 | 8.18E-01 | no | Gm9006 | upstream_gene_variant | MODIFIER |
| 12 | 107779568 | AX-170111033 | 0 | G | A | 0.114 | -4.43E-02 | 1.99E-01 | 2.70E+00 | 2.77E+00 | 8.18E-01 | no | Gm15208 | intron_variant,non_coding_transcript_variant | MODIFIER |
| 12 | 107779778 | AX-170142531 | 0 | G | T | 0.114 | -4.43E-02 | 1.99E-01 | 2.70E+00 | 2.77E+00 | 8.18E-01 | no | Gm15208 | intron_variant,non_coding_transcript_variant | MODIFIER |
| 12 | 107779838 | AX-170260745 | 0 | A | G | 0.114 | -4.43E-02 | 1.99E-01 | 2.70E+00 | 2.77E+00 | 8.18E-01 | no | Gm15208 | intron_variant,non_coding_transcript_variant | MODIFIER |
| 12 | 107787723 | AX-170050126 | 0 | A | G | 0.114 | -4.43E-02 | 1.99E-01 | 2.70E+00 | 2.77E+00 | 8.18E-01 | no | Gm15208 | upstream_gene_variant | MODIFIER |
| 12 | 107793254 | AX-170380627 | 0 | T | C | 0.114 | -4.43E-02 | 1.99E-01 | 2.70E+00 | 2.77E+00 | 8.18E-01 | no | | intergenic_variant | MODIFIER |
| 12 | 107793639 | AX-170246053 | 0 | T | C | 0.114 | -4.43E-02 | 1.99E-01 | 2.70E+00 | 2.77E+00 | 8.18E-01 | no | | intergenic_variant | MODIFIER |
| 12 | 107796535 | AX-170377144 | 0 | C | T | 0.114 | -4.43E-02 | 1.99E-01 | 2.70E+00 | 2.77E+00 | 8.18E-01 | no | | intergenic_variant | MODIFIER |
| 12 | 107797370 | AX-170365083 | 0 | A | G | 0.114 | -4.43E-02 | 1.99E-01 | 2.70E+00 | 2.77E+00 | 8.18E-01 | no | | intergenic_variant | MODIFIER |
| 12 | 107808425 | AX-170007483 | 0 | T | C | 0.114 | -4.43E-02 | 1.99E-01 | 2.70E+00 | 2.77E+00 | 8.18E-01 | no | | regulatory_region_variant | MODIFIER |
| 12 | 107833009 | AX-170187184 | 0 | C | T | 0.114 | -4.43E-02 | 1.99E-01 | 2.70E+00 | 2.77E+00 | 8.18E-01 | no | | intergenic_variant | MODIFIER |
| 12 | 108057825 | AX-170336589 | 0 | G | A | 0.114 | -4.43E-02 | 1.99E-01 | 2.70E+00 | 2.77E+00 | 8.18E-01 | no | | intergenic_variant | MODIFIER |
| 12 | 108220997 | AX-170050132 | 0 | A | G | 0.114 | -4.43E-02 | 1.99E-01 | 2.70E+00 | 2.77E+00 | 8.18E-01 | no | Ccdc85c | intron_variant | MODIFIER |
| 12 | 108343760 | AX-170036817 | 0 | T | C | 0.114 | -4.43E-02 | 1.99E-01 | 2.70E+00 | 2.77E+00 | 8.18E-01 | no | Cyp46a1 | intron_variant | MODIFIER |
| 12 | 108345235 | AX-170391399 | 0 | C | T | 0.114 | -4.43E-02 | 1.99E-01 | 2.70E+00 | 2.77E+00 | 8.18E-01 | no | Cyp46a1 | intron_variant | MODIFIER |
| 12 | 108345340 | AX-170351894 | 0 | C | T | 0.114 | -4.43E-02 | 1.99E-01 | 2.70E+00 | 2.77E+00 | 8.18E-01 | no | Cyp46a1 | intron_variant | MODIFIER |
| 12 | 108348602 | AX-170007486 | 0 | G | A | 0.114 | -4.43E-02 | 1.99E-01 | 2.70E+00 | 2.77E+00 | 8.18E-01 | no | Cyp46a1 | intron_variant | MODIFIER |
| 12 | 108355421 | AX-170270761 | 0 | T | C | 0.114 | -4.43E-02 | 1.99E-01 | 2.70E+00 | 2.77E+00 | 8.18E-01 | no | Cyp46a1 | splice_region_variant,intron_variant | LOW |
| 12 | 108363133 | AX-169999768 | 0 | T | C | 0.114 | -4.43E-02 | 1.99E-01 | 2.70E+00 | 2.77E+00 | 8.18E-01 | no | Cyp46a1 | downstream_gene_variant | MODIFIER |
| 12 | 108371681 | AX-97826833 | 0 | C | T | 0.114 | -4.43E-02 | 1.99E-01 | 2.70E+00 | 2.77E+00 | 8.18E-01 | no | Eml1 | intron_variant,non_coding_transcript_variant | MODIFIER |
| 12 | 108371901 | AX-170007485 | 0 | G | A | 0.114 | -4.43E-02 | 1.99E-01 | 2.70E+00 | 2.77E+00 | 8.18E-01 | no | Eml1 | intron_variant,non_coding_transcript_variant | MODIFIER |
| 12 | 108373658 | AX-170525918 | 0 | C | T | 0.114 | -4.43E-02 | 1.99E-01 | 2.70E+00 | 2.77E+00 | 8.18E-01 | no | Eml1 | intron_variant,non_coding_transcript_variant | MODIFIER |

| | | | | | | | | | | | | | | | |
|---|---|---|---|---|---|---|---|---|---|---|---|---|---|---|---|
| 12 | 108382670 | AX-170373519 | 0 | A | G | 0.114 | -4.43E-02 | 1.99E-01 | 2.70E+00 | 2.77E+00 | 8.18E-01 | no | Eml1 | intron_variant,non_coding_transcript_variant | MODIFIER |
| 12 | 108412106 | AX-170230810 | 0 | T | C | 0.114 | -4.43E-02 | 1.99E-01 | 2.70E+00 | 2.77E+00 | 8.18E-01 | no | Eml1 | intron_variant | MODIFIER |
| 12 | 108415130 | AX-170244396 | 0 | T | C | 0.114 | -4.43E-02 | 1.99E-01 | 2.70E+00 | 2.77E+00 | 8.18E-01 | no | Eml1 | intron_variant | MODIFIER |
| 12 | 108420736 | AX-170476421 | 0 | A | G | 0.114 | -4.43E-02 | 1.99E-01 | 2.70E+00 | 2.77E+00 | 8.18E-01 | no | Eml1 | intron_variant | MODIFIER |
| 12 | 108430161 | AX-170379666 | 0 | A | G | 0.114 | -4.43E-02 | 1.99E-01 | 2.70E+00 | 2.77E+00 | 8.18E-01 | no | Eml1 | intron_variant | MODIFIER |
| 12 | 108624004 | AX-170050141 | 0 | T | A | 0.114 | -4.43E-02 | 1.99E-01 | 2.70E+00 | 2.77E+00 | 8.18E-01 | no | Evl | intron_variant | MODIFIER |
| 1 | 68556628 | AX-170532670 | 0 | G | A | 0.114 | -4.30E-02 | 1.85E-01 | 2.63E+00 | 2.72E+00 | 8.18E-01 | no | Erbb4 | intron_variant | MODIFIER |
| 2 | 92759082 | AX-170219342 | 0 | C | T | 0.053 | -6.80E-02 | 3.00E-01 | 2.72E+00 | 2.75E+00 | 8.18E-01 | no | Gm13816 | non_coding_transcript_exon_variant | MODIFIER |
| 2 | 108053970 | AX-170018615 | 0 | A | C | 0.053 | -6.80E-02 | 3.00E-01 | 2.72E+00 | 2.75E+00 | 8.18E-01 | no | | intergenic_variant | MODIFIER |
| 8 | 29383174 | AX-170330015 | 0 | C | T | 0.175 | 2.49E-02 | 1.11E-01 | 2.67E+00 | 2.76E+00 | 8.18E-01 | no | | intergenic_variant | MODIFIER |
| 4 | 95789014 | AX-97821638 | 1 | G | A | 0.259 | -2.34E-02 | 1.09E-01 | 2.76E+00 | 2.88E+00 | 8.18E-01 | no | Fggy | intron_variant | MODIFIER |
| 5 | 104583304 | AX-170089573 | 1 | A | C | 0.446 | -1.96E-02 | 8.84E-02 | 2.66E+00 | 2.76E+00 | 8.18E-01 | no | | intergenic_variant | MODIFIER |
| 6 | 104832505 | AX-170222613 | 1 | A | G | 0.054 | 6.74E-02 | 2.95E-01 | 2.67E+00 | 2.73E+00 | 8.18E-01 | no | Cntn6 | synonymous_variant | LOW |
| 6 | 104858021 | AX-170493485 | 1 | C | T | 0.054 | 6.74E-02 | 2.95E-01 | 2.67E+00 | 2.73E+00 | 8.18E-01 | no | Cntn6 | intron_variant | MODIFIER |
| 6 | 107492242 | AX-170452318 | 1 | A | G | 0.054 | 6.74E-02 | 2.95E-01 | 2.67E+00 | 2.73E+00 | 8.18E-01 | no | | intergenic_variant | MODIFIER |
| 6 | 112254464 | AX-170130208 | 1 | G | T | 0.054 | 6.74E-02 | 2.95E-01 | 2.67E+00 | 2.73E+00 | 8.18E-01 | no | | intergenic_variant | MODIFIER |
| 9 | 74305373 | AX-97826377 | 0 | T | G | 0.193 | 3.05E-02 | 1.35E-01 | 2.74E+00 | 2.77E+00 | 8.18E-01 | no | | intergenic_variant | MODIFIER |
| 5 | 124943370 | AX-170073388 | 2 | A | T | 0.455 | -2.62E-02 | 1.16E-01 | 2.75E+00 | 2.79E+00 | 8.18E-01 | no | | regulatory_region_variant | MODIFIER |
| 3 | 147365194 | AX-170413077 | 0 | A | G | 0.333 | 1.73E-02 | 7.46E-02 | 2.64E+00 | 2.71E+00 | 8.18E-01 | no | | intergenic_variant | MODIFIER |
| 6 | 118340892 | AX-170326179 | 1 | C | T | 0.054 | 6.75E-02 | 2.96E-01 | 2.67E+00 | 2.73E+00 | 8.19E-01 | no | | intergenic_variant | MODIFIER |
| 10 | 120529136 | AX-170046067 | 1 | G | C | 0.348 | 1.61E-02 | 7.21E-02 | 2.66E+00 | 2.75E+00 | 8.19E-01 | no | | intergenic_variant | MODIFIER |
| 7 | 81894670 | AX-97807862 | 1 | T | G | 0.089 | 4.29E-02 | 1.87E-01 | 2.59E+00 | 2.73E+00 | 8.19E-01 | no | Hdgfl3 | intron_variant | MODIFIER |
| 20 | 134798916 | AX-97783246 | 2 | T | C | 0.118 | 2.67E-02 | 1.17E-01 | 2.64E+00 | 2.73E+00 | 8.19E-01 | no | | intergenic_variant | MODIFIER |
| 6 | 20030712 | AX-97783721 | 0 | G | T | 0.044 | 2.52E-02 | 1.15E-01 | 2.63E+00 | 2.77E+00 | 8.19E-01 | no | | intergenic_variant | MODIFIER |
| 2 | 130115173 | AX-170094411 | 1 | C | T | 0.143 | 5.04E-02 | 2.21E-01 | 2.69E+00 | 2.72E+00 | 8.19E-01 | no | Tgm6 | intron_variant,non_coding_transcript_variant | MODIFIER |
| 2 | 130137820 | AX-97829857 | 1 | C | T | 0.143 | 5.04E-02 | 2.21E-01 | 2.69E+00 | 2.72E+00 | 8.19E-01 | no | Tgm6 | intron_variant | MODIFIER |
| 2 | 137939978 | AX-170445300 | 1 | G | C | 0.143 | 5.04E-02 | 2.21E-01 | 2.69E+00 | 2.72E+00 | 8.19E-01 | no | | regulatory_region_variant | MODIFIER |
| 2 | 145720535 | AX-170213791 | 1 | G | C | 0.143 | 5.04E-02 | 2.21E-01 | 2.69E+00 | 2.72E+00 | 8.19E-01 | no | Rin2 | intron_variant,non_coding_transcript_variant | MODIFIER |
| 2 | 145849871 | AX-170019289 | 1 | G | A | 0.143 | 5.04E-02 | 2.21E-01 | 2.69E+00 | 2.72E+00 | 8.19E-01 | no | Rin2 | intron_variant | MODIFIER |
| 2 | 158550869 | AX-170410480 | 1 | A | C | 0.143 | 5.04E-02 | 2.21E-01 | 2.69E+00 | 2.72E+00 | 8.19E-01 | no | Arhgap40 | downstream_gene_variant | MODIFIER |
| 18 | 64249796 | AX-97839832 | 1 | G | T | 0.42 | 1.87E-02 | 8.45E-02 | 2.74E+00 | 2.81E+00 | 8.19E-01 | no | St8sia3 | upstream_gene_variant | MODIFIER |
| 14 | 50455044 | AX-170053359 | 0 | C | A | 0.044 | -4.30E-02 | 1.91E-01 | 2.63E+00 | 2.74E+00 | 8.19E-01 | no | Olfr740 | downstream_gene_variant | MODIFIER |
| 18 | 49261854 | AX-170416631 | 0 | C | T | 0.061 | -3.50E-02 | 1.58E-01 | 2.59E+00 | 2.75E+00 | 8.19E-01 | no | | intergenic_variant | MODIFIER |
| 8 | 57691159 | AX-170080152 | 0 | A | C | 0.158 | -3.21E-02 | 1.37E-01 | 2.63E+00 | 2.71E+00 | 8.19E-01 | no | | intergenic_variant | MODIFIER |
| 2 | 106503209 | AX-170018584 | 0 | A | G | 0.053 | -6.77E-02 | 3.00E-01 | 2.72E+00 | 2.74E+00 | 8.19E-01 | no | Gm14015 | intron_variant,non_coding_transcript_variant | MODIFIER |
| 2 | 110319710 | AX-168393194 | 1 | T | C | 0.054 | -6.78E-02 | 3.00E-01 | 2.72E+00 | 2.75E+00 | 8.19E-01 | no | | regulatory_region_variant | MODIFIER |
| 5 | 27758837 | AX-97850024 | 1 | A | G | 0.054 | -6.78E-02 | 3.00E-01 | 2.72E+00 | 2.75E+00 | 8.19E-01 | no | Paxip1 | intron_variant | MODIFIER |
| 1 | 68786572 | AX-97813223 | 1 | G | A | 0.116 | 5.20E-02 | 2.30E-01 | 2.68E+00 | 2.74E+00 | 8.19E-01 | no | Erbb4 | intron_variant | MODIFIER |
| 1 | 73392040 | AX-170041652 | 1 | G | C | 0.116 | 5.20E-02 | 2.30E-01 | 2.68E+00 | 2.74E+00 | 8.19E-01 | no | Gm27315 | upstream_gene_variant | MODIFIER |
| 12 | 104045621 | AX-170462844 | 0 | A | G | 0.123 | -4.15E-02 | 1.87E-01 | 2.69E+00 | 2.76E+00 | 8.19E-01 | no | Serpina12 | upstream_gene_variant | MODIFIER |
| 12 | 104131299 | AX-170304071 | 0 | A | T | 0.123 | -4.15E-02 | 1.87E-01 | 2.69E+00 | 2.76E+00 | 8.19E-01 | no | Serpina3b | intron_variant | MODIFIER |
| 12 | 104140474 | AX-170354310 | 0 | C | A | 0.123 | -4.15E-02 | 1.87E-01 | 2.69E+00 | 2.76E+00 | 8.19E-01 | no | Serpina3b | downstream_gene_variant | MODIFIER |
| 12 | 104171469 | AX-170287230 | 0 | T | C | 0.123 | -4.15E-02 | 1.87E-01 | 2.69E+00 | 2.76E+00 | 8.19E-01 | no | Serpina3d-ps | downstream_gene_variant | MODIFIER |
| 12 | 104239776 | AX-97788752 | 0 | G | A | 0.123 | -4.15E-02 | 1.87E-01 | 2.69E+00 | 2.76E+00 | 8.19E-01 | no | Serpina3g | intron_variant | MODIFIER |
| 12 | 104246365 | AX-170108677 | 0 | G | A | 0.123 | -4.15E-02 | 1.87E-01 | 2.69E+00 | 2.76E+00 | 8.19E-01 | no | Serpina3g | downstream_gene_variant | MODIFIER |
| 12 | 104459631 | AX-170244407 | 0 | A | G | 0.123 | -4.15E-02 | 1.87E-01 | 2.69E+00 | 2.76E+00 | 8.19E-01 | no | | regulatory_region_variant | MODIFIER |
| 12 | 104555966 | AX-97831873 | 0 | C | T | 0.123 | -4.15E-02 | 1.87E-01 | 2.69E+00 | 2.76E+00 | 8.19E-01 | no | | intergenic_variant | MODIFIER |
| 12 | 104559127 | AX-170050046 | 0 | G | A | 0.123 | -4.15E-02 | 1.87E-01 | 2.69E+00 | 2.76E+00 | 8.19E-01 | no | | intergenic_variant | MODIFIER |
| 12 | 104565165 | AX-97825720 | 0 | G | A | 0.123 | -4.15E-02 | 1.87E-01 | 2.69E+00 | 2.76E+00 | 8.19E-01 | no | | intergenic_variant | MODIFIER |
| 12 | 104569243 | AX-170280280 | 0 | G | C | 0.123 | -4.15E-02 | 1.87E-01 | 2.69E+00 | 2.76E+00 | 8.19E-01 | no | | intergenic_variant | MODIFIER |
| 12 | 104569350 | AX-170235129 | 0 | G | A | 0.123 | -4.15E-02 | 1.87E-01 | 2.69E+00 | 2.76E+00 | 8.19E-01 | no | | intergenic_variant | MODIFIER |
| 12 | 104577014 | AX-170452969 | 0 | G | A | 0.123 | -4.15E-02 | 1.87E-01 | 2.69E+00 | 2.76E+00 | 8.19E-01 | no | | intergenic_variant | MODIFIER |
| 12 | 104917533 | AX-170391755 | 0 | A | G | 0.123 | -4.15E-02 | 1.87E-01 | 2.69E+00 | 2.76E+00 | 8.19E-01 | no | | intergenic_variant | MODIFIER |
| 12 | 104920151 | AX-170182456 | 0 | G | A | 0.123 | -4.15E-02 | 1.87E-01 | 2.69E+00 | 2.76E+00 | 8.19E-01 | no | | intergenic_variant | MODIFIER |

| | | | | | | | | | | | | | | | |
|---|---|---|---|---|---|---|---|---|---|---|---|---|---|---|---|
| 12 | 104920197 | AX-170471288 | 0 | C | T | 0.123 | -4.15E-02 | 1.87E-01 | 2.69E+00 | 2.76E+00 | 8.19E-01 | no | | intergenic_variant | MODIFIER |
| 12 | 104920272 | AX-170086259 | 0 | G | A | 0.123 | -4.15E-02 | 1.87E-01 | 2.69E+00 | 2.76E+00 | 8.19E-01 | no | | intergenic_variant | MODIFIER |
| 12 | 104946328 | AX-170308640 | 0 | A | C | 0.123 | -4.15E-02 | 1.87E-01 | 2.69E+00 | 2.76E+00 | 8.19E-01 | no | Syne3 | intron_variant | MODIFIER |
| 12 | 104972378 | AX-170159264 | 0 | C | A | 0.123 | -4.15E-02 | 1.87E-01 | 2.69E+00 | 2.76E+00 | 8.19E-01 | no | Syne3 | intron_variant | MODIFIER |
| 12 | 104978854 | AX-170374277 | 0 | G | A | 0.123 | -4.15E-02 | 1.87E-01 | 2.69E+00 | 2.76E+00 | 8.19E-01 | no | Syne3 | intron_variant | MODIFIER |
| 12 | 105051247 | AX-97826808 | 0 | C | T | 0.123 | -4.15E-02 | 1.87E-01 | 2.69E+00 | 2.76E+00 | 8.19E-01 | no | | regulatory_region_variant | MODIFIER |
| 12 | 105366266 | AX-97829980 | 0 | T | C | 0.123 | -4.15E-02 | 1.87E-01 | 2.69E+00 | 2.76E+00 | 8.19E-01 | no | Tunar | intron_variant | MODIFIER |
| 12 | 105458969 | AX-97789231 | 0 | G | A | 0.123 | -4.15E-02 | 1.87E-01 | 2.69E+00 | 2.76E+00 | 8.19E-01 | no | D430019H16Rik | intron_variant | MODIFIER |
| 12 | 106730687 | AX-170007470 | 0 | C | T | 0.123 | -4.15E-02 | 1.87E-01 | 2.69E+00 | 2.76E+00 | 8.19E-01 | no | | intergenic_variant | MODIFIER |
| 12 | 106731240 | AX-170007468 | 0 | T | C | 0.123 | -4.15E-02 | 1.87E-01 | 2.69E+00 | 2.76E+00 | 8.19E-01 | no | | intergenic_variant | MODIFIER |
| 12 | 106744095 | AX-170206341 | 0 | C | T | 0.123 | -4.15E-02 | 1.87E-01 | 2.69E+00 | 2.76E+00 | 8.19E-01 | no | | intergenic_variant | MODIFIER |
| 12 | 106749504 | AX-170457757 | 0 | C | A | 0.123 | -4.15E-02 | 1.87E-01 | 2.69E+00 | 2.76E+00 | 8.19E-01 | no | | intergenic_variant | MODIFIER |
| 12 | 106749612 | AX-170459098 | 0 | G | A | 0.123 | -4.15E-02 | 1.87E-01 | 2.69E+00 | 2.76E+00 | 8.19E-01 | no | | intergenic_variant | MODIFIER |
| 12 | 106754676 | AX-170284034 | 0 | A | G | 0.123 | -4.15E-02 | 1.87E-01 | 2.69E+00 | 2.76E+00 | 8.19E-01 | no | | intergenic_variant | MODIFIER |
| 12 | 106793802 | AX-170007472 | 0 | G | C | 0.123 | -4.15E-02 | 1.87E-01 | 2.69E+00 | 2.76E+00 | 8.19E-01 | no | | intergenic_variant | MODIFIER |
| 12 | 106794174 | AX-97790761 | 0 | G | A | 0.123 | -4.15E-02 | 1.87E-01 | 2.69E+00 | 2.76E+00 | 8.19E-01 | no | | intergenic_variant | MODIFIER |
| 12 | 106794990 | AX-170127239 | 0 | T | C | 0.123 | -4.15E-02 | 1.87E-01 | 2.69E+00 | 2.76E+00 | 8.19E-01 | no | | intergenic_variant | MODIFIER |
| 12 | 106797238 | AX-170304739 | 0 | G | A | 0.123 | -4.15E-02 | 1.87E-01 | 2.69E+00 | 2.76E+00 | 8.19E-01 | no | | regulatory_region_variant | MODIFIER |
| 12 | 106806090 | AX-170396587 | 0 | T | C | 0.123 | -4.15E-02 | 1.87E-01 | 2.69E+00 | 2.76E+00 | 8.19E-01 | no | | intergenic_variant | MODIFIER |
| 12 | 106809790 | AX-170479150 | 0 | A | G | 0.123 | -4.15E-02 | 1.87E-01 | 2.69E+00 | 2.76E+00 | 8.19E-01 | no | | intergenic_variant | MODIFIER |
| 12 | 106815511 | AX-170007467 | 0 | G | A | 0.123 | -4.15E-02 | 1.87E-01 | 2.69E+00 | 2.76E+00 | 8.19E-01 | no | | intergenic_variant | MODIFIER |
| 12 | 106825378 | AX-170050096 | 0 | A | G | 0.123 | -4.15E-02 | 1.87E-01 | 2.69E+00 | 2.76E+00 | 8.19E-01 | no | | intergenic_variant | MODIFIER |
| 12 | 106834630 | AX-170147655 | 0 | A | G | 0.123 | -4.15E-02 | 1.87E-01 | 2.69E+00 | 2.76E+00 | 8.19E-01 | no | 1700013N06Rik | upstream_gene_variant | MODIFIER |
| 12 | 106873102 | AX-170397911 | 0 | G | A | 0.123 | -4.15E-02 | 1.87E-01 | 2.69E+00 | 2.76E+00 | 8.19E-01 | no | 1700013N06Rik | intron_variant,non_coding_transcript_variant | MODIFIER |
| 12 | 106896708 | AX-170261903 | 0 | G | A | 0.123 | -4.15E-02 | 1.87E-01 | 2.69E+00 | 2.76E+00 | 8.19E-01 | no | | intergenic_variant | MODIFIER |
| 12 | 106915720 | AX-170480314 | 0 | T | A | 0.123 | -4.15E-02 | 1.87E-01 | 2.69E+00 | 2.76E+00 | 8.19E-01 | no | | intergenic_variant | MODIFIER |
| 12 | 107010935 | AX-97841330 | 0 | A | G | 0.123 | -4.15E-02 | 1.87E-01 | 2.69E+00 | 2.76E+00 | 8.19E-01 | no | | regulatory_region_variant | MODIFIER |
| 12 | 107075133 | AX-168411650 | 0 | A | G | 0.123 | -4.15E-02 | 1.87E-01 | 2.69E+00 | 2.76E+00 | 8.19E-01 | no | Gm16086 | downstream_gene_variant | MODIFIER |
| 12 | 107093510 | AX-170360665 | 0 | A | G | 0.123 | -4.15E-02 | 1.87E-01 | 2.69E+00 | 2.76E+00 | 8.19E-01 | no | Gm16086 | intron_variant,non_coding_transcript_variant | MODIFIER |
| 12 | 107128471 | AX-170417310 | 0 | A | G | 0.123 | -4.15E-02 | 1.87E-01 | 2.69E+00 | 2.76E+00 | 8.19E-01 | no | Gm16085 | upstream_gene_variant | MODIFIER |
| 12 | 107140688 | AX-170309045 | 0 | G | A | 0.123 | -4.15E-02 | 1.87E-01 | 2.69E+00 | 2.76E+00 | 8.19E-01 | no | | intergenic_variant | MODIFIER |
| 12 | 107169749 | AX-170368568 | 0 | C | A | 0.123 | -4.15E-02 | 1.87E-01 | 2.69E+00 | 2.76E+00 | 8.19E-01 | no | | intergenic_variant | MODIFIER |
| 12 | 107175124 | AX-170453890 | 0 | A | G | 0.123 | -4.15E-02 | 1.87E-01 | 2.69E+00 | 2.76E+00 | 8.19E-01 | no | | intergenic_variant | MODIFIER |
| 12 | 107185533 | AX-170192914 | 0 | C | T | 0.123 | -4.15E-02 | 1.87E-01 | 2.69E+00 | 2.76E+00 | 8.19E-01 | no | Gm2800 | intron_variant,non_coding_transcript_variant | MODIFIER |
| 12 | 107186190 | AX-170164036 | 0 | G | A | 0.123 | -4.15E-02 | 1.87E-01 | 2.69E+00 | 2.76E+00 | 8.19E-01 | no | Gm2800 | intron_variant,non_coding_transcript_variant | MODIFIER |
| 12 | 107189948 | AX-170509985 | 0 | A | G | 0.123 | -4.15E-02 | 1.87E-01 | 2.69E+00 | 2.76E+00 | 8.19E-01 | no | Gm2800 | downstream_gene_variant | MODIFIER |
| 12 | 107204271 | AX-170394661 | 0 | A | C | 0.123 | -4.15E-02 | 1.87E-01 | 2.69E+00 | 2.76E+00 | 8.19E-01 | no | Gm5187 | downstream_gene_variant | MODIFIER |
| 12 | 107273910 | AX-170343769 | 0 | G | A | 0.123 | -4.15E-02 | 1.87E-01 | 2.69E+00 | 2.76E+00 | 8.19E-01 | no | | intergenic_variant | MODIFIER |
| 12 | 107277069 | AX-97843339 | 0 | A | G | 0.123 | -4.15E-02 | 1.87E-01 | 2.69E+00 | 2.76E+00 | 8.19E-01 | no | | intergenic_variant | MODIFIER |
| 12 | 107277146 | AX-170531569 | 0 | G | A | 0.123 | -4.15E-02 | 1.87E-01 | 2.69E+00 | 2.76E+00 | 8.19E-01 | no | | intergenic_variant | MODIFIER |
| 12 | 107278936 | AX-170119268 | 0 | G | A | 0.123 | -4.15E-02 | 1.87E-01 | 2.69E+00 | 2.76E+00 | 8.19E-01 | no | | intergenic_variant | MODIFIER |
| 12 | 107284804 | AX-170395391 | 0 | G | A | 0.123 | -4.15E-02 | 1.87E-01 | 2.69E+00 | 2.76E+00 | 8.19E-01 | no | | intergenic_variant | MODIFIER |
| 12 | 107285264 | AX-170259698 | 0 | T | C | 0.123 | -4.15E-02 | 1.87E-01 | 2.69E+00 | 2.76E+00 | 8.19E-01 | no | | intergenic_variant | MODIFIER |
| 12 | 107286347 | AX-170495808 | 0 | C | T | 0.123 | -4.15E-02 | 1.87E-01 | 2.69E+00 | 2.76E+00 | 8.19E-01 | no | | intergenic_variant | MODIFIER |
| 12 | 107286965 | AX-170050109 | 0 | T | C | 0.123 | -4.15E-02 | 1.87E-01 | 2.69E+00 | 2.76E+00 | 8.19E-01 | no | | intergenic_variant | MODIFIER |
| 12 | 107306132 | AX-170287095 | 0 | C | T | 0.123 | -4.15E-02 | 1.87E-01 | 2.69E+00 | 2.76E+00 | 8.19E-01 | no | | intergenic_variant | MODIFIER |
| 12 | 107308494 | AX-170189656 | 0 | G | T | 0.123 | -4.15E-02 | 1.87E-01 | 2.69E+00 | 2.76E+00 | 8.19E-01 | no | | intergenic_variant | MODIFIER |
| 12 | 107312284 | AX-170353240 | 0 | T | C | 0.123 | -4.15E-02 | 1.87E-01 | 2.69E+00 | 2.76E+00 | 8.19E-01 | no | | intergenic_variant | MODIFIER |
| 12 | 107351950 | AX-170267490 | 0 | T | C | 0.123 | -4.15E-02 | 1.87E-01 | 2.69E+00 | 2.76E+00 | 8.19E-01 | no | | intergenic_variant | MODIFIER |
| 12 | 107385288 | AX-170121043 | 0 | A | T | 0.123 | -4.15E-02 | 1.87E-01 | 2.69E+00 | 2.76E+00 | 8.19E-01 | no | | intergenic_variant | MODIFIER |
| 12 | 107391899 | AX-170197777 | 0 | T | C | 0.123 | -4.15E-02 | 1.87E-01 | 2.69E+00 | 2.76E+00 | 8.19E-01 | no | | intergenic_variant | MODIFIER |
| 12 | 107393056 | AX-170473302 | 0 | C | G | 0.123 | -4.15E-02 | 1.87E-01 | 2.69E+00 | 2.76E+00 | 8.19E-01 | no | | intergenic_variant | MODIFIER |
| 12 | 107417084 | AX-170314629 | 0 | A | C | 0.123 | -4.15E-02 | 1.87E-01 | 2.69E+00 | 2.76E+00 | 8.19E-01 | no | Gm17052 | upstream_gene_variant | MODIFIER |
| 12 | 107425280 | AX-170455129 | 0 | C | T | 0.123 | -4.15E-02 | 1.87E-01 | 2.69E+00 | 2.76E+00 | 8.19E-01 | no | | intergenic_variant | MODIFIER |

| | | | | | | | | | | | | | | | |
|---|---|---|---|---|---|---|---|---|---|---|---|---|---|---|---|
| 12 | 107429815 | AX-170312630 | 0 | G | A | 0.123 | -4.15E-02 | 1.87E-01 | 2.69E+00 | 2.76E+00 | 8.19E-01 | no | | intergenic_variant | MODIFIER |
| 12 | 107437718 | AX-170383400 | 0 | C | G | 0.123 | -4.15E-02 | 1.87E-01 | 2.69E+00 | 2.76E+00 | 8.19E-01 | no | | intergenic_variant | MODIFIER |
| 12 | 107496839 | AX-170441237 | 0 | T | C | 0.123 | -4.15E-02 | 1.87E-01 | 2.69E+00 | 2.76E+00 | 8.19E-01 | no | | intergenic_variant | MODIFIER |
| 12 | 107520931 | AX-170360380 | 0 | C | T | 0.123 | -4.15E-02 | 1.87E-01 | 2.69E+00 | 2.76E+00 | 8.19E-01 | no | | intergenic_variant | MODIFIER |
| 12 | 107548434 | AX-170291995 | 0 | C | T | 0.123 | -4.15E-02 | 1.87E-01 | 2.69E+00 | 2.76E+00 | 8.19E-01 | no | | intergenic_variant | MODIFIER |
| 12 | 107567278 | AX-170258534 | 0 | G | A | 0.123 | -4.15E-02 | 1.87E-01 | 2.69E+00 | 2.76E+00 | 8.19E-01 | no | | intergenic_variant | MODIFIER |
| 12 | 107682048 | AX-170252634 | 0 | A | G | 0.123 | -4.15E-02 | 1.87E-01 | 2.69E+00 | 2.76E+00 | 8.19E-01 | no | | intergenic_variant | MODIFIER |
| 12 | 107735091 | AX-97848863 | 0 | A | G | 0.123 | -4.15E-02 | 1.87E-01 | 2.69E+00 | 2.76E+00 | 8.19E-01 | no | 4930465M20Rik | intron_variant,non_coding_transcript_variant | MODIFIER |
| 12 | 107917326 | AX-170354167 | 0 | C | T | 0.123 | -4.15E-02 | 1.87E-01 | 2.69E+00 | 2.76E+00 | 8.19E-01 | no | Bcl11b | synonymous_variant | LOW |
| 12 | 107923164 | AX-170236890 | 0 | G | A | 0.123 | -4.15E-02 | 1.87E-01 | 2.69E+00 | 2.76E+00 | 8.19E-01 | no | Bcl11b | intron_variant | MODIFIER |
| 12 | 107929269 | AX-170378196 | 0 | G | A | 0.123 | -4.15E-02 | 1.87E-01 | 2.69E+00 | 2.76E+00 | 8.19E-01 | no | Bcl11b | intron_variant | MODIFIER |
| 12 | 107937681 | AX-170510992 | 0 | C | T | 0.123 | -4.15E-02 | 1.87E-01 | 2.69E+00 | 2.76E+00 | 8.19E-01 | no | Bcl11b | intron_variant | MODIFIER |
| 12 | 107967748 | AX-170291891 | 0 | A | G | 0.123 | -4.15E-02 | 1.87E-01 | 2.69E+00 | 2.76E+00 | 8.19E-01 | no | Bcl11b | intron_variant | MODIFIER |
| 12 | 107974829 | AX-170087329 | 0 | A | C | 0.123 | -4.15E-02 | 1.87E-01 | 2.69E+00 | 2.76E+00 | 8.19E-01 | no | Bcl11b | intron_variant | MODIFIER |
| 12 | 108066598 | AX-170150234 | 0 | T | C | 0.123 | -4.15E-02 | 1.87E-01 | 2.69E+00 | 2.76E+00 | 8.19E-01 | no | | intergenic_variant | MODIFIER |
| 12 | 108097851 | AX-170334374 | 0 | C | T | 0.123 | -4.15E-02 | 1.87E-01 | 2.69E+00 | 2.76E+00 | 8.19E-01 | no | | intergenic_variant | MODIFIER |
| 12 | 108098743 | AX-170281838 | 0 | C | T | 0.123 | -4.15E-02 | 1.87E-01 | 2.69E+00 | 2.76E+00 | 8.19E-01 | no | | regulatory_region_variant | MODIFIER |
| 12 | 108113459 | AX-170050130 | 0 | T | A | 0.123 | -4.15E-02 | 1.87E-01 | 2.69E+00 | 2.76E+00 | 8.19E-01 | no | Setd3 | intron_variant | MODIFIER |
| 12 | 108113790 | AX-170245688 | 0 | G | A | 0.123 | -4.15E-02 | 1.87E-01 | 2.69E+00 | 2.76E+00 | 8.19E-01 | no | Setd3 | intron_variant | MODIFIER |
| 12 | 108114007 | AX-170185491 | 0 | T | C | 0.123 | -4.15E-02 | 1.87E-01 | 2.69E+00 | 2.76E+00 | 8.19E-01 | no | Setd3 | intron_variant | MODIFIER |
| 12 | 108121127 | AX-170192568 | 0 | T | C | 0.123 | -4.15E-02 | 1.87E-01 | 2.69E+00 | 2.76E+00 | 8.19E-01 | no | Setd3 | intron_variant | MODIFIER |
| 12 | 108130092 | AX-170196754 | 0 | T | C | 0.123 | -4.15E-02 | 1.87E-01 | 2.69E+00 | 2.76E+00 | 8.19E-01 | no | Setd3 | intron_variant | MODIFIER |
| 12 | 108132165 | AX-168400647 | 0 | A | G | 0.123 | -4.15E-02 | 1.87E-01 | 2.69E+00 | 2.76E+00 | 8.19E-01 | no | Setd3 | intron_variant | MODIFIER |
| 12 | 108155463 | AX-170493352 | 0 | C | T | 0.123 | -4.15E-02 | 1.87E-01 | 2.69E+00 | 2.76E+00 | 8.19E-01 | no | Setd3 | intron_variant | MODIFIER |
| 12 | 108166612 | AX-170335995 | 0 | C | T | 0.123 | -4.15E-02 | 1.87E-01 | 2.69E+00 | 2.76E+00 | 8.19E-01 | no | Setd3 | intron_variant | MODIFIER |
| 12 | 108169457 | AX-170420388 | 0 | G | A | 0.123 | -4.15E-02 | 1.87E-01 | 2.69E+00 | 2.76E+00 | 8.19E-01 | no | Setd3 | intron_variant | MODIFIER |
| 12 | 108172468 | AX-170402216 | 0 | T | C | 0.123 | -4.15E-02 | 1.87E-01 | 2.69E+00 | 2.76E+00 | 8.19E-01 | no | Setd3 | intron_variant | MODIFIER |
| 12 | 108183129 | AX-170337622 | 0 | C | T | 0.123 | -4.15E-02 | 1.87E-01 | 2.69E+00 | 2.76E+00 | 8.19E-01 | no | Setd3 | upstream_gene_variant | MODIFIER |
| 12 | 108186569 | AX-170227899 | 0 | A | G | 0.123 | -4.15E-02 | 1.87E-01 | 2.69E+00 | 2.76E+00 | 8.19E-01 | no | Ccnk | synonymous_variant | LOW |
| 12 | 108218294 | AX-170320594 | 0 | C | G | 0.123 | -4.15E-02 | 1.87E-01 | 2.69E+00 | 2.76E+00 | 8.19E-01 | no | Ccdc85c | intron_variant | MODIFIER |
| 12 | 108232393 | AX-170379654 | 0 | G | A | 0.123 | -4.15E-02 | 1.87E-01 | 2.69E+00 | 2.76E+00 | 8.19E-01 | no | Ccdc85c | intron_variant | MODIFIER |
| 12 | 108305748 | AX-170007484 | 0 | C | A | 0.123 | -4.15E-02 | 1.87E-01 | 2.69E+00 | 2.76E+00 | 8.19E-01 | no | Hhipl1 | upstream_gene_variant | MODIFIER |
| 12 | 108355488 | AX-170507180 | 0 | C | T | 0.123 | -4.15E-02 | 1.87E-01 | 2.69E+00 | 2.76E+00 | 8.19E-01 | no | Cyp46a1 | synonymous_variant | LOW |
| 1 | 115963073 | AX-97805347 | 0 | C | T | 0.167 | 4.60E-02 | 1.96E-01 | 2.62E+00 | 2.70E+00 | 8.19E-01 | no | Cntnap5a | intron_variant | MODIFIER |
| 6 | 104005062 | AX-170144428 | 1 | C | G | 0.054 | 6.73E-02 | 2.97E-01 | 2.67E+00 | 2.73E+00 | 8.19E-01 | no | | intergenic_variant | MODIFIER |
| 2 | 92214718 | AX-97856869 | 0 | G | A | 0.061 | -4.55E-02 | 2.08E-01 | 2.66E+00 | 2.77E+00 | 8.20E-01 | no | Phf21a | intron_variant | MODIFIER |
| 2 | 104831197 | AX-97838290 | 0 | C | T | 0.061 | -4.55E-02 | 2.08E-01 | 2.66E+00 | 2.77E+00 | 8.20E-01 | no | Prrg4 | 3_prime_UTR_variant | MODIFIER |
| 2 | 106127361 | AX-170177577 | 0 | C | T | 0.061 | -4.55E-02 | 2.08E-01 | 2.66E+00 | 2.77E+00 | 8.20E-01 | no | Dcdc5 | intron_variant,non_coding_transcript_variant | MODIFIER |
| 2 | 109732126 | AX-170488390 | 0 | C | T | 0.061 | -4.55E-02 | 2.08E-01 | 2.66E+00 | 2.77E+00 | 8.20E-01 | no | | intergenic_variant | MODIFIER |
| 13 | 54419439 | AX-170363715 | 0 | A | G | 0.518 | -1.43E-02 | 6.75E-02 | 2.69E+00 | 2.80E+00 | 8.20E-01 | no | | regulatory_region_variant | MODIFIER |
| 13 | 54462271 | AX-97852277 | 0 | G | A | 0.518 | -1.43E-02 | 6.75E-02 | 2.69E+00 | 2.80E+00 | 8.20E-01 | no | Thoc3 | intron_variant | MODIFIER |
| 13 | 54547756 | AX-170008340 | 0 | T | C | 0.518 | -1.43E-02 | 6.75E-02 | 2.69E+00 | 2.80E+00 | 8.20E-01 | no | 4833439L19Rik | downstream_gene_variant | MODIFIER |
| 13 | 54582007 | AX-97823481 | 0 | A | C | 0.518 | -1.43E-02 | 6.75E-02 | 2.69E+00 | 2.80E+00 | 8.20E-01 | no | Nop16 | downstream_gene_variant | MODIFIER |
| 13 | 54597355 | AX-97827183 | 0 | A | G | 0.518 | -1.43E-02 | 6.75E-02 | 2.69E+00 | 2.80E+00 | 8.20E-01 | no | Cltb | intron_variant | MODIFIER |
| 13 | 54597618 | AX-170321298 | 0 | C | T | 0.518 | -1.43E-02 | 6.75E-02 | 2.69E+00 | 2.80E+00 | 8.20E-01 | no | Cltb | intron_variant | MODIFIER |
| 13 | 54603324 | AX-170236557 | 0 | G | C | 0.518 | -1.43E-02 | 6.75E-02 | 2.69E+00 | 2.80E+00 | 8.20E-01 | no | Cltb | intron_variant | MODIFIER |
| 13 | 54650656 | AX-170087430 | 0 | C | T | 0.518 | -1.43E-02 | 6.75E-02 | 2.69E+00 | 2.80E+00 | 8.20E-01 | no | Faf2 | intron_variant | MODIFIER |
| 13 | 54651624 | AX-170295105 | 0 | G | A | 0.518 | -1.43E-02 | 6.75E-02 | 2.69E+00 | 2.80E+00 | 8.20E-01 | no | Faf2 | intron_variant | MODIFIER |
| 13 | 54686776 | AX-170269948 | 0 | C | T | 0.518 | -1.43E-02 | 6.75E-02 | 2.69E+00 | 2.80E+00 | 8.20E-01 | no | Rnf44 | intron_variant | MODIFIER |
| 13 | 54709725 | AX-170325428 | 0 | C | T | 0.518 | -1.43E-02 | 6.75E-02 | 2.69E+00 | 2.80E+00 | 8.20E-01 | no | Cdhr2 | intron_variant | MODIFIER |
| 13 | 54715258 | AX-97813628 | 0 | T | C | 0.518 | -1.43E-02 | 6.75E-02 | 2.69E+00 | 2.80E+00 | 8.20E-01 | no | Cdhr2 | intron_variant | MODIFIER |
| 13 | 54726586 | AX-170447461 | 0 | T | C | 0.518 | -1.43E-02 | 6.75E-02 | 2.69E+00 | 2.80E+00 | 8.20E-01 | no | Cdhr2 | missense_variant | MODERATE |
| 13 | 54726628 | AX-170184475 | 0 | A | G | 0.518 | -1.43E-02 | 6.75E-02 | 2.69E+00 | 2.80E+00 | 8.20E-01 | no | Cdhr2 | missense_variant | MODERATE |
| 13 | 54764400 | AX-170283347 | 0 | G | A | 0.518 | -1.43E-02 | 6.75E-02 | 2.69E+00 | 2.80E+00 | 8.20E-01 | no | Sncb | intron_variant | MODIFIER |

| | | | | | | | | | | | | | | | |
|---|---|---|---|---|---|---|---|---|---|---|---|---|---|---|---|
| 13 | 54770462 | AX-97823655 | 0 | A | G | 0.518 | -1.43E-02 | 6.75E-02 | 2.69E+00 | 2.80E+00 | 8.20E-01 | no | Sncb | upstream_gene_variant | MODIFIER |
| 13 | 54824820 | AX-170378195 | 0 | A | C | 0.518 | -1.43E-02 | 6.75E-02 | 2.69E+00 | 2.80E+00 | 8.20E-01 | no | | regulatory_region_variant | MODIFIER |
| 13 | 54837722 | AX-170378984 | 0 | T | C | 0.518 | -1.43E-02 | 6.75E-02 | 2.69E+00 | 2.80E+00 | 8.20E-01 | no | Gm16250 | downstream_gene_variant | MODIFIER |
| 11 | 45273029 | AX-170260018 | 1 | T | G | 0.08 | -4.26E-02 | 1.82E-01 | 2.59E+00 | 2.71E+00 | 8.20E-01 | no | Gm37312 | downstream_gene_variant | MODIFIER |
| 18 | 56053114 | AX-170481773 | 0 | T | A | 0.395 | -2.01E-02 | 8.40E-02 | 2.62E+00 | 2.68E+00 | 8.20E-01 | no | | intergenic_variant | MODIFIER |
| 11 | 33061969 | AX-97793016 | 1 | A | G | 0.107 | 4.31E-02 | 1.87E-01 | 2.64E+00 | 2.72E+00 | 8.20E-01 | no | | intergenic_variant | MODIFIER |
| 20 | 165285571 | AX-170085747 | 0 | A | G | 0.026 | 2.94E-02 | 1.41E-01 | 2.60E+00 | 2.78E+00 | 8.20E-01 | no | Glra2 | intron_variant | MODIFIER |
| 10 | 16175074 | AX-170002810 | 0 | C | T | 0.211 | 2.19E-02 | 1.02E-01 | 2.71E+00 | 2.81E+00 | 8.20E-01 | no | | intergenic_variant | MODIFIER |
| 12 | 73240592 | AX-97807109 | 0 | T | G | 0.07 | 4.52E-02 | 2.06E-01 | 2.71E+00 | 2.78E+00 | 8.20E-01 | no | Mnat1 | intron_variant | MODIFIER |
| 18 | 38166003 | AX-170205930 | 0 | A | C | 0.07 | 4.52E-02 | 2.06E-01 | 2.71E+00 | 2.78E+00 | 8.20E-01 | no | | intergenic_variant | MODIFIER |
| 11 | 33048307 | AX-170004658 | 1 | G | T | 0.107 | 4.32E-02 | 1.89E-01 | 2.65E+00 | 2.72E+00 | 8.20E-01 | no | | intergenic_variant | MODIFIER |
| 7 | 105631014 | AX-97805340 | 1 | A | G | 0.446 | -2.04E-02 | 9.31E-02 | 2.70E+00 | 2.78E+00 | 8.20E-01 | no | Arfip2 | downstream_gene_variant | MODIFIER |
| 1 | 11199836 | AX-170127205 | 1 | C | T | 0.411 | -1.54E-02 | 7.13E-02 | 2.71E+00 | 2.80E+00 | 8.20E-01 | no | Prex2 | splice_region_variant,synonymous_variant | LOW |
| 2 | 146955908 | AX-170019302 | 1 | G | C | 0.134 | 4.63E-02 | 2.07E-01 | 2.70E+00 | 2.74E+00 | 8.20E-01 | no | Kiz | intron_variant | MODIFIER |
| 4 | 139587565 | AX-170531454 | 0 | G | A | 0.158 | -2.12E-02 | 9.81E-02 | 2.70E+00 | 2.79E+00 | 8.21E-01 | no | Iffo2 | intron_variant | MODIFIER |
| 1 | 106511822 | AX-170298298 | 1 | A | G | 0.17 | -3.12E-02 | 1.51E-01 | 2.65E+00 | 2.80E+00 | 8.21E-01 | no | | regulatory_region_variant | MODIFIER |
| 7 | 88931465 | AX-170142015 | 1 | C | T | 0.116 | 4.03E-02 | 1.82E-01 | 2.62E+00 | 2.75E+00 | 8.21E-01 | no | | intergenic_variant | MODIFIER |
| 7 | 90283690 | AX-97840192 | 1 | C | T | 0.125 | 4.39E-02 | 2.01E-01 | 2.67E+00 | 2.76E+00 | 8.21E-01 | no | | intergenic_variant | MODIFIER |
| 8 | 30913654 | AX-170206753 | 0 | A | G | 0.175 | -2.34E-02 | 1.06E-01 | 2.66E+00 | 2.75E+00 | 8.21E-01 | no | | regulatory_region_variant | MODIFIER |
| 16 | 46247443 | AX-170279274 | 1 | C | A | 0.509 | 1.90E-02 | 8.07E-02 | 2.63E+00 | 2.69E+00 | 8.21E-01 | no | | intergenic_variant | MODIFIER |
| 18 | 61413878 | AX-170088355 | 0 | T | G | 0.404 | -1.94E-02 | 8.22E-02 | 2.62E+00 | 2.69E+00 | 8.21E-01 | no | | intergenic_variant | MODIFIER |
| 1 | 3899135 | AX-170485298 | 0 | C | T | 0.518 | 1.63E-02 | 6.76E-02 | 2.60E+00 | 2.68E+00 | 8.21E-01 | no | | intergenic_variant | MODIFIER |
| 5 | 7286412 | AX-170130888 | 0 | G | A | 0.439 | 1.34E-02 | 6.30E-02 | 2.65E+00 | 2.78E+00 | 8.21E-01 | no | Zfp804b | intron_variant | MODIFIER |
| 20 | 167179402 | AX-170369218 | 1 | A | T | 0.188 | 2.74E-02 | 1.17E-01 | 2.61E+00 | 2.70E+00 | 8.21E-01 | no | | regulatory_region_variant | MODIFIER |
| 1 | 36160703 | AX-170086166 | 0 | C | T | 0.456 | 1.72E-02 | 7.75E-02 | 2.67E+00 | 2.75E+00 | 8.21E-01 | no | Uggt1 | intron_variant | MODIFIER |
| 5 | 52581588 | AX-170349688 | 2 | C | A | 0.482 | 2.40E-02 | 9.66E-02 | 2.58E+00 | 2.66E+00 | 8.21E-01 | no | Gm5866 | upstream_gene_variant | MODIFIER |
| 5 | 5855856 | AX-170268857 | 0 | T | C | 0.439 | 1.34E-02 | 6.30E-02 | 2.65E+00 | 2.78E+00 | 8.21E-01 | no | | intergenic_variant | MODIFIER |
| 5 | 5857077 | AX-170158024 | 0 | A | T | 0.439 | 1.34E-02 | 6.30E-02 | 2.65E+00 | 2.78E+00 | 8.21E-01 | no | | intergenic_variant | MODIFIER |
| 5 | 5901404 | AX-170153152 | 0 | C | T | 0.439 | 1.34E-02 | 6.30E-02 | 2.65E+00 | 2.78E+00 | 8.21E-01 | no | | intergenic_variant | MODIFIER |
| 5 | 5923545 | AX-170177965 | 0 | C | T | 0.439 | 1.34E-02 | 6.30E-02 | 2.65E+00 | 2.78E+00 | 8.21E-01 | no | | intergenic_variant | MODIFIER |
| 5 | 6552005 | AX-170512303 | 0 | G | A | 0.439 | 1.34E-02 | 6.30E-02 | 2.65E+00 | 2.78E+00 | 8.21E-01 | no | | intergenic_variant | MODIFIER |
| 5 | 6675298 | AX-170406938 | 0 | T | C | 0.439 | 1.34E-02 | 6.30E-02 | 2.65E+00 | 2.78E+00 | 8.21E-01 | no | | intergenic_variant | MODIFIER |
| 5 | 7067988 | AX-170395212 | 0 | C | T | 0.439 | 1.34E-02 | 6.30E-02 | 2.65E+00 | 2.78E+00 | 8.21E-01 | no | Zfp804b | intron_variant | MODIFIER |
| 5 | 7093173 | AX-170279104 | 0 | A | G | 0.439 | 1.34E-02 | 6.30E-02 | 2.65E+00 | 2.78E+00 | 8.21E-01 | no | Zfp804b | intron_variant | MODIFIER |
| 5 | 7140065 | AX-97853389 | 0 | G | A | 0.439 | 1.34E-02 | 6.30E-02 | 2.65E+00 | 2.78E+00 | 8.21E-01 | no | Zfp804b | intron_variant | MODIFIER |
| 5 | 7180964 | AX-170071234 | 0 | A | G | 0.439 | 1.34E-02 | 6.30E-02 | 2.65E+00 | 2.78E+00 | 8.21E-01 | no | Gm24525 | upstream_gene_variant | MODIFIER |
| 5 | 7251776 | AX-170450858 | 0 | C | T | 0.439 | 1.34E-02 | 6.30E-02 | 2.65E+00 | 2.78E+00 | 8.21E-01 | no | Zfp804b | intron_variant | MODIFIER |
| 5 | 7259168 | AX-170338959 | 0 | G | A | 0.439 | 1.34E-02 | 6.30E-02 | 2.65E+00 | 2.78E+00 | 8.21E-01 | no | Gm42597 | downstream_gene_variant | MODIFIER |
| 9 | 61786130 | AX-170484194 | 0 | C | A | 0.202 | 2.82E-02 | 1.28E-01 | 2.72E+00 | 2.77E+00 | 8.21E-01 | no | | intergenic_variant | MODIFIER |
| 9 | 68588663 | AX-170105300 | 0 | C | T | 0.202 | 2.82E-02 | 1.28E-01 | 2.72E+00 | 2.77E+00 | 8.21E-01 | no | | intergenic_variant | MODIFIER |
| 8 | 23492543 | AX-170382648 | 0 | A | G | 0.167 | 2.48E-02 | 1.14E-01 | 2.69E+00 | 2.77E+00 | 8.21E-01 | no | | intergenic_variant | MODIFIER |
| 15 | 75213041 | AX-170448676 | 0 | T | G | 0.544 | 1.96E-02 | 8.32E-02 | 2.61E+00 | 2.69E+00 | 8.22E-01 | no | BC025446 | upstream_gene_variant | MODIFIER |
| 16 | 76318024 | AX-170103412 | 0 | A | C | 0.465 | 1.77E-02 | 7.33E-02 | 2.59E+00 | 2.68E+00 | 8.22E-01 | no | Nrip1 | intron_variant | MODIFIER |
| 12 | 98184946 | AX-170345662 | 0 | A | G | 0.079 | 4.47E-02 | 2.02E-01 | 2.71E+00 | 2.76E+00 | 8.22E-01 | no | | intergenic_variant | MODIFIER |
| 12 | 98195270 | AX-170524210 | 0 | G | A | 0.079 | 4.47E-02 | 2.02E-01 | 2.71E+00 | 2.76E+00 | 8.22E-01 | no | | intergenic_variant | MODIFIER |
| 12 | 98210073 | AX-170128607 | 0 | G | A | 0.079 | 4.47E-02 | 2.02E-01 | 2.71E+00 | 2.76E+00 | 8.22E-01 | no | Galc | intron_variant | MODIFIER |
| 12 | 98210576 | AX-170510496 | 0 | C | T | 0.079 | 4.47E-02 | 2.02E-01 | 2.71E+00 | 2.76E+00 | 8.22E-01 | no | Galc | intron_variant | MODIFIER |
| 12 | 98210607 | AX-97827640 | 0 | T | G | 0.079 | 4.47E-02 | 2.02E-01 | 2.71E+00 | 2.76E+00 | 8.22E-01 | no | Galc | intron_variant | MODIFIER |
| 12 | 98215003 | AX-170411589 | 0 | T | C | 0.079 | 4.47E-02 | 2.02E-01 | 2.71E+00 | 2.76E+00 | 8.22E-01 | no | Galc | intron_variant | MODIFIER |
| 12 | 98218964 | AX-170305442 | 0 | C | T | 0.079 | 4.47E-02 | 2.02E-01 | 2.71E+00 | 2.76E+00 | 8.22E-01 | no | Galc | intron_variant | MODIFIER |
| 12 | 98239663 | AX-170173641 | 0 | A | G | 0.079 | 4.47E-02 | 2.02E-01 | 2.71E+00 | 2.76E+00 | 8.22E-01 | no | Galc | intron_variant | MODIFIER |
| 12 | 98245050 | AX-170087318 | 0 | G | A | 0.079 | 4.47E-02 | 2.02E-01 | 2.71E+00 | 2.76E+00 | 8.22E-01 | no | Galc | intron_variant | MODIFIER |
| 12 | 98298582 | AX-170410014 | 0 | A | T | 0.079 | 4.47E-02 | 2.02E-01 | 2.71E+00 | 2.76E+00 | 8.22E-01 | no | | intergenic_variant | MODIFIER |
| 12 | 98298791 | AX-170092586 | 0 | C | T | 0.079 | 4.47E-02 | 2.02E-01 | 2.71E+00 | 2.76E+00 | 8.22E-01 | no | | intergenic_variant | MODIFIER |

| | | | | | | | | | | | | | | | |
|---|---|---|---|---|---|---|---|---|---|---|---|---|---|---|---|
| 12 | 98381121 | AX-170007356 | 0 | A | G | 0.079 | 4.47E-02 | 2.02E-01 | 2.71E+00 | 2.76E+00 | 8.22E-01 | no | | intergenic_variant | MODIFIER |
| 12 | 98404492 | AX-97812022 | 0 | C | T | 0.079 | 4.47E-02 | 2.02E-01 | 2.71E+00 | 2.76E+00 | 8.22E-01 | no | | intergenic_variant | MODIFIER |
| 12 | 98418058 | AX-170007357 | 0 | C | T | 0.079 | 4.47E-02 | 2.02E-01 | 2.71E+00 | 2.76E+00 | 8.22E-01 | no | | intergenic_variant | MODIFIER |
| 12 | 98418173 | AX-97822169 | 0 | C | T | 0.079 | 4.47E-02 | 2.02E-01 | 2.71E+00 | 2.76E+00 | 8.22E-01 | no | | intergenic_variant | MODIFIER |
| 12 | 98446398 | AX-170007350 | 0 | A | G | 0.079 | 4.47E-02 | 2.02E-01 | 2.71E+00 | 2.76E+00 | 8.22E-01 | no | Kcnk10 | intron_variant | MODIFIER |
| 12 | 98446947 | AX-97807164 | 0 | G | A | 0.079 | 4.47E-02 | 2.02E-01 | 2.71E+00 | 2.76E+00 | 8.22E-01 | no | Kcnk10 | intron_variant | MODIFIER |
| 12 | 98858683 | AX-170007359 | 0 | T | C | 0.079 | 4.47E-02 | 2.02E-01 | 2.71E+00 | 2.76E+00 | 8.22E-01 | no | Eml5 | intron_variant | MODIFIER |
| 12 | 98869792 | AX-170087321 | 0 | A | C | 0.079 | 4.47E-02 | 2.02E-01 | 2.71E+00 | 2.76E+00 | 8.22E-01 | no | Eml5 | intron_variant | MODIFIER |
| 12 | 98947170 | AX-97790636 | 0 | C | T | 0.079 | 4.47E-02 | 2.02E-01 | 2.71E+00 | 2.76E+00 | 8.22E-01 | no | Ttc8 | intron_variant | MODIFIER |
| 12 | 98978618 | AX-97822141 | 0 | G | A | 0.079 | 4.47E-02 | 2.02E-01 | 2.71E+00 | 2.76E+00 | 8.22E-01 | no | Ttc8 | intron_variant | MODIFIER |
| 12 | 98979502 | AX-170007367 | 0 | G | A | 0.079 | 4.47E-02 | 2.02E-01 | 2.71E+00 | 2.76E+00 | 8.22E-01 | no | Ttc8 | intron_variant | MODIFIER |
| 12 | 98979698 | AX-170221841 | 0 | C | T | 0.079 | 4.47E-02 | 2.02E-01 | 2.71E+00 | 2.76E+00 | 8.22E-01 | no | Ttc8 | intron_variant | MODIFIER |
| 12 | 99014160 | AX-97813710 | 0 | G | A | 0.079 | 4.47E-02 | 2.02E-01 | 2.71E+00 | 2.76E+00 | 8.22E-01 | no | | intergenic_variant | MODIFIER |
| 12 | 99019020 | AX-170178115 | 0 | G | A | 0.079 | 4.47E-02 | 2.02E-01 | 2.71E+00 | 2.76E+00 | 8.22E-01 | no | | intergenic_variant | MODIFIER |
| 12 | 99019171 | AX-170440785 | 0 | T | C | 0.079 | 4.47E-02 | 2.02E-01 | 2.71E+00 | 2.76E+00 | 8.22E-01 | no | | intergenic_variant | MODIFIER |
| 12 | 99158135 | AX-170328812 | 0 | C | T | 0.079 | 4.47E-02 | 2.02E-01 | 2.71E+00 | 2.76E+00 | 8.22E-01 | no | 4930474N09Rik | downstream_gene_variant | MODIFIER |
| 12 | 99166538 | AX-97818982 | 0 | C | T | 0.079 | 4.47E-02 | 2.02E-01 | 2.71E+00 | 2.76E+00 | 8.22E-01 | no | 4930474N09Rik | upstream_gene_variant | MODIFIER |
| 12 | 99201929 | AX-170309126 | 0 | C | T | 0.079 | 4.47E-02 | 2.02E-01 | 2.71E+00 | 2.76E+00 | 8.22E-01 | no | Foxn3 | intron_variant | MODIFIER |
| 2 | 121368872 | AX-170330882 | 1 | G | A | 0.143 | 4.94E-02 | 2.20E-01 | 2.69E+00 | 2.72E+00 | 8.22E-01 | no | Strc | intron_variant | MODIFIER |
| 2 | 123055687 | AX-97838837 | 1 | G | C | 0.143 | 4.94E-02 | 2.20E-01 | 2.69E+00 | 2.72E+00 | 8.22E-01 | no | | intergenic_variant | MODIFIER |
| 10 | 31504949 | AX-170127376 | 0 | C | T | 0.395 | -1.58E-02 | 7.19E-02 | 2.63E+00 | 2.75E+00 | 8.22E-01 | no | Rnf217 | intron_variant | MODIFIER |
| 5 | 127636613 | AX-97816207 | 0 | C | A | 0.412 | 1.90E-02 | 8.17E-02 | 2.60E+00 | 2.71E+00 | 8.22E-01 | no | Glt1d1 | intron_variant | MODIFIER |
| 8 | 90041606 | AX-97781868 | 0 | G | A | 0.202 | -2.43E-02 | 1.03E-01 | 2.59E+00 | 2.70E+00 | 8.22E-01 | no | | intergenic_variant | MODIFIER |
| 7 | 40331205 | AX-170494460 | 0 | T | C | 0.325 | 2.21E-02 | 9.53E-02 | 2.63E+00 | 2.70E+00 | 8.22E-01 | no | | intergenic_variant | MODIFIER |
| 7 | 40334724 | AX-170077126 | 0 | G | A | 0.325 | 2.21E-02 | 9.53E-02 | 2.63E+00 | 2.70E+00 | 8.22E-01 | no | | intergenic_variant | MODIFIER |
| 7 | 40334737 | AX-170077125 | 0 | A | G | 0.325 | 2.21E-02 | 9.53E-02 | 2.63E+00 | 2.70E+00 | 8.22E-01 | no | | intergenic_variant | MODIFIER |
| 7 | 40334978 | AX-170453714 | 0 | A | G | 0.325 | 2.21E-02 | 9.53E-02 | 2.63E+00 | 2.70E+00 | 8.22E-01 | no | | intergenic_variant | MODIFIER |
| 7 | 40335062 | AX-170338779 | 0 | C | G | 0.325 | 2.21E-02 | 9.53E-02 | 2.63E+00 | 2.70E+00 | 8.22E-01 | no | | intergenic_variant | MODIFIER |
| 7 | 40335151 | AX-170186345 | 0 | C | A | 0.325 | 2.21E-02 | 9.53E-02 | 2.63E+00 | 2.70E+00 | 8.22E-01 | no | | intergenic_variant | MODIFIER |
| 7 | 40335184 | AX-170500904 | 0 | C | T | 0.325 | 2.21E-02 | 9.53E-02 | 2.63E+00 | 2.70E+00 | 8.22E-01 | no | | intergenic_variant | MODIFIER |
| 7 | 40358377 | AX-170222451 | 0 | A | G | 0.325 | 2.21E-02 | 9.53E-02 | 2.63E+00 | 2.70E+00 | 8.22E-01 | no | | intergenic_variant | MODIFIER |
| 7 | 40364684 | AX-97834162 | 0 | A | C | 0.325 | 2.21E-02 | 9.53E-02 | 2.63E+00 | 2.70E+00 | 8.22E-01 | no | Gm24910 | downstream_gene_variant | MODIFIER |
| 7 | 40364728 | AX-170253068 | 0 | C | T | 0.325 | 2.21E-02 | 9.53E-02 | 2.63E+00 | 2.70E+00 | 8.22E-01 | no | Gm24910 | downstream_gene_variant | MODIFIER |
| 7 | 40366284 | AX-170246506 | 0 | G | A | 0.325 | 2.21E-02 | 9.53E-02 | 2.63E+00 | 2.70E+00 | 8.22E-01 | no | Gm24910 | downstream_gene_variant | MODIFIER |
| 7 | 40366656 | AX-170114775 | 0 | A | G | 0.325 | 2.21E-02 | 9.53E-02 | 2.63E+00 | 2.70E+00 | 8.22E-01 | no | Gm24910 | downstream_gene_variant | MODIFIER |
| 7 | 40367385 | AX-170501937 | 0 | G | A | 0.325 | 2.21E-02 | 9.53E-02 | 2.63E+00 | 2.70E+00 | 8.22E-01 | no | Gm24910 | downstream_gene_variant | MODIFIER |
| 7 | 40373000 | AX-170306976 | 0 | G | C | 0.325 | 2.21E-02 | 9.53E-02 | 2.63E+00 | 2.70E+00 | 8.22E-01 | no | Gm28807 | downstream_gene_variant | MODIFIER |
| 7 | 40549405 | AX-97784572 | 0 | G | A | 0.325 | 2.21E-02 | 9.53E-02 | 2.63E+00 | 2.70E+00 | 8.22E-01 | no | | intergenic_variant | MODIFIER |
| 7 | 40554154 | AX-170143417 | 0 | C | T | 0.325 | 2.21E-02 | 9.53E-02 | 2.63E+00 | 2.70E+00 | 8.22E-01 | no | | intergenic_variant | MODIFIER |
| 7 | 40569051 | AX-170118801 | 0 | G | A | 0.325 | 2.21E-02 | 9.53E-02 | 2.63E+00 | 2.70E+00 | 8.22E-01 | no | | intergenic_variant | MODIFIER |
| 7 | 40606833 | AX-170239357 | 0 | A | G | 0.325 | 2.21E-02 | 9.53E-02 | 2.63E+00 | 2.70E+00 | 8.22E-01 | no | | intergenic_variant | MODIFIER |
| 7 | 40639200 | AX-170095904 | 0 | C | A | 0.325 | 2.21E-02 | 9.53E-02 | 2.63E+00 | 2.70E+00 | 8.22E-01 | no | | intergenic_variant | MODIFIER |
| 7 | 40681638 | AX-170115963 | 0 | T | C | 0.325 | 2.21E-02 | 9.53E-02 | 2.63E+00 | 2.70E+00 | 8.22E-01 | no | 4933402C06Rik | intron_variant,non_coding_transcript_variant | MODIFIER |
| 7 | 40688411 | AX-170275411 | 0 | T | C | 0.325 | 2.21E-02 | 9.53E-02 | 2.63E+00 | 2.70E+00 | 8.22E-01 | no | 4933402C06Rik | intron_variant,non_coding_transcript_variant | MODIFIER |
| 7 | 40704978 | AX-170446727 | 0 | C | T | 0.325 | 2.21E-02 | 9.53E-02 | 2.63E+00 | 2.70E+00 | 8.22E-01 | no | | intergenic_variant | MODIFIER |
| 7 | 40705265 | AX-170462433 | 0 | A | G | 0.325 | 2.21E-02 | 9.53E-02 | 2.63E+00 | 2.70E+00 | 8.22E-01 | no | | intergenic_variant | MODIFIER |
| 7 | 40723449 | AX-170361522 | 0 | C | T | 0.325 | 2.21E-02 | 9.53E-02 | 2.63E+00 | 2.70E+00 | 8.22E-01 | no | | intergenic_variant | MODIFIER |
| 7 | 44395400 | AX-170174179 | 0 | T | C | 0.325 | 2.21E-02 | 9.53E-02 | 2.63E+00 | 2.70E+00 | 8.22E-01 | no | Syt3 | intron_variant | MODIFIER |
| 7 | 44420820 | AX-170396946 | 0 | T | C | 0.325 | 2.21E-02 | 9.53E-02 | 2.63E+00 | 2.70E+00 | 8.22E-01 | no | | intergenic_variant | MODIFIER |
| 7 | 44422238 | AX-97812048 | 0 | C | G | 0.325 | 2.21E-02 | 9.53E-02 | 2.63E+00 | 2.70E+00 | 8.22E-01 | no | | intergenic_variant | MODIFIER |
| 15 | 101919282 | AX-170261667 | 0 | A | T | 0.114 | -2.95E-02 | 1.34E-01 | 2.65E+00 | 2.75E+00 | 8.22E-01 | no | Krt4 | splice_region_variant,intron_variant | LOW |
| 4 | 154736400 | AX-170469191 | 0 | A | T | 0.114 | -3.45E-02 | 1.57E-01 | 2.64E+00 | 2.75E+00 | 8.22E-01 | no | | intergenic_variant | MODIFIER |
| 8 | 23708523 | AX-170464541 | 1 | C | A | 0.161 | -2.51E-02 | 1.11E-01 | 2.64E+00 | 2.72E+00 | 8.22E-01 | no | Zmat4 | intron_variant | MODIFIER |
| 13 | 46233817 | AX-97794133 | 0 | T | C | 0.491 | -1.33E-02 | 6.24E-02 | 2.67E+00 | 2.78E+00 | 8.22E-01 | no | | intergenic_variant | MODIFIER |

| | | | | | | | | | | | | | | | |
|---|---|---|---|---|---|---|---|---|---|---|---|---|---|---|---|
| 13 | 46257379 | AX-170008258 | 0 | T | C | 0.491 | -1.33E-02 | 6.24E-02 | 2.67E+00 | 2.78E+00 | 8.22E-01 | no | | intergenic_variant | MODIFIER |
| 13 | 46257635 | AX-170008251 | 0 | G | A | 0.491 | -1.33E-02 | 6.24E-02 | 2.67E+00 | 2.78E+00 | 8.22E-01 | no | | regulatory_region_variant | MODIFIER |
| 13 | 46287950 | AX-170008250 | 0 | T | G | 0.491 | -1.33E-02 | 6.24E-02 | 2.67E+00 | 2.78E+00 | 8.22E-01 | no | Stmnd1 | intron_variant | MODIFIER |
| 13 | 46292210 | AX-170208019 | 0 | A | G | 0.491 | -1.33E-02 | 6.24E-02 | 2.67E+00 | 2.78E+00 | 8.22E-01 | no | Stmnd1 | intron_variant | MODIFIER |
| 13 | 46297108 | AX-97823482 | 0 | A | G | 0.491 | -1.33E-02 | 6.24E-02 | 2.67E+00 | 2.78E+00 | 8.22E-01 | no | Stmnd1 | intron_variant | MODIFIER |
| 13 | 46298712 | AX-170507069 | 0 | G | A | 0.491 | -1.33E-02 | 6.24E-02 | 2.67E+00 | 2.78E+00 | 8.22E-01 | no | Stmnd1 | synonymous_variant | LOW |
| 13 | 46298961 | AX-170492342 | 0 | A | G | 0.491 | -1.33E-02 | 6.24E-02 | 2.67E+00 | 2.78E+00 | 8.22E-01 | no | Stmnd1 | intron_variant | MODIFIER |
| 13 | 46302417 | AX-170237526 | 0 | G | A | 0.491 | -1.33E-02 | 6.24E-02 | 2.67E+00 | 2.78E+00 | 8.22E-01 | no | Stmnd1 | downstream_gene_variant | MODIFIER |
| 13 | 46313436 | AX-170008252 | 0 | C | T | 0.491 | -1.33E-02 | 6.24E-02 | 2.67E+00 | 2.78E+00 | 8.22E-01 | no | | intergenic_variant | MODIFIER |
| 13 | 46313637 | AX-170051165 | 0 | T | C | 0.491 | -1.33E-02 | 6.24E-02 | 2.67E+00 | 2.78E+00 | 8.22E-01 | no | | intergenic_variant | MODIFIER |
| 13 | 46315699 | AX-97782930 | 0 | C | T | 0.491 | -1.33E-02 | 6.24E-02 | 2.67E+00 | 2.78E+00 | 8.22E-01 | no | | intergenic_variant | MODIFIER |
| 13 | 46319157 | AX-170295642 | 0 | G | T | 0.491 | -1.33E-02 | 6.24E-02 | 2.67E+00 | 2.78E+00 | 8.22E-01 | no | | intergenic_variant | MODIFIER |
| 13 | 46330840 | AX-170008256 | 0 | T | C | 0.491 | -1.33E-02 | 6.24E-02 | 2.67E+00 | 2.78E+00 | 8.22E-01 | no | | intergenic_variant | MODIFIER |
| 13 | 46334088 | AX-170296539 | 0 | G | A | 0.491 | -1.33E-02 | 6.24E-02 | 2.67E+00 | 2.78E+00 | 8.22E-01 | no | | intergenic_variant | MODIFIER |
| 13 | 46344700 | AX-97810447 | 0 | G | A | 0.491 | -1.33E-02 | 6.24E-02 | 2.67E+00 | 2.78E+00 | 8.22E-01 | no | | intergenic_variant | MODIFIER |
| 13 | 46396441 | AX-97784433 | 0 | A | G | 0.491 | -1.33E-02 | 6.24E-02 | 2.67E+00 | 2.78E+00 | 8.22E-01 | no | | intergenic_variant | MODIFIER |
| 13 | 46396586 | AX-170198041 | 0 | A | G | 0.491 | -1.33E-02 | 6.24E-02 | 2.67E+00 | 2.78E+00 | 8.22E-01 | no | | intergenic_variant | MODIFIER |
| 13 | 46595926 | AX-170338404 | 0 | C | T | 0.491 | -1.33E-02 | 6.24E-02 | 2.67E+00 | 2.78E+00 | 8.22E-01 | no | Cap2 | intron_variant | MODIFIER |
| 13 | 46604703 | AX-170098976 | 0 | G | A | 0.491 | -1.33E-02 | 6.24E-02 | 2.67E+00 | 2.78E+00 | 8.22E-01 | no | Cap2 | intron_variant | MODIFIER |
| 13 | 46627091 | AX-170212379 | 0 | G | A | 0.491 | -1.33E-02 | 6.24E-02 | 2.67E+00 | 2.78E+00 | 8.22E-01 | no | Cap2 | intron_variant | MODIFIER |
| 13 | 46674363 | AX-170144113 | 0 | G | A | 0.491 | -1.33E-02 | 6.24E-02 | 2.67E+00 | 2.78E+00 | 8.22E-01 | no | Fam8a1 | synonymous_variant | LOW |
| 13 | 46725625 | AX-97853282 | 0 | T | C | 0.491 | -1.33E-02 | 6.24E-02 | 2.67E+00 | 2.78E+00 | 8.22E-01 | no | Nup153 | intron_variant | MODIFIER |
| 5 | 126488927 | AX-97804085 | 2 | C | T | 0.464 | 2.06E-02 | 9.23E-02 | 2.65E+00 | 2.73E+00 | 8.22E-01 | no | | intergenic_variant | MODIFIER |
| 1 | 182390846 | AX-170403513 | 0 | C | G | 0.228 | 2.52E-02 | 9.96E-02 | 2.55E+00 | 2.66E+00 | 8.22E-01 | no | Gm5706 | downstream_gene_variant | MODIFIER |
| 5 | 73534761 | AX-170089520 | 1 | G | A | 0.438 | 2.23E-02 | 9.72E-02 | 2.62E+00 | 2.71E+00 | 8.22E-01 | no | Dcun1d4 | intron_variant | MODIFIER |
| 7 | 113852170 | AX-170185121 | 1 | G | A | 0.438 | 2.11E-02 | 9.50E-02 | 2.69E+00 | 2.74E+00 | 8.22E-01 | no | Spon1 | intron_variant | MODIFIER |
| 16 | 97186241 | AX-170170425 | 0 | G | A | 0.474 | 1.62E-02 | 7.68E-02 | 2.68E+00 | 2.79E+00 | 8.22E-01 | no | | intergenic_variant | MODIFIER |
| 4 | 139015806 | AX-97832668 | 0 | G | A | 0.167 | 2.51E-02 | 1.06E-01 | 2.61E+00 | 2.69E+00 | 8.22E-01 | no | Tmco4 | intron_variant | MODIFIER |
| 4 | 95006250 | AX-170386172 | 1 | C | G | 0.304 | 2.39E-02 | 1.04E-01 | 2.62E+00 | 2.71E+00 | 8.23E-01 | no | Gm12694 | intron_variant,non_coding_transcript_variant | MODIFIER |
| 20 | 57097019 | AX-170096774 | 1 | A | T | 0.25 | 1.39E-02 | 6.29E-02 | 2.65E+00 | 2.74E+00 | 8.23E-01 | no | Vgll1 | intron_variant | MODIFIER |
| 8 | 91161933 | AX-170230225 | 1 | A | C | 0.196 | -2.65E-02 | 1.16E-01 | 2.63E+00 | 2.71E+00 | 8.23E-01 | no | Aktip | intron_variant | MODIFIER |
| 1 | 54036069 | AX-170208568 | 1 | A | T | 0.089 | -4.38E-02 | 2.03E-01 | 2.69E+00 | 2.77E+00 | 8.23E-01 | no | Hecw2 | intron_variant | MODIFIER |
| 7 | 93693050 | AX-97798973 | 1 | T | G | 0.116 | 4.68E-02 | 2.11E-01 | 2.65E+00 | 2.73E+00 | 8.23E-01 | no | | intergenic_variant | MODIFIER |
| 8 | 94787648 | AX-170520078 | 1 | A | G | 0.205 | -2.60E-02 | 1.17E-01 | 2.67E+00 | 2.73E+00 | 8.23E-01 | no | | regulatory_region_variant | MODIFIER |
| 5 | 34556740 | AX-170145103 | 0 | A | G | 0.333 | 2.35E-02 | 1.06E-01 | 2.67E+00 | 2.73E+00 | 8.23E-01 | no | Sh3bp2 | intron_variant | MODIFIER |
| 10 | 6555368 | AX-170320607 | 1 | G | A | 0.196 | -2.26E-02 | 1.00E-01 | 2.62E+00 | 2.72E+00 | 8.23E-01 | no | | intergenic_variant | MODIFIER |
| 3 | 32210021 | AX-170479817 | 1 | A | T | 0.152 | -2.20E-02 | 9.89E-02 | 2.66E+00 | 2.73E+00 | 8.23E-01 | no | | intergenic_variant | MODIFIER |
| 7 | 137128962 | AX-170138240 | 0 | A | G | 0.298 | -1.73E-02 | 7.62E-02 | 2.62E+00 | 2.72E+00 | 8.23E-01 | no | Mgmt | 3_prime_UTR_variant | MODIFIER |
| 15 | 91482333 | AX-170225483 | 0 | A | G | 0.14 | 3.15E-02 | 1.38E-01 | 2.64E+00 | 2.71E+00 | 8.23E-01 | no | Slc2a13 | intron_variant | MODIFIER |
| 8 | 30740448 | AX-170162655 | 0 | G | T | 0.175 | 2.52E-02 | 1.14E-01 | 2.66E+00 | 2.73E+00 | 8.23E-01 | no | | intergenic_variant | MODIFIER |
| 8 | 117294956 | AX-170368725 | 0 | A | T | 0.254 | -2.11E-02 | 9.62E-02 | 2.63E+00 | 2.74E+00 | 8.23E-01 | no | Cmip | intron_variant | MODIFIER |
| 7 | 118386014 | AX-170268191 | 0 | G | A | 0.395 | 2.03E-02 | 9.15E-02 | 2.67E+00 | 2.73E+00 | 8.23E-01 | no | Syt17 | intron_variant | MODIFIER |
| 7 | 118647421 | AX-170408989 | 0 | G | A | 0.395 | 2.03E-02 | 9.15E-02 | 2.67E+00 | 2.73E+00 | 8.23E-01 | no | Tmc5 | intron_variant | MODIFIER |
| 20 | 95903369 | AX-97842245 | 0 | A | C | 0.325 | 1.23E-02 | 5.58E-02 | 2.62E+00 | 2.74E+00 | 8.23E-01 | no | Zc3h12b | intron_variant | MODIFIER |
| 4 | 154190894 | AX-97794130 | 1 | A | G | 0.098 | 3.36E-02 | 1.57E-01 | 2.66E+00 | 2.77E+00 | 8.23E-01 | no | Megf6 | intron_variant | MODIFIER |
| 16 | 86161359 | AX-170116100 | 0 | G | T | 0.447 | -1.65E-02 | 7.70E-02 | 2.69E+00 | 2.77E+00 | 8.23E-01 | no | | intergenic_variant | MODIFIER |
| 15 | 90902088 | AX-170226543 | 1 | C | T | 0.134 | 3.56E-02 | 1.57E-01 | 2.64E+00 | 2.70E+00 | 8.24E-01 | no | Spn-ps | upstream_gene_variant | MODIFIER |
| 15 | 94602805 | AX-170408963 | 1 | C | T | 0.134 | 3.56E-02 | 1.57E-01 | 2.64E+00 | 2.70E+00 | 8.24E-01 | no | | intergenic_variant | MODIFIER |
| 7 | 80813838 | AX-170029404 | 0 | C | T | 0.114 | -4.61E-02 | 2.13E-01 | 2.68E+00 | 2.76E+00 | 8.24E-01 | no | Iqgap1 | intron_variant | MODIFIER |
| 7 | 82598670 | AX-170077889 | 0 | G | A | 0.114 | -4.61E-02 | 2.13E-01 | 2.68E+00 | 2.76E+00 | 8.24E-01 | no | Adamtsl3 | intron_variant | MODIFIER |
| 7 | 93575967 | AX-170502752 | 0 | T | A | 0.114 | -4.61E-02 | 2.13E-01 | 2.68E+00 | 2.76E+00 | 8.24E-01 | no | | intergenic_variant | MODIFIER |
| 9 | 88387464 | AX-170255681 | 0 | C | A | 0.132 | 2.72E-02 | 1.28E-01 | 2.70E+00 | 2.78E+00 | 8.24E-01 | no | Snx14 | intron_variant,NMD_transcript_variant | MODIFIER |
| 9 | 89636442 | AX-170032945 | 0 | G | C | 0.132 | 2.72E-02 | 1.28E-01 | 2.70E+00 | 2.78E+00 | 8.24E-01 | no | Gm47409 | upstream_gene_variant | MODIFIER |
| 5 | 99690160 | AX-170270742 | 0 | G | A | 0.421 | 2.23E-02 | 1.00E-01 | 2.65E+00 | 2.73E+00 | 8.24E-01 | no | Rasgef1b | intron_variant,non_coding_transcript_variant | MODIFIER |

| | | | | | | | | | | | | | | | |
|---|---|---|---|---|---|---|---|---|---|---|---|---|---|---|---|
| 18 | 59642637 | AX-170369233 | 0 | A | G | 0.412 | -1.94E-02 | 8.39E-02 | 2.63E+00 | 2.69E+00 | 8.24E-01 | no | | intergenic_variant | MODIFIER |
| 4 | 93202684 | AX-170397130 | 1 | C | G | 0.33 | -2.20E-02 | 1.03E-01 | 2.68E+00 | 2.77E+00 | 8.24E-01 | no | | intergenic_variant | MODIFIER |
| 12 | 68176780 | AX-170413166 | 1 | G | A | 0.054 | -3.17E-02 | 1.53E-01 | 2.63E+00 | 2.78E+00 | 8.24E-01 | no | | intergenic_variant | MODIFIER |
| 4 | 149120028 | AX-170214924 | 1 | G | A | 0.089 | 3.74E-02 | 1.69E-01 | 2.66E+00 | 2.73E+00 | 8.24E-01 | no | Dffa | 3_prime_UTR_variant | MODIFIER |
| 7 | 118031143 | AX-170137647 | 1 | A | G | 0.393 | 2.05E-02 | 9.21E-02 | 2.67E+00 | 2.72E+00 | 8.24E-01 | no | | intergenic_variant | MODIFIER |
| 13 | 54610101 | AX-97817895 | 1 | A | G | 0.518 | -1.39E-02 | 6.73E-02 | 2.69E+00 | 2.80E+00 | 8.24E-01 | no | Cltb | intron_variant | MODIFIER |
| 7 | 82439093 | AX-170467667 | 0 | A | T | 0.114 | 4.64E-02 | 2.11E-01 | 2.65E+00 | 2.74E+00 | 8.24E-01 | no | Adamtsl3 | intron_variant | MODIFIER |
| 1 | 52248476 | AX-170274115 | 1 | T | A | 0.125 | -5.13E-02 | 2.36E-01 | 2.69E+00 | 2.75E+00 | 8.25E-01 | no | | intergenic_variant | MODIFIER |
| 1 | 62882891 | AX-170041434 | 1 | G | A | 0.125 | -5.13E-02 | 2.36E-01 | 2.69E+00 | 2.75E+00 | 8.25E-01 | no | | intergenic_variant | MODIFIER |
| 5 | 110582881 | AX-170372268 | 0 | C | T | 0.456 | 2.55E-02 | 1.14E-01 | 2.67E+00 | 2.71E+00 | 8.25E-01 | no | Galnt9 | intron_variant | MODIFIER |
| 5 | 117168470 | AX-170249814 | 0 | G | A | 0.456 | 2.55E-02 | 1.14E-01 | 2.67E+00 | 2.71E+00 | 8.25E-01 | no | Taok3 | intron_variant | MODIFIER |
| 8 | 102085813 | AX-170229783 | 0 | T | A | 0.228 | 2.40E-02 | 1.10E-01 | 2.67E+00 | 2.74E+00 | 8.25E-01 | no | | intergenic_variant | MODIFIER |
| 3 | 74004272 | AX-170129175 | 1 | C | T | 0.045 | -3.71E-02 | 1.68E-01 | 2.62E+00 | 2.73E+00 | 8.25E-01 | no | Gm20356 | downstream_gene_variant | MODIFIER |
| 10 | 117358856 | AX-170003987 | 1 | C | G | 0.339 | 1.61E-02 | 7.60E-02 | 2.68E+00 | 2.77E+00 | 8.25E-01 | no | Cpsf6 | intron_variant | MODIFIER |
| 1 | 3046097 | AX-169999824 | 1 | A | C | 0.42 | 1.63E-02 | 6.87E-02 | 2.60E+00 | 2.67E+00 | 8.25E-01 | no | | intergenic_variant | MODIFIER |
| 1 | 11272555 | AX-169999943 | 0 | G | A | 0.377 | -1.53E-02 | 7.23E-02 | 2.73E+00 | 2.80E+00 | 8.25E-01 | no | Prex2 | intron_variant | MODIFIER |
| 12 | 116417516 | AX-170387751 | 0 | C | T | 0.14 | -4.13E-02 | 1.90E-01 | 2.67E+00 | 2.74E+00 | 8.25E-01 | no | Ncapg2 | intron_variant | MODIFIER |
| 11 | 116482219 | AX-97828166 | 1 | G | A | 0.054 | -4.14E-02 | 1.96E-01 | 2.65E+00 | 2.77E+00 | 8.25E-01 | no | Prpsap1 | intron_variant | MODIFIER |
| 7 | 90124599 | AX-170029481 | 1 | A | C | 0.134 | -4.58E-02 | 2.12E-01 | 2.67E+00 | 2.75E+00 | 8.25E-01 | no | 2310010J17Rik | downstream_gene_variant | MODIFIER |
| 1 | 170917650 | AX-170462017 | 0 | G | C | 0.211 | 2.97E-02 | 1.32E-01 | 2.65E+00 | 2.71E+00 | 8.25E-01 | no | Fcrla | 3_prime_UTR_variant | MODIFIER |
| 18 | 64258740 | AX-170105581 | 1 | T | C | 0.42 | 1.80E-02 | 8.45E-02 | 2.74E+00 | 2.81E+00 | 8.25E-01 | no | St8sia3 | intron_variant | MODIFIER |
| 1 | 68845656 | AX-170164302 | 1 | C | A | 0.116 | 5.05E-02 | 2.31E-01 | 2.68E+00 | 2.74E+00 | 8.25E-01 | no | Erbb4 | intron_variant | MODIFIER |
| 7 | 40290246 | AX-170028878 | 0 | C | T | 0.333 | 2.17E-02 | 9.60E-02 | 2.63E+00 | 2.70E+00 | 8.26E-01 | no | | intergenic_variant | MODIFIER |
| 7 | 40418691 | AX-170443012 | 0 | T | C | 0.333 | 2.17E-02 | 9.60E-02 | 2.63E+00 | 2.70E+00 | 8.26E-01 | no | Gm28807 | intron_variant,non_coding_transcript_variant | MODIFIER |
| 7 | 40422691 | AX-170077129 | 0 | T | C | 0.333 | 2.17E-02 | 9.60E-02 | 2.63E+00 | 2.70E+00 | 8.26E-01 | no | Gm28807 | intron_variant,non_coding_transcript_variant | MODIFIER |
| 7 | 40447325 | AX-170198885 | 0 | G | A | 0.333 | 2.17E-02 | 9.60E-02 | 2.63E+00 | 2.70E+00 | 8.26E-01 | no | Gm28807 | intron_variant,non_coding_transcript_variant | MODIFIER |
| 7 | 40451361 | AX-97808620 | 0 | G | A | 0.333 | 2.17E-02 | 9.60E-02 | 2.63E+00 | 2.70E+00 | 8.26E-01 | no | Gm28807 | intron_variant,non_coding_transcript_variant | MODIFIER |
| 7 | 40475870 | AX-97825125 | 0 | A | C | 0.333 | 2.17E-02 | 9.60E-02 | 2.63E+00 | 2.70E+00 | 8.26E-01 | no | | intergenic_variant | MODIFIER |
| 7 | 40479069 | AX-170280206 | 0 | C | T | 0.333 | 2.17E-02 | 9.60E-02 | 2.63E+00 | 2.70E+00 | 8.26E-01 | no | | intergenic_variant | MODIFIER |
| 7 | 40483266 | AX-170231047 | 0 | A | G | 0.333 | 2.17E-02 | 9.60E-02 | 2.63E+00 | 2.70E+00 | 8.26E-01 | no | | intergenic_variant | MODIFIER |
| 7 | 40483931 | AX-170423426 | 0 | G | A | 0.333 | 2.17E-02 | 9.60E-02 | 2.63E+00 | 2.70E+00 | 8.26E-01 | no | | intergenic_variant | MODIFIER |
| 7 | 40485819 | AX-170455565 | 0 | T | G | 0.333 | 2.17E-02 | 9.60E-02 | 2.63E+00 | 2.70E+00 | 8.26E-01 | no | | intergenic_variant | MODIFIER |
| 7 | 40494693 | AX-170441239 | 0 | T | C | 0.333 | 2.17E-02 | 9.60E-02 | 2.63E+00 | 2.70E+00 | 8.26E-01 | no | | intergenic_variant | MODIFIER |
| 7 | 40496112 | AX-170118695 | 0 | T | C | 0.333 | 2.17E-02 | 9.60E-02 | 2.63E+00 | 2.70E+00 | 8.26E-01 | no | | intergenic_variant | MODIFIER |
| 7 | 40496312 | AX-170347304 | 0 | C | T | 0.333 | 2.17E-02 | 9.60E-02 | 2.63E+00 | 2.70E+00 | 8.26E-01 | no | | intergenic_variant | MODIFIER |
| 7 | 44446603 | AX-97801960 | 0 | A | G | 0.333 | 2.17E-02 | 9.60E-02 | 2.63E+00 | 2.70E+00 | 8.26E-01 | no | Lrrc4b | intron_variant | MODIFIER |
| 7 | 44447563 | AX-170386023 | 0 | C | T | 0.333 | 2.17E-02 | 9.60E-02 | 2.63E+00 | 2.70E+00 | 8.26E-01 | no | Lrrc4b | intron_variant | MODIFIER |
| 7 | 44485126 | AX-97855740 | 0 | A | C | 0.333 | 2.17E-02 | 9.60E-02 | 2.63E+00 | 2.70E+00 | 8.26E-01 | no | Emc10 | downstream_gene_variant | MODIFIER |
| 7 | 44499007 | AX-170028924 | 0 | G | A | 0.333 | 2.17E-02 | 9.60E-02 | 2.63E+00 | 2.70E+00 | 8.26E-01 | no | Fam71e1 | intron_variant | MODIFIER |
| 7 | 44530309 | AX-97802355 | 0 | G | A | 0.333 | 2.17E-02 | 9.60E-02 | 2.63E+00 | 2.70E+00 | 8.26E-01 | no | Spib | intron_variant | MODIFIER |
| 7 | 44560350 | AX-97805081 | 0 | A | C | 0.333 | 2.17E-02 | 9.60E-02 | 2.63E+00 | 2.70E+00 | 8.26E-01 | no | | intergenic_variant | MODIFIER |
| 7 | 44732803 | AX-170089911 | 0 | C | T | 0.333 | 2.17E-02 | 9.60E-02 | 2.63E+00 | 2.70E+00 | 8.26E-01 | no | Zfp473 | missense_variant | MODERATE |
| 7 | 44770880 | AX-170199696 | 0 | A | G | 0.333 | 2.17E-02 | 9.60E-02 | 2.63E+00 | 2.70E+00 | 8.26E-01 | no | Vrk3 | missense_variant | MODERATE |
| 7 | 44788339 | AX-170487872 | 0 | G | A | 0.333 | 2.17E-02 | 9.60E-02 | 2.63E+00 | 2.70E+00 | 8.26E-01 | no | Gm29791 | downstream_gene_variant | MODIFIER |
| 7 | 44853146 | AX-170212738 | 0 | G | A | 0.333 | 2.17E-02 | 9.60E-02 | 2.63E+00 | 2.70E+00 | 8.26E-01 | no | Pnkp | upstream_gene_variant | MODIFIER |
| 7 | 44862368 | AX-97818074 | 0 | C | T | 0.333 | 2.17E-02 | 9.60E-02 | 2.63E+00 | 2.70E+00 | 8.26E-01 | no | Pnkp | splice_region_variant,synonymous_variant | LOW |
| 7 | 44875971 | AX-170262314 | 0 | G | C | 0.333 | 2.17E-02 | 9.60E-02 | 2.63E+00 | 2.70E+00 | 8.26E-01 | no | Med25 | downstream_gene_variant | MODIFIER |
| 1 | 30539710 | AX-170410959 | 1 | T | C | 0.411 | -1.74E-02 | 8.14E-02 | 2.68E+00 | 2.76E+00 | 8.26E-01 | no | | intergenic_variant | MODIFIER |
| 1 | 30622136 | AX-170160068 | 1 | A | G | 0.411 | -1.74E-02 | 8.14E-02 | 2.68E+00 | 2.76E+00 | 8.26E-01 | no | | intergenic_variant | MODIFIER |
| 1 | 32706155 | AX-170198222 | 1 | C | A | 0.411 | -1.74E-02 | 8.14E-02 | 2.68E+00 | 2.76E+00 | 8.26E-01 | no | Gm37426 | upstream_gene_variant | MODIFIER |
| 8 | 93366490 | AX-170429733 | 0 | G | A | 0.211 | -2.58E-02 | 1.15E-01 | 2.63E+00 | 2.71E+00 | 8.26E-01 | no | Ces1h | intron_variant | MODIFIER |
| 16 | 32291321 | AX-97829726 | 1 | G | A | 0.491 | 1.75E-02 | 7.64E-02 | 2.62E+00 | 2.70E+00 | 8.26E-01 | no | Rnf168 | intron_variant | MODIFIER |
| 8 | 128516281 | AX-170308595 | 0 | G | A | 0.053 | -4.44E-02 | 2.07E-01 | 2.64E+00 | 2.75E+00 | 8.26E-01 | no | | intergenic_variant | MODIFIER |
| 5 | 39045682 | AX-97794034 | 1 | C | T | 0.232 | 2.34E-02 | 1.11E-01 | 2.70E+00 | 2.78E+00 | 8.26E-01 | no | | intergenic_variant | MODIFIER |

| | | | | | | | | | | | | | | | |
|---|---|---|---|---|---|---|---|---|---|---|---|---|---|---|---|
| 4 | 144670678 | AX-170111328 | 1 | G | A | 0.107 | 4.55E-02 | 2.09E-01 | 2.68E+00 | 2.74E+00 | 8.26E-01 | no | Gm436 | synonymous_variant | LOW |
| 5 | 39544084 | AX-170267532 | 0 | C | G | 0.202 | 2.18E-02 | 1.05E-01 | 2.67E+00 | 2.78E+00 | 8.26E-01 | no | 4930513D17Rik | intron_variant,non_coding_transcript_variant | MODIFIER |
| 8 | 85910947 | AX-97845909 | 0 | C | T | 0.202 | -2.67E-02 | 1.26E-01 | 2.69E+00 | 2.76E+00 | 8.26E-01 | no | Phkb | intron_variant | MODIFIER |
| 16 | 92084391 | AX-170232309 | 1 | A | G | 0.455 | -1.70E-02 | 7.92E-02 | 2.67E+00 | 2.75E+00 | 8.26E-01 | no | Mrps6 | intron_variant | MODIFIER |
| 9 | 106097128 | AX-97851089 | 2 | C | T | 0.127 | -2.63E-02 | 1.13E-01 | 2.60E+00 | 2.69E+00 | 8.26E-01 | no | Col6a4 | upstream_gene_variant | MODIFIER |
| 10 | 9609085 | AX-170250544 | 1 | T | C | 0.196 | 2.30E-02 | 1.09E-01 | 2.69E+00 | 2.77E+00 | 8.26E-01 | no | | intergenic_variant | MODIFIER |
| 9 | 85671898 | AX-170384424 | 0 | C | G | 0.14 | -2.88E-02 | 1.27E-01 | 2.62E+00 | 2.69E+00 | 8.26E-01 | no | | intergenic_variant | MODIFIER |
| 9 | 85683304 | AX-170234409 | 0 | C | T | 0.14 | -2.88E-02 | 1.27E-01 | 2.62E+00 | 2.69E+00 | 8.26E-01 | no | Ibtk | downstream_gene_variant | MODIFIER |
| 9 | 85685295 | AX-170185433 | 0 | A | G | 0.14 | -2.88E-02 | 1.27E-01 | 2.62E+00 | 2.69E+00 | 8.26E-01 | no | Ibtk | downstream_gene_variant | MODIFIER |
| 9 | 85689057 | AX-170289518 | 0 | G | A | 0.14 | -2.88E-02 | 1.27E-01 | 2.62E+00 | 2.69E+00 | 8.26E-01 | no | Ibtk | intron_variant | MODIFIER |
| 9 | 85690506 | AX-170366572 | 0 | A | G | 0.14 | -2.88E-02 | 1.27E-01 | 2.62E+00 | 2.69E+00 | 8.26E-01 | no | Ibtk | intron_variant | MODIFIER |
| 9 | 85690643 | AX-170431968 | 0 | T | C | 0.14 | -2.88E-02 | 1.27E-01 | 2.62E+00 | 2.69E+00 | 8.26E-01 | no | Ibtk | intron_variant | MODIFIER |
| 9 | 85691224 | AX-170241842 | 0 | T | C | 0.14 | -2.88E-02 | 1.27E-01 | 2.62E+00 | 2.69E+00 | 8.26E-01 | no | Ibtk | intron_variant | MODIFIER |
| 9 | 85702046 | AX-170096615 | 0 | T | C | 0.14 | -2.88E-02 | 1.27E-01 | 2.62E+00 | 2.69E+00 | 8.26E-01 | no | Ibtk | intron_variant | MODIFIER |
| 9 | 85702108 | AX-170228934 | 0 | T | C | 0.14 | -2.88E-02 | 1.27E-01 | 2.62E+00 | 2.69E+00 | 8.26E-01 | no | Ibtk | intron_variant | MODIFIER |
| 9 | 85718943 | AX-170302060 | 0 | G | A | 0.14 | -2.88E-02 | 1.27E-01 | 2.62E+00 | 2.69E+00 | 8.26E-01 | no | Ibtk | intron_variant | MODIFIER |
| 9 | 85723525 | AX-97801022 | 0 | C | T | 0.14 | -2.88E-02 | 1.27E-01 | 2.62E+00 | 2.69E+00 | 8.26E-01 | no | Ibtk | intron_variant | MODIFIER |
| 9 | 85726637 | AX-170260950 | 0 | A | G | 0.14 | -2.88E-02 | 1.27E-01 | 2.62E+00 | 2.69E+00 | 8.26E-01 | no | Ibtk | synonymous_variant | LOW |
| 9 | 85729748 | AX-170289610 | 0 | T | C | 0.14 | -2.88E-02 | 1.27E-01 | 2.62E+00 | 2.69E+00 | 8.26E-01 | no | Ibtk | intron_variant | MODIFIER |
| 9 | 85732773 | AX-170260566 | 0 | A | G | 0.14 | -2.88E-02 | 1.27E-01 | 2.62E+00 | 2.69E+00 | 8.26E-01 | no | Ibtk | intron_variant | MODIFIER |
| 9 | 85811114 | AX-170262404 | 0 | T | C | 0.14 | -2.88E-02 | 1.27E-01 | 2.62E+00 | 2.69E+00 | 8.26E-01 | no | | intergenic_variant | MODIFIER |
| 9 | 85811318 | AX-170469858 | 0 | A | G | 0.14 | -2.88E-02 | 1.27E-01 | 2.62E+00 | 2.69E+00 | 8.26E-01 | no | | intergenic_variant | MODIFIER |
| 9 | 85812203 | AX-170343314 | 0 | G | A | 0.14 | -2.88E-02 | 1.27E-01 | 2.62E+00 | 2.69E+00 | 8.26E-01 | no | | intergenic_variant | MODIFIER |
| 9 | 85836188 | AX-170454646 | 0 | C | T | 0.14 | -2.88E-02 | 1.27E-01 | 2.62E+00 | 2.69E+00 | 8.26E-01 | no | 9330154J02Rik | intron_variant,non_coding_transcript_variant | MODIFIER |
| 9 | 85848759 | AX-170530407 | 0 | G | A | 0.14 | -2.88E-02 | 1.27E-01 | 2.62E+00 | 2.69E+00 | 8.26E-01 | no | Tpbg | downstream_gene_variant | MODIFIER |
| 9 | 85880209 | AX-97844225 | 0 | A | G | 0.14 | -2.88E-02 | 1.27E-01 | 2.62E+00 | 2.69E+00 | 8.26E-01 | no | | intergenic_variant | MODIFIER |
| 9 | 85886796 | AX-97856444 | 0 | A | G | 0.14 | -2.88E-02 | 1.27E-01 | 2.62E+00 | 2.69E+00 | 8.26E-01 | no | | intergenic_variant | MODIFIER |
| 9 | 85949041 | AX-170347999 | 0 | C | G | 0.14 | -2.88E-02 | 1.27E-01 | 2.62E+00 | 2.69E+00 | 8.26E-01 | no | | regulatory_region_variant | MODIFIER |
| 9 | 85953318 | AX-170307760 | 0 | C | T | 0.14 | -2.88E-02 | 1.27E-01 | 2.62E+00 | 2.69E+00 | 8.26E-01 | no | | intergenic_variant | MODIFIER |
| 9 | 85977999 | AX-170032882 | 0 | C | G | 0.14 | -2.88E-02 | 1.27E-01 | 2.62E+00 | 2.69E+00 | 8.26E-01 | no | | intergenic_variant | MODIFIER |
| 9 | 86009439 | AX-170165958 | 0 | G | A | 0.14 | -2.88E-02 | 1.27E-01 | 2.62E+00 | 2.69E+00 | 8.26E-01 | no | | intergenic_variant | MODIFIER |
| 9 | 86035326 | AX-170168227 | 0 | C | A | 0.14 | -2.88E-02 | 1.27E-01 | 2.62E+00 | 2.69E+00 | 8.26E-01 | no | | intergenic_variant | MODIFIER |
| 9 | 86061668 | AX-170497731 | 0 | C | G | 0.14 | -2.88E-02 | 1.27E-01 | 2.62E+00 | 2.69E+00 | 8.26E-01 | no | | intergenic_variant | MODIFIER |
| 9 | 86062171 | AX-97782696 | 0 | C | T | 0.14 | -2.88E-02 | 1.27E-01 | 2.62E+00 | 2.69E+00 | 8.26E-01 | no | | intergenic_variant | MODIFIER |
| 9 | 86308889 | AX-170192619 | 0 | A | G | 0.14 | -2.88E-02 | 1.27E-01 | 2.62E+00 | 2.69E+00 | 8.26E-01 | no | Ube2cbp | intron_variant | MODIFIER |
| 9 | 86328480 | AX-170487253 | 0 | C | T | 0.14 | -2.88E-02 | 1.27E-01 | 2.62E+00 | 2.69E+00 | 8.26E-01 | no | Ube2cbp | intron_variant | MODIFIER |
| 9 | 86363091 | AX-170083022 | 0 | T | C | 0.14 | -2.88E-02 | 1.27E-01 | 2.62E+00 | 2.69E+00 | 8.26E-01 | no | Ube2cbp | intron_variant | MODIFIER |
| 9 | 86363141 | AX-170083026 | 0 | T | C | 0.14 | -2.88E-02 | 1.27E-01 | 2.62E+00 | 2.69E+00 | 8.26E-01 | no | Ube2cbp | intron_variant | MODIFIER |
| 9 | 86568405 | AX-97796284 | 0 | G | A | 0.14 | -2.88E-02 | 1.27E-01 | 2.62E+00 | 2.69E+00 | 8.26E-01 | no | Rwdd2a | upstream_gene_variant | MODIFIER |
| 9 | 86612708 | AX-170032887 | 0 | C | T | 0.14 | -2.88E-02 | 1.27E-01 | 2.62E+00 | 2.69E+00 | 8.26E-01 | no | Me1 | intron_variant | MODIFIER |
| 9 | 86684002 | AX-97781141 | 0 | G | A | 0.14 | -2.88E-02 | 1.27E-01 | 2.62E+00 | 2.69E+00 | 8.26E-01 | no | Me1 | intron_variant | MODIFIER |
| 9 | 86790310 | AX-170115476 | 0 | T | C | 0.14 | -2.88E-02 | 1.27E-01 | 2.62E+00 | 2.69E+00 | 8.26E-01 | no | Snap91 | intron_variant | MODIFIER |
| 9 | 87300015 | AX-170083041 | 0 | T | A | 0.14 | -2.88E-02 | 1.27E-01 | 2.62E+00 | 2.69E+00 | 8.26E-01 | no | | intergenic_variant | MODIFIER |
| 9 | 87504112 | AX-97805303 | 0 | C | T | 0.14 | -2.88E-02 | 1.27E-01 | 2.62E+00 | 2.69E+00 | 8.26E-01 | no | | intergenic_variant | MODIFIER |
| 9 | 87511373 | AX-170512454 | 0 | T | A | 0.14 | -2.88E-02 | 1.27E-01 | 2.62E+00 | 2.69E+00 | 8.26E-01 | no | | intergenic_variant | MODIFIER |
| 9 | 87530355 | AX-170280103 | 0 | C | T | 0.14 | -2.88E-02 | 1.27E-01 | 2.62E+00 | 2.69E+00 | 8.26E-01 | no | | intergenic_variant | MODIFIER |
| 9 | 87537875 | AX-97835294 | 0 | C | T | 0.14 | -2.88E-02 | 1.27E-01 | 2.62E+00 | 2.69E+00 | 8.26E-01 | no | | intergenic_variant | MODIFIER |
| 9 | 87679514 | AX-170032904 | 0 | C | T | 0.14 | -2.88E-02 | 1.27E-01 | 2.62E+00 | 2.69E+00 | 8.26E-01 | no | | intergenic_variant | MODIFIER |
| 9 | 87679973 | AX-170208227 | 0 | T | C | 0.14 | -2.88E-02 | 1.27E-01 | 2.62E+00 | 2.69E+00 | 8.26E-01 | no | | intergenic_variant | MODIFIER |
| 9 | 87751230 | AX-170470416 | 0 | A | G | 0.14 | -2.88E-02 | 1.27E-01 | 2.62E+00 | 2.69E+00 | 8.26E-01 | no | | intergenic_variant | MODIFIER |
| 9 | 87784629 | AX-170032908 | 0 | T | C | 0.14 | -2.88E-02 | 1.27E-01 | 2.62E+00 | 2.69E+00 | 8.26E-01 | no | | intergenic_variant | MODIFIER |
| 9 | 87784692 | AX-170326688 | 0 | T | C | 0.14 | -2.88E-02 | 1.27E-01 | 2.62E+00 | 2.69E+00 | 8.26E-01 | no | | intergenic_variant | MODIFIER |
| 9 | 87809462 | AX-170175733 | 0 | A | G | 0.14 | -2.88E-02 | 1.27E-01 | 2.62E+00 | 2.69E+00 | 8.26E-01 | no | | intergenic_variant | MODIFIER |
| 9 | 87857692 | AX-170531255 | 0 | C | A | 0.14 | -2.88E-02 | 1.27E-01 | 2.62E+00 | 2.69E+00 | 8.26E-01 | no | | intergenic_variant | MODIFIER |

| | | | | | | | | | | | | | | | |
|---|---|---|---|---|---|---|---|---|---|---|---|---|---|---|---|
| 9 | 87861652 | AX-170314294 | 0 | T | C | 0.14 | -2.88E-02 | 1.27E-01 | 2.62E+00 | 2.69E+00 | 8.26E-01 | no | | intergenic_variant | MODIFIER |
| 9 | 87890420 | AX-170325026 | 0 | C | T | 0.14 | -2.88E-02 | 1.27E-01 | 2.62E+00 | 2.69E+00 | 8.26E-01 | no | Gm25528 | upstream_gene_variant | MODIFIER |
| 9 | 88051291 | AX-170096623 | 0 | G | A | 0.14 | -2.88E-02 | 1.27E-01 | 2.62E+00 | 2.69E+00 | 8.26E-01 | no | | intergenic_variant | MODIFIER |
| 9 | 88094439 | AX-170032921 | 0 | C | T | 0.14 | -2.88E-02 | 1.27E-01 | 2.62E+00 | 2.69E+00 | 8.26E-01 | no | | intergenic_variant | MODIFIER |
| 9 | 88115808 | AX-170032922 | 0 | G | A | 0.14 | -2.88E-02 | 1.27E-01 | 2.62E+00 | 2.69E+00 | 8.26E-01 | no | | intergenic_variant | MODIFIER |
| 9 | 88120046 | AX-170138331 | 0 | C | T | 0.14 | -2.88E-02 | 1.27E-01 | 2.62E+00 | 2.69E+00 | 8.26E-01 | no | Gm2324 | downstream_gene_variant | MODIFIER |
| 9 | 88123936 | AX-97824753 | 0 | A | G | 0.14 | -2.88E-02 | 1.27E-01 | 2.62E+00 | 2.69E+00 | 8.26E-01 | no | Gm2324 | upstream_gene_variant | MODIFIER |
| 9 | 88124990 | AX-170116173 | 0 | G | C | 0.14 | -2.88E-02 | 1.27E-01 | 2.62E+00 | 2.69E+00 | 8.26E-01 | no | Gm2324 | upstream_gene_variant | MODIFIER |
| 9 | 88128044 | AX-170083058 | 0 | G | A | 0.14 | -2.88E-02 | 1.27E-01 | 2.62E+00 | 2.69E+00 | 8.26E-01 | no | | intergenic_variant | MODIFIER |
| 9 | 88153568 | AX-97851087 | 0 | G | A | 0.14 | -2.88E-02 | 1.27E-01 | 2.62E+00 | 2.69E+00 | 8.26E-01 | no | | intergenic_variant | MODIFIER |
| 9 | 88154255 | AX-170473347 | 0 | G | A | 0.14 | -2.88E-02 | 1.27E-01 | 2.62E+00 | 2.69E+00 | 8.26E-01 | no | | intergenic_variant | MODIFIER |
| 9 | 88498812 | AX-97844360 | 0 | T | G | 0.14 | -2.88E-02 | 1.27E-01 | 2.62E+00 | 2.69E+00 | 8.26E-01 | no | | intergenic_variant | MODIFIER |
| 9 | 88536492 | AX-170083062 | 0 | C | T | 0.14 | -2.88E-02 | 1.27E-01 | 2.62E+00 | 2.69E+00 | 8.26E-01 | no | | intergenic_variant | MODIFIER |
| 9 | 88539590 | AX-170083063 | 0 | C | T | 0.14 | -2.88E-02 | 1.27E-01 | 2.62E+00 | 2.69E+00 | 8.26E-01 | no | | regulatory_region_variant | MODIFIER |
| 9 | 88545364 | AX-97782826 | 0 | G | T | 0.14 | -2.88E-02 | 1.27E-01 | 2.62E+00 | 2.69E+00 | 8.26E-01 | no | Zfp949 | upstream_gene_variant | MODIFIER |
| 9 | 88552078 | AX-170411478 | 0 | C | A | 0.14 | -2.88E-02 | 1.27E-01 | 2.62E+00 | 2.69E+00 | 8.26E-01 | no | Zfp949 | intron_variant | MODIFIER |
| 9 | 88552170 | AX-170145127 | 0 | A | C | 0.14 | -2.88E-02 | 1.27E-01 | 2.62E+00 | 2.69E+00 | 8.26E-01 | no | Zfp949 | intron_variant | MODIFIER |
| 9 | 88552249 | AX-170488235 | 0 | T | G | 0.14 | -2.88E-02 | 1.27E-01 | 2.62E+00 | 2.69E+00 | 8.26E-01 | no | Zfp949 | intron_variant | MODIFIER |
| 9 | 88645826 | AX-97788539 | 0 | T | C | 0.14 | -2.88E-02 | 1.27E-01 | 2.62E+00 | 2.69E+00 | 8.26E-01 | no | | intergenic_variant | MODIFIER |
| 9 | 89461812 | AX-170281055 | 0 | A | T | 0.14 | -2.88E-02 | 1.27E-01 | 2.62E+00 | 2.69E+00 | 8.26E-01 | no | | intergenic_variant | MODIFIER |
| 9 | 89467903 | AX-170253937 | 0 | T | G | 0.14 | -2.88E-02 | 1.27E-01 | 2.62E+00 | 2.69E+00 | 8.26E-01 | no | | intergenic_variant | MODIFIER |
| 9 | 89571911 | AX-170274124 | 0 | G | A | 0.14 | -2.88E-02 | 1.27E-01 | 2.62E+00 | 2.69E+00 | 8.26E-01 | no | | intergenic_variant | MODIFIER |
| 9 | 89599694 | AX-170171148 | 0 | G | A | 0.14 | -2.88E-02 | 1.27E-01 | 2.62E+00 | 2.69E+00 | 8.26E-01 | no | Minar1 | intron_variant | MODIFIER |
| 9 | 89605764 | AX-170510630 | 0 | A | C | 0.14 | -2.88E-02 | 1.27E-01 | 2.62E+00 | 2.69E+00 | 8.26E-01 | no | Minar1 | intron_variant | MODIFIER |
| 9 | 89608000 | AX-170384309 | 0 | A | G | 0.14 | -2.88E-02 | 1.27E-01 | 2.62E+00 | 2.69E+00 | 8.26E-01 | no | Minar1 | intron_variant | MODIFIER |
| 9 | 89615505 | AX-170110991 | 0 | T | C | 0.14 | -2.88E-02 | 1.27E-01 | 2.62E+00 | 2.69E+00 | 8.26E-01 | no | Minar1 | intron_variant | MODIFIER |
| 9 | 89946808 | AX-170351700 | 0 | C | T | 0.14 | -2.88E-02 | 1.27E-01 | 2.62E+00 | 2.69E+00 | 8.26E-01 | no | Rasgrf1 | intron_variant | MODIFIER |
| 9 | 89996628 | AX-97838694 | 0 | G | A | 0.14 | -2.88E-02 | 1.27E-01 | 2.62E+00 | 2.69E+00 | 8.26E-01 | no | Rasgrf1 | intron_variant | MODIFIER |
| 9 | 90025864 | AX-170532918 | 0 | A | G | 0.14 | -2.88E-02 | 1.27E-01 | 2.62E+00 | 2.69E+00 | 8.26E-01 | no | Rasgrf1 | intron_variant | MODIFIER |
| 9 | 90039039 | AX-170187725 | 0 | A | G | 0.14 | -2.88E-02 | 1.27E-01 | 2.62E+00 | 2.69E+00 | 8.26E-01 | no | | intergenic_variant | MODIFIER |
| 9 | 90484996 | AX-170083075 | 0 | C | T | 0.14 | -2.88E-02 | 1.27E-01 | 2.62E+00 | 2.69E+00 | 8.26E-01 | no | | intergenic_variant | MODIFIER |
| 18 | 58688466 | AX-170232176 | 0 | C | T | 0.439 | -1.71E-02 | 7.25E-02 | 2.57E+00 | 2.69E+00 | 8.26E-01 | no | | intergenic_variant | MODIFIER |
| 1 | 150730586 | AX-170486287 | 1 | T | C | 0.205 | -2.46E-02 | 1.19E-01 | 2.69E+00 | 2.79E+00 | 8.26E-01 | no | Hmcn1 | intron_variant | MODIFIER |
| 2 | 154730899 | AX-97839634 | 1 | T | C | 0.134 | 4.30E-02 | 1.96E-01 | 2.66E+00 | 2.73E+00 | 8.26E-01 | no | Gm14216 | intron_variant,non_coding_transcript_variant | MODIFIER |
| 15 | 88231331 | AX-170219612 | 1 | A | G | 0.134 | 3.48E-02 | 1.56E-01 | 2.64E+00 | 2.71E+00 | 8.27E-01 | no | | intergenic_variant | MODIFIER |
| 10 | 119507993 | AX-170205803 | 0 | T | C | 0.351 | 1.54E-02 | 7.27E-02 | 2.67E+00 | 2.76E+00 | 8.27E-01 | no | Grip1 | intron_variant | MODIFIER |
| 4 | 145343097 | AX-170323486 | 1 | A | G | 0.107 | 4.43E-02 | 2.04E-01 | 2.67E+00 | 2.73E+00 | 8.27E-01 | no | | intergenic_variant | MODIFIER |
| 20 | 72173682 | AX-170084495 | 0 | G | T | 0.254 | 1.36E-02 | 6.26E-02 | 2.65E+00 | 2.74E+00 | 8.27E-01 | no | | intergenic_variant | MODIFIER |
| 18 | 59348074 | AX-170213494 | 1 | A | G | 0.402 | -1.86E-02 | 8.13E-02 | 2.62E+00 | 2.69E+00 | 8.27E-01 | no | Chsy3 | intron_variant | MODIFIER |
| 18 | 62384398 | AX-170414207 | 1 | C | T | 0.402 | -1.86E-02 | 8.13E-02 | 2.62E+00 | 2.69E+00 | 8.27E-01 | no | Htr4 | intron_variant | MODIFIER |
| 8 | 23330614 | AX-170307517 | 1 | C | T | 0.17 | -2.41E-02 | 1.11E-01 | 2.65E+00 | 2.73E+00 | 8.27E-01 | no | | intergenic_variant | MODIFIER |
| 12 | 101541550 | AX-170385484 | 2 | T | C | 0.082 | -4.11E-02 | 1.93E-01 | 2.67E+00 | 2.75E+00 | 8.27E-01 | no | Catsperb | synonymous_variant | LOW |
| 4 | 127346475 | AX-170467774 | 0 | G | A | 0.298 | -1.20E-02 | 5.91E-02 | 2.72E+00 | 2.83E+00 | 8.27E-01 | no | Gjb4 | downstream_gene_variant | MODIFIER |
| 1 | 13053080 | AX-170040410 | 0 | C | A | 0.377 | 1.72E-02 | 7.67E-02 | 2.64E+00 | 2.70E+00 | 8.27E-01 | no | | intergenic_variant | MODIFIER |
| 1 | 13055719 | AX-170146443 | 0 | G | A | 0.377 | 1.72E-02 | 7.67E-02 | 2.64E+00 | 2.70E+00 | 8.27E-01 | no | Gm37162 | downstream_gene_variant | MODIFIER |
| 1 | 13056361 | AX-170040412 | 0 | A | T | 0.377 | 1.72E-02 | 7.67E-02 | 2.64E+00 | 2.70E+00 | 8.27E-01 | no | Gm37162 | downstream_gene_variant | MODIFIER |
| 1 | 13065985 | AX-170454090 | 0 | G | A | 0.377 | 1.72E-02 | 7.67E-02 | 2.64E+00 | 2.70E+00 | 8.27E-01 | no | Gm5250 | downstream_gene_variant | MODIFIER |
| 1 | 13073341 | AX-170346327 | 0 | T | C | 0.377 | 1.72E-02 | 7.67E-02 | 2.64E+00 | 2.70E+00 | 8.27E-01 | no | Gm29283 | intron_variant,non_coding_transcript_variant | MODIFIER |
| 1 | 13076425 | AX-170130921 | 0 | G | C | 0.377 | 1.72E-02 | 7.67E-02 | 2.64E+00 | 2.70E+00 | 8.27E-01 | no | Gm29283 | non_coding_transcript_exon_variant | MODIFIER |
| 1 | 13105646 | AX-170469300 | 0 | T | C | 0.377 | 1.72E-02 | 7.67E-02 | 2.64E+00 | 2.70E+00 | 8.27E-01 | no | | intergenic_variant | MODIFIER |
| 1 | 13108233 | AX-170403585 | 0 | A | T | 0.377 | 1.72E-02 | 7.67E-02 | 2.64E+00 | 2.70E+00 | 8.27E-01 | no | | intergenic_variant | MODIFIER |
| 1 | 13114284 | AX-170314875 | 0 | G | A | 0.377 | 1.72E-02 | 7.67E-02 | 2.64E+00 | 2.70E+00 | 8.27E-01 | no | Prdm14 | synonymous_variant | LOW |
| 1 | 13122270 | AX-170150764 | 0 | G | A | 0.377 | 1.72E-02 | 7.67E-02 | 2.64E+00 | 2.70E+00 | 8.27E-01 | no | Prdm14 | intron_variant | MODIFIER |
| 1 | 13125617 | AX-170296520 | 0 | A | C | 0.377 | 1.72E-02 | 7.67E-02 | 2.64E+00 | 2.70E+00 | 8.27E-01 | no | Prdm14 | synonymous_variant | LOW |

| | | | | | | | | | | | | | | | |
|---|---|---|---|---|---|---|---|---|---|---|---|---|---|---|---|
| 1 | 13125821 | AX-170429146 | 0 | G | A | 0.377 | 1.72E-02 | 7.67E-02 | 2.64E+00 | 2.70E+00 | 8.27E-01 | no | Prdm14 | synonymous_variant | LOW |
| 1 | 13125845 | AX-170395015 | 0 | A | G | 0.377 | 1.72E-02 | 7.67E-02 | 2.64E+00 | 2.70E+00 | 8.27E-01 | no | Prdm14 | 5_prime_UTR_variant | MODIFIER |
| 1 | 13126179 | AX-170040415 | 0 | G | A | 0.377 | 1.72E-02 | 7.67E-02 | 2.64E+00 | 2.70E+00 | 8.27E-01 | no | Prdm14 | intron_variant | MODIFIER |
| 1 | 13127028 | AX-170040414 | 0 | A | G | 0.377 | 1.72E-02 | 7.67E-02 | 2.64E+00 | 2.70E+00 | 8.27E-01 | no | Prdm14 | intron_variant | MODIFIER |
| 1 | 13163262 | AX-170412034 | 0 | A | G | 0.377 | 1.72E-02 | 7.67E-02 | 2.64E+00 | 2.70E+00 | 8.27E-01 | no | Ncoa2 | intron_variant | MODIFIER |
| 1 | 13182727 | AX-170321216 | 0 | C | T | 0.377 | 1.72E-02 | 7.67E-02 | 2.64E+00 | 2.70E+00 | 8.27E-01 | no | Ncoa2 | intron_variant | MODIFIER |
| 1 | 13191578 | AX-170113664 | 0 | A | C | 0.377 | 1.72E-02 | 7.67E-02 | 2.64E+00 | 2.70E+00 | 8.27E-01 | no | Ncoa2 | intron_variant | MODIFIER |
| 1 | 13192236 | AX-170448069 | 0 | G | T | 0.377 | 1.72E-02 | 7.67E-02 | 2.64E+00 | 2.70E+00 | 8.27E-01 | no | Ncoa2 | intron_variant | MODIFIER |
| 1 | 13195137 | AX-97843211 | 0 | T | C | 0.377 | 1.72E-02 | 7.67E-02 | 2.64E+00 | 2.70E+00 | 8.27E-01 | no | Ncoa2 | intron_variant | MODIFIER |
| 1 | 13201683 | AX-170309253 | 0 | C | T | 0.377 | 1.72E-02 | 7.67E-02 | 2.64E+00 | 2.70E+00 | 8.27E-01 | no | Ncoa2 | intron_variant | MODIFIER |
| 1 | 13202841 | AX-170396626 | 0 | G | A | 0.377 | 1.72E-02 | 7.67E-02 | 2.64E+00 | 2.70E+00 | 8.27E-01 | no | Ncoa2 | intron_variant | MODIFIER |
| 1 | 13203256 | AX-170139384 | 0 | C | T | 0.377 | 1.72E-02 | 7.67E-02 | 2.64E+00 | 2.70E+00 | 8.27E-01 | no | Ncoa2 | intron_variant | MODIFIER |
| 1 | 13204657 | AX-170199705 | 0 | C | G | 0.377 | 1.72E-02 | 7.67E-02 | 2.64E+00 | 2.70E+00 | 8.27E-01 | no | Ncoa2 | intron_variant | MODIFIER |
| 1 | 13204827 | AX-170259105 | 0 | T | C | 0.377 | 1.72E-02 | 7.67E-02 | 2.64E+00 | 2.70E+00 | 8.27E-01 | no | Ncoa2 | intron_variant | MODIFIER |
| 1 | 13214627 | AX-170291621 | 0 | A | G | 0.377 | 1.72E-02 | 7.67E-02 | 2.64E+00 | 2.70E+00 | 8.27E-01 | no | Ncoa2 | intron_variant | MODIFIER |
| 1 | 13226706 | AX-170239876 | 0 | T | C | 0.377 | 1.72E-02 | 7.67E-02 | 2.64E+00 | 2.70E+00 | 8.27E-01 | no | Ncoa2 | intron_variant | MODIFIER |
| 1 | 13234741 | AX-170322715 | 0 | A | C | 0.377 | 1.72E-02 | 7.67E-02 | 2.64E+00 | 2.70E+00 | 8.27E-01 | no | Ncoa2 | intron_variant | MODIFIER |
| 1 | 13235620 | AX-170040417 | 0 | T | A | 0.377 | 1.72E-02 | 7.67E-02 | 2.64E+00 | 2.70E+00 | 8.27E-01 | no | Ncoa2 | intron_variant | MODIFIER |
| 1 | 13237808 | AX-170118501 | 0 | G | A | 0.377 | 1.72E-02 | 7.67E-02 | 2.64E+00 | 2.70E+00 | 8.27E-01 | no | Ncoa2 | intron_variant | MODIFIER |
| 1 | 13240448 | AX-170294490 | 0 | C | T | 0.377 | 1.72E-02 | 7.67E-02 | 2.64E+00 | 2.70E+00 | 8.27E-01 | no | Ncoa2 | intron_variant | MODIFIER |
| 1 | 13241245 | AX-170196464 | 0 | A | C | 0.377 | 1.72E-02 | 7.67E-02 | 2.64E+00 | 2.70E+00 | 8.27E-01 | no | Ncoa2 | intron_variant | MODIFIER |
| 1 | 13251717 | AX-170328719 | 0 | A | G | 0.377 | 1.72E-02 | 7.67E-02 | 2.64E+00 | 2.70E+00 | 8.27E-01 | no | Ncoa2 | intron_variant | MODIFIER |
| 1 | 13252132 | AX-97790943 | 0 | C | T | 0.377 | 1.72E-02 | 7.67E-02 | 2.64E+00 | 2.70E+00 | 8.27E-01 | no | Ncoa2 | intron_variant | MODIFIER |
| 1 | 13259168 | AX-170292458 | 0 | T | C | 0.377 | 1.72E-02 | 7.67E-02 | 2.64E+00 | 2.70E+00 | 8.27E-01 | no | Ncoa2 | intron_variant | MODIFIER |
| 1 | 13275509 | AX-170523298 | 0 | T | C | 0.377 | 1.72E-02 | 7.67E-02 | 2.64E+00 | 2.70E+00 | 8.27E-01 | no | Ncoa2 | intron_variant | MODIFIER |
| 1 | 13298228 | AX-170517314 | 0 | T | G | 0.377 | 1.72E-02 | 7.67E-02 | 2.64E+00 | 2.70E+00 | 8.27E-01 | no | Ncoa2 | intron_variant | MODIFIER |
| 1 | 13301111 | AX-170319953 | 0 | T | A | 0.377 | 1.72E-02 | 7.67E-02 | 2.64E+00 | 2.70E+00 | 8.27E-01 | no | Ncoa2 | intron_variant | MODIFIER |
| 1 | 13316639 | AX-170326533 | 0 | T | C | 0.377 | 1.72E-02 | 7.67E-02 | 2.64E+00 | 2.70E+00 | 8.27E-01 | no | Ncoa2 | intron_variant | MODIFIER |
| 1 | 13342083 | AX-170040421 | 0 | G | C | 0.377 | 1.72E-02 | 7.67E-02 | 2.64E+00 | 2.70E+00 | 8.27E-01 | no | Ncoa2 | intron_variant | MODIFIER |
| 1 | 13343854 | AX-170272324 | 0 | T | C | 0.377 | 1.72E-02 | 7.67E-02 | 2.64E+00 | 2.70E+00 | 8.27E-01 | no | Ncoa2 | intron_variant | MODIFIER |
| 1 | 13346655 | AX-170167904 | 0 | A | G | 0.377 | 1.72E-02 | 7.67E-02 | 2.64E+00 | 2.70E+00 | 8.27E-01 | no | Ncoa2 | intron_variant | MODIFIER |
| 1 | 13349917 | AX-170106935 | 0 | T | C | 0.377 | 1.72E-02 | 7.67E-02 | 2.64E+00 | 2.70E+00 | 8.27E-01 | no | Ncoa2 | intron_variant | MODIFIER |
| 1 | 13356001 | AX-170308212 | 0 | A | G | 0.377 | 1.72E-02 | 7.67E-02 | 2.64E+00 | 2.70E+00 | 8.27E-01 | no | Ncoa2 | intron_variant | MODIFIER |
| 1 | 13365356 | AX-170217928 | 0 | T | G | 0.377 | 1.72E-02 | 7.67E-02 | 2.64E+00 | 2.70E+00 | 8.27E-01 | no | Ncoa2 | intron_variant | MODIFIER |
| 1 | 13367764 | AX-170352456 | 0 | T | C | 0.377 | 1.72E-02 | 7.67E-02 | 2.64E+00 | 2.70E+00 | 8.27E-01 | no | Ncoa2 | intron_variant | MODIFIER |
| 1 | 13368122 | AX-170121730 | 0 | A | C | 0.377 | 1.72E-02 | 7.67E-02 | 2.64E+00 | 2.70E+00 | 8.27E-01 | no | Ncoa2 | intron_variant | MODIFIER |
| 1 | 13382990 | AX-170184015 | 0 | T | C | 0.377 | 1.72E-02 | 7.67E-02 | 2.64E+00 | 2.70E+00 | 8.27E-01 | no | Gm29570 | intron_variant,non_coding_transcript_variant | MODIFIER |
| 1 | 13436010 | AX-169999973 | 0 | T | C | 0.377 | 1.72E-02 | 7.67E-02 | 2.64E+00 | 2.70E+00 | 8.27E-01 | no | | intergenic_variant | MODIFIER |
| 1 | 13455715 | AX-170276784 | 0 | A | G | 0.377 | 1.72E-02 | 7.67E-02 | 2.64E+00 | 2.70E+00 | 8.27E-01 | no | | intergenic_variant | MODIFIER |
| 1 | 13463708 | AX-169999971 | 0 | T | A | 0.377 | 1.72E-02 | 7.67E-02 | 2.64E+00 | 2.70E+00 | 8.27E-01 | no | | intergenic_variant | MODIFIER |
| 1 | 13468192 | AX-170469380 | 0 | C | T | 0.377 | 1.72E-02 | 7.67E-02 | 2.64E+00 | 2.70E+00 | 8.27E-01 | no | | intergenic_variant | MODIFIER |
| 1 | 13500087 | AX-170182688 | 0 | T | C | 0.377 | 1.72E-02 | 7.67E-02 | 2.64E+00 | 2.70E+00 | 8.27E-01 | no | | intergenic_variant | MODIFIER |
| 1 | 13531483 | AX-170256589 | 0 | G | A | 0.377 | 1.72E-02 | 7.67E-02 | 2.64E+00 | 2.70E+00 | 8.27E-01 | no | | regulatory_region_variant | MODIFIER |
| 1 | 13535477 | AX-170353701 | 0 | G | A | 0.377 | 1.72E-02 | 7.67E-02 | 2.64E+00 | 2.70E+00 | 8.27E-01 | no | | intergenic_variant | MODIFIER |
| 1 | 13633693 | AX-170174758 | 0 | G | A | 0.377 | 1.72E-02 | 7.67E-02 | 2.64E+00 | 2.70E+00 | 8.27E-01 | no | Lactb2 | intron_variant | MODIFIER |
| 1 | 13636356 | AX-170175494 | 0 | G | A | 0.377 | 1.72E-02 | 7.67E-02 | 2.64E+00 | 2.70E+00 | 8.27E-01 | no | Lactb2 | intron_variant | MODIFIER |
| 1 | 13636583 | AX-170248224 | 0 | G | T | 0.377 | 1.72E-02 | 7.67E-02 | 2.64E+00 | 2.70E+00 | 8.27E-01 | no | Lactb2 | intron_variant | MODIFIER |
| 1 | 13700937 | AX-170432118 | 0 | G | A | 0.377 | 1.72E-02 | 7.67E-02 | 2.64E+00 | 2.70E+00 | 8.27E-01 | no | Xkr9 | missense_variant | MODERATE |
| 1 | 13707018 | AX-170040429 | 0 | T | C | 0.377 | 1.72E-02 | 7.67E-02 | 2.64E+00 | 2.70E+00 | 8.27E-01 | no | | intergenic_variant | MODIFIER |
| 1 | 13778003 | AX-170124594 | 0 | C | T | 0.377 | 1.72E-02 | 7.67E-02 | 2.64E+00 | 2.70E+00 | 8.27E-01 | no | Gm26273 | upstream_gene_variant | MODIFIER |
| 1 | 13779548 | AX-170105584 | 0 | G | C | 0.377 | 1.72E-02 | 7.67E-02 | 2.64E+00 | 2.70E+00 | 8.27E-01 | no | | intergenic_variant | MODIFIER |
| 1 | 13788101 | AX-170361051 | 0 | T | C | 0.377 | 1.72E-02 | 7.67E-02 | 2.64E+00 | 2.70E+00 | 8.27E-01 | no | Gm5523 | upstream_gene_variant | MODIFIER |
| 1 | 13788138 | AX-170397948 | 0 | A | G | 0.377 | 1.72E-02 | 7.67E-02 | 2.64E+00 | 2.70E+00 | 8.27E-01 | no | Gm5523 | upstream_gene_variant | MODIFIER |
| 1 | 13789723 | AX-170374261 | 0 | T | C | 0.377 | 1.72E-02 | 7.67E-02 | 2.64E+00 | 2.70E+00 | 8.27E-01 | no | Gm5523 | upstream_gene_variant | MODIFIER |

| | | | | | | | | | | | | | | | |
|---|---|---|---|---|---|---|---|---|---|---|---|---|---|---|---|
| 1 | 13810432 | AX-170130863 | 0 | G | A | 0.377 | 1.72E-02 | 7.67E-02 | 2.64E+00 | 2.70E+00 | 8.27E-01 | no | | intergenic_variant | MODIFIER |
| 10 | 119542122 | AX-170307596 | 0 | C | T | 0.342 | -1.60E-02 | 7.44E-02 | 2.66E+00 | 2.74E+00 | 8.27E-01 | no | Grip1 | intron_variant | MODIFIER |
| 9 | 55091631 | AX-170296153 | 1 | G | A | 0.214 | -2.30E-02 | 1.06E-01 | 2.64E+00 | 2.73E+00 | 8.27E-01 | no | Gm7395 | downstream_gene_variant | MODIFIER |
| 7 | 116461381 | AX-170248611 | 2 | A | T | 0.409 | 1.96E-02 | 9.03E-02 | 2.66E+00 | 2.73E+00 | 8.27E-01 | no | | intergenic_variant | MODIFIER |
| 7 | 117933728 | AX-170337706 | 0 | G | A | 0.421 | 2.04E-02 | 9.39E-02 | 2.67E+00 | 2.73E+00 | 8.27E-01 | no | | intergenic_variant | MODIFIER |
| 9 | 113587958 | AX-170160704 | 0 | C | T | 0.132 | -2.27E-02 | 1.05E-01 | 2.64E+00 | 2.74E+00 | 8.27E-01 | no | AU023762 | intron_variant,non_coding_transcript_variant | MODIFIER |
| 9 | 113665081 | AX-97807214 | 0 | T | G | 0.132 | -2.27E-02 | 1.05E-01 | 2.64E+00 | 2.74E+00 | 8.27E-01 | no | Pdcd6ip | intron_variant | MODIFIER |
| 9 | 113691060 | AX-97814049 | 0 | G | A | 0.132 | -2.27E-02 | 1.05E-01 | 2.64E+00 | 2.74E+00 | 8.27E-01 | no | Pdcd6ip | intron_variant | MODIFIER |
| 9 | 113703356 | AX-170033308 | 0 | T | C | 0.132 | -2.27E-02 | 1.05E-01 | 2.64E+00 | 2.74E+00 | 8.27E-01 | no | Pdcd6ip | intron_variant | MODIFIER |
| 9 | 113722288 | AX-170033304 | 0 | G | A | 0.132 | -2.27E-02 | 1.05E-01 | 2.64E+00 | 2.74E+00 | 8.27E-01 | no | | intergenic_variant | MODIFIER |
| 9 | 113743737 | AX-97798449 | 0 | T | C | 0.132 | -2.27E-02 | 1.05E-01 | 2.64E+00 | 2.74E+00 | 8.27E-01 | no | Clasp2 | intron_variant | MODIFIER |
| 9 | 113747938 | AX-170158944 | 0 | C | T | 0.132 | -2.27E-02 | 1.05E-01 | 2.64E+00 | 2.74E+00 | 8.27E-01 | no | Clasp2 | intron_variant | MODIFIER |
| 9 | 113813489 | AX-97817702 | 0 | G | T | 0.132 | -2.27E-02 | 1.05E-01 | 2.64E+00 | 2.74E+00 | 8.27E-01 | no | Clasp2 | intron_variant | MODIFIER |
| 9 | 113814078 | AX-170232338 | 0 | G | A | 0.132 | -2.27E-02 | 1.05E-01 | 2.64E+00 | 2.74E+00 | 8.27E-01 | no | Clasp2 | intron_variant | MODIFIER |
| 9 | 113823401 | AX-170222935 | 0 | A | G | 0.132 | -2.27E-02 | 1.05E-01 | 2.64E+00 | 2.74E+00 | 8.27E-01 | no | Clasp2 | intron_variant | MODIFIER |
| 9 | 113838099 | AX-170149729 | 0 | C | T | 0.132 | -2.27E-02 | 1.05E-01 | 2.64E+00 | 2.74E+00 | 8.27E-01 | no | Clasp2 | intron_variant | MODIFIER |
| 9 | 113839049 | AX-170472780 | 0 | A | G | 0.132 | -2.27E-02 | 1.05E-01 | 2.64E+00 | 2.74E+00 | 8.27E-01 | no | Clasp2 | intron_variant | MODIFIER |
| 9 | 113880303 | AX-97780816 | 0 | T | C | 0.132 | -2.27E-02 | 1.05E-01 | 2.64E+00 | 2.74E+00 | 8.27E-01 | no | Clasp2 | intron_variant | MODIFIER |
| 9 | 113898231 | AX-170311726 | 0 | A | G | 0.132 | -2.27E-02 | 1.05E-01 | 2.64E+00 | 2.74E+00 | 8.27E-01 | no | Clasp2 | intron_variant | MODIFIER |
| 9 | 113898673 | AX-170422968 | 0 | C | A | 0.132 | -2.27E-02 | 1.05E-01 | 2.64E+00 | 2.74E+00 | 8.27E-01 | no | Clasp2 | intron_variant | MODIFIER |
| 9 | 113914595 | AX-170083513 | 0 | A | G | 0.132 | -2.27E-02 | 1.05E-01 | 2.64E+00 | 2.74E+00 | 8.27E-01 | no | Clasp2 | intron_variant | MODIFIER |
| 9 | 113998753 | AX-170334361 | 0 | C | T | 0.132 | -2.27E-02 | 1.05E-01 | 2.64E+00 | 2.74E+00 | 8.27E-01 | no | Fbxl2 | intron_variant | MODIFIER |
| 9 | 113998840 | AX-170501143 | 0 | C | T | 0.132 | -2.27E-02 | 1.05E-01 | 2.64E+00 | 2.74E+00 | 8.27E-01 | no | Fbxl2 | intron_variant | MODIFIER |
| 9 | 114428637 | AX-170033317 | 0 | T | C | 0.132 | -2.27E-02 | 1.05E-01 | 2.64E+00 | 2.74E+00 | 8.27E-01 | no | Glb1 | intron_variant | MODIFIER |
| 9 | 114437747 | AX-170299655 | 0 | T | C | 0.132 | -2.27E-02 | 1.05E-01 | 2.64E+00 | 2.74E+00 | 8.27E-01 | no | Glb1 | synonymous_variant | LOW |
| 9 | 114438002 | AX-170518101 | 0 | A | G | 0.132 | -2.27E-02 | 1.05E-01 | 2.64E+00 | 2.74E+00 | 8.27E-01 | no | Glb1 | intron_variant | MODIFIER |
| 9 | 114439494 | AX-97792706 | 0 | C | T | 0.132 | -2.27E-02 | 1.05E-01 | 2.64E+00 | 2.74E+00 | 8.27E-01 | no | Glb1 | intron_variant | MODIFIER |
| 9 | 114448997 | AX-170033322 | 0 | A | T | 0.132 | -2.27E-02 | 1.05E-01 | 2.64E+00 | 2.74E+00 | 8.27E-01 | no | Glb1 | intron_variant | MODIFIER |
| 9 | 114456514 | AX-170114283 | 0 | T | C | 0.132 | -2.27E-02 | 1.05E-01 | 2.64E+00 | 2.74E+00 | 8.27E-01 | no | Glb1 | synonymous_variant | LOW |
| 9 | 114461788 | AX-170300392 | 0 | A | G | 0.132 | -2.27E-02 | 1.05E-01 | 2.64E+00 | 2.74E+00 | 8.27E-01 | no | Glb1 | intron_variant | MODIFIER |
| 9 | 114462045 | AX-170083522 | 0 | T | C | 0.132 | -2.27E-02 | 1.05E-01 | 2.64E+00 | 2.74E+00 | 8.27E-01 | no | Glb1 | intron_variant | MODIFIER |
| 9 | 114468797 | AX-97827828 | 0 | G | A | 0.132 | -2.27E-02 | 1.05E-01 | 2.64E+00 | 2.74E+00 | 8.27E-01 | no | Glb1 | intron_variant | MODIFIER |
| 13 | 47245639 | AX-170492518 | 1 | G | A | 0.473 | -1.37E-02 | 6.60E-02 | 2.68E+00 | 2.78E+00 | 8.28E-01 | no | Rnf144b | 3_prime_UTR_variant | MODIFIER |
| 13 | 54723908 | AX-170489790 | 1 | G | A | 0.518 | -1.36E-02 | 6.73E-02 | 2.68E+00 | 2.80E+00 | 8.28E-01 | no | Cdhr2 | intron_variant | MODIFIER |
| 8 | 100289426 | AX-170516444 | 0 | T | A | 0.228 | 2.31E-02 | 1.06E-01 | 2.64E+00 | 2.73E+00 | 8.28E-01 | no | | intergenic_variant | MODIFIER |
| 1 | 41984446 | AX-170130999 | 0 | C | A | 0.421 | -1.63E-02 | 7.55E-02 | 2.65E+00 | 2.74E+00 | 8.28E-01 | no | | intergenic_variant | MODIFIER |
| 5 | 118613710 | AX-170073272 | 1 | A | G | 0.455 | -2.47E-02 | 1.16E-01 | 2.75E+00 | 2.79E+00 | 8.28E-01 | no | Med13l | intron_variant | MODIFIER |
| 11 | 34770200 | AX-170437453 | 0 | C | T | 0.096 | -3.92E-02 | 1.88E-01 | 2.68E+00 | 2.77E+00 | 8.28E-01 | no | Dock2 | intron_variant | MODIFIER |
| 9 | 107639250 | AX-170345032 | 1 | C | T | 0.116 | -2.66E-02 | 1.17E-01 | 2.62E+00 | 2.69E+00 | 8.28E-01 | no | Gnai2 | upstream_gene_variant | MODIFIER |
| 4 | 144401055 | AX-170024181 | 1 | T | C | 0.107 | 4.37E-02 | 2.04E-01 | 2.68E+00 | 2.74E+00 | 8.28E-01 | no | Pramef12 | intron_variant | MODIFIER |
| 20 | 87284925 | AX-170090574 | 0 | A | G | 0.272 | 1.26E-02 | 5.86E-02 | 2.63E+00 | 2.74E+00 | 8.28E-01 | no | Il1rapl1 | intron_variant | MODIFIER |
| 11 | 73256869 | AX-170374752 | 1 | C | T | 0.241 | 1.50E-02 | 7.08E-02 | 2.61E+00 | 2.75E+00 | 8.28E-01 | no | Trpv1 | intron_variant | MODIFIER |
| 8 | 95248163 | AX-170263189 | 0 | G | A | 0.228 | 2.39E-02 | 1.14E-01 | 2.69E+00 | 2.77E+00 | 8.28E-01 | no | Cngb1 | intron_variant | MODIFIER |
| 9 | 60128454 | AX-170269072 | 1 | G | A | 0.223 | 2.59E-02 | 1.21E-01 | 2.69E+00 | 2.74E+00 | 8.28E-01 | no | Thsd4 | intron_variant | MODIFIER |
| 2 | 129717929 | AX-170374831 | 0 | A | G | 0.149 | 3.84E-02 | 1.77E-01 | 2.66E+00 | 2.72E+00 | 8.28E-01 | no | | intergenic_variant | MODIFIER |
| 7 | 57215638 | AX-170301254 | 1 | A | T | 0.438 | 1.97E-02 | 9.18E-02 | 2.64E+00 | 2.74E+00 | 8.28E-01 | no | Gabrg3 | intron_variant | MODIFIER |
| 15 | 5310935 | AX-170010939 | 0 | T | C | 0.096 | -3.04E-02 | 9.67E-02 | 2.43E+00 | 2.57E+00 | 8.28E-01 | no | Gm2150 | downstream_gene_variant | MODIFIER |
| 15 | 88081521 | AX-170481216 | 1 | T | C | 0.143 | 3.90E-02 | 1.79E-01 | 2.65E+00 | 2.72E+00 | 8.28E-01 | no | | intergenic_variant | MODIFIER |
| 8 | 75137393 | AX-170300954 | 1 | T | C | 0.196 | -2.81E-02 | 1.31E-01 | 2.67E+00 | 2.74E+00 | 8.28E-01 | no | | intergenic_variant | MODIFIER |
| 9 | 21782326 | AX-170340483 | 0 | G | T | 0.465 | 2.07E-02 | 8.95E-02 | 2.60E+00 | 2.68E+00 | 8.29E-01 | no | Kank2 | intron_variant | MODIFIER |
| 9 | 79110826 | AX-97853036 | 1 | A | G | 0.196 | 2.75E-02 | 1.31E-01 | 2.73E+00 | 2.79E+00 | 8.29E-01 | no | | intergenic_variant | MODIFIER |
| 9 | 112809893 | AX-97843172 | 1 | G | A | 0.125 | -2.31E-02 | 1.06E-01 | 2.64E+00 | 2.73E+00 | 8.29E-01 | no | | intergenic_variant | MODIFIER |
| 9 | 113506208 | AX-168391788 | 1 | G | A | 0.125 | -2.31E-02 | 1.06E-01 | 2.64E+00 | 2.73E+00 | 8.29E-01 | no | AU023762 | intron_variant,non_coding_transcript_variant | MODIFIER |
| 1 | 106498231 | AX-170285949 | 0 | T | G | 0.158 | 3.50E-02 | 1.61E-01 | 2.62E+00 | 2.73E+00 | 8.29E-01 | no | | intergenic_variant | MODIFIER |

| | | | | | | | | | | | | | | | |
|---|---|---|---|---|---|---|---|---|---|---|---|---|---|---|---|
| 1 | 145022802 | AX-170412409 | 0 | G | A | 0.202 | 3.05E-02 | 1.40E-01 | 2.66E+00 | 2.71E+00 | 8.29E-01 | no | | intergenic_variant | MODIFIER |
| 1 | 149575912 | AX-170263483 | 0 | G | A | 0.202 | 3.05E-02 | 1.40E-01 | 2.66E+00 | 2.71E+00 | 8.29E-01 | no | | intergenic_variant | MODIFIER |
| 9 | 113687903 | AX-170033309 | 1 | C | G | 0.134 | -2.23E-02 | 1.05E-01 | 2.65E+00 | 2.74E+00 | 8.29E-01 | no | Pdcd6ip | intron_variant | MODIFIER |
| 6 | 102652974 | AX-170198466 | 1 | A | G | 0.054 | 6.34E-02 | 2.96E-01 | 2.67E+00 | 2.73E+00 | 8.29E-01 | no | | intergenic_variant | MODIFIER |
| 6 | 103502194 | AX-97796217 | 1 | C | G | 0.054 | 6.34E-02 | 2.96E-01 | 2.67E+00 | 2.73E+00 | 8.29E-01 | no | | intergenic_variant | MODIFIER |
| 6 | 103862048 | AX-170299360 | 1 | T | A | 0.054 | 6.34E-02 | 2.96E-01 | 2.67E+00 | 2.73E+00 | 8.29E-01 | no | | intergenic_variant | MODIFIER |
| 6 | 104396624 | AX-170075750 | 1 | C | T | 0.054 | 6.34E-02 | 2.96E-01 | 2.67E+00 | 2.73E+00 | 8.29E-01 | no | | intergenic_variant | MODIFIER |
| 6 | 106959406 | AX-97792443 | 1 | A | T | 0.054 | 6.34E-02 | 2.96E-01 | 2.67E+00 | 2.73E+00 | 8.29E-01 | no | | intergenic_variant | MODIFIER |
| 6 | 107281725 | AX-97786142 | 1 | C | T | 0.054 | 6.34E-02 | 2.96E-01 | 2.67E+00 | 2.73E+00 | 8.29E-01 | no | Gm34933 | upstream_gene_variant | MODIFIER |
| 6 | 110197095 | AX-170075854 | 1 | T | C | 0.054 | 6.34E-02 | 2.96E-01 | 2.67E+00 | 2.73E+00 | 8.29E-01 | no | | intergenic_variant | MODIFIER |
| 6 | 114414629 | AX-170261999 | 1 | T | C | 0.054 | 6.34E-02 | 2.96E-01 | 2.67E+00 | 2.73E+00 | 8.29E-01 | no | Hrh1 | intron_variant | MODIFIER |
| 1 | 9893206 | AX-170425536 | 0 | C | T | 0.412 | -1.46E-02 | 7.11E-02 | 2.72E+00 | 2.80E+00 | 8.29E-01 | no | Sgk3 | intron_variant | MODIFIER |
| 9 | 113862699 | AX-170385567 | 1 | C | T | 0.134 | -2.21E-02 | 1.04E-01 | 2.64E+00 | 2.74E+00 | 8.29E-01 | no | Clasp2 | intron_variant | MODIFIER |
| 6 | 101173194 | AX-170460230 | 0 | A | T | 0.053 | 6.33E-02 | 2.96E-01 | 2.67E+00 | 2.73E+00 | 8.29E-01 | no | Pdzrn3 | intron_variant | MODIFIER |
| 6 | 101516455 | AX-170283245 | 0 | T | A | 0.053 | 6.33E-02 | 2.96E-01 | 2.67E+00 | 2.73E+00 | 8.29E-01 | no | Gm44169 | upstream_gene_variant | MODIFIER |
| 6 | 101516711 | AX-170483498 | 0 | C | T | 0.053 | 6.33E-02 | 2.96E-01 | 2.67E+00 | 2.73E+00 | 8.29E-01 | no | Gm44169 | upstream_gene_variant | MODIFIER |
| 6 | 101516884 | AX-170426568 | 0 | C | T | 0.053 | 6.33E-02 | 2.96E-01 | 2.67E+00 | 2.73E+00 | 8.29E-01 | no | Gm44169 | upstream_gene_variant | MODIFIER |
| 6 | 101517862 | AX-170176896 | 0 | T | C | 0.053 | 6.33E-02 | 2.96E-01 | 2.67E+00 | 2.73E+00 | 8.29E-01 | no | Gm44169 | upstream_gene_variant | MODIFIER |
| 6 | 101521147 | AX-170479621 | 0 | G | A | 0.053 | 6.33E-02 | 2.96E-01 | 2.67E+00 | 2.73E+00 | 8.29E-01 | no | Gm44169 | intron_variant,non_coding_transcript_variant | MODIFIER |
| 6 | 101536301 | AX-170372390 | 0 | A | G | 0.053 | 6.33E-02 | 2.96E-01 | 2.67E+00 | 2.73E+00 | 8.29E-01 | no | Gm44171 | upstream_gene_variant | MODIFIER |
| 6 | 101536447 | AX-170223127 | 0 | G | T | 0.053 | 6.33E-02 | 2.96E-01 | 2.67E+00 | 2.73E+00 | 8.29E-01 | no | Gm44171 | upstream_gene_variant | MODIFIER |
| 6 | 101536963 | AX-170075688 | 0 | T | C | 0.053 | 6.33E-02 | 2.96E-01 | 2.67E+00 | 2.73E+00 | 8.29E-01 | no | Gm44171 | upstream_gene_variant | MODIFIER |
| 6 | 101538022 | AX-170372913 | 0 | A | G | 0.053 | 6.33E-02 | 2.96E-01 | 2.67E+00 | 2.73E+00 | 8.29E-01 | no | Gm44171 | upstream_gene_variant | MODIFIER |
| 6 | 102520938 | AX-170154482 | 0 | T | C | 0.053 | 6.33E-02 | 2.96E-01 | 2.67E+00 | 2.73E+00 | 8.29E-01 | no | Cntn3 | intron_variant | MODIFIER |
| 6 | 102530180 | AX-170027854 | 0 | T | C | 0.053 | 6.33E-02 | 2.96E-01 | 2.67E+00 | 2.73E+00 | 8.29E-01 | no | Cntn3 | intron_variant | MODIFIER |
| 6 | 102644969 | AX-170300209 | 0 | C | A | 0.053 | 6.33E-02 | 2.96E-01 | 2.67E+00 | 2.73E+00 | 8.29E-01 | no | | intergenic_variant | MODIFIER |
| 6 | 102650694 | AX-97812205 | 0 | C | T | 0.053 | 6.33E-02 | 2.96E-01 | 2.67E+00 | 2.73E+00 | 8.29E-01 | no | | intergenic_variant | MODIFIER |
| 6 | 102652778 | AX-170111286 | 0 | A | G | 0.053 | 6.33E-02 | 2.96E-01 | 2.67E+00 | 2.73E+00 | 8.29E-01 | no | | intergenic_variant | MODIFIER |
| 6 | 102706460 | AX-170075714 | 0 | T | G | 0.053 | 6.33E-02 | 2.96E-01 | 2.67E+00 | 2.73E+00 | 8.29E-01 | no | | intergenic_variant | MODIFIER |
| 6 | 102729882 | AX-170323354 | 0 | C | G | 0.053 | 6.33E-02 | 2.96E-01 | 2.67E+00 | 2.73E+00 | 8.29E-01 | no | | intergenic_variant | MODIFIER |
| 6 | 102730256 | AX-170366158 | 0 | G | A | 0.053 | 6.33E-02 | 2.96E-01 | 2.67E+00 | 2.73E+00 | 8.29E-01 | no | | intergenic_variant | MODIFIER |
| 6 | 102774317 | AX-170497658 | 0 | A | G | 0.053 | 6.33E-02 | 2.96E-01 | 2.67E+00 | 2.73E+00 | 8.29E-01 | no | | intergenic_variant | MODIFIER |
| 6 | 102774788 | AX-170144975 | 0 | T | C | 0.053 | 6.33E-02 | 2.96E-01 | 2.67E+00 | 2.73E+00 | 8.29E-01 | no | | intergenic_variant | MODIFIER |
| 6 | 102792199 | AX-170382732 | 0 | G | A | 0.053 | 6.33E-02 | 2.96E-01 | 2.67E+00 | 2.73E+00 | 8.29E-01 | no | Mir6373 | upstream_gene_variant | MODIFIER |
| 6 | 102809960 | AX-170280004 | 0 | A | C | 0.053 | 6.33E-02 | 2.96E-01 | 2.67E+00 | 2.73E+00 | 8.29E-01 | no | | intergenic_variant | MODIFIER |
| 6 | 102882767 | AX-97851046 | 0 | T | C | 0.053 | 6.33E-02 | 2.96E-01 | 2.67E+00 | 2.73E+00 | 8.29E-01 | no | | intergenic_variant | MODIFIER |
| 6 | 103318148 | AX-170284198 | 0 | G | A | 0.053 | 6.33E-02 | 2.96E-01 | 2.67E+00 | 2.73E+00 | 8.29E-01 | no | | regulatory_region_variant | MODIFIER |
| 6 | 103322010 | AX-170181878 | 0 | C | A | 0.053 | 6.33E-02 | 2.96E-01 | 2.67E+00 | 2.73E+00 | 8.29E-01 | no | | intergenic_variant | MODIFIER |
| 6 | 103331119 | AX-170516822 | 0 | T | C | 0.053 | 6.33E-02 | 2.96E-01 | 2.67E+00 | 2.73E+00 | 8.29E-01 | no | | intergenic_variant | MODIFIER |
| 6 | 103363900 | AX-170128298 | 0 | T | C | 0.053 | 6.33E-02 | 2.96E-01 | 2.67E+00 | 2.73E+00 | 8.29E-01 | no | | intergenic_variant | MODIFIER |
| 6 | 103382685 | AX-170400768 | 0 | C | T | 0.053 | 6.33E-02 | 2.96E-01 | 2.67E+00 | 2.73E+00 | 8.29E-01 | no | | regulatory_region_variant | MODIFIER |
| 6 | 103383620 | AX-170420108 | 0 | A | G | 0.053 | 6.33E-02 | 2.96E-01 | 2.67E+00 | 2.73E+00 | 8.29E-01 | no | | intergenic_variant | MODIFIER |
| 6 | 103385733 | AX-170330669 | 0 | G | A | 0.053 | 6.33E-02 | 2.96E-01 | 2.67E+00 | 2.73E+00 | 8.29E-01 | no | | intergenic_variant | MODIFIER |
| 6 | 103391553 | AX-170317991 | 0 | G | A | 0.053 | 6.33E-02 | 2.96E-01 | 2.67E+00 | 2.73E+00 | 8.29E-01 | no | | intergenic_variant | MODIFIER |
| 6 | 103391679 | AX-97789483 | 0 | A | G | 0.053 | 6.33E-02 | 2.96E-01 | 2.67E+00 | 2.73E+00 | 8.29E-01 | no | | intergenic_variant | MODIFIER |
| 6 | 103401425 | AX-170123302 | 0 | G | C | 0.053 | 6.33E-02 | 2.96E-01 | 2.67E+00 | 2.73E+00 | 8.29E-01 | no | | intergenic_variant | MODIFIER |
| 6 | 103418052 | AX-170075729 | 0 | T | C | 0.053 | 6.33E-02 | 2.96E-01 | 2.67E+00 | 2.73E+00 | 8.29E-01 | no | | intergenic_variant | MODIFIER |
| 6 | 103418745 | AX-170225139 | 0 | C | T | 0.053 | 6.33E-02 | 2.96E-01 | 2.67E+00 | 2.73E+00 | 8.29E-01 | no | | intergenic_variant | MODIFIER |
| 6 | 103440239 | AX-170350085 | 0 | C | G | 0.053 | 6.33E-02 | 2.96E-01 | 2.67E+00 | 2.73E+00 | 8.29E-01 | no | | intergenic_variant | MODIFIER |
| 6 | 103477678 | AX-170494317 | 0 | G | C | 0.053 | 6.33E-02 | 2.96E-01 | 2.67E+00 | 2.73E+00 | 8.29E-01 | no | | intergenic_variant | MODIFIER |
| 6 | 103550757 | AX-170273182 | 0 | T | A | 0.053 | 6.33E-02 | 2.96E-01 | 2.67E+00 | 2.73E+00 | 8.29E-01 | no | Chl1 | intron_variant | MODIFIER |
| 6 | 103558064 | AX-170075731 | 0 | T | C | 0.053 | 6.33E-02 | 2.96E-01 | 2.67E+00 | 2.73E+00 | 8.29E-01 | no | Chl1 | intron_variant | MODIFIER |
| 6 | 103562859 | AX-170207725 | 0 | C | T | 0.053 | 6.33E-02 | 2.96E-01 | 2.67E+00 | 2.73E+00 | 8.29E-01 | no | Chl1 | intron_variant | MODIFIER |
| 6 | 103563279 | AX-170496333 | 0 | G | A | 0.053 | 6.33E-02 | 2.96E-01 | 2.67E+00 | 2.73E+00 | 8.29E-01 | no | Chl1 | intron_variant | MODIFIER |

| | | | | | | | | | | | | | | | |
|---|---|---|---|---|---|---|---|---|---|---|---|---|---|---|---|
| 6 | 103565362 | AX-170497671 | 0 | A | G | 0.053 | 6.33E-02 | 2.96E-01 | 2.67E+00 | 2.73E+00 | 8.29E-01 | no | Chl1 | intron_variant | MODIFIER |
| 6 | 103596642 | AX-170452912 | 0 | C | G | 0.053 | 6.33E-02 | 2.96E-01 | 2.67E+00 | 2.73E+00 | 8.29E-01 | no | Chl1 | intron_variant | MODIFIER |
| 6 | 103599715 | AX-170434725 | 0 | A | G | 0.053 | 6.33E-02 | 2.96E-01 | 2.67E+00 | 2.73E+00 | 8.29E-01 | no | Chl1 | intron_variant | MODIFIER |
| 6 | 103599813 | AX-170194302 | 0 | C | T | 0.053 | 6.33E-02 | 2.96E-01 | 2.67E+00 | 2.73E+00 | 8.29E-01 | no | Chl1 | intron_variant | MODIFIER |
| 6 | 103604395 | AX-170027861 | 0 | C | G | 0.053 | 6.33E-02 | 2.96E-01 | 2.67E+00 | 2.73E+00 | 8.29E-01 | no | Chl1 | intron_variant | MODIFIER |
| 6 | 103638789 | AX-170443839 | 0 | G | T | 0.053 | 6.33E-02 | 2.96E-01 | 2.67E+00 | 2.73E+00 | 8.29E-01 | no | Chl1 | intron_variant | MODIFIER |
| 6 | 103674099 | AX-170027859 | 0 | A | T | 0.053 | 6.33E-02 | 2.96E-01 | 2.67E+00 | 2.73E+00 | 8.29E-01 | no | Chl1 | intron_variant | MODIFIER |
| 6 | 103705517 | AX-170155563 | 0 | G | A | 0.053 | 6.33E-02 | 2.96E-01 | 2.67E+00 | 2.73E+00 | 8.29E-01 | no | Chl1 | intron_variant | MODIFIER |
| 6 | 103718529 | AX-170451557 | 0 | A | C | 0.053 | 6.33E-02 | 2.96E-01 | 2.67E+00 | 2.73E+00 | 8.29E-01 | no | Chl1 | intron_variant | MODIFIER |
| 6 | 103726927 | AX-170389557 | 0 | A | G | 0.053 | 6.33E-02 | 2.96E-01 | 2.67E+00 | 2.73E+00 | 8.29E-01 | no | Chl1 | intron_variant | MODIFIER |
| 6 | 103727176 | AX-170472697 | 0 | C | T | 0.053 | 6.33E-02 | 2.96E-01 | 2.67E+00 | 2.73E+00 | 8.29E-01 | no | Chl1 | intron_variant | MODIFIER |
| 6 | 103727444 | AX-170438963 | 0 | T | A | 0.053 | 6.33E-02 | 2.96E-01 | 2.67E+00 | 2.73E+00 | 8.29E-01 | no | Chl1 | intron_variant | MODIFIER |
| 6 | 103727494 | AX-170347684 | 0 | C | T | 0.053 | 6.33E-02 | 2.96E-01 | 2.67E+00 | 2.73E+00 | 8.29E-01 | no | Chl1 | intron_variant | MODIFIER |
| 6 | 103727538 | AX-170390344 | 0 | A | G | 0.053 | 6.33E-02 | 2.96E-01 | 2.67E+00 | 2.73E+00 | 8.29E-01 | no | Chl1 | intron_variant | MODIFIER |
| 6 | 103727757 | AX-170465088 | 0 | T | G | 0.053 | 6.33E-02 | 2.96E-01 | 2.67E+00 | 2.73E+00 | 8.29E-01 | no | Chl1 | intron_variant | MODIFIER |
| 6 | 103727815 | AX-170135696 | 0 | G | A | 0.053 | 6.33E-02 | 2.96E-01 | 2.67E+00 | 2.73E+00 | 8.29E-01 | no | Chl1 | intron_variant | MODIFIER |
| 6 | 103763122 | AX-170481034 | 0 | C | A | 0.053 | 6.33E-02 | 2.96E-01 | 2.67E+00 | 2.73E+00 | 8.29E-01 | no | | intergenic_variant | MODIFIER |
| 6 | 103764461 | AX-170458715 | 0 | T | G | 0.053 | 6.33E-02 | 2.96E-01 | 2.67E+00 | 2.73E+00 | 8.29E-01 | no | | intergenic_variant | MODIFIER |
| 6 | 103768334 | AX-170139341 | 0 | C | T | 0.053 | 6.33E-02 | 2.96E-01 | 2.67E+00 | 2.73E+00 | 8.29E-01 | no | | intergenic_variant | MODIFIER |
| 6 | 103774135 | AX-170132589 | 0 | T | C | 0.053 | 6.33E-02 | 2.96E-01 | 2.67E+00 | 2.73E+00 | 8.29E-01 | no | | intergenic_variant | MODIFIER |
| 6 | 103777991 | AX-170248632 | 0 | G | C | 0.053 | 6.33E-02 | 2.96E-01 | 2.67E+00 | 2.73E+00 | 8.29E-01 | no | | intergenic_variant | MODIFIER |
| 6 | 103778024 | AX-170297509 | 0 | G | C | 0.053 | 6.33E-02 | 2.96E-01 | 2.67E+00 | 2.73E+00 | 8.29E-01 | no | | intergenic_variant | MODIFIER |
| 6 | 103781202 | AX-170075737 | 0 | A | G | 0.053 | 6.33E-02 | 2.96E-01 | 2.67E+00 | 2.73E+00 | 8.29E-01 | no | | intergenic_variant | MODIFIER |
| 6 | 103802170 | AX-170272934 | 0 | T | G | 0.053 | 6.33E-02 | 2.96E-01 | 2.67E+00 | 2.73E+00 | 8.29E-01 | no | | intergenic_variant | MODIFIER |
| 6 | 103808520 | AX-170321927 | 0 | C | T | 0.053 | 6.33E-02 | 2.96E-01 | 2.67E+00 | 2.73E+00 | 8.29E-01 | no | | intergenic_variant | MODIFIER |
| 6 | 103810164 | AX-170193555 | 0 | A | G | 0.053 | 6.33E-02 | 2.96E-01 | 2.67E+00 | 2.73E+00 | 8.29E-01 | no | | regulatory_region_variant | MODIFIER |
| 6 | 103812312 | AX-170377469 | 0 | A | C | 0.053 | 6.33E-02 | 2.96E-01 | 2.67E+00 | 2.73E+00 | 8.29E-01 | no | | intergenic_variant | MODIFIER |
| 6 | 103858780 | AX-97841552 | 0 | G | A | 0.053 | 6.33E-02 | 2.96E-01 | 2.67E+00 | 2.73E+00 | 8.29E-01 | no | | intergenic_variant | MODIFIER |
| 6 | 103859593 | AX-170370766 | 0 | G | C | 0.053 | 6.33E-02 | 2.96E-01 | 2.67E+00 | 2.73E+00 | 8.29E-01 | no | | intergenic_variant | MODIFIER |
| 6 | 103863716 | AX-170104203 | 0 | A | G | 0.053 | 6.33E-02 | 2.96E-01 | 2.67E+00 | 2.73E+00 | 8.29E-01 | no | | intergenic_variant | MODIFIER |
| 6 | 103874889 | AX-170350651 | 0 | G | A | 0.053 | 6.33E-02 | 2.96E-01 | 2.67E+00 | 2.73E+00 | 8.29E-01 | no | | intergenic_variant | MODIFIER |
| 6 | 103877763 | AX-170126395 | 0 | T | C | 0.053 | 6.33E-02 | 2.96E-01 | 2.67E+00 | 2.73E+00 | 8.29E-01 | no | | intergenic_variant | MODIFIER |
| 6 | 103886459 | AX-170444635 | 0 | T | C | 0.053 | 6.33E-02 | 2.96E-01 | 2.67E+00 | 2.73E+00 | 8.29E-01 | no | | intergenic_variant | MODIFIER |
| 6 | 103890202 | AX-97814928 | 0 | C | T | 0.053 | 6.33E-02 | 2.96E-01 | 2.67E+00 | 2.73E+00 | 8.29E-01 | no | | intergenic_variant | MODIFIER |
| 6 | 103891877 | AX-170265377 | 0 | T | G | 0.053 | 6.33E-02 | 2.96E-01 | 2.67E+00 | 2.73E+00 | 8.29E-01 | no | | intergenic_variant | MODIFIER |
| 6 | 103892061 | AX-97814929 | 0 | C | A | 0.053 | 6.33E-02 | 2.96E-01 | 2.67E+00 | 2.73E+00 | 8.29E-01 | no | | intergenic_variant | MODIFIER |
| 6 | 103922295 | AX-170351456 | 0 | C | T | 0.053 | 6.33E-02 | 2.96E-01 | 2.67E+00 | 2.73E+00 | 8.29E-01 | no | | intergenic_variant | MODIFIER |
| 6 | 103923034 | AX-170075742 | 0 | T | C | 0.053 | 6.33E-02 | 2.96E-01 | 2.67E+00 | 2.73E+00 | 8.29E-01 | no | | intergenic_variant | MODIFIER |
| 6 | 103926578 | AX-170433881 | 0 | A | G | 0.053 | 6.33E-02 | 2.96E-01 | 2.67E+00 | 2.73E+00 | 8.29E-01 | no | | intergenic_variant | MODIFIER |
| 6 | 103931101 | AX-170131195 | 0 | A | C | 0.053 | 6.33E-02 | 2.96E-01 | 2.67E+00 | 2.73E+00 | 8.29E-01 | no | | intergenic_variant | MODIFIER |
| 6 | 103998083 | AX-170257580 | 0 | A | G | 0.053 | 6.33E-02 | 2.96E-01 | 2.67E+00 | 2.73E+00 | 8.29E-01 | no | | intergenic_variant | MODIFIER |
| 6 | 103999094 | AX-170214817 | 0 | C | A | 0.053 | 6.33E-02 | 2.96E-01 | 2.67E+00 | 2.73E+00 | 8.29E-01 | no | | intergenic_variant | MODIFIER |
| 6 | 104005001 | AX-170527757 | 0 | A | G | 0.053 | 6.33E-02 | 2.96E-01 | 2.67E+00 | 2.73E+00 | 8.29E-01 | no | | intergenic_variant | MODIFIER |
| 6 | 104011154 | AX-170442527 | 0 | A | G | 0.053 | 6.33E-02 | 2.96E-01 | 2.67E+00 | 2.73E+00 | 8.29E-01 | no | | intergenic_variant | MODIFIER |
| 6 | 104011335 | AX-170095735 | 0 | C | T | 0.053 | 6.33E-02 | 2.96E-01 | 2.67E+00 | 2.73E+00 | 8.29E-01 | no | | intergenic_variant | MODIFIER |
| 6 | 104020754 | AX-170494388 | 0 | C | T | 0.053 | 6.33E-02 | 2.96E-01 | 2.67E+00 | 2.73E+00 | 8.29E-01 | no | | intergenic_variant | MODIFIER |
| 6 | 104029424 | AX-170075745 | 0 | A | C | 0.053 | 6.33E-02 | 2.96E-01 | 2.67E+00 | 2.73E+00 | 8.29E-01 | no | | regulatory_region_variant | MODIFIER |
| 6 | 104030551 | AX-170075744 | 0 | C | T | 0.053 | 6.33E-02 | 2.96E-01 | 2.67E+00 | 2.73E+00 | 8.29E-01 | no | | intergenic_variant | MODIFIER |
| 6 | 104040327 | AX-170308220 | 0 | G | A | 0.053 | 6.33E-02 | 2.96E-01 | 2.67E+00 | 2.73E+00 | 8.29E-01 | no | | intergenic_variant | MODIFIER |
| 6 | 104083446 | AX-170147809 | 0 | T | C | 0.053 | 6.33E-02 | 2.96E-01 | 2.67E+00 | 2.73E+00 | 8.29E-01 | no | | intergenic_variant | MODIFIER |
| 6 | 104084598 | AX-170206270 | 0 | A | C | 0.053 | 6.33E-02 | 2.96E-01 | 2.67E+00 | 2.73E+00 | 8.29E-01 | no | | intergenic_variant | MODIFIER |
| 6 | 104098638 | AX-97810214 | 0 | T | C | 0.053 | 6.33E-02 | 2.96E-01 | 2.67E+00 | 2.73E+00 | 8.29E-01 | no | | intergenic_variant | MODIFIER |
| 6 | 104104299 | AX-170176176 | 0 | A | G | 0.053 | 6.33E-02 | 2.96E-01 | 2.67E+00 | 2.73E+00 | 8.29E-01 | no | | intergenic_variant | MODIFIER |
| 6 | 104117342 | AX-97789922 | 0 | T | C | 0.053 | 6.33E-02 | 2.96E-01 | 2.67E+00 | 2.73E+00 | 8.29E-01 | no | | intergenic_variant | MODIFIER |

| | | | | | | | | | | | | | | | |
|---|---|---|---|---|---|---|---|---|---|---|---|---|---|---|---|
| 6 | 104160429 | AX-170455746 | 0 | G | A | 0.053 | 6.33E-02 | 2.96E-01 | 2.67E+00 | 2.73E+00 | 8.29E-01 | no | | intergenic_variant | MODIFIER |
| 6 | 104161502 | AX-170507526 | 0 | T | C | 0.053 | 6.33E-02 | 2.96E-01 | 2.67E+00 | 2.73E+00 | 8.29E-01 | no | | intergenic_variant | MODIFIER |
| 6 | 104161542 | AX-170516953 | 0 | T | C | 0.053 | 6.33E-02 | 2.96E-01 | 2.67E+00 | 2.73E+00 | 8.29E-01 | no | | intergenic_variant | MODIFIER |
| 6 | 104212622 | AX-170296149 | 0 | T | C | 0.053 | 6.33E-02 | 2.96E-01 | 2.67E+00 | 2.73E+00 | 8.29E-01 | no | | intergenic_variant | MODIFIER |
| 6 | 104212694 | AX-170131119 | 0 | T | C | 0.053 | 6.33E-02 | 2.96E-01 | 2.67E+00 | 2.73E+00 | 8.29E-01 | no | | intergenic_variant | MODIFIER |
| 6 | 104237075 | AX-97802711 | 0 | T | C | 0.053 | 6.33E-02 | 2.96E-01 | 2.67E+00 | 2.73E+00 | 8.29E-01 | no | | intergenic_variant | MODIFIER |
| 6 | 104256919 | AX-170369516 | 0 | C | G | 0.053 | 6.33E-02 | 2.96E-01 | 2.67E+00 | 2.73E+00 | 8.29E-01 | no | | intergenic_variant | MODIFIER |
| 6 | 104279456 | AX-170353435 | 0 | A | C | 0.053 | 6.33E-02 | 2.96E-01 | 2.67E+00 | 2.73E+00 | 8.29E-01 | no | | intergenic_variant | MODIFIER |
| 6 | 104293846 | AX-170456299 | 0 | A | G | 0.053 | 6.33E-02 | 2.96E-01 | 2.67E+00 | 2.73E+00 | 8.29E-01 | no | | intergenic_variant | MODIFIER |
| 6 | 104294035 | AX-170273870 | 0 | G | A | 0.053 | 6.33E-02 | 2.96E-01 | 2.67E+00 | 2.73E+00 | 8.29E-01 | no | | intergenic_variant | MODIFIER |
| 6 | 104299997 | AX-170415502 | 0 | A | G | 0.053 | 6.33E-02 | 2.96E-01 | 2.67E+00 | 2.73E+00 | 8.29E-01 | no | Gm18100 | upstream_gene_variant | MODIFIER |
| 6 | 104303522 | AX-170504776 | 0 | A | G | 0.053 | 6.33E-02 | 2.96E-01 | 2.67E+00 | 2.73E+00 | 8.29E-01 | no | Gm18100 | downstream_gene_variant | MODIFIER |
| 6 | 104307085 | AX-170471750 | 0 | C | G | 0.053 | 6.33E-02 | 2.96E-01 | 2.67E+00 | 2.73E+00 | 8.29E-01 | no | | intergenic_variant | MODIFIER |
| 6 | 104318265 | AX-170512275 | 0 | T | C | 0.053 | 6.33E-02 | 2.96E-01 | 2.67E+00 | 2.73E+00 | 8.29E-01 | no | | intergenic_variant | MODIFIER |
| 6 | 104388801 | AX-97844460 | 0 | C | T | 0.053 | 6.33E-02 | 2.96E-01 | 2.67E+00 | 2.73E+00 | 8.29E-01 | no | | intergenic_variant | MODIFIER |
| 6 | 104445038 | AX-170433893 | 0 | G | T | 0.053 | 6.33E-02 | 2.96E-01 | 2.67E+00 | 2.73E+00 | 8.29E-01 | no | | intergenic_variant | MODIFIER |
| 6 | 104445318 | AX-170167003 | 0 | T | A | 0.053 | 6.33E-02 | 2.96E-01 | 2.67E+00 | 2.73E+00 | 8.29E-01 | no | | intergenic_variant | MODIFIER |
| 6 | 104462172 | AX-170213213 | 0 | T | C | 0.053 | 6.33E-02 | 2.96E-01 | 2.67E+00 | 2.73E+00 | 8.29E-01 | no | | intergenic_variant | MODIFIER |
| 6 | 104492456 | AX-170166799 | 0 | A | G | 0.053 | 6.33E-02 | 2.96E-01 | 2.67E+00 | 2.73E+00 | 8.29E-01 | no | Cntn6 | upstream_gene_variant | MODIFIER |
| 6 | 104514773 | AX-170208277 | 0 | T | C | 0.053 | 6.33E-02 | 2.96E-01 | 2.67E+00 | 2.73E+00 | 8.29E-01 | no | Cntn6 | intron_variant | MODIFIER |
| 6 | 104537917 | AX-170278604 | 0 | T | G | 0.053 | 6.33E-02 | 2.96E-01 | 2.67E+00 | 2.73E+00 | 8.29E-01 | no | Cntn6 | intron_variant | MODIFIER |
| 6 | 104544010 | AX-170444642 | 0 | T | C | 0.053 | 6.33E-02 | 2.96E-01 | 2.67E+00 | 2.73E+00 | 8.29E-01 | no | Cntn6 | intron_variant | MODIFIER |
| 6 | 104563887 | AX-170341588 | 0 | T | C | 0.053 | 6.33E-02 | 2.96E-01 | 2.67E+00 | 2.73E+00 | 8.29E-01 | no | Cntn6 | intron_variant | MODIFIER |
| 6 | 104566145 | AX-170157116 | 0 | A | G | 0.053 | 6.33E-02 | 2.96E-01 | 2.67E+00 | 2.73E+00 | 8.29E-01 | no | Cntn6 | intron_variant | MODIFIER |
| 6 | 104582059 | AX-170075754 | 0 | T | G | 0.053 | 6.33E-02 | 2.96E-01 | 2.67E+00 | 2.73E+00 | 8.29E-01 | no | Cntn6 | intron_variant | MODIFIER |
| 6 | 104607840 | AX-170354674 | 0 | C | A | 0.053 | 6.33E-02 | 2.96E-01 | 2.67E+00 | 2.73E+00 | 8.29E-01 | no | Cntn6 | intron_variant | MODIFIER |
| 6 | 104623737 | AX-170027875 | 0 | G | A | 0.053 | 6.33E-02 | 2.96E-01 | 2.67E+00 | 2.73E+00 | 8.29E-01 | no | Cntn6 | intron_variant | MODIFIER |
| 6 | 104630458 | AX-97842377 | 0 | A | G | 0.053 | 6.33E-02 | 2.96E-01 | 2.67E+00 | 2.73E+00 | 8.29E-01 | no | Cntn6 | intron_variant | MODIFIER |
| 6 | 104680330 | AX-170197878 | 0 | G | C | 0.053 | 6.33E-02 | 2.96E-01 | 2.67E+00 | 2.73E+00 | 8.29E-01 | no | Cntn6 | intron_variant | MODIFIER |
| 6 | 104683960 | AX-170352381 | 0 | T | C | 0.053 | 6.33E-02 | 2.96E-01 | 2.67E+00 | 2.73E+00 | 8.29E-01 | no | Cntn6 | intron_variant | MODIFIER |
| 6 | 104684147 | AX-170245543 | 0 | G | T | 0.053 | 6.33E-02 | 2.96E-01 | 2.67E+00 | 2.73E+00 | 8.29E-01 | no | Cntn6 | intron_variant | MODIFIER |
| 6 | 104733844 | AX-170290048 | 0 | C | T | 0.053 | 6.33E-02 | 2.96E-01 | 2.67E+00 | 2.73E+00 | 8.29E-01 | no | Cntn6 | intron_variant | MODIFIER |
| 6 | 104764310 | AX-170422545 | 0 | T | G | 0.053 | 6.33E-02 | 2.96E-01 | 2.67E+00 | 2.73E+00 | 8.29E-01 | no | Cntn6 | intron_variant | MODIFIER |
| 6 | 104939104 | AX-170397999 | 0 | T | G | 0.053 | 6.33E-02 | 2.96E-01 | 2.67E+00 | 2.73E+00 | 8.29E-01 | no | | intergenic_variant | MODIFIER |
| 6 | 104943827 | AX-170325435 | 0 | T | C | 0.053 | 6.33E-02 | 2.96E-01 | 2.67E+00 | 2.73E+00 | 8.29E-01 | no | | intergenic_variant | MODIFIER |
| 6 | 104978070 | AX-170319927 | 0 | C | T | 0.053 | 6.33E-02 | 2.96E-01 | 2.67E+00 | 2.73E+00 | 8.29E-01 | no | | intergenic_variant | MODIFIER |
| 6 | 104989022 | AX-170173158 | 0 | A | T | 0.053 | 6.33E-02 | 2.96E-01 | 2.67E+00 | 2.73E+00 | 8.29E-01 | no | | intergenic_variant | MODIFIER |
| 6 | 105002109 | AX-170149910 | 0 | C | A | 0.053 | 6.33E-02 | 2.96E-01 | 2.67E+00 | 2.73E+00 | 8.29E-01 | no | | intergenic_variant | MODIFIER |
| 6 | 105065279 | AX-170434303 | 0 | A | G | 0.053 | 6.33E-02 | 2.96E-01 | 2.67E+00 | 2.73E+00 | 8.29E-01 | no | Gm5315 | upstream_gene_variant | MODIFIER |
| 6 | 105252218 | AX-170158062 | 0 | G | A | 0.053 | 6.33E-02 | 2.96E-01 | 2.67E+00 | 2.73E+00 | 8.29E-01 | no | | intergenic_variant | MODIFIER |
| 6 | 105276395 | AX-170337430 | 0 | T | C | 0.053 | 6.33E-02 | 2.96E-01 | 2.67E+00 | 2.73E+00 | 8.29E-01 | no | | intergenic_variant | MODIFIER |
| 6 | 105284424 | AX-170283521 | 0 | C | T | 0.053 | 6.33E-02 | 2.96E-01 | 2.67E+00 | 2.73E+00 | 8.29E-01 | no | | intergenic_variant | MODIFIER |
| 6 | 105309177 | AX-170208285 | 0 | T | G | 0.053 | 6.33E-02 | 2.96E-01 | 2.67E+00 | 2.73E+00 | 8.29E-01 | no | | intergenic_variant | MODIFIER |
| 6 | 105428426 | AX-170075768 | 0 | G | C | 0.053 | 6.33E-02 | 2.96E-01 | 2.67E+00 | 2.73E+00 | 8.29E-01 | no | | intergenic_variant | MODIFIER |
| 6 | 105433962 | AX-170172940 | 0 | C | T | 0.053 | 6.33E-02 | 2.96E-01 | 2.67E+00 | 2.73E+00 | 8.29E-01 | no | | intergenic_variant | MODIFIER |
| 6 | 105434173 | AX-170177256 | 0 | C | G | 0.053 | 6.33E-02 | 2.96E-01 | 2.67E+00 | 2.73E+00 | 8.29E-01 | no | | intergenic_variant | MODIFIER |
| 6 | 105452821 | AX-170257738 | 0 | C | A | 0.053 | 6.33E-02 | 2.96E-01 | 2.67E+00 | 2.73E+00 | 8.29E-01 | no | | intergenic_variant | MODIFIER |
| 6 | 105651361 | AX-170416551 | 0 | T | C | 0.053 | 6.33E-02 | 2.96E-01 | 2.67E+00 | 2.73E+00 | 8.29E-01 | no | | intergenic_variant | MODIFIER |
| 6 | 105651482 | AX-97828914 | 0 | T | C | 0.053 | 6.33E-02 | 2.96E-01 | 2.67E+00 | 2.73E+00 | 8.29E-01 | no | | intergenic_variant | MODIFIER |
| 6 | 105657011 | AX-170444310 | 0 | G | A | 0.053 | 6.33E-02 | 2.96E-01 | 2.67E+00 | 2.73E+00 | 8.29E-01 | no | | intergenic_variant | MODIFIER |
| 6 | 105660258 | AX-170133566 | 0 | C | T | 0.053 | 6.33E-02 | 2.96E-01 | 2.67E+00 | 2.73E+00 | 8.29E-01 | no | | intergenic_variant | MODIFIER |
| 6 | 105670781 | AX-170075772 | 0 | A | G | 0.053 | 6.33E-02 | 2.96E-01 | 2.67E+00 | 2.73E+00 | 8.29E-01 | no | Gm15631 | downstream_gene_variant | MODIFIER |
| 6 | 105703273 | AX-170310189 | 0 | C | T | 0.053 | 6.33E-02 | 2.96E-01 | 2.67E+00 | 2.73E+00 | 8.29E-01 | no | Cntn4 | intron_variant | MODIFIER |
| 6 | 105703358 | AX-170389763 | 0 | C | A | 0.053 | 6.33E-02 | 2.96E-01 | 2.67E+00 | 2.73E+00 | 8.29E-01 | no | Cntn4 | intron_variant | MODIFIER |

| | | | | | | | | | | | | | | | |
|---|---|---|---|---|---|---|---|---|---|---|---|---|---|---|---|
| 6 | 105711178 | AX-170521317 | 0 | T | C | 0.053 | 6.33E-02 | 2.96E-01 | 2.67E+00 | 2.73E+00 | 8.29E-01 | no | Cntn4 | intron_variant | MODIFIER |
| 6 | 105731926 | AX-170469516 | 0 | T | C | 0.053 | 6.33E-02 | 2.96E-01 | 2.67E+00 | 2.73E+00 | 8.29E-01 | no | Cntn4 | intron_variant | MODIFIER |
| 6 | 105737900 | AX-170301052 | 0 | A | G | 0.053 | 6.33E-02 | 2.96E-01 | 2.67E+00 | 2.73E+00 | 8.29E-01 | no | Cntn4 | intron_variant | MODIFIER |
| 6 | 105744839 | AX-170314909 | 0 | G | A | 0.053 | 6.33E-02 | 2.96E-01 | 2.67E+00 | 2.73E+00 | 8.29E-01 | no | Cntn4 | intron_variant | MODIFIER |
| 6 | 105779828 | AX-170370736 | 0 | T | C | 0.053 | 6.33E-02 | 2.96E-01 | 2.67E+00 | 2.73E+00 | 8.29E-01 | no | Cntn4 | intron_variant | MODIFIER |
| 6 | 105861994 | AX-170309151 | 0 | G | T | 0.053 | 6.33E-02 | 2.96E-01 | 2.67E+00 | 2.73E+00 | 8.29E-01 | no | Cntn4 | intron_variant | MODIFIER |
| 6 | 105865809 | AX-170317587 | 0 | A | G | 0.053 | 6.33E-02 | 2.96E-01 | 2.67E+00 | 2.73E+00 | 8.29E-01 | no | Cntn4 | intron_variant | MODIFIER |
| 6 | 105870103 | AX-170075775 | 0 | A | G | 0.053 | 6.33E-02 | 2.96E-01 | 2.67E+00 | 2.73E+00 | 8.29E-01 | no | Cntn4 | intron_variant | MODIFIER |
| 6 | 105893078 | AX-170075777 | 0 | A | G | 0.053 | 6.33E-02 | 2.96E-01 | 2.67E+00 | 2.73E+00 | 8.29E-01 | no | Cntn4 | intron_variant | MODIFIER |
| 6 | 105953148 | AX-170106810 | 0 | A | G | 0.053 | 6.33E-02 | 2.96E-01 | 2.67E+00 | 2.73E+00 | 8.29E-01 | no | Cntn4 | intron_variant | MODIFIER |
| 6 | 105979102 | AX-170286622 | 0 | T | G | 0.053 | 6.33E-02 | 2.96E-01 | 2.67E+00 | 2.73E+00 | 8.29E-01 | no | Cntn4 | intron_variant | MODIFIER |
| 6 | 106113651 | AX-170467071 | 0 | G | A | 0.053 | 6.33E-02 | 2.96E-01 | 2.67E+00 | 2.73E+00 | 8.29E-01 | no | Cntn4 | intron_variant | MODIFIER |
| 6 | 106778169 | AX-170467108 | 0 | T | C | 0.053 | 6.33E-02 | 2.96E-01 | 2.67E+00 | 2.73E+00 | 8.29E-01 | no | Crbn | downstream_gene_variant | MODIFIER |
| 6 | 106779547 | AX-170262221 | 0 | C | T | 0.053 | 6.33E-02 | 2.96E-01 | 2.67E+00 | 2.73E+00 | 8.29E-01 | no | Crbn | downstream_gene_variant | MODIFIER |
| 6 | 106779568 | AX-170103438 | 0 | C | T | 0.053 | 6.33E-02 | 2.96E-01 | 2.67E+00 | 2.73E+00 | 8.29E-01 | no | Crbn | downstream_gene_variant | MODIFIER |
| 6 | 106789513 | AX-170098024 | 0 | T | A | 0.053 | 6.33E-02 | 2.96E-01 | 2.67E+00 | 2.73E+00 | 8.29E-01 | no | Crbn | intron_variant | MODIFIER |
| 6 | 106795855 | AX-170364449 | 0 | T | C | 0.053 | 6.33E-02 | 2.96E-01 | 2.67E+00 | 2.73E+00 | 8.29E-01 | no | Crbn | intron_variant | MODIFIER |
| 6 | 106810922 | AX-170027905 | 0 | T | C | 0.053 | 6.33E-02 | 2.96E-01 | 2.67E+00 | 2.73E+00 | 8.29E-01 | no | | intergenic_variant | MODIFIER |
| 6 | 106860402 | AX-170530838 | 0 | G | A | 0.053 | 6.33E-02 | 2.96E-01 | 2.67E+00 | 2.73E+00 | 8.29E-01 | no | | intergenic_variant | MODIFIER |
| 6 | 106860441 | AX-97801637 | 0 | G | T | 0.053 | 6.33E-02 | 2.96E-01 | 2.67E+00 | 2.73E+00 | 8.29E-01 | no | | intergenic_variant | MODIFIER |
| 6 | 106871892 | AX-170027908 | 0 | A | C | 0.053 | 6.33E-02 | 2.96E-01 | 2.67E+00 | 2.73E+00 | 8.29E-01 | no | | intergenic_variant | MODIFIER |
| 6 | 106874654 | AX-170027904 | 0 | T | G | 0.053 | 6.33E-02 | 2.96E-01 | 2.67E+00 | 2.73E+00 | 8.29E-01 | no | | intergenic_variant | MODIFIER |
| 6 | 106875911 | AX-170150694 | 0 | C | T | 0.053 | 6.33E-02 | 2.96E-01 | 2.67E+00 | 2.73E+00 | 8.29E-01 | no | | intergenic_variant | MODIFIER |
| 6 | 106889603 | AX-170224656 | 0 | T | C | 0.053 | 6.33E-02 | 2.96E-01 | 2.67E+00 | 2.73E+00 | 8.29E-01 | no | | intergenic_variant | MODIFIER |
| 6 | 106904069 | AX-170171542 | 0 | A | G | 0.053 | 6.33E-02 | 2.96E-01 | 2.67E+00 | 2.73E+00 | 8.29E-01 | no | | intergenic_variant | MODIFIER |
| 6 | 106904181 | AX-170307843 | 0 | G | A | 0.053 | 6.33E-02 | 2.96E-01 | 2.67E+00 | 2.73E+00 | 8.29E-01 | no | | intergenic_variant | MODIFIER |
| 6 | 106923416 | AX-97786920 | 0 | A | G | 0.053 | 6.33E-02 | 2.96E-01 | 2.67E+00 | 2.73E+00 | 8.29E-01 | no | | intergenic_variant | MODIFIER |
| 6 | 106947160 | AX-170413399 | 0 | T | C | 0.053 | 6.33E-02 | 2.96E-01 | 2.67E+00 | 2.73E+00 | 8.29E-01 | no | | intergenic_variant | MODIFIER |
| 6 | 106960057 | AX-170119755 | 0 | T | G | 0.053 | 6.33E-02 | 2.96E-01 | 2.67E+00 | 2.73E+00 | 8.29E-01 | no | | intergenic_variant | MODIFIER |
| 6 | 106969253 | AX-170309272 | 0 | A | T | 0.053 | 6.33E-02 | 2.96E-01 | 2.67E+00 | 2.73E+00 | 8.29E-01 | no | | intergenic_variant | MODIFIER |
| 6 | 106971221 | AX-97797693 | 0 | T | A | 0.053 | 6.33E-02 | 2.96E-01 | 2.67E+00 | 2.73E+00 | 8.29E-01 | no | | intergenic_variant | MODIFIER |
| 6 | 106979446 | AX-170261425 | 0 | G | T | 0.053 | 6.33E-02 | 2.96E-01 | 2.67E+00 | 2.73E+00 | 8.29E-01 | no | | intergenic_variant | MODIFIER |
| 6 | 106983957 | AX-170324311 | 0 | C | T | 0.053 | 6.33E-02 | 2.96E-01 | 2.67E+00 | 2.73E+00 | 8.29E-01 | no | | intergenic_variant | MODIFIER |
| 6 | 106992745 | AX-97797834 | 0 | C | T | 0.053 | 6.33E-02 | 2.96E-01 | 2.67E+00 | 2.73E+00 | 8.29E-01 | no | | intergenic_variant | MODIFIER |
| 6 | 107018259 | AX-170318057 | 0 | G | A | 0.053 | 6.33E-02 | 2.96E-01 | 2.67E+00 | 2.73E+00 | 8.29E-01 | no | | intergenic_variant | MODIFIER |
| 6 | 107034051 | AX-170075793 | 0 | T | C | 0.053 | 6.33E-02 | 2.96E-01 | 2.67E+00 | 2.73E+00 | 8.29E-01 | no | | intergenic_variant | MODIFIER |
| 6 | 107034092 | AX-170075796 | 0 | C | T | 0.053 | 6.33E-02 | 2.96E-01 | 2.67E+00 | 2.73E+00 | 8.29E-01 | no | | intergenic_variant | MODIFIER |
| 6 | 107036012 | AX-170483671 | 0 | A | T | 0.053 | 6.33E-02 | 2.96E-01 | 2.67E+00 | 2.73E+00 | 8.29E-01 | no | | intergenic_variant | MODIFIER |
| 6 | 107037185 | AX-170244828 | 0 | C | T | 0.053 | 6.33E-02 | 2.96E-01 | 2.67E+00 | 2.73E+00 | 8.29E-01 | no | | intergenic_variant | MODIFIER |
| 6 | 107037767 | AX-170128215 | 0 | A | G | 0.053 | 6.33E-02 | 2.96E-01 | 2.67E+00 | 2.73E+00 | 8.29E-01 | no | | intergenic_variant | MODIFIER |
| 6 | 107044708 | AX-97800587 | 0 | A | G | 0.053 | 6.33E-02 | 2.96E-01 | 2.67E+00 | 2.73E+00 | 8.29E-01 | no | | intergenic_variant | MODIFIER |
| 6 | 107083575 | AX-170415772 | 0 | T | G | 0.053 | 6.33E-02 | 2.96E-01 | 2.67E+00 | 2.73E+00 | 8.29E-01 | no | | intergenic_variant | MODIFIER |
| 6 | 107092919 | AX-170251887 | 0 | A | G | 0.053 | 6.33E-02 | 2.96E-01 | 2.67E+00 | 2.73E+00 | 8.29E-01 | no | Gm6064 | downstream_gene_variant | MODIFIER |
| 6 | 107094410 | AX-170130045 | 0 | T | G | 0.053 | 6.33E-02 | 2.96E-01 | 2.67E+00 | 2.73E+00 | 8.29E-01 | no | Gm6064 | upstream_gene_variant | MODIFIER |
| 6 | 107138892 | AX-170126026 | 0 | C | T | 0.053 | 6.33E-02 | 2.96E-01 | 2.67E+00 | 2.73E+00 | 8.29E-01 | no | | intergenic_variant | MODIFIER |
| 6 | 107156958 | AX-170252182 | 0 | G | A | 0.053 | 6.33E-02 | 2.96E-01 | 2.67E+00 | 2.73E+00 | 8.29E-01 | no | | intergenic_variant | MODIFIER |
| 6 | 107162385 | AX-170118409 | 0 | A | G | 0.053 | 6.33E-02 | 2.96E-01 | 2.67E+00 | 2.73E+00 | 8.29E-01 | no | | intergenic_variant | MODIFIER |
| 6 | 107163649 | AX-170075801 | 0 | C | T | 0.053 | 6.33E-02 | 2.96E-01 | 2.67E+00 | 2.73E+00 | 8.29E-01 | no | | intergenic_variant | MODIFIER |
| 6 | 107178745 | AX-97835719 | 0 | C | T | 0.053 | 6.33E-02 | 2.96E-01 | 2.67E+00 | 2.73E+00 | 8.29E-01 | no | Gm34933 | downstream_gene_variant | MODIFIER |
| 6 | 107184110 | AX-170415582 | 0 | A | G | 0.053 | 6.33E-02 | 2.96E-01 | 2.67E+00 | 2.73E+00 | 8.29E-01 | no | Gm34933 | intron_variant,non_coding_transcript_variant | MODIFIER |
| 6 | 107205602 | AX-170110404 | 0 | G | A | 0.053 | 6.33E-02 | 2.96E-01 | 2.67E+00 | 2.73E+00 | 8.29E-01 | no | Gm34933 | intron_variant,non_coding_transcript_variant | MODIFIER |
| 6 | 107214748 | AX-170227480 | 0 | G | A | 0.053 | 6.33E-02 | 2.96E-01 | 2.67E+00 | 2.73E+00 | 8.29E-01 | no | Gm34933 | intron_variant,non_coding_transcript_variant | MODIFIER |
| 6 | 107221127 | AX-170410743 | 0 | G | A | 0.053 | 6.33E-02 | 2.96E-01 | 2.67E+00 | 2.73E+00 | 8.29E-01 | no | Gm34933 | intron_variant,non_coding_transcript_variant | MODIFIER |
| 6 | 107222146 | AX-170027911 | 0 | C | T | 0.053 | 6.33E-02 | 2.96E-01 | 2.67E+00 | 2.73E+00 | 8.29E-01 | no | Gm34933 | intron_variant,non_coding_transcript_variant | MODIFIER |

| | | | | | | | | | | | | | | | |
|---|---|---|---|---|---|---|---|---|---|---|---|---|---|---|---|
| 6 | 107230864 | AX-170206303 | 0 | C | T | 0.053 | 6.33E-02 | 2.96E-01 | 2.67E+00 | 2.73E+00 | 8.29E-01 | no | Gm34933 | intron_variant,non_coding_transcript_variant | MODIFIER |
| 6 | 107256464 | AX-170027917 | 0 | A | C | 0.053 | 6.33E-02 | 2.96E-01 | 2.67E+00 | 2.73E+00 | 8.29E-01 | no | Gm34933 | intron_variant,non_coding_transcript_variant | MODIFIER |
| 6 | 107265750 | AX-170309775 | 0 | G | A | 0.053 | 6.33E-02 | 2.96E-01 | 2.67E+00 | 2.73E+00 | 8.29E-01 | no | Gm34933 | intron_variant,non_coding_transcript_variant | MODIFIER |
| 6 | 107270557 | AX-170518736 | 0 | T | A | 0.053 | 6.33E-02 | 2.96E-01 | 2.67E+00 | 2.73E+00 | 8.29E-01 | no | Gm34933 | intron_variant,non_coding_transcript_variant | MODIFIER |
| 6 | 107283234 | AX-170476023 | 0 | T | C | 0.105 | 3.16E-02 | 1.48E-01 | 2.67E+00 | 2.73E+00 | 8.29E-01 | no | | intergenic_variant | MODIFIER |
| 6 | 107343793 | AX-170331823 | 0 | C | T | 0.053 | 6.33E-02 | 2.96E-01 | 2.67E+00 | 2.73E+00 | 8.29E-01 | no | Gm44399 | downstream_gene_variant | MODIFIER |
| 6 | 107344182 | AX-170349009 | 0 | A | T | 0.053 | 6.33E-02 | 2.96E-01 | 2.67E+00 | 2.73E+00 | 8.29E-01 | no | Gm44399 | downstream_gene_variant | MODIFIER |
| 6 | 107361482 | AX-170075802 | 0 | G | A | 0.053 | 6.33E-02 | 2.96E-01 | 2.67E+00 | 2.73E+00 | 8.29E-01 | no | | intergenic_variant | MODIFIER |
| 6 | 107365793 | AX-170145718 | 0 | C | T | 0.053 | 6.33E-02 | 2.96E-01 | 2.67E+00 | 2.73E+00 | 8.29E-01 | no | | intergenic_variant | MODIFIER |
| 6 | 107372700 | AX-97825116 | 0 | T | G | 0.053 | 6.33E-02 | 2.96E-01 | 2.67E+00 | 2.73E+00 | 8.29E-01 | no | | intergenic_variant | MODIFIER |
| 6 | 107423321 | AX-170303518 | 0 | C | T | 0.053 | 6.33E-02 | 2.96E-01 | 2.67E+00 | 2.73E+00 | 8.29E-01 | no | | intergenic_variant | MODIFIER |
| 6 | 107427827 | AX-97805300 | 0 | G | T | 0.053 | 6.33E-02 | 2.96E-01 | 2.67E+00 | 2.73E+00 | 8.29E-01 | no | | intergenic_variant | MODIFIER |
| 6 | 107444306 | AX-97789680 | 0 | A | G | 0.053 | 6.33E-02 | 2.96E-01 | 2.67E+00 | 2.73E+00 | 8.29E-01 | no | Gm27690 | downstream_gene_variant | MODIFIER |
| 6 | 107447584 | AX-170114896 | 0 | T | G | 0.053 | 6.33E-02 | 2.96E-01 | 2.67E+00 | 2.73E+00 | 8.29E-01 | no | Gm27690 | downstream_gene_variant | MODIFIER |
| 6 | 107534174 | AX-97845725 | 0 | G | T | 0.053 | 6.33E-02 | 2.96E-01 | 2.67E+00 | 2.73E+00 | 8.29E-01 | no | Lrrn1 | intron_variant | MODIFIER |
| 6 | 107555540 | AX-170075803 | 0 | T | G | 0.053 | 6.33E-02 | 2.96E-01 | 2.67E+00 | 2.73E+00 | 8.29E-01 | no | Lrrn1 | intron_variant | MODIFIER |
| 6 | 107634991 | AX-170142775 | 0 | G | A | 0.053 | 6.33E-02 | 2.96E-01 | 2.67E+00 | 2.73E+00 | 8.29E-01 | no | 4933431M02Rik | downstream_gene_variant | MODIFIER |
| 6 | 107636570 | AX-170274731 | 0 | A | G | 0.053 | 6.33E-02 | 2.96E-01 | 2.67E+00 | 2.73E+00 | 8.29E-01 | no | 4933431M02Rik | downstream_gene_variant | MODIFIER |
| 6 | 109906225 | AX-170505049 | 0 | G | C | 0.053 | 6.33E-02 | 2.96E-01 | 2.67E+00 | 2.73E+00 | 8.29E-01 | no | | intergenic_variant | MODIFIER |
| 6 | 109906411 | AX-97834003 | 0 | T | C | 0.053 | 6.33E-02 | 2.96E-01 | 2.67E+00 | 2.73E+00 | 8.29E-01 | no | | intergenic_variant | MODIFIER |
| 6 | 109906529 | AX-170075850 | 0 | G | A | 0.053 | 6.33E-02 | 2.96E-01 | 2.67E+00 | 2.73E+00 | 8.29E-01 | no | | intergenic_variant | MODIFIER |
| 6 | 109942948 | AX-170027961 | 0 | T | G | 0.053 | 6.33E-02 | 2.96E-01 | 2.67E+00 | 2.73E+00 | 8.29E-01 | no | | intergenic_variant | MODIFIER |
| 6 | 109950984 | AX-170465377 | 0 | A | G | 0.053 | 6.33E-02 | 2.96E-01 | 2.67E+00 | 2.73E+00 | 8.29E-01 | no | | intergenic_variant | MODIFIER |
| 6 | 109951898 | AX-170351147 | 0 | G | A | 0.053 | 6.33E-02 | 2.96E-01 | 2.67E+00 | 2.73E+00 | 8.29E-01 | no | | intergenic_variant | MODIFIER |
| 6 | 109957636 | AX-170415640 | 0 | G | C | 0.053 | 6.33E-02 | 2.96E-01 | 2.67E+00 | 2.73E+00 | 8.29E-01 | no | | intergenic_variant | MODIFIER |
| 6 | 109977158 | AX-97791159 | 0 | A | G | 0.053 | 6.33E-02 | 2.96E-01 | 2.67E+00 | 2.73E+00 | 8.29E-01 | no | | intergenic_variant | MODIFIER |
| 6 | 110015479 | AX-97815666 | 0 | T | G | 0.053 | 6.33E-02 | 2.96E-01 | 2.67E+00 | 2.73E+00 | 8.29E-01 | no | | intergenic_variant | MODIFIER |
| 6 | 110031206 | AX-170413989 | 0 | T | C | 0.053 | 6.33E-02 | 2.96E-01 | 2.67E+00 | 2.73E+00 | 8.29E-01 | no | Gm44162 | upstream_gene_variant | MODIFIER |
| 6 | 110041936 | AX-170447904 | 0 | C | T | 0.053 | 6.33E-02 | 2.96E-01 | 2.67E+00 | 2.73E+00 | 8.29E-01 | no | | intergenic_variant | MODIFIER |
| 6 | 110047474 | AX-170027956 | 0 | C | G | 0.053 | 6.33E-02 | 2.96E-01 | 2.67E+00 | 2.73E+00 | 8.29E-01 | no | | intergenic_variant | MODIFIER |
| 6 | 110048148 | AX-170135453 | 0 | T | G | 0.053 | 6.33E-02 | 2.96E-01 | 2.67E+00 | 2.73E+00 | 8.29E-01 | no | | intergenic_variant | MODIFIER |
| 6 | 110052798 | AX-170269748 | 0 | A | G | 0.053 | 6.33E-02 | 2.96E-01 | 2.67E+00 | 2.73E+00 | 8.29E-01 | no | | intergenic_variant | MODIFIER |
| 6 | 110085601 | AX-97835439 | 0 | G | A | 0.053 | 6.33E-02 | 2.96E-01 | 2.67E+00 | 2.73E+00 | 8.29E-01 | no | | intergenic_variant | MODIFIER |
| 6 | 110099658 | AX-170208735 | 0 | T | C | 0.053 | 6.33E-02 | 2.96E-01 | 2.67E+00 | 2.73E+00 | 8.29E-01 | no | | intergenic_variant | MODIFIER |
| 6 | 110124755 | AX-170280709 | 0 | A | G | 0.053 | 6.33E-02 | 2.96E-01 | 2.67E+00 | 2.73E+00 | 8.29E-01 | no | | intergenic_variant | MODIFIER |
| 6 | 110130241 | AX-170266235 | 0 | A | G | 0.053 | 6.33E-02 | 2.96E-01 | 2.67E+00 | 2.73E+00 | 8.29E-01 | no | | intergenic_variant | MODIFIER |
| 6 | 110135999 | AX-170344325 | 0 | A | G | 0.053 | 6.33E-02 | 2.96E-01 | 2.67E+00 | 2.73E+00 | 8.29E-01 | no | | intergenic_variant | MODIFIER |
| 6 | 110143071 | AX-170235338 | 0 | A | T | 0.053 | 6.33E-02 | 2.96E-01 | 2.67E+00 | 2.73E+00 | 8.29E-01 | no | | intergenic_variant | MODIFIER |
| 6 | 110163199 | AX-170320225 | 0 | A | C | 0.053 | 6.33E-02 | 2.96E-01 | 2.67E+00 | 2.73E+00 | 8.29E-01 | no | | intergenic_variant | MODIFIER |
| 6 | 110170202 | AX-170525263 | 0 | G | A | 0.053 | 6.33E-02 | 2.96E-01 | 2.67E+00 | 2.73E+00 | 8.29E-01 | no | | intergenic_variant | MODIFIER |
| 6 | 110174917 | AX-170391495 | 0 | C | T | 0.053 | 6.33E-02 | 2.96E-01 | 2.67E+00 | 2.73E+00 | 8.29E-01 | no | | intergenic_variant | MODIFIER |
| 6 | 110202989 | AX-170219230 | 0 | T | C | 0.053 | 6.33E-02 | 2.96E-01 | 2.67E+00 | 2.73E+00 | 8.29E-01 | no | | intergenic_variant | MODIFIER |
| 6 | 110228463 | AX-170095752 | 0 | C | T | 0.053 | 6.33E-02 | 2.96E-01 | 2.67E+00 | 2.73E+00 | 8.29E-01 | no | | regulatory_region_variant | MODIFIER |
| 6 | 110241592 | AX-170126042 | 0 | A | G | 0.053 | 6.33E-02 | 2.96E-01 | 2.67E+00 | 2.73E+00 | 8.29E-01 | no | | intergenic_variant | MODIFIER |
| 6 | 110248625 | AX-170436747 | 0 | A | G | 0.053 | 6.33E-02 | 2.96E-01 | 2.67E+00 | 2.73E+00 | 8.29E-01 | no | | intergenic_variant | MODIFIER |
| 6 | 110892982 | AX-170365016 | 0 | T | C | 0.053 | 6.33E-02 | 2.96E-01 | 2.67E+00 | 2.73E+00 | 8.29E-01 | no | Grm7 | intron_variant | MODIFIER |
| 6 | 110898404 | AX-170442705 | 0 | A | G | 0.053 | 6.33E-02 | 2.96E-01 | 2.67E+00 | 2.73E+00 | 8.29E-01 | no | Grm7 | intron_variant | MODIFIER |
| 6 | 110899179 | AX-170514105 | 0 | C | G | 0.053 | 6.33E-02 | 2.96E-01 | 2.67E+00 | 2.73E+00 | 8.29E-01 | no | Grm7 | intron_variant | MODIFIER |
| 6 | 110900049 | AX-170475856 | 0 | G | T | 0.053 | 6.33E-02 | 2.96E-01 | 2.67E+00 | 2.73E+00 | 8.29E-01 | no | Grm7 | intron_variant | MODIFIER |
| 6 | 110917939 | AX-170216736 | 0 | A | C | 0.053 | 6.33E-02 | 2.96E-01 | 2.67E+00 | 2.73E+00 | 8.29E-01 | no | Grm7 | intron_variant | MODIFIER |
| 6 | 110944130 | AX-170310258 | 0 | T | A | 0.053 | 6.33E-02 | 2.96E-01 | 2.67E+00 | 2.73E+00 | 8.29E-01 | no | Grm7 | intron_variant | MODIFIER |
| 6 | 110966239 | AX-170027983 | 0 | T | G | 0.053 | 6.33E-02 | 2.96E-01 | 2.67E+00 | 2.73E+00 | 8.29E-01 | no | Grm7 | intron_variant | MODIFIER |
| 6 | 111043155 | AX-170027977 | 0 | G | A | 0.053 | 6.33E-02 | 2.96E-01 | 2.67E+00 | 2.73E+00 | 8.29E-01 | no | Grm7 | intron_variant | MODIFIER |
| 6 | 111061586 | AX-170027975 | 0 | C | T | 0.053 | 6.33E-02 | 2.96E-01 | 2.67E+00 | 2.73E+00 | 8.29E-01 | no | Grm7 | intron_variant | MODIFIER |

| | | | | | | | | | | | | | | | |
|---|---|---|---|---|---|---|---|---|---|---|---|---|---|---|---|
| 6 | 111094495 | AX-170089813 | 0 | A | G | 0.053 | 6.33E-02 | 2.96E-01 | 2.67E+00 | 2.73E+00 | 8.29E-01 | no | Grm7 | intron_variant | MODIFIER |
| 6 | 111133554 | AX-170194835 | 0 | A | G | 0.053 | 6.33E-02 | 2.96E-01 | 2.67E+00 | 2.73E+00 | 8.29E-01 | no | Grm7 | intron_variant | MODIFIER |
| 6 | 111133727 | AX-170295206 | 0 | A | G | 0.053 | 6.33E-02 | 2.96E-01 | 2.67E+00 | 2.73E+00 | 8.29E-01 | no | Grm7 | intron_variant | MODIFIER |
| 6 | 111133894 | AX-170278428 | 0 | C | T | 0.053 | 6.33E-02 | 2.96E-01 | 2.67E+00 | 2.73E+00 | 8.29E-01 | no | Grm7 | intron_variant | MODIFIER |
| 6 | 111192452 | AX-170508930 | 0 | C | T | 0.053 | 6.33E-02 | 2.96E-01 | 2.67E+00 | 2.73E+00 | 8.29E-01 | no | Grm7 | intron_variant | MODIFIER |
| 6 | 111192727 | AX-170313059 | 0 | T | C | 0.053 | 6.33E-02 | 2.96E-01 | 2.67E+00 | 2.73E+00 | 8.29E-01 | no | Grm7 | intron_variant | MODIFIER |
| 6 | 111193489 | AX-170438996 | 0 | A | G | 0.053 | 6.33E-02 | 2.96E-01 | 2.67E+00 | 2.73E+00 | 8.29E-01 | no | Grm7 | intron_variant | MODIFIER |
| 6 | 111193743 | AX-170075872 | 0 | T | G | 0.053 | 6.33E-02 | 2.96E-01 | 2.67E+00 | 2.73E+00 | 8.29E-01 | no | Grm7 | intron_variant | MODIFIER |
| 6 | 111274513 | AX-97851806 | 0 | C | T | 0.053 | 6.33E-02 | 2.96E-01 | 2.67E+00 | 2.73E+00 | 8.29E-01 | no | Grm7 | intron_variant | MODIFIER |
| 6 | 111335374 | AX-170273358 | 0 | C | T | 0.053 | 6.33E-02 | 2.96E-01 | 2.67E+00 | 2.73E+00 | 8.29E-01 | no | Grm7 | intron_variant | MODIFIER |
| 6 | 111369028 | AX-97837392 | 0 | A | G | 0.053 | 6.33E-02 | 2.96E-01 | 2.67E+00 | 2.73E+00 | 8.29E-01 | no | Grm7 | intron_variant | MODIFIER |
| 6 | 111433959 | AX-97816745 | 0 | T | G | 0.053 | 6.33E-02 | 2.96E-01 | 2.67E+00 | 2.73E+00 | 8.29E-01 | no | Grm7 | intron_variant | MODIFIER |
| 6 | 111437371 | AX-170209565 | 0 | C | T | 0.053 | 6.33E-02 | 2.96E-01 | 2.67E+00 | 2.73E+00 | 8.29E-01 | no | Grm7 | intron_variant | MODIFIER |
| 6 | 111511082 | AX-170370701 | 0 | C | T | 0.053 | 6.33E-02 | 2.96E-01 | 2.67E+00 | 2.73E+00 | 8.29E-01 | no | Grm7 | intron_variant | MODIFIER |
| 6 | 111527649 | AX-170410522 | 0 | A | G | 0.053 | 6.33E-02 | 2.96E-01 | 2.67E+00 | 2.73E+00 | 8.29E-01 | no | Grm7 | intron_variant | MODIFIER |
| 6 | 111529221 | AX-170232535 | 0 | T | C | 0.053 | 6.33E-02 | 2.96E-01 | 2.67E+00 | 2.73E+00 | 8.29E-01 | no | Grm7 | intron_variant | MODIFIER |
| 6 | 111530012 | AX-170284960 | 0 | T | C | 0.053 | 6.33E-02 | 2.96E-01 | 2.67E+00 | 2.73E+00 | 8.29E-01 | no | Grm7 | intron_variant | MODIFIER |
| 6 | 111530402 | AX-170480819 | 0 | C | T | 0.053 | 6.33E-02 | 2.96E-01 | 2.67E+00 | 2.73E+00 | 8.29E-01 | no | Grm7 | intron_variant | MODIFIER |
| 6 | 111531239 | AX-170453385 | 0 | T | C | 0.053 | 6.33E-02 | 2.96E-01 | 2.67E+00 | 2.73E+00 | 8.29E-01 | no | Grm7 | intron_variant | MODIFIER |
| 6 | 111683593 | AX-170075882 | 0 | C | T | 0.053 | 6.33E-02 | 2.96E-01 | 2.67E+00 | 2.73E+00 | 8.29E-01 | no | | intergenic_variant | MODIFIER |
| 6 | 111688453 | AX-170336899 | 0 | C | T | 0.053 | 6.33E-02 | 2.96E-01 | 2.67E+00 | 2.73E+00 | 8.29E-01 | no | | intergenic_variant | MODIFIER |
| 6 | 111688538 | AX-97801698 | 0 | C | T | 0.053 | 6.33E-02 | 2.96E-01 | 2.67E+00 | 2.73E+00 | 8.29E-01 | no | | intergenic_variant | MODIFIER |
| 6 | 111688935 | AX-170075880 | 0 | T | A | 0.053 | 6.33E-02 | 2.96E-01 | 2.67E+00 | 2.73E+00 | 8.29E-01 | no | | intergenic_variant | MODIFIER |
| 6 | 111733060 | AX-170315928 | 0 | T | G | 0.053 | 6.33E-02 | 2.96E-01 | 2.67E+00 | 2.73E+00 | 8.29E-01 | no | | intergenic_variant | MODIFIER |
| 6 | 111743133 | AX-170415072 | 0 | C | G | 0.053 | 6.33E-02 | 2.96E-01 | 2.67E+00 | 2.73E+00 | 8.29E-01 | no | | intergenic_variant | MODIFIER |
| 6 | 111757265 | AX-170443596 | 0 | G | A | 0.053 | 6.33E-02 | 2.96E-01 | 2.67E+00 | 2.73E+00 | 8.29E-01 | no | | intergenic_variant | MODIFIER |
| 6 | 112215814 | AX-170334949 | 0 | G | A | 0.053 | 6.33E-02 | 2.96E-01 | 2.67E+00 | 2.73E+00 | 8.29E-01 | no | 1700054K19Rik | downstream_gene_variant | MODIFIER |
| 6 | 112224014 | AX-97784596 | 0 | G | A | 0.053 | 6.33E-02 | 2.96E-01 | 2.67E+00 | 2.73E+00 | 8.29E-01 | no | | intergenic_variant | MODIFIER |
| 6 | 112247216 | AX-170382966 | 0 | G | A | 0.053 | 6.33E-02 | 2.96E-01 | 2.67E+00 | 2.73E+00 | 8.29E-01 | no | | intergenic_variant | MODIFIER |
| 6 | 112254629 | AX-170415068 | 0 | G | C | 0.053 | 6.33E-02 | 2.96E-01 | 2.67E+00 | 2.73E+00 | 8.29E-01 | no | | intergenic_variant | MODIFIER |
| 6 | 112255034 | AX-170154854 | 0 | C | A | 0.053 | 6.33E-02 | 2.96E-01 | 2.67E+00 | 2.73E+00 | 8.29E-01 | no | | intergenic_variant | MODIFIER |
| 6 | 112256458 | AX-170207081 | 0 | G | A | 0.053 | 6.33E-02 | 2.96E-01 | 2.67E+00 | 2.73E+00 | 8.29E-01 | no | | regulatory_region_variant | MODIFIER |
| 6 | 112258137 | AX-170095757 | 0 | A | G | 0.053 | 6.33E-02 | 2.96E-01 | 2.67E+00 | 2.73E+00 | 8.29E-01 | no | | intergenic_variant | MODIFIER |
| 6 | 112261537 | AX-170434874 | 0 | T | C | 0.053 | 6.33E-02 | 2.96E-01 | 2.67E+00 | 2.73E+00 | 8.29E-01 | no | | intergenic_variant | MODIFIER |
| 6 | 112272910 | AX-170171235 | 0 | C | T | 0.053 | 6.33E-02 | 2.96E-01 | 2.67E+00 | 2.73E+00 | 8.29E-01 | no | Lmcd1 | upstream_gene_variant | MODIFIER |
| 6 | 112275452 | AX-170445136 | 0 | T | C | 0.053 | 6.33E-02 | 2.96E-01 | 2.67E+00 | 2.73E+00 | 8.29E-01 | no | Lmcd1 | intron_variant | MODIFIER |
| 6 | 112278283 | AX-170095758 | 0 | G | T | 0.053 | 6.33E-02 | 2.96E-01 | 2.67E+00 | 2.73E+00 | 8.29E-01 | no | Lmcd1 | intron_variant | MODIFIER |
| 6 | 112296616 | AX-97847195 | 0 | C | G | 0.053 | 6.33E-02 | 2.96E-01 | 2.67E+00 | 2.73E+00 | 8.29E-01 | no | Lmcd1 | intron_variant | MODIFIER |
| 6 | 112542720 | AX-170322800 | 0 | T | A | 0.053 | 6.33E-02 | 2.96E-01 | 2.67E+00 | 2.73E+00 | 8.29E-01 | no | | intergenic_variant | MODIFIER |
| 6 | 112552811 | AX-170329631 | 0 | A | G | 0.053 | 6.33E-02 | 2.96E-01 | 2.67E+00 | 2.73E+00 | 8.29E-01 | no | | regulatory_region_variant | MODIFIER |
| 6 | 112582341 | AX-97803624 | 0 | G | A | 0.053 | 6.33E-02 | 2.96E-01 | 2.67E+00 | 2.73E+00 | 8.29E-01 | no | | intergenic_variant | MODIFIER |
| 6 | 112582423 | AX-170299387 | 0 | T | A | 0.053 | 6.33E-02 | 2.96E-01 | 2.67E+00 | 2.73E+00 | 8.29E-01 | no | | intergenic_variant | MODIFIER |
| 6 | 112582523 | AX-170027999 | 0 | C | A | 0.053 | 6.33E-02 | 2.96E-01 | 2.67E+00 | 2.73E+00 | 8.29E-01 | no | | intergenic_variant | MODIFIER |
| 6 | 112728122 | AX-170478156 | 0 | C | G | 0.053 | 6.33E-02 | 2.96E-01 | 2.67E+00 | 2.73E+00 | 8.29E-01 | no | Srgap3 | intron_variant | MODIFIER |
| 6 | 112732187 | AX-97808747 | 0 | T | C | 0.053 | 6.33E-02 | 2.96E-01 | 2.67E+00 | 2.73E+00 | 8.29E-01 | no | Srgap3 | intron_variant | MODIFIER |
| 6 | 112755669 | AX-97808398 | 0 | A | G | 0.053 | 6.33E-02 | 2.96E-01 | 2.67E+00 | 2.73E+00 | 8.29E-01 | no | Srgap3 | intron_variant | MODIFIER |
| 6 | 112755769 | AX-170483005 | 0 | C | G | 0.053 | 6.33E-02 | 2.96E-01 | 2.67E+00 | 2.73E+00 | 8.29E-01 | no | Srgap3 | intron_variant | MODIFIER |
| 6 | 112768316 | AX-170277068 | 0 | T | C | 0.053 | 6.33E-02 | 2.96E-01 | 2.67E+00 | 2.73E+00 | 8.29E-01 | no | Srgap3 | intron_variant,non_coding_transcript_variant | MODIFIER |
| 6 | 112774083 | AX-97783172 | 0 | C | T | 0.053 | 6.33E-02 | 2.96E-01 | 2.67E+00 | 2.73E+00 | 8.29E-01 | no | Srgap3 | intron_variant,non_coding_transcript_variant | MODIFIER |
| 6 | 112790550 | AX-170415028 | 0 | G | A | 0.053 | 6.33E-02 | 2.96E-01 | 2.67E+00 | 2.73E+00 | 8.29E-01 | no | Srgap3 | intron_variant,non_coding_transcript_variant | MODIFIER |
| 6 | 112792944 | AX-170436330 | 0 | T | A | 0.053 | 6.33E-02 | 2.96E-01 | 2.67E+00 | 2.73E+00 | 8.29E-01 | no | Srgap3 | intron_variant,non_coding_transcript_variant | MODIFIER |
| 6 | 112796750 | AX-170177746 | 0 | A | G | 0.053 | 6.33E-02 | 2.96E-01 | 2.67E+00 | 2.73E+00 | 8.29E-01 | no | Srgap3 | intron_variant,non_coding_transcript_variant | MODIFIER |
| 6 | 112796812 | AX-170161702 | 0 | G | A | 0.053 | 6.33E-02 | 2.96E-01 | 2.67E+00 | 2.73E+00 | 8.29E-01 | no | Srgap3 | intron_variant,non_coding_transcript_variant | MODIFIER |
| 6 | 112800137 | AX-97822052 | 0 | A | G | 0.053 | 6.33E-02 | 2.96E-01 | 2.67E+00 | 2.73E+00 | 8.29E-01 | no | Srgap3 | intron_variant,non_coding_transcript_variant | MODIFIER |

| | | | | | | | | | | | | | | | |
|---|---|---|---|---|---|---|---|---|---|---|---|---|---|---|---|
| 6 | 112809221 | AX-170278087 | 0 | G | A | 0.053 | 6.33E-02 | 2.96E-01 | 2.67E+00 | 2.73E+00 | 8.29E-01 | no | Srgap3 | intron_variant,non_coding_transcript_variant | MODIFIER |
| 6 | 112922650 | AX-170209475 | 0 | C | G | 0.053 | 6.33E-02 | 2.96E-01 | 2.67E+00 | 2.73E+00 | 8.29E-01 | no | Srgap3 | intron_variant,non_coding_transcript_variant | MODIFIER |
| 6 | 112927300 | AX-97787604 | 0 | C | T | 0.053 | 6.33E-02 | 2.96E-01 | 2.67E+00 | 2.73E+00 | 8.29E-01 | no | Srgap3 | intron_variant,non_coding_transcript_variant | MODIFIER |
| 6 | 112927734 | AX-170243179 | 0 | C | G | 0.053 | 6.33E-02 | 2.96E-01 | 2.67E+00 | 2.73E+00 | 8.29E-01 | no | Srgap3 | intron_variant,non_coding_transcript_variant | MODIFIER |
| 6 | 112929852 | AX-170159952 | 0 | G | A | 0.053 | 6.33E-02 | 2.96E-01 | 2.67E+00 | 2.73E+00 | 8.29E-01 | no | Srgap3 | intron_variant,non_coding_transcript_variant | MODIFIER |
| 6 | 112931383 | AX-170497888 | 0 | T | C | 0.053 | 6.33E-02 | 2.96E-01 | 2.67E+00 | 2.73E+00 | 8.29E-01 | no | Srgap3 | intron_variant,non_coding_transcript_variant | MODIFIER |
| 6 | 112942815 | AX-170520137 | 0 | T | C | 0.053 | 6.33E-02 | 2.96E-01 | 2.67E+00 | 2.73E+00 | 8.29E-01 | no | Srgap3 | intron_variant,non_coding_transcript_variant | MODIFIER |
| 6 | 112942861 | AX-170473583 | 0 | A | G | 0.053 | 6.33E-02 | 2.96E-01 | 2.67E+00 | 2.73E+00 | 8.29E-01 | no | Srgap3 | intron_variant,non_coding_transcript_variant | MODIFIER |
| 6 | 112959451 | AX-97839623 | 0 | C | T | 0.053 | 6.33E-02 | 2.96E-01 | 2.67E+00 | 2.73E+00 | 8.29E-01 | no | | intergenic_variant | MODIFIER |
| 6 | 112959523 | AX-170184870 | 0 | G | A | 0.053 | 6.33E-02 | 2.96E-01 | 2.67E+00 | 2.73E+00 | 8.29E-01 | no | | intergenic_variant | MODIFIER |
| 6 | 112968040 | AX-170324044 | 0 | A | T | 0.053 | 6.33E-02 | 2.96E-01 | 2.67E+00 | 2.73E+00 | 8.29E-01 | no | | intergenic_variant | MODIFIER |
| 6 | 113000595 | AX-170210877 | 0 | T | C | 0.053 | 6.33E-02 | 2.96E-01 | 2.67E+00 | 2.73E+00 | 8.29E-01 | no | | regulatory_region_variant | MODIFIER |
| 6 | 113000627 | AX-170243426 | 0 | G | A | 0.053 | 6.33E-02 | 2.96E-01 | 2.67E+00 | 2.73E+00 | 8.29E-01 | no | | regulatory_region_variant | MODIFIER |
| 6 | 113000754 | AX-170496293 | 0 | T | C | 0.053 | 6.33E-02 | 2.96E-01 | 2.67E+00 | 2.73E+00 | 8.29E-01 | no | | regulatory_region_variant | MODIFIER |
| 6 | 113000850 | AX-170157366 | 0 | A | C | 0.053 | 6.33E-02 | 2.96E-01 | 2.67E+00 | 2.73E+00 | 8.29E-01 | no | | regulatory_region_variant | MODIFIER |
| 6 | 113001329 | AX-170095762 | 0 | G | A | 0.053 | 6.33E-02 | 2.96E-01 | 2.67E+00 | 2.73E+00 | 8.29E-01 | no | | regulatory_region_variant | MODIFIER |
| 6 | 113187870 | AX-170028003 | 0 | C | T | 0.053 | 6.33E-02 | 2.96E-01 | 2.67E+00 | 2.73E+00 | 8.29E-01 | no | Lhfpl4 | intron_variant | MODIFIER |
| 6 | 113211838 | AX-97787275 | 0 | G | T | 0.053 | 6.33E-02 | 2.96E-01 | 2.67E+00 | 2.73E+00 | 8.29E-01 | no | | intergenic_variant | MODIFIER |
| 6 | 113223168 | AX-97821313 | 0 | A | G | 0.053 | 6.33E-02 | 2.96E-01 | 2.67E+00 | 2.73E+00 | 8.29E-01 | no | | intergenic_variant | MODIFIER |
| 6 | 113238731 | AX-170028016 | 0 | G | A | 0.053 | 6.33E-02 | 2.96E-01 | 2.67E+00 | 2.73E+00 | 8.29E-01 | no | Mtmr14 | intron_variant | MODIFIER |
| 6 | 113239370 | AX-170028012 | 0 | T | G | 0.053 | 6.33E-02 | 2.96E-01 | 2.67E+00 | 2.73E+00 | 8.29E-01 | no | Mtmr14 | intron_variant | MODIFIER |
| 6 | 113243591 | AX-170028013 | 0 | G | A | 0.053 | 6.33E-02 | 2.96E-01 | 2.67E+00 | 2.73E+00 | 8.29E-01 | no | Mtmr14 | intron_variant | MODIFIER |
| 6 | 113273276 | AX-170438311 | 0 | G | T | 0.053 | 6.33E-02 | 2.96E-01 | 2.67E+00 | 2.73E+00 | 8.29E-01 | no | Mtmr14 | intron_variant | MODIFIER |
| 6 | 113277730 | AX-170028010 | 0 | A | G | 0.053 | 6.33E-02 | 2.96E-01 | 2.67E+00 | 2.73E+00 | 8.29E-01 | no | Cpne9 | upstream_gene_variant | MODIFIER |
| 6 | 114027405 | AX-97803920 | 0 | C | T | 0.053 | 6.33E-02 | 2.96E-01 | 2.67E+00 | 2.73E+00 | 8.29E-01 | no | Atp2b2 | intron_variant | MODIFIER |
| 6 | 114031043 | AX-170156913 | 0 | C | G | 0.053 | 6.33E-02 | 2.96E-01 | 2.67E+00 | 2.73E+00 | 8.29E-01 | no | Atp2b2 | intron_variant | MODIFIER |
| 6 | 114035202 | AX-170290280 | 0 | C | T | 0.053 | 6.33E-02 | 2.96E-01 | 2.67E+00 | 2.73E+00 | 8.29E-01 | no | Atp2b2 | intron_variant | MODIFIER |
| 6 | 114035407 | AX-170128262 | 0 | G | A | 0.053 | 6.33E-02 | 2.96E-01 | 2.67E+00 | 2.73E+00 | 8.29E-01 | no | Atp2b2 | intron_variant | MODIFIER |
| 6 | 114035900 | AX-170484797 | 0 | A | G | 0.053 | 6.33E-02 | 2.96E-01 | 2.67E+00 | 2.73E+00 | 8.29E-01 | no | Atp2b2 | intron_variant | MODIFIER |
| 6 | 114036106 | AX-170261746 | 0 | G | A | 0.053 | 6.33E-02 | 2.96E-01 | 2.67E+00 | 2.73E+00 | 8.29E-01 | no | Atp2b2 | intron_variant | MODIFIER |
| 6 | 114036243 | AX-170441014 | 0 | T | C | 0.053 | 6.33E-02 | 2.96E-01 | 2.67E+00 | 2.73E+00 | 8.29E-01 | no | Atp2b2 | intron_variant | MODIFIER |
| 6 | 114036271 | AX-170191220 | 0 | T | C | 0.053 | 6.33E-02 | 2.96E-01 | 2.67E+00 | 2.73E+00 | 8.29E-01 | no | Atp2b2 | intron_variant | MODIFIER |
| 6 | 114039273 | AX-170330570 | 0 | C | T | 0.053 | 6.33E-02 | 2.96E-01 | 2.67E+00 | 2.73E+00 | 8.29E-01 | no | Atp2b2 | intron_variant | MODIFIER |
| 6 | 114046835 | AX-97837399 | 0 | A | G | 0.053 | 6.33E-02 | 2.96E-01 | 2.67E+00 | 2.73E+00 | 8.29E-01 | no | Atp2b2 | upstream_gene_variant | MODIFIER |
| 6 | 114099026 | AX-170028027 | 0 | T | C | 0.053 | 6.33E-02 | 2.96E-01 | 2.67E+00 | 2.73E+00 | 8.29E-01 | no | | intergenic_variant | MODIFIER |
| 6 | 114099112 | AX-97839388 | 0 | G | A | 0.053 | 6.33E-02 | 2.96E-01 | 2.67E+00 | 2.73E+00 | 8.29E-01 | no | | intergenic_variant | MODIFIER |
| 6 | 114128449 | AX-97802149 | 0 | A | G | 0.053 | 6.33E-02 | 2.96E-01 | 2.67E+00 | 2.73E+00 | 8.29E-01 | no | Slc6a11 | upstream_gene_variant | MODIFIER |
| 6 | 114161319 | AX-97815822 | 0 | A | G | 0.053 | 6.33E-02 | 2.96E-01 | 2.67E+00 | 2.73E+00 | 8.29E-01 | no | Slc6a11 | intron_variant | MODIFIER |
| 6 | 114166977 | AX-170028022 | 0 | A | G | 0.053 | 6.33E-02 | 2.96E-01 | 2.67E+00 | 2.73E+00 | 8.29E-01 | no | Slc6a11 | intron_variant | MODIFIER |
| 6 | 114245847 | AX-170395493 | 0 | A | T | 0.053 | 6.33E-02 | 2.96E-01 | 2.67E+00 | 2.73E+00 | 8.29E-01 | no | Slc6a11 | intron_variant | MODIFIER |
| 6 | 114246178 | AX-170028029 | 0 | C | T | 0.053 | 6.33E-02 | 2.96E-01 | 2.67E+00 | 2.73E+00 | 8.29E-01 | no | Slc6a11 | intron_variant | MODIFIER |
| 6 | 114248251 | AX-97787551 | 0 | T | C | 0.053 | 6.33E-02 | 2.96E-01 | 2.67E+00 | 2.73E+00 | 8.29E-01 | no | Slc6a11 | 3_prime_UTR_variant | MODIFIER |
| 6 | 114281151 | AX-97829072 | 0 | G | T | 0.053 | 6.33E-02 | 2.96E-01 | 2.67E+00 | 2.73E+00 | 8.29E-01 | no | Slc6a1 | upstream_gene_variant | MODIFIER |
| 6 | 114330193 | AX-170089816 | 0 | A | G | 0.053 | 6.33E-02 | 2.96E-01 | 2.67E+00 | 2.73E+00 | 8.29E-01 | no | | regulatory_region_variant | MODIFIER |
| 6 | 114356944 | AX-170261261 | 0 | G | A | 0.053 | 6.33E-02 | 2.96E-01 | 2.67E+00 | 2.73E+00 | 8.29E-01 | no | | intergenic_variant | MODIFIER |
| 6 | 114366861 | AX-170353417 | 0 | C | T | 0.053 | 6.33E-02 | 2.96E-01 | 2.67E+00 | 2.73E+00 | 8.29E-01 | no | | regulatory_region_variant | MODIFIER |
| 6 | 114372810 | AX-170237541 | 0 | A | G | 0.053 | 6.33E-02 | 2.96E-01 | 2.67E+00 | 2.73E+00 | 8.29E-01 | no | | intergenic_variant | MODIFIER |
| 6 | 114373547 | AX-170119236 | 0 | C | A | 0.053 | 6.33E-02 | 2.96E-01 | 2.67E+00 | 2.73E+00 | 8.29E-01 | no | | intergenic_variant | MODIFIER |
| 6 | 114378013 | AX-170334573 | 0 | G | A | 0.053 | 6.33E-02 | 2.96E-01 | 2.67E+00 | 2.73E+00 | 8.29E-01 | no | | intergenic_variant | MODIFIER |
| 6 | 114381236 | AX-97793701 | 0 | T | C | 0.105 | 3.16E-02 | 1.48E-01 | 2.67E+00 | 2.73E+00 | 8.29E-01 | no | | intergenic_variant | MODIFIER |
| 6 | 114381262 | AX-170108398 | 0 | C | T | 0.053 | 6.33E-02 | 2.96E-01 | 2.67E+00 | 2.73E+00 | 8.29E-01 | no | | intergenic_variant | MODIFIER |
| 6 | 114407245 | AX-170307271 | 0 | A | G | 0.053 | 6.33E-02 | 2.96E-01 | 2.67E+00 | 2.73E+00 | 8.29E-01 | no | Hrh1 | intron_variant | MODIFIER |
| 6 | 114409488 | AX-170501844 | 0 | C | G | 0.053 | 6.33E-02 | 2.96E-01 | 2.67E+00 | 2.73E+00 | 8.29E-01 | no | Hrh1 | intron_variant | MODIFIER |
| 6 | 114423814 | AX-170394816 | 0 | A | G | 0.053 | 6.33E-02 | 2.96E-01 | 2.67E+00 | 2.73E+00 | 8.29E-01 | no | Hrh1 | intron_variant | MODIFIER |

| | | | | | | | | | | | | | | | |
|---|---|---|---|---|---|---|---|---|---|---|---|---|---|---|---|
| 6 | 114437216 | AX-170327477 | 0 | G | T | 0.053 | 6.33E-02 | 2.96E-01 | 2.67E+00 | 2.73E+00 | 8.29E-01 | no | Hrh1 | intron_variant | MODIFIER |
| 6 | 114462312 | AX-170490335 | 0 | A | G | 0.053 | 6.33E-02 | 2.96E-01 | 2.67E+00 | 2.73E+00 | 8.29E-01 | no | Hrh1 | intron_variant | MODIFIER |
| 6 | 114464665 | AX-170235248 | 0 | A | G | 0.053 | 6.33E-02 | 2.96E-01 | 2.67E+00 | 2.73E+00 | 8.29E-01 | no | Hrh1 | intron_variant | MODIFIER |
| 6 | 114475173 | AX-170320042 | 0 | T | C | 0.053 | 6.33E-02 | 2.96E-01 | 2.67E+00 | 2.73E+00 | 8.29E-01 | no | Hrh1 | intron_variant | MODIFIER |
| 6 | 114475394 | AX-170153826 | 0 | G | A | 0.053 | 6.33E-02 | 2.96E-01 | 2.67E+00 | 2.73E+00 | 8.29E-01 | no | Hrh1 | intron_variant | MODIFIER |
| 6 | 114485382 | AX-170470179 | 0 | T | C | 0.053 | 6.33E-02 | 2.96E-01 | 2.67E+00 | 2.73E+00 | 8.29E-01 | no | Hrh1 | downstream_gene_variant | MODIFIER |
| 6 | 114488365 | AX-170508003 | 0 | C | T | 0.053 | 6.33E-02 | 2.96E-01 | 2.67E+00 | 2.73E+00 | 8.29E-01 | no | | intergenic_variant | MODIFIER |
| 6 | 114649595 | AX-170089818 | 0 | T | C | 0.053 | 6.33E-02 | 2.96E-01 | 2.67E+00 | 2.73E+00 | 8.29E-01 | no | Atg7 | intron_variant | MODIFIER |
| 6 | 114662216 | AX-170434020 | 0 | A | G | 0.053 | 6.33E-02 | 2.96E-01 | 2.67E+00 | 2.73E+00 | 8.29E-01 | no | Atg7 | intron_variant | MODIFIER |
| 6 | 114690118 | AX-170159352 | 0 | A | C | 0.053 | 6.33E-02 | 2.96E-01 | 2.67E+00 | 2.73E+00 | 8.29E-01 | no | Atg7 | intron_variant | MODIFIER |
| 6 | 114692996 | AX-170190516 | 0 | A | G | 0.053 | 6.33E-02 | 2.96E-01 | 2.67E+00 | 2.73E+00 | 8.29E-01 | no | Atg7 | intron_variant | MODIFIER |
| 6 | 114699063 | AX-170028031 | 0 | T | C | 0.053 | 6.33E-02 | 2.96E-01 | 2.67E+00 | 2.73E+00 | 8.29E-01 | no | Atg7 | intron_variant | MODIFIER |
| 6 | 114706796 | AX-97831218 | 0 | A | C | 0.053 | 6.33E-02 | 2.96E-01 | 2.67E+00 | 2.73E+00 | 8.29E-01 | no | Atg7 | intron_variant | MODIFIER |
| 6 | 114733071 | AX-170028030 | 0 | A | G | 0.053 | 6.33E-02 | 2.96E-01 | 2.67E+00 | 2.73E+00 | 8.29E-01 | no | Atg7 | intron_variant | MODIFIER |
| 6 | 114768736 | AX-170304005 | 0 | A | G | 0.053 | 6.33E-02 | 2.96E-01 | 2.67E+00 | 2.73E+00 | 8.29E-01 | no | Atg7 | intron_variant | MODIFIER |
| 6 | 114788560 | AX-170284590 | 0 | A | G | 0.053 | 6.33E-02 | 2.96E-01 | 2.67E+00 | 2.73E+00 | 8.29E-01 | no | Atg7 | intron_variant | MODIFIER |
| 6 | 114789169 | AX-170287232 | 0 | A | G | 0.053 | 6.33E-02 | 2.96E-01 | 2.67E+00 | 2.73E+00 | 8.29E-01 | no | Atg7 | intron_variant | MODIFIER |
| 6 | 114801413 | AX-170472475 | 0 | A | G | 0.053 | 6.33E-02 | 2.96E-01 | 2.67E+00 | 2.73E+00 | 8.29E-01 | no | Atg7 | intron_variant | MODIFIER |
| 6 | 114803681 | AX-97797014 | 0 | A | C | 0.053 | 6.33E-02 | 2.96E-01 | 2.67E+00 | 2.73E+00 | 8.29E-01 | no | Atg7 | intron_variant | MODIFIER |
| 6 | 114810431 | AX-170355591 | 0 | T | C | 0.053 | 6.33E-02 | 2.96E-01 | 2.67E+00 | 2.73E+00 | 8.29E-01 | no | Atg7 | intron_variant | MODIFIER |
| 6 | 114882824 | AX-97807906 | 0 | G | C | 0.053 | 6.33E-02 | 2.96E-01 | 2.67E+00 | 2.73E+00 | 8.29E-01 | no | Vgll4 | intron_variant | MODIFIER |
| 6 | 114988619 | AX-170229645 | 0 | A | G | 0.053 | 6.33E-02 | 2.96E-01 | 2.67E+00 | 2.73E+00 | 8.29E-01 | no | | regulatory_region_variant | MODIFIER |
| 6 | 114988732 | AX-170319494 | 0 | A | G | 0.053 | 6.33E-02 | 2.96E-01 | 2.67E+00 | 2.73E+00 | 8.29E-01 | no | | regulatory_region_variant | MODIFIER |
| 6 | 115080528 | AX-170505977 | 0 | T | C | 0.053 | 6.33E-02 | 2.96E-01 | 2.67E+00 | 2.73E+00 | 8.29E-01 | no | | intergenic_variant | MODIFIER |
| 6 | 115342614 | AX-97823738 | 0 | G | C | 0.053 | 6.33E-02 | 2.96E-01 | 2.67E+00 | 2.73E+00 | 8.29E-01 | no | | intergenic_variant | MODIFIER |
| 6 | 116897484 | AX-170028059 | 0 | A | G | 0.053 | 6.33E-02 | 2.96E-01 | 2.67E+00 | 2.73E+00 | 8.29E-01 | no | | intergenic_variant | MODIFIER |
| 6 | 116943087 | AX-170105287 | 0 | G | A | 0.053 | 6.33E-02 | 2.96E-01 | 2.67E+00 | 2.73E+00 | 8.29E-01 | no | | intergenic_variant | MODIFIER |
| 6 | 117100576 | AX-170339935 | 0 | A | C | 0.053 | 6.33E-02 | 2.96E-01 | 2.67E+00 | 2.73E+00 | 8.29E-01 | no | | intergenic_variant | MODIFIER |
| 6 | 117100622 | AX-170395035 | 0 | G | A | 0.053 | 6.33E-02 | 2.96E-01 | 2.67E+00 | 2.73E+00 | 8.29E-01 | no | | intergenic_variant | MODIFIER |
| 6 | 118079307 | AX-170028067 | 0 | G | C | 0.053 | 6.33E-02 | 2.96E-01 | 2.67E+00 | 2.73E+00 | 8.29E-01 | no | Rasgef1a | intron_variant | MODIFIER |
| 6 | 118280460 | AX-170137074 | 0 | T | C | 0.053 | 6.33E-02 | 2.96E-01 | 2.67E+00 | 2.73E+00 | 8.29E-01 | no | | intergenic_variant | MODIFIER |
| 6 | 118284391 | AX-170366932 | 0 | A | G | 0.053 | 6.33E-02 | 2.96E-01 | 2.67E+00 | 2.73E+00 | 8.29E-01 | no | | regulatory_region_variant | MODIFIER |
| 6 | 118291929 | AX-170393844 | 0 | G | T | 0.053 | 6.33E-02 | 2.96E-01 | 2.67E+00 | 2.73E+00 | 8.29E-01 | no | | intergenic_variant | MODIFIER |
| 6 | 118334402 | AX-97844428 | 0 | A | G | 0.053 | 6.33E-02 | 2.96E-01 | 2.67E+00 | 2.73E+00 | 8.29E-01 | no | | intergenic_variant | MODIFIER |
| 6 | 118341005 | AX-170487947 | 0 | G | C | 0.053 | 6.33E-02 | 2.96E-01 | 2.67E+00 | 2.73E+00 | 8.29E-01 | no | | intergenic_variant | MODIFIER |
| 6 | 118466892 | AX-170132429 | 0 | G | A | 0.053 | 6.33E-02 | 2.96E-01 | 2.67E+00 | 2.73E+00 | 8.29E-01 | no | Zfp9 | synonymous_variant | LOW |
| 6 | 118523359 | AX-170513006 | 0 | G | A | 0.053 | 6.33E-02 | 2.96E-01 | 2.67E+00 | 2.73E+00 | 8.29E-01 | no | Ankrd26 | synonymous_variant | LOW |
| 6 | 118527644 | AX-170488414 | 0 | A | G | 0.053 | 6.33E-02 | 2.96E-01 | 2.67E+00 | 2.73E+00 | 8.29E-01 | no | Ankrd26 | intron_variant | MODIFIER |
| 6 | 118541496 | AX-170120625 | 0 | C | T | 0.053 | 6.33E-02 | 2.96E-01 | 2.67E+00 | 2.73E+00 | 8.29E-01 | no | Ankrd26 | intron_variant | MODIFIER |
| 6 | 118548076 | AX-97813747 | 0 | C | T | 0.053 | 6.33E-02 | 2.96E-01 | 2.67E+00 | 2.73E+00 | 8.29E-01 | no | Ankrd26 | intron_variant | MODIFIER |
| 6 | 118643993 | AX-170398048 | 0 | T | G | 0.053 | 6.33E-02 | 2.96E-01 | 2.67E+00 | 2.73E+00 | 8.29E-01 | no | Cacna1c | intron_variant | MODIFIER |
| 6 | 118684670 | AX-170076033 | 0 | G | A | 0.053 | 6.33E-02 | 2.96E-01 | 2.67E+00 | 2.73E+00 | 8.29E-01 | no | Cacna1c | intron_variant | MODIFIER |
| 6 | 118685646 | AX-170270651 | 0 | T | C | 0.053 | 6.33E-02 | 2.96E-01 | 2.67E+00 | 2.73E+00 | 8.29E-01 | no | Cacna1c | intron_variant | MODIFIER |
| 6 | 118723644 | AX-170240794 | 0 | T | C | 0.053 | 6.33E-02 | 2.96E-01 | 2.67E+00 | 2.73E+00 | 8.29E-01 | no | Cacna1c | intron_variant | MODIFIER |
| 6 | 118748869 | AX-170487207 | 0 | T | C | 0.053 | 6.33E-02 | 2.96E-01 | 2.67E+00 | 2.73E+00 | 8.29E-01 | no | Cacna1c | intron_variant | MODIFIER |
| 6 | 118749696 | AX-97828565 | 0 | A | G | 0.053 | 6.33E-02 | 2.96E-01 | 2.67E+00 | 2.73E+00 | 8.29E-01 | no | Cacna1c | intron_variant | MODIFIER |
| 6 | 118772505 | AX-97824748 | 0 | C | T | 0.053 | 6.33E-02 | 2.96E-01 | 2.67E+00 | 2.73E+00 | 8.29E-01 | no | Cacna1c | intron_variant | MODIFIER |
| 6 | 118977380 | AX-170098219 | 0 | A | G | 0.053 | 6.33E-02 | 2.96E-01 | 2.67E+00 | 2.73E+00 | 8.29E-01 | no | Cacna1c | intron_variant | MODIFIER |
| 6 | 119003242 | AX-97795421 | 0 | T | C | 0.053 | 6.33E-02 | 2.96E-01 | 2.67E+00 | 2.73E+00 | 8.29E-01 | no | Cacna1c | intron_variant | MODIFIER |
| 6 | 119065043 | AX-170396262 | 0 | G | A | 0.053 | 6.33E-02 | 2.96E-01 | 2.67E+00 | 2.73E+00 | 8.29E-01 | no | Cacna1c | intron_variant | MODIFIER |
| 6 | 119071011 | AX-170230808 | 0 | A | G | 0.053 | 6.33E-02 | 2.96E-01 | 2.67E+00 | 2.73E+00 | 8.29E-01 | no | Cacna1c | intron_variant | MODIFIER |
| 6 | 119078684 | AX-170266374 | 0 | G | T | 0.053 | 6.33E-02 | 2.96E-01 | 2.67E+00 | 2.73E+00 | 8.29E-01 | no | Cacna1c | intron_variant | MODIFIER |
| 6 | 119080143 | AX-170161313 | 0 | G | A | 0.053 | 6.33E-02 | 2.96E-01 | 2.67E+00 | 2.73E+00 | 8.29E-01 | no | Cacna1c | intron_variant | MODIFIER |
| 6 | 119095519 | AX-170171761 | 0 | A | C | 0.053 | 6.33E-02 | 2.96E-01 | 2.67E+00 | 2.73E+00 | 8.29E-01 | no | Cacna1c | intron_variant | MODIFIER |

| | | | | | | | | | | | | | | | |
|---|---|---|---|---|---|---|---|---|---|---|---|---|---|---|---|
| 6 | 119097921 | AX-170507704 | 0 | T | C | 0.053 | 6.33E-02 | 2.96E-01 | 2.67E+00 | 2.73E+00 | 8.29E-01 | no | Cacna1c | intron_variant | MODIFIER |
| 6 | 119112263 | AX-170028080 | 0 | T | G | 0.053 | 6.33E-02 | 2.96E-01 | 2.67E+00 | 2.73E+00 | 8.29E-01 | no | Cacna1c | upstream_gene_variant | MODIFIER |
| 6 | 119112464 | AX-170028086 | 0 | T | C | 0.053 | 6.33E-02 | 2.96E-01 | 2.67E+00 | 2.73E+00 | 8.29E-01 | no | Cacna1c | upstream_gene_variant | MODIFIER |
| 6 | 119139182 | AX-170295783 | 0 | T | C | 0.053 | 6.33E-02 | 2.96E-01 | 2.67E+00 | 2.73E+00 | 8.29E-01 | no | Cacna1c | intron_variant | MODIFIER |
| 6 | 119139246 | AX-170028083 | 0 | A | C | 0.053 | 6.33E-02 | 2.96E-01 | 2.67E+00 | 2.73E+00 | 8.29E-01 | no | Cacna1c | intron_variant | MODIFIER |
| 6 | 119216438 | AX-170221151 | 0 | T | C | 0.053 | 6.33E-02 | 2.96E-01 | 2.67E+00 | 2.73E+00 | 8.29E-01 | no | Dcp1b | intron_variant | MODIFIER |
| 6 | 119218578 | AX-170133501 | 0 | C | G | 0.053 | 6.33E-02 | 2.96E-01 | 2.67E+00 | 2.73E+00 | 8.29E-01 | no | Dcp1b | intron_variant | MODIFIER |
| 6 | 119357896 | AX-170327260 | 0 | C | T | 0.053 | 6.33E-02 | 2.96E-01 | 2.67E+00 | 2.73E+00 | 8.29E-01 | no | Adipor2 | intron_variant | MODIFIER |
| 6 | 119505374 | AX-97851376 | 0 | T | G | 0.053 | 6.33E-02 | 2.96E-01 | 2.67E+00 | 2.73E+00 | 8.29E-01 | no | Wnt5b | intron_variant | MODIFIER |
| 6 | 119514605 | AX-170243314 | 0 | G | T | 0.053 | 6.33E-02 | 2.96E-01 | 2.67E+00 | 2.73E+00 | 8.29E-01 | no | Wnt5b | intron_variant | MODIFIER |
| 6 | 119515227 | AX-170076050 | 0 | C | G | 0.053 | 6.33E-02 | 2.96E-01 | 2.67E+00 | 2.73E+00 | 8.29E-01 | no | Wnt5b | intron_variant | MODIFIER |
| 6 | 119526384 | AX-170326203 | 0 | T | C | 0.053 | 6.33E-02 | 2.96E-01 | 2.67E+00 | 2.73E+00 | 8.29E-01 | no | Wnt5b | intron_variant | MODIFIER |
| 6 | 119536848 | AX-170286986 | 0 | T | C | 0.053 | 6.33E-02 | 2.96E-01 | 2.67E+00 | 2.73E+00 | 8.29E-01 | no | Wnt5b | intron_variant | MODIFIER |
| 6 | 119537033 | AX-170328882 | 0 | C | T | 0.053 | 6.33E-02 | 2.96E-01 | 2.67E+00 | 2.73E+00 | 8.29E-01 | no | Wnt5b | intron_variant | MODIFIER |
| 6 | 119538417 | AX-170276833 | 0 | A | G | 0.053 | 6.33E-02 | 2.96E-01 | 2.67E+00 | 2.73E+00 | 8.29E-01 | no | Wnt5b | intron_variant | MODIFIER |
| 6 | 119545730 | AX-170269995 | 0 | T | C | 0.053 | 6.33E-02 | 2.96E-01 | 2.67E+00 | 2.73E+00 | 8.29E-01 | no | Wnt5b | upstream_gene_variant | MODIFIER |
| 6 | 119550866 | AX-170095782 | 0 | T | C | 0.053 | 6.33E-02 | 2.96E-01 | 2.67E+00 | 2.73E+00 | 8.29E-01 | no | | intergenic_variant | MODIFIER |
| 6 | 119566176 | AX-170028087 | 0 | T | C | 0.053 | 6.33E-02 | 2.96E-01 | 2.67E+00 | 2.73E+00 | 8.29E-01 | no | Erc1 | downstream_gene_variant | MODIFIER |
| 6 | 119568265 | AX-170367609 | 0 | T | C | 0.053 | 6.33E-02 | 2.96E-01 | 2.67E+00 | 2.73E+00 | 8.29E-01 | no | Erc1 | downstream_gene_variant | MODIFIER |
| 6 | 119595578 | AX-170076053 | 0 | T | C | 0.053 | 6.33E-02 | 2.96E-01 | 2.67E+00 | 2.73E+00 | 8.29E-01 | no | Erc1 | intron_variant | MODIFIER |
| 6 | 119596320 | AX-170076051 | 0 | G | A | 0.053 | 6.33E-02 | 2.96E-01 | 2.67E+00 | 2.73E+00 | 8.29E-01 | no | Erc1 | intron_variant | MODIFIER |
| 6 | 119757213 | AX-170028092 | 0 | A | G | 0.053 | 6.33E-02 | 2.96E-01 | 2.67E+00 | 2.73E+00 | 8.29E-01 | no | Erc1 | intron_variant | MODIFIER |
| 6 | 119761634 | AX-170111406 | 0 | G | C | 0.053 | 6.33E-02 | 2.96E-01 | 2.67E+00 | 2.73E+00 | 8.29E-01 | no | Erc1 | intron_variant | MODIFIER |
| 6 | 120127521 | AX-170221497 | 0 | G | A | 0.053 | 6.33E-02 | 2.96E-01 | 2.67E+00 | 2.73E+00 | 8.29E-01 | no | Ninj2 | intron_variant | MODIFIER |
| 6 | 120226208 | AX-170385719 | 0 | A | C | 0.053 | 6.33E-02 | 2.96E-01 | 2.67E+00 | 2.73E+00 | 8.29E-01 | no | B4galnt3 | intron_variant | MODIFIER |
| 6 | 120369831 | AX-170101804 | 0 | A | C | 0.053 | 6.33E-02 | 2.96E-01 | 2.67E+00 | 2.73E+00 | 8.29E-01 | no | Kdm5a | intron_variant | MODIFIER |
| 6 | 120584545 | AX-170428054 | 0 | A | C | 0.053 | 6.33E-02 | 2.96E-01 | 2.67E+00 | 2.73E+00 | 8.29E-01 | no | Gm44124 | upstream_gene_variant | MODIFIER |
| 6 | 120589097 | AX-170527823 | 0 | A | G | 0.053 | 6.33E-02 | 2.96E-01 | 2.67E+00 | 2.73E+00 | 8.29E-01 | no | | regulatory_region_variant | MODIFIER |
| 6 | 120612585 | AX-170294422 | 0 | T | C | 0.053 | 6.33E-02 | 2.96E-01 | 2.67E+00 | 2.73E+00 | 8.29E-01 | no | | regulatory_region_variant | MODIFIER |
| 6 | 120612853 | AX-170028114 | 0 | G | C | 0.053 | 6.33E-02 | 2.96E-01 | 2.67E+00 | 2.73E+00 | 8.29E-01 | no | | intergenic_variant | MODIFIER |
| 6 | 120707998 | AX-170095784 | 0 | A | G | 0.053 | 6.33E-02 | 2.96E-01 | 2.67E+00 | 2.73E+00 | 8.29E-01 | no | Cecr2 | intron_variant | MODIFIER |
| 6 | 120724281 | AX-168381979 | 0 | T | C | 0.053 | 6.33E-02 | 2.96E-01 | 2.67E+00 | 2.73E+00 | 8.29E-01 | no | Cecr2 | intron_variant | MODIFIER |
| 6 | 120978091 | AX-170119772 | 0 | A | G | 0.053 | 6.33E-02 | 2.96E-01 | 2.67E+00 | 2.73E+00 | 8.29E-01 | no | Mical3 | intron_variant,non_coding_transcript_variant | MODIFIER |
| 6 | 120984406 | AX-170377709 | 0 | A | G | 0.053 | 6.33E-02 | 2.96E-01 | 2.67E+00 | 2.73E+00 | 8.29E-01 | no | Mical3 | intron_variant,non_coding_transcript_variant | MODIFIER |
| 6 | 121040416 | AX-170300334 | 0 | A | G | 0.053 | 6.33E-02 | 2.96E-01 | 2.67E+00 | 2.73E+00 | 8.29E-01 | no | Mical3 | synonymous_variant | LOW |
| 6 | 121057009 | AX-170338064 | 0 | A | G | 0.053 | 6.33E-02 | 2.96E-01 | 2.67E+00 | 2.73E+00 | 8.29E-01 | no | Mical3 | intron_variant | MODIFIER |
| 6 | 121167034 | AX-170076074 | 0 | T | A | 0.053 | 6.33E-02 | 2.96E-01 | 2.67E+00 | 2.73E+00 | 8.29E-01 | no | | regulatory_region_variant | MODIFIER |
| 6 | 121292521 | AX-170302452 | 0 | T | C | 0.053 | 6.33E-02 | 2.96E-01 | 2.67E+00 | 2.73E+00 | 8.29E-01 | no | | intergenic_variant | MODIFIER |
| 6 | 121298338 | AX-170028124 | 0 | T | C | 0.053 | 6.33E-02 | 2.96E-01 | 2.67E+00 | 2.73E+00 | 8.29E-01 | no | Slc6a13 | upstream_gene_variant | MODIFIER |
| 6 | 121302789 | AX-170393507 | 0 | T | C | 0.053 | 6.33E-02 | 2.96E-01 | 2.67E+00 | 2.73E+00 | 8.29E-01 | no | Slc6a13 | synonymous_variant | LOW |
| 6 | 121322559 | AX-170521233 | 0 | A | G | 0.053 | 6.33E-02 | 2.96E-01 | 2.67E+00 | 2.73E+00 | 8.29E-01 | no | Slc6a13 | intron_variant | MODIFIER |
| 6 | 121322580 | AX-170028123 | 0 | C | T | 0.053 | 6.33E-02 | 2.96E-01 | 2.67E+00 | 2.73E+00 | 8.29E-01 | no | Slc6a13 | intron_variant | MODIFIER |
| 6 | 121322678 | AX-170453616 | 0 | C | T | 0.053 | 6.33E-02 | 2.96E-01 | 2.67E+00 | 2.73E+00 | 8.29E-01 | no | Slc6a13 | intron_variant | MODIFIER |
| 6 | 121332846 | AX-170076080 | 0 | C | T | 0.053 | 6.33E-02 | 2.96E-01 | 2.67E+00 | 2.73E+00 | 8.29E-01 | no | Slc6a13 | intron_variant | MODIFIER |
| 6 | 121334301 | AX-170076075 | 0 | T | C | 0.053 | 6.33E-02 | 2.96E-01 | 2.67E+00 | 2.73E+00 | 8.29E-01 | no | Slc6a13 | intron_variant | MODIFIER |
| 6 | 121334997 | AX-170202162 | 0 | G | A | 0.053 | 6.33E-02 | 2.96E-01 | 2.67E+00 | 2.73E+00 | 8.29E-01 | no | Slc6a13 | intron_variant | MODIFIER |
| 6 | 121491432 | AX-170448903 | 0 | C | T | 0.053 | 6.33E-02 | 2.96E-01 | 2.67E+00 | 2.73E+00 | 8.29E-01 | no | | intergenic_variant | MODIFIER |
| 6 | 122131944 | AX-97847109 | 0 | G | A | 0.053 | 6.33E-02 | 2.96E-01 | 2.67E+00 | 2.73E+00 | 8.29E-01 | no | Gm10319 | upstream_gene_variant | MODIFIER |
| 6 | 122262008 | AX-97826824 | 0 | A | G | 0.053 | 6.33E-02 | 2.96E-01 | 2.67E+00 | 2.73E+00 | 8.29E-01 | no | | intergenic_variant | MODIFIER |
| 6 | 122361054 | AX-97834167 | 0 | T | C | 0.053 | 6.33E-02 | 2.96E-01 | 2.67E+00 | 2.73E+00 | 8.29E-01 | no | | intergenic_variant | MODIFIER |
| 6 | 122413424 | AX-97793405 | 0 | G | A | 0.053 | 6.33E-02 | 2.96E-01 | 2.67E+00 | 2.73E+00 | 8.29E-01 | no | Gm19254 | upstream_gene_variant | MODIFIER |
| 6 | 122415564 | AX-170280070 | 0 | C | T | 0.053 | 6.33E-02 | 2.96E-01 | 2.67E+00 | 2.73E+00 | 8.29E-01 | no | Gm19254 | downstream_gene_variant | MODIFIER |
| 6 | 122430456 | AX-170143868 | 0 | T | C | 0.053 | 6.33E-02 | 2.96E-01 | 2.67E+00 | 2.73E+00 | 8.29E-01 | no | | intergenic_variant | MODIFIER |
| 6 | 122432676 | AX-170097141 | 0 | G | A | 0.053 | 6.33E-02 | 2.96E-01 | 2.67E+00 | 2.73E+00 | 8.29E-01 | no | | intergenic_variant | MODIFIER |

| | | | | | | | | | | | | | | | |
|---|---|---|---|---|---|---|---|---|---|---|---|---|---|---|---|
| 6 | 122445295 | AX-97843400 | 0 | G | A | 0.053 | 6.33E-02 | 2.96E-01 | 2.67E+00 | 2.73E+00 | 8.29E-01 | no | Rimklb | downstream_gene_variant | MODIFIER |
| 6 | 122560209 | AX-97843813 | 0 | T | C | 0.053 | 6.33E-02 | 2.96E-01 | 2.67E+00 | 2.73E+00 | 8.29E-01 | no | Aicda | intron_variant | MODIFIER |
| 6 | 122580711 | AX-97801802 | 0 | A | G | 0.053 | 6.33E-02 | 2.96E-01 | 2.67E+00 | 2.73E+00 | 8.29E-01 | no | Apobec1 | synonymous_variant | LOW |
| 6 | 122598974 | AX-170147252 | 0 | G | T | 0.053 | 6.33E-02 | 2.96E-01 | 2.67E+00 | 2.73E+00 | 8.29E-01 | no | Apobec1 | intron_variant | MODIFIER |
| 6 | 122598990 | AX-170262576 | 0 | A | G | 0.053 | 6.33E-02 | 2.96E-01 | 2.67E+00 | 2.73E+00 | 8.29E-01 | no | Apobec1 | intron_variant | MODIFIER |
| 6 | 122599109 | AX-170115139 | 0 | C | G | 0.053 | 6.33E-02 | 2.96E-01 | 2.67E+00 | 2.73E+00 | 8.29E-01 | no | Apobec1 | intron_variant | MODIFIER |
| 6 | 122606813 | AX-97850787 | 0 | A | G | 0.053 | 6.33E-02 | 2.96E-01 | 2.67E+00 | 2.73E+00 | 8.29E-01 | no | Gdf3 | synonymous_variant | LOW |
| 6 | 122624507 | AX-170028146 | 0 | A | G | 0.053 | 6.33E-02 | 2.96E-01 | 2.67E+00 | 2.73E+00 | 8.29E-01 | no | Dppa3 | upstream_gene_variant | MODIFIER |
| 6 | 122632465 | AX-97833314 | 0 | G | T | 0.053 | 6.33E-02 | 2.96E-01 | 2.67E+00 | 2.73E+00 | 8.29E-01 | no | Dppa3 | downstream_gene_variant | MODIFIER |
| 6 | 122632618 | AX-170289298 | 0 | A | T | 0.053 | 6.33E-02 | 2.96E-01 | 2.67E+00 | 2.73E+00 | 8.29E-01 | no | Dppa3 | downstream_gene_variant | MODIFIER |
| 6 | 122644136 | AX-97783687 | 0 | C | T | 0.053 | 6.33E-02 | 2.96E-01 | 2.67E+00 | 2.73E+00 | 8.29E-01 | no | | regulatory_region_variant | MODIFIER |
| 6 | 122649892 | AX-170200564 | 0 | T | C | 0.053 | 6.33E-02 | 2.96E-01 | 2.67E+00 | 2.73E+00 | 8.29E-01 | no | | intergenic_variant | MODIFIER |
| 6 | 122660901 | AX-170076094 | 0 | G | A | 0.053 | 6.33E-02 | 2.96E-01 | 2.67E+00 | 2.73E+00 | 8.29E-01 | no | | regulatory_region_variant | MODIFIER |
| 6 | 122666237 | AX-170028147 | 0 | G | T | 0.053 | 6.33E-02 | 2.96E-01 | 2.67E+00 | 2.73E+00 | 8.29E-01 | no | | regulatory_region_variant | MODIFIER |
| 6 | 122683143 | AX-170293671 | 0 | A | T | 0.053 | 6.33E-02 | 2.96E-01 | 2.67E+00 | 2.73E+00 | 8.29E-01 | no | | regulatory_region_variant | MODIFIER |
| 6 | 122683999 | AX-170242854 | 0 | T | G | 0.053 | 6.33E-02 | 2.96E-01 | 2.67E+00 | 2.73E+00 | 8.29E-01 | no | | intergenic_variant | MODIFIER |
| 6 | 122684532 | AX-170028145 | 0 | T | A | 0.053 | 6.33E-02 | 2.96E-01 | 2.67E+00 | 2.73E+00 | 8.29E-01 | no | | intergenic_variant | MODIFIER |
| 6 | 122736355 | AX-97837560 | 0 | C | A | 0.053 | 6.33E-02 | 2.96E-01 | 2.67E+00 | 2.73E+00 | 8.29E-01 | no | Slc2a3 | intron_variant | MODIFIER |
| 6 | 139148066 | AX-170416191 | 0 | T | A | 0.053 | 6.33E-02 | 2.96E-01 | 2.67E+00 | 2.73E+00 | 8.29E-01 | no | | intergenic_variant | MODIFIER |
| 6 | 139151933 | AX-170445352 | 0 | C | T | 0.053 | 6.33E-02 | 2.96E-01 | 2.67E+00 | 2.73E+00 | 8.29E-01 | no | | intergenic_variant | MODIFIER |
| 6 | 139151988 | AX-97808216 | 0 | T | G | 0.053 | 6.33E-02 | 2.96E-01 | 2.67E+00 | 2.73E+00 | 8.29E-01 | no | | intergenic_variant | MODIFIER |
| 6 | 139164988 | AX-170256887 | 0 | G | C | 0.053 | 6.33E-02 | 2.96E-01 | 2.67E+00 | 2.73E+00 | 8.29E-01 | no | | intergenic_variant | MODIFIER |
| 6 | 139177659 | AX-97789264 | 0 | A | G | 0.053 | 6.33E-02 | 2.96E-01 | 2.67E+00 | 2.73E+00 | 8.29E-01 | no | | intergenic_variant | MODIFIER |
| 6 | 139186793 | AX-170501680 | 0 | C | T | 0.053 | 6.33E-02 | 2.96E-01 | 2.67E+00 | 2.73E+00 | 8.29E-01 | no | | intergenic_variant | MODIFIER |
| 6 | 139186905 | AX-170535035 | 0 | G | A | 0.053 | 6.33E-02 | 2.96E-01 | 2.67E+00 | 2.73E+00 | 8.29E-01 | no | | intergenic_variant | MODIFIER |
| 6 | 139190544 | AX-170184990 | 0 | C | G | 0.053 | 6.33E-02 | 2.96E-01 | 2.67E+00 | 2.73E+00 | 8.29E-01 | no | | intergenic_variant | MODIFIER |
| 6 | 139202480 | AX-170095828 | 0 | T | C | 0.053 | 6.33E-02 | 2.96E-01 | 2.67E+00 | 2.73E+00 | 8.29E-01 | no | | intergenic_variant | MODIFIER |
| 6 | 139223535 | AX-97835918 | 0 | T | C | 0.053 | 6.33E-02 | 2.96E-01 | 2.67E+00 | 2.73E+00 | 8.29E-01 | no | | intergenic_variant | MODIFIER |
| 6 | 139228808 | AX-170285989 | 0 | T | A | 0.053 | 6.33E-02 | 2.96E-01 | 2.67E+00 | 2.73E+00 | 8.29E-01 | no | | intergenic_variant | MODIFIER |
| 6 | 139242215 | AX-170531423 | 0 | G | A | 0.053 | 6.33E-02 | 2.96E-01 | 2.67E+00 | 2.73E+00 | 8.29E-01 | no | | intergenic_variant | MODIFIER |
| 6 | 139247672 | AX-170251503 | 0 | A | T | 0.053 | 6.33E-02 | 2.96E-01 | 2.67E+00 | 2.73E+00 | 8.29E-01 | no | | intergenic_variant | MODIFIER |
| 6 | 139254125 | AX-97789023 | 0 | T | C | 0.053 | 6.33E-02 | 2.96E-01 | 2.67E+00 | 2.73E+00 | 8.29E-01 | no | | intergenic_variant | MODIFIER |
| 6 | 139256650 | AX-170285951 | 0 | G | A | 0.053 | 6.33E-02 | 2.96E-01 | 2.67E+00 | 2.73E+00 | 8.29E-01 | no | | intergenic_variant | MODIFIER |
| 6 | 139263539 | AX-170183274 | 0 | G | C | 0.053 | 6.33E-02 | 2.96E-01 | 2.67E+00 | 2.73E+00 | 8.29E-01 | no | | intergenic_variant | MODIFIER |
| 6 | 139264010 | AX-170076404 | 0 | G | A | 0.053 | 6.33E-02 | 2.96E-01 | 2.67E+00 | 2.73E+00 | 8.29E-01 | no | | intergenic_variant | MODIFIER |
| 6 | 139265461 | AX-170380151 | 0 | A | G | 0.053 | 6.33E-02 | 2.96E-01 | 2.67E+00 | 2.73E+00 | 8.29E-01 | no | | intergenic_variant | MODIFIER |
| 6 | 139280126 | AX-170523610 | 0 | G | C | 0.053 | 6.33E-02 | 2.96E-01 | 2.67E+00 | 2.73E+00 | 8.29E-01 | no | | intergenic_variant | MODIFIER |
| 6 | 139288316 | AX-97790859 | 0 | C | T | 0.053 | 6.33E-02 | 2.96E-01 | 2.67E+00 | 2.73E+00 | 8.29E-01 | no | | intergenic_variant | MODIFIER |
| 6 | 139295411 | AX-170214198 | 0 | C | T | 0.053 | 6.33E-02 | 2.96E-01 | 2.67E+00 | 2.73E+00 | 8.29E-01 | no | | intergenic_variant | MODIFIER |
| 6 | 139312196 | AX-170469391 | 0 | C | T | 0.053 | 6.33E-02 | 2.96E-01 | 2.67E+00 | 2.73E+00 | 8.29E-01 | no | 4922502N22Rik | downstream_gene_variant | MODIFIER |
| 6 | 139314663 | AX-170290561 | 0 | T | C | 0.053 | 6.33E-02 | 2.96E-01 | 2.67E+00 | 2.73E+00 | 8.29E-01 | no | 4922502N22Rik | downstream_gene_variant | MODIFIER |
| 6 | 139315241 | AX-170396568 | 0 | G | A | 0.053 | 6.33E-02 | 2.96E-01 | 2.67E+00 | 2.73E+00 | 8.29E-01 | no | 4922502N22Rik | downstream_gene_variant | MODIFIER |
| 6 | 139316693 | AX-170097294 | 0 | A | G | 0.053 | 6.33E-02 | 2.96E-01 | 2.67E+00 | 2.73E+00 | 8.29E-01 | no | 4922502N22Rik | intron_variant,non_coding_transcript_variant | MODIFIER |
| 6 | 139317103 | AX-170346472 | 0 | T | G | 0.053 | 6.33E-02 | 2.96E-01 | 2.67E+00 | 2.73E+00 | 8.29E-01 | no | 4922502N22Rik | non_coding_transcript_exon_variant | MODIFIER |
| 6 | 139318650 | AX-170218608 | 0 | G | A | 0.053 | 6.33E-02 | 2.96E-01 | 2.67E+00 | 2.73E+00 | 8.29E-01 | no | 4922502N22Rik | intron_variant,non_coding_transcript_variant | MODIFIER |
| 6 | 139322152 | AX-97784149 | 0 | T | G | 0.053 | 6.33E-02 | 2.96E-01 | 2.67E+00 | 2.73E+00 | 8.29E-01 | no | 4922502N22Rik | intron_variant,non_coding_transcript_variant | MODIFIER |
| 6 | 139341575 | AX-170508754 | 0 | A | T | 0.053 | 6.33E-02 | 2.96E-01 | 2.67E+00 | 2.73E+00 | 8.29E-01 | no | | intergenic_variant | MODIFIER |
| 6 | 139368151 | AX-170450028 | 0 | G | A | 0.053 | 6.33E-02 | 2.96E-01 | 2.67E+00 | 2.73E+00 | 8.29E-01 | no | | intergenic_variant | MODIFIER |
| 6 | 139368730 | AX-170178048 | 0 | C | A | 0.053 | 6.33E-02 | 2.96E-01 | 2.67E+00 | 2.73E+00 | 8.29E-01 | no | | intergenic_variant | MODIFIER |
| 6 | 139404122 | AX-170028378 | 0 | A | G | 0.053 | 6.33E-02 | 2.96E-01 | 2.67E+00 | 2.73E+00 | 8.29E-01 | no | | intergenic_variant | MODIFIER |
| 6 | 139420137 | AX-170076405 | 0 | G | A | 0.053 | 6.33E-02 | 2.96E-01 | 2.67E+00 | 2.73E+00 | 8.29E-01 | no | | intergenic_variant | MODIFIER |
| 6 | 139438024 | AX-170226830 | 0 | C | T | 0.053 | 6.33E-02 | 2.96E-01 | 2.67E+00 | 2.73E+00 | 8.29E-01 | no | | intergenic_variant | MODIFIER |
| 6 | 139444936 | AX-170429863 | 0 | A | T | 0.053 | 6.33E-02 | 2.96E-01 | 2.67E+00 | 2.73E+00 | 8.29E-01 | no | | intergenic_variant | MODIFIER |
| 6 | 139446366 | AX-170323778 | 0 | T | C | 0.053 | 6.33E-02 | 2.96E-01 | 2.67E+00 | 2.73E+00 | 8.29E-01 | no | | intergenic_variant | MODIFIER |

| | | | | | | | | | | | | | | | |
|---|---|---|---|---|---|---|---|---|---|---|---|---|---|---|---|
| 6 | 139460644 | AX-97792771 | 0 | G | A | 0.053 | 6.33E-02 | 2.96E-01 | 2.67E+00 | 2.73E+00 | 8.29E-01 | no | | regulatory_region_variant | MODIFIER |
| 6 | 139475645 | AX-170028377 | 0 | T | C | 0.053 | 6.33E-02 | 2.96E-01 | 2.67E+00 | 2.73E+00 | 8.29E-01 | no | | intergenic_variant | MODIFIER |
| 6 | 139492994 | AX-170028380 | 0 | A | C | 0.053 | 6.33E-02 | 2.96E-01 | 2.67E+00 | 2.73E+00 | 8.29E-01 | no | Rergl | 3_prime_UTR_variant | MODIFIER |
| 6 | 139500470 | AX-170288802 | 0 | T | C | 0.053 | 6.33E-02 | 2.96E-01 | 2.67E+00 | 2.73E+00 | 8.29E-01 | no | Rergl | intron_variant | MODIFIER |
| 6 | 139502023 | AX-170264126 | 0 | T | C | 0.053 | 6.33E-02 | 2.96E-01 | 2.67E+00 | 2.73E+00 | 8.29E-01 | no | Rergl | upstream_gene_variant | MODIFIER |
| 6 | 139515928 | AX-170529174 | 0 | G | C | 0.053 | 6.33E-02 | 2.96E-01 | 2.67E+00 | 2.73E+00 | 8.29E-01 | no | | intergenic_variant | MODIFIER |
| 6 | 139526562 | AX-97793270 | 0 | A | G | 0.053 | 6.33E-02 | 2.96E-01 | 2.67E+00 | 2.73E+00 | 8.29E-01 | no | | intergenic_variant | MODIFIER |
| 6 | 139542482 | AX-170028376 | 0 | A | C | 0.053 | 6.33E-02 | 2.96E-01 | 2.67E+00 | 2.73E+00 | 8.29E-01 | no | | intergenic_variant | MODIFIER |
| 6 | 139542521 | AX-170301962 | 0 | C | A | 0.053 | 6.33E-02 | 2.96E-01 | 2.67E+00 | 2.73E+00 | 8.29E-01 | no | | intergenic_variant | MODIFIER |
| 6 | 139542609 | AX-170028383 | 0 | T | C | 0.053 | 6.33E-02 | 2.96E-01 | 2.67E+00 | 2.73E+00 | 8.29E-01 | no | | intergenic_variant | MODIFIER |
| 6 | 139543250 | AX-170286735 | 0 | T | C | 0.053 | 6.33E-02 | 2.96E-01 | 2.67E+00 | 2.73E+00 | 8.29E-01 | no | | intergenic_variant | MODIFIER |
| 6 | 139546519 | AX-97840949 | 0 | C | T | 0.053 | 6.33E-02 | 2.96E-01 | 2.67E+00 | 2.73E+00 | 8.29E-01 | no | | regulatory_region_variant | MODIFIER |
| 6 | 139571464 | AX-170028381 | 0 | A | G | 0.053 | 6.33E-02 | 2.96E-01 | 2.67E+00 | 2.73E+00 | 8.29E-01 | no | 4930443G03Rik | downstream_gene_variant | MODIFIER |
| 6 | 139589317 | AX-170116738 | 0 | T | C | 0.053 | 6.33E-02 | 2.96E-01 | 2.67E+00 | 2.73E+00 | 8.29E-01 | no | Pik3c2g | intron_variant | MODIFIER |
| 6 | 139589576 | AX-170393243 | 0 | C | G | 0.053 | 6.33E-02 | 2.96E-01 | 2.67E+00 | 2.73E+00 | 8.29E-01 | no | Pik3c2g | intron_variant | MODIFIER |
| 6 | 139590118 | AX-170455827 | 0 | A | G | 0.053 | 6.33E-02 | 2.96E-01 | 2.67E+00 | 2.73E+00 | 8.29E-01 | no | Pik3c2g | intron_variant | MODIFIER |
| 6 | 139590780 | AX-170207247 | 0 | C | T | 0.053 | 6.33E-02 | 2.96E-01 | 2.67E+00 | 2.73E+00 | 8.29E-01 | no | Pik3c2g | intron_variant | MODIFIER |
| 6 | 139591567 | AX-170463576 | 0 | A | G | 0.053 | 6.33E-02 | 2.96E-01 | 2.67E+00 | 2.73E+00 | 8.29E-01 | no | Pik3c2g | intron_variant | MODIFIER |
| 6 | 139593164 | AX-170530245 | 0 | C | T | 0.053 | 6.33E-02 | 2.96E-01 | 2.67E+00 | 2.73E+00 | 8.29E-01 | no | Pik3c2g | intron_variant | MODIFIER |
| 6 | 139594735 | AX-170102586 | 0 | T | C | 0.053 | 6.33E-02 | 2.96E-01 | 2.67E+00 | 2.73E+00 | 8.29E-01 | no | Pik3c2g | upstream_gene_variant | MODIFIER |
| 6 | 139595191 | AX-170028382 | 0 | T | A | 0.053 | 6.33E-02 | 2.96E-01 | 2.67E+00 | 2.73E+00 | 8.29E-01 | no | Pik3c2g | upstream_gene_variant | MODIFIER |
| 6 | 139595605 | AX-170152035 | 0 | C | T | 0.053 | 6.33E-02 | 2.96E-01 | 2.67E+00 | 2.73E+00 | 8.29E-01 | no | Pik3c2g | upstream_gene_variant | MODIFIER |
| 6 | 139597573 | AX-170345174 | 0 | A | C | 0.053 | 6.33E-02 | 2.96E-01 | 2.67E+00 | 2.73E+00 | 8.29E-01 | no | Pik3c2g | upstream_gene_variant | MODIFIER |
| 6 | 139597745 | AX-170102121 | 0 | A | G | 0.053 | 6.33E-02 | 2.96E-01 | 2.67E+00 | 2.73E+00 | 8.29E-01 | no | Pik3c2g | upstream_gene_variant | MODIFIER |
| 6 | 139602527 | AX-170519831 | 0 | A | G | 0.053 | 6.33E-02 | 2.96E-01 | 2.67E+00 | 2.73E+00 | 8.29E-01 | no | Pik3c2g | intron_variant,non_coding_transcript_variant | MODIFIER |
| 6 | 139602582 | AX-170413681 | 0 | T | G | 0.053 | 6.33E-02 | 2.96E-01 | 2.67E+00 | 2.73E+00 | 8.29E-01 | no | Pik3c2g | intron_variant,non_coding_transcript_variant | MODIFIER |
| 6 | 139603391 | AX-97803621 | 0 | G | A | 0.053 | 6.33E-02 | 2.96E-01 | 2.67E+00 | 2.73E+00 | 8.29E-01 | no | Pik3c2g | intron_variant,non_coding_transcript_variant | MODIFIER |
| 6 | 139657035 | AX-97843349 | 0 | C | A | 0.053 | 6.33E-02 | 2.96E-01 | 2.67E+00 | 2.73E+00 | 8.29E-01 | no | Pik3c2g | intron_variant | MODIFIER |
| 6 | 139716397 | AX-170169069 | 0 | A | T | 0.053 | 6.33E-02 | 2.96E-01 | 2.67E+00 | 2.73E+00 | 8.29E-01 | no | Pik3c2g | intron_variant | MODIFIER |
| 6 | 139718817 | AX-170251577 | 0 | C | T | 0.053 | 6.33E-02 | 2.96E-01 | 2.67E+00 | 2.73E+00 | 8.29E-01 | no | Pik3c2g | intron_variant | MODIFIER |
| 6 | 139729873 | AX-168392298 | 0 | A | G | 0.053 | 6.33E-02 | 2.96E-01 | 2.67E+00 | 2.73E+00 | 8.29E-01 | no | Pik3c2g | intron_variant | MODIFIER |
| 6 | 139734265 | AX-170089855 | 0 | T | G | 0.053 | 6.33E-02 | 2.96E-01 | 2.67E+00 | 2.73E+00 | 8.29E-01 | no | Pik3c2g | intron_variant | MODIFIER |
| 6 | 139735870 | AX-170028373 | 0 | T | G | 0.053 | 6.33E-02 | 2.96E-01 | 2.67E+00 | 2.73E+00 | 8.29E-01 | no | Pik3c2g | intron_variant | MODIFIER |
| 6 | 139746111 | AX-170385038 | 0 | A | G | 0.053 | 6.33E-02 | 2.96E-01 | 2.67E+00 | 2.73E+00 | 8.29E-01 | no | Pik3c2g | intron_variant | MODIFIER |
| 6 | 139751763 | AX-170161276 | 0 | C | G | 0.053 | 6.33E-02 | 2.96E-01 | 2.67E+00 | 2.73E+00 | 8.29E-01 | no | Pik3c2g | intron_variant | MODIFIER |
| 6 | 139805653 | AX-97801978 | 0 | T | G | 0.053 | 6.33E-02 | 2.96E-01 | 2.67E+00 | 2.73E+00 | 8.29E-01 | no | Pik3c2g | intron_variant | MODIFIER |
| 6 | 139834147 | AX-97810806 | 0 | T | C | 0.053 | 6.33E-02 | 2.96E-01 | 2.67E+00 | 2.73E+00 | 8.29E-01 | no | Pik3c2g | intron_variant | MODIFIER |
| 6 | 139834856 | AX-170141655 | 0 | T | C | 0.053 | 6.33E-02 | 2.96E-01 | 2.67E+00 | 2.73E+00 | 8.29E-01 | no | Pik3c2g | intron_variant | MODIFIER |
| 6 | 139841224 | AX-170399744 | 0 | T | C | 0.053 | 6.33E-02 | 2.96E-01 | 2.67E+00 | 2.73E+00 | 8.29E-01 | no | Pik3c2g | upstream_gene_variant | MODIFIER |
| 6 | 139845269 | AX-170129181 | 0 | G | A | 0.053 | 6.33E-02 | 2.96E-01 | 2.67E+00 | 2.73E+00 | 8.29E-01 | no | Pik3c2g | intron_variant | MODIFIER |
| 6 | 139846482 | AX-170409161 | 0 | C | T | 0.053 | 6.33E-02 | 2.96E-01 | 2.67E+00 | 2.73E+00 | 8.29E-01 | no | Pik3c2g | intron_variant | MODIFIER |
| 6 | 139846645 | AX-170396360 | 0 | T | C | 0.053 | 6.33E-02 | 2.96E-01 | 2.67E+00 | 2.73E+00 | 8.29E-01 | no | Pik3c2g | intron_variant | MODIFIER |
| 6 | 139857209 | AX-170184236 | 0 | T | C | 0.053 | 6.33E-02 | 2.96E-01 | 2.67E+00 | 2.73E+00 | 8.29E-01 | no | Pik3c2g | intron_variant | MODIFIER |
| 6 | 139870909 | AX-97808120 | 0 | C | T | 0.053 | 6.33E-02 | 2.96E-01 | 2.67E+00 | 2.73E+00 | 8.29E-01 | no | Pik3c2g | intron_variant | MODIFIER |
| 6 | 139874071 | AX-170428580 | 0 | C | G | 0.053 | 6.33E-02 | 2.96E-01 | 2.67E+00 | 2.73E+00 | 8.29E-01 | no | Pik3c2g | intron_variant | MODIFIER |
| 6 | 139905751 | AX-97836129 | 0 | T | C | 0.053 | 6.33E-02 | 2.96E-01 | 2.67E+00 | 2.73E+00 | 8.29E-01 | no | Pik3c2g | intron_variant | MODIFIER |
| 6 | 139916084 | AX-97808025 | 0 | A | G | 0.053 | 6.33E-02 | 2.96E-01 | 2.67E+00 | 2.73E+00 | 8.29E-01 | no | Pik3c2g | intron_variant | MODIFIER |
| 6 | 139917252 | AX-168391279 | 0 | T | G | 0.053 | 6.33E-02 | 2.96E-01 | 2.67E+00 | 2.73E+00 | 8.29E-01 | no | Pik3c2g | intron_variant | MODIFIER |
| 6 | 140111627 | AX-170527248 | 0 | T | A | 0.053 | 6.33E-02 | 2.96E-01 | 2.67E+00 | 2.73E+00 | 8.29E-01 | no | | intergenic_variant | MODIFIER |
| 6 | 144031921 | AX-97815060 | 0 | G | A | 0.053 | 6.33E-02 | 2.96E-01 | 2.67E+00 | 2.73E+00 | 8.29E-01 | no | Sox5 | intron_variant | MODIFIER |
| 6 | 144037928 | AX-170453964 | 0 | T | C | 0.053 | 6.33E-02 | 2.96E-01 | 2.67E+00 | 2.73E+00 | 8.29E-01 | no | Sox5 | intron_variant | MODIFIER |
| 6 | 144091739 | AX-97847703 | 0 | A | G | 0.053 | 6.33E-02 | 2.96E-01 | 2.67E+00 | 2.73E+00 | 8.29E-01 | no | Sox5 | intron_variant | MODIFIER |
| 6 | 144093262 | AX-170379149 | 0 | A | G | 0.053 | 6.33E-02 | 2.96E-01 | 2.67E+00 | 2.73E+00 | 8.29E-01 | no | Sox5 | intron_variant | MODIFIER |
| 6 | 144130851 | AX-170099089 | 0 | T | C | 0.053 | 6.33E-02 | 2.96E-01 | 2.67E+00 | 2.73E+00 | 8.29E-01 | no | Sox5 | intron_variant | MODIFIER |

| | | | | | | | | | | | | | | | |
|---|---|---|---|---|---|---|---|---|---|---|---|---|---|---|---|
| 6 | 144134297 | AX-170423566 | 0 | G | C | 0.053 | 6.33E-02 | 2.96E-01 | 2.67E+00 | 2.73E+00 | 8.29E-01 | no | Sox5 | intron_variant | MODIFIER |
| 6 | 144134549 | AX-97786197 | 0 | G | T | 0.053 | 6.33E-02 | 2.96E-01 | 2.67E+00 | 2.73E+00 | 8.29E-01 | no | Sox5 | intron_variant | MODIFIER |
| 6 | 144134942 | AX-170264984 | 0 | A | C | 0.053 | 6.33E-02 | 2.96E-01 | 2.67E+00 | 2.73E+00 | 8.29E-01 | no | Sox5 | intron_variant | MODIFIER |
| 6 | 144299781 | AX-97792281 | 0 | T | C | 0.053 | 6.33E-02 | 2.96E-01 | 2.67E+00 | 2.73E+00 | 8.29E-01 | no | Sox5 | intron_variant,non_coding_transcript_variant | MODIFIER |
| 6 | 144314114 | AX-170028448 | 0 | A | G | 0.053 | 6.33E-02 | 2.96E-01 | 2.67E+00 | 2.73E+00 | 8.29E-01 | no | Sox5 | intron_variant,non_coding_transcript_variant | MODIFIER |
| 6 | 144334636 | AX-170181222 | 0 | C | A | 0.053 | 6.33E-02 | 2.96E-01 | 2.67E+00 | 2.73E+00 | 8.29E-01 | no | Sox5 | intron_variant,non_coding_transcript_variant | MODIFIER |
| 6 | 144335173 | AX-170208276 | 0 | A | C | 0.053 | 6.33E-02 | 2.96E-01 | 2.67E+00 | 2.73E+00 | 8.29E-01 | no | Sox5 | intron_variant,non_coding_transcript_variant | MODIFIER |
| 6 | 144339568 | AX-170028443 | 0 | T | C | 0.053 | 6.33E-02 | 2.96E-01 | 2.67E+00 | 2.73E+00 | 8.29E-01 | no | Sox5 | intron_variant,non_coding_transcript_variant | MODIFIER |
| 6 | 144359655 | AX-97857031 | 0 | A | G | 0.053 | 6.33E-02 | 2.96E-01 | 2.67E+00 | 2.73E+00 | 8.29E-01 | no | Sox5 | intron_variant,non_coding_transcript_variant | MODIFIER |
| 6 | 144368950 | AX-170028446 | 0 | A | G | 0.053 | 6.33E-02 | 2.96E-01 | 2.67E+00 | 2.73E+00 | 8.29E-01 | no | Sox5 | intron_variant,non_coding_transcript_variant | MODIFIER |
| 6 | 144391576 | AX-170076527 | 0 | A | G | 0.053 | 6.33E-02 | 2.96E-01 | 2.67E+00 | 2.73E+00 | 8.29E-01 | no | Sox5 | intron_variant,non_coding_transcript_variant | MODIFIER |
| 6 | 144407881 | AX-170383280 | 0 | T | C | 0.053 | 6.33E-02 | 2.96E-01 | 2.67E+00 | 2.73E+00 | 8.29E-01 | no | Sox5 | intron_variant,non_coding_transcript_variant | MODIFIER |
| 6 | 144412137 | AX-170290046 | 0 | T | G | 0.053 | 6.33E-02 | 2.96E-01 | 2.67E+00 | 2.73E+00 | 8.29E-01 | no | Sox5 | intron_variant,non_coding_transcript_variant | MODIFIER |
| 6 | 144412971 | AX-170028449 | 0 | G | A | 0.053 | 6.33E-02 | 2.96E-01 | 2.67E+00 | 2.73E+00 | 8.29E-01 | no | Sox5 | intron_variant,non_coding_transcript_variant | MODIFIER |
| 6 | 144450053 | AX-170151430 | 0 | A | C | 0.053 | 6.33E-02 | 2.96E-01 | 2.67E+00 | 2.73E+00 | 8.29E-01 | no | Sox5 | intron_variant,non_coding_transcript_variant | MODIFIER |
| 6 | 144457390 | AX-170464095 | 0 | C | T | 0.053 | 6.33E-02 | 2.96E-01 | 2.67E+00 | 2.73E+00 | 8.29E-01 | no | Sox5 | intron_variant,non_coding_transcript_variant | MODIFIER |
| 6 | 144457676 | AX-170223107 | 0 | G | C | 0.053 | 6.33E-02 | 2.96E-01 | 2.67E+00 | 2.73E+00 | 8.29E-01 | no | Sox5 | intron_variant,non_coding_transcript_variant | MODIFIER |
| 6 | 144458410 | AX-170356839 | 0 | A | G | 0.053 | 6.33E-02 | 2.96E-01 | 2.67E+00 | 2.73E+00 | 8.29E-01 | no | Sox5 | intron_variant,non_coding_transcript_variant | MODIFIER |
| 6 | 144459833 | AX-170331428 | 0 | A | G | 0.053 | 6.33E-02 | 2.96E-01 | 2.67E+00 | 2.73E+00 | 8.29E-01 | no | Sox5 | intron_variant,non_coding_transcript_variant | MODIFIER |
| 6 | 144461033 | AX-170114188 | 0 | A | G | 0.053 | 6.33E-02 | 2.96E-01 | 2.67E+00 | 2.73E+00 | 8.29E-01 | no | Sox5 | intron_variant,non_coding_transcript_variant | MODIFIER |
| 6 | 144461695 | AX-170028447 | 0 | A | G | 0.053 | 6.33E-02 | 2.96E-01 | 2.67E+00 | 2.73E+00 | 8.29E-01 | no | Sox5 | intron_variant,non_coding_transcript_variant | MODIFIER |
| 6 | 144461924 | AX-170028441 | 0 | G | A | 0.053 | 6.33E-02 | 2.96E-01 | 2.67E+00 | 2.73E+00 | 8.29E-01 | no | Sox5 | intron_variant,non_coding_transcript_variant | MODIFIER |
| 6 | 144493892 | AX-170076530 | 0 | T | C | 0.053 | 6.33E-02 | 2.96E-01 | 2.67E+00 | 2.73E+00 | 8.29E-01 | no | Sox5 | intron_variant,non_coding_transcript_variant | MODIFIER |
| 6 | 145222272 | AX-97798590 | 0 | G | A | 0.053 | 6.33E-02 | 2.96E-01 | 2.67E+00 | 2.73E+00 | 8.29E-01 | no | Kras | intron_variant | MODIFIER |
| 6 | 146806584 | AX-170028477 | 0 | C | T | 0.053 | 6.33E-02 | 2.96E-01 | 2.67E+00 | 2.73E+00 | 8.29E-01 | no | Arntl2 | intron_variant | MODIFIER |
| 6 | 106874477 | AX-170378839 | 1 | C | T | 0.054 | 6.33E-02 | 2.96E-01 | 2.67E+00 | 2.73E+00 | 8.29E-01 | no | | intergenic_variant | MODIFIER |
| 6 | 110236155 | AX-170100514 | 1 | T | C | 0.054 | 6.33E-02 | 2.96E-01 | 2.67E+00 | 2.73E+00 | 8.29E-01 | no | | intergenic_variant | MODIFIER |
| 7 | 70068238 | AX-170221343 | 0 | A | G | 0.404 | 2.19E-02 | 9.79E-02 | 2.62E+00 | 2.69E+00 | 8.29E-01 | no | | regulatory_region_variant | MODIFIER |
| 8 | 18316447 | AX-170168728 | 0 | C | A | 0.175 | 2.23E-02 | 1.05E-01 | 2.64E+00 | 2.73E+00 | 8.30E-01 | no | 1700014L14Rik | downstream_gene_variant | MODIFIER |
| 8 | 18340062 | AX-170417057 | 0 | C | T | 0.175 | 2.23E-02 | 1.05E-01 | 2.64E+00 | 2.73E+00 | 8.30E-01 | no | | intergenic_variant | MODIFIER |
| 8 | 18419451 | AX-170309488 | 0 | G | A | 0.175 | 2.23E-02 | 1.05E-01 | 2.64E+00 | 2.73E+00 | 8.30E-01 | no | | intergenic_variant | MODIFIER |
| 8 | 18421741 | AX-170252360 | 0 | T | C | 0.175 | 2.23E-02 | 1.05E-01 | 2.64E+00 | 2.73E+00 | 8.30E-01 | no | | intergenic_variant | MODIFIER |
| 8 | 18441284 | AX-170079458 | 0 | T | C | 0.175 | 2.23E-02 | 1.05E-01 | 2.64E+00 | 2.73E+00 | 8.30E-01 | no | | intergenic_variant | MODIFIER |
| 8 | 18442015 | AX-170121320 | 0 | C | T | 0.175 | 2.23E-02 | 1.05E-01 | 2.64E+00 | 2.73E+00 | 8.30E-01 | no | | intergenic_variant | MODIFIER |
| 8 | 18457812 | AX-170197649 | 0 | A | G | 0.175 | 2.23E-02 | 1.05E-01 | 2.64E+00 | 2.73E+00 | 8.30E-01 | no | | intergenic_variant | MODIFIER |
| 8 | 18502837 | AX-170431938 | 0 | T | G | 0.175 | 2.23E-02 | 1.05E-01 | 2.64E+00 | 2.73E+00 | 8.30E-01 | no | | intergenic_variant | MODIFIER |
| 8 | 18523137 | AX-97832032 | 0 | C | T | 0.175 | 2.23E-02 | 1.05E-01 | 2.64E+00 | 2.73E+00 | 8.30E-01 | no | | intergenic_variant | MODIFIER |
| 8 | 18528941 | AX-170481218 | 0 | A | G | 0.175 | 2.23E-02 | 1.05E-01 | 2.64E+00 | 2.73E+00 | 8.30E-01 | no | | intergenic_variant | MODIFIER |
| 8 | 18549952 | AX-170528167 | 0 | A | G | 0.175 | 2.23E-02 | 1.05E-01 | 2.64E+00 | 2.73E+00 | 8.30E-01 | no | | intergenic_variant | MODIFIER |
| 8 | 18550146 | AX-170208116 | 0 | G | A | 0.175 | 2.23E-02 | 1.05E-01 | 2.64E+00 | 2.73E+00 | 8.30E-01 | no | | intergenic_variant | MODIFIER |
| 8 | 18556277 | AX-170361290 | 0 | C | T | 0.175 | 2.23E-02 | 1.05E-01 | 2.64E+00 | 2.73E+00 | 8.30E-01 | no | | intergenic_variant | MODIFIER |
| 8 | 18562703 | AX-170030332 | 0 | C | A | 0.175 | 2.23E-02 | 1.05E-01 | 2.64E+00 | 2.73E+00 | 8.30E-01 | no | | intergenic_variant | MODIFIER |
| 8 | 18581692 | AX-97843102 | 0 | C | T | 0.175 | 2.23E-02 | 1.05E-01 | 2.64E+00 | 2.73E+00 | 8.30E-01 | no | | intergenic_variant | MODIFIER |
| 8 | 18584102 | AX-170030331 | 0 | C | T | 0.175 | 2.23E-02 | 1.05E-01 | 2.64E+00 | 2.73E+00 | 8.30E-01 | no | | intergenic_variant | MODIFIER |
| 8 | 18630825 | AX-170030330 | 0 | G | A | 0.175 | 2.23E-02 | 1.05E-01 | 2.64E+00 | 2.73E+00 | 8.30E-01 | no | Mcph1 | intron_variant | MODIFIER |
| 8 | 18636212 | AX-170337025 | 0 | T | A | 0.175 | 2.23E-02 | 1.05E-01 | 2.64E+00 | 2.73E+00 | 8.30E-01 | no | Mcph1 | intron_variant | MODIFIER |
| 8 | 18636396 | AX-170309283 | 0 | A | C | 0.175 | 2.23E-02 | 1.05E-01 | 2.64E+00 | 2.73E+00 | 8.30E-01 | no | Mcph1 | intron_variant | MODIFIER |
| 8 | 18643119 | AX-170405744 | 0 | T | C | 0.175 | 2.23E-02 | 1.05E-01 | 2.64E+00 | 2.73E+00 | 8.30E-01 | no | Mcph1 | intron_variant | MODIFIER |
| 8 | 18651765 | AX-170224194 | 0 | A | G | 0.175 | 2.23E-02 | 1.05E-01 | 2.64E+00 | 2.73E+00 | 8.30E-01 | no | Mcph1 | intron_variant | MODIFIER |
| 8 | 18671491 | AX-170160809 | 0 | T | C | 0.175 | 2.23E-02 | 1.05E-01 | 2.64E+00 | 2.73E+00 | 8.30E-01 | no | Mcph1 | intron_variant | MODIFIER |
| 8 | 18692327 | AX-170148549 | 0 | A | G | 0.175 | 2.23E-02 | 1.05E-01 | 2.64E+00 | 2.73E+00 | 8.30E-01 | no | Angpt2 | intron_variant | MODIFIER |
| 8 | 18698323 | AX-170167186 | 0 | G | A | 0.175 | 2.23E-02 | 1.05E-01 | 2.64E+00 | 2.73E+00 | 8.30E-01 | no | Angpt2 | intron_variant | MODIFIER |
| 8 | 18700418 | AX-170079465 | 0 | C | T | 0.175 | 2.23E-02 | 1.05E-01 | 2.64E+00 | 2.73E+00 | 8.30E-01 | no | Angpt2 | intron_variant | MODIFIER |
| 8 | 18700541 | AX-170190845 | 0 | T | C | 0.175 | 2.23E-02 | 1.05E-01 | 2.64E+00 | 2.73E+00 | 8.30E-01 | no | Angpt2 | intron_variant | MODIFIER |

| | | | | | | | | | | | | | | | |
|---|---|---|---|---|---|---|---|---|---|---|---|---|---|---|---|
| 8 | 18715842 | AX-170339320 | 0 | A | G | 0.175 | 2.23E-02 | 1.05E-01 | 2.64E+00 | 2.73E+00 | 8.30E-01 | no | Angpt2 | intron_variant | MODIFIER |
| 8 | 18727935 | AX-170472400 | 0 | A | G | 0.175 | 2.23E-02 | 1.05E-01 | 2.64E+00 | 2.73E+00 | 8.30E-01 | no | Angpt2 | intron_variant | MODIFIER |
| 8 | 18742167 | AX-170499311 | 0 | A | G | 0.175 | 2.23E-02 | 1.05E-01 | 2.64E+00 | 2.73E+00 | 8.30E-01 | no | Angpt2 | upstream_gene_variant | MODIFIER |
| 8 | 18760943 | AX-170393561 | 0 | G | A | 0.175 | 2.23E-02 | 1.05E-01 | 2.64E+00 | 2.73E+00 | 8.30E-01 | no | Mcph1 | intron_variant | MODIFIER |
| 8 | 18878939 | AX-170377512 | 0 | G | T | 0.175 | 2.23E-02 | 1.05E-01 | 2.64E+00 | 2.73E+00 | 8.30E-01 | no | Agpat5 | intron_variant | MODIFIER |
| 8 | 18928698 | AX-170179361 | 0 | G | A | 0.175 | 2.23E-02 | 1.05E-01 | 2.64E+00 | 2.73E+00 | 8.30E-01 | no | Xkr5 | downstream_gene_variant | MODIFIER |
| 8 | 18929394 | AX-170375072 | 0 | A | G | 0.175 | 2.23E-02 | 1.05E-01 | 2.64E+00 | 2.73E+00 | 8.30E-01 | no | Xkr5 | downstream_gene_variant | MODIFIER |
| 8 | 18954155 | AX-170445972 | 0 | T | G | 0.175 | 2.23E-02 | 1.05E-01 | 2.64E+00 | 2.73E+00 | 8.30E-01 | no | Xkr5 | upstream_gene_variant | MODIFIER |
| 8 | 18964646 | AX-170171686 | 0 | A | G | 0.175 | 2.23E-02 | 1.05E-01 | 2.64E+00 | 2.73E+00 | 8.30E-01 | no | | intergenic_variant | MODIFIER |
| 8 | 18984332 | AX-170464895 | 0 | G | A | 0.175 | 2.23E-02 | 1.05E-01 | 2.64E+00 | 2.73E+00 | 8.30E-01 | no | Defb37 | downstream_gene_variant | MODIFIER |
| 8 | 18986516 | AX-170389607 | 0 | G | A | 0.175 | 2.23E-02 | 1.05E-01 | 2.64E+00 | 2.73E+00 | 8.30E-01 | no | Defb37 | intron_variant | MODIFIER |
| 8 | 19026403 | AX-170292671 | 0 | C | T | 0.175 | 2.23E-02 | 1.05E-01 | 2.64E+00 | 2.73E+00 | 8.30E-01 | no | Defb38 | intron_variant | MODIFIER |
| 8 | 19026730 | AX-170372425 | 0 | T | C | 0.175 | 2.23E-02 | 1.05E-01 | 2.64E+00 | 2.73E+00 | 8.30E-01 | no | Defb38 | upstream_gene_variant | MODIFIER |
| 8 | 19028948 | AX-170231048 | 0 | A | G | 0.175 | 2.23E-02 | 1.05E-01 | 2.64E+00 | 2.73E+00 | 8.30E-01 | no | Defb38 | upstream_gene_variant | MODIFIER |
| 8 | 19036523 | AX-97795836 | 0 | C | A | 0.175 | 2.23E-02 | 1.05E-01 | 2.64E+00 | 2.73E+00 | 8.30E-01 | no | Gm15366 | downstream_gene_variant | MODIFIER |
| 8 | 19098207 | AX-170295380 | 0 | A | G | 0.175 | 2.23E-02 | 1.05E-01 | 2.64E+00 | 2.73E+00 | 8.30E-01 | no | | intergenic_variant | MODIFIER |
| 8 | 19114162 | AX-170030334 | 0 | C | A | 0.175 | 2.23E-02 | 1.05E-01 | 2.64E+00 | 2.73E+00 | 8.30E-01 | no | Defb12 | intron_variant | MODIFIER |
| 8 | 19138538 | AX-170030333 | 0 | T | G | 0.175 | 2.23E-02 | 1.05E-01 | 2.64E+00 | 2.73E+00 | 8.30E-01 | no | Spag11b | upstream_gene_variant | MODIFIER |
| 8 | 19157721 | AX-170079476 | 0 | T | C | 0.175 | 2.23E-02 | 1.05E-01 | 2.64E+00 | 2.73E+00 | 8.30E-01 | no | Spag11a | upstream_gene_variant | MODIFIER |
| 8 | 19167688 | AX-170275984 | 0 | T | C | 0.175 | 2.23E-02 | 1.05E-01 | 2.64E+00 | 2.73E+00 | 8.30E-01 | no | Gm39139 | upstream_gene_variant | MODIFIER |
| 8 | 19169459 | AX-170453795 | 0 | C | A | 0.175 | 2.23E-02 | 1.05E-01 | 2.64E+00 | 2.73E+00 | 8.30E-01 | no | Gm39139 | upstream_gene_variant | MODIFIER |
| 8 | 19240964 | AX-170447437 | 0 | A | G | 0.175 | 2.23E-02 | 1.05E-01 | 2.64E+00 | 2.73E+00 | 8.30E-01 | no | Defb46 | intron_variant | MODIFIER |
| 8 | 19241299 | AX-170096190 | 0 | C | T | 0.175 | 2.23E-02 | 1.05E-01 | 2.64E+00 | 2.73E+00 | 8.30E-01 | no | Defb46 | intron_variant | MODIFIER |
| 8 | 19242365 | AX-170397084 | 0 | T | C | 0.175 | 2.23E-02 | 1.05E-01 | 2.64E+00 | 2.73E+00 | 8.30E-01 | no | Defb46 | downstream_gene_variant | MODIFIER |
| 8 | 19245226 | AX-170178253 | 0 | T | C | 0.175 | 2.23E-02 | 1.05E-01 | 2.64E+00 | 2.73E+00 | 8.30E-01 | no | Defb46 | downstream_gene_variant | MODIFIER |
| 8 | 19251019 | AX-170284796 | 0 | T | C | 0.175 | 2.23E-02 | 1.05E-01 | 2.64E+00 | 2.73E+00 | 8.30E-01 | no | Defb5 | downstream_gene_variant | MODIFIER |
| 8 | 19278562 | AX-170140920 | 0 | C | T | 0.175 | 2.23E-02 | 1.05E-01 | 2.64E+00 | 2.73E+00 | 8.30E-01 | no | Gm15357 | downstream_gene_variant | MODIFIER |
| 8 | 19281519 | AX-170129480 | 0 | T | C | 0.175 | 2.23E-02 | 1.05E-01 | 2.64E+00 | 2.73E+00 | 8.30E-01 | no | Gm15357 | upstream_gene_variant | MODIFIER |
| 8 | 19281893 | AX-170499894 | 0 | T | G | 0.175 | 2.23E-02 | 1.05E-01 | 2.64E+00 | 2.73E+00 | 8.30E-01 | no | Gm15357 | upstream_gene_variant | MODIFIER |
| 8 | 19322269 | AX-170512984 | 0 | C | A | 0.175 | 2.23E-02 | 1.05E-01 | 2.64E+00 | 2.73E+00 | 8.30E-01 | no | | intergenic_variant | MODIFIER |
| 8 | 19323381 | AX-170203167 | 0 | G | C | 0.175 | 2.23E-02 | 1.05E-01 | 2.64E+00 | 2.73E+00 | 8.30E-01 | no | | intergenic_variant | MODIFIER |
| 8 | 19404580 | AX-170194363 | 0 | C | T | 0.175 | 2.23E-02 | 1.05E-01 | 2.64E+00 | 2.73E+00 | 8.30E-01 | no | | intergenic_variant | MODIFIER |
| 8 | 19408292 | AX-170520604 | 0 | G | A | 0.175 | 2.23E-02 | 1.05E-01 | 2.64E+00 | 2.73E+00 | 8.30E-01 | no | | intergenic_variant | MODIFIER |
| 8 | 19476503 | AX-170410017 | 0 | G | A | 0.175 | 2.23E-02 | 1.05E-01 | 2.64E+00 | 2.73E+00 | 8.30E-01 | no | Defa-ps15 | upstream_gene_variant | MODIFIER |
| 8 | 19479149 | AX-170411332 | 0 | G | T | 0.175 | 2.23E-02 | 1.05E-01 | 2.64E+00 | 2.73E+00 | 8.30E-01 | no | Defa-ps15 | upstream_gene_variant | MODIFIER |
| 8 | 19498211 | AX-170339764 | 0 | T | C | 0.175 | 2.23E-02 | 1.05E-01 | 2.64E+00 | 2.73E+00 | 8.30E-01 | no | Defb7 | downstream_gene_variant | MODIFIER |
| 8 | 19580397 | AX-170219031 | 0 | G | T | 0.175 | 2.23E-02 | 1.05E-01 | 2.64E+00 | 2.73E+00 | 8.30E-01 | no | Selenot-ps | non_coding_transcript_exon_variant | MODIFIER |
| 8 | 20888087 | AX-170375416 | 0 | G | T | 0.175 | 2.23E-02 | 1.05E-01 | 2.64E+00 | 2.73E+00 | 8.30E-01 | no | Defb33 | upstream_gene_variant | MODIFIER |
| 8 | 20941784 | AX-170320860 | 0 | C | A | 0.175 | 2.23E-02 | 1.05E-01 | 2.64E+00 | 2.73E+00 | 8.30E-01 | no | AY761185 | downstream_gene_variant | MODIFIER |
| 8 | 20942022 | AX-170306630 | 0 | T | G | 0.175 | 2.23E-02 | 1.05E-01 | 2.64E+00 | 2.73E+00 | 8.30E-01 | no | AY761185 | downstream_gene_variant | MODIFIER |
| 8 | 20981873 | AX-170503264 | 0 | G | A | 0.175 | 2.23E-02 | 1.05E-01 | 2.64E+00 | 2.73E+00 | 8.30E-01 | no | | intergenic_variant | MODIFIER |
| 8 | 21011929 | AX-170307820 | 0 | A | G | 0.175 | 2.23E-02 | 1.05E-01 | 2.64E+00 | 2.73E+00 | 8.30E-01 | no | | intergenic_variant | MODIFIER |
| 8 | 21091988 | AX-170155526 | 0 | C | G | 0.175 | 2.23E-02 | 1.05E-01 | 2.64E+00 | 2.73E+00 | 8.30E-01 | no | Gm14851 | downstream_gene_variant | MODIFIER |
| 8 | 21747390 | AX-170430690 | 0 | C | G | 0.175 | 2.23E-02 | 1.05E-01 | 2.64E+00 | 2.73E+00 | 8.30E-01 | no | Defa-ps11 | upstream_gene_variant | MODIFIER |
| 8 | 21762894 | AX-97797640 | 0 | A | G | 0.175 | 2.23E-02 | 1.05E-01 | 2.64E+00 | 2.73E+00 | 8.30E-01 | no | | intergenic_variant | MODIFIER |
| 8 | 21768314 | AX-170466930 | 0 | G | A | 0.175 | 2.23E-02 | 1.05E-01 | 2.64E+00 | 2.73E+00 | 8.30E-01 | no | | intergenic_variant | MODIFIER |
| 8 | 21778317 | AX-170515120 | 0 | G | T | 0.175 | 2.23E-02 | 1.05E-01 | 2.64E+00 | 2.73E+00 | 8.30E-01 | no | Defb1 | intron_variant | MODIFIER |
| 8 | 21794847 | AX-170388137 | 0 | A | G | 0.175 | 2.23E-02 | 1.05E-01 | 2.64E+00 | 2.73E+00 | 8.30E-01 | no | Defb1 | 3_prime_UTR_variant | MODIFIER |
| 8 | 21803496 | AX-170030338 | 0 | G | A | 0.175 | 2.23E-02 | 1.05E-01 | 2.64E+00 | 2.73E+00 | 8.30E-01 | no | | intergenic_variant | MODIFIER |
| 8 | 21815172 | AX-170030337 | 0 | G | C | 0.175 | 2.23E-02 | 1.05E-01 | 2.64E+00 | 2.73E+00 | 8.30E-01 | no | Gm15316 | downstream_gene_variant | MODIFIER |
| 8 | 21823733 | AX-170266970 | 0 | A | G | 0.175 | 2.23E-02 | 1.05E-01 | 2.64E+00 | 2.73E+00 | 8.30E-01 | no | Defb50 | intron_variant | MODIFIER |
| 8 | 21856519 | AX-170079489 | 0 | A | G | 0.175 | 2.23E-02 | 1.05E-01 | 2.64E+00 | 2.73E+00 | 8.30E-01 | no | Defb10 | upstream_gene_variant | MODIFIER |
| 8 | 21882203 | AX-170311502 | 0 | C | G | 0.175 | 2.23E-02 | 1.05E-01 | 2.64E+00 | 2.73E+00 | 8.30E-01 | no | Defb9 | intron_variant | MODIFIER |
| 8 | 21889049 | AX-170422750 | 0 | T | C | 0.175 | 2.23E-02 | 1.05E-01 | 2.64E+00 | 2.73E+00 | 8.30E-01 | no | Defb9 | upstream_gene_variant | MODIFIER |

| | | | | | | | | | | | | | | | |
|---|---|---|---|---|---|---|---|---|---|---|---|---|---|---|---|
| 8 | 21890897 | AX-170096193 | 0 | T | A | 0.175 | 2.23E-02 | 1.05E-01 | 2.64E+00 | 2.73E+00 | 8.30E-01 | no | | intergenic_variant | MODIFIER |
| 8 | 101761429 | AX-170327799 | 0 | G | A | 0.219 | 2.41E-02 | 1.15E-01 | 2.68E+00 | 2.75E+00 | 8.30E-01 | no | | intergenic_variant | MODIFIER |
| 8 | 101767798 | AX-170179463 | 0 | T | C | 0.219 | 2.41E-02 | 1.15E-01 | 2.68E+00 | 2.75E+00 | 8.30E-01 | no | | intergenic_variant | MODIFIER |
| 8 | 101786703 | AX-170317572 | 0 | A | G | 0.219 | 2.41E-02 | 1.15E-01 | 2.68E+00 | 2.75E+00 | 8.30E-01 | no | | intergenic_variant | MODIFIER |
| 8 | 101792021 | AX-170119457 | 0 | T | G | 0.219 | 2.41E-02 | 1.15E-01 | 2.68E+00 | 2.75E+00 | 8.30E-01 | no | | intergenic_variant | MODIFIER |
| 8 | 101792269 | AX-97810131 | 0 | C | T | 0.219 | 2.41E-02 | 1.15E-01 | 2.68E+00 | 2.75E+00 | 8.30E-01 | no | | intergenic_variant | MODIFIER |
| 8 | 101842555 | AX-170425312 | 0 | T | C | 0.219 | 2.41E-02 | 1.15E-01 | 2.68E+00 | 2.75E+00 | 8.30E-01 | no | | intergenic_variant | MODIFIER |
| 8 | 101846351 | AX-170183872 | 0 | G | A | 0.219 | 2.41E-02 | 1.15E-01 | 2.68E+00 | 2.75E+00 | 8.30E-01 | no | | intergenic_variant | MODIFIER |
| 8 | 101846503 | AX-170346367 | 0 | T | G | 0.219 | 2.41E-02 | 1.15E-01 | 2.68E+00 | 2.75E+00 | 8.30E-01 | no | | intergenic_variant | MODIFIER |
| 8 | 101861218 | AX-170195708 | 0 | G | A | 0.219 | 2.41E-02 | 1.15E-01 | 2.68E+00 | 2.75E+00 | 8.30E-01 | no | | intergenic_variant | MODIFIER |
| 8 | 101912610 | AX-170345642 | 0 | C | T | 0.219 | 2.41E-02 | 1.15E-01 | 2.68E+00 | 2.75E+00 | 8.30E-01 | no | | intergenic_variant | MODIFIER |
| 8 | 101913374 | AX-170472917 | 0 | A | G | 0.219 | 2.41E-02 | 1.15E-01 | 2.68E+00 | 2.75E+00 | 8.30E-01 | no | | intergenic_variant | MODIFIER |
| 8 | 101916653 | AX-170425986 | 0 | C | T | 0.219 | 2.41E-02 | 1.15E-01 | 2.68E+00 | 2.75E+00 | 8.30E-01 | no | | intergenic_variant | MODIFIER |
| 8 | 101916877 | AX-170483348 | 0 | T | C | 0.219 | 2.41E-02 | 1.15E-01 | 2.68E+00 | 2.75E+00 | 8.30E-01 | no | | intergenic_variant | MODIFIER |
| 8 | 101917552 | AX-170392779 | 0 | T | C | 0.219 | 2.41E-02 | 1.15E-01 | 2.68E+00 | 2.75E+00 | 8.30E-01 | no | | intergenic_variant | MODIFIER |
| 8 | 101917794 | AX-170210392 | 0 | G | A | 0.219 | 2.41E-02 | 1.15E-01 | 2.68E+00 | 2.75E+00 | 8.30E-01 | no | | intergenic_variant | MODIFIER |
| 8 | 101918368 | AX-170498389 | 0 | C | T | 0.219 | 2.41E-02 | 1.15E-01 | 2.68E+00 | 2.75E+00 | 8.30E-01 | no | | intergenic_variant | MODIFIER |
| 8 | 101946084 | AX-97830854 | 0 | G | T | 0.219 | 2.41E-02 | 1.15E-01 | 2.68E+00 | 2.75E+00 | 8.30E-01 | no | | intergenic_variant | MODIFIER |
| 8 | 101955000 | AX-170408929 | 0 | C | T | 0.219 | 2.41E-02 | 1.15E-01 | 2.68E+00 | 2.75E+00 | 8.30E-01 | no | | intergenic_variant | MODIFIER |
| 8 | 101961590 | AX-170264169 | 0 | G | A | 0.219 | 2.41E-02 | 1.15E-01 | 2.68E+00 | 2.75E+00 | 8.30E-01 | no | | intergenic_variant | MODIFIER |
| 8 | 102019472 | AX-170324925 | 0 | C | A | 0.219 | 2.41E-02 | 1.15E-01 | 2.68E+00 | 2.75E+00 | 8.30E-01 | no | | intergenic_variant | MODIFIER |
| 8 | 102031521 | AX-170128729 | 0 | C | T | 0.219 | 2.41E-02 | 1.15E-01 | 2.68E+00 | 2.75E+00 | 8.30E-01 | no | | intergenic_variant | MODIFIER |
| 8 | 102063585 | AX-170184000 | 0 | C | A | 0.219 | 2.41E-02 | 1.15E-01 | 2.68E+00 | 2.75E+00 | 8.30E-01 | no | | intergenic_variant | MODIFIER |
| 8 | 102063875 | AX-170218703 | 0 | G | T | 0.219 | 2.41E-02 | 1.15E-01 | 2.68E+00 | 2.75E+00 | 8.30E-01 | no | | intergenic_variant | MODIFIER |
| 8 | 102090302 | AX-170462395 | 0 | T | C | 0.219 | 2.41E-02 | 1.15E-01 | 2.68E+00 | 2.75E+00 | 8.30E-01 | no | | intergenic_variant | MODIFIER |
| 8 | 102121179 | AX-170533798 | 0 | C | T | 0.219 | 2.41E-02 | 1.15E-01 | 2.68E+00 | 2.75E+00 | 8.30E-01 | no | | intergenic_variant | MODIFIER |
| 8 | 102149683 | AX-97795178 | 0 | C | G | 0.219 | 2.41E-02 | 1.15E-01 | 2.68E+00 | 2.75E+00 | 8.30E-01 | no | | intergenic_variant | MODIFIER |
| 8 | 102159214 | AX-170205781 | 0 | C | G | 0.219 | 2.41E-02 | 1.15E-01 | 2.68E+00 | 2.75E+00 | 8.30E-01 | no | | intergenic_variant | MODIFIER |
| 8 | 102159663 | AX-170500737 | 0 | C | T | 0.219 | 2.41E-02 | 1.15E-01 | 2.68E+00 | 2.75E+00 | 8.30E-01 | no | | intergenic_variant | MODIFIER |
| 8 | 102160003 | AX-170528005 | 0 | G | A | 0.219 | 2.41E-02 | 1.15E-01 | 2.68E+00 | 2.75E+00 | 8.30E-01 | no | | intergenic_variant | MODIFIER |
| 8 | 102335593 | AX-170031401 | 0 | G | A | 0.219 | 2.41E-02 | 1.15E-01 | 2.68E+00 | 2.75E+00 | 8.30E-01 | no | | intergenic_variant | MODIFIER |
| 8 | 102406768 | AX-170343650 | 0 | T | C | 0.219 | 2.41E-02 | 1.15E-01 | 2.68E+00 | 2.75E+00 | 8.30E-01 | no | | intergenic_variant | MODIFIER |
| 8 | 102409593 | AX-170310975 | 0 | C | T | 0.219 | 2.41E-02 | 1.15E-01 | 2.68E+00 | 2.75E+00 | 8.30E-01 | no | | intergenic_variant | MODIFIER |
| 8 | 102411440 | AX-170495443 | 0 | T | G | 0.219 | 2.41E-02 | 1.15E-01 | 2.68E+00 | 2.75E+00 | 8.30E-01 | no | | intergenic_variant | MODIFIER |
| 8 | 102449156 | AX-170251150 | 0 | C | T | 0.219 | 2.41E-02 | 1.15E-01 | 2.68E+00 | 2.75E+00 | 8.30E-01 | no | | intergenic_variant | MODIFIER |
| 8 | 102449317 | AX-170388050 | 0 | C | T | 0.219 | 2.41E-02 | 1.15E-01 | 2.68E+00 | 2.75E+00 | 8.30E-01 | no | | intergenic_variant | MODIFIER |
| 8 | 102485120 | AX-170436636 | 0 | G | A | 0.219 | 2.41E-02 | 1.15E-01 | 2.68E+00 | 2.75E+00 | 8.30E-01 | no | | intergenic_variant | MODIFIER |
| 8 | 102493817 | AX-170429882 | 0 | T | C | 0.219 | 2.41E-02 | 1.15E-01 | 2.68E+00 | 2.75E+00 | 8.30E-01 | no | | intergenic_variant | MODIFIER |
| 8 | 102493999 | AX-97811199 | 0 | C | T | 0.219 | 2.41E-02 | 1.15E-01 | 2.68E+00 | 2.75E+00 | 8.30E-01 | no | | intergenic_variant | MODIFIER |
| 8 | 102512050 | AX-170199970 | 0 | C | T | 0.219 | 2.41E-02 | 1.15E-01 | 2.68E+00 | 2.75E+00 | 8.30E-01 | no | | intergenic_variant | MODIFIER |
| 8 | 102515060 | AX-170473034 | 0 | T | G | 0.219 | 2.41E-02 | 1.15E-01 | 2.68E+00 | 2.75E+00 | 8.30E-01 | no | | intergenic_variant | MODIFIER |
| 8 | 102516259 | AX-170499229 | 0 | A | C | 0.219 | 2.41E-02 | 1.15E-01 | 2.68E+00 | 2.75E+00 | 8.30E-01 | no | | intergenic_variant | MODIFIER |
| 8 | 102586228 | AX-170297150 | 0 | G | A | 0.219 | 2.41E-02 | 1.15E-01 | 2.68E+00 | 2.75E+00 | 8.30E-01 | no | | intergenic_variant | MODIFIER |
| 8 | 102607114 | AX-170522748 | 0 | T | G | 0.219 | 2.41E-02 | 1.15E-01 | 2.68E+00 | 2.75E+00 | 8.30E-01 | no | | intergenic_variant | MODIFIER |
| 8 | 102610314 | AX-170206738 | 0 | T | G | 0.219 | 2.41E-02 | 1.15E-01 | 2.68E+00 | 2.75E+00 | 8.30E-01 | no | | intergenic_variant | MODIFIER |
| 8 | 102610363 | AX-170466208 | 0 | T | C | 0.219 | 2.41E-02 | 1.15E-01 | 2.68E+00 | 2.75E+00 | 8.30E-01 | no | | intergenic_variant | MODIFIER |
| 8 | 102610549 | AX-170081014 | 0 | C | A | 0.219 | 2.41E-02 | 1.15E-01 | 2.68E+00 | 2.75E+00 | 8.30E-01 | no | | intergenic_variant | MODIFIER |
| 8 | 102623692 | AX-170253067 | 0 | C | G | 0.219 | 2.41E-02 | 1.15E-01 | 2.68E+00 | 2.75E+00 | 8.30E-01 | no | | intergenic_variant | MODIFIER |
| 8 | 102640765 | AX-170096391 | 0 | T | G | 0.219 | 2.41E-02 | 1.15E-01 | 2.68E+00 | 2.75E+00 | 8.30E-01 | no | Cdh11 | intron_variant | MODIFIER |
| 8 | 102653705 | AX-170178623 | 0 | C | A | 0.219 | 2.41E-02 | 1.15E-01 | 2.68E+00 | 2.75E+00 | 8.30E-01 | no | Cdh11 | intron_variant | MODIFIER |
| 8 | 102657459 | AX-170526365 | 0 | G | C | 0.219 | 2.41E-02 | 1.15E-01 | 2.68E+00 | 2.75E+00 | 8.30E-01 | no | Cdh11 | intron_variant | MODIFIER |
| 8 | 102716265 | AX-170344127 | 0 | C | T | 0.219 | 2.41E-02 | 1.15E-01 | 2.68E+00 | 2.75E+00 | 8.30E-01 | no | Cdh11 | intron_variant | MODIFIER |
| 8 | 102719758 | AX-97824006 | 0 | T | G | 0.219 | 2.41E-02 | 1.15E-01 | 2.68E+00 | 2.75E+00 | 8.30E-01 | no | Cdh11 | intron_variant | MODIFIER |
| 8 | 102727067 | AX-170174775 | 0 | T | A | 0.219 | 2.41E-02 | 1.15E-01 | 2.68E+00 | 2.75E+00 | 8.30E-01 | no | Cdh11 | intron_variant | MODIFIER |

| | | | | | | | | | | | | | | | |
|---|---|---|---|---|---|---|---|---|---|---|---|---|---|---|---|
| 8 | 102734145 | AX-170114672 | 0 | T | C | 0.219 | 2.41E-02 | 1.15E-01 | 2.68E+00 | 2.75E+00 | 8.30E-01 | no | Cdh11 | intron_variant | MODIFIER |
| 8 | 102737994 | AX-170441184 | 0 | C | T | 0.219 | 2.41E-02 | 1.15E-01 | 2.68E+00 | 2.75E+00 | 8.30E-01 | no | Cdh11 | intron_variant | MODIFIER |
| 18 | 53600358 | AX-170224770 | 0 | T | C | 0.237 | -2.44E-02 | 1.09E-01 | 2.63E+00 | 2.69E+00 | 8.30E-01 | no | | intergenic_variant | MODIFIER |
| 2 | 93788940 | AX-170230698 | 1 | T | C | 0.054 | -6.37E-02 | 3.01E-01 | 2.72E+00 | 2.75E+00 | 8.30E-01 | no | Ext2 | intron_variant | MODIFIER |
| 1 | 55715181 | AX-170041293 | 1 | C | T | 0.08 | -4.11E-02 | 1.96E-01 | 2.68E+00 | 2.75E+00 | 8.30E-01 | no | Plcl1 | intron_variant | MODIFIER |
| 2 | 107869175 | AX-170417705 | 1 | C | T | 0.054 | -6.34E-02 | 3.00E-01 | 2.72E+00 | 2.75E+00 | 8.30E-01 | no | | intergenic_variant | MODIFIER |
| 2 | 110349721 | AX-170018655 | 1 | A | T | 0.054 | -6.34E-02 | 3.00E-01 | 2.72E+00 | 2.75E+00 | 8.30E-01 | no | | intergenic_variant | MODIFIER |
| 9 | 77434165 | AX-170459428 | 1 | C | T | 0.196 | 2.85E-02 | 1.35E-01 | 2.72E+00 | 2.76E+00 | 8.30E-01 | no | Lrrc1 | intron_variant | MODIFIER |
| 9 | 33342080 | AX-170118129 | 1 | C | T | 0.429 | 1.94E-02 | 8.83E-02 | 2.64E+00 | 2.70E+00 | 8.30E-01 | no | | intergenic_variant | MODIFIER |
| 8 | 34927756 | AX-170180936 | 0 | T | C | 0.158 | -2.93E-02 | 1.33E-01 | 2.64E+00 | 2.70E+00 | 8.30E-01 | no | Tnks | intron_variant | MODIFIER |
| 1 | 63641006 | AX-97826485 | 2 | G | T | 0.136 | 5.06E-02 | 2.38E-01 | 2.68E+00 | 2.73E+00 | 8.30E-01 | no | Dytn | intron_variant | MODIFIER |
| 8 | 106649819 | AX-170454745 | 1 | C | G | 0.214 | 2.33E-02 | 1.10E-01 | 2.67E+00 | 2.74E+00 | 8.30E-01 | no | Cdh1 | intron_variant | MODIFIER |
| 8 | 57250071 | AX-170484302 | 1 | G | A | 0.152 | -3.25E-02 | 1.47E-01 | 2.64E+00 | 2.69E+00 | 8.30E-01 | no | | intergenic_variant | MODIFIER |
| 20 | 65651343 | AX-170204498 | 1 | C | A | 0.25 | 1.34E-02 | 6.29E-02 | 2.64E+00 | 2.73E+00 | 8.30E-01 | no | | intergenic_variant | MODIFIER |
| 1 | 153062704 | AX-170483505 | 0 | T | C | 0.211 | 2.57E-02 | 1.17E-01 | 2.63E+00 | 2.71E+00 | 8.30E-01 | no | Nmnat2 | intron_variant | MODIFIER |
| 18 | 69889933 | AX-170382517 | 0 | T | C | 0.263 | -2.13E-02 | 9.87E-02 | 2.66E+00 | 2.72E+00 | 8.30E-01 | no | AC108433.1 | intron_variant,non_coding_transcript_variant | MODIFIER |
| 18 | 69925023 | AX-97835020 | 0 | C | T | 0.263 | -2.13E-02 | 9.87E-02 | 2.66E+00 | 2.72E+00 | 8.30E-01 | no | Ccdc68 | upstream_gene_variant | MODIFIER |
| 18 | 69928744 | AX-170504844 | 0 | A | G | 0.263 | -2.13E-02 | 9.87E-02 | 2.66E+00 | 2.72E+00 | 8.30E-01 | no | Ccdc68 | intron_variant | MODIFIER |
| 18 | 69928808 | AX-170491137 | 0 | T | A | 0.263 | -2.13E-02 | 9.87E-02 | 2.66E+00 | 2.72E+00 | 8.30E-01 | no | Ccdc68 | intron_variant | MODIFIER |
| 18 | 69928865 | AX-170530895 | 0 | T | C | 0.263 | -2.13E-02 | 9.87E-02 | 2.66E+00 | 2.72E+00 | 8.30E-01 | no | Ccdc68 | intron_variant | MODIFIER |
| 18 | 69955489 | AX-170457233 | 0 | T | C | 0.263 | -2.13E-02 | 9.87E-02 | 2.66E+00 | 2.72E+00 | 8.30E-01 | no | Ccdc68 | intron_variant | MODIFIER |
| 18 | 69959393 | AX-170171226 | 0 | G | T | 0.263 | -2.13E-02 | 9.87E-02 | 2.66E+00 | 2.72E+00 | 8.30E-01 | no | Ccdc68 | intron_variant | MODIFIER |
| 18 | 69973299 | AX-170385105 | 0 | C | A | 0.263 | -2.13E-02 | 9.87E-02 | 2.66E+00 | 2.72E+00 | 8.30E-01 | no | Ccdc68 | downstream_gene_variant | MODIFIER |
| 18 | 69977717 | AX-170093939 | 0 | T | G | 0.263 | -2.13E-02 | 9.87E-02 | 2.66E+00 | 2.72E+00 | 8.30E-01 | no | Rab27b | downstream_gene_variant | MODIFIER |
| 18 | 69985350 | AX-170061238 | 0 | A | T | 0.263 | -2.13E-02 | 9.87E-02 | 2.66E+00 | 2.72E+00 | 8.30E-01 | no | Rab27b | synonymous_variant | LOW |
| 18 | 69985510 | AX-170061239 | 0 | G | C | 0.263 | -2.13E-02 | 9.87E-02 | 2.66E+00 | 2.72E+00 | 8.30E-01 | no | Rab27b | intron_variant | MODIFIER |
| 18 | 69987451 | AX-170319910 | 0 | C | T | 0.263 | -2.13E-02 | 9.87E-02 | 2.66E+00 | 2.72E+00 | 8.30E-01 | no | Rab27b | intron_variant | MODIFIER |
| 18 | 69989004 | AX-170253649 | 0 | T | A | 0.263 | -2.13E-02 | 9.87E-02 | 2.66E+00 | 2.72E+00 | 8.30E-01 | no | Rab27b | intron_variant | MODIFIER |
| 18 | 69989205 | AX-170261403 | 0 | A | T | 0.263 | -2.13E-02 | 9.87E-02 | 2.66E+00 | 2.72E+00 | 8.30E-01 | no | Rab27b | intron_variant | MODIFIER |
| 18 | 69990475 | AX-170186463 | 0 | G | T | 0.263 | -2.13E-02 | 9.87E-02 | 2.66E+00 | 2.72E+00 | 8.30E-01 | no | Rab27b | intron_variant | MODIFIER |
| 18 | 69990500 | AX-170093940 | 0 | C | T | 0.263 | -2.13E-02 | 9.87E-02 | 2.66E+00 | 2.72E+00 | 8.30E-01 | no | Rab27b | intron_variant | MODIFIER |
| 18 | 69996033 | AX-170148173 | 0 | T | C | 0.263 | -2.13E-02 | 9.87E-02 | 2.66E+00 | 2.72E+00 | 8.30E-01 | no | Rab27b | intron_variant | MODIFIER |
| 18 | 69996959 | AX-170527580 | 0 | C | T | 0.263 | -2.13E-02 | 9.87E-02 | 2.66E+00 | 2.72E+00 | 8.30E-01 | no | Rab27b | intron_variant | MODIFIER |
| 18 | 69999041 | AX-170463574 | 0 | A | G | 0.263 | -2.13E-02 | 9.87E-02 | 2.66E+00 | 2.72E+00 | 8.30E-01 | no | Rab27b | intron_variant | MODIFIER |
| 18 | 70107833 | AX-170203833 | 0 | T | C | 0.263 | -2.13E-02 | 9.87E-02 | 2.66E+00 | 2.72E+00 | 8.30E-01 | no | Rab27b | intron_variant | MODIFIER |
| 18 | 70649272 | AX-97831592 | 0 | G | A | 0.263 | -2.13E-02 | 9.87E-02 | 2.66E+00 | 2.72E+00 | 8.30E-01 | no | AC166779.2 | upstream_gene_variant | MODIFIER |
| 18 | 70667159 | AX-170369997 | 0 | T | C | 0.263 | -2.13E-02 | 9.87E-02 | 2.66E+00 | 2.72E+00 | 8.30E-01 | no | | intergenic_variant | MODIFIER |
| 18 | 70682667 | AX-170272335 | 0 | T | C | 0.263 | -2.13E-02 | 9.87E-02 | 2.66E+00 | 2.72E+00 | 8.30E-01 | no | | intergenic_variant | MODIFIER |
| 18 | 70708979 | AX-170260380 | 0 | T | A | 0.263 | -2.13E-02 | 9.87E-02 | 2.66E+00 | 2.72E+00 | 8.30E-01 | no | AC166779.4 | upstream_gene_variant | MODIFIER |
| 12 | 117022856 | AX-97816111 | 1 | C | T | 0.125 | -4.03E-02 | 1.95E-01 | 2.69E+00 | 2.77E+00 | 8.30E-01 | no | Ptprn2 | synonymous_variant | LOW |
| 10 | 4676027 | AX-170416866 | 0 | C | T | 0.193 | -2.03E-02 | 9.49E-02 | 2.63E+00 | 2.73E+00 | 8.30E-01 | no | Esr1 | intron_variant | MODIFIER |
| 9 | 54738302 | AX-97838292 | 1 | T | C | 0.205 | -2.48E-02 | 1.18E-01 | 2.69E+00 | 2.75E+00 | 8.31E-01 | no | Wdr61 | upstream_gene_variant | MODIFIER |
| 9 | 119233615 | AX-170102530 | 0 | C | T | 0.202 | -1.75E-02 | 8.23E-02 | 2.62E+00 | 2.73E+00 | 8.31E-01 | no | Oxsr1 | downstream_gene_variant | MODIFIER |
| 9 | 119238047 | AX-170420934 | 0 | A | G | 0.202 | -1.75E-02 | 8.23E-02 | 2.62E+00 | 2.73E+00 | 8.31E-01 | no | Oxsr1 | downstream_gene_variant | MODIFIER |
| 9 | 119242067 | AX-170033401 | 0 | T | C | 0.202 | -1.75E-02 | 8.23E-02 | 2.62E+00 | 2.73E+00 | 8.31E-01 | no | Oxsr1 | intron_variant | MODIFIER |
| 9 | 119251676 | AX-97820923 | 0 | A | G | 0.202 | -1.75E-02 | 8.23E-02 | 2.62E+00 | 2.73E+00 | 8.31E-01 | no | Oxsr1 | intron_variant | MODIFIER |
| 9 | 119251829 | AX-168415572 | 0 | C | T | 0.202 | -1.75E-02 | 8.23E-02 | 2.62E+00 | 2.73E+00 | 8.31E-01 | no | Oxsr1 | intron_variant | MODIFIER |
| 9 | 119255979 | AX-170319014 | 0 | C | T | 0.202 | -1.75E-02 | 8.23E-02 | 2.62E+00 | 2.73E+00 | 8.31E-01 | no | Oxsr1 | intron_variant | MODIFIER |
| 9 | 119261655 | AX-170424775 | 0 | C | T | 0.202 | -1.75E-02 | 8.23E-02 | 2.62E+00 | 2.73E+00 | 8.31E-01 | no | Oxsr1 | intron_variant | MODIFIER |
| 9 | 119272856 | AX-170033400 | 0 | G | C | 0.202 | -1.75E-02 | 8.23E-02 | 2.62E+00 | 2.73E+00 | 8.31E-01 | no | Oxsr1 | intron_variant | MODIFIER |
| 9 | 119338140 | AX-170033406 | 0 | G | A | 0.202 | -1.75E-02 | 8.23E-02 | 2.62E+00 | 2.73E+00 | 8.31E-01 | no | Myd88 | synonymous_variant | LOW |
| 9 | 120307491 | AX-97835101 | 0 | C | A | 0.202 | -1.75E-02 | 8.23E-02 | 2.62E+00 | 2.73E+00 | 8.31E-01 | no | Myrip | intron_variant | MODIFIER |
| 9 | 120314322 | AX-170519995 | 0 | A | G | 0.202 | -1.75E-02 | 8.23E-02 | 2.62E+00 | 2.73E+00 | 8.31E-01 | no | Myrip | intron_variant | MODIFIER |
| 9 | 120324722 | AX-97822159 | 0 | A | G | 0.202 | -1.75E-02 | 8.23E-02 | 2.62E+00 | 2.73E+00 | 8.31E-01 | no | Myrip | intron_variant | MODIFIER |

| | | | | | | | | | | | | | | | |
|---|---|---|---|---|---|---|---|---|---|---|---|---|---|---|---|
| 9 | 120325701 | AX-170033429 | 0 | A | G | 0.202 | -1.75E-02 | 8.23E-02 | 2.62E+00 | 2.73E+00 | 8.31E-01 | no | Myrip | intron_variant | MODIFIER |
| 9 | 120374232 | AX-170230854 | 0 | A | G | 0.202 | -1.75E-02 | 8.23E-02 | 2.62E+00 | 2.73E+00 | 8.31E-01 | no | Myrip | intron_variant | MODIFIER |
| 9 | 120378358 | AX-170273150 | 0 | A | C | 0.202 | -1.75E-02 | 8.23E-02 | 2.62E+00 | 2.73E+00 | 8.31E-01 | no | Myrip | intron_variant | MODIFIER |
| 9 | 120378585 | AX-170175004 | 0 | T | C | 0.202 | -1.75E-02 | 8.23E-02 | 2.62E+00 | 2.73E+00 | 8.31E-01 | no | Myrip | intron_variant | MODIFIER |
| 9 | 120381037 | AX-97792613 | 0 | T | C | 0.202 | -1.75E-02 | 8.23E-02 | 2.62E+00 | 2.73E+00 | 8.31E-01 | no | Myrip | intron_variant | MODIFIER |
| 9 | 120383397 | AX-170165427 | 0 | A | G | 0.202 | -1.75E-02 | 8.23E-02 | 2.62E+00 | 2.73E+00 | 8.31E-01 | no | Myrip | intron_variant | MODIFIER |
| 9 | 120388357 | AX-170262077 | 0 | T | C | 0.202 | -1.75E-02 | 8.23E-02 | 2.62E+00 | 2.73E+00 | 8.31E-01 | no | Myrip | intron_variant | MODIFIER |
| 9 | 120396254 | AX-170090434 | 0 | G | A | 0.202 | -1.75E-02 | 8.23E-02 | 2.62E+00 | 2.73E+00 | 8.31E-01 | no | Myrip | intron_variant | MODIFIER |
| 9 | 120417123 | AX-170374702 | 0 | A | G | 0.202 | -1.75E-02 | 8.23E-02 | 2.62E+00 | 2.73E+00 | 8.31E-01 | no | Myrip | intron_variant | MODIFIER |
| 9 | 120420848 | AX-170472139 | 0 | A | G | 0.202 | -1.75E-02 | 8.23E-02 | 2.62E+00 | 2.73E+00 | 8.31E-01 | no | Myrip | intron_variant | MODIFIER |
| 9 | 120421911 | AX-97834615 | 0 | A | G | 0.202 | -1.75E-02 | 8.23E-02 | 2.62E+00 | 2.73E+00 | 8.31E-01 | no | Myrip | intron_variant | MODIFIER |
| 9 | 120422214 | AX-170377597 | 0 | A | C | 0.202 | -1.75E-02 | 8.23E-02 | 2.62E+00 | 2.73E+00 | 8.31E-01 | no | Myrip | synonymous_variant | LOW |
| 9 | 120423032 | AX-170491166 | 0 | C | T | 0.202 | -1.75E-02 | 8.23E-02 | 2.62E+00 | 2.73E+00 | 8.31E-01 | no | Myrip | intron_variant | MODIFIER |
| 9 | 120423773 | AX-170083638 | 0 | A | G | 0.202 | -1.75E-02 | 8.23E-02 | 2.62E+00 | 2.73E+00 | 8.31E-01 | no | Myrip | intron_variant | MODIFIER |
| 9 | 120424134 | AX-170083637 | 0 | T | C | 0.202 | -1.75E-02 | 8.23E-02 | 2.62E+00 | 2.73E+00 | 8.31E-01 | no | Myrip | intron_variant | MODIFIER |
| 9 | 120426167 | AX-170280935 | 0 | T | C | 0.202 | -1.75E-02 | 8.23E-02 | 2.62E+00 | 2.73E+00 | 8.31E-01 | no | Myrip | intron_variant | MODIFIER |
| 9 | 120527135 | AX-170456723 | 0 | C | T | 0.202 | -1.75E-02 | 8.23E-02 | 2.62E+00 | 2.73E+00 | 8.31E-01 | no | Gm7229 | downstream_gene_variant | MODIFIER |
| 9 | 120531789 | AX-97826119 | 0 | G | A | 0.202 | -1.75E-02 | 8.23E-02 | 2.62E+00 | 2.73E+00 | 8.31E-01 | no | | intergenic_variant | MODIFIER |
| 9 | 120537964 | AX-170391680 | 0 | G | A | 0.202 | -1.75E-02 | 8.23E-02 | 2.62E+00 | 2.73E+00 | 8.31E-01 | no | Entpd3 | upstream_gene_variant | MODIFIER |
| 9 | 120538017 | AX-170344944 | 0 | A | G | 0.202 | -1.75E-02 | 8.23E-02 | 2.62E+00 | 2.73E+00 | 8.31E-01 | no | Entpd3 | upstream_gene_variant | MODIFIER |
| 9 | 120542780 | AX-97847643 | 0 | A | G | 0.202 | -1.75E-02 | 8.23E-02 | 2.62E+00 | 2.73E+00 | 8.31E-01 | no | Entpd3 | intron_variant | MODIFIER |
| 9 | 120543775 | AX-170293220 | 0 | G | A | 0.202 | -1.75E-02 | 8.23E-02 | 2.62E+00 | 2.73E+00 | 8.31E-01 | no | Entpd3 | intron_variant | MODIFIER |
| 9 | 120554398 | AX-170450807 | 0 | T | C | 0.202 | -1.75E-02 | 8.23E-02 | 2.62E+00 | 2.73E+00 | 8.31E-01 | no | Entpd3 | intron_variant | MODIFIER |
| 9 | 120554851 | AX-170334393 | 0 | T | C | 0.202 | -1.75E-02 | 8.23E-02 | 2.62E+00 | 2.73E+00 | 8.31E-01 | no | Entpd3 | intron_variant | MODIFIER |
| 9 | 120561018 | AX-170294877 | 0 | A | G | 0.202 | -1.75E-02 | 8.23E-02 | 2.62E+00 | 2.73E+00 | 8.31E-01 | no | Entpd3 | missense_variant | MODERATE |
| 9 | 120561283 | AX-170173193 | 0 | A | G | 0.202 | -1.75E-02 | 8.23E-02 | 2.62E+00 | 2.73E+00 | 8.31E-01 | no | Entpd3 | intron_variant | MODIFIER |
| 9 | 120697675 | AX-97792995 | 0 | A | C | 0.202 | -1.75E-02 | 8.23E-02 | 2.62E+00 | 2.73E+00 | 8.31E-01 | no | | intergenic_variant | MODIFIER |
| 9 | 120736171 | AX-170149670 | 0 | A | G | 0.202 | -1.75E-02 | 8.23E-02 | 2.62E+00 | 2.73E+00 | 8.31E-01 | no | | regulatory_region_variant | MODIFIER |
| 9 | 120736236 | AX-170474152 | 0 | G | C | 0.202 | -1.75E-02 | 8.23E-02 | 2.62E+00 | 2.73E+00 | 8.31E-01 | no | | regulatory_region_variant | MODIFIER |
| 9 | 120738393 | AX-97829224 | 0 | T | C | 0.202 | -1.75E-02 | 8.23E-02 | 2.62E+00 | 2.73E+00 | 8.31E-01 | no | | regulatory_region_variant | MODIFIER |
| 9 | 120760411 | AX-170354877 | 0 | G | A | 0.202 | -1.75E-02 | 8.23E-02 | 2.62E+00 | 2.73E+00 | 8.31E-01 | no | Gm18101 | upstream_gene_variant | MODIFIER |
| 9 | 120766288 | AX-170090436 | 0 | T | C | 0.202 | -1.75E-02 | 8.23E-02 | 2.62E+00 | 2.73E+00 | 8.31E-01 | no | | intergenic_variant | MODIFIER |
| 9 | 120799707 | AX-97857163 | 0 | C | T | 0.202 | -1.75E-02 | 8.23E-02 | 2.62E+00 | 2.73E+00 | 8.31E-01 | no | Gm47066 | downstream_gene_variant | MODIFIER |
| 9 | 120809774 | AX-170475077 | 0 | A | G | 0.202 | -1.75E-02 | 8.23E-02 | 2.62E+00 | 2.73E+00 | 8.31E-01 | no | | intergenic_variant | MODIFIER |
| 9 | 120817466 | AX-170158929 | 0 | T | C | 0.202 | -1.75E-02 | 8.23E-02 | 2.62E+00 | 2.73E+00 | 8.31E-01 | no | | intergenic_variant | MODIFIER |
| 9 | 120821433 | AX-97789100 | 0 | T | C | 0.202 | -1.75E-02 | 8.23E-02 | 2.62E+00 | 2.73E+00 | 8.31E-01 | no | | intergenic_variant | MODIFIER |
| 9 | 120839614 | AX-170326362 | 0 | T | C | 0.202 | -1.75E-02 | 8.23E-02 | 2.62E+00 | 2.73E+00 | 8.31E-01 | no | Gm39459 | intron_variant,non_coding_transcript_variant | MODIFIER |
| 9 | 120856339 | AX-170361526 | 0 | T | C | 0.202 | -1.75E-02 | 8.23E-02 | 2.62E+00 | 2.73E+00 | 8.31E-01 | no | Gm47069 | downstream_gene_variant | MODIFIER |
| 12 | 95233336 | AX-170209712 | 2 | G | A | 0.045 | 4.49E-02 | 2.11E-01 | 2.64E+00 | 2.73E+00 | 8.31E-01 | no | | intergenic_variant | MODIFIER |
| 11 | 62225425 | AX-97855208 | 1 | C | T | 0.054 | -3.45E-02 | 1.68E-01 | 2.63E+00 | 2.76E+00 | 8.31E-01 | no | Specc1 | downstream_gene_variant | MODIFIER |
| 20 | 157716342 | AX-170381085 | 0 | G | A | 0.132 | 2.54E-02 | 1.20E-01 | 2.65E+00 | 2.73E+00 | 8.31E-01 | no | Smpx | intron_variant,NMD_transcript_variant | MODIFIER |
| 4 | 107371206 | AX-170111850 | 2 | T | A | 0.273 | 2.39E-02 | 1.05E-01 | 2.59E+00 | 2.69E+00 | 8.31E-01 | no | Ndc1 | intron_variant,NMD_transcript_variant | MODIFIER |
| 8 | 46945168 | AX-170272170 | 1 | G | C | 0.152 | 3.07E-02 | 1.48E-01 | 2.72E+00 | 2.78E+00 | 8.31E-01 | no | 1700011L03Rik | downstream_gene_variant | MODIFIER |
| 8 | 48139628 | AX-170517769 | 1 | C | T | 0.152 | 3.07E-02 | 1.48E-01 | 2.72E+00 | 2.78E+00 | 8.31E-01 | no | Dctd | intron_variant | MODIFIER |
| 1 | 54005448 | AX-170520902 | 2 | T | C | 0.1 | 4.38E-02 | 2.08E-01 | 2.65E+00 | 2.74E+00 | 8.31E-01 | no | Hecw2 | intron_variant | MODIFIER |
| 3 | 31759687 | AX-170343642 | 1 | A | T | 0.161 | -1.99E-02 | 9.46E-02 | 2.65E+00 | 2.74E+00 | 8.31E-01 | no | | intergenic_variant | MODIFIER |
| 2 | 103006449 | AX-170099807 | 0 | G | C | 0.053 | -6.31E-02 | 3.00E-01 | 2.72E+00 | 2.74E+00 | 8.31E-01 | no | | intergenic_variant | MODIFIER |
| 4 | 142545914 | AX-170332589 | 1 | C | T | 0.089 | 3.55E-02 | 1.68E-01 | 2.67E+00 | 2.74E+00 | 8.31E-01 | no | | intergenic_variant | MODIFIER |
| 3 | 144090962 | AX-170281563 | 1 | A | G | 0.366 | -1.38E-02 | 6.82E-02 | 2.67E+00 | 2.77E+00 | 8.31E-01 | no | | intergenic_variant | MODIFIER |
| 13 | 47409870 | AX-97823402 | 1 | C | T | 0.5 | 1.35E-02 | 6.29E-02 | 2.62E+00 | 2.73E+00 | 8.31E-01 | no | Gm35733 | intron_variant,non_coding_transcript_variant | MODIFIER |
| 5 | 104158797 | AX-170425187 | 0 | A | G | 0.447 | 2.47E-02 | 1.14E-01 | 2.65E+00 | 2.70E+00 | 8.31E-01 | no | | regulatory_region_variant | MODIFIER |
| 20 | 147334866 | AX-170035186 | 1 | A | C | 0.027 | 2.73E-02 | 1.37E-01 | 2.59E+00 | 2.76E+00 | 8.31E-01 | no | | intergenic_variant | MODIFIER |
| 1 | 76414533 | AX-170442375 | 1 | G | T | 0.143 | 5.15E-02 | 2.44E-01 | 2.68E+00 | 2.73E+00 | 8.32E-01 | no | | intergenic_variant | MODIFIER |
| 1 | 78576010 | AX-170344576 | 1 | G | A | 0.143 | 5.15E-02 | 2.44E-01 | 2.68E+00 | 2.73E+00 | 8.32E-01 | no | | intergenic_variant | MODIFIER |

| | | | | | | | | | | | | | | | |
|---|---|---|---|---|---|---|---|---|---|---|---|---|---|---|---|
| 5 | 98437302 | AX-97808837 | 1 | C | G | 0.438 | 2.21E-02 | 1.04E-01 | 2.65E+00 | 2.72E+00 | 8.32E-01 | no | Cfap299 | intron_variant | MODIFIER |
| 2 | 122292295 | AX-170510990 | 1 | C | T | 0.143 | 4.67E-02 | 2.21E-01 | 2.69E+00 | 2.72E+00 | 8.32E-01 | no | Duox2 | missense_variant | MODERATE |
| 2 | 124859371 | AX-170177017 | 1 | C | T | 0.143 | 4.67E-02 | 2.21E-01 | 2.69E+00 | 2.72E+00 | 8.32E-01 | no | | regulatory_region_variant | MODIFIER |
| 2 | 149261934 | AX-170300116 | 1 | C | T | 0.143 | 4.67E-02 | 2.21E-01 | 2.69E+00 | 2.72E+00 | 8.32E-01 | no | Gm14135 | non_coding_transcript_exon_variant | MODIFIER |
| 15 | 95445108 | AX-170534155 | 0 | A | G | 0.14 | -3.84E-02 | 1.89E-01 | 2.72E+00 | 2.79E+00 | 8.32E-01 | no | Nell2 | intron_variant | MODIFIER |
| 1 | 33127815 | AX-170524566 | 0 | A | T | 0.421 | 1.72E-02 | 8.17E-02 | 2.67E+00 | 2.74E+00 | 8.32E-01 | no | | intergenic_variant | MODIFIER |
| 17 | 52974963 | AX-170014390 | 2 | C | T | 0.064 | 2.53E-02 | 1.25E-01 | 2.60E+00 | 2.76E+00 | 8.32E-01 | no | Kcnh8 | intron_variant | MODIFIER |
| 18 | 57651419 | AX-170501286 | 0 | T | C | 0.412 | 1.74E-02 | 8.52E-02 | 2.74E+00 | 2.81E+00 | 8.32E-01 | no | Ccdc192 | intron_variant | MODIFIER |
| 1 | 78135387 | AX-97801331 | 1 | C | T | 0.134 | 4.82E-02 | 2.28E-01 | 2.67E+00 | 2.73E+00 | 8.32E-01 | no | Pax3 | intron_variant | MODIFIER |
| 4 | 128486593 | AX-97805572 | 0 | A | C | 0.14 | -2.30E-02 | 1.14E-01 | 2.73E+00 | 2.80E+00 | 8.32E-01 | no | Csmd2 | intron_variant,non_coding_transcript_variant | MODIFIER |
| 7 | 55965394 | AX-170130376 | 0 | C | T | 0.421 | 2.09E-02 | 9.80E-02 | 2.64E+00 | 2.72E+00 | 8.32E-01 | no | Nipa2 | upstream_gene_variant | MODIFIER |
| 7 | 57134588 | AX-170356821 | 0 | C | T | 0.421 | 2.09E-02 | 9.80E-02 | 2.64E+00 | 2.72E+00 | 8.32E-01 | no | Gabrg3 | intron_variant | MODIFIER |
| 9 | 35465056 | AX-170359398 | 1 | T | C | 0.438 | -1.81E-02 | 8.93E-02 | 2.73E+00 | 2.79E+00 | 8.32E-01 | no | Cdon | intron_variant | MODIFIER |
| 8 | 83612954 | AX-170336838 | 1 | G | A | 0.205 | 2.69E-02 | 1.29E-01 | 2.67E+00 | 2.74E+00 | 8.33E-01 | no | Dnajb1 | downstream_gene_variant | MODIFIER |
| 3 | 49227752 | AX-170164153 | 0 | C | T | 0.07 | -2.78E-02 | 1.32E-01 | 2.60E+00 | 2.73E+00 | 8.33E-01 | no | | intergenic_variant | MODIFIER |
| 2 | 107657669 | AX-170345626 | 1 | G | T | 0.062 | 4.62E-02 | 2.14E-01 | 2.62E+00 | 2.71E+00 | 8.33E-01 | no | | intergenic_variant | MODIFIER |
| 12 | 107837503 | AX-170406816 | 1 | G | A | 0.116 | -4.10E-02 | 2.01E-01 | 2.70E+00 | 2.77E+00 | 8.33E-01 | no | | intergenic_variant | MODIFIER |
| 9 | 54015775 | AX-170278972 | 1 | T | A | 0.214 | -2.44E-02 | 1.18E-01 | 2.69E+00 | 2.76E+00 | 8.33E-01 | no | | intergenic_variant | MODIFIER |
| 9 | 55027465 | AX-170202199 | 1 | G | T | 0.214 | -2.44E-02 | 1.18E-01 | 2.69E+00 | 2.76E+00 | 8.33E-01 | no | Chrna3 | upstream_gene_variant | MODIFIER |
| 1 | 67235541 | AX-170307372 | 1 | A | G | 0.107 | 4.37E-02 | 2.08E-01 | 2.67E+00 | 2.73E+00 | 8.33E-01 | no | Cps1 | downstream_gene_variant | MODIFIER |
| 16 | 85959239 | AX-170466559 | 1 | T | C | 0.464 | 1.67E-02 | 7.72E-02 | 2.63E+00 | 2.71E+00 | 8.33E-01 | no | | intergenic_variant | MODIFIER |
| 6 | 103567533 | AX-97809616 | 1 | C | T | 0.054 | 6.26E-02 | 2.99E-01 | 2.67E+00 | 2.73E+00 | 8.33E-01 | no | Chl1 | intron_variant | MODIFIER |
| 6 | 144255286 | AX-170387592 | 1 | C | T | 0.054 | 6.26E-02 | 2.99E-01 | 2.67E+00 | 2.73E+00 | 8.33E-01 | no | Sox5 | intron_variant,non_coding_transcript_variant | MODIFIER |
| 7 | 122544572 | AX-170142308 | 0 | T | C | 0.404 | 1.82E-02 | 8.82E-02 | 2.68E+00 | 2.75E+00 | 8.33E-01 | no | Prkcb | intron_variant | MODIFIER |
| 7 | 122568822 | AX-97803060 | 0 | C | T | 0.404 | 1.82E-02 | 8.82E-02 | 2.68E+00 | 2.75E+00 | 8.33E-01 | no | Prkcb | intron_variant | MODIFIER |
| 7 | 122591825 | AX-170276166 | 0 | T | G | 0.404 | 1.82E-02 | 8.82E-02 | 2.68E+00 | 2.75E+00 | 8.33E-01 | no | Prkcb | intron_variant | MODIFIER |
| 7 | 122596520 | AX-170163896 | 0 | T | C | 0.404 | 1.82E-02 | 8.82E-02 | 2.68E+00 | 2.75E+00 | 8.33E-01 | no | Prkcb | intron_variant | MODIFIER |
| 7 | 122596592 | AX-97786699 | 0 | A | G | 0.404 | 1.82E-02 | 8.82E-02 | 2.68E+00 | 2.75E+00 | 8.33E-01 | no | Prkcb | intron_variant | MODIFIER |
| 7 | 122597337 | AX-170078685 | 0 | T | A | 0.404 | 1.82E-02 | 8.82E-02 | 2.68E+00 | 2.75E+00 | 8.33E-01 | no | Prkcb | intron_variant | MODIFIER |
| 7 | 122600993 | AX-170457169 | 0 | A | G | 0.404 | 1.82E-02 | 8.82E-02 | 2.68E+00 | 2.75E+00 | 8.33E-01 | no | Prkcb | intron_variant | MODIFIER |
| 7 | 122615080 | AX-97817758 | 0 | G | A | 0.404 | 1.82E-02 | 8.82E-02 | 2.68E+00 | 2.75E+00 | 8.33E-01 | no | Prkcb | intron_variant | MODIFIER |
| 7 | 122622342 | AX-170029875 | 0 | C | T | 0.404 | 1.82E-02 | 8.82E-02 | 2.68E+00 | 2.75E+00 | 8.33E-01 | no | Prkcb | intron_variant | MODIFIER |
| 13 | 71483335 | AX-170238087 | 0 | C | G | 0.053 | 3.71E-02 | 1.82E-01 | 2.65E+00 | 2.75E+00 | 8.33E-01 | no | 1700112M02Rik | upstream_gene_variant | MODIFIER |
| 7 | 103916262 | AX-170215789 | 0 | C | A | 0.456 | -1.85E-02 | 9.20E-02 | 2.71E+00 | 2.79E+00 | 8.33E-01 | no | | regulatory_region_variant | MODIFIER |
| 10 | 119442980 | AX-97796359 | 1 | A | G | 0.33 | 1.45E-02 | 7.09E-02 | 2.66E+00 | 2.75E+00 | 8.33E-01 | no | Grip1os3 | intron_variant,non_coding_transcript_variant | MODIFIER |
| 1 | 173486431 | AX-170091792 | 0 | C | G | 0.228 | 2.75E-02 | 1.21E-01 | 2.60E+00 | 2.67E+00 | 8.34E-01 | no | Ifi206 | intron_variant | MODIFIER |
| 18 | 69955358 | AX-170186582 | 1 | G | A | 0.268 | -2.06E-02 | 9.81E-02 | 2.66E+00 | 2.73E+00 | 8.34E-01 | no | Ccdc68 | intron_variant | MODIFIER |
| 18 | 69987698 | AX-170259242 | 1 | C | G | 0.268 | -2.06E-02 | 9.81E-02 | 2.66E+00 | 2.73E+00 | 8.34E-01 | no | Rab27b | intron_variant | MODIFIER |
| 1 | 122751666 | AX-170240009 | 0 | G | C | 0.553 | 1.83E-02 | 8.10E-02 | 2.60E+00 | 2.68E+00 | 8.34E-01 | no | | intergenic_variant | MODIFIER |
| 1 | 9631057 | AX-170233998 | 1 | T | A | 0.08 | 3.68E-02 | 1.75E-01 | 2.60E+00 | 2.73E+00 | 8.34E-01 | no | Vxn | downstream_gene_variant | MODIFIER |
| 20 | 92769943 | AX-97807064 | 0 | G | A | 0.272 | 1.27E-02 | 5.95E-02 | 2.62E+00 | 2.72E+00 | 8.34E-01 | no | | intergenic_variant | MODIFIER |
| 12 | 116720275 | AX-170332561 | 0 | G | A | 0.132 | -3.01E-02 | 1.47E-01 | 2.62E+00 | 2.74E+00 | 8.34E-01 | no | Ptprn2 | intron_variant | MODIFIER |
| 9 | 35047364 | AX-170397946 | 1 | C | T | 0.429 | -1.66E-02 | 8.32E-02 | 2.70E+00 | 2.78E+00 | 8.34E-01 | no | St3gal4 | 3_prime_UTR_variant | MODIFIER |
| 13 | 30939920 | AX-170008006 | 1 | A | T | 0.17 | -2.62E-02 | 1.19E-01 | 2.62E+00 | 2.69E+00 | 8.34E-01 | no | Exoc2 | intron_variant | MODIFIER |
| 13 | 32956811 | AX-170244362 | 1 | T | C | 0.17 | -2.62E-02 | 1.19E-01 | 2.62E+00 | 2.69E+00 | 8.34E-01 | no | | regulatory_region_variant | MODIFIER |
| 13 | 36421868 | AX-170008116 | 1 | A | G | 0.17 | -2.62E-02 | 1.19E-01 | 2.62E+00 | 2.69E+00 | 8.34E-01 | no | Fars2 | intron_variant | MODIFIER |
| 13 | 37283381 | AX-170174987 | 1 | A | G | 0.17 | -2.62E-02 | 1.19E-01 | 2.62E+00 | 2.69E+00 | 8.34E-01 | no | | intergenic_variant | MODIFIER |
| 13 | 47314625 | AX-170417970 | 0 | G | A | 0.465 | -1.27E-02 | 6.43E-02 | 2.66E+00 | 2.77E+00 | 8.34E-01 | no | | intergenic_variant | MODIFIER |
| 9 | 88552314 | AX-170158263 | 0 | C | G | 0.149 | -2.62E-02 | 1.22E-01 | 2.63E+00 | 2.71E+00 | 8.34E-01 | no | Zfp949 | intron_variant | MODIFIER |
| 15 | 92351121 | AX-170097741 | 1 | C | A | 0.152 | 3.83E-02 | 1.79E-01 | 2.64E+00 | 2.70E+00 | 8.34E-01 | no | Gm26760 | intron_variant,non_coding_transcript_variant | MODIFIER |
| 18 | 13015351 | AX-170446641 | 0 | C | T | 0.053 | 3.43E-02 | 1.55E-01 | 2.55E+00 | 2.71E+00 | 8.34E-01 | no | Hrh4 | intron_variant | MODIFIER |
| 7 | 94094024 | AX-170212563 | 1 | G | A | 0.089 | -3.38E-02 | 1.69E-01 | 2.62E+00 | 2.76E+00 | 8.34E-01 | no | Gm32647 | intron_variant,non_coding_transcript_variant | MODIFIER |
| 12 | 98296711 | AX-170097521 | 0 | A | T | 0.474 | -5.93E-02 | 2.96E-01 | 2.64E+00 | 2.76E+00 | 8.34E-01 | no | | intergenic_variant | MODIFIER |
| 9 | 33150868 | AX-97848231 | 1 | T | G | 0.429 | 1.87E-02 | 8.74E-02 | 2.64E+00 | 2.70E+00 | 8.34E-01 | no | Gm32171 | upstream_gene_variant | MODIFIER |

| | | | | | | | | | | | | | | | |
|---|---|---|---|---|---|---|---|---|---|---|---|---|---|---|---|
| 1 | 9160619 | AX-170490330 | 0 | T | C | 0.509 | 1.58E-02 | 7.05E-02 | 2.60E+00 | 2.68E+00 | 8.35E-01 | no | Sntg1 | intron_variant,non_coding_transcript_variant | MODIFIER |
| 8 | 102451944 | AX-170320269 | 0 | T | A | 0.228 | 2.24E-02 | 1.09E-01 | 2.66E+00 | 2.74E+00 | 8.35E-01 | no | | intergenic_variant | MODIFIER |
| 8 | 102523104 | AX-97816280 | 0 | G | T | 0.228 | 2.24E-02 | 1.09E-01 | 2.66E+00 | 2.74E+00 | 8.35E-01 | no | Gm24602 | upstream_gene_variant | MODIFIER |
| 8 | 102630112 | AX-170144981 | 0 | G | T | 0.228 | 2.24E-02 | 1.09E-01 | 2.66E+00 | 2.74E+00 | 8.35E-01 | no | Cdh11 | downstream_gene_variant | MODIFIER |
| 2 | 160353530 | AX-170324566 | 2 | G | A | 0.136 | 4.08E-02 | 1.95E-01 | 2.66E+00 | 2.73E+00 | 8.35E-01 | no | | regulatory_region_variant | MODIFIER |
| 8 | 75228417 | AX-170463788 | 1 | A | G | 0.188 | -2.50E-02 | 1.17E-01 | 2.62E+00 | 2.71E+00 | 8.35E-01 | no | Rasd2 | downstream_gene_variant | MODIFIER |
| 5 | 119154123 | AX-170361426 | 0 | G | A | 0.439 | 2.24E-02 | 1.08E-01 | 2.67E+00 | 2.72E+00 | 8.35E-01 | no | | intergenic_variant | MODIFIER |
| 7 | 122158672 | AX-170290447 | 0 | G | C | 0.386 | 1.83E-02 | 8.87E-02 | 2.68E+00 | 2.74E+00 | 8.35E-01 | no | Plk1 | upstream_gene_variant | MODIFIER |
| 7 | 122158704 | AX-97802284 | 0 | T | C | 0.386 | 1.83E-02 | 8.87E-02 | 2.68E+00 | 2.74E+00 | 8.35E-01 | no | Plk1 | upstream_gene_variant | MODIFIER |
| 7 | 122158726 | AX-170078676 | 0 | T | G | 0.386 | 1.83E-02 | 8.87E-02 | 2.68E+00 | 2.74E+00 | 8.35E-01 | no | Plk1 | upstream_gene_variant | MODIFIER |
| 7 | 122161719 | AX-170029869 | 0 | C | T | 0.386 | 1.83E-02 | 8.87E-02 | 2.68E+00 | 2.74E+00 | 8.35E-01 | no | Plk1 | intron_variant | MODIFIER |
| 7 | 122171986 | AX-170386547 | 0 | A | G | 0.386 | 1.83E-02 | 8.87E-02 | 2.68E+00 | 2.74E+00 | 8.35E-01 | no | Ern2 | intron_variant | MODIFIER |
| 7 | 122172095 | AX-170367652 | 0 | G | A | 0.386 | 1.83E-02 | 8.87E-02 | 2.68E+00 | 2.74E+00 | 8.35E-01 | no | Ern2 | intron_variant | MODIFIER |
| 7 | 122178110 | AX-170318825 | 0 | T | C | 0.386 | 1.83E-02 | 8.87E-02 | 2.68E+00 | 2.74E+00 | 8.35E-01 | no | Ern2 | intron_variant | MODIFIER |
| 7 | 122179445 | AX-97791551 | 0 | C | T | 0.386 | 1.83E-02 | 8.87E-02 | 2.68E+00 | 2.74E+00 | 8.35E-01 | no | Ern2 | intron_variant | MODIFIER |
| 7 | 122179543 | AX-170029871 | 0 | T | C | 0.386 | 1.83E-02 | 8.87E-02 | 2.68E+00 | 2.74E+00 | 8.35E-01 | no | Ern2 | intron_variant | MODIFIER |
| 7 | 122179631 | AX-170320983 | 0 | G | C | 0.386 | 1.83E-02 | 8.87E-02 | 2.68E+00 | 2.74E+00 | 8.35E-01 | no | Ern2 | intron_variant | MODIFIER |
| 7 | 122193087 | AX-170309560 | 0 | C | T | 0.386 | 1.83E-02 | 8.87E-02 | 2.68E+00 | 2.74E+00 | 8.35E-01 | no | | intergenic_variant | MODIFIER |
| 7 | 122222195 | AX-170272161 | 0 | C | T | 0.386 | 1.83E-02 | 8.87E-02 | 2.68E+00 | 2.74E+00 | 8.35E-01 | no | Chp2 | 3_prime_UTR_variant | MODIFIER |
| 7 | 122223931 | AX-170507147 | 0 | C | T | 0.386 | 1.83E-02 | 8.87E-02 | 2.68E+00 | 2.74E+00 | 8.35E-01 | no | Chp2 | downstream_gene_variant | MODIFIER |
| 9 | 64032678 | AX-170425049 | 0 | C | T | 0.184 | -2.75E-02 | 1.32E-01 | 2.68E+00 | 2.72E+00 | 8.35E-01 | no | | intergenic_variant | MODIFIER |
| 2 | 123841733 | AX-170189300 | 0 | G | C | 0.14 | -4.52E-02 | 2.21E-01 | 2.74E+00 | 2.77E+00 | 8.35E-01 | no | | intergenic_variant | MODIFIER |
| 2 | 131476013 | AX-97834498 | 0 | G | A | 0.14 | -4.52E-02 | 2.21E-01 | 2.74E+00 | 2.77E+00 | 8.35E-01 | no | | intergenic_variant | MODIFIER |
| 2 | 147242247 | AX-170379127 | 0 | C | T | 0.14 | -4.52E-02 | 2.21E-01 | 2.74E+00 | 2.77E+00 | 8.35E-01 | no | Nkx2-2os | intron_variant,non_coding_transcript_variant | MODIFIER |
| 4 | 142417084 | AX-170420929 | 1 | G | A | 0.098 | 4.15E-02 | 2.00E-01 | 2.67E+00 | 2.73E+00 | 8.35E-01 | no | | intergenic_variant | MODIFIER |
| 9 | 107368413 | AX-170528845 | 1 | C | T | 0.125 | 2.33E-02 | 1.17E-01 | 2.73E+00 | 2.81E+00 | 8.35E-01 | no | | regulatory_region_variant | MODIFIER |
| 1 | 4689883 | AX-170040259 | 1 | G | C | 0.527 | 1.61E-02 | 7.04E-02 | 2.59E+00 | 2.66E+00 | 8.35E-01 | no | Gm6085 | upstream_gene_variant | MODIFIER |
| 9 | 111943461 | AX-170118164 | 0 | C | G | 0.105 | 2.29E-02 | 1.15E-01 | 2.70E+00 | 2.78E+00 | 8.35E-01 | no | | intergenic_variant | MODIFIER |
| 13 | 49007098 | AX-170308178 | 0 | A | G | 0.5 | 1.42E-02 | 6.63E-02 | 2.61E+00 | 2.71E+00 | 8.35E-01 | no | | intergenic_variant | MODIFIER |
| 7 | 60850428 | AX-170029116 | 0 | A | T | 0.342 | -2.07E-02 | 1.04E-01 | 2.66E+00 | 2.76E+00 | 8.35E-01 | no | | intergenic_variant | MODIFIER |
| 12 | 47291376 | AX-97814723 | 0 | A | C | 0.053 | -3.52E-02 | 1.74E-01 | 2.63E+00 | 2.75E+00 | 8.35E-01 | no | | intergenic_variant | MODIFIER |
| 9 | 53898570 | AX-170096550 | 0 | T | C | 0.211 | -2.41E-02 | 1.19E-01 | 2.69E+00 | 2.75E+00 | 8.36E-01 | no | | intergenic_variant | MODIFIER |
| 9 | 53908736 | AX-170032381 | 0 | G | A | 0.211 | -2.41E-02 | 1.19E-01 | 2.69E+00 | 2.75E+00 | 8.36E-01 | no | Elmod1 | downstream_gene_variant | MODIFIER |
| 9 | 53912019 | AX-170201937 | 0 | G | A | 0.211 | -2.41E-02 | 1.19E-01 | 2.69E+00 | 2.75E+00 | 8.36E-01 | no | Elmod1 | 3_prime_UTR_variant | MODIFIER |
| 9 | 53930169 | AX-97821751 | 0 | C | T | 0.211 | -2.41E-02 | 1.19E-01 | 2.69E+00 | 2.75E+00 | 8.36E-01 | no | Elmod1 | intron_variant | MODIFIER |
| 9 | 53967547 | AX-170105773 | 0 | G | C | 0.211 | -2.41E-02 | 1.19E-01 | 2.69E+00 | 2.75E+00 | 8.36E-01 | no | Elmod1 | intron_variant | MODIFIER |
| 9 | 53970932 | AX-170224304 | 0 | C | T | 0.211 | -2.41E-02 | 1.19E-01 | 2.69E+00 | 2.75E+00 | 8.36E-01 | no | Elmod1 | intron_variant | MODIFIER |
| 9 | 53973257 | AX-170350693 | 0 | C | T | 0.211 | -2.41E-02 | 1.19E-01 | 2.69E+00 | 2.75E+00 | 8.36E-01 | no | Elmod1 | intron_variant | MODIFIER |
| 9 | 53973686 | AX-170372079 | 0 | G | A | 0.211 | -2.41E-02 | 1.19E-01 | 2.69E+00 | 2.75E+00 | 8.36E-01 | no | Elmod1 | intron_variant | MODIFIER |
| 9 | 54010307 | AX-170414209 | 0 | A | T | 0.211 | -2.41E-02 | 1.19E-01 | 2.69E+00 | 2.75E+00 | 8.36E-01 | no | | intergenic_variant | MODIFIER |
| 9 | 54014519 | AX-170475651 | 0 | G | C | 0.211 | -2.41E-02 | 1.19E-01 | 2.69E+00 | 2.75E+00 | 8.36E-01 | no | | intergenic_variant | MODIFIER |
| 9 | 54015698 | AX-97784485 | 0 | T | C | 0.211 | -2.41E-02 | 1.19E-01 | 2.69E+00 | 2.75E+00 | 8.36E-01 | no | | intergenic_variant | MODIFIER |
| 9 | 54037440 | AX-170412989 | 0 | G | A | 0.211 | -2.41E-02 | 1.19E-01 | 2.69E+00 | 2.75E+00 | 8.36E-01 | no | Tnfaip8l3 | intron_variant | MODIFIER |
| 9 | 54178688 | AX-170142882 | 0 | G | T | 0.211 | -2.41E-02 | 1.19E-01 | 2.69E+00 | 2.75E+00 | 8.36E-01 | no | Cyp19a1 | intron_variant | MODIFIER |
| 9 | 54181698 | AX-170294768 | 0 | A | G | 0.211 | -2.41E-02 | 1.19E-01 | 2.69E+00 | 2.75E+00 | 8.36E-01 | no | Cyp19a1 | intron_variant | MODIFIER |
| 9 | 54182008 | AX-170327146 | 0 | A | G | 0.211 | -2.41E-02 | 1.19E-01 | 2.69E+00 | 2.75E+00 | 8.36E-01 | no | Cyp19a1 | intron_variant | MODIFIER |
| 9 | 54183252 | AX-170510136 | 0 | A | G | 0.211 | -2.41E-02 | 1.19E-01 | 2.69E+00 | 2.75E+00 | 8.36E-01 | no | Cyp19a1 | intron_variant | MODIFIER |
| 9 | 54209004 | AX-170405957 | 0 | G | C | 0.211 | -2.41E-02 | 1.19E-01 | 2.69E+00 | 2.75E+00 | 8.36E-01 | no | Cyp19a1 | intron_variant,non_coding_transcript_variant | MODIFIER |
| 9 | 54210715 | AX-170389227 | 0 | C | T | 0.211 | -2.41E-02 | 1.19E-01 | 2.69E+00 | 2.75E+00 | 8.36E-01 | no | Cyp19a1 | intron_variant,non_coding_transcript_variant | MODIFIER |
| 9 | 54251551 | AX-170163226 | 0 | T | C | 0.211 | -2.41E-02 | 1.19E-01 | 2.69E+00 | 2.75E+00 | 8.36E-01 | no | Cyp19a1 | intron_variant,non_coding_transcript_variant | MODIFIER |
| 9 | 54251962 | AX-170168219 | 0 | A | G | 0.211 | -2.41E-02 | 1.19E-01 | 2.69E+00 | 2.75E+00 | 8.36E-01 | no | Cyp19a1 | intron_variant,non_coding_transcript_variant | MODIFIER |
| 9 | 54263050 | AX-170173110 | 0 | C | T | 0.211 | -2.41E-02 | 1.19E-01 | 2.69E+00 | 2.75E+00 | 8.36E-01 | no | Cyp19a1 | intron_variant,non_coding_transcript_variant | MODIFIER |
| 9 | 54263675 | AX-170489814 | 0 | T | C | 0.211 | -2.41E-02 | 1.19E-01 | 2.69E+00 | 2.75E+00 | 8.36E-01 | no | Gldnos | downstream_gene_variant | MODIFIER |
| 9 | 54664474 | AX-170217812 | 0 | C | T | 0.211 | -2.41E-02 | 1.19E-01 | 2.69E+00 | 2.75E+00 | 8.36E-01 | no | Acsbg1 | upstream_gene_variant | MODIFIER |

| | | | | | | | | | | | | | | | |
|---|---|---|---|---|---|---|---|---|---|---|---|---|---|---|---|
| 9 | 54677213 | AX-170096553 | 0 | G | A | 0.211 | -2.41E-02 | 1.19E-01 | 2.69E+00 | 2.75E+00 | 8.36E-01 | no | | regulatory_region_variant | MODIFIER |
| 9 | 54894653 | AX-170032397 | 0 | A | G | 0.211 | -2.41E-02 | 1.19E-01 | 2.69E+00 | 2.75E+00 | 8.36E-01 | no | Ireb2 | intron_variant | MODIFIER |
| 9 | 54895551 | AX-170091327 | 0 | A | G | 0.211 | -2.41E-02 | 1.19E-01 | 2.69E+00 | 2.75E+00 | 8.36E-01 | no | Ireb2 | intron_variant | MODIFIER |
| 9 | 54904313 | AX-170315617 | 0 | G | T | 0.211 | -2.41E-02 | 1.19E-01 | 2.69E+00 | 2.75E+00 | 8.36E-01 | no | Ireb2 | intron_variant | MODIFIER |
| 9 | 54921923 | AX-97827526 | 0 | A | G | 0.211 | -2.41E-02 | 1.19E-01 | 2.69E+00 | 2.75E+00 | 8.36E-01 | no | Hykk | intron_variant | MODIFIER |
| 9 | 54932863 | AX-170429999 | 0 | T | C | 0.211 | -2.41E-02 | 1.19E-01 | 2.69E+00 | 2.75E+00 | 8.36E-01 | no | Hykk | intron_variant | MODIFIER |
| 9 | 54941363 | AX-170188031 | 0 | A | G | 0.211 | -2.41E-02 | 1.19E-01 | 2.69E+00 | 2.75E+00 | 8.36E-01 | no | Hykk | intron_variant | MODIFIER |
| 9 | 54976428 | AX-97799622 | 0 | T | G | 0.211 | -2.41E-02 | 1.19E-01 | 2.69E+00 | 2.75E+00 | 8.36E-01 | no | Chrna5 | upstream_gene_variant | MODIFIER |
| 9 | 54994012 | AX-170496102 | 0 | C | T | 0.211 | -2.41E-02 | 1.19E-01 | 2.69E+00 | 2.75E+00 | 8.36E-01 | no | Chrna5 | intron_variant | MODIFIER |
| 9 | 54994800 | AX-170333476 | 0 | T | C | 0.211 | -2.41E-02 | 1.19E-01 | 2.69E+00 | 2.75E+00 | 8.36E-01 | no | Chrna5 | intron_variant | MODIFIER |
| 9 | 54994878 | AX-170389815 | 0 | G | A | 0.211 | -2.41E-02 | 1.19E-01 | 2.69E+00 | 2.75E+00 | 8.36E-01 | no | Chrna5 | intron_variant | MODIFIER |
| 9 | 55023267 | AX-97805044 | 0 | C | A | 0.211 | -2.41E-02 | 1.19E-01 | 2.69E+00 | 2.75E+00 | 8.36E-01 | no | Chrna3 | intron_variant | MODIFIER |
| 9 | 55027128 | AX-170504363 | 0 | C | T | 0.211 | -2.41E-02 | 1.19E-01 | 2.69E+00 | 2.75E+00 | 8.36E-01 | no | Chrna3 | upstream_gene_variant | MODIFIER |
| 9 | 55036433 | AX-170418700 | 0 | A | G | 0.211 | -2.41E-02 | 1.19E-01 | 2.69E+00 | 2.75E+00 | 8.36E-01 | no | Chrnb4 | synonymous_variant | LOW |
| 9 | 55045412 | AX-97849603 | 0 | A | G | 0.211 | -2.41E-02 | 1.19E-01 | 2.69E+00 | 2.75E+00 | 8.36E-01 | no | Chrnb4 | intron_variant | MODIFIER |
| 9 | 55055261 | AX-170176030 | 0 | T | G | 0.211 | -2.41E-02 | 1.19E-01 | 2.69E+00 | 2.75E+00 | 8.36E-01 | no | Gm47886 | downstream_gene_variant | MODIFIER |
| 9 | 55056057 | AX-170449967 | 0 | C | T | 0.211 | -2.41E-02 | 1.19E-01 | 2.69E+00 | 2.75E+00 | 8.36E-01 | no | Gm47886 | downstream_gene_variant | MODIFIER |
| 9 | 55059600 | AX-97810839 | 0 | T | G | 0.211 | -2.41E-02 | 1.19E-01 | 2.69E+00 | 2.75E+00 | 8.36E-01 | no | Gm47886 | downstream_gene_variant | MODIFIER |
| 9 | 55294868 | AX-170118315 | 0 | T | G | 0.211 | -2.41E-02 | 1.19E-01 | 2.69E+00 | 2.75E+00 | 8.36E-01 | no | Nrg4 | intron_variant | MODIFIER |
| 9 | 55307929 | AX-170090305 | 0 | A | G | 0.211 | -2.41E-02 | 1.19E-01 | 2.69E+00 | 2.75E+00 | 8.36E-01 | no | Gm15625 | downstream_gene_variant | MODIFIER |
| 9 | 55314196 | AX-170260561 | 0 | A | T | 0.211 | -2.41E-02 | 1.19E-01 | 2.69E+00 | 2.75E+00 | 8.36E-01 | no | Nrg4 | intron_variant | MODIFIER |
| 9 | 55314710 | AX-170314799 | 0 | C | T | 0.211 | -2.41E-02 | 1.19E-01 | 2.69E+00 | 2.75E+00 | 8.36E-01 | no | Nrg4 | intron_variant | MODIFIER |
| 9 | 55318449 | AX-170204555 | 0 | C | T | 0.211 | -2.41E-02 | 1.19E-01 | 2.69E+00 | 2.75E+00 | 8.36E-01 | no | Nrg4 | intron_variant | MODIFIER |
| 9 | 55319252 | AX-170384304 | 0 | G | A | 0.211 | -2.41E-02 | 1.19E-01 | 2.69E+00 | 2.75E+00 | 8.36E-01 | no | Nrg4 | intron_variant | MODIFIER |
| 9 | 55320658 | AX-97807528 | 0 | A | C | 0.211 | -2.41E-02 | 1.19E-01 | 2.69E+00 | 2.75E+00 | 8.36E-01 | no | Nrg4 | intron_variant | MODIFIER |
| 9 | 55322037 | AX-170401782 | 0 | A | G | 0.211 | -2.41E-02 | 1.19E-01 | 2.69E+00 | 2.75E+00 | 8.36E-01 | no | Tmem266 | upstream_gene_variant | MODIFIER |
| 9 | 55444850 | AX-170032413 | 0 | G | C | 0.211 | -2.41E-02 | 1.19E-01 | 2.69E+00 | 2.75E+00 | 8.36E-01 | no | | intergenic_variant | MODIFIER |
| 9 | 55693875 | AX-97814521 | 0 | G | T | 0.211 | -2.41E-02 | 1.19E-01 | 2.69E+00 | 2.75E+00 | 8.36E-01 | no | Scaper | intron_variant | MODIFIER |
| 8 | 19452131 | AX-170357347 | 0 | A | G | 0.167 | 2.10E-02 | 1.03E-01 | 2.64E+00 | 2.74E+00 | 8.36E-01 | no | Defb8 | upstream_gene_variant | MODIFIER |
| 1 | 25976141 | AX-97849374 | 0 | C | T | 0.404 | -1.63E-02 | 8.21E-02 | 2.69E+00 | 2.78E+00 | 8.36E-01 | no | | intergenic_variant | MODIFIER |
| 1 | 38870403 | AX-170098957 | 0 | T | G | 0.456 | -1.61E-02 | 7.74E-02 | 2.65E+00 | 2.73E+00 | 8.36E-01 | no | Chst10 | intron_variant | MODIFIER |
| 1 | 78228655 | AX-168397080 | 1 | T | C | 0.143 | 4.86E-02 | 2.36E-01 | 2.68E+00 | 2.73E+00 | 8.36E-01 | no | | intergenic_variant | MODIFIER |
| 16 | 97744529 | AX-170293900 | 0 | A | G | 0.465 | -1.67E-02 | 7.65E-02 | 2.59E+00 | 2.70E+00 | 8.36E-01 | no | Ripk4 | intron_variant | MODIFIER |
| 8 | 41824801 | AX-97781917 | 0 | T | C | 0.149 | 2.62E-02 | 1.32E-01 | 2.72E+00 | 2.78E+00 | 8.36E-01 | no | 2810404M03Rik | upstream_gene_variant | MODIFIER |
| 1 | 33213687 | AX-170136102 | 1 | G | T | 0.429 | 1.67E-02 | 8.11E-02 | 2.65E+00 | 2.73E+00 | 8.36E-01 | no | Gm37591 | intron_variant,non_coding_transcript_variant | MODIFIER |
| 4 | 97178257 | AX-170120374 | 0 | G | A | 0.237 | 2.40E-02 | 1.08E-01 | 2.60E+00 | 2.69E+00 | 8.36E-01 | no | Gm10192 | downstream_gene_variant | MODIFIER |
| 1 | 76019807 | AX-170393122 | 1 | A | T | 0.134 | 4.57E-02 | 2.22E-01 | 2.67E+00 | 2.73E+00 | 8.36E-01 | no | Gm29065 | upstream_gene_variant | MODIFIER |
| 10 | 10574023 | AX-170349061 | 2 | T | G | 0.218 | 2.11E-02 | 1.06E-01 | 2.69E+00 | 2.77E+00 | 8.36E-01 | no | | intergenic_variant | MODIFIER |
| 8 | 100460726 | AX-170401904 | 1 | T | C | 0.232 | 2.31E-02 | 1.13E-01 | 2.67E+00 | 2.74E+00 | 8.36E-01 | no | | intergenic_variant | MODIFIER |
| 20 | 144538263 | AX-97818386 | 0 | C | T | 0.149 | -2.57E-02 | 1.22E-01 | 2.61E+00 | 2.72E+00 | 8.36E-01 | no | Trpc5 | intron_variant | MODIFIER |
| 10 | 12562522 | AX-170127266 | 1 | G | A | 0.214 | 2.09E-02 | 1.05E-01 | 2.69E+00 | 2.77E+00 | 8.36E-01 | no | Utrn | intron_variant | MODIFIER |
| 7 | 136586936 | AX-170214218 | 1 | C | T | 0.295 | -1.70E-02 | 8.07E-02 | 2.62E+00 | 2.72E+00 | 8.36E-01 | no | | intergenic_variant | MODIFIER |
| 9 | 85503383 | AX-170032875 | 0 | A | G | 0.202 | -2.49E-02 | 1.23E-01 | 2.68E+00 | 2.74E+00 | 8.36E-01 | no | | intergenic_variant | MODIFIER |
| 9 | 85504895 | AX-170430808 | 0 | T | G | 0.202 | -2.49E-02 | 1.23E-01 | 2.68E+00 | 2.74E+00 | 8.36E-01 | no | | intergenic_variant | MODIFIER |
| 9 | 85517807 | AX-170314600 | 0 | G | C | 0.202 | -2.49E-02 | 1.23E-01 | 2.68E+00 | 2.74E+00 | 8.36E-01 | no | Gm48834 | downstream_gene_variant | MODIFIER |
| 9 | 85518290 | AX-170083009 | 0 | T | A | 0.202 | -2.49E-02 | 1.23E-01 | 2.68E+00 | 2.74E+00 | 8.36E-01 | no | Gm48834 | downstream_gene_variant | MODIFIER |
| 9 | 85518407 | AX-170083006 | 0 | A | C | 0.202 | -2.49E-02 | 1.23E-01 | 2.68E+00 | 2.74E+00 | 8.36E-01 | no | Gm48834 | downstream_gene_variant | MODIFIER |
| 9 | 85600706 | AX-170360771 | 0 | T | C | 0.202 | -2.49E-02 | 1.23E-01 | 2.68E+00 | 2.74E+00 | 8.36E-01 | no | | intergenic_variant | MODIFIER |
| 14 | 45169397 | AX-170169879 | 1 | G | A | 0.116 | -1.80E-02 | 7.47E-02 | 2.50E+00 | 2.68E+00 | 8.37E-01 | no | Txndc16 | intron_variant | MODIFIER |
| 1 | 162179534 | AX-170002151 | 1 | A | C | 0.232 | 2.47E-02 | 1.17E-01 | 2.64E+00 | 2.71E+00 | 8.37E-01 | no | Dnm3 | intron_variant | MODIFIER |
| 16 | 94619287 | AX-97819208 | 1 | T | A | 0.429 | -1.71E-02 | 8.07E-02 | 2.62E+00 | 2.71E+00 | 8.37E-01 | no | Dyrk1a | intron_variant | MODIFIER |
| 12 | 72899878 | AX-170125256 | 1 | A | T | 0.045 | 6.89E-02 | 3.41E-01 | 2.70E+00 | 2.75E+00 | 8.37E-01 | no | 4930447C04Rik | intron_variant | MODIFIER |
| 15 | 94743833 | AX-170377972 | 2 | A | G | 0.136 | 3.41E-02 | 1.62E-01 | 2.65E+00 | 2.71E+00 | 8.37E-01 | no | Tmem117 | intron_variant | MODIFIER |
| 10 | 11936485 | AX-170357937 | 0 | A | G | 0.228 | 1.98E-02 | 9.87E-02 | 2.66E+00 | 2.75E+00 | 8.37E-01 | no | Gm22194 | upstream_gene_variant | MODIFIER |

| | | | | | | | | | | | | | | | |
|---|---|---|---|---|---|---|---|---|---|---|---|---|---|---|---|
| 5 | 107602878 | AX-170202115 | 0 | T | C | 0.456 | 2.38E-02 | 1.14E-01 | 2.67E+00 | 2.71E+00 | 8.37E-01 | no | Rpap2 | intron_variant | MODIFIER |
| 5 | 112981794 | AX-170475185 | 0 | G | A | 0.456 | 2.38E-02 | 1.14E-01 | 2.67E+00 | 2.71E+00 | 8.37E-01 | no | Grk3 | intron_variant | MODIFIER |
| 5 | 113073719 | AX-170098832 | 0 | C | A | 0.456 | 2.38E-02 | 1.14E-01 | 2.67E+00 | 2.71E+00 | 8.37E-01 | no | Crybb2 | upstream_gene_variant | MODIFIER |
| 5 | 115247645 | AX-170240742 | 0 | A | G | 0.456 | 2.38E-02 | 1.14E-01 | 2.67E+00 | 2.71E+00 | 8.37E-01 | no | Rnf10 | intron_variant | MODIFIER |
| 1 | 70889296 | AX-170173599 | 1 | T | C | 0.098 | -3.46E-02 | 1.75E-01 | 2.66E+00 | 2.76E+00 | 8.37E-01 | no | Vwc2l | downstream_gene_variant | MODIFIER |
| 15 | 70925224 | AX-168414965 | 1 | A | G | 0.518 | 1.72E-02 | 7.91E-02 | 2.60E+00 | 2.70E+00 | 8.37E-01 | no | | intergenic_variant | MODIFIER |
| 9 | 104686104 | AX-97817374 | 0 | A | G | 0.123 | 2.34E-02 | 1.19E-01 | 2.72E+00 | 2.80E+00 | 8.37E-01 | no | Cpne4 | intron_variant | MODIFIER |
| 4 | 90077507 | AX-97798882 | 1 | A | C | 0.33 | 2.13E-02 | 1.03E-01 | 2.64E+00 | 2.73E+00 | 8.37E-01 | no | | intergenic_variant | MODIFIER |
| 8 | 54137958 | AX-170516271 | 0 | T | C | 0.158 | 2.77E-02 | 1.42E-01 | 2.73E+00 | 2.81E+00 | 8.37E-01 | no | Vegfc | intron_variant | MODIFIER |
| 1 | 73222449 | AX-170349462 | 1 | A | G | 0.134 | 4.18E-02 | 2.04E-01 | 2.64E+00 | 2.73E+00 | 8.37E-01 | no | | intergenic_variant | MODIFIER |
| 1 | 74445180 | AX-170041682 | 1 | T | C | 0.134 | 4.18E-02 | 2.04E-01 | 2.64E+00 | 2.73E+00 | 8.37E-01 | no | Usp37 | intron_variant | MODIFIER |
| 1 | 77169981 | AX-170452835 | 1 | A | T | 0.134 | 4.18E-02 | 2.04E-01 | 2.64E+00 | 2.73E+00 | 8.37E-01 | no | | intergenic_variant | MODIFIER |
| 1 | 77207656 | AX-170129984 | 1 | T | A | 0.134 | 4.18E-02 | 2.04E-01 | 2.64E+00 | 2.73E+00 | 8.37E-01 | no | | intergenic_variant | MODIFIER |
| 1 | 78576155 | AX-170104897 | 1 | C | T | 0.134 | 4.18E-02 | 2.04E-01 | 2.64E+00 | 2.73E+00 | 8.37E-01 | no | | intergenic_variant | MODIFIER |
| 18 | 28507492 | AX-97793947 | 0 | T | A | 0.044 | 6.88E-02 | 3.42E-01 | 2.70E+00 | 2.75E+00 | 8.37E-01 | no | | intergenic_variant | MODIFIER |
| 18 | 30312984 | AX-170369705 | 0 | G | A | 0.044 | 6.88E-02 | 3.42E-01 | 2.70E+00 | 2.75E+00 | 8.37E-01 | no | Pik3c3 | intron_variant | MODIFIER |
| 18 | 31311339 | AX-97834378 | 0 | G | C | 0.044 | 6.88E-02 | 3.42E-01 | 2.70E+00 | 2.75E+00 | 8.37E-01 | no | Rit2 | intron_variant,NMD_transcript_variant | MODIFIER |
| 18 | 32949240 | AX-170223461 | 0 | C | T | 0.044 | 6.88E-02 | 3.42E-01 | 2.70E+00 | 2.75E+00 | 8.37E-01 | no | Camk4 | intron_variant | MODIFIER |
| 18 | 47192516 | AX-170495271 | 0 | C | T | 0.044 | 6.88E-02 | 3.42E-01 | 2.70E+00 | 2.75E+00 | 8.37E-01 | no | Commd10 | intron_variant,NMD_transcript_variant | MODIFIER |
| 18 | 49954998 | AX-170139209 | 0 | A | T | 0.044 | 6.88E-02 | 3.42E-01 | 2.70E+00 | 2.75E+00 | 8.37E-01 | no | Dmxl1 | intron_variant | MODIFIER |
| 18 | 51577696 | AX-97834767 | 0 | G | A | 0.044 | 6.88E-02 | 3.42E-01 | 2.70E+00 | 2.75E+00 | 8.37E-01 | no | | intergenic_variant | MODIFIER |
| 11 | 74320683 | AX-170409360 | 0 | A | G | 0.272 | 1.40E-02 | 6.88E-02 | 2.61E+00 | 2.74E+00 | 8.37E-01 | no | n-R5s71 | downstream_gene_variant | MODIFIER |
| 20 | 86919754 | AX-170250319 | 0 | C | G | 0.254 | -1.23E-02 | 6.14E-02 | 2.65E+00 | 2.75E+00 | 8.37E-01 | no | Il1rapl1 | intron_variant | MODIFIER |
| 20 | 87167983 | AX-170034349 | 0 | A | T | 0.254 | -1.23E-02 | 6.14E-02 | 2.65E+00 | 2.75E+00 | 8.37E-01 | no | Il1rapl1 | intron_variant | MODIFIER |
| 20 | 87294711 | AX-170034343 | 0 | A | T | 0.254 | -1.23E-02 | 6.14E-02 | 2.65E+00 | 2.75E+00 | 8.37E-01 | no | Il1rapl1 | intron_variant | MODIFIER |
| 20 | 87345977 | AX-170445053 | 0 | G | C | 0.254 | -1.23E-02 | 6.14E-02 | 2.65E+00 | 2.75E+00 | 8.37E-01 | no | Il1rapl1 | intron_variant | MODIFIER |
| 20 | 87362028 | AX-97795417 | 0 | T | C | 0.254 | -1.23E-02 | 6.14E-02 | 2.65E+00 | 2.75E+00 | 8.37E-01 | no | Il1rapl1 | intron_variant | MODIFIER |
| 20 | 87374750 | AX-170110441 | 0 | T | G | 0.254 | -1.23E-02 | 6.14E-02 | 2.65E+00 | 2.75E+00 | 8.37E-01 | no | Il1rapl1 | intron_variant | MODIFIER |
| 20 | 87459296 | AX-170323344 | 0 | G | C | 0.254 | -1.23E-02 | 6.14E-02 | 2.65E+00 | 2.75E+00 | 8.37E-01 | no | Il1rapl1 | intron_variant | MODIFIER |
| 20 | 87503340 | AX-170179576 | 0 | A | C | 0.254 | -1.23E-02 | 6.14E-02 | 2.65E+00 | 2.75E+00 | 8.37E-01 | no | Il1rapl1 | intron_variant | MODIFIER |
| 20 | 87504122 | AX-168407611 | 0 | A | C | 0.254 | -1.23E-02 | 6.14E-02 | 2.65E+00 | 2.75E+00 | 8.37E-01 | no | Il1rapl1 | intron_variant | MODIFIER |
| 10 | 109314675 | AX-97845412 | 0 | G | A | 0.342 | -1.78E-02 | 8.55E-02 | 2.64E+00 | 2.72E+00 | 8.37E-01 | no | | intergenic_variant | MODIFIER |
| 10 | 109527228 | AX-170126800 | 0 | A | C | 0.342 | -1.78E-02 | 8.55E-02 | 2.64E+00 | 2.72E+00 | 8.37E-01 | no | | intergenic_variant | MODIFIER |
| 10 | 109528337 | AX-170045875 | 0 | G | A | 0.342 | -1.78E-02 | 8.55E-02 | 2.64E+00 | 2.72E+00 | 8.37E-01 | no | | intergenic_variant | MODIFIER |
| 10 | 109567746 | AX-170123843 | 0 | G | A | 0.342 | -1.78E-02 | 8.55E-02 | 2.64E+00 | 2.72E+00 | 8.37E-01 | no | | intergenic_variant | MODIFIER |
| 10 | 109573066 | AX-97852953 | 0 | C | T | 0.342 | -1.78E-02 | 8.55E-02 | 2.64E+00 | 2.72E+00 | 8.37E-01 | no | | intergenic_variant | MODIFIER |
| 10 | 109573109 | AX-170163504 | 0 | A | G | 0.342 | -1.78E-02 | 8.55E-02 | 2.64E+00 | 2.72E+00 | 8.37E-01 | no | | intergenic_variant | MODIFIER |
| 10 | 109574182 | AX-170140844 | 0 | A | G | 0.342 | -1.78E-02 | 8.55E-02 | 2.64E+00 | 2.72E+00 | 8.37E-01 | no | | intergenic_variant | MODIFIER |
| 10 | 109641650 | AX-170492939 | 0 | A | G | 0.342 | -1.78E-02 | 8.55E-02 | 2.64E+00 | 2.72E+00 | 8.37E-01 | no | | intergenic_variant | MODIFIER |
| 10 | 109641886 | AX-170188652 | 0 | G | A | 0.342 | -1.78E-02 | 8.55E-02 | 2.64E+00 | 2.72E+00 | 8.37E-01 | no | | intergenic_variant | MODIFIER |
| 10 | 109657415 | AX-170386260 | 0 | T | C | 0.342 | -1.78E-02 | 8.55E-02 | 2.64E+00 | 2.72E+00 | 8.37E-01 | no | | intergenic_variant | MODIFIER |
| 10 | 109687275 | AX-170192174 | 0 | C | G | 0.342 | -1.78E-02 | 8.55E-02 | 2.64E+00 | 2.72E+00 | 8.37E-01 | no | Nav3 | intron_variant | MODIFIER |
| 10 | 109695604 | AX-97846382 | 0 | C | A | 0.342 | -1.78E-02 | 8.55E-02 | 2.64E+00 | 2.72E+00 | 8.37E-01 | no | Nav3 | intron_variant | MODIFIER |
| 10 | 109731513 | AX-170526206 | 0 | A | G | 0.342 | -1.78E-02 | 8.55E-02 | 2.64E+00 | 2.72E+00 | 8.37E-01 | no | Nav3 | intron_variant | MODIFIER |
| 10 | 109731754 | AX-170118764 | 0 | C | T | 0.342 | -1.78E-02 | 8.55E-02 | 2.64E+00 | 2.72E+00 | 8.37E-01 | no | Nav3 | intron_variant | MODIFIER |
| 10 | 109732099 | AX-170217398 | 0 | G | A | 0.342 | -1.78E-02 | 8.55E-02 | 2.64E+00 | 2.72E+00 | 8.37E-01 | no | Nav3 | intron_variant | MODIFIER |
| 10 | 109732544 | AX-170526878 | 0 | G | A | 0.342 | -1.78E-02 | 8.55E-02 | 2.64E+00 | 2.72E+00 | 8.37E-01 | no | Nav3 | intron_variant | MODIFIER |
| 10 | 109732562 | AX-170163393 | 0 | G | A | 0.342 | -1.78E-02 | 8.55E-02 | 2.64E+00 | 2.72E+00 | 8.37E-01 | no | Nav3 | intron_variant | MODIFIER |
| 10 | 109732593 | AX-170456077 | 0 | A | C | 0.342 | -1.78E-02 | 8.55E-02 | 2.64E+00 | 2.72E+00 | 8.37E-01 | no | Nav3 | intron_variant | MODIFIER |
| 10 | 109746669 | AX-170086904 | 0 | A | C | 0.342 | -1.78E-02 | 8.55E-02 | 2.64E+00 | 2.72E+00 | 8.37E-01 | no | Nav3 | intron_variant | MODIFIER |
| 8 | 38065669 | AX-97806952 | 1 | C | T | 0.161 | 2.78E-02 | 1.39E-01 | 2.73E+00 | 2.78E+00 | 8.37E-01 | no | Sgcz | intron_variant | MODIFIER |
| 1 | 30796657 | AX-170344538 | 1 | G | A | 0.411 | 1.66E-02 | 8.14E-02 | 2.67E+00 | 2.74E+00 | 8.37E-01 | no | Gm2914 | upstream_gene_variant | MODIFIER |
| 7 | 57284809 | AX-97846791 | 1 | T | C | 0.42 | 1.97E-02 | 9.40E-02 | 2.64E+00 | 2.71E+00 | 8.37E-01 | no | Gabrg3 | intron_variant | MODIFIER |
| 7 | 57661586 | AX-170077445 | 1 | A | T | 0.42 | 1.97E-02 | 9.40E-02 | 2.64E+00 | 2.71E+00 | 8.37E-01 | no | Gabrb3 | intron_variant | MODIFIER |

| | | | | | | | | | | | | | | | |
|---|---|---|---|---|---|---|---|---|---|---|---|---|---|---|---|
| 2 | 146855725 | AX-170165856 | 0 | G | T | 0.14 | -4.42E-02 | 2.20E-01 | 2.73E+00 | 2.77E+00 | 8.38E-01 | no | Kiz | upstream_gene_variant | MODIFIER |
| 1 | 75769883 | AX-170279345 | 1 | C | T | 0.143 | 4.83E-02 | 2.37E-01 | 2.67E+00 | 2.73E+00 | 8.38E-01 | no | | intergenic_variant | MODIFIER |
| 9 | 61560182 | AX-170165211 | 0 | T | C | 0.193 | 2.79E-02 | 1.39E-01 | 2.73E+00 | 2.76E+00 | 8.38E-01 | no | | intergenic_variant | MODIFIER |
| 9 | 73806180 | AX-170082804 | 0 | G | A | 0.193 | 2.79E-02 | 1.39E-01 | 2.73E+00 | 2.76E+00 | 8.38E-01 | no | Unc13c | intron_variant | MODIFIER |
| 8 | 101281960 | AX-170080986 | 0 | C | T | 0.228 | 2.31E-02 | 1.15E-01 | 2.67E+00 | 2.75E+00 | 8.38E-01 | no | | intergenic_variant | MODIFIER |
| 1 | 28506558 | AX-170488913 | 1 | A | G | 0.402 | -1.58E-02 | 7.90E-02 | 2.66E+00 | 2.75E+00 | 8.38E-01 | no | | intergenic_variant | MODIFIER |
| 9 | 35326617 | AX-97786590 | 1 | T | C | 0.42 | -1.62E-02 | 8.34E-02 | 2.70E+00 | 2.78E+00 | 8.38E-01 | no | | regulatory_region_variant | MODIFIER |
| 8 | 97579763 | AX-97816811 | 1 | A | G | 0.214 | 2.36E-02 | 1.18E-01 | 2.68E+00 | 2.74E+00 | 8.38E-01 | no | | intergenic_variant | MODIFIER |
| 10 | 10549693 | AX-170212966 | 1 | G | A | 0.223 | 2.07E-02 | 1.05E-01 | 2.70E+00 | 2.77E+00 | 8.38E-01 | no | Rab32 | intron_variant | MODIFIER |
| 1 | 118933790 | AX-170293012 | 0 | A | T | 0.158 | 3.99E-02 | 1.88E-01 | 2.62E+00 | 2.70E+00 | 8.38E-01 | no | Gli2 | intron_variant | MODIFIER |
| 7 | 107301477 | AX-170200980 | 0 | T | C | 0.465 | 1.97E-02 | 9.30E-02 | 2.63E+00 | 2.70E+00 | 8.38E-01 | no | Platr28 | downstream_gene_variant | MODIFIER |
| 20 | 130309296 | AX-170415335 | 0 | A | C | 0.325 | 1.10E-02 | 5.29E-02 | 2.60E+00 | 2.72E+00 | 8.38E-01 | no | Diaph2 | intron_variant | MODIFIER |
| 2 | 121700833 | AX-170260516 | 2 | C | T | 0.145 | 4.55E-02 | 2.22E-01 | 2.68E+00 | 2.71E+00 | 8.38E-01 | no | Frmd5 | intron_variant | MODIFIER |
| 4 | 141931552 | AX-170120548 | 1 | C | T | 0.107 | 4.14E-02 | 2.05E-01 | 2.67E+00 | 2.73E+00 | 8.38E-01 | no | Fhad1 | intron_variant | MODIFIER |
| 7 | 106650442 | AX-170410344 | 1 | G | A | 0.482 | 1.93E-02 | 9.23E-02 | 2.63E+00 | 2.71E+00 | 8.39E-01 | no | | regulatory_region_variant | MODIFIER |
| 5 | 47679770 | AX-97780354 | 1 | A | G | 0.491 | 2.27E-02 | 9.77E-02 | 2.57E+00 | 2.65E+00 | 8.39E-01 | no | | intergenic_variant | MODIFIER |
| 6 | 112269793 | AX-170355460 | 1 | A | G | 0.054 | 5.97E-02 | 2.96E-01 | 2.68E+00 | 2.74E+00 | 8.39E-01 | no | Lmcd1 | upstream_gene_variant | MODIFIER |
| 10 | 109566153 | AX-170116438 | 1 | G | A | 0.339 | -1.76E-02 | 8.54E-02 | 2.64E+00 | 2.72E+00 | 8.39E-01 | no | | intergenic_variant | MODIFIER |
| 8 | 38837139 | AX-170168573 | 1 | G | A | 0.143 | -3.05E-02 | 1.46E-01 | 2.65E+00 | 2.70E+00 | 8.39E-01 | no | | intergenic_variant | MODIFIER |
| 15 | 6824981 | AX-170171901 | 0 | C | T | 0.237 | -2.22E-02 | 1.03E-01 | 2.62E+00 | 2.68E+00 | 8.39E-01 | no | Osmr | intron_variant | MODIFIER |
| 9 | 85927351 | AX-170512638 | 1 | T | A | 0.143 | -2.53E-02 | 1.19E-01 | 2.62E+00 | 2.70E+00 | 8.39E-01 | no | | intergenic_variant | MODIFIER |
| 7 | 40009806 | AX-170168211 | 0 | G | C | 0.351 | 1.92E-02 | 9.20E-02 | 2.63E+00 | 2.71E+00 | 8.39E-01 | no | | intergenic_variant | MODIFIER |
| 2 | 114389073 | AX-97837541 | 1 | T | C | 0.143 | 4.54E-02 | 2.24E-01 | 2.69E+00 | 2.72E+00 | 8.39E-01 | no | | intergenic_variant | MODIFIER |
| 2 | 123967122 | AX-170290581 | 1 | G | A | 0.143 | 4.54E-02 | 2.24E-01 | 2.69E+00 | 2.72E+00 | 8.39E-01 | no | | intergenic_variant | MODIFIER |
| 2 | 129925348 | AX-170019012 | 1 | C | G | 0.143 | 4.54E-02 | 2.24E-01 | 2.69E+00 | 2.72E+00 | 8.39E-01 | no | | intergenic_variant | MODIFIER |
| 2 | 75844095 | AX-170305850 | 0 | T | C | 0.053 | -4.93E-02 | 2.39E-01 | 2.64E+00 | 2.72E+00 | 8.39E-01 | no | Agps | intron_variant | MODIFIER |
| 12 | 69117544 | AX-170335568 | 0 | C | T | 0.053 | -4.93E-02 | 2.39E-01 | 2.64E+00 | 2.72E+00 | 8.39E-01 | no | Gm47518 | intron_variant,non_coding_transcript_variant | MODIFIER |
| 8 | 94389350 | AX-170130492 | 0 | A | G | 0.219 | 2.03E-02 | 1.03E-01 | 2.67E+00 | 2.76E+00 | 8.39E-01 | no | Herpud1 | splice_region_variant,intron_variant | LOW |
| 6 | 105578542 | AX-170128420 | 1 | T | A | 0.054 | 5.99E-02 | 2.96E-01 | 2.67E+00 | 2.73E+00 | 8.39E-01 | no | | intergenic_variant | MODIFIER |
| 4 | 70966625 | AX-97803687 | 1 | T | C | 0.223 | -1.90E-02 | 1.02E-01 | 2.70E+00 | 2.81E+00 | 8.39E-01 | no | | intergenic_variant | MODIFIER |
| 16 | 92183784 | AX-170263165 | 0 | T | C | 0.465 | -1.55E-02 | 7.84E-02 | 2.68E+00 | 2.75E+00 | 8.39E-01 | no | Gm41496 | intron_variant,non_coding_transcript_variant | MODIFIER |
| 18 | 52936251 | AX-170122893 | 1 | A | C | 0.241 | -2.32E-02 | 1.09E-01 | 2.62E+00 | 2.69E+00 | 8.39E-01 | no | | intergenic_variant | MODIFIER |
| 18 | 45266319 | AX-170283022 | 0 | C | G | 0.079 | 3.18E-02 | 1.60E-01 | 2.69E+00 | 2.75E+00 | 8.39E-01 | no | Kcnn2 | upstream_gene_variant | MODIFIER |
| 9 | 53910681 | AX-170032376 | 0 | G | T | 0.202 | -2.38E-02 | 1.20E-01 | 2.69E+00 | 2.75E+00 | 8.39E-01 | no | Elmod1 | downstream_gene_variant | MODIFIER |
| 9 | 53954651 | AX-170350546 | 0 | C | A | 0.202 | -2.38E-02 | 1.20E-01 | 2.69E+00 | 2.75E+00 | 8.39E-01 | no | Elmod1 | intron_variant | MODIFIER |
| 9 | 53955296 | AX-170410363 | 0 | T | C | 0.202 | -2.38E-02 | 1.20E-01 | 2.69E+00 | 2.75E+00 | 8.39E-01 | no | Elmod1 | intron_variant | MODIFIER |
| 9 | 53956726 | AX-170508440 | 0 | T | C | 0.202 | -2.38E-02 | 1.20E-01 | 2.69E+00 | 2.75E+00 | 8.39E-01 | no | Elmod1 | intron_variant | MODIFIER |
| 9 | 53966540 | AX-170388210 | 0 | T | C | 0.202 | -2.38E-02 | 1.20E-01 | 2.69E+00 | 2.75E+00 | 8.39E-01 | no | Elmod1 | intron_variant | MODIFIER |
| 9 | 53970734 | AX-170309543 | 0 | G | C | 0.202 | -2.38E-02 | 1.20E-01 | 2.69E+00 | 2.75E+00 | 8.39E-01 | no | Elmod1 | intron_variant | MODIFIER |
| 9 | 53973424 | AX-170356563 | 0 | A | G | 0.202 | -2.38E-02 | 1.20E-01 | 2.69E+00 | 2.75E+00 | 8.39E-01 | no | Elmod1 | intron_variant | MODIFIER |
| 9 | 54030853 | AX-170275985 | 0 | T | C | 0.202 | -2.38E-02 | 1.20E-01 | 2.69E+00 | 2.75E+00 | 8.39E-01 | no | Tnfaip8l3 | intron_variant | MODIFIER |
| 9 | 54030892 | AX-170472624 | 0 | T | G | 0.202 | -2.38E-02 | 1.20E-01 | 2.69E+00 | 2.75E+00 | 8.39E-01 | no | Tnfaip8l3 | intron_variant | MODIFIER |
| 9 | 54061491 | AX-97789498 | 0 | T | G | 0.202 | -2.38E-02 | 1.20E-01 | 2.69E+00 | 2.75E+00 | 8.39E-01 | no | Tnfaip8l3 | intron_variant | MODIFIER |
| 9 | 54101621 | AX-170097623 | 0 | T | C | 0.202 | -2.38E-02 | 1.20E-01 | 2.69E+00 | 2.75E+00 | 8.39E-01 | no | | intergenic_variant | MODIFIER |
| 9 | 54101739 | AX-170082475 | 0 | A | G | 0.202 | -2.38E-02 | 1.20E-01 | 2.69E+00 | 2.75E+00 | 8.39E-01 | no | | intergenic_variant | MODIFIER |
| 9 | 54132613 | AX-97791504 | 0 | A | G | 0.202 | -2.38E-02 | 1.20E-01 | 2.69E+00 | 2.75E+00 | 8.39E-01 | no | | intergenic_variant | MODIFIER |
| 9 | 54139062 | AX-170217767 | 0 | C | T | 0.202 | -2.38E-02 | 1.20E-01 | 2.69E+00 | 2.75E+00 | 8.39E-01 | no | | intergenic_variant | MODIFIER |
| 9 | 54148166 | AX-97805612 | 0 | C | A | 0.202 | -2.38E-02 | 1.20E-01 | 2.69E+00 | 2.75E+00 | 8.39E-01 | no | | intergenic_variant | MODIFIER |
| 9 | 54165596 | AX-170107450 | 0 | G | T | 0.202 | -2.38E-02 | 1.20E-01 | 2.69E+00 | 2.75E+00 | 8.39E-01 | no | Cyp19a1 | downstream_gene_variant | MODIFIER |
| 9 | 54168933 | AX-170101389 | 0 | T | C | 0.202 | -2.38E-02 | 1.20E-01 | 2.69E+00 | 2.75E+00 | 8.39E-01 | no | Cyp19a1 | intron_variant | MODIFIER |
| 9 | 54179342 | AX-170421699 | 0 | T | C | 0.202 | -2.38E-02 | 1.20E-01 | 2.69E+00 | 2.75E+00 | 8.39E-01 | no | Cyp19a1 | intron_variant | MODIFIER |
| 9 | 54197959 | AX-170090303 | 0 | T | C | 0.202 | -2.38E-02 | 1.20E-01 | 2.69E+00 | 2.75E+00 | 8.39E-01 | no | Cyp19a1 | upstream_gene_variant | MODIFIER |
| 9 | 54199488 | AX-170166747 | 0 | T | C | 0.202 | -2.38E-02 | 1.20E-01 | 2.69E+00 | 2.75E+00 | 8.39E-01 | no | Cyp19a1 | intron_variant,non_coding_transcript_variant | MODIFIER |
| 9 | 54212405 | AX-170307587 | 0 | C | T | 0.202 | -2.38E-02 | 1.20E-01 | 2.69E+00 | 2.75E+00 | 8.39E-01 | no | Cyp19a1 | intron_variant,non_coding_transcript_variant | MODIFIER |

| | | | | | | | | | | | | | | | |
|---|---|---|---|---|---|---|---|---|---|---|---|---|---|---|---|
| 9 | 54221440 | AX-170248842 | 0 | A | T | 0.202 | -2.38E-02 | 1.20E-01 | 2.69E+00 | 2.75E+00 | 8.39E-01 | no | Cyp19a1 | intron_variant,non_coding_transcript_variant | MODIFIER |
| 9 | 54222034 | AX-97790374 | 0 | T | C | 0.202 | -2.38E-02 | 1.20E-01 | 2.69E+00 | 2.75E+00 | 8.39E-01 | no | Cyp19a1 | intron_variant,non_coding_transcript_variant | MODIFIER |
| 9 | 54222276 | AX-170531192 | 0 | G | A | 0.202 | -2.38E-02 | 1.20E-01 | 2.69E+00 | 2.75E+00 | 8.39E-01 | no | Cyp19a1 | intron_variant,non_coding_transcript_variant | MODIFIER |
| 9 | 54230444 | AX-170082482 | 0 | C | T | 0.202 | -2.38E-02 | 1.20E-01 | 2.69E+00 | 2.75E+00 | 8.39E-01 | no | Cyp19a1 | intron_variant,non_coding_transcript_variant | MODIFIER |
| 9 | 54281796 | AX-170478555 | 0 | G | A | 0.202 | -2.38E-02 | 1.20E-01 | 2.69E+00 | 2.75E+00 | 8.39E-01 | no | Gldn | upstream_gene_variant | MODIFIER |
| 9 | 54282978 | AX-170414234 | 0 | A | G | 0.202 | -2.38E-02 | 1.20E-01 | 2.69E+00 | 2.75E+00 | 8.39E-01 | no | Gldn | upstream_gene_variant | MODIFIER |
| 9 | 54283444 | AX-170336923 | 0 | G | A | 0.202 | -2.38E-02 | 1.20E-01 | 2.69E+00 | 2.75E+00 | 8.39E-01 | no | Gldn | upstream_gene_variant | MODIFIER |
| 9 | 54297549 | AX-170310665 | 0 | G | A | 0.202 | -2.38E-02 | 1.20E-01 | 2.69E+00 | 2.75E+00 | 8.39E-01 | no | Gldn | intron_variant | MODIFIER |
| 9 | 54308199 | AX-170517483 | 0 | A | G | 0.202 | -2.38E-02 | 1.20E-01 | 2.69E+00 | 2.75E+00 | 8.39E-01 | no | Gldn | intron_variant | MODIFIER |
| 9 | 54309696 | AX-170517507 | 0 | C | T | 0.202 | -2.38E-02 | 1.20E-01 | 2.69E+00 | 2.75E+00 | 8.39E-01 | no | Gldn | intron_variant | MODIFIER |
| 9 | 54313919 | AX-170268408 | 0 | A | T | 0.202 | -2.38E-02 | 1.20E-01 | 2.69E+00 | 2.75E+00 | 8.39E-01 | no | Gldn | intron_variant | MODIFIER |
| 9 | 54315537 | AX-170382988 | 0 | A | G | 0.202 | -2.38E-02 | 1.20E-01 | 2.69E+00 | 2.75E+00 | 8.39E-01 | no | Gldn | intron_variant | MODIFIER |
| 9 | 54333508 | AX-170397726 | 0 | T | C | 0.202 | -2.38E-02 | 1.20E-01 | 2.69E+00 | 2.75E+00 | 8.39E-01 | no | Gldn | intron_variant | MODIFIER |
| 9 | 54470394 | AX-170098920 | 0 | A | G | 0.202 | -2.38E-02 | 1.20E-01 | 2.69E+00 | 2.75E+00 | 8.39E-01 | no | Dmxl2 | intron_variant | MODIFIER |
| 9 | 54492288 | AX-170382003 | 0 | A | G | 0.202 | -2.38E-02 | 1.20E-01 | 2.69E+00 | 2.75E+00 | 8.39E-01 | no | Dmxl2 | intron_variant | MODIFIER |
| 9 | 54531810 | AX-170512295 | 0 | A | G | 0.202 | -2.38E-02 | 1.20E-01 | 2.69E+00 | 2.75E+00 | 8.39E-01 | no | | intergenic_variant | MODIFIER |
| 9 | 54542668 | AX-170176254 | 0 | G | C | 0.202 | -2.38E-02 | 1.20E-01 | 2.69E+00 | 2.75E+00 | 8.39E-01 | no | Cib2 | downstream_gene_variant | MODIFIER |
| 9 | 54575794 | AX-170409433 | 0 | C | A | 0.202 | -2.38E-02 | 1.20E-01 | 2.69E+00 | 2.75E+00 | 8.39E-01 | no | | intergenic_variant | MODIFIER |
| 9 | 54657956 | AX-170159394 | 0 | C | T | 0.202 | -2.38E-02 | 1.20E-01 | 2.69E+00 | 2.75E+00 | 8.39E-01 | no | Acsbg1 | intron_variant | MODIFIER |
| 9 | 54659624 | AX-170312296 | 0 | C | T | 0.202 | -2.38E-02 | 1.20E-01 | 2.69E+00 | 2.75E+00 | 8.39E-01 | no | Acsbg1 | intron_variant | MODIFIER |
| 9 | 54660463 | AX-170274896 | 0 | C | T | 0.202 | -2.38E-02 | 1.20E-01 | 2.69E+00 | 2.75E+00 | 8.39E-01 | no | Acsbg1 | intron_variant | MODIFIER |
| 9 | 54677519 | AX-170409837 | 0 | G | A | 0.202 | -2.38E-02 | 1.20E-01 | 2.69E+00 | 2.75E+00 | 8.39E-01 | no | | intergenic_variant | MODIFIER |
| 9 | 54711379 | AX-170360887 | 0 | T | C | 0.202 | -2.38E-02 | 1.20E-01 | 2.69E+00 | 2.75E+00 | 8.39E-01 | no | Dnaja4 | intron_variant | MODIFIER |
| 9 | 55194655 | AX-97829683 | 0 | C | T | 0.202 | -2.38E-02 | 1.20E-01 | 2.69E+00 | 2.75E+00 | 8.39E-01 | no | Ube2q2 | intron_variant | MODIFIER |
| 9 | 55467340 | AX-97832227 | 0 | T | G | 0.202 | -2.38E-02 | 1.20E-01 | 2.69E+00 | 2.75E+00 | 8.39E-01 | no | Etfa | intron_variant | MODIFIER |
| 9 | 55485435 | AX-97842272 | 0 | T | C | 0.202 | -2.38E-02 | 1.20E-01 | 2.69E+00 | 2.75E+00 | 8.39E-01 | no | Etfa | intron_variant | MODIFIER |
| 9 | 55860262 | AX-97815463 | 0 | C | T | 0.202 | -2.38E-02 | 1.20E-01 | 2.69E+00 | 2.75E+00 | 8.39E-01 | no | Scaper | intron_variant | MODIFIER |
| 9 | 55862546 | AX-170388147 | 0 | T | A | 0.202 | -2.38E-02 | 1.20E-01 | 2.69E+00 | 2.75E+00 | 8.39E-01 | no | Scaper | intron_variant | MODIFIER |
| 9 | 55918864 | AX-97786286 | 0 | G | A | 0.202 | -2.38E-02 | 1.20E-01 | 2.69E+00 | 2.75E+00 | 8.39E-01 | no | Scaper | intron_variant | MODIFIER |
| 9 | 55934818 | AX-170032414 | 0 | G | A | 0.202 | -2.38E-02 | 1.20E-01 | 2.69E+00 | 2.75E+00 | 8.39E-01 | no | Gm7435 | upstream_gene_variant | MODIFIER |
| 9 | 55942673 | AX-170222530 | 0 | T | C | 0.202 | -2.38E-02 | 1.20E-01 | 2.69E+00 | 2.75E+00 | 8.39E-01 | no | Gm7435 | intron_variant,non_coding_transcript_variant | MODIFIER |
| 9 | 55953653 | AX-170162233 | 0 | C | T | 0.202 | -2.38E-02 | 1.20E-01 | 2.69E+00 | 2.75E+00 | 8.39E-01 | no | Gm7435 | downstream_gene_variant | MODIFIER |
| 9 | 55964732 | AX-170296531 | 0 | A | G | 0.202 | -2.38E-02 | 1.20E-01 | 2.69E+00 | 2.75E+00 | 8.39E-01 | no | 4930563M21Rik | intron_variant | MODIFIER |
| 9 | 55973746 | AX-170032421 | 0 | T | A | 0.202 | -2.38E-02 | 1.20E-01 | 2.69E+00 | 2.75E+00 | 8.39E-01 | no | 4930563M21Rik | missense_variant | MODERATE |
| 9 | 55996913 | AX-170242317 | 0 | T | G | 0.202 | -2.38E-02 | 1.20E-01 | 2.69E+00 | 2.75E+00 | 8.39E-01 | no | 4930563M21Rik | intron_variant | MODIFIER |
| 9 | 55997217 | AX-97790713 | 0 | T | C | 0.202 | -2.38E-02 | 1.20E-01 | 2.69E+00 | 2.75E+00 | 8.39E-01 | no | 4930563M21Rik | intron_variant | MODIFIER |
| 9 | 55997319 | AX-170107339 | 0 | A | G | 0.202 | -2.38E-02 | 1.20E-01 | 2.69E+00 | 2.75E+00 | 8.39E-01 | no | 4930563M21Rik | intron_variant | MODIFIER |
| 9 | 56021488 | AX-170032420 | 0 | A | G | 0.202 | -2.38E-02 | 1.20E-01 | 2.69E+00 | 2.75E+00 | 8.39E-01 | no | Gm7439 | non_coding_transcript_exon_variant | MODIFIER |
| 9 | 56022485 | AX-170242747 | 0 | C | G | 0.202 | -2.38E-02 | 1.20E-01 | 2.69E+00 | 2.75E+00 | 8.39E-01 | no | Gm7439 | downstream_gene_variant | MODIFIER |
| 9 | 56044639 | AX-170329607 | 0 | A | G | 0.202 | -2.38E-02 | 1.20E-01 | 2.69E+00 | 2.75E+00 | 8.39E-01 | no | Rcn2 | intron_variant | MODIFIER |
| 9 | 56048677 | AX-170250931 | 0 | G | T | 0.202 | -2.38E-02 | 1.20E-01 | 2.69E+00 | 2.75E+00 | 8.39E-01 | no | Rcn2 | intron_variant | MODIFIER |
| 9 | 56049407 | AX-170032417 | 0 | A | G | 0.202 | -2.38E-02 | 1.20E-01 | 2.69E+00 | 2.75E+00 | 8.39E-01 | no | Rcn2 | intron_variant | MODIFIER |
| 9 | 56070024 | AX-170032422 | 0 | C | T | 0.202 | -2.38E-02 | 1.20E-01 | 2.69E+00 | 2.75E+00 | 8.39E-01 | no | | regulatory_region_variant | MODIFIER |
| 9 | 56093200 | AX-170448758 | 0 | A | G | 0.202 | -2.38E-02 | 1.20E-01 | 2.69E+00 | 2.75E+00 | 8.39E-01 | no | Pstpip1 | intron_variant | MODIFIER |
| 9 | 56093579 | AX-170090310 | 0 | C | T | 0.202 | -2.38E-02 | 1.20E-01 | 2.69E+00 | 2.75E+00 | 8.39E-01 | no | Pstpip1 | intron_variant | MODIFIER |
| 9 | 56096144 | AX-170032419 | 0 | A | G | 0.202 | -2.38E-02 | 1.20E-01 | 2.69E+00 | 2.75E+00 | 8.39E-01 | no | Pstpip1 | intron_variant | MODIFIER |
| 9 | 56096360 | AX-170032418 | 0 | G | A | 0.202 | -2.38E-02 | 1.20E-01 | 2.69E+00 | 2.75E+00 | 8.39E-01 | no | Pstpip1 | intron_variant | MODIFIER |
| 9 | 56130201 | AX-170181539 | 0 | A | G | 0.202 | -2.38E-02 | 1.20E-01 | 2.69E+00 | 2.75E+00 | 8.39E-01 | no | Pstpip1 | downstream_gene_variant | MODIFIER |
| 9 | 56132928 | AX-97809076 | 0 | C | T | 0.202 | -2.38E-02 | 1.20E-01 | 2.69E+00 | 2.75E+00 | 8.39E-01 | no | Tspan3 | downstream_gene_variant | MODIFIER |
| 9 | 56133301 | AX-170032416 | 0 | T | C | 0.202 | -2.38E-02 | 1.20E-01 | 2.69E+00 | 2.75E+00 | 8.39E-01 | no | Tspan3 | downstream_gene_variant | MODIFIER |
| 9 | 56134878 | AX-170393728 | 0 | C | T | 0.202 | -2.38E-02 | 1.20E-01 | 2.69E+00 | 2.75E+00 | 8.39E-01 | no | Tspan3 | downstream_gene_variant | MODIFIER |
| 9 | 56137808 | AX-170280707 | 0 | T | C | 0.202 | -2.38E-02 | 1.20E-01 | 2.69E+00 | 2.75E+00 | 8.39E-01 | no | Tspan3 | intron_variant | MODIFIER |
| 9 | 56215810 | AX-170504703 | 0 | C | T | 0.202 | -2.38E-02 | 1.20E-01 | 2.69E+00 | 2.75E+00 | 8.39E-01 | no | Peak1 | intron_variant | MODIFIER |
| 9 | 56221656 | AX-97786842 | 0 | T | C | 0.202 | -2.38E-02 | 1.20E-01 | 2.69E+00 | 2.75E+00 | 8.39E-01 | no | Peak1 | intron_variant | MODIFIER |

| | | | | | | | | | | | | | | | |
|---|---|---|---|---|---|---|---|---|---|---|---|---|---|---|---|
| 9 | 56231235 | AX-170459052 | 0 | C | G | 0.202 | -2.38E-02 | 1.20E-01 | 2.69E+00 | 2.75E+00 | 8.39E-01 | no | Peak1 | intron_variant | MODIFIER |
| 9 | 56380055 | AX-170115498 | 0 | A | G | 0.202 | -2.38E-02 | 1.20E-01 | 2.69E+00 | 2.75E+00 | 8.39E-01 | no | Peak1 | intron_variant | MODIFIER |
| 9 | 56515607 | AX-170147438 | 0 | A | G | 0.202 | -2.38E-02 | 1.20E-01 | 2.69E+00 | 2.75E+00 | 8.39E-01 | no | | regulatory_region_variant | MODIFIER |
| 9 | 56515722 | AX-170090311 | 0 | C | T | 0.202 | -2.38E-02 | 1.20E-01 | 2.69E+00 | 2.75E+00 | 8.39E-01 | no | | regulatory_region_variant | MODIFIER |
| 9 | 56576098 | AX-170268149 | 0 | A | G | 0.202 | -2.38E-02 | 1.20E-01 | 2.69E+00 | 2.75E+00 | 8.39E-01 | no | | intergenic_variant | MODIFIER |
| 9 | 56577104 | AX-170082506 | 0 | A | C | 0.202 | -2.38E-02 | 1.20E-01 | 2.69E+00 | 2.75E+00 | 8.39E-01 | no | | regulatory_region_variant | MODIFIER |
| 9 | 56579230 | AX-97852843 | 0 | A | G | 0.202 | -2.38E-02 | 1.20E-01 | 2.69E+00 | 2.75E+00 | 8.39E-01 | no | | intergenic_variant | MODIFIER |
| 9 | 56583478 | AX-170032433 | 0 | A | G | 0.202 | -2.38E-02 | 1.20E-01 | 2.69E+00 | 2.75E+00 | 8.39E-01 | no | | regulatory_region_variant | MODIFIER |
| 9 | 56586960 | AX-97804930 | 0 | C | T | 0.202 | -2.38E-02 | 1.20E-01 | 2.69E+00 | 2.75E+00 | 8.39E-01 | no | | intergenic_variant | MODIFIER |
| 9 | 56667017 | AX-170377439 | 0 | A | G | 0.202 | -2.38E-02 | 1.20E-01 | 2.69E+00 | 2.75E+00 | 8.39E-01 | no | Lingo1 | intron_variant | MODIFIER |
| 9 | 56714978 | AX-170032431 | 0 | G | T | 0.202 | -2.38E-02 | 1.20E-01 | 2.69E+00 | 2.75E+00 | 8.39E-01 | no | Lingo1 | intron_variant,non_coding_transcript_variant | MODIFIER |
| 9 | 56799090 | AX-170202987 | 0 | G | C | 0.202 | -2.38E-02 | 1.20E-01 | 2.69E+00 | 2.75E+00 | 8.39E-01 | no | Lingo1 | upstream_gene_variant | MODIFIER |
| 9 | 56799164 | AX-170345449 | 0 | G | A | 0.202 | -2.38E-02 | 1.20E-01 | 2.69E+00 | 2.75E+00 | 8.39E-01 | no | Lingo1 | upstream_gene_variant | MODIFIER |
| 9 | 56802731 | AX-170228103 | 0 | A | C | 0.202 | -2.38E-02 | 1.20E-01 | 2.69E+00 | 2.75E+00 | 8.39E-01 | no | Gm47235 | upstream_gene_variant | MODIFIER |
| 9 | 56803200 | AX-170366779 | 0 | G | A | 0.202 | -2.38E-02 | 1.20E-01 | 2.69E+00 | 2.75E+00 | 8.39E-01 | no | Gm47235 | upstream_gene_variant | MODIFIER |
| 9 | 56805824 | AX-170424648 | 0 | T | C | 0.202 | -2.38E-02 | 1.20E-01 | 2.69E+00 | 2.75E+00 | 8.39E-01 | no | Gm47235 | upstream_gene_variant | MODIFIER |
| 9 | 56806009 | AX-170123893 | 0 | G | A | 0.202 | -2.38E-02 | 1.20E-01 | 2.69E+00 | 2.75E+00 | 8.39E-01 | no | Gm47235 | upstream_gene_variant | MODIFIER |
| 9 | 56808596 | AX-170199750 | 0 | A | C | 0.202 | -2.38E-02 | 1.20E-01 | 2.69E+00 | 2.75E+00 | 8.39E-01 | no | | intergenic_variant | MODIFIER |
| 9 | 56833740 | AX-170204780 | 0 | A | G | 0.202 | -2.38E-02 | 1.20E-01 | 2.69E+00 | 2.75E+00 | 8.39E-01 | no | | intergenic_variant | MODIFIER |
| 9 | 56838202 | AX-170164310 | 0 | G | A | 0.202 | -2.38E-02 | 1.20E-01 | 2.69E+00 | 2.75E+00 | 8.39E-01 | no | | intergenic_variant | MODIFIER |
| 9 | 56849546 | AX-97785310 | 0 | T | C | 0.202 | -2.38E-02 | 1.20E-01 | 2.69E+00 | 2.75E+00 | 8.39E-01 | no | Odf3l1 | intron_variant | MODIFIER |
| 9 | 56849958 | AX-170266311 | 0 | A | G | 0.202 | -2.38E-02 | 1.20E-01 | 2.69E+00 | 2.75E+00 | 8.39E-01 | no | Odf3l1 | synonymous_variant | LOW |
| 12 | 101362004 | AX-170416933 | 1 | T | A | 0.08 | 4.07E-02 | 2.05E-01 | 2.71E+00 | 2.75E+00 | 8.39E-01 | no | Gm18795 | intron_variant,non_coding_transcript_variant | MODIFIER |
| 13 | 8770631 | AX-170353557 | 2 | G | T | 0.209 | -2.62E-02 | 1.14E-01 | 2.57E+00 | 2.66E+00 | 8.39E-01 | no | Adarb2 | downstream_gene_variant | MODIFIER |
| 1 | 187962387 | AX-170364394 | 1 | C | T | 0.241 | -1.59E-02 | 9.12E-02 | 2.68E+00 | 2.83E+00 | 8.40E-01 | no | Esrrg | intron_variant | MODIFIER |
| 8 | 54114286 | AX-170147576 | 0 | A | C | 0.158 | 2.55E-02 | 1.32E-01 | 2.70E+00 | 2.78E+00 | 8.40E-01 | no | Vegfc | intron_variant | MODIFIER |
| 1 | 25737422 | AX-170219649 | 1 | G | A | 0.411 | 1.70E-02 | 8.35E-02 | 2.65E+00 | 2.72E+00 | 8.40E-01 | no | Adgrb3 | intron_variant | MODIFIER |
| 8 | 95313644 | AX-170462859 | 0 | G | A | 0.237 | -2.28E-02 | 1.15E-01 | 2.68E+00 | 2.74E+00 | 8.40E-01 | no | Tepp | intron_variant | MODIFIER |
| 9 | 54031180 | AX-170194708 | 1 | C | T | 0.214 | -2.30E-02 | 1.17E-01 | 2.69E+00 | 2.76E+00 | 8.40E-01 | no | Tnfaip8l3 | intron_variant | MODIFIER |
| 5 | 88998362 | AX-170025548 | 2 | A | T | 0.409 | -1.90E-02 | 9.77E-02 | 2.67E+00 | 2.76E+00 | 8.40E-01 | no | Slc4a4 | intron_variant | MODIFIER |
| 7 | 92412158 | AX-170122899 | 0 | A | T | 0.132 | -4.19E-02 | 2.12E-01 | 2.67E+00 | 2.74E+00 | 8.40E-01 | no | Dlg2 | intron_variant | MODIFIER |
| 7 | 93150569 | AX-170508521 | 0 | G | A | 0.132 | -4.19E-02 | 2.12E-01 | 2.67E+00 | 2.74E+00 | 8.40E-01 | no | Gm25860 | upstream_gene_variant | MODIFIER |
| 8 | 95133196 | AX-170222695 | 0 | C | T | 0.228 | 2.19E-02 | 1.10E-01 | 2.65E+00 | 2.74E+00 | 8.40E-01 | no | Kifc3 | intron_variant | MODIFIER |
| 5 | 101682672 | AX-97839523 | 0 | G | A | 0.412 | 2.18E-02 | 1.09E-01 | 2.69E+00 | 2.74E+00 | 8.40E-01 | no | | intergenic_variant | MODIFIER |
| 5 | 101683254 | AX-170072906 | 0 | C | T | 0.412 | 2.18E-02 | 1.09E-01 | 2.69E+00 | 2.74E+00 | 8.40E-01 | no | | intergenic_variant | MODIFIER |
| 5 | 101683659 | AX-170072907 | 0 | G | A | 0.412 | 2.18E-02 | 1.09E-01 | 2.69E+00 | 2.74E+00 | 8.40E-01 | no | | regulatory_region_variant | MODIFIER |
| 5 | 101684744 | AX-170227112 | 0 | G | A | 0.412 | 2.18E-02 | 1.09E-01 | 2.69E+00 | 2.74E+00 | 8.40E-01 | no | | intergenic_variant | MODIFIER |
| 5 | 101695794 | AX-170114237 | 0 | A | C | 0.412 | 2.18E-02 | 1.09E-01 | 2.69E+00 | 2.74E+00 | 8.40E-01 | no | Gm42133 | downstream_gene_variant | MODIFIER |
| 5 | 101725706 | AX-170182520 | 0 | A | G | 0.412 | 2.18E-02 | 1.09E-01 | 2.69E+00 | 2.74E+00 | 8.40E-01 | no | | intergenic_variant | MODIFIER |
| 5 | 101741915 | AX-170119634 | 0 | T | C | 0.412 | 2.18E-02 | 1.09E-01 | 2.69E+00 | 2.74E+00 | 8.40E-01 | no | | regulatory_region_variant | MODIFIER |
| 5 | 101779623 | AX-97794971 | 0 | A | G | 0.412 | 2.18E-02 | 1.09E-01 | 2.69E+00 | 2.74E+00 | 8.40E-01 | no | Cds1 | intron_variant | MODIFIER |
| 5 | 101795324 | AX-170159836 | 0 | T | A | 0.412 | 2.18E-02 | 1.09E-01 | 2.69E+00 | 2.74E+00 | 8.40E-01 | no | Cds1 | intron_variant | MODIFIER |
| 5 | 101795412 | AX-97844014 | 0 | A | C | 0.412 | 2.18E-02 | 1.09E-01 | 2.69E+00 | 2.74E+00 | 8.40E-01 | no | Cds1 | intron_variant | MODIFIER |
| 5 | 101849714 | AX-170174175 | 0 | C | A | 0.412 | 2.18E-02 | 1.09E-01 | 2.69E+00 | 2.74E+00 | 8.40E-01 | no | Wdfy3 | intron_variant | MODIFIER |
| 5 | 101857366 | AX-170089568 | 0 | G | C | 0.412 | 2.18E-02 | 1.09E-01 | 2.69E+00 | 2.74E+00 | 8.40E-01 | no | Wdfy3 | intron_variant | MODIFIER |
| 5 | 101881996 | AX-97827094 | 0 | G | A | 0.412 | 2.18E-02 | 1.09E-01 | 2.69E+00 | 2.74E+00 | 8.40E-01 | no | Wdfy3 | intron_variant | MODIFIER |
| 5 | 101898993 | AX-97807971 | 0 | A | G | 0.412 | 2.18E-02 | 1.09E-01 | 2.69E+00 | 2.74E+00 | 8.40E-01 | no | Wdfy3 | intron_variant | MODIFIER |
| 5 | 101964105 | AX-170293108 | 0 | T | C | 0.412 | 2.18E-02 | 1.09E-01 | 2.69E+00 | 2.74E+00 | 8.40E-01 | no | Wdfy3 | intron_variant | MODIFIER |
| 5 | 101964324 | AX-170025745 | 0 | C | T | 0.412 | 2.18E-02 | 1.09E-01 | 2.69E+00 | 2.74E+00 | 8.40E-01 | no | Wdfy3 | intron_variant | MODIFIER |
| 5 | 101968751 | AX-170025747 | 0 | G | A | 0.412 | 2.18E-02 | 1.09E-01 | 2.69E+00 | 2.74E+00 | 8.40E-01 | no | Wdfy3 | intron_variant | MODIFIER |
| 5 | 102019320 | AX-170025749 | 0 | C | T | 0.412 | 2.18E-02 | 1.09E-01 | 2.69E+00 | 2.74E+00 | 8.40E-01 | no | Wdfy3 | intron_variant | MODIFIER |
| 5 | 102039105 | AX-170097470 | 0 | A | C | 0.412 | 2.18E-02 | 1.09E-01 | 2.69E+00 | 2.74E+00 | 8.40E-01 | no | Wdfy3 | intron_variant | MODIFIER |
| 9 | 57956304 | AX-170304579 | 1 | G | A | 0.223 | -2.39E-02 | 1.21E-01 | 2.69E+00 | 2.75E+00 | 8.40E-01 | no | Sema7a | intron_variant | MODIFIER |
| 1 | 169125134 | AX-170467457 | 0 | G | A | 0.228 | 2.48E-02 | 1.24E-01 | 2.67E+00 | 2.73E+00 | 8.40E-01 | no | | intergenic_variant | MODIFIER |

| | | | | | | | | | | | | | | | |
|---|---|---|---|---|---|---|---|---|---|---|---|---|---|---|---|
| 9 | 88396620 | AX-97828092 | 0 | G | A | 0.132 | -2.84E-02 | 1.34E-01 | 2.63E+00 | 2.69E+00 | 8.40E-01 | no | Snx14 | intron_variant,NMD_transcript_variant | MODIFIER |
| 4 | 72734212 | AX-170089275 | 0 | C | A | 0.237 | 2.16E-02 | 9.86E-02 | 2.59E+00 | 2.68E+00 | 8.40E-01 | no | | intergenic_variant | MODIFIER |
| 5 | 73432698 | AX-97800219 | 1 | A | G | 0.42 | -1.77E-02 | 9.35E-02 | 2.68E+00 | 2.78E+00 | 8.40E-01 | no | Cwh43 | intron_variant | MODIFIER |
| 1 | 191078143 | AX-170043965 | 1 | C | G | 0.259 | 2.24E-02 | 9.30E-02 | 2.53E+00 | 2.65E+00 | 8.40E-01 | no | Nsl1 | downstream_gene_variant | MODIFIER |
| 4 | 81248410 | AX-97784563 | 1 | A | C | 0.232 | 2.32E-02 | 1.06E-01 | 2.60E+00 | 2.68E+00 | 8.41E-01 | no | | intergenic_variant | MODIFIER |
| 10 | 10745973 | AX-170434119 | 1 | T | C | 0.223 | 2.06E-02 | 1.06E-01 | 2.69E+00 | 2.77E+00 | 8.41E-01 | no | Grm1 | intron_variant | MODIFIER |
| 12 | 78780416 | AX-170007110 | 0 | T | G | 0.044 | -4.39E-02 | 2.05E-01 | 2.59E+00 | 2.69E+00 | 8.41E-01 | no | Mpp5 | intron_variant | MODIFIER |
| 5 | 73430721 | AX-170397353 | 0 | C | T | 0.439 | -1.72E-02 | 9.18E-02 | 2.67E+00 | 2.78E+00 | 8.41E-01 | no | Cwh43 | intron_variant | MODIFIER |
| 11 | 46984269 | AX-170318008 | 0 | G | A | 0.079 | 3.61E-02 | 1.93E-01 | 2.67E+00 | 2.78E+00 | 8.41E-01 | no | Sgcd | intron_variant | MODIFIER |
| 1 | 5009755 | AX-170485315 | 2 | G | A | 0.491 | 1.46E-02 | 6.67E-02 | 2.59E+00 | 2.67E+00 | 8.41E-01 | no | Rgs20 | intron_variant | MODIFIER |
| 12 | 116550301 | AX-170346040 | 1 | G | A | 0.152 | -3.01E-02 | 1.58E-01 | 2.65E+00 | 2.76E+00 | 8.42E-01 | no | Ptprn2 | intron_variant | MODIFIER |
| 9 | 35132257 | AX-170116520 | 0 | T | C | 0.43 | 1.72E-02 | 8.48E-02 | 2.64E+00 | 2.72E+00 | 8.42E-01 | no | Dcps | intron_variant | MODIFIER |
| 18 | 57407712 | AX-170162232 | 0 | T | C | 0.412 | 1.63E-02 | 8.53E-02 | 2.74E+00 | 2.80E+00 | 8.42E-01 | no | 4930511M06Rik | intron_variant,non_coding_transcript_variant | MODIFIER |
| 18 | 57481409 | AX-170265947 | 0 | T | C | 0.412 | 1.63E-02 | 8.53E-02 | 2.74E+00 | 2.80E+00 | 8.42E-01 | no | Ctxn3 | downstream_gene_variant | MODIFIER |
| 18 | 57594527 | AX-170369447 | 0 | T | C | 0.412 | 1.63E-02 | 8.53E-02 | 2.74E+00 | 2.80E+00 | 8.42E-01 | no | Ccdc192 | intron_variant | MODIFIER |
| 18 | 57598802 | AX-170098747 | 0 | A | G | 0.412 | 1.63E-02 | 8.53E-02 | 2.74E+00 | 2.80E+00 | 8.42E-01 | no | Ccdc192 | intron_variant | MODIFIER |
| 18 | 57598978 | AX-170093897 | 0 | G | T | 0.412 | 1.63E-02 | 8.53E-02 | 2.74E+00 | 2.80E+00 | 8.42E-01 | no | Ccdc192 | intron_variant | MODIFIER |
| 18 | 57651379 | AX-97800358 | 0 | G | A | 0.412 | 1.63E-02 | 8.53E-02 | 2.74E+00 | 2.80E+00 | 8.42E-01 | no | Ccdc192 | intron_variant | MODIFIER |
| 18 | 57662331 | AX-170422003 | 0 | A | G | 0.412 | 1.63E-02 | 8.53E-02 | 2.74E+00 | 2.80E+00 | 8.42E-01 | no | Ccdc192 | intron_variant | MODIFIER |
| 18 | 57664400 | AX-170204871 | 0 | T | C | 0.412 | 1.63E-02 | 8.53E-02 | 2.74E+00 | 2.80E+00 | 8.42E-01 | no | Ccdc192 | intron_variant | MODIFIER |
| 18 | 57678374 | AX-170336785 | 0 | A | G | 0.412 | 1.63E-02 | 8.53E-02 | 2.74E+00 | 2.80E+00 | 8.42E-01 | no | Ccdc192 | intron_variant | MODIFIER |
| 18 | 57747247 | AX-170060948 | 0 | A | G | 0.412 | 1.63E-02 | 8.53E-02 | 2.74E+00 | 2.80E+00 | 8.42E-01 | no | AC102232.1 | intron_variant,non_coding_transcript_variant | MODIFIER |
| 18 | 57851122 | AX-97800421 | 0 | G | A | 0.412 | 1.63E-02 | 8.53E-02 | 2.74E+00 | 2.80E+00 | 8.42E-01 | no | | regulatory_region_variant | MODIFIER |
| 18 | 57852246 | AX-170213892 | 0 | T | C | 0.412 | 1.63E-02 | 8.53E-02 | 2.74E+00 | 2.80E+00 | 8.42E-01 | no | | intergenic_variant | MODIFIER |
| 18 | 57860387 | AX-170060951 | 0 | G | A | 0.412 | 1.63E-02 | 8.53E-02 | 2.74E+00 | 2.80E+00 | 8.42E-01 | no | | intergenic_variant | MODIFIER |
| 18 | 57922594 | AX-170183605 | 0 | A | G | 0.412 | 1.63E-02 | 8.53E-02 | 2.74E+00 | 2.80E+00 | 8.42E-01 | no | Slc12a2 | intron_variant | MODIFIER |
| 18 | 57934171 | AX-170320195 | 0 | C | T | 0.412 | 1.63E-02 | 8.53E-02 | 2.74E+00 | 2.80E+00 | 8.42E-01 | no | Slc12a2 | intron_variant | MODIFIER |
| 18 | 57954736 | AX-170439727 | 0 | T | C | 0.412 | 1.63E-02 | 8.53E-02 | 2.74E+00 | 2.80E+00 | 8.42E-01 | no | | intergenic_variant | MODIFIER |
| 18 | 57960443 | AX-170398519 | 0 | A | G | 0.412 | 1.63E-02 | 8.53E-02 | 2.74E+00 | 2.80E+00 | 8.42E-01 | no | | intergenic_variant | MODIFIER |
| 18 | 58047507 | AX-170360505 | 0 | T | A | 0.412 | 1.63E-02 | 8.53E-02 | 2.74E+00 | 2.80E+00 | 8.42E-01 | no | Fbn2 | intron_variant | MODIFIER |
| 18 | 58097440 | AX-170237667 | 0 | C | T | 0.412 | 1.63E-02 | 8.53E-02 | 2.74E+00 | 2.80E+00 | 8.42E-01 | no | Fbn2 | intron_variant | MODIFIER |
| 18 | 58101452 | AX-170319719 | 0 | C | T | 0.412 | 1.63E-02 | 8.53E-02 | 2.74E+00 | 2.80E+00 | 8.42E-01 | no | Fbn2 | intron_variant | MODIFIER |
| 18 | 58105649 | AX-170346735 | 0 | A | G | 0.412 | 1.63E-02 | 8.53E-02 | 2.74E+00 | 2.80E+00 | 8.42E-01 | no | Fbn2 | intron_variant | MODIFIER |
| 18 | 58122539 | AX-97787696 | 0 | T | C | 0.412 | 1.63E-02 | 8.53E-02 | 2.74E+00 | 2.80E+00 | 8.42E-01 | no | Fbn2 | intron_variant | MODIFIER |
| 18 | 58131356 | AX-170391145 | 0 | A | G | 0.412 | 1.63E-02 | 8.53E-02 | 2.74E+00 | 2.80E+00 | 8.42E-01 | no | Fbn2 | intron_variant | MODIFIER |
| 18 | 58153731 | AX-170093900 | 0 | A | G | 0.412 | 1.63E-02 | 8.53E-02 | 2.74E+00 | 2.80E+00 | 8.42E-01 | no | Fbn2 | intron_variant | MODIFIER |
| 18 | 58181280 | AX-170060959 | 0 | A | G | 0.412 | 1.63E-02 | 8.53E-02 | 2.74E+00 | 2.80E+00 | 8.42E-01 | no | Fbn2 | intron_variant | MODIFIER |
| 18 | 58215361 | AX-170015784 | 0 | G | C | 0.412 | 1.63E-02 | 8.53E-02 | 2.74E+00 | 2.80E+00 | 8.42E-01 | no | Fbn2 | upstream_gene_variant | MODIFIER |
| 18 | 58227107 | AX-97837131 | 0 | A | G | 0.412 | 1.63E-02 | 8.53E-02 | 2.74E+00 | 2.80E+00 | 8.42E-01 | no | Gm25660 | downstream_gene_variant | MODIFIER |
| 18 | 58227423 | AX-170346710 | 0 | A | G | 0.412 | 1.63E-02 | 8.53E-02 | 2.74E+00 | 2.80E+00 | 8.42E-01 | no | Gm25660 | downstream_gene_variant | MODIFIER |
| 18 | 58232061 | AX-170253134 | 0 | C | A | 0.412 | 1.63E-02 | 8.53E-02 | 2.74E+00 | 2.80E+00 | 8.42E-01 | no | Gm25660 | upstream_gene_variant | MODIFIER |
| 18 | 58295672 | AX-170238814 | 0 | C | T | 0.412 | 1.63E-02 | 8.53E-02 | 2.74E+00 | 2.80E+00 | 8.42E-01 | no | | intergenic_variant | MODIFIER |
| 18 | 58306422 | AX-170489579 | 0 | A | G | 0.412 | 1.63E-02 | 8.53E-02 | 2.74E+00 | 2.80E+00 | 8.42E-01 | no | | intergenic_variant | MODIFIER |
| 18 | 58323499 | AX-170339624 | 0 | A | G | 0.412 | 1.63E-02 | 8.53E-02 | 2.74E+00 | 2.80E+00 | 8.42E-01 | no | | intergenic_variant | MODIFIER |
| 18 | 58341541 | AX-170529157 | 0 | C | T | 0.412 | 1.63E-02 | 8.53E-02 | 2.74E+00 | 2.80E+00 | 8.42E-01 | no | | intergenic_variant | MODIFIER |
| 18 | 58347513 | AX-168404282 | 0 | T | C | 0.412 | 1.63E-02 | 8.53E-02 | 2.74E+00 | 2.80E+00 | 8.42E-01 | no | | intergenic_variant | MODIFIER |
| 18 | 58351187 | AX-170093901 | 0 | A | G | 0.412 | 1.63E-02 | 8.53E-02 | 2.74E+00 | 2.80E+00 | 8.42E-01 | no | | intergenic_variant | MODIFIER |
| 13 | 37488585 | AX-97794951 | 0 | A | G | 0.193 | -2.41E-02 | 1.15E-01 | 2.62E+00 | 2.69E+00 | 8.42E-01 | no | | intergenic_variant | MODIFIER |
| 13 | 37490293 | AX-170506998 | 0 | C | T | 0.193 | -2.41E-02 | 1.15E-01 | 2.62E+00 | 2.69E+00 | 8.42E-01 | no | | intergenic_variant | MODIFIER |
| 13 | 37525631 | AX-170204004 | 0 | C | A | 0.193 | -2.41E-02 | 1.15E-01 | 2.62E+00 | 2.69E+00 | 8.42E-01 | no | | regulatory_region_variant | MODIFIER |
| 13 | 37537495 | AX-170295257 | 0 | G | A | 0.193 | -2.41E-02 | 1.15E-01 | 2.62E+00 | 2.69E+00 | 8.42E-01 | no | Gm30918 | upstream_gene_variant | MODIFIER |
| 13 | 37537861 | AX-170189692 | 0 | G | C | 0.193 | -2.41E-02 | 1.15E-01 | 2.62E+00 | 2.69E+00 | 8.42E-01 | no | Gm30918 | upstream_gene_variant | MODIFIER |
| 13 | 37539967 | AX-170050989 | 0 | C | T | 0.193 | -2.41E-02 | 1.15E-01 | 2.62E+00 | 2.69E+00 | 8.42E-01 | no | Gm30918 | upstream_gene_variant | MODIFIER |
| 13 | 37543405 | AX-170438986 | 0 | T | C | 0.193 | -2.41E-02 | 1.15E-01 | 2.62E+00 | 2.69E+00 | 8.42E-01 | no | Gm30918 | upstream_gene_variant | MODIFIER |

| | | | | | | | | | | | | | | | |
|---|---|---|---|---|---|---|---|---|---|---|---|---|---|---|---|
| 13 | 37543603 | AX-170343277 | 0 | A | G | 0.193 | -2.41E-02 | 1.15E-01 | 2.62E+00 | 2.69E+00 | 8.42E-01 | no | Gm30918 | upstream_gene_variant | MODIFIER |
| 13 | 37544191 | AX-97851833 | 0 | C | T | 0.193 | -2.41E-02 | 1.15E-01 | 2.62E+00 | 2.69E+00 | 8.42E-01 | no | Gm30918 | upstream_gene_variant | MODIFIER |
| 13 | 37550925 | AX-170167180 | 0 | C | A | 0.193 | -2.41E-02 | 1.15E-01 | 2.62E+00 | 2.69E+00 | 8.42E-01 | no | Gm30918 | downstream_gene_variant | MODIFIER |
| 13 | 37598236 | AX-97794511 | 0 | G | A | 0.193 | -2.41E-02 | 1.15E-01 | 2.62E+00 | 2.69E+00 | 8.42E-01 | no | | regulatory_region_variant | MODIFIER |
| 13 | 37651924 | AX-170214868 | 0 | C | T | 0.193 | -2.41E-02 | 1.15E-01 | 2.62E+00 | 2.69E+00 | 8.42E-01 | no | Gm40916 | upstream_gene_variant | MODIFIER |
| 13 | 37652388 | AX-97846772 | 0 | T | C | 0.193 | -2.41E-02 | 1.15E-01 | 2.62E+00 | 2.69E+00 | 8.42E-01 | no | AI463229 | downstream_gene_variant | MODIFIER |
| 13 | 37656069 | AX-170458732 | 0 | G | A | 0.193 | -2.41E-02 | 1.15E-01 | 2.62E+00 | 2.69E+00 | 8.42E-01 | no | AI463229 | downstream_gene_variant | MODIFIER |
| 13 | 37661688 | AX-97789767 | 0 | C | T | 0.193 | -2.41E-02 | 1.15E-01 | 2.62E+00 | 2.69E+00 | 8.42E-01 | no | AI463229 | intron_variant,non_coding_transcript_variant | MODIFIER |
| 13 | 37850989 | AX-170467151 | 0 | T | C | 0.193 | -2.41E-02 | 1.15E-01 | 2.62E+00 | 2.69E+00 | 8.42E-01 | no | Rreb1 | intron_variant | MODIFIER |
| 13 | 37882820 | AX-170368235 | 0 | T | C | 0.193 | -2.41E-02 | 1.15E-01 | 2.62E+00 | 2.69E+00 | 8.42E-01 | no | Rreb1 | intron_variant | MODIFIER |
| 13 | 37963173 | AX-170442677 | 0 | G | A | 0.193 | -2.41E-02 | 1.15E-01 | 2.62E+00 | 2.69E+00 | 8.42E-01 | no | Ssr1 | downstream_gene_variant | MODIFIER |
| 7 | 136975801 | AX-170232816 | 1 | A | G | 0.295 | -1.47E-02 | 7.16E-02 | 2.60E+00 | 2.71E+00 | 8.42E-01 | no | Mgmt | intron_variant | MODIFIER |
| 7 | 117771064 | AX-170425283 | 0 | C | T | 0.395 | 1.79E-02 | 9.02E-02 | 2.67E+00 | 2.73E+00 | 8.42E-01 | no | | intergenic_variant | MODIFIER |
| 5 | 122693589 | AX-170377927 | 0 | C | A | 0.447 | -2.18E-02 | 1.12E-01 | 2.72E+00 | 2.76E+00 | 8.42E-01 | no | P2rx7 | downstream_gene_variant | MODIFIER |
| 5 | 125332524 | AX-170374765 | 0 | G | T | 0.447 | -2.18E-02 | 1.12E-01 | 2.72E+00 | 2.76E+00 | 8.42E-01 | no | Scarb1 | intron_variant | MODIFIER |
| 5 | 125352527 | AX-97823329 | 0 | A | G | 0.447 | -2.18E-02 | 1.12E-01 | 2.72E+00 | 2.76E+00 | 8.42E-01 | no | | regulatory_region_variant | MODIFIER |
| 5 | 125409819 | AX-97784584 | 0 | T | C | 0.447 | -2.18E-02 | 1.12E-01 | 2.72E+00 | 2.76E+00 | 8.42E-01 | no | Dhx37 | downstream_gene_variant | MODIFIER |
| 5 | 125415631 | AX-170340438 | 0 | A | G | 0.447 | -2.18E-02 | 1.12E-01 | 2.72E+00 | 2.76E+00 | 8.42E-01 | no | Dhx37 | intron_variant | MODIFIER |
| 5 | 125454935 | AX-97793390 | 0 | G | A | 0.447 | -2.18E-02 | 1.12E-01 | 2.72E+00 | 2.76E+00 | 8.42E-01 | no | Bri3bp | 3_prime_UTR_variant | MODIFIER |
| 5 | 125462378 | AX-170026096 | 0 | T | C | 0.447 | -2.18E-02 | 1.12E-01 | 2.72E+00 | 2.76E+00 | 8.42E-01 | no | Bri3bp | downstream_gene_variant | MODIFIER |
| 5 | 125499531 | AX-170026099 | 0 | G | A | 0.447 | -2.18E-02 | 1.12E-01 | 2.72E+00 | 2.76E+00 | 8.42E-01 | no | Aacs | intron_variant | MODIFIER |
| 5 | 125499717 | AX-170386246 | 0 | G | C | 0.447 | -2.18E-02 | 1.12E-01 | 2.72E+00 | 2.76E+00 | 8.42E-01 | no | Aacs | intron_variant | MODIFIER |
| 5 | 125514434 | AX-97821960 | 0 | G | A | 0.447 | -2.18E-02 | 1.12E-01 | 2.72E+00 | 2.76E+00 | 8.42E-01 | no | Aacs | intron_variant | MODIFIER |
| 5 | 125515278 | AX-170026100 | 0 | A | C | 0.447 | -2.18E-02 | 1.12E-01 | 2.72E+00 | 2.76E+00 | 8.42E-01 | no | Aacs | intron_variant | MODIFIER |
| 5 | 125516669 | AX-170481395 | 0 | A | G | 0.447 | -2.18E-02 | 1.12E-01 | 2.72E+00 | 2.76E+00 | 8.42E-01 | no | Aacs | 3_prime_UTR_variant | MODIFIER |
| 5 | 125518001 | AX-170493542 | 0 | A | G | 0.447 | -2.18E-02 | 1.12E-01 | 2.72E+00 | 2.76E+00 | 8.42E-01 | no | Aacs | downstream_gene_variant | MODIFIER |
| 5 | 125617819 | AX-97826520 | 0 | G | A | 0.447 | -2.18E-02 | 1.12E-01 | 2.72E+00 | 2.76E+00 | 8.42E-01 | no | Tmem132b | intron_variant | MODIFIER |
| 5 | 125655192 | AX-97806496 | 0 | C | T | 0.447 | -2.18E-02 | 1.12E-01 | 2.72E+00 | 2.76E+00 | 8.42E-01 | no | Tmem132b | intron_variant | MODIFIER |
| 5 | 125661700 | AX-170026095 | 0 | A | G | 0.447 | -2.18E-02 | 1.12E-01 | 2.72E+00 | 2.76E+00 | 8.42E-01 | no | Tmem132b | intron_variant | MODIFIER |
| 5 | 125672181 | AX-170121580 | 0 | T | C | 0.447 | -2.18E-02 | 1.12E-01 | 2.72E+00 | 2.76E+00 | 8.42E-01 | no | Tmem132b | intron_variant | MODIFIER |
| 5 | 125675136 | AX-170290401 | 0 | A | C | 0.447 | -2.18E-02 | 1.12E-01 | 2.72E+00 | 2.76E+00 | 8.42E-01 | no | Tmem132b | intron_variant | MODIFIER |
| 5 | 125679089 | AX-97850410 | 0 | T | C | 0.447 | -2.18E-02 | 1.12E-01 | 2.72E+00 | 2.76E+00 | 8.42E-01 | no | Tmem132b | intron_variant | MODIFIER |
| 5 | 125682487 | AX-170139930 | 0 | T | C | 0.447 | -2.18E-02 | 1.12E-01 | 2.72E+00 | 2.76E+00 | 8.42E-01 | no | Tmem132b | intron_variant | MODIFIER |
| 5 | 125685208 | AX-170228786 | 0 | C | T | 0.447 | -2.18E-02 | 1.12E-01 | 2.72E+00 | 2.76E+00 | 8.42E-01 | no | Tmem132b | intron_variant | MODIFIER |
| 5 | 125690984 | AX-170350100 | 0 | A | G | 0.447 | -2.18E-02 | 1.12E-01 | 2.72E+00 | 2.76E+00 | 8.42E-01 | no | Tmem132b | intron_variant | MODIFIER |
| 5 | 125691327 | AX-170340509 | 0 | G | A | 0.447 | -2.18E-02 | 1.12E-01 | 2.72E+00 | 2.76E+00 | 8.42E-01 | no | Tmem132b | intron_variant | MODIFIER |
| 5 | 125704062 | AX-170379428 | 0 | T | C | 0.447 | -2.18E-02 | 1.12E-01 | 2.72E+00 | 2.76E+00 | 8.42E-01 | no | Tmem132b | intron_variant | MODIFIER |
| 5 | 125705488 | AX-97813011 | 0 | T | C | 0.447 | -2.18E-02 | 1.12E-01 | 2.72E+00 | 2.76E+00 | 8.42E-01 | no | Tmem132b | intron_variant | MODIFIER |
| 5 | 125705699 | AX-170242056 | 0 | A | G | 0.447 | -2.18E-02 | 1.12E-01 | 2.72E+00 | 2.76E+00 | 8.42E-01 | no | Tmem132b | intron_variant | MODIFIER |
| 5 | 125706290 | AX-170291331 | 0 | T | C | 0.447 | -2.18E-02 | 1.12E-01 | 2.72E+00 | 2.76E+00 | 8.42E-01 | no | Tmem132b | intron_variant | MODIFIER |
| 5 | 125726122 | AX-170073407 | 0 | T | C | 0.447 | -2.18E-02 | 1.12E-01 | 2.72E+00 | 2.76E+00 | 8.42E-01 | no | Tmem132b | intron_variant | MODIFIER |
| 5 | 125731609 | AX-97805584 | 0 | A | G | 0.447 | -2.18E-02 | 1.12E-01 | 2.72E+00 | 2.76E+00 | 8.42E-01 | no | Tmem132b | intron_variant | MODIFIER |
| 5 | 125735567 | AX-170522809 | 0 | T | G | 0.447 | -2.18E-02 | 1.12E-01 | 2.72E+00 | 2.76E+00 | 8.42E-01 | no | Tmem132b | intron_variant | MODIFIER |
| 5 | 125743534 | AX-170360816 | 0 | T | C | 0.447 | -2.18E-02 | 1.12E-01 | 2.72E+00 | 2.76E+00 | 8.42E-01 | no | Tmem132b | intron_variant | MODIFIER |
| 5 | 125744390 | AX-170478913 | 0 | G | A | 0.447 | -2.18E-02 | 1.12E-01 | 2.72E+00 | 2.76E+00 | 8.42E-01 | no | Tmem132b | intron_variant | MODIFIER |
| 5 | 125746038 | AX-170312943 | 0 | C | T | 0.447 | -2.18E-02 | 1.12E-01 | 2.72E+00 | 2.76E+00 | 8.42E-01 | no | Tmem132b | intron_variant | MODIFIER |
| 5 | 125749267 | AX-170244546 | 0 | C | T | 0.447 | -2.18E-02 | 1.12E-01 | 2.72E+00 | 2.76E+00 | 8.42E-01 | no | Tmem132b | synonymous_variant | LOW |
| 5 | 125752259 | AX-170331038 | 0 | G | T | 0.447 | -2.18E-02 | 1.12E-01 | 2.72E+00 | 2.76E+00 | 8.42E-01 | no | Tmem132b | intron_variant | MODIFIER |
| 5 | 125759039 | AX-170522536 | 0 | G | A | 0.447 | -2.18E-02 | 1.12E-01 | 2.72E+00 | 2.76E+00 | 8.42E-01 | no | Tmem132b | intron_variant | MODIFIER |
| 5 | 125760616 | AX-97783606 | 0 | T | C | 0.447 | -2.18E-02 | 1.12E-01 | 2.72E+00 | 2.76E+00 | 8.42E-01 | no | Tmem132b | intron_variant | MODIFIER |
| 5 | 125769954 | AX-170461060 | 0 | A | G | 0.447 | -2.18E-02 | 1.12E-01 | 2.72E+00 | 2.76E+00 | 8.42E-01 | no | Tmem132b | intron_variant | MODIFIER |
| 5 | 125776223 | AX-170212783 | 0 | C | T | 0.447 | -2.18E-02 | 1.12E-01 | 2.72E+00 | 2.76E+00 | 8.42E-01 | no | Tmem132b | intron_variant | MODIFIER |
| 5 | 125781467 | AX-170276234 | 0 | T | G | 0.447 | -2.18E-02 | 1.12E-01 | 2.72E+00 | 2.76E+00 | 8.42E-01 | no | Tmem132b | intron_variant | MODIFIER |
| 5 | 125792327 | AX-170498733 | 0 | A | G | 0.447 | -2.18E-02 | 1.12E-01 | 2.72E+00 | 2.76E+00 | 8.42E-01 | no | Tmem132b | 3_prime_UTR_variant | MODIFIER |

| | | | | | | | | | | | | | | | |
|---|---|---|---|---|---|---|---|---|---|---|---|---|---|---|---|
| 5 | 125811875 | AX-97836541 | 0 | A | T | 0.447 | -2.18E-02 | 1.12E-01 | 2.72E+00 | 2.76E+00 | 8.42E-01 | no | | intergenic_variant | MODIFIER |
| 5 | 125852800 | AX-97814522 | 0 | A | G | 0.447 | -2.18E-02 | 1.12E-01 | 2.72E+00 | 2.76E+00 | 8.42E-01 | no | Gm4868 | upstream_gene_variant | MODIFIER |
| 5 | 125864297 | AX-170449053 | 0 | G | A | 0.447 | -2.18E-02 | 1.12E-01 | 2.72E+00 | 2.76E+00 | 8.42E-01 | no | | intergenic_variant | MODIFIER |
| 5 | 125888832 | AX-97790614 | 0 | A | G | 0.447 | -2.18E-02 | 1.12E-01 | 2.72E+00 | 2.76E+00 | 8.42E-01 | no | | intergenic_variant | MODIFIER |
| 5 | 125888884 | AX-170496118 | 0 | A | G | 0.447 | -2.18E-02 | 1.12E-01 | 2.72E+00 | 2.76E+00 | 8.42E-01 | no | | intergenic_variant | MODIFIER |
| 5 | 125923637 | AX-170218124 | 0 | C | T | 0.447 | -2.18E-02 | 1.12E-01 | 2.72E+00 | 2.76E+00 | 8.42E-01 | no | | intergenic_variant | MODIFIER |
| 5 | 125926974 | AX-97829053 | 0 | A | G | 0.447 | -2.18E-02 | 1.12E-01 | 2.72E+00 | 2.76E+00 | 8.42E-01 | no | 1700048F04Rik | upstream_gene_variant | MODIFIER |
| 5 | 125954473 | AX-97800803 | 0 | T | C | 0.447 | -2.18E-02 | 1.12E-01 | 2.72E+00 | 2.76E+00 | 8.42E-01 | no | Gm42945 | downstream_gene_variant | MODIFIER |
| 5 | 125986084 | AX-170422584 | 0 | A | G | 0.447 | -2.18E-02 | 1.12E-01 | 2.72E+00 | 2.76E+00 | 8.42E-01 | no | Gm42952 | downstream_gene_variant | MODIFIER |
| 5 | 125994131 | AX-168408061 | 0 | G | A | 0.447 | -2.18E-02 | 1.12E-01 | 2.72E+00 | 2.76E+00 | 8.42E-01 | no | | intergenic_variant | MODIFIER |
| 5 | 125999293 | AX-170290778 | 0 | A | C | 0.447 | -2.18E-02 | 1.12E-01 | 2.72E+00 | 2.76E+00 | 8.42E-01 | no | | intergenic_variant | MODIFIER |
| 5 | 126004791 | AX-97833806 | 0 | G | A | 0.447 | -2.18E-02 | 1.12E-01 | 2.72E+00 | 2.76E+00 | 8.42E-01 | no | | intergenic_variant | MODIFIER |
| 5 | 126028383 | AX-170026113 | 0 | A | C | 0.447 | -2.18E-02 | 1.12E-01 | 2.72E+00 | 2.76E+00 | 8.42E-01 | no | Gm7774 | upstream_gene_variant | MODIFIER |
| 5 | 126029377 | AX-97816432 | 0 | G | T | 0.447 | -2.18E-02 | 1.12E-01 | 2.72E+00 | 2.76E+00 | 8.42E-01 | no | Gm7774 | upstream_gene_variant | MODIFIER |
| 5 | 126031590 | AX-170300650 | 0 | C | T | 0.447 | -2.18E-02 | 1.12E-01 | 2.72E+00 | 2.76E+00 | 8.42E-01 | no | | intergenic_variant | MODIFIER |
| 5 | 126031664 | AX-170026108 | 0 | G | A | 0.447 | -2.18E-02 | 1.12E-01 | 2.72E+00 | 2.76E+00 | 8.42E-01 | no | | intergenic_variant | MODIFIER |
| 5 | 126033846 | AX-170469877 | 0 | G | C | 0.447 | -2.18E-02 | 1.12E-01 | 2.72E+00 | 2.76E+00 | 8.42E-01 | no | | intergenic_variant | MODIFIER |
| 5 | 126038186 | AX-170306121 | 0 | A | G | 0.447 | -2.18E-02 | 1.12E-01 | 2.72E+00 | 2.76E+00 | 8.42E-01 | no | | intergenic_variant | MODIFIER |
| 5 | 126039502 | AX-170272614 | 0 | C | T | 0.447 | -2.18E-02 | 1.12E-01 | 2.72E+00 | 2.76E+00 | 8.42E-01 | no | | intergenic_variant | MODIFIER |
| 5 | 126043342 | AX-170312390 | 0 | G | A | 0.447 | -2.18E-02 | 1.12E-01 | 2.72E+00 | 2.76E+00 | 8.42E-01 | no | | intergenic_variant | MODIFIER |
| 5 | 126043358 | AX-170026104 | 0 | G | A | 0.447 | -2.18E-02 | 1.12E-01 | 2.72E+00 | 2.76E+00 | 8.42E-01 | no | | intergenic_variant | MODIFIER |
| 5 | 126044835 | AX-170026112 | 0 | T | G | 0.447 | -2.18E-02 | 1.12E-01 | 2.72E+00 | 2.76E+00 | 8.42E-01 | no | | intergenic_variant | MODIFIER |
| 5 | 126059919 | AX-170273433 | 0 | A | G | 0.447 | -2.18E-02 | 1.12E-01 | 2.72E+00 | 2.76E+00 | 8.42E-01 | no | | intergenic_variant | MODIFIER |
| 5 | 126061234 | AX-170308629 | 0 | C | G | 0.447 | -2.18E-02 | 1.12E-01 | 2.72E+00 | 2.76E+00 | 8.42E-01 | no | | intergenic_variant | MODIFIER |
| 5 | 126063542 | AX-170073416 | 0 | T | C | 0.447 | -2.18E-02 | 1.12E-01 | 2.72E+00 | 2.76E+00 | 8.42E-01 | no | | intergenic_variant | MODIFIER |
| 5 | 126064890 | AX-170073414 | 0 | T | C | 0.447 | -2.18E-02 | 1.12E-01 | 2.72E+00 | 2.76E+00 | 8.42E-01 | no | | intergenic_variant | MODIFIER |
| 5 | 126077131 | AX-97814525 | 0 | T | C | 0.447 | -2.18E-02 | 1.12E-01 | 2.72E+00 | 2.76E+00 | 8.42E-01 | no | | intergenic_variant | MODIFIER |
| 5 | 126113508 | AX-170026105 | 0 | A | G | 0.447 | -2.18E-02 | 1.12E-01 | 2.72E+00 | 2.76E+00 | 8.42E-01 | no | | intergenic_variant | MODIFIER |
| 5 | 126123182 | AX-170340692 | 0 | G | C | 0.447 | -2.18E-02 | 1.12E-01 | 2.72E+00 | 2.76E+00 | 8.42E-01 | no | | intergenic_variant | MODIFIER |
| 5 | 126123324 | AX-170404002 | 0 | A | G | 0.447 | -2.18E-02 | 1.12E-01 | 2.72E+00 | 2.76E+00 | 8.42E-01 | no | | intergenic_variant | MODIFIER |
| 5 | 126123342 | AX-170479554 | 0 | G | A | 0.447 | -2.18E-02 | 1.12E-01 | 2.72E+00 | 2.76E+00 | 8.42E-01 | no | | intergenic_variant | MODIFIER |
| 5 | 126125385 | AX-97837216 | 0 | T | G | 0.447 | -2.18E-02 | 1.12E-01 | 2.72E+00 | 2.76E+00 | 8.42E-01 | no | | intergenic_variant | MODIFIER |
| 5 | 126131609 | AX-170278565 | 0 | T | C | 0.447 | -2.18E-02 | 1.12E-01 | 2.72E+00 | 2.76E+00 | 8.42E-01 | no | | intergenic_variant | MODIFIER |
| 5 | 126157108 | AX-97796208 | 0 | A | C | 0.447 | -2.18E-02 | 1.12E-01 | 2.72E+00 | 2.76E+00 | 8.42E-01 | no | | intergenic_variant | MODIFIER |
| 5 | 126157644 | AX-170073418 | 0 | C | T | 0.447 | -2.18E-02 | 1.12E-01 | 2.72E+00 | 2.76E+00 | 8.42E-01 | no | | intergenic_variant | MODIFIER |
| 5 | 126157928 | AX-170073417 | 0 | A | T | 0.447 | -2.18E-02 | 1.12E-01 | 2.72E+00 | 2.76E+00 | 8.42E-01 | no | | intergenic_variant | MODIFIER |
| 5 | 126158231 | AX-170194247 | 0 | C | A | 0.447 | -2.18E-02 | 1.12E-01 | 2.72E+00 | 2.76E+00 | 8.42E-01 | no | | intergenic_variant | MODIFIER |
| 5 | 126158293 | AX-170168362 | 0 | T | C | 0.447 | -2.18E-02 | 1.12E-01 | 2.72E+00 | 2.76E+00 | 8.42E-01 | no | | intergenic_variant | MODIFIER |
| 5 | 126160936 | AX-170125249 | 0 | T | G | 0.447 | -2.18E-02 | 1.12E-01 | 2.72E+00 | 2.76E+00 | 8.42E-01 | no | | regulatory_region_variant | MODIFIER |
| 5 | 126164611 | AX-170402043 | 0 | C | T | 0.447 | -2.18E-02 | 1.12E-01 | 2.72E+00 | 2.76E+00 | 8.42E-01 | no | | intergenic_variant | MODIFIER |
| 5 | 126190422 | AX-168415646 | 0 | T | C | 0.447 | -2.18E-02 | 1.12E-01 | 2.72E+00 | 2.76E+00 | 8.42E-01 | no | | intergenic_variant | MODIFIER |
| 5 | 126190685 | AX-170026109 | 0 | C | G | 0.447 | -2.18E-02 | 1.12E-01 | 2.72E+00 | 2.76E+00 | 8.42E-01 | no | | intergenic_variant | MODIFIER |
| 5 | 126193201 | AX-170441673 | 0 | C | T | 0.447 | -2.18E-02 | 1.12E-01 | 2.72E+00 | 2.76E+00 | 8.42E-01 | no | | intergenic_variant | MODIFIER |
| 5 | 126203922 | AX-170212573 | 0 | G | A | 0.447 | -2.18E-02 | 1.12E-01 | 2.72E+00 | 2.76E+00 | 8.42E-01 | no | | intergenic_variant | MODIFIER |
| 5 | 126215378 | AX-170480459 | 0 | G | A | 0.447 | -2.18E-02 | 1.12E-01 | 2.72E+00 | 2.76E+00 | 8.42E-01 | no | | intergenic_variant | MODIFIER |
| 5 | 126242581 | AX-170089605 | 0 | T | C | 0.447 | -2.18E-02 | 1.12E-01 | 2.72E+00 | 2.76E+00 | 8.42E-01 | no | | intergenic_variant | MODIFIER |
| 5 | 126249927 | AX-97828807 | 0 | A | G | 0.447 | -2.18E-02 | 1.12E-01 | 2.72E+00 | 2.76E+00 | 8.42E-01 | no | | intergenic_variant | MODIFIER |
| 5 | 126266711 | AX-170469260 | 0 | C | T | 0.447 | -2.18E-02 | 1.12E-01 | 2.72E+00 | 2.76E+00 | 8.42E-01 | no | | intergenic_variant | MODIFIER |
| 5 | 126271521 | AX-170124830 | 0 | C | T | 0.447 | -2.18E-02 | 1.12E-01 | 2.72E+00 | 2.76E+00 | 8.42E-01 | no | | intergenic_variant | MODIFIER |
| 5 | 126271609 | AX-170226494 | 0 | G | A | 0.447 | -2.18E-02 | 1.12E-01 | 2.72E+00 | 2.76E+00 | 8.42E-01 | no | | intergenic_variant | MODIFIER |
| 5 | 126275989 | AX-170143099 | 0 | C | T | 0.447 | -2.18E-02 | 1.12E-01 | 2.72E+00 | 2.76E+00 | 8.42E-01 | no | | intergenic_variant | MODIFIER |
| 5 | 126287132 | AX-97803858 | 0 | G | A | 0.447 | -2.18E-02 | 1.12E-01 | 2.72E+00 | 2.76E+00 | 8.42E-01 | no | | intergenic_variant | MODIFIER |
| 5 | 126327865 | AX-97808131 | 0 | T | C | 0.447 | -2.18E-02 | 1.12E-01 | 2.72E+00 | 2.76E+00 | 8.42E-01 | no | | intergenic_variant | MODIFIER |
| 5 | 126343286 | AX-170273816 | 0 | A | G | 0.447 | -2.18E-02 | 1.12E-01 | 2.72E+00 | 2.76E+00 | 8.42E-01 | no | | intergenic_variant | MODIFIER |

| | | | | | | | | | | | | | | | |
|---|---|---|---|---|---|---|---|---|---|---|---|---|---|---|---|
| 5 | 126343822 | AX-170294704 | 0 | A | G | 0.447 | -2.18E-02 | 1.12E-01 | 2.72E+00 | 2.76E+00 | 8.42E-01 | no | | regulatory_region_variant | MODIFIER |
| 5 | 126344599 | AX-170307322 | 0 | A | G | 0.447 | -2.18E-02 | 1.12E-01 | 2.72E+00 | 2.76E+00 | 8.42E-01 | no | | intergenic_variant | MODIFIER |
| 5 | 126350428 | AX-170531220 | 0 | T | C | 0.447 | -2.18E-02 | 1.12E-01 | 2.72E+00 | 2.76E+00 | 8.42E-01 | no | | intergenic_variant | MODIFIER |
| 5 | 126354447 | AX-97835106 | 0 | T | G | 0.447 | -2.18E-02 | 1.12E-01 | 2.72E+00 | 2.76E+00 | 8.42E-01 | no | | intergenic_variant | MODIFIER |
| 5 | 126369736 | AX-170121220 | 0 | A | G | 0.447 | -2.18E-02 | 1.12E-01 | 2.72E+00 | 2.76E+00 | 8.42E-01 | no | | intergenic_variant | MODIFIER |
| 4 | 95761970 | AX-170297660 | 0 | C | G | 0.254 | -2.09E-02 | 1.14E-01 | 2.76E+00 | 2.88E+00 | 8.42E-01 | no | Fggy | intron_variant | MODIFIER |
| 8 | 95800070 | AX-170031306 | 1 | T | C | 0.223 | 2.26E-02 | 1.15E-01 | 2.68E+00 | 2.74E+00 | 8.42E-01 | no | Cnot1 | intron_variant | MODIFIER |
| 10 | 120050624 | AX-170530080 | 1 | G | T | 0.339 | 1.40E-02 | 7.21E-02 | 2.66E+00 | 2.75E+00 | 8.42E-01 | no | Grip1 | intron_variant | MODIFIER |
| 1 | 64191888 | AX-97810106 | 1 | T | C | 0.116 | 3.31E-02 | 1.66E-01 | 2.64E+00 | 2.73E+00 | 8.42E-01 | no | Gm28981 | intron_variant,non_coding_transcript_variant | MODIFIER |
| 2 | 113381513 | AX-97842946 | 0 | A | G | 0.053 | 4.12E-02 | 2.12E-01 | 2.65E+00 | 2.75E+00 | 8.42E-01 | no | Fmn1 | intron_variant | MODIFIER |
| 7 | 61275576 | AX-170095957 | 1 | T | C | 0.42 | -1.82E-02 | 9.48E-02 | 2.67E+00 | 2.76E+00 | 8.42E-01 | no | A230006K03Rik | intron_variant,non_coding_transcript_variant | MODIFIER |
| 8 | 19141376 | AX-170423665 | 0 | C | T | 0.184 | 1.89E-02 | 9.46E-02 | 2.63E+00 | 2.73E+00 | 8.42E-01 | no | Spag11b | splice_region_variant,synonymous_variant | LOW |
| 13 | 54728808 | AX-170472223 | 1 | T | C | 0.509 | -1.21E-02 | 6.61E-02 | 2.68E+00 | 2.79E+00 | 8.42E-01 | no | Cdhr2 | intron_variant | MODIFIER |
| 5 | 85197493 | AX-170112968 | 0 | C | T | 0.43 | -1.92E-02 | 9.77E-02 | 2.65E+00 | 2.74E+00 | 8.42E-01 | no | | intergenic_variant | MODIFIER |
| 9 | 105453751 | AX-170096663 | 1 | G | A | 0.116 | -2.46E-02 | 1.17E-01 | 2.62E+00 | 2.69E+00 | 8.43E-01 | no | Atp2c1 | intron_variant | MODIFIER |
| 10 | 119056129 | AX-170046041 | 1 | G | T | 0.339 | 1.42E-02 | 7.28E-02 | 2.66E+00 | 2.75E+00 | 8.43E-01 | no | | regulatory_region_variant | MODIFIER |
| 9 | 40475314 | AX-170535992 | 1 | G | A | 0.438 | -1.44E-02 | 7.37E-02 | 2.65E+00 | 2.74E+00 | 8.43E-01 | no | Gramd1b | intron_variant | MODIFIER |
| 6 | 144417025 | AX-97805529 | 0 | G | A | 0.088 | -3.58E-02 | 1.89E-01 | 2.67E+00 | 2.77E+00 | 8.43E-01 | no | Sox5 | intron_variant,non_coding_transcript_variant | MODIFIER |
| 15 | 74369198 | AX-170231956 | 1 | A | G | 0.545 | 1.73E-02 | 8.29E-02 | 2.62E+00 | 2.69E+00 | 8.43E-01 | no | | intergenic_variant | MODIFIER |
| 15 | 75405187 | AX-170506590 | 1 | G | A | 0.545 | 1.73E-02 | 8.29E-02 | 2.62E+00 | 2.69E+00 | 8.43E-01 | no | | intergenic_variant | MODIFIER |
| 1 | 28506529 | AX-170311806 | 0 | C | T | 0.412 | 1.59E-02 | 8.10E-02 | 2.67E+00 | 2.74E+00 | 8.43E-01 | no | | intergenic_variant | MODIFIER |
| 1 | 28805301 | AX-170040722 | 0 | G | A | 0.412 | 1.59E-02 | 8.10E-02 | 2.67E+00 | 2.74E+00 | 8.43E-01 | no | Gm5697 | upstream_gene_variant | MODIFIER |
| 1 | 28943821 | AX-97780226 | 0 | A | G | 0.412 | 1.59E-02 | 8.10E-02 | 2.67E+00 | 2.74E+00 | 8.43E-01 | no | | intergenic_variant | MODIFIER |
| 1 | 28978498 | AX-170253789 | 0 | G | T | 0.412 | 1.59E-02 | 8.10E-02 | 2.67E+00 | 2.74E+00 | 8.43E-01 | no | | intergenic_variant | MODIFIER |
| 2 | 147746545 | AX-170250681 | 1 | T | A | 0.143 | 4.39E-02 | 2.21E-01 | 2.69E+00 | 2.72E+00 | 8.43E-01 | no | | intergenic_variant | MODIFIER |
| 9 | 81235510 | AX-170529097 | 1 | T | C | 0.205 | 2.42E-02 | 1.27E-01 | 2.73E+00 | 2.79E+00 | 8.43E-01 | no | | intergenic_variant | MODIFIER |
| 16 | 19465521 | AX-170012607 | 0 | T | C | 0.263 | 9.50E-03 | 4.83E-02 | 2.61E+00 | 2.74E+00 | 8.43E-01 | no | | intergenic_variant | MODIFIER |
| 3 | 147503021 | AX-170151615 | 1 | T | C | 0.348 | 1.47E-02 | 7.38E-02 | 2.65E+00 | 2.72E+00 | 8.43E-01 | no | | intergenic_variant | MODIFIER |
| 3 | 148012455 | AX-170504007 | 1 | G | A | 0.348 | 1.47E-02 | 7.38E-02 | 2.65E+00 | 2.72E+00 | 8.43E-01 | no | | intergenic_variant | MODIFIER |
| 3 | 148027297 | AX-170409551 | 1 | A | G | 0.348 | 1.47E-02 | 7.38E-02 | 2.65E+00 | 2.72E+00 | 8.43E-01 | no | | intergenic_variant | MODIFIER |
| 9 | 32742651 | AX-170416408 | 0 | T | C | 0.43 | 1.84E-02 | 8.88E-02 | 2.63E+00 | 2.70E+00 | 8.43E-01 | no | Ets1 | intron_variant | MODIFIER |
| 1 | 54466262 | AX-170529636 | 0 | T | A | 0.105 | -4.55E-02 | 2.35E-01 | 2.70E+00 | 2.75E+00 | 8.43E-01 | no | | intergenic_variant | MODIFIER |
| 5 | 123914009 | AX-170369170 | 0 | T | C | 0.447 | -2.01E-02 | 1.07E-01 | 2.72E+00 | 2.79E+00 | 8.43E-01 | no | Denr | intron_variant | MODIFIER |
| 7 | 117733983 | AX-170377493 | 1 | A | G | 0.402 | 1.79E-02 | 9.01E-02 | 2.66E+00 | 2.72E+00 | 8.43E-01 | no | | intergenic_variant | MODIFIER |
| 7 | 118444337 | AX-97813715 | 1 | A | G | 0.402 | 1.79E-02 | 9.01E-02 | 2.66E+00 | 2.72E+00 | 8.43E-01 | no | Syt17 | upstream_gene_variant | MODIFIER |
| 9 | 88384085 | AX-97826844 | 1 | G | C | 0.125 | -2.68E-02 | 1.30E-01 | 2.62E+00 | 2.70E+00 | 8.44E-01 | no | Snx14 | intron_variant,NMD_transcript_variant | MODIFIER |
| 5 | 102546640 | AX-170489639 | 1 | G | A | 0.438 | 2.16E-02 | 1.11E-01 | 2.68E+00 | 2.74E+00 | 8.44E-01 | no | Arhgap24 | intron_variant | MODIFIER |
| 18 | 58394355 | AX-170480398 | 2 | A | C | 0.4 | 2.22E-02 | 9.40E-02 | 2.55E+00 | 2.64E+00 | 8.44E-01 | no | | regulatory_region_variant | MODIFIER |
| 9 | 80720198 | AX-170454375 | 0 | A | C | 0.202 | -2.57E-02 | 1.28E-01 | 2.66E+00 | 2.71E+00 | 8.44E-01 | no | | intergenic_variant | MODIFIER |
| 1 | 32661814 | AX-170040793 | 0 | G | A | 0.43 | 1.59E-02 | 8.10E-02 | 2.65E+00 | 2.73E+00 | 8.44E-01 | no | Khdrbs2 | downstream_gene_variant | MODIFIER |
| 2 | 147592948 | AX-170226300 | 0 | C | T | 0.14 | -4.21E-02 | 2.19E-01 | 2.73E+00 | 2.77E+00 | 8.44E-01 | no | | intergenic_variant | MODIFIER |
| 7 | 81726703 | AX-170185690 | 1 | C | T | 0.116 | 4.14E-02 | 2.11E-01 | 2.65E+00 | 2.73E+00 | 8.44E-01 | no | | regulatory_region_variant | MODIFIER |
| 7 | 82317570 | AX-170438459 | 1 | C | T | 0.116 | 4.14E-02 | 2.11E-01 | 2.65E+00 | 2.73E+00 | 8.44E-01 | no | | intergenic_variant | MODIFIER |
| 18 | 64231256 | AX-170172419 | 1 | C | T | 0.411 | -1.72E-02 | 8.30E-02 | 2.62E+00 | 2.69E+00 | 8.44E-01 | no | St8sia3os | intron_variant,non_coding_transcript_variant | MODIFIER |
| 3 | 146633319 | AX-170175372 | 0 | T | C | 0.342 | 1.46E-02 | 7.34E-02 | 2.65E+00 | 2.72E+00 | 8.44E-01 | no | Uox | downstream_gene_variant | MODIFIER |
| 11 | 10127786 | AX-170046390 | 0 | A | G | 0.044 | 4.57E-02 | 2.40E-01 | 2.63E+00 | 2.75E+00 | 8.44E-01 | no | | intergenic_variant | MODIFIER |
| 11 | 12475950 | AX-170004301 | 0 | T | C | 0.088 | 2.29E-02 | 1.20E-01 | 2.63E+00 | 2.75E+00 | 8.44E-01 | no | | regulatory_region_variant | MODIFIER |
| 11 | 22691728 | AX-170518562 | 0 | C | T | 0.044 | 4.57E-02 | 2.40E-01 | 2.63E+00 | 2.75E+00 | 8.44E-01 | no | | intergenic_variant | MODIFIER |
| 9 | 58669274 | AX-170185582 | 1 | C | A | 0.214 | -2.37E-02 | 1.24E-01 | 2.70E+00 | 2.75E+00 | 8.44E-01 | no | Rec114 | intron_variant | MODIFIER |
| 4 | 74000732 | AX-97845154 | 1 | A | C | 0.241 | 1.97E-02 | 9.36E-02 | 2.60E+00 | 2.69E+00 | 8.44E-01 | no | | intergenic_variant | MODIFIER |
| 8 | 36698071 | AX-170438978 | 0 | C | T | 0.149 | -2.91E-02 | 1.48E-01 | 2.67E+00 | 2.72E+00 | 8.44E-01 | no | Dlc1 | intron_variant | MODIFIER |
| 8 | 37085070 | AX-170458980 | 0 | G | A | 0.149 | -2.91E-02 | 1.48E-01 | 2.67E+00 | 2.72E+00 | 8.44E-01 | no | Gm10683 | downstream_gene_variant | MODIFIER |
| 2 | 117504364 | AX-97803710 | 0 | T | C | 0.132 | 3.82E-02 | 1.97E-01 | 2.69E+00 | 2.74E+00 | 8.44E-01 | no | | intergenic_variant | MODIFIER |
| 9 | 78144081 | AX-170262556 | 1 | C | T | 0.196 | 2.61E-02 | 1.36E-01 | 2.73E+00 | 2.76E+00 | 8.45E-01 | no | Ick | intron_variant | MODIFIER |

| | | | | | | | | | | | | | | | |
|---|---|---|---|---|---|---|---|---|---|---|---|---|---|---|---|
| 7 | 67772572 | AX-170339993 | 0 | G | A | 0.421 | -1.85E-02 | 9.77E-02 | 2.68E+00 | 2.76E+00 | 8.45E-01 | no | | regulatory_region_variant | MODIFIER |
| 1 | 66691892 | AX-170481616 | 1 | C | T | 0.116 | 4.50E-02 | 2.32E-01 | 2.68E+00 | 2.73E+00 | 8.45E-01 | no | Unc80 | intron_variant | MODIFIER |
| 1 | 66923327 | AX-170502144 | 1 | C | T | 0.116 | 4.50E-02 | 2.32E-01 | 2.68E+00 | 2.73E+00 | 8.45E-01 | no | Myl1 | downstream_gene_variant | MODIFIER |
| 1 | 67338318 | AX-97803200 | 1 | G | A | 0.116 | 4.50E-02 | 2.32E-01 | 2.68E+00 | 2.73E+00 | 8.45E-01 | no | | intergenic_variant | MODIFIER |
| 1 | 68750496 | AX-170164705 | 1 | G | A | 0.116 | 4.50E-02 | 2.32E-01 | 2.68E+00 | 2.73E+00 | 8.45E-01 | no | Erbb4 | intron_variant | MODIFIER |
| 5 | 113256497 | AX-170364401 | 1 | C | A | 0.464 | 2.21E-02 | 1.09E-01 | 2.64E+00 | 2.70E+00 | 8.45E-01 | no | Sgsm1 | intron_variant | MODIFIER |
| 5 | 115591013 | AX-170025942 | 1 | A | C | 0.464 | 2.21E-02 | 1.09E-01 | 2.64E+00 | 2.70E+00 | 8.45E-01 | no | Gcn1l1 | synonymous_variant | LOW |
| 4 | 98042636 | AX-97836405 | 1 | C | A | 0.232 | 2.22E-02 | 1.08E-01 | 2.62E+00 | 2.70E+00 | 8.45E-01 | no | Nfia | intron_variant | MODIFIER |
| 1 | 10622401 | AX-170348128 | 0 | G | A | 0.404 | 1.51E-02 | 7.33E-02 | 2.62E+00 | 2.69E+00 | 8.45E-01 | no | Cpa6 | intron_variant | MODIFIER |
| 7 | 24615633 | AX-170506114 | 0 | A | T | 0.404 | 1.72E-02 | 8.41E-02 | 2.59E+00 | 2.71E+00 | 8.45E-01 | no | Phldb3 | intron_variant | MODIFIER |
| 6 | 110891311 | AX-170437677 | 0 | C | T | 0.053 | 5.69E-02 | 2.95E-01 | 2.68E+00 | 2.74E+00 | 8.45E-01 | no | Grm7 | intron_variant | MODIFIER |
| 1 | 73249279 | AX-170302087 | 2 | C | T | 0.127 | 3.78E-02 | 1.88E-01 | 2.61E+00 | 2.72E+00 | 8.45E-01 | no | | intergenic_variant | MODIFIER |
| 12 | 103964136 | AX-170366473 | 0 | G | A | 0.105 | -3.36E-02 | 1.79E-01 | 2.69E+00 | 2.76E+00 | 8.45E-01 | no | | intergenic_variant | MODIFIER |
| 12 | 103978024 | AX-97850306 | 0 | T | C | 0.105 | -3.36E-02 | 1.79E-01 | 2.69E+00 | 2.76E+00 | 8.45E-01 | no | Serpina11 | downstream_gene_variant | MODIFIER |
| 12 | 103980755 | AX-170236880 | 0 | C | G | 0.105 | -3.36E-02 | 1.79E-01 | 2.69E+00 | 2.76E+00 | 8.45E-01 | no | Serpina11 | intron_variant | MODIFIER |
| 8 | 47104266 | AX-170531370 | 1 | C | T | 0.152 | 2.83E-02 | 1.50E-01 | 2.73E+00 | 2.78E+00 | 8.45E-01 | no | | intergenic_variant | MODIFIER |
| 1 | 29085048 | AX-170345240 | 0 | G | A | 0.421 | 1.58E-02 | 8.17E-02 | 2.67E+00 | 2.74E+00 | 8.45E-01 | no | | intergenic_variant | MODIFIER |
| 1 | 29085416 | AX-170113006 | 0 | C | T | 0.421 | 1.58E-02 | 8.17E-02 | 2.67E+00 | 2.74E+00 | 8.45E-01 | no | | intergenic_variant | MODIFIER |
| 1 | 30146334 | AX-170516996 | 0 | A | G | 0.421 | 1.58E-02 | 8.17E-02 | 2.67E+00 | 2.74E+00 | 8.45E-01 | no | | intergenic_variant | MODIFIER |
| 1 | 30207638 | AX-97790616 | 0 | T | G | 0.421 | 1.58E-02 | 8.17E-02 | 2.67E+00 | 2.74E+00 | 8.45E-01 | no | | intergenic_variant | MODIFIER |
| 1 | 30217828 | AX-170330685 | 0 | G | A | 0.421 | 1.58E-02 | 8.17E-02 | 2.67E+00 | 2.74E+00 | 8.45E-01 | no | | intergenic_variant | MODIFIER |
| 1 | 30298803 | AX-170198953 | 0 | A | T | 0.421 | 1.58E-02 | 8.17E-02 | 2.67E+00 | 2.74E+00 | 8.45E-01 | no | | intergenic_variant | MODIFIER |
| 1 | 30358025 | AX-97811158 | 0 | A | G | 0.421 | 1.58E-02 | 8.17E-02 | 2.67E+00 | 2.74E+00 | 8.45E-01 | no | | intergenic_variant | MODIFIER |
| 1 | 30405870 | AX-170492954 | 0 | A | C | 0.421 | 1.58E-02 | 8.17E-02 | 2.67E+00 | 2.74E+00 | 8.45E-01 | no | | intergenic_variant | MODIFIER |
| 1 | 30406299 | AX-97802517 | 0 | T | C | 0.421 | 1.58E-02 | 8.17E-02 | 2.67E+00 | 2.74E+00 | 8.45E-01 | no | | intergenic_variant | MODIFIER |
| 1 | 30488503 | AX-97808229 | 0 | C | T | 0.421 | 1.58E-02 | 8.17E-02 | 2.67E+00 | 2.74E+00 | 8.45E-01 | no | | intergenic_variant | MODIFIER |
| 1 | 30501469 | AX-97814936 | 0 | A | C | 0.421 | 1.58E-02 | 8.17E-02 | 2.67E+00 | 2.74E+00 | 8.45E-01 | no | | intergenic_variant | MODIFIER |
| 1 | 30516107 | AX-170000209 | 0 | G | A | 0.421 | 1.58E-02 | 8.17E-02 | 2.67E+00 | 2.74E+00 | 8.45E-01 | no | | intergenic_variant | MODIFIER |
| 1 | 30541943 | AX-170127566 | 0 | G | A | 0.421 | 1.58E-02 | 8.17E-02 | 2.67E+00 | 2.74E+00 | 8.45E-01 | no | | intergenic_variant | MODIFIER |
| 1 | 30548443 | AX-170223146 | 0 | G | T | 0.421 | 1.58E-02 | 8.17E-02 | 2.67E+00 | 2.74E+00 | 8.45E-01 | no | | intergenic_variant | MODIFIER |
| 1 | 30589059 | AX-170000212 | 0 | C | T | 0.421 | 1.58E-02 | 8.17E-02 | 2.67E+00 | 2.74E+00 | 8.45E-01 | no | | intergenic_variant | MODIFIER |
| 1 | 30589097 | AX-170492219 | 0 | C | T | 0.421 | 1.58E-02 | 8.17E-02 | 2.67E+00 | 2.74E+00 | 8.45E-01 | no | | intergenic_variant | MODIFIER |
| 1 | 30635106 | AX-170422524 | 0 | T | G | 0.421 | 1.58E-02 | 8.17E-02 | 2.67E+00 | 2.74E+00 | 8.45E-01 | no | | intergenic_variant | MODIFIER |
| 1 | 30638123 | AX-170289059 | 0 | T | G | 0.421 | 1.58E-02 | 8.17E-02 | 2.67E+00 | 2.74E+00 | 8.45E-01 | no | | regulatory_region_variant | MODIFIER |
| 1 | 30638248 | AX-170505631 | 0 | C | T | 0.421 | 1.58E-02 | 8.17E-02 | 2.67E+00 | 2.74E+00 | 8.45E-01 | no | | intergenic_variant | MODIFIER |
| 1 | 30639173 | AX-170497298 | 0 | T | C | 0.421 | 1.58E-02 | 8.17E-02 | 2.67E+00 | 2.74E+00 | 8.45E-01 | no | | intergenic_variant | MODIFIER |
| 1 | 30643010 | AX-170110496 | 0 | T | C | 0.421 | 1.58E-02 | 8.17E-02 | 2.67E+00 | 2.74E+00 | 8.45E-01 | no | | intergenic_variant | MODIFIER |
| 1 | 30648873 | AX-97843344 | 0 | T | A | 0.421 | 1.58E-02 | 8.17E-02 | 2.67E+00 | 2.74E+00 | 8.45E-01 | no | | intergenic_variant | MODIFIER |
| 1 | 30677668 | AX-97847710 | 0 | T | C | 0.421 | 1.58E-02 | 8.17E-02 | 2.67E+00 | 2.74E+00 | 8.45E-01 | no | | intergenic_variant | MODIFIER |
| 1 | 30681941 | AX-170478966 | 0 | G | A | 0.421 | 1.58E-02 | 8.17E-02 | 2.67E+00 | 2.74E+00 | 8.45E-01 | no | | intergenic_variant | MODIFIER |
| 1 | 30686017 | AX-170119167 | 0 | G | A | 0.421 | 1.58E-02 | 8.17E-02 | 2.67E+00 | 2.74E+00 | 8.45E-01 | no | | intergenic_variant | MODIFIER |
| 1 | 32350897 | AX-97790097 | 0 | G | T | 0.421 | 1.58E-02 | 8.17E-02 | 2.67E+00 | 2.74E+00 | 8.45E-01 | no | Khdrbs2 | intron_variant | MODIFIER |
| 1 | 32473336 | AX-97803763 | 0 | C | T | 0.421 | 1.58E-02 | 8.17E-02 | 2.67E+00 | 2.74E+00 | 8.45E-01 | no | Khdrbs2 | intron_variant | MODIFIER |
| 1 | 32477193 | AX-170465592 | 0 | C | T | 0.421 | 1.58E-02 | 8.17E-02 | 2.67E+00 | 2.74E+00 | 8.45E-01 | no | Khdrbs2 | intron_variant | MODIFIER |
| 1 | 32673500 | AX-170298555 | 0 | G | A | 0.421 | 1.58E-02 | 8.17E-02 | 2.67E+00 | 2.74E+00 | 8.45E-01 | no | | intergenic_variant | MODIFIER |
| 1 | 32701620 | AX-170133688 | 0 | G | T | 0.421 | 1.58E-02 | 8.17E-02 | 2.67E+00 | 2.74E+00 | 8.45E-01 | no | | intergenic_variant | MODIFIER |
| 1 | 32703290 | AX-170100361 | 0 | G | A | 0.421 | 1.58E-02 | 8.17E-02 | 2.67E+00 | 2.74E+00 | 8.45E-01 | no | | intergenic_variant | MODIFIER |
| 1 | 32987924 | AX-170103269 | 0 | A | G | 0.421 | 1.58E-02 | 8.17E-02 | 2.67E+00 | 2.74E+00 | 8.45E-01 | no | | intergenic_variant | MODIFIER |
| 1 | 32988785 | AX-170322395 | 0 | A | G | 0.421 | 1.58E-02 | 8.17E-02 | 2.67E+00 | 2.74E+00 | 8.45E-01 | no | | intergenic_variant | MODIFIER |
| 1 | 33022749 | AX-170284945 | 0 | T | C | 0.421 | 1.58E-02 | 8.17E-02 | 2.67E+00 | 2.74E+00 | 8.45E-01 | no | | intergenic_variant | MODIFIER |
| 1 | 33022898 | AX-170464755 | 0 | G | T | 0.421 | 1.58E-02 | 8.17E-02 | 2.67E+00 | 2.74E+00 | 8.45E-01 | no | | regulatory_region_variant | MODIFIER |
| 1 | 33060861 | AX-170103289 | 0 | C | G | 0.421 | 1.58E-02 | 8.17E-02 | 2.67E+00 | 2.74E+00 | 8.45E-01 | no | Gm37724 | upstream_gene_variant | MODIFIER |
| 1 | 33062464 | AX-170040807 | 0 | T | C | 0.421 | 1.58E-02 | 8.17E-02 | 2.67E+00 | 2.74E+00 | 8.45E-01 | no | Gm37724 | upstream_gene_variant | MODIFIER |
| 1 | 33086368 | AX-97815077 | 0 | T | C | 0.421 | 1.58E-02 | 8.17E-02 | 2.67E+00 | 2.74E+00 | 8.45E-01 | no | | intergenic_variant | MODIFIER |

| | | | | | | | | | | | | | | | |
|---|---|---|---|---|---|---|---|---|---|---|---|---|---|---|---|
| 1 | 33087600 | AX-170276942 | 0 | T | C | 0.421 | 1.58E-02 | 8.17E-02 | 2.67E+00 | 2.74E+00 | 8.45E-01 | no | | intergenic_variant | MODIFIER |
| 1 | 33122883 | AX-170124164 | 0 | G | A | 0.421 | 1.58E-02 | 8.17E-02 | 2.67E+00 | 2.74E+00 | 8.45E-01 | no | | intergenic_variant | MODIFIER |
| 1 | 33126761 | AX-170527195 | 0 | T | C | 0.421 | 1.58E-02 | 8.17E-02 | 2.67E+00 | 2.74E+00 | 8.45E-01 | no | | intergenic_variant | MODIFIER |
| 1 | 33127633 | AX-97854253 | 0 | T | C | 0.421 | 1.58E-02 | 8.17E-02 | 2.67E+00 | 2.74E+00 | 8.45E-01 | no | | intergenic_variant | MODIFIER |
| 1 | 33131312 | AX-170117947 | 0 | A | G | 0.421 | 1.58E-02 | 8.17E-02 | 2.67E+00 | 2.74E+00 | 8.45E-01 | no | | intergenic_variant | MODIFIER |
| 1 | 33131742 | AX-170126549 | 0 | C | T | 0.421 | 1.58E-02 | 8.17E-02 | 2.67E+00 | 2.74E+00 | 8.45E-01 | no | | intergenic_variant | MODIFIER |
| 1 | 33147755 | AX-170000243 | 0 | C | A | 0.421 | 1.58E-02 | 8.17E-02 | 2.67E+00 | 2.74E+00 | 8.45E-01 | no | | intergenic_variant | MODIFIER |
| 1 | 33154964 | AX-97811256 | 0 | C | T | 0.421 | 1.58E-02 | 8.17E-02 | 2.67E+00 | 2.74E+00 | 8.45E-01 | no | | intergenic_variant | MODIFIER |
| 1 | 33213054 | AX-97840840 | 0 | A | G | 0.421 | 1.58E-02 | 8.17E-02 | 2.67E+00 | 2.74E+00 | 8.45E-01 | no | Gm37591 | non_coding_transcript_exon_variant | MODIFIER |
| 1 | 33213476 | AX-170000240 | 0 | G | A | 0.421 | 1.58E-02 | 8.17E-02 | 2.67E+00 | 2.74E+00 | 8.45E-01 | no | Gm37591 | intron_variant,non_coding_transcript_variant | MODIFIER |
| 1 | 33215368 | AX-170377998 | 0 | T | C | 0.421 | 1.58E-02 | 8.17E-02 | 2.67E+00 | 2.74E+00 | 8.45E-01 | no | Gm37591 | non_coding_transcript_exon_variant | MODIFIER |
| 1 | 33235764 | AX-97847874 | 0 | G | A | 0.421 | 1.58E-02 | 8.17E-02 | 2.67E+00 | 2.74E+00 | 8.45E-01 | no | Gm2987 | upstream_gene_variant | MODIFIER |
| 1 | 33236014 | AX-170374900 | 0 | A | G | 0.421 | 1.58E-02 | 8.17E-02 | 2.67E+00 | 2.74E+00 | 8.45E-01 | no | Gm2987 | non_coding_transcript_exon_variant | MODIFIER |
| 1 | 33241570 | AX-170137622 | 0 | C | G | 0.421 | 1.58E-02 | 8.17E-02 | 2.67E+00 | 2.74E+00 | 8.45E-01 | no | Gm2987 | downstream_gene_variant | MODIFIER |
| 1 | 33241773 | AX-170251079 | 0 | G | C | 0.421 | 1.58E-02 | 8.17E-02 | 2.67E+00 | 2.74E+00 | 8.45E-01 | no | Gm2987 | downstream_gene_variant | MODIFIER |
| 1 | 33247976 | AX-170123343 | 0 | A | G | 0.421 | 1.58E-02 | 8.17E-02 | 2.67E+00 | 2.74E+00 | 8.45E-01 | no | | intergenic_variant | MODIFIER |
| 1 | 33250001 | AX-97787636 | 0 | T | C | 0.421 | 1.58E-02 | 8.17E-02 | 2.67E+00 | 2.74E+00 | 8.45E-01 | no | | intergenic_variant | MODIFIER |
| 1 | 33268453 | AX-170140939 | 0 | G | C | 0.421 | 1.58E-02 | 8.17E-02 | 2.67E+00 | 2.74E+00 | 8.45E-01 | no | | intergenic_variant | MODIFIER |
| 1 | 33268754 | AX-170480055 | 0 | T | C | 0.421 | 1.58E-02 | 8.17E-02 | 2.67E+00 | 2.74E+00 | 8.45E-01 | no | | intergenic_variant | MODIFIER |
| 1 | 33270198 | AX-170194628 | 0 | C | A | 0.421 | 1.58E-02 | 8.17E-02 | 2.67E+00 | 2.74E+00 | 8.45E-01 | no | | intergenic_variant | MODIFIER |
| 1 | 33299039 | AX-170494325 | 0 | A | C | 0.421 | 1.58E-02 | 8.17E-02 | 2.67E+00 | 2.74E+00 | 8.45E-01 | no | | intergenic_variant | MODIFIER |
| 1 | 33326233 | AX-170288551 | 0 | A | G | 0.421 | 1.58E-02 | 8.17E-02 | 2.67E+00 | 2.74E+00 | 8.45E-01 | no | | intergenic_variant | MODIFIER |
| 1 | 33331805 | AX-170404102 | 0 | A | G | 0.421 | 1.58E-02 | 8.17E-02 | 2.67E+00 | 2.74E+00 | 8.45E-01 | no | | intergenic_variant | MODIFIER |
| 1 | 33331840 | AX-97782799 | 0 | C | T | 0.421 | 1.58E-02 | 8.17E-02 | 2.67E+00 | 2.74E+00 | 8.45E-01 | no | | intergenic_variant | MODIFIER |
| 1 | 33452486 | AX-170254749 | 0 | A | G | 0.421 | 1.58E-02 | 8.17E-02 | 2.67E+00 | 2.74E+00 | 8.45E-01 | no | Prim2 | downstream_gene_variant | MODIFIER |
| 1 | 33452912 | AX-170102834 | 0 | A | T | 0.421 | 1.58E-02 | 8.17E-02 | 2.67E+00 | 2.74E+00 | 8.45E-01 | no | Prim2 | downstream_gene_variant | MODIFIER |
| 1 | 33463126 | AX-170349412 | 0 | C | T | 0.421 | 1.58E-02 | 8.17E-02 | 2.67E+00 | 2.74E+00 | 8.45E-01 | no | Prim2 | intron_variant | MODIFIER |
| 1 | 33463300 | AX-170243624 | 0 | A | G | 0.421 | 1.58E-02 | 8.17E-02 | 2.67E+00 | 2.74E+00 | 8.45E-01 | no | Prim2 | intron_variant | MODIFIER |
| 1 | 33494474 | AX-170456309 | 0 | C | T | 0.421 | 1.58E-02 | 8.17E-02 | 2.67E+00 | 2.74E+00 | 8.45E-01 | no | Prim2 | intron_variant | MODIFIER |
| 1 | 33497713 | AX-170199573 | 0 | T | C | 0.421 | 1.58E-02 | 8.17E-02 | 2.67E+00 | 2.74E+00 | 8.45E-01 | no | Prim2 | intron_variant | MODIFIER |
| 1 | 33497959 | AX-170040822 | 0 | C | G | 0.421 | 1.58E-02 | 8.17E-02 | 2.67E+00 | 2.74E+00 | 8.45E-01 | no | Prim2 | intron_variant | MODIFIER |
| 1 | 33515485 | AX-170040823 | 0 | G | A | 0.421 | 1.58E-02 | 8.17E-02 | 2.67E+00 | 2.74E+00 | 8.45E-01 | no | Prim2 | intron_variant | MODIFIER |
| 1 | 33515499 | AX-170040819 | 0 | A | C | 0.421 | 1.58E-02 | 8.17E-02 | 2.67E+00 | 2.74E+00 | 8.45E-01 | no | Prim2 | intron_variant | MODIFIER |
| 1 | 33521748 | AX-170239890 | 0 | A | G | 0.421 | 1.58E-02 | 8.17E-02 | 2.67E+00 | 2.74E+00 | 8.45E-01 | no | Prim2 | intron_variant | MODIFIER |
| 1 | 33546474 | AX-170204222 | 0 | G | A | 0.421 | 1.58E-02 | 8.17E-02 | 2.67E+00 | 2.74E+00 | 8.45E-01 | no | Prim2 | intron_variant | MODIFIER |
| 1 | 33584201 | AX-170161447 | 0 | G | A | 0.421 | 1.58E-02 | 8.17E-02 | 2.67E+00 | 2.74E+00 | 8.45E-01 | no | Prim2 | intron_variant | MODIFIER |
| 1 | 33589287 | AX-170418098 | 0 | A | G | 0.421 | 1.58E-02 | 8.17E-02 | 2.67E+00 | 2.74E+00 | 8.45E-01 | no | Prim2 | intron_variant | MODIFIER |
| 1 | 33614723 | AX-170197920 | 0 | T | C | 0.421 | 1.58E-02 | 8.17E-02 | 2.67E+00 | 2.74E+00 | 8.45E-01 | no | Prim2 | intron_variant | MODIFIER |
| 1 | 33676161 | AX-170271896 | 0 | G | A | 0.421 | 1.58E-02 | 8.17E-02 | 2.67E+00 | 2.74E+00 | 8.45E-01 | no | Gm38173 | upstream_gene_variant | MODIFIER |
| 1 | 33696910 | AX-170515379 | 0 | T | C | 0.421 | 1.58E-02 | 8.17E-02 | 2.67E+00 | 2.74E+00 | 8.45E-01 | no | | intergenic_variant | MODIFIER |
| 1 | 33697154 | AX-170040825 | 0 | C | T | 0.421 | 1.58E-02 | 8.17E-02 | 2.67E+00 | 2.74E+00 | 8.45E-01 | no | | intergenic_variant | MODIFIER |
| 1 | 33746402 | AX-170126368 | 0 | T | C | 0.421 | 1.58E-02 | 8.17E-02 | 2.67E+00 | 2.74E+00 | 8.45E-01 | no | Bag2 | 3_prime_UTR_variant | MODIFIER |
| 1 | 33748248 | AX-97848594 | 0 | T | C | 0.421 | 1.58E-02 | 8.17E-02 | 2.67E+00 | 2.74E+00 | 8.45E-01 | no | Bag2 | synonymous_variant | LOW |
| 8 | 85168015 | AX-170031161 | 0 | C | T | 0.184 | -2.43E-02 | 1.28E-01 | 2.68E+00 | 2.76E+00 | 8.45E-01 | no | Cks1brt | upstream_gene_variant | MODIFIER |
| 8 | 75159575 | AX-170350571 | 0 | T | C | 0.193 | 2.51E-02 | 1.33E-01 | 2.69E+00 | 2.76E+00 | 8.46E-01 | no | | intergenic_variant | MODIFIER |
| 16 | 93087136 | AX-170182603 | 1 | G | A | 0.393 | -1.61E-02 | 7.95E-02 | 2.61E+00 | 2.71E+00 | 8.46E-01 | no | | intergenic_variant | MODIFIER |
| 9 | 64033688 | AX-170148057 | 1 | A | G | 0.196 | 2.63E-02 | 1.37E-01 | 2.74E+00 | 2.77E+00 | 8.46E-01 | no | 1700055C04Rik | downstream_gene_variant | MODIFIER |
| 9 | 64893070 | AX-170409387 | 1 | G | A | 0.196 | 2.63E-02 | 1.37E-01 | 2.74E+00 | 2.77E+00 | 8.46E-01 | no | Dennd4a | intron_variant | MODIFIER |
| 9 | 65293039 | AX-170481468 | 1 | G | A | 0.196 | 2.63E-02 | 1.37E-01 | 2.74E+00 | 2.77E+00 | 8.46E-01 | no | Clpx | upstream_gene_variant | MODIFIER |
| 9 | 65634758 | AX-170032565 | 1 | C | T | 0.196 | 2.63E-02 | 1.37E-01 | 2.74E+00 | 2.77E+00 | 8.46E-01 | no | Rbpms2 | intron_variant | MODIFIER |
| 9 | 68545177 | AX-170481914 | 1 | G | A | 0.196 | 2.63E-02 | 1.37E-01 | 2.74E+00 | 2.77E+00 | 8.46E-01 | no | | intergenic_variant | MODIFIER |
| 8 | 94963339 | AX-170294056 | 1 | T | A | 0.223 | 2.22E-02 | 1.15E-01 | 2.67E+00 | 2.74E+00 | 8.46E-01 | no | Gm10286 | downstream_gene_variant | MODIFIER |
| 8 | 95515176 | AX-170080876 | 1 | T | C | 0.223 | 2.22E-02 | 1.15E-01 | 2.67E+00 | 2.74E+00 | 8.46E-01 | no | | intergenic_variant | MODIFIER |
| 5 | 117838113 | AX-170496709 | 0 | G | A | 0.456 | -2.11E-02 | 1.12E-01 | 2.71E+00 | 2.77E+00 | 8.46E-01 | no | Nos1 | upstream_gene_variant | MODIFIER |

| | | | | | | | | | | | | | | | |
|---|---|---|---|---|---|---|---|---|---|---|---|---|---|---|---|
| 8 | 65866073 | AX-170030923 | 0 | G | A | 0.14 | 2.39E-02 | 1.33E-01 | 2.69E+00 | 2.79E+00 | 8.46E-01 | no | MAR1 | intron_variant | MODIFIER |
| 1 | 30357571 | AX-170411568 | 0 | T | C | 0.412 | 1.56E-02 | 8.17E-02 | 2.67E+00 | 2.74E+00 | 8.46E-01 | no | | intergenic_variant | MODIFIER |
| 1 | 33213398 | AX-170000239 | 0 | G | A | 0.412 | 1.56E-02 | 8.17E-02 | 2.67E+00 | 2.74E+00 | 8.46E-01 | no | Gm37591 | intron_variant,non_coding_transcript_variant | MODIFIER |
| 11 | 47924862 | AX-97823495 | 2 | A | G | 0.082 | 3.59E-02 | 1.97E-01 | 2.67E+00 | 2.78E+00 | 8.46E-01 | no | Sgcd | intron_variant,non_coding_transcript_variant | MODIFIER |
| 8 | 73682750 | AX-170505489 | 1 | A | T | 0.205 | -2.42E-02 | 1.24E-01 | 2.65E+00 | 2.73E+00 | 8.46E-01 | no | | intergenic_variant | MODIFIER |
| 1 | 75645533 | AX-170341554 | 0 | T | C | 0.14 | -3.73E-02 | 1.88E-01 | 2.62E+00 | 2.72E+00 | 8.46E-01 | no | | intergenic_variant | MODIFIER |
| 13 | 47594772 | AX-97823410 | 0 | T | C | 0.509 | 1.30E-02 | 6.46E-02 | 2.61E+00 | 2.71E+00 | 8.46E-01 | no | | intergenic_variant | MODIFIER |
| 13 | 47598718 | AX-170401560 | 0 | T | C | 0.509 | 1.30E-02 | 6.46E-02 | 2.61E+00 | 2.71E+00 | 8.46E-01 | no | | intergenic_variant | MODIFIER |
| 13 | 47612102 | AX-170477851 | 0 | C | A | 0.509 | 1.30E-02 | 6.46E-02 | 2.61E+00 | 2.71E+00 | 8.46E-01 | no | | intergenic_variant | MODIFIER |
| 13 | 47617188 | AX-170334292 | 0 | T | C | 0.509 | 1.30E-02 | 6.46E-02 | 2.61E+00 | 2.71E+00 | 8.46E-01 | no | | intergenic_variant | MODIFIER |
| 13 | 47623412 | AX-170387090 | 0 | A | G | 0.509 | 1.30E-02 | 6.46E-02 | 2.61E+00 | 2.71E+00 | 8.46E-01 | no | | intergenic_variant | MODIFIER |
| 13 | 47659665 | AX-170501101 | 0 | G | A | 0.509 | 1.30E-02 | 6.46E-02 | 2.61E+00 | 2.71E+00 | 8.46E-01 | no | | intergenic_variant | MODIFIER |
| 13 | 47675890 | AX-97815096 | 0 | T | C | 0.509 | 1.30E-02 | 6.46E-02 | 2.61E+00 | 2.71E+00 | 8.46E-01 | no | | intergenic_variant | MODIFIER |
| 13 | 47676180 | AX-170118766 | 0 | G | T | 0.509 | 1.30E-02 | 6.46E-02 | 2.61E+00 | 2.71E+00 | 8.46E-01 | no | | intergenic_variant | MODIFIER |
| 13 | 47704436 | AX-170308240 | 0 | A | G | 0.509 | 1.30E-02 | 6.46E-02 | 2.61E+00 | 2.71E+00 | 8.46E-01 | no | | intergenic_variant | MODIFIER |
| 13 | 47704913 | AX-170189912 | 0 | A | G | 0.509 | 1.30E-02 | 6.46E-02 | 2.61E+00 | 2.71E+00 | 8.46E-01 | no | | intergenic_variant | MODIFIER |
| 13 | 47724486 | AX-168414955 | 0 | A | G | 0.509 | 1.30E-02 | 6.46E-02 | 2.61E+00 | 2.71E+00 | 8.46E-01 | no | | intergenic_variant | MODIFIER |
| 13 | 47725726 | AX-97827162 | 0 | C | A | 0.509 | 1.30E-02 | 6.46E-02 | 2.61E+00 | 2.71E+00 | 8.46E-01 | no | | intergenic_variant | MODIFIER |
| 13 | 47753618 | AX-170411410 | 0 | A | G | 0.509 | 1.30E-02 | 6.46E-02 | 2.61E+00 | 2.71E+00 | 8.46E-01 | no | | intergenic_variant | MODIFIER |
| 13 | 47763178 | AX-168395276 | 0 | G | T | 0.509 | 1.30E-02 | 6.46E-02 | 2.61E+00 | 2.71E+00 | 8.46E-01 | no | | intergenic_variant | MODIFIER |
| 13 | 47764768 | AX-170394943 | 0 | A | G | 0.509 | 1.30E-02 | 6.46E-02 | 2.61E+00 | 2.71E+00 | 8.46E-01 | no | | regulatory_region_variant | MODIFIER |
| 13 | 47767226 | AX-97821561 | 0 | G | T | 0.509 | 1.30E-02 | 6.46E-02 | 2.61E+00 | 2.71E+00 | 8.46E-01 | no | | intergenic_variant | MODIFIER |
| 13 | 47799432 | AX-97790733 | 0 | A | G | 0.509 | 1.30E-02 | 6.46E-02 | 2.61E+00 | 2.71E+00 | 8.46E-01 | no | | intergenic_variant | MODIFIER |
| 9 | 56841985 | AX-170082511 | 0 | C | G | 0.211 | -2.16E-02 | 1.15E-01 | 2.69E+00 | 2.76E+00 | 8.46E-01 | no | | intergenic_variant | MODIFIER |
| 9 | 35034511 | AX-170340123 | 1 | T | G | 0.42 | -1.67E-02 | 8.99E-02 | 2.72E+00 | 2.78E+00 | 8.46E-01 | no | Kirrel3 | intron_variant | MODIFIER |
| 11 | 117237253 | AX-170006012 | 0 | A | G | 0.053 | -3.84E-02 | 2.01E-01 | 2.63E+00 | 2.74E+00 | 8.46E-01 | no | SEP9 | intron_variant | MODIFIER |
| 8 | 32794674 | AX-170101213 | 1 | G | A | 0.143 | -2.51E-02 | 1.29E-01 | 2.66E+00 | 2.72E+00 | 8.46E-01 | no | Nrg1 | intron_variant | MODIFIER |
| 10 | 115665219 | AX-170003967 | 1 | T | G | 0.375 | 1.39E-02 | 7.45E-02 | 2.66E+00 | 2.76E+00 | 8.46E-01 | no | | regulatory_region_variant | MODIFIER |
| 12 | 117404978 | AX-170232155 | 1 | G | T | 0.143 | -3.70E-02 | 1.96E-01 | 2.69E+00 | 2.75E+00 | 8.46E-01 | no | | intergenic_variant | MODIFIER |
| 14 | 105562264 | AX-97797902 | 0 | G | A | 0.044 | -3.14E-02 | 1.60E-01 | 2.59E+00 | 2.73E+00 | 8.46E-01 | no | | intergenic_variant | MODIFIER |
| 4 | 148098267 | AX-170297013 | 1 | T | A | 0.08 | 4.20E-02 | 2.15E-01 | 2.65E+00 | 2.72E+00 | 8.46E-01 | no | Draxin | downstream_gene_variant | MODIFIER |
| 1 | 30579184 | AX-170337336 | 1 | T | C | 0.411 | 1.55E-02 | 8.14E-02 | 2.67E+00 | 2.74E+00 | 8.46E-01 | no | | intergenic_variant | MODIFIER |
| 8 | 93755560 | AX-170125835 | 1 | G | A | 0.214 | -2.26E-02 | 1.18E-01 | 2.68E+00 | 2.74E+00 | 8.46E-01 | no | 4930488L21Rik | intron_variant,non_coding_transcript_variant | MODIFIER |
| 9 | 33551141 | AX-170082055 | 1 | G | C | 0.429 | 1.73E-02 | 8.74E-02 | 2.65E+00 | 2.71E+00 | 8.47E-01 | no | | intergenic_variant | MODIFIER |
| 12 | 71245446 | AX-97789192 | 0 | T | C | 0.061 | -5.78E-02 | 3.01E-01 | 2.71E+00 | 2.73E+00 | 8.47E-01 | no | 2700049A03Rik | downstream_gene_variant | MODIFIER |
| 8 | 32721260 | AX-170419994 | 0 | A | G | 0.14 | -2.44E-02 | 1.26E-01 | 2.65E+00 | 2.73E+00 | 8.47E-01 | no | Nrg1 | intron_variant | MODIFIER |
| 20 | 16176108 | AX-170459480 | 1 | G | A | 0.054 | -1.90E-02 | 1.02E-01 | 2.56E+00 | 2.75E+00 | 8.47E-01 | no | | intergenic_variant | MODIFIER |
| 9 | 89468025 | AX-170445692 | 0 | C | T | 0.149 | -2.30E-02 | 1.15E-01 | 2.62E+00 | 2.71E+00 | 8.47E-01 | no | | intergenic_variant | MODIFIER |
| 12 | 103834438 | AX-170217003 | 2 | G | T | 0.118 | -2.97E-02 | 1.61E-01 | 2.65E+00 | 2.76E+00 | 8.47E-01 | no | Serpina1b | upstream_gene_variant | MODIFIER |
| 7 | 103468334 | AX-97797365 | 1 | C | A | 0.464 | 1.86E-02 | 9.16E-02 | 2.63E+00 | 2.69E+00 | 8.47E-01 | no | Olfr608 | upstream_gene_variant | MODIFIER |
| 1 | 53800122 | AX-170154972 | 0 | C | T | 0.105 | -4.47E-02 | 2.36E-01 | 2.70E+00 | 2.75E+00 | 8.47E-01 | no | | intergenic_variant | MODIFIER |
| 1 | 6854582 | AX-169999875 | 1 | A | C | 0.5 | 1.46E-02 | 7.20E-02 | 2.63E+00 | 2.70E+00 | 8.47E-01 | no | St18 | intron_variant | MODIFIER |
| 8 | 22393816 | AX-97803398 | 0 | G | A | 0.184 | -1.99E-02 | 1.06E-01 | 2.66E+00 | 2.75E+00 | 8.47E-01 | no | Slc25a15 | intron_variant | MODIFIER |
| 8 | 53482305 | AX-170286159 | 0 | T | G | 0.149 | 2.51E-02 | 1.37E-01 | 2.70E+00 | 2.78E+00 | 8.47E-01 | no | Gm45554 | downstream_gene_variant | MODIFIER |
| 18 | 58633033 | AX-170165951 | 0 | A | G | 0.421 | 1.56E-02 | 8.46E-02 | 2.74E+00 | 2.80E+00 | 8.47E-01 | no | | intergenic_variant | MODIFIER |
| 18 | 58634797 | AX-170015789 | 0 | T | C | 0.421 | 1.56E-02 | 8.46E-02 | 2.74E+00 | 2.80E+00 | 8.47E-01 | no | | intergenic_variant | MODIFIER |
| 18 | 58688128 | AX-97825163 | 0 | T | C | 0.421 | 1.56E-02 | 8.46E-02 | 2.74E+00 | 2.80E+00 | 8.47E-01 | no | | intergenic_variant | MODIFIER |
| 18 | 58689116 | AX-170424935 | 0 | A | G | 0.421 | 1.56E-02 | 8.46E-02 | 2.74E+00 | 2.80E+00 | 8.47E-01 | no | | intergenic_variant | MODIFIER |
| 18 | 58690119 | AX-97824280 | 0 | C | T | 0.421 | 1.56E-02 | 8.46E-02 | 2.74E+00 | 2.80E+00 | 8.47E-01 | no | | intergenic_variant | MODIFIER |
| 18 | 58693091 | AX-170508776 | 0 | A | G | 0.421 | 1.56E-02 | 8.46E-02 | 2.74E+00 | 2.80E+00 | 8.47E-01 | no | | intergenic_variant | MODIFIER |
| 18 | 58708776 | AX-170252374 | 0 | T | C | 0.421 | 1.56E-02 | 8.46E-02 | 2.74E+00 | 2.80E+00 | 8.47E-01 | no | | intergenic_variant | MODIFIER |
| 18 | 58735410 | AX-170015791 | 0 | C | T | 0.421 | 1.56E-02 | 8.46E-02 | 2.74E+00 | 2.80E+00 | 8.47E-01 | no | | intergenic_variant | MODIFIER |
| 18 | 58751327 | AX-170060975 | 0 | G | A | 0.421 | 1.56E-02 | 8.46E-02 | 2.74E+00 | 2.80E+00 | 8.47E-01 | no | | intergenic_variant | MODIFIER |
| 18 | 58784914 | AX-97835576 | 0 | C | T | 0.421 | 1.56E-02 | 8.46E-02 | 2.74E+00 | 2.80E+00 | 8.47E-01 | no | AC158924.1 | non_coding_transcript_exon_variant | MODIFIER |

| | | | | | | | | | | | | | | | |
|---|---|---|---|---|---|---|---|---|---|---|---|---|---|---|---|
| 18 | 58818624 | AX-170280432 | 0 | C | A | 0.421 | 1.56E-02 | 8.46E-02 | 2.74E+00 | 2.80E+00 | 8.47E-01 | no | | intergenic_variant | MODIFIER |
| 18 | 58821102 | AX-97835600 | 0 | C | T | 0.421 | 1.56E-02 | 8.46E-02 | 2.74E+00 | 2.80E+00 | 8.47E-01 | no | | intergenic_variant | MODIFIER |
| 18 | 58860655 | AX-170305612 | 0 | T | C | 0.421 | 1.56E-02 | 8.46E-02 | 2.74E+00 | 2.80E+00 | 8.47E-01 | no | Adamts19 | intron_variant | MODIFIER |
| 18 | 58868250 | AX-170103824 | 0 | C | T | 0.421 | 1.56E-02 | 8.46E-02 | 2.74E+00 | 2.80E+00 | 8.47E-01 | no | Adamts19 | intron_variant | MODIFIER |
| 18 | 58881957 | AX-170430475 | 0 | A | G | 0.421 | 1.56E-02 | 8.46E-02 | 2.74E+00 | 2.80E+00 | 8.47E-01 | no | Adamts19 | intron_variant | MODIFIER |
| 18 | 58882319 | AX-170368217 | 0 | C | T | 0.421 | 1.56E-02 | 8.46E-02 | 2.74E+00 | 2.80E+00 | 8.47E-01 | no | Adamts19 | intron_variant | MODIFIER |
| 18 | 58948524 | AX-170420029 | 0 | A | G | 0.421 | 1.56E-02 | 8.46E-02 | 2.74E+00 | 2.80E+00 | 8.47E-01 | no | Adamts19 | intron_variant | MODIFIER |
| 18 | 58970704 | AX-97837756 | 0 | A | G | 0.421 | 1.56E-02 | 8.46E-02 | 2.74E+00 | 2.80E+00 | 8.47E-01 | no | Adamts19 | intron_variant | MODIFIER |
| 18 | 58974495 | AX-170187776 | 0 | G | T | 0.421 | 1.56E-02 | 8.46E-02 | 2.74E+00 | 2.80E+00 | 8.47E-01 | no | Adamts19 | intron_variant | MODIFIER |
| 18 | 58987502 | AX-170428745 | 0 | T | C | 0.421 | 1.56E-02 | 8.46E-02 | 2.74E+00 | 2.80E+00 | 8.47E-01 | no | Adamts19 | intron_variant | MODIFIER |
| 18 | 58990299 | AX-170108566 | 0 | A | G | 0.421 | 1.56E-02 | 8.46E-02 | 2.74E+00 | 2.80E+00 | 8.47E-01 | no | Adamts19 | intron_variant | MODIFIER |
| 18 | 58990324 | AX-170518266 | 0 | A | G | 0.421 | 1.56E-02 | 8.46E-02 | 2.74E+00 | 2.80E+00 | 8.47E-01 | no | Adamts19 | intron_variant | MODIFIER |
| 18 | 58992498 | AX-170060979 | 0 | C | T | 0.421 | 1.56E-02 | 8.46E-02 | 2.74E+00 | 2.80E+00 | 8.47E-01 | no | Adamts19 | intron_variant | MODIFIER |
| 18 | 59048575 | AX-97822807 | 0 | T | C | 0.421 | 1.56E-02 | 8.46E-02 | 2.74E+00 | 2.80E+00 | 8.47E-01 | no | Adamts19 | intron_variant | MODIFIER |
| 18 | 59166586 | AX-170372675 | 0 | A | G | 0.421 | 1.56E-02 | 8.46E-02 | 2.74E+00 | 2.80E+00 | 8.47E-01 | no | | intergenic_variant | MODIFIER |
| 18 | 59193698 | AX-170223143 | 0 | C | T | 0.421 | 1.56E-02 | 8.46E-02 | 2.74E+00 | 2.80E+00 | 8.47E-01 | no | Chsy3 | intron_variant | MODIFIER |
| 18 | 59206529 | AX-97838276 | 0 | A | G | 0.421 | 1.56E-02 | 8.46E-02 | 2.74E+00 | 2.80E+00 | 8.47E-01 | no | Chsy3 | intron_variant | MODIFIER |
| 18 | 59249802 | AX-170315346 | 0 | T | C | 0.421 | 1.56E-02 | 8.46E-02 | 2.74E+00 | 2.80E+00 | 8.47E-01 | no | Chsy3 | intron_variant | MODIFIER |
| 16 | 77785392 | AX-170481669 | 0 | G | A | 0.465 | 1.55E-02 | 7.50E-02 | 2.60E+00 | 2.69E+00 | 8.47E-01 | no | | intergenic_variant | MODIFIER |
| 7 | 51440291 | AX-170138038 | 0 | T | C | 0.421 | 1.90E-02 | 9.61E-02 | 2.64E+00 | 2.71E+00 | 8.47E-01 | no | | intergenic_variant | MODIFIER |
| 4 | 148330899 | AX-170329151 | 1 | G | A | 0.098 | 3.84E-02 | 2.00E-01 | 2.67E+00 | 2.73E+00 | 8.47E-01 | no | | intergenic_variant | MODIFIER |
| 4 | 148355807 | AX-97849740 | 1 | C | T | 0.098 | 3.84E-02 | 2.00E-01 | 2.67E+00 | 2.73E+00 | 8.47E-01 | no | Gm13200 | downstream_gene_variant | MODIFIER |
| 18 | 29592344 | AX-170110164 | 0 | G | C | 0.044 | -4.79E-02 | 2.28E-01 | 2.57E+00 | 2.69E+00 | 8.47E-01 | no | | intergenic_variant | MODIFIER |
| 1 | 8265307 | AX-170514917 | 0 | A | G | 0.518 | -1.25E-02 | 6.98E-02 | 2.72E+00 | 2.81E+00 | 8.47E-01 | no | | intergenic_variant | MODIFIER |
| 1 | 114815896 | AX-97823442 | 1 | C | A | 0.17 | 3.97E-02 | 2.00E-01 | 2.63E+00 | 2.71E+00 | 8.48E-01 | no | | intergenic_variant | MODIFIER |
| 10 | 10620300 | AX-170387579 | 1 | G | A | 0.232 | 1.87E-02 | 1.01E-01 | 2.69E+00 | 2.77E+00 | 8.48E-01 | no | Gm5177 | downstream_gene_variant | MODIFIER |
| 1 | 156438206 | AX-97809534 | 1 | T | C | 0.232 | 2.75E-02 | 1.40E-01 | 2.66E+00 | 2.70E+00 | 8.48E-01 | no | Soat1 | intron_variant | MODIFIER |
| 15 | 86715341 | AX-170318276 | 0 | A | T | 0.5 | 1.48E-02 | 6.84E-02 | 2.57E+00 | 2.67E+00 | 8.48E-01 | no | Gm49464 | downstream_gene_variant | MODIFIER |
| 5 | 122046103 | AX-170506654 | 1 | T | C | 0.455 | 2.10E-02 | 1.08E-01 | 2.65E+00 | 2.71E+00 | 8.48E-01 | no | Cux2 | intron_variant | MODIFIER |
| 4 | 141239656 | AX-170489306 | 0 | C | G | 0.096 | 3.82E-02 | 2.01E-01 | 2.67E+00 | 2.73E+00 | 8.48E-01 | no | Arhgef19 | upstream_gene_variant | MODIFIER |
| 4 | 141246028 | AX-170485946 | 0 | T | C | 0.096 | 3.82E-02 | 2.01E-01 | 2.67E+00 | 2.73E+00 | 8.48E-01 | no | Arhgef19 | intron_variant | MODIFIER |
| 4 | 141252550 | AX-170131543 | 0 | G | C | 0.096 | 3.82E-02 | 2.01E-01 | 2.67E+00 | 2.73E+00 | 8.48E-01 | no | Arhgef19 | intron_variant | MODIFIER |
| 4 | 141275942 | AX-97807671 | 0 | G | A | 0.096 | 3.82E-02 | 2.01E-01 | 2.67E+00 | 2.73E+00 | 8.48E-01 | no | Gm13056 | upstream_gene_variant | MODIFIER |
| 4 | 141302911 | AX-170353406 | 0 | C | T | 0.096 | 3.82E-02 | 2.01E-01 | 2.67E+00 | 2.73E+00 | 8.48E-01 | no | Epha2 | intron_variant | MODIFIER |
| 4 | 141322549 | AX-97788863 | 0 | A | C | 0.096 | 3.82E-02 | 2.01E-01 | 2.67E+00 | 2.73E+00 | 8.48E-01 | no | Epha2 | intron_variant | MODIFIER |
| 4 | 141338439 | AX-97821272 | 0 | T | C | 0.096 | 3.82E-02 | 2.01E-01 | 2.67E+00 | 2.73E+00 | 8.48E-01 | no | | regulatory_region_variant | MODIFIER |
| 4 | 141749374 | AX-97815622 | 0 | T | C | 0.096 | 3.82E-02 | 2.01E-01 | 2.67E+00 | 2.73E+00 | 8.48E-01 | no | Agmat | intron_variant | MODIFIER |
| 4 | 141749718 | AX-170321190 | 0 | G | T | 0.096 | 3.82E-02 | 2.01E-01 | 2.67E+00 | 2.73E+00 | 8.48E-01 | no | Agmat | missense_variant | MODERATE |
| 4 | 141754862 | AX-170367599 | 0 | T | C | 0.096 | 3.82E-02 | 2.01E-01 | 2.67E+00 | 2.73E+00 | 8.48E-01 | no | Agmat | synonymous_variant | LOW |
| 4 | 141762612 | AX-97828382 | 0 | C | T | 0.096 | 3.82E-02 | 2.01E-01 | 2.67E+00 | 2.73E+00 | 8.48E-01 | no | Dnajc16 | 3_prime_UTR_variant | MODIFIER |
| 4 | 141770105 | AX-170130341 | 0 | A | C | 0.096 | 3.82E-02 | 2.01E-01 | 2.67E+00 | 2.73E+00 | 8.48E-01 | no | Dnajc16 | intron_variant | MODIFIER |
| 4 | 141780790 | AX-97836552 | 0 | C | T | 0.096 | 3.82E-02 | 2.01E-01 | 2.67E+00 | 2.73E+00 | 8.48E-01 | no | Dnajc16 | intron_variant | MODIFIER |
| 4 | 141782262 | AX-170317876 | 0 | T | C | 0.096 | 3.82E-02 | 2.01E-01 | 2.67E+00 | 2.73E+00 | 8.48E-01 | no | Dnajc16 | intron_variant | MODIFIER |
| 4 | 141783110 | AX-170429374 | 0 | T | A | 0.096 | 3.82E-02 | 2.01E-01 | 2.67E+00 | 2.73E+00 | 8.48E-01 | no | Dnajc16 | splice_region_variant,intron_variant | LOW |
| 4 | 141831863 | AX-170198810 | 0 | G | A | 0.096 | 3.82E-02 | 2.01E-01 | 2.67E+00 | 2.73E+00 | 8.48E-01 | no | Ctrc | intron_variant,NMD_transcript_variant | MODIFIER |
| 4 | 141872547 | AX-170070897 | 0 | G | A | 0.096 | 3.82E-02 | 2.01E-01 | 2.67E+00 | 2.73E+00 | 8.48E-01 | no | Efhd2 | intron_variant | MODIFIER |
| 4 | 141892481 | AX-97813782 | 0 | T | A | 0.096 | 3.82E-02 | 2.01E-01 | 2.67E+00 | 2.73E+00 | 8.48E-01 | no | Fhad1 | intron_variant | MODIFIER |
| 4 | 141931754 | AX-170107026 | 0 | G | A | 0.096 | 3.82E-02 | 2.01E-01 | 2.67E+00 | 2.73E+00 | 8.48E-01 | no | Fhad1 | intron_variant | MODIFIER |
| 4 | 141932370 | AX-170442975 | 0 | C | A | 0.096 | 3.82E-02 | 2.01E-01 | 2.67E+00 | 2.73E+00 | 8.48E-01 | no | Fhad1 | intron_variant | MODIFIER |
| 4 | 142088816 | AX-170252365 | 0 | T | C | 0.096 | 3.82E-02 | 2.01E-01 | 2.67E+00 | 2.73E+00 | 8.48E-01 | no | Tmem51 | upstream_gene_variant | MODIFIER |
| 4 | 142189805 | AX-170089391 | 0 | C | T | 0.096 | 3.82E-02 | 2.01E-01 | 2.67E+00 | 2.73E+00 | 8.48E-01 | no | Kazn | intron_variant | MODIFIER |
| 4 | 142190136 | AX-170389621 | 0 | C | T | 0.096 | 3.82E-02 | 2.01E-01 | 2.67E+00 | 2.73E+00 | 8.48E-01 | no | Kazn | intron_variant | MODIFIER |
| 4 | 142191070 | AX-170383852 | 0 | T | A | 0.096 | 3.82E-02 | 2.01E-01 | 2.67E+00 | 2.73E+00 | 8.48E-01 | no | Kazn | intron_variant | MODIFIER |
| 4 | 142193552 | AX-170182471 | 0 | C | T | 0.096 | 3.82E-02 | 2.01E-01 | 2.67E+00 | 2.73E+00 | 8.48E-01 | no | Kazn | intron_variant | MODIFIER |

| | | | | | | | | | | | | | | | |
|---|---|---|---|---|---|---|---|---|---|---|---|---|---|---|---|
| 4 | 142227741 | AX-170417390 | 0 | C | T | 0.096 | 3.82E-02 | 2.01E-01 | 2.67E+00 | 2.73E+00 | 8.48E-01 | no | Kazn | intron_variant | MODIFIER |
| 4 | 142231338 | AX-97839871 | 0 | A | G | 0.096 | 3.82E-02 | 2.01E-01 | 2.67E+00 | 2.73E+00 | 8.48E-01 | no | Kazn | intron_variant | MODIFIER |
| 4 | 142231421 | AX-170439101 | 0 | C | T | 0.096 | 3.82E-02 | 2.01E-01 | 2.67E+00 | 2.73E+00 | 8.48E-01 | no | Kazn | intron_variant | MODIFIER |
| 4 | 142231484 | AX-170070904 | 0 | C | A | 0.096 | 3.82E-02 | 2.01E-01 | 2.67E+00 | 2.73E+00 | 8.48E-01 | no | Kazn | intron_variant | MODIFIER |
| 4 | 142234770 | AX-170115238 | 0 | A | G | 0.096 | 3.82E-02 | 2.01E-01 | 2.67E+00 | 2.73E+00 | 8.48E-01 | no | Kazn | intron_variant | MODIFIER |
| 4 | 142400231 | AX-170251629 | 0 | C | A | 0.096 | 3.82E-02 | 2.01E-01 | 2.67E+00 | 2.73E+00 | 8.48E-01 | no | | intergenic_variant | MODIFIER |
| 4 | 142404357 | AX-170344924 | 0 | A | G | 0.096 | 3.82E-02 | 2.01E-01 | 2.67E+00 | 2.73E+00 | 8.48E-01 | no | | intergenic_variant | MODIFIER |
| 4 | 142404380 | AX-170229993 | 0 | A | G | 0.096 | 3.82E-02 | 2.01E-01 | 2.67E+00 | 2.73E+00 | 8.48E-01 | no | | intergenic_variant | MODIFIER |
| 4 | 142412745 | AX-170176516 | 0 | G | A | 0.096 | 3.82E-02 | 2.01E-01 | 2.67E+00 | 2.73E+00 | 8.48E-01 | no | | regulatory_region_variant | MODIFIER |
| 4 | 142413905 | AX-170379522 | 0 | G | C | 0.096 | 3.82E-02 | 2.01E-01 | 2.67E+00 | 2.73E+00 | 8.48E-01 | no | | intergenic_variant | MODIFIER |
| 4 | 142422525 | AX-170070909 | 0 | T | G | 0.096 | 3.82E-02 | 2.01E-01 | 2.67E+00 | 2.73E+00 | 8.48E-01 | no | | intergenic_variant | MODIFIER |
| 4 | 142489389 | AX-170276352 | 0 | T | C | 0.096 | 3.82E-02 | 2.01E-01 | 2.67E+00 | 2.73E+00 | 8.48E-01 | no | | intergenic_variant | MODIFIER |
| 4 | 142503390 | AX-170474655 | 0 | T | G | 0.096 | 3.82E-02 | 2.01E-01 | 2.67E+00 | 2.73E+00 | 8.48E-01 | no | | intergenic_variant | MODIFIER |
| 4 | 142505370 | AX-170402408 | 0 | A | G | 0.096 | 3.82E-02 | 2.01E-01 | 2.67E+00 | 2.73E+00 | 8.48E-01 | no | | intergenic_variant | MODIFIER |
| 4 | 142512214 | AX-170308264 | 0 | A | G | 0.096 | 3.82E-02 | 2.01E-01 | 2.67E+00 | 2.73E+00 | 8.48E-01 | no | | intergenic_variant | MODIFIER |
| 4 | 142514896 | AX-170184058 | 0 | T | C | 0.096 | 3.82E-02 | 2.01E-01 | 2.67E+00 | 2.73E+00 | 8.48E-01 | no | | intergenic_variant | MODIFIER |
| 4 | 142515096 | AX-170373705 | 0 | G | T | 0.096 | 3.82E-02 | 2.01E-01 | 2.67E+00 | 2.73E+00 | 8.48E-01 | no | | regulatory_region_variant | MODIFIER |
| 4 | 142522602 | AX-170400370 | 0 | T | A | 0.096 | 3.82E-02 | 2.01E-01 | 2.67E+00 | 2.73E+00 | 8.48E-01 | no | | intergenic_variant | MODIFIER |
| 4 | 142522740 | AX-170439368 | 0 | G | T | 0.096 | 3.82E-02 | 2.01E-01 | 2.67E+00 | 2.73E+00 | 8.48E-01 | no | | intergenic_variant | MODIFIER |
| 4 | 142541676 | AX-170426462 | 0 | C | T | 0.096 | 3.82E-02 | 2.01E-01 | 2.67E+00 | 2.73E+00 | 8.48E-01 | no | | intergenic_variant | MODIFIER |
| 4 | 142548659 | AX-170476928 | 0 | C | T | 0.096 | 3.82E-02 | 2.01E-01 | 2.67E+00 | 2.73E+00 | 8.48E-01 | no | | intergenic_variant | MODIFIER |
| 4 | 142548879 | AX-170455658 | 0 | T | C | 0.096 | 3.82E-02 | 2.01E-01 | 2.67E+00 | 2.73E+00 | 8.48E-01 | no | | intergenic_variant | MODIFIER |
| 4 | 142840487 | AX-97791237 | 0 | C | T | 0.096 | 3.82E-02 | 2.01E-01 | 2.67E+00 | 2.73E+00 | 8.48E-01 | no | 6330411D24Rik | intron_variant,non_coding_transcript_variant | MODIFIER |
| 4 | 142841614 | AX-170274804 | 0 | T | C | 0.096 | 3.82E-02 | 2.01E-01 | 2.67E+00 | 2.73E+00 | 8.48E-01 | no | 6330411D24Rik | intron_variant,non_coding_transcript_variant | MODIFIER |
| 4 | 142841856 | AX-170391125 | 0 | G | A | 0.096 | 3.82E-02 | 2.01E-01 | 2.67E+00 | 2.73E+00 | 8.48E-01 | no | 6330411D24Rik | intron_variant,non_coding_transcript_variant | MODIFIER |
| 4 | 142888145 | AX-97820029 | 0 | C | T | 0.096 | 3.82E-02 | 2.01E-01 | 2.67E+00 | 2.73E+00 | 8.48E-01 | no | 6330411D24Rik | intron_variant,non_coding_transcript_variant | MODIFIER |
| 4 | 143038292 | AX-97797954 | 0 | T | C | 0.096 | 3.82E-02 | 2.01E-01 | 2.67E+00 | 2.73E+00 | 8.48E-01 | no | 6330411D24Rik | intron_variant,non_coding_transcript_variant | MODIFIER |
| 4 | 143131086 | AX-170199493 | 0 | A | G | 0.096 | 3.82E-02 | 2.01E-01 | 2.67E+00 | 2.73E+00 | 8.48E-01 | no | Prdm2 | intron_variant | MODIFIER |
| 4 | 143136230 | AX-97786415 | 0 | T | C | 0.096 | 3.82E-02 | 2.01E-01 | 2.67E+00 | 2.73E+00 | 8.48E-01 | no | Prdm2 | intron_variant | MODIFIER |
| 4 | 143194943 | AX-97811181 | 0 | C | T | 0.096 | 3.82E-02 | 2.01E-01 | 2.67E+00 | 2.73E+00 | 8.48E-01 | no | Prdm2 | intron_variant | MODIFIER |
| 4 | 143200937 | AX-170070924 | 0 | A | G | 0.096 | 3.82E-02 | 2.01E-01 | 2.67E+00 | 2.73E+00 | 8.48E-01 | no | Prdm2 | intron_variant | MODIFIER |
| 4 | 143206728 | AX-97780352 | 0 | T | G | 0.096 | 3.82E-02 | 2.01E-01 | 2.67E+00 | 2.73E+00 | 8.48E-01 | no | Prdm2 | intron_variant | MODIFIER |
| 4 | 143298103 | AX-97803803 | 0 | T | C | 0.096 | 3.82E-02 | 2.01E-01 | 2.67E+00 | 2.73E+00 | 8.48E-01 | no | Pdpn | intron_variant | MODIFIER |
| 4 | 143894306 | AX-97831705 | 0 | T | G | 0.096 | 3.82E-02 | 2.01E-01 | 2.67E+00 | 2.73E+00 | 8.48E-01 | no | Pramef6 | intron_variant | MODIFIER |
| 4 | 144386367 | AX-97810570 | 0 | A | G | 0.096 | 3.82E-02 | 2.01E-01 | 2.67E+00 | 2.73E+00 | 8.48E-01 | no | | intergenic_variant | MODIFIER |
| 4 | 144516421 | AX-97828815 | 0 | A | G | 0.096 | 3.82E-02 | 2.01E-01 | 2.67E+00 | 2.73E+00 | 8.48E-01 | no | 9430007A20Rik | upstream_gene_variant | MODIFIER |
| 4 | 144640641 | AX-170280541 | 0 | C | T | 0.096 | 3.82E-02 | 2.01E-01 | 2.67E+00 | 2.73E+00 | 8.48E-01 | no | | intergenic_variant | MODIFIER |
| 4 | 144790248 | AX-170476585 | 0 | G | A | 0.096 | 3.82E-02 | 2.01E-01 | 2.67E+00 | 2.73E+00 | 8.48E-01 | no | Gm438 | upstream_gene_variant | MODIFIER |
| 4 | 144861925 | AX-97794171 | 0 | A | G | 0.096 | 3.82E-02 | 2.01E-01 | 2.67E+00 | 2.73E+00 | 8.48E-01 | no | | intergenic_variant | MODIFIER |
| 4 | 144869187 | AX-170474860 | 0 | A | G | 0.096 | 3.82E-02 | 2.01E-01 | 2.67E+00 | 2.73E+00 | 8.48E-01 | no | | intergenic_variant | MODIFIER |
| 4 | 144894584 | AX-170166634 | 0 | G | C | 0.096 | 3.82E-02 | 2.01E-01 | 2.67E+00 | 2.73E+00 | 8.48E-01 | no | Dhrs3 | intron_variant,NMD_transcript_variant | MODIFIER |
| 4 | 145142790 | AX-170129640 | 0 | T | C | 0.096 | 3.82E-02 | 2.01E-01 | 2.67E+00 | 2.73E+00 | 8.48E-01 | no | Vps13d | intron_variant | MODIFIER |
| 4 | 145143848 | AX-170168639 | 0 | A | G | 0.096 | 3.82E-02 | 2.01E-01 | 2.67E+00 | 2.73E+00 | 8.48E-01 | no | Vps13d | intron_variant | MODIFIER |
| 4 | 145236429 | AX-97844316 | 0 | C | G | 0.096 | 3.82E-02 | 2.01E-01 | 2.67E+00 | 2.73E+00 | 8.48E-01 | no | Tnfrsf1b | intron_variant | MODIFIER |
| 4 | 145293120 | AX-170178373 | 0 | T | C | 0.096 | 3.82E-02 | 2.01E-01 | 2.67E+00 | 2.73E+00 | 8.48E-01 | no | Tnfrsf8 | missense_variant | MODERATE |
| 4 | 145336041 | AX-170319385 | 0 | C | T | 0.096 | 3.82E-02 | 2.01E-01 | 2.67E+00 | 2.73E+00 | 8.48E-01 | no | | intergenic_variant | MODIFIER |
| 4 | 145355590 | AX-170511517 | 0 | C | T | 0.096 | 3.82E-02 | 2.01E-01 | 2.67E+00 | 2.73E+00 | 8.48E-01 | no | | intergenic_variant | MODIFIER |
| 4 | 145369249 | AX-97856278 | 0 | A | G | 0.096 | 3.82E-02 | 2.01E-01 | 2.67E+00 | 2.73E+00 | 8.48E-01 | no | | intergenic_variant | MODIFIER |
| 4 | 147885783 | AX-170070977 | 0 | T | C | 0.096 | 3.82E-02 | 2.01E-01 | 2.67E+00 | 2.73E+00 | 8.48E-01 | no | Mfn2 | intron_variant | MODIFIER |
| 4 | 147896432 | AX-170201299 | 0 | G | T | 0.096 | 3.82E-02 | 2.01E-01 | 2.67E+00 | 2.73E+00 | 8.48E-01 | no | Mfn2 | intron_variant | MODIFIER |
| 4 | 148063390 | AX-170228545 | 0 | T | C | 0.096 | 3.82E-02 | 2.01E-01 | 2.67E+00 | 2.73E+00 | 8.48E-01 | no | Mthfr | downstream_gene_variant | MODIFIER |
| 4 | 148080083 | AX-170444809 | 0 | C | T | 0.096 | 3.82E-02 | 2.01E-01 | 2.67E+00 | 2.73E+00 | 8.48E-01 | no | Agtrap | 3_prime_UTR_variant | MODIFIER |
| 4 | 148089161 | AX-170262202 | 0 | A | G | 0.096 | 3.82E-02 | 2.01E-01 | 2.67E+00 | 2.73E+00 | 8.48E-01 | no | Agtrap | upstream_gene_variant | MODIFIER |
| 4 | 148095018 | AX-170347616 | 0 | T | C | 0.096 | 3.82E-02 | 2.01E-01 | 2.67E+00 | 2.73E+00 | 8.48E-01 | no | Draxin | downstream_gene_variant | MODIFIER |

| | | | | | | | | | | | | | | | |
|---|---|---|---|---|---|---|---|---|---|---|---|---|---|---|---|
| 4 | 148098919 | AX-170149875 | 0 | C | T | 0.096 | 3.82E-02 | 2.01E-01 | 2.67E+00 | 2.73E+00 | 8.48E-01 | no | Draxin | 3_prime_UTR_variant | MODIFIER |
| 4 | 148103947 | AX-170465665 | 0 | T | C | 0.096 | 3.82E-02 | 2.01E-01 | 2.67E+00 | 2.73E+00 | 8.48E-01 | no | Draxin | intron_variant | MODIFIER |
| 4 | 148104929 | AX-97856634 | 0 | T | C | 0.096 | 3.82E-02 | 2.01E-01 | 2.67E+00 | 2.73E+00 | 8.48E-01 | no | Draxin | intron_variant | MODIFIER |
| 4 | 148113164 | AX-170218167 | 0 | A | G | 0.096 | 3.82E-02 | 2.01E-01 | 2.67E+00 | 2.73E+00 | 8.48E-01 | no | Draxin | intron_variant | MODIFIER |
| 4 | 148131958 | AX-170461166 | 0 | A | G | 0.096 | 3.82E-02 | 2.01E-01 | 2.67E+00 | 2.73E+00 | 8.48E-01 | no | Draxin | upstream_gene_variant | MODIFIER |
| 4 | 148134916 | AX-170155652 | 0 | T | C | 0.096 | 3.82E-02 | 2.01E-01 | 2.67E+00 | 2.73E+00 | 8.48E-01 | no | Draxin | upstream_gene_variant | MODIFIER |
| 4 | 148135487 | AX-170471912 | 0 | G | A | 0.096 | 3.82E-02 | 2.01E-01 | 2.67E+00 | 2.73E+00 | 8.48E-01 | no | Mad2l2 | upstream_gene_variant | MODIFIER |
| 4 | 148135499 | AX-170473908 | 0 | C | T | 0.096 | 3.82E-02 | 2.01E-01 | 2.67E+00 | 2.73E+00 | 8.48E-01 | no | Mad2l2 | upstream_gene_variant | MODIFIER |
| 4 | 148139300 | AX-97809400 | 0 | C | T | 0.096 | 3.82E-02 | 2.01E-01 | 2.67E+00 | 2.73E+00 | 8.48E-01 | no | Mad2l2 | upstream_gene_variant | MODIFIER |
| 4 | 148145003 | AX-170239925 | 0 | T | C | 0.096 | 3.82E-02 | 2.01E-01 | 2.67E+00 | 2.73E+00 | 8.48E-01 | no | Fbxo6 | downstream_gene_variant | MODIFIER |
| 4 | 148156654 | AX-170376186 | 0 | C | T | 0.096 | 3.82E-02 | 2.01E-01 | 2.67E+00 | 2.73E+00 | 8.48E-01 | no | Fbxo6 | upstream_gene_variant | MODIFIER |
| 4 | 148167303 | AX-170400660 | 0 | G | T | 0.096 | 3.82E-02 | 2.01E-01 | 2.67E+00 | 2.73E+00 | 8.48E-01 | no | Fbxo2 | downstream_gene_variant | MODIFIER |
| 4 | 148177357 | AX-97831037 | 0 | A | G | 0.096 | 3.82E-02 | 2.01E-01 | 2.67E+00 | 2.73E+00 | 8.48E-01 | no | | intergenic_variant | MODIFIER |
| 4 | 148196446 | AX-170216265 | 0 | A | G | 0.096 | 3.82E-02 | 2.01E-01 | 2.67E+00 | 2.73E+00 | 8.48E-01 | no | | intergenic_variant | MODIFIER |
| 4 | 148213478 | AX-170418163 | 0 | T | C | 0.096 | 3.82E-02 | 2.01E-01 | 2.67E+00 | 2.73E+00 | 8.48E-01 | no | | intergenic_variant | MODIFIER |
| 4 | 148213840 | AX-170458430 | 0 | T | C | 0.096 | 3.82E-02 | 2.01E-01 | 2.67E+00 | 2.73E+00 | 8.48E-01 | no | | intergenic_variant | MODIFIER |
| 4 | 148214177 | AX-170457070 | 0 | T | C | 0.096 | 3.82E-02 | 2.01E-01 | 2.67E+00 | 2.73E+00 | 8.48E-01 | no | | regulatory_region_variant | MODIFIER |
| 4 | 148237906 | AX-170280841 | 0 | A | T | 0.096 | 3.82E-02 | 2.01E-01 | 2.67E+00 | 2.73E+00 | 8.48E-01 | no | Disp3 | downstream_gene_variant | MODIFIER |
| 4 | 148241059 | AX-170237775 | 0 | G | C | 0.096 | 3.82E-02 | 2.01E-01 | 2.67E+00 | 2.73E+00 | 8.48E-01 | no | Disp3 | 3_prime_UTR_variant | MODIFIER |
| 4 | 148243065 | AX-170162362 | 0 | G | A | 0.096 | 3.82E-02 | 2.01E-01 | 2.67E+00 | 2.73E+00 | 8.48E-01 | no | Disp3 | intron_variant | MODIFIER |
| 4 | 148243245 | AX-170227305 | 0 | T | C | 0.096 | 3.82E-02 | 2.01E-01 | 2.67E+00 | 2.73E+00 | 8.48E-01 | no | Disp3 | intron_variant | MODIFIER |
| 4 | 148262482 | AX-170486514 | 0 | G | A | 0.096 | 3.82E-02 | 2.01E-01 | 2.67E+00 | 2.73E+00 | 8.48E-01 | no | Disp3 | intron_variant | MODIFIER |
| 4 | 148263962 | AX-170132199 | 0 | G | A | 0.096 | 3.82E-02 | 2.01E-01 | 2.67E+00 | 2.73E+00 | 8.48E-01 | no | Disp3 | intron_variant | MODIFIER |
| 4 | 148274542 | AX-170438558 | 0 | A | C | 0.096 | 3.82E-02 | 2.01E-01 | 2.67E+00 | 2.73E+00 | 8.48E-01 | no | Disp3 | intron_variant | MODIFIER |
| 4 | 148279331 | AX-97825119 | 0 | A | C | 0.096 | 3.82E-02 | 2.01E-01 | 2.67E+00 | 2.73E+00 | 8.48E-01 | no | Disp3 | intron_variant | MODIFIER |
| 4 | 148282356 | AX-170244840 | 0 | A | G | 0.096 | 3.82E-02 | 2.01E-01 | 2.67E+00 | 2.73E+00 | 8.48E-01 | no | Disp3 | intron_variant | MODIFIER |
| 4 | 148288790 | AX-170070989 | 0 | A | G | 0.096 | 3.82E-02 | 2.01E-01 | 2.67E+00 | 2.73E+00 | 8.48E-01 | no | Disp3 | upstream_gene_variant | MODIFIER |
| 4 | 148290110 | AX-170420452 | 0 | T | C | 0.096 | 3.82E-02 | 2.01E-01 | 2.67E+00 | 2.73E+00 | 8.48E-01 | no | Disp3 | upstream_gene_variant | MODIFIER |
| 4 | 148293524 | AX-170070990 | 0 | T | C | 0.096 | 3.82E-02 | 2.01E-01 | 2.67E+00 | 2.73E+00 | 8.48E-01 | no | | regulatory_region_variant | MODIFIER |
| 4 | 148295892 | AX-170377786 | 0 | A | G | 0.096 | 3.82E-02 | 2.01E-01 | 2.67E+00 | 2.73E+00 | 8.48E-01 | no | | intergenic_variant | MODIFIER |
| 4 | 148296558 | AX-170349141 | 0 | T | C | 0.096 | 3.82E-02 | 2.01E-01 | 2.67E+00 | 2.73E+00 | 8.48E-01 | no | | regulatory_region_variant | MODIFIER |
| 4 | 148305971 | AX-170205347 | 0 | A | C | 0.096 | 3.82E-02 | 2.01E-01 | 2.67E+00 | 2.73E+00 | 8.48E-01 | no | Gm13206 | upstream_gene_variant | MODIFIER |
| 4 | 148309265 | AX-170200840 | 0 | A | G | 0.096 | 3.82E-02 | 2.01E-01 | 2.67E+00 | 2.73E+00 | 8.48E-01 | no | Gm13206 | intron_variant,non_coding_transcript_variant | MODIFIER |
| 4 | 148315141 | AX-170357619 | 0 | A | G | 0.096 | 3.82E-02 | 2.01E-01 | 2.67E+00 | 2.73E+00 | 8.48E-01 | no | Gm13206 | intron_variant,non_coding_transcript_variant | MODIFIER |
| 4 | 148325069 | AX-170130315 | 0 | A | G | 0.096 | 3.82E-02 | 2.01E-01 | 2.67E+00 | 2.73E+00 | 8.48E-01 | no | | regulatory_region_variant | MODIFIER |
| 4 | 148337464 | AX-170296386 | 0 | A | T | 0.096 | 3.82E-02 | 2.01E-01 | 2.67E+00 | 2.73E+00 | 8.48E-01 | no | | intergenic_variant | MODIFIER |
| 4 | 148340292 | AX-170263942 | 0 | T | G | 0.096 | 3.82E-02 | 2.01E-01 | 2.67E+00 | 2.73E+00 | 8.48E-01 | no | | regulatory_region_variant | MODIFIER |
| 4 | 148354916 | AX-170458600 | 0 | T | C | 0.096 | 3.82E-02 | 2.01E-01 | 2.67E+00 | 2.73E+00 | 8.48E-01 | no | Gm13200 | downstream_gene_variant | MODIFIER |
| 4 | 148374344 | AX-170318469 | 0 | A | G | 0.096 | 3.82E-02 | 2.01E-01 | 2.67E+00 | 2.73E+00 | 8.48E-01 | no | | intergenic_variant | MODIFIER |
| 4 | 148377203 | AX-170334680 | 0 | G | A | 0.096 | 3.82E-02 | 2.01E-01 | 2.67E+00 | 2.73E+00 | 8.48E-01 | no | | regulatory_region_variant | MODIFIER |
| 4 | 148386552 | AX-170512718 | 0 | C | T | 0.096 | 3.82E-02 | 2.01E-01 | 2.67E+00 | 2.73E+00 | 8.48E-01 | no | | regulatory_region_variant | MODIFIER |
| 4 | 148386961 | AX-170134687 | 0 | T | C | 0.096 | 3.82E-02 | 2.01E-01 | 2.67E+00 | 2.73E+00 | 8.48E-01 | no | | intergenic_variant | MODIFIER |
| 4 | 148393136 | AX-170106948 | 0 | A | G | 0.096 | 3.82E-02 | 2.01E-01 | 2.67E+00 | 2.73E+00 | 8.48E-01 | no | | regulatory_region_variant | MODIFIER |
| 4 | 148417027 | AX-170194830 | 0 | A | T | 0.096 | 3.82E-02 | 2.01E-01 | 2.67E+00 | 2.73E+00 | 8.48E-01 | no | | regulatory_region_variant | MODIFIER |
| 4 | 148419835 | AX-170320142 | 0 | T | C | 0.096 | 3.82E-02 | 2.01E-01 | 2.67E+00 | 2.73E+00 | 8.48E-01 | no | | intergenic_variant | MODIFIER |
| 4 | 148421160 | AX-170070992 | 0 | A | G | 0.096 | 3.82E-02 | 2.01E-01 | 2.67E+00 | 2.73E+00 | 8.48E-01 | no | | intergenic_variant | MODIFIER |
| 4 | 148425842 | AX-170070995 | 0 | T | C | 0.096 | 3.82E-02 | 2.01E-01 | 2.67E+00 | 2.73E+00 | 8.48E-01 | no | | intergenic_variant | MODIFIER |
| 4 | 148426523 | AX-170493120 | 0 | C | T | 0.096 | 3.82E-02 | 2.01E-01 | 2.67E+00 | 2.73E+00 | 8.48E-01 | no | | intergenic_variant | MODIFIER |
| 4 | 148427274 | AX-97792042 | 0 | T | C | 0.096 | 3.82E-02 | 2.01E-01 | 2.67E+00 | 2.73E+00 | 8.48E-01 | no | | regulatory_region_variant | MODIFIER |
| 4 | 148453679 | AX-170156534 | 0 | A | G | 0.096 | 3.82E-02 | 2.01E-01 | 2.67E+00 | 2.73E+00 | 8.48E-01 | no | Mtor | intron_variant | MODIFIER |
| 4 | 148464348 | AX-170156539 | 0 | G | A | 0.096 | 3.82E-02 | 2.01E-01 | 2.67E+00 | 2.73E+00 | 8.48E-01 | no | Mtor | synonymous_variant | LOW |
| 4 | 148467749 | AX-170304300 | 0 | G | A | 0.096 | 3.82E-02 | 2.01E-01 | 2.67E+00 | 2.73E+00 | 8.48E-01 | no | Mtor | intron_variant | MODIFIER |
| 4 | 148471769 | AX-170334529 | 0 | C | G | 0.096 | 3.82E-02 | 2.01E-01 | 2.67E+00 | 2.73E+00 | 8.48E-01 | no | Mtor | intron_variant | MODIFIER |
| 4 | 148479848 | AX-170197901 | 0 | T | C | 0.096 | 3.82E-02 | 2.01E-01 | 2.67E+00 | 2.73E+00 | 8.48E-01 | no | Mtor | intron_variant | MODIFIER |

| | | | | | | | | | | | | | | | |
|---|---|---|---|---|---|---|---|---|---|---|---|---|---|---|---|
| 4 | 148490436 | AX-97844751 | 0 | T | C | 0.096 | 3.82E-02 | 2.01E-01 | 2.67E+00 | 2.73E+00 | 8.48E-01 | no | Angptl7 | downstream_gene_variant | MODIFIER |
| 4 | 148496387 | AX-170181495 | 0 | A | G | 0.096 | 3.82E-02 | 2.01E-01 | 2.67E+00 | 2.73E+00 | 8.48E-01 | no | Angptl7 | intron_variant | MODIFIER |
| 4 | 148500796 | AX-170509876 | 0 | C | T | 0.096 | 3.82E-02 | 2.01E-01 | 2.67E+00 | 2.73E+00 | 8.48E-01 | no | Angptl7 | upstream_gene_variant | MODIFIER |
| 4 | 148507655 | AX-170482305 | 0 | C | G | 0.096 | 3.82E-02 | 2.01E-01 | 2.67E+00 | 2.73E+00 | 8.48E-01 | no | Mtor | intron_variant | MODIFIER |
| 4 | 148512822 | AX-170337163 | 0 | G | T | 0.096 | 3.82E-02 | 2.01E-01 | 2.67E+00 | 2.73E+00 | 8.48E-01 | no | Mtor | intron_variant | MODIFIER |
| 4 | 148539761 | AX-170070996 | 0 | A | G | 0.096 | 3.82E-02 | 2.01E-01 | 2.67E+00 | 2.73E+00 | 8.48E-01 | no | Mtor | intron_variant | MODIFIER |
| 4 | 148549702 | AX-170155509 | 0 | T | A | 0.096 | 3.82E-02 | 2.01E-01 | 2.67E+00 | 2.73E+00 | 8.48E-01 | no | Mtor | intron_variant | MODIFIER |
| 4 | 148550198 | AX-170393092 | 0 | G | A | 0.096 | 3.82E-02 | 2.01E-01 | 2.67E+00 | 2.73E+00 | 8.48E-01 | no | Mtor | synonymous_variant | LOW |
| 4 | 148580944 | AX-170291594 | 0 | T | C | 0.096 | 3.82E-02 | 2.01E-01 | 2.67E+00 | 2.73E+00 | 8.48E-01 | no | Exosc10 | intron_variant | MODIFIER |
| 4 | 148581863 | AX-170024201 | 0 | A | G | 0.096 | 3.82E-02 | 2.01E-01 | 2.67E+00 | 2.73E+00 | 8.48E-01 | no | Exosc10 | intron_variant | MODIFIER |
| 4 | 148611220 | AX-170024202 | 0 | A | C | 0.096 | 3.82E-02 | 2.01E-01 | 2.67E+00 | 2.73E+00 | 8.48E-01 | no | Tardbp | downstream_gene_variant | MODIFIER |
| 4 | 148640375 | AX-170244871 | 0 | C | T | 0.096 | 3.82E-02 | 2.01E-01 | 2.67E+00 | 2.73E+00 | 8.48E-01 | no | Gm572 | upstream_gene_variant | MODIFIER |
| 4 | 148641378 | AX-170534365 | 0 | G | A | 0.096 | 3.82E-02 | 2.01E-01 | 2.67E+00 | 2.73E+00 | 8.48E-01 | no | Gm572 | upstream_gene_variant | MODIFIER |
| 4 | 148644354 | AX-97823207 | 0 | T | C | 0.096 | 3.82E-02 | 2.01E-01 | 2.67E+00 | 2.73E+00 | 8.48E-01 | no | Gm572 | intron_variant | MODIFIER |
| 4 | 148644395 | AX-170024204 | 0 | G | T | 0.096 | 3.82E-02 | 2.01E-01 | 2.67E+00 | 2.73E+00 | 8.48E-01 | no | Gm572 | intron_variant | MODIFIER |
| 4 | 148698170 | AX-170193060 | 0 | C | A | 0.096 | 3.82E-02 | 2.01E-01 | 2.67E+00 | 2.73E+00 | 8.48E-01 | no | | intergenic_variant | MODIFIER |
| 4 | 148700942 | AX-170476191 | 0 | C | A | 0.096 | 3.82E-02 | 2.01E-01 | 2.67E+00 | 2.73E+00 | 8.48E-01 | no | | intergenic_variant | MODIFIER |
| 4 | 148701204 | AX-170228351 | 0 | A | G | 0.096 | 3.82E-02 | 2.01E-01 | 2.67E+00 | 2.73E+00 | 8.48E-01 | no | | intergenic_variant | MODIFIER |
| 4 | 148702177 | AX-170427473 | 0 | A | G | 0.096 | 3.82E-02 | 2.01E-01 | 2.67E+00 | 2.73E+00 | 8.48E-01 | no | | intergenic_variant | MODIFIER |
| 4 | 148713421 | AX-170089394 | 0 | G | A | 0.096 | 3.82E-02 | 2.01E-01 | 2.67E+00 | 2.73E+00 | 8.48E-01 | no | Gm13203 | upstream_gene_variant | MODIFIER |
| 4 | 148728469 | AX-170267552 | 0 | G | A | 0.096 | 3.82E-02 | 2.01E-01 | 2.67E+00 | 2.73E+00 | 8.48E-01 | no | | intergenic_variant | MODIFIER |
| 4 | 148728974 | AX-170377002 | 0 | T | C | 0.096 | 3.82E-02 | 2.01E-01 | 2.67E+00 | 2.73E+00 | 8.48E-01 | no | | intergenic_variant | MODIFIER |
| 4 | 148732546 | AX-97787776 | 0 | T | G | 0.096 | 3.82E-02 | 2.01E-01 | 2.67E+00 | 2.73E+00 | 8.48E-01 | no | | intergenic_variant | MODIFIER |
| 4 | 148734301 | AX-170024207 | 0 | T | C | 0.096 | 3.82E-02 | 2.01E-01 | 2.67E+00 | 2.73E+00 | 8.48E-01 | no | | regulatory_region_variant | MODIFIER |
| 4 | 148769094 | AX-170336696 | 0 | C | G | 0.096 | 3.82E-02 | 2.01E-01 | 2.67E+00 | 2.73E+00 | 8.48E-01 | no | | intergenic_variant | MODIFIER |
| 4 | 148772309 | AX-170349091 | 0 | A | T | 0.096 | 3.82E-02 | 2.01E-01 | 2.67E+00 | 2.73E+00 | 8.48E-01 | no | | intergenic_variant | MODIFIER |
| 4 | 148774276 | AX-170322515 | 0 | C | T | 0.096 | 3.82E-02 | 2.01E-01 | 2.67E+00 | 2.73E+00 | 8.48E-01 | no | | intergenic_variant | MODIFIER |
| 4 | 148774592 | AX-170024205 | 0 | T | C | 0.096 | 3.82E-02 | 2.01E-01 | 2.67E+00 | 2.73E+00 | 8.48E-01 | no | | intergenic_variant | MODIFIER |
| 4 | 148814333 | AX-97846854 | 0 | T | C | 0.096 | 3.82E-02 | 2.01E-01 | 2.67E+00 | 2.73E+00 | 8.48E-01 | no | Casz1 | intron_variant | MODIFIER |
| 4 | 148835046 | AX-170024209 | 0 | G | A | 0.096 | 3.82E-02 | 2.01E-01 | 2.67E+00 | 2.73E+00 | 8.48E-01 | no | Casz1 | intron_variant | MODIFIER |
| 4 | 148840377 | AX-97833974 | 0 | C | T | 0.096 | 3.82E-02 | 2.01E-01 | 2.67E+00 | 2.73E+00 | 8.48E-01 | no | Casz1 | intron_variant | MODIFIER |
| 4 | 148849236 | AX-170441068 | 0 | G | A | 0.096 | 3.82E-02 | 2.01E-01 | 2.67E+00 | 2.73E+00 | 8.48E-01 | no | Casz1 | intron_variant | MODIFIER |
| 4 | 148851183 | AX-170333341 | 0 | G | A | 0.096 | 3.82E-02 | 2.01E-01 | 2.67E+00 | 2.73E+00 | 8.48E-01 | no | Casz1 | intron_variant | MODIFIER |
| 4 | 148855319 | AX-170395788 | 0 | C | T | 0.096 | 3.82E-02 | 2.01E-01 | 2.67E+00 | 2.73E+00 | 8.48E-01 | no | Casz1 | intron_variant | MODIFIER |
| 4 | 148857725 | AX-97808240 | 0 | T | C | 0.096 | 3.82E-02 | 2.01E-01 | 2.67E+00 | 2.73E+00 | 8.48E-01 | no | Casz1 | intron_variant | MODIFIER |
| 4 | 148882237 | AX-170178508 | 0 | T | C | 0.096 | 3.82E-02 | 2.01E-01 | 2.67E+00 | 2.73E+00 | 8.48E-01 | no | Casz1 | intron_variant | MODIFIER |
| 4 | 148886797 | AX-170335369 | 0 | T | G | 0.096 | 3.82E-02 | 2.01E-01 | 2.67E+00 | 2.73E+00 | 8.48E-01 | no | Casz1 | intron_variant | MODIFIER |
| 4 | 148937253 | AX-170245960 | 0 | C | T | 0.096 | 3.82E-02 | 2.01E-01 | 2.67E+00 | 2.73E+00 | 8.48E-01 | no | Casz1 | intron_variant | MODIFIER |
| 4 | 148995833 | AX-170297168 | 0 | G | A | 0.096 | 3.82E-02 | 2.01E-01 | 2.67E+00 | 2.73E+00 | 8.48E-01 | no | Pex14 | intron_variant | MODIFIER |
| 4 | 148997728 | AX-170188088 | 0 | C | A | 0.096 | 3.82E-02 | 2.01E-01 | 2.67E+00 | 2.73E+00 | 8.48E-01 | no | Pex14 | intron_variant | MODIFIER |
| 4 | 148999380 | AX-170515822 | 0 | T | C | 0.096 | 3.82E-02 | 2.01E-01 | 2.67E+00 | 2.73E+00 | 8.48E-01 | no | Pex14 | intron_variant | MODIFIER |
| 4 | 148999775 | AX-170355645 | 0 | G | A | 0.096 | 3.82E-02 | 2.01E-01 | 2.67E+00 | 2.73E+00 | 8.48E-01 | no | Pex14 | intron_variant | MODIFIER |
| 4 | 149002890 | AX-170346511 | 0 | T | C | 0.096 | 3.82E-02 | 2.01E-01 | 2.67E+00 | 2.73E+00 | 8.48E-01 | no | Pex14 | intron_variant | MODIFIER |
| 4 | 149051404 | AX-170250833 | 0 | C | G | 0.096 | 3.82E-02 | 2.01E-01 | 2.67E+00 | 2.73E+00 | 8.48E-01 | no | Pex14 | intron_variant | MODIFIER |
| 4 | 149052195 | AX-170206895 | 0 | C | T | 0.096 | 3.82E-02 | 2.01E-01 | 2.67E+00 | 2.73E+00 | 8.48E-01 | no | Pex14 | intron_variant | MODIFIER |
| 4 | 149071558 | AX-170242295 | 0 | G | A | 0.096 | 3.82E-02 | 2.01E-01 | 2.67E+00 | 2.73E+00 | 8.48E-01 | no | Pex14 | intron_variant | MODIFIER |
| 4 | 149083864 | AX-170024217 | 0 | C | T | 0.096 | 3.82E-02 | 2.01E-01 | 2.67E+00 | 2.73E+00 | 8.48E-01 | no | Pex14 | intron_variant | MODIFIER |
| 4 | 149100342 | AX-170024222 | 0 | C | T | 0.096 | 3.82E-02 | 2.01E-01 | 2.67E+00 | 2.73E+00 | 8.48E-01 | no | Dffa | upstream_gene_variant | MODIFIER |
| 4 | 149128877 | AX-170508605 | 0 | G | C | 0.096 | 3.82E-02 | 2.01E-01 | 2.67E+00 | 2.73E+00 | 8.48E-01 | no | Cenps | missense_variant | MODERATE |
| 4 | 149131966 | AX-170288506 | 0 | C | T | 0.096 | 3.82E-02 | 2.01E-01 | 2.67E+00 | 2.73E+00 | 8.48E-01 | no | Cenps | intron_variant | MODIFIER |
| 4 | 149148449 | AX-97785935 | 0 | T | C | 0.096 | 3.82E-02 | 2.01E-01 | 2.67E+00 | 2.73E+00 | 8.48E-01 | no | Pgd | downstream_gene_variant | MODIFIER |
| 4 | 149159668 | AX-170281633 | 0 | T | C | 0.096 | 3.82E-02 | 2.01E-01 | 2.67E+00 | 2.73E+00 | 8.48E-01 | no | Pgd | intron_variant | MODIFIER |
| 4 | 149221614 | AX-97828396 | 0 | A | G | 0.096 | 3.82E-02 | 2.01E-01 | 2.67E+00 | 2.73E+00 | 8.48E-01 | no | Kif1b | intron_variant | MODIFIER |
| 4 | 149266165 | AX-170024225 | 0 | T | C | 0.096 | 3.82E-02 | 2.01E-01 | 2.67E+00 | 2.73E+00 | 8.48E-01 | no | Kif1b | intron_variant | MODIFIER |

| | | | | | | | | | | | | | | | |
|---|---|---|---|---|---|---|---|---|---|---|---|---|---|---|---|
| 4 | 149274012 | AX-170024219 | 0 | C | A | 0.096 | 3.82E-02 | 2.01E-01 | 2.67E+00 | 2.73E+00 | 8.48E-01 | no | Kif1b | intron_variant | MODIFIER |
| 4 | 149314127 | AX-97828422 | 0 | T | C | 0.096 | 3.82E-02 | 2.01E-01 | 2.67E+00 | 2.73E+00 | 8.48E-01 | no | | regulatory_region_variant | MODIFIER |
| 4 | 149331345 | AX-170253472 | 0 | A | G | 0.096 | 3.82E-02 | 2.01E-01 | 2.67E+00 | 2.73E+00 | 8.48E-01 | no | Ube4b | synonymous_variant | LOW |
| 4 | 149331841 | AX-170409413 | 0 | G | C | 0.096 | 3.82E-02 | 2.01E-01 | 2.67E+00 | 2.73E+00 | 8.48E-01 | no | Ube4b | intron_variant | MODIFIER |
| 4 | 149336787 | AX-170024224 | 0 | G | A | 0.096 | 3.82E-02 | 2.01E-01 | 2.67E+00 | 2.73E+00 | 8.48E-01 | no | Ube4b | intron_variant | MODIFIER |
| 4 | 149338347 | AX-97855439 | 0 | T | C | 0.096 | 3.82E-02 | 2.01E-01 | 2.67E+00 | 2.73E+00 | 8.48E-01 | no | Ube4b | intron_variant | MODIFIER |
| 4 | 149340264 | AX-170131830 | 0 | A | G | 0.096 | 3.82E-02 | 2.01E-01 | 2.67E+00 | 2.73E+00 | 8.48E-01 | no | Ube4b | intron_variant | MODIFIER |
| 4 | 149351613 | AX-170147453 | 0 | C | A | 0.096 | 3.82E-02 | 2.01E-01 | 2.67E+00 | 2.73E+00 | 8.48E-01 | no | Ube4b | intron_variant | MODIFIER |
| 4 | 149352961 | AX-170150728 | 0 | G | A | 0.096 | 3.82E-02 | 2.01E-01 | 2.67E+00 | 2.73E+00 | 8.48E-01 | no | Ube4b | synonymous_variant | LOW |
| 4 | 149358070 | AX-97813152 | 0 | A | G | 0.096 | 3.82E-02 | 2.01E-01 | 2.67E+00 | 2.73E+00 | 8.48E-01 | no | Ube4b | intron_variant | MODIFIER |
| 4 | 149395703 | AX-97816299 | 0 | G | A | 0.096 | 3.82E-02 | 2.01E-01 | 2.67E+00 | 2.73E+00 | 8.48E-01 | no | Ube4b | intron_variant,non_coding_transcript_variant | MODIFIER |
| 4 | 149421889 | AX-97835481 | 0 | A | G | 0.096 | 3.82E-02 | 2.01E-01 | 2.67E+00 | 2.73E+00 | 8.48E-01 | no | Ube4b | intron_variant,non_coding_transcript_variant | MODIFIER |
| 4 | 149463034 | AX-170358146 | 0 | C | T | 0.096 | 3.82E-02 | 2.01E-01 | 2.67E+00 | 2.73E+00 | 8.48E-01 | no | Nmnat1 | downstream_gene_variant | MODIFIER |
| 4 | 149474338 | AX-170275217 | 0 | C | T | 0.096 | 3.82E-02 | 2.01E-01 | 2.67E+00 | 2.73E+00 | 8.48E-01 | no | Nmnat1 | intron_variant | MODIFIER |
| 4 | 149475734 | AX-170306538 | 0 | T | C | 0.096 | 3.82E-02 | 2.01E-01 | 2.67E+00 | 2.73E+00 | 8.48E-01 | no | Nmnat1 | intron_variant | MODIFIER |
| 4 | 149476189 | AX-170334744 | 0 | G | A | 0.096 | 3.82E-02 | 2.01E-01 | 2.67E+00 | 2.73E+00 | 8.48E-01 | no | Nmnat1 | intron_variant | MODIFIER |
| 4 | 149478551 | AX-170139749 | 0 | A | G | 0.096 | 3.82E-02 | 2.01E-01 | 2.67E+00 | 2.73E+00 | 8.48E-01 | no | Nmnat1 | intron_variant | MODIFIER |
| 4 | 149478643 | AX-170368933 | 0 | T | C | 0.096 | 3.82E-02 | 2.01E-01 | 2.67E+00 | 2.73E+00 | 8.48E-01 | no | Nmnat1 | intron_variant | MODIFIER |
| 4 | 149487195 | AX-170181351 | 0 | C | G | 0.096 | 3.82E-02 | 2.01E-01 | 2.67E+00 | 2.73E+00 | 8.48E-01 | no | Lzic | intron_variant | MODIFIER |
| 4 | 149499331 | AX-170467366 | 0 | C | T | 0.096 | 3.82E-02 | 2.01E-01 | 2.67E+00 | 2.73E+00 | 8.48E-01 | no | Lzic | downstream_gene_variant | MODIFIER |
| 4 | 149499863 | AX-170166709 | 0 | T | C | 0.096 | 3.82E-02 | 2.01E-01 | 2.67E+00 | 2.73E+00 | 8.48E-01 | no | Lzic | downstream_gene_variant | MODIFIER |
| 4 | 149509732 | AX-97837559 | 0 | G | T | 0.096 | 3.82E-02 | 2.01E-01 | 2.67E+00 | 2.73E+00 | 8.48E-01 | no | | intergenic_variant | MODIFIER |
| 4 | 149525131 | AX-168384809 | 0 | A | G | 0.096 | 3.82E-02 | 2.01E-01 | 2.67E+00 | 2.73E+00 | 8.48E-01 | no | Ctnnbip1 | intron_variant | MODIFIER |
| 4 | 149532641 | AX-170071011 | 0 | C | T | 0.096 | 3.82E-02 | 2.01E-01 | 2.67E+00 | 2.73E+00 | 8.48E-01 | no | Ctnnbip1 | intron_variant | MODIFIER |
| 4 | 149547669 | AX-170242348 | 0 | C | T | 0.096 | 3.82E-02 | 2.01E-01 | 2.67E+00 | 2.73E+00 | 8.48E-01 | no | Ctnnbip1 | intron_variant | MODIFIER |
| 4 | 149728061 | AX-170024233 | 0 | A | G | 0.096 | 3.82E-02 | 2.01E-01 | 2.67E+00 | 2.73E+00 | 8.48E-01 | no | Tmem201 | missense_variant | MODERATE |
| 4 | 149728557 | AX-97847420 | 0 | A | G | 0.096 | 3.82E-02 | 2.01E-01 | 2.67E+00 | 2.73E+00 | 8.48E-01 | no | Tmem201 | intron_variant | MODIFIER |
| 4 | 149850302 | AX-170024231 | 0 | G | A | 0.096 | 3.82E-02 | 2.01E-01 | 2.67E+00 | 2.73E+00 | 8.48E-01 | no | | intergenic_variant | MODIFIER |
| 4 | 149852563 | AX-170135430 | 0 | T | C | 0.096 | 3.82E-02 | 2.01E-01 | 2.67E+00 | 2.73E+00 | 8.48E-01 | no | | intergenic_variant | MODIFIER |
| 4 | 149858712 | AX-170448680 | 0 | A | G | 0.096 | 3.82E-02 | 2.01E-01 | 2.67E+00 | 2.73E+00 | 8.48E-01 | no | | regulatory_region_variant | MODIFIER |
| 4 | 149865892 | AX-170244839 | 0 | A | C | 0.096 | 3.82E-02 | 2.01E-01 | 2.67E+00 | 2.73E+00 | 8.48E-01 | no | | intergenic_variant | MODIFIER |
| 4 | 149866386 | AX-97787051 | 0 | A | G | 0.096 | 3.82E-02 | 2.01E-01 | 2.67E+00 | 2.73E+00 | 8.48E-01 | no | | intergenic_variant | MODIFIER |
| 4 | 149866453 | AX-170275118 | 0 | C | T | 0.096 | 3.82E-02 | 2.01E-01 | 2.67E+00 | 2.73E+00 | 8.48E-01 | no | | intergenic_variant | MODIFIER |
| 4 | 149886695 | AX-170372985 | 0 | A | G | 0.096 | 3.82E-02 | 2.01E-01 | 2.67E+00 | 2.73E+00 | 8.48E-01 | no | | regulatory_region_variant | MODIFIER |
| 4 | 149950756 | AX-170359991 | 0 | G | C | 0.096 | 3.82E-02 | 2.01E-01 | 2.67E+00 | 2.73E+00 | 8.48E-01 | no | Spsb1 | intron_variant | MODIFIER |
| 4 | 149954162 | AX-170498787 | 0 | C | T | 0.096 | 3.82E-02 | 2.01E-01 | 2.67E+00 | 2.73E+00 | 8.48E-01 | no | Spsb1 | intron_variant | MODIFIER |
| 4 | 149965853 | AX-170401446 | 0 | T | C | 0.096 | 3.82E-02 | 2.01E-01 | 2.67E+00 | 2.73E+00 | 8.48E-01 | no | | regulatory_region_variant | MODIFIER |
| 4 | 149978929 | AX-170154001 | 0 | C | T | 0.096 | 3.82E-02 | 2.01E-01 | 2.67E+00 | 2.73E+00 | 8.48E-01 | no | H6pd | downstream_gene_variant | MODIFIER |
| 4 | 149995056 | AX-170304896 | 0 | T | G | 0.096 | 3.82E-02 | 2.01E-01 | 2.67E+00 | 2.73E+00 | 8.48E-01 | no | H6pd | intron_variant | MODIFIER |
| 4 | 149995224 | AX-170071018 | 0 | G | A | 0.096 | 3.82E-02 | 2.01E-01 | 2.67E+00 | 2.73E+00 | 8.48E-01 | no | H6pd | intron_variant | MODIFIER |
| 4 | 149995418 | AX-170123213 | 0 | C | T | 0.096 | 3.82E-02 | 2.01E-01 | 2.67E+00 | 2.73E+00 | 8.48E-01 | no | H6pd | intron_variant | MODIFIER |
| 4 | 149996046 | AX-170470551 | 0 | T | C | 0.096 | 3.82E-02 | 2.01E-01 | 2.67E+00 | 2.73E+00 | 8.48E-01 | no | H6pd | missense_variant | MODERATE |
| 4 | 149997971 | AX-170471471 | 0 | T | C | 0.096 | 3.82E-02 | 2.01E-01 | 2.67E+00 | 2.73E+00 | 8.48E-01 | no | H6pd | intron_variant | MODIFIER |
| 4 | 149999986 | AX-170377322 | 0 | G | A | 0.096 | 3.82E-02 | 2.01E-01 | 2.67E+00 | 2.73E+00 | 8.48E-01 | no | H6pd | intron_variant | MODIFIER |
| 4 | 150010214 | AX-170470027 | 0 | G | A | 0.096 | 3.82E-02 | 2.01E-01 | 2.67E+00 | 2.73E+00 | 8.48E-01 | no | H6pd | upstream_gene_variant | MODIFIER |
| 4 | 150015262 | AX-170318537 | 0 | G | A | 0.096 | 3.82E-02 | 2.01E-01 | 2.67E+00 | 2.73E+00 | 8.48E-01 | no | | intergenic_variant | MODIFIER |
| 4 | 150100489 | AX-170477637 | 0 | T | C | 0.096 | 3.82E-02 | 2.01E-01 | 2.67E+00 | 2.73E+00 | 8.48E-01 | no | Gpr157 | intron_variant | MODIFIER |
| 4 | 150118397 | AX-170167867 | 0 | T | C | 0.096 | 3.82E-02 | 2.01E-01 | 2.67E+00 | 2.73E+00 | 8.48E-01 | no | Slc2a5 | upstream_gene_variant | MODIFIER |
| 4 | 150179399 | AX-170434153 | 0 | A | C | 0.096 | 3.82E-02 | 2.01E-01 | 2.67E+00 | 2.73E+00 | 8.48E-01 | no | | intergenic_variant | MODIFIER |
| 4 | 150179485 | AX-170071020 | 0 | G | T | 0.096 | 3.82E-02 | 2.01E-01 | 2.67E+00 | 2.73E+00 | 8.48E-01 | no | | intergenic_variant | MODIFIER |
| 4 | 150205615 | AX-170188741 | 0 | G | A | 0.096 | 3.82E-02 | 2.01E-01 | 2.67E+00 | 2.73E+00 | 8.48E-01 | no | Car6 | upstream_gene_variant | MODIFIER |
| 4 | 150220669 | AX-170301773 | 0 | T | C | 0.096 | 3.82E-02 | 2.01E-01 | 2.67E+00 | 2.73E+00 | 8.48E-01 | no | | regulatory_region_variant | MODIFIER |
| 4 | 150238425 | AX-97828385 | 0 | A | G | 0.096 | 3.82E-02 | 2.01E-01 | 2.67E+00 | 2.73E+00 | 8.48E-01 | no | Eno1 | intron_variant | MODIFIER |
| 4 | 150238983 | AX-170294709 | 0 | C | T | 0.096 | 3.82E-02 | 2.01E-01 | 2.67E+00 | 2.73E+00 | 8.48E-01 | no | Eno1 | upstream_gene_variant | MODIFIER |

| | | | | | | | | | | | | | | | |
|---|---|---|---|---|---|---|---|---|---|---|---|---|---|---|---|
| 4 | 150243685 | AX-170416783 | 0 | C | T | 0.096 | 3.82E-02 | 2.01E-01 | 2.67E+00 | 2.73E+00 | 8.48E-01 | no | Eno1 | 5_prime_UTR_variant | MODIFIER |
| 4 | 150243778 | AX-170150930 | 0 | A | G | 0.096 | 3.82E-02 | 2.01E-01 | 2.67E+00 | 2.73E+00 | 8.48E-01 | no | Eno1 | 5_prime_UTR_variant | MODIFIER |
| 4 | 150255391 | AX-170024240 | 0 | A | C | 0.096 | 3.82E-02 | 2.01E-01 | 2.67E+00 | 2.73E+00 | 8.48E-01 | no | | regulatory_region_variant | MODIFIER |
| 4 | 150256152 | AX-170024242 | 0 | T | C | 0.096 | 3.82E-02 | 2.01E-01 | 2.67E+00 | 2.73E+00 | 8.48E-01 | no | | intergenic_variant | MODIFIER |
| 4 | 150262664 | AX-170415871 | 0 | A | T | 0.096 | 3.82E-02 | 2.01E-01 | 2.67E+00 | 2.73E+00 | 8.48E-01 | no | | intergenic_variant | MODIFIER |
| 4 | 150268612 | AX-170359190 | 0 | A | G | 0.096 | 3.82E-02 | 2.01E-01 | 2.67E+00 | 2.73E+00 | 8.48E-01 | no | | regulatory_region_variant | MODIFIER |
| 4 | 150297203 | AX-170024239 | 0 | G | C | 0.096 | 3.82E-02 | 2.01E-01 | 2.67E+00 | 2.73E+00 | 8.48E-01 | no | Rere | intron_variant | MODIFIER |
| 4 | 150299482 | AX-170024241 | 0 | T | A | 0.096 | 3.82E-02 | 2.01E-01 | 2.67E+00 | 2.73E+00 | 8.48E-01 | no | Rere | intron_variant | MODIFIER |
| 4 | 150309421 | AX-170024245 | 0 | T | C | 0.096 | 3.82E-02 | 2.01E-01 | 2.67E+00 | 2.73E+00 | 8.48E-01 | no | Rere | intron_variant | MODIFIER |
| 4 | 150327208 | AX-170024243 | 0 | G | A | 0.096 | 3.82E-02 | 2.01E-01 | 2.67E+00 | 2.73E+00 | 8.48E-01 | no | Rere | intron_variant | MODIFIER |
| 4 | 150369873 | AX-97836533 | 0 | C | T | 0.096 | 3.82E-02 | 2.01E-01 | 2.67E+00 | 2.73E+00 | 8.48E-01 | no | Rere | intron_variant | MODIFIER |
| 4 | 150385590 | AX-97809617 | 0 | A | G | 0.096 | 3.82E-02 | 2.01E-01 | 2.67E+00 | 2.73E+00 | 8.48E-01 | no | Rere | intron_variant | MODIFIER |
| 4 | 150402261 | AX-170071027 | 0 | G | A | 0.096 | 3.82E-02 | 2.01E-01 | 2.67E+00 | 2.73E+00 | 8.48E-01 | no | Rere | intron_variant | MODIFIER |
| 4 | 150451832 | AX-170183870 | 0 | C | T | 0.096 | 3.82E-02 | 2.01E-01 | 2.67E+00 | 2.73E+00 | 8.48E-01 | no | Rere | intron_variant | MODIFIER |
| 4 | 150476349 | AX-170458568 | 0 | G | C | 0.096 | 3.82E-02 | 2.01E-01 | 2.67E+00 | 2.73E+00 | 8.48E-01 | no | Rere | intron_variant | MODIFIER |
| 4 | 150517242 | AX-170494279 | 0 | G | T | 0.096 | 3.82E-02 | 2.01E-01 | 2.67E+00 | 2.73E+00 | 8.48E-01 | no | Rere | intron_variant | MODIFIER |
| 4 | 150549352 | AX-170219582 | 0 | C | T | 0.096 | 3.82E-02 | 2.01E-01 | 2.67E+00 | 2.73E+00 | 8.48E-01 | no | Rere | intron_variant | MODIFIER |
| 1 | 116868790 | AX-97808472 | 0 | C | T | 0.167 | 3.89E-02 | 1.95E-01 | 2.62E+00 | 2.71E+00 | 8.48E-01 | no | | intergenic_variant | MODIFIER |
| 11 | 34734001 | AX-170046763 | 0 | A | G | 0.105 | 3.55E-02 | 1.84E-01 | 2.64E+00 | 2.73E+00 | 8.48E-01 | no | Dock2 | intron_variant | MODIFIER |
| 7 | 60603484 | AX-170116861 | 1 | T | G | 0.42 | 3.41E-02 | 1.85E-01 | 2.62E+00 | 2.75E+00 | 8.48E-01 | no | | intergenic_variant | MODIFIER |
| 10 | 9699600 | AX-168412301 | 1 | C | A | 0.205 | 1.95E-02 | 1.05E-01 | 2.67E+00 | 2.76E+00 | 8.48E-01 | no | | regulatory_region_variant | MODIFIER |
| 8 | 51594307 | AX-170160327 | 0 | G | A | 0.149 | 2.70E-02 | 1.47E-01 | 2.73E+00 | 2.80E+00 | 8.48E-01 | no | | intergenic_variant | MODIFIER |
| 9 | 34966467 | AX-170446488 | 1 | T | C | 0.438 | 1.72E-02 | 8.74E-02 | 2.65E+00 | 2.71E+00 | 8.48E-01 | no | Kirrel3 | intron_variant | MODIFIER |
| 12 | 99575500 | AX-170257287 | 1 | G | A | 0.054 | 3.22E-02 | 1.80E-01 | 2.66E+00 | 2.77E+00 | 8.48E-01 | no | Gm26839 | intron_variant,non_coding_transcript_variant | MODIFIER |
| 13 | 37814540 | AX-97824897 | 1 | G | T | 0.188 | 2.01E-02 | 1.12E-01 | 2.68E+00 | 2.78E+00 | 8.48E-01 | no | Rreb1 | intron_variant | MODIFIER |
| 8 | 95043451 | AX-170321363 | 2 | C | G | 0.227 | 2.15E-02 | 1.14E-01 | 2.66E+00 | 2.74E+00 | 8.48E-01 | no | Adgrg3 | intron_variant | MODIFIER |
| 7 | 98537065 | AX-170266218 | 1 | G | A | 0.134 | -3.96E-02 | 2.12E-01 | 2.67E+00 | 2.75E+00 | 8.48E-01 | no | | regulatory_region_variant | MODIFIER |
| 7 | 82245627 | AX-170077879 | 0 | C | T | 0.114 | -3.91E-02 | 2.12E-01 | 2.67E+00 | 2.76E+00 | 8.48E-01 | no | Sh3gl3 | intron_variant | MODIFIER |
| 3 | 143902283 | AX-170409022 | 1 | T | C | 0.366 | 1.35E-02 | 7.03E-02 | 2.63E+00 | 2.73E+00 | 8.48E-01 | no | Gm42708 | downstream_gene_variant | MODIFIER |
| 1 | 54000308 | AX-97799952 | 0 | G | A | 0.105 | 4.41E-02 | 2.35E-01 | 2.70E+00 | 2.74E+00 | 8.48E-01 | no | Hecw2 | intron_variant | MODIFIER |
| 12 | 109695952 | AX-170449748 | 0 | G | A | 0.14 | -3.60E-02 | 1.93E-01 | 2.68E+00 | 2.75E+00 | 8.49E-01 | no | B830012L14Rik | non_coding_transcript_exon_variant | MODIFIER |
| 12 | 111153394 | AX-170368665 | 0 | G | T | 0.14 | -3.60E-02 | 1.93E-01 | 2.68E+00 | 2.75E+00 | 8.49E-01 | no | | regulatory_region_variant | MODIFIER |
| 12 | 111161692 | AX-97830682 | 0 | C | T | 0.14 | -3.60E-02 | 1.93E-01 | 2.68E+00 | 2.75E+00 | 8.49E-01 | no | Traf3 | upstream_gene_variant | MODIFIER |
| 12 | 111228755 | AX-97854312 | 0 | C | A | 0.14 | -3.60E-02 | 1.93E-01 | 2.68E+00 | 2.75E+00 | 8.49E-01 | no | Traf3 | intron_variant | MODIFIER |
| 12 | 111234870 | AX-170125545 | 0 | C | T | 0.14 | -3.60E-02 | 1.93E-01 | 2.68E+00 | 2.75E+00 | 8.49E-01 | no | Traf3 | intron_variant | MODIFIER |
| 12 | 111238752 | AX-170391545 | 0 | A | G | 0.14 | -3.60E-02 | 1.93E-01 | 2.68E+00 | 2.75E+00 | 8.49E-01 | no | Traf3 | intron_variant | MODIFIER |
| 12 | 111268377 | AX-170347711 | 0 | G | A | 0.14 | -3.60E-02 | 1.93E-01 | 2.68E+00 | 2.75E+00 | 8.49E-01 | no | Traf3 | downstream_gene_variant | MODIFIER |
| 12 | 111283917 | AX-170433878 | 0 | T | C | 0.14 | -3.60E-02 | 1.93E-01 | 2.68E+00 | 2.75E+00 | 8.49E-01 | no | A230087F16Rik | upstream_gene_variant | MODIFIER |
| 12 | 111295920 | AX-170332133 | 0 | C | T | 0.14 | -3.60E-02 | 1.93E-01 | 2.68E+00 | 2.75E+00 | 8.49E-01 | no | Cdc42bpb | intron_variant | MODIFIER |
| 12 | 111295995 | AX-170174188 | 0 | G | A | 0.14 | -3.60E-02 | 1.93E-01 | 2.68E+00 | 2.75E+00 | 8.49E-01 | no | Cdc42bpb | synonymous_variant | LOW |
| 12 | 111308024 | AX-170452525 | 0 | T | C | 0.14 | -3.60E-02 | 1.93E-01 | 2.68E+00 | 2.75E+00 | 8.49E-01 | no | Cdc42bpb | intron_variant | MODIFIER |
| 12 | 111321095 | AX-170151876 | 0 | C | A | 0.14 | -3.60E-02 | 1.93E-01 | 2.68E+00 | 2.75E+00 | 8.49E-01 | no | Cdc42bpb | missense_variant | MODERATE |
| 12 | 111371494 | AX-170332930 | 0 | T | C | 0.14 | -3.60E-02 | 1.93E-01 | 2.68E+00 | 2.75E+00 | 8.49E-01 | no | Cdc42bpb | intron_variant | MODIFIER |
| 12 | 111415238 | AX-170211775 | 0 | T | C | 0.14 | -3.60E-02 | 1.93E-01 | 2.68E+00 | 2.75E+00 | 8.49E-01 | no | Exoc3l4 | upstream_gene_variant | MODIFIER |
| 12 | 111514913 | AX-168415273 | 0 | T | G | 0.14 | -3.60E-02 | 1.93E-01 | 2.68E+00 | 2.75E+00 | 8.49E-01 | no | | regulatory_region_variant | MODIFIER |
| 12 | 111536502 | AX-97826858 | 0 | G | A | 0.14 | -3.60E-02 | 1.93E-01 | 2.68E+00 | 2.75E+00 | 8.49E-01 | no | Eif5 | upstream_gene_variant | MODIFIER |
| 12 | 111544379 | AX-170180326 | 0 | G | A | 0.14 | -3.60E-02 | 1.93E-01 | 2.68E+00 | 2.75E+00 | 8.49E-01 | no | Eif5 | intron_variant | MODIFIER |
| 12 | 111548739 | AX-170425095 | 0 | C | G | 0.14 | -3.60E-02 | 1.93E-01 | 2.68E+00 | 2.75E+00 | 8.49E-01 | no | Eif5 | downstream_gene_variant | MODIFIER |
| 12 | 111583638 | AX-170163036 | 0 | C | T | 0.14 | -3.60E-02 | 1.93E-01 | 2.68E+00 | 2.75E+00 | 8.49E-01 | no | Mark3 | intron_variant | MODIFIER |
| 12 | 111627686 | AX-170007536 | 0 | A | C | 0.14 | -3.60E-02 | 1.93E-01 | 2.68E+00 | 2.75E+00 | 8.49E-01 | no | Mark3 | intron_variant | MODIFIER |
| 12 | 111627786 | AX-97855972 | 0 | G | A | 0.14 | -3.60E-02 | 1.93E-01 | 2.68E+00 | 2.75E+00 | 8.49E-01 | no | Mark3 | intron_variant | MODIFIER |
| 12 | 111650672 | AX-170143245 | 0 | T | C | 0.14 | -3.60E-02 | 1.93E-01 | 2.68E+00 | 2.75E+00 | 8.49E-01 | no | Mark3 | intron_variant | MODIFIER |
| 12 | 111658260 | AX-170471751 | 0 | C | A | 0.14 | -3.60E-02 | 1.93E-01 | 2.68E+00 | 2.75E+00 | 8.49E-01 | no | Mark3 | downstream_gene_variant | MODIFIER |
| 12 | 111667165 | AX-170318998 | 0 | G | A | 0.14 | -3.60E-02 | 1.93E-01 | 2.68E+00 | 2.75E+00 | 8.49E-01 | no | Ckb | downstream_gene_variant | MODIFIER |

| | | | | | | | | | | | | | | | |
|---|---|---|---|---|---|---|---|---|---|---|---|---|---|---|---|
| 12 | 111681976 | AX-97814541 | 0 | A | G | 0.14 | -3.60E-02 | 1.93E-01 | 2.68E+00 | 2.75E+00 | 8.49E-01 | no | Trmt61a | intron_variant | MODIFIER |
| 12 | 111684102 | AX-170007537 | 0 | C | T | 0.14 | -3.60E-02 | 1.93E-01 | 2.68E+00 | 2.75E+00 | 8.49E-01 | no | Trmt61a | downstream_gene_variant | MODIFIER |
| 12 | 111687701 | AX-170007533 | 0 | T | C | 0.14 | -3.60E-02 | 1.93E-01 | 2.68E+00 | 2.75E+00 | 8.49E-01 | no | Trmt61a | downstream_gene_variant | MODIFIER |
| 12 | 111733128 | AX-97833038 | 0 | C | T | 0.14 | -3.60E-02 | 1.93E-01 | 2.68E+00 | 2.75E+00 | 8.49E-01 | no | Apopt1 | intron_variant | MODIFIER |
| 12 | 111747518 | AX-170318416 | 0 | C | G | 0.14 | -3.60E-02 | 1.93E-01 | 2.68E+00 | 2.75E+00 | 8.49E-01 | no | Apopt1 | intron_variant | MODIFIER |
| 12 | 111779022 | AX-170385020 | 0 | G | C | 0.14 | -3.60E-02 | 1.93E-01 | 2.68E+00 | 2.75E+00 | 8.49E-01 | no | Klc1 | intron_variant | MODIFIER |
| 12 | 111804688 | AX-170167307 | 0 | G | A | 0.14 | -3.60E-02 | 1.93E-01 | 2.68E+00 | 2.75E+00 | 8.49E-01 | no | Xrcc3 | intron_variant | MODIFIER |
| 12 | 111821387 | AX-170365454 | 0 | C | G | 0.14 | -3.60E-02 | 1.93E-01 | 2.68E+00 | 2.75E+00 | 8.49E-01 | no | Zfyve21 | intron_variant | MODIFIER |
| 12 | 111824729 | AX-170535354 | 0 | A | T | 0.14 | -3.60E-02 | 1.93E-01 | 2.68E+00 | 2.75E+00 | 8.49E-01 | no | Zfyve21 | intron_variant | MODIFIER |
| 12 | 111824792 | AX-170214002 | 0 | A | G | 0.14 | -3.60E-02 | 1.93E-01 | 2.68E+00 | 2.75E+00 | 8.49E-01 | no | Zfyve21 | intron_variant | MODIFIER |
| 12 | 111824940 | AX-170087334 | 0 | T | C | 0.14 | -3.60E-02 | 1.93E-01 | 2.68E+00 | 2.75E+00 | 8.49E-01 | no | Zfyve21 | intron_variant | MODIFIER |
| 12 | 111825089 | AX-170333595 | 0 | C | A | 0.14 | -3.60E-02 | 1.93E-01 | 2.68E+00 | 2.75E+00 | 8.49E-01 | no | Zfyve21 | synonymous_variant | LOW |
| 12 | 111831321 | AX-170485866 | 0 | T | A | 0.14 | -3.60E-02 | 1.93E-01 | 2.68E+00 | 2.75E+00 | 8.49E-01 | no | Zfyve21 | downstream_gene_variant | MODIFIER |
| 12 | 111832113 | AX-170166401 | 0 | A | G | 0.14 | -3.60E-02 | 1.93E-01 | 2.68E+00 | 2.75E+00 | 8.49E-01 | no | Zfyve21 | downstream_gene_variant | MODIFIER |
| 12 | 111832211 | AX-170304049 | 0 | C | A | 0.14 | -3.60E-02 | 1.93E-01 | 2.68E+00 | 2.75E+00 | 8.49E-01 | no | Zfyve21 | downstream_gene_variant | MODIFIER |
| 12 | 111833110 | AX-170381894 | 0 | C | T | 0.14 | -3.60E-02 | 1.93E-01 | 2.68E+00 | 2.75E+00 | 8.49E-01 | no | Zfyve21 | downstream_gene_variant | MODIFIER |
| 12 | 111840633 | AX-170240599 | 0 | C | T | 0.14 | -3.60E-02 | 1.93E-01 | 2.68E+00 | 2.75E+00 | 8.49E-01 | no | Ppp1r13b | intron_variant | MODIFIER |
| 12 | 111841563 | AX-170331493 | 0 | C | T | 0.14 | -3.60E-02 | 1.93E-01 | 2.68E+00 | 2.75E+00 | 8.49E-01 | no | Ppp1r13b | intron_variant | MODIFIER |
| 12 | 111844508 | AX-170224537 | 0 | C | T | 0.14 | -3.60E-02 | 1.93E-01 | 2.68E+00 | 2.75E+00 | 8.49E-01 | no | Ppp1r13b | intron_variant | MODIFIER |
| 12 | 111844654 | AX-170195800 | 0 | T | C | 0.14 | -3.60E-02 | 1.93E-01 | 2.68E+00 | 2.75E+00 | 8.49E-01 | no | Ppp1r13b | intron_variant | MODIFIER |
| 12 | 111846107 | AX-170233061 | 0 | C | T | 0.14 | -3.60E-02 | 1.93E-01 | 2.68E+00 | 2.75E+00 | 8.49E-01 | no | Ppp1r13b | intron_variant | MODIFIER |
| 12 | 111853636 | AX-170357197 | 0 | T | C | 0.14 | -3.60E-02 | 1.93E-01 | 2.68E+00 | 2.75E+00 | 8.49E-01 | no | Ppp1r13b | intron_variant | MODIFIER |
| 12 | 111859645 | AX-170467440 | 0 | C | A | 0.14 | -3.60E-02 | 1.93E-01 | 2.68E+00 | 2.75E+00 | 8.49E-01 | no | Ppp1r13b | intron_variant | MODIFIER |
| 12 | 111869912 | AX-170312977 | 0 | G | A | 0.14 | -3.60E-02 | 1.93E-01 | 2.68E+00 | 2.75E+00 | 8.49E-01 | no | Ppp1r13b | intron_variant | MODIFIER |
| 12 | 111873420 | AX-170117439 | 0 | T | G | 0.14 | -3.60E-02 | 1.93E-01 | 2.68E+00 | 2.75E+00 | 8.49E-01 | no | Ppp1r13b | intron_variant | MODIFIER |
| 12 | 111896955 | AX-170050220 | 0 | G | A | 0.14 | -3.60E-02 | 1.93E-01 | 2.68E+00 | 2.75E+00 | 8.49E-01 | no | Ppp1r13b | intron_variant | MODIFIER |
| 12 | 111911397 | AX-170225321 | 0 | C | G | 0.14 | -3.60E-02 | 1.93E-01 | 2.68E+00 | 2.75E+00 | 8.49E-01 | no | Ppp1r13b | upstream_gene_variant | MODIFIER |
| 12 | 111945469 | AX-170156398 | 0 | G | T | 0.14 | -3.60E-02 | 1.93E-01 | 2.68E+00 | 2.75E+00 | 8.49E-01 | no | 5033406O09Rik | upstream_gene_variant | MODIFIER |
| 12 | 111946691 | AX-170157210 | 0 | C | T | 0.14 | -3.60E-02 | 1.93E-01 | 2.68E+00 | 2.75E+00 | 8.49E-01 | no | 5033406O09Rik | upstream_gene_variant | MODIFIER |
| 12 | 111947780 | AX-170339358 | 0 | A | G | 0.14 | -3.60E-02 | 1.93E-01 | 2.68E+00 | 2.75E+00 | 8.49E-01 | no | 5033406O09Rik | upstream_gene_variant | MODIFIER |
| 12 | 111951556 | AX-170119261 | 0 | C | T | 0.14 | -3.60E-02 | 1.93E-01 | 2.68E+00 | 2.75E+00 | 8.49E-01 | no | Gm36757 | non_coding_transcript_exon_variant | MODIFIER |
| 12 | 111956822 | AX-170307978 | 0 | A | G | 0.14 | -3.60E-02 | 1.93E-01 | 2.68E+00 | 2.75E+00 | 8.49E-01 | no | Atp5mpl | downstream_gene_variant | MODIFIER |
| 12 | 111958260 | AX-170112316 | 0 | C | T | 0.14 | -3.60E-02 | 1.93E-01 | 2.68E+00 | 2.75E+00 | 8.49E-01 | no | Atp5mpl | downstream_gene_variant | MODIFIER |
| 12 | 111958396 | AX-170161932 | 0 | C | T | 0.14 | -3.60E-02 | 1.93E-01 | 2.68E+00 | 2.75E+00 | 8.49E-01 | no | Atp5mpl | downstream_gene_variant | MODIFIER |
| 12 | 111958525 | AX-170440791 | 0 | C | T | 0.14 | -3.60E-02 | 1.93E-01 | 2.68E+00 | 2.75E+00 | 8.49E-01 | no | Atp5mpl | downstream_gene_variant | MODIFIER |
| 12 | 111958842 | AX-170206559 | 0 | C | T | 0.14 | -3.60E-02 | 1.93E-01 | 2.68E+00 | 2.75E+00 | 8.49E-01 | no | Atp5mpl | downstream_gene_variant | MODIFIER |
| 12 | 111965964 | AX-170388786 | 0 | C | T | 0.14 | -3.60E-02 | 1.93E-01 | 2.68E+00 | 2.75E+00 | 8.49E-01 | no | Atp5mpl | intron_variant | MODIFIER |
| 12 | 111970384 | AX-170375005 | 0 | A | C | 0.14 | -3.60E-02 | 1.93E-01 | 2.68E+00 | 2.75E+00 | 8.49E-01 | no | Atp5mpl | upstream_gene_variant | MODIFIER |
| 12 | 111974452 | AX-170163246 | 0 | C | T | 0.14 | -3.60E-02 | 1.93E-01 | 2.68E+00 | 2.75E+00 | 8.49E-01 | no | Tdrd9 | intron_variant | MODIFIER |
| 12 | 111975797 | AX-97784475 | 0 | G | A | 0.14 | -3.60E-02 | 1.93E-01 | 2.68E+00 | 2.75E+00 | 8.49E-01 | no | Tdrd9 | intron_variant | MODIFIER |
| 12 | 111980218 | AX-170514124 | 0 | A | G | 0.14 | -3.60E-02 | 1.93E-01 | 2.68E+00 | 2.75E+00 | 8.49E-01 | no | Tdrd9 | intron_variant | MODIFIER |
| 12 | 111981718 | AX-170166808 | 0 | G | T | 0.14 | -3.60E-02 | 1.93E-01 | 2.68E+00 | 2.75E+00 | 8.49E-01 | no | Tdrd9 | intron_variant | MODIFIER |
| 12 | 111995984 | AX-97851198 | 0 | C | T | 0.14 | -3.60E-02 | 1.93E-01 | 2.68E+00 | 2.75E+00 | 8.49E-01 | no | Tdrd9 | intron_variant | MODIFIER |
| 12 | 112000291 | AX-170453627 | 0 | G | A | 0.14 | -3.60E-02 | 1.93E-01 | 2.68E+00 | 2.75E+00 | 8.49E-01 | no | Tdrd9 | intron_variant | MODIFIER |
| 12 | 112007839 | AX-170524731 | 0 | T | C | 0.14 | -3.60E-02 | 1.93E-01 | 2.68E+00 | 2.75E+00 | 8.49E-01 | no | Tdrd9 | intron_variant | MODIFIER |
| 12 | 112062827 | AX-170220708 | 0 | A | G | 0.14 | -3.60E-02 | 1.93E-01 | 2.68E+00 | 2.75E+00 | 8.49E-01 | no | Tdrd9 | intron_variant | MODIFIER |
| 12 | 112064284 | AX-170533331 | 0 | G | A | 0.14 | -3.60E-02 | 1.93E-01 | 2.68E+00 | 2.75E+00 | 8.49E-01 | no | Tdrd9 | intron_variant | MODIFIER |
| 12 | 112074960 | AX-170420517 | 0 | A | C | 0.14 | -3.60E-02 | 1.93E-01 | 2.68E+00 | 2.75E+00 | 8.49E-01 | no | | intergenic_variant | MODIFIER |
| 12 | 112103288 | AX-170383101 | 0 | T | C | 0.14 | -3.60E-02 | 1.93E-01 | 2.68E+00 | 2.75E+00 | 8.49E-01 | no | Aspg | upstream_gene_variant | MODIFIER |
| 12 | 112112145 | AX-170368484 | 0 | T | C | 0.14 | -3.60E-02 | 1.93E-01 | 2.68E+00 | 2.75E+00 | 8.49E-01 | no | Aspg | intron_variant | MODIFIER |
| 12 | 112114617 | AX-170302765 | 0 | C | T | 0.14 | -3.60E-02 | 1.93E-01 | 2.68E+00 | 2.75E+00 | 8.49E-01 | no | Aspg | intron_variant | MODIFIER |
| 12 | 112123653 | AX-97829518 | 0 | A | G | 0.14 | -3.60E-02 | 1.93E-01 | 2.68E+00 | 2.75E+00 | 8.49E-01 | no | Aspg | intron_variant | MODIFIER |
| 12 | 112142741 | AX-170007547 | 0 | G | A | 0.14 | -3.60E-02 | 1.93E-01 | 2.68E+00 | 2.75E+00 | 8.49E-01 | no | Kif26a | upstream_gene_variant | MODIFIER |
| 12 | 112156213 | AX-97854531 | 0 | G | A | 0.14 | -3.60E-02 | 1.93E-01 | 2.68E+00 | 2.75E+00 | 8.49E-01 | no | Kif26a | intron_variant | MODIFIER |

| | | | | | | | | | | | | | | | |
|---|---|---|---|---|---|---|---|---|---|---|---|---|---|---|---|
| 12 | 112159819 | AX-170007543 | 0 | T | A | 0.14 | -3.60E-02 | 1.93E-01 | 2.68E+00 | 2.75E+00 | 8.49E-01 | no | Kif26a | intron_variant | MODIFIER |
| 12 | 112162692 | AX-170050225 | 0 | A | G | 0.14 | -3.60E-02 | 1.93E-01 | 2.68E+00 | 2.75E+00 | 8.49E-01 | no | Kif26a | intron_variant | MODIFIER |
| 12 | 112171877 | AX-97822147 | 0 | T | C | 0.14 | -3.60E-02 | 1.93E-01 | 2.68E+00 | 2.75E+00 | 8.49E-01 | no | Kif26a | intron_variant | MODIFIER |
| 12 | 112173069 | AX-170007546 | 0 | C | T | 0.14 | -3.60E-02 | 1.93E-01 | 2.68E+00 | 2.75E+00 | 8.49E-01 | no | Kif26a | intron_variant | MODIFIER |
| 12 | 112173426 | AX-170481330 | 0 | C | T | 0.14 | -3.60E-02 | 1.93E-01 | 2.68E+00 | 2.75E+00 | 8.49E-01 | no | Kif26a | missense_variant | MODERATE |
| 12 | 112192563 | AX-97850426 | 0 | G | T | 0.14 | -3.60E-02 | 1.93E-01 | 2.68E+00 | 2.75E+00 | 8.49E-01 | no | | intergenic_variant | MODIFIER |
| 12 | 112205401 | AX-170507648 | 0 | T | C | 0.14 | -3.60E-02 | 1.93E-01 | 2.68E+00 | 2.75E+00 | 8.49E-01 | no | Gm20368 | upstream_gene_variant | MODIFIER |
| 12 | 112205900 | AX-170007550 | 0 | A | G | 0.14 | -3.60E-02 | 1.93E-01 | 2.68E+00 | 2.75E+00 | 8.49E-01 | no | Gm20368 | upstream_gene_variant | MODIFIER |
| 12 | 112211516 | AX-170007542 | 0 | C | T | 0.14 | -3.60E-02 | 1.93E-01 | 2.68E+00 | 2.75E+00 | 8.49E-01 | no | | regulatory_region_variant | MODIFIER |
| 12 | 112211914 | AX-170243016 | 0 | A | G | 0.14 | -3.60E-02 | 1.93E-01 | 2.68E+00 | 2.75E+00 | 8.49E-01 | no | | intergenic_variant | MODIFIER |
| 12 | 112213920 | AX-170252977 | 0 | C | T | 0.14 | -3.60E-02 | 1.93E-01 | 2.68E+00 | 2.75E+00 | 8.49E-01 | no | | intergenic_variant | MODIFIER |
| 12 | 112215788 | AX-170425585 | 0 | T | C | 0.14 | -3.60E-02 | 1.93E-01 | 2.68E+00 | 2.75E+00 | 8.49E-01 | no | | intergenic_variant | MODIFIER |
| 12 | 112253732 | AX-170007544 | 0 | T | C | 0.14 | -3.60E-02 | 1.93E-01 | 2.68E+00 | 2.75E+00 | 8.49E-01 | no | | regulatory_region_variant | MODIFIER |
| 12 | 112256607 | AX-97840241 | 0 | C | T | 0.14 | -3.60E-02 | 1.93E-01 | 2.68E+00 | 2.75E+00 | 8.49E-01 | no | | intergenic_variant | MODIFIER |
| 12 | 112262894 | AX-97853351 | 0 | A | G | 0.14 | -3.60E-02 | 1.93E-01 | 2.68E+00 | 2.75E+00 | 8.49E-01 | no | | regulatory_region_variant | MODIFIER |
| 12 | 112263013 | AX-170366268 | 0 | T | C | 0.14 | -3.60E-02 | 1.93E-01 | 2.68E+00 | 2.75E+00 | 8.49E-01 | no | | regulatory_region_variant | MODIFIER |
| 12 | 112279776 | AX-170475475 | 0 | A | T | 0.14 | -3.60E-02 | 1.93E-01 | 2.68E+00 | 2.75E+00 | 8.49E-01 | no | | intergenic_variant | MODIFIER |
| 12 | 112279898 | AX-170385140 | 0 | G | A | 0.14 | -3.60E-02 | 1.93E-01 | 2.68E+00 | 2.75E+00 | 8.49E-01 | no | | intergenic_variant | MODIFIER |
| 12 | 112280872 | AX-170403739 | 0 | T | C | 0.14 | -3.60E-02 | 1.93E-01 | 2.68E+00 | 2.75E+00 | 8.49E-01 | no | | intergenic_variant | MODIFIER |
| 12 | 112319184 | AX-170239513 | 0 | A | G | 0.14 | -3.60E-02 | 1.93E-01 | 2.68E+00 | 2.75E+00 | 8.49E-01 | no | | intergenic_variant | MODIFIER |
| 12 | 112322866 | AX-170243037 | 0 | C | T | 0.14 | -3.60E-02 | 1.93E-01 | 2.68E+00 | 2.75E+00 | 8.49E-01 | no | | intergenic_variant | MODIFIER |
| 12 | 112332961 | AX-97796450 | 0 | T | C | 0.14 | -3.60E-02 | 1.93E-01 | 2.68E+00 | 2.75E+00 | 8.49E-01 | no | | regulatory_region_variant | MODIFIER |
| 12 | 112345378 | AX-170050229 | 0 | T | A | 0.14 | -3.60E-02 | 1.93E-01 | 2.68E+00 | 2.75E+00 | 8.49E-01 | no | Gm38123 | downstream_gene_variant | MODIFIER |
| 12 | 112366075 | AX-170087335 | 0 | T | C | 0.14 | -3.60E-02 | 1.93E-01 | 2.68E+00 | 2.75E+00 | 8.49E-01 | no | | intergenic_variant | MODIFIER |
| 12 | 112375820 | AX-170148429 | 0 | C | T | 0.14 | -3.60E-02 | 1.93E-01 | 2.68E+00 | 2.75E+00 | 8.49E-01 | no | | intergenic_variant | MODIFIER |
| 12 | 112378801 | AX-170417882 | 0 | A | G | 0.14 | -3.60E-02 | 1.93E-01 | 2.68E+00 | 2.75E+00 | 8.49E-01 | no | | regulatory_region_variant | MODIFIER |
| 12 | 112379587 | AX-170252848 | 0 | T | C | 0.14 | -3.60E-02 | 1.93E-01 | 2.68E+00 | 2.75E+00 | 8.49E-01 | no | | intergenic_variant | MODIFIER |
| 12 | 112396484 | AX-170207702 | 0 | G | A | 0.14 | -3.60E-02 | 1.93E-01 | 2.68E+00 | 2.75E+00 | 8.49E-01 | no | E330035G20Rik | intron_variant,non_coding_transcript_variant | MODIFIER |
| 12 | 112414960 | AX-170127974 | 0 | T | G | 0.14 | -3.60E-02 | 1.93E-01 | 2.68E+00 | 2.75E+00 | 8.49E-01 | no | A730018C14Rik | non_coding_transcript_exon_variant | MODIFIER |
| 12 | 112421897 | AX-97822466 | 0 | C | T | 0.14 | -3.60E-02 | 1.93E-01 | 2.68E+00 | 2.75E+00 | 8.49E-01 | no | A730018C14Rik | intron_variant,non_coding_transcript_variant | MODIFIER |
| 12 | 112422152 | AX-170527719 | 0 | C | T | 0.14 | -3.60E-02 | 1.93E-01 | 2.68E+00 | 2.75E+00 | 8.49E-01 | no | A730018C14Rik | intron_variant,non_coding_transcript_variant | MODIFIER |
| 12 | 112426346 | AX-170256038 | 0 | G | A | 0.14 | -3.60E-02 | 1.93E-01 | 2.68E+00 | 2.75E+00 | 8.49E-01 | no | A730018C14Rik | upstream_gene_variant | MODIFIER |
| 12 | 112431275 | AX-170348072 | 0 | C | T | 0.14 | -3.60E-02 | 1.93E-01 | 2.68E+00 | 2.75E+00 | 8.49E-01 | no | B020018J22Rik | intron_variant,non_coding_transcript_variant | MODIFIER |
| 12 | 112448438 | AX-97829014 | 0 | C | T | 0.14 | -3.60E-02 | 1.93E-01 | 2.68E+00 | 2.75E+00 | 8.49E-01 | no | | intergenic_variant | MODIFIER |
| 12 | 112452997 | AX-170450566 | 0 | C | G | 0.14 | -3.60E-02 | 1.93E-01 | 2.68E+00 | 2.75E+00 | 8.49E-01 | no | | intergenic_variant | MODIFIER |
| 12 | 112454092 | AX-170050231 | 0 | T | C | 0.14 | -3.60E-02 | 1.93E-01 | 2.68E+00 | 2.75E+00 | 8.49E-01 | no | | intergenic_variant | MODIFIER |
| 12 | 112456477 | AX-170050233 | 0 | A | T | 0.14 | -3.60E-02 | 1.93E-01 | 2.68E+00 | 2.75E+00 | 8.49E-01 | no | | regulatory_region_variant | MODIFIER |
| 12 | 112458926 | AX-170336266 | 0 | C | T | 0.14 | -3.60E-02 | 1.93E-01 | 2.68E+00 | 2.75E+00 | 8.49E-01 | no | | intergenic_variant | MODIFIER |
| 12 | 112538195 | AX-170495501 | 0 | A | G | 0.14 | -3.60E-02 | 1.93E-01 | 2.68E+00 | 2.75E+00 | 8.49E-01 | no | | intergenic_variant | MODIFIER |
| 12 | 112541958 | AX-170326748 | 0 | G | A | 0.14 | -3.60E-02 | 1.93E-01 | 2.68E+00 | 2.75E+00 | 8.49E-01 | no | | regulatory_region_variant | MODIFIER |
| 12 | 112550314 | AX-97830675 | 0 | T | C | 0.14 | -3.60E-02 | 1.93E-01 | 2.68E+00 | 2.75E+00 | 8.49E-01 | no | | intergenic_variant | MODIFIER |
| 12 | 112567644 | AX-97820697 | 0 | C | T | 0.14 | -3.60E-02 | 1.93E-01 | 2.68E+00 | 2.75E+00 | 8.49E-01 | no | | intergenic_variant | MODIFIER |
| 12 | 112574632 | AX-170492591 | 0 | C | T | 0.14 | -3.60E-02 | 1.93E-01 | 2.68E+00 | 2.75E+00 | 8.49E-01 | no | | intergenic_variant | MODIFIER |
| 12 | 112579080 | AX-170352959 | 0 | C | T | 0.14 | -3.60E-02 | 1.93E-01 | 2.68E+00 | 2.75E+00 | 8.49E-01 | no | | regulatory_region_variant | MODIFIER |
| 12 | 112584032 | AX-170273140 | 0 | G | A | 0.14 | -3.60E-02 | 1.93E-01 | 2.68E+00 | 2.75E+00 | 8.49E-01 | no | Inf2 | upstream_gene_variant | MODIFIER |
| 12 | 112602121 | AX-97838093 | 0 | A | T | 0.14 | -3.60E-02 | 1.93E-01 | 2.68E+00 | 2.75E+00 | 8.49E-01 | no | Inf2 | intron_variant | MODIFIER |
| 12 | 112619591 | AX-170203676 | 0 | A | G | 0.14 | -3.60E-02 | 1.93E-01 | 2.68E+00 | 2.75E+00 | 8.49E-01 | no | Adssl1 | upstream_gene_variant | MODIFIER |
| 12 | 112621825 | AX-97798370 | 0 | A | G | 0.14 | -3.60E-02 | 1.93E-01 | 2.68E+00 | 2.75E+00 | 8.49E-01 | no | Adssl1 | intron_variant | MODIFIER |
| 12 | 112624393 | AX-170495191 | 0 | G | C | 0.14 | -3.60E-02 | 1.93E-01 | 2.68E+00 | 2.75E+00 | 8.49E-01 | no | Adssl1 | intron_variant | MODIFIER |
| 12 | 112627161 | AX-170395060 | 0 | A | G | 0.14 | -3.60E-02 | 1.93E-01 | 2.68E+00 | 2.75E+00 | 8.49E-01 | no | Adssl1 | intron_variant | MODIFIER |
| 12 | 112627295 | AX-170235107 | 0 | G | A | 0.14 | -3.60E-02 | 1.93E-01 | 2.68E+00 | 2.75E+00 | 8.49E-01 | no | Adssl1 | intron_variant | MODIFIER |
| 12 | 112637360 | AX-97831763 | 0 | G | T | 0.14 | -3.60E-02 | 1.93E-01 | 2.68E+00 | 2.75E+00 | 8.49E-01 | no | Adssl1 | intron_variant | MODIFIER |
| 12 | 112652156 | AX-170396960 | 0 | C | T | 0.14 | -3.60E-02 | 1.93E-01 | 2.68E+00 | 2.75E+00 | 8.49E-01 | no | Akt1 | downstream_gene_variant | MODIFIER |
| 12 | 113149927 | AX-170112884 | 0 | T | C | 0.14 | -3.60E-02 | 1.93E-01 | 2.68E+00 | 2.75E+00 | 8.49E-01 | no | Crip1 | upstream_gene_variant | MODIFIER |

| | | | | | | | | | | | | | | | |
|---|---|---|---|---|---|---|---|---|---|---|---|---|---|---|---|
| 12 | 113150588 | AX-170507287 | 0 | C | A | 0.14 | -3.60E-02 | 1.93E-01 | 2.68E+00 | 2.75E+00 | 8.49E-01 | no | Crip1 | upstream_gene_variant | MODIFIER |
| 12 | 113155060 | AX-170245865 | 0 | C | T | 0.14 | -3.60E-02 | 1.93E-01 | 2.68E+00 | 2.75E+00 | 8.49E-01 | no | Crip1 | downstream_gene_variant | MODIFIER |
| 12 | 113155076 | AX-170415891 | 0 | T | G | 0.14 | -3.60E-02 | 1.93E-01 | 2.68E+00 | 2.75E+00 | 8.49E-01 | no | Crip1 | downstream_gene_variant | MODIFIER |
| 12 | 113155867 | AX-170050247 | 0 | C | A | 0.14 | -3.60E-02 | 1.93E-01 | 2.68E+00 | 2.75E+00 | 8.49E-01 | no | Crip1 | downstream_gene_variant | MODIFIER |
| 12 | 113161321 | AX-170307001 | 0 | G | A | 0.14 | -3.60E-02 | 1.93E-01 | 2.68E+00 | 2.75E+00 | 8.49E-01 | no | Tedc1 | missense_variant | MODERATE |
| 12 | 113162013 | AX-170514805 | 0 | G | C | 0.14 | -3.60E-02 | 1.93E-01 | 2.68E+00 | 2.75E+00 | 8.49E-01 | no | Tedc1 | intron_variant | MODIFIER |
| 12 | 113165839 | AX-170476223 | 0 | G | A | 0.14 | -3.60E-02 | 1.93E-01 | 2.68E+00 | 2.75E+00 | 8.49E-01 | no | Tedc1 | downstream_gene_variant | MODIFIER |
| 12 | 113192593 | AX-170007560 | 0 | C | T | 0.14 | -3.60E-02 | 1.93E-01 | 2.68E+00 | 2.75E+00 | 8.49E-01 | no | Tmem121 | downstream_gene_variant | MODIFIER |
| 12 | 113215287 | AX-170290532 | 0 | G | T | 0.14 | -3.60E-02 | 1.93E-01 | 2.68E+00 | 2.75E+00 | 8.49E-01 | no | Gm25622 | downstream_gene_variant | MODIFIER |
| 12 | 113215678 | AX-170238431 | 0 | C | T | 0.14 | -3.60E-02 | 1.93E-01 | 2.68E+00 | 2.75E+00 | 8.49E-01 | no | Gm25622 | downstream_gene_variant | MODIFIER |
| 12 | 113221682 | AX-170269904 | 0 | C | T | 0.14 | -3.60E-02 | 1.93E-01 | 2.68E+00 | 2.75E+00 | 8.49E-01 | no | | regulatory_region_variant | MODIFIER |
| 12 | 113226625 | AX-170313309 | 0 | G | A | 0.14 | -3.60E-02 | 1.93E-01 | 2.68E+00 | 2.75E+00 | 8.49E-01 | no | | intergenic_variant | MODIFIER |
| 12 | 113228379 | AX-170171394 | 0 | T | C | 0.14 | -3.60E-02 | 1.93E-01 | 2.68E+00 | 2.75E+00 | 8.49E-01 | no | | regulatory_region_variant | MODIFIER |
| 12 | 113242698 | AX-170309192 | 0 | A | C | 0.14 | -3.60E-02 | 1.93E-01 | 2.68E+00 | 2.75E+00 | 8.49E-01 | no | | intergenic_variant | MODIFIER |
| 12 | 113243265 | AX-170118837 | 0 | C | T | 0.14 | -3.60E-02 | 1.93E-01 | 2.68E+00 | 2.75E+00 | 8.49E-01 | no | | intergenic_variant | MODIFIER |
| 12 | 113244583 | AX-97809009 | 0 | T | C | 0.14 | -3.60E-02 | 1.93E-01 | 2.68E+00 | 2.75E+00 | 8.49E-01 | no | | regulatory_region_variant | MODIFIER |
| 12 | 113244711 | AX-170392482 | 0 | G | T | 0.14 | -3.60E-02 | 1.93E-01 | 2.68E+00 | 2.75E+00 | 8.49E-01 | no | | regulatory_region_variant | MODIFIER |
| 12 | 113251953 | AX-170147159 | 0 | C | T | 0.14 | -3.60E-02 | 1.93E-01 | 2.68E+00 | 2.75E+00 | 8.49E-01 | no | Igha | downstream_gene_variant | MODIFIER |
| 12 | 113251987 | AX-170205400 | 0 | G | A | 0.14 | -3.60E-02 | 1.93E-01 | 2.68E+00 | 2.75E+00 | 8.49E-01 | no | Igha | downstream_gene_variant | MODIFIER |
| 12 | 113258517 | AX-170007565 | 0 | G | A | 0.14 | -3.60E-02 | 1.93E-01 | 2.68E+00 | 2.75E+00 | 8.49E-01 | no | Igha | downstream_gene_variant | MODIFIER |
| 12 | 113270308 | AX-170007564 | 0 | G | A | 0.14 | -3.60E-02 | 1.93E-01 | 2.68E+00 | 2.75E+00 | 8.49E-01 | no | Ighe | downstream_gene_variant | MODIFIER |
| 12 | 113282315 | AX-170465054 | 0 | A | G | 0.14 | -3.60E-02 | 1.93E-01 | 2.68E+00 | 2.75E+00 | 8.49E-01 | no | Ighg2c | downstream_gene_variant | MODIFIER |
| 12 | 113295717 | AX-170527437 | 0 | A | G | 0.14 | -3.60E-02 | 1.93E-01 | 2.68E+00 | 2.75E+00 | 8.49E-01 | no | | intergenic_variant | MODIFIER |
| 12 | 113295934 | AX-170450469 | 0 | T | C | 0.14 | -3.60E-02 | 1.93E-01 | 2.68E+00 | 2.75E+00 | 8.49E-01 | no | | intergenic_variant | MODIFIER |
| 12 | 113297316 | AX-170496157 | 0 | T | C | 0.14 | -3.60E-02 | 1.93E-01 | 2.68E+00 | 2.75E+00 | 8.49E-01 | no | | intergenic_variant | MODIFIER |
| 12 | 113299448 | AX-170166337 | 0 | T | C | 0.14 | -3.60E-02 | 1.93E-01 | 2.68E+00 | 2.75E+00 | 8.49E-01 | no | Ighg2b | downstream_gene_variant | MODIFIER |
| 12 | 113303212 | AX-170228996 | 0 | C | A | 0.14 | -3.60E-02 | 1.93E-01 | 2.68E+00 | 2.75E+00 | 8.49E-01 | no | Ighg2b | downstream_gene_variant | MODIFIER |
| 12 | 113308344 | AX-170300400 | 0 | T | C | 0.14 | -3.60E-02 | 1.93E-01 | 2.68E+00 | 2.75E+00 | 8.49E-01 | no | Ighg2b | upstream_gene_variant | MODIFIER |
| 12 | 113308389 | AX-170153648 | 0 | G | C | 0.14 | -3.60E-02 | 1.93E-01 | 2.68E+00 | 2.75E+00 | 8.49E-01 | no | Ighg2b | upstream_gene_variant | MODIFIER |
| 12 | 113309024 | AX-170514071 | 0 | C | T | 0.14 | -3.60E-02 | 1.93E-01 | 2.68E+00 | 2.75E+00 | 8.49E-01 | no | Ighg2b | upstream_gene_variant | MODIFIER |
| 12 | 113309429 | AX-170234481 | 0 | G | C | 0.14 | -3.60E-02 | 1.93E-01 | 2.68E+00 | 2.75E+00 | 8.49E-01 | no | Ighg2b | upstream_gene_variant | MODIFIER |
| 12 | 113309494 | AX-170425114 | 0 | C | T | 0.14 | -3.60E-02 | 1.93E-01 | 2.68E+00 | 2.75E+00 | 8.49E-01 | no | Ighg2b | upstream_gene_variant | MODIFIER |
| 12 | 113342018 | AX-97784731 | 0 | C | T | 0.14 | -3.60E-02 | 1.93E-01 | 2.68E+00 | 2.75E+00 | 8.49E-01 | no | | intergenic_variant | MODIFIER |
| 12 | 113342205 | AX-170516293 | 0 | T | C | 0.14 | -3.60E-02 | 1.93E-01 | 2.68E+00 | 2.75E+00 | 8.49E-01 | no | | intergenic_variant | MODIFIER |
| 12 | 113342476 | AX-170050254 | 0 | C | T | 0.14 | -3.60E-02 | 1.93E-01 | 2.68E+00 | 2.75E+00 | 8.49E-01 | no | | intergenic_variant | MODIFIER |
| 12 | 113355328 | AX-170050256 | 0 | T | C | 0.14 | -3.60E-02 | 1.93E-01 | 2.68E+00 | 2.75E+00 | 8.49E-01 | no | Ighg3 | downstream_gene_variant | MODIFIER |
| 12 | 113360772 | AX-170128231 | 0 | A | G | 0.14 | -3.60E-02 | 1.93E-01 | 2.68E+00 | 2.75E+00 | 8.49E-01 | no | Ighg3 | intron_variant | MODIFIER |
| 12 | 113365798 | AX-170511523 | 0 | T | C | 0.14 | -3.60E-02 | 1.93E-01 | 2.68E+00 | 2.75E+00 | 8.49E-01 | no | Ighg3 | upstream_gene_variant | MODIFIER |
| 12 | 113366269 | AX-170311947 | 0 | T | C | 0.14 | -3.60E-02 | 1.93E-01 | 2.68E+00 | 2.75E+00 | 8.49E-01 | no | Gm17421 | upstream_gene_variant | MODIFIER |
| 12 | 113367052 | AX-170233936 | 0 | A | G | 0.14 | -3.60E-02 | 1.93E-01 | 2.68E+00 | 2.75E+00 | 8.49E-01 | no | Gm17421 | upstream_gene_variant | MODIFIER |
| 12 | 113369084 | AX-170379264 | 0 | C | T | 0.14 | -3.60E-02 | 1.93E-01 | 2.68E+00 | 2.75E+00 | 8.49E-01 | no | Gm17421 | upstream_gene_variant | MODIFIER |
| 12 | 113369267 | AX-97802848 | 0 | T | C | 0.14 | -3.60E-02 | 1.93E-01 | 2.68E+00 | 2.75E+00 | 8.49E-01 | no | Gm17421 | upstream_gene_variant | MODIFIER |
| 12 | 113369339 | AX-170225655 | 0 | A | G | 0.14 | -3.60E-02 | 1.93E-01 | 2.68E+00 | 2.75E+00 | 8.49E-01 | no | Gm17421 | upstream_gene_variant | MODIFIER |
| 12 | 113370591 | AX-170386570 | 0 | C | G | 0.14 | -3.60E-02 | 1.93E-01 | 2.68E+00 | 2.75E+00 | 8.49E-01 | no | Gm17421 | intron_variant,non_coding_transcript_variant | MODIFIER |
| 12 | 113372187 | AX-170328492 | 0 | T | C | 0.14 | -3.60E-02 | 1.93E-01 | 2.68E+00 | 2.75E+00 | 8.49E-01 | no | Gm17421 | non_coding_transcript_exon_variant | MODIFIER |
| 12 | 113372239 | AX-170181353 | 0 | T | C | 0.14 | -3.60E-02 | 1.93E-01 | 2.68E+00 | 2.75E+00 | 8.49E-01 | no | Gm17421 | downstream_gene_variant | MODIFIER |
| 12 | 113402695 | AX-170489108 | 0 | G | A | 0.14 | -3.60E-02 | 1.93E-01 | 2.68E+00 | 2.75E+00 | 8.49E-01 | no | Ighd | downstream_gene_variant | MODIFIER |
| 12 | 113412924 | AX-170360366 | 0 | A | C | 0.14 | -3.60E-02 | 1.93E-01 | 2.68E+00 | 2.75E+00 | 8.49E-01 | no | Ighd | intron_variant | MODIFIER |
| 12 | 113532514 | AX-97826911 | 0 | A | G | 0.14 | -3.60E-02 | 1.93E-01 | 2.68E+00 | 2.75E+00 | 8.49E-01 | no | Gm16968 | upstream_gene_variant | MODIFIER |
| 12 | 113547384 | AX-170050257 | 0 | C | G | 0.14 | -3.60E-02 | 1.93E-01 | 2.68E+00 | 2.75E+00 | 8.49E-01 | no | Adam6a | downstream_gene_variant | MODIFIER |
| 12 | 113564939 | AX-170515416 | 0 | G | A | 0.14 | -3.60E-02 | 1.93E-01 | 2.68E+00 | 2.75E+00 | 8.49E-01 | no | | intergenic_variant | MODIFIER |
| 12 | 113565396 | AX-168388282 | 0 | G | A | 0.14 | -3.60E-02 | 1.93E-01 | 2.68E+00 | 2.75E+00 | 8.49E-01 | no | | intergenic_variant | MODIFIER |
| 12 | 113574112 | AX-170436879 | 0 | T | C | 0.14 | -3.60E-02 | 1.93E-01 | 2.68E+00 | 2.75E+00 | 8.49E-01 | no | Ighv5-2 | downstream_gene_variant | MODIFIER |
| 12 | 113730450 | AX-170523598 | 0 | C | A | 0.14 | -3.60E-02 | 1.93E-01 | 2.68E+00 | 2.75E+00 | 8.49E-01 | no | | intergenic_variant | MODIFIER |

| | | | | | | | | | | | | | | | |
|---|---|---|---|---|---|---|---|---|---|---|---|---|---|---|---|
| 12 | 113762770 | AX-97824654 | 0 | G | T | 0.14 | -3.60E-02 | 1.93E-01 | 2.68E+00 | 2.75E+00 | 8.49E-01 | no | Ighv5-12-4 | upstream_gene_variant | MODIFIER |
| 12 | 113847903 | AX-170335085 | 0 | G | T | 0.14 | -3.60E-02 | 1.93E-01 | 2.68E+00 | 2.75E+00 | 8.49E-01 | no | | intergenic_variant | MODIFIER |
| 12 | 113860119 | AX-169999769 | 0 | A | C | 0.14 | -3.60E-02 | 1.93E-01 | 2.68E+00 | 2.75E+00 | 8.49E-01 | no | Ighv5-17 | upstream_gene_variant | MODIFIER |
| 12 | 113918569 | AX-170050267 | 0 | C | T | 0.14 | -3.60E-02 | 1.93E-01 | 2.68E+00 | 2.75E+00 | 8.49E-01 | no | | intergenic_variant | MODIFIER |
| 12 | 114091621 | AX-170254090 | 0 | C | T | 0.14 | -3.60E-02 | 1.93E-01 | 2.68E+00 | 2.75E+00 | 8.49E-01 | no | Ighv9-1 | downstream_gene_variant | MODIFIER |
| 12 | 114091824 | AX-170434542 | 0 | C | T | 0.14 | -3.60E-02 | 1.93E-01 | 2.68E+00 | 2.75E+00 | 8.49E-01 | no | Ighv9-1 | downstream_gene_variant | MODIFIER |
| 12 | 114109132 | AX-170450121 | 0 | T | A | 0.14 | -3.60E-02 | 1.93E-01 | 2.68E+00 | 2.75E+00 | 8.49E-01 | no | Ighv9-2 | missense_variant | MODERATE |
| 12 | 114354932 | AX-170440941 | 0 | T | A | 0.14 | -3.60E-02 | 1.93E-01 | 2.68E+00 | 2.75E+00 | 8.49E-01 | no | Ighv13-2 | downstream_gene_variant | MODIFIER |
| 12 | 114355283 | AX-170204117 | 0 | G | C | 0.14 | -3.60E-02 | 1.93E-01 | 2.68E+00 | 2.75E+00 | 8.49E-01 | no | Ighv13-2 | downstream_gene_variant | MODIFIER |
| 12 | 114564130 | AX-170370319 | 0 | A | C | 0.14 | -3.60E-02 | 1.93E-01 | 2.68E+00 | 2.75E+00 | 8.49E-01 | no | Ighv15-2 | downstream_gene_variant | MODIFIER |
| 12 | 114565730 | AX-97782928 | 0 | G | A | 0.14 | -3.60E-02 | 1.93E-01 | 2.68E+00 | 2.75E+00 | 8.49E-01 | no | Ighv15-2 | upstream_gene_variant | MODIFIER |
| 12 | 114630467 | AX-170221238 | 0 | G | C | 0.14 | -3.60E-02 | 1.93E-01 | 2.68E+00 | 2.75E+00 | 8.49E-01 | no | Ighv1-13 | upstream_gene_variant | MODIFIER |
| 12 | 114631285 | AX-170219618 | 0 | A | G | 0.14 | -3.60E-02 | 1.93E-01 | 2.68E+00 | 2.75E+00 | 8.49E-01 | no | Ighv1-13 | downstream_gene_variant | MODIFIER |
| 12 | 114738031 | AX-170007579 | 0 | G | C | 0.14 | -3.60E-02 | 1.93E-01 | 2.68E+00 | 2.75E+00 | 8.49E-01 | no | Ighv1-21 | downstream_gene_variant | MODIFIER |
| 12 | 114837138 | AX-170338723 | 0 | A | C | 0.14 | -3.60E-02 | 1.93E-01 | 2.68E+00 | 2.75E+00 | 8.49E-01 | no | Ighv1-32 | upstream_gene_variant | MODIFIER |
| 12 | 114837608 | AX-170283281 | 0 | G | A | 0.14 | -3.60E-02 | 1.93E-01 | 2.68E+00 | 2.75E+00 | 8.49E-01 | no | Ighv1-32 | upstream_gene_variant | MODIFIER |
| 12 | 114839758 | AX-170050283 | 0 | C | T | 0.14 | -3.60E-02 | 1.93E-01 | 2.68E+00 | 2.75E+00 | 8.49E-01 | no | Ighv1-33 | downstream_gene_variant | MODIFIER |
| 12 | 114917951 | AX-170385480 | 0 | G | A | 0.14 | -3.60E-02 | 1.93E-01 | 2.68E+00 | 2.75E+00 | 8.49E-01 | no | Ighv1-39 | upstream_gene_variant | MODIFIER |
| 12 | 114933308 | AX-97792010 | 0 | A | G | 0.14 | -3.60E-02 | 1.93E-01 | 2.68E+00 | 2.75E+00 | 8.49E-01 | no | Ighv1-42 | downstream_gene_variant | MODIFIER |
| 12 | 114939049 | AX-170218285 | 0 | G | T | 0.14 | -3.60E-02 | 1.93E-01 | 2.68E+00 | 2.75E+00 | 8.49E-01 | no | Ighv1-42 | upstream_gene_variant | MODIFIER |
| 12 | 114979787 | AX-170530733 | 0 | G | A | 0.14 | -3.60E-02 | 1.93E-01 | 2.68E+00 | 2.75E+00 | 8.49E-01 | no | Ighv1-46 | upstream_gene_variant | MODIFIER |
| 12 | 114991870 | AX-97805096 | 0 | A | G | 0.14 | -3.60E-02 | 1.93E-01 | 2.68E+00 | 2.75E+00 | 8.49E-01 | no | Ighv1-47 | upstream_gene_variant | MODIFIER |
| 12 | 115025161 | AX-170440406 | 0 | C | T | 0.14 | -3.60E-02 | 1.93E-01 | 2.68E+00 | 2.75E+00 | 8.49E-01 | no | Ighv8-4 | upstream_gene_variant | MODIFIER |
| 12 | 115051070 | AX-97820820 | 0 | T | C | 0.14 | -3.60E-02 | 1.93E-01 | 2.68E+00 | 2.75E+00 | 8.49E-01 | no | Ighv1-49 | downstream_gene_variant | MODIFIER |
| 12 | 115121609 | AX-170158961 | 0 | T | C | 0.14 | -3.60E-02 | 1.93E-01 | 2.68E+00 | 2.75E+00 | 8.49E-01 | no | Ighv1-50 | upstream_gene_variant | MODIFIER |
| 12 | 115121684 | AX-170407890 | 0 | A | G | 0.14 | -3.60E-02 | 1.93E-01 | 2.68E+00 | 2.75E+00 | 8.49E-01 | no | Ighv1-50 | upstream_gene_variant | MODIFIER |
| 12 | 115131759 | AX-170314839 | 0 | C | T | 0.14 | -3.60E-02 | 1.93E-01 | 2.68E+00 | 2.75E+00 | 8.49E-01 | no | Ighv1-51 | splice_region_variant,non_coding_transcript_exon_variant | LOW |
| 12 | 115150006 | AX-170342320 | 0 | A | G | 0.14 | -3.60E-02 | 1.93E-01 | 2.68E+00 | 2.75E+00 | 8.49E-01 | no | Ighv1-52 | upstream_gene_variant | MODIFIER |
| 12 | 115640192 | AX-97853896 | 0 | G | C | 0.14 | -3.60E-02 | 1.93E-01 | 2.68E+00 | 2.75E+00 | 8.49E-01 | no | | intergenic_variant | MODIFIER |
| 12 | 116029498 | AX-97847521 | 0 | G | T | 0.14 | -3.60E-02 | 1.93E-01 | 2.68E+00 | 2.75E+00 | 8.49E-01 | no | | intergenic_variant | MODIFIER |
| 12 | 116050283 | AX-170007590 | 0 | G | A | 0.14 | -3.60E-02 | 1.93E-01 | 2.68E+00 | 2.75E+00 | 8.49E-01 | no | Zfp386 | intron_variant | MODIFIER |
| 12 | 116050578 | AX-170007588 | 0 | A | G | 0.14 | -3.60E-02 | 1.93E-01 | 2.68E+00 | 2.75E+00 | 8.49E-01 | no | Zfp386 | intron_variant | MODIFIER |
| 12 | 116059105 | AX-170007583 | 0 | G | A | 0.14 | -3.60E-02 | 1.93E-01 | 2.68E+00 | 2.75E+00 | 8.49E-01 | no | Zfp386 | missense_variant | MODERATE |
| 12 | 116093091 | AX-170007581 | 0 | G | C | 0.14 | -3.60E-02 | 1.93E-01 | 2.68E+00 | 2.75E+00 | 8.49E-01 | no | Vipr2 | intron_variant | MODIFIER |
| 12 | 116097322 | AX-170007585 | 0 | C | G | 0.14 | -3.60E-02 | 1.93E-01 | 2.68E+00 | 2.75E+00 | 8.49E-01 | no | Vipr2 | intron_variant | MODIFIER |
| 12 | 116097787 | AX-170527142 | 0 | G | A | 0.14 | -3.60E-02 | 1.93E-01 | 2.68E+00 | 2.75E+00 | 8.49E-01 | no | Vipr2 | intron_variant | MODIFIER |
| 12 | 116103445 | AX-170185968 | 0 | A | G | 0.14 | -3.60E-02 | 1.93E-01 | 2.68E+00 | 2.75E+00 | 8.49E-01 | no | Vipr2 | intron_variant | MODIFIER |
| 12 | 116110279 | AX-170373381 | 0 | C | T | 0.14 | -3.60E-02 | 1.93E-01 | 2.68E+00 | 2.75E+00 | 8.49E-01 | no | Vipr2 | intron_variant | MODIFIER |
| 12 | 116130672 | AX-97794043 | 0 | C | T | 0.14 | -3.60E-02 | 1.93E-01 | 2.68E+00 | 2.75E+00 | 8.49E-01 | no | Vipr2 | intron_variant | MODIFIER |
| 12 | 116137458 | AX-170007587 | 0 | A | G | 0.14 | -3.60E-02 | 1.93E-01 | 2.68E+00 | 2.75E+00 | 8.49E-01 | no | Vipr2 | intron_variant | MODIFIER |
| 12 | 116137808 | AX-170275789 | 0 | C | T | 0.14 | -3.60E-02 | 1.93E-01 | 2.68E+00 | 2.75E+00 | 8.49E-01 | no | Vipr2 | intron_variant | MODIFIER |
| 12 | 116155389 | AX-170050312 | 0 | A | C | 0.14 | -3.60E-02 | 1.93E-01 | 2.68E+00 | 2.75E+00 | 8.49E-01 | no | Gm44372 | upstream_gene_variant | MODIFIER |
| 12 | 116166271 | AX-170050313 | 0 | A | G | 0.14 | -3.60E-02 | 1.93E-01 | 2.68E+00 | 2.75E+00 | 8.49E-01 | no | Gm48620 | upstream_gene_variant | MODIFIER |
| 12 | 116179294 | AX-170007589 | 0 | A | G | 0.14 | -3.60E-02 | 1.93E-01 | 2.68E+00 | 2.75E+00 | 8.49E-01 | no | | intergenic_variant | MODIFIER |
| 12 | 116181254 | AX-170389230 | 0 | G | A | 0.14 | -3.60E-02 | 1.93E-01 | 2.68E+00 | 2.75E+00 | 8.49E-01 | no | | intergenic_variant | MODIFIER |
| 12 | 116194791 | AX-170365738 | 0 | T | A | 0.14 | -3.60E-02 | 1.93E-01 | 2.68E+00 | 2.75E+00 | 8.49E-01 | no | | intergenic_variant | MODIFIER |
| 12 | 116208459 | AX-170259895 | 0 | T | C | 0.14 | -3.60E-02 | 1.93E-01 | 2.68E+00 | 2.75E+00 | 8.49E-01 | no | Wdr60 | intron_variant | MODIFIER |
| 12 | 116226204 | AX-170350868 | 0 | C | T | 0.14 | -3.60E-02 | 1.93E-01 | 2.68E+00 | 2.75E+00 | 8.49E-01 | no | Wdr60 | intron_variant | MODIFIER |
| 12 | 116227961 | AX-97808301 | 0 | G | A | 0.14 | -3.60E-02 | 1.93E-01 | 2.68E+00 | 2.75E+00 | 8.49E-01 | no | Wdr60 | intron_variant | MODIFIER |
| 12 | 116232216 | AX-170209542 | 0 | A | T | 0.14 | -3.60E-02 | 1.93E-01 | 2.68E+00 | 2.75E+00 | 8.49E-01 | no | Wdr60 | intron_variant | MODIFIER |
| 12 | 116233395 | AX-170422282 | 0 | A | T | 0.14 | -3.60E-02 | 1.93E-01 | 2.68E+00 | 2.75E+00 | 8.49E-01 | no | Wdr60 | synonymous_variant | LOW |
| 12 | 116248247 | AX-97852363 | 0 | A | G | 0.14 | -3.60E-02 | 1.93E-01 | 2.68E+00 | 2.75E+00 | 8.49E-01 | no | Wdr60 | synonymous_variant | LOW |
| 12 | 116257981 | AX-170146165 | 0 | C | G | 0.14 | -3.60E-02 | 1.93E-01 | 2.68E+00 | 2.75E+00 | 8.49E-01 | no | Wdr60 | intron_variant | MODIFIER |
| 12 | 116295121 | AX-97856850 | 0 | A | G | 0.14 | -3.60E-02 | 1.93E-01 | 2.68E+00 | 2.75E+00 | 8.49E-01 | no | Esyt2 | intron_variant | MODIFIER |

| | | | | | | | | | | | | | | | |
|---|---|---|---|---|---|---|---|---|---|---|---|---|---|---|---|
| 12 | 116301391 | AX-170431667 | 0 | G | A | 0.14 | -3.60E-02 | 1.93E-01 | 2.68E+00 | 2.75E+00 | 8.49E-01 | no | Esyt2 | intron_variant | MODIFIER |
| 12 | 116312231 | AX-97848156 | 0 | A | G | 0.14 | -3.60E-02 | 1.93E-01 | 2.68E+00 | 2.75E+00 | 8.49E-01 | no | Esyt2 | intron_variant | MODIFIER |
| 12 | 116318783 | AX-170420375 | 0 | C | T | 0.14 | -3.60E-02 | 1.93E-01 | 2.68E+00 | 2.75E+00 | 8.49E-01 | no | Esyt2 | intron_variant | MODIFIER |
| 12 | 116334096 | AX-170306404 | 0 | A | C | 0.14 | -3.60E-02 | 1.93E-01 | 2.68E+00 | 2.75E+00 | 8.49E-01 | no | Esyt2 | intron_variant | MODIFIER |
| 12 | 116338817 | AX-168379517 | 0 | T | C | 0.14 | -3.60E-02 | 1.93E-01 | 2.68E+00 | 2.75E+00 | 8.49E-01 | no | Esyt2 | intron_variant | MODIFIER |
| 12 | 116339700 | AX-170124205 | 0 | A | G | 0.14 | -3.60E-02 | 1.93E-01 | 2.68E+00 | 2.75E+00 | 8.49E-01 | no | Esyt2 | intron_variant | MODIFIER |
| 12 | 116339722 | AX-97822414 | 0 | T | G | 0.14 | -3.60E-02 | 1.93E-01 | 2.68E+00 | 2.75E+00 | 8.49E-01 | no | Esyt2 | intron_variant | MODIFIER |
| 12 | 116339767 | AX-170505009 | 0 | A | C | 0.14 | -3.60E-02 | 1.93E-01 | 2.68E+00 | 2.75E+00 | 8.49E-01 | no | Esyt2 | intron_variant | MODIFIER |
| 12 | 116362980 | AX-170050316 | 0 | G | C | 0.14 | -3.60E-02 | 1.93E-01 | 2.68E+00 | 2.75E+00 | 8.49E-01 | no | Esyt2 | intron_variant | MODIFIER |
| 12 | 116365022 | AX-170395438 | 0 | C | T | 0.14 | -3.60E-02 | 1.93E-01 | 2.68E+00 | 2.75E+00 | 8.49E-01 | no | Esyt2 | intron_variant | MODIFIER |
| 12 | 116394898 | AX-97783573 | 0 | G | A | 0.14 | -3.60E-02 | 1.93E-01 | 2.68E+00 | 2.75E+00 | 8.49E-01 | no | Esyt2 | downstream_gene_variant | MODIFIER |
| 12 | 116408059 | AX-170488921 | 0 | T | C | 0.14 | -3.60E-02 | 1.93E-01 | 2.68E+00 | 2.75E+00 | 8.49E-01 | no | Ncapg2 | intron_variant | MODIFIER |
| 12 | 116409208 | AX-170200205 | 0 | G | T | 0.14 | -3.60E-02 | 1.93E-01 | 2.68E+00 | 2.75E+00 | 8.49E-01 | no | Ncapg2 | intron_variant | MODIFIER |
| 12 | 116410070 | AX-97822522 | 0 | T | G | 0.14 | -3.60E-02 | 1.93E-01 | 2.68E+00 | 2.75E+00 | 8.49E-01 | no | Ncapg2 | intron_variant | MODIFIER |
| 12 | 116439884 | AX-170179743 | 0 | G | T | 0.14 | -3.60E-02 | 1.93E-01 | 2.68E+00 | 2.75E+00 | 8.49E-01 | no | Ncapg2 | intron_variant | MODIFIER |
| 12 | 116447111 | AX-97786210 | 0 | G | A | 0.14 | -3.60E-02 | 1.93E-01 | 2.68E+00 | 2.75E+00 | 8.49E-01 | no | Ncapg2 | intron_variant | MODIFIER |
| 12 | 116450086 | AX-170489781 | 0 | C | T | 0.14 | -3.60E-02 | 1.93E-01 | 2.68E+00 | 2.75E+00 | 8.49E-01 | no | Ncapg2 | intron_variant | MODIFIER |
| 12 | 116450383 | AX-170418267 | 0 | T | G | 0.14 | -3.60E-02 | 1.93E-01 | 2.68E+00 | 2.75E+00 | 8.49E-01 | no | Ncapg2 | missense_variant | MODERATE |
| 12 | 116459139 | AX-170205128 | 0 | G | A | 0.14 | -3.60E-02 | 1.93E-01 | 2.68E+00 | 2.75E+00 | 8.49E-01 | no | Ncapg2 | intron_variant | MODIFIER |
| 12 | 116462442 | AX-170506477 | 0 | T | C | 0.14 | -3.60E-02 | 1.93E-01 | 2.68E+00 | 2.75E+00 | 8.49E-01 | no | Ncapg2 | downstream_gene_variant | MODIFIER |
| 12 | 116467489 | AX-170504540 | 0 | G | C | 0.14 | -3.60E-02 | 1.93E-01 | 2.68E+00 | 2.75E+00 | 8.49E-01 | no | Ncapg2 | downstream_gene_variant | MODIFIER |
| 12 | 116468025 | AX-170449438 | 0 | C | T | 0.14 | -3.60E-02 | 1.93E-01 | 2.68E+00 | 2.75E+00 | 8.49E-01 | no | Ncapg2 | downstream_gene_variant | MODIFIER |
| 12 | 116468084 | AX-170469020 | 0 | T | C | 0.14 | -3.60E-02 | 1.93E-01 | 2.68E+00 | 2.75E+00 | 8.49E-01 | no | Ncapg2 | downstream_gene_variant | MODIFIER |
| 12 | 116523196 | AX-97815746 | 0 | G | A | 0.14 | -3.60E-02 | 1.93E-01 | 2.68E+00 | 2.75E+00 | 8.49E-01 | no | Ptprn2 | intron_variant | MODIFIER |
| 12 | 116524698 | AX-170050319 | 0 | G | A | 0.14 | -3.60E-02 | 1.93E-01 | 2.68E+00 | 2.75E+00 | 8.49E-01 | no | Ptprn2 | intron_variant | MODIFIER |
| 12 | 116531140 | AX-170050318 | 0 | C | T | 0.14 | -3.60E-02 | 1.93E-01 | 2.68E+00 | 2.75E+00 | 8.49E-01 | no | Ptprn2 | intron_variant | MODIFIER |
| 12 | 116531467 | AX-170284055 | 0 | T | G | 0.14 | -3.60E-02 | 1.93E-01 | 2.68E+00 | 2.75E+00 | 8.49E-01 | no | Ptprn2 | intron_variant | MODIFIER |
| 12 | 116532746 | AX-170487461 | 0 | A | G | 0.14 | -3.60E-02 | 1.93E-01 | 2.68E+00 | 2.75E+00 | 8.49E-01 | no | Ptprn2 | intron_variant | MODIFIER |
| 12 | 116534095 | AX-170198133 | 0 | C | T | 0.14 | -3.60E-02 | 1.93E-01 | 2.68E+00 | 2.75E+00 | 8.49E-01 | no | Ptprn2 | intron_variant | MODIFIER |
| 12 | 116550160 | AX-170485650 | 0 | C | A | 0.14 | -3.60E-02 | 1.93E-01 | 2.68E+00 | 2.75E+00 | 8.49E-01 | no | Ptprn2 | intron_variant | MODIFIER |
| 12 | 116551365 | AX-170511294 | 0 | G | A | 0.14 | -3.60E-02 | 1.93E-01 | 2.68E+00 | 2.75E+00 | 8.49E-01 | no | Ptprn2 | intron_variant | MODIFIER |
| 12 | 116552695 | AX-170361187 | 0 | T | G | 0.14 | -3.60E-02 | 1.93E-01 | 2.68E+00 | 2.75E+00 | 8.49E-01 | no | Ptprn2 | intron_variant | MODIFIER |
| 12 | 116564248 | AX-170144947 | 0 | C | T | 0.14 | -3.60E-02 | 1.93E-01 | 2.68E+00 | 2.75E+00 | 8.49E-01 | no | Ptprn2 | intron_variant | MODIFIER |
| 12 | 116579292 | AX-97846274 | 0 | A | G | 0.14 | -3.60E-02 | 1.93E-01 | 2.68E+00 | 2.75E+00 | 8.49E-01 | no | Ptprn2 | intron_variant | MODIFIER |
| 12 | 116581823 | AX-170007593 | 0 | T | A | 0.14 | -3.60E-02 | 1.93E-01 | 2.68E+00 | 2.75E+00 | 8.49E-01 | no | Ptprn2 | intron_variant | MODIFIER |
| 12 | 116582891 | AX-170269272 | 0 | A | G | 0.14 | -3.60E-02 | 1.93E-01 | 2.68E+00 | 2.75E+00 | 8.49E-01 | no | Ptprn2 | intron_variant | MODIFIER |
| 12 | 116608987 | AX-170050323 | 0 | A | C | 0.14 | -3.60E-02 | 1.93E-01 | 2.68E+00 | 2.75E+00 | 8.49E-01 | no | Ptprn2 | intron_variant | MODIFIER |
| 12 | 116615177 | AX-97783294 | 0 | T | C | 0.14 | -3.60E-02 | 1.93E-01 | 2.68E+00 | 2.75E+00 | 8.49E-01 | no | Ptprn2 | intron_variant | MODIFIER |
| 12 | 116618859 | AX-170511536 | 0 | T | G | 0.14 | -3.60E-02 | 1.93E-01 | 2.68E+00 | 2.75E+00 | 8.49E-01 | no | Ptprn2 | intron_variant | MODIFIER |
| 12 | 116651129 | AX-170525054 | 0 | G | A | 0.14 | -3.60E-02 | 1.93E-01 | 2.68E+00 | 2.75E+00 | 8.49E-01 | no | Ptprn2 | intron_variant | MODIFIER |
| 12 | 116661354 | AX-97812769 | 0 | G | A | 0.14 | -3.60E-02 | 1.93E-01 | 2.68E+00 | 2.75E+00 | 8.49E-01 | no | Ptprn2 | intron_variant | MODIFIER |
| 12 | 116677213 | AX-170459519 | 0 | G | C | 0.14 | -3.60E-02 | 1.93E-01 | 2.68E+00 | 2.75E+00 | 8.49E-01 | no | Ptprn2 | intron_variant | MODIFIER |
| 12 | 116688668 | AX-170098897 | 0 | G | T | 0.14 | -3.60E-02 | 1.93E-01 | 2.68E+00 | 2.75E+00 | 8.49E-01 | no | Ptprn2 | intron_variant | MODIFIER |
| 12 | 116688755 | AX-170198070 | 0 | A | G | 0.14 | -3.60E-02 | 1.93E-01 | 2.68E+00 | 2.75E+00 | 8.49E-01 | no | Ptprn2 | intron_variant | MODIFIER |
| 12 | 116693107 | AX-170392989 | 0 | A | G | 0.14 | -3.60E-02 | 1.93E-01 | 2.68E+00 | 2.75E+00 | 8.49E-01 | no | Ptprn2 | intron_variant | MODIFIER |
| 12 | 116707543 | AX-170486783 | 0 | C | A | 0.14 | -3.60E-02 | 1.93E-01 | 2.68E+00 | 2.75E+00 | 8.49E-01 | no | Ptprn2 | intron_variant | MODIFIER |
| 12 | 116707895 | AX-170194277 | 0 | C | T | 0.14 | -3.60E-02 | 1.93E-01 | 2.68E+00 | 2.75E+00 | 8.49E-01 | no | Ptprn2 | intron_variant | MODIFIER |
| 12 | 116739770 | AX-170213444 | 0 | G | A | 0.14 | -3.60E-02 | 1.93E-01 | 2.68E+00 | 2.75E+00 | 8.49E-01 | no | Ptprn2 | intron_variant | MODIFIER |
| 12 | 116741057 | AX-170363988 | 0 | C | T | 0.14 | -3.60E-02 | 1.93E-01 | 2.68E+00 | 2.75E+00 | 8.49E-01 | no | Ptprn2 | intron_variant | MODIFIER |
| 12 | 116745863 | AX-170144928 | 0 | A | G | 0.14 | -3.60E-02 | 1.93E-01 | 2.68E+00 | 2.75E+00 | 8.49E-01 | no | Ptprn2 | intron_variant | MODIFIER |
| 12 | 116746210 | AX-170264618 | 0 | A | G | 0.14 | -3.60E-02 | 1.93E-01 | 2.68E+00 | 2.75E+00 | 8.49E-01 | no | Ptprn2 | intron_variant | MODIFIER |
| 12 | 116748031 | AX-170050326 | 0 | T | C | 0.14 | -3.60E-02 | 1.93E-01 | 2.68E+00 | 2.75E+00 | 8.49E-01 | no | Ptprn2 | intron_variant | MODIFIER |
| 12 | 116751745 | AX-170404353 | 0 | C | T | 0.14 | -3.60E-02 | 1.93E-01 | 2.68E+00 | 2.75E+00 | 8.49E-01 | no | Ptprn2 | intron_variant | MODIFIER |
| 12 | 116769572 | AX-97849591 | 0 | A | G | 0.14 | -3.60E-02 | 1.93E-01 | 2.68E+00 | 2.75E+00 | 8.49E-01 | no | Ptprn2 | intron_variant | MODIFIER |

| | | | | | | | | | | | | | | | |
|---|---|---|---|---|---|---|---|---|---|---|---|---|---|---|---|
| 12 | 116778274 | AX-170386020 | 0 | G | A | 0.14 | -3.60E-02 | 1.93E-01 | 2.68E+00 | 2.75E+00 | 8.49E-01 | no | Ptprn2 | intron_variant | MODIFIER |
| 12 | 116786305 | AX-170502987 | 0 | C | G | 0.14 | -3.60E-02 | 1.93E-01 | 2.68E+00 | 2.75E+00 | 8.49E-01 | no | Ptprn2 | intron_variant | MODIFIER |
| 12 | 116788643 | AX-170328416 | 0 | C | G | 0.14 | -3.60E-02 | 1.93E-01 | 2.68E+00 | 2.75E+00 | 8.49E-01 | no | Ptprn2 | intron_variant | MODIFIER |
| 12 | 116819850 | AX-97830134 | 0 | A | G | 0.14 | -3.60E-02 | 1.93E-01 | 2.68E+00 | 2.75E+00 | 8.49E-01 | no | Ptprn2 | intron_variant | MODIFIER |
| 12 | 116833204 | AX-170050330 | 0 | A | C | 0.14 | -3.60E-02 | 1.93E-01 | 2.68E+00 | 2.75E+00 | 8.49E-01 | no | Ptprn2 | intron_variant | MODIFIER |
| 12 | 116833976 | AX-170467423 | 0 | G | A | 0.14 | -3.60E-02 | 1.93E-01 | 2.68E+00 | 2.75E+00 | 8.49E-01 | no | Ptprn2 | intron_variant | MODIFIER |
| 12 | 116859013 | AX-170173888 | 0 | A | G | 0.14 | -3.60E-02 | 1.93E-01 | 2.68E+00 | 2.75E+00 | 8.49E-01 | no | Ptprn2 | synonymous_variant | LOW |
| 12 | 116859298 | AX-170407569 | 0 | G | A | 0.14 | -3.60E-02 | 1.93E-01 | 2.68E+00 | 2.75E+00 | 8.49E-01 | no | Ptprn2 | intron_variant | MODIFIER |
| 12 | 116859608 | AX-170235559 | 0 | G | A | 0.14 | -3.60E-02 | 1.93E-01 | 2.68E+00 | 2.75E+00 | 8.49E-01 | no | Ptprn2 | intron_variant | MODIFIER |
| 12 | 116861384 | AX-170373745 | 0 | C | T | 0.14 | -3.60E-02 | 1.93E-01 | 2.68E+00 | 2.75E+00 | 8.49E-01 | no | Ptprn2 | intron_variant | MODIFIER |
| 12 | 116862636 | AX-168373900 | 0 | T | C | 0.14 | -3.60E-02 | 1.93E-01 | 2.68E+00 | 2.75E+00 | 8.49E-01 | no | Ptprn2 | intron_variant | MODIFIER |
| 12 | 116862941 | AX-168393158 | 0 | A | G | 0.14 | -3.60E-02 | 1.93E-01 | 2.68E+00 | 2.75E+00 | 8.49E-01 | no | Ptprn2 | intron_variant | MODIFIER |
| 12 | 116863309 | AX-170170644 | 0 | G | A | 0.14 | -3.60E-02 | 1.93E-01 | 2.68E+00 | 2.75E+00 | 8.49E-01 | no | Ptprn2 | intron_variant | MODIFIER |
| 12 | 116890446 | AX-97825102 | 0 | A | G | 0.14 | -3.60E-02 | 1.93E-01 | 2.68E+00 | 2.75E+00 | 8.49E-01 | no | Ptprn2 | intron_variant | MODIFIER |
| 12 | 116893524 | AX-170418818 | 0 | G | A | 0.14 | -3.60E-02 | 1.93E-01 | 2.68E+00 | 2.75E+00 | 8.49E-01 | no | Ptprn2 | intron_variant | MODIFIER |
| 12 | 116894626 | AX-170097358 | 0 | G | A | 0.14 | -3.60E-02 | 1.93E-01 | 2.68E+00 | 2.75E+00 | 8.49E-01 | no | Ptprn2 | intron_variant | MODIFIER |
| 12 | 116897444 | AX-170122188 | 0 | G | A | 0.14 | -3.60E-02 | 1.93E-01 | 2.68E+00 | 2.75E+00 | 8.49E-01 | no | Ptprn2 | intron_variant | MODIFIER |
| 12 | 116930844 | AX-170531410 | 0 | C | A | 0.14 | -3.60E-02 | 1.93E-01 | 2.68E+00 | 2.75E+00 | 8.49E-01 | no | Ptprn2 | intron_variant | MODIFIER |
| 12 | 116934862 | AX-170403203 | 0 | C | A | 0.14 | -3.60E-02 | 1.93E-01 | 2.68E+00 | 2.75E+00 | 8.49E-01 | no | Ptprn2 | intron_variant | MODIFIER |
| 12 | 116941400 | AX-170386843 | 0 | T | G | 0.14 | -3.60E-02 | 1.93E-01 | 2.68E+00 | 2.75E+00 | 8.49E-01 | no | Ptprn2 | intron_variant | MODIFIER |
| 12 | 116949251 | AX-170346255 | 0 | C | T | 0.14 | -3.60E-02 | 1.93E-01 | 2.68E+00 | 2.75E+00 | 8.49E-01 | no | Ptprn2 | intron_variant | MODIFIER |
| 12 | 116967929 | AX-170357065 | 0 | G | A | 0.14 | -3.60E-02 | 1.93E-01 | 2.68E+00 | 2.75E+00 | 8.49E-01 | no | Ptprn2 | intron_variant | MODIFIER |
| 12 | 116985940 | AX-97825812 | 0 | G | A | 0.14 | -3.60E-02 | 1.93E-01 | 2.68E+00 | 2.75E+00 | 8.49E-01 | no | Ptprn2 | intron_variant | MODIFIER |
| 12 | 117039398 | AX-97850079 | 0 | C | T | 0.14 | -3.60E-02 | 1.93E-01 | 2.68E+00 | 2.75E+00 | 8.49E-01 | no | Ptprn2 | intron_variant | MODIFIER |
| 12 | 117040399 | AX-170405262 | 0 | T | C | 0.14 | -3.60E-02 | 1.93E-01 | 2.68E+00 | 2.75E+00 | 8.49E-01 | no | Ptprn2 | intron_variant | MODIFIER |
| 12 | 117041187 | AX-170513821 | 0 | A | C | 0.14 | -3.60E-02 | 1.93E-01 | 2.68E+00 | 2.75E+00 | 8.49E-01 | no | Ptprn2 | intron_variant | MODIFIER |
| 12 | 117042005 | AX-170315774 | 0 | G | A | 0.14 | -3.60E-02 | 1.93E-01 | 2.68E+00 | 2.75E+00 | 8.49E-01 | no | Ptprn2 | intron_variant | MODIFIER |
| 12 | 117042494 | AX-170264583 | 0 | G | A | 0.14 | -3.60E-02 | 1.93E-01 | 2.68E+00 | 2.75E+00 | 8.49E-01 | no | Ptprn2 | intron_variant | MODIFIER |
| 12 | 117042875 | AX-170288038 | 0 | T | G | 0.14 | -3.60E-02 | 1.93E-01 | 2.68E+00 | 2.75E+00 | 8.49E-01 | no | Ptprn2 | intron_variant | MODIFIER |
| 12 | 117043059 | AX-170312700 | 0 | A | G | 0.14 | -3.60E-02 | 1.93E-01 | 2.68E+00 | 2.75E+00 | 8.49E-01 | no | Ptprn2 | intron_variant | MODIFIER |
| 12 | 117045249 | AX-170216201 | 0 | A | C | 0.14 | -3.60E-02 | 1.93E-01 | 2.68E+00 | 2.75E+00 | 8.49E-01 | no | Ptprn2 | intron_variant | MODIFIER |
| 12 | 117048360 | AX-170515904 | 0 | T | G | 0.14 | -3.60E-02 | 1.93E-01 | 2.68E+00 | 2.75E+00 | 8.49E-01 | no | Ptprn2 | intron_variant | MODIFIER |
| 12 | 117049315 | AX-170214718 | 0 | A | G | 0.14 | -3.60E-02 | 1.93E-01 | 2.68E+00 | 2.75E+00 | 8.49E-01 | no | Ptprn2 | intron_variant | MODIFIER |
| 12 | 117057236 | AX-170492892 | 0 | G | A | 0.14 | -3.60E-02 | 1.93E-01 | 2.68E+00 | 2.75E+00 | 8.49E-01 | no | Ptprn2 | intron_variant | MODIFIER |
| 12 | 117057895 | AX-170359990 | 0 | G | T | 0.14 | -3.60E-02 | 1.93E-01 | 2.68E+00 | 2.75E+00 | 8.49E-01 | no | Ptprn2 | intron_variant | MODIFIER |
| 12 | 117058410 | AX-170050340 | 0 | A | G | 0.14 | -3.60E-02 | 1.93E-01 | 2.68E+00 | 2.75E+00 | 8.49E-01 | no | Ptprn2 | intron_variant | MODIFIER |
| 12 | 117184464 | AX-170007598 | 0 | A | C | 0.14 | -3.60E-02 | 1.93E-01 | 2.68E+00 | 2.75E+00 | 8.49E-01 | no | Ptprn2 | intron_variant | MODIFIER |
| 12 | 117184605 | AX-170521830 | 0 | G | A | 0.14 | -3.60E-02 | 1.93E-01 | 2.68E+00 | 2.75E+00 | 8.49E-01 | no | Ptprn2 | missense_variant | MODERATE |
| 12 | 117195523 | AX-170269231 | 0 | A | G | 0.14 | -3.60E-02 | 1.93E-01 | 2.68E+00 | 2.75E+00 | 8.49E-01 | no | Ptprn2 | intron_variant | MODIFIER |
| 12 | 117203738 | AX-170410471 | 0 | G | C | 0.14 | -3.60E-02 | 1.93E-01 | 2.68E+00 | 2.75E+00 | 8.49E-01 | no | Ptprn2 | intron_variant | MODIFIER |
| 12 | 117332911 | AX-170413481 | 0 | G | A | 0.14 | -3.60E-02 | 1.93E-01 | 2.68E+00 | 2.75E+00 | 8.49E-01 | no | Gm5441 | intron_variant,non_coding_transcript_variant | MODIFIER |
| 12 | 117341136 | AX-170473370 | 0 | C | T | 0.14 | -3.60E-02 | 1.93E-01 | 2.68E+00 | 2.75E+00 | 8.49E-01 | no | Gm5441 | upstream_gene_variant | MODIFIER |
| 12 | 117361021 | AX-97822436 | 0 | T | C | 0.14 | -3.60E-02 | 1.93E-01 | 2.68E+00 | 2.75E+00 | 8.49E-01 | no | | intergenic_variant | MODIFIER |
| 12 | 117407168 | AX-170490369 | 0 | A | G | 0.14 | -3.60E-02 | 1.93E-01 | 2.68E+00 | 2.75E+00 | 8.49E-01 | no | | regulatory_region_variant | MODIFIER |
| 12 | 117715891 | AX-170190269 | 0 | A | G | 0.14 | -3.60E-02 | 1.93E-01 | 2.68E+00 | 2.75E+00 | 8.49E-01 | no | Rapgef5 | intron_variant | MODIFIER |
| 12 | 117719114 | AX-170007604 | 0 | C | T | 0.14 | -3.60E-02 | 1.93E-01 | 2.68E+00 | 2.75E+00 | 8.49E-01 | no | Rapgef5 | splice_region_variant,intron_variant | LOW |
| 12 | 117719848 | AX-97826566 | 0 | G | A | 0.14 | -3.60E-02 | 1.93E-01 | 2.68E+00 | 2.75E+00 | 8.49E-01 | no | Rapgef5 | intron_variant | MODIFIER |
| 12 | 117756459 | AX-170050355 | 0 | C | T | 0.14 | -3.60E-02 | 1.93E-01 | 2.68E+00 | 2.75E+00 | 8.49E-01 | no | Rapgef5 | 3_prime_UTR_variant | MODIFIER |
| 12 | 117772929 | AX-170362848 | 0 | C | A | 0.14 | -3.60E-02 | 1.93E-01 | 2.68E+00 | 2.75E+00 | 8.49E-01 | no | | regulatory_region_variant | MODIFIER |
| 12 | 117776514 | AX-170007609 | 0 | G | A | 0.14 | -3.60E-02 | 1.93E-01 | 2.68E+00 | 2.75E+00 | 8.49E-01 | no | | intergenic_variant | MODIFIER |
| 12 | 117778268 | AX-97792487 | 0 | A | G | 0.14 | -3.60E-02 | 1.93E-01 | 2.68E+00 | 2.75E+00 | 8.49E-01 | no | | intergenic_variant | MODIFIER |
| 12 | 117782531 | AX-170308457 | 0 | T | C | 0.14 | -3.60E-02 | 1.93E-01 | 2.68E+00 | 2.75E+00 | 8.49E-01 | no | | intergenic_variant | MODIFIER |
| 12 | 117820385 | AX-170007603 | 0 | A | G | 0.14 | -3.60E-02 | 1.93E-01 | 2.68E+00 | 2.75E+00 | 8.49E-01 | no | Cdca7l | intron_variant,non_coding_transcript_variant | MODIFIER |
| 12 | 117822030 | AX-97822437 | 0 | T | C | 0.14 | -3.60E-02 | 1.93E-01 | 2.68E+00 | 2.75E+00 | 8.49E-01 | no | Cdca7l | intron_variant,non_coding_transcript_variant | MODIFIER |

| | | | | | | | | | | | | | | | |
|---|---|---|---|---|---|---|---|---|---|---|---|---|---|---|---|
| 12 | 117823122 | AX-170408624 | 0 | C | T | 0.14 | -3.60E-02 | 1.93E-01 | 2.68E+00 | 2.75E+00 | 8.49E-01 | no | Cdca7l | intron_variant,non_coding_transcript_variant | MODIFIER |
| 12 | 117829907 | AX-170153068 | 0 | C | T | 0.14 | -3.60E-02 | 1.93E-01 | 2.68E+00 | 2.75E+00 | 8.49E-01 | no | Gm20613 | downstream_gene_variant | MODIFIER |
| 12 | 117865006 | AX-170271895 | 0 | T | G | 0.14 | -3.60E-02 | 1.93E-01 | 2.68E+00 | 2.75E+00 | 8.49E-01 | no | Cdca7l | intron_variant | MODIFIER |
| 12 | 117888534 | AX-170257374 | 0 | C | T | 0.14 | -3.60E-02 | 1.93E-01 | 2.68E+00 | 2.75E+00 | 8.49E-01 | no | Dnah11 | intron_variant | MODIFIER |
| 12 | 117894251 | AX-97780513 | 0 | T | C | 0.14 | -3.60E-02 | 1.93E-01 | 2.68E+00 | 2.75E+00 | 8.49E-01 | no | Dnah11 | intron_variant | MODIFIER |
| 12 | 117894665 | AX-170373153 | 0 | C | T | 0.14 | -3.60E-02 | 1.93E-01 | 2.68E+00 | 2.75E+00 | 8.49E-01 | no | Dnah11 | intron_variant | MODIFIER |
| 12 | 117895113 | AX-170098839 | 0 | A | G | 0.14 | -3.60E-02 | 1.93E-01 | 2.68E+00 | 2.75E+00 | 8.49E-01 | no | Dnah11 | intron_variant | MODIFIER |
| 12 | 117899565 | AX-170154326 | 0 | T | G | 0.14 | -3.60E-02 | 1.93E-01 | 2.68E+00 | 2.75E+00 | 8.49E-01 | no | Dnah11 | intron_variant | MODIFIER |
| 12 | 117902303 | AX-170439277 | 0 | G | A | 0.14 | -3.60E-02 | 1.93E-01 | 2.68E+00 | 2.75E+00 | 8.49E-01 | no | Dnah11 | intron_variant | MODIFIER |
| 12 | 117992258 | AX-170391414 | 0 | C | T | 0.14 | -3.60E-02 | 1.93E-01 | 2.68E+00 | 2.75E+00 | 8.49E-01 | no | Dnah11 | intron_variant | MODIFIER |
| 12 | 118012587 | AX-97822415 | 0 | G | T | 0.14 | -3.60E-02 | 1.93E-01 | 2.68E+00 | 2.75E+00 | 8.49E-01 | no | Dnah11 | intron_variant | MODIFIER |
| 12 | 118028533 | AX-170534125 | 0 | T | C | 0.14 | -3.60E-02 | 1.93E-01 | 2.68E+00 | 2.75E+00 | 8.49E-01 | no | Dnah11 | intron_variant | MODIFIER |
| 12 | 118051033 | AX-170371111 | 0 | C | T | 0.14 | -3.60E-02 | 1.93E-01 | 2.68E+00 | 2.75E+00 | 8.49E-01 | no | Dnah11 | intron_variant | MODIFIER |
| 12 | 118051961 | AX-170514032 | 0 | T | A | 0.14 | -3.60E-02 | 1.93E-01 | 2.68E+00 | 2.75E+00 | 8.49E-01 | no | Dnah11 | intron_variant | MODIFIER |
| 12 | 118075812 | AX-97782088 | 0 | A | C | 0.14 | -3.60E-02 | 1.93E-01 | 2.68E+00 | 2.75E+00 | 8.49E-01 | no | Dnah11 | intron_variant | MODIFIER |
| 12 | 118079368 | AX-170241806 | 0 | A | G | 0.14 | -3.60E-02 | 1.93E-01 | 2.68E+00 | 2.75E+00 | 8.49E-01 | no | Dnah11 | intron_variant | MODIFIER |
| 12 | 118094680 | AX-97848723 | 0 | C | T | 0.14 | -3.60E-02 | 1.93E-01 | 2.68E+00 | 2.75E+00 | 8.49E-01 | no | Dnah11 | intron_variant | MODIFIER |
| 12 | 118095279 | AX-170321129 | 0 | T | C | 0.14 | -3.60E-02 | 1.93E-01 | 2.68E+00 | 2.75E+00 | 8.49E-01 | no | Dnah11 | intron_variant | MODIFIER |
| 12 | 118099584 | AX-170244485 | 0 | T | C | 0.14 | -3.60E-02 | 1.93E-01 | 2.68E+00 | 2.75E+00 | 8.49E-01 | no | Dnah11 | intron_variant | MODIFIER |
| 12 | 118128574 | AX-97822434 | 0 | A | G | 0.14 | -3.60E-02 | 1.93E-01 | 2.68E+00 | 2.75E+00 | 8.49E-01 | no | Dnah11 | intron_variant | MODIFIER |
| 12 | 118128653 | AX-170288354 | 0 | T | A | 0.14 | -3.60E-02 | 1.93E-01 | 2.68E+00 | 2.75E+00 | 8.49E-01 | no | Dnah11 | intron_variant | MODIFIER |
| 12 | 118128975 | AX-170419434 | 0 | T | C | 0.14 | -3.60E-02 | 1.93E-01 | 2.68E+00 | 2.75E+00 | 8.49E-01 | no | Dnah11 | intron_variant | MODIFIER |
| 12 | 118138614 | AX-97805886 | 0 | A | G | 0.14 | -3.60E-02 | 1.93E-01 | 2.68E+00 | 2.75E+00 | 8.49E-01 | no | Dnah11 | intron_variant | MODIFIER |
| 12 | 118174267 | AX-97822435 | 0 | C | T | 0.14 | -3.60E-02 | 1.93E-01 | 2.68E+00 | 2.75E+00 | 8.49E-01 | no | Dnah11 | intron_variant | MODIFIER |
| 12 | 118188575 | AX-170375701 | 0 | A | G | 0.14 | -3.60E-02 | 1.93E-01 | 2.68E+00 | 2.75E+00 | 8.49E-01 | no | Dnah11 | intron_variant | MODIFIER |
| 12 | 118506263 | AX-168383536 | 0 | C | T | 0.14 | -3.60E-02 | 1.93E-01 | 2.68E+00 | 2.75E+00 | 8.49E-01 | no | Gm25577 | upstream_gene_variant | MODIFIER |
| 12 | 118583885 | AX-97799339 | 0 | C | A | 0.14 | -3.60E-02 | 1.93E-01 | 2.68E+00 | 2.75E+00 | 8.49E-01 | no | | regulatory_region_variant | MODIFIER |
| 12 | 118793555 | AX-170374624 | 0 | C | T | 0.14 | -3.60E-02 | 1.93E-01 | 2.68E+00 | 2.75E+00 | 8.49E-01 | no | Gm6171 | non_coding_transcript_exon_variant | MODIFIER |
| 12 | 118798745 | AX-170486695 | 0 | A | C | 0.14 | -3.60E-02 | 1.93E-01 | 2.68E+00 | 2.75E+00 | 8.49E-01 | no | Gm6171 | upstream_gene_variant | MODIFIER |
| 12 | 118799440 | AX-170092634 | 0 | G | A | 0.14 | -3.60E-02 | 1.93E-01 | 2.68E+00 | 2.75E+00 | 8.49E-01 | no | | intergenic_variant | MODIFIER |
| 12 | 118810038 | AX-170007622 | 0 | T | C | 0.14 | -3.60E-02 | 1.93E-01 | 2.68E+00 | 2.75E+00 | 8.49E-01 | no | | intergenic_variant | MODIFIER |
| 12 | 118828070 | AX-97822668 | 0 | G | A | 0.14 | -3.60E-02 | 1.93E-01 | 2.68E+00 | 2.75E+00 | 8.49E-01 | no | | intergenic_variant | MODIFIER |
| 12 | 118829549 | AX-170232703 | 0 | T | C | 0.14 | -3.60E-02 | 1.93E-01 | 2.68E+00 | 2.75E+00 | 8.49E-01 | no | | intergenic_variant | MODIFIER |
| 12 | 118844440 | AX-97856298 | 0 | C | T | 0.14 | -3.60E-02 | 1.93E-01 | 2.68E+00 | 2.75E+00 | 8.49E-01 | no | Sp8 | upstream_gene_variant | MODIFIER |
| 12 | 118858269 | AX-97799395 | 0 | T | C | 0.14 | -3.60E-02 | 1.93E-01 | 2.68E+00 | 2.75E+00 | 8.49E-01 | no | | intergenic_variant | MODIFIER |
| 12 | 118910993 | AX-97822463 | 0 | G | A | 0.14 | -3.60E-02 | 1.93E-01 | 2.68E+00 | 2.75E+00 | 8.49E-01 | no | Abcb5 | intron_variant | MODIFIER |
| 12 | 118911331 | AX-170393815 | 0 | A | G | 0.14 | -3.60E-02 | 1.93E-01 | 2.68E+00 | 2.75E+00 | 8.49E-01 | no | Abcb5 | intron_variant | MODIFIER |
| 12 | 118950258 | AX-97792162 | 0 | G | A | 0.14 | -3.60E-02 | 1.93E-01 | 2.68E+00 | 2.75E+00 | 8.49E-01 | no | Abcb5 | intron_variant | MODIFIER |
| 1 | 75395641 | AX-170292825 | 2 | G | A | 0.127 | 3.96E-02 | 2.09E-01 | 2.68E+00 | 2.73E+00 | 8.49E-01 | no | Speg | intron_variant | MODIFIER |
| 8 | 53968450 | AX-97842200 | 0 | T | C | 0.158 | -2.80E-02 | 1.39E-01 | 2.63E+00 | 2.69E+00 | 8.49E-01 | no | | intergenic_variant | MODIFIER |
| 5 | 6552233 | AX-170221839 | 0 | A | G | 0.421 | 1.10E-02 | 6.35E-02 | 2.64E+00 | 2.78E+00 | 8.49E-01 | no | | intergenic_variant | MODIFIER |
| 6 | 102652962 | AX-170234277 | 1 | G | A | 0.054 | 5.61E-02 | 2.96E-01 | 2.67E+00 | 2.73E+00 | 8.49E-01 | no | | intergenic_variant | MODIFIER |
| 6 | 109932174 | AX-97795343 | 1 | A | G | 0.054 | 5.61E-02 | 2.96E-01 | 2.67E+00 | 2.73E+00 | 8.49E-01 | no | | intergenic_variant | MODIFIER |
| 6 | 120445998 | AX-97785022 | 1 | G | A | 0.054 | 5.61E-02 | 2.96E-01 | 2.67E+00 | 2.73E+00 | 8.49E-01 | no | Kdm5a | downstream_gene_variant | MODIFIER |
| 8 | 57308208 | AX-97830131 | 1 | G | A | 0.143 | 2.60E-02 | 1.44E-01 | 2.73E+00 | 2.81E+00 | 8.49E-01 | no | Hand2os1 | intron_variant,non_coding_transcript_variant | MODIFIER |
| 6 | 101161674 | AX-170426517 | 0 | G | A | 0.07 | -5.19E-02 | 2.76E-01 | 2.70E+00 | 2.74E+00 | 8.49E-01 | no | Pdzrn3 | intron_variant | MODIFIER |
| 6 | 138772300 | AX-170440567 | 0 | A | T | 0.07 | -5.19E-02 | 2.76E-01 | 2.70E+00 | 2.74E+00 | 8.49E-01 | no | | intergenic_variant | MODIFIER |
| 13 | 42655894 | AX-170220609 | 1 | A | G | 0.482 | -1.12E-02 | 6.29E-02 | 2.67E+00 | 2.78E+00 | 8.49E-01 | no | | intergenic_variant | MODIFIER |
| 12 | 103007771 | AX-168382859 | 1 | C | T | 0.134 | -3.57E-02 | 1.93E-01 | 2.68E+00 | 2.75E+00 | 8.49E-01 | no | Unc79 | intron_variant | MODIFIER |
| 12 | 114840273 | AX-170087340 | 1 | G | T | 0.143 | -3.53E-02 | 1.90E-01 | 2.68E+00 | 2.75E+00 | 8.49E-01 | no | Ighv1-33 | downstream_gene_variant | MODIFIER |
| 7 | 57634023 | AX-170326939 | 0 | A | C | 0.43 | -1.78E-02 | 9.81E-02 | 2.69E+00 | 2.77E+00 | 8.49E-01 | no | Gabrb3 | intron_variant | MODIFIER |
| 8 | 82967506 | AX-170414793 | 1 | C | A | 0.205 | 2.41E-02 | 1.28E-01 | 2.67E+00 | 2.74E+00 | 8.49E-01 | no | Rnf150 | intron_variant | MODIFIER |
| 2 | 120921146 | AX-170510767 | 1 | C | T | 0.143 | 4.16E-02 | 2.20E-01 | 2.70E+00 | 2.73E+00 | 8.49E-01 | no | Ubr1 | intron_variant | MODIFIER |
| 9 | 76264932 | AX-97853558 | 1 | T | A | 0.188 | 2.57E-02 | 1.38E-01 | 2.74E+00 | 2.77E+00 | 8.49E-01 | no | Hcrtr2 | intron_variant | MODIFIER |

| | | | | | | | | | | | | | | | |
|---|---|---|---|---|---|---|---|---|---|---|---|---|---|---|---|
| 8 | 94785762 | AX-170322508 | 0 | T | C | 0.219 | 2.05E-02 | 1.11E-01 | 2.68E+00 | 2.76E+00 | 8.49E-01 | no | Cx3cl1 | downstream_gene_variant | MODIFIER |
| 3 | 36876456 | AX-170511952 | 0 | C | T | 0.193 | 1.62E-02 | 8.88E-02 | 2.66E+00 | 2.76E+00 | 8.49E-01 | no | 4932438A13Rik | intron_variant | MODIFIER |
| 1 | 116435350 | AX-170145674 | 1 | A | G | 0.17 | 3.88E-02 | 1.94E-01 | 2.62E+00 | 2.70E+00 | 8.49E-01 | no | Cntnap5a | intron_variant | MODIFIER |
| 9 | 32885420 | AX-170400582 | 0 | G | A | 0.43 | -1.51E-02 | 8.38E-02 | 2.71E+00 | 2.78E+00 | 8.49E-01 | no | | intergenic_variant | MODIFIER |
| 9 | 34458844 | AX-97804678 | 0 | C | T | 0.43 | -1.51E-02 | 8.38E-02 | 2.71E+00 | 2.78E+00 | 8.49E-01 | no | Gm47858 | upstream_gene_variant | MODIFIER |
| 11 | 33031766 | AX-170036461 | 1 | C | G | 0.107 | 3.66E-02 | 1.91E-01 | 2.65E+00 | 2.72E+00 | 8.49E-01 | no | | intergenic_variant | MODIFIER |
| 6 | 104907190 | AX-170528527 | 0 | G | T | 0.061 | 4.22E-02 | 2.30E-01 | 2.68E+00 | 2.76E+00 | 8.49E-01 | no | | intergenic_variant | MODIFIER |
| 6 | 139933642 | AX-170028388 | 0 | T | G | 0.061 | 4.22E-02 | 2.30E-01 | 2.68E+00 | 2.76E+00 | 8.49E-01 | no | Pik3c2g | intron_variant | MODIFIER |
| 2 | 101191321 | AX-170465571 | 0 | T | G | 0.061 | 4.11E-02 | 2.18E-01 | 2.64E+00 | 2.73E+00 | 8.49E-01 | no | | intergenic_variant | MODIFIER |
| 2 | 108221728 | AX-170322123 | 0 | C | G | 0.061 | 4.11E-02 | 2.18E-01 | 2.64E+00 | 2.73E+00 | 8.49E-01 | no | Gm13912 | intron_variant,non_coding_transcript_variant | MODIFIER |
| 2 | 101320529 | AX-170018486 | 0 | T | G | 0.061 | 4.00E-02 | 2.17E-01 | 2.67E+00 | 2.75E+00 | 8.50E-01 | no | | intergenic_variant | MODIFIER |
| 2 | 106231039 | AX-170490932 | 0 | G | A | 0.061 | 4.00E-02 | 2.17E-01 | 2.67E+00 | 2.75E+00 | 8.50E-01 | no | Dcdc5 | intron_variant,non_coding_transcript_variant | MODIFIER |
| 7 | 119043392 | AX-170307541 | 1 | C | T | 0.402 | 1.76E-02 | 9.23E-02 | 2.67E+00 | 2.72E+00 | 8.50E-01 | no | | intergenic_variant | MODIFIER |
| 9 | 113660609 | AX-170315767 | 1 | C | T | 0.134 | -1.95E-02 | 1.04E-01 | 2.65E+00 | 2.74E+00 | 8.50E-01 | no | Pdcd6ip | intron_variant | MODIFIER |
| 11 | 104691933 | AX-170512298 | 0 | A | G | 0.053 | 3.38E-02 | 1.88E-01 | 2.63E+00 | 2.76E+00 | 8.50E-01 | no | Gm11639 | intron_variant | MODIFIER |
| 18 | 57474791 | AX-170504987 | 1 | T | C | 0.411 | 1.54E-02 | 8.51E-02 | 2.73E+00 | 2.80E+00 | 8.50E-01 | no | Ctxn3 | intron_variant | MODIFIER |
| 9 | 80494643 | AX-170335816 | 1 | A | C | 0.196 | 2.39E-02 | 1.30E-01 | 2.72E+00 | 2.77E+00 | 8.50E-01 | no | | intergenic_variant | MODIFIER |
| 13 | 38205492 | AX-170504834 | 0 | G | A | 0.175 | -2.34E-02 | 1.12E-01 | 2.59E+00 | 2.68E+00 | 8.50E-01 | no | Snrnp48 | intron_variant | MODIFIER |
| 13 | 38205564 | AX-170220760 | 0 | T | C | 0.175 | -2.34E-02 | 1.12E-01 | 2.59E+00 | 2.68E+00 | 8.50E-01 | no | Snrnp48 | intron_variant | MODIFIER |
| 13 | 38205903 | AX-97854848 | 0 | C | T | 0.175 | -2.34E-02 | 1.12E-01 | 2.59E+00 | 2.68E+00 | 8.50E-01 | no | Snrnp48 | intron_variant | MODIFIER |
| 13 | 38220021 | AX-170087410 | 0 | C | T | 0.175 | -2.34E-02 | 1.12E-01 | 2.59E+00 | 2.68E+00 | 8.50E-01 | no | Snrnp48 | intron_variant | MODIFIER |
| 13 | 38225826 | AX-170136641 | 0 | C | T | 0.175 | -2.34E-02 | 1.12E-01 | 2.59E+00 | 2.68E+00 | 8.50E-01 | no | Snrnp48 | intron_variant | MODIFIER |
| 13 | 38232399 | AX-170456987 | 0 | T | C | 0.175 | -2.34E-02 | 1.12E-01 | 2.59E+00 | 2.68E+00 | 8.50E-01 | no | Snrnp48 | downstream_gene_variant | MODIFIER |
| 13 | 38244342 | AX-97830469 | 0 | C | T | 0.175 | -2.34E-02 | 1.12E-01 | 2.59E+00 | 2.68E+00 | 8.50E-01 | no | | intergenic_variant | MODIFIER |
| 13 | 38257658 | AX-170430240 | 0 | T | C | 0.175 | -2.34E-02 | 1.12E-01 | 2.59E+00 | 2.68E+00 | 8.50E-01 | no | | intergenic_variant | MODIFIER |
| 13 | 38261165 | AX-170239199 | 0 | G | A | 0.175 | -2.34E-02 | 1.12E-01 | 2.59E+00 | 2.68E+00 | 8.50E-01 | no | | intergenic_variant | MODIFIER |
| 13 | 38282002 | AX-97806158 | 0 | G | A | 0.175 | -2.34E-02 | 1.12E-01 | 2.59E+00 | 2.68E+00 | 8.50E-01 | no | | intergenic_variant | MODIFIER |
| 13 | 38293619 | AX-170175562 | 0 | A | G | 0.175 | -2.34E-02 | 1.12E-01 | 2.59E+00 | 2.68E+00 | 8.50E-01 | no | | intergenic_variant | MODIFIER |
| 13 | 38294878 | AX-97787487 | 0 | G | T | 0.175 | -2.34E-02 | 1.12E-01 | 2.59E+00 | 2.68E+00 | 8.50E-01 | no | | intergenic_variant | MODIFIER |
| 13 | 38524596 | AX-168419935 | 0 | T | C | 0.175 | -2.34E-02 | 1.12E-01 | 2.59E+00 | 2.68E+00 | 8.50E-01 | no | Txndc5 | intron_variant | MODIFIER |
| 13 | 38573148 | AX-97850543 | 0 | T | C | 0.175 | -2.34E-02 | 1.12E-01 | 2.59E+00 | 2.68E+00 | 8.50E-01 | no | | intergenic_variant | MODIFIER |
| 13 | 38622879 | AX-170008147 | 0 | A | T | 0.175 | -2.34E-02 | 1.12E-01 | 2.59E+00 | 2.68E+00 | 8.50E-01 | no | Bloc1s5 | intron_variant | MODIFIER |
| 13 | 38691441 | AX-168391811 | 0 | T | C | 0.175 | -2.34E-02 | 1.12E-01 | 2.59E+00 | 2.68E+00 | 8.50E-01 | no | | intergenic_variant | MODIFIER |
| 13 | 38705643 | AX-170008145 | 0 | T | C | 0.175 | -2.34E-02 | 1.12E-01 | 2.59E+00 | 2.68E+00 | 8.50E-01 | no | Gm46392 | upstream_gene_variant | MODIFIER |
| 13 | 38740194 | AX-170137700 | 0 | C | G | 0.175 | -2.34E-02 | 1.12E-01 | 2.59E+00 | 2.68E+00 | 8.50E-01 | no | Gm46392 | intron_variant,non_coding_transcript_variant | MODIFIER |
| 13 | 38750081 | AX-97823374 | 0 | C | T | 0.175 | -2.34E-02 | 1.12E-01 | 2.59E+00 | 2.68E+00 | 8.50E-01 | no | Gm46392 | intron_variant,non_coding_transcript_variant | MODIFIER |
| 13 | 38761575 | AX-170377935 | 0 | A | G | 0.175 | -2.34E-02 | 1.12E-01 | 2.59E+00 | 2.68E+00 | 8.50E-01 | no | Gm46392 | intron_variant,non_coding_transcript_variant | MODIFIER |
| 13 | 38761850 | AX-170415410 | 0 | C | T | 0.175 | -2.34E-02 | 1.12E-01 | 2.59E+00 | 2.68E+00 | 8.50E-01 | no | Gm46392 | intron_variant,non_coding_transcript_variant | MODIFIER |
| 5 | 97422965 | AX-97783204 | 0 | C | T | 0.439 | 2.00E-02 | 1.06E-01 | 2.67E+00 | 2.73E+00 | 8.50E-01 | no | 4930467D21Rik | intron_variant,non_coding_transcript_variant | MODIFIER |
| 5 | 97433069 | AX-170510592 | 0 | G | A | 0.439 | 2.00E-02 | 1.06E-01 | 2.67E+00 | 2.73E+00 | 8.50E-01 | no | 4930467D21Rik | intron_variant,non_coding_transcript_variant | MODIFIER |
| 16 | 88602783 | AX-170530082 | 0 | T | C | 0.465 | -1.47E-02 | 7.95E-02 | 2.67E+00 | 2.75E+00 | 8.50E-01 | no | | intergenic_variant | MODIFIER |
| 16 | 88602889 | AX-97833476 | 0 | T | C | 0.465 | -1.47E-02 | 7.95E-02 | 2.67E+00 | 2.75E+00 | 8.50E-01 | no | | intergenic_variant | MODIFIER |
| 16 | 88629198 | AX-170267948 | 0 | C | G | 0.465 | -1.47E-02 | 7.95E-02 | 2.67E+00 | 2.75E+00 | 8.50E-01 | no | 2310079G19Rik | upstream_gene_variant | MODIFIER |
| 16 | 88631513 | AX-170460067 | 0 | A | G | 0.465 | -1.47E-02 | 7.95E-02 | 2.67E+00 | 2.75E+00 | 8.50E-01 | no | 2310079G19Rik | upstream_gene_variant | MODIFIER |
| 16 | 88632168 | AX-170115856 | 0 | C | T | 0.465 | -1.47E-02 | 7.95E-02 | 2.67E+00 | 2.75E+00 | 8.50E-01 | no | 2310079G19Rik | upstream_gene_variant | MODIFIER |
| 16 | 88636470 | AX-170093557 | 0 | T | C | 0.465 | -1.47E-02 | 7.95E-02 | 2.67E+00 | 2.75E+00 | 8.50E-01 | no | Gm49688 | intron_variant,non_coding_transcript_variant | MODIFIER |
| 16 | 88648504 | AX-170508218 | 0 | G | A | 0.465 | -1.47E-02 | 7.95E-02 | 2.67E+00 | 2.75E+00 | 8.50E-01 | no | Krtap26-1 | upstream_gene_variant | MODIFIER |
| 16 | 88665064 | AX-97791483 | 0 | G | A | 0.465 | -1.47E-02 | 7.95E-02 | 2.67E+00 | 2.75E+00 | 8.50E-01 | no | | intergenic_variant | MODIFIER |
| 16 | 88665340 | AX-170058051 | 0 | G | A | 0.465 | -1.47E-02 | 7.95E-02 | 2.67E+00 | 2.75E+00 | 8.50E-01 | no | | intergenic_variant | MODIFIER |
| 16 | 88665936 | AX-170432940 | 0 | C | A | 0.465 | -1.47E-02 | 7.95E-02 | 2.67E+00 | 2.75E+00 | 8.50E-01 | no | | intergenic_variant | MODIFIER |
| 16 | 88779391 | AX-170333245 | 0 | T | A | 0.465 | -1.47E-02 | 7.95E-02 | 2.67E+00 | 2.75E+00 | 8.50E-01 | no | Gm5965 | downstream_gene_variant | MODIFIER |
| 16 | 88858560 | AX-170198823 | 0 | G | C | 0.465 | -1.47E-02 | 7.95E-02 | 2.67E+00 | 2.75E+00 | 8.50E-01 | no | | intergenic_variant | MODIFIER |
| 16 | 88865401 | AX-170220775 | 0 | T | G | 0.465 | -1.47E-02 | 7.95E-02 | 2.67E+00 | 2.75E+00 | 8.50E-01 | no | Krtap19-1 | downstream_gene_variant | MODIFIER |
| 16 | 88866615 | AX-170497896 | 0 | A | G | 0.465 | -1.47E-02 | 7.95E-02 | 2.67E+00 | 2.75E+00 | 8.50E-01 | no | Krtap19-1 | downstream_gene_variant | MODIFIER |

| | | | | | | | | | | | | | | | |
|---|---|---|---|---|---|---|---|---|---|---|---|---|---|---|---|
| 16 | 88896900 | AX-170446229 | 0 | G | A | 0.465 | -1.47E-02 | 7.95E-02 | 2.67E+00 | 2.75E+00 | 8.50E-01 | no | Krtap19-5 | upstream_gene_variant | MODIFIER |
| 16 | 88897574 | AX-170426825 | 0 | C | T | 0.465 | -1.47E-02 | 7.95E-02 | 2.67E+00 | 2.75E+00 | 8.50E-01 | no | Krtap19-5 | upstream_gene_variant | MODIFIER |
| 16 | 88914460 | AX-170246528 | 0 | T | C | 0.465 | -1.47E-02 | 7.95E-02 | 2.67E+00 | 2.75E+00 | 8.50E-01 | no | | intergenic_variant | MODIFIER |
| 16 | 88916008 | AX-170206565 | 0 | T | C | 0.465 | -1.47E-02 | 7.95E-02 | 2.67E+00 | 2.75E+00 | 8.50E-01 | no | | intergenic_variant | MODIFIER |
| 16 | 88916393 | AX-97850958 | 0 | A | G | 0.465 | -1.47E-02 | 7.95E-02 | 2.67E+00 | 2.75E+00 | 8.50E-01 | no | | intergenic_variant | MODIFIER |
| 16 | 88919713 | AX-170499891 | 0 | C | T | 0.465 | -1.47E-02 | 7.95E-02 | 2.67E+00 | 2.75E+00 | 8.50E-01 | no | Krtap19-9a | downstream_gene_variant | MODIFIER |
| 16 | 88933809 | AX-170013616 | 0 | G | A | 0.465 | -1.47E-02 | 7.95E-02 | 2.67E+00 | 2.75E+00 | 8.50E-01 | no | Krtap19-9b | upstream_gene_variant | MODIFIER |
| 16 | 88934246 | AX-97815554 | 0 | T | G | 0.465 | -1.47E-02 | 7.95E-02 | 2.67E+00 | 2.75E+00 | 8.50E-01 | no | Krtap19-9b | upstream_gene_variant | MODIFIER |
| 16 | 88944263 | AX-170285639 | 0 | A | T | 0.465 | -1.47E-02 | 7.95E-02 | 2.67E+00 | 2.75E+00 | 8.50E-01 | no | | intergenic_variant | MODIFIER |
| 16 | 88982852 | AX-170013618 | 0 | G | A | 0.465 | -1.47E-02 | 7.95E-02 | 2.67E+00 | 2.75E+00 | 8.50E-01 | no | | intergenic_variant | MODIFIER |
| 16 | 88984710 | AX-170013613 | 0 | G | A | 0.465 | -1.47E-02 | 7.95E-02 | 2.67E+00 | 2.75E+00 | 8.50E-01 | no | | intergenic_variant | MODIFIER |
| 16 | 88991266 | AX-170088094 | 0 | C | T | 0.465 | -1.47E-02 | 7.95E-02 | 2.67E+00 | 2.75E+00 | 8.50E-01 | no | | intergenic_variant | MODIFIER |
| 16 | 88992072 | AX-170219693 | 0 | T | C | 0.465 | -1.47E-02 | 7.95E-02 | 2.67E+00 | 2.75E+00 | 8.50E-01 | no | | intergenic_variant | MODIFIER |
| 16 | 88992577 | AX-170013615 | 0 | C | T | 0.465 | -1.47E-02 | 7.95E-02 | 2.67E+00 | 2.75E+00 | 8.50E-01 | no | | intergenic_variant | MODIFIER |
| 16 | 89057447 | AX-170088095 | 0 | T | G | 0.465 | -1.47E-02 | 7.95E-02 | 2.67E+00 | 2.75E+00 | 8.50E-01 | no | | intergenic_variant | MODIFIER |
| 16 | 89125370 | AX-170013612 | 0 | C | T | 0.465 | -1.47E-02 | 7.95E-02 | 2.67E+00 | 2.75E+00 | 8.50E-01 | no | | intergenic_variant | MODIFIER |
| 16 | 89144713 | AX-170286646 | 0 | A | G | 0.465 | -1.47E-02 | 7.95E-02 | 2.67E+00 | 2.75E+00 | 8.50E-01 | no | Gm6358 | downstream_gene_variant | MODIFIER |
| 16 | 89144865 | AX-170506675 | 0 | C | G | 0.465 | -1.47E-02 | 7.95E-02 | 2.67E+00 | 2.75E+00 | 8.50E-01 | no | Gm6358 | downstream_gene_variant | MODIFIER |
| 16 | 89157777 | AX-170058059 | 0 | A | C | 0.465 | -1.47E-02 | 7.95E-02 | 2.67E+00 | 2.75E+00 | 8.50E-01 | no | Gm9789 | upstream_gene_variant | MODIFIER |
| 16 | 89161545 | AX-170135842 | 0 | G | A | 0.465 | -1.47E-02 | 7.95E-02 | 2.67E+00 | 2.75E+00 | 8.50E-01 | no | Gm9789 | downstream_gene_variant | MODIFIER |
| 16 | 89169439 | AX-170489701 | 0 | T | A | 0.465 | -1.47E-02 | 7.95E-02 | 2.67E+00 | 2.75E+00 | 8.50E-01 | no | Gm7735 | upstream_gene_variant | MODIFIER |
| 16 | 89296309 | AX-170264183 | 0 | G | A | 0.465 | -1.47E-02 | 7.95E-02 | 2.67E+00 | 2.75E+00 | 8.50E-01 | no | | intergenic_variant | MODIFIER |
| 16 | 89296650 | AX-170407605 | 0 | G | A | 0.465 | -1.47E-02 | 7.95E-02 | 2.67E+00 | 2.75E+00 | 8.50E-01 | no | | intergenic_variant | MODIFIER |
| 16 | 89297484 | AX-170341303 | 0 | A | G | 0.465 | -1.47E-02 | 7.95E-02 | 2.67E+00 | 2.75E+00 | 8.50E-01 | no | | intergenic_variant | MODIFIER |
| 16 | 89309376 | AX-170246178 | 0 | C | T | 0.465 | -1.47E-02 | 7.95E-02 | 2.67E+00 | 2.75E+00 | 8.50E-01 | no | | intergenic_variant | MODIFIER |
| 16 | 89309603 | AX-170444093 | 0 | G | A | 0.465 | -1.47E-02 | 7.95E-02 | 2.67E+00 | 2.75E+00 | 8.50E-01 | no | | intergenic_variant | MODIFIER |
| 16 | 89339277 | AX-170486530 | 0 | A | G | 0.465 | -1.47E-02 | 7.95E-02 | 2.67E+00 | 2.75E+00 | 8.50E-01 | no | | intergenic_variant | MODIFIER |
| 16 | 89352505 | AX-97820897 | 0 | G | T | 0.465 | -1.47E-02 | 7.95E-02 | 2.67E+00 | 2.75E+00 | 8.50E-01 | no | | intergenic_variant | MODIFIER |
| 16 | 89359786 | AX-97832731 | 0 | G | A | 0.465 | -1.47E-02 | 7.95E-02 | 2.67E+00 | 2.75E+00 | 8.50E-01 | no | | intergenic_variant | MODIFIER |
| 16 | 89365385 | AX-170524247 | 0 | A | G | 0.465 | -1.47E-02 | 7.95E-02 | 2.67E+00 | 2.75E+00 | 8.50E-01 | no | | intergenic_variant | MODIFIER |
| 16 | 89449861 | AX-170088096 | 0 | C | A | 0.465 | -1.47E-02 | 7.95E-02 | 2.67E+00 | 2.75E+00 | 8.50E-01 | no | Gm23655 | downstream_gene_variant | MODIFIER |
| 16 | 89452923 | AX-170013623 | 0 | T | C | 0.465 | -1.47E-02 | 7.95E-02 | 2.67E+00 | 2.75E+00 | 8.50E-01 | no | Gm23655 | upstream_gene_variant | MODIFIER |
| 16 | 89464376 | AX-170149950 | 0 | C | T | 0.465 | -1.47E-02 | 7.95E-02 | 2.67E+00 | 2.75E+00 | 8.50E-01 | no | | intergenic_variant | MODIFIER |
| 16 | 89487256 | AX-170013625 | 0 | G | C | 0.465 | -1.47E-02 | 7.95E-02 | 2.67E+00 | 2.75E+00 | 8.50E-01 | no | Krtap8-1 | downstream_gene_variant | MODIFIER |
| 16 | 89487274 | AX-170439413 | 0 | T | C | 0.465 | -1.47E-02 | 7.95E-02 | 2.67E+00 | 2.75E+00 | 8.50E-01 | no | Krtap8-1 | downstream_gene_variant | MODIFIER |
| 16 | 89487370 | AX-170013624 | 0 | G | A | 0.465 | -1.47E-02 | 7.95E-02 | 2.67E+00 | 2.75E+00 | 8.50E-01 | no | Krtap8-1 | downstream_gene_variant | MODIFIER |
| 16 | 89497562 | AX-170058066 | 0 | C | T | 0.465 | -1.47E-02 | 7.95E-02 | 2.67E+00 | 2.75E+00 | 8.50E-01 | no | | intergenic_variant | MODIFIER |
| 16 | 89499654 | AX-170058063 | 0 | A | G | 0.465 | -1.47E-02 | 7.95E-02 | 2.67E+00 | 2.75E+00 | 8.50E-01 | no | | intergenic_variant | MODIFIER |
| 16 | 90039513 | AX-170013627 | 0 | C | T | 0.465 | -1.47E-02 | 7.95E-02 | 2.67E+00 | 2.75E+00 | 8.50E-01 | no | Tiam1 | intron_variant | MODIFIER |
| 16 | 90421052 | AX-97828032 | 0 | C | T | 0.465 | -1.47E-02 | 7.95E-02 | 2.67E+00 | 2.75E+00 | 8.50E-01 | no | Hunk | intron_variant | MODIFIER |
| 16 | 90427305 | AX-170527237 | 0 | A | G | 0.465 | -1.47E-02 | 7.95E-02 | 2.67E+00 | 2.75E+00 | 8.50E-01 | no | Hunk | intron_variant | MODIFIER |
| 16 | 90450338 | AX-170458695 | 0 | G | A | 0.465 | -1.47E-02 | 7.95E-02 | 2.67E+00 | 2.75E+00 | 8.50E-01 | no | Hunk | intron_variant | MODIFIER |
| 16 | 90452474 | AX-170464096 | 0 | A | G | 0.465 | -1.47E-02 | 7.95E-02 | 2.67E+00 | 2.75E+00 | 8.50E-01 | no | Hunk | intron_variant | MODIFIER |
| 16 | 90452523 | AX-170404753 | 0 | C | T | 0.465 | -1.47E-02 | 7.95E-02 | 2.67E+00 | 2.75E+00 | 8.50E-01 | no | Hunk | intron_variant | MODIFIER |
| 16 | 90456232 | AX-170531751 | 0 | A | G | 0.465 | -1.47E-02 | 7.95E-02 | 2.67E+00 | 2.75E+00 | 8.50E-01 | no | Hunk | intron_variant | MODIFIER |
| 16 | 90459215 | AX-170147387 | 0 | A | G | 0.465 | -1.47E-02 | 7.95E-02 | 2.67E+00 | 2.75E+00 | 8.50E-01 | no | Hunk | intron_variant | MODIFIER |
| 16 | 90471861 | AX-170058082 | 0 | C | T | 0.465 | -1.47E-02 | 7.95E-02 | 2.67E+00 | 2.75E+00 | 8.50E-01 | no | Hunk | intron_variant | MODIFIER |
| 16 | 90472547 | AX-170303775 | 0 | G | A | 0.465 | -1.47E-02 | 7.95E-02 | 2.67E+00 | 2.75E+00 | 8.50E-01 | no | Hunk | synonymous_variant | LOW |
| 16 | 90502971 | AX-170406680 | 0 | G | A | 0.465 | -1.47E-02 | 7.95E-02 | 2.67E+00 | 2.75E+00 | 8.50E-01 | no | Hunk | downstream_gene_variant | MODIFIER |
| 11 | 31917940 | AX-170125884 | 0 | A | G | 0.158 | 1.84E-02 | 9.68E-02 | 2.63E+00 | 2.73E+00 | 8.50E-01 | no | Cpeb4 | intron_variant | MODIFIER |
| 18 | 57782113 | AX-170341118 | 1 | A | G | 0.411 | 1.54E-02 | 8.53E-02 | 2.73E+00 | 2.80E+00 | 8.50E-01 | no | AC102232.1 | upstream_gene_variant | MODIFIER |
| 1 | 53350061 | AX-170162839 | 1 | T | C | 0.107 | 4.38E-02 | 2.34E-01 | 2.69E+00 | 2.74E+00 | 8.50E-01 | no | Asnsd1 | splice_region_variant,5_prime_UTR_variant | LOW |
| 13 | 46978105 | AX-170245252 | 0 | C | G | 0.482 | -1.15E-02 | 6.41E-02 | 2.66E+00 | 2.76E+00 | 8.50E-01 | no | Gm24769 | upstream_gene_variant | MODIFIER |
| 13 | 46984895 | AX-97817856 | 0 | T | G | 0.482 | -1.15E-02 | 6.41E-02 | 2.66E+00 | 2.76E+00 | 8.50E-01 | no | Gm24769 | downstream_gene_variant | MODIFIER |

| | | | | | | | | | | | | | | | |
|---|---|---|---|---|---|---|---|---|---|---|---|---|---|---|---|
| 13 | 47069751 | AX-170431446 | 0 | C | T | 0.482 | -1.15E-02 | 6.41E-02 | 2.66E+00 | 2.76E+00 | 8.50E-01 | no | Kdm1b | intron_variant | MODIFIER |
| 13 | 47070470 | AX-170395658 | 0 | A | G | 0.482 | -1.15E-02 | 6.41E-02 | 2.66E+00 | 2.76E+00 | 8.50E-01 | no | Kdm1b | intron_variant | MODIFIER |
| 13 | 47074120 | AX-97814652 | 0 | C | T | 0.482 | -1.15E-02 | 6.41E-02 | 2.66E+00 | 2.76E+00 | 8.50E-01 | no | Kdm1b | intron_variant | MODIFIER |
| 13 | 47078896 | AX-170480367 | 0 | G | T | 0.482 | -1.15E-02 | 6.41E-02 | 2.66E+00 | 2.76E+00 | 8.50E-01 | no | Kdm1b | intron_variant | MODIFIER |
| 13 | 47080276 | AX-170221146 | 0 | A | G | 0.482 | -1.15E-02 | 6.41E-02 | 2.66E+00 | 2.76E+00 | 8.50E-01 | no | Dek | downstream_gene_variant | MODIFIER |
| 13 | 47112274 | AX-170484747 | 0 | T | C | 0.482 | -1.15E-02 | 6.41E-02 | 2.66E+00 | 2.76E+00 | 8.50E-01 | no | 1700026N04Rik | downstream_gene_variant | MODIFIER |
| 13 | 47121876 | AX-97823376 | 0 | T | C | 0.482 | -1.15E-02 | 6.41E-02 | 2.66E+00 | 2.76E+00 | 8.50E-01 | no | Rnf144b | upstream_gene_variant | MODIFIER |
| 13 | 47130455 | AX-170454474 | 0 | C | T | 0.482 | -1.15E-02 | 6.41E-02 | 2.66E+00 | 2.76E+00 | 8.50E-01 | no | Rnf144b | intron_variant | MODIFIER |
| 13 | 47132611 | AX-170499237 | 0 | C | T | 0.482 | -1.15E-02 | 6.41E-02 | 2.66E+00 | 2.76E+00 | 8.50E-01 | no | Rnf144b | intron_variant | MODIFIER |
| 13 | 47136110 | AX-97782539 | 0 | C | T | 0.482 | -1.15E-02 | 6.41E-02 | 2.66E+00 | 2.76E+00 | 8.50E-01 | no | Rnf144b | intron_variant | MODIFIER |
| 13 | 47144313 | AX-170399724 | 0 | G | A | 0.482 | -1.15E-02 | 6.41E-02 | 2.66E+00 | 2.76E+00 | 8.50E-01 | no | Rnf144b | intron_variant | MODIFIER |
| 13 | 47232412 | AX-170051185 | 0 | C | A | 0.482 | -1.15E-02 | 6.41E-02 | 2.66E+00 | 2.76E+00 | 8.50E-01 | no | Rnf144b | intron_variant | MODIFIER |
| 13 | 47238480 | AX-97804653 | 0 | T | C | 0.482 | -1.15E-02 | 6.41E-02 | 2.66E+00 | 2.76E+00 | 8.50E-01 | no | Rnf144b | intron_variant | MODIFIER |
| 13 | 47252541 | AX-170008273 | 0 | C | T | 0.482 | -1.15E-02 | 6.41E-02 | 2.66E+00 | 2.76E+00 | 8.50E-01 | no | Rnf144b | downstream_gene_variant | MODIFIER |
| 8 | 35160585 | AX-170435794 | 1 | G | A | 0.152 | -2.77E-02 | 1.43E-01 | 2.65E+00 | 2.71E+00 | 8.50E-01 | no | | regulatory_region_variant | MODIFIER |
| 8 | 30824955 | AX-170373291 | 1 | A | G | 0.17 | 2.16E-02 | 1.16E-01 | 2.67E+00 | 2.74E+00 | 8.50E-01 | no | | intergenic_variant | MODIFIER |
| 15 | 94143526 | AX-170418189 | 0 | G | A | 0.149 | -3.22E-02 | 1.78E-01 | 2.70E+00 | 2.77E+00 | 8.50E-01 | no | | intergenic_variant | MODIFIER |
| 15 | 101688431 | AX-170162308 | 1 | C | T | 0.116 | 2.85E-02 | 1.50E-01 | 2.65E+00 | 2.72E+00 | 8.50E-01 | no | Krt6a | downstream_gene_variant | MODIFIER |
| 8 | 95671651 | AX-170495712 | 0 | T | C | 0.219 | 1.98E-02 | 1.08E-01 | 2.67E+00 | 2.75E+00 | 8.50E-01 | no | Gm6013 | downstream_gene_variant | MODIFIER |
| 13 | 47626957 | AX-170178435 | 1 | A | G | 0.509 | 1.26E-02 | 6.45E-02 | 2.61E+00 | 2.71E+00 | 8.50E-01 | no | | intergenic_variant | MODIFIER |
| 1 | 148439089 | AX-170408917 | 0 | A | G | 0.202 | 2.38E-02 | 1.25E-01 | 2.65E+00 | 2.72E+00 | 8.50E-01 | no | | intergenic_variant | MODIFIER |
| 12 | 111153155 | AX-170147857 | 1 | C | T | 0.143 | -3.52E-02 | 1.92E-01 | 2.68E+00 | 2.75E+00 | 8.50E-01 | no | | regulatory_region_variant | MODIFIER |
| 12 | 116365947 | AX-170287129 | 1 | G | A | 0.143 | -3.52E-02 | 1.92E-01 | 2.68E+00 | 2.75E+00 | 8.50E-01 | no | Esyt2 | intron_variant | MODIFIER |
| 7 | 66391845 | AX-170077584 | 0 | T | A | 0.404 | 1.74E-02 | 9.16E-02 | 2.64E+00 | 2.73E+00 | 8.50E-01 | no | Lrrk1 | upstream_gene_variant | MODIFIER |
| 20 | 145645972 | AX-97851501 | 1 | T | C | 0.125 | 1.96E-02 | 1.03E-01 | 2.61E+00 | 2.73E+00 | 8.51E-01 | no | Gm15060 | upstream_gene_variant | MODIFIER |
| 12 | 107305685 | AX-170438370 | 0 | G | T | 0.132 | -2.83E-02 | 1.59E-01 | 2.66E+00 | 2.77E+00 | 8.51E-01 | no | | intergenic_variant | MODIFIER |
| 12 | 107399534 | AX-170503413 | 0 | C | A | 0.132 | -2.83E-02 | 1.59E-01 | 2.66E+00 | 2.77E+00 | 8.51E-01 | no | | intergenic_variant | MODIFIER |
| 12 | 107682109 | AX-170471931 | 0 | C | A | 0.132 | -2.83E-02 | 1.59E-01 | 2.66E+00 | 2.77E+00 | 8.51E-01 | no | | intergenic_variant | MODIFIER |
| 12 | 108164248 | AX-170116044 | 0 | C | T | 0.132 | -2.83E-02 | 1.59E-01 | 2.66E+00 | 2.77E+00 | 8.51E-01 | no | Setd3 | intron_variant | MODIFIER |
| 5 | 81271742 | AX-170072614 | 0 | C | T | 0.439 | 1.92E-02 | 1.03E-01 | 2.67E+00 | 2.73E+00 | 8.51E-01 | no | Adgrl3 | intron_variant | MODIFIER |
| 1 | 54213844 | AX-97813389 | 0 | C | T | 0.088 | 3.40E-02 | 1.82E-01 | 2.61E+00 | 2.74E+00 | 8.51E-01 | no | | intergenic_variant | MODIFIER |
| 8 | 94391484 | AX-170469187 | 1 | T | C | 0.205 | -2.22E-02 | 1.20E-01 | 2.68E+00 | 2.74E+00 | 8.51E-01 | no | Herpud1 | intron_variant | MODIFIER |
| 6 | 103557019 | AX-170452983 | 1 | T | C | 0.054 | 5.52E-02 | 2.96E-01 | 2.67E+00 | 2.73E+00 | 8.51E-01 | no | Chl1 | intron_variant | MODIFIER |
| 13 | 47230432 | AX-97851548 | 0 | G | A | 0.474 | -1.14E-02 | 6.37E-02 | 2.66E+00 | 2.76E+00 | 8.51E-01 | no | Rnf144b | intron_variant | MODIFIER |
| 6 | 105870390 | AX-170039352 | 1 | C | T | 0.054 | 5.53E-02 | 2.97E-01 | 2.67E+00 | 2.73E+00 | 8.51E-01 | no | Cntn4 | intron_variant | MODIFIER |
| 6 | 110950675 | AX-170027979 | 1 | C | T | 0.054 | 5.53E-02 | 2.97E-01 | 2.67E+00 | 2.73E+00 | 8.51E-01 | no | Grm7 | intron_variant | MODIFIER |
| 2 | 145470724 | AX-170412733 | 0 | G | A | 0.132 | 3.61E-02 | 1.91E-01 | 2.65E+00 | 2.72E+00 | 8.51E-01 | no | Slc24a3 | intron_variant | MODIFIER |
| 18 | 89209816 | AX-170212917 | 1 | C | A | 0.518 | -1.08E-02 | 6.23E-02 | 2.64E+00 | 2.77E+00 | 8.51E-01 | no | Cd226 | intron_variant | MODIFIER |
| 13 | 45911701 | AX-97808710 | 0 | G | T | 0.474 | -1.11E-02 | 6.28E-02 | 2.67E+00 | 2.77E+00 | 8.51E-01 | no | Atxn1 | intron_variant | MODIFIER |
| 13 | 46207070 | AX-170234350 | 0 | A | G | 0.474 | -1.11E-02 | 6.28E-02 | 2.67E+00 | 2.77E+00 | 8.51E-01 | no | | intergenic_variant | MODIFIER |
| 13 | 46207275 | AX-97816031 | 0 | G | A | 0.474 | -1.11E-02 | 6.28E-02 | 2.67E+00 | 2.77E+00 | 8.51E-01 | no | | intergenic_variant | MODIFIER |
| 13 | 46234780 | AX-170117640 | 0 | T | C | 0.474 | -1.11E-02 | 6.28E-02 | 2.67E+00 | 2.77E+00 | 8.51E-01 | no | | intergenic_variant | MODIFIER |
| 13 | 46258063 | AX-170189829 | 0 | T | C | 0.474 | -1.11E-02 | 6.28E-02 | 2.67E+00 | 2.77E+00 | 8.51E-01 | no | | regulatory_region_variant | MODIFIER |
| 13 | 46363804 | AX-170521314 | 0 | A | G | 0.474 | -1.11E-02 | 6.28E-02 | 2.67E+00 | 2.77E+00 | 8.51E-01 | no | | regulatory_region_variant | MODIFIER |
| 13 | 46369428 | AX-97784046 | 0 | G | A | 0.474 | -1.11E-02 | 6.28E-02 | 2.67E+00 | 2.77E+00 | 8.51E-01 | no | | intergenic_variant | MODIFIER |
| 13 | 46390558 | AX-170166936 | 0 | G | A | 0.474 | -1.11E-02 | 6.28E-02 | 2.67E+00 | 2.77E+00 | 8.51E-01 | no | | intergenic_variant | MODIFIER |
| 13 | 46394311 | AX-170403588 | 0 | A | T | 0.474 | -1.11E-02 | 6.28E-02 | 2.67E+00 | 2.77E+00 | 8.51E-01 | no | | intergenic_variant | MODIFIER |
| 13 | 46424795 | AX-97850532 | 0 | G | A | 0.474 | -1.11E-02 | 6.28E-02 | 2.67E+00 | 2.77E+00 | 8.51E-01 | no | Rbm24 | intron_variant | MODIFIER |
| 13 | 46425019 | AX-170491087 | 0 | T | G | 0.474 | -1.11E-02 | 6.28E-02 | 2.67E+00 | 2.77E+00 | 8.51E-01 | no | Rbm24 | intron_variant | MODIFIER |
| 13 | 46432137 | AX-170285971 | 0 | A | C | 0.474 | -1.11E-02 | 6.28E-02 | 2.67E+00 | 2.77E+00 | 8.51E-01 | no | Rbm24 | downstream_gene_variant | MODIFIER |
| 13 | 46442563 | AX-97826666 | 0 | C | A | 0.474 | -1.11E-02 | 6.28E-02 | 2.67E+00 | 2.77E+00 | 8.51E-01 | no | | intergenic_variant | MODIFIER |
| 13 | 46586776 | AX-170397940 | 0 | T | C | 0.474 | -1.11E-02 | 6.28E-02 | 2.67E+00 | 2.77E+00 | 8.51E-01 | no | Cap2 | intron_variant | MODIFIER |
| 13 | 46617981 | AX-97807223 | 0 | A | G | 0.474 | -1.11E-02 | 6.28E-02 | 2.67E+00 | 2.77E+00 | 8.51E-01 | no | Cap2 | intron_variant | MODIFIER |
| 13 | 46619252 | AX-170307776 | 0 | T | A | 0.474 | -1.11E-02 | 6.28E-02 | 2.67E+00 | 2.77E+00 | 8.51E-01 | no | Cap2 | intron_variant | MODIFIER |

| | | | | | | | | | | | | | | | |
|---|---|---|---|---|---|---|---|---|---|---|---|---|---|---|---|
| 13 | 46619951 | AX-170187992 | 0 | A | G | 0.474 | -1.11E-02 | 6.28E-02 | 2.67E+00 | 2.77E+00 | 8.51E-01 | no | Cap2 | intron_variant | MODIFIER |
| 13 | 46659976 | AX-97791471 | 0 | C | T | 0.474 | -1.11E-02 | 6.28E-02 | 2.67E+00 | 2.77E+00 | 8.51E-01 | no | | intergenic_variant | MODIFIER |
| 1 | 52833644 | AX-170510781 | 1 | C | G | 0.107 | 4.30E-02 | 2.33E-01 | 2.69E+00 | 2.74E+00 | 8.51E-01 | no | | intergenic_variant | MODIFIER |
| 1 | 53904479 | AX-170161354 | 1 | T | A | 0.107 | 4.30E-02 | 2.33E-01 | 2.69E+00 | 2.74E+00 | 8.51E-01 | no | Hecw2 | intron_variant | MODIFIER |
| 8 | 83839666 | AX-97809108 | 0 | T | G | 0.219 | 2.06E-02 | 1.09E-01 | 2.65E+00 | 2.73E+00 | 8.52E-01 | no | | intergenic_variant | MODIFIER |
| 12 | 105715530 | AX-170296640 | 0 | C | A | 0.14 | -1.98E-02 | 1.15E-01 | 2.62E+00 | 2.77E+00 | 8.52E-01 | no | Ak7 | intron_variant | MODIFIER |
| 8 | 49703807 | AX-97798581 | 1 | G | A | 0.152 | -2.92E-02 | 1.49E-01 | 2.64E+00 | 2.69E+00 | 8.52E-01 | no | | intergenic_variant | MODIFIER |
| 8 | 52344920 | AX-170090107 | 1 | C | A | 0.152 | -2.92E-02 | 1.49E-01 | 2.64E+00 | 2.69E+00 | 8.52E-01 | no | | intergenic_variant | MODIFIER |
| 8 | 57369897 | AX-170249794 | 1 | C | A | 0.152 | -2.92E-02 | 1.49E-01 | 2.64E+00 | 2.69E+00 | 8.52E-01 | no | | intergenic_variant | MODIFIER |
| 15 | 101085184 | AX-170411039 | 1 | C | T | 0.134 | 3.02E-02 | 1.61E-01 | 2.66E+00 | 2.73E+00 | 8.52E-01 | no | | intergenic_variant | MODIFIER |
| 15 | 101552634 | AX-170056495 | 1 | C | A | 0.134 | 3.02E-02 | 1.61E-01 | 2.66E+00 | 2.73E+00 | 8.52E-01 | no | Krt82 | upstream_gene_variant | MODIFIER |
| 15 | 102331847 | AX-170362830 | 1 | T | C | 0.134 | 3.02E-02 | 1.61E-01 | 2.66E+00 | 2.73E+00 | 8.52E-01 | no | Myg1 | missense_variant | MODERATE |
| 7 | 107272237 | AX-97829771 | 0 | A | C | 0.447 | -1.69E-02 | 9.43E-02 | 2.70E+00 | 2.77E+00 | 8.52E-01 | no | Platr28 | intron_variant,non_coding_transcript_variant | MODIFIER |
| 1 | 136476292 | AX-170481869 | 1 | T | A | 0.205 | 2.13E-02 | 1.12E-01 | 2.62E+00 | 2.72E+00 | 8.52E-01 | no | Kif14 | intron_variant | MODIFIER |
| 16 | 87265754 | AX-170013588 | 0 | A | G | 0.421 | -1.50E-02 | 8.13E-02 | 2.67E+00 | 2.74E+00 | 8.52E-01 | no | | regulatory_region_variant | MODIFIER |
| 13 | 48947743 | AX-97785791 | 0 | A | C | 0.491 | -1.16E-02 | 6.59E-02 | 2.66E+00 | 2.77E+00 | 8.52E-01 | no | Fam120a | intron_variant | MODIFIER |
| 7 | 91789847 | AX-97803365 | 1 | C | T | 0.116 | 3.91E-02 | 2.12E-01 | 2.66E+00 | 2.74E+00 | 8.52E-01 | no | Dlg2 | intron_variant | MODIFIER |
| 7 | 93576050 | AX-170203789 | 1 | T | A | 0.116 | 3.91E-02 | 2.12E-01 | 2.66E+00 | 2.74E+00 | 8.52E-01 | no | | intergenic_variant | MODIFIER |
| 9 | 104999547 | AX-170278838 | 0 | C | T | 0.123 | 2.05E-02 | 1.16E-01 | 2.72E+00 | 2.80E+00 | 8.52E-01 | no | Cpne4 | intron_variant | MODIFIER |
| 1 | 23333837 | AX-97788542 | 1 | C | A | 0.42 | 1.57E-02 | 8.18E-02 | 2.63E+00 | 2.71E+00 | 8.52E-01 | no | | intergenic_variant | MODIFIER |
| 1 | 63969716 | AX-170124894 | 0 | G | A | 0.114 | 4.38E-02 | 2.36E-01 | 2.67E+00 | 2.73E+00 | 8.52E-01 | no | Gm13749 | downstream_gene_variant | MODIFIER |
| 7 | 39691748 | AX-97820387 | 0 | G | T | 0.351 | -1.61E-02 | 9.21E-02 | 2.69E+00 | 2.78E+00 | 8.52E-01 | no | | intergenic_variant | MODIFIER |
| 16 | 46090924 | AX-170012962 | 1 | A | G | 0.482 | -1.61E-02 | 8.15E-02 | 2.63E+00 | 2.68E+00 | 8.52E-01 | no | Cd96 | intron_variant | MODIFIER |
| 5 | 101182430 | AX-170095413 | 0 | T | A | 0.412 | 1.96E-02 | 1.03E-01 | 2.64E+00 | 2.71E+00 | 8.52E-01 | no | | intergenic_variant | MODIFIER |
| 12 | 107525079 | AX-97789234 | 0 | C | T | 0.123 | -2.89E-02 | 1.66E-01 | 2.66E+00 | 2.77E+00 | 8.52E-01 | no | | intergenic_variant | MODIFIER |
| 2 | 119181193 | AX-170124386 | 1 | A | G | 0.134 | 3.77E-02 | 2.01E-01 | 2.66E+00 | 2.72E+00 | 8.52E-01 | no | Dnajc17 | intron_variant | MODIFIER |
| 2 | 121828859 | AX-170158581 | 1 | T | C | 0.134 | 3.77E-02 | 2.01E-01 | 2.66E+00 | 2.72E+00 | 8.52E-01 | no | | intergenic_variant | MODIFIER |
| 2 | 124585557 | AX-170157136 | 1 | T | A | 0.134 | 3.77E-02 | 2.01E-01 | 2.66E+00 | 2.72E+00 | 8.52E-01 | no | Sema6d | intron_variant | MODIFIER |
| 2 | 125212188 | AX-170282072 | 1 | A | T | 0.134 | 3.77E-02 | 2.01E-01 | 2.66E+00 | 2.72E+00 | 8.52E-01 | no | Slc12a1 | intron_variant | MODIFIER |
| 2 | 125789014 | AX-170303791 | 1 | G | A | 0.134 | 3.77E-02 | 2.01E-01 | 2.66E+00 | 2.72E+00 | 8.52E-01 | no | | regulatory_region_variant | MODIFIER |
| 2 | 137374937 | AX-170019144 | 1 | G | T | 0.134 | 3.77E-02 | 2.01E-01 | 2.66E+00 | 2.72E+00 | 8.52E-01 | no | | intergenic_variant | MODIFIER |
| 2 | 143132372 | AX-170503495 | 1 | T | C | 0.134 | 3.77E-02 | 2.01E-01 | 2.66E+00 | 2.72E+00 | 8.52E-01 | no | Gm14083 | upstream_gene_variant | MODIFIER |
| 15 | 95487309 | AX-170203659 | 1 | T | C | 0.143 | -3.30E-02 | 1.88E-01 | 2.71E+00 | 2.79E+00 | 8.52E-01 | no | Nell2 | intron_variant | MODIFIER |
| 6 | 147661013 | AX-170303660 | 0 | C | T | 0.053 | -3.10E-02 | 1.73E-01 | 2.63E+00 | 2.75E+00 | 8.53E-01 | no | | intergenic_variant | MODIFIER |
| 5 | 122675656 | AX-170026054 | 1 | A | T | 0.438 | -1.97E-02 | 1.09E-01 | 2.71E+00 | 2.76E+00 | 8.53E-01 | no | P2rx7 | intron_variant | MODIFIER |
| 5 | 96593051 | AX-97799965 | 0 | C | T | 0.439 | 1.89E-02 | 1.04E-01 | 2.67E+00 | 2.74E+00 | 8.53E-01 | no | Fras1 | intron_variant | MODIFIER |
| 5 | 28571671 | AX-170507900 | 1 | G | A | 0.357 | 1.98E-02 | 1.07E-01 | 2.68E+00 | 2.73E+00 | 8.53E-01 | no | 9530036O11Rik | intron_variant,non_coding_transcript_variant | MODIFIER |
| 5 | 125097215 | AX-170443881 | 2 | C | A | 0.464 | -2.07E-02 | 1.15E-01 | 2.74E+00 | 2.78E+00 | 8.53E-01 | no | Ncor2 | intron_variant | MODIFIER |
| 9 | 56380869 | AX-170153005 | 0 | G | A | 0.211 | -2.01E-02 | 1.12E-01 | 2.68E+00 | 2.76E+00 | 8.53E-01 | no | Peak1 | intron_variant | MODIFIER |
| 4 | 88513304 | AX-97800473 | 0 | G | A | 0.254 | 2.21E-02 | 1.02E-01 | 2.56E+00 | 2.66E+00 | 8.53E-01 | no | | intergenic_variant | MODIFIER |
| 6 | 107264933 | AX-170209087 | 1 | C | T | 0.054 | -2.47E-02 | 1.39E-01 | 2.58E+00 | 2.75E+00 | 8.53E-01 | no | Gm34933 | intron_variant,non_coding_transcript_variant | MODIFIER |
| 2 | 48089918 | AX-170206427 | 1 | G | T | 0.036 | 2.92E-02 | 1.72E-01 | 2.63E+00 | 2.77E+00 | 8.53E-01 | no | Gm13481 | intron_variant,non_coding_transcript_variant | MODIFIER |
| 7 | 91294926 | AX-170208174 | 2 | A | G | 0.118 | -3.71E-02 | 2.00E-01 | 2.62E+00 | 2.73E+00 | 8.53E-01 | no | Dlg2 | intron_variant | MODIFIER |
| 12 | 116781433 | AX-170131009 | 1 | C | T | 0.152 | 2.89E-02 | 1.61E-01 | 2.63E+00 | 2.75E+00 | 8.53E-01 | no | Ptprn2 | intron_variant | MODIFIER |
| 1 | 155753596 | AX-97851851 | 0 | G | A | 0.219 | -2.41E-02 | 1.36E-01 | 2.72E+00 | 2.78E+00 | 8.53E-01 | no | Lhx4 | upstream_gene_variant | MODIFIER |
| 14 | 84282096 | AX-170193694 | 0 | G | T | 0.044 | -2.20E-02 | 1.17E-01 | 2.57E+00 | 2.73E+00 | 8.53E-01 | no | | intergenic_variant | MODIFIER |
| 7 | 82245045 | AX-170077880 | 0 | T | C | 0.123 | 3.27E-02 | 1.73E-01 | 2.61E+00 | 2.72E+00 | 8.53E-01 | no | Sh3gl3 | intron_variant | MODIFIER |
| 18 | 70684124 | AX-170198370 | 1 | T | G | 0.259 | -1.85E-02 | 9.88E-02 | 2.66E+00 | 2.72E+00 | 8.53E-01 | no | | intergenic_variant | MODIFIER |
| 2 | 138496307 | AX-170267020 | 0 | T | A | 0.123 | 3.92E-02 | 2.10E-01 | 2.67E+00 | 2.72E+00 | 8.53E-01 | no | Btbd3 | intron_variant,non_coding_transcript_variant | MODIFIER |
| 12 | 114435509 | AX-170202023 | 1 | G | A | 0.125 | -3.47E-02 | 1.95E-01 | 2.69E+00 | 2.76E+00 | 8.54E-01 | no | Ighv6-6 | upstream_gene_variant | MODIFIER |
| 1 | 12947186 | AX-170202409 | 1 | T | A | 0.375 | -1.29E-02 | 7.38E-02 | 2.71E+00 | 2.79E+00 | 8.54E-01 | no | Slco5a1 | intron_variant | MODIFIER |
| 1 | 5816040 | AX-170303833 | 1 | T | A | 0.089 | 3.00E-02 | 1.64E-01 | 2.62E+00 | 2.74E+00 | 8.54E-01 | no | | intergenic_variant | MODIFIER |
| 1 | 5867649 | AX-170101200 | 1 | G | A | 0.089 | 3.00E-02 | 1.64E-01 | 2.62E+00 | 2.74E+00 | 8.54E-01 | no | | intergenic_variant | MODIFIER |
| 8 | 80980401 | AX-170377706 | 0 | T | C | 0.202 | 2.37E-02 | 1.29E-01 | 2.67E+00 | 2.73E+00 | 8.54E-01 | no | Usp38 | downstream_gene_variant | MODIFIER |

| | | | | | | | | | | | | | | | |
|---|---|---|---|---|---|---|---|---|---|---|---|---|---|---|---|
| 5 | 121568594 | AX-170156867 | 0 | T | G | 0.447 | -1.97E-02 | 1.10E-01 | 2.71E+00 | 2.77E+00 | 8.54E-01 | no | Aldh2 | intron_variant | MODIFIER |
| 5 | 121697719 | AX-170383374 | 0 | G | A | 0.447 | -1.97E-02 | 1.10E-01 | 2.71E+00 | 2.77E+00 | 8.54E-01 | no | | intergenic_variant | MODIFIER |
| 10 | 121200866 | AX-170276216 | 0 | T | C | 0.351 | 1.30E-02 | 7.10E-02 | 2.64E+00 | 2.74E+00 | 8.54E-01 | no | 4930432O09Rik | non_coding_transcript_exon_variant | MODIFIER |
| 2 | 88911708 | AX-97845369 | 0 | C | A | 0.07 | -3.15E-02 | 1.82E-01 | 2.64E+00 | 2.76E+00 | 8.54E-01 | no | Olfr1209 | upstream_gene_variant | MODIFIER |
| 13 | 48462354 | AX-170494295 | 0 | A | T | 0.509 | -1.15E-02 | 6.66E-02 | 2.66E+00 | 2.77E+00 | 8.54E-01 | no | | intergenic_variant | MODIFIER |
| 13 | 49915813 | AX-170202667 | 0 | G | T | 0.509 | -1.15E-02 | 6.66E-02 | 2.66E+00 | 2.77E+00 | 8.54E-01 | no | | intergenic_variant | MODIFIER |
| 13 | 50417384 | AX-97802626 | 0 | T | C | 0.509 | -1.15E-02 | 6.66E-02 | 2.66E+00 | 2.77E+00 | 8.54E-01 | no | Fbxw17 | upstream_gene_variant | MODIFIER |
| 13 | 50437008 | AX-97786918 | 0 | C | G | 0.509 | -1.15E-02 | 6.66E-02 | 2.66E+00 | 2.77E+00 | 8.54E-01 | no | Fbxw17 | downstream_gene_variant | MODIFIER |
| 13 | 50441527 | AX-168412296 | 0 | T | C | 0.509 | -1.15E-02 | 6.66E-02 | 2.66E+00 | 2.77E+00 | 8.54E-01 | no | Gm8739 | intron_variant,non_coding_transcript_variant | MODIFIER |
| 13 | 50980061 | AX-97799973 | 0 | C | T | 0.509 | -1.15E-02 | 6.66E-02 | 2.66E+00 | 2.77E+00 | 8.54E-01 | no | | intergenic_variant | MODIFIER |
| 13 | 51122454 | AX-97851759 | 0 | G | A | 0.509 | -1.15E-02 | 6.66E-02 | 2.66E+00 | 2.77E+00 | 8.54E-01 | no | Spin1 | intron_variant | MODIFIER |
| 13 | 51183867 | AX-170209908 | 0 | A | G | 0.509 | -1.15E-02 | 6.66E-02 | 2.66E+00 | 2.77E+00 | 8.54E-01 | no | | intergenic_variant | MODIFIER |
| 13 | 51221902 | AX-170397919 | 0 | C | T | 0.509 | -1.15E-02 | 6.66E-02 | 2.66E+00 | 2.77E+00 | 8.54E-01 | no | | regulatory_region_variant | MODIFIER |
| 13 | 51301100 | AX-97808975 | 0 | G | A | 0.509 | -1.15E-02 | 6.66E-02 | 2.66E+00 | 2.77E+00 | 8.54E-01 | no | | regulatory_region_variant | MODIFIER |
| 13 | 51323538 | AX-97857157 | 0 | T | C | 0.509 | -1.15E-02 | 6.66E-02 | 2.66E+00 | 2.77E+00 | 8.54E-01 | no | | intergenic_variant | MODIFIER |
| 13 | 51338535 | AX-170274271 | 0 | T | G | 0.509 | -1.15E-02 | 6.66E-02 | 2.66E+00 | 2.77E+00 | 8.54E-01 | no | Gm6056 | upstream_gene_variant | MODIFIER |
| 13 | 51338687 | AX-170522075 | 0 | T | A | 0.509 | -1.15E-02 | 6.66E-02 | 2.66E+00 | 2.77E+00 | 8.54E-01 | no | Gm6056 | upstream_gene_variant | MODIFIER |
| 13 | 51339588 | AX-170520674 | 0 | C | T | 0.509 | -1.15E-02 | 6.66E-02 | 2.66E+00 | 2.77E+00 | 8.54E-01 | no | Gm6056 | non_coding_transcript_exon_variant | MODIFIER |
| 13 | 51345608 | AX-97828636 | 0 | G | A | 0.509 | -1.15E-02 | 6.66E-02 | 2.66E+00 | 2.77E+00 | 8.54E-01 | no | Gm47554 | downstream_gene_variant | MODIFIER |
| 13 | 51349724 | AX-168385739 | 0 | T | C | 0.509 | -1.15E-02 | 6.66E-02 | 2.66E+00 | 2.77E+00 | 8.54E-01 | no | Gm47554 | downstream_gene_variant | MODIFIER |
| 13 | 51351375 | AX-170051261 | 0 | T | C | 0.509 | -1.15E-02 | 6.66E-02 | 2.66E+00 | 2.77E+00 | 8.54E-01 | no | | intergenic_variant | MODIFIER |
| 13 | 51354797 | AX-170051266 | 0 | C | T | 0.509 | -1.15E-02 | 6.66E-02 | 2.66E+00 | 2.77E+00 | 8.54E-01 | no | | intergenic_variant | MODIFIER |
| 13 | 51411777 | AX-97805322 | 0 | G | A | 0.509 | -1.15E-02 | 6.66E-02 | 2.66E+00 | 2.77E+00 | 8.54E-01 | no | S1pr3 | intron_variant | MODIFIER |
| 13 | 51464751 | AX-170271820 | 0 | C | T | 0.509 | -1.15E-02 | 6.66E-02 | 2.66E+00 | 2.77E+00 | 8.54E-01 | no | Shc3 | intron_variant | MODIFIER |
| 13 | 51469575 | AX-97790718 | 0 | A | G | 0.509 | -1.15E-02 | 6.66E-02 | 2.66E+00 | 2.77E+00 | 8.54E-01 | no | Shc3 | intron_variant | MODIFIER |
| 13 | 51489265 | AX-97823456 | 0 | G | A | 0.509 | -1.15E-02 | 6.66E-02 | 2.66E+00 | 2.77E+00 | 8.54E-01 | no | Shc3 | intron_variant | MODIFIER |
| 13 | 51489294 | AX-170272035 | 0 | T | C | 0.509 | -1.15E-02 | 6.66E-02 | 2.66E+00 | 2.77E+00 | 8.54E-01 | no | Shc3 | intron_variant | MODIFIER |
| 13 | 51511485 | AX-170470195 | 0 | T | G | 0.509 | -1.15E-02 | 6.66E-02 | 2.66E+00 | 2.77E+00 | 8.54E-01 | no | Shc3 | intron_variant | MODIFIER |
| 13 | 51516116 | AX-170311487 | 0 | C | A | 0.509 | -1.15E-02 | 6.66E-02 | 2.66E+00 | 2.77E+00 | 8.54E-01 | no | Shc3 | intron_variant | MODIFIER |
| 13 | 51516160 | AX-170456184 | 0 | T | C | 0.509 | -1.15E-02 | 6.66E-02 | 2.66E+00 | 2.77E+00 | 8.54E-01 | no | Shc3 | intron_variant | MODIFIER |
| 13 | 51516228 | AX-97855164 | 0 | T | C | 0.509 | -1.15E-02 | 6.66E-02 | 2.66E+00 | 2.77E+00 | 8.54E-01 | no | Shc3 | intron_variant | MODIFIER |
| 8 | 93491492 | AX-170374454 | 0 | G | A | 0.219 | -2.15E-02 | 1.17E-01 | 2.67E+00 | 2.73E+00 | 8.54E-01 | no | | intergenic_variant | MODIFIER |
| 5 | 122233595 | AX-170026042 | 1 | C | A | 0.464 | -2.00E-02 | 1.12E-01 | 2.71E+00 | 2.76E+00 | 8.54E-01 | no | Hvcn1 | missense_variant | MODERATE |
| 9 | 32742903 | AX-170504695 | 0 | C | T | 0.43 | 1.60E-02 | 8.60E-02 | 2.65E+00 | 2.72E+00 | 8.54E-01 | no | Ets1 | intron_variant | MODIFIER |
| 18 | 88977125 | AX-170381752 | 1 | G | A | 0.509 | -1.06E-02 | 6.20E-02 | 2.63E+00 | 2.76E+00 | 8.54E-01 | no | Rttn | intron_variant | MODIFIER |
| 7 | 116638784 | AX-170529066 | 1 | T | C | 0.429 | 1.74E-02 | 9.43E-02 | 2.67E+00 | 2.72E+00 | 8.54E-01 | no | | intergenic_variant | MODIFIER |
| 4 | 147890848 | AX-170070978 | 1 | G | A | 0.107 | 3.73E-02 | 2.05E-01 | 2.67E+00 | 2.73E+00 | 8.54E-01 | no | Mfn2 | intron_variant | MODIFIER |
| 8 | 78070764 | AX-170125316 | 0 | G | A | 0.202 | -2.44E-02 | 1.33E-01 | 2.67E+00 | 2.73E+00 | 8.54E-01 | no | | intergenic_variant | MODIFIER |
| 8 | 78073687 | AX-170148265 | 0 | T | A | 0.202 | -2.44E-02 | 1.33E-01 | 2.67E+00 | 2.73E+00 | 8.54E-01 | no | | intergenic_variant | MODIFIER |
| 8 | 78089589 | AX-170137913 | 0 | A | G | 0.202 | -2.44E-02 | 1.33E-01 | 2.67E+00 | 2.73E+00 | 8.54E-01 | no | | intergenic_variant | MODIFIER |
| 8 | 78092135 | AX-170421513 | 0 | T | G | 0.202 | -2.44E-02 | 1.33E-01 | 2.67E+00 | 2.73E+00 | 8.54E-01 | no | | intergenic_variant | MODIFIER |
| 8 | 78106918 | AX-170485929 | 0 | G | A | 0.202 | -2.44E-02 | 1.33E-01 | 2.67E+00 | 2.73E+00 | 8.54E-01 | no | Gm45288 | upstream_gene_variant | MODIFIER |
| 8 | 78121734 | AX-170150871 | 0 | T | A | 0.202 | -2.44E-02 | 1.33E-01 | 2.67E+00 | 2.73E+00 | 8.54E-01 | no | | intergenic_variant | MODIFIER |
| 8 | 78122055 | AX-170334263 | 0 | G | A | 0.202 | -2.44E-02 | 1.33E-01 | 2.67E+00 | 2.73E+00 | 8.54E-01 | no | | intergenic_variant | MODIFIER |
| 8 | 78127571 | AX-170466615 | 0 | T | C | 0.202 | -2.44E-02 | 1.33E-01 | 2.67E+00 | 2.73E+00 | 8.54E-01 | no | | intergenic_variant | MODIFIER |
| 8 | 78128277 | AX-170417256 | 0 | A | C | 0.202 | -2.44E-02 | 1.33E-01 | 2.67E+00 | 2.73E+00 | 8.54E-01 | no | | intergenic_variant | MODIFIER |
| 8 | 78132942 | AX-170282496 | 0 | T | G | 0.202 | -2.44E-02 | 1.33E-01 | 2.67E+00 | 2.73E+00 | 8.54E-01 | no | | intergenic_variant | MODIFIER |
| 8 | 78133874 | AX-170489965 | 0 | A | C | 0.202 | -2.44E-02 | 1.33E-01 | 2.67E+00 | 2.73E+00 | 8.54E-01 | no | | intergenic_variant | MODIFIER |
| 8 | 78134070 | AX-170372911 | 0 | C | T | 0.202 | -2.44E-02 | 1.33E-01 | 2.67E+00 | 2.73E+00 | 8.54E-01 | no | | intergenic_variant | MODIFIER |
| 8 | 78134111 | AX-170489730 | 0 | C | G | 0.202 | -2.44E-02 | 1.33E-01 | 2.67E+00 | 2.73E+00 | 8.54E-01 | no | | intergenic_variant | MODIFIER |
| 8 | 78162715 | AX-97788744 | 0 | T | G | 0.202 | -2.44E-02 | 1.33E-01 | 2.67E+00 | 2.73E+00 | 8.54E-01 | no | | intergenic_variant | MODIFIER |
| 8 | 78163971 | AX-170164937 | 0 | C | T | 0.202 | -2.44E-02 | 1.33E-01 | 2.67E+00 | 2.73E+00 | 8.54E-01 | no | | intergenic_variant | MODIFIER |
| 8 | 78164302 | AX-170362945 | 0 | C | A | 0.202 | -2.44E-02 | 1.33E-01 | 2.67E+00 | 2.73E+00 | 8.54E-01 | no | | intergenic_variant | MODIFIER |
| 9 | 66768527 | AX-170480252 | 1 | T | C | 0.196 | -2.30E-02 | 1.25E-01 | 2.67E+00 | 2.73E+00 | 8.54E-01 | no | Car12 | downstream_gene_variant | MODIFIER |

| | | | | | | | | | | | | | | | |
|---|---|---|---|---|---|---|---|---|---|---|---|---|---|---|---|
| 6 | 110515142 | AX-97836026 | 1 | C | T | 0.071 | 4.99E-02 | 2.76E-01 | 2.71E+00 | 2.75E+00 | 8.54E-01 | no | | intergenic_variant | MODIFIER |
| 1 | 172904189 | AX-170448053 | 0 | G | A | 0.219 | 2.33E-02 | 1.15E-01 | 2.60E+00 | 2.68E+00 | 8.54E-01 | no | | intergenic_variant | MODIFIER |
| 8 | 19478769 | AX-170422683 | 1 | T | A | 0.17 | 1.95E-02 | 1.08E-01 | 2.66E+00 | 2.74E+00 | 8.54E-01 | no | Defa-ps15 | upstream_gene_variant | MODIFIER |
| 5 | 6675110 | AX-170295099 | 1 | A | G | 0.42 | 1.05E-02 | 6.32E-02 | 2.64E+00 | 2.78E+00 | 8.54E-01 | no | | intergenic_variant | MODIFIER |
| 9 | 113660710 | AX-170128089 | 1 | G | A | 0.134 | -1.90E-02 | 1.04E-01 | 2.64E+00 | 2.73E+00 | 8.54E-01 | no | Pdcd6ip | intron_variant | MODIFIER |
| 1 | 64150647 | AX-170000665 | 1 | C | T | 0.107 | 3.97E-02 | 2.17E-01 | 2.68E+00 | 2.73E+00 | 8.54E-01 | no | Gm13750 | non_coding_transcript_exon_variant | MODIFIER |
| 9 | 85498871 | AX-170032874 | 1 | C | T | 0.196 | -2.18E-02 | 1.21E-01 | 2.67E+00 | 2.74E+00 | 8.54E-01 | no | | intergenic_variant | MODIFIER |
| 13 | 23276366 | AX-97826979 | 0 | C | A | 0.167 | -2.49E-02 | 1.21E-01 | 2.59E+00 | 2.66E+00 | 8.54E-01 | no | 4930557F10Rik | non_coding_transcript_exon_variant | MODIFIER |
| 5 | 96778278 | AX-170505208 | 0 | A | G | 0.439 | 1.96E-02 | 1.09E-01 | 2.69E+00 | 2.74E+00 | 8.55E-01 | no | Fras1 | intron_variant | MODIFIER |
| 1 | 100479749 | AX-97852612 | 1 | A | G | 0.152 | 3.66E-02 | 1.94E-01 | 2.62E+00 | 2.71E+00 | 8.55E-01 | no | Cntnap5b | intron_variant | MODIFIER |
| 7 | 133007870 | AX-170030017 | 0 | C | G | 0.254 | -1.62E-02 | 8.15E-02 | 2.59E+00 | 2.69E+00 | 8.55E-01 | no | Ctbp2 | intron_variant | MODIFIER |
| 4 | 55713891 | AX-97787186 | 1 | T | C | 0.375 | -1.91E-02 | 1.07E-01 | 2.67E+00 | 2.75E+00 | 8.55E-01 | no | | intergenic_variant | MODIFIER |
| 4 | 56446874 | AX-170207149 | 1 | A | T | 0.375 | -1.91E-02 | 1.07E-01 | 2.67E+00 | 2.75E+00 | 8.55E-01 | no | | intergenic_variant | MODIFIER |
| 8 | 35357783 | AX-170079735 | 0 | T | C | 0.149 | -2.67E-02 | 1.43E-01 | 2.65E+00 | 2.71E+00 | 8.55E-01 | no | | intergenic_variant | MODIFIER |
| 4 | 141415096 | AX-170165023 | 0 | A | G | 0.105 | 3.72E-02 | 2.05E-01 | 2.68E+00 | 2.74E+00 | 8.55E-01 | no | Clcnkb | intron_variant | MODIFIER |
| 4 | 141422307 | AX-170335113 | 0 | C | T | 0.105 | 3.72E-02 | 2.05E-01 | 2.68E+00 | 2.74E+00 | 8.55E-01 | no | Hspb7 | intron_variant | MODIFIER |
| 4 | 141463324 | AX-170281219 | 0 | G | A | 0.105 | 3.72E-02 | 2.05E-01 | 2.68E+00 | 2.74E+00 | 8.55E-01 | no | Zbtb17 | missense_variant | MODERATE |
| 4 | 141922233 | AX-170408199 | 0 | G | C | 0.105 | 3.72E-02 | 2.05E-01 | 2.68E+00 | 2.74E+00 | 8.55E-01 | no | Fhad1 | intron_variant | MODIFIER |
| 4 | 142083138 | AX-168399956 | 0 | G | A | 0.105 | 3.72E-02 | 2.05E-01 | 2.68E+00 | 2.74E+00 | 8.55E-01 | no | Tmem51 | intron_variant | MODIFIER |
| 4 | 142083220 | AX-170223928 | 0 | G | A | 0.105 | 3.72E-02 | 2.05E-01 | 2.68E+00 | 2.74E+00 | 8.55E-01 | no | Tmem51 | intron_variant | MODIFIER |
| 4 | 142083490 | AX-170117092 | 0 | C | T | 0.105 | 3.72E-02 | 2.05E-01 | 2.68E+00 | 2.74E+00 | 8.55E-01 | no | Tmem51 | intron_variant | MODIFIER |
| 4 | 142083529 | AX-170431472 | 0 | A | T | 0.105 | 3.72E-02 | 2.05E-01 | 2.68E+00 | 2.74E+00 | 8.55E-01 | no | Tmem51 | intron_variant | MODIFIER |
| 4 | 142209439 | AX-170391050 | 0 | A | C | 0.105 | 3.72E-02 | 2.05E-01 | 2.68E+00 | 2.74E+00 | 8.55E-01 | no | Kazn | intron_variant | MODIFIER |
| 4 | 142232360 | AX-170275561 | 0 | G | T | 0.105 | 3.72E-02 | 2.05E-01 | 2.68E+00 | 2.74E+00 | 8.55E-01 | no | Kazn | intron_variant | MODIFIER |
| 4 | 142232716 | AX-170352349 | 0 | C | T | 0.105 | 3.72E-02 | 2.05E-01 | 2.68E+00 | 2.74E+00 | 8.55E-01 | no | Kazn | intron_variant | MODIFIER |
| 4 | 142232920 | AX-170264781 | 0 | C | T | 0.105 | 3.72E-02 | 2.05E-01 | 2.68E+00 | 2.74E+00 | 8.55E-01 | no | Kazn | intron_variant | MODIFIER |
| 4 | 142716204 | AX-170526027 | 0 | T | G | 0.105 | 3.72E-02 | 2.05E-01 | 2.68E+00 | 2.74E+00 | 8.55E-01 | no | 6330411D24Rik | intron_variant,non_coding_transcript_variant | MODIFIER |
| 4 | 142995544 | AX-170337947 | 0 | C | A | 0.105 | 3.72E-02 | 2.05E-01 | 2.68E+00 | 2.74E+00 | 8.55E-01 | no | 6330411D24Rik | intron_variant,non_coding_transcript_variant | MODIFIER |
| 4 | 143001070 | AX-170420791 | 0 | C | G | 0.105 | 3.72E-02 | 2.05E-01 | 2.68E+00 | 2.74E+00 | 8.55E-01 | no | 6330411D24Rik | intron_variant,non_coding_transcript_variant | MODIFIER |
| 4 | 143001163 | AX-170258542 | 0 | G | T | 0.105 | 3.72E-02 | 2.05E-01 | 2.68E+00 | 2.74E+00 | 8.55E-01 | no | 6330411D24Rik | intron_variant,non_coding_transcript_variant | MODIFIER |
| 4 | 143013755 | AX-170171491 | 0 | A | G | 0.105 | 3.72E-02 | 2.05E-01 | 2.68E+00 | 2.74E+00 | 8.55E-01 | no | 6330411D24Rik | intron_variant,non_coding_transcript_variant | MODIFIER |
| 4 | 143030450 | AX-170125182 | 0 | C | G | 0.105 | 3.72E-02 | 2.05E-01 | 2.68E+00 | 2.74E+00 | 8.55E-01 | no | 6330411D24Rik | intron_variant,non_coding_transcript_variant | MODIFIER |
| 4 | 143032959 | AX-170178914 | 0 | G | A | 0.105 | 3.72E-02 | 2.05E-01 | 2.68E+00 | 2.74E+00 | 8.55E-01 | no | 6330411D24Rik | intron_variant,non_coding_transcript_variant | MODIFIER |
| 4 | 143034742 | AX-170461811 | 0 | T | C | 0.105 | 3.72E-02 | 2.05E-01 | 2.68E+00 | 2.74E+00 | 8.55E-01 | no | 6330411D24Rik | intron_variant,non_coding_transcript_variant | MODIFIER |
| 4 | 143034768 | AX-170444659 | 0 | C | G | 0.105 | 3.72E-02 | 2.05E-01 | 2.68E+00 | 2.74E+00 | 8.55E-01 | no | 6330411D24Rik | intron_variant,non_coding_transcript_variant | MODIFIER |
| 4 | 143035553 | AX-170469222 | 0 | C | T | 0.105 | 3.72E-02 | 2.05E-01 | 2.68E+00 | 2.74E+00 | 8.55E-01 | no | 6330411D24Rik | intron_variant,non_coding_transcript_variant | MODIFIER |
| 4 | 143056831 | AX-170405020 | 0 | G | A | 0.105 | 3.72E-02 | 2.05E-01 | 2.68E+00 | 2.74E+00 | 8.55E-01 | no | 6330411D24Rik | intron_variant,non_coding_transcript_variant | MODIFIER |
| 4 | 143057035 | AX-170474169 | 0 | T | C | 0.105 | 3.72E-02 | 2.05E-01 | 2.68E+00 | 2.74E+00 | 8.55E-01 | no | 6330411D24Rik | intron_variant,non_coding_transcript_variant | MODIFIER |
| 4 | 143089370 | AX-97829068 | 0 | C | T | 0.105 | 3.72E-02 | 2.05E-01 | 2.68E+00 | 2.74E+00 | 8.55E-01 | no | | intergenic_variant | MODIFIER |
| 4 | 143101359 | AX-170024159 | 0 | A | G | 0.105 | 3.72E-02 | 2.05E-01 | 2.68E+00 | 2.74E+00 | 8.55E-01 | no | | intergenic_variant | MODIFIER |
| 4 | 143131597 | AX-170130042 | 0 | C | T | 0.105 | 3.72E-02 | 2.05E-01 | 2.68E+00 | 2.74E+00 | 8.55E-01 | no | Prdm2 | intron_variant | MODIFIER |
| 4 | 143144067 | AX-170024158 | 0 | T | C | 0.105 | 3.72E-02 | 2.05E-01 | 2.68E+00 | 2.74E+00 | 8.55E-01 | no | Prdm2 | intron_variant | MODIFIER |
| 4 | 143156118 | AX-170024162 | 0 | T | C | 0.105 | 3.72E-02 | 2.05E-01 | 2.68E+00 | 2.74E+00 | 8.55E-01 | no | Prdm2 | intron_variant | MODIFIER |
| 4 | 143174817 | AX-170070920 | 0 | T | C | 0.105 | 3.72E-02 | 2.05E-01 | 2.68E+00 | 2.74E+00 | 8.55E-01 | no | Prdm2 | intron_variant | MODIFIER |
| 4 | 143192585 | AX-170024156 | 0 | G | A | 0.105 | 3.72E-02 | 2.05E-01 | 2.68E+00 | 2.74E+00 | 8.55E-01 | no | Prdm2 | intron_variant | MODIFIER |
| 4 | 143202956 | AX-170070922 | 0 | A | C | 0.105 | 3.72E-02 | 2.05E-01 | 2.68E+00 | 2.74E+00 | 8.55E-01 | no | Prdm2 | intron_variant | MODIFIER |
| 4 | 143229304 | AX-97856859 | 0 | T | C | 0.105 | 3.72E-02 | 2.05E-01 | 2.68E+00 | 2.74E+00 | 8.55E-01 | no | | intergenic_variant | MODIFIER |
| 4 | 143250534 | AX-170086205 | 0 | G | A | 0.105 | 3.72E-02 | 2.05E-01 | 2.68E+00 | 2.74E+00 | 8.55E-01 | no | | intergenic_variant | MODIFIER |
| 4 | 143261926 | AX-97798757 | 0 | C | T | 0.105 | 3.72E-02 | 2.05E-01 | 2.68E+00 | 2.74E+00 | 8.55E-01 | no | | intergenic_variant | MODIFIER |
| 4 | 143281920 | AX-170441480 | 0 | C | T | 0.105 | 3.72E-02 | 2.05E-01 | 2.68E+00 | 2.74E+00 | 8.55E-01 | no | Pdpn | intron_variant | MODIFIER |
| 4 | 143282141 | AX-170114465 | 0 | T | C | 0.105 | 3.72E-02 | 2.05E-01 | 2.68E+00 | 2.74E+00 | 8.55E-01 | no | Pdpn | intron_variant | MODIFIER |
| 4 | 143290189 | AX-170456268 | 0 | T | C | 0.105 | 3.72E-02 | 2.05E-01 | 2.68E+00 | 2.74E+00 | 8.55E-01 | no | Pdpn | intron_variant | MODIFIER |
| 4 | 143290231 | AX-170185300 | 0 | C | T | 0.105 | 3.72E-02 | 2.05E-01 | 2.68E+00 | 2.74E+00 | 8.55E-01 | no | Pdpn | intron_variant | MODIFIER |
| 4 | 143321220 | AX-97793949 | 0 | A | G | 0.105 | 3.72E-02 | 2.05E-01 | 2.68E+00 | 2.74E+00 | 8.55E-01 | no | | intergenic_variant | MODIFIER |

| | | | | | | | | | | | | | | | |
|---|---|---|---|---|---|---|---|---|---|---|---|---|---|---|---|
| 4 | 143321241 | AX-170490458 | 0 | C | T | 0.105 | 3.72E-02 | 2.05E-01 | 2.68E+00 | 2.74E+00 | 8.55E-01 | no | | intergenic_variant | MODIFIER |
| 4 | 143334224 | AX-170373171 | 0 | T | C | 0.105 | 3.72E-02 | 2.05E-01 | 2.68E+00 | 2.74E+00 | 8.55E-01 | no | | regulatory_region_variant | MODIFIER |
| 4 | 143339973 | AX-170370486 | 0 | A | G | 0.105 | 3.72E-02 | 2.05E-01 | 2.68E+00 | 2.74E+00 | 8.55E-01 | no | Gm26313 | upstream_gene_variant | MODIFIER |
| 4 | 143341277 | AX-170161040 | 0 | C | T | 0.105 | 3.72E-02 | 2.05E-01 | 2.68E+00 | 2.74E+00 | 8.55E-01 | no | Gm26313 | downstream_gene_variant | MODIFIER |
| 4 | 143343932 | AX-170377639 | 0 | A | G | 0.105 | 3.72E-02 | 2.05E-01 | 2.68E+00 | 2.74E+00 | 8.55E-01 | no | Gm26313 | downstream_gene_variant | MODIFIER |
| 4 | 143343995 | AX-170292010 | 0 | G | A | 0.105 | 3.72E-02 | 2.05E-01 | 2.68E+00 | 2.74E+00 | 8.55E-01 | no | Gm26313 | downstream_gene_variant | MODIFIER |
| 4 | 143362073 | AX-170181072 | 0 | T | C | 0.105 | 3.72E-02 | 2.05E-01 | 2.68E+00 | 2.74E+00 | 8.55E-01 | no | Lrrc38 | intron_variant | MODIFIER |
| 4 | 143376244 | AX-170384296 | 0 | G | C | 0.105 | 3.72E-02 | 2.05E-01 | 2.68E+00 | 2.74E+00 | 8.55E-01 | no | | intergenic_variant | MODIFIER |
| 4 | 143404813 | AX-170316737 | 0 | C | A | 0.105 | 3.72E-02 | 2.05E-01 | 2.68E+00 | 2.74E+00 | 8.55E-01 | no | Pramel1 | downstream_gene_variant | MODIFIER |
| 4 | 143418257 | AX-170185687 | 0 | C | T | 0.105 | 3.72E-02 | 2.05E-01 | 2.68E+00 | 2.74E+00 | 8.55E-01 | no | Pramef8 | intron_variant | MODIFIER |
| 4 | 143418661 | AX-170240483 | 0 | T | G | 0.105 | 3.72E-02 | 2.05E-01 | 2.68E+00 | 2.74E+00 | 8.55E-01 | no | Pramef8 | intron_variant | MODIFIER |
| 4 | 143420477 | AX-170426160 | 0 | A | G | 0.105 | 3.72E-02 | 2.05E-01 | 2.68E+00 | 2.74E+00 | 8.55E-01 | no | Pramef8 | 3_prime_UTR_variant | MODIFIER |
| 4 | 143429441 | AX-170298652 | 0 | A | G | 0.105 | 3.72E-02 | 2.05E-01 | 2.68E+00 | 2.74E+00 | 8.55E-01 | no | | intergenic_variant | MODIFIER |
| 4 | 143495171 | AX-170070927 | 0 | A | G | 0.105 | 3.72E-02 | 2.05E-01 | 2.68E+00 | 2.74E+00 | 8.55E-01 | no | Gm13041 | upstream_gene_variant | MODIFIER |
| 4 | 143501626 | AX-97806430 | 0 | C | T | 0.105 | 3.72E-02 | 2.05E-01 | 2.68E+00 | 2.74E+00 | 8.55E-01 | no | Gm13041 | non_coding_transcript_exon_variant | MODIFIER |
| 4 | 143502286 | AX-170070929 | 0 | C | T | 0.105 | 3.72E-02 | 2.05E-01 | 2.68E+00 | 2.74E+00 | 8.55E-01 | no | Gm13041 | downstream_gene_variant | MODIFIER |
| 4 | 143502385 | AX-97793788 | 0 | A | G | 0.105 | 3.72E-02 | 2.05E-01 | 2.68E+00 | 2.74E+00 | 8.55E-01 | no | Gm13041 | downstream_gene_variant | MODIFIER |
| 4 | 143571145 | AX-97831561 | 0 | C | T | 0.105 | 3.72E-02 | 2.05E-01 | 2.68E+00 | 2.74E+00 | 8.55E-01 | no | BC080695 | missense_variant | MODERATE |
| 4 | 143691944 | AX-170024173 | 0 | A | C | 0.105 | 3.72E-02 | 2.05E-01 | 2.68E+00 | 2.74E+00 | 8.55E-01 | no | Gm13089 | downstream_gene_variant | MODIFIER |
| 4 | 143692203 | AX-170024170 | 0 | G | T | 0.105 | 3.72E-02 | 2.05E-01 | 2.68E+00 | 2.74E+00 | 8.55E-01 | no | Gm13089 | downstream_gene_variant | MODIFIER |
| 4 | 143719569 | AX-170070928 | 0 | A | G | 0.105 | 3.72E-02 | 2.05E-01 | 2.68E+00 | 2.74E+00 | 8.55E-01 | no | Gm13078 | intron_variant | MODIFIER |
| 4 | 143791713 | AX-97810722 | 0 | C | T | 0.105 | 3.72E-02 | 2.05E-01 | 2.68E+00 | 2.74E+00 | 8.55E-01 | no | Gm13023 | intron_variant | MODIFIER |
| 4 | 143808354 | AX-170121748 | 0 | G | C | 0.105 | 3.72E-02 | 2.05E-01 | 2.68E+00 | 2.74E+00 | 8.55E-01 | no | Gm13084 | downstream_gene_variant | MODIFIER |
| 4 | 143838504 | AX-170024177 | 0 | G | A | 0.105 | 3.72E-02 | 2.05E-01 | 2.68E+00 | 2.74E+00 | 8.55E-01 | no | | intergenic_variant | MODIFIER |
| 4 | 143842414 | AX-170024166 | 0 | C | T | 0.105 | 3.72E-02 | 2.05E-01 | 2.68E+00 | 2.74E+00 | 8.55E-01 | no | Gm13103 | upstream_gene_variant | MODIFIER |
| 4 | 143894350 | AX-170388228 | 0 | G | C | 0.105 | 3.72E-02 | 2.05E-01 | 2.68E+00 | 2.74E+00 | 8.55E-01 | no | Pramef6 | intron_variant | MODIFIER |
| 4 | 143951203 | AX-170024174 | 0 | T | C | 0.105 | 3.72E-02 | 2.05E-01 | 2.68E+00 | 2.74E+00 | 8.55E-01 | no | Pramef25 | upstream_gene_variant | MODIFIER |
| 4 | 144157955 | AX-170286811 | 0 | G | A | 0.105 | 3.72E-02 | 2.05E-01 | 2.68E+00 | 2.74E+00 | 8.55E-01 | no | Oog3 | synonymous_variant | LOW |
| 4 | 144190638 | AX-170400999 | 0 | C | A | 0.105 | 3.72E-02 | 2.05E-01 | 2.68E+00 | 2.74E+00 | 8.55E-01 | no | Oog2 | upstream_gene_variant | MODIFIER |
| 4 | 144240302 | AX-97838642 | 0 | T | C | 0.105 | 3.72E-02 | 2.05E-01 | 2.68E+00 | 2.74E+00 | 8.55E-01 | no | | intergenic_variant | MODIFIER |
| 4 | 144319566 | AX-170288164 | 0 | T | C | 0.105 | 3.72E-02 | 2.05E-01 | 2.68E+00 | 2.74E+00 | 8.55E-01 | no | | intergenic_variant | MODIFIER |
| 4 | 144357060 | AX-170024183 | 0 | A | G | 0.105 | 3.72E-02 | 2.05E-01 | 2.68E+00 | 2.74E+00 | 8.55E-01 | no | Gm13119 | upstream_gene_variant | MODIFIER |
| 4 | 144388010 | AX-170024182 | 0 | T | A | 0.105 | 3.72E-02 | 2.05E-01 | 2.68E+00 | 2.74E+00 | 8.55E-01 | no | Pramef12 | downstream_gene_variant | MODIFIER |
| 4 | 144395714 | AX-170277618 | 0 | T | C | 0.105 | 3.72E-02 | 2.05E-01 | 2.68E+00 | 2.74E+00 | 8.55E-01 | no | Pramef12 | missense_variant | MODERATE |
| 4 | 144401147 | AX-170024180 | 0 | C | T | 0.105 | 3.72E-02 | 2.05E-01 | 2.68E+00 | 2.74E+00 | 8.55E-01 | no | Pramef12 | intron_variant | MODIFIER |
| 4 | 144440181 | AX-170024184 | 0 | T | A | 0.105 | 3.72E-02 | 2.05E-01 | 2.68E+00 | 2.74E+00 | 8.55E-01 | no | 1700012P22Rik | upstream_gene_variant | MODIFIER |
| 4 | 144453362 | AX-170024185 | 0 | G | A | 0.105 | 3.72E-02 | 2.05E-01 | 2.68E+00 | 2.74E+00 | 8.55E-01 | no | Aadacl3 | downstream_gene_variant | MODIFIER |
| 4 | 144465744 | AX-170481161 | 0 | G | C | 0.105 | 3.72E-02 | 2.05E-01 | 2.68E+00 | 2.74E+00 | 8.55E-01 | no | Aadacl3 | upstream_gene_variant | MODIFIER |
| 4 | 144465877 | AX-170287785 | 0 | C | G | 0.105 | 3.72E-02 | 2.05E-01 | 2.68E+00 | 2.74E+00 | 8.55E-01 | no | Aadacl3 | upstream_gene_variant | MODIFIER |
| 4 | 144470192 | AX-97801567 | 0 | C | T | 0.105 | 3.72E-02 | 2.05E-01 | 2.68E+00 | 2.74E+00 | 8.55E-01 | no | | intergenic_variant | MODIFIER |
| 4 | 144486113 | AX-97784179 | 0 | T | G | 0.105 | 3.72E-02 | 2.05E-01 | 2.68E+00 | 2.74E+00 | 8.55E-01 | no | | intergenic_variant | MODIFIER |
| 4 | 144490068 | AX-170406027 | 0 | C | T | 0.105 | 3.72E-02 | 2.05E-01 | 2.68E+00 | 2.74E+00 | 8.55E-01 | no | | intergenic_variant | MODIFIER |
| 4 | 144543906 | AX-170070944 | 0 | G | A | 0.105 | 3.72E-02 | 2.05E-01 | 2.68E+00 | 2.74E+00 | 8.55E-01 | no | Gm13126 | downstream_gene_variant | MODIFIER |
| 4 | 144544111 | AX-97780713 | 0 | G | A | 0.105 | 3.72E-02 | 2.05E-01 | 2.68E+00 | 2.74E+00 | 8.55E-01 | no | Gm13126 | downstream_gene_variant | MODIFIER |
| 4 | 144578984 | AX-97799433 | 0 | A | G | 0.105 | 3.72E-02 | 2.05E-01 | 2.68E+00 | 2.74E+00 | 8.55E-01 | no | | intergenic_variant | MODIFIER |
| 4 | 144610709 | AX-97820611 | 0 | C | T | 0.105 | 3.72E-02 | 2.05E-01 | 2.68E+00 | 2.74E+00 | 8.55E-01 | no | Aadacl4 | upstream_gene_variant | MODIFIER |
| 4 | 144621681 | AX-170150140 | 0 | A | C | 0.105 | 3.72E-02 | 2.05E-01 | 2.68E+00 | 2.74E+00 | 8.55E-01 | no | Aadacl4 | intron_variant | MODIFIER |
| 4 | 144638196 | AX-97813878 | 0 | C | T | 0.105 | 3.72E-02 | 2.05E-01 | 2.68E+00 | 2.74E+00 | 8.55E-01 | no | | intergenic_variant | MODIFIER |
| 4 | 144638245 | AX-170333200 | 0 | A | T | 0.105 | 3.72E-02 | 2.05E-01 | 2.68E+00 | 2.74E+00 | 8.55E-01 | no | | intergenic_variant | MODIFIER |
| 4 | 144640958 | AX-170413108 | 0 | A | T | 0.105 | 3.72E-02 | 2.05E-01 | 2.68E+00 | 2.74E+00 | 8.55E-01 | no | | intergenic_variant | MODIFIER |
| 4 | 144674787 | AX-170122404 | 0 | C | A | 0.105 | 3.72E-02 | 2.05E-01 | 2.68E+00 | 2.74E+00 | 8.55E-01 | no | Gm436 | intron_variant | MODIFIER |
| 4 | 144685481 | AX-97832091 | 0 | T | C | 0.105 | 3.72E-02 | 2.05E-01 | 2.68E+00 | 2.74E+00 | 8.55E-01 | no | Gm436 | intron_variant | MODIFIER |
| 4 | 144715728 | AX-97831197 | 0 | C | T | 0.105 | 3.72E-02 | 2.05E-01 | 2.68E+00 | 2.74E+00 | 8.55E-01 | no | Gm13178 | intron_variant | MODIFIER |
| 4 | 144715847 | AX-170337094 | 0 | G | A | 0.105 | 3.72E-02 | 2.05E-01 | 2.68E+00 | 2.74E+00 | 8.55E-01 | no | Gm13178 | intron_variant | MODIFIER |

| | | | | | | | | | | | | | | | |
|---|---|---|---|---|---|---|---|---|---|---|---|---|---|---|---|
| 4 | 144716034 | AX-170202311 | 0 | A | G | 0.105 | 3.72E-02 | 2.05E-01 | 2.68E+00 | 2.74E+00 | 8.55E-01 | no | Gm13178 | intron_variant | MODIFIER |
| 4 | 144750145 | AX-170146239 | 0 | G | A | 0.105 | 3.72E-02 | 2.05E-01 | 2.68E+00 | 2.74E+00 | 8.55E-01 | no | | intergenic_variant | MODIFIER |
| 4 | 144779199 | AX-170317735 | 0 | G | A | 0.105 | 3.72E-02 | 2.05E-01 | 2.68E+00 | 2.74E+00 | 8.55E-01 | no | Gm438 | intron_variant | MODIFIER |
| 4 | 144779366 | AX-170161454 | 0 | T | C | 0.105 | 3.72E-02 | 2.05E-01 | 2.68E+00 | 2.74E+00 | 8.55E-01 | no | Gm438 | intron_variant | MODIFIER |
| 4 | 144780027 | AX-170178498 | 0 | G | A | 0.105 | 3.72E-02 | 2.05E-01 | 2.68E+00 | 2.74E+00 | 8.55E-01 | no | Gm438 | intron_variant | MODIFIER |
| 4 | 144780561 | AX-170491101 | 0 | G | A | 0.105 | 3.72E-02 | 2.05E-01 | 2.68E+00 | 2.74E+00 | 8.55E-01 | no | Gm438 | missense_variant | MODERATE |
| 4 | 144790614 | AX-170145514 | 0 | A | G | 0.105 | 3.72E-02 | 2.05E-01 | 2.68E+00 | 2.74E+00 | 8.55E-01 | no | Gm438 | upstream_gene_variant | MODIFIER |
| 4 | 144792627 | AX-170121608 | 0 | T | C | 0.105 | 3.72E-02 | 2.05E-01 | 2.68E+00 | 2.74E+00 | 8.55E-01 | no | Prdx2-rs3 | non_coding_transcript_exon_variant | MODIFIER |
| 4 | 144861341 | AX-170070949 | 0 | A | G | 0.105 | 3.72E-02 | 2.05E-01 | 2.68E+00 | 2.74E+00 | 8.55E-01 | no | | intergenic_variant | MODIFIER |
| 4 | 144885692 | AX-97809961 | 0 | A | G | 0.105 | 3.72E-02 | 2.05E-01 | 2.68E+00 | 2.74E+00 | 8.55E-01 | no | | intergenic_variant | MODIFIER |
| 4 | 144900941 | AX-170413638 | 0 | G | A | 0.105 | 3.72E-02 | 2.05E-01 | 2.68E+00 | 2.74E+00 | 8.55E-01 | no | Dhrs3 | intron_variant,NMD_transcript_variant | MODIFIER |
| 4 | 144905035 | AX-170399430 | 0 | T | C | 0.105 | 3.72E-02 | 2.05E-01 | 2.68E+00 | 2.74E+00 | 8.55E-01 | no | Dhrs3 | intron_variant,NMD_transcript_variant | MODIFIER |
| 4 | 144941886 | AX-170237762 | 0 | G | C | 0.105 | 3.72E-02 | 2.05E-01 | 2.68E+00 | 2.74E+00 | 8.55E-01 | no | | regulatory_region_variant | MODIFIER |
| 4 | 144942361 | AX-170383164 | 0 | T | C | 0.105 | 3.72E-02 | 2.05E-01 | 2.68E+00 | 2.74E+00 | 8.55E-01 | no | | intergenic_variant | MODIFIER |
| 4 | 144973265 | AX-170024190 | 0 | T | C | 0.105 | 3.72E-02 | 2.05E-01 | 2.68E+00 | 2.74E+00 | 8.55E-01 | no | Vps13d | 3_prime_UTR_variant | MODIFIER |
| 4 | 144976199 | AX-170124929 | 0 | C | T | 0.105 | 3.72E-02 | 2.05E-01 | 2.68E+00 | 2.74E+00 | 8.55E-01 | no | Vps13d | intron_variant | MODIFIER |
| 4 | 144978939 | AX-170342268 | 0 | G | A | 0.105 | 3.72E-02 | 2.05E-01 | 2.68E+00 | 2.74E+00 | 8.55E-01 | no | Vps13d | intron_variant | MODIFIER |
| 4 | 144982630 | AX-170139875 | 0 | C | T | 0.105 | 3.72E-02 | 2.05E-01 | 2.68E+00 | 2.74E+00 | 8.55E-01 | no | Vps13d | intron_variant | MODIFIER |
| 4 | 144992890 | AX-170024188 | 0 | C | G | 0.105 | 3.72E-02 | 2.05E-01 | 2.68E+00 | 2.74E+00 | 8.55E-01 | no | Vps13d | intron_variant | MODIFIER |
| 4 | 144993302 | AX-170024187 | 0 | T | C | 0.105 | 3.72E-02 | 2.05E-01 | 2.68E+00 | 2.74E+00 | 8.55E-01 | no | Vps13d | intron_variant | MODIFIER |
| 4 | 144994426 | AX-170024189 | 0 | C | T | 0.105 | 3.72E-02 | 2.05E-01 | 2.68E+00 | 2.74E+00 | 8.55E-01 | no | Vps13d | intron_variant | MODIFIER |
| 4 | 145014187 | AX-170070958 | 0 | C | T | 0.105 | 3.72E-02 | 2.05E-01 | 2.68E+00 | 2.74E+00 | 8.55E-01 | no | Vps13d | intron_variant | MODIFIER |
| 4 | 145020147 | AX-170308215 | 0 | G | T | 0.105 | 3.72E-02 | 2.05E-01 | 2.68E+00 | 2.74E+00 | 8.55E-01 | no | Vps13d | synonymous_variant | LOW |
| 4 | 145045487 | AX-97823702 | 0 | A | G | 0.105 | 3.72E-02 | 2.05E-01 | 2.68E+00 | 2.74E+00 | 8.55E-01 | no | Vps13d | intron_variant | MODIFIER |
| 4 | 145075845 | AX-170117547 | 0 | G | A | 0.105 | 3.72E-02 | 2.05E-01 | 2.68E+00 | 2.74E+00 | 8.55E-01 | no | Vps13d | intron_variant | MODIFIER |
| 4 | 145077456 | AX-170342011 | 0 | T | C | 0.105 | 3.72E-02 | 2.05E-01 | 2.68E+00 | 2.74E+00 | 8.55E-01 | no | Vps13d | intron_variant | MODIFIER |
| 4 | 145086329 | AX-170193063 | 0 | G | T | 0.105 | 3.72E-02 | 2.05E-01 | 2.68E+00 | 2.74E+00 | 8.55E-01 | no | Vps13d | intron_variant | MODIFIER |
| 4 | 145105601 | AX-97843780 | 0 | A | G | 0.105 | 3.72E-02 | 2.05E-01 | 2.68E+00 | 2.74E+00 | 8.55E-01 | no | Vps13d | intron_variant | MODIFIER |
| 4 | 145110291 | AX-170183235 | 0 | A | G | 0.105 | 3.72E-02 | 2.05E-01 | 2.68E+00 | 2.74E+00 | 8.55E-01 | no | Vps13d | intron_variant | MODIFIER |
| 4 | 145115672 | AX-170155684 | 0 | T | A | 0.105 | 3.72E-02 | 2.05E-01 | 2.68E+00 | 2.74E+00 | 8.55E-01 | no | Vps13d | intron_variant | MODIFIER |
| 4 | 145144934 | AX-170208561 | 0 | G | A | 0.105 | 3.72E-02 | 2.05E-01 | 2.68E+00 | 2.74E+00 | 8.55E-01 | no | Vps13d | intron_variant | MODIFIER |
| 4 | 145162034 | AX-170193016 | 0 | C | T | 0.105 | 3.72E-02 | 2.05E-01 | 2.68E+00 | 2.74E+00 | 8.55E-01 | no | Vps13d | synonymous_variant | LOW |
| 4 | 145168521 | AX-170451290 | 0 | A | G | 0.105 | 3.72E-02 | 2.05E-01 | 2.68E+00 | 2.74E+00 | 8.55E-01 | no | Vps13d | synonymous_variant | LOW |
| 4 | 145169484 | AX-170166988 | 0 | C | G | 0.105 | 3.72E-02 | 2.05E-01 | 2.68E+00 | 2.74E+00 | 8.55E-01 | no | Vps13d | intron_variant | MODIFIER |
| 4 | 145186864 | AX-170221360 | 0 | T | C | 0.105 | 3.72E-02 | 2.05E-01 | 2.68E+00 | 2.74E+00 | 8.55E-01 | no | Vps13d | intron_variant | MODIFIER |
| 4 | 145197080 | AX-170275605 | 0 | A | G | 0.105 | 3.72E-02 | 2.05E-01 | 2.68E+00 | 2.74E+00 | 8.55E-01 | no | Vps13d | upstream_gene_variant | MODIFIER |
| 4 | 145202687 | AX-97783934 | 0 | G | C | 0.105 | 3.72E-02 | 2.05E-01 | 2.68E+00 | 2.74E+00 | 8.55E-01 | no | | intergenic_variant | MODIFIER |
| 4 | 145209607 | AX-170414998 | 0 | T | C | 0.105 | 3.72E-02 | 2.05E-01 | 2.68E+00 | 2.74E+00 | 8.55E-01 | no | Tnfrsf1b | downstream_gene_variant | MODIFIER |
| 4 | 145229746 | AX-170372886 | 0 | G | A | 0.105 | 3.72E-02 | 2.05E-01 | 2.68E+00 | 2.74E+00 | 8.55E-01 | no | Tnfrsf1b | intron_variant | MODIFIER |
| 4 | 145261624 | AX-170303621 | 0 | G | A | 0.105 | 3.72E-02 | 2.05E-01 | 2.68E+00 | 2.74E+00 | 8.55E-01 | no | | intergenic_variant | MODIFIER |
| 4 | 145309317 | AX-170352041 | 0 | C | T | 0.105 | 3.72E-02 | 2.05E-01 | 2.68E+00 | 2.74E+00 | 8.55E-01 | no | Tnfrsf8 | intron_variant | MODIFIER |
| 4 | 145309867 | AX-170470152 | 0 | G | A | 0.105 | 3.72E-02 | 2.05E-01 | 2.68E+00 | 2.74E+00 | 8.55E-01 | no | Tnfrsf8 | intron_variant | MODIFIER |
| 4 | 145335634 | AX-170358094 | 0 | A | C | 0.105 | 3.72E-02 | 2.05E-01 | 2.68E+00 | 2.74E+00 | 8.55E-01 | no | | intergenic_variant | MODIFIER |
| 4 | 145335678 | AX-170174377 | 0 | A | G | 0.105 | 3.72E-02 | 2.05E-01 | 2.68E+00 | 2.74E+00 | 8.55E-01 | no | | intergenic_variant | MODIFIER |
| 4 | 145336773 | AX-170374851 | 0 | G | A | 0.105 | 3.72E-02 | 2.05E-01 | 2.68E+00 | 2.74E+00 | 8.55E-01 | no | | intergenic_variant | MODIFIER |
| 4 | 145337630 | AX-170328441 | 0 | A | C | 0.105 | 3.72E-02 | 2.05E-01 | 2.68E+00 | 2.74E+00 | 8.55E-01 | no | | intergenic_variant | MODIFIER |
| 4 | 145353306 | AX-170244897 | 0 | T | C | 0.105 | 3.72E-02 | 2.05E-01 | 2.68E+00 | 2.74E+00 | 8.55E-01 | no | | intergenic_variant | MODIFIER |
| 4 | 145391231 | AX-170182787 | 0 | G | A | 0.105 | 3.72E-02 | 2.05E-01 | 2.68E+00 | 2.74E+00 | 8.55E-01 | no | Gm13024 | non_coding_transcript_exon_variant | MODIFIER |
| 4 | 145482703 | AX-170141724 | 0 | C | T | 0.105 | 3.72E-02 | 2.05E-01 | 2.68E+00 | 2.74E+00 | 8.55E-01 | no | | intergenic_variant | MODIFIER |
| 4 | 146533536 | AX-170422172 | 0 | A | C | 0.105 | 3.72E-02 | 2.05E-01 | 2.68E+00 | 2.74E+00 | 8.55E-01 | no | Zfp981 | intron_variant | MODIFIER |
| 4 | 147080345 | AX-170473639 | 0 | T | C | 0.105 | 3.72E-02 | 2.05E-01 | 2.68E+00 | 2.74E+00 | 8.55E-01 | no | C230088H06Rik | intron_variant,non_coding_transcript_variant | MODIFIER |
| 4 | 147420259 | AX-170287811 | 0 | T | C | 0.105 | 3.72E-02 | 2.05E-01 | 2.68E+00 | 2.74E+00 | 8.55E-01 | no | Gm26624 | intron_variant,non_coding_transcript_variant | MODIFIER |
| 4 | 147491717 | AX-170518390 | 0 | A | C | 0.105 | 3.72E-02 | 2.05E-01 | 2.68E+00 | 2.74E+00 | 8.55E-01 | no | Zfp982 | upstream_gene_variant | MODIFIER |
| 4 | 147829708 | AX-170024196 | 0 | A | G | 0.105 | 3.72E-02 | 2.05E-01 | 2.68E+00 | 2.74E+00 | 8.55E-01 | no | Zfp933 | intron_variant | MODIFIER |

| | | | | | | | | | | | | | | | |
|---|---|---|---|---|---|---|---|---|---|---|---|---|---|---|---|
| 4 | 147845974 | AX-97781608 | 0 | T | A | 0.105 | 3.72E-02 | 2.05E-01 | 2.68E+00 | 2.74E+00 | 8.55E-01 | no | Zfp933 | intron_variant | MODIFIER |
| 4 | 147850517 | AX-170024198 | 0 | A | G | 0.105 | 3.72E-02 | 2.05E-01 | 2.68E+00 | 2.74E+00 | 8.55E-01 | no | Zfp933 | upstream_gene_variant | MODIFIER |
| 4 | 147850637 | AX-170024197 | 0 | C | T | 0.105 | 3.72E-02 | 2.05E-01 | 2.68E+00 | 2.74E+00 | 8.55E-01 | no | Zfp933 | upstream_gene_variant | MODIFIER |
| 4 | 147863898 | AX-170336227 | 0 | G | T | 0.105 | 3.72E-02 | 2.05E-01 | 2.68E+00 | 2.74E+00 | 8.55E-01 | no | Miip | intron_variant | MODIFIER |
| 4 | 147885688 | AX-170070979 | 0 | C | T | 0.105 | 3.72E-02 | 2.05E-01 | 2.68E+00 | 2.74E+00 | 8.55E-01 | no | Mfn2 | missense_variant | MODERATE |
| 4 | 147891973 | AX-170387091 | 0 | C | A | 0.105 | 3.72E-02 | 2.05E-01 | 2.68E+00 | 2.74E+00 | 8.55E-01 | no | Mfn2 | intron_variant | MODIFIER |
| 4 | 147895934 | AX-170445848 | 0 | C | T | 0.105 | 3.72E-02 | 2.05E-01 | 2.68E+00 | 2.74E+00 | 8.55E-01 | no | Mfn2 | intron_variant | MODIFIER |
| 4 | 147899754 | AX-170334722 | 0 | C | G | 0.105 | 3.72E-02 | 2.05E-01 | 2.68E+00 | 2.74E+00 | 8.55E-01 | no | Mfn2 | intron_variant | MODIFIER |
| 4 | 147900274 | AX-170335541 | 0 | G | A | 0.105 | 3.72E-02 | 2.05E-01 | 2.68E+00 | 2.74E+00 | 8.55E-01 | no | Mfn2 | intron_variant | MODIFIER |
| 4 | 147918221 | AX-170533791 | 0 | A | C | 0.105 | 3.72E-02 | 2.05E-01 | 2.68E+00 | 2.74E+00 | 8.55E-01 | no | Plod1 | intron_variant | MODIFIER |
| 4 | 147925810 | AX-170180773 | 0 | C | T | 0.105 | 3.72E-02 | 2.05E-01 | 2.68E+00 | 2.74E+00 | 8.55E-01 | no | Plod1 | intron_variant | MODIFIER |
| 4 | 147926520 | AX-170154030 | 0 | A | G | 0.105 | 3.72E-02 | 2.05E-01 | 2.68E+00 | 2.74E+00 | 8.55E-01 | no | Plod1 | intron_variant | MODIFIER |
| 4 | 147940073 | AX-170475572 | 0 | G | A | 0.105 | 3.72E-02 | 2.05E-01 | 2.68E+00 | 2.74E+00 | 8.55E-01 | no | Plod1 | upstream_gene_variant | MODIFIER |
| 4 | 147940091 | AX-170533671 | 0 | C | T | 0.105 | 3.72E-02 | 2.05E-01 | 2.68E+00 | 2.74E+00 | 8.55E-01 | no | Plod1 | upstream_gene_variant | MODIFIER |
| 4 | 147969127 | AX-170300615 | 0 | A | G | 0.105 | 3.72E-02 | 2.05E-01 | 2.68E+00 | 2.74E+00 | 8.55E-01 | no | | regulatory_region_variant | MODIFIER |
| 4 | 147979304 | AX-170461606 | 0 | G | A | 0.105 | 3.72E-02 | 2.05E-01 | 2.68E+00 | 2.74E+00 | 8.55E-01 | no | | intergenic_variant | MODIFIER |
| 4 | 147979865 | AX-170423126 | 0 | C | T | 0.105 | 3.72E-02 | 2.05E-01 | 2.68E+00 | 2.74E+00 | 8.55E-01 | no | | regulatory_region_variant | MODIFIER |
| 4 | 147981217 | AX-170510447 | 0 | T | C | 0.105 | 3.72E-02 | 2.05E-01 | 2.68E+00 | 2.74E+00 | 8.55E-01 | no | Nppb | upstream_gene_variant | MODIFIER |
| 4 | 147986619 | AX-170308976 | 0 | T | C | 0.105 | 3.72E-02 | 2.05E-01 | 2.68E+00 | 2.74E+00 | 8.55E-01 | no | Nppb | intron_variant | MODIFIER |
| 4 | 148004217 | AX-170308874 | 0 | G | T | 0.105 | 3.72E-02 | 2.05E-01 | 2.68E+00 | 2.74E+00 | 8.55E-01 | no | Clcn6 | downstream_gene_variant | MODIFIER |
| 4 | 148012981 | AX-170245350 | 0 | T | C | 0.105 | 3.72E-02 | 2.05E-01 | 2.68E+00 | 2.74E+00 | 8.55E-01 | no | Clcn6 | intron_variant | MODIFIER |
| 4 | 148015508 | AX-170457509 | 0 | T | C | 0.105 | 3.72E-02 | 2.05E-01 | 2.68E+00 | 2.74E+00 | 8.55E-01 | no | Clcn6 | intron_variant | MODIFIER |
| 4 | 148016802 | AX-97804155 | 0 | T | C | 0.105 | 3.72E-02 | 2.05E-01 | 2.68E+00 | 2.74E+00 | 8.55E-01 | no | Clcn6 | intron_variant | MODIFIER |
| 4 | 148017912 | AX-170432122 | 0 | G | A | 0.105 | 3.72E-02 | 2.05E-01 | 2.68E+00 | 2.74E+00 | 8.55E-01 | no | Clcn6 | synonymous_variant | LOW |
| 4 | 148047855 | AX-170070981 | 0 | G | A | 0.105 | 3.72E-02 | 2.05E-01 | 2.68E+00 | 2.74E+00 | 8.55E-01 | no | Mthfr | intron_variant | MODIFIER |
| 4 | 148054011 | AX-170180724 | 0 | G | A | 0.105 | 3.72E-02 | 2.05E-01 | 2.68E+00 | 2.74E+00 | 8.55E-01 | no | Mthfr | intron_variant | MODIFIER |
| 4 | 149524497 | AX-170449592 | 0 | T | G | 0.105 | 3.72E-02 | 2.05E-01 | 2.68E+00 | 2.74E+00 | 8.55E-01 | no | Ctnnbip1 | intron_variant | MODIFIER |
| 4 | 149535724 | AX-170473945 | 0 | C | A | 0.105 | 3.72E-02 | 2.05E-01 | 2.68E+00 | 2.74E+00 | 8.55E-01 | no | Ctnnbip1 | intron_variant | MODIFIER |
| 4 | 149541488 | AX-97821373 | 0 | C | T | 0.105 | 3.72E-02 | 2.05E-01 | 2.68E+00 | 2.74E+00 | 8.55E-01 | no | Ctnnbip1 | intron_variant | MODIFIER |
| 4 | 149623683 | AX-97835498 | 0 | T | C | 0.105 | 3.72E-02 | 2.05E-01 | 2.68E+00 | 2.74E+00 | 8.55E-01 | no | Clstn1 | intron_variant | MODIFIER |
| 4 | 149624490 | AX-170146466 | 0 | G | A | 0.105 | 3.72E-02 | 2.05E-01 | 2.68E+00 | 2.74E+00 | 8.55E-01 | no | Clstn1 | intron_variant | MODIFIER |
| 4 | 149626451 | AX-170024237 | 0 | C | A | 0.105 | 3.72E-02 | 2.05E-01 | 2.68E+00 | 2.74E+00 | 8.55E-01 | no | Clstn1 | intron_variant | MODIFIER |
| 4 | 149691433 | AX-97821405 | 0 | T | C | 0.105 | 3.72E-02 | 2.05E-01 | 2.68E+00 | 2.74E+00 | 8.55E-01 | no | Pik3cd | intron_variant | MODIFIER |
| 4 | 149703136 | AX-170121100 | 0 | G | A | 0.105 | 3.72E-02 | 2.05E-01 | 2.68E+00 | 2.74E+00 | 8.55E-01 | no | Pik3cd | upstream_gene_variant | MODIFIER |
| 4 | 149708499 | AX-97814030 | 0 | T | C | 0.105 | 3.72E-02 | 2.05E-01 | 2.68E+00 | 2.74E+00 | 8.55E-01 | no | | intergenic_variant | MODIFIER |
| 4 | 149720366 | AX-170024236 | 0 | G | A | 0.105 | 3.72E-02 | 2.05E-01 | 2.68E+00 | 2.74E+00 | 8.55E-01 | no | Tmem201 | intron_variant | MODIFIER |
| 4 | 149758784 | AX-170024232 | 0 | C | T | 0.105 | 3.72E-02 | 2.05E-01 | 2.68E+00 | 2.74E+00 | 8.55E-01 | no | Slc25a33 | intron_variant | MODIFIER |
| 4 | 149765138 | AX-170024235 | 0 | T | C | 0.105 | 3.72E-02 | 2.05E-01 | 2.68E+00 | 2.74E+00 | 8.55E-01 | no | Slc25a33 | intron_variant | MODIFIER |
| 4 | 149765324 | AX-170461639 | 0 | T | C | 0.105 | 3.72E-02 | 2.05E-01 | 2.68E+00 | 2.74E+00 | 8.55E-01 | no | Slc25a33 | intron_variant | MODIFIER |
| 4 | 149770777 | AX-97821402 | 0 | G | A | 0.105 | 3.72E-02 | 2.05E-01 | 2.68E+00 | 2.74E+00 | 8.55E-01 | no | Slc25a33 | intron_variant | MODIFIER |
| 4 | 149779843 | AX-170089396 | 0 | C | A | 0.105 | 3.72E-02 | 2.05E-01 | 2.68E+00 | 2.74E+00 | 8.55E-01 | no | Gm13073 | downstream_gene_variant | MODIFIER |
| 4 | 149817187 | AX-97798883 | 0 | A | G | 0.105 | 3.72E-02 | 2.05E-01 | 2.68E+00 | 2.74E+00 | 8.55E-01 | no | n-R5s193 | upstream_gene_variant | MODIFIER |
| 4 | 149848085 | AX-97820198 | 0 | A | C | 0.105 | 3.72E-02 | 2.05E-01 | 2.68E+00 | 2.74E+00 | 8.55E-01 | no | | intergenic_variant | MODIFIER |
| 4 | 149921216 | AX-170276565 | 0 | G | A | 0.105 | 3.72E-02 | 2.05E-01 | 2.68E+00 | 2.74E+00 | 8.55E-01 | no | Spsb1 | intron_variant | MODIFIER |
| 18 | 58873807 | AX-97832294 | 1 | C | T | 0.42 | 1.48E-02 | 8.47E-02 | 2.73E+00 | 2.80E+00 | 8.55E-01 | no | Adamts19 | intron_variant | MODIFIER |
| 10 | 42990995 | AX-170466802 | 0 | C | T | 0.368 | -1.42E-02 | 7.53E-02 | 2.62E+00 | 2.72E+00 | 8.55E-01 | no | | regulatory_region_variant | MODIFIER |
| 7 | 111872629 | AX-170266194 | 1 | A | T | 0.455 | -1.65E-02 | 9.28E-02 | 2.68E+00 | 2.75E+00 | 8.55E-01 | no | | intergenic_variant | MODIFIER |
| 13 | 38026773 | AX-170092726 | 0 | G | A | 0.184 | 1.88E-02 | 1.10E-01 | 2.70E+00 | 2.79E+00 | 8.55E-01 | no | Cage1 | intron_variant | MODIFIER |
| 5 | 6658240 | AX-170435095 | 1 | T | A | 0.438 | 1.07E-02 | 6.31E-02 | 2.64E+00 | 2.77E+00 | 8.55E-01 | no | | intergenic_variant | MODIFIER |
| 1 | 4718067 | AX-170533486 | 2 | C | G | 0.509 | 1.41E-02 | 7.03E-02 | 2.60E+00 | 2.68E+00 | 8.55E-01 | no | | intergenic_variant | MODIFIER |
| 1 | 150657938 | AX-170366548 | 0 | A | T | 0.211 | 2.33E-02 | 1.22E-01 | 2.62E+00 | 2.71E+00 | 8.55E-01 | no | Hmcn1 | intron_variant | MODIFIER |
| 1 | 45488861 | AX-170000404 | 1 | A | G | 0.089 | -3.66E-02 | 2.01E-01 | 2.63E+00 | 2.73E+00 | 8.55E-01 | no | Col5a2 | intron_variant | MODIFIER |
| 1 | 117950520 | AX-170355140 | 0 | C | A | 0.175 | 3.25E-02 | 1.64E-01 | 2.58E+00 | 2.70E+00 | 8.55E-01 | no | Gm28168 | downstream_gene_variant | MODIFIER |
| 2 | 147535613 | AX-170435727 | 0 | A | T | 0.105 | -3.18E-02 | 1.86E-01 | 2.67E+00 | 2.78E+00 | 8.55E-01 | no | | intergenic_variant | MODIFIER |

| | | | | | | | | | | | | | | | |
|---|---|---|---|---|---|---|---|---|---|---|---|---|---|---|---|
| 1 | 185275324 | AX-170272605 | 0 | G | C | 0.228 | -1.82E-02 | 1.10E-01 | 2.70E+00 | 2.81E+00 | 8.55E-01 | no | Rab3gap2 | intron_variant | MODIFIER |
| 9 | 35490066 | AX-97827366 | 1 | T | C | 0.429 | 1.55E-02 | 8.45E-02 | 2.65E+00 | 2.73E+00 | 8.56E-01 | no | Cdon | intron_variant | MODIFIER |
| 18 | 60940006 | AX-170198439 | 0 | C | T | 0.412 | 1.41E-02 | 8.20E-02 | 2.72E+00 | 2.80E+00 | 8.56E-01 | no | Camk2a | intron_variant | MODIFIER |
| 18 | 62126513 | AX-170015833 | 0 | G | A | 0.412 | 1.41E-02 | 8.20E-02 | 2.72E+00 | 2.80E+00 | 8.56E-01 | no | | regulatory_region_variant | MODIFIER |
| 13 | 49172630 | AX-170211723 | 0 | G | A | 0.482 | -1.13E-02 | 6.68E-02 | 2.66E+00 | 2.77E+00 | 8.56E-01 | no | | regulatory_region_variant | MODIFIER |
| 7 | 93652862 | AX-97853195 | 0 | G | A | 0.123 | 4.05E-02 | 2.21E-01 | 2.64E+00 | 2.73E+00 | 8.56E-01 | no | | intergenic_variant | MODIFIER |
| 16 | 19186828 | AX-170147731 | 1 | G | A | 0.134 | -1.71E-02 | 9.56E-02 | 2.61E+00 | 2.74E+00 | 8.56E-01 | no | | intergenic_variant | MODIFIER |
| 8 | 26629616 | AX-170295803 | 2 | C | A | 0.173 | -2.00E-02 | 1.12E-01 | 2.66E+00 | 2.74E+00 | 8.56E-01 | no | Gm32050 | upstream_gene_variant | MODIFIER |
| 15 | 87394040 | AX-97830071 | 1 | G | A | 0.134 | 2.92E-02 | 1.56E-01 | 2.65E+00 | 2.71E+00 | 8.56E-01 | no | | intergenic_variant | MODIFIER |
| 8 | 100503745 | AX-170325725 | 0 | A | T | 0.237 | 1.96E-02 | 1.12E-01 | 2.67E+00 | 2.76E+00 | 8.56E-01 | no | | intergenic_variant | MODIFIER |
| 8 | 103492330 | AX-170210925 | 0 | T | G | 0.219 | -1.97E-02 | 1.10E-01 | 2.67E+00 | 2.74E+00 | 8.56E-01 | no | | regulatory_region_variant | MODIFIER |
| 6 | 105383545 | AX-170443562 | 1 | T | G | 0.062 | 4.15E-02 | 2.37E-01 | 2.69E+00 | 2.76E+00 | 8.56E-01 | no | | intergenic_variant | MODIFIER |
| 5 | 92981459 | AX-170262726 | 1 | A | G | 0.411 | 1.88E-02 | 1.03E-01 | 2.66E+00 | 2.72E+00 | 8.56E-01 | no | | intergenic_variant | MODIFIER |
| 5 | 97639574 | AX-170115347 | 1 | G | A | 0.411 | 1.88E-02 | 1.03E-01 | 2.66E+00 | 2.72E+00 | 8.56E-01 | no | 4930467D21Rik | intron_variant,non_coding_transcript_variant | MODIFIER |
| 10 | 5662192 | AX-170091851 | 0 | C | G | 0.211 | -1.92E-02 | 1.04E-01 | 2.64E+00 | 2.72E+00 | 8.56E-01 | no | | intergenic_variant | MODIFIER |
| 10 | 5662407 | AX-170228803 | 0 | G | A | 0.211 | -1.92E-02 | 1.04E-01 | 2.64E+00 | 2.72E+00 | 8.56E-01 | no | | intergenic_variant | MODIFIER |
| 10 | 5692181 | AX-170515394 | 0 | A | G | 0.211 | -1.92E-02 | 1.04E-01 | 2.64E+00 | 2.72E+00 | 8.56E-01 | no | | intergenic_variant | MODIFIER |
| 10 | 5770927 | AX-170470168 | 0 | T | C | 0.211 | -1.92E-02 | 1.04E-01 | 2.64E+00 | 2.72E+00 | 8.56E-01 | no | | intergenic_variant | MODIFIER |
| 10 | 5788182 | AX-170103354 | 0 | C | A | 0.211 | -1.92E-02 | 1.04E-01 | 2.64E+00 | 2.72E+00 | 8.56E-01 | no | | intergenic_variant | MODIFIER |
| 10 | 5846330 | AX-170208716 | 0 | C | T | 0.211 | -1.92E-02 | 1.04E-01 | 2.64E+00 | 2.72E+00 | 8.56E-01 | no | Rgs17 | intron_variant | MODIFIER |
| 10 | 5861405 | AX-170453854 | 0 | C | A | 0.211 | -1.92E-02 | 1.04E-01 | 2.64E+00 | 2.72E+00 | 8.56E-01 | no | Rgs17 | intron_variant | MODIFIER |
| 10 | 5861934 | AX-170162669 | 0 | G | A | 0.211 | -1.92E-02 | 1.04E-01 | 2.64E+00 | 2.72E+00 | 8.56E-01 | no | Rgs17 | intron_variant | MODIFIER |
| 10 | 5864563 | AX-170269014 | 0 | C | T | 0.211 | -1.92E-02 | 1.04E-01 | 2.64E+00 | 2.72E+00 | 8.56E-01 | no | Rgs17 | intron_variant | MODIFIER |
| 10 | 5927012 | AX-170327890 | 0 | C | T | 0.211 | -1.92E-02 | 1.04E-01 | 2.64E+00 | 2.72E+00 | 8.56E-01 | no | Rgs17 | upstream_gene_variant | MODIFIER |
| 10 | 6028947 | AX-170250128 | 0 | A | G | 0.211 | -1.92E-02 | 1.04E-01 | 2.64E+00 | 2.72E+00 | 8.56E-01 | no | | intergenic_variant | MODIFIER |
| 10 | 6042474 | AX-170288317 | 0 | A | G | 0.211 | -1.92E-02 | 1.04E-01 | 2.64E+00 | 2.72E+00 | 8.56E-01 | no | | intergenic_variant | MODIFIER |
| 10 | 6047136 | AX-170198240 | 0 | G | C | 0.211 | -1.92E-02 | 1.04E-01 | 2.64E+00 | 2.72E+00 | 8.56E-01 | no | | intergenic_variant | MODIFIER |
| 10 | 6121241 | AX-170210930 | 0 | T | C | 0.211 | -1.92E-02 | 1.04E-01 | 2.64E+00 | 2.72E+00 | 8.56E-01 | no | | intergenic_variant | MODIFIER |
| 10 | 6148536 | AX-170259021 | 0 | T | C | 0.211 | -1.92E-02 | 1.04E-01 | 2.64E+00 | 2.72E+00 | 8.56E-01 | no | | intergenic_variant | MODIFIER |
| 10 | 6152302 | AX-170101168 | 0 | G | A | 0.211 | -1.92E-02 | 1.04E-01 | 2.64E+00 | 2.72E+00 | 8.56E-01 | no | | intergenic_variant | MODIFIER |
| 10 | 6154566 | AX-170383202 | 0 | A | G | 0.211 | -1.92E-02 | 1.04E-01 | 2.64E+00 | 2.72E+00 | 8.56E-01 | no | | intergenic_variant | MODIFIER |
| 10 | 6159962 | AX-170002635 | 0 | T | C | 0.211 | -1.92E-02 | 1.04E-01 | 2.64E+00 | 2.72E+00 | 8.56E-01 | no | | intergenic_variant | MODIFIER |
| 10 | 6167797 | AX-170386577 | 0 | A | G | 0.211 | -1.92E-02 | 1.04E-01 | 2.64E+00 | 2.72E+00 | 8.56E-01 | no | | intergenic_variant | MODIFIER |
| 10 | 6168125 | AX-97815351 | 0 | G | C | 0.211 | -1.92E-02 | 1.04E-01 | 2.64E+00 | 2.72E+00 | 8.56E-01 | no | | intergenic_variant | MODIFIER |
| 10 | 6168391 | AX-170471574 | 0 | A | C | 0.211 | -1.92E-02 | 1.04E-01 | 2.64E+00 | 2.72E+00 | 8.56E-01 | no | | intergenic_variant | MODIFIER |
| 10 | 6195353 | AX-97836178 | 0 | T | A | 0.211 | -1.92E-02 | 1.04E-01 | 2.64E+00 | 2.72E+00 | 8.56E-01 | no | | intergenic_variant | MODIFIER |
| 10 | 6209202 | AX-170297408 | 0 | T | C | 0.211 | -1.92E-02 | 1.04E-01 | 2.64E+00 | 2.72E+00 | 8.56E-01 | no | | intergenic_variant | MODIFIER |
| 10 | 6242217 | AX-170320506 | 0 | C | A | 0.211 | -1.92E-02 | 1.04E-01 | 2.64E+00 | 2.72E+00 | 8.56E-01 | no | | intergenic_variant | MODIFIER |
| 10 | 6242556 | AX-170373396 | 0 | G | A | 0.211 | -1.92E-02 | 1.04E-01 | 2.64E+00 | 2.72E+00 | 8.56E-01 | no | | intergenic_variant | MODIFIER |
| 10 | 6243160 | AX-170308313 | 0 | T | C | 0.211 | -1.92E-02 | 1.04E-01 | 2.64E+00 | 2.72E+00 | 8.56E-01 | no | | intergenic_variant | MODIFIER |
| 10 | 6294346 | AX-170187065 | 0 | G | A | 0.211 | -1.92E-02 | 1.04E-01 | 2.64E+00 | 2.72E+00 | 8.56E-01 | no | | intergenic_variant | MODIFIER |
| 10 | 6314039 | AX-170504440 | 0 | A | G | 0.211 | -1.92E-02 | 1.04E-01 | 2.64E+00 | 2.72E+00 | 8.56E-01 | no | | intergenic_variant | MODIFIER |
| 10 | 6314354 | AX-170192960 | 0 | T | G | 0.211 | -1.92E-02 | 1.04E-01 | 2.64E+00 | 2.72E+00 | 8.56E-01 | no | | intergenic_variant | MODIFIER |
| 10 | 6375694 | AX-170248748 | 0 | T | C | 0.211 | -1.92E-02 | 1.04E-01 | 2.64E+00 | 2.72E+00 | 8.56E-01 | no | | intergenic_variant | MODIFIER |
| 10 | 6375799 | AX-170126874 | 0 | C | T | 0.211 | -1.92E-02 | 1.04E-01 | 2.64E+00 | 2.72E+00 | 8.56E-01 | no | | intergenic_variant | MODIFIER |
| 10 | 6375867 | AX-170346602 | 0 | C | T | 0.211 | -1.92E-02 | 1.04E-01 | 2.64E+00 | 2.72E+00 | 8.56E-01 | no | | intergenic_variant | MODIFIER |
| 10 | 6398614 | AX-97844500 | 0 | T | A | 0.211 | -1.92E-02 | 1.04E-01 | 2.64E+00 | 2.72E+00 | 8.56E-01 | no | | intergenic_variant | MODIFIER |
| 10 | 6872733 | AX-170239923 | 0 | C | T | 0.211 | -1.92E-02 | 1.04E-01 | 2.64E+00 | 2.72E+00 | 8.56E-01 | no | Oprm1 | intron_variant | MODIFIER |
| 10 | 6880469 | AX-170451429 | 0 | G | A | 0.211 | -1.92E-02 | 1.04E-01 | 2.64E+00 | 2.72E+00 | 8.56E-01 | no | Oprm1 | intron_variant | MODIFIER |
| 10 | 6919038 | AX-170247293 | 0 | A | T | 0.211 | -1.92E-02 | 1.04E-01 | 2.64E+00 | 2.72E+00 | 8.56E-01 | no | Oprm1 | intron_variant | MODIFIER |
| 10 | 6932297 | AX-170193034 | 0 | T | C | 0.211 | -1.92E-02 | 1.04E-01 | 2.64E+00 | 2.72E+00 | 8.56E-01 | no | Oprm1 | intron_variant | MODIFIER |
| 10 | 7069739 | AX-170476194 | 0 | C | A | 0.211 | -1.92E-02 | 1.04E-01 | 2.64E+00 | 2.72E+00 | 8.56E-01 | no | | intergenic_variant | MODIFIER |
| 10 | 7072743 | AX-170086730 | 0 | G | C | 0.211 | -1.92E-02 | 1.04E-01 | 2.64E+00 | 2.72E+00 | 8.56E-01 | no | | intergenic_variant | MODIFIER |
| 10 | 7114261 | AX-170313012 | 0 | C | T | 0.211 | -1.92E-02 | 1.04E-01 | 2.64E+00 | 2.72E+00 | 8.56E-01 | no | Cnksr3 | downstream_gene_variant | MODIFIER |

| | | | | | | | | | | | | | | | |
|---|---|---|---|---|---|---|---|---|---|---|---|---|---|---|---|
| 10 | 7118210 | AX-170224188 | 0 | T | C | 0.211 | -1.92E-02 | 1.04E-01 | 2.64E+00 | 2.72E+00 | 8.56E-01 | no | Cnksr3 | downstream_gene_variant | MODIFIER |
| 10 | 7120852 | AX-170306744 | 0 | A | G | 0.211 | -1.92E-02 | 1.04E-01 | 2.64E+00 | 2.72E+00 | 8.56E-01 | no | Cnksr3 | intron_variant | MODIFIER |
| 10 | 7128126 | AX-97828969 | 0 | T | A | 0.211 | -1.92E-02 | 1.04E-01 | 2.64E+00 | 2.72E+00 | 8.56E-01 | no | Cnksr3 | intron_variant | MODIFIER |
| 10 | 7128308 | AX-170447821 | 0 | C | A | 0.211 | -1.92E-02 | 1.04E-01 | 2.64E+00 | 2.72E+00 | 8.56E-01 | no | Cnksr3 | intron_variant | MODIFIER |
| 10 | 7315814 | AX-170262928 | 0 | C | T | 0.211 | -1.92E-02 | 1.04E-01 | 2.64E+00 | 2.72E+00 | 8.56E-01 | no | | intergenic_variant | MODIFIER |
| 10 | 7323798 | AX-170002627 | 0 | C | A | 0.211 | -1.92E-02 | 1.04E-01 | 2.64E+00 | 2.72E+00 | 8.56E-01 | no | | intergenic_variant | MODIFIER |
| 10 | 7325127 | AX-170412923 | 0 | A | G | 0.211 | -1.92E-02 | 1.04E-01 | 2.64E+00 | 2.72E+00 | 8.56E-01 | no | | regulatory_region_variant | MODIFIER |
| 10 | 7325385 | AX-168374624 | 0 | T | C | 0.211 | -1.92E-02 | 1.04E-01 | 2.64E+00 | 2.72E+00 | 8.56E-01 | no | | intergenic_variant | MODIFIER |
| 10 | 7337260 | AX-170102621 | 0 | T | C | 0.211 | -1.92E-02 | 1.04E-01 | 2.64E+00 | 2.72E+00 | 8.56E-01 | no | | intergenic_variant | MODIFIER |
| 10 | 7737582 | AX-97800544 | 0 | C | A | 0.211 | -1.92E-02 | 1.04E-01 | 2.64E+00 | 2.72E+00 | 8.56E-01 | no | Katna1 | intron_variant | MODIFIER |
| 10 | 7741272 | AX-170464619 | 0 | G | T | 0.211 | -1.92E-02 | 1.04E-01 | 2.64E+00 | 2.72E+00 | 8.56E-01 | no | Katna1 | intron_variant | MODIFIER |
| 10 | 7776170 | AX-97814708 | 0 | T | C | 0.211 | -1.92E-02 | 1.04E-01 | 2.64E+00 | 2.72E+00 | 8.56E-01 | no | Ginm1 | intron_variant | MODIFIER |
| 10 | 7789177 | AX-97800099 | 0 | C | T | 0.211 | -1.92E-02 | 1.04E-01 | 2.64E+00 | 2.72E+00 | 8.56E-01 | no | Ppil4 | upstream_gene_variant | MODIFIER |
| 10 | 7792734 | AX-170380376 | 0 | C | T | 0.211 | -1.92E-02 | 1.04E-01 | 2.64E+00 | 2.72E+00 | 8.56E-01 | no | Ppil4 | upstream_gene_variant | MODIFIER |
| 10 | 7811239 | AX-170356068 | 0 | C | T | 0.211 | -1.92E-02 | 1.04E-01 | 2.64E+00 | 2.72E+00 | 8.56E-01 | no | Ppil4 | intron_variant | MODIFIER |
| 10 | 7816121 | AX-97825416 | 0 | T | C | 0.211 | -1.92E-02 | 1.04E-01 | 2.64E+00 | 2.72E+00 | 8.56E-01 | no | Ppil4 | intron_variant | MODIFIER |
| 10 | 7816806 | AX-170142664 | 0 | C | T | 0.211 | -1.92E-02 | 1.04E-01 | 2.64E+00 | 2.72E+00 | 8.56E-01 | no | Ppil4 | intron_variant | MODIFIER |
| 10 | 7817724 | AX-170441328 | 0 | T | C | 0.211 | -1.92E-02 | 1.04E-01 | 2.64E+00 | 2.72E+00 | 8.56E-01 | no | Ppil4 | intron_variant | MODIFIER |
| 10 | 7886008 | AX-170002661 | 0 | T | C | 0.211 | -1.92E-02 | 1.04E-01 | 2.64E+00 | 2.72E+00 | 8.56E-01 | no | | intergenic_variant | MODIFIER |
| 10 | 7886169 | AX-170002659 | 0 | C | G | 0.211 | -1.92E-02 | 1.04E-01 | 2.64E+00 | 2.72E+00 | 8.56E-01 | no | | intergenic_variant | MODIFIER |
| 10 | 7886182 | AX-170002660 | 0 | A | G | 0.211 | -1.92E-02 | 1.04E-01 | 2.64E+00 | 2.72E+00 | 8.56E-01 | no | | intergenic_variant | MODIFIER |
| 10 | 8009418 | AX-170534702 | 0 | T | C | 0.211 | -1.92E-02 | 1.04E-01 | 2.64E+00 | 2.72E+00 | 8.56E-01 | no | Gm48614 | intron_variant,non_coding_transcript_variant | MODIFIER |
| 10 | 8028453 | AX-170284322 | 0 | A | G | 0.211 | -1.92E-02 | 1.04E-01 | 2.64E+00 | 2.72E+00 | 8.56E-01 | no | | regulatory_region_variant | MODIFIER |
| 10 | 8118214 | AX-170131432 | 0 | G | T | 0.211 | -1.92E-02 | 1.04E-01 | 2.64E+00 | 2.72E+00 | 8.56E-01 | no | | intergenic_variant | MODIFIER |
| 10 | 8119191 | AX-170509681 | 0 | G | A | 0.211 | -1.92E-02 | 1.04E-01 | 2.64E+00 | 2.72E+00 | 8.56E-01 | no | | intergenic_variant | MODIFIER |
| 10 | 8164176 | AX-170339453 | 0 | A | C | 0.211 | -1.92E-02 | 1.04E-01 | 2.64E+00 | 2.72E+00 | 8.56E-01 | no | | intergenic_variant | MODIFIER |
| 10 | 8164421 | AX-170278245 | 0 | T | C | 0.211 | -1.92E-02 | 1.04E-01 | 2.64E+00 | 2.72E+00 | 8.56E-01 | no | | intergenic_variant | MODIFIER |
| 10 | 8164458 | AX-170175340 | 0 | A | G | 0.211 | -1.92E-02 | 1.04E-01 | 2.64E+00 | 2.72E+00 | 8.56E-01 | no | | intergenic_variant | MODIFIER |
| 10 | 8184077 | AX-170044090 | 0 | G | A | 0.211 | -1.92E-02 | 1.04E-01 | 2.64E+00 | 2.72E+00 | 8.56E-01 | no | Gm30906 | intron_variant,non_coding_transcript_variant | MODIFIER |
| 10 | 8185695 | AX-170044091 | 0 | C | A | 0.211 | -1.92E-02 | 1.04E-01 | 2.64E+00 | 2.72E+00 | 8.56E-01 | no | Gm30906 | intron_variant,non_coding_transcript_variant | MODIFIER |
| 10 | 8199601 | AX-170322590 | 0 | A | G | 0.211 | -1.92E-02 | 1.04E-01 | 2.64E+00 | 2.72E+00 | 8.56E-01 | no | | intergenic_variant | MODIFIER |
| 10 | 8203247 | AX-170123542 | 0 | C | G | 0.211 | -1.92E-02 | 1.04E-01 | 2.64E+00 | 2.72E+00 | 8.56E-01 | no | Ust | downstream_gene_variant | MODIFIER |
| 10 | 8209083 | AX-170285913 | 0 | C | G | 0.211 | -1.92E-02 | 1.04E-01 | 2.64E+00 | 2.72E+00 | 8.56E-01 | no | Ust | intron_variant | MODIFIER |
| 10 | 8209985 | AX-170112066 | 0 | T | C | 0.211 | -1.92E-02 | 1.04E-01 | 2.64E+00 | 2.72E+00 | 8.56E-01 | no | Ust | intron_variant | MODIFIER |
| 10 | 8210100 | AX-97814707 | 0 | T | C | 0.211 | -1.92E-02 | 1.04E-01 | 2.64E+00 | 2.72E+00 | 8.56E-01 | no | Ust | intron_variant | MODIFIER |
| 10 | 8232101 | AX-170445191 | 0 | G | C | 0.211 | -1.92E-02 | 1.04E-01 | 2.64E+00 | 2.72E+00 | 8.56E-01 | no | Ust | intron_variant | MODIFIER |
| 10 | 8247919 | AX-170307324 | 0 | A | T | 0.211 | -1.92E-02 | 1.04E-01 | 2.64E+00 | 2.72E+00 | 8.56E-01 | no | Ust | intron_variant | MODIFIER |
| 10 | 8248831 | AX-170371750 | 0 | G | A | 0.211 | -1.92E-02 | 1.04E-01 | 2.64E+00 | 2.72E+00 | 8.56E-01 | no | Ust | intron_variant | MODIFIER |
| 10 | 8249010 | AX-170151909 | 0 | A | G | 0.211 | -1.92E-02 | 1.04E-01 | 2.64E+00 | 2.72E+00 | 8.56E-01 | no | Ust | intron_variant | MODIFIER |
| 10 | 8257781 | AX-170086734 | 0 | G | C | 0.211 | -1.92E-02 | 1.04E-01 | 2.64E+00 | 2.72E+00 | 8.56E-01 | no | Ust | intron_variant | MODIFIER |
| 10 | 8284436 | AX-170277715 | 0 | G | A | 0.211 | -1.92E-02 | 1.04E-01 | 2.64E+00 | 2.72E+00 | 8.56E-01 | no | Ust | intron_variant | MODIFIER |
| 10 | 8286456 | AX-97855315 | 0 | T | C | 0.211 | -1.92E-02 | 1.04E-01 | 2.64E+00 | 2.72E+00 | 8.56E-01 | no | Ust | intron_variant | MODIFIER |
| 10 | 8321964 | AX-97791081 | 0 | A | G | 0.211 | -1.92E-02 | 1.04E-01 | 2.64E+00 | 2.72E+00 | 8.56E-01 | no | Ust | intron_variant | MODIFIER |
| 10 | 8322691 | AX-170402962 | 0 | G | C | 0.211 | -1.92E-02 | 1.04E-01 | 2.64E+00 | 2.72E+00 | 8.56E-01 | no | Ust | intron_variant | MODIFIER |
| 10 | 8324214 | AX-170444906 | 0 | C | T | 0.211 | -1.92E-02 | 1.04E-01 | 2.64E+00 | 2.72E+00 | 8.56E-01 | no | Ust | intron_variant | MODIFIER |
| 10 | 8333504 | AX-170002670 | 0 | G | A | 0.211 | -1.92E-02 | 1.04E-01 | 2.64E+00 | 2.72E+00 | 8.56E-01 | no | Ust | intron_variant | MODIFIER |
| 10 | 8337632 | AX-97840001 | 0 | G | T | 0.211 | -1.92E-02 | 1.04E-01 | 2.64E+00 | 2.72E+00 | 8.56E-01 | no | Ust | intron_variant | MODIFIER |
| 10 | 8337925 | AX-170323463 | 0 | G | A | 0.211 | -1.92E-02 | 1.04E-01 | 2.64E+00 | 2.72E+00 | 8.56E-01 | no | Ust | intron_variant | MODIFIER |
| 10 | 8338262 | AX-170002672 | 0 | A | T | 0.211 | -1.92E-02 | 1.04E-01 | 2.64E+00 | 2.72E+00 | 8.56E-01 | no | Ust | intron_variant | MODIFIER |
| 10 | 8348459 | AX-170186796 | 0 | C | T | 0.211 | -1.92E-02 | 1.04E-01 | 2.64E+00 | 2.72E+00 | 8.56E-01 | no | Ust | intron_variant | MODIFIER |
| 10 | 8391360 | AX-170044095 | 0 | A | C | 0.211 | -1.92E-02 | 1.04E-01 | 2.64E+00 | 2.72E+00 | 8.56E-01 | no | Ust | intron_variant | MODIFIER |
| 10 | 8396882 | AX-170044096 | 0 | T | C | 0.211 | -1.92E-02 | 1.04E-01 | 2.64E+00 | 2.72E+00 | 8.56E-01 | no | Ust | intron_variant | MODIFIER |
| 10 | 8404421 | AX-170422932 | 0 | G | C | 0.211 | -1.92E-02 | 1.04E-01 | 2.64E+00 | 2.72E+00 | 8.56E-01 | no | Ust | intron_variant | MODIFIER |
| 10 | 8409675 | AX-170002671 | 0 | G | A | 0.211 | -1.92E-02 | 1.04E-01 | 2.64E+00 | 2.72E+00 | 8.56E-01 | no | Ust | intron_variant | MODIFIER |

| | | | | | | | | | | | | | | | |
|---|---|---|---|---|---|---|---|---|---|---|---|---|---|---|---|
| 10 | 8416559 | AX-170002673 | 0 | T | C | 0.211 | -1.92E-02 | 1.04E-01 | 2.64E+00 | 2.72E+00 | 8.56E-01 | no | Ust | intron_variant | MODIFIER |
| 10 | 8416627 | AX-97826825 | 0 | A | G | 0.211 | -1.92E-02 | 1.04E-01 | 2.64E+00 | 2.72E+00 | 8.56E-01 | no | Ust | intron_variant | MODIFIER |
| 10 | 8442560 | AX-170002666 | 0 | G | C | 0.211 | -1.92E-02 | 1.04E-01 | 2.64E+00 | 2.72E+00 | 8.56E-01 | no | Ust | intron_variant | MODIFIER |
| 10 | 8443793 | AX-170002675 | 0 | A | G | 0.211 | -1.92E-02 | 1.04E-01 | 2.64E+00 | 2.72E+00 | 8.56E-01 | no | Ust | intron_variant | MODIFIER |
| 10 | 8469840 | AX-97845863 | 0 | G | T | 0.211 | -1.92E-02 | 1.04E-01 | 2.64E+00 | 2.72E+00 | 8.56E-01 | no | Ust | intron_variant | MODIFIER |
| 10 | 8492885 | AX-170002669 | 0 | T | C | 0.211 | -1.92E-02 | 1.04E-01 | 2.64E+00 | 2.72E+00 | 8.56E-01 | no | Ust | intron_variant | MODIFIER |
| 10 | 8501464 | AX-170519829 | 0 | C | A | 0.211 | -1.92E-02 | 1.04E-01 | 2.64E+00 | 2.72E+00 | 8.56E-01 | no | Ust | intron_variant | MODIFIER |
| 10 | 7034963 | AX-170391994 | 0 | T | C | 0.219 | -1.85E-02 | 1.00E-01 | 2.64E+00 | 2.72E+00 | 8.56E-01 | no | Oprm1 | intron_variant | MODIFIER |
| 10 | 7034994 | AX-170159490 | 0 | T | C | 0.219 | -1.85E-02 | 1.00E-01 | 2.64E+00 | 2.72E+00 | 8.56E-01 | no | Oprm1 | intron_variant | MODIFIER |
| 10 | 7830017 | AX-170162196 | 0 | G | A | 0.219 | -1.85E-02 | 1.00E-01 | 2.64E+00 | 2.72E+00 | 8.56E-01 | no | Zc3h12d | upstream_gene_variant | MODIFIER |
| 10 | 7919058 | AX-170123313 | 0 | T | C | 0.219 | -1.85E-02 | 1.00E-01 | 2.64E+00 | 2.72E+00 | 8.56E-01 | no | Tab2 | downstream_gene_variant | MODIFIER |
| 10 | 8535095 | AX-168400917 | 0 | A | G | 0.219 | -1.85E-02 | 1.00E-01 | 2.64E+00 | 2.72E+00 | 8.56E-01 | no | | intergenic_variant | MODIFIER |
| 10 | 8543586 | AX-170440335 | 0 | G | A | 0.219 | -1.85E-02 | 1.00E-01 | 2.64E+00 | 2.72E+00 | 8.56E-01 | no | | intergenic_variant | MODIFIER |
| 10 | 8564070 | AX-97825469 | 0 | T | C | 0.219 | -1.85E-02 | 1.00E-01 | 2.64E+00 | 2.72E+00 | 8.56E-01 | no | | intergenic_variant | MODIFIER |
| 10 | 8568406 | AX-170292222 | 0 | T | C | 0.219 | -1.85E-02 | 1.00E-01 | 2.64E+00 | 2.72E+00 | 8.56E-01 | no | | intergenic_variant | MODIFIER |
| 10 | 8570175 | AX-170239611 | 0 | G | A | 0.219 | -1.85E-02 | 1.00E-01 | 2.64E+00 | 2.72E+00 | 8.56E-01 | no | | intergenic_variant | MODIFIER |
| 10 | 8572217 | AX-170417553 | 0 | A | T | 0.219 | -1.85E-02 | 1.00E-01 | 2.64E+00 | 2.72E+00 | 8.56E-01 | no | | intergenic_variant | MODIFIER |
| 10 | 8661211 | AX-170125664 | 0 | G | A | 0.219 | -1.85E-02 | 1.00E-01 | 2.64E+00 | 2.72E+00 | 8.56E-01 | no | | intergenic_variant | MODIFIER |
| 10 | 8662843 | AX-170271713 | 0 | A | G | 0.219 | -1.85E-02 | 1.00E-01 | 2.64E+00 | 2.72E+00 | 8.56E-01 | no | | intergenic_variant | MODIFIER |
| 10 | 8903117 | AX-170044103 | 0 | G | A | 0.219 | -1.85E-02 | 1.00E-01 | 2.64E+00 | 2.72E+00 | 8.56E-01 | no | Gm48727 | upstream_gene_variant | MODIFIER |
| 10 | 8979985 | AX-170106754 | 0 | C | T | 0.219 | -1.85E-02 | 1.00E-01 | 2.64E+00 | 2.72E+00 | 8.56E-01 | no | | intergenic_variant | MODIFIER |
| 10 | 8995315 | AX-170354772 | 0 | C | T | 0.219 | -1.85E-02 | 1.00E-01 | 2.64E+00 | 2.72E+00 | 8.56E-01 | no | | regulatory_region_variant | MODIFIER |
| 10 | 8996602 | AX-170518578 | 0 | G | C | 0.219 | -1.85E-02 | 1.00E-01 | 2.64E+00 | 2.72E+00 | 8.56E-01 | no | | intergenic_variant | MODIFIER |
| 12 | 113256803 | AX-170375371 | 0 | C | T | 0.149 | -3.19E-02 | 1.83E-01 | 2.68E+00 | 2.76E+00 | 8.56E-01 | no | Igha | downstream_gene_variant | MODIFIER |
| 13 | 47105045 | AX-97823401 | 1 | T | C | 0.473 | -1.09E-02 | 6.30E-02 | 2.65E+00 | 2.76E+00 | 8.56E-01 | no | Dek | intron_variant | MODIFIER |
| 7 | 117162026 | AX-170515003 | 1 | A | G | 0.402 | 1.64E-02 | 8.99E-02 | 2.66E+00 | 2.72E+00 | 8.57E-01 | no | | intergenic_variant | MODIFIER |
| 4 | 92543272 | AX-97830171 | 0 | G | A | 0.325 | 1.92E-02 | 1.05E-01 | 2.65E+00 | 2.72E+00 | 8.57E-01 | no | Gm12637 | intron_variant,non_coding_transcript_variant | MODIFIER |
| 5 | 34495875 | AX-170154849 | 0 | G | C | 0.342 | -1.62E-02 | 9.45E-02 | 2.69E+00 | 2.77E+00 | 8.57E-01 | no | Tnip2 | downstream_gene_variant | MODIFIER |
| 1 | 36002241 | AX-170480763 | 1 | A | G | 0.455 | 1.39E-02 | 7.91E-02 | 2.68E+00 | 2.75E+00 | 8.57E-01 | no | Gm33222 | downstream_gene_variant | MODIFIER |
| 7 | 40685324 | AX-170519055 | 0 | C | T | 0.325 | 1.74E-02 | 9.37E-02 | 2.63E+00 | 2.71E+00 | 8.57E-01 | no | 4933402C06Rik | intron_variant,non_coding_transcript_variant | MODIFIER |
| 1 | 75533126 | AX-170417185 | 0 | C | T | 0.132 | 4.01E-02 | 2.24E-01 | 2.67E+00 | 2.73E+00 | 8.57E-01 | no | Stk11ip | intron_variant | MODIFIER |
| 1 | 75533214 | AX-170326327 | 0 | T | G | 0.132 | 4.01E-02 | 2.24E-01 | 2.67E+00 | 2.73E+00 | 8.57E-01 | no | Stk11ip | intron_variant | MODIFIER |
| 1 | 75551053 | AX-170134375 | 0 | T | C | 0.132 | 4.01E-02 | 2.24E-01 | 2.67E+00 | 2.73E+00 | 8.57E-01 | no | Slc4a3 | intron_variant,NMD_transcript_variant | MODIFIER |
| 1 | 75553330 | AX-170329223 | 0 | C | T | 0.132 | 4.01E-02 | 2.24E-01 | 2.67E+00 | 2.73E+00 | 8.57E-01 | no | Slc4a3 | 3_prime_UTR_variant,NMD_transcript_variant | MODIFIER |
| 1 | 75567146 | AX-170423064 | 0 | G | A | 0.132 | 4.01E-02 | 2.24E-01 | 2.67E+00 | 2.73E+00 | 8.57E-01 | no | Gm15180 | upstream_gene_variant | MODIFIER |
| 1 | 75588795 | AX-170291162 | 0 | C | T | 0.132 | 4.01E-02 | 2.24E-01 | 2.67E+00 | 2.73E+00 | 8.57E-01 | no | | regulatory_region_variant | MODIFIER |
| 1 | 75588849 | AX-170359212 | 0 | C | A | 0.132 | 4.01E-02 | 2.24E-01 | 2.67E+00 | 2.73E+00 | 8.57E-01 | no | | regulatory_region_variant | MODIFIER |
| 1 | 75596235 | AX-170380300 | 0 | A | C | 0.132 | 4.01E-02 | 2.24E-01 | 2.67E+00 | 2.73E+00 | 8.57E-01 | no | | intergenic_variant | MODIFIER |
| 1 | 75635758 | AX-170229586 | 0 | A | G | 0.132 | 4.01E-02 | 2.24E-01 | 2.67E+00 | 2.73E+00 | 8.57E-01 | no | Gm5257 | upstream_gene_variant | MODIFIER |
| 1 | 75636352 | AX-170449817 | 0 | A | G | 0.132 | 4.01E-02 | 2.24E-01 | 2.67E+00 | 2.73E+00 | 8.57E-01 | no | Gm5257 | upstream_gene_variant | MODIFIER |
| 1 | 75650170 | AX-170465430 | 0 | G | C | 0.132 | 4.01E-02 | 2.24E-01 | 2.67E+00 | 2.73E+00 | 8.57E-01 | no | | intergenic_variant | MODIFIER |
| 1 | 75680663 | AX-170225660 | 0 | T | C | 0.132 | 4.01E-02 | 2.24E-01 | 2.67E+00 | 2.73E+00 | 8.57E-01 | no | | regulatory_region_variant | MODIFIER |
| 1 | 75985395 | AX-97844567 | 0 | C | T | 0.132 | 4.01E-02 | 2.24E-01 | 2.67E+00 | 2.73E+00 | 8.57E-01 | no | | intergenic_variant | MODIFIER |
| 1 | 75993273 | AX-170387403 | 0 | A | G | 0.132 | 4.01E-02 | 2.24E-01 | 2.67E+00 | 2.73E+00 | 8.57E-01 | no | | intergenic_variant | MODIFIER |
| 1 | 75995094 | AX-170183086 | 0 | G | A | 0.132 | 4.01E-02 | 2.24E-01 | 2.67E+00 | 2.73E+00 | 8.57E-01 | no | | intergenic_variant | MODIFIER |
| 1 | 75998730 | AX-170399971 | 0 | T | C | 0.132 | 4.01E-02 | 2.24E-01 | 2.67E+00 | 2.73E+00 | 8.57E-01 | no | | intergenic_variant | MODIFIER |
| 1 | 76001715 | AX-170183516 | 0 | A | T | 0.132 | 4.01E-02 | 2.24E-01 | 2.67E+00 | 2.73E+00 | 8.57E-01 | no | | regulatory_region_variant | MODIFIER |
| 1 | 76002840 | AX-97815857 | 0 | A | G | 0.132 | 4.01E-02 | 2.24E-01 | 2.67E+00 | 2.73E+00 | 8.57E-01 | no | | intergenic_variant | MODIFIER |
| 1 | 76013696 | AX-170454539 | 0 | A | C | 0.132 | 4.01E-02 | 2.24E-01 | 2.67E+00 | 2.73E+00 | 8.57E-01 | no | Gm29065 | downstream_gene_variant | MODIFIER |
| 1 | 76019543 | AX-170318981 | 0 | A | G | 0.132 | 4.01E-02 | 2.24E-01 | 2.67E+00 | 2.73E+00 | 8.57E-01 | no | Gm29065 | upstream_gene_variant | MODIFIER |
| 1 | 76019783 | AX-170334612 | 0 | C | T | 0.132 | 4.01E-02 | 2.24E-01 | 2.67E+00 | 2.73E+00 | 8.57E-01 | no | Gm29065 | upstream_gene_variant | MODIFIER |
| 1 | 76043364 | AX-170125800 | 0 | C | T | 0.132 | 4.01E-02 | 2.24E-01 | 2.67E+00 | 2.73E+00 | 8.57E-01 | no | | intergenic_variant | MODIFIER |
| 1 | 76046078 | AX-170197591 | 0 | G | A | 0.132 | 4.01E-02 | 2.24E-01 | 2.67E+00 | 2.73E+00 | 8.57E-01 | no | | intergenic_variant | MODIFIER |
| 1 | 76049197 | AX-170319761 | 0 | T | C | 0.132 | 4.01E-02 | 2.24E-01 | 2.67E+00 | 2.73E+00 | 8.57E-01 | no | | regulatory_region_variant | MODIFIER |

| | | | | | | | | | | | | | | | |
|---|---|---|---|---|---|---|---|---|---|---|---|---|---|---|---|
| 1 | 76081545 | AX-170441753 | 0 | C | T | 0.132 | 4.01E-02 | 2.24E-01 | 2.67E+00 | 2.73E+00 | 8.57E-01 | no | | intergenic_variant | MODIFIER |
| 1 | 76083418 | AX-170512397 | 0 | A | G | 0.132 | 4.01E-02 | 2.24E-01 | 2.67E+00 | 2.73E+00 | 8.57E-01 | no | | regulatory_region_variant | MODIFIER |
| 1 | 76084476 | AX-170108780 | 0 | A | G | 0.132 | 4.01E-02 | 2.24E-01 | 2.67E+00 | 2.73E+00 | 8.57E-01 | no | | regulatory_region_variant | MODIFIER |
| 1 | 76084578 | AX-170478511 | 0 | A | G | 0.132 | 4.01E-02 | 2.24E-01 | 2.67E+00 | 2.73E+00 | 8.57E-01 | no | | regulatory_region_variant | MODIFIER |
| 1 | 76094885 | AX-170239222 | 0 | T | C | 0.132 | 4.01E-02 | 2.24E-01 | 2.67E+00 | 2.73E+00 | 8.57E-01 | no | | intergenic_variant | MODIFIER |
| 1 | 76094969 | AX-97841568 | 0 | A | G | 0.132 | 4.01E-02 | 2.24E-01 | 2.67E+00 | 2.73E+00 | 8.57E-01 | no | | intergenic_variant | MODIFIER |
| 1 | 76095017 | AX-170467063 | 0 | A | C | 0.132 | 4.01E-02 | 2.24E-01 | 2.67E+00 | 2.73E+00 | 8.57E-01 | no | | intergenic_variant | MODIFIER |
| 1 | 76112183 | AX-170332104 | 0 | G | A | 0.132 | 4.01E-02 | 2.24E-01 | 2.67E+00 | 2.73E+00 | 8.57E-01 | no | Gm29068 | upstream_gene_variant | MODIFIER |
| 1 | 76122266 | AX-170478440 | 0 | G | C | 0.132 | 4.01E-02 | 2.24E-01 | 2.67E+00 | 2.73E+00 | 8.57E-01 | no | | intergenic_variant | MODIFIER |
| 1 | 76122336 | AX-170213154 | 0 | C | G | 0.132 | 4.01E-02 | 2.24E-01 | 2.67E+00 | 2.73E+00 | 8.57E-01 | no | | intergenic_variant | MODIFIER |
| 1 | 76139841 | AX-170281906 | 0 | T | C | 0.132 | 4.01E-02 | 2.24E-01 | 2.67E+00 | 2.73E+00 | 8.57E-01 | no | | intergenic_variant | MODIFIER |
| 1 | 76142789 | AX-170336127 | 0 | C | T | 0.132 | 4.01E-02 | 2.24E-01 | 2.67E+00 | 2.73E+00 | 8.57E-01 | no | | intergenic_variant | MODIFIER |
| 1 | 76143743 | AX-170155129 | 0 | T | C | 0.132 | 4.01E-02 | 2.24E-01 | 2.67E+00 | 2.73E+00 | 8.57E-01 | no | | regulatory_region_variant | MODIFIER |
| 1 | 76149367 | AX-170352974 | 0 | C | T | 0.132 | 4.01E-02 | 2.24E-01 | 2.67E+00 | 2.73E+00 | 8.57E-01 | no | | intergenic_variant | MODIFIER |
| 1 | 76173003 | AX-97843384 | 0 | G | A | 0.132 | 4.01E-02 | 2.24E-01 | 2.67E+00 | 2.73E+00 | 8.57E-01 | no | | intergenic_variant | MODIFIER |
| 1 | 76173094 | AX-170385946 | 0 | G | T | 0.132 | 4.01E-02 | 2.24E-01 | 2.67E+00 | 2.73E+00 | 8.57E-01 | no | | intergenic_variant | MODIFIER |
| 1 | 76173581 | AX-170437152 | 0 | A | G | 0.132 | 4.01E-02 | 2.24E-01 | 2.67E+00 | 2.73E+00 | 8.57E-01 | no | | intergenic_variant | MODIFIER |
| 1 | 76201114 | AX-170000839 | 0 | G | C | 0.132 | 4.01E-02 | 2.24E-01 | 2.67E+00 | 2.73E+00 | 8.57E-01 | no | | intergenic_variant | MODIFIER |
| 1 | 76301940 | AX-170000842 | 0 | A | G | 0.132 | 4.01E-02 | 2.24E-01 | 2.67E+00 | 2.73E+00 | 8.57E-01 | no | | intergenic_variant | MODIFIER |
| 1 | 76340473 | AX-170000840 | 0 | A | T | 0.132 | 4.01E-02 | 2.24E-01 | 2.67E+00 | 2.73E+00 | 8.57E-01 | no | | intergenic_variant | MODIFIER |
| 1 | 76771783 | AX-170000849 | 0 | C | T | 0.132 | 4.01E-02 | 2.24E-01 | 2.67E+00 | 2.73E+00 | 8.57E-01 | no | | intergenic_variant | MODIFIER |
| 1 | 76812900 | AX-170113049 | 0 | T | C | 0.132 | 4.01E-02 | 2.24E-01 | 2.67E+00 | 2.73E+00 | 8.57E-01 | no | | intergenic_variant | MODIFIER |
| 1 | 76914832 | AX-170109587 | 0 | A | G | 0.132 | 4.01E-02 | 2.24E-01 | 2.67E+00 | 2.73E+00 | 8.57E-01 | no | Gm816 | intron_variant,non_coding_transcript_variant | MODIFIER |
| 1 | 76917586 | AX-170463531 | 0 | C | T | 0.132 | 4.01E-02 | 2.24E-01 | 2.67E+00 | 2.73E+00 | 8.57E-01 | no | Gm816 | intron_variant,non_coding_transcript_variant | MODIFIER |
| 1 | 76918532 | AX-97793389 | 0 | T | G | 0.132 | 4.01E-02 | 2.24E-01 | 2.67E+00 | 2.73E+00 | 8.57E-01 | no | Gm816 | intron_variant,non_coding_transcript_variant | MODIFIER |
| 1 | 76989446 | AX-170222735 | 0 | T | C | 0.132 | 4.01E-02 | 2.24E-01 | 2.67E+00 | 2.73E+00 | 8.57E-01 | no | Gm816 | intron_variant,non_coding_transcript_variant | MODIFIER |
| 1 | 76998261 | AX-170324715 | 0 | C | G | 0.132 | 4.01E-02 | 2.24E-01 | 2.67E+00 | 2.73E+00 | 8.57E-01 | no | Gm816 | intron_variant,non_coding_transcript_variant | MODIFIER |
| 1 | 77046591 | AX-170497672 | 0 | G | A | 0.132 | 4.01E-02 | 2.24E-01 | 2.67E+00 | 2.73E+00 | 8.57E-01 | no | | intergenic_variant | MODIFIER |
| 1 | 77047761 | AX-170305377 | 0 | C | T | 0.132 | 4.01E-02 | 2.24E-01 | 2.67E+00 | 2.73E+00 | 8.57E-01 | no | | intergenic_variant | MODIFIER |
| 1 | 77047938 | AX-168406298 | 0 | A | G | 0.132 | 4.01E-02 | 2.24E-01 | 2.67E+00 | 2.73E+00 | 8.57E-01 | no | | intergenic_variant | MODIFIER |
| 1 | 77052082 | AX-170041727 | 0 | G | A | 0.132 | 4.01E-02 | 2.24E-01 | 2.67E+00 | 2.73E+00 | 8.57E-01 | no | | intergenic_variant | MODIFIER |
| 1 | 77053923 | AX-170041726 | 0 | T | C | 0.132 | 4.01E-02 | 2.24E-01 | 2.67E+00 | 2.73E+00 | 8.57E-01 | no | | intergenic_variant | MODIFIER |
| 1 | 77054242 | AX-170480860 | 0 | A | G | 0.132 | 4.01E-02 | 2.24E-01 | 2.67E+00 | 2.73E+00 | 8.57E-01 | no | | intergenic_variant | MODIFIER |
| 1 | 77054438 | AX-170311054 | 0 | G | A | 0.132 | 4.01E-02 | 2.24E-01 | 2.67E+00 | 2.73E+00 | 8.57E-01 | no | | intergenic_variant | MODIFIER |
| 1 | 77055114 | AX-170527764 | 0 | C | T | 0.132 | 4.01E-02 | 2.24E-01 | 2.67E+00 | 2.73E+00 | 8.57E-01 | no | | intergenic_variant | MODIFIER |
| 1 | 77140997 | AX-97853476 | 0 | A | G | 0.132 | 4.01E-02 | 2.24E-01 | 2.67E+00 | 2.73E+00 | 8.57E-01 | no | Gm25974 | upstream_gene_variant | MODIFIER |
| 1 | 77152051 | AX-170525151 | 0 | G | A | 0.132 | 4.01E-02 | 2.24E-01 | 2.67E+00 | 2.73E+00 | 8.57E-01 | no | | intergenic_variant | MODIFIER |
| 1 | 77156732 | AX-170525195 | 0 | G | A | 0.132 | 4.01E-02 | 2.24E-01 | 2.67E+00 | 2.73E+00 | 8.57E-01 | no | | intergenic_variant | MODIFIER |
| 1 | 77170108 | AX-170213428 | 0 | T | C | 0.132 | 4.01E-02 | 2.24E-01 | 2.67E+00 | 2.73E+00 | 8.57E-01 | no | | intergenic_variant | MODIFIER |
| 1 | 77254336 | AX-170140034 | 0 | G | A | 0.132 | 4.01E-02 | 2.24E-01 | 2.67E+00 | 2.73E+00 | 8.57E-01 | no | | intergenic_variant | MODIFIER |
| 1 | 77323200 | AX-170515899 | 0 | A | G | 0.132 | 4.01E-02 | 2.24E-01 | 2.67E+00 | 2.73E+00 | 8.57E-01 | no | | intergenic_variant | MODIFIER |
| 1 | 77370673 | AX-170000852 | 0 | C | T | 0.132 | 4.01E-02 | 2.24E-01 | 2.67E+00 | 2.73E+00 | 8.57E-01 | no | Epha4 | intron_variant | MODIFIER |
| 1 | 77371067 | AX-170246472 | 0 | T | C | 0.132 | 4.01E-02 | 2.24E-01 | 2.67E+00 | 2.73E+00 | 8.57E-01 | no | Epha4 | intron_variant | MODIFIER |
| 1 | 77387396 | AX-170324841 | 0 | T | C | 0.132 | 4.01E-02 | 2.24E-01 | 2.67E+00 | 2.73E+00 | 8.57E-01 | no | Epha4 | intron_variant | MODIFIER |
| 1 | 77431390 | AX-97812152 | 0 | G | A | 0.132 | 4.01E-02 | 2.24E-01 | 2.67E+00 | 2.73E+00 | 8.57E-01 | no | Epha4 | intron_variant | MODIFIER |
| 1 | 77445677 | AX-170293077 | 0 | C | T | 0.132 | 4.01E-02 | 2.24E-01 | 2.67E+00 | 2.73E+00 | 8.57E-01 | no | Epha4 | intron_variant | MODIFIER |
| 1 | 77454349 | AX-170136325 | 0 | G | A | 0.132 | 4.01E-02 | 2.24E-01 | 2.67E+00 | 2.73E+00 | 8.57E-01 | no | Epha4 | intron_variant | MODIFIER |
| 1 | 77456107 | AX-170207340 | 0 | T | C | 0.132 | 4.01E-02 | 2.24E-01 | 2.67E+00 | 2.73E+00 | 8.57E-01 | no | Epha4 | intron_variant | MODIFIER |
| 1 | 77460216 | AX-97832158 | 0 | G | A | 0.132 | 4.01E-02 | 2.24E-01 | 2.67E+00 | 2.73E+00 | 8.57E-01 | no | Epha4 | intron_variant | MODIFIER |
| 1 | 77463156 | AX-170137091 | 0 | G | C | 0.132 | 4.01E-02 | 2.24E-01 | 2.67E+00 | 2.73E+00 | 8.57E-01 | no | Epha4 | intron_variant | MODIFIER |
| 1 | 77525764 | AX-170493405 | 0 | A | G | 0.132 | 4.01E-02 | 2.24E-01 | 2.67E+00 | 2.73E+00 | 8.57E-01 | no | | intergenic_variant | MODIFIER |
| 1 | 77567300 | AX-170041740 | 0 | C | A | 0.132 | 4.01E-02 | 2.24E-01 | 2.67E+00 | 2.73E+00 | 8.57E-01 | no | Gm28385 | upstream_gene_variant | MODIFIER |
| 1 | 77587676 | AX-170463884 | 0 | T | C | 0.132 | 4.01E-02 | 2.24E-01 | 2.67E+00 | 2.73E+00 | 8.57E-01 | no | | intergenic_variant | MODIFIER |
| 1 | 77591470 | AX-170282633 | 0 | G | A | 0.132 | 4.01E-02 | 2.24E-01 | 2.67E+00 | 2.73E+00 | 8.57E-01 | no | | intergenic_variant | MODIFIER |

| | | | | | | | | | | | | | | | |
|---|---|---|---|---|---|---|---|---|---|---|---|---|---|---|---|
| 1 | 77633733 | AX-170502533 | 0 | C | T | 0.132 | 4.01E-02 | 2.24E-01 | 2.67E+00 | 2.73E+00 | 8.57E-01 | no | | intergenic_variant | MODIFIER |
| 1 | 77639149 | AX-170367556 | 0 | C | T | 0.132 | 4.01E-02 | 2.24E-01 | 2.67E+00 | 2.73E+00 | 8.57E-01 | no | | intergenic_variant | MODIFIER |
| 1 | 77645904 | AX-170103605 | 0 | A | C | 0.132 | 4.01E-02 | 2.24E-01 | 2.67E+00 | 2.73E+00 | 8.57E-01 | no | | intergenic_variant | MODIFIER |
| 1 | 77647635 | AX-170443559 | 0 | G | A | 0.132 | 4.01E-02 | 2.24E-01 | 2.67E+00 | 2.73E+00 | 8.57E-01 | no | | intergenic_variant | MODIFIER |
| 1 | 77663449 | AX-170260219 | 0 | A | G | 0.132 | 4.01E-02 | 2.24E-01 | 2.67E+00 | 2.73E+00 | 8.57E-01 | no | | intergenic_variant | MODIFIER |
| 1 | 77664522 | AX-170098182 | 0 | T | C | 0.132 | 4.01E-02 | 2.24E-01 | 2.67E+00 | 2.73E+00 | 8.57E-01 | no | | intergenic_variant | MODIFIER |
| 1 | 77673263 | AX-170487602 | 0 | G | A | 0.132 | 4.01E-02 | 2.24E-01 | 2.67E+00 | 2.73E+00 | 8.57E-01 | no | | intergenic_variant | MODIFIER |
| 1 | 77673352 | AX-170041745 | 0 | T | C | 0.132 | 4.01E-02 | 2.24E-01 | 2.67E+00 | 2.73E+00 | 8.57E-01 | no | | intergenic_variant | MODIFIER |
| 1 | 77673370 | AX-170041746 | 0 | A | G | 0.132 | 4.01E-02 | 2.24E-01 | 2.67E+00 | 2.73E+00 | 8.57E-01 | no | | intergenic_variant | MODIFIER |
| 1 | 77715227 | AX-170492128 | 0 | A | T | 0.132 | 4.01E-02 | 2.24E-01 | 2.67E+00 | 2.73E+00 | 8.57E-01 | no | | intergenic_variant | MODIFIER |
| 1 | 77738175 | AX-170168915 | 0 | G | A | 0.132 | 4.01E-02 | 2.24E-01 | 2.67E+00 | 2.73E+00 | 8.57E-01 | no | | intergenic_variant | MODIFIER |
| 1 | 77738221 | AX-170176942 | 0 | G | A | 0.132 | 4.01E-02 | 2.24E-01 | 2.67E+00 | 2.73E+00 | 8.57E-01 | no | | intergenic_variant | MODIFIER |
| 1 | 77823405 | AX-170000865 | 0 | A | G | 0.132 | 4.01E-02 | 2.24E-01 | 2.67E+00 | 2.73E+00 | 8.57E-01 | no | | intergenic_variant | MODIFIER |
| 1 | 77866604 | AX-170086505 | 0 | C | T | 0.132 | 4.01E-02 | 2.24E-01 | 2.67E+00 | 2.73E+00 | 8.57E-01 | no | | intergenic_variant | MODIFIER |
| 1 | 78021969 | AX-170525738 | 0 | T | C | 0.132 | 4.01E-02 | 2.24E-01 | 2.67E+00 | 2.73E+00 | 8.57E-01 | no | | regulatory_region_variant | MODIFIER |
| 1 | 78025801 | AX-170214737 | 0 | C | T | 0.132 | 4.01E-02 | 2.24E-01 | 2.67E+00 | 2.73E+00 | 8.57E-01 | no | | intergenic_variant | MODIFIER |
| 1 | 78025888 | AX-170041749 | 0 | A | G | 0.132 | 4.01E-02 | 2.24E-01 | 2.67E+00 | 2.73E+00 | 8.57E-01 | no | | intergenic_variant | MODIFIER |
| 1 | 78025979 | AX-170041751 | 0 | A | G | 0.132 | 4.01E-02 | 2.24E-01 | 2.67E+00 | 2.73E+00 | 8.57E-01 | no | | intergenic_variant | MODIFIER |
| 1 | 78026038 | AX-170041750 | 0 | G | C | 0.132 | 4.01E-02 | 2.24E-01 | 2.67E+00 | 2.73E+00 | 8.57E-01 | no | | intergenic_variant | MODIFIER |
| 1 | 78027216 | AX-170501052 | 0 | A | G | 0.132 | 4.01E-02 | 2.24E-01 | 2.67E+00 | 2.73E+00 | 8.57E-01 | no | | intergenic_variant | MODIFIER |
| 1 | 78027304 | AX-97811512 | 0 | G | A | 0.132 | 4.01E-02 | 2.24E-01 | 2.67E+00 | 2.73E+00 | 8.57E-01 | no | | intergenic_variant | MODIFIER |
| 1 | 78040590 | AX-170199415 | 0 | A | C | 0.132 | 4.01E-02 | 2.24E-01 | 2.67E+00 | 2.73E+00 | 8.57E-01 | no | | intergenic_variant | MODIFIER |
| 1 | 78055310 | AX-97785087 | 0 | T | C | 0.132 | 4.01E-02 | 2.24E-01 | 2.67E+00 | 2.73E+00 | 8.57E-01 | no | | intergenic_variant | MODIFIER |
| 1 | 78088535 | AX-97805315 | 0 | C | A | 0.132 | 4.01E-02 | 2.24E-01 | 2.67E+00 | 2.73E+00 | 8.57E-01 | no | | intergenic_variant | MODIFIER |
| 1 | 78090426 | AX-170267185 | 0 | G | A | 0.132 | 4.01E-02 | 2.24E-01 | 2.67E+00 | 2.73E+00 | 8.57E-01 | no | | intergenic_variant | MODIFIER |
| 1 | 78135561 | AX-170217640 | 0 | C | G | 0.132 | 4.01E-02 | 2.24E-01 | 2.67E+00 | 2.73E+00 | 8.57E-01 | no | Pax3 | intron_variant | MODIFIER |
| 1 | 78135574 | AX-170407823 | 0 | A | G | 0.132 | 4.01E-02 | 2.24E-01 | 2.67E+00 | 2.73E+00 | 8.57E-01 | no | Pax3 | intron_variant | MODIFIER |
| 1 | 78137701 | AX-170232439 | 0 | C | G | 0.132 | 4.01E-02 | 2.24E-01 | 2.67E+00 | 2.73E+00 | 8.57E-01 | no | Pax3 | intron_variant | MODIFIER |
| 1 | 78138407 | AX-170534886 | 0 | A | G | 0.132 | 4.01E-02 | 2.24E-01 | 2.67E+00 | 2.73E+00 | 8.57E-01 | no | Pax3 | intron_variant | MODIFIER |
| 1 | 78142482 | AX-170264387 | 0 | C | T | 0.132 | 4.01E-02 | 2.24E-01 | 2.67E+00 | 2.73E+00 | 8.57E-01 | no | Pax3 | intron_variant | MODIFIER |
| 1 | 78144748 | AX-170041754 | 0 | C | T | 0.132 | 4.01E-02 | 2.24E-01 | 2.67E+00 | 2.73E+00 | 8.57E-01 | no | Pax3 | intron_variant | MODIFIER |
| 1 | 78220687 | AX-97812124 | 0 | G | A | 0.132 | 4.01E-02 | 2.24E-01 | 2.67E+00 | 2.73E+00 | 8.57E-01 | no | | intergenic_variant | MODIFIER |
| 1 | 78594124 | AX-170125539 | 0 | T | C | 0.132 | 4.01E-02 | 2.24E-01 | 2.67E+00 | 2.73E+00 | 8.57E-01 | no | 5730419F03Rik | downstream_gene_variant | MODIFIER |
| 1 | 78647672 | AX-170380128 | 0 | G | T | 0.132 | 4.01E-02 | 2.24E-01 | 2.67E+00 | 2.73E+00 | 8.57E-01 | no | | regulatory_region_variant | MODIFIER |
| 1 | 78694050 | AX-170120985 | 0 | A | G | 0.132 | 4.01E-02 | 2.24E-01 | 2.67E+00 | 2.73E+00 | 8.57E-01 | no | Acsl3 | intron_variant | MODIFIER |
| 1 | 78705246 | AX-170126471 | 0 | T | C | 0.132 | 4.01E-02 | 2.24E-01 | 2.67E+00 | 2.73E+00 | 8.57E-01 | no | Acsl3 | synonymous_variant | LOW |
| 1 | 78732151 | AX-170138940 | 0 | A | G | 0.132 | 4.01E-02 | 2.24E-01 | 2.67E+00 | 2.73E+00 | 8.57E-01 | no | | intergenic_variant | MODIFIER |
| 1 | 78820060 | AX-170162128 | 0 | C | T | 0.132 | 4.01E-02 | 2.24E-01 | 2.67E+00 | 2.73E+00 | 8.57E-01 | no | Kcne4 | downstream_gene_variant | MODIFIER |
| 11 | 13465937 | AX-170215493 | 0 | A | G | 0.07 | -3.34E-02 | 1.89E-01 | 2.63E+00 | 2.74E+00 | 8.57E-01 | no | | intergenic_variant | MODIFIER |
| 8 | 57458348 | AX-170080146 | 2 | G | A | 0.145 | 2.38E-02 | 1.41E-01 | 2.72E+00 | 2.80E+00 | 8.57E-01 | no | Scrg1 | intron_variant | MODIFIER |
| 5 | 32782192 | AX-97786226 | 0 | G | C | 0.351 | 1.87E-02 | 1.04E-01 | 2.67E+00 | 2.73E+00 | 8.57E-01 | no | Pisd | intron_variant | MODIFIER |
| 9 | 58972169 | AX-170141501 | 0 | G | A | 0.211 | 2.19E-02 | 1.22E-01 | 2.68E+00 | 2.73E+00 | 8.57E-01 | no | Neo1 | intron_variant | MODIFIER |
| 14 | 64996040 | AX-97826559 | 1 | G | A | 0.054 | 3.34E-02 | 1.92E-01 | 2.65E+00 | 2.75E+00 | 8.57E-01 | no | Ints9 | intron_variant | MODIFIER |
| 7 | 109309932 | AX-170029749 | 0 | C | T | 0.465 | -1.57E-02 | 9.22E-02 | 2.71E+00 | 2.78E+00 | 8.58E-01 | no | Stk33 | intron_variant | MODIFIER |
| 8 | 52437150 | AX-97836821 | 0 | A | G | 0.149 | 2.55E-02 | 1.48E-01 | 2.74E+00 | 2.79E+00 | 8.58E-01 | no | | intergenic_variant | MODIFIER |
| 4 | 72594620 | AX-170069667 | 1 | C | T | 0.232 | 2.03E-02 | 1.04E-01 | 2.61E+00 | 2.69E+00 | 8.58E-01 | no | | intergenic_variant | MODIFIER |
| 10 | 8395702 | AX-97823205 | 0 | G | T | 0.219 | -1.86E-02 | 1.03E-01 | 2.64E+00 | 2.73E+00 | 8.58E-01 | no | Ust | intron_variant | MODIFIER |
| 1 | 36478840 | AX-97814722 | 1 | A | G | 0.455 | 1.38E-02 | 7.93E-02 | 2.68E+00 | 2.75E+00 | 8.58E-01 | no | Cnnm4 | intron_variant | MODIFIER |
| 1 | 37021943 | AX-170389490 | 1 | G | A | 0.455 | 1.38E-02 | 7.93E-02 | 2.68E+00 | 2.75E+00 | 8.58E-01 | no | Vwa3b | upstream_gene_variant | MODIFIER |
| 10 | 114815250 | AX-170374311 | 1 | A | G | 0.366 | 1.34E-02 | 7.75E-02 | 2.67E+00 | 2.75E+00 | 8.58E-01 | no | Gm15723 | intron_variant,non_coding_transcript_variant | MODIFIER |
| 3 | 55346347 | AX-170338241 | 1 | T | C | 0.045 | -2.22E-02 | 1.25E-01 | 2.57E+00 | 2.74E+00 | 8.58E-01 | no | Dclk1 | intron_variant | MODIFIER |
| 5 | 127266405 | AX-97832419 | 0 | A | G | 0.421 | 1.61E-02 | 8.87E-02 | 2.63E+00 | 2.72E+00 | 8.58E-01 | no | Tmem132c | intron_variant | MODIFIER |
| 6 | 104029710 | AX-170502078 | 1 | G | A | 0.054 | 5.31E-02 | 2.98E-01 | 2.67E+00 | 2.73E+00 | 8.58E-01 | no | | intergenic_variant | MODIFIER |
| 6 | 107495982 | AX-170075806 | 1 | G | C | 0.054 | 5.31E-02 | 2.98E-01 | 2.67E+00 | 2.73E+00 | 8.58E-01 | no | | intergenic_variant | MODIFIER |

| | | | | | | | | | | | | | | | |
|---|---|---|---|---|---|---|---|---|---|---|---|---|---|---|---|
| 6 | 122658870 | AX-170175704 | 1 | C | T | 0.054 | 5.31E-02 | 2.98E-01 | 2.67E+00 | 2.73E+00 | 8.58E-01 | no | | intergenic_variant | MODIFIER |
| 5 | 93031109 | AX-170349958 | 0 | G | A | 0.421 | 1.88E-02 | 1.06E-01 | 2.67E+00 | 2.73E+00 | 8.58E-01 | no | | regulatory_region_variant | MODIFIER |
| 8 | 94371546 | AX-170090181 | 0 | C | A | 0.211 | -2.13E-02 | 1.18E-01 | 2.66E+00 | 2.72E+00 | 8.58E-01 | no | | intergenic_variant | MODIFIER |
| 8 | 94549804 | AX-97844802 | 0 | C | T | 0.211 | -2.13E-02 | 1.18E-01 | 2.66E+00 | 2.72E+00 | 8.58E-01 | no | Cpne2 | intron_variant | MODIFIER |
| 8 | 94571773 | AX-170411086 | 0 | C | A | 0.211 | -2.13E-02 | 1.18E-01 | 2.66E+00 | 2.72E+00 | 8.58E-01 | no | Fam192a | downstream_gene_variant | MODIFIER |
| 8 | 94677449 | AX-170365362 | 0 | C | T | 0.211 | -2.13E-02 | 1.18E-01 | 2.66E+00 | 2.72E+00 | 8.58E-01 | no | Pllp | intron_variant | MODIFIER |
| 1 | 10743552 | AX-169999936 | 2 | T | C | 0.373 | -1.21E-02 | 7.24E-02 | 2.69E+00 | 2.78E+00 | 8.58E-01 | no | | intergenic_variant | MODIFIER |
| 7 | 87688773 | AX-170366685 | 1 | T | G | 0.107 | 3.61E-02 | 2.02E-01 | 2.63E+00 | 2.73E+00 | 8.58E-01 | no | Grm5 | intron_variant | MODIFIER |
| 10 | 117877339 | AX-170469186 | 0 | A | G | 0.333 | 1.28E-02 | 7.41E-02 | 2.66E+00 | 2.75E+00 | 8.58E-01 | no | | intergenic_variant | MODIFIER |
| 10 | 117882898 | AX-170423156 | 0 | A | C | 0.333 | 1.28E-02 | 7.41E-02 | 2.66E+00 | 2.75E+00 | 8.58E-01 | no | | intergenic_variant | MODIFIER |
| 10 | 117886215 | AX-170500535 | 0 | G | A | 0.333 | 1.28E-02 | 7.41E-02 | 2.66E+00 | 2.75E+00 | 8.58E-01 | no | | intergenic_variant | MODIFIER |
| 10 | 117887026 | AX-170194272 | 0 | G | A | 0.333 | 1.28E-02 | 7.41E-02 | 2.66E+00 | 2.75E+00 | 8.58E-01 | no | | intergenic_variant | MODIFIER |
| 10 | 117888400 | AX-170092128 | 0 | G | A | 0.333 | 1.28E-02 | 7.41E-02 | 2.66E+00 | 2.75E+00 | 8.58E-01 | no | | intergenic_variant | MODIFIER |
| 15 | 69907381 | AX-170473461 | 1 | G | A | 0.018 | -2.51E-02 | 1.38E-01 | 2.55E+00 | 2.73E+00 | 8.58E-01 | no | | intergenic_variant | MODIFIER |
| 7 | 7850686 | AX-170095858 | 0 | C | A | 0.175 | -1.53E-02 | 7.92E-02 | 2.57E+00 | 2.71E+00 | 8.58E-01 | no | Vmn2r-ps40 | intron_variant,non_coding_transcript_variant | MODIFIER |
| 12 | 72453469 | AX-170126876 | 0 | G | A | 0.061 | -4.01E-02 | 2.16E-01 | 2.62E+00 | 2.71E+00 | 8.59E-01 | no | Lrrc9 | intron_variant,non_coding_transcript_variant | MODIFIER |
| 18 | 34380327 | AX-170233826 | 0 | A | C | 0.061 | -4.01E-02 | 2.16E-01 | 2.62E+00 | 2.71E+00 | 8.59E-01 | no | | regulatory_region_variant | MODIFIER |
| 18 | 38922575 | AX-170250399 | 0 | C | T | 0.061 | -4.01E-02 | 2.16E-01 | 2.62E+00 | 2.71E+00 | 8.59E-01 | no | Fgf1 | upstream_gene_variant | MODIFIER |
| 18 | 38942846 | AX-170254839 | 0 | C | A | 0.061 | -4.01E-02 | 2.16E-01 | 2.62E+00 | 2.71E+00 | 8.59E-01 | no | AC110730.1 | intron_variant,non_coding_transcript_variant | MODIFIER |
| 4 | 144033961 | AX-170360700 | 1 | T | G | 0.08 | -3.87E-02 | 2.17E-01 | 2.64E+00 | 2.73E+00 | 8.59E-01 | no | Gm13108 | non_coding_transcript_exon_variant | MODIFIER |
| 8 | 51797043 | AX-170030686 | 2 | T | A | 0.155 | 2.01E-02 | 1.28E-01 | 2.69E+00 | 2.81E+00 | 8.59E-01 | no | | intergenic_variant | MODIFIER |
| 8 | 72685173 | AX-170455256 | 0 | T | C | 0.193 | -2.40E-02 | 1.35E-01 | 2.67E+00 | 2.73E+00 | 8.59E-01 | no | Nwd1 | intron_variant | MODIFIER |
| 8 | 72697549 | AX-170080398 | 0 | T | C | 0.193 | -2.40E-02 | 1.35E-01 | 2.67E+00 | 2.73E+00 | 8.59E-01 | no | Nwd1 | intron_variant | MODIFIER |
| 8 | 72697858 | AX-170322969 | 0 | T | G | 0.193 | -2.40E-02 | 1.35E-01 | 2.67E+00 | 2.73E+00 | 8.59E-01 | no | Nwd1 | missense_variant | MODERATE |
| 8 | 72697994 | AX-170449906 | 0 | C | A | 0.193 | -2.40E-02 | 1.35E-01 | 2.67E+00 | 2.73E+00 | 8.59E-01 | no | Nwd1 | intron_variant | MODIFIER |
| 8 | 72699796 | AX-170369796 | 0 | C | T | 0.193 | -2.40E-02 | 1.35E-01 | 2.67E+00 | 2.73E+00 | 8.59E-01 | no | Nwd1 | intron_variant | MODIFIER |
| 8 | 72702615 | AX-170148987 | 0 | A | G | 0.193 | -2.40E-02 | 1.35E-01 | 2.67E+00 | 2.73E+00 | 8.59E-01 | no | Nwd1 | intron_variant | MODIFIER |
| 8 | 72719326 | AX-170473488 | 0 | G | A | 0.193 | -2.40E-02 | 1.35E-01 | 2.67E+00 | 2.73E+00 | 8.59E-01 | no | Sin3b | upstream_gene_variant | MODIFIER |
| 8 | 72748871 | AX-170254159 | 0 | T | C | 0.193 | -2.40E-02 | 1.35E-01 | 2.67E+00 | 2.73E+00 | 8.59E-01 | no | Sin3b | intron_variant | MODIFIER |
| 8 | 72749784 | AX-170356318 | 0 | T | C | 0.193 | -2.40E-02 | 1.35E-01 | 2.67E+00 | 2.73E+00 | 8.59E-01 | no | Sin3b | synonymous_variant | LOW |
| 8 | 72756599 | AX-170167609 | 0 | C | T | 0.193 | -2.40E-02 | 1.35E-01 | 2.67E+00 | 2.73E+00 | 8.59E-01 | no | Sin3b | intron_variant | MODIFIER |
| 8 | 72762186 | AX-170237891 | 0 | C | T | 0.193 | -2.40E-02 | 1.35E-01 | 2.67E+00 | 2.73E+00 | 8.59E-01 | no | Sin3b | downstream_gene_variant | MODIFIER |
| 8 | 72764426 | AX-170260657 | 0 | A | G | 0.193 | -2.40E-02 | 1.35E-01 | 2.67E+00 | 2.73E+00 | 8.59E-01 | no | F2rl3 | downstream_gene_variant | MODIFIER |
| 8 | 72780961 | AX-97822924 | 0 | G | C | 0.193 | -2.40E-02 | 1.35E-01 | 2.67E+00 | 2.73E+00 | 8.59E-01 | no | Gm45704 | intron_variant,non_coding_transcript_variant | MODIFIER |
| 8 | 72781115 | AX-170358988 | 0 | A | G | 0.193 | -2.40E-02 | 1.35E-01 | 2.67E+00 | 2.73E+00 | 8.59E-01 | no | Gm45704 | intron_variant,non_coding_transcript_variant | MODIFIER |
| 8 | 72822440 | AX-170291749 | 0 | G | A | 0.193 | -2.40E-02 | 1.35E-01 | 2.67E+00 | 2.73E+00 | 8.59E-01 | no | Large1 | intron_variant | MODIFIER |
| 8 | 72825069 | AX-170373983 | 0 | C | T | 0.193 | -2.40E-02 | 1.35E-01 | 2.67E+00 | 2.73E+00 | 8.59E-01 | no | Large1 | intron_variant | MODIFIER |
| 8 | 72825382 | AX-97802194 | 0 | G | A | 0.193 | -2.40E-02 | 1.35E-01 | 2.67E+00 | 2.73E+00 | 8.59E-01 | no | Large1 | intron_variant | MODIFIER |
| 8 | 72833230 | AX-170492090 | 0 | G | A | 0.193 | -2.40E-02 | 1.35E-01 | 2.67E+00 | 2.73E+00 | 8.59E-01 | no | Large1 | intron_variant | MODIFIER |
| 8 | 72836566 | AX-97813312 | 0 | G | T | 0.193 | -2.40E-02 | 1.35E-01 | 2.67E+00 | 2.73E+00 | 8.59E-01 | no | Large1 | intron_variant | MODIFIER |
| 8 | 72873517 | AX-170080402 | 0 | C | T | 0.193 | -2.40E-02 | 1.35E-01 | 2.67E+00 | 2.73E+00 | 8.59E-01 | no | Large1 | intron_variant | MODIFIER |
| 8 | 72900644 | AX-170098649 | 0 | T | C | 0.193 | -2.40E-02 | 1.35E-01 | 2.67E+00 | 2.73E+00 | 8.59E-01 | no | Large1 | intron_variant | MODIFIER |
| 8 | 72917682 | AX-170525093 | 0 | A | G | 0.193 | -2.40E-02 | 1.35E-01 | 2.67E+00 | 2.73E+00 | 8.59E-01 | no | Large1 | intron_variant | MODIFIER |
| 8 | 72917745 | AX-170385245 | 0 | A | G | 0.193 | -2.40E-02 | 1.35E-01 | 2.67E+00 | 2.73E+00 | 8.59E-01 | no | Large1 | intron_variant | MODIFIER |
| 8 | 72934300 | AX-170158556 | 0 | A | C | 0.193 | -2.40E-02 | 1.35E-01 | 2.67E+00 | 2.73E+00 | 8.59E-01 | no | Large1 | intron_variant | MODIFIER |
| 8 | 72954020 | AX-170126792 | 0 | G | T | 0.193 | -2.40E-02 | 1.35E-01 | 2.67E+00 | 2.73E+00 | 8.59E-01 | no | Large1 | intron_variant | MODIFIER |
| 8 | 72985700 | AX-170361996 | 0 | T | C | 0.193 | -2.40E-02 | 1.35E-01 | 2.67E+00 | 2.73E+00 | 8.59E-01 | no | Large1 | intron_variant | MODIFIER |
| 8 | 72996785 | AX-97812735 | 0 | C | T | 0.193 | -2.40E-02 | 1.35E-01 | 2.67E+00 | 2.73E+00 | 8.59E-01 | no | Large1 | intron_variant | MODIFIER |
| 8 | 72997121 | AX-170125774 | 0 | T | C | 0.193 | -2.40E-02 | 1.35E-01 | 2.67E+00 | 2.73E+00 | 8.59E-01 | no | Large1 | intron_variant | MODIFIER |
| 8 | 73005218 | AX-170377097 | 0 | C | T | 0.193 | -2.40E-02 | 1.35E-01 | 2.67E+00 | 2.73E+00 | 8.59E-01 | no | Large1 | intron_variant | MODIFIER |
| 8 | 73027399 | AX-97847607 | 0 | T | C | 0.193 | -2.40E-02 | 1.35E-01 | 2.67E+00 | 2.73E+00 | 8.59E-01 | no | Large1 | intron_variant | MODIFIER |
| 8 | 73031019 | AX-170211597 | 0 | C | T | 0.193 | -2.40E-02 | 1.35E-01 | 2.67E+00 | 2.73E+00 | 8.59E-01 | no | Large1 | intron_variant | MODIFIER |
| 8 | 73031052 | AX-170236424 | 0 | C | T | 0.193 | -2.40E-02 | 1.35E-01 | 2.67E+00 | 2.73E+00 | 8.59E-01 | no | Large1 | intron_variant | MODIFIER |
| 8 | 73058800 | AX-170467342 | 0 | A | G | 0.193 | -2.40E-02 | 1.35E-01 | 2.67E+00 | 2.73E+00 | 8.59E-01 | no | Large1 | intron_variant | MODIFIER |

| | | | | | | | | | | | | | | | |
|---|---|---|---|---|---|---|---|---|---|---|---|---|---|---|---|
| 8 | 73067789 | AX-170185781 | 0 | G | C | 0.193 | -2.40E-02 | 1.35E-01 | 2.67E+00 | 2.73E+00 | 8.59E-01 | no | Large1 | intron_variant | MODIFIER |
| 8 | 73091671 | AX-170080407 | 0 | A | G | 0.193 | -2.40E-02 | 1.35E-01 | 2.67E+00 | 2.73E+00 | 8.59E-01 | no | Large1 | intron_variant | MODIFIER |
| 8 | 73092776 | AX-170306106 | 0 | T | C | 0.193 | -2.40E-02 | 1.35E-01 | 2.67E+00 | 2.73E+00 | 8.59E-01 | no | Large1 | intron_variant | MODIFIER |
| 8 | 73093286 | AX-170090146 | 0 | G | T | 0.193 | -2.40E-02 | 1.35E-01 | 2.67E+00 | 2.73E+00 | 8.59E-01 | no | Large1 | intron_variant | MODIFIER |
| 8 | 73103434 | AX-170080406 | 0 | T | G | 0.193 | -2.40E-02 | 1.35E-01 | 2.67E+00 | 2.73E+00 | 8.59E-01 | no | Large1 | intron_variant | MODIFIER |
| 8 | 73121063 | AX-97780665 | 0 | A | T | 0.193 | -2.40E-02 | 1.35E-01 | 2.67E+00 | 2.73E+00 | 8.59E-01 | no | Large1 | intron_variant | MODIFIER |
| 8 | 73121284 | AX-170030995 | 0 | A | C | 0.193 | -2.40E-02 | 1.35E-01 | 2.67E+00 | 2.73E+00 | 8.59E-01 | no | Large1 | intron_variant | MODIFIER |
| 8 | 73129730 | AX-170425834 | 0 | A | T | 0.193 | -2.40E-02 | 1.35E-01 | 2.67E+00 | 2.73E+00 | 8.59E-01 | no | Large1 | intron_variant | MODIFIER |
| 8 | 73134581 | AX-170467263 | 0 | T | C | 0.193 | -2.40E-02 | 1.35E-01 | 2.67E+00 | 2.73E+00 | 8.59E-01 | no | Large1 | intron_variant | MODIFIER |
| 8 | 73136569 | AX-170339114 | 0 | C | T | 0.193 | -2.40E-02 | 1.35E-01 | 2.67E+00 | 2.73E+00 | 8.59E-01 | no | Large1 | intron_variant | MODIFIER |
| 8 | 73158454 | AX-170133508 | 0 | C | T | 0.193 | -2.40E-02 | 1.35E-01 | 2.67E+00 | 2.73E+00 | 8.59E-01 | no | Large1 | intron_variant | MODIFIER |
| 8 | 73158562 | AX-170301391 | 0 | T | C | 0.193 | -2.40E-02 | 1.35E-01 | 2.67E+00 | 2.73E+00 | 8.59E-01 | no | Large1 | intron_variant | MODIFIER |
| 8 | 73158854 | AX-170108293 | 0 | C | T | 0.193 | -2.40E-02 | 1.35E-01 | 2.67E+00 | 2.73E+00 | 8.59E-01 | no | Large1 | intron_variant | MODIFIER |
| 8 | 73177955 | AX-170096319 | 0 | A | G | 0.193 | -2.40E-02 | 1.35E-01 | 2.67E+00 | 2.73E+00 | 8.59E-01 | no | Large1 | intron_variant | MODIFIER |
| 8 | 73198944 | AX-170287266 | 0 | C | T | 0.193 | -2.40E-02 | 1.35E-01 | 2.67E+00 | 2.73E+00 | 8.59E-01 | no | Large1 | intron_variant | MODIFIER |
| 8 | 73200257 | AX-170430807 | 0 | C | A | 0.193 | -2.40E-02 | 1.35E-01 | 2.67E+00 | 2.73E+00 | 8.59E-01 | no | Large1 | intron_variant | MODIFIER |
| 8 | 73215230 | AX-170498490 | 0 | G | A | 0.193 | -2.40E-02 | 1.35E-01 | 2.67E+00 | 2.73E+00 | 8.59E-01 | no | Large1 | intron_variant | MODIFIER |
| 8 | 73216951 | AX-97814236 | 0 | T | G | 0.193 | -2.40E-02 | 1.35E-01 | 2.67E+00 | 2.73E+00 | 8.59E-01 | no | Large1 | intron_variant | MODIFIER |
| 8 | 73217105 | AX-170123175 | 0 | G | A | 0.193 | -2.40E-02 | 1.35E-01 | 2.67E+00 | 2.73E+00 | 8.59E-01 | no | Large1 | intron_variant | MODIFIER |
| 8 | 73230609 | AX-170535065 | 0 | A | G | 0.193 | -2.40E-02 | 1.35E-01 | 2.67E+00 | 2.73E+00 | 8.59E-01 | no | Large1 | intron_variant | MODIFIER |
| 8 | 73243831 | AX-97796855 | 0 | A | G | 0.193 | -2.40E-02 | 1.35E-01 | 2.67E+00 | 2.73E+00 | 8.59E-01 | no | Large1 | intron_variant | MODIFIER |
| 8 | 73279864 | AX-170177911 | 0 | C | T | 0.193 | -2.40E-02 | 1.35E-01 | 2.67E+00 | 2.73E+00 | 8.59E-01 | no | Large1 | intron_variant | MODIFIER |
| 8 | 73280258 | AX-170475875 | 0 | C | T | 0.193 | -2.40E-02 | 1.35E-01 | 2.67E+00 | 2.73E+00 | 8.59E-01 | no | Large1 | intron_variant | MODIFIER |
| 8 | 73290781 | AX-170146988 | 0 | C | G | 0.193 | -2.40E-02 | 1.35E-01 | 2.67E+00 | 2.73E+00 | 8.59E-01 | no | Large1 | intron_variant | MODIFIER |
| 8 | 73324534 | AX-170285822 | 0 | A | T | 0.193 | -2.40E-02 | 1.35E-01 | 2.67E+00 | 2.73E+00 | 8.59E-01 | no | Large1 | intron_variant | MODIFIER |
| 8 | 73337147 | AX-170080415 | 0 | G | A | 0.193 | -2.40E-02 | 1.35E-01 | 2.67E+00 | 2.73E+00 | 8.59E-01 | no | Large1 | intron_variant | MODIFIER |
| 8 | 73337252 | AX-170080414 | 0 | C | G | 0.193 | -2.40E-02 | 1.35E-01 | 2.67E+00 | 2.73E+00 | 8.59E-01 | no | Large1 | intron_variant | MODIFIER |
| 8 | 73354029 | AX-170270050 | 0 | T | C | 0.193 | -2.40E-02 | 1.35E-01 | 2.67E+00 | 2.73E+00 | 8.59E-01 | no | Large1 | upstream_gene_variant | MODIFIER |
| 8 | 73376165 | AX-170201969 | 0 | G | A | 0.193 | -2.40E-02 | 1.35E-01 | 2.67E+00 | 2.73E+00 | 8.59E-01 | no | | intergenic_variant | MODIFIER |
| 8 | 73380701 | AX-170267848 | 0 | C | T | 0.193 | -2.40E-02 | 1.35E-01 | 2.67E+00 | 2.73E+00 | 8.59E-01 | no | | intergenic_variant | MODIFIER |
| 8 | 73387188 | AX-97788128 | 0 | C | T | 0.193 | -2.40E-02 | 1.35E-01 | 2.67E+00 | 2.73E+00 | 8.59E-01 | no | | intergenic_variant | MODIFIER |
| 8 | 73387686 | AX-170474890 | 0 | T | C | 0.193 | -2.40E-02 | 1.35E-01 | 2.67E+00 | 2.73E+00 | 8.59E-01 | no | | intergenic_variant | MODIFIER |
| 8 | 73387774 | AX-170289896 | 0 | T | C | 0.193 | -2.40E-02 | 1.35E-01 | 2.67E+00 | 2.73E+00 | 8.59E-01 | no | | intergenic_variant | MODIFIER |
| 8 | 73389865 | AX-170442202 | 0 | T | G | 0.193 | -2.40E-02 | 1.35E-01 | 2.67E+00 | 2.73E+00 | 8.59E-01 | no | | intergenic_variant | MODIFIER |
| 8 | 73404600 | AX-170080416 | 0 | G | A | 0.193 | -2.40E-02 | 1.35E-01 | 2.67E+00 | 2.73E+00 | 8.59E-01 | no | | intergenic_variant | MODIFIER |
| 8 | 73404636 | AX-170271269 | 0 | T | C | 0.193 | -2.40E-02 | 1.35E-01 | 2.67E+00 | 2.73E+00 | 8.59E-01 | no | | intergenic_variant | MODIFIER |
| 8 | 73405593 | AX-170531616 | 0 | A | G | 0.193 | -2.40E-02 | 1.35E-01 | 2.67E+00 | 2.73E+00 | 8.59E-01 | no | | intergenic_variant | MODIFIER |
| 8 | 73408006 | AX-170343361 | 0 | T | G | 0.193 | -2.40E-02 | 1.35E-01 | 2.67E+00 | 2.73E+00 | 8.59E-01 | no | | intergenic_variant | MODIFIER |
| 8 | 73459202 | AX-170091307 | 0 | A | T | 0.193 | -2.40E-02 | 1.35E-01 | 2.67E+00 | 2.73E+00 | 8.59E-01 | no | | intergenic_variant | MODIFIER |
| 8 | 73459522 | AX-170223418 | 0 | A | G | 0.193 | -2.40E-02 | 1.35E-01 | 2.67E+00 | 2.73E+00 | 8.59E-01 | no | | intergenic_variant | MODIFIER |
| 8 | 73487871 | AX-170445232 | 0 | A | G | 0.193 | -2.40E-02 | 1.35E-01 | 2.67E+00 | 2.73E+00 | 8.59E-01 | no | | intergenic_variant | MODIFIER |
| 8 | 73487962 | AX-170490252 | 0 | G | A | 0.193 | -2.40E-02 | 1.35E-01 | 2.67E+00 | 2.73E+00 | 8.59E-01 | no | | intergenic_variant | MODIFIER |
| 8 | 73488478 | AX-170428631 | 0 | T | G | 0.193 | -2.40E-02 | 1.35E-01 | 2.67E+00 | 2.73E+00 | 8.59E-01 | no | | intergenic_variant | MODIFIER |
| 8 | 73499880 | AX-170399771 | 0 | T | G | 0.193 | -2.40E-02 | 1.35E-01 | 2.67E+00 | 2.73E+00 | 8.59E-01 | no | | intergenic_variant | MODIFIER |
| 8 | 73512849 | AX-97805169 | 0 | A | G | 0.193 | -2.40E-02 | 1.35E-01 | 2.67E+00 | 2.73E+00 | 8.59E-01 | no | | intergenic_variant | MODIFIER |
| 8 | 73513619 | AX-170291546 | 0 | A | G | 0.193 | -2.40E-02 | 1.35E-01 | 2.67E+00 | 2.73E+00 | 8.59E-01 | no | | intergenic_variant | MODIFIER |
| 8 | 73514146 | AX-170098345 | 0 | A | C | 0.193 | -2.40E-02 | 1.35E-01 | 2.67E+00 | 2.73E+00 | 8.59E-01 | no | | intergenic_variant | MODIFIER |
| 8 | 73572182 | AX-170410736 | 0 | C | A | 0.193 | -2.40E-02 | 1.35E-01 | 2.67E+00 | 2.73E+00 | 8.59E-01 | no | | intergenic_variant | MODIFIER |
| 8 | 73595329 | AX-170430637 | 0 | G | A | 0.193 | -2.40E-02 | 1.35E-01 | 2.67E+00 | 2.73E+00 | 8.59E-01 | no | | intergenic_variant | MODIFIER |
| 8 | 73599205 | AX-97808541 | 0 | T | G | 0.193 | -2.40E-02 | 1.35E-01 | 2.67E+00 | 2.73E+00 | 8.59E-01 | no | | intergenic_variant | MODIFIER |
| 8 | 73607820 | AX-170031005 | 0 | A | G | 0.193 | -2.40E-02 | 1.35E-01 | 2.67E+00 | 2.73E+00 | 8.59E-01 | no | | intergenic_variant | MODIFIER |
| 8 | 73612948 | AX-170031004 | 0 | A | G | 0.193 | -2.40E-02 | 1.35E-01 | 2.67E+00 | 2.73E+00 | 8.59E-01 | no | | intergenic_variant | MODIFIER |
| 8 | 73613924 | AX-170080420 | 0 | G | C | 0.193 | -2.40E-02 | 1.35E-01 | 2.67E+00 | 2.73E+00 | 8.59E-01 | no | | intergenic_variant | MODIFIER |
| 8 | 73616584 | AX-170080421 | 0 | A | C | 0.193 | -2.40E-02 | 1.35E-01 | 2.67E+00 | 2.73E+00 | 8.59E-01 | no | | intergenic_variant | MODIFIER |

| | | | | | | | | | | | | | | | |
|---|---|---|---|---|---|---|---|---|---|---|---|---|---|---|---|
| 8 | 73627137 | AX-170080417 | 0 | T | C | 0.193 | -2.40E-02 | 1.35E-01 | 2.67E+00 | 2.73E+00 | 8.59E-01 | no | | intergenic_variant | MODIFIER |
| 8 | 73642241 | AX-170199775 | 0 | A | T | 0.193 | -2.40E-02 | 1.35E-01 | 2.67E+00 | 2.73E+00 | 8.59E-01 | no | Gm23255 | upstream_gene_variant | MODIFIER |
| 8 | 73660407 | AX-97790227 | 0 | A | G | 0.193 | -2.40E-02 | 1.35E-01 | 2.67E+00 | 2.73E+00 | 8.59E-01 | no | | intergenic_variant | MODIFIER |
| 8 | 73689700 | AX-170253809 | 0 | G | A | 0.193 | -2.40E-02 | 1.35E-01 | 2.67E+00 | 2.73E+00 | 8.59E-01 | no | | intergenic_variant | MODIFIER |
| 8 | 73694525 | AX-97848006 | 0 | T | C | 0.193 | -2.40E-02 | 1.35E-01 | 2.67E+00 | 2.73E+00 | 8.59E-01 | no | | intergenic_variant | MODIFIER |
| 8 | 73711160 | AX-170315368 | 0 | C | T | 0.193 | -2.40E-02 | 1.35E-01 | 2.67E+00 | 2.73E+00 | 8.59E-01 | no | | intergenic_variant | MODIFIER |
| 8 | 73716085 | AX-170535382 | 0 | T | A | 0.193 | -2.40E-02 | 1.35E-01 | 2.67E+00 | 2.73E+00 | 8.59E-01 | no | | intergenic_variant | MODIFIER |
| 8 | 73721112 | AX-170476060 | 0 | A | G | 0.193 | -2.40E-02 | 1.35E-01 | 2.67E+00 | 2.73E+00 | 8.59E-01 | no | | intergenic_variant | MODIFIER |
| 8 | 73721480 | AX-170443887 | 0 | G | A | 0.193 | -2.40E-02 | 1.35E-01 | 2.67E+00 | 2.73E+00 | 8.59E-01 | no | | intergenic_variant | MODIFIER |
| 8 | 73727332 | AX-170137270 | 0 | A | G | 0.193 | -2.40E-02 | 1.35E-01 | 2.67E+00 | 2.73E+00 | 8.59E-01 | no | | intergenic_variant | MODIFIER |
| 8 | 73749056 | AX-170275860 | 0 | G | A | 0.193 | -2.40E-02 | 1.35E-01 | 2.67E+00 | 2.73E+00 | 8.59E-01 | no | Crry-ps | downstream_gene_variant | MODIFIER |
| 8 | 73749795 | AX-170201493 | 0 | C | G | 0.193 | -2.40E-02 | 1.35E-01 | 2.67E+00 | 2.73E+00 | 8.59E-01 | no | Crry-ps | non_coding_transcript_exon_variant | MODIFIER |
| 8 | 73761039 | AX-170304142 | 0 | G | A | 0.193 | -2.40E-02 | 1.35E-01 | 2.67E+00 | 2.73E+00 | 8.59E-01 | no | | intergenic_variant | MODIFIER |
| 8 | 73761105 | AX-170424116 | 0 | G | A | 0.193 | -2.40E-02 | 1.35E-01 | 2.67E+00 | 2.73E+00 | 8.59E-01 | no | | intergenic_variant | MODIFIER |
| 8 | 73767876 | AX-170186261 | 0 | C | T | 0.193 | -2.40E-02 | 1.35E-01 | 2.67E+00 | 2.73E+00 | 8.59E-01 | no | | intergenic_variant | MODIFIER |
| 8 | 73779599 | AX-170506156 | 0 | A | G | 0.193 | -2.40E-02 | 1.35E-01 | 2.67E+00 | 2.73E+00 | 8.59E-01 | no | | intergenic_variant | MODIFIER |
| 8 | 73802451 | AX-97806809 | 0 | G | A | 0.193 | -2.40E-02 | 1.35E-01 | 2.67E+00 | 2.73E+00 | 8.59E-01 | no | | intergenic_variant | MODIFIER |
| 8 | 73811745 | AX-170080422 | 0 | G | A | 0.193 | -2.40E-02 | 1.35E-01 | 2.67E+00 | 2.73E+00 | 8.59E-01 | no | | intergenic_variant | MODIFIER |
| 8 | 73814402 | AX-170423920 | 0 | C | T | 0.193 | -2.40E-02 | 1.35E-01 | 2.67E+00 | 2.73E+00 | 8.59E-01 | no | | intergenic_variant | MODIFIER |
| 8 | 73843070 | AX-170080424 | 0 | A | G | 0.193 | -2.40E-02 | 1.35E-01 | 2.67E+00 | 2.73E+00 | 8.59E-01 | no | | intergenic_variant | MODIFIER |
| 8 | 73860417 | AX-170207302 | 0 | C | T | 0.193 | -2.40E-02 | 1.35E-01 | 2.67E+00 | 2.73E+00 | 8.59E-01 | no | | intergenic_variant | MODIFIER |
| 8 | 73879861 | AX-170080425 | 0 | G | A | 0.193 | -2.40E-02 | 1.35E-01 | 2.67E+00 | 2.73E+00 | 8.59E-01 | no | | intergenic_variant | MODIFIER |
| 8 | 73880594 | AX-170333463 | 0 | G | T | 0.193 | -2.40E-02 | 1.35E-01 | 2.67E+00 | 2.73E+00 | 8.59E-01 | no | | intergenic_variant | MODIFIER |
| 8 | 73904076 | AX-170361511 | 0 | C | T | 0.193 | -2.40E-02 | 1.35E-01 | 2.67E+00 | 2.73E+00 | 8.59E-01 | no | | regulatory_region_variant | MODIFIER |
| 8 | 73904155 | AX-170195782 | 0 | G | A | 0.193 | -2.40E-02 | 1.35E-01 | 2.67E+00 | 2.73E+00 | 8.59E-01 | no | | regulatory_region_variant | MODIFIER |
| 8 | 73911282 | AX-170298711 | 0 | G | A | 0.193 | -2.40E-02 | 1.35E-01 | 2.67E+00 | 2.73E+00 | 8.59E-01 | no | | intergenic_variant | MODIFIER |
| 8 | 73927977 | AX-170169376 | 0 | T | G | 0.193 | -2.40E-02 | 1.35E-01 | 2.67E+00 | 2.73E+00 | 8.59E-01 | no | | intergenic_variant | MODIFIER |
| 8 | 73932513 | AX-170123013 | 0 | G | A | 0.193 | -2.40E-02 | 1.35E-01 | 2.67E+00 | 2.73E+00 | 8.59E-01 | no | | intergenic_variant | MODIFIER |
| 8 | 73983163 | AX-170430178 | 0 | C | T | 0.193 | -2.40E-02 | 1.35E-01 | 2.67E+00 | 2.73E+00 | 8.59E-01 | no | | intergenic_variant | MODIFIER |
| 8 | 73983488 | AX-170451005 | 0 | G | T | 0.193 | -2.40E-02 | 1.35E-01 | 2.67E+00 | 2.73E+00 | 8.59E-01 | no | | intergenic_variant | MODIFIER |
| 8 | 74085791 | AX-170138892 | 0 | A | G | 0.193 | -2.40E-02 | 1.35E-01 | 2.67E+00 | 2.73E+00 | 8.59E-01 | no | Gm45821 | intron_variant,non_coding_transcript_variant | MODIFIER |
| 8 | 74089512 | AX-170424423 | 0 | T | A | 0.193 | -2.40E-02 | 1.35E-01 | 2.67E+00 | 2.73E+00 | 8.59E-01 | no | Gm45821 | intron_variant,non_coding_transcript_variant | MODIFIER |
| 8 | 74089583 | AX-170230043 | 0 | C | A | 0.193 | -2.40E-02 | 1.35E-01 | 2.67E+00 | 2.73E+00 | 8.59E-01 | no | Gm45821 | intron_variant,non_coding_transcript_variant | MODIFIER |
| 8 | 74168780 | AX-97814893 | 0 | C | A | 0.193 | -2.40E-02 | 1.35E-01 | 2.67E+00 | 2.73E+00 | 8.59E-01 | no | | intergenic_variant | MODIFIER |
| 8 | 74228914 | AX-170425048 | 0 | C | A | 0.193 | -2.40E-02 | 1.35E-01 | 2.67E+00 | 2.73E+00 | 8.59E-01 | no | | intergenic_variant | MODIFIER |
| 8 | 74242942 | AX-170263825 | 0 | T | C | 0.193 | -2.40E-02 | 1.35E-01 | 2.67E+00 | 2.73E+00 | 8.59E-01 | no | | intergenic_variant | MODIFIER |
| 8 | 74242994 | AX-97815919 | 0 | G | C | 0.193 | -2.40E-02 | 1.35E-01 | 2.67E+00 | 2.73E+00 | 8.59E-01 | no | | intergenic_variant | MODIFIER |
| 8 | 74298365 | AX-97855655 | 0 | C | T | 0.193 | -2.40E-02 | 1.35E-01 | 2.67E+00 | 2.73E+00 | 8.59E-01 | no | | intergenic_variant | MODIFIER |
| 8 | 74298955 | AX-170198786 | 0 | G | A | 0.193 | -2.40E-02 | 1.35E-01 | 2.67E+00 | 2.73E+00 | 8.59E-01 | no | | intergenic_variant | MODIFIER |
| 8 | 74300609 | AX-170031018 | 0 | T | G | 0.193 | -2.40E-02 | 1.35E-01 | 2.67E+00 | 2.73E+00 | 8.59E-01 | no | | intergenic_variant | MODIFIER |
| 8 | 74313964 | AX-170031016 | 0 | A | G | 0.193 | -2.40E-02 | 1.35E-01 | 2.67E+00 | 2.73E+00 | 8.59E-01 | no | | intergenic_variant | MODIFIER |
| 8 | 74314328 | AX-97844671 | 0 | T | C | 0.193 | -2.40E-02 | 1.35E-01 | 2.67E+00 | 2.73E+00 | 8.59E-01 | no | | intergenic_variant | MODIFIER |
| 8 | 74362496 | AX-170031009 | 0 | C | A | 0.193 | -2.40E-02 | 1.35E-01 | 2.67E+00 | 2.73E+00 | 8.59E-01 | no | | intergenic_variant | MODIFIER |
| 8 | 74376659 | AX-170349881 | 0 | C | T | 0.193 | -2.40E-02 | 1.35E-01 | 2.67E+00 | 2.73E+00 | 8.59E-01 | no | | intergenic_variant | MODIFIER |
| 8 | 74377300 | AX-170513171 | 0 | C | A | 0.193 | -2.40E-02 | 1.35E-01 | 2.67E+00 | 2.73E+00 | 8.59E-01 | no | | intergenic_variant | MODIFIER |
| 8 | 74391574 | AX-170534791 | 0 | G | C | 0.193 | -2.40E-02 | 1.35E-01 | 2.67E+00 | 2.73E+00 | 8.59E-01 | no | | intergenic_variant | MODIFIER |
| 8 | 74440362 | AX-97795595 | 0 | A | G | 0.193 | -2.40E-02 | 1.35E-01 | 2.67E+00 | 2.73E+00 | 8.59E-01 | no | | intergenic_variant | MODIFIER |
| 8 | 74446886 | AX-170031012 | 0 | G | A | 0.193 | -2.40E-02 | 1.35E-01 | 2.67E+00 | 2.73E+00 | 8.59E-01 | no | | intergenic_variant | MODIFIER |
| 8 | 74471837 | AX-97791510 | 0 | G | A | 0.193 | -2.40E-02 | 1.35E-01 | 2.67E+00 | 2.73E+00 | 8.59E-01 | no | | intergenic_variant | MODIFIER |
| 8 | 74472015 | AX-170031008 | 0 | A | G | 0.193 | -2.40E-02 | 1.35E-01 | 2.67E+00 | 2.73E+00 | 8.59E-01 | no | | intergenic_variant | MODIFIER |
| 8 | 74473883 | AX-170031013 | 0 | A | G | 0.193 | -2.40E-02 | 1.35E-01 | 2.67E+00 | 2.73E+00 | 8.59E-01 | no | | intergenic_variant | MODIFIER |
| 8 | 74495377 | AX-170272774 | 0 | G | A | 0.193 | -2.40E-02 | 1.35E-01 | 2.67E+00 | 2.73E+00 | 8.59E-01 | no | | intergenic_variant | MODIFIER |
| 8 | 74550416 | AX-170096322 | 0 | T | G | 0.193 | -2.40E-02 | 1.35E-01 | 2.67E+00 | 2.73E+00 | 8.59E-01 | no | | intergenic_variant | MODIFIER |
| 8 | 74561176 | AX-170090147 | 0 | T | C | 0.193 | -2.40E-02 | 1.35E-01 | 2.67E+00 | 2.73E+00 | 8.59E-01 | no | | intergenic_variant | MODIFIER |

| | | | | | | | | | | | | | | | |
|---|---|---|---|---|---|---|---|---|---|---|---|---|---|---|---|
| 8 | 74598253 | AX-97782358 | 0 | C | A | 0.193 | -2.40E-02 | 1.35E-01 | 2.67E+00 | 2.73E+00 | 8.59E-01 | no | Gm11033 | intron_variant,non_coding_transcript_variant | MODIFIER |
| 8 | 74602390 | AX-170299262 | 0 | C | T | 0.193 | -2.40E-02 | 1.35E-01 | 2.67E+00 | 2.73E+00 | 8.59E-01 | no | Gm11033 | intron_variant,non_coding_transcript_variant | MODIFIER |
| 8 | 74613382 | AX-170387866 | 0 | G | A | 0.193 | -2.40E-02 | 1.35E-01 | 2.67E+00 | 2.73E+00 | 8.59E-01 | no | Gm11033 | intron_variant,non_coding_transcript_variant | MODIFIER |
| 8 | 74613577 | AX-170100073 | 0 | A | G | 0.193 | -2.40E-02 | 1.35E-01 | 2.67E+00 | 2.73E+00 | 8.59E-01 | no | Gm11033 | intron_variant,non_coding_transcript_variant | MODIFIER |
| 8 | 74619659 | AX-170266764 | 0 | G | A | 0.193 | -2.40E-02 | 1.35E-01 | 2.67E+00 | 2.73E+00 | 8.59E-01 | no | Gm11033 | intron_variant,non_coding_transcript_variant | MODIFIER |
| 8 | 74621161 | AX-170420408 | 0 | C | A | 0.193 | -2.40E-02 | 1.35E-01 | 2.67E+00 | 2.73E+00 | 8.59E-01 | no | Gm11033 | intron_variant,non_coding_transcript_variant | MODIFIER |
| 8 | 74636760 | AX-170114230 | 0 | G | C | 0.193 | -2.40E-02 | 1.35E-01 | 2.67E+00 | 2.73E+00 | 8.59E-01 | no | Gm11033 | intron_variant,non_coding_transcript_variant | MODIFIER |
| 8 | 74643050 | AX-170326209 | 0 | T | C | 0.193 | -2.40E-02 | 1.35E-01 | 2.67E+00 | 2.73E+00 | 8.59E-01 | no | Gm11033 | intron_variant,non_coding_transcript_variant | MODIFIER |
| 8 | 74670364 | AX-170105395 | 0 | A | C | 0.193 | -2.40E-02 | 1.35E-01 | 2.67E+00 | 2.73E+00 | 8.59E-01 | no | Gm11033 | intron_variant,non_coding_transcript_variant | MODIFIER |
| 8 | 74698571 | AX-170459676 | 0 | C | T | 0.193 | -2.40E-02 | 1.35E-01 | 2.67E+00 | 2.73E+00 | 8.59E-01 | no | Gm11033 | intron_variant,non_coding_transcript_variant | MODIFIER |
| 8 | 74734728 | AX-97815656 | 0 | A | G | 0.193 | -2.40E-02 | 1.35E-01 | 2.67E+00 | 2.73E+00 | 8.59E-01 | no | Gm11033 | intron_variant,non_coding_transcript_variant | MODIFIER |
| 8 | 74758568 | AX-170031025 | 0 | A | G | 0.193 | -2.40E-02 | 1.35E-01 | 2.67E+00 | 2.73E+00 | 8.59E-01 | no | Gm11033 | intron_variant,non_coding_transcript_variant | MODIFIER |
| 8 | 74762289 | AX-170408961 | 0 | G | A | 0.193 | -2.40E-02 | 1.35E-01 | 2.67E+00 | 2.73E+00 | 8.59E-01 | no | Gm11033 | intron_variant,non_coding_transcript_variant | MODIFIER |
| 8 | 74764134 | AX-170505148 | 0 | C | A | 0.193 | -2.40E-02 | 1.35E-01 | 2.67E+00 | 2.73E+00 | 8.59E-01 | no | Gm11033 | intron_variant,non_coding_transcript_variant | MODIFIER |
| 8 | 74764485 | AX-170327182 | 0 | A | G | 0.193 | -2.40E-02 | 1.35E-01 | 2.67E+00 | 2.73E+00 | 8.59E-01 | no | Gm11033 | intron_variant,non_coding_transcript_variant | MODIFIER |
| 8 | 74770857 | AX-170291153 | 0 | G | C | 0.193 | -2.40E-02 | 1.35E-01 | 2.67E+00 | 2.73E+00 | 8.59E-01 | no | Gm11033 | intron_variant,non_coding_transcript_variant | MODIFIER |
| 8 | 74825568 | AX-170367494 | 0 | C | A | 0.193 | -2.40E-02 | 1.35E-01 | 2.67E+00 | 2.73E+00 | 8.59E-01 | no | Gm11033 | intron_variant,non_coding_transcript_variant | MODIFIER |
| 8 | 74826246 | AX-170496878 | 0 | A | G | 0.193 | -2.40E-02 | 1.35E-01 | 2.67E+00 | 2.73E+00 | 8.59E-01 | no | Gm11033 | intron_variant,non_coding_transcript_variant | MODIFIER |
| 8 | 74850182 | AX-97804494 | 0 | A | G | 0.193 | -2.40E-02 | 1.35E-01 | 2.67E+00 | 2.73E+00 | 8.59E-01 | no | | intergenic_variant | MODIFIER |
| 8 | 74851441 | AX-170031020 | 0 | T | C | 0.193 | -2.40E-02 | 1.35E-01 | 2.67E+00 | 2.73E+00 | 8.59E-01 | no | | intergenic_variant | MODIFIER |
| 8 | 74864976 | AX-170240485 | 0 | A | G | 0.193 | -2.40E-02 | 1.35E-01 | 2.67E+00 | 2.73E+00 | 8.59E-01 | no | | intergenic_variant | MODIFIER |
| 8 | 74868116 | AX-170326920 | 0 | C | A | 0.193 | -2.40E-02 | 1.35E-01 | 2.67E+00 | 2.73E+00 | 8.59E-01 | no | | intergenic_variant | MODIFIER |
| 8 | 74887988 | AX-170080437 | 0 | C | G | 0.193 | -2.40E-02 | 1.35E-01 | 2.67E+00 | 2.73E+00 | 8.59E-01 | no | Isx | intron_variant | MODIFIER |
| 8 | 74899763 | AX-170305114 | 0 | A | G | 0.193 | -2.40E-02 | 1.35E-01 | 2.67E+00 | 2.73E+00 | 8.59E-01 | no | | intergenic_variant | MODIFIER |
| 8 | 74901200 | AX-97856020 | 0 | A | G | 0.193 | -2.40E-02 | 1.35E-01 | 2.67E+00 | 2.73E+00 | 8.59E-01 | no | | intergenic_variant | MODIFIER |
| 8 | 74933587 | AX-170336793 | 0 | T | C | 0.193 | -2.40E-02 | 1.35E-01 | 2.67E+00 | 2.73E+00 | 8.59E-01 | no | | intergenic_variant | MODIFIER |
| 8 | 74953357 | AX-170201913 | 0 | T | A | 0.193 | -2.40E-02 | 1.35E-01 | 2.67E+00 | 2.73E+00 | 8.59E-01 | no | | intergenic_variant | MODIFIER |
| 8 | 74962141 | AX-170369912 | 0 | G | A | 0.193 | -2.40E-02 | 1.35E-01 | 2.67E+00 | 2.73E+00 | 8.59E-01 | no | | intergenic_variant | MODIFIER |
| 8 | 74966798 | AX-170258669 | 0 | T | A | 0.193 | -2.40E-02 | 1.35E-01 | 2.67E+00 | 2.73E+00 | 8.59E-01 | no | | intergenic_variant | MODIFIER |
| 8 | 75004351 | AX-170389653 | 0 | A | G | 0.193 | -2.40E-02 | 1.35E-01 | 2.67E+00 | 2.73E+00 | 8.59E-01 | no | Hmgxb4 | intron_variant | MODIFIER |
| 8 | 75019021 | AX-170438356 | 0 | T | C | 0.193 | -2.40E-02 | 1.35E-01 | 2.67E+00 | 2.73E+00 | 8.59E-01 | no | Hmgxb4 | intron_variant | MODIFIER |
| 8 | 75019045 | AX-170485672 | 0 | C | G | 0.193 | -2.40E-02 | 1.35E-01 | 2.67E+00 | 2.73E+00 | 8.59E-01 | no | Hmgxb4 | intron_variant | MODIFIER |
| 8 | 75027911 | AX-170284139 | 0 | G | A | 0.193 | -2.40E-02 | 1.35E-01 | 2.67E+00 | 2.73E+00 | 8.59E-01 | no | Hmgxb4 | intron_variant | MODIFIER |
| 8 | 75038153 | AX-170147304 | 0 | C | T | 0.193 | -2.40E-02 | 1.35E-01 | 2.67E+00 | 2.73E+00 | 8.59E-01 | no | Tom1 | intron_variant | MODIFIER |
| 8 | 75042414 | AX-170142117 | 0 | T | C | 0.193 | -2.40E-02 | 1.35E-01 | 2.67E+00 | 2.73E+00 | 8.59E-01 | no | Tom1 | intron_variant | MODIFIER |
| 8 | 75042837 | AX-170245319 | 0 | G | A | 0.193 | -2.40E-02 | 1.35E-01 | 2.67E+00 | 2.73E+00 | 8.59E-01 | no | Tom1 | intron_variant | MODIFIER |
| 8 | 75044223 | AX-170327387 | 0 | A | C | 0.193 | -2.40E-02 | 1.35E-01 | 2.67E+00 | 2.73E+00 | 8.59E-01 | no | Tom1 | intron_variant | MODIFIER |
| 8 | 75044380 | AX-170228574 | 0 | T | C | 0.193 | -2.40E-02 | 1.35E-01 | 2.67E+00 | 2.73E+00 | 8.59E-01 | no | Tom1 | intron_variant | MODIFIER |
| 8 | 75056710 | AX-170345665 | 0 | G | A | 0.193 | -2.40E-02 | 1.35E-01 | 2.67E+00 | 2.73E+00 | 8.59E-01 | no | Tom1 | intron_variant | MODIFIER |
| 8 | 75060261 | AX-170368634 | 0 | A | G | 0.193 | -2.40E-02 | 1.35E-01 | 2.67E+00 | 2.73E+00 | 8.59E-01 | no | Tom1 | intron_variant | MODIFIER |
| 8 | 75063663 | AX-170396160 | 0 | T | C | 0.193 | -2.40E-02 | 1.35E-01 | 2.67E+00 | 2.73E+00 | 8.59E-01 | no | Tom1 | intron_variant | MODIFIER |
| 8 | 75070900 | AX-170235781 | 0 | C | G | 0.193 | -2.40E-02 | 1.35E-01 | 2.67E+00 | 2.73E+00 | 8.59E-01 | no | Tom1 | downstream_gene_variant | MODIFIER |
| 8 | 75101669 | AX-170126371 | 0 | C | T | 0.193 | -2.40E-02 | 1.35E-01 | 2.67E+00 | 2.73E+00 | 8.59E-01 | no | Hmox1 | downstream_gene_variant | MODIFIER |
| 8 | 75114094 | AX-170355599 | 0 | C | T | 0.193 | -2.40E-02 | 1.35E-01 | 2.67E+00 | 2.73E+00 | 8.59E-01 | no | Mcm5 | intron_variant | MODIFIER |
| 8 | 75133889 | AX-170112849 | 0 | C | T | 0.193 | -2.40E-02 | 1.35E-01 | 2.67E+00 | 2.73E+00 | 8.59E-01 | no | | intergenic_variant | MODIFIER |
| 8 | 75137902 | AX-170207703 | 0 | T | C | 0.193 | -2.40E-02 | 1.35E-01 | 2.67E+00 | 2.73E+00 | 8.59E-01 | no | | intergenic_variant | MODIFIER |
| 8 | 75147867 | AX-170250346 | 0 | A | G | 0.193 | -2.40E-02 | 1.35E-01 | 2.67E+00 | 2.73E+00 | 8.59E-01 | no | | intergenic_variant | MODIFIER |
| 8 | 75149232 | AX-170525807 | 0 | T | C | 0.193 | -2.40E-02 | 1.35E-01 | 2.67E+00 | 2.73E+00 | 8.59E-01 | no | | intergenic_variant | MODIFIER |
| 8 | 75158610 | AX-170233796 | 0 | A | G | 0.193 | -2.40E-02 | 1.35E-01 | 2.67E+00 | 2.73E+00 | 8.59E-01 | no | | intergenic_variant | MODIFIER |
| 8 | 75159727 | AX-170489121 | 0 | C | T | 0.193 | -2.40E-02 | 1.35E-01 | 2.67E+00 | 2.73E+00 | 8.59E-01 | no | | intergenic_variant | MODIFIER |
| 8 | 75190216 | AX-170136717 | 0 | T | C | 0.193 | -2.40E-02 | 1.35E-01 | 2.67E+00 | 2.73E+00 | 8.59E-01 | no | | intergenic_variant | MODIFIER |
| 8 | 75191436 | AX-170368094 | 0 | C | A | 0.193 | -2.40E-02 | 1.35E-01 | 2.67E+00 | 2.73E+00 | 8.59E-01 | no | | intergenic_variant | MODIFIER |
| 8 | 75192494 | AX-170498969 | 0 | A | T | 0.193 | -2.40E-02 | 1.35E-01 | 2.67E+00 | 2.73E+00 | 8.59E-01 | no | | intergenic_variant | MODIFIER |
| 8 | 75198121 | AX-170234301 | 0 | T | C | 0.193 | -2.40E-02 | 1.35E-01 | 2.67E+00 | 2.73E+00 | 8.59E-01 | no | Gm45903 | upstream_gene_variant | MODIFIER |

| | | | | | | | | | | | | | | | |
|---|---|---|---|---|---|---|---|---|---|---|---|---|---|---|---|
| 8 | 75205033 | AX-170245857 | 0 | C | A | 0.193 | -2.40E-02 | 1.35E-01 | 2.67E+00 | 2.73E+00 | 8.59E-01 | no | Gm45903 | intron_variant,non_coding_transcript_variant | MODIFIER |
| 8 | 75210595 | AX-170485609 | 0 | A | G | 0.193 | -2.40E-02 | 1.35E-01 | 2.67E+00 | 2.73E+00 | 8.59E-01 | no | Rasd2 | upstream_gene_variant | MODIFIER |
| 8 | 75215461 | AX-170153898 | 0 | T | C | 0.193 | -2.40E-02 | 1.35E-01 | 2.67E+00 | 2.73E+00 | 8.59E-01 | no | Rasd2 | intron_variant | MODIFIER |
| 8 | 75221366 | AX-170080441 | 0 | C | T | 0.193 | -2.40E-02 | 1.35E-01 | 2.67E+00 | 2.73E+00 | 8.59E-01 | no | Rasd2 | intron_variant | MODIFIER |
| 8 | 75221987 | AX-170096325 | 0 | T | C | 0.193 | -2.40E-02 | 1.35E-01 | 2.67E+00 | 2.73E+00 | 8.59E-01 | no | Rasd2 | synonymous_variant | LOW |
| 8 | 75224607 | AX-170080444 | 0 | C | G | 0.193 | -2.40E-02 | 1.35E-01 | 2.67E+00 | 2.73E+00 | 8.59E-01 | no | Rasd2 | downstream_gene_variant | MODIFIER |
| 8 | 75265304 | AX-170478781 | 0 | G | A | 0.193 | -2.40E-02 | 1.35E-01 | 2.67E+00 | 2.73E+00 | 8.59E-01 | no | | intergenic_variant | MODIFIER |
| 8 | 75294221 | AX-97799847 | 0 | C | T | 0.193 | -2.40E-02 | 1.35E-01 | 2.67E+00 | 2.73E+00 | 8.59E-01 | no | Gm45907 | upstream_gene_variant | MODIFIER |
| 8 | 75310718 | AX-170031030 | 0 | A | C | 0.193 | -2.40E-02 | 1.35E-01 | 2.67E+00 | 2.73E+00 | 8.59E-01 | no | | intergenic_variant | MODIFIER |
| 8 | 75317731 | AX-170266173 | 0 | C | T | 0.193 | -2.40E-02 | 1.35E-01 | 2.67E+00 | 2.73E+00 | 8.59E-01 | no | | intergenic_variant | MODIFIER |
| 8 | 75375187 | AX-170301630 | 0 | C | T | 0.193 | -2.40E-02 | 1.35E-01 | 2.67E+00 | 2.73E+00 | 8.59E-01 | no | | intergenic_variant | MODIFIER |
| 8 | 75376189 | AX-170402642 | 0 | A | G | 0.193 | -2.40E-02 | 1.35E-01 | 2.67E+00 | 2.73E+00 | 8.59E-01 | no | | intergenic_variant | MODIFIER |
| 8 | 75381594 | AX-170239839 | 0 | G | A | 0.193 | -2.40E-02 | 1.35E-01 | 2.67E+00 | 2.73E+00 | 8.59E-01 | no | Umpk-ps | upstream_gene_variant | MODIFIER |
| 8 | 75394679 | AX-170423500 | 0 | A | T | 0.193 | -2.40E-02 | 1.35E-01 | 2.67E+00 | 2.73E+00 | 8.59E-01 | no | Gm7984 | upstream_gene_variant | MODIFIER |
| 8 | 75408924 | AX-170130201 | 0 | C | T | 0.193 | -2.40E-02 | 1.35E-01 | 2.67E+00 | 2.73E+00 | 8.59E-01 | no | Gm10358 | upstream_gene_variant | MODIFIER |
| 8 | 75412561 | AX-170357737 | 0 | A | G | 0.193 | -2.40E-02 | 1.35E-01 | 2.67E+00 | 2.73E+00 | 8.59E-01 | no | | intergenic_variant | MODIFIER |
| 8 | 75456855 | AX-170096326 | 0 | G | A | 0.193 | -2.40E-02 | 1.35E-01 | 2.67E+00 | 2.73E+00 | 8.59E-01 | no | Iqcm | intron_variant | MODIFIER |
| 8 | 75474323 | AX-97817348 | 0 | A | G | 0.193 | -2.40E-02 | 1.35E-01 | 2.67E+00 | 2.73E+00 | 8.59E-01 | no | Iqcm | intron_variant | MODIFIER |
| 8 | 75475738 | AX-170175182 | 0 | A | G | 0.193 | -2.40E-02 | 1.35E-01 | 2.67E+00 | 2.73E+00 | 8.59E-01 | no | Iqcm | intron_variant | MODIFIER |
| 8 | 75477005 | AX-170519556 | 0 | T | C | 0.193 | -2.40E-02 | 1.35E-01 | 2.67E+00 | 2.73E+00 | 8.59E-01 | no | Iqcm | intron_variant | MODIFIER |
| 8 | 75477398 | AX-170121015 | 0 | G | A | 0.193 | -2.40E-02 | 1.35E-01 | 2.67E+00 | 2.73E+00 | 8.59E-01 | no | Iqcm | intron_variant | MODIFIER |
| 8 | 75478586 | AX-170145975 | 0 | A | T | 0.193 | -2.40E-02 | 1.35E-01 | 2.67E+00 | 2.73E+00 | 8.59E-01 | no | Iqcm | intron_variant | MODIFIER |
| 8 | 75509555 | AX-170299235 | 0 | T | A | 0.193 | -2.40E-02 | 1.35E-01 | 2.67E+00 | 2.73E+00 | 8.59E-01 | no | Iqcm | intron_variant | MODIFIER |
| 8 | 75531414 | AX-170103448 | 0 | G | T | 0.193 | -2.40E-02 | 1.35E-01 | 2.67E+00 | 2.73E+00 | 8.59E-01 | no | Iqcm | intron_variant | MODIFIER |
| 8 | 75531917 | AX-170449747 | 0 | T | C | 0.193 | -2.40E-02 | 1.35E-01 | 2.67E+00 | 2.73E+00 | 8.59E-01 | no | Iqcm | intron_variant | MODIFIER |
| 8 | 75537727 | AX-170212995 | 0 | T | A | 0.193 | -2.40E-02 | 1.35E-01 | 2.67E+00 | 2.73E+00 | 8.59E-01 | no | Iqcm | intron_variant | MODIFIER |
| 8 | 75542484 | AX-170236261 | 0 | G | A | 0.193 | -2.40E-02 | 1.35E-01 | 2.67E+00 | 2.73E+00 | 8.59E-01 | no | Iqcm | intron_variant | MODIFIER |
| 8 | 75542827 | AX-170091308 | 0 | A | C | 0.193 | -2.40E-02 | 1.35E-01 | 2.67E+00 | 2.73E+00 | 8.59E-01 | no | Iqcm | intron_variant | MODIFIER |
| 8 | 75542928 | AX-170429380 | 0 | G | A | 0.193 | -2.40E-02 | 1.35E-01 | 2.67E+00 | 2.73E+00 | 8.59E-01 | no | Iqcm | intron_variant | MODIFIER |
| 8 | 75579374 | AX-170117431 | 0 | C | T | 0.193 | -2.40E-02 | 1.35E-01 | 2.67E+00 | 2.73E+00 | 8.59E-01 | no | Iqcm | intron_variant | MODIFIER |
| 8 | 75579842 | AX-170379554 | 0 | C | A | 0.193 | -2.40E-02 | 1.35E-01 | 2.67E+00 | 2.73E+00 | 8.59E-01 | no | Iqcm | intron_variant | MODIFIER |
| 8 | 75594862 | AX-170311293 | 0 | G | A | 0.193 | -2.40E-02 | 1.35E-01 | 2.67E+00 | 2.73E+00 | 8.59E-01 | no | Iqcm | intron_variant | MODIFIER |
| 8 | 75616825 | AX-170178260 | 0 | T | A | 0.193 | -2.40E-02 | 1.35E-01 | 2.67E+00 | 2.73E+00 | 8.59E-01 | no | Iqcm | intron_variant | MODIFIER |
| 8 | 75644716 | AX-170080450 | 0 | C | A | 0.193 | -2.40E-02 | 1.35E-01 | 2.67E+00 | 2.73E+00 | 8.59E-01 | no | Iqcm | intron_variant | MODIFIER |
| 8 | 75645432 | AX-170416956 | 0 | G | T | 0.193 | -2.40E-02 | 1.35E-01 | 2.67E+00 | 2.73E+00 | 8.59E-01 | no | Iqcm | intron_variant | MODIFIER |
| 8 | 75660877 | AX-170255293 | 0 | G | A | 0.193 | -2.40E-02 | 1.35E-01 | 2.67E+00 | 2.73E+00 | 8.59E-01 | no | Iqcm | intron_variant | MODIFIER |
| 8 | 75670352 | AX-170098509 | 0 | T | C | 0.193 | -2.40E-02 | 1.35E-01 | 2.67E+00 | 2.73E+00 | 8.59E-01 | no | Iqcm | intron_variant | MODIFIER |
| 8 | 75678619 | AX-170424694 | 0 | C | A | 0.193 | -2.40E-02 | 1.35E-01 | 2.67E+00 | 2.73E+00 | 8.59E-01 | no | Iqcm | intron_variant | MODIFIER |
| 8 | 75682392 | AX-170296953 | 0 | C | T | 0.193 | -2.40E-02 | 1.35E-01 | 2.67E+00 | 2.73E+00 | 8.59E-01 | no | Iqcm | intron_variant | MODIFIER |
| 8 | 75713876 | AX-170186839 | 0 | C | A | 0.193 | -2.40E-02 | 1.35E-01 | 2.67E+00 | 2.73E+00 | 8.59E-01 | no | Iqcm | intron_variant | MODIFIER |
| 8 | 75727408 | AX-170141358 | 0 | T | C | 0.193 | -2.40E-02 | 1.35E-01 | 2.67E+00 | 2.73E+00 | 8.59E-01 | no | Iqcm | intron_variant | MODIFIER |
| 8 | 75743871 | AX-97838037 | 0 | A | T | 0.193 | -2.40E-02 | 1.35E-01 | 2.67E+00 | 2.73E+00 | 8.59E-01 | no | Iqcm | intron_variant | MODIFIER |
| 8 | 75828628 | AX-170321404 | 0 | C | T | 0.193 | -2.40E-02 | 1.35E-01 | 2.67E+00 | 2.73E+00 | 8.59E-01 | no | Iqcm | intron_variant | MODIFIER |
| 8 | 75921774 | AX-97840470 | 0 | G | A | 0.193 | -2.40E-02 | 1.35E-01 | 2.67E+00 | 2.73E+00 | 8.59E-01 | no | Iqcm | intron_variant | MODIFIER |
| 8 | 76267233 | AX-170104133 | 0 | T | A | 0.193 | -2.40E-02 | 1.35E-01 | 2.67E+00 | 2.73E+00 | 8.59E-01 | no | Gm45740 | intron_variant,non_coding_transcript_variant | MODIFIER |
| 8 | 76277245 | AX-170231700 | 0 | C | T | 0.193 | -2.40E-02 | 1.35E-01 | 2.67E+00 | 2.73E+00 | 8.59E-01 | no | Gm22509 | upstream_gene_variant | MODIFIER |
| 8 | 76380100 | AX-170395163 | 0 | T | C | 0.193 | -2.40E-02 | 1.35E-01 | 2.67E+00 | 2.73E+00 | 8.59E-01 | no | Gm45740 | intron_variant,non_coding_transcript_variant | MODIFIER |
| 8 | 76402478 | AX-170031051 | 0 | G | T | 0.193 | -2.40E-02 | 1.35E-01 | 2.67E+00 | 2.73E+00 | 8.59E-01 | no | Gm45740 | intron_variant,non_coding_transcript_variant | MODIFIER |
| 8 | 76406396 | AX-170031044 | 0 | A | G | 0.193 | -2.40E-02 | 1.35E-01 | 2.67E+00 | 2.73E+00 | 8.59E-01 | no | Gm45740 | intron_variant,non_coding_transcript_variant | MODIFIER |
| 8 | 76441623 | AX-170080468 | 0 | T | C | 0.193 | -2.40E-02 | 1.35E-01 | 2.67E+00 | 2.73E+00 | 8.59E-01 | no | | intergenic_variant | MODIFIER |
| 8 | 76441754 | AX-170031048 | 0 | A | G | 0.193 | -2.40E-02 | 1.35E-01 | 2.67E+00 | 2.73E+00 | 8.59E-01 | no | | intergenic_variant | MODIFIER |
| 8 | 76445389 | AX-170457414 | 0 | C | T | 0.193 | -2.40E-02 | 1.35E-01 | 2.67E+00 | 2.73E+00 | 8.59E-01 | no | | intergenic_variant | MODIFIER |
| 8 | 76446790 | AX-170526437 | 0 | T | C | 0.193 | -2.40E-02 | 1.35E-01 | 2.67E+00 | 2.73E+00 | 8.59E-01 | no | | intergenic_variant | MODIFIER |
| 8 | 76450367 | AX-170511381 | 0 | T | G | 0.193 | -2.40E-02 | 1.35E-01 | 2.67E+00 | 2.73E+00 | 8.59E-01 | no | | intergenic_variant | MODIFIER |

| | | | | | | | | | | | | | | | |
|---|---|---|---|---|---|---|---|---|---|---|---|---|---|---|---|
| 8 | 76452699 | AX-170031049 | 0 | C | T | 0.193 | -2.40E-02 | 1.35E-01 | 2.67E+00 | 2.73E+00 | 8.59E-01 | no | | intergenic_variant | MODIFIER |
| 8 | 76579969 | AX-170031050 | 0 | A | T | 0.193 | -2.40E-02 | 1.35E-01 | 2.67E+00 | 2.73E+00 | 8.59E-01 | no | | intergenic_variant | MODIFIER |
| 8 | 77699437 | AX-97807024 | 0 | A | G | 0.193 | -2.40E-02 | 1.35E-01 | 2.67E+00 | 2.73E+00 | 8.59E-01 | no | Ednra | intron_variant | MODIFIER |
| 8 | 77719832 | AX-97788855 | 0 | T | C | 0.193 | -2.40E-02 | 1.35E-01 | 2.67E+00 | 2.73E+00 | 8.59E-01 | no | Ednra | intron_variant | MODIFIER |
| 8 | 77762095 | AX-170128822 | 0 | G | C | 0.193 | -2.40E-02 | 1.35E-01 | 2.67E+00 | 2.73E+00 | 8.59E-01 | no | 4933431K23Rik | upstream_gene_variant | MODIFIER |
| 8 | 77762586 | AX-170111717 | 0 | T | C | 0.193 | -2.40E-02 | 1.35E-01 | 2.67E+00 | 2.73E+00 | 8.59E-01 | no | 4933431K23Rik | upstream_gene_variant | MODIFIER |
| 8 | 77876932 | AX-170147739 | 0 | G | A | 0.193 | -2.40E-02 | 1.35E-01 | 2.67E+00 | 2.73E+00 | 8.59E-01 | no | | intergenic_variant | MODIFIER |
| 8 | 77877728 | AX-170372975 | 0 | T | G | 0.193 | -2.40E-02 | 1.35E-01 | 2.67E+00 | 2.73E+00 | 8.59E-01 | no | | intergenic_variant | MODIFIER |
| 8 | 77881312 | AX-170144439 | 0 | G | A | 0.193 | -2.40E-02 | 1.35E-01 | 2.67E+00 | 2.73E+00 | 8.59E-01 | no | | intergenic_variant | MODIFIER |
| 8 | 77905648 | AX-170244921 | 0 | T | G | 0.193 | -2.40E-02 | 1.35E-01 | 2.67E+00 | 2.73E+00 | 8.59E-01 | no | | intergenic_variant | MODIFIER |
| 8 | 77905938 | AX-97803937 | 0 | C | A | 0.193 | -2.40E-02 | 1.35E-01 | 2.67E+00 | 2.73E+00 | 8.59E-01 | no | | intergenic_variant | MODIFIER |
| 8 | 77905974 | AX-170202306 | 0 | A | C | 0.193 | -2.40E-02 | 1.35E-01 | 2.67E+00 | 2.73E+00 | 8.59E-01 | no | | intergenic_variant | MODIFIER |
| 8 | 77936341 | AX-170280710 | 0 | C | T | 0.193 | -2.40E-02 | 1.35E-01 | 2.67E+00 | 2.73E+00 | 8.59E-01 | no | | intergenic_variant | MODIFIER |
| 8 | 77936540 | AX-170212972 | 0 | A | G | 0.193 | -2.40E-02 | 1.35E-01 | 2.67E+00 | 2.73E+00 | 8.59E-01 | no | | intergenic_variant | MODIFIER |
| 8 | 77949006 | AX-170431264 | 0 | G | A | 0.193 | -2.40E-02 | 1.35E-01 | 2.67E+00 | 2.73E+00 | 8.59E-01 | no | | intergenic_variant | MODIFIER |
| 8 | 77957193 | AX-170031078 | 0 | C | T | 0.193 | -2.40E-02 | 1.35E-01 | 2.67E+00 | 2.73E+00 | 8.59E-01 | no | | intergenic_variant | MODIFIER |
| 8 | 77969943 | AX-170343005 | 0 | C | T | 0.193 | -2.40E-02 | 1.35E-01 | 2.67E+00 | 2.73E+00 | 8.59E-01 | no | | regulatory_region_variant | MODIFIER |
| 8 | 77970264 | AX-97786486 | 0 | C | T | 0.193 | -2.40E-02 | 1.35E-01 | 2.67E+00 | 2.73E+00 | 8.59E-01 | no | | regulatory_region_variant | MODIFIER |
| 8 | 77972129 | AX-170248758 | 0 | G | A | 0.193 | -2.40E-02 | 1.35E-01 | 2.67E+00 | 2.73E+00 | 8.59E-01 | no | | intergenic_variant | MODIFIER |
| 8 | 77973408 | AX-170427651 | 0 | T | A | 0.193 | -2.40E-02 | 1.35E-01 | 2.67E+00 | 2.73E+00 | 8.59E-01 | no | | regulatory_region_variant | MODIFIER |
| 8 | 77976951 | AX-170183816 | 0 | G | A | 0.193 | -2.40E-02 | 1.35E-01 | 2.67E+00 | 2.73E+00 | 8.59E-01 | no | | intergenic_variant | MODIFIER |
| 8 | 77979610 | AX-170080495 | 0 | A | G | 0.193 | -2.40E-02 | 1.35E-01 | 2.67E+00 | 2.73E+00 | 8.59E-01 | no | | intergenic_variant | MODIFIER |
| 8 | 77985611 | AX-97834777 | 0 | T | C | 0.193 | -2.40E-02 | 1.35E-01 | 2.67E+00 | 2.73E+00 | 8.59E-01 | no | | intergenic_variant | MODIFIER |
| 8 | 78025451 | AX-170464648 | 0 | A | G | 0.193 | -2.40E-02 | 1.35E-01 | 2.67E+00 | 2.73E+00 | 8.59E-01 | no | | regulatory_region_variant | MODIFIER |
| 8 | 78025702 | AX-170282697 | 0 | A | C | 0.193 | -2.40E-02 | 1.35E-01 | 2.67E+00 | 2.73E+00 | 8.59E-01 | no | | regulatory_region_variant | MODIFIER |
| 8 | 78025950 | AX-170466830 | 0 | G | C | 0.193 | -2.40E-02 | 1.35E-01 | 2.67E+00 | 2.73E+00 | 8.59E-01 | no | | regulatory_region_variant | MODIFIER |
| 8 | 78026972 | AX-170453327 | 0 | T | A | 0.193 | -2.40E-02 | 1.35E-01 | 2.67E+00 | 2.73E+00 | 8.59E-01 | no | | intergenic_variant | MODIFIER |
| 8 | 78027966 | AX-170031075 | 0 | A | G | 0.193 | -2.40E-02 | 1.35E-01 | 2.67E+00 | 2.73E+00 | 8.59E-01 | no | | intergenic_variant | MODIFIER |
| 8 | 78028011 | AX-170031077 | 0 | G | A | 0.193 | -2.40E-02 | 1.35E-01 | 2.67E+00 | 2.73E+00 | 8.59E-01 | no | | intergenic_variant | MODIFIER |
| 8 | 78028322 | AX-170137407 | 0 | A | C | 0.193 | -2.40E-02 | 1.35E-01 | 2.67E+00 | 2.73E+00 | 8.59E-01 | no | | intergenic_variant | MODIFIER |
| 8 | 78029274 | AX-97847437 | 0 | C | A | 0.193 | -2.40E-02 | 1.35E-01 | 2.67E+00 | 2.73E+00 | 8.59E-01 | no | | intergenic_variant | MODIFIER |
| 8 | 78050970 | AX-170449797 | 0 | T | C | 0.193 | -2.40E-02 | 1.35E-01 | 2.67E+00 | 2.73E+00 | 8.59E-01 | no | Gm29895 | non_coding_transcript_exon_variant | MODIFIER |
| 8 | 78070818 | AX-97849487 | 0 | C | T | 0.193 | -2.40E-02 | 1.35E-01 | 2.67E+00 | 2.73E+00 | 8.59E-01 | no | | intergenic_variant | MODIFIER |
| 8 | 78079150 | AX-170439293 | 0 | T | C | 0.193 | -2.40E-02 | 1.35E-01 | 2.67E+00 | 2.73E+00 | 8.59E-01 | no | | intergenic_variant | MODIFIER |
| 8 | 78081745 | AX-170344893 | 0 | T | C | 0.193 | -2.40E-02 | 1.35E-01 | 2.67E+00 | 2.73E+00 | 8.59E-01 | no | | intergenic_variant | MODIFIER |
| 8 | 78081873 | AX-170416677 | 0 | G | A | 0.193 | -2.40E-02 | 1.35E-01 | 2.67E+00 | 2.73E+00 | 8.59E-01 | no | | intergenic_variant | MODIFIER |
| 8 | 78090854 | AX-170212354 | 0 | T | C | 0.193 | -2.40E-02 | 1.35E-01 | 2.67E+00 | 2.73E+00 | 8.59E-01 | no | | regulatory_region_variant | MODIFIER |
| 8 | 78136662 | AX-97810541 | 0 | C | T | 0.193 | -2.40E-02 | 1.35E-01 | 2.67E+00 | 2.73E+00 | 8.59E-01 | no | | intergenic_variant | MODIFIER |
| 8 | 78147692 | AX-170364651 | 0 | A | G | 0.193 | -2.40E-02 | 1.35E-01 | 2.67E+00 | 2.73E+00 | 8.59E-01 | no | | intergenic_variant | MODIFIER |
| 5 | 38031415 | AX-97846068 | 0 | G | A | 0.342 | -1.78E-02 | 1.05E-01 | 2.67E+00 | 2.76E+00 | 8.59E-01 | no | Gm43762 | upstream_gene_variant | MODIFIER |
| 13 | 47619580 | AX-97794742 | 2 | A | G | 0.5 | 1.17E-02 | 6.40E-02 | 2.62E+00 | 2.72E+00 | 8.59E-01 | no | | intergenic_variant | MODIFIER |
| 12 | 68576153 | AX-170501206 | 2 | G | A | 0.073 | -4.00E-02 | 2.29E-01 | 2.66E+00 | 2.74E+00 | 8.59E-01 | no | | intergenic_variant | MODIFIER |
| 11 | 46984969 | AX-170393550 | 1 | T | A | 0.08 | 3.21E-02 | 1.94E-01 | 2.67E+00 | 2.78E+00 | 8.59E-01 | no | Sgcd | intron_variant | MODIFIER |
| 11 | 47593779 | AX-170232948 | 1 | A | G | 0.08 | 3.21E-02 | 1.94E-01 | 2.67E+00 | 2.78E+00 | 8.59E-01 | no | Sgcd | intron_variant,non_coding_transcript_variant | MODIFIER |
| 2 | 159715174 | AX-170393234 | 1 | T | C | 0.134 | -3.03E-02 | 1.68E-01 | 2.61E+00 | 2.73E+00 | 8.59E-01 | no | | regulatory_region_variant | MODIFIER |
| 10 | 8208541 | AX-170360673 | 1 | C | T | 0.214 | -1.86E-02 | 1.04E-01 | 2.64E+00 | 2.73E+00 | 8.59E-01 | no | Ust | intron_variant | MODIFIER |
| 12 | 114341034 | AX-170226475 | 0 | T | A | 0.123 | -3.34E-02 | 1.95E-01 | 2.68E+00 | 2.76E+00 | 8.59E-01 | no | | intergenic_variant | MODIFIER |
| 2 | 142266624 | AX-170019222 | 1 | G | A | 0.143 | 3.99E-02 | 2.25E-01 | 2.69E+00 | 2.72E+00 | 8.59E-01 | no | Macrod2 | intron_variant | MODIFIER |
| 2 | 149449592 | AX-170065237 | 1 | G | C | 0.143 | 3.99E-02 | 2.25E-01 | 2.69E+00 | 2.72E+00 | 8.59E-01 | no | | intergenic_variant | MODIFIER |
| 2 | 156524137 | AX-170065371 | 1 | T | C | 0.143 | 3.99E-02 | 2.25E-01 | 2.69E+00 | 2.72E+00 | 8.59E-01 | no | Epb41l1 | intron_variant | MODIFIER |
| 10 | 33653023 | AX-170003023 | 0 | C | T | 0.386 | -1.35E-02 | 7.35E-02 | 2.61E+00 | 2.72E+00 | 8.59E-01 | no | Gm15939 | intron_variant,non_coding_transcript_variant | MODIFIER |
| 10 | 43049605 | AX-170229166 | 0 | G | A | 0.368 | -1.37E-02 | 7.53E-02 | 2.62E+00 | 2.72E+00 | 8.59E-01 | no | Sobp | intron_variant | MODIFIER |
| 1 | 5907589 | AX-170158029 | 1 | C | T | 0.098 | 3.07E-02 | 1.72E-01 | 2.62E+00 | 2.73E+00 | 8.59E-01 | no | | intergenic_variant | MODIFIER |
| 1 | 65792098 | AX-97805606 | 1 | T | C | 0.143 | 4.18E-02 | 2.37E-01 | 2.68E+00 | 2.73E+00 | 8.59E-01 | no | | intergenic_variant | MODIFIER |

| | | | | | | | | | | | | | | | |
|---|---|---|---|---|---|---|---|---|---|---|---|---|---|---|---|
| 1 | 70244590 | AX-170419132 | 1 | C | T | 0.143 | 4.18E-02 | 2.37E-01 | 2.68E+00 | 2.73E+00 | 8.59E-01 | no | Spag16 | intron_variant | MODIFIER |
| 14 | 41279246 | AX-170309686 | 0 | T | G | 0.491 | 5.24E-02 | 2.81E-01 | 2.59E+00 | 2.71E+00 | 8.60E-01 | no | Gm2832 | intron_variant | MODIFIER |
| 5 | 33970431 | AX-170071749 | 0 | C | T | 0.36 | 1.88E-02 | 1.08E-01 | 2.68E+00 | 2.74E+00 | 8.60E-01 | no | | intergenic_variant | MODIFIER |
| 15 | 97564421 | AX-170407736 | 0 | C | T | 0.158 | -2.52E-02 | 1.50E-01 | 2.69E+00 | 2.77E+00 | 8.60E-01 | no | | intergenic_variant | MODIFIER |
| 8 | 90289716 | AX-97812520 | 0 | C | G | 0.202 | -2.12E-02 | 1.18E-01 | 2.64E+00 | 2.72E+00 | 8.60E-01 | no | Tox3 | intron_variant | MODIFIER |
| 11 | 34153171 | AX-170138437 | 0 | C | T | 0.096 | -3.18E-02 | 1.87E-01 | 2.67E+00 | 2.76E+00 | 8.60E-01 | no | | intergenic_variant | MODIFIER |
| 8 | 101858995 | AX-170463648 | 1 | G | T | 0.223 | 1.95E-02 | 1.13E-01 | 2.68E+00 | 2.75E+00 | 8.60E-01 | no | | intergenic_variant | MODIFIER |
| 2 | 122006471 | AX-170244806 | 1 | C | A | 0.134 | 3.59E-02 | 2.05E-01 | 2.69E+00 | 2.74E+00 | 8.60E-01 | no | Ctdspl2 | intron_variant | MODIFIER |
| 7 | 81710687 | AX-170303947 | 1 | C | T | 0.125 | 3.43E-02 | 1.84E-01 | 2.61E+00 | 2.71E+00 | 8.60E-01 | no | Homer2 | upstream_gene_variant | MODIFIER |
| 16 | 45612613 | AX-170099786 | 2 | A | G | 0.455 | -1.45E-02 | 7.67E-02 | 2.62E+00 | 2.70E+00 | 8.60E-01 | no | Gcsam | intron_variant | MODIFIER |
| 1 | 65505631 | AX-170392970 | 0 | T | C | 0.14 | 4.18E-02 | 2.39E-01 | 2.68E+00 | 2.74E+00 | 8.60E-01 | no | | intergenic_variant | MODIFIER |
| 1 | 65515601 | AX-170335208 | 0 | G | A | 0.14 | 4.18E-02 | 2.39E-01 | 2.68E+00 | 2.74E+00 | 8.60E-01 | no | | intergenic_variant | MODIFIER |
| 1 | 65543957 | AX-97802661 | 0 | T | C | 0.14 | 4.18E-02 | 2.39E-01 | 2.68E+00 | 2.74E+00 | 8.60E-01 | no | | intergenic_variant | MODIFIER |
| 1 | 65565267 | AX-97834332 | 0 | C | T | 0.14 | 4.18E-02 | 2.39E-01 | 2.68E+00 | 2.74E+00 | 8.60E-01 | no | | intergenic_variant | MODIFIER |
| 1 | 65565475 | AX-170456592 | 0 | A | G | 0.14 | 4.18E-02 | 2.39E-01 | 2.68E+00 | 2.74E+00 | 8.60E-01 | no | | intergenic_variant | MODIFIER |
| 1 | 65590741 | AX-97835119 | 0 | A | G | 0.14 | 4.18E-02 | 2.39E-01 | 2.68E+00 | 2.74E+00 | 8.60E-01 | no | | intergenic_variant | MODIFIER |
| 1 | 65615912 | AX-170157003 | 0 | A | G | 0.14 | 4.18E-02 | 2.39E-01 | 2.68E+00 | 2.74E+00 | 8.60E-01 | no | | intergenic_variant | MODIFIER |
| 1 | 65633031 | AX-97796347 | 0 | A | G | 0.14 | 4.18E-02 | 2.39E-01 | 2.68E+00 | 2.74E+00 | 8.60E-01 | no | | intergenic_variant | MODIFIER |
| 1 | 65642057 | AX-170253668 | 0 | G | T | 0.14 | 4.18E-02 | 2.39E-01 | 2.68E+00 | 2.74E+00 | 8.60E-01 | no | | intergenic_variant | MODIFIER |
| 1 | 65648821 | AX-97794573 | 0 | T | G | 0.14 | 4.18E-02 | 2.39E-01 | 2.68E+00 | 2.74E+00 | 8.60E-01 | no | | regulatory_region_variant | MODIFIER |
| 1 | 65675399 | AX-170252442 | 0 | T | C | 0.14 | 4.18E-02 | 2.39E-01 | 2.68E+00 | 2.74E+00 | 8.60E-01 | no | | intergenic_variant | MODIFIER |
| 1 | 65742148 | AX-170340837 | 0 | C | G | 0.14 | 4.18E-02 | 2.39E-01 | 2.68E+00 | 2.74E+00 | 8.60E-01 | no | | intergenic_variant | MODIFIER |
| 1 | 65755857 | AX-170330717 | 0 | G | C | 0.14 | 4.18E-02 | 2.39E-01 | 2.68E+00 | 2.74E+00 | 8.60E-01 | no | | intergenic_variant | MODIFIER |
| 1 | 65755954 | AX-97811812 | 0 | T | C | 0.14 | 4.18E-02 | 2.39E-01 | 2.68E+00 | 2.74E+00 | 8.60E-01 | no | | intergenic_variant | MODIFIER |
| 1 | 65811875 | AX-170000696 | 0 | T | A | 0.14 | 4.18E-02 | 2.39E-01 | 2.68E+00 | 2.74E+00 | 8.60E-01 | no | | intergenic_variant | MODIFIER |
| 1 | 65838598 | AX-170097286 | 0 | C | T | 0.14 | 4.18E-02 | 2.39E-01 | 2.68E+00 | 2.74E+00 | 8.60E-01 | no | | intergenic_variant | MODIFIER |
| 1 | 65838801 | AX-170000693 | 0 | C | T | 0.14 | 4.18E-02 | 2.39E-01 | 2.68E+00 | 2.74E+00 | 8.60E-01 | no | | intergenic_variant | MODIFIER |
| 1 | 65839220 | AX-170000694 | 0 | A | G | 0.14 | 4.18E-02 | 2.39E-01 | 2.68E+00 | 2.74E+00 | 8.60E-01 | no | | intergenic_variant | MODIFIER |
| 1 | 65851153 | AX-97801155 | 0 | T | C | 0.14 | 4.18E-02 | 2.39E-01 | 2.68E+00 | 2.74E+00 | 8.60E-01 | no | | intergenic_variant | MODIFIER |
| 1 | 65856525 | AX-170295637 | 0 | C | T | 0.14 | 4.18E-02 | 2.39E-01 | 2.68E+00 | 2.74E+00 | 8.60E-01 | no | | intergenic_variant | MODIFIER |
| 1 | 65893356 | AX-97829763 | 0 | G | T | 0.14 | 4.18E-02 | 2.39E-01 | 2.68E+00 | 2.74E+00 | 8.60E-01 | no | | intergenic_variant | MODIFIER |
| 1 | 65912285 | AX-170234322 | 0 | G | T | 0.14 | 4.18E-02 | 2.39E-01 | 2.68E+00 | 2.74E+00 | 8.60E-01 | no | | intergenic_variant | MODIFIER |
| 1 | 65914629 | AX-170299618 | 0 | T | C | 0.14 | 4.18E-02 | 2.39E-01 | 2.68E+00 | 2.74E+00 | 8.60E-01 | no | | intergenic_variant | MODIFIER |
| 1 | 65919422 | AX-97820352 | 0 | A | G | 0.14 | 4.18E-02 | 2.39E-01 | 2.68E+00 | 2.74E+00 | 8.60E-01 | no | | intergenic_variant | MODIFIER |
| 1 | 65933910 | AX-170041483 | 0 | G | A | 0.14 | 4.18E-02 | 2.39E-01 | 2.68E+00 | 2.74E+00 | 8.60E-01 | no | | intergenic_variant | MODIFIER |
| 1 | 65934142 | AX-170091538 | 0 | C | A | 0.14 | 4.18E-02 | 2.39E-01 | 2.68E+00 | 2.74E+00 | 8.60E-01 | no | | intergenic_variant | MODIFIER |
| 1 | 65955925 | AX-170000697 | 0 | G | A | 0.14 | 4.18E-02 | 2.39E-01 | 2.68E+00 | 2.74E+00 | 8.60E-01 | no | | intergenic_variant | MODIFIER |
| 1 | 65956043 | AX-170086485 | 0 | A | G | 0.14 | 4.18E-02 | 2.39E-01 | 2.68E+00 | 2.74E+00 | 8.60E-01 | no | | intergenic_variant | MODIFIER |
| 1 | 65964778 | AX-170000698 | 0 | C | A | 0.14 | 4.18E-02 | 2.39E-01 | 2.68E+00 | 2.74E+00 | 8.60E-01 | no | | intergenic_variant | MODIFIER |
| 1 | 65981104 | AX-170220887 | 0 | G | T | 0.14 | 4.18E-02 | 2.39E-01 | 2.68E+00 | 2.74E+00 | 8.60E-01 | no | | intergenic_variant | MODIFIER |
| 1 | 65982194 | AX-170526330 | 0 | G | C | 0.14 | 4.18E-02 | 2.39E-01 | 2.68E+00 | 2.74E+00 | 8.60E-01 | no | | intergenic_variant | MODIFIER |
| 1 | 65982392 | AX-170480942 | 0 | G | C | 0.14 | 4.18E-02 | 2.39E-01 | 2.68E+00 | 2.74E+00 | 8.60E-01 | no | | intergenic_variant | MODIFIER |
| 1 | 65982486 | AX-97835388 | 0 | C | A | 0.14 | 4.18E-02 | 2.39E-01 | 2.68E+00 | 2.74E+00 | 8.60E-01 | no | | intergenic_variant | MODIFIER |
| 1 | 66000391 | AX-97841509 | 0 | G | A | 0.14 | 4.18E-02 | 2.39E-01 | 2.68E+00 | 2.74E+00 | 8.60E-01 | no | | regulatory_region_variant | MODIFIER |
| 1 | 66004184 | AX-170000702 | 0 | A | G | 0.14 | 4.18E-02 | 2.39E-01 | 2.68E+00 | 2.74E+00 | 8.60E-01 | no | | regulatory_region_variant | MODIFIER |
| 1 | 66043843 | AX-97854748 | 0 | T | C | 0.14 | 4.18E-02 | 2.39E-01 | 2.68E+00 | 2.74E+00 | 8.60E-01 | no | | intergenic_variant | MODIFIER |
| 1 | 66043902 | AX-170443188 | 0 | A | C | 0.14 | 4.18E-02 | 2.39E-01 | 2.68E+00 | 2.74E+00 | 8.60E-01 | no | | intergenic_variant | MODIFIER |
| 1 | 66096311 | AX-170149587 | 0 | G | A | 0.14 | 4.18E-02 | 2.39E-01 | 2.68E+00 | 2.74E+00 | 8.60E-01 | no | | intergenic_variant | MODIFIER |
| 1 | 66129862 | AX-170505834 | 0 | C | A | 0.14 | 4.18E-02 | 2.39E-01 | 2.68E+00 | 2.74E+00 | 8.60E-01 | no | | intergenic_variant | MODIFIER |
| 1 | 66129918 | AX-170222388 | 0 | T | C | 0.14 | 4.18E-02 | 2.39E-01 | 2.68E+00 | 2.74E+00 | 8.60E-01 | no | | intergenic_variant | MODIFIER |
| 1 | 66131132 | AX-170278142 | 0 | C | T | 0.14 | 4.18E-02 | 2.39E-01 | 2.68E+00 | 2.74E+00 | 8.60E-01 | no | | intergenic_variant | MODIFIER |
| 1 | 66135039 | AX-170422010 | 0 | T | C | 0.14 | 4.18E-02 | 2.39E-01 | 2.68E+00 | 2.74E+00 | 8.60E-01 | no | | intergenic_variant | MODIFIER |
| 1 | 66136284 | AX-170318683 | 0 | C | T | 0.14 | 4.18E-02 | 2.39E-01 | 2.68E+00 | 2.74E+00 | 8.60E-01 | no | | intergenic_variant | MODIFIER |
| 1 | 66136728 | AX-170035613 | 0 | C | T | 0.14 | 4.18E-02 | 2.39E-01 | 2.68E+00 | 2.74E+00 | 8.60E-01 | no | | intergenic_variant | MODIFIER |

| | | | | | | | | | | | | | | | |
|---|---|---|---|---|---|---|---|---|---|---|---|---|---|---|---|
| 1 | 66139742 | AX-170041488 | 0 | A | G | 0.14 | 4.18E-02 | 2.39E-01 | 2.68E+00 | 2.74E+00 | 8.60E-01 | no | | intergenic_variant | MODIFIER |
| 1 | 66158651 | AX-170212853 | 0 | T | C | 0.14 | 4.18E-02 | 2.39E-01 | 2.68E+00 | 2.74E+00 | 8.60E-01 | no | | intergenic_variant | MODIFIER |
| 1 | 66168474 | AX-97810963 | 0 | A | G | 0.14 | 4.18E-02 | 2.39E-01 | 2.68E+00 | 2.74E+00 | 8.60E-01 | no | | intergenic_variant | MODIFIER |
| 1 | 68944658 | AX-170254688 | 0 | C | T | 0.14 | 4.18E-02 | 2.39E-01 | 2.68E+00 | 2.74E+00 | 8.60E-01 | no | Erbb4 | intron_variant | MODIFIER |
| 1 | 68958745 | AX-170157383 | 0 | A | G | 0.14 | 4.18E-02 | 2.39E-01 | 2.68E+00 | 2.74E+00 | 8.60E-01 | no | Erbb4 | intron_variant | MODIFIER |
| 1 | 68973622 | AX-170531536 | 0 | T | C | 0.14 | 4.18E-02 | 2.39E-01 | 2.68E+00 | 2.74E+00 | 8.60E-01 | no | Erbb4 | intron_variant | MODIFIER |
| 1 | 68995621 | AX-170419287 | 0 | C | G | 0.14 | 4.18E-02 | 2.39E-01 | 2.68E+00 | 2.74E+00 | 8.60E-01 | no | Erbb4 | intron_variant | MODIFIER |
| 1 | 69012065 | AX-170317121 | 0 | G | T | 0.14 | 4.18E-02 | 2.39E-01 | 2.68E+00 | 2.74E+00 | 8.60E-01 | no | Erbb4 | intron_variant | MODIFIER |
| 1 | 69019958 | AX-170126794 | 0 | T | C | 0.14 | 4.18E-02 | 2.39E-01 | 2.68E+00 | 2.74E+00 | 8.60E-01 | no | Erbb4 | intron_variant | MODIFIER |
| 1 | 69049308 | AX-97811901 | 0 | A | C | 0.14 | 4.18E-02 | 2.39E-01 | 2.68E+00 | 2.74E+00 | 8.60E-01 | no | Erbb4 | intron_variant | MODIFIER |
| 1 | 69049901 | AX-170041553 | 0 | C | T | 0.14 | 4.18E-02 | 2.39E-01 | 2.68E+00 | 2.74E+00 | 8.60E-01 | no | Erbb4 | intron_variant | MODIFIER |
| 1 | 69058689 | AX-170294059 | 0 | G | A | 0.14 | 4.18E-02 | 2.39E-01 | 2.68E+00 | 2.74E+00 | 8.60E-01 | no | Erbb4 | intron_variant | MODIFIER |
| 1 | 69076396 | AX-97857302 | 0 | T | C | 0.14 | 4.18E-02 | 2.39E-01 | 2.68E+00 | 2.74E+00 | 8.60E-01 | no | Erbb4 | intron_variant | MODIFIER |
| 1 | 69116979 | AX-97800791 | 0 | A | C | 0.14 | 4.18E-02 | 2.39E-01 | 2.68E+00 | 2.74E+00 | 8.60E-01 | no | | regulatory_region_variant | MODIFIER |
| 1 | 69131421 | AX-170364852 | 0 | C | T | 0.14 | 4.18E-02 | 2.39E-01 | 2.68E+00 | 2.74E+00 | 8.60E-01 | no | Gm8840 | upstream_gene_variant | MODIFIER |
| 1 | 69131833 | AX-170165792 | 0 | G | A | 0.14 | 4.18E-02 | 2.39E-01 | 2.68E+00 | 2.74E+00 | 8.60E-01 | no | Gm8840 | upstream_gene_variant | MODIFIER |
| 1 | 69148778 | AX-170377229 | 0 | C | T | 0.14 | 4.18E-02 | 2.39E-01 | 2.68E+00 | 2.74E+00 | 8.60E-01 | no | | intergenic_variant | MODIFIER |
| 1 | 69157191 | AX-170000746 | 0 | G | A | 0.14 | 4.18E-02 | 2.39E-01 | 2.68E+00 | 2.74E+00 | 8.60E-01 | no | | intergenic_variant | MODIFIER |
| 1 | 69250019 | AX-170135098 | 0 | C | T | 0.14 | 4.18E-02 | 2.39E-01 | 2.68E+00 | 2.74E+00 | 8.60E-01 | no | | intergenic_variant | MODIFIER |
| 1 | 69311434 | AX-170145026 | 0 | G | T | 0.14 | 4.18E-02 | 2.39E-01 | 2.68E+00 | 2.74E+00 | 8.60E-01 | no | | intergenic_variant | MODIFIER |
| 1 | 69311511 | AX-170284745 | 0 | T | G | 0.14 | 4.18E-02 | 2.39E-01 | 2.68E+00 | 2.74E+00 | 8.60E-01 | no | | intergenic_variant | MODIFIER |
| 1 | 69311553 | AX-170325846 | 0 | T | G | 0.14 | 4.18E-02 | 2.39E-01 | 2.68E+00 | 2.74E+00 | 8.60E-01 | no | | intergenic_variant | MODIFIER |
| 1 | 69434218 | AX-97856596 | 0 | A | G | 0.14 | 4.18E-02 | 2.39E-01 | 2.68E+00 | 2.74E+00 | 8.60E-01 | no | | regulatory_region_variant | MODIFIER |
| 1 | 69457935 | AX-97839237 | 0 | C | A | 0.14 | 4.18E-02 | 2.39E-01 | 2.68E+00 | 2.74E+00 | 8.60E-01 | no | | intergenic_variant | MODIFIER |
| 1 | 69489592 | AX-170086491 | 0 | A | G | 0.14 | 4.18E-02 | 2.39E-01 | 2.68E+00 | 2.74E+00 | 8.60E-01 | no | Gm29113 | downstream_gene_variant | MODIFIER |
| 1 | 69505400 | AX-170368378 | 0 | C | T | 0.14 | 4.18E-02 | 2.39E-01 | 2.68E+00 | 2.74E+00 | 8.60E-01 | no | | intergenic_variant | MODIFIER |
| 1 | 69508101 | AX-97800258 | 0 | A | C | 0.14 | 4.18E-02 | 2.39E-01 | 2.68E+00 | 2.74E+00 | 8.60E-01 | no | | regulatory_region_variant | MODIFIER |
| 1 | 69512271 | AX-170296303 | 0 | C | T | 0.14 | 4.18E-02 | 2.39E-01 | 2.68E+00 | 2.74E+00 | 8.60E-01 | no | Gm29114 | upstream_gene_variant | MODIFIER |
| 1 | 69639337 | AX-170091546 | 0 | C | A | 0.14 | 4.18E-02 | 2.39E-01 | 2.68E+00 | 2.74E+00 | 8.60E-01 | no | Ikzf2 | intron_variant | MODIFIER |
| 1 | 69711252 | AX-97822307 | 0 | C | T | 0.14 | 4.18E-02 | 2.39E-01 | 2.68E+00 | 2.74E+00 | 8.60E-01 | no | | intergenic_variant | MODIFIER |
| 1 | 69739121 | AX-170533880 | 0 | C | G | 0.14 | 4.18E-02 | 2.39E-01 | 2.68E+00 | 2.74E+00 | 8.60E-01 | no | | intergenic_variant | MODIFIER |
| 1 | 69767081 | AX-170000757 | 0 | T | A | 0.14 | 4.18E-02 | 2.39E-01 | 2.68E+00 | 2.74E+00 | 8.60E-01 | no | Gm37930 | upstream_gene_variant | MODIFIER |
| 1 | 69870612 | AX-97792345 | 0 | C | T | 0.14 | 4.18E-02 | 2.39E-01 | 2.68E+00 | 2.74E+00 | 8.60E-01 | no | Spag16 | intron_variant | MODIFIER |
| 1 | 69886235 | AX-97811928 | 0 | C | T | 0.14 | 4.18E-02 | 2.39E-01 | 2.68E+00 | 2.74E+00 | 8.60E-01 | no | Spag16 | intron_variant | MODIFIER |
| 1 | 69886267 | AX-170204863 | 0 | A | G | 0.14 | 4.18E-02 | 2.39E-01 | 2.68E+00 | 2.74E+00 | 8.60E-01 | no | Spag16 | intron_variant | MODIFIER |
| 1 | 69905865 | AX-170145152 | 0 | G | A | 0.14 | 4.18E-02 | 2.39E-01 | 2.68E+00 | 2.74E+00 | 8.60E-01 | no | Spag16 | intron_variant | MODIFIER |
| 1 | 69976666 | AX-170391109 | 0 | T | C | 0.14 | 4.18E-02 | 2.39E-01 | 2.68E+00 | 2.74E+00 | 8.60E-01 | no | Spag16 | intron_variant | MODIFIER |
| 1 | 70009939 | AX-97800092 | 0 | C | T | 0.14 | 4.18E-02 | 2.39E-01 | 2.68E+00 | 2.74E+00 | 8.60E-01 | no | Spag16 | intron_variant | MODIFIER |
| 1 | 70024402 | AX-170176598 | 0 | G | T | 0.14 | 4.18E-02 | 2.39E-01 | 2.68E+00 | 2.74E+00 | 8.60E-01 | no | Spag16 | intron_variant | MODIFIER |
| 1 | 70029042 | AX-170530678 | 0 | G | T | 0.14 | 4.18E-02 | 2.39E-01 | 2.68E+00 | 2.74E+00 | 8.60E-01 | no | Spag16 | intron_variant | MODIFIER |
| 1 | 70100065 | AX-97810165 | 0 | T | C | 0.14 | 4.18E-02 | 2.39E-01 | 2.68E+00 | 2.74E+00 | 8.60E-01 | no | Spag16 | intron_variant | MODIFIER |
| 1 | 70125519 | AX-97841228 | 0 | T | C | 0.14 | 4.18E-02 | 2.39E-01 | 2.68E+00 | 2.74E+00 | 8.60E-01 | no | Spag16 | intron_variant | MODIFIER |
| 1 | 70165775 | AX-97811925 | 0 | G | T | 0.14 | 4.18E-02 | 2.39E-01 | 2.68E+00 | 2.74E+00 | 8.60E-01 | no | Spag16 | intron_variant | MODIFIER |
| 1 | 70207365 | AX-97802878 | 0 | C | T | 0.14 | 4.18E-02 | 2.39E-01 | 2.68E+00 | 2.74E+00 | 8.60E-01 | no | Spag16 | intron_variant | MODIFIER |
| 1 | 70244727 | AX-170214355 | 0 | A | G | 0.14 | 4.18E-02 | 2.39E-01 | 2.68E+00 | 2.74E+00 | 8.60E-01 | no | Spag16 | intron_variant | MODIFIER |
| 1 | 70316665 | AX-170272629 | 0 | A | G | 0.14 | 4.18E-02 | 2.39E-01 | 2.68E+00 | 2.74E+00 | 8.60E-01 | no | Spag16 | intron_variant | MODIFIER |
| 1 | 70327567 | AX-170000764 | 0 | A | G | 0.14 | 4.18E-02 | 2.39E-01 | 2.68E+00 | 2.74E+00 | 8.60E-01 | no | Spag16 | intron_variant | MODIFIER |
| 1 | 70339696 | AX-168410165 | 0 | T | C | 0.14 | 4.18E-02 | 2.39E-01 | 2.68E+00 | 2.74E+00 | 8.60E-01 | no | Spag16 | intron_variant | MODIFIER |
| 1 | 70343560 | AX-170277952 | 0 | T | C | 0.14 | 4.18E-02 | 2.39E-01 | 2.68E+00 | 2.74E+00 | 8.60E-01 | no | Spag16 | intron_variant | MODIFIER |
| 1 | 70511422 | AX-97856399 | 0 | T | G | 0.14 | 4.18E-02 | 2.39E-01 | 2.68E+00 | 2.74E+00 | 8.60E-01 | no | Spag16 | intron_variant | MODIFIER |
| 1 | 70759517 | AX-97825089 | 0 | G | C | 0.14 | 4.18E-02 | 2.39E-01 | 2.68E+00 | 2.74E+00 | 8.60E-01 | no | Vwc2l | intron_variant | MODIFIER |
| 1 | 70881889 | AX-170419992 | 0 | G | C | 0.14 | 4.18E-02 | 2.39E-01 | 2.68E+00 | 2.74E+00 | 8.60E-01 | no | Vwc2l | intron_variant | MODIFIER |
| 1 | 70886639 | AX-170000769 | 0 | C | A | 0.14 | 4.18E-02 | 2.39E-01 | 2.68E+00 | 2.74E+00 | 8.60E-01 | no | Vwc2l | downstream_gene_variant | MODIFIER |
| 1 | 70887307 | AX-170298897 | 0 | G | A | 0.14 | 4.18E-02 | 2.39E-01 | 2.68E+00 | 2.74E+00 | 8.60E-01 | no | Vwc2l | downstream_gene_variant | MODIFIER |

| | | | | | | | | | | | | | | | |
|---|---|---|---|---|---|---|---|---|---|---|---|---|---|---|---|
| 1 | 70956019 | AX-97791636 | 0 | C | T | 0.14 | 4.18E-02 | 2.39E-01 | 2.68E+00 | 2.74E+00 | 8.60E-01 | no | | intergenic_variant | MODIFIER |
| 1 | 71000540 | AX-170481434 | 0 | A | G | 0.14 | 4.18E-02 | 2.39E-01 | 2.68E+00 | 2.74E+00 | 8.60E-01 | no | | intergenic_variant | MODIFIER |
| 1 | 71064371 | AX-170200522 | 0 | G | T | 0.14 | 4.18E-02 | 2.39E-01 | 2.68E+00 | 2.74E+00 | 8.60E-01 | no | Bard1 | intron_variant | MODIFIER |
| 1 | 71079458 | AX-170041596 | 0 | C | T | 0.14 | 4.18E-02 | 2.39E-01 | 2.68E+00 | 2.74E+00 | 8.60E-01 | no | Bard1 | intron_variant | MODIFIER |
| 1 | 71087412 | AX-97811952 | 0 | T | C | 0.14 | 4.18E-02 | 2.39E-01 | 2.68E+00 | 2.74E+00 | 8.60E-01 | no | Bard1 | intron_variant | MODIFIER |
| 1 | 71090002 | AX-170041595 | 0 | T | G | 0.14 | 4.18E-02 | 2.39E-01 | 2.68E+00 | 2.74E+00 | 8.60E-01 | no | Bard1 | intron_variant | MODIFIER |
| 1 | 71097350 | AX-97842175 | 0 | G | A | 0.14 | 4.18E-02 | 2.39E-01 | 2.68E+00 | 2.74E+00 | 8.60E-01 | no | Bard1 | intron_variant | MODIFIER |
| 1 | 71135703 | AX-170502851 | 0 | G | A | 0.14 | 4.18E-02 | 2.39E-01 | 2.68E+00 | 2.74E+00 | 8.60E-01 | no | | intergenic_variant | MODIFIER |
| 1 | 71136257 | AX-170313025 | 0 | G | A | 0.14 | 4.18E-02 | 2.39E-01 | 2.68E+00 | 2.74E+00 | 8.60E-01 | no | | intergenic_variant | MODIFIER |
| 1 | 71154319 | AX-170091551 | 0 | C | T | 0.14 | 4.18E-02 | 2.39E-01 | 2.68E+00 | 2.74E+00 | 8.60E-01 | no | Gm8870 | downstream_gene_variant | MODIFIER |
| 1 | 71154590 | AX-170390278 | 0 | A | T | 0.14 | 4.18E-02 | 2.39E-01 | 2.68E+00 | 2.74E+00 | 8.60E-01 | no | Gm8870 | downstream_gene_variant | MODIFIER |
| 1 | 71196641 | AX-170349432 | 0 | G | T | 0.14 | 4.18E-02 | 2.39E-01 | 2.68E+00 | 2.74E+00 | 8.60E-01 | no | | intergenic_variant | MODIFIER |
| 1 | 71210977 | AX-97818740 | 0 | A | G | 0.14 | 4.18E-02 | 2.39E-01 | 2.68E+00 | 2.74E+00 | 8.60E-01 | no | | intergenic_variant | MODIFIER |
| 1 | 71243858 | AX-97822755 | 0 | C | A | 0.14 | 4.18E-02 | 2.39E-01 | 2.68E+00 | 2.74E+00 | 8.60E-01 | no | Abca12 | intron_variant | MODIFIER |
| 1 | 71244795 | AX-170110711 | 0 | T | C | 0.14 | 4.18E-02 | 2.39E-01 | 2.68E+00 | 2.74E+00 | 8.60E-01 | no | Abca12 | synonymous_variant | LOW |
| 1 | 71258730 | AX-170425480 | 0 | T | G | 0.14 | 4.18E-02 | 2.39E-01 | 2.68E+00 | 2.74E+00 | 8.60E-01 | no | Abca12 | intron_variant | MODIFIER |
| 1 | 71259181 | AX-170116199 | 0 | T | C | 0.14 | 4.18E-02 | 2.39E-01 | 2.68E+00 | 2.74E+00 | 8.60E-01 | no | Abca12 | intron_variant | MODIFIER |
| 1 | 71263765 | AX-170304639 | 0 | G | A | 0.14 | 4.18E-02 | 2.39E-01 | 2.68E+00 | 2.74E+00 | 8.60E-01 | no | Abca12 | intron_variant | MODIFIER |
| 1 | 71305825 | AX-170041600 | 0 | T | C | 0.14 | 4.18E-02 | 2.39E-01 | 2.68E+00 | 2.74E+00 | 8.60E-01 | no | Abca12 | intron_variant | MODIFIER |
| 1 | 72503279 | AX-170190170 | 0 | C | T | 0.14 | 4.18E-02 | 2.39E-01 | 2.68E+00 | 2.74E+00 | 8.60E-01 | no | MAR4 | intron_variant | MODIFIER |
| 1 | 72504565 | AX-170332459 | 0 | A | G | 0.14 | 4.18E-02 | 2.39E-01 | 2.68E+00 | 2.74E+00 | 8.60E-01 | no | MAR4 | intron_variant | MODIFIER |
| 1 | 72505423 | AX-170397225 | 0 | T | C | 0.14 | 4.18E-02 | 2.39E-01 | 2.68E+00 | 2.74E+00 | 8.60E-01 | no | MAR4 | intron_variant | MODIFIER |
| 1 | 72505877 | AX-170142606 | 0 | C | A | 0.14 | 4.18E-02 | 2.39E-01 | 2.68E+00 | 2.74E+00 | 8.60E-01 | no | MAR4 | intron_variant | MODIFIER |
| 1 | 72506057 | AX-170123155 | 0 | G | A | 0.14 | 4.18E-02 | 2.39E-01 | 2.68E+00 | 2.74E+00 | 8.60E-01 | no | MAR4 | intron_variant | MODIFIER |
| 1 | 72507987 | AX-97785251 | 0 | C | T | 0.14 | 4.18E-02 | 2.39E-01 | 2.68E+00 | 2.74E+00 | 8.60E-01 | no | MAR4 | intron_variant | MODIFIER |
| 1 | 72509393 | AX-170373923 | 0 | T | G | 0.14 | 4.18E-02 | 2.39E-01 | 2.68E+00 | 2.74E+00 | 8.60E-01 | no | MAR4 | intron_variant | MODIFIER |
| 1 | 72509896 | AX-170152234 | 0 | A | G | 0.14 | 4.18E-02 | 2.39E-01 | 2.68E+00 | 2.74E+00 | 8.60E-01 | no | MAR4 | intron_variant | MODIFIER |
| 1 | 72520851 | AX-170209563 | 0 | A | C | 0.14 | 4.18E-02 | 2.39E-01 | 2.68E+00 | 2.74E+00 | 8.60E-01 | no | MAR4 | intron_variant | MODIFIER |
| 1 | 72527419 | AX-170000788 | 0 | C | G | 0.14 | 4.18E-02 | 2.39E-01 | 2.68E+00 | 2.74E+00 | 8.60E-01 | no | MAR4 | intron_variant | MODIFIER |
| 1 | 72530458 | AX-170478299 | 0 | T | C | 0.14 | 4.18E-02 | 2.39E-01 | 2.68E+00 | 2.74E+00 | 8.60E-01 | no | MAR4 | intron_variant | MODIFIER |
| 1 | 72533575 | AX-97806935 | 0 | C | A | 0.14 | 4.18E-02 | 2.39E-01 | 2.68E+00 | 2.74E+00 | 8.60E-01 | no | MAR4 | intron_variant | MODIFIER |
| 1 | 72537988 | AX-170100628 | 0 | C | T | 0.14 | 4.18E-02 | 2.39E-01 | 2.68E+00 | 2.74E+00 | 8.60E-01 | no | MAR4 | upstream_gene_variant | MODIFIER |
| 1 | 72538500 | AX-170224362 | 0 | G | T | 0.14 | 4.18E-02 | 2.39E-01 | 2.68E+00 | 2.74E+00 | 8.60E-01 | no | MAR4 | upstream_gene_variant | MODIFIER |
| 1 | 72550212 | AX-170357570 | 0 | T | C | 0.14 | 4.18E-02 | 2.39E-01 | 2.68E+00 | 2.74E+00 | 8.60E-01 | no | | intergenic_variant | MODIFIER |
| 1 | 72554024 | AX-97833189 | 0 | T | C | 0.14 | 4.18E-02 | 2.39E-01 | 2.68E+00 | 2.74E+00 | 8.60E-01 | no | | intergenic_variant | MODIFIER |
| 1 | 72559618 | AX-170177079 | 0 | C | T | 0.14 | 4.18E-02 | 2.39E-01 | 2.68E+00 | 2.74E+00 | 8.60E-01 | no | | intergenic_variant | MODIFIER |
| 1 | 72592746 | AX-97808851 | 0 | A | G | 0.14 | 4.18E-02 | 2.39E-01 | 2.68E+00 | 2.74E+00 | 8.60E-01 | no | Smarcal1 | intron_variant | MODIFIER |
| 1 | 72598113 | AX-170522200 | 0 | T | C | 0.14 | 4.18E-02 | 2.39E-01 | 2.68E+00 | 2.74E+00 | 8.60E-01 | no | Smarcal1 | intron_variant | MODIFIER |
| 1 | 72604838 | AX-170144259 | 0 | G | A | 0.14 | 4.18E-02 | 2.39E-01 | 2.68E+00 | 2.74E+00 | 8.60E-01 | no | Smarcal1 | intron_variant | MODIFIER |
| 1 | 72608273 | AX-170091554 | 0 | A | G | 0.14 | 4.18E-02 | 2.39E-01 | 2.68E+00 | 2.74E+00 | 8.60E-01 | no | Smarcal1 | intron_variant | MODIFIER |
| 1 | 72610882 | AX-170041630 | 0 | G | A | 0.14 | 4.18E-02 | 2.39E-01 | 2.68E+00 | 2.74E+00 | 8.60E-01 | no | Smarcal1 | intron_variant | MODIFIER |
| 1 | 72617843 | AX-170350280 | 0 | A | G | 0.14 | 4.18E-02 | 2.39E-01 | 2.68E+00 | 2.74E+00 | 8.60E-01 | no | Smarcal1 | intron_variant | MODIFIER |
| 1 | 72632626 | AX-97812010 | 0 | A | G | 0.14 | 4.18E-02 | 2.39E-01 | 2.68E+00 | 2.74E+00 | 8.60E-01 | no | Smarcal1 | intron_variant | MODIFIER |
| 1 | 72650582 | AX-170086497 | 0 | T | C | 0.14 | 4.18E-02 | 2.39E-01 | 2.68E+00 | 2.74E+00 | 8.60E-01 | no | Ankar | intron_variant | MODIFIER |
| 1 | 72718764 | AX-170466897 | 0 | A | G | 0.14 | 4.18E-02 | 2.39E-01 | 2.68E+00 | 2.74E+00 | 8.60E-01 | no | Rpl37a | downstream_gene_variant | MODIFIER |
| 1 | 72737055 | AX-97785816 | 0 | G | A | 0.14 | 4.18E-02 | 2.39E-01 | 2.68E+00 | 2.74E+00 | 8.60E-01 | no | | regulatory_region_variant | MODIFIER |
| 1 | 72737086 | AX-170161135 | 0 | C | T | 0.14 | 4.18E-02 | 2.39E-01 | 2.68E+00 | 2.74E+00 | 8.60E-01 | no | | regulatory_region_variant | MODIFIER |
| 1 | 72741185 | AX-170330722 | 0 | A | G | 0.14 | 4.18E-02 | 2.39E-01 | 2.68E+00 | 2.74E+00 | 8.60E-01 | no | | intergenic_variant | MODIFIER |
| 1 | 72741829 | AX-170480106 | 0 | A | C | 0.14 | 4.18E-02 | 2.39E-01 | 2.68E+00 | 2.74E+00 | 8.60E-01 | no | | intergenic_variant | MODIFIER |
| 1 | 72744075 | AX-170447498 | 0 | A | G | 0.14 | 4.18E-02 | 2.39E-01 | 2.68E+00 | 2.74E+00 | 8.60E-01 | no | | intergenic_variant | MODIFIER |
| 1 | 72744291 | AX-170272317 | 0 | T | G | 0.14 | 4.18E-02 | 2.39E-01 | 2.68E+00 | 2.74E+00 | 8.60E-01 | no | | intergenic_variant | MODIFIER |
| 1 | 72744751 | AX-170275034 | 0 | A | G | 0.14 | 4.18E-02 | 2.39E-01 | 2.68E+00 | 2.74E+00 | 8.60E-01 | no | | intergenic_variant | MODIFIER |
| 1 | 72748241 | AX-170041634 | 0 | C | T | 0.14 | 4.18E-02 | 2.39E-01 | 2.68E+00 | 2.74E+00 | 8.60E-01 | no | | intergenic_variant | MODIFIER |
| 1 | 72748282 | AX-168401833 | 0 | T | C | 0.14 | 4.18E-02 | 2.39E-01 | 2.68E+00 | 2.74E+00 | 8.60E-01 | no | | intergenic_variant | MODIFIER |

| | | | | | | | | | | | | | | | | |
|---|---|---|---|---|---|---|---|---|---|---|---|---|---|---|---|---|
| 1 | 72748736 | AX-170041635 | 0 | A | G | 0.14 | 4.18E-02 | 2.39E-01 | 2.68E+00 | 2.74E+00 | 8.60E-01 | no | | | intergenic_variant | MODIFIER |
| 1 | 72748795 | AX-170444273 | 0 | A | G | 0.14 | 4.18E-02 | 2.39E-01 | 2.68E+00 | 2.74E+00 | 8.60E-01 | no | | | intergenic_variant | MODIFIER |
| 1 | 72750182 | AX-170041633 | 0 | T | C | 0.14 | 4.18E-02 | 2.39E-01 | 2.68E+00 | 2.74E+00 | 8.60E-01 | no | | | intergenic_variant | MODIFIER |
| 1 | 72751731 | AX-170425242 | 0 | A | G | 0.14 | 4.18E-02 | 2.39E-01 | 2.68E+00 | 2.74E+00 | 8.60E-01 | no | | | regulatory_region_variant | MODIFIER |
| 1 | 73220264 | AX-170342814 | 0 | G | A | 0.14 | 4.18E-02 | 2.39E-01 | 2.68E+00 | 2.74E+00 | 8.60E-01 | no | | | intergenic_variant | MODIFIER |
| 1 | 73224026 | AX-170439309 | 0 | A | G | 0.14 | 4.18E-02 | 2.39E-01 | 2.68E+00 | 2.74E+00 | 8.60E-01 | no | | | intergenic_variant | MODIFIER |
| 1 | 73224311 | AX-170195894 | 0 | T | C | 0.14 | 4.18E-02 | 2.39E-01 | 2.68E+00 | 2.74E+00 | 8.60E-01 | no | | | intergenic_variant | MODIFIER |
| 1 | 73226630 | AX-170363691 | 0 | C | T | 0.14 | 4.18E-02 | 2.39E-01 | 2.68E+00 | 2.74E+00 | 8.60E-01 | no | | | intergenic_variant | MODIFIER |
| 1 | 73226906 | AX-170453085 | 0 | G | T | 0.14 | 4.18E-02 | 2.39E-01 | 2.68E+00 | 2.74E+00 | 8.60E-01 | no | | | intergenic_variant | MODIFIER |
| 1 | 74058376 | AX-170041672 | 0 | G | C | 0.14 | 4.18E-02 | 2.39E-01 | 2.68E+00 | 2.74E+00 | 8.60E-01 | no | Tns1 | | intron_variant | MODIFIER |
| 1 | 74063711 | AX-170041670 | 0 | A | T | 0.14 | 4.18E-02 | 2.39E-01 | 2.68E+00 | 2.74E+00 | 8.60E-01 | no | Tns1 | | intron_variant | MODIFIER |
| 1 | 74429219 | AX-170178391 | 0 | T | C | 0.14 | 4.18E-02 | 2.39E-01 | 2.68E+00 | 2.74E+00 | 8.60E-01 | no | Vil1 | | intron_variant | MODIFIER |
| 1 | 74432447 | AX-97856658 | 0 | C | A | 0.14 | 4.18E-02 | 2.39E-01 | 2.68E+00 | 2.74E+00 | 8.60E-01 | no | Vil1 | | intron_variant | MODIFIER |
| 1 | 74436973 | AX-170399799 | 0 | G | A | 0.14 | 4.18E-02 | 2.39E-01 | 2.68E+00 | 2.74E+00 | 8.60E-01 | no | Vil1 | | downstream_gene_variant | MODIFIER |
| 1 | 74436995 | AX-170435486 | 0 | T | C | 0.14 | 4.18E-02 | 2.39E-01 | 2.68E+00 | 2.74E+00 | 8.60E-01 | no | Vil1 | | downstream_gene_variant | MODIFIER |
| 1 | 74443505 | AX-170041681 | 0 | C | T | 0.14 | 4.18E-02 | 2.39E-01 | 2.68E+00 | 2.74E+00 | 8.60E-01 | no | Usp37 | | intron_variant | MODIFIER |
| 1 | 74476943 | AX-170252363 | 0 | G | A | 0.14 | 4.18E-02 | 2.39E-01 | 2.68E+00 | 2.74E+00 | 8.60E-01 | no | Usp37 | | intron_variant | MODIFIER |
| 1 | 74477367 | AX-170305316 | 0 | C | T | 0.14 | 4.18E-02 | 2.39E-01 | 2.68E+00 | 2.74E+00 | 8.60E-01 | no | Usp37 | | intron_variant | MODIFIER |
| 1 | 74486201 | AX-97812008 | 0 | A | G | 0.14 | 4.18E-02 | 2.39E-01 | 2.68E+00 | 2.74E+00 | 8.60E-01 | no | Usp37 | | intron_variant | MODIFIER |
| 1 | 74504679 | AX-97806366 | 0 | T | C | 0.14 | 4.18E-02 | 2.39E-01 | 2.68E+00 | 2.74E+00 | 8.60E-01 | no | Usp37 | | intron_variant | MODIFIER |
| 1 | 74631605 | AX-170476677 | 0 | G | A | 0.14 | 4.18E-02 | 2.39E-01 | 2.68E+00 | 2.74E+00 | 8.60E-01 | no | Stk36 | | missense_variant | MODERATE |
| 1 | 74824824 | AX-97811035 | 0 | C | T | 0.14 | 4.18E-02 | 2.39E-01 | 2.68E+00 | 2.74E+00 | 8.60E-01 | no | | | regulatory_region_variant | MODIFIER |
| 1 | 74901757 | AX-170512315 | 0 | T | C | 0.14 | 4.18E-02 | 2.39E-01 | 2.68E+00 | 2.74E+00 | 8.60E-01 | no | Mir375 | | upstream_gene_variant | MODIFIER |
| 1 | 74902086 | AX-170364305 | 0 | A | T | 0.14 | 4.18E-02 | 2.39E-01 | 2.68E+00 | 2.74E+00 | 8.60E-01 | no | Mir375 | | upstream_gene_variant | MODIFIER |
| 1 | 74962595 | AX-170449390 | 0 | C | G | 0.14 | 4.18E-02 | 2.39E-01 | 2.68E+00 | 2.74E+00 | 8.60E-01 | no | Nhej1 | | downstream_gene_variant | MODIFIER |
| 1 | 74963849 | AX-97797356 | 0 | C | T | 0.14 | 4.18E-02 | 2.39E-01 | 2.68E+00 | 2.74E+00 | 8.60E-01 | no | Nhej1 | | downstream_gene_variant | MODIFIER |
| 1 | 75010584 | AX-170086498 | 0 | C | T | 0.14 | 4.18E-02 | 2.39E-01 | 2.68E+00 | 2.74E+00 | 8.60E-01 | no | Nhej1 | | intron_variant | MODIFIER |
| 1 | 75128344 | AX-97812037 | 0 | C | A | 0.14 | 4.18E-02 | 2.39E-01 | 2.68E+00 | 2.74E+00 | 8.60E-01 | no | Slc23a3 | | intron_variant | MODIFIER |
| 1 | 75131797 | AX-170000817 | 0 | A | T | 0.14 | 4.18E-02 | 2.39E-01 | 2.68E+00 | 2.74E+00 | 8.60E-01 | no | Slc23a3 | | intron_variant | MODIFIER |
| 1 | 75272502 | AX-170510218 | 0 | T | C | 0.14 | 4.18E-02 | 2.39E-01 | 2.68E+00 | 2.74E+00 | 8.60E-01 | no | Resp18 | | intron_variant | MODIFIER |
| 1 | 75276050 | AX-170262102 | 0 | G | A | 0.14 | 4.18E-02 | 2.39E-01 | 2.68E+00 | 2.74E+00 | 8.60E-01 | no | Resp18 | | intron_variant | MODIFIER |
| 1 | 75427690 | AX-170434756 | 0 | T | C | 0.14 | 4.18E-02 | 2.39E-01 | 2.68E+00 | 2.74E+00 | 8.60E-01 | no | Speg | | synonymous_variant | LOW |
| 1 | 75434838 | AX-170293631 | 0 | C | T | 0.14 | 4.18E-02 | 2.39E-01 | 2.68E+00 | 2.74E+00 | 8.60E-01 | no | Gmppa | | upstream_gene_variant | MODIFIER |
| 1 | 75442944 | AX-97841127 | 0 | G | A | 0.14 | 4.18E-02 | 2.39E-01 | 2.68E+00 | 2.74E+00 | 8.60E-01 | no | Gmppa | | synonymous_variant | LOW |
| 1 | 75464653 | AX-170372743 | 0 | T | A | 0.14 | 4.18E-02 | 2.39E-01 | 2.68E+00 | 2.74E+00 | 8.60E-01 | no | Asic4 | | intron_variant | MODIFIER |
| 1 | 75472884 | AX-170527853 | 0 | G | A | 0.14 | 4.18E-02 | 2.39E-01 | 2.68E+00 | 2.74E+00 | 8.60E-01 | no | Asic4 | | missense_variant | MODERATE |
| 1 | 75497230 | AX-169999762 | 0 | T | C | 0.14 | 4.18E-02 | 2.39E-01 | 2.68E+00 | 2.74E+00 | 8.60E-01 | no | Obsl1 | | missense_variant | MODERATE |
| 1 | 75498183 | AX-170374692 | 0 | C | T | 0.14 | 4.18E-02 | 2.39E-01 | 2.68E+00 | 2.74E+00 | 8.60E-01 | no | Obsl1 | | missense_variant | MODERATE |
| 1 | 75587888 | AX-170139454 | 0 | T | C | 0.14 | 4.18E-02 | 2.39E-01 | 2.68E+00 | 2.74E+00 | 8.60E-01 | no | | | regulatory_region_variant | MODIFIER |
| 1 | 75598826 | AX-170343402 | 0 | A | G | 0.14 | 4.18E-02 | 2.39E-01 | 2.68E+00 | 2.74E+00 | 8.60E-01 | no | | | intergenic_variant | MODIFIER |
| 1 | 75631043 | AX-170000827 | 0 | T | C | 0.14 | 4.18E-02 | 2.39E-01 | 2.68E+00 | 2.74E+00 | 8.60E-01 | no | Gm15182 | | downstream_gene_variant | MODIFIER |
| 1 | 75642470 | AX-170480424 | 0 | A | G | 0.14 | 4.18E-02 | 2.39E-01 | 2.68E+00 | 2.74E+00 | 8.60E-01 | no | | | intergenic_variant | MODIFIER |
| 1 | 75645915 | AX-170465631 | 0 | C | T | 0.14 | 4.18E-02 | 2.39E-01 | 2.68E+00 | 2.74E+00 | 8.60E-01 | no | | | intergenic_variant | MODIFIER |
| 1 | 75679304 | AX-170434300 | 0 | G | A | 0.14 | 4.18E-02 | 2.39E-01 | 2.68E+00 | 2.74E+00 | 8.60E-01 | no | | | intergenic_variant | MODIFIER |
| 1 | 75685339 | AX-170000828 | 0 | T | C | 0.14 | 4.18E-02 | 2.39E-01 | 2.68E+00 | 2.74E+00 | 8.60E-01 | no | | | intergenic_variant | MODIFIER |
| 1 | 75705008 | AX-97813610 | 0 | G | A | 0.14 | 4.18E-02 | 2.39E-01 | 2.68E+00 | 2.74E+00 | 8.60E-01 | no | | | regulatory_region_variant | MODIFIER |
| 1 | 75705184 | AX-170362997 | 0 | G | A | 0.14 | 4.18E-02 | 2.39E-01 | 2.68E+00 | 2.74E+00 | 8.60E-01 | no | | | regulatory_region_variant | MODIFIER |
| 1 | 76064853 | AX-97812091 | 0 | T | C | 0.14 | 4.18E-02 | 2.39E-01 | 2.68E+00 | 2.74E+00 | 8.60E-01 | no | | | intergenic_variant | MODIFIER |
| 1 | 76263454 | AX-170532276 | 0 | G | A | 0.14 | 4.18E-02 | 2.39E-01 | 2.68E+00 | 2.74E+00 | 8.60E-01 | no | | | intergenic_variant | MODIFIER |
| 1 | 76319215 | AX-170319430 | 0 | G | A | 0.14 | 4.18E-02 | 2.39E-01 | 2.68E+00 | 2.74E+00 | 8.60E-01 | no | | | intergenic_variant | MODIFIER |
| 1 | 76339925 | AX-170534322 | 0 | G | A | 0.14 | 4.18E-02 | 2.39E-01 | 2.68E+00 | 2.74E+00 | 8.60E-01 | no | | | intergenic_variant | MODIFIER |
| 1 | 76394666 | AX-97812039 | 0 | C | T | 0.14 | 4.18E-02 | 2.39E-01 | 2.68E+00 | 2.74E+00 | 8.60E-01 | no | | | intergenic_variant | MODIFIER |
| 1 | 76400480 | AX-170041712 | 0 | T | C | 0.14 | 4.18E-02 | 2.39E-01 | 2.68E+00 | 2.74E+00 | 8.60E-01 | no | | | intergenic_variant | MODIFIER |
| 1 | 76401702 | AX-170000837 | 0 | C | T | 0.14 | 4.18E-02 | 2.39E-01 | 2.68E+00 | 2.74E+00 | 8.60E-01 | no | | | intergenic_variant | MODIFIER |

| | | | | | | | | | | | | | | | |
|---|---|---|---|---|---|---|---|---|---|---|---|---|---|---|---|
| 1 | 76410562 | AX-170091565 | 0 | A | C | 0.14 | 4.18E-02 | 2.39E-01 | 2.68E+00 | 2.74E+00 | 8.60E-01 | no | | intergenic_variant | MODIFIER |
| 1 | 76451132 | AX-170438638 | 0 | C | G | 0.14 | 4.18E-02 | 2.39E-01 | 2.68E+00 | 2.74E+00 | 8.60E-01 | no | Gm29069 | intron_variant,non_coding_transcript_variant | MODIFIER |
| 1 | 76452425 | AX-170409250 | 0 | C | T | 0.14 | 4.18E-02 | 2.39E-01 | 2.68E+00 | 2.74E+00 | 8.60E-01 | no | Gm29069 | intron_variant,non_coding_transcript_variant | MODIFIER |
| 1 | 76458821 | AX-170153204 | 0 | C | T | 0.14 | 4.18E-02 | 2.39E-01 | 2.68E+00 | 2.74E+00 | 8.60E-01 | no | Gm29069 | intron_variant,non_coding_transcript_variant | MODIFIER |
| 1 | 76459471 | AX-97794751 | 0 | A | G | 0.14 | 4.18E-02 | 2.39E-01 | 2.68E+00 | 2.74E+00 | 8.60E-01 | no | Gm29069 | intron_variant,non_coding_transcript_variant | MODIFIER |
| 1 | 76492320 | AX-170515136 | 0 | G | C | 0.14 | 4.18E-02 | 2.39E-01 | 2.68E+00 | 2.74E+00 | 8.60E-01 | no | Gm29069 | intron_variant,non_coding_transcript_variant | MODIFIER |
| 1 | 76501043 | AX-168417731 | 0 | T | C | 0.14 | 4.18E-02 | 2.39E-01 | 2.68E+00 | 2.74E+00 | 8.60E-01 | no | Gm29069 | intron_variant,non_coding_transcript_variant | MODIFIER |
| 1 | 76501497 | AX-170086502 | 0 | G | T | 0.14 | 4.18E-02 | 2.39E-01 | 2.68E+00 | 2.74E+00 | 8.60E-01 | no | Gm29069 | intron_variant,non_coding_transcript_variant | MODIFIER |
| 1 | 76503770 | AX-170247770 | 0 | G | A | 0.14 | 4.18E-02 | 2.39E-01 | 2.68E+00 | 2.74E+00 | 8.60E-01 | no | Gm29069 | intron_variant,non_coding_transcript_variant | MODIFIER |
| 1 | 76507200 | AX-170151106 | 0 | C | T | 0.14 | 4.18E-02 | 2.39E-01 | 2.68E+00 | 2.74E+00 | 8.60E-01 | no | Mir6343 | upstream_gene_variant | MODIFIER |
| 1 | 76522497 | AX-170041717 | 0 | A | T | 0.14 | 4.18E-02 | 2.39E-01 | 2.68E+00 | 2.74E+00 | 8.60E-01 | no | Gm29069 | intron_variant,non_coding_transcript_variant | MODIFIER |
| 1 | 76523232 | AX-170136708 | 0 | C | T | 0.14 | 4.18E-02 | 2.39E-01 | 2.68E+00 | 2.74E+00 | 8.60E-01 | no | Gm29069 | intron_variant,non_coding_transcript_variant | MODIFIER |
| 1 | 76642407 | AX-170195169 | 0 | G | A | 0.14 | 4.18E-02 | 2.39E-01 | 2.68E+00 | 2.74E+00 | 8.60E-01 | no | | intergenic_variant | MODIFIER |
| 1 | 76642924 | AX-170161375 | 0 | T | A | 0.14 | 4.18E-02 | 2.39E-01 | 2.68E+00 | 2.74E+00 | 8.60E-01 | no | | intergenic_variant | MODIFIER |
| 1 | 76643667 | AX-170183704 | 0 | C | T | 0.14 | 4.18E-02 | 2.39E-01 | 2.68E+00 | 2.74E+00 | 8.60E-01 | no | | intergenic_variant | MODIFIER |
| 1 | 76644867 | AX-170528195 | 0 | T | C | 0.14 | 4.18E-02 | 2.39E-01 | 2.68E+00 | 2.74E+00 | 8.60E-01 | no | | intergenic_variant | MODIFIER |
| 1 | 76675647 | AX-170361367 | 0 | T | G | 0.14 | 4.18E-02 | 2.39E-01 | 2.68E+00 | 2.74E+00 | 8.60E-01 | no | | intergenic_variant | MODIFIER |
| 1 | 76677164 | AX-97820366 | 0 | C | T | 0.14 | 4.18E-02 | 2.39E-01 | 2.68E+00 | 2.74E+00 | 8.60E-01 | no | | regulatory_region_variant | MODIFIER |
| 1 | 76935849 | AX-170041723 | 0 | G | A | 0.14 | 4.18E-02 | 2.39E-01 | 2.68E+00 | 2.74E+00 | 8.60E-01 | no | Gm816 | intron_variant,non_coding_transcript_variant | MODIFIER |
| 1 | 76935985 | AX-170041725 | 0 | T | C | 0.14 | 4.18E-02 | 2.39E-01 | 2.68E+00 | 2.74E+00 | 8.60E-01 | no | Gm816 | intron_variant,non_coding_transcript_variant | MODIFIER |
| 1 | 76936608 | AX-170171977 | 0 | A | G | 0.14 | 4.18E-02 | 2.39E-01 | 2.68E+00 | 2.74E+00 | 8.60E-01 | no | Gm816 | intron_variant,non_coding_transcript_variant | MODIFIER |
| 1 | 76937264 | AX-170484955 | 0 | T | C | 0.14 | 4.18E-02 | 2.39E-01 | 2.68E+00 | 2.74E+00 | 8.60E-01 | no | Gm816 | intron_variant,non_coding_transcript_variant | MODIFIER |
| 1 | 76937911 | AX-170417373 | 0 | T | C | 0.14 | 4.18E-02 | 2.39E-01 | 2.68E+00 | 2.74E+00 | 8.60E-01 | no | Gm816 | intron_variant,non_coding_transcript_variant | MODIFIER |
| 1 | 76944834 | AX-170297659 | 0 | C | T | 0.14 | 4.18E-02 | 2.39E-01 | 2.68E+00 | 2.74E+00 | 8.60E-01 | no | Gm816 | intron_variant,non_coding_transcript_variant | MODIFIER |
| 1 | 76949329 | AX-170334770 | 0 | G | A | 0.14 | 4.18E-02 | 2.39E-01 | 2.68E+00 | 2.74E+00 | 8.60E-01 | no | Gm816 | intron_variant,non_coding_transcript_variant | MODIFIER |
| 1 | 76962804 | AX-97790270 | 0 | A | G | 0.14 | 4.18E-02 | 2.39E-01 | 2.68E+00 | 2.74E+00 | 8.60E-01 | no | Gm816 | intron_variant,non_coding_transcript_variant | MODIFIER |
| 1 | 76977058 | AX-170216069 | 0 | A | C | 0.14 | 4.18E-02 | 2.39E-01 | 2.68E+00 | 2.74E+00 | 8.60E-01 | no | Gm816 | intron_variant,non_coding_transcript_variant | MODIFIER |
| 1 | 76982496 | AX-170203817 | 0 | T | C | 0.14 | 4.18E-02 | 2.39E-01 | 2.68E+00 | 2.74E+00 | 8.60E-01 | no | Gm816 | intron_variant,non_coding_transcript_variant | MODIFIER |
| 1 | 76985187 | AX-170525993 | 0 | T | C | 0.14 | 4.18E-02 | 2.39E-01 | 2.68E+00 | 2.74E+00 | 8.60E-01 | no | Gm816 | intron_variant,non_coding_transcript_variant | MODIFIER |
| 1 | 76991042 | AX-170468427 | 0 | C | T | 0.14 | 4.18E-02 | 2.39E-01 | 2.68E+00 | 2.74E+00 | 8.60E-01 | no | Gm816 | intron_variant,non_coding_transcript_variant | MODIFIER |
| 1 | 76991201 | AX-170091567 | 0 | C | T | 0.14 | 4.18E-02 | 2.39E-01 | 2.68E+00 | 2.74E+00 | 8.60E-01 | no | Gm816 | intron_variant,non_coding_transcript_variant | MODIFIER |
| 1 | 77028085 | AX-97786996 | 0 | G | A | 0.14 | 4.18E-02 | 2.39E-01 | 2.68E+00 | 2.74E+00 | 8.60E-01 | no | Gm816 | upstream_gene_variant | MODIFIER |
| 1 | 77100851 | AX-170352442 | 0 | A | G | 0.14 | 4.18E-02 | 2.39E-01 | 2.68E+00 | 2.74E+00 | 8.60E-01 | no | | intergenic_variant | MODIFIER |
| 1 | 77102645 | AX-170379916 | 0 | A | G | 0.14 | 4.18E-02 | 2.39E-01 | 2.68E+00 | 2.74E+00 | 8.60E-01 | no | | regulatory_region_variant | MODIFIER |
| 1 | 77107692 | AX-170287150 | 0 | T | C | 0.14 | 4.18E-02 | 2.39E-01 | 2.68E+00 | 2.74E+00 | 8.60E-01 | no | | intergenic_variant | MODIFIER |
| 1 | 77108675 | AX-97855607 | 0 | G | A | 0.14 | 4.18E-02 | 2.39E-01 | 2.68E+00 | 2.74E+00 | 8.60E-01 | no | | intergenic_variant | MODIFIER |
| 1 | 77127179 | AX-170394319 | 0 | T | G | 0.14 | 4.18E-02 | 2.39E-01 | 2.68E+00 | 2.74E+00 | 8.60E-01 | no | | intergenic_variant | MODIFIER |
| 1 | 77157936 | AX-97812065 | 0 | G | T | 0.14 | 4.18E-02 | 2.39E-01 | 2.68E+00 | 2.74E+00 | 8.60E-01 | no | | intergenic_variant | MODIFIER |
| 1 | 77168619 | AX-170135450 | 0 | A | G | 0.14 | 4.18E-02 | 2.39E-01 | 2.68E+00 | 2.74E+00 | 8.60E-01 | no | | regulatory_region_variant | MODIFIER |
| 1 | 77187288 | AX-170091568 | 0 | A | G | 0.14 | 4.18E-02 | 2.39E-01 | 2.68E+00 | 2.74E+00 | 8.60E-01 | no | | intergenic_variant | MODIFIER |
| 1 | 77206252 | AX-170139650 | 0 | G | T | 0.14 | 4.18E-02 | 2.39E-01 | 2.68E+00 | 2.74E+00 | 8.60E-01 | no | | regulatory_region_variant | MODIFIER |
| 1 | 77207689 | AX-97790596 | 0 | G | A | 0.14 | 4.18E-02 | 2.39E-01 | 2.68E+00 | 2.74E+00 | 8.60E-01 | no | | intergenic_variant | MODIFIER |
| 1 | 77209947 | AX-170130104 | 0 | C | A | 0.14 | 4.18E-02 | 2.39E-01 | 2.68E+00 | 2.74E+00 | 8.60E-01 | no | | intergenic_variant | MODIFIER |
| 1 | 77248239 | AX-170300567 | 0 | G | A | 0.14 | 4.18E-02 | 2.39E-01 | 2.68E+00 | 2.74E+00 | 8.60E-01 | no | | regulatory_region_variant | MODIFIER |
| 1 | 77252024 | AX-170482497 | 0 | T | C | 0.14 | 4.18E-02 | 2.39E-01 | 2.68E+00 | 2.74E+00 | 8.60E-01 | no | | intergenic_variant | MODIFIER |
| 1 | 77255816 | AX-170331330 | 0 | T | C | 0.14 | 4.18E-02 | 2.39E-01 | 2.68E+00 | 2.74E+00 | 8.60E-01 | no | | intergenic_variant | MODIFIER |
| 1 | 77255846 | AX-97825044 | 0 | C | T | 0.14 | 4.18E-02 | 2.39E-01 | 2.68E+00 | 2.74E+00 | 8.60E-01 | no | | intergenic_variant | MODIFIER |
| 1 | 77257994 | AX-170000853 | 0 | A | C | 0.14 | 4.18E-02 | 2.39E-01 | 2.68E+00 | 2.74E+00 | 8.60E-01 | no | | regulatory_region_variant | MODIFIER |
| 1 | 77264466 | AX-170000850 | 0 | G | A | 0.14 | 4.18E-02 | 2.39E-01 | 2.68E+00 | 2.74E+00 | 8.60E-01 | no | | intergenic_variant | MODIFIER |
| 1 | 77264668 | AX-170293843 | 0 | G | T | 0.14 | 4.18E-02 | 2.39E-01 | 2.68E+00 | 2.74E+00 | 8.60E-01 | no | | intergenic_variant | MODIFIER |
| 1 | 77272773 | AX-170000855 | 0 | A | G | 0.14 | 4.18E-02 | 2.39E-01 | 2.68E+00 | 2.74E+00 | 8.60E-01 | no | | intergenic_variant | MODIFIER |
| 1 | 77289214 | AX-97811231 | 0 | G | A | 0.14 | 4.18E-02 | 2.39E-01 | 2.68E+00 | 2.74E+00 | 8.60E-01 | no | | intergenic_variant | MODIFIER |
| 1 | 77293884 | AX-170000854 | 0 | G | A | 0.14 | 4.18E-02 | 2.39E-01 | 2.68E+00 | 2.74E+00 | 8.60E-01 | no | | intergenic_variant | MODIFIER |
| 1 | 77295117 | AX-170152996 | 0 | G | A | 0.14 | 4.18E-02 | 2.39E-01 | 2.68E+00 | 2.74E+00 | 8.60E-01 | no | | intergenic_variant | MODIFIER |

| | | | | | | | | | | | | | | | |
|---|---|---|---|---|---|---|---|---|---|---|---|---|---|---|---|
| 1 | 77295927 | AX-170471939 | 0 | A | G | 0.14 | 4.18E-02 | 2.39E-01 | 2.68E+00 | 2.74E+00 | 8.60E-01 | no | | intergenic_variant | MODIFIER |
| 1 | 77306641 | AX-170000856 | 0 | C | T | 0.14 | 4.18E-02 | 2.39E-01 | 2.68E+00 | 2.74E+00 | 8.60E-01 | no | | intergenic_variant | MODIFIER |
| 1 | 77310411 | AX-97793966 | 0 | A | G | 0.14 | 4.18E-02 | 2.39E-01 | 2.68E+00 | 2.74E+00 | 8.60E-01 | no | | intergenic_variant | MODIFIER |
| 1 | 77329338 | AX-170248139 | 0 | A | G | 0.14 | 4.18E-02 | 2.39E-01 | 2.68E+00 | 2.74E+00 | 8.60E-01 | no | | intergenic_variant | MODIFIER |
| 1 | 77337784 | AX-97824587 | 0 | G | A | 0.14 | 4.18E-02 | 2.39E-01 | 2.68E+00 | 2.74E+00 | 8.60E-01 | no | | regulatory_region_variant | MODIFIER |
| 1 | 77379895 | AX-170272057 | 0 | T | C | 0.14 | 4.18E-02 | 2.39E-01 | 2.68E+00 | 2.74E+00 | 8.60E-01 | no | Epha4 | intron_variant | MODIFIER |
| 1 | 77382675 | AX-97836320 | 0 | C | A | 0.14 | 4.18E-02 | 2.39E-01 | 2.68E+00 | 2.74E+00 | 8.60E-01 | no | Epha4 | intron_variant | MODIFIER |
| 1 | 77398680 | AX-170228407 | 0 | T | A | 0.14 | 4.18E-02 | 2.39E-01 | 2.68E+00 | 2.74E+00 | 8.60E-01 | no | Epha4 | intron_variant | MODIFIER |
| 1 | 77413348 | AX-97811306 | 0 | T | C | 0.14 | 4.18E-02 | 2.39E-01 | 2.68E+00 | 2.74E+00 | 8.60E-01 | no | Epha4 | intron_variant | MODIFIER |
| 1 | 77491440 | AX-97832028 | 0 | G | A | 0.14 | 4.18E-02 | 2.39E-01 | 2.68E+00 | 2.74E+00 | 8.60E-01 | no | Epha4 | intron_variant | MODIFIER |
| 1 | 77500574 | AX-170333344 | 0 | A | G | 0.14 | 4.18E-02 | 2.39E-01 | 2.68E+00 | 2.74E+00 | 8.60E-01 | no | Epha4 | intron_variant | MODIFIER |
| 1 | 77525402 | AX-170102421 | 0 | A | G | 0.14 | 4.18E-02 | 2.39E-01 | 2.68E+00 | 2.74E+00 | 8.60E-01 | no | | intergenic_variant | MODIFIER |
| 1 | 77525500 | AX-97835259 | 0 | T | C | 0.14 | 4.18E-02 | 2.39E-01 | 2.68E+00 | 2.74E+00 | 8.60E-01 | no | | intergenic_variant | MODIFIER |
| 1 | 77526224 | AX-170247895 | 0 | T | C | 0.14 | 4.18E-02 | 2.39E-01 | 2.68E+00 | 2.74E+00 | 8.60E-01 | no | | intergenic_variant | MODIFIER |
| 1 | 77527385 | AX-170112791 | 0 | A | T | 0.14 | 4.18E-02 | 2.39E-01 | 2.68E+00 | 2.74E+00 | 8.60E-01 | no | | intergenic_variant | MODIFIER |
| 1 | 77566776 | AX-97808790 | 0 | A | G | 0.14 | 4.18E-02 | 2.39E-01 | 2.68E+00 | 2.74E+00 | 8.60E-01 | no | Gm28385 | upstream_gene_variant | MODIFIER |
| 1 | 77619130 | AX-97821761 | 0 | G | A | 0.14 | 4.18E-02 | 2.39E-01 | 2.68E+00 | 2.74E+00 | 8.60E-01 | no | | intergenic_variant | MODIFIER |
| 1 | 77622050 | AX-170464081 | 0 | A | G | 0.14 | 4.18E-02 | 2.39E-01 | 2.68E+00 | 2.74E+00 | 8.60E-01 | no | | intergenic_variant | MODIFIER |
| 1 | 77659216 | AX-170123484 | 0 | G | A | 0.14 | 4.18E-02 | 2.39E-01 | 2.68E+00 | 2.74E+00 | 8.60E-01 | no | | intergenic_variant | MODIFIER |
| 1 | 77666042 | AX-170456556 | 0 | A | G | 0.14 | 4.18E-02 | 2.39E-01 | 2.68E+00 | 2.74E+00 | 8.60E-01 | no | | intergenic_variant | MODIFIER |
| 1 | 77666747 | AX-97853425 | 0 | G | T | 0.14 | 4.18E-02 | 2.39E-01 | 2.68E+00 | 2.74E+00 | 8.60E-01 | no | | intergenic_variant | MODIFIER |
| 1 | 77666972 | AX-170503024 | 0 | A | G | 0.14 | 4.18E-02 | 2.39E-01 | 2.68E+00 | 2.74E+00 | 8.60E-01 | no | | intergenic_variant | MODIFIER |
| 1 | 77672214 | AX-170279487 | 0 | C | T | 0.14 | 4.18E-02 | 2.39E-01 | 2.68E+00 | 2.74E+00 | 8.60E-01 | no | | intergenic_variant | MODIFIER |
| 1 | 77672307 | AX-170403637 | 0 | G | T | 0.14 | 4.18E-02 | 2.39E-01 | 2.68E+00 | 2.74E+00 | 8.60E-01 | no | | intergenic_variant | MODIFIER |
| 1 | 77673148 | AX-170529469 | 0 | G | T | 0.14 | 4.18E-02 | 2.39E-01 | 2.68E+00 | 2.74E+00 | 8.60E-01 | no | | intergenic_variant | MODIFIER |
| 1 | 77711836 | AX-170424236 | 0 | G | A | 0.14 | 4.18E-02 | 2.39E-01 | 2.68E+00 | 2.74E+00 | 8.60E-01 | no | | intergenic_variant | MODIFIER |
| 1 | 77719347 | AX-97824705 | 0 | T | C | 0.14 | 4.18E-02 | 2.39E-01 | 2.68E+00 | 2.74E+00 | 8.60E-01 | no | | intergenic_variant | MODIFIER |
| 1 | 77728931 | AX-97801737 | 0 | G | T | 0.14 | 4.18E-02 | 2.39E-01 | 2.68E+00 | 2.74E+00 | 8.60E-01 | no | | intergenic_variant | MODIFIER |
| 1 | 77739994 | AX-170302971 | 0 | G | T | 0.14 | 4.18E-02 | 2.39E-01 | 2.68E+00 | 2.74E+00 | 8.60E-01 | no | | intergenic_variant | MODIFIER |
| 1 | 77746968 | AX-170000860 | 0 | C | T | 0.14 | 4.18E-02 | 2.39E-01 | 2.68E+00 | 2.74E+00 | 8.60E-01 | no | | intergenic_variant | MODIFIER |
| 1 | 78022754 | AX-170334498 | 0 | C | T | 0.14 | 4.18E-02 | 2.39E-01 | 2.68E+00 | 2.74E+00 | 8.60E-01 | no | | regulatory_region_variant | MODIFIER |
| 1 | 78025681 | AX-170301002 | 0 | T | C | 0.14 | 4.18E-02 | 2.39E-01 | 2.68E+00 | 2.74E+00 | 8.60E-01 | no | | intergenic_variant | MODIFIER |
| 1 | 78026779 | AX-170214421 | 0 | A | G | 0.14 | 4.18E-02 | 2.39E-01 | 2.68E+00 | 2.74E+00 | 8.60E-01 | no | | intergenic_variant | MODIFIER |
| 1 | 78029752 | AX-170256949 | 0 | T | C | 0.14 | 4.18E-02 | 2.39E-01 | 2.68E+00 | 2.74E+00 | 8.60E-01 | no | | intergenic_variant | MODIFIER |
| 1 | 78059764 | AX-170469596 | 0 | A | G | 0.14 | 4.18E-02 | 2.39E-01 | 2.68E+00 | 2.74E+00 | 8.60E-01 | no | | intergenic_variant | MODIFIER |
| 1 | 78146666 | AX-170041755 | 0 | G | A | 0.14 | 4.18E-02 | 2.39E-01 | 2.68E+00 | 2.74E+00 | 8.60E-01 | no | Pax3 | intron_variant | MODIFIER |
| 1 | 78163884 | AX-170507918 | 0 | G | A | 0.14 | 4.18E-02 | 2.39E-01 | 2.68E+00 | 2.74E+00 | 8.60E-01 | no | Pax3 | intron_variant | MODIFIER |
| 1 | 78165835 | AX-170282389 | 0 | T | C | 0.14 | 4.18E-02 | 2.39E-01 | 2.68E+00 | 2.74E+00 | 8.60E-01 | no | Pax3 | intron_variant | MODIFIER |
| 1 | 78165871 | AX-170372215 | 0 | A | G | 0.14 | 4.18E-02 | 2.39E-01 | 2.68E+00 | 2.74E+00 | 8.60E-01 | no | Pax3 | intron_variant | MODIFIER |
| 1 | 78170634 | AX-170441095 | 0 | A | G | 0.14 | 4.18E-02 | 2.39E-01 | 2.68E+00 | 2.74E+00 | 8.60E-01 | no | Pax3 | intron_variant | MODIFIER |
| 1 | 78170990 | AX-170506976 | 0 | T | G | 0.14 | 4.18E-02 | 2.39E-01 | 2.68E+00 | 2.74E+00 | 8.60E-01 | no | Pax3 | intron_variant | MODIFIER |
| 1 | 78223818 | AX-170000870 | 0 | T | C | 0.14 | 4.18E-02 | 2.39E-01 | 2.68E+00 | 2.74E+00 | 8.60E-01 | no | | intergenic_variant | MODIFIER |
| 1 | 78230522 | AX-170428559 | 0 | T | G | 0.14 | 4.18E-02 | 2.39E-01 | 2.68E+00 | 2.74E+00 | 8.60E-01 | no | | intergenic_variant | MODIFIER |
| 1 | 78232397 | AX-170205598 | 0 | A | T | 0.14 | 4.18E-02 | 2.39E-01 | 2.68E+00 | 2.74E+00 | 8.60E-01 | no | | regulatory_region_variant | MODIFIER |
| 1 | 78235090 | AX-97857456 | 0 | T | G | 0.14 | 4.18E-02 | 2.39E-01 | 2.68E+00 | 2.74E+00 | 8.60E-01 | no | | intergenic_variant | MODIFIER |
| 1 | 78237396 | AX-170360093 | 0 | G | A | 0.14 | 4.18E-02 | 2.39E-01 | 2.68E+00 | 2.74E+00 | 8.60E-01 | no | | intergenic_variant | MODIFIER |
| 1 | 78237518 | AX-170340347 | 0 | G | A | 0.14 | 4.18E-02 | 2.39E-01 | 2.68E+00 | 2.74E+00 | 8.60E-01 | no | | intergenic_variant | MODIFIER |
| 1 | 78250369 | AX-170000869 | 0 | C | T | 0.14 | 4.18E-02 | 2.39E-01 | 2.68E+00 | 2.74E+00 | 8.60E-01 | no | | intergenic_variant | MODIFIER |
| 1 | 78263300 | AX-170405103 | 0 | T | C | 0.14 | 4.18E-02 | 2.39E-01 | 2.68E+00 | 2.74E+00 | 8.60E-01 | no | | intergenic_variant | MODIFIER |
| 1 | 78294980 | AX-168409860 | 0 | T | C | 0.14 | 4.18E-02 | 2.39E-01 | 2.68E+00 | 2.74E+00 | 8.60E-01 | no | | intergenic_variant | MODIFIER |
| 1 | 78295279 | AX-170000866 | 0 | C | T | 0.14 | 4.18E-02 | 2.39E-01 | 2.68E+00 | 2.74E+00 | 8.60E-01 | no | | intergenic_variant | MODIFIER |
| 1 | 78315951 | AX-170000871 | 0 | A | G | 0.14 | 4.18E-02 | 2.39E-01 | 2.68E+00 | 2.74E+00 | 8.60E-01 | no | Sgpp2 | intron_variant | MODIFIER |
| 1 | 78318493 | AX-170170385 | 0 | A | G | 0.14 | 4.18E-02 | 2.39E-01 | 2.68E+00 | 2.74E+00 | 8.60E-01 | no | Sgpp2 | intron_variant | MODIFIER |
| 1 | 78333494 | AX-170392326 | 0 | A | G | 0.14 | 4.18E-02 | 2.39E-01 | 2.68E+00 | 2.74E+00 | 8.60E-01 | no | Sgpp2 | intron_variant | MODIFIER |

| | | | | | | | | | | | | | | | |
|---|---|---|---|---|---|---|---|---|---|---|---|---|---|---|---|
| 1 | 78335293 | AX-97856088 | 0 | C | T | 0.14 | 4.18E-02 | 2.39E-01 | 2.68E+00 | 2.74E+00 | 8.60E-01 | no | Sgpp2 | intron_variant | MODIFIER |
| 1 | 78349263 | AX-97812123 | 0 | C | T | 0.14 | 4.18E-02 | 2.39E-01 | 2.68E+00 | 2.74E+00 | 8.60E-01 | no | Sgpp2 | intron_variant | MODIFIER |
| 1 | 78359695 | AX-170357026 | 0 | C | G | 0.14 | 4.18E-02 | 2.39E-01 | 2.68E+00 | 2.74E+00 | 8.60E-01 | no | Sgpp2 | intron_variant | MODIFIER |
| 1 | 78363523 | AX-170244459 | 0 | G | A | 0.14 | 4.18E-02 | 2.39E-01 | 2.68E+00 | 2.74E+00 | 8.60E-01 | no | Sgpp2 | intron_variant | MODIFIER |
| 1 | 78376961 | AX-170323516 | 0 | T | G | 0.14 | 4.18E-02 | 2.39E-01 | 2.68E+00 | 2.74E+00 | 8.60E-01 | no | Sgpp2 | intron_variant | MODIFIER |
| 1 | 78392098 | AX-97788611 | 0 | C | T | 0.14 | 4.18E-02 | 2.39E-01 | 2.68E+00 | 2.74E+00 | 8.60E-01 | no | Sgpp2 | intron_variant | MODIFIER |
| 1 | 78392850 | AX-170244872 | 0 | C | T | 0.14 | 4.18E-02 | 2.39E-01 | 2.68E+00 | 2.74E+00 | 8.60E-01 | no | Sgpp2 | intron_variant | MODIFIER |
| 1 | 78399059 | AX-170330419 | 0 | G | T | 0.14 | 4.18E-02 | 2.39E-01 | 2.68E+00 | 2.74E+00 | 8.60E-01 | no | Sgpp2 | intron_variant | MODIFIER |
| 1 | 78399790 | AX-170041758 | 0 | G | A | 0.14 | 4.18E-02 | 2.39E-01 | 2.68E+00 | 2.74E+00 | 8.60E-01 | no | Sgpp2 | intron_variant | MODIFIER |
| 1 | 78401940 | AX-170140287 | 0 | T | C | 0.14 | 4.18E-02 | 2.39E-01 | 2.68E+00 | 2.74E+00 | 8.60E-01 | no | Sgpp2 | intron_variant | MODIFIER |
| 1 | 78401971 | AX-170530037 | 0 | G | A | 0.14 | 4.18E-02 | 2.39E-01 | 2.68E+00 | 2.74E+00 | 8.60E-01 | no | Sgpp2 | intron_variant | MODIFIER |
| 1 | 78402055 | AX-170343974 | 0 | C | T | 0.14 | 4.18E-02 | 2.39E-01 | 2.68E+00 | 2.74E+00 | 8.60E-01 | no | Sgpp2 | intron_variant | MODIFIER |
| 1 | 78403452 | AX-97846603 | 0 | T | C | 0.14 | 4.18E-02 | 2.39E-01 | 2.68E+00 | 2.74E+00 | 8.60E-01 | no | Sgpp2 | intron_variant | MODIFIER |
| 1 | 78420861 | AX-170245186 | 0 | C | G | 0.14 | 4.18E-02 | 2.39E-01 | 2.68E+00 | 2.74E+00 | 8.60E-01 | no | Sgpp2 | downstream_gene_variant | MODIFIER |
| 1 | 78460000 | AX-170420965 | 0 | A | G | 0.14 | 4.18E-02 | 2.39E-01 | 2.68E+00 | 2.74E+00 | 8.60E-01 | no | Farsb | intron_variant | MODIFIER |
| 1 | 78513810 | AX-97812150 | 0 | T | G | 0.14 | 4.18E-02 | 2.39E-01 | 2.68E+00 | 2.74E+00 | 8.60E-01 | no | Mogat1 | intron_variant | MODIFIER |
| 1 | 78513827 | AX-170300162 | 0 | T | C | 0.14 | 4.18E-02 | 2.39E-01 | 2.68E+00 | 2.74E+00 | 8.60E-01 | no | Mogat1 | intron_variant | MODIFIER |
| 1 | 78542646 | AX-170360445 | 0 | A | G | 0.14 | 4.18E-02 | 2.39E-01 | 2.68E+00 | 2.74E+00 | 8.60E-01 | no | Mogat1 | downstream_gene_variant | MODIFIER |
| 1 | 78550546 | AX-170041761 | 0 | T | C | 0.14 | 4.18E-02 | 2.39E-01 | 2.68E+00 | 2.74E+00 | 8.60E-01 | no | | intergenic_variant | MODIFIER |
| 1 | 78550793 | AX-170041760 | 0 | A | C | 0.14 | 4.18E-02 | 2.39E-01 | 2.68E+00 | 2.74E+00 | 8.60E-01 | no | | intergenic_variant | MODIFIER |
| 1 | 78576277 | AX-170304870 | 0 | C | T | 0.14 | 4.18E-02 | 2.39E-01 | 2.68E+00 | 2.74E+00 | 8.60E-01 | no | | intergenic_variant | MODIFIER |
| 1 | 78576802 | AX-170438640 | 0 | T | A | 0.14 | 4.18E-02 | 2.39E-01 | 2.68E+00 | 2.74E+00 | 8.60E-01 | no | | intergenic_variant | MODIFIER |
| 1 | 78584042 | AX-97818264 | 0 | G | A | 0.14 | 4.18E-02 | 2.39E-01 | 2.68E+00 | 2.74E+00 | 8.60E-01 | no | | intergenic_variant | MODIFIER |
| 1 | 78584391 | AX-170335264 | 0 | T | C | 0.14 | 4.18E-02 | 2.39E-01 | 2.68E+00 | 2.74E+00 | 8.60E-01 | no | | intergenic_variant | MODIFIER |
| 1 | 78605181 | AX-170458315 | 0 | G | A | 0.14 | 4.18E-02 | 2.39E-01 | 2.68E+00 | 2.74E+00 | 8.60E-01 | no | 5730419F03Rik | intron_variant,non_coding_transcript_variant | MODIFIER |
| 1 | 78621353 | AX-97799292 | 0 | G | A | 0.14 | 4.18E-02 | 2.39E-01 | 2.68E+00 | 2.74E+00 | 8.60E-01 | no | Gm29611 | downstream_gene_variant | MODIFIER |
| 1 | 78627874 | AX-170372600 | 0 | G | A | 0.14 | 4.18E-02 | 2.39E-01 | 2.68E+00 | 2.74E+00 | 8.60E-01 | no | | intergenic_variant | MODIFIER |
| 1 | 78647578 | AX-170271816 | 0 | A | C | 0.14 | 4.18E-02 | 2.39E-01 | 2.68E+00 | 2.74E+00 | 8.60E-01 | no | | regulatory_region_variant | MODIFIER |
| 1 | 78687715 | AX-97812121 | 0 | A | G | 0.14 | 4.18E-02 | 2.39E-01 | 2.68E+00 | 2.74E+00 | 8.60E-01 | no | Acsl3 | intron_variant | MODIFIER |
| 1 | 78722177 | AX-170436553 | 0 | C | T | 0.14 | 4.18E-02 | 2.39E-01 | 2.68E+00 | 2.74E+00 | 8.60E-01 | no | | regulatory_region_variant | MODIFIER |
| 1 | 78773963 | AX-170000875 | 0 | A | T | 0.14 | 4.18E-02 | 2.39E-01 | 2.68E+00 | 2.74E+00 | 8.60E-01 | no | | intergenic_variant | MODIFIER |
| 1 | 78774937 | AX-170362127 | 0 | T | G | 0.14 | 4.18E-02 | 2.39E-01 | 2.68E+00 | 2.74E+00 | 8.60E-01 | no | | intergenic_variant | MODIFIER |
| 1 | 78775038 | AX-170156849 | 0 | G | A | 0.14 | 4.18E-02 | 2.39E-01 | 2.68E+00 | 2.74E+00 | 8.60E-01 | no | | intergenic_variant | MODIFIER |
| 12 | 59699284 | AX-170506604 | 0 | C | T | 0.053 | 4.20E-02 | 2.50E-01 | 2.70E+00 | 2.77E+00 | 8.60E-01 | no | | intergenic_variant | MODIFIER |
| 12 | 61459333 | AX-170323828 | 0 | G | T | 0.053 | 4.20E-02 | 2.50E-01 | 2.70E+00 | 2.77E+00 | 8.60E-01 | no | Nanog-ps2 | downstream_gene_variant | MODIFIER |
| 1 | 74973257 | AX-97847305 | 1 | G | A | 0.143 | 4.19E-02 | 2.39E-01 | 2.68E+00 | 2.73E+00 | 8.60E-01 | no | Nhej1 | intron_variant | MODIFIER |
| 20 | 56959736 | AX-170211329 | 0 | C | T | 0.246 | 1.12E-02 | 6.35E-02 | 2.64E+00 | 2.73E+00 | 8.60E-01 | no | Adgrg4 | upstream_gene_variant | MODIFIER |
| 17 | 80592047 | AX-170475120 | 0 | T | C | 0.053 | -2.44E-02 | 1.43E-01 | 2.63E+00 | 2.75E+00 | 8.60E-01 | no | Map4k3 | intron_variant | MODIFIER |
| 10 | 117918176 | AX-97827295 | 1 | T | A | 0.348 | 1.26E-02 | 7.38E-02 | 2.66E+00 | 2.75E+00 | 8.60E-01 | no | | regulatory_region_variant | MODIFIER |
| 1 | 78486797 | AX-170232184 | 1 | C | T | 0.143 | 4.18E-02 | 2.38E-01 | 2.68E+00 | 2.73E+00 | 8.60E-01 | no | Farsb | intron_variant | MODIFIER |
| 9 | 32901686 | AX-170246413 | 0 | G | A | 0.439 | 1.52E-02 | 8.53E-02 | 2.65E+00 | 2.72E+00 | 8.60E-01 | no | Gm27162 | downstream_gene_variant | MODIFIER |
| 16 | 94337370 | AX-170194595 | 1 | G | C | 0.42 | -1.48E-02 | 8.01E-02 | 2.61E+00 | 2.71E+00 | 8.60E-01 | no | Ripply3 | downstream_gene_variant | MODIFIER |
| 9 | 33301507 | AX-170252061 | 1 | T | C | 0.455 | 1.47E-02 | 8.14E-02 | 2.64E+00 | 2.72E+00 | 8.60E-01 | no | Gm47775 | intron_variant,non_coding_transcript_variant | MODIFIER |
| 10 | 4845114 | AX-168412563 | 1 | T | G | 0.205 | 1.64E-02 | 9.67E-02 | 2.64E+00 | 2.75E+00 | 8.60E-01 | no | Esr1 | intron_variant | MODIFIER |
| 9 | 34744446 | AX-170141907 | 0 | A | T | 0.421 | 1.52E-02 | 8.46E-02 | 2.64E+00 | 2.72E+00 | 8.60E-01 | no | Kirrel3 | intron_variant | MODIFIER |
| 5 | 36094626 | AX-170219931 | 0 | A | G | 0.325 | 1.91E-02 | 1.08E-01 | 2.66E+00 | 2.73E+00 | 8.61E-01 | no | Sorcs2 | intron_variant | MODIFIER |
| 18 | 66658519 | AX-170289464 | 0 | C | G | 0.088 | -2.84E-02 | 1.73E-01 | 2.65E+00 | 2.76E+00 | 8.61E-01 | no | AC133607.2 | intron_variant,non_coding_transcript_variant | MODIFIER |
| 10 | 56177641 | AX-170185883 | 1 | T | C | 0.295 | 1.26E-02 | 8.00E-02 | 2.69E+00 | 2.80E+00 | 8.61E-01 | no | Tbc1d32 | intron_variant | MODIFIER |
| 10 | 57811607 | AX-170358896 | 1 | C | A | 0.295 | 1.26E-02 | 8.00E-02 | 2.69E+00 | 2.80E+00 | 8.61E-01 | no | Smpdl3a | 3_prime_UTR_variant | MODIFIER |
| 1 | 30642336 | AX-97811235 | 1 | A | G | 0.42 | 1.42E-02 | 8.17E-02 | 2.67E+00 | 2.74E+00 | 8.61E-01 | no | | intergenic_variant | MODIFIER |
| 1 | 41512185 | AX-170097053 | 0 | T | C | 0.421 | -1.28E-02 | 7.37E-02 | 2.65E+00 | 2.74E+00 | 8.61E-01 | no | | intergenic_variant | MODIFIER |
| 10 | 117812355 | AX-170115574 | 0 | T | A | 0.342 | 1.26E-02 | 7.48E-02 | 2.67E+00 | 2.75E+00 | 8.61E-01 | no | Rap1b | downstream_gene_variant | MODIFIER |
| 10 | 117825438 | AX-170321668 | 0 | C | T | 0.342 | 1.26E-02 | 7.48E-02 | 2.67E+00 | 2.75E+00 | 8.61E-01 | no | Rap1b | intron_variant | MODIFIER |
| 10 | 117925322 | AX-170003998 | 0 | G | A | 0.342 | 1.26E-02 | 7.48E-02 | 2.67E+00 | 2.75E+00 | 8.61E-01 | no | 4933411E08Rik | upstream_gene_variant | MODIFIER |

| | | | | | | | | | | | | | | | |
|---|---|---|---|---|---|---|---|---|---|---|---|---|---|---|---|
| 10 | 117936857 | AX-170003993 | 0 | A | T | 0.342 | 1.26E-02 | 7.48E-02 | 2.67E+00 | 2.75E+00 | 8.61E-01 | no | 4933411E08Rik | intron_variant,non_coding_transcript_variant | MODIFIER |
| 18 | 86815166 | AX-170428141 | 0 | A | C | 0.167 | -1.99E-02 | 1.03E-01 | 2.58E+00 | 2.69E+00 | 8.61E-01 | no | | intergenic_variant | MODIFIER |
| 7 | 96572757 | AX-170267367 | 0 | C | T | 0.123 | 3.36E-02 | 1.83E-01 | 2.61E+00 | 2.71E+00 | 8.61E-01 | no | Tenm4 | intron_variant | MODIFIER |
| 13 | 32016712 | AX-170008029 | 0 | G | A | 0.184 | -2.15E-02 | 1.16E-01 | 2.63E+00 | 2.70E+00 | 8.61E-01 | no | Gmds | intron_variant | MODIFIER |
| 4 | 142195568 | AX-170427188 | 1 | A | G | 0.089 | 3.01E-02 | 1.73E-01 | 2.66E+00 | 2.73E+00 | 8.61E-01 | no | Kazn | intron_variant | MODIFIER |
| 4 | 143278003 | AX-170227414 | 1 | G | A | 0.089 | 3.01E-02 | 1.73E-01 | 2.66E+00 | 2.73E+00 | 8.61E-01 | no | Pdpn | intron_variant | MODIFIER |
| 4 | 147461196 | AX-170070973 | 1 | T | C | 0.089 | 3.01E-02 | 1.73E-01 | 2.66E+00 | 2.73E+00 | 8.61E-01 | no | Gm26624 | downstream_gene_variant | MODIFIER |
| 15 | 92168850 | AX-170200959 | 0 | C | A | 0.14 | 3.21E-02 | 1.79E-01 | 2.65E+00 | 2.71E+00 | 8.61E-01 | no | Cntn1 | intron_variant | MODIFIER |
| 9 | 69577432 | AX-170511805 | 1 | G | T | 0.196 | 2.17E-02 | 1.28E-01 | 2.72E+00 | 2.77E+00 | 8.61E-01 | no | | intergenic_variant | MODIFIER |
| 1 | 41235338 | AX-170177098 | 1 | T | C | 0.411 | -1.32E-02 | 7.50E-02 | 2.64E+00 | 2.73E+00 | 8.61E-01 | no | | intergenic_variant | MODIFIER |
| 9 | 59232596 | AX-170032469 | 1 | C | A | 0.214 | -2.05E-02 | 1.21E-01 | 2.70E+00 | 2.76E+00 | 8.61E-01 | no | Gm33053 | intron_variant,non_coding_transcript_variant | MODIFIER |
| 2 | 155183460 | AX-97849143 | 1 | C | T | 0.143 | 3.84E-02 | 2.20E-01 | 2.69E+00 | 2.72E+00 | 8.61E-01 | no | Itch | intron_variant | MODIFIER |
| 9 | 71437503 | AX-170276205 | 0 | G | A | 0.211 | 2.21E-02 | 1.30E-01 | 2.72E+00 | 2.77E+00 | 8.61E-01 | no | | intergenic_variant | MODIFIER |
| 11 | 46424089 | AX-170100762 | 0 | G | A | 0.088 | -3.26E-02 | 1.82E-01 | 2.62E+00 | 2.72E+00 | 8.62E-01 | no | | intergenic_variant | MODIFIER |
| 4 | 149523977 | AX-168386485 | 0 | T | C | 0.096 | -2.93E-02 | 1.69E-01 | 2.64E+00 | 2.73E+00 | 8.62E-01 | no | Ctnnbip1 | intron_variant | MODIFIER |
| 20 | 66887719 | AX-97852857 | 1 | T | C | 0.232 | 1.11E-02 | 6.28E-02 | 2.63E+00 | 2.73E+00 | 8.62E-01 | no | | intergenic_variant | MODIFIER |
| 5 | 43181434 | AX-170071916 | 0 | G | C | 0.211 | -1.92E-02 | 1.04E-01 | 2.61E+00 | 2.71E+00 | 8.62E-01 | no | Gm7854 | intron_variant,non_coding_transcript_variant | MODIFIER |
| 15 | 79916121 | AX-170012070 | 1 | A | G | 0.554 | 1.40E-02 | 7.77E-02 | 2.62E+00 | 2.71E+00 | 8.62E-01 | no | Cbx7 | 3_prime_UTR_variant | MODIFIER |
| 8 | 93781141 | AX-170458728 | 0 | C | T | 0.219 | -1.84E-02 | 1.09E-01 | 2.66E+00 | 2.75E+00 | 8.62E-01 | no | 4930488L21Rik | intron_variant,non_coding_transcript_variant | MODIFIER |
| 12 | 111506803 | AX-170107448 | 0 | G | A | 0.14 | -3.24E-02 | 1.91E-01 | 2.67E+00 | 2.75E+00 | 8.62E-01 | no | | intergenic_variant | MODIFIER |
| 5 | 99692740 | AX-170184859 | 0 | T | C | 0.421 | 1.78E-02 | 1.01E-01 | 2.65E+00 | 2.72E+00 | 8.62E-01 | no | Rasgef1b | intron_variant,non_coding_transcript_variant | MODIFIER |
| 1 | 53896238 | AX-170516306 | 0 | C | A | 0.105 | -3.31E-02 | 1.84E-01 | 2.62E+00 | 2.71E+00 | 8.62E-01 | no | Hecw2 | intron_variant | MODIFIER |
| 1 | 36971069 | AX-170122476 | 1 | G | A | 0.455 | 1.33E-02 | 7.90E-02 | 2.68E+00 | 2.75E+00 | 8.62E-01 | no | | intergenic_variant | MODIFIER |
| 10 | 31025668 | AX-170256388 | 0 | C | T | 0.404 | -1.34E-02 | 7.54E-02 | 2.61E+00 | 2.72E+00 | 8.62E-01 | no | Gm30676 | intron_variant,non_coding_transcript_variant | MODIFIER |
| 3 | 147102668 | AX-170121869 | 0 | A | G | 0.351 | 1.28E-02 | 7.30E-02 | 2.65E+00 | 2.72E+00 | 8.62E-01 | no | Gm29771 | intron_variant,non_coding_transcript_variant | MODIFIER |
| 2 | 118565271 | AX-170018816 | 1 | A | G | 0.134 | 2.79E-02 | 1.66E-01 | 2.67E+00 | 2.75E+00 | 8.62E-01 | no | | intergenic_variant | MODIFIER |
| 9 | 57814622 | AX-170082517 | 0 | C | T | 0.219 | -2.07E-02 | 1.21E-01 | 2.68E+00 | 2.74E+00 | 8.62E-01 | no | Arid3b | intron_variant | MODIFIER |
| 1 | 164815313 | AX-170188686 | 0 | G | A | 0.219 | 2.13E-02 | 1.17E-01 | 2.61E+00 | 2.71E+00 | 8.62E-01 | no | Dpt | intron_variant | MODIFIER |
| 8 | 73748774 | AX-97798220 | 0 | G | A | 0.202 | 2.05E-02 | 1.24E-01 | 2.67E+00 | 2.76E+00 | 8.62E-01 | no | Crry-ps | downstream_gene_variant | MODIFIER |
| 12 | 104016798 | AX-170337655 | 1 | G | A | 0.134 | -3.28E-02 | 1.94E-01 | 2.68E+00 | 2.75E+00 | 8.62E-01 | no | Serpina9 | upstream_gene_variant | MODIFIER |
| 12 | 104305114 | AX-170050038 | 1 | G | A | 0.134 | -3.28E-02 | 1.94E-01 | 2.68E+00 | 2.75E+00 | 8.62E-01 | no | | intergenic_variant | MODIFIER |
| 13 | 46629834 | AX-170100632 | 0 | T | C | 0.474 | -1.02E-02 | 6.29E-02 | 2.66E+00 | 2.77E+00 | 8.62E-01 | no | Cap2 | intron_variant | MODIFIER |
| 2 | 155923026 | AX-170019446 | 2 | C | T | 0.145 | 3.83E-02 | 2.21E-01 | 2.69E+00 | 2.72E+00 | 8.63E-01 | no | Uqcc1 | intron_variant | MODIFIER |
| 16 | 93268595 | AX-170392334 | 0 | G | A | 0.386 | 1.25E-02 | 7.87E-02 | 2.68E+00 | 2.78E+00 | 8.63E-01 | no | | intergenic_variant | MODIFIER |
| 15 | 99050270 | AX-170012367 | 1 | G | A | 0.107 | 2.50E-02 | 1.43E-01 | 2.64E+00 | 2.73E+00 | 8.63E-01 | no | Prph | upstream_gene_variant | MODIFIER |
| 9 | 34859608 | AX-170337169 | 0 | G | C | 0.43 | 1.53E-02 | 8.66E-02 | 2.65E+00 | 2.72E+00 | 8.63E-01 | no | Kirrel3 | intron_variant | MODIFIER |
| 3 | 147750878 | AX-170525563 | 1 | C | A | 0.348 | 1.30E-02 | 7.45E-02 | 2.65E+00 | 2.72E+00 | 8.63E-01 | no | | regulatory_region_variant | MODIFIER |
| 12 | 69524233 | AX-170166979 | 0 | G | C | 0.07 | -3.97E-02 | 2.27E-01 | 2.62E+00 | 2.73E+00 | 8.63E-01 | no | | regulatory_region_variant | MODIFIER |
| 4 | 149526253 | AX-170246297 | 1 | G | A | 0.089 | 2.91E-02 | 1.69E-01 | 2.66E+00 | 2.73E+00 | 8.63E-01 | no | Ctnnbip1 | intron_variant | MODIFIER |
| 4 | 151258400 | AX-170481693 | 1 | G | T | 0.089 | 2.91E-02 | 1.69E-01 | 2.66E+00 | 2.73E+00 | 8.63E-01 | no | Camta1 | intron_variant | MODIFIER |
| 8 | 42323887 | AX-97784546 | 1 | A | G | 0.152 | -2.27E-02 | 1.33E-01 | 2.65E+00 | 2.74E+00 | 8.63E-01 | no | | intergenic_variant | MODIFIER |
| 9 | 77263596 | AX-170310019 | 1 | C | T | 0.205 | 2.22E-02 | 1.32E-01 | 2.72E+00 | 2.77E+00 | 8.63E-01 | no | Mlip | intron_variant | MODIFIER |
| 9 | 27922454 | AX-170175989 | 0 | G | T | 0.456 | 1.62E-02 | 9.03E-02 | 2.63E+00 | 2.71E+00 | 8.63E-01 | no | Opcml | intron_variant | MODIFIER |
| 11 | 32921195 | AX-170090882 | 0 | C | T | 0.105 | 3.31E-02 | 1.89E-01 | 2.65E+00 | 2.72E+00 | 8.63E-01 | no | Gm12111 | intron_variant,non_coding_transcript_variant | MODIFIER |
| 11 | 32928887 | AX-170392463 | 0 | C | T | 0.105 | 3.31E-02 | 1.89E-01 | 2.65E+00 | 2.72E+00 | 8.63E-01 | no | Gm12111 | upstream_gene_variant | MODIFIER |
| 11 | 32930554 | AX-170222077 | 0 | G | A | 0.105 | 3.31E-02 | 1.89E-01 | 2.65E+00 | 2.72E+00 | 8.63E-01 | no | Gm12111 | upstream_gene_variant | MODIFIER |
| 11 | 32960938 | AX-170124300 | 0 | G | A | 0.105 | 3.31E-02 | 1.89E-01 | 2.65E+00 | 2.72E+00 | 8.63E-01 | no | | intergenic_variant | MODIFIER |
| 11 | 32978625 | AX-97853554 | 0 | C | T | 0.105 | 3.31E-02 | 1.89E-01 | 2.65E+00 | 2.72E+00 | 8.63E-01 | no | | intergenic_variant | MODIFIER |
| 11 | 32982848 | AX-170179621 | 0 | T | A | 0.105 | 3.31E-02 | 1.89E-01 | 2.65E+00 | 2.72E+00 | 8.63E-01 | no | Gm26052 | downstream_gene_variant | MODIFIER |
| 11 | 32990398 | AX-170166697 | 0 | A | G | 0.105 | 3.31E-02 | 1.89E-01 | 2.65E+00 | 2.72E+00 | 8.63E-01 | no | | intergenic_variant | MODIFIER |
| 11 | 32994257 | AX-170123518 | 0 | C | G | 0.105 | 3.31E-02 | 1.89E-01 | 2.65E+00 | 2.72E+00 | 8.63E-01 | no | | intergenic_variant | MODIFIER |
| 11 | 33000799 | AX-97817680 | 0 | C | A | 0.105 | 3.31E-02 | 1.89E-01 | 2.65E+00 | 2.72E+00 | 8.63E-01 | no | | intergenic_variant | MODIFIER |
| 11 | 33001242 | AX-170046733 | 0 | T | C | 0.105 | 3.31E-02 | 1.89E-01 | 2.65E+00 | 2.72E+00 | 8.63E-01 | no | | intergenic_variant | MODIFIER |
| 11 | 33001424 | AX-170370790 | 0 | C | G | 0.105 | 3.31E-02 | 1.89E-01 | 2.65E+00 | 2.72E+00 | 8.63E-01 | no | | intergenic_variant | MODIFIER |

| | | | | | | | | | | | | | | | |
|---|---|---|---|---|---|---|---|---|---|---|---|---|---|---|---|
| 11 | 33021572 | AX-170046734 | 0 | A | G | 0.105 | 3.31E-02 | 1.89E-01 | 2.65E+00 | 2.72E+00 | 8.63E-01 | no | | intergenic_variant | MODIFIER |
| 11 | 33048082 | AX-170284218 | 0 | A | G | 0.105 | 3.31E-02 | 1.89E-01 | 2.65E+00 | 2.72E+00 | 8.63E-01 | no | | intergenic_variant | MODIFIER |
| 11 | 33050825 | AX-170098186 | 0 | G | C | 0.105 | 3.31E-02 | 1.89E-01 | 2.65E+00 | 2.72E+00 | 8.63E-01 | no | | intergenic_variant | MODIFIER |
| 11 | 33055926 | AX-170251786 | 0 | G | C | 0.105 | 3.31E-02 | 1.89E-01 | 2.65E+00 | 2.72E+00 | 8.63E-01 | no | | intergenic_variant | MODIFIER |
| 11 | 33057801 | AX-170490463 | 0 | G | T | 0.105 | 3.31E-02 | 1.89E-01 | 2.65E+00 | 2.72E+00 | 8.63E-01 | no | | intergenic_variant | MODIFIER |
| 11 | 33070397 | AX-170157963 | 0 | G | A | 0.105 | 3.31E-02 | 1.89E-01 | 2.65E+00 | 2.72E+00 | 8.63E-01 | no | Gm12114 | intron_variant,non_coding_transcript_variant | MODIFIER |
| 11 | 33070487 | AX-170320784 | 0 | A | T | 0.105 | 3.31E-02 | 1.89E-01 | 2.65E+00 | 2.72E+00 | 8.63E-01 | no | Gm12114 | intron_variant,non_coding_transcript_variant | MODIFIER |
| 11 | 33072632 | AX-170328332 | 0 | C | A | 0.105 | 3.31E-02 | 1.89E-01 | 2.65E+00 | 2.72E+00 | 8.63E-01 | no | Gm12114 | intron_variant,non_coding_transcript_variant | MODIFIER |
| 11 | 33072650 | AX-170004661 | 0 | C | T | 0.105 | 3.31E-02 | 1.89E-01 | 2.65E+00 | 2.72E+00 | 8.63E-01 | no | Gm12114 | intron_variant,non_coding_transcript_variant | MODIFIER |
| 11 | 33109868 | AX-170036463 | 0 | G | C | 0.105 | 3.31E-02 | 1.89E-01 | 2.65E+00 | 2.72E+00 | 8.63E-01 | no | | intergenic_variant | MODIFIER |
| 11 | 33111681 | AX-170451657 | 0 | A | G | 0.105 | 3.31E-02 | 1.89E-01 | 2.65E+00 | 2.72E+00 | 8.63E-01 | no | | intergenic_variant | MODIFIER |
| 11 | 33112276 | AX-97816907 | 0 | C | T | 0.105 | 3.31E-02 | 1.89E-01 | 2.65E+00 | 2.72E+00 | 8.63E-01 | no | | regulatory_region_variant | MODIFIER |
| 11 | 33116460 | AX-170240046 | 0 | C | T | 0.105 | 3.31E-02 | 1.89E-01 | 2.65E+00 | 2.72E+00 | 8.63E-01 | no | Fgf18 | downstream_gene_variant | MODIFIER |
| 11 | 33120984 | AX-170004659 | 0 | C | A | 0.105 | 3.31E-02 | 1.89E-01 | 2.65E+00 | 2.72E+00 | 8.63E-01 | no | Fgf18 | intron_variant | MODIFIER |
| 11 | 33124469 | AX-170004657 | 0 | G | A | 0.105 | 3.31E-02 | 1.89E-01 | 2.65E+00 | 2.72E+00 | 8.63E-01 | no | Fgf18 | intron_variant | MODIFIER |
| 11 | 33134249 | AX-170272427 | 0 | A | G | 0.105 | 3.31E-02 | 1.89E-01 | 2.65E+00 | 2.72E+00 | 8.63E-01 | no | Fgf18 | synonymous_variant | LOW |
| 11 | 33140031 | AX-170004656 | 0 | T | C | 0.105 | 3.31E-02 | 1.89E-01 | 2.65E+00 | 2.72E+00 | 8.63E-01 | no | Fgf18 | intron_variant | MODIFIER |
| 11 | 33140899 | AX-97802010 | 0 | A | G | 0.105 | 3.31E-02 | 1.89E-01 | 2.65E+00 | 2.72E+00 | 8.63E-01 | no | Fgf18 | intron_variant | MODIFIER |
| 11 | 33167831 | AX-97856301 | 0 | A | G | 0.105 | 3.31E-02 | 1.89E-01 | 2.65E+00 | 2.72E+00 | 8.63E-01 | no | Npm1 | upstream_gene_variant | MODIFIER |
| 11 | 33178928 | AX-170174816 | 0 | T | A | 0.105 | 3.31E-02 | 1.89E-01 | 2.65E+00 | 2.72E+00 | 8.63E-01 | no | | intergenic_variant | MODIFIER |
| 11 | 33195992 | AX-97809380 | 0 | T | C | 0.105 | 3.31E-02 | 1.89E-01 | 2.65E+00 | 2.72E+00 | 8.63E-01 | no | Tlx3 | downstream_gene_variant | MODIFIER |
| 11 | 33196209 | AX-170121721 | 0 | G | C | 0.105 | 3.31E-02 | 1.89E-01 | 2.65E+00 | 2.72E+00 | 8.63E-01 | no | Tlx3 | downstream_gene_variant | MODIFIER |
| 11 | 33198849 | AX-170349850 | 0 | A | G | 0.105 | 3.31E-02 | 1.89E-01 | 2.65E+00 | 2.72E+00 | 8.63E-01 | no | Tlx3 | downstream_gene_variant | MODIFIER |
| 11 | 33217751 | AX-170521443 | 0 | A | G | 0.105 | 3.31E-02 | 1.89E-01 | 2.65E+00 | 2.72E+00 | 8.63E-01 | no | Ranbp17 | intron_variant | MODIFIER |
| 11 | 33218087 | AX-170493727 | 0 | G | A | 0.105 | 3.31E-02 | 1.89E-01 | 2.65E+00 | 2.72E+00 | 8.63E-01 | no | Ranbp17 | intron_variant | MODIFIER |
| 11 | 33242131 | AX-97802708 | 0 | G | A | 0.105 | 3.31E-02 | 1.89E-01 | 2.65E+00 | 2.72E+00 | 8.63E-01 | no | Ranbp17 | intron_variant | MODIFIER |
| 11 | 33264834 | AX-97848274 | 0 | C | T | 0.105 | 3.31E-02 | 1.89E-01 | 2.65E+00 | 2.72E+00 | 8.63E-01 | no | Ranbp17 | intron_variant | MODIFIER |
| 11 | 33265121 | AX-170355620 | 0 | G | A | 0.105 | 3.31E-02 | 1.89E-01 | 2.65E+00 | 2.72E+00 | 8.63E-01 | no | Ranbp17 | intron_variant | MODIFIER |
| 11 | 33317129 | AX-170046739 | 0 | C | T | 0.105 | 3.31E-02 | 1.89E-01 | 2.65E+00 | 2.72E+00 | 8.63E-01 | no | Ranbp17 | intron_variant | MODIFIER |
| 11 | 33331676 | AX-170046740 | 0 | G | A | 0.105 | 3.31E-02 | 1.89E-01 | 2.65E+00 | 2.72E+00 | 8.63E-01 | no | Ranbp17 | intron_variant | MODIFIER |
| 11 | 33332017 | AX-170046735 | 0 | G | A | 0.105 | 3.31E-02 | 1.89E-01 | 2.65E+00 | 2.72E+00 | 8.63E-01 | no | Ranbp17 | intron_variant | MODIFIER |
| 11 | 33348725 | AX-97780389 | 0 | A | G | 0.105 | 3.31E-02 | 1.89E-01 | 2.65E+00 | 2.72E+00 | 8.63E-01 | no | Ranbp17 | intron_variant | MODIFIER |
| 11 | 33356820 | AX-170137792 | 0 | C | T | 0.105 | 3.31E-02 | 1.89E-01 | 2.65E+00 | 2.72E+00 | 8.63E-01 | no | Ranbp17 | intron_variant | MODIFIER |
| 11 | 33383785 | AX-170036456 | 0 | G | A | 0.105 | 3.31E-02 | 1.89E-01 | 2.65E+00 | 2.72E+00 | 8.63E-01 | no | Ranbp17 | intron_variant | MODIFIER |
| 11 | 33394158 | AX-97793013 | 0 | G | A | 0.105 | 3.31E-02 | 1.89E-01 | 2.65E+00 | 2.72E+00 | 8.63E-01 | no | Ranbp17 | intron_variant | MODIFIER |
| 11 | 33415245 | AX-97852945 | 0 | C | T | 0.105 | 3.31E-02 | 1.89E-01 | 2.65E+00 | 2.72E+00 | 8.63E-01 | no | Ranbp17 | intron_variant | MODIFIER |
| 11 | 33421844 | AX-170258361 | 0 | A | G | 0.105 | 3.31E-02 | 1.89E-01 | 2.65E+00 | 2.72E+00 | 8.63E-01 | no | Ranbp17 | intron_variant | MODIFIER |
| 11 | 33455345 | AX-170231341 | 0 | C | T | 0.105 | 3.31E-02 | 1.89E-01 | 2.65E+00 | 2.72E+00 | 8.63E-01 | no | Ranbp17 | intron_variant | MODIFIER |
| 11 | 33459372 | AX-170117917 | 0 | A | T | 0.105 | 3.31E-02 | 1.89E-01 | 2.65E+00 | 2.72E+00 | 8.63E-01 | no | Ranbp17 | intron_variant | MODIFIER |
| 11 | 33480600 | AX-97828448 | 0 | C | A | 0.105 | 3.31E-02 | 1.89E-01 | 2.65E+00 | 2.72E+00 | 8.63E-01 | no | Ranbp17 | intron_variant | MODIFIER |
| 11 | 33480777 | AX-170345400 | 0 | A | G | 0.105 | 3.31E-02 | 1.89E-01 | 2.65E+00 | 2.72E+00 | 8.63E-01 | no | Ranbp17 | intron_variant | MODIFIER |
| 11 | 33484475 | AX-170535230 | 0 | A | G | 0.105 | 3.31E-02 | 1.89E-01 | 2.65E+00 | 2.72E+00 | 8.63E-01 | no | Ranbp17 | intron_variant | MODIFIER |
| 11 | 33487895 | AX-170118936 | 0 | A | G | 0.105 | 3.31E-02 | 1.89E-01 | 2.65E+00 | 2.72E+00 | 8.63E-01 | no | Ranbp17 | intron_variant | MODIFIER |
| 11 | 33511728 | AX-170397383 | 0 | T | C | 0.105 | 3.31E-02 | 1.89E-01 | 2.65E+00 | 2.72E+00 | 8.63E-01 | no | Ranbp17 | intron_variant | MODIFIER |
| 11 | 33514168 | AX-170254980 | 0 | G | A | 0.105 | 3.31E-02 | 1.89E-01 | 2.65E+00 | 2.72E+00 | 8.63E-01 | no | Ranbp17 | upstream_gene_variant | MODIFIER |
| 11 | 33525214 | AX-97855233 | 0 | T | C | 0.105 | 3.31E-02 | 1.89E-01 | 2.65E+00 | 2.72E+00 | 8.63E-01 | no | | regulatory_region_variant | MODIFIER |
| 11 | 33531079 | AX-170432170 | 0 | A | G | 0.105 | 3.31E-02 | 1.89E-01 | 2.65E+00 | 2.72E+00 | 8.63E-01 | no | | intergenic_variant | MODIFIER |
| 11 | 33541379 | AX-170450930 | 0 | C | A | 0.105 | 3.31E-02 | 1.89E-01 | 2.65E+00 | 2.72E+00 | 8.63E-01 | no | | intergenic_variant | MODIFIER |
| 11 | 33553648 | AX-170450751 | 0 | T | C | 0.105 | 3.31E-02 | 1.89E-01 | 2.65E+00 | 2.72E+00 | 8.63E-01 | no | Gabrp | intron_variant | MODIFIER |
| 11 | 33563520 | AX-170004669 | 0 | T | C | 0.105 | 3.31E-02 | 1.89E-01 | 2.65E+00 | 2.72E+00 | 8.63E-01 | no | Gabrp | intron_variant | MODIFIER |
| 11 | 33565766 | AX-170004662 | 0 | C | G | 0.105 | 3.31E-02 | 1.89E-01 | 2.65E+00 | 2.72E+00 | 8.63E-01 | no | Gabrp | intron_variant | MODIFIER |
| 11 | 33570755 | AX-170178681 | 0 | A | G | 0.105 | 3.31E-02 | 1.89E-01 | 2.65E+00 | 2.72E+00 | 8.63E-01 | no | Gabrp | intron_variant | MODIFIER |
| 11 | 33577192 | AX-170467216 | 0 | C | T | 0.105 | 3.31E-02 | 1.89E-01 | 2.65E+00 | 2.72E+00 | 8.63E-01 | no | Gabrp | intron_variant | MODIFIER |
| 11 | 33606539 | AX-170046743 | 0 | A | G | 0.105 | 3.31E-02 | 1.89E-01 | 2.65E+00 | 2.72E+00 | 8.63E-01 | no | D130052B06Rik | intron_variant | MODIFIER |

| | | | | | | | | | | | | | | | |
|---|---|---|---|---|---|---|---|---|---|---|---|---|---|---|---|
| 11 | 33607105 | AX-97818151 | 0 | G | C | 0.105 | 3.31E-02 | 1.89E-01 | 2.65E+00 | 2.72E+00 | 8.63E-01 | no | D130052B06Rik | intron_variant | MODIFIER |
| 11 | 33615832 | AX-170386790 | 0 | T | C | 0.105 | 3.31E-02 | 1.89E-01 | 2.65E+00 | 2.72E+00 | 8.63E-01 | no | D130052B06Rik | intron_variant | MODIFIER |
| 11 | 33627466 | AX-170436390 | 0 | T | C | 0.105 | 3.31E-02 | 1.89E-01 | 2.65E+00 | 2.72E+00 | 8.63E-01 | no | Kcnip1 | downstream_gene_variant | MODIFIER |
| 11 | 33627793 | AX-170004664 | 0 | A | G | 0.105 | 3.31E-02 | 1.89E-01 | 2.65E+00 | 2.72E+00 | 8.63E-01 | no | Kcnip1 | downstream_gene_variant | MODIFIER |
| 11 | 33628121 | AX-170004665 | 0 | C | G | 0.105 | 3.31E-02 | 1.89E-01 | 2.65E+00 | 2.72E+00 | 8.63E-01 | no | Kcnip1 | downstream_gene_variant | MODIFIER |
| 11 | 33638252 | AX-170004666 | 0 | A | G | 0.105 | 3.31E-02 | 1.89E-01 | 2.65E+00 | 2.72E+00 | 8.63E-01 | no | Kcnip1 | intron_variant | MODIFIER |
| 11 | 33642745 | AX-170286496 | 0 | A | G | 0.105 | 3.31E-02 | 1.89E-01 | 2.65E+00 | 2.72E+00 | 8.63E-01 | no | Kcnip1 | intron_variant | MODIFIER |
| 11 | 33680651 | AX-170004667 | 0 | C | T | 0.105 | 3.31E-02 | 1.89E-01 | 2.65E+00 | 2.72E+00 | 8.63E-01 | no | Kcnip1 | intron_variant | MODIFIER |
| 11 | 33684428 | AX-170158389 | 0 | G | A | 0.105 | 3.31E-02 | 1.89E-01 | 2.65E+00 | 2.72E+00 | 8.63E-01 | no | Kcnip1 | intron_variant | MODIFIER |
| 11 | 33685637 | AX-97783628 | 0 | A | G | 0.105 | 3.31E-02 | 1.89E-01 | 2.65E+00 | 2.72E+00 | 8.63E-01 | no | Kcnip1 | intron_variant | MODIFIER |
| 11 | 33703655 | AX-97817989 | 0 | G | A | 0.105 | 3.31E-02 | 1.89E-01 | 2.65E+00 | 2.72E+00 | 8.63E-01 | no | Kcnip1 | intron_variant | MODIFIER |
| 11 | 33721240 | AX-97857458 | 0 | C | T | 0.105 | 3.31E-02 | 1.89E-01 | 2.65E+00 | 2.72E+00 | 8.63E-01 | no | Kcnip1 | intron_variant | MODIFIER |
| 11 | 33721650 | AX-170526654 | 0 | T | C | 0.105 | 3.31E-02 | 1.89E-01 | 2.65E+00 | 2.72E+00 | 8.63E-01 | no | Kcnip1 | intron_variant | MODIFIER |
| 11 | 33729549 | AX-168408939 | 0 | G | A | 0.105 | 3.31E-02 | 1.89E-01 | 2.65E+00 | 2.72E+00 | 8.63E-01 | no | Kcnip1 | intron_variant | MODIFIER |
| 11 | 33749999 | AX-170451637 | 0 | T | A | 0.105 | 3.31E-02 | 1.89E-01 | 2.65E+00 | 2.72E+00 | 8.63E-01 | no | Kcnip1 | intron_variant | MODIFIER |
| 11 | 33752249 | AX-170270508 | 0 | G | A | 0.105 | 3.31E-02 | 1.89E-01 | 2.65E+00 | 2.72E+00 | 8.63E-01 | no | Kcnip1 | intron_variant | MODIFIER |
| 11 | 33755473 | AX-170490761 | 0 | T | A | 0.105 | 3.31E-02 | 1.89E-01 | 2.65E+00 | 2.72E+00 | 8.63E-01 | no | Kcnip1 | intron_variant | MODIFIER |
| 11 | 33756100 | AX-97855241 | 0 | T | C | 0.105 | 3.31E-02 | 1.89E-01 | 2.65E+00 | 2.72E+00 | 8.63E-01 | no | Kcnip1 | intron_variant | MODIFIER |
| 11 | 33784894 | AX-97781748 | 0 | G | C | 0.105 | 3.31E-02 | 1.89E-01 | 2.65E+00 | 2.72E+00 | 8.63E-01 | no | Kcnip1 | intron_variant | MODIFIER |
| 11 | 33784930 | AX-170036458 | 0 | A | C | 0.105 | 3.31E-02 | 1.89E-01 | 2.65E+00 | 2.72E+00 | 8.63E-01 | no | Kcnip1 | intron_variant | MODIFIER |
| 11 | 33814962 | AX-170292029 | 0 | T | C | 0.105 | 3.31E-02 | 1.89E-01 | 2.65E+00 | 2.72E+00 | 8.63E-01 | no | Kcnip1 | intron_variant | MODIFIER |
| 11 | 33815889 | AX-170427597 | 0 | G | A | 0.105 | 3.31E-02 | 1.89E-01 | 2.65E+00 | 2.72E+00 | 8.63E-01 | no | Kcnip1 | intron_variant | MODIFIER |
| 11 | 33824664 | AX-170303910 | 0 | C | G | 0.105 | 3.31E-02 | 1.89E-01 | 2.65E+00 | 2.72E+00 | 8.63E-01 | no | Kcnip1 | intron_variant | MODIFIER |
| 11 | 33837738 | AX-170201768 | 0 | C | T | 0.105 | 3.31E-02 | 1.89E-01 | 2.65E+00 | 2.72E+00 | 8.63E-01 | no | Kcnip1 | intron_variant | MODIFIER |
| 11 | 33837863 | AX-170394958 | 0 | G | C | 0.105 | 3.31E-02 | 1.89E-01 | 2.65E+00 | 2.72E+00 | 8.63E-01 | no | Kcnip1 | intron_variant | MODIFIER |
| 11 | 33839407 | AX-170470511 | 0 | G | A | 0.105 | 3.31E-02 | 1.89E-01 | 2.65E+00 | 2.72E+00 | 8.63E-01 | no | Kcnip1 | intron_variant | MODIFIER |
| 11 | 33839694 | AX-97818181 | 0 | T | G | 0.105 | 3.31E-02 | 1.89E-01 | 2.65E+00 | 2.72E+00 | 8.63E-01 | no | Kcnip1 | intron_variant | MODIFIER |
| 11 | 33849310 | AX-170036459 | 0 | G | A | 0.105 | 3.31E-02 | 1.89E-01 | 2.65E+00 | 2.72E+00 | 8.63E-01 | no | Kcnip1 | intron_variant,non_coding_transcript_variant | MODIFIER |
| 11 | 33866882 | AX-97782337 | 0 | A | G | 0.105 | 3.31E-02 | 1.89E-01 | 2.65E+00 | 2.72E+00 | 8.63E-01 | no | Kcnip1 | intron_variant,non_coding_transcript_variant | MODIFIER |
| 11 | 33869634 | AX-170243836 | 0 | T | C | 0.105 | 3.31E-02 | 1.89E-01 | 2.65E+00 | 2.72E+00 | 8.63E-01 | no | Kcnip1 | intron_variant,non_coding_transcript_variant | MODIFIER |
| 11 | 33870778 | AX-170203359 | 0 | A | G | 0.105 | 3.31E-02 | 1.89E-01 | 2.65E+00 | 2.72E+00 | 8.63E-01 | no | Kcnip1 | intron_variant,non_coding_transcript_variant | MODIFIER |
| 11 | 33873828 | AX-170211583 | 0 | G | A | 0.105 | 3.31E-02 | 1.89E-01 | 2.65E+00 | 2.72E+00 | 8.63E-01 | no | Kcnip1 | intron_variant,non_coding_transcript_variant | MODIFIER |
| 11 | 33908235 | AX-97832188 | 0 | A | C | 0.105 | 3.31E-02 | 1.89E-01 | 2.65E+00 | 2.72E+00 | 8.63E-01 | no | Kcnip1 | intron_variant,non_coding_transcript_variant | MODIFIER |
| 11 | 33914324 | AX-97795418 | 0 | A | G | 0.105 | 3.31E-02 | 1.89E-01 | 2.65E+00 | 2.72E+00 | 8.63E-01 | no | Kcnip1 | intron_variant,non_coding_transcript_variant | MODIFIER |
| 11 | 33916501 | AX-170162881 | 0 | A | C | 0.105 | 3.31E-02 | 1.89E-01 | 2.65E+00 | 2.72E+00 | 8.63E-01 | no | Kcnip1 | intron_variant,non_coding_transcript_variant | MODIFIER |
| 11 | 33916856 | AX-170349099 | 0 | A | G | 0.105 | 3.31E-02 | 1.89E-01 | 2.65E+00 | 2.72E+00 | 8.63E-01 | no | Kcnip1 | intron_variant,non_coding_transcript_variant | MODIFIER |
| 9 | 32857290 | AX-170223247 | 1 | G | T | 0.438 | 1.54E-02 | 8.64E-02 | 2.65E+00 | 2.71E+00 | 8.63E-01 | no | | intergenic_variant | MODIFIER |
| 4 | 93297044 | AX-170023420 | 0 | T | A | 0.333 | 1.87E-02 | 1.07E-01 | 2.65E+00 | 2.72E+00 | 8.63E-01 | no | | intergenic_variant | MODIFIER |
| 4 | 93619036 | AX-170383586 | 0 | C | T | 0.333 | 1.87E-02 | 1.07E-01 | 2.65E+00 | 2.72E+00 | 8.63E-01 | no | Gm12649 | intron_variant,non_coding_transcript_variant | MODIFIER |
| 6 | 105110958 | AX-170207514 | 1 | C | T | 0.071 | -3.60E-02 | 2.05E-01 | 2.63E+00 | 2.72E+00 | 8.63E-01 | no | | intergenic_variant | MODIFIER |
| 5 | 118149797 | AX-170370528 | 1 | T | C | 0.455 | -1.94E-02 | 1.15E-01 | 2.73E+00 | 2.77E+00 | 8.63E-01 | no | Fbxw8 | intron_variant | MODIFIER |
| 7 | 94518181 | AX-170134713 | 0 | A | T | 0.123 | -3.10E-02 | 1.90E-01 | 2.67E+00 | 2.76E+00 | 8.63E-01 | no | Gm32647 | intron_variant,non_coding_transcript_variant | MODIFIER |
| 4 | 142970783 | AX-170125350 | 1 | G | A | 0.089 | 2.90E-02 | 1.69E-01 | 2.66E+00 | 2.73E+00 | 8.63E-01 | no | 6330411D24Rik | intron_variant,non_coding_transcript_variant | MODIFIER |
| 6 | 147028249 | AX-97835495 | 2 | G | A | 0.055 | 4.28E-02 | 2.60E-01 | 2.70E+00 | 2.77E+00 | 8.63E-01 | no | Ppfibp1 | intron_variant | MODIFIER |
| 7 | 90356459 | AX-170334540 | 1 | T | C | 0.125 | -3.19E-02 | 1.90E-01 | 2.64E+00 | 2.74E+00 | 8.63E-01 | no | Sytl2 | intron_variant | MODIFIER |
| 10 | 13832673 | AX-170348213 | 0 | C | T | 0.202 | -1.88E-02 | 1.08E-01 | 2.64E+00 | 2.73E+00 | 8.63E-01 | no | Aig1 | intron_variant | MODIFIER |
| 8 | 74764054 | AX-97790113 | 1 | G | A | 0.196 | -2.27E-02 | 1.33E-01 | 2.67E+00 | 2.74E+00 | 8.63E-01 | no | Gm11033 | intron_variant,non_coding_transcript_variant | MODIFIER |
| 4 | 140911309 | AX-170024122 | 0 | G | A | 0.088 | 2.89E-02 | 1.69E-01 | 2.66E+00 | 2.73E+00 | 8.63E-01 | no | Padi2 | intron_variant | MODIFIER |
| 4 | 140944118 | AX-170244302 | 0 | C | T | 0.088 | 2.89E-02 | 1.69E-01 | 2.66E+00 | 2.73E+00 | 8.63E-01 | no | Padi2 | intron_variant | MODIFIER |
| 4 | 140953259 | AX-170070887 | 0 | C | G | 0.088 | 2.89E-02 | 1.69E-01 | 2.66E+00 | 2.73E+00 | 8.63E-01 | no | Padi2 | downstream_gene_variant | MODIFIER |
| 4 | 140959253 | AX-170095176 | 0 | A | G | 0.088 | 2.89E-02 | 1.69E-01 | 2.66E+00 | 2.73E+00 | 8.63E-01 | no | Sdhb | upstream_gene_variant | MODIFIER |
| 4 | 140960797 | AX-170509441 | 0 | T | C | 0.088 | 2.89E-02 | 1.69E-01 | 2.66E+00 | 2.73E+00 | 8.63E-01 | no | Sdhb | upstream_gene_variant | MODIFIER |
| 4 | 140975669 | AX-170070886 | 0 | G | T | 0.088 | 2.89E-02 | 1.69E-01 | 2.66E+00 | 2.73E+00 | 8.63E-01 | no | Sdhb | intron_variant | MODIFIER |
| 4 | 140980957 | AX-170125305 | 0 | A | T | 0.088 | 2.89E-02 | 1.69E-01 | 2.66E+00 | 2.73E+00 | 8.63E-01 | no | Sdhb | downstream_gene_variant | MODIFIER |

| | | | | | | | | | | | | | | | |
|---|---|---|---|---|---|---|---|---|---|---|---|---|---|---|---|
| 4 | 140989988 | AX-170333236 | 0 | G | A | 0.088 | 2.89E-02 | 1.69E-01 | 2.66E+00 | 2.73E+00 | 8.63E-01 | no | Atp13a2 | intron_variant | MODIFIER |
| 4 | 140995966 | AX-170161247 | 0 | T | G | 0.088 | 2.89E-02 | 1.69E-01 | 2.66E+00 | 2.73E+00 | 8.63E-01 | no | Atp13a2 | intron_variant | MODIFIER |
| 4 | 140996029 | AX-170259913 | 0 | A | G | 0.088 | 2.89E-02 | 1.69E-01 | 2.66E+00 | 2.73E+00 | 8.63E-01 | no | Atp13a2 | intron_variant | MODIFIER |
| 4 | 141026628 | AX-97811547 | 0 | T | G | 0.088 | 2.89E-02 | 1.69E-01 | 2.66E+00 | 2.73E+00 | 8.63E-01 | no | Crocc | intron_variant | MODIFIER |
| 4 | 141027135 | AX-170241335 | 0 | C | A | 0.088 | 2.89E-02 | 1.69E-01 | 2.66E+00 | 2.73E+00 | 8.63E-01 | no | Crocc | intron_variant | MODIFIER |
| 4 | 141049162 | AX-170260423 | 0 | C | T | 0.088 | 2.89E-02 | 1.69E-01 | 2.66E+00 | 2.73E+00 | 8.63E-01 | no | Crocc | intron_variant | MODIFIER |
| 4 | 141061174 | AX-170351131 | 0 | A | C | 0.088 | 2.89E-02 | 1.69E-01 | 2.66E+00 | 2.73E+00 | 8.63E-01 | no | Crocc | upstream_gene_variant | MODIFIER |
| 4 | 141066363 | AX-170314069 | 0 | A | G | 0.088 | 2.89E-02 | 1.69E-01 | 2.66E+00 | 2.73E+00 | 8.63E-01 | no | Necap2 | downstream_gene_variant | MODIFIER |
| 4 | 141105565 | AX-170377737 | 0 | G | A | 0.088 | 2.89E-02 | 1.69E-01 | 2.66E+00 | 2.73E+00 | 8.63E-01 | no | Spata21 | intron_variant | MODIFIER |
| 4 | 141110936 | AX-97827743 | 0 | C | T | 0.088 | 2.89E-02 | 1.69E-01 | 2.66E+00 | 2.73E+00 | 8.63E-01 | no | Szrd1 | downstream_gene_variant | MODIFIER |
| 4 | 141124951 | AX-170294212 | 0 | A | G | 0.088 | 2.89E-02 | 1.69E-01 | 2.66E+00 | 2.73E+00 | 8.63E-01 | no | Szrd1 | intron_variant,non_coding_transcript_variant | MODIFIER |
| 4 | 141157871 | AX-170478675 | 0 | C | G | 0.088 | 2.89E-02 | 1.69E-01 | 2.66E+00 | 2.73E+00 | 8.63E-01 | no | Fbxo42 | intron_variant | MODIFIER |
| 4 | 141163871 | AX-170095177 | 0 | A | G | 0.088 | 2.89E-02 | 1.69E-01 | 2.66E+00 | 2.73E+00 | 8.63E-01 | no | Fbxo42 | intron_variant | MODIFIER |
| 4 | 141166604 | AX-170390770 | 0 | C | G | 0.088 | 2.89E-02 | 1.69E-01 | 2.66E+00 | 2.73E+00 | 8.63E-01 | no | Fbxo42 | intron_variant | MODIFIER |
| 4 | 141171798 | AX-170070888 | 0 | G | C | 0.088 | 2.89E-02 | 1.69E-01 | 2.66E+00 | 2.73E+00 | 8.63E-01 | no | Fbxo42 | intron_variant | MODIFIER |
| 4 | 141174258 | AX-170295964 | 0 | T | C | 0.088 | 2.89E-02 | 1.69E-01 | 2.66E+00 | 2.73E+00 | 8.63E-01 | no | Fbxo42 | intron_variant | MODIFIER |
| 4 | 141181000 | AX-170070889 | 0 | A | G | 0.088 | 2.89E-02 | 1.69E-01 | 2.66E+00 | 2.73E+00 | 8.63E-01 | no | Fbxo42 | intron_variant | MODIFIER |
| 4 | 141184419 | AX-170396796 | 0 | T | C | 0.088 | 2.89E-02 | 1.69E-01 | 2.66E+00 | 2.73E+00 | 8.63E-01 | no | Fbxo42 | intron_variant | MODIFIER |
| 4 | 141232362 | AX-170504961 | 0 | A | G | 0.088 | 2.89E-02 | 1.69E-01 | 2.66E+00 | 2.73E+00 | 8.63E-01 | no | | intergenic_variant | MODIFIER |
| 4 | 141861586 | AX-170274933 | 0 | T | G | 0.088 | 2.89E-02 | 1.69E-01 | 2.66E+00 | 2.73E+00 | 8.63E-01 | no | Efhd2 | intron_variant | MODIFIER |
| 4 | 141918988 | AX-170272561 | 0 | A | G | 0.088 | 2.89E-02 | 1.69E-01 | 2.66E+00 | 2.73E+00 | 8.63E-01 | no | Fhad1 | intron_variant | MODIFIER |
| 4 | 141931362 | AX-170532973 | 0 | A | C | 0.088 | 2.89E-02 | 1.69E-01 | 2.66E+00 | 2.73E+00 | 8.63E-01 | no | Fhad1 | intron_variant | MODIFIER |
| 4 | 142136810 | AX-170488983 | 0 | T | C | 0.088 | 2.89E-02 | 1.69E-01 | 2.66E+00 | 2.73E+00 | 8.63E-01 | no | Kazn | downstream_gene_variant | MODIFIER |
| 4 | 142259822 | AX-170346492 | 0 | T | C | 0.088 | 2.89E-02 | 1.69E-01 | 2.66E+00 | 2.73E+00 | 8.63E-01 | no | | intergenic_variant | MODIFIER |
| 4 | 142277929 | AX-170351508 | 0 | T | C | 0.088 | 2.89E-02 | 1.69E-01 | 2.66E+00 | 2.73E+00 | 8.63E-01 | no | | regulatory_region_variant | MODIFIER |
| 4 | 142283271 | AX-97832923 | 0 | C | T | 0.088 | 2.89E-02 | 1.69E-01 | 2.66E+00 | 2.73E+00 | 8.63E-01 | no | | intergenic_variant | MODIFIER |
| 4 | 142302638 | AX-170303538 | 0 | C | T | 0.088 | 2.89E-02 | 1.69E-01 | 2.66E+00 | 2.73E+00 | 8.63E-01 | no | | regulatory_region_variant | MODIFIER |
| 4 | 142326541 | AX-97848259 | 0 | C | T | 0.088 | 2.89E-02 | 1.69E-01 | 2.66E+00 | 2.73E+00 | 8.63E-01 | no | Gm13052 | intron_variant,non_coding_transcript_variant | MODIFIER |
| 4 | 142329833 | AX-170151080 | 0 | A | G | 0.088 | 2.89E-02 | 1.69E-01 | 2.66E+00 | 2.73E+00 | 8.63E-01 | no | Gm13052 | intron_variant,non_coding_transcript_variant | MODIFIER |
| 4 | 142407934 | AX-170342307 | 0 | G | A | 0.088 | 2.89E-02 | 1.69E-01 | 2.66E+00 | 2.73E+00 | 8.63E-01 | no | | intergenic_variant | MODIFIER |
| 4 | 142542418 | AX-170334370 | 0 | G | A | 0.088 | 2.89E-02 | 1.69E-01 | 2.66E+00 | 2.73E+00 | 8.63E-01 | no | | intergenic_variant | MODIFIER |
| 4 | 142545715 | AX-170330093 | 0 | A | G | 0.088 | 2.89E-02 | 1.69E-01 | 2.66E+00 | 2.73E+00 | 8.63E-01 | no | | intergenic_variant | MODIFIER |
| 4 | 142545965 | AX-170231764 | 0 | G | T | 0.088 | 2.89E-02 | 1.69E-01 | 2.66E+00 | 2.73E+00 | 8.63E-01 | no | | intergenic_variant | MODIFIER |
| 4 | 142547007 | AX-170248487 | 0 | C | T | 0.088 | 2.89E-02 | 1.69E-01 | 2.66E+00 | 2.73E+00 | 8.63E-01 | no | | intergenic_variant | MODIFIER |
| 4 | 142912003 | AX-170334749 | 0 | C | A | 0.088 | 2.89E-02 | 1.69E-01 | 2.66E+00 | 2.73E+00 | 8.63E-01 | no | 6330411D24Rik | intron_variant,non_coding_transcript_variant | MODIFIER |
| 4 | 142945932 | AX-170477772 | 0 | C | A | 0.088 | 2.89E-02 | 1.69E-01 | 2.66E+00 | 2.73E+00 | 8.63E-01 | no | 6330411D24Rik | intron_variant,non_coding_transcript_variant | MODIFIER |
| 4 | 142957094 | AX-170494205 | 0 | G | A | 0.088 | 2.89E-02 | 1.69E-01 | 2.66E+00 | 2.73E+00 | 8.63E-01 | no | 6330411D24Rik | intron_variant,non_coding_transcript_variant | MODIFIER |
| 4 | 142958719 | AX-170141108 | 0 | A | G | 0.088 | 2.89E-02 | 1.69E-01 | 2.66E+00 | 2.73E+00 | 8.63E-01 | no | 6330411D24Rik | intron_variant,non_coding_transcript_variant | MODIFIER |
| 4 | 142960993 | AX-170505864 | 0 | G | A | 0.088 | 2.89E-02 | 1.69E-01 | 2.66E+00 | 2.73E+00 | 8.63E-01 | no | 6330411D24Rik | intron_variant,non_coding_transcript_variant | MODIFIER |
| 4 | 142963468 | AX-170264790 | 0 | G | A | 0.088 | 2.89E-02 | 1.69E-01 | 2.66E+00 | 2.73E+00 | 8.63E-01 | no | 6330411D24Rik | intron_variant,non_coding_transcript_variant | MODIFIER |
| 4 | 142968184 | AX-170278330 | 0 | T | A | 0.088 | 2.89E-02 | 1.69E-01 | 2.66E+00 | 2.73E+00 | 8.63E-01 | no | 6330411D24Rik | intron_variant,non_coding_transcript_variant | MODIFIER |
| 4 | 142970101 | AX-170499332 | 0 | A | T | 0.088 | 2.89E-02 | 1.69E-01 | 2.66E+00 | 2.73E+00 | 8.63E-01 | no | 6330411D24Rik | intron_variant,non_coding_transcript_variant | MODIFIER |
| 4 | 142972075 | AX-170521873 | 0 | G | A | 0.088 | 2.89E-02 | 1.69E-01 | 2.66E+00 | 2.73E+00 | 8.63E-01 | no | 6330411D24Rik | intron_variant,non_coding_transcript_variant | MODIFIER |
| 4 | 142973945 | AX-170149253 | 0 | C | T | 0.088 | 2.89E-02 | 1.69E-01 | 2.66E+00 | 2.73E+00 | 8.63E-01 | no | 6330411D24Rik | intron_variant,non_coding_transcript_variant | MODIFIER |
| 4 | 143033245 | AX-170405770 | 0 | A | G | 0.088 | 2.89E-02 | 1.69E-01 | 2.66E+00 | 2.73E+00 | 8.63E-01 | no | 6330411D24Rik | intron_variant,non_coding_transcript_variant | MODIFIER |
| 4 | 143037130 | AX-170222030 | 0 | T | C | 0.088 | 2.89E-02 | 1.69E-01 | 2.66E+00 | 2.73E+00 | 8.63E-01 | no | 6330411D24Rik | intron_variant,non_coding_transcript_variant | MODIFIER |
| 4 | 143059144 | AX-170220268 | 0 | A | G | 0.088 | 2.89E-02 | 1.69E-01 | 2.66E+00 | 2.73E+00 | 8.63E-01 | no | 6330411D24Rik | intron_variant,non_coding_transcript_variant | MODIFIER |
| 4 | 143066333 | AX-97781926 | 0 | A | G | 0.088 | 2.89E-02 | 1.69E-01 | 2.66E+00 | 2.73E+00 | 8.63E-01 | no | 6330411D24Rik | intron_variant,non_coding_transcript_variant | MODIFIER |
| 4 | 143692271 | AX-97821296 | 0 | A | T | 0.088 | 2.89E-02 | 1.69E-01 | 2.66E+00 | 2.73E+00 | 8.63E-01 | no | Gm13089 | downstream_gene_variant | MODIFIER |
| 4 | 143896325 | AX-170070930 | 0 | C | T | 0.088 | 2.89E-02 | 1.69E-01 | 2.66E+00 | 2.73E+00 | 8.63E-01 | no | Pramef6 | intron_variant | MODIFIER |
| 4 | 144978994 | AX-170366336 | 0 | T | C | 0.088 | 2.89E-02 | 1.69E-01 | 2.66E+00 | 2.73E+00 | 8.63E-01 | no | Vps13d | intron_variant | MODIFIER |
| 4 | 145071011 | AX-170485200 | 0 | T | C | 0.088 | 2.89E-02 | 1.69E-01 | 2.66E+00 | 2.73E+00 | 8.63E-01 | no | Vps13d | intron_variant | MODIFIER |
| 4 | 145085317 | AX-170249124 | 0 | C | A | 0.088 | 2.89E-02 | 1.69E-01 | 2.66E+00 | 2.73E+00 | 8.63E-01 | no | Vps13d | missense_variant | MODERATE |
| 4 | 145140311 | AX-97840052 | 0 | A | G | 0.088 | 2.89E-02 | 1.69E-01 | 2.66E+00 | 2.73E+00 | 8.63E-01 | no | Vps13d | intron_variant | MODIFIER |

| | | | | | | | | | | | | | | | |
|---|---|---|---|---|---|---|---|---|---|---|---|---|---|---|---|
| 4 | 145292211 | AX-170511869 | 0 | C | A | 0.088 | 2.89E-02 | 1.69E-01 | 2.66E+00 | 2.73E+00 | 8.63E-01 | no | Tnfrsf8 | intron_variant | MODIFIER |
| 4 | 145292609 | AX-170244143 | 0 | A | G | 0.088 | 2.89E-02 | 1.69E-01 | 2.66E+00 | 2.73E+00 | 8.63E-01 | no | Tnfrsf8 | missense_variant | MODERATE |
| 4 | 145299848 | AX-97851909 | 0 | T | C | 0.088 | 2.89E-02 | 1.69E-01 | 2.66E+00 | 2.73E+00 | 8.63E-01 | no | Tnfrsf8 | intron_variant | MODIFIER |
| 4 | 145302652 | AX-170451776 | 0 | A | G | 0.088 | 2.89E-02 | 1.69E-01 | 2.66E+00 | 2.73E+00 | 8.63E-01 | no | Tnfrsf8 | intron_variant | MODIFIER |
| 4 | 145308207 | AX-97788464 | 0 | T | C | 0.088 | 2.89E-02 | 1.69E-01 | 2.66E+00 | 2.73E+00 | 8.63E-01 | no | Tnfrsf8 | intron_variant | MODIFIER |
| 4 | 147933474 | AX-97856547 | 0 | A | C | 0.088 | 2.89E-02 | 1.69E-01 | 2.66E+00 | 2.73E+00 | 8.63E-01 | no | Plod1 | intron_variant | MODIFIER |
| 4 | 148059827 | AX-97808524 | 0 | C | T | 0.088 | 2.89E-02 | 1.69E-01 | 2.66E+00 | 2.73E+00 | 8.63E-01 | no | Mthfr | downstream_gene_variant | MODIFIER |
| 4 | 148890444 | AX-170024214 | 0 | G | A | 0.088 | 2.89E-02 | 1.69E-01 | 2.66E+00 | 2.73E+00 | 8.63E-01 | no | Casz1 | intron_variant | MODIFIER |
| 4 | 148986977 | AX-97838307 | 0 | T | C | 0.088 | 2.89E-02 | 1.69E-01 | 2.66E+00 | 2.73E+00 | 8.63E-01 | no | Pex14 | intron_variant | MODIFIER |
| 4 | 149005307 | AX-170406140 | 0 | A | G | 0.088 | 2.89E-02 | 1.69E-01 | 2.66E+00 | 2.73E+00 | 8.63E-01 | no | Pex14 | intron_variant | MODIFIER |
| 4 | 149005376 | AX-170399752 | 0 | C | T | 0.088 | 2.89E-02 | 1.69E-01 | 2.66E+00 | 2.73E+00 | 8.63E-01 | no | Pex14 | intron_variant | MODIFIER |
| 4 | 149086102 | AX-97783964 | 0 | A | G | 0.088 | 2.89E-02 | 1.69E-01 | 2.66E+00 | 2.73E+00 | 8.63E-01 | no | Pex14 | intron_variant | MODIFIER |
| 4 | 149124050 | AX-170199015 | 0 | A | G | 0.088 | 2.89E-02 | 1.69E-01 | 2.66E+00 | 2.73E+00 | 8.63E-01 | no | Cenps | downstream_gene_variant | MODIFIER |
| 4 | 149174201 | AX-170024218 | 0 | A | G | 0.088 | 2.89E-02 | 1.69E-01 | 2.66E+00 | 2.73E+00 | 8.63E-01 | no | Kif1b | downstream_gene_variant | MODIFIER |
| 4 | 149236099 | AX-170024216 | 0 | G | A | 0.088 | 2.89E-02 | 1.69E-01 | 2.66E+00 | 2.73E+00 | 8.63E-01 | no | Kif1b | 3_prime_UTR_variant | MODIFIER |
| 4 | 149242112 | AX-97799727 | 0 | G | A | 0.088 | 2.89E-02 | 1.69E-01 | 2.66E+00 | 2.73E+00 | 8.63E-01 | no | Kif1b | intron_variant | MODIFIER |
| 4 | 149302265 | AX-168400648 | 0 | A | G | 0.088 | 2.89E-02 | 1.69E-01 | 2.66E+00 | 2.73E+00 | 8.63E-01 | no | Kif1b | intron_variant | MODIFIER |
| 4 | 149334866 | AX-170295732 | 0 | A | G | 0.088 | 2.89E-02 | 1.69E-01 | 2.66E+00 | 2.73E+00 | 8.63E-01 | no | Ube4b | intron_variant | MODIFIER |
| 4 | 149529631 | AX-170457402 | 0 | A | G | 0.088 | 2.89E-02 | 1.69E-01 | 2.66E+00 | 2.73E+00 | 8.63E-01 | no | Ctnnbip1 | intron_variant | MODIFIER |
| 4 | 149530229 | AX-170248238 | 0 | C | G | 0.088 | 2.89E-02 | 1.69E-01 | 2.66E+00 | 2.73E+00 | 8.63E-01 | no | Ctnnbip1 | intron_variant | MODIFIER |
| 4 | 149530257 | AX-170192781 | 0 | A | G | 0.088 | 2.89E-02 | 1.69E-01 | 2.66E+00 | 2.73E+00 | 8.63E-01 | no | Ctnnbip1 | intron_variant | MODIFIER |
| 4 | 149532550 | AX-168415381 | 0 | C | T | 0.088 | 2.89E-02 | 1.69E-01 | 2.66E+00 | 2.73E+00 | 8.63E-01 | no | Ctnnbip1 | intron_variant | MODIFIER |
| 4 | 149541697 | AX-170443071 | 0 | A | G | 0.088 | 2.89E-02 | 1.69E-01 | 2.66E+00 | 2.73E+00 | 8.63E-01 | no | Ctnnbip1 | intron_variant | MODIFIER |
| 4 | 149558923 | AX-170201787 | 0 | T | C | 0.088 | 2.89E-02 | 1.69E-01 | 2.66E+00 | 2.73E+00 | 8.63E-01 | no | Ctnnbip1 | intron_variant | MODIFIER |
| 4 | 149559860 | AX-97844425 | 0 | T | C | 0.088 | 2.89E-02 | 1.69E-01 | 2.66E+00 | 2.73E+00 | 8.63E-01 | no | Ctnnbip1 | intron_variant | MODIFIER |
| 4 | 149568112 | AX-170173143 | 0 | A | G | 0.088 | 2.89E-02 | 1.69E-01 | 2.66E+00 | 2.73E+00 | 8.63E-01 | no | Ctnnbip1 | downstream_gene_variant | MODIFIER |
| 4 | 149571046 | AX-170474458 | 0 | A | G | 0.088 | 2.89E-02 | 1.69E-01 | 2.66E+00 | 2.73E+00 | 8.63E-01 | no | Ctnnbip1 | downstream_gene_variant | MODIFIER |
| 4 | 149723067 | AX-170257758 | 0 | T | C | 0.088 | 2.89E-02 | 1.69E-01 | 2.66E+00 | 2.73E+00 | 8.63E-01 | no | Tmem201 | intron_variant | MODIFIER |
| 4 | 149948856 | AX-170184161 | 0 | C | T | 0.088 | 2.89E-02 | 1.69E-01 | 2.66E+00 | 2.73E+00 | 8.63E-01 | no | Spsb1 | intron_variant | MODIFIER |
| 4 | 149958787 | AX-170360158 | 0 | G | T | 0.088 | 2.89E-02 | 1.69E-01 | 2.66E+00 | 2.73E+00 | 8.63E-01 | no | Spsb1 | upstream_gene_variant | MODIFIER |
| 4 | 149994494 | AX-97851723 | 0 | A | G | 0.088 | 2.89E-02 | 1.69E-01 | 2.66E+00 | 2.73E+00 | 8.63E-01 | no | H6pd | synonymous_variant | LOW |
| 4 | 149998990 | AX-170122774 | 0 | A | G | 0.088 | 2.89E-02 | 1.69E-01 | 2.66E+00 | 2.73E+00 | 8.63E-01 | no | H6pd | intron_variant | MODIFIER |
| 4 | 150018079 | AX-170183685 | 0 | C | G | 0.088 | 2.89E-02 | 1.69E-01 | 2.66E+00 | 2.73E+00 | 8.63E-01 | no | | intergenic_variant | MODIFIER |
| 4 | 150018788 | AX-97809912 | 0 | G | A | 0.088 | 2.89E-02 | 1.69E-01 | 2.66E+00 | 2.73E+00 | 8.63E-01 | no | | intergenic_variant | MODIFIER |
| 4 | 150020177 | AX-170150486 | 0 | T | C | 0.088 | 2.89E-02 | 1.69E-01 | 2.66E+00 | 2.73E+00 | 8.63E-01 | no | | regulatory_region_variant | MODIFIER |
| 4 | 150024545 | AX-170530101 | 0 | A | G | 0.088 | 2.89E-02 | 1.69E-01 | 2.66E+00 | 2.73E+00 | 8.63E-01 | no | | intergenic_variant | MODIFIER |
| 4 | 150055273 | AX-97791742 | 0 | C | T | 0.088 | 2.89E-02 | 1.69E-01 | 2.66E+00 | 2.73E+00 | 8.63E-01 | no | Gm13067 | intron_variant,non_coding_transcript_variant | MODIFIER |
| 4 | 150120241 | AX-97803769 | 0 | A | G | 0.088 | 2.89E-02 | 1.69E-01 | 2.66E+00 | 2.73E+00 | 8.63E-01 | no | Slc2a5 | intron_variant | MODIFIER |
| 4 | 150389482 | AX-170143810 | 0 | A | G | 0.088 | 2.89E-02 | 1.69E-01 | 2.66E+00 | 2.73E+00 | 8.63E-01 | no | Rere | intron_variant | MODIFIER |
| 4 | 150392140 | AX-170325486 | 0 | T | A | 0.088 | 2.89E-02 | 1.69E-01 | 2.66E+00 | 2.73E+00 | 8.63E-01 | no | Rere | intron_variant | MODIFIER |
| 4 | 150445981 | AX-170143279 | 0 | A | G | 0.088 | 2.89E-02 | 1.69E-01 | 2.66E+00 | 2.73E+00 | 8.63E-01 | no | Rere | intron_variant | MODIFIER |
| 4 | 150476763 | AX-170510820 | 0 | G | A | 0.088 | 2.89E-02 | 1.69E-01 | 2.66E+00 | 2.73E+00 | 8.63E-01 | no | Rere | intron_variant | MODIFIER |
| 4 | 150484287 | AX-170339014 | 0 | C | T | 0.088 | 2.89E-02 | 1.69E-01 | 2.66E+00 | 2.73E+00 | 8.63E-01 | no | Rere | upstream_gene_variant | MODIFIER |
| 4 | 150576493 | AX-170512749 | 0 | T | G | 0.088 | 2.89E-02 | 1.69E-01 | 2.66E+00 | 2.73E+00 | 8.63E-01 | no | Rere | intron_variant | MODIFIER |
| 4 | 150599737 | AX-170477845 | 0 | T | C | 0.088 | 2.89E-02 | 1.69E-01 | 2.66E+00 | 2.73E+00 | 8.63E-01 | no | Rere | intron_variant | MODIFIER |
| 4 | 150607961 | AX-170352264 | 0 | T | C | 0.088 | 2.89E-02 | 1.69E-01 | 2.66E+00 | 2.73E+00 | 8.63E-01 | no | Rere | intron_variant | MODIFIER |
| 4 | 151257419 | AX-170480894 | 0 | A | G | 0.088 | 2.89E-02 | 1.69E-01 | 2.66E+00 | 2.73E+00 | 8.63E-01 | no | Camta1 | intron_variant | MODIFIER |
| 4 | 151257669 | AX-170484965 | 0 | C | T | 0.088 | 2.89E-02 | 1.69E-01 | 2.66E+00 | 2.73E+00 | 8.63E-01 | no | Camta1 | intron_variant | MODIFIER |
| 4 | 149421926 | AX-170166750 | 1 | C | T | 0.089 | 2.89E-02 | 1.68E-01 | 2.66E+00 | 2.73E+00 | 8.63E-01 | no | Ube4b | intron_variant,non_coding_transcript_variant | MODIFIER |
| 1 | 187279380 | AX-170243556 | 0 | A | T | 0.228 | -1.59E-02 | 1.04E-01 | 2.71E+00 | 2.82E+00 | 8.63E-01 | no | Gpatch2 | intron_variant | MODIFIER |
| 7 | 117674955 | AX-170484681 | 1 | T | A | 0.384 | 1.54E-02 | 8.92E-02 | 2.67E+00 | 2.73E+00 | 8.64E-01 | no | Xylt1 | downstream_gene_variant | MODIFIER |
| 7 | 118392480 | AX-170263228 | 1 | T | A | 0.384 | 1.54E-02 | 8.92E-02 | 2.67E+00 | 2.73E+00 | 8.64E-01 | no | Syt17 | intron_variant | MODIFIER |
| 7 | 119071016 | AX-170535308 | 1 | A | G | 0.384 | 1.54E-02 | 8.92E-02 | 2.67E+00 | 2.73E+00 | 8.64E-01 | no | | intergenic_variant | MODIFIER |
| 6 | 100873685 | AX-170027823 | 0 | T | C | 0.079 | 3.73E-02 | 2.12E-01 | 2.63E+00 | 2.72E+00 | 8.64E-01 | no | Ppp4r2 | downstream_gene_variant | MODIFIER |

| | | | | | | | | | | | | | | | |
|---|---|---|---|---|---|---|---|---|---|---|---|---|---|---|---|
| 2 | 115549863 | AX-170471098 | 1 | G | A | 0.143 | 3.78E-02 | 2.20E-01 | 2.69E+00 | 2.72E+00 | 8.64E-01 | no | | intergenic_variant | MODIFIER |
| 2 | 142350855 | AX-170019227 | 1 | C | T | 0.143 | 3.78E-02 | 2.20E-01 | 2.69E+00 | 2.72E+00 | 8.64E-01 | no | Macrod2 | intron_variant | MODIFIER |
| 2 | 119018092 | AX-170018824 | 0 | C | G | 0.14 | -3.73E-02 | 2.22E-01 | 2.74E+00 | 2.77E+00 | 8.64E-01 | no | Ccdc32 | 3_prime_UTR_variant | MODIFIER |
| 2 | 122285154 | AX-170064770 | 0 | C | T | 0.14 | -3.73E-02 | 2.22E-01 | 2.74E+00 | 2.77E+00 | 8.64E-01 | no | Duox2 | splice_region_variant,intron_variant | LOW |
| 2 | 139165582 | AX-170088767 | 0 | T | C | 0.14 | -3.73E-02 | 2.22E-01 | 2.74E+00 | 2.77E+00 | 8.64E-01 | no | | intergenic_variant | MODIFIER |
| 7 | 103733399 | AX-97843401 | 0 | G | A | 0.447 | -1.47E-02 | 9.10E-02 | 2.71E+00 | 2.78E+00 | 8.64E-01 | no | Olfr628 | downstream_gene_variant | MODIFIER |
| 2 | 160013034 | AX-170019515 | 0 | G | T | 0.149 | 3.61E-02 | 2.14E-01 | 2.69E+00 | 2.74E+00 | 8.64E-01 | no | 9130015L21Rik | upstream_gene_variant | MODIFIER |
| 7 | 91471788 | AX-170137587 | 0 | C | T | 0.132 | -3.56E-02 | 2.11E-01 | 2.66E+00 | 2.74E+00 | 8.64E-01 | no | Dlg2 | intron_variant | MODIFIER |
| 7 | 97402842 | AX-170248731 | 0 | G | A | 0.132 | -3.56E-02 | 2.11E-01 | 2.66E+00 | 2.74E+00 | 8.64E-01 | no | Ndufc2 | intron_variant | MODIFIER |
| 10 | 31973768 | AX-170483186 | 1 | C | T | 0.384 | -1.28E-02 | 7.36E-02 | 2.61E+00 | 2.73E+00 | 8.64E-01 | no | Nkain2 | intron_variant | MODIFIER |
| 6 | 111457553 | AX-170075878 | 0 | C | G | 0.061 | 4.37E-02 | 2.58E-01 | 2.68E+00 | 2.74E+00 | 8.64E-01 | no | Grm7 | intron_variant | MODIFIER |
| 6 | 112793959 | AX-170440512 | 0 | A | G | 0.061 | 4.37E-02 | 2.58E-01 | 2.68E+00 | 2.74E+00 | 8.64E-01 | no | Srgap3 | intron_variant,non_coding_transcript_variant | MODIFIER |
| 6 | 113170385 | AX-97855068 | 0 | A | G | 0.061 | 4.37E-02 | 2.58E-01 | 2.68E+00 | 2.74E+00 | 8.64E-01 | no | Lhfpl4 | 3_prime_UTR_variant | MODIFIER |
| 18 | 58418655 | AX-170215285 | 0 | T | A | 0.404 | 1.79E-02 | 9.27E-02 | 2.60E+00 | 2.67E+00 | 8.64E-01 | no | | intergenic_variant | MODIFIER |
| 13 | 47283927 | AX-170008270 | 0 | G | C | 0.465 | 1.12E-02 | 6.43E-02 | 2.62E+00 | 2.73E+00 | 8.64E-01 | no | | intergenic_variant | MODIFIER |
| 1 | 11184476 | AX-97855436 | 1 | T | C | 0.384 | -1.16E-02 | 7.18E-02 | 2.70E+00 | 2.78E+00 | 8.64E-01 | no | Prex2 | synonymous_variant | LOW |
| 5 | 96573171 | AX-170472057 | 0 | G | A | 0.43 | 1.74E-02 | 1.01E-01 | 2.65E+00 | 2.72E+00 | 8.64E-01 | no | Fras1 | intron_variant | MODIFIER |
| 2 | 159887526 | AX-170019521 | 0 | G | A | 0.149 | 2.90E-02 | 1.76E-01 | 2.67E+00 | 2.75E+00 | 8.64E-01 | no | | intergenic_variant | MODIFIER |
| 5 | 71585308 | AX-170216131 | 1 | T | A | 0.438 | -1.49E-02 | 9.28E-02 | 2.67E+00 | 2.77E+00 | 8.65E-01 | no | Gabra4 | intron_variant | MODIFIER |
| 9 | 117748191 | AX-170102856 | 0 | C | T | 0.123 | -1.68E-02 | 9.26E-02 | 2.58E+00 | 2.71E+00 | 8.65E-01 | no | Gm45881 | intron_variant,non_coding_transcript_variant | MODIFIER |
| 6 | 102763351 | AX-170400293 | 0 | G | A | 0.053 | 5.01E-02 | 2.96E-01 | 2.67E+00 | 2.74E+00 | 8.65E-01 | no | | intergenic_variant | MODIFIER |
| 6 | 105763806 | AX-170244358 | 0 | C | A | 0.053 | 5.01E-02 | 2.96E-01 | 2.67E+00 | 2.74E+00 | 8.65E-01 | no | Cntn4 | intron_variant | MODIFIER |
| 6 | 107055670 | AX-170236973 | 0 | A | C | 0.053 | 5.01E-02 | 2.96E-01 | 2.67E+00 | 2.74E+00 | 8.65E-01 | no | | intergenic_variant | MODIFIER |
| 7 | 40469883 | AX-170182991 | 0 | A | T | 0.333 | 1.66E-02 | 9.53E-02 | 2.64E+00 | 2.72E+00 | 8.65E-01 | no | Gm28807 | upstream_gene_variant | MODIFIER |
| 4 | 150335912 | AX-97785464 | 1 | G | T | 0.089 | 2.86E-02 | 1.68E-01 | 2.66E+00 | 2.73E+00 | 8.65E-01 | no | Rere | intron_variant | MODIFIER |
| 12 | 53204199 | AX-170390659 | 1 | C | T | 0.054 | -2.50E-02 | 1.28E-01 | 2.55E+00 | 2.69E+00 | 8.65E-01 | no | | intergenic_variant | MODIFIER |
| 4 | 141475118 | AX-170070895 | 0 | C | T | 0.096 | 3.38E-02 | 2.00E-01 | 2.67E+00 | 2.74E+00 | 8.65E-01 | no | Spen | missense_variant | MODERATE |
| 4 | 142195694 | AX-170366923 | 0 | C | T | 0.096 | 3.38E-02 | 2.00E-01 | 2.67E+00 | 2.74E+00 | 8.65E-01 | no | Kazn | intron_variant | MODIFIER |
| 4 | 149358740 | AX-170497498 | 0 | G | A | 0.096 | 3.38E-02 | 2.00E-01 | 2.67E+00 | 2.74E+00 | 8.65E-01 | no | Ube4b | intron_variant | MODIFIER |
| 4 | 137133749 | AX-170372531 | 0 | A | G | 0.167 | -1.53E-02 | 1.02E-01 | 2.69E+00 | 2.81E+00 | 8.65E-01 | no | | regulatory_region_variant | MODIFIER |
| 12 | 117058494 | AX-170050339 | 0 | C | T | 0.149 | 2.79E-02 | 1.67E-01 | 2.64E+00 | 2.74E+00 | 8.65E-01 | no | Ptprn2 | intron_variant | MODIFIER |
| 10 | 115002729 | AX-170289952 | 1 | A | G | 0.375 | 1.27E-02 | 7.57E-02 | 2.65E+00 | 2.74E+00 | 8.65E-01 | no | | intergenic_variant | MODIFIER |
| 4 | 108914771 | AX-170147154 | 1 | T | A | 0.25 | 2.15E-02 | 1.11E-01 | 2.58E+00 | 2.68E+00 | 8.65E-01 | no | Rab3b | intron_variant | MODIFIER |
| 12 | 73084886 | AX-170266528 | 0 | C | T | 0.07 | 3.60E-02 | 2.22E-01 | 2.67E+00 | 2.76E+00 | 8.65E-01 | no | | intergenic_variant | MODIFIER |
| 4 | 142505281 | AX-170163721 | 0 | C | T | 0.105 | 3.05E-02 | 1.77E-01 | 2.65E+00 | 2.72E+00 | 8.65E-01 | no | | intergenic_variant | MODIFIER |
| 4 | 148143664 | AX-170518697 | 0 | G | C | 0.105 | 3.05E-02 | 1.77E-01 | 2.65E+00 | 2.72E+00 | 8.65E-01 | no | Fbxo6 | downstream_gene_variant | MODIFIER |
| 1 | 101357315 | AX-170292923 | 0 | T | C | 0.14 | 2.78E-02 | 1.62E-01 | 2.62E+00 | 2.73E+00 | 8.65E-01 | no | | intergenic_variant | MODIFIER |
| 7 | 123070242 | AX-170283157 | 0 | G | A | 0.404 | 1.39E-02 | 8.41E-02 | 2.67E+00 | 2.75E+00 | 8.65E-01 | no | Gm45847 | downstream_gene_variant | MODIFIER |
| 15 | 95365230 | AX-170275987 | 1 | G | T | 0.143 | 2.85E-02 | 1.66E-01 | 2.65E+00 | 2.72E+00 | 8.65E-01 | no | Nell2 | intron_variant | MODIFIER |
| 2 | 124912126 | AX-170249838 | 1 | G | A | 0.125 | 3.60E-02 | 2.09E-01 | 2.67E+00 | 2.71E+00 | 8.65E-01 | no | | intergenic_variant | MODIFIER |
| 12 | 109649654 | AX-170340117 | 0 | G | T | 0.132 | -3.02E-02 | 1.83E-01 | 2.67E+00 | 2.75E+00 | 8.65E-01 | no | DQ267101 | upstream_gene_variant | MODIFIER |
| 12 | 109673412 | AX-170316455 | 0 | T | C | 0.132 | -3.02E-02 | 1.83E-01 | 2.67E+00 | 2.75E+00 | 8.65E-01 | no | Gm23347 | upstream_gene_variant | MODIFIER |
| 12 | 109749013 | AX-170493272 | 0 | G | A | 0.132 | -3.02E-02 | 1.83E-01 | 2.67E+00 | 2.75E+00 | 8.65E-01 | no | Mir3072 | downstream_gene_variant | MODIFIER |
| 12 | 109752127 | AX-170447069 | 0 | T | C | 0.132 | -3.02E-02 | 1.83E-01 | 2.67E+00 | 2.75E+00 | 8.65E-01 | no | Mir3072 | downstream_gene_variant | MODIFIER |
| 12 | 109785497 | AX-170482757 | 0 | C | T | 0.132 | -3.02E-02 | 1.83E-01 | 2.67E+00 | 2.75E+00 | 8.65E-01 | no | | intergenic_variant | MODIFIER |
| 12 | 109830395 | AX-170245167 | 0 | T | C | 0.132 | -3.02E-02 | 1.83E-01 | 2.67E+00 | 2.75E+00 | 8.65E-01 | no | Gm9661 | downstream_gene_variant | MODIFIER |
| 12 | 109830722 | AX-170370662 | 0 | C | T | 0.132 | -3.02E-02 | 1.83E-01 | 2.67E+00 | 2.75E+00 | 8.65E-01 | no | Gm9661 | downstream_gene_variant | MODIFIER |
| 12 | 109831902 | AX-170337398 | 0 | A | G | 0.132 | -3.02E-02 | 1.83E-01 | 2.67E+00 | 2.75E+00 | 8.65E-01 | no | Gm9661 | downstream_gene_variant | MODIFIER |
| 12 | 109832644 | AX-170483754 | 0 | T | C | 0.132 | -3.02E-02 | 1.83E-01 | 2.67E+00 | 2.75E+00 | 8.65E-01 | no | Gm9661 | downstream_gene_variant | MODIFIER |
| 12 | 109848800 | AX-170319094 | 0 | T | G | 0.132 | -3.02E-02 | 1.83E-01 | 2.67E+00 | 2.75E+00 | 8.65E-01 | no | | intergenic_variant | MODIFIER |
| 12 | 110873042 | AX-170171485 | 0 | T | C | 0.132 | -3.02E-02 | 1.83E-01 | 2.67E+00 | 2.75E+00 | 8.65E-01 | no | Cinp | 3_prime_UTR_variant | MODIFIER |
| 12 | 110877381 | AX-170461039 | 0 | A | G | 0.132 | -3.02E-02 | 1.83E-01 | 2.67E+00 | 2.75E+00 | 8.65E-01 | no | Cinp | intron_variant | MODIFIER |
| 12 | 110903406 | AX-97796525 | 0 | A | G | 0.132 | -3.02E-02 | 1.83E-01 | 2.67E+00 | 2.75E+00 | 8.65E-01 | no | Tecpr2 | intron_variant | MODIFIER |
| 12 | 110905868 | AX-170298797 | 0 | G | A | 0.132 | -3.02E-02 | 1.83E-01 | 2.67E+00 | 2.75E+00 | 8.65E-01 | no | Tecpr2 | intron_variant | MODIFIER |

| | | | | | | | | | | | | | | | |
|---|---|---|---|---|---|---|---|---|---|---|---|---|---|---|---|
| 12 | 110908730 | AX-170166365 | 0 | A | G | 0.132 | -3.02E-02 | 1.83E-01 | 2.67E+00 | 2.75E+00 | 8.65E-01 | no | Tecpr2 | intron_variant | MODIFIER |
| 12 | 110909912 | AX-170409379 | 0 | C | T | 0.132 | -3.02E-02 | 1.83E-01 | 2.67E+00 | 2.75E+00 | 8.65E-01 | no | Tecpr2 | intron_variant | MODIFIER |
| 12 | 114195765 | AX-170401775 | 0 | A | G | 0.132 | -3.02E-02 | 1.83E-01 | 2.67E+00 | 2.75E+00 | 8.65E-01 | no | Ighv3-3 | downstream_gene_variant | MODIFIER |
| 12 | 115121920 | AX-97803606 | 0 | A | C | 0.132 | -3.02E-02 | 1.83E-01 | 2.67E+00 | 2.75E+00 | 8.65E-01 | no | Ighv1-50 | upstream_gene_variant | MODIFIER |
| 12 | 117715934 | AX-170150270 | 0 | G | A | 0.132 | -3.02E-02 | 1.83E-01 | 2.67E+00 | 2.75E+00 | 8.65E-01 | no | Rapgef5 | intron_variant | MODIFIER |
| 12 | 117873375 | AX-168388740 | 0 | A | C | 0.132 | -3.02E-02 | 1.83E-01 | 2.67E+00 | 2.75E+00 | 8.65E-01 | no | Cdca7l | intron_variant | MODIFIER |
| 12 | 117950868 | AX-97831203 | 0 | A | G | 0.132 | -3.02E-02 | 1.83E-01 | 2.67E+00 | 2.75E+00 | 8.65E-01 | no | Dnah11 | intron_variant | MODIFIER |
| 12 | 118454088 | AX-97853129 | 0 | A | T | 0.132 | -3.02E-02 | 1.83E-01 | 2.67E+00 | 2.75E+00 | 8.65E-01 | no | | intergenic_variant | MODIFIER |
| 12 | 118458673 | AX-170323696 | 0 | T | G | 0.132 | -3.02E-02 | 1.83E-01 | 2.67E+00 | 2.75E+00 | 8.65E-01 | no | | intergenic_variant | MODIFIER |
| 12 | 118517970 | AX-170301739 | 0 | T | C | 0.132 | -3.02E-02 | 1.83E-01 | 2.67E+00 | 2.75E+00 | 8.65E-01 | no | D230030E09Rik | downstream_gene_variant | MODIFIER |
| 8 | 50192630 | AX-170507866 | 0 | G | A | 0.158 | -2.56E-02 | 1.42E-01 | 2.64E+00 | 2.69E+00 | 8.65E-01 | no | Gm45341 | intron_variant,non_coding_transcript_variant | MODIFIER |
| 18 | 57806396 | AX-170344706 | 1 | A | G | 0.411 | 1.37E-02 | 8.52E-02 | 2.73E+00 | 2.79E+00 | 8.65E-01 | no | | intergenic_variant | MODIFIER |
| 3 | 63341342 | AX-170435563 | 1 | G | A | 0.054 | 2.33E-02 | 1.52E-01 | 2.63E+00 | 2.77E+00 | 8.66E-01 | no | Mme | intron_variant | MODIFIER |
| 2 | 121365011 | AX-170264593 | 0 | C | T | 0.14 | -3.65E-02 | 2.21E-01 | 2.73E+00 | 2.76E+00 | 8.66E-01 | no | Ckmt1 | downstream_gene_variant | MODIFIER |
| 2 | 129721240 | AX-97789045 | 0 | C | T | 0.14 | -3.65E-02 | 2.21E-01 | 2.73E+00 | 2.76E+00 | 8.66E-01 | no | | intergenic_variant | MODIFIER |
| 5 | 89356569 | AX-97853188 | 0 | G | T | 0.421 | 1.83E-02 | 1.10E-01 | 2.69E+00 | 2.74E+00 | 8.66E-01 | no | | intergenic_variant | MODIFIER |
| 8 | 75579959 | AX-170428800 | 1 | C | A | 0.196 | 2.15E-02 | 1.32E-01 | 2.69E+00 | 2.76E+00 | 8.66E-01 | no | Iqcm | intron_variant | MODIFIER |
| 4 | 143170211 | AX-170024165 | 1 | T | G | 0.098 | 3.17E-02 | 1.92E-01 | 2.67E+00 | 2.74E+00 | 8.66E-01 | no | Prdm2 | intron_variant | MODIFIER |
| 1 | 51383379 | AX-170250953 | 0 | A | G | 0.105 | -3.89E-02 | 2.35E-01 | 2.70E+00 | 2.75E+00 | 8.66E-01 | no | | intergenic_variant | MODIFIER |
| 20 | 168917037 | AX-170035456 | 0 | A | G | 0.184 | 2.07E-02 | 1.18E-01 | 2.62E+00 | 2.71E+00 | 8.66E-01 | no | Arhgap6 | intron_variant | MODIFIER |
| 9 | 78869977 | AX-170032777 | 0 | A | G | 0.202 | 2.15E-02 | 1.31E-01 | 2.73E+00 | 2.77E+00 | 8.66E-01 | no | | intergenic_variant | MODIFIER |
| 9 | 78877623 | AX-170209099 | 0 | G | A | 0.202 | 2.15E-02 | 1.31E-01 | 2.73E+00 | 2.77E+00 | 8.66E-01 | no | | intergenic_variant | MODIFIER |
| 9 | 78878364 | AX-170220571 | 0 | C | T | 0.202 | 2.15E-02 | 1.31E-01 | 2.73E+00 | 2.77E+00 | 8.66E-01 | no | | intergenic_variant | MODIFIER |
| 9 | 78878526 | AX-170454788 | 0 | G | T | 0.202 | 2.15E-02 | 1.31E-01 | 2.73E+00 | 2.77E+00 | 8.66E-01 | no | | intergenic_variant | MODIFIER |
| 9 | 78880012 | AX-170393286 | 0 | A | G | 0.202 | 2.15E-02 | 1.31E-01 | 2.73E+00 | 2.77E+00 | 8.66E-01 | no | | regulatory_region_variant | MODIFIER |
| 9 | 78880227 | AX-170442075 | 0 | A | G | 0.202 | 2.15E-02 | 1.31E-01 | 2.73E+00 | 2.77E+00 | 8.66E-01 | no | | regulatory_region_variant | MODIFIER |
| 9 | 78924660 | AX-170131699 | 0 | G | A | 0.202 | 2.15E-02 | 1.31E-01 | 2.73E+00 | 2.77E+00 | 8.66E-01 | no | | intergenic_variant | MODIFIER |
| 9 | 78955501 | AX-170152964 | 0 | A | G | 0.202 | 2.15E-02 | 1.31E-01 | 2.73E+00 | 2.77E+00 | 8.66E-01 | no | | intergenic_variant | MODIFIER |
| 9 | 79019116 | AX-97825649 | 0 | T | C | 0.202 | 2.15E-02 | 1.31E-01 | 2.73E+00 | 2.77E+00 | 8.66E-01 | no | | intergenic_variant | MODIFIER |
| 9 | 79034983 | AX-170481352 | 0 | C | T | 0.202 | 2.15E-02 | 1.31E-01 | 2.73E+00 | 2.77E+00 | 8.66E-01 | no | | intergenic_variant | MODIFIER |
| 9 | 79045493 | AX-170096598 | 0 | T | G | 0.202 | 2.15E-02 | 1.31E-01 | 2.73E+00 | 2.77E+00 | 8.66E-01 | no | | regulatory_region_variant | MODIFIER |
| 9 | 79107941 | AX-97822201 | 0 | T | G | 0.202 | 2.15E-02 | 1.31E-01 | 2.73E+00 | 2.77E+00 | 8.66E-01 | no | | intergenic_variant | MODIFIER |
| 9 | 79153220 | AX-170253671 | 0 | C | T | 0.202 | 2.15E-02 | 1.31E-01 | 2.73E+00 | 2.77E+00 | 8.66E-01 | no | | intergenic_variant | MODIFIER |
| 9 | 79167566 | AX-170395586 | 0 | A | G | 0.202 | 2.15E-02 | 1.31E-01 | 2.73E+00 | 2.77E+00 | 8.66E-01 | no | Gm47499 | intron_variant,non_coding_transcript_variant | MODIFIER |
| 9 | 79173188 | AX-170343240 | 0 | A | G | 0.202 | 2.15E-02 | 1.31E-01 | 2.73E+00 | 2.77E+00 | 8.66E-01 | no | Gm47499 | intron_variant,non_coding_transcript_variant | MODIFIER |
| 9 | 79201864 | AX-170368651 | 0 | T | C | 0.202 | 2.15E-02 | 1.31E-01 | 2.73E+00 | 2.77E+00 | 8.66E-01 | no | Gm47499 | splice_region_variant,non_coding_transcript_exon_variant | LOW |
| 9 | 79221121 | AX-97810544 | 0 | T | C | 0.202 | 2.15E-02 | 1.31E-01 | 2.73E+00 | 2.77E+00 | 8.66E-01 | no | Gm47501 | intron_variant,non_coding_transcript_variant | MODIFIER |
| 9 | 79235969 | AX-170395299 | 0 | T | C | 0.202 | 2.15E-02 | 1.31E-01 | 2.73E+00 | 2.77E+00 | 8.66E-01 | no | Gm28620 | upstream_gene_variant | MODIFIER |
| 9 | 79251363 | AX-170145948 | 0 | T | C | 0.202 | 2.15E-02 | 1.31E-01 | 2.73E+00 | 2.77E+00 | 8.66E-01 | no | Gm47501 | intron_variant,non_coding_transcript_variant | MODIFIER |
| 9 | 79268365 | AX-170082899 | 0 | A | G | 0.202 | 2.15E-02 | 1.31E-01 | 2.73E+00 | 2.77E+00 | 8.66E-01 | no | Gm47500 | downstream_gene_variant | MODIFIER |
| 9 | 79270898 | AX-97827543 | 0 | A | C | 0.202 | 2.15E-02 | 1.31E-01 | 2.73E+00 | 2.77E+00 | 8.66E-01 | no | Gm47501 | upstream_gene_variant | MODIFIER |
| 9 | 79316070 | AX-170479359 | 0 | G | C | 0.202 | 2.15E-02 | 1.31E-01 | 2.73E+00 | 2.77E+00 | 8.66E-01 | no | | intergenic_variant | MODIFIER |
| 9 | 79317378 | AX-97839686 | 0 | T | C | 0.202 | 2.15E-02 | 1.31E-01 | 2.73E+00 | 2.77E+00 | 8.66E-01 | no | | regulatory_region_variant | MODIFIER |
| 9 | 79342182 | AX-170445125 | 0 | T | C | 0.202 | 2.15E-02 | 1.31E-01 | 2.73E+00 | 2.77E+00 | 8.66E-01 | no | Gm49264 | downstream_gene_variant | MODIFIER |
| 9 | 79435514 | AX-170032783 | 0 | C | T | 0.202 | 2.15E-02 | 1.31E-01 | 2.73E+00 | 2.77E+00 | 8.66E-01 | no | | intergenic_variant | MODIFIER |
| 9 | 79452620 | AX-97798872 | 0 | G | A | 0.202 | 2.15E-02 | 1.31E-01 | 2.73E+00 | 2.77E+00 | 8.66E-01 | no | Gm10635 | intron_variant,non_coding_transcript_variant | MODIFIER |
| 9 | 79476090 | AX-170286332 | 0 | G | A | 0.202 | 2.15E-02 | 1.31E-01 | 2.73E+00 | 2.77E+00 | 8.66E-01 | no | Gm10635 | intron_variant,non_coding_transcript_variant | MODIFIER |
| 9 | 79479304 | AX-170381811 | 0 | G | A | 0.202 | 2.15E-02 | 1.31E-01 | 2.73E+00 | 2.77E+00 | 8.66E-01 | no | Gm10635 | intron_variant,non_coding_transcript_variant | MODIFIER |
| 9 | 79479743 | AX-170166288 | 0 | G | A | 0.202 | 2.15E-02 | 1.31E-01 | 2.73E+00 | 2.77E+00 | 8.66E-01 | no | Gm10635 | intron_variant,non_coding_transcript_variant | MODIFIER |
| 9 | 79515628 | AX-170214671 | 0 | G | A | 0.202 | 2.15E-02 | 1.31E-01 | 2.73E+00 | 2.77E+00 | 8.66E-01 | no | Gm10635 | intron_variant,non_coding_transcript_variant | MODIFIER |
| 9 | 79515652 | AX-97819056 | 0 | T | C | 0.202 | 2.15E-02 | 1.31E-01 | 2.73E+00 | 2.77E+00 | 8.66E-01 | no | Gm10635 | intron_variant,non_coding_transcript_variant | MODIFIER |
| 9 | 79520055 | AX-170156158 | 0 | A | G | 0.202 | 2.15E-02 | 1.31E-01 | 2.73E+00 | 2.77E+00 | 8.66E-01 | no | Gm10635 | upstream_gene_variant | MODIFIER |
| 9 | 79562745 | AX-170098039 | 0 | C | T | 0.202 | 2.15E-02 | 1.31E-01 | 2.73E+00 | 2.77E+00 | 8.66E-01 | no | | intergenic_variant | MODIFIER |
| 9 | 79575960 | AX-97809057 | 0 | T | A | 0.202 | 2.15E-02 | 1.31E-01 | 2.73E+00 | 2.77E+00 | 8.66E-01 | no | Gm8125 | downstream_gene_variant | MODIFIER |

| | | | | | | | | | | | | | | | |
|---|---|---|---|---|---|---|---|---|---|---|---|---|---|---|---|
| 9 | 79600440 | AX-170354432 | 0 | G | A | 0.202 | 2.15E-02 | 1.31E-01 | 2.73E+00 | 2.77E+00 | 8.66E-01 | no | Col12a1 | 3_prime_UTR_variant | MODIFIER |
| 9 | 79608946 | AX-170462387 | 0 | A | C | 0.202 | 2.15E-02 | 1.31E-01 | 2.73E+00 | 2.77E+00 | 8.66E-01 | no | Col12a1 | intron_variant | MODIFIER |
| 9 | 79659499 | AX-170136397 | 0 | A | G | 0.202 | 2.15E-02 | 1.31E-01 | 2.73E+00 | 2.77E+00 | 8.66E-01 | no | Col12a1 | intron_variant | MODIFIER |
| 9 | 79671969 | AX-97786839 | 0 | A | G | 0.202 | 2.15E-02 | 1.31E-01 | 2.73E+00 | 2.77E+00 | 8.66E-01 | no | Col12a1 | intron_variant | MODIFIER |
| 9 | 79722964 | AX-170368272 | 0 | A | G | 0.202 | 2.15E-02 | 1.31E-01 | 2.73E+00 | 2.77E+00 | 8.66E-01 | no | Col12a1 | upstream_gene_variant | MODIFIER |
| 9 | 79769947 | AX-97795179 | 0 | G | A | 0.202 | 2.15E-02 | 1.31E-01 | 2.73E+00 | 2.77E+00 | 8.66E-01 | no | Tmem30a | 3_prime_UTR_variant | MODIFIER |
| 9 | 79898390 | AX-170032796 | 0 | A | G | 0.202 | 2.15E-02 | 1.31E-01 | 2.73E+00 | 2.77E+00 | 8.66E-01 | no | Filip1 | synonymous_variant | LOW |
| 9 | 79992290 | AX-170301711 | 0 | A | T | 0.202 | 2.15E-02 | 1.31E-01 | 2.73E+00 | 2.77E+00 | 8.66E-01 | no | Filip1 | intron_variant | MODIFIER |
| 9 | 80036565 | AX-97794830 | 0 | T | G | 0.202 | 2.15E-02 | 1.31E-01 | 2.73E+00 | 2.77E+00 | 8.66E-01 | no | Gm25433 | upstream_gene_variant | MODIFIER |
| 9 | 80077829 | AX-170321513 | 0 | G | A | 0.202 | 2.15E-02 | 1.31E-01 | 2.73E+00 | 2.77E+00 | 8.66E-01 | no | Senp6 | intron_variant | MODIFIER |
| 9 | 80078325 | AX-97809554 | 0 | C | A | 0.202 | 2.15E-02 | 1.31E-01 | 2.73E+00 | 2.77E+00 | 8.66E-01 | no | Senp6 | intron_variant | MODIFIER |
| 9 | 80103584 | AX-170195031 | 0 | C | T | 0.202 | 2.15E-02 | 1.31E-01 | 2.73E+00 | 2.77E+00 | 8.66E-01 | no | Senp6 | intron_variant | MODIFIER |
| 9 | 80146973 | AX-170260365 | 0 | G | A | 0.202 | 2.15E-02 | 1.31E-01 | 2.73E+00 | 2.77E+00 | 8.66E-01 | no | Senp6 | downstream_gene_variant | MODIFIER |
| 9 | 80147073 | AX-170116568 | 0 | C | G | 0.202 | 2.15E-02 | 1.31E-01 | 2.73E+00 | 2.77E+00 | 8.66E-01 | no | Senp6 | downstream_gene_variant | MODIFIER |
| 9 | 80152274 | AX-170090345 | 0 | A | G | 0.202 | 2.15E-02 | 1.31E-01 | 2.73E+00 | 2.77E+00 | 8.66E-01 | no | | intergenic_variant | MODIFIER |
| 9 | 80189097 | AX-170205217 | 0 | T | C | 0.202 | 2.15E-02 | 1.31E-01 | 2.73E+00 | 2.77E+00 | 8.66E-01 | no | Myo6 | intron_variant | MODIFIER |
| 9 | 80198000 | AX-170501973 | 0 | C | T | 0.202 | 2.15E-02 | 1.31E-01 | 2.73E+00 | 2.77E+00 | 8.66E-01 | no | Myo6 | intron_variant | MODIFIER |
| 9 | 80200125 | AX-97845024 | 0 | A | G | 0.202 | 2.15E-02 | 1.31E-01 | 2.73E+00 | 2.77E+00 | 8.66E-01 | no | Myo6 | intron_variant | MODIFIER |
| 9 | 80208868 | AX-170391534 | 0 | C | T | 0.202 | 2.15E-02 | 1.31E-01 | 2.73E+00 | 2.77E+00 | 8.66E-01 | no | Myo6 | intron_variant | MODIFIER |
| 9 | 80220249 | AX-170090346 | 0 | G | A | 0.202 | 2.15E-02 | 1.31E-01 | 2.73E+00 | 2.77E+00 | 8.66E-01 | no | Myo6 | intron_variant | MODIFIER |
| 9 | 80220281 | AX-170295155 | 0 | A | G | 0.202 | 2.15E-02 | 1.31E-01 | 2.73E+00 | 2.77E+00 | 8.66E-01 | no | Myo6 | intron_variant | MODIFIER |
| 9 | 80253461 | AX-170082917 | 0 | G | A | 0.202 | 2.15E-02 | 1.31E-01 | 2.73E+00 | 2.77E+00 | 8.66E-01 | no | Myo6 | intron_variant | MODIFIER |
| 9 | 80254266 | AX-170082916 | 0 | C | A | 0.202 | 2.15E-02 | 1.31E-01 | 2.73E+00 | 2.77E+00 | 8.66E-01 | no | Myo6 | intron_variant | MODIFIER |
| 9 | 80280538 | AX-170329491 | 0 | A | T | 0.202 | 2.15E-02 | 1.31E-01 | 2.73E+00 | 2.77E+00 | 8.66E-01 | no | Myo6 | intron_variant | MODIFIER |
| 9 | 80280642 | AX-170518148 | 0 | G | A | 0.202 | 2.15E-02 | 1.31E-01 | 2.73E+00 | 2.77E+00 | 8.66E-01 | no | Myo6 | intron_variant | MODIFIER |
| 9 | 80294519 | AX-170176671 | 0 | T | C | 0.202 | 2.15E-02 | 1.31E-01 | 2.73E+00 | 2.77E+00 | 8.66E-01 | no | Myo6 | intron_variant | MODIFIER |
| 9 | 80518689 | AX-170032799 | 0 | C | T | 0.202 | 2.15E-02 | 1.31E-01 | 2.73E+00 | 2.77E+00 | 8.66E-01 | no | | intergenic_variant | MODIFIER |
| 9 | 80528791 | AX-170096603 | 0 | A | G | 0.202 | 2.15E-02 | 1.31E-01 | 2.73E+00 | 2.77E+00 | 8.66E-01 | no | | intergenic_variant | MODIFIER |
| 9 | 80528873 | AX-170032798 | 0 | G | T | 0.202 | 2.15E-02 | 1.31E-01 | 2.73E+00 | 2.77E+00 | 8.66E-01 | no | | intergenic_variant | MODIFIER |
| 9 | 80538057 | AX-170442994 | 0 | G | T | 0.202 | 2.15E-02 | 1.31E-01 | 2.73E+00 | 2.77E+00 | 8.66E-01 | no | | intergenic_variant | MODIFIER |
| 9 | 80538099 | AX-170167712 | 0 | A | G | 0.202 | 2.15E-02 | 1.31E-01 | 2.73E+00 | 2.77E+00 | 8.66E-01 | no | | intergenic_variant | MODIFIER |
| 9 | 80588222 | AX-170244060 | 0 | C | T | 0.202 | 2.15E-02 | 1.31E-01 | 2.73E+00 | 2.77E+00 | 8.66E-01 | no | Gm48387 | intron_variant,non_coding_transcript_variant | MODIFIER |
| 9 | 80596188 | AX-170090347 | 0 | G | A | 0.202 | 2.15E-02 | 1.31E-01 | 2.73E+00 | 2.77E+00 | 8.66E-01 | no | Gm48387 | intron_variant,non_coding_transcript_variant | MODIFIER |
| 9 | 80609447 | AX-170217465 | 0 | A | C | 0.202 | 2.15E-02 | 1.31E-01 | 2.73E+00 | 2.77E+00 | 8.66E-01 | no | | intergenic_variant | MODIFIER |
| 9 | 80714429 | AX-170136720 | 0 | T | C | 0.202 | 2.15E-02 | 1.31E-01 | 2.73E+00 | 2.77E+00 | 8.66E-01 | no | Gm39380 | downstream_gene_variant | MODIFIER |
| 9 | 80746079 | AX-170035813 | 0 | A | G | 0.202 | 2.15E-02 | 1.31E-01 | 2.73E+00 | 2.77E+00 | 8.66E-01 | no | | intergenic_variant | MODIFIER |
| 9 | 80746520 | AX-170125561 | 0 | T | G | 0.202 | 2.15E-02 | 1.31E-01 | 2.73E+00 | 2.77E+00 | 8.66E-01 | no | | intergenic_variant | MODIFIER |
| 9 | 80751877 | AX-170035812 | 0 | A | G | 0.202 | 2.15E-02 | 1.31E-01 | 2.73E+00 | 2.77E+00 | 8.66E-01 | no | | intergenic_variant | MODIFIER |
| 9 | 80770538 | AX-170433647 | 0 | A | T | 0.202 | 2.15E-02 | 1.31E-01 | 2.73E+00 | 2.77E+00 | 8.66E-01 | no | | intergenic_variant | MODIFIER |
| 9 | 80777095 | AX-170082931 | 0 | G | A | 0.202 | 2.15E-02 | 1.31E-01 | 2.73E+00 | 2.77E+00 | 8.66E-01 | no | | intergenic_variant | MODIFIER |
| 9 | 80781907 | AX-170413594 | 0 | T | C | 0.202 | 2.15E-02 | 1.31E-01 | 2.73E+00 | 2.77E+00 | 8.66E-01 | no | | intergenic_variant | MODIFIER |
| 9 | 80800895 | AX-170082934 | 0 | C | G | 0.202 | 2.15E-02 | 1.31E-01 | 2.73E+00 | 2.77E+00 | 8.66E-01 | no | Gm27224 | upstream_gene_variant | MODIFIER |
| 9 | 80815142 | AX-170198965 | 0 | G | A | 0.202 | 2.15E-02 | 1.31E-01 | 2.73E+00 | 2.77E+00 | 8.66E-01 | no | Gm27224 | intron_variant,non_coding_transcript_variant | MODIFIER |
| 9 | 80819282 | AX-170305968 | 0 | G | C | 0.202 | 2.15E-02 | 1.31E-01 | 2.73E+00 | 2.77E+00 | 8.66E-01 | no | Gm27224 | intron_variant,non_coding_transcript_variant | MODIFIER |
| 9 | 80901956 | AX-170105198 | 0 | G | A | 0.202 | 2.15E-02 | 1.31E-01 | 2.73E+00 | 2.77E+00 | 8.66E-01 | no | | intergenic_variant | MODIFIER |
| 9 | 80909335 | AX-170348048 | 0 | G | A | 0.202 | 2.15E-02 | 1.31E-01 | 2.73E+00 | 2.77E+00 | 8.66E-01 | no | | intergenic_variant | MODIFIER |
| 9 | 80947307 | AX-170419928 | 0 | A | G | 0.202 | 2.15E-02 | 1.31E-01 | 2.73E+00 | 2.77E+00 | 8.66E-01 | no | | intergenic_variant | MODIFIER |
| 9 | 80947737 | AX-170208380 | 0 | T | C | 0.202 | 2.15E-02 | 1.31E-01 | 2.73E+00 | 2.77E+00 | 8.66E-01 | no | | intergenic_variant | MODIFIER |
| 9 | 80986884 | AX-170205377 | 0 | C | A | 0.202 | 2.15E-02 | 1.31E-01 | 2.73E+00 | 2.77E+00 | 8.66E-01 | no | Gm26713 | intron_variant,non_coding_transcript_variant | MODIFIER |
| 9 | 80996960 | AX-170410952 | 0 | G | A | 0.202 | 2.15E-02 | 1.31E-01 | 2.73E+00 | 2.77E+00 | 8.66E-01 | no | Gm26713 | intron_variant,non_coding_transcript_variant | MODIFIER |
| 9 | 81007904 | AX-170484200 | 0 | G | A | 0.202 | 2.15E-02 | 1.31E-01 | 2.73E+00 | 2.77E+00 | 8.66E-01 | no | Gm26713 | intron_variant,non_coding_transcript_variant | MODIFIER |
| 9 | 81008787 | AX-170446021 | 0 | A | G | 0.202 | 2.15E-02 | 1.31E-01 | 2.73E+00 | 2.77E+00 | 8.66E-01 | no | Gm26713 | intron_variant,non_coding_transcript_variant | MODIFIER |
| 9 | 81010618 | AX-170238491 | 0 | G | A | 0.202 | 2.15E-02 | 1.31E-01 | 2.73E+00 | 2.77E+00 | 8.66E-01 | no | Gm26713 | intron_variant,non_coding_transcript_variant | MODIFIER |
| 9 | 81012192 | AX-170135042 | 0 | C | G | 0.202 | 2.15E-02 | 1.31E-01 | 2.73E+00 | 2.77E+00 | 8.66E-01 | no | Gm26713 | intron_variant,non_coding_transcript_variant | MODIFIER |

| | | | | | | | | | | | | | | | |
|---|---|---|---|---|---|---|---|---|---|---|---|---|---|---|---|
| 9 | 81059791 | AX-170183752 | 0 | C | A | 0.202 | 2.15E-02 | 1.31E-01 | 2.73E+00 | 2.77E+00 | 8.66E-01 | no | Gm26713 | intron_variant,non_coding_transcript_variant | MODIFIER |
| 9 | 81263153 | AX-170320667 | 0 | T | C | 0.202 | 2.15E-02 | 1.31E-01 | 2.73E+00 | 2.77E+00 | 8.66E-01 | no | | intergenic_variant | MODIFIER |
| 9 | 81275600 | AX-170194068 | 0 | T | C | 0.202 | 2.15E-02 | 1.31E-01 | 2.73E+00 | 2.77E+00 | 8.66E-01 | no | | intergenic_variant | MODIFIER |
| 9 | 81276467 | AX-170221158 | 0 | C | T | 0.202 | 2.15E-02 | 1.31E-01 | 2.73E+00 | 2.77E+00 | 8.66E-01 | no | | intergenic_variant | MODIFIER |
| 9 | 81407724 | AX-170517233 | 0 | A | G | 0.202 | 2.15E-02 | 1.31E-01 | 2.73E+00 | 2.77E+00 | 8.66E-01 | no | | intergenic_variant | MODIFIER |
| 9 | 83040245 | AX-170291323 | 0 | T | G | 0.202 | 2.15E-02 | 1.31E-01 | 2.73E+00 | 2.77E+00 | 8.66E-01 | no | Gm39383 | intron_variant,non_coding_transcript_variant | MODIFIER |
| 9 | 83042143 | AX-170534387 | 0 | T | A | 0.202 | 2.15E-02 | 1.31E-01 | 2.73E+00 | 2.77E+00 | 8.66E-01 | no | Gm39383 | non_coding_transcript_exon_variant | MODIFIER |
| 9 | 83068213 | AX-170349783 | 0 | T | C | 0.202 | 2.15E-02 | 1.31E-01 | 2.73E+00 | 2.77E+00 | 8.66E-01 | no | | intergenic_variant | MODIFIER |
| 9 | 83072854 | AX-170284861 | 0 | A | G | 0.202 | 2.15E-02 | 1.31E-01 | 2.73E+00 | 2.77E+00 | 8.66E-01 | no | Gm27248 | downstream_gene_variant | MODIFIER |
| 9 | 83110483 | AX-170365274 | 0 | A | T | 0.202 | 2.15E-02 | 1.31E-01 | 2.73E+00 | 2.77E+00 | 8.66E-01 | no | Hmgn3 | intron_variant | MODIFIER |
| 9 | 60460437 | AX-170369711 | 0 | G | T | 0.202 | -2.26E-02 | 1.34E-01 | 2.69E+00 | 2.72E+00 | 8.66E-01 | no | Thsd4 | intron_variant | MODIFIER |
| 5 | 123615072 | AX-170073359 | 1 | C | T | 0.464 | -1.80E-02 | 1.10E-01 | 2.70E+00 | 2.76E+00 | 8.66E-01 | no | Clip1 | intron_variant | MODIFIER |
| 1 | 63477494 | AX-170245900 | 0 | T | A | 0.123 | 3.11E-02 | 1.90E-01 | 2.66E+00 | 2.75E+00 | 8.66E-01 | no | Adam23 | intron_variant | MODIFIER |
| 1 | 13161985 | AX-170436944 | 0 | C | T | 0.377 | 1.34E-02 | 7.72E-02 | 2.65E+00 | 2.71E+00 | 8.66E-01 | no | Ncoa2 | intron_variant | MODIFIER |
| 15 | 100783283 | AX-170145785 | 1 | C | A | 0.116 | 2.55E-02 | 1.50E-01 | 2.65E+00 | 2.72E+00 | 8.66E-01 | no | Slc4a8 | intron_variant | MODIFIER |
| 10 | 10834235 | AX-170091860 | 1 | G | A | 0.232 | 1.63E-02 | 1.01E-01 | 2.68E+00 | 2.76E+00 | 8.66E-01 | no | Grm1 | intron_variant | MODIFIER |
| 11 | 49097091 | AX-97851545 | 0 | T | C | 0.088 | -3.02E-02 | 1.61E-01 | 2.58E+00 | 2.70E+00 | 8.66E-01 | no | Ifi47 | downstream_gene_variant | MODIFIER |
| 4 | 108924357 | AX-170099833 | 0 | A | T | 0.254 | 2.15E-02 | 1.12E-01 | 2.59E+00 | 2.68E+00 | 8.66E-01 | no | Rab3b | intron_variant | MODIFIER |
| 18 | 58098796 | AX-170423816 | 0 | T | C | 0.404 | 1.86E-02 | 9.63E-02 | 2.60E+00 | 2.66E+00 | 8.66E-01 | no | Fbn2 | intron_variant | MODIFIER |
| 1 | 50203045 | AX-97823744 | 0 | G | C | 0.105 | -3.91E-02 | 2.36E-01 | 2.70E+00 | 2.74E+00 | 8.66E-01 | no | | intergenic_variant | MODIFIER |
| 13 | 34238702 | AX-170008071 | 1 | C | G | 0.188 | -2.08E-02 | 1.16E-01 | 2.62E+00 | 2.70E+00 | 8.66E-01 | no | Slc22a23 | intron_variant | MODIFIER |
| 14 | 33371044 | AX-170369055 | 1 | A | G | 0.045 | 3.00E-02 | 1.77E-01 | 2.61E+00 | 2.73E+00 | 8.66E-01 | no | Arhgap22 | downstream_gene_variant | MODIFIER |
| 12 | 68849296 | AX-170191359 | 1 | G | T | 0.054 | 4.12E-02 | 2.57E-01 | 2.70E+00 | 2.77E+00 | 8.66E-01 | no | | intergenic_variant | MODIFIER |
| 18 | 34347204 | AX-170015510 | 1 | A | G | 0.054 | 4.12E-02 | 2.57E-01 | 2.70E+00 | 2.77E+00 | 8.66E-01 | no | Reep5 | intron_variant | MODIFIER |
| 18 | 43642840 | AX-170532142 | 1 | T | A | 0.054 | 4.12E-02 | 2.57E-01 | 2.70E+00 | 2.77E+00 | 8.66E-01 | no | Jakmip2 | intron_variant | MODIFIER |
| 18 | 46160048 | AX-170060749 | 1 | G | A | 0.054 | 4.12E-02 | 2.57E-01 | 2.70E+00 | 2.77E+00 | 8.66E-01 | no | | intergenic_variant | MODIFIER |
| 1 | 107319125 | AX-170322223 | 1 | A | G | 0.161 | 3.43E-02 | 1.90E-01 | 2.62E+00 | 2.70E+00 | 8.66E-01 | no | | intergenic_variant | MODIFIER |
| 4 | 144296758 | AX-170070941 | 0 | G | A | 0.105 | 2.89E-02 | 1.68E-01 | 2.62E+00 | 2.72E+00 | 8.67E-01 | no | Gm13120 | non_coding_transcript_exon_variant | MODIFIER |
| 4 | 148489771 | AX-170211589 | 0 | T | A | 0.105 | 2.89E-02 | 1.68E-01 | 2.62E+00 | 2.72E+00 | 8.67E-01 | no | Mtor | intron_variant | MODIFIER |
| 4 | 149164031 | AX-97804632 | 0 | C | T | 0.105 | 2.89E-02 | 1.68E-01 | 2.62E+00 | 2.72E+00 | 8.67E-01 | no | Pgd | intron_variant | MODIFIER |
| 6 | 144744986 | AX-170336982 | 0 | C | T | 0.07 | -4.57E-02 | 2.77E-01 | 2.71E+00 | 2.74E+00 | 8.67E-01 | no | Sox5 | intron_variant,non_coding_transcript_variant | MODIFIER |
| 6 | 145344467 | AX-170254088 | 0 | G | A | 0.07 | -4.57E-02 | 2.77E-01 | 2.71E+00 | 2.74E+00 | 8.67E-01 | no | | intergenic_variant | MODIFIER |
| 1 | 170206374 | AX-97812535 | 0 | C | A | 0.237 | 2.21E-02 | 1.31E-01 | 2.68E+00 | 2.72E+00 | 8.67E-01 | no | Uhmk1 | intron_variant | MODIFIER |
| 1 | 35995551 | AX-170332873 | 0 | T | C | 0.456 | 1.29E-02 | 7.91E-02 | 2.67E+00 | 2.75E+00 | 8.67E-01 | no | | intergenic_variant | MODIFIER |
| 1 | 35995739 | AX-170296881 | 0 | T | C | 0.456 | 1.29E-02 | 7.91E-02 | 2.67E+00 | 2.75E+00 | 8.67E-01 | no | | intergenic_variant | MODIFIER |
| 1 | 35999939 | AX-170173394 | 0 | C | A | 0.456 | 1.29E-02 | 7.91E-02 | 2.67E+00 | 2.75E+00 | 8.67E-01 | no | Gm33222 | downstream_gene_variant | MODIFIER |
| 1 | 36000002 | AX-170527676 | 0 | G | T | 0.456 | 1.29E-02 | 7.91E-02 | 2.67E+00 | 2.75E+00 | 8.67E-01 | no | Gm33222 | downstream_gene_variant | MODIFIER |
| 1 | 36000382 | AX-170299506 | 0 | G | A | 0.456 | 1.29E-02 | 7.91E-02 | 2.67E+00 | 2.75E+00 | 8.67E-01 | no | Gm33222 | downstream_gene_variant | MODIFIER |
| 1 | 36001484 | AX-170366892 | 0 | C | T | 0.456 | 1.29E-02 | 7.91E-02 | 2.67E+00 | 2.75E+00 | 8.67E-01 | no | Gm33222 | downstream_gene_variant | MODIFIER |
| 1 | 36002647 | AX-170331683 | 0 | A | G | 0.456 | 1.29E-02 | 7.91E-02 | 2.67E+00 | 2.75E+00 | 8.67E-01 | no | Gm33222 | downstream_gene_variant | MODIFIER |
| 1 | 36003242 | AX-170187623 | 0 | T | C | 0.456 | 1.29E-02 | 7.91E-02 | 2.67E+00 | 2.75E+00 | 8.67E-01 | no | Gm33222 | non_coding_transcript_exon_variant | MODIFIER |
| 1 | 36004285 | AX-170289797 | 0 | G | A | 0.456 | 1.29E-02 | 7.91E-02 | 2.67E+00 | 2.75E+00 | 8.67E-01 | no | Gm33222 | intron_variant,non_coding_transcript_variant | MODIFIER |
| 1 | 36007178 | AX-170091467 | 0 | T | C | 0.456 | 1.29E-02 | 7.91E-02 | 2.67E+00 | 2.75E+00 | 8.67E-01 | no | 4930535G08Rik | upstream_gene_variant | MODIFIER |
| 1 | 36030844 | AX-170137034 | 0 | C | T | 0.456 | 1.29E-02 | 7.91E-02 | 2.67E+00 | 2.75E+00 | 8.67E-01 | no | Gm38049 | non_coding_transcript_exon_variant | MODIFIER |
| 1 | 36038377 | AX-97856676 | 0 | G | A | 0.456 | 1.29E-02 | 7.91E-02 | 2.67E+00 | 2.75E+00 | 8.67E-01 | no | | intergenic_variant | MODIFIER |
| 1 | 36039251 | AX-170498294 | 0 | C | T | 0.456 | 1.29E-02 | 7.91E-02 | 2.67E+00 | 2.75E+00 | 8.67E-01 | no | | intergenic_variant | MODIFIER |
| 1 | 36041144 | AX-170147981 | 0 | G | A | 0.456 | 1.29E-02 | 7.91E-02 | 2.67E+00 | 2.75E+00 | 8.67E-01 | no | | intergenic_variant | MODIFIER |
| 1 | 36087174 | AX-170492173 | 0 | T | C | 0.456 | 1.29E-02 | 7.91E-02 | 2.67E+00 | 2.75E+00 | 8.67E-01 | no | Hs6st1 | intron_variant | MODIFIER |
| 1 | 36087259 | AX-170283616 | 0 | G | T | 0.456 | 1.29E-02 | 7.91E-02 | 2.67E+00 | 2.75E+00 | 8.67E-01 | no | Hs6st1 | intron_variant | MODIFIER |
| 1 | 36089666 | AX-170457331 | 0 | A | G | 0.456 | 1.29E-02 | 7.91E-02 | 2.67E+00 | 2.75E+00 | 8.67E-01 | no | Hs6st1 | intron_variant | MODIFIER |
| 1 | 36093675 | AX-170223553 | 0 | C | T | 0.456 | 1.29E-02 | 7.91E-02 | 2.67E+00 | 2.75E+00 | 8.67E-01 | no | Hs6st1 | intron_variant | MODIFIER |
| 1 | 36136825 | AX-170123331 | 0 | G | A | 0.456 | 1.29E-02 | 7.91E-02 | 2.67E+00 | 2.75E+00 | 8.67E-01 | no | Uggt1 | downstream_gene_variant | MODIFIER |
| 1 | 36144277 | AX-170513396 | 0 | G | A | 0.456 | 1.29E-02 | 7.91E-02 | 2.67E+00 | 2.75E+00 | 8.67E-01 | no | Uggt1 | intron_variant | MODIFIER |
| 1 | 36146661 | AX-170040878 | 0 | G | A | 0.456 | 1.29E-02 | 7.91E-02 | 2.67E+00 | 2.75E+00 | 8.67E-01 | no | Uggt1 | synonymous_variant | LOW |

| | | | | | | | | | | | | | | | |
|---|---|---|---|---|---|---|---|---|---|---|---|---|---|---|---|
| 1 | 36146688 | AX-170040880 | 0 | C | T | 0.456 | 1.29E-02 | 7.91E-02 | 2.67E+00 | 2.75E+00 | 8.67E-01 | no | Uggt1 | synonymous_variant | LOW |
| 1 | 36157797 | AX-97793429 | 0 | A | C | 0.456 | 1.29E-02 | 7.91E-02 | 2.67E+00 | 2.75E+00 | 8.67E-01 | no | Uggt1 | intron_variant | MODIFIER |
| 1 | 36161474 | AX-170289609 | 0 | G | A | 0.456 | 1.29E-02 | 7.91E-02 | 2.67E+00 | 2.75E+00 | 8.67E-01 | no | Uggt1 | intron_variant | MODIFIER |
| 1 | 36162596 | AX-170208509 | 0 | T | C | 0.456 | 1.29E-02 | 7.91E-02 | 2.67E+00 | 2.75E+00 | 8.67E-01 | no | Uggt1 | intron_variant | MODIFIER |
| 1 | 36177295 | AX-170230216 | 0 | A | G | 0.456 | 1.29E-02 | 7.91E-02 | 2.67E+00 | 2.75E+00 | 8.67E-01 | no | Uggt1 | intron_variant | MODIFIER |
| 1 | 36192422 | AX-170214392 | 0 | G | A | 0.456 | 1.29E-02 | 7.91E-02 | 2.67E+00 | 2.75E+00 | 8.67E-01 | no | Uggt1 | synonymous_variant | LOW |
| 1 | 36217911 | AX-170413398 | 0 | A | G | 0.456 | 1.29E-02 | 7.91E-02 | 2.67E+00 | 2.75E+00 | 8.67E-01 | no | Uggt1 | intron_variant | MODIFIER |
| 1 | 36225970 | AX-170254085 | 0 | C | A | 0.456 | 1.29E-02 | 7.91E-02 | 2.67E+00 | 2.75E+00 | 8.67E-01 | no | Uggt1 | intron_variant | MODIFIER |
| 1 | 36229951 | AX-170427923 | 0 | G | T | 0.456 | 1.29E-02 | 7.91E-02 | 2.67E+00 | 2.75E+00 | 8.67E-01 | no | Uggt1 | intron_variant | MODIFIER |
| 1 | 36246033 | AX-170256902 | 0 | A | G | 0.456 | 1.29E-02 | 7.91E-02 | 2.67E+00 | 2.75E+00 | 8.67E-01 | no | Uggt1 | upstream_gene_variant | MODIFIER |
| 1 | 36253398 | AX-170178721 | 0 | A | G | 0.456 | 1.29E-02 | 7.91E-02 | 2.67E+00 | 2.75E+00 | 8.67E-01 | no | | regulatory_region_variant | MODIFIER |
| 1 | 36265540 | AX-170237551 | 0 | G | A | 0.456 | 1.29E-02 | 7.91E-02 | 2.67E+00 | 2.75E+00 | 8.67E-01 | no | Neurl3 | 3_prime_UTR_variant | MODIFIER |
| 1 | 36269029 | AX-170000286 | 0 | G | T | 0.456 | 1.29E-02 | 7.91E-02 | 2.67E+00 | 2.75E+00 | 8.67E-01 | no | Neurl3 | intron_variant | MODIFIER |
| 1 | 36323160 | AX-170000280 | 0 | A | C | 0.456 | 1.29E-02 | 7.91E-02 | 2.67E+00 | 2.75E+00 | 8.67E-01 | no | Arid5a | downstream_gene_variant | MODIFIER |
| 1 | 36323320 | AX-170000283 | 0 | A | G | 0.456 | 1.29E-02 | 7.91E-02 | 2.67E+00 | 2.75E+00 | 8.67E-01 | no | Arid5a | downstream_gene_variant | MODIFIER |
| 1 | 36327373 | AX-170156992 | 0 | C | T | 0.456 | 1.29E-02 | 7.91E-02 | 2.67E+00 | 2.75E+00 | 8.67E-01 | no | Arid5a | downstream_gene_variant | MODIFIER |
| 1 | 36333045 | AX-97802507 | 0 | G | A | 0.456 | 1.29E-02 | 7.91E-02 | 2.67E+00 | 2.75E+00 | 8.67E-01 | no | Kansl3 | downstream_gene_variant | MODIFIER |
| 1 | 36338128 | AX-170393378 | 0 | G | A | 0.456 | 1.29E-02 | 7.91E-02 | 2.67E+00 | 2.75E+00 | 8.67E-01 | no | Kansl3 | intron_variant | MODIFIER |
| 1 | 36345072 | AX-170161737 | 0 | C | T | 0.456 | 1.29E-02 | 7.91E-02 | 2.67E+00 | 2.75E+00 | 8.67E-01 | no | Kansl3 | intron_variant | MODIFIER |
| 1 | 36345771 | AX-170147640 | 0 | A | G | 0.456 | 1.29E-02 | 7.91E-02 | 2.67E+00 | 2.75E+00 | 8.67E-01 | no | Kansl3 | intron_variant | MODIFIER |
| 1 | 36353912 | AX-170000279 | 0 | G | A | 0.456 | 1.29E-02 | 7.91E-02 | 2.67E+00 | 2.75E+00 | 8.67E-01 | no | Kansl3 | intron_variant | MODIFIER |
| 1 | 36365077 | AX-170292951 | 0 | T | C | 0.456 | 1.29E-02 | 7.91E-02 | 2.67E+00 | 2.75E+00 | 8.67E-01 | no | Kansl3 | intron_variant | MODIFIER |
| 1 | 36412948 | AX-170458658 | 0 | T | C | 0.456 | 1.29E-02 | 7.91E-02 | 2.67E+00 | 2.75E+00 | 8.67E-01 | no | Fer1l5 | upstream_gene_variant | MODIFIER |
| 1 | 36435251 | AX-170214151 | 0 | T | C | 0.456 | 1.29E-02 | 7.91E-02 | 2.67E+00 | 2.75E+00 | 8.67E-01 | no | Lman2l | intron_variant,NMD_transcript_variant | MODIFIER |
| 1 | 36521897 | AX-97853569 | 0 | G | A | 0.456 | 1.29E-02 | 7.91E-02 | 2.67E+00 | 2.75E+00 | 8.67E-01 | no | Cnnm3 | intron_variant | MODIFIER |
| 1 | 36769934 | AX-170000291 | 0 | A | G | 0.456 | 1.29E-02 | 7.91E-02 | 2.67E+00 | 2.75E+00 | 8.67E-01 | no | Zap70 | intron_variant | MODIFIER |
| 1 | 36787065 | AX-170241147 | 0 | C | T | 0.456 | 1.29E-02 | 7.91E-02 | 2.67E+00 | 2.75E+00 | 8.67E-01 | no | Zap70 | downstream_gene_variant | MODIFIER |
| 1 | 36791983 | AX-97824364 | 0 | C | T | 0.456 | 1.29E-02 | 7.91E-02 | 2.67E+00 | 2.75E+00 | 8.67E-01 | no | Tmem131 | downstream_gene_variant | MODIFIER |
| 1 | 36795115 | AX-170352980 | 0 | A | G | 0.456 | 1.29E-02 | 7.91E-02 | 2.67E+00 | 2.75E+00 | 8.67E-01 | no | Tmem131 | synonymous_variant | LOW |
| 1 | 36795475 | AX-170269147 | 0 | T | G | 0.456 | 1.29E-02 | 7.91E-02 | 2.67E+00 | 2.75E+00 | 8.67E-01 | no | Tmem131 | intron_variant | MODIFIER |
| 1 | 36798457 | AX-170355592 | 0 | A | G | 0.456 | 1.29E-02 | 7.91E-02 | 2.67E+00 | 2.75E+00 | 8.67E-01 | no | Tmem131 | intron_variant | MODIFIER |
| 1 | 36816825 | AX-170234580 | 0 | C | T | 0.456 | 1.29E-02 | 7.91E-02 | 2.67E+00 | 2.75E+00 | 8.67E-01 | no | Tmem131 | intron_variant | MODIFIER |
| 1 | 36818212 | AX-170262933 | 0 | T | C | 0.456 | 1.29E-02 | 7.91E-02 | 2.67E+00 | 2.75E+00 | 8.67E-01 | no | Tmem131 | intron_variant | MODIFIER |
| 1 | 36818672 | AX-170325055 | 0 | C | T | 0.456 | 1.29E-02 | 7.91E-02 | 2.67E+00 | 2.75E+00 | 8.67E-01 | no | Tmem131 | synonymous_variant | LOW |
| 1 | 36824590 | AX-170228687 | 0 | C | T | 0.456 | 1.29E-02 | 7.91E-02 | 2.67E+00 | 2.75E+00 | 8.67E-01 | no | Tmem131 | intron_variant | MODIFIER |
| 1 | 36825260 | AX-170370854 | 0 | G | A | 0.456 | 1.29E-02 | 7.91E-02 | 2.67E+00 | 2.75E+00 | 8.67E-01 | no | Tmem131 | synonymous_variant | LOW |
| 1 | 36826618 | AX-170000292 | 0 | A | C | 0.456 | 1.29E-02 | 7.91E-02 | 2.67E+00 | 2.75E+00 | 8.67E-01 | no | Tmem131 | intron_variant | MODIFIER |
| 1 | 36826721 | AX-170000294 | 0 | T | C | 0.456 | 1.29E-02 | 7.91E-02 | 2.67E+00 | 2.75E+00 | 8.67E-01 | no | Tmem131 | intron_variant | MODIFIER |
| 1 | 36827331 | AX-170000287 | 0 | A | C | 0.456 | 1.29E-02 | 7.91E-02 | 2.67E+00 | 2.75E+00 | 8.67E-01 | no | Tmem131 | synonymous_variant | LOW |
| 1 | 36834717 | AX-170117841 | 0 | A | G | 0.456 | 1.29E-02 | 7.91E-02 | 2.67E+00 | 2.75E+00 | 8.67E-01 | no | Tmem131 | intron_variant | MODIFIER |
| 1 | 36835360 | AX-170222959 | 0 | A | G | 0.456 | 1.29E-02 | 7.91E-02 | 2.67E+00 | 2.75E+00 | 8.67E-01 | no | Tmem131 | intron_variant | MODIFIER |
| 1 | 36835381 | AX-170512702 | 0 | G | T | 0.456 | 1.29E-02 | 7.91E-02 | 2.67E+00 | 2.75E+00 | 8.67E-01 | no | Tmem131 | intron_variant | MODIFIER |
| 1 | 36843246 | AX-168414371 | 0 | A | G | 0.456 | 1.29E-02 | 7.91E-02 | 2.67E+00 | 2.75E+00 | 8.67E-01 | no | Tmem131 | intron_variant | MODIFIER |
| 1 | 36856905 | AX-97824356 | 0 | G | A | 0.456 | 1.29E-02 | 7.91E-02 | 2.67E+00 | 2.75E+00 | 8.67E-01 | no | Tmem131 | intron_variant | MODIFIER |
| 1 | 36857065 | AX-170000293 | 0 | C | T | 0.456 | 1.29E-02 | 7.91E-02 | 2.67E+00 | 2.75E+00 | 8.67E-01 | no | Tmem131 | intron_variant | MODIFIER |
| 1 | 36859738 | AX-170040894 | 0 | C | T | 0.456 | 1.29E-02 | 7.91E-02 | 2.67E+00 | 2.75E+00 | 8.67E-01 | no | Tmem131 | intron_variant | MODIFIER |
| 1 | 36859988 | AX-170000288 | 0 | C | T | 0.456 | 1.29E-02 | 7.91E-02 | 2.67E+00 | 2.75E+00 | 8.67E-01 | no | Tmem131 | intron_variant | MODIFIER |
| 1 | 36895177 | AX-97825250 | 0 | G | A | 0.456 | 1.29E-02 | 7.91E-02 | 2.67E+00 | 2.75E+00 | 8.67E-01 | no | Tmem131 | intron_variant | MODIFIER |
| 1 | 36901397 | AX-170387351 | 0 | G | T | 0.456 | 1.29E-02 | 7.91E-02 | 2.67E+00 | 2.75E+00 | 8.67E-01 | no | Tmem131 | intron_variant | MODIFIER |
| 1 | 36904828 | AX-170336148 | 0 | C | T | 0.456 | 1.29E-02 | 7.91E-02 | 2.67E+00 | 2.75E+00 | 8.67E-01 | no | Tmem131 | intron_variant | MODIFIER |
| 1 | 36962193 | AX-170241122 | 0 | C | G | 0.456 | 1.29E-02 | 7.91E-02 | 2.67E+00 | 2.75E+00 | 8.67E-01 | no | Rpl12-ps1 | upstream_gene_variant | MODIFIER |
| 1 | 36965008 | AX-97855688 | 0 | A | G | 0.456 | 1.29E-02 | 7.91E-02 | 2.67E+00 | 2.75E+00 | 8.67E-01 | no | | intergenic_variant | MODIFIER |
| 1 | 36966019 | AX-170277706 | 0 | G | C | 0.456 | 1.29E-02 | 7.91E-02 | 2.67E+00 | 2.75E+00 | 8.67E-01 | no | | intergenic_variant | MODIFIER |
| 1 | 36966736 | AX-170367097 | 0 | A | G | 0.456 | 1.29E-02 | 7.91E-02 | 2.67E+00 | 2.75E+00 | 8.67E-01 | no | | intergenic_variant | MODIFIER |

| | | | | | | | | | | | | | | | |
|---|---|---|---|---|---|---|---|---|---|---|---|---|---|---|---|
| 1 | 36976902 | AX-170091469 | 0 | G | A | 0.456 | 1.29E-02 | 7.91E-02 | 2.67E+00 | 2.75E+00 | 8.67E-01 | no | | regulatory_region_variant | MODIFIER |
| 1 | 36986104 | AX-170040900 | 0 | C | T | 0.456 | 1.29E-02 | 7.91E-02 | 2.67E+00 | 2.75E+00 | 8.67E-01 | no | | regulatory_region_variant | MODIFIER |
| 1 | 37016554 | AX-170171641 | 0 | G | A | 0.456 | 1.29E-02 | 7.91E-02 | 2.67E+00 | 2.75E+00 | 8.67E-01 | no | | intergenic_variant | MODIFIER |
| 1 | 37020521 | AX-170460838 | 0 | T | C | 0.456 | 1.29E-02 | 7.91E-02 | 2.67E+00 | 2.75E+00 | 8.67E-01 | no | | intergenic_variant | MODIFIER |
| 1 | 37027094 | AX-170419843 | 0 | T | C | 0.456 | 1.29E-02 | 7.91E-02 | 2.67E+00 | 2.75E+00 | 8.67E-01 | no | Vwa3b | upstream_gene_variant | MODIFIER |
| 1 | 37030925 | AX-170160225 | 0 | C | T | 0.456 | 1.29E-02 | 7.91E-02 | 2.67E+00 | 2.75E+00 | 8.67E-01 | no | Vwa3b | upstream_gene_variant | MODIFIER |
| 1 | 37031361 | AX-170164131 | 0 | C | T | 0.456 | 1.29E-02 | 7.91E-02 | 2.67E+00 | 2.75E+00 | 8.67E-01 | no | Vwa3b | upstream_gene_variant | MODIFIER |
| 1 | 37031952 | AX-168417872 | 0 | C | T | 0.456 | 1.29E-02 | 7.91E-02 | 2.67E+00 | 2.75E+00 | 8.67E-01 | no | Vwa3b | upstream_gene_variant | MODIFIER |
| 1 | 37034497 | AX-170251681 | 0 | G | A | 0.456 | 1.29E-02 | 7.91E-02 | 2.67E+00 | 2.75E+00 | 8.67E-01 | no | Vwa3b | upstream_gene_variant | MODIFIER |
| 1 | 37042068 | AX-170236314 | 0 | G | A | 0.456 | 1.29E-02 | 7.91E-02 | 2.67E+00 | 2.75E+00 | 8.67E-01 | no | Vwa3b | intron_variant,NMD_transcript_variant | MODIFIER |
| 1 | 37058146 | AX-97846824 | 0 | C | T | 0.456 | 1.29E-02 | 7.91E-02 | 2.67E+00 | 2.75E+00 | 8.67E-01 | no | Vwa3b | intron_variant,NMD_transcript_variant | MODIFIER |
| 1 | 37065703 | AX-170146059 | 0 | C | T | 0.456 | 1.29E-02 | 7.91E-02 | 2.67E+00 | 2.75E+00 | 8.67E-01 | no | Vwa3b | intron_variant,NMD_transcript_variant | MODIFIER |
| 1 | 37082315 | AX-170206078 | 0 | A | G | 0.456 | 1.29E-02 | 7.91E-02 | 2.67E+00 | 2.75E+00 | 8.67E-01 | no | Vwa3b | intron_variant,NMD_transcript_variant | MODIFIER |
| 1 | 37085147 | AX-97833225 | 0 | G | A | 0.456 | 1.29E-02 | 7.91E-02 | 2.67E+00 | 2.75E+00 | 8.67E-01 | no | Vwa3b | intron_variant,NMD_transcript_variant | MODIFIER |
| 1 | 37087471 | AX-170040901 | 0 | A | G | 0.456 | 1.29E-02 | 7.91E-02 | 2.67E+00 | 2.75E+00 | 8.67E-01 | no | Vwa3b | intron_variant,NMD_transcript_variant | MODIFIER |
| 1 | 37094204 | AX-170211511 | 0 | C | T | 0.456 | 1.29E-02 | 7.91E-02 | 2.67E+00 | 2.75E+00 | 8.67E-01 | no | Vwa3b | intron_variant,NMD_transcript_variant | MODIFIER |
| 1 | 37107897 | AX-170478018 | 0 | T | C | 0.456 | 1.29E-02 | 7.91E-02 | 2.67E+00 | 2.75E+00 | 8.67E-01 | no | Vwa3b | intron_variant,NMD_transcript_variant | MODIFIER |
| 1 | 37109118 | AX-170201157 | 0 | G | A | 0.456 | 1.29E-02 | 7.91E-02 | 2.67E+00 | 2.75E+00 | 8.67E-01 | no | Vwa3b | synonymous_variant,NMD_transcript_variant | LOW |
| 1 | 37155712 | AX-170395971 | 0 | G | A | 0.456 | 1.29E-02 | 7.91E-02 | 2.67E+00 | 2.75E+00 | 8.67E-01 | no | Vwa3b | intron_variant,NMD_transcript_variant | MODIFIER |
| 1 | 37157465 | AX-170180763 | 0 | A | G | 0.456 | 1.29E-02 | 7.91E-02 | 2.67E+00 | 2.75E+00 | 8.67E-01 | no | Vwa3b | intron_variant,NMD_transcript_variant | MODIFIER |
| 1 | 37157836 | AX-97827567 | 0 | T | C | 0.456 | 1.29E-02 | 7.91E-02 | 2.67E+00 | 2.75E+00 | 8.67E-01 | no | Vwa3b | intron_variant,NMD_transcript_variant | MODIFIER |
| 1 | 37180323 | AX-170533405 | 0 | C | G | 0.456 | 1.29E-02 | 7.91E-02 | 2.67E+00 | 2.75E+00 | 8.67E-01 | no | Vwa3b | intron_variant,NMD_transcript_variant | MODIFIER |
| 1 | 37182077 | AX-170430463 | 0 | T | C | 0.456 | 1.29E-02 | 7.91E-02 | 2.67E+00 | 2.75E+00 | 8.67E-01 | no | Vwa3b | intron_variant,NMD_transcript_variant | MODIFIER |
| 1 | 37183892 | AX-97852429 | 0 | C | T | 0.456 | 1.29E-02 | 7.91E-02 | 2.67E+00 | 2.75E+00 | 8.67E-01 | no | Vwa3b | intron_variant,NMD_transcript_variant | MODIFIER |
| 1 | 37185205 | AX-170164040 | 0 | G | A | 0.456 | 1.29E-02 | 7.91E-02 | 2.67E+00 | 2.75E+00 | 8.67E-01 | no | Vwa3b | intron_variant,NMD_transcript_variant | MODIFIER |
| 1 | 37221902 | AX-170444124 | 0 | C | A | 0.456 | 1.29E-02 | 7.91E-02 | 2.67E+00 | 2.75E+00 | 8.67E-01 | no | Cnga3 | intron_variant | MODIFIER |
| 1 | 37221999 | AX-170503641 | 0 | C | T | 0.456 | 1.29E-02 | 7.91E-02 | 2.67E+00 | 2.75E+00 | 8.67E-01 | no | Cnga3 | intron_variant | MODIFIER |
| 1 | 37225738 | AX-170510375 | 0 | A | G | 0.456 | 1.29E-02 | 7.91E-02 | 2.67E+00 | 2.75E+00 | 8.67E-01 | no | Cnga3 | intron_variant | MODIFIER |
| 1 | 37230879 | AX-170483374 | 0 | T | G | 0.456 | 1.29E-02 | 7.91E-02 | 2.67E+00 | 2.75E+00 | 8.67E-01 | no | Cnga3 | intron_variant | MODIFIER |
| 1 | 37231237 | AX-170386470 | 0 | C | T | 0.456 | 1.29E-02 | 7.91E-02 | 2.67E+00 | 2.75E+00 | 8.67E-01 | no | Cnga3 | intron_variant | MODIFIER |
| 1 | 37231448 | AX-170333874 | 0 | T | C | 0.456 | 1.29E-02 | 7.91E-02 | 2.67E+00 | 2.75E+00 | 8.67E-01 | no | Cnga3 | intron_variant | MODIFIER |
| 1 | 37238223 | AX-170423952 | 0 | G | A | 0.456 | 1.29E-02 | 7.91E-02 | 2.67E+00 | 2.75E+00 | 8.67E-01 | no | Cnga3 | intron_variant | MODIFIER |
| 1 | 37268793 | AX-170399668 | 0 | T | C | 0.456 | 1.29E-02 | 7.91E-02 | 2.67E+00 | 2.75E+00 | 8.67E-01 | no | | intergenic_variant | MODIFIER |
| 1 | 37306247 | AX-97852970 | 0 | T | C | 0.456 | 1.29E-02 | 7.91E-02 | 2.67E+00 | 2.75E+00 | 8.67E-01 | no | Inpp4a | intron_variant | MODIFIER |
| 1 | 37330814 | AX-97824382 | 0 | G | A | 0.456 | 1.29E-02 | 7.91E-02 | 2.67E+00 | 2.75E+00 | 8.67E-01 | no | Inpp4a | intron_variant | MODIFIER |
| 1 | 37341663 | AX-170394047 | 0 | A | G | 0.456 | 1.29E-02 | 7.91E-02 | 2.67E+00 | 2.75E+00 | 8.67E-01 | no | Inpp4a | intron_variant | MODIFIER |
| 1 | 37380943 | AX-97808357 | 0 | G | A | 0.456 | 1.29E-02 | 7.91E-02 | 2.67E+00 | 2.75E+00 | 8.67E-01 | no | Inpp4a | intron_variant | MODIFIER |
| 1 | 37517073 | AX-170361907 | 0 | G | A | 0.456 | 1.29E-02 | 7.91E-02 | 2.67E+00 | 2.75E+00 | 8.67E-01 | no | Mgat4a | intron_variant | MODIFIER |
| 1 | 37518051 | AX-170383458 | 0 | C | T | 0.456 | 1.29E-02 | 7.91E-02 | 2.67E+00 | 2.75E+00 | 8.67E-01 | no | Mgat4a | intron_variant | MODIFIER |
| 13 | 45916934 | AX-170204204 | 0 | A | T | 0.474 | 1.10E-02 | 6.24E-02 | 2.60E+00 | 2.72E+00 | 8.67E-01 | no | Atxn1 | intron_variant | MODIFIER |
| 8 | 35349172 | AX-170263000 | 1 | T | A | 0.152 | -2.49E-02 | 1.43E-01 | 2.65E+00 | 2.70E+00 | 8.67E-01 | no | | regulatory_region_variant | MODIFIER |
| 9 | 88288622 | AX-170509408 | 0 | A | C | 0.149 | 1.86E-02 | 1.14E-01 | 2.64E+00 | 2.75E+00 | 8.67E-01 | no | Gm47620 | downstream_gene_variant | MODIFIER |
| 8 | 22026440 | AX-170274885 | 0 | C | T | 0.175 | -1.86E-02 | 1.13E-01 | 2.66E+00 | 2.74E+00 | 8.67E-01 | no | Atp7b | intron_variant | MODIFIER |
| 1 | 63657115 | AX-170234017 | 2 | G | A | 0.127 | 3.52E-02 | 2.12E-01 | 2.65E+00 | 2.74E+00 | 8.67E-01 | no | Dytn | intron_variant | MODIFIER |
| 7 | 117827076 | AX-170438523 | 1 | T | A | 0.366 | -1.41E-02 | 8.65E-02 | 2.66E+00 | 2.75E+00 | 8.67E-01 | no | | intergenic_variant | MODIFIER |
| 6 | 145306820 | AX-170305828 | 1 | C | T | 0.071 | 4.56E-02 | 2.77E-01 | 2.71E+00 | 2.75E+00 | 8.67E-01 | no | 2010013B24Rik | intron_variant,non_coding_transcript_variant | MODIFIER |
| 9 | 57900701 | AX-170442262 | 0 | G | C | 0.211 | -2.04E-02 | 1.23E-01 | 2.68E+00 | 2.74E+00 | 8.67E-01 | no | | regulatory_region_variant | MODIFIER |
| 4 | 149480761 | AX-170095186 | 0 | C | T | 0.096 | 3.31E-02 | 2.00E-01 | 2.67E+00 | 2.74E+00 | 8.67E-01 | no | Lzic | upstream_gene_variant | MODIFIER |
| 15 | 93390997 | AX-170190560 | 1 | A | G | 0.152 | 3.15E-02 | 1.83E-01 | 2.65E+00 | 2.71E+00 | 8.67E-01 | no | Zcrb1 | intron_variant | MODIFIER |
| 15 | 94540716 | AX-97826204 | 1 | C | T | 0.152 | 3.15E-02 | 1.83E-01 | 2.65E+00 | 2.71E+00 | 8.67E-01 | no | Pus7l | missense_variant | MODERATE |
| 15 | 95325349 | AX-170128192 | 1 | C | G | 0.152 | 3.15E-02 | 1.83E-01 | 2.65E+00 | 2.71E+00 | 8.67E-01 | no | Nell2 | intron_variant | MODIFIER |
| 4 | 144058612 | AX-170280912 | 1 | A | T | 0.062 | 3.59E-02 | 2.02E-01 | 2.59E+00 | 2.71E+00 | 8.67E-01 | no | Pramel4 | upstream_gene_variant | MODIFIER |
| 1 | 36355412 | AX-170000281 | 1 | A | G | 0.455 | 1.28E-02 | 7.90E-02 | 2.67E+00 | 2.75E+00 | 8.67E-01 | no | Kansl3 | intron_variant | MODIFIER |
| 1 | 7940716 | AX-170396146 | 0 | T | A | 0.421 | 1.26E-02 | 7.17E-02 | 2.63E+00 | 2.70E+00 | 8.67E-01 | no | | intergenic_variant | MODIFIER |

| | | | | | | | | | | | | | | | |
|---|---|---|---|---|---|---|---|---|---|---|---|---|---|---|---|
| 18 | 53014561 | AX-170257327 | 1 | G | C | 0.241 | -1.92E-02 | 1.08E-01 | 2.63E+00 | 2.69E+00 | 8.68E-01 | no | | intergenic_variant | MODIFIER |
| 6 | 146481871 | AX-170171429 | 0 | G | A | 0.07 | -4.54E-02 | 2.76E-01 | 2.70E+00 | 2.74E+00 | 8.68E-01 | no | Itpr2 | intron_variant | MODIFIER |
| 20 | 59453315 | AX-170201572 | 2 | A | T | 0.236 | 1.06E-02 | 6.37E-02 | 2.63E+00 | 2.73E+00 | 8.68E-01 | no | Fgf13 | intron_variant | MODIFIER |
| 1 | 74974397 | AX-170511760 | 1 | C | A | 0.125 | 3.41E-02 | 2.07E-01 | 2.68E+00 | 2.74E+00 | 8.68E-01 | no | Nhej1 | intron_variant | MODIFIER |
| 9 | 34952185 | AX-170332795 | 1 | C | T | 0.446 | 1.52E-02 | 8.79E-02 | 2.64E+00 | 2.71E+00 | 8.68E-01 | no | Kirrel3 | intron_variant | MODIFIER |
| 15 | 88118096 | AX-170149199 | 1 | A | G | 0.134 | 2.76E-02 | 1.62E-01 | 2.65E+00 | 2.71E+00 | 8.68E-01 | no | | intergenic_variant | MODIFIER |
| 15 | 88237828 | AX-170295990 | 1 | T | G | 0.134 | 2.76E-02 | 1.62E-01 | 2.65E+00 | 2.71E+00 | 8.68E-01 | no | | intergenic_variant | MODIFIER |
| 15 | 91529190 | AX-170249185 | 1 | T | C | 0.134 | 2.76E-02 | 1.62E-01 | 2.65E+00 | 2.71E+00 | 8.68E-01 | no | Slc2a13 | intron_variant | MODIFIER |
| 15 | 94230971 | AX-170131940 | 1 | A | T | 0.134 | 2.76E-02 | 1.62E-01 | 2.65E+00 | 2.71E+00 | 8.68E-01 | no | | intergenic_variant | MODIFIER |
| 9 | 117106081 | AX-170160343 | 0 | C | A | 0.132 | -1.65E-02 | 9.53E-02 | 2.60E+00 | 2.72E+00 | 8.68E-01 | no | Rbms3 | intron_variant | MODIFIER |
| 4 | 144785160 | AX-97802765 | 0 | A | G | 0.114 | -2.87E-02 | 1.82E-01 | 2.67E+00 | 2.76E+00 | 8.68E-01 | no | Gm438 | intron_variant | MODIFIER |
| 8 | 102123632 | AX-170511844 | 0 | G | A | 0.219 | 1.87E-02 | 1.16E-01 | 2.68E+00 | 2.75E+00 | 8.68E-01 | no | | intergenic_variant | MODIFIER |
| 8 | 47001738 | AX-170176158 | 0 | G | T | 0.158 | -2.29E-02 | 1.37E-01 | 2.66E+00 | 2.73E+00 | 8.68E-01 | no | Enpp6 | intron_variant | MODIFIER |
| 8 | 72802541 | AX-170466955 | 0 | C | T | 0.202 | -2.07E-02 | 1.26E-01 | 2.66E+00 | 2.74E+00 | 8.68E-01 | no | | intergenic_variant | MODIFIER |
| 1 | 148435643 | AX-97786934 | 1 | C | A | 0.196 | 2.14E-02 | 1.25E-01 | 2.64E+00 | 2.71E+00 | 8.68E-01 | no | | intergenic_variant | MODIFIER |
| 18 | 90549862 | AX-97852428 | 0 | A | T | 0.518 | -9.73E-03 | 6.32E-02 | 2.63E+00 | 2.76E+00 | 8.68E-01 | no | | intergenic_variant | MODIFIER |
| 6 | 134931542 | AX-170191942 | 0 | C | A | 0.57 | 1.48E-02 | 8.46E-02 | 2.63E+00 | 2.70E+00 | 8.68E-01 | no | Lockd | intron_variant,non_coding_transcript_variant | MODIFIER |
| 16 | 18383430 | AX-170474982 | 0 | C | T | 0.132 | -1.56E-02 | 9.62E-02 | 2.61E+00 | 2.74E+00 | 8.69E-01 | no | Arvcf | intron_variant | MODIFIER |
| 16 | 18464860 | AX-170012595 | 0 | C | T | 0.132 | -1.56E-02 | 9.62E-02 | 2.61E+00 | 2.74E+00 | 8.69E-01 | no | Txnrd2 | intron_variant,NMD_transcript_variant | MODIFIER |
| 16 | 18907614 | AX-170012603 | 0 | C | T | 0.132 | -1.56E-02 | 9.62E-02 | 2.61E+00 | 2.74E+00 | 8.69E-01 | no | Hira | intron_variant | MODIFIER |
| 16 | 19153172 | AX-97831130 | 0 | C | T | 0.132 | -1.56E-02 | 9.62E-02 | 2.61E+00 | 2.74E+00 | 8.69E-01 | no | | regulatory_region_variant | MODIFIER |
| 16 | 19513046 | AX-170307340 | 0 | G | A | 0.132 | -1.56E-02 | 9.62E-02 | 2.61E+00 | 2.74E+00 | 8.69E-01 | no | Olfr167 | downstream_gene_variant | MODIFIER |
| 16 | 19612343 | AX-170314851 | 0 | A | G | 0.132 | -1.56E-02 | 9.62E-02 | 2.61E+00 | 2.74E+00 | 8.69E-01 | no | Olfr170 | upstream_gene_variant | MODIFIER |
| 6 | 145276590 | AX-170243573 | 0 | C | T | 0.07 | 4.49E-02 | 2.76E-01 | 2.71E+00 | 2.75E+00 | 8.69E-01 | no | | regulatory_region_variant | MODIFIER |
| 1 | 145655145 | AX-170381384 | 2 | T | C | 0.2 | 2.18E-02 | 1.27E-01 | 2.63E+00 | 2.72E+00 | 8.69E-01 | no | | intergenic_variant | MODIFIER |
| 1 | 47310528 | AX-170381718 | 0 | C | A | 0.105 | 3.83E-02 | 2.35E-01 | 2.69E+00 | 2.74E+00 | 8.69E-01 | no | | intergenic_variant | MODIFIER |
| 1 | 49360171 | AX-170267156 | 0 | C | T | 0.105 | 3.83E-02 | 2.35E-01 | 2.69E+00 | 2.74E+00 | 8.69E-01 | no | | intergenic_variant | MODIFIER |
| 1 | 55393497 | AX-97810939 | 0 | T | C | 0.105 | 3.83E-02 | 2.35E-01 | 2.69E+00 | 2.74E+00 | 8.69E-01 | no | | intergenic_variant | MODIFIER |
| 1 | 60821360 | AX-170535534 | 0 | T | G | 0.105 | 3.83E-02 | 2.35E-01 | 2.69E+00 | 2.74E+00 | 8.69E-01 | no | | regulatory_region_variant | MODIFIER |
| 16 | 93160044 | AX-170164569 | 1 | G | A | 0.375 | -1.39E-02 | 7.93E-02 | 2.62E+00 | 2.71E+00 | 8.69E-01 | no | | intergenic_variant | MODIFIER |
| 10 | 57811130 | AX-170003293 | 0 | A | G | 0.289 | 1.20E-02 | 8.18E-02 | 2.70E+00 | 2.81E+00 | 8.69E-01 | no | Smpdl3a | intron_variant | MODIFIER |
| 10 | 118983084 | AX-170246051 | 1 | C | T | 0.348 | 1.18E-02 | 7.34E-02 | 2.66E+00 | 2.75E+00 | 8.69E-01 | no | | intergenic_variant | MODIFIER |
| 10 | 119539685 | AX-97817740 | 1 | C | A | 0.348 | 1.18E-02 | 7.34E-02 | 2.66E+00 | 2.75E+00 | 8.69E-01 | no | Grip1 | intron_variant | MODIFIER |
| 20 | 56968022 | AX-97853070 | 1 | C | A | 0.232 | -1.00E-02 | 6.33E-02 | 2.65E+00 | 2.75E+00 | 8.69E-01 | no | Adgrg4 | intron_variant | MODIFIER |
| 8 | 97815227 | AX-170031347 | 2 | C | T | 0.209 | 1.91E-02 | 1.18E-01 | 2.68E+00 | 2.75E+00 | 8.69E-01 | no | | intergenic_variant | MODIFIER |
| 9 | 83449420 | AX-170217917 | 0 | G | C | 0.211 | 2.05E-02 | 1.28E-01 | 2.71E+00 | 2.76E+00 | 8.69E-01 | no | Gm2087 | intron_variant,non_coding_transcript_variant | MODIFIER |
| 9 | 83520558 | AX-168378964 | 0 | G | A | 0.211 | 2.05E-02 | 1.28E-01 | 2.71E+00 | 2.76E+00 | 8.69E-01 | no | | intergenic_variant | MODIFIER |
| 9 | 83523450 | AX-168390366 | 0 | C | T | 0.211 | 2.05E-02 | 1.28E-01 | 2.71E+00 | 2.76E+00 | 8.69E-01 | no | | intergenic_variant | MODIFIER |
| 9 | 83539602 | AX-170415715 | 0 | A | G | 0.211 | 2.05E-02 | 1.28E-01 | 2.71E+00 | 2.76E+00 | 8.69E-01 | no | | regulatory_region_variant | MODIFIER |
| 9 | 83551276 | AX-170131939 | 0 | G | A | 0.211 | 2.05E-02 | 1.28E-01 | 2.71E+00 | 2.76E+00 | 8.69E-01 | no | Sh3bgrl2 | intron_variant | MODIFIER |
| 9 | 83563949 | AX-170244832 | 0 | C | T | 0.211 | 2.05E-02 | 1.28E-01 | 2.71E+00 | 2.76E+00 | 8.69E-01 | no | Sh3bgrl2 | intron_variant | MODIFIER |
| 9 | 83568274 | AX-170167970 | 0 | T | G | 0.211 | 2.05E-02 | 1.28E-01 | 2.71E+00 | 2.76E+00 | 8.69E-01 | no | Sh3bgrl2 | intron_variant | MODIFIER |
| 9 | 83568903 | AX-170482505 | 0 | A | G | 0.211 | 2.05E-02 | 1.28E-01 | 2.71E+00 | 2.76E+00 | 8.69E-01 | no | Sh3bgrl2 | intron_variant | MODIFIER |
| 9 | 83574893 | AX-170321535 | 0 | C | T | 0.211 | 2.05E-02 | 1.28E-01 | 2.71E+00 | 2.76E+00 | 8.69E-01 | no | Sh3bgrl2 | intron_variant | MODIFIER |
| 9 | 83578483 | AX-170352677 | 0 | G | A | 0.211 | 2.05E-02 | 1.28E-01 | 2.71E+00 | 2.76E+00 | 8.69E-01 | no | Sh3bgrl2 | intron_variant | MODIFIER |
| 9 | 83578524 | AX-170364300 | 0 | G | A | 0.211 | 2.05E-02 | 1.28E-01 | 2.71E+00 | 2.76E+00 | 8.69E-01 | no | Sh3bgrl2 | intron_variant | MODIFIER |
| 9 | 83580072 | AX-170247807 | 0 | T | C | 0.211 | 2.05E-02 | 1.28E-01 | 2.71E+00 | 2.76E+00 | 8.69E-01 | no | Sh3bgrl2 | intron_variant | MODIFIER |
| 9 | 83583035 | AX-170276837 | 0 | G | A | 0.211 | 2.05E-02 | 1.28E-01 | 2.71E+00 | 2.76E+00 | 8.69E-01 | no | Sh3bgrl2 | intron_variant | MODIFIER |
| 9 | 83599426 | AX-170167569 | 0 | G | A | 0.211 | 2.05E-02 | 1.28E-01 | 2.71E+00 | 2.76E+00 | 8.69E-01 | no | Sh3bgrl2 | 3_prime_UTR_variant | MODIFIER |
| 9 | 83701532 | AX-97816105 | 0 | C | T | 0.211 | 2.05E-02 | 1.28E-01 | 2.71E+00 | 2.76E+00 | 8.69E-01 | no | | intergenic_variant | MODIFIER |
| 9 | 83730938 | AX-170209164 | 0 | C | T | 0.211 | 2.05E-02 | 1.28E-01 | 2.71E+00 | 2.76E+00 | 8.69E-01 | no | | intergenic_variant | MODIFIER |
| 9 | 84133206 | AX-170186547 | 0 | T | C | 0.211 | 2.05E-02 | 1.28E-01 | 2.71E+00 | 2.76E+00 | 8.69E-01 | no | | intergenic_variant | MODIFIER |
| 9 | 84145719 | AX-170283681 | 0 | C | A | 0.211 | 2.05E-02 | 1.28E-01 | 2.71E+00 | 2.76E+00 | 8.69E-01 | no | | intergenic_variant | MODIFIER |
| 9 | 84146164 | AX-97815982 | 0 | A | C | 0.211 | 2.05E-02 | 1.28E-01 | 2.71E+00 | 2.76E+00 | 8.69E-01 | no | | intergenic_variant | MODIFIER |

| | | | | | | | | | | | | | | | |
|---|---|---|---|---|---|---|---|---|---|---|---|---|---|---|---|
| 9 | 84149119 | AX-170348650 | 0 | T | C | 0.211 | 2.05E-02 | 1.28E-01 | 2.71E+00 | 2.76E+00 | 8.69E-01 | no | | intergenic_variant | MODIFIER |
| 9 | 84186664 | AX-170166477 | 0 | G | A | 0.211 | 2.05E-02 | 1.28E-01 | 2.71E+00 | 2.76E+00 | 8.69E-01 | no | Gm26146 | upstream_gene_variant | MODIFIER |
| 9 | 84188797 | AX-170292581 | 0 | C | G | 0.211 | 2.05E-02 | 1.28E-01 | 2.71E+00 | 2.76E+00 | 8.69E-01 | no | Gm26146 | downstream_gene_variant | MODIFIER |
| 9 | 84189701 | AX-170340703 | 0 | G | A | 0.211 | 2.05E-02 | 1.28E-01 | 2.71E+00 | 2.76E+00 | 8.69E-01 | no | Gm26146 | downstream_gene_variant | MODIFIER |
| 9 | 84200445 | AX-97800557 | 0 | A | G | 0.211 | 2.05E-02 | 1.28E-01 | 2.71E+00 | 2.76E+00 | 8.69E-01 | no | | intergenic_variant | MODIFIER |
| 9 | 84232884 | AX-170082986 | 0 | T | C | 0.211 | 2.05E-02 | 1.28E-01 | 2.71E+00 | 2.76E+00 | 8.69E-01 | no | | intergenic_variant | MODIFIER |
| 9 | 84234036 | AX-170514911 | 0 | T | C | 0.211 | 2.05E-02 | 1.28E-01 | 2.71E+00 | 2.76E+00 | 8.69E-01 | no | | intergenic_variant | MODIFIER |
| 9 | 84239979 | AX-97823472 | 0 | G | A | 0.211 | 2.05E-02 | 1.28E-01 | 2.71E+00 | 2.76E+00 | 8.69E-01 | no | | intergenic_variant | MODIFIER |
| 9 | 84250806 | AX-170497707 | 0 | C | T | 0.211 | 2.05E-02 | 1.28E-01 | 2.71E+00 | 2.76E+00 | 8.69E-01 | no | | intergenic_variant | MODIFIER |
| 9 | 84257626 | AX-170127666 | 0 | C | A | 0.211 | 2.05E-02 | 1.28E-01 | 2.71E+00 | 2.76E+00 | 8.69E-01 | no | | intergenic_variant | MODIFIER |
| 9 | 84288189 | AX-170105167 | 0 | T | C | 0.211 | 2.05E-02 | 1.28E-01 | 2.71E+00 | 2.76E+00 | 8.69E-01 | no | | intergenic_variant | MODIFIER |
| 9 | 84310793 | AX-170404944 | 0 | T | A | 0.211 | 2.05E-02 | 1.28E-01 | 2.71E+00 | 2.76E+00 | 8.69E-01 | no | | intergenic_variant | MODIFIER |
| 9 | 84312736 | AX-170486051 | 0 | G | A | 0.211 | 2.05E-02 | 1.28E-01 | 2.71E+00 | 2.76E+00 | 8.69E-01 | no | | intergenic_variant | MODIFIER |
| 9 | 84316365 | AX-170250643 | 0 | A | G | 0.211 | 2.05E-02 | 1.28E-01 | 2.71E+00 | 2.76E+00 | 8.69E-01 | no | | intergenic_variant | MODIFIER |
| 9 | 84319644 | AX-97805926 | 0 | A | G | 0.211 | 2.05E-02 | 1.28E-01 | 2.71E+00 | 2.76E+00 | 8.69E-01 | no | | intergenic_variant | MODIFIER |
| 9 | 84324404 | AX-170111710 | 0 | T | C | 0.211 | 2.05E-02 | 1.28E-01 | 2.71E+00 | 2.76E+00 | 8.69E-01 | no | | intergenic_variant | MODIFIER |
| 9 | 84352597 | AX-97782789 | 0 | T | G | 0.211 | 2.05E-02 | 1.28E-01 | 2.71E+00 | 2.76E+00 | 8.69E-01 | no | | intergenic_variant | MODIFIER |
| 9 | 84392937 | AX-97790958 | 0 | G | A | 0.211 | 2.05E-02 | 1.28E-01 | 2.71E+00 | 2.76E+00 | 8.69E-01 | no | | intergenic_variant | MODIFIER |
| 9 | 84394675 | AX-170032862 | 0 | C | T | 0.211 | 2.05E-02 | 1.28E-01 | 2.71E+00 | 2.76E+00 | 8.69E-01 | no | | regulatory_region_variant | MODIFIER |
| 9 | 84435505 | AX-170187474 | 0 | G | A | 0.211 | 2.05E-02 | 1.28E-01 | 2.71E+00 | 2.76E+00 | 8.69E-01 | no | | intergenic_variant | MODIFIER |
| 9 | 84437772 | AX-170508596 | 0 | T | C | 0.211 | 2.05E-02 | 1.28E-01 | 2.71E+00 | 2.76E+00 | 8.69E-01 | no | | intergenic_variant | MODIFIER |
| 9 | 84439703 | AX-170394998 | 0 | T | A | 0.211 | 2.05E-02 | 1.28E-01 | 2.71E+00 | 2.76E+00 | 8.69E-01 | no | | intergenic_variant | MODIFIER |
| 9 | 84454377 | AX-97803129 | 0 | C | T | 0.211 | 2.05E-02 | 1.28E-01 | 2.71E+00 | 2.76E+00 | 8.69E-01 | no | | intergenic_variant | MODIFIER |
| 9 | 84495988 | AX-97824500 | 0 | T | C | 0.211 | 2.05E-02 | 1.28E-01 | 2.71E+00 | 2.76E+00 | 8.69E-01 | no | | intergenic_variant | MODIFIER |
| 9 | 84510241 | AX-170379137 | 0 | A | G | 0.211 | 2.05E-02 | 1.28E-01 | 2.71E+00 | 2.76E+00 | 8.69E-01 | no | | intergenic_variant | MODIFIER |
| 9 | 84510514 | AX-170130068 | 0 | A | T | 0.211 | 2.05E-02 | 1.28E-01 | 2.71E+00 | 2.76E+00 | 8.69E-01 | no | | intergenic_variant | MODIFIER |
| 9 | 84511361 | AX-170168569 | 0 | G | A | 0.211 | 2.05E-02 | 1.28E-01 | 2.71E+00 | 2.76E+00 | 8.69E-01 | no | | intergenic_variant | MODIFIER |
| 9 | 84559613 | AX-170032859 | 0 | C | A | 0.211 | 2.05E-02 | 1.28E-01 | 2.71E+00 | 2.76E+00 | 8.69E-01 | no | | intergenic_variant | MODIFIER |
| 9 | 84635440 | AX-97791714 | 0 | T | C | 0.211 | 2.05E-02 | 1.28E-01 | 2.71E+00 | 2.76E+00 | 8.69E-01 | no | | intergenic_variant | MODIFIER |
| 9 | 84641958 | AX-170082994 | 0 | G | C | 0.211 | 2.05E-02 | 1.28E-01 | 2.71E+00 | 2.76E+00 | 8.69E-01 | no | | intergenic_variant | MODIFIER |
| 9 | 84831997 | AX-170090360 | 0 | T | C | 0.211 | 2.05E-02 | 1.28E-01 | 2.71E+00 | 2.76E+00 | 8.69E-01 | no | 4930554C24Rik | intron_variant,non_coding_transcript_variant | MODIFIER |
| 9 | 84950001 | AX-97786340 | 0 | C | T | 0.211 | 2.05E-02 | 1.28E-01 | 2.71E+00 | 2.76E+00 | 8.69E-01 | no | | intergenic_variant | MODIFIER |
| 9 | 84959627 | AX-170437830 | 0 | C | T | 0.211 | 2.05E-02 | 1.28E-01 | 2.71E+00 | 2.76E+00 | 8.69E-01 | no | | intergenic_variant | MODIFIER |
| 9 | 84961565 | AX-170314957 | 0 | A | C | 0.211 | 2.05E-02 | 1.28E-01 | 2.71E+00 | 2.76E+00 | 8.69E-01 | no | | intergenic_variant | MODIFIER |
| 9 | 84961689 | AX-170173485 | 0 | A | G | 0.211 | 2.05E-02 | 1.28E-01 | 2.71E+00 | 2.76E+00 | 8.69E-01 | no | | intergenic_variant | MODIFIER |
| 9 | 85079555 | AX-170521341 | 0 | A | C | 0.211 | 2.05E-02 | 1.28E-01 | 2.71E+00 | 2.76E+00 | 8.69E-01 | no | | intergenic_variant | MODIFIER |
| 10 | 119026556 | AX-170371981 | 0 | G | A | 0.333 | 1.15E-02 | 7.21E-02 | 2.66E+00 | 2.75E+00 | 8.69E-01 | no | Gm47461 | upstream_gene_variant | MODIFIER |
| 10 | 119030309 | AX-170481864 | 0 | G | A | 0.333 | 1.15E-02 | 7.21E-02 | 2.66E+00 | 2.75E+00 | 8.69E-01 | no | Gm33677 | non_coding_transcript_exon_variant | MODIFIER |
| 10 | 119031115 | AX-170298457 | 0 | A | G | 0.333 | 1.15E-02 | 7.21E-02 | 2.66E+00 | 2.75E+00 | 8.69E-01 | no | Gm33677 | intron_variant,non_coding_transcript_variant | MODIFIER |
| 10 | 119031794 | AX-170157921 | 0 | A | C | 0.333 | 1.15E-02 | 7.21E-02 | 2.66E+00 | 2.75E+00 | 8.69E-01 | no | Gm33677 | intron_variant,non_coding_transcript_variant | MODIFIER |
| 10 | 119033855 | AX-97838589 | 0 | C | T | 0.333 | 1.15E-02 | 7.21E-02 | 2.66E+00 | 2.75E+00 | 8.69E-01 | no | Gm33677 | intron_variant,non_coding_transcript_variant | MODIFIER |
| 10 | 119042161 | AX-170119613 | 0 | C | T | 0.333 | 1.15E-02 | 7.21E-02 | 2.66E+00 | 2.75E+00 | 8.69E-01 | no | Gm33677 | intron_variant,non_coding_transcript_variant | MODIFIER |
| 10 | 119049813 | AX-170107920 | 0 | G | A | 0.333 | 1.15E-02 | 7.21E-02 | 2.66E+00 | 2.75E+00 | 8.69E-01 | no | Gm33677 | intron_variant,non_coding_transcript_variant | MODIFIER |
| 10 | 119049850 | AX-170119512 | 0 | T | C | 0.333 | 1.15E-02 | 7.21E-02 | 2.66E+00 | 2.75E+00 | 8.69E-01 | no | Gm33677 | intron_variant,non_coding_transcript_variant | MODIFIER |
| 10 | 119052723 | AX-170046042 | 0 | G | A | 0.333 | 1.15E-02 | 7.21E-02 | 2.66E+00 | 2.75E+00 | 8.69E-01 | no | Gm33677 | upstream_gene_variant | MODIFIER |
| 10 | 119056234 | AX-170046043 | 0 | C | T | 0.333 | 1.15E-02 | 7.21E-02 | 2.66E+00 | 2.75E+00 | 8.69E-01 | no | | regulatory_region_variant | MODIFIER |
| 10 | 119056719 | AX-170046040 | 0 | C | T | 0.333 | 1.15E-02 | 7.21E-02 | 2.66E+00 | 2.75E+00 | 8.69E-01 | no | | regulatory_region_variant | MODIFIER |
| 10 | 119209272 | AX-170245784 | 0 | C | G | 0.333 | 1.15E-02 | 7.21E-02 | 2.66E+00 | 2.75E+00 | 8.69E-01 | no | Cand1 | intron_variant | MODIFIER |
| 10 | 119332091 | AX-97826245 | 0 | T | C | 0.333 | 1.15E-02 | 7.21E-02 | 2.66E+00 | 2.75E+00 | 8.69E-01 | no | 4930477N07Rik | downstream_gene_variant | MODIFIER |
| 10 | 119362673 | AX-170004016 | 0 | T | C | 0.333 | 1.15E-02 | 7.21E-02 | 2.66E+00 | 2.75E+00 | 8.69E-01 | no | | intergenic_variant | MODIFIER |
| 10 | 119362719 | AX-170272094 | 0 | G | A | 0.333 | 1.15E-02 | 7.21E-02 | 2.66E+00 | 2.75E+00 | 8.69E-01 | no | | intergenic_variant | MODIFIER |
| 10 | 119363034 | AX-97816982 | 0 | T | C | 0.333 | 1.15E-02 | 7.21E-02 | 2.66E+00 | 2.75E+00 | 8.69E-01 | no | | intergenic_variant | MODIFIER |
| 10 | 119370493 | AX-170173000 | 0 | A | G | 0.333 | 1.15E-02 | 7.21E-02 | 2.66E+00 | 2.75E+00 | 8.69E-01 | no | | intergenic_variant | MODIFIER |
| 10 | 119406985 | AX-170435702 | 0 | A | G | 0.333 | 1.15E-02 | 7.21E-02 | 2.66E+00 | 2.75E+00 | 8.69E-01 | no | Gm40778 | downstream_gene_variant | MODIFIER |

| | | | | | | | | | | | | | | | |
|---|---|---|---|---|---|---|---|---|---|---|---|---|---|---|---|
| 10 | 119414809 | AX-170320596 | 0 | C | G | 0.333 | 1.15E-02 | 7.21E-02 | 2.66E+00 | 2.75E+00 | 8.69E-01 | no | 1700025F24Rik | intron_variant,non_coding_transcript_variant | MODIFIER |
| 10 | 119419692 | AX-97839487 | 0 | C | T | 0.333 | 1.15E-02 | 7.21E-02 | 2.66E+00 | 2.75E+00 | 8.69E-01 | no | 1700025F24Rik | intron_variant,non_coding_transcript_variant | MODIFIER |
| 10 | 119574590 | AX-97825557 | 0 | T | C | 0.333 | 1.15E-02 | 7.21E-02 | 2.66E+00 | 2.75E+00 | 8.69E-01 | no | Grip1 | intron_variant | MODIFIER |
| 10 | 119588658 | AX-170288613 | 0 | C | A | 0.333 | 1.15E-02 | 7.21E-02 | 2.66E+00 | 2.75E+00 | 8.69E-01 | no | Grip1 | intron_variant | MODIFIER |
| 10 | 119591148 | AX-97793263 | 0 | A | G | 0.333 | 1.15E-02 | 7.21E-02 | 2.66E+00 | 2.75E+00 | 8.69E-01 | no | Grip1 | intron_variant | MODIFIER |
| 10 | 119591680 | AX-170359552 | 0 | G | T | 0.333 | 1.15E-02 | 7.21E-02 | 2.66E+00 | 2.75E+00 | 8.69E-01 | no | Grip1 | intron_variant | MODIFIER |
| 10 | 119592532 | AX-170389160 | 0 | C | G | 0.333 | 1.15E-02 | 7.21E-02 | 2.66E+00 | 2.75E+00 | 8.69E-01 | no | Grip1 | intron_variant | MODIFIER |
| 10 | 119592563 | AX-170521995 | 0 | G | C | 0.333 | 1.15E-02 | 7.21E-02 | 2.66E+00 | 2.75E+00 | 8.69E-01 | no | Grip1 | intron_variant | MODIFIER |
| 10 | 119595049 | AX-170159879 | 0 | T | C | 0.333 | 1.15E-02 | 7.21E-02 | 2.66E+00 | 2.75E+00 | 8.69E-01 | no | Grip1 | intron_variant | MODIFIER |
| 10 | 119597450 | AX-170159457 | 0 | T | G | 0.333 | 1.15E-02 | 7.21E-02 | 2.66E+00 | 2.75E+00 | 8.69E-01 | no | Grip1 | intron_variant | MODIFIER |
| 10 | 119600441 | AX-170148622 | 0 | A | G | 0.333 | 1.15E-02 | 7.21E-02 | 2.66E+00 | 2.75E+00 | 8.69E-01 | no | Grip1 | intron_variant | MODIFIER |
| 10 | 119600750 | AX-170383793 | 0 | A | G | 0.333 | 1.15E-02 | 7.21E-02 | 2.66E+00 | 2.75E+00 | 8.69E-01 | no | Grip1 | intron_variant | MODIFIER |
| 10 | 119600890 | AX-170373092 | 0 | A | T | 0.333 | 1.15E-02 | 7.21E-02 | 2.66E+00 | 2.75E+00 | 8.69E-01 | no | Grip1 | intron_variant | MODIFIER |
| 10 | 119615267 | AX-97825559 | 0 | C | T | 0.333 | 1.15E-02 | 7.21E-02 | 2.66E+00 | 2.75E+00 | 8.69E-01 | no | Grip1 | intron_variant | MODIFIER |
| 10 | 119616843 | AX-170427040 | 0 | G | A | 0.333 | 1.15E-02 | 7.21E-02 | 2.66E+00 | 2.75E+00 | 8.69E-01 | no | Grip1 | intron_variant | MODIFIER |
| 10 | 119617021 | AX-170377083 | 0 | A | G | 0.333 | 1.15E-02 | 7.21E-02 | 2.66E+00 | 2.75E+00 | 8.69E-01 | no | Grip1 | intron_variant | MODIFIER |
| 10 | 119618185 | AX-170113994 | 0 | C | T | 0.333 | 1.15E-02 | 7.21E-02 | 2.66E+00 | 2.75E+00 | 8.69E-01 | no | Grip1 | intron_variant | MODIFIER |
| 10 | 119618236 | AX-170304075 | 0 | C | T | 0.333 | 1.15E-02 | 7.21E-02 | 2.66E+00 | 2.75E+00 | 8.69E-01 | no | Grip1 | intron_variant | MODIFIER |
| 10 | 119637670 | AX-97845796 | 0 | G | A | 0.333 | 1.15E-02 | 7.21E-02 | 2.66E+00 | 2.75E+00 | 8.69E-01 | no | Grip1 | intron_variant | MODIFIER |
| 10 | 119671545 | AX-170446350 | 0 | G | T | 0.333 | 1.15E-02 | 7.21E-02 | 2.66E+00 | 2.75E+00 | 8.69E-01 | no | Grip1 | intron_variant | MODIFIER |
| 10 | 119691473 | AX-97824331 | 0 | A | G | 0.333 | 1.15E-02 | 7.21E-02 | 2.66E+00 | 2.75E+00 | 8.69E-01 | no | Grip1 | intron_variant | MODIFIER |
| 10 | 119691493 | AX-170234812 | 0 | T | C | 0.333 | 1.15E-02 | 7.21E-02 | 2.66E+00 | 2.75E+00 | 8.69E-01 | no | Grip1 | intron_variant | MODIFIER |
| 10 | 119696044 | AX-170004028 | 0 | T | A | 0.333 | 1.15E-02 | 7.21E-02 | 2.66E+00 | 2.75E+00 | 8.69E-01 | no | Grip1 | intron_variant | MODIFIER |
| 10 | 119706511 | AX-170369355 | 0 | T | C | 0.333 | 1.15E-02 | 7.21E-02 | 2.66E+00 | 2.75E+00 | 8.69E-01 | no | Grip1 | intron_variant | MODIFIER |
| 10 | 120066345 | AX-170455653 | 0 | T | C | 0.333 | 1.15E-02 | 7.21E-02 | 2.66E+00 | 2.75E+00 | 8.69E-01 | no | Grip1 | intron_variant | MODIFIER |
| 10 | 120066562 | AX-97832518 | 0 | A | G | 0.333 | 1.15E-02 | 7.21E-02 | 2.66E+00 | 2.75E+00 | 8.69E-01 | no | Grip1 | intron_variant | MODIFIER |
| 10 | 120142306 | AX-170478520 | 0 | G | A | 0.333 | 1.15E-02 | 7.21E-02 | 2.66E+00 | 2.75E+00 | 8.69E-01 | no | Irak3 | 3_prime_UTR_variant | MODIFIER |
| 10 | 120230098 | AX-97817739 | 0 | T | C | 0.333 | 1.15E-02 | 7.21E-02 | 2.66E+00 | 2.75E+00 | 8.69E-01 | no | Llph | intron_variant | MODIFIER |
| 10 | 120271313 | AX-170046055 | 0 | A | G | 0.333 | 1.15E-02 | 7.21E-02 | 2.66E+00 | 2.75E+00 | 8.69E-01 | no | | regulatory_region_variant | MODIFIER |
| 10 | 120280580 | AX-170359255 | 0 | T | G | 0.333 | 1.15E-02 | 7.21E-02 | 2.66E+00 | 2.75E+00 | 8.69E-01 | no | Gm48115 | upstream_gene_variant | MODIFIER |
| 10 | 120284626 | AX-170004032 | 0 | G | T | 0.333 | 1.15E-02 | 7.21E-02 | 2.66E+00 | 2.75E+00 | 8.69E-01 | no | Gm48118 | downstream_gene_variant | MODIFIER |
| 10 | 120339332 | AX-97800411 | 0 | A | C | 0.333 | 1.15E-02 | 7.21E-02 | 2.66E+00 | 2.75E+00 | 8.69E-01 | no | | regulatory_region_variant | MODIFIER |
| 10 | 120372447 | AX-97790981 | 0 | A | G | 0.333 | 1.15E-02 | 7.21E-02 | 2.66E+00 | 2.75E+00 | 8.69E-01 | no | 1700006J14Rik | intron_variant,non_coding_transcript_variant | MODIFIER |
| 10 | 120453383 | AX-170430402 | 0 | T | C | 0.333 | 1.15E-02 | 7.21E-02 | 2.66E+00 | 2.75E+00 | 8.69E-01 | no | Hmga2 | intron_variant | MODIFIER |
| 10 | 120528510 | AX-170424405 | 0 | G | C | 0.333 | 1.15E-02 | 7.21E-02 | 2.66E+00 | 2.75E+00 | 8.69E-01 | no | | intergenic_variant | MODIFIER |
| 10 | 120582576 | AX-170521767 | 0 | C | T | 0.333 | 1.15E-02 | 7.21E-02 | 2.66E+00 | 2.75E+00 | 8.69E-01 | no | | regulatory_region_variant | MODIFIER |
| 10 | 120588113 | AX-97854737 | 0 | A | G | 0.333 | 1.15E-02 | 7.21E-02 | 2.66E+00 | 2.75E+00 | 8.69E-01 | no | | regulatory_region_variant | MODIFIER |
| 9 | 58431208 | AX-170447755 | 0 | C | T | 0.219 | 1.96E-02 | 1.20E-01 | 2.68E+00 | 2.74E+00 | 8.69E-01 | no | | intergenic_variant | MODIFIER |
| 20 | 114769987 | AX-170249162 | 0 | A | G | 0.316 | 8.91E-03 | 5.35E-02 | 2.61E+00 | 2.73E+00 | 8.69E-01 | no | Gm5942 | downstream_gene_variant | MODIFIER |
| 1 | 36790915 | AX-170287947 | 1 | A | T | 0.446 | 1.23E-02 | 7.77E-02 | 2.68E+00 | 2.76E+00 | 8.69E-01 | no | Tmem131 | downstream_gene_variant | MODIFIER |
| 20 | 130720464 | AX-97806876 | 1 | A | C | 0.143 | 2.11E-02 | 1.32E-01 | 2.64E+00 | 2.74E+00 | 8.70E-01 | no | | intergenic_variant | MODIFIER |
| 20 | 139827968 | AX-170296250 | 1 | G | A | 0.143 | 2.11E-02 | 1.32E-01 | 2.64E+00 | 2.74E+00 | 8.70E-01 | no | Morc4 | intron_variant | MODIFIER |
| 20 | 140675268 | AX-170035092 | 1 | A | T | 0.143 | 2.11E-02 | 1.32E-01 | 2.64E+00 | 2.74E+00 | 8.70E-01 | no | Mid2 | intron_variant | MODIFIER |
| 15 | 97987550 | AX-170012345 | 1 | C | T | 0.134 | 2.65E-02 | 1.57E-01 | 2.65E+00 | 2.71E+00 | 8.70E-01 | no | Col2a1 | missense_variant | MODERATE |
| 18 | 89085446 | AX-170061644 | 0 | A | G | 0.509 | -9.45E-03 | 6.20E-02 | 2.62E+00 | 2.76E+00 | 8.70E-01 | no | Rttn | intron_variant | MODIFIER |
| 18 | 56493751 | AX-170205709 | 1 | C | T | 0.411 | 1.31E-02 | 8.57E-02 | 2.71E+00 | 2.79E+00 | 8.70E-01 | no | Gramd3 | intron_variant | MODIFIER |
| 12 | 68505965 | AX-170289496 | 0 | A | C | 0.105 | -2.07E-02 | 1.30E-01 | 2.62E+00 | 2.74E+00 | 8.70E-01 | no | Gm19176 | upstream_gene_variant | MODIFIER |
| 6 | 114349926 | AX-170404845 | 0 | C | A | 0.053 | 4.82E-02 | 2.96E-01 | 2.67E+00 | 2.73E+00 | 8.70E-01 | no | | intergenic_variant | MODIFIER |
| 15 | 86858263 | AX-170035554 | 0 | G | A | 0.149 | 3.03E-02 | 1.80E-01 | 2.65E+00 | 2.71E+00 | 8.70E-01 | no | | intergenic_variant | MODIFIER |
| 15 | 87909514 | AX-170279027 | 0 | T | C | 0.149 | 3.03E-02 | 1.80E-01 | 2.65E+00 | 2.71E+00 | 8.70E-01 | no | | intergenic_variant | MODIFIER |
| 15 | 87919307 | AX-170144448 | 0 | C | T | 0.149 | 3.03E-02 | 1.80E-01 | 2.65E+00 | 2.71E+00 | 8.70E-01 | no | | intergenic_variant | MODIFIER |
| 15 | 87941606 | AX-170035553 | 0 | T | C | 0.149 | 3.03E-02 | 1.80E-01 | 2.65E+00 | 2.71E+00 | 8.70E-01 | no | | intergenic_variant | MODIFIER |
| 15 | 88081305 | AX-97855369 | 0 | T | C | 0.149 | 3.03E-02 | 1.80E-01 | 2.65E+00 | 2.71E+00 | 8.70E-01 | no | | intergenic_variant | MODIFIER |
| 15 | 88085548 | AX-170137753 | 0 | C | T | 0.149 | 3.03E-02 | 1.80E-01 | 2.65E+00 | 2.71E+00 | 8.70E-01 | no | | intergenic_variant | MODIFIER |

| | | | | | | | | | | | | | | | |
|---|---|---|---|---|---|---|---|---|---|---|---|---|---|---|---|
| 15 | 88085828 | AX-170144605 | 0 | G | A | 0.149 | 3.03E-02 | 1.80E-01 | 2.65E+00 | 2.71E+00 | 8.70E-01 | no | | intergenic_variant | MODIFIER |
| 15 | 88086044 | AX-170381695 | 0 | T | C | 0.149 | 3.03E-02 | 1.80E-01 | 2.65E+00 | 2.71E+00 | 8.70E-01 | no | | intergenic_variant | MODIFIER |
| 15 | 88086461 | AX-97831311 | 0 | T | C | 0.149 | 3.03E-02 | 1.80E-01 | 2.65E+00 | 2.71E+00 | 8.70E-01 | no | | intergenic_variant | MODIFIER |
| 15 | 88086949 | AX-170056232 | 0 | G | C | 0.149 | 3.03E-02 | 1.80E-01 | 2.65E+00 | 2.71E+00 | 8.70E-01 | no | | intergenic_variant | MODIFIER |
| 15 | 88114482 | AX-170035552 | 0 | C | T | 0.149 | 3.03E-02 | 1.80E-01 | 2.65E+00 | 2.71E+00 | 8.70E-01 | no | | intergenic_variant | MODIFIER |
| 15 | 88117632 | AX-170150233 | 0 | T | G | 0.149 | 3.03E-02 | 1.80E-01 | 2.65E+00 | 2.71E+00 | 8.70E-01 | no | | intergenic_variant | MODIFIER |
| 15 | 88132285 | AX-170183377 | 0 | C | T | 0.149 | 3.03E-02 | 1.80E-01 | 2.65E+00 | 2.71E+00 | 8.70E-01 | no | | intergenic_variant | MODIFIER |
| 15 | 88164353 | AX-97856704 | 0 | G | A | 0.149 | 3.03E-02 | 1.80E-01 | 2.65E+00 | 2.71E+00 | 8.70E-01 | no | | intergenic_variant | MODIFIER |
| 15 | 88204761 | AX-97842436 | 0 | A | G | 0.149 | 3.03E-02 | 1.80E-01 | 2.65E+00 | 2.71E+00 | 8.70E-01 | no | | intergenic_variant | MODIFIER |
| 15 | 88218779 | AX-170335651 | 0 | T | G | 0.149 | 3.03E-02 | 1.80E-01 | 2.65E+00 | 2.71E+00 | 8.70E-01 | no | | intergenic_variant | MODIFIER |
| 15 | 88231166 | AX-170449867 | 0 | T | A | 0.149 | 3.03E-02 | 1.80E-01 | 2.65E+00 | 2.71E+00 | 8.70E-01 | no | | intergenic_variant | MODIFIER |
| 15 | 88236055 | AX-170315960 | 0 | T | A | 0.149 | 3.03E-02 | 1.80E-01 | 2.65E+00 | 2.71E+00 | 8.70E-01 | no | | intergenic_variant | MODIFIER |
| 15 | 88244433 | AX-170167715 | 0 | C | T | 0.149 | 3.03E-02 | 1.80E-01 | 2.65E+00 | 2.71E+00 | 8.70E-01 | no | | intergenic_variant | MODIFIER |
| 15 | 88250485 | AX-170056235 | 0 | A | G | 0.149 | 3.03E-02 | 1.80E-01 | 2.65E+00 | 2.71E+00 | 8.70E-01 | no | | intergenic_variant | MODIFIER |
| 15 | 88420204 | AX-170012215 | 0 | A | G | 0.149 | 3.03E-02 | 1.80E-01 | 2.65E+00 | 2.71E+00 | 8.70E-01 | no | | intergenic_variant | MODIFIER |
| 15 | 88420693 | AX-170273308 | 0 | T | G | 0.149 | 3.03E-02 | 1.80E-01 | 2.65E+00 | 2.71E+00 | 8.70E-01 | no | | intergenic_variant | MODIFIER |
| 15 | 89552751 | AX-170462784 | 0 | C | T | 0.149 | 3.03E-02 | 1.80E-01 | 2.65E+00 | 2.71E+00 | 8.70E-01 | no | Shank3 | intron_variant | MODIFIER |
| 15 | 89564697 | AX-170311452 | 0 | C | A | 0.149 | 3.03E-02 | 1.80E-01 | 2.65E+00 | 2.71E+00 | 8.70E-01 | no | Acr | upstream_gene_variant | MODIFIER |
| 15 | 89574454 | AX-170265508 | 0 | C | T | 0.149 | 3.03E-02 | 1.80E-01 | 2.65E+00 | 2.71E+00 | 8.70E-01 | no | Acr | 3_prime_UTR_variant | MODIFIER |
| 15 | 89590536 | AX-97854460 | 0 | G | A | 0.149 | 3.03E-02 | 1.80E-01 | 2.65E+00 | 2.71E+00 | 8.70E-01 | no | Rabl2 | intron_variant | MODIFIER |
| 15 | 89746269 | AX-170221205 | 0 | T | G | 0.149 | 3.03E-02 | 1.80E-01 | 2.65E+00 | 2.71E+00 | 8.70E-01 | no | | intergenic_variant | MODIFIER |
| 15 | 89845145 | AX-168382548 | 0 | A | G | 0.149 | 3.03E-02 | 1.80E-01 | 2.65E+00 | 2.71E+00 | 8.70E-01 | no | Syt10 | upstream_gene_variant | MODIFIER |
| 15 | 90306335 | AX-170110679 | 0 | T | A | 0.149 | 3.03E-02 | 1.80E-01 | 2.65E+00 | 2.71E+00 | 8.70E-01 | no | | intergenic_variant | MODIFIER |
| 15 | 90434304 | AX-170380593 | 0 | T | C | 0.149 | 3.03E-02 | 1.80E-01 | 2.65E+00 | 2.71E+00 | 8.70E-01 | no | | intergenic_variant | MODIFIER |
| 15 | 90451339 | AX-170056281 | 0 | G | T | 0.149 | 3.03E-02 | 1.80E-01 | 2.65E+00 | 2.71E+00 | 8.70E-01 | no | | intergenic_variant | MODIFIER |
| 15 | 90453400 | AX-170129625 | 0 | T | C | 0.149 | 3.03E-02 | 1.80E-01 | 2.65E+00 | 2.71E+00 | 8.70E-01 | no | | intergenic_variant | MODIFIER |
| 15 | 90460709 | AX-170012243 | 0 | C | T | 0.149 | 3.03E-02 | 1.80E-01 | 2.65E+00 | 2.71E+00 | 8.70E-01 | no | | intergenic_variant | MODIFIER |
| 15 | 90486105 | AX-170012239 | 0 | A | G | 0.149 | 3.03E-02 | 1.80E-01 | 2.65E+00 | 2.71E+00 | 8.70E-01 | no | Cpne8 | downstream_gene_variant | MODIFIER |
| 15 | 90493369 | AX-97782135 | 0 | A | G | 0.149 | 3.03E-02 | 1.80E-01 | 2.65E+00 | 2.71E+00 | 8.70E-01 | no | Cpne8 | intron_variant | MODIFIER |
| 15 | 90507814 | AX-170012238 | 0 | G | A | 0.149 | 3.03E-02 | 1.80E-01 | 2.65E+00 | 2.71E+00 | 8.70E-01 | no | Cpne8 | intron_variant | MODIFIER |
| 15 | 90517838 | AX-97847076 | 0 | T | C | 0.149 | 3.03E-02 | 1.80E-01 | 2.65E+00 | 2.71E+00 | 8.70E-01 | no | Cpne8 | intron_variant | MODIFIER |
| 15 | 90545941 | AX-170012236 | 0 | A | T | 0.149 | 3.03E-02 | 1.80E-01 | 2.65E+00 | 2.71E+00 | 8.70E-01 | no | Cpne8 | intron_variant | MODIFIER |
| 15 | 90564719 | AX-170093334 | 0 | G | A | 0.149 | 3.03E-02 | 1.80E-01 | 2.65E+00 | 2.71E+00 | 8.70E-01 | no | Cpne8 | intron_variant | MODIFIER |
| 15 | 90565607 | AX-170156437 | 0 | T | C | 0.149 | 3.03E-02 | 1.80E-01 | 2.65E+00 | 2.71E+00 | 8.70E-01 | no | Cpne8 | intron_variant | MODIFIER |
| 15 | 90599438 | AX-97830124 | 0 | A | G | 0.149 | 3.03E-02 | 1.80E-01 | 2.65E+00 | 2.71E+00 | 8.70E-01 | no | Cpne8 | intron_variant | MODIFIER |
| 15 | 90599461 | AX-170423851 | 0 | G | A | 0.149 | 3.03E-02 | 1.80E-01 | 2.65E+00 | 2.71E+00 | 8.70E-01 | no | Cpne8 | intron_variant | MODIFIER |
| 15 | 90604904 | AX-170140413 | 0 | G | C | 0.149 | 3.03E-02 | 1.80E-01 | 2.65E+00 | 2.71E+00 | 8.70E-01 | no | Cpne8 | intron_variant | MODIFIER |
| 15 | 90920940 | AX-97807695 | 0 | T | C | 0.149 | 3.03E-02 | 1.80E-01 | 2.65E+00 | 2.71E+00 | 8.70E-01 | no | Gm36618 | intron_variant,non_coding_transcript_variant | MODIFIER |
| 15 | 90928982 | AX-170012247 | 0 | G | A | 0.149 | 3.03E-02 | 1.80E-01 | 2.65E+00 | 2.71E+00 | 8.70E-01 | no | Kif21a | downstream_gene_variant | MODIFIER |
| 15 | 90932282 | AX-170342769 | 0 | C | A | 0.149 | 3.03E-02 | 1.80E-01 | 2.65E+00 | 2.71E+00 | 8.70E-01 | no | Kif21a | downstream_gene_variant | MODIFIER |
| 15 | 90980397 | AX-170056293 | 0 | G | C | 0.149 | 3.03E-02 | 1.80E-01 | 2.65E+00 | 2.71E+00 | 8.70E-01 | no | Kif21a | intron_variant | MODIFIER |
| 15 | 91010785 | AX-170012244 | 0 | T | A | 0.149 | 3.03E-02 | 1.80E-01 | 2.65E+00 | 2.71E+00 | 8.70E-01 | no | Kif21a | intron_variant | MODIFIER |
| 15 | 91013674 | AX-170453523 | 0 | A | C | 0.149 | 3.03E-02 | 1.80E-01 | 2.65E+00 | 2.71E+00 | 8.70E-01 | no | Kif21a | intron_variant | MODIFIER |
| 15 | 91015957 | AX-170012245 | 0 | T | C | 0.149 | 3.03E-02 | 1.80E-01 | 2.65E+00 | 2.71E+00 | 8.70E-01 | no | Kif21a | intron_variant | MODIFIER |
| 15 | 91061619 | AX-170167629 | 0 | A | G | 0.149 | 3.03E-02 | 1.80E-01 | 2.65E+00 | 2.71E+00 | 8.70E-01 | no | | intergenic_variant | MODIFIER |
| 15 | 91062357 | AX-170163791 | 0 | G | C | 0.149 | 3.03E-02 | 1.80E-01 | 2.65E+00 | 2.71E+00 | 8.70E-01 | no | | intergenic_variant | MODIFIER |
| 15 | 91089840 | AX-97786853 | 0 | G | A | 0.149 | 3.03E-02 | 1.80E-01 | 2.65E+00 | 2.71E+00 | 8.70E-01 | no | | intergenic_variant | MODIFIER |
| 15 | 91105276 | AX-170308783 | 0 | G | C | 0.149 | 3.03E-02 | 1.80E-01 | 2.65E+00 | 2.71E+00 | 8.70E-01 | no | | regulatory_region_variant | MODIFIER |
| 15 | 91113001 | AX-170175495 | 0 | C | G | 0.149 | 3.03E-02 | 1.80E-01 | 2.65E+00 | 2.71E+00 | 8.70E-01 | no | | regulatory_region_variant | MODIFIER |
| 15 | 91114705 | AX-170530616 | 0 | A | G | 0.149 | 3.03E-02 | 1.80E-01 | 2.65E+00 | 2.71E+00 | 8.70E-01 | no | | intergenic_variant | MODIFIER |
| 15 | 91138250 | AX-170457247 | 0 | T | C | 0.149 | 3.03E-02 | 1.80E-01 | 2.65E+00 | 2.71E+00 | 8.70E-01 | no | | intergenic_variant | MODIFIER |
| 15 | 91159429 | AX-170316386 | 0 | A | C | 0.149 | 3.03E-02 | 1.80E-01 | 2.65E+00 | 2.71E+00 | 8.70E-01 | no | Abcd2 | intron_variant | MODIFIER |
| 15 | 91184917 | AX-170455681 | 0 | G | A | 0.149 | 3.03E-02 | 1.80E-01 | 2.65E+00 | 2.71E+00 | 8.70E-01 | no | Abcd2 | intron_variant | MODIFIER |
| 15 | 91329932 | AX-170415408 | 0 | G | T | 0.149 | 3.03E-02 | 1.80E-01 | 2.65E+00 | 2.71E+00 | 8.70E-01 | no | Slc2a13 | intron_variant | MODIFIER |

| | | | | | | | | | | | | | | | |
|---|---|---|---|---|---|---|---|---|---|---|---|---|---|---|---|
| 15 | 91340027 | AX-170293500 | 0 | T | C | 0.149 | 3.03E-02 | 1.80E-01 | 2.65E+00 | 2.71E+00 | 8.70E-01 | no | Slc2a13 | intron_variant | MODIFIER |
| 15 | 91357863 | AX-170202352 | 0 | A | C | 0.149 | 3.03E-02 | 1.80E-01 | 2.65E+00 | 2.71E+00 | 8.70E-01 | no | Slc2a13 | intron_variant | MODIFIER |
| 15 | 91358521 | AX-170456761 | 0 | G | A | 0.149 | 3.03E-02 | 1.80E-01 | 2.65E+00 | 2.71E+00 | 8.70E-01 | no | Slc2a13 | intron_variant | MODIFIER |
| 15 | 91358992 | AX-170510065 | 0 | T | C | 0.149 | 3.03E-02 | 1.80E-01 | 2.65E+00 | 2.71E+00 | 8.70E-01 | no | Slc2a13 | intron_variant | MODIFIER |
| 15 | 91359205 | AX-170056296 | 0 | T | C | 0.149 | 3.03E-02 | 1.80E-01 | 2.65E+00 | 2.71E+00 | 8.70E-01 | no | Slc2a13 | intron_variant | MODIFIER |
| 15 | 91360137 | AX-170480753 | 0 | G | A | 0.149 | 3.03E-02 | 1.80E-01 | 2.65E+00 | 2.71E+00 | 8.70E-01 | no | Slc2a13 | intron_variant | MODIFIER |
| 15 | 91380769 | AX-170416385 | 0 | A | G | 0.149 | 3.03E-02 | 1.80E-01 | 2.65E+00 | 2.71E+00 | 8.70E-01 | no | Slc2a13 | intron_variant | MODIFIER |
| 15 | 91383261 | AX-97848798 | 0 | C | T | 0.149 | 3.03E-02 | 1.80E-01 | 2.65E+00 | 2.71E+00 | 8.70E-01 | no | Slc2a13 | intron_variant | MODIFIER |
| 15 | 91468801 | AX-97830194 | 0 | A | G | 0.149 | 3.03E-02 | 1.80E-01 | 2.65E+00 | 2.71E+00 | 8.70E-01 | no | Slc2a13 | intron_variant | MODIFIER |
| 15 | 91731553 | AX-170309839 | 0 | C | G | 0.149 | 3.03E-02 | 1.80E-01 | 2.65E+00 | 2.71E+00 | 8.70E-01 | no | Lrrk2 | missense_variant | MODERATE |
| 15 | 91760574 | AX-170056301 | 0 | T | C | 0.149 | 3.03E-02 | 1.80E-01 | 2.65E+00 | 2.71E+00 | 8.70E-01 | no | Lrrk2 | intron_variant | MODIFIER |
| 15 | 91762000 | AX-170142327 | 0 | T | C | 0.149 | 3.03E-02 | 1.80E-01 | 2.65E+00 | 2.71E+00 | 8.70E-01 | no | Lrrk2 | intron_variant | MODIFIER |
| 15 | 91768326 | AX-170056302 | 0 | G | C | 0.149 | 3.03E-02 | 1.80E-01 | 2.65E+00 | 2.71E+00 | 8.70E-01 | no | Lrrk2 | intron_variant | MODIFIER |
| 15 | 91810058 | AX-170535660 | 0 | C | T | 0.149 | 3.03E-02 | 1.80E-01 | 2.65E+00 | 2.71E+00 | 8.70E-01 | no | Lrrk2 | intron_variant | MODIFIER |
| 15 | 91820685 | AX-97794396 | 0 | A | G | 0.149 | 3.03E-02 | 1.80E-01 | 2.65E+00 | 2.71E+00 | 8.70E-01 | no | Lrrk2 | downstream_gene_variant | MODIFIER |
| 15 | 91825371 | AX-170252234 | 0 | C | T | 0.149 | 3.03E-02 | 1.80E-01 | 2.65E+00 | 2.71E+00 | 8.70E-01 | no | Gm15382 | downstream_gene_variant | MODIFIER |
| 15 | 91827325 | AX-170428441 | 0 | A | G | 0.149 | 3.03E-02 | 1.80E-01 | 2.65E+00 | 2.71E+00 | 8.70E-01 | no | Gm15382 | downstream_gene_variant | MODIFIER |
| 15 | 91833322 | AX-170240395 | 0 | T | G | 0.149 | 3.03E-02 | 1.80E-01 | 2.65E+00 | 2.71E+00 | 8.70E-01 | no | Smgc | upstream_gene_variant | MODIFIER |
| 15 | 91838543 | AX-170339633 | 0 | G | A | 0.149 | 3.03E-02 | 1.80E-01 | 2.65E+00 | 2.71E+00 | 8.70E-01 | no | Smgc | intron_variant | MODIFIER |
| 15 | 91838975 | AX-97798359 | 0 | C | T | 0.149 | 3.03E-02 | 1.80E-01 | 2.65E+00 | 2.71E+00 | 8.70E-01 | no | Smgc | intron_variant | MODIFIER |
| 15 | 91855471 | AX-170189650 | 0 | T | C | 0.149 | 3.03E-02 | 1.80E-01 | 2.65E+00 | 2.71E+00 | 8.70E-01 | no | Smgc | intron_variant | MODIFIER |
| 15 | 91871339 | AX-170240089 | 0 | A | G | 0.149 | 3.03E-02 | 1.80E-01 | 2.65E+00 | 2.71E+00 | 8.70E-01 | no | Muc19 | intron_variant,non_coding_transcript_variant | MODIFIER |
| 15 | 91871613 | AX-97825800 | 0 | A | C | 0.149 | 3.03E-02 | 1.80E-01 | 2.65E+00 | 2.71E+00 | 8.70E-01 | no | Muc19 | intron_variant,non_coding_transcript_variant | MODIFIER |
| 15 | 91882175 | AX-170470190 | 0 | C | T | 0.149 | 3.03E-02 | 1.80E-01 | 2.65E+00 | 2.71E+00 | 8.70E-01 | no | Muc19 | intron_variant,non_coding_transcript_variant | MODIFIER |
| 15 | 91892619 | AX-170217118 | 0 | C | T | 0.149 | 3.03E-02 | 1.80E-01 | 2.65E+00 | 2.71E+00 | 8.70E-01 | no | Muc19 | intron_variant,non_coding_transcript_variant | MODIFIER |
| 15 | 91892918 | AX-170192936 | 0 | T | C | 0.149 | 3.03E-02 | 1.80E-01 | 2.65E+00 | 2.71E+00 | 8.70E-01 | no | Muc19 | non_coding_transcript_exon_variant | MODIFIER |
| 15 | 91895055 | AX-170430988 | 0 | C | T | 0.149 | 3.03E-02 | 1.80E-01 | 2.65E+00 | 2.71E+00 | 8.70E-01 | no | Muc19 | intron_variant,non_coding_transcript_variant | MODIFIER |
| 15 | 91897876 | AX-170293419 | 0 | C | T | 0.149 | 3.03E-02 | 1.80E-01 | 2.65E+00 | 2.71E+00 | 8.70E-01 | no | Muc19 | intron_variant,non_coding_transcript_variant | MODIFIER |
| 15 | 91934336 | AX-170171393 | 0 | T | C | 0.149 | 3.03E-02 | 1.80E-01 | 2.65E+00 | 2.71E+00 | 8.70E-01 | no | Muc19 | non_coding_transcript_exon_variant | MODIFIER |
| 15 | 91938107 | AX-168415742 | 0 | C | T | 0.149 | 3.03E-02 | 1.80E-01 | 2.65E+00 | 2.71E+00 | 8.70E-01 | no | Muc19 | downstream_gene_variant | MODIFIER |
| 15 | 91959119 | AX-170429284 | 0 | C | T | 0.149 | 3.03E-02 | 1.80E-01 | 2.65E+00 | 2.71E+00 | 8.70E-01 | no | Gm18979 | upstream_gene_variant | MODIFIER |
| 15 | 91997215 | AX-97806843 | 0 | C | T | 0.149 | 3.03E-02 | 1.80E-01 | 2.65E+00 | 2.71E+00 | 8.70E-01 | no | | intergenic_variant | MODIFIER |
| 15 | 92050237 | AX-170012265 | 0 | A | C | 0.149 | 3.03E-02 | 1.80E-01 | 2.65E+00 | 2.71E+00 | 8.70E-01 | no | Cntn1 | upstream_gene_variant | MODIFIER |
| 15 | 92053056 | AX-170501806 | 0 | C | A | 0.149 | 3.03E-02 | 1.80E-01 | 2.65E+00 | 2.71E+00 | 8.70E-01 | no | Cntn1 | intron_variant,non_coding_transcript_variant | MODIFIER |
| 15 | 92053461 | AX-170145761 | 0 | G | A | 0.149 | 3.03E-02 | 1.80E-01 | 2.65E+00 | 2.71E+00 | 8.70E-01 | no | Cntn1 | intron_variant,non_coding_transcript_variant | MODIFIER |
| 15 | 92078512 | AX-170012260 | 0 | A | G | 0.149 | 3.03E-02 | 1.80E-01 | 2.65E+00 | 2.71E+00 | 8.70E-01 | no | Cntn1 | intron_variant,non_coding_transcript_variant | MODIFIER |
| 15 | 92093219 | AX-170335081 | 0 | G | A | 0.149 | 3.03E-02 | 1.80E-01 | 2.65E+00 | 2.71E+00 | 8.70E-01 | no | Cntn1 | intron_variant,non_coding_transcript_variant | MODIFIER |
| 15 | 92094143 | AX-97790226 | 0 | G | A | 0.149 | 3.03E-02 | 1.80E-01 | 2.65E+00 | 2.71E+00 | 8.70E-01 | no | Cntn1 | intron_variant,non_coding_transcript_variant | MODIFIER |
| 15 | 92098157 | AX-170371176 | 0 | T | C | 0.149 | 3.03E-02 | 1.80E-01 | 2.65E+00 | 2.71E+00 | 8.70E-01 | no | Cntn1 | intron_variant,non_coding_transcript_variant | MODIFIER |
| 15 | 92102497 | AX-170012264 | 0 | T | C | 0.149 | 3.03E-02 | 1.80E-01 | 2.65E+00 | 2.71E+00 | 8.70E-01 | no | Cntn1 | intron_variant,non_coding_transcript_variant | MODIFIER |
| 15 | 92186629 | AX-170296115 | 0 | A | G | 0.149 | 3.03E-02 | 1.80E-01 | 2.65E+00 | 2.71E+00 | 8.70E-01 | no | Cntn1 | intron_variant | MODIFIER |
| 15 | 92187317 | AX-170157232 | 0 | C | G | 0.149 | 3.03E-02 | 1.80E-01 | 2.65E+00 | 2.71E+00 | 8.70E-01 | no | Cntn1 | intron_variant | MODIFIER |
| 15 | 92293867 | AX-97821418 | 0 | G | A | 0.149 | 3.03E-02 | 1.80E-01 | 2.65E+00 | 2.71E+00 | 8.70E-01 | no | Cntn1 | intron_variant | MODIFIER |
| 15 | 92318051 | AX-170337176 | 0 | T | C | 0.149 | 3.03E-02 | 1.80E-01 | 2.65E+00 | 2.71E+00 | 8.70E-01 | no | Cntn1 | synonymous_variant | LOW |
| 15 | 92318107 | AX-97794058 | 0 | C | T | 0.149 | 3.03E-02 | 1.80E-01 | 2.65E+00 | 2.71E+00 | 8.70E-01 | no | Cntn1 | intron_variant | MODIFIER |
| 15 | 92329746 | AX-170469991 | 0 | G | T | 0.149 | 3.03E-02 | 1.80E-01 | 2.65E+00 | 2.71E+00 | 8.70E-01 | no | Cntn1 | intron_variant | MODIFIER |
| 15 | 92333268 | AX-170439163 | 0 | T | C | 0.149 | 3.03E-02 | 1.80E-01 | 2.65E+00 | 2.71E+00 | 8.70E-01 | no | Cntn1 | intron_variant | MODIFIER |
| 15 | 92342108 | AX-170389371 | 0 | C | T | 0.149 | 3.03E-02 | 1.80E-01 | 2.65E+00 | 2.71E+00 | 8.70E-01 | no | Cntn1 | downstream_gene_variant | MODIFIER |
| 15 | 92346905 | AX-170334917 | 0 | C | G | 0.149 | 3.03E-02 | 1.80E-01 | 2.65E+00 | 2.71E+00 | 8.70E-01 | no | Cntn1 | downstream_gene_variant | MODIFIER |
| 15 | 92349828 | AX-97790145 | 0 | C | T | 0.149 | 3.03E-02 | 1.80E-01 | 2.65E+00 | 2.71E+00 | 8.70E-01 | no | Gm26760 | intron_variant,non_coding_transcript_variant | MODIFIER |
| 15 | 92389681 | AX-170365378 | 0 | C | T | 0.149 | 3.03E-02 | 1.80E-01 | 2.65E+00 | 2.71E+00 | 8.70E-01 | no | | intergenic_variant | MODIFIER |
| 15 | 92393404 | AX-97803117 | 0 | G | A | 0.149 | 3.03E-02 | 1.80E-01 | 2.65E+00 | 2.71E+00 | 8.70E-01 | no | Pdzrn4 | upstream_gene_variant | MODIFIER |
| 15 | 92393537 | AX-170106920 | 0 | T | C | 0.149 | 3.03E-02 | 1.80E-01 | 2.65E+00 | 2.71E+00 | 8.70E-01 | no | Pdzrn4 | upstream_gene_variant | MODIFIER |
| 15 | 92449449 | AX-97799074 | 0 | C | T | 0.149 | 3.03E-02 | 1.80E-01 | 2.65E+00 | 2.71E+00 | 8.70E-01 | no | Pdzrn4 | intron_variant | MODIFIER |

| | | | | | | | | | | | | | | | |
|---|---|---|---|---|---|---|---|---|---|---|---|---|---|---|---|
| 15 | 92457520 | AX-170391773 | 0 | A | G | 0.149 | 3.03E-02 | 1.80E-01 | 2.65E+00 | 2.71E+00 | 8.70E-01 | no | Pdzrn4 | intron_variant | MODIFIER |
| 15 | 92458172 | AX-170468392 | 0 | A | T | 0.149 | 3.03E-02 | 1.80E-01 | 2.65E+00 | 2.71E+00 | 8.70E-01 | no | Pdzrn4 | intron_variant | MODIFIER |
| 15 | 92458982 | AX-170056319 | 0 | G | A | 0.149 | 3.03E-02 | 1.80E-01 | 2.65E+00 | 2.71E+00 | 8.70E-01 | no | Pdzrn4 | intron_variant | MODIFIER |
| 15 | 92459113 | AX-170056320 | 0 | A | C | 0.149 | 3.03E-02 | 1.80E-01 | 2.65E+00 | 2.71E+00 | 8.70E-01 | no | Pdzrn4 | intron_variant | MODIFIER |
| 15 | 92459777 | AX-170520917 | 0 | G | A | 0.149 | 3.03E-02 | 1.80E-01 | 2.65E+00 | 2.71E+00 | 8.70E-01 | no | Pdzrn4 | intron_variant | MODIFIER |
| 15 | 92626307 | AX-170211797 | 0 | G | A | 0.149 | 3.03E-02 | 1.80E-01 | 2.65E+00 | 2.71E+00 | 8.70E-01 | no | Pdzrn4 | intron_variant | MODIFIER |
| 15 | 92626501 | AX-170366285 | 0 | A | G | 0.149 | 3.03E-02 | 1.80E-01 | 2.65E+00 | 2.71E+00 | 8.70E-01 | no | Pdzrn4 | intron_variant | MODIFIER |
| 15 | 92628122 | AX-170272261 | 0 | G | T | 0.149 | 3.03E-02 | 1.80E-01 | 2.65E+00 | 2.71E+00 | 8.70E-01 | no | Pdzrn4 | intron_variant | MODIFIER |
| 15 | 92628322 | AX-170012275 | 0 | A | G | 0.149 | 3.03E-02 | 1.80E-01 | 2.65E+00 | 2.71E+00 | 8.70E-01 | no | Pdzrn4 | intron_variant | MODIFIER |
| 15 | 92661753 | AX-170012269 | 0 | T | C | 0.149 | 3.03E-02 | 1.80E-01 | 2.65E+00 | 2.71E+00 | 8.70E-01 | no | Pdzrn4 | intron_variant | MODIFIER |
| 15 | 92667320 | AX-97830228 | 0 | A | G | 0.149 | 3.03E-02 | 1.80E-01 | 2.65E+00 | 2.71E+00 | 8.70E-01 | no | Pdzrn4 | intron_variant | MODIFIER |
| 15 | 92761546 | AX-97835288 | 0 | A | G | 0.149 | 3.03E-02 | 1.80E-01 | 2.65E+00 | 2.71E+00 | 8.70E-01 | no | Pdzrn4 | intron_variant | MODIFIER |
| 15 | 92794001 | AX-97849003 | 0 | G | T | 0.149 | 3.03E-02 | 1.80E-01 | 2.65E+00 | 2.71E+00 | 8.70E-01 | no | | intergenic_variant | MODIFIER |
| 15 | 92908288 | AX-97794084 | 0 | A | G | 0.149 | 3.03E-02 | 1.80E-01 | 2.65E+00 | 2.71E+00 | 8.70E-01 | no | | intergenic_variant | MODIFIER |
| 15 | 92929678 | AX-170151435 | 0 | A | G | 0.149 | 3.03E-02 | 1.80E-01 | 2.65E+00 | 2.71E+00 | 8.70E-01 | no | Gm49418 | upstream_gene_variant | MODIFIER |
| 15 | 92942800 | AX-97846885 | 0 | T | C | 0.149 | 3.03E-02 | 1.80E-01 | 2.65E+00 | 2.71E+00 | 8.70E-01 | no | Gm49418 | intron_variant,non_coding_transcript_variant | MODIFIER |
| 15 | 92955563 | AX-170292559 | 0 | A | G | 0.149 | 3.03E-02 | 1.80E-01 | 2.65E+00 | 2.71E+00 | 8.70E-01 | no | | intergenic_variant | MODIFIER |
| 15 | 92995415 | AX-97835212 | 0 | T | G | 0.149 | 3.03E-02 | 1.80E-01 | 2.65E+00 | 2.71E+00 | 8.70E-01 | no | | intergenic_variant | MODIFIER |
| 15 | 93008861 | AX-170131810 | 0 | C | A | 0.149 | 3.03E-02 | 1.80E-01 | 2.65E+00 | 2.71E+00 | 8.70E-01 | no | Gm30085 | downstream_gene_variant | MODIFIER |
| 15 | 93009038 | AX-170170926 | 0 | C | T | 0.149 | 3.03E-02 | 1.80E-01 | 2.65E+00 | 2.71E+00 | 8.70E-01 | no | Gm30085 | downstream_gene_variant | MODIFIER |
| 15 | 93012900 | AX-170336478 | 0 | G | C | 0.149 | 3.03E-02 | 1.80E-01 | 2.65E+00 | 2.71E+00 | 8.70E-01 | no | | intergenic_variant | MODIFIER |
| 15 | 93016576 | AX-97838460 | 0 | G | T | 0.149 | 3.03E-02 | 1.80E-01 | 2.65E+00 | 2.71E+00 | 8.70E-01 | no | | intergenic_variant | MODIFIER |
| 15 | 93026040 | AX-170479388 | 0 | G | A | 0.149 | 3.03E-02 | 1.80E-01 | 2.65E+00 | 2.71E+00 | 8.70E-01 | no | | regulatory_region_variant | MODIFIER |
| 15 | 93031148 | AX-170447399 | 0 | T | C | 0.149 | 3.03E-02 | 1.80E-01 | 2.65E+00 | 2.71E+00 | 8.70E-01 | no | | intergenic_variant | MODIFIER |
| 15 | 93034728 | AX-170426262 | 0 | C | A | 0.149 | 3.03E-02 | 1.80E-01 | 2.65E+00 | 2.71E+00 | 8.70E-01 | no | | intergenic_variant | MODIFIER |
| 15 | 93040980 | AX-97782998 | 0 | A | G | 0.149 | 3.03E-02 | 1.80E-01 | 2.65E+00 | 2.71E+00 | 8.70E-01 | no | | intergenic_variant | MODIFIER |
| 15 | 93064415 | AX-97816539 | 0 | A | C | 0.149 | 3.03E-02 | 1.80E-01 | 2.65E+00 | 2.71E+00 | 8.70E-01 | no | | intergenic_variant | MODIFIER |
| 15 | 93105959 | AX-97829900 | 0 | C | T | 0.149 | 3.03E-02 | 1.80E-01 | 2.65E+00 | 2.71E+00 | 8.70E-01 | no | | regulatory_region_variant | MODIFIER |
| 15 | 93121022 | AX-97829904 | 0 | G | A | 0.149 | 3.03E-02 | 1.80E-01 | 2.65E+00 | 2.71E+00 | 8.70E-01 | no | | intergenic_variant | MODIFIER |
| 15 | 93127013 | AX-170012284 | 0 | C | T | 0.149 | 3.03E-02 | 1.80E-01 | 2.65E+00 | 2.71E+00 | 8.70E-01 | no | | intergenic_variant | MODIFIER |
| 15 | 93144077 | AX-97785545 | 0 | G | A | 0.149 | 3.03E-02 | 1.80E-01 | 2.65E+00 | 2.71E+00 | 8.70E-01 | no | | intergenic_variant | MODIFIER |
| 15 | 93151616 | AX-170267607 | 0 | C | G | 0.149 | 3.03E-02 | 1.80E-01 | 2.65E+00 | 2.71E+00 | 8.70E-01 | no | | intergenic_variant | MODIFIER |
| 15 | 93152784 | AX-170473227 | 0 | T | C | 0.149 | 3.03E-02 | 1.80E-01 | 2.65E+00 | 2.71E+00 | 8.70E-01 | no | | intergenic_variant | MODIFIER |
| 15 | 93158303 | AX-170012285 | 0 | G | T | 0.149 | 3.03E-02 | 1.80E-01 | 2.65E+00 | 2.71E+00 | 8.70E-01 | no | | intergenic_variant | MODIFIER |
| 15 | 93158696 | AX-97789178 | 0 | T | C | 0.149 | 3.03E-02 | 1.80E-01 | 2.65E+00 | 2.71E+00 | 8.70E-01 | no | | intergenic_variant | MODIFIER |
| 15 | 93158784 | AX-170117192 | 0 | C | A | 0.149 | 3.03E-02 | 1.80E-01 | 2.65E+00 | 2.71E+00 | 8.70E-01 | no | | intergenic_variant | MODIFIER |
| 15 | 93226313 | AX-97829897 | 0 | T | C | 0.149 | 3.03E-02 | 1.80E-01 | 2.65E+00 | 2.71E+00 | 8.70E-01 | no | | intergenic_variant | MODIFIER |
| 15 | 93241198 | AX-97798639 | 0 | C | A | 0.149 | 3.03E-02 | 1.80E-01 | 2.65E+00 | 2.71E+00 | 8.70E-01 | no | Gxylt1 | 3_prime_UTR_variant | MODIFIER |
| 15 | 93241867 | AX-170165137 | 0 | A | G | 0.149 | 3.03E-02 | 1.80E-01 | 2.65E+00 | 2.71E+00 | 8.70E-01 | no | Gxylt1 | 3_prime_UTR_variant | MODIFIER |
| 15 | 93278217 | AX-170087940 | 0 | A | G | 0.149 | 3.03E-02 | 1.80E-01 | 2.65E+00 | 2.71E+00 | 8.70E-01 | no | Gxylt1 | upstream_gene_variant | MODIFIER |
| 15 | 93281821 | AX-170012288 | 0 | A | G | 0.149 | 3.03E-02 | 1.80E-01 | 2.65E+00 | 2.71E+00 | 8.70E-01 | no | Yaf2 | downstream_gene_variant | MODIFIER |
| 15 | 93287676 | AX-97854606 | 0 | C | T | 0.149 | 3.03E-02 | 1.80E-01 | 2.65E+00 | 2.71E+00 | 8.70E-01 | no | Yaf2 | intron_variant | MODIFIER |
| 15 | 93297522 | AX-170170622 | 0 | C | A | 0.149 | 3.03E-02 | 1.80E-01 | 2.65E+00 | 2.71E+00 | 8.70E-01 | no | Yaf2 | intron_variant | MODIFIER |
| 15 | 93303877 | AX-170012279 | 0 | T | G | 0.149 | 3.03E-02 | 1.80E-01 | 2.65E+00 | 2.71E+00 | 8.70E-01 | no | Yaf2 | intron_variant | MODIFIER |
| 15 | 93304423 | AX-170012281 | 0 | G | A | 0.149 | 3.03E-02 | 1.80E-01 | 2.65E+00 | 2.71E+00 | 8.70E-01 | no | Yaf2 | intron_variant | MODIFIER |
| 15 | 93381010 | AX-97830732 | 0 | T | C | 0.149 | 3.03E-02 | 1.80E-01 | 2.65E+00 | 2.71E+00 | 8.70E-01 | no | | intergenic_variant | MODIFIER |
| 15 | 93395365 | AX-97830762 | 0 | C | T | 0.149 | 3.03E-02 | 1.80E-01 | 2.65E+00 | 2.71E+00 | 8.70E-01 | no | Pphln1 | upstream_gene_variant | MODIFIER |
| 15 | 93430710 | AX-97829951 | 0 | C | A | 0.149 | 3.03E-02 | 1.80E-01 | 2.65E+00 | 2.71E+00 | 8.70E-01 | no | Pphln1 | intron_variant | MODIFIER |
| 15 | 93445727 | AX-97829950 | 0 | A | G | 0.149 | 3.03E-02 | 1.80E-01 | 2.65E+00 | 2.71E+00 | 8.70E-01 | no | Pphln1 | intron_variant | MODIFIER |
| 15 | 93449566 | AX-170176329 | 0 | T | C | 0.149 | 3.03E-02 | 1.80E-01 | 2.65E+00 | 2.71E+00 | 8.70E-01 | no | Pphln1 | intron_variant | MODIFIER |
| 15 | 93449601 | AX-170310595 | 0 | C | A | 0.149 | 3.03E-02 | 1.80E-01 | 2.65E+00 | 2.71E+00 | 8.70E-01 | no | Pphln1 | intron_variant | MODIFIER |
| 15 | 93449690 | AX-170379882 | 0 | C | T | 0.149 | 3.03E-02 | 1.80E-01 | 2.65E+00 | 2.71E+00 | 8.70E-01 | no | Pphln1 | intron_variant | MODIFIER |
| 15 | 93449877 | AX-170405362 | 0 | T | G | 0.149 | 3.03E-02 | 1.80E-01 | 2.65E+00 | 2.71E+00 | 8.70E-01 | no | Pphln1 | intron_variant | MODIFIER |
| 15 | 93472430 | AX-97826630 | 0 | T | C | 0.149 | 3.03E-02 | 1.80E-01 | 2.65E+00 | 2.71E+00 | 8.70E-01 | no | Pphln1 | intron_variant | MODIFIER |

| | | | | | | | | | | | | | | | |
|---|---|---|---|---|---|---|---|---|---|---|---|---|---|---|---|
| 15 | 93481025 | AX-170435967 | 0 | C | A | 0.149 | 3.03E-02 | 1.80E-01 | 2.65E+00 | 2.71E+00 | 8.70E-01 | no | Pphln1 | intron_variant | MODIFIER |
| 15 | 93482648 | AX-170216460 | 0 | C | A | 0.149 | 3.03E-02 | 1.80E-01 | 2.65E+00 | 2.71E+00 | 8.70E-01 | no | Pphln1 | intron_variant | MODIFIER |
| 15 | 93483801 | AX-97795333 | 0 | A | G | 0.149 | 3.03E-02 | 1.80E-01 | 2.65E+00 | 2.71E+00 | 8.70E-01 | no | Pphln1 | intron_variant | MODIFIER |
| 15 | 93485939 | AX-170184839 | 0 | T | A | 0.149 | 3.03E-02 | 1.80E-01 | 2.65E+00 | 2.71E+00 | 8.70E-01 | no | Pphln1 | intron_variant | MODIFIER |
| 15 | 93511025 | AX-170489621 | 0 | T | G | 0.149 | 3.03E-02 | 1.80E-01 | 2.65E+00 | 2.71E+00 | 8.70E-01 | no | Prickle1 | intron_variant | MODIFIER |
| 15 | 93518432 | AX-170461103 | 0 | T | A | 0.149 | 3.03E-02 | 1.80E-01 | 2.65E+00 | 2.71E+00 | 8.70E-01 | no | Prickle1 | intron_variant | MODIFIER |
| 15 | 93525040 | AX-170056332 | 0 | A | G | 0.149 | 3.03E-02 | 1.80E-01 | 2.65E+00 | 2.71E+00 | 8.70E-01 | no | Prickle1 | intron_variant | MODIFIER |
| 15 | 93528946 | AX-170093341 | 0 | A | G | 0.149 | 3.03E-02 | 1.80E-01 | 2.65E+00 | 2.71E+00 | 8.70E-01 | no | Prickle1 | intron_variant | MODIFIER |
| 15 | 93530601 | AX-170364641 | 0 | A | G | 0.149 | 3.03E-02 | 1.80E-01 | 2.65E+00 | 2.71E+00 | 8.70E-01 | no | Prickle1 | intron_variant | MODIFIER |
| 15 | 93530782 | AX-170470542 | 0 | G | C | 0.149 | 3.03E-02 | 1.80E-01 | 2.65E+00 | 2.71E+00 | 8.70E-01 | no | Prickle1 | intron_variant | MODIFIER |
| 15 | 93531014 | AX-170498857 | 0 | G | A | 0.149 | 3.03E-02 | 1.80E-01 | 2.65E+00 | 2.71E+00 | 8.70E-01 | no | Prickle1 | intron_variant | MODIFIER |
| 15 | 93532776 | AX-170108729 | 0 | T | C | 0.149 | 3.03E-02 | 1.80E-01 | 2.65E+00 | 2.71E+00 | 8.70E-01 | no | Prickle1 | intron_variant | MODIFIER |
| 15 | 93539203 | AX-170282463 | 0 | T | C | 0.149 | 3.03E-02 | 1.80E-01 | 2.65E+00 | 2.71E+00 | 8.70E-01 | no | Prickle1 | intron_variant | MODIFIER |
| 15 | 93557104 | AX-97855950 | 0 | A | G | 0.149 | 3.03E-02 | 1.80E-01 | 2.65E+00 | 2.71E+00 | 8.70E-01 | no | Prickle1 | intron_variant | MODIFIER |
| 15 | 93608910 | AX-170012295 | 0 | A | G | 0.149 | 3.03E-02 | 1.80E-01 | 2.65E+00 | 2.71E+00 | 8.70E-01 | no | | intergenic_variant | MODIFIER |
| 15 | 93609456 | AX-170012294 | 0 | G | A | 0.149 | 3.03E-02 | 1.80E-01 | 2.65E+00 | 2.71E+00 | 8.70E-01 | no | | intergenic_variant | MODIFIER |
| 15 | 93616221 | AX-170012292 | 0 | G | T | 0.149 | 3.03E-02 | 1.80E-01 | 2.65E+00 | 2.71E+00 | 8.70E-01 | no | | intergenic_variant | MODIFIER |
| 15 | 93619283 | AX-170250409 | 0 | T | C | 0.149 | 3.03E-02 | 1.80E-01 | 2.65E+00 | 2.71E+00 | 8.70E-01 | no | | intergenic_variant | MODIFIER |
| 15 | 93646658 | AX-97839944 | 0 | G | A | 0.149 | 3.03E-02 | 1.80E-01 | 2.65E+00 | 2.71E+00 | 8.70E-01 | no | | intergenic_variant | MODIFIER |
| 15 | 93650879 | AX-170260084 | 0 | A | G | 0.149 | 3.03E-02 | 1.80E-01 | 2.65E+00 | 2.71E+00 | 8.70E-01 | no | | regulatory_region_variant | MODIFIER |
| 15 | 93653393 | AX-170382938 | 0 | C | T | 0.149 | 3.03E-02 | 1.80E-01 | 2.65E+00 | 2.71E+00 | 8.70E-01 | no | | intergenic_variant | MODIFIER |
| 15 | 93672763 | AX-170157371 | 0 | G | A | 0.149 | 3.03E-02 | 1.80E-01 | 2.65E+00 | 2.71E+00 | 8.70E-01 | no | | intergenic_variant | MODIFIER |
| 15 | 93676533 | AX-170056334 | 0 | C | T | 0.149 | 3.03E-02 | 1.80E-01 | 2.65E+00 | 2.71E+00 | 8.70E-01 | no | Zfp42-ps1 | downstream_gene_variant | MODIFIER |
| 15 | 93677555 | AX-170056333 | 0 | A | G | 0.149 | 3.03E-02 | 1.80E-01 | 2.65E+00 | 2.71E+00 | 8.70E-01 | no | Zfp42-ps1 | downstream_gene_variant | MODIFIER |
| 15 | 93683762 | AX-170473739 | 0 | G | A | 0.149 | 3.03E-02 | 1.80E-01 | 2.65E+00 | 2.71E+00 | 8.70E-01 | no | Zfp42-ps1 | upstream_gene_variant | MODIFIER |
| 15 | 93687797 | AX-170122777 | 0 | A | G | 0.149 | 3.03E-02 | 1.80E-01 | 2.65E+00 | 2.71E+00 | 8.70E-01 | no | | intergenic_variant | MODIFIER |
| 15 | 93688215 | AX-170507234 | 0 | C | T | 0.149 | 3.03E-02 | 1.80E-01 | 2.65E+00 | 2.71E+00 | 8.70E-01 | no | | intergenic_variant | MODIFIER |
| 15 | 93688261 | AX-97843845 | 0 | A | G | 0.149 | 3.03E-02 | 1.80E-01 | 2.65E+00 | 2.71E+00 | 8.70E-01 | no | | intergenic_variant | MODIFIER |
| 15 | 93688755 | AX-170491796 | 0 | G | A | 0.149 | 3.03E-02 | 1.80E-01 | 2.65E+00 | 2.71E+00 | 8.70E-01 | no | | intergenic_variant | MODIFIER |
| 15 | 93703750 | AX-170104542 | 0 | T | C | 0.149 | 3.03E-02 | 1.80E-01 | 2.65E+00 | 2.71E+00 | 8.70E-01 | no | Gm41386 | downstream_gene_variant | MODIFIER |
| 15 | 93708623 | AX-170261366 | 0 | C | T | 0.149 | 3.03E-02 | 1.80E-01 | 2.65E+00 | 2.71E+00 | 8.70E-01 | no | Gm41386 | intron_variant,non_coding_transcript_variant | MODIFIER |
| 15 | 93709923 | AX-170168240 | 0 | A | G | 0.149 | 3.03E-02 | 1.80E-01 | 2.65E+00 | 2.71E+00 | 8.70E-01 | no | Gm41386 | intron_variant,non_coding_transcript_variant | MODIFIER |
| 15 | 93710280 | AX-170279705 | 0 | T | C | 0.149 | 3.03E-02 | 1.80E-01 | 2.65E+00 | 2.71E+00 | 8.70E-01 | no | Gm41386 | intron_variant,non_coding_transcript_variant | MODIFIER |
| 15 | 93711405 | AX-170254832 | 0 | T | C | 0.149 | 3.03E-02 | 1.80E-01 | 2.65E+00 | 2.71E+00 | 8.70E-01 | no | Gm41386 | intron_variant,non_coding_transcript_variant | MODIFIER |
| 15 | 93712266 | AX-170477323 | 0 | T | C | 0.149 | 3.03E-02 | 1.80E-01 | 2.65E+00 | 2.71E+00 | 8.70E-01 | no | Gm41386 | intron_variant,non_coding_transcript_variant | MODIFIER |
| 15 | 93712472 | AX-170179534 | 0 | A | C | 0.149 | 3.03E-02 | 1.80E-01 | 2.65E+00 | 2.71E+00 | 8.70E-01 | no | Gm41386 | intron_variant,non_coding_transcript_variant | MODIFIER |
| 15 | 93718379 | AX-97812011 | 0 | C | T | 0.149 | 3.03E-02 | 1.80E-01 | 2.65E+00 | 2.71E+00 | 8.70E-01 | no | | intergenic_variant | MODIFIER |
| 15 | 93718944 | AX-170131255 | 0 | C | T | 0.149 | 3.03E-02 | 1.80E-01 | 2.65E+00 | 2.71E+00 | 8.70E-01 | no | | intergenic_variant | MODIFIER |
| 15 | 93730536 | AX-170353049 | 0 | T | C | 0.149 | 3.03E-02 | 1.80E-01 | 2.65E+00 | 2.71E+00 | 8.70E-01 | no | Gm30339 | upstream_gene_variant | MODIFIER |
| 15 | 93748160 | AX-97844085 | 0 | T | C | 0.149 | 3.03E-02 | 1.80E-01 | 2.65E+00 | 2.71E+00 | 8.70E-01 | no | | intergenic_variant | MODIFIER |
| 15 | 93752306 | AX-170152022 | 0 | G | C | 0.149 | 3.03E-02 | 1.80E-01 | 2.65E+00 | 2.71E+00 | 8.70E-01 | no | | intergenic_variant | MODIFIER |
| 15 | 93758443 | AX-170335166 | 0 | G | C | 0.149 | 3.03E-02 | 1.80E-01 | 2.65E+00 | 2.71E+00 | 8.70E-01 | no | | intergenic_variant | MODIFIER |
| 15 | 93763272 | AX-170527860 | 0 | A | G | 0.149 | 3.03E-02 | 1.80E-01 | 2.65E+00 | 2.71E+00 | 8.70E-01 | no | | intergenic_variant | MODIFIER |
| 15 | 93772883 | AX-97801487 | 0 | C | T | 0.149 | 3.03E-02 | 1.80E-01 | 2.65E+00 | 2.71E+00 | 8.70E-01 | no | | intergenic_variant | MODIFIER |
| 15 | 93778572 | AX-170297592 | 0 | G | A | 0.149 | 3.03E-02 | 1.80E-01 | 2.65E+00 | 2.71E+00 | 8.70E-01 | no | | intergenic_variant | MODIFIER |
| 15 | 93778677 | AX-170133533 | 0 | C | T | 0.149 | 3.03E-02 | 1.80E-01 | 2.65E+00 | 2.71E+00 | 8.70E-01 | no | | intergenic_variant | MODIFIER |
| 15 | 93779260 | AX-170353853 | 0 | T | C | 0.149 | 3.03E-02 | 1.80E-01 | 2.65E+00 | 2.71E+00 | 8.70E-01 | no | | intergenic_variant | MODIFIER |
| 15 | 93779714 | AX-170288565 | 0 | A | G | 0.149 | 3.03E-02 | 1.80E-01 | 2.65E+00 | 2.71E+00 | 8.70E-01 | no | | intergenic_variant | MODIFIER |
| 15 | 93786853 | AX-170384889 | 0 | C | G | 0.149 | 3.03E-02 | 1.80E-01 | 2.65E+00 | 2.71E+00 | 8.70E-01 | no | | regulatory_region_variant | MODIFIER |
| 15 | 93787144 | AX-97806406 | 0 | G | A | 0.149 | 3.03E-02 | 1.80E-01 | 2.65E+00 | 2.71E+00 | 8.70E-01 | no | | intergenic_variant | MODIFIER |
| 15 | 93806056 | AX-170442384 | 0 | G | A | 0.149 | 3.03E-02 | 1.80E-01 | 2.65E+00 | 2.71E+00 | 8.70E-01 | no | | intergenic_variant | MODIFIER |
| 15 | 93835367 | AX-170528027 | 0 | C | T | 0.149 | 3.03E-02 | 1.80E-01 | 2.65E+00 | 2.71E+00 | 8.70E-01 | no | Gm49445 | intron_variant,non_coding_transcript_variant | MODIFIER |
| 15 | 93836705 | AX-170104892 | 0 | C | T | 0.149 | 3.03E-02 | 1.80E-01 | 2.65E+00 | 2.71E+00 | 8.70E-01 | no | Gm49445 | intron_variant,non_coding_transcript_variant | MODIFIER |
| 15 | 93856211 | AX-170436468 | 0 | C | T | 0.149 | 3.03E-02 | 1.80E-01 | 2.65E+00 | 2.71E+00 | 8.70E-01 | no | | intergenic_variant | MODIFIER |

| | | | | | | | | | | | | | | | |
|---|---|---|---|---|---|---|---|---|---|---|---|---|---|---|---|
| 15 | 93857116 | AX-170277612 | 0 | G | A | 0.149 | 3.03E-02 | 1.80E-01 | 2.65E+00 | 2.71E+00 | 8.70E-01 | no | | regulatory_region_variant | MODIFIER |
| 15 | 93876311 | AX-170141387 | 0 | T | C | 0.149 | 3.03E-02 | 1.80E-01 | 2.65E+00 | 2.71E+00 | 8.70E-01 | no | | intergenic_variant | MODIFIER |
| 15 | 93877687 | AX-170151402 | 0 | T | C | 0.149 | 3.03E-02 | 1.80E-01 | 2.65E+00 | 2.71E+00 | 8.70E-01 | no | | intergenic_variant | MODIFIER |
| 15 | 93888015 | AX-170297213 | 0 | C | A | 0.149 | 3.03E-02 | 1.80E-01 | 2.65E+00 | 2.71E+00 | 8.70E-01 | no | | intergenic_variant | MODIFIER |
| 15 | 93903613 | AX-170293754 | 0 | T | A | 0.149 | 3.03E-02 | 1.80E-01 | 2.65E+00 | 2.71E+00 | 8.70E-01 | no | 9430014N10Rik | downstream_gene_variant | MODIFIER |
| 15 | 93971122 | AX-170056341 | 0 | C | T | 0.149 | 3.03E-02 | 1.80E-01 | 2.65E+00 | 2.71E+00 | 8.70E-01 | no | | intergenic_variant | MODIFIER |
| 15 | 93971501 | AX-170056342 | 0 | C | A | 0.149 | 3.03E-02 | 1.80E-01 | 2.65E+00 | 2.71E+00 | 8.70E-01 | no | | intergenic_variant | MODIFIER |
| 15 | 93980601 | AX-170217748 | 0 | T | G | 0.149 | 3.03E-02 | 1.80E-01 | 2.65E+00 | 2.71E+00 | 8.70E-01 | no | | intergenic_variant | MODIFIER |
| 15 | 93987254 | AX-170341844 | 0 | G | A | 0.149 | 3.03E-02 | 1.80E-01 | 2.65E+00 | 2.71E+00 | 8.70E-01 | no | | intergenic_variant | MODIFIER |
| 15 | 93987522 | AX-170248774 | 0 | A | G | 0.149 | 3.03E-02 | 1.80E-01 | 2.65E+00 | 2.71E+00 | 8.70E-01 | no | | intergenic_variant | MODIFIER |
| 15 | 93992820 | AX-170466631 | 0 | A | G | 0.149 | 3.03E-02 | 1.80E-01 | 2.65E+00 | 2.71E+00 | 8.70E-01 | no | | intergenic_variant | MODIFIER |
| 15 | 93993102 | AX-170427437 | 0 | T | C | 0.149 | 3.03E-02 | 1.80E-01 | 2.65E+00 | 2.71E+00 | 8.70E-01 | no | | intergenic_variant | MODIFIER |
| 15 | 93993762 | AX-170300900 | 0 | T | C | 0.149 | 3.03E-02 | 1.80E-01 | 2.65E+00 | 2.71E+00 | 8.70E-01 | no | | intergenic_variant | MODIFIER |
| 15 | 93993803 | AX-170313901 | 0 | A | G | 0.149 | 3.03E-02 | 1.80E-01 | 2.65E+00 | 2.71E+00 | 8.70E-01 | no | | intergenic_variant | MODIFIER |
| 15 | 94007911 | AX-170473036 | 0 | G | A | 0.149 | 3.03E-02 | 1.80E-01 | 2.65E+00 | 2.71E+00 | 8.70E-01 | no | | regulatory_region_variant | MODIFIER |
| 15 | 94021483 | AX-170454215 | 0 | C | T | 0.149 | 3.03E-02 | 1.80E-01 | 2.65E+00 | 2.71E+00 | 8.70E-01 | no | | intergenic_variant | MODIFIER |
| 15 | 94022584 | AX-170464387 | 0 | T | C | 0.149 | 3.03E-02 | 1.80E-01 | 2.65E+00 | 2.71E+00 | 8.70E-01 | no | | intergenic_variant | MODIFIER |
| 15 | 94022816 | AX-170402213 | 0 | C | T | 0.149 | 3.03E-02 | 1.80E-01 | 2.65E+00 | 2.71E+00 | 8.70E-01 | no | | intergenic_variant | MODIFIER |
| 15 | 94097850 | AX-170412120 | 0 | A | G | 0.149 | 3.03E-02 | 1.80E-01 | 2.65E+00 | 2.71E+00 | 8.70E-01 | no | | intergenic_variant | MODIFIER |
| 15 | 94114898 | AX-97782461 | 0 | G | A | 0.149 | 3.03E-02 | 1.80E-01 | 2.65E+00 | 2.71E+00 | 8.70E-01 | no | | intergenic_variant | MODIFIER |
| 15 | 94141142 | AX-170316383 | 0 | T | C | 0.149 | 3.03E-02 | 1.80E-01 | 2.65E+00 | 2.71E+00 | 8.70E-01 | no | | intergenic_variant | MODIFIER |
| 15 | 94141216 | AX-170377526 | 0 | T | C | 0.149 | 3.03E-02 | 1.80E-01 | 2.65E+00 | 2.71E+00 | 8.70E-01 | no | | intergenic_variant | MODIFIER |
| 15 | 94143190 | AX-170012297 | 0 | G | A | 0.149 | 3.03E-02 | 1.80E-01 | 2.65E+00 | 2.71E+00 | 8.70E-01 | no | | intergenic_variant | MODIFIER |
| 15 | 94147835 | AX-170418660 | 0 | A | G | 0.149 | 3.03E-02 | 1.80E-01 | 2.65E+00 | 2.71E+00 | 8.70E-01 | no | | intergenic_variant | MODIFIER |
| 15 | 94150314 | AX-170407040 | 0 | A | G | 0.149 | 3.03E-02 | 1.80E-01 | 2.65E+00 | 2.71E+00 | 8.70E-01 | no | | intergenic_variant | MODIFIER |
| 15 | 94174405 | AX-170325758 | 0 | A | C | 0.149 | 3.03E-02 | 1.80E-01 | 2.65E+00 | 2.71E+00 | 8.70E-01 | no | | regulatory_region_variant | MODIFIER |
| 15 | 94181401 | AX-97816952 | 0 | G | A | 0.149 | 3.03E-02 | 1.80E-01 | 2.65E+00 | 2.71E+00 | 8.70E-01 | no | | intergenic_variant | MODIFIER |
| 15 | 94187688 | AX-170193030 | 0 | A | T | 0.149 | 3.03E-02 | 1.80E-01 | 2.65E+00 | 2.71E+00 | 8.70E-01 | no | | intergenic_variant | MODIFIER |
| 15 | 94189780 | AX-170087941 | 0 | T | C | 0.149 | 3.03E-02 | 1.80E-01 | 2.65E+00 | 2.71E+00 | 8.70E-01 | no | | regulatory_region_variant | MODIFIER |
| 15 | 94202624 | AX-170357446 | 0 | A | G | 0.149 | 3.03E-02 | 1.80E-01 | 2.65E+00 | 2.71E+00 | 8.70E-01 | no | | intergenic_variant | MODIFIER |
| 15 | 94209139 | AX-170056348 | 0 | G | A | 0.149 | 3.03E-02 | 1.80E-01 | 2.65E+00 | 2.71E+00 | 8.70E-01 | no | | regulatory_region_variant | MODIFIER |
| 15 | 94214298 | AX-170302949 | 0 | A | C | 0.149 | 3.03E-02 | 1.80E-01 | 2.65E+00 | 2.71E+00 | 8.70E-01 | no | | regulatory_region_variant | MODIFIER |
| 15 | 94214849 | AX-170056350 | 0 | G | A | 0.149 | 3.03E-02 | 1.80E-01 | 2.65E+00 | 2.71E+00 | 8.70E-01 | no | | regulatory_region_variant | MODIFIER |
| 15 | 94218755 | AX-97782203 | 0 | A | G | 0.149 | 3.03E-02 | 1.80E-01 | 2.65E+00 | 2.71E+00 | 8.70E-01 | no | | intergenic_variant | MODIFIER |
| 15 | 94236877 | AX-170130262 | 0 | T | G | 0.149 | 3.03E-02 | 1.80E-01 | 2.65E+00 | 2.71E+00 | 8.70E-01 | no | | regulatory_region_variant | MODIFIER |
| 15 | 94246217 | AX-97845873 | 0 | T | C | 0.149 | 3.03E-02 | 1.80E-01 | 2.65E+00 | 2.71E+00 | 8.70E-01 | no | | regulatory_region_variant | MODIFIER |
| 15 | 94248854 | AX-170145565 | 0 | C | T | 0.149 | 3.03E-02 | 1.80E-01 | 2.65E+00 | 2.71E+00 | 8.70E-01 | no | | regulatory_region_variant | MODIFIER |
| 15 | 94264171 | AX-97780925 | 0 | T | C | 0.149 | 3.03E-02 | 1.80E-01 | 2.65E+00 | 2.71E+00 | 8.70E-01 | no | | regulatory_region_variant | MODIFIER |
| 15 | 94265791 | AX-170277092 | 0 | G | T | 0.149 | 3.03E-02 | 1.80E-01 | 2.65E+00 | 2.71E+00 | 8.70E-01 | no | Adamts20 | downstream_gene_variant | MODIFIER |
| 15 | 94270770 | AX-170435426 | 0 | A | G | 0.149 | 3.03E-02 | 1.80E-01 | 2.65E+00 | 2.71E+00 | 8.70E-01 | no | Adamts20 | intron_variant | MODIFIER |
| 15 | 94276382 | AX-170146133 | 0 | A | G | 0.149 | 3.03E-02 | 1.80E-01 | 2.65E+00 | 2.71E+00 | 8.70E-01 | no | Adamts20 | intron_variant | MODIFIER |
| 15 | 94281114 | AX-170526602 | 0 | A | T | 0.149 | 3.03E-02 | 1.80E-01 | 2.65E+00 | 2.71E+00 | 8.70E-01 | no | Adamts20 | intron_variant | MODIFIER |
| 15 | 94294586 | AX-170534659 | 0 | C | G | 0.149 | 3.03E-02 | 1.80E-01 | 2.65E+00 | 2.71E+00 | 8.70E-01 | no | Adamts20 | intron_variant | MODIFIER |
| 15 | 94303075 | AX-97803752 | 0 | A | G | 0.149 | 3.03E-02 | 1.80E-01 | 2.65E+00 | 2.71E+00 | 8.70E-01 | no | Adamts20 | intron_variant | MODIFIER |
| 15 | 94303337 | AX-170488856 | 0 | G | T | 0.149 | 3.03E-02 | 1.80E-01 | 2.65E+00 | 2.71E+00 | 8.70E-01 | no | Adamts20 | intron_variant | MODIFIER |
| 15 | 94312336 | AX-170151954 | 0 | A | G | 0.149 | 3.03E-02 | 1.80E-01 | 2.65E+00 | 2.71E+00 | 8.70E-01 | no | Adamts20 | intron_variant | MODIFIER |
| 15 | 94312415 | AX-170130515 | 0 | T | A | 0.149 | 3.03E-02 | 1.80E-01 | 2.65E+00 | 2.71E+00 | 8.70E-01 | no | Adamts20 | intron_variant | MODIFIER |
| 15 | 94315646 | AX-97816209 | 0 | G | A | 0.149 | 3.03E-02 | 1.80E-01 | 2.65E+00 | 2.71E+00 | 8.70E-01 | no | Adamts20 | intron_variant | MODIFIER |
| 15 | 94439467 | AX-97805790 | 0 | A | T | 0.149 | 3.03E-02 | 1.80E-01 | 2.65E+00 | 2.71E+00 | 8.70E-01 | no | Adamts20 | intron_variant,non_coding_transcript_variant | MODIFIER |
| 15 | 94462699 | AX-170314120 | 0 | C | T | 0.149 | 3.03E-02 | 1.80E-01 | 2.65E+00 | 2.71E+00 | 8.70E-01 | no | Adamts20 | intron_variant,non_coding_transcript_variant | MODIFIER |
| 15 | 94560266 | AX-170167460 | 0 | A | T | 0.149 | 3.03E-02 | 1.80E-01 | 2.65E+00 | 2.71E+00 | 8.70E-01 | no | Irak4 | intron_variant | MODIFIER |
| 15 | 94570797 | AX-97831249 | 0 | G | A | 0.149 | 3.03E-02 | 1.80E-01 | 2.65E+00 | 2.71E+00 | 8.70E-01 | no | Irak4 | downstream_gene_variant | MODIFIER |
| 15 | 94588449 | AX-170309693 | 0 | A | G | 0.149 | 3.03E-02 | 1.80E-01 | 2.65E+00 | 2.71E+00 | 8.70E-01 | no | Twf1 | intron_variant | MODIFIER |
| 15 | 94684298 | AX-170453463 | 0 | C | T | 0.149 | 3.03E-02 | 1.80E-01 | 2.65E+00 | 2.71E+00 | 8.70E-01 | no | Tmem117 | intron_variant | MODIFIER |

| | | | | | | | | | | | | | | | |
|---|---|---|---|---|---|---|---|---|---|---|---|---|---|---|---|
| 15 | 94689305 | AX-170287576 | 0 | G | C | 0.149 | 3.03E-02 | 1.80E-01 | 2.65E+00 | 2.71E+00 | 8.70E-01 | no | Tmem117 | intron_variant | MODIFIER |
| 15 | 94701211 | AX-170412114 | 0 | T | G | 0.149 | 3.03E-02 | 1.80E-01 | 2.65E+00 | 2.71E+00 | 8.70E-01 | no | Tmem117 | intron_variant | MODIFIER |
| 15 | 94701735 | AX-170292013 | 0 | G | C | 0.149 | 3.03E-02 | 1.80E-01 | 2.65E+00 | 2.71E+00 | 8.70E-01 | no | Tmem117 | intron_variant | MODIFIER |
| 15 | 94736603 | AX-170143052 | 0 | A | G | 0.149 | 3.03E-02 | 1.80E-01 | 2.65E+00 | 2.71E+00 | 8.70E-01 | no | Tmem117 | intron_variant | MODIFIER |
| 15 | 94738706 | AX-170434102 | 0 | C | T | 0.149 | 3.03E-02 | 1.80E-01 | 2.65E+00 | 2.71E+00 | 8.70E-01 | no | Tmem117 | intron_variant | MODIFIER |
| 15 | 94743792 | AX-170248202 | 0 | T | C | 0.149 | 3.03E-02 | 1.80E-01 | 2.65E+00 | 2.71E+00 | 8.70E-01 | no | Tmem117 | intron_variant | MODIFIER |
| 15 | 94760102 | AX-170329904 | 0 | T | C | 0.149 | 3.03E-02 | 1.80E-01 | 2.65E+00 | 2.71E+00 | 8.70E-01 | no | Tmem117 | intron_variant | MODIFIER |
| 15 | 94760456 | AX-170012304 | 0 | C | T | 0.149 | 3.03E-02 | 1.80E-01 | 2.65E+00 | 2.71E+00 | 8.70E-01 | no | Tmem117 | intron_variant | MODIFIER |
| 15 | 94760949 | AX-170387654 | 0 | G | A | 0.149 | 3.03E-02 | 1.80E-01 | 2.65E+00 | 2.71E+00 | 8.70E-01 | no | Tmem117 | intron_variant | MODIFIER |
| 15 | 94762419 | AX-170308111 | 0 | C | T | 0.149 | 3.03E-02 | 1.80E-01 | 2.65E+00 | 2.71E+00 | 8.70E-01 | no | Tmem117 | intron_variant | MODIFIER |
| 15 | 94763868 | AX-170227427 | 0 | C | A | 0.149 | 3.03E-02 | 1.80E-01 | 2.65E+00 | 2.71E+00 | 8.70E-01 | no | Tmem117 | intron_variant | MODIFIER |
| 15 | 94765733 | AX-170317779 | 0 | G | T | 0.149 | 3.03E-02 | 1.80E-01 | 2.65E+00 | 2.71E+00 | 8.70E-01 | no | Tmem117 | intron_variant | MODIFIER |
| 15 | 94768854 | AX-97830760 | 0 | T | C | 0.149 | 3.03E-02 | 1.80E-01 | 2.65E+00 | 2.71E+00 | 8.70E-01 | no | Tmem117 | intron_variant | MODIFIER |
| 15 | 94777000 | AX-170197258 | 0 | C | T | 0.149 | 3.03E-02 | 1.80E-01 | 2.65E+00 | 2.71E+00 | 8.70E-01 | no | Tmem117 | intron_variant | MODIFIER |
| 15 | 94780280 | AX-170457294 | 0 | A | G | 0.149 | 3.03E-02 | 1.80E-01 | 2.65E+00 | 2.71E+00 | 8.70E-01 | no | Tmem117 | intron_variant | MODIFIER |
| 15 | 94791845 | AX-170213103 | 0 | G | C | 0.149 | 3.03E-02 | 1.80E-01 | 2.65E+00 | 2.71E+00 | 8.70E-01 | no | Tmem117 | intron_variant | MODIFIER |
| 15 | 94792306 | AX-170220528 | 0 | A | T | 0.149 | 3.03E-02 | 1.80E-01 | 2.65E+00 | 2.71E+00 | 8.70E-01 | no | Tmem117 | intron_variant | MODIFIER |
| 15 | 94798114 | AX-170303288 | 0 | G | A | 0.149 | 3.03E-02 | 1.80E-01 | 2.65E+00 | 2.71E+00 | 8.70E-01 | no | Tmem117 | intron_variant | MODIFIER |
| 15 | 94800334 | AX-170056362 | 0 | C | T | 0.149 | 3.03E-02 | 1.80E-01 | 2.65E+00 | 2.71E+00 | 8.70E-01 | no | Tmem117 | intron_variant | MODIFIER |
| 15 | 94802450 | AX-170012308 | 0 | T | C | 0.149 | 3.03E-02 | 1.80E-01 | 2.65E+00 | 2.71E+00 | 8.70E-01 | no | Tmem117 | intron_variant | MODIFIER |
| 15 | 94803523 | AX-170402752 | 0 | A | T | 0.149 | 3.03E-02 | 1.80E-01 | 2.65E+00 | 2.71E+00 | 8.70E-01 | no | Tmem117 | intron_variant | MODIFIER |
| 15 | 94804101 | AX-170167196 | 0 | T | C | 0.149 | 3.03E-02 | 1.80E-01 | 2.65E+00 | 2.71E+00 | 8.70E-01 | no | Tmem117 | intron_variant | MODIFIER |
| 15 | 94811534 | AX-97789255 | 0 | T | A | 0.149 | 3.03E-02 | 1.80E-01 | 2.65E+00 | 2.71E+00 | 8.70E-01 | no | Tmem117 | intron_variant | MODIFIER |
| 15 | 94833570 | AX-97842118 | 0 | C | A | 0.149 | 3.03E-02 | 1.80E-01 | 2.65E+00 | 2.71E+00 | 8.70E-01 | no | Tmem117 | intron_variant | MODIFIER |
| 15 | 94872212 | AX-97840872 | 0 | A | G | 0.149 | 3.03E-02 | 1.80E-01 | 2.65E+00 | 2.71E+00 | 8.70E-01 | no | Tmem117 | intron_variant | MODIFIER |
| 15 | 94876286 | AX-170456905 | 0 | T | C | 0.149 | 3.03E-02 | 1.80E-01 | 2.65E+00 | 2.71E+00 | 8.70E-01 | no | Tmem117 | intron_variant | MODIFIER |
| 15 | 94881039 | AX-170110477 | 0 | T | A | 0.149 | 3.03E-02 | 1.80E-01 | 2.65E+00 | 2.71E+00 | 8.70E-01 | no | Tmem117 | intron_variant | MODIFIER |
| 15 | 94882339 | AX-170289262 | 0 | G | A | 0.149 | 3.03E-02 | 1.80E-01 | 2.65E+00 | 2.71E+00 | 8.70E-01 | no | Tmem117 | intron_variant | MODIFIER |
| 15 | 94882870 | AX-170386215 | 0 | G | A | 0.149 | 3.03E-02 | 1.80E-01 | 2.65E+00 | 2.71E+00 | 8.70E-01 | no | Tmem117 | intron_variant | MODIFIER |
| 15 | 94883533 | AX-170455204 | 0 | C | T | 0.149 | 3.03E-02 | 1.80E-01 | 2.65E+00 | 2.71E+00 | 8.70E-01 | no | Tmem117 | intron_variant | MODIFIER |
| 15 | 94888095 | AX-170488096 | 0 | C | T | 0.149 | 3.03E-02 | 1.80E-01 | 2.65E+00 | 2.71E+00 | 8.70E-01 | no | Tmem117 | intron_variant | MODIFIER |
| 15 | 95137244 | AX-170056370 | 0 | T | A | 0.149 | 3.03E-02 | 1.80E-01 | 2.65E+00 | 2.71E+00 | 8.70E-01 | no | 1700129L04Rik | intron_variant,non_coding_transcript_variant | MODIFIER |
| 15 | 95140674 | AX-168382914 | 0 | G | A | 0.149 | 3.03E-02 | 1.80E-01 | 2.65E+00 | 2.71E+00 | 8.70E-01 | no | 1700129L04Rik | intron_variant,non_coding_transcript_variant | MODIFIER |
| 15 | 95141119 | AX-170389031 | 0 | A | G | 0.149 | 3.03E-02 | 1.80E-01 | 2.65E+00 | 2.71E+00 | 8.70E-01 | no | 1700129L04Rik | intron_variant,non_coding_transcript_variant | MODIFIER |
| 15 | 95142048 | AX-170227527 | 0 | G | C | 0.149 | 3.03E-02 | 1.80E-01 | 2.65E+00 | 2.71E+00 | 8.70E-01 | no | 1700129L04Rik | intron_variant,non_coding_transcript_variant | MODIFIER |
| 15 | 95145030 | AX-97827572 | 0 | T | G | 0.149 | 3.03E-02 | 1.80E-01 | 2.65E+00 | 2.71E+00 | 8.70E-01 | no | 1700129L04Rik | intron_variant,non_coding_transcript_variant | MODIFIER |
| 15 | 95145996 | AX-170231549 | 0 | G | A | 0.149 | 3.03E-02 | 1.80E-01 | 2.65E+00 | 2.71E+00 | 8.70E-01 | no | 1700129L04Rik | intron_variant,non_coding_transcript_variant | MODIFIER |
| 15 | 95173667 | AX-170087944 | 0 | G | A | 0.149 | 3.03E-02 | 1.80E-01 | 2.65E+00 | 2.71E+00 | 8.70E-01 | no | Nell2 | intron_variant | MODIFIER |
| 15 | 95174712 | AX-170012311 | 0 | C | T | 0.149 | 3.03E-02 | 1.80E-01 | 2.65E+00 | 2.71E+00 | 8.70E-01 | no | Nell2 | intron_variant | MODIFIER |
| 15 | 95203696 | AX-170156048 | 0 | A | T | 0.149 | 3.03E-02 | 1.80E-01 | 2.65E+00 | 2.71E+00 | 8.70E-01 | no | Nell2 | intron_variant | MODIFIER |
| 15 | 95204394 | AX-170166441 | 0 | A | G | 0.149 | 3.03E-02 | 1.80E-01 | 2.65E+00 | 2.71E+00 | 8.70E-01 | no | Nell2 | intron_variant | MODIFIER |
| 15 | 95221939 | AX-97830283 | 0 | G | A | 0.149 | 3.03E-02 | 1.80E-01 | 2.65E+00 | 2.71E+00 | 8.70E-01 | no | Nell2 | intron_variant | MODIFIER |
| 15 | 95235905 | AX-170470876 | 0 | T | C | 0.149 | 3.03E-02 | 1.80E-01 | 2.65E+00 | 2.71E+00 | 8.70E-01 | no | Nell2 | intron_variant | MODIFIER |
| 15 | 95237562 | AX-170012309 | 0 | C | G | 0.149 | 3.03E-02 | 1.80E-01 | 2.65E+00 | 2.71E+00 | 8.70E-01 | no | Nell2 | intron_variant | MODIFIER |
| 15 | 95237797 | AX-170433464 | 0 | A | G | 0.149 | 3.03E-02 | 1.80E-01 | 2.65E+00 | 2.71E+00 | 8.70E-01 | no | Nell2 | intron_variant | MODIFIER |
| 15 | 95237815 | AX-170012317 | 0 | A | T | 0.149 | 3.03E-02 | 1.80E-01 | 2.65E+00 | 2.71E+00 | 8.70E-01 | no | Nell2 | intron_variant | MODIFIER |
| 15 | 95239353 | AX-170515351 | 0 | C | A | 0.149 | 3.03E-02 | 1.80E-01 | 2.65E+00 | 2.71E+00 | 8.70E-01 | no | Nell2 | intron_variant | MODIFIER |
| 15 | 95242460 | AX-170118257 | 0 | A | G | 0.149 | 3.03E-02 | 1.80E-01 | 2.65E+00 | 2.71E+00 | 8.70E-01 | no | Nell2 | intron_variant | MODIFIER |
| 15 | 95242674 | AX-170123219 | 0 | C | G | 0.149 | 3.03E-02 | 1.80E-01 | 2.65E+00 | 2.71E+00 | 8.70E-01 | no | Nell2 | intron_variant | MODIFIER |
| 15 | 95242773 | AX-170056371 | 0 | G | C | 0.149 | 3.03E-02 | 1.80E-01 | 2.65E+00 | 2.71E+00 | 8.70E-01 | no | Nell2 | intron_variant | MODIFIER |
| 15 | 95243020 | AX-170056373 | 0 | G | A | 0.149 | 3.03E-02 | 1.80E-01 | 2.65E+00 | 2.71E+00 | 8.70E-01 | no | Nell2 | intron_variant | MODIFIER |
| 15 | 95244447 | AX-170528840 | 0 | A | T | 0.149 | 3.03E-02 | 1.80E-01 | 2.65E+00 | 2.71E+00 | 8.70E-01 | no | Nell2 | intron_variant | MODIFIER |
| 15 | 95245094 | AX-170432339 | 0 | C | T | 0.149 | 3.03E-02 | 1.80E-01 | 2.65E+00 | 2.71E+00 | 8.70E-01 | no | Nell2 | intron_variant | MODIFIER |
| 15 | 95246307 | AX-170093345 | 0 | G | T | 0.149 | 3.03E-02 | 1.80E-01 | 2.65E+00 | 2.71E+00 | 8.70E-01 | no | Nell2 | intron_variant | MODIFIER |

| | | | | | | | | | | | | | | | |
|---|---|---|---|---|---|---|---|---|---|---|---|---|---|---|---|
| 15 | 95255480 | AX-170012312 | 0 | A | G | 0.149 | 3.03E-02 | 1.80E-01 | 2.65E+00 | 2.71E+00 | 8.70E-01 | no | Nell2 | intron_variant | MODIFIER |
| 15 | 95264162 | AX-97811908 | 0 | G | A | 0.149 | 3.03E-02 | 1.80E-01 | 2.65E+00 | 2.71E+00 | 8.70E-01 | no | Nell2 | intron_variant | MODIFIER |
| 15 | 95281743 | AX-170012310 | 0 | G | A | 0.149 | 3.03E-02 | 1.80E-01 | 2.65E+00 | 2.71E+00 | 8.70E-01 | no | Nell2 | intron_variant | MODIFIER |
| 15 | 95281768 | AX-170346277 | 0 | A | G | 0.149 | 3.03E-02 | 1.80E-01 | 2.65E+00 | 2.71E+00 | 8.70E-01 | no | Nell2 | intron_variant | MODIFIER |
| 15 | 95283174 | AX-170322159 | 0 | G | A | 0.149 | 3.03E-02 | 1.80E-01 | 2.65E+00 | 2.71E+00 | 8.70E-01 | no | Nell2 | intron_variant | MODIFIER |
| 15 | 95286828 | AX-170393074 | 0 | A | G | 0.149 | 3.03E-02 | 1.80E-01 | 2.65E+00 | 2.71E+00 | 8.70E-01 | no | Nell2 | intron_variant | MODIFIER |
| 15 | 95298521 | AX-170448373 | 0 | T | C | 0.149 | 3.03E-02 | 1.80E-01 | 2.65E+00 | 2.71E+00 | 8.70E-01 | no | Nell2 | intron_variant | MODIFIER |
| 15 | 95300613 | AX-170012316 | 0 | T | C | 0.149 | 3.03E-02 | 1.80E-01 | 2.65E+00 | 2.71E+00 | 8.70E-01 | no | Nell2 | intron_variant | MODIFIER |
| 15 | 95300649 | AX-170426345 | 0 | T | A | 0.149 | 3.03E-02 | 1.80E-01 | 2.65E+00 | 2.71E+00 | 8.70E-01 | no | Nell2 | intron_variant | MODIFIER |
| 15 | 95314132 | AX-170012315 | 0 | A | G | 0.149 | 3.03E-02 | 1.80E-01 | 2.65E+00 | 2.71E+00 | 8.70E-01 | no | Nell2 | intron_variant | MODIFIER |
| 15 | 95314546 | AX-170363403 | 0 | G | A | 0.149 | 3.03E-02 | 1.80E-01 | 2.65E+00 | 2.71E+00 | 8.70E-01 | no | Nell2 | intron_variant | MODIFIER |
| 15 | 95325426 | AX-170233690 | 0 | C | T | 0.149 | 3.03E-02 | 1.80E-01 | 2.65E+00 | 2.71E+00 | 8.70E-01 | no | Nell2 | intron_variant | MODIFIER |
| 15 | 95325906 | AX-97793965 | 0 | A | C | 0.149 | 3.03E-02 | 1.80E-01 | 2.65E+00 | 2.71E+00 | 8.70E-01 | no | Nell2 | intron_variant | MODIFIER |
| 15 | 95329331 | AX-170288407 | 0 | A | G | 0.149 | 3.03E-02 | 1.80E-01 | 2.65E+00 | 2.71E+00 | 8.70E-01 | no | Nell2 | intron_variant | MODIFIER |
| 15 | 95329613 | AX-170199492 | 0 | T | G | 0.149 | 3.03E-02 | 1.80E-01 | 2.65E+00 | 2.71E+00 | 8.70E-01 | no | Nell2 | intron_variant | MODIFIER |
| 15 | 95339358 | AX-170496950 | 0 | T | A | 0.149 | 3.03E-02 | 1.80E-01 | 2.65E+00 | 2.71E+00 | 8.70E-01 | no | Nell2 | intron_variant | MODIFIER |
| 15 | 95346220 | AX-97849098 | 0 | C | T | 0.149 | 3.03E-02 | 1.80E-01 | 2.65E+00 | 2.71E+00 | 8.70E-01 | no | Nell2 | intron_variant | MODIFIER |
| 15 | 95354568 | AX-170201853 | 0 | G | T | 0.149 | 3.03E-02 | 1.80E-01 | 2.65E+00 | 2.71E+00 | 8.70E-01 | no | Nell2 | intron_variant | MODIFIER |
| 15 | 95371365 | AX-97810499 | 0 | T | C | 0.149 | 3.03E-02 | 1.80E-01 | 2.65E+00 | 2.71E+00 | 8.70E-01 | no | Nell2 | intron_variant | MODIFIER |
| 15 | 95382633 | AX-170507436 | 0 | G | T | 0.149 | 3.03E-02 | 1.80E-01 | 2.65E+00 | 2.71E+00 | 8.70E-01 | no | Nell2 | intron_variant | MODIFIER |
| 15 | 95382699 | AX-170152529 | 0 | C | T | 0.149 | 3.03E-02 | 1.80E-01 | 2.65E+00 | 2.71E+00 | 8.70E-01 | no | Nell2 | intron_variant | MODIFIER |
| 15 | 95388547 | AX-170399507 | 0 | A | T | 0.149 | 3.03E-02 | 1.80E-01 | 2.65E+00 | 2.71E+00 | 8.70E-01 | no | Nell2 | intron_variant | MODIFIER |
| 15 | 95390289 | AX-170111997 | 0 | A | G | 0.149 | 3.03E-02 | 1.80E-01 | 2.65E+00 | 2.71E+00 | 8.70E-01 | no | Nell2 | intron_variant | MODIFIER |
| 15 | 95400012 | AX-97789477 | 0 | A | C | 0.149 | 3.03E-02 | 1.80E-01 | 2.65E+00 | 2.71E+00 | 8.70E-01 | no | Nell2 | intron_variant | MODIFIER |
| 15 | 95431998 | AX-170526113 | 0 | G | C | 0.149 | 3.03E-02 | 1.80E-01 | 2.65E+00 | 2.71E+00 | 8.70E-01 | no | Nell2 | intron_variant | MODIFIER |
| 15 | 95432100 | AX-97796043 | 0 | A | G | 0.149 | 3.03E-02 | 1.80E-01 | 2.65E+00 | 2.71E+00 | 8.70E-01 | no | Nell2 | intron_variant | MODIFIER |
| 15 | 95433575 | AX-170363367 | 0 | C | T | 0.149 | 3.03E-02 | 1.80E-01 | 2.65E+00 | 2.71E+00 | 8.70E-01 | no | Nell2 | intron_variant | MODIFIER |
| 15 | 95468722 | AX-170221482 | 0 | C | T | 0.149 | 3.03E-02 | 1.80E-01 | 2.65E+00 | 2.71E+00 | 8.70E-01 | no | Nell2 | intron_variant | MODIFIER |
| 15 | 95475935 | AX-170235223 | 0 | T | C | 0.149 | 3.03E-02 | 1.80E-01 | 2.65E+00 | 2.71E+00 | 8.70E-01 | no | Nell2 | intron_variant | MODIFIER |
| 15 | 95486739 | AX-97801981 | 0 | G | A | 0.149 | 3.03E-02 | 1.80E-01 | 2.65E+00 | 2.71E+00 | 8.70E-01 | no | Nell2 | intron_variant | MODIFIER |
| 15 | 95486815 | AX-170494909 | 0 | A | G | 0.149 | 3.03E-02 | 1.80E-01 | 2.65E+00 | 2.71E+00 | 8.70E-01 | no | Nell2 | intron_variant | MODIFIER |
| 15 | 95487383 | AX-170142902 | 0 | C | A | 0.149 | 3.03E-02 | 1.80E-01 | 2.65E+00 | 2.71E+00 | 8.70E-01 | no | Nell2 | intron_variant | MODIFIER |
| 15 | 95488704 | AX-170161207 | 0 | C | T | 0.149 | 3.03E-02 | 1.80E-01 | 2.65E+00 | 2.71E+00 | 8.70E-01 | no | Nell2 | intron_variant | MODIFIER |
| 15 | 95489383 | AX-170056380 | 0 | T | C | 0.149 | 3.03E-02 | 1.80E-01 | 2.65E+00 | 2.71E+00 | 8.70E-01 | no | Nell2 | intron_variant | MODIFIER |
| 15 | 95495326 | AX-170221200 | 0 | G | C | 0.149 | 3.03E-02 | 1.80E-01 | 2.65E+00 | 2.71E+00 | 8.70E-01 | no | Nell2 | intron_variant | MODIFIER |
| 15 | 95530050 | AX-97856836 | 0 | G | A | 0.149 | 3.03E-02 | 1.80E-01 | 2.65E+00 | 2.71E+00 | 8.70E-01 | no | Nell2 | upstream_gene_variant | MODIFIER |
| 15 | 95536742 | AX-170197270 | 0 | T | G | 0.149 | 3.03E-02 | 1.80E-01 | 2.65E+00 | 2.71E+00 | 8.70E-01 | no | | intergenic_variant | MODIFIER |
| 15 | 95541486 | AX-170176195 | 0 | C | T | 0.149 | 3.03E-02 | 1.80E-01 | 2.65E+00 | 2.71E+00 | 8.70E-01 | no | | intergenic_variant | MODIFIER |
| 15 | 95544249 | AX-97799042 | 0 | A | C | 0.149 | 3.03E-02 | 1.80E-01 | 2.65E+00 | 2.71E+00 | 8.70E-01 | no | | intergenic_variant | MODIFIER |
| 15 | 95556960 | AX-170110875 | 0 | G | A | 0.149 | 3.03E-02 | 1.80E-01 | 2.65E+00 | 2.71E+00 | 8.70E-01 | no | | intergenic_variant | MODIFIER |
| 15 | 95561098 | AX-170210602 | 0 | T | C | 0.149 | 3.03E-02 | 1.80E-01 | 2.65E+00 | 2.71E+00 | 8.70E-01 | no | | intergenic_variant | MODIFIER |
| 15 | 95563565 | AX-170284928 | 0 | G | A | 0.149 | 3.03E-02 | 1.80E-01 | 2.65E+00 | 2.71E+00 | 8.70E-01 | no | | intergenic_variant | MODIFIER |
| 15 | 95566823 | AX-170227042 | 0 | T | C | 0.149 | 3.03E-02 | 1.80E-01 | 2.65E+00 | 2.71E+00 | 8.70E-01 | no | | intergenic_variant | MODIFIER |
| 15 | 95567353 | AX-170141120 | 0 | G | A | 0.149 | 3.03E-02 | 1.80E-01 | 2.65E+00 | 2.71E+00 | 8.70E-01 | no | | intergenic_variant | MODIFIER |
| 15 | 95567605 | AX-170207449 | 0 | T | C | 0.149 | 3.03E-02 | 1.80E-01 | 2.65E+00 | 2.71E+00 | 8.70E-01 | no | | intergenic_variant | MODIFIER |
| 15 | 95579922 | AX-170316753 | 0 | G | A | 0.149 | 3.03E-02 | 1.80E-01 | 2.65E+00 | 2.71E+00 | 8.70E-01 | no | | intergenic_variant | MODIFIER |
| 15 | 95582026 | AX-97830285 | 0 | C | T | 0.149 | 3.03E-02 | 1.80E-01 | 2.65E+00 | 2.71E+00 | 8.70E-01 | no | | intergenic_variant | MODIFIER |
| 15 | 95589546 | AX-170403229 | 0 | G | A | 0.149 | 3.03E-02 | 1.80E-01 | 2.65E+00 | 2.71E+00 | 8.70E-01 | no | | intergenic_variant | MODIFIER |
| 15 | 95592703 | AX-170256721 | 0 | C | T | 0.149 | 3.03E-02 | 1.80E-01 | 2.65E+00 | 2.71E+00 | 8.70E-01 | no | | intergenic_variant | MODIFIER |
| 15 | 95595349 | AX-170093346 | 0 | A | G | 0.149 | 3.03E-02 | 1.80E-01 | 2.65E+00 | 2.71E+00 | 8.70E-01 | no | | intergenic_variant | MODIFIER |
| 15 | 95624162 | AX-97814716 | 0 | G | A | 0.149 | 3.03E-02 | 1.80E-01 | 2.65E+00 | 2.71E+00 | 8.70E-01 | no | Dbx2 | 3_prime_UTR_variant | MODIFIER |
| 15 | 95634739 | AX-170012320 | 0 | T | C | 0.149 | 3.03E-02 | 1.80E-01 | 2.65E+00 | 2.71E+00 | 8.70E-01 | no | Dbx2 | intron_variant | MODIFIER |
| 15 | 95639968 | AX-170374876 | 0 | A | C | 0.149 | 3.03E-02 | 1.80E-01 | 2.65E+00 | 2.71E+00 | 8.70E-01 | no | Dbx2 | intron_variant | MODIFIER |
| 15 | 95643389 | AX-97797097 | 0 | C | T | 0.149 | 3.03E-02 | 1.80E-01 | 2.65E+00 | 2.71E+00 | 8.70E-01 | no | Dbx2 | intron_variant | MODIFIER |

| | | | | | | | | | | | | | | | |
|---|---|---|---|---|---|---|---|---|---|---|---|---|---|---|---|
| 15 | 95683004 | AX-97793643 | 0 | C | T | 0.149 | 3.03E-02 | 1.80E-01 | 2.65E+00 | 2.71E+00 | 8.70E-01 | no | | intergenic_variant | MODIFIER |
| 15 | 95683343 | AX-170012319 | 0 | G | A | 0.149 | 3.03E-02 | 1.80E-01 | 2.65E+00 | 2.71E+00 | 8.70E-01 | no | | intergenic_variant | MODIFIER |
| 15 | 95702048 | AX-97853696 | 0 | G | T | 0.149 | 3.03E-02 | 1.80E-01 | 2.65E+00 | 2.71E+00 | 8.70E-01 | no | | regulatory_region_variant | MODIFIER |
| 15 | 95702930 | AX-170012322 | 0 | G | A | 0.149 | 3.03E-02 | 1.80E-01 | 2.65E+00 | 2.71E+00 | 8.70E-01 | no | | intergenic_variant | MODIFIER |
| 15 | 95714319 | AX-168389752 | 0 | G | A | 0.149 | 3.03E-02 | 1.80E-01 | 2.65E+00 | 2.71E+00 | 8.70E-01 | no | Gm8843 | upstream_gene_variant | MODIFIER |
| 15 | 95714460 | AX-97854072 | 0 | T | G | 0.149 | 3.03E-02 | 1.80E-01 | 2.65E+00 | 2.71E+00 | 8.70E-01 | no | Gm8843 | upstream_gene_variant | MODIFIER |
| 15 | 95726242 | AX-170377773 | 0 | A | C | 0.149 | 3.03E-02 | 1.80E-01 | 2.65E+00 | 2.71E+00 | 8.70E-01 | no | | regulatory_region_variant | MODIFIER |
| 15 | 95750967 | AX-97830048 | 0 | A | G | 0.149 | 3.03E-02 | 1.80E-01 | 2.65E+00 | 2.71E+00 | 8.70E-01 | no | Gm6961 | intron_variant,non_coding_transcript_variant | MODIFIER |
| 15 | 95756855 | AX-170012323 | 0 | G | A | 0.149 | 3.03E-02 | 1.80E-01 | 2.65E+00 | 2.71E+00 | 8.70E-01 | no | Gm49169 | upstream_gene_variant | MODIFIER |
| 15 | 95757157 | AX-170012318 | 0 | G | A | 0.149 | 3.03E-02 | 1.80E-01 | 2.65E+00 | 2.71E+00 | 8.70E-01 | no | Gm49169 | upstream_gene_variant | MODIFIER |
| 15 | 95758133 | AX-170439232 | 0 | G | A | 0.149 | 3.03E-02 | 1.80E-01 | 2.65E+00 | 2.71E+00 | 8.70E-01 | no | Gm49169 | upstream_gene_variant | MODIFIER |
| 15 | 95776695 | AX-170012321 | 0 | A | G | 0.149 | 3.03E-02 | 1.80E-01 | 2.65E+00 | 2.71E+00 | 8.70E-01 | no | Gm8849 | downstream_gene_variant | MODIFIER |
| 15 | 95787554 | AX-97856894 | 0 | C | T | 0.149 | 3.03E-02 | 1.80E-01 | 2.65E+00 | 2.71E+00 | 8.70E-01 | no | Ano6 | upstream_gene_variant | MODIFIER |
| 15 | 95794945 | AX-170462773 | 0 | T | A | 0.149 | 3.03E-02 | 1.80E-01 | 2.65E+00 | 2.71E+00 | 8.70E-01 | no | Ano6 | intron_variant | MODIFIER |
| 15 | 95812263 | AX-170235449 | 0 | T | C | 0.149 | 3.03E-02 | 1.80E-01 | 2.65E+00 | 2.71E+00 | 8.70E-01 | no | Ano6 | intron_variant | MODIFIER |
| 15 | 95813074 | AX-170188998 | 0 | T | C | 0.149 | 3.03E-02 | 1.80E-01 | 2.65E+00 | 2.71E+00 | 8.70E-01 | no | Ano6 | intron_variant | MODIFIER |
| 15 | 95813100 | AX-97833413 | 0 | C | T | 0.149 | 3.03E-02 | 1.80E-01 | 2.65E+00 | 2.71E+00 | 8.70E-01 | no | Ano6 | intron_variant | MODIFIER |
| 15 | 95813199 | AX-170469019 | 0 | T | C | 0.149 | 3.03E-02 | 1.80E-01 | 2.65E+00 | 2.71E+00 | 8.70E-01 | no | Ano6 | intron_variant | MODIFIER |
| 15 | 95815665 | AX-170394541 | 0 | T | C | 0.149 | 3.03E-02 | 1.80E-01 | 2.65E+00 | 2.71E+00 | 8.70E-01 | no | Ano6 | intron_variant | MODIFIER |
| 15 | 95817233 | AX-170499167 | 0 | C | T | 0.149 | 3.03E-02 | 1.80E-01 | 2.65E+00 | 2.71E+00 | 8.70E-01 | no | Ano6 | intron_variant | MODIFIER |
| 15 | 95827212 | AX-170375743 | 0 | G | A | 0.149 | 3.03E-02 | 1.80E-01 | 2.65E+00 | 2.71E+00 | 8.70E-01 | no | Ano6 | intron_variant | MODIFIER |
| 15 | 95827409 | AX-170142023 | 0 | C | T | 0.149 | 3.03E-02 | 1.80E-01 | 2.65E+00 | 2.71E+00 | 8.70E-01 | no | Ano6 | intron_variant | MODIFIER |
| 15 | 95871753 | AX-170354854 | 0 | A | C | 0.149 | 3.03E-02 | 1.80E-01 | 2.65E+00 | 2.71E+00 | 8.70E-01 | no | Ano6 | intron_variant | MODIFIER |
| 15 | 95874677 | AX-170242606 | 0 | C | T | 0.149 | 3.03E-02 | 1.80E-01 | 2.65E+00 | 2.71E+00 | 8.70E-01 | no | Ano6 | intron_variant | MODIFIER |
| 15 | 95890369 | AX-170162748 | 0 | G | A | 0.149 | 3.03E-02 | 1.80E-01 | 2.65E+00 | 2.71E+00 | 8.70E-01 | no | Ano6 | intron_variant | MODIFIER |
| 15 | 95909925 | AX-170056388 | 0 | T | C | 0.149 | 3.03E-02 | 1.80E-01 | 2.65E+00 | 2.71E+00 | 8.70E-01 | no | Ano6 | intron_variant | MODIFIER |
| 15 | 95910050 | AX-170434215 | 0 | A | G | 0.149 | 3.03E-02 | 1.80E-01 | 2.65E+00 | 2.71E+00 | 8.70E-01 | no | Ano6 | intron_variant | MODIFIER |
| 15 | 95910442 | AX-170456641 | 0 | G | A | 0.149 | 3.03E-02 | 1.80E-01 | 2.65E+00 | 2.71E+00 | 8.70E-01 | no | Ano6 | intron_variant | MODIFIER |
| 15 | 95911102 | AX-170438275 | 0 | A | T | 0.149 | 3.03E-02 | 1.80E-01 | 2.65E+00 | 2.71E+00 | 8.70E-01 | no | Ano6 | intron_variant | MODIFIER |
| 15 | 95919952 | AX-170322076 | 0 | C | T | 0.149 | 3.03E-02 | 1.80E-01 | 2.65E+00 | 2.71E+00 | 8.70E-01 | no | Ano6 | intron_variant | MODIFIER |
| 15 | 95921682 | AX-170273270 | 0 | G | C | 0.149 | 3.03E-02 | 1.80E-01 | 2.65E+00 | 2.71E+00 | 8.70E-01 | no | Ano6 | intron_variant | MODIFIER |
| 15 | 95924780 | AX-170409201 | 0 | C | T | 0.149 | 3.03E-02 | 1.80E-01 | 2.65E+00 | 2.71E+00 | 8.70E-01 | no | Ano6 | intron_variant | MODIFIER |
| 15 | 95925529 | AX-170237725 | 0 | C | T | 0.149 | 3.03E-02 | 1.80E-01 | 2.65E+00 | 2.71E+00 | 8.70E-01 | no | Ano6 | intron_variant | MODIFIER |
| 15 | 95934843 | AX-170335830 | 0 | A | G | 0.149 | 3.03E-02 | 1.80E-01 | 2.65E+00 | 2.71E+00 | 8.70E-01 | no | Ano6 | intron_variant | MODIFIER |
| 15 | 95949152 | AX-170175287 | 0 | A | G | 0.149 | 3.03E-02 | 1.80E-01 | 2.65E+00 | 2.71E+00 | 8.70E-01 | no | Ano6 | intron_variant | MODIFIER |
| 15 | 95949676 | AX-170344759 | 0 | G | A | 0.149 | 3.03E-02 | 1.80E-01 | 2.65E+00 | 2.71E+00 | 8.70E-01 | no | Ano6 | intron_variant | MODIFIER |
| 15 | 95953082 | AX-170250016 | 0 | T | G | 0.149 | 3.03E-02 | 1.80E-01 | 2.65E+00 | 2.71E+00 | 8.70E-01 | no | Ano6 | intron_variant | MODIFIER |
| 15 | 95953108 | AX-170487622 | 0 | A | G | 0.149 | 3.03E-02 | 1.80E-01 | 2.65E+00 | 2.71E+00 | 8.70E-01 | no | Ano6 | intron_variant | MODIFIER |
| 15 | 95958795 | AX-170235068 | 0 | A | C | 0.149 | 3.03E-02 | 1.80E-01 | 2.65E+00 | 2.71E+00 | 8.70E-01 | no | Ano6 | intron_variant | MODIFIER |
| 15 | 95959054 | AX-97844468 | 0 | G | A | 0.149 | 3.03E-02 | 1.80E-01 | 2.65E+00 | 2.71E+00 | 8.70E-01 | no | Ano6 | intron_variant | MODIFIER |
| 15 | 95981605 | AX-170056390 | 0 | C | G | 0.149 | 3.03E-02 | 1.80E-01 | 2.65E+00 | 2.71E+00 | 8.70E-01 | no | Gm49172 | intron_variant,non_coding_transcript_variant | MODIFIER |
| 15 | 95984021 | AX-170209370 | 0 | C | T | 0.149 | 3.03E-02 | 1.80E-01 | 2.65E+00 | 2.71E+00 | 8.70E-01 | no | Gm49172 | intron_variant,non_coding_transcript_variant | MODIFIER |
| 15 | 95987341 | AX-170327982 | 0 | A | G | 0.149 | 3.03E-02 | 1.80E-01 | 2.65E+00 | 2.71E+00 | 8.70E-01 | no | Gm49172 | intron_variant,non_coding_transcript_variant | MODIFIER |
| 15 | 95991518 | AX-170185939 | 0 | T | C | 0.149 | 3.03E-02 | 1.80E-01 | 2.65E+00 | 2.71E+00 | 8.70E-01 | no | Gm49172 | intron_variant,non_coding_transcript_variant | MODIFIER |
| 15 | 95991947 | AX-97848583 | 0 | C | A | 0.149 | 3.03E-02 | 1.80E-01 | 2.65E+00 | 2.71E+00 | 8.70E-01 | no | Gm49172 | intron_variant,non_coding_transcript_variant | MODIFIER |
| 15 | 95992189 | AX-170400339 | 0 | T | G | 0.149 | 3.03E-02 | 1.80E-01 | 2.65E+00 | 2.71E+00 | 8.70E-01 | no | Gm49172 | intron_variant,non_coding_transcript_variant | MODIFIER |
| 15 | 96181065 | AX-170456283 | 0 | C | T | 0.149 | 3.03E-02 | 1.80E-01 | 2.65E+00 | 2.71E+00 | 8.70E-01 | no | 2610037D02Rik | intron_variant,non_coding_transcript_variant | MODIFIER |
| 15 | 96199437 | AX-170128958 | 0 | G | A | 0.149 | 3.03E-02 | 1.80E-01 | 2.65E+00 | 2.71E+00 | 8.70E-01 | no | 2610037D02Rik | intron_variant,non_coding_transcript_variant | MODIFIER |
| 15 | 96200660 | AX-97830095 | 0 | C | T | 0.149 | 3.03E-02 | 1.80E-01 | 2.65E+00 | 2.71E+00 | 8.70E-01 | no | 2610037D02Rik | intron_variant,non_coding_transcript_variant | MODIFIER |
| 15 | 96200934 | AX-170012325 | 0 | T | C | 0.149 | 3.03E-02 | 1.80E-01 | 2.65E+00 | 2.71E+00 | 8.70E-01 | no | 2610037D02Rik | intron_variant,non_coding_transcript_variant | MODIFIER |
| 15 | 96201118 | AX-170387947 | 0 | C | T | 0.149 | 3.03E-02 | 1.80E-01 | 2.65E+00 | 2.71E+00 | 8.70E-01 | no | 2610037D02Rik | intron_variant,non_coding_transcript_variant | MODIFIER |
| 15 | 96209171 | AX-170100048 | 0 | G | A | 0.149 | 3.03E-02 | 1.80E-01 | 2.65E+00 | 2.71E+00 | 8.70E-01 | no | 2610037D02Rik | intron_variant,non_coding_transcript_variant | MODIFIER |
| 15 | 96211528 | AX-170381130 | 0 | C | T | 0.149 | 3.03E-02 | 1.80E-01 | 2.65E+00 | 2.71E+00 | 8.70E-01 | no | 2610037D02Rik | intron_variant,non_coding_transcript_variant | MODIFIER |
| 15 | 96221471 | AX-170056393 | 0 | G | A | 0.149 | 3.03E-02 | 1.80E-01 | 2.65E+00 | 2.71E+00 | 8.70E-01 | no | 2610037D02Rik | intron_variant,non_coding_transcript_variant | MODIFIER |

| | | | | | | | | | | | | | | | |
|---|---|---|---|---|---|---|---|---|---|---|---|---|---|---|---|
| 15 | 96227138 | AX-97805840 | 0 | A | G | 0.149 | 3.03E-02 | 1.80E-01 | 2.65E+00 | 2.71E+00 | 8.70E-01 | no | 2610037D02Rik | intron_variant,non_coding_transcript_variant | MODIFIER |
| 15 | 96246468 | AX-97853317 | 0 | G | A | 0.149 | 3.03E-02 | 1.80E-01 | 2.65E+00 | 2.71E+00 | 8.70E-01 | no | 2610037D02Rik | intron_variant,non_coding_transcript_variant | MODIFIER |
| 15 | 96246543 | AX-170012327 | 0 | C | T | 0.149 | 3.03E-02 | 1.80E-01 | 2.65E+00 | 2.71E+00 | 8.70E-01 | no | 2610037D02Rik | intron_variant,non_coding_transcript_variant | MODIFIER |
| 15 | 96255509 | AX-170282308 | 0 | C | G | 0.149 | 3.03E-02 | 1.80E-01 | 2.65E+00 | 2.71E+00 | 8.70E-01 | no | 2610037D02Rik | intron_variant,non_coding_transcript_variant | MODIFIER |
| 15 | 96255578 | AX-170262535 | 0 | G | T | 0.149 | 3.03E-02 | 1.80E-01 | 2.65E+00 | 2.71E+00 | 8.70E-01 | no | 2610037D02Rik | intron_variant,non_coding_transcript_variant | MODIFIER |
| 15 | 96341145 | AX-97839698 | 0 | A | G | 0.149 | 3.03E-02 | 1.80E-01 | 2.65E+00 | 2.71E+00 | 8.70E-01 | no | Arid2 | intron_variant | MODIFIER |
| 15 | 96470607 | AX-97829148 | 0 | C | T | 0.149 | 3.03E-02 | 1.80E-01 | 2.65E+00 | 2.71E+00 | 8.70E-01 | no | | intergenic_variant | MODIFIER |
| 15 | 96482739 | AX-170231444 | 0 | A | G | 0.149 | 3.03E-02 | 1.80E-01 | 2.65E+00 | 2.71E+00 | 8.70E-01 | no | | intergenic_variant | MODIFIER |
| 15 | 96550112 | AX-170469912 | 0 | A | G | 0.149 | 3.03E-02 | 1.80E-01 | 2.65E+00 | 2.71E+00 | 8.70E-01 | no | | intergenic_variant | MODIFIER |
| 15 | 96559860 | AX-170492671 | 0 | A | G | 0.149 | 3.03E-02 | 1.80E-01 | 2.65E+00 | 2.71E+00 | 8.70E-01 | no | | intergenic_variant | MODIFIER |
| 15 | 96560074 | AX-170134562 | 0 | C | T | 0.149 | 3.03E-02 | 1.80E-01 | 2.65E+00 | 2.71E+00 | 8.70E-01 | no | | intergenic_variant | MODIFIER |
| 15 | 96575559 | AX-97833257 | 0 | A | G | 0.149 | 3.03E-02 | 1.80E-01 | 2.65E+00 | 2.71E+00 | 8.70E-01 | no | Slc38a1 | 3_prime_UTR_variant | MODIFIER |
| 15 | 96632729 | AX-170499489 | 0 | T | A | 0.149 | 3.03E-02 | 1.80E-01 | 2.65E+00 | 2.71E+00 | 8.70E-01 | no | Slc38a1 | intron_variant | MODIFIER |
| 15 | 96632858 | AX-170171259 | 0 | T | A | 0.149 | 3.03E-02 | 1.80E-01 | 2.65E+00 | 2.71E+00 | 8.70E-01 | no | Slc38a1 | intron_variant | MODIFIER |
| 15 | 96634907 | AX-170308333 | 0 | T | C | 0.149 | 3.03E-02 | 1.80E-01 | 2.65E+00 | 2.71E+00 | 8.70E-01 | no | Slc38a1 | intron_variant | MODIFIER |
| 15 | 96635214 | AX-170463652 | 0 | C | T | 0.149 | 3.03E-02 | 1.80E-01 | 2.65E+00 | 2.71E+00 | 8.70E-01 | no | Slc38a1 | intron_variant | MODIFIER |
| 15 | 96705968 | AX-97852943 | 0 | T | G | 0.149 | 3.03E-02 | 1.80E-01 | 2.65E+00 | 2.71E+00 | 8.70E-01 | no | | intergenic_variant | MODIFIER |
| 15 | 96772632 | AX-170315859 | 0 | C | G | 0.149 | 3.03E-02 | 1.80E-01 | 2.65E+00 | 2.71E+00 | 8.70E-01 | no | | regulatory_region_variant | MODIFIER |
| 15 | 96783419 | AX-170093352 | 0 | C | T | 0.149 | 3.03E-02 | 1.80E-01 | 2.65E+00 | 2.71E+00 | 8.70E-01 | no | | intergenic_variant | MODIFIER |
| 15 | 96791689 | AX-97849555 | 0 | T | C | 0.149 | 3.03E-02 | 1.80E-01 | 2.65E+00 | 2.71E+00 | 8.70E-01 | no | | intergenic_variant | MODIFIER |
| 15 | 96927365 | AX-97830195 | 0 | A | G | 0.149 | 3.03E-02 | 1.80E-01 | 2.65E+00 | 2.71E+00 | 8.70E-01 | no | Rpl10a-ps3 | upstream_gene_variant | MODIFIER |
| 15 | 96941028 | AX-97855887 | 0 | A | G | 0.149 | 3.03E-02 | 1.80E-01 | 2.65E+00 | 2.71E+00 | 8.70E-01 | no | | intergenic_variant | MODIFIER |
| 15 | 96946992 | AX-170301285 | 0 | G | A | 0.149 | 3.03E-02 | 1.80E-01 | 2.65E+00 | 2.71E+00 | 8.70E-01 | no | | intergenic_variant | MODIFIER |
| 15 | 96953370 | AX-170269521 | 0 | T | C | 0.149 | 3.03E-02 | 1.80E-01 | 2.65E+00 | 2.71E+00 | 8.70E-01 | no | | regulatory_region_variant | MODIFIER |
| 15 | 96963847 | AX-97808418 | 0 | T | C | 0.149 | 3.03E-02 | 1.80E-01 | 2.65E+00 | 2.71E+00 | 8.70E-01 | no | | regulatory_region_variant | MODIFIER |
| 15 | 97013203 | AX-97829636 | 0 | A | C | 0.149 | 3.03E-02 | 1.80E-01 | 2.65E+00 | 2.71E+00 | 8.70E-01 | no | Slc38a4 | intron_variant | MODIFIER |
| 15 | 97127535 | AX-170105353 | 0 | C | A | 0.149 | 3.03E-02 | 1.80E-01 | 2.65E+00 | 2.71E+00 | 8.70E-01 | no | | intergenic_variant | MODIFIER |
| 15 | 97128307 | AX-170407765 | 0 | A | G | 0.149 | 3.03E-02 | 1.80E-01 | 2.65E+00 | 2.71E+00 | 8.70E-01 | no | | intergenic_variant | MODIFIER |
| 15 | 97128330 | AX-170171665 | 0 | A | C | 0.149 | 3.03E-02 | 1.80E-01 | 2.65E+00 | 2.71E+00 | 8.70E-01 | no | | intergenic_variant | MODIFIER |
| 15 | 97143191 | AX-170369262 | 0 | G | C | 0.149 | 3.03E-02 | 1.80E-01 | 2.65E+00 | 2.71E+00 | 8.70E-01 | no | | intergenic_variant | MODIFIER |
| 15 | 97144577 | AX-170182316 | 0 | T | A | 0.149 | 3.03E-02 | 1.80E-01 | 2.65E+00 | 2.71E+00 | 8.70E-01 | no | | intergenic_variant | MODIFIER |
| 15 | 97436658 | AX-170242754 | 0 | G | T | 0.149 | 3.03E-02 | 1.80E-01 | 2.65E+00 | 2.71E+00 | 8.70E-01 | no | | intergenic_variant | MODIFIER |
| 15 | 97442867 | AX-170056425 | 0 | T | C | 0.149 | 3.03E-02 | 1.80E-01 | 2.65E+00 | 2.71E+00 | 8.70E-01 | no | | intergenic_variant | MODIFIER |
| 15 | 97444366 | AX-97830277 | 0 | C | T | 0.149 | 3.03E-02 | 1.80E-01 | 2.65E+00 | 2.71E+00 | 8.70E-01 | no | | intergenic_variant | MODIFIER |
| 15 | 97455617 | AX-170269783 | 0 | T | A | 0.149 | 3.03E-02 | 1.80E-01 | 2.65E+00 | 2.71E+00 | 8.70E-01 | no | | regulatory_region_variant | MODIFIER |
| 15 | 97456320 | AX-170395354 | 0 | G | A | 0.149 | 3.03E-02 | 1.80E-01 | 2.65E+00 | 2.71E+00 | 8.70E-01 | no | | regulatory_region_variant | MODIFIER |
| 15 | 97470137 | AX-170404642 | 0 | T | C | 0.149 | 3.03E-02 | 1.80E-01 | 2.65E+00 | 2.71E+00 | 8.70E-01 | no | | regulatory_region_variant | MODIFIER |
| 15 | 97479246 | AX-170254148 | 0 | C | A | 0.149 | 3.03E-02 | 1.80E-01 | 2.65E+00 | 2.71E+00 | 8.70E-01 | no | | intergenic_variant | MODIFIER |
| 15 | 97514433 | AX-170105051 | 0 | G | A | 0.149 | 3.03E-02 | 1.80E-01 | 2.65E+00 | 2.71E+00 | 8.70E-01 | no | | intergenic_variant | MODIFIER |
| 15 | 97515398 | AX-170517302 | 0 | C | T | 0.149 | 3.03E-02 | 1.80E-01 | 2.65E+00 | 2.71E+00 | 8.70E-01 | no | | intergenic_variant | MODIFIER |
| 15 | 97526218 | AX-170093356 | 0 | A | G | 0.149 | 3.03E-02 | 1.80E-01 | 2.65E+00 | 2.71E+00 | 8.70E-01 | no | | intergenic_variant | MODIFIER |
| 15 | 97531999 | AX-170387788 | 0 | C | T | 0.149 | 3.03E-02 | 1.80E-01 | 2.65E+00 | 2.71E+00 | 8.70E-01 | no | | intergenic_variant | MODIFIER |
| 15 | 97532412 | AX-170158554 | 0 | C | T | 0.149 | 3.03E-02 | 1.80E-01 | 2.65E+00 | 2.71E+00 | 8.70E-01 | no | | intergenic_variant | MODIFIER |
| 15 | 97534809 | AX-170334621 | 0 | G | A | 0.149 | 3.03E-02 | 1.80E-01 | 2.65E+00 | 2.71E+00 | 8.70E-01 | no | | regulatory_region_variant | MODIFIER |
| 15 | 97540788 | AX-97806984 | 0 | G | T | 0.149 | 3.03E-02 | 1.80E-01 | 2.65E+00 | 2.71E+00 | 8.70E-01 | no | | intergenic_variant | MODIFIER |
| 15 | 97548066 | AX-170481071 | 0 | T | C | 0.149 | 3.03E-02 | 1.80E-01 | 2.65E+00 | 2.71E+00 | 8.70E-01 | no | | intergenic_variant | MODIFIER |
| 15 | 97571013 | AX-97817578 | 0 | G | A | 0.149 | 3.03E-02 | 1.80E-01 | 2.65E+00 | 2.71E+00 | 8.70E-01 | no | | intergenic_variant | MODIFIER |
| 15 | 97576133 | AX-170123763 | 0 | G | T | 0.149 | 3.03E-02 | 1.80E-01 | 2.65E+00 | 2.71E+00 | 8.70E-01 | no | | intergenic_variant | MODIFIER |
| 15 | 97578434 | AX-170427388 | 0 | T | C | 0.149 | 3.03E-02 | 1.80E-01 | 2.65E+00 | 2.71E+00 | 8.70E-01 | no | | intergenic_variant | MODIFIER |
| 15 | 97579002 | AX-112530150 | 0 | A | G | 0.149 | 3.03E-02 | 1.80E-01 | 2.65E+00 | 2.71E+00 | 8.70E-01 | no | | intergenic_variant | MODIFIER |
| 15 | 97594740 | AX-97830284 | 0 | T | C | 0.149 | 3.03E-02 | 1.80E-01 | 2.65E+00 | 2.71E+00 | 8.70E-01 | no | | regulatory_region_variant | MODIFIER |
| 15 | 97602291 | AX-170114181 | 0 | T | C | 0.149 | 3.03E-02 | 1.80E-01 | 2.65E+00 | 2.71E+00 | 8.70E-01 | no | | intergenic_variant | MODIFIER |
| 1 | 52429872 | AX-170326593 | 0 | G | A | 0.132 | -3.91E-02 | 2.42E-01 | 2.70E+00 | 2.74E+00 | 8.70E-01 | no | | intergenic_variant | MODIFIER |
| 11 | 33232125 | AX-170389142 | 1 | A | T | 0.08 | 3.44E-02 | 2.05E-01 | 2.64E+00 | 2.72E+00 | 8.70E-01 | no | Ranbp17 | intron_variant | MODIFIER |

| | | | | | | | | | | | | | | | |
|---|---|---|---|---|---|---|---|---|---|---|---|---|---|---|---|
| 9 | 53899882 | AX-170501748 | 1 | G | A | 0.205 | -1.91E-02 | 1.20E-01 | 2.69E+00 | 2.76E+00 | 8.70E-01 | no | | regulatory_region_variant | MODIFIER |
| 15 | 95624221 | AX-170154408 | 1 | G | A | 0.152 | 3.03E-02 | 1.79E-01 | 2.65E+00 | 2.71E+00 | 8.70E-01 | no | Dbx2 | 3_prime_UTR_variant | MODIFIER |
| 1 | 8133749 | AX-170091383 | 2 | G | T | 0.445 | -1.03E-02 | 6.78E-02 | 2.70E+00 | 2.79E+00 | 8.70E-01 | no | | intergenic_variant | MODIFIER |
| 13 | 47231308 | AX-170051186 | 2 | T | C | 0.473 | -9.93E-03 | 6.44E-02 | 2.66E+00 | 2.76E+00 | 8.70E-01 | no | Rnf144b | intron_variant | MODIFIER |
| 9 | 53076870 | AX-170321459 | 0 | T | C | 0.228 | 1.79E-02 | 1.10E-01 | 2.66E+00 | 2.73E+00 | 8.70E-01 | no | | intergenic_variant | MODIFIER |
| 9 | 57550888 | AX-97854613 | 1 | C | G | 0.205 | -1.99E-02 | 1.24E-01 | 2.69E+00 | 2.75E+00 | 8.70E-01 | no | Mpi | intron_variant | MODIFIER |
| 15 | 92154058 | AX-170213875 | 1 | G | C | 0.134 | 2.64E-02 | 1.57E-01 | 2.65E+00 | 2.71E+00 | 8.70E-01 | no | Cntn1 | intron_variant | MODIFIER |
| 5 | 123650123 | AX-170211977 | 1 | C | G | 0.455 | -1.77E-02 | 1.11E-01 | 2.70E+00 | 2.76E+00 | 8.70E-01 | no | Clip1 | intron_variant | MODIFIER |
| 7 | 68622748 | AX-170468413 | 0 | G | C | 0.404 | 1.69E-02 | 9.98E-02 | 2.65E+00 | 2.71E+00 | 8.70E-01 | no | Gm5334 | downstream_gene_variant | MODIFIER |
| 8 | 97846912 | AX-170031349 | 0 | G | T | 0.219 | -1.77E-02 | 1.12E-01 | 2.66E+00 | 2.75E+00 | 8.70E-01 | no | Gm39231 | intron_variant,non_coding_transcript_variant | MODIFIER |
| 15 | 100692616 | AX-170352181 | 1 | T | A | 0.098 | 2.73E-02 | 1.65E-01 | 2.64E+00 | 2.72E+00 | 8.70E-01 | no | Cela1 | upstream_gene_variant | MODIFIER |
| 1 | 9594506 | AX-170328945 | 1 | C | T | 0.098 | 2.86E-02 | 1.72E-01 | 2.62E+00 | 2.73E+00 | 8.70E-01 | no | 2610203C22Rik | intron_variant,non_coding_transcript_variant | MODIFIER |
| 9 | 57938101 | AX-170337998 | 1 | C | T | 0.223 | 1.96E-02 | 1.21E-01 | 2.68E+00 | 2.74E+00 | 8.70E-01 | no | Sema7a | upstream_gene_variant | MODIFIER |
| 18 | 89714843 | AX-170388213 | 1 | G | A | 0.518 | -9.38E-03 | 6.24E-02 | 2.63E+00 | 2.76E+00 | 8.70E-01 | no | Dok6 | intron_variant | MODIFIER |
| 15 | 86724109 | AX-170012191 | 0 | A | C | 0.132 | 2.63E-02 | 1.57E-01 | 2.65E+00 | 2.72E+00 | 8.70E-01 | no | Gm49465 | upstream_gene_variant | MODIFIER |
| 15 | 86732895 | AX-170141244 | 0 | C | T | 0.132 | 2.63E-02 | 1.57E-01 | 2.65E+00 | 2.72E+00 | 8.70E-01 | no | 4930543I03Rik | upstream_gene_variant | MODIFIER |
| 15 | 86733750 | AX-170376433 | 0 | G | A | 0.132 | 2.63E-02 | 1.57E-01 | 2.65E+00 | 2.72E+00 | 8.70E-01 | no | 4930543I03Rik | upstream_gene_variant | MODIFIER |
| 15 | 86735077 | AX-170393491 | 0 | C | T | 0.132 | 2.63E-02 | 1.57E-01 | 2.65E+00 | 2.72E+00 | 8.70E-01 | no | 4930543I03Rik | upstream_gene_variant | MODIFIER |
| 15 | 86735674 | AX-170384659 | 0 | G | A | 0.132 | 2.63E-02 | 1.57E-01 | 2.65E+00 | 2.72E+00 | 8.70E-01 | no | 4930543I03Rik | upstream_gene_variant | MODIFIER |
| 15 | 86740967 | AX-97827998 | 0 | A | G | 0.132 | 2.63E-02 | 1.57E-01 | 2.65E+00 | 2.72E+00 | 8.70E-01 | no | 4930543I03Rik | non_coding_transcript_exon_variant | MODIFIER |
| 15 | 86750730 | AX-170012197 | 0 | T | C | 0.132 | 2.63E-02 | 1.57E-01 | 2.65E+00 | 2.72E+00 | 8.70E-01 | no | 4930543I03Rik | intron_variant,non_coding_transcript_variant | MODIFIER |
| 15 | 86754021 | AX-170012192 | 0 | C | A | 0.132 | 2.63E-02 | 1.57E-01 | 2.65E+00 | 2.72E+00 | 8.70E-01 | no | 4930543I03Rik | intron_variant,non_coding_transcript_variant | MODIFIER |
| 15 | 86772082 | AX-97830070 | 0 | T | G | 0.132 | 2.63E-02 | 1.57E-01 | 2.65E+00 | 2.72E+00 | 8.70E-01 | no | | intergenic_variant | MODIFIER |
| 15 | 86781286 | AX-170130079 | 0 | A | G | 0.132 | 2.63E-02 | 1.57E-01 | 2.65E+00 | 2.72E+00 | 8.70E-01 | no | | intergenic_variant | MODIFIER |
| 15 | 86786759 | AX-170263668 | 0 | C | T | 0.132 | 2.63E-02 | 1.57E-01 | 2.65E+00 | 2.72E+00 | 8.70E-01 | no | | intergenic_variant | MODIFIER |
| 15 | 86797877 | AX-170087932 | 0 | C | T | 0.132 | 2.63E-02 | 1.57E-01 | 2.65E+00 | 2.72E+00 | 8.70E-01 | no | | intergenic_variant | MODIFIER |
| 15 | 86801214 | AX-170012195 | 0 | G | C | 0.132 | 2.63E-02 | 1.57E-01 | 2.65E+00 | 2.72E+00 | 8.70E-01 | no | | intergenic_variant | MODIFIER |
| 15 | 86801327 | AX-170012196 | 0 | G | T | 0.132 | 2.63E-02 | 1.57E-01 | 2.65E+00 | 2.72E+00 | 8.70E-01 | no | | intergenic_variant | MODIFIER |
| 15 | 86802185 | AX-170056203 | 0 | G | C | 0.132 | 2.63E-02 | 1.57E-01 | 2.65E+00 | 2.72E+00 | 8.70E-01 | no | | intergenic_variant | MODIFIER |
| 15 | 86802838 | AX-170131931 | 0 | T | C | 0.132 | 2.63E-02 | 1.57E-01 | 2.65E+00 | 2.72E+00 | 8.70E-01 | no | | intergenic_variant | MODIFIER |
| 15 | 86804651 | AX-170366139 | 0 | T | C | 0.132 | 2.63E-02 | 1.57E-01 | 2.65E+00 | 2.72E+00 | 8.70E-01 | no | | intergenic_variant | MODIFIER |
| 15 | 86805078 | AX-170415534 | 0 | A | G | 0.132 | 2.63E-02 | 1.57E-01 | 2.65E+00 | 2.72E+00 | 8.70E-01 | no | | intergenic_variant | MODIFIER |
| 15 | 86806378 | AX-170181326 | 0 | T | C | 0.132 | 2.63E-02 | 1.57E-01 | 2.65E+00 | 2.72E+00 | 8.70E-01 | no | | intergenic_variant | MODIFIER |
| 15 | 86809049 | AX-170483243 | 0 | A | G | 0.132 | 2.63E-02 | 1.57E-01 | 2.65E+00 | 2.72E+00 | 8.70E-01 | no | | intergenic_variant | MODIFIER |
| 15 | 86810776 | AX-170272208 | 0 | A | G | 0.132 | 2.63E-02 | 1.57E-01 | 2.65E+00 | 2.72E+00 | 8.70E-01 | no | | intergenic_variant | MODIFIER |
| 15 | 86815232 | AX-170427254 | 0 | G | A | 0.132 | 2.63E-02 | 1.57E-01 | 2.65E+00 | 2.72E+00 | 8.70E-01 | no | | intergenic_variant | MODIFIER |
| 15 | 86818197 | AX-97799407 | 0 | C | T | 0.132 | 2.63E-02 | 1.57E-01 | 2.65E+00 | 2.72E+00 | 8.70E-01 | no | | intergenic_variant | MODIFIER |
| 15 | 86819327 | AX-170201952 | 0 | T | C | 0.132 | 2.63E-02 | 1.57E-01 | 2.65E+00 | 2.72E+00 | 8.70E-01 | no | | intergenic_variant | MODIFIER |
| 15 | 86821405 | AX-170525292 | 0 | A | G | 0.132 | 2.63E-02 | 1.57E-01 | 2.65E+00 | 2.72E+00 | 8.70E-01 | no | | intergenic_variant | MODIFIER |
| 15 | 86821722 | AX-170178565 | 0 | G | A | 0.132 | 2.63E-02 | 1.57E-01 | 2.65E+00 | 2.72E+00 | 8.70E-01 | no | | intergenic_variant | MODIFIER |
| 15 | 86825737 | AX-170223613 | 0 | A | G | 0.132 | 2.63E-02 | 1.57E-01 | 2.65E+00 | 2.72E+00 | 8.70E-01 | no | | intergenic_variant | MODIFIER |
| 15 | 86826600 | AX-170436671 | 0 | T | A | 0.132 | 2.63E-02 | 1.57E-01 | 2.65E+00 | 2.72E+00 | 8.70E-01 | no | | intergenic_variant | MODIFIER |
| 15 | 86827300 | AX-170203379 | 0 | C | A | 0.132 | 2.63E-02 | 1.57E-01 | 2.65E+00 | 2.72E+00 | 8.70E-01 | no | | intergenic_variant | MODIFIER |
| 15 | 86828277 | AX-170012189 | 0 | A | T | 0.132 | 2.63E-02 | 1.57E-01 | 2.65E+00 | 2.72E+00 | 8.70E-01 | no | | intergenic_variant | MODIFIER |
| 15 | 86835054 | AX-170286025 | 0 | A | G | 0.132 | 2.63E-02 | 1.57E-01 | 2.65E+00 | 2.72E+00 | 8.70E-01 | no | | intergenic_variant | MODIFIER |
| 15 | 86835645 | AX-170162739 | 0 | G | A | 0.132 | 2.63E-02 | 1.57E-01 | 2.65E+00 | 2.72E+00 | 8.70E-01 | no | | regulatory_region_variant | MODIFIER |
| 15 | 86835754 | AX-170136713 | 0 | T | C | 0.132 | 2.63E-02 | 1.57E-01 | 2.65E+00 | 2.72E+00 | 8.70E-01 | no | | regulatory_region_variant | MODIFIER |
| 15 | 86836142 | AX-170461716 | 0 | T | G | 0.132 | 2.63E-02 | 1.57E-01 | 2.65E+00 | 2.72E+00 | 8.70E-01 | no | | regulatory_region_variant | MODIFIER |
| 15 | 86836243 | AX-170241761 | 0 | T | C | 0.132 | 2.63E-02 | 1.57E-01 | 2.65E+00 | 2.72E+00 | 8.70E-01 | no | | regulatory_region_variant | MODIFIER |
| 15 | 86843762 | AX-97830046 | 0 | G | A | 0.132 | 2.63E-02 | 1.57E-01 | 2.65E+00 | 2.72E+00 | 8.70E-01 | no | | intergenic_variant | MODIFIER |
| 15 | 87058862 | AX-97830052 | 0 | A | G | 0.132 | 2.63E-02 | 1.57E-01 | 2.65E+00 | 2.72E+00 | 8.70E-01 | no | | intergenic_variant | MODIFIER |
| 15 | 87074939 | AX-170461200 | 0 | G | C | 0.132 | 2.63E-02 | 1.57E-01 | 2.65E+00 | 2.72E+00 | 8.70E-01 | no | | regulatory_region_variant | MODIFIER |
| 15 | 87095096 | AX-170447288 | 0 | T | C | 0.132 | 2.63E-02 | 1.57E-01 | 2.65E+00 | 2.72E+00 | 8.70E-01 | no | Gm49469 | upstream_gene_variant | MODIFIER |
| 15 | 87100502 | AX-97795426 | 0 | C | T | 0.132 | 2.63E-02 | 1.57E-01 | 2.65E+00 | 2.72E+00 | 8.70E-01 | no | Gm49469 | downstream_gene_variant | MODIFIER |

| | | | | | | | | | | | | | | | |
|---|---|---|---|---|---|---|---|---|---|---|---|---|---|---|---|
| 15 | 87100807 | AX-170401369 | 0 | T | G | 0.132 | 2.63E-02 | 1.57E-01 | 2.65E+00 | 2.72E+00 | 8.70E-01 | no | Gm49469 | downstream_gene_variant | MODIFIER |
| 15 | 87111731 | AX-170330866 | 0 | C | T | 0.132 | 2.63E-02 | 1.57E-01 | 2.65E+00 | 2.72E+00 | 8.70E-01 | no | | intergenic_variant | MODIFIER |
| 15 | 87111769 | AX-170383728 | 0 | C | G | 0.132 | 2.63E-02 | 1.57E-01 | 2.65E+00 | 2.72E+00 | 8.70E-01 | no | | intergenic_variant | MODIFIER |
| 15 | 87125012 | AX-97830045 | 0 | C | T | 0.132 | 2.63E-02 | 1.57E-01 | 2.65E+00 | 2.72E+00 | 8.70E-01 | no | | intergenic_variant | MODIFIER |
| 15 | 87171477 | AX-97855327 | 0 | C | T | 0.132 | 2.63E-02 | 1.57E-01 | 2.65E+00 | 2.72E+00 | 8.70E-01 | no | n-R5s42 | downstream_gene_variant | MODIFIER |
| 15 | 87171527 | AX-170423826 | 0 | C | T | 0.132 | 2.63E-02 | 1.57E-01 | 2.65E+00 | 2.72E+00 | 8.70E-01 | no | n-R5s42 | downstream_gene_variant | MODIFIER |
| 15 | 87172603 | AX-170354550 | 0 | G | A | 0.132 | 2.63E-02 | 1.57E-01 | 2.65E+00 | 2.72E+00 | 8.70E-01 | no | n-R5s42 | downstream_gene_variant | MODIFIER |
| 15 | 87255453 | AX-97789862 | 0 | G | A | 0.132 | 2.63E-02 | 1.57E-01 | 2.65E+00 | 2.72E+00 | 8.70E-01 | no | A930027H12Rik | downstream_gene_variant | MODIFIER |
| 15 | 87282853 | AX-170099932 | 0 | T | C | 0.132 | 2.63E-02 | 1.57E-01 | 2.65E+00 | 2.72E+00 | 8.70E-01 | no | | intergenic_variant | MODIFIER |
| 15 | 87313057 | AX-170303519 | 0 | A | C | 0.132 | 2.63E-02 | 1.57E-01 | 2.65E+00 | 2.72E+00 | 8.70E-01 | no | | intergenic_variant | MODIFIER |
| 15 | 87313786 | AX-170179699 | 0 | G | C | 0.132 | 2.63E-02 | 1.57E-01 | 2.65E+00 | 2.72E+00 | 8.70E-01 | no | | intergenic_variant | MODIFIER |
| 15 | 87316322 | AX-170365424 | 0 | C | T | 0.132 | 2.63E-02 | 1.57E-01 | 2.65E+00 | 2.72E+00 | 8.70E-01 | no | | intergenic_variant | MODIFIER |
| 15 | 87316720 | AX-170294138 | 0 | C | T | 0.132 | 2.63E-02 | 1.57E-01 | 2.65E+00 | 2.72E+00 | 8.70E-01 | no | | intergenic_variant | MODIFIER |
| 15 | 87319304 | AX-170012204 | 0 | T | C | 0.132 | 2.63E-02 | 1.57E-01 | 2.65E+00 | 2.72E+00 | 8.70E-01 | no | | intergenic_variant | MODIFIER |
| 15 | 87319411 | AX-170219771 | 0 | C | T | 0.132 | 2.63E-02 | 1.57E-01 | 2.65E+00 | 2.72E+00 | 8.70E-01 | no | | intergenic_variant | MODIFIER |
| 15 | 87321392 | AX-170360118 | 0 | A | C | 0.132 | 2.63E-02 | 1.57E-01 | 2.65E+00 | 2.72E+00 | 8.70E-01 | no | | regulatory_region_variant | MODIFIER |
| 15 | 87327094 | AX-170133379 | 0 | C | T | 0.132 | 2.63E-02 | 1.57E-01 | 2.65E+00 | 2.72E+00 | 8.70E-01 | no | | intergenic_variant | MODIFIER |
| 15 | 87328586 | AX-170012198 | 0 | C | T | 0.132 | 2.63E-02 | 1.57E-01 | 2.65E+00 | 2.72E+00 | 8.70E-01 | no | | intergenic_variant | MODIFIER |
| 15 | 87336507 | AX-170328091 | 0 | T | C | 0.132 | 2.63E-02 | 1.57E-01 | 2.65E+00 | 2.72E+00 | 8.70E-01 | no | | intergenic_variant | MODIFIER |
| 15 | 87336550 | AX-170143624 | 0 | A | T | 0.132 | 2.63E-02 | 1.57E-01 | 2.65E+00 | 2.72E+00 | 8.70E-01 | no | | intergenic_variant | MODIFIER |
| 15 | 87350521 | AX-170012200 | 0 | T | C | 0.132 | 2.63E-02 | 1.57E-01 | 2.65E+00 | 2.72E+00 | 8.70E-01 | no | | regulatory_region_variant | MODIFIER |
| 15 | 87350568 | AX-97845895 | 0 | T | G | 0.132 | 2.63E-02 | 1.57E-01 | 2.65E+00 | 2.72E+00 | 8.70E-01 | no | | regulatory_region_variant | MODIFIER |
| 15 | 87350706 | AX-170240844 | 0 | C | A | 0.132 | 2.63E-02 | 1.57E-01 | 2.65E+00 | 2.72E+00 | 8.70E-01 | no | | intergenic_variant | MODIFIER |
| 15 | 87359914 | AX-170012202 | 0 | A | G | 0.132 | 2.63E-02 | 1.57E-01 | 2.65E+00 | 2.72E+00 | 8.70E-01 | no | | intergenic_variant | MODIFIER |
| 15 | 87359960 | AX-170502449 | 0 | T | C | 0.132 | 2.63E-02 | 1.57E-01 | 2.65E+00 | 2.72E+00 | 8.70E-01 | no | | intergenic_variant | MODIFIER |
| 15 | 87362337 | AX-97785655 | 0 | A | C | 0.132 | 2.63E-02 | 1.57E-01 | 2.65E+00 | 2.72E+00 | 8.70E-01 | no | | intergenic_variant | MODIFIER |
| 15 | 87383602 | AX-170468595 | 0 | A | G | 0.132 | 2.63E-02 | 1.57E-01 | 2.65E+00 | 2.72E+00 | 8.70E-01 | no | | intergenic_variant | MODIFIER |
| 15 | 87385474 | AX-170391547 | 0 | G | A | 0.132 | 2.63E-02 | 1.57E-01 | 2.65E+00 | 2.72E+00 | 8.70E-01 | no | | intergenic_variant | MODIFIER |
| 15 | 87386029 | AX-170056215 | 0 | G | A | 0.132 | 2.63E-02 | 1.57E-01 | 2.65E+00 | 2.72E+00 | 8.70E-01 | no | | intergenic_variant | MODIFIER |
| 15 | 87386541 | AX-170212049 | 0 | A | G | 0.132 | 2.63E-02 | 1.57E-01 | 2.65E+00 | 2.72E+00 | 8.70E-01 | no | | regulatory_region_variant | MODIFIER |
| 15 | 87386803 | AX-170150900 | 0 | G | A | 0.132 | 2.63E-02 | 1.57E-01 | 2.65E+00 | 2.72E+00 | 8.70E-01 | no | | regulatory_region_variant | MODIFIER |
| 15 | 87405696 | AX-170180638 | 0 | C | T | 0.132 | 2.63E-02 | 1.57E-01 | 2.65E+00 | 2.72E+00 | 8.70E-01 | no | | intergenic_variant | MODIFIER |
| 15 | 87407471 | AX-170447414 | 0 | G | A | 0.132 | 2.63E-02 | 1.57E-01 | 2.65E+00 | 2.72E+00 | 8.70E-01 | no | | intergenic_variant | MODIFIER |
| 15 | 87407959 | AX-168416334 | 0 | G | A | 0.132 | 2.63E-02 | 1.57E-01 | 2.65E+00 | 2.72E+00 | 8.70E-01 | no | | intergenic_variant | MODIFIER |
| 15 | 87414214 | AX-170341176 | 0 | A | G | 0.132 | 2.63E-02 | 1.57E-01 | 2.65E+00 | 2.72E+00 | 8.70E-01 | no | | intergenic_variant | MODIFIER |
| 15 | 87415110 | AX-97804498 | 0 | T | C | 0.132 | 2.63E-02 | 1.57E-01 | 2.65E+00 | 2.72E+00 | 8.70E-01 | no | | intergenic_variant | MODIFIER |
| 15 | 87426308 | AX-170399064 | 0 | A | G | 0.132 | 2.63E-02 | 1.57E-01 | 2.65E+00 | 2.72E+00 | 8.70E-01 | no | | intergenic_variant | MODIFIER |
| 15 | 87430499 | AX-170460209 | 0 | A | G | 0.132 | 2.63E-02 | 1.57E-01 | 2.65E+00 | 2.72E+00 | 8.70E-01 | no | | intergenic_variant | MODIFIER |
| 15 | 87454260 | AX-97823767 | 0 | T | G | 0.132 | 2.63E-02 | 1.57E-01 | 2.65E+00 | 2.72E+00 | 8.70E-01 | no | | intergenic_variant | MODIFIER |
| 15 | 87455002 | AX-170056219 | 0 | A | G | 0.132 | 2.63E-02 | 1.57E-01 | 2.65E+00 | 2.72E+00 | 8.70E-01 | no | | intergenic_variant | MODIFIER |
| 15 | 87464720 | AX-170056216 | 0 | C | A | 0.132 | 2.63E-02 | 1.57E-01 | 2.65E+00 | 2.72E+00 | 8.70E-01 | no | | intergenic_variant | MODIFIER |
| 15 | 87466177 | AX-170227486 | 0 | C | T | 0.132 | 2.63E-02 | 1.57E-01 | 2.65E+00 | 2.72E+00 | 8.70E-01 | no | | intergenic_variant | MODIFIER |
| 15 | 87470400 | AX-170349424 | 0 | C | G | 0.132 | 2.63E-02 | 1.57E-01 | 2.65E+00 | 2.72E+00 | 8.70E-01 | no | | intergenic_variant | MODIFIER |
| 15 | 87470586 | AX-170365944 | 0 | C | T | 0.132 | 2.63E-02 | 1.57E-01 | 2.65E+00 | 2.72E+00 | 8.70E-01 | no | | intergenic_variant | MODIFIER |
| 15 | 87472053 | AX-170260187 | 0 | T | C | 0.132 | 2.63E-02 | 1.57E-01 | 2.65E+00 | 2.72E+00 | 8.70E-01 | no | | intergenic_variant | MODIFIER |
| 15 | 87475831 | AX-97802343 | 0 | G | T | 0.132 | 2.63E-02 | 1.57E-01 | 2.65E+00 | 2.72E+00 | 8.70E-01 | no | | intergenic_variant | MODIFIER |
| 15 | 87494261 | AX-170376996 | 0 | T | C | 0.132 | 2.63E-02 | 1.57E-01 | 2.65E+00 | 2.72E+00 | 8.70E-01 | no | | intergenic_variant | MODIFIER |
| 15 | 87496687 | AX-170466178 | 0 | C | T | 0.132 | 2.63E-02 | 1.57E-01 | 2.65E+00 | 2.72E+00 | 8.70E-01 | no | | intergenic_variant | MODIFIER |
| 15 | 87497464 | AX-170528462 | 0 | C | T | 0.132 | 2.63E-02 | 1.57E-01 | 2.65E+00 | 2.72E+00 | 8.70E-01 | no | | intergenic_variant | MODIFIER |
| 15 | 87497696 | AX-170399341 | 0 | A | G | 0.132 | 2.63E-02 | 1.57E-01 | 2.65E+00 | 2.72E+00 | 8.70E-01 | no | | intergenic_variant | MODIFIER |
| 15 | 87499463 | AX-170314587 | 0 | C | A | 0.132 | 2.63E-02 | 1.57E-01 | 2.65E+00 | 2.72E+00 | 8.70E-01 | no | | intergenic_variant | MODIFIER |
| 15 | 87500044 | AX-170515100 | 0 | G | A | 0.132 | 2.63E-02 | 1.57E-01 | 2.65E+00 | 2.72E+00 | 8.70E-01 | no | | intergenic_variant | MODIFIER |
| 15 | 87500679 | AX-170299474 | 0 | C | A | 0.132 | 2.63E-02 | 1.57E-01 | 2.65E+00 | 2.72E+00 | 8.70E-01 | no | | intergenic_variant | MODIFIER |
| 15 | 87501174 | AX-170148396 | 0 | A | G | 0.132 | 2.63E-02 | 1.57E-01 | 2.65E+00 | 2.72E+00 | 8.70E-01 | no | | intergenic_variant | MODIFIER |

| | | | | | | | | | | | | | | | |
|---|---|---|---|---|---|---|---|---|---|---|---|---|---|---|---|
| 15 | 87503909 | AX-97815321 | 0 | G | A | 0.132 | 2.63E-02 | 1.57E-01 | 2.65E+00 | 2.72E+00 | 8.70E-01 | no | | intergenic_variant | MODIFIER |
| 15 | 87505780 | AX-170208755 | 0 | T | C | 0.132 | 2.63E-02 | 1.57E-01 | 2.65E+00 | 2.72E+00 | 8.70E-01 | no | | intergenic_variant | MODIFIER |
| 15 | 87509035 | AX-170367084 | 0 | C | T | 0.132 | 2.63E-02 | 1.57E-01 | 2.65E+00 | 2.72E+00 | 8.70E-01 | no | | intergenic_variant | MODIFIER |
| 15 | 87509059 | AX-170130518 | 0 | A | T | 0.132 | 2.63E-02 | 1.57E-01 | 2.65E+00 | 2.72E+00 | 8.70E-01 | no | | intergenic_variant | MODIFIER |
| 15 | 87512068 | AX-97839116 | 0 | T | C | 0.132 | 2.63E-02 | 1.57E-01 | 2.65E+00 | 2.72E+00 | 8.70E-01 | no | | intergenic_variant | MODIFIER |
| 15 | 87512785 | AX-170500106 | 0 | C | T | 0.132 | 2.63E-02 | 1.57E-01 | 2.65E+00 | 2.72E+00 | 8.70E-01 | no | | intergenic_variant | MODIFIER |
| 15 | 87627359 | AX-97801165 | 0 | T | C | 0.132 | 2.63E-02 | 1.57E-01 | 2.65E+00 | 2.72E+00 | 8.70E-01 | no | Fam19a5 | intron_variant | MODIFIER |
| 15 | 87657255 | AX-97800960 | 0 | T | C | 0.132 | 2.63E-02 | 1.57E-01 | 2.65E+00 | 2.72E+00 | 8.70E-01 | no | Fam19a5 | intron_variant | MODIFIER |
| 15 | 87672134 | AX-97812202 | 0 | A | G | 0.132 | 2.63E-02 | 1.57E-01 | 2.65E+00 | 2.72E+00 | 8.70E-01 | no | Fam19a5 | intron_variant | MODIFIER |
| 15 | 87672846 | AX-170503089 | 0 | A | C | 0.132 | 2.63E-02 | 1.57E-01 | 2.65E+00 | 2.72E+00 | 8.70E-01 | no | Fam19a5 | intron_variant | MODIFIER |
| 15 | 87674648 | AX-170392006 | 0 | T | C | 0.132 | 2.63E-02 | 1.57E-01 | 2.65E+00 | 2.72E+00 | 8.70E-01 | no | Fam19a5 | intron_variant | MODIFIER |
| 15 | 87678526 | AX-170502079 | 0 | C | T | 0.132 | 2.63E-02 | 1.57E-01 | 2.65E+00 | 2.72E+00 | 8.70E-01 | no | Fam19a5 | intron_variant | MODIFIER |
| 15 | 87704846 | AX-170386443 | 0 | A | G | 0.132 | 2.63E-02 | 1.57E-01 | 2.65E+00 | 2.72E+00 | 8.70E-01 | no | Fam19a5 | intron_variant | MODIFIER |
| 15 | 87819907 | AX-97820586 | 0 | T | C | 0.132 | 2.63E-02 | 1.57E-01 | 2.65E+00 | 2.72E+00 | 8.70E-01 | no | | intergenic_variant | MODIFIER |
| 15 | 87863033 | AX-170273147 | 0 | T | C | 0.132 | 2.63E-02 | 1.57E-01 | 2.65E+00 | 2.72E+00 | 8.70E-01 | no | | intergenic_variant | MODIFIER |
| 15 | 87879971 | AX-97830687 | 0 | A | G | 0.132 | 2.63E-02 | 1.57E-01 | 2.65E+00 | 2.72E+00 | 8.70E-01 | no | | intergenic_variant | MODIFIER |
| 15 | 87886967 | AX-170112885 | 0 | T | A | 0.132 | 2.63E-02 | 1.57E-01 | 2.65E+00 | 2.72E+00 | 8.70E-01 | no | | intergenic_variant | MODIFIER |
| 15 | 87887878 | AX-170306677 | 0 | A | C | 0.132 | 2.63E-02 | 1.57E-01 | 2.65E+00 | 2.72E+00 | 8.70E-01 | no | | intergenic_variant | MODIFIER |
| 15 | 87887921 | AX-170172385 | 0 | C | T | 0.132 | 2.63E-02 | 1.57E-01 | 2.65E+00 | 2.72E+00 | 8.70E-01 | no | | intergenic_variant | MODIFIER |
| 15 | 87896587 | AX-97830713 | 0 | C | T | 0.132 | 2.63E-02 | 1.57E-01 | 2.65E+00 | 2.72E+00 | 8.70E-01 | no | | intergenic_variant | MODIFIER |
| 15 | 87904747 | AX-170252258 | 0 | T | C | 0.132 | 2.63E-02 | 1.57E-01 | 2.65E+00 | 2.72E+00 | 8.70E-01 | no | | intergenic_variant | MODIFIER |
| 15 | 87988948 | AX-97812201 | 0 | A | G | 0.132 | 2.63E-02 | 1.57E-01 | 2.65E+00 | 2.72E+00 | 8.70E-01 | no | Gm49472 | downstream_gene_variant | MODIFIER |
| 15 | 88109983 | AX-170183292 | 0 | C | T | 0.132 | 2.63E-02 | 1.57E-01 | 2.65E+00 | 2.72E+00 | 8.70E-01 | no | | intergenic_variant | MODIFIER |
| 15 | 88112068 | AX-170262060 | 0 | T | C | 0.132 | 2.63E-02 | 1.57E-01 | 2.65E+00 | 2.72E+00 | 8.70E-01 | no | | intergenic_variant | MODIFIER |
| 15 | 88112239 | AX-170416323 | 0 | G | A | 0.132 | 2.63E-02 | 1.57E-01 | 2.65E+00 | 2.72E+00 | 8.70E-01 | no | | intergenic_variant | MODIFIER |
| 15 | 88113520 | AX-170126655 | 0 | T | G | 0.132 | 2.63E-02 | 1.57E-01 | 2.65E+00 | 2.72E+00 | 8.70E-01 | no | | intergenic_variant | MODIFIER |
| 15 | 88117279 | AX-170355420 | 0 | T | G | 0.132 | 2.63E-02 | 1.57E-01 | 2.65E+00 | 2.72E+00 | 8.70E-01 | no | | intergenic_variant | MODIFIER |
| 15 | 88124954 | AX-170364067 | 0 | A | G | 0.132 | 2.63E-02 | 1.57E-01 | 2.65E+00 | 2.72E+00 | 8.70E-01 | no | | intergenic_variant | MODIFIER |
| 15 | 88128831 | AX-170274929 | 0 | G | A | 0.132 | 2.63E-02 | 1.57E-01 | 2.65E+00 | 2.72E+00 | 8.70E-01 | no | | intergenic_variant | MODIFIER |
| 15 | 88134065 | AX-170230146 | 0 | G | A | 0.132 | 2.63E-02 | 1.57E-01 | 2.65E+00 | 2.72E+00 | 8.70E-01 | no | | intergenic_variant | MODIFIER |
| 15 | 88136580 | AX-170165143 | 0 | T | C | 0.132 | 2.63E-02 | 1.57E-01 | 2.65E+00 | 2.72E+00 | 8.70E-01 | no | | intergenic_variant | MODIFIER |
| 15 | 88187560 | AX-170056234 | 0 | T | C | 0.132 | 2.63E-02 | 1.57E-01 | 2.65E+00 | 2.72E+00 | 8.70E-01 | no | | intergenic_variant | MODIFIER |
| 15 | 88204023 | AX-170424792 | 0 | A | G | 0.132 | 2.63E-02 | 1.57E-01 | 2.65E+00 | 2.72E+00 | 8.70E-01 | no | | intergenic_variant | MODIFIER |
| 15 | 88219017 | AX-170444092 | 0 | G | T | 0.132 | 2.63E-02 | 1.57E-01 | 2.65E+00 | 2.72E+00 | 8.70E-01 | no | | intergenic_variant | MODIFIER |
| 15 | 88228875 | AX-170510011 | 0 | A | G | 0.132 | 2.63E-02 | 1.57E-01 | 2.65E+00 | 2.72E+00 | 8.70E-01 | no | | intergenic_variant | MODIFIER |
| 15 | 88232706 | AX-170261395 | 0 | T | C | 0.132 | 2.63E-02 | 1.57E-01 | 2.65E+00 | 2.72E+00 | 8.70E-01 | no | | intergenic_variant | MODIFIER |
| 15 | 88245734 | AX-170436760 | 0 | G | T | 0.132 | 2.63E-02 | 1.57E-01 | 2.65E+00 | 2.72E+00 | 8.70E-01 | no | | intergenic_variant | MODIFIER |
| 15 | 88250846 | AX-170056236 | 0 | T | C | 0.132 | 2.63E-02 | 1.57E-01 | 2.65E+00 | 2.72E+00 | 8.70E-01 | no | | intergenic_variant | MODIFIER |
| 15 | 88305145 | AX-168412619 | 0 | T | C | 0.132 | 2.63E-02 | 1.57E-01 | 2.65E+00 | 2.72E+00 | 8.70E-01 | no | 4930445N06Rik | intron_variant,non_coding_transcript_variant | MODIFIER |
| 15 | 88416742 | AX-170175353 | 0 | C | T | 0.132 | 2.63E-02 | 1.57E-01 | 2.65E+00 | 2.72E+00 | 8.70E-01 | no | | intergenic_variant | MODIFIER |
| 15 | 88417691 | AX-168419964 | 0 | C | T | 0.132 | 2.63E-02 | 1.57E-01 | 2.65E+00 | 2.72E+00 | 8.70E-01 | no | | intergenic_variant | MODIFIER |
| 15 | 88469498 | AX-170379430 | 0 | T | G | 0.132 | 2.63E-02 | 1.57E-01 | 2.65E+00 | 2.72E+00 | 8.70E-01 | no | | intergenic_variant | MODIFIER |
| 15 | 88470832 | AX-97799716 | 0 | G | A | 0.132 | 2.63E-02 | 1.57E-01 | 2.65E+00 | 2.72E+00 | 8.70E-01 | no | | intergenic_variant | MODIFIER |
| 15 | 88489222 | AX-97844006 | 0 | T | G | 0.132 | 2.63E-02 | 1.57E-01 | 2.65E+00 | 2.72E+00 | 8.70E-01 | no | | intergenic_variant | MODIFIER |
| 15 | 89810721 | AX-170398326 | 0 | C | T | 0.132 | 2.63E-02 | 1.57E-01 | 2.65E+00 | 2.72E+00 | 8.70E-01 | no | Syt10 | intron_variant | MODIFIER |
| 15 | 89824648 | AX-170182178 | 0 | C | T | 0.132 | 2.63E-02 | 1.57E-01 | 2.65E+00 | 2.72E+00 | 8.70E-01 | no | Syt10 | intron_variant | MODIFIER |
| 15 | 89841154 | AX-170012233 | 0 | G | T | 0.132 | 2.63E-02 | 1.57E-01 | 2.65E+00 | 2.72E+00 | 8.70E-01 | no | Syt10 | intron_variant | MODIFIER |
| 15 | 90786915 | AX-170308504 | 0 | T | A | 0.132 | 2.63E-02 | 1.57E-01 | 2.65E+00 | 2.72E+00 | 8.70E-01 | no | | intergenic_variant | MODIFIER |
| 15 | 90788481 | AX-97837336 | 0 | G | A | 0.132 | 2.63E-02 | 1.57E-01 | 2.65E+00 | 2.72E+00 | 8.70E-01 | no | | intergenic_variant | MODIFIER |
| 15 | 90788857 | AX-170163534 | 0 | G | C | 0.132 | 2.63E-02 | 1.57E-01 | 2.65E+00 | 2.72E+00 | 8.70E-01 | no | | intergenic_variant | MODIFIER |
| 15 | 90791234 | AX-170224439 | 0 | C | T | 0.132 | 2.63E-02 | 1.57E-01 | 2.65E+00 | 2.72E+00 | 8.70E-01 | no | | intergenic_variant | MODIFIER |
| 15 | 90814306 | AX-170479445 | 0 | A | G | 0.132 | 2.63E-02 | 1.57E-01 | 2.65E+00 | 2.72E+00 | 8.70E-01 | no | | intergenic_variant | MODIFIER |
| 15 | 90815064 | AX-170525324 | 0 | G | A | 0.132 | 2.63E-02 | 1.57E-01 | 2.65E+00 | 2.72E+00 | 8.70E-01 | no | | intergenic_variant | MODIFIER |
| 15 | 90816617 | AX-170093335 | 0 | C | T | 0.132 | 2.63E-02 | 1.57E-01 | 2.65E+00 | 2.72E+00 | 8.70E-01 | no | | intergenic_variant | MODIFIER |

| | | | | | | | | | | | | | | | |
|---|---|---|---|---|---|---|---|---|---|---|---|---|---|---|---|
| 15 | 90817641 | AX-170410701 | 0 | A | G | 0.132 | 2.63E-02 | 1.57E-01 | 2.65E+00 | 2.72E+00 | 8.70E-01 | no | | intergenic_variant | MODIFIER |
| 15 | 90847004 | AX-97799616 | 0 | A | C | 0.132 | 2.63E-02 | 1.57E-01 | 2.65E+00 | 2.72E+00 | 8.70E-01 | no | | intergenic_variant | MODIFIER |
| 15 | 90847058 | AX-170247859 | 0 | G | A | 0.132 | 2.63E-02 | 1.57E-01 | 2.65E+00 | 2.72E+00 | 8.70E-01 | no | | intergenic_variant | MODIFIER |
| 15 | 90887004 | AX-170263477 | 0 | G | A | 0.132 | 2.63E-02 | 1.57E-01 | 2.65E+00 | 2.72E+00 | 8.70E-01 | no | Tcea1-ps1 | downstream_gene_variant | MODIFIER |
| 15 | 90888106 | AX-170130477 | 0 | A | G | 0.132 | 2.63E-02 | 1.57E-01 | 2.65E+00 | 2.72E+00 | 8.70E-01 | no | Tcea1-ps1 | downstream_gene_variant | MODIFIER |
| 15 | 90888262 | AX-170510455 | 0 | A | G | 0.132 | 2.63E-02 | 1.57E-01 | 2.65E+00 | 2.72E+00 | 8.70E-01 | no | Tcea1-ps1 | downstream_gene_variant | MODIFIER |
| 15 | 90898452 | AX-97831185 | 0 | C | A | 0.132 | 2.63E-02 | 1.57E-01 | 2.65E+00 | 2.72E+00 | 8.70E-01 | no | Spn-ps | non_coding_transcript_exon_variant | MODIFIER |
| 15 | 90900590 | AX-170127665 | 0 | A | G | 0.132 | 2.63E-02 | 1.57E-01 | 2.65E+00 | 2.72E+00 | 8.70E-01 | no | Spn-ps | upstream_gene_variant | MODIFIER |
| 15 | 90949749 | AX-97832081 | 0 | C | A | 0.132 | 2.63E-02 | 1.57E-01 | 2.65E+00 | 2.72E+00 | 8.70E-01 | no | Kif21a | intron_variant | MODIFIER |
| 15 | 90956254 | AX-170133780 | 0 | T | C | 0.132 | 2.63E-02 | 1.57E-01 | 2.65E+00 | 2.72E+00 | 8.70E-01 | no | Kif21a | intron_variant | MODIFIER |
| 15 | 90957311 | AX-170012249 | 0 | A | G | 0.132 | 2.63E-02 | 1.57E-01 | 2.65E+00 | 2.72E+00 | 8.70E-01 | no | Kif21a | intron_variant | MODIFIER |
| 15 | 90959639 | AX-170140968 | 0 | A | G | 0.132 | 2.63E-02 | 1.57E-01 | 2.65E+00 | 2.72E+00 | 8.70E-01 | no | Kif21a | intron_variant | MODIFIER |
| 15 | 90992503 | AX-170056290 | 0 | A | G | 0.132 | 2.63E-02 | 1.57E-01 | 2.65E+00 | 2.72E+00 | 8.70E-01 | no | Kif21a | intron_variant | MODIFIER |
| 15 | 90997851 | AX-97803241 | 0 | A | G | 0.132 | 2.63E-02 | 1.57E-01 | 2.65E+00 | 2.72E+00 | 8.70E-01 | no | Kif21a | synonymous_variant | LOW |
| 15 | 91057841 | AX-97815567 | 0 | A | G | 0.132 | 2.63E-02 | 1.57E-01 | 2.65E+00 | 2.72E+00 | 8.70E-01 | no | | regulatory_region_variant | MODIFIER |
| 15 | 91061721 | AX-170404322 | 0 | C | T | 0.132 | 2.63E-02 | 1.57E-01 | 2.65E+00 | 2.72E+00 | 8.70E-01 | no | | intergenic_variant | MODIFIER |
| 15 | 91061752 | AX-170468523 | 0 | T | C | 0.132 | 2.63E-02 | 1.57E-01 | 2.65E+00 | 2.72E+00 | 8.70E-01 | no | | intergenic_variant | MODIFIER |
| 15 | 91183082 | AX-170056295 | 0 | G | C | 0.132 | 2.63E-02 | 1.57E-01 | 2.65E+00 | 2.72E+00 | 8.70E-01 | no | Abcd2 | intron_variant | MODIFIER |
| 15 | 91208287 | AX-170276158 | 0 | C | G | 0.132 | 2.63E-02 | 1.57E-01 | 2.65E+00 | 2.72E+00 | 8.70E-01 | no | CN725425 | intron_variant | MODIFIER |
| 15 | 91314049 | AX-170359558 | 0 | A | C | 0.132 | 2.63E-02 | 1.57E-01 | 2.65E+00 | 2.72E+00 | 8.70E-01 | no | Slc2a13 | intron_variant | MODIFIER |
| 15 | 91457359 | AX-170473137 | 0 | G | A | 0.132 | 2.63E-02 | 1.57E-01 | 2.65E+00 | 2.72E+00 | 8.70E-01 | no | Slc2a13 | intron_variant | MODIFIER |
| 15 | 91463379 | AX-170514915 | 0 | C | T | 0.132 | 2.63E-02 | 1.57E-01 | 2.65E+00 | 2.72E+00 | 8.70E-01 | no | Slc2a13 | intron_variant | MODIFIER |
| 15 | 91464339 | AX-170482582 | 0 | G | A | 0.132 | 2.63E-02 | 1.57E-01 | 2.65E+00 | 2.72E+00 | 8.70E-01 | no | Slc2a13 | intron_variant | MODIFIER |
| 15 | 91482067 | AX-170264546 | 0 | A | G | 0.132 | 2.63E-02 | 1.57E-01 | 2.65E+00 | 2.72E+00 | 8.70E-01 | no | Slc2a13 | intron_variant | MODIFIER |
| 15 | 91482313 | AX-170187436 | 0 | T | A | 0.132 | 2.63E-02 | 1.57E-01 | 2.65E+00 | 2.72E+00 | 8.70E-01 | no | Slc2a13 | intron_variant | MODIFIER |
| 15 | 91491454 | AX-170218449 | 0 | T | C | 0.132 | 2.63E-02 | 1.57E-01 | 2.65E+00 | 2.72E+00 | 8.70E-01 | no | Slc2a13 | intron_variant | MODIFIER |
| 15 | 91493310 | AX-170317739 | 0 | T | C | 0.132 | 2.63E-02 | 1.57E-01 | 2.65E+00 | 2.72E+00 | 8.70E-01 | no | Slc2a13 | intron_variant | MODIFIER |
| 15 | 91493500 | AX-97799114 | 0 | G | A | 0.132 | 2.63E-02 | 1.57E-01 | 2.65E+00 | 2.72E+00 | 8.70E-01 | no | Slc2a13 | intron_variant | MODIFIER |
| 15 | 91494559 | AX-170134647 | 0 | C | A | 0.132 | 2.63E-02 | 1.57E-01 | 2.65E+00 | 2.72E+00 | 8.70E-01 | no | Slc2a13 | intron_variant | MODIFIER |
| 15 | 91497571 | AX-170453654 | 0 | G | A | 0.132 | 2.63E-02 | 1.57E-01 | 2.65E+00 | 2.72E+00 | 8.70E-01 | no | Slc2a13 | synonymous_variant | LOW |
| 15 | 91497772 | AX-170401529 | 0 | C | T | 0.132 | 2.63E-02 | 1.57E-01 | 2.65E+00 | 2.72E+00 | 8.70E-01 | no | Slc2a13 | intron_variant | MODIFIER |
| 15 | 91518225 | AX-170278888 | 0 | G | C | 0.132 | 2.63E-02 | 1.57E-01 | 2.65E+00 | 2.72E+00 | 8.70E-01 | no | Slc2a13 | intron_variant | MODIFIER |
| 15 | 91529394 | AX-97782838 | 0 | T | G | 0.132 | 2.63E-02 | 1.57E-01 | 2.65E+00 | 2.72E+00 | 8.70E-01 | no | Slc2a13 | intron_variant | MODIFIER |
| 15 | 91530953 | AX-170251814 | 0 | A | G | 0.132 | 2.63E-02 | 1.57E-01 | 2.65E+00 | 2.72E+00 | 8.70E-01 | no | Slc2a13 | intron_variant | MODIFIER |
| 15 | 91531836 | AX-170365193 | 0 | A | G | 0.132 | 2.63E-02 | 1.57E-01 | 2.65E+00 | 2.72E+00 | 8.70E-01 | no | Slc2a13 | intron_variant | MODIFIER |
| 15 | 91825561 | AX-170442708 | 0 | A | G | 0.132 | 2.63E-02 | 1.57E-01 | 2.65E+00 | 2.72E+00 | 8.70E-01 | no | Gm15382 | downstream_gene_variant | MODIFIER |
| 15 | 91857524 | AX-170289657 | 0 | A | G | 0.132 | 2.63E-02 | 1.57E-01 | 2.65E+00 | 2.72E+00 | 8.70E-01 | no | Smgc | intron_variant | MODIFIER |
| 15 | 91943355 | AX-170056306 | 0 | C | T | 0.132 | 2.63E-02 | 1.57E-01 | 2.65E+00 | 2.72E+00 | 8.70E-01 | no | | intergenic_variant | MODIFIER |
| 15 | 91946971 | AX-170365949 | 0 | T | C | 0.132 | 2.63E-02 | 1.57E-01 | 2.65E+00 | 2.72E+00 | 8.70E-01 | no | | intergenic_variant | MODIFIER |
| 15 | 92050683 | AX-170403948 | 0 | G | C | 0.132 | 2.63E-02 | 1.57E-01 | 2.65E+00 | 2.72E+00 | 8.70E-01 | no | Cntn1 | upstream_gene_variant | MODIFIER |
| 15 | 92052789 | AX-170105944 | 0 | A | G | 0.132 | 2.63E-02 | 1.57E-01 | 2.65E+00 | 2.72E+00 | 8.70E-01 | no | Cntn1 | intron_variant,non_coding_transcript_variant | MODIFIER |
| 15 | 92052846 | AX-170198150 | 0 | C | T | 0.132 | 2.63E-02 | 1.57E-01 | 2.65E+00 | 2.72E+00 | 8.70E-01 | no | Cntn1 | intron_variant,non_coding_transcript_variant | MODIFIER |
| 15 | 92053343 | AX-170482699 | 0 | G | A | 0.132 | 2.63E-02 | 1.57E-01 | 2.65E+00 | 2.72E+00 | 8.70E-01 | no | Cntn1 | intron_variant,non_coding_transcript_variant | MODIFIER |
| 15 | 92057604 | AX-170514540 | 0 | C | T | 0.132 | 2.63E-02 | 1.57E-01 | 2.65E+00 | 2.72E+00 | 8.70E-01 | no | Cntn1 | intron_variant,non_coding_transcript_variant | MODIFIER |
| 15 | 92069459 | AX-170531665 | 0 | C | G | 0.132 | 2.63E-02 | 1.57E-01 | 2.65E+00 | 2.72E+00 | 8.70E-01 | no | Cntn1 | intron_variant,non_coding_transcript_variant | MODIFIER |
| 15 | 92095820 | AX-170471280 | 0 | G | A | 0.132 | 2.63E-02 | 1.57E-01 | 2.65E+00 | 2.72E+00 | 8.70E-01 | no | Cntn1 | intron_variant,non_coding_transcript_variant | MODIFIER |
| 15 | 92095933 | AX-170149908 | 0 | A | C | 0.132 | 2.63E-02 | 1.57E-01 | 2.65E+00 | 2.72E+00 | 8.70E-01 | no | Cntn1 | intron_variant,non_coding_transcript_variant | MODIFIER |
| 15 | 92132102 | AX-170359697 | 0 | A | T | 0.132 | 2.63E-02 | 1.57E-01 | 2.65E+00 | 2.72E+00 | 8.70E-01 | no | Cntn1 | intron_variant | MODIFIER |
| 15 | 92144301 | AX-97802722 | 0 | C | A | 0.132 | 2.63E-02 | 1.57E-01 | 2.65E+00 | 2.72E+00 | 8.70E-01 | no | Cntn1 | intron_variant | MODIFIER |
| 15 | 92153429 | AX-170400509 | 0 | T | A | 0.132 | 2.63E-02 | 1.57E-01 | 2.65E+00 | 2.72E+00 | 8.70E-01 | no | Cntn1 | intron_variant | MODIFIER |
| 15 | 92154264 | AX-170218940 | 0 | T | C | 0.132 | 2.63E-02 | 1.57E-01 | 2.65E+00 | 2.72E+00 | 8.70E-01 | no | Cntn1 | intron_variant | MODIFIER |
| 15 | 92154855 | AX-170056314 | 0 | G | A | 0.132 | 2.63E-02 | 1.57E-01 | 2.65E+00 | 2.72E+00 | 8.70E-01 | no | Cntn1 | intron_variant | MODIFIER |
| 15 | 92189655 | AX-170254728 | 0 | A | G | 0.132 | 2.63E-02 | 1.57E-01 | 2.65E+00 | 2.72E+00 | 8.70E-01 | no | Cntn1 | intron_variant | MODIFIER |
| 15 | 92190482 | AX-170149943 | 0 | T | C | 0.132 | 2.63E-02 | 1.57E-01 | 2.65E+00 | 2.72E+00 | 8.70E-01 | no | Cntn1 | intron_variant | MODIFIER |

| | | | | | | | | | | | | | | | |
|---|---|---|---|---|---|---|---|---|---|---|---|---|---|---|---|
| 15 | 92190993 | AX-170119092 | 0 | G | A | 0.132 | 2.63E-02 | 1.57E-01 | 2.65E+00 | 2.72E+00 | 8.70E-01 | no | Cntn1 | intron_variant | MODIFIER |
| 15 | 92197672 | AX-170407571 | 0 | C | T | 0.132 | 2.63E-02 | 1.57E-01 | 2.65E+00 | 2.72E+00 | 8.70E-01 | no | Cntn1 | intron_variant | MODIFIER |
| 15 | 92289176 | AX-170056317 | 0 | C | A | 0.132 | 2.63E-02 | 1.57E-01 | 2.65E+00 | 2.72E+00 | 8.70E-01 | no | Cntn1 | intron_variant | MODIFIER |
| 15 | 92290703 | AX-170513515 | 0 | T | C | 0.132 | 2.63E-02 | 1.57E-01 | 2.65E+00 | 2.72E+00 | 8.70E-01 | no | Cntn1 | intron_variant | MODIFIER |
| 15 | 92329104 | AX-170297412 | 0 | C | A | 0.132 | 2.63E-02 | 1.57E-01 | 2.65E+00 | 2.72E+00 | 8.70E-01 | no | Cntn1 | intron_variant | MODIFIER |
| 15 | 92518960 | AX-170146621 | 0 | A | G | 0.132 | 2.63E-02 | 1.57E-01 | 2.65E+00 | 2.72E+00 | 8.70E-01 | no | Pdzrn4 | intron_variant | MODIFIER |
| 15 | 92525292 | AX-170372589 | 0 | G | A | 0.132 | 2.63E-02 | 1.57E-01 | 2.65E+00 | 2.72E+00 | 8.70E-01 | no | Pdzrn4 | intron_variant | MODIFIER |
| 15 | 92530024 | AX-170380282 | 0 | C | T | 0.132 | 2.63E-02 | 1.57E-01 | 2.65E+00 | 2.72E+00 | 8.70E-01 | no | Pdzrn4 | intron_variant | MODIFIER |
| 15 | 92634520 | AX-170518398 | 0 | C | A | 0.132 | 2.63E-02 | 1.57E-01 | 2.65E+00 | 2.72E+00 | 8.70E-01 | no | Pdzrn4 | intron_variant | MODIFIER |
| 15 | 93902460 | AX-97821967 | 0 | A | G | 0.132 | 2.63E-02 | 1.57E-01 | 2.65E+00 | 2.72E+00 | 8.70E-01 | no | 9430014N10Rik | downstream_gene_variant | MODIFIER |
| 15 | 93903946 | AX-170525201 | 0 | A | C | 0.132 | 2.63E-02 | 1.57E-01 | 2.65E+00 | 2.72E+00 | 8.70E-01 | no | 9430014N10Rik | downstream_gene_variant | MODIFIER |
| 15 | 93904853 | AX-170139139 | 0 | T | C | 0.132 | 2.63E-02 | 1.57E-01 | 2.65E+00 | 2.72E+00 | 8.70E-01 | no | 9430014N10Rik | downstream_gene_variant | MODIFIER |
| 15 | 94215705 | AX-170186755 | 0 | A | G | 0.132 | 2.63E-02 | 1.57E-01 | 2.65E+00 | 2.72E+00 | 8.70E-01 | no | | intergenic_variant | MODIFIER |
| 15 | 94569179 | AX-170167202 | 0 | A | G | 0.132 | 2.63E-02 | 1.57E-01 | 2.65E+00 | 2.72E+00 | 8.70E-01 | no | Irak4 | downstream_gene_variant | MODIFIER |
| 15 | 94577807 | AX-170159250 | 0 | T | C | 0.132 | 2.63E-02 | 1.57E-01 | 2.65E+00 | 2.72E+00 | 8.70E-01 | no | Twf1 | downstream_gene_variant | MODIFIER |
| 15 | 94616045 | AX-170190479 | 0 | C | T | 0.132 | 2.63E-02 | 1.57E-01 | 2.65E+00 | 2.72E+00 | 8.70E-01 | no | | intergenic_variant | MODIFIER |
| 15 | 94616083 | AX-170056357 | 0 | T | C | 0.132 | 2.63E-02 | 1.57E-01 | 2.65E+00 | 2.72E+00 | 8.70E-01 | no | | intergenic_variant | MODIFIER |
| 15 | 94616804 | AX-170056358 | 0 | A | G | 0.132 | 2.63E-02 | 1.57E-01 | 2.65E+00 | 2.72E+00 | 8.70E-01 | no | | intergenic_variant | MODIFIER |
| 15 | 94616864 | AX-170056356 | 0 | C | A | 0.132 | 2.63E-02 | 1.57E-01 | 2.65E+00 | 2.72E+00 | 8.70E-01 | no | | intergenic_variant | MODIFIER |
| 15 | 94616995 | AX-170402379 | 0 | C | G | 0.132 | 2.63E-02 | 1.57E-01 | 2.65E+00 | 2.72E+00 | 8.70E-01 | no | | intergenic_variant | MODIFIER |
| 15 | 94618772 | AX-97835500 | 0 | T | C | 0.132 | 2.63E-02 | 1.57E-01 | 2.65E+00 | 2.72E+00 | 8.70E-01 | no | | intergenic_variant | MODIFIER |
| 15 | 94640063 | AX-170012303 | 0 | A | G | 0.132 | 2.63E-02 | 1.57E-01 | 2.65E+00 | 2.72E+00 | 8.70E-01 | no | Tmem117 | intron_variant | MODIFIER |
| 15 | 94651759 | AX-97851175 | 0 | G | A | 0.132 | 2.63E-02 | 1.57E-01 | 2.65E+00 | 2.72E+00 | 8.70E-01 | no | Tmem117 | intron_variant | MODIFIER |
| 15 | 94664982 | AX-170012307 | 0 | A | G | 0.132 | 2.63E-02 | 1.57E-01 | 2.65E+00 | 2.72E+00 | 8.70E-01 | no | Tmem117 | intron_variant | MODIFIER |
| 15 | 94671054 | AX-97830250 | 0 | T | C | 0.132 | 2.63E-02 | 1.57E-01 | 2.65E+00 | 2.72E+00 | 8.70E-01 | no | Tmem117 | intron_variant | MODIFIER |
| 15 | 94744244 | AX-170407874 | 0 | A | T | 0.132 | 2.63E-02 | 1.57E-01 | 2.65E+00 | 2.72E+00 | 8.70E-01 | no | Tmem117 | intron_variant | MODIFIER |
| 15 | 96306935 | AX-170238824 | 0 | G | A | 0.132 | 2.63E-02 | 1.57E-01 | 2.65E+00 | 2.72E+00 | 8.70E-01 | no | Arid2 | intron_variant | MODIFIER |
| 15 | 96333573 | AX-170123975 | 0 | T | G | 0.132 | 2.63E-02 | 1.57E-01 | 2.65E+00 | 2.72E+00 | 8.70E-01 | no | Arid2 | intron_variant | MODIFIER |
| 15 | 96333628 | AX-170333901 | 0 | T | C | 0.132 | 2.63E-02 | 1.57E-01 | 2.65E+00 | 2.72E+00 | 8.70E-01 | no | Arid2 | intron_variant | MODIFIER |
| 15 | 96500649 | AX-170318814 | 0 | T | C | 0.132 | 2.63E-02 | 1.57E-01 | 2.65E+00 | 2.72E+00 | 8.70E-01 | no | Gm41392 | downstream_gene_variant | MODIFIER |
| 15 | 96518068 | AX-170473661 | 0 | T | C | 0.132 | 2.63E-02 | 1.57E-01 | 2.65E+00 | 2.72E+00 | 8.70E-01 | no | | intergenic_variant | MODIFIER |
| 15 | 96533306 | AX-170420896 | 0 | T | C | 0.132 | 2.63E-02 | 1.57E-01 | 2.65E+00 | 2.72E+00 | 8.70E-01 | no | | intergenic_variant | MODIFIER |
| 15 | 96534031 | AX-170428067 | 0 | T | C | 0.132 | 2.63E-02 | 1.57E-01 | 2.65E+00 | 2.72E+00 | 8.70E-01 | no | | regulatory_region_variant | MODIFIER |
| 15 | 96548284 | AX-170101736 | 0 | A | G | 0.132 | 2.63E-02 | 1.57E-01 | 2.65E+00 | 2.72E+00 | 8.70E-01 | no | | regulatory_region_variant | MODIFIER |
| 15 | 96552018 | AX-170121394 | 0 | T | G | 0.132 | 2.63E-02 | 1.57E-01 | 2.65E+00 | 2.72E+00 | 8.70E-01 | no | | regulatory_region_variant | MODIFIER |
| 15 | 96565579 | AX-170312720 | 0 | T | C | 0.132 | 2.63E-02 | 1.57E-01 | 2.65E+00 | 2.72E+00 | 8.70E-01 | no | | intergenic_variant | MODIFIER |
| 15 | 96570526 | AX-170380141 | 0 | A | G | 0.132 | 2.63E-02 | 1.57E-01 | 2.65E+00 | 2.72E+00 | 8.70E-01 | no | Slc38a1 | downstream_gene_variant | MODIFIER |
| 15 | 96576353 | AX-170514584 | 0 | A | G | 0.132 | 2.63E-02 | 1.57E-01 | 2.65E+00 | 2.72E+00 | 8.70E-01 | no | Slc38a1 | 3_prime_UTR_variant | MODIFIER |
| 15 | 96601039 | AX-170461515 | 0 | T | G | 0.132 | 2.63E-02 | 1.57E-01 | 2.65E+00 | 2.72E+00 | 8.70E-01 | no | Slc38a1 | intron_variant | MODIFIER |
| 15 | 96605821 | AX-170509026 | 0 | A | G | 0.132 | 2.63E-02 | 1.57E-01 | 2.65E+00 | 2.72E+00 | 8.70E-01 | no | Slc38a1 | intron_variant | MODIFIER |
| 15 | 96605875 | AX-170416212 | 0 | A | G | 0.132 | 2.63E-02 | 1.57E-01 | 2.65E+00 | 2.72E+00 | 8.70E-01 | no | Slc38a1 | intron_variant | MODIFIER |
| 15 | 96626503 | AX-170524787 | 0 | T | C | 0.132 | 2.63E-02 | 1.57E-01 | 2.65E+00 | 2.72E+00 | 8.70E-01 | no | Slc38a1 | intron_variant | MODIFIER |
| 15 | 96638221 | AX-170247497 | 0 | G | A | 0.132 | 2.63E-02 | 1.57E-01 | 2.65E+00 | 2.72E+00 | 8.70E-01 | no | Slc38a1 | intron_variant | MODIFIER |
| 15 | 96638321 | AX-170354534 | 0 | A | G | 0.132 | 2.63E-02 | 1.57E-01 | 2.65E+00 | 2.72E+00 | 8.70E-01 | no | Slc38a1 | intron_variant | MODIFIER |
| 15 | 96676595 | AX-170012334 | 0 | T | A | 0.132 | 2.63E-02 | 1.57E-01 | 2.65E+00 | 2.72E+00 | 8.70E-01 | no | Gm22045 | downstream_gene_variant | MODIFIER |
| 15 | 96682732 | AX-170487964 | 0 | A | C | 0.132 | 2.63E-02 | 1.57E-01 | 2.65E+00 | 2.72E+00 | 8.70E-01 | no | Slc38a2 | downstream_gene_variant | MODIFIER |
| 15 | 96704932 | AX-170439180 | 0 | A | G | 0.132 | 2.63E-02 | 1.57E-01 | 2.65E+00 | 2.72E+00 | 8.70E-01 | no | Gm38144 | downstream_gene_variant | MODIFIER |
| 15 | 96707866 | AX-170183214 | 0 | T | C | 0.132 | 2.63E-02 | 1.57E-01 | 2.65E+00 | 2.72E+00 | 8.70E-01 | no | | regulatory_region_variant | MODIFIER |
| 15 | 96711804 | AX-170012332 | 0 | C | G | 0.132 | 2.63E-02 | 1.57E-01 | 2.65E+00 | 2.72E+00 | 8.70E-01 | no | | regulatory_region_variant | MODIFIER |
| 15 | 96742884 | AX-170012333 | 0 | C | G | 0.132 | 2.63E-02 | 1.57E-01 | 2.65E+00 | 2.72E+00 | 8.70E-01 | no | | regulatory_region_variant | MODIFIER |
| 15 | 96742928 | AX-170012336 | 0 | G | A | 0.132 | 2.63E-02 | 1.57E-01 | 2.65E+00 | 2.72E+00 | 8.70E-01 | no | | regulatory_region_variant | MODIFIER |
| 15 | 96753416 | AX-170461163 | 0 | G | A | 0.132 | 2.63E-02 | 1.57E-01 | 2.65E+00 | 2.72E+00 | 8.70E-01 | no | | intergenic_variant | MODIFIER |
| 15 | 96794400 | AX-170229523 | 0 | A | G | 0.132 | 2.63E-02 | 1.57E-01 | 2.65E+00 | 2.72E+00 | 8.70E-01 | no | | regulatory_region_variant | MODIFIER |
| 15 | 96829680 | AX-170205011 | 0 | A | C | 0.132 | 2.63E-02 | 1.57E-01 | 2.65E+00 | 2.72E+00 | 8.70E-01 | no | | intergenic_variant | MODIFIER |

| | | | | | | | | | | | | | | | |
|---|---|---|---|---|---|---|---|---|---|---|---|---|---|---|---|
| 15 | 96861473 | AX-170476278 | 0 | T | C | 0.132 | 2.63E-02 | 1.57E-01 | 2.65E+00 | 2.72E+00 | 8.70E-01 | no | | intergenic_variant | MODIFIER |
| 15 | 97657322 | AX-170162796 | 0 | G | T | 0.132 | 2.63E-02 | 1.57E-01 | 2.65E+00 | 2.72E+00 | 8.70E-01 | no | | intergenic_variant | MODIFIER |
| 15 | 97657757 | AX-170211454 | 0 | C | T | 0.132 | 2.63E-02 | 1.57E-01 | 2.65E+00 | 2.72E+00 | 8.70E-01 | no | | regulatory_region_variant | MODIFIER |
| 15 | 97657819 | AX-170446064 | 0 | T | A | 0.132 | 2.63E-02 | 1.57E-01 | 2.65E+00 | 2.72E+00 | 8.70E-01 | no | | regulatory_region_variant | MODIFIER |
| 15 | 97659916 | AX-170178462 | 0 | A | G | 0.132 | 2.63E-02 | 1.57E-01 | 2.65E+00 | 2.72E+00 | 8.70E-01 | no | | intergenic_variant | MODIFIER |
| 15 | 97660168 | AX-170162802 | 0 | A | G | 0.132 | 2.63E-02 | 1.57E-01 | 2.65E+00 | 2.72E+00 | 8.70E-01 | no | | intergenic_variant | MODIFIER |
| 15 | 97663626 | AX-170056434 | 0 | G | C | 0.132 | 2.63E-02 | 1.57E-01 | 2.65E+00 | 2.72E+00 | 8.70E-01 | no | | regulatory_region_variant | MODIFIER |
| 15 | 97677800 | AX-170087946 | 0 | G | T | 0.132 | 2.63E-02 | 1.57E-01 | 2.65E+00 | 2.72E+00 | 8.70E-01 | no | Rpap3 | intron_variant | MODIFIER |
| 15 | 97711195 | AX-170393904 | 0 | T | C | 0.132 | 2.63E-02 | 1.57E-01 | 2.65E+00 | 2.72E+00 | 8.70E-01 | no | Endou | 3_prime_UTR_variant | MODIFIER |
| 15 | 97711349 | AX-170365729 | 0 | T | C | 0.132 | 2.63E-02 | 1.57E-01 | 2.65E+00 | 2.72E+00 | 8.70E-01 | no | Endou | 3_prime_UTR_variant | MODIFIER |
| 15 | 97711406 | AX-170171872 | 0 | T | C | 0.132 | 2.63E-02 | 1.57E-01 | 2.65E+00 | 2.72E+00 | 8.70E-01 | no | Endou | 3_prime_UTR_variant | MODIFIER |
| 15 | 97712343 | AX-170227655 | 0 | G | A | 0.132 | 2.63E-02 | 1.57E-01 | 2.65E+00 | 2.72E+00 | 8.70E-01 | no | Endou | intron_variant | MODIFIER |
| 15 | 97718451 | AX-170283985 | 0 | T | C | 0.132 | 2.63E-02 | 1.57E-01 | 2.65E+00 | 2.72E+00 | 8.70E-01 | no | Endou | intron_variant | MODIFIER |
| 15 | 97723921 | AX-170268766 | 0 | A | G | 0.132 | 2.63E-02 | 1.57E-01 | 2.65E+00 | 2.72E+00 | 8.70E-01 | no | Endou | upstream_gene_variant | MODIFIER |
| 15 | 97739516 | AX-170233652 | 0 | G | A | 0.132 | 2.63E-02 | 1.57E-01 | 2.65E+00 | 2.72E+00 | 8.70E-01 | no | | intergenic_variant | MODIFIER |
| 15 | 97739532 | AX-170471109 | 0 | G | A | 0.132 | 2.63E-02 | 1.57E-01 | 2.65E+00 | 2.72E+00 | 8.70E-01 | no | | intergenic_variant | MODIFIER |
| 15 | 97739969 | AX-170371650 | 0 | C | T | 0.132 | 2.63E-02 | 1.57E-01 | 2.65E+00 | 2.72E+00 | 8.70E-01 | no | Rapgef3 | downstream_gene_variant | MODIFIER |
| 15 | 97742134 | AX-97845062 | 0 | C | T | 0.132 | 2.63E-02 | 1.57E-01 | 2.65E+00 | 2.72E+00 | 8.70E-01 | no | Rapgef3 | downstream_gene_variant | MODIFIER |
| 15 | 97753467 | AX-170012346 | 0 | C | T | 0.132 | 2.63E-02 | 1.57E-01 | 2.65E+00 | 2.72E+00 | 8.70E-01 | no | Rapgef3 | intron_variant,non_coding_transcript_variant | MODIFIER |
| 15 | 97766728 | AX-170303664 | 0 | T | C | 0.132 | 2.63E-02 | 1.57E-01 | 2.65E+00 | 2.72E+00 | 8.70E-01 | no | Rapgef3 | intron_variant,non_coding_transcript_variant | MODIFIER |
| 15 | 97800471 | AX-97857029 | 0 | A | G | 0.132 | 2.63E-02 | 1.57E-01 | 2.65E+00 | 2.72E+00 | 8.70E-01 | no | Hdac7 | intron_variant | MODIFIER |
| 15 | 97823924 | AX-170012343 | 0 | A | C | 0.132 | 2.63E-02 | 1.57E-01 | 2.65E+00 | 2.72E+00 | 8.70E-01 | no | Hdac7 | intron_variant | MODIFIER |
| 15 | 97826351 | AX-97830281 | 0 | G | A | 0.132 | 2.63E-02 | 1.57E-01 | 2.65E+00 | 2.72E+00 | 8.70E-01 | no | Hdac7 | intron_variant | MODIFIER |
| 15 | 97826924 | AX-170012352 | 0 | G | A | 0.132 | 2.63E-02 | 1.57E-01 | 2.65E+00 | 2.72E+00 | 8.70E-01 | no | Hdac7 | intron_variant | MODIFIER |
| 15 | 97844678 | AX-170461123 | 0 | A | G | 0.132 | 2.63E-02 | 1.57E-01 | 2.65E+00 | 2.72E+00 | 8.70E-01 | no | Hdac7 | upstream_gene_variant | MODIFIER |
| 15 | 97845363 | AX-97789814 | 0 | A | G | 0.132 | 2.63E-02 | 1.57E-01 | 2.65E+00 | 2.72E+00 | 8.70E-01 | no | Hdac7 | upstream_gene_variant | MODIFIER |
| 15 | 97848587 | AX-170442575 | 0 | C | A | 0.132 | 2.63E-02 | 1.57E-01 | 2.65E+00 | 2.72E+00 | 8.70E-01 | no | Hdac7 | upstream_gene_variant | MODIFIER |
| 15 | 97850825 | AX-170457266 | 0 | C | T | 0.132 | 2.63E-02 | 1.57E-01 | 2.65E+00 | 2.72E+00 | 8.70E-01 | no | Vdr | downstream_gene_variant | MODIFIER |
| 15 | 97856150 | AX-170449856 | 0 | T | C | 0.132 | 2.63E-02 | 1.57E-01 | 2.65E+00 | 2.72E+00 | 8.70E-01 | no | Vdr | 3_prime_UTR_variant | MODIFIER |
| 15 | 97863331 | AX-170265678 | 0 | A | T | 0.132 | 2.63E-02 | 1.57E-01 | 2.65E+00 | 2.72E+00 | 8.70E-01 | no | Vdr | intron_variant | MODIFIER |
| 15 | 97863979 | AX-170240335 | 0 | G | C | 0.132 | 2.63E-02 | 1.57E-01 | 2.65E+00 | 2.72E+00 | 8.70E-01 | no | Vdr | intron_variant | MODIFIER |
| 15 | 97864294 | AX-170106006 | 0 | G | A | 0.132 | 2.63E-02 | 1.57E-01 | 2.65E+00 | 2.72E+00 | 8.70E-01 | no | Vdr | intron_variant | MODIFIER |
| 15 | 97864415 | AX-170133456 | 0 | T | G | 0.132 | 2.63E-02 | 1.57E-01 | 2.65E+00 | 2.72E+00 | 8.70E-01 | no | Vdr | intron_variant | MODIFIER |
| 15 | 97873946 | AX-170012347 | 0 | C | T | 0.132 | 2.63E-02 | 1.57E-01 | 2.65E+00 | 2.72E+00 | 8.70E-01 | no | Vdr | intron_variant | MODIFIER |
| 15 | 97873981 | AX-97849110 | 0 | A | G | 0.132 | 2.63E-02 | 1.57E-01 | 2.65E+00 | 2.72E+00 | 8.70E-01 | no | Vdr | intron_variant | MODIFIER |
| 15 | 97875565 | AX-170335668 | 0 | T | C | 0.132 | 2.63E-02 | 1.57E-01 | 2.65E+00 | 2.72E+00 | 8.70E-01 | no | Vdr | intron_variant | MODIFIER |
| 15 | 97876860 | AX-170396722 | 0 | A | G | 0.132 | 2.63E-02 | 1.57E-01 | 2.65E+00 | 2.72E+00 | 8.70E-01 | no | Vdr | splice_region_variant,intron_variant | LOW |
| 15 | 97885407 | AX-170177817 | 0 | T | C | 0.132 | 2.63E-02 | 1.57E-01 | 2.65E+00 | 2.72E+00 | 8.70E-01 | no | Vdr | intron_variant | MODIFIER |
| 15 | 97886118 | AX-170373636 | 0 | C | T | 0.132 | 2.63E-02 | 1.57E-01 | 2.65E+00 | 2.72E+00 | 8.70E-01 | no | Vdr | intron_variant | MODIFIER |
| 15 | 97890489 | AX-97830328 | 0 | C | T | 0.132 | 2.63E-02 | 1.57E-01 | 2.65E+00 | 2.72E+00 | 8.70E-01 | no | Vdr | intron_variant | MODIFIER |
| 15 | 97891007 | AX-170262491 | 0 | A | G | 0.132 | 2.63E-02 | 1.57E-01 | 2.65E+00 | 2.72E+00 | 8.70E-01 | no | Vdr | intron_variant | MODIFIER |
| 15 | 97891227 | AX-170496042 | 0 | G | C | 0.132 | 2.63E-02 | 1.57E-01 | 2.65E+00 | 2.72E+00 | 8.70E-01 | no | Vdr | intron_variant | MODIFIER |
| 15 | 97901061 | AX-170115039 | 0 | A | G | 0.132 | 2.63E-02 | 1.57E-01 | 2.65E+00 | 2.72E+00 | 8.70E-01 | no | Vdr | intron_variant | MODIFIER |
| 15 | 97918246 | AX-170192105 | 0 | A | C | 0.132 | 2.63E-02 | 1.57E-01 | 2.65E+00 | 2.72E+00 | 8.70E-01 | no | | intergenic_variant | MODIFIER |
| 15 | 97919590 | AX-97819634 | 0 | C | T | 0.132 | 2.63E-02 | 1.57E-01 | 2.65E+00 | 2.72E+00 | 8.70E-01 | no | | intergenic_variant | MODIFIER |
| 15 | 97951801 | AX-97830330 | 0 | C | T | 0.132 | 2.63E-02 | 1.57E-01 | 2.65E+00 | 2.72E+00 | 8.70E-01 | no | | intergenic_variant | MODIFIER |
| 15 | 97952210 | AX-170123089 | 0 | C | G | 0.132 | 2.63E-02 | 1.57E-01 | 2.65E+00 | 2.72E+00 | 8.70E-01 | no | | regulatory_region_variant | MODIFIER |
| 15 | 98025567 | AX-97830332 | 0 | C | T | 0.132 | 2.63E-02 | 1.57E-01 | 2.65E+00 | 2.72E+00 | 8.70E-01 | no | | regulatory_region_variant | MODIFIER |
| 15 | 98062894 | AX-170202758 | 0 | G | A | 0.132 | 2.63E-02 | 1.57E-01 | 2.65E+00 | 2.72E+00 | 8.70E-01 | no | Senp1 | intron_variant | MODIFIER |
| 15 | 98103903 | AX-97800418 | 0 | G | A | 0.132 | 2.63E-02 | 1.57E-01 | 2.65E+00 | 2.72E+00 | 8.70E-01 | no | Pfkm | upstream_gene_variant | MODIFIER |
| 15 | 98117818 | AX-170225060 | 0 | T | A | 0.132 | 2.63E-02 | 1.57E-01 | 2.65E+00 | 2.72E+00 | 8.70E-01 | no | Pfkm | intron_variant | MODIFIER |
| 15 | 98118103 | AX-170356065 | 0 | C | T | 0.132 | 2.63E-02 | 1.57E-01 | 2.65E+00 | 2.72E+00 | 8.70E-01 | no | Pfkm | intron_variant | MODIFIER |
| 15 | 98123767 | AX-170367817 | 0 | T | C | 0.132 | 2.63E-02 | 1.57E-01 | 2.65E+00 | 2.72E+00 | 8.70E-01 | no | Pfkm | intron_variant | MODIFIER |
| 15 | 98126562 | AX-170272444 | 0 | G | A | 0.132 | 2.63E-02 | 1.57E-01 | 2.65E+00 | 2.72E+00 | 8.70E-01 | no | Pfkm | intron_variant | MODIFIER |

| | | | | | | | | | | | | | | | |
|---|---|---|---|---|---|---|---|---|---|---|---|---|---|---|---|
| 15 | 98130818 | AX-97857666 | 0 | G | A | 0.132 | 2.63E-02 | 1.57E-01 | 2.65E+00 | 2.72E+00 | 8.70E-01 | no | Pfkm | intron_variant | MODIFIER |
| 15 | 98162530 | AX-170347903 | 0 | T | C | 0.132 | 2.63E-02 | 1.57E-01 | 2.65E+00 | 2.72E+00 | 8.70E-01 | no | Ccdc184 | upstream_gene_variant | MODIFIER |
| 15 | 98319350 | AX-97830820 | 0 | C | T | 0.132 | 2.63E-02 | 1.57E-01 | 2.65E+00 | 2.72E+00 | 8.70E-01 | no | Olfr285 | upstream_gene_variant | MODIFIER |
| 15 | 98337503 | AX-170409543 | 0 | T | C | 0.132 | 2.63E-02 | 1.57E-01 | 2.65E+00 | 2.72E+00 | 8.70E-01 | no | Olfr284 | downstream_gene_variant | MODIFIER |
| 15 | 98341944 | AX-170012358 | 0 | A | G | 0.132 | 2.63E-02 | 1.57E-01 | 2.65E+00 | 2.72E+00 | 8.70E-01 | no | Olfr284 | upstream_gene_variant | MODIFIER |
| 15 | 98342080 | AX-170413756 | 0 | T | C | 0.132 | 2.63E-02 | 1.57E-01 | 2.65E+00 | 2.72E+00 | 8.70E-01 | no | Olfr284 | upstream_gene_variant | MODIFIER |
| 15 | 98342921 | AX-97815911 | 0 | G | A | 0.132 | 2.63E-02 | 1.57E-01 | 2.65E+00 | 2.72E+00 | 8.70E-01 | no | Olfr284 | upstream_gene_variant | MODIFIER |
| 15 | 98360232 | AX-170012354 | 0 | T | G | 0.132 | 2.63E-02 | 1.57E-01 | 2.65E+00 | 2.72E+00 | 8.70E-01 | no | | intergenic_variant | MODIFIER |
| 15 | 98362447 | AX-170213169 | 0 | G | A | 0.132 | 2.63E-02 | 1.57E-01 | 2.65E+00 | 2.72E+00 | 8.70E-01 | no | | intergenic_variant | MODIFIER |
| 15 | 98363150 | AX-168396080 | 0 | C | T | 0.132 | 2.63E-02 | 1.57E-01 | 2.65E+00 | 2.72E+00 | 8.70E-01 | no | | intergenic_variant | MODIFIER |
| 15 | 98376463 | AX-97815413 | 0 | G | A | 0.132 | 2.63E-02 | 1.57E-01 | 2.65E+00 | 2.72E+00 | 8.70E-01 | no | Olfr283 | downstream_gene_variant | MODIFIER |
| 15 | 98381929 | AX-170519602 | 0 | T | C | 0.132 | 2.63E-02 | 1.57E-01 | 2.65E+00 | 2.72E+00 | 8.70E-01 | no | Olfr283 | upstream_gene_variant | MODIFIER |
| 15 | 98382037 | AX-170437701 | 0 | G | A | 0.132 | 2.63E-02 | 1.57E-01 | 2.65E+00 | 2.72E+00 | 8.70E-01 | no | Olfr283 | upstream_gene_variant | MODIFIER |
| 15 | 98397543 | AX-170012359 | 0 | G | A | 0.132 | 2.63E-02 | 1.57E-01 | 2.65E+00 | 2.72E+00 | 8.70E-01 | no | 1700031M16Rik | upstream_gene_variant | MODIFIER |
| 15 | 98446079 | AX-170012360 | 0 | T | G | 0.132 | 2.63E-02 | 1.57E-01 | 2.65E+00 | 2.72E+00 | 8.70E-01 | no | Gm6979 | upstream_gene_variant | MODIFIER |
| 15 | 98446136 | AX-170529810 | 0 | T | C | 0.132 | 2.63E-02 | 1.57E-01 | 2.65E+00 | 2.72E+00 | 8.70E-01 | no | Gm6979 | upstream_gene_variant | MODIFIER |
| 15 | 98446307 | AX-170520477 | 0 | G | T | 0.132 | 2.63E-02 | 1.57E-01 | 2.65E+00 | 2.72E+00 | 8.70E-01 | no | Gm6979 | upstream_gene_variant | MODIFIER |
| 15 | 98452501 | AX-170267585 | 0 | A | C | 0.132 | 2.63E-02 | 1.57E-01 | 2.65E+00 | 2.72E+00 | 8.70E-01 | no | Olfr281 | upstream_gene_variant | MODIFIER |
| 6 | 110938598 | AX-170027976 | 0 | T | A | 0.053 | 4.78E-02 | 2.96E-01 | 2.67E+00 | 2.74E+00 | 8.70E-01 | no | Grm7 | intron_variant | MODIFIER |
| 9 | 108611475 | AX-170405992 | 0 | G | A | 0.123 | 1.77E-02 | 1.16E-01 | 2.71E+00 | 2.78E+00 | 8.70E-01 | no | Arih2 | intron_variant | MODIFIER |
| 18 | 88484905 | AX-170330079 | 0 | G | A | 0.518 | -9.34E-03 | 6.23E-02 | 2.63E+00 | 2.76E+00 | 8.71E-01 | no | | intergenic_variant | MODIFIER |
| 18 | 88485252 | AX-170527221 | 0 | G | C | 0.518 | -9.34E-03 | 6.23E-02 | 2.63E+00 | 2.76E+00 | 8.71E-01 | no | | intergenic_variant | MODIFIER |
| 18 | 88485997 | AX-170422221 | 0 | A | G | 0.518 | -9.34E-03 | 6.23E-02 | 2.63E+00 | 2.76E+00 | 8.71E-01 | no | | intergenic_variant | MODIFIER |
| 18 | 88975823 | AX-170472498 | 0 | A | G | 0.518 | -9.34E-03 | 6.23E-02 | 2.63E+00 | 2.76E+00 | 8.71E-01 | no | Rttn | intron_variant | MODIFIER |
| 18 | 89046692 | AX-170423445 | 0 | T | C | 0.518 | -9.34E-03 | 6.23E-02 | 2.63E+00 | 2.76E+00 | 8.71E-01 | no | Rttn | intron_variant | MODIFIER |
| 18 | 89047919 | AX-170471744 | 0 | G | A | 0.518 | -9.34E-03 | 6.23E-02 | 2.63E+00 | 2.76E+00 | 8.71E-01 | no | Rttn | intron_variant | MODIFIER |
| 18 | 89090050 | AX-170061647 | 0 | C | T | 0.518 | -9.34E-03 | 6.23E-02 | 2.63E+00 | 2.76E+00 | 8.71E-01 | no | Rttn | intron_variant | MODIFIER |
| 18 | 89108657 | AX-170061645 | 0 | C | A | 0.518 | -9.34E-03 | 6.23E-02 | 2.63E+00 | 2.76E+00 | 8.71E-01 | no | Rttn | intron_variant | MODIFIER |
| 18 | 89160258 | AX-170513756 | 0 | G | C | 0.518 | -9.34E-03 | 6.23E-02 | 2.63E+00 | 2.76E+00 | 8.71E-01 | no | | intergenic_variant | MODIFIER |
| 18 | 89206605 | AX-170446217 | 0 | T | G | 0.518 | -9.34E-03 | 6.23E-02 | 2.63E+00 | 2.76E+00 | 8.71E-01 | no | Cd226 | missense_variant | MODERATE |
| 18 | 89207638 | AX-170410138 | 0 | G | A | 0.518 | -9.34E-03 | 6.23E-02 | 2.63E+00 | 2.76E+00 | 8.71E-01 | no | Cd226 | intron_variant | MODIFIER |
| 18 | 89240028 | AX-170490112 | 0 | C | G | 0.518 | -9.34E-03 | 6.23E-02 | 2.63E+00 | 2.76E+00 | 8.71E-01 | no | Cd226 | intron_variant | MODIFIER |
| 18 | 89241772 | AX-168405856 | 0 | G | A | 0.518 | -9.34E-03 | 6.23E-02 | 2.63E+00 | 2.76E+00 | 8.71E-01 | no | Cd226 | intron_variant | MODIFIER |
| 18 | 89242087 | AX-97856750 | 0 | G | A | 0.518 | -9.34E-03 | 6.23E-02 | 2.63E+00 | 2.76E+00 | 8.71E-01 | no | Cd226 | intron_variant | MODIFIER |
| 18 | 89242166 | AX-170419407 | 0 | G | A | 0.518 | -9.34E-03 | 6.23E-02 | 2.63E+00 | 2.76E+00 | 8.71E-01 | no | Cd226 | intron_variant | MODIFIER |
| 18 | 89257657 | AX-170182765 | 0 | C | T | 0.518 | -9.34E-03 | 6.23E-02 | 2.63E+00 | 2.76E+00 | 8.71E-01 | no | Cd226 | intron_variant | MODIFIER |
| 18 | 89675369 | AX-170097566 | 0 | G | A | 0.518 | -9.34E-03 | 6.23E-02 | 2.63E+00 | 2.76E+00 | 8.71E-01 | no | Dok6 | intron_variant | MODIFIER |
| 18 | 89675724 | AX-170488986 | 0 | C | T | 0.518 | -9.34E-03 | 6.23E-02 | 2.63E+00 | 2.76E+00 | 8.71E-01 | no | Dok6 | intron_variant | MODIFIER |
| 18 | 89676406 | AX-97849158 | 0 | C | T | 0.518 | -9.34E-03 | 6.23E-02 | 2.63E+00 | 2.76E+00 | 8.71E-01 | no | Dok6 | intron_variant | MODIFIER |
| 18 | 89691225 | AX-170196707 | 0 | A | T | 0.518 | -9.34E-03 | 6.23E-02 | 2.63E+00 | 2.76E+00 | 8.71E-01 | no | Dok6 | intron_variant | MODIFIER |
| 18 | 89692423 | AX-170504966 | 0 | A | G | 0.518 | -9.34E-03 | 6.23E-02 | 2.63E+00 | 2.76E+00 | 8.71E-01 | no | Dok6 | intron_variant | MODIFIER |
| 18 | 89728169 | AX-170102884 | 0 | T | C | 0.518 | -9.34E-03 | 6.23E-02 | 2.63E+00 | 2.76E+00 | 8.71E-01 | no | Dok6 | intron_variant | MODIFIER |
| 18 | 89756640 | AX-97788984 | 0 | A | G | 0.518 | -9.34E-03 | 6.23E-02 | 2.63E+00 | 2.76E+00 | 8.71E-01 | no | Dok6 | intron_variant | MODIFIER |
| 18 | 89852583 | AX-170166775 | 0 | C | T | 0.518 | -9.34E-03 | 6.23E-02 | 2.63E+00 | 2.76E+00 | 8.71E-01 | no | | intergenic_variant | MODIFIER |
| 18 | 89877475 | AX-170061663 | 0 | C | G | 0.518 | -9.34E-03 | 6.23E-02 | 2.63E+00 | 2.76E+00 | 8.71E-01 | no | | intergenic_variant | MODIFIER |
| 18 | 89943551 | AX-170451072 | 0 | C | G | 0.518 | -9.34E-03 | 6.23E-02 | 2.63E+00 | 2.76E+00 | 8.71E-01 | no | Gm27037 | non_coding_transcript_exon_variant | MODIFIER |
| 18 | 89953855 | AX-170482650 | 0 | A | G | 0.518 | -9.34E-03 | 6.23E-02 | 2.63E+00 | 2.76E+00 | 8.71E-01 | no | | intergenic_variant | MODIFIER |
| 18 | 89965568 | AX-97796156 | 0 | T | C | 0.518 | -9.34E-03 | 6.23E-02 | 2.63E+00 | 2.76E+00 | 8.71E-01 | no | | intergenic_variant | MODIFIER |
| 18 | 90004771 | AX-170016174 | 0 | C | A | 0.518 | -9.34E-03 | 6.23E-02 | 2.63E+00 | 2.76E+00 | 8.71E-01 | no | | intergenic_variant | MODIFIER |
| 18 | 90026854 | AX-170016178 | 0 | T | C | 0.518 | -9.34E-03 | 6.23E-02 | 2.63E+00 | 2.76E+00 | 8.71E-01 | no | | intergenic_variant | MODIFIER |
| 15 | 93622379 | AX-97803869 | 0 | A | G | 0.158 | 2.68E-02 | 1.54E-01 | 2.60E+00 | 2.71E+00 | 8.71E-01 | no | | intergenic_variant | MODIFIER |
| 7 | 116731637 | AX-170504837 | 0 | C | T | 0.43 | 1.50E-02 | 9.26E-02 | 2.68E+00 | 2.73E+00 | 8.71E-01 | no | | intergenic_variant | MODIFIER |
| 4 | 72130011 | AX-170365986 | 0 | C | A | 0.219 | 1.76E-02 | 9.97E-02 | 2.61E+00 | 2.70E+00 | 8.71E-01 | no | Tle1 | intron_variant | MODIFIER |
| 18 | 56040379 | AX-97783553 | 0 | C | A | 0.404 | -1.50E-02 | 8.70E-02 | 2.64E+00 | 2.70E+00 | 8.71E-01 | no | | regulatory_region_variant | MODIFIER |

| | | | | | | | | | | | | | | | |
|---|---|---|---|---|---|---|---|---|---|---|---|---|---|---|---|
| 18 | 56040829 | AX-170414601 | 0 | G | C | 0.404 | -1.50E-02 | 8.70E-02 | 2.64E+00 | 2.70E+00 | 8.71E-01 | no | | regulatory_region_variant | MODIFIER |
| 18 | 56041002 | AX-170374246 | 0 | T | C | 0.404 | -1.50E-02 | 8.70E-02 | 2.64E+00 | 2.70E+00 | 8.71E-01 | no | | regulatory_region_variant | MODIFIER |
| 18 | 56047795 | AX-170060915 | 0 | T | C | 0.404 | -1.50E-02 | 8.70E-02 | 2.64E+00 | 2.70E+00 | 8.71E-01 | no | | regulatory_region_variant | MODIFIER |
| 18 | 56048497 | AX-170060913 | 0 | T | C | 0.404 | -1.50E-02 | 8.70E-02 | 2.64E+00 | 2.70E+00 | 8.71E-01 | no | | regulatory_region_variant | MODIFIER |
| 18 | 56062250 | AX-170249413 | 0 | T | C | 0.404 | -1.50E-02 | 8.70E-02 | 2.64E+00 | 2.70E+00 | 8.71E-01 | no | | intergenic_variant | MODIFIER |
| 18 | 56070824 | AX-170233244 | 0 | C | T | 0.404 | -1.50E-02 | 8.70E-02 | 2.64E+00 | 2.70E+00 | 8.71E-01 | no | | regulatory_region_variant | MODIFIER |
| 18 | 56070850 | AX-170179668 | 0 | C | T | 0.404 | -1.50E-02 | 8.70E-02 | 2.64E+00 | 2.70E+00 | 8.71E-01 | no | | regulatory_region_variant | MODIFIER |
| 18 | 56434053 | AX-170472060 | 0 | T | C | 0.404 | -1.50E-02 | 8.70E-02 | 2.64E+00 | 2.70E+00 | 8.71E-01 | no | Gramd3 | intron_variant | MODIFIER |
| 18 | 56485469 | AX-170203964 | 0 | T | G | 0.404 | -1.50E-02 | 8.70E-02 | 2.64E+00 | 2.70E+00 | 8.71E-01 | no | Gramd3 | intron_variant | MODIFIER |
| 18 | 56487402 | AX-170140004 | 0 | C | T | 0.404 | -1.50E-02 | 8.70E-02 | 2.64E+00 | 2.70E+00 | 8.71E-01 | no | Gramd3 | intron_variant | MODIFIER |
| 18 | 56489483 | AX-170257323 | 0 | A | C | 0.404 | -1.50E-02 | 8.70E-02 | 2.64E+00 | 2.70E+00 | 8.71E-01 | no | Gramd3 | intron_variant | MODIFIER |
| 18 | 56489574 | AX-170236771 | 0 | A | G | 0.404 | -1.50E-02 | 8.70E-02 | 2.64E+00 | 2.70E+00 | 8.71E-01 | no | Gramd3 | intron_variant | MODIFIER |
| 18 | 56603348 | AX-170318229 | 0 | G | C | 0.404 | -1.50E-02 | 8.70E-02 | 2.64E+00 | 2.70E+00 | 8.71E-01 | no | AC162035.1 | upstream_gene_variant | MODIFIER |
| 18 | 56637053 | AX-170358045 | 0 | T | A | 0.404 | -1.50E-02 | 8.70E-02 | 2.64E+00 | 2.70E+00 | 8.71E-01 | no | | intergenic_variant | MODIFIER |
| 18 | 56660608 | AX-170494559 | 0 | A | T | 0.404 | -1.50E-02 | 8.70E-02 | 2.64E+00 | 2.70E+00 | 8.71E-01 | no | | intergenic_variant | MODIFIER |
| 18 | 56665797 | AX-170060921 | 0 | C | T | 0.404 | -1.50E-02 | 8.70E-02 | 2.64E+00 | 2.70E+00 | 8.71E-01 | no | AC158763.1 | upstream_gene_variant | MODIFIER |
| 18 | 56666322 | AX-170060920 | 0 | G | C | 0.404 | -1.50E-02 | 8.70E-02 | 2.64E+00 | 2.70E+00 | 8.71E-01 | no | AC158763.1 | upstream_gene_variant | MODIFIER |
| 18 | 56669510 | AX-170060923 | 0 | A | C | 0.404 | -1.50E-02 | 8.70E-02 | 2.64E+00 | 2.70E+00 | 8.71E-01 | no | AC158763.1 | downstream_gene_variant | MODIFIER |
| 18 | 56717456 | AX-97850188 | 0 | T | G | 0.404 | -1.50E-02 | 8.70E-02 | 2.64E+00 | 2.70E+00 | 8.71E-01 | no | Lmnb1 | intron_variant | MODIFIER |
| 18 | 56772804 | AX-97790300 | 0 | A | G | 0.404 | -1.50E-02 | 8.70E-02 | 2.64E+00 | 2.70E+00 | 8.71E-01 | no | MAR3 | downstream_gene_variant | MODIFIER |
| 18 | 56837680 | AX-170060924 | 0 | G | C | 0.404 | -1.50E-02 | 8.70E-02 | 2.64E+00 | 2.70E+00 | 8.71E-01 | no | MAR3 | intron_variant | MODIFIER |
| 18 | 56841400 | AX-170262268 | 0 | C | T | 0.404 | -1.50E-02 | 8.70E-02 | 2.64E+00 | 2.70E+00 | 8.71E-01 | no | MAR3 | intron_variant | MODIFIER |
| 18 | 56841759 | AX-97780870 | 0 | C | T | 0.404 | -1.50E-02 | 8.70E-02 | 2.64E+00 | 2.70E+00 | 8.71E-01 | no | MAR3 | intron_variant | MODIFIER |
| 18 | 56842015 | AX-170472185 | 0 | A | G | 0.404 | -1.50E-02 | 8.70E-02 | 2.64E+00 | 2.70E+00 | 8.71E-01 | no | MAR3 | intron_variant | MODIFIER |
| 18 | 56862443 | AX-170105474 | 0 | C | T | 0.404 | -1.50E-02 | 8.70E-02 | 2.64E+00 | 2.70E+00 | 8.71E-01 | no | MAR3 | intron_variant | MODIFIER |
| 18 | 57059452 | AX-170322645 | 0 | A | G | 0.404 | -1.50E-02 | 8.70E-02 | 2.64E+00 | 2.70E+00 | 8.71E-01 | no | | regulatory_region_variant | MODIFIER |
| 5 | 98037482 | AX-170185695 | 0 | G | A | 0.412 | 1.81E-02 | 1.06E-01 | 2.64E+00 | 2.70E+00 | 8.71E-01 | no | Gm42759 | downstream_gene_variant | MODIFIER |
| 6 | 140088995 | AX-170076419 | 0 | C | G | 0.07 | -3.53E-02 | 2.02E-01 | 2.60E+00 | 2.71E+00 | 8.71E-01 | no | | intergenic_variant | MODIFIER |
| 18 | 47856801 | AX-170015660 | 0 | C | A | 0.044 | 5.41E-02 | 3.41E-01 | 2.70E+00 | 2.75E+00 | 8.71E-01 | no | | intergenic_variant | MODIFIER |
| 9 | 104544744 | AX-170245759 | 0 | G | C | 0.132 | 1.68E-02 | 1.11E-01 | 2.69E+00 | 2.78E+00 | 8.71E-01 | no | Cpne4 | upstream_gene_variant | MODIFIER |
| 2 | 158297026 | AX-170370147 | 1 | C | A | 0.143 | 3.56E-02 | 2.20E-01 | 2.69E+00 | 2.73E+00 | 8.71E-01 | no | | intergenic_variant | MODIFIER |
| 1 | 69173536 | AX-170117653 | 0 | T | A | 0.14 | -3.72E-02 | 2.37E-01 | 2.70E+00 | 2.76E+00 | 8.71E-01 | no | | intergenic_variant | MODIFIER |
| 7 | 117298633 | AX-170269133 | 0 | G | A | 0.404 | 1.45E-02 | 8.98E-02 | 2.67E+00 | 2.73E+00 | 8.71E-01 | no | | intergenic_variant | MODIFIER |
| 10 | 118987968 | AX-97790975 | 0 | C | T | 0.342 | 1.14E-02 | 7.28E-02 | 2.66E+00 | 2.75E+00 | 8.71E-01 | no | | intergenic_variant | MODIFIER |
| 10 | 118988224 | AX-170317596 | 0 | T | C | 0.342 | 1.14E-02 | 7.28E-02 | 2.66E+00 | 2.75E+00 | 8.71E-01 | no | | regulatory_region_variant | MODIFIER |
| 10 | 119058246 | AX-170374936 | 0 | A | G | 0.342 | 1.14E-02 | 7.28E-02 | 2.66E+00 | 2.75E+00 | 8.71E-01 | no | | regulatory_region_variant | MODIFIER |
| 10 | 119058748 | AX-170169156 | 0 | T | C | 0.342 | 1.14E-02 | 7.28E-02 | 2.66E+00 | 2.75E+00 | 8.71E-01 | no | | regulatory_region_variant | MODIFIER |
| 10 | 119058790 | AX-170197181 | 0 | C | T | 0.342 | 1.14E-02 | 7.28E-02 | 2.66E+00 | 2.75E+00 | 8.71E-01 | no | | regulatory_region_variant | MODIFIER |
| 10 | 119058934 | AX-97792743 | 0 | C | T | 0.342 | 1.14E-02 | 7.28E-02 | 2.66E+00 | 2.75E+00 | 8.71E-01 | no | | regulatory_region_variant | MODIFIER |
| 10 | 119059369 | AX-170097235 | 0 | G | A | 0.342 | 1.14E-02 | 7.28E-02 | 2.66E+00 | 2.75E+00 | 8.71E-01 | no | | intergenic_variant | MODIFIER |
| 10 | 119106447 | AX-97790919 | 0 | G | A | 0.342 | 1.14E-02 | 7.28E-02 | 2.66E+00 | 2.75E+00 | 8.71E-01 | no | 1700064J06Rik | downstream_gene_variant | MODIFIER |
| 10 | 119115711 | AX-170508753 | 0 | C | T | 0.342 | 1.14E-02 | 7.28E-02 | 2.66E+00 | 2.75E+00 | 8.71E-01 | no | Gm47480 | intron_variant,non_coding_transcript_variant | MODIFIER |
| 10 | 119159989 | AX-97825526 | 0 | G | A | 0.342 | 1.14E-02 | 7.28E-02 | 2.66E+00 | 2.75E+00 | 8.71E-01 | no | Gm34045 | intron_variant,non_coding_transcript_variant | MODIFIER |
| 10 | 119288080 | AX-170004011 | 0 | T | G | 0.342 | 1.14E-02 | 7.28E-02 | 2.66E+00 | 2.75E+00 | 8.71E-01 | no | | intergenic_variant | MODIFIER |
| 10 | 119288611 | AX-97816987 | 0 | T | G | 0.342 | 1.14E-02 | 7.28E-02 | 2.66E+00 | 2.75E+00 | 8.71E-01 | no | | intergenic_variant | MODIFIER |
| 10 | 119430667 | AX-170106796 | 0 | C | T | 0.342 | 1.14E-02 | 7.28E-02 | 2.66E+00 | 2.75E+00 | 8.71E-01 | no | Grip1os3 | intron_variant,non_coding_transcript_variant | MODIFIER |
| 10 | 119463188 | AX-97825528 | 0 | T | C | 0.342 | 1.14E-02 | 7.28E-02 | 2.66E+00 | 2.75E+00 | 8.71E-01 | no | Grip1 | intron_variant | MODIFIER |
| 10 | 119468139 | AX-170149309 | 0 | T | C | 0.342 | 1.14E-02 | 7.28E-02 | 2.66E+00 | 2.75E+00 | 8.71E-01 | no | Grip1 | intron_variant | MODIFIER |
| 10 | 119476409 | AX-170361837 | 0 | C | G | 0.342 | 1.14E-02 | 7.28E-02 | 2.66E+00 | 2.75E+00 | 8.71E-01 | no | Grip1 | intron_variant | MODIFIER |
| 10 | 119479830 | AX-170232077 | 0 | G | T | 0.342 | 1.14E-02 | 7.28E-02 | 2.66E+00 | 2.75E+00 | 8.71E-01 | no | Grip1 | intron_variant | MODIFIER |
| 10 | 119498757 | AX-97838990 | 0 | T | C | 0.342 | 1.14E-02 | 7.28E-02 | 2.66E+00 | 2.75E+00 | 8.71E-01 | no | Grip1 | intron_variant | MODIFIER |
| 10 | 119500601 | AX-170154896 | 0 | T | C | 0.342 | 1.14E-02 | 7.28E-02 | 2.66E+00 | 2.75E+00 | 8.71E-01 | no | Grip1 | intron_variant | MODIFIER |
| 10 | 119519342 | AX-97813656 | 0 | C | A | 0.342 | 1.14E-02 | 7.28E-02 | 2.66E+00 | 2.75E+00 | 8.71E-01 | no | Grip1 | intron_variant | MODIFIER |
| 10 | 119530656 | AX-170192212 | 0 | G | T | 0.342 | 1.14E-02 | 7.28E-02 | 2.66E+00 | 2.75E+00 | 8.71E-01 | no | Grip1 | intron_variant | MODIFIER |

| | | | | | | | | | | | | | | | |
|---|---|---|---|---|---|---|---|---|---|---|---|---|---|---|---|
| 10 | 119535002 | AX-168411909 | 0 | C | T | 0.342 | 1.14E-02 | 7.28E-02 | 2.66E+00 | 2.75E+00 | 8.71E-01 | no | Grip1 | intron_variant | MODIFIER |
| 10 | 119535109 | AX-170046047 | 0 | T | C | 0.342 | 1.14E-02 | 7.28E-02 | 2.66E+00 | 2.75E+00 | 8.71E-01 | no | Grip1 | intron_variant | MODIFIER |
| 10 | 119537887 | AX-170456943 | 0 | T | C | 0.342 | 1.14E-02 | 7.28E-02 | 2.66E+00 | 2.75E+00 | 8.71E-01 | no | Grip1 | intron_variant | MODIFIER |
| 10 | 119540340 | AX-170517331 | 0 | C | T | 0.342 | 1.14E-02 | 7.28E-02 | 2.66E+00 | 2.75E+00 | 8.71E-01 | no | Grip1 | intron_variant | MODIFIER |
| 10 | 119542835 | AX-170286357 | 0 | A | G | 0.342 | 1.14E-02 | 7.28E-02 | 2.66E+00 | 2.75E+00 | 8.71E-01 | no | Grip1 | intron_variant | MODIFIER |
| 10 | 119937377 | AX-170004023 | 0 | A | T | 0.342 | 1.14E-02 | 7.28E-02 | 2.66E+00 | 2.75E+00 | 8.71E-01 | no | Grip1 | intron_variant | MODIFIER |
| 10 | 119942009 | AX-97785751 | 0 | T | C | 0.342 | 1.14E-02 | 7.28E-02 | 2.66E+00 | 2.75E+00 | 8.71E-01 | no | Grip1 | intron_variant | MODIFIER |
| 10 | 120031288 | AX-170103232 | 0 | A | G | 0.342 | 1.14E-02 | 7.28E-02 | 2.66E+00 | 2.75E+00 | 8.71E-01 | no | Grip1 | intron_variant | MODIFIER |
| 10 | 120031583 | AX-97793590 | 0 | A | G | 0.342 | 1.14E-02 | 7.28E-02 | 2.66E+00 | 2.75E+00 | 8.71E-01 | no | Grip1 | intron_variant | MODIFIER |
| 10 | 120270351 | AX-170046058 | 0 | C | T | 0.342 | 1.14E-02 | 7.28E-02 | 2.66E+00 | 2.75E+00 | 8.71E-01 | no | | regulatory_region_variant | MODIFIER |
| 10 | 120270691 | AX-97826137 | 0 | T | C | 0.342 | 1.14E-02 | 7.28E-02 | 2.66E+00 | 2.75E+00 | 8.71E-01 | no | | regulatory_region_variant | MODIFIER |
| 10 | 120345854 | AX-170380687 | 0 | C | T | 0.342 | 1.14E-02 | 7.28E-02 | 2.66E+00 | 2.75E+00 | 8.71E-01 | no | | intergenic_variant | MODIFIER |
| 10 | 120372504 | AX-170460117 | 0 | C | T | 0.342 | 1.14E-02 | 7.28E-02 | 2.66E+00 | 2.75E+00 | 8.71E-01 | no | 1700006J14Rik | intron_variant,non_coding_transcript_variant | MODIFIER |
| 9 | 58619876 | AX-170032461 | 0 | G | A | 0.211 | -1.96E-02 | 1.23E-01 | 2.69E+00 | 2.74E+00 | 8.72E-01 | no | Nptn | intron_variant | MODIFIER |
| 1 | 128662040 | AX-170334990 | 0 | T | C | 0.535 | 1.46E-02 | 8.53E-02 | 2.63E+00 | 2.70E+00 | 8.72E-01 | no | | regulatory_region_variant | MODIFIER |
| 6 | 104387919 | AX-170075752 | 0 | G | A | 0.061 | 3.53E-02 | 2.10E-01 | 2.60E+00 | 2.72E+00 | 8.72E-01 | no | | intergenic_variant | MODIFIER |
| 6 | 120528093 | AX-170102779 | 0 | G | C | 0.061 | 3.53E-02 | 2.10E-01 | 2.60E+00 | 2.72E+00 | 8.72E-01 | no | Hdhd5 | intron_variant | MODIFIER |
| 4 | 93829261 | AX-170133173 | 0 | C | G | 0.316 | 1.70E-02 | 1.03E-01 | 2.64E+00 | 2.72E+00 | 8.72E-01 | no | | intergenic_variant | MODIFIER |
| 7 | 118097407 | AX-97789047 | 0 | G | A | 0.395 | 1.49E-02 | 9.27E-02 | 2.68E+00 | 2.73E+00 | 8.72E-01 | no | | intergenic_variant | MODIFIER |
| 6 | 105381919 | AX-97812184 | 1 | A | G | 0.071 | 4.37E-02 | 2.76E-01 | 2.71E+00 | 2.75E+00 | 8.72E-01 | no | | intergenic_variant | MODIFIER |
| 6 | 145376754 | AX-170291415 | 1 | C | T | 0.071 | 4.37E-02 | 2.76E-01 | 2.71E+00 | 2.75E+00 | 8.72E-01 | no | Lmntd1 | intron_variant,NMD_transcript_variant | MODIFIER |
| 13 | 38026880 | AX-170138025 | 1 | G | T | 0.179 | -1.99E-02 | 1.17E-01 | 2.63E+00 | 2.70E+00 | 8.72E-01 | no | Cage1 | intron_variant | MODIFIER |
| 3 | 37542656 | AX-97838876 | 0 | C | T | 0.202 | -1.44E-02 | 9.28E-02 | 2.66E+00 | 2.75E+00 | 8.72E-01 | no | Spata5 | intron_variant | MODIFIER |
| 1 | 69189215 | AX-170492457 | 0 | C | A | 0.105 | 3.03E-02 | 1.87E-01 | 2.62E+00 | 2.73E+00 | 8.72E-01 | no | | intergenic_variant | MODIFIER |
| 13 | 52738584 | AX-170482561 | 0 | G | C | 0.509 | 1.13E-02 | 6.67E-02 | 2.61E+00 | 2.71E+00 | 8.72E-01 | no | BB123696 | upstream_gene_variant | MODIFIER |
| 4 | 94328833 | AX-170238743 | 1 | G | A | 0.339 | 1.77E-02 | 1.07E-01 | 2.64E+00 | 2.72E+00 | 8.72E-01 | no | Gm12648 | intron_variant,non_coding_transcript_variant | MODIFIER |
| 1 | 176570429 | AX-170002352 | 0 | G | A | 0.219 | -1.68E-02 | 1.14E-01 | 2.70E+00 | 2.79E+00 | 8.72E-01 | no | | intergenic_variant | MODIFIER |
| 4 | 92790945 | AX-170106012 | 1 | T | C | 0.321 | -1.48E-02 | 9.67E-02 | 2.66E+00 | 2.76E+00 | 8.72E-01 | no | | intergenic_variant | MODIFIER |
| 11 | 13080026 | AX-170004307 | 0 | A | G | 0.053 | 3.30E-02 | 2.12E-01 | 2.65E+00 | 2.75E+00 | 8.72E-01 | no | | intergenic_variant | MODIFIER |
| 11 | 24156979 | AX-170334585 | 0 | T | C | 0.053 | 3.30E-02 | 2.12E-01 | 2.65E+00 | 2.75E+00 | 8.72E-01 | no | Bcl11a | intron_variant | MODIFIER |
| 9 | 55953159 | AX-170090308 | 1 | G | A | 0.205 | -1.95E-02 | 1.24E-01 | 2.70E+00 | 2.75E+00 | 8.72E-01 | no | Gm7435 | downstream_gene_variant | MODIFIER |
| 10 | 13490972 | AX-170086748 | 2 | A | T | 0.209 | -1.72E-02 | 1.04E-01 | 2.64E+00 | 2.72E+00 | 8.72E-01 | no | | intergenic_variant | MODIFIER |
| 15 | 93683050 | AX-170379310 | 0 | A | G | 0.158 | 2.95E-02 | 1.74E-01 | 2.63E+00 | 2.71E+00 | 8.72E-01 | no | Zfp42-ps1 | upstream_gene_variant | MODIFIER |
| 16 | 94381658 | AX-170237164 | 0 | A | G | 0.404 | -1.45E-02 | 8.33E-02 | 2.61E+00 | 2.70E+00 | 8.73E-01 | no | Ttc3 | intron_variant | MODIFIER |
| 1 | 78401717 | AX-170221722 | 0 | C | A | 0.14 | -3.69E-02 | 2.36E-01 | 2.70E+00 | 2.75E+00 | 8.73E-01 | no | Sgpp2 | intron_variant | MODIFIER |
| 5 | 101742227 | AX-170391977 | 0 | G | A | 0.43 | 1.77E-02 | 1.12E-01 | 2.69E+00 | 2.74E+00 | 8.73E-01 | no | | regulatory_region_variant | MODIFIER |
| 1 | 42034691 | AX-170131766 | 0 | G | A | 0.439 | 1.12E-02 | 7.38E-02 | 2.67E+00 | 2.76E+00 | 8.73E-01 | no | | intergenic_variant | MODIFIER |
| 7 | 40576837 | AX-170349428 | 0 | G | A | 0.325 | -1.28E-02 | 8.46E-02 | 2.66E+00 | 2.76E+00 | 8.73E-01 | no | | intergenic_variant | MODIFIER |
| 8 | 55107308 | AX-170118137 | 1 | T | C | 0.143 | 2.33E-02 | 1.52E-01 | 2.74E+00 | 2.80E+00 | 8.73E-01 | no | | intergenic_variant | MODIFIER |
| 8 | 57330521 | AX-97813363 | 1 | G | A | 0.143 | 2.33E-02 | 1.52E-01 | 2.74E+00 | 2.80E+00 | 8.73E-01 | no | 5033428I22Rik | upstream_gene_variant | MODIFIER |
| 10 | 30655197 | AX-170044486 | 0 | T | C | 0.193 | 1.43E-02 | 9.65E-02 | 2.67E+00 | 2.77E+00 | 8.73E-01 | no | Ncoa7 | intron_variant | MODIFIER |
| 7 | 60909224 | AX-97821672 | 1 | A | C | 0.429 | -1.50E-02 | 9.80E-02 | 2.68E+00 | 2.76E+00 | 8.73E-01 | no | | intergenic_variant | MODIFIER |
| 1 | 55510450 | AX-170292646 | 1 | C | T | 0.107 | -3.65E-02 | 2.34E-01 | 2.70E+00 | 2.75E+00 | 8.73E-01 | no | Plcl1 | intron_variant | MODIFIER |
| 5 | 69833508 | AX-170436514 | 1 | A | G | 0.446 | 1.59E-02 | 9.46E-02 | 2.62E+00 | 2.71E+00 | 8.73E-01 | no | | intergenic_variant | MODIFIER |
| 1 | 151140618 | AX-170457586 | 0 | T | C | 0.018 | 2.12E-02 | 1.67E-01 | 2.62E+00 | 2.80E+00 | 8.73E-01 | no | Gm8941 | downstream_gene_variant | MODIFIER |
| 6 | 107270468 | AX-170320939 | 0 | G | A | 0.053 | 4.69E-02 | 2.96E-01 | 2.67E+00 | 2.74E+00 | 8.73E-01 | no | Gm34933 | intron_variant,non_coding_transcript_variant | MODIFIER |
| 9 | 58767288 | AX-170035810 | 1 | A | G | 0.223 | 1.89E-02 | 1.18E-01 | 2.66E+00 | 2.73E+00 | 8.73E-01 | no | | regulatory_region_variant | MODIFIER |
| 12 | 103691202 | AX-170491236 | 1 | A | T | 0.125 | -3.08E-02 | 1.96E-01 | 2.66E+00 | 2.74E+00 | 8.73E-01 | no | Serpina1f | intron_variant | MODIFIER |
| 12 | 103766893 | AX-170340266 | 1 | C | T | 0.125 | -3.08E-02 | 1.96E-01 | 2.66E+00 | 2.74E+00 | 8.73E-01 | no | Serpina1d | intron_variant | MODIFIER |
| 12 | 104147733 | AX-170349792 | 1 | G | A | 0.125 | -3.08E-02 | 1.96E-01 | 2.66E+00 | 2.74E+00 | 8.73E-01 | no | Serpina3c | intron_variant | MODIFIER |
| 2 | 119707810 | AX-170406553 | 2 | T | A | 0.145 | 3.66E-02 | 2.29E-01 | 2.70E+00 | 2.72E+00 | 8.73E-01 | no | Rtf1 | intron_variant | MODIFIER |
| 8 | 72898402 | AX-170512774 | 0 | G | T | 0.202 | -1.97E-02 | 1.25E-01 | 2.66E+00 | 2.74E+00 | 8.73E-01 | no | Large1 | intron_variant | MODIFIER |
| 8 | 73052197 | AX-170096318 | 0 | T | C | 0.202 | -1.97E-02 | 1.25E-01 | 2.66E+00 | 2.74E+00 | 8.73E-01 | no | Large1 | intron_variant | MODIFIER |
| 8 | 73268085 | AX-170425577 | 0 | C | A | 0.202 | -1.97E-02 | 1.25E-01 | 2.66E+00 | 2.74E+00 | 8.73E-01 | no | Large1 | intron_variant | MODIFIER |

| | | | | | | | | | | | | | | | |
|---|---|---|---|---|---|---|---|---|---|---|---|---|---|---|---|
| 8 | 74123936 | AX-170307410 | 0 | C | T | 0.202 | -1.97E-02 | 1.25E-01 | 2.66E+00 | 2.74E+00 | 8.73E-01 | no | | intergenic_variant | MODIFIER |
| 4 | 145115593 | AX-170362759 | 0 | C | T | 0.114 | 2.78E-02 | 1.71E-01 | 2.63E+00 | 2.72E+00 | 8.73E-01 | no | Vps13d | synonymous_variant | LOW |
| 4 | 148013574 | AX-170284839 | 0 | G | A | 0.114 | 2.78E-02 | 1.71E-01 | 2.63E+00 | 2.72E+00 | 8.73E-01 | no | Clcn6 | intron_variant | MODIFIER |
| 10 | 16201038 | AX-97802845 | 0 | G | C | 0.193 | 1.47E-02 | 1.02E-01 | 2.68E+00 | 2.78E+00 | 8.74E-01 | no | | intergenic_variant | MODIFIER |
| 8 | 18885519 | AX-170196837 | 2 | A | T | 0.155 | -1.60E-02 | 1.01E-01 | 2.62E+00 | 2.74E+00 | 8.74E-01 | no | Agpat5 | downstream_gene_variant | MODIFIER |
| 8 | 90102996 | AX-170331951 | 1 | A | G | 0.196 | -1.90E-02 | 1.17E-01 | 2.64E+00 | 2.72E+00 | 8.74E-01 | no | | intergenic_variant | MODIFIER |
| 9 | 84564246 | AX-170427210 | 0 | G | T | 0.219 | 1.85E-02 | 1.20E-01 | 2.68E+00 | 2.75E+00 | 8.74E-01 | no | | intergenic_variant | MODIFIER |
| 7 | 135310004 | AX-97839677 | 0 | C | T | 0.263 | -1.34E-02 | 7.91E-02 | 2.60E+00 | 2.71E+00 | 8.74E-01 | no | | intergenic_variant | MODIFIER |
| 1 | 8361179 | AX-170124667 | 1 | T | C | 0.527 | 1.24E-02 | 7.24E-02 | 2.62E+00 | 2.70E+00 | 8.74E-01 | no | Sntg1 | downstream_gene_variant | MODIFIER |
| 9 | 67847453 | AX-170181450 | 0 | T | C | 0.193 | -2.17E-02 | 1.37E-01 | 2.70E+00 | 2.73E+00 | 8.74E-01 | no | Vps13c | intron_variant | MODIFIER |
| 11 | 46636808 | AX-170380736 | 1 | A | G | 0.08 | -3.01E-02 | 1.82E-01 | 2.62E+00 | 2.72E+00 | 8.74E-01 | no | Gm4926 | non_coding_transcript_exon_variant | MODIFIER |
| 11 | 47328119 | AX-170046977 | 1 | T | C | 0.08 | -3.01E-02 | 1.82E-01 | 2.62E+00 | 2.72E+00 | 8.74E-01 | no | Sgcd | intron_variant | MODIFIER |
| 7 | 91822500 | AX-97780737 | 0 | G | A | 0.132 | -3.29E-02 | 2.11E-01 | 2.66E+00 | 2.74E+00 | 8.74E-01 | no | Dlg2 | intron_variant | MODIFIER |
| 16 | 94337118 | AX-170387679 | 0 | T | C | 0.412 | 1.13E-02 | 7.92E-02 | 2.68E+00 | 2.78E+00 | 8.74E-01 | no | Ripply3 | downstream_gene_variant | MODIFIER |
| 1 | 37210169 | AX-170040907 | 0 | A | G | 0.447 | -1.24E-02 | 7.94E-02 | 2.66E+00 | 2.74E+00 | 8.74E-01 | no | Cnga3 | upstream_gene_variant | MODIFIER |
| 1 | 55740857 | AX-170433270 | 1 | C | T | 0.107 | 3.69E-02 | 2.36E-01 | 2.69E+00 | 2.74E+00 | 8.74E-01 | no | Plcl1 | intron_variant | MODIFIER |
| 1 | 60761603 | AX-170507135 | 1 | G | C | 0.107 | 3.69E-02 | 2.36E-01 | 2.69E+00 | 2.74E+00 | 8.74E-01 | no | Cd28 | intron_variant | MODIFIER |
| 1 | 39304953 | AX-170091475 | 0 | T | C | 0.395 | -1.22E-02 | 7.29E-02 | 2.63E+00 | 2.71E+00 | 8.74E-01 | no | Npas2 | intron_variant | MODIFIER |
| 7 | 81099333 | AX-170190280 | 0 | C | T | 0.114 | 2.52E-02 | 1.57E-01 | 2.60E+00 | 2.73E+00 | 8.74E-01 | no | Alpk3 | intron_variant | MODIFIER |
| 1 | 82300095 | AX-170368221 | 0 | C | T | 0.158 | 3.34E-02 | 2.06E-01 | 2.65E+00 | 2.72E+00 | 8.74E-01 | no | | intergenic_variant | MODIFIER |
| 8 | 90608213 | AX-170314324 | 0 | T | C | 0.193 | 1.76E-02 | 1.18E-01 | 2.69E+00 | 2.77E+00 | 8.74E-01 | no | | intergenic_variant | MODIFIER |
| 9 | 79730767 | AX-97823596 | 0 | G | A | 0.202 | -2.03E-02 | 1.24E-01 | 2.65E+00 | 2.71E+00 | 8.74E-01 | no | | intergenic_variant | MODIFIER |
| 9 | 80311447 | AX-170437741 | 0 | C | T | 0.202 | -2.03E-02 | 1.24E-01 | 2.65E+00 | 2.71E+00 | 8.74E-01 | no | Myo6 | 3_prime_UTR_variant | MODIFIER |
| 9 | 68589601 | AX-170082731 | 1 | C | T | 0.196 | 2.13E-02 | 1.38E-01 | 2.73E+00 | 2.76E+00 | 8.74E-01 | no | | intergenic_variant | MODIFIER |
| 2 | 42524578 | AX-170250310 | 0 | T | C | 0.044 | 3.05E-02 | 2.05E-01 | 2.68E+00 | 2.76E+00 | 8.75E-01 | no | Lrp1b | intron_variant | MODIFIER |
| 17 | 89549516 | AX-97847117 | 0 | T | C | 0.044 | 3.05E-02 | 2.05E-01 | 2.68E+00 | 2.76E+00 | 8.75E-01 | no | | intergenic_variant | MODIFIER |
| 4 | 93617820 | AX-170209021 | 0 | G | T | 0.325 | 1.69E-02 | 1.05E-01 | 2.64E+00 | 2.72E+00 | 8.75E-01 | no | Gm12649 | intron_variant,non_coding_transcript_variant | MODIFIER |
| 1 | 42150522 | AX-170000358 | 1 | C | T | 0.446 | -1.17E-02 | 7.43E-02 | 2.64E+00 | 2.73E+00 | 8.75E-01 | no | | intergenic_variant | MODIFIER |
| 1 | 186114719 | AX-97801333 | 0 | T | C | 0.026 | 1.91E-02 | 1.23E-01 | 2.55E+00 | 2.74E+00 | 8.75E-01 | no | Lyplal1 | intron_variant | MODIFIER |
| 20 | 109971316 | AX-97807354 | 1 | G | A | 0.304 | 8.22E-03 | 5.33E-02 | 2.62E+00 | 2.74E+00 | 8.75E-01 | no | | intergenic_variant | MODIFIER |
| 1 | 61105536 | AX-170000624 | 0 | C | T | 0.14 | 3.12E-02 | 1.90E-01 | 2.62E+00 | 2.72E+00 | 8.75E-01 | no | | intergenic_variant | MODIFIER |
| 1 | 4026064 | AX-170433541 | 1 | G | A | 0.527 | -9.75E-03 | 6.90E-02 | 2.72E+00 | 2.81E+00 | 8.75E-01 | no | Rp1 | intron_variant | MODIFIER |
| 5 | 43564391 | AX-170375073 | 0 | T | A | 0.184 | -1.71E-02 | 1.04E-01 | 2.62E+00 | 2.72E+00 | 8.75E-01 | no | C1qtnf7 | intron_variant | MODIFIER |
| 8 | 90816546 | AX-170173792 | 2 | A | T | 0.191 | -1.94E-02 | 1.14E-01 | 2.61E+00 | 2.70E+00 | 8.75E-01 | no | Gm19935 | 3_prime_UTR_variant | MODIFIER |
| 5 | 37523874 | AX-170506505 | 0 | G | A | 0.342 | 1.64E-02 | 1.08E-01 | 2.68E+00 | 2.75E+00 | 8.75E-01 | no | Stk32b | intron_variant | MODIFIER |
| 20 | 140715241 | AX-170035089 | 0 | G | A | 0.149 | 2.13E-02 | 1.35E-01 | 2.64E+00 | 2.73E+00 | 8.75E-01 | no | Mid2 | intron_variant | MODIFIER |
| 8 | 47746312 | AX-97848286 | 1 | C | T | 0.152 | 2.23E-02 | 1.47E-01 | 2.72E+00 | 2.77E+00 | 8.75E-01 | no | Gm31786 | intron_variant,non_coding_transcript_variant | MODIFIER |
| 4 | 148195034 | AX-170521609 | 2 | T | C | 0.064 | 3.20E-02 | 2.18E-01 | 2.67E+00 | 2.76E+00 | 8.75E-01 | no | | intergenic_variant | MODIFIER |
| 9 | 123036384 | AX-97850039 | 0 | G | A | 0.149 | -1.34E-02 | 8.78E-02 | 2.63E+00 | 2.74E+00 | 8.76E-01 | no | Tgm4 | intron_variant | MODIFIER |
| 5 | 33588179 | AX-170447564 | 0 | G | A | 0.342 | 1.64E-02 | 1.03E-01 | 2.65E+00 | 2.72E+00 | 8.76E-01 | no | | intergenic_variant | MODIFIER |
| 6 | 147117708 | AX-97852520 | 1 | C | T | 0.08 | 3.34E-02 | 2.05E-01 | 2.62E+00 | 2.72E+00 | 8.76E-01 | no | Klhl42 | downstream_gene_variant | MODIFIER |
| 1 | 38741436 | AX-170172768 | 1 | C | T | 0.464 | 1.16E-02 | 7.71E-02 | 2.67E+00 | 2.76E+00 | 8.76E-01 | no | | intergenic_variant | MODIFIER |
| 7 | 109511900 | AX-97828527 | 1 | G | A | 0.482 | -1.40E-02 | 9.50E-02 | 2.70E+00 | 2.78E+00 | 8.76E-01 | no | Trim66 | upstream_gene_variant | MODIFIER |
| 8 | 93747679 | AX-170418201 | 1 | G | A | 0.214 | -1.85E-02 | 1.19E-01 | 2.67E+00 | 2.73E+00 | 8.76E-01 | no | 4930488L21Rik | downstream_gene_variant | MODIFIER |
| 1 | 70926271 | AX-170206656 | 0 | G | A | 0.132 | 3.43E-02 | 2.22E-01 | 2.67E+00 | 2.74E+00 | 8.76E-01 | no | | intergenic_variant | MODIFIER |
| 1 | 10029129 | AX-170271678 | 1 | G | A | 0.098 | 2.73E-02 | 1.72E-01 | 2.62E+00 | 2.73E+00 | 8.76E-01 | no | Cops5 | intron_variant | MODIFIER |
| 10 | 9608714 | AX-170456702 | 0 | T | C | 0.202 | 1.61E-02 | 1.08E-01 | 2.68E+00 | 2.76E+00 | 8.76E-01 | no | | intergenic_variant | MODIFIER |
| 10 | 9619697 | AX-97812307 | 0 | A | C | 0.202 | 1.61E-02 | 1.08E-01 | 2.68E+00 | 2.76E+00 | 8.76E-01 | no | Samd5 | downstream_gene_variant | MODIFIER |
| 10 | 9622698 | AX-170227490 | 0 | C | G | 0.202 | 1.61E-02 | 1.08E-01 | 2.68E+00 | 2.76E+00 | 8.76E-01 | no | Samd5 | downstream_gene_variant | MODIFIER |
| 10 | 10525356 | AX-170002711 | 0 | C | G | 0.202 | 1.61E-02 | 1.08E-01 | 2.68E+00 | 2.76E+00 | 8.76E-01 | no | | intergenic_variant | MODIFIER |
| 10 | 10748972 | AX-170446121 | 0 | T | C | 0.202 | 1.61E-02 | 1.08E-01 | 2.68E+00 | 2.76E+00 | 8.76E-01 | no | Grm1 | intron_variant | MODIFIER |
| 10 | 11098520 | AX-97814727 | 0 | A | G | 0.202 | 1.61E-02 | 1.08E-01 | 2.68E+00 | 2.76E+00 | 8.76E-01 | no | | intergenic_variant | MODIFIER |
| 10 | 12222567 | AX-97804704 | 0 | G | A | 0.202 | 1.61E-02 | 1.08E-01 | 2.68E+00 | 2.76E+00 | 8.76E-01 | no | | intergenic_variant | MODIFIER |
| 10 | 12283723 | AX-170436243 | 0 | C | T | 0.202 | 1.61E-02 | 1.08E-01 | 2.68E+00 | 2.76E+00 | 8.76E-01 | no | | intergenic_variant | MODIFIER |

| | | | | | | | | | | | | | | | |
|---|---|---|---|---|---|---|---|---|---|---|---|---|---|---|---|
| 10 | 12289392 | AX-170246289 | 0 | A | G | 0.202 | 1.61E-02 | 1.08E-01 | 2.68E+00 | 2.76E+00 | 8.76E-01 | no | | intergenic_variant | MODIFIER |
| 10 | 12289854 | AX-170289369 | 0 | T | C | 0.202 | 1.61E-02 | 1.08E-01 | 2.68E+00 | 2.76E+00 | 8.76E-01 | no | | intergenic_variant | MODIFIER |
| 10 | 12293499 | AX-170140954 | 0 | G | C | 0.202 | 1.61E-02 | 1.08E-01 | 2.68E+00 | 2.76E+00 | 8.76E-01 | no | | intergenic_variant | MODIFIER |
| 9 | 52563391 | AX-170129399 | 0 | A | G | 0.228 | 1.72E-02 | 1.13E-01 | 2.68E+00 | 2.75E+00 | 8.76E-01 | no | | intergenic_variant | MODIFIER |
| 7 | 88454640 | AX-170311651 | 1 | A | G | 0.125 | -2.99E-02 | 1.99E-01 | 2.66E+00 | 2.75E+00 | 8.76E-01 | no | Rab38 | intron_variant | MODIFIER |
| 6 | 133197395 | AX-170505678 | 2 | C | G | 0.527 | 1.51E-02 | 8.55E-02 | 2.60E+00 | 2.68E+00 | 8.76E-01 | no | Gm25019 | downstream_gene_variant | MODIFIER |
| 12 | 71147134 | AX-170303200 | 0 | G | A | 0.07 | 3.57E-02 | 2.28E-01 | 2.66E+00 | 2.73E+00 | 8.76E-01 | no | 2700049A03Rik | intron_variant | MODIFIER |
| 18 | 41545832 | AX-170334874 | 0 | G | A | 0.07 | 3.57E-02 | 2.28E-01 | 2.66E+00 | 2.73E+00 | 8.76E-01 | no | | intergenic_variant | MODIFIER |
| 18 | 41828859 | AX-170484741 | 0 | C | T | 0.07 | 3.57E-02 | 2.28E-01 | 2.66E+00 | 2.73E+00 | 8.76E-01 | no | AC154172.2 | intron_variant,non_coding_transcript_variant | MODIFIER |
| 13 | 54329290 | AX-170087429 | 0 | A | T | 0.526 | 1.20E-02 | 6.94E-02 | 2.59E+00 | 2.70E+00 | 8.76E-01 | no | Gm48622 | upstream_gene_variant | MODIFIER |
| 5 | 73608401 | AX-170342393 | 1 | T | C | 0.429 | 1.58E-02 | 9.82E-02 | 2.64E+00 | 2.72E+00 | 8.76E-01 | no | Lrrc66 | missense_variant | MODERATE |
| 13 | 22242969 | AX-170007904 | 1 | T | A | 0.161 | -2.04E-02 | 1.14E-01 | 2.59E+00 | 2.68E+00 | 8.76E-01 | no | Vmn1r194 | upstream_gene_variant | MODIFIER |
| 7 | 106444834 | AX-170110838 | 0 | G | A | 0.482 | -1.38E-02 | 9.47E-02 | 2.70E+00 | 2.78E+00 | 8.76E-01 | no | | intergenic_variant | MODIFIER |
| 10 | 114932482 | AX-170397436 | 0 | T | C | 0.368 | 1.14E-02 | 7.53E-02 | 2.66E+00 | 2.75E+00 | 8.76E-01 | no | | intergenic_variant | MODIFIER |
| 9 | 104590110 | AX-170281679 | 0 | C | T | 0.14 | 1.49E-02 | 1.03E-01 | 2.67E+00 | 2.77E+00 | 8.76E-01 | no | Cpne4 | intron_variant | MODIFIER |
| 6 | 139748768 | AX-170412038 | 0 | A | G | 0.053 | 3.24E-02 | 2.17E-01 | 2.63E+00 | 2.75E+00 | 8.77E-01 | no | Pik3c2g | intron_variant | MODIFIER |
| 1 | 37117183 | AX-170327469 | 1 | T | C | 0.446 | 1.17E-02 | 7.84E-02 | 2.67E+00 | 2.75E+00 | 8.77E-01 | no | Vwa3b | intron_variant,NMD_transcript_variant | MODIFIER |
| 9 | 85690991 | AX-170247380 | 0 | T | C | 0.132 | -2.17E-02 | 1.33E-01 | 2.64E+00 | 2.71E+00 | 8.77E-01 | no | Ibtk | intron_variant | MODIFIER |
| 9 | 85788796 | AX-170211062 | 0 | G | C | 0.132 | -2.17E-02 | 1.33E-01 | 2.64E+00 | 2.71E+00 | 8.77E-01 | no | Gm18766 | downstream_gene_variant | MODIFIER |
| 9 | 85800819 | AX-170412836 | 0 | A | G | 0.132 | -2.17E-02 | 1.33E-01 | 2.64E+00 | 2.71E+00 | 8.77E-01 | no | | intergenic_variant | MODIFIER |
| 9 | 85836772 | AX-170266497 | 0 | G | A | 0.132 | -2.17E-02 | 1.33E-01 | 2.64E+00 | 2.71E+00 | 8.77E-01 | no | 9330154J02Rik | intron_variant,non_coding_transcript_variant | MODIFIER |
| 9 | 85844981 | AX-170338851 | 0 | A | G | 0.132 | -2.17E-02 | 1.33E-01 | 2.64E+00 | 2.71E+00 | 8.77E-01 | no | Tpbg | synonymous_variant | LOW |
| 9 | 85949286 | AX-170481390 | 0 | A | G | 0.132 | -2.17E-02 | 1.33E-01 | 2.64E+00 | 2.71E+00 | 8.77E-01 | no | | intergenic_variant | MODIFIER |
| 9 | 85995347 | AX-170391381 | 0 | A | G | 0.132 | -2.17E-02 | 1.33E-01 | 2.64E+00 | 2.71E+00 | 8.77E-01 | no | | intergenic_variant | MODIFIER |
| 9 | 86037213 | AX-170032880 | 0 | C | T | 0.132 | -2.17E-02 | 1.33E-01 | 2.64E+00 | 2.71E+00 | 8.77E-01 | no | | intergenic_variant | MODIFIER |
| 9 | 86037302 | AX-170032879 | 0 | A | G | 0.132 | -2.17E-02 | 1.33E-01 | 2.64E+00 | 2.71E+00 | 8.77E-01 | no | | intergenic_variant | MODIFIER |
| 9 | 86037483 | AX-170439919 | 0 | A | C | 0.132 | -2.17E-02 | 1.33E-01 | 2.64E+00 | 2.71E+00 | 8.77E-01 | no | | intergenic_variant | MODIFIER |
| 9 | 86037796 | AX-170483471 | 0 | T | G | 0.132 | -2.17E-02 | 1.33E-01 | 2.64E+00 | 2.71E+00 | 8.77E-01 | no | | intergenic_variant | MODIFIER |
| 9 | 86053212 | AX-170264651 | 0 | C | G | 0.132 | -2.17E-02 | 1.33E-01 | 2.64E+00 | 2.71E+00 | 8.77E-01 | no | | intergenic_variant | MODIFIER |
| 9 | 86053499 | AX-170139258 | 0 | A | G | 0.132 | -2.17E-02 | 1.33E-01 | 2.64E+00 | 2.71E+00 | 8.77E-01 | no | | intergenic_variant | MODIFIER |
| 9 | 86061515 | AX-170032884 | 0 | T | C | 0.132 | -2.17E-02 | 1.33E-01 | 2.64E+00 | 2.71E+00 | 8.77E-01 | no | | intergenic_variant | MODIFIER |
| 9 | 86061936 | AX-170032881 | 0 | T | C | 0.132 | -2.17E-02 | 1.33E-01 | 2.64E+00 | 2.71E+00 | 8.77E-01 | no | | intergenic_variant | MODIFIER |
| 9 | 86070474 | AX-170171837 | 0 | A | G | 0.132 | -2.17E-02 | 1.33E-01 | 2.64E+00 | 2.71E+00 | 8.77E-01 | no | | intergenic_variant | MODIFIER |
| 9 | 86095365 | AX-170032878 | 0 | A | G | 0.132 | -2.17E-02 | 1.33E-01 | 2.64E+00 | 2.71E+00 | 8.77E-01 | no | | intergenic_variant | MODIFIER |
| 9 | 86133183 | AX-170144899 | 0 | A | G | 0.132 | -2.17E-02 | 1.33E-01 | 2.64E+00 | 2.71E+00 | 8.77E-01 | no | | intergenic_variant | MODIFIER |
| 9 | 86133252 | AX-170459229 | 0 | A | G | 0.132 | -2.17E-02 | 1.33E-01 | 2.64E+00 | 2.71E+00 | 8.77E-01 | no | | intergenic_variant | MODIFIER |
| 9 | 86319502 | AX-170482369 | 0 | T | C | 0.132 | -2.17E-02 | 1.33E-01 | 2.64E+00 | 2.71E+00 | 8.77E-01 | no | Ube2cbp | intron_variant | MODIFIER |
| 9 | 86320501 | AX-170249778 | 0 | T | G | 0.132 | -2.17E-02 | 1.33E-01 | 2.64E+00 | 2.71E+00 | 8.77E-01 | no | Ube2cbp | intron_variant | MODIFIER |
| 9 | 86328178 | AX-170246417 | 0 | G | C | 0.132 | -2.17E-02 | 1.33E-01 | 2.64E+00 | 2.71E+00 | 8.77E-01 | no | Ube2cbp | intron_variant | MODIFIER |
| 9 | 86336672 | AX-170350330 | 0 | T | C | 0.132 | -2.17E-02 | 1.33E-01 | 2.64E+00 | 2.71E+00 | 8.77E-01 | no | Ube2cbp | intron_variant | MODIFIER |
| 9 | 86377827 | AX-170189447 | 0 | A | G | 0.132 | -2.17E-02 | 1.33E-01 | 2.64E+00 | 2.71E+00 | 8.77E-01 | no | Ube2cbp | intron_variant | MODIFIER |
| 9 | 86463729 | AX-170224689 | 0 | A | C | 0.132 | -2.17E-02 | 1.33E-01 | 2.64E+00 | 2.71E+00 | 8.77E-01 | no | Ube2cbp | intron_variant | MODIFIER |
| 9 | 86550916 | AX-170149157 | 0 | A | G | 0.132 | -2.17E-02 | 1.33E-01 | 2.64E+00 | 2.71E+00 | 8.77E-01 | no | Dop1a | intron_variant | MODIFIER |
| 9 | 86596203 | AX-97849192 | 0 | A | G | 0.132 | -2.17E-02 | 1.33E-01 | 2.64E+00 | 2.71E+00 | 8.77E-01 | no | Me1 | splice_region_variant,intron_variant | LOW |
| 9 | 86713209 | AX-170330044 | 0 | A | G | 0.132 | -2.17E-02 | 1.33E-01 | 2.64E+00 | 2.71E+00 | 8.77E-01 | no | Gm37484 | non_coding_transcript_exon_variant | MODIFIER |
| 9 | 86780902 | AX-170381093 | 0 | T | C | 0.132 | -2.17E-02 | 1.33E-01 | 2.64E+00 | 2.71E+00 | 8.77E-01 | no | Snap91 | intron_variant | MODIFIER |
| 9 | 86808887 | AX-170437380 | 0 | T | C | 0.132 | -2.17E-02 | 1.33E-01 | 2.64E+00 | 2.71E+00 | 8.77E-01 | no | Snap91 | intron_variant | MODIFIER |
| 9 | 86810289 | AX-170163472 | 0 | A | G | 0.132 | -2.17E-02 | 1.33E-01 | 2.64E+00 | 2.71E+00 | 8.77E-01 | no | Snap91 | intron_variant | MODIFIER |
| 9 | 86867907 | AX-97830946 | 0 | A | G | 0.132 | -2.17E-02 | 1.33E-01 | 2.64E+00 | 2.71E+00 | 8.77E-01 | no | Snap91 | intron_variant | MODIFIER |
| 9 | 86868681 | AX-170358639 | 0 | A | G | 0.132 | -2.17E-02 | 1.33E-01 | 2.64E+00 | 2.71E+00 | 8.77E-01 | no | Snap91 | intron_variant | MODIFIER |
| 9 | 86901562 | AX-170239596 | 0 | T | G | 0.132 | -2.17E-02 | 1.33E-01 | 2.64E+00 | 2.71E+00 | 8.77E-01 | no | | intergenic_variant | MODIFIER |
| 9 | 86901800 | AX-97783690 | 0 | C | T | 0.132 | -2.17E-02 | 1.33E-01 | 2.64E+00 | 2.71E+00 | 8.77E-01 | no | | intergenic_variant | MODIFIER |
| 9 | 86904920 | AX-170083034 | 0 | T | C | 0.132 | -2.17E-02 | 1.33E-01 | 2.64E+00 | 2.71E+00 | 8.77E-01 | no | | intergenic_variant | MODIFIER |
| 9 | 86905452 | AX-170083032 | 0 | A | G | 0.132 | -2.17E-02 | 1.33E-01 | 2.64E+00 | 2.71E+00 | 8.77E-01 | no | | intergenic_variant | MODIFIER |

| | | | | | | | | | | | | | | | |
|---|---|---|---|---|---|---|---|---|---|---|---|---|---|---|---|
| 9 | 86905514 | AX-170174949 | 0 | A | G | 0.132 | -2.17E-02 | 1.33E-01 | 2.64E+00 | 2.71E+00 | 8.77E-01 | no | | intergenic_variant | MODIFIER |
| 9 | 86951655 | AX-170083033 | 0 | T | G | 0.132 | -2.17E-02 | 1.33E-01 | 2.64E+00 | 2.71E+00 | 8.77E-01 | no | | regulatory_region_variant | MODIFIER |
| 9 | 86952110 | AX-170340002 | 0 | A | T | 0.132 | -2.17E-02 | 1.33E-01 | 2.64E+00 | 2.71E+00 | 8.77E-01 | no | | regulatory_region_variant | MODIFIER |
| 9 | 86952460 | AX-170358721 | 0 | A | G | 0.132 | -2.17E-02 | 1.33E-01 | 2.64E+00 | 2.71E+00 | 8.77E-01 | no | | regulatory_region_variant | MODIFIER |
| 9 | 86953001 | AX-170228148 | 0 | A | G | 0.132 | -2.17E-02 | 1.33E-01 | 2.64E+00 | 2.71E+00 | 8.77E-01 | no | | regulatory_region_variant | MODIFIER |
| 9 | 86964443 | AX-97792702 | 0 | A | G | 0.132 | -2.17E-02 | 1.33E-01 | 2.64E+00 | 2.71E+00 | 8.77E-01 | no | | intergenic_variant | MODIFIER |
| 9 | 86966620 | AX-170283927 | 0 | C | G | 0.132 | -2.17E-02 | 1.33E-01 | 2.64E+00 | 2.71E+00 | 8.77E-01 | no | | intergenic_variant | MODIFIER |
| 9 | 87013087 | AX-170126948 | 0 | C | T | 0.132 | -2.17E-02 | 1.33E-01 | 2.64E+00 | 2.71E+00 | 8.77E-01 | no | Ripply2 | upstream_gene_variant | MODIFIER |
| 9 | 87018228 | AX-170456302 | 0 | A | G | 0.132 | -2.17E-02 | 1.33E-01 | 2.64E+00 | 2.71E+00 | 8.77E-01 | no | Ripply2 | intron_variant | MODIFIER |
| 9 | 87026688 | AX-97809761 | 0 | T | C | 0.132 | -2.17E-02 | 1.33E-01 | 2.64E+00 | 2.71E+00 | 8.77E-01 | no | Cyb5r4 | intron_variant | MODIFIER |
| 9 | 87045691 | AX-97848908 | 0 | C | T | 0.132 | -2.17E-02 | 1.33E-01 | 2.64E+00 | 2.71E+00 | 8.77E-01 | no | Cyb5r4 | intron_variant | MODIFIER |
| 9 | 87054613 | AX-170273974 | 0 | A | G | 0.132 | -2.17E-02 | 1.33E-01 | 2.64E+00 | 2.71E+00 | 8.77E-01 | no | Cyb5r4 | intron_variant | MODIFIER |
| 9 | 87056338 | AX-170141755 | 0 | A | G | 0.132 | -2.17E-02 | 1.33E-01 | 2.64E+00 | 2.71E+00 | 8.77E-01 | no | Cyb5r4 | intron_variant | MODIFIER |
| 9 | 87066697 | AX-170096619 | 0 | C | A | 0.132 | -2.17E-02 | 1.33E-01 | 2.64E+00 | 2.71E+00 | 8.77E-01 | no | Cyb5r4 | synonymous_variant | LOW |
| 9 | 87066980 | AX-170032901 | 0 | A | G | 0.132 | -2.17E-02 | 1.33E-01 | 2.64E+00 | 2.71E+00 | 8.77E-01 | no | Cyb5r4 | intron_variant | MODIFIER |
| 9 | 87068242 | AX-170198154 | 0 | A | C | 0.132 | -2.17E-02 | 1.33E-01 | 2.64E+00 | 2.71E+00 | 8.77E-01 | no | Cyb5r4 | intron_variant | MODIFIER |
| 9 | 87078572 | AX-170261509 | 0 | T | C | 0.132 | -2.17E-02 | 1.33E-01 | 2.64E+00 | 2.71E+00 | 8.77E-01 | no | Cyb5r4 | downstream_gene_variant | MODIFIER |
| 9 | 87138424 | AX-170167139 | 0 | T | C | 0.132 | -2.17E-02 | 1.33E-01 | 2.64E+00 | 2.71E+00 | 8.77E-01 | no | | intergenic_variant | MODIFIER |
| 9 | 87139056 | AX-170032899 | 0 | C | A | 0.132 | -2.17E-02 | 1.33E-01 | 2.64E+00 | 2.71E+00 | 8.77E-01 | no | | intergenic_variant | MODIFIER |
| 9 | 87140116 | AX-170517880 | 0 | A | G | 0.132 | -2.17E-02 | 1.33E-01 | 2.64E+00 | 2.71E+00 | 8.77E-01 | no | Mrap2 | upstream_gene_variant | MODIFIER |
| 9 | 87178526 | AX-170235723 | 0 | C | T | 0.132 | -2.17E-02 | 1.33E-01 | 2.64E+00 | 2.71E+00 | 8.77E-01 | no | Mrap2 | intron_variant | MODIFIER |
| 9 | 87240607 | AX-97795545 | 0 | G | T | 0.132 | -2.17E-02 | 1.33E-01 | 2.64E+00 | 2.71E+00 | 8.77E-01 | no | Cep162 | intron_variant | MODIFIER |
| 9 | 87300051 | AX-170083040 | 0 | G | A | 0.132 | -2.17E-02 | 1.33E-01 | 2.64E+00 | 2.71E+00 | 8.77E-01 | no | | intergenic_variant | MODIFIER |
| 9 | 87302371 | AX-170110272 | 0 | G | C | 0.132 | -2.17E-02 | 1.33E-01 | 2.64E+00 | 2.71E+00 | 8.77E-01 | no | | intergenic_variant | MODIFIER |
| 9 | 87303286 | AX-170414002 | 0 | G | T | 0.132 | -2.17E-02 | 1.33E-01 | 2.64E+00 | 2.71E+00 | 8.77E-01 | no | | intergenic_variant | MODIFIER |
| 9 | 87330521 | AX-170096620 | 0 | C | T | 0.132 | -2.17E-02 | 1.33E-01 | 2.64E+00 | 2.71E+00 | 8.77E-01 | no | | intergenic_variant | MODIFIER |
| 9 | 87331668 | AX-170468733 | 0 | T | C | 0.132 | -2.17E-02 | 1.33E-01 | 2.64E+00 | 2.71E+00 | 8.77E-01 | no | | intergenic_variant | MODIFIER |
| 9 | 87344582 | AX-170220860 | 0 | T | C | 0.132 | -2.17E-02 | 1.33E-01 | 2.64E+00 | 2.71E+00 | 8.77E-01 | no | | intergenic_variant | MODIFIER |
| 9 | 87346823 | AX-170244833 | 0 | G | A | 0.132 | -2.17E-02 | 1.33E-01 | 2.64E+00 | 2.71E+00 | 8.77E-01 | no | | intergenic_variant | MODIFIER |
| 9 | 87354426 | AX-170139124 | 0 | A | T | 0.132 | -2.17E-02 | 1.33E-01 | 2.64E+00 | 2.71E+00 | 8.77E-01 | no | | intergenic_variant | MODIFIER |
| 9 | 87354753 | AX-170511711 | 0 | A | G | 0.132 | -2.17E-02 | 1.33E-01 | 2.64E+00 | 2.71E+00 | 8.77E-01 | no | | intergenic_variant | MODIFIER |
| 9 | 87360160 | AX-170319860 | 0 | T | C | 0.132 | -2.17E-02 | 1.33E-01 | 2.64E+00 | 2.71E+00 | 8.77E-01 | no | | intergenic_variant | MODIFIER |
| 9 | 87370281 | AX-170465274 | 0 | T | C | 0.132 | -2.17E-02 | 1.33E-01 | 2.64E+00 | 2.71E+00 | 8.77E-01 | no | | intergenic_variant | MODIFIER |
| 9 | 87373432 | AX-97817425 | 0 | T | C | 0.132 | -2.17E-02 | 1.33E-01 | 2.64E+00 | 2.71E+00 | 8.77E-01 | no | | intergenic_variant | MODIFIER |
| 9 | 87378355 | AX-170222327 | 0 | A | G | 0.132 | -2.17E-02 | 1.33E-01 | 2.64E+00 | 2.71E+00 | 8.77E-01 | no | | intergenic_variant | MODIFIER |
| 9 | 87379862 | AX-170126931 | 0 | G | C | 0.132 | -2.17E-02 | 1.33E-01 | 2.64E+00 | 2.71E+00 | 8.77E-01 | no | | regulatory_region_variant | MODIFIER |
| 9 | 87410045 | AX-170100920 | 0 | G | A | 0.132 | -2.17E-02 | 1.33E-01 | 2.64E+00 | 2.71E+00 | 8.77E-01 | no | | intergenic_variant | MODIFIER |
| 9 | 87411071 | AX-170357887 | 0 | A | G | 0.132 | -2.17E-02 | 1.33E-01 | 2.64E+00 | 2.71E+00 | 8.77E-01 | no | | intergenic_variant | MODIFIER |
| 9 | 87411604 | AX-170183467 | 0 | C | T | 0.132 | -2.17E-02 | 1.33E-01 | 2.64E+00 | 2.71E+00 | 8.77E-01 | no | | intergenic_variant | MODIFIER |
| 9 | 87419093 | AX-170096621 | 0 | C | T | 0.132 | -2.17E-02 | 1.33E-01 | 2.64E+00 | 2.71E+00 | 8.77E-01 | no | | intergenic_variant | MODIFIER |
| 9 | 87420942 | AX-170083042 | 0 | C | T | 0.132 | -2.17E-02 | 1.33E-01 | 2.64E+00 | 2.71E+00 | 8.77E-01 | no | | intergenic_variant | MODIFIER |
| 9 | 87425433 | AX-170507249 | 0 | C | T | 0.132 | -2.17E-02 | 1.33E-01 | 2.64E+00 | 2.71E+00 | 8.77E-01 | no | | regulatory_region_variant | MODIFIER |
| 9 | 87429556 | AX-170503334 | 0 | A | T | 0.132 | -2.17E-02 | 1.33E-01 | 2.64E+00 | 2.71E+00 | 8.77E-01 | no | | intergenic_variant | MODIFIER |
| 9 | 87429667 | AX-170226103 | 0 | G | A | 0.132 | -2.17E-02 | 1.33E-01 | 2.64E+00 | 2.71E+00 | 8.77E-01 | no | | intergenic_variant | MODIFIER |
| 9 | 87430260 | AX-170206381 | 0 | A | G | 0.132 | -2.17E-02 | 1.33E-01 | 2.64E+00 | 2.71E+00 | 8.77E-01 | no | | intergenic_variant | MODIFIER |
| 9 | 87437270 | AX-170323112 | 0 | A | G | 0.132 | -2.17E-02 | 1.33E-01 | 2.64E+00 | 2.71E+00 | 8.77E-01 | no | | intergenic_variant | MODIFIER |
| 9 | 87441244 | AX-170361441 | 0 | G | A | 0.132 | -2.17E-02 | 1.33E-01 | 2.64E+00 | 2.71E+00 | 8.77E-01 | no | | intergenic_variant | MODIFIER |
| 9 | 87441837 | AX-170262113 | 0 | C | T | 0.132 | -2.17E-02 | 1.33E-01 | 2.64E+00 | 2.71E+00 | 8.77E-01 | no | | intergenic_variant | MODIFIER |
| 9 | 87448229 | AX-97807793 | 0 | A | G | 0.132 | -2.17E-02 | 1.33E-01 | 2.64E+00 | 2.71E+00 | 8.77E-01 | no | | intergenic_variant | MODIFIER |
| 9 | 87452724 | AX-170314552 | 0 | G | T | 0.132 | -2.17E-02 | 1.33E-01 | 2.64E+00 | 2.71E+00 | 8.77E-01 | no | | intergenic_variant | MODIFIER |
| 9 | 87456997 | AX-170356514 | 0 | T | G | 0.132 | -2.17E-02 | 1.33E-01 | 2.64E+00 | 2.71E+00 | 8.77E-01 | no | | intergenic_variant | MODIFIER |
| 9 | 87572001 | AX-170380375 | 0 | T | G | 0.132 | -2.17E-02 | 1.33E-01 | 2.64E+00 | 2.71E+00 | 8.77E-01 | no | | intergenic_variant | MODIFIER |
| 9 | 87684917 | AX-170251208 | 0 | A | G | 0.132 | -2.17E-02 | 1.33E-01 | 2.64E+00 | 2.71E+00 | 8.77E-01 | no | | intergenic_variant | MODIFIER |
| 9 | 87723685 | AX-170032903 | 0 | T | C | 0.132 | -2.17E-02 | 1.33E-01 | 2.64E+00 | 2.71E+00 | 8.77E-01 | no | Tbx18 | intron_variant | MODIFIER |

| | | | | | | | | | | | | | | | |
|---|---|---|---|---|---|---|---|---|---|---|---|---|---|---|---|
| 9 | 87739648 | AX-170274032 | 0 | T | C | 0.132 | -2.17E-02 | 1.33E-01 | 2.64E+00 | 2.71E+00 | 8.77E-01 | no | Gm37374 | non_coding_transcript_exon_variant | MODIFIER |
| 9 | 87746288 | AX-97795504 | 0 | T | G | 0.132 | -2.17E-02 | 1.33E-01 | 2.64E+00 | 2.71E+00 | 8.77E-01 | no | | intergenic_variant | MODIFIER |
| 9 | 87748625 | AX-170413061 | 0 | A | G | 0.132 | -2.17E-02 | 1.33E-01 | 2.64E+00 | 2.71E+00 | 8.77E-01 | no | | intergenic_variant | MODIFIER |
| 9 | 88038832 | AX-170083053 | 0 | G | T | 0.132 | -2.17E-02 | 1.33E-01 | 2.64E+00 | 2.71E+00 | 8.77E-01 | no | Gm28877 | downstream_gene_variant | MODIFIER |
| 9 | 88074492 | AX-170376976 | 0 | C | T | 0.132 | -2.17E-02 | 1.33E-01 | 2.64E+00 | 2.71E+00 | 8.77E-01 | no | | intergenic_variant | MODIFIER |
| 9 | 88166781 | AX-170032914 | 0 | T | C | 0.132 | -2.17E-02 | 1.33E-01 | 2.64E+00 | 2.71E+00 | 8.77E-01 | no | | intergenic_variant | MODIFIER |
| 9 | 88173687 | AX-170090366 | 0 | G | A | 0.132 | -2.17E-02 | 1.33E-01 | 2.64E+00 | 2.71E+00 | 8.77E-01 | no | | intergenic_variant | MODIFIER |
| 9 | 88179278 | AX-170376143 | 0 | G | C | 0.132 | -2.17E-02 | 1.33E-01 | 2.64E+00 | 2.71E+00 | 8.77E-01 | no | | intergenic_variant | MODIFIER |
| 9 | 88196062 | AX-170032913 | 0 | C | T | 0.132 | -2.17E-02 | 1.33E-01 | 2.64E+00 | 2.71E+00 | 8.77E-01 | no | | intergenic_variant | MODIFIER |
| 9 | 88209902 | AX-170430376 | 0 | G | A | 0.132 | -2.17E-02 | 1.33E-01 | 2.64E+00 | 2.71E+00 | 8.77E-01 | no | | intergenic_variant | MODIFIER |
| 9 | 88219921 | AX-170032918 | 0 | C | G | 0.132 | -2.17E-02 | 1.33E-01 | 2.64E+00 | 2.71E+00 | 8.77E-01 | no | | intergenic_variant | MODIFIER |
| 9 | 88221847 | AX-97819833 | 0 | G | A | 0.132 | -2.17E-02 | 1.33E-01 | 2.64E+00 | 2.71E+00 | 8.77E-01 | no | | regulatory_region_variant | MODIFIER |
| 9 | 88260279 | AX-97854318 | 0 | T | C | 0.132 | -2.17E-02 | 1.33E-01 | 2.64E+00 | 2.71E+00 | 8.77E-01 | no | Gm28231 | non_coding_transcript_exon_variant | MODIFIER |
| 9 | 88293145 | AX-170488841 | 0 | T | C | 0.132 | -2.17E-02 | 1.33E-01 | 2.64E+00 | 2.71E+00 | 8.77E-01 | no | | intergenic_variant | MODIFIER |
| 9 | 88298072 | AX-170248167 | 0 | G | A | 0.132 | -2.17E-02 | 1.33E-01 | 2.64E+00 | 2.71E+00 | 8.77E-01 | no | | regulatory_region_variant | MODIFIER |
| 9 | 88298360 | AX-170090367 | 0 | C | A | 0.132 | -2.17E-02 | 1.33E-01 | 2.64E+00 | 2.71E+00 | 8.77E-01 | no | | regulatory_region_variant | MODIFIER |
| 9 | 88341638 | AX-97793719 | 0 | A | G | 0.132 | -2.17E-02 | 1.33E-01 | 2.64E+00 | 2.71E+00 | 8.77E-01 | no | Nt5e | intron_variant | MODIFIER |
| 9 | 88350654 | AX-170368439 | 0 | G | C | 0.132 | -2.17E-02 | 1.33E-01 | 2.64E+00 | 2.71E+00 | 8.77E-01 | no | Nt5e | intron_variant | MODIFIER |
| 9 | 88435433 | AX-170467225 | 0 | T | C | 0.132 | -2.17E-02 | 1.33E-01 | 2.64E+00 | 2.71E+00 | 8.77E-01 | no | Snx14 | intron_variant,NMD_transcript_variant | MODIFIER |
| 9 | 88450807 | AX-97846502 | 0 | G | A | 0.132 | -2.17E-02 | 1.33E-01 | 2.64E+00 | 2.71E+00 | 8.77E-01 | no | Syncrip | 3_prime_UTR_variant | MODIFIER |
| 9 | 88484548 | AX-170251882 | 0 | G | C | 0.132 | -2.17E-02 | 1.33E-01 | 2.64E+00 | 2.71E+00 | 8.77E-01 | no | Syncrip | upstream_gene_variant | MODIFIER |
| 9 | 89313132 | AX-170326637 | 0 | G | T | 0.132 | -2.17E-02 | 1.33E-01 | 2.64E+00 | 2.71E+00 | 8.77E-01 | no | Mthfs | intron_variant,non_coding_transcript_variant | MODIFIER |
| 9 | 89313681 | AX-97800301 | 0 | T | C | 0.132 | -2.17E-02 | 1.33E-01 | 2.64E+00 | 2.71E+00 | 8.77E-01 | no | Mthfs | intron_variant,non_coding_transcript_variant | MODIFIER |
| 9 | 89332732 | AX-170386979 | 0 | A | G | 0.132 | -2.17E-02 | 1.33E-01 | 2.64E+00 | 2.71E+00 | 8.77E-01 | no | Mthfs | intron_variant,non_coding_transcript_variant | MODIFIER |
| 9 | 89333440 | AX-170415283 | 0 | A | G | 0.132 | -2.17E-02 | 1.33E-01 | 2.64E+00 | 2.71E+00 | 8.77E-01 | no | Mthfs | intron_variant,non_coding_transcript_variant | MODIFIER |
| 9 | 89401654 | AX-170417706 | 0 | G | A | 0.132 | -2.17E-02 | 1.33E-01 | 2.64E+00 | 2.71E+00 | 8.77E-01 | no | | intergenic_variant | MODIFIER |
| 9 | 89403108 | AX-170349683 | 0 | A | G | 0.132 | -2.17E-02 | 1.33E-01 | 2.64E+00 | 2.71E+00 | 8.77E-01 | no | | intergenic_variant | MODIFIER |
| 9 | 89414291 | AX-170202471 | 0 | G | A | 0.132 | -2.17E-02 | 1.33E-01 | 2.64E+00 | 2.71E+00 | 8.77E-01 | no | | intergenic_variant | MODIFIER |
| 9 | 89416903 | AX-97790154 | 0 | T | C | 0.132 | -2.17E-02 | 1.33E-01 | 2.64E+00 | 2.71E+00 | 8.77E-01 | no | | intergenic_variant | MODIFIER |
| 9 | 89438767 | AX-170120843 | 0 | A | G | 0.132 | -2.17E-02 | 1.33E-01 | 2.64E+00 | 2.71E+00 | 8.77E-01 | no | | intergenic_variant | MODIFIER |
| 9 | 89452888 | AX-170391980 | 0 | A | C | 0.132 | -2.17E-02 | 1.33E-01 | 2.64E+00 | 2.71E+00 | 8.77E-01 | no | | intergenic_variant | MODIFIER |
| 9 | 89453056 | AX-170032942 | 0 | C | T | 0.132 | -2.17E-02 | 1.33E-01 | 2.64E+00 | 2.71E+00 | 8.77E-01 | no | | intergenic_variant | MODIFIER |
| 9 | 89521117 | AX-97822834 | 0 | G | A | 0.132 | -2.17E-02 | 1.33E-01 | 2.64E+00 | 2.71E+00 | 8.77E-01 | no | | intergenic_variant | MODIFIER |
| 9 | 89529788 | AX-170032936 | 0 | C | A | 0.132 | -2.17E-02 | 1.33E-01 | 2.64E+00 | 2.71E+00 | 8.77E-01 | no | | intergenic_variant | MODIFIER |
| 9 | 89559566 | AX-170032935 | 0 | C | A | 0.132 | -2.17E-02 | 1.33E-01 | 2.64E+00 | 2.71E+00 | 8.77E-01 | no | Gm47403 | intron_variant,non_coding_transcript_variant | MODIFIER |
| 9 | 89571788 | AX-170032943 | 0 | T | C | 0.132 | -2.17E-02 | 1.33E-01 | 2.64E+00 | 2.71E+00 | 8.77E-01 | no | | intergenic_variant | MODIFIER |
| 9 | 89637755 | AX-170032939 | 0 | A | G | 0.132 | -2.17E-02 | 1.33E-01 | 2.64E+00 | 2.71E+00 | 8.77E-01 | no | Gm47409 | upstream_gene_variant | MODIFIER |
| 9 | 89738640 | AX-97811333 | 0 | T | C | 0.132 | -2.17E-02 | 1.33E-01 | 2.64E+00 | 2.71E+00 | 8.77E-01 | no | Ankrd34c | upstream_gene_variant | MODIFIER |
| 9 | 89765729 | AX-97852826 | 0 | A | G | 0.132 | -2.17E-02 | 1.33E-01 | 2.64E+00 | 2.71E+00 | 8.77E-01 | no | | intergenic_variant | MODIFIER |
| 9 | 89785150 | AX-170509599 | 0 | C | T | 0.132 | -2.17E-02 | 1.33E-01 | 2.64E+00 | 2.71E+00 | 8.77E-01 | no | | intergenic_variant | MODIFIER |
| 9 | 89803110 | AX-97842351 | 0 | A | G | 0.132 | -2.17E-02 | 1.33E-01 | 2.64E+00 | 2.71E+00 | 8.77E-01 | no | Mir184 | upstream_gene_variant | MODIFIER |
| 9 | 89806784 | AX-170483690 | 0 | A | G | 0.132 | -2.17E-02 | 1.33E-01 | 2.64E+00 | 2.71E+00 | 8.77E-01 | no | Mir184 | upstream_gene_variant | MODIFIER |
| 9 | 89806818 | AX-170329816 | 0 | C | T | 0.132 | -2.17E-02 | 1.33E-01 | 2.64E+00 | 2.71E+00 | 8.77E-01 | no | Mir184 | upstream_gene_variant | MODIFIER |
| 9 | 89812924 | AX-170170607 | 0 | G | A | 0.132 | -2.17E-02 | 1.33E-01 | 2.64E+00 | 2.71E+00 | 8.77E-01 | no | | intergenic_variant | MODIFIER |
| 9 | 89825184 | AX-170194169 | 0 | C | A | 0.132 | -2.17E-02 | 1.33E-01 | 2.64E+00 | 2.71E+00 | 8.77E-01 | no | 4930524O08Rik | upstream_gene_variant | MODIFIER |
| 9 | 89825694 | AX-170211612 | 0 | G | C | 0.132 | -2.17E-02 | 1.33E-01 | 2.64E+00 | 2.71E+00 | 8.77E-01 | no | 4930524O08Rik | upstream_gene_variant | MODIFIER |
| 9 | 89825921 | AX-170106893 | 0 | G | A | 0.132 | -2.17E-02 | 1.33E-01 | 2.64E+00 | 2.71E+00 | 8.77E-01 | no | 4930524O08Rik | upstream_gene_variant | MODIFIER |
| 9 | 89838482 | AX-97850018 | 0 | T | C | 0.132 | -2.17E-02 | 1.33E-01 | 2.64E+00 | 2.71E+00 | 8.77E-01 | no | 4930524O08Rik | intron_variant,non_coding_transcript_variant | MODIFIER |
| 9 | 89864331 | AX-170341609 | 0 | A | G | 0.132 | -2.17E-02 | 1.33E-01 | 2.64E+00 | 2.71E+00 | 8.77E-01 | no | 4930524O08Rik | downstream_gene_variant | MODIFIER |
| 9 | 89928702 | AX-97853468 | 0 | A | G | 0.132 | -2.17E-02 | 1.33E-01 | 2.64E+00 | 2.71E+00 | 8.77E-01 | no | Rasgrf1 | intron_variant | MODIFIER |
| 9 | 89967032 | AX-170382557 | 0 | G | A | 0.132 | -2.17E-02 | 1.33E-01 | 2.64E+00 | 2.71E+00 | 8.77E-01 | no | Rasgrf1 | intron_variant | MODIFIER |
| 9 | 89971583 | AX-97836774 | 0 | C | T | 0.132 | -2.17E-02 | 1.33E-01 | 2.64E+00 | 2.71E+00 | 8.77E-01 | no | Rasgrf1 | intron_variant | MODIFIER |
| 9 | 89991556 | AX-170170508 | 0 | C | T | 0.132 | -2.17E-02 | 1.33E-01 | 2.64E+00 | 2.71E+00 | 8.77E-01 | no | Rasgrf1 | synonymous_variant | LOW |
| 9 | 89991646 | AX-170172975 | 0 | T | C | 0.132 | -2.17E-02 | 1.33E-01 | 2.64E+00 | 2.71E+00 | 8.77E-01 | no | Rasgrf1 | synonymous_variant | LOW |

| | | | | | | | | | | | | | | | |
|---|---|---|---|---|---|---|---|---|---|---|---|---|---|---|---|
| 9 | 89996059 | AX-170510668 | 0 | A | G | 0.132 | -2.17E-02 | 1.33E-01 | 2.64E+00 | 2.71E+00 | 8.77E-01 | no | Rasgrf1 | intron_variant | MODIFIER |
| 9 | 90025369 | AX-170142412 | 0 | T | G | 0.132 | -2.17E-02 | 1.33E-01 | 2.64E+00 | 2.71E+00 | 8.77E-01 | no | Rasgrf1 | intron_variant | MODIFIER |
| 9 | 90025825 | AX-170171458 | 0 | A | G | 0.132 | -2.17E-02 | 1.33E-01 | 2.64E+00 | 2.71E+00 | 8.77E-01 | no | Rasgrf1 | intron_variant | MODIFIER |
| 9 | 90031276 | AX-170333669 | 0 | C | T | 0.132 | -2.17E-02 | 1.33E-01 | 2.64E+00 | 2.71E+00 | 8.77E-01 | no | Rasgrf1 | downstream_gene_variant | MODIFIER |
| 7 | 91727881 | AX-170029503 | 2 | A | G | 0.118 | 3.40E-02 | 2.19E-01 | 2.65E+00 | 2.73E+00 | 8.77E-01 | no | Dlg2 | intron_variant | MODIFIER |
| 12 | 111092157 | AX-170007530 | 0 | C | T | 0.123 | -2.93E-02 | 1.96E-01 | 2.68E+00 | 2.75E+00 | 8.77E-01 | no | Rcor1 | intron_variant | MODIFIER |
| 13 | 39887909 | AX-170428151 | 0 | G | T | 0.167 | -1.91E-02 | 1.08E-01 | 2.59E+00 | 2.69E+00 | 8.77E-01 | no | Gm47316 | upstream_gene_variant | MODIFIER |
| 4 | 148420112 | AX-97804635 | 0 | C | T | 0.096 | -3.00E-02 | 1.99E-01 | 2.68E+00 | 2.75E+00 | 8.77E-01 | no | | intergenic_variant | MODIFIER |
| 20 | 72959715 | AX-170176554 | 1 | A | T | 0.241 | 9.87E-03 | 6.31E-02 | 2.64E+00 | 2.73E+00 | 8.77E-01 | no | Nsdhl | downstream_gene_variant | MODIFIER |
| 8 | 108529544 | AX-170081122 | 0 | G | A | 0.219 | 1.68E-02 | 1.11E-01 | 2.67E+00 | 2.74E+00 | 8.77E-01 | no | Gm39244 | intron_variant,non_coding_transcript_variant | MODIFIER |
| 12 | 112275345 | AX-170380264 | 0 | A | G | 0.149 | -2.38E-02 | 1.63E-01 | 2.65E+00 | 2.76E+00 | 8.77E-01 | no | | regulatory_region_variant | MODIFIER |
| 1 | 68290610 | AX-170281975 | 2 | G | A | 0.118 | 3.56E-02 | 2.30E-01 | 2.68E+00 | 2.73E+00 | 8.77E-01 | no | Gm15670 | downstream_gene_variant | MODIFIER |
| 16 | 93047255 | AX-170268426 | 0 | T | C | 0.386 | -1.31E-02 | 7.96E-02 | 2.62E+00 | 2.71E+00 | 8.77E-01 | no | | intergenic_variant | MODIFIER |
| 16 | 93048523 | AX-170181212 | 0 | C | G | 0.386 | -1.31E-02 | 7.96E-02 | 2.62E+00 | 2.71E+00 | 8.77E-01 | no | | intergenic_variant | MODIFIER |
| 16 | 93052184 | AX-170121127 | 0 | C | T | 0.386 | -1.31E-02 | 7.96E-02 | 2.62E+00 | 2.71E+00 | 8.77E-01 | no | | intergenic_variant | MODIFIER |
| 16 | 93056807 | AX-170477194 | 0 | T | C | 0.386 | -1.31E-02 | 7.96E-02 | 2.62E+00 | 2.71E+00 | 8.77E-01 | no | | intergenic_variant | MODIFIER |
| 16 | 93062370 | AX-170388626 | 0 | T | C | 0.386 | -1.31E-02 | 7.96E-02 | 2.62E+00 | 2.71E+00 | 8.77E-01 | no | | intergenic_variant | MODIFIER |
| 16 | 93080026 | AX-170289734 | 0 | A | G | 0.386 | -1.31E-02 | 7.96E-02 | 2.62E+00 | 2.71E+00 | 8.77E-01 | no | | regulatory_region_variant | MODIFIER |
| 16 | 93081558 | AX-170440685 | 0 | G | A | 0.386 | -1.31E-02 | 7.96E-02 | 2.62E+00 | 2.71E+00 | 8.77E-01 | no | | intergenic_variant | MODIFIER |
| 16 | 93090728 | AX-170239795 | 0 | G | A | 0.386 | -1.31E-02 | 7.96E-02 | 2.62E+00 | 2.71E+00 | 8.77E-01 | no | | intergenic_variant | MODIFIER |
| 16 | 93114749 | AX-170287949 | 0 | C | T | 0.386 | -1.31E-02 | 7.96E-02 | 2.62E+00 | 2.71E+00 | 8.77E-01 | no | | intergenic_variant | MODIFIER |
| 16 | 93127900 | AX-170141786 | 0 | G | A | 0.386 | -1.31E-02 | 7.96E-02 | 2.62E+00 | 2.71E+00 | 8.77E-01 | no | | regulatory_region_variant | MODIFIER |
| 16 | 93132005 | AX-170439069 | 0 | C | T | 0.386 | -1.31E-02 | 7.96E-02 | 2.62E+00 | 2.71E+00 | 8.77E-01 | no | | regulatory_region_variant | MODIFIER |
| 16 | 93156563 | AX-170058134 | 0 | A | G | 0.386 | -1.31E-02 | 7.96E-02 | 2.62E+00 | 2.71E+00 | 8.77E-01 | no | | intergenic_variant | MODIFIER |
| 16 | 93158160 | AX-170383709 | 0 | G | A | 0.386 | -1.31E-02 | 7.96E-02 | 2.62E+00 | 2.71E+00 | 8.77E-01 | no | | intergenic_variant | MODIFIER |
| 16 | 93160250 | AX-170459191 | 0 | G | A | 0.386 | -1.31E-02 | 7.96E-02 | 2.62E+00 | 2.71E+00 | 8.77E-01 | no | | intergenic_variant | MODIFIER |
| 16 | 93258670 | AX-170013680 | 0 | A | T | 0.386 | -1.31E-02 | 7.96E-02 | 2.62E+00 | 2.71E+00 | 8.77E-01 | no | | intergenic_variant | MODIFIER |
| 16 | 93261134 | AX-170013677 | 0 | C | A | 0.386 | -1.31E-02 | 7.96E-02 | 2.62E+00 | 2.71E+00 | 8.77E-01 | no | | intergenic_variant | MODIFIER |
| 16 | 93261462 | AX-170013679 | 0 | G | T | 0.386 | -1.31E-02 | 7.96E-02 | 2.62E+00 | 2.71E+00 | 8.77E-01 | no | | intergenic_variant | MODIFIER |
| 16 | 93275037 | AX-170013676 | 0 | T | C | 0.386 | -1.31E-02 | 7.96E-02 | 2.62E+00 | 2.71E+00 | 8.77E-01 | no | | intergenic_variant | MODIFIER |
| 16 | 93285611 | AX-170446097 | 0 | C | T | 0.386 | -1.31E-02 | 7.96E-02 | 2.62E+00 | 2.71E+00 | 8.77E-01 | no | | intergenic_variant | MODIFIER |
| 16 | 93301822 | AX-170159965 | 0 | C | A | 0.386 | -1.31E-02 | 7.96E-02 | 2.62E+00 | 2.71E+00 | 8.77E-01 | no | | intergenic_variant | MODIFIER |
| 16 | 93318212 | AX-170152952 | 0 | A | G | 0.386 | -1.31E-02 | 7.96E-02 | 2.62E+00 | 2.71E+00 | 8.77E-01 | no | | intergenic_variant | MODIFIER |
| 16 | 93319331 | AX-170192539 | 0 | G | A | 0.386 | -1.31E-02 | 7.96E-02 | 2.62E+00 | 2.71E+00 | 8.77E-01 | no | | intergenic_variant | MODIFIER |
| 16 | 93319379 | AX-97781913 | 0 | C | T | 0.386 | -1.31E-02 | 7.96E-02 | 2.62E+00 | 2.71E+00 | 8.77E-01 | no | | intergenic_variant | MODIFIER |
| 16 | 93321050 | AX-170288350 | 0 | T | A | 0.386 | -1.31E-02 | 7.96E-02 | 2.62E+00 | 2.71E+00 | 8.77E-01 | no | | intergenic_variant | MODIFIER |
| 16 | 93321700 | AX-170371783 | 0 | A | G | 0.386 | -1.31E-02 | 7.96E-02 | 2.62E+00 | 2.71E+00 | 8.77E-01 | no | | intergenic_variant | MODIFIER |
| 16 | 93321791 | AX-170262283 | 0 | A | C | 0.386 | -1.31E-02 | 7.96E-02 | 2.62E+00 | 2.71E+00 | 8.77E-01 | no | | intergenic_variant | MODIFIER |
| 16 | 93324515 | AX-170503823 | 0 | A | G | 0.386 | -1.31E-02 | 7.96E-02 | 2.62E+00 | 2.71E+00 | 8.77E-01 | no | | intergenic_variant | MODIFIER |
| 16 | 93331389 | AX-170357011 | 0 | G | A | 0.386 | -1.31E-02 | 7.96E-02 | 2.62E+00 | 2.71E+00 | 8.77E-01 | no | | intergenic_variant | MODIFIER |
| 16 | 93350291 | AX-170306791 | 0 | T | G | 0.386 | -1.31E-02 | 7.96E-02 | 2.62E+00 | 2.71E+00 | 8.77E-01 | no | 1810053B23Rik | intron_variant,non_coding_transcript_variant | MODIFIER |
| 16 | 93353068 | AX-170501437 | 0 | C | T | 0.386 | -1.31E-02 | 7.96E-02 | 2.62E+00 | 2.71E+00 | 8.77E-01 | no | 1810053B23Rik | intron_variant,non_coding_transcript_variant | MODIFIER |
| 16 | 93353086 | AX-170355813 | 0 | T | C | 0.386 | -1.31E-02 | 7.96E-02 | 2.62E+00 | 2.71E+00 | 8.77E-01 | no | 1810053B23Rik | intron_variant,non_coding_transcript_variant | MODIFIER |
| 16 | 93365222 | AX-170232729 | 0 | G | A | 0.386 | -1.31E-02 | 7.96E-02 | 2.62E+00 | 2.71E+00 | 8.77E-01 | no | Mir802 | upstream_gene_variant | MODIFIER |
| 16 | 93393097 | AX-170354376 | 0 | G | T | 0.386 | -1.31E-02 | 7.96E-02 | 2.62E+00 | 2.71E+00 | 8.77E-01 | no | 1700029J03Rik | downstream_gene_variant | MODIFIER |
| 16 | 93395929 | AX-170293097 | 0 | G | A | 0.386 | -1.31E-02 | 7.96E-02 | 2.62E+00 | 2.71E+00 | 8.77E-01 | no | 1700029J03Rik | downstream_gene_variant | MODIFIER |
| 16 | 93405669 | AX-170088101 | 0 | C | T | 0.386 | -1.31E-02 | 7.96E-02 | 2.62E+00 | 2.71E+00 | 8.77E-01 | no | 1700029J03Rik | intron_variant,non_coding_transcript_variant | MODIFIER |
| 16 | 93413443 | AX-170453840 | 0 | G | A | 0.386 | -1.31E-02 | 7.96E-02 | 2.62E+00 | 2.71E+00 | 8.77E-01 | no | 1700029J03Rik | intron_variant,non_coding_transcript_variant | MODIFIER |
| 10 | 117551756 | AX-170173090 | 0 | G | A | 0.342 | 1.13E-02 | 7.72E-02 | 2.68E+00 | 2.76E+00 | 8.77E-01 | no | | intergenic_variant | MODIFIER |
| 1 | 147855166 | AX-170277158 | 0 | T | C | 0.202 | -1.57E-02 | 1.16E-01 | 2.67E+00 | 2.79E+00 | 8.77E-01 | no | | intergenic_variant | MODIFIER |
| 11 | 74197136 | AX-170133431 | 0 | G | A | 0.263 | 1.06E-02 | 7.19E-02 | 2.61E+00 | 2.75E+00 | 8.77E-01 | no | Olfr403 | downstream_gene_variant | MODIFIER |
| 1 | 47824732 | AX-170322425 | 2 | A | T | 0.091 | -2.96E-02 | 1.99E-01 | 2.65E+00 | 2.75E+00 | 8.77E-01 | no | | intergenic_variant | MODIFIER |
| 1 | 162126272 | AX-170441011 | 1 | C | T | 0.214 | 2.20E-02 | 1.38E-01 | 2.66E+00 | 2.71E+00 | 8.77E-01 | no | Dnm3 | intron_variant | MODIFIER |
| 2 | 55895328 | AX-170063716 | 0 | C | T | 0.044 | 2.98E-02 | 2.01E-01 | 2.65E+00 | 2.75E+00 | 8.78E-01 | no | | intergenic_variant | MODIFIER |

| | | | | | | | | | | | | | | | |
|---|---|---|---|---|---|---|---|---|---|---|---|---|---|---|---|
| 12 | 12144108 | AX-170481026 | 0 | A | C | 0.044 | 2.98E-02 | 2.01E-01 | 2.65E+00 | 2.75E+00 | 8.78E-01 | no | | intergenic_variant | MODIFIER |
| 1 | 54512315 | AX-170443013 | 1 | T | C | 0.107 | 2.87E-02 | 1.84E-01 | 2.62E+00 | 2.73E+00 | 8.78E-01 | no | Pgap1 | intron_variant | MODIFIER |
| 1 | 52466121 | AX-170403841 | 1 | G | A | 0.125 | 3.00E-02 | 2.00E-01 | 2.67E+00 | 2.75E+00 | 8.78E-01 | no | Nab1 | intron_variant | MODIFIER |
| 5 | 122024253 | AX-170351107 | 1 | G | A | 0.455 | 1.72E-02 | 1.10E-01 | 2.67E+00 | 2.72E+00 | 8.78E-01 | no | Cux2 | intron_variant | MODIFIER |
| 8 | 28356721 | AX-170335476 | 0 | G | A | 0.167 | 1.77E-02 | 1.17E-01 | 2.68E+00 | 2.74E+00 | 8.78E-01 | no | | intergenic_variant | MODIFIER |
| 8 | 28449828 | AX-97830153 | 0 | C | T | 0.167 | 1.77E-02 | 1.17E-01 | 2.68E+00 | 2.74E+00 | 8.78E-01 | no | | intergenic_variant | MODIFIER |
| 8 | 28515709 | AX-97827801 | 0 | G | A | 0.167 | 1.77E-02 | 1.17E-01 | 2.68E+00 | 2.74E+00 | 8.78E-01 | no | Gm8110 | upstream_gene_variant | MODIFIER |
| 8 | 29097075 | AX-170333788 | 0 | T | C | 0.167 | 1.77E-02 | 1.17E-01 | 2.68E+00 | 2.74E+00 | 8.78E-01 | no | Unc5d | intron_variant | MODIFIER |
| 8 | 29369568 | AX-170030424 | 0 | T | C | 0.167 | 1.77E-02 | 1.17E-01 | 2.68E+00 | 2.74E+00 | 8.78E-01 | no | | intergenic_variant | MODIFIER |
| 8 | 29373192 | AX-170519555 | 0 | A | G | 0.167 | 1.77E-02 | 1.17E-01 | 2.68E+00 | 2.74E+00 | 8.78E-01 | no | | intergenic_variant | MODIFIER |
| 8 | 29375421 | AX-170245100 | 0 | T | G | 0.167 | 1.77E-02 | 1.17E-01 | 2.68E+00 | 2.74E+00 | 8.78E-01 | no | | intergenic_variant | MODIFIER |
| 8 | 29376336 | AX-170079616 | 0 | A | G | 0.167 | 1.77E-02 | 1.17E-01 | 2.68E+00 | 2.74E+00 | 8.78E-01 | no | | intergenic_variant | MODIFIER |
| 8 | 29376618 | AX-170507161 | 0 | G | A | 0.167 | 1.77E-02 | 1.17E-01 | 2.68E+00 | 2.74E+00 | 8.78E-01 | no | | intergenic_variant | MODIFIER |
| 8 | 29533229 | AX-170188698 | 0 | T | G | 0.167 | 1.77E-02 | 1.17E-01 | 2.68E+00 | 2.74E+00 | 8.78E-01 | no | | intergenic_variant | MODIFIER |
| 8 | 29537346 | AX-170339689 | 0 | G | A | 0.167 | 1.77E-02 | 1.17E-01 | 2.68E+00 | 2.74E+00 | 8.78E-01 | no | | intergenic_variant | MODIFIER |
| 8 | 29546574 | AX-170162968 | 0 | A | G | 0.167 | 1.77E-02 | 1.17E-01 | 2.68E+00 | 2.74E+00 | 8.78E-01 | no | | intergenic_variant | MODIFIER |
| 8 | 29546893 | AX-97785381 | 0 | G | A | 0.167 | 1.77E-02 | 1.17E-01 | 2.68E+00 | 2.74E+00 | 8.78E-01 | no | | intergenic_variant | MODIFIER |
| 8 | 29560787 | AX-170154770 | 0 | C | T | 0.167 | 1.77E-02 | 1.17E-01 | 2.68E+00 | 2.74E+00 | 8.78E-01 | no | | intergenic_variant | MODIFIER |
| 8 | 29583735 | AX-170151003 | 0 | C | T | 0.167 | 1.77E-02 | 1.17E-01 | 2.68E+00 | 2.74E+00 | 8.78E-01 | no | | intergenic_variant | MODIFIER |
| 8 | 29583829 | AX-97801133 | 0 | C | T | 0.167 | 1.77E-02 | 1.17E-01 | 2.68E+00 | 2.74E+00 | 8.78E-01 | no | | intergenic_variant | MODIFIER |
| 8 | 29587747 | AX-170103748 | 0 | G | T | 0.167 | 1.77E-02 | 1.17E-01 | 2.68E+00 | 2.74E+00 | 8.78E-01 | no | | intergenic_variant | MODIFIER |
| 8 | 29595450 | AX-170499055 | 0 | C | T | 0.167 | 1.77E-02 | 1.17E-01 | 2.68E+00 | 2.74E+00 | 8.78E-01 | no | | intergenic_variant | MODIFIER |
| 8 | 29597798 | AX-170292411 | 0 | T | C | 0.167 | 1.77E-02 | 1.17E-01 | 2.68E+00 | 2.74E+00 | 8.78E-01 | no | | intergenic_variant | MODIFIER |
| 8 | 29597989 | AX-170143765 | 0 | G | A | 0.167 | 1.77E-02 | 1.17E-01 | 2.68E+00 | 2.74E+00 | 8.78E-01 | no | | intergenic_variant | MODIFIER |
| 8 | 29602134 | AX-170451198 | 0 | A | G | 0.167 | 1.77E-02 | 1.17E-01 | 2.68E+00 | 2.74E+00 | 8.78E-01 | no | | intergenic_variant | MODIFIER |
| 8 | 29613342 | AX-170148811 | 0 | C | G | 0.167 | 1.77E-02 | 1.17E-01 | 2.68E+00 | 2.74E+00 | 8.78E-01 | no | | intergenic_variant | MODIFIER |
| 8 | 29614248 | AX-170122122 | 0 | C | G | 0.167 | 1.77E-02 | 1.17E-01 | 2.68E+00 | 2.74E+00 | 8.78E-01 | no | | intergenic_variant | MODIFIER |
| 8 | 29678897 | AX-97846459 | 0 | A | C | 0.167 | 1.77E-02 | 1.17E-01 | 2.68E+00 | 2.74E+00 | 8.78E-01 | no | | intergenic_variant | MODIFIER |
| 8 | 29703522 | AX-170261265 | 0 | C | A | 0.167 | 1.77E-02 | 1.17E-01 | 2.68E+00 | 2.74E+00 | 8.78E-01 | no | | intergenic_variant | MODIFIER |
| 8 | 29703613 | AX-170275511 | 0 | T | C | 0.167 | 1.77E-02 | 1.17E-01 | 2.68E+00 | 2.74E+00 | 8.78E-01 | no | | intergenic_variant | MODIFIER |
| 8 | 29705775 | AX-170216294 | 0 | C | T | 0.167 | 1.77E-02 | 1.17E-01 | 2.68E+00 | 2.74E+00 | 8.78E-01 | no | | intergenic_variant | MODIFIER |
| 8 | 29714733 | AX-170183096 | 0 | T | G | 0.167 | 1.77E-02 | 1.17E-01 | 2.68E+00 | 2.74E+00 | 8.78E-01 | no | | intergenic_variant | MODIFIER |
| 8 | 29740638 | AX-170111583 | 0 | T | C | 0.167 | 1.77E-02 | 1.17E-01 | 2.68E+00 | 2.74E+00 | 8.78E-01 | no | | intergenic_variant | MODIFIER |
| 8 | 29747329 | AX-97803993 | 0 | C | T | 0.167 | 1.77E-02 | 1.17E-01 | 2.68E+00 | 2.74E+00 | 8.78E-01 | no | | intergenic_variant | MODIFIER |
| 8 | 29749447 | AX-170230424 | 0 | G | A | 0.167 | 1.77E-02 | 1.17E-01 | 2.68E+00 | 2.74E+00 | 8.78E-01 | no | | intergenic_variant | MODIFIER |
| 8 | 29749482 | AX-170341444 | 0 | G | A | 0.167 | 1.77E-02 | 1.17E-01 | 2.68E+00 | 2.74E+00 | 8.78E-01 | no | | intergenic_variant | MODIFIER |
| 8 | 29781747 | AX-170142154 | 0 | C | G | 0.167 | 1.77E-02 | 1.17E-01 | 2.68E+00 | 2.74E+00 | 8.78E-01 | no | | intergenic_variant | MODIFIER |
| 8 | 29833282 | AX-97856490 | 0 | T | C | 0.167 | 1.77E-02 | 1.17E-01 | 2.68E+00 | 2.74E+00 | 8.78E-01 | no | | intergenic_variant | MODIFIER |
| 8 | 29833317 | AX-170466478 | 0 | A | T | 0.167 | 1.77E-02 | 1.17E-01 | 2.68E+00 | 2.74E+00 | 8.78E-01 | no | | intergenic_variant | MODIFIER |
| 8 | 29838535 | AX-170264877 | 0 | T | C | 0.167 | 1.77E-02 | 1.17E-01 | 2.68E+00 | 2.74E+00 | 8.78E-01 | no | | intergenic_variant | MODIFIER |
| 8 | 29875632 | AX-97818005 | 0 | T | C | 0.167 | 1.77E-02 | 1.17E-01 | 2.68E+00 | 2.74E+00 | 8.78E-01 | no | | intergenic_variant | MODIFIER |
| 8 | 29877894 | AX-170358001 | 0 | G | T | 0.167 | 1.77E-02 | 1.17E-01 | 2.68E+00 | 2.74E+00 | 8.78E-01 | no | | intergenic_variant | MODIFIER |
| 8 | 29883235 | AX-170030429 | 0 | T | C | 0.167 | 1.77E-02 | 1.17E-01 | 2.68E+00 | 2.74E+00 | 8.78E-01 | no | | intergenic_variant | MODIFIER |
| 8 | 29908030 | AX-97822623 | 0 | C | A | 0.167 | 1.77E-02 | 1.17E-01 | 2.68E+00 | 2.74E+00 | 8.78E-01 | no | | intergenic_variant | MODIFIER |
| 8 | 29922796 | AX-170305001 | 0 | T | C | 0.167 | 1.77E-02 | 1.17E-01 | 2.68E+00 | 2.74E+00 | 8.78E-01 | no | | intergenic_variant | MODIFIER |
| 8 | 29928152 | AX-97827240 | 0 | T | G | 0.167 | 1.77E-02 | 1.17E-01 | 2.68E+00 | 2.74E+00 | 8.78E-01 | no | | regulatory_region_variant | MODIFIER |
| 8 | 29966284 | AX-170030431 | 0 | G | A | 0.167 | 1.77E-02 | 1.17E-01 | 2.68E+00 | 2.74E+00 | 8.78E-01 | no | | intergenic_variant | MODIFIER |
| 8 | 29968349 | AX-170492099 | 0 | G | A | 0.167 | 1.77E-02 | 1.17E-01 | 2.68E+00 | 2.74E+00 | 8.78E-01 | no | | intergenic_variant | MODIFIER |
| 8 | 29974639 | AX-170193022 | 0 | T | G | 0.167 | 1.77E-02 | 1.17E-01 | 2.68E+00 | 2.74E+00 | 8.78E-01 | no | | intergenic_variant | MODIFIER |
| 8 | 29978789 | AX-170397559 | 0 | A | G | 0.167 | 1.77E-02 | 1.17E-01 | 2.68E+00 | 2.74E+00 | 8.78E-01 | no | | regulatory_region_variant | MODIFIER |
| 8 | 29999717 | AX-170475318 | 0 | A | C | 0.167 | 1.77E-02 | 1.17E-01 | 2.68E+00 | 2.74E+00 | 8.78E-01 | no | | intergenic_variant | MODIFIER |
| 8 | 29999740 | AX-97794904 | 0 | G | T | 0.167 | 1.77E-02 | 1.17E-01 | 2.68E+00 | 2.74E+00 | 8.78E-01 | no | | intergenic_variant | MODIFIER |
| 8 | 30062360 | AX-170420250 | 0 | C | A | 0.167 | 1.77E-02 | 1.17E-01 | 2.68E+00 | 2.74E+00 | 8.78E-01 | no | | intergenic_variant | MODIFIER |
| 8 | 30075466 | AX-170158189 | 0 | T | C | 0.167 | 1.77E-02 | 1.17E-01 | 2.68E+00 | 2.74E+00 | 8.78E-01 | no | | intergenic_variant | MODIFIER |

| | | | | | | | | | | | | | | | |
|---|---|---|---|---|---|---|---|---|---|---|---|---|---|---|---|
| 8 | 30077253 | AX-170260907 | 0 | G | A | 0.167 | 1.77E-02 | 1.17E-01 | 2.68E+00 | 2.74E+00 | 8.78E-01 | no | Gm45492 | downstream_gene_variant | MODIFIER |
| 8 | 30105367 | AX-170079627 | 0 | C | T | 0.167 | 1.77E-02 | 1.17E-01 | 2.68E+00 | 2.74E+00 | 8.78E-01 | no | | intergenic_variant | MODIFIER |
| 8 | 30114244 | AX-170079630 | 0 | A | C | 0.167 | 1.77E-02 | 1.17E-01 | 2.68E+00 | 2.74E+00 | 8.78E-01 | no | Gm39154 | intron_variant,non_coding_transcript_variant | MODIFIER |
| 8 | 30115101 | AX-170431487 | 0 | C | T | 0.167 | 1.77E-02 | 1.17E-01 | 2.68E+00 | 2.74E+00 | 8.78E-01 | no | Gm39154 | intron_variant,non_coding_transcript_variant | MODIFIER |
| 8 | 30124744 | AX-170256541 | 0 | G | A | 0.167 | 1.77E-02 | 1.17E-01 | 2.68E+00 | 2.74E+00 | 8.78E-01 | no | Gm39154 | upstream_gene_variant | MODIFIER |
| 8 | 30185501 | AX-170205190 | 0 | G | C | 0.167 | 1.77E-02 | 1.17E-01 | 2.68E+00 | 2.74E+00 | 8.78E-01 | no | Gm6853 | upstream_gene_variant | MODIFIER |
| 8 | 30232526 | AX-170285685 | 0 | C | G | 0.167 | 1.77E-02 | 1.17E-01 | 2.68E+00 | 2.74E+00 | 8.78E-01 | no | | intergenic_variant | MODIFIER |
| 8 | 30249386 | AX-170096212 | 0 | G | C | 0.167 | 1.77E-02 | 1.17E-01 | 2.68E+00 | 2.74E+00 | 8.78E-01 | no | | intergenic_variant | MODIFIER |
| 8 | 30262052 | AX-170464878 | 0 | A | C | 0.167 | 1.77E-02 | 1.17E-01 | 2.68E+00 | 2.74E+00 | 8.78E-01 | no | | intergenic_variant | MODIFIER |
| 8 | 30264332 | AX-170232557 | 0 | A | C | 0.167 | 1.77E-02 | 1.17E-01 | 2.68E+00 | 2.74E+00 | 8.78E-01 | no | | intergenic_variant | MODIFIER |
| 8 | 30275864 | AX-170285124 | 0 | T | C | 0.167 | 1.77E-02 | 1.17E-01 | 2.68E+00 | 2.74E+00 | 8.78E-01 | no | | intergenic_variant | MODIFIER |
| 8 | 30307523 | AX-97841862 | 0 | C | A | 0.167 | 1.77E-02 | 1.17E-01 | 2.68E+00 | 2.74E+00 | 8.78E-01 | no | | intergenic_variant | MODIFIER |
| 8 | 30313840 | AX-170161007 | 0 | T | A | 0.167 | 1.77E-02 | 1.17E-01 | 2.68E+00 | 2.74E+00 | 8.78E-01 | no | | intergenic_variant | MODIFIER |
| 8 | 30346016 | AX-170217726 | 0 | T | C | 0.167 | 1.77E-02 | 1.17E-01 | 2.68E+00 | 2.74E+00 | 8.78E-01 | no | | intergenic_variant | MODIFIER |
| 8 | 30357494 | AX-170253520 | 0 | C | T | 0.167 | 1.77E-02 | 1.17E-01 | 2.68E+00 | 2.74E+00 | 8.78E-01 | no | | intergenic_variant | MODIFIER |
| 8 | 30359711 | AX-170266788 | 0 | T | C | 0.167 | 1.77E-02 | 1.17E-01 | 2.68E+00 | 2.74E+00 | 8.78E-01 | no | | intergenic_variant | MODIFIER |
| 8 | 30387949 | AX-170330416 | 0 | A | G | 0.167 | 1.77E-02 | 1.17E-01 | 2.68E+00 | 2.74E+00 | 8.78E-01 | no | | intergenic_variant | MODIFIER |
| 8 | 30389622 | AX-170227344 | 0 | G | A | 0.167 | 1.77E-02 | 1.17E-01 | 2.68E+00 | 2.74E+00 | 8.78E-01 | no | | intergenic_variant | MODIFIER |
| 8 | 30421443 | AX-170203043 | 0 | C | A | 0.167 | 1.77E-02 | 1.17E-01 | 2.68E+00 | 2.74E+00 | 8.78E-01 | no | | intergenic_variant | MODIFIER |
| 8 | 30434042 | AX-170264208 | 0 | C | T | 0.167 | 1.77E-02 | 1.17E-01 | 2.68E+00 | 2.74E+00 | 8.78E-01 | no | | intergenic_variant | MODIFIER |
| 8 | 30438131 | AX-170158267 | 0 | A | G | 0.167 | 1.77E-02 | 1.17E-01 | 2.68E+00 | 2.74E+00 | 8.78E-01 | no | Gm45490 | downstream_gene_variant | MODIFIER |
| 8 | 30481050 | AX-97853470 | 0 | A | G | 0.167 | 1.77E-02 | 1.17E-01 | 2.68E+00 | 2.74E+00 | 8.78E-01 | no | Gm45489 | downstream_gene_variant | MODIFIER |
| 8 | 30490470 | AX-170079635 | 0 | C | T | 0.167 | 1.77E-02 | 1.17E-01 | 2.68E+00 | 2.74E+00 | 8.78E-01 | no | Gm45489 | intron_variant,non_coding_transcript_variant | MODIFIER |
| 8 | 30502141 | AX-170030437 | 0 | G | T | 0.167 | 1.77E-02 | 1.17E-01 | 2.68E+00 | 2.74E+00 | 8.78E-01 | no | Gm45489 | intron_variant,non_coding_transcript_variant | MODIFIER |
| 8 | 30532727 | AX-170427282 | 0 | A | G | 0.167 | 1.77E-02 | 1.17E-01 | 2.68E+00 | 2.74E+00 | 8.78E-01 | no | | regulatory_region_variant | MODIFIER |
| 8 | 30544272 | AX-170244174 | 0 | A | G | 0.167 | 1.77E-02 | 1.17E-01 | 2.68E+00 | 2.74E+00 | 8.78E-01 | no | | intergenic_variant | MODIFIER |
| 8 | 30544349 | AX-170427928 | 0 | T | C | 0.167 | 1.77E-02 | 1.17E-01 | 2.68E+00 | 2.74E+00 | 8.78E-01 | no | | intergenic_variant | MODIFIER |
| 8 | 30550989 | AX-170446963 | 0 | A | G | 0.167 | 1.77E-02 | 1.17E-01 | 2.68E+00 | 2.74E+00 | 8.78E-01 | no | | intergenic_variant | MODIFIER |
| 8 | 30551201 | AX-170207774 | 0 | T | C | 0.167 | 1.77E-02 | 1.17E-01 | 2.68E+00 | 2.74E+00 | 8.78E-01 | no | | intergenic_variant | MODIFIER |
| 8 | 30554924 | AX-170253425 | 0 | T | C | 0.167 | 1.77E-02 | 1.17E-01 | 2.68E+00 | 2.74E+00 | 8.78E-01 | no | | intergenic_variant | MODIFIER |
| 8 | 30555412 | AX-170140017 | 0 | C | T | 0.167 | 1.77E-02 | 1.17E-01 | 2.68E+00 | 2.74E+00 | 8.78E-01 | no | | intergenic_variant | MODIFIER |
| 8 | 30561205 | AX-170231378 | 0 | A | G | 0.167 | 1.77E-02 | 1.17E-01 | 2.68E+00 | 2.74E+00 | 8.78E-01 | no | | intergenic_variant | MODIFIER |
| 8 | 30561254 | AX-97854220 | 0 | T | C | 0.167 | 1.77E-02 | 1.17E-01 | 2.68E+00 | 2.74E+00 | 8.78E-01 | no | | intergenic_variant | MODIFIER |
| 8 | 30584531 | AX-170134575 | 0 | T | C | 0.167 | 1.77E-02 | 1.17E-01 | 2.68E+00 | 2.74E+00 | 8.78E-01 | no | | intergenic_variant | MODIFIER |
| 8 | 30603704 | AX-170187181 | 0 | A | C | 0.167 | 1.77E-02 | 1.17E-01 | 2.68E+00 | 2.74E+00 | 8.78E-01 | no | | intergenic_variant | MODIFIER |
| 8 | 30622826 | AX-170096213 | 0 | T | A | 0.167 | 1.77E-02 | 1.17E-01 | 2.68E+00 | 2.74E+00 | 8.78E-01 | no | | intergenic_variant | MODIFIER |
| 8 | 30630365 | AX-170485092 | 0 | A | G | 0.167 | 1.77E-02 | 1.17E-01 | 2.68E+00 | 2.74E+00 | 8.78E-01 | no | | regulatory_region_variant | MODIFIER |
| 8 | 30632121 | AX-170479905 | 0 | G | A | 0.167 | 1.77E-02 | 1.17E-01 | 2.68E+00 | 2.74E+00 | 8.78E-01 | no | | intergenic_variant | MODIFIER |
| 8 | 30660245 | AX-170424465 | 0 | C | T | 0.167 | 1.77E-02 | 1.17E-01 | 2.68E+00 | 2.74E+00 | 8.78E-01 | no | | intergenic_variant | MODIFIER |
| 8 | 30662204 | AX-97801585 | 0 | A | G | 0.167 | 1.77E-02 | 1.17E-01 | 2.68E+00 | 2.74E+00 | 8.78E-01 | no | | intergenic_variant | MODIFIER |
| 8 | 30681070 | AX-170256610 | 0 | C | G | 0.167 | 1.77E-02 | 1.17E-01 | 2.68E+00 | 2.74E+00 | 8.78E-01 | no | | intergenic_variant | MODIFIER |
| 8 | 30683730 | AX-170164655 | 0 | T | C | 0.167 | 1.77E-02 | 1.17E-01 | 2.68E+00 | 2.74E+00 | 8.78E-01 | no | | intergenic_variant | MODIFIER |
| 8 | 30686112 | AX-170227970 | 0 | G | A | 0.167 | 1.77E-02 | 1.17E-01 | 2.68E+00 | 2.74E+00 | 8.78E-01 | no | | intergenic_variant | MODIFIER |
| 8 | 30702900 | AX-170249446 | 0 | A | G | 0.167 | 1.77E-02 | 1.17E-01 | 2.68E+00 | 2.74E+00 | 8.78E-01 | no | | intergenic_variant | MODIFIER |
| 8 | 30703565 | AX-170461201 | 0 | A | C | 0.167 | 1.77E-02 | 1.17E-01 | 2.68E+00 | 2.74E+00 | 8.78E-01 | no | | intergenic_variant | MODIFIER |
| 8 | 30715642 | AX-170229519 | 0 | A | C | 0.167 | 1.77E-02 | 1.17E-01 | 2.68E+00 | 2.74E+00 | 8.78E-01 | no | | intergenic_variant | MODIFIER |
| 8 | 30717262 | AX-170419649 | 0 | G | C | 0.167 | 1.77E-02 | 1.17E-01 | 2.68E+00 | 2.74E+00 | 8.78E-01 | no | | intergenic_variant | MODIFIER |
| 8 | 30727761 | AX-170242617 | 0 | G | A | 0.167 | 1.77E-02 | 1.17E-01 | 2.68E+00 | 2.74E+00 | 8.78E-01 | no | | intergenic_variant | MODIFIER |
| 8 | 30741145 | AX-97791878 | 0 | T | C | 0.167 | 1.77E-02 | 1.17E-01 | 2.68E+00 | 2.74E+00 | 8.78E-01 | no | | intergenic_variant | MODIFIER |
| 8 | 30748300 | AX-170246832 | 0 | A | G | 0.167 | 1.77E-02 | 1.17E-01 | 2.68E+00 | 2.74E+00 | 8.78E-01 | no | | intergenic_variant | MODIFIER |
| 8 | 30749150 | AX-170105044 | 0 | A | G | 0.167 | 1.77E-02 | 1.17E-01 | 2.68E+00 | 2.74E+00 | 8.78E-01 | no | | intergenic_variant | MODIFIER |
| 8 | 30778229 | AX-170079642 | 0 | T | C | 0.167 | 1.77E-02 | 1.17E-01 | 2.68E+00 | 2.74E+00 | 8.78E-01 | no | | intergenic_variant | MODIFIER |
| 8 | 30798049 | AX-170238089 | 0 | C | T | 0.167 | 1.77E-02 | 1.17E-01 | 2.68E+00 | 2.74E+00 | 8.78E-01 | no | | intergenic_variant | MODIFIER |
| 8 | 30816330 | AX-170315633 | 0 | T | C | 0.167 | 1.77E-02 | 1.17E-01 | 2.68E+00 | 2.74E+00 | 8.78E-01 | no | | intergenic_variant | MODIFIER |

| | | | | | | | | | | | | | | | |
|---|---|---|---|---|---|---|---|---|---|---|---|---|---|---|---|
| 8 | 30823806 | AX-97797342 | 0 | G | A | 0.167 | 1.77E-02 | 1.17E-01 | 2.68E+00 | 2.74E+00 | 8.78E-01 | no | | regulatory_region_variant | MODIFIER |
| 8 | 30825187 | AX-170511087 | 0 | G | A | 0.167 | 1.77E-02 | 1.17E-01 | 2.68E+00 | 2.74E+00 | 8.78E-01 | no | | intergenic_variant | MODIFIER |
| 8 | 30830141 | AX-170374145 | 0 | T | A | 0.167 | 1.77E-02 | 1.17E-01 | 2.68E+00 | 2.74E+00 | 8.78E-01 | no | | intergenic_variant | MODIFIER |
| 8 | 30851596 | AX-170184776 | 0 | G | A | 0.167 | 1.77E-02 | 1.17E-01 | 2.68E+00 | 2.74E+00 | 8.78E-01 | no | | intergenic_variant | MODIFIER |
| 8 | 30851742 | AX-170339414 | 0 | A | T | 0.167 | 1.77E-02 | 1.17E-01 | 2.68E+00 | 2.74E+00 | 8.78E-01 | no | | intergenic_variant | MODIFIER |
| 8 | 30859591 | AX-170329646 | 0 | C | T | 0.167 | 1.77E-02 | 1.17E-01 | 2.68E+00 | 2.74E+00 | 8.78E-01 | no | | intergenic_variant | MODIFIER |
| 8 | 30862718 | AX-170527588 | 0 | G | A | 0.167 | 1.77E-02 | 1.17E-01 | 2.68E+00 | 2.74E+00 | 8.78E-01 | no | | intergenic_variant | MODIFIER |
| 8 | 30862748 | AX-170167423 | 0 | C | A | 0.167 | 1.77E-02 | 1.17E-01 | 2.68E+00 | 2.74E+00 | 8.78E-01 | no | | intergenic_variant | MODIFIER |
| 8 | 30864073 | AX-170124833 | 0 | G | A | 0.167 | 1.77E-02 | 1.17E-01 | 2.68E+00 | 2.74E+00 | 8.78E-01 | no | | intergenic_variant | MODIFIER |
| 8 | 30864728 | AX-170481171 | 0 | A | G | 0.167 | 1.77E-02 | 1.17E-01 | 2.68E+00 | 2.74E+00 | 8.78E-01 | no | | intergenic_variant | MODIFIER |
| 8 | 30894848 | AX-170079641 | 0 | T | C | 0.167 | 1.77E-02 | 1.17E-01 | 2.68E+00 | 2.74E+00 | 8.78E-01 | no | | intergenic_variant | MODIFIER |
| 8 | 30904688 | AX-97791920 | 0 | A | G | 0.167 | 1.77E-02 | 1.17E-01 | 2.68E+00 | 2.74E+00 | 8.78E-01 | no | | intergenic_variant | MODIFIER |
| 8 | 30909235 | AX-170520926 | 0 | C | G | 0.167 | 1.77E-02 | 1.17E-01 | 2.68E+00 | 2.74E+00 | 8.78E-01 | no | | intergenic_variant | MODIFIER |
| 8 | 30913630 | AX-170337445 | 0 | C | T | 0.167 | 1.77E-02 | 1.17E-01 | 2.68E+00 | 2.74E+00 | 8.78E-01 | no | | regulatory_region_variant | MODIFIER |
| 8 | 30915418 | AX-170235992 | 0 | T | C | 0.167 | 1.77E-02 | 1.17E-01 | 2.68E+00 | 2.74E+00 | 8.78E-01 | no | | regulatory_region_variant | MODIFIER |
| 8 | 30927494 | AX-170152030 | 0 | G | T | 0.167 | 1.77E-02 | 1.17E-01 | 2.68E+00 | 2.74E+00 | 8.78E-01 | no | | intergenic_variant | MODIFIER |
| 8 | 30951972 | AX-170394101 | 0 | G | A | 0.167 | 1.77E-02 | 1.17E-01 | 2.68E+00 | 2.74E+00 | 8.78E-01 | no | | intergenic_variant | MODIFIER |
| 8 | 30953229 | AX-97852305 | 0 | G | T | 0.167 | 1.77E-02 | 1.17E-01 | 2.68E+00 | 2.74E+00 | 8.78E-01 | no | | intergenic_variant | MODIFIER |
| 8 | 30956621 | AX-170079643 | 0 | A | G | 0.167 | 1.77E-02 | 1.17E-01 | 2.68E+00 | 2.74E+00 | 8.78E-01 | no | | intergenic_variant | MODIFIER |
| 8 | 30956823 | AX-170079644 | 0 | G | A | 0.167 | 1.77E-02 | 1.17E-01 | 2.68E+00 | 2.74E+00 | 8.78E-01 | no | | intergenic_variant | MODIFIER |
| 8 | 30967537 | AX-170030442 | 0 | A | T | 0.167 | 1.77E-02 | 1.17E-01 | 2.68E+00 | 2.74E+00 | 8.78E-01 | no | | intergenic_variant | MODIFIER |
| 8 | 30967678 | AX-97789081 | 0 | C | T | 0.167 | 1.77E-02 | 1.17E-01 | 2.68E+00 | 2.74E+00 | 8.78E-01 | no | | intergenic_variant | MODIFIER |
| 2 | 122986616 | AX-170064776 | 1 | T | G | 0.116 | 3.03E-02 | 1.93E-01 | 2.65E+00 | 2.72E+00 | 8.78E-01 | no | | intergenic_variant | MODIFIER |
| 6 | 105923140 | AX-170223068 | 0 | G | A | 0.07 | -4.18E-02 | 2.76E-01 | 2.70E+00 | 2.74E+00 | 8.78E-01 | no | Cntn4 | intron_variant | MODIFIER |
| 7 | 89610090 | AX-170029480 | 0 | G | A | 0.132 | -3.19E-02 | 2.11E-01 | 2.66E+00 | 2.74E+00 | 8.78E-01 | no | Gm2546 | downstream_gene_variant | MODIFIER |
| 7 | 116374333 | AX-170090013 | 0 | C | T | 0.412 | -1.36E-02 | 9.17E-02 | 2.71E+00 | 2.76E+00 | 8.78E-01 | no | Pik3c2a | intron_variant | MODIFIER |
| 1 | 13559338 | AX-170255015 | 1 | T | C | 0.366 | 1.21E-02 | 7.35E-02 | 2.63E+00 | 2.70E+00 | 8.78E-01 | no | | intergenic_variant | MODIFIER |
| 11 | 20140741 | AX-170381560 | 1 | A | C | 0.054 | 2.91E-02 | 1.89E-01 | 2.59E+00 | 2.73E+00 | 8.78E-01 | no | Gm12030 | upstream_gene_variant | MODIFIER |
| 18 | 63488476 | AX-97825220 | 1 | C | T | 0.402 | 1.62E-02 | 9.39E-02 | 2.62E+00 | 2.68E+00 | 8.78E-01 | no | | intergenic_variant | MODIFIER |
| 5 | 39149069 | AX-170357942 | 0 | G | A | 0.228 | 1.59E-02 | 1.11E-01 | 2.69E+00 | 2.77E+00 | 8.78E-01 | no | | intergenic_variant | MODIFIER |
| 18 | 65709407 | AX-170205203 | 1 | T | A | 0.411 | 1.13E-02 | 7.98E-02 | 2.71E+00 | 2.79E+00 | 8.78E-01 | no | Oacyl | intron_variant | MODIFIER |
| 13 | 47625056 | AX-170514377 | 0 | G | A | 0.526 | 1.03E-02 | 6.43E-02 | 2.62E+00 | 2.72E+00 | 8.78E-01 | no | | intergenic_variant | MODIFIER |
| 13 | 47625797 | AX-170289772 | 0 | C | T | 0.526 | 1.03E-02 | 6.43E-02 | 2.62E+00 | 2.72E+00 | 8.78E-01 | no | | intergenic_variant | MODIFIER |
| 13 | 47659416 | AX-170266910 | 0 | G | T | 0.526 | 1.03E-02 | 6.43E-02 | 2.62E+00 | 2.72E+00 | 8.78E-01 | no | | intergenic_variant | MODIFIER |
| 13 | 47685842 | AX-97841754 | 0 | T | C | 0.526 | 1.03E-02 | 6.43E-02 | 2.62E+00 | 2.72E+00 | 8.78E-01 | no | | intergenic_variant | MODIFIER |
| 13 | 47805340 | AX-170221702 | 0 | A | G | 0.526 | 1.03E-02 | 6.43E-02 | 2.62E+00 | 2.72E+00 | 8.78E-01 | no | | intergenic_variant | MODIFIER |
| 13 | 47810344 | AX-168412925 | 0 | A | G | 0.526 | 1.03E-02 | 6.43E-02 | 2.62E+00 | 2.72E+00 | 8.78E-01 | no | | intergenic_variant | MODIFIER |
| 13 | 47813050 | AX-170008280 | 0 | C | T | 0.526 | 1.03E-02 | 6.43E-02 | 2.62E+00 | 2.72E+00 | 8.78E-01 | no | | intergenic_variant | MODIFIER |
| 5 | 119042800 | AX-97801964 | 1 | G | A | 0.464 | 1.73E-02 | 1.12E-01 | 2.67E+00 | 2.72E+00 | 8.79E-01 | no | | regulatory_region_variant | MODIFIER |
| 9 | 89808276 | AX-170117706 | 1 | G | A | 0.134 | -2.10E-02 | 1.31E-01 | 2.64E+00 | 2.71E+00 | 8.79E-01 | no | 1700017I07Rik | downstream_gene_variant | MODIFIER |
| 2 | 121824051 | AX-170527405 | 1 | A | G | 0.125 | 3.25E-02 | 2.14E-01 | 2.68E+00 | 2.73E+00 | 8.79E-01 | no | | intergenic_variant | MODIFIER |
| 1 | 6891095 | AX-170206116 | 0 | C | T | 0.518 | 1.17E-02 | 7.11E-02 | 2.63E+00 | 2.70E+00 | 8.79E-01 | no | | intergenic_variant | MODIFIER |
| 10 | 109719688 | AX-170092095 | 0 | C | T | 0.342 | 1.21E-02 | 8.45E-02 | 2.68E+00 | 2.77E+00 | 8.79E-01 | no | Nav3 | intron_variant | MODIFIER |
| 5 | 97503030 | AX-170162476 | 2 | T | A | 0.427 | 1.60E-02 | 1.05E-01 | 2.66E+00 | 2.73E+00 | 8.79E-01 | no | 4930467D21Rik | intron_variant,non_coding_transcript_variant | MODIFIER |
| 4 | 148660876 | AX-170429807 | 1 | T | A | 0.098 | 2.51E-02 | 1.63E-01 | 2.62E+00 | 2.73E+00 | 8.79E-01 | no | Gm572 | intron_variant | MODIFIER |
| 1 | 64146458 | AX-170456071 | 0 | A | G | 0.123 | 3.18E-02 | 2.10E-01 | 2.66E+00 | 2.73E+00 | 8.79E-01 | no | Gm13750 | upstream_gene_variant | MODIFIER |
| 15 | 97731249 | AX-170262309 | 0 | G | A | 0.14 | 2.27E-02 | 1.40E-01 | 2.62E+00 | 2.71E+00 | 8.79E-01 | no | Endou | synonymous_variant | LOW |
| 1 | 45689329 | AX-170215360 | 0 | T | C | 0.105 | -2.47E-02 | 1.73E-01 | 2.64E+00 | 2.76E+00 | 8.79E-01 | no | Gm23216 | upstream_gene_variant | MODIFIER |
| 10 | 7783438 | AX-170097528 | 0 | C | T | 0.211 | 1.43E-02 | 9.95E-02 | 2.66E+00 | 2.76E+00 | 8.79E-01 | no | Ginm1 | upstream_gene_variant | MODIFIER |
| 18 | 66932185 | AX-170404459 | 0 | T | C | 0.088 | 2.87E-02 | 1.81E-01 | 2.61E+00 | 2.72E+00 | 8.79E-01 | no | AC127234.1 | intron_variant,non_coding_transcript_variant | MODIFIER |
| 10 | 6167277 | AX-170216367 | 1 | C | A | 0.214 | -1.50E-02 | 1.01E-01 | 2.64E+00 | 2.74E+00 | 8.79E-01 | no | | intergenic_variant | MODIFIER |
| 11 | 18370243 | AX-170158350 | 2 | G | A | 0.064 | 3.09E-02 | 2.14E-01 | 2.66E+00 | 2.75E+00 | 8.79E-01 | no | | intergenic_variant | MODIFIER |
| 5 | 5401889 | AX-97792688 | 0 | G | A | 0.43 | -1.04E-02 | 6.44E-02 | 2.59E+00 | 2.72E+00 | 8.79E-01 | no | Cdk14 | intron_variant | MODIFIER |
| 5 | 5421966 | AX-170334225 | 0 | G | A | 0.43 | -1.04E-02 | 6.44E-02 | 2.59E+00 | 2.72E+00 | 8.79E-01 | no | Cdk14 | upstream_gene_variant | MODIFIER |

| | | | | | | | | | | | | | | | |
|---|---|---|---|---|---|---|---|---|---|---|---|---|---|---|---|
| 5 | 5422424 | AX-170139944 | 0 | T | C | 0.43 | -1.04E-02 | 6.44E-02 | 2.59E+00 | 2.72E+00 | 8.79E-01 | no | Cdk14 | upstream_gene_variant | MODIFIER |
| 5 | 5435228 | AX-170387577 | 0 | T | A | 0.43 | -1.04E-02 | 6.44E-02 | 2.59E+00 | 2.72E+00 | 8.79E-01 | no | 1700015F17Rik | downstream_gene_variant | MODIFIER |
| 5 | 5436429 | AX-170174293 | 0 | T | C | 0.43 | -1.04E-02 | 6.44E-02 | 2.59E+00 | 2.72E+00 | 8.79E-01 | no | 1700015F17Rik | downstream_gene_variant | MODIFIER |
| 5 | 5477491 | AX-170336943 | 0 | G | T | 0.43 | -1.04E-02 | 6.44E-02 | 2.59E+00 | 2.72E+00 | 8.79E-01 | no | 1700015F17Rik | intron_variant | MODIFIER |
| 7 | 137403599 | AX-170224765 | 1 | A | C | 0.304 | -1.23E-02 | 7.80E-02 | 2.62E+00 | 2.72E+00 | 8.80E-01 | no | 9430038I01Rik | intron_variant | MODIFIER |
| 16 | 92114059 | AX-97854051 | 1 | C | T | 0.455 | -1.19E-02 | 7.96E-02 | 2.66E+00 | 2.74E+00 | 8.80E-01 | no | Mrps6 | downstream_gene_variant | MODIFIER |
| 5 | 125980209 | AX-97820116 | 0 | T | C | 0.447 | -1.64E-02 | 1.12E-01 | 2.71E+00 | 2.76E+00 | 8.80E-01 | no | Gm42952 | intron_variant,non_coding_transcript_variant | MODIFIER |
| 12 | 108394821 | AX-97855623 | 0 | A | C | 0.237 | -1.47E-02 | 1.02E-01 | 2.68E+00 | 2.76E+00 | 8.80E-01 | no | Eml1 | intron_variant,non_coding_transcript_variant | MODIFIER |
| 11 | 43358462 | AX-170245849 | 0 | G | A | 0.105 | -3.14E-02 | 2.10E-01 | 2.66E+00 | 2.74E+00 | 8.80E-01 | no | | regulatory_region_variant | MODIFIER |
| 3 | 145298901 | AX-170089097 | 1 | A | G | 0.348 | -1.05E-02 | 7.19E-02 | 2.66E+00 | 2.75E+00 | 8.80E-01 | no | Col24a1 | intron_variant | MODIFIER |
| 1 | 75314241 | AX-170041698 | 1 | G | A | 0.134 | 3.12E-02 | 2.08E-01 | 2.65E+00 | 2.74E+00 | 8.80E-01 | no | Dnpep | intron_variant | MODIFIER |
| 5 | 33565983 | AX-97823444 | 0 | C | T | 0.36 | 1.60E-02 | 1.06E-01 | 2.67E+00 | 2.73E+00 | 8.80E-01 | no | | intergenic_variant | MODIFIER |
| 10 | 8247704 | AX-170194414 | 1 | C | G | 0.214 | -1.60E-02 | 1.02E-01 | 2.63E+00 | 2.72E+00 | 8.80E-01 | no | Ust | intron_variant | MODIFIER |
| 8 | 30323385 | AX-97791965 | 1 | G | T | 0.17 | 1.71E-02 | 1.15E-01 | 2.67E+00 | 2.74E+00 | 8.80E-01 | no | | intergenic_variant | MODIFIER |
| 7 | 116964371 | AX-170289866 | 1 | G | T | 0.402 | 1.42E-02 | 9.18E-02 | 2.66E+00 | 2.71E+00 | 8.80E-01 | no | | intergenic_variant | MODIFIER |
| 11 | 17988602 | AX-170046501 | 0 | G | A | 0.07 | 2.65E-02 | 1.85E-01 | 2.64E+00 | 2.75E+00 | 8.80E-01 | no | | intergenic_variant | MODIFIER |
| 7 | 122018869 | AX-170183421 | 0 | G | C | 0.412 | 1.35E-02 | 9.07E-02 | 2.68E+00 | 2.74E+00 | 8.80E-01 | no | Gga2 | intron_variant | MODIFIER |
| 10 | 8284729 | AX-170413730 | 1 | G | C | 0.214 | -1.63E-02 | 1.06E-01 | 2.64E+00 | 2.72E+00 | 8.80E-01 | no | Ust | intron_variant | MODIFIER |
| 7 | 116339651 | AX-170429155 | 0 | T | C | 0.395 | -1.21E-02 | 8.60E-02 | 2.68E+00 | 2.77E+00 | 8.80E-01 | no | Pik3c2a | 3_prime_UTR_variant | MODIFIER |
| 12 | 101696000 | AX-170007398 | 0 | T | C | 0.061 | 2.87E-02 | 1.94E-01 | 2.65E+00 | 2.74E+00 | 8.80E-01 | no | Tc2n | intron_variant,NMD_transcript_variant | MODIFIER |
| 8 | 68515352 | AX-170188018 | 1 | G | A | 0.179 | -2.04E-02 | 1.31E-01 | 2.65E+00 | 2.72E+00 | 8.80E-01 | no | Csgalnact1 | intron_variant | MODIFIER |
| 12 | 63490322 | AX-97785459 | 0 | G | A | 0.07 | -2.83E-02 | 1.85E-01 | 2.62E+00 | 2.73E+00 | 8.80E-01 | no | | intergenic_variant | MODIFIER |
| 13 | 53817037 | AX-170368946 | 0 | C | T | 0.509 | -9.23E-03 | 6.83E-02 | 2.66E+00 | 2.78E+00 | 8.80E-01 | no | | intergenic_variant | MODIFIER |
| 1 | 175514926 | AX-170113563 | 0 | G | A | 0.228 | 1.93E-02 | 1.16E-01 | 2.61E+00 | 2.69E+00 | 8.80E-01 | no | | intergenic_variant | MODIFIER |
| 20 | 98252044 | AX-170519806 | 0 | G | A | 0.316 | 8.45E-03 | 5.63E-02 | 2.62E+00 | 2.74E+00 | 8.80E-01 | no | Ar | intron_variant | MODIFIER |
| 20 | 101002111 | AX-170084873 | 0 | T | G | 0.316 | 8.45E-03 | 5.63E-02 | 2.62E+00 | 2.74E+00 | 8.80E-01 | no | Tex11 | intron_variant | MODIFIER |
| 20 | 102223507 | AX-170295730 | 0 | A | G | 0.316 | 8.45E-03 | 5.63E-02 | 2.62E+00 | 2.74E+00 | 8.80E-01 | no | | intergenic_variant | MODIFIER |
| 20 | 102424804 | AX-170520800 | 0 | G | A | 0.316 | 8.45E-03 | 5.63E-02 | 2.62E+00 | 2.74E+00 | 8.80E-01 | no | Hdac8 | intron_variant | MODIFIER |
| 20 | 102436440 | AX-170356925 | 0 | C | T | 0.316 | 8.45E-03 | 5.63E-02 | 2.62E+00 | 2.74E+00 | 8.80E-01 | no | Hdac8 | intron_variant | MODIFIER |
| 20 | 102442813 | AX-170452704 | 0 | G | C | 0.316 | 8.45E-03 | 5.63E-02 | 2.62E+00 | 2.74E+00 | 8.80E-01 | no | Hdac8 | intron_variant | MODIFIER |
| 20 | 102471581 | AX-170146208 | 0 | T | C | 0.316 | 8.45E-03 | 5.63E-02 | 2.62E+00 | 2.74E+00 | 8.80E-01 | no | Hdac8 | intron_variant | MODIFIER |
| 20 | 102516351 | AX-97788603 | 0 | G | A | 0.316 | 8.45E-03 | 5.63E-02 | 2.62E+00 | 2.74E+00 | 8.80E-01 | no | Phka1 | intron_variant | MODIFIER |
| 20 | 102535820 | AX-170034596 | 0 | T | A | 0.316 | 8.45E-03 | 5.63E-02 | 2.62E+00 | 2.74E+00 | 8.80E-01 | no | Phka1 | intron_variant | MODIFIER |
| 20 | 103157085 | AX-170437489 | 0 | A | G | 0.316 | 8.45E-03 | 5.63E-02 | 2.62E+00 | 2.74E+00 | 8.80E-01 | no | | intergenic_variant | MODIFIER |
| 20 | 103161767 | AX-170131877 | 0 | A | G | 0.316 | 8.45E-03 | 5.63E-02 | 2.62E+00 | 2.74E+00 | 8.80E-01 | no | Gm6206 | downstream_gene_variant | MODIFIER |
| 20 | 103182659 | AX-170217945 | 0 | A | G | 0.316 | 8.45E-03 | 5.63E-02 | 2.62E+00 | 2.74E+00 | 8.80E-01 | no | 4930519F16Rik | downstream_gene_variant | MODIFIER |
| 20 | 104175509 | AX-168387551 | 0 | A | G | 0.316 | 8.45E-03 | 5.63E-02 | 2.62E+00 | 2.74E+00 | 8.80E-01 | no | Nexmif | intron_variant | MODIFIER |
| 20 | 104392869 | AX-97801284 | 0 | A | G | 0.316 | 8.45E-03 | 5.63E-02 | 2.62E+00 | 2.74E+00 | 8.80E-01 | no | Abcb7 | intron_variant | MODIFIER |
| 13 | 46961792 | AX-97802821 | 0 | A | G | 0.465 | -9.09E-03 | 6.47E-02 | 2.66E+00 | 2.76E+00 | 8.81E-01 | no | | regulatory_region_variant | MODIFIER |
| 13 | 47017456 | AX-97840228 | 0 | T | C | 0.465 | -9.09E-03 | 6.47E-02 | 2.66E+00 | 2.76E+00 | 8.81E-01 | no | Nhlrc1 | upstream_gene_variant | MODIFIER |
| 13 | 47037917 | AX-97857247 | 0 | T | C | 0.465 | -9.09E-03 | 6.47E-02 | 2.66E+00 | 2.76E+00 | 8.81E-01 | no | Tpmt | intron_variant | MODIFIER |
| 13 | 47049524 | AX-170166260 | 0 | A | G | 0.465 | -9.09E-03 | 6.47E-02 | 2.66E+00 | 2.76E+00 | 8.81E-01 | no | Kdm1b | intron_variant | MODIFIER |
| 13 | 47241018 | AX-170417768 | 0 | T | C | 0.465 | -9.09E-03 | 6.47E-02 | 2.66E+00 | 2.76E+00 | 8.81E-01 | no | Rnf144b | intron_variant | MODIFIER |
| 13 | 47280082 | AX-170442633 | 0 | A | G | 0.465 | -9.09E-03 | 6.47E-02 | 2.66E+00 | 2.76E+00 | 8.81E-01 | no | | intergenic_variant | MODIFIER |
| 13 | 47281529 | AX-170402603 | 0 | T | G | 0.465 | -9.09E-03 | 6.47E-02 | 2.66E+00 | 2.76E+00 | 8.81E-01 | no | | intergenic_variant | MODIFIER |
| 13 | 47281559 | AX-170386993 | 0 | C | T | 0.465 | -9.09E-03 | 6.47E-02 | 2.66E+00 | 2.76E+00 | 8.81E-01 | no | | intergenic_variant | MODIFIER |
| 13 | 47281939 | AX-170097744 | 0 | G | A | 0.465 | -9.09E-03 | 6.47E-02 | 2.66E+00 | 2.76E+00 | 8.81E-01 | no | | intergenic_variant | MODIFIER |
| 13 | 47311405 | AX-170495058 | 0 | G | T | 0.465 | -9.09E-03 | 6.47E-02 | 2.66E+00 | 2.76E+00 | 8.81E-01 | no | | intergenic_variant | MODIFIER |
| 13 | 47313814 | AX-170227759 | 0 | C | T | 0.465 | -9.09E-03 | 6.47E-02 | 2.66E+00 | 2.76E+00 | 8.81E-01 | no | | intergenic_variant | MODIFIER |
| 13 | 47319107 | AX-97852509 | 0 | C | T | 0.465 | -9.09E-03 | 6.47E-02 | 2.66E+00 | 2.76E+00 | 8.81E-01 | no | | regulatory_region_variant | MODIFIER |
| 13 | 47333433 | AX-170008269 | 0 | T | A | 0.465 | -9.09E-03 | 6.47E-02 | 2.66E+00 | 2.76E+00 | 8.81E-01 | no | Gm48612 | downstream_gene_variant | MODIFIER |
| 13 | 47345492 | AX-170092743 | 0 | G | T | 0.465 | -9.09E-03 | 6.47E-02 | 2.66E+00 | 2.76E+00 | 8.81E-01 | no | Gm48612 | upstream_gene_variant | MODIFIER |
| 13 | 47345588 | AX-170310158 | 0 | C | T | 0.465 | -9.09E-03 | 6.47E-02 | 2.66E+00 | 2.76E+00 | 8.81E-01 | no | Gm48612 | upstream_gene_variant | MODIFIER |
| 13 | 47348995 | AX-170051191 | 0 | C | T | 0.465 | -9.09E-03 | 6.47E-02 | 2.66E+00 | 2.76E+00 | 8.81E-01 | no | | regulatory_region_variant | MODIFIER |

| | | | | | | | | | | | | | | | |
|---|---|---|---|---|---|---|---|---|---|---|---|---|---|---|---|
| 13 | 47351872 | AX-170008267 | 0 | A | G | 0.465 | -9.09E-03 | 6.47E-02 | 2.66E+00 | 2.76E+00 | 8.81E-01 | no | | intergenic_variant | MODIFIER |
| 13 | 47429826 | AX-170263820 | 0 | T | C | 0.465 | -9.09E-03 | 6.47E-02 | 2.66E+00 | 2.76E+00 | 8.81E-01 | no | Gm35733 | intron_variant,non_coding_transcript_variant | MODIFIER |
| 13 | 47442784 | AX-97838094 | 0 | T | C | 0.465 | -9.09E-03 | 6.47E-02 | 2.66E+00 | 2.76E+00 | 8.81E-01 | no | Gm35733 | downstream_gene_variant | MODIFIER |
| 13 | 47452791 | AX-170008274 | 0 | T | C | 0.465 | -9.09E-03 | 6.47E-02 | 2.66E+00 | 2.76E+00 | 8.81E-01 | no | | regulatory_region_variant | MODIFIER |
| 13 | 47452806 | AX-170475287 | 0 | C | A | 0.465 | -9.09E-03 | 6.47E-02 | 2.66E+00 | 2.76E+00 | 8.81E-01 | no | | intergenic_variant | MODIFIER |
| 13 | 47460186 | AX-97826280 | 0 | T | C | 0.465 | -9.09E-03 | 6.47E-02 | 2.66E+00 | 2.76E+00 | 8.81E-01 | no | | intergenic_variant | MODIFIER |
| 12 | 111943790 | AX-170487412 | 0 | A | G | 0.149 | 2.43E-02 | 1.60E-01 | 2.62E+00 | 2.73E+00 | 8.81E-01 | no | 5033406O09Rik | intron_variant,non_coding_transcript_variant | MODIFIER |
| 10 | 6548865 | AX-170257207 | 1 | G | A | 0.205 | 1.44E-02 | 1.02E-01 | 2.66E+00 | 2.76E+00 | 8.81E-01 | no | | intergenic_variant | MODIFIER |
| 18 | 56812610 | AX-97820355 | 0 | C | T | 0.412 | -1.31E-02 | 8.26E-02 | 2.63E+00 | 2.71E+00 | 8.81E-01 | no | MAR3 | upstream_gene_variant | MODIFIER |
| 8 | 68834214 | AX-170303748 | 0 | T | C | 0.184 | -1.85E-02 | 1.23E-01 | 2.65E+00 | 2.73E+00 | 8.81E-01 | no | Ints10 | downstream_gene_variant | MODIFIER |
| 10 | 59101306 | AX-170177970 | 0 | C | G | 0.272 | 9.77E-03 | 7.60E-02 | 2.68E+00 | 2.80E+00 | 8.81E-01 | no | Sh3rf3 | intron_variant,non_coding_transcript_variant | MODIFIER |
| 10 | 14282988 | AX-170285483 | 1 | A | T | 0.205 | 1.46E-02 | 1.05E-01 | 2.69E+00 | 2.77E+00 | 8.81E-01 | no | | intergenic_variant | MODIFIER |
| 9 | 87785298 | AX-97841859 | 1 | T | C | 0.134 | -2.05E-02 | 1.31E-01 | 2.64E+00 | 2.71E+00 | 8.81E-01 | no | | intergenic_variant | MODIFIER |
| 5 | 106378215 | AX-170267591 | 0 | C | T | 0.447 | 1.78E-02 | 1.17E-01 | 2.68E+00 | 2.71E+00 | 8.81E-01 | no | | intergenic_variant | MODIFIER |
| 15 | 86737864 | AX-170337004 | 0 | T | C | 0.132 | 2.40E-02 | 1.57E-01 | 2.66E+00 | 2.72E+00 | 8.81E-01 | no | 4930543I03Rik | non_coding_transcript_exon_variant | MODIFIER |
| 15 | 97719794 | AX-97830280 | 0 | G | A | 0.132 | 2.40E-02 | 1.57E-01 | 2.66E+00 | 2.72E+00 | 8.81E-01 | no | Endou | intron_variant | MODIFIER |
| 4 | 115902908 | AX-170160160 | 0 | A | G | 0.211 | 1.82E-02 | 1.12E-01 | 2.61E+00 | 2.70E+00 | 8.81E-01 | no | 6430628N08Rik | downstream_gene_variant | MODIFIER |
| 8 | 116317696 | AX-170081273 | 0 | T | C | 0.254 | -1.44E-02 | 9.62E-02 | 2.62E+00 | 2.73E+00 | 8.81E-01 | no | | intergenic_variant | MODIFIER |
| 8 | 117079241 | AX-170427446 | 0 | T | C | 0.254 | -1.44E-02 | 9.62E-02 | 2.62E+00 | 2.73E+00 | 8.81E-01 | no | Pkd1l2 | intron_variant | MODIFIER |
| 1 | 179870692 | AX-170280116 | 0 | A | T | 0.237 | 1.78E-02 | 1.01E-01 | 2.57E+00 | 2.69E+00 | 8.81E-01 | no | | regulatory_region_variant | MODIFIER |
| 1 | 180221967 | AX-170002406 | 0 | A | G | 0.237 | 1.78E-02 | 1.01E-01 | 2.57E+00 | 2.69E+00 | 8.81E-01 | no | | intergenic_variant | MODIFIER |
| 2 | 121761942 | AX-170064763 | 0 | C | A | 0.14 | -3.23E-02 | 2.22E-01 | 2.73E+00 | 2.76E+00 | 8.81E-01 | no | Frmd5 | intron_variant | MODIFIER |
| 2 | 122805762 | AX-170141589 | 0 | C | T | 0.14 | -3.23E-02 | 2.22E-01 | 2.73E+00 | 2.76E+00 | 8.81E-01 | no | Sqor | intron_variant | MODIFIER |
| 12 | 58203745 | AX-170384427 | 0 | A | G | 0.053 | 4.01E-02 | 2.77E-01 | 2.69E+00 | 2.75E+00 | 8.82E-01 | no | | intergenic_variant | MODIFIER |
| 12 | 59712028 | AX-170502739 | 0 | A | G | 0.053 | 4.01E-02 | 2.77E-01 | 2.69E+00 | 2.75E+00 | 8.82E-01 | no | | intergenic_variant | MODIFIER |
| 18 | 34003070 | AX-170060523 | 0 | A | T | 0.053 | 4.01E-02 | 2.77E-01 | 2.69E+00 | 2.75E+00 | 8.82E-01 | no | Epb41l4a | intron_variant | MODIFIER |
| 18 | 40322420 | AX-170470887 | 0 | T | C | 0.053 | 4.01E-02 | 2.77E-01 | 2.69E+00 | 2.75E+00 | 8.82E-01 | no | Kctd16 | intron_variant | MODIFIER |
| 18 | 43546178 | AX-170328555 | 0 | A | G | 0.053 | 4.01E-02 | 2.77E-01 | 2.69E+00 | 2.75E+00 | 8.82E-01 | no | Jakmip2 | intron_variant | MODIFIER |
| 18 | 46294422 | AX-170015647 | 0 | A | G | 0.053 | 4.01E-02 | 2.77E-01 | 2.69E+00 | 2.75E+00 | 8.82E-01 | no | Ccdc112 | intron_variant | MODIFIER |
| 1 | 66602433 | AX-170200655 | 1 | C | T | 0.116 | 3.43E-02 | 2.30E-01 | 2.68E+00 | 2.73E+00 | 8.82E-01 | no | Unc80 | intron_variant | MODIFIER |
| 15 | 91328085 | AX-170298738 | 0 | C | T | 0.132 | -1.95E-02 | 1.38E-01 | 2.66E+00 | 2.76E+00 | 8.82E-01 | no | Slc2a13 | intron_variant | MODIFIER |
| 13 | 117409659 | AX-170052561 | 0 | C | T | 0.026 | -2.59E-02 | 1.75E-01 | 2.57E+00 | 2.74E+00 | 8.82E-01 | no | | intergenic_variant | MODIFIER |
| 7 | 117865259 | AX-170204109 | 2 | C | A | 0.391 | 1.36E-02 | 9.06E-02 | 2.67E+00 | 2.73E+00 | 8.82E-01 | no | | intergenic_variant | MODIFIER |
| 2 | 122448893 | AX-170412657 | 0 | T | C | 0.132 | 3.10E-02 | 2.07E-01 | 2.68E+00 | 2.72E+00 | 8.82E-01 | no | Slc28a2 | intron_variant | MODIFIER |
| 2 | 124808672 | AX-97795006 | 0 | G | C | 0.132 | 3.10E-02 | 2.07E-01 | 2.68E+00 | 2.72E+00 | 8.82E-01 | no | | intergenic_variant | MODIFIER |
| 10 | 6558884 | AX-170280418 | 0 | C | T | 0.202 | -1.60E-02 | 1.04E-01 | 2.63E+00 | 2.72E+00 | 8.82E-01 | no | | intergenic_variant | MODIFIER |
| 15 | 91826861 | AX-170225307 | 0 | G | T | 0.158 | 2.48E-02 | 1.55E-01 | 2.61E+00 | 2.71E+00 | 8.82E-01 | no | Gm15382 | downstream_gene_variant | MODIFIER |
| 15 | 93527615 | AX-170496918 | 0 | C | G | 0.158 | 2.48E-02 | 1.55E-01 | 2.61E+00 | 2.71E+00 | 8.82E-01 | no | Prickle1 | intron_variant | MODIFIER |
| 15 | 93991755 | AX-170365135 | 0 | G | A | 0.158 | 2.48E-02 | 1.55E-01 | 2.61E+00 | 2.71E+00 | 8.82E-01 | no | | intergenic_variant | MODIFIER |
| 5 | 97979045 | AX-170354936 | 0 | C | T | 0.43 | -1.55E-02 | 1.08E-01 | 2.70E+00 | 2.76E+00 | 8.82E-01 | no | Antxr2 | intron_variant | MODIFIER |
| 5 | 98053952 | AX-170194447 | 0 | T | C | 0.43 | -1.55E-02 | 1.08E-01 | 2.70E+00 | 2.76E+00 | 8.82E-01 | no | Gm42760 | upstream_gene_variant | MODIFIER |
| 5 | 98054871 | AX-170142662 | 0 | T | C | 0.43 | -1.55E-02 | 1.08E-01 | 2.70E+00 | 2.76E+00 | 8.82E-01 | no | Gm42760 | upstream_gene_variant | MODIFIER |
| 5 | 98066092 | AX-170385790 | 0 | C | T | 0.43 | -1.55E-02 | 1.08E-01 | 2.70E+00 | 2.76E+00 | 8.82E-01 | no | | intergenic_variant | MODIFIER |
| 5 | 98074134 | AX-170285196 | 0 | C | T | 0.43 | -1.55E-02 | 1.08E-01 | 2.70E+00 | 2.76E+00 | 8.82E-01 | no | Gm8041 | downstream_gene_variant | MODIFIER |
| 5 | 98112891 | AX-170072840 | 0 | T | C | 0.43 | -1.55E-02 | 1.08E-01 | 2.70E+00 | 2.76E+00 | 8.82E-01 | no | | intergenic_variant | MODIFIER |
| 5 | 98174638 | AX-170363891 | 0 | T | C | 0.43 | -1.55E-02 | 1.08E-01 | 2.70E+00 | 2.76E+00 | 8.82E-01 | no | Prdm8 | intron_variant | MODIFIER |
| 5 | 98175756 | AX-170246989 | 0 | A | G | 0.43 | -1.55E-02 | 1.08E-01 | 2.70E+00 | 2.76E+00 | 8.82E-01 | no | Prdm8 | intron_variant | MODIFIER |
| 5 | 98176822 | AX-170248117 | 0 | G | T | 0.43 | -1.55E-02 | 1.08E-01 | 2.70E+00 | 2.76E+00 | 8.82E-01 | no | Prdm8 | upstream_gene_variant | MODIFIER |
| 5 | 98193827 | AX-170025687 | 0 | T | C | 0.43 | -1.55E-02 | 1.08E-01 | 2.70E+00 | 2.76E+00 | 8.82E-01 | no | Prdm8 | downstream_gene_variant | MODIFIER |
| 5 | 98220182 | AX-170025683 | 0 | A | G | 0.43 | -1.55E-02 | 1.08E-01 | 2.70E+00 | 2.76E+00 | 8.82E-01 | no | | intergenic_variant | MODIFIER |
| 5 | 98228750 | AX-170457224 | 0 | C | T | 0.43 | -1.55E-02 | 1.08E-01 | 2.70E+00 | 2.76E+00 | 8.82E-01 | no | | intergenic_variant | MODIFIER |
| 5 | 98233230 | AX-170477492 | 0 | G | A | 0.43 | -1.55E-02 | 1.08E-01 | 2.70E+00 | 2.76E+00 | 8.82E-01 | no | | intergenic_variant | MODIFIER |
| 5 | 98370551 | AX-170460095 | 0 | T | C | 0.43 | -1.55E-02 | 1.08E-01 | 2.70E+00 | 2.76E+00 | 8.82E-01 | no | Cfap299 | intron_variant | MODIFIER |
| 5 | 98378313 | AX-170072841 | 0 | G | T | 0.43 | -1.55E-02 | 1.08E-01 | 2.70E+00 | 2.76E+00 | 8.82E-01 | no | Cfap299 | intron_variant | MODIFIER |

| | | | | | | | | | | | | | | | |
|---|---|---|---|---|---|---|---|---|---|---|---|---|---|---|---|
| 5 | 98384061 | AX-170072842 | 0 | T | C | 0.43 | -1.55E-02 | 1.08E-01 | 2.70E+00 | 2.76E+00 | 8.82E-01 | no | Cfap299 | intron_variant | MODIFIER |
| 5 | 98387658 | AX-170072846 | 0 | A | G | 0.43 | -1.55E-02 | 1.08E-01 | 2.70E+00 | 2.76E+00 | 8.82E-01 | no | Cfap299 | intron_variant | MODIFIER |
| 5 | 98461720 | AX-97822597 | 0 | G | A | 0.43 | -1.55E-02 | 1.08E-01 | 2.70E+00 | 2.76E+00 | 8.82E-01 | no | Cfap299 | intron_variant | MODIFIER |
| 5 | 99011515 | AX-97807949 | 0 | C | A | 0.43 | -1.55E-02 | 1.08E-01 | 2.70E+00 | 2.76E+00 | 8.82E-01 | no | Prkg2 | intron_variant | MODIFIER |
| 5 | 99117913 | AX-97828655 | 0 | C | G | 0.43 | -1.55E-02 | 1.08E-01 | 2.70E+00 | 2.76E+00 | 8.82E-01 | no | Gm43251 | intron_variant,non_coding_transcript_variant | MODIFIER |
| 5 | 99236142 | AX-170025702 | 0 | G | T | 0.43 | -1.55E-02 | 1.08E-01 | 2.70E+00 | 2.76E+00 | 8.82E-01 | no | Rasgef1b | intron_variant | MODIFIER |
| 5 | 99257175 | AX-97800782 | 0 | G | A | 0.43 | -1.55E-02 | 1.08E-01 | 2.70E+00 | 2.76E+00 | 8.82E-01 | no | Rasgef1b | upstream_gene_variant | MODIFIER |
| 5 | 99272501 | AX-170025705 | 0 | G | A | 0.43 | -1.55E-02 | 1.08E-01 | 2.70E+00 | 2.76E+00 | 8.82E-01 | no | Gm35394 | upstream_gene_variant | MODIFIER |
| 5 | 99277580 | AX-97825620 | 0 | G | A | 0.43 | -1.55E-02 | 1.08E-01 | 2.70E+00 | 2.76E+00 | 8.82E-01 | no | Gm35394 | non_coding_transcript_exon_variant | MODIFIER |
| 5 | 99295456 | AX-170415665 | 0 | C | A | 0.43 | -1.55E-02 | 1.08E-01 | 2.70E+00 | 2.76E+00 | 8.82E-01 | no | Rasgef1b | downstream_gene_variant | MODIFIER |
| 5 | 99295493 | AX-170356574 | 0 | A | G | 0.43 | -1.55E-02 | 1.08E-01 | 2.70E+00 | 2.76E+00 | 8.82E-01 | no | Rasgef1b | downstream_gene_variant | MODIFIER |
| 5 | 99299507 | AX-170025701 | 0 | C | T | 0.43 | -1.55E-02 | 1.08E-01 | 2.70E+00 | 2.76E+00 | 8.82E-01 | no | Rasgef1b | 3_prime_UTR_variant | MODIFIER |
| 5 | 99299829 | AX-170025707 | 0 | C | G | 0.43 | -1.55E-02 | 1.08E-01 | 2.70E+00 | 2.76E+00 | 8.82E-01 | no | Rasgef1b | 3_prime_UTR_variant | MODIFIER |
| 5 | 99300319 | AX-97804060 | 0 | A | G | 0.43 | -1.55E-02 | 1.08E-01 | 2.70E+00 | 2.76E+00 | 8.82E-01 | no | Rasgef1b | intron_variant | MODIFIER |
| 5 | 99320002 | AX-170025700 | 0 | G | A | 0.43 | -1.55E-02 | 1.08E-01 | 2.70E+00 | 2.76E+00 | 8.82E-01 | no | Rasgef1b | intron_variant | MODIFIER |
| 5 | 99320544 | AX-170319174 | 0 | G | A | 0.43 | -1.55E-02 | 1.08E-01 | 2.70E+00 | 2.76E+00 | 8.82E-01 | no | Rasgef1b | intron_variant | MODIFIER |
| 5 | 99322557 | AX-97849593 | 0 | C | T | 0.43 | -1.55E-02 | 1.08E-01 | 2.70E+00 | 2.76E+00 | 8.82E-01 | no | Rasgef1b | intron_variant | MODIFIER |
| 5 | 99477304 | AX-170211410 | 0 | C | T | 0.43 | -1.55E-02 | 1.08E-01 | 2.70E+00 | 2.76E+00 | 8.82E-01 | no | Rasgef1b | intron_variant | MODIFIER |
| 5 | 99478360 | AX-170325564 | 0 | C | A | 0.43 | -1.55E-02 | 1.08E-01 | 2.70E+00 | 2.76E+00 | 8.82E-01 | no | Rasgef1b | intron_variant | MODIFIER |
| 5 | 99484625 | AX-170211518 | 0 | G | A | 0.43 | -1.55E-02 | 1.08E-01 | 2.70E+00 | 2.76E+00 | 8.82E-01 | no | Rasgef1b | intron_variant | MODIFIER |
| 5 | 99494171 | AX-170141533 | 0 | G | A | 0.43 | -1.55E-02 | 1.08E-01 | 2.70E+00 | 2.76E+00 | 8.82E-01 | no | Rasgef1b | intron_variant | MODIFIER |
| 5 | 99494198 | AX-97808291 | 0 | C | G | 0.43 | -1.55E-02 | 1.08E-01 | 2.70E+00 | 2.76E+00 | 8.82E-01 | no | Rasgef1b | intron_variant | MODIFIER |
| 5 | 99494222 | AX-170145724 | 0 | G | A | 0.43 | -1.55E-02 | 1.08E-01 | 2.70E+00 | 2.76E+00 | 8.82E-01 | no | Rasgef1b | intron_variant | MODIFIER |
| 5 | 99494736 | AX-97833293 | 0 | A | G | 0.43 | -1.55E-02 | 1.08E-01 | 2.70E+00 | 2.76E+00 | 8.82E-01 | no | Rasgef1b | intron_variant | MODIFIER |
| 5 | 99503749 | AX-170305348 | 0 | T | A | 0.43 | -1.55E-02 | 1.08E-01 | 2.70E+00 | 2.76E+00 | 8.82E-01 | no | Rasgef1b | intron_variant | MODIFIER |
| 5 | 99508155 | AX-170129897 | 0 | G | A | 0.43 | -1.55E-02 | 1.08E-01 | 2.70E+00 | 2.76E+00 | 8.82E-01 | no | Rasgef1b | intron_variant | MODIFIER |
| 5 | 99691314 | AX-170382386 | 0 | T | A | 0.43 | -1.55E-02 | 1.08E-01 | 2.70E+00 | 2.76E+00 | 8.82E-01 | no | Rasgef1b | intron_variant,non_coding_transcript_variant | MODIFIER |
| 5 | 99801563 | AX-97783478 | 0 | G | A | 0.43 | -1.55E-02 | 1.08E-01 | 2.70E+00 | 2.76E+00 | 8.82E-01 | no | | intergenic_variant | MODIFIER |
| 5 | 100061904 | AX-170204808 | 0 | C | T | 0.43 | -1.55E-02 | 1.08E-01 | 2.70E+00 | 2.76E+00 | 8.82E-01 | no | Enoph1 | intron_variant | MODIFIER |
| 5 | 100078121 | AX-170165397 | 0 | G | A | 0.43 | -1.55E-02 | 1.08E-01 | 2.70E+00 | 2.76E+00 | 8.82E-01 | no | Tmem150c | 3_prime_UTR_variant | MODIFIER |
| 5 | 100092406 | AX-170072870 | 0 | A | G | 0.43 | -1.55E-02 | 1.08E-01 | 2.70E+00 | 2.76E+00 | 8.82E-01 | no | Tmem150c | intron_variant | MODIFIER |
| 5 | 100092498 | AX-170072869 | 0 | T | C | 0.43 | -1.55E-02 | 1.08E-01 | 2.70E+00 | 2.76E+00 | 8.82E-01 | no | Tmem150c | intron_variant | MODIFIER |
| 5 | 100106744 | AX-170290884 | 0 | T | C | 0.43 | -1.55E-02 | 1.08E-01 | 2.70E+00 | 2.76E+00 | 8.82E-01 | no | Tmem150c | intron_variant | MODIFIER |
| 5 | 100109126 | AX-170439671 | 0 | C | G | 0.43 | -1.55E-02 | 1.08E-01 | 2.70E+00 | 2.76E+00 | 8.82E-01 | no | Tmem150c | intron_variant | MODIFIER |
| 5 | 100109623 | AX-170429179 | 0 | G | A | 0.43 | -1.55E-02 | 1.08E-01 | 2.70E+00 | 2.76E+00 | 8.82E-01 | no | Tmem150c | intron_variant | MODIFIER |
| 5 | 100109689 | AX-170108961 | 0 | T | C | 0.43 | -1.55E-02 | 1.08E-01 | 2.70E+00 | 2.76E+00 | 8.82E-01 | no | Tmem150c | intron_variant | MODIFIER |
| 5 | 100109720 | AX-170150111 | 0 | C | T | 0.43 | -1.55E-02 | 1.08E-01 | 2.70E+00 | 2.76E+00 | 8.82E-01 | no | Tmem150c | intron_variant | MODIFIER |
| 5 | 100125775 | AX-170367589 | 0 | A | G | 0.43 | -1.55E-02 | 1.08E-01 | 2.70E+00 | 2.76E+00 | 8.82E-01 | no | Tmem150c | intron_variant | MODIFIER |
| 5 | 100129377 | AX-170481633 | 0 | G | A | 0.43 | -1.55E-02 | 1.08E-01 | 2.70E+00 | 2.76E+00 | 8.82E-01 | no | Tmem150c | intron_variant | MODIFIER |
| 5 | 100144521 | AX-170268148 | 0 | A | G | 0.43 | -1.55E-02 | 1.08E-01 | 2.70E+00 | 2.76E+00 | 8.82E-01 | no | Tmem150c | intron_variant | MODIFIER |
| 5 | 100164242 | AX-170106318 | 0 | C | T | 0.43 | -1.55E-02 | 1.08E-01 | 2.70E+00 | 2.76E+00 | 8.82E-01 | no | Tmem150c | upstream_gene_variant | MODIFIER |
| 5 | 100168868 | AX-170472944 | 0 | C | T | 0.43 | -1.55E-02 | 1.08E-01 | 2.70E+00 | 2.76E+00 | 8.82E-01 | no | | intergenic_variant | MODIFIER |
| 5 | 100168922 | AX-97813981 | 0 | T | C | 0.43 | -1.55E-02 | 1.08E-01 | 2.70E+00 | 2.76E+00 | 8.82E-01 | no | | intergenic_variant | MODIFIER |
| 5 | 100174553 | AX-170214353 | 0 | T | C | 0.43 | -1.55E-02 | 1.08E-01 | 2.70E+00 | 2.76E+00 | 8.82E-01 | no | | intergenic_variant | MODIFIER |
| 5 | 100198841 | AX-170280874 | 0 | G | T | 0.43 | -1.55E-02 | 1.08E-01 | 2.70E+00 | 2.76E+00 | 8.82E-01 | no | Gm9932 | non_coding_transcript_exon_variant | MODIFIER |
| 5 | 100200863 | AX-97820300 | 0 | A | G | 0.43 | -1.55E-02 | 1.08E-01 | 2.70E+00 | 2.76E+00 | 8.82E-01 | no | Gm9932 | downstream_gene_variant | MODIFIER |
| 5 | 100218893 | AX-170342142 | 0 | T | C | 0.43 | -1.55E-02 | 1.08E-01 | 2.70E+00 | 2.76E+00 | 8.82E-01 | no | Gm8091 | downstream_gene_variant | MODIFIER |
| 5 | 100220947 | AX-170510196 | 0 | C | T | 0.43 | -1.55E-02 | 1.08E-01 | 2.70E+00 | 2.76E+00 | 8.82E-01 | no | Gm8091 | intron_variant,non_coding_transcript_variant | MODIFIER |
| 5 | 100238698 | AX-170072873 | 0 | G | A | 0.43 | -1.55E-02 | 1.08E-01 | 2.70E+00 | 2.76E+00 | 8.82E-01 | no | Gm8091 | intron_variant,non_coding_transcript_variant | MODIFIER |
| 5 | 100239888 | AX-170072874 | 0 | G | A | 0.43 | -1.55E-02 | 1.08E-01 | 2.70E+00 | 2.76E+00 | 8.82E-01 | no | Gm8091 | intron_variant,non_coding_transcript_variant | MODIFIER |
| 5 | 100258634 | AX-170366650 | 0 | G | A | 0.43 | -1.55E-02 | 1.08E-01 | 2.70E+00 | 2.76E+00 | 8.82E-01 | no | Gm8091 | non_coding_transcript_exon_variant | MODIFIER |
| 5 | 100265897 | AX-170497123 | 0 | A | T | 0.43 | -1.55E-02 | 1.08E-01 | 2.70E+00 | 2.76E+00 | 8.82E-01 | no | Gm8091 | intron_variant,non_coding_transcript_variant | MODIFIER |
| 5 | 100606050 | AX-170072876 | 0 | T | C | 0.43 | -1.55E-02 | 1.08E-01 | 2.70E+00 | 2.76E+00 | 8.82E-01 | no | | regulatory_region_variant | MODIFIER |
| 5 | 100606289 | AX-170359282 | 0 | T | A | 0.43 | -1.55E-02 | 1.08E-01 | 2.70E+00 | 2.76E+00 | 8.82E-01 | no | | regulatory_region_variant | MODIFIER |

| | | | | | | | | | | | | | | | |
|---|---|---|---|---|---|---|---|---|---|---|---|---|---|---|---|
| 5 | 100614238 | AX-97807355 | 0 | G | A | 0.43 | -1.55E-02 | 1.08E-01 | 2.70E+00 | 2.76E+00 | 8.82E-01 | no | | regulatory_region_variant | MODIFIER |
| 5 | 100652030 | AX-170513828 | 0 | C | T | 0.43 | -1.55E-02 | 1.08E-01 | 2.70E+00 | 2.76E+00 | 8.82E-01 | no | Coq2 | downstream_gene_variant | MODIFIER |
| 5 | 100654571 | AX-170328790 | 0 | T | C | 0.43 | -1.55E-02 | 1.08E-01 | 2.70E+00 | 2.76E+00 | 8.82E-01 | no | Coq2 | downstream_gene_variant | MODIFIER |
| 5 | 100654902 | AX-170248924 | 0 | T | C | 0.43 | -1.55E-02 | 1.08E-01 | 2.70E+00 | 2.76E+00 | 8.82E-01 | no | Coq2 | 3_prime_UTR_variant | MODIFIER |
| 5 | 100655631 | AX-170450545 | 0 | G | A | 0.43 | -1.55E-02 | 1.08E-01 | 2.70E+00 | 2.76E+00 | 8.82E-01 | no | Coq2 | intron_variant | MODIFIER |
| 5 | 100656922 | AX-97835915 | 0 | T | C | 0.43 | -1.55E-02 | 1.08E-01 | 2.70E+00 | 2.76E+00 | 8.82E-01 | no | Coq2 | intron_variant | MODIFIER |
| 5 | 100658163 | AX-170457891 | 0 | T | C | 0.43 | -1.55E-02 | 1.08E-01 | 2.70E+00 | 2.76E+00 | 8.82E-01 | no | Coq2 | intron_variant | MODIFIER |
| 5 | 100659648 | AX-170485286 | 0 | T | C | 0.43 | -1.55E-02 | 1.08E-01 | 2.70E+00 | 2.76E+00 | 8.82E-01 | no | Coq2 | intron_variant | MODIFIER |
| 5 | 100667655 | AX-170326868 | 0 | C | T | 0.43 | -1.55E-02 | 1.08E-01 | 2.70E+00 | 2.76E+00 | 8.82E-01 | no | Coq2 | intron_variant | MODIFIER |
| 5 | 100680184 | AX-170344645 | 0 | A | C | 0.43 | -1.55E-02 | 1.08E-01 | 2.70E+00 | 2.76E+00 | 8.82E-01 | no | Hpse | 3_prime_UTR_variant | MODIFIER |
| 5 | 100682547 | AX-170448353 | 0 | C | T | 0.43 | -1.55E-02 | 1.08E-01 | 2.70E+00 | 2.76E+00 | 8.82E-01 | no | Hpse | intron_variant | MODIFIER |
| 5 | 100727246 | AX-170522255 | 0 | G | T | 0.43 | -1.55E-02 | 1.08E-01 | 2.70E+00 | 2.76E+00 | 8.82E-01 | no | | intergenic_variant | MODIFIER |
| 5 | 100728240 | AX-170282670 | 0 | A | C | 0.43 | -1.55E-02 | 1.08E-01 | 2.70E+00 | 2.76E+00 | 8.82E-01 | no | | intergenic_variant | MODIFIER |
| 5 | 100729072 | AX-170072883 | 0 | A | G | 0.43 | -1.55E-02 | 1.08E-01 | 2.70E+00 | 2.76E+00 | 8.82E-01 | no | | regulatory_region_variant | MODIFIER |
| 5 | 100756560 | AX-170072882 | 0 | G | A | 0.43 | -1.55E-02 | 1.08E-01 | 2.70E+00 | 2.76E+00 | 8.82E-01 | no | | intergenic_variant | MODIFIER |
| 5 | 100756602 | AX-170072884 | 0 | C | A | 0.43 | -1.55E-02 | 1.08E-01 | 2.70E+00 | 2.76E+00 | 8.82E-01 | no | | intergenic_variant | MODIFIER |
| 5 | 100756801 | AX-97836186 | 0 | A | C | 0.43 | -1.55E-02 | 1.08E-01 | 2.70E+00 | 2.76E+00 | 8.82E-01 | no | | regulatory_region_variant | MODIFIER |
| 5 | 100767601 | AX-170419166 | 0 | A | C | 0.43 | -1.55E-02 | 1.08E-01 | 2.70E+00 | 2.76E+00 | 8.82E-01 | no | Helq | intron_variant | MODIFIER |
| 5 | 100768086 | AX-170320597 | 0 | A | G | 0.43 | -1.55E-02 | 1.08E-01 | 2.70E+00 | 2.76E+00 | 8.82E-01 | no | Helq | intron_variant | MODIFIER |
| 5 | 100778204 | AX-170288157 | 0 | A | G | 0.43 | -1.55E-02 | 1.08E-01 | 2.70E+00 | 2.76E+00 | 8.82E-01 | no | Helq | intron_variant | MODIFIER |
| 5 | 100785076 | AX-170373253 | 0 | T | C | 0.43 | -1.55E-02 | 1.08E-01 | 2.70E+00 | 2.76E+00 | 8.82E-01 | no | Helq | intron_variant | MODIFIER |
| 5 | 100788566 | AX-170421060 | 0 | T | C | 0.43 | -1.55E-02 | 1.08E-01 | 2.70E+00 | 2.76E+00 | 8.82E-01 | no | Helq | intron_variant | MODIFIER |
| 5 | 100794970 | AX-170273725 | 0 | C | G | 0.43 | -1.55E-02 | 1.08E-01 | 2.70E+00 | 2.76E+00 | 8.82E-01 | no | Mrps18c | upstream_gene_variant | MODIFIER |
| 5 | 100796121 | AX-170247901 | 0 | G | A | 0.43 | -1.55E-02 | 1.08E-01 | 2.70E+00 | 2.76E+00 | 8.82E-01 | no | Mrps18c | upstream_gene_variant | MODIFIER |
| 5 | 100798973 | AX-97820202 | 0 | C | T | 0.43 | -1.55E-02 | 1.08E-01 | 2.70E+00 | 2.76E+00 | 8.82E-01 | no | Mrps18c | intron_variant | MODIFIER |
| 5 | 100943013 | AX-170517610 | 0 | A | G | 0.43 | -1.55E-02 | 1.08E-01 | 2.70E+00 | 2.76E+00 | 8.82E-01 | no | | intergenic_variant | MODIFIER |
| 5 | 100943073 | AX-170204826 | 0 | C | T | 0.43 | -1.55E-02 | 1.08E-01 | 2.70E+00 | 2.76E+00 | 8.82E-01 | no | | intergenic_variant | MODIFIER |
| 5 | 100943531 | AX-97808441 | 0 | C | G | 0.43 | -1.55E-02 | 1.08E-01 | 2.70E+00 | 2.76E+00 | 8.82E-01 | no | | intergenic_variant | MODIFIER |
| 5 | 100948116 | AX-170072889 | 0 | A | G | 0.43 | -1.55E-02 | 1.08E-01 | 2.70E+00 | 2.76E+00 | 8.82E-01 | no | | intergenic_variant | MODIFIER |
| 5 | 100949603 | AX-170137258 | 0 | G | A | 0.43 | -1.55E-02 | 1.08E-01 | 2.70E+00 | 2.76E+00 | 8.82E-01 | no | | intergenic_variant | MODIFIER |
| 5 | 100949771 | AX-170095410 | 0 | A | C | 0.43 | -1.55E-02 | 1.08E-01 | 2.70E+00 | 2.76E+00 | 8.82E-01 | no | | intergenic_variant | MODIFIER |
| 5 | 100958086 | AX-170433496 | 0 | T | C | 0.43 | -1.55E-02 | 1.08E-01 | 2.70E+00 | 2.76E+00 | 8.82E-01 | no | | intergenic_variant | MODIFIER |
| 5 | 100978150 | AX-170325667 | 0 | A | G | 0.43 | -1.55E-02 | 1.08E-01 | 2.70E+00 | 2.76E+00 | 8.82E-01 | no | | intergenic_variant | MODIFIER |
| 5 | 101357331 | AX-97846965 | 0 | C | G | 0.43 | -1.55E-02 | 1.08E-01 | 2.70E+00 | 2.76E+00 | 8.82E-01 | no | | intergenic_variant | MODIFIER |
| 5 | 101375505 | AX-170143206 | 0 | T | C | 0.43 | -1.55E-02 | 1.08E-01 | 2.70E+00 | 2.76E+00 | 8.82E-01 | no | | intergenic_variant | MODIFIER |
| 5 | 101383799 | AX-170025735 | 0 | C | A | 0.43 | -1.55E-02 | 1.08E-01 | 2.70E+00 | 2.76E+00 | 8.82E-01 | no | | intergenic_variant | MODIFIER |
| 5 | 101383876 | AX-170025734 | 0 | A | G | 0.43 | -1.55E-02 | 1.08E-01 | 2.70E+00 | 2.76E+00 | 8.82E-01 | no | | intergenic_variant | MODIFIER |
| 5 | 101383918 | AX-170025743 | 0 | C | T | 0.43 | -1.55E-02 | 1.08E-01 | 2.70E+00 | 2.76E+00 | 8.82E-01 | no | | intergenic_variant | MODIFIER |
| 5 | 101451688 | AX-170218671 | 0 | A | G | 0.43 | -1.55E-02 | 1.08E-01 | 2.70E+00 | 2.76E+00 | 8.82E-01 | no | | intergenic_variant | MODIFIER |
| 5 | 101494842 | AX-170095414 | 0 | G | A | 0.43 | -1.55E-02 | 1.08E-01 | 2.70E+00 | 2.76E+00 | 8.82E-01 | no | | intergenic_variant | MODIFIER |
| 5 | 101496139 | AX-170475445 | 0 | C | G | 0.43 | -1.55E-02 | 1.08E-01 | 2.70E+00 | 2.76E+00 | 8.82E-01 | no | | intergenic_variant | MODIFIER |
| 5 | 101498866 | AX-170025738 | 0 | C | G | 0.43 | -1.55E-02 | 1.08E-01 | 2.70E+00 | 2.76E+00 | 8.82E-01 | no | | intergenic_variant | MODIFIER |
| 5 | 101548474 | AX-170025732 | 0 | A | G | 0.43 | -1.55E-02 | 1.08E-01 | 2.70E+00 | 2.76E+00 | 8.82E-01 | no | | intergenic_variant | MODIFIER |
| 5 | 101554690 | AX-97800517 | 0 | G | T | 0.43 | -1.55E-02 | 1.08E-01 | 2.70E+00 | 2.76E+00 | 8.82E-01 | no | | intergenic_variant | MODIFIER |
| 5 | 101557130 | AX-170025740 | 0 | G | A | 0.43 | -1.55E-02 | 1.08E-01 | 2.70E+00 | 2.76E+00 | 8.82E-01 | no | | intergenic_variant | MODIFIER |
| 5 | 101558583 | AX-170025741 | 0 | C | T | 0.43 | -1.55E-02 | 1.08E-01 | 2.70E+00 | 2.76E+00 | 8.82E-01 | no | | regulatory_region_variant | MODIFIER |
| 10 | 15717736 | AX-170110307 | 0 | A | G | 0.184 | 1.49E-02 | 1.07E-01 | 2.68E+00 | 2.77E+00 | 8.82E-01 | no | Gm40604 | intron_variant,non_coding_transcript_variant | MODIFIER |
| 9 | 111814446 | AX-170105042 | 2 | C | T | 0.109 | 1.61E-02 | 1.16E-01 | 2.69E+00 | 2.77E+00 | 8.82E-01 | no | | intergenic_variant | MODIFIER |
| 11 | 49117638 | AX-170092239 | 0 | C | T | 0.088 | -2.54E-02 | 1.59E-01 | 2.60E+00 | 2.71E+00 | 8.82E-01 | no | Olfr56 | 5_prime_UTR_variant | MODIFIER |
| 7 | 66449479 | AX-170247515 | 0 | A | G | 0.404 | -1.32E-02 | 9.44E-02 | 2.68E+00 | 2.76E+00 | 8.82E-01 | no | | regulatory_region_variant | MODIFIER |
| 18 | 31701223 | AX-170448188 | 0 | C | G | 0.053 | 3.64E-02 | 2.45E-01 | 2.64E+00 | 2.73E+00 | 8.82E-01 | no | Sap130 | intron_variant | MODIFIER |
| 7 | 57530273 | AX-170172083 | 1 | T | A | 0.429 | -1.37E-02 | 9.79E-02 | 2.68E+00 | 2.76E+00 | 8.82E-01 | no | Gabrb3 | intron_variant | MODIFIER |
| 13 | 48758742 | AX-170433664 | 1 | G | A | 0.5 | 1.05E-02 | 6.75E-02 | 2.62E+00 | 2.72E+00 | 8.82E-01 | no | Rpl17-ps6 | downstream_gene_variant | MODIFIER |
| 1 | 54699608 | AX-170130116 | 0 | C | A | 0.096 | 3.12E-02 | 2.17E-01 | 2.69E+00 | 2.75E+00 | 8.82E-01 | no | Ankrd44 | intron_variant | MODIFIER |

| | | | | | | | | | | | | | | | |
|---|---|---|---|---|---|---|---|---|---|---|---|---|---|---|---|
| 1 | 59902147 | AX-170199311 | 0 | C | T | 0.096 | 3.12E-02 | 2.17E-01 | 2.69E+00 | 2.75E+00 | 8.82E-01 | no | | regulatory_region_variant | MODIFIER |
| 1 | 155854952 | AX-170216706 | 2 | A | G | 0.209 | 2.12E-02 | 1.38E-01 | 2.66E+00 | 2.71E+00 | 8.83E-01 | no | Cep350 | intron_variant,non_coding_transcript_variant | MODIFIER |
| 7 | 70044312 | AX-170219345 | 2 | G | T | 0.391 | -1.31E-02 | 9.56E-02 | 2.68E+00 | 2.77E+00 | 8.83E-01 | no | | intergenic_variant | MODIFIER |
| 6 | 139716601 | AX-170262532 | 1 | C | T | 0.071 | 3.99E-02 | 2.76E-01 | 2.71E+00 | 2.75E+00 | 8.83E-01 | no | Pik3c2g | intron_variant | MODIFIER |
| 8 | 94149943 | AX-170303987 | 0 | T | C | 0.219 | 1.58E-02 | 1.09E-01 | 2.66E+00 | 2.74E+00 | 8.83E-01 | no | Mt3 | upstream_gene_variant | MODIFIER |
| 11 | 9716435 | AX-170417792 | 1 | T | C | 0.062 | 2.99E-02 | 2.13E-01 | 2.66E+00 | 2.75E+00 | 8.83E-01 | no | | intergenic_variant | MODIFIER |
| 5 | 104943993 | AX-170394285 | 0 | A | G | 0.456 | 1.69E-02 | 1.12E-01 | 2.67E+00 | 2.72E+00 | 8.83E-01 | no | Abcg3 | intron_variant | MODIFIER |
| 11 | 33365035 | AX-97798272 | 0 | A | G | 0.105 | 2.39E-02 | 1.64E-01 | 2.63E+00 | 2.74E+00 | 8.83E-01 | no | Ranbp17 | intron_variant | MODIFIER |
| 6 | 147330116 | AX-170076582 | 1 | A | C | 0.062 | -3.31E-02 | 2.25E-01 | 2.65E+00 | 2.73E+00 | 8.83E-01 | no | | intergenic_variant | MODIFIER |
| 7 | 51887343 | AX-170155111 | 2 | C | T | 0.418 | 1.51E-02 | 1.01E-01 | 2.65E+00 | 2.73E+00 | 8.83E-01 | no | Gas2 | intron_variant | MODIFIER |
| 1 | 65532759 | AX-170041480 | 0 | G | A | 0.149 | 2.85E-02 | 1.88E-01 | 2.62E+00 | 2.72E+00 | 8.83E-01 | no | | intergenic_variant | MODIFIER |
| 1 | 72687754 | AX-97789519 | 0 | C | T | 0.149 | 2.85E-02 | 1.88E-01 | 2.62E+00 | 2.72E+00 | 8.83E-01 | no | Ankar | synonymous_variant | LOW |
| 1 | 76278637 | AX-170000841 | 0 | A | T | 0.149 | 2.85E-02 | 1.88E-01 | 2.62E+00 | 2.72E+00 | 8.83E-01 | no | | intergenic_variant | MODIFIER |
| 1 | 77025272 | AX-170145070 | 0 | C | T | 0.149 | 2.85E-02 | 1.88E-01 | 2.62E+00 | 2.72E+00 | 8.83E-01 | no | Gm816 | intron_variant,non_coding_transcript_variant | MODIFIER |
| 1 | 77673103 | AX-170342261 | 0 | G | A | 0.149 | 2.85E-02 | 1.88E-01 | 2.62E+00 | 2.72E+00 | 8.83E-01 | no | | intergenic_variant | MODIFIER |
| 1 | 78376816 | AX-170000872 | 0 | T | A | 0.149 | 2.85E-02 | 1.88E-01 | 2.62E+00 | 2.72E+00 | 8.83E-01 | no | Sgpp2 | intron_variant | MODIFIER |
| 9 | 66002999 | AX-170032576 | 1 | G | A | 0.196 | 1.94E-02 | 1.35E-01 | 2.73E+00 | 2.76E+00 | 8.83E-01 | no | Csnk1g1 | intron_variant | MODIFIER |
| 2 | 119047333 | AX-170123910 | 1 | G | T | 0.143 | 3.25E-02 | 2.21E-01 | 2.69E+00 | 2.72E+00 | 8.83E-01 | no | Knl1 | 5_prime_UTR_variant | MODIFIER |
| 2 | 121136496 | AX-170490156 | 1 | C | T | 0.143 | 3.25E-02 | 2.21E-01 | 2.69E+00 | 2.72E+00 | 8.83E-01 | no | Adal | upstream_gene_variant | MODIFIER |
| 2 | 123299816 | AX-170103954 | 1 | G | A | 0.143 | 3.25E-02 | 2.21E-01 | 2.69E+00 | 2.72E+00 | 8.83E-01 | no | | intergenic_variant | MODIFIER |
| 2 | 125985169 | AX-170480782 | 1 | G | A | 0.143 | 3.25E-02 | 2.21E-01 | 2.69E+00 | 2.72E+00 | 8.83E-01 | no | Galk2 | downstream_gene_variant | MODIFIER |
| 2 | 129264556 | AX-170192091 | 1 | A | C | 0.143 | 3.25E-02 | 2.21E-01 | 2.69E+00 | 2.72E+00 | 8.83E-01 | no | Ckap2l | downstream_gene_variant | MODIFIER |
| 2 | 129609818 | AX-97851455 | 1 | G | T | 0.143 | 3.25E-02 | 2.21E-01 | 2.69E+00 | 2.72E+00 | 8.83E-01 | no | Sirpa | intron_variant | MODIFIER |
| 2 | 130058622 | AX-170206871 | 1 | G | A | 0.143 | 3.25E-02 | 2.21E-01 | 2.69E+00 | 2.72E+00 | 8.83E-01 | no | | regulatory_region_variant | MODIFIER |
| 2 | 139603591 | AX-97795517 | 1 | G | T | 0.143 | 3.25E-02 | 2.21E-01 | 2.69E+00 | 2.72E+00 | 8.83E-01 | no | Sptlc3 | intron_variant | MODIFIER |
| 2 | 143588517 | AX-97842346 | 1 | C | T | 0.143 | 3.25E-02 | 2.21E-01 | 2.69E+00 | 2.72E+00 | 8.83E-01 | no | Pcsk2 | intron_variant | MODIFIER |
| 2 | 147236399 | AX-170127095 | 1 | C | T | 0.143 | 3.25E-02 | 2.21E-01 | 2.69E+00 | 2.72E+00 | 8.83E-01 | no | Nkx2-2os | intron_variant,non_coding_transcript_variant | MODIFIER |
| 2 | 156278536 | AX-97834809 | 1 | A | G | 0.143 | 3.25E-02 | 2.21E-01 | 2.69E+00 | 2.72E+00 | 8.83E-01 | no | Phf20 | intron_variant | MODIFIER |
| 2 | 157697578 | AX-170401267 | 1 | C | T | 0.143 | 3.25E-02 | 2.21E-01 | 2.69E+00 | 2.72E+00 | 8.83E-01 | no | | regulatory_region_variant | MODIFIER |
| 1 | 45753369 | AX-170106292 | 1 | G | A | 0.446 | -1.09E-02 | 7.43E-02 | 2.64E+00 | 2.73E+00 | 8.83E-01 | no | | intergenic_variant | MODIFIER |
| 3 | 25544286 | AX-170381791 | 0 | T | C | 0.246 | -9.70E-03 | 7.06E-02 | 2.59E+00 | 2.75E+00 | 8.83E-01 | no | Nlgn1 | intron_variant | MODIFIER |
| 7 | 75647057 | AX-168405470 | 0 | G | A | 0.36 | -1.30E-02 | 9.20E-02 | 2.64E+00 | 2.75E+00 | 8.84E-01 | no | Akap13 | intron_variant | MODIFIER |
| 5 | 73187456 | AX-170482297 | 0 | C | A | 0.456 | 1.36E-02 | 9.03E-02 | 2.62E+00 | 2.73E+00 | 8.84E-01 | no | Fryl | intron_variant | MODIFIER |
| 16 | 44351445 | AX-170534039 | 1 | T | C | 0.473 | -1.30E-02 | 8.00E-02 | 2.62E+00 | 2.69E+00 | 8.84E-01 | no | Spice1 | intron_variant | MODIFIER |
| 12 | 70852610 | AX-170529570 | 2 | C | T | 0.055 | 3.57E-02 | 2.57E-01 | 2.69E+00 | 2.77E+00 | 8.84E-01 | no | Frmd6 | intron_variant | MODIFIER |
| 7 | 137163477 | AX-170078987 | 1 | C | T | 0.295 | -1.22E-02 | 8.00E-02 | 2.62E+00 | 2.72E+00 | 8.84E-01 | no | Gm45499 | upstream_gene_variant | MODIFIER |
| 10 | 118716929 | AX-97805777 | 1 | T | C | 0.348 | 1.05E-02 | 7.34E-02 | 2.65E+00 | 2.74E+00 | 8.84E-01 | no | Gm33337 | upstream_gene_variant | MODIFIER |
| 8 | 68184603 | AX-170386481 | 0 | T | C | 0.149 | -2.25E-02 | 1.45E-01 | 2.64E+00 | 2.71E+00 | 8.84E-01 | no | Psd3 | intron_variant | MODIFIER |
| 15 | 93054630 | AX-170217581 | 0 | G | A | 0.149 | 2.67E-02 | 1.79E-01 | 2.66E+00 | 2.72E+00 | 8.84E-01 | no | | intergenic_variant | MODIFIER |
| 1 | 71516455 | AX-170504309 | 1 | C | T | 0.116 | 3.39E-02 | 2.34E-01 | 2.68E+00 | 2.73E+00 | 8.84E-01 | no | | intergenic_variant | MODIFIER |
| 1 | 33245775 | AX-170508962 | 0 | T | G | 0.404 | -1.19E-02 | 8.34E-02 | 2.67E+00 | 2.74E+00 | 8.84E-01 | no | | intergenic_variant | MODIFIER |
| 1 | 56904715 | AX-97843802 | 1 | C | T | 0.152 | -2.33E-02 | 1.52E-01 | 2.66E+00 | 2.70E+00 | 8.84E-01 | no | | intergenic_variant | MODIFIER |
| 8 | 57282427 | AX-170253888 | 1 | G | A | 0.152 | -2.33E-02 | 1.52E-01 | 2.66E+00 | 2.70E+00 | 8.84E-01 | no | Hand2os1 | intron_variant,non_coding_transcript_variant | MODIFIER |
| 7 | 92086123 | AX-170324399 | 0 | C | A | 0.132 | -3.05E-02 | 2.11E-01 | 2.66E+00 | 2.74E+00 | 8.84E-01 | no | Dlg2 | intron_variant | MODIFIER |
| 1 | 70244299 | AX-170299606 | 1 | C | T | 0.143 | 3.44E-02 | 2.37E-01 | 2.68E+00 | 2.73E+00 | 8.84E-01 | no | Spag16 | intron_variant | MODIFIER |
| 2 | 113636146 | AX-170483771 | 0 | T | G | 0.14 | 3.24E-02 | 2.23E-01 | 2.70E+00 | 2.73E+00 | 8.84E-01 | no | Fmn1 | intron_variant | MODIFIER |
| 2 | 113653694 | AX-170235541 | 0 | G | A | 0.14 | 3.24E-02 | 2.23E-01 | 2.70E+00 | 2.73E+00 | 8.84E-01 | no | Fmn1 | intron_variant | MODIFIER |
| 2 | 113658875 | AX-97839603 | 0 | C | T | 0.14 | 3.24E-02 | 2.23E-01 | 2.70E+00 | 2.73E+00 | 8.84E-01 | no | Fmn1 | intron_variant | MODIFIER |
| 2 | 113665157 | AX-170477470 | 0 | T | C | 0.14 | 3.24E-02 | 2.23E-01 | 2.70E+00 | 2.73E+00 | 8.84E-01 | no | Fmn1 | intron_variant | MODIFIER |
| 2 | 113683850 | AX-170396520 | 0 | G | A | 0.14 | 3.24E-02 | 2.23E-01 | 2.70E+00 | 2.73E+00 | 8.84E-01 | no | Fmn1 | intron_variant | MODIFIER |
| 2 | 113689229 | AX-170124819 | 0 | T | C | 0.14 | 3.24E-02 | 2.23E-01 | 2.70E+00 | 2.73E+00 | 8.84E-01 | no | Fmn1 | intron_variant | MODIFIER |
| 2 | 113690619 | AX-170303423 | 0 | T | C | 0.14 | 3.24E-02 | 2.23E-01 | 2.70E+00 | 2.73E+00 | 8.84E-01 | no | Fmn1 | intron_variant | MODIFIER |
| 2 | 113690823 | AX-170228427 | 0 | G | A | 0.14 | 3.24E-02 | 2.23E-01 | 2.70E+00 | 2.73E+00 | 8.84E-01 | no | Fmn1 | intron_variant | MODIFIER |
| 2 | 113704051 | AX-170119737 | 0 | C | T | 0.14 | 3.24E-02 | 2.23E-01 | 2.70E+00 | 2.73E+00 | 8.84E-01 | no | Fmn1 | intron_variant | MODIFIER |

| | | | | | | | | | | | | | | | |
|---|---|---|---|---|---|---|---|---|---|---|---|---|---|---|---|
| 2 | 113712616 | AX-170104742 | 0 | T | A | 0.14 | 3.24E-02 | 2.23E-01 | 2.70E+00 | 2.73E+00 | 8.84E-01 | no | Fmn1 | downstream_gene_variant | MODIFIER |
| 2 | 113723268 | AX-170425249 | 0 | G | A | 0.14 | 3.24E-02 | 2.23E-01 | 2.70E+00 | 2.73E+00 | 8.84E-01 | no | | intergenic_variant | MODIFIER |
| 2 | 113796787 | AX-170226926 | 0 | G | C | 0.14 | 3.24E-02 | 2.23E-01 | 2.70E+00 | 2.73E+00 | 8.84E-01 | no | Scg5 | intron_variant | MODIFIER |
| 2 | 113797351 | AX-170313109 | 0 | A | G | 0.14 | 3.24E-02 | 2.23E-01 | 2.70E+00 | 2.73E+00 | 8.84E-01 | no | Scg5 | intron_variant | MODIFIER |
| 2 | 113800645 | AX-170094376 | 0 | A | G | 0.14 | 3.24E-02 | 2.23E-01 | 2.70E+00 | 2.73E+00 | 8.84E-01 | no | Scg5 | intron_variant | MODIFIER |
| 2 | 113862920 | AX-170506438 | 0 | C | T | 0.14 | 3.24E-02 | 2.23E-01 | 2.70E+00 | 2.73E+00 | 8.84E-01 | no | | intergenic_variant | MODIFIER |
| 2 | 114008229 | AX-97839609 | 0 | G | A | 0.14 | 3.24E-02 | 2.23E-01 | 2.70E+00 | 2.73E+00 | 8.84E-01 | no | Gjd2 | downstream_gene_variant | MODIFIER |
| 2 | 114010459 | AX-170018728 | 0 | C | T | 0.14 | 3.24E-02 | 2.23E-01 | 2.70E+00 | 2.73E+00 | 8.84E-01 | no | Gjd2 | 3_prime_UTR_variant | MODIFIER |
| 2 | 114044450 | AX-170497275 | 0 | A | T | 0.14 | 3.24E-02 | 2.23E-01 | 2.70E+00 | 2.73E+00 | 8.84E-01 | no | Actc1 | downstream_gene_variant | MODIFIER |
| 2 | 114048491 | AX-170337809 | 0 | A | G | 0.14 | 3.24E-02 | 2.23E-01 | 2.70E+00 | 2.73E+00 | 8.84E-01 | no | Actc1 | intron_variant | MODIFIER |
| 2 | 114048541 | AX-170163547 | 0 | C | T | 0.14 | 3.24E-02 | 2.23E-01 | 2.70E+00 | 2.73E+00 | 8.84E-01 | no | Actc1 | intron_variant | MODIFIER |
| 2 | 114048734 | AX-170018720 | 0 | T | C | 0.14 | 3.24E-02 | 2.23E-01 | 2.70E+00 | 2.73E+00 | 8.84E-01 | no | Actc1 | intron_variant | MODIFIER |
| 2 | 114050044 | AX-170018726 | 0 | G | A | 0.14 | 3.24E-02 | 2.23E-01 | 2.70E+00 | 2.73E+00 | 8.84E-01 | no | Actc1 | intron_variant | MODIFIER |
| 2 | 114050127 | AX-97847673 | 0 | A | G | 0.14 | 3.24E-02 | 2.23E-01 | 2.70E+00 | 2.73E+00 | 8.84E-01 | no | Actc1 | intron_variant | MODIFIER |
| 2 | 114064405 | AX-170304440 | 0 | A | C | 0.14 | 3.24E-02 | 2.23E-01 | 2.70E+00 | 2.73E+00 | 8.84E-01 | no | A530058N18Rik | intron_variant,non_coding_transcript_variant | MODIFIER |
| 2 | 114071721 | AX-97785276 | 0 | C | A | 0.14 | 3.24E-02 | 2.23E-01 | 2.70E+00 | 2.73E+00 | 8.84E-01 | no | A530058N18Rik | downstream_gene_variant | MODIFIER |
| 2 | 114084646 | AX-170064638 | 0 | C | A | 0.14 | 3.24E-02 | 2.23E-01 | 2.70E+00 | 2.73E+00 | 8.84E-01 | no | | intergenic_variant | MODIFIER |
| 2 | 114095133 | AX-170018722 | 0 | G | A | 0.14 | 3.24E-02 | 2.23E-01 | 2.70E+00 | 2.73E+00 | 8.84E-01 | no | | intergenic_variant | MODIFIER |
| 2 | 114099940 | AX-170018723 | 0 | T | C | 0.14 | 3.24E-02 | 2.23E-01 | 2.70E+00 | 2.73E+00 | 8.84E-01 | no | Aqr | downstream_gene_variant | MODIFIER |
| 2 | 114100218 | AX-170301654 | 0 | A | C | 0.14 | 3.24E-02 | 2.23E-01 | 2.70E+00 | 2.73E+00 | 8.84E-01 | no | Aqr | downstream_gene_variant | MODIFIER |
| 2 | 114100673 | AX-170241106 | 0 | G | T | 0.14 | 3.24E-02 | 2.23E-01 | 2.70E+00 | 2.73E+00 | 8.84E-01 | no | Aqr | downstream_gene_variant | MODIFIER |
| 2 | 114101776 | AX-97796062 | 0 | C | T | 0.14 | 3.24E-02 | 2.23E-01 | 2.70E+00 | 2.73E+00 | 8.84E-01 | no | Aqr | intron_variant | MODIFIER |
| 2 | 114104862 | AX-170340291 | 0 | A | G | 0.14 | 3.24E-02 | 2.23E-01 | 2.70E+00 | 2.73E+00 | 8.84E-01 | no | Aqr | intron_variant | MODIFIER |
| 2 | 114127389 | AX-170094377 | 0 | A | G | 0.14 | 3.24E-02 | 2.23E-01 | 2.70E+00 | 2.73E+00 | 8.84E-01 | no | Aqr | intron_variant | MODIFIER |
| 2 | 114139076 | AX-170450322 | 0 | C | T | 0.14 | 3.24E-02 | 2.23E-01 | 2.70E+00 | 2.73E+00 | 8.84E-01 | no | Aqr | intron_variant | MODIFIER |
| 2 | 114144498 | AX-170171223 | 0 | C | G | 0.14 | 3.24E-02 | 2.23E-01 | 2.70E+00 | 2.73E+00 | 8.84E-01 | no | Aqr | intron_variant | MODIFIER |
| 2 | 114146971 | AX-170475539 | 0 | A | G | 0.14 | 3.24E-02 | 2.23E-01 | 2.70E+00 | 2.73E+00 | 8.84E-01 | no | Aqr | missense_variant | MODERATE |
| 2 | 114148268 | AX-170364018 | 0 | C | T | 0.14 | 3.24E-02 | 2.23E-01 | 2.70E+00 | 2.73E+00 | 8.84E-01 | no | Aqr | intron_variant | MODIFIER |
| 2 | 114150378 | AX-170344894 | 0 | C | T | 0.14 | 3.24E-02 | 2.23E-01 | 2.70E+00 | 2.73E+00 | 8.84E-01 | no | Aqr | intron_variant | MODIFIER |
| 2 | 114151255 | AX-170018721 | 0 | T | A | 0.14 | 3.24E-02 | 2.23E-01 | 2.70E+00 | 2.73E+00 | 8.84E-01 | no | Aqr | intron_variant | MODIFIER |
| 2 | 114160728 | AX-97797592 | 0 | T | C | 0.14 | 3.24E-02 | 2.23E-01 | 2.70E+00 | 2.73E+00 | 8.84E-01 | no | Aqr | intron_variant | MODIFIER |
| 2 | 114172855 | AX-170461869 | 0 | G | A | 0.14 | 3.24E-02 | 2.23E-01 | 2.70E+00 | 2.73E+00 | 8.84E-01 | no | Aqr | intron_variant | MODIFIER |
| 2 | 114174527 | AX-170522926 | 0 | A | G | 0.14 | 3.24E-02 | 2.23E-01 | 2.70E+00 | 2.73E+00 | 8.84E-01 | no | Aqr | intron_variant | MODIFIER |
| 2 | 114176810 | AX-170459531 | 0 | A | G | 0.14 | 3.24E-02 | 2.23E-01 | 2.70E+00 | 2.73E+00 | 8.84E-01 | no | Aqr | upstream_gene_variant | MODIFIER |
| 2 | 114188626 | AX-170094378 | 0 | A | G | 0.14 | 3.24E-02 | 2.23E-01 | 2.70E+00 | 2.73E+00 | 8.84E-01 | no | Zfp770 | downstream_gene_variant | MODIFIER |
| 2 | 114192269 | AX-170159134 | 0 | G | A | 0.14 | 3.24E-02 | 2.23E-01 | 2.70E+00 | 2.73E+00 | 8.84E-01 | no | Zfp770 | downstream_gene_variant | MODIFIER |
| 2 | 114203013 | AX-97838903 | 0 | G | C | 0.14 | 3.24E-02 | 2.23E-01 | 2.70E+00 | 2.73E+00 | 8.84E-01 | no | Zfp770 | upstream_gene_variant | MODIFIER |
| 2 | 114261165 | AX-170099716 | 0 | G | C | 0.14 | 3.24E-02 | 2.23E-01 | 2.70E+00 | 2.73E+00 | 8.84E-01 | no | | regulatory_region_variant | MODIFIER |
| 2 | 114272220 | AX-170408761 | 0 | T | A | 0.14 | 3.24E-02 | 2.23E-01 | 2.70E+00 | 2.73E+00 | 8.84E-01 | no | Gm25103 | downstream_gene_variant | MODIFIER |
| 2 | 114316841 | AX-170145894 | 0 | T | C | 0.14 | 3.24E-02 | 2.23E-01 | 2.70E+00 | 2.73E+00 | 8.84E-01 | no | | intergenic_variant | MODIFIER |
| 2 | 114325350 | AX-97791856 | 0 | A | G | 0.14 | 3.24E-02 | 2.23E-01 | 2.70E+00 | 2.73E+00 | 8.84E-01 | no | | intergenic_variant | MODIFIER |
| 2 | 114334543 | AX-170214242 | 0 | T | C | 0.14 | 3.24E-02 | 2.23E-01 | 2.70E+00 | 2.73E+00 | 8.84E-01 | no | | intergenic_variant | MODIFIER |
| 2 | 114353313 | AX-170064645 | 0 | A | G | 0.14 | 3.24E-02 | 2.23E-01 | 2.70E+00 | 2.73E+00 | 8.84E-01 | no | | intergenic_variant | MODIFIER |
| 2 | 114354486 | AX-170116533 | 0 | C | T | 0.14 | 3.24E-02 | 2.23E-01 | 2.70E+00 | 2.73E+00 | 8.84E-01 | no | | intergenic_variant | MODIFIER |
| 2 | 114363779 | AX-170239295 | 0 | T | G | 0.14 | 3.24E-02 | 2.23E-01 | 2.70E+00 | 2.73E+00 | 8.84E-01 | no | | intergenic_variant | MODIFIER |
| 2 | 114382395 | AX-170296010 | 0 | G | A | 0.14 | 3.24E-02 | 2.23E-01 | 2.70E+00 | 2.73E+00 | 8.84E-01 | no | | intergenic_variant | MODIFIER |
| 2 | 114410310 | AX-170354426 | 0 | T | C | 0.14 | 3.24E-02 | 2.23E-01 | 2.70E+00 | 2.73E+00 | 8.84E-01 | no | | intergenic_variant | MODIFIER |
| 2 | 114417967 | AX-97854972 | 0 | A | C | 0.14 | 3.24E-02 | 2.23E-01 | 2.70E+00 | 2.73E+00 | 8.84E-01 | no | | intergenic_variant | MODIFIER |
| 2 | 114425746 | AX-170386960 | 0 | T | C | 0.14 | 3.24E-02 | 2.23E-01 | 2.70E+00 | 2.73E+00 | 8.84E-01 | no | | intergenic_variant | MODIFIER |
| 2 | 114436562 | AX-97827626 | 0 | G | A | 0.14 | 3.24E-02 | 2.23E-01 | 2.70E+00 | 2.73E+00 | 8.84E-01 | no | | intergenic_variant | MODIFIER |
| 2 | 114436708 | AX-170451661 | 0 | C | G | 0.14 | 3.24E-02 | 2.23E-01 | 2.70E+00 | 2.73E+00 | 8.84E-01 | no | | intergenic_variant | MODIFIER |
| 2 | 114436946 | AX-170337377 | 0 | T | G | 0.14 | 3.24E-02 | 2.23E-01 | 2.70E+00 | 2.73E+00 | 8.84E-01 | no | | intergenic_variant | MODIFIER |
| 2 | 114475113 | AX-97838605 | 0 | A | G | 0.14 | 3.24E-02 | 2.23E-01 | 2.70E+00 | 2.73E+00 | 8.84E-01 | no | | intergenic_variant | MODIFIER |
| 2 | 114490170 | AX-170430293 | 0 | G | T | 0.14 | 3.24E-02 | 2.23E-01 | 2.70E+00 | 2.73E+00 | 8.84E-01 | no | | intergenic_variant | MODIFIER |

| | | | | | | | | | | | | | | | |
|---|---|---|---|---|---|---|---|---|---|---|---|---|---|---|---|
| 2 | 114490832 | AX-170171687 | 0 | C | G | 0.14 | 3.24E-02 | 2.23E-01 | 2.70E+00 | 2.73E+00 | 8.84E-01 | no | | intergenic_variant | MODIFIER |
| 2 | 114527612 | AX-170254854 | 0 | A | C | 0.14 | 3.24E-02 | 2.23E-01 | 2.70E+00 | 2.73E+00 | 8.84E-01 | no | Dph6 | intron_variant | MODIFIER |
| 2 | 114535278 | AX-97827014 | 0 | C | T | 0.14 | 3.24E-02 | 2.23E-01 | 2.70E+00 | 2.73E+00 | 8.84E-01 | no | Dph6 | intron_variant | MODIFIER |
| 2 | 114577939 | AX-170114915 | 0 | A | G | 0.14 | 3.24E-02 | 2.23E-01 | 2.70E+00 | 2.73E+00 | 8.84E-01 | no | Dph6 | intron_variant | MODIFIER |
| 2 | 114590808 | AX-170018743 | 0 | G | A | 0.14 | 3.24E-02 | 2.23E-01 | 2.70E+00 | 2.73E+00 | 8.84E-01 | no | Dph6 | intron_variant | MODIFIER |
| 2 | 114636589 | AX-170094379 | 0 | G | T | 0.14 | 3.24E-02 | 2.23E-01 | 2.70E+00 | 2.73E+00 | 8.84E-01 | no | Dph6 | intron_variant | MODIFIER |
| 2 | 114658084 | AX-170018736 | 0 | A | C | 0.14 | 3.24E-02 | 2.23E-01 | 2.70E+00 | 2.73E+00 | 8.84E-01 | no | Dph6 | upstream_gene_variant | MODIFIER |
| 2 | 114736029 | AX-170396465 | 0 | G | A | 0.14 | 3.24E-02 | 2.23E-01 | 2.70E+00 | 2.73E+00 | 8.84E-01 | no | | intergenic_variant | MODIFIER |
| 2 | 114772004 | AX-97833725 | 0 | T | C | 0.14 | 3.24E-02 | 2.23E-01 | 2.70E+00 | 2.73E+00 | 8.84E-01 | no | | intergenic_variant | MODIFIER |
| 2 | 114811420 | AX-97838632 | 0 | C | T | 0.14 | 3.24E-02 | 2.23E-01 | 2.70E+00 | 2.73E+00 | 8.84E-01 | no | | intergenic_variant | MODIFIER |
| 2 | 115100069 | AX-170088715 | 0 | C | A | 0.14 | 3.24E-02 | 2.23E-01 | 2.70E+00 | 2.73E+00 | 8.84E-01 | no | Gm28494 | intron_variant,non_coding_transcript_variant | MODIFIER |
| 2 | 115122925 | AX-97838668 | 0 | A | G | 0.14 | 3.24E-02 | 2.23E-01 | 2.70E+00 | 2.73E+00 | 8.84E-01 | no | Gm28494 | intron_variant,non_coding_transcript_variant | MODIFIER |
| 2 | 115124735 | AX-170182039 | 0 | A | T | 0.14 | 3.24E-02 | 2.23E-01 | 2.70E+00 | 2.73E+00 | 8.84E-01 | no | Gm28494 | intron_variant,non_coding_transcript_variant | MODIFIER |
| 2 | 115352172 | AX-97812060 | 0 | C | A | 0.14 | 3.24E-02 | 2.23E-01 | 2.70E+00 | 2.73E+00 | 8.84E-01 | no | | intergenic_variant | MODIFIER |
| 2 | 115527658 | AX-97804796 | 0 | T | C | 0.14 | 3.24E-02 | 2.23E-01 | 2.70E+00 | 2.73E+00 | 8.84E-01 | no | 3110099E03Rik | intron_variant,non_coding_transcript_variant | MODIFIER |
| 2 | 115538143 | AX-170362457 | 0 | T | C | 0.14 | 3.24E-02 | 2.23E-01 | 2.70E+00 | 2.73E+00 | 8.84E-01 | no | | regulatory_region_variant | MODIFIER |
| 2 | 115560409 | AX-170235200 | 0 | T | C | 0.14 | 3.24E-02 | 2.23E-01 | 2.70E+00 | 2.73E+00 | 8.84E-01 | no | Gm13977 | non_coding_transcript_exon_variant | MODIFIER |
| 2 | 115562322 | AX-97793176 | 0 | T | C | 0.14 | 3.24E-02 | 2.23E-01 | 2.70E+00 | 2.73E+00 | 8.84E-01 | no | Gm13977 | intron_variant,non_coding_transcript_variant | MODIFIER |
| 2 | 115565844 | AX-170488711 | 0 | G | A | 0.14 | 3.24E-02 | 2.23E-01 | 2.70E+00 | 2.73E+00 | 8.84E-01 | no | Gm13977 | intron_variant,non_coding_transcript_variant | MODIFIER |
| 2 | 115606903 | AX-97822534 | 0 | A | C | 0.14 | 3.24E-02 | 2.23E-01 | 2.70E+00 | 2.73E+00 | 8.84E-01 | no | BC052040 | intron_variant | MODIFIER |
| 2 | 115613027 | AX-170130545 | 0 | G | A | 0.14 | 3.24E-02 | 2.23E-01 | 2.70E+00 | 2.73E+00 | 8.84E-01 | no | BC052040 | intron_variant | MODIFIER |
| 2 | 115632581 | AX-97787542 | 0 | A | C | 0.14 | 3.24E-02 | 2.23E-01 | 2.70E+00 | 2.73E+00 | 8.84E-01 | no | BC052040 | intron_variant | MODIFIER |
| 2 | 115638444 | AX-170130776 | 0 | C | G | 0.14 | 3.24E-02 | 2.23E-01 | 2.70E+00 | 2.73E+00 | 8.84E-01 | no | BC052040 | intron_variant | MODIFIER |
| 2 | 115723017 | AX-170127159 | 0 | G | C | 0.14 | 3.24E-02 | 2.23E-01 | 2.70E+00 | 2.73E+00 | 8.84E-01 | no | BC052040 | intron_variant | MODIFIER |
| 2 | 115728707 | AX-170319288 | 0 | A | G | 0.14 | 3.24E-02 | 2.23E-01 | 2.70E+00 | 2.73E+00 | 8.84E-01 | no | BC052040 | intron_variant | MODIFIER |
| 2 | 115728806 | AX-170263309 | 0 | C | T | 0.14 | 3.24E-02 | 2.23E-01 | 2.70E+00 | 2.73E+00 | 8.84E-01 | no | BC052040 | intron_variant | MODIFIER |
| 2 | 115729084 | AX-97828005 | 0 | T | C | 0.14 | 3.24E-02 | 2.23E-01 | 2.70E+00 | 2.73E+00 | 8.84E-01 | no | BC052040 | intron_variant | MODIFIER |
| 2 | 115740796 | AX-170018754 | 0 | T | C | 0.14 | 3.24E-02 | 2.23E-01 | 2.70E+00 | 2.73E+00 | 8.84E-01 | no | BC052040 | intron_variant | MODIFIER |
| 2 | 115768347 | AX-97835971 | 0 | C | T | 0.14 | 3.24E-02 | 2.23E-01 | 2.70E+00 | 2.73E+00 | 8.84E-01 | no | BC052040 | intron_variant | MODIFIER |
| 2 | 115772538 | AX-170018759 | 0 | G | A | 0.14 | 3.24E-02 | 2.23E-01 | 2.70E+00 | 2.73E+00 | 8.84E-01 | no | BC052040 | intron_variant | MODIFIER |
| 2 | 115804011 | AX-170018755 | 0 | A | T | 0.14 | 3.24E-02 | 2.23E-01 | 2.70E+00 | 2.73E+00 | 8.84E-01 | no | | intergenic_variant | MODIFIER |
| 2 | 115818013 | AX-97833014 | 0 | T | C | 0.14 | 3.24E-02 | 2.23E-01 | 2.70E+00 | 2.73E+00 | 8.84E-01 | no | | regulatory_region_variant | MODIFIER |
| 2 | 115847423 | AX-97795404 | 0 | G | A | 0.14 | 3.24E-02 | 2.23E-01 | 2.70E+00 | 2.73E+00 | 8.84E-01 | no | | intergenic_variant | MODIFIER |
| 2 | 115867104 | AX-170226333 | 0 | C | A | 0.14 | 3.24E-02 | 2.23E-01 | 2.70E+00 | 2.73E+00 | 8.84E-01 | no | Meis2 | intron_variant | MODIFIER |
| 2 | 115873719 | AX-170018756 | 0 | G | T | 0.14 | 3.24E-02 | 2.23E-01 | 2.70E+00 | 2.73E+00 | 8.84E-01 | no | Meis2 | intron_variant | MODIFIER |
| 2 | 115890915 | AX-170245982 | 0 | C | T | 0.14 | 3.24E-02 | 2.23E-01 | 2.70E+00 | 2.73E+00 | 8.84E-01 | no | Meis2 | intron_variant | MODIFIER |
| 2 | 115891051 | AX-170188537 | 0 | A | G | 0.14 | 3.24E-02 | 2.23E-01 | 2.70E+00 | 2.73E+00 | 8.84E-01 | no | Meis2 | intron_variant | MODIFIER |
| 2 | 115891123 | AX-170302519 | 0 | T | C | 0.14 | 3.24E-02 | 2.23E-01 | 2.70E+00 | 2.73E+00 | 8.84E-01 | no | Meis2 | intron_variant | MODIFIER |
| 2 | 115900351 | AX-97826759 | 0 | T | C | 0.14 | 3.24E-02 | 2.23E-01 | 2.70E+00 | 2.73E+00 | 8.84E-01 | no | Meis2 | intron_variant | MODIFIER |
| 2 | 115901822 | AX-170064661 | 0 | T | C | 0.14 | 3.24E-02 | 2.23E-01 | 2.70E+00 | 2.73E+00 | 8.84E-01 | no | Meis2 | intron_variant | MODIFIER |
| 2 | 116011321 | AX-97794984 | 0 | T | A | 0.14 | 3.24E-02 | 2.23E-01 | 2.70E+00 | 2.73E+00 | 8.84E-01 | no | Meis2 | intron_variant | MODIFIER |
| 2 | 116108206 | AX-97838733 | 0 | G | T | 0.14 | 3.24E-02 | 2.23E-01 | 2.70E+00 | 2.73E+00 | 8.84E-01 | no | | intergenic_variant | MODIFIER |
| 2 | 116122030 | AX-97850089 | 0 | A | G | 0.14 | 3.24E-02 | 2.23E-01 | 2.70E+00 | 2.73E+00 | 8.84E-01 | no | | intergenic_variant | MODIFIER |
| 2 | 116232894 | AX-97853686 | 0 | C | T | 0.14 | 3.24E-02 | 2.23E-01 | 2.70E+00 | 2.73E+00 | 8.84E-01 | no | | intergenic_variant | MODIFIER |
| 2 | 116341597 | AX-170514388 | 0 | A | G | 0.14 | 3.24E-02 | 2.23E-01 | 2.70E+00 | 2.73E+00 | 8.84E-01 | no | | intergenic_variant | MODIFIER |
| 2 | 116562911 | AX-170064672 | 0 | G | A | 0.14 | 3.24E-02 | 2.23E-01 | 2.70E+00 | 2.73E+00 | 8.84E-01 | no | | intergenic_variant | MODIFIER |
| 2 | 116564862 | AX-97832315 | 0 | A | G | 0.14 | 3.24E-02 | 2.23E-01 | 2.70E+00 | 2.73E+00 | 8.84E-01 | no | | intergenic_variant | MODIFIER |
| 2 | 116567491 | AX-170305512 | 0 | T | C | 0.14 | 3.24E-02 | 2.23E-01 | 2.70E+00 | 2.73E+00 | 8.84E-01 | no | | intergenic_variant | MODIFIER |
| 2 | 116567992 | AX-170154292 | 0 | G | A | 0.14 | 3.24E-02 | 2.23E-01 | 2.70E+00 | 2.73E+00 | 8.84E-01 | no | | intergenic_variant | MODIFIER |
| 2 | 116570547 | AX-168419967 | 0 | C | A | 0.14 | 3.24E-02 | 2.23E-01 | 2.70E+00 | 2.73E+00 | 8.84E-01 | no | | intergenic_variant | MODIFIER |
| 2 | 116606522 | AX-168377052 | 0 | G | A | 0.14 | 3.24E-02 | 2.23E-01 | 2.70E+00 | 2.73E+00 | 8.84E-01 | no | | intergenic_variant | MODIFIER |
| 2 | 116607365 | AX-97815629 | 0 | A | C | 0.14 | 3.24E-02 | 2.23E-01 | 2.70E+00 | 2.73E+00 | 8.84E-01 | no | | intergenic_variant | MODIFIER |
| 2 | 118874478 | AX-170113857 | 0 | C | G | 0.14 | 3.24E-02 | 2.23E-01 | 2.70E+00 | 2.73E+00 | 8.84E-01 | no | Ivd | intron_variant | MODIFIER |
| 2 | 118875945 | AX-97826733 | 0 | G | A | 0.14 | 3.24E-02 | 2.23E-01 | 2.70E+00 | 2.73E+00 | 8.84E-01 | no | Ivd | intron_variant | MODIFIER |

| | | | | | | | | | | | | | | | |
|---|---|---|---|---|---|---|---|---|---|---|---|---|---|---|---|
| 2 | 118891691 | AX-170496070 | 0 | T | C | 0.14 | 3.24E-02 | 2.23E-01 | 2.70E+00 | 2.73E+00 | 8.84E-01 | no | | intergenic_variant | MODIFIER |
| 2 | 118892840 | AX-170064707 | 0 | C | T | 0.14 | 3.24E-02 | 2.23E-01 | 2.70E+00 | 2.73E+00 | 8.84E-01 | no | | intergenic_variant | MODIFIER |
| 2 | 118898297 | AX-170064709 | 0 | G | T | 0.14 | 3.24E-02 | 2.23E-01 | 2.70E+00 | 2.73E+00 | 8.84E-01 | no | Bahd1 | upstream_gene_variant | MODIFIER |
| 2 | 118931125 | AX-170018820 | 0 | G | A | 0.14 | 3.24E-02 | 2.23E-01 | 2.70E+00 | 2.73E+00 | 8.84E-01 | no | Chst14 | downstream_gene_variant | MODIFIER |
| 2 | 118966485 | AX-170018826 | 0 | C | T | 0.14 | 3.24E-02 | 2.23E-01 | 2.70E+00 | 2.73E+00 | 8.84E-01 | no | | intergenic_variant | MODIFIER |
| 2 | 118967167 | AX-97826730 | 0 | G | T | 0.14 | 3.24E-02 | 2.23E-01 | 2.70E+00 | 2.73E+00 | 8.84E-01 | no | | intergenic_variant | MODIFIER |
| 2 | 118970642 | AX-170018823 | 0 | T | C | 0.14 | 3.24E-02 | 2.23E-01 | 2.70E+00 | 2.73E+00 | 8.84E-01 | no | | intergenic_variant | MODIFIER |
| 2 | 118979390 | AX-170277386 | 0 | T | C | 0.14 | 3.24E-02 | 2.23E-01 | 2.70E+00 | 2.73E+00 | 8.84E-01 | no | | intergenic_variant | MODIFIER |
| 2 | 118988136 | AX-170018819 | 0 | A | G | 0.14 | 3.24E-02 | 2.23E-01 | 2.70E+00 | 2.73E+00 | 8.84E-01 | no | | intergenic_variant | MODIFIER |
| 2 | 118988363 | AX-170518831 | 0 | C | T | 0.14 | 3.24E-02 | 2.23E-01 | 2.70E+00 | 2.73E+00 | 8.84E-01 | no | | intergenic_variant | MODIFIER |
| 2 | 119007007 | AX-170409770 | 0 | G | A | 0.14 | 3.24E-02 | 2.23E-01 | 2.70E+00 | 2.73E+00 | 8.84E-01 | no | Uqcrh-ps2 | upstream_gene_variant | MODIFIER |
| 2 | 119017385 | AX-97812721 | 0 | T | C | 0.14 | 3.24E-02 | 2.23E-01 | 2.70E+00 | 2.73E+00 | 8.84E-01 | no | Ccdc32 | downstream_gene_variant | MODIFIER |
| 2 | 119046940 | AX-170265837 | 0 | T | C | 0.14 | 3.24E-02 | 2.23E-01 | 2.70E+00 | 2.73E+00 | 8.84E-01 | no | Knl1 | upstream_gene_variant | MODIFIER |
| 2 | 119047020 | AX-170329232 | 0 | A | G | 0.14 | 3.24E-02 | 2.23E-01 | 2.70E+00 | 2.73E+00 | 8.84E-01 | no | Knl1 | upstream_gene_variant | MODIFIER |
| 2 | 119047608 | AX-170328847 | 0 | T | C | 0.14 | 3.24E-02 | 2.23E-01 | 2.70E+00 | 2.73E+00 | 8.84E-01 | no | Knl1 | intron_variant | MODIFIER |
| 2 | 119109172 | AX-170260042 | 0 | T | C | 0.14 | 3.24E-02 | 2.23E-01 | 2.70E+00 | 2.73E+00 | 8.84E-01 | no | Rad51 | upstream_gene_variant | MODIFIER |
| 2 | 119112528 | AX-170233237 | 0 | G | C | 0.14 | 3.24E-02 | 2.23E-01 | 2.70E+00 | 2.73E+00 | 8.84E-01 | no | Rad51 | upstream_gene_variant | MODIFIER |
| 2 | 119113341 | AX-170064713 | 0 | T | G | 0.14 | 3.24E-02 | 2.23E-01 | 2.70E+00 | 2.73E+00 | 8.84E-01 | no | Rad51 | intron_variant | MODIFIER |
| 2 | 119174164 | AX-97837583 | 0 | A | C | 0.14 | 3.24E-02 | 2.23E-01 | 2.70E+00 | 2.73E+00 | 8.84E-01 | no | Dnajc17 | intron_variant | MODIFIER |
| 2 | 119200084 | AX-170099096 | 0 | C | T | 0.14 | 3.24E-02 | 2.23E-01 | 2.70E+00 | 2.73E+00 | 8.84E-01 | no | Dnajc17 | intron_variant | MODIFIER |
| 2 | 119203290 | AX-170446120 | 0 | T | C | 0.14 | 3.24E-02 | 2.23E-01 | 2.70E+00 | 2.73E+00 | 8.84E-01 | no | Dnajc17 | intron_variant | MODIFIER |
| 2 | 119239456 | AX-170251551 | 0 | A | G | 0.14 | 3.24E-02 | 2.23E-01 | 2.70E+00 | 2.73E+00 | 8.84E-01 | no | Spint1 | intron_variant | MODIFIER |
| 2 | 119242044 | AX-170196605 | 0 | T | C | 0.14 | 3.24E-02 | 2.23E-01 | 2.70E+00 | 2.73E+00 | 8.84E-01 | no | Spint1 | intron_variant | MODIFIER |
| 2 | 119242268 | AX-170188084 | 0 | G | C | 0.14 | 3.24E-02 | 2.23E-01 | 2.70E+00 | 2.73E+00 | 8.84E-01 | no | Spint1 | intron_variant | MODIFIER |
| 2 | 119264445 | AX-97839741 | 0 | A | G | 0.14 | 3.24E-02 | 2.23E-01 | 2.70E+00 | 2.73E+00 | 8.84E-01 | no | Rhov | downstream_gene_variant | MODIFIER |
| 2 | 119288487 | AX-97838754 | 0 | C | T | 0.14 | 3.24E-02 | 2.23E-01 | 2.70E+00 | 2.73E+00 | 8.84E-01 | no | Vps18 | upstream_gene_variant | MODIFIER |
| 2 | 119291128 | AX-170281444 | 0 | C | A | 0.14 | 3.24E-02 | 2.23E-01 | 2.70E+00 | 2.73E+00 | 8.84E-01 | no | Vps18 | intron_variant | MODIFIER |
| 2 | 119305636 | AX-170286469 | 0 | G | A | 0.14 | 3.24E-02 | 2.23E-01 | 2.70E+00 | 2.73E+00 | 8.84E-01 | no | | regulatory_region_variant | MODIFIER |
| 2 | 119309729 | AX-170229285 | 0 | C | T | 0.14 | 3.24E-02 | 2.23E-01 | 2.70E+00 | 2.73E+00 | 8.84E-01 | no | | regulatory_region_variant | MODIFIER |
| 2 | 119330593 | AX-170471002 | 0 | G | C | 0.14 | 3.24E-02 | 2.23E-01 | 2.70E+00 | 2.73E+00 | 8.84E-01 | no | Dll4 | intron_variant | MODIFIER |
| 2 | 119347880 | AX-97806505 | 0 | G | T | 0.14 | 3.24E-02 | 2.23E-01 | 2.70E+00 | 2.73E+00 | 8.84E-01 | no | Chac1 | upstream_gene_variant | MODIFIER |
| 2 | 119375808 | AX-170064714 | 0 | C | T | 0.14 | 3.24E-02 | 2.23E-01 | 2.70E+00 | 2.73E+00 | 8.84E-01 | no | Ino80 | intron_variant | MODIFIER |
| 2 | 119377039 | AX-170064718 | 0 | G | A | 0.14 | 3.24E-02 | 2.23E-01 | 2.70E+00 | 2.73E+00 | 8.84E-01 | no | Ino80 | intron_variant | MODIFIER |
| 2 | 119383323 | AX-170283287 | 0 | T | C | 0.14 | 3.24E-02 | 2.23E-01 | 2.70E+00 | 2.73E+00 | 8.84E-01 | no | Ino80 | intron_variant | MODIFIER |
| 2 | 119399346 | AX-170286310 | 0 | C | G | 0.14 | 3.24E-02 | 2.23E-01 | 2.70E+00 | 2.73E+00 | 8.84E-01 | no | Ino80 | intron_variant | MODIFIER |
| 2 | 119400995 | AX-170221908 | 0 | G | T | 0.14 | 3.24E-02 | 2.23E-01 | 2.70E+00 | 2.73E+00 | 8.84E-01 | no | Ino80 | intron_variant | MODIFIER |
| 2 | 119410968 | AX-170374531 | 0 | G | C | 0.14 | 3.24E-02 | 2.23E-01 | 2.70E+00 | 2.73E+00 | 8.84E-01 | no | Ino80 | intron_variant | MODIFIER |
| 2 | 119447456 | AX-170209083 | 0 | T | C | 0.14 | 3.24E-02 | 2.23E-01 | 2.70E+00 | 2.73E+00 | 8.84E-01 | no | Ino80 | intron_variant | MODIFIER |
| 2 | 119448739 | AX-170479558 | 0 | G | C | 0.14 | 3.24E-02 | 2.23E-01 | 2.70E+00 | 2.73E+00 | 8.84E-01 | no | Ino80 | intron_variant | MODIFIER |
| 2 | 119448961 | AX-97786966 | 0 | G | C | 0.14 | 3.24E-02 | 2.23E-01 | 2.70E+00 | 2.73E+00 | 8.84E-01 | no | Ino80 | intron_variant | MODIFIER |
| 2 | 119493070 | AX-170018834 | 0 | A | G | 0.14 | 3.24E-02 | 2.23E-01 | 2.70E+00 | 2.73E+00 | 8.84E-01 | no | | intergenic_variant | MODIFIER |
| 2 | 119516015 | AX-170235157 | 0 | G | A | 0.14 | 3.24E-02 | 2.23E-01 | 2.70E+00 | 2.73E+00 | 8.84E-01 | no | Exd1 | downstream_gene_variant | MODIFIER |
| 2 | 119518456 | AX-170473123 | 0 | A | C | 0.14 | 3.24E-02 | 2.23E-01 | 2.70E+00 | 2.73E+00 | 8.84E-01 | no | Exd1 | 3_prime_UTR_variant | MODIFIER |
| 2 | 119524263 | AX-97856374 | 0 | A | G | 0.14 | 3.24E-02 | 2.23E-01 | 2.70E+00 | 2.73E+00 | 8.84E-01 | no | Exd1 | intron_variant | MODIFIER |
| 2 | 119546195 | AX-170064719 | 0 | C | A | 0.14 | 3.24E-02 | 2.23E-01 | 2.70E+00 | 2.73E+00 | 8.84E-01 | no | Chp1 | upstream_gene_variant | MODIFIER |
| 2 | 119570396 | AX-97856245 | 0 | G | A | 0.14 | 3.24E-02 | 2.23E-01 | 2.70E+00 | 2.73E+00 | 8.84E-01 | no | Chp1 | intron_variant | MODIFIER |
| 2 | 119599650 | AX-97828494 | 0 | C | T | 0.14 | 3.24E-02 | 2.23E-01 | 2.70E+00 | 2.73E+00 | 8.84E-01 | no | 1700020I14Rik | non_coding_transcript_exon_variant | MODIFIER |
| 2 | 119600825 | AX-170064721 | 0 | A | C | 0.14 | 3.24E-02 | 2.23E-01 | 2.70E+00 | 2.73E+00 | 8.84E-01 | no | 1700020I14Rik | downstream_gene_variant | MODIFIER |
| 2 | 119621787 | AX-170064720 | 0 | A | G | 0.14 | 3.24E-02 | 2.23E-01 | 2.70E+00 | 2.73E+00 | 8.84E-01 | no | Nusap1 | intron_variant | MODIFIER |
| 2 | 119627702 | AX-170147601 | 0 | A | G | 0.14 | 3.24E-02 | 2.23E-01 | 2.70E+00 | 2.73E+00 | 8.84E-01 | no | Nusap1 | intron_variant | MODIFIER |
| 2 | 119641656 | AX-170441172 | 0 | G | A | 0.14 | 3.24E-02 | 2.23E-01 | 2.70E+00 | 2.73E+00 | 8.84E-01 | no | Nusap1 | intron_variant | MODIFIER |
| 2 | 119727974 | AX-170018829 | 0 | T | G | 0.14 | 3.24E-02 | 2.23E-01 | 2.70E+00 | 2.73E+00 | 8.84E-01 | no | Rtf1 | intron_variant | MODIFIER |
| 2 | 119737097 | AX-170408621 | 0 | A | G | 0.14 | 3.24E-02 | 2.23E-01 | 2.70E+00 | 2.73E+00 | 8.84E-01 | no | Rtf1 | downstream_gene_variant | MODIFIER |
| 2 | 119766175 | AX-170431413 | 0 | C | G | 0.14 | 3.24E-02 | 2.23E-01 | 2.70E+00 | 2.73E+00 | 8.84E-01 | no | Rpap1 | intron_variant | MODIFIER |

| | | | | | | | | | | | | | | | |
|---|---|---|---|---|---|---|---|---|---|---|---|---|---|---|---|
| 2 | 119776810 | AX-170181358 | 0 | C | T | 0.14 | 3.24E-02 | 2.23E-01 | 2.70E+00 | 2.73E+00 | 8.84E-01 | no | Rpap1 | missense_variant | MODERATE |
| 2 | 119777580 | AX-97809156 | 0 | A | G | 0.14 | 3.24E-02 | 2.23E-01 | 2.70E+00 | 2.73E+00 | 8.84E-01 | no | Rpap1 | intron_variant | MODIFIER |
| 2 | 119806611 | AX-170400884 | 0 | T | C | 0.14 | 3.24E-02 | 2.23E-01 | 2.70E+00 | 2.73E+00 | 8.84E-01 | no | Tyro3 | intron_variant | MODIFIER |
| 2 | 119811883 | AX-170103777 | 0 | T | A | 0.14 | 3.24E-02 | 2.23E-01 | 2.70E+00 | 2.73E+00 | 8.84E-01 | no | Tyro3 | intron_variant | MODIFIER |
| 2 | 119812556 | AX-170344261 | 0 | G | C | 0.14 | 3.24E-02 | 2.23E-01 | 2.70E+00 | 2.73E+00 | 8.84E-01 | no | Tyro3 | synonymous_variant | LOW |
| 2 | 119818926 | AX-170298167 | 0 | C | T | 0.14 | 3.24E-02 | 2.23E-01 | 2.70E+00 | 2.73E+00 | 8.84E-01 | no | Tyro3 | downstream_gene_variant | MODIFIER |
| 2 | 119857816 | AX-170064724 | 0 | T | C | 0.14 | 3.24E-02 | 2.23E-01 | 2.70E+00 | 2.73E+00 | 8.84E-01 | no | | intergenic_variant | MODIFIER |
| 2 | 119857948 | AX-170420453 | 0 | C | G | 0.14 | 3.24E-02 | 2.23E-01 | 2.70E+00 | 2.73E+00 | 8.84E-01 | no | | intergenic_variant | MODIFIER |
| 2 | 119858038 | AX-170064726 | 0 | G | A | 0.14 | 3.24E-02 | 2.23E-01 | 2.70E+00 | 2.73E+00 | 8.84E-01 | no | | intergenic_variant | MODIFIER |
| 2 | 119858604 | AX-170355116 | 0 | A | C | 0.14 | 3.24E-02 | 2.23E-01 | 2.70E+00 | 2.73E+00 | 8.84E-01 | no | | regulatory_region_variant | MODIFIER |
| 2 | 119869634 | AX-170141432 | 0 | C | T | 0.14 | 3.24E-02 | 2.23E-01 | 2.70E+00 | 2.73E+00 | 8.84E-01 | no | | intergenic_variant | MODIFIER |
| 2 | 119944814 | AX-170460695 | 0 | G | C | 0.14 | 3.24E-02 | 2.23E-01 | 2.70E+00 | 2.73E+00 | 8.84E-01 | no | Mga | intron_variant | MODIFIER |
| 2 | 119972205 | AX-170123377 | 0 | G | A | 0.14 | 3.24E-02 | 2.23E-01 | 2.70E+00 | 2.73E+00 | 8.84E-01 | no | Mga | downstream_gene_variant | MODIFIER |
| 2 | 119973413 | AX-97832603 | 0 | A | G | 0.14 | 3.24E-02 | 2.23E-01 | 2.70E+00 | 2.73E+00 | 8.84E-01 | no | Mapkbp1 | intron_variant | MODIFIER |
| 2 | 119975164 | AX-170296096 | 0 | C | T | 0.14 | 3.24E-02 | 2.23E-01 | 2.70E+00 | 2.73E+00 | 8.84E-01 | no | Mapkbp1 | intron_variant | MODIFIER |
| 2 | 120007476 | AX-170180243 | 0 | A | T | 0.14 | 3.24E-02 | 2.23E-01 | 2.70E+00 | 2.73E+00 | 8.84E-01 | no | Mapkbp1 | intron_variant | MODIFIER |
| 2 | 120014138 | AX-170398269 | 0 | T | C | 0.14 | 3.24E-02 | 2.23E-01 | 2.70E+00 | 2.73E+00 | 8.84E-01 | no | Mapkbp1 | intron_variant | MODIFIER |
| 2 | 120026396 | AX-170390933 | 0 | A | G | 0.14 | 3.24E-02 | 2.23E-01 | 2.70E+00 | 2.73E+00 | 8.84E-01 | no | Jmjd7 | upstream_gene_variant | MODIFIER |
| 2 | 120032981 | AX-170284179 | 0 | G | A | 0.14 | 3.24E-02 | 2.23E-01 | 2.70E+00 | 2.73E+00 | 8.84E-01 | no | Jmjd7 | downstream_gene_variant | MODIFIER |
| 2 | 120047283 | AX-170064730 | 0 | C | T | 0.14 | 3.24E-02 | 2.23E-01 | 2.70E+00 | 2.73E+00 | 8.84E-01 | no | Gm28042 | downstream_gene_variant | MODIFIER |
| 2 | 120047807 | AX-170018843 | 0 | T | C | 0.14 | 3.24E-02 | 2.23E-01 | 2.70E+00 | 2.73E+00 | 8.84E-01 | no | Gm28042 | downstream_gene_variant | MODIFIER |
| 2 | 120062544 | AX-170284419 | 0 | C | T | 0.14 | 3.24E-02 | 2.23E-01 | 2.70E+00 | 2.73E+00 | 8.84E-01 | no | Sptbn5 | non_coding_transcript_exon_variant | MODIFIER |
| 2 | 120066508 | AX-170447580 | 0 | G | A | 0.14 | 3.24E-02 | 2.23E-01 | 2.70E+00 | 2.73E+00 | 8.84E-01 | no | Sptbn5 | non_coding_transcript_exon_variant | MODIFIER |
| 2 | 120066567 | AX-170520276 | 0 | C | T | 0.14 | 3.24E-02 | 2.23E-01 | 2.70E+00 | 2.73E+00 | 8.84E-01 | no | Sptbn5 | splice_acceptor_variant,non_coding_transcript_variant | HIGH |
| 2 | 120069541 | AX-170018841 | 0 | G | A | 0.14 | 3.24E-02 | 2.23E-01 | 2.70E+00 | 2.73E+00 | 8.84E-01 | no | Sptbn5 | downstream_gene_variant | MODIFIER |
| 2 | 120072979 | AX-170238218 | 0 | A | G | 0.14 | 3.24E-02 | 2.23E-01 | 2.70E+00 | 2.73E+00 | 8.84E-01 | no | Sptbn5 | downstream_gene_variant | MODIFIER |
| 2 | 120074533 | AX-97794950 | 0 | G | A | 0.14 | 3.24E-02 | 2.23E-01 | 2.70E+00 | 2.73E+00 | 8.84E-01 | no | Sptbn5 | intron_variant,non_coding_transcript_variant | MODIFIER |
| 2 | 120074606 | AX-170018842 | 0 | C | T | 0.14 | 3.24E-02 | 2.23E-01 | 2.70E+00 | 2.73E+00 | 8.84E-01 | no | Sptbn5 | intron_variant,non_coding_transcript_variant | MODIFIER |
| 2 | 120092565 | AX-170100297 | 0 | G | A | 0.14 | 3.24E-02 | 2.23E-01 | 2.70E+00 | 2.73E+00 | 8.84E-01 | no | Ehd4 | intron_variant | MODIFIER |
| 2 | 120102769 | AX-170372651 | 0 | A | G | 0.14 | 3.24E-02 | 2.23E-01 | 2.70E+00 | 2.73E+00 | 8.84E-01 | no | Ehd4 | intron_variant | MODIFIER |
| 2 | 120103107 | AX-170191326 | 0 | A | G | 0.14 | 3.24E-02 | 2.23E-01 | 2.70E+00 | 2.73E+00 | 8.84E-01 | no | Ehd4 | intron_variant | MODIFIER |
| 2 | 120106983 | AX-170462303 | 0 | C | T | 0.14 | 3.24E-02 | 2.23E-01 | 2.70E+00 | 2.73E+00 | 8.84E-01 | no | Ehd4 | intron_variant | MODIFIER |
| 2 | 120117636 | AX-170437997 | 0 | G | A | 0.14 | 3.24E-02 | 2.23E-01 | 2.70E+00 | 2.73E+00 | 8.84E-01 | no | Ehd4 | intron_variant | MODIFIER |
| 2 | 120120492 | AX-170154196 | 0 | A | G | 0.14 | 3.24E-02 | 2.23E-01 | 2.70E+00 | 2.73E+00 | 8.84E-01 | no | Ehd4 | intron_variant | MODIFIER |
| 2 | 120133468 | AX-170179813 | 0 | A | G | 0.14 | 3.24E-02 | 2.23E-01 | 2.70E+00 | 2.73E+00 | 8.84E-01 | no | Ehd4 | intron_variant | MODIFIER |
| 2 | 120133815 | AX-97848743 | 0 | G | A | 0.14 | 3.24E-02 | 2.23E-01 | 2.70E+00 | 2.73E+00 | 8.84E-01 | no | Ehd4 | intron_variant | MODIFIER |
| 2 | 120136884 | AX-170259539 | 0 | T | C | 0.14 | 3.24E-02 | 2.23E-01 | 2.70E+00 | 2.73E+00 | 8.84E-01 | no | Ehd4 | intron_variant | MODIFIER |
| 2 | 120170358 | AX-170293743 | 0 | C | T | 0.14 | 3.24E-02 | 2.23E-01 | 2.70E+00 | 2.73E+00 | 8.84E-01 | no | Pla2g4e | intron_variant | MODIFIER |
| 2 | 120183147 | AX-170512878 | 0 | T | C | 0.14 | 3.24E-02 | 2.23E-01 | 2.70E+00 | 2.73E+00 | 8.84E-01 | no | Pla2g4e | intron_variant | MODIFIER |
| 2 | 120190125 | AX-170064733 | 0 | T | C | 0.14 | 3.24E-02 | 2.23E-01 | 2.70E+00 | 2.73E+00 | 8.84E-01 | no | Pla2g4e | intron_variant | MODIFIER |
| 2 | 120214819 | AX-170064735 | 0 | T | C | 0.14 | 3.24E-02 | 2.23E-01 | 2.70E+00 | 2.73E+00 | 8.84E-01 | no | Pla2g4e | intron_variant | MODIFIER |
| 2 | 120231224 | AX-170361228 | 0 | C | T | 0.14 | 3.24E-02 | 2.23E-01 | 2.70E+00 | 2.73E+00 | 8.84E-01 | no | Pla2g4e | intron_variant | MODIFIER |
| 2 | 120239702 | AX-170251610 | 0 | G | A | 0.14 | 3.24E-02 | 2.23E-01 | 2.70E+00 | 2.73E+00 | 8.84E-01 | no | Pla2g4e | intron_variant | MODIFIER |
| 2 | 120249517 | AX-97790808 | 0 | C | T | 0.14 | 3.24E-02 | 2.23E-01 | 2.70E+00 | 2.73E+00 | 8.84E-01 | no | Pla2g4e | upstream_gene_variant | MODIFIER |
| 2 | 120262757 | AX-170172279 | 0 | T | A | 0.14 | 3.24E-02 | 2.23E-01 | 2.70E+00 | 2.73E+00 | 8.84E-01 | no | Pla2g4d | downstream_gene_variant | MODIFIER |
| 2 | 120279164 | AX-170405999 | 0 | C | T | 0.14 | 3.24E-02 | 2.23E-01 | 2.70E+00 | 2.73E+00 | 8.84E-01 | no | Pla2g4d | intron_variant | MODIFIER |
| 2 | 120288114 | AX-97832605 | 0 | T | C | 0.14 | 3.24E-02 | 2.23E-01 | 2.70E+00 | 2.73E+00 | 8.84E-01 | no | Pla2g4d | intron_variant | MODIFIER |
| 2 | 120291615 | AX-170328378 | 0 | G | A | 0.14 | 3.24E-02 | 2.23E-01 | 2.70E+00 | 2.73E+00 | 8.84E-01 | no | Pla2g4d | upstream_gene_variant | MODIFIER |
| 2 | 120317779 | AX-170294959 | 0 | C | T | 0.14 | 3.24E-02 | 2.23E-01 | 2.70E+00 | 2.73E+00 | 8.84E-01 | no | Vps39 | 3_prime_UTR_variant | MODIFIER |
| 2 | 120324932 | AX-170227240 | 0 | A | G | 0.14 | 3.24E-02 | 2.23E-01 | 2.70E+00 | 2.73E+00 | 8.84E-01 | no | Vps39 | intron_variant | MODIFIER |
| 2 | 120325350 | AX-170213692 | 0 | C | T | 0.14 | 3.24E-02 | 2.23E-01 | 2.70E+00 | 2.73E+00 | 8.84E-01 | no | Vps39 | intron_variant | MODIFIER |
| 2 | 120337682 | AX-97833138 | 0 | G | A | 0.14 | 3.24E-02 | 2.23E-01 | 2.70E+00 | 2.73E+00 | 8.84E-01 | no | Vps39 | intron_variant | MODIFIER |
| 2 | 120338618 | AX-170471972 | 0 | T | A | 0.14 | 3.24E-02 | 2.23E-01 | 2.70E+00 | 2.73E+00 | 8.84E-01 | no | Vps39 | intron_variant | MODIFIER |
| 2 | 120344351 | AX-170305559 | 0 | C | T | 0.14 | 3.24E-02 | 2.23E-01 | 2.70E+00 | 2.73E+00 | 8.84E-01 | no | Vps39 | intron_variant | MODIFIER |

| | | | | | | | | | | | | | | | |
|---|---|---|---|---|---|---|---|---|---|---|---|---|---|---|---|
| 2 | 120365866 | AX-170529677 | 0 | A | G | 0.14 | 3.24E-02 | 2.23E-01 | 2.70E+00 | 2.73E+00 | 8.84E-01 | no | Tmem87a | intron_variant | MODIFIER |
| 2 | 120377253 | AX-170379345 | 0 | A | G | 0.14 | 3.24E-02 | 2.23E-01 | 2.70E+00 | 2.73E+00 | 8.84E-01 | no | Tmem87a | intron_variant | MODIFIER |
| 2 | 120380019 | AX-97804488 | 0 | T | C | 0.14 | 3.24E-02 | 2.23E-01 | 2.70E+00 | 2.73E+00 | 8.84E-01 | no | Tmem87a | intron_variant | MODIFIER |
| 2 | 120454338 | AX-170064737 | 0 | T | C | 0.14 | 3.24E-02 | 2.23E-01 | 2.70E+00 | 2.73E+00 | 8.84E-01 | no | Ganc | intron_variant,non_coding_transcript_variant | MODIFIER |
| 2 | 120455032 | AX-170208009 | 0 | T | A | 0.14 | 3.24E-02 | 2.23E-01 | 2.70E+00 | 2.73E+00 | 8.84E-01 | no | Ganc | intron_variant,non_coding_transcript_variant | MODIFIER |
| 2 | 120460067 | AX-97801012 | 0 | G | A | 0.14 | 3.24E-02 | 2.23E-01 | 2.70E+00 | 2.73E+00 | 8.84E-01 | no | Capn3 | upstream_gene_variant | MODIFIER |
| 2 | 120465322 | AX-170278035 | 0 | T | C | 0.14 | 3.24E-02 | 2.23E-01 | 2.70E+00 | 2.73E+00 | 8.84E-01 | no | Capn3 | intron_variant | MODIFIER |
| 2 | 120467686 | AX-170516534 | 0 | T | C | 0.14 | 3.24E-02 | 2.23E-01 | 2.70E+00 | 2.73E+00 | 8.84E-01 | no | Capn3 | intron_variant | MODIFIER |
| 2 | 120481638 | AX-170018849 | 0 | C | T | 0.14 | 3.24E-02 | 2.23E-01 | 2.70E+00 | 2.73E+00 | 8.84E-01 | no | Capn3 | intron_variant | MODIFIER |
| 2 | 120496644 | AX-170161069 | 0 | T | C | 0.14 | 3.24E-02 | 2.23E-01 | 2.70E+00 | 2.73E+00 | 8.84E-01 | no | Capn3 | intron_variant | MODIFIER |
| 2 | 120500751 | AX-170291871 | 0 | T | A | 0.14 | 3.24E-02 | 2.23E-01 | 2.70E+00 | 2.73E+00 | 8.84E-01 | no | Capn3 | intron_variant | MODIFIER |
| 2 | 120501154 | AX-170360619 | 0 | A | G | 0.14 | 3.24E-02 | 2.23E-01 | 2.70E+00 | 2.73E+00 | 8.84E-01 | no | Capn3 | intron_variant | MODIFIER |
| 2 | 120512941 | AX-170150661 | 0 | A | G | 0.14 | 3.24E-02 | 2.23E-01 | 2.70E+00 | 2.73E+00 | 8.84E-01 | no | Zfp106 | intron_variant | MODIFIER |
| 2 | 120525952 | AX-97795483 | 0 | A | G | 0.14 | 3.24E-02 | 2.23E-01 | 2.70E+00 | 2.73E+00 | 8.84E-01 | no | Zfp106 | intron_variant | MODIFIER |
| 2 | 120581674 | AX-170018850 | 0 | G | A | 0.14 | 3.24E-02 | 2.23E-01 | 2.70E+00 | 2.73E+00 | 8.84E-01 | no | Snap23 | intron_variant | MODIFIER |
| 2 | 120581822 | AX-170018848 | 0 | G | C | 0.14 | 3.24E-02 | 2.23E-01 | 2.70E+00 | 2.73E+00 | 8.84E-01 | no | Snap23 | intron_variant | MODIFIER |
| 2 | 120584231 | AX-170159141 | 0 | C | A | 0.14 | 3.24E-02 | 2.23E-01 | 2.70E+00 | 2.73E+00 | 8.84E-01 | no | Snap23 | intron_variant | MODIFIER |
| 2 | 120596813 | AX-170233352 | 0 | C | A | 0.14 | 3.24E-02 | 2.23E-01 | 2.70E+00 | 2.73E+00 | 8.84E-01 | no | Snap23 | intron_variant | MODIFIER |
| 2 | 120606234 | AX-97818002 | 0 | A | G | 0.14 | 3.24E-02 | 2.23E-01 | 2.70E+00 | 2.73E+00 | 8.84E-01 | no | Haus2 | upstream_gene_variant | MODIFIER |
| 2 | 120606835 | AX-170018853 | 0 | A | G | 0.14 | 3.24E-02 | 2.23E-01 | 2.70E+00 | 2.73E+00 | 8.84E-01 | no | Haus2 | upstream_gene_variant | MODIFIER |
| 2 | 120606878 | AX-170018845 | 0 | T | A | 0.14 | 3.24E-02 | 2.23E-01 | 2.70E+00 | 2.73E+00 | 8.84E-01 | no | Haus2 | upstream_gene_variant | MODIFIER |
| 2 | 120633538 | AX-170351652 | 0 | T | A | 0.14 | 3.24E-02 | 2.23E-01 | 2.70E+00 | 2.73E+00 | 8.84E-01 | no | Stard9 | intron_variant,NMD_transcript_variant | MODIFIER |
| 2 | 120639347 | AX-97798587 | 0 | G | A | 0.14 | 3.24E-02 | 2.23E-01 | 2.70E+00 | 2.73E+00 | 8.84E-01 | no | n-R5s205 | downstream_gene_variant | MODIFIER |
| 2 | 120653229 | AX-170189167 | 0 | T | A | 0.14 | 3.24E-02 | 2.23E-01 | 2.70E+00 | 2.73E+00 | 8.84E-01 | no | Stard9 | intron_variant,NMD_transcript_variant | MODIFIER |
| 2 | 120666471 | AX-97809504 | 0 | A | G | 0.14 | 3.24E-02 | 2.23E-01 | 2.70E+00 | 2.73E+00 | 8.84E-01 | no | Stard9 | intron_variant,NMD_transcript_variant | MODIFIER |
| 2 | 120682820 | AX-170478359 | 0 | C | G | 0.14 | 3.24E-02 | 2.23E-01 | 2.70E+00 | 2.73E+00 | 8.84E-01 | no | Stard9 | intron_variant,NMD_transcript_variant | MODIFIER |
| 2 | 120682912 | AX-97799689 | 0 | T | C | 0.14 | 3.24E-02 | 2.23E-01 | 2.70E+00 | 2.73E+00 | 8.84E-01 | no | Stard9 | intron_variant,NMD_transcript_variant | MODIFIER |
| 2 | 120684206 | AX-170064743 | 0 | T | G | 0.14 | 3.24E-02 | 2.23E-01 | 2.70E+00 | 2.73E+00 | 8.84E-01 | no | Stard9 | intron_variant,NMD_transcript_variant | MODIFIER |
| 2 | 120691151 | AX-170064748 | 0 | G | A | 0.14 | 3.24E-02 | 2.23E-01 | 2.70E+00 | 2.73E+00 | 8.84E-01 | no | Stard9 | intron_variant,NMD_transcript_variant | MODIFIER |
| 2 | 120699874 | AX-170064745 | 0 | T | C | 0.14 | 3.24E-02 | 2.23E-01 | 2.70E+00 | 2.73E+00 | 8.84E-01 | no | Stard9 | upstream_gene_variant | MODIFIER |
| 2 | 120702754 | AX-170064746 | 0 | G | A | 0.14 | 3.24E-02 | 2.23E-01 | 2.70E+00 | 2.73E+00 | 8.84E-01 | no | Stard9 | missense_variant,NMD_transcript_variant | MODERATE |
| 2 | 120719118 | AX-170156940 | 0 | G | C | 0.14 | 3.24E-02 | 2.23E-01 | 2.70E+00 | 2.73E+00 | 8.84E-01 | no | Stard9 | downstream_gene_variant | MODIFIER |
| 2 | 120724557 | AX-170251222 | 0 | C | A | 0.14 | 3.24E-02 | 2.23E-01 | 2.70E+00 | 2.73E+00 | 8.84E-01 | no | Cdan1 | intron_variant | MODIFIER |
| 2 | 120754295 | AX-97824876 | 0 | G | C | 0.14 | 3.24E-02 | 2.23E-01 | 2.70E+00 | 2.73E+00 | 8.84E-01 | no | Ttbk2 | intron_variant | MODIFIER |
| 2 | 120767502 | AX-170420654 | 0 | A | G | 0.14 | 3.24E-02 | 2.23E-01 | 2.70E+00 | 2.73E+00 | 8.84E-01 | no | Ttbk2 | intron_variant | MODIFIER |
| 2 | 120773222 | AX-97822509 | 0 | T | C | 0.14 | 3.24E-02 | 2.23E-01 | 2.70E+00 | 2.73E+00 | 8.84E-01 | no | Ttbk2 | splice_region_variant,intron_variant | LOW |
| 2 | 120790775 | AX-170144632 | 0 | A | G | 0.14 | 3.24E-02 | 2.23E-01 | 2.70E+00 | 2.73E+00 | 8.84E-01 | no | Ttbk2 | intron_variant | MODIFIER |
| 2 | 120794173 | AX-170514421 | 0 | T | C | 0.14 | 3.24E-02 | 2.23E-01 | 2.70E+00 | 2.73E+00 | 8.84E-01 | no | Ttbk2 | intron_variant | MODIFIER |
| 2 | 120794431 | AX-170535256 | 0 | T | G | 0.14 | 3.24E-02 | 2.23E-01 | 2.70E+00 | 2.73E+00 | 8.84E-01 | no | Ttbk2 | intron_variant | MODIFIER |
| 2 | 120835085 | AX-97843192 | 0 | A | G | 0.14 | 3.24E-02 | 2.23E-01 | 2.70E+00 | 2.73E+00 | 8.84E-01 | no | Ttbk2 | intron_variant | MODIFIER |
| 2 | 120848277 | AX-170184821 | 0 | C | A | 0.14 | 3.24E-02 | 2.23E-01 | 2.70E+00 | 2.73E+00 | 8.84E-01 | no | Ttbk2 | intron_variant | MODIFIER |
| 2 | 120848298 | AX-170430518 | 0 | A | G | 0.14 | 3.24E-02 | 2.23E-01 | 2.70E+00 | 2.73E+00 | 8.84E-01 | no | Ttbk2 | intron_variant | MODIFIER |
| 2 | 120877802 | AX-170234910 | 0 | A | G | 0.14 | 3.24E-02 | 2.23E-01 | 2.70E+00 | 2.73E+00 | 8.84E-01 | no | Ubr1 | intron_variant | MODIFIER |
| 2 | 120882242 | AX-170422515 | 0 | A | G | 0.14 | 3.24E-02 | 2.23E-01 | 2.70E+00 | 2.73E+00 | 8.84E-01 | no | Ubr1 | synonymous_variant | LOW |
| 2 | 120897161 | AX-170292496 | 0 | C | T | 0.14 | 3.24E-02 | 2.23E-01 | 2.70E+00 | 2.73E+00 | 8.84E-01 | no | Ubr1 | intron_variant | MODIFIER |
| 2 | 120909436 | AX-170145693 | 0 | T | C | 0.14 | 3.24E-02 | 2.23E-01 | 2.70E+00 | 2.73E+00 | 8.84E-01 | no | Ubr1 | missense_variant | MODERATE |
| 2 | 120914510 | AX-170119898 | 0 | A | G | 0.14 | 3.24E-02 | 2.23E-01 | 2.70E+00 | 2.73E+00 | 8.84E-01 | no | Ubr1 | intron_variant | MODIFIER |
| 2 | 120924297 | AX-170473534 | 0 | G | C | 0.14 | 3.24E-02 | 2.23E-01 | 2.70E+00 | 2.73E+00 | 8.84E-01 | no | Ubr1 | intron_variant | MODIFIER |
| 2 | 120942858 | AX-170108574 | 0 | C | T | 0.14 | 3.24E-02 | 2.23E-01 | 2.70E+00 | 2.73E+00 | 8.84E-01 | no | Ubr1 | intron_variant | MODIFIER |
| 2 | 120969052 | AX-170018854 | 0 | T | A | 0.14 | 3.24E-02 | 2.23E-01 | 2.70E+00 | 2.73E+00 | 8.84E-01 | no | Ubr1 | intron_variant | MODIFIER |
| 2 | 120969613 | AX-170018859 | 0 | A | G | 0.14 | 3.24E-02 | 2.23E-01 | 2.70E+00 | 2.73E+00 | 8.84E-01 | no | Ubr1 | intron_variant | MODIFIER |
| 2 | 120969859 | AX-170018862 | 0 | A | C | 0.14 | 3.24E-02 | 2.23E-01 | 2.70E+00 | 2.73E+00 | 8.84E-01 | no | Ubr1 | intron_variant | MODIFIER |
| 2 | 120980505 | AX-170425204 | 0 | A | G | 0.14 | 3.24E-02 | 2.23E-01 | 2.70E+00 | 2.73E+00 | 8.84E-01 | no | Tmem62 | missense_variant | MODERATE |
| 2 | 121001741 | AX-170064749 | 0 | C | T | 0.14 | 3.24E-02 | 2.23E-01 | 2.70E+00 | 2.73E+00 | 8.84E-01 | no | Tmem62 | intron_variant | MODIFIER |

| | | | | | | | | | | | | | | | |
|---|---|---|---|---|---|---|---|---|---|---|---|---|---|---|---|
| 2 | 121004885 | AX-170142242 | 0 | G | C | 0.14 | 3.24E-02 | 2.23E-01 | 2.70E+00 | 2.73E+00 | 8.84E-01 | no | Ccndbp1 | upstream_gene_variant | MODIFIER |
| 2 | 121005673 | AX-97832781 | 0 | A | G | 0.14 | 3.24E-02 | 2.23E-01 | 2.70E+00 | 2.73E+00 | 8.84E-01 | no | Ccndbp1 | upstream_gene_variant | MODIFIER |
| 2 | 121011456 | AX-170018864 | 0 | C | A | 0.14 | 3.24E-02 | 2.23E-01 | 2.70E+00 | 2.73E+00 | 8.84E-01 | no | Ccndbp1 | synonymous_variant | LOW |
| 2 | 121011477 | AX-170018860 | 0 | T | C | 0.14 | 3.24E-02 | 2.23E-01 | 2.70E+00 | 2.73E+00 | 8.84E-01 | no | Ccndbp1 | synonymous_variant | LOW |
| 2 | 121027079 | AX-170094388 | 0 | T | C | 0.14 | 3.24E-02 | 2.23E-01 | 2.70E+00 | 2.73E+00 | 8.84E-01 | no | Epb42 | splice_region_variant,intron_variant | LOW |
| 2 | 121046059 | AX-170018857 | 0 | C | T | 0.14 | 3.24E-02 | 2.23E-01 | 2.70E+00 | 2.73E+00 | 8.84E-01 | no | Tgm5 | downstream_gene_variant | MODIFIER |
| 2 | 121048696 | AX-170105195 | 0 | C | T | 0.14 | 3.24E-02 | 2.23E-01 | 2.70E+00 | 2.73E+00 | 8.84E-01 | no | Tgm5 | intron_variant | MODIFIER |
| 2 | 121048892 | AX-170402091 | 0 | G | A | 0.14 | 3.24E-02 | 2.23E-01 | 2.70E+00 | 2.73E+00 | 8.84E-01 | no | Tgm5 | synonymous_variant | LOW |
| 2 | 121058507 | AX-97838603 | 0 | A | G | 0.14 | 3.24E-02 | 2.23E-01 | 2.70E+00 | 2.73E+00 | 8.84E-01 | no | Tgm5 | intron_variant | MODIFIER |
| 2 | 121067339 | AX-170288420 | 0 | G | T | 0.14 | 3.24E-02 | 2.23E-01 | 2.70E+00 | 2.73E+00 | 8.84E-01 | no | Tgm5 | intron_variant | MODIFIER |
| 2 | 121096705 | AX-170018856 | 0 | C | A | 0.14 | 3.24E-02 | 2.23E-01 | 2.70E+00 | 2.73E+00 | 8.84E-01 | no | Tgm7 | intron_variant | MODIFIER |
| 2 | 121103278 | AX-170276506 | 0 | T | G | 0.14 | 3.24E-02 | 2.23E-01 | 2.70E+00 | 2.73E+00 | 8.84E-01 | no | Tgm7 | intron_variant | MODIFIER |
| 2 | 121103777 | AX-170018861 | 0 | T | C | 0.14 | 3.24E-02 | 2.23E-01 | 2.70E+00 | 2.73E+00 | 8.84E-01 | no | Tgm7 | intron_variant | MODIFIER |
| 2 | 121108602 | AX-170340790 | 0 | A | C | 0.14 | 3.24E-02 | 2.23E-01 | 2.70E+00 | 2.73E+00 | 8.84E-01 | no | Tgm7 | missense_variant | MODERATE |
| 2 | 121115796 | AX-170196219 | 0 | T | G | 0.14 | 3.24E-02 | 2.23E-01 | 2.70E+00 | 2.73E+00 | 8.84E-01 | no | | regulatory_region_variant | MODIFIER |
| 2 | 121119866 | AX-170309632 | 0 | T | C | 0.14 | 3.24E-02 | 2.23E-01 | 2.70E+00 | 2.73E+00 | 8.84E-01 | no | | intergenic_variant | MODIFIER |
| 2 | 121120731 | AX-97839779 | 0 | A | G | 0.14 | 3.24E-02 | 2.23E-01 | 2.70E+00 | 2.73E+00 | 8.84E-01 | no | | intergenic_variant | MODIFIER |
| 2 | 121122344 | AX-170253304 | 0 | G | C | 0.14 | 3.24E-02 | 2.23E-01 | 2.70E+00 | 2.73E+00 | 8.84E-01 | no | | intergenic_variant | MODIFIER |
| 2 | 121148590 | AX-170499919 | 0 | C | A | 0.14 | 3.24E-02 | 2.23E-01 | 2.70E+00 | 2.73E+00 | 8.84E-01 | no | Adal | intron_variant | MODIFIER |
| 2 | 121188409 | AX-170064753 | 0 | C | T | 0.14 | 3.24E-02 | 2.23E-01 | 2.70E+00 | 2.73E+00 | 8.84E-01 | no | Tubgcp4 | intron_variant | MODIFIER |
| 2 | 121190683 | AX-170491680 | 0 | G | A | 0.14 | 3.24E-02 | 2.23E-01 | 2.70E+00 | 2.73E+00 | 8.84E-01 | no | Tubgcp4 | intron_variant | MODIFIER |
| 2 | 121193723 | AX-170064751 | 0 | G | A | 0.14 | 3.24E-02 | 2.23E-01 | 2.70E+00 | 2.73E+00 | 8.84E-01 | no | Tubgcp4 | intron_variant | MODIFIER |
| 2 | 121214005 | AX-170216638 | 0 | G | A | 0.14 | 3.24E-02 | 2.23E-01 | 2.70E+00 | 2.73E+00 | 8.84E-01 | no | Trp53bp1 | intron_variant | MODIFIER |
| 2 | 121214024 | AX-170481734 | 0 | C | T | 0.14 | 3.24E-02 | 2.23E-01 | 2.70E+00 | 2.73E+00 | 8.84E-01 | no | Trp53bp1 | intron_variant | MODIFIER |
| 2 | 121214319 | AX-170064752 | 0 | G | A | 0.14 | 3.24E-02 | 2.23E-01 | 2.70E+00 | 2.73E+00 | 8.84E-01 | no | Trp53bp1 | intron_variant | MODIFIER |
| 2 | 121225491 | AX-170295601 | 0 | G | A | 0.14 | 3.24E-02 | 2.23E-01 | 2.70E+00 | 2.73E+00 | 8.84E-01 | no | Trp53bp1 | intron_variant | MODIFIER |
| 2 | 121242158 | AX-170118555 | 0 | G | A | 0.14 | 3.24E-02 | 2.23E-01 | 2.70E+00 | 2.73E+00 | 8.84E-01 | no | Trp53bp1 | intron_variant | MODIFIER |
| 2 | 121244929 | AX-170164426 | 0 | G | A | 0.14 | 3.24E-02 | 2.23E-01 | 2.70E+00 | 2.73E+00 | 8.84E-01 | no | Trp53bp1 | intron_variant | MODIFIER |
| 2 | 121245007 | AX-170128218 | 0 | G | A | 0.14 | 3.24E-02 | 2.23E-01 | 2.70E+00 | 2.73E+00 | 8.84E-01 | no | Trp53bp1 | intron_variant | MODIFIER |
| 2 | 121293468 | AX-170180309 | 0 | C | T | 0.14 | 3.24E-02 | 2.23E-01 | 2.70E+00 | 2.73E+00 | 8.84E-01 | no | Map1a | intron_variant | MODIFIER |
| 2 | 121297809 | AX-170274710 | 0 | C | T | 0.14 | 3.24E-02 | 2.23E-01 | 2.70E+00 | 2.73E+00 | 8.84E-01 | no | Map1a | intron_variant | MODIFIER |
| 2 | 121303324 | AX-170367733 | 0 | G | A | 0.14 | 3.24E-02 | 2.23E-01 | 2.70E+00 | 2.73E+00 | 8.84E-01 | no | Map1a | synonymous_variant | LOW |
| 2 | 121315901 | AX-97837121 | 0 | T | A | 0.14 | 3.24E-02 | 2.23E-01 | 2.70E+00 | 2.73E+00 | 8.84E-01 | no | Ppip5k1 | intron_variant | MODIFIER |
| 2 | 121321907 | AX-170299550 | 0 | A | G | 0.14 | 3.24E-02 | 2.23E-01 | 2.70E+00 | 2.73E+00 | 8.84E-01 | no | Ppip5k1 | intron_variant | MODIFIER |
| 2 | 121324067 | AX-97839805 | 0 | G | A | 0.14 | 3.24E-02 | 2.23E-01 | 2.70E+00 | 2.73E+00 | 8.84E-01 | no | Ppip5k1 | intron_variant | MODIFIER |
| 2 | 121366748 | AX-170347499 | 0 | C | T | 0.14 | 3.24E-02 | 2.23E-01 | 2.70E+00 | 2.73E+00 | 8.84E-01 | no | Ckmt1 | downstream_gene_variant | MODIFIER |
| 2 | 121374729 | AX-170524563 | 0 | A | C | 0.14 | 3.24E-02 | 2.23E-01 | 2.70E+00 | 2.73E+00 | 8.84E-01 | no | Strc | synonymous_variant | LOW |
| 2 | 121399205 | AX-97812851 | 0 | G | C | 0.14 | 3.24E-02 | 2.23E-01 | 2.70E+00 | 2.73E+00 | 8.84E-01 | no | Catsper2 | intron_variant | MODIFIER |
| 2 | 121413111 | AX-170100210 | 0 | C | T | 0.14 | 3.24E-02 | 2.23E-01 | 2.70E+00 | 2.73E+00 | 8.84E-01 | no | Pdia3 | upstream_gene_variant | MODIFIER |
| 2 | 121418967 | AX-170437527 | 0 | G | C | 0.14 | 3.24E-02 | 2.23E-01 | 2.70E+00 | 2.73E+00 | 8.84E-01 | no | Pdia3 | intron_variant | MODIFIER |
| 2 | 121451852 | AX-170018870 | 0 | C | T | 0.14 | 3.24E-02 | 2.23E-01 | 2.70E+00 | 2.73E+00 | 8.84E-01 | no | Serinc4 | 3_prime_UTR_variant | MODIFIER |
| 2 | 121507746 | AX-170018865 | 0 | T | C | 0.14 | 3.24E-02 | 2.23E-01 | 2.70E+00 | 2.73E+00 | 8.84E-01 | no | Wdr76 | intron_variant | MODIFIER |
| 2 | 121507788 | AX-170018866 | 0 | T | G | 0.14 | 3.24E-02 | 2.23E-01 | 2.70E+00 | 2.73E+00 | 8.84E-01 | no | Wdr76 | intron_variant | MODIFIER |
| 2 | 121508014 | AX-170018869 | 0 | T | C | 0.14 | 3.24E-02 | 2.23E-01 | 2.70E+00 | 2.73E+00 | 8.84E-01 | no | Wdr76 | intron_variant | MODIFIER |
| 2 | 121518474 | AX-170018868 | 0 | T | G | 0.14 | 3.24E-02 | 2.23E-01 | 2.70E+00 | 2.73E+00 | 8.84E-01 | no | Wdr76 | intron_variant | MODIFIER |
| 2 | 121535583 | AX-170018867 | 0 | C | G | 0.14 | 3.24E-02 | 2.23E-01 | 2.70E+00 | 2.73E+00 | 8.84E-01 | no | Wdr76 | splice_region_variant,intron_variant | LOW |
| 2 | 121536936 | AX-170064760 | 0 | T | C | 0.14 | 3.24E-02 | 2.23E-01 | 2.70E+00 | 2.73E+00 | 8.84E-01 | no | Wdr76 | intron_variant | MODIFIER |
| 2 | 121547159 | AX-170018873 | 0 | T | A | 0.14 | 3.24E-02 | 2.23E-01 | 2.70E+00 | 2.73E+00 | 8.84E-01 | no | Wdr76 | downstream_gene_variant | MODIFIER |
| 2 | 121564260 | AX-170348629 | 0 | A | G | 0.14 | 3.24E-02 | 2.23E-01 | 2.70E+00 | 2.73E+00 | 8.84E-01 | no | Frmd5 | intron_variant | MODIFIER |
| 2 | 121566721 | AX-170367180 | 0 | T | C | 0.14 | 3.24E-02 | 2.23E-01 | 2.70E+00 | 2.73E+00 | 8.84E-01 | no | Frmd5 | intron_variant | MODIFIER |
| 2 | 121568220 | AX-170094389 | 0 | G | A | 0.14 | 3.24E-02 | 2.23E-01 | 2.70E+00 | 2.73E+00 | 8.84E-01 | no | Frmd5 | intron_variant | MODIFIER |
| 2 | 121609496 | AX-170521492 | 0 | T | C | 0.14 | 3.24E-02 | 2.23E-01 | 2.70E+00 | 2.73E+00 | 8.84E-01 | no | Frmd5 | intron_variant | MODIFIER |
| 2 | 121611769 | AX-170411788 | 0 | T | A | 0.14 | 3.24E-02 | 2.23E-01 | 2.70E+00 | 2.73E+00 | 8.84E-01 | no | Frmd5 | intron_variant | MODIFIER |
| 2 | 121612142 | AX-170194146 | 0 | G | A | 0.14 | 3.24E-02 | 2.23E-01 | 2.70E+00 | 2.73E+00 | 8.84E-01 | no | Frmd5 | intron_variant | MODIFIER |

| | | | | | | | | | | | | | | | |
|---|---|---|---|---|---|---|---|---|---|---|---|---|---|---|---|
| 2 | 121616241 | AX-97788988 | 0 | C | T | 0.14 | 3.24E-02 | 2.23E-01 | 2.70E+00 | 2.73E+00 | 8.84E-01 | no | Frmd5 | intron_variant | MODIFIER |
| 2 | 121630289 | AX-170437641 | 0 | G | A | 0.14 | 3.24E-02 | 2.23E-01 | 2.70E+00 | 2.73E+00 | 8.84E-01 | no | Frmd5 | intron_variant | MODIFIER |
| 2 | 121644141 | AX-170204286 | 0 | G | A | 0.14 | 3.24E-02 | 2.23E-01 | 2.70E+00 | 2.73E+00 | 8.84E-01 | no | Frmd5 | intron_variant | MODIFIER |
| 2 | 121665382 | AX-170338062 | 0 | A | G | 0.14 | 3.24E-02 | 2.23E-01 | 2.70E+00 | 2.73E+00 | 8.84E-01 | no | Frmd5 | intron_variant | MODIFIER |
| 2 | 121666726 | AX-170418397 | 0 | C | T | 0.14 | 3.24E-02 | 2.23E-01 | 2.70E+00 | 2.73E+00 | 8.84E-01 | no | Frmd5 | intron_variant | MODIFIER |
| 2 | 121689352 | AX-97784711 | 0 | G | A | 0.14 | 3.24E-02 | 2.23E-01 | 2.70E+00 | 2.73E+00 | 8.84E-01 | no | Frmd5 | intron_variant | MODIFIER |
| 2 | 121723401 | AX-170160824 | 0 | T | G | 0.14 | 3.24E-02 | 2.23E-01 | 2.70E+00 | 2.73E+00 | 8.84E-01 | no | Frmd5 | intron_variant | MODIFIER |
| 2 | 121725649 | AX-170508833 | 0 | T | C | 0.14 | 3.24E-02 | 2.23E-01 | 2.70E+00 | 2.73E+00 | 8.84E-01 | no | Frmd5 | intron_variant | MODIFIER |
| 2 | 121726648 | AX-170263689 | 0 | G | A | 0.14 | 3.24E-02 | 2.23E-01 | 2.70E+00 | 2.73E+00 | 8.84E-01 | no | Frmd5 | intron_variant | MODIFIER |
| 2 | 121764857 | AX-170386121 | 0 | T | C | 0.14 | 3.24E-02 | 2.23E-01 | 2.70E+00 | 2.73E+00 | 8.84E-01 | no | Frmd5 | intron_variant | MODIFIER |
| 2 | 121783086 | AX-170116798 | 0 | T | G | 0.14 | 3.24E-02 | 2.23E-01 | 2.70E+00 | 2.73E+00 | 8.84E-01 | no | Frmd5 | intron_variant | MODIFIER |
| 2 | 121787432 | AX-170154446 | 0 | G | A | 0.14 | 3.24E-02 | 2.23E-01 | 2.70E+00 | 2.73E+00 | 8.84E-01 | no | Frmd5 | intron_variant | MODIFIER |
| 2 | 121790201 | AX-170502804 | 0 | A | G | 0.14 | 3.24E-02 | 2.23E-01 | 2.70E+00 | 2.73E+00 | 8.84E-01 | no | Frmd5 | intron_variant | MODIFIER |
| 2 | 121824788 | AX-170521568 | 0 | A | G | 0.14 | 3.24E-02 | 2.23E-01 | 2.70E+00 | 2.73E+00 | 8.84E-01 | no | | regulatory_region_variant | MODIFIER |
| 2 | 121843653 | AX-97853154 | 0 | T | C | 0.14 | 3.24E-02 | 2.23E-01 | 2.70E+00 | 2.73E+00 | 8.84E-01 | no | | intergenic_variant | MODIFIER |
| 2 | 121847611 | AX-170264366 | 0 | T | A | 0.14 | 3.24E-02 | 2.23E-01 | 2.70E+00 | 2.73E+00 | 8.84E-01 | no | | intergenic_variant | MODIFIER |
| 2 | 121903371 | AX-97797095 | 0 | A | G | 0.14 | 3.24E-02 | 2.23E-01 | 2.70E+00 | 2.73E+00 | 8.84E-01 | no | Casc4 | intron_variant | MODIFIER |
| 2 | 121931713 | AX-170117690 | 0 | A | C | 0.14 | 3.24E-02 | 2.23E-01 | 2.70E+00 | 2.73E+00 | 8.84E-01 | no | Casc4 | intron_variant | MODIFIER |
| 2 | 121932408 | AX-170468873 | 0 | G | C | 0.14 | 3.24E-02 | 2.23E-01 | 2.70E+00 | 2.73E+00 | 8.84E-01 | no | Casc4 | intron_variant | MODIFIER |
| 2 | 121956288 | AX-170094390 | 0 | T | C | 0.14 | 3.24E-02 | 2.23E-01 | 2.70E+00 | 2.73E+00 | 8.84E-01 | no | Ctdspl2 | 5_prime_UTR_variant | MODIFIER |
| 2 | 121968674 | AX-170431382 | 0 | G | A | 0.14 | 3.24E-02 | 2.23E-01 | 2.70E+00 | 2.73E+00 | 8.84E-01 | no | Ctdspl2 | intron_variant | MODIFIER |
| 2 | 121969177 | AX-170139175 | 0 | G | A | 0.14 | 3.24E-02 | 2.23E-01 | 2.70E+00 | 2.73E+00 | 8.84E-01 | no | Ctdspl2 | synonymous_variant | LOW |
| 2 | 121974594 | AX-170064765 | 0 | G | A | 0.14 | 3.24E-02 | 2.23E-01 | 2.70E+00 | 2.73E+00 | 8.84E-01 | no | Ctdspl2 | intron_variant | MODIFIER |
| 2 | 121975560 | AX-170018879 | 0 | T | G | 0.14 | 3.24E-02 | 2.23E-01 | 2.70E+00 | 2.73E+00 | 8.84E-01 | no | Ctdspl2 | intron_variant | MODIFIER |
| 2 | 121984980 | AX-170094391 | 0 | C | T | 0.14 | 3.24E-02 | 2.23E-01 | 2.70E+00 | 2.73E+00 | 8.84E-01 | no | Ctdspl2 | intron_variant | MODIFIER |
| 2 | 121988198 | AX-170420820 | 0 | C | T | 0.14 | 3.24E-02 | 2.23E-01 | 2.70E+00 | 2.73E+00 | 8.84E-01 | no | Ctdspl2 | intron_variant | MODIFIER |
| 2 | 122007673 | AX-97846063 | 0 | T | C | 0.14 | 3.24E-02 | 2.23E-01 | 2.70E+00 | 2.73E+00 | 8.84E-01 | no | Ctdspl2 | intron_variant | MODIFIER |
| 2 | 122015867 | AX-170122023 | 0 | A | G | 0.14 | 3.24E-02 | 2.23E-01 | 2.70E+00 | 2.73E+00 | 8.84E-01 | no | Ctdspl2 | downstream_gene_variant | MODIFIER |
| 2 | 122083373 | AX-170513473 | 0 | G | T | 0.14 | 3.24E-02 | 2.23E-01 | 2.70E+00 | 2.73E+00 | 8.84E-01 | no | Spg11 | intron_variant | MODIFIER |
| 2 | 122097209 | AX-97826380 | 0 | T | C | 0.14 | 3.24E-02 | 2.23E-01 | 2.70E+00 | 2.73E+00 | 8.84E-01 | no | Spg11 | intron_variant | MODIFIER |
| 2 | 122123313 | AX-170205381 | 0 | G | A | 0.14 | 3.24E-02 | 2.23E-01 | 2.70E+00 | 2.73E+00 | 8.84E-01 | no | Patl2 | intron_variant | MODIFIER |
| 2 | 122124929 | AX-170305690 | 0 | T | C | 0.14 | 3.24E-02 | 2.23E-01 | 2.70E+00 | 2.73E+00 | 8.84E-01 | no | Patl2 | intron_variant | MODIFIER |
| 2 | 122125337 | AX-170018878 | 0 | C | T | 0.14 | 3.24E-02 | 2.23E-01 | 2.70E+00 | 2.73E+00 | 8.84E-01 | no | Patl2 | synonymous_variant | LOW |
| 2 | 122126701 | AX-170018874 | 0 | T | C | 0.14 | 3.24E-02 | 2.23E-01 | 2.70E+00 | 2.73E+00 | 8.84E-01 | no | Patl2 | synonymous_variant | LOW |
| 2 | 122129286 | AX-170109020 | 0 | G | A | 0.14 | 3.24E-02 | 2.23E-01 | 2.70E+00 | 2.73E+00 | 8.84E-01 | no | Patl2 | intron_variant | MODIFIER |
| 2 | 122130623 | AX-97838783 | 0 | T | C | 0.14 | 3.24E-02 | 2.23E-01 | 2.70E+00 | 2.73E+00 | 8.84E-01 | no | Patl2 | intron_variant | MODIFIER |
| 2 | 122141541 | AX-170156563 | 0 | G | A | 0.14 | 3.24E-02 | 2.23E-01 | 2.70E+00 | 2.73E+00 | 8.84E-01 | no | Patl2 | intron_variant | MODIFIER |
| 2 | 122186766 | AX-170471201 | 0 | C | T | 0.14 | 3.24E-02 | 2.23E-01 | 2.70E+00 | 2.73E+00 | 8.84E-01 | no | Terb2 | intron_variant | MODIFIER |
| 2 | 122186884 | AX-97838781 | 0 | T | C | 0.14 | 3.24E-02 | 2.23E-01 | 2.70E+00 | 2.73E+00 | 8.84E-01 | no | Terb2 | intron_variant | MODIFIER |
| 2 | 122193870 | AX-170018883 | 0 | C | A | 0.14 | 3.24E-02 | 2.23E-01 | 2.70E+00 | 2.73E+00 | 8.84E-01 | no | Terb2 | intron_variant | MODIFIER |
| 2 | 122205986 | AX-170214950 | 0 | C | T | 0.14 | 3.24E-02 | 2.23E-01 | 2.70E+00 | 2.73E+00 | 8.84E-01 | no | Terb2 | 3_prime_UTR_variant | MODIFIER |
| 2 | 122230022 | AX-170371974 | 0 | G | A | 0.14 | 3.24E-02 | 2.23E-01 | 2.70E+00 | 2.73E+00 | 8.84E-01 | no | Sord | upstream_gene_variant | MODIFIER |
| 2 | 122232617 | AX-170465950 | 0 | T | C | 0.14 | 3.24E-02 | 2.23E-01 | 2.70E+00 | 2.73E+00 | 8.84E-01 | no | Sord | upstream_gene_variant | MODIFIER |
| 2 | 122246087 | AX-170377564 | 0 | A | G | 0.14 | 3.24E-02 | 2.23E-01 | 2.70E+00 | 2.73E+00 | 8.84E-01 | no | Sord | intron_variant | MODIFIER |
| 2 | 122278164 | AX-170431362 | 0 | G | C | 0.14 | 3.24E-02 | 2.23E-01 | 2.70E+00 | 2.73E+00 | 8.84E-01 | no | Duox2 | downstream_gene_variant | MODIFIER |
| 2 | 122287339 | AX-170064766 | 0 | G | A | 0.14 | 3.24E-02 | 2.23E-01 | 2.70E+00 | 2.73E+00 | 8.84E-01 | no | Duox2 | intron_variant | MODIFIER |
| 2 | 122292115 | AX-170064769 | 0 | C | T | 0.14 | 3.24E-02 | 2.23E-01 | 2.70E+00 | 2.73E+00 | 8.84E-01 | no | Duox2 | intron_variant | MODIFIER |
| 2 | 122293756 | AX-170064768 | 0 | A | G | 0.14 | 3.24E-02 | 2.23E-01 | 2.70E+00 | 2.73E+00 | 8.84E-01 | no | Duox2 | intron_variant | MODIFIER |
| 2 | 122294092 | AX-170156042 | 0 | C | T | 0.14 | 3.24E-02 | 2.23E-01 | 2.70E+00 | 2.73E+00 | 8.84E-01 | no | Duoxa2 | upstream_gene_variant | MODIFIER |
| 2 | 122318583 | AX-170315694 | 0 | A | T | 0.14 | 3.24E-02 | 2.23E-01 | 2.70E+00 | 2.73E+00 | 8.84E-01 | no | Duox1 | intron_variant | MODIFIER |
| 2 | 122329518 | AX-170265264 | 0 | T | C | 0.14 | 3.24E-02 | 2.23E-01 | 2.70E+00 | 2.73E+00 | 8.84E-01 | no | Duox1 | intron_variant | MODIFIER |
| 2 | 122329926 | AX-170094393 | 0 | G | A | 0.14 | 3.24E-02 | 2.23E-01 | 2.70E+00 | 2.73E+00 | 8.84E-01 | no | Duox1 | intron_variant | MODIFIER |
| 2 | 122360216 | AX-170494095 | 0 | C | A | 0.14 | 3.24E-02 | 2.23E-01 | 2.70E+00 | 2.73E+00 | 8.84E-01 | no | Shf | intron_variant | MODIFIER |
| 2 | 122364679 | AX-170157457 | 0 | T | G | 0.14 | 3.24E-02 | 2.23E-01 | 2.70E+00 | 2.73E+00 | 8.84E-01 | no | Shf | intron_variant | MODIFIER |

| | | | | | | | | | | | | | | | |
|---|---|---|---|---|---|---|---|---|---|---|---|---|---|---|---|
| 2 | 122389305 | AX-97838954 | 0 | A | C | 0.14 | 3.24E-02 | 2.23E-01 | 2.70E+00 | 2.73E+00 | 8.84E-01 | no | Gm24409 | upstream_gene_variant | MODIFIER |
| 2 | 122396830 | AX-170429339 | 0 | T | C | 0.14 | 3.24E-02 | 2.23E-01 | 2.70E+00 | 2.73E+00 | 8.84E-01 | no | | intergenic_variant | MODIFIER |
| 2 | 122397065 | AX-170088734 | 0 | T | C | 0.14 | 3.24E-02 | 2.23E-01 | 2.70E+00 | 2.73E+00 | 8.84E-01 | no | | intergenic_variant | MODIFIER |
| 2 | 122458764 | AX-170324574 | 0 | C | T | 0.14 | 3.24E-02 | 2.23E-01 | 2.70E+00 | 2.73E+00 | 8.84E-01 | no | Slc28a2 | intron_variant | MODIFIER |
| 2 | 122469172 | AX-170018894 | 0 | C | T | 0.14 | 3.24E-02 | 2.23E-01 | 2.70E+00 | 2.73E+00 | 8.84E-01 | no | Bambi-ps1 | upstream_gene_variant | MODIFIER |
| 2 | 122489172 | AX-170018888 | 0 | G | C | 0.14 | 3.24E-02 | 2.23E-01 | 2.70E+00 | 2.73E+00 | 8.84E-01 | no | Gm14085 | intron_variant | MODIFIER |
| 2 | 122519184 | AX-170249663 | 0 | C | T | 0.14 | 3.24E-02 | 2.23E-01 | 2.70E+00 | 2.73E+00 | 8.84E-01 | no | Gm14085 | intron_variant | MODIFIER |
| 2 | 122558071 | AX-170088736 | 0 | A | G | 0.14 | 3.24E-02 | 2.23E-01 | 2.70E+00 | 2.73E+00 | 8.84E-01 | no | | intergenic_variant | MODIFIER |
| 2 | 122571462 | AX-170018893 | 0 | A | G | 0.14 | 3.24E-02 | 2.23E-01 | 2.70E+00 | 2.73E+00 | 8.84E-01 | no | | intergenic_variant | MODIFIER |
| 2 | 122585457 | AX-170109254 | 0 | C | G | 0.14 | 3.24E-02 | 2.23E-01 | 2.70E+00 | 2.73E+00 | 8.84E-01 | no | | intergenic_variant | MODIFIER |
| 2 | 122598057 | AX-170288650 | 0 | G | A | 0.14 | 3.24E-02 | 2.23E-01 | 2.70E+00 | 2.73E+00 | 8.84E-01 | no | Gatm | intron_variant | MODIFIER |
| 2 | 122615277 | AX-97825459 | 0 | G | A | 0.14 | 3.24E-02 | 2.23E-01 | 2.70E+00 | 2.73E+00 | 8.84E-01 | no | Gatm | upstream_gene_variant | MODIFIER |
| 2 | 122620402 | AX-170258072 | 0 | A | G | 0.14 | 3.24E-02 | 2.23E-01 | 2.70E+00 | 2.73E+00 | 8.84E-01 | no | | intergenic_variant | MODIFIER |
| 2 | 122655758 | AX-170018887 | 0 | C | T | 0.14 | 3.24E-02 | 2.23E-01 | 2.70E+00 | 2.73E+00 | 8.84E-01 | no | | intergenic_variant | MODIFIER |
| 2 | 122674143 | AX-170064773 | 0 | T | C | 0.14 | 3.24E-02 | 2.23E-01 | 2.70E+00 | 2.73E+00 | 8.84E-01 | no | | intergenic_variant | MODIFIER |
| 2 | 122685091 | AX-170018892 | 0 | A | G | 0.14 | 3.24E-02 | 2.23E-01 | 2.70E+00 | 2.73E+00 | 8.84E-01 | no | Slc30a4 | 3_prime_UTR_variant | MODIFIER |
| 2 | 122685194 | AX-170018890 | 0 | C | G | 0.14 | 3.24E-02 | 2.23E-01 | 2.70E+00 | 2.73E+00 | 8.84E-01 | no | Slc30a4 | missense_variant | MODERATE |
| 2 | 122685340 | AX-170088737 | 0 | T | C | 0.14 | 3.24E-02 | 2.23E-01 | 2.70E+00 | 2.73E+00 | 8.84E-01 | no | Slc30a4 | splice_region_variant,intron_variant | LOW |
| 2 | 122688224 | AX-170064771 | 0 | C | T | 0.14 | 3.24E-02 | 2.23E-01 | 2.70E+00 | 2.73E+00 | 8.84E-01 | no | Slc30a4 | intron_variant | MODIFIER |
| 2 | 122712140 | AX-170384710 | 0 | G | T | 0.14 | 3.24E-02 | 2.23E-01 | 2.70E+00 | 2.73E+00 | 8.84E-01 | no | 4930417H01Rik | intron_variant,non_coding_transcript_variant | MODIFIER |
| 2 | 122712648 | AX-170064774 | 0 | A | G | 0.14 | 3.24E-02 | 2.23E-01 | 2.70E+00 | 2.73E+00 | 8.84E-01 | no | 4930417H01Rik | intron_variant,non_coding_transcript_variant | MODIFIER |
| 2 | 122716955 | AX-170064772 | 0 | A | G | 0.14 | 3.24E-02 | 2.23E-01 | 2.70E+00 | 2.73E+00 | 8.84E-01 | no | 4930417H01Rik | intron_variant,non_coding_transcript_variant | MODIFIER |
| 2 | 122719180 | AX-97802053 | 0 | T | C | 0.14 | 3.24E-02 | 2.23E-01 | 2.70E+00 | 2.73E+00 | 8.84E-01 | no | 4930417H01Rik | intron_variant,non_coding_transcript_variant | MODIFIER |
| 2 | 122723524 | AX-170222505 | 0 | C | A | 0.14 | 3.24E-02 | 2.23E-01 | 2.70E+00 | 2.73E+00 | 8.84E-01 | no | 4930417H01Rik | downstream_gene_variant | MODIFIER |
| 2 | 122725235 | AX-170425648 | 0 | A | G | 0.14 | 3.24E-02 | 2.23E-01 | 2.70E+00 | 2.73E+00 | 8.84E-01 | no | 4930417H01Rik | downstream_gene_variant | MODIFIER |
| 2 | 122730165 | AX-170358613 | 0 | G | A | 0.14 | 3.24E-02 | 2.23E-01 | 2.70E+00 | 2.73E+00 | 8.84E-01 | no | Bloc1s6os | downstream_gene_variant | MODIFIER |
| 2 | 122776827 | AX-97817501 | 0 | C | T | 0.14 | 3.24E-02 | 2.23E-01 | 2.70E+00 | 2.73E+00 | 8.84E-01 | no | Sqor | intron_variant | MODIFIER |
| 2 | 122784976 | AX-170426116 | 0 | G | A | 0.14 | 3.24E-02 | 2.23E-01 | 2.70E+00 | 2.73E+00 | 8.84E-01 | no | Sqor | missense_variant | MODERATE |
| 2 | 122790231 | AX-170393471 | 0 | T | C | 0.14 | 3.24E-02 | 2.23E-01 | 2.70E+00 | 2.73E+00 | 8.84E-01 | no | Sqor | intron_variant | MODIFIER |
| 2 | 122808602 | AX-170333227 | 0 | C | T | 0.14 | 3.24E-02 | 2.23E-01 | 2.70E+00 | 2.73E+00 | 8.84E-01 | no | Sqor | intron_variant | MODIFIER |
| 2 | 122808703 | AX-97795513 | 0 | A | G | 0.14 | 3.24E-02 | 2.23E-01 | 2.70E+00 | 2.73E+00 | 8.84E-01 | no | Sqor | intron_variant | MODIFIER |
| 2 | 122824249 | AX-170204321 | 0 | T | C | 0.14 | 3.24E-02 | 2.23E-01 | 2.70E+00 | 2.73E+00 | 8.84E-01 | no | | intergenic_variant | MODIFIER |
| 2 | 122843410 | AX-97840149 | 0 | C | G | 0.14 | 3.24E-02 | 2.23E-01 | 2.70E+00 | 2.73E+00 | 8.84E-01 | no | | intergenic_variant | MODIFIER |
| 2 | 122848730 | AX-170487085 | 0 | A | G | 0.14 | 3.24E-02 | 2.23E-01 | 2.70E+00 | 2.73E+00 | 8.84E-01 | no | | intergenic_variant | MODIFIER |
| 2 | 122857959 | AX-170132056 | 0 | C | T | 0.14 | 3.24E-02 | 2.23E-01 | 2.70E+00 | 2.73E+00 | 8.84E-01 | no | | intergenic_variant | MODIFIER |
| 2 | 122869553 | AX-170112338 | 0 | G | C | 0.14 | 3.24E-02 | 2.23E-01 | 2.70E+00 | 2.73E+00 | 8.84E-01 | no | | intergenic_variant | MODIFIER |
| 2 | 122887041 | AX-170392754 | 0 | A | G | 0.14 | 3.24E-02 | 2.23E-01 | 2.70E+00 | 2.73E+00 | 8.84E-01 | no | | intergenic_variant | MODIFIER |
| 2 | 122896820 | AX-170241118 | 0 | C | A | 0.14 | 3.24E-02 | 2.23E-01 | 2.70E+00 | 2.73E+00 | 8.84E-01 | no | | intergenic_variant | MODIFIER |
| 2 | 122896865 | AX-170221655 | 0 | T | C | 0.14 | 3.24E-02 | 2.23E-01 | 2.70E+00 | 2.73E+00 | 8.84E-01 | no | | intergenic_variant | MODIFIER |
| 2 | 122896908 | AX-170367320 | 0 | G | A | 0.14 | 3.24E-02 | 2.23E-01 | 2.70E+00 | 2.73E+00 | 8.84E-01 | no | | intergenic_variant | MODIFIER |
| 2 | 122897987 | AX-170521132 | 0 | G | C | 0.14 | 3.24E-02 | 2.23E-01 | 2.70E+00 | 2.73E+00 | 8.84E-01 | no | | intergenic_variant | MODIFIER |
| 2 | 122921575 | AX-170270308 | 0 | C | A | 0.14 | 3.24E-02 | 2.23E-01 | 2.70E+00 | 2.73E+00 | 8.84E-01 | no | | intergenic_variant | MODIFIER |
| 2 | 122957406 | AX-170178953 | 0 | C | T | 0.14 | 3.24E-02 | 2.23E-01 | 2.70E+00 | 2.73E+00 | 8.84E-01 | no | | intergenic_variant | MODIFIER |
| 2 | 122957760 | AX-97817274 | 0 | A | C | 0.14 | 3.24E-02 | 2.23E-01 | 2.70E+00 | 2.73E+00 | 8.84E-01 | no | | intergenic_variant | MODIFIER |
| 2 | 122983683 | AX-170064779 | 0 | T | A | 0.14 | 3.24E-02 | 2.23E-01 | 2.70E+00 | 2.73E+00 | 8.84E-01 | no | | intergenic_variant | MODIFIER |
| 2 | 123016565 | AX-170164978 | 0 | C | A | 0.14 | 3.24E-02 | 2.23E-01 | 2.70E+00 | 2.73E+00 | 8.84E-01 | no | | intergenic_variant | MODIFIER |
| 2 | 123025700 | AX-97801895 | 0 | G | A | 0.14 | 3.24E-02 | 2.23E-01 | 2.70E+00 | 2.73E+00 | 8.84E-01 | no | | intergenic_variant | MODIFIER |
| 2 | 123043042 | AX-170110112 | 0 | T | C | 0.14 | 3.24E-02 | 2.23E-01 | 2.70E+00 | 2.73E+00 | 8.84E-01 | no | | intergenic_variant | MODIFIER |
| 2 | 123062477 | AX-170516567 | 0 | G | A | 0.14 | 3.24E-02 | 2.23E-01 | 2.70E+00 | 2.73E+00 | 8.84E-01 | no | | intergenic_variant | MODIFIER |
| 2 | 123104261 | AX-170494477 | 0 | C | G | 0.14 | 3.24E-02 | 2.23E-01 | 2.70E+00 | 2.73E+00 | 8.84E-01 | no | | intergenic_variant | MODIFIER |
| 2 | 123133890 | AX-170018898 | 0 | G | A | 0.14 | 3.24E-02 | 2.23E-01 | 2.70E+00 | 2.73E+00 | 8.84E-01 | no | | intergenic_variant | MODIFIER |
| 2 | 123170829 | AX-170018899 | 0 | T | C | 0.14 | 3.24E-02 | 2.23E-01 | 2.70E+00 | 2.73E+00 | 8.84E-01 | no | | regulatory_region_variant | MODIFIER |
| 2 | 123172537 | AX-170018901 | 0 | A | G | 0.14 | 3.24E-02 | 2.23E-01 | 2.70E+00 | 2.73E+00 | 8.84E-01 | no | | intergenic_variant | MODIFIER |
| 2 | 123173617 | AX-97838842 | 0 | T | C | 0.14 | 3.24E-02 | 2.23E-01 | 2.70E+00 | 2.73E+00 | 8.84E-01 | no | | intergenic_variant | MODIFIER |

| | | | | | | | | | | | | | | | |
|---|---|---|---|---|---|---|---|---|---|---|---|---|---|---|---|
| 2 | 123178377 | AX-170373260 | 0 | C | T | 0.14 | 3.24E-02 | 2.23E-01 | 2.70E+00 | 2.73E+00 | 8.84E-01 | no | | intergenic_variant | MODIFIER |
| 2 | 123200385 | AX-170391054 | 0 | T | C | 0.14 | 3.24E-02 | 2.23E-01 | 2.70E+00 | 2.73E+00 | 8.84E-01 | no | | intergenic_variant | MODIFIER |
| 2 | 123204743 | AX-97799373 | 0 | C | T | 0.14 | 3.24E-02 | 2.23E-01 | 2.70E+00 | 2.73E+00 | 8.84E-01 | no | | intergenic_variant | MODIFIER |
| 2 | 123220528 | AX-170018897 | 0 | G | T | 0.14 | 3.24E-02 | 2.23E-01 | 2.70E+00 | 2.73E+00 | 8.84E-01 | no | | intergenic_variant | MODIFIER |
| 2 | 123243658 | AX-170239523 | 0 | C | A | 0.14 | 3.24E-02 | 2.23E-01 | 2.70E+00 | 2.73E+00 | 8.84E-01 | no | | intergenic_variant | MODIFIER |
| 2 | 123244446 | AX-170339772 | 0 | A | C | 0.14 | 3.24E-02 | 2.23E-01 | 2.70E+00 | 2.73E+00 | 8.84E-01 | no | | intergenic_variant | MODIFIER |
| 2 | 123245158 | AX-170235187 | 0 | G | C | 0.14 | 3.24E-02 | 2.23E-01 | 2.70E+00 | 2.73E+00 | 8.84E-01 | no | | intergenic_variant | MODIFIER |
| 2 | 123245705 | AX-170197084 | 0 | G | A | 0.14 | 3.24E-02 | 2.23E-01 | 2.70E+00 | 2.73E+00 | 8.84E-01 | no | | intergenic_variant | MODIFIER |
| 2 | 123265398 | AX-170064784 | 0 | A | C | 0.14 | 3.24E-02 | 2.23E-01 | 2.70E+00 | 2.73E+00 | 8.84E-01 | no | | intergenic_variant | MODIFIER |
| 2 | 123265428 | AX-170344548 | 0 | A | G | 0.14 | 3.24E-02 | 2.23E-01 | 2.70E+00 | 2.73E+00 | 8.84E-01 | no | | intergenic_variant | MODIFIER |
| 2 | 123277371 | AX-170064781 | 0 | G | T | 0.14 | 3.24E-02 | 2.23E-01 | 2.70E+00 | 2.73E+00 | 8.84E-01 | no | Gm13988 | downstream_gene_variant | MODIFIER |
| 2 | 123297130 | AX-170446572 | 0 | T | A | 0.14 | 3.24E-02 | 2.23E-01 | 2.70E+00 | 2.73E+00 | 8.84E-01 | no | | intergenic_variant | MODIFIER |
| 2 | 123298343 | AX-170064783 | 0 | C | T | 0.14 | 3.24E-02 | 2.23E-01 | 2.70E+00 | 2.73E+00 | 8.84E-01 | no | | intergenic_variant | MODIFIER |
| 2 | 123299567 | AX-170358056 | 0 | A | C | 0.14 | 3.24E-02 | 2.23E-01 | 2.70E+00 | 2.73E+00 | 8.84E-01 | no | | intergenic_variant | MODIFIER |
| 2 | 123301629 | AX-170220701 | 0 | G | A | 0.14 | 3.24E-02 | 2.23E-01 | 2.70E+00 | 2.73E+00 | 8.84E-01 | no | | intergenic_variant | MODIFIER |
| 2 | 123302612 | AX-170247685 | 0 | A | T | 0.14 | 3.24E-02 | 2.23E-01 | 2.70E+00 | 2.73E+00 | 8.84E-01 | no | | intergenic_variant | MODIFIER |
| 2 | 123311617 | AX-170446190 | 0 | T | C | 0.14 | 3.24E-02 | 2.23E-01 | 2.70E+00 | 2.73E+00 | 8.84E-01 | no | | intergenic_variant | MODIFIER |
| 2 | 123320365 | AX-170116474 | 0 | G | A | 0.14 | 3.24E-02 | 2.23E-01 | 2.70E+00 | 2.73E+00 | 8.84E-01 | no | | intergenic_variant | MODIFIER |
| 2 | 123320881 | AX-170482181 | 0 | T | G | 0.14 | 3.24E-02 | 2.23E-01 | 2.70E+00 | 2.73E+00 | 8.84E-01 | no | | intergenic_variant | MODIFIER |
| 2 | 123324988 | AX-170109847 | 0 | C | A | 0.14 | 3.24E-02 | 2.23E-01 | 2.70E+00 | 2.73E+00 | 8.84E-01 | no | | intergenic_variant | MODIFIER |
| 2 | 123334007 | AX-97801015 | 0 | T | C | 0.14 | 3.24E-02 | 2.23E-01 | 2.70E+00 | 2.73E+00 | 8.84E-01 | no | | intergenic_variant | MODIFIER |
| 2 | 123346568 | AX-170094396 | 0 | T | C | 0.14 | 3.24E-02 | 2.23E-01 | 2.70E+00 | 2.73E+00 | 8.84E-01 | no | | intergenic_variant | MODIFIER |
| 2 | 123352516 | AX-170381710 | 0 | T | G | 0.14 | 3.24E-02 | 2.23E-01 | 2.70E+00 | 2.73E+00 | 8.84E-01 | no | | intergenic_variant | MODIFIER |
| 2 | 123366405 | AX-170228367 | 0 | A | G | 0.14 | 3.24E-02 | 2.23E-01 | 2.70E+00 | 2.73E+00 | 8.84E-01 | no | | intergenic_variant | MODIFIER |
| 2 | 123366679 | AX-170517406 | 0 | T | A | 0.14 | 3.24E-02 | 2.23E-01 | 2.70E+00 | 2.73E+00 | 8.84E-01 | no | | intergenic_variant | MODIFIER |
| 2 | 123371622 | AX-97795359 | 0 | C | T | 0.14 | 3.24E-02 | 2.23E-01 | 2.70E+00 | 2.73E+00 | 8.84E-01 | no | | intergenic_variant | MODIFIER |
| 2 | 123398970 | AX-170290484 | 0 | A | G | 0.14 | 3.24E-02 | 2.23E-01 | 2.70E+00 | 2.73E+00 | 8.84E-01 | no | | intergenic_variant | MODIFIER |
| 2 | 123414644 | AX-170094397 | 0 | C | T | 0.14 | 3.24E-02 | 2.23E-01 | 2.70E+00 | 2.73E+00 | 8.84E-01 | no | | intergenic_variant | MODIFIER |
| 2 | 123416643 | AX-170422915 | 0 | A | G | 0.14 | 3.24E-02 | 2.23E-01 | 2.70E+00 | 2.73E+00 | 8.84E-01 | no | | intergenic_variant | MODIFIER |
| 2 | 123416808 | AX-170171134 | 0 | C | T | 0.14 | 3.24E-02 | 2.23E-01 | 2.70E+00 | 2.73E+00 | 8.84E-01 | no | | intergenic_variant | MODIFIER |
| 2 | 123423101 | AX-170234249 | 0 | T | C | 0.14 | 3.24E-02 | 2.23E-01 | 2.70E+00 | 2.73E+00 | 8.84E-01 | no | | intergenic_variant | MODIFIER |
| 2 | 123426614 | AX-97847578 | 0 | G | A | 0.14 | 3.24E-02 | 2.23E-01 | 2.70E+00 | 2.73E+00 | 8.84E-01 | no | | intergenic_variant | MODIFIER |
| 2 | 123428356 | AX-170221173 | 0 | A | G | 0.14 | 3.24E-02 | 2.23E-01 | 2.70E+00 | 2.73E+00 | 8.84E-01 | no | | intergenic_variant | MODIFIER |
| 2 | 123428431 | AX-170197044 | 0 | G | A | 0.14 | 3.24E-02 | 2.23E-01 | 2.70E+00 | 2.73E+00 | 8.84E-01 | no | | intergenic_variant | MODIFIER |
| 2 | 123428646 | AX-170143369 | 0 | A | G | 0.14 | 3.24E-02 | 2.23E-01 | 2.70E+00 | 2.73E+00 | 8.84E-01 | no | | intergenic_variant | MODIFIER |
| 2 | 123436602 | AX-170131517 | 0 | C | T | 0.14 | 3.24E-02 | 2.23E-01 | 2.70E+00 | 2.73E+00 | 8.84E-01 | no | | intergenic_variant | MODIFIER |
| 2 | 123452484 | AX-170286251 | 0 | C | G | 0.14 | 3.24E-02 | 2.23E-01 | 2.70E+00 | 2.73E+00 | 8.84E-01 | no | | intergenic_variant | MODIFIER |
| 2 | 123458476 | AX-97821506 | 0 | T | C | 0.14 | 3.24E-02 | 2.23E-01 | 2.70E+00 | 2.73E+00 | 8.84E-01 | no | | regulatory_region_variant | MODIFIER |
| 2 | 123469465 | AX-170405733 | 0 | G | A | 0.14 | 3.24E-02 | 2.23E-01 | 2.70E+00 | 2.73E+00 | 8.84E-01 | no | | intergenic_variant | MODIFIER |
| 2 | 123470663 | AX-170064788 | 0 | A | G | 0.14 | 3.24E-02 | 2.23E-01 | 2.70E+00 | 2.73E+00 | 8.84E-01 | no | | intergenic_variant | MODIFIER |
| 2 | 123471719 | AX-170256741 | 0 | C | T | 0.14 | 3.24E-02 | 2.23E-01 | 2.70E+00 | 2.73E+00 | 8.84E-01 | no | | intergenic_variant | MODIFIER |
| 2 | 123481064 | AX-170387361 | 0 | A | G | 0.14 | 3.24E-02 | 2.23E-01 | 2.70E+00 | 2.73E+00 | 8.84E-01 | no | | intergenic_variant | MODIFIER |
| 2 | 123487070 | AX-170346932 | 0 | T | C | 0.14 | 3.24E-02 | 2.23E-01 | 2.70E+00 | 2.73E+00 | 8.84E-01 | no | | intergenic_variant | MODIFIER |
| 2 | 123526165 | AX-97839851 | 0 | G | A | 0.14 | 3.24E-02 | 2.23E-01 | 2.70E+00 | 2.73E+00 | 8.84E-01 | no | | intergenic_variant | MODIFIER |
| 2 | 123538658 | AX-170296093 | 0 | A | G | 0.14 | 3.24E-02 | 2.23E-01 | 2.70E+00 | 2.73E+00 | 8.84E-01 | no | | intergenic_variant | MODIFIER |
| 2 | 123574233 | AX-170309397 | 0 | C | G | 0.14 | 3.24E-02 | 2.23E-01 | 2.70E+00 | 2.73E+00 | 8.84E-01 | no | | intergenic_variant | MODIFIER |
| 2 | 123616176 | AX-170213698 | 0 | G | A | 0.14 | 3.24E-02 | 2.23E-01 | 2.70E+00 | 2.73E+00 | 8.84E-01 | no | | intergenic_variant | MODIFIER |
| 2 | 123618664 | AX-170018908 | 0 | G | A | 0.14 | 3.24E-02 | 2.23E-01 | 2.70E+00 | 2.73E+00 | 8.84E-01 | no | | intergenic_variant | MODIFIER |
| 2 | 123621106 | AX-170273811 | 0 | A | T | 0.14 | 3.24E-02 | 2.23E-01 | 2.70E+00 | 2.73E+00 | 8.84E-01 | no | | intergenic_variant | MODIFIER |
| 2 | 123636402 | AX-97789197 | 0 | T | C | 0.14 | 3.24E-02 | 2.23E-01 | 2.70E+00 | 2.73E+00 | 8.84E-01 | no | | intergenic_variant | MODIFIER |
| 2 | 123684884 | AX-170018912 | 0 | T | C | 0.14 | 3.24E-02 | 2.23E-01 | 2.70E+00 | 2.73E+00 | 8.84E-01 | no | | intergenic_variant | MODIFIER |
| 2 | 123708740 | AX-170247637 | 0 | G | T | 0.14 | 3.24E-02 | 2.23E-01 | 2.70E+00 | 2.73E+00 | 8.84E-01 | no | | intergenic_variant | MODIFIER |
| 2 | 123719419 | AX-170402206 | 0 | C | T | 0.14 | 3.24E-02 | 2.23E-01 | 2.70E+00 | 2.73E+00 | 8.84E-01 | no | | intergenic_variant | MODIFIER |
| 2 | 123722175 | AX-97841590 | 0 | A | G | 0.14 | 3.24E-02 | 2.23E-01 | 2.70E+00 | 2.73E+00 | 8.84E-01 | no | | intergenic_variant | MODIFIER |

| | | | | | | | | | | | | | | | |
|---|---|---|---|---|---|---|---|---|---|---|---|---|---|---|---|
| 2 | 123768099 | AX-170407217 | 0 | C | T | 0.14 | 3.24E-02 | 2.23E-01 | 2.70E+00 | 2.73E+00 | 8.84E-01 | no | | intergenic_variant | MODIFIER |
| 2 | 123774732 | AX-170218764 | 0 | A | G | 0.14 | 3.24E-02 | 2.23E-01 | 2.70E+00 | 2.73E+00 | 8.84E-01 | no | | intergenic_variant | MODIFIER |
| 2 | 123782478 | AX-97793481 | 0 | C | A | 0.14 | 3.24E-02 | 2.23E-01 | 2.70E+00 | 2.73E+00 | 8.84E-01 | no | | intergenic_variant | MODIFIER |
| 2 | 123783083 | AX-170064791 | 0 | A | G | 0.14 | 3.24E-02 | 2.23E-01 | 2.70E+00 | 2.73E+00 | 8.84E-01 | no | | intergenic_variant | MODIFIER |
| 2 | 123801255 | AX-170475563 | 0 | A | G | 0.14 | 3.24E-02 | 2.23E-01 | 2.70E+00 | 2.73E+00 | 8.84E-01 | no | | intergenic_variant | MODIFIER |
| 2 | 123802908 | AX-170155132 | 0 | C | T | 0.14 | 3.24E-02 | 2.23E-01 | 2.70E+00 | 2.73E+00 | 8.84E-01 | no | | intergenic_variant | MODIFIER |
| 2 | 123802933 | AX-170378249 | 0 | C | G | 0.14 | 3.24E-02 | 2.23E-01 | 2.70E+00 | 2.73E+00 | 8.84E-01 | no | | intergenic_variant | MODIFIER |
| 2 | 123808932 | AX-170417075 | 0 | T | G | 0.14 | 3.24E-02 | 2.23E-01 | 2.70E+00 | 2.73E+00 | 8.84E-01 | no | | intergenic_variant | MODIFIER |
| 2 | 123814211 | AX-170088739 | 0 | C | T | 0.14 | 3.24E-02 | 2.23E-01 | 2.70E+00 | 2.73E+00 | 8.84E-01 | no | | regulatory_region_variant | MODIFIER |
| 2 | 123814278 | AX-170279791 | 0 | G | C | 0.14 | 3.24E-02 | 2.23E-01 | 2.70E+00 | 2.73E+00 | 8.84E-01 | no | | regulatory_region_variant | MODIFIER |
| 2 | 123820802 | AX-170462652 | 0 | G | A | 0.14 | 3.24E-02 | 2.23E-01 | 2.70E+00 | 2.73E+00 | 8.84E-01 | no | | intergenic_variant | MODIFIER |
| 2 | 123822100 | AX-170517885 | 0 | A | G | 0.14 | 3.24E-02 | 2.23E-01 | 2.70E+00 | 2.73E+00 | 8.84E-01 | no | | intergenic_variant | MODIFIER |
| 2 | 123823057 | AX-170306699 | 0 | T | C | 0.14 | 3.24E-02 | 2.23E-01 | 2.70E+00 | 2.73E+00 | 8.84E-01 | no | | intergenic_variant | MODIFIER |
| 2 | 123832351 | AX-170504390 | 0 | A | G | 0.14 | 3.24E-02 | 2.23E-01 | 2.70E+00 | 2.73E+00 | 8.84E-01 | no | | intergenic_variant | MODIFIER |
| 2 | 123832653 | AX-170443489 | 0 | A | C | 0.14 | 3.24E-02 | 2.23E-01 | 2.70E+00 | 2.73E+00 | 8.84E-01 | no | | intergenic_variant | MODIFIER |
| 2 | 123841519 | AX-170239609 | 0 | A | T | 0.14 | 3.24E-02 | 2.23E-01 | 2.70E+00 | 2.73E+00 | 8.84E-01 | no | | intergenic_variant | MODIFIER |
| 2 | 123866617 | AX-170367543 | 0 | G | T | 0.14 | 3.24E-02 | 2.23E-01 | 2.70E+00 | 2.73E+00 | 8.84E-01 | no | | intergenic_variant | MODIFIER |
| 2 | 123874220 | AX-97839850 | 0 | C | T | 0.14 | 3.24E-02 | 2.23E-01 | 2.70E+00 | 2.73E+00 | 8.84E-01 | no | | intergenic_variant | MODIFIER |
| 2 | 123876266 | AX-170205065 | 0 | A | T | 0.14 | 3.24E-02 | 2.23E-01 | 2.70E+00 | 2.73E+00 | 8.84E-01 | no | | intergenic_variant | MODIFIER |
| 2 | 123919444 | AX-170094399 | 0 | A | G | 0.14 | 3.24E-02 | 2.23E-01 | 2.70E+00 | 2.73E+00 | 8.84E-01 | no | | intergenic_variant | MODIFIER |
| 2 | 123927257 | AX-170352452 | 0 | C | A | 0.14 | 3.24E-02 | 2.23E-01 | 2.70E+00 | 2.73E+00 | 8.84E-01 | no | | intergenic_variant | MODIFIER |
| 2 | 123927735 | AX-170160745 | 0 | G | A | 0.14 | 3.24E-02 | 2.23E-01 | 2.70E+00 | 2.73E+00 | 8.84E-01 | no | | intergenic_variant | MODIFIER |
| 2 | 123935409 | AX-97834180 | 0 | C | T | 0.14 | 3.24E-02 | 2.23E-01 | 2.70E+00 | 2.73E+00 | 8.84E-01 | no | | intergenic_variant | MODIFIER |
| 2 | 123939604 | AX-170281732 | 0 | A | G | 0.14 | 3.24E-02 | 2.23E-01 | 2.70E+00 | 2.73E+00 | 8.84E-01 | no | | intergenic_variant | MODIFIER |
| 2 | 123942206 | AX-170458740 | 0 | G | A | 0.14 | 3.24E-02 | 2.23E-01 | 2.70E+00 | 2.73E+00 | 8.84E-01 | no | | intergenic_variant | MODIFIER |
| 2 | 123967011 | AX-170118253 | 0 | G | A | 0.14 | 3.24E-02 | 2.23E-01 | 2.70E+00 | 2.73E+00 | 8.84E-01 | no | | intergenic_variant | MODIFIER |
| 2 | 123978391 | AX-97820407 | 0 | G | A | 0.14 | 3.24E-02 | 2.23E-01 | 2.70E+00 | 2.73E+00 | 8.84E-01 | no | | intergenic_variant | MODIFIER |
| 2 | 123980488 | AX-170494496 | 0 | A | C | 0.14 | 3.24E-02 | 2.23E-01 | 2.70E+00 | 2.73E+00 | 8.84E-01 | no | | intergenic_variant | MODIFIER |
| 2 | 123991958 | AX-170479074 | 0 | A | G | 0.14 | 3.24E-02 | 2.23E-01 | 2.70E+00 | 2.73E+00 | 8.84E-01 | no | | intergenic_variant | MODIFIER |
| 2 | 123992073 | AX-170371619 | 0 | A | G | 0.14 | 3.24E-02 | 2.23E-01 | 2.70E+00 | 2.73E+00 | 8.84E-01 | no | | intergenic_variant | MODIFIER |
| 2 | 124008494 | AX-97837896 | 0 | C | T | 0.14 | 3.24E-02 | 2.23E-01 | 2.70E+00 | 2.73E+00 | 8.84E-01 | no | | intergenic_variant | MODIFIER |
| 2 | 124009003 | AX-170497055 | 0 | T | C | 0.14 | 3.24E-02 | 2.23E-01 | 2.70E+00 | 2.73E+00 | 8.84E-01 | no | | intergenic_variant | MODIFIER |
| 2 | 124025498 | AX-97796858 | 0 | T | G | 0.14 | 3.24E-02 | 2.23E-01 | 2.70E+00 | 2.73E+00 | 8.84E-01 | no | | intergenic_variant | MODIFIER |
| 2 | 124048969 | AX-170396056 | 0 | C | T | 0.14 | 3.24E-02 | 2.23E-01 | 2.70E+00 | 2.73E+00 | 8.84E-01 | no | | intergenic_variant | MODIFIER |
| 2 | 124049000 | AX-170372147 | 0 | G | A | 0.14 | 3.24E-02 | 2.23E-01 | 2.70E+00 | 2.73E+00 | 8.84E-01 | no | | intergenic_variant | MODIFIER |
| 2 | 124050531 | AX-170106800 | 0 | T | C | 0.14 | 3.24E-02 | 2.23E-01 | 2.70E+00 | 2.73E+00 | 8.84E-01 | no | | intergenic_variant | MODIFIER |
| 2 | 124061654 | AX-97855438 | 0 | T | C | 0.14 | 3.24E-02 | 2.23E-01 | 2.70E+00 | 2.73E+00 | 8.84E-01 | no | | intergenic_variant | MODIFIER |
| 2 | 124064502 | AX-170407120 | 0 | T | C | 0.14 | 3.24E-02 | 2.23E-01 | 2.70E+00 | 2.73E+00 | 8.84E-01 | no | | intergenic_variant | MODIFIER |
| 2 | 124064781 | AX-170438430 | 0 | T | C | 0.14 | 3.24E-02 | 2.23E-01 | 2.70E+00 | 2.73E+00 | 8.84E-01 | no | | intergenic_variant | MODIFIER |
| 2 | 124070568 | AX-97824020 | 0 | T | G | 0.14 | 3.24E-02 | 2.23E-01 | 2.70E+00 | 2.73E+00 | 8.84E-01 | no | | intergenic_variant | MODIFIER |
| 2 | 124097430 | AX-170064798 | 0 | A | C | 0.14 | 3.24E-02 | 2.23E-01 | 2.70E+00 | 2.73E+00 | 8.84E-01 | no | Sema6d | intron_variant | MODIFIER |
| 2 | 124101985 | AX-170064799 | 0 | T | C | 0.14 | 3.24E-02 | 2.23E-01 | 2.70E+00 | 2.73E+00 | 8.84E-01 | no | Sema6d | intron_variant | MODIFIER |
| 2 | 124102440 | AX-170445961 | 0 | A | G | 0.14 | 3.24E-02 | 2.23E-01 | 2.70E+00 | 2.73E+00 | 8.84E-01 | no | Sema6d | intron_variant | MODIFIER |
| 2 | 124114126 | AX-170212250 | 0 | A | T | 0.14 | 3.24E-02 | 2.23E-01 | 2.70E+00 | 2.73E+00 | 8.84E-01 | no | Sema6d | intron_variant | MODIFIER |
| 2 | 124123146 | AX-170274673 | 0 | C | A | 0.14 | 3.24E-02 | 2.23E-01 | 2.70E+00 | 2.73E+00 | 8.84E-01 | no | Sema6d | intron_variant | MODIFIER |
| 2 | 124127225 | AX-170018917 | 0 | A | G | 0.14 | 3.24E-02 | 2.23E-01 | 2.70E+00 | 2.73E+00 | 8.84E-01 | no | Sema6d | intron_variant | MODIFIER |
| 2 | 124129935 | AX-170018913 | 0 | A | C | 0.14 | 3.24E-02 | 2.23E-01 | 2.70E+00 | 2.73E+00 | 8.84E-01 | no | Sema6d | intron_variant | MODIFIER |
| 2 | 124132374 | AX-170367675 | 0 | A | G | 0.14 | 3.24E-02 | 2.23E-01 | 2.70E+00 | 2.73E+00 | 8.84E-01 | no | Sema6d | intron_variant | MODIFIER |
| 2 | 124180321 | AX-170444179 | 0 | C | T | 0.14 | 3.24E-02 | 2.23E-01 | 2.70E+00 | 2.73E+00 | 8.84E-01 | no | Sema6d | intron_variant | MODIFIER |
| 2 | 124184232 | AX-170018916 | 0 | C | T | 0.14 | 3.24E-02 | 2.23E-01 | 2.70E+00 | 2.73E+00 | 8.84E-01 | no | Sema6d | intron_variant | MODIFIER |
| 2 | 124186048 | AX-170193710 | 0 | C | G | 0.14 | 3.24E-02 | 2.23E-01 | 2.70E+00 | 2.73E+00 | 8.84E-01 | no | Sema6d | intron_variant | MODIFIER |
| 2 | 124198232 | AX-170370087 | 0 | G | A | 0.14 | 3.24E-02 | 2.23E-01 | 2.70E+00 | 2.73E+00 | 8.84E-01 | no | Sema6d | intron_variant | MODIFIER |
| 2 | 124207480 | AX-97843884 | 0 | T | C | 0.14 | 3.24E-02 | 2.23E-01 | 2.70E+00 | 2.73E+00 | 8.84E-01 | no | Sema6d | intron_variant | MODIFIER |
| 2 | 124208829 | AX-170178753 | 0 | C | T | 0.14 | 3.24E-02 | 2.23E-01 | 2.70E+00 | 2.73E+00 | 8.84E-01 | no | Sema6d | intron_variant | MODIFIER |

| | | | | | | | | | | | | | | | |
|---|---|---|---|---|---|---|---|---|---|---|---|---|---|---|---|
| 2 | 124234423 | AX-170419614 | 0 | T | G | 0.14 | 3.24E-02 | 2.23E-01 | 2.70E+00 | 2.73E+00 | 8.84E-01 | no | Sema6d | intron_variant | MODIFIER |
| 2 | 124236010 | AX-170475165 | 0 | C | T | 0.14 | 3.24E-02 | 2.23E-01 | 2.70E+00 | 2.73E+00 | 8.84E-01 | no | Sema6d | intron_variant | MODIFIER |
| 2 | 124237467 | AX-170136096 | 0 | A | G | 0.14 | 3.24E-02 | 2.23E-01 | 2.70E+00 | 2.73E+00 | 8.84E-01 | no | Sema6d | intron_variant | MODIFIER |
| 2 | 124246775 | AX-170156244 | 0 | A | G | 0.14 | 3.24E-02 | 2.23E-01 | 2.70E+00 | 2.73E+00 | 8.84E-01 | no | Sema6d | intron_variant | MODIFIER |
| 2 | 124246911 | AX-97791185 | 0 | C | A | 0.14 | 3.24E-02 | 2.23E-01 | 2.70E+00 | 2.73E+00 | 8.84E-01 | no | Sema6d | intron_variant | MODIFIER |
| 2 | 124267893 | AX-170348749 | 0 | G | T | 0.14 | 3.24E-02 | 2.23E-01 | 2.70E+00 | 2.73E+00 | 8.84E-01 | no | Sema6d | intron_variant | MODIFIER |
| 2 | 124283556 | AX-170064802 | 0 | T | C | 0.14 | 3.24E-02 | 2.23E-01 | 2.70E+00 | 2.73E+00 | 8.84E-01 | no | Sema6d | intron_variant | MODIFIER |
| 2 | 124285063 | AX-97846131 | 0 | A | G | 0.14 | 3.24E-02 | 2.23E-01 | 2.70E+00 | 2.73E+00 | 8.84E-01 | no | Sema6d | intron_variant | MODIFIER |
| 2 | 124287712 | AX-170337309 | 0 | C | T | 0.14 | 3.24E-02 | 2.23E-01 | 2.70E+00 | 2.73E+00 | 8.84E-01 | no | Sema6d | intron_variant | MODIFIER |
| 2 | 124287891 | AX-170179722 | 0 | C | T | 0.14 | 3.24E-02 | 2.23E-01 | 2.70E+00 | 2.73E+00 | 8.84E-01 | no | Sema6d | intron_variant | MODIFIER |
| 2 | 124308117 | AX-170110392 | 0 | C | T | 0.14 | 3.24E-02 | 2.23E-01 | 2.70E+00 | 2.73E+00 | 8.84E-01 | no | Sema6d | intron_variant | MODIFIER |
| 2 | 124323484 | AX-170151265 | 0 | G | C | 0.14 | 3.24E-02 | 2.23E-01 | 2.70E+00 | 2.73E+00 | 8.84E-01 | no | Sema6d | intron_variant | MODIFIER |
| 2 | 124327319 | AX-170271216 | 0 | T | C | 0.14 | 3.24E-02 | 2.23E-01 | 2.70E+00 | 2.73E+00 | 8.84E-01 | no | Sema6d | intron_variant | MODIFIER |
| 2 | 124337428 | AX-170341430 | 0 | G | A | 0.14 | 3.24E-02 | 2.23E-01 | 2.70E+00 | 2.73E+00 | 8.84E-01 | no | Sema6d | intron_variant | MODIFIER |
| 2 | 124360938 | AX-170512606 | 0 | T | C | 0.14 | 3.24E-02 | 2.23E-01 | 2.70E+00 | 2.73E+00 | 8.84E-01 | no | Sema6d | intron_variant | MODIFIER |
| 2 | 124373893 | AX-170194220 | 0 | T | A | 0.14 | 3.24E-02 | 2.23E-01 | 2.70E+00 | 2.73E+00 | 8.84E-01 | no | Sema6d | intron_variant | MODIFIER |
| 2 | 124393237 | AX-170448316 | 0 | G | A | 0.14 | 3.24E-02 | 2.23E-01 | 2.70E+00 | 2.73E+00 | 8.84E-01 | no | Sema6d | intron_variant | MODIFIER |
| 2 | 124393435 | AX-170320110 | 0 | A | G | 0.14 | 3.24E-02 | 2.23E-01 | 2.70E+00 | 2.73E+00 | 8.84E-01 | no | Sema6d | intron_variant | MODIFIER |
| 2 | 124399472 | AX-170516948 | 0 | C | T | 0.14 | 3.24E-02 | 2.23E-01 | 2.70E+00 | 2.73E+00 | 8.84E-01 | no | Sema6d | intron_variant | MODIFIER |
| 2 | 124404499 | AX-170282667 | 0 | G | A | 0.14 | 3.24E-02 | 2.23E-01 | 2.70E+00 | 2.73E+00 | 8.84E-01 | no | Sema6d | intron_variant | MODIFIER |
| 2 | 124408616 | AX-170399017 | 0 | C | T | 0.14 | 3.24E-02 | 2.23E-01 | 2.70E+00 | 2.73E+00 | 8.84E-01 | no | Sema6d | intron_variant | MODIFIER |
| 2 | 124409286 | AX-170188828 | 0 | A | G | 0.14 | 3.24E-02 | 2.23E-01 | 2.70E+00 | 2.73E+00 | 8.84E-01 | no | Sema6d | intron_variant | MODIFIER |
| 2 | 124412165 | AX-170398976 | 0 | G | A | 0.14 | 3.24E-02 | 2.23E-01 | 2.70E+00 | 2.73E+00 | 8.84E-01 | no | Sema6d | intron_variant | MODIFIER |
| 2 | 124413474 | AX-170064804 | 0 | G | C | 0.14 | 3.24E-02 | 2.23E-01 | 2.70E+00 | 2.73E+00 | 8.84E-01 | no | Sema6d | intron_variant | MODIFIER |
| 2 | 124413834 | AX-170449492 | 0 | C | T | 0.14 | 3.24E-02 | 2.23E-01 | 2.70E+00 | 2.73E+00 | 8.84E-01 | no | Sema6d | intron_variant | MODIFIER |
| 2 | 124429973 | AX-170119419 | 0 | C | T | 0.14 | 3.24E-02 | 2.23E-01 | 2.70E+00 | 2.73E+00 | 8.84E-01 | no | Sema6d | intron_variant | MODIFIER |
| 2 | 124449153 | AX-170188997 | 0 | T | C | 0.14 | 3.24E-02 | 2.23E-01 | 2.70E+00 | 2.73E+00 | 8.84E-01 | no | Sema6d | intron_variant | MODIFIER |
| 2 | 124461889 | AX-170330632 | 0 | C | T | 0.14 | 3.24E-02 | 2.23E-01 | 2.70E+00 | 2.73E+00 | 8.84E-01 | no | Sema6d | intron_variant | MODIFIER |
| 2 | 124467257 | AX-170321754 | 0 | G | C | 0.14 | 3.24E-02 | 2.23E-01 | 2.70E+00 | 2.73E+00 | 8.84E-01 | no | Sema6d | intron_variant | MODIFIER |
| 2 | 124472081 | AX-170448209 | 0 | T | C | 0.14 | 3.24E-02 | 2.23E-01 | 2.70E+00 | 2.73E+00 | 8.84E-01 | no | Sema6d | intron_variant | MODIFIER |
| 2 | 124486758 | AX-170476390 | 0 | G | A | 0.14 | 3.24E-02 | 2.23E-01 | 2.70E+00 | 2.73E+00 | 8.84E-01 | no | Sema6d | intron_variant | MODIFIER |
| 2 | 124514224 | AX-170244023 | 0 | A | C | 0.14 | 3.24E-02 | 2.23E-01 | 2.70E+00 | 2.73E+00 | 8.84E-01 | no | Sema6d | intron_variant | MODIFIER |
| 2 | 124520293 | AX-170296526 | 0 | G | A | 0.14 | 3.24E-02 | 2.23E-01 | 2.70E+00 | 2.73E+00 | 8.84E-01 | no | Sema6d | intron_variant | MODIFIER |
| 2 | 124520470 | AX-170197192 | 0 | G | A | 0.14 | 3.24E-02 | 2.23E-01 | 2.70E+00 | 2.73E+00 | 8.84E-01 | no | Sema6d | intron_variant | MODIFIER |
| 2 | 124522760 | AX-170397644 | 0 | G | A | 0.14 | 3.24E-02 | 2.23E-01 | 2.70E+00 | 2.73E+00 | 8.84E-01 | no | Sema6d | intron_variant | MODIFIER |
| 2 | 124524315 | AX-170307507 | 0 | G | A | 0.14 | 3.24E-02 | 2.23E-01 | 2.70E+00 | 2.73E+00 | 8.84E-01 | no | Sema6d | intron_variant | MODIFIER |
| 2 | 124529362 | AX-170458160 | 0 | A | G | 0.14 | 3.24E-02 | 2.23E-01 | 2.70E+00 | 2.73E+00 | 8.84E-01 | no | Sema6d | intron_variant | MODIFIER |
| 2 | 124538655 | AX-170424602 | 0 | A | G | 0.14 | 3.24E-02 | 2.23E-01 | 2.70E+00 | 2.73E+00 | 8.84E-01 | no | Sema6d | intron_variant | MODIFIER |
| 2 | 124538859 | AX-170134056 | 0 | C | T | 0.14 | 3.24E-02 | 2.23E-01 | 2.70E+00 | 2.73E+00 | 8.84E-01 | no | Sema6d | intron_variant | MODIFIER |
| 2 | 124539704 | AX-170217054 | 0 | A | C | 0.14 | 3.24E-02 | 2.23E-01 | 2.70E+00 | 2.73E+00 | 8.84E-01 | no | Sema6d | intron_variant | MODIFIER |
| 2 | 124541918 | AX-170461780 | 0 | G | A | 0.14 | 3.24E-02 | 2.23E-01 | 2.70E+00 | 2.73E+00 | 8.84E-01 | no | Sema6d | intron_variant | MODIFIER |
| 2 | 124560467 | AX-170486263 | 0 | G | A | 0.14 | 3.24E-02 | 2.23E-01 | 2.70E+00 | 2.73E+00 | 8.84E-01 | no | Sema6d | intron_variant | MODIFIER |
| 2 | 124564038 | AX-170364201 | 0 | T | C | 0.14 | 3.24E-02 | 2.23E-01 | 2.70E+00 | 2.73E+00 | 8.84E-01 | no | Sema6d | intron_variant | MODIFIER |
| 2 | 124564189 | AX-170136688 | 0 | T | C | 0.14 | 3.24E-02 | 2.23E-01 | 2.70E+00 | 2.73E+00 | 8.84E-01 | no | Sema6d | intron_variant | MODIFIER |
| 2 | 124579894 | AX-97809985 | 0 | A | C | 0.14 | 3.24E-02 | 2.23E-01 | 2.70E+00 | 2.73E+00 | 8.84E-01 | no | Sema6d | intron_variant | MODIFIER |
| 2 | 124582074 | AX-170064807 | 0 | T | C | 0.14 | 3.24E-02 | 2.23E-01 | 2.70E+00 | 2.73E+00 | 8.84E-01 | no | Sema6d | intron_variant | MODIFIER |
| 2 | 124589601 | AX-170368994 | 0 | G | A | 0.14 | 3.24E-02 | 2.23E-01 | 2.70E+00 | 2.73E+00 | 8.84E-01 | no | Sema6d | intron_variant | MODIFIER |
| 2 | 124598560 | AX-170385999 | 0 | G | A | 0.14 | 3.24E-02 | 2.23E-01 | 2.70E+00 | 2.73E+00 | 8.84E-01 | no | Sema6d | intron_variant | MODIFIER |
| 2 | 124614130 | AX-170277789 | 0 | T | G | 0.14 | 3.24E-02 | 2.23E-01 | 2.70E+00 | 2.73E+00 | 8.84E-01 | no | Sema6d | intron_variant | MODIFIER |
| 2 | 124615844 | AX-170088740 | 0 | T | A | 0.14 | 3.24E-02 | 2.23E-01 | 2.70E+00 | 2.73E+00 | 8.84E-01 | no | Sema6d | intron_variant | MODIFIER |
| 2 | 124636167 | AX-170455722 | 0 | A | C | 0.14 | 3.24E-02 | 2.23E-01 | 2.70E+00 | 2.73E+00 | 8.84E-01 | no | Sema6d | intron_variant | MODIFIER |
| 2 | 124644869 | AX-170018924 | 0 | C | T | 0.14 | 3.24E-02 | 2.23E-01 | 2.70E+00 | 2.73E+00 | 8.84E-01 | no | Sema6d | intron_variant | MODIFIER |
| 2 | 124650891 | AX-170018926 | 0 | C | T | 0.14 | 3.24E-02 | 2.23E-01 | 2.70E+00 | 2.73E+00 | 8.84E-01 | no | Sema6d | intron_variant | MODIFIER |
| 2 | 124660154 | AX-170180221 | 0 | C | T | 0.14 | 3.24E-02 | 2.23E-01 | 2.70E+00 | 2.73E+00 | 8.84E-01 | no | Sema6d | intron_variant | MODIFIER |

| | | | | | | | | | | | | | | | |
|---|---|---|---|---|---|---|---|---|---|---|---|---|---|---|---|
| 2 | 124661381 | AX-170383467 | 0 | G | A | 0.14 | 3.24E-02 | 2.23E-01 | 2.70E+00 | 2.73E+00 | 8.84E-01 | no | Sema6d | intron_variant | MODIFIER |
| 2 | 124664727 | AX-170235976 | 0 | C | T | 0.14 | 3.24E-02 | 2.23E-01 | 2.70E+00 | 2.73E+00 | 8.84E-01 | no | Sema6d | synonymous_variant | LOW |
| 2 | 124668623 | AX-170426138 | 0 | T | G | 0.14 | 3.24E-02 | 2.23E-01 | 2.70E+00 | 2.73E+00 | 8.84E-01 | no | Sema6d | downstream_gene_variant | MODIFIER |
| 2 | 124697202 | AX-168406723 | 0 | C | T | 0.14 | 3.24E-02 | 2.23E-01 | 2.70E+00 | 2.73E+00 | 8.84E-01 | no | | intergenic_variant | MODIFIER |
| 2 | 124704213 | AX-170513038 | 0 | G | A | 0.14 | 3.24E-02 | 2.23E-01 | 2.70E+00 | 2.73E+00 | 8.84E-01 | no | Gm13994 | upstream_gene_variant | MODIFIER |
| 2 | 124708979 | AX-170018925 | 0 | G | A | 0.14 | 3.24E-02 | 2.23E-01 | 2.70E+00 | 2.73E+00 | 8.84E-01 | no | Gm13994 | non_coding_transcript_exon_variant | MODIFIER |
| 2 | 124719656 | AX-170285742 | 0 | A | G | 0.14 | 3.24E-02 | 2.23E-01 | 2.70E+00 | 2.73E+00 | 8.84E-01 | no | | regulatory_region_variant | MODIFIER |
| 2 | 124721125 | AX-170088741 | 0 | A | T | 0.14 | 3.24E-02 | 2.23E-01 | 2.70E+00 | 2.73E+00 | 8.84E-01 | no | | regulatory_region_variant | MODIFIER |
| 2 | 124721137 | AX-170115190 | 0 | C | T | 0.14 | 3.24E-02 | 2.23E-01 | 2.70E+00 | 2.73E+00 | 8.84E-01 | no | | regulatory_region_variant | MODIFIER |
| 2 | 124722561 | AX-170344592 | 0 | T | C | 0.14 | 3.24E-02 | 2.23E-01 | 2.70E+00 | 2.73E+00 | 8.84E-01 | no | | intergenic_variant | MODIFIER |
| 2 | 124768241 | AX-170018919 | 0 | C | T | 0.14 | 3.24E-02 | 2.23E-01 | 2.70E+00 | 2.73E+00 | 8.84E-01 | no | | intergenic_variant | MODIFIER |
| 2 | 124770315 | AX-170018920 | 0 | T | C | 0.14 | 3.24E-02 | 2.23E-01 | 2.70E+00 | 2.73E+00 | 8.84E-01 | no | | intergenic_variant | MODIFIER |
| 2 | 124772615 | AX-97782633 | 0 | A | G | 0.14 | 3.24E-02 | 2.23E-01 | 2.70E+00 | 2.73E+00 | 8.84E-01 | no | | regulatory_region_variant | MODIFIER |
| 2 | 124789943 | AX-170222493 | 0 | T | C | 0.14 | 3.24E-02 | 2.23E-01 | 2.70E+00 | 2.73E+00 | 8.84E-01 | no | | intergenic_variant | MODIFIER |
| 2 | 124790090 | AX-170419168 | 0 | G | T | 0.14 | 3.24E-02 | 2.23E-01 | 2.70E+00 | 2.73E+00 | 8.84E-01 | no | | intergenic_variant | MODIFIER |
| 2 | 124790102 | AX-170221913 | 0 | A | G | 0.14 | 3.24E-02 | 2.23E-01 | 2.70E+00 | 2.73E+00 | 8.84E-01 | no | | intergenic_variant | MODIFIER |
| 2 | 124790238 | AX-170018927 | 0 | C | T | 0.14 | 3.24E-02 | 2.23E-01 | 2.70E+00 | 2.73E+00 | 8.84E-01 | no | | intergenic_variant | MODIFIER |
| 2 | 124794034 | AX-170064808 | 0 | A | G | 0.14 | 3.24E-02 | 2.23E-01 | 2.70E+00 | 2.73E+00 | 8.84E-01 | no | | intergenic_variant | MODIFIER |
| 2 | 124796647 | AX-170354266 | 0 | G | A | 0.14 | 3.24E-02 | 2.23E-01 | 2.70E+00 | 2.73E+00 | 8.84E-01 | no | | intergenic_variant | MODIFIER |
| 2 | 124798124 | AX-170161926 | 0 | A | G | 0.14 | 3.24E-02 | 2.23E-01 | 2.70E+00 | 2.73E+00 | 8.84E-01 | no | | intergenic_variant | MODIFIER |
| 2 | 124812217 | AX-170465919 | 0 | A | G | 0.14 | 3.24E-02 | 2.23E-01 | 2.70E+00 | 2.73E+00 | 8.84E-01 | no | | intergenic_variant | MODIFIER |
| 2 | 124812363 | AX-170258704 | 0 | T | C | 0.14 | 3.24E-02 | 2.23E-01 | 2.70E+00 | 2.73E+00 | 8.84E-01 | no | | intergenic_variant | MODIFIER |
| 2 | 124847764 | AX-170018929 | 0 | A | G | 0.14 | 3.24E-02 | 2.23E-01 | 2.70E+00 | 2.73E+00 | 8.84E-01 | no | | intergenic_variant | MODIFIER |
| 2 | 124856238 | AX-170111670 | 0 | A | G | 0.14 | 3.24E-02 | 2.23E-01 | 2.70E+00 | 2.73E+00 | 8.84E-01 | no | | intergenic_variant | MODIFIER |
| 2 | 124889223 | AX-170443573 | 0 | C | T | 0.14 | 3.24E-02 | 2.23E-01 | 2.70E+00 | 2.73E+00 | 8.84E-01 | no | | intergenic_variant | MODIFIER |
| 2 | 124903638 | AX-170113767 | 0 | T | C | 0.14 | 3.24E-02 | 2.23E-01 | 2.70E+00 | 2.73E+00 | 8.84E-01 | no | | intergenic_variant | MODIFIER |
| 2 | 124905666 | AX-170018922 | 0 | C | T | 0.14 | 3.24E-02 | 2.23E-01 | 2.70E+00 | 2.73E+00 | 8.84E-01 | no | | intergenic_variant | MODIFIER |
| 2 | 124921105 | AX-170126083 | 0 | C | A | 0.14 | 3.24E-02 | 2.23E-01 | 2.70E+00 | 2.73E+00 | 8.84E-01 | no | | regulatory_region_variant | MODIFIER |
| 2 | 124924748 | AX-170294299 | 0 | T | G | 0.14 | 3.24E-02 | 2.23E-01 | 2.70E+00 | 2.73E+00 | 8.84E-01 | no | | intergenic_variant | MODIFIER |
| 2 | 124930153 | AX-97842578 | 0 | T | C | 0.14 | 3.24E-02 | 2.23E-01 | 2.70E+00 | 2.73E+00 | 8.84E-01 | no | | intergenic_variant | MODIFIER |
| 2 | 124930205 | AX-170487283 | 0 | C | T | 0.14 | 3.24E-02 | 2.23E-01 | 2.70E+00 | 2.73E+00 | 8.84E-01 | no | | intergenic_variant | MODIFIER |
| 2 | 124930936 | AX-170362846 | 0 | G | T | 0.14 | 3.24E-02 | 2.23E-01 | 2.70E+00 | 2.73E+00 | 8.84E-01 | no | | intergenic_variant | MODIFIER |
| 2 | 124933794 | AX-170101365 | 0 | T | C | 0.14 | 3.24E-02 | 2.23E-01 | 2.70E+00 | 2.73E+00 | 8.84E-01 | no | | intergenic_variant | MODIFIER |
| 2 | 124933930 | AX-170529005 | 0 | G | A | 0.14 | 3.24E-02 | 2.23E-01 | 2.70E+00 | 2.73E+00 | 8.84E-01 | no | | intergenic_variant | MODIFIER |
| 2 | 124936229 | AX-170454520 | 0 | T | C | 0.14 | 3.24E-02 | 2.23E-01 | 2.70E+00 | 2.73E+00 | 8.84E-01 | no | | intergenic_variant | MODIFIER |
| 2 | 124937180 | AX-170398922 | 0 | T | C | 0.14 | 3.24E-02 | 2.23E-01 | 2.70E+00 | 2.73E+00 | 8.84E-01 | no | | intergenic_variant | MODIFIER |
| 2 | 124952716 | AX-170018928 | 0 | G | A | 0.14 | 3.24E-02 | 2.23E-01 | 2.70E+00 | 2.73E+00 | 8.84E-01 | no | | intergenic_variant | MODIFIER |
| 2 | 124959982 | AX-170226064 | 0 | A | G | 0.14 | 3.24E-02 | 2.23E-01 | 2.70E+00 | 2.73E+00 | 8.84E-01 | no | | intergenic_variant | MODIFIER |
| 2 | 124969195 | AX-170217143 | 0 | C | T | 0.14 | 3.24E-02 | 2.23E-01 | 2.70E+00 | 2.73E+00 | 8.84E-01 | no | | intergenic_variant | MODIFIER |
| 2 | 124969487 | AX-97840518 | 0 | G | T | 0.14 | 3.24E-02 | 2.23E-01 | 2.70E+00 | 2.73E+00 | 8.84E-01 | no | | intergenic_variant | MODIFIER |
| 2 | 124973549 | AX-170064813 | 0 | G | A | 0.14 | 3.24E-02 | 2.23E-01 | 2.70E+00 | 2.73E+00 | 8.84E-01 | no | | intergenic_variant | MODIFIER |
| 2 | 124974064 | AX-168377350 | 0 | C | T | 0.14 | 3.24E-02 | 2.23E-01 | 2.70E+00 | 2.73E+00 | 8.84E-01 | no | | intergenic_variant | MODIFIER |
| 2 | 124974448 | AX-170064810 | 0 | C | T | 0.14 | 3.24E-02 | 2.23E-01 | 2.70E+00 | 2.73E+00 | 8.84E-01 | no | | intergenic_variant | MODIFIER |
| 2 | 124975735 | AX-168392062 | 0 | G | T | 0.14 | 3.24E-02 | 2.23E-01 | 2.70E+00 | 2.73E+00 | 8.84E-01 | no | | intergenic_variant | MODIFIER |
| 2 | 125046569 | AX-170162927 | 0 | A | G | 0.14 | 3.24E-02 | 2.23E-01 | 2.70E+00 | 2.73E+00 | 8.84E-01 | no | | intergenic_variant | MODIFIER |
| 2 | 125047724 | AX-170331614 | 0 | A | C | 0.14 | 3.24E-02 | 2.23E-01 | 2.70E+00 | 2.73E+00 | 8.84E-01 | no | | intergenic_variant | MODIFIER |
| 2 | 125048109 | AX-170137680 | 0 | A | G | 0.14 | 3.24E-02 | 2.23E-01 | 2.70E+00 | 2.73E+00 | 8.84E-01 | no | | intergenic_variant | MODIFIER |
| 2 | 125052078 | AX-170099535 | 0 | G | A | 0.14 | 3.24E-02 | 2.23E-01 | 2.70E+00 | 2.73E+00 | 8.84E-01 | no | | intergenic_variant | MODIFIER |
| 2 | 125063148 | AX-97845528 | 0 | G | A | 0.14 | 3.24E-02 | 2.23E-01 | 2.70E+00 | 2.73E+00 | 8.84E-01 | no | Slc24a5 | upstream_gene_variant | MODIFIER |
| 2 | 125068654 | AX-170319524 | 0 | T | A | 0.14 | 3.24E-02 | 2.23E-01 | 2.70E+00 | 2.73E+00 | 8.84E-01 | no | Slc24a5 | intron_variant | MODIFIER |
| 2 | 125071105 | AX-170155005 | 0 | C | A | 0.14 | 3.24E-02 | 2.23E-01 | 2.70E+00 | 2.73E+00 | 8.84E-01 | no | Slc24a5 | intron_variant | MODIFIER |
| 2 | 125077497 | AX-170303416 | 0 | C | T | 0.14 | 3.24E-02 | 2.23E-01 | 2.70E+00 | 2.73E+00 | 8.84E-01 | no | Slc24a5 | intron_variant | MODIFIER |
| 2 | 125077542 | AX-170202035 | 0 | C | A | 0.14 | 3.24E-02 | 2.23E-01 | 2.70E+00 | 2.73E+00 | 8.84E-01 | no | Slc24a5 | intron_variant | MODIFIER |
| 2 | 125078642 | AX-170261451 | 0 | T | G | 0.14 | 3.24E-02 | 2.23E-01 | 2.70E+00 | 2.73E+00 | 8.84E-01 | no | Slc24a5 | intron_variant | MODIFIER |

| | | | | | | | | | | | | | | | |
|---|---|---|---|---|---|---|---|---|---|---|---|---|---|---|---|
| 2 | 125083949 | AX-170173039 | 0 | A | G | 0.14 | 3.24E-02 | 2.23E-01 | 2.70E+00 | 2.73E+00 | 8.84E-01 | no | Myef2 | downstream_gene_variant | MODIFIER |
| 2 | 125099403 | AX-170243159 | 0 | T | A | 0.14 | 3.24E-02 | 2.23E-01 | 2.70E+00 | 2.73E+00 | 8.84E-01 | no | Myef2 | intron_variant | MODIFIER |
| 2 | 125107287 | AX-170272356 | 0 | T | C | 0.14 | 3.24E-02 | 2.23E-01 | 2.70E+00 | 2.73E+00 | 8.84E-01 | no | Myef2 | intron_variant | MODIFIER |
| 2 | 125109113 | AX-97812290 | 0 | A | G | 0.14 | 3.24E-02 | 2.23E-01 | 2.70E+00 | 2.73E+00 | 8.84E-01 | no | Myef2 | intron_variant | MODIFIER |
| 2 | 125119856 | AX-170457898 | 0 | G | C | 0.14 | 3.24E-02 | 2.23E-01 | 2.70E+00 | 2.73E+00 | 8.84E-01 | no | Myef2 | intron_variant | MODIFIER |
| 2 | 125126773 | AX-170166199 | 0 | A | T | 0.14 | 3.24E-02 | 2.23E-01 | 2.70E+00 | 2.73E+00 | 8.84E-01 | no | Myef2 | upstream_gene_variant | MODIFIER |
| 2 | 125130796 | AX-170356699 | 0 | C | T | 0.14 | 3.24E-02 | 2.23E-01 | 2.70E+00 | 2.73E+00 | 8.84E-01 | no | Gm14004 | intron_variant,non_coding_transcript_variant | MODIFIER |
| 2 | 125130892 | AX-97832830 | 0 | C | T | 0.14 | 3.24E-02 | 2.23E-01 | 2.70E+00 | 2.73E+00 | 8.84E-01 | no | Gm14004 | intron_variant,non_coding_transcript_variant | MODIFIER |
| 2 | 125132332 | AX-170342241 | 0 | G | A | 0.14 | 3.24E-02 | 2.23E-01 | 2.70E+00 | 2.73E+00 | 8.84E-01 | no | Ctxn2 | upstream_gene_variant | MODIFIER |
| 2 | 125135637 | AX-170136789 | 0 | A | G | 0.14 | 3.24E-02 | 2.23E-01 | 2.70E+00 | 2.73E+00 | 8.84E-01 | no | Ctxn2 | upstream_gene_variant | MODIFIER |
| 2 | 125171378 | AX-170215803 | 0 | T | C | 0.14 | 3.24E-02 | 2.23E-01 | 2.70E+00 | 2.73E+00 | 8.84E-01 | no | Slc12a1 | intron_variant | MODIFIER |
| 2 | 125176289 | AX-170018941 | 0 | A | C | 0.14 | 3.24E-02 | 2.23E-01 | 2.70E+00 | 2.73E+00 | 8.84E-01 | no | Slc12a1 | intron_variant | MODIFIER |
| 2 | 125176623 | AX-170018937 | 0 | G | A | 0.14 | 3.24E-02 | 2.23E-01 | 2.70E+00 | 2.73E+00 | 8.84E-01 | no | Slc12a1 | intron_variant | MODIFIER |
| 2 | 125188778 | AX-170486983 | 0 | A | C | 0.14 | 3.24E-02 | 2.23E-01 | 2.70E+00 | 2.73E+00 | 8.84E-01 | no | Slc12a1 | intron_variant | MODIFIER |
| 2 | 125206952 | AX-170018932 | 0 | T | C | 0.14 | 3.24E-02 | 2.23E-01 | 2.70E+00 | 2.73E+00 | 8.84E-01 | no | Slc12a1 | intron_variant | MODIFIER |
| 2 | 125207012 | AX-170213907 | 0 | T | C | 0.14 | 3.24E-02 | 2.23E-01 | 2.70E+00 | 2.73E+00 | 8.84E-01 | no | Slc12a1 | intron_variant | MODIFIER |
| 2 | 125212766 | AX-170098223 | 0 | G | A | 0.14 | 3.24E-02 | 2.23E-01 | 2.70E+00 | 2.73E+00 | 8.84E-01 | no | Slc12a1 | intron_variant | MODIFIER |
| 2 | 125213311 | AX-97829445 | 0 | T | C | 0.14 | 3.24E-02 | 2.23E-01 | 2.70E+00 | 2.73E+00 | 8.84E-01 | no | Slc12a1 | intron_variant | MODIFIER |
| 2 | 125216011 | AX-170368846 | 0 | A | G | 0.14 | 3.24E-02 | 2.23E-01 | 2.70E+00 | 2.73E+00 | 8.84E-01 | no | Slc12a1 | intron_variant | MODIFIER |
| 2 | 125259468 | AX-170225975 | 0 | A | G | 0.14 | 3.24E-02 | 2.23E-01 | 2.70E+00 | 2.73E+00 | 8.84E-01 | no | Dut | downstream_gene_variant | MODIFIER |
| 2 | 125260366 | AX-97857318 | 0 | T | G | 0.14 | 3.24E-02 | 2.23E-01 | 2.70E+00 | 2.73E+00 | 8.84E-01 | no | Dut | downstream_gene_variant | MODIFIER |
| 2 | 125261138 | AX-170154670 | 0 | T | C | 0.14 | 3.24E-02 | 2.23E-01 | 2.70E+00 | 2.73E+00 | 8.84E-01 | no | Dut | downstream_gene_variant | MODIFIER |
| 2 | 125264016 | AX-170506151 | 0 | A | T | 0.14 | 3.24E-02 | 2.23E-01 | 2.70E+00 | 2.73E+00 | 8.84E-01 | no | | regulatory_region_variant | MODIFIER |
| 2 | 125267938 | AX-170377388 | 0 | C | A | 0.14 | 3.24E-02 | 2.23E-01 | 2.70E+00 | 2.73E+00 | 8.84E-01 | no | | intergenic_variant | MODIFIER |
| 2 | 125288667 | AX-97795428 | 0 | C | T | 0.14 | 3.24E-02 | 2.23E-01 | 2.70E+00 | 2.73E+00 | 8.84E-01 | no | Gm14003 | upstream_gene_variant | MODIFIER |
| 2 | 125308390 | AX-170332540 | 0 | A | G | 0.14 | 3.24E-02 | 2.23E-01 | 2.70E+00 | 2.73E+00 | 8.84E-01 | no | Fbn1 | intron_variant | MODIFIER |
| 2 | 125560241 | AX-97839876 | 0 | A | G | 0.14 | 3.24E-02 | 2.23E-01 | 2.70E+00 | 2.73E+00 | 8.84E-01 | no | Cep152 | downstream_gene_variant | MODIFIER |
| 2 | 125694884 | AX-170100956 | 0 | C | T | 0.14 | 3.24E-02 | 2.23E-01 | 2.70E+00 | 2.73E+00 | 8.84E-01 | no | Shc4 | missense_variant | MODERATE |
| 2 | 125697053 | AX-170355256 | 0 | T | C | 0.14 | 3.24E-02 | 2.23E-01 | 2.70E+00 | 2.73E+00 | 8.84E-01 | no | Shc4 | intron_variant | MODIFIER |
| 2 | 125698737 | AX-170283435 | 0 | G | A | 0.14 | 3.24E-02 | 2.23E-01 | 2.70E+00 | 2.73E+00 | 8.84E-01 | no | Shc4 | intron_variant | MODIFIER |
| 2 | 125707738 | AX-170346810 | 0 | G | A | 0.14 | 3.24E-02 | 2.23E-01 | 2.70E+00 | 2.73E+00 | 8.84E-01 | no | Shc4 | intron_variant | MODIFIER |
| 2 | 125757762 | AX-170064825 | 0 | A | G | 0.14 | 3.24E-02 | 2.23E-01 | 2.70E+00 | 2.73E+00 | 8.84E-01 | no | Secisbp2l | intron_variant | MODIFIER |
| 2 | 125787290 | AX-170064828 | 0 | C | G | 0.14 | 3.24E-02 | 2.23E-01 | 2.70E+00 | 2.73E+00 | 8.84E-01 | no | Secisbp2l | upstream_gene_variant | MODIFIER |
| 2 | 125809286 | AX-170267330 | 0 | T | G | 0.14 | 3.24E-02 | 2.23E-01 | 2.70E+00 | 2.73E+00 | 8.84E-01 | no | | intergenic_variant | MODIFIER |
| 2 | 125823099 | AX-170122821 | 0 | T | C | 0.14 | 3.24E-02 | 2.23E-01 | 2.70E+00 | 2.73E+00 | 8.84E-01 | no | | intergenic_variant | MODIFIER |
| 2 | 125835176 | AX-170378369 | 0 | A | G | 0.14 | 3.24E-02 | 2.23E-01 | 2.70E+00 | 2.73E+00 | 8.84E-01 | no | Cops2 | intron_variant | MODIFIER |
| 2 | 125839552 | AX-170117791 | 0 | T | G | 0.14 | 3.24E-02 | 2.23E-01 | 2.70E+00 | 2.73E+00 | 8.84E-01 | no | Cops2 | intron_variant | MODIFIER |
| 2 | 125970467 | AX-170232564 | 0 | C | T | 0.14 | 3.24E-02 | 2.23E-01 | 2.70E+00 | 2.73E+00 | 8.84E-01 | no | Galk2 | intron_variant | MODIFIER |
| 2 | 125973245 | AX-170359295 | 0 | C | A | 0.14 | 3.24E-02 | 2.23E-01 | 2.70E+00 | 2.73E+00 | 8.84E-01 | no | Galk2 | intron_variant | MODIFIER |
| 2 | 125988290 | AX-97834134 | 0 | C | T | 0.14 | 3.24E-02 | 2.23E-01 | 2.70E+00 | 2.73E+00 | 8.84E-01 | no | Galk2 | downstream_gene_variant | MODIFIER |
| 2 | 125988341 | AX-170064830 | 0 | G | C | 0.14 | 3.24E-02 | 2.23E-01 | 2.70E+00 | 2.73E+00 | 8.84E-01 | no | Galk2 | downstream_gene_variant | MODIFIER |
| 2 | 125989020 | AX-170064831 | 0 | A | G | 0.14 | 3.24E-02 | 2.23E-01 | 2.70E+00 | 2.73E+00 | 8.84E-01 | no | Galk2 | downstream_gene_variant | MODIFIER |
| 2 | 125990198 | AX-170064832 | 0 | C | T | 0.14 | 3.24E-02 | 2.23E-01 | 2.70E+00 | 2.73E+00 | 8.84E-01 | no | Fam227b | intron_variant | MODIFIER |
| 2 | 126035538 | AX-170198424 | 0 | C | G | 0.14 | 3.24E-02 | 2.23E-01 | 2.70E+00 | 2.73E+00 | 8.84E-01 | no | Fgf7 | 5_prime_UTR_variant | MODIFIER |
| 2 | 126035762 | AX-170196288 | 0 | G | A | 0.14 | 3.24E-02 | 2.23E-01 | 2.70E+00 | 2.73E+00 | 8.84E-01 | no | Fgf7 | synonymous_variant | LOW |
| 2 | 126036225 | AX-170450996 | 0 | C | A | 0.14 | 3.24E-02 | 2.23E-01 | 2.70E+00 | 2.73E+00 | 8.84E-01 | no | Fgf7 | intron_variant | MODIFIER |
| 2 | 126041717 | AX-170341847 | 0 | T | C | 0.14 | 3.24E-02 | 2.23E-01 | 2.70E+00 | 2.73E+00 | 8.84E-01 | no | Fgf7 | intron_variant | MODIFIER |
| 2 | 126054418 | AX-170088742 | 0 | G | A | 0.14 | 3.24E-02 | 2.23E-01 | 2.70E+00 | 2.73E+00 | 8.84E-01 | no | Fgf7 | intron_variant | MODIFIER |
| 2 | 126117886 | AX-97781201 | 0 | C | T | 0.14 | 3.24E-02 | 2.23E-01 | 2.70E+00 | 2.73E+00 | 8.84E-01 | no | Fam227b | intron_variant | MODIFIER |
| 2 | 126169633 | AX-97823788 | 0 | T | C | 0.14 | 3.24E-02 | 2.23E-01 | 2.70E+00 | 2.73E+00 | 8.84E-01 | no | Dtwd1 | downstream_gene_variant | MODIFIER |
| 2 | 126173728 | AX-170470247 | 0 | C | T | 0.14 | 3.24E-02 | 2.23E-01 | 2.70E+00 | 2.73E+00 | 8.84E-01 | no | | intergenic_variant | MODIFIER |
| 2 | 126930973 | AX-170462091 | 0 | T | C | 0.14 | 3.24E-02 | 2.23E-01 | 2.70E+00 | 2.73E+00 | 8.84E-01 | no | Sppl2a | intron_variant | MODIFIER |
| 2 | 126932984 | AX-170394001 | 0 | A | G | 0.14 | 3.24E-02 | 2.23E-01 | 2.70E+00 | 2.73E+00 | 8.84E-01 | no | Sppl2a | missense_variant | MODERATE |
| 2 | 126964961 | AX-170444725 | 0 | T | A | 0.14 | 3.24E-02 | 2.23E-01 | 2.70E+00 | 2.73E+00 | 8.84E-01 | no | | intergenic_variant | MODIFIER |

| | | | | | | | | | | | | | | | |
|---|---|---|---|---|---|---|---|---|---|---|---|---|---|---|---|
| 2 | 126972370 | AX-170270538 | 0 | T | G | 0.14 | 3.24E-02 | 2.23E-01 | 2.70E+00 | 2.73E+00 | 8.84E-01 | no | | intergenic_variant | MODIFIER |
| 2 | 126998316 | AX-170064844 | 0 | A | G | 0.14 | 3.24E-02 | 2.23E-01 | 2.70E+00 | 2.73E+00 | 8.84E-01 | no | | intergenic_variant | MODIFIER |
| 2 | 126998455 | AX-170064846 | 0 | T | C | 0.14 | 3.24E-02 | 2.23E-01 | 2.70E+00 | 2.73E+00 | 8.84E-01 | no | | intergenic_variant | MODIFIER |
| 2 | 127027532 | AX-170288497 | 0 | G | A | 0.14 | 3.24E-02 | 2.23E-01 | 2.70E+00 | 2.73E+00 | 8.84E-01 | no | Ap4e1 | intron_variant | MODIFIER |
| 2 | 127027936 | AX-170141292 | 0 | C | A | 0.14 | 3.24E-02 | 2.23E-01 | 2.70E+00 | 2.73E+00 | 8.84E-01 | no | Ap4e1 | intron_variant | MODIFIER |
| 2 | 127049815 | AX-170493784 | 0 | A | G | 0.14 | 3.24E-02 | 2.23E-01 | 2.70E+00 | 2.73E+00 | 8.84E-01 | no | Ap4e1 | intron_variant | MODIFIER |
| 2 | 127059667 | AX-170140350 | 0 | G | A | 0.14 | 3.24E-02 | 2.23E-01 | 2.70E+00 | 2.73E+00 | 8.84E-01 | no | Ap4e1 | intron_variant | MODIFIER |
| 2 | 127129455 | AX-97838947 | 0 | G | A | 0.14 | 3.24E-02 | 2.23E-01 | 2.70E+00 | 2.73E+00 | 8.84E-01 | no | Ncaph | intron_variant | MODIFIER |
| 2 | 127180342 | AX-97849867 | 0 | T | C | 0.14 | 3.24E-02 | 2.23E-01 | 2.70E+00 | 2.73E+00 | 8.84E-01 | no | | intergenic_variant | MODIFIER |
| 2 | 127195529 | AX-170344785 | 0 | A | G | 0.14 | 3.24E-02 | 2.23E-01 | 2.70E+00 | 2.73E+00 | 8.84E-01 | no | 1810024B03Rik | intron_variant | MODIFIER |
| 2 | 127195655 | AX-170482328 | 0 | C | T | 0.14 | 3.24E-02 | 2.23E-01 | 2.70E+00 | 2.73E+00 | 8.84E-01 | no | 1810024B03Rik | intron_variant | MODIFIER |
| 2 | 127204336 | AX-170191292 | 0 | T | C | 0.14 | 3.24E-02 | 2.23E-01 | 2.70E+00 | 2.73E+00 | 8.84E-01 | no | 1810024B03Rik | intron_variant | MODIFIER |
| 2 | 127204450 | AX-170109559 | 0 | G | A | 0.14 | 3.24E-02 | 2.23E-01 | 2.70E+00 | 2.73E+00 | 8.84E-01 | no | 1810024B03Rik | intron_variant | MODIFIER |
| 2 | 127216865 | AX-170018976 | 0 | G | T | 0.14 | 3.24E-02 | 2.23E-01 | 2.70E+00 | 2.73E+00 | 8.84E-01 | no | Snrnp200 | intron_variant | MODIFIER |
| 2 | 127226083 | AX-170276616 | 0 | A | G | 0.14 | 3.24E-02 | 2.23E-01 | 2.70E+00 | 2.73E+00 | 8.84E-01 | no | Snrnp200 | synonymous_variant | LOW |
| 2 | 127226541 | AX-170176810 | 0 | G | A | 0.14 | 3.24E-02 | 2.23E-01 | 2.70E+00 | 2.73E+00 | 8.84E-01 | no | Snrnp200 | intron_variant | MODIFIER |
| 2 | 127228861 | AX-170424583 | 0 | A | G | 0.14 | 3.24E-02 | 2.23E-01 | 2.70E+00 | 2.73E+00 | 8.84E-01 | no | Snrnp200 | intron_variant | MODIFIER |
| 2 | 127254102 | AX-170474945 | 0 | A | G | 0.14 | 3.24E-02 | 2.23E-01 | 2.70E+00 | 2.73E+00 | 8.84E-01 | no | Tmem127 | intron_variant | MODIFIER |
| 2 | 127254352 | AX-170371119 | 0 | G | C | 0.14 | 3.24E-02 | 2.23E-01 | 2.70E+00 | 2.73E+00 | 8.84E-01 | no | Tmem127 | intron_variant | MODIFIER |
| 2 | 127411623 | AX-170479431 | 0 | T | A | 0.14 | 3.24E-02 | 2.23E-01 | 2.70E+00 | 2.73E+00 | 8.84E-01 | no | | intergenic_variant | MODIFIER |
| 2 | 127413135 | AX-170324457 | 0 | C | T | 0.14 | 3.24E-02 | 2.23E-01 | 2.70E+00 | 2.73E+00 | 8.84E-01 | no | | intergenic_variant | MODIFIER |
| 2 | 127420407 | AX-170064853 | 0 | T | C | 0.14 | 3.24E-02 | 2.23E-01 | 2.70E+00 | 2.73E+00 | 8.84E-01 | no | Gpat2 | upstream_gene_variant | MODIFIER |
| 2 | 127427990 | AX-170035648 | 0 | A | T | 0.14 | 3.24E-02 | 2.23E-01 | 2.70E+00 | 2.73E+00 | 8.84E-01 | no | Gpat2 | intron_variant | MODIFIER |
| 2 | 127436936 | AX-170105812 | 0 | C | T | 0.14 | 3.24E-02 | 2.23E-01 | 2.70E+00 | 2.73E+00 | 8.84E-01 | no | Fahd2a | intron_variant | MODIFIER |
| 2 | 127437782 | AX-97814232 | 0 | G | A | 0.14 | 3.24E-02 | 2.23E-01 | 2.70E+00 | 2.73E+00 | 8.84E-01 | no | Fahd2a | intron_variant | MODIFIER |
| 2 | 127443170 | AX-170362241 | 0 | T | C | 0.14 | 3.24E-02 | 2.23E-01 | 2.70E+00 | 2.73E+00 | 8.84E-01 | no | Fahd2a | intron_variant | MODIFIER |
| 2 | 127456469 | AX-97832835 | 0 | C | T | 0.14 | 3.24E-02 | 2.23E-01 | 2.70E+00 | 2.73E+00 | 8.84E-01 | no | Kcnip3 | downstream_gene_variant | MODIFIER |
| 2 | 127499938 | AX-170278317 | 0 | A | G | 0.14 | 3.24E-02 | 2.23E-01 | 2.70E+00 | 2.73E+00 | 8.84E-01 | no | Kcnip3 | intron_variant | MODIFIER |
| 2 | 127508204 | AX-97808684 | 0 | A | G | 0.14 | 3.24E-02 | 2.23E-01 | 2.70E+00 | 2.73E+00 | 8.84E-01 | no | Kcnip3 | intron_variant | MODIFIER |
| 2 | 127694393 | AX-97805185 | 0 | T | C | 0.14 | 3.24E-02 | 2.23E-01 | 2.70E+00 | 2.73E+00 | 8.84E-01 | no | Gm14245 | downstream_gene_variant | MODIFIER |
| 2 | 127694408 | AX-170018978 | 0 | T | C | 0.14 | 3.24E-02 | 2.23E-01 | 2.70E+00 | 2.73E+00 | 8.84E-01 | no | Gm14245 | downstream_gene_variant | MODIFIER |
| 2 | 127721275 | AX-97796862 | 0 | G | A | 0.14 | 3.24E-02 | 2.23E-01 | 2.70E+00 | 2.73E+00 | 8.84E-01 | no | Mall | intron_variant | MODIFIER |
| 2 | 127724870 | AX-170365395 | 0 | C | T | 0.14 | 3.24E-02 | 2.23E-01 | 2.70E+00 | 2.73E+00 | 8.84E-01 | no | Mall | intron_variant | MODIFIER |
| 2 | 127724934 | AX-170275515 | 0 | T | G | 0.14 | 3.24E-02 | 2.23E-01 | 2.70E+00 | 2.73E+00 | 8.84E-01 | no | Mall | intron_variant | MODIFIER |
| 2 | 127725726 | AX-170188406 | 0 | C | A | 0.14 | 3.24E-02 | 2.23E-01 | 2.70E+00 | 2.73E+00 | 8.84E-01 | no | Mall | intron_variant | MODIFIER |
| 2 | 127727565 | AX-170491384 | 0 | T | C | 0.14 | 3.24E-02 | 2.23E-01 | 2.70E+00 | 2.73E+00 | 8.84E-01 | no | Mall | intron_variant | MODIFIER |
| 2 | 127740192 | AX-170018981 | 0 | G | A | 0.14 | 3.24E-02 | 2.23E-01 | 2.70E+00 | 2.73E+00 | 8.84E-01 | no | Nphp1 | downstream_gene_variant | MODIFIER |
| 2 | 127772698 | AX-97840038 | 0 | G | C | 0.14 | 3.24E-02 | 2.23E-01 | 2.70E+00 | 2.73E+00 | 8.84E-01 | no | Nphp1 | intron_variant | MODIFIER |
| 2 | 127818965 | AX-170018979 | 0 | T | C | 0.14 | 3.24E-02 | 2.23E-01 | 2.70E+00 | 2.73E+00 | 8.84E-01 | no | Bub1 | intron_variant | MODIFIER |
| 2 | 127961142 | AX-170410302 | 0 | G | C | 0.14 | 3.24E-02 | 2.23E-01 | 2.70E+00 | 2.73E+00 | 8.84E-01 | no | Acoxl | intron_variant | MODIFIER |
| 2 | 128261981 | AX-170338692 | 0 | T | C | 0.14 | 3.24E-02 | 2.23E-01 | 2.70E+00 | 2.73E+00 | 8.84E-01 | no | Gm14005 | intron_variant,non_coding_transcript_variant | MODIFIER |
| 2 | 128309783 | AX-170064865 | 0 | T | C | 0.14 | 3.24E-02 | 2.23E-01 | 2.70E+00 | 2.73E+00 | 8.84E-01 | no | Gm14005 | intron_variant,non_coding_transcript_variant | MODIFIER |
| 2 | 128322083 | AX-170523271 | 0 | T | C | 0.14 | 3.24E-02 | 2.23E-01 | 2.70E+00 | 2.73E+00 | 8.84E-01 | no | Gm14005 | intron_variant,non_coding_transcript_variant | MODIFIER |
| 2 | 128419254 | AX-97838090 | 0 | A | C | 0.14 | 3.24E-02 | 2.23E-01 | 2.70E+00 | 2.73E+00 | 8.84E-01 | no | Gm14005 | intron_variant,non_coding_transcript_variant | MODIFIER |
| 2 | 128424375 | AX-170310971 | 0 | G | A | 0.14 | 3.24E-02 | 2.23E-01 | 2.70E+00 | 2.73E+00 | 8.84E-01 | no | Gm14005 | intron_variant,non_coding_transcript_variant | MODIFIER |
| 2 | 128427931 | AX-170355199 | 0 | A | G | 0.14 | 3.24E-02 | 2.23E-01 | 2.70E+00 | 2.73E+00 | 8.84E-01 | no | Gm14005 | intron_variant,non_coding_transcript_variant | MODIFIER |
| 2 | 128431803 | AX-97795062 | 0 | T | G | 0.14 | 3.24E-02 | 2.23E-01 | 2.70E+00 | 2.73E+00 | 8.84E-01 | no | Gm14005 | upstream_gene_variant | MODIFIER |
| 2 | 128453497 | AX-97857134 | 0 | T | C | 0.14 | 3.24E-02 | 2.23E-01 | 2.70E+00 | 2.73E+00 | 8.84E-01 | no | Gm14005 | intron_variant,non_coding_transcript_variant | MODIFIER |
| 2 | 128466213 | AX-170433525 | 0 | G | T | 0.14 | 3.24E-02 | 2.23E-01 | 2.70E+00 | 2.73E+00 | 8.84E-01 | no | Gm14005 | intron_variant,non_coding_transcript_variant | MODIFIER |
| 2 | 128480318 | AX-170238828 | 0 | A | G | 0.14 | 3.24E-02 | 2.23E-01 | 2.70E+00 | 2.73E+00 | 8.84E-01 | no | Gm14005 | intron_variant,non_coding_transcript_variant | MODIFIER |
| 2 | 128501144 | AX-170291700 | 0 | C | G | 0.14 | 3.24E-02 | 2.23E-01 | 2.70E+00 | 2.73E+00 | 8.84E-01 | no | Gm14005 | intron_variant,non_coding_transcript_variant | MODIFIER |
| 2 | 128512831 | AX-170316206 | 0 | A | G | 0.14 | 3.24E-02 | 2.23E-01 | 2.70E+00 | 2.73E+00 | 8.84E-01 | no | | regulatory_region_variant | MODIFIER |
| 2 | 128512962 | AX-170484756 | 0 | G | A | 0.14 | 3.24E-02 | 2.23E-01 | 2.70E+00 | 2.73E+00 | 8.84E-01 | no | | regulatory_region_variant | MODIFIER |
| 2 | 128515103 | AX-170113607 | 0 | C | G | 0.14 | 3.24E-02 | 2.23E-01 | 2.70E+00 | 2.73E+00 | 8.84E-01 | no | | regulatory_region_variant | MODIFIER |

| | | | | | | | | | | | | | | | |
|---|---|---|---|---|---|---|---|---|---|---|---|---|---|---|---|
| 2 | 128516568 | AX-170458406 | 0 | A | G | 0.14 | 3.24E-02 | 2.23E-01 | 2.70E+00 | 2.73E+00 | 8.84E-01 | no | | regulatory_region_variant | MODIFIER |
| 2 | 128568919 | AX-97818490 | 0 | A | G | 0.14 | 3.24E-02 | 2.23E-01 | 2.70E+00 | 2.73E+00 | 8.84E-01 | no | | regulatory_region_variant | MODIFIER |
| 2 | 128593732 | AX-170439114 | 0 | C | A | 0.14 | 3.24E-02 | 2.23E-01 | 2.70E+00 | 2.73E+00 | 8.84E-01 | no | Spdye4c | intron_variant,NMD_transcript_variant | MODIFIER |
| 2 | 128746011 | AX-170493628 | 0 | T | C | 0.14 | 3.24E-02 | 2.23E-01 | 2.70E+00 | 2.73E+00 | 8.84E-01 | no | Mertk | intron_variant | MODIFIER |
| 2 | 128884370 | AX-97801408 | 0 | C | T | 0.14 | 3.24E-02 | 2.23E-01 | 2.70E+00 | 2.73E+00 | 8.84E-01 | no | Fbln7 | intron_variant | MODIFIER |
| 2 | 129009780 | AX-97834095 | 0 | A | G | 0.14 | 3.24E-02 | 2.23E-01 | 2.70E+00 | 2.73E+00 | 8.84E-01 | no | Zc3h6 | intron_variant | MODIFIER |
| 2 | 129038340 | AX-97838972 | 0 | A | G | 0.14 | 3.24E-02 | 2.23E-01 | 2.70E+00 | 2.73E+00 | 8.84E-01 | no | Vinac1 | synonymous_variant | LOW |
| 2 | 129045123 | AX-170431366 | 0 | G | A | 0.14 | 3.24E-02 | 2.23E-01 | 2.70E+00 | 2.73E+00 | 8.84E-01 | no | Vinac1 | intron_variant | MODIFIER |
| 2 | 129080555 | AX-170413153 | 0 | A | T | 0.14 | 3.24E-02 | 2.23E-01 | 2.70E+00 | 2.73E+00 | 8.84E-01 | no | Ttl | intron_variant | MODIFIER |
| 2 | 129088497 | AX-97814080 | 0 | C | T | 0.14 | 3.24E-02 | 2.23E-01 | 2.70E+00 | 2.73E+00 | 8.84E-01 | no | Ttl | intron_variant | MODIFIER |
| 2 | 129103798 | AX-97839121 | 0 | G | A | 0.14 | 3.24E-02 | 2.23E-01 | 2.70E+00 | 2.73E+00 | 8.84E-01 | no | Polr1b | intron_variant | MODIFIER |
| 2 | 129129144 | AX-170323255 | 0 | G | A | 0.14 | 3.24E-02 | 2.23E-01 | 2.70E+00 | 2.73E+00 | 8.84E-01 | no | Chchd5 | upstream_gene_variant | MODIFIER |
| 2 | 129131498 | AX-97781393 | 0 | A | G | 0.14 | 3.24E-02 | 2.23E-01 | 2.70E+00 | 2.73E+00 | 8.84E-01 | no | Chchd5 | intron_variant | MODIFIER |
| 2 | 129133730 | AX-170298624 | 0 | C | T | 0.14 | 3.24E-02 | 2.23E-01 | 2.70E+00 | 2.73E+00 | 8.84E-01 | no | Chchd5 | 3_prime_UTR_variant | MODIFIER |
| 2 | 129150807 | AX-97795908 | 0 | T | C | 0.14 | 3.24E-02 | 2.23E-01 | 2.70E+00 | 2.73E+00 | 8.84E-01 | no | | intergenic_variant | MODIFIER |
| 2 | 129167301 | AX-170395129 | 0 | C | T | 0.14 | 3.24E-02 | 2.23E-01 | 2.70E+00 | 2.73E+00 | 8.84E-01 | no | Gm14026 | upstream_gene_variant | MODIFIER |
| 2 | 129193279 | AX-170376719 | 0 | T | A | 0.14 | 3.24E-02 | 2.23E-01 | 2.70E+00 | 2.73E+00 | 8.84E-01 | no | Gm4430 | downstream_gene_variant | MODIFIER |
| 2 | 129218811 | AX-97835499 | 0 | C | A | 0.14 | 3.24E-02 | 2.23E-01 | 2.70E+00 | 2.73E+00 | 8.84E-01 | no | 9830144P21Rik | intron_variant,non_coding_transcript_variant | MODIFIER |
| 2 | 129222961 | AX-97791889 | 0 | C | A | 0.14 | 3.24E-02 | 2.23E-01 | 2.70E+00 | 2.73E+00 | 8.84E-01 | no | 9830144P21Rik | downstream_gene_variant | MODIFIER |
| 2 | 129252671 | AX-170019003 | 0 | A | G | 0.14 | 3.24E-02 | 2.23E-01 | 2.70E+00 | 2.73E+00 | 8.84E-01 | no | Gm14024 | intron_variant,non_coding_transcript_variant | MODIFIER |
| 2 | 129255752 | AX-97857555 | 0 | A | G | 0.14 | 3.24E-02 | 2.23E-01 | 2.70E+00 | 2.73E+00 | 8.84E-01 | no | Gm14024 | intron_variant,non_coding_transcript_variant | MODIFIER |
| 2 | 129265263 | AX-170342566 | 0 | T | C | 0.14 | 3.24E-02 | 2.23E-01 | 2.70E+00 | 2.73E+00 | 8.84E-01 | no | Ckap2l | downstream_gene_variant | MODIFIER |
| 2 | 129270635 | AX-97854786 | 0 | T | C | 0.14 | 3.24E-02 | 2.23E-01 | 2.70E+00 | 2.73E+00 | 8.84E-01 | no | Ckap2l | missense_variant | MODERATE |
| 2 | 129295242 | AX-170371399 | 0 | T | C | 0.14 | 3.24E-02 | 2.23E-01 | 2.70E+00 | 2.73E+00 | 8.84E-01 | no | Il1a | downstream_gene_variant | MODIFIER |
| 2 | 129295438 | AX-170368446 | 0 | C | A | 0.14 | 3.24E-02 | 2.23E-01 | 2.70E+00 | 2.73E+00 | 8.84E-01 | no | Il1a | downstream_gene_variant | MODIFIER |
| 2 | 129296106 | AX-170157541 | 0 | C | T | 0.14 | 3.24E-02 | 2.23E-01 | 2.70E+00 | 2.73E+00 | 8.84E-01 | no | Il1a | downstream_gene_variant | MODIFIER |
| 2 | 129296548 | AX-170064890 | 0 | G | C | 0.14 | 3.24E-02 | 2.23E-01 | 2.70E+00 | 2.73E+00 | 8.84E-01 | no | Il1a | downstream_gene_variant | MODIFIER |
| 2 | 129393620 | AX-97806589 | 0 | T | C | 0.14 | 3.24E-02 | 2.23E-01 | 2.70E+00 | 2.73E+00 | 8.84E-01 | no | | regulatory_region_variant | MODIFIER |
| 2 | 129430990 | AX-97805269 | 0 | C | T | 0.14 | 3.24E-02 | 2.23E-01 | 2.70E+00 | 2.73E+00 | 8.84E-01 | no | | intergenic_variant | MODIFIER |
| 2 | 129475176 | AX-97856905 | 0 | C | T | 0.14 | 3.24E-02 | 2.23E-01 | 2.70E+00 | 2.73E+00 | 8.84E-01 | no | F830045P16Rik | intron_variant | MODIFIER |
| 2 | 129490565 | AX-170165274 | 0 | C | T | 0.14 | 3.24E-02 | 2.23E-01 | 2.70E+00 | 2.73E+00 | 8.84E-01 | no | F830045P16Rik | intron_variant | MODIFIER |
| 2 | 129498662 | AX-97794977 | 0 | C | T | 0.14 | 3.24E-02 | 2.23E-01 | 2.70E+00 | 2.73E+00 | 8.84E-01 | no | F830045P16Rik | intron_variant | MODIFIER |
| 2 | 129520040 | AX-97802471 | 0 | G | A | 0.14 | 3.24E-02 | 2.23E-01 | 2.70E+00 | 2.73E+00 | 8.84E-01 | no | F830045P16Rik | intron_variant | MODIFIER |
| 2 | 129573053 | AX-170377692 | 0 | A | C | 0.14 | 3.24E-02 | 2.23E-01 | 2.70E+00 | 2.73E+00 | 8.84E-01 | no | | intergenic_variant | MODIFIER |
| 2 | 129574971 | AX-97838971 | 0 | C | T | 0.14 | 3.24E-02 | 2.23E-01 | 2.70E+00 | 2.73E+00 | 8.84E-01 | no | | regulatory_region_variant | MODIFIER |
| 2 | 129575897 | AX-170139941 | 0 | A | G | 0.14 | 3.24E-02 | 2.23E-01 | 2.70E+00 | 2.73E+00 | 8.84E-01 | no | | regulatory_region_variant | MODIFIER |
| 2 | 129598024 | AX-170433609 | 0 | A | G | 0.14 | 3.24E-02 | 2.23E-01 | 2.70E+00 | 2.73E+00 | 8.84E-01 | no | Sirpa | intron_variant | MODIFIER |
| 2 | 129602752 | AX-170384925 | 0 | C | A | 0.14 | 3.24E-02 | 2.23E-01 | 2.70E+00 | 2.73E+00 | 8.84E-01 | no | Sirpa | intron_variant | MODIFIER |
| 2 | 129603024 | AX-170520141 | 0 | T | C | 0.14 | 3.24E-02 | 2.23E-01 | 2.70E+00 | 2.73E+00 | 8.84E-01 | no | Sirpa | intron_variant | MODIFIER |
| 2 | 129608217 | AX-170200456 | 0 | G | A | 0.14 | 3.24E-02 | 2.23E-01 | 2.70E+00 | 2.73E+00 | 8.84E-01 | no | Sirpa | intron_variant | MODIFIER |
| 2 | 129611410 | AX-170173580 | 0 | T | A | 0.14 | 3.24E-02 | 2.23E-01 | 2.70E+00 | 2.73E+00 | 8.84E-01 | no | Sirpa | intron_variant | MODIFIER |
| 2 | 129614905 | AX-170177889 | 0 | A | G | 0.14 | 3.24E-02 | 2.23E-01 | 2.70E+00 | 2.73E+00 | 8.84E-01 | no | Sirpa | intron_variant | MODIFIER |
| 2 | 129616737 | AX-170349313 | 0 | T | C | 0.14 | 3.24E-02 | 2.23E-01 | 2.70E+00 | 2.73E+00 | 8.84E-01 | no | Sirpa | intron_variant | MODIFIER |
| 2 | 129619481 | AX-170448820 | 0 | G | A | 0.14 | 3.24E-02 | 2.23E-01 | 2.70E+00 | 2.73E+00 | 8.84E-01 | no | Sirpa | intron_variant | MODIFIER |
| 2 | 129619640 | AX-170514139 | 0 | C | G | 0.14 | 3.24E-02 | 2.23E-01 | 2.70E+00 | 2.73E+00 | 8.84E-01 | no | Sirpa | intron_variant | MODIFIER |
| 2 | 129624758 | AX-170381171 | 0 | C | A | 0.14 | 3.24E-02 | 2.23E-01 | 2.70E+00 | 2.73E+00 | 8.84E-01 | no | Sirpa | intron_variant | MODIFIER |
| 2 | 129625609 | AX-170128479 | 0 | A | G | 0.14 | 3.24E-02 | 2.23E-01 | 2.70E+00 | 2.73E+00 | 8.84E-01 | no | Sirpa | intron_variant | MODIFIER |
| 2 | 129631820 | AX-97828012 | 0 | G | T | 0.14 | 3.24E-02 | 2.23E-01 | 2.70E+00 | 2.73E+00 | 8.84E-01 | no | Sirpa | 3_prime_UTR_variant | MODIFIER |
| 2 | 129645442 | AX-170417741 | 0 | G | A | 0.14 | 3.24E-02 | 2.23E-01 | 2.70E+00 | 2.73E+00 | 8.84E-01 | no | | intergenic_variant | MODIFIER |
| 2 | 129658593 | AX-97818004 | 0 | A | C | 0.14 | 3.24E-02 | 2.23E-01 | 2.70E+00 | 2.73E+00 | 8.84E-01 | no | | intergenic_variant | MODIFIER |
| 2 | 129675314 | AX-170140379 | 0 | T | C | 0.14 | 3.24E-02 | 2.23E-01 | 2.70E+00 | 2.73E+00 | 8.84E-01 | no | | regulatory_region_variant | MODIFIER |
| 2 | 129686991 | AX-170436612 | 0 | G | T | 0.14 | 3.24E-02 | 2.23E-01 | 2.70E+00 | 2.73E+00 | 8.84E-01 | no | Pdyn | 3_prime_UTR_variant | MODIFIER |
| 2 | 129687692 | AX-97841780 | 0 | C | A | 0.14 | 3.24E-02 | 2.23E-01 | 2.70E+00 | 2.73E+00 | 8.84E-01 | no | Pdyn | 3_prime_UTR_variant | MODIFIER |
| 2 | 129689244 | AX-170337289 | 0 | C | T | 0.14 | 3.24E-02 | 2.23E-01 | 2.70E+00 | 2.73E+00 | 8.84E-01 | no | Pdyn | intron_variant | MODIFIER |

| | | | | | | | | | | | | | | | |
|---|---|---|---|---|---|---|---|---|---|---|---|---|---|---|---|
| 2 | 129689464 | AX-170476877 | 0 | T | C | 0.14 | 3.24E-02 | 2.23E-01 | 2.70E+00 | 2.73E+00 | 8.84E-01 | no | Pdyn | intron_variant | MODIFIER |
| 2 | 129712861 | AX-170353519 | 0 | T | G | 0.14 | 3.24E-02 | 2.23E-01 | 2.70E+00 | 2.73E+00 | 8.84E-01 | no | | intergenic_variant | MODIFIER |
| 2 | 129717409 | AX-170402931 | 0 | T | C | 0.14 | 3.24E-02 | 2.23E-01 | 2.70E+00 | 2.73E+00 | 8.84E-01 | no | | intergenic_variant | MODIFIER |
| 2 | 129717501 | AX-170277320 | 0 | C | T | 0.14 | 3.24E-02 | 2.23E-01 | 2.70E+00 | 2.73E+00 | 8.84E-01 | no | | intergenic_variant | MODIFIER |
| 2 | 129803380 | AX-170019019 | 0 | C | T | 0.14 | 3.24E-02 | 2.23E-01 | 2.70E+00 | 2.73E+00 | 8.84E-01 | no | Stk35 | intron_variant | MODIFIER |
| 2 | 129808434 | AX-170019013 | 0 | C | T | 0.14 | 3.24E-02 | 2.23E-01 | 2.70E+00 | 2.73E+00 | 8.84E-01 | no | Stk35 | intron_variant | MODIFIER |
| 2 | 129816891 | AX-170019016 | 0 | C | T | 0.14 | 3.24E-02 | 2.23E-01 | 2.70E+00 | 2.73E+00 | 8.84E-01 | no | Stk35 | intron_variant | MODIFIER |
| 2 | 129818264 | AX-170323607 | 0 | A | C | 0.14 | 3.24E-02 | 2.23E-01 | 2.70E+00 | 2.73E+00 | 8.84E-01 | no | Stk35 | intron_variant | MODIFIER |
| 2 | 129833109 | AX-97840883 | 0 | T | C | 0.14 | 3.24E-02 | 2.23E-01 | 2.70E+00 | 2.73E+00 | 8.84E-01 | no | Stk35 | downstream_gene_variant | MODIFIER |
| 2 | 129836327 | AX-170019017 | 0 | G | T | 0.14 | 3.24E-02 | 2.23E-01 | 2.70E+00 | 2.73E+00 | 8.84E-01 | no | Stk35 | downstream_gene_variant | MODIFIER |
| 2 | 129839595 | AX-170318525 | 0 | C | T | 0.14 | 3.24E-02 | 2.23E-01 | 2.70E+00 | 2.73E+00 | 8.84E-01 | no | | intergenic_variant | MODIFIER |
| 2 | 129864558 | AX-97814311 | 0 | C | T | 0.14 | 3.24E-02 | 2.23E-01 | 2.70E+00 | 2.73E+00 | 8.84E-01 | no | | intergenic_variant | MODIFIER |
| 2 | 129874793 | AX-170517704 | 0 | G | A | 0.14 | 3.24E-02 | 2.23E-01 | 2.70E+00 | 2.73E+00 | 8.84E-01 | no | | intergenic_variant | MODIFIER |
| 2 | 129877696 | AX-170440366 | 0 | T | C | 0.14 | 3.24E-02 | 2.23E-01 | 2.70E+00 | 2.73E+00 | 8.84E-01 | no | | intergenic_variant | MODIFIER |
| 2 | 129878563 | AX-170342130 | 0 | A | T | 0.14 | 3.24E-02 | 2.23E-01 | 2.70E+00 | 2.73E+00 | 8.84E-01 | no | | intergenic_variant | MODIFIER |
| 2 | 129879994 | AX-170477107 | 0 | T | C | 0.14 | 3.24E-02 | 2.23E-01 | 2.70E+00 | 2.73E+00 | 8.84E-01 | no | | regulatory_region_variant | MODIFIER |
| 2 | 129881423 | AX-97852659 | 0 | G | A | 0.14 | 3.24E-02 | 2.23E-01 | 2.70E+00 | 2.73E+00 | 8.84E-01 | no | | intergenic_variant | MODIFIER |
| 2 | 129897030 | AX-97829512 | 0 | T | C | 0.14 | 3.24E-02 | 2.23E-01 | 2.70E+00 | 2.73E+00 | 8.84E-01 | no | | regulatory_region_variant | MODIFIER |
| 2 | 129897401 | AX-170019011 | 0 | C | T | 0.14 | 3.24E-02 | 2.23E-01 | 2.70E+00 | 2.73E+00 | 8.84E-01 | no | | intergenic_variant | MODIFIER |
| 2 | 129925169 | AX-170019014 | 0 | C | T | 0.14 | 3.24E-02 | 2.23E-01 | 2.70E+00 | 2.73E+00 | 8.84E-01 | no | | intergenic_variant | MODIFIER |
| 2 | 129925374 | AX-97838945 | 0 | C | T | 0.14 | 3.24E-02 | 2.23E-01 | 2.70E+00 | 2.73E+00 | 8.84E-01 | no | | intergenic_variant | MODIFIER |
| 2 | 129943299 | AX-170064905 | 0 | C | T | 0.14 | 3.24E-02 | 2.23E-01 | 2.70E+00 | 2.73E+00 | 8.84E-01 | no | | intergenic_variant | MODIFIER |
| 2 | 129943838 | AX-170235997 | 0 | T | A | 0.14 | 3.24E-02 | 2.23E-01 | 2.70E+00 | 2.73E+00 | 8.84E-01 | no | Gm28196 | upstream_gene_variant | MODIFIER |
| 2 | 129952310 | AX-170167152 | 0 | G | A | 0.14 | 3.24E-02 | 2.23E-01 | 2.70E+00 | 2.73E+00 | 8.84E-01 | no | Gm28196 | downstream_gene_variant | MODIFIER |
| 2 | 129952879 | AX-170356408 | 0 | G | A | 0.14 | 3.24E-02 | 2.23E-01 | 2.70E+00 | 2.73E+00 | 8.84E-01 | no | Gm28196 | downstream_gene_variant | MODIFIER |
| 2 | 129954870 | AX-170167821 | 0 | G | A | 0.14 | 3.24E-02 | 2.23E-01 | 2.70E+00 | 2.73E+00 | 8.84E-01 | no | | regulatory_region_variant | MODIFIER |
| 2 | 129955262 | AX-170185544 | 0 | T | G | 0.14 | 3.24E-02 | 2.23E-01 | 2.70E+00 | 2.73E+00 | 8.84E-01 | no | | regulatory_region_variant | MODIFIER |
| 2 | 129956057 | AX-97848557 | 0 | T | C | 0.14 | 3.24E-02 | 2.23E-01 | 2.70E+00 | 2.73E+00 | 8.84E-01 | no | | regulatory_region_variant | MODIFIER |
| 2 | 129960350 | AX-170529071 | 0 | A | G | 0.14 | 3.24E-02 | 2.23E-01 | 2.70E+00 | 2.73E+00 | 8.84E-01 | no | | intergenic_variant | MODIFIER |
| 2 | 129978800 | AX-97791453 | 0 | T | C | 0.14 | 3.24E-02 | 2.23E-01 | 2.70E+00 | 2.73E+00 | 8.84E-01 | no | | intergenic_variant | MODIFIER |
| 2 | 129980693 | AX-170260039 | 0 | G | C | 0.14 | 3.24E-02 | 2.23E-01 | 2.70E+00 | 2.73E+00 | 8.84E-01 | no | | intergenic_variant | MODIFIER |
| 2 | 129981179 | AX-170339386 | 0 | T | C | 0.14 | 3.24E-02 | 2.23E-01 | 2.70E+00 | 2.73E+00 | 8.84E-01 | no | | intergenic_variant | MODIFIER |
| 2 | 129988991 | AX-170357135 | 0 | A | C | 0.14 | 3.24E-02 | 2.23E-01 | 2.70E+00 | 2.73E+00 | 8.84E-01 | no | | intergenic_variant | MODIFIER |
| 2 | 129995253 | AX-170479922 | 0 | T | C | 0.14 | 3.24E-02 | 2.23E-01 | 2.70E+00 | 2.73E+00 | 8.84E-01 | no | | intergenic_variant | MODIFIER |
| 2 | 130000374 | AX-170248383 | 0 | C | T | 0.14 | 3.24E-02 | 2.23E-01 | 2.70E+00 | 2.73E+00 | 8.84E-01 | no | | intergenic_variant | MODIFIER |
| 2 | 130001194 | AX-170403137 | 0 | A | C | 0.14 | 3.24E-02 | 2.23E-01 | 2.70E+00 | 2.73E+00 | 8.84E-01 | no | | intergenic_variant | MODIFIER |
| 2 | 130001678 | AX-168417261 | 0 | A | G | 0.14 | 3.24E-02 | 2.23E-01 | 2.70E+00 | 2.73E+00 | 8.84E-01 | no | | intergenic_variant | MODIFIER |
| 2 | 130005339 | AX-97815526 | 0 | T | C | 0.14 | 3.24E-02 | 2.23E-01 | 2.70E+00 | 2.73E+00 | 8.84E-01 | no | | intergenic_variant | MODIFIER |
| 2 | 130006568 | AX-170400121 | 0 | A | G | 0.14 | 3.24E-02 | 2.23E-01 | 2.70E+00 | 2.73E+00 | 8.84E-01 | no | | intergenic_variant | MODIFIER |
| 2 | 130027780 | AX-97805436 | 0 | T | C | 0.14 | 3.24E-02 | 2.23E-01 | 2.70E+00 | 2.73E+00 | 8.84E-01 | no | Tgm3 | intron_variant | MODIFIER |
| 2 | 130033432 | AX-170266186 | 0 | T | C | 0.14 | 3.24E-02 | 2.23E-01 | 2.70E+00 | 2.73E+00 | 8.84E-01 | no | Tgm3 | intron_variant | MODIFIER |
| 2 | 130035074 | AX-170392990 | 0 | C | T | 0.14 | 3.24E-02 | 2.23E-01 | 2.70E+00 | 2.73E+00 | 8.84E-01 | no | Tgm3 | intron_variant | MODIFIER |
| 2 | 130041640 | AX-170328824 | 0 | G | A | 0.14 | 3.24E-02 | 2.23E-01 | 2.70E+00 | 2.73E+00 | 8.84E-01 | no | Tgm3 | intron_variant | MODIFIER |
| 2 | 130041741 | AX-170199089 | 0 | T | C | 0.14 | 3.24E-02 | 2.23E-01 | 2.70E+00 | 2.73E+00 | 8.84E-01 | no | Tgm3 | intron_variant | MODIFIER |
| 2 | 130041965 | AX-170300495 | 0 | G | A | 0.14 | 3.24E-02 | 2.23E-01 | 2.70E+00 | 2.73E+00 | 8.84E-01 | no | Tgm3 | synonymous_variant | LOW |
| 2 | 130042224 | AX-170265137 | 0 | T | C | 0.14 | 3.24E-02 | 2.23E-01 | 2.70E+00 | 2.73E+00 | 8.84E-01 | no | Tgm3 | intron_variant | MODIFIER |
| 2 | 130042259 | AX-170302518 | 0 | T | G | 0.14 | 3.24E-02 | 2.23E-01 | 2.70E+00 | 2.73E+00 | 8.84E-01 | no | Tgm3 | intron_variant | MODIFIER |
| 2 | 130042447 | AX-170228892 | 0 | A | G | 0.14 | 3.24E-02 | 2.23E-01 | 2.70E+00 | 2.73E+00 | 8.84E-01 | no | Tgm3 | intron_variant | MODIFIER |
| 2 | 130042659 | AX-170188431 | 0 | C | T | 0.14 | 3.24E-02 | 2.23E-01 | 2.70E+00 | 2.73E+00 | 8.84E-01 | no | Tgm3 | intron_variant | MODIFIER |
| 2 | 130055308 | AX-170420021 | 0 | G | C | 0.14 | 3.24E-02 | 2.23E-01 | 2.70E+00 | 2.73E+00 | 8.84E-01 | no | Tgm3 | downstream_gene_variant | MODIFIER |
| 2 | 130058370 | AX-170332035 | 0 | G | A | 0.14 | 3.24E-02 | 2.23E-01 | 2.70E+00 | 2.73E+00 | 8.84E-01 | no | | regulatory_region_variant | MODIFIER |
| 2 | 130068459 | AX-170336406 | 0 | G | T | 0.14 | 3.24E-02 | 2.23E-01 | 2.70E+00 | 2.73E+00 | 8.84E-01 | no | Gm22424 | upstream_gene_variant | MODIFIER |
| 2 | 130085522 | AX-97817244 | 0 | G | A | 0.14 | 3.24E-02 | 2.23E-01 | 2.70E+00 | 2.73E+00 | 8.84E-01 | no | Gm24665 | upstream_gene_variant | MODIFIER |
| 2 | 130121322 | AX-170343744 | 0 | A | G | 0.14 | 3.24E-02 | 2.23E-01 | 2.70E+00 | 2.73E+00 | 8.84E-01 | no | Tgm6 | upstream_gene_variant | MODIFIER |

| | | | | | | | | | | | | | | | |
|---|---|---|---|---|---|---|---|---|---|---|---|---|---|---|---|
| 2 | 130121997 | AX-170064910 | 0 | C | T | 0.14 | 3.24E-02 | 2.23E-01 | 2.70E+00 | 2.73E+00 | 8.84E-01 | no | Tgm6 | upstream_gene_variant | MODIFIER |
| 2 | 130123317 | AX-170508814 | 0 | T | A | 0.14 | 3.24E-02 | 2.23E-01 | 2.70E+00 | 2.73E+00 | 8.84E-01 | no | Tgm6 | 5_prime_UTR_variant | MODIFIER |
| 2 | 130127112 | AX-170112739 | 0 | T | G | 0.14 | 3.24E-02 | 2.23E-01 | 2.70E+00 | 2.73E+00 | 8.84E-01 | no | Tgm6 | intron_variant | MODIFIER |
| 2 | 130151275 | AX-170471212 | 0 | G | A | 0.14 | 3.24E-02 | 2.23E-01 | 2.70E+00 | 2.73E+00 | 8.84E-01 | no | Tgm6 | splice_region_variant,synonymous_variant | LOW |
| 2 | 130151857 | AX-170307691 | 0 | A | G | 0.14 | 3.24E-02 | 2.23E-01 | 2.70E+00 | 2.73E+00 | 8.84E-01 | no | Tgm6 | intron_variant | MODIFIER |
| 2 | 130158299 | AX-97795633 | 0 | C | A | 0.14 | 3.24E-02 | 2.23E-01 | 2.70E+00 | 2.73E+00 | 8.84E-01 | no | Tgm6 | downstream_gene_variant | MODIFIER |
| 2 | 130158355 | AX-170391203 | 0 | A | G | 0.14 | 3.24E-02 | 2.23E-01 | 2.70E+00 | 2.73E+00 | 8.84E-01 | no | Tgm6 | downstream_gene_variant | MODIFIER |
| 2 | 130170861 | AX-168389471 | 0 | T | C | 0.14 | 3.24E-02 | 2.23E-01 | 2.70E+00 | 2.73E+00 | 8.84E-01 | no | Snrpb | downstream_gene_variant | MODIFIER |
| 2 | 130177430 | AX-97856319 | 0 | A | G | 0.14 | 3.24E-02 | 2.23E-01 | 2.70E+00 | 2.73E+00 | 8.84E-01 | no | Snrpb | intron_variant | MODIFIER |
| 2 | 130414818 | AX-170088748 | 0 | C | T | 0.14 | 3.24E-02 | 2.23E-01 | 2.70E+00 | 2.73E+00 | 8.84E-01 | no | Pced1a | downstream_gene_variant | MODIFIER |
| 2 | 130416734 | AX-170019021 | 0 | T | A | 0.14 | 3.24E-02 | 2.23E-01 | 2.70E+00 | 2.73E+00 | 8.84E-01 | no | Pced1a | downstream_gene_variant | MODIFIER |
| 2 | 130416796 | AX-170019024 | 0 | A | G | 0.14 | 3.24E-02 | 2.23E-01 | 2.70E+00 | 2.73E+00 | 8.84E-01 | no | Pced1a | downstream_gene_variant | MODIFIER |
| 2 | 130438571 | AX-97845947 | 0 | A | C | 0.14 | 3.24E-02 | 2.23E-01 | 2.70E+00 | 2.73E+00 | 8.84E-01 | no | Vps16 | missense_variant,splice_region_variant | MODERATE |
| 2 | 130439594 | AX-170181860 | 0 | T | C | 0.14 | 3.24E-02 | 2.23E-01 | 2.70E+00 | 2.73E+00 | 8.84E-01 | no | Vps16 | intron_variant | MODIFIER |
| 2 | 130459862 | AX-170088749 | 0 | A | G | 0.14 | 3.24E-02 | 2.23E-01 | 2.70E+00 | 2.73E+00 | 8.84E-01 | no | Ptpra | intron_variant | MODIFIER |
| 2 | 130467733 | AX-170019027 | 0 | A | G | 0.14 | 3.24E-02 | 2.23E-01 | 2.70E+00 | 2.73E+00 | 8.84E-01 | no | Ptpra | intron_variant | MODIFIER |
| 2 | 130484482 | AX-97838977 | 0 | G | A | 0.14 | 3.24E-02 | 2.23E-01 | 2.70E+00 | 2.73E+00 | 8.84E-01 | no | Ptpra | intron_variant | MODIFIER |
| 2 | 130495451 | AX-170225888 | 0 | T | C | 0.14 | 3.24E-02 | 2.23E-01 | 2.70E+00 | 2.73E+00 | 8.84E-01 | no | Ptpra | intron_variant | MODIFIER |
| 2 | 130500793 | AX-97848959 | 0 | G | A | 0.14 | 3.24E-02 | 2.23E-01 | 2.70E+00 | 2.73E+00 | 8.84E-01 | no | Ptpra | intron_variant | MODIFIER |
| 2 | 130522746 | AX-170482599 | 0 | A | G | 0.14 | 3.24E-02 | 2.23E-01 | 2.70E+00 | 2.73E+00 | 8.84E-01 | no | Ptpra | intron_variant | MODIFIER |
| 2 | 130539605 | AX-168395885 | 0 | A | G | 0.14 | 3.24E-02 | 2.23E-01 | 2.70E+00 | 2.73E+00 | 8.84E-01 | no | Ptpra | intron_variant | MODIFIER |
| 2 | 130541379 | AX-170327893 | 0 | T | C | 0.14 | 3.24E-02 | 2.23E-01 | 2.70E+00 | 2.73E+00 | 8.84E-01 | no | Ptpra | synonymous_variant | LOW |
| 2 | 130952088 | AX-97807633 | 0 | C | T | 0.14 | 3.24E-02 | 2.23E-01 | 2.70E+00 | 2.73E+00 | 8.84E-01 | no | Atrn | intron_variant | MODIFIER |
| 2 | 131415043 | AX-170019045 | 0 | G | A | 0.14 | 3.24E-02 | 2.23E-01 | 2.70E+00 | 2.73E+00 | 8.84E-01 | no | Gm14283 | non_coding_transcript_exon_variant | MODIFIER |
| 2 | 131451617 | AX-170505536 | 0 | C | T | 0.14 | 3.24E-02 | 2.23E-01 | 2.70E+00 | 2.73E+00 | 8.84E-01 | no | Gm14304 | downstream_gene_variant | MODIFIER |
| 2 | 131466103 | AX-170448969 | 0 | C | T | 0.14 | 3.24E-02 | 2.23E-01 | 2.70E+00 | 2.73E+00 | 8.84E-01 | no | | intergenic_variant | MODIFIER |
| 2 | 131475988 | AX-170470489 | 0 | A | C | 0.14 | 3.24E-02 | 2.23E-01 | 2.70E+00 | 2.73E+00 | 8.84E-01 | no | | intergenic_variant | MODIFIER |
| 2 | 131486151 | AX-170168273 | 0 | A | G | 0.14 | 3.24E-02 | 2.23E-01 | 2.70E+00 | 2.73E+00 | 8.84E-01 | no | | intergenic_variant | MODIFIER |
| 2 | 131499098 | AX-170120934 | 0 | T | A | 0.14 | 3.24E-02 | 2.23E-01 | 2.70E+00 | 2.73E+00 | 8.84E-01 | no | Smox | intron_variant | MODIFIER |
| 2 | 131525639 | AX-170315523 | 0 | C | G | 0.14 | 3.24E-02 | 2.23E-01 | 2.70E+00 | 2.73E+00 | 8.84E-01 | no | Smox | downstream_gene_variant | MODIFIER |
| 2 | 131558438 | AX-170290058 | 0 | C | G | 0.14 | 3.24E-02 | 2.23E-01 | 2.70E+00 | 2.73E+00 | 8.84E-01 | no | Adra1d | intron_variant | MODIFIER |
| 2 | 131681262 | AX-97822938 | 0 | A | G | 0.14 | 3.24E-02 | 2.23E-01 | 2.70E+00 | 2.73E+00 | 8.84E-01 | no | Gm14280 | intron_variant,non_coding_transcript_variant | MODIFIER |
| 2 | 131749425 | AX-170240489 | 0 | T | A | 0.14 | 3.24E-02 | 2.23E-01 | 2.70E+00 | 2.73E+00 | 8.84E-01 | no | | regulatory_region_variant | MODIFIER |
| 2 | 131750862 | AX-170379637 | 0 | A | G | 0.14 | 3.24E-02 | 2.23E-01 | 2.70E+00 | 2.73E+00 | 8.84E-01 | no | | intergenic_variant | MODIFIER |
| 2 | 131758400 | AX-170122547 | 0 | G | A | 0.14 | 3.24E-02 | 2.23E-01 | 2.70E+00 | 2.73E+00 | 8.84E-01 | no | | intergenic_variant | MODIFIER |
| 2 | 131776708 | AX-170302647 | 0 | T | C | 0.14 | 3.24E-02 | 2.23E-01 | 2.70E+00 | 2.73E+00 | 8.84E-01 | no | | intergenic_variant | MODIFIER |
| 2 | 131805014 | AX-170189399 | 0 | T | C | 0.14 | 3.24E-02 | 2.23E-01 | 2.70E+00 | 2.73E+00 | 8.84E-01 | no | | intergenic_variant | MODIFIER |
| 2 | 131807035 | AX-170140672 | 0 | A | C | 0.14 | 3.24E-02 | 2.23E-01 | 2.70E+00 | 2.73E+00 | 8.84E-01 | no | | intergenic_variant | MODIFIER |
| 2 | 131807214 | AX-170271955 | 0 | C | T | 0.14 | 3.24E-02 | 2.23E-01 | 2.70E+00 | 2.73E+00 | 8.84E-01 | no | | intergenic_variant | MODIFIER |
| 2 | 131853680 | AX-170019053 | 0 | A | G | 0.14 | 3.24E-02 | 2.23E-01 | 2.70E+00 | 2.73E+00 | 8.84E-01 | no | Erv3 | 3_prime_UTR_variant | MODIFIER |
| 2 | 131856651 | AX-97840158 | 0 | A | G | 0.14 | 3.24E-02 | 2.23E-01 | 2.70E+00 | 2.73E+00 | 8.84E-01 | no | Erv3 | 5_prime_UTR_variant | MODIFIER |
| 2 | 132540231 | AX-97852835 | 0 | T | C | 0.14 | 3.24E-02 | 2.23E-01 | 2.70E+00 | 2.73E+00 | 8.84E-01 | no | Gpcpd1 | intron_variant | MODIFIER |
| 2 | 132584525 | AX-97839522 | 0 | C | T | 0.14 | 3.24E-02 | 2.23E-01 | 2.70E+00 | 2.73E+00 | 8.84E-01 | no | Gpcpd1 | intron_variant | MODIFIER |
| 2 | 132584569 | AX-170280623 | 0 | A | G | 0.14 | 3.24E-02 | 2.23E-01 | 2.70E+00 | 2.73E+00 | 8.84E-01 | no | Gpcpd1 | intron_variant | MODIFIER |
| 2 | 132584637 | AX-170203604 | 0 | C | T | 0.14 | 3.24E-02 | 2.23E-01 | 2.70E+00 | 2.73E+00 | 8.84E-01 | no | Gpcpd1 | intron_variant | MODIFIER |
| 2 | 132584790 | AX-170273980 | 0 | C | A | 0.14 | 3.24E-02 | 2.23E-01 | 2.70E+00 | 2.73E+00 | 8.84E-01 | no | Gpcpd1 | intron_variant | MODIFIER |
| 2 | 132588590 | AX-170126808 | 0 | G | A | 0.14 | 3.24E-02 | 2.23E-01 | 2.70E+00 | 2.73E+00 | 8.84E-01 | no | Gpcpd1 | upstream_gene_variant | MODIFIER |
| 2 | 132603282 | AX-170345839 | 0 | G | A | 0.14 | 3.24E-02 | 2.23E-01 | 2.70E+00 | 2.73E+00 | 8.84E-01 | no | AU019990 | intron_variant,non_coding_transcript_variant | MODIFIER |
| 2 | 132610146 | AX-97802040 | 0 | T | C | 0.14 | 3.24E-02 | 2.23E-01 | 2.70E+00 | 2.73E+00 | 8.84E-01 | no | AU019990 | intron_variant,non_coding_transcript_variant | MODIFIER |
| 2 | 132638502 | AX-170159583 | 0 | T | A | 0.14 | 3.24E-02 | 2.23E-01 | 2.70E+00 | 2.73E+00 | 8.84E-01 | no | AU019990 | intron_variant,non_coding_transcript_variant | MODIFIER |
| 2 | 132695042 | AX-170094415 | 0 | A | G | 0.14 | 3.24E-02 | 2.23E-01 | 2.70E+00 | 2.73E+00 | 8.84E-01 | no | Shld1 | intron_variant | MODIFIER |
| 2 | 132800710 | AX-170304189 | 0 | A | G | 0.14 | 3.24E-02 | 2.23E-01 | 2.70E+00 | 2.73E+00 | 8.84E-01 | no | Trmt6 | downstream_gene_variant | MODIFIER |
| 2 | 132813404 | AX-97821419 | 0 | C | T | 0.14 | 3.24E-02 | 2.23E-01 | 2.70E+00 | 2.73E+00 | 8.84E-01 | no | Mcm8 | upstream_gene_variant | MODIFIER |
| 2 | 132817992 | AX-170105106 | 0 | A | T | 0.14 | 3.24E-02 | 2.23E-01 | 2.70E+00 | 2.73E+00 | 8.84E-01 | no | Mcm8 | intron_variant | MODIFIER |

| | | | | | | | | | | | | | | | |
|---|---|---|---|---|---|---|---|---|---|---|---|---|---|---|---|
| 2 | 132820744 | AX-170148995 | 0 | T | C | 0.14 | 3.24E-02 | 2.23E-01 | 2.70E+00 | 2.73E+00 | 8.84E-01 | no | Mcm8 | intron_variant | MODIFIER |
| 2 | 132828719 | AX-170396364 | 0 | T | C | 0.14 | 3.24E-02 | 2.23E-01 | 2.70E+00 | 2.73E+00 | 8.84E-01 | no | Mcm8 | intron_variant | MODIFIER |
| 2 | 132829350 | AX-170210976 | 0 | T | G | 0.14 | 3.24E-02 | 2.23E-01 | 2.70E+00 | 2.73E+00 | 8.84E-01 | no | Mcm8 | intron_variant | MODIFIER |
| 2 | 132829516 | AX-170088754 | 0 | A | G | 0.14 | 3.24E-02 | 2.23E-01 | 2.70E+00 | 2.73E+00 | 8.84E-01 | no | Mcm8 | intron_variant | MODIFIER |
| 2 | 132843809 | AX-170223420 | 0 | A | T | 0.14 | 3.24E-02 | 2.23E-01 | 2.70E+00 | 2.73E+00 | 8.84E-01 | no | Mcm8 | 3_prime_UTR_variant | MODIFIER |
| 2 | 133486093 | AX-170019075 | 0 | T | C | 0.14 | 3.24E-02 | 2.23E-01 | 2.70E+00 | 2.73E+00 | 8.84E-01 | no | | intergenic_variant | MODIFIER |
| 2 | 133505740 | AX-170245135 | 0 | T | C | 0.14 | 3.24E-02 | 2.23E-01 | 2.70E+00 | 2.73E+00 | 8.84E-01 | no | | intergenic_variant | MODIFIER |
| 2 | 133507358 | AX-97805430 | 0 | G | A | 0.14 | 3.24E-02 | 2.23E-01 | 2.70E+00 | 2.73E+00 | 8.84E-01 | no | | intergenic_variant | MODIFIER |
| 2 | 133509615 | AX-170219114 | 0 | A | G | 0.14 | 3.24E-02 | 2.23E-01 | 2.70E+00 | 2.73E+00 | 8.84E-01 | no | | intergenic_variant | MODIFIER |
| 2 | 133520532 | AX-170019082 | 0 | A | G | 0.14 | 3.24E-02 | 2.23E-01 | 2.70E+00 | 2.73E+00 | 8.84E-01 | no | | intergenic_variant | MODIFIER |
| 2 | 133540905 | AX-97849826 | 0 | C | T | 0.14 | 3.24E-02 | 2.23E-01 | 2.70E+00 | 2.73E+00 | 8.84E-01 | no | | intergenic_variant | MODIFIER |
| 2 | 133649399 | AX-97841199 | 0 | G | C | 0.14 | 3.24E-02 | 2.23E-01 | 2.70E+00 | 2.73E+00 | 8.84E-01 | no | | intergenic_variant | MODIFIER |
| 2 | 133680246 | AX-170177985 | 0 | C | A | 0.14 | 3.24E-02 | 2.23E-01 | 2.70E+00 | 2.73E+00 | 8.84E-01 | no | | intergenic_variant | MODIFIER |
| 2 | 133683381 | AX-97787800 | 0 | A | G | 0.14 | 3.24E-02 | 2.23E-01 | 2.70E+00 | 2.73E+00 | 8.84E-01 | no | | intergenic_variant | MODIFIER |
| 2 | 133689227 | AX-170506195 | 0 | G | T | 0.14 | 3.24E-02 | 2.23E-01 | 2.70E+00 | 2.73E+00 | 8.84E-01 | no | | intergenic_variant | MODIFIER |
| 2 | 133695297 | AX-170285837 | 0 | A | T | 0.14 | 3.24E-02 | 2.23E-01 | 2.70E+00 | 2.73E+00 | 8.84E-01 | no | | intergenic_variant | MODIFIER |
| 2 | 133709346 | AX-97799811 | 0 | A | G | 0.14 | 3.24E-02 | 2.23E-01 | 2.70E+00 | 2.73E+00 | 8.84E-01 | no | | intergenic_variant | MODIFIER |
| 2 | 136767956 | AX-170019141 | 0 | C | T | 0.14 | 3.24E-02 | 2.23E-01 | 2.70E+00 | 2.73E+00 | 8.84E-01 | no | Snap25 | intron_variant | MODIFIER |
| 2 | 136768182 | AX-170019143 | 0 | G | A | 0.14 | 3.24E-02 | 2.23E-01 | 2.70E+00 | 2.73E+00 | 8.84E-01 | no | Snap25 | intron_variant | MODIFIER |
| 2 | 136771825 | AX-170019140 | 0 | C | T | 0.14 | 3.24E-02 | 2.23E-01 | 2.70E+00 | 2.73E+00 | 8.84E-01 | no | Snap25 | intron_variant | MODIFIER |
| 2 | 136803682 | AX-97851700 | 0 | T | C | 0.14 | 3.24E-02 | 2.23E-01 | 2.70E+00 | 2.73E+00 | 8.84E-01 | no | | intergenic_variant | MODIFIER |
| 2 | 136808045 | AX-170019142 | 0 | C | T | 0.14 | 3.24E-02 | 2.23E-01 | 2.70E+00 | 2.73E+00 | 8.84E-01 | no | | regulatory_region_variant | MODIFIER |
| 2 | 136808611 | AX-170459084 | 0 | G | A | 0.14 | 3.24E-02 | 2.23E-01 | 2.70E+00 | 2.73E+00 | 8.84E-01 | no | | intergenic_variant | MODIFIER |
| 2 | 136839602 | AX-170523070 | 0 | T | C | 0.14 | 3.24E-02 | 2.23E-01 | 2.70E+00 | 2.73E+00 | 8.84E-01 | no | | intergenic_variant | MODIFIER |
| 2 | 136844374 | AX-170490505 | 0 | T | C | 0.14 | 3.24E-02 | 2.23E-01 | 2.70E+00 | 2.73E+00 | 8.84E-01 | no | | intergenic_variant | MODIFIER |
| 2 | 136848453 | AX-170065009 | 0 | G | A | 0.14 | 3.24E-02 | 2.23E-01 | 2.70E+00 | 2.73E+00 | 8.84E-01 | no | | intergenic_variant | MODIFIER |
| 2 | 136848525 | AX-170065011 | 0 | C | T | 0.14 | 3.24E-02 | 2.23E-01 | 2.70E+00 | 2.73E+00 | 8.84E-01 | no | | intergenic_variant | MODIFIER |
| 2 | 136848693 | AX-170327187 | 0 | C | T | 0.14 | 3.24E-02 | 2.23E-01 | 2.70E+00 | 2.73E+00 | 8.84E-01 | no | | intergenic_variant | MODIFIER |
| 2 | 136853480 | AX-97840306 | 0 | A | G | 0.14 | 3.24E-02 | 2.23E-01 | 2.70E+00 | 2.73E+00 | 8.84E-01 | no | | regulatory_region_variant | MODIFIER |
| 2 | 136878453 | AX-97818642 | 0 | A | G | 0.14 | 3.24E-02 | 2.23E-01 | 2.70E+00 | 2.73E+00 | 8.84E-01 | no | Mkks | intron_variant | MODIFIER |
| 2 | 137025830 | AX-170457103 | 0 | A | G | 0.14 | 3.24E-02 | 2.23E-01 | 2.70E+00 | 2.73E+00 | 8.84E-01 | no | Slx4ip | intron_variant | MODIFIER |
| 2 | 137072527 | AX-170212117 | 0 | T | C | 0.14 | 3.24E-02 | 2.23E-01 | 2.70E+00 | 2.73E+00 | 8.84E-01 | no | Slx4ip | downstream_gene_variant | MODIFIER |
| 2 | 137083517 | AX-170302882 | 0 | A | G | 0.14 | 3.24E-02 | 2.23E-01 | 2.70E+00 | 2.73E+00 | 8.84E-01 | no | Jag1 | intron_variant | MODIFIER |
| 2 | 137083707 | AX-170400127 | 0 | T | C | 0.14 | 3.24E-02 | 2.23E-01 | 2.70E+00 | 2.73E+00 | 8.84E-01 | no | Jag1 | intron_variant | MODIFIER |
| 2 | 137086126 | AX-170099905 | 0 | C | T | 0.14 | 3.24E-02 | 2.23E-01 | 2.70E+00 | 2.73E+00 | 8.84E-01 | no | Jag1 | intron_variant | MODIFIER |
| 2 | 137087753 | AX-170411006 | 0 | G | A | 0.14 | 3.24E-02 | 2.23E-01 | 2.70E+00 | 2.73E+00 | 8.84E-01 | no | Jag1 | intron_variant | MODIFIER |
| 2 | 137087804 | AX-170365156 | 0 | C | T | 0.14 | 3.24E-02 | 2.23E-01 | 2.70E+00 | 2.73E+00 | 8.84E-01 | no | Jag1 | intron_variant | MODIFIER |
| 2 | 137142178 | AX-170466381 | 0 | T | C | 0.14 | 3.24E-02 | 2.23E-01 | 2.70E+00 | 2.73E+00 | 8.84E-01 | no | | intergenic_variant | MODIFIER |
| 2 | 137142596 | AX-170385892 | 0 | T | C | 0.14 | 3.24E-02 | 2.23E-01 | 2.70E+00 | 2.73E+00 | 8.84E-01 | no | | intergenic_variant | MODIFIER |
| 2 | 137189817 | AX-170019147 | 0 | A | G | 0.14 | 3.24E-02 | 2.23E-01 | 2.70E+00 | 2.73E+00 | 8.84E-01 | no | | regulatory_region_variant | MODIFIER |
| 2 | 137191218 | AX-170273343 | 0 | A | G | 0.14 | 3.24E-02 | 2.23E-01 | 2.70E+00 | 2.73E+00 | 8.84E-01 | no | | intergenic_variant | MODIFIER |
| 2 | 137193951 | AX-170422872 | 0 | T | C | 0.14 | 3.24E-02 | 2.23E-01 | 2.70E+00 | 2.73E+00 | 8.84E-01 | no | | intergenic_variant | MODIFIER |
| 2 | 137203833 | AX-170019149 | 0 | T | C | 0.14 | 3.24E-02 | 2.23E-01 | 2.70E+00 | 2.73E+00 | 8.84E-01 | no | Gm28214 | downstream_gene_variant | MODIFIER |
| 2 | 137216521 | AX-168413454 | 0 | A | G | 0.14 | 3.24E-02 | 2.23E-01 | 2.70E+00 | 2.73E+00 | 8.84E-01 | no | | intergenic_variant | MODIFIER |
| 2 | 137230535 | AX-170019146 | 0 | G | A | 0.14 | 3.24E-02 | 2.23E-01 | 2.70E+00 | 2.73E+00 | 8.84E-01 | no | | regulatory_region_variant | MODIFIER |
| 2 | 137247355 | AX-170019150 | 0 | T | C | 0.14 | 3.24E-02 | 2.23E-01 | 2.70E+00 | 2.73E+00 | 8.84E-01 | no | | intergenic_variant | MODIFIER |
| 2 | 137322167 | AX-170019145 | 0 | G | A | 0.14 | 3.24E-02 | 2.23E-01 | 2.70E+00 | 2.73E+00 | 8.84E-01 | no | Gm14055 | intron_variant,non_coding_transcript_variant | MODIFIER |
| 2 | 137331634 | AX-97785372 | 0 | T | G | 0.14 | 3.24E-02 | 2.23E-01 | 2.70E+00 | 2.73E+00 | 8.84E-01 | no | Gm14055 | downstream_gene_variant | MODIFIER |
| 2 | 137335565 | AX-170019154 | 0 | C | A | 0.14 | 3.24E-02 | 2.23E-01 | 2.70E+00 | 2.73E+00 | 8.84E-01 | no | | intergenic_variant | MODIFIER |
| 2 | 137342281 | AX-170019148 | 0 | A | G | 0.14 | 3.24E-02 | 2.23E-01 | 2.70E+00 | 2.73E+00 | 8.84E-01 | no | | intergenic_variant | MODIFIER |
| 2 | 137357280 | AX-170019151 | 0 | G | C | 0.14 | 3.24E-02 | 2.23E-01 | 2.70E+00 | 2.73E+00 | 8.84E-01 | no | | regulatory_region_variant | MODIFIER |
| 2 | 137370923 | AX-170247456 | 0 | C | T | 0.14 | 3.24E-02 | 2.23E-01 | 2.70E+00 | 2.73E+00 | 8.84E-01 | no | | intergenic_variant | MODIFIER |
| 2 | 137375079 | AX-170148229 | 0 | T | C | 0.14 | 3.24E-02 | 2.23E-01 | 2.70E+00 | 2.73E+00 | 8.84E-01 | no | | intergenic_variant | MODIFIER |
| 2 | 137376462 | AX-170065016 | 0 | C | T | 0.14 | 3.24E-02 | 2.23E-01 | 2.70E+00 | 2.73E+00 | 8.84E-01 | no | | intergenic_variant | MODIFIER |

| | | | | | | | | | | | | | | | |
|---|---|---|---|---|---|---|---|---|---|---|---|---|---|---|---|
| 2 | 137587677 | AX-97841448 | 0 | T | C | 0.14 | 3.24E-02 | 2.23E-01 | 2.70E+00 | 2.73E+00 | 8.84E-01 | no | | intergenic_variant | MODIFIER |
| 2 | 137614780 | AX-97786206 | 0 | C | T | 0.14 | 3.24E-02 | 2.23E-01 | 2.70E+00 | 2.73E+00 | 8.84E-01 | no | | intergenic_variant | MODIFIER |
| 2 | 137692062 | AX-97841269 | 0 | C | G | 0.14 | 3.24E-02 | 2.23E-01 | 2.70E+00 | 2.73E+00 | 8.84E-01 | no | Gm14064 | intron_variant,non_coding_transcript_variant | MODIFIER |
| 2 | 137789017 | AX-170019160 | 0 | A | G | 0.14 | 3.24E-02 | 2.23E-01 | 2.70E+00 | 2.73E+00 | 8.84E-01 | no | Gm14064 | intron_variant,non_coding_transcript_variant | MODIFIER |
| 2 | 137821929 | AX-170450776 | 0 | A | G | 0.14 | 3.24E-02 | 2.23E-01 | 2.70E+00 | 2.73E+00 | 8.84E-01 | no | Gm14064 | intron_variant,non_coding_transcript_variant | MODIFIER |
| 2 | 137822203 | AX-97804252 | 0 | T | G | 0.14 | 3.24E-02 | 2.23E-01 | 2.70E+00 | 2.73E+00 | 8.84E-01 | no | Gm14064 | intron_variant,non_coding_transcript_variant | MODIFIER |
| 2 | 137823207 | AX-170019162 | 0 | G | A | 0.14 | 3.24E-02 | 2.23E-01 | 2.70E+00 | 2.73E+00 | 8.84E-01 | no | Gm14064 | intron_variant,non_coding_transcript_variant | MODIFIER |
| 2 | 137907962 | AX-170289723 | 0 | A | T | 0.14 | 3.24E-02 | 2.23E-01 | 2.70E+00 | 2.73E+00 | 8.84E-01 | no | | intergenic_variant | MODIFIER |
| 2 | 137908215 | AX-97839188 | 0 | C | T | 0.14 | 3.24E-02 | 2.23E-01 | 2.70E+00 | 2.73E+00 | 8.84E-01 | no | | intergenic_variant | MODIFIER |
| 2 | 137931574 | AX-170534698 | 0 | C | T | 0.14 | 3.24E-02 | 2.23E-01 | 2.70E+00 | 2.73E+00 | 8.84E-01 | no | | intergenic_variant | MODIFIER |
| 2 | 137931648 | AX-170436038 | 0 | C | T | 0.14 | 3.24E-02 | 2.23E-01 | 2.70E+00 | 2.73E+00 | 8.84E-01 | no | | intergenic_variant | MODIFIER |
| 2 | 137931704 | AX-97857151 | 0 | C | T | 0.14 | 3.24E-02 | 2.23E-01 | 2.70E+00 | 2.73E+00 | 8.84E-01 | no | | intergenic_variant | MODIFIER |
| 2 | 137932197 | AX-170169441 | 0 | A | C | 0.14 | 3.24E-02 | 2.23E-01 | 2.70E+00 | 2.73E+00 | 8.84E-01 | no | | intergenic_variant | MODIFIER |
| 2 | 137942970 | AX-170019161 | 0 | A | G | 0.14 | 3.24E-02 | 2.23E-01 | 2.70E+00 | 2.73E+00 | 8.84E-01 | no | | regulatory_region_variant | MODIFIER |
| 2 | 137942993 | AX-170019159 | 0 | A | G | 0.14 | 3.24E-02 | 2.23E-01 | 2.70E+00 | 2.73E+00 | 8.84E-01 | no | | regulatory_region_variant | MODIFIER |
| 2 | 137943298 | AX-170421229 | 0 | G | A | 0.14 | 3.24E-02 | 2.23E-01 | 2.70E+00 | 2.73E+00 | 8.84E-01 | no | | regulatory_region_variant | MODIFIER |
| 2 | 137943480 | AX-170173417 | 0 | A | G | 0.14 | 3.24E-02 | 2.23E-01 | 2.70E+00 | 2.73E+00 | 8.84E-01 | no | | intergenic_variant | MODIFIER |
| 2 | 137945972 | AX-97839505 | 0 | G | A | 0.14 | 3.24E-02 | 2.23E-01 | 2.70E+00 | 2.73E+00 | 8.84E-01 | no | | intergenic_variant | MODIFIER |
| 2 | 138073216 | AX-170478914 | 0 | A | T | 0.14 | 3.24E-02 | 2.23E-01 | 2.70E+00 | 2.73E+00 | 8.84E-01 | no | | intergenic_variant | MODIFIER |
| 2 | 138073299 | AX-97797038 | 0 | G | T | 0.14 | 3.24E-02 | 2.23E-01 | 2.70E+00 | 2.73E+00 | 8.84E-01 | no | | intergenic_variant | MODIFIER |
| 2 | 138096354 | AX-170500879 | 0 | C | T | 0.14 | 3.24E-02 | 2.23E-01 | 2.70E+00 | 2.73E+00 | 8.84E-01 | no | | intergenic_variant | MODIFIER |
| 2 | 138097324 | AX-170418636 | 0 | A | C | 0.14 | 3.24E-02 | 2.23E-01 | 2.70E+00 | 2.73E+00 | 8.84E-01 | no | | intergenic_variant | MODIFIER |
| 2 | 138097514 | AX-170132701 | 0 | C | T | 0.14 | 3.24E-02 | 2.23E-01 | 2.70E+00 | 2.73E+00 | 8.84E-01 | no | | intergenic_variant | MODIFIER |
| 2 | 138109578 | AX-97817312 | 0 | A | G | 0.14 | 3.24E-02 | 2.23E-01 | 2.70E+00 | 2.73E+00 | 8.84E-01 | no | | intergenic_variant | MODIFIER |
| 2 | 138114630 | AX-170188964 | 0 | G | A | 0.14 | 3.24E-02 | 2.23E-01 | 2.70E+00 | 2.73E+00 | 8.84E-01 | no | | regulatory_region_variant | MODIFIER |
| 2 | 138117456 | AX-170448070 | 0 | A | G | 0.14 | 3.24E-02 | 2.23E-01 | 2.70E+00 | 2.73E+00 | 8.84E-01 | no | | intergenic_variant | MODIFIER |
| 2 | 138139532 | AX-170199524 | 0 | A | G | 0.14 | 3.24E-02 | 2.23E-01 | 2.70E+00 | 2.73E+00 | 8.84E-01 | no | | intergenic_variant | MODIFIER |
| 2 | 138206484 | AX-97786062 | 0 | C | T | 0.14 | 3.24E-02 | 2.23E-01 | 2.70E+00 | 2.73E+00 | 8.84E-01 | no | | intergenic_variant | MODIFIER |
| 2 | 138220769 | AX-170530869 | 0 | A | T | 0.14 | 3.24E-02 | 2.23E-01 | 2.70E+00 | 2.73E+00 | 8.84E-01 | no | | intergenic_variant | MODIFIER |
| 2 | 138220906 | AX-170316723 | 0 | A | G | 0.14 | 3.24E-02 | 2.23E-01 | 2.70E+00 | 2.73E+00 | 8.84E-01 | no | | intergenic_variant | MODIFIER |
| 2 | 138221689 | AX-170258235 | 0 | T | C | 0.14 | 3.24E-02 | 2.23E-01 | 2.70E+00 | 2.73E+00 | 8.84E-01 | no | | intergenic_variant | MODIFIER |
| 2 | 138221719 | AX-170449274 | 0 | T | G | 0.14 | 3.24E-02 | 2.23E-01 | 2.70E+00 | 2.73E+00 | 8.84E-01 | no | | intergenic_variant | MODIFIER |
| 2 | 138223501 | AX-97797230 | 0 | A | G | 0.14 | 3.24E-02 | 2.23E-01 | 2.70E+00 | 2.73E+00 | 8.84E-01 | no | | intergenic_variant | MODIFIER |
| 2 | 138239791 | AX-170149230 | 0 | C | T | 0.14 | 3.24E-02 | 2.23E-01 | 2.70E+00 | 2.73E+00 | 8.84E-01 | no | | intergenic_variant | MODIFIER |
| 2 | 138310135 | AX-170489136 | 0 | G | T | 0.14 | 3.24E-02 | 2.23E-01 | 2.70E+00 | 2.73E+00 | 8.84E-01 | no | Btbd3 | intron_variant,non_coding_transcript_variant | MODIFIER |
| 2 | 138318791 | AX-170396277 | 0 | A | G | 0.14 | 3.24E-02 | 2.23E-01 | 2.70E+00 | 2.73E+00 | 8.84E-01 | no | Btbd3 | intron_variant,non_coding_transcript_variant | MODIFIER |
| 2 | 138364870 | AX-170216438 | 0 | A | G | 0.14 | 3.24E-02 | 2.23E-01 | 2.70E+00 | 2.73E+00 | 8.84E-01 | no | Btbd3 | intron_variant,non_coding_transcript_variant | MODIFIER |
| 2 | 138365160 | AX-170285521 | 0 | G | T | 0.14 | 3.24E-02 | 2.23E-01 | 2.70E+00 | 2.73E+00 | 8.84E-01 | no | Btbd3 | intron_variant,non_coding_transcript_variant | MODIFIER |
| 2 | 138368481 | AX-170119561 | 0 | C | T | 0.14 | 3.24E-02 | 2.23E-01 | 2.70E+00 | 2.73E+00 | 8.84E-01 | no | Btbd3 | intron_variant,non_coding_transcript_variant | MODIFIER |
| 2 | 138369698 | AX-170409142 | 0 | G | C | 0.14 | 3.24E-02 | 2.23E-01 | 2.70E+00 | 2.73E+00 | 8.84E-01 | no | Btbd3 | intron_variant,non_coding_transcript_variant | MODIFIER |
| 2 | 138398808 | AX-170209603 | 0 | A | G | 0.14 | 3.24E-02 | 2.23E-01 | 2.70E+00 | 2.73E+00 | 8.84E-01 | no | Btbd3 | intron_variant,non_coding_transcript_variant | MODIFIER |
| 2 | 138438085 | AX-97854295 | 0 | A | G | 0.14 | 3.24E-02 | 2.23E-01 | 2.70E+00 | 2.73E+00 | 8.84E-01 | no | Btbd3 | intron_variant,non_coding_transcript_variant | MODIFIER |
| 2 | 138446149 | AX-170019167 | 0 | G | A | 0.14 | 3.24E-02 | 2.23E-01 | 2.70E+00 | 2.73E+00 | 8.84E-01 | no | Btbd3 | intron_variant,non_coding_transcript_variant | MODIFIER |
| 2 | 138447823 | AX-170019163 | 0 | A | G | 0.14 | 3.24E-02 | 2.23E-01 | 2.70E+00 | 2.73E+00 | 8.84E-01 | no | Btbd3 | intron_variant,non_coding_transcript_variant | MODIFIER |
| 2 | 138448644 | AX-170019165 | 0 | C | T | 0.14 | 3.24E-02 | 2.23E-01 | 2.70E+00 | 2.73E+00 | 8.84E-01 | no | Btbd3 | intron_variant,non_coding_transcript_variant | MODIFIER |
| 2 | 138460515 | AX-97846135 | 0 | A | G | 0.14 | 3.24E-02 | 2.23E-01 | 2.70E+00 | 2.73E+00 | 8.84E-01 | no | Btbd3 | intron_variant,non_coding_transcript_variant | MODIFIER |
| 2 | 138488919 | AX-170019166 | 0 | A | T | 0.14 | 3.24E-02 | 2.23E-01 | 2.70E+00 | 2.73E+00 | 8.84E-01 | no | Btbd3 | intron_variant,non_coding_transcript_variant | MODIFIER |
| 2 | 138548184 | AX-97790272 | 0 | T | C | 0.14 | 3.24E-02 | 2.23E-01 | 2.70E+00 | 2.73E+00 | 8.84E-01 | no | Btbd3 | intron_variant,non_coding_transcript_variant | MODIFIER |
| 2 | 138548450 | AX-170217451 | 0 | C | G | 0.14 | 3.24E-02 | 2.23E-01 | 2.70E+00 | 2.73E+00 | 8.84E-01 | no | Btbd3 | intron_variant,non_coding_transcript_variant | MODIFIER |
| 2 | 138549173 | AX-170516297 | 0 | C | G | 0.14 | 3.24E-02 | 2.23E-01 | 2.70E+00 | 2.73E+00 | 8.84E-01 | no | Btbd3 | intron_variant,non_coding_transcript_variant | MODIFIER |
| 2 | 138650758 | AX-170065035 | 0 | T | C | 0.14 | 3.24E-02 | 2.23E-01 | 2.70E+00 | 2.73E+00 | 8.84E-01 | no | Gm14061 | downstream_gene_variant | MODIFIER |
| 2 | 138658964 | AX-170489883 | 0 | C | T | 0.14 | 3.24E-02 | 2.23E-01 | 2.70E+00 | 2.73E+00 | 8.84E-01 | no | | intergenic_variant | MODIFIER |
| 2 | 138665918 | AX-97790445 | 0 | A | G | 0.14 | 3.24E-02 | 2.23E-01 | 2.70E+00 | 2.73E+00 | 8.84E-01 | no | | intergenic_variant | MODIFIER |
| 2 | 138666128 | AX-170361146 | 0 | G | A | 0.14 | 3.24E-02 | 2.23E-01 | 2.70E+00 | 2.73E+00 | 8.84E-01 | no | | intergenic_variant | MODIFIER |

| | | | | | | | | | | | | | | | |
|---|---|---|---|---|---|---|---|---|---|---|---|---|---|---|---|
| 2 | 138679788 | AX-97798262 | 0 | C | T | 0.14 | 3.24E-02 | 2.23E-01 | 2.70E+00 | 2.73E+00 | 8.84E-01 | no | | intergenic_variant | MODIFIER |
| 2 | 138688344 | AX-170260581 | 0 | A | G | 0.14 | 3.24E-02 | 2.23E-01 | 2.70E+00 | 2.73E+00 | 8.84E-01 | no | | intergenic_variant | MODIFIER |
| 2 | 138730220 | AX-170439045 | 0 | T | C | 0.14 | 3.24E-02 | 2.23E-01 | 2.70E+00 | 2.73E+00 | 8.84E-01 | no | | intergenic_variant | MODIFIER |
| 2 | 138741085 | AX-170314705 | 0 | G | T | 0.14 | 3.24E-02 | 2.23E-01 | 2.70E+00 | 2.73E+00 | 8.84E-01 | no | | intergenic_variant | MODIFIER |
| 2 | 138742420 | AX-97811118 | 0 | A | G | 0.14 | 3.24E-02 | 2.23E-01 | 2.70E+00 | 2.73E+00 | 8.84E-01 | no | | intergenic_variant | MODIFIER |
| 2 | 138771131 | AX-170393178 | 0 | G | A | 0.14 | 3.24E-02 | 2.23E-01 | 2.70E+00 | 2.73E+00 | 8.84E-01 | no | | intergenic_variant | MODIFIER |
| 2 | 138780144 | AX-170232766 | 0 | G | A | 0.14 | 3.24E-02 | 2.23E-01 | 2.70E+00 | 2.73E+00 | 8.84E-01 | no | | intergenic_variant | MODIFIER |
| 2 | 138780462 | AX-170333266 | 0 | T | C | 0.14 | 3.24E-02 | 2.23E-01 | 2.70E+00 | 2.73E+00 | 8.84E-01 | no | | intergenic_variant | MODIFIER |
| 2 | 138780601 | AX-170400478 | 0 | T | C | 0.14 | 3.24E-02 | 2.23E-01 | 2.70E+00 | 2.73E+00 | 8.84E-01 | no | | intergenic_variant | MODIFIER |
| 2 | 138815870 | AX-97855850 | 0 | A | G | 0.14 | 3.24E-02 | 2.23E-01 | 2.70E+00 | 2.73E+00 | 8.84E-01 | no | Mir1952 | upstream_gene_variant | MODIFIER |
| 2 | 138818576 | AX-170496040 | 0 | C | G | 0.14 | 3.24E-02 | 2.23E-01 | 2.70E+00 | 2.73E+00 | 8.84E-01 | no | Mir1952 | upstream_gene_variant | MODIFIER |
| 2 | 138820798 | AX-97798890 | 0 | T | C | 0.14 | 3.24E-02 | 2.23E-01 | 2.70E+00 | 2.73E+00 | 8.84E-01 | no | Mir1952 | downstream_gene_variant | MODIFIER |
| 2 | 138838901 | AX-170175154 | 0 | A | C | 0.14 | 3.24E-02 | 2.23E-01 | 2.70E+00 | 2.73E+00 | 8.84E-01 | no | | intergenic_variant | MODIFIER |
| 2 | 138839780 | AX-170529112 | 0 | G | C | 0.14 | 3.24E-02 | 2.23E-01 | 2.70E+00 | 2.73E+00 | 8.84E-01 | no | | intergenic_variant | MODIFIER |
| 2 | 138841648 | AX-170389041 | 0 | T | A | 0.14 | 3.24E-02 | 2.23E-01 | 2.70E+00 | 2.73E+00 | 8.84E-01 | no | | intergenic_variant | MODIFIER |
| 2 | 138883423 | AX-170100350 | 0 | G | A | 0.14 | 3.24E-02 | 2.23E-01 | 2.70E+00 | 2.73E+00 | 8.84E-01 | no | | intergenic_variant | MODIFIER |
| 2 | 138901019 | AX-170100429 | 0 | C | A | 0.14 | 3.24E-02 | 2.23E-01 | 2.70E+00 | 2.73E+00 | 8.84E-01 | no | | intergenic_variant | MODIFIER |
| 2 | 138975489 | AX-170442549 | 0 | G | A | 0.14 | 3.24E-02 | 2.23E-01 | 2.70E+00 | 2.73E+00 | 8.84E-01 | no | | regulatory_region_variant | MODIFIER |
| 2 | 138975591 | AX-170019171 | 0 | T | C | 0.14 | 3.24E-02 | 2.23E-01 | 2.70E+00 | 2.73E+00 | 8.84E-01 | no | | regulatory_region_variant | MODIFIER |
| 2 | 138990445 | AX-170335295 | 0 | G | T | 0.14 | 3.24E-02 | 2.23E-01 | 2.70E+00 | 2.73E+00 | 8.84E-01 | no | | intergenic_variant | MODIFIER |
| 2 | 138992217 | AX-170019168 | 0 | C | T | 0.14 | 3.24E-02 | 2.23E-01 | 2.70E+00 | 2.73E+00 | 8.84E-01 | no | | intergenic_variant | MODIFIER |
| 2 | 139014718 | AX-97822555 | 0 | G | T | 0.14 | 3.24E-02 | 2.23E-01 | 2.70E+00 | 2.73E+00 | 8.84E-01 | no | | intergenic_variant | MODIFIER |
| 2 | 139054693 | AX-97818116 | 0 | A | G | 0.14 | 3.24E-02 | 2.23E-01 | 2.70E+00 | 2.73E+00 | 8.84E-01 | no | | intergenic_variant | MODIFIER |
| 2 | 139071713 | AX-170019172 | 0 | A | T | 0.14 | 3.24E-02 | 2.23E-01 | 2.70E+00 | 2.73E+00 | 8.84E-01 | no | | intergenic_variant | MODIFIER |
| 2 | 139071823 | AX-170019173 | 0 | A | C | 0.14 | 3.24E-02 | 2.23E-01 | 2.70E+00 | 2.73E+00 | 8.84E-01 | no | | intergenic_variant | MODIFIER |
| 2 | 139089013 | AX-97800638 | 0 | G | A | 0.14 | 3.24E-02 | 2.23E-01 | 2.70E+00 | 2.73E+00 | 8.84E-01 | no | | intergenic_variant | MODIFIER |
| 2 | 139102707 | AX-97827897 | 0 | G | A | 0.14 | 3.24E-02 | 2.23E-01 | 2.70E+00 | 2.73E+00 | 8.84E-01 | no | | intergenic_variant | MODIFIER |
| 2 | 139102834 | AX-170107962 | 0 | G | A | 0.14 | 3.24E-02 | 2.23E-01 | 2.70E+00 | 2.73E+00 | 8.84E-01 | no | | intergenic_variant | MODIFIER |
| 2 | 139119559 | AX-170019174 | 0 | G | T | 0.14 | 3.24E-02 | 2.23E-01 | 2.70E+00 | 2.73E+00 | 8.84E-01 | no | | intergenic_variant | MODIFIER |
| 2 | 139121228 | AX-97797821 | 0 | A | G | 0.14 | 3.24E-02 | 2.23E-01 | 2.70E+00 | 2.73E+00 | 8.84E-01 | no | | intergenic_variant | MODIFIER |
| 2 | 139139764 | AX-170065048 | 0 | G | A | 0.14 | 3.24E-02 | 2.23E-01 | 2.70E+00 | 2.73E+00 | 8.84E-01 | no | | intergenic_variant | MODIFIER |
| 2 | 139208008 | AX-97800213 | 0 | C | T | 0.14 | 3.24E-02 | 2.23E-01 | 2.70E+00 | 2.73E+00 | 8.84E-01 | no | | intergenic_variant | MODIFIER |
| 2 | 139412196 | AX-97807253 | 0 | C | A | 0.14 | 3.24E-02 | 2.23E-01 | 2.70E+00 | 2.73E+00 | 8.84E-01 | no | Gm14066 | intron_variant,non_coding_transcript_variant | MODIFIER |
| 2 | 139563907 | AX-97857291 | 0 | G | T | 0.14 | 3.24E-02 | 2.23E-01 | 2.70E+00 | 2.73E+00 | 8.84E-01 | no | Sptlc3 | intron_variant | MODIFIER |
| 2 | 139624911 | AX-97845371 | 0 | C | T | 0.14 | 3.24E-02 | 2.23E-01 | 2.70E+00 | 2.73E+00 | 8.84E-01 | no | Sptlc3 | intron_variant | MODIFIER |
| 2 | 139725399 | AX-170168549 | 0 | G | A | 0.14 | 3.24E-02 | 2.23E-01 | 2.70E+00 | 2.73E+00 | 8.84E-01 | no | Ism1 | intron_variant | MODIFIER |
| 2 | 139727632 | AX-170019181 | 0 | T | C | 0.14 | 3.24E-02 | 2.23E-01 | 2.70E+00 | 2.73E+00 | 8.84E-01 | no | Ism1 | intron_variant | MODIFIER |
| 2 | 139728004 | AX-170314853 | 0 | G | A | 0.14 | 3.24E-02 | 2.23E-01 | 2.70E+00 | 2.73E+00 | 8.84E-01 | no | Ism1 | intron_variant | MODIFIER |
| 2 | 139728146 | AX-97822982 | 0 | A | G | 0.14 | 3.24E-02 | 2.23E-01 | 2.70E+00 | 2.73E+00 | 8.84E-01 | no | Ism1 | intron_variant | MODIFIER |
| 2 | 139729485 | AX-170307152 | 0 | T | C | 0.14 | 3.24E-02 | 2.23E-01 | 2.70E+00 | 2.73E+00 | 8.84E-01 | no | Ism1 | intron_variant | MODIFIER |
| 2 | 139755111 | AX-97822873 | 0 | C | G | 0.14 | 3.24E-02 | 2.23E-01 | 2.70E+00 | 2.73E+00 | 8.84E-01 | no | Ism1 | intron_variant | MODIFIER |
| 2 | 139865608 | AX-97819263 | 0 | T | C | 0.14 | 3.24E-02 | 2.23E-01 | 2.70E+00 | 2.73E+00 | 8.84E-01 | no | Tasp1 | intron_variant | MODIFIER |
| 2 | 139908823 | AX-170165116 | 0 | G | A | 0.14 | 3.24E-02 | 2.23E-01 | 2.70E+00 | 2.73E+00 | 8.84E-01 | no | Tasp1 | intron_variant | MODIFIER |
| 2 | 139962528 | AX-97806454 | 0 | G | A | 0.14 | 3.24E-02 | 2.23E-01 | 2.70E+00 | 2.73E+00 | 8.84E-01 | no | Tasp1 | intron_variant | MODIFIER |
| 2 | 139965044 | AX-170148345 | 0 | G | A | 0.14 | 3.24E-02 | 2.23E-01 | 2.70E+00 | 2.73E+00 | 8.84E-01 | no | Tasp1 | intron_variant | MODIFIER |
| 2 | 139980202 | AX-170262226 | 0 | T | C | 0.14 | 3.24E-02 | 2.23E-01 | 2.70E+00 | 2.73E+00 | 8.84E-01 | no | Tasp1 | intron_variant | MODIFIER |
| 2 | 139980885 | AX-170094429 | 0 | G | C | 0.14 | 3.24E-02 | 2.23E-01 | 2.70E+00 | 2.73E+00 | 8.84E-01 | no | Tasp1 | intron_variant | MODIFIER |
| 2 | 140267906 | AX-170019189 | 0 | A | G | 0.14 | 3.24E-02 | 2.23E-01 | 2.70E+00 | 2.73E+00 | 8.84E-01 | no | Sel1l2 | intron_variant | MODIFIER |
| 2 | 140290677 | AX-170286920 | 0 | T | C | 0.14 | 3.24E-02 | 2.23E-01 | 2.70E+00 | 2.73E+00 | 8.84E-01 | no | Sel1l2 | downstream_gene_variant | MODIFIER |
| 2 | 140291159 | AX-170019191 | 0 | G | A | 0.14 | 3.24E-02 | 2.23E-01 | 2.70E+00 | 2.73E+00 | 8.84E-01 | no | Sel1l2 | downstream_gene_variant | MODIFIER |
| 2 | 140342423 | AX-170335753 | 0 | A | T | 0.14 | 3.24E-02 | 2.23E-01 | 2.70E+00 | 2.73E+00 | 8.84E-01 | no | Sel1l2 | intron_variant | MODIFIER |
| 2 | 140658879 | AX-97840347 | 0 | G | A | 0.14 | 3.24E-02 | 2.23E-01 | 2.70E+00 | 2.73E+00 | 8.84E-01 | no | Flrt3 | downstream_gene_variant | MODIFIER |
| 2 | 140989512 | AX-97794057 | 0 | A | G | 0.14 | 3.24E-02 | 2.23E-01 | 2.70E+00 | 2.73E+00 | 8.84E-01 | no | Macrod2 | intron_variant | MODIFIER |
| 2 | 141002113 | AX-170065078 | 0 | A | G | 0.14 | 3.24E-02 | 2.23E-01 | 2.70E+00 | 2.73E+00 | 8.84E-01 | no | Macrod2 | intron_variant | MODIFIER |

| | | | | | | | | | | | | | | | |
|---|---|---|---|---|---|---|---|---|---|---|---|---|---|---|---|
| 2 | 141016735 | AX-170220212 | 0 | C | T | 0.14 | 3.24E-02 | 2.23E-01 | 2.70E+00 | 2.73E+00 | 8.84E-01 | no | Macrod2 | intron_variant | MODIFIER |
| 2 | 141036211 | AX-170161067 | 0 | G | A | 0.14 | 3.24E-02 | 2.23E-01 | 2.70E+00 | 2.73E+00 | 8.84E-01 | no | Macrod2 | intron_variant | MODIFIER |
| 2 | 141053021 | AX-170088772 | 0 | C | T | 0.14 | 3.24E-02 | 2.23E-01 | 2.70E+00 | 2.73E+00 | 8.84E-01 | no | Macrod2 | intron_variant | MODIFIER |
| 2 | 141059024 | AX-170219901 | 0 | T | C | 0.14 | 3.24E-02 | 2.23E-01 | 2.70E+00 | 2.73E+00 | 8.84E-01 | no | Macrod2 | intron_variant | MODIFIER |
| 2 | 141087452 | AX-170177273 | 0 | G | C | 0.14 | 3.24E-02 | 2.23E-01 | 2.70E+00 | 2.73E+00 | 8.84E-01 | no | Macrod2 | intron_variant | MODIFIER |
| 2 | 141099479 | AX-170306957 | 0 | T | C | 0.14 | 3.24E-02 | 2.23E-01 | 2.70E+00 | 2.73E+00 | 8.84E-01 | no | Macrod2 | intron_variant | MODIFIER |
| 2 | 141114285 | AX-97843502 | 0 | C | T | 0.14 | 3.24E-02 | 2.23E-01 | 2.70E+00 | 2.73E+00 | 8.84E-01 | no | Macrod2 | intron_variant | MODIFIER |
| 2 | 141146765 | AX-170065082 | 0 | G | A | 0.14 | 3.24E-02 | 2.23E-01 | 2.70E+00 | 2.73E+00 | 8.84E-01 | no | Macrod2 | intron_variant | MODIFIER |
| 2 | 141147673 | AX-170065085 | 0 | C | G | 0.14 | 3.24E-02 | 2.23E-01 | 2.70E+00 | 2.73E+00 | 8.84E-01 | no | Macrod2 | intron_variant | MODIFIER |
| 2 | 141159193 | AX-97840378 | 0 | C | A | 0.14 | 3.24E-02 | 2.23E-01 | 2.70E+00 | 2.73E+00 | 8.84E-01 | no | Macrod2 | intron_variant | MODIFIER |
| 2 | 141207746 | AX-170280507 | 0 | T | C | 0.14 | 3.24E-02 | 2.23E-01 | 2.70E+00 | 2.73E+00 | 8.84E-01 | no | Macrod2 | intron_variant | MODIFIER |
| 2 | 141207863 | AX-97832160 | 0 | T | G | 0.14 | 3.24E-02 | 2.23E-01 | 2.70E+00 | 2.73E+00 | 8.84E-01 | no | Macrod2 | intron_variant | MODIFIER |
| 2 | 141232848 | AX-170019208 | 0 | G | C | 0.14 | 3.24E-02 | 2.23E-01 | 2.70E+00 | 2.73E+00 | 8.84E-01 | no | Macrod2 | intron_variant | MODIFIER |
| 2 | 141276613 | AX-170507398 | 0 | G | A | 0.14 | 3.24E-02 | 2.23E-01 | 2.70E+00 | 2.73E+00 | 8.84E-01 | no | Macrod2 | intron_variant | MODIFIER |
| 2 | 141283684 | AX-97793857 | 0 | C | T | 0.14 | 3.24E-02 | 2.23E-01 | 2.70E+00 | 2.73E+00 | 8.84E-01 | no | Macrod2 | intron_variant | MODIFIER |
| 2 | 142020442 | AX-170221228 | 0 | T | C | 0.14 | 3.24E-02 | 2.23E-01 | 2.70E+00 | 2.73E+00 | 8.84E-01 | no | Macrod2 | intron_variant | MODIFIER |
| 2 | 142025928 | AX-170496246 | 0 | T | G | 0.14 | 3.24E-02 | 2.23E-01 | 2.70E+00 | 2.73E+00 | 8.84E-01 | no | Macrod2 | intron_variant | MODIFIER |
| 2 | 142027386 | AX-170168662 | 0 | T | C | 0.14 | 3.24E-02 | 2.23E-01 | 2.70E+00 | 2.73E+00 | 8.84E-01 | no | Macrod2 | intron_variant | MODIFIER |
| 2 | 142107375 | AX-170088775 | 0 | T | C | 0.14 | 3.24E-02 | 2.23E-01 | 2.70E+00 | 2.73E+00 | 8.84E-01 | no | Macrod2 | intron_variant | MODIFIER |
| 2 | 142121108 | AX-170506141 | 0 | G | A | 0.14 | 3.24E-02 | 2.23E-01 | 2.70E+00 | 2.73E+00 | 8.84E-01 | no | Macrod2 | intron_variant | MODIFIER |
| 2 | 142122148 | AX-170294057 | 0 | T | C | 0.14 | 3.24E-02 | 2.23E-01 | 2.70E+00 | 2.73E+00 | 8.84E-01 | no | Macrod2 | intron_variant | MODIFIER |
| 2 | 142123358 | AX-170340414 | 0 | G | C | 0.14 | 3.24E-02 | 2.23E-01 | 2.70E+00 | 2.73E+00 | 8.84E-01 | no | Macrod2 | intron_variant | MODIFIER |
| 2 | 142123500 | AX-97788867 | 0 | T | C | 0.14 | 3.24E-02 | 2.23E-01 | 2.70E+00 | 2.73E+00 | 8.84E-01 | no | Macrod2 | intron_variant | MODIFIER |
| 2 | 142156023 | AX-97830442 | 0 | C | T | 0.14 | 3.24E-02 | 2.23E-01 | 2.70E+00 | 2.73E+00 | 8.84E-01 | no | Macrod2 | intron_variant | MODIFIER |
| 2 | 142169562 | AX-170019230 | 0 | G | A | 0.14 | 3.24E-02 | 2.23E-01 | 2.70E+00 | 2.73E+00 | 8.84E-01 | no | Macrod2 | intron_variant | MODIFIER |
| 2 | 142188987 | AX-170019226 | 0 | C | T | 0.14 | 3.24E-02 | 2.23E-01 | 2.70E+00 | 2.73E+00 | 8.84E-01 | no | Macrod2 | intron_variant | MODIFIER |
| 2 | 142189556 | AX-97840434 | 0 | G | A | 0.14 | 3.24E-02 | 2.23E-01 | 2.70E+00 | 2.73E+00 | 8.84E-01 | no | Macrod2 | intron_variant | MODIFIER |
| 2 | 142193980 | AX-170411595 | 0 | G | A | 0.14 | 3.24E-02 | 2.23E-01 | 2.70E+00 | 2.73E+00 | 8.84E-01 | no | Macrod2 | intron_variant | MODIFIER |
| 2 | 142250852 | AX-170148801 | 0 | T | C | 0.14 | 3.24E-02 | 2.23E-01 | 2.70E+00 | 2.73E+00 | 8.84E-01 | no | Macrod2 | intron_variant | MODIFIER |
| 2 | 142253412 | AX-97809093 | 0 | G | T | 0.14 | 3.24E-02 | 2.23E-01 | 2.70E+00 | 2.73E+00 | 8.84E-01 | no | Macrod2 | intron_variant | MODIFIER |
| 2 | 142253642 | AX-170232795 | 0 | G | A | 0.14 | 3.24E-02 | 2.23E-01 | 2.70E+00 | 2.73E+00 | 8.84E-01 | no | Macrod2 | intron_variant | MODIFIER |
| 2 | 142370317 | AX-170019231 | 0 | G | A | 0.14 | 3.24E-02 | 2.23E-01 | 2.70E+00 | 2.73E+00 | 8.84E-01 | no | Macrod2 | intron_variant | MODIFIER |
| 2 | 142388696 | AX-170088777 | 0 | G | T | 0.14 | 3.24E-02 | 2.23E-01 | 2.70E+00 | 2.73E+00 | 8.84E-01 | no | Macrod2 | downstream_gene_variant | MODIFIER |
| 2 | 142400726 | AX-97803615 | 0 | A | G | 0.14 | 3.24E-02 | 2.23E-01 | 2.70E+00 | 2.73E+00 | 8.84E-01 | no | | intergenic_variant | MODIFIER |
| 2 | 142439093 | AX-170251589 | 0 | T | C | 0.14 | 3.24E-02 | 2.23E-01 | 2.70E+00 | 2.73E+00 | 8.84E-01 | no | | intergenic_variant | MODIFIER |
| 2 | 142447865 | AX-97788138 | 0 | A | C | 0.14 | 3.24E-02 | 2.23E-01 | 2.70E+00 | 2.73E+00 | 8.84E-01 | no | | intergenic_variant | MODIFIER |
| 2 | 142490697 | AX-170173156 | 0 | A | G | 0.14 | 3.24E-02 | 2.23E-01 | 2.70E+00 | 2.73E+00 | 8.84E-01 | no | | intergenic_variant | MODIFIER |
| 2 | 142491077 | AX-97836027 | 0 | G | A | 0.14 | 3.24E-02 | 2.23E-01 | 2.70E+00 | 2.73E+00 | 8.84E-01 | no | | intergenic_variant | MODIFIER |
| 2 | 142504311 | AX-97798161 | 0 | C | T | 0.14 | 3.24E-02 | 2.23E-01 | 2.70E+00 | 2.73E+00 | 8.84E-01 | no | | intergenic_variant | MODIFIER |
| 2 | 142517946 | AX-170452546 | 0 | A | G | 0.14 | 3.24E-02 | 2.23E-01 | 2.70E+00 | 2.73E+00 | 8.84E-01 | no | | intergenic_variant | MODIFIER |
| 2 | 142579175 | AX-97842788 | 0 | C | T | 0.14 | 3.24E-02 | 2.23E-01 | 2.70E+00 | 2.73E+00 | 8.84E-01 | no | Rbmx2-ps | downstream_gene_variant | MODIFIER |
| 2 | 142598018 | AX-170094433 | 0 | G | A | 0.14 | 3.24E-02 | 2.23E-01 | 2.70E+00 | 2.73E+00 | 8.84E-01 | no | | intergenic_variant | MODIFIER |
| 2 | 142598277 | AX-170430243 | 0 | T | C | 0.14 | 3.24E-02 | 2.23E-01 | 2.70E+00 | 2.73E+00 | 8.84E-01 | no | | intergenic_variant | MODIFIER |
| 2 | 142696413 | AX-97848158 | 0 | T | C | 0.14 | 3.24E-02 | 2.23E-01 | 2.70E+00 | 2.73E+00 | 8.84E-01 | no | Kif16b | intron_variant | MODIFIER |
| 2 | 142700579 | AX-170065110 | 0 | A | C | 0.14 | 3.24E-02 | 2.23E-01 | 2.70E+00 | 2.73E+00 | 8.84E-01 | no | Kif16b | intron_variant | MODIFIER |
| 2 | 142700671 | AX-170065112 | 0 | T | A | 0.14 | 3.24E-02 | 2.23E-01 | 2.70E+00 | 2.73E+00 | 8.84E-01 | no | Kif16b | intron_variant | MODIFIER |
| 2 | 142714726 | AX-170192669 | 0 | T | C | 0.14 | 3.24E-02 | 2.23E-01 | 2.70E+00 | 2.73E+00 | 8.84E-01 | no | Kif16b | synonymous_variant | LOW |
| 2 | 142770424 | AX-170019238 | 0 | G | A | 0.14 | 3.24E-02 | 2.23E-01 | 2.70E+00 | 2.73E+00 | 8.84E-01 | no | Kif16b | intron_variant | MODIFIER |
| 2 | 142785813 | AX-170356184 | 0 | T | A | 0.14 | 3.24E-02 | 2.23E-01 | 2.70E+00 | 2.73E+00 | 8.84E-01 | no | Kif16b | intron_variant | MODIFIER |
| 2 | 142786877 | AX-97806590 | 0 | A | G | 0.14 | 3.24E-02 | 2.23E-01 | 2.70E+00 | 2.73E+00 | 8.84E-01 | no | Kif16b | intron_variant | MODIFIER |
| 2 | 142820786 | AX-97813566 | 0 | G | A | 0.14 | 3.24E-02 | 2.23E-01 | 2.70E+00 | 2.73E+00 | 8.84E-01 | no | Kif16b | intron_variant | MODIFIER |
| 2 | 142829469 | AX-170019233 | 0 | G | A | 0.14 | 3.24E-02 | 2.23E-01 | 2.70E+00 | 2.73E+00 | 8.84E-01 | no | Kif16b | intron_variant | MODIFIER |
| 2 | 142860709 | AX-97785910 | 0 | C | T | 0.14 | 3.24E-02 | 2.23E-01 | 2.70E+00 | 2.73E+00 | 8.84E-01 | no | Kif16b | intron_variant | MODIFIER |
| 2 | 142861311 | AX-170019235 | 0 | G | A | 0.14 | 3.24E-02 | 2.23E-01 | 2.70E+00 | 2.73E+00 | 8.84E-01 | no | Kif16b | intron_variant | MODIFIER |

| | | | | | | | | | | | | | | | |
|---|---|---|---|---|---|---|---|---|---|---|---|---|---|---|---|
| 2 | 142891952 | AX-170154997 | 0 | C | T | 0.14 | 3.24E-02 | 2.23E-01 | 2.70E+00 | 2.73E+00 | 8.84E-01 | no | Kif16b | intron_variant | MODIFIER |
| 2 | 142896817 | AX-170019240 | 0 | T | G | 0.14 | 3.24E-02 | 2.23E-01 | 2.70E+00 | 2.73E+00 | 8.84E-01 | no | Kif16b | intron_variant | MODIFIER |
| 2 | 142962570 | AX-170311454 | 0 | A | C | 0.14 | 3.24E-02 | 2.23E-01 | 2.70E+00 | 2.73E+00 | 8.84E-01 | no | | intergenic_variant | MODIFIER |
| 2 | 142972207 | AX-170528013 | 0 | C | T | 0.14 | 3.24E-02 | 2.23E-01 | 2.70E+00 | 2.73E+00 | 8.84E-01 | no | | intergenic_variant | MODIFIER |
| 2 | 143007717 | AX-170088778 | 0 | G | T | 0.14 | 3.24E-02 | 2.23E-01 | 2.70E+00 | 2.73E+00 | 8.84E-01 | no | | intergenic_variant | MODIFIER |
| 2 | 143069500 | AX-170065116 | 0 | C | A | 0.14 | 3.24E-02 | 2.23E-01 | 2.70E+00 | 2.73E+00 | 8.84E-01 | no | Snrpb2 | intron_variant | MODIFIER |
| 2 | 143076100 | AX-170525515 | 0 | C | T | 0.14 | 3.24E-02 | 2.23E-01 | 2.70E+00 | 2.73E+00 | 8.84E-01 | no | Snrpb2 | downstream_gene_variant | MODIFIER |
| 2 | 143079422 | AX-170297979 | 0 | A | G | 0.14 | 3.24E-02 | 2.23E-01 | 2.70E+00 | 2.73E+00 | 8.84E-01 | no | Otor | intron_variant | MODIFIER |
| 2 | 143079901 | AX-170338683 | 0 | G | A | 0.14 | 3.24E-02 | 2.23E-01 | 2.70E+00 | 2.73E+00 | 8.84E-01 | no | Otor | intron_variant | MODIFIER |
| 2 | 143085856 | AX-97817981 | 0 | T | G | 0.14 | 3.24E-02 | 2.23E-01 | 2.70E+00 | 2.73E+00 | 8.84E-01 | no | Otor | downstream_gene_variant | MODIFIER |
| 2 | 143097995 | AX-170173706 | 0 | T | A | 0.14 | 3.24E-02 | 2.23E-01 | 2.70E+00 | 2.73E+00 | 8.84E-01 | no | | intergenic_variant | MODIFIER |
| 2 | 143099072 | AX-170226658 | 0 | A | G | 0.14 | 3.24E-02 | 2.23E-01 | 2.70E+00 | 2.73E+00 | 8.84E-01 | no | | intergenic_variant | MODIFIER |
| 2 | 143099393 | AX-170175156 | 0 | T | C | 0.14 | 3.24E-02 | 2.23E-01 | 2.70E+00 | 2.73E+00 | 8.84E-01 | no | | intergenic_variant | MODIFIER |
| 2 | 143103487 | AX-170372089 | 0 | C | A | 0.14 | 3.24E-02 | 2.23E-01 | 2.70E+00 | 2.73E+00 | 8.84E-01 | no | | intergenic_variant | MODIFIER |
| 2 | 143207675 | AX-170109146 | 0 | C | G | 0.14 | 3.24E-02 | 2.23E-01 | 2.70E+00 | 2.73E+00 | 8.84E-01 | no | | intergenic_variant | MODIFIER |
| 2 | 143214065 | AX-170408957 | 0 | A | T | 0.14 | 3.24E-02 | 2.23E-01 | 2.70E+00 | 2.73E+00 | 8.84E-01 | no | | intergenic_variant | MODIFIER |
| 2 | 143215047 | AX-97839277 | 0 | T | G | 0.14 | 3.24E-02 | 2.23E-01 | 2.70E+00 | 2.73E+00 | 8.84E-01 | no | | intergenic_variant | MODIFIER |
| 2 | 143306591 | AX-170019248 | 0 | C | T | 0.14 | 3.24E-02 | 2.23E-01 | 2.70E+00 | 2.73E+00 | 8.84E-01 | no | | regulatory_region_variant | MODIFIER |
| 2 | 143319584 | AX-97854223 | 0 | T | C | 0.14 | 3.24E-02 | 2.23E-01 | 2.70E+00 | 2.73E+00 | 8.84E-01 | no | | intergenic_variant | MODIFIER |
| 2 | 143334598 | AX-97856514 | 0 | T | C | 0.14 | 3.24E-02 | 2.23E-01 | 2.70E+00 | 2.73E+00 | 8.84E-01 | no | | intergenic_variant | MODIFIER |
| 2 | 143366956 | AX-168419929 | 0 | C | T | 0.14 | 3.24E-02 | 2.23E-01 | 2.70E+00 | 2.73E+00 | 8.84E-01 | no | | intergenic_variant | MODIFIER |
| 2 | 143370537 | AX-170088780 | 0 | T | G | 0.14 | 3.24E-02 | 2.23E-01 | 2.70E+00 | 2.73E+00 | 8.84E-01 | no | | intergenic_variant | MODIFIER |
| 2 | 143370881 | AX-170019244 | 0 | C | A | 0.14 | 3.24E-02 | 2.23E-01 | 2.70E+00 | 2.73E+00 | 8.84E-01 | no | | intergenic_variant | MODIFIER |
| 2 | 143409233 | AX-97795685 | 0 | C | T | 0.14 | 3.24E-02 | 2.23E-01 | 2.70E+00 | 2.73E+00 | 8.84E-01 | no | | intergenic_variant | MODIFIER |
| 2 | 143420108 | AX-170019247 | 0 | C | T | 0.14 | 3.24E-02 | 2.23E-01 | 2.70E+00 | 2.73E+00 | 8.84E-01 | no | | intergenic_variant | MODIFIER |
| 2 | 143437710 | AX-97854521 | 0 | G | A | 0.14 | 3.24E-02 | 2.23E-01 | 2.70E+00 | 2.73E+00 | 8.84E-01 | no | | intergenic_variant | MODIFIER |
| 2 | 143452323 | AX-170105334 | 0 | G | A | 0.14 | 3.24E-02 | 2.23E-01 | 2.70E+00 | 2.73E+00 | 8.84E-01 | no | | regulatory_region_variant | MODIFIER |
| 2 | 143459482 | AX-170019251 | 0 | C | T | 0.14 | 3.24E-02 | 2.23E-01 | 2.70E+00 | 2.73E+00 | 8.84E-01 | no | | intergenic_variant | MODIFIER |
| 2 | 143495391 | AX-97825417 | 0 | G | A | 0.14 | 3.24E-02 | 2.23E-01 | 2.70E+00 | 2.73E+00 | 8.84E-01 | no | | intergenic_variant | MODIFIER |
| 2 | 143495903 | AX-170521551 | 0 | G | A | 0.14 | 3.24E-02 | 2.23E-01 | 2.70E+00 | 2.73E+00 | 8.84E-01 | no | | intergenic_variant | MODIFIER |
| 2 | 143523444 | AX-170279444 | 0 | C | G | 0.14 | 3.24E-02 | 2.23E-01 | 2.70E+00 | 2.73E+00 | 8.84E-01 | no | | intergenic_variant | MODIFIER |
| 2 | 143534658 | AX-170191231 | 0 | G | A | 0.14 | 3.24E-02 | 2.23E-01 | 2.70E+00 | 2.73E+00 | 8.84E-01 | no | Pcsk2os1 | downstream_gene_variant | MODIFIER |
| 2 | 143552678 | AX-170467631 | 0 | G | A | 0.14 | 3.24E-02 | 2.23E-01 | 2.70E+00 | 2.73E+00 | 8.84E-01 | no | Pcsk2 | intron_variant | MODIFIER |
| 2 | 143565352 | AX-97795225 | 0 | A | G | 0.14 | 3.24E-02 | 2.23E-01 | 2.70E+00 | 2.73E+00 | 8.84E-01 | no | Pcsk2 | intron_variant | MODIFIER |
| 2 | 143653717 | AX-170483023 | 0 | G | A | 0.14 | 3.24E-02 | 2.23E-01 | 2.70E+00 | 2.73E+00 | 8.84E-01 | no | Pcsk2 | intron_variant | MODIFIER |
| 2 | 143665191 | AX-170523784 | 0 | T | C | 0.14 | 3.24E-02 | 2.23E-01 | 2.70E+00 | 2.73E+00 | 8.84E-01 | no | Pcsk2 | intron_variant | MODIFIER |
| 2 | 143680428 | AX-97839274 | 0 | A | G | 0.14 | 3.24E-02 | 2.23E-01 | 2.70E+00 | 2.73E+00 | 8.84E-01 | no | Pcsk2 | intron_variant | MODIFIER |
| 2 | 143780068 | AX-170151070 | 0 | T | C | 0.14 | 3.24E-02 | 2.23E-01 | 2.70E+00 | 2.73E+00 | 8.84E-01 | no | Pcsk2 | intron_variant | MODIFIER |
| 2 | 143924535 | AX-170445213 | 0 | T | C | 0.14 | 3.24E-02 | 2.23E-01 | 2.70E+00 | 2.73E+00 | 8.84E-01 | no | Dstn | intron_variant | MODIFIER |
| 2 | 143956409 | AX-97842899 | 0 | C | T | 0.14 | 3.24E-02 | 2.23E-01 | 2.70E+00 | 2.73E+00 | 8.84E-01 | no | Rrbp1 | intron_variant | MODIFIER |
| 2 | 143983182 | AX-170279691 | 0 | T | C | 0.14 | 3.24E-02 | 2.23E-01 | 2.70E+00 | 2.73E+00 | 8.84E-01 | no | Rrbp1 | intron_variant | MODIFIER |
| 2 | 143991724 | AX-170509665 | 0 | G | A | 0.14 | 3.24E-02 | 2.23E-01 | 2.70E+00 | 2.73E+00 | 8.84E-01 | no | Rrbp1 | intron_variant | MODIFIER |
| 2 | 144013451 | AX-170487744 | 0 | A | C | 0.14 | 3.24E-02 | 2.23E-01 | 2.70E+00 | 2.73E+00 | 8.84E-01 | no | Rrbp1 | upstream_gene_variant | MODIFIER |
| 2 | 144014572 | AX-170112148 | 0 | G | A | 0.14 | 3.24E-02 | 2.23E-01 | 2.70E+00 | 2.73E+00 | 8.84E-01 | no | Rrbp1 | upstream_gene_variant | MODIFIER |
| 2 | 144016485 | AX-170259515 | 0 | A | G | 0.14 | 3.24E-02 | 2.23E-01 | 2.70E+00 | 2.73E+00 | 8.84E-01 | no | | intergenic_variant | MODIFIER |
| 2 | 144025686 | AX-170065133 | 0 | A | G | 0.14 | 3.24E-02 | 2.23E-01 | 2.70E+00 | 2.73E+00 | 8.84E-01 | no | | regulatory_region_variant | MODIFIER |
| 2 | 144038148 | AX-97800094 | 0 | T | C | 0.14 | 3.24E-02 | 2.23E-01 | 2.70E+00 | 2.73E+00 | 8.84E-01 | no | Banf2 | intron_variant | MODIFIER |
| 2 | 144040457 | AX-168411548 | 0 | C | T | 0.14 | 3.24E-02 | 2.23E-01 | 2.70E+00 | 2.73E+00 | 8.84E-01 | no | Banf2 | intron_variant | MODIFIER |
| 2 | 144051069 | AX-170146626 | 0 | A | G | 0.14 | 3.24E-02 | 2.23E-01 | 2.70E+00 | 2.73E+00 | 8.84E-01 | no | Banf2 | intron_variant | MODIFIER |
| 2 | 144094000 | AX-170397173 | 0 | G | A | 0.14 | 3.24E-02 | 2.23E-01 | 2.70E+00 | 2.73E+00 | 8.84E-01 | no | Gm5535 | downstream_gene_variant | MODIFIER |
| 2 | 144094362 | AX-170451292 | 0 | T | C | 0.14 | 3.24E-02 | 2.23E-01 | 2.70E+00 | 2.73E+00 | 8.84E-01 | no | Gm5535 | downstream_gene_variant | MODIFIER |
| 2 | 144121646 | AX-170432054 | 0 | G | A | 0.14 | 3.24E-02 | 2.23E-01 | 2.70E+00 | 2.73E+00 | 8.84E-01 | no | Gm5535 | intron_variant,non_coding_transcript_variant | MODIFIER |
| 2 | 144133086 | AX-97791900 | 0 | A | G | 0.14 | 3.24E-02 | 2.23E-01 | 2.70E+00 | 2.73E+00 | 8.84E-01 | no | Gm5535 | intron_variant,non_coding_transcript_variant | MODIFIER |
| 2 | 144133972 | AX-170238226 | 0 | T | C | 0.14 | 3.24E-02 | 2.23E-01 | 2.70E+00 | 2.73E+00 | 8.84E-01 | no | Gm5535 | intron_variant,non_coding_transcript_variant | MODIFIER |

| | | | | | | | | | | | | | | | |
|---|---|---|---|---|---|---|---|---|---|---|---|---|---|---|---|
| 2 | 144198971 | AX-170149265 | 0 | C | G | 0.14 | 3.24E-02 | 2.23E-01 | 2.70E+00 | 2.73E+00 | 8.84E-01 | no | Gm5535 | intron_variant,non_coding_transcript_variant | MODIFIER |
| 2 | 144207339 | AX-97795252 | 0 | G | A | 0.14 | 3.24E-02 | 2.23E-01 | 2.70E+00 | 2.73E+00 | 8.84E-01 | no | Gm5535 | intron_variant,non_coding_transcript_variant | MODIFIER |
| 2 | 144604063 | AX-170227301 | 0 | T | C | 0.14 | 3.24E-02 | 2.23E-01 | 2.70E+00 | 2.73E+00 | 8.84E-01 | no | Dtd1 | intron_variant | MODIFIER |
| 2 | 144604186 | AX-170220183 | 0 | A | G | 0.14 | 3.24E-02 | 2.23E-01 | 2.70E+00 | 2.73E+00 | 8.84E-01 | no | Dtd1 | intron_variant | MODIFIER |
| 2 | 144666057 | AX-170178346 | 0 | C | T | 0.14 | 3.24E-02 | 2.23E-01 | 2.70E+00 | 2.73E+00 | 8.84E-01 | no | Dtd1 | intron_variant | MODIFIER |
| 2 | 144843586 | AX-97846022 | 0 | A | C | 0.14 | 3.24E-02 | 2.23E-01 | 2.70E+00 | 2.73E+00 | 8.84E-01 | no | | intergenic_variant | MODIFIER |
| 2 | 145052306 | AX-170094435 | 0 | G | A | 0.14 | 3.24E-02 | 2.23E-01 | 2.70E+00 | 2.73E+00 | 8.84E-01 | no | | intergenic_variant | MODIFIER |
| 2 | 145460287 | AX-97822227 | 0 | A | G | 0.14 | 3.24E-02 | 2.23E-01 | 2.70E+00 | 2.73E+00 | 8.84E-01 | no | Slc24a3 | intron_variant | MODIFIER |
| 2 | 145460559 | AX-170065164 | 0 | C | T | 0.14 | 3.24E-02 | 2.23E-01 | 2.70E+00 | 2.73E+00 | 8.84E-01 | no | Slc24a3 | intron_variant | MODIFIER |
| 2 | 145460963 | AX-170261320 | 0 | A | T | 0.14 | 3.24E-02 | 2.23E-01 | 2.70E+00 | 2.73E+00 | 8.84E-01 | no | Slc24a3 | intron_variant | MODIFIER |
| 2 | 145473959 | AX-170181995 | 0 | T | C | 0.14 | 3.24E-02 | 2.23E-01 | 2.70E+00 | 2.73E+00 | 8.84E-01 | no | Slc24a3 | intron_variant | MODIFIER |
| 2 | 145489781 | AX-97835614 | 0 | T | A | 0.14 | 3.24E-02 | 2.23E-01 | 2.70E+00 | 2.73E+00 | 8.84E-01 | no | Slc24a3 | intron_variant | MODIFIER |
| 2 | 145513361 | AX-170403241 | 0 | T | C | 0.14 | 3.24E-02 | 2.23E-01 | 2.70E+00 | 2.73E+00 | 8.84E-01 | no | Slc24a3 | intron_variant | MODIFIER |
| 2 | 145513458 | AX-170431809 | 0 | A | C | 0.14 | 3.24E-02 | 2.23E-01 | 2.70E+00 | 2.73E+00 | 8.84E-01 | no | Slc24a3 | intron_variant | MODIFIER |
| 2 | 145518318 | AX-170330153 | 0 | T | C | 0.14 | 3.24E-02 | 2.23E-01 | 2.70E+00 | 2.73E+00 | 8.84E-01 | no | Slc24a3 | intron_variant | MODIFIER |
| 2 | 145519918 | AX-170312252 | 0 | C | T | 0.14 | 3.24E-02 | 2.23E-01 | 2.70E+00 | 2.73E+00 | 8.84E-01 | no | Slc24a3 | intron_variant | MODIFIER |
| 2 | 145525795 | AX-170150918 | 0 | A | G | 0.14 | 3.24E-02 | 2.23E-01 | 2.70E+00 | 2.73E+00 | 8.84E-01 | no | Slc24a3 | intron_variant | MODIFIER |
| 2 | 145525848 | AX-170304957 | 0 | T | G | 0.14 | 3.24E-02 | 2.23E-01 | 2.70E+00 | 2.73E+00 | 8.84E-01 | no | Slc24a3 | intron_variant | MODIFIER |
| 2 | 145527966 | AX-170464883 | 0 | T | G | 0.14 | 3.24E-02 | 2.23E-01 | 2.70E+00 | 2.73E+00 | 8.84E-01 | no | Slc24a3 | intron_variant | MODIFIER |
| 2 | 145529092 | AX-97787507 | 0 | C | T | 0.14 | 3.24E-02 | 2.23E-01 | 2.70E+00 | 2.73E+00 | 8.84E-01 | no | Slc24a3 | intron_variant | MODIFIER |
| 2 | 145554507 | AX-97823828 | 0 | T | C | 0.14 | 3.24E-02 | 2.23E-01 | 2.70E+00 | 2.73E+00 | 8.84E-01 | no | Slc24a3 | intron_variant | MODIFIER |
| 2 | 145554870 | AX-170382079 | 0 | G | C | 0.14 | 3.24E-02 | 2.23E-01 | 2.70E+00 | 2.73E+00 | 8.84E-01 | no | Slc24a3 | intron_variant | MODIFIER |
| 2 | 145570054 | AX-170065166 | 0 | T | C | 0.14 | 3.24E-02 | 2.23E-01 | 2.70E+00 | 2.73E+00 | 8.84E-01 | no | Slc24a3 | intron_variant | MODIFIER |
| 2 | 145584469 | AX-170451392 | 0 | A | C | 0.14 | 3.24E-02 | 2.23E-01 | 2.70E+00 | 2.73E+00 | 8.84E-01 | no | Slc24a3 | intron_variant | MODIFIER |
| 2 | 145584503 | AX-170138365 | 0 | G | C | 0.14 | 3.24E-02 | 2.23E-01 | 2.70E+00 | 2.73E+00 | 8.84E-01 | no | Slc24a3 | intron_variant | MODIFIER |
| 2 | 145600151 | AX-97846710 | 0 | T | C | 0.14 | 3.24E-02 | 2.23E-01 | 2.70E+00 | 2.73E+00 | 8.84E-01 | no | Slc24a3 | intron_variant | MODIFIER |
| 2 | 145607855 | AX-170182934 | 0 | G | A | 0.14 | 3.24E-02 | 2.23E-01 | 2.70E+00 | 2.73E+00 | 8.84E-01 | no | Slc24a3 | intron_variant | MODIFIER |
| 2 | 145630679 | AX-170212434 | 0 | A | G | 0.14 | 3.24E-02 | 2.23E-01 | 2.70E+00 | 2.73E+00 | 8.84E-01 | no | Slc24a3 | intron_variant | MODIFIER |
| 2 | 145631647 | AX-170189589 | 0 | C | T | 0.14 | 3.24E-02 | 2.23E-01 | 2.70E+00 | 2.73E+00 | 8.84E-01 | no | Slc24a3 | intron_variant | MODIFIER |
| 2 | 145631713 | AX-170142912 | 0 | C | T | 0.14 | 3.24E-02 | 2.23E-01 | 2.70E+00 | 2.73E+00 | 8.84E-01 | no | Slc24a3 | intron_variant | MODIFIER |
| 2 | 145631741 | AX-170134126 | 0 | A | C | 0.14 | 3.24E-02 | 2.23E-01 | 2.70E+00 | 2.73E+00 | 8.84E-01 | no | Slc24a3 | intron_variant | MODIFIER |
| 2 | 145631857 | AX-97791675 | 0 | T | C | 0.14 | 3.24E-02 | 2.23E-01 | 2.70E+00 | 2.73E+00 | 8.84E-01 | no | Slc24a3 | intron_variant | MODIFIER |
| 2 | 145636222 | AX-170194968 | 0 | T | C | 0.14 | 3.24E-02 | 2.23E-01 | 2.70E+00 | 2.73E+00 | 8.84E-01 | no | Slc24a3 | intron_variant | MODIFIER |
| 2 | 145636400 | AX-170107885 | 0 | A | G | 0.14 | 3.24E-02 | 2.23E-01 | 2.70E+00 | 2.73E+00 | 8.84E-01 | no | Slc24a3 | intron_variant | MODIFIER |
| 2 | 145662797 | AX-97802224 | 0 | T | C | 0.14 | 3.24E-02 | 2.23E-01 | 2.70E+00 | 2.73E+00 | 8.84E-01 | no | | intergenic_variant | MODIFIER |
| 2 | 145676975 | AX-170434222 | 0 | G | A | 0.14 | 3.24E-02 | 2.23E-01 | 2.70E+00 | 2.73E+00 | 8.84E-01 | no | Rin2 | intron_variant,non_coding_transcript_variant | MODIFIER |
| 2 | 145689034 | AX-170457016 | 0 | T | C | 0.14 | 3.24E-02 | 2.23E-01 | 2.70E+00 | 2.73E+00 | 8.84E-01 | no | Rin2 | intron_variant,non_coding_transcript_variant | MODIFIER |
| 2 | 145706152 | AX-97806421 | 0 | C | A | 0.14 | 3.24E-02 | 2.23E-01 | 2.70E+00 | 2.73E+00 | 8.84E-01 | no | Gm11763 | upstream_gene_variant | MODIFIER |
| 2 | 145720337 | AX-170460304 | 0 | T | C | 0.14 | 3.24E-02 | 2.23E-01 | 2.70E+00 | 2.73E+00 | 8.84E-01 | no | Rin2 | intron_variant,non_coding_transcript_variant | MODIFIER |
| 2 | 145724177 | AX-97847942 | 0 | C | T | 0.14 | 3.24E-02 | 2.23E-01 | 2.70E+00 | 2.73E+00 | 8.84E-01 | no | Rin2 | intron_variant,non_coding_transcript_variant | MODIFIER |
| 2 | 145728166 | AX-170065172 | 0 | C | T | 0.14 | 3.24E-02 | 2.23E-01 | 2.70E+00 | 2.73E+00 | 8.84E-01 | no | Rin2 | intron_variant,non_coding_transcript_variant | MODIFIER |
| 2 | 145728756 | AX-170207682 | 0 | T | A | 0.14 | 3.24E-02 | 2.23E-01 | 2.70E+00 | 2.73E+00 | 8.84E-01 | no | Rin2 | intron_variant,non_coding_transcript_variant | MODIFIER |
| 2 | 145732079 | AX-170246980 | 0 | C | T | 0.14 | 3.24E-02 | 2.23E-01 | 2.70E+00 | 2.73E+00 | 8.84E-01 | no | Rin2 | intron_variant,non_coding_transcript_variant | MODIFIER |
| 2 | 145735643 | AX-170448595 | 0 | G | T | 0.14 | 3.24E-02 | 2.23E-01 | 2.70E+00 | 2.73E+00 | 8.84E-01 | no | Rin2 | intron_variant,non_coding_transcript_variant | MODIFIER |
| 2 | 145764453 | AX-170181041 | 0 | A | G | 0.14 | 3.24E-02 | 2.23E-01 | 2.70E+00 | 2.73E+00 | 8.84E-01 | no | Rin2 | intron_variant,non_coding_transcript_variant | MODIFIER |
| 2 | 145768617 | AX-97855833 | 0 | A | C | 0.14 | 3.24E-02 | 2.23E-01 | 2.70E+00 | 2.73E+00 | 8.84E-01 | no | Rin2 | intron_variant,non_coding_transcript_variant | MODIFIER |
| 2 | 145801273 | AX-170181030 | 0 | T | C | 0.14 | 3.24E-02 | 2.23E-01 | 2.70E+00 | 2.73E+00 | 8.84E-01 | no | Rin2 | intron_variant | MODIFIER |
| 2 | 145808248 | AX-97848994 | 0 | G | A | 0.14 | 3.24E-02 | 2.23E-01 | 2.70E+00 | 2.73E+00 | 8.84E-01 | no | Rin2 | intron_variant | MODIFIER |
| 2 | 145813219 | AX-170487640 | 0 | A | G | 0.14 | 3.24E-02 | 2.23E-01 | 2.70E+00 | 2.73E+00 | 8.84E-01 | no | Rin2 | intron_variant | MODIFIER |
| 2 | 145824946 | AX-170311660 | 0 | G | A | 0.14 | 3.24E-02 | 2.23E-01 | 2.70E+00 | 2.73E+00 | 8.84E-01 | no | Rin2 | intron_variant | MODIFIER |
| 2 | 145825922 | AX-170377199 | 0 | C | A | 0.14 | 3.24E-02 | 2.23E-01 | 2.70E+00 | 2.73E+00 | 8.84E-01 | no | Rin2 | intron_variant | MODIFIER |
| 2 | 145831220 | AX-170179186 | 0 | A | G | 0.14 | 3.24E-02 | 2.23E-01 | 2.70E+00 | 2.73E+00 | 8.84E-01 | no | Rin2 | intron_variant | MODIFIER |
| 2 | 145832099 | AX-170514486 | 0 | A | G | 0.14 | 3.24E-02 | 2.23E-01 | 2.70E+00 | 2.73E+00 | 8.84E-01 | no | Rin2 | intron_variant | MODIFIER |
| 2 | 145840546 | AX-170353187 | 0 | G | A | 0.14 | 3.24E-02 | 2.23E-01 | 2.70E+00 | 2.73E+00 | 8.84E-01 | no | Rin2 | intron_variant | MODIFIER |

| | | | | | | | | | | | | | | | |
|---|---|---|---|---|---|---|---|---|---|---|---|---|---|---|---|
| 2 | 145847008 | AX-170400453 | 0 | C | G | 0.14 | 3.24E-02 | 2.23E-01 | 2.70E+00 | 2.73E+00 | 8.84E-01 | no | Rin2 | intron_variant | MODIFIER |
| 2 | 145849963 | AX-170019283 | 0 | G | A | 0.14 | 3.24E-02 | 2.23E-01 | 2.70E+00 | 2.73E+00 | 8.84E-01 | no | Rin2 | intron_variant | MODIFIER |
| 2 | 145850168 | AX-97839347 | 0 | A | G | 0.14 | 3.24E-02 | 2.23E-01 | 2.70E+00 | 2.73E+00 | 8.84E-01 | no | Rin2 | intron_variant | MODIFIER |
| 2 | 146556734 | AX-97786052 | 0 | T | C | 0.14 | 3.24E-02 | 2.23E-01 | 2.70E+00 | 2.73E+00 | 8.84E-01 | no | | intergenic_variant | MODIFIER |
| 2 | 146563324 | AX-170431567 | 0 | G | T | 0.14 | 3.24E-02 | 2.23E-01 | 2.70E+00 | 2.73E+00 | 8.84E-01 | no | | intergenic_variant | MODIFIER |
| 2 | 146565698 | AX-170299281 | 0 | T | C | 0.14 | 3.24E-02 | 2.23E-01 | 2.70E+00 | 2.73E+00 | 8.84E-01 | no | | intergenic_variant | MODIFIER |
| 2 | 146590557 | AX-170167653 | 0 | T | C | 0.14 | 3.24E-02 | 2.23E-01 | 2.70E+00 | 2.73E+00 | 8.84E-01 | no | | intergenic_variant | MODIFIER |
| 2 | 146591146 | AX-97840263 | 0 | C | T | 0.14 | 3.24E-02 | 2.23E-01 | 2.70E+00 | 2.73E+00 | 8.84E-01 | no | | intergenic_variant | MODIFIER |
| 2 | 146591200 | AX-170143157 | 0 | T | C | 0.14 | 3.24E-02 | 2.23E-01 | 2.70E+00 | 2.73E+00 | 8.84E-01 | no | | intergenic_variant | MODIFIER |
| 2 | 146593886 | AX-170356577 | 0 | C | T | 0.14 | 3.24E-02 | 2.23E-01 | 2.70E+00 | 2.73E+00 | 8.84E-01 | no | | intergenic_variant | MODIFIER |
| 2 | 146611225 | AX-170351536 | 0 | A | T | 0.14 | 3.24E-02 | 2.23E-01 | 2.70E+00 | 2.73E+00 | 8.84E-01 | no | | intergenic_variant | MODIFIER |
| 2 | 146611436 | AX-170389732 | 0 | C | T | 0.14 | 3.24E-02 | 2.23E-01 | 2.70E+00 | 2.73E+00 | 8.84E-01 | no | | intergenic_variant | MODIFIER |
| 2 | 146611502 | AX-170170612 | 0 | G | A | 0.14 | 3.24E-02 | 2.23E-01 | 2.70E+00 | 2.73E+00 | 8.84E-01 | no | | intergenic_variant | MODIFIER |
| 2 | 146649620 | AX-170202332 | 0 | A | G | 0.14 | 3.24E-02 | 2.23E-01 | 2.70E+00 | 2.73E+00 | 8.84E-01 | no | | intergenic_variant | MODIFIER |
| 2 | 146658089 | AX-97819686 | 0 | C | T | 0.14 | 3.24E-02 | 2.23E-01 | 2.70E+00 | 2.73E+00 | 8.84E-01 | no | | intergenic_variant | MODIFIER |
| 2 | 146658194 | AX-170312280 | 0 | A | G | 0.14 | 3.24E-02 | 2.23E-01 | 2.70E+00 | 2.73E+00 | 8.84E-01 | no | | intergenic_variant | MODIFIER |
| 2 | 146662401 | AX-170163322 | 0 | C | G | 0.14 | 3.24E-02 | 2.23E-01 | 2.70E+00 | 2.73E+00 | 8.84E-01 | no | | intergenic_variant | MODIFIER |
| 2 | 146662460 | AX-170483724 | 0 | T | C | 0.14 | 3.24E-02 | 2.23E-01 | 2.70E+00 | 2.73E+00 | 8.84E-01 | no | | intergenic_variant | MODIFIER |
| 2 | 146695738 | AX-170487168 | 0 | T | C | 0.14 | 3.24E-02 | 2.23E-01 | 2.70E+00 | 2.73E+00 | 8.84E-01 | no | | intergenic_variant | MODIFIER |
| 2 | 146696481 | AX-170112160 | 0 | A | G | 0.14 | 3.24E-02 | 2.23E-01 | 2.70E+00 | 2.73E+00 | 8.84E-01 | no | | intergenic_variant | MODIFIER |
| 2 | 146708245 | AX-170098416 | 0 | A | G | 0.14 | 3.24E-02 | 2.23E-01 | 2.70E+00 | 2.73E+00 | 8.84E-01 | no | | intergenic_variant | MODIFIER |
| 2 | 146708633 | AX-170413096 | 0 | T | C | 0.14 | 3.24E-02 | 2.23E-01 | 2.70E+00 | 2.73E+00 | 8.84E-01 | no | | regulatory_region_variant | MODIFIER |
| 2 | 146712463 | AX-170360469 | 0 | T | A | 0.14 | 3.24E-02 | 2.23E-01 | 2.70E+00 | 2.73E+00 | 8.84E-01 | no | | intergenic_variant | MODIFIER |
| 2 | 146724842 | AX-170512614 | 0 | G | A | 0.14 | 3.24E-02 | 2.23E-01 | 2.70E+00 | 2.73E+00 | 8.84E-01 | no | | regulatory_region_variant | MODIFIER |
| 2 | 146726906 | AX-97840504 | 0 | A | G | 0.14 | 3.24E-02 | 2.23E-01 | 2.70E+00 | 2.73E+00 | 8.84E-01 | no | | intergenic_variant | MODIFIER |
| 2 | 146734795 | AX-170065190 | 0 | G | A | 0.14 | 3.24E-02 | 2.23E-01 | 2.70E+00 | 2.73E+00 | 8.84E-01 | no | | intergenic_variant | MODIFIER |
| 2 | 146761475 | AX-97836942 | 0 | A | G | 0.14 | 3.24E-02 | 2.23E-01 | 2.70E+00 | 2.73E+00 | 8.84E-01 | no | | intergenic_variant | MODIFIER |
| 2 | 146772660 | AX-97829145 | 0 | T | C | 0.14 | 3.24E-02 | 2.23E-01 | 2.70E+00 | 2.73E+00 | 8.84E-01 | no | | intergenic_variant | MODIFIER |
| 2 | 146782509 | AX-170427975 | 0 | G | A | 0.14 | 3.24E-02 | 2.23E-01 | 2.70E+00 | 2.73E+00 | 8.84E-01 | no | | intergenic_variant | MODIFIER |
| 2 | 146800747 | AX-170126585 | 0 | G | A | 0.14 | 3.24E-02 | 2.23E-01 | 2.70E+00 | 2.73E+00 | 8.84E-01 | no | | intergenic_variant | MODIFIER |
| 2 | 146801930 | AX-97813520 | 0 | C | T | 0.14 | 3.24E-02 | 2.23E-01 | 2.70E+00 | 2.73E+00 | 8.84E-01 | no | | intergenic_variant | MODIFIER |
| 2 | 146851830 | AX-170222296 | 0 | A | T | 0.14 | 3.24E-02 | 2.23E-01 | 2.70E+00 | 2.73E+00 | 8.84E-01 | no | Kiz | upstream_gene_variant | MODIFIER |
| 2 | 146852429 | AX-97812172 | 0 | A | C | 0.14 | 3.24E-02 | 2.23E-01 | 2.70E+00 | 2.73E+00 | 8.84E-01 | no | Kiz | upstream_gene_variant | MODIFIER |
| 2 | 146857971 | AX-170116586 | 0 | C | T | 0.14 | 3.24E-02 | 2.23E-01 | 2.70E+00 | 2.73E+00 | 8.84E-01 | no | Kiz | intron_variant | MODIFIER |
| 2 | 146860522 | AX-170019301 | 0 | A | G | 0.14 | 3.24E-02 | 2.23E-01 | 2.70E+00 | 2.73E+00 | 8.84E-01 | no | Kiz | intron_variant | MODIFIER |
| 2 | 146860964 | AX-170299983 | 0 | G | A | 0.14 | 3.24E-02 | 2.23E-01 | 2.70E+00 | 2.73E+00 | 8.84E-01 | no | Kiz | intron_variant | MODIFIER |
| 2 | 146868696 | AX-170254017 | 0 | A | G | 0.14 | 3.24E-02 | 2.23E-01 | 2.70E+00 | 2.73E+00 | 8.84E-01 | no | Kiz | intron_variant | MODIFIER |
| 2 | 146869703 | AX-170449868 | 0 | T | C | 0.14 | 3.24E-02 | 2.23E-01 | 2.70E+00 | 2.73E+00 | 8.84E-01 | no | Kiz | intron_variant | MODIFIER |
| 2 | 146897839 | AX-97829789 | 0 | T | G | 0.14 | 3.24E-02 | 2.23E-01 | 2.70E+00 | 2.73E+00 | 8.84E-01 | no | Kiz | intron_variant | MODIFIER |
| 2 | 146931068 | AX-170019306 | 0 | T | C | 0.14 | 3.24E-02 | 2.23E-01 | 2.70E+00 | 2.73E+00 | 8.84E-01 | no | Kiz | intron_variant | MODIFIER |
| 2 | 146935461 | AX-170019304 | 0 | G | A | 0.14 | 3.24E-02 | 2.23E-01 | 2.70E+00 | 2.73E+00 | 8.84E-01 | no | Kiz | intron_variant | MODIFIER |
| 2 | 146951213 | AX-97803124 | 0 | A | G | 0.14 | 3.24E-02 | 2.23E-01 | 2.70E+00 | 2.73E+00 | 8.84E-01 | no | Kiz | intron_variant | MODIFIER |
| 2 | 146955573 | AX-170019305 | 0 | G | A | 0.14 | 3.24E-02 | 2.23E-01 | 2.70E+00 | 2.73E+00 | 8.84E-01 | no | Kiz | intron_variant | MODIFIER |
| 2 | 146956091 | AX-170458179 | 0 | G | A | 0.14 | 3.24E-02 | 2.23E-01 | 2.70E+00 | 2.73E+00 | 8.84E-01 | no | Kiz | intron_variant | MODIFIER |
| 2 | 146956511 | AX-170019303 | 0 | G | C | 0.14 | 3.24E-02 | 2.23E-01 | 2.70E+00 | 2.73E+00 | 8.84E-01 | no | Kiz | intron_variant | MODIFIER |
| 2 | 146989263 | AX-170397580 | 0 | G | A | 0.14 | 3.24E-02 | 2.23E-01 | 2.70E+00 | 2.73E+00 | 8.84E-01 | no | | intergenic_variant | MODIFIER |
| 2 | 147012288 | AX-97840505 | 0 | T | C | 0.14 | 3.24E-02 | 2.23E-01 | 2.70E+00 | 2.73E+00 | 8.84E-01 | no | Xrn2 | upstream_gene_variant | MODIFIER |
| 2 | 147027134 | AX-170502568 | 0 | C | G | 0.14 | 3.24E-02 | 2.23E-01 | 2.70E+00 | 2.73E+00 | 8.84E-01 | no | Xrn2 | intron_variant | MODIFIER |
| 2 | 147055788 | AX-170429324 | 0 | C | T | 0.14 | 3.24E-02 | 2.23E-01 | 2.70E+00 | 2.73E+00 | 8.84E-01 | no | Xrn2 | intron_variant | MODIFIER |
| 2 | 147056019 | AX-170426648 | 0 | G | A | 0.14 | 3.24E-02 | 2.23E-01 | 2.70E+00 | 2.73E+00 | 8.84E-01 | no | Xrn2 | intron_variant | MODIFIER |
| 2 | 147060149 | AX-170492935 | 0 | G | A | 0.14 | 3.24E-02 | 2.23E-01 | 2.70E+00 | 2.73E+00 | 8.84E-01 | no | Xrn2 | intron_variant | MODIFIER |
| 2 | 147096805 | AX-170193038 | 0 | C | T | 0.14 | 3.24E-02 | 2.23E-01 | 2.70E+00 | 2.73E+00 | 8.84E-01 | no | | intergenic_variant | MODIFIER |
| 2 | 147100240 | AX-170319482 | 0 | T | C | 0.14 | 3.24E-02 | 2.23E-01 | 2.70E+00 | 2.73E+00 | 8.84E-01 | no | | intergenic_variant | MODIFIER |
| 2 | 147106284 | AX-170415276 | 0 | G | A | 0.14 | 3.24E-02 | 2.23E-01 | 2.70E+00 | 2.73E+00 | 8.84E-01 | no | | intergenic_variant | MODIFIER |

| | | | | | | | | | | | | | | |
|---|---|---|---|---|---|---|---|---|---|---|---|---|---|---|
| 2 | 147119105 | AX-97840529 | 0 | G | A | 0.14 | 3.24E-02 | 2.23E-01 | 2.70E+00 | 2.73E+00 | 8.84E-01 | no | | intergenic_variant | MODIFIER |
| 2 | 147130502 | AX-170426046 | 0 | C | T | 0.14 | 3.24E-02 | 2.23E-01 | 2.70E+00 | 2.73E+00 | 8.84E-01 | no | | intergenic_variant | MODIFIER |
| 2 | 147132953 | AX-170389065 | 0 | A | G | 0.14 | 3.24E-02 | 2.23E-01 | 2.70E+00 | 2.73E+00 | 8.84E-01 | no | | intergenic_variant | MODIFIER |
| 2 | 147134008 | AX-170378316 | 0 | A | G | 0.14 | 3.24E-02 | 2.23E-01 | 2.70E+00 | 2.73E+00 | 8.84E-01 | no | | intergenic_variant | MODIFIER |
| 2 | 147162224 | AX-170118791 | 0 | T | C | 0.14 | 3.24E-02 | 2.23E-01 | 2.70E+00 | 2.73E+00 | 8.84E-01 | no | | intergenic_variant | MODIFIER |
| 2 | 147169536 | AX-97841324 | 0 | A | G | 0.14 | 3.24E-02 | 2.23E-01 | 2.70E+00 | 2.73E+00 | 8.84E-01 | no | | intergenic_variant | MODIFIER |
| 2 | 147174980 | AX-170414519 | 0 | G | A | 0.14 | 3.24E-02 | 2.23E-01 | 2.70E+00 | 2.73E+00 | 8.84E-01 | no | Nkx2-2 | downstream_gene_variant | MODIFIER |
| 2 | 147189412 | AX-97839371 | 0 | C | G | 0.14 | 3.24E-02 | 2.23E-01 | 2.70E+00 | 2.73E+00 | 8.84E-01 | no | Nkx2-2 | upstream_gene_variant | MODIFIER |
| 2 | 147213749 | AX-97811695 | 0 | T | G | 0.14 | 3.24E-02 | 2.23E-01 | 2.70E+00 | 2.73E+00 | 8.84E-01 | no | Nkx2-2os | intron_variant,non_coding_transcript_variant | MODIFIER |
| 2 | 147218124 | AX-170173014 | 0 | T | C | 0.14 | 3.24E-02 | 2.23E-01 | 2.70E+00 | 2.73E+00 | 8.84E-01 | no | Nkx2-2os | intron_variant,non_coding_transcript_variant | MODIFIER |
| 2 | 147232198 | AX-97816055 | 0 | A | G | 0.14 | 3.24E-02 | 2.23E-01 | 2.70E+00 | 2.73E+00 | 8.84E-01 | no | Nkx2-2os | intron_variant,non_coding_transcript_variant | MODIFIER |
| 2 | 147236286 | AX-170260731 | 0 | G | A | 0.14 | 3.24E-02 | 2.23E-01 | 2.70E+00 | 2.73E+00 | 8.84E-01 | no | Nkx2-2os | intron_variant,non_coding_transcript_variant | MODIFIER |
| 2 | 147236537 | AX-170297242 | 0 | T | C | 0.14 | 3.24E-02 | 2.23E-01 | 2.70E+00 | 2.73E+00 | 8.84E-01 | no | Nkx2-2os | intron_variant,non_coding_transcript_variant | MODIFIER |
| 2 | 147236944 | AX-170252784 | 0 | A | G | 0.14 | 3.24E-02 | 2.23E-01 | 2.70E+00 | 2.73E+00 | 8.84E-01 | no | Nkx2-2os | intron_variant,non_coding_transcript_variant | MODIFIER |
| 2 | 147257743 | AX-170065197 | 0 | T | C | 0.14 | 3.24E-02 | 2.23E-01 | 2.70E+00 | 2.73E+00 | 8.84E-01 | no | Nkx2-2os | intron_variant,non_coding_transcript_variant | MODIFIER |
| 2 | 147258367 | AX-170065199 | 0 | A | G | 0.14 | 3.24E-02 | 2.23E-01 | 2.70E+00 | 2.73E+00 | 8.84E-01 | no | Nkx2-2os | intron_variant,non_coding_transcript_variant | MODIFIER |
| 2 | 147259106 | AX-170296889 | 0 | C | T | 0.14 | 3.24E-02 | 2.23E-01 | 2.70E+00 | 2.73E+00 | 8.84E-01 | no | Nkx2-2os | intron_variant,non_coding_transcript_variant | MODIFIER |
| 2 | 147259371 | AX-170148202 | 0 | A | T | 0.14 | 3.24E-02 | 2.23E-01 | 2.70E+00 | 2.73E+00 | 8.84E-01 | no | Nkx2-2os | intron_variant,non_coding_transcript_variant | MODIFIER |
| 2 | 147261306 | AX-170250092 | 0 | G | A | 0.14 | 3.24E-02 | 2.23E-01 | 2.70E+00 | 2.73E+00 | 8.84E-01 | no | Nkx2-2os | intron_variant,non_coding_transcript_variant | MODIFIER |
| 2 | 147263060 | AX-170319872 | 0 | A | G | 0.14 | 3.24E-02 | 2.23E-01 | 2.70E+00 | 2.73E+00 | 8.84E-01 | no | Nkx2-2os | intron_variant,non_coding_transcript_variant | MODIFIER |
| 2 | 147285449 | AX-170329734 | 0 | T | C | 0.14 | 3.24E-02 | 2.23E-01 | 2.70E+00 | 2.73E+00 | 8.84E-01 | no | Gm14115 | upstream_gene_variant | MODIFIER |
| 2 | 147302972 | AX-170454384 | 0 | C | T | 0.14 | 3.24E-02 | 2.23E-01 | 2.70E+00 | 2.73E+00 | 8.84E-01 | no | Nkx2-2os | intron_variant,non_coding_transcript_variant | MODIFIER |
| 2 | 147303666 | AX-170094440 | 0 | A | G | 0.14 | 3.24E-02 | 2.23E-01 | 2.70E+00 | 2.73E+00 | 8.84E-01 | no | Nkx2-2os | non_coding_transcript_exon_variant | MODIFIER |
| 2 | 147305584 | AX-170465300 | 0 | G | T | 0.14 | 3.24E-02 | 2.23E-01 | 2.70E+00 | 2.73E+00 | 8.84E-01 | no | Nkx2-2os | non_coding_transcript_exon_variant | MODIFIER |
| 2 | 147327462 | AX-170110619 | 0 | A | C | 0.14 | 3.24E-02 | 2.23E-01 | 2.70E+00 | 2.73E+00 | 8.84E-01 | no | Gm14110 | upstream_gene_variant | MODIFIER |
| 2 | 147327569 | AX-170019314 | 0 | A | C | 0.14 | 3.24E-02 | 2.23E-01 | 2.70E+00 | 2.73E+00 | 8.84E-01 | no | Gm14110 | upstream_gene_variant | MODIFIER |
| 2 | 147333576 | AX-170277303 | 0 | A | G | 0.14 | 3.24E-02 | 2.23E-01 | 2.70E+00 | 2.73E+00 | 8.84E-01 | no | Nkx2-2os | downstream_gene_variant | MODIFIER |
| 2 | 147343739 | AX-170215597 | 0 | G | A | 0.14 | 3.24E-02 | 2.23E-01 | 2.70E+00 | 2.73E+00 | 8.84E-01 | no | | intergenic_variant | MODIFIER |
| 2 | 147370310 | AX-97835233 | 0 | G | A | 0.14 | 3.24E-02 | 2.23E-01 | 2.70E+00 | 2.73E+00 | 8.84E-01 | no | Pax1 | intron_variant | MODIFIER |
| 2 | 147378823 | AX-170019313 | 0 | G | A | 0.14 | 3.24E-02 | 2.23E-01 | 2.70E+00 | 2.73E+00 | 8.84E-01 | no | Pax1 | downstream_gene_variant | MODIFIER |
| 2 | 147402131 | AX-97856541 | 0 | T | C | 0.14 | 3.24E-02 | 2.23E-01 | 2.70E+00 | 2.73E+00 | 8.84E-01 | no | Gm14109 | downstream_gene_variant | MODIFIER |
| 2 | 147410003 | AX-170019307 | 0 | T | C | 0.14 | 3.24E-02 | 2.23E-01 | 2.70E+00 | 2.73E+00 | 8.84E-01 | no | | intergenic_variant | MODIFIER |
| 2 | 147415123 | AX-170309280 | 0 | A | G | 0.14 | 3.24E-02 | 2.23E-01 | 2.70E+00 | 2.73E+00 | 8.84E-01 | no | | regulatory_region_variant | MODIFIER |
| 2 | 147415153 | AX-170111932 | 0 | T | C | 0.14 | 3.24E-02 | 2.23E-01 | 2.70E+00 | 2.73E+00 | 8.84E-01 | no | | regulatory_region_variant | MODIFIER |
| 2 | 147423722 | AX-170065201 | 0 | C | T | 0.14 | 3.24E-02 | 2.23E-01 | 2.70E+00 | 2.73E+00 | 8.84E-01 | no | | intergenic_variant | MODIFIER |
| 2 | 147431919 | AX-170019315 | 0 | A | G | 0.14 | 3.24E-02 | 2.23E-01 | 2.70E+00 | 2.73E+00 | 8.84E-01 | no | | intergenic_variant | MODIFIER |
| 2 | 147449207 | AX-97839394 | 0 | A | T | 0.14 | 3.24E-02 | 2.23E-01 | 2.70E+00 | 2.73E+00 | 8.84E-01 | no | | intergenic_variant | MODIFIER |
| 2 | 147485193 | AX-97839405 | 0 | T | C | 0.14 | 3.24E-02 | 2.23E-01 | 2.70E+00 | 2.73E+00 | 8.84E-01 | no | | intergenic_variant | MODIFIER |
| 2 | 147488850 | AX-170019312 | 0 | A | G | 0.14 | 3.24E-02 | 2.23E-01 | 2.70E+00 | 2.73E+00 | 8.84E-01 | no | | intergenic_variant | MODIFIER |
| 2 | 147494812 | AX-170102205 | 0 | G | A | 0.14 | 3.24E-02 | 2.23E-01 | 2.70E+00 | 2.73E+00 | 8.84E-01 | no | | intergenic_variant | MODIFIER |
| 2 | 147495036 | AX-170065202 | 0 | T | C | 0.14 | 3.24E-02 | 2.23E-01 | 2.70E+00 | 2.73E+00 | 8.84E-01 | no | | intergenic_variant | MODIFIER |
| 2 | 147515349 | AX-170438515 | 0 | T | G | 0.14 | 3.24E-02 | 2.23E-01 | 2.70E+00 | 2.73E+00 | 8.84E-01 | no | | intergenic_variant | MODIFIER |
| 2 | 147530123 | AX-170019316 | 0 | C | T | 0.14 | 3.24E-02 | 2.23E-01 | 2.70E+00 | 2.73E+00 | 8.84E-01 | no | | intergenic_variant | MODIFIER |
| 2 | 147530339 | AX-170456472 | 0 | G | C | 0.14 | 3.24E-02 | 2.23E-01 | 2.70E+00 | 2.73E+00 | 8.84E-01 | no | | intergenic_variant | MODIFIER |
| 2 | 147530851 | AX-170154108 | 0 | A | G | 0.14 | 3.24E-02 | 2.23E-01 | 2.70E+00 | 2.73E+00 | 8.84E-01 | no | | intergenic_variant | MODIFIER |
| 2 | 147560606 | AX-170019310 | 0 | G | C | 0.14 | 3.24E-02 | 2.23E-01 | 2.70E+00 | 2.73E+00 | 8.84E-01 | no | | intergenic_variant | MODIFIER |
| 2 | 147560844 | AX-170520685 | 0 | C | T | 0.14 | 3.24E-02 | 2.23E-01 | 2.70E+00 | 2.73E+00 | 8.84E-01 | no | | regulatory_region_variant | MODIFIER |
| 2 | 147565497 | AX-170269688 | 0 | C | T | 0.14 | 3.24E-02 | 2.23E-01 | 2.70E+00 | 2.73E+00 | 8.84E-01 | no | | intergenic_variant | MODIFIER |
| 2 | 147575422 | AX-97793743 | 0 | C | T | 0.14 | 3.24E-02 | 2.23E-01 | 2.70E+00 | 2.73E+00 | 8.84E-01 | no | | intergenic_variant | MODIFIER |
| 2 | 147581774 | AX-170150483 | 0 | A | C | 0.14 | 3.24E-02 | 2.23E-01 | 2.70E+00 | 2.73E+00 | 8.84E-01 | no | | intergenic_variant | MODIFIER |
| 2 | 147586362 | AX-170376776 | 0 | C | G | 0.14 | 3.24E-02 | 2.23E-01 | 2.70E+00 | 2.73E+00 | 8.84E-01 | no | | intergenic_variant | MODIFIER |
| 2 | 147586748 | AX-170125877 | 0 | T | G | 0.14 | 3.24E-02 | 2.23E-01 | 2.70E+00 | 2.73E+00 | 8.84E-01 | no | | intergenic_variant | MODIFIER |
| 2 | 147587373 | AX-170150927 | 0 | A | C | 0.14 | 3.24E-02 | 2.23E-01 | 2.70E+00 | 2.73E+00 | 8.84E-01 | no | | intergenic_variant | MODIFIER |
| 2 | 147591884 | AX-170328938 | 0 | G | T | 0.14 | 3.24E-02 | 2.23E-01 | 2.70E+00 | 2.73E+00 | 8.84E-01 | no | | intergenic_variant | MODIFIER |

| | | | | | | | | | | | | | | | |
|---|---|---|---|---|---|---|---|---|---|---|---|---|---|---|---|
| 2 | 147592110 | AX-170124465 | 0 | G | A | 0.14 | 3.24E-02 | 2.23E-01 | 2.70E+00 | 2.73E+00 | 8.84E-01 | no | | intergenic_variant | MODIFIER |
| 2 | 147597784 | AX-170305785 | 0 | C | T | 0.14 | 3.24E-02 | 2.23E-01 | 2.70E+00 | 2.73E+00 | 8.84E-01 | no | | intergenic_variant | MODIFIER |
| 2 | 147600998 | AX-170372858 | 0 | T | C | 0.14 | 3.24E-02 | 2.23E-01 | 2.70E+00 | 2.73E+00 | 8.84E-01 | no | | intergenic_variant | MODIFIER |
| 2 | 147613741 | AX-170161477 | 0 | T | G | 0.14 | 3.24E-02 | 2.23E-01 | 2.70E+00 | 2.73E+00 | 8.84E-01 | no | | intergenic_variant | MODIFIER |
| 2 | 147623268 | AX-170375825 | 0 | C | T | 0.14 | 3.24E-02 | 2.23E-01 | 2.70E+00 | 2.73E+00 | 8.84E-01 | no | | intergenic_variant | MODIFIER |
| 2 | 147623573 | AX-170065206 | 0 | C | T | 0.14 | 3.24E-02 | 2.23E-01 | 2.70E+00 | 2.73E+00 | 8.84E-01 | no | | intergenic_variant | MODIFIER |
| 2 | 147626484 | AX-170065205 | 0 | C | T | 0.14 | 3.24E-02 | 2.23E-01 | 2.70E+00 | 2.73E+00 | 8.84E-01 | no | | intergenic_variant | MODIFIER |
| 2 | 147634075 | AX-170434825 | 0 | T | C | 0.14 | 3.24E-02 | 2.23E-01 | 2.70E+00 | 2.73E+00 | 8.84E-01 | no | | intergenic_variant | MODIFIER |
| 2 | 147635195 | AX-170065207 | 0 | A | G | 0.14 | 3.24E-02 | 2.23E-01 | 2.70E+00 | 2.73E+00 | 8.84E-01 | no | | intergenic_variant | MODIFIER |
| 2 | 147636862 | AX-170156890 | 0 | C | T | 0.14 | 3.24E-02 | 2.23E-01 | 2.70E+00 | 2.73E+00 | 8.84E-01 | no | | intergenic_variant | MODIFIER |
| 2 | 147636921 | AX-170150218 | 0 | C | T | 0.14 | 3.24E-02 | 2.23E-01 | 2.70E+00 | 2.73E+00 | 8.84E-01 | no | | intergenic_variant | MODIFIER |
| 2 | 147639898 | AX-170456182 | 0 | G | A | 0.14 | 3.24E-02 | 2.23E-01 | 2.70E+00 | 2.73E+00 | 8.84E-01 | no | Gm14107 | downstream_gene_variant | MODIFIER |
| 2 | 147642298 | AX-170413788 | 0 | A | G | 0.14 | 3.24E-02 | 2.23E-01 | 2.70E+00 | 2.73E+00 | 8.84E-01 | no | Gm14107 | downstream_gene_variant | MODIFIER |
| 2 | 147645589 | AX-170275631 | 0 | A | G | 0.14 | 3.24E-02 | 2.23E-01 | 2.70E+00 | 2.73E+00 | 8.84E-01 | no | Gm14107 | upstream_gene_variant | MODIFIER |
| 2 | 147648744 | AX-97847029 | 0 | G | A | 0.14 | 3.24E-02 | 2.23E-01 | 2.70E+00 | 2.73E+00 | 8.84E-01 | no | | intergenic_variant | MODIFIER |
| 2 | 147683534 | AX-170088787 | 0 | C | T | 0.14 | 3.24E-02 | 2.23E-01 | 2.70E+00 | 2.73E+00 | 8.84E-01 | no | | intergenic_variant | MODIFIER |
| 2 | 147688802 | AX-170108593 | 0 | A | G | 0.14 | 3.24E-02 | 2.23E-01 | 2.70E+00 | 2.73E+00 | 8.84E-01 | no | | regulatory_region_variant | MODIFIER |
| 2 | 147697318 | AX-170038283 | 0 | G | T | 0.14 | 3.24E-02 | 2.23E-01 | 2.70E+00 | 2.73E+00 | 8.84E-01 | no | | intergenic_variant | MODIFIER |
| 2 | 147699727 | AX-97848757 | 0 | A | G | 0.14 | 3.24E-02 | 2.23E-01 | 2.70E+00 | 2.73E+00 | 8.84E-01 | no | | regulatory_region_variant | MODIFIER |
| 2 | 147739664 | AX-97842104 | 0 | C | G | 0.14 | 3.24E-02 | 2.23E-01 | 2.70E+00 | 2.73E+00 | 8.84E-01 | no | | intergenic_variant | MODIFIER |
| 2 | 147740205 | AX-170519392 | 0 | G | T | 0.14 | 3.24E-02 | 2.23E-01 | 2.70E+00 | 2.73E+00 | 8.84E-01 | no | | intergenic_variant | MODIFIER |
| 2 | 147744620 | AX-170482102 | 0 | G | A | 0.14 | 3.24E-02 | 2.23E-01 | 2.70E+00 | 2.73E+00 | 8.84E-01 | no | | intergenic_variant | MODIFIER |
| 2 | 147745602 | AX-170432693 | 0 | C | T | 0.14 | 3.24E-02 | 2.23E-01 | 2.70E+00 | 2.73E+00 | 8.84E-01 | no | | intergenic_variant | MODIFIER |
| 2 | 147747175 | AX-170442318 | 0 | T | C | 0.14 | 3.24E-02 | 2.23E-01 | 2.70E+00 | 2.73E+00 | 8.84E-01 | no | | intergenic_variant | MODIFIER |
| 2 | 147765007 | AX-170270246 | 0 | A | C | 0.14 | 3.24E-02 | 2.23E-01 | 2.70E+00 | 2.73E+00 | 8.84E-01 | no | | intergenic_variant | MODIFIER |
| 2 | 147786747 | AX-170163930 | 0 | T | C | 0.14 | 3.24E-02 | 2.23E-01 | 2.70E+00 | 2.73E+00 | 8.84E-01 | no | | intergenic_variant | MODIFIER |
| 2 | 147793333 | AX-170065211 | 0 | C | T | 0.14 | 3.24E-02 | 2.23E-01 | 2.70E+00 | 2.73E+00 | 8.84E-01 | no | | intergenic_variant | MODIFIER |
| 2 | 147793465 | AX-170065208 | 0 | G | C | 0.14 | 3.24E-02 | 2.23E-01 | 2.70E+00 | 2.73E+00 | 8.84E-01 | no | | intergenic_variant | MODIFIER |
| 2 | 148067287 | AX-170219841 | 0 | G | A | 0.14 | 3.24E-02 | 2.23E-01 | 2.70E+00 | 2.73E+00 | 8.84E-01 | no | Gm14108 | downstream_gene_variant | MODIFIER |
| 2 | 148266645 | AX-170065220 | 0 | A | C | 0.14 | 3.24E-02 | 2.23E-01 | 2.70E+00 | 2.73E+00 | 8.84E-01 | no | | intergenic_variant | MODIFIER |
| 2 | 148267202 | AX-170065218 | 0 | A | C | 0.14 | 3.24E-02 | 2.23E-01 | 2.70E+00 | 2.73E+00 | 8.84E-01 | no | | intergenic_variant | MODIFIER |
| 2 | 148271434 | AX-168373561 | 0 | A | G | 0.14 | 3.24E-02 | 2.23E-01 | 2.70E+00 | 2.73E+00 | 8.84E-01 | no | | intergenic_variant | MODIFIER |
| 2 | 148278631 | AX-170530880 | 0 | G | A | 0.14 | 3.24E-02 | 2.23E-01 | 2.70E+00 | 2.73E+00 | 8.84E-01 | no | | intergenic_variant | MODIFIER |
| 2 | 148282316 | AX-170065216 | 0 | A | G | 0.14 | 3.24E-02 | 2.23E-01 | 2.70E+00 | 2.73E+00 | 8.84E-01 | no | | intergenic_variant | MODIFIER |
| 2 | 148294449 | AX-97856845 | 0 | T | C | 0.14 | 3.24E-02 | 2.23E-01 | 2.70E+00 | 2.73E+00 | 8.84E-01 | no | | intergenic_variant | MODIFIER |
| 2 | 148516688 | AX-97842640 | 0 | C | T | 0.14 | 3.24E-02 | 2.23E-01 | 2.70E+00 | 2.73E+00 | 8.84E-01 | no | | intergenic_variant | MODIFIER |
| 2 | 148534573 | AX-97840562 | 0 | C | T | 0.14 | 3.24E-02 | 2.23E-01 | 2.70E+00 | 2.73E+00 | 8.84E-01 | no | | intergenic_variant | MODIFIER |
| 2 | 148546348 | AX-170311923 | 0 | G | C | 0.14 | 3.24E-02 | 2.23E-01 | 2.70E+00 | 2.73E+00 | 8.84E-01 | no | | intergenic_variant | MODIFIER |
| 2 | 148551627 | AX-97804723 | 0 | C | T | 0.14 | 3.24E-02 | 2.23E-01 | 2.70E+00 | 2.73E+00 | 8.84E-01 | no | | intergenic_variant | MODIFIER |
| 2 | 148575248 | AX-170019337 | 0 | C | T | 0.14 | 3.24E-02 | 2.23E-01 | 2.70E+00 | 2.73E+00 | 8.84E-01 | no | | intergenic_variant | MODIFIER |
| 2 | 148603259 | AX-170019329 | 0 | G | A | 0.14 | 3.24E-02 | 2.23E-01 | 2.70E+00 | 2.73E+00 | 8.84E-01 | no | | intergenic_variant | MODIFIER |
| 2 | 148603981 | AX-170345493 | 0 | A | G | 0.14 | 3.24E-02 | 2.23E-01 | 2.70E+00 | 2.73E+00 | 8.84E-01 | no | | intergenic_variant | MODIFIER |
| 2 | 148604875 | AX-97801865 | 0 | G | A | 0.14 | 3.24E-02 | 2.23E-01 | 2.70E+00 | 2.73E+00 | 8.84E-01 | no | | intergenic_variant | MODIFIER |
| 2 | 148605109 | AX-170289067 | 0 | G | C | 0.14 | 3.24E-02 | 2.23E-01 | 2.70E+00 | 2.73E+00 | 8.84E-01 | no | | intergenic_variant | MODIFIER |
| 2 | 148622970 | AX-97783621 | 0 | T | C | 0.14 | 3.24E-02 | 2.23E-01 | 2.70E+00 | 2.73E+00 | 8.84E-01 | no | Gm25866 | upstream_gene_variant | MODIFIER |
| 2 | 148629099 | AX-170065223 | 0 | C | A | 0.14 | 3.24E-02 | 2.23E-01 | 2.70E+00 | 2.73E+00 | 8.84E-01 | no | Gm25866 | downstream_gene_variant | MODIFIER |
| 2 | 148705848 | AX-170222148 | 0 | T | C | 0.14 | 3.24E-02 | 2.23E-01 | 2.70E+00 | 2.73E+00 | 8.84E-01 | no | Napb | intron_variant | MODIFIER |
| 2 | 148721339 | AX-170422120 | 0 | T | C | 0.14 | 3.24E-02 | 2.23E-01 | 2.70E+00 | 2.73E+00 | 8.84E-01 | no | Napb | intron_variant | MODIFIER |
| 2 | 148751217 | AX-170232234 | 0 | G | A | 0.14 | 3.24E-02 | 2.23E-01 | 2.70E+00 | 2.73E+00 | 8.84E-01 | no | Cstl1 | intron_variant | MODIFIER |
| 2 | 148753356 | AX-170307568 | 0 | T | C | 0.14 | 3.24E-02 | 2.23E-01 | 2.70E+00 | 2.73E+00 | 8.84E-01 | no | Cstl1 | intron_variant | MODIFIER |
| 2 | 148754952 | AX-170434447 | 0 | G | A | 0.14 | 3.24E-02 | 2.23E-01 | 2.70E+00 | 2.73E+00 | 8.84E-01 | no | Cstl1 | intron_variant | MODIFIER |
| 2 | 148756246 | AX-170254279 | 0 | G | T | 0.14 | 3.24E-02 | 2.23E-01 | 2.70E+00 | 2.73E+00 | 8.84E-01 | no | Cstl1 | downstream_gene_variant | MODIFIER |
| 2 | 148779329 | AX-170496170 | 0 | A | G | 0.14 | 3.24E-02 | 2.23E-01 | 2.70E+00 | 2.73E+00 | 8.84E-01 | no | 8030411F24Rik | upstream_gene_variant | MODIFIER |
| 2 | 148781702 | AX-97795311 | 0 | C | T | 0.14 | 3.24E-02 | 2.23E-01 | 2.70E+00 | 2.73E+00 | 8.84E-01 | no | 8030411F24Rik | upstream_gene_variant | MODIFIER |

| | | | | | | | | | | | | | | | |
|---|---|---|---|---|---|---|---|---|---|---|---|---|---|---|---|
| 2 | 148840198 | AX-170494480 | 0 | G | A | 0.14 | 3.24E-02 | 2.23E-01 | 2.70E+00 | 2.73E+00 | 8.84E-01 | no | Cst9 | downstream_gene_variant | MODIFIER |
| 2 | 148847959 | AX-97795242 | 0 | G | A | 0.14 | 3.24E-02 | 2.23E-01 | 2.70E+00 | 2.73E+00 | 8.84E-01 | no | 9230104L09Rik | synonymous_variant | LOW |
| 2 | 148850734 | AX-170201213 | 0 | A | G | 0.14 | 3.24E-02 | 2.23E-01 | 2.70E+00 | 2.73E+00 | 8.84E-01 | no | 9230104L09Rik | synonymous_variant | LOW |
| 2 | 148872595 | AX-170214975 | 0 | A | G | 0.14 | 3.24E-02 | 2.23E-01 | 2.70E+00 | 2.73E+00 | 8.84E-01 | no | Cst3 | intron_variant | MODIFIER |
| 2 | 148873708 | AX-170359383 | 0 | C | T | 0.14 | 3.24E-02 | 2.23E-01 | 2.70E+00 | 2.73E+00 | 8.84E-01 | no | Cst3 | intron_variant | MODIFIER |
| 2 | 148873748 | AX-170105973 | 0 | C | T | 0.14 | 3.24E-02 | 2.23E-01 | 2.70E+00 | 2.73E+00 | 8.84E-01 | no | Cst3 | intron_variant | MODIFIER |
| 2 | 148931368 | AX-97806667 | 0 | A | G | 0.14 | 3.24E-02 | 2.23E-01 | 2.70E+00 | 2.73E+00 | 8.84E-01 | no | Gm14129 | intron_variant,non_coding_transcript_variant | MODIFIER |
| 2 | 148946042 | AX-170065233 | 0 | A | G | 0.14 | 3.24E-02 | 2.23E-01 | 2.70E+00 | 2.73E+00 | 8.84E-01 | no | | intergenic_variant | MODIFIER |
| 2 | 148999102 | AX-170065228 | 0 | C | T | 0.14 | 3.24E-02 | 2.23E-01 | 2.70E+00 | 2.73E+00 | 8.84E-01 | no | Gm1330 | intron_variant | MODIFIER |
| 2 | 149001977 | AX-170065230 | 0 | A | G | 0.14 | 3.24E-02 | 2.23E-01 | 2.70E+00 | 2.73E+00 | 8.84E-01 | no | Gm1330 | intron_variant | MODIFIER |
| 2 | 149002141 | AX-170019340 | 0 | G | C | 0.14 | 3.24E-02 | 2.23E-01 | 2.70E+00 | 2.73E+00 | 8.84E-01 | no | Gm1330 | intron_variant | MODIFIER |
| 2 | 149057026 | AX-170411242 | 0 | G | T | 0.14 | 3.24E-02 | 2.23E-01 | 2.70E+00 | 2.73E+00 | 8.84E-01 | no | | intergenic_variant | MODIFIER |
| 2 | 149087763 | AX-170019341 | 0 | G | A | 0.14 | 3.24E-02 | 2.23E-01 | 2.70E+00 | 2.73E+00 | 8.84E-01 | no | | intergenic_variant | MODIFIER |
| 2 | 149147589 | AX-170019338 | 0 | T | C | 0.14 | 3.24E-02 | 2.23E-01 | 2.70E+00 | 2.73E+00 | 8.84E-01 | no | | intergenic_variant | MODIFIER |
| 2 | 149149965 | AX-170019345 | 0 | C | T | 0.14 | 3.24E-02 | 2.23E-01 | 2.70E+00 | 2.73E+00 | 8.84E-01 | no | | intergenic_variant | MODIFIER |
| 2 | 149175480 | AX-170019339 | 0 | T | C | 0.14 | 3.24E-02 | 2.23E-01 | 2.70E+00 | 2.73E+00 | 8.84E-01 | no | | intergenic_variant | MODIFIER |
| 2 | 149215341 | AX-170522931 | 0 | G | C | 0.14 | 3.24E-02 | 2.23E-01 | 2.70E+00 | 2.73E+00 | 8.84E-01 | no | | intergenic_variant | MODIFIER |
| 2 | 149217614 | AX-97833807 | 0 | C | T | 0.14 | 3.24E-02 | 2.23E-01 | 2.70E+00 | 2.73E+00 | 8.84E-01 | no | | intergenic_variant | MODIFIER |
| 2 | 149262294 | AX-170442766 | 0 | T | C | 0.14 | 3.24E-02 | 2.23E-01 | 2.70E+00 | 2.73E+00 | 8.84E-01 | no | Gm14135 | downstream_gene_variant | MODIFIER |
| 2 | 149262395 | AX-97827318 | 0 | T | G | 0.14 | 3.24E-02 | 2.23E-01 | 2.70E+00 | 2.73E+00 | 8.84E-01 | no | Gm14135 | downstream_gene_variant | MODIFIER |
| 2 | 149311396 | AX-170509742 | 0 | T | A | 0.14 | 3.24E-02 | 2.23E-01 | 2.70E+00 | 2.73E+00 | 8.84E-01 | no | | intergenic_variant | MODIFIER |
| 2 | 149333926 | AX-170311190 | 0 | T | C | 0.14 | 3.24E-02 | 2.23E-01 | 2.70E+00 | 2.73E+00 | 8.84E-01 | no | Gm14132 | non_coding_transcript_exon_variant | MODIFIER |
| 2 | 149344839 | AX-170333296 | 0 | G | A | 0.14 | 3.24E-02 | 2.23E-01 | 2.70E+00 | 2.73E+00 | 8.84E-01 | no | | intergenic_variant | MODIFIER |
| 2 | 149350278 | AX-170233169 | 0 | T | G | 0.14 | 3.24E-02 | 2.23E-01 | 2.70E+00 | 2.73E+00 | 8.84E-01 | no | | regulatory_region_variant | MODIFIER |
| 2 | 149358327 | AX-170266522 | 0 | G | A | 0.14 | 3.24E-02 | 2.23E-01 | 2.70E+00 | 2.73E+00 | 8.84E-01 | no | Gm14131 | downstream_gene_variant | MODIFIER |
| 2 | 149374908 | AX-170103493 | 0 | A | G | 0.14 | 3.24E-02 | 2.23E-01 | 2.70E+00 | 2.73E+00 | 8.84E-01 | no | | intergenic_variant | MODIFIER |
| 2 | 149375164 | AX-97786371 | 0 | C | T | 0.14 | 3.24E-02 | 2.23E-01 | 2.70E+00 | 2.73E+00 | 8.84E-01 | no | | intergenic_variant | MODIFIER |
| 2 | 149410882 | AX-170274755 | 0 | A | G | 0.14 | 3.24E-02 | 2.23E-01 | 2.70E+00 | 2.73E+00 | 8.84E-01 | no | Cst10 | downstream_gene_variant | MODIFIER |
| 2 | 149422036 | AX-170297576 | 0 | A | C | 0.14 | 3.24E-02 | 2.23E-01 | 2.70E+00 | 2.73E+00 | 8.84E-01 | no | | intergenic_variant | MODIFIER |
| 2 | 149437762 | AX-170234723 | 0 | C | G | 0.14 | 3.24E-02 | 2.23E-01 | 2.70E+00 | 2.73E+00 | 8.84E-01 | no | | intergenic_variant | MODIFIER |
| 2 | 149635305 | AX-170019348 | 0 | T | C | 0.14 | 3.24E-02 | 2.23E-01 | 2.70E+00 | 2.73E+00 | 8.84E-01 | no | | intergenic_variant | MODIFIER |
| 2 | 149637022 | AX-170345049 | 0 | T | C | 0.14 | 3.24E-02 | 2.23E-01 | 2.70E+00 | 2.73E+00 | 8.84E-01 | no | | intergenic_variant | MODIFIER |
| 2 | 149735474 | AX-170035646 | 0 | C | A | 0.14 | 3.24E-02 | 2.23E-01 | 2.70E+00 | 2.73E+00 | 8.84E-01 | no | | intergenic_variant | MODIFIER |
| 2 | 149815560 | AX-170114505 | 0 | G | A | 0.14 | 3.24E-02 | 2.23E-01 | 2.70E+00 | 2.73E+00 | 8.84E-01 | no | | intergenic_variant | MODIFIER |
| 2 | 149817331 | AX-170223044 | 0 | A | T | 0.14 | 3.24E-02 | 2.23E-01 | 2.70E+00 | 2.73E+00 | 8.84E-01 | no | | intergenic_variant | MODIFIER |
| 2 | 149819049 | AX-170141825 | 0 | A | G | 0.14 | 3.24E-02 | 2.23E-01 | 2.70E+00 | 2.73E+00 | 8.84E-01 | no | | intergenic_variant | MODIFIER |
| 2 | 149836228 | AX-170382163 | 0 | C | T | 0.14 | 3.24E-02 | 2.23E-01 | 2.70E+00 | 2.73E+00 | 8.84E-01 | no | Syndig1 | intron_variant | MODIFIER |
| 2 | 149943533 | AX-97788950 | 0 | G | A | 0.14 | 3.24E-02 | 2.23E-01 | 2.70E+00 | 2.73E+00 | 8.84E-01 | no | Syndig1 | intron_variant | MODIFIER |
| 2 | 149979013 | AX-170242742 | 0 | A | G | 0.14 | 3.24E-02 | 2.23E-01 | 2.70E+00 | 2.73E+00 | 8.84E-01 | no | Syndig1 | intron_variant | MODIFIER |
| 2 | 149983304 | AX-97806492 | 0 | C | T | 0.14 | 3.24E-02 | 2.23E-01 | 2.70E+00 | 2.73E+00 | 8.84E-01 | no | Syndig1 | intron_variant | MODIFIER |
| 2 | 150045040 | AX-97839530 | 0 | C | A | 0.14 | 3.24E-02 | 2.23E-01 | 2.70E+00 | 2.73E+00 | 8.84E-01 | no | | intergenic_variant | MODIFIER |
| 2 | 150091383 | AX-97853295 | 0 | G | C | 0.14 | 3.24E-02 | 2.23E-01 | 2.70E+00 | 2.73E+00 | 8.84E-01 | no | | intergenic_variant | MODIFIER |
| 2 | 150105738 | AX-97832185 | 0 | T | C | 0.14 | 3.24E-02 | 2.23E-01 | 2.70E+00 | 2.73E+00 | 8.84E-01 | no | Gm14140 | downstream_gene_variant | MODIFIER |
| 2 | 154595219 | AX-170223833 | 0 | G | A | 0.14 | 3.24E-02 | 2.23E-01 | 2.70E+00 | 2.73E+00 | 8.84E-01 | no | Pxmp4 | intron_variant | MODIFIER |
| 2 | 154596133 | AX-97796473 | 0 | C | A | 0.14 | 3.24E-02 | 2.23E-01 | 2.70E+00 | 2.73E+00 | 8.84E-01 | no | Pxmp4 | intron_variant | MODIFIER |
| 2 | 154605071 | AX-168373843 | 0 | T | C | 0.14 | 3.24E-02 | 2.23E-01 | 2.70E+00 | 2.73E+00 | 8.84E-01 | no | Pxmp4 | upstream_gene_variant | MODIFIER |
| 2 | 154644014 | AX-97804874 | 0 | A | G | 0.14 | 3.24E-02 | 2.23E-01 | 2.70E+00 | 2.73E+00 | 8.84E-01 | no | Zfp341 | intron_variant | MODIFIER |
| 2 | 154646353 | AX-170348210 | 0 | C | T | 0.14 | 3.24E-02 | 2.23E-01 | 2.70E+00 | 2.73E+00 | 8.84E-01 | no | Zfp341 | 3_prime_UTR_variant | MODIFIER |
| 2 | 154647772 | AX-97857270 | 0 | G | T | 0.14 | 3.24E-02 | 2.23E-01 | 2.70E+00 | 2.73E+00 | 8.84E-01 | no | Zfp341 | downstream_gene_variant | MODIFIER |
| 2 | 154685192 | AX-170156928 | 0 | A | G | 0.14 | 3.24E-02 | 2.23E-01 | 2.70E+00 | 2.73E+00 | 8.84E-01 | no | Chmp4b | intron_variant | MODIFIER |
| 2 | 154702225 | AX-170430847 | 0 | A | G | 0.14 | 3.24E-02 | 2.23E-01 | 2.70E+00 | 2.73E+00 | 8.84E-01 | no | Rpl5-ps2 | upstream_gene_variant | MODIFIER |
| 2 | 154703339 | AX-170258224 | 0 | T | C | 0.14 | 3.24E-02 | 2.23E-01 | 2.70E+00 | 2.73E+00 | 8.84E-01 | no | Rpl5-ps2 | upstream_gene_variant | MODIFIER |
| 2 | 154706895 | AX-170300935 | 0 | A | G | 0.14 | 3.24E-02 | 2.23E-01 | 2.70E+00 | 2.73E+00 | 8.84E-01 | no | Rpl5-ps2 | downstream_gene_variant | MODIFIER |
| 2 | 154743556 | AX-170270510 | 0 | C | T | 0.14 | 3.24E-02 | 2.23E-01 | 2.70E+00 | 2.73E+00 | 8.84E-01 | no | | regulatory_region_variant | MODIFIER |

| | | | | | | | | | | | | | | | |
|---|---|---|---|---|---|---|---|---|---|---|---|---|---|---|---|
| 2 | 154743606 | AX-170131332 | 0 | C | T | 0.14 | 3.24E-02 | 2.23E-01 | 2.70E+00 | 2.73E+00 | 8.84E-01 | no | | regulatory_region_variant | MODIFIER |
| 2 | 154747179 | AX-170357683 | 0 | A | G | 0.14 | 3.24E-02 | 2.23E-01 | 2.70E+00 | 2.73E+00 | 8.84E-01 | no | | intergenic_variant | MODIFIER |
| 2 | 154751217 | AX-170259993 | 0 | G | A | 0.14 | 3.24E-02 | 2.23E-01 | 2.70E+00 | 2.73E+00 | 8.84E-01 | no | | intergenic_variant | MODIFIER |
| 2 | 154757824 | AX-97809079 | 0 | G | A | 0.14 | 3.24E-02 | 2.23E-01 | 2.70E+00 | 2.73E+00 | 8.84E-01 | no | | regulatory_region_variant | MODIFIER |
| 2 | 154771086 | AX-170019417 | 0 | G | A | 0.14 | 3.24E-02 | 2.23E-01 | 2.70E+00 | 2.73E+00 | 8.84E-01 | no | Gm14214 | upstream_gene_variant | MODIFIER |
| 2 | 154773017 | AX-97820749 | 0 | A | G | 0.14 | 3.24E-02 | 2.23E-01 | 2.70E+00 | 2.73E+00 | 8.84E-01 | no | Gm14214 | upstream_gene_variant | MODIFIER |
| 2 | 154786904 | AX-170225746 | 0 | A | G | 0.14 | 3.24E-02 | 2.23E-01 | 2.70E+00 | 2.73E+00 | 8.84E-01 | no | Raly | upstream_gene_variant | MODIFIER |
| 2 | 154808327 | AX-170207164 | 0 | A | G | 0.14 | 3.24E-02 | 2.23E-01 | 2.70E+00 | 2.73E+00 | 8.84E-01 | no | Raly | intron_variant | MODIFIER |
| 2 | 154819305 | AX-170260094 | 0 | G | A | 0.14 | 3.24E-02 | 2.23E-01 | 2.70E+00 | 2.73E+00 | 8.84E-01 | no | Raly | intron_variant | MODIFIER |
| 2 | 154821969 | AX-170289707 | 0 | G | A | 0.14 | 3.24E-02 | 2.23E-01 | 2.70E+00 | 2.73E+00 | 8.84E-01 | no | Raly | intron_variant | MODIFIER |
| 2 | 154846713 | AX-170164892 | 0 | G | A | 0.14 | 3.24E-02 | 2.23E-01 | 2.70E+00 | 2.73E+00 | 8.84E-01 | no | Raly | intron_variant | MODIFIER |
| 2 | 154852944 | AX-170515665 | 0 | T | C | 0.14 | 3.24E-02 | 2.23E-01 | 2.70E+00 | 2.73E+00 | 8.84E-01 | no | Raly | intron_variant | MODIFIER |
| 2 | 154864899 | AX-170533376 | 0 | G | T | 0.14 | 3.24E-02 | 2.23E-01 | 2.70E+00 | 2.73E+00 | 8.84E-01 | no | Raly | intron_variant | MODIFIER |
| 2 | 154871658 | AX-97835555 | 0 | T | C | 0.14 | 3.24E-02 | 2.23E-01 | 2.70E+00 | 2.73E+00 | 8.84E-01 | no | Raly | downstream_gene_variant | MODIFIER |
| 2 | 154875711 | AX-170233454 | 0 | A | G | 0.14 | 3.24E-02 | 2.23E-01 | 2.70E+00 | 2.73E+00 | 8.84E-01 | no | Eif2s2 | intron_variant | MODIFIER |
| 2 | 154886660 | AX-170309220 | 0 | G | T | 0.14 | 3.24E-02 | 2.23E-01 | 2.70E+00 | 2.73E+00 | 8.84E-01 | no | Eif2s2 | intron_variant | MODIFIER |
| 2 | 154940160 | AX-170088797 | 0 | A | C | 0.14 | 3.24E-02 | 2.23E-01 | 2.70E+00 | 2.73E+00 | 8.84E-01 | no | a | intron_variant | MODIFIER |
| 2 | 154944251 | AX-170177979 | 0 | A | T | 0.14 | 3.24E-02 | 2.23E-01 | 2.70E+00 | 2.73E+00 | 8.84E-01 | no | a | intron_variant | MODIFIER |
| 2 | 154956385 | AX-170268705 | 0 | A | T | 0.14 | 3.24E-02 | 2.23E-01 | 2.70E+00 | 2.73E+00 | 8.84E-01 | no | a | intron_variant | MODIFIER |
| 2 | 154957126 | AX-170159675 | 0 | T | C | 0.14 | 3.24E-02 | 2.23E-01 | 2.70E+00 | 2.73E+00 | 8.84E-01 | no | a | intron_variant | MODIFIER |
| 2 | 154966737 | AX-97845093 | 0 | G | A | 0.14 | 3.24E-02 | 2.23E-01 | 2.70E+00 | 2.73E+00 | 8.84E-01 | no | a | intron_variant | MODIFIER |
| 2 | 154997482 | AX-97851157 | 0 | G | A | 0.14 | 3.24E-02 | 2.23E-01 | 2.70E+00 | 2.73E+00 | 8.84E-01 | no | a | intron_variant | MODIFIER |
| 2 | 155013625 | AX-170460195 | 0 | A | G | 0.14 | 3.24E-02 | 2.23E-01 | 2.70E+00 | 2.73E+00 | 8.84E-01 | no | a | 5_prime_UTR_variant | MODIFIER |
| 2 | 155035247 | AX-170094454 | 0 | C | T | 0.14 | 3.24E-02 | 2.23E-01 | 2.70E+00 | 2.73E+00 | 8.84E-01 | no | a | intron_variant | MODIFIER |
| 2 | 155036336 | AX-170065350 | 0 | C | T | 0.14 | 3.24E-02 | 2.23E-01 | 2.70E+00 | 2.73E+00 | 8.84E-01 | no | a | intron_variant | MODIFIER |
| 2 | 155064613 | AX-97839628 | 0 | C | T | 0.14 | 3.24E-02 | 2.23E-01 | 2.70E+00 | 2.73E+00 | 8.84E-01 | no | Ahcy | intron_variant | MODIFIER |
| 2 | 155066919 | AX-170204354 | 0 | T | C | 0.14 | 3.24E-02 | 2.23E-01 | 2.70E+00 | 2.73E+00 | 8.84E-01 | no | Ahcy | intron_variant | MODIFIER |
| 2 | 155066954 | AX-170281901 | 0 | C | T | 0.14 | 3.24E-02 | 2.23E-01 | 2.70E+00 | 2.73E+00 | 8.84E-01 | no | Ahcy | intron_variant | MODIFIER |
| 2 | 155073099 | AX-170286437 | 0 | T | C | 0.14 | 3.24E-02 | 2.23E-01 | 2.70E+00 | 2.73E+00 | 8.84E-01 | no | Ahcy | intron_variant | MODIFIER |
| 2 | 155080299 | AX-170094455 | 0 | G | A | 0.14 | 3.24E-02 | 2.23E-01 | 2.70E+00 | 2.73E+00 | 8.84E-01 | no | Gm45609 | intron_variant,non_coding_transcript_variant | MODIFIER |
| 2 | 155086768 | AX-170476583 | 0 | C | T | 0.14 | 3.24E-02 | 2.23E-01 | 2.70E+00 | 2.73E+00 | 8.84E-01 | no | Gm45609 | intron_variant,non_coding_transcript_variant | MODIFIER |
| 2 | 155112325 | AX-170384872 | 0 | G | A | 0.14 | 3.24E-02 | 2.23E-01 | 2.70E+00 | 2.73E+00 | 8.84E-01 | no | | intergenic_variant | MODIFIER |
| 2 | 155143810 | AX-97843404 | 0 | A | G | 0.14 | 3.24E-02 | 2.23E-01 | 2.70E+00 | 2.73E+00 | 8.84E-01 | no | Itch | intron_variant | MODIFIER |
| 2 | 155144383 | AX-170533071 | 0 | G | A | 0.14 | 3.24E-02 | 2.23E-01 | 2.70E+00 | 2.73E+00 | 8.84E-01 | no | Itch | intron_variant | MODIFIER |
| 2 | 155178698 | AX-170019436 | 0 | C | T | 0.14 | 3.24E-02 | 2.23E-01 | 2.70E+00 | 2.73E+00 | 8.84E-01 | no | Itch | intron_variant | MODIFIER |
| 2 | 155179807 | AX-170216956 | 0 | A | G | 0.14 | 3.24E-02 | 2.23E-01 | 2.70E+00 | 2.73E+00 | 8.84E-01 | no | Itch | intron_variant | MODIFIER |
| 2 | 155183936 | AX-170247331 | 0 | C | T | 0.14 | 3.24E-02 | 2.23E-01 | 2.70E+00 | 2.73E+00 | 8.84E-01 | no | Itch | intron_variant | MODIFIER |
| 2 | 155208116 | AX-170019425 | 0 | C | A | 0.14 | 3.24E-02 | 2.23E-01 | 2.70E+00 | 2.73E+00 | 8.84E-01 | no | Itch | intron_variant | MODIFIER |
| 2 | 155210214 | AX-170088799 | 0 | C | T | 0.14 | 3.24E-02 | 2.23E-01 | 2.70E+00 | 2.73E+00 | 8.84E-01 | no | Itch | intron_variant | MODIFIER |
| 2 | 155240876 | AX-170435518 | 0 | A | C | 0.14 | 3.24E-02 | 2.23E-01 | 2.70E+00 | 2.73E+00 | 8.84E-01 | no | Dynlrb1 | intron_variant,non_coding_transcript_variant | MODIFIER |
| 2 | 155250469 | AX-170019430 | 0 | C | A | 0.14 | 3.24E-02 | 2.23E-01 | 2.70E+00 | 2.73E+00 | 8.84E-01 | no | Dynlrb1 | downstream_gene_variant | MODIFIER |
| 2 | 155258494 | AX-170403465 | 0 | A | C | 0.14 | 3.24E-02 | 2.23E-01 | 2.70E+00 | 2.73E+00 | 8.84E-01 | no | | regulatory_region_variant | MODIFIER |
| 2 | 155267354 | AX-97795943 | 0 | G | A | 0.14 | 3.24E-02 | 2.23E-01 | 2.70E+00 | 2.73E+00 | 8.84E-01 | no | | intergenic_variant | MODIFIER |
| 2 | 155287448 | AX-97832990 | 0 | T | C | 0.14 | 3.24E-02 | 2.23E-01 | 2.70E+00 | 2.73E+00 | 8.84E-01 | no | Pigu | intron_variant | MODIFIER |
| 2 | 155299663 | AX-170019426 | 0 | A | G | 0.14 | 3.24E-02 | 2.23E-01 | 2.70E+00 | 2.73E+00 | 8.84E-01 | no | Pigu | intron_variant | MODIFIER |
| 2 | 155301188 | AX-97853403 | 0 | A | G | 0.14 | 3.24E-02 | 2.23E-01 | 2.70E+00 | 2.73E+00 | 8.84E-01 | no | Pigu | intron_variant | MODIFIER |
| 2 | 155308742 | AX-170065352 | 0 | T | C | 0.14 | 3.24E-02 | 2.23E-01 | 2.70E+00 | 2.73E+00 | 8.84E-01 | no | Pigu | intron_variant | MODIFIER |
| 2 | 155309596 | AX-170491782 | 0 | T | C | 0.14 | 3.24E-02 | 2.23E-01 | 2.70E+00 | 2.73E+00 | 8.84E-01 | no | Pigu | intron_variant | MODIFIER |
| 2 | 155327071 | AX-170019428 | 0 | A | T | 0.14 | 3.24E-02 | 2.23E-01 | 2.70E+00 | 2.73E+00 | 8.84E-01 | no | Pigu | intron_variant | MODIFIER |
| 2 | 155333279 | AX-97807660 | 0 | G | A | 0.14 | 3.24E-02 | 2.23E-01 | 2.70E+00 | 2.73E+00 | 8.84E-01 | no | Pigu | intron_variant | MODIFIER |
| 2 | 155333704 | AX-170019429 | 0 | G | A | 0.14 | 3.24E-02 | 2.23E-01 | 2.70E+00 | 2.73E+00 | 8.84E-01 | no | Pigu | intron_variant | MODIFIER |
| 2 | 155358881 | AX-170019431 | 0 | T | C | 0.14 | 3.24E-02 | 2.23E-01 | 2.70E+00 | 2.73E+00 | 8.84E-01 | no | Pigu | upstream_gene_variant | MODIFIER |
| 2 | 155359836 | AX-97851153 | 0 | C | A | 0.14 | 3.24E-02 | 2.23E-01 | 2.70E+00 | 2.73E+00 | 8.84E-01 | no | Pigu | upstream_gene_variant | MODIFIER |
| 2 | 155412055 | AX-170019435 | 0 | A | G | 0.14 | 3.24E-02 | 2.23E-01 | 2.70E+00 | 2.73E+00 | 8.84E-01 | no | Ncoa6 | intron_variant | MODIFIER |

| | | | | | | | | | | | | | | | |
|---|---|---|---|---|---|---|---|---|---|---|---|---|---|---|---|
| 2 | 155418181 | AX-97802983 | 0 | T | C | 0.14 | 3.24E-02 | 2.23E-01 | 2.70E+00 | 2.73E+00 | 8.84E-01 | no | Ncoa6 | intron_variant | MODIFIER |
| 2 | 155422872 | AX-170304715 | 0 | T | C | 0.14 | 3.24E-02 | 2.23E-01 | 2.70E+00 | 2.73E+00 | 8.84E-01 | no | Ncoa6 | missense_variant | MODERATE |
| 2 | 155449795 | AX-170364868 | 0 | T | C | 0.14 | 3.24E-02 | 2.23E-01 | 2.70E+00 | 2.73E+00 | 8.84E-01 | no | Ncoa6 | intron_variant | MODIFIER |
| 2 | 155454052 | AX-170478619 | 0 | A | G | 0.14 | 3.24E-02 | 2.23E-01 | 2.70E+00 | 2.73E+00 | 8.84E-01 | no | Ncoa6 | intron_variant | MODIFIER |
| 2 | 155479428 | AX-170019434 | 0 | C | T | 0.14 | 3.24E-02 | 2.23E-01 | 2.70E+00 | 2.73E+00 | 8.84E-01 | no | | intergenic_variant | MODIFIER |
| 2 | 155498289 | AX-97811079 | 0 | T | C | 0.14 | 3.24E-02 | 2.23E-01 | 2.70E+00 | 2.73E+00 | 8.84E-01 | no | Ggt7 | intron_variant | MODIFIER |
| 2 | 155533807 | AX-170397968 | 0 | A | G | 0.14 | 3.24E-02 | 2.23E-01 | 2.70E+00 | 2.73E+00 | 8.84E-01 | no | Acss2 | intron_variant | MODIFIER |
| 2 | 155535376 | AX-170295625 | 0 | A | G | 0.14 | 3.24E-02 | 2.23E-01 | 2.70E+00 | 2.73E+00 | 8.84E-01 | no | Acss2 | intron_variant | MODIFIER |
| 2 | 155549877 | AX-170065356 | 0 | T | C | 0.14 | 3.24E-02 | 2.23E-01 | 2.70E+00 | 2.73E+00 | 8.84E-01 | no | Acss2 | splice_region_variant,intron_variant | LOW |
| 2 | 155553809 | AX-168392131 | 0 | C | T | 0.14 | 3.24E-02 | 2.23E-01 | 2.70E+00 | 2.73E+00 | 8.84E-01 | no | Acss2 | intron_variant | MODIFIER |
| 2 | 155559225 | AX-97799366 | 0 | T | C | 0.14 | 3.24E-02 | 2.23E-01 | 2.70E+00 | 2.73E+00 | 8.84E-01 | no | Acss2 | intron_variant | MODIFIER |
| 2 | 155561262 | AX-170065360 | 0 | A | G | 0.14 | 3.24E-02 | 2.23E-01 | 2.70E+00 | 2.73E+00 | 8.84E-01 | no | Acss2 | intron_variant | MODIFIER |
| 2 | 155564572 | AX-170065358 | 0 | A | G | 0.14 | 3.24E-02 | 2.23E-01 | 2.70E+00 | 2.73E+00 | 8.84E-01 | no | Acss2 | downstream_gene_variant | MODIFIER |
| 2 | 155568601 | AX-170415467 | 0 | C | T | 0.14 | 3.24E-02 | 2.23E-01 | 2.70E+00 | 2.73E+00 | 8.84E-01 | no | Acss2 | intron_variant | MODIFIER |
| 2 | 155571915 | AX-97800605 | 0 | T | A | 0.14 | 3.24E-02 | 2.23E-01 | 2.70E+00 | 2.73E+00 | 8.84E-01 | no | Acss2 | intron_variant | MODIFIER |
| 2 | 155602591 | AX-170280348 | 0 | T | C | 0.14 | 3.24E-02 | 2.23E-01 | 2.70E+00 | 2.73E+00 | 8.84E-01 | no | | intergenic_variant | MODIFIER |
| 2 | 155602650 | AX-170100682 | 0 | C | T | 0.14 | 3.24E-02 | 2.23E-01 | 2.70E+00 | 2.73E+00 | 8.84E-01 | no | | intergenic_variant | MODIFIER |
| 2 | 155603209 | AX-170312495 | 0 | G | A | 0.14 | 3.24E-02 | 2.23E-01 | 2.70E+00 | 2.73E+00 | 8.84E-01 | no | | regulatory_region_variant | MODIFIER |
| 2 | 155620413 | AX-170286668 | 0 | C | T | 0.14 | 3.24E-02 | 2.23E-01 | 2.70E+00 | 2.73E+00 | 8.84E-01 | no | Myh7b | synonymous_variant | LOW |
| 2 | 155621257 | AX-97855496 | 0 | T | C | 0.14 | 3.24E-02 | 2.23E-01 | 2.70E+00 | 2.73E+00 | 8.84E-01 | no | Myh7b | intron_variant | MODIFIER |
| 2 | 155670420 | AX-170139955 | 0 | G | A | 0.14 | 3.24E-02 | 2.23E-01 | 2.70E+00 | 2.73E+00 | 8.84E-01 | no | Trpc4ap | intron_variant | MODIFIER |
| 2 | 155679992 | AX-170352527 | 0 | G | A | 0.14 | 3.24E-02 | 2.23E-01 | 2.70E+00 | 2.73E+00 | 8.84E-01 | no | Trpc4ap | intron_variant | MODIFIER |
| 2 | 155680021 | AX-97802182 | 0 | G | T | 0.14 | 3.24E-02 | 2.23E-01 | 2.70E+00 | 2.73E+00 | 8.84E-01 | no | Trpc4ap | intron_variant | MODIFIER |
| 2 | 155708661 | AX-97848833 | 0 | T | C | 0.14 | 3.24E-02 | 2.23E-01 | 2.70E+00 | 2.73E+00 | 8.84E-01 | no | Edem2 | intron_variant | MODIFIER |
| 2 | 155726866 | AX-170019444 | 0 | A | G | 0.14 | 3.24E-02 | 2.23E-01 | 2.70E+00 | 2.73E+00 | 8.84E-01 | no | Edem2 | intron_variant | MODIFIER |
| 2 | 155727002 | AX-170019450 | 0 | T | G | 0.14 | 3.24E-02 | 2.23E-01 | 2.70E+00 | 2.73E+00 | 8.84E-01 | no | Edem2 | intron_variant | MODIFIER |
| 2 | 155727021 | AX-97832992 | 0 | C | T | 0.14 | 3.24E-02 | 2.23E-01 | 2.70E+00 | 2.73E+00 | 8.84E-01 | no | Edem2 | intron_variant | MODIFIER |
| 2 | 155760341 | AX-170019439 | 0 | G | A | 0.14 | 3.24E-02 | 2.23E-01 | 2.70E+00 | 2.73E+00 | 8.84E-01 | no | Procr | downstream_gene_variant | MODIFIER |
| 2 | 155803841 | AX-170019447 | 0 | T | C | 0.14 | 3.24E-02 | 2.23E-01 | 2.70E+00 | 2.73E+00 | 8.84E-01 | no | Mmp24 | intron_variant | MODIFIER |
| 2 | 155828819 | AX-97804251 | 0 | T | C | 0.14 | 3.24E-02 | 2.23E-01 | 2.70E+00 | 2.73E+00 | 8.84E-01 | no | Eif6 | upstream_gene_variant | MODIFIER |
| 2 | 155828921 | AX-170019440 | 0 | G | A | 0.14 | 3.24E-02 | 2.23E-01 | 2.70E+00 | 2.73E+00 | 8.84E-01 | no | Eif6 | upstream_gene_variant | MODIFIER |
| 2 | 155879957 | AX-170214514 | 0 | T | C | 0.14 | 3.24E-02 | 2.23E-01 | 2.70E+00 | 2.73E+00 | 8.84E-01 | no | Uqcc1 | intron_variant | MODIFIER |
| 2 | 155885867 | AX-170019443 | 0 | G | A | 0.14 | 3.24E-02 | 2.23E-01 | 2.70E+00 | 2.73E+00 | 8.84E-01 | no | Uqcc1 | intron_variant | MODIFIER |
| 2 | 155886901 | AX-170019442 | 0 | A | G | 0.14 | 3.24E-02 | 2.23E-01 | 2.70E+00 | 2.73E+00 | 8.84E-01 | no | Uqcc1 | intron_variant | MODIFIER |
| 2 | 155898173 | AX-170517547 | 0 | C | T | 0.14 | 3.24E-02 | 2.23E-01 | 2.70E+00 | 2.73E+00 | 8.84E-01 | no | Uqcc1 | intron_variant | MODIFIER |
| 2 | 155899912 | AX-170530052 | 0 | C | T | 0.14 | 3.24E-02 | 2.23E-01 | 2.70E+00 | 2.73E+00 | 8.84E-01 | no | Uqcc1 | intron_variant | MODIFIER |
| 2 | 155906756 | AX-170186827 | 0 | C | A | 0.14 | 3.24E-02 | 2.23E-01 | 2.70E+00 | 2.73E+00 | 8.84E-01 | no | Uqcc1 | intron_variant | MODIFIER |
| 2 | 155914434 | AX-170359665 | 0 | T | C | 0.14 | 3.24E-02 | 2.23E-01 | 2.70E+00 | 2.73E+00 | 8.84E-01 | no | Uqcc1 | intron_variant | MODIFIER |
| 2 | 155923067 | AX-170019445 | 0 | C | T | 0.14 | 3.24E-02 | 2.23E-01 | 2.70E+00 | 2.73E+00 | 8.84E-01 | no | Uqcc1 | intron_variant | MODIFIER |
| 2 | 155928608 | AX-170088800 | 0 | G | A | 0.14 | 3.24E-02 | 2.23E-01 | 2.70E+00 | 2.73E+00 | 8.84E-01 | no | Uqcc1 | intron_variant | MODIFIER |
| 2 | 155944764 | AX-170455439 | 0 | A | G | 0.14 | 3.24E-02 | 2.23E-01 | 2.70E+00 | 2.73E+00 | 8.84E-01 | no | Gdf5 | missense_variant | MODERATE |
| 2 | 155952436 | AX-170519060 | 0 | A | G | 0.14 | 3.24E-02 | 2.23E-01 | 2.70E+00 | 2.73E+00 | 8.84E-01 | no | Cep250 | upstream_gene_variant | MODIFIER |
| 2 | 155961221 | AX-97852837 | 0 | T | C | 0.14 | 3.24E-02 | 2.23E-01 | 2.70E+00 | 2.73E+00 | 8.84E-01 | no | Cep250 | intron_variant | MODIFIER |
| 2 | 155961238 | AX-170019441 | 0 | G | A | 0.14 | 3.24E-02 | 2.23E-01 | 2.70E+00 | 2.73E+00 | 8.84E-01 | no | Cep250 | intron_variant | MODIFIER |
| 2 | 155988224 | AX-97832876 | 0 | A | G | 0.14 | 3.24E-02 | 2.23E-01 | 2.70E+00 | 2.73E+00 | 8.84E-01 | no | Cep250 | intron_variant | MODIFIER |
| 2 | 155993516 | AX-170243264 | 0 | A | G | 0.14 | 3.24E-02 | 2.23E-01 | 2.70E+00 | 2.73E+00 | 8.84E-01 | no | Cep250 | intron_variant | MODIFIER |
| 2 | 155998118 | AX-170065366 | 0 | A | G | 0.14 | 3.24E-02 | 2.23E-01 | 2.70E+00 | 2.73E+00 | 8.84E-01 | no | Cep250 | intron_variant | MODIFIER |
| 2 | 156009063 | AX-97855889 | 0 | T | C | 0.14 | 3.24E-02 | 2.23E-01 | 2.70E+00 | 2.73E+00 | 8.84E-01 | no | Ergic3 | intron_variant | MODIFIER |
| 2 | 156009422 | AX-170521959 | 0 | C | T | 0.14 | 3.24E-02 | 2.23E-01 | 2.70E+00 | 2.73E+00 | 8.84E-01 | no | Ergic3 | intron_variant | MODIFIER |
| 2 | 156014945 | AX-170394227 | 0 | A | G | 0.14 | 3.24E-02 | 2.23E-01 | 2.70E+00 | 2.73E+00 | 8.84E-01 | no | Ergic3 | intron_variant | MODIFIER |
| 2 | 156018431 | AX-170307010 | 0 | C | A | 0.14 | 3.24E-02 | 2.23E-01 | 2.70E+00 | 2.73E+00 | 8.84E-01 | no | Ergic3 | downstream_gene_variant | MODIFIER |
| 2 | 156022441 | AX-97832985 | 0 | A | G | 0.14 | 3.24E-02 | 2.23E-01 | 2.70E+00 | 2.73E+00 | 8.84E-01 | no | Ergic3 | downstream_gene_variant | MODIFIER |
| 2 | 156022526 | AX-170509431 | 0 | A | G | 0.14 | 3.24E-02 | 2.23E-01 | 2.70E+00 | 2.73E+00 | 8.84E-01 | no | Ergic3 | downstream_gene_variant | MODIFIER |
| 2 | 156049619 | AX-170460371 | 0 | A | G | 0.14 | 3.24E-02 | 2.23E-01 | 2.70E+00 | 2.73E+00 | 8.84E-01 | no | Fer1l4 | splice_region_variant,intron_variant | LOW |

| | | | | | | | | | | | | | | | |
|---|---|---|---|---|---|---|---|---|---|---|---|---|---|---|---|
| 2 | 156063437 | AX-97812884 | 0 | C | T | 0.14 | 3.24E-02 | 2.23E-01 | 2.70E+00 | 2.73E+00 | 8.84E-01 | no | Spag4 | upstream_gene_variant | MODIFIER |
| 2 | 156075892 | AX-97828490 | 0 | A | G | 0.14 | 3.24E-02 | 2.23E-01 | 2.70E+00 | 2.73E+00 | 8.84E-01 | no | Cpne1 | intron_variant | MODIFIER |
| 2 | 156091255 | AX-170144699 | 0 | A | G | 0.14 | 3.24E-02 | 2.23E-01 | 2.70E+00 | 2.73E+00 | 8.84E-01 | no | Rbm12 | downstream_gene_variant | MODIFIER |
| 2 | 156093514 | AX-170257547 | 0 | G | A | 0.14 | 3.24E-02 | 2.23E-01 | 2.70E+00 | 2.73E+00 | 8.84E-01 | no | Rbm12 | 3_prime_UTR_variant | MODIFIER |
| 2 | 156097364 | AX-170281120 | 0 | A | G | 0.14 | 3.24E-02 | 2.23E-01 | 2.70E+00 | 2.73E+00 | 8.84E-01 | no | Rbm12 | synonymous_variant | LOW |
| 2 | 156098099 | AX-97851327 | 0 | C | T | 0.14 | 3.24E-02 | 2.23E-01 | 2.70E+00 | 2.73E+00 | 8.84E-01 | no | Rbm12 | synonymous_variant | LOW |
| 2 | 156122268 | AX-170433387 | 0 | T | C | 0.14 | 3.24E-02 | 2.23E-01 | 2.70E+00 | 2.73E+00 | 8.84E-01 | no | Nfs1 | downstream_gene_variant | MODIFIER |
| 2 | 156133684 | AX-97855200 | 0 | C | T | 0.14 | 3.24E-02 | 2.23E-01 | 2.70E+00 | 2.73E+00 | 8.84E-01 | no | Nfs1 | intron_variant | MODIFIER |
| 2 | 156138784 | AX-170189814 | 0 | C | T | 0.14 | 3.24E-02 | 2.23E-01 | 2.70E+00 | 2.73E+00 | 8.84E-01 | no | Nfs1 | intron_variant | MODIFIER |
| 2 | 156139648 | AX-170356871 | 0 | C | T | 0.14 | 3.24E-02 | 2.23E-01 | 2.70E+00 | 2.73E+00 | 8.84E-01 | no | Nfs1 | intron_variant | MODIFIER |
| 2 | 156146781 | AX-170384899 | 0 | A | G | 0.14 | 3.24E-02 | 2.23E-01 | 2.70E+00 | 2.73E+00 | 8.84E-01 | no | Nfs1 | upstream_gene_variant | MODIFIER |
| 2 | 156149470 | AX-170088801 | 0 | A | G | 0.14 | 3.24E-02 | 2.23E-01 | 2.70E+00 | 2.73E+00 | 8.84E-01 | no | Rbm39 | intron_variant | MODIFIER |
| 2 | 156149552 | AX-170185436 | 0 | A | T | 0.14 | 3.24E-02 | 2.23E-01 | 2.70E+00 | 2.73E+00 | 8.84E-01 | no | Rbm39 | intron_variant | MODIFIER |
| 2 | 156173353 | AX-97833046 | 0 | G | T | 0.14 | 3.24E-02 | 2.23E-01 | 2.70E+00 | 2.73E+00 | 8.84E-01 | no | Rbm39 | intron_variant | MODIFIER |
| 2 | 156179057 | AX-170230460 | 0 | A | T | 0.14 | 3.24E-02 | 2.23E-01 | 2.70E+00 | 2.73E+00 | 8.84E-01 | no | Rbm39 | intron_variant | MODIFIER |
| 2 | 156224920 | AX-170529998 | 0 | T | C | 0.14 | 3.24E-02 | 2.23E-01 | 2.70E+00 | 2.73E+00 | 8.84E-01 | no | Phf20 | intron_variant | MODIFIER |
| 2 | 156249211 | AX-170065368 | 0 | A | G | 0.14 | 3.24E-02 | 2.23E-01 | 2.70E+00 | 2.73E+00 | 8.84E-01 | no | Phf20 | intron_variant | MODIFIER |
| 2 | 156249515 | AX-170409200 | 0 | G | A | 0.14 | 3.24E-02 | 2.23E-01 | 2.70E+00 | 2.73E+00 | 8.84E-01 | no | Phf20 | intron_variant | MODIFIER |
| 2 | 156250873 | AX-170296837 | 0 | T | G | 0.14 | 3.24E-02 | 2.23E-01 | 2.70E+00 | 2.73E+00 | 8.84E-01 | no | Phf20 | intron_variant | MODIFIER |
| 2 | 156254422 | AX-170019464 | 0 | G | A | 0.14 | 3.24E-02 | 2.23E-01 | 2.70E+00 | 2.73E+00 | 8.84E-01 | no | Phf20 | intron_variant | MODIFIER |
| 2 | 156256281 | AX-170421844 | 0 | T | G | 0.14 | 3.24E-02 | 2.23E-01 | 2.70E+00 | 2.73E+00 | 8.84E-01 | no | Phf20 | intron_variant | MODIFIER |
| 2 | 156257897 | AX-97839654 | 0 | C | T | 0.14 | 3.24E-02 | 2.23E-01 | 2.70E+00 | 2.73E+00 | 8.84E-01 | no | Phf20 | intron_variant | MODIFIER |
| 2 | 156286036 | AX-170019456 | 0 | T | C | 0.14 | 3.24E-02 | 2.23E-01 | 2.70E+00 | 2.73E+00 | 8.84E-01 | no | Phf20 | intron_variant | MODIFIER |
| 2 | 156287162 | AX-170465931 | 0 | A | T | 0.14 | 3.24E-02 | 2.23E-01 | 2.70E+00 | 2.73E+00 | 8.84E-01 | no | Phf20 | intron_variant | MODIFIER |
| 2 | 156287254 | AX-170385938 | 0 | T | C | 0.14 | 3.24E-02 | 2.23E-01 | 2.70E+00 | 2.73E+00 | 8.84E-01 | no | Phf20 | intron_variant | MODIFIER |
| 2 | 156289929 | AX-170356898 | 0 | T | G | 0.14 | 3.24E-02 | 2.23E-01 | 2.70E+00 | 2.73E+00 | 8.84E-01 | no | Phf20 | intron_variant | MODIFIER |
| 2 | 156298029 | AX-97795402 | 0 | C | G | 0.14 | 3.24E-02 | 2.23E-01 | 2.70E+00 | 2.73E+00 | 8.84E-01 | no | Phf20 | intron_variant | MODIFIER |
| 2 | 156328545 | AX-170019453 | 0 | A | G | 0.14 | 3.24E-02 | 2.23E-01 | 2.70E+00 | 2.73E+00 | 8.84E-01 | no | Cnbd2 | intron_variant | MODIFIER |
| 2 | 156337952 | AX-97813433 | 0 | C | T | 0.14 | 3.24E-02 | 2.23E-01 | 2.70E+00 | 2.73E+00 | 8.84E-01 | no | Cnbd2 | intron_variant | MODIFIER |
| 2 | 156340188 | AX-170468443 | 0 | A | G | 0.14 | 3.24E-02 | 2.23E-01 | 2.70E+00 | 2.73E+00 | 8.84E-01 | no | Cnbd2 | intron_variant | MODIFIER |
| 2 | 156341015 | AX-170103969 | 0 | G | A | 0.14 | 3.24E-02 | 2.23E-01 | 2.70E+00 | 2.73E+00 | 8.84E-01 | no | Cnbd2 | intron_variant | MODIFIER |
| 2 | 156341764 | AX-170525598 | 0 | A | G | 0.14 | 3.24E-02 | 2.23E-01 | 2.70E+00 | 2.73E+00 | 8.84E-01 | no | Cnbd2 | intron_variant | MODIFIER |
| 2 | 156349338 | AX-170124676 | 0 | A | G | 0.14 | 3.24E-02 | 2.23E-01 | 2.70E+00 | 2.73E+00 | 8.84E-01 | no | Cnbd2 | intron_variant | MODIFIER |
| 2 | 156349823 | AX-170172651 | 0 | C | T | 0.14 | 3.24E-02 | 2.23E-01 | 2.70E+00 | 2.73E+00 | 8.84E-01 | no | Cnbd2 | intron_variant | MODIFIER |
| 2 | 156351174 | AX-170019462 | 0 | A | T | 0.14 | 3.24E-02 | 2.23E-01 | 2.70E+00 | 2.73E+00 | 8.84E-01 | no | Cnbd2 | intron_variant | MODIFIER |
| 2 | 156368434 | AX-170019452 | 0 | C | T | 0.14 | 3.24E-02 | 2.23E-01 | 2.70E+00 | 2.73E+00 | 8.84E-01 | no | Cnbd2 | intron_variant | MODIFIER |
| 2 | 156403991 | AX-170019460 | 0 | C | A | 0.14 | 3.24E-02 | 2.23E-01 | 2.70E+00 | 2.73E+00 | 8.84E-01 | no | | intergenic_variant | MODIFIER |
| 2 | 156404158 | AX-170019457 | 0 | T | C | 0.14 | 3.24E-02 | 2.23E-01 | 2.70E+00 | 2.73E+00 | 8.84E-01 | no | | intergenic_variant | MODIFIER |
| 2 | 156420295 | AX-170094458 | 0 | A | G | 0.14 | 3.24E-02 | 2.23E-01 | 2.70E+00 | 2.73E+00 | 8.84E-01 | no | Epb41l1 | upstream_gene_variant | MODIFIER |
| 2 | 156449855 | AX-170019458 | 0 | T | C | 0.14 | 3.24E-02 | 2.23E-01 | 2.70E+00 | 2.73E+00 | 8.84E-01 | no | Epb41l1 | intron_variant | MODIFIER |
| 2 | 156451042 | AX-170065373 | 0 | T | C | 0.14 | 3.24E-02 | 2.23E-01 | 2.70E+00 | 2.73E+00 | 8.84E-01 | no | Epb41l1 | intron_variant | MODIFIER |
| 2 | 156453983 | AX-97794860 | 0 | C | T | 0.14 | 3.24E-02 | 2.23E-01 | 2.70E+00 | 2.73E+00 | 8.84E-01 | no | Epb41l1 | intron_variant | MODIFIER |
| 2 | 156494722 | AX-170019463 | 0 | C | T | 0.14 | 3.24E-02 | 2.23E-01 | 2.70E+00 | 2.73E+00 | 8.84E-01 | no | Epb41l1 | intron_variant | MODIFIER |
| 2 | 156513504 | AX-170019459 | 0 | A | T | 0.14 | 3.24E-02 | 2.23E-01 | 2.70E+00 | 2.73E+00 | 8.84E-01 | no | Epb41l1 | intron_variant | MODIFIER |
| 2 | 156526643 | AX-170389785 | 0 | C | T | 0.14 | 3.24E-02 | 2.23E-01 | 2.70E+00 | 2.73E+00 | 8.84E-01 | no | Epb41l1 | intron_variant | MODIFIER |
| 2 | 156530706 | AX-170339378 | 0 | G | C | 0.14 | 3.24E-02 | 2.23E-01 | 2.70E+00 | 2.73E+00 | 8.84E-01 | no | Epb41l1 | intron_variant | MODIFIER |
| 2 | 156532892 | AX-170389869 | 0 | A | G | 0.14 | 3.24E-02 | 2.23E-01 | 2.70E+00 | 2.73E+00 | 8.84E-01 | no | Epb41l1 | intron_variant | MODIFIER |
| 2 | 156532939 | AX-170333437 | 0 | A | G | 0.14 | 3.24E-02 | 2.23E-01 | 2.70E+00 | 2.73E+00 | 8.84E-01 | no | Epb41l1 | intron_variant | MODIFIER |
| 2 | 156533270 | AX-170420555 | 0 | A | G | 0.14 | 3.24E-02 | 2.23E-01 | 2.70E+00 | 2.73E+00 | 8.84E-01 | no | Epb41l1 | intron_variant | MODIFIER |
| 2 | 156536672 | AX-170516773 | 0 | G | A | 0.14 | 3.24E-02 | 2.23E-01 | 2.70E+00 | 2.73E+00 | 8.84E-01 | no | Epb41l1 | intron_variant | MODIFIER |
| 2 | 156541718 | AX-170019454 | 0 | C | T | 0.14 | 3.24E-02 | 2.23E-01 | 2.70E+00 | 2.73E+00 | 8.84E-01 | no | Epb41l1 | 3_prime_UTR_variant | MODIFIER |
| 2 | 156599460 | AX-170498201 | 0 | G | C | 0.14 | 3.24E-02 | 2.23E-01 | 2.70E+00 | 2.73E+00 | 8.84E-01 | no | | intergenic_variant | MODIFIER |
| 2 | 157253092 | AX-170379855 | 0 | A | C | 0.14 | 3.24E-02 | 2.23E-01 | 2.70E+00 | 2.73E+00 | 8.84E-01 | no | Mroh8 | intron_variant | MODIFIER |
| 2 | 157289378 | AX-170019484 | 0 | G | A | 0.14 | 3.24E-02 | 2.23E-01 | 2.70E+00 | 2.73E+00 | 8.84E-01 | no | Rpn2 | intron_variant | MODIFIER |

| | | | | | | | | | | | | | | | |
|---|---|---|---|---|---|---|---|---|---|---|---|---|---|---|---|
| 2 | 157354408 | AX-170371481 | 0 | C | T | 0.14 | 3.24E-02 | 2.23E-01 | 2.70E+00 | 2.73E+00 | 8.84E-01 | no | | regulatory_region_variant | MODIFIER |
| 2 | 157354983 | AX-97806334 | 0 | G | A | 0.14 | 3.24E-02 | 2.23E-01 | 2.70E+00 | 2.73E+00 | 8.84E-01 | no | | intergenic_variant | MODIFIER |
| 2 | 157362347 | AX-170419713 | 0 | C | G | 0.14 | 3.24E-02 | 2.23E-01 | 2.70E+00 | 2.73E+00 | 8.84E-01 | no | | regulatory_region_variant | MODIFIER |
| 2 | 157362764 | AX-170359894 | 0 | A | G | 0.14 | 3.24E-02 | 2.23E-01 | 2.70E+00 | 2.73E+00 | 8.84E-01 | no | Manbal | upstream_gene_variant | MODIFIER |
| 2 | 157368123 | AX-170019478 | 0 | C | T | 0.14 | 3.24E-02 | 2.23E-01 | 2.70E+00 | 2.73E+00 | 8.84E-01 | no | Manbal | intron_variant | MODIFIER |
| 2 | 157385699 | AX-97805885 | 0 | A | G | 0.14 | 3.24E-02 | 2.23E-01 | 2.70E+00 | 2.73E+00 | 8.84E-01 | no | Manbal | intron_variant | MODIFIER |
| 2 | 157386804 | AX-170276228 | 0 | G | A | 0.14 | 3.24E-02 | 2.23E-01 | 2.70E+00 | 2.73E+00 | 8.84E-01 | no | Manbal | intron_variant | MODIFIER |
| 2 | 157388500 | AX-170174806 | 0 | A | G | 0.14 | 3.24E-02 | 2.23E-01 | 2.70E+00 | 2.73E+00 | 8.84E-01 | no | Manbal | intron_variant | MODIFIER |
| 2 | 157391498 | AX-170346363 | 0 | G | A | 0.14 | 3.24E-02 | 2.23E-01 | 2.70E+00 | 2.73E+00 | 8.84E-01 | no | Manbal | intron_variant | MODIFIER |
| 2 | 157432908 | AX-170380497 | 0 | T | C | 0.14 | 3.24E-02 | 2.23E-01 | 2.70E+00 | 2.73E+00 | 8.84E-01 | no | Src | intron_variant | MODIFIER |
| 2 | 157434128 | AX-170520182 | 0 | C | G | 0.14 | 3.24E-02 | 2.23E-01 | 2.70E+00 | 2.73E+00 | 8.84E-01 | no | Src | intron_variant | MODIFIER |
| 2 | 157436368 | AX-170336704 | 0 | T | C | 0.14 | 3.24E-02 | 2.23E-01 | 2.70E+00 | 2.73E+00 | 8.84E-01 | no | Src | intron_variant | MODIFIER |
| 2 | 157475278 | AX-170065388 | 0 | T | C | 0.14 | 3.24E-02 | 2.23E-01 | 2.70E+00 | 2.73E+00 | 8.84E-01 | no | Src | downstream_gene_variant | MODIFIER |
| 2 | 157475714 | AX-170065385 | 0 | C | T | 0.14 | 3.24E-02 | 2.23E-01 | 2.70E+00 | 2.73E+00 | 8.84E-01 | no | Src | downstream_gene_variant | MODIFIER |
| 2 | 157478571 | AX-170065386 | 0 | G | A | 0.14 | 3.24E-02 | 2.23E-01 | 2.70E+00 | 2.73E+00 | 8.84E-01 | no | | regulatory_region_variant | MODIFIER |
| 2 | 157556140 | AX-170442654 | 0 | C | T | 0.14 | 3.24E-02 | 2.23E-01 | 2.70E+00 | 2.73E+00 | 8.84E-01 | no | Nnat | upstream_gene_variant | MODIFIER |
| 2 | 157673935 | AX-170210693 | 0 | A | G | 0.14 | 3.24E-02 | 2.23E-01 | 2.70E+00 | 2.73E+00 | 8.84E-01 | no | | intergenic_variant | MODIFIER |
| 2 | 157678215 | AX-170354666 | 0 | G | A | 0.14 | 3.24E-02 | 2.23E-01 | 2.70E+00 | 2.73E+00 | 8.84E-01 | no | | intergenic_variant | MODIFIER |
| 2 | 157688383 | AX-170202219 | 0 | A | C | 0.14 | 3.24E-02 | 2.23E-01 | 2.70E+00 | 2.73E+00 | 8.84E-01 | no | | regulatory_region_variant | MODIFIER |
| 2 | 157695837 | AX-170336067 | 0 | C | T | 0.14 | 3.24E-02 | 2.23E-01 | 2.70E+00 | 2.73E+00 | 8.84E-01 | no | | regulatory_region_variant | MODIFIER |
| 2 | 157706175 | AX-170516396 | 0 | A | G | 0.14 | 3.24E-02 | 2.23E-01 | 2.70E+00 | 2.73E+00 | 8.84E-01 | no | | regulatory_region_variant | MODIFIER |
| 2 | 157714008 | AX-170312649 | 0 | G | A | 0.14 | 3.24E-02 | 2.23E-01 | 2.70E+00 | 2.73E+00 | 8.84E-01 | no | Gm25407 | upstream_gene_variant | MODIFIER |
| 2 | 157743892 | AX-170088811 | 0 | G | C | 0.14 | 3.24E-02 | 2.23E-01 | 2.70E+00 | 2.73E+00 | 8.84E-01 | no | Ctnnbl1 | intron_variant | MODIFIER |
| 2 | 157763021 | AX-97803339 | 0 | G | T | 0.14 | 3.24E-02 | 2.23E-01 | 2.70E+00 | 2.73E+00 | 8.84E-01 | no | Ctnnbl1 | intron_variant | MODIFIER |
| 2 | 157765589 | AX-170182641 | 0 | C | T | 0.14 | 3.24E-02 | 2.23E-01 | 2.70E+00 | 2.73E+00 | 8.84E-01 | no | Ctnnbl1 | intron_variant | MODIFIER |
| 2 | 157769362 | AX-170336056 | 0 | C | T | 0.14 | 3.24E-02 | 2.23E-01 | 2.70E+00 | 2.73E+00 | 8.84E-01 | no | Ctnnbl1 | intron_variant | MODIFIER |
| 2 | 157770351 | AX-170447792 | 0 | A | G | 0.14 | 3.24E-02 | 2.23E-01 | 2.70E+00 | 2.73E+00 | 8.84E-01 | no | Ctnnbl1 | intron_variant | MODIFIER |
| 2 | 157789168 | AX-170019494 | 0 | G | A | 0.14 | 3.24E-02 | 2.23E-01 | 2.70E+00 | 2.73E+00 | 8.84E-01 | no | Ctnnbl1 | intron_variant | MODIFIER |
| 2 | 157789766 | AX-170441551 | 0 | A | G | 0.14 | 3.24E-02 | 2.23E-01 | 2.70E+00 | 2.73E+00 | 8.84E-01 | no | Ctnnbl1 | intron_variant | MODIFIER |
| 2 | 157806691 | AX-170019496 | 0 | T | C | 0.14 | 3.24E-02 | 2.23E-01 | 2.70E+00 | 2.73E+00 | 8.84E-01 | no | Ctnnbl1 | missense_variant | MODERATE |
| 2 | 157839020 | AX-97795967 | 0 | C | T | 0.14 | 3.24E-02 | 2.23E-01 | 2.70E+00 | 2.73E+00 | 8.84E-01 | no | Ctnnbl1 | intron_variant | MODIFIER |
| 2 | 157876844 | AX-170229384 | 0 | A | T | 0.14 | 3.24E-02 | 2.23E-01 | 2.70E+00 | 2.73E+00 | 8.84E-01 | no | Ctnnbl1 | intron_variant | MODIFIER |
| 2 | 157887275 | AX-170181824 | 0 | A | T | 0.14 | 3.24E-02 | 2.23E-01 | 2.70E+00 | 2.73E+00 | 8.84E-01 | no | Ctnnbl1 | intron_variant | MODIFIER |
| 2 | 157957443 | AX-170191691 | 0 | G | A | 0.14 | 3.24E-02 | 2.23E-01 | 2.70E+00 | 2.73E+00 | 8.84E-01 | no | | intergenic_variant | MODIFIER |
| 2 | 157979580 | AX-170019488 | 0 | G | C | 0.14 | 3.24E-02 | 2.23E-01 | 2.70E+00 | 2.73E+00 | 8.84E-01 | no | Tti1 | downstream_gene_variant | MODIFIER |
| 2 | 158013869 | AX-170019495 | 0 | C | T | 0.14 | 3.24E-02 | 2.23E-01 | 2.70E+00 | 2.73E+00 | 8.84E-01 | no | Tti1 | intron_variant | MODIFIER |
| 2 | 158132040 | AX-170366657 | 0 | C | T | 0.14 | 3.24E-02 | 2.23E-01 | 2.70E+00 | 2.73E+00 | 8.84E-01 | no | Tgm2 | intron_variant | MODIFIER |
| 2 | 158243352 | AX-170146916 | 0 | C | A | 0.14 | 3.24E-02 | 2.23E-01 | 2.70E+00 | 2.73E+00 | 8.84E-01 | no | Mir7685 | downstream_gene_variant | MODIFIER |
| 2 | 158297280 | AX-97795423 | 0 | A | G | 0.14 | 3.24E-02 | 2.23E-01 | 2.70E+00 | 2.73E+00 | 8.84E-01 | no | | intergenic_variant | MODIFIER |
| 2 | 158305019 | AX-170451223 | 0 | C | T | 0.14 | 3.24E-02 | 2.23E-01 | 2.70E+00 | 2.73E+00 | 8.84E-01 | no | Lbp | upstream_gene_variant | MODIFIER |
| 2 | 158305492 | AX-170302030 | 0 | T | C | 0.14 | 3.24E-02 | 2.23E-01 | 2.70E+00 | 2.73E+00 | 8.84E-01 | no | Lbp | upstream_gene_variant | MODIFIER |
| 2 | 158317871 | AX-170476756 | 0 | C | A | 0.14 | 3.24E-02 | 2.23E-01 | 2.70E+00 | 2.73E+00 | 8.84E-01 | no | Lbp | intron_variant | MODIFIER |
| 2 | 158320369 | AX-170142784 | 0 | C | T | 0.14 | 3.24E-02 | 2.23E-01 | 2.70E+00 | 2.73E+00 | 8.84E-01 | no | Lbp | missense_variant | MODERATE |
| 2 | 158320513 | AX-170483469 | 0 | G | C | 0.14 | 3.24E-02 | 2.23E-01 | 2.70E+00 | 2.73E+00 | 8.84E-01 | no | Lbp | intron_variant | MODIFIER |
| 2 | 158321624 | AX-170254197 | 0 | G | A | 0.14 | 3.24E-02 | 2.23E-01 | 2.70E+00 | 2.73E+00 | 8.84E-01 | no | Lbp | intron_variant | MODIFIER |
| 2 | 158332889 | AX-170390130 | 0 | A | G | 0.14 | 3.24E-02 | 2.23E-01 | 2.70E+00 | 2.73E+00 | 8.84E-01 | no | Lbp | downstream_gene_variant | MODIFIER |
| 2 | 158461726 | AX-170349243 | 0 | T | C | 0.14 | 3.24E-02 | 2.23E-01 | 2.70E+00 | 2.73E+00 | 8.84E-01 | no | Ralgapb | intron_variant | MODIFIER |
| 2 | 158495425 | AX-170396237 | 0 | T | C | 0.14 | 3.24E-02 | 2.23E-01 | 2.70E+00 | 2.73E+00 | 8.84E-01 | no | Ralgapb | intron_variant | MODIFIER |
| 2 | 158498214 | AX-170303771 | 0 | A | G | 0.14 | 3.24E-02 | 2.23E-01 | 2.70E+00 | 2.73E+00 | 8.84E-01 | no | Ralgapb | 3_prime_UTR_variant | MODIFIER |
| 2 | 158498330 | AX-170382143 | 0 | A | G | 0.14 | 3.24E-02 | 2.23E-01 | 2.70E+00 | 2.73E+00 | 8.84E-01 | no | Ralgapb | 3_prime_UTR_variant | MODIFIER |
| 2 | 158517508 | AX-170065402 | 0 | T | C | 0.14 | 3.24E-02 | 2.23E-01 | 2.70E+00 | 2.73E+00 | 8.84E-01 | no | Arhgap40 | intron_variant | MODIFIER |
| 2 | 158518492 | AX-170065404 | 0 | A | G | 0.14 | 3.24E-02 | 2.23E-01 | 2.70E+00 | 2.73E+00 | 8.84E-01 | no | Arhgap40 | intron_variant | MODIFIER |
| 2 | 158518637 | AX-170296774 | 0 | A | C | 0.14 | 3.24E-02 | 2.23E-01 | 2.70E+00 | 2.73E+00 | 8.84E-01 | no | Arhgap40 | intron_variant | MODIFIER |
| 2 | 158518964 | AX-170320554 | 0 | A | G | 0.14 | 3.24E-02 | 2.23E-01 | 2.70E+00 | 2.73E+00 | 8.84E-01 | no | Arhgap40 | intron_variant | MODIFIER |

| | | | | | | | | | | | | | | | |
|---|---|---|---|---|---|---|---|---|---|---|---|---|---|---|---|
| 2 | 158523085 | AX-170326910 | 0 | G | A | 0.14 | 3.24E-02 | 2.23E-01 | 2.70E+00 | 2.73E+00 | 8.84E-01 | no | Arhgap40 | intron_variant | MODIFIER |
| 2 | 158524752 | AX-97809491 | 0 | A | G | 0.14 | 3.24E-02 | 2.23E-01 | 2.70E+00 | 2.73E+00 | 8.84E-01 | no | Arhgap40 | intron_variant | MODIFIER |
| 2 | 158526638 | AX-170288279 | 0 | G | A | 0.14 | 3.24E-02 | 2.23E-01 | 2.70E+00 | 2.73E+00 | 8.84E-01 | no | Arhgap40 | intron_variant | MODIFIER |
| 2 | 158528294 | AX-170288283 | 0 | C | T | 0.14 | 3.24E-02 | 2.23E-01 | 2.70E+00 | 2.73E+00 | 8.84E-01 | no | Arhgap40 | intron_variant | MODIFIER |
| 2 | 158543975 | AX-170387189 | 0 | C | T | 0.14 | 3.24E-02 | 2.23E-01 | 2.70E+00 | 2.73E+00 | 8.84E-01 | no | Arhgap40 | intron_variant | MODIFIER |
| 2 | 158566835 | AX-170330225 | 0 | T | A | 0.14 | 3.24E-02 | 2.23E-01 | 2.70E+00 | 2.73E+00 | 8.84E-01 | no | | intergenic_variant | MODIFIER |
| 2 | 158568654 | AX-170177658 | 0 | C | T | 0.14 | 3.24E-02 | 2.23E-01 | 2.70E+00 | 2.73E+00 | 8.84E-01 | no | | intergenic_variant | MODIFIER |
| 2 | 158569080 | AX-170261050 | 0 | A | C | 0.14 | 3.24E-02 | 2.23E-01 | 2.70E+00 | 2.73E+00 | 8.84E-01 | no | | intergenic_variant | MODIFIER |
| 2 | 158571111 | AX-170439760 | 0 | A | G | 0.14 | 3.24E-02 | 2.23E-01 | 2.70E+00 | 2.73E+00 | 8.84E-01 | no | | regulatory_region_variant | MODIFIER |
| 2 | 158571438 | AX-170482529 | 0 | G | A | 0.14 | 3.24E-02 | 2.23E-01 | 2.70E+00 | 2.73E+00 | 8.84E-01 | no | | intergenic_variant | MODIFIER |
| 2 | 158580414 | AX-170385679 | 0 | G | A | 0.14 | 3.24E-02 | 2.23E-01 | 2.70E+00 | 2.73E+00 | 8.84E-01 | no | | intergenic_variant | MODIFIER |
| 2 | 158582962 | AX-170358405 | 0 | G | A | 0.14 | 3.24E-02 | 2.23E-01 | 2.70E+00 | 2.73E+00 | 8.84E-01 | no | | intergenic_variant | MODIFIER |
| 2 | 158584141 | AX-97811424 | 0 | A | G | 0.14 | 3.24E-02 | 2.23E-01 | 2.70E+00 | 2.73E+00 | 8.84E-01 | no | | intergenic_variant | MODIFIER |
| 2 | 158818336 | AX-170236703 | 0 | T | C | 0.14 | 3.24E-02 | 2.23E-01 | 2.70E+00 | 2.73E+00 | 8.84E-01 | no | Dhx35 | splice_region_variant,intron_variant | LOW |
| 2 | 159093069 | AX-97810523 | 0 | A | G | 0.14 | 3.24E-02 | 2.23E-01 | 2.70E+00 | 2.73E+00 | 8.84E-01 | no | | intergenic_variant | MODIFIER |
| 2 | 159093575 | AX-170226911 | 0 | G | A | 0.14 | 3.24E-02 | 2.23E-01 | 2.70E+00 | 2.73E+00 | 8.84E-01 | no | | intergenic_variant | MODIFIER |
| 1 | 21047365 | AX-170442841 | 1 | A | C | 0.393 | 1.33E-02 | 8.57E-02 | 2.65E+00 | 2.70E+00 | 8.84E-01 | no | Tram2 | intron_variant | MODIFIER |
| 12 | 102978214 | AX-97836907 | 0 | C | T | 0.079 | -2.88E-02 | 1.95E-01 | 2.65E+00 | 2.73E+00 | 8.84E-01 | no | Unc79 | intron_variant | MODIFIER |
| 12 | 102992869 | AX-170207096 | 0 | T | C | 0.079 | -2.88E-02 | 1.95E-01 | 2.65E+00 | 2.73E+00 | 8.84E-01 | no | Unc79 | intron_variant | MODIFIER |
| 12 | 102995814 | AX-170415963 | 0 | G | A | 0.079 | -2.88E-02 | 1.95E-01 | 2.65E+00 | 2.73E+00 | 8.84E-01 | no | Unc79 | intron_variant | MODIFIER |
| 16 | 76749492 | AX-170511374 | 0 | T | C | 0.456 | -9.85E-03 | 7.50E-02 | 2.69E+00 | 2.78E+00 | 8.84E-01 | no | Gm30790 | downstream_gene_variant | MODIFIER |
| 4 | 103804530 | AX-170323570 | 0 | A | G | 0.474 | 9.28E-03 | 5.67E-02 | 2.60E+00 | 2.69E+00 | 8.84E-01 | no | Dab1 | intron_variant | MODIFIER |
| 4 | 103870302 | AX-170321097 | 0 | G | A | 0.237 | 1.86E-02 | 1.13E-01 | 2.60E+00 | 2.69E+00 | 8.84E-01 | no | Dab1 | intron_variant | MODIFIER |
| 4 | 104970248 | AX-170263100 | 0 | A | G | 0.474 | 9.28E-03 | 5.67E-02 | 2.60E+00 | 2.69E+00 | 8.84E-01 | no | Fyb2 | intron_variant | MODIFIER |
| 7 | 67387188 | AX-97782694 | 0 | T | C | 0.421 | -1.38E-02 | 9.99E-02 | 2.69E+00 | 2.76E+00 | 8.84E-01 | no | | intergenic_variant | MODIFIER |
| 7 | 67397053 | AX-170029197 | 0 | G | A | 0.421 | -1.38E-02 | 9.99E-02 | 2.69E+00 | 2.76E+00 | 8.84E-01 | no | | intergenic_variant | MODIFIER |
| 7 | 67410357 | AX-170077611 | 0 | G | T | 0.421 | -1.38E-02 | 9.99E-02 | 2.69E+00 | 2.76E+00 | 8.84E-01 | no | | intergenic_variant | MODIFIER |
| 7 | 67436361 | AX-170029196 | 0 | C | A | 0.421 | -1.38E-02 | 9.99E-02 | 2.69E+00 | 2.76E+00 | 8.84E-01 | no | Gm33926 | intron_variant,non_coding_transcript_variant | MODIFIER |
| 7 | 67439900 | AX-170077612 | 0 | A | G | 0.421 | -1.38E-02 | 9.99E-02 | 2.69E+00 | 2.76E+00 | 8.84E-01 | no | Gm33926 | intron_variant,non_coding_transcript_variant | MODIFIER |
| 7 | 67461815 | AX-170514145 | 0 | C | G | 0.421 | -1.38E-02 | 9.99E-02 | 2.69E+00 | 2.76E+00 | 8.84E-01 | no | | regulatory_region_variant | MODIFIER |
| 7 | 67462563 | AX-170394711 | 0 | T | C | 0.421 | -1.38E-02 | 9.99E-02 | 2.69E+00 | 2.76E+00 | 8.84E-01 | no | | regulatory_region_variant | MODIFIER |
| 7 | 67463487 | AX-170532688 | 0 | T | G | 0.421 | -1.38E-02 | 9.99E-02 | 2.69E+00 | 2.76E+00 | 8.84E-01 | no | | regulatory_region_variant | MODIFIER |
| 7 | 67469207 | AX-170247190 | 0 | G | C | 0.421 | -1.38E-02 | 9.99E-02 | 2.69E+00 | 2.76E+00 | 8.84E-01 | no | | regulatory_region_variant | MODIFIER |
| 7 | 67475181 | AX-170447329 | 0 | C | G | 0.421 | -1.38E-02 | 9.99E-02 | 2.69E+00 | 2.76E+00 | 8.84E-01 | no | | regulatory_region_variant | MODIFIER |
| 7 | 67500207 | AX-170291912 | 0 | A | G | 0.421 | -1.38E-02 | 9.99E-02 | 2.69E+00 | 2.76E+00 | 8.84E-01 | no | | regulatory_region_variant | MODIFIER |
| 7 | 67512023 | AX-170117472 | 0 | A | C | 0.421 | -1.38E-02 | 9.99E-02 | 2.69E+00 | 2.76E+00 | 8.84E-01 | no | Lrrc28 | downstream_gene_variant | MODIFIER |
| 7 | 67512089 | AX-170371137 | 0 | C | G | 0.421 | -1.38E-02 | 9.99E-02 | 2.69E+00 | 2.76E+00 | 8.84E-01 | no | Lrrc28 | downstream_gene_variant | MODIFIER |
| 7 | 67512295 | AX-170338746 | 0 | T | A | 0.421 | -1.38E-02 | 9.99E-02 | 2.69E+00 | 2.76E+00 | 8.84E-01 | no | Lrrc28 | downstream_gene_variant | MODIFIER |
| 7 | 67512422 | AX-168382837 | 0 | A | G | 0.421 | -1.38E-02 | 9.99E-02 | 2.69E+00 | 2.76E+00 | 8.84E-01 | no | Lrrc28 | downstream_gene_variant | MODIFIER |
| 7 | 67513399 | AX-170302536 | 0 | A | G | 0.421 | -1.38E-02 | 9.99E-02 | 2.69E+00 | 2.76E+00 | 8.84E-01 | no | Lrrc28 | downstream_gene_variant | MODIFIER |
| 7 | 67542847 | AX-170135254 | 0 | A | G | 0.421 | -1.38E-02 | 9.99E-02 | 2.69E+00 | 2.76E+00 | 8.84E-01 | no | Lrrc28 | intron_variant | MODIFIER |
| 7 | 67559538 | AX-170077616 | 0 | T | C | 0.421 | -1.38E-02 | 9.99E-02 | 2.69E+00 | 2.76E+00 | 8.84E-01 | no | Lrrc28 | intron_variant | MODIFIER |
| 7 | 67569591 | AX-170399334 | 0 | G | A | 0.421 | -1.38E-02 | 9.99E-02 | 2.69E+00 | 2.76E+00 | 8.84E-01 | no | Lrrc28 | intron_variant | MODIFIER |
| 7 | 67611248 | AX-170412186 | 0 | C | T | 0.421 | -1.38E-02 | 9.99E-02 | 2.69E+00 | 2.76E+00 | 8.84E-01 | no | Lrrc28 | intron_variant | MODIFIER |
| 7 | 67626270 | AX-170480921 | 0 | G | A | 0.421 | -1.38E-02 | 9.99E-02 | 2.69E+00 | 2.76E+00 | 8.84E-01 | no | Lrrc28 | intron_variant | MODIFIER |
| 7 | 67633241 | AX-170357797 | 0 | G | A | 0.421 | -1.38E-02 | 9.99E-02 | 2.69E+00 | 2.76E+00 | 8.84E-01 | no | Lrrc28 | intron_variant | MODIFIER |
| 7 | 67637437 | AX-170417219 | 0 | C | T | 0.421 | -1.38E-02 | 9.99E-02 | 2.69E+00 | 2.76E+00 | 8.84E-01 | no | Lrrc28 | intron_variant | MODIFIER |
| 7 | 67651848 | AX-170317219 | 0 | T | C | 0.421 | -1.38E-02 | 9.99E-02 | 2.69E+00 | 2.76E+00 | 8.84E-01 | no | Ttc23 | intron_variant | MODIFIER |
| 7 | 67668026 | AX-97798813 | 0 | C | T | 0.421 | -1.38E-02 | 9.99E-02 | 2.69E+00 | 2.76E+00 | 8.84E-01 | no | Ttc23 | intron_variant | MODIFIER |
| 7 | 67754990 | AX-170295566 | 0 | C | T | 0.421 | -1.38E-02 | 9.99E-02 | 2.69E+00 | 2.76E+00 | 8.84E-01 | no | Synm | intron_variant | MODIFIER |
| 7 | 67772473 | AX-170330130 | 0 | C | T | 0.421 | -1.38E-02 | 9.99E-02 | 2.69E+00 | 2.76E+00 | 8.84E-01 | no | | regulatory_region_variant | MODIFIER |
| 7 | 68321710 | AX-170208722 | 0 | C | T | 0.421 | -1.38E-02 | 9.99E-02 | 2.69E+00 | 2.76E+00 | 8.84E-01 | no | Fam169b | intron_variant | MODIFIER |
| 7 | 68392307 | AX-170412277 | 0 | C | T | 0.421 | -1.38E-02 | 9.99E-02 | 2.69E+00 | 2.76E+00 | 8.84E-01 | no | Gm44889 | downstream_gene_variant | MODIFIER |
| 7 | 68469230 | AX-170029207 | 0 | T | G | 0.421 | -1.38E-02 | 9.99E-02 | 2.69E+00 | 2.76E+00 | 8.84E-01 | no | Gm34549 | intron_variant,non_coding_transcript_variant | MODIFIER |

| | | | | | | | | | | | | | | | |
|---|---|---|---|---|---|---|---|---|---|---|---|---|---|---|---|
| 7 | 68490138 | AX-170029208 | 0 | G | A | 0.421 | -1.38E-02 | 9.99E-02 | 2.69E+00 | 2.76E+00 | 8.84E-01 | no | Gm34549 | intron_variant,non_coding_transcript_variant | MODIFIER |
| 7 | 68509197 | AX-170089946 | 0 | T | G | 0.421 | -1.38E-02 | 9.99E-02 | 2.69E+00 | 2.76E+00 | 8.84E-01 | no | | intergenic_variant | MODIFIER |
| 7 | 68546236 | AX-170196744 | 0 | G | A | 0.421 | -1.38E-02 | 9.99E-02 | 2.69E+00 | 2.76E+00 | 8.84E-01 | no | | intergenic_variant | MODIFIER |
| 7 | 68546587 | AX-170029214 | 0 | T | A | 0.421 | -1.38E-02 | 9.99E-02 | 2.69E+00 | 2.76E+00 | 8.84E-01 | no | | intergenic_variant | MODIFIER |
| 7 | 68554817 | AX-97834192 | 0 | T | G | 0.421 | -1.38E-02 | 9.99E-02 | 2.69E+00 | 2.76E+00 | 8.84E-01 | no | | intergenic_variant | MODIFIER |
| 7 | 68555222 | AX-170267129 | 0 | C | G | 0.421 | -1.38E-02 | 9.99E-02 | 2.69E+00 | 2.76E+00 | 8.84E-01 | no | | intergenic_variant | MODIFIER |
| 7 | 68903779 | AX-170324281 | 0 | C | T | 0.421 | -1.38E-02 | 9.99E-02 | 2.69E+00 | 2.76E+00 | 8.84E-01 | no | Gm34664 | intron_variant,non_coding_transcript_variant | MODIFIER |
| 7 | 69400993 | AX-170299130 | 0 | G | A | 0.421 | -1.38E-02 | 9.99E-02 | 2.69E+00 | 2.76E+00 | 8.84E-01 | no | Gm34838 | upstream_gene_variant | MODIFIER |
| 9 | 85878260 | AX-170530263 | 1 | C | T | 0.143 | -1.94E-02 | 1.27E-01 | 2.64E+00 | 2.71E+00 | 8.85E-01 | no | | intergenic_variant | MODIFIER |
| 9 | 86319873 | AX-170128547 | 1 | G | T | 0.143 | -1.94E-02 | 1.27E-01 | 2.64E+00 | 2.71E+00 | 8.85E-01 | no | Ube2cbp | intron_variant | MODIFIER |
| 2 | 124561764 | AX-97847080 | 0 | C | A | 0.149 | -2.38E-02 | 1.77E-01 | 2.67E+00 | 2.77E+00 | 8.85E-01 | no | Sema6d | intron_variant | MODIFIER |
| 9 | 57177416 | AX-170140400 | 2 | T | C | 0.218 | 1.78E-02 | 1.23E-01 | 2.68E+00 | 2.73E+00 | 8.85E-01 | no | 1700041C23Rik | intron_variant,non_coding_transcript_variant | MODIFIER |
| 8 | 81791023 | AX-170512928 | 1 | C | T | 0.205 | 1.88E-02 | 1.31E-01 | 2.68E+00 | 2.74E+00 | 8.85E-01 | no | Inpp4b | intron_variant | MODIFIER |
| 8 | 82993988 | AX-170361463 | 1 | G | A | 0.205 | 1.88E-02 | 1.31E-01 | 2.68E+00 | 2.74E+00 | 8.85E-01 | no | Rnf150 | intron_variant | MODIFIER |
| 7 | 51246644 | AX-170447503 | 1 | T | C | 0.42 | 1.49E-02 | 1.01E-01 | 2.65E+00 | 2.73E+00 | 8.85E-01 | no | | intergenic_variant | MODIFIER |
| 7 | 57268505 | AX-97805245 | 1 | A | C | 0.42 | 1.49E-02 | 1.01E-01 | 2.65E+00 | 2.73E+00 | 8.85E-01 | no | Gabrg3 | intron_variant | MODIFIER |
| 2 | 128417223 | AX-170064869 | 1 | T | C | 0.143 | 3.21E-02 | 2.21E-01 | 2.69E+00 | 2.72E+00 | 8.85E-01 | no | Gm14005 | intron_variant,non_coding_transcript_variant | MODIFIER |
| 6 | 139317013 | AX-170035738 | 1 | C | T | 0.071 | 3.94E-02 | 2.76E-01 | 2.71E+00 | 2.74E+00 | 8.85E-01 | no | 4922502N22Rik | non_coding_transcript_exon_variant | MODIFIER |
| 1 | 24315139 | AX-170371022 | 1 | T | C | 0.429 | 1.23E-02 | 8.19E-02 | 2.64E+00 | 2.72E+00 | 8.85E-01 | no | Col19a1 | intron_variant | MODIFIER |
| 9 | 105199253 | AX-170511333 | 1 | C | T | 0.116 | -1.84E-02 | 1.16E-01 | 2.62E+00 | 2.70E+00 | 8.85E-01 | no | Nek11 | intron_variant | MODIFIER |
| 10 | 10768947 | AX-170300278 | 0 | A | G | 0.211 | 1.44E-02 | 1.04E-01 | 2.68E+00 | 2.76E+00 | 8.85E-01 | no | Grm1 | intron_variant | MODIFIER |
| 10 | 10771810 | AX-170506538 | 0 | T | G | 0.211 | 1.44E-02 | 1.04E-01 | 2.68E+00 | 2.76E+00 | 8.85E-01 | no | Grm1 | intron_variant | MODIFIER |
| 10 | 10782343 | AX-170090753 | 0 | G | A | 0.211 | 1.44E-02 | 1.04E-01 | 2.68E+00 | 2.76E+00 | 8.85E-01 | no | Grm1 | intron_variant | MODIFIER |
| 10 | 10803878 | AX-170200374 | 0 | T | C | 0.211 | 1.44E-02 | 1.04E-01 | 2.68E+00 | 2.76E+00 | 8.85E-01 | no | Grm1 | intron_variant | MODIFIER |
| 10 | 10817156 | AX-170112335 | 0 | A | G | 0.211 | 1.44E-02 | 1.04E-01 | 2.68E+00 | 2.76E+00 | 8.85E-01 | no | Grm1 | intron_variant | MODIFIER |
| 10 | 10842196 | AX-170109754 | 0 | A | G | 0.211 | 1.44E-02 | 1.04E-01 | 2.68E+00 | 2.76E+00 | 8.85E-01 | no | Grm1 | intron_variant | MODIFIER |
| 10 | 11839961 | AX-170117831 | 0 | T | G | 0.211 | 1.44E-02 | 1.04E-01 | 2.68E+00 | 2.76E+00 | 8.85E-01 | no | Gm48697 | intron_variant,non_coding_transcript_variant | MODIFIER |
| 10 | 12146479 | AX-168395274 | 0 | G | A | 0.211 | 1.44E-02 | 1.04E-01 | 2.68E+00 | 2.76E+00 | 8.85E-01 | no | | intergenic_variant | MODIFIER |
| 7 | 87769097 | AX-170148712 | 0 | C | A | 0.123 | -2.26E-02 | 1.54E-01 | 2.60E+00 | 2.73E+00 | 8.85E-01 | no | Grm5 | intron_variant | MODIFIER |
| 15 | 97732023 | AX-170416073 | 1 | G | A | 0.134 | 2.34E-02 | 1.56E-01 | 2.65E+00 | 2.72E+00 | 8.85E-01 | no | Endou | upstream_gene_variant | MODIFIER |
| 9 | 110897466 | AX-170385054 | 0 | A | G | 0.105 | 1.44E-02 | 1.08E-01 | 2.68E+00 | 2.77E+00 | 8.85E-01 | no | Als2cl | intron_variant | MODIFIER |
| 20 | 53089031 | AX-97834206 | 0 | C | A | 0.254 | -8.89E-03 | 6.36E-02 | 2.66E+00 | 2.75E+00 | 8.85E-01 | no | Plac1 | intron_variant | MODIFIER |
| 5 | 97715453 | AX-170400892 | 0 | T | A | 0.421 | -1.51E-02 | 1.07E-01 | 2.70E+00 | 2.75E+00 | 8.85E-01 | no | | intergenic_variant | MODIFIER |
| 7 | 50743458 | AX-170341837 | 0 | C | A | 0.43 | 1.49E-02 | 1.00E-01 | 2.65E+00 | 2.72E+00 | 8.85E-01 | no | Nell1 | intron_variant | MODIFIER |
| 7 | 50822680 | AX-170357731 | 0 | C | T | 0.43 | 1.49E-02 | 1.00E-01 | 2.65E+00 | 2.72E+00 | 8.85E-01 | no | Nell1 | intron_variant | MODIFIER |
| 7 | 50823312 | AX-170302532 | 0 | G | C | 0.43 | 1.49E-02 | 1.00E-01 | 2.65E+00 | 2.72E+00 | 8.85E-01 | no | Nell1 | intron_variant | MODIFIER |
| 7 | 50824814 | AX-170226430 | 0 | G | A | 0.43 | 1.49E-02 | 1.00E-01 | 2.65E+00 | 2.72E+00 | 8.85E-01 | no | Nell1 | intron_variant | MODIFIER |
| 7 | 50825227 | AX-97808438 | 0 | C | T | 0.43 | 1.49E-02 | 1.00E-01 | 2.65E+00 | 2.72E+00 | 8.85E-01 | no | Nell1 | intron_variant | MODIFIER |
| 7 | 50826978 | AX-170349162 | 0 | C | G | 0.43 | 1.49E-02 | 1.00E-01 | 2.65E+00 | 2.72E+00 | 8.85E-01 | no | Nell1 | intron_variant | MODIFIER |
| 7 | 50866176 | AX-170248891 | 0 | C | T | 0.43 | 1.49E-02 | 1.00E-01 | 2.65E+00 | 2.72E+00 | 8.85E-01 | no | Nell1 | 3_prime_UTR_variant | MODIFIER |
| 7 | 50866247 | AX-170101315 | 0 | T | C | 0.43 | 1.49E-02 | 1.00E-01 | 2.65E+00 | 2.72E+00 | 8.85E-01 | no | Nell1 | 3_prime_UTR_variant | MODIFIER |
| 7 | 50907013 | AX-170418017 | 0 | G | A | 0.43 | 1.49E-02 | 1.00E-01 | 2.65E+00 | 2.72E+00 | 8.85E-01 | no | | intergenic_variant | MODIFIER |
| 7 | 50908539 | AX-170383404 | 0 | C | T | 0.43 | 1.49E-02 | 1.00E-01 | 2.65E+00 | 2.72E+00 | 8.85E-01 | no | | intergenic_variant | MODIFIER |
| 7 | 50910306 | AX-170405370 | 0 | T | A | 0.43 | 1.49E-02 | 1.00E-01 | 2.65E+00 | 2.72E+00 | 8.85E-01 | no | | intergenic_variant | MODIFIER |
| 7 | 50910517 | AX-170288249 | 0 | A | C | 0.43 | 1.49E-02 | 1.00E-01 | 2.65E+00 | 2.72E+00 | 8.85E-01 | no | | intergenic_variant | MODIFIER |
| 7 | 50910534 | AX-97816205 | 0 | T | C | 0.43 | 1.49E-02 | 1.00E-01 | 2.65E+00 | 2.72E+00 | 8.85E-01 | no | | intergenic_variant | MODIFIER |
| 7 | 50915357 | AX-170238242 | 0 | T | C | 0.43 | 1.49E-02 | 1.00E-01 | 2.65E+00 | 2.72E+00 | 8.85E-01 | no | | intergenic_variant | MODIFIER |
| 7 | 51011181 | AX-170028990 | 0 | C | A | 0.43 | 1.49E-02 | 1.00E-01 | 2.65E+00 | 2.72E+00 | 8.85E-01 | no | | intergenic_variant | MODIFIER |
| 7 | 51016396 | AX-170077316 | 0 | C | G | 0.43 | 1.49E-02 | 1.00E-01 | 2.65E+00 | 2.72E+00 | 8.85E-01 | no | | intergenic_variant | MODIFIER |
| 7 | 51017000 | AX-170077318 | 0 | T | A | 0.43 | 1.49E-02 | 1.00E-01 | 2.65E+00 | 2.72E+00 | 8.85E-01 | no | | intergenic_variant | MODIFIER |
| 7 | 51018601 | AX-170028989 | 0 | A | G | 0.43 | 1.49E-02 | 1.00E-01 | 2.65E+00 | 2.72E+00 | 8.85E-01 | no | | intergenic_variant | MODIFIER |
| 7 | 51020741 | AX-170526436 | 0 | G | A | 0.43 | 1.49E-02 | 1.00E-01 | 2.65E+00 | 2.72E+00 | 8.85E-01 | no | | intergenic_variant | MODIFIER |
| 7 | 51021588 | AX-170495731 | 0 | T | C | 0.43 | 1.49E-02 | 1.00E-01 | 2.65E+00 | 2.72E+00 | 8.85E-01 | no | | intergenic_variant | MODIFIER |
| 7 | 51036893 | AX-170028994 | 0 | T | A | 0.43 | 1.49E-02 | 1.00E-01 | 2.65E+00 | 2.72E+00 | 8.85E-01 | no | | intergenic_variant | MODIFIER |

| | | | | | | | | | | | | | | | |
|---|---|---|---|---|---|---|---|---|---|---|---|---|---|---|---|
| 7 | 51042073 | AX-170194663 | 0 | A | G | 0.43 | 1.49E-02 | 1.00E-01 | 2.65E+00 | 2.72E+00 | 8.85E-01 | no | | intergenic_variant | MODIFIER |
| 7 | 51043816 | AX-97857389 | 0 | C | T | 0.43 | 1.49E-02 | 1.00E-01 | 2.65E+00 | 2.72E+00 | 8.85E-01 | no | | intergenic_variant | MODIFIER |
| 7 | 51065719 | AX-97804183 | 0 | C | T | 0.43 | 1.49E-02 | 1.00E-01 | 2.65E+00 | 2.72E+00 | 8.85E-01 | no | | intergenic_variant | MODIFIER |
| 7 | 51083114 | AX-97802415 | 0 | T | A | 0.43 | 1.49E-02 | 1.00E-01 | 2.65E+00 | 2.72E+00 | 8.85E-01 | no | | intergenic_variant | MODIFIER |
| 7 | 51174708 | AX-97819194 | 0 | A | G | 0.43 | 1.49E-02 | 1.00E-01 | 2.65E+00 | 2.72E+00 | 8.85E-01 | no | Gm16478 | non_coding_transcript_exon_variant | MODIFIER |
| 7 | 51192769 | AX-170493594 | 0 | C | T | 0.43 | 1.49E-02 | 1.00E-01 | 2.65E+00 | 2.72E+00 | 8.85E-01 | no | | intergenic_variant | MODIFIER |
| 7 | 51204316 | AX-170144375 | 0 | G | A | 0.43 | 1.49E-02 | 1.00E-01 | 2.65E+00 | 2.72E+00 | 8.85E-01 | no | | intergenic_variant | MODIFIER |
| 7 | 51215402 | AX-170095927 | 0 | A | G | 0.43 | 1.49E-02 | 1.00E-01 | 2.65E+00 | 2.72E+00 | 8.85E-01 | no | | intergenic_variant | MODIFIER |
| 7 | 51224880 | AX-170495516 | 0 | C | T | 0.43 | 1.49E-02 | 1.00E-01 | 2.65E+00 | 2.72E+00 | 8.85E-01 | no | | intergenic_variant | MODIFIER |
| 7 | 51225638 | AX-170410602 | 0 | A | T | 0.43 | 1.49E-02 | 1.00E-01 | 2.65E+00 | 2.72E+00 | 8.85E-01 | no | | intergenic_variant | MODIFIER |
| 7 | 51233914 | AX-170503835 | 0 | G | C | 0.43 | 1.49E-02 | 1.00E-01 | 2.65E+00 | 2.72E+00 | 8.85E-01 | no | | intergenic_variant | MODIFIER |
| 7 | 51242028 | AX-97841652 | 0 | C | A | 0.43 | 1.49E-02 | 1.00E-01 | 2.65E+00 | 2.72E+00 | 8.85E-01 | no | | intergenic_variant | MODIFIER |
| 7 | 51243951 | AX-170482044 | 0 | G | A | 0.43 | 1.49E-02 | 1.00E-01 | 2.65E+00 | 2.72E+00 | 8.85E-01 | no | | intergenic_variant | MODIFIER |
| 7 | 51244954 | AX-170338676 | 0 | C | A | 0.43 | 1.49E-02 | 1.00E-01 | 2.65E+00 | 2.72E+00 | 8.85E-01 | no | | intergenic_variant | MODIFIER |
| 7 | 51245064 | AX-170325703 | 0 | G | T | 0.43 | 1.49E-02 | 1.00E-01 | 2.65E+00 | 2.72E+00 | 8.85E-01 | no | | intergenic_variant | MODIFIER |
| 7 | 51247612 | AX-170035753 | 0 | C | T | 0.43 | 1.49E-02 | 1.00E-01 | 2.65E+00 | 2.72E+00 | 8.85E-01 | no | | intergenic_variant | MODIFIER |
| 7 | 51258807 | AX-170077322 | 0 | T | A | 0.43 | 1.49E-02 | 1.00E-01 | 2.65E+00 | 2.72E+00 | 8.85E-01 | no | | intergenic_variant | MODIFIER |
| 7 | 51386543 | AX-170192913 | 0 | T | C | 0.43 | 1.49E-02 | 1.00E-01 | 2.65E+00 | 2.72E+00 | 8.85E-01 | no | | intergenic_variant | MODIFIER |
| 7 | 51397841 | AX-170398406 | 0 | C | T | 0.43 | 1.49E-02 | 1.00E-01 | 2.65E+00 | 2.72E+00 | 8.85E-01 | no | Gm45002 | downstream_gene_variant | MODIFIER |
| 7 | 51405742 | AX-170222644 | 0 | T | C | 0.43 | 1.49E-02 | 1.00E-01 | 2.65E+00 | 2.72E+00 | 8.85E-01 | no | | intergenic_variant | MODIFIER |
| 7 | 51417085 | AX-170450192 | 0 | T | A | 0.43 | 1.49E-02 | 1.00E-01 | 2.65E+00 | 2.72E+00 | 8.85E-01 | no | | intergenic_variant | MODIFIER |
| 7 | 51448974 | AX-170077325 | 0 | G | A | 0.43 | 1.49E-02 | 1.00E-01 | 2.65E+00 | 2.72E+00 | 8.85E-01 | no | | intergenic_variant | MODIFIER |
| 7 | 51449696 | AX-170193820 | 0 | G | A | 0.43 | 1.49E-02 | 1.00E-01 | 2.65E+00 | 2.72E+00 | 8.85E-01 | no | | intergenic_variant | MODIFIER |
| 7 | 51450061 | AX-170324256 | 0 | G | A | 0.43 | 1.49E-02 | 1.00E-01 | 2.65E+00 | 2.72E+00 | 8.85E-01 | no | | intergenic_variant | MODIFIER |
| 7 | 51464798 | AX-170029000 | 0 | C | T | 0.43 | 1.49E-02 | 1.00E-01 | 2.65E+00 | 2.72E+00 | 8.85E-01 | no | | regulatory_region_variant | MODIFIER |
| 7 | 51497089 | AX-170452992 | 0 | C | T | 0.43 | 1.49E-02 | 1.00E-01 | 2.65E+00 | 2.72E+00 | 8.85E-01 | no | | intergenic_variant | MODIFIER |
| 7 | 51587584 | AX-170445194 | 0 | T | C | 0.43 | 1.49E-02 | 1.00E-01 | 2.65E+00 | 2.72E+00 | 8.85E-01 | no | Ano5 | intron_variant | MODIFIER |
| 7 | 51598948 | AX-97841978 | 0 | T | C | 0.43 | 1.49E-02 | 1.00E-01 | 2.65E+00 | 2.72E+00 | 8.85E-01 | no | Ano5 | downstream_gene_variant | MODIFIER |
| 7 | 51640473 | AX-170505310 | 0 | G | A | 0.43 | 1.49E-02 | 1.00E-01 | 2.65E+00 | 2.72E+00 | 8.85E-01 | no | Slc17a6 | intron_variant | MODIFIER |
| 7 | 51640521 | AX-170357925 | 0 | A | G | 0.43 | 1.49E-02 | 1.00E-01 | 2.65E+00 | 2.72E+00 | 8.85E-01 | no | Slc17a6 | intron_variant | MODIFIER |
| 7 | 51641344 | AX-170494816 | 0 | G | A | 0.43 | 1.49E-02 | 1.00E-01 | 2.65E+00 | 2.72E+00 | 8.85E-01 | no | Slc17a6 | intron_variant | MODIFIER |
| 7 | 51644208 | AX-170166127 | 0 | T | A | 0.43 | 1.49E-02 | 1.00E-01 | 2.65E+00 | 2.72E+00 | 8.85E-01 | no | Slc17a6 | intron_variant | MODIFIER |
| 7 | 51665931 | AX-170077329 | 0 | C | T | 0.43 | 1.49E-02 | 1.00E-01 | 2.65E+00 | 2.72E+00 | 8.85E-01 | no | Slc17a6 | intron_variant | MODIFIER |
| 7 | 51760733 | AX-170370800 | 0 | A | T | 0.43 | 1.49E-02 | 1.00E-01 | 2.65E+00 | 2.72E+00 | 8.85E-01 | no | Gm29296 | intron_variant,non_coding_transcript_variant | MODIFIER |
| 7 | 51770566 | AX-170323396 | 0 | T | C | 0.43 | 1.49E-02 | 1.00E-01 | 2.65E+00 | 2.72E+00 | 8.85E-01 | no | Gm29296 | intron_variant,non_coding_transcript_variant | MODIFIER |
| 7 | 51778846 | AX-97789440 | 0 | C | A | 0.43 | 1.49E-02 | 1.00E-01 | 2.65E+00 | 2.72E+00 | 8.85E-01 | no | | intergenic_variant | MODIFIER |
| 7 | 51787622 | AX-170461761 | 0 | A | G | 0.43 | 1.49E-02 | 1.00E-01 | 2.65E+00 | 2.72E+00 | 8.85E-01 | no | | intergenic_variant | MODIFIER |
| 7 | 51813439 | AX-97850470 | 0 | C | G | 0.43 | 1.49E-02 | 1.00E-01 | 2.65E+00 | 2.72E+00 | 8.85E-01 | no | | intergenic_variant | MODIFIER |
| 7 | 51830870 | AX-170309396 | 0 | G | T | 0.43 | 1.49E-02 | 1.00E-01 | 2.65E+00 | 2.72E+00 | 8.85E-01 | no | | intergenic_variant | MODIFIER |
| 7 | 51844465 | AX-170191823 | 0 | C | T | 0.43 | 1.49E-02 | 1.00E-01 | 2.65E+00 | 2.72E+00 | 8.85E-01 | no | | intergenic_variant | MODIFIER |
| 7 | 51845161 | AX-170392447 | 0 | G | A | 0.43 | 1.49E-02 | 1.00E-01 | 2.65E+00 | 2.72E+00 | 8.85E-01 | no | | intergenic_variant | MODIFIER |
| 7 | 51845978 | AX-97794233 | 0 | C | T | 0.43 | 1.49E-02 | 1.00E-01 | 2.65E+00 | 2.72E+00 | 8.85E-01 | no | Gm45001 | upstream_gene_variant | MODIFIER |
| 7 | 51849210 | AX-97781571 | 0 | A | C | 0.43 | 1.49E-02 | 1.00E-01 | 2.65E+00 | 2.72E+00 | 8.85E-01 | no | Gm45001 | upstream_gene_variant | MODIFIER |
| 7 | 51864727 | AX-170253650 | 0 | A | C | 0.43 | 1.49E-02 | 1.00E-01 | 2.65E+00 | 2.72E+00 | 8.85E-01 | no | Gas2 | intron_variant,non_coding_transcript_variant | MODIFIER |
| 7 | 51886140 | AX-170095930 | 0 | C | T | 0.43 | 1.49E-02 | 1.00E-01 | 2.65E+00 | 2.72E+00 | 8.85E-01 | no | Gas2 | intron_variant | MODIFIER |
| 7 | 51887266 | AX-170104923 | 0 | T | C | 0.43 | 1.49E-02 | 1.00E-01 | 2.65E+00 | 2.72E+00 | 8.85E-01 | no | Gas2 | intron_variant | MODIFIER |
| 7 | 51908547 | AX-170252889 | 0 | C | A | 0.43 | 1.49E-02 | 1.00E-01 | 2.65E+00 | 2.72E+00 | 8.85E-01 | no | Gas2 | intron_variant | MODIFIER |
| 7 | 51999196 | AX-170209558 | 0 | C | T | 0.43 | 1.49E-02 | 1.00E-01 | 2.65E+00 | 2.72E+00 | 8.85E-01 | no | Gas2 | downstream_gene_variant | MODIFIER |
| 7 | 52001610 | AX-170523850 | 0 | C | A | 0.43 | 1.49E-02 | 1.00E-01 | 2.65E+00 | 2.72E+00 | 8.85E-01 | no | Svip | intron_variant | MODIFIER |
| 7 | 52014090 | AX-170077341 | 0 | A | G | 0.43 | 1.49E-02 | 1.00E-01 | 2.65E+00 | 2.72E+00 | 8.85E-01 | no | 1700015G11Rik | intron_variant | MODIFIER |
| 7 | 52014117 | AX-170077340 | 0 | G | A | 0.43 | 1.49E-02 | 1.00E-01 | 2.65E+00 | 2.72E+00 | 8.85E-01 | no | 1700015G11Rik | intron_variant | MODIFIER |
| 7 | 52035559 | AX-170277039 | 0 | T | C | 0.43 | 1.49E-02 | 1.00E-01 | 2.65E+00 | 2.72E+00 | 8.85E-01 | no | | intergenic_variant | MODIFIER |
| 7 | 52038873 | AX-170162659 | 0 | T | G | 0.43 | 1.49E-02 | 1.00E-01 | 2.65E+00 | 2.72E+00 | 8.85E-01 | no | | intergenic_variant | MODIFIER |
| 7 | 52074098 | AX-170504113 | 0 | T | A | 0.43 | 1.49E-02 | 1.00E-01 | 2.65E+00 | 2.72E+00 | 8.85E-01 | no | Gm22211 | downstream_gene_variant | MODIFIER |

| | | | | | | | | | | | | | | | |
|---|---|---|---|---|---|---|---|---|---|---|---|---|---|---|---|
| 7 | 52087254 | AX-170242427 | 0 | T | C | 0.43 | 1.49E-02 | 1.00E-01 | 2.65E+00 | 2.72E+00 | 8.85E-01 | no | | intergenic_variant | MODIFIER |
| 7 | 52089590 | AX-97800454 | 0 | T | C | 0.43 | 1.49E-02 | 1.00E-01 | 2.65E+00 | 2.72E+00 | 8.85E-01 | no | | intergenic_variant | MODIFIER |
| 7 | 52122552 | AX-97812312 | 0 | C | G | 0.43 | 1.49E-02 | 1.00E-01 | 2.65E+00 | 2.72E+00 | 8.85E-01 | no | | intergenic_variant | MODIFIER |
| 7 | 52148701 | AX-170029006 | 0 | A | G | 0.43 | 1.49E-02 | 1.00E-01 | 2.65E+00 | 2.72E+00 | 8.85E-01 | no | | intergenic_variant | MODIFIER |
| 7 | 52148940 | AX-170029003 | 0 | C | G | 0.43 | 1.49E-02 | 1.00E-01 | 2.65E+00 | 2.72E+00 | 8.85E-01 | no | | intergenic_variant | MODIFIER |
| 7 | 52149135 | AX-170287126 | 0 | T | A | 0.43 | 1.49E-02 | 1.00E-01 | 2.65E+00 | 2.72E+00 | 8.85E-01 | no | | intergenic_variant | MODIFIER |
| 7 | 52161717 | AX-97849442 | 0 | T | C | 0.43 | 1.49E-02 | 1.00E-01 | 2.65E+00 | 2.72E+00 | 8.85E-01 | no | | intergenic_variant | MODIFIER |
| 7 | 52170753 | AX-170444867 | 0 | A | G | 0.43 | 1.49E-02 | 1.00E-01 | 2.65E+00 | 2.72E+00 | 8.85E-01 | no | | intergenic_variant | MODIFIER |
| 7 | 52171540 | AX-170189205 | 0 | G | T | 0.43 | 1.49E-02 | 1.00E-01 | 2.65E+00 | 2.72E+00 | 8.85E-01 | no | | intergenic_variant | MODIFIER |
| 7 | 52184310 | AX-170214402 | 0 | C | G | 0.43 | 1.49E-02 | 1.00E-01 | 2.65E+00 | 2.72E+00 | 8.85E-01 | no | | intergenic_variant | MODIFIER |
| 7 | 52185673 | AX-170413530 | 0 | A | C | 0.43 | 1.49E-02 | 1.00E-01 | 2.65E+00 | 2.72E+00 | 8.85E-01 | no | | intergenic_variant | MODIFIER |
| 7 | 52200691 | AX-170077346 | 0 | T | G | 0.43 | 1.49E-02 | 1.00E-01 | 2.65E+00 | 2.72E+00 | 8.85E-01 | no | | intergenic_variant | MODIFIER |
| 7 | 52239043 | AX-170456362 | 0 | G | A | 0.43 | 1.49E-02 | 1.00E-01 | 2.65E+00 | 2.72E+00 | 8.85E-01 | no | | intergenic_variant | MODIFIER |
| 7 | 52239161 | AX-170508310 | 0 | A | G | 0.43 | 1.49E-02 | 1.00E-01 | 2.65E+00 | 2.72E+00 | 8.85E-01 | no | | intergenic_variant | MODIFIER |
| 7 | 52243748 | AX-97786650 | 0 | A | G | 0.43 | 1.49E-02 | 1.00E-01 | 2.65E+00 | 2.72E+00 | 8.85E-01 | no | | intergenic_variant | MODIFIER |
| 7 | 52261583 | AX-170293236 | 0 | G | C | 0.43 | 1.49E-02 | 1.00E-01 | 2.65E+00 | 2.72E+00 | 8.85E-01 | no | | intergenic_variant | MODIFIER |
| 7 | 52304000 | AX-170319857 | 0 | T | C | 0.43 | 1.49E-02 | 1.00E-01 | 2.65E+00 | 2.72E+00 | 8.85E-01 | no | | intergenic_variant | MODIFIER |
| 7 | 52304248 | AX-97818904 | 0 | T | C | 0.43 | 1.49E-02 | 1.00E-01 | 2.65E+00 | 2.72E+00 | 8.85E-01 | no | | intergenic_variant | MODIFIER |
| 7 | 52308621 | AX-170283065 | 0 | C | T | 0.43 | 1.49E-02 | 1.00E-01 | 2.65E+00 | 2.72E+00 | 8.85E-01 | no | | intergenic_variant | MODIFIER |
| 7 | 52321109 | AX-170177828 | 0 | T | C | 0.43 | 1.49E-02 | 1.00E-01 | 2.65E+00 | 2.72E+00 | 8.85E-01 | no | | intergenic_variant | MODIFIER |
| 7 | 52362820 | AX-97782409 | 0 | A | G | 0.43 | 1.49E-02 | 1.00E-01 | 2.65E+00 | 2.72E+00 | 8.85E-01 | no | | intergenic_variant | MODIFIER |
| 7 | 52405300 | AX-170411077 | 0 | C | T | 0.43 | 1.49E-02 | 1.00E-01 | 2.65E+00 | 2.72E+00 | 8.85E-01 | no | | intergenic_variant | MODIFIER |
| 7 | 52413565 | AX-170259294 | 0 | A | G | 0.43 | 1.49E-02 | 1.00E-01 | 2.65E+00 | 2.72E+00 | 8.85E-01 | no | Gm20074 | downstream_gene_variant | MODIFIER |
| 7 | 52553245 | AX-170295846 | 0 | T | G | 0.43 | 1.49E-02 | 1.00E-01 | 2.65E+00 | 2.72E+00 | 8.85E-01 | no | | intergenic_variant | MODIFIER |
| 7 | 52656229 | AX-170196543 | 0 | A | G | 0.43 | 1.49E-02 | 1.00E-01 | 2.65E+00 | 2.72E+00 | 8.85E-01 | no | | intergenic_variant | MODIFIER |
| 7 | 52659422 | AX-170029013 | 0 | C | T | 0.43 | 1.49E-02 | 1.00E-01 | 2.65E+00 | 2.72E+00 | 8.85E-01 | no | | intergenic_variant | MODIFIER |
| 7 | 52690091 | AX-170507961 | 0 | T | G | 0.43 | 1.49E-02 | 1.00E-01 | 2.65E+00 | 2.72E+00 | 8.85E-01 | no | | intergenic_variant | MODIFIER |
| 7 | 52734450 | AX-170029011 | 0 | G | C | 0.43 | 1.49E-02 | 1.00E-01 | 2.65E+00 | 2.72E+00 | 8.85E-01 | no | | intergenic_variant | MODIFIER |
| 7 | 52734980 | AX-170029008 | 0 | G | C | 0.43 | 1.49E-02 | 1.00E-01 | 2.65E+00 | 2.72E+00 | 8.85E-01 | no | | intergenic_variant | MODIFIER |
| 7 | 52768893 | AX-170077357 | 0 | C | T | 0.43 | 1.49E-02 | 1.00E-01 | 2.65E+00 | 2.72E+00 | 8.85E-01 | no | | intergenic_variant | MODIFIER |
| 7 | 52846493 | AX-97780903 | 0 | C | T | 0.43 | 1.49E-02 | 1.00E-01 | 2.65E+00 | 2.72E+00 | 8.85E-01 | no | | intergenic_variant | MODIFIER |
| 7 | 52846580 | AX-170367959 | 0 | A | G | 0.43 | 1.49E-02 | 1.00E-01 | 2.65E+00 | 2.72E+00 | 8.85E-01 | no | | intergenic_variant | MODIFIER |
| 7 | 52847323 | AX-170347108 | 0 | T | A | 0.43 | 1.49E-02 | 1.00E-01 | 2.65E+00 | 2.72E+00 | 8.85E-01 | no | | intergenic_variant | MODIFIER |
| 7 | 52868778 | AX-170471609 | 0 | G | A | 0.43 | 1.49E-02 | 1.00E-01 | 2.65E+00 | 2.72E+00 | 8.85E-01 | no | | intergenic_variant | MODIFIER |
| 7 | 52869144 | AX-170151344 | 0 | C | T | 0.43 | 1.49E-02 | 1.00E-01 | 2.65E+00 | 2.72E+00 | 8.85E-01 | no | | intergenic_variant | MODIFIER |
| 7 | 52869686 | AX-170333417 | 0 | T | C | 0.43 | 1.49E-02 | 1.00E-01 | 2.65E+00 | 2.72E+00 | 8.85E-01 | no | | intergenic_variant | MODIFIER |
| 7 | 52872488 | AX-170089917 | 0 | T | C | 0.43 | 1.49E-02 | 1.00E-01 | 2.65E+00 | 2.72E+00 | 8.85E-01 | no | | intergenic_variant | MODIFIER |
| 7 | 52983423 | AX-170429248 | 0 | T | A | 0.43 | 1.49E-02 | 1.00E-01 | 2.65E+00 | 2.72E+00 | 8.85E-01 | no | | intergenic_variant | MODIFIER |
| 7 | 52996852 | AX-170135621 | 0 | C | A | 0.43 | 1.49E-02 | 1.00E-01 | 2.65E+00 | 2.72E+00 | 8.85E-01 | no | | intergenic_variant | MODIFIER |
| 7 | 52997298 | AX-170233521 | 0 | G | C | 0.43 | 1.49E-02 | 1.00E-01 | 2.65E+00 | 2.72E+00 | 8.85E-01 | no | | intergenic_variant | MODIFIER |
| 7 | 53014123 | AX-170534329 | 0 | T | A | 0.43 | 1.49E-02 | 1.00E-01 | 2.65E+00 | 2.72E+00 | 8.85E-01 | no | | intergenic_variant | MODIFIER |
| 7 | 53031893 | AX-170213713 | 0 | G | C | 0.43 | 1.49E-02 | 1.00E-01 | 2.65E+00 | 2.72E+00 | 8.85E-01 | no | | intergenic_variant | MODIFIER |
| 7 | 53045120 | AX-170095936 | 0 | C | T | 0.43 | 1.49E-02 | 1.00E-01 | 2.65E+00 | 2.72E+00 | 8.85E-01 | no | | intergenic_variant | MODIFIER |
| 7 | 53122273 | AX-170382439 | 0 | G | T | 0.43 | 1.49E-02 | 1.00E-01 | 2.65E+00 | 2.72E+00 | 8.85E-01 | no | | intergenic_variant | MODIFIER |
| 7 | 53157183 | AX-170130378 | 0 | T | C | 0.43 | 1.49E-02 | 1.00E-01 | 2.65E+00 | 2.72E+00 | 8.85E-01 | no | | intergenic_variant | MODIFIER |
| 7 | 53744175 | AX-170077370 | 0 | A | C | 0.43 | 1.49E-02 | 1.00E-01 | 2.65E+00 | 2.72E+00 | 8.85E-01 | no | | intergenic_variant | MODIFIER |
| 7 | 53821300 | AX-170242633 | 0 | A | C | 0.43 | 1.49E-02 | 1.00E-01 | 2.65E+00 | 2.72E+00 | 8.85E-01 | no | | intergenic_variant | MODIFIER |
| 7 | 53854704 | AX-170173285 | 0 | A | T | 0.43 | 1.49E-02 | 1.00E-01 | 2.65E+00 | 2.72E+00 | 8.85E-01 | no | | intergenic_variant | MODIFIER |
| 7 | 53879225 | AX-170178917 | 0 | G | C | 0.43 | 1.49E-02 | 1.00E-01 | 2.65E+00 | 2.72E+00 | 8.85E-01 | no | | intergenic_variant | MODIFIER |
| 7 | 53892001 | AX-170103830 | 0 | G | A | 0.43 | 1.49E-02 | 1.00E-01 | 2.65E+00 | 2.72E+00 | 8.85E-01 | no | Mir6238 | upstream_gene_variant | MODIFIER |
| 7 | 53909931 | AX-170459136 | 0 | G | A | 0.43 | 1.49E-02 | 1.00E-01 | 2.65E+00 | 2.72E+00 | 8.85E-01 | no | | intergenic_variant | MODIFIER |
| 7 | 53912332 | AX-170089920 | 0 | C | A | 0.43 | 1.49E-02 | 1.00E-01 | 2.65E+00 | 2.72E+00 | 8.85E-01 | no | | intergenic_variant | MODIFIER |
| 7 | 53913845 | AX-170515158 | 0 | T | C | 0.43 | 1.49E-02 | 1.00E-01 | 2.65E+00 | 2.72E+00 | 8.85E-01 | no | | intergenic_variant | MODIFIER |
| 7 | 53923214 | AX-170077376 | 0 | C | T | 0.43 | 1.49E-02 | 1.00E-01 | 2.65E+00 | 2.72E+00 | 8.85E-01 | no | | intergenic_variant | MODIFIER |

| | | | | | | | | | | | | | | | |
|---|---|---|---|---|---|---|---|---|---|---|---|---|---|---|---|
| 7 | 53939998 | AX-170488608 | 0 | C | T | 0.43 | 1.49E-02 | 1.00E-01 | 2.65E+00 | 2.72E+00 | 8.85E-01 | no | | intergenic_variant | MODIFIER |
| 7 | 53995962 | AX-170029027 | 0 | A | G | 0.43 | 1.49E-02 | 1.00E-01 | 2.65E+00 | 2.72E+00 | 8.85E-01 | no | Gm44682 | non_coding_transcript_exon_variant | MODIFIER |
| 7 | 53998027 | AX-97790114 | 0 | A | G | 0.43 | 1.49E-02 | 1.00E-01 | 2.65E+00 | 2.72E+00 | 8.85E-01 | no | Gm44682 | non_coding_transcript_exon_variant | MODIFIER |
| 7 | 54005661 | AX-97819514 | 0 | T | C | 0.43 | 1.49E-02 | 1.00E-01 | 2.65E+00 | 2.72E+00 | 8.85E-01 | no | Gm27921 | downstream_gene_variant | MODIFIER |
| 7 | 54010265 | AX-170189160 | 0 | A | G | 0.43 | 1.49E-02 | 1.00E-01 | 2.65E+00 | 2.72E+00 | 8.85E-01 | no | Gm27921 | upstream_gene_variant | MODIFIER |
| 7 | 54043824 | AX-97798150 | 0 | A | T | 0.43 | 1.49E-02 | 1.00E-01 | 2.65E+00 | 2.72E+00 | 8.85E-01 | no | | intergenic_variant | MODIFIER |
| 7 | 54062396 | AX-97833782 | 0 | A | C | 0.43 | 1.49E-02 | 1.00E-01 | 2.65E+00 | 2.72E+00 | 8.85E-01 | no | | intergenic_variant | MODIFIER |
| 7 | 54092857 | AX-170489663 | 0 | C | G | 0.43 | 1.49E-02 | 1.00E-01 | 2.65E+00 | 2.72E+00 | 8.85E-01 | no | | intergenic_variant | MODIFIER |
| 7 | 54092965 | AX-170237357 | 0 | A | G | 0.43 | 1.49E-02 | 1.00E-01 | 2.65E+00 | 2.72E+00 | 8.85E-01 | no | | intergenic_variant | MODIFIER |
| 7 | 54097067 | AX-170310059 | 0 | A | T | 0.43 | 1.49E-02 | 1.00E-01 | 2.65E+00 | 2.72E+00 | 8.85E-01 | no | | intergenic_variant | MODIFIER |
| 7 | 54099174 | AX-97787702 | 0 | C | T | 0.43 | 1.49E-02 | 1.00E-01 | 2.65E+00 | 2.72E+00 | 8.85E-01 | no | | intergenic_variant | MODIFIER |
| 7 | 54267876 | AX-170446557 | 0 | C | T | 0.43 | 1.49E-02 | 1.00E-01 | 2.65E+00 | 2.72E+00 | 8.85E-01 | no | | intergenic_variant | MODIFIER |
| 7 | 54272906 | AX-97825755 | 0 | C | T | 0.43 | 1.49E-02 | 1.00E-01 | 2.65E+00 | 2.72E+00 | 8.85E-01 | no | | intergenic_variant | MODIFIER |
| 7 | 54319087 | AX-170510836 | 0 | G | A | 0.43 | 1.49E-02 | 1.00E-01 | 2.65E+00 | 2.72E+00 | 8.85E-01 | no | | intergenic_variant | MODIFIER |
| 7 | 54485664 | AX-168410570 | 0 | A | G | 0.43 | 1.49E-02 | 1.00E-01 | 2.65E+00 | 2.72E+00 | 8.85E-01 | no | | intergenic_variant | MODIFIER |
| 7 | 54641112 | AX-170029038 | 0 | A | G | 0.43 | 1.49E-02 | 1.00E-01 | 2.65E+00 | 2.72E+00 | 8.85E-01 | no | | intergenic_variant | MODIFIER |
| 7 | 54641178 | AX-170288100 | 0 | T | C | 0.43 | 1.49E-02 | 1.00E-01 | 2.65E+00 | 2.72E+00 | 8.85E-01 | no | | intergenic_variant | MODIFIER |
| 7 | 54641982 | AX-170221217 | 0 | T | C | 0.43 | 1.49E-02 | 1.00E-01 | 2.65E+00 | 2.72E+00 | 8.85E-01 | no | | intergenic_variant | MODIFIER |
| 7 | 54657072 | AX-97802862 | 0 | A | G | 0.43 | 1.49E-02 | 1.00E-01 | 2.65E+00 | 2.72E+00 | 8.85E-01 | no | | intergenic_variant | MODIFIER |
| 7 | 54746246 | AX-170512243 | 0 | T | C | 0.43 | 1.49E-02 | 1.00E-01 | 2.65E+00 | 2.72E+00 | 8.85E-01 | no | | intergenic_variant | MODIFIER |
| 7 | 54746555 | AX-170511820 | 0 | T | C | 0.43 | 1.49E-02 | 1.00E-01 | 2.65E+00 | 2.72E+00 | 8.85E-01 | no | | intergenic_variant | MODIFIER |
| 7 | 54796811 | AX-97822282 | 0 | T | G | 0.43 | 1.49E-02 | 1.00E-01 | 2.65E+00 | 2.72E+00 | 8.85E-01 | no | | intergenic_variant | MODIFIER |
| 7 | 54830859 | AX-170401523 | 0 | C | T | 0.43 | 1.49E-02 | 1.00E-01 | 2.65E+00 | 2.72E+00 | 8.85E-01 | no | Luzp2 | upstream_gene_variant | MODIFIER |
| 7 | 54831961 | AX-170077389 | 0 | C | T | 0.43 | 1.49E-02 | 1.00E-01 | 2.65E+00 | 2.72E+00 | 8.85E-01 | no | Luzp2 | upstream_gene_variant | MODIFIER |
| 7 | 54838649 | AX-170516299 | 0 | C | T | 0.43 | 1.49E-02 | 1.00E-01 | 2.65E+00 | 2.72E+00 | 8.85E-01 | no | Luzp2 | intron_variant | MODIFIER |
| 7 | 54839076 | AX-170320906 | 0 | T | C | 0.43 | 1.49E-02 | 1.00E-01 | 2.65E+00 | 2.72E+00 | 8.85E-01 | no | Luzp2 | intron_variant | MODIFIER |
| 7 | 54841213 | AX-170299429 | 0 | T | C | 0.43 | 1.49E-02 | 1.00E-01 | 2.65E+00 | 2.72E+00 | 8.85E-01 | no | Luzp2 | intron_variant | MODIFIER |
| 7 | 54841664 | AX-170197445 | 0 | G | C | 0.43 | 1.49E-02 | 1.00E-01 | 2.65E+00 | 2.72E+00 | 8.85E-01 | no | Luzp2 | intron_variant | MODIFIER |
| 7 | 54843140 | AX-170255036 | 0 | A | G | 0.43 | 1.49E-02 | 1.00E-01 | 2.65E+00 | 2.72E+00 | 8.85E-01 | no | Luzp2 | intron_variant | MODIFIER |
| 7 | 54844662 | AX-170314066 | 0 | T | G | 0.43 | 1.49E-02 | 1.00E-01 | 2.65E+00 | 2.72E+00 | 8.85E-01 | no | Luzp2 | intron_variant | MODIFIER |
| 7 | 54847364 | AX-170226390 | 0 | T | G | 0.43 | 1.49E-02 | 1.00E-01 | 2.65E+00 | 2.72E+00 | 8.85E-01 | no | Luzp2 | intron_variant | MODIFIER |
| 7 | 54848497 | AX-170274491 | 0 | G | A | 0.43 | 1.49E-02 | 1.00E-01 | 2.65E+00 | 2.72E+00 | 8.85E-01 | no | Luzp2 | intron_variant | MODIFIER |
| 7 | 54980581 | AX-170462722 | 0 | C | T | 0.43 | 1.49E-02 | 1.00E-01 | 2.65E+00 | 2.72E+00 | 8.85E-01 | no | Luzp2 | intron_variant | MODIFIER |
| 7 | 54985808 | AX-168380789 | 0 | T | C | 0.43 | 1.49E-02 | 1.00E-01 | 2.65E+00 | 2.72E+00 | 8.85E-01 | no | Luzp2 | intron_variant | MODIFIER |
| 7 | 54997038 | AX-170077392 | 0 | A | C | 0.43 | 1.49E-02 | 1.00E-01 | 2.65E+00 | 2.72E+00 | 8.85E-01 | no | Luzp2 | intron_variant | MODIFIER |
| 7 | 55502641 | AX-170402987 | 0 | C | A | 0.43 | 1.49E-02 | 1.00E-01 | 2.65E+00 | 2.72E+00 | 8.85E-01 | no | | intergenic_variant | MODIFIER |
| 7 | 55548989 | AX-170089925 | 0 | T | C | 0.43 | 1.49E-02 | 1.00E-01 | 2.65E+00 | 2.72E+00 | 8.85E-01 | no | | intergenic_variant | MODIFIER |
| 7 | 55555056 | AX-170029051 | 0 | C | T | 0.43 | 1.49E-02 | 1.00E-01 | 2.65E+00 | 2.72E+00 | 8.85E-01 | no | | intergenic_variant | MODIFIER |
| 7 | 55557796 | AX-170186680 | 0 | T | C | 0.43 | 1.49E-02 | 1.00E-01 | 2.65E+00 | 2.72E+00 | 8.85E-01 | no | | intergenic_variant | MODIFIER |
| 7 | 55588992 | AX-97788504 | 0 | G | A | 0.43 | 1.49E-02 | 1.00E-01 | 2.65E+00 | 2.72E+00 | 8.85E-01 | no | | intergenic_variant | MODIFIER |
| 7 | 55797102 | AX-170231797 | 0 | C | T | 0.43 | 1.49E-02 | 1.00E-01 | 2.65E+00 | 2.72E+00 | 8.85E-01 | no | Tubgcp5 | intron_variant | MODIFIER |
| 7 | 55797563 | AX-170112229 | 0 | T | C | 0.43 | 1.49E-02 | 1.00E-01 | 2.65E+00 | 2.72E+00 | 8.85E-01 | no | Tubgcp5 | intron_variant | MODIFIER |
| 7 | 55797893 | AX-97802442 | 0 | T | C | 0.43 | 1.49E-02 | 1.00E-01 | 2.65E+00 | 2.72E+00 | 8.85E-01 | no | Tubgcp5 | intron_variant | MODIFIER |
| 7 | 55871098 | AX-170176520 | 0 | T | C | 0.43 | 1.49E-02 | 1.00E-01 | 2.65E+00 | 2.72E+00 | 8.85E-01 | no | Cyfip1 | intron_variant | MODIFIER |
| 7 | 55875163 | AX-170336539 | 0 | C | T | 0.43 | 1.49E-02 | 1.00E-01 | 2.65E+00 | 2.72E+00 | 8.85E-01 | no | Cyfip1 | intron_variant | MODIFIER |
| 7 | 55899521 | AX-170332271 | 0 | T | C | 0.43 | 1.49E-02 | 1.00E-01 | 2.65E+00 | 2.72E+00 | 8.85E-01 | no | Cyfip1 | intron_variant | MODIFIER |
| 7 | 55904118 | AX-170101904 | 0 | G | A | 0.43 | 1.49E-02 | 1.00E-01 | 2.65E+00 | 2.72E+00 | 8.85E-01 | no | Cyfip1 | intron_variant | MODIFIER |
| 7 | 55913386 | AX-170524245 | 0 | C | A | 0.43 | 1.49E-02 | 1.00E-01 | 2.65E+00 | 2.72E+00 | 8.85E-01 | no | Cyfip1 | intron_variant | MODIFIER |
| 7 | 55917232 | AX-170077412 | 0 | A | G | 0.43 | 1.49E-02 | 1.00E-01 | 2.65E+00 | 2.72E+00 | 8.85E-01 | no | Cyfip1 | intron_variant | MODIFIER |
| 7 | 55926755 | AX-170179946 | 0 | T | C | 0.43 | 1.49E-02 | 1.00E-01 | 2.65E+00 | 2.72E+00 | 8.85E-01 | no | Cyfip1 | intron_variant | MODIFIER |
| 7 | 55938662 | AX-97832563 | 0 | G | T | 0.43 | 1.49E-02 | 1.00E-01 | 2.65E+00 | 2.72E+00 | 8.85E-01 | no | Nipa2 | intron_variant | MODIFIER |
| 7 | 55974060 | AX-170313885 | 0 | C | T | 0.43 | 1.49E-02 | 1.00E-01 | 2.65E+00 | 2.72E+00 | 8.85E-01 | no | Nipa1 | downstream_gene_variant | MODIFIER |
| 7 | 55974164 | AX-170135551 | 0 | G | T | 0.43 | 1.49E-02 | 1.00E-01 | 2.65E+00 | 2.72E+00 | 8.85E-01 | no | Nipa1 | downstream_gene_variant | MODIFIER |
| 7 | 56000966 | AX-170114693 | 0 | G | C | 0.43 | 1.49E-02 | 1.00E-01 | 2.65E+00 | 2.72E+00 | 8.85E-01 | no | Nipa1 | intron_variant | MODIFIER |

| | | | | | | | | | | | | | | | |
|---|---|---|---|---|---|---|---|---|---|---|---|---|---|---|---|
| 7 | 56024446 | AX-170444172 | 0 | A | G | 0.43 | 1.49E-02 | 1.00E-01 | 2.65E+00 | 2.72E+00 | 8.85E-01 | no | Nipa1 | upstream_gene_variant | MODIFIER |
| 7 | 56041722 | AX-170113314 | 0 | C | G | 0.43 | 1.49E-02 | 1.00E-01 | 2.65E+00 | 2.72E+00 | 8.85E-01 | no | Gm34121 | intron_variant,non_coding_transcript_variant | MODIFIER |
| 7 | 56044669 | AX-97848412 | 0 | C | T | 0.43 | 1.49E-02 | 1.00E-01 | 2.65E+00 | 2.72E+00 | 8.85E-01 | no | Gm34121 | intron_variant,non_coding_transcript_variant | MODIFIER |
| 7 | 56049425 | AX-170529258 | 0 | C | T | 0.43 | 1.49E-02 | 1.00E-01 | 2.65E+00 | 2.72E+00 | 8.85E-01 | no | Herc2 | upstream_gene_variant | MODIFIER |
| 7 | 56052720 | AX-170293432 | 0 | C | T | 0.43 | 1.49E-02 | 1.00E-01 | 2.65E+00 | 2.72E+00 | 8.85E-01 | no | Herc2 | intron_variant | MODIFIER |
| 7 | 56068278 | AX-170434397 | 0 | G | C | 0.43 | 1.49E-02 | 1.00E-01 | 2.65E+00 | 2.72E+00 | 8.85E-01 | no | Herc2 | intron_variant | MODIFIER |
| 7 | 56132741 | AX-97816223 | 0 | C | G | 0.43 | 1.49E-02 | 1.00E-01 | 2.65E+00 | 2.72E+00 | 8.85E-01 | no | Herc2 | intron_variant | MODIFIER |
| 7 | 56133958 | AX-170376334 | 0 | C | G | 0.43 | 1.49E-02 | 1.00E-01 | 2.65E+00 | 2.72E+00 | 8.85E-01 | no | Herc2 | intron_variant | MODIFIER |
| 7 | 56146266 | AX-170405255 | 0 | C | T | 0.43 | 1.49E-02 | 1.00E-01 | 2.65E+00 | 2.72E+00 | 8.85E-01 | no | Herc2 | intron_variant | MODIFIER |
| 7 | 56165350 | AX-170029056 | 0 | G | A | 0.43 | 1.49E-02 | 1.00E-01 | 2.65E+00 | 2.72E+00 | 8.85E-01 | no | Herc2 | intron_variant | MODIFIER |
| 7 | 56178598 | AX-170422862 | 0 | A | G | 0.43 | 1.49E-02 | 1.00E-01 | 2.65E+00 | 2.72E+00 | 8.85E-01 | no | Herc2 | intron_variant | MODIFIER |
| 7 | 56178985 | AX-170029054 | 0 | A | C | 0.43 | 1.49E-02 | 1.00E-01 | 2.65E+00 | 2.72E+00 | 8.85E-01 | no | Herc2 | intron_variant | MODIFIER |
| 7 | 56190627 | AX-170340846 | 0 | C | T | 0.43 | 1.49E-02 | 1.00E-01 | 2.65E+00 | 2.72E+00 | 8.85E-01 | no | Herc2 | intron_variant | MODIFIER |
| 7 | 56225997 | AX-170271158 | 0 | T | C | 0.43 | 1.49E-02 | 1.00E-01 | 2.65E+00 | 2.72E+00 | 8.85E-01 | no | Herc2 | intron_variant | MODIFIER |
| 7 | 56384262 | AX-170163165 | 0 | C | A | 0.43 | 1.49E-02 | 1.00E-01 | 2.65E+00 | 2.72E+00 | 8.85E-01 | no | Oca2 | intron_variant | MODIFIER |
| 7 | 56428312 | AX-170185124 | 0 | G | A | 0.43 | 1.49E-02 | 1.00E-01 | 2.65E+00 | 2.72E+00 | 8.85E-01 | no | Oca2 | intron_variant | MODIFIER |
| 7 | 56429368 | AX-97806475 | 0 | C | A | 0.43 | 1.49E-02 | 1.00E-01 | 2.65E+00 | 2.72E+00 | 8.85E-01 | no | Oca2 | missense_variant | MODERATE |
| 7 | 56666104 | AX-97783515 | 0 | A | G | 0.43 | 1.49E-02 | 1.00E-01 | 2.65E+00 | 2.72E+00 | 8.85E-01 | no | | intergenic_variant | MODIFIER |
| 7 | 56731459 | AX-97841517 | 0 | T | G | 0.43 | 1.49E-02 | 1.00E-01 | 2.65E+00 | 2.72E+00 | 8.85E-01 | no | Gabrg3 | intron_variant | MODIFIER |
| 7 | 56768839 | AX-170077428 | 0 | G | A | 0.43 | 1.49E-02 | 1.00E-01 | 2.65E+00 | 2.72E+00 | 8.85E-01 | no | Gabrg3 | intron_variant | MODIFIER |
| 7 | 56774916 | AX-97835586 | 0 | T | C | 0.43 | 1.49E-02 | 1.00E-01 | 2.65E+00 | 2.72E+00 | 8.85E-01 | no | Gabrg3 | intron_variant | MODIFIER |
| 7 | 56779196 | AX-170289203 | 0 | G | A | 0.43 | 1.49E-02 | 1.00E-01 | 2.65E+00 | 2.72E+00 | 8.85E-01 | no | Gabrg3 | intron_variant | MODIFIER |
| 7 | 56795385 | AX-170381893 | 0 | A | G | 0.43 | 1.49E-02 | 1.00E-01 | 2.65E+00 | 2.72E+00 | 8.85E-01 | no | Gabrg3 | intron_variant | MODIFIER |
| 7 | 56895565 | AX-170366303 | 0 | T | C | 0.43 | 1.49E-02 | 1.00E-01 | 2.65E+00 | 2.72E+00 | 8.85E-01 | no | Gabrg3 | intron_variant | MODIFIER |
| 7 | 56915187 | AX-170173276 | 0 | G | C | 0.43 | 1.49E-02 | 1.00E-01 | 2.65E+00 | 2.72E+00 | 8.85E-01 | no | Gabrg3 | intron_variant | MODIFIER |
| 7 | 57124314 | AX-170193104 | 0 | T | C | 0.43 | 1.49E-02 | 1.00E-01 | 2.65E+00 | 2.72E+00 | 8.85E-01 | no | Gabrg3 | intron_variant | MODIFIER |
| 7 | 57126079 | AX-97805881 | 0 | A | G | 0.43 | 1.49E-02 | 1.00E-01 | 2.65E+00 | 2.72E+00 | 8.85E-01 | no | Gabrg3 | intron_variant | MODIFIER |
| 7 | 57132999 | AX-170126855 | 0 | C | A | 0.43 | 1.49E-02 | 1.00E-01 | 2.65E+00 | 2.72E+00 | 8.85E-01 | no | Gabrg3 | intron_variant | MODIFIER |
| 7 | 57138211 | AX-170371630 | 0 | A | T | 0.43 | 1.49E-02 | 1.00E-01 | 2.65E+00 | 2.72E+00 | 8.85E-01 | no | Gabrg3 | intron_variant | MODIFIER |
| 7 | 57211016 | AX-170029078 | 0 | G | A | 0.43 | 1.49E-02 | 1.00E-01 | 2.65E+00 | 2.72E+00 | 8.85E-01 | no | Gabrg3 | intron_variant | MODIFIER |
| 7 | 57219827 | AX-170132808 | 0 | A | G | 0.43 | 1.49E-02 | 1.00E-01 | 2.65E+00 | 2.72E+00 | 8.85E-01 | no | Gabrg3 | intron_variant | MODIFIER |
| 7 | 57244529 | AX-97823750 | 0 | T | C | 0.43 | 1.49E-02 | 1.00E-01 | 2.65E+00 | 2.72E+00 | 8.85E-01 | no | Gabrg3 | intron_variant | MODIFIER |
| 7 | 57244927 | AX-170176986 | 0 | G | A | 0.43 | 1.49E-02 | 1.00E-01 | 2.65E+00 | 2.72E+00 | 8.85E-01 | no | Gabrg3 | intron_variant | MODIFIER |
| 7 | 57246777 | AX-170029075 | 0 | C | T | 0.43 | 1.49E-02 | 1.00E-01 | 2.65E+00 | 2.72E+00 | 8.85E-01 | no | Gabrg3 | intron_variant | MODIFIER |
| 7 | 57246948 | AX-170077436 | 0 | G | A | 0.43 | 1.49E-02 | 1.00E-01 | 2.65E+00 | 2.72E+00 | 8.85E-01 | no | Gabrg3 | intron_variant | MODIFIER |
| 7 | 57247758 | AX-170491553 | 0 | T | C | 0.43 | 1.49E-02 | 1.00E-01 | 2.65E+00 | 2.72E+00 | 8.85E-01 | no | Gabrg3 | intron_variant | MODIFIER |
| 7 | 57269606 | AX-170376055 | 0 | G | C | 0.43 | 1.49E-02 | 1.00E-01 | 2.65E+00 | 2.72E+00 | 8.85E-01 | no | Gabrg3 | intron_variant | MODIFIER |
| 7 | 57300102 | AX-170089929 | 0 | C | T | 0.43 | 1.49E-02 | 1.00E-01 | 2.65E+00 | 2.72E+00 | 8.85E-01 | no | Gabrg3 | intron_variant | MODIFIER |
| 7 | 57304688 | AX-170029077 | 0 | T | C | 0.43 | 1.49E-02 | 1.00E-01 | 2.65E+00 | 2.72E+00 | 8.85E-01 | no | Gabrg3 | intron_variant | MODIFIER |
| 7 | 57306313 | AX-170478336 | 0 | T | C | 0.43 | 1.49E-02 | 1.00E-01 | 2.65E+00 | 2.72E+00 | 8.85E-01 | no | Gabrg3 | intron_variant | MODIFIER |
| 7 | 57345504 | AX-170390921 | 0 | T | C | 0.43 | 1.49E-02 | 1.00E-01 | 2.65E+00 | 2.72E+00 | 8.85E-01 | no | Gabrg3 | intron_variant | MODIFIER |
| 7 | 57347443 | AX-170289932 | 0 | C | T | 0.43 | 1.49E-02 | 1.00E-01 | 2.65E+00 | 2.72E+00 | 8.85E-01 | no | Gabrg3 | intron_variant | MODIFIER |
| 7 | 57389293 | AX-170235900 | 0 | G | C | 0.43 | 1.49E-02 | 1.00E-01 | 2.65E+00 | 2.72E+00 | 8.85E-01 | no | Gabrg3 | upstream_gene_variant | MODIFIER |
| 7 | 57422223 | AX-170218538 | 0 | A | G | 0.43 | 1.49E-02 | 1.00E-01 | 2.65E+00 | 2.72E+00 | 8.85E-01 | no | Gabra5 | intron_variant | MODIFIER |
| 7 | 57455043 | AX-170077442 | 0 | C | G | 0.43 | 1.49E-02 | 1.00E-01 | 2.65E+00 | 2.72E+00 | 8.85E-01 | no | Gabra5 | intron_variant | MODIFIER |
| 7 | 57487870 | AX-170330789 | 0 | A | C | 0.43 | 1.49E-02 | 1.00E-01 | 2.65E+00 | 2.72E+00 | 8.85E-01 | no | Gabra5 | intron_variant | MODIFIER |
| 7 | 57494742 | AX-170463517 | 0 | T | C | 0.43 | 1.49E-02 | 1.00E-01 | 2.65E+00 | 2.72E+00 | 8.85E-01 | no | Gabra5 | intron_variant | MODIFIER |
| 7 | 57503755 | AX-170109829 | 0 | C | A | 0.43 | 1.49E-02 | 1.00E-01 | 2.65E+00 | 2.72E+00 | 8.85E-01 | no | Gabra5 | intron_variant | MODIFIER |
| 7 | 57504163 | AX-97798023 | 0 | A | C | 0.43 | 1.49E-02 | 1.00E-01 | 2.65E+00 | 2.72E+00 | 8.85E-01 | no | Gabra5 | intron_variant | MODIFIER |
| 7 | 57507382 | AX-170189018 | 0 | G | A | 0.43 | 1.49E-02 | 1.00E-01 | 2.65E+00 | 2.72E+00 | 8.85E-01 | no | Gabra5 | intron_variant | MODIFIER |
| 7 | 57519382 | AX-170348138 | 0 | C | A | 0.43 | 1.49E-02 | 1.00E-01 | 2.65E+00 | 2.72E+00 | 8.85E-01 | no | Gabrb3 | intron_variant | MODIFIER |
| 7 | 57530107 | AX-170318269 | 0 | G | A | 0.43 | 1.49E-02 | 1.00E-01 | 2.65E+00 | 2.72E+00 | 8.85E-01 | no | Gabrb3 | intron_variant | MODIFIER |
| 7 | 57530215 | AX-170156277 | 0 | C | G | 0.43 | 1.49E-02 | 1.00E-01 | 2.65E+00 | 2.72E+00 | 8.85E-01 | no | Gabrb3 | intron_variant | MODIFIER |
| 7 | 57538108 | AX-170464451 | 0 | A | G | 0.43 | 1.49E-02 | 1.00E-01 | 2.65E+00 | 2.72E+00 | 8.85E-01 | no | Gabrb3 | intron_variant | MODIFIER |

| | | | | | | | | | | | | | | | |
|---|---|---|---|---|---|---|---|---|---|---|---|---|---|---|---|
| 7 | 57539520 | AX-170493115 | 0 | C | A | 0.43 | 1.49E-02 | 1.00E-01 | 2.65E+00 | 2.72E+00 | 8.85E-01 | no | Gabrb3 | intron_variant | MODIFIER |
| 7 | 57539673 | AX-170392506 | 0 | C | T | 0.43 | 1.49E-02 | 1.00E-01 | 2.65E+00 | 2.72E+00 | 8.85E-01 | no | Gabrb3 | intron_variant | MODIFIER |
| 7 | 57550331 | AX-170350843 | 0 | C | T | 0.43 | 1.49E-02 | 1.00E-01 | 2.65E+00 | 2.72E+00 | 8.85E-01 | no | Gabrb3 | intron_variant | MODIFIER |
| 7 | 57582362 | AX-170320960 | 0 | C | T | 0.43 | 1.49E-02 | 1.00E-01 | 2.65E+00 | 2.72E+00 | 8.85E-01 | no | Gabrb3 | intron_variant | MODIFIER |
| 7 | 57588693 | AX-97797675 | 0 | A | C | 0.43 | 1.49E-02 | 1.00E-01 | 2.65E+00 | 2.72E+00 | 8.85E-01 | no | Gabrb3 | upstream_gene_variant | MODIFIER |
| 7 | 57615435 | AX-170416020 | 0 | C | T | 0.43 | 1.49E-02 | 1.00E-01 | 2.65E+00 | 2.72E+00 | 8.85E-01 | no | Gabrb3 | intron_variant | MODIFIER |
| 7 | 57621809 | AX-170424078 | 0 | G | A | 0.43 | 1.49E-02 | 1.00E-01 | 2.65E+00 | 2.72E+00 | 8.85E-01 | no | Gabrb3 | intron_variant | MODIFIER |
| 7 | 57629968 | AX-170355910 | 0 | T | C | 0.43 | 1.49E-02 | 1.00E-01 | 2.65E+00 | 2.72E+00 | 8.85E-01 | no | Gabrb3 | intron_variant | MODIFIER |
| 7 | 57630241 | AX-170275289 | 0 | C | G | 0.43 | 1.49E-02 | 1.00E-01 | 2.65E+00 | 2.72E+00 | 8.85E-01 | no | Gabrb3 | intron_variant | MODIFIER |
| 7 | 57630427 | AX-170308804 | 0 | A | G | 0.43 | 1.49E-02 | 1.00E-01 | 2.65E+00 | 2.72E+00 | 8.85E-01 | no | Gabrb3 | intron_variant | MODIFIER |
| 7 | 57630574 | AX-170317056 | 0 | A | C | 0.43 | 1.49E-02 | 1.00E-01 | 2.65E+00 | 2.72E+00 | 8.85E-01 | no | Gabrb3 | intron_variant | MODIFIER |
| 7 | 57633824 | AX-170199689 | 0 | C | T | 0.43 | 1.49E-02 | 1.00E-01 | 2.65E+00 | 2.72E+00 | 8.85E-01 | no | Gabrb3 | intron_variant | MODIFIER |
| 7 | 57634909 | AX-170508211 | 0 | G | A | 0.43 | 1.49E-02 | 1.00E-01 | 2.65E+00 | 2.72E+00 | 8.85E-01 | no | Gabrb3 | intron_variant | MODIFIER |
| 7 | 57640587 | AX-97839222 | 0 | A | C | 0.43 | 1.49E-02 | 1.00E-01 | 2.65E+00 | 2.72E+00 | 8.85E-01 | no | Gabrb3 | intron_variant | MODIFIER |
| 7 | 57671288 | AX-170398023 | 0 | T | C | 0.43 | 1.49E-02 | 1.00E-01 | 2.65E+00 | 2.72E+00 | 8.85E-01 | no | Gabrb3 | intron_variant | MODIFIER |
| 7 | 57672618 | AX-170424460 | 0 | C | T | 0.43 | 1.49E-02 | 1.00E-01 | 2.65E+00 | 2.72E+00 | 8.85E-01 | no | Gabrb3 | intron_variant | MODIFIER |
| 7 | 57672849 | AX-170531671 | 0 | T | C | 0.43 | 1.49E-02 | 1.00E-01 | 2.65E+00 | 2.72E+00 | 8.85E-01 | no | Gabrb3 | intron_variant | MODIFIER |
| 7 | 57674035 | AX-97804548 | 0 | T | G | 0.43 | 1.49E-02 | 1.00E-01 | 2.65E+00 | 2.72E+00 | 8.85E-01 | no | Gabrb3 | intron_variant | MODIFIER |
| 7 | 57679794 | AX-170381436 | 0 | C | T | 0.43 | 1.49E-02 | 1.00E-01 | 2.65E+00 | 2.72E+00 | 8.85E-01 | no | Gabrb3 | intron_variant | MODIFIER |
| 12 | 68906558 | AX-97811104 | 0 | T | C | 0.061 | 4.24E-02 | 3.01E-01 | 2.74E+00 | 2.76E+00 | 8.86E-01 | no | Gm47515 | intron_variant,non_coding_transcript_variant | MODIFIER |
| 10 | 114870084 | AX-170445410 | 0 | G | A | 0.368 | -1.10E-02 | 7.69E-02 | 2.65E+00 | 2.74E+00 | 8.86E-01 | no | Gm31182 | upstream_gene_variant | MODIFIER |
| 1 | 65682697 | AX-170271550 | 1 | G | T | 0.152 | 2.80E-02 | 1.87E-01 | 2.61E+00 | 2.72E+00 | 8.86E-01 | no | | intergenic_variant | MODIFIER |
| 1 | 20267903 | AX-170421606 | 0 | T | C | 0.404 | 1.26E-02 | 8.13E-02 | 2.63E+00 | 2.70E+00 | 8.86E-01 | no | Pkhd1 | intron_variant | MODIFIER |
| 8 | 93668454 | AX-97856922 | 0 | A | G | 0.211 | 1.66E-02 | 1.19E-01 | 2.69E+00 | 2.75E+00 | 8.86E-01 | no | Gm26843 | intron_variant,non_coding_transcript_variant | MODIFIER |
| 8 | 18512935 | AX-170086035 | 0 | A | G | 0.167 | 1.52E-02 | 1.08E-01 | 2.64E+00 | 2.74E+00 | 8.86E-01 | no | Gm24469 | upstream_gene_variant | MODIFIER |
| 6 | 110135286 | AX-170488814 | 0 | G | C | 0.061 | -3.18E-02 | 2.33E-01 | 2.65E+00 | 2.75E+00 | 8.86E-01 | no | | intergenic_variant | MODIFIER |
| 18 | 30360147 | AX-170324649 | 0 | G | T | 0.053 | 3.27E-02 | 2.24E-01 | 2.60E+00 | 2.73E+00 | 8.86E-01 | no | | intergenic_variant | MODIFIER |
| 18 | 31559157 | AX-170122313 | 0 | G | T | 0.053 | 3.27E-02 | 2.24E-01 | 2.60E+00 | 2.73E+00 | 8.86E-01 | no | Slc25a46 | intron_variant,NMD_transcript_variant | MODIFIER |
| 12 | 69389979 | AX-170413366 | 0 | G | A | 0.061 | -4.31E-02 | 3.01E-01 | 2.70E+00 | 2.73E+00 | 8.86E-01 | no | | regulatory_region_variant | MODIFIER |
| 9 | 104405699 | AX-170083335 | 0 | C | T | 0.123 | -1.83E-02 | 1.18E-01 | 2.63E+00 | 2.70E+00 | 8.86E-01 | no | | intergenic_variant | MODIFIER |
| 9 | 105187708 | AX-170421050 | 0 | G | A | 0.123 | -1.83E-02 | 1.18E-01 | 2.63E+00 | 2.70E+00 | 8.86E-01 | no | Nek11 | intron_variant | MODIFIER |
| 9 | 105195034 | AX-170521406 | 0 | C | A | 0.123 | -1.83E-02 | 1.18E-01 | 2.63E+00 | 2.70E+00 | 8.86E-01 | no | Nek11 | intron_variant | MODIFIER |
| 9 | 105200329 | AX-170098168 | 0 | G | A | 0.123 | -1.83E-02 | 1.18E-01 | 2.63E+00 | 2.70E+00 | 8.86E-01 | no | Nek11 | intron_variant | MODIFIER |
| 9 | 107912841 | AX-170306450 | 0 | G | A | 0.123 | -1.83E-02 | 1.18E-01 | 2.63E+00 | 2.70E+00 | 8.86E-01 | no | Mst1r | intron_variant | MODIFIER |
| 2 | 118767821 | AX-170471948 | 1 | A | T | 0.125 | 2.52E-02 | 1.77E-01 | 2.65E+00 | 2.74E+00 | 8.86E-01 | no | Phgr1 | upstream_gene_variant | MODIFIER |
| 9 | 74671335 | AX-170247357 | 2 | T | A | 0.191 | 1.72E-02 | 1.26E-01 | 2.70E+00 | 2.77E+00 | 8.86E-01 | no | | regulatory_region_variant | MODIFIER |
| 4 | 80683727 | AX-170298210 | 0 | A | G | 0.237 | 1.62E-02 | 1.04E-01 | 2.62E+00 | 2.70E+00 | 8.86E-01 | no | | intergenic_variant | MODIFIER |
| 13 | 48119885 | AX-170207645 | 0 | T | C | 0.491 | -8.81E-03 | 6.68E-02 | 2.66E+00 | 2.76E+00 | 8.86E-01 | no | A330033J07Rik | intron_variant,non_coding_transcript_variant | MODIFIER |
| 13 | 48380363 | AX-97807480 | 0 | A | G | 0.491 | -8.81E-03 | 6.68E-02 | 2.66E+00 | 2.76E+00 | 8.86E-01 | no | Gm48648 | intron_variant,non_coding_transcript_variant | MODIFIER |
| 13 | 48614102 | AX-97798236 | 0 | A | G | 0.491 | -8.81E-03 | 6.68E-02 | 2.66E+00 | 2.76E+00 | 8.86E-01 | no | Ptpdc1 | intron_variant | MODIFIER |
| 13 | 48736441 | AX-170303728 | 0 | C | T | 0.491 | -8.81E-03 | 6.68E-02 | 2.66E+00 | 2.76E+00 | 8.86E-01 | no | | intergenic_variant | MODIFIER |
| 13 | 48736486 | AX-170380507 | 0 | T | C | 0.491 | -8.81E-03 | 6.68E-02 | 2.66E+00 | 2.76E+00 | 8.86E-01 | no | | intergenic_variant | MODIFIER |
| 13 | 48736770 | AX-170097843 | 0 | G | A | 0.491 | -8.81E-03 | 6.68E-02 | 2.66E+00 | 2.76E+00 | 8.86E-01 | no | | intergenic_variant | MODIFIER |
| 13 | 48737140 | AX-170254317 | 0 | A | G | 0.491 | -8.81E-03 | 6.68E-02 | 2.66E+00 | 2.76E+00 | 8.86E-01 | no | | intergenic_variant | MODIFIER |
| 13 | 48760877 | AX-170274172 | 0 | C | T | 0.491 | -8.81E-03 | 6.68E-02 | 2.66E+00 | 2.76E+00 | 8.86E-01 | no | Rpl17-ps6 | downstream_gene_variant | MODIFIER |
| 13 | 48761518 | AX-170317556 | 0 | C | A | 0.491 | -8.81E-03 | 6.68E-02 | 2.66E+00 | 2.76E+00 | 8.86E-01 | no | Rpl17-ps6 | downstream_gene_variant | MODIFIER |
| 13 | 48762951 | AX-170051220 | 0 | G | A | 0.491 | -8.81E-03 | 6.68E-02 | 2.66E+00 | 2.76E+00 | 8.86E-01 | no | Rpl17-ps6 | downstream_gene_variant | MODIFIER |
| 13 | 48815846 | AX-97784247 | 0 | G | A | 0.491 | -8.81E-03 | 6.68E-02 | 2.66E+00 | 2.76E+00 | 8.86E-01 | no | Phf2 | intron_variant | MODIFIER |
| 13 | 48835331 | AX-170302869 | 0 | C | T | 0.491 | -8.81E-03 | 6.68E-02 | 2.66E+00 | 2.76E+00 | 8.86E-01 | no | Phf2 | intron_variant | MODIFIER |
| 13 | 48838575 | AX-97793524 | 0 | T | C | 0.491 | -8.81E-03 | 6.68E-02 | 2.66E+00 | 2.76E+00 | 8.86E-01 | no | Phf2 | intron_variant | MODIFIER |
| 13 | 48872966 | AX-170104306 | 0 | G | A | 0.491 | -8.81E-03 | 6.68E-02 | 2.66E+00 | 2.76E+00 | 8.86E-01 | no | Phf2 | upstream_gene_variant | MODIFIER |
| 13 | 48882005 | AX-97803261 | 0 | A | G | 0.491 | -8.81E-03 | 6.68E-02 | 2.66E+00 | 2.76E+00 | 8.86E-01 | no | Fam120a | intron_variant | MODIFIER |
| 13 | 48882158 | AX-170348794 | 0 | A | G | 0.491 | -8.81E-03 | 6.68E-02 | 2.66E+00 | 2.76E+00 | 8.86E-01 | no | Fam120a | intron_variant | MODIFIER |
| 13 | 48895247 | AX-97793351 | 0 | T | G | 0.491 | -8.81E-03 | 6.68E-02 | 2.66E+00 | 2.76E+00 | 8.86E-01 | no | Fam120a | intron_variant | MODIFIER |

| | | | | | | | | | | | | | | | |
|---|---|---|---|---|---|---|---|---|---|---|---|---|---|---|---|
| 13 | 48978132 | AX-97823454 | 0 | C | T | 0.491 | -8.81E-03 | 6.68E-02 | 2.66E+00 | 2.76E+00 | 8.86E-01 | no | | regulatory_region_variant | MODIFIER |
| 13 | 48999130 | AX-170409859 | 0 | C | T | 0.491 | -8.81E-03 | 6.68E-02 | 2.66E+00 | 2.76E+00 | 8.86E-01 | no | | regulatory_region_variant | MODIFIER |
| 13 | 49005879 | AX-170345776 | 0 | T | A | 0.491 | -8.81E-03 | 6.68E-02 | 2.66E+00 | 2.76E+00 | 8.86E-01 | no | | intergenic_variant | MODIFIER |
| 13 | 49006683 | AX-170535001 | 0 | A | G | 0.491 | -8.81E-03 | 6.68E-02 | 2.66E+00 | 2.76E+00 | 8.86E-01 | no | | intergenic_variant | MODIFIER |
| 13 | 49045585 | AX-97784348 | 0 | A | C | 0.491 | -8.81E-03 | 6.68E-02 | 2.66E+00 | 2.76E+00 | 8.86E-01 | no | Wnk2 | intron_variant | MODIFIER |
| 13 | 49060328 | AX-170051225 | 0 | T | C | 0.491 | -8.81E-03 | 6.68E-02 | 2.66E+00 | 2.76E+00 | 8.86E-01 | no | Wnk2 | intron_variant | MODIFIER |
| 13 | 49066971 | AX-97788380 | 0 | G | A | 0.491 | -8.81E-03 | 6.68E-02 | 2.66E+00 | 2.76E+00 | 8.86E-01 | no | Wnk2 | intron_variant | MODIFIER |
| 13 | 49098707 | AX-97816861 | 0 | G | T | 0.491 | -8.81E-03 | 6.68E-02 | 2.66E+00 | 2.76E+00 | 8.86E-01 | no | Wnk2 | intron_variant | MODIFIER |
| 13 | 49169060 | AX-170403093 | 0 | A | T | 0.491 | -8.81E-03 | 6.68E-02 | 2.66E+00 | 2.76E+00 | 8.86E-01 | no | | intergenic_variant | MODIFIER |
| 13 | 49171584 | AX-97782765 | 0 | C | T | 0.491 | -8.81E-03 | 6.68E-02 | 2.66E+00 | 2.76E+00 | 8.86E-01 | no | | intergenic_variant | MODIFIER |
| 13 | 49294519 | AX-97846260 | 0 | G | A | 0.491 | -8.81E-03 | 6.68E-02 | 2.66E+00 | 2.76E+00 | 8.86E-01 | no | Fgd3 | intron_variant | MODIFIER |
| 13 | 49405613 | AX-97835343 | 0 | T | C | 0.491 | -8.81E-03 | 6.68E-02 | 2.66E+00 | 2.76E+00 | 8.86E-01 | no | | intergenic_variant | MODIFIER |
| 13 | 49483206 | AX-97823458 | 0 | T | C | 0.491 | -8.81E-03 | 6.68E-02 | 2.66E+00 | 2.76E+00 | 8.86E-01 | no | Cenpp | intron_variant | MODIFIER |
| 13 | 49725349 | AX-97784257 | 0 | C | T | 0.491 | -8.81E-03 | 6.68E-02 | 2.66E+00 | 2.76E+00 | 8.86E-01 | no | Iars | intron_variant | MODIFIER |
| 13 | 51231302 | AX-97837959 | 0 | A | G | 0.491 | -8.81E-03 | 6.68E-02 | 2.66E+00 | 2.76E+00 | 8.86E-01 | no | | intergenic_variant | MODIFIER |
| 1 | 28997242 | AX-170091444 | 1 | C | T | 0.42 | 1.13E-02 | 7.96E-02 | 2.66E+00 | 2.74E+00 | 8.86E-01 | no | | intergenic_variant | MODIFIER |
| 1 | 52632969 | AX-170394670 | 0 | C | T | 0.114 | 3.15E-02 | 2.13E-01 | 2.65E+00 | 2.72E+00 | 8.86E-01 | no | Nemp2 | intron_variant | MODIFIER |
| 1 | 59760063 | AX-170389145 | 0 | C | T | 0.114 | 3.15E-02 | 2.13E-01 | 2.65E+00 | 2.72E+00 | 8.86E-01 | no | Bmpr2 | upstream_gene_variant | MODIFIER |
| 1 | 59953335 | AX-170535659 | 0 | A | G | 0.114 | 3.15E-02 | 2.13E-01 | 2.65E+00 | 2.72E+00 | 8.86E-01 | no | Fam117b | intron_variant | MODIFIER |
| 1 | 60176233 | AX-170298340 | 0 | T | A | 0.114 | 3.15E-02 | 2.13E-01 | 2.65E+00 | 2.72E+00 | 8.86E-01 | no | Nbeal1 | upstream_gene_variant | MODIFIER |
| 1 | 60479396 | AX-170000614 | 0 | C | A | 0.114 | 3.15E-02 | 2.13E-01 | 2.65E+00 | 2.72E+00 | 8.86E-01 | no | Raph1 | downstream_gene_variant | MODIFIER |
| 1 | 60724174 | AX-170291010 | 0 | C | T | 0.114 | 3.15E-02 | 2.13E-01 | 2.65E+00 | 2.72E+00 | 8.86E-01 | no | Cd28 | intron_variant,non_coding_transcript_variant | MODIFIER |
| 6 | 104019266 | AX-170246807 | 0 | G | A | 0.053 | 4.21E-02 | 2.96E-01 | 2.67E+00 | 2.73E+00 | 8.86E-01 | no | | intergenic_variant | MODIFIER |
| 9 | 38002508 | AX-170273371 | 1 | G | A | 0.509 | -1.06E-02 | 7.75E-02 | 2.66E+00 | 2.75E+00 | 8.86E-01 | no | Olfr882-ps1 | upstream_gene_variant | MODIFIER |
| 7 | 81926416 | AX-170412509 | 1 | C | T | 0.116 | 3.04E-02 | 2.13E-01 | 2.66E+00 | 2.73E+00 | 8.87E-01 | no | Hdgfl3 | intron_variant | MODIFIER |
| 5 | 128473031 | AX-97842263 | 0 | T | C | 0.158 | -1.51E-02 | 8.28E-02 | 2.55E+00 | 2.67E+00 | 8.87E-01 | no | | intergenic_variant | MODIFIER |
| 1 | 74154686 | AX-170136317 | 0 | G | A | 0.123 | -2.87E-02 | 2.07E-01 | 2.69E+00 | 2.75E+00 | 8.87E-01 | no | Cxcr2 | intron_variant | MODIFIER |
| 12 | 112414925 | AX-170355292 | 1 | T | A | 0.116 | -2.41E-02 | 1.81E-01 | 2.67E+00 | 2.76E+00 | 8.87E-01 | no | A730018C14Rik | non_coding_transcript_exon_variant | MODIFIER |
| 12 | 113223108 | AX-170484675 | 1 | A | G | 0.116 | -2.41E-02 | 1.81E-01 | 2.67E+00 | 2.76E+00 | 8.87E-01 | no | | regulatory_region_variant | MODIFIER |
| 18 | 43658364 | AX-170326108 | 1 | A | G | 0.062 | 4.19E-02 | 3.01E-01 | 2.73E+00 | 2.76E+00 | 8.87E-01 | no | Jakmip2 | intron_variant | MODIFIER |
| 9 | 88073588 | AX-170099008 | 0 | T | A | 0.14 | 1.61E-02 | 1.23E-01 | 2.69E+00 | 2.77E+00 | 8.87E-01 | no | | regulatory_region_variant | MODIFIER |
| 5 | 125197762 | AX-170421518 | 1 | T | C | 0.455 | -1.60E-02 | 1.16E-01 | 2.73E+00 | 2.76E+00 | 8.87E-01 | no | | intergenic_variant | MODIFIER |
| 10 | 121201392 | AX-170173423 | 1 | C | T | 0.348 | 1.02E-02 | 7.35E-02 | 2.65E+00 | 2.74E+00 | 8.87E-01 | no | 4930432O09Rik | intron_variant,non_coding_transcript_variant | MODIFIER |
| 2 | 146822809 | AX-170376079 | 1 | G | T | 0.143 | 3.13E-02 | 2.20E-01 | 2.69E+00 | 2.72E+00 | 8.87E-01 | no | | intergenic_variant | MODIFIER |
| 2 | 156338343 | AX-170484715 | 1 | T | A | 0.143 | 3.13E-02 | 2.20E-01 | 2.69E+00 | 2.72E+00 | 8.87E-01 | no | Cnbd2 | intron_variant | MODIFIER |
| 12 | 68750640 | AX-97795617 | 1 | C | T | 0.062 | 4.18E-02 | 3.01E-01 | 2.74E+00 | 2.76E+00 | 8.87E-01 | no | | intergenic_variant | MODIFIER |
| 12 | 71139792 | AX-170007017 | 1 | G | A | 0.062 | 4.18E-02 | 3.01E-01 | 2.74E+00 | 2.76E+00 | 8.87E-01 | no | Timm9 | upstream_gene_variant | MODIFIER |
| 7 | 135303444 | AX-170030039 | 0 | A | G | 0.263 | 9.82E-03 | 7.78E-02 | 2.65E+00 | 2.77E+00 | 8.87E-01 | no | | intergenic_variant | MODIFIER |
| 10 | 109726318 | AX-170444241 | 0 | G | A | 0.351 | -1.19E-02 | 8.36E-02 | 2.65E+00 | 2.73E+00 | 8.87E-01 | no | Nav3 | intron_variant | MODIFIER |
| 5 | 106818866 | AX-170147645 | 1 | C | G | 0.429 | 1.59E-02 | 1.05E-01 | 2.64E+00 | 2.71E+00 | 8.87E-01 | no | | intergenic_variant | MODIFIER |
| 16 | 45103305 | AX-170012945 | 0 | A | G | 0.482 | -1.24E-02 | 8.17E-02 | 2.64E+00 | 2.70E+00 | 8.87E-01 | no | Ccdc80 | intron_variant | MODIFIER |
| 1 | 52062301 | AX-170465354 | 0 | T | A | 0.132 | -2.85E-02 | 1.91E-01 | 2.63E+00 | 2.72E+00 | 8.88E-01 | no | Stat4 | intron_variant | MODIFIER |
| 1 | 162141168 | AX-170527921 | 0 | A | G | 0.219 | -1.96E-02 | 1.44E-01 | 2.73E+00 | 2.77E+00 | 8.88E-01 | no | Dnm3 | intron_variant | MODIFIER |
| 15 | 92302862 | AX-170407474 | 0 | G | T | 0.14 | 2.48E-02 | 1.69E-01 | 2.63E+00 | 2.72E+00 | 8.88E-01 | no | Cntn1 | intron_variant | MODIFIER |
| 7 | 10516759 | AX-97795853 | 1 | G | A | 0.17 | -1.36E-02 | 8.12E-02 | 2.56E+00 | 2.69E+00 | 8.88E-01 | no | Vmn1r68 | downstream_gene_variant | MODIFIER |
| 8 | 101780413 | AX-170118388 | 1 | G | C | 0.214 | 1.60E-02 | 1.16E-01 | 2.68E+00 | 2.74E+00 | 8.88E-01 | no | | intergenic_variant | MODIFIER |
| 10 | 14530021 | AX-97836658 | 0 | A | G | 0.211 | -1.59E-02 | 1.07E-01 | 2.64E+00 | 2.71E+00 | 8.88E-01 | no | Adgrg6 | intron_variant | MODIFIER |
| 10 | 15091507 | AX-170002797 | 0 | T | G | 0.211 | -1.59E-02 | 1.07E-01 | 2.64E+00 | 2.71E+00 | 8.88E-01 | no | | intergenic_variant | MODIFIER |
| 8 | 39960298 | AX-170403415 | 0 | G | T | 0.158 | 1.88E-02 | 1.39E-01 | 2.70E+00 | 2.76E+00 | 8.88E-01 | no | | intergenic_variant | MODIFIER |
| 8 | 40070689 | AX-170136366 | 0 | G | A | 0.158 | 1.88E-02 | 1.39E-01 | 2.70E+00 | 2.76E+00 | 8.88E-01 | no | | intergenic_variant | MODIFIER |
| 15 | 103293055 | AX-170093362 | 1 | T | C | 0.116 | 2.18E-02 | 1.53E-01 | 2.66E+00 | 2.73E+00 | 8.88E-01 | no | Copz1 | intron_variant | MODIFIER |
| 5 | 27906583 | AX-97807400 | 0 | G | A | 0.333 | -1.40E-02 | 1.05E-01 | 2.69E+00 | 2.76E+00 | 8.88E-01 | no | | intergenic_variant | MODIFIER |
| 7 | 111947630 | AX-170330115 | 0 | A | T | 0.465 | -1.25E-02 | 9.23E-02 | 2.69E+00 | 2.75E+00 | 8.88E-01 | no | Gm23345 | upstream_gene_variant | MODIFIER |
| 12 | 29813130 | AX-170435759 | 0 | T | C | 0.044 | -2.95E-02 | 2.09E-01 | 2.64E+00 | 2.73E+00 | 8.88E-01 | no | Myt1l | intron_variant | MODIFIER |

| | | | | | | | | | | | | | | | |
|---|---|---|---|---|---|---|---|---|---|---|---|---|---|---|---|
| 4 | 143003485 | AX-170244461 | 0 | A | T | 0.114 | 2.40E-02 | 1.68E-01 | 2.63E+00 | 2.73E+00 | 8.88E-01 | no | 6330411D24Rik | intron_variant,non_coding_transcript_variant | MODIFIER |
| 7 | 60556916 | AX-170495540 | 1 | C | A | 0.42 | -1.29E-02 | 9.45E-02 | 2.66E+00 | 2.74E+00 | 8.88E-01 | no | | intergenic_variant | MODIFIER |
| 10 | 114398052 | AX-170455171 | 0 | A | G | 0.368 | -1.14E-02 | 8.08E-02 | 2.66E+00 | 2.73E+00 | 8.89E-01 | no | Trhde | downstream_gene_variant | MODIFIER |
| 10 | 114449061 | AX-170242285 | 0 | G | C | 0.368 | -1.14E-02 | 8.08E-02 | 2.66E+00 | 2.73E+00 | 8.89E-01 | no | Trhde | intron_variant | MODIFIER |
| 10 | 114491277 | AX-170478513 | 0 | T | C | 0.368 | -1.14E-02 | 8.08E-02 | 2.66E+00 | 2.73E+00 | 8.89E-01 | no | Trhde | intron_variant | MODIFIER |
| 10 | 114502216 | AX-97852586 | 0 | G | A | 0.368 | -1.14E-02 | 8.08E-02 | 2.66E+00 | 2.73E+00 | 8.89E-01 | no | Trhde | intron_variant | MODIFIER |
| 10 | 114504738 | AX-170210050 | 0 | T | C | 0.368 | -1.14E-02 | 8.08E-02 | 2.66E+00 | 2.73E+00 | 8.89E-01 | no | Trhde | intron_variant | MODIFIER |
| 10 | 114519399 | AX-170324524 | 0 | C | G | 0.368 | -1.14E-02 | 8.08E-02 | 2.66E+00 | 2.73E+00 | 8.89E-01 | no | Trhde | intron_variant | MODIFIER |
| 10 | 114523094 | AX-170308969 | 0 | G | A | 0.368 | -1.14E-02 | 8.08E-02 | 2.66E+00 | 2.73E+00 | 8.89E-01 | no | Trhde | intron_variant | MODIFIER |
| 10 | 114572531 | AX-97819423 | 0 | C | G | 0.368 | -1.14E-02 | 8.08E-02 | 2.66E+00 | 2.73E+00 | 8.89E-01 | no | Trhde | intron_variant | MODIFIER |
| 20 | 104497371 | AX-97806611 | 0 | T | C | 0.333 | 7.66E-03 | 5.50E-02 | 2.62E+00 | 2.74E+00 | 8.89E-01 | no | Uprt | intron_variant | MODIFIER |
| 20 | 104611487 | AX-97853581 | 0 | A | G | 0.333 | 7.66E-03 | 5.50E-02 | 2.62E+00 | 2.74E+00 | 8.89E-01 | no | Zdhhc15 | intron_variant | MODIFIER |
| 10 | 5484572 | AX-170320230 | 2 | A | G | 0.209 | 1.29E-02 | 9.63E-02 | 2.64E+00 | 2.75E+00 | 8.89E-01 | no | Syne1 | intron_variant | MODIFIER |
| 1 | 44262980 | AX-97825307 | 0 | T | C | 0.105 | 2.45E-02 | 1.75E-01 | 2.62E+00 | 2.74E+00 | 8.89E-01 | no | | intergenic_variant | MODIFIER |
| 8 | 78165019 | AX-170499803 | 1 | G | A | 0.205 | -1.80E-02 | 1.30E-01 | 2.67E+00 | 2.74E+00 | 8.89E-01 | no | | intergenic_variant | MODIFIER |
| 15 | 98738191 | AX-97851311 | 1 | G | A | 0.116 | 2.12E-02 | 1.50E-01 | 2.65E+00 | 2.73E+00 | 8.89E-01 | no | Arf3 | 3_prime_UTR_variant | MODIFIER |
| 5 | 89931445 | AX-170284149 | 1 | A | T | 0.42 | -1.38E-02 | 1.02E-01 | 2.66E+00 | 2.74E+00 | 8.89E-01 | no | | intergenic_variant | MODIFIER |
| 1 | 71246656 | AX-170282705 | 0 | A | G | 0.14 | -3.24E-02 | 2.39E-01 | 2.70E+00 | 2.75E+00 | 8.89E-01 | no | Abca12 | intron_variant | MODIFIER |
| 7 | 116795802 | AX-170255680 | 0 | C | A | 0.412 | 1.30E-02 | 9.21E-02 | 2.67E+00 | 2.73E+00 | 8.89E-01 | no | | intergenic_variant | MODIFIER |
| 1 | 47985350 | AX-97820234 | 1 | T | C | 0.107 | 3.25E-02 | 2.35E-01 | 2.69E+00 | 2.74E+00 | 8.89E-01 | no | | intergenic_variant | MODIFIER |
| 1 | 49152938 | AX-170460714 | 1 | G | T | 0.107 | 3.25E-02 | 2.35E-01 | 2.69E+00 | 2.74E+00 | 8.89E-01 | no | | intergenic_variant | MODIFIER |
| 12 | 114932873 | AX-170442907 | 1 | G | A | 0.125 | -2.15E-02 | 1.60E-01 | 2.66E+00 | 2.75E+00 | 8.89E-01 | no | Ighv1-42 | downstream_gene_variant | MODIFIER |
| 1 | 70852613 | AX-170280059 | 1 | A | G | 0.134 | 2.67E-02 | 1.96E-01 | 2.65E+00 | 2.74E+00 | 8.89E-01 | no | Vwc2l | intron_variant | MODIFIER |
| 15 | 94857780 | AX-170308440 | 1 | A | T | 0.152 | 2.58E-02 | 1.79E-01 | 2.65E+00 | 2.72E+00 | 8.89E-01 | no | Tmem117 | intron_variant | MODIFIER |
| 15 | 98565481 | AX-170326345 | 0 | A | G | 0.114 | 2.11E-02 | 1.50E-01 | 2.66E+00 | 2.73E+00 | 8.89E-01 | no | Ccnt1 | intron_variant | MODIFIER |
| 15 | 98581982 | AX-170468300 | 0 | T | G | 0.114 | 2.11E-02 | 1.50E-01 | 2.66E+00 | 2.73E+00 | 8.89E-01 | no | 4930415O20Rik | intron_variant | MODIFIER |
| 15 | 98582094 | AX-170411599 | 0 | G | A | 0.114 | 2.11E-02 | 1.50E-01 | 2.66E+00 | 2.73E+00 | 8.89E-01 | no | 4930415O20Rik | intron_variant | MODIFIER |
| 15 | 98582352 | AX-170056448 | 0 | C | T | 0.114 | 2.11E-02 | 1.50E-01 | 2.66E+00 | 2.73E+00 | 8.89E-01 | no | 4930415O20Rik | intron_variant | MODIFIER |
| 15 | 98582590 | AX-170056447 | 0 | A | G | 0.114 | 2.11E-02 | 1.50E-01 | 2.66E+00 | 2.73E+00 | 8.89E-01 | no | 4930415O20Rik | intron_variant | MODIFIER |
| 15 | 98600974 | AX-170174184 | 0 | A | G | 0.114 | 2.11E-02 | 1.50E-01 | 2.66E+00 | 2.73E+00 | 8.89E-01 | no | Adcy6 | synonymous_variant | LOW |
| 15 | 98601987 | AX-97781727 | 0 | C | T | 0.114 | 2.11E-02 | 1.50E-01 | 2.66E+00 | 2.73E+00 | 8.89E-01 | no | Adcy6 | intron_variant | MODIFIER |
| 15 | 98606000 | AX-170423740 | 0 | C | T | 0.114 | 2.11E-02 | 1.50E-01 | 2.66E+00 | 2.73E+00 | 8.89E-01 | no | Adcy6 | intron_variant | MODIFIER |
| 15 | 98688790 | AX-170373937 | 0 | G | A | 0.114 | 2.11E-02 | 1.50E-01 | 2.66E+00 | 2.73E+00 | 8.89E-01 | no | Gm34152 | downstream_gene_variant | MODIFIER |
| 15 | 98702546 | AX-170437420 | 0 | G | A | 0.114 | 2.11E-02 | 1.50E-01 | 2.66E+00 | 2.73E+00 | 8.89E-01 | no | | intergenic_variant | MODIFIER |
| 15 | 98708341 | AX-97851611 | 0 | C | T | 0.114 | 2.11E-02 | 1.50E-01 | 2.66E+00 | 2.73E+00 | 8.89E-01 | no | Ccdc65 | 5_prime_UTR_variant | MODIFIER |
| 15 | 98709882 | AX-170181543 | 0 | T | C | 0.114 | 2.11E-02 | 1.50E-01 | 2.66E+00 | 2.73E+00 | 8.89E-01 | no | Ccdc65 | intron_variant | MODIFIER |
| 15 | 98710465 | AX-97830378 | 0 | T | G | 0.114 | 2.11E-02 | 1.50E-01 | 2.66E+00 | 2.73E+00 | 8.89E-01 | no | Ccdc65 | intron_variant | MODIFIER |
| 15 | 98734754 | AX-170321471 | 0 | G | A | 0.114 | 2.11E-02 | 1.50E-01 | 2.66E+00 | 2.73E+00 | 8.89E-01 | no | Arf3 | downstream_gene_variant | MODIFIER |
| 15 | 98734841 | AX-170121435 | 0 | T | C | 0.114 | 2.11E-02 | 1.50E-01 | 2.66E+00 | 2.73E+00 | 8.89E-01 | no | Arf3 | downstream_gene_variant | MODIFIER |
| 15 | 98842291 | AX-170256819 | 0 | T | C | 0.114 | 2.11E-02 | 1.50E-01 | 2.66E+00 | 2.73E+00 | 8.89E-01 | no | Kmt2d | intron_variant | MODIFIER |
| 15 | 98842387 | AX-170187163 | 0 | G | A | 0.114 | 2.11E-02 | 1.50E-01 | 2.66E+00 | 2.73E+00 | 8.89E-01 | no | Kmt2d | intron_variant | MODIFIER |
| 15 | 98925013 | AX-97799413 | 0 | T | A | 0.114 | 2.11E-02 | 1.50E-01 | 2.66E+00 | 2.73E+00 | 8.89E-01 | no | | regulatory_region_variant | MODIFIER |
| 15 | 98955670 | AX-170012363 | 0 | A | G | 0.114 | 2.11E-02 | 1.50E-01 | 2.66E+00 | 2.73E+00 | 8.89E-01 | no | Tuba1a | upstream_gene_variant | MODIFIER |
| 15 | 98956189 | AX-170012361 | 0 | T | C | 0.114 | 2.11E-02 | 1.50E-01 | 2.66E+00 | 2.73E+00 | 8.89E-01 | no | Tuba1a | upstream_gene_variant | MODIFIER |
| 15 | 98982038 | AX-97836047 | 0 | C | A | 0.114 | 2.11E-02 | 1.50E-01 | 2.66E+00 | 2.73E+00 | 8.89E-01 | no | 4930578M01Rik | upstream_gene_variant | MODIFIER |
| 15 | 98983601 | AX-97830380 | 0 | T | C | 0.114 | 2.11E-02 | 1.50E-01 | 2.66E+00 | 2.73E+00 | 8.89E-01 | no | 4930578M01Rik | upstream_gene_variant | MODIFIER |
| 15 | 98983737 | AX-170012368 | 0 | C | T | 0.114 | 2.11E-02 | 1.50E-01 | 2.66E+00 | 2.73E+00 | 8.89E-01 | no | 4930578M01Rik | upstream_gene_variant | MODIFIER |
| 15 | 99004730 | AX-170314170 | 0 | C | T | 0.114 | 2.11E-02 | 1.50E-01 | 2.66E+00 | 2.73E+00 | 8.89E-01 | no | Gm8973 | upstream_gene_variant | MODIFIER |
| 15 | 100271510 | AX-170151807 | 0 | A | G | 0.114 | 2.11E-02 | 1.50E-01 | 2.66E+00 | 2.73E+00 | 8.89E-01 | no | | intergenic_variant | MODIFIER |
| 15 | 100317821 | AX-170119331 | 0 | C | T | 0.114 | 2.11E-02 | 1.50E-01 | 2.66E+00 | 2.73E+00 | 8.89E-01 | no | Mettl7a1 | downstream_gene_variant | MODIFIER |
| 15 | 100330322 | AX-170525059 | 0 | A | C | 0.114 | 2.11E-02 | 1.50E-01 | 2.66E+00 | 2.73E+00 | 8.89E-01 | no | Mettl7a3 | upstream_gene_variant | MODIFIER |
| 15 | 100384666 | AX-170513540 | 0 | C | T | 0.114 | 2.11E-02 | 1.50E-01 | 2.66E+00 | 2.73E+00 | 8.89E-01 | no | Slc11a2 | downstream_gene_variant | MODIFIER |
| 15 | 100385962 | AX-170516477 | 0 | G | A | 0.114 | 2.11E-02 | 1.50E-01 | 2.66E+00 | 2.73E+00 | 8.89E-01 | no | Slc11a2 | downstream_gene_variant | MODIFIER |
| 15 | 100396389 | AX-170368431 | 0 | C | T | 0.114 | 2.11E-02 | 1.50E-01 | 2.66E+00 | 2.73E+00 | 8.89E-01 | no | Slc11a2 | intron_variant | MODIFIER |

| | | | | | | | | | | | | | | | |
|---|---|---|---|---|---|---|---|---|---|---|---|---|---|---|---|
| 15 | 100450700 | AX-170152502 | 0 | A | C | 0.114 | 2.11E-02 | 1.50E-01 | 2.66E+00 | 2.73E+00 | 8.89E-01 | no | n-R5s43 | downstream_gene_variant | MODIFIER |
| 15 | 100451375 | AX-170313102 | 0 | T | C | 0.114 | 2.11E-02 | 1.50E-01 | 2.66E+00 | 2.73E+00 | 8.89E-01 | no | n-R5s43 | downstream_gene_variant | MODIFIER |
| 15 | 100453254 | AX-168397842 | 0 | A | C | 0.114 | 2.11E-02 | 1.50E-01 | 2.66E+00 | 2.73E+00 | 8.89E-01 | no | n-R5s43 | downstream_gene_variant | MODIFIER |
| 15 | 100458316 | AX-170440403 | 0 | T | C | 0.114 | 2.11E-02 | 1.50E-01 | 2.66E+00 | 2.73E+00 | 8.89E-01 | no | 5330439K02Rik | downstream_gene_variant | MODIFIER |
| 15 | 100463229 | AX-170262721 | 0 | G | C | 0.114 | 2.11E-02 | 1.50E-01 | 2.66E+00 | 2.73E+00 | 8.89E-01 | no | 5330439K02Rik | intron_variant,non_coding_transcript_variant | MODIFIER |
| 15 | 100484037 | AX-170012384 | 0 | T | C | 0.114 | 2.11E-02 | 1.50E-01 | 2.66E+00 | 2.73E+00 | 8.89E-01 | no | Letmd1 | downstream_gene_variant | MODIFIER |
| 15 | 100508265 | AX-170012385 | 0 | C | T | 0.114 | 2.11E-02 | 1.50E-01 | 2.66E+00 | 2.73E+00 | 8.89E-01 | no | Tfcp2 | intron_variant | MODIFIER |
| 15 | 100583651 | AX-170012383 | 0 | T | C | 0.114 | 2.11E-02 | 1.50E-01 | 2.66E+00 | 2.73E+00 | 8.89E-01 | no | Pou6f1 | intron_variant | MODIFIER |
| 15 | 100583707 | AX-170175441 | 0 | T | C | 0.114 | 2.11E-02 | 1.50E-01 | 2.66E+00 | 2.73E+00 | 8.89E-01 | no | Pou6f1 | intron_variant | MODIFIER |
| 15 | 100692698 | AX-170319930 | 0 | C | T | 0.114 | 2.11E-02 | 1.50E-01 | 2.66E+00 | 2.73E+00 | 8.89E-01 | no | Cela1 | upstream_gene_variant | MODIFIER |
| 15 | 100693213 | AX-170104151 | 0 | G | A | 0.114 | 2.11E-02 | 1.50E-01 | 2.66E+00 | 2.73E+00 | 8.89E-01 | no | Galnt6 | 3_prime_UTR_variant | MODIFIER |
| 15 | 100693558 | AX-170319102 | 0 | A | G | 0.114 | 2.11E-02 | 1.50E-01 | 2.66E+00 | 2.73E+00 | 8.89E-01 | no | Galnt6 | intron_variant | MODIFIER |
| 15 | 100694226 | AX-170186196 | 0 | T | C | 0.114 | 2.11E-02 | 1.50E-01 | 2.66E+00 | 2.73E+00 | 8.89E-01 | no | Galnt6 | intron_variant | MODIFIER |
| 15 | 100694248 | AX-170233748 | 0 | A | G | 0.114 | 2.11E-02 | 1.50E-01 | 2.66E+00 | 2.73E+00 | 8.89E-01 | no | Galnt6 | intron_variant | MODIFIER |
| 15 | 100694853 | AX-170056481 | 0 | T | C | 0.114 | 2.11E-02 | 1.50E-01 | 2.66E+00 | 2.73E+00 | 8.89E-01 | no | Galnt6 | intron_variant | MODIFIER |
| 15 | 100694879 | AX-170056480 | 0 | C | T | 0.114 | 2.11E-02 | 1.50E-01 | 2.66E+00 | 2.73E+00 | 8.89E-01 | no | Galnt6 | intron_variant | MODIFIER |
| 15 | 100701422 | AX-170274964 | 0 | T | C | 0.114 | 2.11E-02 | 1.50E-01 | 2.66E+00 | 2.73E+00 | 8.89E-01 | no | Galnt6 | intron_variant | MODIFIER |
| 15 | 100706155 | AX-170158272 | 0 | G | A | 0.114 | 2.11E-02 | 1.50E-01 | 2.66E+00 | 2.73E+00 | 8.89E-01 | no | Galnt6 | intron_variant | MODIFIER |
| 15 | 100706773 | AX-170327395 | 0 | G | T | 0.114 | 2.11E-02 | 1.50E-01 | 2.66E+00 | 2.73E+00 | 8.89E-01 | no | Galnt6 | intron_variant | MODIFIER |
| 15 | 100707719 | AX-170149332 | 0 | C | T | 0.114 | 2.11E-02 | 1.50E-01 | 2.66E+00 | 2.73E+00 | 8.89E-01 | no | Galnt6 | intron_variant | MODIFIER |
| 15 | 100708861 | AX-170495016 | 0 | T | C | 0.114 | 2.11E-02 | 1.50E-01 | 2.66E+00 | 2.73E+00 | 8.89E-01 | no | Galnt6 | intron_variant | MODIFIER |
| 15 | 100709172 | AX-170119388 | 0 | T | C | 0.114 | 2.11E-02 | 1.50E-01 | 2.66E+00 | 2.73E+00 | 8.89E-01 | no | Galnt6 | intron_variant | MODIFIER |
| 15 | 100716658 | AX-170476468 | 0 | T | C | 0.114 | 2.11E-02 | 1.50E-01 | 2.66E+00 | 2.73E+00 | 8.89E-01 | no | Galnt6 | intron_variant | MODIFIER |
| 15 | 100716748 | AX-170100494 | 0 | T | G | 0.114 | 2.11E-02 | 1.50E-01 | 2.66E+00 | 2.73E+00 | 8.89E-01 | no | Galnt6 | intron_variant | MODIFIER |
| 15 | 100717021 | AX-170335957 | 0 | G | A | 0.114 | 2.11E-02 | 1.50E-01 | 2.66E+00 | 2.73E+00 | 8.89E-01 | no | Galnt6 | intron_variant | MODIFIER |
| 15 | 100718426 | AX-170352729 | 0 | T | C | 0.114 | 2.11E-02 | 1.50E-01 | 2.66E+00 | 2.73E+00 | 8.89E-01 | no | Galnt6 | intron_variant | MODIFIER |
| 15 | 100720119 | AX-170521123 | 0 | T | C | 0.114 | 2.11E-02 | 1.50E-01 | 2.66E+00 | 2.73E+00 | 8.89E-01 | no | Galnt6 | 5_prime_UTR_variant | MODIFIER |
| 15 | 100721788 | AX-170298727 | 0 | G | C | 0.114 | 2.11E-02 | 1.50E-01 | 2.66E+00 | 2.73E+00 | 8.89E-01 | no | Galnt6 | intron_variant | MODIFIER |
| 15 | 100764729 | AX-170468697 | 0 | T | C | 0.114 | 2.11E-02 | 1.50E-01 | 2.66E+00 | 2.73E+00 | 8.89E-01 | no | Slc4a8 | intron_variant | MODIFIER |
| 15 | 100768286 | AX-170161677 | 0 | T | C | 0.114 | 2.11E-02 | 1.50E-01 | 2.66E+00 | 2.73E+00 | 8.89E-01 | no | Slc4a8 | intron_variant | MODIFIER |
| 15 | 100768301 | AX-170516828 | 0 | C | T | 0.114 | 2.11E-02 | 1.50E-01 | 2.66E+00 | 2.73E+00 | 8.89E-01 | no | Slc4a8 | intron_variant | MODIFIER |
| 15 | 100774561 | AX-170460415 | 0 | G | A | 0.114 | 2.11E-02 | 1.50E-01 | 2.66E+00 | 2.73E+00 | 8.89E-01 | no | Slc4a8 | intron_variant | MODIFIER |
| 15 | 100783202 | AX-170249924 | 0 | A | T | 0.114 | 2.11E-02 | 1.50E-01 | 2.66E+00 | 2.73E+00 | 8.89E-01 | no | Slc4a8 | intron_variant | MODIFIER |
| 15 | 100785563 | AX-170056482 | 0 | G | C | 0.114 | 2.11E-02 | 1.50E-01 | 2.66E+00 | 2.73E+00 | 8.89E-01 | no | Slc4a8 | intron_variant | MODIFIER |
| 15 | 100801358 | AX-170093357 | 0 | G | A | 0.114 | 2.11E-02 | 1.50E-01 | 2.66E+00 | 2.73E+00 | 8.89E-01 | no | Slc4a8 | intron_variant | MODIFIER |
| 15 | 100802261 | AX-170317841 | 0 | T | C | 0.114 | 2.11E-02 | 1.50E-01 | 2.66E+00 | 2.73E+00 | 8.89E-01 | no | Slc4a8 | intron_variant | MODIFIER |
| 15 | 100803597 | AX-170240365 | 0 | T | C | 0.114 | 2.11E-02 | 1.50E-01 | 2.66E+00 | 2.73E+00 | 8.89E-01 | no | Slc4a8 | intron_variant | MODIFIER |
| 15 | 100812555 | AX-170232786 | 0 | G | A | 0.114 | 2.11E-02 | 1.50E-01 | 2.66E+00 | 2.73E+00 | 8.89E-01 | no | Slc4a8 | intron_variant | MODIFIER |
| 15 | 100817680 | AX-170313810 | 0 | A | G | 0.114 | 2.11E-02 | 1.50E-01 | 2.66E+00 | 2.73E+00 | 8.89E-01 | no | Slc4a8 | 3_prime_UTR_variant | MODIFIER |
| 15 | 101020420 | AX-170468586 | 0 | C | G | 0.114 | 2.11E-02 | 1.50E-01 | 2.66E+00 | 2.73E+00 | 8.89E-01 | no | Scn8a | intron_variant | MODIFIER |
| 15 | 101020713 | AX-170134588 | 0 | C | T | 0.114 | 2.11E-02 | 1.50E-01 | 2.66E+00 | 2.73E+00 | 8.89E-01 | no | Scn8a | intron_variant | MODIFIER |
| 15 | 101027438 | AX-170056489 | 0 | A | G | 0.114 | 2.11E-02 | 1.50E-01 | 2.66E+00 | 2.73E+00 | 8.89E-01 | no | Scn8a | intron_variant | MODIFIER |
| 15 | 101055513 | AX-170378508 | 0 | A | G | 0.114 | 2.11E-02 | 1.50E-01 | 2.66E+00 | 2.73E+00 | 8.89E-01 | no | Fignl2 | upstream_gene_variant | MODIFIER |
| 15 | 101056358 | AX-170452752 | 0 | G | C | 0.114 | 2.11E-02 | 1.50E-01 | 2.66E+00 | 2.73E+00 | 8.89E-01 | no | Fignl2 | upstream_gene_variant | MODIFIER |
| 15 | 101056836 | AX-170296660 | 0 | G | A | 0.114 | 2.11E-02 | 1.50E-01 | 2.66E+00 | 2.73E+00 | 8.89E-01 | no | Fignl2 | upstream_gene_variant | MODIFIER |
| 15 | 101097863 | AX-170160738 | 0 | C | T | 0.114 | 2.11E-02 | 1.50E-01 | 2.66E+00 | 2.73E+00 | 8.89E-01 | no | | regulatory_region_variant | MODIFIER |
| 15 | 101115732 | AX-170267150 | 0 | C | T | 0.114 | 2.11E-02 | 1.50E-01 | 2.66E+00 | 2.73E+00 | 8.89E-01 | no | Ankrd33 | upstream_gene_variant | MODIFIER |
| 15 | 101115951 | AX-170211420 | 0 | G | A | 0.114 | 2.11E-02 | 1.50E-01 | 2.66E+00 | 2.73E+00 | 8.89E-01 | no | Ankrd33 | intron_variant | MODIFIER |
| 15 | 101282904 | AX-97838233 | 0 | C | T | 0.114 | 2.11E-02 | 1.50E-01 | 2.66E+00 | 2.73E+00 | 8.89E-01 | no | Atg101 | upstream_gene_variant | MODIFIER |
| 15 | 101295654 | AX-170415507 | 0 | G | A | 0.114 | 2.11E-02 | 1.50E-01 | 2.66E+00 | 2.73E+00 | 8.89E-01 | no | Atg101 | downstream_gene_variant | MODIFIER |
| 15 | 101453456 | AX-170375267 | 0 | A | G | 0.114 | 2.11E-02 | 1.50E-01 | 2.66E+00 | 2.73E+00 | 8.89E-01 | no | Krt88 | intron_variant | MODIFIER |
| 15 | 101470270 | AX-170131500 | 0 | T | C | 0.114 | 2.11E-02 | 1.50E-01 | 2.66E+00 | 2.73E+00 | 8.89E-01 | no | Krt86 | upstream_gene_variant | MODIFIER |
| 15 | 101480064 | AX-97817541 | 0 | A | G | 0.114 | 2.11E-02 | 1.50E-01 | 2.66E+00 | 2.73E+00 | 8.89E-01 | no | Krt86 | downstream_gene_variant | MODIFIER |
| 15 | 101486764 | AX-170460182 | 0 | T | G | 0.114 | 2.11E-02 | 1.50E-01 | 2.66E+00 | 2.73E+00 | 8.89E-01 | no | Krt83 | intron_variant | MODIFIER |

| | | | | | | | | | | | | | | | |
|---|---|---|---|---|---|---|---|---|---|---|---|---|---|---|---|
| 15 | 101490371 | AX-170483594 | 0 | A | G | 0.114 | 2.11E-02 | 1.50E-01 | 2.66E+00 | 2.73E+00 | 8.89E-01 | no | Krt83 | intron_variant | MODIFIER |
| 15 | 101529782 | AX-170333265 | 0 | T | C | 0.114 | 2.11E-02 | 1.50E-01 | 2.66E+00 | 2.73E+00 | 8.89E-01 | no | Krt84 | intron_variant | MODIFIER |
| 15 | 101531830 | AX-170398142 | 0 | C | T | 0.114 | 2.11E-02 | 1.50E-01 | 2.66E+00 | 2.73E+00 | 8.89E-01 | no | Krt84 | intron_variant | MODIFIER |
| 15 | 101556903 | AX-170056498 | 0 | A | G | 0.114 | 2.11E-02 | 1.50E-01 | 2.66E+00 | 2.73E+00 | 8.89E-01 | no | Krt90 | intron_variant | MODIFIER |
| 15 | 101562706 | AX-170102859 | 0 | A | G | 0.114 | 2.11E-02 | 1.50E-01 | 2.66E+00 | 2.73E+00 | 8.89E-01 | no | Krt90 | synonymous_variant | LOW |
| 15 | 101676816 | AX-97784056 | 0 | C | A | 0.114 | 2.11E-02 | 1.50E-01 | 2.66E+00 | 2.73E+00 | 8.89E-01 | no | Krt6b | intron_variant | MODIFIER |
| 15 | 101678239 | AX-170395965 | 0 | C | T | 0.114 | 2.11E-02 | 1.50E-01 | 2.66E+00 | 2.73E+00 | 8.89E-01 | no | Krt6b | intron_variant | MODIFIER |
| 15 | 101688475 | AX-170489037 | 0 | A | G | 0.114 | 2.11E-02 | 1.50E-01 | 2.66E+00 | 2.73E+00 | 8.89E-01 | no | Krt6a | downstream_gene_variant | MODIFIER |
| 15 | 101737069 | AX-170439767 | 0 | C | T | 0.114 | 2.11E-02 | 1.50E-01 | 2.66E+00 | 2.73E+00 | 8.89E-01 | no | Krt71 | intron_variant | MODIFIER |
| 15 | 101738191 | AX-170244789 | 0 | T | C | 0.114 | 2.11E-02 | 1.50E-01 | 2.66E+00 | 2.73E+00 | 8.89E-01 | no | Krt71 | intron_variant | MODIFIER |
| 15 | 101753691 | AX-170397069 | 0 | A | C | 0.114 | 2.11E-02 | 1.50E-01 | 2.66E+00 | 2.73E+00 | 8.89E-01 | no | Krt74 | downstream_gene_variant | MODIFIER |
| 15 | 101761363 | AX-170164535 | 0 | A | G | 0.114 | 2.11E-02 | 1.50E-01 | 2.66E+00 | 2.73E+00 | 8.89E-01 | no | Krt74 | intron_variant,non_coding_transcript_variant | MODIFIER |
| 15 | 101771229 | AX-170378422 | 0 | T | C | 0.114 | 2.11E-02 | 1.50E-01 | 2.66E+00 | 2.73E+00 | 8.89E-01 | no | Krt72 | downstream_gene_variant | MODIFIER |
| 15 | 101882956 | AX-170122484 | 0 | G | A | 0.114 | 2.11E-02 | 1.50E-01 | 2.66E+00 | 2.73E+00 | 8.89E-01 | no | Krt76 | downstream_gene_variant | MODIFIER |
| 15 | 101904290 | AX-170278016 | 0 | T | C | 0.114 | 2.11E-02 | 1.50E-01 | 2.66E+00 | 2.73E+00 | 8.89E-01 | no | Gm49434 | downstream_gene_variant | MODIFIER |
| 15 | 101909485 | AX-170112981 | 0 | C | T | 0.114 | 2.11E-02 | 1.50E-01 | 2.66E+00 | 2.73E+00 | 8.89E-01 | no | Gm49434 | intron_variant,non_coding_transcript_variant | MODIFIER |
| 15 | 101915022 | AX-170319140 | 0 | C | G | 0.114 | 2.11E-02 | 1.50E-01 | 2.66E+00 | 2.73E+00 | 8.89E-01 | no | Krt4 | downstream_gene_variant | MODIFIER |
| 15 | 101918369 | AX-170533402 | 0 | T | C | 0.114 | 2.11E-02 | 1.50E-01 | 2.66E+00 | 2.73E+00 | 8.89E-01 | no | Krt4 | downstream_gene_variant | MODIFIER |
| 15 | 101918834 | AX-170379964 | 0 | A | G | 0.114 | 2.11E-02 | 1.50E-01 | 2.66E+00 | 2.73E+00 | 8.89E-01 | no | Krt4 | 3_prime_UTR_variant | MODIFIER |
| 15 | 101920013 | AX-170394744 | 0 | A | G | 0.114 | 2.11E-02 | 1.50E-01 | 2.66E+00 | 2.73E+00 | 8.89E-01 | no | Krt4 | intron_variant | MODIFIER |
| 15 | 102269210 | AX-97839544 | 0 | G | A | 0.114 | 2.11E-02 | 1.50E-01 | 2.66E+00 | 2.73E+00 | 8.89E-01 | no | Gm9918 | upstream_gene_variant | MODIFIER |
| 15 | 102306017 | AX-170244433 | 0 | T | C | 0.114 | 2.11E-02 | 1.50E-01 | 2.66E+00 | 2.73E+00 | 8.89E-01 | no | Espl1 | synonymous_variant | LOW |
| 15 | 102309052 | AX-97822058 | 0 | A | G | 0.114 | 2.11E-02 | 1.50E-01 | 2.66E+00 | 2.73E+00 | 8.89E-01 | no | Espl1 | intron_variant | MODIFIER |
| 15 | 102408133 | AX-170173066 | 0 | T | C | 0.114 | 2.11E-02 | 1.50E-01 | 2.66E+00 | 2.73E+00 | 8.89E-01 | no | Sp1 | intron_variant | MODIFIER |
| 15 | 102428745 | AX-170418928 | 0 | G | A | 0.114 | 2.11E-02 | 1.50E-01 | 2.66E+00 | 2.73E+00 | 8.89E-01 | no | Sp1 | intron_variant | MODIFIER |
| 15 | 102428868 | AX-170328916 | 0 | C | G | 0.114 | 2.11E-02 | 1.50E-01 | 2.66E+00 | 2.73E+00 | 8.89E-01 | no | Sp1 | intron_variant | MODIFIER |
| 15 | 102714816 | AX-170395407 | 0 | C | T | 0.114 | 2.11E-02 | 1.50E-01 | 2.66E+00 | 2.73E+00 | 8.89E-01 | no | Calcoco1 | intron_variant | MODIFIER |
| 15 | 102786274 | AX-170394713 | 0 | G | A | 0.114 | 2.11E-02 | 1.50E-01 | 2.66E+00 | 2.73E+00 | 8.89E-01 | no | | intergenic_variant | MODIFIER |
| 15 | 102795189 | AX-170305072 | 0 | A | G | 0.114 | 2.11E-02 | 1.50E-01 | 2.66E+00 | 2.73E+00 | 8.89E-01 | no | Gm49473 | non_coding_transcript_exon_variant | MODIFIER |
| 15 | 102800433 | AX-170270817 | 0 | T | C | 0.114 | 2.11E-02 | 1.50E-01 | 2.66E+00 | 2.73E+00 | 8.89E-01 | no | Gm36560 | intron_variant,non_coding_transcript_variant | MODIFIER |
| 15 | 102825193 | AX-170187491 | 0 | A | G | 0.114 | 2.11E-02 | 1.50E-01 | 2.66E+00 | 2.73E+00 | 8.89E-01 | no | | intergenic_variant | MODIFIER |
| 15 | 102841498 | AX-170195787 | 0 | A | G | 0.114 | 2.11E-02 | 1.50E-01 | 2.66E+00 | 2.73E+00 | 8.89E-01 | no | | intergenic_variant | MODIFIER |
| 15 | 102847660 | AX-170329246 | 0 | A | G | 0.114 | 2.11E-02 | 1.50E-01 | 2.66E+00 | 2.73E+00 | 8.89E-01 | no | | intergenic_variant | MODIFIER |
| 15 | 102848170 | AX-170448772 | 0 | C | T | 0.114 | 2.11E-02 | 1.50E-01 | 2.66E+00 | 2.73E+00 | 8.89E-01 | no | | intergenic_variant | MODIFIER |
| 15 | 103272771 | AX-170012425 | 0 | C | T | 0.114 | 2.11E-02 | 1.50E-01 | 2.66E+00 | 2.73E+00 | 8.89E-01 | no | Copz1 | upstream_gene_variant | MODIFIER |
| 4 | 144837205 | AX-170342180 | 2 | A | T | 0.082 | 2.86E-02 | 2.02E-01 | 2.63E+00 | 2.73E+00 | 8.89E-01 | no | | intergenic_variant | MODIFIER |
| 1 | 76771602 | AX-170000845 | 0 | T | C | 0.14 | 3.25E-02 | 2.37E-01 | 2.68E+00 | 2.74E+00 | 8.89E-01 | no | | intergenic_variant | MODIFIER |
| 1 | 104938662 | AX-170001266 | 0 | A | G | 0.149 | -2.13E-02 | 1.51E-01 | 2.62E+00 | 2.73E+00 | 8.90E-01 | no | Cdh20 | intron_variant | MODIFIER |
| 15 | 101316658 | AX-97841988 | 0 | C | T | 0.114 | 2.10E-02 | 1.50E-01 | 2.66E+00 | 2.73E+00 | 8.90E-01 | no | Gm49499 | non_coding_transcript_exon_variant | MODIFIER |
| 13 | 53134456 | AX-170397062 | 0 | A | T | 0.518 | -8.23E-03 | 6.63E-02 | 2.66E+00 | 2.77E+00 | 8.90E-01 | no | Ror2 | intron_variant | MODIFIER |
| 1 | 53330815 | AX-170249059 | 2 | T | C | 0.091 | 2.89E-02 | 2.13E-01 | 2.69E+00 | 2.74E+00 | 8.90E-01 | no | Osgepl1 | downstream_gene_variant | MODIFIER |
| 9 | 109647195 | AX-170168160 | 0 | T | C | 0.132 | -1.57E-02 | 1.06E-01 | 2.63E+00 | 2.71E+00 | 8.90E-01 | no | Fbxw25 | intron_variant | MODIFIER |
| 15 | 81194206 | AX-97834580 | 1 | G | A | 0.545 | 1.23E-02 | 7.94E-02 | 2.61E+00 | 2.70E+00 | 8.90E-01 | no | Mrtfa | upstream_gene_variant | MODIFIER |
| 7 | 81525372 | AX-170203164 | 1 | T | C | 0.116 | 2.90E-02 | 2.10E-01 | 2.65E+00 | 2.74E+00 | 8.90E-01 | no | Mir1839 | upstream_gene_variant | MODIFIER |
| 8 | 30255597 | AX-170311193 | 0 | G | A | 0.175 | 1.41E-02 | 1.03E-01 | 2.65E+00 | 2.74E+00 | 8.90E-01 | no | | intergenic_variant | MODIFIER |
| 12 | 116833294 | AX-97854542 | 1 | C | A | 0.143 | -2.57E-02 | 1.91E-01 | 2.68E+00 | 2.75E+00 | 8.90E-01 | no | Ptprn2 | intron_variant | MODIFIER |
| 8 | 18055929 | AX-97833063 | 0 | C | A | 0.175 | -1.50E-02 | 1.10E-01 | 2.65E+00 | 2.74E+00 | 8.90E-01 | no | | intergenic_variant | MODIFIER |
| 13 | 49040657 | AX-170051223 | 0 | G | A | 0.482 | -8.49E-03 | 6.65E-02 | 2.65E+00 | 2.76E+00 | 8.90E-01 | no | Wnk2 | intron_variant | MODIFIER |
| 8 | 109198915 | AX-170110176 | 0 | C | T | 0.237 | 1.46E-02 | 1.08E-01 | 2.66E+00 | 2.74E+00 | 8.90E-01 | no | D030068K23Rik | intron_variant,non_coding_transcript_variant | MODIFIER |
| 9 | 113219007 | AX-170033303 | 0 | G | A | 0.132 | -1.44E-02 | 1.03E-01 | 2.64E+00 | 2.73E+00 | 8.90E-01 | no | | intergenic_variant | MODIFIER |
| 8 | 81491991 | AX-170517210 | 1 | A | T | 0.205 | 1.77E-02 | 1.28E-01 | 2.67E+00 | 2.73E+00 | 8.90E-01 | no | Inpp4b | intron_variant,NMD_transcript_variant | MODIFIER |
| 8 | 49353453 | AX-97810220 | 0 | A | C | 0.158 | 1.74E-02 | 1.34E-01 | 2.70E+00 | 2.77E+00 | 8.90E-01 | no | | intergenic_variant | MODIFIER |
| 7 | 104574779 | AX-170255622 | 1 | G | A | 0.482 | -1.23E-02 | 9.53E-02 | 2.70E+00 | 2.77E+00 | 8.90E-01 | no | Olfr653 | upstream_gene_variant | MODIFIER |
| 10 | 9276073 | AX-170446195 | 0 | T | A | 0.202 | -1.46E-02 | 9.85E-02 | 2.62E+00 | 2.71E+00 | 8.90E-01 | no | | intergenic_variant | MODIFIER |

| | | | | | | | | | | | | | | | |
|---|---|---|---|---|---|---|---|---|---|---|---|---|---|---|---|
| 9 | 63947244 | AX-170267182 | 1 | T | C | 0.196 | -1.87E-02 | 1.37E-01 | 2.71E+00 | 2.73E+00 | 8.91E-01 | no | | regulatory_region_variant | MODIFIER |
| 16 | 75757024 | AX-170231299 | 1 | G | T | 0.5 | 1.20E-02 | 7.52E-02 | 2.60E+00 | 2.69E+00 | 8.91E-01 | no | Hspa13 | 3_prime_UTR_variant | MODIFIER |
| 11 | 33922274 | AX-170188494 | 1 | G | A | 0.107 | 2.67E-02 | 1.90E-01 | 2.65E+00 | 2.72E+00 | 8.91E-01 | no | Kcnip1 | intron_variant,non_coding_transcript_variant | MODIFIER |
| 8 | 84792522 | AX-170360222 | 1 | G | A | 0.196 | 1.76E-02 | 1.30E-01 | 2.68E+00 | 2.74E+00 | 8.91E-01 | no | Nfix | intron_variant | MODIFIER |
| 9 | 105243581 | AX-170357182 | 1 | T | A | 0.125 | 1.49E-02 | 1.18E-01 | 2.71E+00 | 2.78E+00 | 8.91E-01 | no | Nek11 | intron_variant | MODIFIER |
| 7 | 116268221 | AX-170489139 | 1 | C | G | 0.411 | 1.30E-02 | 9.34E-02 | 2.67E+00 | 2.72E+00 | 8.91E-01 | no | Plekha7 | intron_variant | MODIFIER |
| 13 | 48925472 | AX-97812516 | 1 | C | T | 0.491 | -8.42E-03 | 6.69E-02 | 2.66E+00 | 2.76E+00 | 8.91E-01 | no | Fam120a | intron_variant | MODIFIER |
| 3 | 147674502 | AX-170530191 | 0 | T | C | 0.342 | -9.61E-03 | 7.35E-02 | 2.68E+00 | 2.75E+00 | 8.91E-01 | no | | intergenic_variant | MODIFIER |
| 1 | 146878124 | AX-97824551 | 0 | A | T | 0.202 | -1.62E-02 | 1.30E-01 | 2.69E+00 | 2.78E+00 | 8.91E-01 | no | Brinp3 | intron_variant | MODIFIER |
| 1 | 152015971 | AX-170240101 | 0 | T | A | 0.202 | -1.62E-02 | 1.30E-01 | 2.69E+00 | 2.78E+00 | 8.91E-01 | no | 1700025G04Rik | intron_variant | MODIFIER |
| 1 | 152327496 | AX-170201305 | 0 | T | G | 0.202 | -1.62E-02 | 1.30E-01 | 2.69E+00 | 2.78E+00 | 8.91E-01 | no | | intergenic_variant | MODIFIER |
| 15 | 92144687 | AX-170322018 | 0 | G | A | 0.132 | 2.19E-02 | 1.56E-01 | 2.66E+00 | 2.72E+00 | 8.91E-01 | no | Cntn1 | intron_variant | MODIFIER |
| 15 | 97731038 | AX-170410178 | 0 | C | T | 0.132 | 2.19E-02 | 1.56E-01 | 2.66E+00 | 2.72E+00 | 8.91E-01 | no | Endou | intron_variant | MODIFIER |
| 2 | 72370727 | AX-170266503 | 0 | A | G | 0.053 | 2.35E-02 | 1.83E-01 | 2.62E+00 | 2.75E+00 | 8.91E-01 | no | Map3k20 | intron_variant | MODIFIER |
| 5 | 97974428 | AX-170480149 | 2 | T | C | 0.391 | -1.16E-02 | 9.18E-02 | 2.66E+00 | 2.76E+00 | 8.91E-01 | no | Antxr2 | intron_variant | MODIFIER |
| 1 | 46085378 | AX-97811409 | 0 | T | A | 0.333 | 9.27E-03 | 7.26E-02 | 2.66E+00 | 2.76E+00 | 8.91E-01 | no | Dnah7b | intron_variant | MODIFIER |
| 7 | 117517713 | AX-170393395 | 1 | C | T | 0.402 | -1.21E-02 | 9.17E-02 | 2.69E+00 | 2.75E+00 | 8.91E-01 | no | Xylt1 | intron_variant | MODIFIER |
| 9 | 76063155 | AX-170188440 | 1 | G | T | 0.196 | -1.83E-02 | 1.34E-01 | 2.69E+00 | 2.73E+00 | 8.91E-01 | no | Hmgcll1 | intron_variant | MODIFIER |
| 5 | 98387923 | AX-170072845 | 1 | C | G | 0.42 | -1.38E-02 | 1.05E-01 | 2.70E+00 | 2.76E+00 | 8.91E-01 | no | Cfap299 | intron_variant | MODIFIER |
| 6 | 104349079 | AX-170097927 | 0 | G | T | 0.079 | 2.02E-02 | 1.64E-01 | 2.61E+00 | 2.76E+00 | 8.91E-01 | no | | intergenic_variant | MODIFIER |
| 6 | 110604220 | AX-170213653 | 1 | C | A | 0.071 | 3.68E-02 | 2.76E-01 | 2.71E+00 | 2.75E+00 | 8.92E-01 | no | | intergenic_variant | MODIFIER |
| 1 | 89460455 | AX-97828724 | 0 | T | G | 0.158 | 2.37E-02 | 1.73E-01 | 2.64E+00 | 2.73E+00 | 8.92E-01 | no | Agap1 | intron_variant | MODIFIER |
| 5 | 101727839 | AX-97810349 | 0 | A | G | 0.43 | 1.49E-02 | 1.11E-01 | 2.69E+00 | 2.74E+00 | 8.92E-01 | no | | intergenic_variant | MODIFIER |
| 5 | 101935232 | AX-97854115 | 0 | G | T | 0.43 | 1.49E-02 | 1.11E-01 | 2.69E+00 | 2.74E+00 | 8.92E-01 | no | Wdfy3 | intron_variant | MODIFIER |
| 5 | 101960986 | AX-97810637 | 0 | T | C | 0.43 | 1.49E-02 | 1.11E-01 | 2.69E+00 | 2.74E+00 | 8.92E-01 | no | Wdfy3 | intron_variant | MODIFIER |
| 5 | 102021837 | AX-170025750 | 0 | G | A | 0.43 | 1.49E-02 | 1.11E-01 | 2.69E+00 | 2.74E+00 | 8.92E-01 | no | Wdfy3 | intron_variant | MODIFIER |
| 5 | 102022321 | AX-170532560 | 0 | G | A | 0.43 | 1.49E-02 | 1.11E-01 | 2.69E+00 | 2.74E+00 | 8.92E-01 | no | Wdfy3 | intron_variant | MODIFIER |
| 5 | 102027350 | AX-170429161 | 0 | C | T | 0.43 | 1.49E-02 | 1.11E-01 | 2.69E+00 | 2.74E+00 | 8.92E-01 | no | Wdfy3 | intron_variant | MODIFIER |
| 5 | 102027595 | AX-170163766 | 0 | G | A | 0.43 | 1.49E-02 | 1.11E-01 | 2.69E+00 | 2.74E+00 | 8.92E-01 | no | Wdfy3 | intron_variant | MODIFIER |
| 5 | 102028881 | AX-97830370 | 0 | T | C | 0.43 | 1.49E-02 | 1.11E-01 | 2.69E+00 | 2.74E+00 | 8.92E-01 | no | Wdfy3 | intron_variant | MODIFIER |
| 5 | 102029203 | AX-170353490 | 0 | T | C | 0.43 | 1.49E-02 | 1.11E-01 | 2.69E+00 | 2.74E+00 | 8.92E-01 | no | Wdfy3 | intron_variant | MODIFIER |
| 5 | 102037559 | AX-170039073 | 0 | G | C | 0.43 | 1.49E-02 | 1.11E-01 | 2.69E+00 | 2.74E+00 | 8.92E-01 | no | Wdfy3 | intron_variant | MODIFIER |
| 5 | 102132303 | AX-170072912 | 0 | A | G | 0.43 | 1.49E-02 | 1.11E-01 | 2.69E+00 | 2.74E+00 | 8.92E-01 | no | | intergenic_variant | MODIFIER |
| 5 | 102135624 | AX-170140617 | 0 | G | A | 0.43 | 1.49E-02 | 1.11E-01 | 2.69E+00 | 2.74E+00 | 8.92E-01 | no | | intergenic_variant | MODIFIER |
| 5 | 102159605 | AX-170526615 | 0 | A | G | 0.43 | 1.49E-02 | 1.11E-01 | 2.69E+00 | 2.74E+00 | 8.92E-01 | no | Gm17978 | non_coding_transcript_exon_variant | MODIFIER |
| 5 | 102206029 | AX-170362880 | 0 | T | C | 0.43 | 1.49E-02 | 1.11E-01 | 2.69E+00 | 2.74E+00 | 8.92E-01 | no | | intergenic_variant | MODIFIER |
| 5 | 102207534 | AX-170404096 | 0 | A | G | 0.43 | 1.49E-02 | 1.11E-01 | 2.69E+00 | 2.74E+00 | 8.92E-01 | no | | regulatory_region_variant | MODIFIER |
| 5 | 102209087 | AX-170095416 | 0 | G | A | 0.43 | 1.49E-02 | 1.11E-01 | 2.69E+00 | 2.74E+00 | 8.92E-01 | no | | intergenic_variant | MODIFIER |
| 5 | 102211100 | AX-170311850 | 0 | C | T | 0.43 | 1.49E-02 | 1.11E-01 | 2.69E+00 | 2.74E+00 | 8.92E-01 | no | | intergenic_variant | MODIFIER |
| 5 | 102211204 | AX-170311872 | 0 | T | C | 0.43 | 1.49E-02 | 1.11E-01 | 2.69E+00 | 2.74E+00 | 8.92E-01 | no | | intergenic_variant | MODIFIER |
| 5 | 102229924 | AX-170395088 | 0 | T | C | 0.43 | 1.49E-02 | 1.11E-01 | 2.69E+00 | 2.74E+00 | 8.92E-01 | no | | intergenic_variant | MODIFIER |
| 5 | 102240947 | AX-170508393 | 0 | T | A | 0.43 | 1.49E-02 | 1.11E-01 | 2.69E+00 | 2.74E+00 | 8.92E-01 | no | | intergenic_variant | MODIFIER |
| 5 | 102267062 | AX-170464103 | 0 | A | C | 0.43 | 1.49E-02 | 1.11E-01 | 2.69E+00 | 2.74E+00 | 8.92E-01 | no | | regulatory_region_variant | MODIFIER |
| 5 | 102285854 | AX-170487086 | 0 | T | A | 0.43 | 1.49E-02 | 1.11E-01 | 2.69E+00 | 2.74E+00 | 8.92E-01 | no | | intergenic_variant | MODIFIER |
| 5 | 102301433 | AX-170187454 | 0 | G | C | 0.43 | 1.49E-02 | 1.11E-01 | 2.69E+00 | 2.74E+00 | 8.92E-01 | no | | intergenic_variant | MODIFIER |
| 5 | 102303211 | AX-170513225 | 0 | T | A | 0.43 | 1.49E-02 | 1.11E-01 | 2.69E+00 | 2.74E+00 | 8.92E-01 | no | | intergenic_variant | MODIFIER |
| 5 | 102313082 | AX-170072918 | 0 | T | A | 0.43 | 1.49E-02 | 1.11E-01 | 2.69E+00 | 2.74E+00 | 8.92E-01 | no | | intergenic_variant | MODIFIER |
| 5 | 102315901 | AX-170072916 | 0 | A | G | 0.43 | 1.49E-02 | 1.11E-01 | 2.69E+00 | 2.74E+00 | 8.92E-01 | no | | intergenic_variant | MODIFIER |
| 5 | 102322627 | AX-170115196 | 0 | A | G | 0.43 | 1.49E-02 | 1.11E-01 | 2.69E+00 | 2.74E+00 | 8.92E-01 | no | | intergenic_variant | MODIFIER |
| 5 | 102328071 | AX-170089569 | 0 | C | A | 0.43 | 1.49E-02 | 1.11E-01 | 2.69E+00 | 2.74E+00 | 8.92E-01 | no | | intergenic_variant | MODIFIER |
| 5 | 102541246 | AX-97818006 | 0 | C | A | 0.43 | 1.49E-02 | 1.11E-01 | 2.69E+00 | 2.74E+00 | 8.92E-01 | no | Arhgap24 | intron_variant | MODIFIER |
| 5 | 102546534 | AX-97813369 | 0 | A | G | 0.43 | 1.49E-02 | 1.11E-01 | 2.69E+00 | 2.74E+00 | 8.92E-01 | no | Arhgap24 | intron_variant | MODIFIER |
| 5 | 102550521 | AX-170072919 | 0 | G | C | 0.43 | 1.49E-02 | 1.11E-01 | 2.69E+00 | 2.74E+00 | 8.92E-01 | no | Arhgap24 | intron_variant | MODIFIER |
| 5 | 102550608 | AX-170484338 | 0 | T | C | 0.43 | 1.49E-02 | 1.11E-01 | 2.69E+00 | 2.74E+00 | 8.92E-01 | no | Arhgap24 | intron_variant | MODIFIER |

| | | | | | | | | | | | | | | | |
|---|---|---|---|---|---|---|---|---|---|---|---|---|---|---|---|
| 5 | 102550763 | AX-170119117 | 0 | C | T | 0.43 | 1.49E-02 | 1.11E-01 | 2.69E+00 | 2.74E+00 | 8.92E-01 | no | Arhgap24 | intron_variant | MODIFIER |
| 5 | 102553414 | AX-170273993 | 0 | A | G | 0.43 | 1.49E-02 | 1.11E-01 | 2.69E+00 | 2.74E+00 | 8.92E-01 | no | Arhgap24 | intron_variant | MODIFIER |
| 5 | 102626446 | AX-170025757 | 0 | T | C | 0.43 | 1.49E-02 | 1.11E-01 | 2.69E+00 | 2.74E+00 | 8.92E-01 | no | Arhgap24 | intron_variant | MODIFIER |
| 5 | 102630089 | AX-170025754 | 0 | C | A | 0.43 | 1.49E-02 | 1.11E-01 | 2.69E+00 | 2.74E+00 | 8.92E-01 | no | Arhgap24 | intron_variant | MODIFIER |
| 5 | 102630336 | AX-97844714 | 0 | C | T | 0.43 | 1.49E-02 | 1.11E-01 | 2.69E+00 | 2.74E+00 | 8.92E-01 | no | Arhgap24 | intron_variant | MODIFIER |
| 5 | 102634172 | AX-168373928 | 0 | T | G | 0.43 | 1.49E-02 | 1.11E-01 | 2.69E+00 | 2.74E+00 | 8.92E-01 | no | Arhgap24 | intron_variant | MODIFIER |
| 5 | 102643400 | AX-170343244 | 0 | C | A | 0.43 | 1.49E-02 | 1.11E-01 | 2.69E+00 | 2.74E+00 | 8.92E-01 | no | Arhgap24 | intron_variant | MODIFIER |
| 5 | 102643548 | AX-170106894 | 0 | A | G | 0.43 | 1.49E-02 | 1.11E-01 | 2.69E+00 | 2.74E+00 | 8.92E-01 | no | Arhgap24 | intron_variant | MODIFIER |
| 5 | 102663782 | AX-97782046 | 0 | T | C | 0.43 | 1.49E-02 | 1.11E-01 | 2.69E+00 | 2.74E+00 | 8.92E-01 | no | Arhgap24 | intron_variant | MODIFIER |
| 5 | 102673794 | AX-170025753 | 0 | C | T | 0.43 | 1.49E-02 | 1.11E-01 | 2.69E+00 | 2.74E+00 | 8.92E-01 | no | Arhgap24 | intron_variant | MODIFIER |
| 4 | 148513227 | AX-170209847 | 0 | G | T | 0.096 | 2.69E-02 | 2.00E-01 | 2.67E+00 | 2.74E+00 | 8.92E-01 | no | Mtor | intron_variant | MODIFIER |
| 5 | 127843760 | AX-170313770 | 0 | G | A | 0.412 | 1.18E-02 | 8.20E-02 | 2.61E+00 | 2.72E+00 | 8.92E-01 | no | Tmem132d | intron_variant | MODIFIER |
| 11 | 45023305 | AX-170498523 | 1 | C | T | 0.089 | 2.27E-02 | 1.82E-01 | 2.65E+00 | 2.76E+00 | 8.92E-01 | no | | intergenic_variant | MODIFIER |
| 4 | 77491155 | AX-170172891 | 1 | A | G | 0.321 | 1.53E-02 | 1.09E-01 | 2.64E+00 | 2.72E+00 | 8.92E-01 | no | Ptprd | intron_variant,non_coding_transcript_variant | MODIFIER |
| 9 | 76253463 | AX-170380969 | 1 | C | T | 0.196 | 1.83E-02 | 1.38E-01 | 2.74E+00 | 2.76E+00 | 8.92E-01 | no | Hcrtr2 | intron_variant | MODIFIER |
| 1 | 5780946 | AX-169999861 | 0 | C | T | 0.421 | 1.03E-02 | 6.99E-02 | 2.63E+00 | 2.70E+00 | 8.92E-01 | no | | intergenic_variant | MODIFIER |
| 1 | 6702692 | AX-97839840 | 0 | T | C | 0.421 | 1.03E-02 | 6.99E-02 | 2.63E+00 | 2.70E+00 | 8.92E-01 | no | St18 | intron_variant | MODIFIER |
| 1 | 6815425 | AX-97857148 | 0 | A | G | 0.421 | 1.03E-02 | 6.99E-02 | 2.63E+00 | 2.70E+00 | 8.92E-01 | no | St18 | intron_variant | MODIFIER |
| 7 | 26364928 | AX-170312997 | 0 | C | T | 0.395 | 1.29E-02 | 8.83E-02 | 2.60E+00 | 2.72E+00 | 8.92E-01 | no | Nlrp9c | intron_variant | MODIFIER |
| 7 | 26384144 | AX-170385409 | 0 | C | T | 0.395 | 1.29E-02 | 8.83E-02 | 2.60E+00 | 2.72E+00 | 8.92E-01 | no | Nlrp9c | intron_variant | MODIFIER |
| 7 | 26414486 | AX-97840023 | 0 | T | C | 0.395 | 1.29E-02 | 8.83E-02 | 2.60E+00 | 2.72E+00 | 8.92E-01 | no | | intergenic_variant | MODIFIER |
| 7 | 26801467 | AX-170236302 | 0 | G | T | 0.395 | 1.29E-02 | 8.83E-02 | 2.60E+00 | 2.72E+00 | 8.92E-01 | no | | intergenic_variant | MODIFIER |
| 7 | 26813949 | AX-97855666 | 0 | A | G | 0.395 | 1.29E-02 | 8.83E-02 | 2.60E+00 | 2.72E+00 | 8.92E-01 | no | Cyp2g1 | intron_variant | MODIFIER |
| 7 | 26814245 | AX-170281143 | 0 | A | G | 0.395 | 1.29E-02 | 8.83E-02 | 2.60E+00 | 2.72E+00 | 8.92E-01 | no | Cyp2g1 | missense_variant | MODERATE |
| 7 | 26815812 | AX-170391621 | 0 | T | C | 0.395 | 1.29E-02 | 8.83E-02 | 2.60E+00 | 2.72E+00 | 8.92E-01 | no | Cyp2g1 | intron_variant | MODIFIER |
| 7 | 26832129 | AX-97813928 | 0 | T | C | 0.395 | 1.29E-02 | 8.83E-02 | 2.60E+00 | 2.72E+00 | 8.92E-01 | no | Cyp2a5 | upstream_gene_variant | MODIFIER |
| 7 | 27046879 | AX-170161994 | 0 | C | A | 0.395 | 1.29E-02 | 8.83E-02 | 2.60E+00 | 2.72E+00 | 8.92E-01 | no | Gm4290 | upstream_gene_variant | MODIFIER |
| 7 | 27049027 | AX-170230989 | 0 | C | T | 0.395 | 1.29E-02 | 8.83E-02 | 2.60E+00 | 2.72E+00 | 8.92E-01 | no | Gm4290 | downstream_gene_variant | MODIFIER |
| 7 | 27074594 | AX-170184513 | 0 | A | G | 0.395 | 1.29E-02 | 8.83E-02 | 2.60E+00 | 2.72E+00 | 8.92E-01 | no | | intergenic_variant | MODIFIER |
| 7 | 27092950 | AX-97842754 | 0 | C | A | 0.395 | 1.29E-02 | 8.83E-02 | 2.60E+00 | 2.72E+00 | 8.92E-01 | no | | regulatory_region_variant | MODIFIER |
| 7 | 27119665 | AX-97803879 | 0 | C | T | 0.395 | 1.29E-02 | 8.83E-02 | 2.60E+00 | 2.72E+00 | 8.92E-01 | no | Cyp2f2 | upstream_gene_variant | MODIFIER |
| 7 | 27124208 | AX-97790517 | 0 | G | A | 0.395 | 1.29E-02 | 8.83E-02 | 2.60E+00 | 2.72E+00 | 8.92E-01 | no | Cyp2f2 | intron_variant | MODIFIER |
| 7 | 27129771 | AX-170402112 | 0 | G | T | 0.395 | 1.29E-02 | 8.83E-02 | 2.60E+00 | 2.72E+00 | 8.92E-01 | no | Cyp2f2 | missense_variant | MODERATE |
| 7 | 27130966 | AX-170239226 | 0 | C | T | 0.395 | 1.29E-02 | 8.83E-02 | 2.60E+00 | 2.72E+00 | 8.92E-01 | no | Cyp2f2 | intron_variant | MODIFIER |
| 7 | 27132518 | AX-170392927 | 0 | C | G | 0.395 | 1.29E-02 | 8.83E-02 | 2.60E+00 | 2.72E+00 | 8.92E-01 | no | Cyp2f2 | synonymous_variant | LOW |
| 7 | 27133700 | AX-170432484 | 0 | A | C | 0.395 | 1.29E-02 | 8.83E-02 | 2.60E+00 | 2.72E+00 | 8.92E-01 | no | Cyp2f2 | downstream_gene_variant | MODIFIER |
| 7 | 27145113 | AX-170439202 | 0 | G | A | 0.395 | 1.29E-02 | 8.83E-02 | 2.60E+00 | 2.72E+00 | 8.92E-01 | no | Gm44640 | downstream_gene_variant | MODIFIER |
| 7 | 27319404 | AX-97814091 | 0 | T | C | 0.395 | 1.29E-02 | 8.83E-02 | 2.60E+00 | 2.72E+00 | 8.92E-01 | no | Ltbp4 | splice_region_variant,intron_variant | LOW |
| 7 | 27510402 | AX-170028726 | 0 | G | A | 0.395 | 1.29E-02 | 8.83E-02 | 2.60E+00 | 2.72E+00 | 8.92E-01 | no | Prx | intron_variant | MODIFIER |
| 7 | 27520538 | AX-170235440 | 0 | C | T | 0.395 | 1.29E-02 | 8.83E-02 | 2.60E+00 | 2.72E+00 | 8.92E-01 | no | Prx | downstream_gene_variant | MODIFIER |
| 7 | 27525045 | AX-170371687 | 0 | A | G | 0.395 | 1.29E-02 | 8.83E-02 | 2.60E+00 | 2.72E+00 | 8.92E-01 | no | Hipk4 | upstream_gene_variant | MODIFIER |
| 7 | 27606064 | AX-170146709 | 0 | C | T | 0.395 | 1.29E-02 | 8.83E-02 | 2.60E+00 | 2.72E+00 | 8.92E-01 | no | Akt2 | intron_variant | MODIFIER |
| 18 | 89105363 | AX-170264414 | 0 | A | G | 0.518 | -7.52E-03 | 6.07E-02 | 2.62E+00 | 2.76E+00 | 8.92E-01 | no | Rttn | intron_variant | MODIFIER |
| 8 | 82498888 | AX-97797090 | 0 | A | G | 0.211 | 1.62E-02 | 1.19E-01 | 2.66E+00 | 2.73E+00 | 8.93E-01 | no | | intergenic_variant | MODIFIER |
| 8 | 84032100 | AX-170283422 | 0 | A | G | 0.211 | 1.62E-02 | 1.19E-01 | 2.66E+00 | 2.73E+00 | 8.93E-01 | no | Il27ra | missense_variant | MODERATE |
| 8 | 100647956 | AX-170031381 | 0 | T | C | 0.237 | -1.39E-02 | 1.08E-01 | 2.66E+00 | 2.75E+00 | 8.93E-01 | no | | intergenic_variant | MODIFIER |
| 9 | 86019996 | AX-97783044 | 0 | G | A | 0.14 | -1.70E-02 | 1.21E-01 | 2.63E+00 | 2.72E+00 | 8.93E-01 | no | | intergenic_variant | MODIFIER |
| 11 | 44897623 | AX-170436564 | 2 | A | T | 0.091 | 2.30E-02 | 1.82E-01 | 2.65E+00 | 2.75E+00 | 8.93E-01 | no | Ebf1 | intron_variant | MODIFIER |
| 10 | 6829788 | AX-170091847 | 1 | A | T | 0.196 | -1.44E-02 | 1.05E-01 | 2.64E+00 | 2.73E+00 | 8.93E-01 | no | Oprm1 | intron_variant | MODIFIER |
| 2 | 120240219 | AX-170162159 | 1 | A | G | 0.125 | 2.37E-02 | 1.74E-01 | 2.67E+00 | 2.73E+00 | 8.93E-01 | no | Pla2g4e | intron_variant | MODIFIER |
| 2 | 121315647 | AX-170236876 | 1 | T | C | 0.125 | 2.37E-02 | 1.74E-01 | 2.67E+00 | 2.73E+00 | 8.93E-01 | no | Ppip5k1 | intron_variant | MODIFIER |
| 2 | 121383533 | AX-170396010 | 1 | T | C | 0.125 | 2.37E-02 | 1.74E-01 | 2.67E+00 | 2.73E+00 | 8.93E-01 | no | Strc | upstream_gene_variant | MODIFIER |
| 2 | 142121422 | AX-170239490 | 1 | A | G | 0.125 | 2.37E-02 | 1.74E-01 | 2.67E+00 | 2.73E+00 | 8.93E-01 | no | Macrod2 | intron_variant | MODIFIER |
| 2 | 142432717 | AX-170224195 | 1 | G | C | 0.125 | 2.37E-02 | 1.74E-01 | 2.67E+00 | 2.73E+00 | 8.93E-01 | no | | intergenic_variant | MODIFIER |

| | | | | | | | | | | | | | | | |
|---|---|---|---|---|---|---|---|---|---|---|---|---|---|---|---|
| 2 | 145567315 | AX-170337342 | 1 | A | T | 0.125 | 2.37E-02 | 1.74E-01 | 2.67E+00 | 2.73E+00 | 8.93E-01 | no | Slc24a3 | intron_variant | MODIFIER |
| 20 | 95066181 | AX-170034471 | 0 | G | C | 0.272 | 8.21E-03 | 5.94E-02 | 2.63E+00 | 2.73E+00 | 8.93E-01 | no | Arhgef9 | intron_variant | MODIFIER |
| 18 | 69818580 | AX-170449969 | 0 | A | G | 0.105 | 2.30E-02 | 1.53E-01 | 2.59E+00 | 2.71E+00 | 8.93E-01 | no | | intergenic_variant | MODIFIER |
| 8 | 35806527 | AX-170318623 | 0 | A | G | 0.158 | -1.87E-02 | 1.34E-01 | 2.65E+00 | 2.72E+00 | 8.93E-01 | no | Gm19410 | intron_variant | MODIFIER |
| 2 | 120513302 | AX-170209233 | 0 | G | A | 0.149 | -2.46E-02 | 1.93E-01 | 2.69E+00 | 2.76E+00 | 8.93E-01 | no | Zfp106 | intron_variant | MODIFIER |
| 2 | 121957890 | AX-170237433 | 0 | G | A | 0.149 | -2.46E-02 | 1.93E-01 | 2.69E+00 | 2.76E+00 | 8.93E-01 | no | Ctdspl2 | intron_variant | MODIFIER |
| 2 | 124167169 | AX-170018915 | 0 | G | A | 0.149 | -2.46E-02 | 1.93E-01 | 2.69E+00 | 2.76E+00 | 8.93E-01 | no | Sema6d | intron_variant | MODIFIER |
| 2 | 143769352 | AX-97809218 | 0 | C | T | 0.149 | -2.46E-02 | 1.93E-01 | 2.69E+00 | 2.76E+00 | 8.93E-01 | no | Pcsk2 | intron_variant | MODIFIER |
| 2 | 147325555 | AX-170160197 | 0 | G | A | 0.149 | -2.46E-02 | 1.93E-01 | 2.69E+00 | 2.76E+00 | 8.93E-01 | no | Gm14110 | upstream_gene_variant | MODIFIER |
| 2 | 156243880 | AX-170484486 | 0 | G | A | 0.149 | -2.46E-02 | 1.93E-01 | 2.69E+00 | 2.76E+00 | 8.93E-01 | no | Phf20 | intron_variant | MODIFIER |
| 2 | 156342860 | AX-170181278 | 0 | G | A | 0.149 | -2.46E-02 | 1.93E-01 | 2.69E+00 | 2.76E+00 | 8.93E-01 | no | Cnbd2 | intron_variant | MODIFIER |
| 1 | 114943609 | AX-170363462 | 1 | G | A | 0.161 | 2.73E-02 | 1.89E-01 | 2.62E+00 | 2.71E+00 | 8.93E-01 | no | | intergenic_variant | MODIFIER |
| 18 | 41762109 | AX-170060664 | 1 | G | A | 0.062 | -3.27E-02 | 2.15E-01 | 2.60E+00 | 2.70E+00 | 8.93E-01 | no | AC154172.2 | intron_variant,non_coding_transcript_variant | MODIFIER |
| 1 | 13255895 | AX-170439291 | 1 | C | G | 0.384 | 1.08E-02 | 7.65E-02 | 2.65E+00 | 2.71E+00 | 8.93E-01 | no | Ncoa2 | intron_variant | MODIFIER |
| 8 | 22162516 | AX-170303862 | 0 | G | A | 0.193 | 1.34E-02 | 9.93E-02 | 2.63E+00 | 2.73E+00 | 8.94E-01 | no | Nek3 | intron_variant | MODIFIER |
| 1 | 66814875 | AX-170356084 | 1 | G | T | 0.107 | -2.85E-02 | 2.22E-01 | 2.66E+00 | 2.75E+00 | 8.94E-01 | no | Kansl1l | intron_variant | MODIFIER |
| 1 | 69174115 | AX-97852829 | 1 | G | A | 0.107 | -2.85E-02 | 2.22E-01 | 2.66E+00 | 2.75E+00 | 8.94E-01 | no | | intergenic_variant | MODIFIER |
| 15 | 90447941 | AX-170056282 | 1 | G | A | 0.143 | 2.26E-02 | 1.66E-01 | 2.65E+00 | 2.73E+00 | 8.94E-01 | no | | intergenic_variant | MODIFIER |
| 8 | 32624082 | AX-170475523 | 0 | G | A | 0.149 | -1.62E-02 | 1.19E-01 | 2.65E+00 | 2.73E+00 | 8.94E-01 | no | Nrg1 | intron_variant | MODIFIER |
| 3 | 148531702 | AX-170021893 | 0 | T | G | 0.36 | 9.71E-03 | 6.99E-02 | 2.64E+00 | 2.72E+00 | 8.94E-01 | no | | intergenic_variant | MODIFIER |
| 8 | 23048842 | AX-97808315 | 1 | C | T | 0.161 | -1.56E-02 | 1.15E-01 | 2.65E+00 | 2.73E+00 | 8.94E-01 | no | Ank1 | intron_variant | MODIFIER |
| 10 | 12968313 | AX-170465653 | 0 | G | A | 0.202 | -1.49E-02 | 1.06E-01 | 2.64E+00 | 2.72E+00 | 8.94E-01 | no | Stx11 | upstream_gene_variant | MODIFIER |
| 10 | 13013594 | AX-170430639 | 0 | C | T | 0.202 | -1.49E-02 | 1.06E-01 | 2.64E+00 | 2.72E+00 | 8.94E-01 | no | Sf3b5 | downstream_gene_variant | MODIFIER |
| 10 | 13013811 | AX-170284564 | 0 | C | T | 0.202 | -1.49E-02 | 1.06E-01 | 2.64E+00 | 2.72E+00 | 8.94E-01 | no | Sf3b5 | downstream_gene_variant | MODIFIER |
| 10 | 13186253 | AX-170002766 | 0 | A | T | 0.202 | -1.49E-02 | 1.06E-01 | 2.64E+00 | 2.72E+00 | 8.94E-01 | no | Ltv1 | intron_variant | MODIFIER |
| 10 | 13194246 | AX-170086747 | 0 | A | C | 0.202 | -1.49E-02 | 1.06E-01 | 2.64E+00 | 2.72E+00 | 8.94E-01 | no | Ltv1 | upstream_gene_variant | MODIFIER |
| 10 | 13201897 | AX-170188176 | 0 | A | G | 0.202 | -1.49E-02 | 1.06E-01 | 2.64E+00 | 2.72E+00 | 8.94E-01 | no | Gm47402 | upstream_gene_variant | MODIFIER |
| 10 | 13218484 | AX-170002765 | 0 | C | T | 0.202 | -1.49E-02 | 1.06E-01 | 2.64E+00 | 2.72E+00 | 8.94E-01 | no | Phactr2 | intron_variant | MODIFIER |
| 10 | 13227548 | AX-170442531 | 0 | A | G | 0.202 | -1.49E-02 | 1.06E-01 | 2.64E+00 | 2.72E+00 | 8.94E-01 | no | Phactr2 | intron_variant | MODIFIER |
| 10 | 13231235 | AX-97818633 | 0 | C | T | 0.202 | -1.49E-02 | 1.06E-01 | 2.64E+00 | 2.72E+00 | 8.94E-01 | no | Phactr2 | intron_variant | MODIFIER |
| 10 | 13240952 | AX-170472069 | 0 | C | G | 0.202 | -1.49E-02 | 1.06E-01 | 2.64E+00 | 2.72E+00 | 8.94E-01 | no | Phactr2 | intron_variant | MODIFIER |
| 10 | 13243054 | AX-170154558 | 0 | C | A | 0.202 | -1.49E-02 | 1.06E-01 | 2.64E+00 | 2.72E+00 | 8.94E-01 | no | Phactr2 | intron_variant | MODIFIER |
| 10 | 13285229 | AX-170002760 | 0 | G | T | 0.202 | -1.49E-02 | 1.06E-01 | 2.64E+00 | 2.72E+00 | 8.94E-01 | no | Phactr2 | intron_variant | MODIFIER |
| 10 | 13289893 | AX-170002761 | 0 | C | T | 0.202 | -1.49E-02 | 1.06E-01 | 2.64E+00 | 2.72E+00 | 8.94E-01 | no | Phactr2 | intron_variant | MODIFIER |
| 10 | 13313585 | AX-170141570 | 0 | A | G | 0.202 | -1.49E-02 | 1.06E-01 | 2.64E+00 | 2.72E+00 | 8.94E-01 | no | Phactr2 | intron_variant | MODIFIER |
| 10 | 13317568 | AX-97817067 | 0 | G | A | 0.202 | -1.49E-02 | 1.06E-01 | 2.64E+00 | 2.72E+00 | 8.94E-01 | no | Phactr2 | intron_variant | MODIFIER |
| 10 | 13418473 | AX-97789130 | 0 | C | T | 0.202 | -1.49E-02 | 1.06E-01 | 2.64E+00 | 2.72E+00 | 8.94E-01 | no | Phactr2 | intron_variant | MODIFIER |
| 10 | 13436558 | AX-170520709 | 0 | A | G | 0.202 | -1.49E-02 | 1.06E-01 | 2.64E+00 | 2.72E+00 | 8.94E-01 | no | Phactr2 | intron_variant | MODIFIER |
| 10 | 13438786 | AX-170335573 | 0 | T | C | 0.202 | -1.49E-02 | 1.06E-01 | 2.64E+00 | 2.72E+00 | 8.94E-01 | no | Phactr2 | intron_variant | MODIFIER |
| 10 | 13438981 | AX-97850316 | 0 | A | G | 0.202 | -1.49E-02 | 1.06E-01 | 2.64E+00 | 2.72E+00 | 8.94E-01 | no | Phactr2 | intron_variant | MODIFIER |
| 10 | 13466312 | AX-97820733 | 0 | A | G | 0.202 | -1.49E-02 | 1.06E-01 | 2.64E+00 | 2.72E+00 | 8.94E-01 | no | Phactr2 | intron_variant | MODIFIER |
| 10 | 13658822 | AX-170002772 | 0 | A | G | 0.202 | -1.49E-02 | 1.06E-01 | 2.64E+00 | 2.72E+00 | 8.94E-01 | no | Aig1 | intron_variant | MODIFIER |
| 10 | 13668011 | AX-170002770 | 0 | T | A | 0.202 | -1.49E-02 | 1.06E-01 | 2.64E+00 | 2.72E+00 | 8.94E-01 | no | Aig1 | intron_variant | MODIFIER |
| 10 | 13668122 | AX-97820101 | 0 | T | C | 0.202 | -1.49E-02 | 1.06E-01 | 2.64E+00 | 2.72E+00 | 8.94E-01 | no | Aig1 | intron_variant | MODIFIER |
| 10 | 14097083 | AX-170002783 | 0 | A | G | 0.202 | -1.49E-02 | 1.06E-01 | 2.64E+00 | 2.72E+00 | 8.94E-01 | no | Hivep2 | intron_variant | MODIFIER |
| 10 | 14102302 | AX-97782289 | 0 | A | C | 0.202 | -1.49E-02 | 1.06E-01 | 2.64E+00 | 2.72E+00 | 8.94E-01 | no | Hivep2 | intron_variant | MODIFIER |
| 10 | 14130071 | AX-170398105 | 0 | T | C | 0.202 | -1.49E-02 | 1.06E-01 | 2.64E+00 | 2.72E+00 | 8.94E-01 | no | Hivep2 | synonymous_variant | LOW |
| 10 | 14135197 | AX-170420220 | 0 | T | C | 0.202 | -1.49E-02 | 1.06E-01 | 2.64E+00 | 2.72E+00 | 8.94E-01 | no | Hivep2 | intron_variant | MODIFIER |
| 10 | 14139214 | AX-170002781 | 0 | G | A | 0.202 | -1.49E-02 | 1.06E-01 | 2.64E+00 | 2.72E+00 | 8.94E-01 | no | Hivep2 | intron_variant | MODIFIER |
| 10 | 14139249 | AX-97825520 | 0 | G | A | 0.202 | -1.49E-02 | 1.06E-01 | 2.64E+00 | 2.72E+00 | 8.94E-01 | no | Hivep2 | intron_variant | MODIFIER |
| 10 | 14154021 | AX-170002784 | 0 | A | G | 0.202 | -1.49E-02 | 1.06E-01 | 2.64E+00 | 2.72E+00 | 8.94E-01 | no | Hivep2 | downstream_gene_variant | MODIFIER |
| 10 | 14176132 | AX-97819061 | 0 | C | A | 0.202 | -1.49E-02 | 1.06E-01 | 2.64E+00 | 2.72E+00 | 8.94E-01 | no | | intergenic_variant | MODIFIER |
| 10 | 14188703 | AX-170164058 | 0 | G | T | 0.202 | -1.49E-02 | 1.06E-01 | 2.64E+00 | 2.72E+00 | 8.94E-01 | no | | intergenic_variant | MODIFIER |
| 10 | 14188800 | AX-170497094 | 0 | A | C | 0.202 | -1.49E-02 | 1.06E-01 | 2.64E+00 | 2.72E+00 | 8.94E-01 | no | | intergenic_variant | MODIFIER |

| | | | | | | | | | | | | | | | |
|---|---|---|---|---|---|---|---|---|---|---|---|---|---|---|---|
| 10 | 14189160 | AX-170408331 | 0 | T | C | 0.202 | -1.49E-02 | 1.06E-01 | 2.64E+00 | 2.72E+00 | 8.94E-01 | no | | intergenic_variant | MODIFIER |
| 10 | 14189539 | AX-170091868 | 0 | G | A | 0.202 | -1.49E-02 | 1.06E-01 | 2.64E+00 | 2.72E+00 | 8.94E-01 | no | | regulatory_region_variant | MODIFIER |
| 10 | 14190047 | AX-170535205 | 0 | G | T | 0.202 | -1.49E-02 | 1.06E-01 | 2.64E+00 | 2.72E+00 | 8.94E-01 | no | | intergenic_variant | MODIFIER |
| 10 | 14190091 | AX-170421497 | 0 | A | T | 0.202 | -1.49E-02 | 1.06E-01 | 2.64E+00 | 2.72E+00 | 8.94E-01 | no | | intergenic_variant | MODIFIER |
| 10 | 14193374 | AX-170399639 | 0 | T | C | 0.202 | -1.49E-02 | 1.06E-01 | 2.64E+00 | 2.72E+00 | 8.94E-01 | no | Gm48843 | upstream_gene_variant | MODIFIER |
| 10 | 14193864 | AX-170475923 | 0 | A | C | 0.202 | -1.49E-02 | 1.06E-01 | 2.64E+00 | 2.72E+00 | 8.94E-01 | no | Gm48843 | upstream_gene_variant | MODIFIER |
| 10 | 14209875 | AX-97814824 | 0 | T | C | 0.202 | -1.49E-02 | 1.06E-01 | 2.64E+00 | 2.72E+00 | 8.94E-01 | no | Gm48843 | downstream_gene_variant | MODIFIER |
| 10 | 14238044 | AX-97826798 | 0 | A | G | 0.202 | -1.49E-02 | 1.06E-01 | 2.64E+00 | 2.72E+00 | 8.94E-01 | no | Gm24343 | downstream_gene_variant | MODIFIER |
| 10 | 14240279 | AX-170109405 | 0 | C | G | 0.202 | -1.49E-02 | 1.06E-01 | 2.64E+00 | 2.72E+00 | 8.94E-01 | no | Gm48845 | intron_variant,non_coding_transcript_variant | MODIFIER |
| 10 | 14268017 | AX-170086749 | 0 | C | T | 0.202 | -1.49E-02 | 1.06E-01 | 2.64E+00 | 2.72E+00 | 8.94E-01 | no | Gm48845 | downstream_gene_variant | MODIFIER |
| 10 | 14282876 | AX-170185698 | 0 | C | G | 0.202 | -1.49E-02 | 1.06E-01 | 2.64E+00 | 2.72E+00 | 8.94E-01 | no | | intergenic_variant | MODIFIER |
| 10 | 14336221 | AX-170123952 | 0 | G | A | 0.202 | -1.49E-02 | 1.06E-01 | 2.64E+00 | 2.72E+00 | 8.94E-01 | no | Gm48854 | intron_variant,non_coding_transcript_variant | MODIFIER |
| 10 | 14343742 | AX-97850227 | 0 | C | A | 0.202 | -1.49E-02 | 1.06E-01 | 2.64E+00 | 2.72E+00 | 8.94E-01 | no | | intergenic_variant | MODIFIER |
| 10 | 14347898 | AX-170114388 | 0 | T | C | 0.202 | -1.49E-02 | 1.06E-01 | 2.64E+00 | 2.72E+00 | 8.94E-01 | no | | intergenic_variant | MODIFIER |
| 10 | 14364186 | AX-170431036 | 0 | G | C | 0.202 | -1.49E-02 | 1.06E-01 | 2.64E+00 | 2.72E+00 | 8.94E-01 | no | | intergenic_variant | MODIFIER |
| 10 | 14404507 | AX-170405617 | 0 | T | C | 0.202 | -1.49E-02 | 1.06E-01 | 2.64E+00 | 2.72E+00 | 8.94E-01 | no | Adgrg6 | 3_prime_UTR_variant | MODIFIER |
| 10 | 14404585 | AX-170374731 | 0 | T | C | 0.202 | -1.49E-02 | 1.06E-01 | 2.64E+00 | 2.72E+00 | 8.94E-01 | no | Adgrg6 | 3_prime_UTR_variant | MODIFIER |
| 10 | 14407396 | AX-170091869 | 0 | C | A | 0.202 | -1.49E-02 | 1.06E-01 | 2.64E+00 | 2.72E+00 | 8.94E-01 | no | Adgrg6 | intron_variant | MODIFIER |
| 10 | 14408160 | AX-170100777 | 0 | T | C | 0.202 | -1.49E-02 | 1.06E-01 | 2.64E+00 | 2.72E+00 | 8.94E-01 | no | Adgrg6 | intron_variant | MODIFIER |
| 10 | 14733127 | AX-170323663 | 0 | C | T | 0.202 | -1.49E-02 | 1.06E-01 | 2.64E+00 | 2.72E+00 | 8.94E-01 | no | Nmbr | intron_variant,non_coding_transcript_variant | MODIFIER |
| 10 | 14743233 | AX-170220977 | 0 | T | G | 0.202 | -1.49E-02 | 1.06E-01 | 2.64E+00 | 2.72E+00 | 8.94E-01 | no | Gm8355 | downstream_gene_variant | MODIFIER |
| 10 | 14755087 | AX-97814011 | 0 | T | G | 0.202 | -1.49E-02 | 1.06E-01 | 2.64E+00 | 2.72E+00 | 8.94E-01 | no | B230364G03Rik | downstream_gene_variant | MODIFIER |
| 10 | 14756577 | AX-170273117 | 0 | A | G | 0.202 | -1.49E-02 | 1.06E-01 | 2.64E+00 | 2.72E+00 | 8.94E-01 | no | Nmbr | upstream_gene_variant | MODIFIER |
| 10 | 14758420 | AX-170384775 | 0 | A | G | 0.202 | -1.49E-02 | 1.06E-01 | 2.64E+00 | 2.72E+00 | 8.94E-01 | no | Nmbr | upstream_gene_variant | MODIFIER |
| 3 | 146641922 | AX-170021856 | 0 | C | T | 0.342 | 1.00E-02 | 7.40E-02 | 2.66E+00 | 2.73E+00 | 8.94E-01 | no | 4930503B20Rik | downstream_gene_variant | MODIFIER |
| 3 | 146648433 | AX-97821240 | 0 | T | C | 0.342 | 1.00E-02 | 7.40E-02 | 2.66E+00 | 2.73E+00 | 8.94E-01 | no | 4930503B20Rik | intron_variant | MODIFIER |
| 3 | 146677782 | AX-97797610 | 0 | A | T | 0.342 | 1.00E-02 | 7.40E-02 | 2.66E+00 | 2.73E+00 | 8.94E-01 | no | Samd13 | intron_variant | MODIFIER |
| 3 | 146708632 | AX-97783401 | 0 | G | A | 0.342 | 1.00E-02 | 7.40E-02 | 2.66E+00 | 2.73E+00 | 8.94E-01 | no | | intergenic_variant | MODIFIER |
| 3 | 146741102 | AX-170021851 | 0 | T | C | 0.342 | 1.00E-02 | 7.40E-02 | 2.66E+00 | 2.73E+00 | 8.94E-01 | no | Prkacb | intron_variant | MODIFIER |
| 3 | 146759575 | AX-170504126 | 0 | T | C | 0.342 | 1.00E-02 | 7.40E-02 | 2.66E+00 | 2.73E+00 | 8.94E-01 | no | Prkacb | intron_variant | MODIFIER |
| 3 | 146766404 | AX-170390662 | 0 | A | C | 0.342 | 1.00E-02 | 7.40E-02 | 2.66E+00 | 2.73E+00 | 8.94E-01 | no | Prkacb | intron_variant | MODIFIER |
| 3 | 146766674 | AX-170374264 | 0 | A | G | 0.342 | 1.00E-02 | 7.40E-02 | 2.66E+00 | 2.73E+00 | 8.94E-01 | no | Prkacb | intron_variant | MODIFIER |
| 3 | 146774411 | AX-97854435 | 0 | G | A | 0.342 | 1.00E-02 | 7.40E-02 | 2.66E+00 | 2.73E+00 | 8.94E-01 | no | Prkacb | intron_variant | MODIFIER |
| 3 | 146803983 | AX-170382784 | 0 | A | G | 0.342 | 1.00E-02 | 7.40E-02 | 2.66E+00 | 2.73E+00 | 8.94E-01 | no | Prkacb | intron_variant | MODIFIER |
| 3 | 146804725 | AX-170406637 | 0 | C | T | 0.342 | 1.00E-02 | 7.40E-02 | 2.66E+00 | 2.73E+00 | 8.94E-01 | no | Prkacb | intron_variant | MODIFIER |
| 3 | 146845062 | AX-97806850 | 0 | A | G | 0.342 | 1.00E-02 | 7.40E-02 | 2.66E+00 | 2.73E+00 | 8.94E-01 | no | | intergenic_variant | MODIFIER |
| 3 | 146850819 | AX-170150787 | 0 | C | T | 0.342 | 1.00E-02 | 7.40E-02 | 2.66E+00 | 2.73E+00 | 8.94E-01 | no | Ttll7 | upstream_gene_variant | MODIFIER |
| 3 | 146852320 | AX-170293371 | 0 | T | C | 0.342 | 1.00E-02 | 7.40E-02 | 2.66E+00 | 2.73E+00 | 8.94E-01 | no | Ttll7 | upstream_gene_variant | MODIFIER |
| 3 | 146853327 | AX-170258244 | 0 | T | G | 0.342 | 1.00E-02 | 7.40E-02 | 2.66E+00 | 2.73E+00 | 8.94E-01 | no | Ttll7 | intron_variant | MODIFIER |
| 3 | 146857667 | AX-170068368 | 0 | C | A | 0.342 | 1.00E-02 | 7.40E-02 | 2.66E+00 | 2.73E+00 | 8.94E-01 | no | Ttll7 | intron_variant | MODIFIER |
| 3 | 146857760 | AX-170068367 | 0 | T | C | 0.342 | 1.00E-02 | 7.40E-02 | 2.66E+00 | 2.73E+00 | 8.94E-01 | no | Ttll7 | intron_variant | MODIFIER |
| 3 | 147136059 | AX-170159556 | 0 | T | C | 0.342 | 1.00E-02 | 7.40E-02 | 2.66E+00 | 2.73E+00 | 8.94E-01 | no | | intergenic_variant | MODIFIER |
| 3 | 147176275 | AX-170155642 | 0 | A | C | 0.342 | 1.00E-02 | 7.40E-02 | 2.66E+00 | 2.73E+00 | 8.94E-01 | no | | intergenic_variant | MODIFIER |
| 3 | 147207051 | AX-170382173 | 0 | C | T | 0.342 | 1.00E-02 | 7.40E-02 | 2.66E+00 | 2.73E+00 | 8.94E-01 | no | | intergenic_variant | MODIFIER |
| 3 | 147211244 | AX-170299609 | 0 | G | A | 0.342 | 1.00E-02 | 7.40E-02 | 2.66E+00 | 2.73E+00 | 8.94E-01 | no | | regulatory_region_variant | MODIFIER |
| 3 | 147211633 | AX-170240868 | 0 | A | G | 0.342 | 1.00E-02 | 7.40E-02 | 2.66E+00 | 2.73E+00 | 8.94E-01 | no | | regulatory_region_variant | MODIFIER |
| 3 | 147242640 | AX-97805920 | 0 | C | T | 0.342 | 1.00E-02 | 7.40E-02 | 2.66E+00 | 2.73E+00 | 8.94E-01 | no | | intergenic_variant | MODIFIER |
| 3 | 147244593 | AX-170173154 | 0 | A | G | 0.342 | 1.00E-02 | 7.40E-02 | 2.66E+00 | 2.73E+00 | 8.94E-01 | no | | intergenic_variant | MODIFIER |
| 3 | 147249739 | AX-170266964 | 0 | A | C | 0.342 | 1.00E-02 | 7.40E-02 | 2.66E+00 | 2.73E+00 | 8.94E-01 | no | | intergenic_variant | MODIFIER |
| 3 | 147287936 | AX-170402731 | 0 | C | T | 0.342 | 1.00E-02 | 7.40E-02 | 2.66E+00 | 2.73E+00 | 8.94E-01 | no | | regulatory_region_variant | MODIFIER |
| 3 | 147341745 | AX-97790595 | 0 | C | T | 0.342 | 1.00E-02 | 7.40E-02 | 2.66E+00 | 2.73E+00 | 8.94E-01 | no | | intergenic_variant | MODIFIER |
| 3 | 147354715 | AX-170162378 | 0 | A | G | 0.342 | 1.00E-02 | 7.40E-02 | 2.66E+00 | 2.73E+00 | 8.94E-01 | no | | intergenic_variant | MODIFIER |
| 3 | 147387547 | AX-170181572 | 0 | T | C | 0.342 | 1.00E-02 | 7.40E-02 | 2.66E+00 | 2.73E+00 | 8.94E-01 | no | Gm36831 | upstream_gene_variant | MODIFIER |
| 3 | 147387787 | AX-170250819 | 0 | A | G | 0.342 | 1.00E-02 | 7.40E-02 | 2.66E+00 | 2.73E+00 | 8.94E-01 | no | Gm36831 | upstream_gene_variant | MODIFIER |

| | | | | | | | | | | | | | | | |
|---|---|---|---|---|---|---|---|---|---|---|---|---|---|---|---|
| 3 | 147388056 | AX-170419573 | 0 | G | A | 0.342 | 1.00E-02 | 7.40E-02 | 2.66E+00 | 2.73E+00 | 8.94E-01 | no | Gm36831 | upstream_gene_variant | MODIFIER |
| 3 | 147388137 | AX-170526226 | 0 | A | G | 0.342 | 1.00E-02 | 7.40E-02 | 2.66E+00 | 2.73E+00 | 8.94E-01 | no | Gm36831 | upstream_gene_variant | MODIFIER |
| 3 | 147398279 | AX-170365996 | 0 | T | A | 0.342 | 1.00E-02 | 7.40E-02 | 2.66E+00 | 2.73E+00 | 8.94E-01 | no | Gm36831 | intron_variant,non_coding_transcript_variant | MODIFIER |
| 3 | 147421101 | AX-170035666 | 0 | G | A | 0.342 | 1.00E-02 | 7.40E-02 | 2.66E+00 | 2.73E+00 | 8.94E-01 | no | | intergenic_variant | MODIFIER |
| 3 | 147424606 | AX-97853329 | 0 | C | T | 0.342 | 1.00E-02 | 7.40E-02 | 2.66E+00 | 2.73E+00 | 8.94E-01 | no | Gm37041 | upstream_gene_variant | MODIFIER |
| 3 | 147434339 | AX-170143124 | 0 | G | C | 0.342 | 1.00E-02 | 7.40E-02 | 2.66E+00 | 2.73E+00 | 8.94E-01 | no | Gm37041 | downstream_gene_variant | MODIFIER |
| 3 | 147477092 | AX-170408799 | 0 | A | G | 0.342 | 1.00E-02 | 7.40E-02 | 2.66E+00 | 2.73E+00 | 8.94E-01 | no | | intergenic_variant | MODIFIER |
| 3 | 147477595 | AX-97831640 | 0 | T | C | 0.342 | 1.00E-02 | 7.40E-02 | 2.66E+00 | 2.73E+00 | 8.94E-01 | no | | intergenic_variant | MODIFIER |
| 3 | 147479663 | AX-170128478 | 0 | G | C | 0.342 | 1.00E-02 | 7.40E-02 | 2.66E+00 | 2.73E+00 | 8.94E-01 | no | | intergenic_variant | MODIFIER |
| 3 | 147480872 | AX-170520987 | 0 | C | G | 0.342 | 1.00E-02 | 7.40E-02 | 2.66E+00 | 2.73E+00 | 8.94E-01 | no | | intergenic_variant | MODIFIER |
| 3 | 147515450 | AX-170500294 | 0 | T | A | 0.342 | 1.00E-02 | 7.40E-02 | 2.66E+00 | 2.73E+00 | 8.94E-01 | no | | intergenic_variant | MODIFIER |
| 3 | 147525870 | AX-170136682 | 0 | A | G | 0.342 | 1.00E-02 | 7.40E-02 | 2.66E+00 | 2.73E+00 | 8.94E-01 | no | | intergenic_variant | MODIFIER |
| 3 | 147543351 | AX-170021869 | 0 | A | G | 0.342 | 1.00E-02 | 7.40E-02 | 2.66E+00 | 2.73E+00 | 8.94E-01 | no | | intergenic_variant | MODIFIER |
| 3 | 147590209 | AX-97852880 | 0 | A | G | 0.342 | 1.00E-02 | 7.40E-02 | 2.66E+00 | 2.73E+00 | 8.94E-01 | no | | intergenic_variant | MODIFIER |
| 3 | 147607166 | AX-170021871 | 0 | T | C | 0.342 | 1.00E-02 | 7.40E-02 | 2.66E+00 | 2.73E+00 | 8.94E-01 | no | | intergenic_variant | MODIFIER |
| 3 | 147673339 | AX-97823172 | 0 | T | C | 0.342 | 1.00E-02 | 7.40E-02 | 2.66E+00 | 2.73E+00 | 8.94E-01 | no | | intergenic_variant | MODIFIER |
| 3 | 147676006 | AX-170153395 | 0 | T | C | 0.342 | 1.00E-02 | 7.40E-02 | 2.66E+00 | 2.73E+00 | 8.94E-01 | no | | intergenic_variant | MODIFIER |
| 3 | 147678570 | AX-170298338 | 0 | T | C | 0.342 | 1.00E-02 | 7.40E-02 | 2.66E+00 | 2.73E+00 | 8.94E-01 | no | | intergenic_variant | MODIFIER |
| 3 | 147706850 | AX-170515939 | 0 | T | A | 0.342 | 1.00E-02 | 7.40E-02 | 2.66E+00 | 2.73E+00 | 8.94E-01 | no | | regulatory_region_variant | MODIFIER |
| 3 | 147722435 | AX-97800211 | 0 | G | T | 0.342 | 1.00E-02 | 7.40E-02 | 2.66E+00 | 2.73E+00 | 8.94E-01 | no | | intergenic_variant | MODIFIER |
| 3 | 147728952 | AX-170522182 | 0 | C | T | 0.342 | 1.00E-02 | 7.40E-02 | 2.66E+00 | 2.73E+00 | 8.94E-01 | no | | intergenic_variant | MODIFIER |
| 3 | 147747048 | AX-170507009 | 0 | G | A | 0.342 | 1.00E-02 | 7.40E-02 | 2.66E+00 | 2.73E+00 | 8.94E-01 | no | | intergenic_variant | MODIFIER |
| 3 | 147785676 | AX-170068386 | 0 | G | A | 0.342 | 1.00E-02 | 7.40E-02 | 2.66E+00 | 2.73E+00 | 8.94E-01 | no | | intergenic_variant | MODIFIER |
| 3 | 147791611 | AX-97790840 | 0 | G | A | 0.342 | 1.00E-02 | 7.40E-02 | 2.66E+00 | 2.73E+00 | 8.94E-01 | no | | intergenic_variant | MODIFIER |
| 3 | 147794728 | AX-170397490 | 0 | T | G | 0.342 | 1.00E-02 | 7.40E-02 | 2.66E+00 | 2.73E+00 | 8.94E-01 | no | | intergenic_variant | MODIFIER |
| 3 | 147896625 | AX-170213570 | 0 | T | A | 0.342 | 1.00E-02 | 7.40E-02 | 2.66E+00 | 2.73E+00 | 8.94E-01 | no | | intergenic_variant | MODIFIER |
| 3 | 147961791 | AX-170068391 | 0 | G | C | 0.342 | 1.00E-02 | 7.40E-02 | 2.66E+00 | 2.73E+00 | 8.94E-01 | no | | intergenic_variant | MODIFIER |
| 3 | 147966362 | AX-170068390 | 0 | T | C | 0.342 | 1.00E-02 | 7.40E-02 | 2.66E+00 | 2.73E+00 | 8.94E-01 | no | | regulatory_region_variant | MODIFIER |
| 3 | 147981546 | AX-97802028 | 0 | G | T | 0.342 | 1.00E-02 | 7.40E-02 | 2.66E+00 | 2.73E+00 | 8.94E-01 | no | | intergenic_variant | MODIFIER |
| 3 | 147983129 | AX-170237043 | 0 | A | G | 0.342 | 1.00E-02 | 7.40E-02 | 2.66E+00 | 2.73E+00 | 8.94E-01 | no | | intergenic_variant | MODIFIER |
| 3 | 148015258 | AX-170021884 | 0 | T | C | 0.342 | 1.00E-02 | 7.40E-02 | 2.66E+00 | 2.73E+00 | 8.94E-01 | no | | intergenic_variant | MODIFIER |
| 3 | 148031026 | AX-170102311 | 0 | G | A | 0.342 | 1.00E-02 | 7.40E-02 | 2.66E+00 | 2.73E+00 | 8.94E-01 | no | | intergenic_variant | MODIFIER |
| 3 | 148038522 | AX-170409941 | 0 | A | C | 0.342 | 1.00E-02 | 7.40E-02 | 2.66E+00 | 2.73E+00 | 8.94E-01 | no | | intergenic_variant | MODIFIER |
| 3 | 148040635 | AX-170021876 | 0 | A | G | 0.342 | 1.00E-02 | 7.40E-02 | 2.66E+00 | 2.73E+00 | 8.94E-01 | no | | intergenic_variant | MODIFIER |
| 3 | 148042383 | AX-97794768 | 0 | A | G | 0.342 | 1.00E-02 | 7.40E-02 | 2.66E+00 | 2.73E+00 | 8.94E-01 | no | | intergenic_variant | MODIFIER |
| 3 | 148043983 | AX-170159027 | 0 | T | C | 0.342 | 1.00E-02 | 7.40E-02 | 2.66E+00 | 2.73E+00 | 8.94E-01 | no | | intergenic_variant | MODIFIER |
| 1 | 41629826 | AX-97799877 | 2 | G | A | 0.418 | -1.00E-02 | 7.58E-02 | 2.65E+00 | 2.74E+00 | 8.94E-01 | no | | intergenic_variant | MODIFIER |
| 12 | 101054122 | AX-97822194 | 0 | T | C | 0.061 | 2.75E-02 | 2.05E-01 | 2.65E+00 | 2.73E+00 | 8.94E-01 | no | Ppp4r3a | intron_variant | MODIFIER |
| 12 | 101629804 | AX-97853372 | 0 | T | C | 0.061 | 2.75E-02 | 2.05E-01 | 2.65E+00 | 2.73E+00 | 8.94E-01 | no | Catsperb | downstream_gene_variant | MODIFIER |
| 12 | 101769400 | AX-170245668 | 0 | G | A | 0.061 | 2.75E-02 | 2.05E-01 | 2.65E+00 | 2.73E+00 | 8.94E-01 | no | Fbln5 | intron_variant | MODIFIER |
| 12 | 101924382 | AX-170243929 | 0 | G | A | 0.061 | 2.75E-02 | 2.05E-01 | 2.65E+00 | 2.73E+00 | 8.94E-01 | no | Atxn3 | intron_variant | MODIFIER |
| 12 | 101942027 | AX-170141762 | 0 | A | G | 0.061 | 2.75E-02 | 2.05E-01 | 2.65E+00 | 2.73E+00 | 8.94E-01 | no | Atxn3 | intron_variant | MODIFIER |
| 12 | 101991855 | AX-170049985 | 0 | A | G | 0.061 | 2.75E-02 | 2.05E-01 | 2.65E+00 | 2.73E+00 | 8.94E-01 | no | Cpsf2 | intron_variant | MODIFIER |
| 12 | 101992115 | AX-97793260 | 0 | T | C | 0.061 | 2.75E-02 | 2.05E-01 | 2.65E+00 | 2.73E+00 | 8.94E-01 | no | Cpsf2 | intron_variant | MODIFIER |
| 12 | 101992482 | AX-170049983 | 0 | C | T | 0.061 | 2.75E-02 | 2.05E-01 | 2.65E+00 | 2.73E+00 | 8.94E-01 | no | Cpsf2 | intron_variant | MODIFIER |
| 12 | 101994976 | AX-170190053 | 0 | G | A | 0.061 | 2.75E-02 | 2.05E-01 | 2.65E+00 | 2.73E+00 | 8.94E-01 | no | Cpsf2 | intron_variant | MODIFIER |
| 12 | 101995063 | AX-170290575 | 0 | C | T | 0.061 | 2.75E-02 | 2.05E-01 | 2.65E+00 | 2.73E+00 | 8.94E-01 | no | Cpsf2 | intron_variant | MODIFIER |
| 12 | 101997895 | AX-170156932 | 0 | T | A | 0.061 | 2.75E-02 | 2.05E-01 | 2.65E+00 | 2.73E+00 | 8.94E-01 | no | Cpsf2 | intron_variant | MODIFIER |
| 12 | 101999224 | AX-170506136 | 0 | G | A | 0.061 | 2.75E-02 | 2.05E-01 | 2.65E+00 | 2.73E+00 | 8.94E-01 | no | Cpsf2 | intron_variant | MODIFIER |
| 12 | 102216427 | AX-97809558 | 0 | A | G | 0.061 | 2.75E-02 | 2.05E-01 | 2.65E+00 | 2.73E+00 | 8.94E-01 | no | Slc24a4 | intron_variant | MODIFIER |
| 12 | 102222778 | AX-168380870 | 0 | A | G | 0.061 | 2.75E-02 | 2.05E-01 | 2.65E+00 | 2.73E+00 | 8.94E-01 | no | Slc24a4 | synonymous_variant | LOW |
| 12 | 102229211 | AX-170049992 | 0 | C | T | 0.061 | 2.75E-02 | 2.05E-01 | 2.65E+00 | 2.73E+00 | 8.94E-01 | no | Slc24a4 | intron_variant | MODIFIER |
| 12 | 102229360 | AX-170403313 | 0 | G | A | 0.061 | 2.75E-02 | 2.05E-01 | 2.65E+00 | 2.73E+00 | 8.94E-01 | no | Slc24a4 | intron_variant | MODIFIER |
| 12 | 102249815 | AX-97836295 | 0 | T | C | 0.061 | 2.75E-02 | 2.05E-01 | 2.65E+00 | 2.73E+00 | 8.94E-01 | no | Slc24a4 | intron_variant | MODIFIER |

| | | | | | | | | | | | | | | | |
|---|---|---|---|---|---|---|---|---|---|---|---|---|---|---|---|
| 12 | 102418004 | AX-170155797 | 0 | A | T | 0.061 | 2.75E-02 | 2.05E-01 | 2.65E+00 | 2.73E+00 | 8.94E-01 | no | Lgmn | intron_variant | MODIFIER |
| 10 | 8646949 | AX-170521678 | 1 | C | T | 0.232 | 1.20E-02 | 9.65E-02 | 2.66E+00 | 2.76E+00 | 8.94E-01 | no | Gm24374 | downstream_gene_variant | MODIFIER |
| 16 | 89455949 | AX-97807473 | 1 | C | T | 0.473 | -1.03E-02 | 7.93E-02 | 2.67E+00 | 2.74E+00 | 8.95E-01 | no | Gm23655 | upstream_gene_variant | MODIFIER |
| 4 | 128420622 | AX-170495568 | 0 | A | G | 0.158 | 1.56E-02 | 1.08E-01 | 2.62E+00 | 2.71E+00 | 8.95E-01 | no | Csmd2 | downstream_gene_variant | MODIFIER |
| 4 | 128818723 | AX-97799323 | 0 | A | G | 0.158 | 1.56E-02 | 1.08E-01 | 2.62E+00 | 2.71E+00 | 8.95E-01 | no | | intergenic_variant | MODIFIER |
| 7 | 138067968 | AX-170278115 | 0 | G | T | 0.289 | -1.09E-02 | 7.97E-02 | 2.62E+00 | 2.73E+00 | 8.95E-01 | no | | intergenic_variant | MODIFIER |
| 18 | 35989633 | AX-170015531 | 0 | G | A | 0.07 | 2.82E-02 | 2.09E-01 | 2.62E+00 | 2.73E+00 | 8.95E-01 | no | Psd2 | intron_variant | MODIFIER |
| 18 | 40416715 | AX-170060650 | 0 | C | T | 0.07 | 2.82E-02 | 2.09E-01 | 2.62E+00 | 2.73E+00 | 8.95E-01 | no | Kctd16 | intron_variant | MODIFIER |
| 18 | 43089474 | AX-170015614 | 0 | G | A | 0.07 | 2.82E-02 | 2.09E-01 | 2.62E+00 | 2.73E+00 | 8.95E-01 | no | AC099605.1 | intron_variant,non_coding_transcript_variant | MODIFIER |
| 1 | 25881990 | AX-170439818 | 1 | A | G | 0.393 | -1.04E-02 | 8.30E-02 | 2.67E+00 | 2.76E+00 | 8.95E-01 | no | | intergenic_variant | MODIFIER |
| 8 | 51798946 | AX-170030687 | 1 | C | T | 0.152 | -2.08E-02 | 1.50E-01 | 2.66E+00 | 2.71E+00 | 8.95E-01 | no | | intergenic_variant | MODIFIER |
| 1 | 5815878 | AX-97803635 | 0 | T | C | 0.088 | 2.21E-02 | 1.61E-01 | 2.60E+00 | 2.73E+00 | 8.95E-01 | no | | intergenic_variant | MODIFIER |
| 8 | 53259143 | AX-170030730 | 1 | T | A | 0.152 | -2.11E-02 | 1.50E-01 | 2.66E+00 | 2.70E+00 | 8.95E-01 | no | | intergenic_variant | MODIFIER |
| 6 | 147161229 | AX-170207526 | 0 | C | A | 0.079 | -2.96E-02 | 2.25E-01 | 2.62E+00 | 2.74E+00 | 8.95E-01 | no | | intergenic_variant | MODIFIER |
| 5 | 102609195 | AX-170286793 | 1 | G | C | 0.429 | 1.44E-02 | 1.11E-01 | 2.69E+00 | 2.74E+00 | 8.95E-01 | no | Arhgap24 | intron_variant | MODIFIER |
| 2 | 161881441 | AX-97849738 | 0 | A | G | 0.158 | 2.12E-02 | 1.65E-01 | 2.63E+00 | 2.74E+00 | 8.95E-01 | no | Ptprt | intron_variant | MODIFIER |
| 9 | 58703876 | AX-170114210 | 0 | G | A | 0.219 | 1.55E-02 | 1.19E-01 | 2.68E+00 | 2.74E+00 | 8.95E-01 | no | Rec114 | intron_variant | MODIFIER |
| 8 | 102535246 | AX-170138405 | 2 | A | G | 0.218 | 1.45E-02 | 1.13E-01 | 2.67E+00 | 2.74E+00 | 8.95E-01 | no | Gm45422 | downstream_gene_variant | MODIFIER |
| 15 | 101718322 | AX-170237493 | 1 | C | G | 0.134 | 2.17E-02 | 1.64E-01 | 2.67E+00 | 2.73E+00 | 8.95E-01 | no | | intergenic_variant | MODIFIER |
| 18 | 80137959 | AX-170159544 | 1 | G | A | 0.393 | -1.14E-02 | 8.49E-02 | 2.64E+00 | 2.73E+00 | 8.96E-01 | no | Adnp2 | intron_variant | MODIFIER |
| 13 | 42519037 | AX-97834589 | 0 | A | G | 0.491 | 9.07E-03 | 6.28E-02 | 2.60E+00 | 2.71E+00 | 8.96E-01 | no | | intergenic_variant | MODIFIER |
| 10 | 10531025 | AX-170389988 | 1 | G | A | 0.223 | 1.32E-02 | 1.07E-01 | 2.68E+00 | 2.76E+00 | 8.96E-01 | no | | intergenic_variant | MODIFIER |
| 2 | 99273156 | AX-170497773 | 2 | A | G | 0.055 | -3.85E-02 | 2.99E-01 | 2.71E+00 | 2.74E+00 | 8.96E-01 | no | | intergenic_variant | MODIFIER |
| 20 | 143836063 | AX-97815653 | 0 | G | A | 0.149 | 1.76E-02 | 1.36E-01 | 2.65E+00 | 2.74E+00 | 8.96E-01 | no | | intergenic_variant | MODIFIER |
| 20 | 115474221 | AX-170085077 | 0 | C | T | 0.316 | 7.29E-03 | 5.34E-02 | 2.61E+00 | 2.73E+00 | 8.96E-01 | no | | intergenic_variant | MODIFIER |
| 4 | 83432135 | AX-170216412 | 0 | A | T | 0.237 | 1.37E-02 | 9.46E-02 | 2.61E+00 | 2.71E+00 | 8.96E-01 | no | Snapc3 | intron_variant | MODIFIER |
| 3 | 148721156 | AX-170403624 | 0 | G | A | 0.333 | 9.50E-03 | 7.12E-02 | 2.64E+00 | 2.73E+00 | 8.96E-01 | no | | intergenic_variant | MODIFIER |
| 13 | 37795545 | AX-170454532 | 1 | G | C | 0.179 | 1.38E-02 | 1.16E-01 | 2.70E+00 | 2.78E+00 | 8.96E-01 | no | Rreb1 | intron_variant | MODIFIER |
| 20 | 87425610 | AX-170125014 | 1 | T | C | 0.259 | -7.76E-03 | 6.14E-02 | 2.64E+00 | 2.74E+00 | 8.96E-01 | no | Il1rapl1 | intron_variant | MODIFIER |
| 1 | 174560886 | AX-170129357 | 1 | T | C | 0.446 | -7.44E-03 | 6.48E-02 | 2.71E+00 | 2.80E+00 | 8.96E-01 | no | Fmn2 | intron_variant | MODIFIER |
| 20 | 111521980 | AX-170085020 | 0 | T | A | 0.307 | -6.81E-03 | 5.34E-02 | 2.63E+00 | 2.74E+00 | 8.96E-01 | no | Rps6ka6 | intron_variant | MODIFIER |
| 15 | 82921731 | AX-97804851 | 0 | G | A | 0.553 | 1.19E-02 | 8.18E-02 | 2.63E+00 | 2.70E+00 | 8.96E-01 | no | Tcf20 | intron_variant | MODIFIER |
| 6 | 118549600 | AX-170139453 | 0 | A | G | 0.079 | -2.66E-02 | 2.11E-01 | 2.68E+00 | 2.75E+00 | 8.96E-01 | no | Ankrd26 | intron_variant | MODIFIER |
| 6 | 145175378 | AX-170513809 | 0 | A | T | 0.079 | -2.66E-02 | 2.11E-01 | 2.68E+00 | 2.75E+00 | 8.96E-01 | no | Lrmp | downstream_gene_variant | MODIFIER |
| 15 | 101113046 | AX-170012396 | 1 | T | C | 0.134 | 2.11E-02 | 1.61E-01 | 2.66E+00 | 2.73E+00 | 8.96E-01 | no | Ankrd33 | upstream_gene_variant | MODIFIER |
| 18 | 65029759 | AX-97809033 | 0 | A | C | 0.412 | 1.00E-02 | 8.26E-02 | 2.71E+00 | 2.77E+00 | 8.96E-01 | no | Nedd4l | intron_variant | MODIFIER |
| 18 | 65033110 | AX-170385802 | 0 | A | G | 0.412 | 1.00E-02 | 8.26E-02 | 2.71E+00 | 2.77E+00 | 8.96E-01 | no | Nedd4l | intron_variant | MODIFIER |
| 18 | 65057402 | AX-170088361 | 0 | A | C | 0.412 | 1.00E-02 | 8.26E-02 | 2.71E+00 | 2.77E+00 | 8.96E-01 | no | Nedd4l | intron_variant | MODIFIER |
| 18 | 65057447 | AX-170455592 | 0 | G | A | 0.412 | 1.00E-02 | 8.26E-02 | 2.71E+00 | 2.77E+00 | 8.96E-01 | no | Nedd4l | intron_variant | MODIFIER |
| 18 | 65058014 | AX-170151079 | 0 | C | T | 0.412 | 1.00E-02 | 8.26E-02 | 2.71E+00 | 2.77E+00 | 8.96E-01 | no | Nedd4l | intron_variant | MODIFIER |
| 18 | 65061975 | AX-170511708 | 0 | A | G | 0.412 | 1.00E-02 | 8.26E-02 | 2.71E+00 | 2.77E+00 | 8.96E-01 | no | Nedd4l | intron_variant | MODIFIER |
| 18 | 65076445 | AX-170093916 | 0 | T | C | 0.412 | 1.00E-02 | 8.26E-02 | 2.71E+00 | 2.77E+00 | 8.96E-01 | no | Nedd4l | intron_variant | MODIFIER |
| 18 | 65084596 | AX-170322033 | 0 | A | C | 0.412 | 1.00E-02 | 8.26E-02 | 2.71E+00 | 2.77E+00 | 8.96E-01 | no | Nedd4l | intron_variant | MODIFIER |
| 18 | 65084899 | AX-170517624 | 0 | A | G | 0.412 | 1.00E-02 | 8.26E-02 | 2.71E+00 | 2.77E+00 | 8.96E-01 | no | Nedd4l | intron_variant | MODIFIER |
| 18 | 65126850 | AX-170093917 | 0 | G | T | 0.412 | 1.00E-02 | 8.26E-02 | 2.71E+00 | 2.77E+00 | 8.96E-01 | no | Nedd4l | intron_variant | MODIFIER |
| 8 | 81068852 | AX-170297846 | 1 | G | C | 0.205 | 1.66E-02 | 1.30E-01 | 2.68E+00 | 2.74E+00 | 8.97E-01 | no | Gm45538 | intron_variant,non_coding_transcript_variant | MODIFIER |
| 1 | 107215172 | AX-97812678 | 0 | G | A | 0.167 | 2.69E-02 | 1.94E-01 | 2.63E+00 | 2.71E+00 | 8.97E-01 | no | Serpinb3b | intron_variant | MODIFIER |
| 1 | 116056741 | AX-170086584 | 0 | G | A | 0.167 | 2.69E-02 | 1.94E-01 | 2.63E+00 | 2.71E+00 | 8.97E-01 | no | Cntnap5a | intron_variant | MODIFIER |
| 18 | 57727224 | AX-170015780 | 0 | G | C | 0.351 | 1.12E-02 | 9.61E-02 | 2.64E+00 | 2.76E+00 | 8.97E-01 | no | Ccdc192 | intron_variant | MODIFIER |
| 18 | 57727862 | AX-170015776 | 0 | A | G | 0.351 | 1.12E-02 | 9.61E-02 | 2.64E+00 | 2.76E+00 | 8.97E-01 | no | Ccdc192 | intron_variant | MODIFIER |
| 8 | 90100706 | AX-170372543 | 0 | G | A | 0.193 | -1.59E-02 | 1.19E-01 | 2.65E+00 | 2.72E+00 | 8.97E-01 | no | | regulatory_region_variant | MODIFIER |
| 12 | 104565239 | AX-170050045 | 0 | C | T | 0.132 | 2.12E-02 | 1.63E-01 | 2.64E+00 | 2.74E+00 | 8.97E-01 | no | | intergenic_variant | MODIFIER |
| 9 | 59418808 | AX-170411905 | 0 | A | G | 0.211 | 1.41E-02 | 1.10E-01 | 2.64E+00 | 2.74E+00 | 8.97E-01 | no | Arih1 | intron_variant,NMD_transcript_variant | MODIFIER |
| 15 | 100967159 | AX-97830873 | 0 | A | G | 0.132 | 2.11E-02 | 1.62E-01 | 2.67E+00 | 2.73E+00 | 8.97E-01 | no | Scn8a | intron_variant | MODIFIER |

| | | | | | | | | | | | | | | | |
|---|---|---|---|---|---|---|---|---|---|---|---|---|---|---|---|
| 15 | 100999564 | AX-97830870 | 0 | G | A | 0.132 | 2.11E-02 | 1.62E-01 | 2.67E+00 | 2.73E+00 | 8.97E-01 | no | Scn8a | intron_variant | MODIFIER |
| 15 | 101000367 | AX-170401365 | 0 | G | C | 0.132 | 2.11E-02 | 1.62E-01 | 2.67E+00 | 2.73E+00 | 8.97E-01 | no | Scn8a | intron_variant | MODIFIER |
| 15 | 101002265 | AX-170141802 | 0 | G | C | 0.132 | 2.11E-02 | 1.62E-01 | 2.67E+00 | 2.73E+00 | 8.97E-01 | no | Scn8a | intron_variant | MODIFIER |
| 15 | 101027651 | AX-170056487 | 0 | C | T | 0.132 | 2.11E-02 | 1.62E-01 | 2.67E+00 | 2.73E+00 | 8.97E-01 | no | Scn8a | intron_variant | MODIFIER |
| 15 | 101027949 | AX-170474661 | 0 | T | C | 0.132 | 2.11E-02 | 1.62E-01 | 2.67E+00 | 2.73E+00 | 8.97E-01 | no | Scn8a | intron_variant | MODIFIER |
| 15 | 101070035 | AX-97845767 | 0 | G | A | 0.132 | 2.11E-02 | 1.62E-01 | 2.67E+00 | 2.73E+00 | 8.97E-01 | no | Fignl2 | intron_variant | MODIFIER |
| 15 | 101122346 | AX-97851096 | 0 | C | T | 0.132 | 2.11E-02 | 1.62E-01 | 2.67E+00 | 2.73E+00 | 8.97E-01 | no | Ankrd33 | downstream_gene_variant | MODIFIER |
| 15 | 101250858 | AX-97830360 | 0 | T | C | 0.132 | 2.11E-02 | 1.62E-01 | 2.67E+00 | 2.73E+00 | 8.97E-01 | no | Nr4a1 | upstream_gene_variant | MODIFIER |
| 15 | 101400613 | AX-170087951 | 0 | C | T | 0.132 | 2.11E-02 | 1.62E-01 | 2.67E+00 | 2.73E+00 | 8.97E-01 | no | Gm49500 | downstream_gene_variant | MODIFIER |
| 15 | 101423296 | AX-97855713 | 0 | G | A | 0.132 | 2.11E-02 | 1.62E-01 | 2.67E+00 | 2.73E+00 | 8.97E-01 | no | Krt7 | synonymous_variant | LOW |
| 15 | 101544240 | AX-97789620 | 0 | C | A | 0.132 | 2.11E-02 | 1.62E-01 | 2.67E+00 | 2.73E+00 | 8.97E-01 | no | Krt82 | intron_variant | MODIFIER |
| 15 | 101545863 | AX-170343178 | 0 | T | C | 0.132 | 2.11E-02 | 1.62E-01 | 2.67E+00 | 2.73E+00 | 8.97E-01 | no | Krt82 | intron_variant | MODIFIER |
| 15 | 101550547 | AX-170428839 | 0 | T | C | 0.132 | 2.11E-02 | 1.62E-01 | 2.67E+00 | 2.73E+00 | 8.97E-01 | no | Krt82 | synonymous_variant | LOW |
| 15 | 101558229 | AX-170520161 | 0 | G | A | 0.132 | 2.11E-02 | 1.62E-01 | 2.67E+00 | 2.73E+00 | 8.97E-01 | no | Krt90 | intron_variant | MODIFIER |
| 15 | 101701404 | AX-97824561 | 0 | C | T | 0.132 | 2.11E-02 | 1.62E-01 | 2.67E+00 | 2.73E+00 | 8.97E-01 | no | | intergenic_variant | MODIFIER |
| 15 | 101702294 | AX-170012404 | 0 | A | G | 0.132 | 2.11E-02 | 1.62E-01 | 2.67E+00 | 2.73E+00 | 8.97E-01 | no | Krt5 | downstream_gene_variant | MODIFIER |
| 15 | 101707394 | AX-170012403 | 0 | T | C | 0.132 | 2.11E-02 | 1.62E-01 | 2.67E+00 | 2.73E+00 | 8.97E-01 | no | Krt5 | 3_prime_UTR_variant | MODIFIER |
| 15 | 101753527 | AX-97823506 | 0 | A | G | 0.132 | 2.11E-02 | 1.62E-01 | 2.67E+00 | 2.73E+00 | 8.97E-01 | no | Krt74 | downstream_gene_variant | MODIFIER |
| 15 | 101768965 | AX-170012402 | 0 | G | A | 0.132 | 2.11E-02 | 1.62E-01 | 2.67E+00 | 2.73E+00 | 8.97E-01 | no | | intergenic_variant | MODIFIER |
| 15 | 101771307 | AX-170173411 | 0 | T | C | 0.132 | 2.11E-02 | 1.62E-01 | 2.67E+00 | 2.73E+00 | 8.97E-01 | no | Krt72 | downstream_gene_variant | MODIFIER |
| 15 | 101773356 | AX-170341754 | 0 | A | G | 0.132 | 2.11E-02 | 1.62E-01 | 2.67E+00 | 2.73E+00 | 8.97E-01 | no | Krt72 | downstream_gene_variant | MODIFIER |
| 15 | 101775649 | AX-170404167 | 0 | C | T | 0.132 | 2.11E-02 | 1.62E-01 | 2.67E+00 | 2.73E+00 | 8.97E-01 | no | Krt72 | downstream_gene_variant | MODIFIER |
| 15 | 101785486 | AX-97856579 | 0 | A | G | 0.132 | 2.11E-02 | 1.62E-01 | 2.67E+00 | 2.73E+00 | 8.97E-01 | no | Krt72 | intron_variant | MODIFIER |
| 15 | 101855241 | AX-97836158 | 0 | C | A | 0.132 | 2.11E-02 | 1.62E-01 | 2.67E+00 | 2.73E+00 | 8.97E-01 | no | Krt1 | upstream_gene_variant | MODIFIER |
| 15 | 101867275 | AX-170355040 | 0 | A | G | 0.132 | 2.11E-02 | 1.62E-01 | 2.67E+00 | 2.73E+00 | 8.97E-01 | no | Krt77 | intron_variant | MODIFIER |
| 15 | 101870782 | AX-170517979 | 0 | G | A | 0.132 | 2.11E-02 | 1.62E-01 | 2.67E+00 | 2.73E+00 | 8.97E-01 | no | Krt77 | upstream_gene_variant | MODIFIER |
| 15 | 101871056 | AX-97785648 | 0 | T | C | 0.132 | 2.11E-02 | 1.62E-01 | 2.67E+00 | 2.73E+00 | 8.97E-01 | no | Krt77 | upstream_gene_variant | MODIFIER |
| 15 | 102081839 | AX-170093360 | 0 | T | C | 0.132 | 2.11E-02 | 1.62E-01 | 2.67E+00 | 2.73E+00 | 8.97E-01 | no | Eif4b | intron_variant | MODIFIER |
| 15 | 102299530 | AX-170522293 | 0 | C | T | 0.132 | 2.11E-02 | 1.62E-01 | 2.67E+00 | 2.73E+00 | 8.97E-01 | no | Espl1 | splice_region_variant,intron_variant | LOW |
| 15 | 102325447 | AX-170149316 | 0 | A | G | 0.132 | 2.11E-02 | 1.62E-01 | 2.67E+00 | 2.73E+00 | 8.97E-01 | no | Pfdn5 | upstream_gene_variant | MODIFIER |
| 15 | 102332284 | AX-170525179 | 0 | C | T | 0.132 | 2.11E-02 | 1.62E-01 | 2.67E+00 | 2.73E+00 | 8.97E-01 | no | Myg1 | intron_variant | MODIFIER |
| 15 | 102336789 | AX-170111883 | 0 | A | G | 0.132 | 2.11E-02 | 1.62E-01 | 2.67E+00 | 2.73E+00 | 8.97E-01 | no | Myg1 | intron_variant | MODIFIER |
| 15 | 102338143 | AX-97840045 | 0 | T | C | 0.132 | 2.11E-02 | 1.62E-01 | 2.67E+00 | 2.73E+00 | 8.97E-01 | no | Myg1 | downstream_gene_variant | MODIFIER |
| 15 | 102339326 | AX-170310287 | 0 | G | T | 0.132 | 2.11E-02 | 1.62E-01 | 2.67E+00 | 2.73E+00 | 8.97E-01 | no | Myg1 | downstream_gene_variant | MODIFIER |
| 15 | 102358134 | AX-170119483 | 0 | G | A | 0.132 | 2.11E-02 | 1.62E-01 | 2.67E+00 | 2.73E+00 | 8.97E-01 | no | Sp7 | 3_prime_UTR_variant | MODIFIER |
| 15 | 102359751 | AX-97783225 | 0 | G | A | 0.132 | 2.11E-02 | 1.62E-01 | 2.67E+00 | 2.73E+00 | 8.97E-01 | no | Sp7 | intron_variant | MODIFIER |
| 15 | 102391221 | AX-170385062 | 0 | G | A | 0.132 | 2.11E-02 | 1.62E-01 | 2.67E+00 | 2.73E+00 | 8.97E-01 | no | | regulatory_region_variant | MODIFIER |
| 15 | 102433276 | AX-170196105 | 0 | G | A | 0.132 | 2.11E-02 | 1.62E-01 | 2.67E+00 | 2.73E+00 | 8.97E-01 | no | Sp1 | 3_prime_UTR_variant | MODIFIER |
| 15 | 102461170 | AX-170155995 | 0 | C | T | 0.132 | 2.11E-02 | 1.62E-01 | 2.67E+00 | 2.73E+00 | 8.97E-01 | no | Prr13 | intron_variant | MODIFIER |
| 15 | 102465453 | AX-97815132 | 0 | G | A | 0.132 | 2.11E-02 | 1.62E-01 | 2.67E+00 | 2.73E+00 | 8.97E-01 | no | Prr13 | downstream_gene_variant | MODIFIER |
| 15 | 102488182 | AX-170312496 | 0 | C | T | 0.132 | 2.11E-02 | 1.62E-01 | 2.67E+00 | 2.73E+00 | 8.97E-01 | no | Pcbp2 | intron_variant | MODIFIER |
| 15 | 102500631 | AX-170271135 | 0 | C | T | 0.132 | 2.11E-02 | 1.62E-01 | 2.67E+00 | 2.73E+00 | 8.97E-01 | no | Map3k12 | synonymous_variant | LOW |
| 15 | 102502623 | AX-170301548 | 0 | T | C | 0.132 | 2.11E-02 | 1.62E-01 | 2.67E+00 | 2.73E+00 | 8.97E-01 | no | Map3k12 | intron_variant | MODIFIER |
| 15 | 102713967 | AX-170012421 | 0 | T | C | 0.132 | 2.11E-02 | 1.62E-01 | 2.67E+00 | 2.73E+00 | 8.97E-01 | no | Calcoco1 | intron_variant | MODIFIER |
| 15 | 102749054 | AX-170012415 | 0 | C | T | 0.132 | 2.11E-02 | 1.62E-01 | 2.67E+00 | 2.73E+00 | 8.97E-01 | no | | intergenic_variant | MODIFIER |
| 15 | 102795874 | AX-97830361 | 0 | G | A | 0.132 | 2.11E-02 | 1.62E-01 | 2.67E+00 | 2.73E+00 | 8.97E-01 | no | Gm49473 | upstream_gene_variant | MODIFIER |
| 15 | 102830525 | AX-170012424 | 0 | T | C | 0.132 | 2.11E-02 | 1.62E-01 | 2.67E+00 | 2.73E+00 | 8.97E-01 | no | | intergenic_variant | MODIFIER |
| 15 | 102874306 | AX-170012413 | 0 | C | T | 0.132 | 2.11E-02 | 1.62E-01 | 2.67E+00 | 2.73E+00 | 8.97E-01 | no | | intergenic_variant | MODIFIER |
| 15 | 102952665 | AX-170012412 | 0 | C | T | 0.132 | 2.11E-02 | 1.62E-01 | 2.67E+00 | 2.73E+00 | 8.97E-01 | no | Hoxc11 | upstream_gene_variant | MODIFIER |
| 15 | 102963993 | AX-170012418 | 0 | C | G | 0.132 | 2.11E-02 | 1.62E-01 | 2.67E+00 | 2.73E+00 | 8.97E-01 | no | Hoxc10 | upstream_gene_variant | MODIFIER |
| 15 | 102976567 | AX-97802672 | 0 | C | T | 0.132 | 2.11E-02 | 1.62E-01 | 2.67E+00 | 2.73E+00 | 8.97E-01 | no | Hoxc10 | downstream_gene_variant | MODIFIER |
| 15 | 103005645 | AX-170489676 | 0 | G | T | 0.132 | 2.11E-02 | 1.62E-01 | 2.67E+00 | 2.73E+00 | 8.97E-01 | no | Hoxc6 | upstream_gene_variant | MODIFIER |
| 15 | 103005825 | AX-97844252 | 0 | G | A | 0.132 | 2.11E-02 | 1.62E-01 | 2.67E+00 | 2.73E+00 | 8.97E-01 | no | Hoxc6 | upstream_gene_variant | MODIFIER |
| 15 | 103313148 | AX-170012429 | 0 | G | A | 0.132 | 2.11E-02 | 1.62E-01 | 2.67E+00 | 2.73E+00 | 8.97E-01 | no | Gpr84 | upstream_gene_variant | MODIFIER |

| | | | | | | | | | | | | | | | |
|---|---|---|---|---|---|---|---|---|---|---|---|---|---|---|---|
| 15 | 103313546 | AX-97814377 | 0 | C | T | 0.132 | 2.11E-02 | 1.62E-01 | 2.67E+00 | 2.73E+00 | 8.97E-01 | no | Gpr84 | upstream_gene_variant | MODIFIER |
| 15 | 103316787 | AX-170283165 | 0 | C | A | 0.132 | 2.11E-02 | 1.62E-01 | 2.67E+00 | 2.73E+00 | 8.97E-01 | no | Zfp385a | intron_variant | MODIFIER |
| 15 | 103334045 | AX-97829594 | 0 | G | A | 0.132 | 2.11E-02 | 1.62E-01 | 2.67E+00 | 2.73E+00 | 8.97E-01 | no | Zfp385a | intron_variant | MODIFIER |
| 15 | 103334218 | AX-170012432 | 0 | G | A | 0.132 | 2.11E-02 | 1.62E-01 | 2.67E+00 | 2.73E+00 | 8.97E-01 | no | Zfp385a | intron_variant | MODIFIER |
| 15 | 103335822 | AX-170012428 | 0 | G | A | 0.132 | 2.11E-02 | 1.62E-01 | 2.67E+00 | 2.73E+00 | 8.97E-01 | no | Zfp385a | intron_variant | MODIFIER |
| 15 | 103350886 | AX-170012427 | 0 | A | G | 0.132 | 2.11E-02 | 1.62E-01 | 2.67E+00 | 2.73E+00 | 8.97E-01 | no | Itga5 | intron_variant | MODIFIER |
| 15 | 103359595 | AX-97836680 | 0 | T | C | 0.132 | 2.11E-02 | 1.62E-01 | 2.67E+00 | 2.73E+00 | 8.97E-01 | no | Itga5 | intron_variant | MODIFIER |
| 15 | 103392585 | AX-170012426 | 0 | T | C | 0.132 | 2.11E-02 | 1.62E-01 | 2.67E+00 | 2.73E+00 | 8.97E-01 | no | | intergenic_variant | MODIFIER |
| 15 | 103394890 | AX-97855026 | 0 | A | G | 0.132 | 2.11E-02 | 1.62E-01 | 2.67E+00 | 2.73E+00 | 8.97E-01 | no | | intergenic_variant | MODIFIER |
| 15 | 103400849 | AX-170166923 | 0 | A | G | 0.132 | 2.11E-02 | 1.62E-01 | 2.67E+00 | 2.73E+00 | 8.97E-01 | no | Gtsf1 | downstream_gene_variant | MODIFIER |
| 15 | 103448405 | AX-170473481 | 0 | G | C | 0.132 | 2.11E-02 | 1.62E-01 | 2.67E+00 | 2.73E+00 | 8.97E-01 | no | Gtsf2 | 5_prime_UTR_variant | MODIFIER |
| 10 | 16120426 | AX-170492779 | 1 | C | T | 0.196 | 1.24E-02 | 1.04E-01 | 2.68E+00 | 2.77E+00 | 8.97E-01 | no | | intergenic_variant | MODIFIER |
| 13 | 37331507 | AX-170365215 | 0 | T | C | 0.175 | -1.63E-02 | 1.17E-01 | 2.64E+00 | 2.71E+00 | 8.97E-01 | no | Gm47711 | downstream_gene_variant | MODIFIER |
| 13 | 37350858 | AX-170184972 | 0 | A | G | 0.175 | -1.63E-02 | 1.17E-01 | 2.64E+00 | 2.71E+00 | 8.97E-01 | no | Ly86 | intron_variant | MODIFIER |
| 13 | 37372240 | AX-170469095 | 0 | T | C | 0.175 | -1.63E-02 | 1.17E-01 | 2.64E+00 | 2.71E+00 | 8.97E-01 | no | Ly86 | intron_variant | MODIFIER |
| 13 | 37389777 | AX-170513315 | 0 | T | C | 0.175 | -1.63E-02 | 1.17E-01 | 2.64E+00 | 2.71E+00 | 8.97E-01 | no | Ly86 | intron_variant | MODIFIER |
| 13 | 37389932 | AX-97839614 | 0 | C | T | 0.175 | -1.63E-02 | 1.17E-01 | 2.64E+00 | 2.71E+00 | 8.97E-01 | no | Ly86 | intron_variant | MODIFIER |
| 13 | 37390566 | AX-170337878 | 0 | T | C | 0.175 | -1.63E-02 | 1.17E-01 | 2.64E+00 | 2.71E+00 | 8.97E-01 | no | Ly86 | intron_variant | MODIFIER |
| 13 | 37390955 | AX-170292995 | 0 | T | C | 0.175 | -1.63E-02 | 1.17E-01 | 2.64E+00 | 2.71E+00 | 8.97E-01 | no | Ly86 | intron_variant | MODIFIER |
| 13 | 37393650 | AX-170487305 | 0 | C | T | 0.175 | -1.63E-02 | 1.17E-01 | 2.64E+00 | 2.71E+00 | 8.97E-01 | no | Ly86 | intron_variant | MODIFIER |
| 13 | 37413161 | AX-170389766 | 0 | T | G | 0.175 | -1.63E-02 | 1.17E-01 | 2.64E+00 | 2.71E+00 | 8.97E-01 | no | Ly86 | intron_variant | MODIFIER |
| 13 | 37441586 | AX-170303110 | 0 | T | C | 0.175 | -1.63E-02 | 1.17E-01 | 2.64E+00 | 2.71E+00 | 8.97E-01 | no | Gm47730 | upstream_gene_variant | MODIFIER |
| 13 | 37489267 | AX-170508762 | 0 | C | T | 0.175 | -1.63E-02 | 1.17E-01 | 2.64E+00 | 2.71E+00 | 8.97E-01 | no | | intergenic_variant | MODIFIER |
| 13 | 37493432 | AX-170353736 | 0 | T | C | 0.175 | -1.63E-02 | 1.17E-01 | 2.64E+00 | 2.71E+00 | 8.97E-01 | no | | intergenic_variant | MODIFIER |
| 13 | 37494549 | AX-170382856 | 0 | T | C | 0.175 | -1.63E-02 | 1.17E-01 | 2.64E+00 | 2.71E+00 | 8.97E-01 | no | | intergenic_variant | MODIFIER |
| 13 | 37524277 | AX-170008129 | 0 | T | A | 0.175 | -1.63E-02 | 1.17E-01 | 2.64E+00 | 2.71E+00 | 8.97E-01 | no | | intergenic_variant | MODIFIER |
| 13 | 37524746 | AX-170008132 | 0 | C | T | 0.175 | -1.63E-02 | 1.17E-01 | 2.64E+00 | 2.71E+00 | 8.97E-01 | no | | intergenic_variant | MODIFIER |
| 13 | 37525027 | AX-170008130 | 0 | A | G | 0.175 | -1.63E-02 | 1.17E-01 | 2.64E+00 | 2.71E+00 | 8.97E-01 | no | | intergenic_variant | MODIFIER |
| 13 | 37525422 | AX-170410839 | 0 | T | C | 0.175 | -1.63E-02 | 1.17E-01 | 2.64E+00 | 2.71E+00 | 8.97E-01 | no | | intergenic_variant | MODIFIER |
| 13 | 37561900 | AX-170404728 | 0 | T | C | 0.175 | -1.63E-02 | 1.17E-01 | 2.64E+00 | 2.71E+00 | 8.97E-01 | no | Gm47751 | downstream_gene_variant | MODIFIER |
| 13 | 37571282 | AX-170008127 | 0 | T | C | 0.175 | -1.63E-02 | 1.17E-01 | 2.64E+00 | 2.71E+00 | 8.97E-01 | no | | intergenic_variant | MODIFIER |
| 13 | 37589546 | AX-170171637 | 0 | A | G | 0.175 | -1.63E-02 | 1.17E-01 | 2.64E+00 | 2.71E+00 | 8.97E-01 | no | Gm47754 | downstream_gene_variant | MODIFIER |
| 13 | 37593626 | AX-170160403 | 0 | T | C | 0.175 | -1.63E-02 | 1.17E-01 | 2.64E+00 | 2.71E+00 | 8.97E-01 | no | Gm47754 | downstream_gene_variant | MODIFIER |
| 13 | 37686361 | AX-170473367 | 0 | C | A | 0.175 | -1.63E-02 | 1.17E-01 | 2.64E+00 | 2.71E+00 | 8.97E-01 | no | Gm49350 | non_coding_transcript_exon_variant | MODIFIER |
| 13 | 37714357 | AX-170238008 | 0 | C | G | 0.175 | -1.63E-02 | 1.17E-01 | 2.64E+00 | 2.71E+00 | 8.97E-01 | no | Gm40918 | intron_variant,non_coding_transcript_variant | MODIFIER |
| 13 | 37714572 | AX-170137084 | 0 | G | A | 0.175 | -1.63E-02 | 1.17E-01 | 2.64E+00 | 2.71E+00 | 8.97E-01 | no | Gm40918 | intron_variant,non_coding_transcript_variant | MODIFIER |
| 13 | 37714698 | AX-170092724 | 0 | G | A | 0.175 | -1.63E-02 | 1.17E-01 | 2.64E+00 | 2.71E+00 | 8.97E-01 | no | Gm40918 | intron_variant,non_coding_transcript_variant | MODIFIER |
| 13 | 37732134 | AX-170237993 | 0 | A | G | 0.175 | -1.63E-02 | 1.17E-01 | 2.64E+00 | 2.71E+00 | 8.97E-01 | no | | regulatory_region_variant | MODIFIER |
| 13 | 37742554 | AX-97812435 | 0 | G | A | 0.175 | -1.63E-02 | 1.17E-01 | 2.64E+00 | 2.71E+00 | 8.97E-01 | no | | intergenic_variant | MODIFIER |
| 13 | 37749802 | AX-170310818 | 0 | T | C | 0.175 | -1.63E-02 | 1.17E-01 | 2.64E+00 | 2.71E+00 | 8.97E-01 | no | | regulatory_region_variant | MODIFIER |
| 13 | 37754243 | AX-168397926 | 0 | T | C | 0.175 | -1.63E-02 | 1.17E-01 | 2.64E+00 | 2.71E+00 | 8.97E-01 | no | | intergenic_variant | MODIFIER |
| 13 | 37759289 | AX-170213758 | 0 | T | C | 0.175 | -1.63E-02 | 1.17E-01 | 2.64E+00 | 2.71E+00 | 8.97E-01 | no | Gm31600 | upstream_gene_variant | MODIFIER |
| 13 | 37790477 | AX-97786826 | 0 | G | A | 0.175 | -1.63E-02 | 1.17E-01 | 2.64E+00 | 2.71E+00 | 8.97E-01 | no | Rreb1 | intron_variant | MODIFIER |
| 13 | 37841710 | AX-170461191 | 0 | T | C | 0.175 | -1.63E-02 | 1.17E-01 | 2.64E+00 | 2.71E+00 | 8.97E-01 | no | Rreb1 | intron_variant | MODIFIER |
| 13 | 37879178 | AX-170385211 | 0 | C | T | 0.175 | -1.63E-02 | 1.17E-01 | 2.64E+00 | 2.71E+00 | 8.97E-01 | no | Rreb1 | intron_variant | MODIFIER |
| 13 | 37926739 | AX-170199377 | 0 | A | G | 0.175 | -1.63E-02 | 1.17E-01 | 2.64E+00 | 2.71E+00 | 8.97E-01 | no | Rreb1 | intron_variant | MODIFIER |
| 13 | 37948347 | AX-170387611 | 0 | A | G | 0.175 | -1.63E-02 | 1.17E-01 | 2.64E+00 | 2.71E+00 | 8.97E-01 | no | Rreb1 | intron_variant | MODIFIER |
| 13 | 37978763 | AX-97781394 | 0 | G | A | 0.175 | -1.63E-02 | 1.17E-01 | 2.64E+00 | 2.71E+00 | 8.97E-01 | no | Ssr1 | 3_prime_UTR_variant | MODIFIER |
| 13 | 37989466 | AX-97781896 | 0 | G | A | 0.175 | -1.63E-02 | 1.17E-01 | 2.64E+00 | 2.71E+00 | 8.97E-01 | no | Ssr1 | intron_variant | MODIFIER |
| 13 | 37990157 | AX-170366346 | 0 | A | G | 0.175 | -1.63E-02 | 1.17E-01 | 2.64E+00 | 2.71E+00 | 8.97E-01 | no | Ssr1 | intron_variant | MODIFIER |
| 13 | 38010418 | AX-97832484 | 0 | C | T | 0.175 | -1.63E-02 | 1.17E-01 | 2.64E+00 | 2.71E+00 | 8.97E-01 | no | Cage1 | intron_variant | MODIFIER |
| 13 | 38023863 | AX-170433997 | 0 | T | C | 0.175 | -1.63E-02 | 1.17E-01 | 2.64E+00 | 2.71E+00 | 8.97E-01 | no | Cage1 | intron_variant | MODIFIER |
| 13 | 38028304 | AX-170408923 | 0 | C | A | 0.175 | -1.63E-02 | 1.17E-01 | 2.64E+00 | 2.71E+00 | 8.97E-01 | no | Cage1 | intron_variant | MODIFIER |
| 13 | 38028329 | AX-170419694 | 0 | C | T | 0.175 | -1.63E-02 | 1.17E-01 | 2.64E+00 | 2.71E+00 | 8.97E-01 | no | Cage1 | intron_variant | MODIFIER |

| | | | | | | | | | | | | | | | |
|---|---|---|---|---|---|---|---|---|---|---|---|---|---|---|---|
| 13 | 38029084 | AX-170467364 | 0 | C | T | 0.175 | -1.63E-02 | 1.17E-01 | 2.64E+00 | 2.71E+00 | 8.97E-01 | no | Cage1 | intron_variant | MODIFIER |
| 13 | 38045407 | AX-170092727 | 0 | C | T | 0.175 | -1.63E-02 | 1.17E-01 | 2.64E+00 | 2.71E+00 | 8.97E-01 | no | Riok1 | intron_variant | MODIFIER |
| 13 | 38050326 | AX-170508937 | 0 | A | G | 0.175 | -1.63E-02 | 1.17E-01 | 2.64E+00 | 2.71E+00 | 8.97E-01 | no | Riok1 | intron_variant | MODIFIER |
| 13 | 38050607 | AX-170309607 | 0 | C | T | 0.175 | -1.63E-02 | 1.17E-01 | 2.64E+00 | 2.71E+00 | 8.97E-01 | no | Riok1 | intron_variant | MODIFIER |
| 13 | 38050656 | AX-170008139 | 0 | A | G | 0.175 | -1.63E-02 | 1.17E-01 | 2.64E+00 | 2.71E+00 | 8.97E-01 | no | Riok1 | intron_variant | MODIFIER |
| 13 | 38053074 | AX-170333039 | 0 | A | G | 0.175 | -1.63E-02 | 1.17E-01 | 2.64E+00 | 2.71E+00 | 8.97E-01 | no | Riok1 | intron_variant | MODIFIER |
| 13 | 38058093 | AX-170008140 | 0 | A | G | 0.175 | -1.63E-02 | 1.17E-01 | 2.64E+00 | 2.71E+00 | 8.97E-01 | no | Riok1 | intron_variant | MODIFIER |
| 13 | 38058931 | AX-170379845 | 0 | C | T | 0.175 | -1.63E-02 | 1.17E-01 | 2.64E+00 | 2.71E+00 | 8.97E-01 | no | Riok1 | intron_variant | MODIFIER |
| 13 | 38060931 | AX-170276016 | 0 | T | C | 0.175 | -1.63E-02 | 1.17E-01 | 2.64E+00 | 2.71E+00 | 8.97E-01 | no | Riok1 | 3_prime_UTR_variant | MODIFIER |
| 13 | 38063095 | AX-170501768 | 0 | G | A | 0.175 | -1.63E-02 | 1.17E-01 | 2.64E+00 | 2.71E+00 | 8.97E-01 | no | Riok1 | downstream_gene_variant | MODIFIER |
| 13 | 38063889 | AX-170362786 | 0 | C | T | 0.175 | -1.63E-02 | 1.17E-01 | 2.64E+00 | 2.71E+00 | 8.97E-01 | no | Riok1 | downstream_gene_variant | MODIFIER |
| 13 | 38085895 | AX-170008134 | 0 | G | C | 0.175 | -1.63E-02 | 1.17E-01 | 2.64E+00 | 2.71E+00 | 8.97E-01 | no | Gm27387 | upstream_gene_variant | MODIFIER |
| 13 | 38096053 | AX-170008135 | 0 | C | T | 0.175 | -1.63E-02 | 1.17E-01 | 2.64E+00 | 2.71E+00 | 8.97E-01 | no | | intergenic_variant | MODIFIER |
| 13 | 38116787 | AX-170008141 | 0 | A | C | 0.175 | -1.63E-02 | 1.17E-01 | 2.64E+00 | 2.71E+00 | 8.97E-01 | no | | intergenic_variant | MODIFIER |
| 13 | 38119329 | AX-97850260 | 0 | G | A | 0.175 | -1.63E-02 | 1.17E-01 | 2.64E+00 | 2.71E+00 | 8.97E-01 | no | | intergenic_variant | MODIFIER |
| 13 | 38130621 | AX-170008138 | 0 | C | A | 0.175 | -1.63E-02 | 1.17E-01 | 2.64E+00 | 2.71E+00 | 8.97E-01 | no | | intergenic_variant | MODIFIER |
| 13 | 38135450 | AX-97785813 | 0 | T | C | 0.175 | -1.63E-02 | 1.17E-01 | 2.64E+00 | 2.71E+00 | 8.97E-01 | no | | intergenic_variant | MODIFIER |
| 13 | 38154816 | AX-170008136 | 0 | T | C | 0.175 | -1.63E-02 | 1.17E-01 | 2.64E+00 | 2.71E+00 | 8.97E-01 | no | Dsp | intron_variant | MODIFIER |
| 13 | 38178846 | AX-170484035 | 0 | A | T | 0.175 | -1.63E-02 | 1.17E-01 | 2.64E+00 | 2.71E+00 | 8.97E-01 | no | Dsp | intron_variant | MODIFIER |
| 10 | 128153496 | AX-170269533 | 1 | A | C | 0.33 | -9.77E-03 | 7.31E-02 | 2.62E+00 | 2.73E+00 | 8.97E-01 | no | Rbms2 | intron_variant | MODIFIER |
| 9 | 74890189 | AX-170519441 | 0 | G | A | 0.211 | 1.60E-02 | 1.26E-01 | 2.69E+00 | 2.74E+00 | 8.97E-01 | no | Onecut1 | 3_prime_UTR_variant | MODIFIER |
| 1 | 68607002 | AX-170101047 | 0 | A | C | 0.123 | -2.55E-02 | 2.08E-01 | 2.69E+00 | 2.76E+00 | 8.97E-01 | no | Erbb4 | intron_variant | MODIFIER |
| 1 | 71943112 | AX-97811951 | 0 | C | T | 0.123 | -2.55E-02 | 2.08E-01 | 2.69E+00 | 2.76E+00 | 8.97E-01 | no | | intergenic_variant | MODIFIER |
| 1 | 72363236 | AX-170357545 | 0 | G | T | 0.123 | -2.55E-02 | 2.08E-01 | 2.69E+00 | 2.76E+00 | 8.97E-01 | no | Xrcc5 | intron_variant | MODIFIER |
| 10 | 126872997 | AX-170046208 | 1 | T | C | 0.33 | -1.01E-02 | 7.44E-02 | 2.62E+00 | 2.72E+00 | 8.97E-01 | no | Atp23 | intron_variant | MODIFIER |
| 10 | 128176082 | AX-170153558 | 1 | T | G | 0.33 | -1.01E-02 | 7.44E-02 | 2.62E+00 | 2.72E+00 | 8.97E-01 | no | Rbms2 | intron_variant | MODIFIER |
| 3 | 147074267 | AX-170410671 | 1 | C | T | 0.348 | 9.72E-03 | 7.33E-02 | 2.65E+00 | 2.72E+00 | 8.97E-01 | no | | intergenic_variant | MODIFIER |
| 10 | 31054186 | AX-170309183 | 0 | A | G | 0.377 | -1.03E-02 | 7.65E-02 | 2.61E+00 | 2.73E+00 | 8.97E-01 | no | Gm30676 | intron_variant,non_coding_transcript_variant | MODIFIER |
| 8 | 22457824 | AX-170252546 | 0 | G | A | 0.167 | -1.47E-02 | 1.13E-01 | 2.65E+00 | 2.73E+00 | 8.98E-01 | no | Smim19 | downstream_gene_variant | MODIFIER |
| 8 | 22458048 | AX-170320172 | 0 | C | T | 0.167 | -1.47E-02 | 1.13E-01 | 2.65E+00 | 2.73E+00 | 8.98E-01 | no | Smim19 | downstream_gene_variant | MODIFIER |
| 8 | 22549056 | AX-170079506 | 0 | G | C | 0.167 | -1.47E-02 | 1.13E-01 | 2.65E+00 | 2.73E+00 | 8.98E-01 | no | Slc20a2 | intron_variant | MODIFIER |
| 8 | 22551047 | AX-170079503 | 0 | T | C | 0.167 | -1.47E-02 | 1.13E-01 | 2.65E+00 | 2.73E+00 | 8.98E-01 | no | Slc20a2 | intron_variant | MODIFIER |
| 8 | 22561110 | AX-97782754 | 0 | C | T | 0.167 | -1.47E-02 | 1.13E-01 | 2.65E+00 | 2.73E+00 | 8.98E-01 | no | Slc20a2 | synonymous_variant | LOW |
| 8 | 22719101 | AX-170152863 | 0 | T | C | 0.167 | -1.47E-02 | 1.13E-01 | 2.65E+00 | 2.73E+00 | 8.98E-01 | no | | intergenic_variant | MODIFIER |
| 8 | 22745092 | AX-97806364 | 0 | A | G | 0.167 | -1.47E-02 | 1.13E-01 | 2.65E+00 | 2.73E+00 | 8.98E-01 | no | | intergenic_variant | MODIFIER |
| 8 | 22746933 | AX-170239758 | 0 | C | T | 0.167 | -1.47E-02 | 1.13E-01 | 2.65E+00 | 2.73E+00 | 8.98E-01 | no | | intergenic_variant | MODIFIER |
| 8 | 22761842 | AX-170090062 | 0 | T | C | 0.167 | -1.47E-02 | 1.13E-01 | 2.65E+00 | 2.73E+00 | 8.98E-01 | no | Plat | intron_variant | MODIFIER |
| 8 | 22791187 | AX-97823748 | 0 | T | C | 0.167 | -1.47E-02 | 1.13E-01 | 2.65E+00 | 2.73E+00 | 8.98E-01 | no | Ap3m2 | intron_variant | MODIFIER |
| 8 | 22816034 | AX-170412978 | 0 | C | A | 0.167 | -1.47E-02 | 1.13E-01 | 2.65E+00 | 2.73E+00 | 8.98E-01 | no | | intergenic_variant | MODIFIER |
| 8 | 22961784 | AX-170030346 | 0 | A | C | 0.167 | -1.47E-02 | 1.13E-01 | 2.65E+00 | 2.73E+00 | 8.98E-01 | no | Gm10043 | upstream_gene_variant | MODIFIER |
| 8 | 22962959 | AX-170096197 | 0 | C | G | 0.167 | -1.47E-02 | 1.13E-01 | 2.65E+00 | 2.73E+00 | 8.98E-01 | no | Gm10043 | upstream_gene_variant | MODIFIER |
| 8 | 22964204 | AX-170030347 | 0 | T | C | 0.167 | -1.47E-02 | 1.13E-01 | 2.65E+00 | 2.73E+00 | 8.98E-01 | no | | intergenic_variant | MODIFIER |
| 8 | 22979827 | AX-170176613 | 0 | C | A | 0.167 | -1.47E-02 | 1.13E-01 | 2.65E+00 | 2.73E+00 | 8.98E-01 | no | Ank1 | intron_variant | MODIFIER |
| 8 | 22981837 | AX-170440191 | 0 | C | T | 0.167 | -1.47E-02 | 1.13E-01 | 2.65E+00 | 2.73E+00 | 8.98E-01 | no | Ank1 | intron_variant | MODIFIER |
| 8 | 22984035 | AX-97838361 | 0 | G | A | 0.167 | -1.47E-02 | 1.13E-01 | 2.65E+00 | 2.73E+00 | 8.98E-01 | no | Ank1 | intron_variant | MODIFIER |
| 8 | 22984199 | AX-170109109 | 0 | G | A | 0.167 | -1.47E-02 | 1.13E-01 | 2.65E+00 | 2.73E+00 | 8.98E-01 | no | Ank1 | intron_variant | MODIFIER |
| 8 | 22985515 | AX-170030344 | 0 | G | A | 0.167 | -1.47E-02 | 1.13E-01 | 2.65E+00 | 2.73E+00 | 8.98E-01 | no | Ank1 | intron_variant | MODIFIER |
| 8 | 23007911 | AX-170314506 | 0 | C | T | 0.167 | -1.47E-02 | 1.13E-01 | 2.65E+00 | 2.73E+00 | 8.98E-01 | no | Ank1 | intron_variant | MODIFIER |
| 8 | 23008395 | AX-170344614 | 0 | T | G | 0.167 | -1.47E-02 | 1.13E-01 | 2.65E+00 | 2.73E+00 | 8.98E-01 | no | Ank1 | intron_variant | MODIFIER |
| 8 | 23011388 | AX-97787633 | 0 | T | C | 0.167 | -1.47E-02 | 1.13E-01 | 2.65E+00 | 2.73E+00 | 8.98E-01 | no | Ank1 | intron_variant | MODIFIER |
| 8 | 23014922 | AX-170358108 | 0 | G | T | 0.167 | -1.47E-02 | 1.13E-01 | 2.65E+00 | 2.73E+00 | 8.98E-01 | no | Ank1 | intron_variant | MODIFIER |
| 8 | 23030087 | AX-170193235 | 0 | T | C | 0.167 | -1.47E-02 | 1.13E-01 | 2.65E+00 | 2.73E+00 | 8.98E-01 | no | Ank1 | intron_variant | MODIFIER |
| 8 | 23033591 | AX-170333360 | 0 | C | T | 0.167 | -1.47E-02 | 1.13E-01 | 2.65E+00 | 2.73E+00 | 8.98E-01 | no | Ank1 | intron_variant | MODIFIER |
| 8 | 23222079 | AX-170090051 | 0 | A | G | 0.167 | -1.47E-02 | 1.13E-01 | 2.65E+00 | 2.73E+00 | 8.98E-01 | no | Gins4 | downstream_gene_variant | MODIFIER |

| | | | | | | | | | | | | | | | |
|---|---|---|---|---|---|---|---|---|---|---|---|---|---|---|---|
| 8 | 23230036 | AX-170373451 | 0 | C | T | 0.167 | -1.47E-02 | 1.13E-01 | 2.65E+00 | 2.73E+00 | 8.98E-01 | no | Gins4 | intron_variant | MODIFIER |
| 8 | 23232860 | AX-170409523 | 0 | C | T | 0.167 | -1.47E-02 | 1.13E-01 | 2.65E+00 | 2.73E+00 | 8.98E-01 | no | Gins4 | intron_variant | MODIFIER |
| 8 | 23240006 | AX-97791301 | 0 | G | A | 0.167 | -1.47E-02 | 1.13E-01 | 2.65E+00 | 2.73E+00 | 8.98E-01 | no | Gins4 | upstream_gene_variant | MODIFIER |
| 8 | 23287870 | AX-170464113 | 0 | C | T | 0.167 | -1.47E-02 | 1.13E-01 | 2.65E+00 | 2.73E+00 | 8.98E-01 | no | Gm24335 | downstream_gene_variant | MODIFIER |
| 8 | 23296737 | AX-97823522 | 0 | C | T | 0.167 | -1.47E-02 | 1.13E-01 | 2.65E+00 | 2.73E+00 | 8.98E-01 | no | | intergenic_variant | MODIFIER |
| 8 | 23299947 | AX-170344779 | 0 | G | C | 0.167 | -1.47E-02 | 1.13E-01 | 2.65E+00 | 2.73E+00 | 8.98E-01 | no | | regulatory_region_variant | MODIFIER |
| 8 | 23302491 | AX-170280255 | 0 | A | T | 0.167 | -1.47E-02 | 1.13E-01 | 2.65E+00 | 2.73E+00 | 8.98E-01 | no | | intergenic_variant | MODIFIER |
| 8 | 23318224 | AX-170293245 | 0 | G | C | 0.167 | -1.47E-02 | 1.13E-01 | 2.65E+00 | 2.73E+00 | 8.98E-01 | no | | intergenic_variant | MODIFIER |
| 8 | 23324431 | AX-170164187 | 0 | A | G | 0.167 | -1.47E-02 | 1.13E-01 | 2.65E+00 | 2.73E+00 | 8.98E-01 | no | | intergenic_variant | MODIFIER |
| 8 | 23330799 | AX-170227451 | 0 | A | T | 0.167 | -1.47E-02 | 1.13E-01 | 2.65E+00 | 2.73E+00 | 8.98E-01 | no | | intergenic_variant | MODIFIER |
| 8 | 23353524 | AX-97828656 | 0 | T | C | 0.167 | -1.47E-02 | 1.13E-01 | 2.65E+00 | 2.73E+00 | 8.98E-01 | no | | intergenic_variant | MODIFIER |
| 8 | 23361598 | AX-97792527 | 0 | T | G | 0.167 | -1.47E-02 | 1.13E-01 | 2.65E+00 | 2.73E+00 | 8.98E-01 | no | | regulatory_region_variant | MODIFIER |
| 8 | 23400020 | AX-97830638 | 0 | T | C | 0.167 | -1.47E-02 | 1.13E-01 | 2.65E+00 | 2.73E+00 | 8.98E-01 | no | | intergenic_variant | MODIFIER |
| 8 | 23407612 | AX-170418736 | 0 | T | C | 0.167 | -1.47E-02 | 1.13E-01 | 2.65E+00 | 2.73E+00 | 8.98E-01 | no | Sfrp1 | upstream_gene_variant | MODIFIER |
| 8 | 23422704 | AX-170030355 | 0 | G | A | 0.167 | -1.47E-02 | 1.13E-01 | 2.65E+00 | 2.73E+00 | 8.98E-01 | no | Sfrp1 | intron_variant | MODIFIER |
| 8 | 23428239 | AX-170079519 | 0 | C | G | 0.167 | -1.47E-02 | 1.13E-01 | 2.65E+00 | 2.73E+00 | 8.98E-01 | no | Sfrp1 | intron_variant | MODIFIER |
| 8 | 23433500 | AX-170030357 | 0 | C | T | 0.167 | -1.47E-02 | 1.13E-01 | 2.65E+00 | 2.73E+00 | 8.98E-01 | no | Sfrp1 | intron_variant | MODIFIER |
| 8 | 23439682 | AX-170090063 | 0 | T | C | 0.167 | -1.47E-02 | 1.13E-01 | 2.65E+00 | 2.73E+00 | 8.98E-01 | no | Sfrp1 | intron_variant | MODIFIER |
| 8 | 23441005 | AX-170282783 | 0 | C | T | 0.167 | -1.47E-02 | 1.13E-01 | 2.65E+00 | 2.73E+00 | 8.98E-01 | no | Sfrp1 | intron_variant | MODIFIER |
| 8 | 23466725 | AX-170258373 | 0 | C | T | 0.167 | -1.47E-02 | 1.13E-01 | 2.65E+00 | 2.73E+00 | 8.98E-01 | no | | intergenic_variant | MODIFIER |
| 8 | 23470741 | AX-97845453 | 0 | T | C | 0.167 | -1.47E-02 | 1.13E-01 | 2.65E+00 | 2.73E+00 | 8.98E-01 | no | | regulatory_region_variant | MODIFIER |
| 8 | 23476272 | AX-170401490 | 0 | A | C | 0.167 | -1.47E-02 | 1.13E-01 | 2.65E+00 | 2.73E+00 | 8.98E-01 | no | | regulatory_region_variant | MODIFIER |
| 8 | 23477380 | AX-170030354 | 0 | T | C | 0.167 | -1.47E-02 | 1.13E-01 | 2.65E+00 | 2.73E+00 | 8.98E-01 | no | | intergenic_variant | MODIFIER |
| 8 | 23489908 | AX-170256253 | 0 | A | G | 0.167 | -1.47E-02 | 1.13E-01 | 2.65E+00 | 2.73E+00 | 8.98E-01 | no | | intergenic_variant | MODIFIER |
| 8 | 23491626 | AX-97849105 | 0 | G | A | 0.167 | -1.47E-02 | 1.13E-01 | 2.65E+00 | 2.73E+00 | 8.98E-01 | no | | intergenic_variant | MODIFIER |
| 8 | 23525303 | AX-97849437 | 0 | T | C | 0.167 | -1.47E-02 | 1.13E-01 | 2.65E+00 | 2.73E+00 | 8.98E-01 | no | | intergenic_variant | MODIFIER |
| 8 | 23525363 | AX-170030352 | 0 | T | C | 0.167 | -1.47E-02 | 1.13E-01 | 2.65E+00 | 2.73E+00 | 8.98E-01 | no | | intergenic_variant | MODIFIER |
| 8 | 23543148 | AX-97802097 | 0 | A | G | 0.167 | -1.47E-02 | 1.13E-01 | 2.65E+00 | 2.73E+00 | 8.98E-01 | no | | intergenic_variant | MODIFIER |
| 8 | 23555286 | AX-170030351 | 0 | A | C | 0.167 | -1.47E-02 | 1.13E-01 | 2.65E+00 | 2.73E+00 | 8.98E-01 | no | | intergenic_variant | MODIFIER |
| 8 | 23580561 | AX-97824943 | 0 | G | A | 0.167 | -1.47E-02 | 1.13E-01 | 2.65E+00 | 2.73E+00 | 8.98E-01 | no | | intergenic_variant | MODIFIER |
| 8 | 23593980 | AX-97836278 | 0 | C | T | 0.167 | -1.47E-02 | 1.13E-01 | 2.65E+00 | 2.73E+00 | 8.98E-01 | no | | intergenic_variant | MODIFIER |
| 8 | 23595349 | AX-170343637 | 0 | T | C | 0.167 | -1.47E-02 | 1.13E-01 | 2.65E+00 | 2.73E+00 | 8.98E-01 | no | | intergenic_variant | MODIFIER |
| 8 | 23600868 | AX-170030349 | 0 | G | A | 0.167 | -1.47E-02 | 1.13E-01 | 2.65E+00 | 2.73E+00 | 8.98E-01 | no | | intergenic_variant | MODIFIER |
| 8 | 23601562 | AX-170030358 | 0 | C | G | 0.167 | -1.47E-02 | 1.13E-01 | 2.65E+00 | 2.73E+00 | 8.98E-01 | no | | intergenic_variant | MODIFIER |
| 8 | 23602090 | AX-170211281 | 0 | T | C | 0.167 | -1.47E-02 | 1.13E-01 | 2.65E+00 | 2.73E+00 | 8.98E-01 | no | | intergenic_variant | MODIFIER |
| 8 | 23605700 | AX-170174381 | 0 | G | C | 0.167 | -1.47E-02 | 1.13E-01 | 2.65E+00 | 2.73E+00 | 8.98E-01 | no | | regulatory_region_variant | MODIFIER |
| 8 | 23633815 | AX-170440496 | 0 | C | G | 0.167 | -1.47E-02 | 1.13E-01 | 2.65E+00 | 2.73E+00 | 8.98E-01 | no | Zmat4 | upstream_gene_variant | MODIFIER |
| 8 | 23645238 | AX-170483148 | 0 | A | G | 0.167 | -1.47E-02 | 1.13E-01 | 2.65E+00 | 2.73E+00 | 8.98E-01 | no | Zmat4 | intron_variant | MODIFIER |
| 8 | 23645712 | AX-97808748 | 0 | T | C | 0.167 | -1.47E-02 | 1.13E-01 | 2.65E+00 | 2.73E+00 | 8.98E-01 | no | Zmat4 | intron_variant | MODIFIER |
| 8 | 23647713 | AX-170260434 | 0 | A | G | 0.167 | -1.47E-02 | 1.13E-01 | 2.65E+00 | 2.73E+00 | 8.98E-01 | no | Zmat4 | intron_variant | MODIFIER |
| 8 | 23649205 | AX-170354267 | 0 | G | T | 0.167 | -1.47E-02 | 1.13E-01 | 2.65E+00 | 2.73E+00 | 8.98E-01 | no | Zmat4 | intron_variant | MODIFIER |
| 8 | 23657054 | AX-170266845 | 0 | T | C | 0.167 | -1.47E-02 | 1.13E-01 | 2.65E+00 | 2.73E+00 | 8.98E-01 | no | Zmat4 | intron_variant | MODIFIER |
| 8 | 23670357 | AX-170270468 | 0 | C | G | 0.167 | -1.47E-02 | 1.13E-01 | 2.65E+00 | 2.73E+00 | 8.98E-01 | no | Zmat4 | intron_variant | MODIFIER |
| 8 | 23684204 | AX-170211774 | 0 | A | C | 0.167 | -1.47E-02 | 1.13E-01 | 2.65E+00 | 2.73E+00 | 8.98E-01 | no | Zmat4 | intron_variant | MODIFIER |
| 8 | 23684371 | AX-170341893 | 0 | G | T | 0.167 | -1.47E-02 | 1.13E-01 | 2.65E+00 | 2.73E+00 | 8.98E-01 | no | Zmat4 | intron_variant | MODIFIER |
| 8 | 23687281 | AX-170526077 | 0 | C | T | 0.167 | -1.47E-02 | 1.13E-01 | 2.65E+00 | 2.73E+00 | 8.98E-01 | no | Zmat4 | intron_variant | MODIFIER |
| 8 | 23691696 | AX-97793081 | 0 | A | G | 0.167 | -1.47E-02 | 1.13E-01 | 2.65E+00 | 2.73E+00 | 8.98E-01 | no | Zmat4 | intron_variant | MODIFIER |
| 8 | 23701307 | AX-170472065 | 0 | C | T | 0.167 | -1.47E-02 | 1.13E-01 | 2.65E+00 | 2.73E+00 | 8.98E-01 | no | Zmat4 | intron_variant | MODIFIER |
| 8 | 23728863 | AX-170432333 | 0 | G | A | 0.167 | -1.47E-02 | 1.13E-01 | 2.65E+00 | 2.73E+00 | 8.98E-01 | no | Zmat4 | intron_variant | MODIFIER |
| 8 | 23800941 | AX-170475486 | 0 | G | A | 0.167 | -1.47E-02 | 1.13E-01 | 2.65E+00 | 2.73E+00 | 8.98E-01 | no | Zmat4 | intron_variant | MODIFIER |
| 8 | 23804153 | AX-97806118 | 0 | G | A | 0.167 | -1.47E-02 | 1.13E-01 | 2.65E+00 | 2.73E+00 | 8.98E-01 | no | Zmat4 | intron_variant | MODIFIER |
| 8 | 23806026 | AX-170285295 | 0 | T | C | 0.167 | -1.47E-02 | 1.13E-01 | 2.65E+00 | 2.73E+00 | 8.98E-01 | no | Zmat4 | intron_variant | MODIFIER |
| 8 | 23838585 | AX-170518083 | 0 | A | G | 0.167 | -1.47E-02 | 1.13E-01 | 2.65E+00 | 2.73E+00 | 8.98E-01 | no | Zmat4 | intron_variant | MODIFIER |
| 8 | 26556335 | AX-170272085 | 0 | C | T | 0.167 | -1.47E-02 | 1.13E-01 | 2.65E+00 | 2.73E+00 | 8.98E-01 | no | Gm39149 | downstream_gene_variant | MODIFIER |

| | | | | | | | | | | | | | | | |
|---|---|---|---|---|---|---|---|---|---|---|---|---|---|---|---|
| 8 | 26563714 | AX-170480627 | 0 | C | G | 0.167 | -1.47E-02 | 1.13E-01 | 2.65E+00 | 2.73E+00 | 8.98E-01 | no | Gm39149 | intron_variant,non_coding_transcript_variant | MODIFIER |
| 8 | 26564414 | AX-170079573 | 0 | T | G | 0.167 | -1.47E-02 | 1.13E-01 | 2.65E+00 | 2.73E+00 | 8.98E-01 | no | Gm39149 | intron_variant,non_coding_transcript_variant | MODIFIER |
| 8 | 26567215 | AX-170420545 | 0 | A | G | 0.167 | -1.47E-02 | 1.13E-01 | 2.65E+00 | 2.73E+00 | 8.98E-01 | no | Gm39149 | intron_variant,non_coding_transcript_variant | MODIFIER |
| 8 | 26570236 | AX-170107205 | 0 | T | C | 0.167 | -1.47E-02 | 1.13E-01 | 2.65E+00 | 2.73E+00 | 8.98E-01 | no | Gm39149 | intron_variant,non_coding_transcript_variant | MODIFIER |
| 8 | 26597877 | AX-170475696 | 0 | A | G | 0.167 | -1.47E-02 | 1.13E-01 | 2.65E+00 | 2.73E+00 | 8.98E-01 | no | | intergenic_variant | MODIFIER |
| 8 | 26598875 | AX-170030385 | 0 | C | T | 0.167 | -1.47E-02 | 1.13E-01 | 2.65E+00 | 2.73E+00 | 8.98E-01 | no | | intergenic_variant | MODIFIER |
| 8 | 26598896 | AX-170030380 | 0 | C | A | 0.167 | -1.47E-02 | 1.13E-01 | 2.65E+00 | 2.73E+00 | 8.98E-01 | no | | intergenic_variant | MODIFIER |
| 8 | 26606105 | AX-170447520 | 0 | A | G | 0.167 | -1.47E-02 | 1.13E-01 | 2.65E+00 | 2.73E+00 | 8.98E-01 | no | | intergenic_variant | MODIFIER |
| 8 | 26629259 | AX-170030384 | 0 | T | G | 0.167 | -1.47E-02 | 1.13E-01 | 2.65E+00 | 2.73E+00 | 8.98E-01 | no | Gm32050 | upstream_gene_variant | MODIFIER |
| 8 | 26633066 | AX-170030379 | 0 | A | G | 0.167 | -1.47E-02 | 1.13E-01 | 2.65E+00 | 2.73E+00 | 8.98E-01 | no | Gm45584 | upstream_gene_variant | MODIFIER |
| 8 | 26634237 | AX-170030378 | 0 | T | C | 0.167 | -1.47E-02 | 1.13E-01 | 2.65E+00 | 2.73E+00 | 8.98E-01 | no | Gm45584 | upstream_gene_variant | MODIFIER |
| 8 | 26636433 | AX-170532033 | 0 | G | A | 0.167 | -1.47E-02 | 1.13E-01 | 2.65E+00 | 2.73E+00 | 8.98E-01 | no | Gm45584 | upstream_gene_variant | MODIFIER |
| 8 | 26677296 | AX-170449790 | 0 | T | C | 0.167 | -1.47E-02 | 1.13E-01 | 2.65E+00 | 2.73E+00 | 8.98E-01 | no | | regulatory_region_variant | MODIFIER |
| 8 | 26690823 | AX-170098497 | 0 | T | C | 0.167 | -1.47E-02 | 1.13E-01 | 2.65E+00 | 2.73E+00 | 8.98E-01 | no | | intergenic_variant | MODIFIER |
| 8 | 26690941 | AX-170418713 | 0 | A | C | 0.167 | -1.47E-02 | 1.13E-01 | 2.65E+00 | 2.73E+00 | 8.98E-01 | no | | intergenic_variant | MODIFIER |
| 8 | 26691073 | AX-170030382 | 0 | T | C | 0.167 | -1.47E-02 | 1.13E-01 | 2.65E+00 | 2.73E+00 | 8.98E-01 | no | | intergenic_variant | MODIFIER |
| 8 | 26709200 | AX-170167801 | 0 | T | C | 0.167 | -1.47E-02 | 1.13E-01 | 2.65E+00 | 2.73E+00 | 8.98E-01 | no | | intergenic_variant | MODIFIER |
| 8 | 26717938 | AX-170030387 | 0 | T | C | 0.167 | -1.47E-02 | 1.13E-01 | 2.65E+00 | 2.73E+00 | 8.98E-01 | no | | intergenic_variant | MODIFIER |
| 8 | 26717996 | AX-170030383 | 0 | G | A | 0.167 | -1.47E-02 | 1.13E-01 | 2.65E+00 | 2.73E+00 | 8.98E-01 | no | | intergenic_variant | MODIFIER |
| 8 | 26731049 | AX-97817603 | 0 | T | C | 0.167 | -1.47E-02 | 1.13E-01 | 2.65E+00 | 2.73E+00 | 8.98E-01 | no | | intergenic_variant | MODIFIER |
| 8 | 26804978 | AX-170261036 | 0 | A | G | 0.167 | -1.47E-02 | 1.13E-01 | 2.65E+00 | 2.73E+00 | 8.98E-01 | no | | regulatory_region_variant | MODIFIER |
| 7 | 54043927 | AX-170029028 | 1 | G | C | 0.411 | 1.31E-02 | 9.52E-02 | 2.64E+00 | 2.71E+00 | 8.98E-01 | no | | intergenic_variant | MODIFIER |
| 8 | 88352964 | AX-168398423 | 0 | G | T | 0.193 | -1.55E-02 | 1.21E-01 | 2.65E+00 | 2.74E+00 | 8.98E-01 | no | Brd7 | intron_variant | MODIFIER |
| 1 | 52660090 | AX-170526652 | 2 | C | T | 0.109 | 3.01E-02 | 2.36E-01 | 2.69E+00 | 2.74E+00 | 8.98E-01 | no | Mfsd6 | 3_prime_UTR_variant | MODIFIER |
| 12 | 71327936 | AX-97807339 | 1 | C | T | 0.062 | -3.83E-02 | 2.99E-01 | 2.71E+00 | 2.73E+00 | 8.98E-01 | no | | intergenic_variant | MODIFIER |
| 20 | 156642934 | AX-97801261 | 0 | C | T | 0.132 | 1.42E-02 | 1.12E-01 | 2.65E+00 | 2.74E+00 | 8.98E-01 | no | | intergenic_variant | MODIFIER |
| 8 | 72685116 | AX-170278895 | 0 | C | A | 0.193 | -1.77E-02 | 1.39E-01 | 2.68E+00 | 2.73E+00 | 8.98E-01 | no | Nwd1 | intron_variant | MODIFIER |
| 8 | 77767378 | AX-170244795 | 0 | C | T | 0.193 | -1.77E-02 | 1.39E-01 | 2.68E+00 | 2.73E+00 | 8.98E-01 | no | 4933431K23Rik | non_coding_transcript_exon_variant | MODIFIER |
| 8 | 37987931 | AX-170030525 | 2 | T | C | 0.145 | 1.59E-02 | 1.32E-01 | 2.69E+00 | 2.76E+00 | 8.98E-01 | no | Sgcz | intron_variant | MODIFIER |
| 7 | 106865138 | AX-170345834 | 0 | G | A | 0.474 | -1.14E-02 | 9.35E-02 | 2.68E+00 | 2.75E+00 | 8.98E-01 | no | Olfr704 | missense_variant | MODERATE |
| 9 | 69115117 | AX-170498141 | 0 | A | T | 0.202 | 1.58E-02 | 1.26E-01 | 2.72E+00 | 2.75E+00 | 8.98E-01 | no | Rora | intron_variant | MODIFIER |
| 5 | 73059301 | AX-97799676 | 1 | G | T | 0.438 | 1.32E-02 | 9.60E-02 | 2.63E+00 | 2.72E+00 | 8.98E-01 | no | Fryl | intron_variant | MODIFIER |
| 15 | 100813210 | AX-170451966 | 0 | G | T | 0.114 | 1.93E-02 | 1.50E-01 | 2.66E+00 | 2.73E+00 | 8.98E-01 | no | Slc4a8 | intron_variant | MODIFIER |
| 3 | 148041139 | AX-170021872 | 2 | C | T | 0.364 | -8.35E-03 | 7.07E-02 | 2.67E+00 | 2.76E+00 | 8.98E-01 | no | | intergenic_variant | MODIFIER |
| 8 | 103245920 | AX-170272062 | 0 | C | A | 0.219 | -1.30E-02 | 1.06E-01 | 2.66E+00 | 2.75E+00 | 8.98E-01 | no | | regulatory_region_variant | MODIFIER |
| 8 | 103904021 | AX-170102581 | 0 | A | G | 0.219 | -1.30E-02 | 1.06E-01 | 2.66E+00 | 2.75E+00 | 8.98E-01 | no | | regulatory_region_variant | MODIFIER |
| 10 | 13971139 | AX-170044183 | 0 | G | A | 0.219 | 1.12E-02 | 9.63E-02 | 2.66E+00 | 2.76E+00 | 8.99E-01 | no | Hivep2 | intron_variant | MODIFIER |
| 5 | 73050649 | AX-170475244 | 0 | C | T | 0.43 | 1.31E-02 | 9.82E-02 | 2.64E+00 | 2.72E+00 | 8.99E-01 | no | Fryl | intron_variant | MODIFIER |
| 5 | 73050835 | AX-170137457 | 0 | T | C | 0.43 | 1.31E-02 | 9.82E-02 | 2.64E+00 | 2.72E+00 | 8.99E-01 | no | Fryl | intron_variant | MODIFIER |
| 5 | 73053177 | AX-170124406 | 0 | C | T | 0.43 | 1.31E-02 | 9.82E-02 | 2.64E+00 | 2.72E+00 | 8.99E-01 | no | Fryl | intron_variant | MODIFIER |
| 5 | 73133263 | AX-170072456 | 0 | A | C | 0.43 | 1.31E-02 | 9.82E-02 | 2.64E+00 | 2.72E+00 | 8.99E-01 | no | Fryl | synonymous_variant | LOW |
| 5 | 73137710 | AX-170072454 | 0 | G | A | 0.43 | 1.31E-02 | 9.82E-02 | 2.64E+00 | 2.72E+00 | 8.99E-01 | no | Fryl | intron_variant | MODIFIER |
| 5 | 73148833 | AX-170354105 | 0 | A | G | 0.43 | 1.31E-02 | 9.82E-02 | 2.64E+00 | 2.72E+00 | 8.99E-01 | no | Fryl | upstream_gene_variant | MODIFIER |
| 5 | 73161943 | AX-170292798 | 0 | T | C | 0.43 | 1.31E-02 | 9.82E-02 | 2.64E+00 | 2.72E+00 | 8.99E-01 | no | Fryl | intron_variant | MODIFIER |
| 5 | 73167022 | AX-170499160 | 0 | G | A | 0.43 | 1.31E-02 | 9.82E-02 | 2.64E+00 | 2.72E+00 | 8.99E-01 | no | Fryl | intron_variant | MODIFIER |
| 5 | 73232724 | AX-170176759 | 0 | A | G | 0.43 | 1.31E-02 | 9.82E-02 | 2.64E+00 | 2.72E+00 | 8.99E-01 | no | Fryl | intron_variant | MODIFIER |
| 5 | 73235401 | AX-170423629 | 0 | G | A | 0.43 | 1.31E-02 | 9.82E-02 | 2.64E+00 | 2.72E+00 | 8.99E-01 | no | Fryl | intron_variant | MODIFIER |
| 5 | 73269787 | AX-170101556 | 0 | A | G | 0.43 | 1.31E-02 | 9.82E-02 | 2.64E+00 | 2.72E+00 | 8.99E-01 | no | 1700025M24Rik | intron_variant,non_coding_transcript_variant | MODIFIER |
| 5 | 73283985 | AX-97808581 | 0 | G | A | 0.43 | 1.31E-02 | 9.82E-02 | 2.64E+00 | 2.72E+00 | 8.99E-01 | no | 1700025M24Rik | intron_variant,non_coding_transcript_variant | MODIFIER |
| 5 | 73286142 | AX-170375577 | 0 | A | G | 0.43 | 1.31E-02 | 9.82E-02 | 2.64E+00 | 2.72E+00 | 8.99E-01 | no | 1700025M24Rik | upstream_gene_variant | MODIFIER |
| 5 | 73288908 | AX-170072459 | 0 | T | C | 0.43 | 1.31E-02 | 9.82E-02 | 2.64E+00 | 2.72E+00 | 8.99E-01 | no | Ociad1 | upstream_gene_variant | MODIFIER |
| 5 | 73291709 | AX-170072460 | 0 | T | G | 0.43 | 1.31E-02 | 9.82E-02 | 2.64E+00 | 2.72E+00 | 8.99E-01 | no | Ociad1 | upstream_gene_variant | MODIFIER |
| 5 | 73313363 | AX-97832841 | 0 | T | C | 0.43 | 1.31E-02 | 9.82E-02 | 2.64E+00 | 2.72E+00 | 8.99E-01 | no | Ociad1 | intron_variant | MODIFIER |
| 5 | 73313531 | AX-170366803 | 0 | T | C | 0.43 | 1.31E-02 | 9.82E-02 | 2.64E+00 | 2.72E+00 | 8.99E-01 | no | Ociad1 | 3_prime_UTR_variant | MODIFIER |

| | | | | | | | | | | | | | | | |
|---|---|---|---|---|---|---|---|---|---|---|---|---|---|---|---|
| 5 | 73335938 | AX-170512398 | 0 | C | T | 0.43 | 1.31E-02 | 9.82E-02 | 2.64E+00 | 2.72E+00 | 8.99E-01 | no | Ociad2 | 5_prime_UTR_variant | MODIFIER |
| 5 | 73336480 | AX-170138962 | 0 | G | C | 0.43 | 1.31E-02 | 9.82E-02 | 2.64E+00 | 2.72E+00 | 8.99E-01 | no | Ociad2 | intron_variant | MODIFIER |
| 5 | 73337295 | AX-170273627 | 0 | G | A | 0.43 | 1.31E-02 | 9.82E-02 | 2.64E+00 | 2.72E+00 | 8.99E-01 | no | Ociad2 | intron_variant | MODIFIER |
| 5 | 73340435 | AX-170319508 | 0 | C | T | 0.43 | 1.31E-02 | 9.82E-02 | 2.64E+00 | 2.72E+00 | 8.99E-01 | no | Ociad2 | upstream_gene_variant | MODIFIER |
| 5 | 73340930 | AX-170350855 | 0 | T | C | 0.43 | 1.31E-02 | 9.82E-02 | 2.64E+00 | 2.72E+00 | 8.99E-01 | no | Ociad2 | upstream_gene_variant | MODIFIER |
| 5 | 73409882 | AX-170272448 | 0 | A | G | 0.43 | 1.31E-02 | 9.82E-02 | 2.64E+00 | 2.72E+00 | 8.99E-01 | no | Cwh43 | intron_variant | MODIFIER |
| 5 | 73413118 | AX-170114118 | 0 | G | A | 0.43 | 1.31E-02 | 9.82E-02 | 2.64E+00 | 2.72E+00 | 8.99E-01 | no | Cwh43 | synonymous_variant | LOW |
| 5 | 73434323 | AX-170025280 | 0 | T | C | 0.43 | 1.31E-02 | 9.82E-02 | 2.64E+00 | 2.72E+00 | 8.99E-01 | no | Cwh43 | synonymous_variant | LOW |
| 5 | 73458076 | AX-170025281 | 0 | C | T | 0.43 | 1.31E-02 | 9.82E-02 | 2.64E+00 | 2.72E+00 | 8.99E-01 | no | Cwh43 | downstream_gene_variant | MODIFIER |
| 5 | 73501988 | AX-170025283 | 0 | A | G | 0.43 | 1.31E-02 | 9.82E-02 | 2.64E+00 | 2.72E+00 | 8.99E-01 | no | Dcun1d4 | intron_variant | MODIFIER |
| 5 | 73504638 | AX-170414007 | 0 | C | T | 0.43 | 1.31E-02 | 9.82E-02 | 2.64E+00 | 2.72E+00 | 8.99E-01 | no | Dcun1d4 | intron_variant | MODIFIER |
| 5 | 73524436 | AX-170140194 | 0 | A | G | 0.43 | 1.31E-02 | 9.82E-02 | 2.64E+00 | 2.72E+00 | 8.99E-01 | no | Dcun1d4 | intron_variant | MODIFIER |
| 5 | 73541350 | AX-170476762 | 0 | T | C | 0.43 | 1.31E-02 | 9.82E-02 | 2.64E+00 | 2.72E+00 | 8.99E-01 | no | Dcun1d4 | intron_variant | MODIFIER |
| 5 | 73553705 | AX-170188397 | 0 | A | G | 0.43 | 1.31E-02 | 9.82E-02 | 2.64E+00 | 2.72E+00 | 8.99E-01 | no | Dcun1d4 | intron_variant | MODIFIER |
| 5 | 73580493 | AX-170233051 | 0 | G | A | 0.43 | 1.31E-02 | 9.82E-02 | 2.64E+00 | 2.72E+00 | 8.99E-01 | no | | intergenic_variant | MODIFIER |
| 5 | 73581391 | AX-170333467 | 0 | T | G | 0.43 | 1.31E-02 | 9.82E-02 | 2.64E+00 | 2.72E+00 | 8.99E-01 | no | | intergenic_variant | MODIFIER |
| 5 | 73607599 | AX-170072469 | 0 | A | G | 0.43 | 1.31E-02 | 9.82E-02 | 2.64E+00 | 2.72E+00 | 8.99E-01 | no | Lrrc66 | synonymous_variant | LOW |
| 5 | 73608571 | AX-170267315 | 0 | T | C | 0.43 | 1.31E-02 | 9.82E-02 | 2.64E+00 | 2.72E+00 | 8.99E-01 | no | Lrrc66 | synonymous_variant | LOW |
| 2 | 118594385 | AX-97826728 | 1 | C | T | 0.134 | 2.49E-02 | 1.97E-01 | 2.69E+00 | 2.74E+00 | 8.99E-01 | no | Bub1b | upstream_gene_variant | MODIFIER |
| 5 | 96740873 | AX-170402028 | 0 | T | C | 0.439 | 1.30E-02 | 1.04E-01 | 2.68E+00 | 2.74E+00 | 8.99E-01 | no | Fras1 | synonymous_variant | LOW |
| 8 | 36266432 | AX-97806515 | 2 | A | G | 0.145 | 1.72E-02 | 1.42E-01 | 2.70E+00 | 2.76E+00 | 8.99E-01 | no | | intergenic_variant | MODIFIER |
| 6 | 131422873 | AX-97853610 | 0 | T | C | 0.07 | 2.80E-02 | 2.18E-01 | 2.66E+00 | 2.73E+00 | 8.99E-01 | no | | intergenic_variant | MODIFIER |
| 12 | 101288087 | AX-170473101 | 0 | A | G | 0.07 | 2.80E-02 | 2.18E-01 | 2.66E+00 | 2.73E+00 | 8.99E-01 | no | Gm18795 | intron_variant,non_coding_transcript_variant | MODIFIER |
| 12 | 101288105 | AX-97826865 | 0 | G | A | 0.07 | 2.80E-02 | 2.18E-01 | 2.66E+00 | 2.73E+00 | 8.99E-01 | no | Gm18795 | intron_variant,non_coding_transcript_variant | MODIFIER |
| 7 | 70046288 | AX-170213236 | 1 | T | C | 0.393 | 1.25E-02 | 9.00E-02 | 2.62E+00 | 2.71E+00 | 8.99E-01 | no | | intergenic_variant | MODIFIER |
| 20 | 130288103 | AX-170161086 | 0 | A | G | 0.316 | 7.06E-03 | 5.29E-02 | 2.60E+00 | 2.72E+00 | 8.99E-01 | no | Diaph2 | intron_variant | MODIFIER |
| 20 | 130295704 | AX-170103212 | 0 | C | G | 0.316 | 7.06E-03 | 5.29E-02 | 2.60E+00 | 2.72E+00 | 8.99E-01 | no | Diaph2 | intron_variant | MODIFIER |
| 20 | 130310151 | AX-97838550 | 0 | T | C | 0.316 | 7.06E-03 | 5.29E-02 | 2.60E+00 | 2.72E+00 | 8.99E-01 | no | Diaph2 | intron_variant | MODIFIER |
| 20 | 130351805 | AX-170483794 | 0 | G | A | 0.316 | 7.06E-03 | 5.29E-02 | 2.60E+00 | 2.72E+00 | 8.99E-01 | no | Diaph2 | intron_variant | MODIFIER |
| 20 | 130361901 | AX-170237175 | 0 | A | G | 0.316 | 7.06E-03 | 5.29E-02 | 2.60E+00 | 2.72E+00 | 8.99E-01 | no | Diaph2 | intron_variant | MODIFIER |
| 20 | 130381004 | AX-170238969 | 0 | A | G | 0.316 | 7.06E-03 | 5.29E-02 | 2.60E+00 | 2.72E+00 | 8.99E-01 | no | Diaph2 | intron_variant | MODIFIER |
| 20 | 130385353 | AX-170359060 | 0 | T | C | 0.316 | 7.06E-03 | 5.29E-02 | 2.60E+00 | 2.72E+00 | 8.99E-01 | no | Diaph2 | intron_variant | MODIFIER |
| 20 | 130389869 | AX-97784239 | 0 | C | T | 0.316 | 7.06E-03 | 5.29E-02 | 2.60E+00 | 2.72E+00 | 8.99E-01 | no | Diaph2 | intron_variant | MODIFIER |
| 20 | 130390562 | AX-170181239 | 0 | T | C | 0.316 | 7.06E-03 | 5.29E-02 | 2.60E+00 | 2.72E+00 | 8.99E-01 | no | Diaph2 | intron_variant | MODIFIER |
| 20 | 130448615 | AX-170338890 | 0 | C | A | 0.316 | 7.06E-03 | 5.29E-02 | 2.60E+00 | 2.72E+00 | 8.99E-01 | no | Diaph2 | intron_variant | MODIFIER |
| 20 | 130474985 | AX-170289123 | 0 | T | C | 0.316 | 7.06E-03 | 5.29E-02 | 2.60E+00 | 2.72E+00 | 8.99E-01 | no | | intergenic_variant | MODIFIER |
| 20 | 130644183 | AX-170034935 | 0 | T | C | 0.316 | 7.06E-03 | 5.29E-02 | 2.60E+00 | 2.72E+00 | 8.99E-01 | no | | intergenic_variant | MODIFIER |
| 20 | 130653334 | AX-170034934 | 0 | A | G | 0.316 | 7.06E-03 | 5.29E-02 | 2.60E+00 | 2.72E+00 | 8.99E-01 | no | | intergenic_variant | MODIFIER |
| 20 | 130662803 | AX-170034937 | 0 | A | C | 0.316 | 7.06E-03 | 5.29E-02 | 2.60E+00 | 2.72E+00 | 8.99E-01 | no | | intergenic_variant | MODIFIER |
| 20 | 130666200 | AX-170085246 | 0 | C | A | 0.316 | 7.06E-03 | 5.29E-02 | 2.60E+00 | 2.72E+00 | 8.99E-01 | no | | intergenic_variant | MODIFIER |
| 20 | 130666337 | AX-170085241 | 0 | T | C | 0.316 | 7.06E-03 | 5.29E-02 | 2.60E+00 | 2.72E+00 | 8.99E-01 | no | | intergenic_variant | MODIFIER |
| 20 | 130673345 | AX-170034936 | 0 | T | C | 0.316 | 7.06E-03 | 5.29E-02 | 2.60E+00 | 2.72E+00 | 8.99E-01 | no | | intergenic_variant | MODIFIER |
| 20 | 130674119 | AX-170034938 | 0 | T | C | 0.316 | 7.06E-03 | 5.29E-02 | 2.60E+00 | 2.72E+00 | 8.99E-01 | no | | intergenic_variant | MODIFIER |
| 20 | 130677220 | AX-170223206 | 0 | G | A | 0.316 | 7.06E-03 | 5.29E-02 | 2.60E+00 | 2.72E+00 | 8.99E-01 | no | | intergenic_variant | MODIFIER |
| 4 | 142279078 | AX-170162090 | 2 | T | C | 0.1 | -1.63E-02 | 1.25E-01 | 2.61E+00 | 2.73E+00 | 8.99E-01 | no | | intergenic_variant | MODIFIER |
| 1 | 53985448 | AX-170113339 | 1 | T | C | 0.098 | 2.62E-02 | 2.09E-01 | 2.64E+00 | 2.74E+00 | 8.99E-01 | no | Hecw2 | intron_variant | MODIFIER |
| 18 | 66519486 | AX-170520312 | 1 | C | T | 0.089 | -2.04E-02 | 1.75E-01 | 2.64E+00 | 2.76E+00 | 8.99E-01 | no | AC100212.4 | upstream_gene_variant | MODIFIER |
| 6 | 146993063 | AX-169999806 | 1 | A | T | 0.054 | -2.67E-02 | 2.11E-01 | 2.62E+00 | 2.74E+00 | 8.99E-01 | no | Ppfibp1 | intron_variant | MODIFIER |
| 18 | 39340467 | AX-170060629 | 1 | G | A | 0.062 | 3.74E-02 | 3.02E-01 | 2.73E+00 | 2.76E+00 | 8.99E-01 | no | Arhgap26 | intron_variant | MODIFIER |
| 1 | 109220266 | AX-97812730 | 1 | T | C | 0.152 | 2.65E-02 | 1.92E-01 | 2.62E+00 | 2.71E+00 | 8.99E-01 | no | | regulatory_region_variant | MODIFIER |
| 5 | 117529239 | AX-170292166 | 0 | C | T | 0.456 | -1.42E-02 | 1.16E-01 | 2.73E+00 | 2.76E+00 | 9.00E-01 | no | Ksr2 | intron_variant,non_coding_transcript_variant | MODIFIER |
| 5 | 117530605 | AX-170336329 | 0 | T | C | 0.456 | -1.42E-02 | 1.16E-01 | 2.73E+00 | 2.76E+00 | 9.00E-01 | no | Ksr2 | intron_variant,non_coding_transcript_variant | MODIFIER |
| 5 | 117534809 | AX-170322701 | 0 | G | A | 0.456 | -1.42E-02 | 1.16E-01 | 2.73E+00 | 2.76E+00 | 9.00E-01 | no | Ksr2 | intron_variant,non_coding_transcript_variant | MODIFIER |
| 5 | 117544287 | AX-170165776 | 0 | A | G | 0.456 | -1.42E-02 | 1.16E-01 | 2.73E+00 | 2.76E+00 | 9.00E-01 | no | Ksr2 | intron_variant,non_coding_transcript_variant | MODIFIER |

| | | | | | | | | | | | | | | | |
|---|---|---|---|---|---|---|---|---|---|---|---|---|---|---|---|
| 5 | 117551313 | AX-97813356 | 0 | A | C | 0.456 | -1.42E-02 | 1.16E-01 | 2.73E+00 | 2.76E+00 | 9.00E-01 | no | Ksr2 | intron_variant,non_coding_transcript_variant | MODIFIER |
| 5 | 117567679 | AX-170025968 | 0 | C | T | 0.456 | -1.42E-02 | 1.16E-01 | 2.73E+00 | 2.76E+00 | 9.00E-01 | no | Ksr2 | intron_variant | MODIFIER |
| 5 | 117570284 | AX-168373263 | 0 | C | T | 0.456 | -1.42E-02 | 1.16E-01 | 2.73E+00 | 2.76E+00 | 9.00E-01 | no | Ksr2 | intron_variant | MODIFIER |
| 5 | 117583117 | AX-170440698 | 0 | G | A | 0.456 | -1.42E-02 | 1.16E-01 | 2.73E+00 | 2.76E+00 | 9.00E-01 | no | Ksr2 | intron_variant | MODIFIER |
| 5 | 117597871 | AX-170497058 | 0 | A | T | 0.456 | -1.42E-02 | 1.16E-01 | 2.73E+00 | 2.76E+00 | 9.00E-01 | no | Ksr2 | intron_variant | MODIFIER |
| 5 | 117600573 | AX-170480981 | 0 | C | T | 0.456 | -1.42E-02 | 1.16E-01 | 2.73E+00 | 2.76E+00 | 9.00E-01 | no | Ksr2 | intron_variant | MODIFIER |
| 5 | 117600872 | AX-170466675 | 0 | T | C | 0.456 | -1.42E-02 | 1.16E-01 | 2.73E+00 | 2.76E+00 | 9.00E-01 | no | Ksr2 | intron_variant | MODIFIER |
| 5 | 117605247 | AX-170394203 | 0 | G | T | 0.456 | -1.42E-02 | 1.16E-01 | 2.73E+00 | 2.76E+00 | 9.00E-01 | no | Ksr2 | intron_variant | MODIFIER |
| 5 | 117606097 | AX-170427521 | 0 | A | T | 0.456 | -1.42E-02 | 1.16E-01 | 2.73E+00 | 2.76E+00 | 9.00E-01 | no | Ksr2 | intron_variant | MODIFIER |
| 5 | 117606321 | AX-170308543 | 0 | G | A | 0.456 | -1.42E-02 | 1.16E-01 | 2.73E+00 | 2.76E+00 | 9.00E-01 | no | Ksr2 | intron_variant | MODIFIER |
| 5 | 117618242 | AX-170521565 | 0 | T | G | 0.456 | -1.42E-02 | 1.16E-01 | 2.73E+00 | 2.76E+00 | 9.00E-01 | no | Ksr2 | intron_variant | MODIFIER |
| 5 | 117619088 | AX-170275074 | 0 | T | C | 0.456 | -1.42E-02 | 1.16E-01 | 2.73E+00 | 2.76E+00 | 9.00E-01 | no | Ksr2 | intron_variant | MODIFIER |
| 5 | 117627124 | AX-97857449 | 0 | G | T | 0.456 | -1.42E-02 | 1.16E-01 | 2.73E+00 | 2.76E+00 | 9.00E-01 | no | Ksr2 | intron_variant | MODIFIER |
| 5 | 117627715 | AX-170226437 | 0 | A | G | 0.456 | -1.42E-02 | 1.16E-01 | 2.73E+00 | 2.76E+00 | 9.00E-01 | no | Ksr2 | intron_variant | MODIFIER |
| 5 | 117648884 | AX-97825034 | 0 | G | A | 0.456 | -1.42E-02 | 1.16E-01 | 2.73E+00 | 2.76E+00 | 9.00E-01 | no | Ksr2 | intron_variant | MODIFIER |
| 5 | 117682955 | AX-97791241 | 0 | C | T | 0.456 | -1.42E-02 | 1.16E-01 | 2.73E+00 | 2.76E+00 | 9.00E-01 | no | Ksr2 | intron_variant | MODIFIER |
| 5 | 117716928 | AX-97834311 | 0 | G | A | 0.456 | -1.42E-02 | 1.16E-01 | 2.73E+00 | 2.76E+00 | 9.00E-01 | no | Ksr2 | intron_variant | MODIFIER |
| 5 | 117720759 | AX-97833746 | 0 | A | G | 0.456 | -1.42E-02 | 1.16E-01 | 2.73E+00 | 2.76E+00 | 9.00E-01 | no | Ksr2 | intron_variant | MODIFIER |
| 5 | 117748450 | AX-97786524 | 0 | T | G | 0.456 | -1.42E-02 | 1.16E-01 | 2.73E+00 | 2.76E+00 | 9.00E-01 | no | Ksr2 | intron_variant | MODIFIER |
| 5 | 117765720 | AX-170307144 | 0 | T | C | 0.456 | -1.42E-02 | 1.16E-01 | 2.73E+00 | 2.76E+00 | 9.00E-01 | no | Ksr2 | 3_prime_UTR_variant | MODIFIER |
| 5 | 117766548 | AX-170494311 | 0 | T | C | 0.456 | -1.42E-02 | 1.16E-01 | 2.73E+00 | 2.76E+00 | 9.00E-01 | no | Ksr2 | 3_prime_UTR_variant | MODIFIER |
| 5 | 117776459 | AX-170255907 | 0 | G | A | 0.456 | -1.42E-02 | 1.16E-01 | 2.73E+00 | 2.76E+00 | 9.00E-01 | no | Nos1 | upstream_gene_variant | MODIFIER |
| 5 | 117782832 | AX-170218567 | 0 | G | A | 0.456 | -1.42E-02 | 1.16E-01 | 2.73E+00 | 2.76E+00 | 9.00E-01 | no | Nos1 | intron_variant,non_coding_transcript_variant | MODIFIER |
| 5 | 117816540 | AX-97814152 | 0 | A | G | 0.456 | -1.42E-02 | 1.16E-01 | 2.73E+00 | 2.76E+00 | 9.00E-01 | no | Nos1 | intron_variant,non_coding_transcript_variant | MODIFIER |
| 5 | 117818938 | AX-170305941 | 0 | G | A | 0.456 | -1.42E-02 | 1.16E-01 | 2.73E+00 | 2.76E+00 | 9.00E-01 | no | Nos1 | intron_variant,non_coding_transcript_variant | MODIFIER |
| 5 | 117850529 | AX-170025979 | 0 | C | T | 0.456 | -1.42E-02 | 1.16E-01 | 2.73E+00 | 2.76E+00 | 9.00E-01 | no | Nos1 | intron_variant,NMD_transcript_variant | MODIFIER |
| 5 | 117870420 | AX-170025975 | 0 | A | G | 0.456 | -1.42E-02 | 1.16E-01 | 2.73E+00 | 2.76E+00 | 9.00E-01 | no | Nos1 | intron_variant,NMD_transcript_variant | MODIFIER |
| 5 | 117929120 | AX-170255776 | 0 | C | T | 0.456 | -1.42E-02 | 1.16E-01 | 2.73E+00 | 2.76E+00 | 9.00E-01 | no | Nos1 | intron_variant,NMD_transcript_variant | MODIFIER |
| 5 | 117929149 | AX-170201389 | 0 | A | G | 0.456 | -1.42E-02 | 1.16E-01 | 2.73E+00 | 2.76E+00 | 9.00E-01 | no | Nos1 | intron_variant,NMD_transcript_variant | MODIFIER |
| 5 | 117955608 | AX-97812694 | 0 | A | G | 0.456 | -1.42E-02 | 1.16E-01 | 2.73E+00 | 2.76E+00 | 9.00E-01 | no | Nos1 | downstream_gene_variant | MODIFIER |
| 5 | 117994241 | AX-97821640 | 0 | A | G | 0.456 | -1.42E-02 | 1.16E-01 | 2.73E+00 | 2.76E+00 | 9.00E-01 | no | Fbxo21 | intron_variant | MODIFIER |
| 5 | 118003435 | AX-169999794 | 0 | G | A | 0.456 | -1.42E-02 | 1.16E-01 | 2.73E+00 | 2.76E+00 | 9.00E-01 | no | Fbxo21 | intron_variant | MODIFIER |
| 5 | 118003947 | AX-170025978 | 0 | T | C | 0.456 | -1.42E-02 | 1.16E-01 | 2.73E+00 | 2.76E+00 | 9.00E-01 | no | Fbxo21 | intron_variant | MODIFIER |
| 5 | 118004994 | AX-170348468 | 0 | A | G | 0.456 | -1.42E-02 | 1.16E-01 | 2.73E+00 | 2.76E+00 | 9.00E-01 | no | Fbxo21 | intron_variant | MODIFIER |
| 5 | 118005157 | AX-170453676 | 0 | C | T | 0.456 | -1.42E-02 | 1.16E-01 | 2.73E+00 | 2.76E+00 | 9.00E-01 | no | Fbxo21 | intron_variant | MODIFIER |
| 5 | 118006914 | AX-170385108 | 0 | G | A | 0.456 | -1.42E-02 | 1.16E-01 | 2.73E+00 | 2.76E+00 | 9.00E-01 | no | Fbxo21 | intron_variant | MODIFIER |
| 5 | 118011450 | AX-170103201 | 0 | C | A | 0.456 | -1.42E-02 | 1.16E-01 | 2.73E+00 | 2.76E+00 | 9.00E-01 | no | Fbxo21 | downstream_gene_variant | MODIFIER |
| 5 | 118053656 | AX-170114290 | 0 | T | C | 0.456 | -1.42E-02 | 1.16E-01 | 2.73E+00 | 2.76E+00 | 9.00E-01 | no | Tesc | synonymous_variant | LOW |
| 5 | 118055626 | AX-97831710 | 0 | G | A | 0.456 | -1.42E-02 | 1.16E-01 | 2.73E+00 | 2.76E+00 | 9.00E-01 | no | Tesc | intron_variant | MODIFIER |
| 5 | 118074854 | AX-97806111 | 0 | C | T | 0.456 | -1.42E-02 | 1.16E-01 | 2.73E+00 | 2.76E+00 | 9.00E-01 | no | Fbxw8 | intron_variant | MODIFIER |
| 5 | 118076296 | AX-170152244 | 0 | C | T | 0.456 | -1.42E-02 | 1.16E-01 | 2.73E+00 | 2.76E+00 | 9.00E-01 | no | Fbxw8 | intron_variant | MODIFIER |
| 5 | 118076785 | AX-170385549 | 0 | C | T | 0.456 | -1.42E-02 | 1.16E-01 | 2.73E+00 | 2.76E+00 | 9.00E-01 | no | Fbxw8 | intron_variant | MODIFIER |
| 5 | 118114783 | AX-170180738 | 0 | C | T | 0.456 | -1.42E-02 | 1.16E-01 | 2.73E+00 | 2.76E+00 | 9.00E-01 | no | Fbxw8 | intron_variant | MODIFIER |
| 5 | 118134678 | AX-170253030 | 0 | T | C | 0.456 | -1.42E-02 | 1.16E-01 | 2.73E+00 | 2.76E+00 | 9.00E-01 | no | Fbxw8 | intron_variant | MODIFIER |
| 5 | 118140758 | AX-170281100 | 0 | T | C | 0.456 | -1.42E-02 | 1.16E-01 | 2.73E+00 | 2.76E+00 | 9.00E-01 | no | Fbxw8 | intron_variant | MODIFIER |
| 5 | 118147656 | AX-170365209 | 0 | A | G | 0.456 | -1.42E-02 | 1.16E-01 | 2.73E+00 | 2.76E+00 | 9.00E-01 | no | Fbxw8 | intron_variant | MODIFIER |
| 5 | 118150377 | AX-170181902 | 0 | T | C | 0.456 | -1.42E-02 | 1.16E-01 | 2.73E+00 | 2.76E+00 | 9.00E-01 | no | Fbxw8 | intron_variant | MODIFIER |
| 5 | 118194514 | AX-170457208 | 0 | G | A | 0.456 | -1.42E-02 | 1.16E-01 | 2.73E+00 | 2.76E+00 | 9.00E-01 | no | Rnft2 | intron_variant | MODIFIER |
| 5 | 118204602 | AX-97803313 | 0 | C | G | 0.456 | -1.42E-02 | 1.16E-01 | 2.73E+00 | 2.76E+00 | 9.00E-01 | no | Rnft2 | intron_variant | MODIFIER |
| 5 | 118239553 | AX-170204108 | 0 | C | T | 0.456 | -1.42E-02 | 1.16E-01 | 2.73E+00 | 2.76E+00 | 9.00E-01 | no | Rnft2 | intron_variant | MODIFIER |
| 5 | 118239782 | AX-97810832 | 0 | T | C | 0.456 | -1.42E-02 | 1.16E-01 | 2.73E+00 | 2.76E+00 | 9.00E-01 | no | Rnft2 | intron_variant | MODIFIER |
| 5 | 118254960 | AX-170332990 | 0 | G | A | 0.456 | -1.42E-02 | 1.16E-01 | 2.73E+00 | 2.76E+00 | 9.00E-01 | no | 2410131K14Rik | intron_variant | MODIFIER |
| 5 | 118269516 | AX-170352589 | 0 | A | G | 0.456 | -1.42E-02 | 1.16E-01 | 2.73E+00 | 2.76E+00 | 9.00E-01 | no | 2410131K14Rik | downstream_gene_variant | MODIFIER |
| 5 | 118273039 | AX-170393898 | 0 | A | G | 0.456 | -1.42E-02 | 1.16E-01 | 2.73E+00 | 2.76E+00 | 9.00E-01 | no | | regulatory_region_variant | MODIFIER |

| | | | | | | | | | | | | | | | |
|---|---|---|---|---|---|---|---|---|---|---|---|---|---|---|---|
| 5 | 118273885 | AX-170282865 | 0 | A | G | 0.456 | -1.42E-02 | 1.16E-01 | 2.73E+00 | 2.76E+00 | 9.00E-01 | no | | regulatory_region_variant | MODIFIER |
| 5 | 118565216 | AX-170521366 | 0 | G | A | 0.456 | -1.42E-02 | 1.16E-01 | 2.73E+00 | 2.76E+00 | 9.00E-01 | no | Med13l | intron_variant | MODIFIER |
| 5 | 118570474 | AX-97811619 | 0 | A | G | 0.456 | -1.42E-02 | 1.16E-01 | 2.73E+00 | 2.76E+00 | 9.00E-01 | no | Med13l | intron_variant | MODIFIER |
| 5 | 118592186 | AX-170263699 | 0 | C | A | 0.456 | -1.42E-02 | 1.16E-01 | 2.73E+00 | 2.76E+00 | 9.00E-01 | no | Med13l | intron_variant | MODIFIER |
| 5 | 118597696 | AX-170434096 | 0 | A | G | 0.456 | -1.42E-02 | 1.16E-01 | 2.73E+00 | 2.76E+00 | 9.00E-01 | no | Med13l | intron_variant | MODIFIER |
| 5 | 118599137 | AX-170386501 | 0 | C | G | 0.456 | -1.42E-02 | 1.16E-01 | 2.73E+00 | 2.76E+00 | 9.00E-01 | no | Med13l | intron_variant | MODIFIER |
| 5 | 118607895 | AX-170073274 | 0 | G | A | 0.456 | -1.42E-02 | 1.16E-01 | 2.73E+00 | 2.76E+00 | 9.00E-01 | no | Med13l | intron_variant | MODIFIER |
| 5 | 118616484 | AX-168385999 | 0 | T | C | 0.456 | -1.42E-02 | 1.16E-01 | 2.73E+00 | 2.76E+00 | 9.00E-01 | no | Med13l | intron_variant | MODIFIER |
| 5 | 118677451 | AX-170291980 | 0 | C | G | 0.456 | -1.42E-02 | 1.16E-01 | 2.73E+00 | 2.76E+00 | 9.00E-01 | no | Med13l | intron_variant | MODIFIER |
| 5 | 118679155 | AX-97805215 | 0 | G | A | 0.456 | -1.42E-02 | 1.16E-01 | 2.73E+00 | 2.76E+00 | 9.00E-01 | no | Med13l | intron_variant | MODIFIER |
| 5 | 118681730 | AX-170353696 | 0 | A | G | 0.456 | -1.42E-02 | 1.16E-01 | 2.73E+00 | 2.76E+00 | 9.00E-01 | no | Med13l | intron_variant | MODIFIER |
| 5 | 118688455 | AX-170427948 | 0 | C | G | 0.456 | -1.42E-02 | 1.16E-01 | 2.73E+00 | 2.76E+00 | 9.00E-01 | no | Med13l | intron_variant | MODIFIER |
| 5 | 118691616 | AX-170229701 | 0 | T | C | 0.456 | -1.42E-02 | 1.16E-01 | 2.73E+00 | 2.76E+00 | 9.00E-01 | no | Med13l | intron_variant | MODIFIER |
| 5 | 118705067 | AX-170392662 | 0 | G | A | 0.456 | -1.42E-02 | 1.16E-01 | 2.73E+00 | 2.76E+00 | 9.00E-01 | no | Med13l | intron_variant | MODIFIER |
| 5 | 118707290 | AX-170496707 | 0 | C | A | 0.456 | -1.42E-02 | 1.16E-01 | 2.73E+00 | 2.76E+00 | 9.00E-01 | no | Med13l | intron_variant | MODIFIER |
| 5 | 118716687 | AX-170297284 | 0 | G | A | 0.456 | -1.42E-02 | 1.16E-01 | 2.73E+00 | 2.76E+00 | 9.00E-01 | no | Med13l | intron_variant | MODIFIER |
| 5 | 118731440 | AX-97804002 | 0 | C | A | 0.456 | -1.42E-02 | 1.16E-01 | 2.73E+00 | 2.76E+00 | 9.00E-01 | no | Med13l | missense_variant | MODERATE |
| 5 | 118744663 | AX-170383471 | 0 | A | G | 0.456 | -1.42E-02 | 1.16E-01 | 2.73E+00 | 2.76E+00 | 9.00E-01 | no | Med13l | intron_variant | MODIFIER |
| 5 | 118754698 | AX-170499900 | 0 | A | G | 0.456 | -1.42E-02 | 1.16E-01 | 2.73E+00 | 2.76E+00 | 9.00E-01 | no | Med13l | intron_variant | MODIFIER |
| 5 | 118757573 | AX-97792534 | 0 | A | G | 0.456 | -1.42E-02 | 1.16E-01 | 2.73E+00 | 2.76E+00 | 9.00E-01 | no | Med13l | intron_variant | MODIFIER |
| 5 | 118772406 | AX-170073280 | 0 | A | G | 0.456 | -1.42E-02 | 1.16E-01 | 2.73E+00 | 2.76E+00 | 9.00E-01 | no | | intergenic_variant | MODIFIER |
| 5 | 118785503 | AX-170255546 | 0 | G | A | 0.456 | -1.42E-02 | 1.16E-01 | 2.73E+00 | 2.76E+00 | 9.00E-01 | no | | intergenic_variant | MODIFIER |
| 5 | 118789493 | AX-97837453 | 0 | G | A | 0.456 | -1.42E-02 | 1.16E-01 | 2.73E+00 | 2.76E+00 | 9.00E-01 | no | | intergenic_variant | MODIFIER |
| 5 | 118807647 | AX-97843386 | 0 | T | C | 0.456 | -1.42E-02 | 1.16E-01 | 2.73E+00 | 2.76E+00 | 9.00E-01 | no | | intergenic_variant | MODIFIER |
| 5 | 118934819 | AX-97819550 | 0 | A | G | 0.456 | -1.42E-02 | 1.16E-01 | 2.73E+00 | 2.76E+00 | 9.00E-01 | no | Gm43342 | intron_variant,non_coding_transcript_variant | MODIFIER |
| 5 | 118935266 | AX-170442247 | 0 | G | A | 0.456 | -1.42E-02 | 1.16E-01 | 2.73E+00 | 2.76E+00 | 9.00E-01 | no | Gm43342 | intron_variant,non_coding_transcript_variant | MODIFIER |
| 5 | 118946299 | AX-170247564 | 0 | T | C | 0.456 | -1.42E-02 | 1.16E-01 | 2.73E+00 | 2.76E+00 | 9.00E-01 | no | | regulatory_region_variant | MODIFIER |
| 5 | 118947865 | AX-170396265 | 0 | G | A | 0.456 | -1.42E-02 | 1.16E-01 | 2.73E+00 | 2.76E+00 | 9.00E-01 | no | 4930413E15Rik | upstream_gene_variant | MODIFIER |
| 5 | 118958341 | AX-170025990 | 0 | T | G | 0.456 | -1.42E-02 | 1.16E-01 | 2.73E+00 | 2.76E+00 | 9.00E-01 | no | 4930413E15Rik | intron_variant,non_coding_transcript_variant | MODIFIER |
| 5 | 118985914 | AX-170025993 | 0 | G | A | 0.456 | -1.42E-02 | 1.16E-01 | 2.73E+00 | 2.76E+00 | 9.00E-01 | no | | intergenic_variant | MODIFIER |
| 5 | 119008183 | AX-170333124 | 0 | G | C | 0.456 | -1.42E-02 | 1.16E-01 | 2.73E+00 | 2.76E+00 | 9.00E-01 | no | | intergenic_variant | MODIFIER |
| 5 | 119013191 | AX-97788581 | 0 | T | G | 0.456 | -1.42E-02 | 1.16E-01 | 2.73E+00 | 2.76E+00 | 9.00E-01 | no | | intergenic_variant | MODIFIER |
| 5 | 119035032 | AX-170025991 | 0 | G | T | 0.456 | -1.42E-02 | 1.16E-01 | 2.73E+00 | 2.76E+00 | 9.00E-01 | no | | intergenic_variant | MODIFIER |
| 5 | 119047812 | AX-97829486 | 0 | A | G | 0.456 | -1.42E-02 | 1.16E-01 | 2.73E+00 | 2.76E+00 | 9.00E-01 | no | | intergenic_variant | MODIFIER |
| 5 | 119049829 | AX-170387303 | 0 | G | C | 0.456 | -1.42E-02 | 1.16E-01 | 2.73E+00 | 2.76E+00 | 9.00E-01 | no | | intergenic_variant | MODIFIER |
| 5 | 122079287 | AX-97828246 | 0 | G | A | 0.456 | -1.42E-02 | 1.16E-01 | 2.73E+00 | 2.76E+00 | 9.00E-01 | no | | intergenic_variant | MODIFIER |
| 5 | 122139240 | AX-170026038 | 0 | G | A | 0.456 | -1.42E-02 | 1.16E-01 | 2.73E+00 | 2.76E+00 | 9.00E-01 | no | Ccdc63 | upstream_gene_variant | MODIFIER |
| 5 | 122170512 | AX-97820252 | 0 | G | A | 0.456 | -1.42E-02 | 1.16E-01 | 2.73E+00 | 2.76E+00 | 9.00E-01 | no | Ppp1cc | intron_variant | MODIFIER |
| 5 | 122239087 | AX-170026047 | 0 | A | G | 0.456 | -1.42E-02 | 1.16E-01 | 2.73E+00 | 2.76E+00 | 9.00E-01 | no | Hvcn1 | intron_variant | MODIFIER |
| 5 | 122373865 | AX-97822885 | 0 | C | T | 0.456 | -1.42E-02 | 1.16E-01 | 2.73E+00 | 2.76E+00 | 9.00E-01 | no | Fam216a | upstream_gene_variant | MODIFIER |
| 5 | 122519419 | AX-170378939 | 0 | A | G | 0.456 | -1.42E-02 | 1.16E-01 | 2.73E+00 | 2.76E+00 | 9.00E-01 | no | | regulatory_region_variant | MODIFIER |
| 5 | 122519943 | AX-170193206 | 0 | G | A | 0.456 | -1.42E-02 | 1.16E-01 | 2.73E+00 | 2.76E+00 | 9.00E-01 | no | | regulatory_region_variant | MODIFIER |
| 5 | 122525588 | AX-97836498 | 0 | G | A | 0.456 | -1.42E-02 | 1.16E-01 | 2.73E+00 | 2.76E+00 | 9.00E-01 | no | | regulatory_region_variant | MODIFIER |
| 5 | 122554752 | AX-97799822 | 0 | C | T | 0.456 | -1.42E-02 | 1.16E-01 | 2.73E+00 | 2.76E+00 | 9.00E-01 | no | Ift81 | intron_variant | MODIFIER |
| 5 | 122565758 | AX-170391714 | 0 | G | C | 0.456 | -1.42E-02 | 1.16E-01 | 2.73E+00 | 2.76E+00 | 9.00E-01 | no | Ift81 | intron_variant | MODIFIER |
| 5 | 122566323 | AX-170421236 | 0 | A | G | 0.456 | -1.42E-02 | 1.16E-01 | 2.73E+00 | 2.76E+00 | 9.00E-01 | no | Ift81 | intron_variant | MODIFIER |
| 5 | 122569152 | AX-170466570 | 0 | T | C | 0.456 | -1.42E-02 | 1.16E-01 | 2.73E+00 | 2.76E+00 | 9.00E-01 | no | Ift81 | intron_variant | MODIFIER |
| 5 | 122569617 | AX-170484913 | 0 | T | C | 0.456 | -1.42E-02 | 1.16E-01 | 2.73E+00 | 2.76E+00 | 9.00E-01 | no | Ift81 | intron_variant | MODIFIER |
| 5 | 122572386 | AX-97786707 | 0 | A | C | 0.456 | -1.42E-02 | 1.16E-01 | 2.73E+00 | 2.76E+00 | 9.00E-01 | no | Ift81 | intron_variant | MODIFIER |
| 5 | 122586172 | AX-170250531 | 0 | A | T | 0.456 | -1.42E-02 | 1.16E-01 | 2.73E+00 | 2.76E+00 | 9.00E-01 | no | Ift81 | intron_variant | MODIFIER |
| 5 | 122625512 | AX-170354814 | 0 | A | G | 0.456 | -1.42E-02 | 1.16E-01 | 2.73E+00 | 2.76E+00 | 9.00E-01 | no | | regulatory_region_variant | MODIFIER |
| 5 | 122625552 | AX-170095455 | 0 | G | A | 0.456 | -1.42E-02 | 1.16E-01 | 2.73E+00 | 2.76E+00 | 9.00E-01 | no | | regulatory_region_variant | MODIFIER |
| 5 | 122657740 | AX-97804751 | 0 | G | A | 0.456 | -1.42E-02 | 1.16E-01 | 2.73E+00 | 2.76E+00 | 9.00E-01 | no | P2rx7 | intron_variant | MODIFIER |
| 5 | 122666716 | AX-170515247 | 0 | C | T | 0.456 | -1.42E-02 | 1.16E-01 | 2.73E+00 | 2.76E+00 | 9.00E-01 | no | P2rx7 | intron_variant | MODIFIER |

| | | | | | | | | | | | | | | | |
|---|---|---|---|---|---|---|---|---|---|---|---|---|---|---|---|
| 5 | 122694162 | AX-97800776 | 0 | C | T | 0.456 | -1.42E-02 | 1.16E-01 | 2.73E+00 | 2.76E+00 | 9.00E-01 | no | P2rx7 | downstream_gene_variant | MODIFIER |
| 5 | 122797063 | AX-97803401 | 0 | G | A | 0.456 | -1.42E-02 | 1.16E-01 | 2.73E+00 | 2.76E+00 | 9.00E-01 | no | Anapc5 | intron_variant | MODIFIER |
| 5 | 123165952 | AX-97822952 | 0 | C | T | 0.456 | -1.42E-02 | 1.16E-01 | 2.73E+00 | 2.76E+00 | 9.00E-01 | no | Setd1b | synonymous_variant | LOW |
| 5 | 123195589 | AX-97849930 | 0 | C | T | 0.456 | -1.42E-02 | 1.16E-01 | 2.73E+00 | 2.76E+00 | 9.00E-01 | no | Gm26745 | intron_variant,non_coding_transcript_variant | MODIFIER |
| 5 | 123232422 | AX-97782121 | 0 | C | T | 0.456 | -1.42E-02 | 1.16E-01 | 2.73E+00 | 2.76E+00 | 9.00E-01 | no | Psmd9 | intron_variant | MODIFIER |
| 5 | 123243967 | AX-170026056 | 0 | C | T | 0.456 | -1.42E-02 | 1.16E-01 | 2.73E+00 | 2.76E+00 | 9.00E-01 | no | Psmd9 | intron_variant | MODIFIER |
| 5 | 123271172 | AX-170477355 | 0 | G | A | 0.456 | -1.42E-02 | 1.16E-01 | 2.73E+00 | 2.76E+00 | 9.00E-01 | no | Wdr66 | intron_variant | MODIFIER |
| 5 | 123475726 | AX-170364015 | 0 | A | G | 0.456 | -1.42E-02 | 1.16E-01 | 2.73E+00 | 2.76E+00 | 9.00E-01 | no | Il31 | downstream_gene_variant | MODIFIER |
| 5 | 123476537 | AX-97786820 | 0 | G | A | 0.456 | -1.42E-02 | 1.16E-01 | 2.73E+00 | 2.76E+00 | 9.00E-01 | no | Il31 | downstream_gene_variant | MODIFIER |
| 5 | 123477990 | AX-170073357 | 0 | G | A | 0.456 | -1.42E-02 | 1.16E-01 | 2.73E+00 | 2.76E+00 | 9.00E-01 | no | Il31 | downstream_gene_variant | MODIFIER |
| 5 | 123509387 | AX-97850940 | 0 | G | A | 0.456 | -1.42E-02 | 1.16E-01 | 2.73E+00 | 2.76E+00 | 9.00E-01 | no | B3gnt4 | upstream_gene_variant | MODIFIER |
| 5 | 123513076 | AX-170264897 | 0 | A | G | 0.456 | -1.42E-02 | 1.16E-01 | 2.73E+00 | 2.76E+00 | 9.00E-01 | no | B3gnt4 | downstream_gene_variant | MODIFIER |
| 5 | 123513589 | AX-170534286 | 0 | G | A | 0.456 | -1.42E-02 | 1.16E-01 | 2.73E+00 | 2.76E+00 | 9.00E-01 | no | B3gnt4 | downstream_gene_variant | MODIFIER |
| 5 | 123513927 | AX-170411572 | 0 | C | T | 0.456 | -1.42E-02 | 1.16E-01 | 2.73E+00 | 2.76E+00 | 9.00E-01 | no | B3gnt4 | downstream_gene_variant | MODIFIER |
| 5 | 123569597 | AX-170136211 | 0 | T | C | 0.456 | -1.42E-02 | 1.16E-01 | 2.73E+00 | 2.76E+00 | 9.00E-01 | no | Vps33a | synonymous_variant | LOW |
| 5 | 123577177 | AX-97794861 | 0 | T | C | 0.456 | -1.42E-02 | 1.16E-01 | 2.73E+00 | 2.76E+00 | 9.00E-01 | no | Clip1 | downstream_gene_variant | MODIFIER |
| 5 | 123623766 | AX-170089601 | 0 | A | C | 0.456 | -1.42E-02 | 1.16E-01 | 2.73E+00 | 2.76E+00 | 9.00E-01 | no | Clip1 | intron_variant | MODIFIER |
| 5 | 123852740 | AX-170073369 | 0 | A | C | 0.456 | -1.42E-02 | 1.16E-01 | 2.73E+00 | 2.76E+00 | 9.00E-01 | no | | intergenic_variant | MODIFIER |
| 5 | 124059525 | AX-97804367 | 0 | T | C | 0.456 | -1.42E-02 | 1.16E-01 | 2.73E+00 | 2.76E+00 | 9.00E-01 | no | Abcb9 | downstream_gene_variant | MODIFIER |
| 5 | 124276367 | AX-170233219 | 0 | A | C | 0.456 | -1.42E-02 | 1.16E-01 | 2.73E+00 | 2.76E+00 | 9.00E-01 | no | Mphosph9 | intron_variant | MODIFIER |
| 5 | 124276385 | AX-97805752 | 0 | G | A | 0.456 | -1.42E-02 | 1.16E-01 | 2.73E+00 | 2.76E+00 | 9.00E-01 | no | Mphosph9 | intron_variant | MODIFIER |
| 5 | 124311472 | AX-170177015 | 0 | T | C | 0.456 | -1.42E-02 | 1.16E-01 | 2.73E+00 | 2.76E+00 | 9.00E-01 | no | Mphosph9 | intron_variant | MODIFIER |
| 5 | 124459250 | AX-97841148 | 0 | C | A | 0.456 | -1.42E-02 | 1.16E-01 | 2.73E+00 | 2.76E+00 | 9.00E-01 | no | Rilpl2 | downstream_gene_variant | MODIFIER |
| 5 | 124786014 | AX-170165279 | 0 | A | T | 0.456 | -1.42E-02 | 1.16E-01 | 2.73E+00 | 2.76E+00 | 9.00E-01 | no | Dnah10 | intron_variant | MODIFIER |
| 5 | 124790585 | AX-170254522 | 0 | G | T | 0.456 | -1.42E-02 | 1.16E-01 | 2.73E+00 | 2.76E+00 | 9.00E-01 | no | Dnah10 | intron_variant | MODIFIER |
| 5 | 124793502 | AX-170444002 | 0 | C | T | 0.456 | -1.42E-02 | 1.16E-01 | 2.73E+00 | 2.76E+00 | 9.00E-01 | no | Dnah10 | intron_variant | MODIFIER |
| 5 | 124806614 | AX-170513990 | 0 | C | A | 0.456 | -1.42E-02 | 1.16E-01 | 2.73E+00 | 2.76E+00 | 9.00E-01 | no | Dnah10 | intron_variant | MODIFIER |
| 5 | 124808951 | AX-97807733 | 0 | G | A | 0.456 | -1.42E-02 | 1.16E-01 | 2.73E+00 | 2.76E+00 | 9.00E-01 | no | Dnah10 | intron_variant | MODIFIER |
| 5 | 124809609 | AX-170510652 | 0 | A | G | 0.456 | -1.42E-02 | 1.16E-01 | 2.73E+00 | 2.76E+00 | 9.00E-01 | no | Dnah10 | intron_variant | MODIFIER |
| 5 | 124809849 | AX-170135306 | 0 | G | A | 0.456 | -1.42E-02 | 1.16E-01 | 2.73E+00 | 2.76E+00 | 9.00E-01 | no | Dnah10 | intron_variant | MODIFIER |
| 5 | 124809874 | AX-170484164 | 0 | C | T | 0.456 | -1.42E-02 | 1.16E-01 | 2.73E+00 | 2.76E+00 | 9.00E-01 | no | Dnah10 | intron_variant | MODIFIER |
| 5 | 124815803 | AX-170026086 | 0 | A | G | 0.456 | -1.42E-02 | 1.16E-01 | 2.73E+00 | 2.76E+00 | 9.00E-01 | no | Dnah10 | intron_variant | MODIFIER |
| 5 | 124836226 | AX-170510979 | 0 | T | C | 0.456 | -1.42E-02 | 1.16E-01 | 2.73E+00 | 2.76E+00 | 9.00E-01 | no | Ccdc92 | missense_variant | MODERATE |
| 5 | 124837725 | AX-97847861 | 0 | G | A | 0.456 | -1.42E-02 | 1.16E-01 | 2.73E+00 | 2.76E+00 | 9.00E-01 | no | Ccdc92 | intron_variant | MODIFIER |
| 5 | 124864394 | AX-170217126 | 0 | T | C | 0.456 | -1.42E-02 | 1.16E-01 | 2.73E+00 | 2.76E+00 | 9.00E-01 | no | Ccdc92 | upstream_gene_variant | MODIFIER |
| 5 | 124897711 | AX-170294132 | 0 | G | A | 0.456 | -1.42E-02 | 1.16E-01 | 2.73E+00 | 2.76E+00 | 9.00E-01 | no | Zfp664 | intron_variant,non_coding_transcript_variant | MODIFIER |
| 5 | 124902248 | AX-170170587 | 0 | C | A | 0.456 | -1.42E-02 | 1.16E-01 | 2.73E+00 | 2.76E+00 | 9.00E-01 | no | Zfp664 | non_coding_transcript_exon_variant | MODIFIER |
| 5 | 124902551 | AX-170026092 | 0 | G | A | 0.456 | -1.42E-02 | 1.16E-01 | 2.73E+00 | 2.76E+00 | 9.00E-01 | no | Zfp664 | non_coding_transcript_exon_variant | MODIFIER |
| 5 | 124917020 | AX-97816822 | 0 | T | C | 0.456 | -1.42E-02 | 1.16E-01 | 2.73E+00 | 2.76E+00 | 9.00E-01 | no | | intergenic_variant | MODIFIER |
| 5 | 124943800 | AX-170073389 | 0 | T | C | 0.456 | -1.42E-02 | 1.16E-01 | 2.73E+00 | 2.76E+00 | 9.00E-01 | no | | regulatory_region_variant | MODIFIER |
| 5 | 124944061 | AX-170073387 | 0 | T | C | 0.456 | -1.42E-02 | 1.16E-01 | 2.73E+00 | 2.76E+00 | 9.00E-01 | no | | regulatory_region_variant | MODIFIER |
| 5 | 124944460 | AX-170073386 | 0 | T | C | 0.456 | -1.42E-02 | 1.16E-01 | 2.73E+00 | 2.76E+00 | 9.00E-01 | no | | regulatory_region_variant | MODIFIER |
| 5 | 124945949 | AX-97808493 | 0 | C | T | 0.456 | -1.42E-02 | 1.16E-01 | 2.73E+00 | 2.76E+00 | 9.00E-01 | no | | intergenic_variant | MODIFIER |
| 5 | 124960430 | AX-170457338 | 0 | C | T | 0.456 | -1.42E-02 | 1.16E-01 | 2.73E+00 | 2.76E+00 | 9.00E-01 | no | | regulatory_region_variant | MODIFIER |
| 5 | 124980751 | AX-170026087 | 0 | C | T | 0.456 | -1.42E-02 | 1.16E-01 | 2.73E+00 | 2.76E+00 | 9.00E-01 | no | | intergenic_variant | MODIFIER |
| 5 | 125016581 | AX-97840387 | 0 | T | C | 0.456 | -1.42E-02 | 1.16E-01 | 2.73E+00 | 2.76E+00 | 9.00E-01 | no | Rflna | downstream_gene_variant | MODIFIER |
| 5 | 125016874 | AX-170026084 | 0 | G | A | 0.456 | -1.42E-02 | 1.16E-01 | 2.73E+00 | 2.76E+00 | 9.00E-01 | no | Rflna | downstream_gene_variant | MODIFIER |
| 5 | 125031286 | AX-170121535 | 0 | G | C | 0.456 | -1.42E-02 | 1.16E-01 | 2.73E+00 | 2.76E+00 | 9.00E-01 | no | Ncor2 | intron_variant | MODIFIER |
| 5 | 125032244 | AX-97814717 | 0 | A | G | 0.456 | -1.42E-02 | 1.16E-01 | 2.73E+00 | 2.76E+00 | 9.00E-01 | no | Ncor2 | synonymous_variant | LOW |
| 5 | 125049724 | AX-97805644 | 0 | T | C | 0.456 | -1.42E-02 | 1.16E-01 | 2.73E+00 | 2.76E+00 | 9.00E-01 | no | Ncor2 | intron_variant | MODIFIER |
| 5 | 125096488 | AX-170468426 | 0 | T | C | 0.456 | -1.42E-02 | 1.16E-01 | 2.73E+00 | 2.76E+00 | 9.00E-01 | no | Ncor2 | intron_variant | MODIFIER |
| 5 | 125096517 | AX-170120468 | 0 | G | A | 0.456 | -1.42E-02 | 1.16E-01 | 2.73E+00 | 2.76E+00 | 9.00E-01 | no | Ncor2 | intron_variant | MODIFIER |
| 5 | 125097251 | AX-170313674 | 0 | T | A | 0.456 | -1.42E-02 | 1.16E-01 | 2.73E+00 | 2.76E+00 | 9.00E-01 | no | Ncor2 | intron_variant | MODIFIER |
| 5 | 125100866 | AX-97807619 | 0 | G | A | 0.456 | -1.42E-02 | 1.16E-01 | 2.73E+00 | 2.76E+00 | 9.00E-01 | no | Ncor2 | intron_variant | MODIFIER |

| | | | | | | | | | | | | | | | |
|---|---|---|---|---|---|---|---|---|---|---|---|---|---|---|---|
| 5 | 125122121 | AX-170073394 | 0 | A | G | 0.456 | -1.42E-02 | 1.16E-01 | 2.73E+00 | 2.76E+00 | 9.00E-01 | no | Ncor2 | intron_variant | MODIFIER |
| 5 | 125133706 | AX-170178459 | 0 | G | A | 0.456 | -1.42E-02 | 1.16E-01 | 2.73E+00 | 2.76E+00 | 9.00E-01 | no | Ncor2 | intron_variant | MODIFIER |
| 5 | 125154224 | AX-97817321 | 0 | C | T | 0.456 | -1.42E-02 | 1.16E-01 | 2.73E+00 | 2.76E+00 | 9.00E-01 | no | Ncor2 | intron_variant | MODIFIER |
| 5 | 125155593 | AX-170347296 | 0 | A | G | 0.456 | -1.42E-02 | 1.16E-01 | 2.73E+00 | 2.76E+00 | 9.00E-01 | no | Ncor2 | intron_variant | MODIFIER |
| 5 | 125158232 | AX-170195925 | 0 | A | T | 0.456 | -1.42E-02 | 1.16E-01 | 2.73E+00 | 2.76E+00 | 9.00E-01 | no | Ncor2 | intron_variant | MODIFIER |
| 5 | 125190225 | AX-97792851 | 0 | T | C | 0.456 | -1.42E-02 | 1.16E-01 | 2.73E+00 | 2.76E+00 | 9.00E-01 | no | | intergenic_variant | MODIFIER |
| 5 | 125266659 | AX-170174004 | 0 | T | G | 0.456 | -1.42E-02 | 1.16E-01 | 2.73E+00 | 2.76E+00 | 9.00E-01 | no | Gm32585 | intron_variant,non_coding_transcript_variant | MODIFIER |
| 5 | 125291036 | AX-170371944 | 0 | A | G | 0.456 | -1.42E-02 | 1.16E-01 | 2.73E+00 | 2.76E+00 | 9.00E-01 | no | Scarb1 | intron_variant | MODIFIER |
| 5 | 125300083 | AX-97834783 | 0 | G | A | 0.456 | -1.42E-02 | 1.16E-01 | 2.73E+00 | 2.76E+00 | 9.00E-01 | no | Scarb1 | intron_variant | MODIFIER |
| 5 | 125322171 | AX-97782935 | 0 | A | G | 0.456 | -1.42E-02 | 1.16E-01 | 2.73E+00 | 2.76E+00 | 9.00E-01 | no | Scarb1 | intron_variant | MODIFIER |
| 5 | 125324209 | AX-170320519 | 0 | C | A | 0.456 | -1.42E-02 | 1.16E-01 | 2.73E+00 | 2.76E+00 | 9.00E-01 | no | Scarb1 | intron_variant | MODIFIER |
| 5 | 125327578 | AX-170105236 | 0 | A | G | 0.456 | -1.42E-02 | 1.16E-01 | 2.73E+00 | 2.76E+00 | 9.00E-01 | no | Scarb1 | intron_variant | MODIFIER |
| 5 | 125329475 | AX-170375623 | 0 | A | G | 0.456 | -1.42E-02 | 1.16E-01 | 2.73E+00 | 2.76E+00 | 9.00E-01 | no | Scarb1 | intron_variant | MODIFIER |
| 8 | 84010956 | AX-170115744 | 0 | G | A | 0.211 | 1.41E-02 | 1.13E-01 | 2.64E+00 | 2.74E+00 | 9.00E-01 | no | 2210011C24Rik | intron_variant | MODIFIER |
| 15 | 95778238 | AX-97795056 | 0 | C | T | 0.149 | 2.33E-02 | 1.79E-01 | 2.66E+00 | 2.72E+00 | 9.00E-01 | no | Gm8849 | downstream_gene_variant | MODIFIER |
| 2 | 70215435 | AX-97808820 | 0 | C | G | 0.053 | 2.26E-02 | 1.69E-01 | 2.60E+00 | 2.72E+00 | 9.00E-01 | no | Myo3b | intron_variant | MODIFIER |
| 1 | 72053593 | AX-170235939 | 0 | C | T | 0.114 | -2.84E-02 | 2.32E-01 | 2.70E+00 | 2.75E+00 | 9.00E-01 | no | | intergenic_variant | MODIFIER |
| 12 | 68549171 | AX-97821622 | 0 | C | T | 0.061 | -3.77E-02 | 3.00E-01 | 2.71E+00 | 2.73E+00 | 9.00E-01 | no | Gm47490 | downstream_gene_variant | MODIFIER |
| 15 | 74335134 | AX-170227877 | 0 | G | T | 0.553 | 1.12E-02 | 8.16E-02 | 2.63E+00 | 2.71E+00 | 9.00E-01 | no | | intergenic_variant | MODIFIER |
| 11 | 46887910 | AX-170379234 | 0 | C | T | 0.079 | -2.38E-02 | 1.79E-01 | 2.60E+00 | 2.72E+00 | 9.00E-01 | no | | intergenic_variant | MODIFIER |
| 2 | 115943970 | AX-97806646 | 0 | G | T | 0.149 | 2.39E-02 | 1.80E-01 | 2.63E+00 | 2.72E+00 | 9.00E-01 | no | Meis2 | intron_variant | MODIFIER |
| 2 | 124465573 | AX-170388359 | 0 | G | T | 0.149 | 2.39E-02 | 1.80E-01 | 2.63E+00 | 2.72E+00 | 9.00E-01 | no | Sema6d | intron_variant | MODIFIER |
| 2 | 127532301 | AX-97839439 | 0 | C | T | 0.149 | 2.39E-02 | 1.80E-01 | 2.63E+00 | 2.72E+00 | 9.00E-01 | no | Prom2 | intron_variant | MODIFIER |
| 2 | 129052122 | AX-97849092 | 0 | C | T | 0.149 | 2.39E-02 | 1.80E-01 | 2.63E+00 | 2.72E+00 | 9.00E-01 | no | Vinac1 | upstream_gene_variant | MODIFIER |
| 2 | 129131922 | AX-170157980 | 0 | G | A | 0.149 | 2.39E-02 | 1.80E-01 | 2.63E+00 | 2.72E+00 | 9.00E-01 | no | Chchd5 | intron_variant | MODIFIER |
| 2 | 136851731 | AX-170302214 | 0 | C | A | 0.149 | 2.39E-02 | 1.80E-01 | 2.63E+00 | 2.72E+00 | 9.00E-01 | no | | intergenic_variant | MODIFIER |
| 2 | 144038186 | AX-170498546 | 0 | C | T | 0.149 | 2.39E-02 | 1.80E-01 | 2.63E+00 | 2.72E+00 | 9.00E-01 | no | Banf2 | intron_variant | MODIFIER |
| 2 | 145706405 | AX-170065169 | 0 | G | A | 0.149 | 2.39E-02 | 1.80E-01 | 2.63E+00 | 2.72E+00 | 9.00E-01 | no | Gm11763 | upstream_gene_variant | MODIFIER |
| 2 | 154695797 | AX-97821080 | 0 | G | T | 0.149 | 2.39E-02 | 1.80E-01 | 2.63E+00 | 2.72E+00 | 9.00E-01 | no | Chmp4b | downstream_gene_variant | MODIFIER |
| 2 | 157792225 | AX-170278638 | 0 | G | A | 0.149 | 2.39E-02 | 1.80E-01 | 2.63E+00 | 2.72E+00 | 9.00E-01 | no | Ctnnbl1 | intron_variant | MODIFIER |
| 2 | 157830603 | AX-170229075 | 0 | C | A | 0.149 | 2.39E-02 | 1.80E-01 | 2.63E+00 | 2.72E+00 | 9.00E-01 | no | Ctnnbl1 | intron_variant | MODIFIER |
| 20 | 92797636 | AX-97810878 | 0 | T | C | 0.272 | 7.85E-03 | 6.07E-02 | 2.63E+00 | 2.73E+00 | 9.00E-01 | no | | intergenic_variant | MODIFIER |
| 1 | 66499022 | AX-170170046 | 1 | G | A | 0.116 | 2.92E-02 | 2.32E-01 | 2.68E+00 | 2.73E+00 | 9.00E-01 | no | Unc80 | intron_variant | MODIFIER |
| 1 | 71632231 | AX-170494180 | 1 | C | T | 0.116 | 2.92E-02 | 2.32E-01 | 2.68E+00 | 2.73E+00 | 9.00E-01 | no | Fn1 | intron_variant | MODIFIER |
| 1 | 65726439 | AX-170330615 | 1 | C | G | 0.143 | 3.03E-02 | 2.42E-01 | 2.68E+00 | 2.73E+00 | 9.00E-01 | no | | intergenic_variant | MODIFIER |
| 1 | 51826435 | AX-97781184 | 0 | G | A | 0.053 | 1.43E-02 | 1.18E-01 | 2.57E+00 | 2.74E+00 | 9.00E-01 | no | Myo1b | intron_variant | MODIFIER |
| 12 | 106657696 | AX-170367867 | 0 | C | T | 0.14 | 2.31E-02 | 1.85E-01 | 2.65E+00 | 2.74E+00 | 9.00E-01 | no | | intergenic_variant | MODIFIER |
| 1 | 152794105 | AX-170218424 | 0 | T | A | 0.202 | -1.47E-02 | 1.23E-01 | 2.67E+00 | 2.75E+00 | 9.00E-01 | no | | intergenic_variant | MODIFIER |
| 18 | 69929943 | AX-170414557 | 0 | A | G | 0.263 | -1.24E-02 | 9.74E-02 | 2.66E+00 | 2.73E+00 | 9.00E-01 | no | Ccdc68 | intron_variant | MODIFIER |
| 9 | 80561480 | AX-170082926 | 0 | A | G | 0.202 | 1.51E-02 | 1.28E-01 | 2.71E+00 | 2.77E+00 | 9.00E-01 | no | Gm48387 | intron_variant,non_coding_transcript_variant | MODIFIER |
| 5 | 128010962 | AX-97845794 | 1 | C | T | 0.402 | -9.67E-03 | 8.28E-02 | 2.65E+00 | 2.75E+00 | 9.00E-01 | no | Tmem132d | intron_variant | MODIFIER |
| 7 | 4096542 | AX-170281299 | 0 | A | T | 0.193 | 8.98E-03 | 7.31E-02 | 2.59E+00 | 2.74E+00 | 9.00E-01 | no | 9430041J12Rik | intron_variant,non_coding_transcript_variant | MODIFIER |
| 10 | 31035666 | AX-170124566 | 0 | C | T | 0.386 | -9.54E-03 | 7.45E-02 | 2.62E+00 | 2.73E+00 | 9.01E-01 | no | Gm30676 | intron_variant,non_coding_transcript_variant | MODIFIER |
| 10 | 31051346 | AX-97831589 | 0 | G | A | 0.386 | -9.54E-03 | 7.45E-02 | 2.62E+00 | 2.73E+00 | 9.01E-01 | no | Gm30676 | intron_variant,non_coding_transcript_variant | MODIFIER |
| 10 | 31341562 | AX-170256431 | 0 | T | C | 0.386 | -9.54E-03 | 7.45E-02 | 2.62E+00 | 2.73E+00 | 9.01E-01 | no | Tpd52l1 | intron_variant | MODIFIER |
| 10 | 31343129 | AX-170493930 | 0 | A | G | 0.386 | -9.54E-03 | 7.45E-02 | 2.62E+00 | 2.73E+00 | 9.01E-01 | no | Tpd52l1 | intron_variant | MODIFIER |
| 10 | 31344881 | AX-170469253 | 0 | A | G | 0.386 | -9.54E-03 | 7.45E-02 | 2.62E+00 | 2.73E+00 | 9.01E-01 | no | Tpd52l1 | intron_variant | MODIFIER |
| 10 | 31347764 | AX-170142090 | 0 | G | C | 0.386 | -9.54E-03 | 7.45E-02 | 2.62E+00 | 2.73E+00 | 9.01E-01 | no | Tpd52l1 | intron_variant | MODIFIER |
| 10 | 31396108 | AX-170333051 | 0 | T | C | 0.386 | -9.54E-03 | 7.45E-02 | 2.62E+00 | 2.73E+00 | 9.01E-01 | no | Tpd52l1 | intron_variant | MODIFIER |
| 10 | 31396411 | AX-170504714 | 0 | G | A | 0.386 | -9.54E-03 | 7.45E-02 | 2.62E+00 | 2.73E+00 | 9.01E-01 | no | Tpd52l1 | intron_variant | MODIFIER |
| 10 | 31402393 | AX-170044500 | 0 | G | A | 0.386 | -9.54E-03 | 7.45E-02 | 2.62E+00 | 2.73E+00 | 9.01E-01 | no | Tpd52l1 | intron_variant | MODIFIER |
| 10 | 31404224 | AX-97809561 | 0 | A | T | 0.386 | -9.54E-03 | 7.45E-02 | 2.62E+00 | 2.73E+00 | 9.01E-01 | no | Tpd52l1 | intron_variant | MODIFIER |
| 10 | 31438897 | AX-97854142 | 0 | G | C | 0.386 | -9.54E-03 | 7.45E-02 | 2.62E+00 | 2.73E+00 | 9.01E-01 | no | Tpd52l1 | intron_variant | MODIFIER |
| 10 | 31480085 | AX-170124343 | 0 | T | G | 0.386 | -9.54E-03 | 7.45E-02 | 2.62E+00 | 2.73E+00 | 9.01E-01 | no | | intergenic_variant | MODIFIER |

| | | | | | | | | | | | | | | | |
|---|---|---|---|---|---|---|---|---|---|---|---|---|---|---|---|
| 10 | 31491551 | AX-170322742 | 0 | A | G | 0.386 | -9.54E-03 | 7.45E-02 | 2.62E+00 | 2.73E+00 | 9.01E-01 | no | Rnf217 | downstream_gene_variant | MODIFIER |
| 10 | 31491720 | AX-170263897 | 0 | C | T | 0.386 | -9.54E-03 | 7.45E-02 | 2.62E+00 | 2.73E+00 | 9.01E-01 | no | Rnf217 | downstream_gene_variant | MODIFIER |
| 10 | 31492093 | AX-170501799 | 0 | A | G | 0.386 | -9.54E-03 | 7.45E-02 | 2.62E+00 | 2.73E+00 | 9.01E-01 | no | Rnf217 | downstream_gene_variant | MODIFIER |
| 10 | 31503721 | AX-170488912 | 0 | T | A | 0.386 | -9.54E-03 | 7.45E-02 | 2.62E+00 | 2.73E+00 | 9.01E-01 | no | Rnf217 | intron_variant | MODIFIER |
| 10 | 31531011 | AX-170177304 | 0 | A | G | 0.386 | -9.54E-03 | 7.45E-02 | 2.62E+00 | 2.73E+00 | 9.01E-01 | no | Rnf217 | intron_variant | MODIFIER |
| 10 | 31531613 | AX-97815200 | 0 | C | T | 0.386 | -9.54E-03 | 7.45E-02 | 2.62E+00 | 2.73E+00 | 9.01E-01 | no | Rnf217 | intron_variant | MODIFIER |
| 10 | 31720479 | AX-170326570 | 0 | G | A | 0.386 | -9.54E-03 | 7.45E-02 | 2.62E+00 | 2.73E+00 | 9.01E-01 | no | Nkain2 | intron_variant | MODIFIER |
| 10 | 31780311 | AX-170203295 | 0 | C | T | 0.386 | -9.54E-03 | 7.45E-02 | 2.62E+00 | 2.73E+00 | 9.01E-01 | no | Nkain2 | intron_variant | MODIFIER |
| 10 | 31868344 | AX-170248963 | 0 | C | T | 0.386 | -9.54E-03 | 7.45E-02 | 2.62E+00 | 2.73E+00 | 9.01E-01 | no | Nkain2 | intron_variant | MODIFIER |
| 10 | 31868462 | AX-170423068 | 0 | G | A | 0.386 | -9.54E-03 | 7.45E-02 | 2.62E+00 | 2.73E+00 | 9.01E-01 | no | Nkain2 | intron_variant | MODIFIER |
| 10 | 31874916 | AX-170335463 | 0 | T | C | 0.386 | -9.54E-03 | 7.45E-02 | 2.62E+00 | 2.73E+00 | 9.01E-01 | no | Nkain2 | intron_variant | MODIFIER |
| 10 | 31937066 | AX-170222035 | 0 | T | G | 0.386 | -9.54E-03 | 7.45E-02 | 2.62E+00 | 2.73E+00 | 9.01E-01 | no | Nkain2 | intron_variant | MODIFIER |
| 10 | 31970772 | AX-97848555 | 0 | C | A | 0.386 | -9.54E-03 | 7.45E-02 | 2.62E+00 | 2.73E+00 | 9.01E-01 | no | Nkain2 | intron_variant | MODIFIER |
| 10 | 32020148 | AX-170481049 | 0 | A | G | 0.386 | -9.54E-03 | 7.45E-02 | 2.62E+00 | 2.73E+00 | 9.01E-01 | no | Nkain2 | intron_variant | MODIFIER |
| 10 | 32091206 | AX-170104821 | 0 | T | C | 0.386 | -9.54E-03 | 7.45E-02 | 2.62E+00 | 2.73E+00 | 9.01E-01 | no | Nkain2 | intron_variant | MODIFIER |
| 10 | 32100412 | AX-170287517 | 0 | C | G | 0.386 | -9.54E-03 | 7.45E-02 | 2.62E+00 | 2.73E+00 | 9.01E-01 | no | Nkain2 | intron_variant | MODIFIER |
| 10 | 32107726 | AX-97857192 | 0 | T | C | 0.386 | -9.54E-03 | 7.45E-02 | 2.62E+00 | 2.73E+00 | 9.01E-01 | no | Nkain2 | intron_variant | MODIFIER |
| 10 | 32142463 | AX-170091911 | 0 | G | A | 0.386 | -9.54E-03 | 7.45E-02 | 2.62E+00 | 2.73E+00 | 9.01E-01 | no | Nkain2 | intron_variant | MODIFIER |
| 10 | 32280194 | AX-170135353 | 0 | A | G | 0.386 | -9.54E-03 | 7.45E-02 | 2.62E+00 | 2.73E+00 | 9.01E-01 | no | Nkain2 | intron_variant | MODIFIER |
| 10 | 32291781 | AX-170479140 | 0 | C | T | 0.386 | -9.54E-03 | 7.45E-02 | 2.62E+00 | 2.73E+00 | 9.01E-01 | no | Nkain2 | intron_variant | MODIFIER |
| 10 | 32396320 | AX-170175080 | 0 | A | G | 0.386 | -9.54E-03 | 7.45E-02 | 2.62E+00 | 2.73E+00 | 9.01E-01 | no | Nkain2 | intron_variant | MODIFIER |
| 10 | 32469174 | AX-170402267 | 0 | A | G | 0.386 | -9.54E-03 | 7.45E-02 | 2.62E+00 | 2.73E+00 | 9.01E-01 | no | Nkain2 | intron_variant | MODIFIER |
| 10 | 32535281 | AX-170003015 | 0 | T | C | 0.386 | -9.54E-03 | 7.45E-02 | 2.62E+00 | 2.73E+00 | 9.01E-01 | no | Nkain2 | intron_variant | MODIFIER |
| 10 | 32536316 | AX-97794038 | 0 | C | T | 0.386 | -9.54E-03 | 7.45E-02 | 2.62E+00 | 2.73E+00 | 9.01E-01 | no | Nkain2 | intron_variant | MODIFIER |
| 10 | 32545654 | AX-170169262 | 0 | A | C | 0.386 | -9.54E-03 | 7.45E-02 | 2.62E+00 | 2.73E+00 | 9.01E-01 | no | Nkain2 | intron_variant | MODIFIER |
| 10 | 32570938 | AX-170274542 | 0 | A | G | 0.386 | -9.54E-03 | 7.45E-02 | 2.62E+00 | 2.73E+00 | 9.01E-01 | no | Nkain2 | intron_variant | MODIFIER |
| 10 | 32571954 | AX-170226396 | 0 | A | G | 0.386 | -9.54E-03 | 7.45E-02 | 2.62E+00 | 2.73E+00 | 9.01E-01 | no | Nkain2 | intron_variant | MODIFIER |
| 10 | 32574108 | AX-170351941 | 0 | G | A | 0.386 | -9.54E-03 | 7.45E-02 | 2.62E+00 | 2.73E+00 | 9.01E-01 | no | Nkain2 | intron_variant | MODIFIER |
| 10 | 32574265 | AX-170344072 | 0 | T | A | 0.386 | -9.54E-03 | 7.45E-02 | 2.62E+00 | 2.73E+00 | 9.01E-01 | no | Nkain2 | intron_variant | MODIFIER |
| 10 | 32607423 | AX-170086775 | 0 | A | G | 0.386 | -9.54E-03 | 7.45E-02 | 2.62E+00 | 2.73E+00 | 9.01E-01 | no | Nkain2 | intron_variant | MODIFIER |
| 10 | 32659865 | AX-170467816 | 0 | T | G | 0.386 | -9.54E-03 | 7.45E-02 | 2.62E+00 | 2.73E+00 | 9.01E-01 | no | Nkain2 | intron_variant | MODIFIER |
| 10 | 32694874 | AX-170317696 | 0 | A | G | 0.386 | -9.54E-03 | 7.45E-02 | 2.62E+00 | 2.73E+00 | 9.01E-01 | no | Nkain2 | intron_variant | MODIFIER |
| 10 | 32699214 | AX-170176167 | 0 | C | T | 0.386 | -9.54E-03 | 7.45E-02 | 2.62E+00 | 2.73E+00 | 9.01E-01 | no | Nkain2 | intron_variant | MODIFIER |
| 12 | 111974266 | AX-170319656 | 1 | T | C | 0.134 | -2.08E-02 | 1.73E-01 | 2.65E+00 | 2.75E+00 | 9.01E-01 | no | Tdrd9 | intron_variant | MODIFIER |
| 7 | 79579545 | AX-170488749 | 1 | C | G | 0.134 | -2.61E-02 | 2.15E-01 | 2.67E+00 | 2.74E+00 | 9.01E-01 | no | Gm45168 | upstream_gene_variant | MODIFIER |
| 7 | 94284657 | AX-170107900 | 1 | A | T | 0.134 | -2.61E-02 | 2.15E-01 | 2.67E+00 | 2.74E+00 | 9.01E-01 | no | Gm32647 | intron_variant,non_coding_transcript_variant | MODIFIER |
| 18 | 41267159 | AX-170250676 | 0 | A | T | 0.061 | 3.67E-02 | 3.01E-01 | 2.73E+00 | 2.76E+00 | 9.01E-01 | no | | intergenic_variant | MODIFIER |
| 18 | 41660293 | AX-170375226 | 0 | G | A | 0.061 | 3.67E-02 | 3.01E-01 | 2.73E+00 | 2.76E+00 | 9.01E-01 | no | | intergenic_variant | MODIFIER |
| 18 | 41761868 | AX-170414999 | 0 | C | T | 0.061 | 3.67E-02 | 3.01E-01 | 2.73E+00 | 2.76E+00 | 9.01E-01 | no | AC154172.2 | intron_variant,non_coding_transcript_variant | MODIFIER |
| 7 | 113232154 | AX-170442763 | 2 | T | C | 0.436 | 1.18E-02 | 9.50E-02 | 2.68E+00 | 2.73E+00 | 9.01E-01 | no | Arntl | intron_variant | MODIFIER |
| 1 | 66197457 | AX-170154288 | 0 | A | G | 0.114 | 2.89E-02 | 2.33E-01 | 2.69E+00 | 2.73E+00 | 9.01E-01 | no | Map2 | intron_variant | MODIFIER |
| 1 | 66200608 | AX-170226484 | 0 | A | T | 0.114 | 2.89E-02 | 2.33E-01 | 2.69E+00 | 2.73E+00 | 9.01E-01 | no | Map2 | intron_variant | MODIFIER |
| 1 | 66232333 | AX-97853136 | 0 | T | G | 0.114 | 2.89E-02 | 2.33E-01 | 2.69E+00 | 2.73E+00 | 9.01E-01 | no | Map2 | intron_variant | MODIFIER |
| 1 | 66248911 | AX-170437533 | 0 | G | A | 0.114 | 2.89E-02 | 2.33E-01 | 2.69E+00 | 2.73E+00 | 9.01E-01 | no | Map2 | upstream_gene_variant | MODIFIER |
| 1 | 66251282 | AX-170148349 | 0 | A | G | 0.114 | 2.89E-02 | 2.33E-01 | 2.69E+00 | 2.73E+00 | 9.01E-01 | no | Map2 | upstream_gene_variant | MODIFIER |
| 1 | 66263370 | AX-170182864 | 0 | G | A | 0.114 | 2.89E-02 | 2.33E-01 | 2.69E+00 | 2.73E+00 | 9.01E-01 | no | Map2 | intron_variant | MODIFIER |
| 1 | 66263414 | AX-170248833 | 0 | A | T | 0.114 | 2.89E-02 | 2.33E-01 | 2.69E+00 | 2.73E+00 | 9.01E-01 | no | Map2 | intron_variant | MODIFIER |
| 1 | 66298872 | AX-170415889 | 0 | G | C | 0.114 | 2.89E-02 | 2.33E-01 | 2.69E+00 | 2.73E+00 | 9.01E-01 | no | Map2 | intron_variant | MODIFIER |
| 1 | 66300366 | AX-170200487 | 0 | A | T | 0.114 | 2.89E-02 | 2.33E-01 | 2.69E+00 | 2.73E+00 | 9.01E-01 | no | Map2 | intron_variant | MODIFIER |
| 1 | 66319545 | AX-170302959 | 0 | A | G | 0.114 | 2.89E-02 | 2.33E-01 | 2.69E+00 | 2.73E+00 | 9.01E-01 | no | Map2 | intron_variant | MODIFIER |
| 1 | 66328761 | AX-170161684 | 0 | G | A | 0.114 | 2.89E-02 | 2.33E-01 | 2.69E+00 | 2.73E+00 | 9.01E-01 | no | Map2 | intron_variant | MODIFIER |
| 1 | 66351846 | AX-170376045 | 0 | G | T | 0.114 | 2.89E-02 | 2.33E-01 | 2.69E+00 | 2.73E+00 | 9.01E-01 | no | Map2 | intron_variant | MODIFIER |
| 1 | 66351872 | AX-170304048 | 0 | G | C | 0.114 | 2.89E-02 | 2.33E-01 | 2.69E+00 | 2.73E+00 | 9.01E-01 | no | Map2 | intron_variant | MODIFIER |
| 1 | 66355362 | AX-170143012 | 0 | C | A | 0.114 | 2.89E-02 | 2.33E-01 | 2.69E+00 | 2.73E+00 | 9.01E-01 | no | Map2 | intron_variant | MODIFIER |

| | | | | | | | | | | | | | | | |
|---|---|---|---|---|---|---|---|---|---|---|---|---|---|---|---|
| 1 | 66372464 | AX-170000703 | 0 | T | C | 0.114 | 2.89E-02 | 2.33E-01 | 2.69E+00 | 2.73E+00 | 9.01E-01 | no | Map2 | intron_variant | MODIFIER |
| 1 | 66379081 | AX-170399779 | 0 | T | C | 0.114 | 2.89E-02 | 2.33E-01 | 2.69E+00 | 2.73E+00 | 9.01E-01 | no | Map2 | intron_variant | MODIFIER |
| 1 | 66379134 | AX-170000706 | 0 | T | A | 0.114 | 2.89E-02 | 2.33E-01 | 2.69E+00 | 2.73E+00 | 9.01E-01 | no | Map2 | intron_variant | MODIFIER |
| 1 | 66381290 | AX-97787035 | 0 | T | G | 0.114 | 2.89E-02 | 2.33E-01 | 2.69E+00 | 2.73E+00 | 9.01E-01 | no | Map2 | intron_variant | MODIFIER |
| 1 | 66384223 | AX-170041492 | 0 | G | T | 0.114 | 2.89E-02 | 2.33E-01 | 2.69E+00 | 2.73E+00 | 9.01E-01 | no | Map2 | intron_variant | MODIFIER |
| 1 | 66406815 | AX-170000707 | 0 | T | C | 0.114 | 2.89E-02 | 2.33E-01 | 2.69E+00 | 2.73E+00 | 9.01E-01 | no | Map2 | intron_variant | MODIFIER |
| 1 | 66406849 | AX-170000710 | 0 | A | C | 0.114 | 2.89E-02 | 2.33E-01 | 2.69E+00 | 2.73E+00 | 9.01E-01 | no | Map2 | intron_variant | MODIFIER |
| 1 | 66408474 | AX-170000708 | 0 | A | G | 0.114 | 2.89E-02 | 2.33E-01 | 2.69E+00 | 2.73E+00 | 9.01E-01 | no | Map2 | intron_variant | MODIFIER |
| 1 | 66416169 | AX-170041495 | 0 | T | C | 0.114 | 2.89E-02 | 2.33E-01 | 2.69E+00 | 2.73E+00 | 9.01E-01 | no | Map2 | intron_variant | MODIFIER |
| 1 | 66442875 | AX-170000705 | 0 | G | C | 0.114 | 2.89E-02 | 2.33E-01 | 2.69E+00 | 2.73E+00 | 9.01E-01 | no | Map2 | downstream_gene_variant | MODIFIER |
| 1 | 66442985 | AX-170000704 | 0 | A | T | 0.114 | 2.89E-02 | 2.33E-01 | 2.69E+00 | 2.73E+00 | 9.01E-01 | no | Map2 | downstream_gene_variant | MODIFIER |
| 1 | 66449503 | AX-170091540 | 0 | A | G | 0.114 | 2.89E-02 | 2.33E-01 | 2.69E+00 | 2.73E+00 | 9.01E-01 | no | | intergenic_variant | MODIFIER |
| 1 | 66485119 | AX-170118574 | 0 | T | G | 0.114 | 2.89E-02 | 2.33E-01 | 2.69E+00 | 2.73E+00 | 9.01E-01 | no | Unc80 | intron_variant | MODIFIER |
| 1 | 66509948 | AX-170466117 | 0 | C | T | 0.114 | 2.89E-02 | 2.33E-01 | 2.69E+00 | 2.73E+00 | 9.01E-01 | no | Unc80 | intron_variant | MODIFIER |
| 1 | 66511752 | AX-170190981 | 0 | C | G | 0.114 | 2.89E-02 | 2.33E-01 | 2.69E+00 | 2.73E+00 | 9.01E-01 | no | Unc80 | intron_variant | MODIFIER |
| 1 | 66526999 | AX-97853872 | 0 | A | G | 0.114 | 2.89E-02 | 2.33E-01 | 2.69E+00 | 2.73E+00 | 9.01E-01 | no | Unc80 | intron_variant | MODIFIER |
| 1 | 66565827 | AX-97855487 | 0 | A | C | 0.114 | 2.89E-02 | 2.33E-01 | 2.69E+00 | 2.73E+00 | 9.01E-01 | no | Unc80 | intron_variant | MODIFIER |
| 1 | 66593572 | AX-97800558 | 0 | C | T | 0.114 | 2.89E-02 | 2.33E-01 | 2.69E+00 | 2.73E+00 | 9.01E-01 | no | Unc80 | intron_variant | MODIFIER |
| 1 | 66593819 | AX-170155900 | 0 | A | G | 0.114 | 2.89E-02 | 2.33E-01 | 2.69E+00 | 2.73E+00 | 9.01E-01 | no | Unc80 | intron_variant | MODIFIER |
| 1 | 66602245 | AX-170492669 | 0 | C | T | 0.114 | 2.89E-02 | 2.33E-01 | 2.69E+00 | 2.73E+00 | 9.01E-01 | no | Unc80 | intron_variant | MODIFIER |
| 1 | 66602275 | AX-170455430 | 0 | C | G | 0.114 | 2.89E-02 | 2.33E-01 | 2.69E+00 | 2.73E+00 | 9.01E-01 | no | Unc80 | intron_variant | MODIFIER |
| 1 | 66602657 | AX-170309156 | 0 | A | G | 0.114 | 2.89E-02 | 2.33E-01 | 2.69E+00 | 2.73E+00 | 9.01E-01 | no | Unc80 | intron_variant | MODIFIER |
| 1 | 66637230 | AX-170438241 | 0 | C | A | 0.114 | 2.89E-02 | 2.33E-01 | 2.69E+00 | 2.73E+00 | 9.01E-01 | no | Unc80 | intron_variant | MODIFIER |
| 1 | 66644056 | AX-170041502 | 0 | T | C | 0.114 | 2.89E-02 | 2.33E-01 | 2.69E+00 | 2.73E+00 | 9.01E-01 | no | Unc80 | intron_variant | MODIFIER |
| 1 | 66646286 | AX-170086487 | 0 | T | C | 0.114 | 2.89E-02 | 2.33E-01 | 2.69E+00 | 2.73E+00 | 9.01E-01 | no | Unc80 | intron_variant | MODIFIER |
| 1 | 66687742 | AX-97801470 | 0 | T | G | 0.114 | 2.89E-02 | 2.33E-01 | 2.69E+00 | 2.73E+00 | 9.01E-01 | no | Unc80 | intron_variant | MODIFIER |
| 1 | 66688824 | AX-170117163 | 0 | T | G | 0.114 | 2.89E-02 | 2.33E-01 | 2.69E+00 | 2.73E+00 | 9.01E-01 | no | Unc80 | intron_variant | MODIFIER |
| 1 | 66688951 | AX-170186634 | 0 | T | C | 0.114 | 2.89E-02 | 2.33E-01 | 2.69E+00 | 2.73E+00 | 9.01E-01 | no | Unc80 | intron_variant | MODIFIER |
| 1 | 66692765 | AX-170371051 | 0 | T | C | 0.114 | 2.89E-02 | 2.33E-01 | 2.69E+00 | 2.73E+00 | 9.01E-01 | no | Unc80 | intron_variant | MODIFIER |
| 1 | 66709013 | AX-170404725 | 0 | T | C | 0.114 | 2.89E-02 | 2.33E-01 | 2.69E+00 | 2.73E+00 | 9.01E-01 | no | Rpe | intron_variant | MODIFIER |
| 1 | 66734046 | AX-170388254 | 0 | C | T | 0.114 | 2.89E-02 | 2.33E-01 | 2.69E+00 | 2.73E+00 | 9.01E-01 | no | Kansl1l | intron_variant | MODIFIER |
| 1 | 66734099 | AX-170178359 | 0 | A | G | 0.114 | 2.89E-02 | 2.33E-01 | 2.69E+00 | 2.73E+00 | 9.01E-01 | no | Kansl1l | intron_variant | MODIFIER |
| 1 | 66749456 | AX-170441647 | 0 | T | G | 0.114 | 2.89E-02 | 2.33E-01 | 2.69E+00 | 2.73E+00 | 9.01E-01 | no | Kansl1l | intron_variant | MODIFIER |
| 1 | 66761538 | AX-170385764 | 0 | G | A | 0.114 | 2.89E-02 | 2.33E-01 | 2.69E+00 | 2.73E+00 | 9.01E-01 | no | Kansl1l | intron_variant | MODIFIER |
| 1 | 66770768 | AX-97854990 | 0 | A | G | 0.114 | 2.89E-02 | 2.33E-01 | 2.69E+00 | 2.73E+00 | 9.01E-01 | no | Kansl1l | intron_variant | MODIFIER |
| 1 | 66772802 | AX-170410810 | 0 | A | G | 0.114 | 2.89E-02 | 2.33E-01 | 2.69E+00 | 2.73E+00 | 9.01E-01 | no | Kansl1l | intron_variant | MODIFIER |
| 1 | 66809176 | AX-170222265 | 0 | C | T | 0.114 | 2.89E-02 | 2.33E-01 | 2.69E+00 | 2.73E+00 | 9.01E-01 | no | Kansl1l | intron_variant | MODIFIER |
| 1 | 66811198 | AX-97785739 | 0 | G | A | 0.114 | 2.89E-02 | 2.33E-01 | 2.69E+00 | 2.73E+00 | 9.01E-01 | no | Kansl1l | intron_variant | MODIFIER |
| 1 | 66814971 | AX-170171262 | 0 | A | G | 0.114 | 2.89E-02 | 2.33E-01 | 2.69E+00 | 2.73E+00 | 9.01E-01 | no | Kansl1l | intron_variant | MODIFIER |
| 1 | 66843392 | AX-170384576 | 0 | T | C | 0.114 | 2.89E-02 | 2.33E-01 | 2.69E+00 | 2.73E+00 | 9.01E-01 | no | Acadl | intron_variant | MODIFIER |
| 1 | 66847562 | AX-170000716 | 0 | G | T | 0.114 | 2.89E-02 | 2.33E-01 | 2.69E+00 | 2.73E+00 | 9.01E-01 | no | Acadl | intron_variant | MODIFIER |
| 1 | 66848174 | AX-170086294 | 0 | T | G | 0.114 | 2.89E-02 | 2.33E-01 | 2.69E+00 | 2.73E+00 | 9.01E-01 | no | Acadl | intron_variant | MODIFIER |
| 1 | 66852569 | AX-170041506 | 0 | A | G | 0.114 | 2.89E-02 | 2.33E-01 | 2.69E+00 | 2.73E+00 | 9.01E-01 | no | Acadl | intron_variant | MODIFIER |
| 1 | 66853174 | AX-170000717 | 0 | T | C | 0.114 | 2.89E-02 | 2.33E-01 | 2.69E+00 | 2.73E+00 | 9.01E-01 | no | Acadl | synonymous_variant | LOW |
| 1 | 66882488 | AX-170000711 | 0 | C | T | 0.114 | 2.89E-02 | 2.33E-01 | 2.69E+00 | 2.73E+00 | 9.01E-01 | no | | intergenic_variant | MODIFIER |
| 1 | 66887341 | AX-170257760 | 0 | G | A | 0.114 | 2.89E-02 | 2.33E-01 | 2.69E+00 | 2.73E+00 | 9.01E-01 | no | | intergenic_variant | MODIFIER |
| 1 | 66887371 | AX-170147502 | 0 | T | C | 0.114 | 2.89E-02 | 2.33E-01 | 2.69E+00 | 2.73E+00 | 9.01E-01 | no | | intergenic_variant | MODIFIER |
| 1 | 66905704 | AX-170000713 | 0 | G | A | 0.114 | 2.89E-02 | 2.33E-01 | 2.69E+00 | 2.73E+00 | 9.01E-01 | no | | intergenic_variant | MODIFIER |
| 1 | 66917731 | AX-170282706 | 0 | T | C | 0.114 | 2.89E-02 | 2.33E-01 | 2.69E+00 | 2.73E+00 | 9.01E-01 | no | | intergenic_variant | MODIFIER |
| 1 | 66917842 | AX-170321455 | 0 | C | T | 0.114 | 2.89E-02 | 2.33E-01 | 2.69E+00 | 2.73E+00 | 9.01E-01 | no | | intergenic_variant | MODIFIER |
| 1 | 66919474 | AX-97803547 | 0 | T | C | 0.114 | 2.89E-02 | 2.33E-01 | 2.69E+00 | 2.73E+00 | 9.01E-01 | no | Myl1 | downstream_gene_variant | MODIFIER |
| 1 | 66921406 | AX-97784625 | 0 | C | T | 0.114 | 2.89E-02 | 2.33E-01 | 2.69E+00 | 2.73E+00 | 9.01E-01 | no | Myl1 | downstream_gene_variant | MODIFIER |
| 1 | 66926519 | AX-170000714 | 0 | A | C | 0.114 | 2.89E-02 | 2.33E-01 | 2.69E+00 | 2.73E+00 | 9.01E-01 | no | Myl1 | intron_variant | MODIFIER |
| 1 | 66953840 | AX-97810112 | 0 | C | T | 0.114 | 2.89E-02 | 2.33E-01 | 2.69E+00 | 2.73E+00 | 9.01E-01 | no | Gm15826 | downstream_gene_variant | MODIFIER |

| | | | | | | | | | | | | | | | |
|---|---|---|---|---|---|---|---|---|---|---|---|---|---|---|---|
| 1 | 66964549 | AX-170500029 | 0 | C | A | 0.114 | 2.89E-02 | 2.33E-01 | 2.69E+00 | 2.73E+00 | 9.01E-01 | no | | intergenic_variant | MODIFIER |
| 1 | 66964702 | AX-170305147 | 0 | A | G | 0.114 | 2.89E-02 | 2.33E-01 | 2.69E+00 | 2.73E+00 | 9.01E-01 | no | | intergenic_variant | MODIFIER |
| 1 | 66977232 | AX-170350510 | 0 | T | G | 0.114 | 2.89E-02 | 2.33E-01 | 2.69E+00 | 2.73E+00 | 9.01E-01 | no | | intergenic_variant | MODIFIER |
| 1 | 66978116 | AX-170421898 | 0 | G | A | 0.114 | 2.89E-02 | 2.33E-01 | 2.69E+00 | 2.73E+00 | 9.01E-01 | no | | intergenic_variant | MODIFIER |
| 1 | 67003987 | AX-170152865 | 0 | G | A | 0.114 | 2.89E-02 | 2.33E-01 | 2.69E+00 | 2.73E+00 | 9.01E-01 | no | Lancl1 | intron_variant | MODIFIER |
| 1 | 67010207 | AX-97792648 | 0 | T | C | 0.114 | 2.89E-02 | 2.33E-01 | 2.69E+00 | 2.73E+00 | 9.01E-01 | no | Lancl1 | intron_variant | MODIFIER |
| 1 | 67021771 | AX-97781707 | 0 | C | A | 0.114 | 2.89E-02 | 2.33E-01 | 2.69E+00 | 2.73E+00 | 9.01E-01 | no | Lancl1 | intron_variant | MODIFIER |
| 1 | 67031799 | AX-170305336 | 0 | G | A | 0.114 | 2.89E-02 | 2.33E-01 | 2.69E+00 | 2.73E+00 | 9.01E-01 | no | Lancl1 | intron_variant | MODIFIER |
| 1 | 67062554 | AX-170422839 | 0 | G | A | 0.114 | 2.89E-02 | 2.33E-01 | 2.69E+00 | 2.73E+00 | 9.01E-01 | no | | intergenic_variant | MODIFIER |
| 1 | 67093537 | AX-97846618 | 0 | G | A | 0.114 | 2.89E-02 | 2.33E-01 | 2.69E+00 | 2.73E+00 | 9.01E-01 | no | | intergenic_variant | MODIFIER |
| 1 | 67093635 | AX-170263009 | 0 | C | T | 0.114 | 2.89E-02 | 2.33E-01 | 2.69E+00 | 2.73E+00 | 9.01E-01 | no | | intergenic_variant | MODIFIER |
| 1 | 67125928 | AX-170435611 | 0 | C | T | 0.114 | 2.89E-02 | 2.33E-01 | 2.69E+00 | 2.73E+00 | 9.01E-01 | no | Cps1 | intron_variant | MODIFIER |
| 1 | 67126517 | AX-97786085 | 0 | C | T | 0.114 | 2.89E-02 | 2.33E-01 | 2.69E+00 | 2.73E+00 | 9.01E-01 | no | Cps1 | intron_variant | MODIFIER |
| 1 | 67134526 | AX-170455079 | 0 | T | C | 0.114 | 2.89E-02 | 2.33E-01 | 2.69E+00 | 2.73E+00 | 9.01E-01 | no | Cps1 | intron_variant | MODIFIER |
| 1 | 67142244 | AX-170372411 | 0 | C | T | 0.114 | 2.89E-02 | 2.33E-01 | 2.69E+00 | 2.73E+00 | 9.01E-01 | no | Cps1 | intron_variant | MODIFIER |
| 1 | 67148158 | AX-170468083 | 0 | G | A | 0.114 | 2.89E-02 | 2.33E-01 | 2.69E+00 | 2.73E+00 | 9.01E-01 | no | Cps1 | intron_variant | MODIFIER |
| 1 | 67149941 | AX-170201777 | 0 | T | G | 0.114 | 2.89E-02 | 2.33E-01 | 2.69E+00 | 2.73E+00 | 9.01E-01 | no | Cps1 | intron_variant | MODIFIER |
| 1 | 67172574 | AX-170500453 | 0 | A | C | 0.114 | 2.89E-02 | 2.33E-01 | 2.69E+00 | 2.73E+00 | 9.01E-01 | no | Cps1 | intron_variant | MODIFIER |
| 1 | 67178305 | AX-170443080 | 0 | T | C | 0.114 | 2.89E-02 | 2.33E-01 | 2.69E+00 | 2.73E+00 | 9.01E-01 | no | Cps1 | intron_variant | MODIFIER |
| 1 | 67181980 | AX-170507446 | 0 | T | C | 0.114 | 2.89E-02 | 2.33E-01 | 2.69E+00 | 2.73E+00 | 9.01E-01 | no | Cps1 | intron_variant | MODIFIER |
| 1 | 67195418 | AX-170041517 | 0 | T | C | 0.114 | 2.89E-02 | 2.33E-01 | 2.69E+00 | 2.73E+00 | 9.01E-01 | no | Cps1 | intron_variant | MODIFIER |
| 1 | 67195821 | AX-170041519 | 0 | G | A | 0.114 | 2.89E-02 | 2.33E-01 | 2.69E+00 | 2.73E+00 | 9.01E-01 | no | Cps1 | intron_variant | MODIFIER |
| 1 | 67208752 | AX-170086488 | 0 | C | A | 0.114 | 2.89E-02 | 2.33E-01 | 2.69E+00 | 2.73E+00 | 9.01E-01 | no | Cps1 | intron_variant | MODIFIER |
| 1 | 67224169 | AX-97815209 | 0 | G | A | 0.114 | 2.89E-02 | 2.33E-01 | 2.69E+00 | 2.73E+00 | 9.01E-01 | no | Cps1 | intron_variant | MODIFIER |
| 1 | 67261057 | AX-97847608 | 0 | C | T | 0.114 | 2.89E-02 | 2.33E-01 | 2.69E+00 | 2.73E+00 | 9.01E-01 | no | Gm26342 | downstream_gene_variant | MODIFIER |
| 1 | 67285632 | AX-97811907 | 0 | T | C | 0.114 | 2.89E-02 | 2.33E-01 | 2.69E+00 | 2.73E+00 | 9.01E-01 | no | Gm15668 | intron_variant,non_coding_transcript_variant | MODIFIER |
| 1 | 67285683 | AX-170315768 | 0 | T | C | 0.114 | 2.89E-02 | 2.33E-01 | 2.69E+00 | 2.73E+00 | 9.01E-01 | no | Gm15668 | intron_variant,non_coding_transcript_variant | MODIFIER |
| 1 | 67326302 | AX-170517921 | 0 | A | G | 0.114 | 2.89E-02 | 2.33E-01 | 2.69E+00 | 2.73E+00 | 9.01E-01 | no | Gm15668 | downstream_gene_variant | MODIFIER |
| 1 | 67360462 | AX-97800412 | 0 | A | G | 0.114 | 2.89E-02 | 2.33E-01 | 2.69E+00 | 2.73E+00 | 9.01E-01 | no | | intergenic_variant | MODIFIER |
| 1 | 67390530 | AX-170490759 | 0 | A | C | 0.114 | 2.89E-02 | 2.33E-01 | 2.69E+00 | 2.73E+00 | 9.01E-01 | no | | intergenic_variant | MODIFIER |
| 1 | 67397029 | AX-170202163 | 0 | A | T | 0.114 | 2.89E-02 | 2.33E-01 | 2.69E+00 | 2.73E+00 | 9.01E-01 | no | | intergenic_variant | MODIFIER |
| 1 | 67397099 | AX-170351902 | 0 | C | T | 0.114 | 2.89E-02 | 2.33E-01 | 2.69E+00 | 2.73E+00 | 9.01E-01 | no | | intergenic_variant | MODIFIER |
| 1 | 67402248 | AX-97796494 | 0 | A | C | 0.114 | 2.89E-02 | 2.33E-01 | 2.69E+00 | 2.73E+00 | 9.01E-01 | no | | intergenic_variant | MODIFIER |
| 1 | 67403122 | AX-170206802 | 0 | G | A | 0.114 | 2.89E-02 | 2.33E-01 | 2.69E+00 | 2.73E+00 | 9.01E-01 | no | | intergenic_variant | MODIFIER |
| 1 | 67444910 | AX-170335550 | 0 | G | A | 0.114 | 2.89E-02 | 2.33E-01 | 2.69E+00 | 2.73E+00 | 9.01E-01 | no | | intergenic_variant | MODIFIER |
| 1 | 67446388 | AX-170099492 | 0 | T | C | 0.114 | 2.89E-02 | 2.33E-01 | 2.69E+00 | 2.73E+00 | 9.01E-01 | no | | intergenic_variant | MODIFIER |
| 1 | 67457324 | AX-170193485 | 0 | C | T | 0.114 | 2.89E-02 | 2.33E-01 | 2.69E+00 | 2.73E+00 | 9.01E-01 | no | | intergenic_variant | MODIFIER |
| 1 | 67462702 | AX-170135156 | 0 | T | C | 0.114 | 2.89E-02 | 2.33E-01 | 2.69E+00 | 2.73E+00 | 9.01E-01 | no | | intergenic_variant | MODIFIER |
| 1 | 67499694 | AX-170187650 | 0 | T | C | 0.114 | 2.89E-02 | 2.33E-01 | 2.69E+00 | 2.73E+00 | 9.01E-01 | no | | intergenic_variant | MODIFIER |
| 1 | 67500110 | AX-170169582 | 0 | G | C | 0.114 | 2.89E-02 | 2.33E-01 | 2.69E+00 | 2.73E+00 | 9.01E-01 | no | | intergenic_variant | MODIFIER |
| 1 | 67507738 | AX-97840333 | 0 | A | G | 0.114 | 2.89E-02 | 2.33E-01 | 2.69E+00 | 2.73E+00 | 9.01E-01 | no | | intergenic_variant | MODIFIER |
| 1 | 67516195 | AX-170000722 | 0 | G | C | 0.114 | 2.89E-02 | 2.33E-01 | 2.69E+00 | 2.73E+00 | 9.01E-01 | no | | intergenic_variant | MODIFIER |
| 1 | 67516227 | AX-170000728 | 0 | T | G | 0.114 | 2.89E-02 | 2.33E-01 | 2.69E+00 | 2.73E+00 | 9.01E-01 | no | | intergenic_variant | MODIFIER |
| 1 | 67557995 | AX-170041520 | 0 | G | A | 0.114 | 2.89E-02 | 2.33E-01 | 2.69E+00 | 2.73E+00 | 9.01E-01 | no | | intergenic_variant | MODIFIER |
| 1 | 67558529 | AX-97811655 | 0 | G | A | 0.114 | 2.89E-02 | 2.33E-01 | 2.69E+00 | 2.73E+00 | 9.01E-01 | no | | intergenic_variant | MODIFIER |
| 1 | 67559547 | AX-170086249 | 0 | A | G | 0.114 | 2.89E-02 | 2.33E-01 | 2.69E+00 | 2.73E+00 | 9.01E-01 | no | | regulatory_region_variant | MODIFIER |
| 1 | 67574694 | AX-97811904 | 0 | G | A | 0.114 | 2.89E-02 | 2.33E-01 | 2.69E+00 | 2.73E+00 | 9.01E-01 | no | | intergenic_variant | MODIFIER |
| 1 | 67574890 | AX-170000725 | 0 | A | G | 0.114 | 2.89E-02 | 2.33E-01 | 2.69E+00 | 2.73E+00 | 9.01E-01 | no | | intergenic_variant | MODIFIER |
| 1 | 67579383 | AX-170041521 | 0 | A | T | 0.114 | 2.89E-02 | 2.33E-01 | 2.69E+00 | 2.73E+00 | 9.01E-01 | no | | intergenic_variant | MODIFIER |
| 1 | 67586987 | AX-170172261 | 0 | G | A | 0.114 | 2.89E-02 | 2.33E-01 | 2.69E+00 | 2.73E+00 | 9.01E-01 | no | Kif22-ps | non_coding_transcript_exon_variant | MODIFIER |
| 1 | 67640998 | AX-170358487 | 0 | G | A | 0.114 | 2.89E-02 | 2.33E-01 | 2.69E+00 | 2.73E+00 | 9.01E-01 | no | | regulatory_region_variant | MODIFIER |
| 1 | 67641526 | AX-170000723 | 0 | A | G | 0.114 | 2.89E-02 | 2.33E-01 | 2.69E+00 | 2.73E+00 | 9.01E-01 | no | | intergenic_variant | MODIFIER |
| 1 | 67663769 | AX-170231122 | 0 | A | T | 0.114 | 2.89E-02 | 2.33E-01 | 2.69E+00 | 2.73E+00 | 9.01E-01 | no | | intergenic_variant | MODIFIER |
| 1 | 67722259 | AX-170000719 | 0 | G | A | 0.114 | 2.89E-02 | 2.33E-01 | 2.69E+00 | 2.73E+00 | 9.01E-01 | no | | intergenic_variant | MODIFIER |

| | | | | | | | | | | | | | | | |
|---|---|---|---|---|---|---|---|---|---|---|---|---|---|---|---|
| 1 | 67723365 | AX-97795583 | 0 | T | C | 0.114 | 2.89E-02 | 2.33E-01 | 2.69E+00 | 2.73E+00 | 9.01E-01 | no | | intergenic_variant | MODIFIER |
| 1 | 67749123 | AX-170320078 | 0 | A | G | 0.114 | 2.89E-02 | 2.33E-01 | 2.69E+00 | 2.73E+00 | 9.01E-01 | no | | intergenic_variant | MODIFIER |
| 1 | 67762530 | AX-170526855 | 0 | C | T | 0.114 | 2.89E-02 | 2.33E-01 | 2.69E+00 | 2.73E+00 | 9.01E-01 | no | | intergenic_variant | MODIFIER |
| 1 | 67777023 | AX-170086490 | 0 | A | G | 0.114 | 2.89E-02 | 2.33E-01 | 2.69E+00 | 2.73E+00 | 9.01E-01 | no | | regulatory_region_variant | MODIFIER |
| 1 | 67788086 | AX-170506869 | 0 | T | A | 0.114 | 2.89E-02 | 2.33E-01 | 2.69E+00 | 2.73E+00 | 9.01E-01 | no | Gm15669 | upstream_gene_variant | MODIFIER |
| 1 | 67812248 | AX-97798052 | 0 | C | T | 0.114 | 2.89E-02 | 2.33E-01 | 2.69E+00 | 2.73E+00 | 9.01E-01 | no | | intergenic_variant | MODIFIER |
| 1 | 67841029 | AX-170507384 | 0 | A | T | 0.114 | 2.89E-02 | 2.33E-01 | 2.69E+00 | 2.73E+00 | 9.01E-01 | no | | intergenic_variant | MODIFIER |
| 1 | 67876911 | AX-170146441 | 0 | C | A | 0.114 | 2.89E-02 | 2.33E-01 | 2.69E+00 | 2.73E+00 | 9.01E-01 | no | | intergenic_variant | MODIFIER |
| 1 | 67897084 | AX-170041529 | 0 | C | T | 0.114 | 2.89E-02 | 2.33E-01 | 2.69E+00 | 2.73E+00 | 9.01E-01 | no | | intergenic_variant | MODIFIER |
| 1 | 67902479 | AX-97811658 | 0 | T | C | 0.114 | 2.89E-02 | 2.33E-01 | 2.69E+00 | 2.73E+00 | 9.01E-01 | no | Gm37957 | upstream_gene_variant | MODIFIER |
| 1 | 67929116 | AX-170457530 | 0 | T | C | 0.114 | 2.89E-02 | 2.33E-01 | 2.69E+00 | 2.73E+00 | 9.01E-01 | no | | intergenic_variant | MODIFIER |
| 1 | 67932824 | AX-97785567 | 0 | C | A | 0.114 | 2.89E-02 | 2.33E-01 | 2.69E+00 | 2.73E+00 | 9.01E-01 | no | | intergenic_variant | MODIFIER |
| 1 | 67933324 | AX-170532305 | 0 | T | A | 0.114 | 2.89E-02 | 2.33E-01 | 2.69E+00 | 2.73E+00 | 9.01E-01 | no | | intergenic_variant | MODIFIER |
| 1 | 67939690 | AX-170136351 | 0 | T | C | 0.114 | 2.89E-02 | 2.33E-01 | 2.69E+00 | 2.73E+00 | 9.01E-01 | no | | intergenic_variant | MODIFIER |
| 1 | 67943046 | AX-170104333 | 0 | A | T | 0.114 | 2.89E-02 | 2.33E-01 | 2.69E+00 | 2.73E+00 | 9.01E-01 | no | | intergenic_variant | MODIFIER |
| 1 | 67962747 | AX-170421747 | 0 | G | A | 0.114 | 2.89E-02 | 2.33E-01 | 2.69E+00 | 2.73E+00 | 9.01E-01 | no | | intergenic_variant | MODIFIER |
| 1 | 67984929 | AX-170518695 | 0 | G | C | 0.114 | 2.89E-02 | 2.33E-01 | 2.69E+00 | 2.73E+00 | 9.01E-01 | no | | intergenic_variant | MODIFIER |
| 1 | 67985238 | AX-170391158 | 0 | C | G | 0.114 | 2.89E-02 | 2.33E-01 | 2.69E+00 | 2.73E+00 | 9.01E-01 | no | | intergenic_variant | MODIFIER |
| 1 | 67985340 | AX-97829537 | 0 | A | G | 0.114 | 2.89E-02 | 2.33E-01 | 2.69E+00 | 2.73E+00 | 9.01E-01 | no | | intergenic_variant | MODIFIER |
| 1 | 67985509 | AX-170497787 | 0 | T | C | 0.114 | 2.89E-02 | 2.33E-01 | 2.69E+00 | 2.73E+00 | 9.01E-01 | no | | intergenic_variant | MODIFIER |
| 1 | 68028372 | AX-170325665 | 0 | C | T | 0.114 | 2.89E-02 | 2.33E-01 | 2.69E+00 | 2.73E+00 | 9.01E-01 | no | Erbb4 | downstream_gene_variant | MODIFIER |
| 1 | 68035855 | AX-170138294 | 0 | C | G | 0.114 | 2.89E-02 | 2.33E-01 | 2.69E+00 | 2.73E+00 | 9.01E-01 | no | Erbb4 | 3_prime_UTR_variant | MODIFIER |
| 1 | 68039843 | AX-170114183 | 0 | C | T | 0.114 | 2.89E-02 | 2.33E-01 | 2.69E+00 | 2.73E+00 | 9.01E-01 | no | Erbb4 | 3_prime_UTR_variant | MODIFIER |
| 1 | 68040804 | AX-97852950 | 0 | A | G | 0.114 | 2.89E-02 | 2.33E-01 | 2.69E+00 | 2.73E+00 | 9.01E-01 | no | Erbb4 | intron_variant | MODIFIER |
| 1 | 68041445 | AX-170257174 | 0 | T | A | 0.114 | 2.89E-02 | 2.33E-01 | 2.69E+00 | 2.73E+00 | 9.01E-01 | no | Erbb4 | intron_variant | MODIFIER |
| 1 | 68116024 | AX-170041532 | 0 | G | A | 0.114 | 2.89E-02 | 2.33E-01 | 2.69E+00 | 2.73E+00 | 9.01E-01 | no | Erbb4 | intron_variant | MODIFIER |
| 1 | 68118369 | AX-170041530 | 0 | C | T | 0.114 | 2.89E-02 | 2.33E-01 | 2.69E+00 | 2.73E+00 | 9.01E-01 | no | Erbb4 | intron_variant | MODIFIER |
| 1 | 68120419 | AX-170150716 | 0 | A | G | 0.114 | 2.89E-02 | 2.33E-01 | 2.69E+00 | 2.73E+00 | 9.01E-01 | no | Erbb4 | intron_variant | MODIFIER |
| 1 | 68120891 | AX-170138001 | 0 | A | G | 0.114 | 2.89E-02 | 2.33E-01 | 2.69E+00 | 2.73E+00 | 9.01E-01 | no | Erbb4 | intron_variant | MODIFIER |
| 1 | 68121006 | AX-170179988 | 0 | A | C | 0.114 | 2.89E-02 | 2.33E-01 | 2.69E+00 | 2.73E+00 | 9.01E-01 | no | Erbb4 | intron_variant | MODIFIER |
| 1 | 68122817 | AX-170000734 | 0 | A | G | 0.114 | 2.89E-02 | 2.33E-01 | 2.69E+00 | 2.73E+00 | 9.01E-01 | no | Erbb4 | intron_variant | MODIFIER |
| 1 | 68153724 | AX-170367249 | 0 | C | A | 0.114 | 2.89E-02 | 2.33E-01 | 2.69E+00 | 2.73E+00 | 9.01E-01 | no | Erbb4 | intron_variant | MODIFIER |
| 1 | 68163713 | AX-170000729 | 0 | A | G | 0.114 | 2.89E-02 | 2.33E-01 | 2.69E+00 | 2.73E+00 | 9.01E-01 | no | Erbb4 | intron_variant | MODIFIER |
| 1 | 68163839 | AX-170488459 | 0 | G | A | 0.114 | 2.89E-02 | 2.33E-01 | 2.69E+00 | 2.73E+00 | 9.01E-01 | no | Erbb4 | intron_variant | MODIFIER |
| 1 | 68169789 | AX-170327026 | 0 | C | T | 0.114 | 2.89E-02 | 2.33E-01 | 2.69E+00 | 2.73E+00 | 9.01E-01 | no | Erbb4 | intron_variant | MODIFIER |
| 1 | 68171443 | AX-170473628 | 0 | T | C | 0.114 | 2.89E-02 | 2.33E-01 | 2.69E+00 | 2.73E+00 | 9.01E-01 | no | Erbb4 | intron_variant | MODIFIER |
| 1 | 68178921 | AX-97800883 | 0 | G | A | 0.114 | 2.89E-02 | 2.33E-01 | 2.69E+00 | 2.73E+00 | 9.01E-01 | no | Erbb4 | intron_variant | MODIFIER |
| 1 | 68218282 | AX-170150090 | 0 | A | G | 0.114 | 2.89E-02 | 2.33E-01 | 2.69E+00 | 2.73E+00 | 9.01E-01 | no | Erbb4 | intron_variant | MODIFIER |
| 1 | 68258326 | AX-170162991 | 0 | C | G | 0.114 | 2.89E-02 | 2.33E-01 | 2.69E+00 | 2.73E+00 | 9.01E-01 | no | Erbb4 | intron_variant | MODIFIER |
| 1 | 68258400 | AX-97850158 | 0 | C | T | 0.114 | 2.89E-02 | 2.33E-01 | 2.69E+00 | 2.73E+00 | 9.01E-01 | no | Erbb4 | intron_variant | MODIFIER |
| 1 | 68288254 | AX-170212280 | 0 | G | A | 0.114 | 2.89E-02 | 2.33E-01 | 2.69E+00 | 2.73E+00 | 9.01E-01 | no | Erbb4 | intron_variant | MODIFIER |
| 1 | 68290576 | AX-170112231 | 0 | A | G | 0.114 | 2.89E-02 | 2.33E-01 | 2.69E+00 | 2.73E+00 | 9.01E-01 | no | Gm15670 | downstream_gene_variant | MODIFIER |
| 1 | 68305390 | AX-170128220 | 0 | G | T | 0.114 | 2.89E-02 | 2.33E-01 | 2.69E+00 | 2.73E+00 | 9.01E-01 | no | Erbb4 | intron_variant | MODIFIER |
| 1 | 68339431 | AX-170041538 | 0 | C | T | 0.114 | 2.89E-02 | 2.33E-01 | 2.69E+00 | 2.73E+00 | 9.01E-01 | no | Erbb4 | synonymous_variant | LOW |
| 1 | 68346646 | AX-170124533 | 0 | C | T | 0.114 | 2.89E-02 | 2.33E-01 | 2.69E+00 | 2.73E+00 | 9.01E-01 | no | Erbb4 | synonymous_variant | LOW |
| 1 | 68356316 | AX-170091544 | 0 | C | T | 0.114 | 2.89E-02 | 2.33E-01 | 2.69E+00 | 2.73E+00 | 9.01E-01 | no | Erbb4 | intron_variant | MODIFIER |
| 1 | 68361376 | AX-170041541 | 0 | A | G | 0.114 | 2.89E-02 | 2.33E-01 | 2.69E+00 | 2.73E+00 | 9.01E-01 | no | Erbb4 | intron_variant | MODIFIER |
| 1 | 68362958 | AX-170041539 | 0 | C | A | 0.114 | 2.89E-02 | 2.33E-01 | 2.69E+00 | 2.73E+00 | 9.01E-01 | no | Erbb4 | intron_variant | MODIFIER |
| 1 | 68430981 | AX-170103296 | 0 | T | C | 0.114 | 2.89E-02 | 2.33E-01 | 2.69E+00 | 2.73E+00 | 9.01E-01 | no | Erbb4 | intron_variant | MODIFIER |
| 1 | 68432681 | AX-170169855 | 0 | T | C | 0.114 | 2.89E-02 | 2.33E-01 | 2.69E+00 | 2.73E+00 | 9.01E-01 | no | Erbb4 | intron_variant | MODIFIER |
| 1 | 68441994 | AX-170145483 | 0 | C | G | 0.114 | 2.89E-02 | 2.33E-01 | 2.69E+00 | 2.73E+00 | 9.01E-01 | no | Erbb4 | intron_variant | MODIFIER |
| 1 | 68442508 | AX-170167349 | 0 | G | A | 0.114 | 2.89E-02 | 2.33E-01 | 2.69E+00 | 2.73E+00 | 9.01E-01 | no | Erbb4 | intron_variant | MODIFIER |
| 1 | 68442843 | AX-97811872 | 0 | G | T | 0.114 | 2.89E-02 | 2.33E-01 | 2.69E+00 | 2.73E+00 | 9.01E-01 | no | Erbb4 | intron_variant | MODIFIER |
| 1 | 68492845 | AX-170526233 | 0 | T | C | 0.114 | 2.89E-02 | 2.33E-01 | 2.69E+00 | 2.73E+00 | 9.01E-01 | no | Erbb4 | intron_variant | MODIFIER |

| | | | | | | | | | | | | | | | |
|---|---|---|---|---|---|---|---|---|---|---|---|---|---|---|---|
| 1 | 68494264 | AX-170488022 | 0 | C | G | 0.114 | 2.89E-02 | 2.33E-01 | 2.69E+00 | 2.73E+00 | 9.01E-01 | no | Erbb4 | intron_variant | MODIFIER |
| 1 | 68504430 | AX-97829992 | 0 | A | G | 0.114 | 2.89E-02 | 2.33E-01 | 2.69E+00 | 2.73E+00 | 9.01E-01 | no | Erbb4 | intron_variant | MODIFIER |
| 1 | 68537582 | AX-170121044 | 0 | C | T | 0.114 | 2.89E-02 | 2.33E-01 | 2.69E+00 | 2.73E+00 | 9.01E-01 | no | Erbb4 | intron_variant | MODIFIER |
| 1 | 68557046 | AX-97802189 | 0 | A | G | 0.114 | 2.89E-02 | 2.33E-01 | 2.69E+00 | 2.73E+00 | 9.01E-01 | no | Erbb4 | intron_variant | MODIFIER |
| 1 | 68572133 | AX-170135864 | 0 | G | C | 0.114 | 2.89E-02 | 2.33E-01 | 2.69E+00 | 2.73E+00 | 9.01E-01 | no | Erbb4 | intron_variant | MODIFIER |
| 1 | 68605671 | AX-170228762 | 0 | C | T | 0.114 | 2.89E-02 | 2.33E-01 | 2.69E+00 | 2.73E+00 | 9.01E-01 | no | Erbb4 | intron_variant | MODIFIER |
| 1 | 68618775 | AX-170131367 | 0 | A | G | 0.114 | 2.89E-02 | 2.33E-01 | 2.69E+00 | 2.73E+00 | 9.01E-01 | no | Erbb4 | intron_variant | MODIFIER |
| 1 | 68636933 | AX-170532935 | 0 | C | T | 0.114 | 2.89E-02 | 2.33E-01 | 2.69E+00 | 2.73E+00 | 9.01E-01 | no | Erbb4 | intron_variant | MODIFIER |
| 1 | 68650490 | AX-170190664 | 0 | A | G | 0.114 | 2.89E-02 | 2.33E-01 | 2.69E+00 | 2.73E+00 | 9.01E-01 | no | Erbb4 | intron_variant | MODIFIER |
| 1 | 68751845 | AX-170517312 | 0 | C | G | 0.114 | 2.89E-02 | 2.33E-01 | 2.69E+00 | 2.73E+00 | 9.01E-01 | no | Erbb4 | intron_variant | MODIFIER |
| 1 | 68756082 | AX-170423707 | 0 | T | G | 0.114 | 2.89E-02 | 2.33E-01 | 2.69E+00 | 2.73E+00 | 9.01E-01 | no | Erbb4 | intron_variant | MODIFIER |
| 1 | 68772089 | AX-170000744 | 0 | G | C | 0.114 | 2.89E-02 | 2.33E-01 | 2.69E+00 | 2.73E+00 | 9.01E-01 | no | Erbb4 | intron_variant | MODIFIER |
| 1 | 68775871 | AX-170228508 | 0 | T | C | 0.114 | 2.89E-02 | 2.33E-01 | 2.69E+00 | 2.73E+00 | 9.01E-01 | no | Erbb4 | intron_variant | MODIFIER |
| 1 | 68780137 | AX-170191008 | 0 | G | A | 0.114 | 2.89E-02 | 2.33E-01 | 2.69E+00 | 2.73E+00 | 9.01E-01 | no | Erbb4 | intron_variant | MODIFIER |
| 1 | 68781688 | AX-170231461 | 0 | A | G | 0.114 | 2.89E-02 | 2.33E-01 | 2.69E+00 | 2.73E+00 | 9.01E-01 | no | Erbb4 | intron_variant | MODIFIER |
| 1 | 68800262 | AX-97811926 | 0 | T | C | 0.114 | 2.89E-02 | 2.33E-01 | 2.69E+00 | 2.73E+00 | 9.01E-01 | no | Erbb4 | intron_variant | MODIFIER |
| 1 | 68825431 | AX-97803337 | 0 | A | G | 0.114 | 2.89E-02 | 2.33E-01 | 2.69E+00 | 2.73E+00 | 9.01E-01 | no | Erbb4 | intron_variant | MODIFIER |
| 1 | 68845576 | AX-170000742 | 0 | T | G | 0.114 | 2.89E-02 | 2.33E-01 | 2.69E+00 | 2.73E+00 | 9.01E-01 | no | Erbb4 | intron_variant | MODIFIER |
| 1 | 68847324 | AX-97821481 | 0 | A | G | 0.114 | 2.89E-02 | 2.33E-01 | 2.69E+00 | 2.73E+00 | 9.01E-01 | no | Erbb4 | intron_variant | MODIFIER |
| 1 | 68848217 | AX-170000738 | 0 | T | C | 0.114 | 2.89E-02 | 2.33E-01 | 2.69E+00 | 2.73E+00 | 9.01E-01 | no | Erbb4 | intron_variant | MODIFIER |
| 1 | 68860187 | AX-170041548 | 0 | C | G | 0.114 | 2.89E-02 | 2.33E-01 | 2.69E+00 | 2.73E+00 | 9.01E-01 | no | Erbb4 | intron_variant | MODIFIER |
| 1 | 68891652 | AX-170108548 | 0 | A | G | 0.114 | 2.89E-02 | 2.33E-01 | 2.69E+00 | 2.73E+00 | 9.01E-01 | no | Erbb4 | intron_variant | MODIFIER |
| 1 | 68891932 | AX-170283615 | 0 | G | C | 0.114 | 2.89E-02 | 2.33E-01 | 2.69E+00 | 2.73E+00 | 9.01E-01 | no | Erbb4 | intron_variant | MODIFIER |
| 1 | 68911551 | AX-170217066 | 0 | G | A | 0.114 | 2.89E-02 | 2.33E-01 | 2.69E+00 | 2.73E+00 | 9.01E-01 | no | Erbb4 | intron_variant | MODIFIER |
| 1 | 68917938 | AX-97822533 | 0 | A | G | 0.114 | 2.89E-02 | 2.33E-01 | 2.69E+00 | 2.73E+00 | 9.01E-01 | no | Erbb4 | intron_variant | MODIFIER |
| 1 | 68927120 | AX-170388661 | 0 | T | A | 0.114 | 2.89E-02 | 2.33E-01 | 2.69E+00 | 2.73E+00 | 9.01E-01 | no | Erbb4 | intron_variant | MODIFIER |
| 1 | 68986206 | AX-97783930 | 0 | G | A | 0.114 | 2.89E-02 | 2.33E-01 | 2.69E+00 | 2.73E+00 | 9.01E-01 | no | Erbb4 | intron_variant | MODIFIER |
| 1 | 69002487 | AX-170432774 | 0 | T | C | 0.114 | 2.89E-02 | 2.33E-01 | 2.69E+00 | 2.73E+00 | 9.01E-01 | no | Erbb4 | intron_variant | MODIFIER |
| 1 | 69134915 | AX-97837291 | 0 | G | A | 0.114 | 2.89E-02 | 2.33E-01 | 2.69E+00 | 2.73E+00 | 9.01E-01 | no | Gm8840 | upstream_gene_variant | MODIFIER |
| 1 | 69165419 | AX-170502521 | 0 | C | T | 0.114 | 2.89E-02 | 2.33E-01 | 2.69E+00 | 2.73E+00 | 9.01E-01 | no | | regulatory_region_variant | MODIFIER |
| 1 | 69255783 | AX-170214241 | 0 | A | G | 0.114 | 2.89E-02 | 2.33E-01 | 2.69E+00 | 2.73E+00 | 9.01E-01 | no | | intergenic_variant | MODIFIER |
| 1 | 69304249 | AX-97780548 | 0 | T | C | 0.114 | 2.89E-02 | 2.33E-01 | 2.69E+00 | 2.73E+00 | 9.01E-01 | no | | intergenic_variant | MODIFIER |
| 1 | 69333658 | AX-170305879 | 0 | C | T | 0.114 | 2.89E-02 | 2.33E-01 | 2.69E+00 | 2.73E+00 | 9.01E-01 | no | | intergenic_variant | MODIFIER |
| 1 | 69334006 | AX-170339475 | 0 | G | A | 0.114 | 2.89E-02 | 2.33E-01 | 2.69E+00 | 2.73E+00 | 9.01E-01 | no | | intergenic_variant | MODIFIER |
| 1 | 69334479 | AX-97822098 | 0 | G | A | 0.114 | 2.89E-02 | 2.33E-01 | 2.69E+00 | 2.73E+00 | 9.01E-01 | no | | intergenic_variant | MODIFIER |
| 1 | 69362812 | AX-170473918 | 0 | G | C | 0.114 | 2.89E-02 | 2.33E-01 | 2.69E+00 | 2.73E+00 | 9.01E-01 | no | | intergenic_variant | MODIFIER |
| 1 | 69369747 | AX-97812054 | 0 | T | C | 0.114 | 2.89E-02 | 2.33E-01 | 2.69E+00 | 2.73E+00 | 9.01E-01 | no | | intergenic_variant | MODIFIER |
| 1 | 69393724 | AX-170041559 | 0 | T | C | 0.114 | 2.89E-02 | 2.33E-01 | 2.69E+00 | 2.73E+00 | 9.01E-01 | no | | intergenic_variant | MODIFIER |
| 1 | 69393873 | AX-97822207 | 0 | A | G | 0.114 | 2.89E-02 | 2.33E-01 | 2.69E+00 | 2.73E+00 | 9.01E-01 | no | | intergenic_variant | MODIFIER |
| 1 | 69408031 | AX-170259079 | 0 | A | T | 0.114 | 2.89E-02 | 2.33E-01 | 2.69E+00 | 2.73E+00 | 9.01E-01 | no | | intergenic_variant | MODIFIER |
| 1 | 69408488 | AX-97811924 | 0 | T | C | 0.114 | 2.89E-02 | 2.33E-01 | 2.69E+00 | 2.73E+00 | 9.01E-01 | no | | intergenic_variant | MODIFIER |
| 1 | 69527743 | AX-170335912 | 0 | G | A | 0.114 | 2.89E-02 | 2.33E-01 | 2.69E+00 | 2.73E+00 | 9.01E-01 | no | Ikzf2 | downstream_gene_variant | MODIFIER |
| 1 | 69558089 | AX-170501164 | 0 | G | A | 0.114 | 2.89E-02 | 2.33E-01 | 2.69E+00 | 2.73E+00 | 9.01E-01 | no | Ikzf2 | intron_variant | MODIFIER |
| 1 | 69558739 | AX-97846276 | 0 | A | G | 0.114 | 2.89E-02 | 2.33E-01 | 2.69E+00 | 2.73E+00 | 9.01E-01 | no | Ikzf2 | intron_variant | MODIFIER |
| 1 | 69567600 | AX-170148845 | 0 | T | C | 0.114 | 2.89E-02 | 2.33E-01 | 2.69E+00 | 2.73E+00 | 9.01E-01 | no | Ikzf2 | intron_variant | MODIFIER |
| 1 | 69568301 | AX-170421998 | 0 | C | A | 0.114 | 2.89E-02 | 2.33E-01 | 2.69E+00 | 2.73E+00 | 9.01E-01 | no | Ikzf2 | intron_variant | MODIFIER |
| 1 | 69568699 | AX-170471070 | 0 | T | C | 0.114 | 2.89E-02 | 2.33E-01 | 2.69E+00 | 2.73E+00 | 9.01E-01 | no | Ikzf2 | intron_variant | MODIFIER |
| 1 | 69578087 | AX-97825535 | 0 | C | T | 0.114 | 2.89E-02 | 2.33E-01 | 2.69E+00 | 2.73E+00 | 9.01E-01 | no | Ikzf2 | intron_variant | MODIFIER |
| 1 | 69643859 | AX-170041568 | 0 | G | A | 0.114 | 2.89E-02 | 2.33E-01 | 2.69E+00 | 2.73E+00 | 9.01E-01 | no | Ikzf2 | intron_variant | MODIFIER |
| 1 | 69647545 | AX-97814998 | 0 | G | T | 0.114 | 2.89E-02 | 2.33E-01 | 2.69E+00 | 2.73E+00 | 9.01E-01 | no | Ikzf2 | intron_variant | MODIFIER |
| 1 | 69697186 | AX-170216836 | 0 | A | G | 0.114 | 2.89E-02 | 2.33E-01 | 2.69E+00 | 2.73E+00 | 9.01E-01 | no | | intergenic_variant | MODIFIER |
| 1 | 69717898 | AX-170441491 | 0 | T | C | 0.114 | 2.89E-02 | 2.33E-01 | 2.69E+00 | 2.73E+00 | 9.01E-01 | no | | intergenic_variant | MODIFIER |
| 1 | 69726122 | AX-170000760 | 0 | A | G | 0.114 | 2.89E-02 | 2.33E-01 | 2.69E+00 | 2.73E+00 | 9.01E-01 | no | | intergenic_variant | MODIFIER |
| 1 | 69824235 | AX-97792246 | 0 | T | C | 0.114 | 2.89E-02 | 2.33E-01 | 2.69E+00 | 2.73E+00 | 9.01E-01 | no | Spag16 | upstream_gene_variant | MODIFIER |

| | | | | | | | | | | | | | | | |
|---|---|---|---|---|---|---|---|---|---|---|---|---|---|---|---|
| 1 | 70084922 | AX-170179465 | 0 | T | C | 0.114 | 2.89E-02 | 2.33E-01 | 2.69E+00 | 2.73E+00 | 9.01E-01 | no | Spag16 | intron_variant | MODIFIER |
| 1 | 70084954 | AX-170353545 | 0 | A | G | 0.114 | 2.89E-02 | 2.33E-01 | 2.69E+00 | 2.73E+00 | 9.01E-01 | no | Spag16 | intron_variant | MODIFIER |
| 1 | 70191972 | AX-170134211 | 0 | G | A | 0.114 | 2.89E-02 | 2.33E-01 | 2.69E+00 | 2.73E+00 | 9.01E-01 | no | Spag16 | intron_variant | MODIFIER |
| 1 | 70304172 | AX-97786115 | 0 | G | A | 0.114 | 2.89E-02 | 2.33E-01 | 2.69E+00 | 2.73E+00 | 9.01E-01 | no | Spag16 | intron_variant | MODIFIER |
| 1 | 70316469 | AX-170041582 | 0 | C | T | 0.114 | 2.89E-02 | 2.33E-01 | 2.69E+00 | 2.73E+00 | 9.01E-01 | no | Spag16 | intron_variant | MODIFIER |
| 1 | 70328315 | AX-170000762 | 0 | T | G | 0.114 | 2.89E-02 | 2.33E-01 | 2.69E+00 | 2.73E+00 | 9.01E-01 | no | Spag16 | intron_variant | MODIFIER |
| 1 | 70397722 | AX-170297131 | 0 | G | T | 0.114 | 2.89E-02 | 2.33E-01 | 2.69E+00 | 2.73E+00 | 9.01E-01 | no | Spag16 | intron_variant | MODIFIER |
| 1 | 70646492 | AX-170254371 | 0 | A | C | 0.114 | 2.89E-02 | 2.33E-01 | 2.69E+00 | 2.73E+00 | 9.01E-01 | no | Spag16 | intron_variant | MODIFIER |
| 1 | 70657713 | AX-170159061 | 0 | A | T | 0.114 | 2.89E-02 | 2.33E-01 | 2.69E+00 | 2.73E+00 | 9.01E-01 | no | Spag16 | intron_variant | MODIFIER |
| 1 | 70657799 | AX-170516873 | 0 | A | C | 0.114 | 2.89E-02 | 2.33E-01 | 2.69E+00 | 2.73E+00 | 9.01E-01 | no | Spag16 | intron_variant | MODIFIER |
| 1 | 70657870 | AX-170495321 | 0 | A | G | 0.114 | 2.89E-02 | 2.33E-01 | 2.69E+00 | 2.73E+00 | 9.01E-01 | no | Spag16 | intron_variant | MODIFIER |
| 1 | 70659384 | AX-170448928 | 0 | G | A | 0.114 | 2.89E-02 | 2.33E-01 | 2.69E+00 | 2.73E+00 | 9.01E-01 | no | Spag16 | intron_variant | MODIFIER |
| 1 | 70659463 | AX-170238120 | 0 | T | C | 0.114 | 2.89E-02 | 2.33E-01 | 2.69E+00 | 2.73E+00 | 9.01E-01 | no | Spag16 | intron_variant | MODIFIER |
| 1 | 70662582 | AX-97834516 | 0 | G | T | 0.114 | 2.89E-02 | 2.33E-01 | 2.69E+00 | 2.73E+00 | 9.01E-01 | no | Spag16 | intron_variant | MODIFIER |
| 1 | 70692165 | AX-97820926 | 0 | G | A | 0.114 | 2.89E-02 | 2.33E-01 | 2.69E+00 | 2.73E+00 | 9.01E-01 | no | Spag16 | intron_variant | MODIFIER |
| 1 | 70699363 | AX-97819861 | 0 | G | A | 0.114 | 2.89E-02 | 2.33E-01 | 2.69E+00 | 2.73E+00 | 9.01E-01 | no | Spag16 | intron_variant | MODIFIER |
| 1 | 70728215 | AX-170305480 | 0 | A | G | 0.114 | 2.89E-02 | 2.33E-01 | 2.69E+00 | 2.73E+00 | 9.01E-01 | no | Vwc2l | intron_variant | MODIFIER |
| 1 | 70764819 | AX-170169246 | 0 | G | C | 0.114 | 2.89E-02 | 2.33E-01 | 2.69E+00 | 2.73E+00 | 9.01E-01 | no | Vwc2l | intron_variant | MODIFIER |
| 1 | 71356496 | AX-170489863 | 0 | G | A | 0.114 | 2.89E-02 | 2.33E-01 | 2.69E+00 | 2.73E+00 | 9.01E-01 | no | Abca12 | intron_variant | MODIFIER |
| 1 | 71489641 | AX-170155241 | 0 | C | G | 0.114 | 2.89E-02 | 2.33E-01 | 2.69E+00 | 2.73E+00 | 9.01E-01 | no | Gm6947 | downstream_gene_variant | MODIFIER |
| 1 | 71494771 | AX-97834038 | 0 | T | G | 0.114 | 2.89E-02 | 2.33E-01 | 2.69E+00 | 2.73E+00 | 9.01E-01 | no | Gm6947 | upstream_gene_variant | MODIFIER |
| 1 | 71497226 | AX-170041608 | 0 | A | G | 0.114 | 2.89E-02 | 2.33E-01 | 2.69E+00 | 2.73E+00 | 9.01E-01 | no | Gm6947 | upstream_gene_variant | MODIFIER |
| 1 | 71501811 | AX-170041609 | 0 | T | C | 0.114 | 2.89E-02 | 2.33E-01 | 2.69E+00 | 2.73E+00 | 9.01E-01 | no | | intergenic_variant | MODIFIER |
| 1 | 71515770 | AX-170488799 | 0 | T | C | 0.114 | 2.89E-02 | 2.33E-01 | 2.69E+00 | 2.73E+00 | 9.01E-01 | no | | intergenic_variant | MODIFIER |
| 1 | 71525194 | AX-170236218 | 0 | G | A | 0.114 | 2.89E-02 | 2.33E-01 | 2.69E+00 | 2.73E+00 | 9.01E-01 | no | | intergenic_variant | MODIFIER |
| 1 | 71528671 | AX-170458560 | 0 | C | T | 0.114 | 2.89E-02 | 2.33E-01 | 2.69E+00 | 2.73E+00 | 9.01E-01 | no | | intergenic_variant | MODIFIER |
| 1 | 71566097 | AX-170403676 | 0 | C | T | 0.114 | 2.89E-02 | 2.33E-01 | 2.69E+00 | 2.73E+00 | 9.01E-01 | no | Atic | intron_variant | MODIFIER |
| 1 | 71567714 | AX-170086494 | 0 | G | T | 0.114 | 2.89E-02 | 2.33E-01 | 2.69E+00 | 2.73E+00 | 9.01E-01 | no | Atic | intron_variant | MODIFIER |
| 1 | 71579842 | AX-170358524 | 0 | C | T | 0.114 | 2.89E-02 | 2.33E-01 | 2.69E+00 | 2.73E+00 | 9.01E-01 | no | Atic | downstream_gene_variant | MODIFIER |
| 1 | 71591455 | AX-97790458 | 0 | A | G | 0.114 | 2.89E-02 | 2.33E-01 | 2.69E+00 | 2.73E+00 | 9.01E-01 | no | Fn1 | intron_variant | MODIFIER |
| 1 | 71595585 | AX-170497099 | 0 | G | A | 0.114 | 2.89E-02 | 2.33E-01 | 2.69E+00 | 2.73E+00 | 9.01E-01 | no | Fn1 | intron_variant | MODIFIER |
| 1 | 71609525 | AX-170269676 | 0 | T | C | 0.114 | 2.89E-02 | 2.33E-01 | 2.69E+00 | 2.73E+00 | 9.01E-01 | no | Fn1 | synonymous_variant | LOW |
| 1 | 71609770 | AX-97806561 | 0 | C | A | 0.114 | 2.89E-02 | 2.33E-01 | 2.69E+00 | 2.73E+00 | 9.01E-01 | no | Fn1 | intron_variant | MODIFIER |
| 1 | 71632254 | AX-170253867 | 0 | A | G | 0.114 | 2.89E-02 | 2.33E-01 | 2.69E+00 | 2.73E+00 | 9.01E-01 | no | Fn1 | intron_variant | MODIFIER |
| 1 | 71646128 | AX-170310825 | 0 | G | A | 0.114 | 2.89E-02 | 2.33E-01 | 2.69E+00 | 2.73E+00 | 9.01E-01 | no | Fn1 | intron_variant | MODIFIER |
| 1 | 71647183 | AX-170371130 | 0 | A | G | 0.114 | 2.89E-02 | 2.33E-01 | 2.69E+00 | 2.73E+00 | 9.01E-01 | no | Fn1 | intron_variant | MODIFIER |
| 1 | 71647307 | AX-170300626 | 0 | G | A | 0.114 | 2.89E-02 | 2.33E-01 | 2.69E+00 | 2.73E+00 | 9.01E-01 | no | Fn1 | intron_variant | MODIFIER |
| 1 | 71671717 | AX-170225516 | 0 | A | C | 0.114 | 2.89E-02 | 2.33E-01 | 2.69E+00 | 2.73E+00 | 9.01E-01 | no | | intergenic_variant | MODIFIER |
| 1 | 71689928 | AX-170179874 | 0 | A | G | 0.114 | 2.89E-02 | 2.33E-01 | 2.69E+00 | 2.73E+00 | 9.01E-01 | no | | intergenic_variant | MODIFIER |
| 1 | 71690411 | AX-170270375 | 0 | C | T | 0.114 | 2.89E-02 | 2.33E-01 | 2.69E+00 | 2.73E+00 | 9.01E-01 | no | | intergenic_variant | MODIFIER |
| 1 | 71700534 | AX-170218181 | 0 | A | G | 0.114 | 2.89E-02 | 2.33E-01 | 2.69E+00 | 2.73E+00 | 9.01E-01 | no | | intergenic_variant | MODIFIER |
| 1 | 71765695 | AX-170198691 | 0 | A | T | 0.114 | 2.89E-02 | 2.33E-01 | 2.69E+00 | 2.73E+00 | 9.01E-01 | no | | intergenic_variant | MODIFIER |
| 1 | 71768369 | AX-97811954 | 0 | T | G | 0.114 | 2.89E-02 | 2.33E-01 | 2.69E+00 | 2.73E+00 | 9.01E-01 | no | | intergenic_variant | MODIFIER |
| 1 | 71776898 | AX-170514111 | 0 | A | G | 0.114 | 2.89E-02 | 2.33E-01 | 2.69E+00 | 2.73E+00 | 9.01E-01 | no | | regulatory_region_variant | MODIFIER |
| 1 | 71777524 | AX-170280853 | 0 | A | C | 0.114 | 2.89E-02 | 2.33E-01 | 2.69E+00 | 2.73E+00 | 9.01E-01 | no | | intergenic_variant | MODIFIER |
| 1 | 71779087 | AX-170041614 | 0 | T | C | 0.114 | 2.89E-02 | 2.33E-01 | 2.69E+00 | 2.73E+00 | 9.01E-01 | no | | intergenic_variant | MODIFIER |
| 1 | 71792941 | AX-170105939 | 0 | G | A | 0.114 | 2.89E-02 | 2.33E-01 | 2.69E+00 | 2.73E+00 | 9.01E-01 | no | | intergenic_variant | MODIFIER |
| 1 | 71793156 | AX-170204314 | 0 | T | C | 0.114 | 2.89E-02 | 2.33E-01 | 2.69E+00 | 2.73E+00 | 9.01E-01 | no | | intergenic_variant | MODIFIER |
| 1 | 71793264 | AX-170107775 | 0 | C | T | 0.114 | 2.89E-02 | 2.33E-01 | 2.69E+00 | 2.73E+00 | 9.01E-01 | no | | intergenic_variant | MODIFIER |
| 1 | 71793822 | AX-170409698 | 0 | G | A | 0.114 | 2.89E-02 | 2.33E-01 | 2.69E+00 | 2.73E+00 | 9.01E-01 | no | | intergenic_variant | MODIFIER |
| 1 | 71797482 | AX-170328488 | 0 | T | C | 0.114 | 2.89E-02 | 2.33E-01 | 2.69E+00 | 2.73E+00 | 9.01E-01 | no | | intergenic_variant | MODIFIER |
| 1 | 71799558 | AX-170295861 | 0 | A | G | 0.114 | 2.89E-02 | 2.33E-01 | 2.69E+00 | 2.73E+00 | 9.01E-01 | no | | regulatory_region_variant | MODIFIER |
| 1 | 71814499 | AX-170201014 | 0 | C | T | 0.114 | 2.89E-02 | 2.33E-01 | 2.69E+00 | 2.73E+00 | 9.01E-01 | no | | intergenic_variant | MODIFIER |
| 1 | 71817017 | AX-170414215 | 0 | A | C | 0.114 | 2.89E-02 | 2.33E-01 | 2.69E+00 | 2.73E+00 | 9.01E-01 | no | | intergenic_variant | MODIFIER |

| | | | | | | | | | | | | | | | |
|---|---|---|---|---|---|---|---|---|---|---|---|---|---|---|---|
| 1 | 71820053 | AX-170369559 | 0 | G | A | 0.114 | 2.89E-02 | 2.33E-01 | 2.69E+00 | 2.73E+00 | 9.01E-01 | no | | intergenic_variant | MODIFIER |
| 1 | 71820458 | AX-97804008 | 0 | T | C | 0.114 | 2.89E-02 | 2.33E-01 | 2.69E+00 | 2.73E+00 | 9.01E-01 | no | | intergenic_variant | MODIFIER |
| 1 | 71890542 | AX-170041615 | 0 | G | C | 0.114 | 2.89E-02 | 2.33E-01 | 2.69E+00 | 2.73E+00 | 9.01E-01 | no | Gm28818 | downstream_gene_variant | MODIFIER |
| 1 | 71920480 | AX-170000780 | 0 | G | C | 0.114 | 2.89E-02 | 2.33E-01 | 2.69E+00 | 2.73E+00 | 9.01E-01 | no | Gm28818 | upstream_gene_variant | MODIFIER |
| 1 | 71954212 | AX-97853975 | 0 | G | T | 0.114 | 2.89E-02 | 2.33E-01 | 2.69E+00 | 2.73E+00 | 9.01E-01 | no | 4933417E11Rik | downstream_gene_variant | MODIFIER |
| 1 | 71963541 | AX-170312520 | 0 | A | G | 0.114 | 2.89E-02 | 2.33E-01 | 2.69E+00 | 2.73E+00 | 9.01E-01 | no | 4933417E11Rik | non_coding_transcript_exon_variant | MODIFIER |
| 1 | 71964665 | AX-170242771 | 0 | A | G | 0.114 | 2.89E-02 | 2.33E-01 | 2.69E+00 | 2.73E+00 | 9.01E-01 | no | 4933417E11Rik | intron_variant,non_coding_transcript_variant | MODIFIER |
| 1 | 71968437 | AX-170168157 | 0 | G | A | 0.114 | 2.89E-02 | 2.33E-01 | 2.69E+00 | 2.73E+00 | 9.01E-01 | no | 4933417E11Rik | intron_variant,non_coding_transcript_variant | MODIFIER |
| 1 | 71974247 | AX-97811978 | 0 | A | G | 0.114 | 2.89E-02 | 2.33E-01 | 2.69E+00 | 2.73E+00 | 9.01E-01 | no | 4933417E11Rik | intron_variant,non_coding_transcript_variant | MODIFIER |
| 1 | 71995710 | AX-170512873 | 0 | A | G | 0.114 | 2.89E-02 | 2.33E-01 | 2.69E+00 | 2.73E+00 | 9.01E-01 | no | 4930556G22Rik | intron_variant,non_coding_transcript_variant | MODIFIER |
| 1 | 72005288 | AX-170000778 | 0 | G | A | 0.114 | 2.89E-02 | 2.33E-01 | 2.69E+00 | 2.73E+00 | 9.01E-01 | no | Gm5528 | downstream_gene_variant | MODIFIER |
| 1 | 72006268 | AX-97855632 | 0 | T | C | 0.114 | 2.89E-02 | 2.33E-01 | 2.69E+00 | 2.73E+00 | 9.01E-01 | no | Gm5528 | downstream_gene_variant | MODIFIER |
| 1 | 72007975 | AX-170000776 | 0 | G | C | 0.114 | 2.89E-02 | 2.33E-01 | 2.69E+00 | 2.73E+00 | 9.01E-01 | no | Gm5528 | downstream_gene_variant | MODIFIER |
| 1 | 72051902 | AX-170000781 | 0 | T | C | 0.114 | 2.89E-02 | 2.33E-01 | 2.69E+00 | 2.73E+00 | 9.01E-01 | no | | regulatory_region_variant | MODIFIER |
| 1 | 72053021 | AX-97825997 | 0 | T | C | 0.114 | 2.89E-02 | 2.33E-01 | 2.69E+00 | 2.73E+00 | 9.01E-01 | no | | intergenic_variant | MODIFIER |
| 1 | 72053120 | AX-170292968 | 0 | C | T | 0.114 | 2.89E-02 | 2.33E-01 | 2.69E+00 | 2.73E+00 | 9.01E-01 | no | | intergenic_variant | MODIFIER |
| 1 | 72078329 | AX-97854733 | 0 | A | C | 0.114 | 2.89E-02 | 2.33E-01 | 2.69E+00 | 2.73E+00 | 9.01E-01 | no | | intergenic_variant | MODIFIER |
| 1 | 72164327 | AX-170183060 | 0 | C | T | 0.114 | 2.89E-02 | 2.33E-01 | 2.69E+00 | 2.73E+00 | 9.01E-01 | no | Mreg | intron_variant | MODIFIER |
| 1 | 72164373 | AX-170407074 | 0 | A | C | 0.114 | 2.89E-02 | 2.33E-01 | 2.69E+00 | 2.73E+00 | 9.01E-01 | no | Mreg | intron_variant | MODIFIER |
| 1 | 72170169 | AX-170374364 | 0 | C | T | 0.114 | 2.89E-02 | 2.33E-01 | 2.69E+00 | 2.73E+00 | 9.01E-01 | no | Mreg | intron_variant | MODIFIER |
| 1 | 72170262 | AX-170143474 | 0 | G | A | 0.114 | 2.89E-02 | 2.33E-01 | 2.69E+00 | 2.73E+00 | 9.01E-01 | no | Mreg | intron_variant | MODIFIER |
| 1 | 72290411 | AX-97783254 | 0 | C | G | 0.114 | 2.89E-02 | 2.33E-01 | 2.69E+00 | 2.73E+00 | 9.01E-01 | no | Tmem169 | intron_variant | MODIFIER |
| 1 | 72297298 | AX-170160250 | 0 | C | T | 0.114 | 2.89E-02 | 2.33E-01 | 2.69E+00 | 2.73E+00 | 9.01E-01 | no | Tmem169 | intron_variant | MODIFIER |
| 1 | 72301297 | AX-170446167 | 0 | C | T | 0.114 | 2.89E-02 | 2.33E-01 | 2.69E+00 | 2.73E+00 | 9.01E-01 | no | Tmem169 | synonymous_variant | LOW |
| 1 | 72301401 | AX-170191607 | 0 | A | T | 0.114 | 2.89E-02 | 2.33E-01 | 2.69E+00 | 2.73E+00 | 9.01E-01 | no | Tmem169 | 3_prime_UTR_variant | MODIFIER |
| 1 | 72307869 | AX-170086495 | 0 | G | T | 0.114 | 2.89E-02 | 2.33E-01 | 2.69E+00 | 2.73E+00 | 9.01E-01 | no | Xrcc5 | intron_variant | MODIFIER |
| 1 | 72325191 | AX-97783684 | 0 | G | A | 0.114 | 2.89E-02 | 2.33E-01 | 2.69E+00 | 2.73E+00 | 9.01E-01 | no | Xrcc5 | splice_region_variant,intron_variant | LOW |
| 1 | 72326417 | AX-170258576 | 0 | T | C | 0.114 | 2.89E-02 | 2.33E-01 | 2.69E+00 | 2.73E+00 | 9.01E-01 | no | Xrcc5 | intron_variant | MODIFIER |
| 1 | 72338856 | AX-170150523 | 0 | T | C | 0.114 | 2.89E-02 | 2.33E-01 | 2.69E+00 | 2.73E+00 | 9.01E-01 | no | Xrcc5 | intron_variant | MODIFIER |
| 1 | 72343763 | AX-170426263 | 0 | A | G | 0.114 | 2.89E-02 | 2.33E-01 | 2.69E+00 | 2.73E+00 | 9.01E-01 | no | Xrcc5 | intron_variant | MODIFIER |
| 1 | 72369056 | AX-170436752 | 0 | T | C | 0.114 | 2.89E-02 | 2.33E-01 | 2.69E+00 | 2.73E+00 | 9.01E-01 | no | Xrcc5 | intron_variant | MODIFIER |
| 1 | 72384320 | AX-170156995 | 0 | A | G | 0.114 | 2.89E-02 | 2.33E-01 | 2.69E+00 | 2.73E+00 | 9.01E-01 | no | Xrcc5 | intron_variant | MODIFIER |
| 1 | 72394297 | AX-170000783 | 0 | A | G | 0.114 | 2.89E-02 | 2.33E-01 | 2.69E+00 | 2.73E+00 | 9.01E-01 | no | Xrcc5 | intron_variant | MODIFIER |
| 1 | 72407779 | AX-97785531 | 0 | G | A | 0.114 | 2.89E-02 | 2.33E-01 | 2.69E+00 | 2.73E+00 | 9.01E-01 | no | | intergenic_variant | MODIFIER |
| 1 | 72418197 | AX-170000789 | 0 | A | G | 0.114 | 2.89E-02 | 2.33E-01 | 2.69E+00 | 2.73E+00 | 9.01E-01 | no | | intergenic_variant | MODIFIER |
| 1 | 72427382 | AX-170000787 | 0 | T | C | 0.114 | 2.89E-02 | 2.33E-01 | 2.69E+00 | 2.73E+00 | 9.01E-01 | no | MAR4 | 3_prime_UTR_variant | MODIFIER |
| 1 | 72445078 | AX-170000785 | 0 | G | A | 0.114 | 2.89E-02 | 2.33E-01 | 2.69E+00 | 2.73E+00 | 9.01E-01 | no | MAR4 | intron_variant | MODIFIER |
| 1 | 72445243 | AX-170086496 | 0 | T | G | 0.114 | 2.89E-02 | 2.33E-01 | 2.69E+00 | 2.73E+00 | 9.01E-01 | no | MAR4 | intron_variant | MODIFIER |
| 1 | 72445790 | AX-170446818 | 0 | T | C | 0.114 | 2.89E-02 | 2.33E-01 | 2.69E+00 | 2.73E+00 | 9.01E-01 | no | MAR4 | intron_variant | MODIFIER |
| 1 | 72452682 | AX-170041626 | 0 | C | T | 0.114 | 2.89E-02 | 2.33E-01 | 2.69E+00 | 2.73E+00 | 9.01E-01 | no | MAR4 | intron_variant | MODIFIER |
| 1 | 72473309 | AX-170000790 | 0 | G | A | 0.114 | 2.89E-02 | 2.33E-01 | 2.69E+00 | 2.73E+00 | 9.01E-01 | no | MAR4 | intron_variant | MODIFIER |
| 1 | 72473348 | AX-170000786 | 0 | C | T | 0.114 | 2.89E-02 | 2.33E-01 | 2.69E+00 | 2.73E+00 | 9.01E-01 | no | MAR4 | intron_variant | MODIFIER |
| 1 | 72475931 | AX-170041628 | 0 | C | T | 0.114 | 2.89E-02 | 2.33E-01 | 2.69E+00 | 2.73E+00 | 9.01E-01 | no | MAR4 | intron_variant | MODIFIER |
| 1 | 72476889 | AX-97831625 | 0 | A | C | 0.114 | 2.89E-02 | 2.33E-01 | 2.69E+00 | 2.73E+00 | 9.01E-01 | no | MAR4 | intron_variant | MODIFIER |
| 1 | 72855558 | AX-170464133 | 0 | A | G | 0.114 | 2.89E-02 | 2.33E-01 | 2.69E+00 | 2.73E+00 | 9.01E-01 | no | Igfbp5 | downstream_gene_variant | MODIFIER |
| 1 | 72930378 | AX-170505892 | 0 | T | C | 0.114 | 2.89E-02 | 2.33E-01 | 2.69E+00 | 2.73E+00 | 9.01E-01 | no | | regulatory_region_variant | MODIFIER |
| 1 | 72930564 | AX-170273978 | 0 | T | C | 0.114 | 2.89E-02 | 2.33E-01 | 2.69E+00 | 2.73E+00 | 9.01E-01 | no | | intergenic_variant | MODIFIER |
| 1 | 73012521 | AX-170493243 | 0 | G | T | 0.114 | 2.89E-02 | 2.33E-01 | 2.69E+00 | 2.73E+00 | 9.01E-01 | no | Tnp1 | downstream_gene_variant | MODIFIER |
| 1 | 73015159 | AX-170000791 | 0 | T | A | 0.114 | 2.89E-02 | 2.33E-01 | 2.69E+00 | 2.73E+00 | 9.01E-01 | no | Tnp1 | 3_prime_UTR_variant | MODIFIER |
| 1 | 73195361 | AX-170156642 | 0 | A | G | 0.114 | 2.89E-02 | 2.33E-01 | 2.69E+00 | 2.73E+00 | 9.01E-01 | no | | intergenic_variant | MODIFIER |
| 1 | 73235212 | AX-97781148 | 0 | G | T | 0.114 | 2.89E-02 | 2.33E-01 | 2.69E+00 | 2.73E+00 | 9.01E-01 | no | | intergenic_variant | MODIFIER |
| 1 | 73235403 | AX-170218631 | 0 | G | C | 0.114 | 2.89E-02 | 2.33E-01 | 2.69E+00 | 2.73E+00 | 9.01E-01 | no | | intergenic_variant | MODIFIER |
| 1 | 73236118 | AX-170248871 | 0 | T | A | 0.114 | 2.89E-02 | 2.33E-01 | 2.69E+00 | 2.73E+00 | 9.01E-01 | no | | intergenic_variant | MODIFIER |
| 1 | 73239778 | AX-170427163 | 0 | A | G | 0.114 | 2.89E-02 | 2.33E-01 | 2.69E+00 | 2.73E+00 | 9.01E-01 | no | | intergenic_variant | MODIFIER |

| | | | | | | | | | | | | | | | |
|---|---|---|---|---|---|---|---|---|---|---|---|---|---|---|---|
| 1 | 73249086 | AX-170493787 | 0 | A | C | 0.114 | 2.89E-02 | 2.33E-01 | 2.69E+00 | 2.73E+00 | 9.01E-01 | no | | intergenic_variant | MODIFIER |
| 1 | 73249484 | AX-170140334 | 0 | A | T | 0.114 | 2.89E-02 | 2.33E-01 | 2.69E+00 | 2.73E+00 | 9.01E-01 | no | | intergenic_variant | MODIFIER |
| 1 | 73266491 | AX-97848661 | 0 | A | G | 0.114 | 2.89E-02 | 2.33E-01 | 2.69E+00 | 2.73E+00 | 9.01E-01 | no | Gm29185 | upstream_gene_variant | MODIFIER |
| 1 | 73277498 | AX-170235500 | 0 | C | T | 0.114 | 2.89E-02 | 2.33E-01 | 2.69E+00 | 2.73E+00 | 9.01E-01 | no | Gm29185 | intron_variant,non_coding_transcript_variant | MODIFIER |
| 1 | 73283117 | AX-97804333 | 0 | A | G | 0.114 | 2.89E-02 | 2.33E-01 | 2.69E+00 | 2.73E+00 | 9.01E-01 | no | Gm29185 | intron_variant,non_coding_transcript_variant | MODIFIER |
| 1 | 73283159 | AX-170041650 | 0 | C | T | 0.114 | 2.89E-02 | 2.33E-01 | 2.69E+00 | 2.73E+00 | 9.01E-01 | no | Gm29185 | intron_variant,non_coding_transcript_variant | MODIFIER |
| 1 | 73283280 | AX-170041647 | 0 | T | C | 0.114 | 2.89E-02 | 2.33E-01 | 2.69E+00 | 2.73E+00 | 9.01E-01 | no | Gm29185 | intron_variant,non_coding_transcript_variant | MODIFIER |
| 1 | 73285406 | AX-170401683 | 0 | A | G | 0.114 | 2.89E-02 | 2.33E-01 | 2.69E+00 | 2.73E+00 | 9.01E-01 | no | Gm29185 | intron_variant,non_coding_transcript_variant | MODIFIER |
| 1 | 73301666 | AX-170226352 | 0 | C | A | 0.114 | 2.89E-02 | 2.33E-01 | 2.69E+00 | 2.73E+00 | 9.01E-01 | no | Gm29185 | intron_variant,non_coding_transcript_variant | MODIFIER |
| 1 | 73305292 | AX-97816281 | 0 | A | G | 0.114 | 2.89E-02 | 2.33E-01 | 2.69E+00 | 2.73E+00 | 9.01E-01 | no | Gm29185 | intron_variant,non_coding_transcript_variant | MODIFIER |
| 1 | 73305414 | AX-170153802 | 0 | C | G | 0.114 | 2.89E-02 | 2.33E-01 | 2.69E+00 | 2.73E+00 | 9.01E-01 | no | Gm29185 | intron_variant,non_coding_transcript_variant | MODIFIER |
| 1 | 73345641 | AX-97824899 | 0 | C | A | 0.114 | 2.89E-02 | 2.33E-01 | 2.69E+00 | 2.73E+00 | 9.01E-01 | no | | intergenic_variant | MODIFIER |
| 1 | 73363192 | AX-97781297 | 0 | C | G | 0.114 | 2.89E-02 | 2.33E-01 | 2.69E+00 | 2.73E+00 | 9.01E-01 | no | | regulatory_region_variant | MODIFIER |
| 1 | 73367526 | AX-170415559 | 0 | G | C | 0.114 | 2.89E-02 | 2.33E-01 | 2.69E+00 | 2.73E+00 | 9.01E-01 | no | | regulatory_region_variant | MODIFIER |
| 1 | 73369314 | AX-170388367 | 0 | A | G | 0.114 | 2.89E-02 | 2.33E-01 | 2.69E+00 | 2.73E+00 | 9.01E-01 | no | | intergenic_variant | MODIFIER |
| 1 | 73370681 | AX-170500017 | 0 | G | A | 0.114 | 2.89E-02 | 2.33E-01 | 2.69E+00 | 2.73E+00 | 9.01E-01 | no | | intergenic_variant | MODIFIER |
| 1 | 73377024 | AX-170356955 | 0 | C | T | 0.114 | 2.89E-02 | 2.33E-01 | 2.69E+00 | 2.73E+00 | 9.01E-01 | no | | intergenic_variant | MODIFIER |
| 1 | 73379031 | AX-170304376 | 0 | G | A | 0.114 | 2.89E-02 | 2.33E-01 | 2.69E+00 | 2.73E+00 | 9.01E-01 | no | | intergenic_variant | MODIFIER |
| 1 | 73388587 | AX-170277676 | 0 | T | C | 0.114 | 2.89E-02 | 2.33E-01 | 2.69E+00 | 2.73E+00 | 9.01E-01 | no | Gm27315 | upstream_gene_variant | MODIFIER |
| 1 | 73390314 | AX-170174414 | 0 | T | C | 0.114 | 2.89E-02 | 2.33E-01 | 2.69E+00 | 2.73E+00 | 9.01E-01 | no | Gm27315 | upstream_gene_variant | MODIFIER |
| 1 | 73391940 | AX-97852635 | 0 | T | C | 0.114 | 2.89E-02 | 2.33E-01 | 2.69E+00 | 2.73E+00 | 9.01E-01 | no | Gm27315 | upstream_gene_variant | MODIFIER |
| 1 | 73392417 | AX-170041651 | 0 | A | T | 0.114 | 2.89E-02 | 2.33E-01 | 2.69E+00 | 2.73E+00 | 9.01E-01 | no | Gm27315 | downstream_gene_variant | MODIFIER |
| 1 | 73392658 | AX-170512229 | 0 | T | C | 0.114 | 2.89E-02 | 2.33E-01 | 2.69E+00 | 2.73E+00 | 9.01E-01 | no | Gm27315 | downstream_gene_variant | MODIFIER |
| 1 | 74971179 | AX-170512865 | 0 | A | T | 0.114 | 2.89E-02 | 2.33E-01 | 2.69E+00 | 2.73E+00 | 9.01E-01 | no | Nhej1 | intron_variant | MODIFIER |
| 1 | 75290116 | AX-170287125 | 0 | T | A | 0.114 | 2.89E-02 | 2.33E-01 | 2.69E+00 | 2.73E+00 | 9.01E-01 | no | | regulatory_region_variant | MODIFIER |
| 1 | 174583037 | AX-170425534 | 0 | A | G | 0.219 | 1.60E-02 | 1.18E-01 | 2.63E+00 | 2.71E+00 | 9.01E-01 | no | Fmn2 | intron_variant | MODIFIER |
| 8 | 90541957 | AX-170380513 | 1 | C | T | 0.196 | -1.54E-02 | 1.19E-01 | 2.65E+00 | 2.72E+00 | 9.01E-01 | no | | intergenic_variant | MODIFIER |
| 6 | 104040547 | AX-170097288 | 2 | C | A | 0.064 | -2.63E-02 | 2.27E-01 | 2.66E+00 | 2.76E+00 | 9.01E-01 | no | | intergenic_variant | MODIFIER |
| 15 | 99017994 | AX-97816492 | 1 | C | T | 0.116 | 1.92E-02 | 1.52E-01 | 2.66E+00 | 2.73E+00 | 9.01E-01 | no | | regulatory_region_variant | MODIFIER |
| 15 | 98559820 | AX-170192901 | 1 | A | G | 0.107 | -1.82E-02 | 1.55E-01 | 2.68E+00 | 2.75E+00 | 9.01E-01 | no | Ccnt1 | intron_variant | MODIFIER |
| 15 | 100817629 | AX-170347218 | 1 | G | A | 0.107 | -1.82E-02 | 1.55E-01 | 2.68E+00 | 2.75E+00 | 9.01E-01 | no | Slc4a8 | 3_prime_UTR_variant | MODIFIER |
| 20 | 59476327 | AX-170033949 | 1 | T | A | 0.25 | 7.87E-03 | 6.29E-02 | 2.64E+00 | 2.73E+00 | 9.01E-01 | no | Fgf13 | intron_variant | MODIFIER |
| 4 | 145027619 | AX-170420843 | 0 | G | A | 0.105 | 2.50E-02 | 2.04E-01 | 2.68E+00 | 2.74E+00 | 9.01E-01 | no | Vps13d | intron_variant | MODIFIER |
| 9 | 34077952 | AX-97801309 | 1 | C | T | 0.438 | -1.00E-02 | 8.63E-02 | 2.69E+00 | 2.76E+00 | 9.01E-01 | no | | intergenic_variant | MODIFIER |
| 12 | 101304716 | AX-170392100 | 1 | G | A | 0.098 | 2.02E-02 | 1.60E-01 | 2.63E+00 | 2.73E+00 | 9.02E-01 | no | Gm18795 | intron_variant,non_coding_transcript_variant | MODIFIER |
| 7 | 52148990 | AX-170398727 | 1 | T | C | 0.42 | -1.14E-02 | 9.99E-02 | 2.68E+00 | 2.76E+00 | 9.02E-01 | no | | intergenic_variant | MODIFIER |
| 1 | 30146077 | AX-170222557 | 0 | A | C | 0.421 | 9.40E-03 | 7.78E-02 | 2.66E+00 | 2.74E+00 | 9.02E-01 | no | | regulatory_region_variant | MODIFIER |
| 9 | 70272613 | AX-97834808 | 0 | C | T | 0.211 | 1.45E-02 | 1.21E-01 | 2.68E+00 | 2.74E+00 | 9.02E-01 | no | Myo1e | intron_variant | MODIFIER |
| 10 | 122431221 | AX-170263384 | 0 | G | A | 0.333 | 8.66E-03 | 7.29E-02 | 2.64E+00 | 2.74E+00 | 9.02E-01 | no | Gm48877 | intron_variant,non_coding_transcript_variant | MODIFIER |
| 5 | 117810543 | AX-168374447 | 0 | C | T | 0.465 | -1.34E-02 | 1.14E-01 | 2.72E+00 | 2.77E+00 | 9.02E-01 | no | Nos1 | intron_variant,non_coding_transcript_variant | MODIFIER |
| 5 | 124921806 | AX-170134759 | 0 | C | T | 0.465 | -1.34E-02 | 1.14E-01 | 2.72E+00 | 2.77E+00 | 9.02E-01 | no | | intergenic_variant | MODIFIER |
| 18 | 59112454 | AX-170215627 | 0 | G | T | 0.421 | 9.04E-03 | 8.24E-02 | 2.71E+00 | 2.79E+00 | 9.02E-01 | no | | intergenic_variant | MODIFIER |
| 1 | 60281516 | AX-170492820 | 0 | A | C | 0.105 | -2.83E-02 | 2.35E-01 | 2.70E+00 | 2.74E+00 | 9.02E-01 | no | Nbeal1 | intron_variant | MODIFIER |
| 7 | 67561997 | AX-170297926 | 0 | A | T | 0.421 | 1.25E-02 | 9.90E-02 | 2.65E+00 | 2.73E+00 | 9.02E-01 | no | Lrrc28 | intron_variant | MODIFIER |
| 1 | 108439379 | AX-170415230 | 1 | T | A | 0.161 | 2.71E-02 | 2.02E-01 | 2.63E+00 | 2.71E+00 | 9.02E-01 | no | | intergenic_variant | MODIFIER |
| 9 | 85603905 | AX-170333063 | 1 | A | T | 0.205 | -1.46E-02 | 1.20E-01 | 2.67E+00 | 2.74E+00 | 9.02E-01 | no | | intergenic_variant | MODIFIER |
| 11 | 5784111 | AX-97857164 | 0 | C | T | 0.07 | 2.03E-02 | 1.68E-01 | 2.61E+00 | 2.74E+00 | 9.02E-01 | no | Dbnl | upstream_gene_variant | MODIFIER |
| 5 | 113704969 | AX-170378973 | 0 | G | A | 0.456 | -1.33E-02 | 1.14E-01 | 2.72E+00 | 2.77E+00 | 9.02E-01 | no | 1700069L16Rik | intron_variant | MODIFIER |
| 7 | 116892734 | AX-170103651 | 0 | A | G | 0.404 | 1.17E-02 | 9.13E-02 | 2.66E+00 | 2.72E+00 | 9.02E-01 | no | | intergenic_variant | MODIFIER |
| 8 | 18886956 | AX-170377602 | 1 | G | T | 0.179 | 1.27E-02 | 1.03E-01 | 2.64E+00 | 2.73E+00 | 9.02E-01 | no | Agpat5 | downstream_gene_variant | MODIFIER |
| 11 | 22494354 | AX-170408142 | 1 | C | T | 0.054 | 2.29E-02 | 1.83E-01 | 2.59E+00 | 2.73E+00 | 9.02E-01 | no | Gm12051 | downstream_gene_variant | MODIFIER |
| 18 | 41676532 | AX-170235076 | 0 | T | A | 0.061 | -2.92E-02 | 2.12E-01 | 2.61E+00 | 2.71E+00 | 9.02E-01 | no | | intergenic_variant | MODIFIER |
| 8 | 102018348 | AX-170216452 | 0 | G | A | 0.219 | 1.33E-02 | 1.14E-01 | 2.67E+00 | 2.75E+00 | 9.02E-01 | no | | intergenic_variant | MODIFIER |
| 20 | 56911232 | AX-97800712 | 0 | A | G | 0.246 | 7.72E-03 | 6.29E-02 | 2.64E+00 | 2.73E+00 | 9.02E-01 | no | Adgrg4 | intron_variant | MODIFIER |

| | | | | | | | | | | | | | | | |
|---|---|---|---|---|---|---|---|---|---|---|---|---|---|---|---|
| 20 | 56953784 | AX-97819354 | 0 | G | A | 0.246 | 7.72E-03 | 6.29E-02 | 2.64E+00 | 2.73E+00 | 9.02E-01 | no | Adgrg4 | intron_variant | MODIFIER |
| 20 | 56955571 | AX-170033906 | 0 | A | G | 0.246 | 7.72E-03 | 6.29E-02 | 2.64E+00 | 2.73E+00 | 9.02E-01 | no | Adgrg4 | intron_variant | MODIFIER |
| 20 | 56963780 | AX-170347102 | 0 | C | T | 0.246 | 7.72E-03 | 6.29E-02 | 2.64E+00 | 2.73E+00 | 9.02E-01 | no | Adgrg4 | intron_variant | MODIFIER |
| 20 | 56967187 | AX-170469811 | 0 | A | T | 0.246 | 7.72E-03 | 6.29E-02 | 2.64E+00 | 2.73E+00 | 9.02E-01 | no | Adgrg4 | intron_variant | MODIFIER |
| 20 | 56973681 | AX-170033912 | 0 | G | A | 0.246 | 7.72E-03 | 6.29E-02 | 2.64E+00 | 2.73E+00 | 9.02E-01 | no | Adgrg4 | intron_variant | MODIFIER |
| 20 | 56982142 | AX-170033907 | 0 | G | A | 0.246 | 7.72E-03 | 6.29E-02 | 2.64E+00 | 2.73E+00 | 9.02E-01 | no | Adgrg4 | downstream_gene_variant | MODIFIER |
| 20 | 56997883 | AX-97853387 | 0 | C | T | 0.246 | 7.72E-03 | 6.29E-02 | 2.64E+00 | 2.73E+00 | 9.02E-01 | no | Adgrg4 | intron_variant | MODIFIER |
| 20 | 57035241 | AX-170033908 | 0 | T | G | 0.246 | 7.72E-03 | 6.29E-02 | 2.64E+00 | 2.73E+00 | 9.02E-01 | no | | intergenic_variant | MODIFIER |
| 20 | 57036492 | AX-97792572 | 0 | T | C | 0.246 | 7.72E-03 | 6.29E-02 | 2.64E+00 | 2.73E+00 | 9.02E-01 | no | | intergenic_variant | MODIFIER |
| 20 | 57133793 | AX-97849031 | 0 | C | T | 0.246 | 7.72E-03 | 6.29E-02 | 2.64E+00 | 2.73E+00 | 9.02E-01 | no | | intergenic_variant | MODIFIER |
| 20 | 57177390 | AX-97836069 | 0 | T | C | 0.246 | 7.72E-03 | 6.29E-02 | 2.64E+00 | 2.73E+00 | 9.02E-01 | no | | regulatory_region_variant | MODIFIER |
| 20 | 57236974 | AX-97812820 | 0 | T | C | 0.246 | 7.72E-03 | 6.29E-02 | 2.64E+00 | 2.73E+00 | 9.02E-01 | no | Arhgef6 | intron_variant | MODIFIER |
| 20 | 57263302 | AX-97844734 | 0 | G | A | 0.246 | 7.72E-03 | 6.29E-02 | 2.64E+00 | 2.73E+00 | 9.02E-01 | no | Arhgef6 | intron_variant | MODIFIER |
| 20 | 57354247 | AX-97851679 | 0 | A | G | 0.246 | 7.72E-03 | 6.29E-02 | 2.64E+00 | 2.73E+00 | 9.02E-01 | no | | intergenic_variant | MODIFIER |
| 20 | 57523767 | AX-97797599 | 0 | G | A | 0.246 | 7.72E-03 | 6.29E-02 | 2.64E+00 | 2.73E+00 | 9.02E-01 | no | | intergenic_variant | MODIFIER |
| 20 | 57524627 | AX-170376892 | 0 | T | C | 0.246 | 7.72E-03 | 6.29E-02 | 2.64E+00 | 2.73E+00 | 9.02E-01 | no | | intergenic_variant | MODIFIER |
| 20 | 57529431 | AX-170195536 | 0 | C | T | 0.246 | 7.72E-03 | 6.29E-02 | 2.64E+00 | 2.73E+00 | 9.02E-01 | no | | intergenic_variant | MODIFIER |
| 20 | 57534512 | AX-97818519 | 0 | C | A | 0.246 | 7.72E-03 | 6.29E-02 | 2.64E+00 | 2.73E+00 | 9.02E-01 | no | | intergenic_variant | MODIFIER |
| 20 | 57637776 | AX-97799352 | 0 | G | A | 0.246 | 7.72E-03 | 6.29E-02 | 2.64E+00 | 2.73E+00 | 9.02E-01 | no | | intergenic_variant | MODIFIER |
| 20 | 57659901 | AX-97791955 | 0 | A | G | 0.246 | 7.72E-03 | 6.29E-02 | 2.64E+00 | 2.73E+00 | 9.02E-01 | no | Gm5389 | non_coding_transcript_exon_variant | MODIFIER |
| 20 | 57761839 | AX-168408756 | 0 | A | G | 0.246 | 7.72E-03 | 6.29E-02 | 2.64E+00 | 2.73E+00 | 9.02E-01 | no | | intergenic_variant | MODIFIER |
| 20 | 57763163 | AX-97780843 | 0 | G | T | 0.246 | 7.72E-03 | 6.29E-02 | 2.64E+00 | 2.73E+00 | 9.02E-01 | no | | intergenic_variant | MODIFIER |
| 20 | 57867627 | AX-170035819 | 0 | A | G | 0.246 | 7.72E-03 | 6.29E-02 | 2.64E+00 | 2.73E+00 | 9.02E-01 | no | | intergenic_variant | MODIFIER |
| 20 | 59235498 | AX-97837872 | 0 | A | G | 0.246 | 7.72E-03 | 6.29E-02 | 2.64E+00 | 2.73E+00 | 9.02E-01 | no | Fgf13 | intron_variant | MODIFIER |
| 20 | 59320697 | AX-97819526 | 0 | A | C | 0.246 | 7.72E-03 | 6.29E-02 | 2.64E+00 | 2.73E+00 | 9.02E-01 | no | Fgf13 | intron_variant | MODIFIER |
| 20 | 59432194 | AX-170033946 | 0 | G | A | 0.246 | 7.72E-03 | 6.29E-02 | 2.64E+00 | 2.73E+00 | 9.02E-01 | no | Fgf13 | intron_variant | MODIFIER |
| 20 | 59442756 | AX-97838487 | 0 | A | C | 0.246 | 7.72E-03 | 6.29E-02 | 2.64E+00 | 2.73E+00 | 9.02E-01 | no | Fgf13 | intron_variant | MODIFIER |
| 20 | 59453022 | AX-170033948 | 0 | T | C | 0.246 | 7.72E-03 | 6.29E-02 | 2.64E+00 | 2.73E+00 | 9.02E-01 | no | Fgf13 | intron_variant | MODIFIER |
| 20 | 59477553 | AX-168420025 | 0 | C | T | 0.246 | 7.72E-03 | 6.29E-02 | 2.64E+00 | 2.73E+00 | 9.02E-01 | no | Fgf13 | intron_variant | MODIFIER |
| 20 | 59485115 | AX-97820893 | 0 | G | A | 0.246 | 7.72E-03 | 6.29E-02 | 2.64E+00 | 2.73E+00 | 9.02E-01 | no | Fgf13 | intron_variant | MODIFIER |
| 20 | 59563619 | AX-97794336 | 0 | T | C | 0.246 | 7.72E-03 | 6.29E-02 | 2.64E+00 | 2.73E+00 | 9.02E-01 | no | Fgf13 | intron_variant | MODIFIER |
| 20 | 59815332 | AX-97821784 | 0 | C | A | 0.246 | 7.72E-03 | 6.29E-02 | 2.64E+00 | 2.73E+00 | 9.02E-01 | no | | intergenic_variant | MODIFIER |
| 20 | 60715477 | AX-170033968 | 0 | C | T | 0.246 | 7.72E-03 | 6.29E-02 | 2.64E+00 | 2.73E+00 | 9.02E-01 | no | Gm14661 | intron_variant,non_coding_transcript_variant | MODIFIER |
| 20 | 60937611 | AX-97810677 | 0 | C | T | 0.246 | 7.72E-03 | 6.29E-02 | 2.64E+00 | 2.73E+00 | 9.02E-01 | no | | intergenic_variant | MODIFIER |
| 20 | 61154759 | AX-168420029 | 0 | A | G | 0.246 | 7.72E-03 | 6.29E-02 | 2.64E+00 | 2.73E+00 | 9.02E-01 | no | C230004F18Rik | intron_variant,non_coding_transcript_variant | MODIFIER |
| 20 | 61412532 | AX-97823749 | 0 | T | C | 0.246 | 7.72E-03 | 6.29E-02 | 2.64E+00 | 2.73E+00 | 9.02E-01 | no | | intergenic_variant | MODIFIER |
| 20 | 61502945 | AX-97786282 | 0 | T | G | 0.246 | 7.72E-03 | 6.29E-02 | 2.64E+00 | 2.73E+00 | 9.02E-01 | no | | intergenic_variant | MODIFIER |
| 20 | 61586251 | AX-170316871 | 0 | C | T | 0.246 | 7.72E-03 | 6.29E-02 | 2.64E+00 | 2.73E+00 | 9.02E-01 | no | | intergenic_variant | MODIFIER |
| 20 | 61652361 | AX-170255078 | 0 | C | A | 0.246 | 7.72E-03 | 6.29E-02 | 2.64E+00 | 2.73E+00 | 9.02E-01 | no | | intergenic_variant | MODIFIER |
| 20 | 62862435 | AX-170191907 | 0 | G | T | 0.246 | 7.72E-03 | 6.29E-02 | 2.64E+00 | 2.73E+00 | 9.02E-01 | no | | intergenic_variant | MODIFIER |
| 20 | 63658205 | AX-97813193 | 0 | T | G | 0.246 | 7.72E-03 | 6.29E-02 | 2.64E+00 | 2.73E+00 | 9.02E-01 | no | | intergenic_variant | MODIFIER |
| 20 | 64161107 | AX-97806131 | 0 | T | C | 0.246 | 7.72E-03 | 6.29E-02 | 2.64E+00 | 2.73E+00 | 9.02E-01 | no | | intergenic_variant | MODIFIER |
| 20 | 64615642 | AX-170034011 | 0 | A | C | 0.246 | 7.72E-03 | 6.29E-02 | 2.64E+00 | 2.73E+00 | 9.02E-01 | no | | intergenic_variant | MODIFIER |
| 20 | 65294562 | AX-97806157 | 0 | G | T | 0.246 | 7.72E-03 | 6.29E-02 | 2.64E+00 | 2.73E+00 | 9.02E-01 | no | | intergenic_variant | MODIFIER |
| 20 | 65315485 | AX-170034022 | 0 | T | C | 0.246 | 7.72E-03 | 6.29E-02 | 2.64E+00 | 2.73E+00 | 9.02E-01 | no | | intergenic_variant | MODIFIER |
| 20 | 65351858 | AX-170099241 | 0 | A | C | 0.246 | 7.72E-03 | 6.29E-02 | 2.64E+00 | 2.73E+00 | 9.02E-01 | no | | intergenic_variant | MODIFIER |
| 20 | 65370371 | AX-170162637 | 0 | G | T | 0.246 | 7.72E-03 | 6.29E-02 | 2.64E+00 | 2.73E+00 | 9.02E-01 | no | | intergenic_variant | MODIFIER |
| 20 | 65370853 | AX-170471052 | 0 | C | T | 0.246 | 7.72E-03 | 6.29E-02 | 2.64E+00 | 2.73E+00 | 9.02E-01 | no | | intergenic_variant | MODIFIER |
| 20 | 65402709 | AX-170433386 | 0 | G | A | 0.246 | 7.72E-03 | 6.29E-02 | 2.64E+00 | 2.73E+00 | 9.02E-01 | no | | intergenic_variant | MODIFIER |
| 20 | 65438256 | AX-170514605 | 0 | G | A | 0.246 | 7.72E-03 | 6.29E-02 | 2.64E+00 | 2.73E+00 | 9.02E-01 | no | | intergenic_variant | MODIFIER |
| 20 | 65439922 | AX-170104758 | 0 | G | C | 0.246 | 7.72E-03 | 6.29E-02 | 2.64E+00 | 2.73E+00 | 9.02E-01 | no | | intergenic_variant | MODIFIER |
| 20 | 65450873 | AX-97846584 | 0 | C | T | 0.246 | 7.72E-03 | 6.29E-02 | 2.64E+00 | 2.73E+00 | 9.02E-01 | no | | intergenic_variant | MODIFIER |
| 20 | 65512158 | AX-97803128 | 0 | G | A | 0.246 | 7.72E-03 | 6.29E-02 | 2.64E+00 | 2.73E+00 | 9.02E-01 | no | | intergenic_variant | MODIFIER |
| 20 | 65561149 | AX-170414193 | 0 | C | A | 0.246 | 7.72E-03 | 6.29E-02 | 2.64E+00 | 2.73E+00 | 9.02E-01 | no | | intergenic_variant | MODIFIER |

| | | | | | | | | | | | | | | | |
|---|---|---|---|---|---|---|---|---|---|---|---|---|---|---|---|
| 20 | 65600275 | AX-97818065 | 0 | G | T | 0.246 | 7.72E-03 | 6.29E-02 | 2.64E+00 | 2.73E+00 | 9.02E-01 | no | | intergenic_variant | MODIFIER |
| 20 | 65651765 | AX-170165926 | 0 | G | C | 0.246 | 7.72E-03 | 6.29E-02 | 2.64E+00 | 2.73E+00 | 9.02E-01 | no | | intergenic_variant | MODIFIER |
| 20 | 65652202 | AX-170305365 | 0 | T | G | 0.246 | 7.72E-03 | 6.29E-02 | 2.64E+00 | 2.73E+00 | 9.02E-01 | no | | intergenic_variant | MODIFIER |
| 20 | 66297099 | AX-97806040 | 0 | T | A | 0.246 | 7.72E-03 | 6.29E-02 | 2.64E+00 | 2.73E+00 | 9.02E-01 | no | | intergenic_variant | MODIFIER |
| 20 | 66417635 | AX-170034041 | 0 | A | T | 0.246 | 7.72E-03 | 6.29E-02 | 2.64E+00 | 2.73E+00 | 9.02E-01 | no | | intergenic_variant | MODIFIER |
| 20 | 66417830 | AX-170090529 | 0 | A | T | 0.246 | 7.72E-03 | 6.29E-02 | 2.64E+00 | 2.73E+00 | 9.02E-01 | no | | intergenic_variant | MODIFIER |
| 20 | 66449371 | AX-97845499 | 0 | G | A | 0.246 | 7.72E-03 | 6.29E-02 | 2.64E+00 | 2.73E+00 | 9.02E-01 | no | Gm23028 | downstream_gene_variant | MODIFIER |
| 20 | 66504125 | AX-170034037 | 0 | G | A | 0.246 | 7.72E-03 | 6.29E-02 | 2.64E+00 | 2.73E+00 | 9.02E-01 | no | | intergenic_variant | MODIFIER |
| 5 | 85834291 | AX-170436339 | 0 | T | C | 0.43 | -1.31E-02 | 1.08E-01 | 2.69E+00 | 2.74E+00 | 9.03E-01 | no | Gm24626 | downstream_gene_variant | MODIFIER |
| 10 | 36198040 | AX-170241116 | 0 | C | T | 0.377 | -9.86E-03 | 7.53E-02 | 2.61E+00 | 2.72E+00 | 9.03E-01 | no | | intergenic_variant | MODIFIER |
| 10 | 38965551 | AX-170319214 | 0 | T | C | 0.377 | -9.86E-03 | 7.53E-02 | 2.61E+00 | 2.72E+00 | 9.03E-01 | no | Lama4 | 5_prime_UTR_variant | MODIFIER |
| 16 | 92595213 | AX-170476988 | 0 | C | A | 0.474 | -9.75E-03 | 7.96E-02 | 2.65E+00 | 2.73E+00 | 9.03E-01 | no | | intergenic_variant | MODIFIER |
| 9 | 87300138 | AX-170320107 | 1 | T | C | 0.125 | 1.39E-02 | 1.27E-01 | 2.68E+00 | 2.77E+00 | 9.03E-01 | no | | intergenic_variant | MODIFIER |
| 1 | 66372434 | AX-170000709 | 1 | G | A | 0.116 | 2.82E-02 | 2.32E-01 | 2.68E+00 | 2.73E+00 | 9.03E-01 | no | Map2 | intron_variant | MODIFIER |
| 10 | 6579366 | AX-170336018 | 0 | T | C | 0.202 | 1.15E-02 | 9.72E-02 | 2.64E+00 | 2.74E+00 | 9.03E-01 | no | | intergenic_variant | MODIFIER |
| 4 | 96300392 | AX-170345671 | 0 | T | C | 0.237 | 1.51E-02 | 9.71E-02 | 2.56E+00 | 2.69E+00 | 9.03E-01 | no | Cyp2j11 | intron_variant | MODIFIER |
| 18 | 29535363 | AX-170423691 | 0 | C | A | 0.053 | 2.19E-02 | 1.87E-01 | 2.61E+00 | 2.74E+00 | 9.03E-01 | no | | intergenic_variant | MODIFIER |
| 10 | 6541840 | AX-170412931 | 0 | A | C | 0.202 | -1.29E-02 | 1.03E-01 | 2.64E+00 | 2.73E+00 | 9.03E-01 | no | | intergenic_variant | MODIFIER |
| 10 | 6552175 | AX-170231245 | 0 | G | A | 0.202 | -1.29E-02 | 1.03E-01 | 2.64E+00 | 2.73E+00 | 9.03E-01 | no | | intergenic_variant | MODIFIER |
| 10 | 6552188 | AX-170195537 | 0 | A | G | 0.202 | -1.29E-02 | 1.03E-01 | 2.64E+00 | 2.73E+00 | 9.03E-01 | no | | intergenic_variant | MODIFIER |
| 10 | 6553834 | AX-170461609 | 0 | C | A | 0.202 | -1.29E-02 | 1.03E-01 | 2.64E+00 | 2.73E+00 | 9.03E-01 | no | | intergenic_variant | MODIFIER |
| 10 | 6559946 | AX-170475496 | 0 | A | G | 0.202 | -1.29E-02 | 1.03E-01 | 2.64E+00 | 2.73E+00 | 9.03E-01 | no | | intergenic_variant | MODIFIER |
| 10 | 6652755 | AX-170108041 | 0 | C | A | 0.202 | -1.29E-02 | 1.03E-01 | 2.64E+00 | 2.73E+00 | 9.03E-01 | no | | intergenic_variant | MODIFIER |
| 10 | 6677696 | AX-170274404 | 0 | A | G | 0.202 | -1.29E-02 | 1.03E-01 | 2.64E+00 | 2.73E+00 | 9.03E-01 | no | | intergenic_variant | MODIFIER |
| 10 | 6677735 | AX-170155933 | 0 | C | T | 0.202 | -1.29E-02 | 1.03E-01 | 2.64E+00 | 2.73E+00 | 9.03E-01 | no | | intergenic_variant | MODIFIER |
| 10 | 6677851 | AX-170161898 | 0 | G | A | 0.202 | -1.29E-02 | 1.03E-01 | 2.64E+00 | 2.73E+00 | 9.03E-01 | no | | intergenic_variant | MODIFIER |
| 10 | 6678161 | AX-170091848 | 0 | C | G | 0.202 | -1.29E-02 | 1.03E-01 | 2.64E+00 | 2.73E+00 | 9.03E-01 | no | | intergenic_variant | MODIFIER |
| 10 | 6690143 | AX-170002632 | 0 | G | A | 0.202 | -1.29E-02 | 1.03E-01 | 2.64E+00 | 2.73E+00 | 9.03E-01 | no | | intergenic_variant | MODIFIER |
| 10 | 6774597 | AX-170002629 | 0 | T | G | 0.202 | -1.29E-02 | 1.03E-01 | 2.64E+00 | 2.73E+00 | 9.03E-01 | no | Oprm1 | intron_variant | MODIFIER |
| 10 | 7206731 | AX-170439452 | 0 | G | A | 0.202 | -1.29E-02 | 1.03E-01 | 2.64E+00 | 2.73E+00 | 9.03E-01 | no | Cnksr3 | intron_variant | MODIFIER |
| 10 | 7212665 | AX-170297689 | 0 | T | C | 0.202 | -1.29E-02 | 1.03E-01 | 2.64E+00 | 2.73E+00 | 9.03E-01 | no | Cnksr3 | upstream_gene_variant | MODIFIER |
| 10 | 7293854 | AX-170002624 | 0 | A | G | 0.202 | -1.29E-02 | 1.03E-01 | 2.64E+00 | 2.73E+00 | 9.03E-01 | no | | intergenic_variant | MODIFIER |
| 10 | 7294117 | AX-97834677 | 0 | C | G | 0.202 | -1.29E-02 | 1.03E-01 | 2.64E+00 | 2.73E+00 | 9.03E-01 | no | | intergenic_variant | MODIFIER |
| 10 | 8968958 | AX-97842418 | 0 | T | C | 0.202 | -1.29E-02 | 1.03E-01 | 2.64E+00 | 2.73E+00 | 9.03E-01 | no | Gm48728 | downstream_gene_variant | MODIFIER |
| 10 | 8969386 | AX-170002680 | 0 | A | G | 0.202 | -1.29E-02 | 1.03E-01 | 2.64E+00 | 2.73E+00 | 9.03E-01 | no | Gm48728 | downstream_gene_variant | MODIFIER |
| 10 | 8970276 | AX-170215488 | 0 | A | T | 0.202 | -1.29E-02 | 1.03E-01 | 2.64E+00 | 2.73E+00 | 9.03E-01 | no | Gm48728 | downstream_gene_variant | MODIFIER |
| 10 | 8972256 | AX-170439741 | 0 | G | A | 0.202 | -1.29E-02 | 1.03E-01 | 2.64E+00 | 2.73E+00 | 9.03E-01 | no | | regulatory_region_variant | MODIFIER |
| 10 | 9017189 | AX-97826227 | 0 | C | T | 0.202 | -1.29E-02 | 1.03E-01 | 2.64E+00 | 2.73E+00 | 9.03E-01 | no | | intergenic_variant | MODIFIER |
| 10 | 9227350 | AX-97814757 | 0 | T | C | 0.202 | -1.29E-02 | 1.03E-01 | 2.64E+00 | 2.73E+00 | 9.03E-01 | no | Gm48744 | upstream_gene_variant | MODIFIER |
| 10 | 9239854 | AX-170357693 | 0 | C | T | 0.202 | -1.29E-02 | 1.03E-01 | 2.64E+00 | 2.73E+00 | 9.03E-01 | no | | regulatory_region_variant | MODIFIER |
| 15 | 79389526 | AX-170355085 | 0 | T | A | 0.561 | 1.02E-02 | 7.82E-02 | 2.64E+00 | 2.71E+00 | 9.03E-01 | no | Tmem184b | intron_variant | MODIFIER |
| 12 | 59498107 | AX-170190885 | 0 | C | T | 0.044 | -2.14E-02 | 1.68E-01 | 2.59E+00 | 2.73E+00 | 9.03E-01 | no | | intergenic_variant | MODIFIER |
| 6 | 134704557 | AX-170140259 | 0 | G | A | 0.57 | 1.23E-02 | 8.67E-02 | 2.62E+00 | 2.69E+00 | 9.03E-01 | no | Borcs5 | intron_variant | MODIFIER |
| 4 | 72047934 | AX-170472396 | 0 | A | T | 0.228 | 1.45E-02 | 1.05E-01 | 2.62E+00 | 2.70E+00 | 9.04E-01 | no | | intergenic_variant | MODIFIER |
| 2 | 154722192 | AX-170441562 | 0 | G | A | 0.14 | 2.66E-02 | 2.21E-01 | 2.70E+00 | 2.73E+00 | 9.04E-01 | no | Gm14216 | downstream_gene_variant | MODIFIER |
| 18 | 61012106 | AX-170260370 | 1 | G | A | 0.42 | 8.93E-03 | 8.17E-02 | 2.70E+00 | 2.78E+00 | 9.04E-01 | no | Slc6a7 | intron_variant | MODIFIER |
| 11 | 5389940 | AX-170205432 | 1 | T | C | 0.062 | 2.45E-02 | 2.13E-01 | 2.66E+00 | 2.75E+00 | 9.04E-01 | no | Znrf3 | intron_variant | MODIFIER |
| 11 | 11152439 | AX-170290920 | 1 | C | T | 0.062 | 2.45E-02 | 2.13E-01 | 2.66E+00 | 2.75E+00 | 9.04E-01 | no | Vwc2 | intron_variant | MODIFIER |
| 2 | 129267166 | AX-170186342 | 1 | G | A | 0.143 | 2.71E-02 | 2.22E-01 | 2.69E+00 | 2.72E+00 | 9.04E-01 | no | Ckap2l | downstream_gene_variant | MODIFIER |
| 2 | 139103041 | AX-170521760 | 1 | C | T | 0.143 | 2.71E-02 | 2.22E-01 | 2.69E+00 | 2.72E+00 | 9.04E-01 | no | | intergenic_variant | MODIFIER |
| 4 | 154831949 | AX-170235282 | 0 | G | A | 0.114 | 1.89E-02 | 1.64E-01 | 2.65E+00 | 2.75E+00 | 9.04E-01 | no | | intergenic_variant | MODIFIER |
| 18 | 66016076 | AX-170015875 | 0 | C | T | 0.202 | -1.41E-02 | 1.15E-01 | 2.66E+00 | 2.73E+00 | 9.04E-01 | no | Lman1 | intron_variant | MODIFIER |
| 4 | 140998873 | AX-170394671 | 0 | G | A | 0.088 | 2.03E-02 | 1.68E-01 | 2.66E+00 | 2.73E+00 | 9.04E-01 | no | Atp13a2 | intron_variant | MODIFIER |
| 10 | 32131372 | AX-170477956 | 0 | C | T | 0.395 | -9.00E-03 | 7.38E-02 | 2.62E+00 | 2.73E+00 | 9.04E-01 | no | Nkain2 | intron_variant | MODIFIER |

| | | | | | | | | | | | | | | | |
|---|---|---|---|---|---|---|---|---|---|---|---|---|---|---|---|
| 7 | 79561975 | AX-170101818 | 2 | A | G | 0.091 | 2.16E-02 | 1.83E-01 | 2.62E+00 | 2.74E+00 | 9.04E-01 | no | Gm35040 | downstream_gene_variant | MODIFIER |
| 4 | 92571845 | AX-170196577 | 0 | C | T | 0.325 | 1.28E-02 | 1.03E-01 | 2.65E+00 | 2.73E+00 | 9.04E-01 | no | | intergenic_variant | MODIFIER |
| 10 | 120372680 | AX-170102741 | 1 | G | C | 0.33 | 8.41E-03 | 7.22E-02 | 2.66E+00 | 2.74E+00 | 9.04E-01 | no | 1700006J14Rik | intron_variant,non_coding_transcript_variant | MODIFIER |
| 9 | 110896578 | AX-170271477 | 0 | A | G | 0.096 | 1.29E-02 | 1.18E-01 | 2.69E+00 | 2.77E+00 | 9.04E-01 | no | Als2cl | intron_variant | MODIFIER |
| 1 | 68955564 | AX-170496818 | 0 | T | C | 0.132 | 2.29E-02 | 1.96E-01 | 2.66E+00 | 2.74E+00 | 9.04E-01 | no | Erbb4 | intron_variant | MODIFIER |
| 18 | 45147474 | AX-170374063 | 1 | G | A | 0.062 | 3.52E-02 | 3.00E-01 | 2.73E+00 | 2.76E+00 | 9.04E-01 | no | AC134458.3 | upstream_gene_variant | MODIFIER |
| 7 | 82760104 | AX-170358051 | 1 | G | A | 0.116 | 2.58E-02 | 2.15E-01 | 2.65E+00 | 2.73E+00 | 9.04E-01 | no | Efl1 | intron_variant | MODIFIER |
| 7 | 93914700 | AX-170260931 | 1 | A | T | 0.116 | 2.58E-02 | 2.15E-01 | 2.65E+00 | 2.73E+00 | 9.04E-01 | no | | intergenic_variant | MODIFIER |
| 9 | 106661105 | AX-170142334 | 2 | C | T | 0.136 | -1.47E-02 | 1.04E-01 | 2.59E+00 | 2.70E+00 | 9.04E-01 | no | Tex264 | intron_variant | MODIFIER |
| 15 | 101771785 | AX-170173449 | 0 | A | G | 0.114 | 1.81E-02 | 1.49E-01 | 2.66E+00 | 2.73E+00 | 9.05E-01 | no | Krt72 | downstream_gene_variant | MODIFIER |
| 8 | 78332751 | AX-170126621 | 1 | C | A | 0.205 | -1.52E-02 | 1.30E-01 | 2.68E+00 | 2.74E+00 | 9.05E-01 | no | Ttc29 | intron_variant | MODIFIER |
| 9 | 89331520 | AX-170143392 | 1 | T | C | 0.134 | 1.44E-02 | 1.32E-01 | 2.69E+00 | 2.77E+00 | 9.05E-01 | no | Mthfs | intron_variant,non_coding_transcript_variant | MODIFIER |
| 1 | 191020011 | AX-97844905 | 0 | T | G | 0.254 | -8.01E-03 | 9.03E-02 | 2.66E+00 | 2.80E+00 | 9.05E-01 | no | Flvcr1 | intron_variant | MODIFIER |
| 8 | 93149240 | AX-97844913 | 0 | T | C | 0.211 | -1.41E-02 | 1.18E-01 | 2.68E+00 | 2.74E+00 | 9.05E-01 | no | Gm45910 | intron_variant,non_coding_transcript_variant | MODIFIER |
| 8 | 93158294 | AX-170188795 | 0 | C | T | 0.211 | -1.41E-02 | 1.18E-01 | 2.68E+00 | 2.74E+00 | 9.05E-01 | no | Gm45910 | upstream_gene_variant | MODIFIER |
| 8 | 93163932 | AX-170188507 | 0 | T | G | 0.211 | -1.41E-02 | 1.18E-01 | 2.68E+00 | 2.74E+00 | 9.05E-01 | no | Ces1d | downstream_gene_variant | MODIFIER |
| 8 | 93168946 | AX-170031267 | 0 | T | C | 0.211 | -1.41E-02 | 1.18E-01 | 2.68E+00 | 2.74E+00 | 9.05E-01 | no | Ces1d | intron_variant | MODIFIER |
| 8 | 93172948 | AX-170428572 | 0 | A | G | 0.211 | -1.41E-02 | 1.18E-01 | 2.68E+00 | 2.74E+00 | 9.05E-01 | no | Ces1d | intron_variant | MODIFIER |
| 8 | 93175115 | AX-170080835 | 0 | T | C | 0.211 | -1.41E-02 | 1.18E-01 | 2.68E+00 | 2.74E+00 | 9.05E-01 | no | Ces1d | missense_variant | MODERATE |
| 8 | 93206369 | AX-170440655 | 0 | T | C | 0.211 | -1.41E-02 | 1.18E-01 | 2.68E+00 | 2.74E+00 | 9.05E-01 | no | Ces1e | intron_variant | MODIFIER |
| 8 | 93206399 | AX-97853812 | 0 | C | T | 0.211 | -1.41E-02 | 1.18E-01 | 2.68E+00 | 2.74E+00 | 9.05E-01 | no | Ces1e | intron_variant | MODIFIER |
| 8 | 93208283 | AX-170309071 | 0 | T | C | 0.211 | -1.41E-02 | 1.18E-01 | 2.68E+00 | 2.74E+00 | 9.05E-01 | no | Ces1e | intron_variant | MODIFIER |
| 8 | 93208369 | AX-170189059 | 0 | C | G | 0.211 | -1.41E-02 | 1.18E-01 | 2.68E+00 | 2.74E+00 | 9.05E-01 | no | Ces1e | intron_variant | MODIFIER |
| 8 | 93210251 | AX-170126379 | 0 | A | G | 0.211 | -1.41E-02 | 1.18E-01 | 2.68E+00 | 2.74E+00 | 9.05E-01 | no | Ces1e | intron_variant | MODIFIER |
| 8 | 93216535 | AX-170253905 | 0 | T | A | 0.211 | -1.41E-02 | 1.18E-01 | 2.68E+00 | 2.74E+00 | 9.05E-01 | no | Ces1e | intron_variant | MODIFIER |
| 8 | 93219122 | AX-170170947 | 0 | T | G | 0.211 | -1.41E-02 | 1.18E-01 | 2.68E+00 | 2.74E+00 | 9.05E-01 | no | Ces1e | intron_variant | MODIFIER |
| 8 | 93219277 | AX-97821391 | 0 | G | C | 0.211 | -1.41E-02 | 1.18E-01 | 2.68E+00 | 2.74E+00 | 9.05E-01 | no | Ces1e | intron_variant | MODIFIER |
| 8 | 93227953 | AX-170385185 | 0 | T | C | 0.211 | -1.41E-02 | 1.18E-01 | 2.68E+00 | 2.74E+00 | 9.05E-01 | no | Ces1e | intron_variant | MODIFIER |
| 8 | 93234189 | AX-170364108 | 0 | A | C | 0.211 | -1.41E-02 | 1.18E-01 | 2.68E+00 | 2.74E+00 | 9.05E-01 | no | Ces1e | upstream_gene_variant | MODIFIER |
| 8 | 93258475 | AX-97845106 | 0 | G | C | 0.211 | -1.41E-02 | 1.18E-01 | 2.68E+00 | 2.74E+00 | 9.05E-01 | no | Ces1f | intron_variant | MODIFIER |
| 8 | 93260094 | AX-170401935 | 0 | C | T | 0.211 | -1.41E-02 | 1.18E-01 | 2.68E+00 | 2.74E+00 | 9.05E-01 | no | Ces1f | intron_variant | MODIFIER |
| 8 | 93276368 | AX-97824898 | 0 | T | G | 0.211 | -1.41E-02 | 1.18E-01 | 2.68E+00 | 2.74E+00 | 9.05E-01 | no | Ces1f | intron_variant | MODIFIER |
| 8 | 93276415 | AX-170385502 | 0 | A | G | 0.211 | -1.41E-02 | 1.18E-01 | 2.68E+00 | 2.74E+00 | 9.05E-01 | no | Ces1f | intron_variant | MODIFIER |
| 8 | 93287373 | AX-170489099 | 0 | G | A | 0.211 | -1.41E-02 | 1.18E-01 | 2.68E+00 | 2.74E+00 | 9.05E-01 | no | | intergenic_variant | MODIFIER |
| 8 | 93300673 | AX-97789237 | 0 | A | G | 0.211 | -1.41E-02 | 1.18E-01 | 2.68E+00 | 2.74E+00 | 9.05E-01 | no | Ces1g | downstream_gene_variant | MODIFIER |
| 8 | 93303360 | AX-170218362 | 0 | T | C | 0.211 | -1.41E-02 | 1.18E-01 | 2.68E+00 | 2.74E+00 | 9.05E-01 | no | Ces1g | intron_variant | MODIFIER |
| 8 | 93303542 | AX-170080837 | 0 | T | C | 0.211 | -1.41E-02 | 1.18E-01 | 2.68E+00 | 2.74E+00 | 9.05E-01 | no | Ces1g | intron_variant | MODIFIER |
| 8 | 93356675 | AX-170260795 | 0 | T | C | 0.211 | -1.41E-02 | 1.18E-01 | 2.68E+00 | 2.74E+00 | 9.05E-01 | no | Ces1h | intron_variant | MODIFIER |
| 8 | 93361196 | AX-170192231 | 0 | A | G | 0.211 | -1.41E-02 | 1.18E-01 | 2.68E+00 | 2.74E+00 | 9.05E-01 | no | Ces1h | intron_variant | MODIFIER |
| 8 | 93361325 | AX-170173707 | 0 | C | G | 0.211 | -1.41E-02 | 1.18E-01 | 2.68E+00 | 2.74E+00 | 9.05E-01 | no | Ces1h | intron_variant | MODIFIER |
| 8 | 93391565 | AX-170475071 | 0 | T | C | 0.211 | -1.41E-02 | 1.18E-01 | 2.68E+00 | 2.74E+00 | 9.05E-01 | no | Gm45765 | intron_variant,non_coding_transcript_variant | MODIFIER |
| 8 | 93398993 | AX-170479008 | 0 | A | C | 0.211 | -1.41E-02 | 1.18E-01 | 2.68E+00 | 2.74E+00 | 9.05E-01 | no | Gm45765 | intron_variant,non_coding_transcript_variant | MODIFIER |
| 8 | 93399027 | AX-170340016 | 0 | C | T | 0.211 | -1.41E-02 | 1.18E-01 | 2.68E+00 | 2.74E+00 | 9.05E-01 | no | Gm45765 | intron_variant,non_coding_transcript_variant | MODIFIER |
| 8 | 93399188 | AX-170170646 | 0 | C | G | 0.211 | -1.41E-02 | 1.18E-01 | 2.68E+00 | 2.74E+00 | 9.05E-01 | no | Gm45765 | intron_variant,non_coding_transcript_variant | MODIFIER |
| 8 | 93408358 | AX-170122495 | 0 | C | G | 0.211 | -1.41E-02 | 1.18E-01 | 2.68E+00 | 2.74E+00 | 9.05E-01 | no | Gm45765 | intron_variant,non_coding_transcript_variant | MODIFIER |
| 8 | 93410813 | AX-170357210 | 0 | C | A | 0.211 | -1.41E-02 | 1.18E-01 | 2.68E+00 | 2.74E+00 | 9.05E-01 | no | Gm45765 | intron_variant,non_coding_transcript_variant | MODIFIER |
| 8 | 93413438 | AX-170172276 | 0 | C | A | 0.211 | -1.41E-02 | 1.18E-01 | 2.68E+00 | 2.74E+00 | 9.05E-01 | no | Gm45765 | intron_variant,non_coding_transcript_variant | MODIFIER |
| 8 | 93417403 | AX-170336312 | 0 | A | G | 0.211 | -1.41E-02 | 1.18E-01 | 2.68E+00 | 2.74E+00 | 9.05E-01 | no | Gm45765 | intron_variant,non_coding_transcript_variant | MODIFIER |
| 8 | 93417797 | AX-170504218 | 0 | A | G | 0.211 | -1.41E-02 | 1.18E-01 | 2.68E+00 | 2.74E+00 | 9.05E-01 | no | Gm45765 | intron_variant,non_coding_transcript_variant | MODIFIER |
| 8 | 93419735 | AX-97784899 | 0 | G | A | 0.211 | -1.41E-02 | 1.18E-01 | 2.68E+00 | 2.74E+00 | 9.05E-01 | no | Gm45765 | intron_variant,non_coding_transcript_variant | MODIFIER |
| 8 | 93429265 | AX-170119903 | 0 | C | T | 0.211 | -1.41E-02 | 1.18E-01 | 2.68E+00 | 2.74E+00 | 9.05E-01 | no | Gm45766 | downstream_gene_variant | MODIFIER |
| 8 | 93429402 | AX-170414788 | 0 | C | T | 0.211 | -1.41E-02 | 1.18E-01 | 2.68E+00 | 2.74E+00 | 9.05E-01 | no | Gm45766 | downstream_gene_variant | MODIFIER |
| 8 | 93429519 | AX-170258466 | 0 | A | G | 0.211 | -1.41E-02 | 1.18E-01 | 2.68E+00 | 2.74E+00 | 9.05E-01 | no | Gm45766 | downstream_gene_variant | MODIFIER |
| 8 | 93429726 | AX-170315670 | 0 | T | C | 0.211 | -1.41E-02 | 1.18E-01 | 2.68E+00 | 2.74E+00 | 9.05E-01 | no | Gm45766 | downstream_gene_variant | MODIFIER |

| | | | | | | | | | | | | | | | |
|---|---|---|---|---|---|---|---|---|---|---|---|---|---|---|---|
| 8 | 93435149 | AX-170295806 | 0 | T | C | 0.211 | -1.41E-02 | 1.18E-01 | 2.68E+00 | 2.74E+00 | 9.05E-01 | no | Gm45766 | intron_variant,non_coding_transcript_variant | MODIFIER |
| 8 | 93474048 | AX-97790598 | 0 | A | G | 0.211 | -1.41E-02 | 1.18E-01 | 2.68E+00 | 2.74E+00 | 9.05E-01 | no | | intergenic_variant | MODIFIER |
| 8 | 93474163 | AX-170434929 | 0 | A | G | 0.211 | -1.41E-02 | 1.18E-01 | 2.68E+00 | 2.74E+00 | 9.05E-01 | no | | intergenic_variant | MODIFIER |
| 8 | 93490846 | AX-170353330 | 0 | C | T | 0.211 | -1.41E-02 | 1.18E-01 | 2.68E+00 | 2.74E+00 | 9.05E-01 | no | | intergenic_variant | MODIFIER |
| 8 | 93498595 | AX-170419645 | 0 | A | G | 0.211 | -1.41E-02 | 1.18E-01 | 2.68E+00 | 2.74E+00 | 9.05E-01 | no | Ces5a | downstream_gene_variant | MODIFIER |
| 8 | 93521528 | AX-170243658 | 0 | T | C | 0.211 | -1.41E-02 | 1.18E-01 | 2.68E+00 | 2.74E+00 | 9.05E-01 | no | Ces5a | intron_variant | MODIFIER |
| 8 | 93531514 | AX-170513583 | 0 | T | C | 0.211 | -1.41E-02 | 1.18E-01 | 2.68E+00 | 2.74E+00 | 9.05E-01 | no | Ces5a | intron_variant | MODIFIER |
| 8 | 93537058 | AX-170526554 | 0 | A | G | 0.211 | -1.41E-02 | 1.18E-01 | 2.68E+00 | 2.74E+00 | 9.05E-01 | no | Ces5a | upstream_gene_variant | MODIFIER |
| 8 | 93546456 | AX-170258861 | 0 | A | C | 0.211 | -1.41E-02 | 1.18E-01 | 2.68E+00 | 2.74E+00 | 9.05E-01 | no | | intergenic_variant | MODIFIER |
| 8 | 93551107 | AX-170532088 | 0 | C | T | 0.211 | -1.41E-02 | 1.18E-01 | 2.68E+00 | 2.74E+00 | 9.05E-01 | no | | intergenic_variant | MODIFIER |
| 8 | 93552379 | AX-170337459 | 0 | A | C | 0.211 | -1.41E-02 | 1.18E-01 | 2.68E+00 | 2.74E+00 | 9.05E-01 | no | | intergenic_variant | MODIFIER |
| 8 | 93552491 | AX-170466302 | 0 | T | C | 0.211 | -1.41E-02 | 1.18E-01 | 2.68E+00 | 2.74E+00 | 9.05E-01 | no | | intergenic_variant | MODIFIER |
| 8 | 93557601 | AX-170492368 | 0 | G | T | 0.211 | -1.41E-02 | 1.18E-01 | 2.68E+00 | 2.74E+00 | 9.05E-01 | no | | intergenic_variant | MODIFIER |
| 8 | 93557696 | AX-170100555 | 0 | A | G | 0.211 | -1.41E-02 | 1.18E-01 | 2.68E+00 | 2.74E+00 | 9.05E-01 | no | | intergenic_variant | MODIFIER |
| 8 | 93570007 | AX-170394970 | 0 | C | T | 0.211 | -1.41E-02 | 1.18E-01 | 2.68E+00 | 2.74E+00 | 9.05E-01 | no | Gm45708 | intron_variant,non_coding_transcript_variant | MODIFIER |
| 8 | 93570566 | AX-170080843 | 0 | C | T | 0.211 | -1.41E-02 | 1.18E-01 | 2.68E+00 | 2.74E+00 | 9.05E-01 | no | Gm45708 | intron_variant,non_coding_transcript_variant | MODIFIER |
| 8 | 93571287 | AX-170406090 | 0 | C | T | 0.211 | -1.41E-02 | 1.18E-01 | 2.68E+00 | 2.74E+00 | 9.05E-01 | no | Gm45708 | intron_variant,non_coding_transcript_variant | MODIFIER |
| 8 | 93601018 | AX-170240325 | 0 | C | G | 0.211 | -1.41E-02 | 1.18E-01 | 2.68E+00 | 2.74E+00 | 9.05E-01 | no | Gm45708 | intron_variant,non_coding_transcript_variant | MODIFIER |
| 8 | 93605989 | AX-170117853 | 0 | G | A | 0.211 | -1.41E-02 | 1.18E-01 | 2.68E+00 | 2.74E+00 | 9.05E-01 | no | Gm45708 | intron_variant,non_coding_transcript_variant | MODIFIER |
| 8 | 93610274 | AX-170374067 | 0 | T | C | 0.211 | -1.41E-02 | 1.18E-01 | 2.68E+00 | 2.74E+00 | 9.05E-01 | no | Gm45708 | intron_variant,non_coding_transcript_variant | MODIFIER |
| 8 | 93620610 | AX-97791211 | 0 | T | C | 0.211 | -1.41E-02 | 1.18E-01 | 2.68E+00 | 2.74E+00 | 9.05E-01 | no | Gm45708 | intron_variant,non_coding_transcript_variant | MODIFIER |
| 8 | 93633297 | AX-170518618 | 0 | C | T | 0.211 | -1.41E-02 | 1.18E-01 | 2.68E+00 | 2.74E+00 | 9.05E-01 | no | Gm45708 | upstream_gene_variant | MODIFIER |
| 8 | 93640462 | AX-170467388 | 0 | G | A | 0.211 | -1.41E-02 | 1.18E-01 | 2.68E+00 | 2.74E+00 | 9.05E-01 | no | | intergenic_variant | MODIFIER |
| 8 | 93667261 | AX-170202684 | 0 | A | G | 0.211 | -1.41E-02 | 1.18E-01 | 2.68E+00 | 2.74E+00 | 9.05E-01 | no | Gm26843 | intron_variant,non_coding_transcript_variant | MODIFIER |
| 8 | 93675881 | AX-170080846 | 0 | A | G | 0.211 | -1.41E-02 | 1.18E-01 | 2.68E+00 | 2.74E+00 | 9.05E-01 | no | Gm26843 | intron_variant,non_coding_transcript_variant | MODIFIER |
| 8 | 93685268 | AX-170080847 | 0 | A | T | 0.211 | -1.41E-02 | 1.18E-01 | 2.68E+00 | 2.74E+00 | 9.05E-01 | no | Gm26843 | intron_variant,non_coding_transcript_variant | MODIFIER |
| 8 | 93706589 | AX-170161055 | 0 | A | G | 0.211 | -1.41E-02 | 1.18E-01 | 2.68E+00 | 2.74E+00 | 9.05E-01 | no | Gm30606 | downstream_gene_variant | MODIFIER |
| 8 | 93728152 | AX-97793345 | 0 | T | C | 0.211 | -1.41E-02 | 1.18E-01 | 2.68E+00 | 2.74E+00 | 9.05E-01 | no | | intergenic_variant | MODIFIER |
| 8 | 93752467 | AX-97848669 | 0 | T | C | 0.211 | -1.41E-02 | 1.18E-01 | 2.68E+00 | 2.74E+00 | 9.05E-01 | no | 4930488L21Rik | non_coding_transcript_exon_variant | MODIFIER |
| 8 | 93755248 | AX-170031270 | 0 | G | A | 0.211 | -1.41E-02 | 1.18E-01 | 2.68E+00 | 2.74E+00 | 9.05E-01 | no | 4930488L21Rik | intron_variant,non_coding_transcript_variant | MODIFIER |
| 8 | 93756009 | AX-170390910 | 0 | C | T | 0.211 | -1.41E-02 | 1.18E-01 | 2.68E+00 | 2.74E+00 | 9.05E-01 | no | 4930488L21Rik | intron_variant,non_coding_transcript_variant | MODIFIER |
| 8 | 93774344 | AX-97839047 | 0 | A | G | 0.211 | -1.41E-02 | 1.18E-01 | 2.68E+00 | 2.74E+00 | 9.05E-01 | no | 4930488L21Rik | intron_variant,non_coding_transcript_variant | MODIFIER |
| 8 | 93781431 | AX-170139945 | 0 | G | A | 0.211 | -1.41E-02 | 1.18E-01 | 2.68E+00 | 2.74E+00 | 9.05E-01 | no | 4930488L21Rik | intron_variant,non_coding_transcript_variant | MODIFIER |
| 8 | 93781950 | AX-170031271 | 0 | A | G | 0.211 | -1.41E-02 | 1.18E-01 | 2.68E+00 | 2.74E+00 | 9.05E-01 | no | 4930488L21Rik | intron_variant,non_coding_transcript_variant | MODIFIER |
| 8 | 93795140 | AX-97846744 | 0 | G | C | 0.211 | -1.41E-02 | 1.18E-01 | 2.68E+00 | 2.74E+00 | 9.05E-01 | no | 4930488L21Rik | intron_variant,non_coding_transcript_variant | MODIFIER |
| 8 | 93819021 | AX-170031269 | 0 | A | G | 0.211 | -1.41E-02 | 1.18E-01 | 2.68E+00 | 2.74E+00 | 9.05E-01 | no | Gnao1 | intron_variant | MODIFIER |
| 8 | 93822727 | AX-97834461 | 0 | A | G | 0.211 | -1.41E-02 | 1.18E-01 | 2.68E+00 | 2.74E+00 | 9.05E-01 | no | Gnao1 | intron_variant | MODIFIER |
| 8 | 93841852 | AX-170080848 | 0 | C | T | 0.211 | -1.41E-02 | 1.18E-01 | 2.68E+00 | 2.74E+00 | 9.05E-01 | no | Gnao1 | intron_variant | MODIFIER |
| 8 | 93842140 | AX-170080849 | 0 | T | G | 0.211 | -1.41E-02 | 1.18E-01 | 2.68E+00 | 2.74E+00 | 9.05E-01 | no | Gnao1 | intron_variant | MODIFIER |
| 8 | 93843202 | AX-97793925 | 0 | A | G | 0.211 | -1.41E-02 | 1.18E-01 | 2.68E+00 | 2.74E+00 | 9.05E-01 | no | Gnao1 | intron_variant | MODIFIER |
| 8 | 93843374 | AX-170235358 | 0 | A | C | 0.211 | -1.41E-02 | 1.18E-01 | 2.68E+00 | 2.74E+00 | 9.05E-01 | no | Gnao1 | intron_variant | MODIFIER |
| 8 | 93843396 | AX-170496450 | 0 | C | T | 0.211 | -1.41E-02 | 1.18E-01 | 2.68E+00 | 2.74E+00 | 9.05E-01 | no | Gnao1 | intron_variant | MODIFIER |
| 8 | 93851291 | AX-170222236 | 0 | G | A | 0.211 | -1.41E-02 | 1.18E-01 | 2.68E+00 | 2.74E+00 | 9.05E-01 | no | Gnao1 | intron_variant | MODIFIER |
| 8 | 93861050 | AX-170178355 | 0 | G | A | 0.211 | -1.41E-02 | 1.18E-01 | 2.68E+00 | 2.74E+00 | 9.05E-01 | no | Gnao1 | intron_variant | MODIFIER |
| 8 | 93887616 | AX-170392058 | 0 | C | T | 0.211 | -1.41E-02 | 1.18E-01 | 2.68E+00 | 2.74E+00 | 9.05E-01 | no | Gnao1 | intron_variant | MODIFIER |
| 8 | 93892825 | AX-170457597 | 0 | A | G | 0.211 | -1.41E-02 | 1.18E-01 | 2.68E+00 | 2.74E+00 | 9.05E-01 | no | Gnao1 | intron_variant | MODIFIER |
| 8 | 93899042 | AX-170483235 | 0 | G | C | 0.211 | -1.41E-02 | 1.18E-01 | 2.68E+00 | 2.74E+00 | 9.05E-01 | no | Gnao1 | intron_variant | MODIFIER |
| 8 | 93902888 | AX-97815929 | 0 | T | C | 0.211 | -1.41E-02 | 1.18E-01 | 2.68E+00 | 2.74E+00 | 9.05E-01 | no | Gnao1 | intron_variant | MODIFIER |
| 8 | 93920542 | AX-170086147 | 0 | C | T | 0.211 | -1.41E-02 | 1.18E-01 | 2.68E+00 | 2.74E+00 | 9.05E-01 | no | Gnao1 | intron_variant | MODIFIER |
| 8 | 93934334 | AX-97810279 | 0 | T | C | 0.211 | -1.41E-02 | 1.18E-01 | 2.68E+00 | 2.74E+00 | 9.05E-01 | no | Gnao1 | intron_variant | MODIFIER |
| 8 | 93935722 | AX-170242490 | 0 | G | A | 0.211 | -1.41E-02 | 1.18E-01 | 2.68E+00 | 2.74E+00 | 9.05E-01 | no | Gnao1 | intron_variant | MODIFIER |
| 8 | 93949131 | AX-170410961 | 0 | C | T | 0.211 | -1.41E-02 | 1.18E-01 | 2.68E+00 | 2.74E+00 | 9.05E-01 | no | Gnao1 | intron_variant | MODIFIER |
| 8 | 93981402 | AX-170339748 | 0 | A | G | 0.211 | -1.41E-02 | 1.18E-01 | 2.68E+00 | 2.74E+00 | 9.05E-01 | no | Amfr | intron_variant | MODIFIER |
| 8 | 93987476 | AX-170419128 | 0 | G | A | 0.211 | -1.41E-02 | 1.18E-01 | 2.68E+00 | 2.74E+00 | 9.05E-01 | no | Amfr | synonymous_variant | LOW |

| | | | | | | | | | | | | | | | |
|---|---|---|---|---|---|---|---|---|---|---|---|---|---|---|---|
| 8 | 93990256 | AX-97795346 | 0 | G | A | 0.211 | -1.41E-02 | 1.18E-01 | 2.68E+00 | 2.74E+00 | 9.05E-01 | no | Amfr | intron_variant | MODIFIER |
| 8 | 94014762 | AX-170315985 | 0 | A | T | 0.211 | -1.41E-02 | 1.18E-01 | 2.68E+00 | 2.74E+00 | 9.05E-01 | no | Nudt21 | downstream_gene_variant | MODIFIER |
| 8 | 94019713 | AX-170366013 | 0 | A | T | 0.211 | -1.41E-02 | 1.18E-01 | 2.68E+00 | 2.74E+00 | 9.05E-01 | no | Nudt21 | intron_variant | MODIFIER |
| 8 | 94039489 | AX-170080853 | 0 | T | C | 0.211 | -1.41E-02 | 1.18E-01 | 2.68E+00 | 2.74E+00 | 9.05E-01 | no | Nudt21 | upstream_gene_variant | MODIFIER |
| 8 | 94049792 | AX-97816951 | 0 | G | A | 0.211 | -1.41E-02 | 1.18E-01 | 2.68E+00 | 2.74E+00 | 9.05E-01 | no | Ogfod1 | intron_variant | MODIFIER |
| 8 | 94050519 | AX-170080854 | 0 | G | C | 0.211 | -1.41E-02 | 1.18E-01 | 2.68E+00 | 2.74E+00 | 9.05E-01 | no | Ogfod1 | intron_variant | MODIFIER |
| 8 | 94074423 | AX-170325139 | 0 | A | C | 0.211 | -1.41E-02 | 1.18E-01 | 2.68E+00 | 2.74E+00 | 9.05E-01 | no | Bbs2 | intron_variant | MODIFIER |
| 8 | 94085738 | AX-170326471 | 0 | G | A | 0.211 | -1.41E-02 | 1.18E-01 | 2.68E+00 | 2.74E+00 | 9.05E-01 | no | Bbs2 | intron_variant | MODIFIER |
| 8 | 94089084 | AX-170437244 | 0 | G | T | 0.211 | -1.41E-02 | 1.18E-01 | 2.68E+00 | 2.74E+00 | 9.05E-01 | no | Bbs2 | intron_variant | MODIFIER |
| 8 | 94112205 | AX-170091311 | 0 | A | G | 0.211 | -1.41E-02 | 1.18E-01 | 2.68E+00 | 2.74E+00 | 9.05E-01 | no | | regulatory_region_variant | MODIFIER |
| 8 | 94116052 | AX-170190519 | 0 | G | A | 0.211 | -1.41E-02 | 1.18E-01 | 2.68E+00 | 2.74E+00 | 9.05E-01 | no | | intergenic_variant | MODIFIER |
| 8 | 94136156 | AX-170460661 | 0 | T | G | 0.211 | -1.41E-02 | 1.18E-01 | 2.68E+00 | 2.74E+00 | 9.05E-01 | no | Mt4 | upstream_gene_variant | MODIFIER |
| 8 | 94136651 | AX-170211708 | 0 | G | T | 0.211 | -1.41E-02 | 1.18E-01 | 2.68E+00 | 2.74E+00 | 9.05E-01 | no | Mt4 | upstream_gene_variant | MODIFIER |
| 8 | 94138312 | AX-170305246 | 0 | T | A | 0.211 | -1.41E-02 | 1.18E-01 | 2.68E+00 | 2.74E+00 | 9.05E-01 | no | Mt4 | intron_variant | MODIFIER |
| 8 | 94138997 | AX-170494805 | 0 | A | G | 0.211 | -1.41E-02 | 1.18E-01 | 2.68E+00 | 2.74E+00 | 9.05E-01 | no | Mt4 | 3_prime_UTR_variant | MODIFIER |
| 8 | 94172897 | AX-170031284 | 0 | A | G | 0.211 | -1.41E-02 | 1.18E-01 | 2.68E+00 | 2.74E+00 | 9.05E-01 | no | Mt2 | intron_variant | MODIFIER |
| 8 | 94193771 | AX-170080855 | 0 | G | A | 0.211 | -1.41E-02 | 1.18E-01 | 2.68E+00 | 2.74E+00 | 9.05E-01 | no | | intergenic_variant | MODIFIER |
| 8 | 94224969 | AX-97791648 | 0 | A | C | 0.211 | -1.41E-02 | 1.18E-01 | 2.68E+00 | 2.74E+00 | 9.05E-01 | no | Nup93 | intron_variant | MODIFIER |
| 8 | 94238990 | AX-97813431 | 0 | A | G | 0.211 | -1.41E-02 | 1.18E-01 | 2.68E+00 | 2.74E+00 | 9.05E-01 | no | Nup93 | intron_variant | MODIFIER |
| 8 | 94239948 | AX-170196619 | 0 | T | C | 0.211 | -1.41E-02 | 1.18E-01 | 2.68E+00 | 2.74E+00 | 9.05E-01 | no | Nup93 | intron_variant | MODIFIER |
| 8 | 94248807 | AX-170409748 | 0 | G | A | 0.211 | -1.41E-02 | 1.18E-01 | 2.68E+00 | 2.74E+00 | 9.05E-01 | no | Nup93 | intron_variant | MODIFIER |
| 8 | 94261133 | AX-97855762 | 0 | C | T | 0.211 | -1.41E-02 | 1.18E-01 | 2.68E+00 | 2.74E+00 | 9.05E-01 | no | Nup93 | intron_variant | MODIFIER |
| 8 | 94261434 | AX-170031287 | 0 | T | G | 0.211 | -1.41E-02 | 1.18E-01 | 2.68E+00 | 2.74E+00 | 9.05E-01 | no | Nup93 | intron_variant | MODIFIER |
| 8 | 94274970 | AX-170505205 | 0 | A | T | 0.211 | -1.41E-02 | 1.18E-01 | 2.68E+00 | 2.74E+00 | 9.05E-01 | no | Nup93 | intron_variant | MODIFIER |
| 8 | 94275405 | AX-170386035 | 0 | G | A | 0.211 | -1.41E-02 | 1.18E-01 | 2.68E+00 | 2.74E+00 | 9.05E-01 | no | Nup93 | intron_variant | MODIFIER |
| 8 | 94281422 | AX-170229856 | 0 | A | C | 0.211 | -1.41E-02 | 1.18E-01 | 2.68E+00 | 2.74E+00 | 9.05E-01 | no | Nup93 | intron_variant | MODIFIER |
| 8 | 94290835 | AX-170141403 | 0 | G | A | 0.211 | -1.41E-02 | 1.18E-01 | 2.68E+00 | 2.74E+00 | 9.05E-01 | no | Nup93 | intron_variant | MODIFIER |
| 8 | 94293645 | AX-170031280 | 0 | A | G | 0.211 | -1.41E-02 | 1.18E-01 | 2.68E+00 | 2.74E+00 | 9.05E-01 | no | Nup93 | intron_variant | MODIFIER |
| 8 | 94302946 | AX-170517020 | 0 | C | G | 0.211 | -1.41E-02 | 1.18E-01 | 2.68E+00 | 2.74E+00 | 9.05E-01 | no | Nup93 | intron_variant | MODIFIER |
| 8 | 94305165 | AX-97850477 | 0 | T | C | 0.211 | -1.41E-02 | 1.18E-01 | 2.68E+00 | 2.74E+00 | 9.05E-01 | no | Nup93 | intron_variant | MODIFIER |
| 8 | 94307992 | AX-170381770 | 0 | C | T | 0.211 | -1.41E-02 | 1.18E-01 | 2.68E+00 | 2.74E+00 | 9.05E-01 | no | Nup93 | intron_variant | MODIFIER |
| 8 | 94308049 | AX-170175975 | 0 | A | G | 0.211 | -1.41E-02 | 1.18E-01 | 2.68E+00 | 2.74E+00 | 9.05E-01 | no | Nup93 | intron_variant | MODIFIER |
| 8 | 94320926 | AX-170496095 | 0 | T | C | 0.211 | -1.41E-02 | 1.18E-01 | 2.68E+00 | 2.74E+00 | 9.05E-01 | no | Mir138-2 | upstream_gene_variant | MODIFIER |
| 8 | 94330056 | AX-170319574 | 0 | G | A | 0.211 | -1.41E-02 | 1.18E-01 | 2.68E+00 | 2.74E+00 | 9.05E-01 | no | Slc12a3 | intron_variant | MODIFIER |
| 8 | 94337945 | AX-170080859 | 0 | T | C | 0.211 | -1.41E-02 | 1.18E-01 | 2.68E+00 | 2.74E+00 | 9.05E-01 | no | Slc12a3 | intron_variant | MODIFIER |
| 8 | 94348911 | AX-170080858 | 0 | C | A | 0.211 | -1.41E-02 | 1.18E-01 | 2.68E+00 | 2.74E+00 | 9.05E-01 | no | Slc12a3 | intron_variant | MODIFIER |
| 8 | 94368514 | AX-170152059 | 0 | T | C | 0.211 | -1.41E-02 | 1.18E-01 | 2.68E+00 | 2.74E+00 | 9.05E-01 | no | Slc12a3 | downstream_gene_variant | MODIFIER |
| 8 | 94388806 | AX-170446463 | 0 | G | C | 0.211 | -1.41E-02 | 1.18E-01 | 2.68E+00 | 2.74E+00 | 9.05E-01 | no | Herpud1 | intron_variant | MODIFIER |
| 8 | 94415996 | AX-170031282 | 0 | G | A | 0.211 | -1.41E-02 | 1.18E-01 | 2.68E+00 | 2.74E+00 | 9.05E-01 | no | | intergenic_variant | MODIFIER |
| 8 | 94417835 | AX-170091312 | 0 | C | T | 0.211 | -1.41E-02 | 1.18E-01 | 2.68E+00 | 2.74E+00 | 9.05E-01 | no | | regulatory_region_variant | MODIFIER |
| 8 | 94417900 | AX-170039798 | 0 | C | T | 0.211 | -1.41E-02 | 1.18E-01 | 2.68E+00 | 2.74E+00 | 9.05E-01 | no | 9330175E14Rik | downstream_gene_variant | MODIFIER |
| 8 | 94430857 | AX-170098339 | 0 | C | T | 0.211 | -1.41E-02 | 1.18E-01 | 2.68E+00 | 2.74E+00 | 9.05E-01 | no | 9330175E14Rik | non_coding_transcript_exon_variant | MODIFIER |
| 8 | 94445175 | AX-170112977 | 0 | A | G | 0.211 | -1.41E-02 | 1.18E-01 | 2.68E+00 | 2.74E+00 | 9.05E-01 | no | Nlrc5 | intron_variant | MODIFIER |
| 8 | 94456189 | AX-170383425 | 0 | C | T | 0.211 | -1.41E-02 | 1.18E-01 | 2.68E+00 | 2.74E+00 | 9.05E-01 | no | Nlrc5 | intron_variant | MODIFIER |
| 8 | 94456674 | AX-170239175 | 0 | A | G | 0.211 | -1.41E-02 | 1.18E-01 | 2.68E+00 | 2.74E+00 | 9.05E-01 | no | Nlrc5 | intron_variant | MODIFIER |
| 8 | 94517200 | AX-170442257 | 0 | A | G | 0.211 | -1.41E-02 | 1.18E-01 | 2.68E+00 | 2.74E+00 | 9.05E-01 | no | Nlrc5 | intron_variant | MODIFIER |
| 8 | 94556127 | AX-170080860 | 0 | G | A | 0.211 | -1.41E-02 | 1.18E-01 | 2.68E+00 | 2.74E+00 | 9.05E-01 | no | Cpne2 | intron_variant | MODIFIER |
| 8 | 94562244 | AX-170266941 | 0 | T | C | 0.211 | -1.41E-02 | 1.18E-01 | 2.68E+00 | 2.74E+00 | 9.05E-01 | no | Cpne2 | intron_variant | MODIFIER |
| 8 | 94585380 | AX-170500899 | 0 | T | C | 0.211 | -1.41E-02 | 1.18E-01 | 2.68E+00 | 2.74E+00 | 9.05E-01 | no | Fam192a | intron_variant | MODIFIER |
| 8 | 94586344 | AX-170111333 | 0 | C | A | 0.211 | -1.41E-02 | 1.18E-01 | 2.68E+00 | 2.74E+00 | 9.05E-01 | no | Fam192a | intron_variant | MODIFIER |
| 8 | 94587249 | AX-170129759 | 0 | C | G | 0.211 | -1.41E-02 | 1.18E-01 | 2.68E+00 | 2.74E+00 | 9.05E-01 | no | Fam192a | intron_variant | MODIFIER |
| 8 | 94587276 | AX-170298905 | 0 | T | G | 0.211 | -1.41E-02 | 1.18E-01 | 2.68E+00 | 2.74E+00 | 9.05E-01 | no | Fam192a | intron_variant | MODIFIER |
| 8 | 94607644 | AX-170352478 | 0 | T | C | 0.211 | -1.41E-02 | 1.18E-01 | 2.68E+00 | 2.74E+00 | 9.05E-01 | no | Rspry1 | intron_variant | MODIFIER |
| 8 | 94607909 | AX-170403765 | 0 | C | A | 0.211 | -1.41E-02 | 1.18E-01 | 2.68E+00 | 2.74E+00 | 9.05E-01 | no | Rspry1 | intron_variant | MODIFIER |

| | | | | | | | | | | | | | | | |
|---|---|---|---|---|---|---|---|---|---|---|---|---|---|---|---|
| 8 | 94652110 | AX-170287584 | 0 | A | G | 0.211 | -1.41E-02 | 1.18E-01 | 2.68E+00 | 2.74E+00 | 9.05E-01 | no | Rspry1 | intron_variant | MODIFIER |
| 8 | 94654164 | AX-170400101 | 0 | A | G | 0.211 | -1.41E-02 | 1.18E-01 | 2.68E+00 | 2.74E+00 | 9.05E-01 | no | Rspry1 | intron_variant | MODIFIER |
| 8 | 94661725 | AX-170277246 | 0 | T | C | 0.211 | -1.41E-02 | 1.18E-01 | 2.68E+00 | 2.74E+00 | 9.05E-01 | no | Arl2bp | upstream_gene_variant | MODIFIER |
| 8 | 94683798 | AX-170460382 | 0 | A | G | 0.211 | -1.41E-02 | 1.18E-01 | 2.68E+00 | 2.74E+00 | 9.05E-01 | no | Pllp | intron_variant | MODIFIER |
| 8 | 94716152 | AX-170270982 | 0 | G | A | 0.211 | -1.41E-02 | 1.18E-01 | 2.68E+00 | 2.74E+00 | 9.05E-01 | no | | intergenic_variant | MODIFIER |
| 8 | 94716196 | AX-170244366 | 0 | G | A | 0.211 | -1.41E-02 | 1.18E-01 | 2.68E+00 | 2.74E+00 | 9.05E-01 | no | | intergenic_variant | MODIFIER |
| 8 | 94726280 | AX-170031296 | 0 | T | C | 0.211 | -1.41E-02 | 1.18E-01 | 2.68E+00 | 2.74E+00 | 9.05E-01 | no | | regulatory_region_variant | MODIFIER |
| 8 | 94759919 | AX-170180091 | 0 | G | A | 0.211 | -1.41E-02 | 1.18E-01 | 2.68E+00 | 2.74E+00 | 9.05E-01 | no | Gnb1-ps3 | upstream_gene_variant | MODIFIER |
| 8 | 94770277 | AX-97838294 | 0 | T | G | 0.211 | -1.41E-02 | 1.18E-01 | 2.68E+00 | 2.74E+00 | 9.05E-01 | no | Cx3cl1 | upstream_gene_variant | MODIFIER |
| 8 | 94776657 | AX-170208742 | 0 | T | G | 0.211 | -1.41E-02 | 1.18E-01 | 2.68E+00 | 2.74E+00 | 9.05E-01 | no | Cx3cl1 | intron_variant | MODIFIER |
| 8 | 94778733 | AX-170174229 | 0 | C | G | 0.211 | -1.41E-02 | 1.18E-01 | 2.68E+00 | 2.74E+00 | 9.05E-01 | no | Cx3cl1 | intron_variant | MODIFIER |
| 8 | 94780000 | AX-170451885 | 0 | C | A | 0.211 | -1.41E-02 | 1.18E-01 | 2.68E+00 | 2.74E+00 | 9.05E-01 | no | Cx3cl1 | missense_variant | MODERATE |
| 8 | 94781785 | AX-170241229 | 0 | C | T | 0.211 | -1.41E-02 | 1.18E-01 | 2.68E+00 | 2.74E+00 | 9.05E-01 | no | Cx3cl1 | 3_prime_UTR_variant | MODIFIER |
| 8 | 94806270 | AX-170150139 | 0 | C | G | 0.211 | -1.41E-02 | 1.18E-01 | 2.68E+00 | 2.74E+00 | 9.05E-01 | no | Ccl17 | upstream_gene_variant | MODIFIER |
| 8 | 94806288 | AX-170400783 | 0 | T | G | 0.211 | -1.41E-02 | 1.18E-01 | 2.68E+00 | 2.74E+00 | 9.05E-01 | no | Ccl17 | upstream_gene_variant | MODIFIER |
| 8 | 94823326 | AX-170464905 | 0 | G | A | 0.211 | -1.41E-02 | 1.18E-01 | 2.68E+00 | 2.74E+00 | 9.05E-01 | no | Ciapin1 | intron_variant | MODIFIER |
| 8 | 94895563 | AX-170290192 | 0 | C | T | 0.211 | -1.41E-02 | 1.18E-01 | 2.68E+00 | 2.74E+00 | 9.05E-01 | no | | regulatory_region_variant | MODIFIER |
| 8 | 94934589 | AX-170436492 | 0 | G | C | 0.211 | -1.41E-02 | 1.18E-01 | 2.68E+00 | 2.74E+00 | 9.05E-01 | no | Adgrg5 | intron_variant | MODIFIER |
| 8 | 37167381 | AX-170281358 | 0 | A | G | 0.14 | 1.49E-02 | 1.33E-01 | 2.67E+00 | 2.76E+00 | 9.05E-01 | no | Gm45524 | intron_variant,non_coding_transcript_variant | MODIFIER |
| 15 | 100445103 | AX-170087950 | 1 | C | A | 0.107 | 1.74E-02 | 1.46E-01 | 2.66E+00 | 2.73E+00 | 9.05E-01 | no | n-R5s43 | upstream_gene_variant | MODIFIER |
| 12 | 104301707 | AX-170225911 | 1 | A | G | 0.116 | -1.81E-02 | 1.60E-01 | 2.67E+00 | 2.75E+00 | 9.05E-01 | no | | intergenic_variant | MODIFIER |
| 12 | 106660468 | AX-170487413 | 1 | A | G | 0.116 | -1.81E-02 | 1.60E-01 | 2.67E+00 | 2.75E+00 | 9.05E-01 | no | | intergenic_variant | MODIFIER |
| 18 | 64786578 | AX-170277299 | 0 | G | A | 0.421 | 9.13E-03 | 8.41E-02 | 2.71E+00 | 2.78E+00 | 9.05E-01 | no | Gm24504 | upstream_gene_variant | MODIFIER |
| 1 | 169066654 | AX-170043497 | 0 | C | G | 0.219 | -1.21E-02 | 1.19E-01 | 2.66E+00 | 2.77E+00 | 9.05E-01 | no | | intergenic_variant | MODIFIER |
| 7 | 116587228 | AX-170383781 | 0 | C | T | 0.439 | 1.12E-02 | 9.16E-02 | 2.66E+00 | 2.72E+00 | 9.05E-01 | no | | intergenic_variant | MODIFIER |
| 13 | 37328285 | AX-97780251 | 0 | G | A | 0.184 | -1.42E-02 | 1.13E-01 | 2.64E+00 | 2.72E+00 | 9.05E-01 | no | Gm47711 | downstream_gene_variant | MODIFIER |
| 13 | 37846589 | AX-170087409 | 0 | G | T | 0.184 | -1.42E-02 | 1.13E-01 | 2.64E+00 | 2.72E+00 | 9.05E-01 | no | Rreb1 | intron_variant | MODIFIER |
| 10 | 127237461 | AX-170367822 | 0 | C | T | 0.325 | -9.13E-03 | 7.29E-02 | 2.62E+00 | 2.72E+00 | 9.05E-01 | no | Kif5a | intron_variant | MODIFIER |
| 2 | 114778976 | AX-170126524 | 1 | G | A | 0.134 | -2.47E-02 | 2.17E-01 | 2.71E+00 | 2.76E+00 | 9.05E-01 | no | | intergenic_variant | MODIFIER |
| 2 | 129045173 | AX-170305991 | 1 | T | A | 0.134 | -2.47E-02 | 2.17E-01 | 2.71E+00 | 2.76E+00 | 9.05E-01 | no | Vinac1 | intron_variant | MODIFIER |
| 2 | 142594856 | AX-170065107 | 1 | G | A | 0.134 | -2.47E-02 | 2.17E-01 | 2.71E+00 | 2.76E+00 | 9.05E-01 | no | | intergenic_variant | MODIFIER |
| 2 | 146838040 | AX-170519372 | 1 | A | G | 0.134 | -2.47E-02 | 2.17E-01 | 2.71E+00 | 2.76E+00 | 9.05E-01 | no | Gm14114 | intron_variant,non_coding_transcript_variant | MODIFIER |
| 1 | 169621386 | AX-170151800 | 0 | T | C | 0.237 | -1.35E-02 | 1.22E-01 | 2.69E+00 | 2.76E+00 | 9.06E-01 | no | | intergenic_variant | MODIFIER |
| 1 | 170464542 | AX-170115100 | 0 | A | G | 0.237 | -1.35E-02 | 1.22E-01 | 2.69E+00 | 2.76E+00 | 9.06E-01 | no | Nos1ap | intron_variant,non_coding_transcript_variant | MODIFIER |
| 1 | 52009667 | AX-170370601 | 1 | A | T | 0.107 | -2.71E-02 | 2.34E-01 | 2.70E+00 | 2.74E+00 | 9.06E-01 | no | Stat4 | intron_variant | MODIFIER |
| 9 | 66120975 | AX-170455425 | 0 | G | A | 0.202 | -1.55E-02 | 1.30E-01 | 2.69E+00 | 2.73E+00 | 9.06E-01 | no | Snx1 | intron_variant | MODIFIER |
| 9 | 66736845 | AX-170424239 | 0 | C | T | 0.202 | -1.55E-02 | 1.30E-01 | 2.69E+00 | 2.73E+00 | 9.06E-01 | no | Car12 | intron_variant | MODIFIER |
| 13 | 52714328 | AX-168393486 | 0 | C | T | 0.526 | 8.41E-03 | 6.65E-02 | 2.62E+00 | 2.72E+00 | 9.06E-01 | no | | intergenic_variant | MODIFIER |
| 5 | 99956371 | AX-170446718 | 1 | C | G | 0.429 | -1.23E-02 | 1.08E-01 | 2.70E+00 | 2.75E+00 | 9.06E-01 | no | Hnrnpd | downstream_gene_variant | MODIFIER |
| 7 | 54083470 | AX-97791198 | 1 | T | C | 0.429 | 1.22E-02 | 9.91E-02 | 2.65E+00 | 2.72E+00 | 9.06E-01 | no | | intergenic_variant | MODIFIER |
| 7 | 94043690 | AX-170335214 | 1 | A | G | 0.116 | 2.53E-02 | 2.12E-01 | 2.65E+00 | 2.73E+00 | 9.06E-01 | no | Gm32647 | intron_variant,non_coding_transcript_variant | MODIFIER |
| 5 | 73329875 | AX-170506602 | 0 | A | T | 0.439 | 1.20E-02 | 9.62E-02 | 2.64E+00 | 2.72E+00 | 9.06E-01 | no | Ociad2 | intron_variant | MODIFIER |
| 20 | 145128036 | AX-97788567 | 0 | C | T | 0.149 | 1.60E-02 | 1.35E-01 | 2.64E+00 | 2.74E+00 | 9.06E-01 | no | | intergenic_variant | MODIFIER |
| 5 | 117529221 | AX-170380031 | 1 | G | A | 0.464 | -1.26E-02 | 1.12E-01 | 2.71E+00 | 2.77E+00 | 9.06E-01 | no | Ksr2 | intron_variant,non_coding_transcript_variant | MODIFIER |
| 5 | 118948509 | AX-170398671 | 1 | C | T | 0.464 | -1.26E-02 | 1.12E-01 | 2.71E+00 | 2.77E+00 | 9.06E-01 | no | 4930413E15Rik | upstream_gene_variant | MODIFIER |
| 5 | 122684117 | AX-170514141 | 1 | G | C | 0.464 | -1.26E-02 | 1.12E-01 | 2.71E+00 | 2.77E+00 | 9.06E-01 | no | P2rx7 | intron_variant | MODIFIER |
| 1 | 60075579 | AX-170485872 | 0 | A | G | 0.105 | -2.74E-02 | 2.36E-01 | 2.70E+00 | 2.74E+00 | 9.06E-01 | no | Wdr12 | 3_prime_UTR_variant | MODIFIER |
| 20 | 130512497 | AX-170515431 | 0 | A | G | 0.333 | 6.44E-03 | 5.20E-02 | 2.61E+00 | 2.73E+00 | 9.06E-01 | no | Gm14987 | downstream_gene_variant | MODIFIER |
| 1 | 150494336 | AX-170467153 | 1 | G | T | 0.205 | 1.46E-02 | 1.20E-01 | 2.64E+00 | 2.73E+00 | 9.06E-01 | no | | intergenic_variant | MODIFIER |
| 2 | 124974657 | AX-170416999 | 1 | C | A | 0.125 | 2.38E-02 | 2.00E-01 | 2.65E+00 | 2.73E+00 | 9.06E-01 | no | | intergenic_variant | MODIFIER |
| 8 | 93968154 | AX-170173976 | 1 | C | T | 0.214 | -1.37E-02 | 1.17E-01 | 2.67E+00 | 2.74E+00 | 9.06E-01 | no | Gnao1 | 3_prime_UTR_variant | MODIFIER |
| 8 | 94187639 | AX-170517292 | 1 | C | T | 0.214 | -1.37E-02 | 1.17E-01 | 2.67E+00 | 2.74E+00 | 9.06E-01 | no | Gm39228 | downstream_gene_variant | MODIFIER |
| 7 | 40358123 | AX-170214896 | 0 | T | C | 0.316 | 1.23E-02 | 9.52E-02 | 2.63E+00 | 2.71E+00 | 9.06E-01 | no | | intergenic_variant | MODIFIER |
| 8 | 94288277 | AX-170111568 | 1 | T | C | 0.205 | -1.41E-02 | 1.20E-01 | 2.68E+00 | 2.74E+00 | 9.06E-01 | no | Nup93 | intron_variant | MODIFIER |

| | | | | | | | | | | | | | | | |
|---|---|---|---|---|---|---|---|---|---|---|---|---|---|---|---|
| 1 | 77054280 | AX-170183979 | 0 | C | T | 0.132 | -2.50E-02 | 2.22E-01 | 2.68E+00 | 2.75E+00 | 9.06E-01 | no | | intergenic_variant | MODIFIER |
| 1 | 48918262 | AX-170000442 | 2 | T | C | 0.1 | -2.54E-02 | 2.22E-01 | 2.66E+00 | 2.74E+00 | 9.06E-01 | no | | intergenic_variant | MODIFIER |
| 3 | 87348954 | AX-97794128 | 0 | A | G | 0.061 | -1.57E-02 | 1.34E-01 | 2.59E+00 | 2.74E+00 | 9.06E-01 | no | | intergenic_variant | MODIFIER |
| 1 | 148978648 | AX-170043081 | 0 | G | A | 0.202 | 1.68E-02 | 1.38E-01 | 2.67E+00 | 2.72E+00 | 9.06E-01 | no | Gm5703 | non_coding_transcript_exon_variant | MODIFIER |
| 1 | 152790899 | AX-170218075 | 0 | C | A | 0.202 | 1.68E-02 | 1.38E-01 | 2.67E+00 | 2.72E+00 | 9.06E-01 | no | | regulatory_region_variant | MODIFIER |
| 10 | 5806208 | AX-170519727 | 0 | T | C | 0.193 | -1.27E-02 | 1.06E-01 | 2.64E+00 | 2.73E+00 | 9.06E-01 | no | Fbxo5 | upstream_gene_variant | MODIFIER |
| 10 | 5829661 | AX-170420455 | 0 | T | C | 0.193 | -1.27E-02 | 1.06E-01 | 2.64E+00 | 2.73E+00 | 9.06E-01 | no | Rgs17 | downstream_gene_variant | MODIFIER |
| 10 | 5854657 | AX-170166027 | 0 | G | A | 0.193 | -1.27E-02 | 1.06E-01 | 2.64E+00 | 2.73E+00 | 9.06E-01 | no | Rgs17 | intron_variant | MODIFIER |
| 10 | 5871830 | AX-170532952 | 0 | T | C | 0.193 | -1.27E-02 | 1.06E-01 | 2.64E+00 | 2.73E+00 | 9.06E-01 | no | Rgs17 | intron_variant | MODIFIER |
| 10 | 5871961 | AX-170100657 | 0 | G | A | 0.193 | -1.27E-02 | 1.06E-01 | 2.64E+00 | 2.73E+00 | 9.06E-01 | no | Rgs17 | intron_variant | MODIFIER |
| 10 | 5916242 | AX-97821520 | 0 | G | C | 0.193 | -1.27E-02 | 1.06E-01 | 2.64E+00 | 2.73E+00 | 9.06E-01 | no | Rgs17 | intron_variant | MODIFIER |
| 10 | 5944550 | AX-170197887 | 0 | C | T | 0.193 | -1.27E-02 | 1.06E-01 | 2.64E+00 | 2.73E+00 | 9.06E-01 | no | | intergenic_variant | MODIFIER |
| 10 | 5946402 | AX-170147308 | 0 | T | G | 0.193 | -1.27E-02 | 1.06E-01 | 2.64E+00 | 2.73E+00 | 9.06E-01 | no | | intergenic_variant | MODIFIER |
| 10 | 5950777 | AX-170330613 | 0 | T | C | 0.193 | -1.27E-02 | 1.06E-01 | 2.64E+00 | 2.73E+00 | 9.06E-01 | no | | intergenic_variant | MODIFIER |
| 10 | 6420912 | AX-170477660 | 0 | A | C | 0.193 | -1.27E-02 | 1.06E-01 | 2.64E+00 | 2.73E+00 | 9.06E-01 | no | Gm47755 | upstream_gene_variant | MODIFIER |
| 10 | 6490661 | AX-170368942 | 0 | G | A | 0.193 | -1.27E-02 | 1.06E-01 | 2.64E+00 | 2.73E+00 | 9.06E-01 | no | | intergenic_variant | MODIFIER |
| 10 | 6494996 | AX-170298743 | 0 | A | C | 0.193 | -1.27E-02 | 1.06E-01 | 2.64E+00 | 2.73E+00 | 9.06E-01 | no | | intergenic_variant | MODIFIER |
| 10 | 6502044 | AX-170336435 | 0 | G | T | 0.193 | -1.27E-02 | 1.06E-01 | 2.64E+00 | 2.73E+00 | 9.06E-01 | no | | intergenic_variant | MODIFIER |
| 10 | 6552893 | AX-170044059 | 0 | T | C | 0.193 | -1.27E-02 | 1.06E-01 | 2.64E+00 | 2.73E+00 | 9.06E-01 | no | | intergenic_variant | MODIFIER |
| 10 | 6554555 | AX-170128940 | 0 | T | C | 0.193 | -1.27E-02 | 1.06E-01 | 2.64E+00 | 2.73E+00 | 9.06E-01 | no | | intergenic_variant | MODIFIER |
| 10 | 6672464 | AX-170167502 | 0 | T | C | 0.193 | -1.27E-02 | 1.06E-01 | 2.64E+00 | 2.73E+00 | 9.06E-01 | no | | intergenic_variant | MODIFIER |
| 10 | 6707085 | AX-170468870 | 0 | A | G | 0.193 | -1.27E-02 | 1.06E-01 | 2.64E+00 | 2.73E+00 | 9.06E-01 | no | | intergenic_variant | MODIFIER |
| 10 | 6721045 | AX-170477762 | 0 | T | A | 0.193 | -1.27E-02 | 1.06E-01 | 2.64E+00 | 2.73E+00 | 9.06E-01 | no | | intergenic_variant | MODIFIER |
| 10 | 6827971 | AX-170450415 | 0 | C | T | 0.193 | -1.27E-02 | 1.06E-01 | 2.64E+00 | 2.73E+00 | 9.06E-01 | no | Oprm1 | intron_variant | MODIFIER |
| 10 | 6831927 | AX-170529617 | 0 | C | T | 0.193 | -1.27E-02 | 1.06E-01 | 2.64E+00 | 2.73E+00 | 9.06E-01 | no | Oprm1 | intron_variant | MODIFIER |
| 10 | 6872843 | AX-170287528 | 0 | A | C | 0.193 | -1.27E-02 | 1.06E-01 | 2.64E+00 | 2.73E+00 | 9.06E-01 | no | Oprm1 | intron_variant | MODIFIER |
| 10 | 6928396 | AX-170478509 | 0 | T | C | 0.193 | -1.27E-02 | 1.06E-01 | 2.64E+00 | 2.73E+00 | 9.06E-01 | no | Oprm1 | intron_variant | MODIFIER |
| 10 | 6997249 | AX-170300412 | 0 | A | G | 0.193 | -1.27E-02 | 1.06E-01 | 2.64E+00 | 2.73E+00 | 9.06E-01 | no | Oprm1 | intron_variant | MODIFIER |
| 10 | 7003541 | AX-170470382 | 0 | C | T | 0.193 | -1.27E-02 | 1.06E-01 | 2.64E+00 | 2.73E+00 | 9.06E-01 | no | Oprm1 | intron_variant | MODIFIER |
| 10 | 7118376 | AX-170368556 | 0 | G | A | 0.193 | -1.27E-02 | 1.06E-01 | 2.64E+00 | 2.73E+00 | 9.06E-01 | no | Cnksr3 | downstream_gene_variant | MODIFIER |
| 10 | 7130757 | AX-170174305 | 0 | A | T | 0.193 | -1.27E-02 | 1.06E-01 | 2.64E+00 | 2.73E+00 | 9.06E-01 | no | Cnksr3 | intron_variant | MODIFIER |
| 10 | 7166209 | AX-170369709 | 0 | C | T | 0.193 | -1.27E-02 | 1.06E-01 | 2.64E+00 | 2.73E+00 | 9.06E-01 | no | Cnksr3 | intron_variant | MODIFIER |
| 10 | 7166728 | AX-170113865 | 0 | G | A | 0.193 | -1.27E-02 | 1.06E-01 | 2.64E+00 | 2.73E+00 | 9.06E-01 | no | Cnksr3 | intron_variant | MODIFIER |
| 10 | 7192162 | AX-170269595 | 0 | A | G | 0.193 | -1.27E-02 | 1.06E-01 | 2.64E+00 | 2.73E+00 | 9.06E-01 | no | Cnksr3 | intron_variant | MODIFIER |
| 10 | 7195606 | AX-170153041 | 0 | C | T | 0.193 | -1.27E-02 | 1.06E-01 | 2.64E+00 | 2.73E+00 | 9.06E-01 | no | Cnksr3 | intron_variant | MODIFIER |
| 10 | 7202758 | AX-170143371 | 0 | G | A | 0.193 | -1.27E-02 | 1.06E-01 | 2.64E+00 | 2.73E+00 | 9.06E-01 | no | Cnksr3 | intron_variant | MODIFIER |
| 10 | 7204685 | AX-170121223 | 0 | A | G | 0.193 | -1.27E-02 | 1.06E-01 | 2.64E+00 | 2.73E+00 | 9.06E-01 | no | Cnksr3 | intron_variant | MODIFIER |
| 10 | 7205039 | AX-170306446 | 0 | G | A | 0.193 | -1.27E-02 | 1.06E-01 | 2.64E+00 | 2.73E+00 | 9.06E-01 | no | Cnksr3 | intron_variant | MODIFIER |
| 10 | 7229520 | AX-170425670 | 0 | A | G | 0.193 | -1.27E-02 | 1.06E-01 | 2.64E+00 | 2.73E+00 | 9.06E-01 | no | | intergenic_variant | MODIFIER |
| 10 | 7251127 | AX-170478015 | 0 | C | T | 0.193 | -1.27E-02 | 1.06E-01 | 2.64E+00 | 2.73E+00 | 9.06E-01 | no | | intergenic_variant | MODIFIER |
| 10 | 7298352 | AX-170271585 | 0 | T | C | 0.193 | -1.27E-02 | 1.06E-01 | 2.64E+00 | 2.73E+00 | 9.06E-01 | no | | regulatory_region_variant | MODIFIER |
| 10 | 7595344 | AX-170241268 | 0 | T | C | 0.193 | -1.27E-02 | 1.06E-01 | 2.64E+00 | 2.73E+00 | 9.06E-01 | no | Lrp11 | intron_variant | MODIFIER |
| 10 | 7826604 | AX-170269143 | 0 | A | G | 0.193 | -1.27E-02 | 1.06E-01 | 2.64E+00 | 2.73E+00 | 9.06E-01 | no | Ppil4 | downstream_gene_variant | MODIFIER |
| 10 | 8614052 | AX-170208678 | 0 | A | G | 0.193 | -1.27E-02 | 1.06E-01 | 2.64E+00 | 2.73E+00 | 9.06E-01 | no | | intergenic_variant | MODIFIER |
| 10 | 9097279 | AX-97831498 | 0 | C | A | 0.193 | -1.27E-02 | 1.06E-01 | 2.64E+00 | 2.73E+00 | 9.06E-01 | no | | intergenic_variant | MODIFIER |
| 10 | 9099986 | AX-170449141 | 0 | A | G | 0.193 | -1.27E-02 | 1.06E-01 | 2.64E+00 | 2.73E+00 | 9.06E-01 | no | | intergenic_variant | MODIFIER |
| 10 | 9127809 | AX-170433202 | 0 | T | A | 0.193 | -1.27E-02 | 1.06E-01 | 2.64E+00 | 2.73E+00 | 9.06E-01 | no | | intergenic_variant | MODIFIER |
| 10 | 9128248 | AX-97850574 | 0 | G | T | 0.193 | -1.27E-02 | 1.06E-01 | 2.64E+00 | 2.73E+00 | 9.06E-01 | no | | intergenic_variant | MODIFIER |
| 10 | 9158727 | AX-170250860 | 0 | A | G | 0.193 | -1.27E-02 | 1.06E-01 | 2.64E+00 | 2.73E+00 | 9.06E-01 | no | | intergenic_variant | MODIFIER |
| 18 | 78284822 | AX-97843519 | 0 | T | C | 0.175 | 1.22E-02 | 1.16E-01 | 2.70E+00 | 2.78E+00 | 9.06E-01 | no | Slc14a2 | intron_variant | MODIFIER |
| 1 | 101955425 | AX-170042208 | 1 | G | T | 0.152 | 1.60E-02 | 1.38E-01 | 2.61E+00 | 2.74E+00 | 9.06E-01 | no | | intergenic_variant | MODIFIER |
| 8 | 91014911 | AX-170395305 | 2 | G | T | 0.191 | -1.45E-02 | 1.19E-01 | 2.64E+00 | 2.73E+00 | 9.06E-01 | no | Chd9 | intron_variant | MODIFIER |
| 15 | 101879006 | AX-170302513 | 0 | G | T | 0.114 | 1.77E-02 | 1.49E-01 | 2.66E+00 | 2.73E+00 | 9.06E-01 | no | | intergenic_variant | MODIFIER |
| 15 | 5721422 | AX-170111371 | 0 | C | A | 0.254 | -1.30E-02 | 9.90E-02 | 2.62E+00 | 2.71E+00 | 9.06E-01 | no | | intergenic_variant | MODIFIER |

| | | | | | | | | | | | | | | | |
|---|---|---|---|---|---|---|---|---|---|---|---|---|---|---|---|
| 5 | 88439497 | AX-170270280 | 2 | G | T | 0.445 | 1.26E-02 | 1.07E-01 | 2.67E+00 | 2.73E+00 | 9.06E-01 | no | | intergenic_variant | MODIFIER |
| 9 | 57132114 | AX-97805632 | 0 | C | T | 0.219 | -1.39E-02 | 1.22E-01 | 2.69E+00 | 2.74E+00 | 9.07E-01 | no | Man2c1 | intron_variant | MODIFIER |
| 9 | 57252260 | AX-97850326 | 0 | C | T | 0.219 | -1.39E-02 | 1.22E-01 | 2.69E+00 | 2.74E+00 | 9.07E-01 | no | 1700017B05Rik | downstream_gene_variant | MODIFIER |
| 9 | 57380612 | AX-170430705 | 0 | G | C | 0.219 | -1.39E-02 | 1.22E-01 | 2.69E+00 | 2.74E+00 | 9.07E-01 | no | Ppcdc | downstream_gene_variant | MODIFIER |
| 9 | 57402258 | AX-170121383 | 0 | G | C | 0.219 | -1.39E-02 | 1.22E-01 | 2.69E+00 | 2.74E+00 | 9.07E-01 | no | Ppcdc | intron_variant | MODIFIER |
| 9 | 57415204 | AX-170397551 | 0 | G | A | 0.219 | -1.39E-02 | 1.22E-01 | 2.69E+00 | 2.74E+00 | 9.07E-01 | no | Ppcdc | synonymous_variant | LOW |
| 9 | 57422116 | AX-170474913 | 0 | T | C | 0.219 | -1.39E-02 | 1.22E-01 | 2.69E+00 | 2.74E+00 | 9.07E-01 | no | Ppcdc | intron_variant | MODIFIER |
| 9 | 57424073 | AX-170155197 | 0 | T | C | 0.219 | -1.39E-02 | 1.22E-01 | 2.69E+00 | 2.74E+00 | 9.07E-01 | no | Ppcdc | intron_variant | MODIFIER |
| 9 | 57424140 | AX-170484434 | 0 | G | A | 0.219 | -1.39E-02 | 1.22E-01 | 2.69E+00 | 2.74E+00 | 9.07E-01 | no | Ppcdc | intron_variant | MODIFIER |
| 9 | 57453299 | AX-97815555 | 0 | G | A | 0.219 | -1.39E-02 | 1.22E-01 | 2.69E+00 | 2.74E+00 | 9.07E-01 | no | Scamp5 | intron_variant | MODIFIER |
| 9 | 57726660 | AX-170187855 | 0 | A | G | 0.219 | -1.39E-02 | 1.22E-01 | 2.69E+00 | 2.74E+00 | 9.07E-01 | no | Edc3 | intron_variant | MODIFIER |
| 9 | 57740083 | AX-170402194 | 0 | C | T | 0.219 | -1.39E-02 | 1.22E-01 | 2.69E+00 | 2.74E+00 | 9.07E-01 | no | Edc3 | intron_variant | MODIFIER |
| 9 | 57793475 | AX-170179539 | 0 | G | A | 0.219 | -1.39E-02 | 1.22E-01 | 2.69E+00 | 2.74E+00 | 9.07E-01 | no | Arid3b | intron_variant | MODIFIER |
| 9 | 57810984 | AX-170264294 | 0 | A | T | 0.219 | -1.39E-02 | 1.22E-01 | 2.69E+00 | 2.74E+00 | 9.07E-01 | no | Arid3b | intron_variant | MODIFIER |
| 9 | 57811480 | AX-170103248 | 0 | G | A | 0.219 | -1.39E-02 | 1.22E-01 | 2.69E+00 | 2.74E+00 | 9.07E-01 | no | Arid3b | intron_variant | MODIFIER |
| 9 | 57961169 | AX-170236008 | 0 | A | G | 0.219 | -1.39E-02 | 1.22E-01 | 2.69E+00 | 2.74E+00 | 9.07E-01 | no | Sema7a | intron_variant | MODIFIER |
| 9 | 57978235 | AX-170385036 | 0 | G | A | 0.219 | -1.39E-02 | 1.22E-01 | 2.69E+00 | 2.74E+00 | 9.07E-01 | no | | regulatory_region_variant | MODIFIER |
| 9 | 57987324 | AX-170403048 | 0 | A | G | 0.219 | -1.39E-02 | 1.22E-01 | 2.69E+00 | 2.74E+00 | 9.07E-01 | no | | regulatory_region_variant | MODIFIER |
| 9 | 58296260 | AX-97832492 | 0 | C | T | 0.219 | -1.39E-02 | 1.22E-01 | 2.69E+00 | 2.74E+00 | 9.07E-01 | no | Loxl1 | intron_variant | MODIFIER |
| 9 | 58459538 | AX-170303192 | 0 | C | T | 0.219 | -1.39E-02 | 1.22E-01 | 2.69E+00 | 2.74E+00 | 9.07E-01 | no | 4930461G14Rik | intron_variant,non_coding_transcript_variant | MODIFIER |
| 9 | 58468462 | AX-97822267 | 0 | C | T | 0.219 | -1.39E-02 | 1.22E-01 | 2.69E+00 | 2.74E+00 | 9.07E-01 | no | 4930461G14Rik | intron_variant,non_coding_transcript_variant | MODIFIER |
| 9 | 60091229 | AX-170166537 | 0 | C | T | 0.219 | -1.39E-02 | 1.22E-01 | 2.69E+00 | 2.74E+00 | 9.07E-01 | no | Thsd4 | intron_variant | MODIFIER |
| 9 | 60092490 | AX-170499078 | 0 | A | C | 0.219 | -1.39E-02 | 1.22E-01 | 2.69E+00 | 2.74E+00 | 9.07E-01 | no | Thsd4 | intron_variant | MODIFIER |
| 9 | 60094268 | AX-170176118 | 0 | T | C | 0.219 | -1.39E-02 | 1.22E-01 | 2.69E+00 | 2.74E+00 | 9.07E-01 | no | Thsd4 | intron_variant | MODIFIER |
| 9 | 60095762 | AX-170156959 | 0 | C | T | 0.219 | -1.39E-02 | 1.22E-01 | 2.69E+00 | 2.74E+00 | 9.07E-01 | no | Thsd4 | intron_variant | MODIFIER |
| 9 | 60096093 | AX-170302728 | 0 | T | C | 0.219 | -1.39E-02 | 1.22E-01 | 2.69E+00 | 2.74E+00 | 9.07E-01 | no | Thsd4 | intron_variant | MODIFIER |
| 9 | 60098095 | AX-97787402 | 0 | A | G | 0.219 | -1.39E-02 | 1.22E-01 | 2.69E+00 | 2.74E+00 | 9.07E-01 | no | Thsd4 | intron_variant | MODIFIER |
| 9 | 60098873 | AX-170221555 | 0 | T | C | 0.219 | -1.39E-02 | 1.22E-01 | 2.69E+00 | 2.74E+00 | 9.07E-01 | no | Thsd4 | intron_variant | MODIFIER |
| 9 | 60109949 | AX-170456819 | 0 | T | C | 0.219 | -1.39E-02 | 1.22E-01 | 2.69E+00 | 2.74E+00 | 9.07E-01 | no | Thsd4 | intron_variant | MODIFIER |
| 9 | 60115276 | AX-97795551 | 0 | C | T | 0.219 | -1.39E-02 | 1.22E-01 | 2.69E+00 | 2.74E+00 | 9.07E-01 | no | Thsd4 | intron_variant | MODIFIER |
| 9 | 60115507 | AX-170495796 | 0 | A | G | 0.219 | -1.39E-02 | 1.22E-01 | 2.69E+00 | 2.74E+00 | 9.07E-01 | no | Thsd4 | intron_variant | MODIFIER |
| 9 | 60118411 | AX-170195846 | 0 | T | A | 0.219 | -1.39E-02 | 1.22E-01 | 2.69E+00 | 2.74E+00 | 9.07E-01 | no | Thsd4 | intron_variant | MODIFIER |
| 9 | 60130215 | AX-170274205 | 0 | A | G | 0.219 | -1.39E-02 | 1.22E-01 | 2.69E+00 | 2.74E+00 | 9.07E-01 | no | Thsd4 | intron_variant | MODIFIER |
| 9 | 60138847 | AX-97821372 | 0 | T | G | 0.219 | -1.39E-02 | 1.22E-01 | 2.69E+00 | 2.74E+00 | 9.07E-01 | no | Thsd4 | intron_variant | MODIFIER |
| 4 | 58587314 | AX-170491551 | 1 | G | A | 0.384 | -1.19E-02 | 1.08E-01 | 2.68E+00 | 2.76E+00 | 9.07E-01 | no | Gm12578 | upstream_gene_variant | MODIFIER |
| 7 | 112316348 | AX-170078444 | 1 | T | C | 0.446 | 1.10E-02 | 9.28E-02 | 2.67E+00 | 2.73E+00 | 9.07E-01 | no | Mical2 | intron_variant | MODIFIER |
| 18 | 42420738 | AX-170060680 | 0 | A | G | 0.07 | 2.62E-02 | 2.46E-01 | 2.67E+00 | 2.76E+00 | 9.07E-01 | no | | intergenic_variant | MODIFIER |
| 9 | 79268622 | AX-170082898 | 1 | G | A | 0.205 | 1.44E-02 | 1.29E-01 | 2.71E+00 | 2.76E+00 | 9.07E-01 | no | Gm47500 | downstream_gene_variant | MODIFIER |
| 7 | 89775125 | AX-170209404 | 0 | G | A | 0.132 | 2.42E-02 | 2.11E-01 | 2.66E+00 | 2.74E+00 | 9.07E-01 | no | Me3 | intron_variant | MODIFIER |
| 5 | 118888527 | AX-170089592 | 0 | A | T | 0.447 | -1.29E-02 | 1.14E-01 | 2.72E+00 | 2.76E+00 | 9.07E-01 | no | | intergenic_variant | MODIFIER |
| 5 | 118896270 | AX-170120769 | 0 | T | C | 0.447 | -1.29E-02 | 1.14E-01 | 2.72E+00 | 2.76E+00 | 9.07E-01 | no | | intergenic_variant | MODIFIER |
| 5 | 119074345 | AX-170240674 | 0 | C | T | 0.447 | -1.29E-02 | 1.14E-01 | 2.72E+00 | 2.76E+00 | 9.07E-01 | no | | intergenic_variant | MODIFIER |
| 5 | 119076653 | AX-170229158 | 0 | C | T | 0.447 | -1.29E-02 | 1.14E-01 | 2.72E+00 | 2.76E+00 | 9.07E-01 | no | | intergenic_variant | MODIFIER |
| 5 | 119080443 | AX-170339492 | 0 | T | C | 0.447 | -1.29E-02 | 1.14E-01 | 2.72E+00 | 2.76E+00 | 9.07E-01 | no | | intergenic_variant | MODIFIER |
| 5 | 119080929 | AX-97805980 | 0 | C | T | 0.447 | -1.29E-02 | 1.14E-01 | 2.72E+00 | 2.76E+00 | 9.07E-01 | no | | intergenic_variant | MODIFIER |
| 5 | 119106354 | AX-170025996 | 0 | C | G | 0.447 | -1.29E-02 | 1.14E-01 | 2.72E+00 | 2.76E+00 | 9.07E-01 | no | 1700081H04Rik | upstream_gene_variant | MODIFIER |
| 5 | 119110201 | AX-170346081 | 0 | C | G | 0.447 | -1.29E-02 | 1.14E-01 | 2.72E+00 | 2.76E+00 | 9.07E-01 | no | 1700081H04Rik | intron_variant,non_coding_transcript_variant | MODIFIER |
| 5 | 119134537 | AX-97790047 | 0 | G | A | 0.447 | -1.29E-02 | 1.14E-01 | 2.72E+00 | 2.76E+00 | 9.07E-01 | no | | intergenic_variant | MODIFIER |
| 5 | 119145614 | AX-170459373 | 0 | T | C | 0.447 | -1.29E-02 | 1.14E-01 | 2.72E+00 | 2.76E+00 | 9.07E-01 | no | | intergenic_variant | MODIFIER |
| 5 | 119145752 | AX-170525806 | 0 | G | A | 0.447 | -1.29E-02 | 1.14E-01 | 2.72E+00 | 2.76E+00 | 9.07E-01 | no | | intergenic_variant | MODIFIER |
| 5 | 119151794 | AX-170171724 | 0 | T | C | 0.447 | -1.29E-02 | 1.14E-01 | 2.72E+00 | 2.76E+00 | 9.07E-01 | no | | intergenic_variant | MODIFIER |
| 5 | 119152527 | AX-170218641 | 0 | T | C | 0.447 | -1.29E-02 | 1.14E-01 | 2.72E+00 | 2.76E+00 | 9.07E-01 | no | | intergenic_variant | MODIFIER |
| 5 | 119153290 | AX-170451462 | 0 | T | C | 0.447 | -1.29E-02 | 1.14E-01 | 2.72E+00 | 2.76E+00 | 9.07E-01 | no | | regulatory_region_variant | MODIFIER |
| 5 | 119192038 | AX-97815775 | 0 | T | C | 0.447 | -1.29E-02 | 1.14E-01 | 2.72E+00 | 2.76E+00 | 9.07E-01 | no | Gm7538 | downstream_gene_variant | MODIFIER |

| | | | | | | | | | | | | | | | |
|---|---|---|---|---|---|---|---|---|---|---|---|---|---|---|---|
| 5 | 119192134 | AX-170456397 | 0 | T | A | 0.447 | -1.29E-02 | 1.14E-01 | 2.72E+00 | 2.76E+00 | 9.07E-01 | no | Gm7538 | downstream_gene_variant | MODIFIER |
| 5 | 119192198 | AX-170181277 | 0 | T | C | 0.447 | -1.29E-02 | 1.14E-01 | 2.72E+00 | 2.76E+00 | 9.07E-01 | no | Gm7538 | downstream_gene_variant | MODIFIER |
| 5 | 119207047 | AX-170089593 | 0 | C | T | 0.447 | -1.29E-02 | 1.14E-01 | 2.72E+00 | 2.76E+00 | 9.07E-01 | no | | intergenic_variant | MODIFIER |
| 5 | 119231816 | AX-170303303 | 0 | A | G | 0.447 | -1.29E-02 | 1.14E-01 | 2.72E+00 | 2.76E+00 | 9.07E-01 | no | | intergenic_variant | MODIFIER |
| 5 | 119237664 | AX-97782645 | 0 | T | G | 0.447 | -1.29E-02 | 1.14E-01 | 2.72E+00 | 2.76E+00 | 9.07E-01 | no | | intergenic_variant | MODIFIER |
| 5 | 119241205 | AX-170495956 | 0 | C | A | 0.447 | -1.29E-02 | 1.14E-01 | 2.72E+00 | 2.76E+00 | 9.07E-01 | no | | intergenic_variant | MODIFIER |
| 5 | 119277019 | AX-97790214 | 0 | G | A | 0.447 | -1.29E-02 | 1.14E-01 | 2.72E+00 | 2.76E+00 | 9.07E-01 | no | | intergenic_variant | MODIFIER |
| 5 | 119299666 | AX-170368754 | 0 | T | C | 0.447 | -1.29E-02 | 1.14E-01 | 2.72E+00 | 2.76E+00 | 9.07E-01 | no | n-R5s175 | downstream_gene_variant | MODIFIER |
| 5 | 119302221 | AX-170123406 | 0 | A | C | 0.447 | -1.29E-02 | 1.14E-01 | 2.72E+00 | 2.76E+00 | 9.07E-01 | no | n-R5s175 | downstream_gene_variant | MODIFIER |
| 5 | 119304923 | AX-170297114 | 0 | A | T | 0.447 | -1.29E-02 | 1.14E-01 | 2.72E+00 | 2.76E+00 | 9.07E-01 | no | | intergenic_variant | MODIFIER |
| 5 | 119324154 | AX-170433221 | 0 | G | T | 0.447 | -1.29E-02 | 1.14E-01 | 2.72E+00 | 2.76E+00 | 9.07E-01 | no | | intergenic_variant | MODIFIER |
| 5 | 119428033 | AX-170132686 | 0 | G | A | 0.447 | -1.29E-02 | 1.14E-01 | 2.72E+00 | 2.76E+00 | 9.07E-01 | no | | regulatory_region_variant | MODIFIER |
| 5 | 119563696 | AX-170378251 | 0 | T | G | 0.447 | -1.29E-02 | 1.14E-01 | 2.72E+00 | 2.76E+00 | 9.07E-01 | no | | regulatory_region_variant | MODIFIER |
| 5 | 119580870 | AX-97821335 | 0 | A | G | 0.447 | -1.29E-02 | 1.14E-01 | 2.72E+00 | 2.76E+00 | 9.07E-01 | no | Tbx3os1 | intron_variant,non_coding_transcript_variant | MODIFIER |
| 5 | 119644016 | AX-170245292 | 0 | G | A | 0.447 | -1.29E-02 | 1.14E-01 | 2.72E+00 | 2.76E+00 | 9.07E-01 | no | Tbx3os1 | intron_variant,non_coding_transcript_variant | MODIFIER |
| 5 | 119654026 | AX-170288492 | 0 | T | G | 0.447 | -1.29E-02 | 1.14E-01 | 2.72E+00 | 2.76E+00 | 9.07E-01 | no | Tbx3os1 | intron_variant,non_coding_transcript_variant | MODIFIER |
| 5 | 119846847 | AX-170488960 | 0 | C | T | 0.447 | -1.29E-02 | 1.14E-01 | 2.72E+00 | 2.76E+00 | 9.07E-01 | no | Tbx5 | intron_variant | MODIFIER |
| 5 | 119847440 | AX-170364061 | 0 | C | T | 0.447 | -1.29E-02 | 1.14E-01 | 2.72E+00 | 2.76E+00 | 9.07E-01 | no | Tbx5 | intron_variant | MODIFIER |
| 5 | 119847995 | AX-97822140 | 0 | T | G | 0.447 | -1.29E-02 | 1.14E-01 | 2.72E+00 | 2.76E+00 | 9.07E-01 | no | Tbx5 | intron_variant | MODIFIER |
| 5 | 119869426 | AX-170222773 | 0 | T | C | 0.447 | -1.29E-02 | 1.14E-01 | 2.72E+00 | 2.76E+00 | 9.07E-01 | no | Tbx5 | intron_variant | MODIFIER |
| 5 | 119869488 | AX-170358319 | 0 | G | A | 0.447 | -1.29E-02 | 1.14E-01 | 2.72E+00 | 2.76E+00 | 9.07E-01 | no | Tbx5 | intron_variant | MODIFIER |
| 5 | 119873381 | AX-170332975 | 0 | C | T | 0.447 | -1.29E-02 | 1.14E-01 | 2.72E+00 | 2.76E+00 | 9.07E-01 | no | Tbx5 | intron_variant | MODIFIER |
| 5 | 119914324 | AX-97813857 | 0 | A | G | 0.447 | -1.29E-02 | 1.14E-01 | 2.72E+00 | 2.76E+00 | 9.07E-01 | no | Gm43269 | upstream_gene_variant | MODIFIER |
| 5 | 119920872 | AX-170507347 | 0 | G | A | 0.447 | -1.29E-02 | 1.14E-01 | 2.72E+00 | 2.76E+00 | 9.07E-01 | no | Gm43269 | downstream_gene_variant | MODIFIER |
| 5 | 119921890 | AX-170347770 | 0 | T | C | 0.447 | -1.29E-02 | 1.14E-01 | 2.72E+00 | 2.76E+00 | 9.07E-01 | no | | intergenic_variant | MODIFIER |
| 5 | 119923560 | AX-97848146 | 0 | A | C | 0.447 | -1.29E-02 | 1.14E-01 | 2.72E+00 | 2.76E+00 | 9.07E-01 | no | | intergenic_variant | MODIFIER |
| 5 | 119960513 | AX-170494523 | 0 | G | A | 0.447 | -1.29E-02 | 1.14E-01 | 2.72E+00 | 2.76E+00 | 9.07E-01 | no | Gm38554 | intron_variant,non_coding_transcript_variant | MODIFIER |
| 5 | 119971322 | AX-170299807 | 0 | C | T | 0.447 | -1.29E-02 | 1.14E-01 | 2.72E+00 | 2.76E+00 | 9.07E-01 | no | | intergenic_variant | MODIFIER |
| 5 | 119977440 | AX-170116889 | 0 | C | T | 0.447 | -1.29E-02 | 1.14E-01 | 2.72E+00 | 2.76E+00 | 9.07E-01 | no | | intergenic_variant | MODIFIER |
| 5 | 119979175 | AX-170465849 | 0 | T | C | 0.447 | -1.29E-02 | 1.14E-01 | 2.72E+00 | 2.76E+00 | 9.07E-01 | no | | intergenic_variant | MODIFIER |
| 5 | 119982362 | AX-170130255 | 0 | A | G | 0.447 | -1.29E-02 | 1.14E-01 | 2.72E+00 | 2.76E+00 | 9.07E-01 | no | | intergenic_variant | MODIFIER |
| 5 | 119988952 | AX-170186741 | 0 | C | T | 0.447 | -1.29E-02 | 1.14E-01 | 2.72E+00 | 2.76E+00 | 9.07E-01 | no | | intergenic_variant | MODIFIER |
| 5 | 119993341 | AX-170073299 | 0 | C | T | 0.447 | -1.29E-02 | 1.14E-01 | 2.72E+00 | 2.76E+00 | 9.07E-01 | no | | intergenic_variant | MODIFIER |
| 5 | 119993386 | AX-170073297 | 0 | T | A | 0.447 | -1.29E-02 | 1.14E-01 | 2.72E+00 | 2.76E+00 | 9.07E-01 | no | | intergenic_variant | MODIFIER |
| 5 | 119994643 | AX-170073298 | 0 | G | A | 0.447 | -1.29E-02 | 1.14E-01 | 2.72E+00 | 2.76E+00 | 9.07E-01 | no | | intergenic_variant | MODIFIER |
| 5 | 119994694 | AX-170073301 | 0 | A | G | 0.447 | -1.29E-02 | 1.14E-01 | 2.72E+00 | 2.76E+00 | 9.07E-01 | no | | intergenic_variant | MODIFIER |
| 5 | 119995876 | AX-170145503 | 0 | A | C | 0.447 | -1.29E-02 | 1.14E-01 | 2.72E+00 | 2.76E+00 | 9.07E-01 | no | | intergenic_variant | MODIFIER |
| 5 | 119997641 | AX-170125625 | 0 | T | C | 0.447 | -1.29E-02 | 1.14E-01 | 2.72E+00 | 2.76E+00 | 9.07E-01 | no | | intergenic_variant | MODIFIER |
| 5 | 119997762 | AX-170433034 | 0 | A | G | 0.447 | -1.29E-02 | 1.14E-01 | 2.72E+00 | 2.76E+00 | 9.07E-01 | no | | intergenic_variant | MODIFIER |
| 5 | 120012782 | AX-170475981 | 0 | C | T | 0.447 | -1.29E-02 | 1.14E-01 | 2.72E+00 | 2.76E+00 | 9.07E-01 | no | | regulatory_region_variant | MODIFIER |
| 5 | 120041346 | AX-170423476 | 0 | A | T | 0.447 | -1.29E-02 | 1.14E-01 | 2.72E+00 | 2.76E+00 | 9.07E-01 | no | | intergenic_variant | MODIFIER |
| 5 | 120042428 | AX-168375966 | 0 | T | C | 0.447 | -1.29E-02 | 1.14E-01 | 2.72E+00 | 2.76E+00 | 9.07E-01 | no | | intergenic_variant | MODIFIER |
| 5 | 120044389 | AX-168397613 | 0 | G | A | 0.447 | -1.29E-02 | 1.14E-01 | 2.72E+00 | 2.76E+00 | 9.07E-01 | no | | intergenic_variant | MODIFIER |
| 5 | 120065073 | AX-170147179 | 0 | C | T | 0.447 | -1.29E-02 | 1.14E-01 | 2.72E+00 | 2.76E+00 | 9.07E-01 | no | n-R5s176 | downstream_gene_variant | MODIFIER |
| 5 | 120247756 | AX-170142354 | 0 | G | A | 0.447 | -1.29E-02 | 1.14E-01 | 2.72E+00 | 2.76E+00 | 9.07E-01 | no | | intergenic_variant | MODIFIER |
| 5 | 120248221 | AX-170026008 | 0 | C | T | 0.447 | -1.29E-02 | 1.14E-01 | 2.72E+00 | 2.76E+00 | 9.07E-01 | no | | intergenic_variant | MODIFIER |
| 5 | 120770199 | AX-97814028 | 0 | A | T | 0.447 | -1.29E-02 | 1.14E-01 | 2.72E+00 | 2.76E+00 | 9.07E-01 | no | Oas3 | intron_variant | MODIFIER |
| 5 | 120961807 | AX-97824862 | 0 | A | C | 0.447 | -1.29E-02 | 1.14E-01 | 2.72E+00 | 2.76E+00 | 9.07E-01 | no | Rph3a | intron_variant | MODIFIER |
| 5 | 120962870 | AX-170026019 | 0 | C | T | 0.447 | -1.29E-02 | 1.14E-01 | 2.72E+00 | 2.76E+00 | 9.07E-01 | no | Rph3a | intron_variant | MODIFIER |
| 5 | 120968746 | AX-170348670 | 0 | T | C | 0.447 | -1.29E-02 | 1.14E-01 | 2.72E+00 | 2.76E+00 | 9.07E-01 | no | Rph3a | intron_variant | MODIFIER |
| 5 | 120987022 | AX-97845456 | 0 | G | A | 0.447 | -1.29E-02 | 1.14E-01 | 2.72E+00 | 2.76E+00 | 9.07E-01 | no | Rph3a | intron_variant | MODIFIER |
| 5 | 120994117 | AX-170026027 | 0 | C | T | 0.447 | -1.29E-02 | 1.14E-01 | 2.72E+00 | 2.76E+00 | 9.07E-01 | no | Rph3a | intron_variant | MODIFIER |
| 5 | 121020821 | AX-170482886 | 0 | G | C | 0.447 | -1.29E-02 | 1.14E-01 | 2.72E+00 | 2.76E+00 | 9.07E-01 | no | | intergenic_variant | MODIFIER |
| 5 | 121022189 | AX-170116235 | 0 | A | G | 0.447 | -1.29E-02 | 1.14E-01 | 2.72E+00 | 2.76E+00 | 9.07E-01 | no | | intergenic_variant | MODIFIER |

| | | | | | | | | | | | | | | | |
|---|---|---|---|---|---|---|---|---|---|---|---|---|---|---|---|
| 5 | 121023093 | AX-170099985 | 0 | T | C | 0.447 | -1.29E-02 | 1.14E-01 | 2.72E+00 | 2.76E+00 | 9.07E-01 | no | | intergenic_variant | MODIFIER |
| 5 | 121034628 | AX-170095453 | 0 | G | T | 0.447 | -1.29E-02 | 1.14E-01 | 2.72E+00 | 2.76E+00 | 9.07E-01 | no | | intergenic_variant | MODIFIER |
| 5 | 121049198 | AX-170026018 | 0 | T | A | 0.447 | -1.29E-02 | 1.14E-01 | 2.72E+00 | 2.76E+00 | 9.07E-01 | no | | intergenic_variant | MODIFIER |
| 5 | 121049454 | AX-170026026 | 0 | T | C | 0.447 | -1.29E-02 | 1.14E-01 | 2.72E+00 | 2.76E+00 | 9.07E-01 | no | | intergenic_variant | MODIFIER |
| 5 | 121050460 | AX-97822640 | 0 | C | T | 0.447 | -1.29E-02 | 1.14E-01 | 2.72E+00 | 2.76E+00 | 9.07E-01 | no | | intergenic_variant | MODIFIER |
| 5 | 121078302 | AX-170026023 | 0 | T | G | 0.447 | -1.29E-02 | 1.14E-01 | 2.72E+00 | 2.76E+00 | 9.07E-01 | no | | intergenic_variant | MODIFIER |
| 5 | 121082821 | AX-170086122 | 0 | G | A | 0.447 | -1.29E-02 | 1.14E-01 | 2.72E+00 | 2.76E+00 | 9.07E-01 | no | | intergenic_variant | MODIFIER |
| 5 | 121097550 | AX-170157912 | 0 | G | A | 0.447 | -1.29E-02 | 1.14E-01 | 2.72E+00 | 2.76E+00 | 9.07E-01 | no | | intergenic_variant | MODIFIER |
| 5 | 121097731 | AX-97833614 | 0 | C | T | 0.447 | -1.29E-02 | 1.14E-01 | 2.72E+00 | 2.76E+00 | 9.07E-01 | no | | intergenic_variant | MODIFIER |
| 5 | 121097945 | AX-170437510 | 0 | T | C | 0.447 | -1.29E-02 | 1.14E-01 | 2.72E+00 | 2.76E+00 | 9.07E-01 | no | | intergenic_variant | MODIFIER |
| 5 | 121113696 | AX-170336404 | 0 | T | G | 0.447 | -1.29E-02 | 1.14E-01 | 2.72E+00 | 2.76E+00 | 9.07E-01 | no | | regulatory_region_variant | MODIFIER |
| 5 | 121113840 | AX-170174424 | 0 | T | C | 0.447 | -1.29E-02 | 1.14E-01 | 2.72E+00 | 2.76E+00 | 9.07E-01 | no | | regulatory_region_variant | MODIFIER |
| 5 | 121129606 | AX-170122644 | 0 | A | C | 0.447 | -1.29E-02 | 1.14E-01 | 2.72E+00 | 2.76E+00 | 9.07E-01 | no | Ptpn11 | downstream_gene_variant | MODIFIER |
| 5 | 121134649 | AX-97801887 | 0 | T | C | 0.447 | -1.29E-02 | 1.14E-01 | 2.72E+00 | 2.76E+00 | 9.07E-01 | no | Ptpn11 | intron_variant | MODIFIER |
| 5 | 121158347 | AX-97809910 | 0 | G | A | 0.447 | -1.29E-02 | 1.14E-01 | 2.72E+00 | 2.76E+00 | 9.07E-01 | no | Ptpn11 | intron_variant | MODIFIER |
| 5 | 121181629 | AX-97836176 | 0 | A | C | 0.447 | -1.29E-02 | 1.14E-01 | 2.72E+00 | 2.76E+00 | 9.07E-01 | no | Ptpn11 | intron_variant | MODIFIER |
| 5 | 121202482 | AX-170109771 | 0 | G | T | 0.447 | -1.29E-02 | 1.14E-01 | 2.72E+00 | 2.76E+00 | 9.07E-01 | no | Rpl6 | upstream_gene_variant | MODIFIER |
| 5 | 121208753 | AX-170089596 | 0 | G | A | 0.447 | -1.29E-02 | 1.14E-01 | 2.72E+00 | 2.76E+00 | 9.07E-01 | no | Rpl6 | intron_variant | MODIFIER |
| 5 | 121211521 | AX-170131805 | 0 | C | T | 0.447 | -1.29E-02 | 1.14E-01 | 2.72E+00 | 2.76E+00 | 9.07E-01 | no | Rpl6 | downstream_gene_variant | MODIFIER |
| 5 | 121227511 | AX-170325299 | 0 | T | C | 0.447 | -1.29E-02 | 1.14E-01 | 2.72E+00 | 2.76E+00 | 9.07E-01 | no | Hectd4 | intron_variant | MODIFIER |
| 5 | 121229389 | AX-168402711 | 0 | A | G | 0.447 | -1.29E-02 | 1.14E-01 | 2.72E+00 | 2.76E+00 | 9.07E-01 | no | Hectd4 | intron_variant | MODIFIER |
| 5 | 121229439 | AX-170472478 | 0 | C | T | 0.447 | -1.29E-02 | 1.14E-01 | 2.72E+00 | 2.76E+00 | 9.07E-01 | no | Hectd4 | intron_variant | MODIFIER |
| 5 | 121230977 | AX-97784551 | 0 | A | G | 0.447 | -1.29E-02 | 1.14E-01 | 2.72E+00 | 2.76E+00 | 9.07E-01 | no | Hectd4 | intron_variant | MODIFIER |
| 5 | 121252226 | AX-170316449 | 0 | G | A | 0.447 | -1.29E-02 | 1.14E-01 | 2.72E+00 | 2.76E+00 | 9.07E-01 | no | Hectd4 | intron_variant | MODIFIER |
| 5 | 121340942 | AX-97839128 | 0 | A | G | 0.447 | -1.29E-02 | 1.14E-01 | 2.72E+00 | 2.76E+00 | 9.07E-01 | no | Hectd4 | intron_variant | MODIFIER |
| 5 | 121342162 | AX-170186256 | 0 | G | T | 0.447 | -1.29E-02 | 1.14E-01 | 2.72E+00 | 2.76E+00 | 9.07E-01 | no | Hectd4 | intron_variant | MODIFIER |
| 5 | 121349563 | AX-170254456 | 0 | C | G | 0.447 | -1.29E-02 | 1.14E-01 | 2.72E+00 | 2.76E+00 | 9.07E-01 | no | Hectd4 | synonymous_variant | LOW |
| 5 | 121529173 | AX-97793512 | 0 | C | T | 0.447 | -1.29E-02 | 1.14E-01 | 2.72E+00 | 2.76E+00 | 9.07E-01 | no | Mapkapk5 | intron_variant | MODIFIER |
| 5 | 121644656 | AX-170377569 | 0 | T | C | 0.447 | -1.29E-02 | 1.14E-01 | 2.72E+00 | 2.76E+00 | 9.07E-01 | no | Acad10 | intron_variant | MODIFIER |
| 5 | 121646599 | AX-170210785 | 0 | T | C | 0.447 | -1.29E-02 | 1.14E-01 | 2.72E+00 | 2.76E+00 | 9.07E-01 | no | Acad10 | intron_variant | MODIFIER |
| 5 | 121647828 | AX-170089598 | 0 | A | G | 0.447 | -1.29E-02 | 1.14E-01 | 2.72E+00 | 2.76E+00 | 9.07E-01 | no | Acad10 | splice_region_variant,intron_variant | LOW |
| 5 | 121683972 | AX-170089599 | 0 | T | C | 0.447 | -1.29E-02 | 1.14E-01 | 2.72E+00 | 2.76E+00 | 9.07E-01 | no | Brap | intron_variant | MODIFIER |
| 5 | 121686488 | AX-170026030 | 0 | A | G | 0.447 | -1.29E-02 | 1.14E-01 | 2.72E+00 | 2.76E+00 | 9.07E-01 | no | Brap | 3_prime_UTR_variant | MODIFIER |
| 5 | 121692940 | AX-170108902 | 0 | T | A | 0.447 | -1.29E-02 | 1.14E-01 | 2.72E+00 | 2.76E+00 | 9.07E-01 | no | | intergenic_variant | MODIFIER |
| 5 | 121760010 | AX-97822041 | 0 | C | T | 0.447 | -1.29E-02 | 1.14E-01 | 2.72E+00 | 2.76E+00 | 9.07E-01 | no | Atxn2 | intron_variant | MODIFIER |
| 5 | 121766071 | AX-170026032 | 0 | A | G | 0.447 | -1.29E-02 | 1.14E-01 | 2.72E+00 | 2.76E+00 | 9.07E-01 | no | Atxn2 | intron_variant | MODIFIER |
| 5 | 121805149 | AX-170194067 | 0 | A | G | 0.447 | -1.29E-02 | 1.14E-01 | 2.72E+00 | 2.76E+00 | 9.07E-01 | no | Atxn2 | intron_variant | MODIFIER |
| 5 | 121805901 | AX-97849504 | 0 | T | C | 0.447 | -1.29E-02 | 1.14E-01 | 2.72E+00 | 2.76E+00 | 9.07E-01 | no | Atxn2 | intron_variant | MODIFIER |
| 5 | 121810127 | AX-170172337 | 0 | T | C | 0.447 | -1.29E-02 | 1.14E-01 | 2.72E+00 | 2.76E+00 | 9.07E-01 | no | Atxn2 | intron_variant | MODIFIER |
| 5 | 121917835 | AX-170122697 | 0 | A | G | 0.447 | -1.29E-02 | 1.14E-01 | 2.72E+00 | 2.76E+00 | 9.07E-01 | no | Cux2 | intron_variant | MODIFIER |
| 5 | 121937563 | AX-170311366 | 0 | A | G | 0.447 | -1.29E-02 | 1.14E-01 | 2.72E+00 | 2.76E+00 | 9.07E-01 | no | Cux2 | intron_variant | MODIFIER |
| 5 | 121938474 | AX-170156470 | 0 | T | C | 0.447 | -1.29E-02 | 1.14E-01 | 2.72E+00 | 2.76E+00 | 9.07E-01 | no | Cux2 | intron_variant | MODIFIER |
| 5 | 121943697 | AX-170317857 | 0 | T | A | 0.447 | -1.29E-02 | 1.14E-01 | 2.72E+00 | 2.76E+00 | 9.07E-01 | no | Cux2 | intron_variant | MODIFIER |
| 5 | 121959702 | AX-170339344 | 0 | T | C | 0.447 | -1.29E-02 | 1.14E-01 | 2.72E+00 | 2.76E+00 | 9.07E-01 | no | Cux2 | intron_variant | MODIFIER |
| 5 | 121960407 | AX-97855961 | 0 | A | G | 0.447 | -1.29E-02 | 1.14E-01 | 2.72E+00 | 2.76E+00 | 9.07E-01 | no | Cux2 | intron_variant | MODIFIER |
| 5 | 121960450 | AX-170413887 | 0 | C | T | 0.447 | -1.29E-02 | 1.14E-01 | 2.72E+00 | 2.76E+00 | 9.07E-01 | no | Cux2 | intron_variant | MODIFIER |
| 5 | 121960498 | AX-170366912 | 0 | C | G | 0.447 | -1.29E-02 | 1.14E-01 | 2.72E+00 | 2.76E+00 | 9.07E-01 | no | Cux2 | intron_variant | MODIFIER |
| 5 | 121961042 | AX-170358738 | 0 | G | A | 0.447 | -1.29E-02 | 1.14E-01 | 2.72E+00 | 2.76E+00 | 9.07E-01 | no | Cux2 | intron_variant | MODIFIER |
| 5 | 121962113 | AX-170293540 | 0 | T | C | 0.447 | -1.29E-02 | 1.14E-01 | 2.72E+00 | 2.76E+00 | 9.07E-01 | no | Cux2 | intron_variant | MODIFIER |
| 5 | 121986078 | AX-170141096 | 0 | A | C | 0.447 | -1.29E-02 | 1.14E-01 | 2.72E+00 | 2.76E+00 | 9.07E-01 | no | Cux2 | intron_variant | MODIFIER |
| 5 | 121991630 | AX-170182439 | 0 | T | C | 0.447 | -1.29E-02 | 1.14E-01 | 2.72E+00 | 2.76E+00 | 9.07E-01 | no | Cux2 | intron_variant | MODIFIER |
| 5 | 122003880 | AX-170073334 | 0 | T | C | 0.447 | -1.29E-02 | 1.14E-01 | 2.72E+00 | 2.76E+00 | 9.07E-01 | no | Cux2 | intron_variant | MODIFIER |
| 5 | 122004784 | AX-170073332 | 0 | A | C | 0.447 | -1.29E-02 | 1.14E-01 | 2.72E+00 | 2.76E+00 | 9.07E-01 | no | Cux2 | intron_variant | MODIFIER |
| 5 | 122026587 | AX-170387148 | 0 | C | A | 0.447 | -1.29E-02 | 1.14E-01 | 2.72E+00 | 2.76E+00 | 9.07E-01 | no | Cux2 | intron_variant | MODIFIER |

| | | | | | | | | | | | | | | | |
|---|---|---|---|---|---|---|---|---|---|---|---|---|---|---|---|
| 5 | 122039104 | AX-170267882 | 0 | T | A | 0.447 | -1.29E-02 | 1.14E-01 | 2.72E+00 | 2.76E+00 | 9.07E-01 | no | Cux2 | intron_variant | MODIFIER |
| 5 | 122092337 | AX-170218345 | 0 | G | A | 0.447 | -1.29E-02 | 1.14E-01 | 2.72E+00 | 2.76E+00 | 9.07E-01 | no | | regulatory_region_variant | MODIFIER |
| 5 | 122104821 | AX-170294308 | 0 | G | T | 0.447 | -1.29E-02 | 1.14E-01 | 2.72E+00 | 2.76E+00 | 9.07E-01 | no | Myl2 | splice_region_variant,intron_variant | LOW |
| 5 | 122457945 | AX-168379195 | 0 | A | G | 0.447 | -1.29E-02 | 1.14E-01 | 2.72E+00 | 2.76E+00 | 9.07E-01 | no | Atp2a2 | intron_variant | MODIFIER |
| 5 | 122801309 | AX-170456179 | 0 | T | G | 0.447 | -1.29E-02 | 1.14E-01 | 2.72E+00 | 2.76E+00 | 9.07E-01 | no | Anapc5 | intron_variant | MODIFIER |
| 5 | 122813195 | AX-170026048 | 0 | C | T | 0.447 | -1.29E-02 | 1.14E-01 | 2.72E+00 | 2.76E+00 | 9.07E-01 | no | Anapc5 | intron_variant | MODIFIER |
| 5 | 122862168 | AX-97829801 | 0 | G | T | 0.447 | -1.29E-02 | 1.14E-01 | 2.72E+00 | 2.76E+00 | 9.07E-01 | no | Rnf34 | intron_variant | MODIFIER |
| 5 | 122867806 | AX-170485047 | 0 | C | T | 0.447 | -1.29E-02 | 1.14E-01 | 2.72E+00 | 2.76E+00 | 9.07E-01 | no | Rnf34 | intron_variant | MODIFIER |
| 5 | 122874759 | AX-170505060 | 0 | C | T | 0.447 | -1.29E-02 | 1.14E-01 | 2.72E+00 | 2.76E+00 | 9.07E-01 | no | Rnf34 | downstream_gene_variant | MODIFIER |
| 5 | 122890352 | AX-170478819 | 0 | T | C | 0.447 | -1.29E-02 | 1.14E-01 | 2.72E+00 | 2.76E+00 | 9.07E-01 | no | Kdm2b | intron_variant | MODIFIER |
| 5 | 122894599 | AX-170181143 | 0 | G | A | 0.447 | -1.29E-02 | 1.14E-01 | 2.72E+00 | 2.76E+00 | 9.07E-01 | no | Kdm2b | intron_variant | MODIFIER |
| 5 | 122899042 | AX-97788230 | 0 | G | A | 0.447 | -1.29E-02 | 1.14E-01 | 2.72E+00 | 2.76E+00 | 9.07E-01 | no | Kdm2b | intron_variant | MODIFIER |
| 5 | 122912034 | AX-170142748 | 0 | A | G | 0.447 | -1.29E-02 | 1.14E-01 | 2.72E+00 | 2.76E+00 | 9.07E-01 | no | Kdm2b | intron_variant | MODIFIER |
| 5 | 122916136 | AX-170164238 | 0 | A | G | 0.447 | -1.29E-02 | 1.14E-01 | 2.72E+00 | 2.76E+00 | 9.07E-01 | no | Kdm2b | intron_variant | MODIFIER |
| 5 | 122918685 | AX-170307303 | 0 | C | G | 0.447 | -1.29E-02 | 1.14E-01 | 2.72E+00 | 2.76E+00 | 9.07E-01 | no | Kdm2b | intron_variant | MODIFIER |
| 5 | 122924132 | AX-170267120 | 0 | G | A | 0.447 | -1.29E-02 | 1.14E-01 | 2.72E+00 | 2.76E+00 | 9.07E-01 | no | Kdm2b | intron_variant | MODIFIER |
| 5 | 122925636 | AX-170277886 | 0 | T | C | 0.447 | -1.29E-02 | 1.14E-01 | 2.72E+00 | 2.76E+00 | 9.07E-01 | no | Kdm2b | intron_variant | MODIFIER |
| 5 | 122927081 | AX-170137005 | 0 | A | T | 0.447 | -1.29E-02 | 1.14E-01 | 2.72E+00 | 2.76E+00 | 9.07E-01 | no | Kdm2b | intron_variant | MODIFIER |
| 5 | 122930098 | AX-170398893 | 0 | C | A | 0.447 | -1.29E-02 | 1.14E-01 | 2.72E+00 | 2.76E+00 | 9.07E-01 | no | Kdm2b | intron_variant | MODIFIER |
| 5 | 122933018 | AX-97848762 | 0 | C | T | 0.447 | -1.29E-02 | 1.14E-01 | 2.72E+00 | 2.76E+00 | 9.07E-01 | no | Kdm2b | intron_variant | MODIFIER |
| 5 | 122947728 | AX-170269626 | 0 | T | G | 0.447 | -1.29E-02 | 1.14E-01 | 2.72E+00 | 2.76E+00 | 9.07E-01 | no | Kdm2b | splice_region_variant,synonymous_variant | LOW |
| 5 | 122950671 | AX-170161520 | 0 | T | C | 0.447 | -1.29E-02 | 1.14E-01 | 2.72E+00 | 2.76E+00 | 9.07E-01 | no | Kdm2b | intron_variant | MODIFIER |
| 5 | 122961228 | AX-170341685 | 0 | T | C | 0.447 | -1.29E-02 | 1.14E-01 | 2.72E+00 | 2.76E+00 | 9.07E-01 | no | Kdm2b | intron_variant | MODIFIER |
| 5 | 122976909 | AX-170443770 | 0 | T | C | 0.447 | -1.29E-02 | 1.14E-01 | 2.72E+00 | 2.76E+00 | 9.07E-01 | no | Kdm2b | intron_variant | MODIFIER |
| 5 | 122987009 | AX-97843940 | 0 | C | T | 0.447 | -1.29E-02 | 1.14E-01 | 2.72E+00 | 2.76E+00 | 9.07E-01 | no | Kdm2b | intron_variant | MODIFIER |
| 5 | 122987275 | AX-170409268 | 0 | G | A | 0.447 | -1.29E-02 | 1.14E-01 | 2.72E+00 | 2.76E+00 | 9.07E-01 | no | Kdm2b | intron_variant | MODIFIER |
| 5 | 122989634 | AX-170511241 | 0 | T | C | 0.447 | -1.29E-02 | 1.14E-01 | 2.72E+00 | 2.76E+00 | 9.07E-01 | no | Kdm2b | upstream_gene_variant | MODIFIER |
| 5 | 122995457 | AX-170141763 | 0 | G | A | 0.447 | -1.29E-02 | 1.14E-01 | 2.72E+00 | 2.76E+00 | 9.07E-01 | no | A930024E05Rik | intron_variant,non_coding_transcript_variant | MODIFIER |
| 5 | 122995794 | AX-170368429 | 0 | T | C | 0.447 | -1.29E-02 | 1.14E-01 | 2.72E+00 | 2.76E+00 | 9.07E-01 | no | A930024E05Rik | intron_variant,non_coding_transcript_variant | MODIFIER |
| 5 | 123001894 | AX-170202911 | 0 | T | C | 0.447 | -1.29E-02 | 1.14E-01 | 2.72E+00 | 2.76E+00 | 9.07E-01 | no | A930024E05Rik | downstream_gene_variant | MODIFIER |
| 5 | 123003374 | AX-170152972 | 0 | G | C | 0.447 | -1.29E-02 | 1.14E-01 | 2.72E+00 | 2.76E+00 | 9.07E-01 | no | Gm44574 | intron_variant,non_coding_transcript_variant | MODIFIER |
| 5 | 123039321 | AX-170521673 | 0 | A | G | 0.447 | -1.29E-02 | 1.14E-01 | 2.72E+00 | 2.76E+00 | 9.07E-01 | no | Morn3 | synonymous_variant | LOW |
| 5 | 123042627 | AX-170089600 | 0 | C | T | 0.447 | -1.29E-02 | 1.14E-01 | 2.72E+00 | 2.76E+00 | 9.07E-01 | no | Morn3 | intron_variant | MODIFIER |
| 5 | 123042849 | AX-170420952 | 0 | C | T | 0.447 | -1.29E-02 | 1.14E-01 | 2.72E+00 | 2.76E+00 | 9.07E-01 | no | Morn3 | intron_variant | MODIFIER |
| 5 | 123044436 | AX-170360964 | 0 | C | G | 0.447 | -1.29E-02 | 1.14E-01 | 2.72E+00 | 2.76E+00 | 9.07E-01 | no | Morn3 | intron_variant | MODIFIER |
| 5 | 123054927 | AX-97834157 | 0 | C | T | 0.447 | -1.29E-02 | 1.14E-01 | 2.72E+00 | 2.76E+00 | 9.07E-01 | no | Gm26745 | intron_variant,non_coding_transcript_variant | MODIFIER |
| 5 | 123151156 | AX-170476492 | 0 | C | A | 0.447 | -1.29E-02 | 1.14E-01 | 2.72E+00 | 2.76E+00 | 9.07E-01 | no | Setd1b | intron_variant | MODIFIER |
| 5 | 123241089 | AX-170103365 | 0 | G | T | 0.447 | -1.29E-02 | 1.14E-01 | 2.72E+00 | 2.76E+00 | 9.07E-01 | no | Psmd9 | intron_variant | MODIFIER |
| 5 | 123263535 | AX-170026066 | 0 | G | A | 0.447 | -1.29E-02 | 1.14E-01 | 2.72E+00 | 2.76E+00 | 9.07E-01 | no | Wdr66 | intron_variant | MODIFIER |
| 5 | 123265801 | AX-170211348 | 0 | T | C | 0.447 | -1.29E-02 | 1.14E-01 | 2.72E+00 | 2.76E+00 | 9.07E-01 | no | Wdr66 | intron_variant | MODIFIER |
| 5 | 123266214 | AX-170360755 | 0 | C | T | 0.447 | -1.29E-02 | 1.14E-01 | 2.72E+00 | 2.76E+00 | 9.07E-01 | no | Wdr66 | intron_variant | MODIFIER |
| 5 | 123266315 | AX-170478004 | 0 | A | G | 0.447 | -1.29E-02 | 1.14E-01 | 2.72E+00 | 2.76E+00 | 9.07E-01 | no | Wdr66 | intron_variant | MODIFIER |
| 5 | 123529601 | AX-170348281 | 0 | T | A | 0.447 | -1.29E-02 | 1.14E-01 | 2.72E+00 | 2.76E+00 | 9.07E-01 | no | Vps33a | 3_prime_UTR_variant | MODIFIER |
| 5 | 123532027 | AX-170513963 | 0 | C | A | 0.447 | -1.29E-02 | 1.14E-01 | 2.72E+00 | 2.76E+00 | 9.07E-01 | no | Vps33a | intron_variant | MODIFIER |
| 5 | 123566066 | AX-170496239 | 0 | T | G | 0.447 | -1.29E-02 | 1.14E-01 | 2.72E+00 | 2.76E+00 | 9.07E-01 | no | Vps33a | intron_variant | MODIFIER |
| 5 | 123570847 | AX-170531925 | 0 | T | C | 0.447 | -1.29E-02 | 1.14E-01 | 2.72E+00 | 2.76E+00 | 9.07E-01 | no | Vps33a | splice_region_variant,intron_variant | LOW |
| 5 | 123585683 | AX-170304141 | 0 | C | A | 0.447 | -1.29E-02 | 1.14E-01 | 2.72E+00 | 2.76E+00 | 9.07E-01 | no | Clip1 | intron_variant | MODIFIER |
| 5 | 123589387 | AX-170385998 | 0 | A | G | 0.447 | -1.29E-02 | 1.14E-01 | 2.72E+00 | 2.76E+00 | 9.07E-01 | no | Clip1 | intron_variant | MODIFIER |
| 5 | 123610890 | AX-170245808 | 0 | C | T | 0.447 | -1.29E-02 | 1.14E-01 | 2.72E+00 | 2.76E+00 | 9.07E-01 | no | Clip1 | intron_variant | MODIFIER |
| 5 | 123687510 | AX-170345228 | 0 | G | A | 0.447 | -1.29E-02 | 1.14E-01 | 2.72E+00 | 2.76E+00 | 9.07E-01 | no | Clip1 | upstream_gene_variant | MODIFIER |
| 5 | 123735660 | AX-170112284 | 0 | C | T | 0.447 | -1.29E-02 | 1.14E-01 | 2.72E+00 | 2.76E+00 | 9.07E-01 | no | Rsrc2 | intron_variant | MODIFIER |
| 5 | 123782993 | AX-170368891 | 0 | A | G | 0.447 | -1.29E-02 | 1.14E-01 | 2.72E+00 | 2.76E+00 | 9.07E-01 | no | Kntc1 | intron_variant | MODIFIER |
| 5 | 123791054 | AX-170026072 | 0 | A | C | 0.447 | -1.29E-02 | 1.14E-01 | 2.72E+00 | 2.76E+00 | 9.07E-01 | no | Kntc1 | missense_variant,splice_region_variant | MODERATE |
| 5 | 123802571 | AX-170026074 | 0 | G | A | 0.447 | -1.29E-02 | 1.14E-01 | 2.72E+00 | 2.76E+00 | 9.07E-01 | no | Kntc1 | intron_variant | MODIFIER |

| | | | | | | | | | | | | | | | |
|---|---|---|---|---|---|---|---|---|---|---|---|---|---|---|---|
| 5 | 123805103 | AX-97798067 | 0 | G | A | 0.447 | -1.29E-02 | 1.14E-01 | 2.72E+00 | 2.76E+00 | 9.07E-01 | no | Kntc1 | intron_variant | MODIFIER |
| 5 | 123810733 | AX-170124766 | 0 | T | A | 0.447 | -1.29E-02 | 1.14E-01 | 2.72E+00 | 2.76E+00 | 9.07E-01 | no | Kntc1 | intron_variant | MODIFIER |
| 5 | 123811029 | AX-170173574 | 0 | A | G | 0.447 | -1.29E-02 | 1.14E-01 | 2.72E+00 | 2.76E+00 | 9.07E-01 | no | Kntc1 | intron_variant | MODIFIER |
| 5 | 123815091 | AX-170511400 | 0 | A | G | 0.447 | -1.29E-02 | 1.14E-01 | 2.72E+00 | 2.76E+00 | 9.07E-01 | no | Kntc1 | intron_variant | MODIFIER |
| 5 | 123824159 | AX-97847127 | 0 | T | C | 0.447 | -1.29E-02 | 1.14E-01 | 2.72E+00 | 2.76E+00 | 9.07E-01 | no | Kntc1 | downstream_gene_variant | MODIFIER |
| 5 | 123837426 | AX-170026070 | 0 | A | G | 0.447 | -1.29E-02 | 1.14E-01 | 2.72E+00 | 2.76E+00 | 9.07E-01 | no | | regulatory_region_variant | MODIFIER |
| 5 | 123867747 | AX-170073367 | 0 | A | G | 0.447 | -1.29E-02 | 1.14E-01 | 2.72E+00 | 2.76E+00 | 9.07E-01 | no | Hcar2 | upstream_gene_variant | MODIFIER |
| 5 | 123869880 | AX-170464508 | 0 | A | G | 0.447 | -1.29E-02 | 1.14E-01 | 2.72E+00 | 2.76E+00 | 9.07E-01 | no | Hcar2 | upstream_gene_variant | MODIFIER |
| 5 | 123885553 | AX-170280779 | 0 | A | G | 0.447 | -1.29E-02 | 1.14E-01 | 2.72E+00 | 2.76E+00 | 9.07E-01 | no | | intergenic_variant | MODIFIER |
| 5 | 123894755 | AX-170494887 | 0 | G | A | 0.447 | -1.29E-02 | 1.14E-01 | 2.72E+00 | 2.76E+00 | 9.07E-01 | no | | intergenic_variant | MODIFIER |
| 5 | 123929223 | AX-97797673 | 0 | G | A | 0.447 | -1.29E-02 | 1.14E-01 | 2.72E+00 | 2.76E+00 | 9.07E-01 | no | Denr | downstream_gene_variant | MODIFIER |
| 5 | 123947524 | AX-170450207 | 0 | A | G | 0.447 | -1.29E-02 | 1.14E-01 | 2.72E+00 | 2.76E+00 | 9.07E-01 | no | Ccdc62 | intron_variant | MODIFIER |
| 5 | 123970078 | AX-170160378 | 0 | G | A | 0.447 | -1.29E-02 | 1.14E-01 | 2.72E+00 | 2.76E+00 | 9.07E-01 | no | Hip1r | upstream_gene_variant | MODIFIER |
| 5 | 123984609 | AX-170391107 | 0 | C | T | 0.447 | -1.29E-02 | 1.14E-01 | 2.72E+00 | 2.76E+00 | 9.07E-01 | no | Hip1r | intron_variant | MODIFIER |
| 5 | 123987317 | AX-170215735 | 0 | G | A | 0.447 | -1.29E-02 | 1.14E-01 | 2.72E+00 | 2.76E+00 | 9.07E-01 | no | Hip1r | intron_variant | MODIFIER |
| 5 | 124006411 | AX-170246006 | 0 | T | C | 0.447 | -1.29E-02 | 1.14E-01 | 2.72E+00 | 2.76E+00 | 9.07E-01 | no | Hip1r | downstream_gene_variant | MODIFIER |
| 5 | 124007729 | AX-170363777 | 0 | A | G | 0.447 | -1.29E-02 | 1.14E-01 | 2.72E+00 | 2.76E+00 | 9.07E-01 | no | Hip1r | downstream_gene_variant | MODIFIER |
| 5 | 124034612 | AX-170174713 | 0 | C | T | 0.447 | -1.29E-02 | 1.14E-01 | 2.72E+00 | 2.76E+00 | 9.07E-01 | no | Vps37b | upstream_gene_variant | MODIFIER |
| 5 | 124041169 | AX-170303702 | 0 | T | C | 0.447 | -1.29E-02 | 1.14E-01 | 2.72E+00 | 2.76E+00 | 9.07E-01 | no | Gm34086 | intron_variant,non_coding_transcript_variant | MODIFIER |
| 5 | 124223381 | AX-170453944 | 0 | A | G | 0.447 | -1.29E-02 | 1.14E-01 | 2.72E+00 | 2.76E+00 | 9.07E-01 | no | Pitpnm2 | intron_variant | MODIFIER |
| 5 | 124242923 | AX-170255904 | 0 | T | C | 0.447 | -1.29E-02 | 1.14E-01 | 2.72E+00 | 2.76E+00 | 9.07E-01 | no | Pitpnm2 | intron_variant | MODIFIER |
| 5 | 124294261 | AX-170026081 | 0 | A | G | 0.447 | -1.29E-02 | 1.14E-01 | 2.72E+00 | 2.76E+00 | 9.07E-01 | no | Mphosph9 | intron_variant | MODIFIER |
| 5 | 124296631 | AX-170356987 | 0 | C | T | 0.447 | -1.29E-02 | 1.14E-01 | 2.72E+00 | 2.76E+00 | 9.07E-01 | no | Mphosph9 | intron_variant | MODIFIER |
| 5 | 124308660 | AX-97813518 | 0 | C | T | 0.447 | -1.29E-02 | 1.14E-01 | 2.72E+00 | 2.76E+00 | 9.07E-01 | no | Mphosph9 | intron_variant | MODIFIER |
| 5 | 124321346 | AX-170026079 | 0 | T | G | 0.447 | -1.29E-02 | 1.14E-01 | 2.72E+00 | 2.76E+00 | 9.07E-01 | no | Mphosph9 | intron_variant | MODIFIER |
| 5 | 124324799 | AX-170026082 | 0 | C | G | 0.447 | -1.29E-02 | 1.14E-01 | 2.72E+00 | 2.76E+00 | 9.07E-01 | no | Mphosph9 | intron_variant | MODIFIER |
| 5 | 124507624 | AX-170151006 | 0 | A | C | 0.447 | -1.29E-02 | 1.14E-01 | 2.72E+00 | 2.76E+00 | 9.07E-01 | no | Rilpl1 | intron_variant | MODIFIER |
| 5 | 124507784 | AX-170267506 | 0 | T | C | 0.447 | -1.29E-02 | 1.14E-01 | 2.72E+00 | 2.76E+00 | 9.07E-01 | no | Rilpl1 | intron_variant | MODIFIER |
| 5 | 124508112 | AX-170531346 | 0 | A | G | 0.447 | -1.29E-02 | 1.14E-01 | 2.72E+00 | 2.76E+00 | 9.07E-01 | no | Rilpl1 | intron_variant | MODIFIER |
| 5 | 124556707 | AX-170470273 | 0 | T | C | 0.447 | -1.29E-02 | 1.14E-01 | 2.72E+00 | 2.76E+00 | 9.07E-01 | no | Ddx55 | intron_variant | MODIFIER |
| 5 | 124557910 | AX-170457114 | 0 | T | C | 0.447 | -1.29E-02 | 1.14E-01 | 2.72E+00 | 2.76E+00 | 9.07E-01 | no | Ddx55 | intron_variant | MODIFIER |
| 5 | 124572644 | AX-170196613 | 0 | G | T | 0.447 | -1.29E-02 | 1.14E-01 | 2.72E+00 | 2.76E+00 | 9.07E-01 | no | Eif2b1 | intron_variant | MODIFIER |
| 5 | 124578462 | AX-170271408 | 0 | T | C | 0.447 | -1.29E-02 | 1.14E-01 | 2.72E+00 | 2.76E+00 | 9.07E-01 | no | Gtf2h3 | upstream_gene_variant | MODIFIER |
| 5 | 124578583 | AX-170336673 | 0 | G | A | 0.447 | -1.29E-02 | 1.14E-01 | 2.72E+00 | 2.76E+00 | 9.07E-01 | no | Gtf2h3 | upstream_gene_variant | MODIFIER |
| 5 | 124598678 | AX-97788390 | 0 | C | T | 0.447 | -1.29E-02 | 1.14E-01 | 2.72E+00 | 2.76E+00 | 9.07E-01 | no | Gtf2h3 | downstream_gene_variant | MODIFIER |
| 5 | 124613974 | AX-170491500 | 0 | A | G | 0.447 | -1.29E-02 | 1.14E-01 | 2.72E+00 | 2.76E+00 | 9.07E-01 | no | Tctn2 | intron_variant,non_coding_transcript_variant | MODIFIER |
| 5 | 124724309 | AX-170149425 | 0 | C | T | 0.447 | -1.29E-02 | 1.14E-01 | 2.72E+00 | 2.76E+00 | 9.07E-01 | no | Atp6v0a2 | 3_prime_UTR_variant | MODIFIER |
| 1 | 19705037 | AX-170145143 | 1 | A | T | 0.402 | -8.76E-03 | 8.26E-02 | 2.71E+00 | 2.78E+00 | 9.07E-01 | no | | intergenic_variant | MODIFIER |
| 12 | 112167003 | AX-170376026 | 1 | T | C | 0.143 | -2.17E-02 | 1.92E-01 | 2.67E+00 | 2.74E+00 | 9.07E-01 | no | Kif26a | intron_variant | MODIFIER |
| 12 | 116777083 | AX-170222248 | 1 | G | A | 0.143 | -2.17E-02 | 1.92E-01 | 2.67E+00 | 2.74E+00 | 9.07E-01 | no | Ptprn2 | intron_variant | MODIFIER |
| 12 | 117822559 | AX-170533348 | 1 | A | G | 0.143 | -2.17E-02 | 1.92E-01 | 2.67E+00 | 2.74E+00 | 9.07E-01 | no | Cdca7l | intron_variant,non_coding_transcript_variant | MODIFIER |
| 8 | 78316011 | AX-170080506 | 1 | G | A | 0.205 | 1.43E-02 | 1.27E-01 | 2.68E+00 | 2.75E+00 | 9.07E-01 | no | Ttc29 | intron_variant | MODIFIER |
| 20 | 156877998 | AX-170035304 | 0 | A | G | 0.044 | -1.30E-02 | 1.39E-01 | 2.57E+00 | 2.76E+00 | 9.08E-01 | no | | intergenic_variant | MODIFIER |
| 16 | 92598562 | AX-170148099 | 0 | T | C | 0.456 | -8.76E-03 | 7.74E-02 | 2.65E+00 | 2.74E+00 | 9.08E-01 | no | Runx1 | downstream_gene_variant | MODIFIER |
| 12 | 112173503 | AX-170509124 | 0 | C | T | 0.123 | -2.16E-02 | 1.95E-01 | 2.68E+00 | 2.75E+00 | 9.08E-01 | no | Kif26a | intron_variant | MODIFIER |
| 9 | 57059934 | AX-170032439 | 1 | C | T | 0.223 | -1.35E-02 | 1.21E-01 | 2.69E+00 | 2.75E+00 | 9.08E-01 | no | Ptpn9 | synonymous_variant | LOW |
| 1 | 25349795 | AX-170040651 | 1 | A | C | 0.411 | -8.75E-03 | 8.09E-02 | 2.69E+00 | 2.76E+00 | 9.08E-01 | no | Adgrb3 | intron_variant | MODIFIER |
| 15 | 85688023 | AX-170496174 | 0 | C | T | 0.518 | 9.86E-03 | 7.09E-02 | 2.59E+00 | 2.70E+00 | 9.08E-01 | no | Lncppara | intron_variant,non_coding_transcript_variant | MODIFIER |
| 13 | 47393996 | AX-97850204 | 1 | C | T | 0.491 | -6.90E-03 | 6.43E-02 | 2.65E+00 | 2.75E+00 | 9.08E-01 | no | Gm35733 | intron_variant,non_coding_transcript_variant | MODIFIER |
| 4 | 93092599 | AX-170187201 | 0 | T | A | 0.333 | 1.25E-02 | 1.05E-01 | 2.65E+00 | 2.73E+00 | 9.08E-01 | no | | intergenic_variant | MODIFIER |
| 9 | 68618020 | AX-170353116 | 0 | C | G | 0.193 | -1.60E-02 | 1.37E-01 | 2.70E+00 | 2.73E+00 | 9.08E-01 | no | | intergenic_variant | MODIFIER |
| 15 | 78410374 | AX-170284006 | 0 | G | A | 0.553 | 9.28E-03 | 7.65E-02 | 2.63E+00 | 2.73E+00 | 9.08E-01 | no | Mpst | missense_variant | MODERATE |
| 10 | 117457547 | AX-170129747 | 0 | T | G | 0.333 | 8.21E-03 | 7.64E-02 | 2.67E+00 | 2.76E+00 | 9.08E-01 | no | | regulatory_region_variant | MODIFIER |
| 10 | 117459903 | AX-170333332 | 0 | T | C | 0.333 | 8.21E-03 | 7.64E-02 | 2.67E+00 | 2.76E+00 | 9.08E-01 | no | | intergenic_variant | MODIFIER |

| | | | | | | | | | | | | | | | |
|---|---|---|---|---|---|---|---|---|---|---|---|---|---|---|---|
| 10 | 117460605 | AX-170138578 | 0 | A | G | 0.333 | 8.21E-03 | 7.64E-02 | 2.67E+00 | 2.76E+00 | 9.08E-01 | no | | intergenic_variant | MODIFIER |
| 10 | 117460697 | AX-170337107 | 0 | A | G | 0.333 | 8.21E-03 | 7.64E-02 | 2.67E+00 | 2.76E+00 | 9.08E-01 | no | | intergenic_variant | MODIFIER |
| 10 | 117572424 | AX-97786119 | 0 | A | G | 0.333 | 8.21E-03 | 7.64E-02 | 2.67E+00 | 2.76E+00 | 9.08E-01 | no | | regulatory_region_variant | MODIFIER |
| 10 | 117572737 | AX-170046015 | 0 | A | G | 0.333 | 8.21E-03 | 7.64E-02 | 2.67E+00 | 2.76E+00 | 9.08E-01 | no | | regulatory_region_variant | MODIFIER |
| 10 | 117647450 | AX-97797901 | 0 | A | G | 0.333 | 8.21E-03 | 7.64E-02 | 2.67E+00 | 2.76E+00 | 9.08E-01 | no | Cpm | intron_variant | MODIFIER |
| 1 | 52831064 | AX-170227936 | 0 | G | A | 0.114 | -1.95E-02 | 1.87E-01 | 2.66E+00 | 2.76E+00 | 9.08E-01 | no | | intergenic_variant | MODIFIER |
| 1 | 54300499 | AX-170000525 | 0 | T | C | 0.114 | -1.95E-02 | 1.87E-01 | 2.66E+00 | 2.76E+00 | 9.08E-01 | no | Ccdc150 | intron_variant | MODIFIER |
| 18 | 45651883 | AX-170277006 | 1 | T | G | 0.062 | -3.46E-02 | 3.00E-01 | 2.71E+00 | 2.73E+00 | 9.08E-01 | no | Kcnn2 | intron_variant | MODIFIER |
| 10 | 23619743 | AX-170002919 | 0 | C | T | 0.579 | -6.05E-03 | 6.88E-02 | 2.64E+00 | 2.78E+00 | 9.08E-01 | no | 4930520K02Rik | intron_variant,non_coding_transcript_variant | MODIFIER |
| 12 | 117389499 | AX-170481531 | 1 | C | T | 0.143 | -2.14E-02 | 1.89E-01 | 2.67E+00 | 2.74E+00 | 9.08E-01 | no | | regulatory_region_variant | MODIFIER |
| 16 | 89489570 | AX-97856201 | 0 | C | T | 0.465 | 9.18E-03 | 7.98E-02 | 2.66E+00 | 2.74E+00 | 9.08E-01 | no | Krtap8-1 | upstream_gene_variant | MODIFIER |
| 5 | 122939614 | AX-170313064 | 1 | C | A | 0.455 | -1.24E-02 | 1.13E-01 | 2.72E+00 | 2.76E+00 | 9.08E-01 | no | Kdm2b | intron_variant | MODIFIER |
| 9 | 58777785 | AX-170082546 | 1 | C | T | 0.214 | -1.40E-02 | 1.24E-01 | 2.69E+00 | 2.74E+00 | 9.08E-01 | no | | intergenic_variant | MODIFIER |
| 9 | 59330238 | AX-170270127 | 1 | C | G | 0.214 | -1.40E-02 | 1.24E-01 | 2.69E+00 | 2.74E+00 | 9.08E-01 | no | Bbs4 | intron_variant | MODIFIER |
| 9 | 105601591 | AX-170184241 | 0 | T | C | 0.123 | -1.53E-02 | 1.17E-01 | 2.62E+00 | 2.70E+00 | 9.08E-01 | no | | intergenic_variant | MODIFIER |
| 1 | 165816564 | AX-170494199 | 0 | T | G | 0.228 | -1.15E-02 | 1.16E-01 | 2.66E+00 | 2.77E+00 | 9.08E-01 | no | Cd247 | intron_variant | MODIFIER |
| 5 | 124887186 | AX-170026091 | 0 | G | A | 0.465 | -1.19E-02 | 1.10E-01 | 2.70E+00 | 2.76E+00 | 9.08E-01 | no | Zfp664 | 3_prime_UTR_variant | MODIFIER |
| 6 | 121165707 | AX-170095786 | 0 | G | A | 0.07 | -3.17E-02 | 2.77E-01 | 2.70E+00 | 2.74E+00 | 9.08E-01 | no | | regulatory_region_variant | MODIFIER |
| 10 | 60436452 | AX-170003314 | 0 | A | G | 0.298 | 8.11E-03 | 8.38E-02 | 2.69E+00 | 2.79E+00 | 9.08E-01 | no | Cdh23 | intron_variant | MODIFIER |
| 5 | 123523164 | AX-170026057 | 0 | C | T | 0.456 | -1.28E-02 | 1.16E-01 | 2.72E+00 | 2.76E+00 | 9.08E-01 | no | Diablo | intron_variant | MODIFIER |
| 11 | 74629548 | AX-97793393 | 0 | T | A | 0.228 | 8.40E-03 | 7.46E-02 | 2.61E+00 | 2.74E+00 | 9.08E-01 | no | Ccdc92b | intron_variant | MODIFIER |
| 11 | 5466787 | AX-170230191 | 1 | T | A | 0.054 | -2.41E-02 | 2.02E-01 | 2.63E+00 | 2.73E+00 | 9.08E-01 | no | | intergenic_variant | MODIFIER |
| 10 | 7741513 | AX-170424263 | 1 | T | C | 0.196 | -1.23E-02 | 1.05E-01 | 2.64E+00 | 2.73E+00 | 9.08E-01 | no | Katna1 | missense_variant | MODERATE |
| 16 | 92584344 | AX-170236859 | 1 | C | A | 0.473 | -8.87E-03 | 7.88E-02 | 2.66E+00 | 2.74E+00 | 9.08E-01 | no | | intergenic_variant | MODIFIER |
| 12 | 52604667 | AX-97785685 | 0 | C | T | 0.053 | -1.98E-02 | 1.76E-01 | 2.62E+00 | 2.74E+00 | 9.08E-01 | no | Gm23296 | upstream_gene_variant | MODIFIER |
| 15 | 102433844 | AX-170165790 | 0 | A | G | 0.105 | 1.65E-02 | 1.43E-01 | 2.65E+00 | 2.74E+00 | 9.08E-01 | no | Sp1 | 3_prime_UTR_variant | MODIFIER |
| 1 | 11116817 | AX-170305721 | 1 | C | G | 0.098 | 2.04E-02 | 1.72E-01 | 2.62E+00 | 2.73E+00 | 9.08E-01 | no | Prex2 | intron_variant | MODIFIER |
| 10 | 31793599 | AX-170288264 | 0 | G | A | 0.395 | 8.11E-03 | 7.59E-02 | 2.64E+00 | 2.75E+00 | 9.09E-01 | no | Nkain2 | intron_variant | MODIFIER |
| 1 | 3493708 | AX-170330603 | 0 | G | A | 0.518 | 8.96E-03 | 7.06E-02 | 2.64E+00 | 2.71E+00 | 9.09E-01 | no | Xkr4 | intron_variant | MODIFIER |
| 7 | 10988824 | AX-97851881 | 2 | C | A | 0.191 | -9.82E-03 | 7.97E-02 | 2.58E+00 | 2.73E+00 | 9.09E-01 | no | Gm7789 | intron_variant,non_coding_transcript_variant | MODIFIER |
| 18 | 89086353 | AX-170430766 | 1 | G | A | 0.509 | -6.54E-03 | 6.34E-02 | 2.62E+00 | 2.75E+00 | 9.09E-01 | no | Rttn | intron_variant | MODIFIER |
| 18 | 89944276 | AX-170474217 | 1 | G | A | 0.509 | -6.54E-03 | 6.34E-02 | 2.62E+00 | 2.75E+00 | 9.09E-01 | no | Gm27037 | non_coding_transcript_exon_variant | MODIFIER |
| 12 | 77957707 | AX-170418356 | 0 | G | A | 0.044 | -3.16E-02 | 2.74E-01 | 2.68E+00 | 2.73E+00 | 9.09E-01 | no | | intergenic_variant | MODIFIER |
| 10 | 43344658 | AX-170432455 | 0 | G | C | 0.368 | 8.00E-03 | 7.50E-02 | 2.65E+00 | 2.75E+00 | 9.09E-01 | no | Pdss2 | intron_variant | MODIFIER |
| 13 | 54910907 | AX-168386640 | 0 | A | G | 0.509 | 8.62E-03 | 6.53E-02 | 2.59E+00 | 2.71E+00 | 9.09E-01 | no | 5330429C05Rik | upstream_gene_variant | MODIFIER |
| 13 | 54911354 | AX-97799106 | 0 | T | C | 0.509 | 8.62E-03 | 6.53E-02 | 2.59E+00 | 2.71E+00 | 9.09E-01 | no | 5330429C05Rik | upstream_gene_variant | MODIFIER |
| 13 | 55063173 | AX-97841550 | 0 | T | C | 0.509 | 8.62E-03 | 6.53E-02 | 2.59E+00 | 2.71E+00 | 9.09E-01 | no | Uimc1 | intron_variant | MODIFIER |
| 13 | 56224688 | AX-170481475 | 0 | T | C | 0.509 | 8.62E-03 | 6.53E-02 | 2.59E+00 | 2.71E+00 | 9.09E-01 | no | 4930550C17Rik | intron_variant,non_coding_transcript_variant | MODIFIER |
| 13 | 56227215 | AX-170376384 | 0 | T | C | 0.509 | 8.62E-03 | 6.53E-02 | 2.59E+00 | 2.71E+00 | 9.09E-01 | no | 4930550C17Rik | intron_variant,non_coding_transcript_variant | MODIFIER |
| 13 | 56227244 | AX-170378570 | 0 | A | G | 0.509 | 8.62E-03 | 6.53E-02 | 2.59E+00 | 2.71E+00 | 9.09E-01 | no | 4930550C17Rik | intron_variant,non_coding_transcript_variant | MODIFIER |
| 7 | 67593913 | AX-97816828 | 0 | T | C | 0.351 | -1.11E-02 | 1.05E-01 | 2.65E+00 | 2.75E+00 | 9.09E-01 | no | Lrrc28 | downstream_gene_variant | MODIFIER |
| 5 | 123650849 | AX-170182980 | 1 | T | C | 0.446 | -1.24E-02 | 1.13E-01 | 2.72E+00 | 2.76E+00 | 9.09E-01 | no | Clip1 | intron_variant | MODIFIER |
| 5 | 118606122 | AX-97802215 | 0 | G | A | 0.456 | 1.30E-02 | 1.09E-01 | 2.66E+00 | 2.72E+00 | 9.09E-01 | no | Med13l | intron_variant | MODIFIER |
| 5 | 124594268 | AX-170519805 | 0 | C | T | 0.456 | -1.20E-02 | 1.12E-01 | 2.71E+00 | 2.76E+00 | 9.09E-01 | no | Gtf2h3 | intron_variant | MODIFIER |
| 8 | 93324923 | AX-170134574 | 0 | T | G | 0.202 | -1.36E-02 | 1.20E-01 | 2.68E+00 | 2.74E+00 | 9.09E-01 | no | Ces1g | intron_variant | MODIFIER |
| 8 | 93385230 | AX-97805778 | 0 | T | C | 0.202 | -1.36E-02 | 1.20E-01 | 2.68E+00 | 2.74E+00 | 9.09E-01 | no | Gm45765 | intron_variant,non_coding_transcript_variant | MODIFIER |
| 8 | 94391599 | AX-170205195 | 0 | T | C | 0.202 | -1.36E-02 | 1.20E-01 | 2.68E+00 | 2.74E+00 | 9.09E-01 | no | Herpud1 | intron_variant | MODIFIER |
| 8 | 94479237 | AX-170258245 | 0 | G | A | 0.202 | -1.36E-02 | 1.20E-01 | 2.68E+00 | 2.74E+00 | 9.09E-01 | no | Nlrc5 | intron_variant | MODIFIER |
| 8 | 94486876 | AX-170296285 | 0 | T | C | 0.202 | -1.36E-02 | 1.20E-01 | 2.68E+00 | 2.74E+00 | 9.09E-01 | no | Nlrc5 | intron_variant | MODIFIER |
| 8 | 94491256 | AX-170449496 | 0 | T | C | 0.202 | -1.36E-02 | 1.20E-01 | 2.68E+00 | 2.74E+00 | 9.09E-01 | no | Nlrc5 | intron_variant | MODIFIER |
| 8 | 94890125 | AX-170371472 | 0 | A | G | 0.202 | -1.36E-02 | 1.20E-01 | 2.68E+00 | 2.74E+00 | 9.09E-01 | no | | intergenic_variant | MODIFIER |
| 8 | 94895818 | AX-170097444 | 0 | G | A | 0.202 | -1.36E-02 | 1.20E-01 | 2.68E+00 | 2.74E+00 | 9.09E-01 | no | | intergenic_variant | MODIFIER |
| 1 | 8059728 | AX-170456718 | 0 | A | G | 0.412 | 8.77E-03 | 7.10E-02 | 2.64E+00 | 2.71E+00 | 9.09E-01 | no | | intergenic_variant | MODIFIER |
| 2 | 138927435 | AX-170065043 | 1 | C | T | 0.134 | 2.34E-02 | 2.07E-01 | 2.69E+00 | 2.73E+00 | 9.09E-01 | no | | intergenic_variant | MODIFIER |

| | | | | | | | | | | | | | | | |
|---|---|---|---|---|---|---|---|---|---|---|---|---|---|---|---|
| 2 | 156489250 | AX-97795403 | 1 | G | A | 0.134 | 2.34E-02 | 2.07E-01 | 2.69E+00 | 2.73E+00 | 9.09E-01 | no | Epb41l1 | intron_variant | MODIFIER |
| 20 | 61119998 | AX-170142892 | 0 | A | T | 0.044 | -1.58E-02 | 1.13E-01 | 2.53E+00 | 2.71E+00 | 9.10E-01 | no | C230004F18Rik | intron_variant,non_coding_transcript_variant | MODIFIER |
| 7 | 118485912 | AX-170179169 | 0 | A | G | 0.395 | 1.04E-02 | 9.19E-02 | 2.68E+00 | 2.74E+00 | 9.10E-01 | no | Itpripl2 | 3_prime_UTR_variant | MODIFIER |
| 6 | 114353335 | AX-97853174 | 0 | A | G | 0.053 | 3.34E-02 | 2.95E-01 | 2.67E+00 | 2.73E+00 | 9.10E-01 | no | | intergenic_variant | MODIFIER |
| 5 | 32969598 | AX-170369544 | 0 | G | A | 0.36 | 1.20E-02 | 1.07E-01 | 2.68E+00 | 2.74E+00 | 9.10E-01 | no | Depdc5 | intron_variant | MODIFIER |
| 15 | 74963884 | AX-170011999 | 0 | G | C | 0.544 | -8.31E-03 | 8.01E-02 | 2.67E+00 | 2.76E+00 | 9.10E-01 | no | Ly6e | downstream_gene_variant | MODIFIER |
| 8 | 48787200 | AX-170338911 | 1 | G | A | 0.152 | -1.81E-02 | 1.47E-01 | 2.65E+00 | 2.71E+00 | 9.10E-01 | no | Tenm3 | intron_variant | MODIFIER |
| 9 | 57793149 | AX-170181343 | 1 | A | G | 0.223 | -1.31E-02 | 1.20E-01 | 2.69E+00 | 2.75E+00 | 9.10E-01 | no | Arid3b | intron_variant | MODIFIER |
| 9 | 60130621 | AX-170500600 | 1 | G | A | 0.223 | -1.31E-02 | 1.20E-01 | 2.69E+00 | 2.75E+00 | 9.10E-01 | no | Thsd4 | intron_variant | MODIFIER |
| 10 | 6491434 | AX-170395469 | 0 | C | T | 0.211 | -1.14E-02 | 9.97E-02 | 2.63E+00 | 2.73E+00 | 9.10E-01 | no | | intergenic_variant | MODIFIER |
| 9 | 83592745 | AX-170119317 | 1 | G | C | 0.214 | 1.36E-02 | 1.26E-01 | 2.69E+00 | 2.75E+00 | 9.10E-01 | no | Sh3bgrl2 | intron_variant | MODIFIER |
| 18 | 57463535 | AX-170289494 | 1 | G | A | 0.42 | 8.61E-03 | 8.38E-02 | 2.70E+00 | 2.77E+00 | 9.10E-01 | no | Ctxn3 | upstream_gene_variant | MODIFIER |
| 9 | 83609595 | AX-170447182 | 0 | C | T | 0.211 | 1.40E-02 | 1.30E-01 | 2.71E+00 | 2.76E+00 | 9.10E-01 | no | Sh3bgrl2 | intron_variant | MODIFIER |
| 11 | 23149106 | AX-170275021 | 0 | C | T | 0.053 | -2.16E-02 | 2.00E-01 | 2.61E+00 | 2.74E+00 | 9.10E-01 | no | 1700061J23Rik | upstream_gene_variant | MODIFIER |
| 8 | 47209405 | AX-170334216 | 0 | G | A | 0.158 | -1.65E-02 | 1.45E-01 | 2.68E+00 | 2.73E+00 | 9.10E-01 | no | Stox2 | intron_variant | MODIFIER |
| 8 | 48143253 | AX-97802781 | 0 | G | T | 0.158 | -1.65E-02 | 1.45E-01 | 2.68E+00 | 2.73E+00 | 9.10E-01 | no | Dctd | downstream_gene_variant | MODIFIER |
| 1 | 67168862 | AX-170193107 | 1 | A | C | 0.116 | 2.10E-02 | 1.92E-01 | 2.67E+00 | 2.74E+00 | 9.10E-01 | no | Cps1 | intron_variant | MODIFIER |
| 19 | 22519533 | AX-170327620 | 0 | C | T | 0.044 | -2.25E-02 | 2.08E-01 | 2.65E+00 | 2.74E+00 | 9.10E-01 | no | Trpm3 | intron_variant | MODIFIER |
| 15 | 90485803 | AX-170124462 | 1 | C | T | 0.161 | 1.93E-02 | 1.54E-01 | 2.61E+00 | 2.71E+00 | 9.11E-01 | no | Cpne8 | downstream_gene_variant | MODIFIER |
| 2 | 106384975 | AX-170182223 | 1 | C | T | 0.071 | 2.05E-02 | 1.69E-01 | 2.59E+00 | 2.72E+00 | 9.11E-01 | no | Dcdc5 | intron_variant,non_coding_transcript_variant | MODIFIER |
| 4 | 103444389 | AX-170023572 | 1 | A | T | 0.473 | 7.64E-03 | 5.64E-02 | 2.60E+00 | 2.70E+00 | 9.11E-01 | no | Gm12718 | intron_variant,non_coding_transcript_variant | MODIFIER |
| 2 | 148873219 | AX-170461041 | 2 | A | T | 0.118 | 1.90E-02 | 1.69E-01 | 2.67E+00 | 2.73E+00 | 9.11E-01 | no | Cst3 | intron_variant | MODIFIER |
| 7 | 91418852 | AX-170515242 | 0 | C | A | 0.132 | -2.34E-02 | 2.10E-01 | 2.66E+00 | 2.74E+00 | 9.11E-01 | no | Dlg2 | intron_variant | MODIFIER |
| 7 | 93064322 | AX-170518807 | 0 | C | A | 0.132 | -2.34E-02 | 2.10E-01 | 2.66E+00 | 2.74E+00 | 9.11E-01 | no | Gm9934 | intron_variant,non_coding_transcript_variant | MODIFIER |
| 1 | 39918352 | AX-170475533 | 1 | A | G | 0.375 | 7.71E-03 | 7.49E-02 | 2.68E+00 | 2.76E+00 | 9.11E-01 | no | Map4k4 | intron_variant | MODIFIER |
| 1 | 39935771 | AX-170255859 | 1 | A | C | 0.375 | 7.71E-03 | 7.49E-02 | 2.68E+00 | 2.76E+00 | 9.11E-01 | no | Map4k4 | intron_variant | MODIFIER |
| 19 | 43561781 | AX-170245573 | 0 | G | A | 0.053 | 2.06E-02 | 1.65E-01 | 2.59E+00 | 2.72E+00 | 9.11E-01 | no | AC138790.2 | intron_variant,non_coding_transcript_variant | MODIFIER |
| 4 | 144541668 | AX-170220566 | 0 | C | T | 0.105 | 2.27E-02 | 2.04E-01 | 2.68E+00 | 2.74E+00 | 9.11E-01 | no | Gm13126 | downstream_gene_variant | MODIFIER |
| 4 | 144547374 | AX-170301077 | 0 | G | A | 0.105 | 2.27E-02 | 2.04E-01 | 2.68E+00 | 2.74E+00 | 9.11E-01 | no | Gm13126 | non_coding_transcript_exon_variant | MODIFIER |
| 4 | 145104739 | AX-170488515 | 0 | A | G | 0.105 | 2.27E-02 | 2.04E-01 | 2.68E+00 | 2.74E+00 | 9.11E-01 | no | Vps13d | intron_variant | MODIFIER |
| 10 | 116134022 | AX-170003972 | 0 | C | T | 0.342 | 7.98E-03 | 7.72E-02 | 2.67E+00 | 2.76E+00 | 9.11E-01 | no | Ptprr | intron_variant | MODIFIER |
| 10 | 117298586 | AX-170296787 | 0 | T | C | 0.342 | 7.98E-03 | 7.72E-02 | 2.67E+00 | 2.76E+00 | 9.11E-01 | no | | intergenic_variant | MODIFIER |
| 10 | 117366312 | AX-170003990 | 0 | T | C | 0.342 | 7.98E-03 | 7.72E-02 | 2.67E+00 | 2.76E+00 | 9.11E-01 | no | Cpsf6 | intron_variant | MODIFIER |
| 10 | 117367274 | AX-170299204 | 0 | A | G | 0.342 | 7.98E-03 | 7.72E-02 | 2.67E+00 | 2.76E+00 | 9.11E-01 | no | Cpsf6 | intron_variant | MODIFIER |
| 10 | 117576082 | AX-170250972 | 0 | C | T | 0.342 | 7.98E-03 | 7.72E-02 | 2.67E+00 | 2.76E+00 | 9.11E-01 | no | | regulatory_region_variant | MODIFIER |
| 10 | 117616755 | AX-170104795 | 0 | G | A | 0.342 | 7.98E-03 | 7.72E-02 | 2.67E+00 | 2.76E+00 | 9.11E-01 | no | Gm32365 | non_coding_transcript_exon_variant | MODIFIER |
| 9 | 58788004 | AX-170478628 | 1 | T | C | 0.214 | -1.35E-02 | 1.24E-01 | 2.69E+00 | 2.75E+00 | 9.11E-01 | no | | intergenic_variant | MODIFIER |
| 9 | 55917900 | AX-170462087 | 0 | G | A | 0.211 | -1.35E-02 | 1.24E-01 | 2.69E+00 | 2.74E+00 | 9.11E-01 | no | Scaper | intron_variant | MODIFIER |
| 9 | 56945814 | AX-170180556 | 0 | A | G | 0.211 | -1.35E-02 | 1.24E-01 | 2.69E+00 | 2.74E+00 | 9.11E-01 | no | | intergenic_variant | MODIFIER |
| 9 | 57092620 | AX-170032441 | 0 | T | C | 0.211 | -1.35E-02 | 1.24E-01 | 2.69E+00 | 2.74E+00 | 9.11E-01 | no | Sin3a | intron_variant | MODIFIER |
| 9 | 57126055 | AX-170304162 | 0 | G | C | 0.211 | -1.35E-02 | 1.24E-01 | 2.69E+00 | 2.74E+00 | 9.11E-01 | no | Man2c1 | upstream_gene_variant | MODIFIER |
| 9 | 57137925 | AX-170450794 | 0 | G | T | 0.211 | -1.35E-02 | 1.24E-01 | 2.69E+00 | 2.74E+00 | 9.11E-01 | no | Man2c1 | synonymous_variant | LOW |
| 9 | 57193795 | AX-170202649 | 0 | T | C | 0.211 | -1.35E-02 | 1.24E-01 | 2.69E+00 | 2.74E+00 | 9.11E-01 | no | | intergenic_variant | MODIFIER |
| 9 | 57197674 | AX-170426969 | 0 | T | G | 0.211 | -1.35E-02 | 1.24E-01 | 2.69E+00 | 2.74E+00 | 9.11E-01 | no | | intergenic_variant | MODIFIER |
| 9 | 57202613 | AX-170277139 | 0 | A | C | 0.211 | -1.35E-02 | 1.24E-01 | 2.69E+00 | 2.74E+00 | 9.11E-01 | no | | intergenic_variant | MODIFIER |
| 9 | 57228753 | AX-170488384 | 0 | T | C | 0.211 | -1.35E-02 | 1.24E-01 | 2.69E+00 | 2.74E+00 | 9.11E-01 | no | | intergenic_variant | MODIFIER |
| 9 | 57229629 | AX-170202005 | 0 | T | C | 0.211 | -1.35E-02 | 1.24E-01 | 2.69E+00 | 2.74E+00 | 9.11E-01 | no | | intergenic_variant | MODIFIER |
| 9 | 57232629 | AX-170246753 | 0 | G | A | 0.211 | -1.35E-02 | 1.24E-01 | 2.69E+00 | 2.74E+00 | 9.11E-01 | no | Trcg1 | upstream_gene_variant | MODIFIER |
| 9 | 57247133 | AX-170082513 | 0 | T | G | 0.211 | -1.35E-02 | 1.24E-01 | 2.69E+00 | 2.74E+00 | 9.11E-01 | no | Trcg1 | intron_variant | MODIFIER |
| 9 | 57249928 | AX-170379434 | 0 | A | G | 0.211 | -1.35E-02 | 1.24E-01 | 2.69E+00 | 2.74E+00 | 9.11E-01 | no | 1700017B05Rik | downstream_gene_variant | MODIFIER |
| 9 | 57252006 | AX-170299494 | 0 | A | G | 0.211 | -1.35E-02 | 1.24E-01 | 2.69E+00 | 2.74E+00 | 9.11E-01 | no | 1700017B05Rik | downstream_gene_variant | MODIFIER |
| 9 | 57263708 | AX-170383736 | 0 | T | G | 0.211 | -1.35E-02 | 1.24E-01 | 2.69E+00 | 2.74E+00 | 9.11E-01 | no | 1700017B05Rik | upstream_gene_variant | MODIFIER |
| 9 | 57276405 | AX-170476816 | 0 | A | G | 0.211 | -1.35E-02 | 1.24E-01 | 2.69E+00 | 2.74E+00 | 9.11E-01 | no | | regulatory_region_variant | MODIFIER |
| 9 | 57276628 | AX-170348503 | 0 | T | C | 0.211 | -1.35E-02 | 1.24E-01 | 2.69E+00 | 2.74E+00 | 9.11E-01 | no | | intergenic_variant | MODIFIER |

| | | | | | | | | | | | | | | | |
|---|---|---|---|---|---|---|---|---|---|---|---|---|---|---|---|
| 9 | 57288526 | AX-97805633 | 0 | C | T | 0.211 | -1.35E-02 | 1.24E-01 | 2.69E+00 | 2.74E+00 | 9.11E-01 | no | | intergenic_variant | MODIFIER |
| 9 | 57299833 | AX-170193716 | 0 | T | C | 0.211 | -1.35E-02 | 1.24E-01 | 2.69E+00 | 2.74E+00 | 9.11E-01 | no | | regulatory_region_variant | MODIFIER |
| 9 | 57305212 | AX-170154760 | 0 | A | G | 0.211 | -1.35E-02 | 1.24E-01 | 2.69E+00 | 2.74E+00 | 9.11E-01 | no | | intergenic_variant | MODIFIER |
| 9 | 57350996 | AX-170242133 | 0 | T | C | 0.211 | -1.35E-02 | 1.24E-01 | 2.69E+00 | 2.74E+00 | 9.11E-01 | no | | intergenic_variant | MODIFIER |
| 9 | 57352864 | AX-170312697 | 0 | G | A | 0.211 | -1.35E-02 | 1.24E-01 | 2.69E+00 | 2.74E+00 | 9.11E-01 | no | | intergenic_variant | MODIFIER |
| 9 | 57362923 | AX-170444117 | 0 | T | C | 0.211 | -1.35E-02 | 1.24E-01 | 2.69E+00 | 2.74E+00 | 9.11E-01 | no | | intergenic_variant | MODIFIER |
| 9 | 57363240 | AX-170342062 | 0 | G | A | 0.211 | -1.35E-02 | 1.24E-01 | 2.69E+00 | 2.74E+00 | 9.11E-01 | no | | intergenic_variant | MODIFIER |
| 9 | 57363421 | AX-170333336 | 0 | G | A | 0.211 | -1.35E-02 | 1.24E-01 | 2.69E+00 | 2.74E+00 | 9.11E-01 | no | | intergenic_variant | MODIFIER |
| 9 | 57460454 | AX-170032444 | 0 | T | G | 0.211 | -1.35E-02 | 1.24E-01 | 2.69E+00 | 2.74E+00 | 9.11E-01 | no | Scamp5 | intron_variant | MODIFIER |
| 9 | 57472630 | AX-170032451 | 0 | C | T | 0.211 | -1.35E-02 | 1.24E-01 | 2.69E+00 | 2.74E+00 | 9.11E-01 | no | Scamp5 | upstream_gene_variant | MODIFIER |
| 9 | 57506995 | AX-97795074 | 0 | A | G | 0.211 | -1.35E-02 | 1.24E-01 | 2.69E+00 | 2.74E+00 | 9.11E-01 | no | Rpp25 | downstream_gene_variant | MODIFIER |
| 9 | 57562034 | AX-170510802 | 0 | T | C | 0.211 | -1.35E-02 | 1.24E-01 | 2.69E+00 | 2.74E+00 | 9.11E-01 | no | Scamp2 | intron_variant | MODIFIER |
| 9 | 57800766 | AX-97844918 | 0 | A | G | 0.211 | -1.35E-02 | 1.24E-01 | 2.69E+00 | 2.74E+00 | 9.11E-01 | no | Arid3b | intron_variant | MODIFIER |
| 9 | 57811845 | AX-170240318 | 0 | T | C | 0.211 | -1.35E-02 | 1.24E-01 | 2.69E+00 | 2.74E+00 | 9.11E-01 | no | Arid3b | intron_variant | MODIFIER |
| 9 | 57910843 | AX-170388221 | 0 | G | C | 0.211 | -1.35E-02 | 1.24E-01 | 2.69E+00 | 2.74E+00 | 9.11E-01 | no | Ubl7 | upstream_gene_variant | MODIFIER |
| 9 | 57931554 | AX-170371587 | 0 | G | A | 0.211 | -1.35E-02 | 1.24E-01 | 2.69E+00 | 2.74E+00 | 9.11E-01 | no | Ubl7 | downstream_gene_variant | MODIFIER |
| 9 | 57933832 | AX-170279539 | 0 | A | G | 0.211 | -1.35E-02 | 1.24E-01 | 2.69E+00 | 2.74E+00 | 9.11E-01 | no | Ubl7 | downstream_gene_variant | MODIFIER |
| 9 | 58262383 | AX-170138345 | 0 | C | T | 0.211 | -1.35E-02 | 1.24E-01 | 2.69E+00 | 2.74E+00 | 9.11E-01 | no | Stoml1 | 3_prime_UTR_variant | MODIFIER |
| 9 | 58340485 | AX-170082532 | 0 | A | G | 0.211 | -1.35E-02 | 1.24E-01 | 2.69E+00 | 2.74E+00 | 9.11E-01 | no | | intergenic_variant | MODIFIER |
| 9 | 58473529 | AX-170098248 | 0 | A | G | 0.211 | -1.35E-02 | 1.24E-01 | 2.69E+00 | 2.74E+00 | 9.11E-01 | no | 4930461G14Rik | downstream_gene_variant | MODIFIER |
| 9 | 58496268 | AX-170082538 | 0 | C | T | 0.211 | -1.35E-02 | 1.24E-01 | 2.69E+00 | 2.74E+00 | 9.11E-01 | no | Insyn1 | intron_variant | MODIFIER |
| 9 | 58501346 | AX-97851504 | 0 | T | C | 0.211 | -1.35E-02 | 1.24E-01 | 2.69E+00 | 2.74E+00 | 9.11E-01 | no | Insyn1 | downstream_gene_variant | MODIFIER |
| 9 | 58504006 | AX-170412819 | 0 | G | A | 0.211 | -1.35E-02 | 1.24E-01 | 2.69E+00 | 2.74E+00 | 9.11E-01 | no | Insyn1 | downstream_gene_variant | MODIFIER |
| 9 | 58504094 | AX-170392556 | 0 | G | A | 0.211 | -1.35E-02 | 1.24E-01 | 2.69E+00 | 2.74E+00 | 9.11E-01 | no | Insyn1 | downstream_gene_variant | MODIFIER |
| 9 | 58505877 | AX-170293028 | 0 | C | T | 0.211 | -1.35E-02 | 1.24E-01 | 2.69E+00 | 2.74E+00 | 9.11E-01 | no | | intergenic_variant | MODIFIER |
| 9 | 58514209 | AX-170278041 | 0 | A | G | 0.211 | -1.35E-02 | 1.24E-01 | 2.69E+00 | 2.74E+00 | 9.11E-01 | no | | regulatory_region_variant | MODIFIER |
| 9 | 58519109 | AX-170466726 | 0 | T | A | 0.211 | -1.35E-02 | 1.24E-01 | 2.69E+00 | 2.74E+00 | 9.11E-01 | no | | regulatory_region_variant | MODIFIER |
| 9 | 58521801 | AX-170144028 | 0 | T | C | 0.211 | -1.35E-02 | 1.24E-01 | 2.69E+00 | 2.74E+00 | 9.11E-01 | no | Cd276 | downstream_gene_variant | MODIFIER |
| 9 | 58531002 | AX-170237825 | 0 | G | T | 0.211 | -1.35E-02 | 1.24E-01 | 2.69E+00 | 2.74E+00 | 9.11E-01 | no | Cd276 | intron_variant | MODIFIER |
| 9 | 58534687 | AX-170183457 | 0 | T | C | 0.211 | -1.35E-02 | 1.24E-01 | 2.69E+00 | 2.74E+00 | 9.11E-01 | no | Cd276 | intron_variant | MODIFIER |
| 9 | 58562576 | AX-168380931 | 0 | G | A | 0.211 | -1.35E-02 | 1.24E-01 | 2.69E+00 | 2.74E+00 | 9.11E-01 | no | | intergenic_variant | MODIFIER |
| 9 | 58570763 | AX-170032460 | 0 | T | C | 0.211 | -1.35E-02 | 1.24E-01 | 2.69E+00 | 2.74E+00 | 9.11E-01 | no | | intergenic_variant | MODIFIER |
| 9 | 58574076 | AX-97830805 | 0 | G | A | 0.211 | -1.35E-02 | 1.24E-01 | 2.69E+00 | 2.74E+00 | 9.11E-01 | no | | intergenic_variant | MODIFIER |
| 9 | 58630730 | AX-170032459 | 0 | T | A | 0.211 | -1.35E-02 | 1.24E-01 | 2.69E+00 | 2.74E+00 | 9.11E-01 | no | Nptn | intron_variant | MODIFIER |
| 9 | 58657626 | AX-170082543 | 0 | C | T | 0.211 | -1.35E-02 | 1.24E-01 | 2.69E+00 | 2.74E+00 | 9.11E-01 | no | Nptn | downstream_gene_variant | MODIFIER |
| 9 | 58661091 | AX-170032458 | 0 | G | A | 0.211 | -1.35E-02 | 1.24E-01 | 2.69E+00 | 2.74E+00 | 9.11E-01 | no | Rec114 | intron_variant | MODIFIER |
| 9 | 58665993 | AX-170082542 | 0 | A | G | 0.211 | -1.35E-02 | 1.24E-01 | 2.69E+00 | 2.74E+00 | 9.11E-01 | no | Rec114 | intron_variant | MODIFIER |
| 9 | 58668820 | AX-170082540 | 0 | T | C | 0.211 | -1.35E-02 | 1.24E-01 | 2.69E+00 | 2.74E+00 | 9.11E-01 | no | Rec114 | intron_variant | MODIFIER |
| 9 | 58671419 | AX-170113885 | 0 | G | A | 0.211 | -1.35E-02 | 1.24E-01 | 2.69E+00 | 2.74E+00 | 9.11E-01 | no | Rec114 | intron_variant | MODIFIER |
| 9 | 58674356 | AX-170390323 | 0 | A | G | 0.211 | -1.35E-02 | 1.24E-01 | 2.69E+00 | 2.74E+00 | 9.11E-01 | no | Rec114 | intron_variant | MODIFIER |
| 9 | 58700701 | AX-170535075 | 0 | G | A | 0.211 | -1.35E-02 | 1.24E-01 | 2.69E+00 | 2.74E+00 | 9.11E-01 | no | Rec114 | intron_variant | MODIFIER |
| 9 | 58703341 | AX-170114430 | 0 | T | C | 0.211 | -1.35E-02 | 1.24E-01 | 2.69E+00 | 2.74E+00 | 9.11E-01 | no | Rec114 | intron_variant | MODIFIER |
| 9 | 58722994 | AX-170353846 | 0 | G | A | 0.211 | -1.35E-02 | 1.24E-01 | 2.69E+00 | 2.74E+00 | 9.11E-01 | no | Rec114 | intron_variant | MODIFIER |
| 9 | 58725807 | AX-170333679 | 0 | A | G | 0.211 | -1.35E-02 | 1.24E-01 | 2.69E+00 | 2.74E+00 | 9.11E-01 | no | Rec114 | intron_variant | MODIFIER |
| 9 | 58733758 | AX-170238624 | 0 | T | C | 0.211 | -1.35E-02 | 1.24E-01 | 2.69E+00 | 2.74E+00 | 9.11E-01 | no | Rec114 | intron_variant | MODIFIER |
| 9 | 58739574 | AX-97828588 | 0 | A | G | 0.211 | -1.35E-02 | 1.24E-01 | 2.69E+00 | 2.74E+00 | 9.11E-01 | no | Rec114 | intron_variant | MODIFIER |
| 9 | 58754084 | AX-170536040 | 0 | G | C | 0.211 | -1.35E-02 | 1.24E-01 | 2.69E+00 | 2.74E+00 | 9.11E-01 | no | | intergenic_variant | MODIFIER |
| 9 | 58756559 | AX-170474681 | 0 | G | C | 0.211 | -1.35E-02 | 1.24E-01 | 2.69E+00 | 2.74E+00 | 9.11E-01 | no | | regulatory_region_variant | MODIFIER |
| 9 | 58758359 | AX-170533993 | 0 | G | A | 0.211 | -1.35E-02 | 1.24E-01 | 2.69E+00 | 2.74E+00 | 9.11E-01 | no | | intergenic_variant | MODIFIER |
| 9 | 58758792 | AX-170476573 | 0 | G | T | 0.211 | -1.35E-02 | 1.24E-01 | 2.69E+00 | 2.74E+00 | 9.11E-01 | no | | regulatory_region_variant | MODIFIER |
| 9 | 58759577 | AX-170281620 | 0 | G | A | 0.211 | -1.35E-02 | 1.24E-01 | 2.69E+00 | 2.74E+00 | 9.11E-01 | no | | intergenic_variant | MODIFIER |
| 9 | 58759963 | AX-170271449 | 0 | A | T | 0.211 | -1.35E-02 | 1.24E-01 | 2.69E+00 | 2.74E+00 | 9.11E-01 | no | | intergenic_variant | MODIFIER |
| 9 | 58764116 | AX-170346499 | 0 | A | G | 0.211 | -1.35E-02 | 1.24E-01 | 2.69E+00 | 2.74E+00 | 9.11E-01 | no | | intergenic_variant | MODIFIER |
| 9 | 58764440 | AX-170162063 | 0 | G | C | 0.211 | -1.35E-02 | 1.24E-01 | 2.69E+00 | 2.74E+00 | 9.11E-01 | no | | intergenic_variant | MODIFIER |

| | | | | | | | | | | | | | | | |
|---|---|---|---|---|---|---|---|---|---|---|---|---|---|---|---|
| 9 | 58771343 | AX-170299822 | 0 | A | G | 0.211 | -1.35E-02 | 1.24E-01 | 2.69E+00 | 2.74E+00 | 9.11E-01 | no | | intergenic_variant | MODIFIER |
| 9 | 58786791 | AX-170429272 | 0 | G | A | 0.211 | -1.35E-02 | 1.24E-01 | 2.69E+00 | 2.74E+00 | 9.11E-01 | no | | intergenic_variant | MODIFIER |
| 9 | 58788278 | AX-170200495 | 0 | C | T | 0.211 | -1.35E-02 | 1.24E-01 | 2.69E+00 | 2.74E+00 | 9.11E-01 | no | | intergenic_variant | MODIFIER |
| 9 | 58791620 | AX-170395921 | 0 | C | T | 0.211 | -1.35E-02 | 1.24E-01 | 2.69E+00 | 2.74E+00 | 9.11E-01 | no | | intergenic_variant | MODIFIER |
| 9 | 58803936 | AX-97856569 | 0 | T | C | 0.211 | -1.35E-02 | 1.24E-01 | 2.69E+00 | 2.74E+00 | 9.11E-01 | no | | regulatory_region_variant | MODIFIER |
| 9 | 58853179 | AX-170269705 | 0 | A | G | 0.211 | -1.35E-02 | 1.24E-01 | 2.69E+00 | 2.74E+00 | 9.11E-01 | no | Hcn4 | intron_variant | MODIFIER |
| 9 | 58890871 | AX-170123314 | 0 | A | G | 0.211 | -1.35E-02 | 1.24E-01 | 2.69E+00 | 2.74E+00 | 9.11E-01 | no | Neo1 | intron_variant | MODIFIER |
| 9 | 58909438 | AX-170449713 | 0 | C | G | 0.211 | -1.35E-02 | 1.24E-01 | 2.69E+00 | 2.74E+00 | 9.11E-01 | no | Neo1 | intron_variant | MODIFIER |
| 9 | 58917207 | AX-170410747 | 0 | T | C | 0.211 | -1.35E-02 | 1.24E-01 | 2.69E+00 | 2.74E+00 | 9.11E-01 | no | Neo1 | intron_variant | MODIFIER |
| 9 | 58920445 | AX-170399562 | 0 | A | C | 0.211 | -1.35E-02 | 1.24E-01 | 2.69E+00 | 2.74E+00 | 9.11E-01 | no | Neo1 | intron_variant | MODIFIER |
| 9 | 58933134 | AX-170453942 | 0 | A | C | 0.211 | -1.35E-02 | 1.24E-01 | 2.69E+00 | 2.74E+00 | 9.11E-01 | no | Neo1 | intron_variant | MODIFIER |
| 9 | 58935326 | AX-97808570 | 0 | A | G | 0.211 | -1.35E-02 | 1.24E-01 | 2.69E+00 | 2.74E+00 | 9.11E-01 | no | Neo1 | intron_variant | MODIFIER |
| 9 | 58935608 | AX-170438669 | 0 | C | T | 0.211 | -1.35E-02 | 1.24E-01 | 2.69E+00 | 2.74E+00 | 9.11E-01 | no | Neo1 | intron_variant | MODIFIER |
| 9 | 59067785 | AX-170288292 | 0 | T | C | 0.211 | -1.35E-02 | 1.24E-01 | 2.69E+00 | 2.74E+00 | 9.11E-01 | no | | intergenic_variant | MODIFIER |
| 9 | 59071883 | AX-170263829 | 0 | G | T | 0.211 | -1.35E-02 | 1.24E-01 | 2.69E+00 | 2.74E+00 | 9.11E-01 | no | | regulatory_region_variant | MODIFIER |
| 9 | 59084239 | AX-170205037 | 0 | A | G | 0.211 | -1.35E-02 | 1.24E-01 | 2.69E+00 | 2.74E+00 | 9.11E-01 | no | | intergenic_variant | MODIFIER |
| 9 | 59088359 | AX-170216659 | 0 | G | C | 0.211 | -1.35E-02 | 1.24E-01 | 2.69E+00 | 2.74E+00 | 9.11E-01 | no | | intergenic_variant | MODIFIER |
| 9 | 59090612 | AX-170449667 | 0 | A | G | 0.211 | -1.35E-02 | 1.24E-01 | 2.69E+00 | 2.74E+00 | 9.11E-01 | no | | intergenic_variant | MODIFIER |
| 9 | 59133752 | AX-170197666 | 0 | C | T | 0.211 | -1.35E-02 | 1.24E-01 | 2.69E+00 | 2.74E+00 | 9.11E-01 | no | | regulatory_region_variant | MODIFIER |
| 9 | 59144543 | AX-170370975 | 0 | G | T | 0.211 | -1.35E-02 | 1.24E-01 | 2.69E+00 | 2.74E+00 | 9.11E-01 | no | Gm7589 | downstream_gene_variant | MODIFIER |
| 9 | 59177268 | AX-170238740 | 0 | A | G | 0.211 | -1.35E-02 | 1.24E-01 | 2.69E+00 | 2.74E+00 | 9.11E-01 | no | | intergenic_variant | MODIFIER |
| 9 | 59191078 | AX-97815100 | 0 | C | T | 0.211 | -1.35E-02 | 1.24E-01 | 2.69E+00 | 2.74E+00 | 9.11E-01 | no | Gm33053 | downstream_gene_variant | MODIFIER |
| 9 | 59191465 | AX-170082552 | 0 | A | G | 0.211 | -1.35E-02 | 1.24E-01 | 2.69E+00 | 2.74E+00 | 9.11E-01 | no | Gm33053 | non_coding_transcript_exon_variant | MODIFIER |
| 9 | 59198312 | AX-170422892 | 0 | G | A | 0.211 | -1.35E-02 | 1.24E-01 | 2.69E+00 | 2.74E+00 | 9.11E-01 | no | Gm33053 | intron_variant,non_coding_transcript_variant | MODIFIER |
| 9 | 59199751 | AX-170032464 | 0 | T | C | 0.211 | -1.35E-02 | 1.24E-01 | 2.69E+00 | 2.74E+00 | 9.11E-01 | no | Gm33053 | intron_variant,non_coding_transcript_variant | MODIFIER |
| 9 | 59209220 | AX-170209403 | 0 | A | G | 0.211 | -1.35E-02 | 1.24E-01 | 2.69E+00 | 2.74E+00 | 9.11E-01 | no | Gm33053 | intron_variant,non_coding_transcript_variant | MODIFIER |
| 9 | 59210493 | AX-170405041 | 0 | T | G | 0.211 | -1.35E-02 | 1.24E-01 | 2.69E+00 | 2.74E+00 | 9.11E-01 | no | Gm33053 | non_coding_transcript_exon_variant | MODIFIER |
| 9 | 59248693 | AX-170032468 | 0 | T | C | 0.211 | -1.35E-02 | 1.24E-01 | 2.69E+00 | 2.74E+00 | 9.11E-01 | no | | intergenic_variant | MODIFIER |
| 9 | 59279708 | AX-170110541 | 0 | C | T | 0.211 | -1.35E-02 | 1.24E-01 | 2.69E+00 | 2.74E+00 | 9.11E-01 | no | | intergenic_variant | MODIFIER |
| 9 | 59301522 | AX-170032466 | 0 | A | G | 0.211 | -1.35E-02 | 1.24E-01 | 2.69E+00 | 2.74E+00 | 9.11E-01 | no | Adpgk | intron_variant | MODIFIER |
| 9 | 59314087 | AX-170206902 | 0 | C | T | 0.211 | -1.35E-02 | 1.24E-01 | 2.69E+00 | 2.74E+00 | 9.11E-01 | no | Adpgk | intron_variant | MODIFIER |
| 9 | 59321936 | AX-170233164 | 0 | A | G | 0.211 | -1.35E-02 | 1.24E-01 | 2.69E+00 | 2.74E+00 | 9.11E-01 | no | Bbs4 | downstream_gene_variant | MODIFIER |
| 9 | 59346419 | AX-170400661 | 0 | G | C | 0.211 | -1.35E-02 | 1.24E-01 | 2.69E+00 | 2.74E+00 | 9.11E-01 | no | Bbs4 | intron_variant | MODIFIER |
| 9 | 59391873 | AX-97793107 | 0 | C | A | 0.211 | -1.35E-02 | 1.24E-01 | 2.69E+00 | 2.74E+00 | 9.11E-01 | no | Arih1 | downstream_gene_variant | MODIFIER |
| 9 | 59434886 | AX-170189391 | 0 | C | T | 0.211 | -1.35E-02 | 1.24E-01 | 2.69E+00 | 2.74E+00 | 9.11E-01 | no | Arih1 | intron_variant,NMD_transcript_variant | MODIFIER |
| 9 | 59440464 | AX-170082556 | 0 | G | A | 0.211 | -1.35E-02 | 1.24E-01 | 2.69E+00 | 2.74E+00 | 9.11E-01 | no | Arih1 | intron_variant,NMD_transcript_variant | MODIFIER |
| 9 | 59460466 | AX-170370276 | 0 | T | A | 0.211 | -1.35E-02 | 1.24E-01 | 2.69E+00 | 2.74E+00 | 9.11E-01 | no | Arih1 | intron_variant,NMD_transcript_variant | MODIFIER |
| 9 | 59511160 | AX-170174929 | 0 | C | T | 0.211 | -1.35E-02 | 1.24E-01 | 2.69E+00 | 2.74E+00 | 9.11E-01 | no | Gm33180 | intron_variant,non_coding_transcript_variant | MODIFIER |
| 9 | 60052845 | AX-97804957 | 0 | G | A | 0.211 | -1.35E-02 | 1.24E-01 | 2.69E+00 | 2.74E+00 | 9.11E-01 | no | Thsd4 | intron_variant | MODIFIER |
| 1 | 3373734 | AX-97810602 | 0 | C | A | 0.518 | 8.79E-03 | 6.84E-02 | 2.62E+00 | 2.70E+00 | 9.11E-01 | no | Xkr4 | intron_variant | MODIFIER |
| 8 | 95479947 | AX-170149844 | 0 | A | C | 0.228 | 1.26E-02 | 1.16E-01 | 2.68E+00 | 2.74E+00 | 9.11E-01 | no | Csnk2a2 | intron_variant | MODIFIER |
| 13 | 52959744 | AX-170184795 | 1 | G | T | 0.5 | 8.37E-03 | 6.62E-02 | 2.60E+00 | 2.71E+00 | 9.11E-01 | no | n-R5s54 | downstream_gene_variant | MODIFIER |
| 18 | 59657698 | AX-97853167 | 0 | G | C | 0.412 | 1.11E-02 | 8.60E-02 | 2.61E+00 | 2.71E+00 | 9.11E-01 | no | | intergenic_variant | MODIFIER |
| 9 | 105736586 | AX-170083361 | 0 | G | A | 0.123 | -1.46E-02 | 1.18E-01 | 2.64E+00 | 2.71E+00 | 9.11E-01 | no | Col6a6 | intron_variant | MODIFIER |
| 15 | 15546847 | AX-97789391 | 0 | T | C | 0.237 | -1.26E-02 | 1.06E-01 | 2.65E+00 | 2.71E+00 | 9.12E-01 | no | | intergenic_variant | MODIFIER |
| 7 | 91594629 | AX-170317875 | 1 | T | C | 0.134 | 2.29E-02 | 2.11E-01 | 2.66E+00 | 2.74E+00 | 9.12E-01 | no | Dlg2 | intron_variant | MODIFIER |
| 20 | 95100033 | AX-168417090 | 0 | G | A | 0.289 | -5.79E-03 | 5.69E-02 | 2.64E+00 | 2.75E+00 | 9.12E-01 | no | Arhgef9 | intron_variant | MODIFIER |
| 9 | 87344356 | AX-170434618 | 0 | T | C | 0.123 | 1.29E-02 | 1.29E-01 | 2.69E+00 | 2.77E+00 | 9.12E-01 | no | | intergenic_variant | MODIFIER |
| 2 | 119006975 | AX-170455095 | 0 | C | T | 0.14 | 2.45E-02 | 2.21E-01 | 2.70E+00 | 2.73E+00 | 9.12E-01 | no | Uqcrh-ps2 | upstream_gene_variant | MODIFIER |
| 2 | 122915285 | AX-97784419 | 0 | G | A | 0.14 | 2.45E-02 | 2.21E-01 | 2.70E+00 | 2.73E+00 | 9.12E-01 | no | | intergenic_variant | MODIFIER |
| 2 | 137333461 | AX-170441897 | 0 | A | T | 0.14 | 2.45E-02 | 2.21E-01 | 2.70E+00 | 2.73E+00 | 9.12E-01 | no | Gm14055 | downstream_gene_variant | MODIFIER |
| 2 | 142447785 | AX-170019223 | 0 | C | T | 0.14 | 2.45E-02 | 2.21E-01 | 2.70E+00 | 2.73E+00 | 9.12E-01 | no | | intergenic_variant | MODIFIER |
| 2 | 154846641 | AX-170272875 | 0 | G | A | 0.14 | 2.45E-02 | 2.21E-01 | 2.70E+00 | 2.73E+00 | 9.12E-01 | no | Raly | intron_variant | MODIFIER |
| 9 | 67330784 | AX-170377820 | 0 | C | T | 0.193 | -1.53E-02 | 1.37E-01 | 2.70E+00 | 2.73E+00 | 9.12E-01 | no | Tln2 | intron_variant | MODIFIER |

| | | | | | | | | | | | | | | | |
|---|---|---|---|---|---|---|---|---|---|---|---|---|---|---|---|
| 1 | 190065535 | AX-170508947 | 0 | A | C | 0.254 | 1.32E-02 | 9.02E-02 | 2.57E+00 | 2.69E+00 | 9.12E-01 | no | | intergenic_variant | MODIFIER |
| 5 | 93399319 | AX-170025634 | 0 | C | T | 0.43 | -1.05E-02 | 1.02E-01 | 2.68E+00 | 2.75E+00 | 9.12E-01 | no | | intergenic_variant | MODIFIER |
| 15 | 95589605 | AX-170148041 | 0 | A | G | 0.149 | -1.85E-02 | 1.80E-01 | 2.70E+00 | 2.76E+00 | 9.12E-01 | no | | intergenic_variant | MODIFIER |
| 15 | 97512220 | AX-170259706 | 0 | G | C | 0.149 | -1.85E-02 | 1.80E-01 | 2.70E+00 | 2.76E+00 | 9.12E-01 | no | | intergenic_variant | MODIFIER |
| 11 | 46584730 | AX-170398839 | 0 | C | T | 0.079 | 1.77E-02 | 1.78E-01 | 2.63E+00 | 2.75E+00 | 9.13E-01 | no | BC053393 | intron_variant | MODIFIER |
| 16 | 92328110 | AX-97832210 | 1 | T | C | 0.473 | -8.44E-03 | 7.86E-02 | 2.66E+00 | 2.74E+00 | 9.13E-01 | no | Smim34 | downstream_gene_variant | MODIFIER |
| 8 | 54521568 | AX-170453337 | 2 | G | A | 0.155 | 1.54E-02 | 1.51E-01 | 2.72E+00 | 2.77E+00 | 9.13E-01 | no | Spcs3 | 3_prime_UTR_variant | MODIFIER |
| 9 | 37306191 | AX-170247332 | 0 | G | A | 0.509 | -8.32E-03 | 8.11E-02 | 2.67E+00 | 2.75E+00 | 9.13E-01 | no | Ccdc15 | intron_variant | MODIFIER |
| 11 | 5106402 | AX-170178421 | 0 | T | A | 0.07 | -2.27E-02 | 2.02E-01 | 2.62E+00 | 2.73E+00 | 9.13E-01 | no | Rhbdd3 | downstream_gene_variant | MODIFIER |
| 11 | 13609984 | AX-170450037 | 0 | T | C | 0.07 | -2.27E-02 | 2.02E-01 | 2.62E+00 | 2.73E+00 | 9.13E-01 | no | | intergenic_variant | MODIFIER |
| 5 | 118753711 | AX-170495968 | 1 | T | C | 0.455 | -1.22E-02 | 1.16E-01 | 2.72E+00 | 2.76E+00 | 9.13E-01 | no | Med13l | intron_variant | MODIFIER |
| 5 | 125085057 | AX-170026088 | 1 | G | A | 0.455 | -1.22E-02 | 1.16E-01 | 2.72E+00 | 2.76E+00 | 9.13E-01 | no | Ncor2 | splice_region_variant,synonymous_variant | LOW |
| 10 | 6769412 | AX-97813151 | 0 | A | T | 0.193 | 1.11E-02 | 1.02E-01 | 2.64E+00 | 2.74E+00 | 9.13E-01 | no | Oprm1 | intron_variant | MODIFIER |
| 5 | 84242263 | AX-170025461 | 1 | C | T | 0.438 | -1.15E-02 | 1.07E-01 | 2.68E+00 | 2.74E+00 | 9.13E-01 | no | Epha5 | intron_variant | MODIFIER |
| 12 | 101046319 | AX-170154818 | 1 | G | A | 0.08 | 1.90E-02 | 1.76E-01 | 2.64E+00 | 2.74E+00 | 9.13E-01 | no | Ppp4r3a | intron_variant | MODIFIER |
| 1 | 26111379 | AX-170219781 | 1 | A | G | 0.393 | 9.34E-03 | 8.34E-02 | 2.65E+00 | 2.73E+00 | 9.13E-01 | no | | intergenic_variant | MODIFIER |
| 8 | 94391256 | AX-170459013 | 1 | C | G | 0.205 | -1.27E-02 | 1.18E-01 | 2.67E+00 | 2.74E+00 | 9.13E-01 | no | Herpud1 | intron_variant | MODIFIER |
| 1 | 78401333 | AX-170406728 | 0 | G | A | 0.149 | -1.87E-02 | 1.80E-01 | 2.67E+00 | 2.75E+00 | 9.13E-01 | no | Sgpp2 | intron_variant | MODIFIER |
| 12 | 64520071 | AX-170006912 | 1 | G | A | 0.062 | -3.27E-02 | 3.01E-01 | 2.71E+00 | 2.73E+00 | 9.13E-01 | no | | intergenic_variant | MODIFIER |
| 7 | 79218897 | AX-97829471 | 0 | T | C | 0.123 | -1.67E-02 | 1.73E-01 | 2.64E+00 | 2.76E+00 | 9.13E-01 | no | | intergenic_variant | MODIFIER |
| 1 | 8265328 | AX-170458128 | 1 | C | T | 0.518 | 8.89E-03 | 7.16E-02 | 2.63E+00 | 2.70E+00 | 9.13E-01 | no | | intergenic_variant | MODIFIER |
| 1 | 9750481 | AX-97842847 | 1 | A | G | 0.518 | 8.89E-03 | 7.16E-02 | 2.63E+00 | 2.70E+00 | 9.13E-01 | no | Vcpip1 | upstream_gene_variant | MODIFIER |
| 5 | 37752667 | AX-170233644 | 0 | A | G | 0.342 | 1.18E-02 | 1.09E-01 | 2.66E+00 | 2.73E+00 | 9.14E-01 | no | | regulatory_region_variant | MODIFIER |
| 13 | 31441261 | AX-170139265 | 1 | T | C | 0.179 | -1.45E-02 | 1.22E-01 | 2.64E+00 | 2.71E+00 | 9.14E-01 | no | | intergenic_variant | MODIFIER |
| 13 | 32055800 | AX-170143050 | 1 | G | C | 0.179 | -1.45E-02 | 1.22E-01 | 2.64E+00 | 2.71E+00 | 9.14E-01 | no | Gmds | intron_variant | MODIFIER |
| 13 | 33762935 | AX-97832056 | 1 | T | C | 0.179 | -1.45E-02 | 1.22E-01 | 2.64E+00 | 2.71E+00 | 9.14E-01 | no | Vmn2r-ps94 | upstream_gene_variant | MODIFIER |
| 13 | 36210161 | AX-170087405 | 1 | C | A | 0.179 | -1.45E-02 | 1.22E-01 | 2.64E+00 | 2.71E+00 | 9.14E-01 | no | Fars2 | intron_variant | MODIFIER |
| 1 | 51242336 | AX-170414563 | 1 | C | T | 0.107 | 2.53E-02 | 2.35E-01 | 2.69E+00 | 2.74E+00 | 9.14E-01 | no | 9330175M20Rik | intron_variant,non_coding_transcript_variant | MODIFIER |
| 1 | 51258618 | AX-170106264 | 1 | G | A | 0.107 | 2.53E-02 | 2.35E-01 | 2.69E+00 | 2.74E+00 | 9.14E-01 | no | 9330175M20Rik | intron_variant,non_coding_transcript_variant | MODIFIER |
| 13 | 32224878 | AX-97801630 | 0 | T | C | 0.184 | -1.38E-02 | 1.19E-01 | 2.64E+00 | 2.72E+00 | 9.14E-01 | no | Gmds | intron_variant | MODIFIER |
| 12 | 102176535 | AX-170007414 | 2 | T | C | 0.064 | -1.82E-02 | 1.68E-01 | 2.64E+00 | 2.73E+00 | 9.14E-01 | no | Slc24a4 | intron_variant | MODIFIER |
| 4 | 141456317 | AX-170282155 | 0 | T | G | 0.105 | 2.19E-02 | 2.04E-01 | 2.67E+00 | 2.74E+00 | 9.14E-01 | no | Zbtb17 | intron_variant | MODIFIER |
| 15 | 94021586 | AX-170162851 | 0 | G | A | 0.149 | -1.79E-02 | 1.78E-01 | 2.69E+00 | 2.76E+00 | 9.14E-01 | no | | intergenic_variant | MODIFIER |
| 8 | 82965643 | AX-170315949 | 1 | G | A | 0.205 | 1.39E-02 | 1.30E-01 | 2.68E+00 | 2.74E+00 | 9.14E-01 | no | Rnf150 | intron_variant | MODIFIER |
| 9 | 79241470 | AX-97852389 | 0 | G | A | 0.202 | 1.31E-02 | 1.30E-01 | 2.71E+00 | 2.76E+00 | 9.14E-01 | no | Gm28620 | downstream_gene_variant | MODIFIER |
| 7 | 54641277 | AX-170414371 | 0 | C | T | 0.43 | -9.79E-03 | 9.70E-02 | 2.67E+00 | 2.75E+00 | 9.14E-01 | no | | intergenic_variant | MODIFIER |
| 7 | 55349392 | AX-170116408 | 0 | C | T | 0.43 | -9.79E-03 | 9.70E-02 | 2.67E+00 | 2.75E+00 | 9.14E-01 | no | | intergenic_variant | MODIFIER |
| 18 | 58689090 | AX-170404713 | 1 | A | G | 0.42 | 8.32E-03 | 8.48E-02 | 2.71E+00 | 2.77E+00 | 9.14E-01 | no | | intergenic_variant | MODIFIER |
| 2 | 127059860 | AX-170189384 | 0 | T | G | 0.149 | 1.81E-02 | 1.62E-01 | 2.66E+00 | 2.72E+00 | 9.15E-01 | no | Ap4e1 | intron_variant | MODIFIER |
| 2 | 147787451 | AX-170190348 | 0 | A | T | 0.149 | 1.81E-02 | 1.62E-01 | 2.66E+00 | 2.72E+00 | 9.15E-01 | no | | intergenic_variant | MODIFIER |
| 7 | 109304395 | AX-170029751 | 0 | C | T | 0.474 | -9.39E-03 | 9.41E-02 | 2.68E+00 | 2.76E+00 | 9.15E-01 | no | Stk33 | intron_variant | MODIFIER |
| 8 | 81981912 | AX-170185360 | 0 | T | A | 0.202 | -1.38E-02 | 1.30E-01 | 2.68E+00 | 2.74E+00 | 9.15E-01 | no | Inpp4b | intron_variant | MODIFIER |
| 18 | 33506147 | AX-170367998 | 1 | A | C | 0.062 | 3.14E-02 | 3.01E-01 | 2.73E+00 | 2.75E+00 | 9.15E-01 | no | | intergenic_variant | MODIFIER |
| 8 | 94832338 | AX-170323922 | 0 | C | A | 0.219 | -1.16E-02 | 1.10E-01 | 2.66E+00 | 2.74E+00 | 9.15E-01 | no | Ciapin1 | intron_variant | MODIFIER |
| 9 | 69922186 | AX-170320925 | 0 | C | T | 0.193 | -1.51E-02 | 1.40E-01 | 2.71E+00 | 2.73E+00 | 9.15E-01 | no | | intergenic_variant | MODIFIER |
| 8 | 98335039 | AX-170031353 | 1 | T | C | 0.223 | -1.19E-02 | 1.15E-01 | 2.67E+00 | 2.74E+00 | 9.15E-01 | no | | intergenic_variant | MODIFIER |
| 15 | 92190309 | AX-170301590 | 1 | T | G | 0.143 | -1.34E-02 | 1.25E-01 | 2.63E+00 | 2.73E+00 | 9.15E-01 | no | Cntn1 | intron_variant | MODIFIER |
| 10 | 120066279 | AX-170234333 | 1 | C | T | 0.339 | 7.32E-03 | 7.11E-02 | 2.65E+00 | 2.74E+00 | 9.15E-01 | no | Grip1 | intron_variant | MODIFIER |
| 11 | 18675932 | AX-170190396 | 0 | C | T | 0.061 | -2.33E-02 | 2.14E-01 | 2.64E+00 | 2.73E+00 | 9.15E-01 | no | Gm12023 | intron_variant,non_coding_transcript_variant | MODIFIER |
| 2 | 123832092 | AX-170198698 | 1 | C | T | 0.125 | -1.86E-02 | 1.86E-01 | 2.65E+00 | 2.75E+00 | 9.15E-01 | no | | intergenic_variant | MODIFIER |
| 15 | 98406740 | AX-97833755 | 1 | G | T | 0.125 | -1.61E-02 | 1.64E-01 | 2.69E+00 | 2.76E+00 | 9.15E-01 | no | 1700031M16Rik | intron_variant,non_coding_transcript_variant | MODIFIER |
| 2 | 121942931 | AX-170325735 | 0 | A | G | 0.149 | 2.00E-02 | 1.78E-01 | 2.64E+00 | 2.72E+00 | 9.15E-01 | no | | intergenic_variant | MODIFIER |
| 1 | 39091389 | AX-170000325 | 0 | C | T | 0.465 | 7.62E-03 | 7.53E-02 | 2.65E+00 | 2.75E+00 | 9.15E-01 | no | | intergenic_variant | MODIFIER |
| 2 | 114658038 | AX-97851805 | 0 | G | A | 0.14 | 2.32E-02 | 2.20E-01 | 2.70E+00 | 2.73E+00 | 9.16E-01 | no | Dph6 | upstream_gene_variant | MODIFIER |

| | | | | | | | | | | | | | | | |
|---|---|---|---|---|---|---|---|---|---|---|---|---|---|---|---|
| 2 | 123050473 | AX-170513148 | 0 | C | T | 0.14 | 2.32E-02 | 2.20E-01 | 2.70E+00 | 2.73E+00 | 9.16E-01 | no | | intergenic_variant | MODIFIER |
| 2 | 123199754 | AX-170018902 | 0 | T | C | 0.14 | 2.32E-02 | 2.20E-01 | 2.70E+00 | 2.73E+00 | 9.16E-01 | no | | intergenic_variant | MODIFIER |
| 2 | 155770022 | AX-170019438 | 0 | C | T | 0.14 | 2.32E-02 | 2.20E-01 | 2.70E+00 | 2.73E+00 | 9.16E-01 | no | | intergenic_variant | MODIFIER |
| 2 | 156238026 | AX-170224284 | 0 | G | A | 0.14 | 2.32E-02 | 2.20E-01 | 2.70E+00 | 2.73E+00 | 9.16E-01 | no | Phf20 | intron_variant | MODIFIER |
| 5 | 87844705 | AX-170186667 | 0 | C | T | 0.43 | 1.09E-02 | 1.07E-01 | 2.68E+00 | 2.74E+00 | 9.16E-01 | no | Prr27 | intron_variant | MODIFIER |
| 10 | 92260097 | AX-170427205 | 0 | G | A | 0.307 | 7.78E-03 | 8.21E-02 | 2.69E+00 | 2.77E+00 | 9.16E-01 | no | Gm20757 | intron_variant,non_coding_transcript_variant | MODIFIER |
| 10 | 114870032 | AX-170462002 | 2 | G | T | 0.364 | 7.93E-03 | 7.87E-02 | 2.66E+00 | 2.75E+00 | 9.16E-01 | no | Gm31182 | upstream_gene_variant | MODIFIER |
| 4 | 101360130 | AX-170474028 | 0 | C | T | 0.342 | -9.52E-03 | 9.71E-02 | 2.66E+00 | 2.75E+00 | 9.16E-01 | no | Gm12795 | downstream_gene_variant | MODIFIER |
| 18 | 65588623 | AX-170272848 | 0 | T | A | 0.412 | -9.24E-03 | 7.96E-02 | 2.64E+00 | 2.71E+00 | 9.16E-01 | no | Zfp532 | intron_variant | MODIFIER |
| 13 | 31914127 | AX-170175585 | 0 | A | C | 0.184 | 1.08E-02 | 1.14E-01 | 2.68E+00 | 2.76E+00 | 9.16E-01 | no | Gmds | intron_variant | MODIFIER |
| 10 | 119030046 | AX-170437190 | 0 | A | G | 0.325 | 7.28E-03 | 7.24E-02 | 2.66E+00 | 2.75E+00 | 9.16E-01 | no | Gm33677 | non_coding_transcript_exon_variant | MODIFIER |
| 5 | 125614045 | AX-170026098 | 1 | T | C | 0.438 | -1.09E-02 | 1.07E-01 | 2.68E+00 | 2.74E+00 | 9.16E-01 | no | Tmem132b | intron_variant | MODIFIER |
| 5 | 128011076 | AX-170268041 | 1 | C | T | 0.411 | -7.99E-03 | 8.33E-02 | 2.65E+00 | 2.75E+00 | 9.16E-01 | no | Tmem132d | intron_variant | MODIFIER |
| 5 | 126206210 | AX-97780781 | 1 | C | T | 0.464 | 1.08E-02 | 1.00E-01 | 2.65E+00 | 2.73E+00 | 9.16E-01 | no | | intergenic_variant | MODIFIER |
| 11 | 20401560 | AX-170234599 | 0 | G | A | 0.053 | -2.08E-02 | 2.02E-01 | 2.61E+00 | 2.74E+00 | 9.16E-01 | no | | regulatory_region_variant | MODIFIER |
| 5 | 93717612 | AX-97840119 | 0 | G | T | 0.421 | -1.05E-02 | 1.05E-01 | 2.68E+00 | 2.75E+00 | 9.16E-01 | no | | intergenic_variant | MODIFIER |
| 1 | 7269313 | AX-97836024 | 1 | C | T | 0.08 | 1.90E-02 | 1.71E-01 | 2.61E+00 | 2.73E+00 | 9.16E-01 | no | Gm23274 | downstream_gene_variant | MODIFIER |
| 7 | 135003001 | AX-170225044 | 0 | G | T | 0.254 | -9.44E-03 | 8.11E-02 | 2.62E+00 | 2.72E+00 | 9.16E-01 | no | Dock1 | intron_variant | MODIFIER |
| 7 | 82076923 | AX-170263385 | 1 | C | T | 0.116 | 2.25E-02 | 2.11E-01 | 2.65E+00 | 2.73E+00 | 9.16E-01 | no | Gm20744 | intron_variant,non_coding_transcript_variant | MODIFIER |
| 3 | 36474653 | AX-97793898 | 0 | A | G | 0.193 | 8.92E-03 | 8.79E-02 | 2.64E+00 | 2.74E+00 | 9.16E-01 | no | Anxa5 | intron_variant | MODIFIER |
| 3 | 36685072 | AX-97849104 | 0 | T | C | 0.193 | 8.92E-03 | 8.79E-02 | 2.64E+00 | 2.74E+00 | 9.16E-01 | no | Trpc3 | intron_variant | MODIFIER |
| 1 | 40282099 | AX-170439462 | 0 | C | T | 0.412 | -8.44E-03 | 7.65E-02 | 2.65E+00 | 2.72E+00 | 9.16E-01 | no | Il1r1 | intron_variant | MODIFIER |
| 1 | 40436268 | AX-170514995 | 0 | G | A | 0.412 | -8.44E-03 | 7.65E-02 | 2.65E+00 | 2.72E+00 | 9.16E-01 | no | Il1rl1 | upstream_gene_variant | MODIFIER |
| 10 | 114742109 | AX-170086906 | 0 | A | G | 0.36 | 7.84E-03 | 7.81E-02 | 2.66E+00 | 2.75E+00 | 9.16E-01 | no | Trhde | intron_variant | MODIFIER |
| 10 | 114756436 | AX-170139740 | 0 | G | A | 0.36 | 7.84E-03 | 7.81E-02 | 2.66E+00 | 2.75E+00 | 9.16E-01 | no | Trhde | intron_variant | MODIFIER |
| 10 | 114786166 | AX-170494470 | 0 | C | T | 0.36 | 7.84E-03 | 7.81E-02 | 2.66E+00 | 2.75E+00 | 9.16E-01 | no | Trhde | intron_variant | MODIFIER |
| 10 | 114786845 | AX-170362021 | 0 | G | A | 0.36 | 7.84E-03 | 7.81E-02 | 2.66E+00 | 2.75E+00 | 9.16E-01 | no | Trhde | intron_variant | MODIFIER |
| 10 | 114788471 | AX-170106563 | 0 | C | A | 0.36 | 7.84E-03 | 7.81E-02 | 2.66E+00 | 2.75E+00 | 9.16E-01 | no | Trhde | intron_variant | MODIFIER |
| 10 | 114811740 | AX-170098945 | 0 | A | C | 0.36 | 7.84E-03 | 7.81E-02 | 2.66E+00 | 2.75E+00 | 9.16E-01 | no | Gm15723 | intron_variant,non_coding_transcript_variant | MODIFIER |
| 10 | 114811878 | AX-170174508 | 0 | A | T | 0.36 | 7.84E-03 | 7.81E-02 | 2.66E+00 | 2.75E+00 | 9.16E-01 | no | Gm15723 | intron_variant,non_coding_transcript_variant | MODIFIER |
| 10 | 114869460 | AX-170177499 | 0 | A | C | 0.36 | 7.84E-03 | 7.81E-02 | 2.66E+00 | 2.75E+00 | 9.16E-01 | no | | intergenic_variant | MODIFIER |
| 10 | 114869601 | AX-170470538 | 0 | C | T | 0.36 | 7.84E-03 | 7.81E-02 | 2.66E+00 | 2.75E+00 | 9.16E-01 | no | | intergenic_variant | MODIFIER |
| 10 | 114875518 | AX-170376462 | 0 | T | C | 0.36 | 7.84E-03 | 7.81E-02 | 2.66E+00 | 2.75E+00 | 9.16E-01 | no | Gm31182 | intron_variant,non_coding_transcript_variant | MODIFIER |
| 10 | 114892869 | AX-170152584 | 0 | C | A | 0.36 | 7.84E-03 | 7.81E-02 | 2.66E+00 | 2.75E+00 | 9.16E-01 | no | | intergenic_variant | MODIFIER |
| 10 | 114899403 | AX-170151268 | 0 | T | A | 0.36 | 7.84E-03 | 7.81E-02 | 2.66E+00 | 2.75E+00 | 9.16E-01 | no | | intergenic_variant | MODIFIER |
| 10 | 114938643 | AX-170086907 | 0 | G | T | 0.36 | 7.84E-03 | 7.81E-02 | 2.66E+00 | 2.75E+00 | 9.16E-01 | no | | intergenic_variant | MODIFIER |
| 10 | 115001917 | AX-170259267 | 0 | G | A | 0.36 | 7.84E-03 | 7.81E-02 | 2.66E+00 | 2.75E+00 | 9.16E-01 | no | | intergenic_variant | MODIFIER |
| 10 | 115004293 | AX-170443480 | 0 | A | G | 0.36 | 7.84E-03 | 7.81E-02 | 2.66E+00 | 2.75E+00 | 9.16E-01 | no | | intergenic_variant | MODIFIER |
| 10 | 115004884 | AX-170325855 | 0 | G | C | 0.36 | 7.84E-03 | 7.81E-02 | 2.66E+00 | 2.75E+00 | 9.16E-01 | no | | intergenic_variant | MODIFIER |
| 10 | 115016005 | AX-170282967 | 0 | C | G | 0.36 | 7.84E-03 | 7.81E-02 | 2.66E+00 | 2.75E+00 | 9.16E-01 | no | | intergenic_variant | MODIFIER |
| 2 | 118023462 | AX-170357463 | 0 | G | A | 0.132 | 2.06E-02 | 1.99E-01 | 2.69E+00 | 2.74E+00 | 9.16E-01 | no | Gm13986 | intron_variant,non_coding_transcript_variant | MODIFIER |
| 2 | 100929906 | AX-97819347 | 1 | G | A | 0.062 | 2.32E-02 | 2.26E-01 | 2.65E+00 | 2.74E+00 | 9.17E-01 | no | | intergenic_variant | MODIFIER |
| 16 | 93997905 | AX-170330575 | 1 | G | A | 0.429 | -8.98E-03 | 7.88E-02 | 2.63E+00 | 2.72E+00 | 9.17E-01 | no | Cldn14 | intron_variant | MODIFIER |
| 9 | 66699401 | AX-97838426 | 1 | G | A | 0.196 | 1.38E-02 | 1.35E-01 | 2.72E+00 | 2.75E+00 | 9.17E-01 | no | | regulatory_region_variant | MODIFIER |
| 9 | 69027181 | AX-170142577 | 1 | T | C | 0.196 | 1.38E-02 | 1.35E-01 | 2.72E+00 | 2.75E+00 | 9.17E-01 | no | Rora | intron_variant | MODIFIER |
| 8 | 75604927 | AX-170304176 | 1 | C | A | 0.196 | -1.34E-02 | 1.23E-01 | 2.63E+00 | 2.73E+00 | 9.17E-01 | no | Iqcm | intron_variant | MODIFIER |
| 7 | 137219737 | AX-170221980 | 0 | C | G | 0.289 | -8.92E-03 | 8.00E-02 | 2.62E+00 | 2.72E+00 | 9.17E-01 | no | Ebf3 | intron_variant | MODIFIER |
| 3 | 148612301 | AX-168394901 | 0 | C | T | 0.325 | 8.19E-03 | 7.58E-02 | 2.65E+00 | 2.73E+00 | 9.17E-01 | no | | intergenic_variant | MODIFIER |
| 7 | 94323428 | AX-170192415 | 0 | G | A | 0.114 | -2.12E-02 | 2.12E-01 | 2.67E+00 | 2.75E+00 | 9.17E-01 | no | Gm32647 | intron_variant,non_coding_transcript_variant | MODIFIER |
| 2 | 130541361 | AX-97837859 | 1 | G | A | 0.143 | -1.82E-02 | 1.83E-01 | 2.67E+00 | 2.75E+00 | 9.17E-01 | no | Ptpra | synonymous_variant | LOW |
| 7 | 89715622 | AX-170497148 | 1 | G | T | 0.116 | 1.67E-02 | 1.59E-01 | 2.63E+00 | 2.73E+00 | 9.17E-01 | no | Me3 | intron_variant | MODIFIER |
| 7 | 90212883 | AX-170406792 | 1 | A | G | 0.116 | 1.67E-02 | 1.59E-01 | 2.63E+00 | 2.73E+00 | 9.17E-01 | no | Picalm | downstream_gene_variant | MODIFIER |
| 7 | 90249971 | AX-170259865 | 1 | T | C | 0.116 | 1.67E-02 | 1.59E-01 | 2.63E+00 | 2.73E+00 | 9.17E-01 | no | Ccdc83 | intron_variant | MODIFIER |
| 1 | 174674981 | AX-170240820 | 0 | C | A | 0.228 | 1.53E-02 | 1.26E-01 | 2.64E+00 | 2.70E+00 | 9.17E-01 | no | Fmn2 | intron_variant | MODIFIER |

| | | | | | | | | | | | | | | | |
|---|---|---|---|---|---|---|---|---|---|---|---|---|---|---|---|
| 1 | 174873613 | AX-170346563 | 0 | C | G | 0.228 | 1.53E-02 | 1.26E-01 | 2.64E+00 | 2.70E+00 | 9.17E-01 | no | Grem2 | intron_variant | MODIFIER |
| 6 | 138802495 | AX-170516024 | 1 | A | T | 0.071 | 2.82E-02 | 2.76E-01 | 2.70E+00 | 2.74E+00 | 9.17E-01 | no | | intergenic_variant | MODIFIER |
| 7 | 116825968 | AX-170295946 | 0 | C | T | 0.43 | 9.87E-03 | 9.48E-02 | 2.68E+00 | 2.73E+00 | 9.17E-01 | no | Gm4366 | upstream_gene_variant | MODIFIER |
| 7 | 40438002 | AX-170400026 | 0 | C | G | 0.333 | -8.54E-03 | 8.97E-02 | 2.68E+00 | 2.76E+00 | 9.17E-01 | no | Gm28807 | intron_variant,non_coding_transcript_variant | MODIFIER |
| 16 | 44874561 | AX-170320360 | 1 | C | T | 0.482 | 7.51E-03 | 8.08E-02 | 2.71E+00 | 2.78E+00 | 9.18E-01 | no | Cd200r2 | intron_variant | MODIFIER |
| 10 | 127408311 | AX-170004125 | 2 | C | T | 0.336 | -7.79E-03 | 7.34E-02 | 2.62E+00 | 2.73E+00 | 9.18E-01 | no | R3hdm2 | intron_variant | MODIFIER |
| 8 | 83621446 | AX-170351232 | 0 | G | A | 0.211 | 1.27E-02 | 1.17E-01 | 2.64E+00 | 2.73E+00 | 9.18E-01 | no | | regulatory_region_variant | MODIFIER |
| 9 | 62554655 | AX-170180109 | 1 | C | A | 0.196 | 1.39E-02 | 1.37E-01 | 2.72E+00 | 2.75E+00 | 9.18E-01 | no | | regulatory_region_variant | MODIFIER |
| 7 | 116490255 | AX-170515187 | 0 | G | T | 0.412 | -9.24E-03 | 9.41E-02 | 2.70E+00 | 2.75E+00 | 9.18E-01 | no | Gm45031 | downstream_gene_variant | MODIFIER |
| 8 | 23698512 | AX-170079522 | 1 | A | G | 0.17 | 9.95E-03 | 1.06E-01 | 2.65E+00 | 2.75E+00 | 9.18E-01 | no | Zmat4 | intron_variant | MODIFIER |
| 11 | 35026131 | AX-170036465 | 2 | C | T | 0.1 | 2.00E-02 | 1.88E-01 | 2.65E+00 | 2.73E+00 | 9.18E-01 | no | | intergenic_variant | MODIFIER |
| 1 | 67390371 | AX-97798356 | 1 | T | A | 0.116 | 2.40E-02 | 2.31E-01 | 2.68E+00 | 2.73E+00 | 9.18E-01 | no | | intergenic_variant | MODIFIER |
| 1 | 6943721 | AX-170488760 | 1 | T | C | 0.5 | 8.21E-03 | 7.01E-02 | 2.62E+00 | 2.71E+00 | 9.18E-01 | no | | intergenic_variant | MODIFIER |
| 12 | 101973395 | AX-170420598 | 0 | A | C | 0.061 | 2.12E-02 | 2.05E-01 | 2.66E+00 | 2.73E+00 | 9.19E-01 | no | Cpsf2 | upstream_gene_variant | MODIFIER |
| 9 | 62813075 | AX-170126492 | 1 | G | A | 0.205 | -1.30E-02 | 1.26E-01 | 2.68E+00 | 2.73E+00 | 9.19E-01 | no | Fem1b | upstream_gene_variant | MODIFIER |
| 4 | 69053911 | AX-170534132 | 0 | A | T | 0.325 | 1.22E-02 | 1.16E-01 | 2.66E+00 | 2.73E+00 | 9.19E-01 | no | | intergenic_variant | MODIFIER |
| 1 | 6193125 | AX-170107586 | 0 | C | T | 0.088 | 1.97E-02 | 1.83E-01 | 2.61E+00 | 2.73E+00 | 9.19E-01 | no | | intergenic_variant | MODIFIER |
| 1 | 9592074 | AX-170347358 | 0 | A | T | 0.088 | 1.97E-02 | 1.83E-01 | 2.61E+00 | 2.73E+00 | 9.19E-01 | no | 2610203C22Rik | intron_variant,non_coding_transcript_variant | MODIFIER |
| 1 | 10229505 | AX-169999924 | 0 | T | G | 0.088 | 1.97E-02 | 1.83E-01 | 2.61E+00 | 2.73E+00 | 9.19E-01 | no | Arfgef1 | intron_variant | MODIFIER |
| 2 | 110993845 | AX-170088708 | 0 | C | T | 0.061 | -1.91E-02 | 2.10E-01 | 2.65E+00 | 2.76E+00 | 9.19E-01 | no | | intergenic_variant | MODIFIER |
| 20 | 92842959 | AX-170527917 | 0 | A | G | 0.272 | -5.74E-03 | 5.96E-02 | 2.65E+00 | 2.75E+00 | 9.19E-01 | no | | intergenic_variant | MODIFIER |
| 1 | 48914355 | AX-170317860 | 1 | C | A | 0.107 | 2.37E-02 | 2.35E-01 | 2.69E+00 | 2.74E+00 | 9.19E-01 | no | | intergenic_variant | MODIFIER |
| 10 | 22918905 | AX-97815031 | 0 | A | G | 0.57 | -5.01E-03 | 7.18E-02 | 2.64E+00 | 2.79E+00 | 9.19E-01 | no | | intergenic_variant | MODIFIER |
| 12 | 99023698 | AX-170176011 | 0 | C | T | 0.079 | 1.99E-02 | 2.01E-01 | 2.69E+00 | 2.74E+00 | 9.19E-01 | no | | intergenic_variant | MODIFIER |
| 13 | 52769858 | AX-97788429 | 0 | C | T | 0.5 | 7.73E-03 | 6.63E-02 | 2.61E+00 | 2.71E+00 | 9.19E-01 | no | | regulatory_region_variant | MODIFIER |
| 13 | 52770016 | AX-170051304 | 0 | G | A | 0.5 | 7.73E-03 | 6.63E-02 | 2.61E+00 | 2.71E+00 | 9.19E-01 | no | | regulatory_region_variant | MODIFIER |
| 13 | 52770289 | AX-170051306 | 0 | A | G | 0.5 | 7.73E-03 | 6.63E-02 | 2.61E+00 | 2.71E+00 | 9.19E-01 | no | | regulatory_region_variant | MODIFIER |
| 13 | 52919400 | AX-170471967 | 0 | A | T | 0.5 | 7.73E-03 | 6.63E-02 | 2.61E+00 | 2.71E+00 | 9.19E-01 | no | Auh | intron_variant | MODIFIER |
| 13 | 52954441 | AX-170051307 | 0 | T | C | 0.5 | 7.73E-03 | 6.63E-02 | 2.61E+00 | 2.71E+00 | 9.19E-01 | no | | intergenic_variant | MODIFIER |
| 13 | 52959969 | AX-170203201 | 0 | G | T | 0.5 | 7.73E-03 | 6.63E-02 | 2.61E+00 | 2.71E+00 | 9.19E-01 | no | n-R5s54 | downstream_gene_variant | MODIFIER |
| 13 | 52960736 | AX-170460879 | 0 | A | G | 0.5 | 7.73E-03 | 6.63E-02 | 2.61E+00 | 2.71E+00 | 9.19E-01 | no | n-R5s54 | downstream_gene_variant | MODIFIER |
| 13 | 52964828 | AX-170226752 | 0 | G | A | 0.5 | 7.73E-03 | 6.63E-02 | 2.61E+00 | 2.71E+00 | 9.19E-01 | no | Nfil3 | downstream_gene_variant | MODIFIER |
| 13 | 52965370 | AX-170136472 | 0 | G | A | 0.5 | 7.73E-03 | 6.63E-02 | 2.61E+00 | 2.71E+00 | 9.19E-01 | no | Nfil3 | downstream_gene_variant | MODIFIER |
| 13 | 52969301 | AX-170293197 | 0 | C | T | 0.5 | 7.73E-03 | 6.63E-02 | 2.61E+00 | 2.71E+00 | 9.19E-01 | no | Nfil3 | intron_variant | MODIFIER |
| 13 | 52969522 | AX-170109094 | 0 | T | C | 0.5 | 7.73E-03 | 6.63E-02 | 2.61E+00 | 2.71E+00 | 9.19E-01 | no | Nfil3 | intron_variant | MODIFIER |
| 13 | 52973713 | AX-97812129 | 0 | G | A | 0.5 | 7.73E-03 | 6.63E-02 | 2.61E+00 | 2.71E+00 | 9.19E-01 | no | Nfil3 | intron_variant | MODIFIER |
| 13 | 53002646 | AX-97828692 | 0 | C | T | 0.5 | 7.73E-03 | 6.63E-02 | 2.61E+00 | 2.71E+00 | 9.19E-01 | no | | intergenic_variant | MODIFIER |
| 13 | 53007228 | AX-170159529 | 0 | A | C | 0.5 | 7.73E-03 | 6.63E-02 | 2.61E+00 | 2.71E+00 | 9.19E-01 | no | | intergenic_variant | MODIFIER |
| 13 | 53012441 | AX-170290388 | 0 | A | G | 0.5 | 7.73E-03 | 6.63E-02 | 2.61E+00 | 2.71E+00 | 9.19E-01 | no | Gm33424 | upstream_gene_variant | MODIFIER |
| 13 | 53016748 | AX-170142124 | 0 | C | T | 0.5 | 7.73E-03 | 6.63E-02 | 2.61E+00 | 2.71E+00 | 9.19E-01 | no | Gm33424 | intron_variant,non_coding_transcript_variant | MODIFIER |
| 13 | 53035732 | AX-170239428 | 0 | T | C | 0.5 | 7.73E-03 | 6.63E-02 | 2.61E+00 | 2.71E+00 | 9.19E-01 | no | Gm48336 | downstream_gene_variant | MODIFIER |
| 13 | 53049227 | AX-170333439 | 0 | T | C | 0.5 | 7.73E-03 | 6.63E-02 | 2.61E+00 | 2.71E+00 | 9.19E-01 | no | | intergenic_variant | MODIFIER |
| 13 | 53049436 | AX-170051309 | 0 | G | A | 0.5 | 7.73E-03 | 6.63E-02 | 2.61E+00 | 2.71E+00 | 9.19E-01 | no | | intergenic_variant | MODIFIER |
| 13 | 53116708 | AX-170326730 | 0 | T | C | 0.5 | 7.73E-03 | 6.63E-02 | 2.61E+00 | 2.71E+00 | 9.19E-01 | no | Ror2 | intron_variant | MODIFIER |
| 13 | 53118463 | AX-97789579 | 0 | C | T | 0.5 | 7.73E-03 | 6.63E-02 | 2.61E+00 | 2.71E+00 | 9.19E-01 | no | Ror2 | intron_variant | MODIFIER |
| 13 | 53130508 | AX-170312377 | 0 | T | C | 0.5 | 7.73E-03 | 6.63E-02 | 2.61E+00 | 2.71E+00 | 9.19E-01 | no | Ror2 | intron_variant | MODIFIER |
| 13 | 53174623 | AX-97844376 | 0 | G | A | 0.5 | 7.73E-03 | 6.63E-02 | 2.61E+00 | 2.71E+00 | 9.19E-01 | no | Ror2 | intron_variant | MODIFIER |
| 13 | 53174910 | AX-170496967 | 0 | A | C | 0.5 | 7.73E-03 | 6.63E-02 | 2.61E+00 | 2.71E+00 | 9.19E-01 | no | Ror2 | intron_variant | MODIFIER |
| 13 | 53331094 | AX-170102556 | 0 | T | A | 0.5 | 7.73E-03 | 6.63E-02 | 2.61E+00 | 2.71E+00 | 9.19E-01 | no | Sptlc1 | downstream_gene_variant | MODIFIER |
| 13 | 53342408 | AX-170277571 | 0 | T | C | 0.5 | 7.73E-03 | 6.63E-02 | 2.61E+00 | 2.71E+00 | 9.19E-01 | no | Sptlc1 | intron_variant | MODIFIER |
| 13 | 53356280 | AX-170106811 | 0 | G | C | 0.5 | 7.73E-03 | 6.63E-02 | 2.61E+00 | 2.71E+00 | 9.19E-01 | no | Sptlc1 | intron_variant | MODIFIER |
| 13 | 53356565 | AX-170264548 | 0 | C | T | 0.5 | 7.73E-03 | 6.63E-02 | 2.61E+00 | 2.71E+00 | 9.19E-01 | no | Sptlc1 | intron_variant | MODIFIER |
| 13 | 53365272 | AX-170370963 | 0 | C | T | 0.5 | 7.73E-03 | 6.63E-02 | 2.61E+00 | 2.71E+00 | 9.19E-01 | no | Sptlc1 | intron_variant | MODIFIER |
| 13 | 53372789 | AX-170337920 | 0 | T | C | 0.5 | 7.73E-03 | 6.63E-02 | 2.61E+00 | 2.71E+00 | 9.19E-01 | no | Sptlc1 | intron_variant | MODIFIER |

| | | | | | | | | | | | | | | | |
|---|---|---|---|---|---|---|---|---|---|---|---|---|---|---|---|
| 13 | 53373936 | AX-170051316 | 0 | T | C | 0.5 | 7.73E-03 | 6.63E-02 | 2.61E+00 | 2.71E+00 | 9.19E-01 | no | Sptlc1 | intron_variant | MODIFIER |
| 13 | 53389809 | AX-170133187 | 0 | G | T | 0.5 | 7.73E-03 | 6.63E-02 | 2.61E+00 | 2.71E+00 | 9.19E-01 | no | Gm2762 | upstream_gene_variant | MODIFIER |
| 13 | 53398100 | AX-170531360 | 0 | T | C | 0.5 | 7.73E-03 | 6.63E-02 | 2.61E+00 | 2.71E+00 | 9.19E-01 | no | Gm2762 | intron_variant,non_coding_transcript_variant | MODIFIER |
| 13 | 53411514 | AX-170182104 | 0 | A | G | 0.5 | 7.73E-03 | 6.63E-02 | 2.61E+00 | 2.71E+00 | 9.19E-01 | no | Gm2762 | intron_variant,non_coding_transcript_variant | MODIFIER |
| 7 | 40367198 | AX-170490065 | 0 | A | T | 0.333 | -7.91E-03 | 8.99E-02 | 2.66E+00 | 2.76E+00 | 9.19E-01 | no | Gm24910 | downstream_gene_variant | MODIFIER |
| 1 | 7222064 | AX-170513994 | 1 | T | G | 0.098 | 1.82E-02 | 1.72E-01 | 2.62E+00 | 2.73E+00 | 9.19E-01 | no | | intergenic_variant | MODIFIER |
| 2 | 155482323 | AX-97832993 | 0 | T | G | 0.149 | -1.71E-02 | 1.81E-01 | 2.69E+00 | 2.75E+00 | 9.20E-01 | no | Gm14238 | downstream_gene_variant | MODIFIER |
| 6 | 133234350 | AX-170240713 | 2 | T | C | 0.418 | 1.35E-02 | 1.10E-01 | 2.59E+00 | 2.71E+00 | 9.20E-01 | no | | intergenic_variant | MODIFIER |
| 8 | 50279012 | AX-170112206 | 0 | G | A | 0.158 | 1.22E-02 | 1.36E-01 | 2.69E+00 | 2.77E+00 | 9.20E-01 | no | Gm45455 | intron_variant,non_coding_transcript_variant | MODIFIER |
| 8 | 55572207 | AX-170260856 | 0 | C | T | 0.158 | 1.22E-02 | 1.36E-01 | 2.69E+00 | 2.77E+00 | 9.20E-01 | no | | intergenic_variant | MODIFIER |
| 16 | 18787799 | AX-170298770 | 0 | G | A | 0.14 | 9.14E-03 | 9.77E-02 | 2.63E+00 | 2.75E+00 | 9.20E-01 | no | Cdc45 | intron_variant | MODIFIER |
| 7 | 137946129 | AX-170161636 | 1 | T | A | 0.312 | 7.38E-03 | 7.88E-02 | 2.64E+00 | 2.75E+00 | 9.20E-01 | no | | intergenic_variant | MODIFIER |
| 7 | 88306563 | AX-170445624 | 1 | T | G | 0.134 | -2.05E-02 | 2.11E-01 | 2.66E+00 | 2.74E+00 | 9.20E-01 | no | Ctsc | intron_variant | MODIFIER |
| 20 | 134876281 | AX-170035006 | 0 | A | G | 0.14 | 1.20E-02 | 1.12E-01 | 2.63E+00 | 2.72E+00 | 9.20E-01 | no | | intergenic_variant | MODIFIER |
| 20 | 88943204 | AX-170418205 | 0 | T | C | 0.272 | 6.34E-03 | 6.05E-02 | 2.63E+00 | 2.73E+00 | 9.20E-01 | no | | intergenic_variant | MODIFIER |
| 7 | 58870789 | AX-170029098 | 1 | C | T | 0.438 | -9.18E-03 | 9.91E-02 | 2.67E+00 | 2.75E+00 | 9.20E-01 | no | | regulatory_region_variant | MODIFIER |
| 12 | 101087056 | AX-97822217 | 0 | G | A | 0.096 | 1.74E-02 | 1.62E-01 | 2.62E+00 | 2.73E+00 | 9.20E-01 | no | Ppp4r3a | upstream_gene_variant | MODIFIER |
| 7 | 58766946 | AX-170456805 | 1 | G | T | 0.42 | -9.44E-03 | 9.84E-02 | 2.66E+00 | 2.74E+00 | 9.20E-01 | no | Atp10a | intron_variant | MODIFIER |
| 1 | 41888536 | AX-170393917 | 1 | C | T | 0.446 | -7.73E-03 | 7.57E-02 | 2.65E+00 | 2.73E+00 | 9.20E-01 | no | Gm29260 | intron_variant,non_coding_transcript_variant | MODIFIER |
| 8 | 68990228 | AX-170460089 | 1 | G | A | 0.188 | -1.39E-02 | 1.34E-01 | 2.66E+00 | 2.73E+00 | 9.20E-01 | no | Gm33103 | intron_variant,non_coding_transcript_variant | MODIFIER |
| 13 | 42752962 | AX-170231103 | 0 | T | C | 0.491 | 7.25E-03 | 6.23E-02 | 2.60E+00 | 2.71E+00 | 9.20E-01 | no | Phactr1 | intron_variant | MODIFIER |
| 9 | 58506006 | AX-170345469 | 0 | G | A | 0.219 | -1.10E-02 | 1.16E-01 | 2.68E+00 | 2.75E+00 | 9.20E-01 | no | | intergenic_variant | MODIFIER |
| 2 | 128577787 | AX-170310644 | 1 | C | A | 0.143 | 2.23E-02 | 2.20E-01 | 2.69E+00 | 2.73E+00 | 9.21E-01 | no | | intergenic_variant | MODIFIER |
| 10 | 4634647 | AX-97799388 | 0 | G | C | 0.193 | 9.50E-03 | 9.99E-02 | 2.64E+00 | 2.74E+00 | 9.21E-01 | no | Esr1 | intron_variant | MODIFIER |
| 10 | 4648513 | AX-170338936 | 0 | G | T | 0.193 | 9.50E-03 | 9.99E-02 | 2.64E+00 | 2.74E+00 | 9.21E-01 | no | Esr1 | intron_variant | MODIFIER |
| 10 | 4653180 | AX-170410636 | 0 | A | G | 0.193 | 9.50E-03 | 9.99E-02 | 2.64E+00 | 2.74E+00 | 9.21E-01 | no | Esr1 | intron_variant | MODIFIER |
| 10 | 4654118 | AX-170326071 | 0 | A | G | 0.193 | 9.50E-03 | 9.99E-02 | 2.64E+00 | 2.74E+00 | 9.21E-01 | no | Esr1 | intron_variant | MODIFIER |
| 10 | 4682391 | AX-170296525 | 0 | T | C | 0.193 | 9.50E-03 | 9.99E-02 | 2.64E+00 | 2.74E+00 | 9.21E-01 | no | Esr1 | intron_variant | MODIFIER |
| 10 | 4700000 | AX-170212132 | 0 | G | A | 0.193 | 9.50E-03 | 9.99E-02 | 2.64E+00 | 2.74E+00 | 9.21E-01 | no | Esr1 | intron_variant | MODIFIER |
| 10 | 4785733 | AX-97830806 | 0 | C | G | 0.193 | 9.50E-03 | 9.99E-02 | 2.64E+00 | 2.74E+00 | 9.21E-01 | no | Esr1 | intron_variant | MODIFIER |
| 10 | 4857822 | AX-170229968 | 0 | C | T | 0.193 | 9.50E-03 | 9.99E-02 | 2.64E+00 | 2.74E+00 | 9.21E-01 | no | Esr1 | intron_variant | MODIFIER |
| 10 | 4877382 | AX-170086733 | 0 | C | G | 0.193 | 9.50E-03 | 9.99E-02 | 2.64E+00 | 2.74E+00 | 9.21E-01 | no | Esr1 | intron_variant | MODIFIER |
| 10 | 4908958 | AX-170159750 | 0 | T | C | 0.193 | 9.50E-03 | 9.99E-02 | 2.64E+00 | 2.74E+00 | 9.21E-01 | no | Esr1 | intron_variant | MODIFIER |
| 10 | 4909068 | AX-170253900 | 0 | T | C | 0.193 | 9.50E-03 | 9.99E-02 | 2.64E+00 | 2.74E+00 | 9.21E-01 | no | Esr1 | intron_variant | MODIFIER |
| 10 | 4959380 | AX-170333160 | 0 | A | G | 0.193 | 9.50E-03 | 9.99E-02 | 2.64E+00 | 2.74E+00 | 9.21E-01 | no | Esr1 | intron_variant | MODIFIER |
| 10 | 4961093 | AX-97814686 | 0 | G | C | 0.193 | 9.50E-03 | 9.99E-02 | 2.64E+00 | 2.74E+00 | 9.21E-01 | no | Esr1 | intron_variant | MODIFIER |
| 10 | 5059090 | AX-170002641 | 0 | T | C | 0.193 | 9.50E-03 | 9.99E-02 | 2.64E+00 | 2.74E+00 | 9.21E-01 | no | Syne1 | intron_variant | MODIFIER |
| 10 | 5067237 | AX-170129935 | 0 | A | G | 0.193 | 9.50E-03 | 9.99E-02 | 2.64E+00 | 2.74E+00 | 9.21E-01 | no | Syne1 | intron_variant | MODIFIER |
| 10 | 5596691 | AX-170002639 | 0 | T | C | 0.193 | 9.50E-03 | 9.99E-02 | 2.64E+00 | 2.74E+00 | 9.21E-01 | no | Myct1 | intron_variant | MODIFIER |
| 7 | 109496281 | AX-170312878 | 0 | G | A | 0.474 | 1.01E-02 | 9.60E-02 | 2.66E+00 | 2.72E+00 | 9.21E-01 | no | Trim66 | intron_variant | MODIFIER |
| 9 | 59086421 | AX-170423102 | 1 | C | T | 0.205 | 1.18E-02 | 1.21E-01 | 2.68E+00 | 2.74E+00 | 9.21E-01 | no | | intergenic_variant | MODIFIER |
| 5 | 81850093 | AX-97809303 | 1 | G | A | 0.429 | 1.04E-02 | 1.01E-01 | 2.65E+00 | 2.73E+00 | 9.21E-01 | no | | intergenic_variant | MODIFIER |
| 12 | 116140851 | AX-170375574 | 1 | G | A | 0.125 | 1.97E-02 | 1.96E-01 | 2.66E+00 | 2.73E+00 | 9.21E-01 | no | Vipr2 | intron_variant | MODIFIER |
| 1 | 7954253 | AX-169999893 | 1 | T | A | 0.42 | 8.03E-03 | 7.09E-02 | 2.63E+00 | 2.71E+00 | 9.21E-01 | no | | intergenic_variant | MODIFIER |
| 4 | 144528723 | AX-170024179 | 0 | A | C | 0.123 | -1.11E-02 | 1.18E-01 | 2.62E+00 | 2.74E+00 | 9.21E-01 | no | 9430007A20Rik | missense_variant | MODERATE |
| 1 | 52167184 | AX-170185058 | 0 | G | A | 0.132 | -2.33E-02 | 2.40E-01 | 2.69E+00 | 2.74E+00 | 9.21E-01 | no | Gls | intron_variant | MODIFIER |
| 7 | 58758813 | AX-170227773 | 0 | A | C | 0.43 | -9.12E-03 | 9.77E-02 | 2.67E+00 | 2.75E+00 | 9.21E-01 | no | Atp10a | intron_variant | MODIFIER |
| 7 | 58762678 | AX-170268021 | 0 | C | T | 0.43 | -9.12E-03 | 9.77E-02 | 2.67E+00 | 2.75E+00 | 9.21E-01 | no | Atp10a | intron_variant | MODIFIER |
| 7 | 58771268 | AX-170110861 | 0 | G | A | 0.43 | -9.12E-03 | 9.77E-02 | 2.67E+00 | 2.75E+00 | 9.21E-01 | no | Atp10a | intron_variant | MODIFIER |
| 7 | 58771618 | AX-170473800 | 0 | T | A | 0.43 | -9.12E-03 | 9.77E-02 | 2.67E+00 | 2.75E+00 | 9.21E-01 | no | Atp10a | intron_variant | MODIFIER |
| 7 | 58771758 | AX-170456909 | 0 | G | A | 0.43 | -9.12E-03 | 9.77E-02 | 2.67E+00 | 2.75E+00 | 9.21E-01 | no | Atp10a | intron_variant | MODIFIER |
| 7 | 58774153 | AX-170113255 | 0 | G | T | 0.43 | -9.12E-03 | 9.77E-02 | 2.67E+00 | 2.75E+00 | 9.21E-01 | no | Atp10a | intron_variant | MODIFIER |
| 7 | 58774619 | AX-170382490 | 0 | T | G | 0.43 | -9.12E-03 | 9.77E-02 | 2.67E+00 | 2.75E+00 | 9.21E-01 | no | Atp10a | intron_variant | MODIFIER |
| 7 | 58776453 | AX-170470639 | 0 | A | G | 0.43 | -9.12E-03 | 9.77E-02 | 2.67E+00 | 2.75E+00 | 9.21E-01 | no | Atp10a | intron_variant | MODIFIER |

| | | | | | | | | | | | | | | | |
|---|---|---|---|---|---|---|---|---|---|---|---|---|---|---|---|
| 7 | 58777429 | AX-170492162 | 0 | C | T | 0.43 | -9.12E-03 | 9.77E-02 | 2.67E+00 | 2.75E+00 | 9.21E-01 | no | Atp10a | intron_variant | MODIFIER |
| 7 | 58779943 | AX-97856744 | 0 | G | A | 0.43 | -9.12E-03 | 9.77E-02 | 2.67E+00 | 2.75E+00 | 9.21E-01 | no | Atp10a | intron_variant | MODIFIER |
| 7 | 58791422 | AX-170298250 | 0 | C | T | 0.43 | -9.12E-03 | 9.77E-02 | 2.67E+00 | 2.75E+00 | 9.21E-01 | no | Atp10a | intron_variant | MODIFIER |
| 7 | 58796244 | AX-170512544 | 0 | T | C | 0.43 | -9.12E-03 | 9.77E-02 | 2.67E+00 | 2.75E+00 | 9.21E-01 | no | Atp10a | intron_variant | MODIFIER |
| 7 | 58796869 | AX-170104911 | 0 | G | A | 0.43 | -9.12E-03 | 9.77E-02 | 2.67E+00 | 2.75E+00 | 9.21E-01 | no | Atp10a | intron_variant | MODIFIER |
| 7 | 58800539 | AX-170189352 | 0 | T | C | 0.43 | -9.12E-03 | 9.77E-02 | 2.67E+00 | 2.75E+00 | 9.21E-01 | no | Atp10a | intron_variant | MODIFIER |
| 7 | 58800943 | AX-170323935 | 0 | C | T | 0.43 | -9.12E-03 | 9.77E-02 | 2.67E+00 | 2.75E+00 | 9.21E-01 | no | Atp10a | intron_variant | MODIFIER |
| 7 | 58814902 | AX-170029095 | 0 | T | C | 0.43 | -9.12E-03 | 9.77E-02 | 2.67E+00 | 2.75E+00 | 9.21E-01 | no | Atp10a | intron_variant | MODIFIER |
| 7 | 58854342 | AX-97797586 | 0 | T | C | 0.43 | -9.12E-03 | 9.77E-02 | 2.67E+00 | 2.75E+00 | 9.21E-01 | no | | intergenic_variant | MODIFIER |
| 7 | 58870522 | AX-170029097 | 0 | G | A | 0.43 | -9.12E-03 | 9.77E-02 | 2.67E+00 | 2.75E+00 | 9.21E-01 | no | | intergenic_variant | MODIFIER |
| 7 | 58921281 | AX-170029099 | 0 | T | C | 0.43 | -9.12E-03 | 9.77E-02 | 2.67E+00 | 2.75E+00 | 9.21E-01 | no | | intergenic_variant | MODIFIER |
| 7 | 58924961 | AX-170029094 | 0 | T | C | 0.43 | -9.12E-03 | 9.77E-02 | 2.67E+00 | 2.75E+00 | 9.21E-01 | no | Gm44938 | upstream_gene_variant | MODIFIER |
| 7 | 58938769 | AX-170212654 | 0 | A | T | 0.43 | -9.12E-03 | 9.77E-02 | 2.67E+00 | 2.75E+00 | 9.21E-01 | no | | intergenic_variant | MODIFIER |
| 7 | 59149430 | AX-170533703 | 0 | C | T | 0.43 | -9.12E-03 | 9.77E-02 | 2.67E+00 | 2.75E+00 | 9.21E-01 | no | | intergenic_variant | MODIFIER |
| 7 | 59340730 | AX-170216103 | 0 | T | C | 0.43 | -9.12E-03 | 9.77E-02 | 2.67E+00 | 2.75E+00 | 9.21E-01 | no | Gm24966 | downstream_gene_variant | MODIFIER |
| 7 | 59625956 | AX-170089932 | 0 | G | A | 0.43 | -9.12E-03 | 9.77E-02 | 2.67E+00 | 2.75E+00 | 9.21E-01 | no | Snhg14 | intron_variant,non_coding_transcript_variant | MODIFIER |
| 7 | 59649682 | AX-97780308 | 0 | T | C | 0.43 | -9.12E-03 | 9.77E-02 | 2.67E+00 | 2.75E+00 | 9.21E-01 | no | Snhg14 | intron_variant,non_coding_transcript_variant | MODIFIER |
| 7 | 59672197 | AX-170029105 | 0 | G | T | 0.43 | -9.12E-03 | 9.77E-02 | 2.67E+00 | 2.75E+00 | 9.21E-01 | no | Gm26504 | downstream_gene_variant | MODIFIER |
| 7 | 59926119 | AX-97819143 | 0 | T | C | 0.43 | -9.12E-03 | 9.77E-02 | 2.67E+00 | 2.75E+00 | 9.21E-01 | no | Gm22373 | upstream_gene_variant | MODIFIER |
| 7 | 59945053 | AX-170029103 | 0 | C | T | 0.43 | -9.12E-03 | 9.77E-02 | 2.67E+00 | 2.75E+00 | 9.21E-01 | no | Snhg14 | downstream_gene_variant | MODIFIER |
| 7 | 59982433 | AX-170322967 | 0 | T | C | 0.43 | -9.12E-03 | 9.77E-02 | 2.67E+00 | 2.75E+00 | 9.21E-01 | no | Snrpn | downstream_gene_variant | MODIFIER |
| 7 | 59982856 | AX-170232778 | 0 | A | G | 0.43 | -9.12E-03 | 9.77E-02 | 2.67E+00 | 2.75E+00 | 9.21E-01 | no | Snrpn | 3_prime_UTR_variant | MODIFIER |
| 7 | 60023139 | AX-97835070 | 0 | A | T | 0.43 | -9.12E-03 | 9.77E-02 | 2.67E+00 | 2.75E+00 | 9.21E-01 | no | Snrpn | intron_variant | MODIFIER |
| 7 | 60312612 | AX-170077482 | 0 | C | G | 0.43 | -9.12E-03 | 9.77E-02 | 2.67E+00 | 2.75E+00 | 9.21E-01 | no | | intergenic_variant | MODIFIER |
| 7 | 60586764 | AX-97839311 | 0 | A | G | 0.43 | -9.12E-03 | 9.77E-02 | 2.67E+00 | 2.75E+00 | 9.21E-01 | no | | intergenic_variant | MODIFIER |
| 7 | 60798694 | AX-97853279 | 0 | G | A | 0.43 | -9.12E-03 | 9.77E-02 | 2.67E+00 | 2.75E+00 | 9.21E-01 | no | | intergenic_variant | MODIFIER |
| 7 | 60833249 | AX-97797786 | 0 | T | C | 0.43 | -9.12E-03 | 9.77E-02 | 2.67E+00 | 2.75E+00 | 9.21E-01 | no | | intergenic_variant | MODIFIER |
| 7 | 60863572 | AX-170095955 | 0 | T | C | 0.43 | -9.12E-03 | 9.77E-02 | 2.67E+00 | 2.75E+00 | 9.21E-01 | no | | intergenic_variant | MODIFIER |
| 7 | 60863821 | AX-170417956 | 0 | G | A | 0.43 | -9.12E-03 | 9.77E-02 | 2.67E+00 | 2.75E+00 | 9.21E-01 | no | | intergenic_variant | MODIFIER |
| 7 | 60875163 | AX-170120969 | 0 | G | A | 0.43 | -9.12E-03 | 9.77E-02 | 2.67E+00 | 2.75E+00 | 9.21E-01 | no | | intergenic_variant | MODIFIER |
| 7 | 60880791 | AX-97780624 | 0 | T | G | 0.43 | -9.12E-03 | 9.77E-02 | 2.67E+00 | 2.75E+00 | 9.21E-01 | no | | intergenic_variant | MODIFIER |
| 7 | 60909188 | AX-170029113 | 0 | T | G | 0.43 | -9.12E-03 | 9.77E-02 | 2.67E+00 | 2.75E+00 | 9.21E-01 | no | | intergenic_variant | MODIFIER |
| 7 | 60933240 | AX-170029114 | 0 | T | C | 0.43 | -9.12E-03 | 9.77E-02 | 2.67E+00 | 2.75E+00 | 9.21E-01 | no | Gm44643 | intron_variant,non_coding_transcript_variant | MODIFIER |
| 7 | 60933313 | AX-170029111 | 0 | G | T | 0.43 | -9.12E-03 | 9.77E-02 | 2.67E+00 | 2.75E+00 | 9.21E-01 | no | Gm44643 | intron_variant,non_coding_transcript_variant | MODIFIER |
| 7 | 60961116 | AX-97807754 | 0 | A | G | 0.43 | -9.12E-03 | 9.77E-02 | 2.67E+00 | 2.75E+00 | 9.21E-01 | no | Gm44643 | intron_variant,non_coding_transcript_variant | MODIFIER |
| 7 | 60977480 | AX-97804459 | 0 | G | T | 0.43 | -9.12E-03 | 9.77E-02 | 2.67E+00 | 2.75E+00 | 9.21E-01 | no | | intergenic_variant | MODIFIER |
| 7 | 61030158 | AX-170342352 | 0 | G | A | 0.43 | -9.12E-03 | 9.77E-02 | 2.67E+00 | 2.75E+00 | 9.21E-01 | no | | regulatory_region_variant | MODIFIER |
| 7 | 61030373 | AX-170166530 | 0 | A | G | 0.43 | -9.12E-03 | 9.77E-02 | 2.67E+00 | 2.75E+00 | 9.21E-01 | no | | regulatory_region_variant | MODIFIER |
| 7 | 61981281 | AX-170029133 | 0 | G | A | 0.43 | -9.12E-03 | 9.77E-02 | 2.67E+00 | 2.75E+00 | 9.21E-01 | no | Mir344g | downstream_gene_variant | MODIFIER |
| 7 | 61985477 | AX-97807837 | 0 | T | C | 0.43 | -9.12E-03 | 9.77E-02 | 2.67E+00 | 2.75E+00 | 9.21E-01 | no | Mir344g | upstream_gene_variant | MODIFIER |
| 7 | 62290838 | AX-97800896 | 0 | G | A | 0.43 | -9.12E-03 | 9.77E-02 | 2.67E+00 | 2.75E+00 | 9.21E-01 | no | | intergenic_variant | MODIFIER |
| 7 | 62320979 | AX-97807017 | 0 | A | G | 0.43 | -9.12E-03 | 9.77E-02 | 2.67E+00 | 2.75E+00 | 9.21E-01 | no | | intergenic_variant | MODIFIER |
| 7 | 62431707 | AX-170429517 | 0 | C | T | 0.43 | -9.12E-03 | 9.77E-02 | 2.67E+00 | 2.75E+00 | 9.21E-01 | no | | intergenic_variant | MODIFIER |
| 7 | 62436155 | AX-97782007 | 0 | C | T | 0.43 | -9.12E-03 | 9.77E-02 | 2.67E+00 | 2.75E+00 | 9.21E-01 | no | | intergenic_variant | MODIFIER |
| 7 | 62717634 | AX-97799150 | 0 | C | T | 0.43 | -9.12E-03 | 9.77E-02 | 2.67E+00 | 2.75E+00 | 9.21E-01 | no | | intergenic_variant | MODIFIER |
| 7 | 63147913 | AX-97802977 | 0 | T | C | 0.43 | -9.12E-03 | 9.77E-02 | 2.67E+00 | 2.75E+00 | 9.21E-01 | no | Chrna7 | intron_variant | MODIFIER |
| 12 | 100484252 | AX-97844394 | 0 | A | G | 0.079 | 1.90E-02 | 1.91E-01 | 2.66E+00 | 2.73E+00 | 9.21E-01 | no | Ttc7b | intron_variant | MODIFIER |
| 12 | 100489064 | AX-170188393 | 0 | T | A | 0.079 | 1.90E-02 | 1.91E-01 | 2.66E+00 | 2.73E+00 | 9.21E-01 | no | Ttc7b | intron_variant | MODIFIER |
| 12 | 100489106 | AX-170049954 | 0 | T | A | 0.079 | 1.90E-02 | 1.91E-01 | 2.66E+00 | 2.73E+00 | 9.21E-01 | no | Ttc7b | intron_variant | MODIFIER |
| 2 | 122776856 | AX-170332708 | 0 | A | C | 0.14 | 1.82E-02 | 1.68E-01 | 2.62E+00 | 2.72E+00 | 9.21E-01 | no | Sqor | intron_variant | MODIFIER |
| 5 | 88212034 | AX-97847193 | 1 | C | T | 0.42 | -1.02E-02 | 1.05E-01 | 2.68E+00 | 2.74E+00 | 9.21E-01 | no | Gm42164 | downstream_gene_variant | MODIFIER |
| 12 | 43355333 | AX-97781131 | 1 | G | T | 0.045 | -1.35E-02 | 1.35E-01 | 2.56E+00 | 2.74E+00 | 9.21E-01 | no | Gm47030 | intron_variant,non_coding_transcript_variant | MODIFIER |
| 18 | 69213808 | AX-170426627 | 1 | G | T | 0.277 | 8.78E-03 | 9.69E-02 | 2.68E+00 | 2.76E+00 | 9.21E-01 | no | AC121263.2 | upstream_gene_variant | MODIFIER |
| 3 | 147083836 | AX-170498796 | 1 | G | A | 0.348 | 7.70E-03 | 7.49E-02 | 2.65E+00 | 2.73E+00 | 9.21E-01 | no | | regulatory_region_variant | MODIFIER |

| | | | | | | | | | | | | | | | |
|---|---|---|---|---|---|---|---|---|---|---|---|---|---|---|---|
| 3 | 147232058 | AX-170341578 | 1 | C | A | 0.348 | 7.70E-03 | 7.49E-02 | 2.65E+00 | 2.73E+00 | 9.21E-01 | no | | intergenic_variant | MODIFIER |
| 3 | 147698288 | AX-97841673 | 1 | A | C | 0.348 | 7.70E-03 | 7.49E-02 | 2.65E+00 | 2.73E+00 | 9.21E-01 | no | | intergenic_variant | MODIFIER |
| 9 | 77279567 | AX-170032752 | 0 | G | A | 0.211 | -1.26E-02 | 1.27E-01 | 2.68E+00 | 2.73E+00 | 9.22E-01 | no | Mlip | intron_variant | MODIFIER |
| 8 | 55722643 | AX-170238981 | 1 | G | A | 0.152 | -1.63E-02 | 1.52E-01 | 2.66E+00 | 2.71E+00 | 9.22E-01 | no | | intergenic_variant | MODIFIER |
| 15 | 100809603 | AX-170362671 | 0 | T | G | 0.114 | 1.49E-02 | 1.50E-01 | 2.66E+00 | 2.73E+00 | 9.22E-01 | no | Slc4a8 | intron_variant | MODIFIER |
| 18 | 53033626 | AX-170478460 | 0 | T | C | 0.246 | 9.02E-03 | 1.06E-01 | 2.69E+00 | 2.77E+00 | 9.22E-01 | no | | intergenic_variant | MODIFIER |
| 9 | 29513325 | AX-170315218 | 1 | A | C | 0.438 | 9.10E-03 | 8.71E-02 | 2.66E+00 | 2.72E+00 | 9.22E-01 | no | Ntm | intron_variant | MODIFIER |
| 20 | 145646806 | AX-170198799 | 1 | A | G | 0.143 | 1.32E-02 | 1.32E-01 | 2.64E+00 | 2.73E+00 | 9.22E-01 | no | Gm15060 | non_coding_transcript_exon_variant | MODIFIER |
| 11 | 35369546 | AX-170519139 | 0 | G | A | 0.096 | -1.71E-02 | 1.87E-01 | 2.66E+00 | 2.75E+00 | 9.22E-01 | no | Slit3 | intron_variant | MODIFIER |
| 13 | 37350710 | AX-170332218 | 1 | G | T | 0.179 | -1.28E-02 | 1.20E-01 | 2.65E+00 | 2.72E+00 | 9.22E-01 | no | Ly86 | intron_variant | MODIFIER |
| 10 | 88649761 | AX-170313911 | 0 | C | T | 0.482 | -4.97E-03 | 1.12E-01 | 2.62E+00 | 2.81E+00 | 9.22E-01 | no | Gm48106 | downstream_gene_variant | MODIFIER |
| 7 | 79411342 | AX-170104312 | 1 | C | A | 0.116 | 2.09E-02 | 2.11E-01 | 2.65E+00 | 2.73E+00 | 9.22E-01 | no | Fanci | intron_variant | MODIFIER |
| 11 | 31974946 | AX-170004646 | 1 | T | C | 0.08 | -1.55E-02 | 1.69E-01 | 2.64E+00 | 2.75E+00 | 9.22E-01 | no | 4930524B15Rik | intron_variant | MODIFIER |
| 20 | 146271556 | AX-97786614 | 1 | G | T | 0.125 | 1.16E-02 | 1.16E-01 | 2.64E+00 | 2.73E+00 | 9.22E-01 | no | | intergenic_variant | MODIFIER |
| 9 | 106323434 | AX-170033217 | 1 | C | T | 0.125 | 1.01E-02 | 1.17E-01 | 2.70E+00 | 2.77E+00 | 9.22E-01 | no | Poc1a | intron_variant | MODIFIER |
| 9 | 106830510 | AX-97793344 | 1 | G | T | 0.125 | 1.01E-02 | 1.17E-01 | 2.70E+00 | 2.77E+00 | 9.22E-01 | no | Dcaf1 | intron_variant | MODIFIER |
| 9 | 113446926 | AX-97846298 | 1 | C | T | 0.125 | -1.08E-02 | 1.07E-01 | 2.65E+00 | 2.73E+00 | 9.22E-01 | no | | intergenic_variant | MODIFIER |
| 1 | 74078996 | AX-97844426 | 0 | C | T | 0.123 | 2.01E-02 | 2.07E-01 | 2.68E+00 | 2.74E+00 | 9.22E-01 | no | Tns1 | intron_variant | MODIFIER |
| 1 | 74141355 | AX-97812040 | 0 | A | C | 0.123 | 2.01E-02 | 2.07E-01 | 2.68E+00 | 2.74E+00 | 9.22E-01 | no | Rufy4 | intron_variant | MODIFIER |
| 1 | 74148175 | AX-170457395 | 0 | A | G | 0.123 | 2.01E-02 | 2.07E-01 | 2.68E+00 | 2.74E+00 | 9.22E-01 | no | Rufy4 | 3_prime_UTR_variant | MODIFIER |
| 1 | 74148716 | AX-170289942 | 0 | G | A | 0.123 | 2.01E-02 | 2.07E-01 | 2.68E+00 | 2.74E+00 | 9.22E-01 | no | Rufy4 | downstream_gene_variant | MODIFIER |
| 1 | 74543791 | AX-170000811 | 0 | G | C | 0.123 | 2.01E-02 | 2.07E-01 | 2.68E+00 | 2.74E+00 | 9.22E-01 | no | Plcd4 | upstream_gene_variant | MODIFIER |
| 1 | 74543966 | AX-97837934 | 0 | A | G | 0.123 | 2.01E-02 | 2.07E-01 | 2.68E+00 | 2.74E+00 | 9.22E-01 | no | Plcd4 | upstream_gene_variant | MODIFIER |
| 1 | 74560154 | AX-97783035 | 0 | G | C | 0.123 | 2.01E-02 | 2.07E-01 | 2.68E+00 | 2.74E+00 | 9.22E-01 | no | Plcd4 | intron_variant | MODIFIER |
| 1 | 74573805 | AX-170000810 | 0 | C | T | 0.123 | 2.01E-02 | 2.07E-01 | 2.68E+00 | 2.74E+00 | 9.22E-01 | no | Zfp142 | intron_variant | MODIFIER |
| 1 | 74574747 | AX-170110786 | 0 | T | C | 0.123 | 2.01E-02 | 2.07E-01 | 2.68E+00 | 2.74E+00 | 9.22E-01 | no | Zfp142 | intron_variant | MODIFIER |
| 1 | 74577782 | AX-170341151 | 0 | T | C | 0.123 | 2.01E-02 | 2.07E-01 | 2.68E+00 | 2.74E+00 | 9.22E-01 | no | Zfp142 | intron_variant | MODIFIER |
| 1 | 74578090 | AX-170109347 | 0 | C | T | 0.123 | 2.01E-02 | 2.07E-01 | 2.68E+00 | 2.74E+00 | 9.22E-01 | no | Zfp142 | intron_variant | MODIFIER |
| 1 | 74581083 | AX-97849230 | 0 | C | T | 0.123 | 2.01E-02 | 2.07E-01 | 2.68E+00 | 2.74E+00 | 9.22E-01 | no | Zfp142 | intron_variant | MODIFIER |
| 1 | 74907623 | AX-170455340 | 0 | A | G | 0.123 | 2.01E-02 | 2.07E-01 | 2.68E+00 | 2.74E+00 | 9.22E-01 | no | Cfap65 | synonymous_variant | LOW |
| 1 | 74967194 | AX-170313709 | 0 | G | T | 0.123 | 2.01E-02 | 2.07E-01 | 2.68E+00 | 2.74E+00 | 9.22E-01 | no | Nhej1 | 3_prime_UTR_variant | MODIFIER |
| 1 | 74972525 | AX-170399708 | 0 | G | C | 0.123 | 2.01E-02 | 2.07E-01 | 2.68E+00 | 2.74E+00 | 9.22E-01 | no | Nhej1 | intron_variant | MODIFIER |
| 1 | 74973137 | AX-170236414 | 0 | A | C | 0.123 | 2.01E-02 | 2.07E-01 | 2.68E+00 | 2.74E+00 | 9.22E-01 | no | Nhej1 | intron_variant | MODIFIER |
| 1 | 75001007 | AX-170377171 | 0 | G | C | 0.123 | 2.01E-02 | 2.07E-01 | 2.68E+00 | 2.74E+00 | 9.22E-01 | no | Nhej1 | intron_variant | MODIFIER |
| 1 | 75001807 | AX-170363073 | 0 | C | A | 0.123 | 2.01E-02 | 2.07E-01 | 2.68E+00 | 2.74E+00 | 9.22E-01 | no | Nhej1 | intron_variant | MODIFIER |
| 1 | 75001954 | AX-170341531 | 0 | A | G | 0.123 | 2.01E-02 | 2.07E-01 | 2.68E+00 | 2.74E+00 | 9.22E-01 | no | Nhej1 | intron_variant | MODIFIER |
| 1 | 75036793 | AX-97805282 | 0 | T | C | 0.123 | 2.01E-02 | 2.07E-01 | 2.68E+00 | 2.74E+00 | 9.22E-01 | no | Nhej1 | intron_variant | MODIFIER |
| 1 | 75038417 | AX-170323606 | 0 | T | C | 0.123 | 2.01E-02 | 2.07E-01 | 2.68E+00 | 2.74E+00 | 9.22E-01 | no | Nhej1 | intron_variant | MODIFIER |
| 1 | 75044042 | AX-170214420 | 0 | G | A | 0.123 | 2.01E-02 | 2.07E-01 | 2.68E+00 | 2.74E+00 | 9.22E-01 | no | Nhej1 | intron_variant | MODIFIER |
| 1 | 75044898 | AX-170000823 | 0 | G | A | 0.123 | 2.01E-02 | 2.07E-01 | 2.68E+00 | 2.74E+00 | 9.22E-01 | no | Nhej1 | intron_variant | MODIFIER |
| 1 | 75171886 | AX-170315967 | 0 | T | C | 0.123 | 2.01E-02 | 2.07E-01 | 2.68E+00 | 2.74E+00 | 9.22E-01 | no | Zfand2b | downstream_gene_variant | MODIFIER |
| 1 | 75172546 | AX-170389759 | 0 | G | C | 0.123 | 2.01E-02 | 2.07E-01 | 2.68E+00 | 2.74E+00 | 9.22E-01 | no | Zfand2b | downstream_gene_variant | MODIFIER |
| 1 | 75174439 | AX-170118718 | 0 | A | G | 0.123 | 2.01E-02 | 2.07E-01 | 2.68E+00 | 2.74E+00 | 9.22E-01 | no | Zfand2b | downstream_gene_variant | MODIFIER |
| 1 | 75174935 | AX-170000818 | 0 | T | G | 0.123 | 2.01E-02 | 2.07E-01 | 2.68E+00 | 2.74E+00 | 9.22E-01 | no | Zfand2b | downstream_gene_variant | MODIFIER |
| 1 | 75174959 | AX-170000813 | 0 | G | A | 0.123 | 2.01E-02 | 2.07E-01 | 2.68E+00 | 2.74E+00 | 9.22E-01 | no | Zfand2b | downstream_gene_variant | MODIFIER |
| 1 | 75221007 | AX-170000814 | 0 | G | A | 0.123 | 2.01E-02 | 2.07E-01 | 2.68E+00 | 2.74E+00 | 9.22E-01 | no | Tuba4a | upstream_gene_variant | MODIFIER |
| 1 | 75245260 | AX-170347213 | 0 | C | T | 0.123 | 2.01E-02 | 2.07E-01 | 2.68E+00 | 2.74E+00 | 9.22E-01 | no | Ptprn | downstream_gene_variant | MODIFIER |
| 1 | 75272738 | AX-170360245 | 0 | A | T | 0.123 | 2.01E-02 | 2.07E-01 | 2.68E+00 | 2.74E+00 | 9.22E-01 | no | Resp18 | intron_variant | MODIFIER |
| 1 | 75286989 | AX-170146140 | 0 | G | A | 0.123 | 2.01E-02 | 2.07E-01 | 2.68E+00 | 2.74E+00 | 9.22E-01 | no | | intergenic_variant | MODIFIER |
| 1 | 75299007 | AX-170193859 | 0 | C | T | 0.123 | 2.01E-02 | 2.07E-01 | 2.68E+00 | 2.74E+00 | 9.22E-01 | no | | intergenic_variant | MODIFIER |
| 1 | 75303472 | AX-170368098 | 0 | G | T | 0.123 | 2.01E-02 | 2.07E-01 | 2.68E+00 | 2.74E+00 | 9.22E-01 | no | Dnpep | downstream_gene_variant | MODIFIER |
| 1 | 75303973 | AX-170041696 | 0 | A | G | 0.123 | 2.01E-02 | 2.07E-01 | 2.68E+00 | 2.74E+00 | 9.22E-01 | no | Dnpep | downstream_gene_variant | MODIFIER |
| 1 | 75339738 | AX-170205276 | 0 | T | C | 0.123 | 2.01E-02 | 2.07E-01 | 2.68E+00 | 2.74E+00 | 9.22E-01 | no | | regulatory_region_variant | MODIFIER |
| 1 | 75357209 | AX-170379322 | 0 | A | G | 0.123 | 2.01E-02 | 2.07E-01 | 2.68E+00 | 2.74E+00 | 9.22E-01 | no | Des | upstream_gene_variant | MODIFIER |

| | | | | | | | | | | | | | | | |
|---|---|---|---|---|---|---|---|---|---|---|---|---|---|---|---|
| 11 | 48055578 | AX-170150424 | 1 | A | G | 0.08 | 1.59E-02 | 1.82E-01 | 2.63E+00 | 2.75E+00 | 9.22E-01 | no | | intergenic_variant | MODIFIER |
| 12 | 79045957 | AX-170426899 | 0 | T | G | 0.053 | -1.96E-02 | 2.05E-01 | 2.63E+00 | 2.74E+00 | 9.22E-01 | no | Plekhh1 | intron_variant | MODIFIER |
| 13 | 21346415 | AX-170227101 | 0 | T | C | 0.175 | -1.38E-02 | 1.16E-01 | 2.61E+00 | 2.70E+00 | 9.23E-01 | no | Olfr1367 | upstream_gene_variant | MODIFIER |
| 15 | 74359563 | AX-170486371 | 0 | G | A | 0.553 | 8.84E-03 | 8.13E-02 | 2.63E+00 | 2.72E+00 | 9.23E-01 | no | | intergenic_variant | MODIFIER |
| 1 | 152259587 | AX-97798404 | 0 | A | G | 0.202 | -1.25E-02 | 1.37E-01 | 2.70E+00 | 2.75E+00 | 9.23E-01 | no | | intergenic_variant | MODIFIER |
| 7 | 51495238 | AX-97802824 | 1 | A | T | 0.438 | -8.55E-03 | 9.73E-02 | 2.67E+00 | 2.76E+00 | 9.23E-01 | no | | intergenic_variant | MODIFIER |
| 9 | 69966457 | AX-170490276 | 2 | C | T | 0.2 | 1.26E-02 | 1.34E-01 | 2.71E+00 | 2.75E+00 | 9.23E-01 | no | | intergenic_variant | MODIFIER |
| 20 | 72517631 | AX-170433734 | 2 | T | C | 0.245 | 6.08E-03 | 6.23E-02 | 2.64E+00 | 2.74E+00 | 9.23E-01 | no | Gabra3 | intron_variant | MODIFIER |
| 7 | 92437163 | AX-170271488 | 0 | G | A | 0.132 | -2.04E-02 | 2.12E-01 | 2.66E+00 | 2.74E+00 | 9.23E-01 | no | Dlg2 | intron_variant | MODIFIER |
| 2 | 88498658 | AX-170375133 | 0 | A | C | 0.061 | 2.03E-02 | 2.03E-01 | 2.62E+00 | 2.73E+00 | 9.23E-01 | no | Olfr1185-ps1 | intron_variant | MODIFIER |
| 5 | 99350478 | AX-97799983 | 1 | T | C | 0.438 | 1.02E-02 | 1.04E-01 | 2.66E+00 | 2.73E+00 | 9.23E-01 | no | Rasgef1b | intron_variant | MODIFIER |
| 1 | 54907924 | AX-170169307 | 0 | C | A | 0.105 | -2.24E-02 | 2.36E-01 | 2.70E+00 | 2.74E+00 | 9.23E-01 | no | Ankrd44 | intron_variant | MODIFIER |
| 1 | 150016195 | AX-97854632 | 0 | T | G | 0.202 | -1.10E-02 | 1.26E-01 | 2.68E+00 | 2.76E+00 | 9.23E-01 | no | | intergenic_variant | MODIFIER |
| 5 | 83230680 | AX-170072644 | 0 | A | G | 0.026 | 1.08E-02 | 1.31E-01 | 2.57E+00 | 2.75E+00 | 9.23E-01 | no | | intergenic_variant | MODIFIER |
| 2 | 146984677 | AX-170065196 | 0 | T | C | 0.14 | -2.05E-02 | 2.19E-01 | 2.72E+00 | 2.75E+00 | 9.23E-01 | no | | intergenic_variant | MODIFIER |
| 7 | 103216004 | AX-170385355 | 2 | G | C | 0.473 | 9.49E-03 | 9.25E-02 | 2.65E+00 | 2.72E+00 | 9.23E-01 | no | Olfr594 | upstream_gene_variant | MODIFIER |
| 1 | 43173925 | AX-97784406 | 0 | A | G | 0.342 | -7.29E-03 | 7.35E-02 | 2.64E+00 | 2.73E+00 | 9.23E-01 | no | Fhl2 | intron_variant | MODIFIER |
| 15 | 92929642 | AX-97804061 | 0 | C | A | 0.14 | -1.49E-02 | 1.73E-01 | 2.67E+00 | 2.76E+00 | 9.23E-01 | no | Gm49418 | upstream_gene_variant | MODIFIER |
| 16 | 92205595 | AX-170339049 | 1 | C | T | 0.473 | -7.45E-03 | 7.91E-02 | 2.66E+00 | 2.74E+00 | 9.23E-01 | no | Gm29880 | intron_variant,non_coding_transcript_variant | MODIFIER |
| 12 | 109909353 | AX-170335798 | 0 | G | C | 0.132 | -1.69E-02 | 1.82E-01 | 2.67E+00 | 2.74E+00 | 9.23E-01 | no | 4930511J24Rik | upstream_gene_variant | MODIFIER |
| 9 | 32859130 | AX-170288281 | 0 | A | T | 0.421 | 8.75E-03 | 8.70E-02 | 2.66E+00 | 2.73E+00 | 9.23E-01 | no | | intergenic_variant | MODIFIER |
| 8 | 30230108 | AX-170257350 | 0 | A | G | 0.167 | -1.16E-02 | 1.20E-01 | 2.67E+00 | 2.73E+00 | 9.23E-01 | no | | intergenic_variant | MODIFIER |
| 6 | 100991165 | AX-170262073 | 0 | T | A | 0.07 | -2.64E-02 | 2.77E-01 | 2.70E+00 | 2.74E+00 | 9.24E-01 | no | | intergenic_variant | MODIFIER |
| 6 | 144809318 | AX-97785854 | 0 | G | A | 0.07 | -2.64E-02 | 2.77E-01 | 2.70E+00 | 2.74E+00 | 9.24E-01 | no | | intergenic_variant | MODIFIER |
| 3 | 147674106 | AX-170528927 | 1 | T | C | 0.348 | 7.31E-03 | 7.37E-02 | 2.66E+00 | 2.73E+00 | 9.24E-01 | no | | intergenic_variant | MODIFIER |
| 14 | 67697348 | AX-170135971 | 0 | G | A | 0.053 | 1.70E-02 | 1.82E-01 | 2.62E+00 | 2.74E+00 | 9.24E-01 | no | Cdca2 | downstream_gene_variant | MODIFIER |
| 18 | 59253465 | AX-97822373 | 2 | T | G | 0.418 | 7.24E-03 | 8.44E-02 | 2.70E+00 | 2.77E+00 | 9.24E-01 | no | Chsy3 | intron_variant | MODIFIER |
| 2 | 124972113 | AX-170064812 | 0 | T | C | 0.14 | -2.06E-02 | 2.22E-01 | 2.72E+00 | 2.75E+00 | 9.24E-01 | no | | regulatory_region_variant | MODIFIER |
| 8 | 41253701 | AX-170516680 | 1 | C | T | 0.143 | -1.44E-02 | 1.48E-01 | 2.68E+00 | 2.73E+00 | 9.24E-01 | no | Pcm1 | intron_variant | MODIFIER |
| 3 | 146741889 | AX-170295519 | 1 | C | G | 0.348 | 7.17E-03 | 7.03E-02 | 2.64E+00 | 2.72E+00 | 9.24E-01 | no | Prkacb | intron_variant | MODIFIER |
| 2 | 148929036 | AX-170019344 | 1 | A | G | 0.17 | 1.37E-02 | 1.32E-01 | 2.60E+00 | 2.73E+00 | 9.24E-01 | no | Gm14129 | intron_variant,non_coding_transcript_variant | MODIFIER |
| 1 | 99839818 | AX-97826839 | 2 | G | A | 0.136 | 2.10E-02 | 1.82E-01 | 2.60E+00 | 2.71E+00 | 9.24E-01 | no | Cntnap5b | intron_variant | MODIFIER |
| 11 | 17633860 | AX-170271438 | 1 | G | A | 0.062 | 1.89E-02 | 2.13E-01 | 2.65E+00 | 2.75E+00 | 9.24E-01 | no | Gm12016 | downstream_gene_variant | MODIFIER |
| 15 | 88178912 | AX-170221102 | 0 | G | A | 0.132 | 1.56E-02 | 1.56E-01 | 2.66E+00 | 2.72E+00 | 9.24E-01 | no | | intergenic_variant | MODIFIER |
| 5 | 128104479 | AX-97807033 | 0 | T | C | 0.421 | -6.60E-03 | 8.01E-02 | 2.65E+00 | 2.76E+00 | 9.24E-01 | no | Tmem132d | intron_variant | MODIFIER |
| 1 | 78049203 | AX-170462463 | 0 | G | A | 0.132 | 2.11E-02 | 2.23E-01 | 2.67E+00 | 2.74E+00 | 9.24E-01 | no | | intergenic_variant | MODIFIER |
| 7 | 67463735 | AX-170337100 | 1 | G | A | 0.429 | 9.93E-03 | 9.82E-02 | 2.64E+00 | 2.72E+00 | 9.25E-01 | no | | regulatory_region_variant | MODIFIER |
| 7 | 102548760 | AX-170201507 | 0 | C | T | 0.465 | -7.92E-03 | 9.33E-02 | 2.69E+00 | 2.76E+00 | 9.25E-01 | no | Olfr548-ps1 | downstream_gene_variant | MODIFIER |
| 8 | 85384460 | AX-170080641 | 0 | G | A | 0.202 | 1.20E-02 | 1.28E-01 | 2.67E+00 | 2.74E+00 | 9.25E-01 | no | Mylk3 | upstream_gene_variant | MODIFIER |
| 7 | 97423923 | AX-170522716 | 2 | C | T | 0.091 | 1.62E-02 | 1.78E-01 | 2.63E+00 | 2.74E+00 | 9.25E-01 | no | | regulatory_region_variant | MODIFIER |
| 6 | 139011763 | AX-170028365 | 1 | C | T | 0.071 | 2.57E-02 | 2.79E-01 | 2.71E+00 | 2.74E+00 | 9.25E-01 | no | | intergenic_variant | MODIFIER |
| 7 | 69518348 | AX-170264113 | 0 | G | T | 0.404 | 1.01E-02 | 9.80E-02 | 2.65E+00 | 2.72E+00 | 9.25E-01 | no | 4930402F11Rik | non_coding_transcript_exon_variant | MODIFIER |
| 7 | 69524807 | AX-170502518 | 0 | C | A | 0.404 | 1.01E-02 | 9.80E-02 | 2.65E+00 | 2.72E+00 | 9.25E-01 | no | 4930402F11Rik | intron_variant,non_coding_transcript_variant | MODIFIER |
| 7 | 69545678 | AX-170315761 | 0 | T | C | 0.404 | 1.01E-02 | 9.80E-02 | 2.65E+00 | 2.72E+00 | 9.25E-01 | no | 4930402F11Rik | intron_variant,non_coding_transcript_variant | MODIFIER |
| 7 | 69737812 | AX-170386437 | 0 | A | G | 0.404 | 1.01E-02 | 9.80E-02 | 2.65E+00 | 2.72E+00 | 9.25E-01 | no | | intergenic_variant | MODIFIER |
| 7 | 69752619 | AX-97797105 | 0 | G | C | 0.404 | 1.01E-02 | 9.80E-02 | 2.65E+00 | 2.72E+00 | 9.25E-01 | no | | intergenic_variant | MODIFIER |
| 7 | 69758322 | AX-170343827 | 0 | T | A | 0.404 | 1.01E-02 | 9.80E-02 | 2.65E+00 | 2.72E+00 | 9.25E-01 | no | | intergenic_variant | MODIFIER |
| 7 | 69758886 | AX-170397414 | 0 | C | T | 0.404 | 1.01E-02 | 9.80E-02 | 2.65E+00 | 2.72E+00 | 9.25E-01 | no | | intergenic_variant | MODIFIER |
| 7 | 69758912 | AX-170207196 | 0 | C | T | 0.404 | 1.01E-02 | 9.80E-02 | 2.65E+00 | 2.72E+00 | 9.25E-01 | no | | intergenic_variant | MODIFIER |
| 7 | 69783141 | AX-170077654 | 0 | G | C | 0.404 | 1.01E-02 | 9.80E-02 | 2.65E+00 | 2.72E+00 | 9.25E-01 | no | | intergenic_variant | MODIFIER |
| 7 | 69796926 | AX-170077653 | 0 | A | G | 0.404 | 1.01E-02 | 9.80E-02 | 2.65E+00 | 2.72E+00 | 9.25E-01 | no | | intergenic_variant | MODIFIER |
| 7 | 69826226 | AX-170466214 | 0 | T | C | 0.404 | 1.01E-02 | 9.80E-02 | 2.65E+00 | 2.72E+00 | 9.25E-01 | no | | intergenic_variant | MODIFIER |
| 7 | 69826653 | AX-97797506 | 0 | A | G | 0.404 | 1.01E-02 | 9.80E-02 | 2.65E+00 | 2.72E+00 | 9.25E-01 | no | | intergenic_variant | MODIFIER |
| 7 | 69871735 | AX-97817817 | 0 | G | A | 0.404 | 1.01E-02 | 9.80E-02 | 2.65E+00 | 2.72E+00 | 9.25E-01 | no | | intergenic_variant | MODIFIER |

| | | | | | | | | | | | | | | | |
|---|---|---|---|---|---|---|---|---|---|---|---|---|---|---|---|
| 7 | 69876913 | AX-97836214 | 0 | C | T | 0.404 | 1.01E-02 | 9.80E-02 | 2.65E+00 | 2.72E+00 | 9.25E-01 | no | | intergenic_variant | MODIFIER |
| 7 | 69888694 | AX-170138250 | 0 | C | T | 0.404 | 1.01E-02 | 9.80E-02 | 2.65E+00 | 2.72E+00 | 9.25E-01 | no | Gm44690 | upstream_gene_variant | MODIFIER |
| 7 | 69930745 | AX-170191501 | 0 | T | C | 0.404 | 1.01E-02 | 9.80E-02 | 2.65E+00 | 2.72E+00 | 9.25E-01 | no | | intergenic_variant | MODIFIER |
| 7 | 69932778 | AX-170345283 | 0 | G | A | 0.404 | 1.01E-02 | 9.80E-02 | 2.65E+00 | 2.72E+00 | 9.25E-01 | no | | intergenic_variant | MODIFIER |
| 7 | 69949862 | AX-170513710 | 0 | T | C | 0.404 | 1.01E-02 | 9.80E-02 | 2.65E+00 | 2.72E+00 | 9.25E-01 | no | 4933423L19Rik | non_coding_transcript_exon_variant | MODIFIER |
| 7 | 69953022 | AX-170362899 | 0 | T | G | 0.404 | 1.01E-02 | 9.80E-02 | 2.65E+00 | 2.72E+00 | 9.25E-01 | no | Gm24120 | downstream_gene_variant | MODIFIER |
| 7 | 69967209 | AX-170077659 | 0 | T | C | 0.404 | 1.01E-02 | 9.80E-02 | 2.65E+00 | 2.72E+00 | 9.25E-01 | no | | intergenic_variant | MODIFIER |
| 7 | 69991321 | AX-170475345 | 0 | T | C | 0.404 | 1.01E-02 | 9.80E-02 | 2.65E+00 | 2.72E+00 | 9.25E-01 | no | | intergenic_variant | MODIFIER |
| 7 | 70037266 | AX-170439450 | 0 | G | A | 0.404 | 1.01E-02 | 9.80E-02 | 2.65E+00 | 2.72E+00 | 9.25E-01 | no | | intergenic_variant | MODIFIER |
| 7 | 70041254 | AX-170211073 | 0 | T | C | 0.404 | 1.01E-02 | 9.80E-02 | 2.65E+00 | 2.72E+00 | 9.25E-01 | no | | intergenic_variant | MODIFIER |
| 7 | 70044403 | AX-170134059 | 0 | C | T | 0.404 | 1.01E-02 | 9.80E-02 | 2.65E+00 | 2.72E+00 | 9.25E-01 | no | | intergenic_variant | MODIFIER |
| 7 | 70045183 | AX-170269836 | 0 | A | G | 0.404 | 1.01E-02 | 9.80E-02 | 2.65E+00 | 2.72E+00 | 9.25E-01 | no | | intergenic_variant | MODIFIER |
| 7 | 70049057 | AX-170127014 | 0 | A | T | 0.404 | 1.01E-02 | 9.80E-02 | 2.65E+00 | 2.72E+00 | 9.25E-01 | no | | intergenic_variant | MODIFIER |
| 7 | 70053305 | AX-170288326 | 0 | A | C | 0.404 | 1.01E-02 | 9.80E-02 | 2.65E+00 | 2.72E+00 | 9.25E-01 | no | | intergenic_variant | MODIFIER |
| 7 | 70053910 | AX-170372979 | 0 | A | G | 0.404 | 1.01E-02 | 9.80E-02 | 2.65E+00 | 2.72E+00 | 9.25E-01 | no | | intergenic_variant | MODIFIER |
| 7 | 70054882 | AX-170207706 | 0 | C | T | 0.404 | 1.01E-02 | 9.80E-02 | 2.65E+00 | 2.72E+00 | 9.25E-01 | no | | intergenic_variant | MODIFIER |
| 7 | 70080073 | AX-170373027 | 0 | T | C | 0.404 | 1.01E-02 | 9.80E-02 | 2.65E+00 | 2.72E+00 | 9.25E-01 | no | | intergenic_variant | MODIFIER |
| 7 | 70096041 | AX-170401882 | 0 | G | T | 0.404 | 1.01E-02 | 9.80E-02 | 2.65E+00 | 2.72E+00 | 9.25E-01 | no | | intergenic_variant | MODIFIER |
| 7 | 70102511 | AX-170322314 | 0 | C | G | 0.404 | 1.01E-02 | 9.80E-02 | 2.65E+00 | 2.72E+00 | 9.25E-01 | no | | intergenic_variant | MODIFIER |
| 4 | 147335621 | AX-97792483 | 1 | C | T | 0.071 | -1.97E-02 | 2.10E-01 | 2.63E+00 | 2.74E+00 | 9.25E-01 | no | Zfp988 | downstream_gene_variant | MODIFIER |
| 8 | 48536388 | AX-97792839 | 0 | G | A | 0.149 | -1.56E-02 | 1.52E-01 | 2.67E+00 | 2.71E+00 | 9.25E-01 | no | Tenm3 | intron_variant | MODIFIER |
| 8 | 48594494 | AX-97852004 | 0 | G | A | 0.149 | -1.56E-02 | 1.52E-01 | 2.67E+00 | 2.71E+00 | 9.25E-01 | no | Tenm3 | intron_variant | MODIFIER |
| 8 | 48594882 | AX-170359904 | 0 | A | G | 0.149 | -1.56E-02 | 1.52E-01 | 2.67E+00 | 2.71E+00 | 9.25E-01 | no | Tenm3 | intron_variant | MODIFIER |
| 8 | 48599519 | AX-170129116 | 0 | A | T | 0.149 | -1.56E-02 | 1.52E-01 | 2.67E+00 | 2.71E+00 | 9.25E-01 | no | Tenm3 | intron_variant | MODIFIER |
| 8 | 48600300 | AX-170409052 | 0 | T | C | 0.149 | -1.56E-02 | 1.52E-01 | 2.67E+00 | 2.71E+00 | 9.25E-01 | no | Tenm3 | intron_variant | MODIFIER |
| 8 | 48603491 | AX-170519869 | 0 | G | A | 0.149 | -1.56E-02 | 1.52E-01 | 2.67E+00 | 2.71E+00 | 9.25E-01 | no | Tenm3 | intron_variant | MODIFIER |
| 8 | 48639276 | AX-170514890 | 0 | G | A | 0.149 | -1.56E-02 | 1.52E-01 | 2.67E+00 | 2.71E+00 | 9.25E-01 | no | Tenm3 | intron_variant | MODIFIER |
| 8 | 48642472 | AX-170221179 | 0 | A | G | 0.149 | -1.56E-02 | 1.52E-01 | 2.67E+00 | 2.71E+00 | 9.25E-01 | no | Tenm3 | intron_variant | MODIFIER |
| 8 | 48642533 | AX-170201179 | 0 | G | C | 0.149 | -1.56E-02 | 1.52E-01 | 2.67E+00 | 2.71E+00 | 9.25E-01 | no | Tenm3 | intron_variant | MODIFIER |
| 8 | 48649281 | AX-170505161 | 0 | A | G | 0.149 | -1.56E-02 | 1.52E-01 | 2.67E+00 | 2.71E+00 | 9.25E-01 | no | Tenm3 | intron_variant | MODIFIER |
| 8 | 48669374 | AX-170080018 | 0 | G | A | 0.149 | -1.56E-02 | 1.52E-01 | 2.67E+00 | 2.71E+00 | 9.25E-01 | no | Tenm3 | intron_variant | MODIFIER |
| 8 | 48671460 | AX-170080019 | 0 | A | G | 0.149 | -1.56E-02 | 1.52E-01 | 2.67E+00 | 2.71E+00 | 9.25E-01 | no | Tenm3 | intron_variant | MODIFIER |
| 8 | 48671828 | AX-170105405 | 0 | A | G | 0.149 | -1.56E-02 | 1.52E-01 | 2.67E+00 | 2.71E+00 | 9.25E-01 | no | Tenm3 | intron_variant | MODIFIER |
| 8 | 48679090 | AX-97806944 | 0 | G | A | 0.149 | -1.56E-02 | 1.52E-01 | 2.67E+00 | 2.71E+00 | 9.25E-01 | no | Tenm3 | upstream_gene_variant | MODIFIER |
| 8 | 48681867 | AX-170192436 | 0 | A | G | 0.149 | -1.56E-02 | 1.52E-01 | 2.67E+00 | 2.71E+00 | 9.25E-01 | no | Tenm3 | intron_variant | MODIFIER |
| 8 | 48685940 | AX-170323951 | 0 | T | C | 0.149 | -1.56E-02 | 1.52E-01 | 2.67E+00 | 2.71E+00 | 9.25E-01 | no | Tenm3 | intron_variant | MODIFIER |
| 8 | 48689127 | AX-170233380 | 0 | A | G | 0.149 | -1.56E-02 | 1.52E-01 | 2.67E+00 | 2.71E+00 | 9.25E-01 | no | Tenm3 | intron_variant | MODIFIER |
| 8 | 48705654 | AX-170124056 | 0 | T | C | 0.149 | -1.56E-02 | 1.52E-01 | 2.67E+00 | 2.71E+00 | 9.25E-01 | no | Tenm3 | intron_variant | MODIFIER |
| 8 | 48714545 | AX-170104640 | 0 | T | G | 0.149 | -1.56E-02 | 1.52E-01 | 2.67E+00 | 2.71E+00 | 9.25E-01 | no | Tenm3 | intron_variant | MODIFIER |
| 8 | 48752844 | AX-97834612 | 0 | G | A | 0.149 | -1.56E-02 | 1.52E-01 | 2.67E+00 | 2.71E+00 | 9.25E-01 | no | Tenm3 | intron_variant | MODIFIER |
| 8 | 48753465 | AX-170341715 | 0 | G | A | 0.149 | -1.56E-02 | 1.52E-01 | 2.67E+00 | 2.71E+00 | 9.25E-01 | no | Tenm3 | intron_variant | MODIFIER |
| 8 | 48759546 | AX-170086236 | 0 | A | T | 0.149 | -1.56E-02 | 1.52E-01 | 2.67E+00 | 2.71E+00 | 9.25E-01 | no | Tenm3 | intron_variant | MODIFIER |
| 8 | 48929180 | AX-170030635 | 0 | T | C | 0.149 | -1.56E-02 | 1.52E-01 | 2.67E+00 | 2.71E+00 | 9.25E-01 | no | | intergenic_variant | MODIFIER |
| 8 | 48930146 | AX-170030634 | 0 | T | C | 0.149 | -1.56E-02 | 1.52E-01 | 2.67E+00 | 2.71E+00 | 9.25E-01 | no | | intergenic_variant | MODIFIER |
| 8 | 48955584 | AX-170030642 | 0 | C | A | 0.149 | -1.56E-02 | 1.52E-01 | 2.67E+00 | 2.71E+00 | 9.25E-01 | no | Gm25830 | downstream_gene_variant | MODIFIER |
| 8 | 48957805 | AX-170030633 | 0 | A | G | 0.149 | -1.56E-02 | 1.52E-01 | 2.67E+00 | 2.71E+00 | 9.25E-01 | no | Gm25830 | downstream_gene_variant | MODIFIER |
| 8 | 48993542 | AX-97834856 | 0 | A | C | 0.149 | -1.56E-02 | 1.52E-01 | 2.67E+00 | 2.71E+00 | 9.25E-01 | no | | intergenic_variant | MODIFIER |
| 8 | 49017065 | AX-97855857 | 0 | G | A | 0.149 | -1.56E-02 | 1.52E-01 | 2.67E+00 | 2.71E+00 | 9.25E-01 | no | | intergenic_variant | MODIFIER |
| 8 | 49017888 | AX-170190938 | 0 | T | C | 0.149 | -1.56E-02 | 1.52E-01 | 2.67E+00 | 2.71E+00 | 9.25E-01 | no | | intergenic_variant | MODIFIER |
| 8 | 49055109 | AX-170213993 | 0 | T | C | 0.149 | -1.56E-02 | 1.52E-01 | 2.67E+00 | 2.71E+00 | 9.25E-01 | no | | intergenic_variant | MODIFIER |
| 8 | 49056985 | AX-170481464 | 0 | C | G | 0.149 | -1.56E-02 | 1.52E-01 | 2.67E+00 | 2.71E+00 | 9.25E-01 | no | | intergenic_variant | MODIFIER |
| 8 | 49057364 | AX-170090094 | 0 | C | T | 0.149 | -1.56E-02 | 1.52E-01 | 2.67E+00 | 2.71E+00 | 9.25E-01 | no | | intergenic_variant | MODIFIER |
| 8 | 49081183 | AX-170212881 | 0 | A | T | 0.149 | -1.56E-02 | 1.52E-01 | 2.67E+00 | 2.71E+00 | 9.25E-01 | no | | intergenic_variant | MODIFIER |
| 8 | 49082512 | AX-97854436 | 0 | C | A | 0.149 | -1.56E-02 | 1.52E-01 | 2.67E+00 | 2.71E+00 | 9.25E-01 | no | | intergenic_variant | MODIFIER |

| | | | | | | | | | | | | | | | |
|---|---|---|---|---|---|---|---|---|---|---|---|---|---|---|---|
| 8 | 49082630 | AX-170196279 | 0 | C | T | 0.149 | -1.56E-02 | 1.52E-01 | 2.67E+00 | 2.71E+00 | 9.25E-01 | no | | intergenic_variant | MODIFIER |
| 8 | 49101434 | AX-97856708 | 0 | G | A | 0.149 | -1.56E-02 | 1.52E-01 | 2.67E+00 | 2.71E+00 | 9.25E-01 | no | | intergenic_variant | MODIFIER |
| 8 | 49113077 | AX-170305794 | 0 | A | G | 0.149 | -1.56E-02 | 1.52E-01 | 2.67E+00 | 2.71E+00 | 9.25E-01 | no | | intergenic_variant | MODIFIER |
| 8 | 49115818 | AX-170423666 | 0 | C | T | 0.149 | -1.56E-02 | 1.52E-01 | 2.67E+00 | 2.71E+00 | 9.25E-01 | no | | intergenic_variant | MODIFIER |
| 8 | 49116162 | AX-170119787 | 0 | G | A | 0.149 | -1.56E-02 | 1.52E-01 | 2.67E+00 | 2.71E+00 | 9.25E-01 | no | | intergenic_variant | MODIFIER |
| 8 | 49117978 | AX-97808658 | 0 | A | G | 0.149 | -1.56E-02 | 1.52E-01 | 2.67E+00 | 2.71E+00 | 9.25E-01 | no | | intergenic_variant | MODIFIER |
| 8 | 49174785 | AX-97843332 | 0 | A | G | 0.149 | -1.56E-02 | 1.52E-01 | 2.67E+00 | 2.71E+00 | 9.25E-01 | no | | intergenic_variant | MODIFIER |
| 8 | 49175116 | AX-170152565 | 0 | A | G | 0.149 | -1.56E-02 | 1.52E-01 | 2.67E+00 | 2.71E+00 | 9.25E-01 | no | | intergenic_variant | MODIFIER |
| 8 | 49177880 | AX-170490934 | 0 | G | A | 0.149 | -1.56E-02 | 1.52E-01 | 2.67E+00 | 2.71E+00 | 9.25E-01 | no | | intergenic_variant | MODIFIER |
| 8 | 49180865 | AX-170080022 | 0 | A | C | 0.149 | -1.56E-02 | 1.52E-01 | 2.67E+00 | 2.71E+00 | 9.25E-01 | no | | intergenic_variant | MODIFIER |
| 8 | 49180892 | AX-170080021 | 0 | A | C | 0.149 | -1.56E-02 | 1.52E-01 | 2.67E+00 | 2.71E+00 | 9.25E-01 | no | | intergenic_variant | MODIFIER |
| 8 | 49188588 | AX-170119866 | 0 | A | C | 0.149 | -1.56E-02 | 1.52E-01 | 2.67E+00 | 2.71E+00 | 9.25E-01 | no | | intergenic_variant | MODIFIER |
| 8 | 49188970 | AX-170364359 | 0 | A | T | 0.149 | -1.56E-02 | 1.52E-01 | 2.67E+00 | 2.71E+00 | 9.25E-01 | no | | intergenic_variant | MODIFIER |
| 8 | 49189426 | AX-97786200 | 0 | T | G | 0.149 | -1.56E-02 | 1.52E-01 | 2.67E+00 | 2.71E+00 | 9.25E-01 | no | | intergenic_variant | MODIFIER |
| 8 | 49190194 | AX-170203807 | 0 | G | C | 0.149 | -1.56E-02 | 1.52E-01 | 2.67E+00 | 2.71E+00 | 9.25E-01 | no | | intergenic_variant | MODIFIER |
| 8 | 49191311 | AX-170155373 | 0 | A | T | 0.149 | -1.56E-02 | 1.52E-01 | 2.67E+00 | 2.71E+00 | 9.25E-01 | no | | intergenic_variant | MODIFIER |
| 8 | 49219254 | AX-97789028 | 0 | C | T | 0.149 | -1.56E-02 | 1.52E-01 | 2.67E+00 | 2.71E+00 | 9.25E-01 | no | | intergenic_variant | MODIFIER |
| 8 | 49236473 | AX-170199924 | 0 | C | T | 0.149 | -1.56E-02 | 1.52E-01 | 2.67E+00 | 2.71E+00 | 9.25E-01 | no | Gm45832 | upstream_gene_variant | MODIFIER |
| 8 | 49248241 | AX-170239714 | 0 | A | G | 0.149 | -1.56E-02 | 1.52E-01 | 2.67E+00 | 2.71E+00 | 9.25E-01 | no | | intergenic_variant | MODIFIER |
| 8 | 49276758 | AX-170090095 | 0 | A | G | 0.149 | -1.56E-02 | 1.52E-01 | 2.67E+00 | 2.71E+00 | 9.25E-01 | no | | intergenic_variant | MODIFIER |
| 8 | 49296464 | AX-97794044 | 0 | G | A | 0.149 | -1.56E-02 | 1.52E-01 | 2.67E+00 | 2.71E+00 | 9.25E-01 | no | | intergenic_variant | MODIFIER |
| 8 | 49312562 | AX-170030646 | 0 | C | T | 0.149 | -1.56E-02 | 1.52E-01 | 2.67E+00 | 2.71E+00 | 9.25E-01 | no | | intergenic_variant | MODIFIER |
| 8 | 49312980 | AX-97800784 | 0 | T | C | 0.149 | -1.56E-02 | 1.52E-01 | 2.67E+00 | 2.71E+00 | 9.25E-01 | no | | intergenic_variant | MODIFIER |
| 8 | 49315235 | AX-170105676 | 0 | T | C | 0.149 | -1.56E-02 | 1.52E-01 | 2.67E+00 | 2.71E+00 | 9.25E-01 | no | | intergenic_variant | MODIFIER |
| 8 | 49335416 | AX-170030650 | 0 | C | T | 0.149 | -1.56E-02 | 1.52E-01 | 2.67E+00 | 2.71E+00 | 9.25E-01 | no | | intergenic_variant | MODIFIER |
| 8 | 49336915 | AX-170030652 | 0 | A | G | 0.149 | -1.56E-02 | 1.52E-01 | 2.67E+00 | 2.71E+00 | 9.25E-01 | no | | intergenic_variant | MODIFIER |
| 8 | 49369204 | AX-170030648 | 0 | G | T | 0.149 | -1.56E-02 | 1.52E-01 | 2.67E+00 | 2.71E+00 | 9.25E-01 | no | 4930555F03Rik | upstream_gene_variant | MODIFIER |
| 8 | 49381980 | AX-170030651 | 0 | G | A | 0.149 | -1.56E-02 | 1.52E-01 | 2.67E+00 | 2.71E+00 | 9.25E-01 | no | 4930555F03Rik | intron_variant,non_coding_transcript_variant | MODIFIER |
| 8 | 49382970 | AX-170395025 | 0 | C | T | 0.149 | -1.56E-02 | 1.52E-01 | 2.67E+00 | 2.71E+00 | 9.25E-01 | no | 4930555F03Rik | intron_variant,non_coding_transcript_variant | MODIFIER |
| 8 | 49383603 | AX-170344378 | 0 | G | A | 0.149 | -1.56E-02 | 1.52E-01 | 2.67E+00 | 2.71E+00 | 9.25E-01 | no | 4930555F03Rik | intron_variant,non_coding_transcript_variant | MODIFIER |
| 8 | 49405005 | AX-170030653 | 0 | A | G | 0.149 | -1.56E-02 | 1.52E-01 | 2.67E+00 | 2.71E+00 | 9.25E-01 | no | 4930555F03Rik | intron_variant,non_coding_transcript_variant | MODIFIER |
| 8 | 49405040 | AX-97784124 | 0 | C | T | 0.149 | -1.56E-02 | 1.52E-01 | 2.67E+00 | 2.71E+00 | 9.25E-01 | no | 4930555F03Rik | intron_variant,non_coding_transcript_variant | MODIFIER |
| 8 | 49409571 | AX-170150453 | 0 | T | C | 0.149 | -1.56E-02 | 1.52E-01 | 2.67E+00 | 2.71E+00 | 9.25E-01 | no | 4930555F03Rik | intron_variant,non_coding_transcript_variant | MODIFIER |
| 8 | 49409719 | AX-97840396 | 0 | A | G | 0.149 | -1.56E-02 | 1.52E-01 | 2.67E+00 | 2.71E+00 | 9.25E-01 | no | 4930555F03Rik | intron_variant,non_coding_transcript_variant | MODIFIER |
| 8 | 49526632 | AX-97847171 | 0 | C | T | 0.149 | -1.56E-02 | 1.52E-01 | 2.67E+00 | 2.71E+00 | 9.25E-01 | no | Gm45696 | downstream_gene_variant | MODIFIER |
| 8 | 49588693 | AX-97787008 | 0 | C | T | 0.149 | -1.56E-02 | 1.52E-01 | 2.67E+00 | 2.71E+00 | 9.25E-01 | no | | intergenic_variant | MODIFIER |
| 8 | 49618927 | AX-97850523 | 0 | C | T | 0.149 | -1.56E-02 | 1.52E-01 | 2.67E+00 | 2.71E+00 | 9.25E-01 | no | | intergenic_variant | MODIFIER |
| 8 | 49648690 | AX-170359400 | 0 | T | A | 0.149 | -1.56E-02 | 1.52E-01 | 2.67E+00 | 2.71E+00 | 9.25E-01 | no | | intergenic_variant | MODIFIER |
| 8 | 49675956 | AX-170090097 | 0 | C | T | 0.149 | -1.56E-02 | 1.52E-01 | 2.67E+00 | 2.71E+00 | 9.25E-01 | no | | intergenic_variant | MODIFIER |
| 8 | 49689244 | AX-170425294 | 0 | A | G | 0.149 | -1.56E-02 | 1.52E-01 | 2.67E+00 | 2.71E+00 | 9.25E-01 | no | | intergenic_variant | MODIFIER |
| 8 | 49705039 | AX-170480729 | 0 | A | C | 0.149 | -1.56E-02 | 1.52E-01 | 2.67E+00 | 2.71E+00 | 9.25E-01 | no | | intergenic_variant | MODIFIER |
| 8 | 49706944 | AX-170153235 | 0 | T | C | 0.149 | -1.56E-02 | 1.52E-01 | 2.67E+00 | 2.71E+00 | 9.25E-01 | no | | intergenic_variant | MODIFIER |
| 8 | 49719375 | AX-97827315 | 0 | A | G | 0.149 | -1.56E-02 | 1.52E-01 | 2.67E+00 | 2.71E+00 | 9.25E-01 | no | | intergenic_variant | MODIFIER |
| 8 | 49782826 | AX-97847960 | 0 | C | T | 0.149 | -1.56E-02 | 1.52E-01 | 2.67E+00 | 2.71E+00 | 9.25E-01 | no | | intergenic_variant | MODIFIER |
| 8 | 49810169 | AX-170096261 | 0 | T | C | 0.149 | -1.56E-02 | 1.52E-01 | 2.67E+00 | 2.71E+00 | 9.25E-01 | no | | intergenic_variant | MODIFIER |
| 8 | 49842763 | AX-170336292 | 0 | A | C | 0.149 | -1.56E-02 | 1.52E-01 | 2.67E+00 | 2.71E+00 | 9.25E-01 | no | Gm45321 | intron_variant,non_coding_transcript_variant | MODIFIER |
| 8 | 49842985 | AX-170242526 | 0 | A | G | 0.149 | -1.56E-02 | 1.52E-01 | 2.67E+00 | 2.71E+00 | 9.25E-01 | no | Gm45321 | intron_variant,non_coding_transcript_variant | MODIFIER |
| 8 | 49846554 | AX-97836753 | 0 | A | G | 0.149 | -1.56E-02 | 1.52E-01 | 2.67E+00 | 2.71E+00 | 9.25E-01 | no | Gm45321 | intron_variant,non_coding_transcript_variant | MODIFIER |
| 8 | 49850045 | AX-170513102 | 0 | T | C | 0.149 | -1.56E-02 | 1.52E-01 | 2.67E+00 | 2.71E+00 | 9.25E-01 | no | Gm45321 | intron_variant,non_coding_transcript_variant | MODIFIER |
| 8 | 49853520 | AX-170030660 | 0 | T | C | 0.149 | -1.56E-02 | 1.52E-01 | 2.67E+00 | 2.71E+00 | 9.25E-01 | no | Gm45321 | intron_variant,non_coding_transcript_variant | MODIFIER |
| 8 | 49880020 | AX-170030655 | 0 | G | T | 0.149 | -1.56E-02 | 1.52E-01 | 2.67E+00 | 2.71E+00 | 9.25E-01 | no | Gm45321 | intron_variant,non_coding_transcript_variant | MODIFIER |
| 8 | 49884054 | AX-170303366 | 0 | A | G | 0.149 | -1.56E-02 | 1.52E-01 | 2.67E+00 | 2.71E+00 | 9.25E-01 | no | Gm45321 | intron_variant,non_coding_transcript_variant | MODIFIER |
| 8 | 49884834 | AX-170504634 | 0 | C | T | 0.149 | -1.56E-02 | 1.52E-01 | 2.67E+00 | 2.71E+00 | 9.25E-01 | no | Gm45321 | intron_variant,non_coding_transcript_variant | MODIFIER |
| 8 | 49884865 | AX-97826494 | 0 | G | T | 0.149 | -1.56E-02 | 1.52E-01 | 2.67E+00 | 2.71E+00 | 9.25E-01 | no | Gm45321 | intron_variant,non_coding_transcript_variant | MODIFIER |

| | | | | | | | | | | | | | | | |
|---|---|---|---|---|---|---|---|---|---|---|---|---|---|---|---|
| 8 | 49913194 | AX-170178845 | 0 | G | T | 0.149 | -1.56E-02 | 1.52E-01 | 2.67E+00 | 2.71E+00 | 9.25E-01 | no | Gm45321 | intron_variant,non_coding_transcript_variant | MODIFIER |
| 8 | 49915404 | AX-170030658 | 0 | G | T | 0.149 | -1.56E-02 | 1.52E-01 | 2.67E+00 | 2.71E+00 | 9.25E-01 | no | Gm45321 | intron_variant,non_coding_transcript_variant | MODIFIER |
| 8 | 50139545 | AX-170096263 | 0 | G | A | 0.149 | -1.56E-02 | 1.52E-01 | 2.67E+00 | 2.71E+00 | 9.25E-01 | no | Gm45341 | intron_variant,non_coding_transcript_variant | MODIFIER |
| 8 | 50157697 | AX-170425920 | 0 | C | T | 0.149 | -1.56E-02 | 1.52E-01 | 2.67E+00 | 2.71E+00 | 9.25E-01 | no | Gm45341 | intron_variant,non_coding_transcript_variant | MODIFIER |
| 8 | 50237139 | AX-170252713 | 0 | A | G | 0.149 | -1.56E-02 | 1.52E-01 | 2.67E+00 | 2.71E+00 | 9.25E-01 | no | Gm45455 | intron_variant,non_coding_transcript_variant | MODIFIER |
| 8 | 50289339 | AX-97845377 | 0 | A | G | 0.149 | -1.56E-02 | 1.52E-01 | 2.67E+00 | 2.71E+00 | 9.25E-01 | no | Gm45455 | intron_variant,non_coding_transcript_variant | MODIFIER |
| 8 | 50293662 | AX-170515750 | 0 | A | T | 0.149 | -1.56E-02 | 1.52E-01 | 2.67E+00 | 2.71E+00 | 9.25E-01 | no | Gm45455 | intron_variant,non_coding_transcript_variant | MODIFIER |
| 8 | 50293963 | AX-170282327 | 0 | G | A | 0.149 | -1.56E-02 | 1.52E-01 | 2.67E+00 | 2.71E+00 | 9.25E-01 | no | Gm45455 | intron_variant,non_coding_transcript_variant | MODIFIER |
| 8 | 50304660 | AX-170453507 | 0 | G | A | 0.149 | -1.56E-02 | 1.52E-01 | 2.67E+00 | 2.71E+00 | 9.25E-01 | no | Gm45455 | intron_variant,non_coding_transcript_variant | MODIFIER |
| 8 | 50305534 | AX-170336426 | 0 | G | A | 0.149 | -1.56E-02 | 1.52E-01 | 2.67E+00 | 2.71E+00 | 9.25E-01 | no | Gm45455 | intron_variant,non_coding_transcript_variant | MODIFIER |
| 8 | 50323919 | AX-97842720 | 0 | G | A | 0.149 | -1.56E-02 | 1.52E-01 | 2.67E+00 | 2.71E+00 | 9.25E-01 | no | Gm45455 | intron_variant,non_coding_transcript_variant | MODIFIER |
| 8 | 50331001 | AX-170265627 | 0 | T | C | 0.149 | -1.56E-02 | 1.52E-01 | 2.67E+00 | 2.71E+00 | 9.25E-01 | no | Gm45455 | intron_variant,non_coding_transcript_variant | MODIFIER |
| 8 | 50346738 | AX-170226359 | 0 | G | T | 0.149 | -1.56E-02 | 1.52E-01 | 2.67E+00 | 2.71E+00 | 9.25E-01 | no | Gm45455 | intron_variant,non_coding_transcript_variant | MODIFIER |
| 8 | 50352346 | AX-170213613 | 0 | A | G | 0.149 | -1.56E-02 | 1.52E-01 | 2.67E+00 | 2.71E+00 | 9.25E-01 | no | Gm45455 | intron_variant,non_coding_transcript_variant | MODIFIER |
| 8 | 50358245 | AX-170080047 | 0 | G | C | 0.149 | -1.56E-02 | 1.52E-01 | 2.67E+00 | 2.71E+00 | 9.25E-01 | no | Gm45455 | intron_variant,non_coding_transcript_variant | MODIFIER |
| 8 | 50552060 | AX-168392473 | 0 | A | G | 0.149 | -1.56E-02 | 1.52E-01 | 2.67E+00 | 2.71E+00 | 9.25E-01 | no | | intergenic_variant | MODIFIER |
| 8 | 50647994 | AX-97812931 | 0 | A | G | 0.149 | -1.56E-02 | 1.52E-01 | 2.67E+00 | 2.71E+00 | 9.25E-01 | no | | intergenic_variant | MODIFIER |
| 8 | 50750364 | AX-170101854 | 0 | G | A | 0.149 | -1.56E-02 | 1.52E-01 | 2.67E+00 | 2.71E+00 | 9.25E-01 | no | C130073E24Rik | intron_variant,non_coding_transcript_variant | MODIFIER |
| 8 | 50771460 | AX-170090099 | 0 | A | G | 0.149 | -1.56E-02 | 1.52E-01 | 2.67E+00 | 2.71E+00 | 9.25E-01 | no | C130073E24Rik | intron_variant,non_coding_transcript_variant | MODIFIER |
| 8 | 50830009 | AX-97786661 | 0 | G | A | 0.149 | -1.56E-02 | 1.52E-01 | 2.67E+00 | 2.71E+00 | 9.25E-01 | no | C130073E24Rik | intron_variant,non_coding_transcript_variant | MODIFIER |
| 8 | 50837091 | AX-170498883 | 0 | T | C | 0.149 | -1.56E-02 | 1.52E-01 | 2.67E+00 | 2.71E+00 | 9.25E-01 | no | C130073E24Rik | intron_variant,non_coding_transcript_variant | MODIFIER |
| 8 | 50844393 | AX-97803647 | 0 | G | A | 0.149 | -1.56E-02 | 1.52E-01 | 2.67E+00 | 2.71E+00 | 9.25E-01 | no | C130073E24Rik | intron_variant,non_coding_transcript_variant | MODIFIER |
| 8 | 50874407 | AX-97795350 | 0 | G | T | 0.149 | -1.56E-02 | 1.52E-01 | 2.67E+00 | 2.71E+00 | 9.25E-01 | no | C130073E24Rik | intron_variant,non_coding_transcript_variant | MODIFIER |
| 8 | 50875562 | AX-170244696 | 0 | C | A | 0.149 | -1.56E-02 | 1.52E-01 | 2.67E+00 | 2.71E+00 | 9.25E-01 | no | C130073E24Rik | intron_variant,non_coding_transcript_variant | MODIFIER |
| 8 | 51050187 | AX-170090100 | 0 | A | G | 0.149 | -1.56E-02 | 1.52E-01 | 2.67E+00 | 2.71E+00 | 9.25E-01 | no | | intergenic_variant | MODIFIER |
| 8 | 51363297 | AX-97810561 | 0 | T | C | 0.149 | -1.56E-02 | 1.52E-01 | 2.67E+00 | 2.71E+00 | 9.25E-01 | no | | intergenic_variant | MODIFIER |
| 8 | 51480844 | AX-170209943 | 0 | G | T | 0.149 | -1.56E-02 | 1.52E-01 | 2.67E+00 | 2.71E+00 | 9.25E-01 | no | | intergenic_variant | MODIFIER |
| 8 | 51552810 | AX-97794429 | 0 | T | G | 0.149 | -1.56E-02 | 1.52E-01 | 2.67E+00 | 2.71E+00 | 9.25E-01 | no | | intergenic_variant | MODIFIER |
| 8 | 51555604 | AX-170317367 | 0 | T | C | 0.149 | -1.56E-02 | 1.52E-01 | 2.67E+00 | 2.71E+00 | 9.25E-01 | no | | intergenic_variant | MODIFIER |
| 8 | 51595079 | AX-97810083 | 0 | T | C | 0.149 | -1.56E-02 | 1.52E-01 | 2.67E+00 | 2.71E+00 | 9.25E-01 | no | | intergenic_variant | MODIFIER |
| 8 | 51626159 | AX-170096266 | 0 | T | C | 0.149 | -1.56E-02 | 1.52E-01 | 2.67E+00 | 2.71E+00 | 9.25E-01 | no | | intergenic_variant | MODIFIER |
| 8 | 51630014 | AX-170030694 | 0 | T | A | 0.149 | -1.56E-02 | 1.52E-01 | 2.67E+00 | 2.71E+00 | 9.25E-01 | no | | intergenic_variant | MODIFIER |
| 8 | 51640832 | AX-97804214 | 0 | G | A | 0.149 | -1.56E-02 | 1.52E-01 | 2.67E+00 | 2.71E+00 | 9.25E-01 | no | | intergenic_variant | MODIFIER |
| 8 | 51704591 | AX-170192472 | 0 | T | G | 0.149 | -1.56E-02 | 1.52E-01 | 2.67E+00 | 2.71E+00 | 9.25E-01 | no | | intergenic_variant | MODIFIER |
| 8 | 51704727 | AX-170335814 | 0 | T | G | 0.149 | -1.56E-02 | 1.52E-01 | 2.67E+00 | 2.71E+00 | 9.25E-01 | no | | intergenic_variant | MODIFIER |
| 8 | 51711271 | AX-170030695 | 0 | G | T | 0.149 | -1.56E-02 | 1.52E-01 | 2.67E+00 | 2.71E+00 | 9.25E-01 | no | | intergenic_variant | MODIFIER |
| 8 | 51718984 | AX-97844885 | 0 | A | G | 0.149 | -1.56E-02 | 1.52E-01 | 2.67E+00 | 2.71E+00 | 9.25E-01 | no | | intergenic_variant | MODIFIER |
| 8 | 51738632 | AX-170461805 | 0 | T | C | 0.149 | -1.56E-02 | 1.52E-01 | 2.67E+00 | 2.71E+00 | 9.25E-01 | no | | intergenic_variant | MODIFIER |
| 8 | 51742665 | AX-170030693 | 0 | C | A | 0.149 | -1.56E-02 | 1.52E-01 | 2.67E+00 | 2.71E+00 | 9.25E-01 | no | | intergenic_variant | MODIFIER |
| 8 | 51785532 | AX-97852505 | 0 | A | G | 0.149 | -1.56E-02 | 1.52E-01 | 2.67E+00 | 2.71E+00 | 9.25E-01 | no | | intergenic_variant | MODIFIER |
| 8 | 51794402 | AX-170391335 | 0 | C | T | 0.149 | -1.56E-02 | 1.52E-01 | 2.67E+00 | 2.71E+00 | 9.25E-01 | no | | intergenic_variant | MODIFIER |
| 8 | 51794671 | AX-170172492 | 0 | A | G | 0.149 | -1.56E-02 | 1.52E-01 | 2.67E+00 | 2.71E+00 | 9.25E-01 | no | | intergenic_variant | MODIFIER |
| 8 | 51854824 | AX-170030692 | 0 | G | A | 0.149 | -1.56E-02 | 1.52E-01 | 2.67E+00 | 2.71E+00 | 9.25E-01 | no | | intergenic_variant | MODIFIER |
| 8 | 51883626 | AX-170030699 | 0 | A | G | 0.149 | -1.56E-02 | 1.52E-01 | 2.67E+00 | 2.71E+00 | 9.25E-01 | no | Gm6458 | downstream_gene_variant | MODIFIER |
| 8 | 51891687 | AX-170030696 | 0 | A | G | 0.149 | -1.56E-02 | 1.52E-01 | 2.67E+00 | 2.71E+00 | 9.25E-01 | no | | intergenic_variant | MODIFIER |
| 8 | 51891718 | AX-170030690 | 0 | A | G | 0.149 | -1.56E-02 | 1.52E-01 | 2.67E+00 | 2.71E+00 | 9.25E-01 | no | | intergenic_variant | MODIFIER |
| 8 | 51906840 | AX-97836769 | 0 | A | G | 0.149 | -1.56E-02 | 1.52E-01 | 2.67E+00 | 2.71E+00 | 9.25E-01 | no | | regulatory_region_variant | MODIFIER |
| 8 | 51968587 | AX-170526190 | 0 | A | T | 0.149 | -1.56E-02 | 1.52E-01 | 2.67E+00 | 2.71E+00 | 9.25E-01 | no | | intergenic_variant | MODIFIER |
| 8 | 52027930 | AX-97805126 | 0 | T | C | 0.149 | -1.56E-02 | 1.52E-01 | 2.67E+00 | 2.71E+00 | 9.25E-01 | no | Gm45389 | intron_variant,non_coding_transcript_variant | MODIFIER |
| 8 | 52078465 | AX-97786502 | 0 | C | A | 0.149 | -1.56E-02 | 1.52E-01 | 2.67E+00 | 2.71E+00 | 9.25E-01 | no | Gm24929 | upstream_gene_variant | MODIFIER |
| 8 | 52100415 | AX-170495082 | 0 | G | C | 0.149 | -1.56E-02 | 1.52E-01 | 2.67E+00 | 2.71E+00 | 9.25E-01 | no | Gm45388 | downstream_gene_variant | MODIFIER |
| 8 | 52111363 | AX-170438902 | 0 | G | T | 0.149 | -1.56E-02 | 1.52E-01 | 2.67E+00 | 2.71E+00 | 9.25E-01 | no | | intergenic_variant | MODIFIER |
| 8 | 52149068 | AX-97793996 | 0 | C | T | 0.149 | -1.56E-02 | 1.52E-01 | 2.67E+00 | 2.71E+00 | 9.25E-01 | no | | intergenic_variant | MODIFIER |
| 8 | 52149265 | AX-170119015 | 0 | A | T | 0.149 | -1.56E-02 | 1.52E-01 | 2.67E+00 | 2.71E+00 | 9.25E-01 | no | | intergenic_variant | MODIFIER |

| | | | | | | | | | | | | | | | |
|---|---|---|---|---|---|---|---|---|---|---|---|---|---|---|---|
| 8 | 52170623 | AX-97815403 | 0 | G | A | 0.149 | -1.56E-02 | 1.52E-01 | 2.67E+00 | 2.71E+00 | 9.25E-01 | no | | intergenic_variant | MODIFIER |
| 8 | 52192907 | AX-170268504 | 0 | A | G | 0.149 | -1.56E-02 | 1.52E-01 | 2.67E+00 | 2.71E+00 | 9.25E-01 | no | Gm9892 | downstream_gene_variant | MODIFIER |
| 8 | 52192951 | AX-97791614 | 0 | T | C | 0.149 | -1.56E-02 | 1.52E-01 | 2.67E+00 | 2.71E+00 | 9.25E-01 | no | Gm9892 | downstream_gene_variant | MODIFIER |
| 8 | 52199640 | AX-170030708 | 0 | A | C | 0.149 | -1.56E-02 | 1.52E-01 | 2.67E+00 | 2.71E+00 | 9.25E-01 | no | Gm9892 | upstream_gene_variant | MODIFIER |
| 8 | 52216454 | AX-97852172 | 0 | G | A | 0.149 | -1.56E-02 | 1.52E-01 | 2.67E+00 | 2.71E+00 | 9.25E-01 | no | | intergenic_variant | MODIFIER |
| 8 | 52315687 | AX-170030703 | 0 | T | C | 0.149 | -1.56E-02 | 1.52E-01 | 2.67E+00 | 2.71E+00 | 9.25E-01 | no | | intergenic_variant | MODIFIER |
| 8 | 52341658 | AX-170384973 | 0 | G | T | 0.149 | -1.56E-02 | 1.52E-01 | 2.67E+00 | 2.71E+00 | 9.25E-01 | no | Gm25813 | upstream_gene_variant | MODIFIER |
| 8 | 52344500 | AX-170030701 | 0 | C | T | 0.149 | -1.56E-02 | 1.52E-01 | 2.67E+00 | 2.71E+00 | 9.25E-01 | no | | intergenic_variant | MODIFIER |
| 8 | 52408531 | AX-97809409 | 0 | T | C | 0.149 | -1.56E-02 | 1.52E-01 | 2.67E+00 | 2.71E+00 | 9.25E-01 | no | | intergenic_variant | MODIFIER |
| 8 | 52414512 | AX-97807489 | 0 | C | G | 0.149 | -1.56E-02 | 1.52E-01 | 2.67E+00 | 2.71E+00 | 9.25E-01 | no | | intergenic_variant | MODIFIER |
| 8 | 52479964 | AX-170030711 | 0 | G | A | 0.149 | -1.56E-02 | 1.52E-01 | 2.67E+00 | 2.71E+00 | 9.25E-01 | no | | intergenic_variant | MODIFIER |
| 8 | 52525569 | AX-97803679 | 0 | A | G | 0.149 | -1.56E-02 | 1.52E-01 | 2.67E+00 | 2.71E+00 | 9.25E-01 | no | | intergenic_variant | MODIFIER |
| 8 | 52525584 | AX-170347224 | 0 | T | C | 0.149 | -1.56E-02 | 1.52E-01 | 2.67E+00 | 2.71E+00 | 9.25E-01 | no | | intergenic_variant | MODIFIER |
| 8 | 52678732 | AX-97856545 | 0 | A | G | 0.149 | -1.56E-02 | 1.52E-01 | 2.67E+00 | 2.71E+00 | 9.25E-01 | no | | intergenic_variant | MODIFIER |
| 8 | 52690270 | AX-97800438 | 0 | A | C | 0.149 | -1.56E-02 | 1.52E-01 | 2.67E+00 | 2.71E+00 | 9.25E-01 | no | | intergenic_variant | MODIFIER |
| 8 | 52764380 | AX-170332188 | 0 | C | T | 0.149 | -1.56E-02 | 1.52E-01 | 2.67E+00 | 2.71E+00 | 9.25E-01 | no | | intergenic_variant | MODIFIER |
| 8 | 52777659 | AX-97803680 | 0 | T | A | 0.149 | -1.56E-02 | 1.52E-01 | 2.67E+00 | 2.71E+00 | 9.25E-01 | no | Gm9685 | downstream_gene_variant | MODIFIER |
| 8 | 52809932 | AX-170030719 | 0 | G | C | 0.149 | -1.56E-02 | 1.52E-01 | 2.67E+00 | 2.71E+00 | 9.25E-01 | no | | intergenic_variant | MODIFIER |
| 8 | 52810028 | AX-170030716 | 0 | G | A | 0.149 | -1.56E-02 | 1.52E-01 | 2.67E+00 | 2.71E+00 | 9.25E-01 | no | | intergenic_variant | MODIFIER |
| 8 | 52810079 | AX-170030718 | 0 | C | T | 0.149 | -1.56E-02 | 1.52E-01 | 2.67E+00 | 2.71E+00 | 9.25E-01 | no | | intergenic_variant | MODIFIER |
| 8 | 52844437 | AX-170482590 | 0 | C | A | 0.149 | -1.56E-02 | 1.52E-01 | 2.67E+00 | 2.71E+00 | 9.25E-01 | no | | intergenic_variant | MODIFIER |
| 8 | 52856303 | AX-170411366 | 0 | T | C | 0.149 | -1.56E-02 | 1.52E-01 | 2.67E+00 | 2.71E+00 | 9.25E-01 | no | | intergenic_variant | MODIFIER |
| 8 | 52901926 | AX-97800670 | 0 | C | A | 0.149 | -1.56E-02 | 1.52E-01 | 2.67E+00 | 2.71E+00 | 9.25E-01 | no | | intergenic_variant | MODIFIER |
| 8 | 53013264 | AX-170188249 | 0 | C | A | 0.149 | -1.56E-02 | 1.52E-01 | 2.67E+00 | 2.71E+00 | 9.25E-01 | no | | intergenic_variant | MODIFIER |
| 8 | 53127883 | AX-170080081 | 0 | T | A | 0.149 | -1.56E-02 | 1.52E-01 | 2.67E+00 | 2.71E+00 | 9.25E-01 | no | | intergenic_variant | MODIFIER |
| 8 | 53139576 | AX-170080080 | 0 | A | G | 0.149 | -1.56E-02 | 1.52E-01 | 2.67E+00 | 2.71E+00 | 9.25E-01 | no | | intergenic_variant | MODIFIER |
| 8 | 53150575 | AX-97837853 | 0 | C | T | 0.149 | -1.56E-02 | 1.52E-01 | 2.67E+00 | 2.71E+00 | 9.25E-01 | no | | intergenic_variant | MODIFIER |
| 8 | 53176464 | AX-170080084 | 0 | A | G | 0.149 | -1.56E-02 | 1.52E-01 | 2.67E+00 | 2.71E+00 | 9.25E-01 | no | | intergenic_variant | MODIFIER |
| 8 | 53273879 | AX-170030723 | 0 | T | A | 0.149 | -1.56E-02 | 1.52E-01 | 2.67E+00 | 2.71E+00 | 9.25E-01 | no | | intergenic_variant | MODIFIER |
| 8 | 53337363 | AX-170030724 | 0 | T | G | 0.149 | -1.56E-02 | 1.52E-01 | 2.67E+00 | 2.71E+00 | 9.25E-01 | no | | intergenic_variant | MODIFIER |
| 8 | 53349639 | AX-170030727 | 0 | T | C | 0.149 | -1.56E-02 | 1.52E-01 | 2.67E+00 | 2.71E+00 | 9.25E-01 | no | | intergenic_variant | MODIFIER |
| 8 | 53407568 | AX-170132512 | 0 | G | C | 0.149 | -1.56E-02 | 1.52E-01 | 2.67E+00 | 2.71E+00 | 9.25E-01 | no | | intergenic_variant | MODIFIER |
| 8 | 53421289 | AX-170374205 | 0 | C | T | 0.149 | -1.56E-02 | 1.52E-01 | 2.67E+00 | 2.71E+00 | 9.25E-01 | no | | intergenic_variant | MODIFIER |
| 8 | 53425714 | AX-97790380 | 0 | A | G | 0.149 | -1.56E-02 | 1.52E-01 | 2.67E+00 | 2.71E+00 | 9.25E-01 | no | | intergenic_variant | MODIFIER |
| 8 | 53446789 | AX-97801939 | 0 | G | T | 0.149 | -1.56E-02 | 1.52E-01 | 2.67E+00 | 2.71E+00 | 9.25E-01 | no | Gm45554 | intron_variant,non_coding_transcript_variant | MODIFIER |
| 8 | 53485572 | AX-170080090 | 0 | A | G | 0.149 | -1.56E-02 | 1.52E-01 | 2.67E+00 | 2.71E+00 | 9.25E-01 | no | Gm45554 | downstream_gene_variant | MODIFIER |
| 8 | 53486119 | AX-97846699 | 0 | A | G | 0.149 | -1.56E-02 | 1.52E-01 | 2.67E+00 | 2.71E+00 | 9.25E-01 | no | | intergenic_variant | MODIFIER |
| 8 | 53510741 | AX-170432647 | 0 | G | A | 0.149 | -1.56E-02 | 1.52E-01 | 2.67E+00 | 2.71E+00 | 9.25E-01 | no | Aga | upstream_gene_variant | MODIFIER |
| 8 | 53511702 | AX-170090114 | 0 | G | A | 0.149 | -1.56E-02 | 1.52E-01 | 2.67E+00 | 2.71E+00 | 9.25E-01 | no | Aga | upstream_gene_variant | MODIFIER |
| 8 | 53511726 | AX-170123731 | 0 | G | C | 0.149 | -1.56E-02 | 1.52E-01 | 2.67E+00 | 2.71E+00 | 9.25E-01 | no | Aga | upstream_gene_variant | MODIFIER |
| 8 | 53572846 | AX-170265086 | 0 | A | G | 0.149 | -1.56E-02 | 1.52E-01 | 2.67E+00 | 2.71E+00 | 9.25E-01 | no | | intergenic_variant | MODIFIER |
| 8 | 53610286 | AX-170429614 | 0 | A | G | 0.149 | -1.56E-02 | 1.52E-01 | 2.67E+00 | 2.71E+00 | 9.25E-01 | no | Neil3 | intron_variant | MODIFIER |
| 8 | 53773250 | AX-170165934 | 0 | G | A | 0.149 | -1.56E-02 | 1.52E-01 | 2.67E+00 | 2.71E+00 | 9.25E-01 | no | | intergenic_variant | MODIFIER |
| 8 | 53821365 | AX-97794593 | 0 | T | C | 0.149 | -1.56E-02 | 1.52E-01 | 2.67E+00 | 2.71E+00 | 9.25E-01 | no | | intergenic_variant | MODIFIER |
| 8 | 53923224 | AX-97852786 | 0 | A | G | 0.149 | -1.56E-02 | 1.52E-01 | 2.67E+00 | 2.71E+00 | 9.25E-01 | no | | intergenic_variant | MODIFIER |
| 8 | 54016496 | AX-170458059 | 0 | G | C | 0.149 | -1.56E-02 | 1.52E-01 | 2.67E+00 | 2.71E+00 | 9.25E-01 | no | | intergenic_variant | MODIFIER |
| 8 | 54054262 | AX-170198365 | 0 | C | G | 0.149 | -1.56E-02 | 1.52E-01 | 2.67E+00 | 2.71E+00 | 9.25E-01 | no | | intergenic_variant | MODIFIER |
| 8 | 54085930 | AX-170294984 | 0 | G | C | 0.149 | -1.56E-02 | 1.52E-01 | 2.67E+00 | 2.71E+00 | 9.25E-01 | no | Vegfc | intron_variant | MODIFIER |
| 8 | 54133863 | AX-170122795 | 0 | C | A | 0.149 | -1.56E-02 | 1.52E-01 | 2.67E+00 | 2.71E+00 | 9.25E-01 | no | Vegfc | intron_variant | MODIFIER |
| 8 | 54134227 | AX-170420291 | 0 | T | C | 0.149 | -1.56E-02 | 1.52E-01 | 2.67E+00 | 2.71E+00 | 9.25E-01 | no | Vegfc | intron_variant | MODIFIER |
| 8 | 54137919 | AX-170352314 | 0 | G | A | 0.149 | -1.56E-02 | 1.52E-01 | 2.67E+00 | 2.71E+00 | 9.25E-01 | no | Vegfc | intron_variant | MODIFIER |
| 8 | 54137993 | AX-170080102 | 0 | C | G | 0.149 | -1.56E-02 | 1.52E-01 | 2.67E+00 | 2.71E+00 | 9.25E-01 | no | Vegfc | intron_variant | MODIFIER |
| 8 | 54141762 | AX-97824346 | 0 | T | C | 0.149 | -1.56E-02 | 1.52E-01 | 2.67E+00 | 2.71E+00 | 9.25E-01 | no | Vegfc | intron_variant | MODIFIER |
| 8 | 54153765 | AX-170151002 | 0 | T | C | 0.149 | -1.56E-02 | 1.52E-01 | 2.67E+00 | 2.71E+00 | 9.25E-01 | no | Vegfc | intron_variant | MODIFIER |

| | | | | | | | | | | | | | | | |
|---|---|---|---|---|---|---|---|---|---|---|---|---|---|---|---|
| 8 | 54199626 | AX-170155443 | 0 | G | A | 0.149 | -1.56E-02 | 1.52E-01 | 2.67E+00 | 2.71E+00 | 9.25E-01 | no | | intergenic_variant | MODIFIER |
| 8 | 54199849 | AX-170233054 | 0 | A | G | 0.149 | -1.56E-02 | 1.52E-01 | 2.67E+00 | 2.71E+00 | 9.25E-01 | no | | intergenic_variant | MODIFIER |
| 8 | 54218096 | AX-170411230 | 0 | T | C | 0.149 | -1.56E-02 | 1.52E-01 | 2.67E+00 | 2.71E+00 | 9.25E-01 | no | | intergenic_variant | MODIFIER |
| 8 | 54223370 | AX-97840104 | 0 | C | T | 0.149 | -1.56E-02 | 1.52E-01 | 2.67E+00 | 2.71E+00 | 9.25E-01 | no | | intergenic_variant | MODIFIER |
| 8 | 54288215 | AX-170446336 | 0 | T | A | 0.149 | -1.56E-02 | 1.52E-01 | 2.67E+00 | 2.71E+00 | 9.25E-01 | no | | intergenic_variant | MODIFIER |
| 8 | 54288467 | AX-170391696 | 0 | T | C | 0.149 | -1.56E-02 | 1.52E-01 | 2.67E+00 | 2.71E+00 | 9.25E-01 | no | | intergenic_variant | MODIFIER |
| 8 | 54292168 | AX-170096274 | 0 | C | A | 0.149 | -1.56E-02 | 1.52E-01 | 2.67E+00 | 2.71E+00 | 9.25E-01 | no | | intergenic_variant | MODIFIER |
| 8 | 54331012 | AX-170112743 | 0 | G | A | 0.149 | -1.56E-02 | 1.52E-01 | 2.67E+00 | 2.71E+00 | 9.25E-01 | no | | intergenic_variant | MODIFIER |
| 8 | 54388789 | AX-97837369 | 0 | G | T | 0.149 | -1.56E-02 | 1.52E-01 | 2.67E+00 | 2.71E+00 | 9.25E-01 | no | | intergenic_variant | MODIFIER |
| 8 | 54389084 | AX-170030742 | 0 | A | G | 0.149 | -1.56E-02 | 1.52E-01 | 2.67E+00 | 2.71E+00 | 9.25E-01 | no | | intergenic_variant | MODIFIER |
| 8 | 54389144 | AX-170080104 | 0 | A | G | 0.149 | -1.56E-02 | 1.52E-01 | 2.67E+00 | 2.71E+00 | 9.25E-01 | no | | intergenic_variant | MODIFIER |
| 8 | 54406530 | AX-170030744 | 0 | G | A | 0.149 | -1.56E-02 | 1.52E-01 | 2.67E+00 | 2.71E+00 | 9.25E-01 | no | | intergenic_variant | MODIFIER |
| 8 | 54487450 | AX-97798125 | 0 | T | C | 0.149 | -1.56E-02 | 1.52E-01 | 2.67E+00 | 2.71E+00 | 9.25E-01 | no | | regulatory_region_variant | MODIFIER |
| 8 | 54490523 | AX-170417021 | 0 | T | A | 0.149 | -1.56E-02 | 1.52E-01 | 2.67E+00 | 2.71E+00 | 9.25E-01 | no | | intergenic_variant | MODIFIER |
| 8 | 54517622 | AX-97838593 | 0 | G | A | 0.149 | -1.56E-02 | 1.52E-01 | 2.67E+00 | 2.71E+00 | 9.25E-01 | no | Spcs3 | downstream_gene_variant | MODIFIER |
| 8 | 54521145 | AX-170261077 | 0 | A | G | 0.149 | -1.56E-02 | 1.52E-01 | 2.67E+00 | 2.71E+00 | 9.25E-01 | no | Spcs3 | 3_prime_UTR_variant | MODIFIER |
| 8 | 54521175 | AX-170299496 | 0 | A | T | 0.149 | -1.56E-02 | 1.52E-01 | 2.67E+00 | 2.71E+00 | 9.25E-01 | no | Spcs3 | 3_prime_UTR_variant | MODIFIER |
| 8 | 54548815 | AX-97823336 | 0 | A | G | 0.149 | -1.56E-02 | 1.52E-01 | 2.67E+00 | 2.71E+00 | 9.25E-01 | no | Asb5 | upstream_gene_variant | MODIFIER |
| 8 | 54553784 | AX-170307732 | 0 | G | A | 0.149 | -1.56E-02 | 1.52E-01 | 2.67E+00 | 2.71E+00 | 9.25E-01 | no | Asb5 | intron_variant | MODIFIER |
| 8 | 54562640 | AX-170441468 | 0 | A | C | 0.149 | -1.56E-02 | 1.52E-01 | 2.67E+00 | 2.71E+00 | 9.25E-01 | no | Asb5 | intron_variant | MODIFIER |
| 8 | 54562667 | AX-97802609 | 0 | A | G | 0.149 | -1.56E-02 | 1.52E-01 | 2.67E+00 | 2.71E+00 | 9.25E-01 | no | Asb5 | intron_variant | MODIFIER |
| 8 | 54651395 | AX-97847815 | 0 | G | T | 0.149 | -1.56E-02 | 1.52E-01 | 2.67E+00 | 2.71E+00 | 9.25E-01 | no | Wdr17 | intron_variant | MODIFIER |
| 8 | 54659328 | AX-170158262 | 0 | T | C | 0.149 | -1.56E-02 | 1.52E-01 | 2.67E+00 | 2.71E+00 | 9.25E-01 | no | Wdr17 | intron_variant | MODIFIER |
| 8 | 55021625 | AX-170030754 | 0 | A | G | 0.149 | -1.56E-02 | 1.52E-01 | 2.67E+00 | 2.71E+00 | 9.25E-01 | no | Gpm6a | intron_variant | MODIFIER |
| 8 | 55031630 | AX-170533861 | 0 | T | C | 0.149 | -1.56E-02 | 1.52E-01 | 2.67E+00 | 2.71E+00 | 9.25E-01 | no | Gpm6a | intron_variant | MODIFIER |
| 8 | 55041546 | AX-97796428 | 0 | T | C | 0.149 | -1.56E-02 | 1.52E-01 | 2.67E+00 | 2.71E+00 | 9.25E-01 | no | Gpm6a | intron_variant | MODIFIER |
| 8 | 55041796 | AX-170396830 | 0 | C | T | 0.149 | -1.56E-02 | 1.52E-01 | 2.67E+00 | 2.71E+00 | 9.25E-01 | no | Gpm6a | intron_variant | MODIFIER |
| 8 | 55042804 | AX-170428237 | 0 | T | C | 0.149 | -1.56E-02 | 1.52E-01 | 2.67E+00 | 2.71E+00 | 9.25E-01 | no | Gpm6a | intron_variant | MODIFIER |
| 8 | 55049244 | AX-170080114 | 0 | T | G | 0.149 | -1.56E-02 | 1.52E-01 | 2.67E+00 | 2.71E+00 | 9.25E-01 | no | Gpm6a | intron_variant | MODIFIER |
| 8 | 55052692 | AX-170536003 | 0 | T | C | 0.149 | -1.56E-02 | 1.52E-01 | 2.67E+00 | 2.71E+00 | 9.25E-01 | no | Gpm6a | intron_variant | MODIFIER |
| 8 | 55081643 | AX-170030758 | 0 | A | G | 0.149 | -1.56E-02 | 1.52E-01 | 2.67E+00 | 2.71E+00 | 9.25E-01 | no | 9330121K16Rik | downstream_gene_variant | MODIFIER |
| 8 | 55106591 | AX-170141796 | 0 | A | C | 0.149 | -1.56E-02 | 1.52E-01 | 2.67E+00 | 2.71E+00 | 9.25E-01 | no | | intergenic_variant | MODIFIER |
| 8 | 55430703 | AX-97792718 | 0 | T | C | 0.149 | -1.56E-02 | 1.52E-01 | 2.67E+00 | 2.71E+00 | 9.25E-01 | no | | intergenic_variant | MODIFIER |
| 8 | 55575739 | AX-170143980 | 0 | A | C | 0.149 | -1.56E-02 | 1.52E-01 | 2.67E+00 | 2.71E+00 | 9.25E-01 | no | | intergenic_variant | MODIFIER |
| 8 | 55659538 | AX-97828125 | 0 | A | G | 0.149 | -1.56E-02 | 1.52E-01 | 2.67E+00 | 2.71E+00 | 9.25E-01 | no | Gm18320 | non_coding_transcript_exon_variant | MODIFIER |
| 8 | 55680438 | AX-170536177 | 0 | C | G | 0.149 | -1.56E-02 | 1.52E-01 | 2.67E+00 | 2.71E+00 | 9.25E-01 | no | | intergenic_variant | MODIFIER |
| 8 | 55717498 | AX-97837815 | 0 | C | A | 0.149 | -1.56E-02 | 1.52E-01 | 2.67E+00 | 2.71E+00 | 9.25E-01 | no | | intergenic_variant | MODIFIER |
| 8 | 55723000 | AX-170132766 | 0 | G | C | 0.149 | -1.56E-02 | 1.52E-01 | 2.67E+00 | 2.71E+00 | 9.25E-01 | no | | intergenic_variant | MODIFIER |
| 8 | 55805796 | AX-170295572 | 0 | A | G | 0.149 | -1.56E-02 | 1.52E-01 | 2.67E+00 | 2.71E+00 | 9.25E-01 | no | | intergenic_variant | MODIFIER |
| 8 | 55813314 | AX-97785013 | 0 | T | C | 0.149 | -1.56E-02 | 1.52E-01 | 2.67E+00 | 2.71E+00 | 9.25E-01 | no | | intergenic_variant | MODIFIER |
| 8 | 56073941 | AX-97781751 | 0 | A | G | 0.149 | -1.56E-02 | 1.52E-01 | 2.67E+00 | 2.71E+00 | 9.25E-01 | no | Glra3 | intron_variant | MODIFIER |
| 8 | 56094368 | AX-170218236 | 0 | A | G | 0.149 | -1.56E-02 | 1.52E-01 | 2.67E+00 | 2.71E+00 | 9.25E-01 | no | Glra3 | intron_variant | MODIFIER |
| 8 | 56097380 | AX-170198703 | 0 | A | G | 0.149 | -1.56E-02 | 1.52E-01 | 2.67E+00 | 2.71E+00 | 9.25E-01 | no | Glra3 | intron_variant | MODIFIER |
| 8 | 56296737 | AX-97821691 | 0 | C | A | 0.149 | -1.56E-02 | 1.52E-01 | 2.67E+00 | 2.71E+00 | 9.25E-01 | no | Hpgd | intron_variant | MODIFIER |
| 8 | 56307197 | AX-170488755 | 0 | G | A | 0.149 | -1.56E-02 | 1.52E-01 | 2.67E+00 | 2.71E+00 | 9.25E-01 | no | Hpgd | intron_variant | MODIFIER |
| 8 | 56312263 | AX-170208463 | 0 | G | T | 0.149 | -1.56E-02 | 1.52E-01 | 2.67E+00 | 2.71E+00 | 9.25E-01 | no | Hpgd | intron_variant | MODIFIER |
| 8 | 56324425 | AX-170323615 | 0 | A | G | 0.149 | -1.56E-02 | 1.52E-01 | 2.67E+00 | 2.71E+00 | 9.25E-01 | no | Hpgd | downstream_gene_variant | MODIFIER |
| 8 | 56364070 | AX-170030767 | 0 | C | T | 0.149 | -1.56E-02 | 1.52E-01 | 2.67E+00 | 2.71E+00 | 9.25E-01 | no | | intergenic_variant | MODIFIER |
| 8 | 56378672 | AX-170030771 | 0 | G | C | 0.149 | -1.56E-02 | 1.52E-01 | 2.67E+00 | 2.71E+00 | 9.25E-01 | no | | intergenic_variant | MODIFIER |
| 8 | 56408003 | AX-170106650 | 0 | G | C | 0.149 | -1.56E-02 | 1.52E-01 | 2.67E+00 | 2.71E+00 | 9.25E-01 | no | | regulatory_region_variant | MODIFIER |
| 8 | 56475448 | AX-170030775 | 0 | A | G | 0.149 | -1.56E-02 | 1.52E-01 | 2.67E+00 | 2.71E+00 | 9.25E-01 | no | 4930518J21Rik | intron_variant,non_coding_transcript_variant | MODIFIER |
| 8 | 56477886 | AX-170280898 | 0 | G | A | 0.149 | -1.56E-02 | 1.52E-01 | 2.67E+00 | 2.71E+00 | 9.25E-01 | no | 4930518J21Rik | intron_variant,non_coding_transcript_variant | MODIFIER |
| 8 | 56477989 | AX-170356037 | 0 | T | C | 0.149 | -1.56E-02 | 1.52E-01 | 2.67E+00 | 2.71E+00 | 9.25E-01 | no | 4930518J21Rik | intron_variant,non_coding_transcript_variant | MODIFIER |
| 8 | 56486097 | AX-170030772 | 0 | G | T | 0.149 | -1.56E-02 | 1.52E-01 | 2.67E+00 | 2.71E+00 | 9.25E-01 | no | 4930518J21Rik | upstream_gene_variant | MODIFIER |

| | | | | | | | | | | | | | | | |
|---|---|---|---|---|---|---|---|---|---|---|---|---|---|---|---|
| 8 | 56517606 | AX-97801346 | 0 | T | C | 0.149 | -1.56E-02 | 1.52E-01 | 2.67E+00 | 2.71E+00 | 9.25E-01 | no | Gm24375 | downstream_gene_variant | MODIFIER |
| 8 | 56528444 | AX-170030774 | 0 | G | A | 0.149 | -1.56E-02 | 1.52E-01 | 2.67E+00 | 2.71E+00 | 9.25E-01 | no | Cep44 | downstream_gene_variant | MODIFIER |
| 8 | 56531682 | AX-170186346 | 0 | T | C | 0.149 | -1.56E-02 | 1.52E-01 | 2.67E+00 | 2.71E+00 | 9.25E-01 | no | Cep44 | 3_prime_UTR_variant | MODIFIER |
| 8 | 56534851 | AX-170312286 | 0 | A | G | 0.149 | -1.56E-02 | 1.52E-01 | 2.67E+00 | 2.71E+00 | 9.25E-01 | no | Cep44 | intron_variant | MODIFIER |
| 8 | 56550911 | AX-97857005 | 0 | C | T | 0.149 | -1.56E-02 | 1.52E-01 | 2.67E+00 | 2.71E+00 | 9.25E-01 | no | Fbxo8 | upstream_gene_variant | MODIFIER |
| 8 | 56550941 | AX-170030769 | 0 | T | G | 0.149 | -1.56E-02 | 1.52E-01 | 2.67E+00 | 2.71E+00 | 9.25E-01 | no | Fbxo8 | upstream_gene_variant | MODIFIER |
| 8 | 56575906 | AX-170030770 | 0 | A | T | 0.149 | -1.56E-02 | 1.52E-01 | 2.67E+00 | 2.71E+00 | 9.25E-01 | no | Fbxo8 | intron_variant | MODIFIER |
| 8 | 56587867 | AX-170096278 | 0 | G | A | 0.149 | -1.56E-02 | 1.52E-01 | 2.67E+00 | 2.71E+00 | 9.25E-01 | no | Fbxo8 | intron_variant | MODIFIER |
| 8 | 56600657 | AX-170030773 | 0 | T | C | 0.149 | -1.56E-02 | 1.52E-01 | 2.67E+00 | 2.71E+00 | 9.25E-01 | no | | intergenic_variant | MODIFIER |
| 8 | 56613267 | AX-97838341 | 0 | A | C | 0.149 | -1.56E-02 | 1.52E-01 | 2.67E+00 | 2.71E+00 | 9.25E-01 | no | | intergenic_variant | MODIFIER |
| 8 | 56639773 | AX-170251622 | 0 | C | T | 0.149 | -1.56E-02 | 1.52E-01 | 2.67E+00 | 2.71E+00 | 9.25E-01 | no | | intergenic_variant | MODIFIER |
| 8 | 56675058 | AX-170219543 | 0 | T | C | 0.149 | -1.56E-02 | 1.52E-01 | 2.67E+00 | 2.71E+00 | 9.25E-01 | no | | intergenic_variant | MODIFIER |
| 8 | 56694146 | AX-170244922 | 0 | T | C | 0.149 | -1.56E-02 | 1.52E-01 | 2.67E+00 | 2.71E+00 | 9.25E-01 | no | | intergenic_variant | MODIFIER |
| 8 | 56695801 | AX-97806292 | 0 | A | G | 0.149 | -1.56E-02 | 1.52E-01 | 2.67E+00 | 2.71E+00 | 9.25E-01 | no | | intergenic_variant | MODIFIER |
| 8 | 56724515 | AX-170215003 | 0 | A | G | 0.149 | -1.56E-02 | 1.52E-01 | 2.67E+00 | 2.71E+00 | 9.25E-01 | no | | intergenic_variant | MODIFIER |
| 8 | 56828942 | AX-170466911 | 0 | C | T | 0.149 | -1.56E-02 | 1.52E-01 | 2.67E+00 | 2.71E+00 | 9.25E-01 | no | | intergenic_variant | MODIFIER |
| 8 | 56878305 | AX-97782184 | 0 | T | C | 0.149 | -1.56E-02 | 1.52E-01 | 2.67E+00 | 2.71E+00 | 9.25E-01 | no | Gm33501 | intron_variant,non_coding_transcript_variant | MODIFIER |
| 8 | 56905124 | AX-170030781 | 0 | G | A | 0.149 | -1.56E-02 | 1.52E-01 | 2.67E+00 | 2.71E+00 | 9.25E-01 | no | | intergenic_variant | MODIFIER |
| 8 | 56919304 | AX-97827337 | 0 | C | A | 0.149 | -1.56E-02 | 1.52E-01 | 2.67E+00 | 2.71E+00 | 9.25E-01 | no | | intergenic_variant | MODIFIER |
| 8 | 57163003 | AX-97856284 | 0 | G | A | 0.149 | -1.56E-02 | 1.52E-01 | 2.67E+00 | 2.71E+00 | 9.25E-01 | no | | intergenic_variant | MODIFIER |
| 8 | 57163519 | AX-170222162 | 0 | T | C | 0.149 | -1.56E-02 | 1.52E-01 | 2.67E+00 | 2.71E+00 | 9.25E-01 | no | | intergenic_variant | MODIFIER |
| 8 | 57163919 | AX-170403223 | 0 | C | T | 0.149 | -1.56E-02 | 1.52E-01 | 2.67E+00 | 2.71E+00 | 9.25E-01 | no | | intergenic_variant | MODIFIER |
| 8 | 57180360 | AX-170030784 | 0 | G | C | 0.149 | -1.56E-02 | 1.52E-01 | 2.67E+00 | 2.71E+00 | 9.25E-01 | no | | intergenic_variant | MODIFIER |
| 8 | 57181110 | AX-170264031 | 0 | G | A | 0.149 | -1.56E-02 | 1.52E-01 | 2.67E+00 | 2.71E+00 | 9.25E-01 | no | | intergenic_variant | MODIFIER |
| 8 | 57192822 | AX-170030790 | 0 | A | C | 0.149 | -1.56E-02 | 1.52E-01 | 2.67E+00 | 2.71E+00 | 9.25E-01 | no | | regulatory_region_variant | MODIFIER |
| 8 | 57192886 | AX-170030785 | 0 | C | T | 0.149 | -1.56E-02 | 1.52E-01 | 2.67E+00 | 2.71E+00 | 9.25E-01 | no | | regulatory_region_variant | MODIFIER |
| 8 | 57280470 | AX-97797339 | 0 | A | C | 0.149 | -1.56E-02 | 1.52E-01 | 2.67E+00 | 2.71E+00 | 9.25E-01 | no | Hand2os1 | downstream_gene_variant | MODIFIER |
| 8 | 57292917 | AX-170359743 | 0 | A | G | 0.149 | -1.56E-02 | 1.52E-01 | 2.67E+00 | 2.71E+00 | 9.25E-01 | no | Gm17994 | upstream_gene_variant | MODIFIER |
| 8 | 57312292 | AX-170129997 | 0 | A | G | 0.149 | -1.56E-02 | 1.52E-01 | 2.67E+00 | 2.71E+00 | 9.25E-01 | no | Hand2os1 | downstream_gene_variant | MODIFIER |
| 8 | 57326897 | AX-170132525 | 0 | C | T | 0.149 | -1.56E-02 | 1.52E-01 | 2.67E+00 | 2.71E+00 | 9.25E-01 | no | Hand2 | downstream_gene_variant | MODIFIER |
| 8 | 57348102 | AX-170330486 | 0 | T | C | 0.149 | -1.56E-02 | 1.52E-01 | 2.67E+00 | 2.71E+00 | 9.25E-01 | no | 5033428I22Rik | intron_variant,non_coding_transcript_variant | MODIFIER |
| 8 | 57353347 | AX-170524388 | 0 | C | G | 0.149 | -1.56E-02 | 1.52E-01 | 2.67E+00 | 2.71E+00 | 9.25E-01 | no | 5033428I22Rik | intron_variant,non_coding_transcript_variant | MODIFIER |
| 8 | 57361205 | AX-170177689 | 0 | G | A | 0.149 | -1.56E-02 | 1.52E-01 | 2.67E+00 | 2.71E+00 | 9.25E-01 | no | | regulatory_region_variant | MODIFIER |
| 8 | 57365883 | AX-170186212 | 0 | T | C | 0.149 | -1.56E-02 | 1.52E-01 | 2.67E+00 | 2.71E+00 | 9.25E-01 | no | | intergenic_variant | MODIFIER |
| 8 | 57366003 | AX-170523364 | 0 | A | G | 0.149 | -1.56E-02 | 1.52E-01 | 2.67E+00 | 2.71E+00 | 9.25E-01 | no | | intergenic_variant | MODIFIER |
| 8 | 57367800 | AX-170209104 | 0 | C | T | 0.149 | -1.56E-02 | 1.52E-01 | 2.67E+00 | 2.71E+00 | 9.25E-01 | no | | intergenic_variant | MODIFIER |
| 8 | 57453526 | AX-170229830 | 0 | A | G | 0.149 | -1.56E-02 | 1.52E-01 | 2.67E+00 | 2.71E+00 | 9.25E-01 | no | Scrg1 | upstream_gene_variant | MODIFIER |
| 8 | 57456874 | AX-97803728 | 0 | A | C | 0.149 | -1.56E-02 | 1.52E-01 | 2.67E+00 | 2.71E+00 | 9.25E-01 | no | Scrg1 | intron_variant | MODIFIER |
| 8 | 57457463 | AX-170105540 | 0 | G | C | 0.149 | -1.56E-02 | 1.52E-01 | 2.67E+00 | 2.71E+00 | 9.25E-01 | no | Scrg1 | intron_variant | MODIFIER |
| 8 | 57458779 | AX-97814964 | 0 | T | C | 0.149 | -1.56E-02 | 1.52E-01 | 2.67E+00 | 2.71E+00 | 9.25E-01 | no | Scrg1 | intron_variant | MODIFIER |
| 8 | 57499932 | AX-170388163 | 0 | T | C | 0.149 | -1.56E-02 | 1.52E-01 | 2.67E+00 | 2.71E+00 | 9.25E-01 | no | 2500002B13Rik | intron_variant,non_coding_transcript_variant | MODIFIER |
| 8 | 57504272 | AX-170030799 | 0 | A | T | 0.149 | -1.56E-02 | 1.52E-01 | 2.67E+00 | 2.71E+00 | 9.25E-01 | no | 2500002B13Rik | intron_variant,non_coding_transcript_variant | MODIFIER |
| 8 | 57506656 | AX-168375394 | 0 | G | A | 0.149 | -1.56E-02 | 1.52E-01 | 2.67E+00 | 2.71E+00 | 9.25E-01 | no | 2500002B13Rik | non_coding_transcript_exon_variant | MODIFIER |
| 8 | 57659526 | AX-170111036 | 0 | C | T | 0.149 | -1.56E-02 | 1.52E-01 | 2.67E+00 | 2.71E+00 | 9.25E-01 | no | Gm45359 | downstream_gene_variant | MODIFIER |
| 8 | 57663129 | AX-170030798 | 0 | A | G | 0.149 | -1.56E-02 | 1.52E-01 | 2.67E+00 | 2.71E+00 | 9.25E-01 | no | Gm45359 | downstream_gene_variant | MODIFIER |
| 8 | 57665155 | AX-170126714 | 0 | T | C | 0.149 | -1.56E-02 | 1.52E-01 | 2.67E+00 | 2.71E+00 | 9.25E-01 | no | AW046200 | downstream_gene_variant | MODIFIER |
| 8 | 57667032 | AX-97839895 | 0 | A | G | 0.149 | -1.56E-02 | 1.52E-01 | 2.67E+00 | 2.71E+00 | 9.25E-01 | no | AW046200 | downstream_gene_variant | MODIFIER |
| 8 | 57688873 | AX-170489229 | 0 | A | G | 0.149 | -1.56E-02 | 1.52E-01 | 2.67E+00 | 2.71E+00 | 9.25E-01 | no | | intergenic_variant | MODIFIER |
| 8 | 57689873 | AX-170277082 | 0 | G | T | 0.149 | -1.56E-02 | 1.52E-01 | 2.67E+00 | 2.71E+00 | 9.25E-01 | no | | intergenic_variant | MODIFIER |
| 8 | 57696462 | AX-97788595 | 0 | G | A | 0.149 | -1.56E-02 | 1.52E-01 | 2.67E+00 | 2.71E+00 | 9.25E-01 | no | Gm45356 | upstream_gene_variant | MODIFIER |
| 8 | 58602922 | AX-170464368 | 0 | T | A | 0.149 | -1.56E-02 | 1.52E-01 | 2.67E+00 | 2.71E+00 | 9.25E-01 | no | Galntl6 | intron_variant,non_coding_transcript_variant | MODIFIER |
| 9 | 81023800 | AX-170166271 | 0 | A | G | 0.211 | 1.14E-02 | 1.29E-01 | 2.71E+00 | 2.76E+00 | 9.25E-01 | no | Gm26713 | intron_variant,non_coding_transcript_variant | MODIFIER |
| 9 | 81061095 | AX-170032801 | 0 | A | G | 0.211 | 1.14E-02 | 1.29E-01 | 2.71E+00 | 2.76E+00 | 9.25E-01 | no | Gm26713 | intron_variant,non_coding_transcript_variant | MODIFIER |
| 9 | 81076442 | AX-170032804 | 0 | T | A | 0.211 | 1.14E-02 | 1.29E-01 | 2.71E+00 | 2.76E+00 | 9.25E-01 | no | Gm26713 | intron_variant,non_coding_transcript_variant | MODIFIER |

| | | | | | | | | | | | | | | | |
|---|---|---|---|---|---|---|---|---|---|---|---|---|---|---|---|
| 9 | 81139744 | AX-170032805 | 0 | G | A | 0.211 | 1.14E-02 | 1.29E-01 | 2.71E+00 | 2.76E+00 | 9.25E-01 | no | Gm26713 | intron_variant,non_coding_transcript_variant | MODIFIER |
| 9 | 81148736 | AX-170032802 | 0 | A | G | 0.211 | 1.14E-02 | 1.29E-01 | 2.71E+00 | 2.76E+00 | 9.25E-01 | no | Gm26713 | intron_variant,non_coding_transcript_variant | MODIFIER |
| 9 | 81199986 | AX-170215212 | 0 | A | T | 0.211 | 1.14E-02 | 1.29E-01 | 2.71E+00 | 2.76E+00 | 9.25E-01 | no | Gm26713 | intron_variant,non_coding_transcript_variant | MODIFIER |
| 9 | 81292702 | AX-170221147 | 0 | T | C | 0.211 | 1.14E-02 | 1.29E-01 | 2.71E+00 | 2.76E+00 | 9.25E-01 | no | | intergenic_variant | MODIFIER |
| 9 | 81691016 | AX-97790094 | 0 | C | T | 0.211 | 1.14E-02 | 1.29E-01 | 2.71E+00 | 2.76E+00 | 9.25E-01 | no | | intergenic_variant | MODIFIER |
| 9 | 82564589 | AX-170289659 | 0 | C | T | 0.211 | 1.14E-02 | 1.29E-01 | 2.71E+00 | 2.76E+00 | 9.25E-01 | no | | intergenic_variant | MODIFIER |
| 9 | 82797228 | AX-170416043 | 0 | C | T | 0.211 | 1.14E-02 | 1.29E-01 | 2.71E+00 | 2.76E+00 | 9.25E-01 | no | | intergenic_variant | MODIFIER |
| 9 | 82809108 | AX-170032836 | 0 | C | T | 0.211 | 1.14E-02 | 1.29E-01 | 2.71E+00 | 2.76E+00 | 9.25E-01 | no | | intergenic_variant | MODIFIER |
| 9 | 82993006 | AX-97785914 | 0 | C | T | 0.211 | 1.14E-02 | 1.29E-01 | 2.71E+00 | 2.76E+00 | 9.25E-01 | no | | intergenic_variant | MODIFIER |
| 9 | 83007916 | AX-170473219 | 0 | G | C | 0.211 | 1.14E-02 | 1.29E-01 | 2.71E+00 | 2.76E+00 | 9.25E-01 | no | Gm39383 | upstream_gene_variant | MODIFIER |
| 9 | 83020739 | AX-170481139 | 0 | C | T | 0.211 | 1.14E-02 | 1.29E-01 | 2.71E+00 | 2.76E+00 | 9.25E-01 | no | Gm39383 | intron_variant,non_coding_transcript_variant | MODIFIER |
| 9 | 83024045 | AX-170506190 | 0 | T | C | 0.211 | 1.14E-02 | 1.29E-01 | 2.71E+00 | 2.76E+00 | 9.25E-01 | no | Gm39383 | intron_variant,non_coding_transcript_variant | MODIFIER |
| 9 | 83101587 | AX-97813002 | 0 | C | T | 0.211 | 1.14E-02 | 1.29E-01 | 2.71E+00 | 2.76E+00 | 9.25E-01 | no | Gm38398 | intron_variant,non_coding_transcript_variant | MODIFIER |
| 9 | 83110694 | AX-170513635 | 0 | C | T | 0.211 | 1.14E-02 | 1.29E-01 | 2.71E+00 | 2.76E+00 | 9.25E-01 | no | Hmgn3 | intron_variant | MODIFIER |
| 9 | 83117018 | AX-170467901 | 0 | C | A | 0.211 | 1.14E-02 | 1.29E-01 | 2.71E+00 | 2.76E+00 | 9.25E-01 | no | Hmgn3 | intron_variant | MODIFIER |
| 9 | 83117241 | AX-170419322 | 0 | T | C | 0.211 | 1.14E-02 | 1.29E-01 | 2.71E+00 | 2.76E+00 | 9.25E-01 | no | Hmgn3 | intron_variant | MODIFIER |
| 10 | 12156413 | AX-170086745 | 0 | G | C | 0.211 | 9.18E-03 | 1.05E-01 | 2.68E+00 | 2.75E+00 | 9.25E-01 | no | | intergenic_variant | MODIFIER |
| 10 | 12451498 | AX-170383135 | 0 | A | G | 0.211 | 9.18E-03 | 1.05E-01 | 2.68E+00 | 2.75E+00 | 9.25E-01 | no | Utrn | intron_variant | MODIFIER |
| 10 | 12504205 | AX-170185062 | 0 | T | A | 0.211 | 9.18E-03 | 1.05E-01 | 2.68E+00 | 2.75E+00 | 9.25E-01 | no | Utrn | intron_variant | MODIFIER |
| 10 | 12507335 | AX-170091864 | 0 | T | C | 0.211 | 9.18E-03 | 1.05E-01 | 2.68E+00 | 2.75E+00 | 9.25E-01 | no | Utrn | intron_variant | MODIFIER |
| 10 | 12517623 | AX-170476431 | 0 | C | G | 0.211 | 9.18E-03 | 1.05E-01 | 2.68E+00 | 2.75E+00 | 9.25E-01 | no | Utrn | intron_variant | MODIFIER |
| 10 | 12534089 | AX-170520793 | 0 | G | A | 0.211 | 9.18E-03 | 1.05E-01 | 2.68E+00 | 2.75E+00 | 9.25E-01 | no | Utrn | intron_variant | MODIFIER |
| 10 | 12534486 | AX-170492506 | 0 | G | A | 0.211 | 9.18E-03 | 1.05E-01 | 2.68E+00 | 2.75E+00 | 9.25E-01 | no | Utrn | intron_variant | MODIFIER |
| 10 | 12543692 | AX-170108931 | 0 | C | G | 0.211 | 9.18E-03 | 1.05E-01 | 2.68E+00 | 2.75E+00 | 9.25E-01 | no | Utrn | intron_variant | MODIFIER |
| 10 | 12550090 | AX-170141050 | 0 | C | A | 0.211 | 9.18E-03 | 1.05E-01 | 2.68E+00 | 2.75E+00 | 9.25E-01 | no | Utrn | intron_variant | MODIFIER |
| 10 | 12564770 | AX-170002746 | 0 | T | G | 0.211 | 9.18E-03 | 1.05E-01 | 2.68E+00 | 2.75E+00 | 9.25E-01 | no | Utrn | intron_variant | MODIFIER |
| 10 | 12564831 | AX-97824942 | 0 | G | A | 0.211 | 9.18E-03 | 1.05E-01 | 2.68E+00 | 2.75E+00 | 9.25E-01 | no | Utrn | intron_variant | MODIFIER |
| 1 | 188040862 | AX-170176170 | 1 | C | T | 0.25 | 1.25E-02 | 9.53E-02 | 2.57E+00 | 2.70E+00 | 9.25E-01 | no | Esrrg | intron_variant | MODIFIER |
| 7 | 58814500 | AX-97833109 | 0 | G | T | 0.439 | -8.22E-03 | 9.61E-02 | 2.66E+00 | 2.75E+00 | 9.25E-01 | no | Atp10a | intron_variant | MODIFIER |
| 7 | 61030280 | AX-170508857 | 0 | G | T | 0.439 | -8.22E-03 | 9.61E-02 | 2.66E+00 | 2.75E+00 | 9.25E-01 | no | | regulatory_region_variant | MODIFIER |
| 1 | 54478908 | AX-97782551 | 0 | G | A | 0.114 | -1.61E-02 | 1.87E-01 | 2.64E+00 | 2.75E+00 | 9.25E-01 | no | Pgap1 | 3_prime_UTR_variant | MODIFIER |
| 12 | 116940550 | AX-170437099 | 1 | C | A | 0.125 | -1.74E-02 | 1.96E-01 | 2.68E+00 | 2.75E+00 | 9.25E-01 | no | Ptprn2 | intron_variant | MODIFIER |
| 11 | 45179473 | AX-170471015 | 0 | T | G | 0.088 | -1.82E-02 | 1.82E-01 | 2.62E+00 | 2.73E+00 | 9.25E-01 | no | Gm12160 | intron_variant,non_coding_transcript_variant | MODIFIER |
| 8 | 73068014 | AX-97856630 | 1 | C | T | 0.205 | 1.10E-02 | 1.24E-01 | 2.66E+00 | 2.75E+00 | 9.26E-01 | no | Large1 | intron_variant | MODIFIER |
| 1 | 8189226 | AX-170226876 | 0 | C | G | 0.447 | -5.61E-03 | 6.84E-02 | 2.69E+00 | 2.77E+00 | 9.26E-01 | no | | intergenic_variant | MODIFIER |
| 2 | 129610139 | AX-170276738 | 0 | C | T | 0.14 | -2.01E-02 | 2.22E-01 | 2.72E+00 | 2.75E+00 | 9.26E-01 | no | Sirpa | intron_variant | MODIFIER |
| 2 | 146935057 | AX-97817708 | 0 | G | A | 0.14 | -2.01E-02 | 2.22E-01 | 2.72E+00 | 2.75E+00 | 9.26E-01 | no | Kiz | intron_variant | MODIFIER |
| 10 | 12493733 | AX-170186081 | 0 | G | A | 0.219 | 8.83E-03 | 1.00E-01 | 2.66E+00 | 2.75E+00 | 9.26E-01 | no | Utrn | intron_variant | MODIFIER |
| 5 | 89945185 | AX-170342629 | 1 | C | T | 0.411 | 9.63E-03 | 1.04E-01 | 2.68E+00 | 2.74E+00 | 9.26E-01 | no | | intergenic_variant | MODIFIER |
| 12 | 118060490 | AX-170050360 | 0 | C | T | 0.149 | -1.14E-02 | 1.46E-01 | 2.63E+00 | 2.76E+00 | 9.26E-01 | no | Dnah11 | intron_variant | MODIFIER |
| 1 | 151303810 | AX-170519986 | 0 | C | T | 0.211 | 1.31E-02 | 1.35E-01 | 2.67E+00 | 2.73E+00 | 9.26E-01 | no | | intergenic_variant | MODIFIER |
| 4 | 93292819 | AX-170023414 | 0 | C | A | 0.333 | -9.37E-03 | 1.09E-01 | 2.69E+00 | 2.76E+00 | 9.26E-01 | no | | intergenic_variant | MODIFIER |
| 20 | 114961824 | AX-97783109 | 0 | T | C | 0.307 | 5.18E-03 | 5.34E-02 | 2.61E+00 | 2.73E+00 | 9.26E-01 | no | | intergenic_variant | MODIFIER |
| 20 | 114962056 | AX-170355997 | 0 | C | T | 0.307 | 5.18E-03 | 5.34E-02 | 2.61E+00 | 2.73E+00 | 9.26E-01 | no | | intergenic_variant | MODIFIER |
| 20 | 115202685 | AX-170034772 | 0 | T | C | 0.307 | 5.18E-03 | 5.34E-02 | 2.61E+00 | 2.73E+00 | 9.26E-01 | no | | intergenic_variant | MODIFIER |
| 20 | 115203322 | AX-170034771 | 0 | G | T | 0.307 | 5.18E-03 | 5.34E-02 | 2.61E+00 | 2.73E+00 | 9.26E-01 | no | | intergenic_variant | MODIFIER |
| 20 | 115236450 | AX-170034770 | 0 | C | A | 0.307 | 5.18E-03 | 5.34E-02 | 2.61E+00 | 2.73E+00 | 9.26E-01 | no | | intergenic_variant | MODIFIER |
| 20 | 115300970 | AX-170096897 | 0 | A | G | 0.307 | 5.18E-03 | 5.34E-02 | 2.61E+00 | 2.73E+00 | 9.26E-01 | no | | intergenic_variant | MODIFIER |
| 20 | 115381271 | AX-170085076 | 0 | A | G | 0.307 | 5.18E-03 | 5.34E-02 | 2.61E+00 | 2.73E+00 | 9.26E-01 | no | | intergenic_variant | MODIFIER |
| 20 | 115397053 | AX-170090644 | 0 | T | A | 0.307 | 5.18E-03 | 5.34E-02 | 2.61E+00 | 2.73E+00 | 9.26E-01 | no | Gm27414 | upstream_gene_variant | MODIFIER |
| 20 | 115490043 | AX-170190211 | 0 | C | T | 0.307 | 5.18E-03 | 5.34E-02 | 2.61E+00 | 2.73E+00 | 9.26E-01 | no | | intergenic_variant | MODIFIER |
| 20 | 115491586 | AX-170307287 | 0 | T | C | 0.307 | 5.18E-03 | 5.34E-02 | 2.61E+00 | 2.73E+00 | 9.26E-01 | no | | intergenic_variant | MODIFIER |
| 20 | 115976169 | AX-170085083 | 0 | C | T | 0.307 | 5.18E-03 | 5.34E-02 | 2.61E+00 | 2.73E+00 | 9.26E-01 | no | | intergenic_variant | MODIFIER |
| 20 | 115987986 | AX-170488351 | 0 | T | C | 0.307 | 5.18E-03 | 5.34E-02 | 2.61E+00 | 2.73E+00 | 9.26E-01 | no | | intergenic_variant | MODIFIER |

| | | | | | | | | | | | | | | | |
|---|---|---|---|---|---|---|---|---|---|---|---|---|---|---|---|
| 20 | 116063659 | AX-170320282 | 0 | G | T | 0.307 | 5.18E-03 | 5.34E-02 | 2.61E+00 | 2.73E+00 | 9.26E-01 | no | | intergenic_variant | MODIFIER |
| 20 | 116135241 | AX-170520442 | 0 | T | C | 0.307 | 5.18E-03 | 5.34E-02 | 2.61E+00 | 2.73E+00 | 9.26E-01 | no | | intergenic_variant | MODIFIER |
| 20 | 116871170 | AX-170090649 | 0 | A | G | 0.307 | 5.18E-03 | 5.34E-02 | 2.61E+00 | 2.73E+00 | 9.26E-01 | no | | intergenic_variant | MODIFIER |
| 20 | 116963450 | AX-170034796 | 0 | C | T | 0.307 | 5.18E-03 | 5.34E-02 | 2.61E+00 | 2.73E+00 | 9.26E-01 | no | | intergenic_variant | MODIFIER |
| 20 | 117037280 | AX-170085097 | 0 | A | G | 0.307 | 5.18E-03 | 5.34E-02 | 2.61E+00 | 2.73E+00 | 9.26E-01 | no | | intergenic_variant | MODIFIER |
| 20 | 117052276 | AX-170265933 | 0 | C | T | 0.307 | 5.18E-03 | 5.34E-02 | 2.61E+00 | 2.73E+00 | 9.26E-01 | no | | intergenic_variant | MODIFIER |
| 20 | 127288992 | AX-170169883 | 0 | A | G | 0.307 | 5.18E-03 | 5.34E-02 | 2.61E+00 | 2.73E+00 | 9.26E-01 | no | | intergenic_variant | MODIFIER |
| 18 | 68307234 | AX-170271735 | 0 | T | G | 0.281 | -9.15E-03 | 9.39E-02 | 2.66E+00 | 2.73E+00 | 9.26E-01 | no | Rnmt | intron_variant | MODIFIER |
| 1 | 8145598 | AX-170115218 | 1 | T | C | 0.446 | -5.59E-03 | 6.85E-02 | 2.69E+00 | 2.77E+00 | 9.26E-01 | no | | intergenic_variant | MODIFIER |
| 1 | 8169752 | AX-170384398 | 1 | T | G | 0.446 | -5.59E-03 | 6.85E-02 | 2.69E+00 | 2.77E+00 | 9.26E-01 | no | | intergenic_variant | MODIFIER |
| 9 | 61588327 | AX-170403151 | 1 | T | C | 0.188 | -1.29E-02 | 1.36E-01 | 2.70E+00 | 2.73E+00 | 9.26E-01 | no | | intergenic_variant | MODIFIER |
| 13 | 37391012 | AX-170342960 | 1 | C | A | 0.17 | -1.23E-02 | 1.16E-01 | 2.64E+00 | 2.71E+00 | 9.26E-01 | no | Ly86 | intron_variant | MODIFIER |
| 16 | 97767014 | AX-170300011 | 0 | G | A | 0.465 | -7.93E-03 | 7.51E-02 | 2.61E+00 | 2.72E+00 | 9.26E-01 | no | Ripk4 | upstream_gene_variant | MODIFIER |
| 12 | 116433334 | AX-170164988 | 0 | G | A | 0.105 | 1.20E-02 | 1.38E-01 | 2.61E+00 | 2.74E+00 | 9.26E-01 | no | Ncapg2 | intron_variant | MODIFIER |
| 8 | 81227145 | AX-170251182 | 0 | C | G | 0.211 | -1.13E-02 | 1.28E-01 | 2.68E+00 | 2.75E+00 | 9.26E-01 | no | | intergenic_variant | MODIFIER |
| 20 | 60705900 | AX-170033962 | 0 | C | T | 0.246 | 5.80E-03 | 6.28E-02 | 2.64E+00 | 2.74E+00 | 9.26E-01 | no | Gm4909 | downstream_gene_variant | MODIFIER |
| 4 | 150054208 | AX-170179042 | 0 | T | C | 0.096 | 1.24E-02 | 1.34E-01 | 2.63E+00 | 2.74E+00 | 9.26E-01 | no | Gm13067 | intron_variant,non_coding_transcript_variant | MODIFIER |
| 5 | 86630537 | AX-170118614 | 1 | A | G | 0.42 | 9.73E-03 | 1.07E-01 | 2.68E+00 | 2.74E+00 | 9.27E-01 | no | Tmprss11f | intron_variant | MODIFIER |
| 5 | 87783202 | AX-170025529 | 1 | T | G | 0.42 | 9.73E-03 | 1.07E-01 | 2.68E+00 | 2.74E+00 | 9.27E-01 | no | Csn1s2a | intron_variant | MODIFIER |
| 5 | 90592269 | AX-170025578 | 1 | T | C | 0.42 | 9.73E-03 | 1.07E-01 | 2.68E+00 | 2.74E+00 | 9.27E-01 | no | 5830473C10Rik | intron_variant | MODIFIER |
| 18 | 34242128 | AX-170015511 | 0 | T | C | 0.061 | -2.79E-02 | 3.01E-01 | 2.71E+00 | 2.73E+00 | 9.27E-01 | no | Apc | intron_variant | MODIFIER |
| 18 | 41545916 | AX-170060662 | 0 | A | C | 0.061 | -2.79E-02 | 3.01E-01 | 2.71E+00 | 2.73E+00 | 9.27E-01 | no | | intergenic_variant | MODIFIER |
| 18 | 43647791 | AX-170277388 | 0 | A | T | 0.061 | -2.79E-02 | 3.01E-01 | 2.71E+00 | 2.73E+00 | 9.27E-01 | no | Jakmip2 | intron_variant | MODIFIER |
| 2 | 127032764 | AX-170206169 | 1 | G | A | 0.143 | -1.59E-02 | 1.82E-01 | 2.69E+00 | 2.75E+00 | 9.27E-01 | no | Ap4e1 | intron_variant | MODIFIER |
| 9 | 83865152 | AX-170327976 | 0 | G | A | 0.219 | 1.09E-02 | 1.26E-01 | 2.69E+00 | 2.75E+00 | 9.27E-01 | no | Ttk | synonymous_variant | LOW |
| 9 | 83867364 | AX-170300573 | 0 | G | A | 0.219 | 1.09E-02 | 1.26E-01 | 2.69E+00 | 2.75E+00 | 9.27E-01 | no | Ttk | intron_variant | MODIFIER |
| 9 | 83868770 | AX-170224410 | 0 | T | C | 0.219 | 1.09E-02 | 1.26E-01 | 2.69E+00 | 2.75E+00 | 9.27E-01 | no | Ttk | intron_variant | MODIFIER |
| 9 | 83869430 | AX-170219505 | 0 | A | G | 0.219 | 1.09E-02 | 1.26E-01 | 2.69E+00 | 2.75E+00 | 9.27E-01 | no | Ttk | intron_variant | MODIFIER |
| 11 | 44916536 | AX-170046934 | 0 | A | T | 0.07 | 1.56E-02 | 1.85E-01 | 2.63E+00 | 2.75E+00 | 9.27E-01 | no | Ebf1 | intron_variant | MODIFIER |
| 11 | 45154349 | AX-170222709 | 0 | G | A | 0.07 | 1.56E-02 | 1.85E-01 | 2.63E+00 | 2.75E+00 | 9.27E-01 | no | | regulatory_region_variant | MODIFIER |
| 11 | 46316872 | AX-170104518 | 0 | A | T | 0.07 | 1.56E-02 | 1.85E-01 | 2.63E+00 | 2.75E+00 | 9.27E-01 | no | Cyfip2 | upstream_gene_variant | MODIFIER |
| 9 | 81116556 | AX-170032806 | 0 | G | A | 0.219 | 1.02E-02 | 1.20E-01 | 2.68E+00 | 2.75E+00 | 9.27E-01 | no | Gm26713 | intron_variant,non_coding_transcript_variant | MODIFIER |
| 1 | 55702040 | AX-170451425 | 0 | C | T | 0.105 | -2.12E-02 | 2.37E-01 | 2.70E+00 | 2.74E+00 | 9.27E-01 | no | Plcl1 | synonymous_variant | LOW |
| 9 | 34653334 | AX-170229024 | 0 | T | A | 0.43 | -6.58E-03 | 8.31E-02 | 2.67E+00 | 2.76E+00 | 9.27E-01 | no | Kirrel3 | intron_variant | MODIFIER |
| 10 | 8468630 | AX-170535462 | 0 | C | A | 0.211 | 8.29E-03 | 9.68E-02 | 2.64E+00 | 2.74E+00 | 9.27E-01 | no | Ust | intron_variant | MODIFIER |
| 8 | 93528969 | AX-170107516 | 1 | T | C | 0.223 | 9.80E-03 | 1.07E-01 | 2.66E+00 | 2.74E+00 | 9.27E-01 | no | Ces5a | intron_variant | MODIFIER |
| 13 | 35178342 | AX-170365103 | 0 | G | A | 0.184 | -1.19E-02 | 1.13E-01 | 2.63E+00 | 2.71E+00 | 9.27E-01 | no | 1700011B04Rik | upstream_gene_variant | MODIFIER |
| 8 | 56310325 | AX-170187352 | 1 | G | A | 0.152 | -1.47E-02 | 1.48E-01 | 2.66E+00 | 2.72E+00 | 9.28E-01 | no | Hpgd | intron_variant | MODIFIER |
| 8 | 56409104 | AX-170214231 | 1 | A | T | 0.152 | -1.47E-02 | 1.48E-01 | 2.66E+00 | 2.72E+00 | 9.28E-01 | no | | regulatory_region_variant | MODIFIER |
| 7 | 94280412 | AX-170166352 | 1 | T | C | 0.125 | -1.74E-02 | 1.90E-01 | 2.64E+00 | 2.74E+00 | 9.28E-01 | no | Gm32647 | intron_variant,non_coding_transcript_variant | MODIFIER |
| 1 | 72055108 | AX-170211791 | 0 | C | G | 0.114 | 2.07E-02 | 2.30E-01 | 2.69E+00 | 2.74E+00 | 9.28E-01 | no | | intergenic_variant | MODIFIER |
| 13 | 44485120 | AX-170378580 | 0 | G | C | 0.491 | -4.89E-03 | 6.12E-02 | 2.65E+00 | 2.75E+00 | 9.28E-01 | no | Gm34084 | non_coding_transcript_exon_variant | MODIFIER |
| 13 | 45441678 | AX-170434024 | 0 | G | A | 0.491 | -4.89E-03 | 6.12E-02 | 2.65E+00 | 2.75E+00 | 9.28E-01 | no | | intergenic_variant | MODIFIER |
| 13 | 45446421 | AX-97785990 | 0 | T | G | 0.491 | -4.89E-03 | 6.12E-02 | 2.65E+00 | 2.75E+00 | 9.28E-01 | no | | regulatory_region_variant | MODIFIER |
| 13 | 45453687 | AX-170403435 | 0 | G | A | 0.491 | -4.89E-03 | 6.12E-02 | 2.65E+00 | 2.75E+00 | 9.28E-01 | no | | regulatory_region_variant | MODIFIER |
| 13 | 45475947 | AX-170428079 | 0 | C | T | 0.491 | -4.89E-03 | 6.12E-02 | 2.65E+00 | 2.75E+00 | 9.28E-01 | no | | intergenic_variant | MODIFIER |
| 13 | 45483722 | AX-170139467 | 0 | C | T | 0.491 | -4.89E-03 | 6.12E-02 | 2.65E+00 | 2.75E+00 | 9.28E-01 | no | Gm23387 | downstream_gene_variant | MODIFIER |
| 13 | 45488719 | AX-170160278 | 0 | A | C | 0.491 | -4.89E-03 | 6.12E-02 | 2.65E+00 | 2.75E+00 | 9.28E-01 | no | Gm23387 | upstream_gene_variant | MODIFIER |
| 13 | 45512992 | AX-97794680 | 0 | C | A | 0.491 | -4.89E-03 | 6.12E-02 | 2.65E+00 | 2.75E+00 | 9.28E-01 | no | Gmpr | intron_variant | MODIFIER |
| 13 | 45526645 | AX-170510467 | 0 | G | C | 0.491 | -4.89E-03 | 6.12E-02 | 2.65E+00 | 2.75E+00 | 9.28E-01 | no | Gmpr | intron_variant | MODIFIER |
| 13 | 45526723 | AX-170401498 | 0 | T | C | 0.491 | -4.89E-03 | 6.12E-02 | 2.65E+00 | 2.75E+00 | 9.28E-01 | no | Gmpr | intron_variant | MODIFIER |
| 13 | 45529288 | AX-170265614 | 0 | T | C | 0.491 | -4.89E-03 | 6.12E-02 | 2.65E+00 | 2.75E+00 | 9.28E-01 | no | Gmpr | intron_variant | MODIFIER |
| 12 | 106477405 | AX-170117282 | 2 | G | C | 0.1 | 1.69E-02 | 1.85E-01 | 2.63E+00 | 2.74E+00 | 9.28E-01 | no | | regulatory_region_variant | MODIFIER |
| 8 | 109400219 | AX-170236667 | 0 | T | C | 0.246 | 9.53E-03 | 1.01E-01 | 2.63E+00 | 2.73E+00 | 9.28E-01 | no | | regulatory_region_variant | MODIFIER |

| | | | | | | | | | | | | | | | |
|---|---|---|---|---|---|---|---|---|---|---|---|---|---|---|---|
| 10 | 14343963 | AX-170527730 | 0 | A | G | 0.202 | -1.02E-02 | 1.06E-01 | 2.65E+00 | 2.72E+00 | 9.28E-01 | no | | intergenic_variant | MODIFIER |
| 4 | 134727981 | AX-97854218 | 1 | T | C | 0.152 | 1.04E-02 | 9.86E-02 | 2.60E+00 | 2.72E+00 | 9.28E-01 | no | | intergenic_variant | MODIFIER |
| 7 | 103441985 | AX-170029671 | 1 | G | A | 0.473 | -7.98E-03 | 9.58E-02 | 2.68E+00 | 2.75E+00 | 9.28E-01 | no | Olfr605 | downstream_gene_variant | MODIFIER |
| 9 | 59197350 | AX-170194065 | 1 | C | T | 0.205 | -1.07E-02 | 1.22E-01 | 2.69E+00 | 2.74E+00 | 9.28E-01 | no | Gm33053 | intron_variant,non_coding_transcript_variant | MODIFIER |
| 12 | 112452836 | AX-170350196 | 1 | C | T | 0.134 | 1.60E-02 | 1.85E-01 | 2.66E+00 | 2.74E+00 | 9.28E-01 | no | | intergenic_variant | MODIFIER |
| 12 | 116208296 | AX-170309771 | 1 | A | G | 0.134 | 1.60E-02 | 1.85E-01 | 2.66E+00 | 2.74E+00 | 9.28E-01 | no | Wdr60 | intron_variant | MODIFIER |
| 8 | 74675809 | AX-170080432 | 1 | A | G | 0.196 | -1.09E-02 | 1.21E-01 | 2.65E+00 | 2.74E+00 | 9.28E-01 | no | Gm11033 | intron_variant,non_coding_transcript_variant | MODIFIER |
| 2 | 143578916 | AX-170152719 | 1 | T | A | 0.134 | 1.85E-02 | 2.00E-01 | 2.67E+00 | 2.73E+00 | 9.28E-01 | no | Pcsk2 | intron_variant | MODIFIER |
| 1 | 144772462 | AX-170318411 | 0 | A | G | 0.219 | 1.10E-02 | 1.15E-01 | 2.65E+00 | 2.73E+00 | 9.28E-01 | no | Rgs18 | intron_variant | MODIFIER |
| 11 | 13093089 | AX-97801027 | 0 | G | A | 0.061 | -1.98E-02 | 2.14E-01 | 2.64E+00 | 2.73E+00 | 9.28E-01 | no | | intergenic_variant | MODIFIER |
| 2 | 155789139 | AX-170285448 | 0 | T | C | 0.149 | -1.52E-02 | 1.79E-01 | 2.70E+00 | 2.75E+00 | 9.28E-01 | no | Mmp24 | intron_variant | MODIFIER |
| 10 | 5846384 | AX-170520215 | 0 | C | T | 0.202 | 8.23E-03 | 9.99E-02 | 2.65E+00 | 2.75E+00 | 9.28E-01 | no | Rgs17 | intron_variant | MODIFIER |
| 5 | 114365151 | AX-97809792 | 2 | T | C | 0.436 | 9.83E-03 | 1.02E-01 | 2.65E+00 | 2.72E+00 | 9.28E-01 | no | Kctd10 | downstream_gene_variant | MODIFIER |
| 15 | 91078273 | AX-170260225 | 1 | T | G | 0.116 | -1.41E-02 | 1.78E-01 | 2.69E+00 | 2.76E+00 | 9.28E-01 | no | Gm18814 | downstream_gene_variant | MODIFIER |
| 5 | 97938225 | AX-170114752 | 1 | T | C | 0.411 | 9.55E-03 | 1.03E-01 | 2.66E+00 | 2.73E+00 | 9.28E-01 | no | Antxr2 | intron_variant | MODIFIER |
| 1 | 13821707 | AX-170423833 | 0 | A | G | 0.368 | 7.14E-03 | 7.44E-02 | 2.66E+00 | 2.72E+00 | 9.28E-01 | no | | intergenic_variant | MODIFIER |
| 9 | 105946042 | AX-97851697 | 1 | T | C | 0.116 | -1.23E-02 | 1.15E-01 | 2.63E+00 | 2.71E+00 | 9.29E-01 | no | Col6a5 | missense_variant | MODERATE |
| 1 | 66511701 | AX-170153704 | 1 | T | A | 0.107 | 1.84E-02 | 2.10E-01 | 2.67E+00 | 2.74E+00 | 9.29E-01 | no | Unc80 | intron_variant | MODIFIER |
| 1 | 67181823 | AX-170106527 | 1 | A | G | 0.107 | 1.84E-02 | 2.10E-01 | 2.67E+00 | 2.74E+00 | 9.29E-01 | no | Cps1 | intron_variant | MODIFIER |
| 7 | 103713804 | AX-170519243 | 0 | C | T | 0.456 | -7.58E-03 | 9.39E-02 | 2.69E+00 | 2.76E+00 | 9.29E-01 | no | Olfr243 | upstream_gene_variant | MODIFIER |
| 10 | 6681596 | AX-170409539 | 1 | T | C | 0.196 | -9.79E-03 | 1.04E-01 | 2.64E+00 | 2.73E+00 | 9.29E-01 | no | | intergenic_variant | MODIFIER |
| 12 | 109735677 | AX-170050165 | 0 | C | T | 0.123 | -1.67E-02 | 1.95E-01 | 2.67E+00 | 2.74E+00 | 9.29E-01 | no | Mir134 | downstream_gene_variant | MODIFIER |
| 14 | 95235617 | AX-170010456 | 1 | G | C | 0.045 | 1.78E-02 | 2.17E-01 | 2.67E+00 | 2.75E+00 | 9.29E-01 | no | | intergenic_variant | MODIFIER |
| 1 | 8443904 | AX-170288058 | 1 | G | A | 0.518 | -5.55E-03 | 7.22E-02 | 2.69E+00 | 2.77E+00 | 9.29E-01 | no | Sntg1 | intron_variant,non_coding_transcript_variant | MODIFIER |
| 1 | 48952164 | AX-170327136 | 0 | G | A | 0.105 | -2.04E-02 | 2.34E-01 | 2.70E+00 | 2.74E+00 | 9.29E-01 | no | | intergenic_variant | MODIFIER |
| 15 | 78537001 | AX-170322713 | 0 | C | A | 0.553 | 8.32E-03 | 8.27E-02 | 2.64E+00 | 2.72E+00 | 9.29E-01 | no | Sstr3 | downstream_gene_variant | MODIFIER |
| 1 | 11196329 | AX-170496588 | 1 | A | T | 0.393 | -5.83E-03 | 7.28E-02 | 2.69E+00 | 2.76E+00 | 9.29E-01 | no | Prex2 | intron_variant | MODIFIER |
| 8 | 90704188 | AX-170400110 | 1 | C | T | 0.205 | -1.08E-02 | 1.14E-01 | 2.64E+00 | 2.73E+00 | 9.29E-01 | no | | intergenic_variant | MODIFIER |
| 8 | 48603289 | AX-170527361 | 1 | T | G | 0.152 | -1.45E-02 | 1.50E-01 | 2.66E+00 | 2.72E+00 | 9.29E-01 | no | Tenm3 | intron_variant | MODIFIER |
| 15 | 78316891 | AX-170250829 | 0 | G | T | 0.544 | 8.03E-03 | 7.93E-02 | 2.63E+00 | 2.72E+00 | 9.29E-01 | no | | intergenic_variant | MODIFIER |
| 15 | 79202707 | AX-170012068 | 2 | T | C | 0.536 | 7.99E-03 | 7.97E-02 | 2.63E+00 | 2.72E+00 | 9.29E-01 | no | Gm10863 | intron_variant,non_coding_transcript_variant | MODIFIER |
| 15 | 86733464 | AX-170244183 | 0 | C | A | 0.132 | 1.46E-02 | 1.56E-01 | 2.66E+00 | 2.73E+00 | 9.29E-01 | no | 4930543I03Rik | upstream_gene_variant | MODIFIER |
| 9 | 85586904 | AX-170534695 | 0 | A | G | 0.202 | 1.06E-02 | 1.20E-01 | 2.67E+00 | 2.74E+00 | 9.29E-01 | no | | intergenic_variant | MODIFIER |
| 2 | 158453748 | AX-170019498 | 1 | T | A | 0.143 | 2.00E-02 | 2.23E-01 | 2.70E+00 | 2.73E+00 | 9.29E-01 | no | Ralgapb | intron_variant | MODIFIER |
| 8 | 37588466 | AX-170509216 | 1 | T | C | 0.143 | -1.29E-02 | 1.36E-01 | 2.66E+00 | 2.72E+00 | 9.29E-01 | no | Sgcz | intron_variant | MODIFIER |
| 4 | 82272495 | AX-170427269 | 0 | G | A | 0.246 | 1.10E-02 | 1.05E-01 | 2.63E+00 | 2.71E+00 | 9.29E-01 | no | | regulatory_region_variant | MODIFIER |
| 10 | 113660237 | AX-170218992 | 0 | T | A | 0.351 | -7.11E-03 | 7.96E-02 | 2.66E+00 | 2.73E+00 | 9.29E-01 | no | | intergenic_variant | MODIFIER |
| 10 | 113689506 | AX-170441096 | 0 | T | A | 0.351 | -7.11E-03 | 7.96E-02 | 2.66E+00 | 2.73E+00 | 9.29E-01 | no | | intergenic_variant | MODIFIER |
| 10 | 113768124 | AX-170045953 | 0 | T | C | 0.351 | -7.11E-03 | 7.96E-02 | 2.66E+00 | 2.73E+00 | 9.29E-01 | no | | intergenic_variant | MODIFIER |
| 10 | 113777384 | AX-97808733 | 0 | G | A | 0.351 | -7.11E-03 | 7.96E-02 | 2.66E+00 | 2.73E+00 | 9.29E-01 | no | | intergenic_variant | MODIFIER |
| 10 | 113815301 | AX-170401965 | 0 | T | C | 0.351 | -7.11E-03 | 7.96E-02 | 2.66E+00 | 2.73E+00 | 9.29E-01 | no | | intergenic_variant | MODIFIER |
| 10 | 114038926 | AX-170261563 | 0 | G | A | 0.351 | -7.11E-03 | 7.96E-02 | 2.66E+00 | 2.73E+00 | 9.29E-01 | no | | intergenic_variant | MODIFIER |
| 10 | 114038986 | AX-170163685 | 0 | G | A | 0.351 | -7.11E-03 | 7.96E-02 | 2.66E+00 | 2.73E+00 | 9.29E-01 | no | | intergenic_variant | MODIFIER |
| 10 | 114117457 | AX-170523700 | 0 | C | A | 0.351 | -7.11E-03 | 7.96E-02 | 2.66E+00 | 2.73E+00 | 9.29E-01 | no | | intergenic_variant | MODIFIER |
| 10 | 114154907 | AX-97848174 | 0 | A | T | 0.351 | -7.11E-03 | 7.96E-02 | 2.66E+00 | 2.73E+00 | 9.29E-01 | no | | intergenic_variant | MODIFIER |
| 10 | 114154956 | AX-170327680 | 0 | C | A | 0.351 | -7.11E-03 | 7.96E-02 | 2.66E+00 | 2.73E+00 | 9.29E-01 | no | | intergenic_variant | MODIFIER |
| 10 | 114210897 | AX-170003941 | 0 | G | A | 0.351 | -7.11E-03 | 7.96E-02 | 2.66E+00 | 2.73E+00 | 9.29E-01 | no | | intergenic_variant | MODIFIER |
| 10 | 114210988 | AX-97824725 | 0 | A | G | 0.351 | -7.11E-03 | 7.96E-02 | 2.66E+00 | 2.73E+00 | 9.29E-01 | no | | intergenic_variant | MODIFIER |
| 10 | 114399893 | AX-170045961 | 0 | G | C | 0.351 | -7.11E-03 | 7.96E-02 | 2.66E+00 | 2.73E+00 | 9.29E-01 | no | Trhde | 3_prime_UTR_variant | MODIFIER |
| 10 | 114401218 | AX-170312643 | 0 | G | A | 0.351 | -7.11E-03 | 7.96E-02 | 2.66E+00 | 2.73E+00 | 9.29E-01 | no | Trhde | 3_prime_UTR_variant | MODIFIER |
| 10 | 114401473 | AX-170486191 | 0 | G | A | 0.351 | -7.11E-03 | 7.96E-02 | 2.66E+00 | 2.73E+00 | 9.29E-01 | no | Trhde | synonymous_variant | LOW |
| 10 | 114402394 | AX-170424523 | 0 | T | G | 0.351 | -7.11E-03 | 7.96E-02 | 2.66E+00 | 2.73E+00 | 9.29E-01 | no | Trhde | intron_variant | MODIFIER |
| 10 | 114402644 | AX-170098082 | 0 | C | G | 0.351 | -7.11E-03 | 7.96E-02 | 2.66E+00 | 2.73E+00 | 9.29E-01 | no | Trhde | intron_variant | MODIFIER |
| 10 | 114417581 | AX-170448241 | 0 | A | C | 0.351 | -7.11E-03 | 7.96E-02 | 2.66E+00 | 2.73E+00 | 9.29E-01 | no | Trhde | intron_variant | MODIFIER |

| | | | | | | | | | | | | | | | |
|---|---|---|---|---|---|---|---|---|---|---|---|---|---|---|---|
| 10 | 114593619 | AX-170231447 | 0 | A | C | 0.351 | -7.11E-03 | 7.96E-02 | 2.66E+00 | 2.73E+00 | 9.29E-01 | no | Trhde | intron_variant | MODIFIER |
| 10 | 114656190 | AX-97839834 | 0 | T | G | 0.351 | -7.11E-03 | 7.96E-02 | 2.66E+00 | 2.73E+00 | 9.29E-01 | no | Trhde | intron_variant | MODIFIER |
| 10 | 114688336 | AX-170147481 | 0 | T | C | 0.351 | -7.11E-03 | 7.96E-02 | 2.66E+00 | 2.73E+00 | 9.29E-01 | no | Trhde | intron_variant | MODIFIER |
| 10 | 114696772 | AX-170003950 | 0 | C | A | 0.351 | -7.11E-03 | 7.96E-02 | 2.66E+00 | 2.73E+00 | 9.29E-01 | no | Trhde | intron_variant | MODIFIER |
| 10 | 114723715 | AX-170349317 | 0 | T | A | 0.351 | -7.11E-03 | 7.96E-02 | 2.66E+00 | 2.73E+00 | 9.29E-01 | no | Trhde | intron_variant | MODIFIER |
| 10 | 114731182 | AX-170311938 | 0 | C | T | 0.351 | -7.11E-03 | 7.96E-02 | 2.66E+00 | 2.73E+00 | 9.29E-01 | no | Trhde | intron_variant | MODIFIER |
| 5 | 128075925 | AX-170484339 | 0 | G | A | 0.412 | -6.63E-03 | 8.22E-02 | 2.64E+00 | 2.75E+00 | 9.30E-01 | no | Tmem132d | intron_variant | MODIFIER |
| 1 | 76456936 | AX-170372777 | 0 | G | C | 0.149 | -1.74E-02 | 2.12E-01 | 2.67E+00 | 2.75E+00 | 9.30E-01 | no | Gm29069 | intron_variant,non_coding_transcript_variant | MODIFIER |
| 1 | 77525247 | AX-170475937 | 0 | G | A | 0.149 | -1.74E-02 | 2.12E-01 | 2.67E+00 | 2.75E+00 | 9.30E-01 | no | | intergenic_variant | MODIFIER |
| 1 | 25765742 | AX-170000135 | 0 | A | G | 0.404 | 7.47E-03 | 8.24E-02 | 2.65E+00 | 2.73E+00 | 9.30E-01 | no | Adgrb3 | intron_variant | MODIFIER |
| 12 | 107168842 | AX-97825854 | 0 | C | A | 0.123 | -1.56E-02 | 1.85E-01 | 2.67E+00 | 2.74E+00 | 9.30E-01 | no | | intergenic_variant | MODIFIER |
| 1 | 21305488 | AX-97856823 | 0 | A | C | 0.395 | 7.85E-03 | 7.87E-02 | 2.64E+00 | 2.72E+00 | 9.30E-01 | no | | regulatory_region_variant | MODIFIER |
| 13 | 54752164 | AX-97829719 | 0 | G | A | 0.526 | -4.82E-03 | 6.65E-02 | 2.65E+00 | 2.76E+00 | 9.30E-01 | no | Gprin1 | upstream_gene_variant | MODIFIER |
| 10 | 10532321 | AX-97824483 | 0 | G | A | 0.219 | 8.59E-03 | 1.06E-01 | 2.68E+00 | 2.75E+00 | 9.30E-01 | no | | intergenic_variant | MODIFIER |
| 10 | 10535150 | AX-170002706 | 0 | C | T | 0.219 | 8.59E-03 | 1.06E-01 | 2.68E+00 | 2.75E+00 | 9.30E-01 | no | | intergenic_variant | MODIFIER |
| 10 | 10554399 | AX-170390842 | 0 | A | G | 0.219 | 8.59E-03 | 1.06E-01 | 2.68E+00 | 2.75E+00 | 9.30E-01 | no | Rab32 | intron_variant | MODIFIER |
| 10 | 10555362 | AX-97791587 | 0 | A | G | 0.219 | 8.59E-03 | 1.06E-01 | 2.68E+00 | 2.75E+00 | 9.30E-01 | no | Rab32 | intron_variant | MODIFIER |
| 10 | 10731552 | AX-170091859 | 0 | A | G | 0.219 | 8.59E-03 | 1.06E-01 | 2.68E+00 | 2.75E+00 | 9.30E-01 | no | Grm1 | intron_variant | MODIFIER |
| 10 | 10746003 | AX-170109913 | 0 | G | A | 0.219 | 8.59E-03 | 1.06E-01 | 2.68E+00 | 2.75E+00 | 9.30E-01 | no | Grm1 | intron_variant | MODIFIER |
| 10 | 11940366 | AX-170415302 | 0 | G | A | 0.219 | 8.59E-03 | 1.06E-01 | 2.68E+00 | 2.75E+00 | 9.30E-01 | no | Gm22194 | upstream_gene_variant | MODIFIER |
| 10 | 11960680 | AX-97856736 | 0 | T | C | 0.219 | 8.59E-03 | 1.06E-01 | 2.68E+00 | 2.75E+00 | 9.30E-01 | no | Gm25011 | downstream_gene_variant | MODIFIER |
| 10 | 11965904 | AX-170231783 | 0 | T | C | 0.219 | 8.59E-03 | 1.06E-01 | 2.68E+00 | 2.75E+00 | 9.30E-01 | no | | intergenic_variant | MODIFIER |
| 10 | 12145973 | AX-170044156 | 0 | G | A | 0.219 | 8.59E-03 | 1.06E-01 | 2.68E+00 | 2.75E+00 | 9.30E-01 | no | | intergenic_variant | MODIFIER |
| 20 | 140704428 | AX-170258086 | 1 | T | C | 0.143 | 1.24E-02 | 1.37E-01 | 2.64E+00 | 2.73E+00 | 9.30E-01 | no | Mid2 | intron_variant | MODIFIER |
| 12 | 57909633 | AX-170225281 | 0 | C | A | 0.061 | -2.64E-02 | 3.00E-01 | 2.71E+00 | 2.73E+00 | 9.30E-01 | no | | intergenic_variant | MODIFIER |
| 12 | 73084749 | AX-170172484 | 0 | C | T | 0.061 | -2.64E-02 | 3.00E-01 | 2.71E+00 | 2.73E+00 | 9.30E-01 | no | | intergenic_variant | MODIFIER |
| 1 | 13017116 | AX-170320733 | 0 | C | A | 0.395 | 7.05E-03 | 7.23E-02 | 2.65E+00 | 2.72E+00 | 9.30E-01 | no | | intergenic_variant | MODIFIER |
| 13 | 54717001 | AX-170301516 | 1 | G | A | 0.509 | -4.85E-03 | 6.64E-02 | 2.64E+00 | 2.76E+00 | 9.30E-01 | no | Cdhr2 | synonymous_variant | LOW |
| 5 | 89373654 | AX-170025559 | 0 | G | A | 0.43 | 8.91E-03 | 1.06E-01 | 2.68E+00 | 2.74E+00 | 9.30E-01 | no | | intergenic_variant | MODIFIER |
| 4 | 134368988 | AX-170070758 | 1 | T | C | 0.152 | -5.67E-03 | 1.01E-01 | 2.65E+00 | 2.79E+00 | 9.30E-01 | no | Extl1 | intron_variant | MODIFIER |
| 1 | 99745513 | AX-97812492 | 0 | T | C | 0.149 | 1.49E-02 | 1.69E-01 | 2.63E+00 | 2.74E+00 | 9.30E-01 | no | | intergenic_variant | MODIFIER |
| 18 | 52300313 | AX-170509703 | 0 | C | G | 0.044 | -2.20E-02 | 2.29E-01 | 2.61E+00 | 2.73E+00 | 9.30E-01 | no | | intergenic_variant | MODIFIER |
| 5 | 121692616 | AX-170101136 | 1 | A | G | 0.464 | 1.02E-02 | 1.06E-01 | 2.65E+00 | 2.72E+00 | 9.30E-01 | no | | intergenic_variant | MODIFIER |
| 10 | 114899877 | AX-170192589 | 0 | G | A | 0.36 | -6.80E-03 | 7.63E-02 | 2.64E+00 | 2.73E+00 | 9.31E-01 | no | | intergenic_variant | MODIFIER |
| 15 | 68573695 | AX-170492332 | 0 | T | C | 0.351 | 4.62E-03 | 7.35E-02 | 2.67E+00 | 2.78E+00 | 9.31E-01 | no | | intergenic_variant | MODIFIER |
| 9 | 61853558 | AX-170440855 | 0 | A | G | 0.193 | 1.10E-02 | 1.33E-01 | 2.70E+00 | 2.75E+00 | 9.31E-01 | no | Gm19208 | downstream_gene_variant | MODIFIER |
| 1 | 3135620 | AX-169999822 | 0 | C | A | 0.518 | -4.73E-03 | 6.82E-02 | 2.68E+00 | 2.77E+00 | 9.31E-01 | no | | intergenic_variant | MODIFIER |
| 2 | 139040386 | AX-97851934 | 0 | C | T | 0.123 | -1.40E-02 | 1.81E-01 | 2.65E+00 | 2.75E+00 | 9.31E-01 | no | | intergenic_variant | MODIFIER |
| 11 | 8964973 | AX-170004240 | 1 | C | T | 0.062 | 1.73E-02 | 2.14E-01 | 2.65E+00 | 2.75E+00 | 9.31E-01 | no | Pkd1l1 | intron_variant | MODIFIER |
| 1 | 52207461 | AX-170316769 | 1 | T | A | 0.134 | -2.02E-02 | 2.41E-01 | 2.70E+00 | 2.74E+00 | 9.31E-01 | no | Gls | intron_variant | MODIFIER |
| 9 | 106085055 | AX-170491224 | 0 | A | G | 0.132 | 8.76E-03 | 1.13E-01 | 2.68E+00 | 2.76E+00 | 9.31E-01 | no | Col6a4 | intron_variant | MODIFIER |
| 2 | 143271983 | AX-170172937 | 0 | T | C | 0.14 | -1.84E-02 | 2.21E-01 | 2.72E+00 | 2.75E+00 | 9.31E-01 | no | | intergenic_variant | MODIFIER |
| 4 | 149401937 | AX-170505641 | 1 | A | G | 0.089 | 1.67E-02 | 1.92E-01 | 2.65E+00 | 2.74E+00 | 9.31E-01 | no | Ube4b | intron_variant,non_coding_transcript_variant | MODIFIER |
| 7 | 75116300 | AX-97809835 | 0 | G | A | 0.368 | -7.76E-03 | 9.49E-02 | 2.64E+00 | 2.74E+00 | 9.31E-01 | no | Sv2b | 3_prime_UTR_variant | MODIFIER |
| 7 | 76546636 | AX-170029334 | 0 | A | C | 0.368 | -7.76E-03 | 9.49E-02 | 2.64E+00 | 2.74E+00 | 9.31E-01 | no | Agbl1 | intron_variant | MODIFIER |
| 1 | 4915812 | AX-170284874 | 0 | C | A | 0.421 | 6.75E-03 | 7.02E-02 | 2.65E+00 | 2.72E+00 | 9.32E-01 | no | Rgs20 | intron_variant | MODIFIER |
| 1 | 28959399 | AX-170000186 | 0 | T | C | 0.386 | -6.56E-03 | 8.00E-02 | 2.65E+00 | 2.74E+00 | 9.32E-01 | no | | intergenic_variant | MODIFIER |
| 15 | 92339256 | AX-170507328 | 0 | G | A | 0.149 | 1.61E-02 | 1.78E-01 | 2.66E+00 | 2.73E+00 | 9.32E-01 | no | Cntn1 | intron_variant | MODIFIER |
| 9 | 35026327 | AX-170196491 | 0 | G | A | 0.43 | 7.86E-03 | 8.65E-02 | 2.66E+00 | 2.73E+00 | 9.32E-01 | no | Kirrel3 | intron_variant | MODIFIER |
| 9 | 35400942 | AX-170273622 | 0 | T | C | 0.43 | 7.86E-03 | 8.65E-02 | 2.66E+00 | 2.73E+00 | 9.32E-01 | no | | regulatory_region_variant | MODIFIER |
| 9 | 35517010 | AX-170292976 | 0 | G | A | 0.43 | 7.86E-03 | 8.65E-02 | 2.66E+00 | 2.73E+00 | 9.32E-01 | no | Gm1113 | synonymous_variant | LOW |
| 4 | 149762577 | AX-170281786 | 1 | G | A | 0.098 | 1.66E-02 | 1.84E-01 | 2.63E+00 | 2.73E+00 | 9.32E-01 | no | Slc25a33 | intron_variant | MODIFIER |
| 1 | 66513040 | AX-170144998 | 1 | T | C | 0.116 | 2.00E-02 | 2.30E-01 | 2.68E+00 | 2.73E+00 | 9.32E-01 | no | Unc80 | intron_variant | MODIFIER |
| 20 | 112905960 | AX-170508768 | 0 | T | C | 0.289 | -4.41E-03 | 5.36E-02 | 2.62E+00 | 2.74E+00 | 9.32E-01 | no | | intergenic_variant | MODIFIER |

| | | | | | | | | | | | | | | | |
|---|---|---|---|---|---|---|---|---|---|---|---|---|---|---|---|
| 20 | 113081775 | AX-97842271 | 0 | G | A | 0.289 | -4.41E-03 | 5.36E-02 | 2.62E+00 | 2.74E+00 | 9.32E-01 | no | Chm | intron_variant | MODIFIER |
| 4 | 137158506 | AX-170024038 | 1 | T | C | 0.161 | 1.10E-02 | 1.12E-01 | 2.64E+00 | 2.71E+00 | 9.32E-01 | no | Gm13003 | downstream_gene_variant | MODIFIER |
| 10 | 8969546 | AX-170188112 | 0 | C | T | 0.211 | 8.11E-03 | 1.02E-01 | 2.65E+00 | 2.75E+00 | 9.32E-01 | no | Gm48728 | downstream_gene_variant | MODIFIER |
| 1 | 4923356 | AX-170035830 | 0 | A | G | 0.096 | 1.53E-02 | 1.72E-01 | 2.62E+00 | 2.73E+00 | 9.32E-01 | no | Rgs20 | intron_variant | MODIFIER |
| 1 | 5177304 | AX-170040271 | 0 | G | A | 0.096 | 1.53E-02 | 1.72E-01 | 2.62E+00 | 2.73E+00 | 9.32E-01 | no | | intergenic_variant | MODIFIER |
| 1 | 5208497 | AX-170040269 | 0 | C | T | 0.096 | 1.53E-02 | 1.72E-01 | 2.62E+00 | 2.73E+00 | 9.32E-01 | no | | intergenic_variant | MODIFIER |
| 1 | 5515863 | AX-97783375 | 0 | A | T | 0.096 | 1.53E-02 | 1.72E-01 | 2.62E+00 | 2.73E+00 | 9.32E-01 | no | | intergenic_variant | MODIFIER |
| 1 | 5781130 | AX-170127412 | 0 | T | G | 0.096 | 1.53E-02 | 1.72E-01 | 2.62E+00 | 2.73E+00 | 9.32E-01 | no | | intergenic_variant | MODIFIER |
| 1 | 5784429 | AX-170490435 | 0 | A | T | 0.096 | 1.53E-02 | 1.72E-01 | 2.62E+00 | 2.73E+00 | 9.32E-01 | no | | intergenic_variant | MODIFIER |
| 1 | 5784460 | AX-97815024 | 0 | A | G | 0.096 | 1.53E-02 | 1.72E-01 | 2.62E+00 | 2.73E+00 | 9.32E-01 | no | | intergenic_variant | MODIFIER |
| 1 | 5794253 | AX-170040283 | 0 | C | T | 0.096 | 1.53E-02 | 1.72E-01 | 2.62E+00 | 2.73E+00 | 9.32E-01 | no | | intergenic_variant | MODIFIER |
| 1 | 5814590 | AX-170500064 | 0 | G | A | 0.096 | 1.53E-02 | 1.72E-01 | 2.62E+00 | 2.73E+00 | 9.32E-01 | no | | intergenic_variant | MODIFIER |
| 1 | 5815807 | AX-170524255 | 0 | A | G | 0.096 | 1.53E-02 | 1.72E-01 | 2.62E+00 | 2.73E+00 | 9.32E-01 | no | | intergenic_variant | MODIFIER |
| 1 | 5830704 | AX-170086400 | 0 | G | T | 0.096 | 1.53E-02 | 1.72E-01 | 2.62E+00 | 2.73E+00 | 9.32E-01 | no | | intergenic_variant | MODIFIER |
| 1 | 5856524 | AX-170100664 | 0 | A | T | 0.096 | 1.53E-02 | 1.72E-01 | 2.62E+00 | 2.73E+00 | 9.32E-01 | no | | intergenic_variant | MODIFIER |
| 1 | 5866956 | AX-170119083 | 0 | T | G | 0.096 | 1.53E-02 | 1.72E-01 | 2.62E+00 | 2.73E+00 | 9.32E-01 | no | | intergenic_variant | MODIFIER |
| 1 | 5880896 | AX-97819484 | 0 | G | A | 0.096 | 1.53E-02 | 1.72E-01 | 2.62E+00 | 2.73E+00 | 9.32E-01 | no | | intergenic_variant | MODIFIER |
| 1 | 5958611 | AX-170241144 | 0 | G | T | 0.096 | 1.53E-02 | 1.72E-01 | 2.62E+00 | 2.73E+00 | 9.32E-01 | no | | intergenic_variant | MODIFIER |
| 1 | 6036788 | AX-170334046 | 0 | C | A | 0.096 | 1.53E-02 | 1.72E-01 | 2.62E+00 | 2.73E+00 | 9.32E-01 | no | | intergenic_variant | MODIFIER |
| 1 | 6079319 | AX-170040288 | 0 | T | C | 0.096 | 1.53E-02 | 1.72E-01 | 2.62E+00 | 2.73E+00 | 9.32E-01 | no | | intergenic_variant | MODIFIER |
| 1 | 6079479 | AX-170431356 | 0 | T | C | 0.096 | 1.53E-02 | 1.72E-01 | 2.62E+00 | 2.73E+00 | 9.32E-01 | no | | intergenic_variant | MODIFIER |
| 1 | 6079953 | AX-170040289 | 0 | G | A | 0.096 | 1.53E-02 | 1.72E-01 | 2.62E+00 | 2.73E+00 | 9.32E-01 | no | | intergenic_variant | MODIFIER |
| 1 | 6092224 | AX-170415974 | 0 | A | G | 0.096 | 1.53E-02 | 1.72E-01 | 2.62E+00 | 2.73E+00 | 9.32E-01 | no | | intergenic_variant | MODIFIER |
| 1 | 6102884 | AX-170369648 | 0 | G | C | 0.096 | 1.53E-02 | 1.72E-01 | 2.62E+00 | 2.73E+00 | 9.32E-01 | no | | intergenic_variant | MODIFIER |
| 1 | 6154188 | AX-170187255 | 0 | A | G | 0.096 | 1.53E-02 | 1.72E-01 | 2.62E+00 | 2.73E+00 | 9.32E-01 | no | | intergenic_variant | MODIFIER |
| 1 | 6183007 | AX-97810621 | 0 | T | G | 0.096 | 1.53E-02 | 1.72E-01 | 2.62E+00 | 2.73E+00 | 9.32E-01 | no | | intergenic_variant | MODIFIER |
| 1 | 6192704 | AX-170157941 | 0 | A | G | 0.096 | 1.53E-02 | 1.72E-01 | 2.62E+00 | 2.73E+00 | 9.32E-01 | no | | intergenic_variant | MODIFIER |
| 1 | 6237609 | AX-170091379 | 0 | A | G | 0.096 | 1.53E-02 | 1.72E-01 | 2.62E+00 | 2.73E+00 | 9.32E-01 | no | Rb1cc1 | intron_variant | MODIFIER |
| 1 | 6575099 | AX-170493398 | 0 | T | C | 0.096 | 1.53E-02 | 1.72E-01 | 2.62E+00 | 2.73E+00 | 9.32E-01 | no | St18 | intron_variant | MODIFIER |
| 1 | 6980263 | AX-170187458 | 0 | A | G | 0.096 | 1.53E-02 | 1.72E-01 | 2.62E+00 | 2.73E+00 | 9.32E-01 | no | | intergenic_variant | MODIFIER |
| 1 | 6980387 | AX-97819183 | 0 | C | A | 0.096 | 1.53E-02 | 1.72E-01 | 2.62E+00 | 2.73E+00 | 9.32E-01 | no | | intergenic_variant | MODIFIER |
| 1 | 7002118 | AX-170146327 | 0 | A | G | 0.096 | 1.53E-02 | 1.72E-01 | 2.62E+00 | 2.73E+00 | 9.32E-01 | no | Gm37225 | downstream_gene_variant | MODIFIER |
| 1 | 7008006 | AX-97835058 | 0 | T | C | 0.096 | 1.53E-02 | 1.72E-01 | 2.62E+00 | 2.73E+00 | 9.32E-01 | no | | intergenic_variant | MODIFIER |
| 1 | 7010895 | AX-170525061 | 0 | C | T | 0.096 | 1.53E-02 | 1.72E-01 | 2.62E+00 | 2.73E+00 | 9.32E-01 | no | Gm37489 | upstream_gene_variant | MODIFIER |
| 1 | 7027302 | AX-97854081 | 0 | T | G | 0.096 | 1.53E-02 | 1.72E-01 | 2.62E+00 | 2.73E+00 | 9.32E-01 | no | | intergenic_variant | MODIFIER |
| 1 | 7059737 | AX-170142634 | 0 | G | T | 0.096 | 1.53E-02 | 1.72E-01 | 2.62E+00 | 2.73E+00 | 9.32E-01 | no | | intergenic_variant | MODIFIER |
| 1 | 7102666 | AX-170519951 | 0 | A | G | 0.096 | 1.53E-02 | 1.72E-01 | 2.62E+00 | 2.73E+00 | 9.32E-01 | no | Pcmtd1 | intron_variant | MODIFIER |
| 1 | 7102867 | AX-170299209 | 0 | T | C | 0.096 | 1.53E-02 | 1.72E-01 | 2.62E+00 | 2.73E+00 | 9.32E-01 | no | Pcmtd1 | intron_variant | MODIFIER |
| 1 | 7211931 | AX-97791010 | 0 | T | C | 0.096 | 1.53E-02 | 1.72E-01 | 2.62E+00 | 2.73E+00 | 9.32E-01 | no | | intergenic_variant | MODIFIER |
| 1 | 7221702 | AX-170337784 | 0 | A | G | 0.096 | 1.53E-02 | 1.72E-01 | 2.62E+00 | 2.73E+00 | 9.32E-01 | no | | intergenic_variant | MODIFIER |
| 1 | 7221952 | AX-170285556 | 0 | T | C | 0.096 | 1.53E-02 | 1.72E-01 | 2.62E+00 | 2.73E+00 | 9.32E-01 | no | | intergenic_variant | MODIFIER |
| 1 | 7230081 | AX-97844676 | 0 | G | A | 0.096 | 1.53E-02 | 1.72E-01 | 2.62E+00 | 2.73E+00 | 9.32E-01 | no | | intergenic_variant | MODIFIER |
| 1 | 7251684 | AX-169999884 | 0 | T | C | 0.096 | 1.53E-02 | 1.72E-01 | 2.62E+00 | 2.73E+00 | 9.32E-01 | no | | intergenic_variant | MODIFIER |
| 1 | 7311185 | AX-169999882 | 0 | A | G | 0.096 | 1.53E-02 | 1.72E-01 | 2.62E+00 | 2.73E+00 | 9.32E-01 | no | Gm19002 | upstream_gene_variant | MODIFIER |
| 1 | 8083912 | AX-170161561 | 0 | G | A | 0.096 | 1.53E-02 | 1.72E-01 | 2.62E+00 | 2.73E+00 | 9.32E-01 | no | | intergenic_variant | MODIFIER |
| 1 | 8100226 | AX-170035837 | 0 | C | G | 0.096 | 1.53E-02 | 1.72E-01 | 2.62E+00 | 2.73E+00 | 9.32E-01 | no | | intergenic_variant | MODIFIER |
| 1 | 9514170 | AX-170347181 | 0 | T | C | 0.096 | 1.53E-02 | 1.72E-01 | 2.62E+00 | 2.73E+00 | 9.32E-01 | no | | intergenic_variant | MODIFIER |
| 1 | 9515427 | AX-97827370 | 0 | C | T | 0.096 | 1.53E-02 | 1.72E-01 | 2.62E+00 | 2.73E+00 | 9.32E-01 | no | | intergenic_variant | MODIFIER |
| 1 | 9564127 | AX-170434202 | 0 | G | T | 0.096 | 1.53E-02 | 1.72E-01 | 2.62E+00 | 2.73E+00 | 9.32E-01 | no | Adhfe1 | intron_variant,NMD_transcript_variant | MODIFIER |
| 1 | 9567529 | AX-170530812 | 0 | G | A | 0.096 | 1.53E-02 | 1.72E-01 | 2.62E+00 | 2.73E+00 | 9.32E-01 | no | Adhfe1 | intron_variant,NMD_transcript_variant | MODIFIER |
| 1 | 9584165 | AX-170241837 | 0 | G | A | 0.096 | 1.53E-02 | 1.72E-01 | 2.62E+00 | 2.73E+00 | 9.32E-01 | no | 2610203C22Rik | intron_variant,non_coding_transcript_variant | MODIFIER |
| 1 | 9584295 | AX-170121170 | 0 | A | C | 0.096 | 1.53E-02 | 1.72E-01 | 2.62E+00 | 2.73E+00 | 9.32E-01 | no | 2610203C22Rik | intron_variant,non_coding_transcript_variant | MODIFIER |
| 1 | 9588807 | AX-170530335 | 0 | G | A | 0.096 | 1.53E-02 | 1.72E-01 | 2.62E+00 | 2.73E+00 | 9.32E-01 | no | 2610203C22Rik | intron_variant,non_coding_transcript_variant | MODIFIER |
| 1 | 9590800 | AX-170243121 | 0 | G | A | 0.096 | 1.53E-02 | 1.72E-01 | 2.62E+00 | 2.73E+00 | 9.32E-01 | no | 2610203C22Rik | intron_variant,non_coding_transcript_variant | MODIFIER |

| | | | | | | | | | | | | | | | |
|---|---|---|---|---|---|---|---|---|---|---|---|---|---|---|---|
| 1 | 9621851 | AX-170111940 | 0 | C | T | 0.096 | 1.53E-02 | 1.72E-01 | 2.62E+00 | 2.73E+00 | 9.32E-01 | no | Vxn | intron_variant | MODIFIER |
| 1 | 9627567 | AX-170439263 | 0 | G | A | 0.096 | 1.53E-02 | 1.72E-01 | 2.62E+00 | 2.73E+00 | 9.32E-01 | no | Vxn | downstream_gene_variant | MODIFIER |
| 1 | 9633187 | AX-170197534 | 0 | C | T | 0.096 | 1.53E-02 | 1.72E-01 | 2.62E+00 | 2.73E+00 | 9.32E-01 | no | 2610203C22Rik | upstream_gene_variant | MODIFIER |
| 1 | 9642553 | AX-97815036 | 0 | G | A | 0.096 | 1.53E-02 | 1.72E-01 | 2.62E+00 | 2.73E+00 | 9.32E-01 | no | Gm29520 | upstream_gene_variant | MODIFIER |
| 1 | 9672171 | AX-170228648 | 0 | C | T | 0.096 | 1.53E-02 | 1.72E-01 | 2.62E+00 | 2.73E+00 | 9.32E-01 | no | Mybl1 | intron_variant | MODIFIER |
| 1 | 9705396 | AX-170167635 | 0 | A | G | 0.096 | 1.53E-02 | 1.72E-01 | 2.62E+00 | 2.73E+00 | 9.32E-01 | no | | intergenic_variant | MODIFIER |
| 1 | 9776004 | AX-170040349 | 0 | T | C | 0.096 | 1.53E-02 | 1.72E-01 | 2.62E+00 | 2.73E+00 | 9.32E-01 | no | 1700034P13Rik | intron_variant,non_coding_transcript_variant | MODIFIER |
| 1 | 9918246 | AX-170343698 | 0 | T | G | 0.096 | 1.53E-02 | 1.72E-01 | 2.62E+00 | 2.73E+00 | 9.32E-01 | no | Mcmdc2 | intron_variant | MODIFIER |
| 1 | 9918429 | AX-97852812 | 0 | C | T | 0.096 | 1.53E-02 | 1.72E-01 | 2.62E+00 | 2.73E+00 | 9.32E-01 | no | Mcmdc2 | intron_variant | MODIFIER |
| 1 | 9940386 | AX-170220692 | 0 | G | C | 0.096 | 1.53E-02 | 1.72E-01 | 2.62E+00 | 2.73E+00 | 9.32E-01 | no | Mcmdc2 | missense_variant | MODERATE |
| 1 | 9953187 | AX-170281891 | 0 | T | C | 0.096 | 1.53E-02 | 1.72E-01 | 2.62E+00 | 2.73E+00 | 9.32E-01 | no | | regulatory_region_variant | MODIFIER |
| 1 | 9962375 | AX-170156123 | 0 | A | G | 0.096 | 1.53E-02 | 1.72E-01 | 2.62E+00 | 2.73E+00 | 9.32E-01 | no | Tcf24 | 5_prime_UTR_variant | MODIFIER |
| 1 | 9972901 | AX-97848268 | 0 | A | G | 0.096 | 1.53E-02 | 1.72E-01 | 2.62E+00 | 2.73E+00 | 9.32E-01 | no | Ppp1r42 | intron_variant | MODIFIER |
| 1 | 9978791 | AX-170040354 | 0 | G | A | 0.096 | 1.53E-02 | 1.72E-01 | 2.62E+00 | 2.73E+00 | 9.32E-01 | no | Ppp1r42 | intron_variant | MODIFIER |
| 1 | 10004435 | AX-170234624 | 0 | A | G | 0.096 | 1.53E-02 | 1.72E-01 | 2.62E+00 | 2.73E+00 | 9.32E-01 | no | Ppp1r42 | intron_variant | MODIFIER |
| 1 | 10004484 | AX-170476044 | 0 | C | A | 0.096 | 1.53E-02 | 1.72E-01 | 2.62E+00 | 2.73E+00 | 9.32E-01 | no | Ppp1r42 | intron_variant | MODIFIER |
| 1 | 10005845 | AX-170305839 | 0 | C | T | 0.096 | 1.53E-02 | 1.72E-01 | 2.62E+00 | 2.73E+00 | 9.32E-01 | no | Ppp1r42 | intron_variant | MODIFIER |
| 1 | 10025331 | AX-170418423 | 0 | C | T | 0.096 | 1.53E-02 | 1.72E-01 | 2.62E+00 | 2.73E+00 | 9.32E-01 | no | Cops5 | intron_variant | MODIFIER |
| 1 | 10028816 | AX-170459416 | 0 | C | T | 0.096 | 1.53E-02 | 1.72E-01 | 2.62E+00 | 2.73E+00 | 9.32E-01 | no | Cops5 | intron_variant | MODIFIER |
| 1 | 10061395 | AX-170316978 | 0 | T | C | 0.096 | 1.53E-02 | 1.72E-01 | 2.62E+00 | 2.73E+00 | 9.32E-01 | no | Cspp1 | intron_variant | MODIFIER |
| 1 | 10114953 | AX-170308136 | 0 | C | T | 0.096 | 1.53E-02 | 1.72E-01 | 2.62E+00 | 2.73E+00 | 9.32E-01 | no | Cspp1 | intron_variant | MODIFIER |
| 1 | 10135944 | AX-97822419 | 0 | C | T | 0.096 | 1.53E-02 | 1.72E-01 | 2.62E+00 | 2.73E+00 | 9.32E-01 | no | Cspp1 | synonymous_variant | LOW |
| 1 | 10149240 | AX-170197079 | 0 | T | G | 0.096 | 1.53E-02 | 1.72E-01 | 2.62E+00 | 2.73E+00 | 9.32E-01 | no | Arfgef1 | intron_variant | MODIFIER |
| 1 | 10153471 | AX-170202973 | 0 | A | G | 0.096 | 1.53E-02 | 1.72E-01 | 2.62E+00 | 2.73E+00 | 9.32E-01 | no | Arfgef1 | synonymous_variant | LOW |
| 1 | 10153548 | AX-170156464 | 0 | T | C | 0.096 | 1.53E-02 | 1.72E-01 | 2.62E+00 | 2.73E+00 | 9.32E-01 | no | Arfgef1 | intron_variant | MODIFIER |
| 1 | 10155224 | AX-170414807 | 0 | T | C | 0.096 | 1.53E-02 | 1.72E-01 | 2.62E+00 | 2.73E+00 | 9.32E-01 | no | Arfgef1 | intron_variant | MODIFIER |
| 1 | 10210223 | AX-170040359 | 0 | T | C | 0.096 | 1.53E-02 | 1.72E-01 | 2.62E+00 | 2.73E+00 | 9.32E-01 | no | Arfgef1 | intron_variant | MODIFIER |
| 1 | 10212909 | AX-170349535 | 0 | G | T | 0.096 | 1.53E-02 | 1.72E-01 | 2.62E+00 | 2.73E+00 | 9.32E-01 | no | Arfgef1 | intron_variant | MODIFIER |
| 1 | 10213368 | AX-170091396 | 0 | T | G | 0.096 | 1.53E-02 | 1.72E-01 | 2.62E+00 | 2.73E+00 | 9.32E-01 | no | Arfgef1 | intron_variant | MODIFIER |
| 1 | 10218984 | AX-170101479 | 0 | G | T | 0.096 | 1.53E-02 | 1.72E-01 | 2.62E+00 | 2.73E+00 | 9.32E-01 | no | Arfgef1 | intron_variant | MODIFIER |
| 1 | 10257088 | AX-169999926 | 0 | C | A | 0.096 | 1.53E-02 | 1.72E-01 | 2.62E+00 | 2.73E+00 | 9.32E-01 | no | | regulatory_region_variant | MODIFIER |
| 1 | 10259828 | AX-170477843 | 0 | A | G | 0.096 | 1.53E-02 | 1.72E-01 | 2.62E+00 | 2.73E+00 | 9.32E-01 | no | | intergenic_variant | MODIFIER |
| 1 | 10259922 | AX-170152345 | 0 | G | A | 0.096 | 1.53E-02 | 1.72E-01 | 2.62E+00 | 2.73E+00 | 9.32E-01 | no | | intergenic_variant | MODIFIER |
| 1 | 10260830 | AX-170148669 | 0 | T | C | 0.096 | 1.53E-02 | 1.72E-01 | 2.62E+00 | 2.73E+00 | 9.32E-01 | no | | regulatory_region_variant | MODIFIER |
| 1 | 10267631 | AX-170204202 | 0 | C | G | 0.096 | 1.53E-02 | 1.72E-01 | 2.62E+00 | 2.73E+00 | 9.32E-01 | no | | intergenic_variant | MODIFIER |
| 1 | 10276733 | AX-170170265 | 0 | C | T | 0.096 | 1.53E-02 | 1.72E-01 | 2.62E+00 | 2.73E+00 | 9.32E-01 | no | | intergenic_variant | MODIFIER |
| 1 | 10279703 | AX-170360172 | 0 | A | C | 0.096 | 1.53E-02 | 1.72E-01 | 2.62E+00 | 2.73E+00 | 9.32E-01 | no | | intergenic_variant | MODIFIER |
| 1 | 10280180 | AX-170518356 | 0 | C | G | 0.096 | 1.53E-02 | 1.72E-01 | 2.62E+00 | 2.73E+00 | 9.32E-01 | no | | regulatory_region_variant | MODIFIER |
| 1 | 10314058 | AX-170384862 | 0 | T | C | 0.096 | 1.53E-02 | 1.72E-01 | 2.62E+00 | 2.73E+00 | 9.32E-01 | no | | intergenic_variant | MODIFIER |
| 1 | 10327023 | AX-170434620 | 0 | T | A | 0.096 | 1.53E-02 | 1.72E-01 | 2.62E+00 | 2.73E+00 | 9.32E-01 | no | Cpa6 | intron_variant | MODIFIER |
| 1 | 10328584 | AX-170461531 | 0 | C | T | 0.096 | 1.53E-02 | 1.72E-01 | 2.62E+00 | 2.73E+00 | 9.32E-01 | no | Cpa6 | intron_variant | MODIFIER |
| 1 | 11029205 | AX-170040369 | 0 | A | C | 0.096 | 1.53E-02 | 1.72E-01 | 2.62E+00 | 2.73E+00 | 9.32E-01 | no | Prex2 | intron_variant | MODIFIER |
| 1 | 11038063 | AX-170385050 | 0 | A | T | 0.096 | 1.53E-02 | 1.72E-01 | 2.62E+00 | 2.73E+00 | 9.32E-01 | no | Prex2 | intron_variant | MODIFIER |
| 1 | 11048021 | AX-170333147 | 0 | G | A | 0.096 | 1.53E-02 | 1.72E-01 | 2.62E+00 | 2.73E+00 | 9.32E-01 | no | Prex2 | intron_variant | MODIFIER |
| 1 | 11053838 | AX-170330521 | 0 | C | G | 0.096 | 1.53E-02 | 1.72E-01 | 2.62E+00 | 2.73E+00 | 9.32E-01 | no | Prex2 | intron_variant | MODIFIER |
| 1 | 11084542 | AX-170111243 | 0 | T | G | 0.096 | 1.53E-02 | 1.72E-01 | 2.62E+00 | 2.73E+00 | 9.32E-01 | no | Prex2 | intron_variant | MODIFIER |
| 1 | 11118188 | AX-170505661 | 0 | T | C | 0.096 | 1.53E-02 | 1.72E-01 | 2.62E+00 | 2.73E+00 | 9.32E-01 | no | Prex2 | intron_variant | MODIFIER |
| 1 | 11136295 | AX-170208840 | 0 | T | C | 0.096 | 1.53E-02 | 1.72E-01 | 2.62E+00 | 2.73E+00 | 9.32E-01 | no | Prex2 | intron_variant | MODIFIER |
| 1 | 11139652 | AX-170278326 | 0 | C | G | 0.096 | 1.53E-02 | 1.72E-01 | 2.62E+00 | 2.73E+00 | 9.32E-01 | no | Prex2 | intron_variant | MODIFIER |
| 6 | 139764980 | AX-170522286 | 1 | C | T | 0.071 | 2.32E-02 | 2.77E-01 | 2.70E+00 | 2.74E+00 | 9.32E-01 | no | Pik3c2g | intron_variant | MODIFIER |
| 20 | 113740182 | AX-170465002 | 0 | T | C | 0.325 | 4.64E-03 | 5.26E-02 | 2.62E+00 | 2.73E+00 | 9.32E-01 | no | Dach2 | intron_variant | MODIFIER |
| 20 | 114276123 | AX-170090642 | 0 | A | G | 0.325 | 4.64E-03 | 5.26E-02 | 2.62E+00 | 2.73E+00 | 9.32E-01 | no | | intergenic_variant | MODIFIER |
| 20 | 114376069 | AX-170142584 | 0 | C | T | 0.325 | 4.64E-03 | 5.26E-02 | 2.62E+00 | 2.73E+00 | 9.32E-01 | no | | intergenic_variant | MODIFIER |
| 10 | 9424683 | AX-170090752 | 0 | T | C | 0.193 | 8.03E-03 | 1.06E-01 | 2.67E+00 | 2.76E+00 | 9.32E-01 | no | | intergenic_variant | MODIFIER |

| | | | | | | | | | | | | | | | |
|---|---|---|---|---|---|---|---|---|---|---|---|---|---|---|---|
| 2 | 89660670 | AX-97816028 | 0 | C | A | 0.061 | 1.19E-02 | 1.48E-01 | 2.58E+00 | 2.74E+00 | 9.32E-01 | no | Olfr1250 | upstream_gene_variant | MODIFIER |
| 5 | 102629787 | AX-170319233 | 0 | G | C | 0.43 | -9.42E-03 | 1.11E-01 | 2.68E+00 | 2.73E+00 | 9.32E-01 | no | Arhgap24 | intron_variant | MODIFIER |
| 1 | 55558404 | AX-170086471 | 2 | G | A | 0.109 | 2.03E-02 | 2.39E-01 | 2.69E+00 | 2.73E+00 | 9.32E-01 | no | Plcl1 | intron_variant | MODIFIER |
| 10 | 119011110 | AX-170488160 | 0 | T | C | 0.342 | -6.13E-03 | 7.06E-02 | 2.64E+00 | 2.73E+00 | 9.32E-01 | no | | regulatory_region_variant | MODIFIER |
| 10 | 28410666 | AX-170129982 | 2 | C | T | 0.173 | 6.73E-03 | 9.04E-02 | 2.62E+00 | 2.75E+00 | 9.32E-01 | no | Ptprk | intron_variant | MODIFIER |
| 1 | 68527025 | AX-170376493 | 0 | G | A | 0.123 | 1.65E-02 | 1.84E-01 | 2.64E+00 | 2.73E+00 | 9.32E-01 | no | Erbb4 | intron_variant | MODIFIER |
| 1 | 149411662 | AX-170091741 | 1 | A | G | 0.196 | -9.76E-03 | 1.29E-01 | 2.67E+00 | 2.75E+00 | 9.32E-01 | no | Gm29398 | downstream_gene_variant | MODIFIER |
| 3 | 156095280 | AX-170068534 | 0 | A | G | 0.325 | 6.11E-03 | 7.46E-02 | 2.66E+00 | 2.74E+00 | 9.32E-01 | no | | intergenic_variant | MODIFIER |
| 20 | 116117465 | AX-170480007 | 0 | G | A | 0.307 | -4.23E-03 | 5.35E-02 | 2.62E+00 | 2.74E+00 | 9.33E-01 | no | | intergenic_variant | MODIFIER |
| 5 | 100680886 | AX-97836436 | 0 | A | G | 0.421 | -8.23E-03 | 1.04E-01 | 2.69E+00 | 2.75E+00 | 9.33E-01 | no | Hpse | synonymous_variant | LOW |
| 8 | 30490775 | AX-97845360 | 0 | C | T | 0.175 | 9.20E-03 | 1.12E-01 | 2.68E+00 | 2.74E+00 | 9.33E-01 | no | Gm45489 | intron_variant,non_coding_transcript_variant | MODIFIER |
| 1 | 151771045 | AX-170043145 | 1 | C | T | 0.205 | 1.20E-02 | 1.28E-01 | 2.64E+00 | 2.72E+00 | 9.33E-01 | no | Edem3 | intron_variant | MODIFIER |
| 7 | 112320278 | AX-170078445 | 0 | A | G | 0.447 | 7.96E-03 | 9.26E-02 | 2.66E+00 | 2.73E+00 | 9.33E-01 | no | Mical2 | intron_variant | MODIFIER |
| 7 | 51425944 | AX-170429409 | 0 | C | A | 0.43 | 8.54E-03 | 9.65E-02 | 2.65E+00 | 2.73E+00 | 9.33E-01 | no | | intergenic_variant | MODIFIER |
| 7 | 52310406 | AX-170250338 | 0 | A | T | 0.43 | 8.54E-03 | 9.65E-02 | 2.65E+00 | 2.73E+00 | 9.33E-01 | no | | intergenic_variant | MODIFIER |
| 11 | 33824508 | AX-97815065 | 0 | C | T | 0.105 | 1.65E-02 | 1.88E-01 | 2.65E+00 | 2.73E+00 | 9.33E-01 | no | Kcnip1 | intron_variant | MODIFIER |
| 8 | 73067983 | AX-170304761 | 1 | C | T | 0.196 | -1.15E-02 | 1.35E-01 | 2.67E+00 | 2.73E+00 | 9.33E-01 | no | Large1 | intron_variant | MODIFIER |
| 1 | 66228544 | AX-170516987 | 1 | C | T | 0.116 | 1.98E-02 | 2.35E-01 | 2.69E+00 | 2.73E+00 | 9.33E-01 | no | Map2 | intron_variant | MODIFIER |
| 1 | 67032234 | AX-170201643 | 1 | A | G | 0.116 | 1.98E-02 | 2.35E-01 | 2.69E+00 | 2.73E+00 | 9.33E-01 | no | Lancl1 | intron_variant | MODIFIER |
| 1 | 46184913 | AX-170444144 | 1 | G | A | 0.348 | 5.68E-03 | 7.45E-02 | 2.65E+00 | 2.75E+00 | 9.33E-01 | no | Dnah7b | intron_variant | MODIFIER |
| 8 | 50195331 | AX-170383322 | 0 | C | T | 0.158 | -1.27E-02 | 1.38E-01 | 2.65E+00 | 2.72E+00 | 9.33E-01 | no | Gm45341 | intron_variant,non_coding_transcript_variant | MODIFIER |
| 16 | 88933666 | AX-170013620 | 1 | C | T | 0.473 | -6.42E-03 | 7.88E-02 | 2.66E+00 | 2.74E+00 | 9.33E-01 | no | Krtap19-9b | upstream_gene_variant | MODIFIER |
| 16 | 95121560 | AX-97798373 | 0 | T | C | 0.439 | 5.28E-03 | 7.53E-02 | 2.66E+00 | 2.76E+00 | 9.33E-01 | no | | intergenic_variant | MODIFIER |
| 5 | 6897013 | AX-97850408 | 0 | T | G | 0.43 | -6.08E-03 | 6.70E-02 | 2.60E+00 | 2.73E+00 | 9.33E-01 | no | Zfp804b | intron_variant | MODIFIER |
| 3 | 145237338 | AX-170428949 | 1 | T | G | 0.366 | -5.30E-03 | 6.94E-02 | 2.66E+00 | 2.75E+00 | 9.34E-01 | no | | intergenic_variant | MODIFIER |
| 7 | 27251194 | AX-170178386 | 2 | G | A | 0.391 | 8.68E-03 | 8.85E-02 | 2.61E+00 | 2.72E+00 | 9.34E-01 | no | Coq8b | intron_variant | MODIFIER |
| 1 | 60600687 | AX-170132176 | 2 | G | A | 0.109 | -1.88E-02 | 2.34E-01 | 2.69E+00 | 2.74E+00 | 9.34E-01 | no | Gm38388 | upstream_gene_variant | MODIFIER |
| 10 | 114873875 | AX-170367820 | 0 | A | T | 0.351 | 5.73E-03 | 7.39E-02 | 2.64E+00 | 2.74E+00 | 9.34E-01 | no | Gm31182 | upstream_gene_variant | MODIFIER |
| 18 | 78643981 | AX-170359641 | 1 | A | G | 0.402 | -7.80E-03 | 8.70E-02 | 2.64E+00 | 2.73E+00 | 9.34E-01 | no | Gm23895 | upstream_gene_variant | MODIFIER |
| 16 | 26747261 | AX-170219715 | 0 | T | C | 0.246 | -6.77E-03 | 8.64E-02 | 2.61E+00 | 2.74E+00 | 9.34E-01 | no | Gm20319 | intron_variant,non_coding_transcript_variant | MODIFIER |
| 2 | 107617341 | AX-170262590 | 1 | A | G | 0.054 | -1.67E-02 | 1.88E-01 | 2.61E+00 | 2.73E+00 | 9.34E-01 | no | | intergenic_variant | MODIFIER |
| 6 | 104886037 | AX-170267091 | 2 | G | A | 0.073 | 2.26E-02 | 2.77E-01 | 2.70E+00 | 2.74E+00 | 9.34E-01 | no | | intergenic_variant | MODIFIER |
| 10 | 19370883 | AX-170339487 | 0 | T | A | 0.412 | -7.22E-03 | 6.72E-02 | 2.58E+00 | 2.71E+00 | 9.34E-01 | no | | intergenic_variant | MODIFIER |
| 5 | 117757243 | AX-168417250 | 1 | C | T | 0.464 | -8.36E-03 | 1.07E-01 | 2.68E+00 | 2.75E+00 | 9.34E-01 | no | Ksr2 | intron_variant | MODIFIER |
| 1 | 71310835 | AX-97782537 | 1 | A | C | 0.134 | -1.87E-02 | 2.36E-01 | 2.67E+00 | 2.74E+00 | 9.34E-01 | no | Abca12 | intron_variant | MODIFIER |
| 1 | 77673575 | AX-170175580 | 1 | A | G | 0.134 | -1.87E-02 | 2.36E-01 | 2.67E+00 | 2.74E+00 | 9.34E-01 | no | | intergenic_variant | MODIFIER |
| 12 | 75356803 | AX-170444429 | 1 | C | T | 0.045 | -2.30E-02 | 2.74E-01 | 2.68E+00 | 2.73E+00 | 9.34E-01 | no | Rhoj | intron_variant | MODIFIER |
| 4 | 149581390 | AX-97845520 | 0 | C | T | 0.088 | 1.57E-02 | 1.77E-01 | 2.63E+00 | 2.73E+00 | 9.34E-01 | no | | intergenic_variant | MODIFIER |
| 4 | 137040661 | AX-170086118 | 0 | A | T | 0.158 | 1.07E-02 | 1.09E-01 | 2.64E+00 | 2.71E+00 | 9.34E-01 | no | Zbtb40 | intron_variant | MODIFIER |
| 9 | 76254702 | AX-170032738 | 1 | A | C | 0.188 | 1.09E-02 | 1.36E-01 | 2.72E+00 | 2.75E+00 | 9.34E-01 | no | Hcrtr2 | splice_region_variant,synonymous_variant | LOW |
| 9 | 78199783 | AX-170533763 | 1 | G | A | 0.188 | 1.09E-02 | 1.36E-01 | 2.72E+00 | 2.75E+00 | 9.34E-01 | no | Gsta4 | intron_variant | MODIFIER |
| 7 | 118790953 | AX-170531018 | 0 | A | G | 0.395 | -7.16E-03 | 9.12E-02 | 2.69E+00 | 2.75E+00 | 9.34E-01 | no | Vps35l | intron_variant | MODIFIER |
| 7 | 53044542 | AX-170428136 | 0 | T | G | 0.412 | 8.49E-03 | 9.34E-02 | 2.64E+00 | 2.72E+00 | 9.34E-01 | no | | intergenic_variant | MODIFIER |
| 12 | 109841834 | AX-170182674 | 0 | C | T | 0.149 | 1.16E-02 | 1.42E-01 | 2.60E+00 | 2.74E+00 | 9.34E-01 | no | | regulatory_region_variant | MODIFIER |
| 1 | 54838251 | AX-170451809 | 1 | T | C | 0.107 | 1.92E-02 | 2.35E-01 | 2.69E+00 | 2.74E+00 | 9.34E-01 | no | Ankrd44 | intron_variant | MODIFIER |
| 13 | 37732034 | AX-170396996 | 1 | C | T | 0.179 | -1.07E-02 | 1.15E-01 | 2.64E+00 | 2.72E+00 | 9.34E-01 | no | | regulatory_region_variant | MODIFIER |
| 7 | 57506827 | AX-170451965 | 1 | T | C | 0.429 | 8.80E-03 | 1.01E-01 | 2.66E+00 | 2.73E+00 | 9.34E-01 | no | Gabra5 | intron_variant | MODIFIER |
| 5 | 105992935 | AX-170025814 | 0 | T | A | 0.439 | 9.96E-03 | 1.13E-01 | 2.67E+00 | 2.72E+00 | 9.35E-01 | no | | intergenic_variant | MODIFIER |
| 16 | 85952386 | AX-170351049 | 1 | G | A | 0.455 | -5.91E-03 | 7.62E-02 | 2.66E+00 | 2.74E+00 | 9.35E-01 | no | | intergenic_variant | MODIFIER |
| 8 | 48955550 | AX-97803618 | 1 | C | T | 0.152 | 1.10E-02 | 1.49E-01 | 2.71E+00 | 2.76E+00 | 9.35E-01 | no | Gm25830 | downstream_gene_variant | MODIFIER |
| 6 | 110577946 | AX-97852254 | 1 | G | A | 0.071 | 2.23E-02 | 2.77E-01 | 2.70E+00 | 2.74E+00 | 9.35E-01 | no | | intergenic_variant | MODIFIER |
| 6 | 145137423 | AX-97839064 | 1 | G | A | 0.071 | 2.23E-02 | 2.77E-01 | 2.70E+00 | 2.74E+00 | 9.35E-01 | no | Lrmp | intron_variant | MODIFIER |
| 6 | 147348935 | AX-170127952 | 1 | A | G | 0.071 | 2.23E-02 | 2.77E-01 | 2.70E+00 | 2.74E+00 | 9.35E-01 | no | | intergenic_variant | MODIFIER |
| 13 | 40672873 | AX-170247607 | 0 | G | A | 0.167 | 6.81E-03 | 1.08E-01 | 2.66E+00 | 2.77E+00 | 9.35E-01 | no | | intergenic_variant | MODIFIER |

| | | | | | | | | | | | | | | | |
|---|---|---|---|---|---|---|---|---|---|---|---|---|---|---|---|
| 2 | 142021943 | AX-170250630 | 1 | G | T | 0.125 | 1.56E-02 | 1.92E-01 | 2.69E+00 | 2.74E+00 | 9.35E-01 | no | Macrod2 | intron_variant | MODIFIER |
| 1 | 172663616 | AX-170509639 | 0 | T | C | 0.228 | -7.53E-03 | 1.17E-01 | 2.68E+00 | 2.77E+00 | 9.35E-01 | no | | intergenic_variant | MODIFIER |
| 1 | 174666538 | AX-170232293 | 0 | A | G | 0.228 | -7.53E-03 | 1.17E-01 | 2.68E+00 | 2.77E+00 | 9.35E-01 | no | Fmn2 | intron_variant | MODIFIER |
| 6 | 147370540 | AX-170154552 | 1 | C | G | 0.071 | 2.22E-02 | 2.77E-01 | 2.70E+00 | 2.74E+00 | 9.35E-01 | no | | intergenic_variant | MODIFIER |
| 16 | 97985584 | AX-170532822 | 1 | T | C | 0.446 | -7.88E-03 | 7.74E-02 | 2.60E+00 | 2.72E+00 | 9.35E-01 | no | | regulatory_region_variant | MODIFIER |
| 18 | 31262619 | AX-170152592 | 2 | A | C | 0.064 | 2.40E-02 | 3.02E-01 | 2.72E+00 | 2.75E+00 | 9.35E-01 | no | Rit2 | intron_variant,NMD_transcript_variant | MODIFIER |
| 16 | 75616291 | AX-170259154 | 0 | C | T | 0.491 | -4.66E-03 | 7.48E-02 | 2.67E+00 | 2.77E+00 | 9.35E-01 | no | | intergenic_variant | MODIFIER |
| 12 | 70000277 | AX-170315103 | 1 | C | T | 0.062 | 2.41E-02 | 3.03E-01 | 2.73E+00 | 2.75E+00 | 9.35E-01 | no | | intergenic_variant | MODIFIER |
| 5 | 126042147 | AX-170382254 | 2 | T | A | 0.455 | -8.43E-03 | 1.08E-01 | 2.69E+00 | 2.75E+00 | 9.35E-01 | no | | intergenic_variant | MODIFIER |
| 9 | 54677004 | AX-170400061 | 1 | C | G | 0.205 | -9.52E-03 | 1.20E-01 | 2.68E+00 | 2.74E+00 | 9.35E-01 | no | | regulatory_region_variant | MODIFIER |
| 15 | 78731594 | AX-97829824 | 1 | G | T | 0.562 | 8.03E-03 | 8.17E-02 | 2.63E+00 | 2.71E+00 | 9.35E-01 | no | | intergenic_variant | MODIFIER |
| 8 | 74299597 | AX-170485265 | 0 | T | C | 0.202 | 8.89E-03 | 1.17E-01 | 2.67E+00 | 2.75E+00 | 9.35E-01 | no | | intergenic_variant | MODIFIER |
| 7 | 61013302 | AX-170089934 | 1 | T | C | 0.42 | -7.05E-03 | 9.39E-02 | 2.67E+00 | 2.75E+00 | 9.35E-01 | no | | intergenic_variant | MODIFIER |
| 3 | 145767747 | AX-170503200 | 1 | T | C | 0.357 | -5.32E-03 | 7.02E-02 | 2.66E+00 | 2.75E+00 | 9.35E-01 | no | Ddah1 | intron_variant | MODIFIER |
| 5 | 93162358 | AX-170346637 | 0 | C | A | 0.412 | 8.10E-03 | 1.00E-01 | 2.67E+00 | 2.74E+00 | 9.35E-01 | no | SEP11 | intron_variant | MODIFIER |
| 4 | 143059775 | AX-170024160 | 1 | G | A | 0.107 | 1.67E-02 | 2.04E-01 | 2.67E+00 | 2.74E+00 | 9.35E-01 | no | 6330411D24Rik | intron_variant,non_coding_transcript_variant | MODIFIER |
| 1 | 74425572 | AX-170371764 | 0 | A | G | 0.14 | -1.85E-02 | 2.37E-01 | 2.69E+00 | 2.74E+00 | 9.35E-01 | no | Vil1 | synonymous_variant | LOW |
| 8 | 82679936 | AX-170227992 | 1 | A | G | 0.196 | -1.03E-02 | 1.28E-01 | 2.67E+00 | 2.74E+00 | 9.35E-01 | no | | intergenic_variant | MODIFIER |
| 3 | 144599564 | AX-170421781 | 0 | G | A | 0.36 | -5.27E-03 | 7.01E-02 | 2.66E+00 | 2.75E+00 | 9.35E-01 | no | Selenof | downstream_gene_variant | MODIFIER |
| 3 | 144672147 | AX-170021823 | 0 | A | G | 0.36 | -5.27E-03 | 7.01E-02 | 2.66E+00 | 2.75E+00 | 9.35E-01 | no | | regulatory_region_variant | MODIFIER |
| 3 | 144672930 | AX-170068334 | 0 | T | C | 0.36 | -5.27E-03 | 7.01E-02 | 2.66E+00 | 2.75E+00 | 9.35E-01 | no | | intergenic_variant | MODIFIER |
| 3 | 144673107 | AX-170068337 | 0 | A | C | 0.36 | -5.27E-03 | 7.01E-02 | 2.66E+00 | 2.75E+00 | 9.35E-01 | no | | intergenic_variant | MODIFIER |
| 3 | 145192842 | AX-97805424 | 0 | T | C | 0.36 | -5.27E-03 | 7.01E-02 | 2.66E+00 | 2.75E+00 | 9.35E-01 | no | | intergenic_variant | MODIFIER |
| 3 | 145203907 | AX-170280794 | 0 | C | G | 0.36 | -5.27E-03 | 7.01E-02 | 2.66E+00 | 2.75E+00 | 9.35E-01 | no | | intergenic_variant | MODIFIER |
| 3 | 145203953 | AX-170218123 | 0 | T | C | 0.36 | -5.27E-03 | 7.01E-02 | 2.66E+00 | 2.75E+00 | 9.35E-01 | no | | intergenic_variant | MODIFIER |
| 3 | 145212431 | AX-170116573 | 0 | G | A | 0.36 | -5.27E-03 | 7.01E-02 | 2.66E+00 | 2.75E+00 | 9.35E-01 | no | | intergenic_variant | MODIFIER |
| 3 | 145218173 | AX-170140192 | 0 | G | C | 0.36 | -5.27E-03 | 7.01E-02 | 2.66E+00 | 2.75E+00 | 9.35E-01 | no | | intergenic_variant | MODIFIER |
| 3 | 145223394 | AX-170516404 | 0 | T | C | 0.36 | -5.27E-03 | 7.01E-02 | 2.66E+00 | 2.75E+00 | 9.35E-01 | no | | intergenic_variant | MODIFIER |
| 3 | 145225111 | AX-170333496 | 0 | A | G | 0.36 | -5.27E-03 | 7.01E-02 | 2.66E+00 | 2.75E+00 | 9.35E-01 | no | | intergenic_variant | MODIFIER |
| 3 | 145245089 | AX-170235657 | 0 | T | C | 0.36 | -5.27E-03 | 7.01E-02 | 2.66E+00 | 2.75E+00 | 9.35E-01 | no | | intergenic_variant | MODIFIER |
| 3 | 145278578 | AX-97782656 | 0 | G | A | 0.36 | -5.27E-03 | 7.01E-02 | 2.66E+00 | 2.75E+00 | 9.35E-01 | no | | intergenic_variant | MODIFIER |
| 3 | 145286639 | AX-170428991 | 0 | C | T | 0.36 | -5.27E-03 | 7.01E-02 | 2.66E+00 | 2.75E+00 | 9.35E-01 | no | | intergenic_variant | MODIFIER |
| 3 | 145287173 | AX-170518754 | 0 | T | C | 0.36 | -5.27E-03 | 7.01E-02 | 2.66E+00 | 2.75E+00 | 9.35E-01 | no | | intergenic_variant | MODIFIER |
| 3 | 145298773 | AX-170281983 | 0 | A | G | 0.36 | -5.27E-03 | 7.01E-02 | 2.66E+00 | 2.75E+00 | 9.35E-01 | no | Col24a1 | intron_variant | MODIFIER |
| 3 | 145307627 | AX-170255606 | 0 | A | G | 0.36 | -5.27E-03 | 7.01E-02 | 2.66E+00 | 2.75E+00 | 9.35E-01 | no | Col24a1 | intron_variant | MODIFIER |
| 3 | 145317719 | AX-97828163 | 0 | A | G | 0.36 | -5.27E-03 | 7.01E-02 | 2.66E+00 | 2.75E+00 | 9.35E-01 | no | Col24a1 | intron_variant | MODIFIER |
| 3 | 145403067 | AX-170021833 | 0 | A | C | 0.36 | -5.27E-03 | 7.01E-02 | 2.66E+00 | 2.75E+00 | 9.35E-01 | no | Col24a1 | intron_variant | MODIFIER |
| 3 | 145405058 | AX-170021827 | 0 | C | A | 0.36 | -5.27E-03 | 7.01E-02 | 2.66E+00 | 2.75E+00 | 9.35E-01 | no | Col24a1 | synonymous_variant | LOW |
| 3 | 145411293 | AX-170355710 | 0 | G | A | 0.36 | -5.27E-03 | 7.01E-02 | 2.66E+00 | 2.75E+00 | 9.35E-01 | no | Col24a1 | intron_variant | MODIFIER |
| 3 | 145412028 | AX-170021826 | 0 | C | T | 0.36 | -5.27E-03 | 7.01E-02 | 2.66E+00 | 2.75E+00 | 9.35E-01 | no | Col24a1 | intron_variant | MODIFIER |
| 3 | 145416903 | AX-170068353 | 0 | C | G | 0.36 | -5.27E-03 | 7.01E-02 | 2.66E+00 | 2.75E+00 | 9.35E-01 | no | Col24a1 | intron_variant | MODIFIER |
| 3 | 145417030 | AX-97812575 | 0 | C | T | 0.36 | -5.27E-03 | 7.01E-02 | 2.66E+00 | 2.75E+00 | 9.35E-01 | no | Col24a1 | intron_variant | MODIFIER |
| 3 | 145432461 | AX-170021830 | 0 | T | A | 0.36 | -5.27E-03 | 7.01E-02 | 2.66E+00 | 2.75E+00 | 9.35E-01 | no | Col24a1 | intron_variant | MODIFIER |
| 3 | 145464713 | AX-97804491 | 0 | C | T | 0.36 | -5.27E-03 | 7.01E-02 | 2.66E+00 | 2.75E+00 | 9.35E-01 | no | Col24a1 | intron_variant | MODIFIER |
| 3 | 145471588 | AX-170411739 | 0 | A | G | 0.36 | -5.27E-03 | 7.01E-02 | 2.66E+00 | 2.75E+00 | 9.35E-01 | no | Col24a1 | intron_variant | MODIFIER |
| 3 | 145485250 | AX-97824687 | 0 | A | G | 0.36 | -5.27E-03 | 7.01E-02 | 2.66E+00 | 2.75E+00 | 9.35E-01 | no | Col24a1 | intron_variant | MODIFIER |
| 3 | 145533138 | AX-97807262 | 0 | G | A | 0.36 | -5.27E-03 | 7.01E-02 | 2.66E+00 | 2.75E+00 | 9.35E-01 | no | Col24a1 | intron_variant | MODIFIER |
| 3 | 145635735 | AX-170288781 | 0 | A | G | 0.36 | -5.27E-03 | 7.01E-02 | 2.66E+00 | 2.75E+00 | 9.35E-01 | no | | intergenic_variant | MODIFIER |
| 3 | 145635944 | AX-97845636 | 0 | A | G | 0.36 | -5.27E-03 | 7.01E-02 | 2.66E+00 | 2.75E+00 | 9.35E-01 | no | | intergenic_variant | MODIFIER |
| 3 | 145638192 | AX-170439632 | 0 | A | G | 0.36 | -5.27E-03 | 7.01E-02 | 2.66E+00 | 2.75E+00 | 9.35E-01 | no | | intergenic_variant | MODIFIER |
| 3 | 145638579 | AX-170184791 | 0 | A | G | 0.36 | -5.27E-03 | 7.01E-02 | 2.66E+00 | 2.75E+00 | 9.35E-01 | no | | intergenic_variant | MODIFIER |
| 3 | 145768250 | AX-170137547 | 0 | C | T | 0.36 | -5.27E-03 | 7.01E-02 | 2.66E+00 | 2.75E+00 | 9.35E-01 | no | Ddah1 | intron_variant | MODIFIER |
| 3 | 145770455 | AX-97807453 | 0 | T | C | 0.36 | -5.27E-03 | 7.01E-02 | 2.66E+00 | 2.75E+00 | 9.35E-01 | no | Ddah1 | intron_variant | MODIFIER |
| 3 | 145771549 | AX-170207625 | 0 | C | T | 0.36 | -5.27E-03 | 7.01E-02 | 2.66E+00 | 2.75E+00 | 9.35E-01 | no | Ddah1 | intron_variant | MODIFIER |

| | | | | | | | | | | | | | | | |
|---|---|---|---|---|---|---|---|---|---|---|---|---|---|---|---|
| 3 | 145771754 | AX-170098871 | 0 | T | C | 0.36 | -5.27E-03 | 7.01E-02 | 2.66E+00 | 2.75E+00 | 9.35E-01 | no | Ddah1 | intron_variant | MODIFIER |
| 4 | 98055424 | AX-170023477 | 0 | G | C | 0.228 | -7.52E-03 | 1.09E-01 | 2.68E+00 | 2.76E+00 | 9.35E-01 | no | Nfia | intron_variant | MODIFIER |
| 4 | 141373150 | AX-170123416 | 0 | A | G | 0.105 | -1.61E-02 | 2.05E-01 | 2.68E+00 | 2.74E+00 | 9.36E-01 | no | Fam131c | intron_variant | MODIFIER |
| 16 | 92994347 | AX-170360308 | 0 | C | T | 0.482 | -6.16E-03 | 7.94E-02 | 2.66E+00 | 2.74E+00 | 9.36E-01 | no | | intergenic_variant | MODIFIER |
| 7 | 67703465 | AX-170409593 | 0 | G | T | 0.404 | -7.17E-03 | 9.73E-02 | 2.68E+00 | 2.75E+00 | 9.36E-01 | no | Ttc23 | intron_variant | MODIFIER |
| 7 | 67708203 | AX-170400108 | 0 | G | T | 0.404 | -7.17E-03 | 9.73E-02 | 2.68E+00 | 2.75E+00 | 9.36E-01 | no | Ttc23 | intron_variant | MODIFIER |
| 7 | 68549793 | AX-170196986 | 0 | T | C | 0.404 | -7.17E-03 | 9.73E-02 | 2.68E+00 | 2.75E+00 | 9.36E-01 | no | | regulatory_region_variant | MODIFIER |
| 7 | 68583451 | AX-97793417 | 0 | T | C | 0.404 | -7.17E-03 | 9.73E-02 | 2.68E+00 | 2.75E+00 | 9.36E-01 | no | | intergenic_variant | MODIFIER |
| 7 | 68589085 | AX-170029211 | 0 | C | T | 0.404 | -7.17E-03 | 9.73E-02 | 2.68E+00 | 2.75E+00 | 9.36E-01 | no | | intergenic_variant | MODIFIER |
| 7 | 68617188 | AX-170254718 | 0 | G | A | 0.404 | -7.17E-03 | 9.73E-02 | 2.68E+00 | 2.75E+00 | 9.36E-01 | no | Gm5334 | upstream_gene_variant | MODIFIER |
| 7 | 68617337 | AX-170372167 | 0 | A | T | 0.404 | -7.17E-03 | 9.73E-02 | 2.68E+00 | 2.75E+00 | 9.36E-01 | no | Gm5334 | upstream_gene_variant | MODIFIER |
| 7 | 68619885 | AX-97840424 | 0 | G | A | 0.404 | -7.17E-03 | 9.73E-02 | 2.68E+00 | 2.75E+00 | 9.36E-01 | no | Gm5334 | downstream_gene_variant | MODIFIER |
| 7 | 68620071 | AX-170029212 | 0 | G | A | 0.404 | -7.17E-03 | 9.73E-02 | 2.68E+00 | 2.75E+00 | 9.36E-01 | no | Gm5334 | downstream_gene_variant | MODIFIER |
| 7 | 68622560 | AX-170029209 | 0 | G | A | 0.404 | -7.17E-03 | 9.73E-02 | 2.68E+00 | 2.75E+00 | 9.36E-01 | no | Gm5334 | downstream_gene_variant | MODIFIER |
| 7 | 68650342 | AX-170077635 | 0 | G | A | 0.404 | -7.17E-03 | 9.73E-02 | 2.68E+00 | 2.75E+00 | 9.36E-01 | no | | intergenic_variant | MODIFIER |
| 7 | 68666377 | AX-170167555 | 0 | G | A | 0.404 | -7.17E-03 | 9.73E-02 | 2.68E+00 | 2.75E+00 | 9.36E-01 | no | | intergenic_variant | MODIFIER |
| 7 | 68666481 | AX-170137392 | 0 | A | G | 0.404 | -7.17E-03 | 9.73E-02 | 2.68E+00 | 2.75E+00 | 9.36E-01 | no | | intergenic_variant | MODIFIER |
| 7 | 68680663 | AX-97846405 | 0 | G | A | 0.404 | -7.17E-03 | 9.73E-02 | 2.68E+00 | 2.75E+00 | 9.36E-01 | no | | regulatory_region_variant | MODIFIER |
| 7 | 68684007 | AX-170357538 | 0 | A | G | 0.404 | -7.17E-03 | 9.73E-02 | 2.68E+00 | 2.75E+00 | 9.36E-01 | no | | intergenic_variant | MODIFIER |
| 7 | 68944648 | AX-170303790 | 0 | A | G | 0.404 | -7.17E-03 | 9.73E-02 | 2.68E+00 | 2.75E+00 | 9.36E-01 | no | | intergenic_variant | MODIFIER |
| 7 | 68981491 | AX-170493418 | 0 | C | T | 0.404 | -7.17E-03 | 9.73E-02 | 2.68E+00 | 2.75E+00 | 9.36E-01 | no | | regulatory_region_variant | MODIFIER |
| 7 | 68982107 | AX-170305439 | 0 | C | T | 0.404 | -7.17E-03 | 9.73E-02 | 2.68E+00 | 2.75E+00 | 9.36E-01 | no | | intergenic_variant | MODIFIER |
| 7 | 68983580 | AX-170377967 | 0 | C | T | 0.404 | -7.17E-03 | 9.73E-02 | 2.68E+00 | 2.75E+00 | 9.36E-01 | no | | regulatory_region_variant | MODIFIER |
| 7 | 68985228 | AX-170384300 | 0 | T | C | 0.404 | -7.17E-03 | 9.73E-02 | 2.68E+00 | 2.75E+00 | 9.36E-01 | no | Gm44691 | upstream_gene_variant | MODIFIER |
| 7 | 69018615 | AX-170217225 | 0 | G | A | 0.404 | -7.17E-03 | 9.73E-02 | 2.68E+00 | 2.75E+00 | 9.36E-01 | no | | intergenic_variant | MODIFIER |
| 7 | 69039227 | AX-170521659 | 0 | G | T | 0.404 | -7.17E-03 | 9.73E-02 | 2.68E+00 | 2.75E+00 | 9.36E-01 | no | | intergenic_variant | MODIFIER |
| 7 | 69044737 | AX-97795626 | 0 | C | T | 0.404 | -7.17E-03 | 9.73E-02 | 2.68E+00 | 2.75E+00 | 9.36E-01 | no | | intergenic_variant | MODIFIER |
| 7 | 69059047 | AX-170029219 | 0 | A | T | 0.404 | -7.17E-03 | 9.73E-02 | 2.68E+00 | 2.75E+00 | 9.36E-01 | no | | intergenic_variant | MODIFIER |
| 7 | 69215523 | AX-170524840 | 0 | G | C | 0.404 | -7.17E-03 | 9.73E-02 | 2.68E+00 | 2.75E+00 | 9.36E-01 | no | Gm5342 | downstream_gene_variant | MODIFIER |
| 11 | 8576860 | AX-170377705 | 0 | C | G | 0.07 | -1.40E-02 | 1.79E-01 | 2.62E+00 | 2.74E+00 | 9.36E-01 | no | Tns3 | intron_variant | MODIFIER |
| 16 | 92261954 | AX-97829360 | 0 | G | A | 0.474 | -6.18E-03 | 7.87E-02 | 2.66E+00 | 2.74E+00 | 9.36E-01 | no | | intergenic_variant | MODIFIER |
| 16 | 92268277 | AX-170120721 | 0 | C | T | 0.474 | -6.18E-03 | 7.87E-02 | 2.66E+00 | 2.74E+00 | 9.36E-01 | no | | regulatory_region_variant | MODIFIER |
| 16 | 92296856 | AX-170013663 | 0 | G | A | 0.474 | -6.18E-03 | 7.87E-02 | 2.66E+00 | 2.74E+00 | 9.36E-01 | no | Kcne2 | synonymous_variant | LOW |
| 16 | 92320778 | AX-170125338 | 0 | G | A | 0.474 | -6.18E-03 | 7.87E-02 | 2.66E+00 | 2.74E+00 | 9.36E-01 | no | Fam243 | synonymous_variant | LOW |
| 16 | 92321836 | AX-170454892 | 0 | T | C | 0.474 | -6.18E-03 | 7.87E-02 | 2.66E+00 | 2.74E+00 | 9.36E-01 | no | Fam243 | upstream_gene_variant | MODIFIER |
| 16 | 92325728 | AX-170211170 | 0 | G | A | 0.474 | -6.18E-03 | 7.87E-02 | 2.66E+00 | 2.74E+00 | 9.36E-01 | no | Fam243 | upstream_gene_variant | MODIFIER |
| 16 | 92332353 | AX-97791539 | 0 | C | A | 0.474 | -6.18E-03 | 7.87E-02 | 2.66E+00 | 2.74E+00 | 9.36E-01 | no | Smim34 | upstream_gene_variant | MODIFIER |
| 16 | 92332418 | AX-170013660 | 0 | G | A | 0.474 | -6.18E-03 | 7.87E-02 | 2.66E+00 | 2.74E+00 | 9.36E-01 | no | Smim34 | upstream_gene_variant | MODIFIER |
| 16 | 92336182 | AX-170233134 | 0 | C | A | 0.474 | -6.18E-03 | 7.87E-02 | 2.66E+00 | 2.74E+00 | 9.36E-01 | no | | regulatory_region_variant | MODIFIER |
| 16 | 92336314 | AX-170370916 | 0 | C | T | 0.474 | -6.18E-03 | 7.87E-02 | 2.66E+00 | 2.74E+00 | 9.36E-01 | no | | regulatory_region_variant | MODIFIER |
| 16 | 92336390 | AX-170325450 | 0 | A | G | 0.474 | -6.18E-03 | 7.87E-02 | 2.66E+00 | 2.74E+00 | 9.36E-01 | no | | regulatory_region_variant | MODIFIER |
| 16 | 92341086 | AX-170118918 | 0 | G | A | 0.474 | -6.18E-03 | 7.87E-02 | 2.66E+00 | 2.74E+00 | 9.36E-01 | no | Kcne1 | downstream_gene_variant | MODIFIER |
| 16 | 92359268 | AX-170525603 | 0 | G | T | 0.474 | -6.18E-03 | 7.87E-02 | 2.66E+00 | 2.74E+00 | 9.36E-01 | no | Kcne1 | intron_variant | MODIFIER |
| 16 | 92374437 | AX-170013665 | 0 | G | A | 0.474 | -6.18E-03 | 7.87E-02 | 2.66E+00 | 2.74E+00 | 9.36E-01 | no | Gm29805 | downstream_gene_variant | MODIFIER |
| 16 | 92374740 | AX-170491292 | 0 | C | T | 0.474 | -6.18E-03 | 7.87E-02 | 2.66E+00 | 2.74E+00 | 9.36E-01 | no | Gm29805 | downstream_gene_variant | MODIFIER |
| 16 | 92375307 | AX-170462381 | 0 | C | T | 0.474 | -6.18E-03 | 7.87E-02 | 2.66E+00 | 2.74E+00 | 9.36E-01 | no | Gm29805 | downstream_gene_variant | MODIFIER |
| 16 | 92375825 | AX-170337263 | 0 | C | G | 0.474 | -6.18E-03 | 7.87E-02 | 2.66E+00 | 2.74E+00 | 9.36E-01 | no | Gm29805 | downstream_gene_variant | MODIFIER |
| 16 | 92409417 | AX-170013667 | 0 | A | G | 0.474 | -6.18E-03 | 7.87E-02 | 2.66E+00 | 2.74E+00 | 9.36E-01 | no | Rcan1 | intron_variant | MODIFIER |
| 16 | 92411638 | AX-170502435 | 0 | A | G | 0.474 | -6.18E-03 | 7.87E-02 | 2.66E+00 | 2.74E+00 | 9.36E-01 | no | Rcan1 | intron_variant | MODIFIER |
| 16 | 92420145 | AX-170266583 | 0 | G | A | 0.474 | -6.18E-03 | 7.87E-02 | 2.66E+00 | 2.74E+00 | 9.36E-01 | no | Rcan1 | intron_variant | MODIFIER |
| 16 | 92424416 | AX-170268850 | 0 | A | G | 0.474 | -6.18E-03 | 7.87E-02 | 2.66E+00 | 2.74E+00 | 9.36E-01 | no | Rcan1 | intron_variant | MODIFIER |
| 16 | 92426884 | AX-170353278 | 0 | C | G | 0.474 | -6.18E-03 | 7.87E-02 | 2.66E+00 | 2.74E+00 | 9.36E-01 | no | Rcan1 | intron_variant | MODIFIER |
| 16 | 92439457 | AX-170387662 | 0 | C | T | 0.474 | -6.18E-03 | 7.87E-02 | 2.66E+00 | 2.74E+00 | 9.36E-01 | no | Rcan1 | intron_variant | MODIFIER |
| 16 | 92449352 | AX-170322156 | 0 | T | C | 0.474 | -6.18E-03 | 7.87E-02 | 2.66E+00 | 2.74E+00 | 9.36E-01 | no | Rcan1 | intron_variant | MODIFIER |

| | | | | | | | | | | | | | | | |
|---|---|---|---|---|---|---|---|---|---|---|---|---|---|---|---|
| 16 | 92453793 | AX-170252273 | 0 | A | G | 0.474 | -6.18E-03 | 7.87E-02 | 2.66E+00 | 2.74E+00 | 9.36E-01 | no | Rcan1 | intron_variant | MODIFIER |
| 16 | 92453892 | AX-170284226 | 0 | A | C | 0.474 | -6.18E-03 | 7.87E-02 | 2.66E+00 | 2.74E+00 | 9.36E-01 | no | Rcan1 | intron_variant | MODIFIER |
| 16 | 92457426 | AX-170434027 | 0 | C | T | 0.474 | -6.18E-03 | 7.87E-02 | 2.66E+00 | 2.74E+00 | 9.36E-01 | no | Rcan1 | intron_variant | MODIFIER |
| 16 | 92485395 | AX-170138965 | 0 | G | A | 0.474 | -6.18E-03 | 7.87E-02 | 2.66E+00 | 2.74E+00 | 9.36E-01 | no | 2410124H12Rik | intron_variant,non_coding_transcript_variant | MODIFIER |
| 16 | 92487504 | AX-170245856 | 0 | A | G | 0.474 | -6.18E-03 | 7.87E-02 | 2.66E+00 | 2.74E+00 | 9.36E-01 | no | 2410124H12Rik | intron_variant,non_coding_transcript_variant | MODIFIER |
| 16 | 92506688 | AX-170410989 | 0 | C | T | 0.474 | -6.18E-03 | 7.87E-02 | 2.66E+00 | 2.74E+00 | 9.36E-01 | no | Clic6 | intron_variant | MODIFIER |
| 16 | 92510092 | AX-170441065 | 0 | T | G | 0.474 | -6.18E-03 | 7.87E-02 | 2.66E+00 | 2.74E+00 | 9.36E-01 | no | Clic6 | intron_variant | MODIFIER |
| 16 | 92543209 | AX-170058122 | 0 | A | G | 0.474 | -6.18E-03 | 7.87E-02 | 2.66E+00 | 2.74E+00 | 9.36E-01 | no | Clic6 | downstream_gene_variant | MODIFIER |
| 16 | 92544565 | AX-170058120 | 0 | C | T | 0.474 | -6.18E-03 | 7.87E-02 | 2.66E+00 | 2.74E+00 | 9.36E-01 | no | Clic6 | downstream_gene_variant | MODIFIER |
| 16 | 92546949 | AX-170058121 | 0 | A | G | 0.474 | -6.18E-03 | 7.87E-02 | 2.66E+00 | 2.74E+00 | 9.36E-01 | no | Gm49722 | downstream_gene_variant | MODIFIER |
| 16 | 92546974 | AX-97784296 | 0 | A | G | 0.474 | -6.18E-03 | 7.87E-02 | 2.66E+00 | 2.74E+00 | 9.36E-01 | no | Gm49722 | downstream_gene_variant | MODIFIER |
| 16 | 92554501 | AX-170486895 | 0 | T | C | 0.474 | -6.18E-03 | 7.87E-02 | 2.66E+00 | 2.74E+00 | 9.36E-01 | no | Gm49722 | upstream_gene_variant | MODIFIER |
| 16 | 92558955 | AX-170291468 | 0 | C | T | 0.474 | -6.18E-03 | 7.87E-02 | 2.66E+00 | 2.74E+00 | 9.36E-01 | no | | intergenic_variant | MODIFIER |
| 16 | 92562851 | AX-170326194 | 0 | C | T | 0.474 | -6.18E-03 | 7.87E-02 | 2.66E+00 | 2.74E+00 | 9.36E-01 | no | | regulatory_region_variant | MODIFIER |
| 16 | 92564335 | AX-170384655 | 0 | G | C | 0.474 | -6.18E-03 | 7.87E-02 | 2.66E+00 | 2.74E+00 | 9.36E-01 | no | | intergenic_variant | MODIFIER |
| 16 | 92588123 | AX-170318504 | 0 | A | T | 0.474 | -6.18E-03 | 7.87E-02 | 2.66E+00 | 2.74E+00 | 9.36E-01 | no | | regulatory_region_variant | MODIFIER |
| 16 | 92592647 | AX-170369040 | 0 | A | T | 0.474 | -6.18E-03 | 7.87E-02 | 2.66E+00 | 2.74E+00 | 9.36E-01 | no | | regulatory_region_variant | MODIFIER |
| 16 | 92616566 | AX-170362156 | 0 | C | T | 0.474 | -6.18E-03 | 7.87E-02 | 2.66E+00 | 2.74E+00 | 9.36E-01 | no | Runx1 | intron_variant | MODIFIER |
| 16 | 92622763 | AX-170093566 | 0 | G | A | 0.474 | -6.18E-03 | 7.87E-02 | 2.66E+00 | 2.74E+00 | 9.36E-01 | no | Runx1 | intron_variant | MODIFIER |
| 16 | 92626249 | AX-170165736 | 0 | T | C | 0.474 | -6.18E-03 | 7.87E-02 | 2.66E+00 | 2.74E+00 | 9.36E-01 | no | Runx1 | intron_variant | MODIFIER |
| 16 | 92640984 | AX-170173752 | 0 | A | G | 0.474 | -6.18E-03 | 7.87E-02 | 2.66E+00 | 2.74E+00 | 9.36E-01 | no | Runx1 | intron_variant | MODIFIER |
| 16 | 92652285 | AX-170468274 | 0 | C | T | 0.474 | -6.18E-03 | 7.87E-02 | 2.66E+00 | 2.74E+00 | 9.36E-01 | no | Runx1 | intron_variant | MODIFIER |
| 16 | 92652994 | AX-170058125 | 0 | G | A | 0.474 | -6.18E-03 | 7.87E-02 | 2.66E+00 | 2.74E+00 | 9.36E-01 | no | Runx1 | intron_variant | MODIFIER |
| 16 | 92667537 | AX-170466628 | 0 | T | C | 0.474 | -6.18E-03 | 7.87E-02 | 2.66E+00 | 2.74E+00 | 9.36E-01 | no | Runx1 | intron_variant | MODIFIER |
| 16 | 92670479 | AX-170175341 | 0 | A | T | 0.474 | -6.18E-03 | 7.87E-02 | 2.66E+00 | 2.74E+00 | 9.36E-01 | no | Runx1 | intron_variant | MODIFIER |
| 16 | 92671008 | AX-170141233 | 0 | G | T | 0.474 | -6.18E-03 | 7.87E-02 | 2.66E+00 | 2.74E+00 | 9.36E-01 | no | Runx1 | intron_variant | MODIFIER |
| 16 | 92671906 | AX-170175272 | 0 | C | T | 0.474 | -6.18E-03 | 7.87E-02 | 2.66E+00 | 2.74E+00 | 9.36E-01 | no | Runx1 | intron_variant | MODIFIER |
| 16 | 92671988 | AX-170426376 | 0 | G | A | 0.474 | -6.18E-03 | 7.87E-02 | 2.66E+00 | 2.74E+00 | 9.36E-01 | no | Runx1 | intron_variant | MODIFIER |
| 16 | 92672509 | AX-170264326 | 0 | C | T | 0.474 | -6.18E-03 | 7.87E-02 | 2.66E+00 | 2.74E+00 | 9.36E-01 | no | Runx1 | intron_variant | MODIFIER |
| 16 | 92682466 | AX-170037542 | 0 | T | C | 0.474 | -6.18E-03 | 7.87E-02 | 2.66E+00 | 2.74E+00 | 9.36E-01 | no | Runx1 | intron_variant | MODIFIER |
| 16 | 92682651 | AX-170284367 | 0 | A | G | 0.474 | -6.18E-03 | 7.87E-02 | 2.66E+00 | 2.74E+00 | 9.36E-01 | no | Runx1 | intron_variant | MODIFIER |
| 16 | 92692556 | AX-97832132 | 0 | G | A | 0.474 | -6.18E-03 | 7.87E-02 | 2.66E+00 | 2.74E+00 | 9.36E-01 | no | Runx1 | intron_variant | MODIFIER |
| 16 | 92722254 | AX-97791540 | 0 | T | C | 0.474 | -6.18E-03 | 7.87E-02 | 2.66E+00 | 2.74E+00 | 9.36E-01 | no | Runx1 | intron_variant | MODIFIER |
| 16 | 92727304 | AX-170013670 | 0 | G | A | 0.474 | -6.18E-03 | 7.87E-02 | 2.66E+00 | 2.74E+00 | 9.36E-01 | no | Runx1 | intron_variant | MODIFIER |
| 16 | 92728072 | AX-170265921 | 0 | A | C | 0.474 | -6.18E-03 | 7.87E-02 | 2.66E+00 | 2.74E+00 | 9.36E-01 | no | Runx1 | intron_variant | MODIFIER |
| 16 | 92742364 | AX-97848178 | 0 | A | G | 0.474 | -6.18E-03 | 7.87E-02 | 2.66E+00 | 2.74E+00 | 9.36E-01 | no | Runx1 | intron_variant | MODIFIER |
| 16 | 92744198 | AX-170228903 | 0 | A | G | 0.474 | -6.18E-03 | 7.87E-02 | 2.66E+00 | 2.74E+00 | 9.36E-01 | no | Runx1 | intron_variant | MODIFIER |
| 16 | 92747938 | AX-170473777 | 0 | G | C | 0.474 | -6.18E-03 | 7.87E-02 | 2.66E+00 | 2.74E+00 | 9.36E-01 | no | Runx1 | intron_variant | MODIFIER |
| 16 | 92760923 | AX-170535607 | 0 | C | T | 0.474 | -6.18E-03 | 7.87E-02 | 2.66E+00 | 2.74E+00 | 9.36E-01 | no | Runx1 | intron_variant | MODIFIER |
| 16 | 92765227 | AX-170502475 | 0 | C | T | 0.474 | -6.18E-03 | 7.87E-02 | 2.66E+00 | 2.74E+00 | 9.36E-01 | no | Runx1 | intron_variant | MODIFIER |
| 16 | 92767277 | AX-170527009 | 0 | C | T | 0.474 | -6.18E-03 | 7.87E-02 | 2.66E+00 | 2.74E+00 | 9.36E-01 | no | Runx1 | intron_variant | MODIFIER |
| 16 | 92770501 | AX-170058128 | 0 | C | A | 0.474 | -6.18E-03 | 7.87E-02 | 2.66E+00 | 2.74E+00 | 9.36E-01 | no | Runx1 | intron_variant | MODIFIER |
| 16 | 92789987 | AX-170392299 | 0 | T | C | 0.474 | -6.18E-03 | 7.87E-02 | 2.66E+00 | 2.74E+00 | 9.36E-01 | no | Runx1 | intron_variant | MODIFIER |
| 16 | 92790065 | AX-170403656 | 0 | G | T | 0.474 | -6.18E-03 | 7.87E-02 | 2.66E+00 | 2.74E+00 | 9.36E-01 | no | Runx1 | intron_variant | MODIFIER |
| 16 | 92795916 | AX-170058129 | 0 | T | C | 0.474 | -6.18E-03 | 7.87E-02 | 2.66E+00 | 2.74E+00 | 9.36E-01 | no | Runx1 | intron_variant | MODIFIER |
| 16 | 92816299 | AX-170342157 | 0 | T | C | 0.474 | -6.18E-03 | 7.87E-02 | 2.66E+00 | 2.74E+00 | 9.36E-01 | no | Runx1 | intron_variant | MODIFIER |
| 16 | 92836679 | AX-97782075 | 0 | C | T | 0.474 | -6.18E-03 | 7.87E-02 | 2.66E+00 | 2.74E+00 | 9.36E-01 | no | | intergenic_variant | MODIFIER |
| 16 | 92839696 | AX-170217595 | 0 | A | G | 0.474 | -6.18E-03 | 7.87E-02 | 2.66E+00 | 2.74E+00 | 9.36E-01 | no | | intergenic_variant | MODIFIER |
| 16 | 92850367 | AX-170478920 | 0 | G | T | 0.474 | -6.18E-03 | 7.87E-02 | 2.66E+00 | 2.74E+00 | 9.36E-01 | no | | intergenic_variant | MODIFIER |
| 16 | 92854094 | AX-170274368 | 0 | C | T | 0.474 | -6.18E-03 | 7.87E-02 | 2.66E+00 | 2.74E+00 | 9.36E-01 | no | | intergenic_variant | MODIFIER |
| 16 | 92889720 | AX-170353377 | 0 | C | T | 0.474 | -6.18E-03 | 7.87E-02 | 2.66E+00 | 2.74E+00 | 9.36E-01 | no | | intergenic_variant | MODIFIER |
| 16 | 92889876 | AX-170518195 | 0 | A | G | 0.474 | -6.18E-03 | 7.87E-02 | 2.66E+00 | 2.74E+00 | 9.36E-01 | no | | intergenic_variant | MODIFIER |
| 16 | 92890169 | AX-170515852 | 0 | G | A | 0.474 | -6.18E-03 | 7.87E-02 | 2.66E+00 | 2.74E+00 | 9.36E-01 | no | | intergenic_variant | MODIFIER |
| 16 | 92892756 | AX-170037541 | 0 | G | A | 0.474 | -6.18E-03 | 7.87E-02 | 2.66E+00 | 2.74E+00 | 9.36E-01 | no | | intergenic_variant | MODIFIER |

| | | | | | | | | | | | | | | | |
|---|---|---|---|---|---|---|---|---|---|---|---|---|---|---|---|
| 16 | 92895260 | AX-170345258 | 0 | C | T | 0.474 | -6.18E-03 | 7.87E-02 | 2.66E+00 | 2.74E+00 | 9.36E-01 | no | | intergenic_variant | MODIFIER |
| 16 | 92902133 | AX-170379692 | 0 | G | A | 0.474 | -6.18E-03 | 7.87E-02 | 2.66E+00 | 2.74E+00 | 9.36E-01 | no | | intergenic_variant | MODIFIER |
| 16 | 92908865 | AX-170499979 | 0 | G | A | 0.474 | -6.18E-03 | 7.87E-02 | 2.66E+00 | 2.74E+00 | 9.36E-01 | no | | intergenic_variant | MODIFIER |
| 16 | 92915870 | AX-97829046 | 0 | G | T | 0.474 | -6.18E-03 | 7.87E-02 | 2.66E+00 | 2.74E+00 | 9.36E-01 | no | | intergenic_variant | MODIFIER |
| 16 | 92925313 | AX-170193168 | 0 | C | T | 0.474 | -6.18E-03 | 7.87E-02 | 2.66E+00 | 2.74E+00 | 9.36E-01 | no | | intergenic_variant | MODIFIER |
| 16 | 92926022 | AX-170193211 | 0 | G | T | 0.474 | -6.18E-03 | 7.87E-02 | 2.66E+00 | 2.74E+00 | 9.36E-01 | no | | intergenic_variant | MODIFIER |
| 16 | 92953664 | AX-170462106 | 0 | G | A | 0.474 | -6.18E-03 | 7.87E-02 | 2.66E+00 | 2.74E+00 | 9.36E-01 | no | | regulatory_region_variant | MODIFIER |
| 16 | 92976240 | AX-170121951 | 0 | A | T | 0.474 | -6.18E-03 | 7.87E-02 | 2.66E+00 | 2.74E+00 | 9.36E-01 | no | | regulatory_region_variant | MODIFIER |
| 16 | 92976714 | AX-170517963 | 0 | G | T | 0.474 | -6.18E-03 | 7.87E-02 | 2.66E+00 | 2.74E+00 | 9.36E-01 | no | | intergenic_variant | MODIFIER |
| 16 | 92987955 | AX-170130257 | 0 | C | T | 0.474 | -6.18E-03 | 7.87E-02 | 2.66E+00 | 2.74E+00 | 9.36E-01 | no | | regulatory_region_variant | MODIFIER |
| 16 | 92987981 | AX-170501408 | 0 | T | C | 0.474 | -6.18E-03 | 7.87E-02 | 2.66E+00 | 2.74E+00 | 9.36E-01 | no | | regulatory_region_variant | MODIFIER |
| 16 | 92988054 | AX-170478718 | 0 | C | A | 0.474 | -6.18E-03 | 7.87E-02 | 2.66E+00 | 2.74E+00 | 9.36E-01 | no | | regulatory_region_variant | MODIFIER |
| 16 | 92996468 | AX-170323142 | 0 | A | G | 0.474 | -6.18E-03 | 7.87E-02 | 2.66E+00 | 2.74E+00 | 9.36E-01 | no | | intergenic_variant | MODIFIER |
| 16 | 93000312 | AX-170504842 | 0 | T | C | 0.474 | -6.18E-03 | 7.87E-02 | 2.66E+00 | 2.74E+00 | 9.36E-01 | no | | intergenic_variant | MODIFIER |
| 16 | 93006644 | AX-170247111 | 0 | C | A | 0.474 | -6.18E-03 | 7.87E-02 | 2.66E+00 | 2.74E+00 | 9.36E-01 | no | | regulatory_region_variant | MODIFIER |
| 16 | 93008570 | AX-170232632 | 0 | C | A | 0.474 | -6.18E-03 | 7.87E-02 | 2.66E+00 | 2.74E+00 | 9.36E-01 | no | | intergenic_variant | MODIFIER |
| 7 | 55978434 | AX-170351648 | 0 | T | C | 0.404 | 8.16E-03 | 9.41E-02 | 2.63E+00 | 2.73E+00 | 9.36E-01 | no | Nipa1 | 3_prime_UTR_variant | MODIFIER |
| 15 | 97711920 | AX-170535694 | 0 | A | G | 0.132 | 1.16E-02 | 1.42E-01 | 2.65E+00 | 2.73E+00 | 9.36E-01 | no | Endou | 3_prime_UTR_variant | MODIFIER |
| 7 | 91756142 | AX-170029508 | 1 | C | T | 0.134 | -1.68E-02 | 2.15E-01 | 2.66E+00 | 2.74E+00 | 9.36E-01 | no | Dlg2 | intron_variant | MODIFIER |
| 15 | 97713402 | AX-170268083 | 1 | G | T | 0.134 | 1.36E-02 | 1.57E-01 | 2.66E+00 | 2.72E+00 | 9.36E-01 | no | Endou | intron_variant | MODIFIER |
| 20 | 65615174 | AX-97817653 | 1 | T | C | 0.241 | 4.67E-03 | 5.93E-02 | 2.64E+00 | 2.74E+00 | 9.36E-01 | no | | intergenic_variant | MODIFIER |
| 10 | 13196176 | AX-170194550 | 0 | G | A | 0.211 | 7.19E-03 | 1.02E-01 | 2.68E+00 | 2.76E+00 | 9.36E-01 | no | Ltv1 | upstream_gene_variant | MODIFIER |
| 13 | 49056258 | AX-170051224 | 1 | G | A | 0.5 | 6.16E-03 | 6.74E-02 | 2.62E+00 | 2.72E+00 | 9.36E-01 | no | Wnk2 | intron_variant | MODIFIER |
| 1 | 5988177 | AX-170534149 | 0 | C | T | 0.105 | 1.32E-02 | 1.49E-01 | 2.60E+00 | 2.73E+00 | 9.36E-01 | no | | intergenic_variant | MODIFIER |
| 1 | 11084478 | AX-170169076 | 0 | G | A | 0.105 | 1.32E-02 | 1.49E-01 | 2.60E+00 | 2.73E+00 | 9.36E-01 | no | Prex2 | intron_variant | MODIFIER |
| 1 | 28871505 | AX-170419335 | 1 | G | A | 0.402 | -6.05E-03 | 8.02E-02 | 2.66E+00 | 2.74E+00 | 9.36E-01 | no | | intergenic_variant | MODIFIER |
| 12 | 63365596 | AX-170204031 | 1 | C | T | 0.062 | -2.40E-02 | 3.00E-01 | 2.71E+00 | 2.73E+00 | 9.36E-01 | no | | intergenic_variant | MODIFIER |
| 8 | 78812509 | AX-97808220 | 0 | G | A | 0.202 | -1.06E-02 | 1.29E-01 | 2.67E+00 | 2.73E+00 | 9.36E-01 | no | Lsm6 | intron_variant | MODIFIER |
| 15 | 100484324 | AX-170163561 | 0 | A | G | 0.114 | 1.22E-02 | 1.49E-01 | 2.66E+00 | 2.73E+00 | 9.36E-01 | no | Csrnp2 | intron_variant | MODIFIER |
| 9 | 114464563 | AX-170482549 | 0 | A | G | 0.132 | -8.23E-03 | 1.04E-01 | 2.64E+00 | 2.74E+00 | 9.36E-01 | no | Glb1 | intron_variant | MODIFIER |
| 16 | 94390565 | AX-170391356 | 2 | T | C | 0.382 | -7.51E-03 | 8.14E-02 | 2.63E+00 | 2.72E+00 | 9.36E-01 | no | Ttc3 | intron_variant | MODIFIER |
| 5 | 107568827 | AX-170380244 | 0 | T | A | 0.456 | 9.61E-03 | 1.15E-01 | 2.68E+00 | 2.72E+00 | 9.36E-01 | no | Glmn | intron_variant | MODIFIER |
| 5 | 113967944 | AX-170176316 | 0 | G | A | 0.456 | 9.61E-03 | 1.15E-01 | 2.68E+00 | 2.72E+00 | 9.36E-01 | no | Ssh1 | intron_variant | MODIFIER |
| 11 | 33045074 | AX-170422670 | 0 | G | T | 0.105 | 1.57E-02 | 1.88E-01 | 2.65E+00 | 2.73E+00 | 9.36E-01 | no | | intergenic_variant | MODIFIER |
| 5 | 97956157 | AX-170190049 | 0 | G | A | 0.412 | -7.93E-03 | 1.06E-01 | 2.69E+00 | 2.75E+00 | 9.36E-01 | no | Antxr2 | intron_variant | MODIFIER |
| 5 | 97956955 | AX-170204139 | 0 | A | C | 0.412 | -7.93E-03 | 1.06E-01 | 2.69E+00 | 2.75E+00 | 9.36E-01 | no | Antxr2 | intron_variant | MODIFIER |
| 5 | 97957731 | AX-170072837 | 0 | T | C | 0.412 | -7.93E-03 | 1.06E-01 | 2.69E+00 | 2.75E+00 | 9.36E-01 | no | Antxr2 | intron_variant | MODIFIER |
| 5 | 97960200 | AX-170284364 | 0 | A | G | 0.412 | -7.93E-03 | 1.06E-01 | 2.69E+00 | 2.75E+00 | 9.36E-01 | no | Antxr2 | intron_variant | MODIFIER |
| 5 | 97974018 | AX-170504931 | 0 | C | T | 0.412 | -7.93E-03 | 1.06E-01 | 2.69E+00 | 2.75E+00 | 9.36E-01 | no | Antxr2 | intron_variant | MODIFIER |
| 5 | 97975004 | AX-170429586 | 0 | C | T | 0.412 | -7.93E-03 | 1.06E-01 | 2.69E+00 | 2.75E+00 | 9.36E-01 | no | Antxr2 | intron_variant | MODIFIER |
| 5 | 97980151 | AX-170154657 | 0 | C | T | 0.412 | -7.93E-03 | 1.06E-01 | 2.69E+00 | 2.75E+00 | 9.36E-01 | no | Antxr2 | intron_variant | MODIFIER |
| 5 | 98000979 | AX-97849943 | 0 | T | C | 0.412 | -7.93E-03 | 1.06E-01 | 2.69E+00 | 2.75E+00 | 9.36E-01 | no | Antxr2 | intron_variant | MODIFIER |
| 5 | 98035707 | AX-97837644 | 0 | A | G | 0.412 | -7.93E-03 | 1.06E-01 | 2.69E+00 | 2.75E+00 | 9.36E-01 | no | Antxr2 | upstream_gene_variant | MODIFIER |
| 5 | 98036033 | AX-170439073 | 0 | A | G | 0.412 | -7.93E-03 | 1.06E-01 | 2.69E+00 | 2.75E+00 | 9.36E-01 | no | Gm42759 | downstream_gene_variant | MODIFIER |
| 5 | 98113379 | AX-170221948 | 0 | T | G | 0.412 | -7.93E-03 | 1.06E-01 | 2.69E+00 | 2.75E+00 | 9.36E-01 | no | | intergenic_variant | MODIFIER |
| 5 | 98346827 | AX-170147477 | 0 | A | C | 0.412 | -7.93E-03 | 1.06E-01 | 2.69E+00 | 2.75E+00 | 9.36E-01 | no | Cfap299 | intron_variant | MODIFIER |
| 5 | 98353353 | AX-170380068 | 0 | T | G | 0.412 | -7.93E-03 | 1.06E-01 | 2.69E+00 | 2.75E+00 | 9.36E-01 | no | Cfap299 | intron_variant | MODIFIER |
| 5 | 98565661 | AX-170484685 | 0 | G | C | 0.412 | -7.93E-03 | 1.06E-01 | 2.69E+00 | 2.75E+00 | 9.36E-01 | no | Cfap299 | intron_variant | MODIFIER |
| 5 | 99002935 | AX-170439770 | 0 | A | G | 0.412 | -7.93E-03 | 1.06E-01 | 2.69E+00 | 2.75E+00 | 9.36E-01 | no | Prkg2 | intron_variant | MODIFIER |
| 5 | 99227083 | AX-170025699 | 0 | A | G | 0.412 | -7.93E-03 | 1.06E-01 | 2.69E+00 | 2.75E+00 | 9.36E-01 | no | Rasgef1b | intron_variant | MODIFIER |
| 5 | 99592532 | AX-97784065 | 0 | G | A | 0.412 | -7.93E-03 | 1.06E-01 | 2.69E+00 | 2.75E+00 | 9.36E-01 | no | Rasgef1b | intron_variant,non_coding_transcript_variant | MODIFIER |
| 5 | 99727188 | AX-97827571 | 0 | C | T | 0.412 | -7.93E-03 | 1.06E-01 | 2.69E+00 | 2.75E+00 | 9.36E-01 | no | Rasgef1b | intron_variant,non_coding_transcript_variant | MODIFIER |
| 5 | 99747799 | AX-170226616 | 0 | T | C | 0.412 | -7.93E-03 | 1.06E-01 | 2.69E+00 | 2.75E+00 | 9.36E-01 | no | Gm38413 | intron_variant,non_coding_transcript_variant | MODIFIER |
| 5 | 99954534 | AX-97827695 | 0 | T | C | 0.412 | -7.93E-03 | 1.06E-01 | 2.69E+00 | 2.75E+00 | 9.36E-01 | no | Hnrnpd | downstream_gene_variant | MODIFIER |

| | | | | | | | | | | | | | | | |
|---|---|---|---|---|---|---|---|---|---|---|---|---|---|---|---|
| 5 | 100047564 | AX-170277014 | 0 | A | G | 0.412 | -7.93E-03 | 1.06E-01 | 2.69E+00 | 2.75E+00 | 9.36E-01 | no | Enoph1 | intron_variant | MODIFIER |
| 5 | 100047632 | AX-170353014 | 0 | G | A | 0.412 | -7.93E-03 | 1.06E-01 | 2.69E+00 | 2.75E+00 | 9.36E-01 | no | Enoph1 | intron_variant | MODIFIER |
| 5 | 100047700 | AX-170479808 | 0 | C | T | 0.412 | -7.93E-03 | 1.06E-01 | 2.69E+00 | 2.75E+00 | 9.36E-01 | no | Enoph1 | intron_variant | MODIFIER |
| 5 | 100051070 | AX-170342324 | 0 | A | G | 0.412 | -7.93E-03 | 1.06E-01 | 2.69E+00 | 2.75E+00 | 9.36E-01 | no | Enoph1 | intron_variant | MODIFIER |
| 5 | 100054679 | AX-170354213 | 0 | G | A | 0.412 | -7.93E-03 | 1.06E-01 | 2.69E+00 | 2.75E+00 | 9.36E-01 | no | Enoph1 | intron_variant | MODIFIER |
| 5 | 100058036 | AX-97812051 | 0 | T | C | 0.412 | -7.93E-03 | 1.06E-01 | 2.69E+00 | 2.75E+00 | 9.36E-01 | no | Enoph1 | intron_variant | MODIFIER |
| 5 | 100061008 | AX-170373188 | 0 | T | C | 0.412 | -7.93E-03 | 1.06E-01 | 2.69E+00 | 2.75E+00 | 9.36E-01 | no | Enoph1 | missense_variant | MODERATE |
| 5 | 100062009 | AX-170125470 | 0 | A | C | 0.412 | -7.93E-03 | 1.06E-01 | 2.69E+00 | 2.75E+00 | 9.36E-01 | no | Enoph1 | intron_variant | MODIFIER |
| 5 | 100067739 | AX-170263717 | 0 | C | T | 0.412 | -7.93E-03 | 1.06E-01 | 2.69E+00 | 2.75E+00 | 9.36E-01 | no | Enoph1 | intron_variant | MODIFIER |
| 5 | 100092539 | AX-97811763 | 0 | A | G | 0.412 | -7.93E-03 | 1.06E-01 | 2.69E+00 | 2.75E+00 | 9.36E-01 | no | Tmem150c | intron_variant | MODIFIER |
| 5 | 100109460 | AX-97783689 | 0 | T | C | 0.412 | -7.93E-03 | 1.06E-01 | 2.69E+00 | 2.75E+00 | 9.36E-01 | no | Tmem150c | intron_variant | MODIFIER |
| 5 | 100258340 | AX-97810512 | 0 | T | C | 0.412 | -7.93E-03 | 1.06E-01 | 2.69E+00 | 2.75E+00 | 9.36E-01 | no | Gm8091 | intron_variant,non_coding_transcript_variant | MODIFIER |
| 5 | 100274378 | AX-97811664 | 0 | T | C | 0.412 | -7.93E-03 | 1.06E-01 | 2.69E+00 | 2.75E+00 | 9.36E-01 | no | Gm8091 | intron_variant,non_coding_transcript_variant | MODIFIER |
| 5 | 100281008 | AX-170243777 | 0 | C | G | 0.412 | -7.93E-03 | 1.06E-01 | 2.69E+00 | 2.75E+00 | 9.36E-01 | no | Gm8091 | intron_variant,non_coding_transcript_variant | MODIFIER |
| 5 | 101040427 | AX-170270409 | 0 | C | A | 0.412 | -7.93E-03 | 1.06E-01 | 2.69E+00 | 2.75E+00 | 9.36E-01 | no | | regulatory_region_variant | MODIFIER |
| 5 | 101053206 | AX-170294069 | 0 | C | T | 0.412 | -7.93E-03 | 1.06E-01 | 2.69E+00 | 2.75E+00 | 9.36E-01 | no | | intergenic_variant | MODIFIER |
| 5 | 101061149 | AX-170196444 | 0 | G | A | 0.412 | -7.93E-03 | 1.06E-01 | 2.69E+00 | 2.75E+00 | 9.36E-01 | no | | intergenic_variant | MODIFIER |
| 5 | 101064637 | AX-170446335 | 0 | C | T | 0.412 | -7.93E-03 | 1.06E-01 | 2.69E+00 | 2.75E+00 | 9.36E-01 | no | | intergenic_variant | MODIFIER |
| 5 | 101068557 | AX-170072893 | 0 | T | A | 0.412 | -7.93E-03 | 1.06E-01 | 2.69E+00 | 2.75E+00 | 9.36E-01 | no | | intergenic_variant | MODIFIER |
| 5 | 101080673 | AX-170478367 | 0 | G | A | 0.412 | -7.93E-03 | 1.06E-01 | 2.69E+00 | 2.75E+00 | 9.36E-01 | no | | intergenic_variant | MODIFIER |
| 5 | 101080711 | AX-170072891 | 0 | A | T | 0.412 | -7.93E-03 | 1.06E-01 | 2.69E+00 | 2.75E+00 | 9.36E-01 | no | | intergenic_variant | MODIFIER |
| 5 | 101093517 | AX-170117540 | 0 | A | G | 0.412 | -7.93E-03 | 1.06E-01 | 2.69E+00 | 2.75E+00 | 9.36E-01 | no | | intergenic_variant | MODIFIER |
| 5 | 101115775 | AX-97843477 | 0 | C | T | 0.412 | -7.93E-03 | 1.06E-01 | 2.69E+00 | 2.75E+00 | 9.36E-01 | no | | intergenic_variant | MODIFIER |
| 5 | 101138541 | AX-170492387 | 0 | G | A | 0.412 | -7.93E-03 | 1.06E-01 | 2.69E+00 | 2.75E+00 | 9.36E-01 | no | | intergenic_variant | MODIFIER |
| 5 | 101143060 | AX-170207726 | 0 | A | T | 0.412 | -7.93E-03 | 1.06E-01 | 2.69E+00 | 2.75E+00 | 9.36E-01 | no | | intergenic_variant | MODIFIER |
| 5 | 101170201 | AX-170241856 | 0 | G | C | 0.412 | -7.93E-03 | 1.06E-01 | 2.69E+00 | 2.75E+00 | 9.36E-01 | no | | intergenic_variant | MODIFIER |
| 5 | 101170238 | AX-170497897 | 0 | G | A | 0.412 | -7.93E-03 | 1.06E-01 | 2.69E+00 | 2.75E+00 | 9.36E-01 | no | | intergenic_variant | MODIFIER |
| 5 | 101172387 | AX-170501062 | 0 | G | A | 0.412 | -7.93E-03 | 1.06E-01 | 2.69E+00 | 2.75E+00 | 9.36E-01 | no | | intergenic_variant | MODIFIER |
| 5 | 101191190 | AX-170452717 | 0 | T | C | 0.412 | -7.93E-03 | 1.06E-01 | 2.69E+00 | 2.75E+00 | 9.36E-01 | no | | intergenic_variant | MODIFIER |
| 5 | 101203115 | AX-170144670 | 0 | A | G | 0.412 | -7.93E-03 | 1.06E-01 | 2.69E+00 | 2.75E+00 | 9.36E-01 | no | | intergenic_variant | MODIFIER |
| 5 | 101208723 | AX-170216443 | 0 | A | T | 0.412 | -7.93E-03 | 1.06E-01 | 2.69E+00 | 2.75E+00 | 9.36E-01 | no | | regulatory_region_variant | MODIFIER |
| 5 | 101208873 | AX-170072897 | 0 | C | G | 0.412 | -7.93E-03 | 1.06E-01 | 2.69E+00 | 2.75E+00 | 9.36E-01 | no | | intergenic_variant | MODIFIER |
| 5 | 101214565 | AX-170072898 | 0 | G | A | 0.412 | -7.93E-03 | 1.06E-01 | 2.69E+00 | 2.75E+00 | 9.36E-01 | no | | intergenic_variant | MODIFIER |
| 5 | 101215441 | AX-170491938 | 0 | C | T | 0.412 | -7.93E-03 | 1.06E-01 | 2.69E+00 | 2.75E+00 | 9.36E-01 | no | | intergenic_variant | MODIFIER |
| 5 | 101223750 | AX-170362308 | 0 | C | G | 0.412 | -7.93E-03 | 1.06E-01 | 2.69E+00 | 2.75E+00 | 9.36E-01 | no | | intergenic_variant | MODIFIER |
| 5 | 101227759 | AX-170390249 | 0 | G | C | 0.412 | -7.93E-03 | 1.06E-01 | 2.69E+00 | 2.75E+00 | 9.36E-01 | no | | intergenic_variant | MODIFIER |
| 5 | 101229041 | AX-97785876 | 0 | A | G | 0.412 | -7.93E-03 | 1.06E-01 | 2.69E+00 | 2.75E+00 | 9.36E-01 | no | | intergenic_variant | MODIFIER |
| 5 | 101234204 | AX-170508208 | 0 | T | C | 0.412 | -7.93E-03 | 1.06E-01 | 2.69E+00 | 2.75E+00 | 9.36E-01 | no | | intergenic_variant | MODIFIER |
| 5 | 101246621 | AX-97807123 | 0 | C | A | 0.412 | -7.93E-03 | 1.06E-01 | 2.69E+00 | 2.75E+00 | 9.36E-01 | no | | intergenic_variant | MODIFIER |
| 5 | 101289433 | AX-170276243 | 0 | A | G | 0.412 | -7.93E-03 | 1.06E-01 | 2.69E+00 | 2.75E+00 | 9.36E-01 | no | | intergenic_variant | MODIFIER |
| 5 | 101295979 | AX-170468241 | 0 | T | G | 0.412 | -7.93E-03 | 1.06E-01 | 2.69E+00 | 2.75E+00 | 9.36E-01 | no | | regulatory_region_variant | MODIFIER |
| 5 | 101296321 | AX-170197701 | 0 | T | C | 0.412 | -7.93E-03 | 1.06E-01 | 2.69E+00 | 2.75E+00 | 9.36E-01 | no | | regulatory_region_variant | MODIFIER |
| 5 | 101296400 | AX-170238746 | 0 | G | A | 0.412 | -7.93E-03 | 1.06E-01 | 2.69E+00 | 2.75E+00 | 9.36E-01 | no | | regulatory_region_variant | MODIFIER |
| 5 | 101320338 | AX-170229248 | 0 | T | A | 0.412 | -7.93E-03 | 1.06E-01 | 2.69E+00 | 2.75E+00 | 9.36E-01 | no | | intergenic_variant | MODIFIER |
| 5 | 101450130 | AX-170072905 | 0 | T | C | 0.412 | -7.93E-03 | 1.06E-01 | 2.69E+00 | 2.75E+00 | 9.36E-01 | no | | intergenic_variant | MODIFIER |
| 5 | 101492352 | AX-170072904 | 0 | A | C | 0.412 | -7.93E-03 | 1.06E-01 | 2.69E+00 | 2.75E+00 | 9.36E-01 | no | | intergenic_variant | MODIFIER |
| 5 | 101551792 | AX-170362611 | 0 | A | G | 0.412 | -7.93E-03 | 1.06E-01 | 2.69E+00 | 2.75E+00 | 9.36E-01 | no | | intergenic_variant | MODIFIER |
| 18 | 34227798 | AX-97841935 | 0 | A | T | 0.07 | -1.93E-02 | 2.29E-01 | 2.63E+00 | 2.73E+00 | 9.36E-01 | no | Apc | intron_variant | MODIFIER |
| 18 | 39127221 | AX-170374668 | 0 | A | C | 0.07 | -1.93E-02 | 2.29E-01 | 2.63E+00 | 2.73E+00 | 9.36E-01 | no | Arhgap26 | intron_variant | MODIFIER |
| 7 | 119034543 | AX-170534885 | 0 | G | T | 0.395 | 7.23E-03 | 8.88E-02 | 2.67E+00 | 2.73E+00 | 9.37E-01 | no | | regulatory_region_variant | MODIFIER |
| 10 | 119444909 | AX-170407752 | 0 | G | A | 0.333 | -5.70E-03 | 7.00E-02 | 2.64E+00 | 2.73E+00 | 9.37E-01 | no | Grip1os3 | intron_variant,non_coding_transcript_variant | MODIFIER |
| 9 | 107260311 | AX-170033226 | 1 | G | A | 0.125 | 8.12E-03 | 1.18E-01 | 2.69E+00 | 2.76E+00 | 9.37E-01 | no | Mapkapk3 | intron_variant | MODIFIER |
| 7 | 78507976 | AX-97815599 | 0 | A | G | 0.132 | -1.66E-02 | 2.13E-01 | 2.66E+00 | 2.74E+00 | 9.37E-01 | no | Ntrk3 | intron_variant | MODIFIER |
| 7 | 78724253 | AX-170029370 | 0 | C | T | 0.132 | -1.66E-02 | 2.13E-01 | 2.66E+00 | 2.74E+00 | 9.37E-01 | no | Ntrk3 | downstream_gene_variant | MODIFIER |

| | | | | | | | | | | | | | | | |
|---|---|---|---|---|---|---|---|---|---|---|---|---|---|---|---|
| 7 | 78936507 | AX-170148007 | 0 | A | G | 0.132 | -1.66E-02 | 2.13E-01 | 2.66E+00 | 2.74E+00 | 9.37E-01 | no | | intergenic_variant | MODIFIER |
| 7 | 78936559 | AX-170171925 | 0 | T | A | 0.132 | -1.66E-02 | 2.13E-01 | 2.66E+00 | 2.74E+00 | 9.37E-01 | no | | intergenic_variant | MODIFIER |
| 7 | 78937739 | AX-97850445 | 0 | C | T | 0.132 | -1.66E-02 | 2.13E-01 | 2.66E+00 | 2.74E+00 | 9.37E-01 | no | | intergenic_variant | MODIFIER |
| 7 | 78942469 | AX-170281798 | 0 | G | A | 0.132 | -1.66E-02 | 2.13E-01 | 2.66E+00 | 2.74E+00 | 9.37E-01 | no | | intergenic_variant | MODIFIER |
| 7 | 78952173 | AX-170419162 | 0 | A | C | 0.132 | -1.66E-02 | 2.13E-01 | 2.66E+00 | 2.74E+00 | 9.37E-01 | no | | intergenic_variant | MODIFIER |
| 7 | 79018876 | AX-170263361 | 0 | T | C | 0.132 | -1.66E-02 | 2.13E-01 | 2.66E+00 | 2.74E+00 | 9.37E-01 | no | Gm26633 | non_coding_transcript_exon_variant | MODIFIER |
| 7 | 79026759 | AX-170405529 | 0 | A | G | 0.132 | -1.66E-02 | 2.13E-01 | 2.66E+00 | 2.74E+00 | 9.37E-01 | no | Gm26633 | intron_variant,non_coding_transcript_variant | MODIFIER |
| 7 | 79026907 | AX-170095994 | 0 | T | C | 0.132 | -1.66E-02 | 2.13E-01 | 2.66E+00 | 2.74E+00 | 9.37E-01 | no | Gm26633 | intron_variant,non_coding_transcript_variant | MODIFIER |
| 7 | 79110409 | AX-97844446 | 0 | A | G | 0.132 | -1.66E-02 | 2.13E-01 | 2.66E+00 | 2.74E+00 | 9.37E-01 | no | Acan | intron_variant | MODIFIER |
| 7 | 79111527 | AX-170029380 | 0 | A | T | 0.132 | -1.66E-02 | 2.13E-01 | 2.66E+00 | 2.74E+00 | 9.37E-01 | no | Acan | intron_variant | MODIFIER |
| 7 | 79137393 | AX-170510670 | 0 | C | G | 0.132 | -1.66E-02 | 2.13E-01 | 2.66E+00 | 2.74E+00 | 9.37E-01 | no | Mfge8 | intron_variant | MODIFIER |
| 7 | 79138431 | AX-170077821 | 0 | A | G | 0.132 | -1.66E-02 | 2.13E-01 | 2.66E+00 | 2.74E+00 | 9.37E-01 | no | Mfge8 | intron_variant | MODIFIER |
| 7 | 79140281 | AX-170329603 | 0 | G | A | 0.132 | -1.66E-02 | 2.13E-01 | 2.66E+00 | 2.74E+00 | 9.37E-01 | no | Mfge8 | intron_variant | MODIFIER |
| 7 | 79141296 | AX-97811706 | 0 | A | G | 0.132 | -1.66E-02 | 2.13E-01 | 2.66E+00 | 2.74E+00 | 9.37E-01 | no | Mfge8 | intron_variant | MODIFIER |
| 7 | 79141309 | AX-170289834 | 0 | T | A | 0.132 | -1.66E-02 | 2.13E-01 | 2.66E+00 | 2.74E+00 | 9.37E-01 | no | Mfge8 | intron_variant | MODIFIER |
| 7 | 79141713 | AX-170029381 | 0 | C | T | 0.132 | -1.66E-02 | 2.13E-01 | 2.66E+00 | 2.74E+00 | 9.37E-01 | no | Mfge8 | missense_variant | MODERATE |
| 7 | 79143096 | AX-170029379 | 0 | A | G | 0.132 | -1.66E-02 | 2.13E-01 | 2.66E+00 | 2.74E+00 | 9.37E-01 | no | Mfge8 | intron_variant | MODIFIER |
| 7 | 79159088 | AX-170434861 | 0 | T | G | 0.132 | -1.66E-02 | 2.13E-01 | 2.66E+00 | 2.74E+00 | 9.37E-01 | no | | intergenic_variant | MODIFIER |
| 7 | 79176146 | AX-170129893 | 0 | A | G | 0.132 | -1.66E-02 | 2.13E-01 | 2.66E+00 | 2.74E+00 | 9.37E-01 | no | | intergenic_variant | MODIFIER |
| 7 | 79177038 | AX-170428985 | 0 | C | T | 0.132 | -1.66E-02 | 2.13E-01 | 2.66E+00 | 2.74E+00 | 9.37E-01 | no | | intergenic_variant | MODIFIER |
| 7 | 79203008 | AX-170351814 | 0 | A | G | 0.132 | -1.66E-02 | 2.13E-01 | 2.66E+00 | 2.74E+00 | 9.37E-01 | no | | regulatory_region_variant | MODIFIER |
| 7 | 79203062 | AX-170139014 | 0 | A | G | 0.132 | -1.66E-02 | 2.13E-01 | 2.66E+00 | 2.74E+00 | 9.37E-01 | no | | regulatory_region_variant | MODIFIER |
| 7 | 79435276 | AX-170262442 | 0 | G | A | 0.132 | -1.66E-02 | 2.13E-01 | 2.66E+00 | 2.74E+00 | 9.37E-01 | no | Fanci | missense_variant | MODERATE |
| 7 | 79658194 | AX-170508462 | 0 | T | C | 0.132 | -1.66E-02 | 2.13E-01 | 2.66E+00 | 2.74E+00 | 9.37E-01 | no | Ticrr | upstream_gene_variant | MODIFIER |
| 7 | 79667524 | AX-170417516 | 0 | T | C | 0.132 | -1.66E-02 | 2.13E-01 | 2.66E+00 | 2.74E+00 | 9.37E-01 | no | Ticrr | intron_variant | MODIFIER |
| 7 | 79684228 | AX-170292573 | 0 | A | C | 0.132 | -1.66E-02 | 2.13E-01 | 2.66E+00 | 2.74E+00 | 9.37E-01 | no | Ticrr | missense_variant | MODERATE |
| 7 | 79688904 | AX-170077831 | 0 | G | C | 0.132 | -1.66E-02 | 2.13E-01 | 2.66E+00 | 2.74E+00 | 9.37E-01 | no | Ticrr | intron_variant | MODIFIER |
| 7 | 79699649 | AX-170077833 | 0 | G | A | 0.132 | -1.66E-02 | 2.13E-01 | 2.66E+00 | 2.74E+00 | 9.37E-01 | no | Ticrr | downstream_gene_variant | MODIFIER |
| 7 | 88296239 | AX-170485173 | 0 | G | A | 0.132 | -1.66E-02 | 2.13E-01 | 2.66E+00 | 2.74E+00 | 9.37E-01 | no | Ctsc | intron_variant | MODIFIER |
| 7 | 88316222 | AX-170301812 | 0 | A | G | 0.132 | -1.66E-02 | 2.13E-01 | 2.66E+00 | 2.74E+00 | 9.37E-01 | no | Gm44751 | downstream_gene_variant | MODIFIER |
| 7 | 88318248 | AX-170122916 | 0 | G | A | 0.132 | -1.66E-02 | 2.13E-01 | 2.66E+00 | 2.74E+00 | 9.37E-01 | no | Gm44751 | downstream_gene_variant | MODIFIER |
| 7 | 88332096 | AX-170179843 | 0 | G | T | 0.132 | -1.66E-02 | 2.13E-01 | 2.66E+00 | 2.74E+00 | 9.37E-01 | no | | intergenic_variant | MODIFIER |
| 7 | 88524775 | AX-170029473 | 0 | C | T | 0.132 | -1.66E-02 | 2.13E-01 | 2.66E+00 | 2.74E+00 | 9.37E-01 | no | | intergenic_variant | MODIFIER |
| 7 | 88947234 | AX-170335473 | 0 | A | G | 0.132 | -1.66E-02 | 2.13E-01 | 2.66E+00 | 2.74E+00 | 9.37E-01 | no | | intergenic_variant | MODIFIER |
| 7 | 89412396 | AX-170078015 | 0 | T | C | 0.132 | -1.66E-02 | 2.13E-01 | 2.66E+00 | 2.74E+00 | 9.37E-01 | no | Fzd4 | 3_prime_UTR_variant | MODIFIER |
| 7 | 89413995 | AX-170078017 | 0 | G | A | 0.132 | -1.66E-02 | 2.13E-01 | 2.66E+00 | 2.74E+00 | 9.37E-01 | no | Fzd4 | downstream_gene_variant | MODIFIER |
| 7 | 89416450 | AX-170115516 | 0 | G | C | 0.132 | -1.66E-02 | 2.13E-01 | 2.66E+00 | 2.74E+00 | 9.37E-01 | no | Fzd4 | downstream_gene_variant | MODIFIER |
| 7 | 89592185 | AX-97857269 | 0 | A | G | 0.132 | -1.66E-02 | 2.13E-01 | 2.66E+00 | 2.74E+00 | 9.37E-01 | no | | intergenic_variant | MODIFIER |
| 7 | 89603538 | AX-170166608 | 0 | C | A | 0.132 | -1.66E-02 | 2.13E-01 | 2.66E+00 | 2.74E+00 | 9.37E-01 | no | Gm2546 | upstream_gene_variant | MODIFIER |
| 7 | 89640341 | AX-170435352 | 0 | T | C | 0.132 | -1.66E-02 | 2.13E-01 | 2.66E+00 | 2.74E+00 | 9.37E-01 | no | Me3 | intron_variant | MODIFIER |
| 7 | 89657339 | AX-170386346 | 0 | A | G | 0.132 | -1.66E-02 | 2.13E-01 | 2.66E+00 | 2.74E+00 | 9.37E-01 | no | Me3 | intron_variant | MODIFIER |
| 7 | 89697475 | AX-97836023 | 0 | T | G | 0.132 | -1.66E-02 | 2.13E-01 | 2.66E+00 | 2.74E+00 | 9.37E-01 | no | Me3 | intron_variant | MODIFIER |
| 7 | 89720469 | AX-170078020 | 0 | C | T | 0.132 | -1.66E-02 | 2.13E-01 | 2.66E+00 | 2.74E+00 | 9.37E-01 | no | Me3 | intron_variant | MODIFIER |
| 7 | 89724126 | AX-170089973 | 0 | G | A | 0.132 | -1.66E-02 | 2.13E-01 | 2.66E+00 | 2.74E+00 | 9.37E-01 | no | Me3 | intron_variant | MODIFIER |
| 7 | 89725586 | AX-170275006 | 0 | T | C | 0.132 | -1.66E-02 | 2.13E-01 | 2.66E+00 | 2.74E+00 | 9.37E-01 | no | Me3 | intron_variant | MODIFIER |
| 7 | 89728122 | AX-170170207 | 0 | A | G | 0.132 | -1.66E-02 | 2.13E-01 | 2.66E+00 | 2.74E+00 | 9.37E-01 | no | Me3 | intron_variant | MODIFIER |
| 7 | 89737828 | AX-170078021 | 0 | A | C | 0.132 | -1.66E-02 | 2.13E-01 | 2.66E+00 | 2.74E+00 | 9.37E-01 | no | Me3 | intron_variant | MODIFIER |
| 7 | 89743082 | AX-170363716 | 0 | C | T | 0.132 | -1.66E-02 | 2.13E-01 | 2.66E+00 | 2.74E+00 | 9.37E-01 | no | Me3 | intron_variant | MODIFIER |
| 7 | 89775556 | AX-170309862 | 0 | T | C | 0.132 | -1.66E-02 | 2.13E-01 | 2.66E+00 | 2.74E+00 | 9.37E-01 | no | Me3 | intron_variant | MODIFIER |
| 7 | 89792999 | AX-170327687 | 0 | A | G | 0.132 | -1.66E-02 | 2.13E-01 | 2.66E+00 | 2.74E+00 | 9.37E-01 | no | Me3 | intron_variant | MODIFIER |
| 7 | 89838989 | AX-170196876 | 0 | C | T | 0.132 | -1.66E-02 | 2.13E-01 | 2.66E+00 | 2.74E+00 | 9.37E-01 | no | Me3 | intron_variant,non_coding_transcript_variant | MODIFIER |
| 7 | 89839410 | AX-170455058 | 0 | A | G | 0.132 | -1.66E-02 | 2.13E-01 | 2.66E+00 | 2.74E+00 | 9.37E-01 | no | Me3 | intron_variant,non_coding_transcript_variant | MODIFIER |
| 7 | 89840721 | AX-170280913 | 0 | T | C | 0.132 | -1.66E-02 | 2.13E-01 | 2.66E+00 | 2.74E+00 | 9.37E-01 | no | Me3 | intron_variant,non_coding_transcript_variant | MODIFIER |
| 7 | 89850817 | AX-170154899 | 0 | G | A | 0.132 | -1.66E-02 | 2.13E-01 | 2.66E+00 | 2.74E+00 | 9.37E-01 | no | Me3 | intron_variant,non_coding_transcript_variant | MODIFIER |

| | | | | | | | | | | | | | | | |
|---|---|---|---|---|---|---|---|---|---|---|---|---|---|---|---|
| 7 | 89856745 | AX-170148596 | 0 | A | C | 0.132 | -1.66E-02 | 2.13E-01 | 2.66E+00 | 2.74E+00 | 9.37E-01 | no | Me3 | downstream_gene_variant | MODIFIER |
| 7 | 89856821 | AX-170121308 | 0 | A | C | 0.132 | -1.66E-02 | 2.13E-01 | 2.66E+00 | 2.74E+00 | 9.37E-01 | no | Me3 | downstream_gene_variant | MODIFIER |
| 7 | 89856890 | AX-170197430 | 0 | G | T | 0.132 | -1.66E-02 | 2.13E-01 | 2.66E+00 | 2.74E+00 | 9.37E-01 | no | Me3 | downstream_gene_variant | MODIFIER |
| 7 | 89871974 | AX-170311512 | 0 | T | C | 0.132 | -1.66E-02 | 2.13E-01 | 2.66E+00 | 2.74E+00 | 9.37E-01 | no | Ccdc81 | intron_variant | MODIFIER |
| 7 | 89881033 | AX-170353928 | 0 | A | G | 0.132 | -1.66E-02 | 2.13E-01 | 2.66E+00 | 2.74E+00 | 9.37E-01 | no | Ccdc81 | intron_variant | MODIFIER |
| 7 | 89971323 | AX-170134577 | 0 | A | G | 0.132 | -1.66E-02 | 2.13E-01 | 2.66E+00 | 2.74E+00 | 9.37E-01 | no | Eed | intron_variant | MODIFIER |
| 7 | 89982521 | AX-170119605 | 0 | T | C | 0.132 | -1.66E-02 | 2.13E-01 | 2.66E+00 | 2.74E+00 | 9.37E-01 | no | Eed | upstream_gene_variant | MODIFIER |
| 7 | 90041534 | AX-170174142 | 0 | C | T | 0.132 | -1.66E-02 | 2.13E-01 | 2.66E+00 | 2.74E+00 | 9.37E-01 | no | E230029C05Rik | downstream_gene_variant | MODIFIER |
| 7 | 90041593 | AX-170279827 | 0 | T | C | 0.132 | -1.66E-02 | 2.13E-01 | 2.66E+00 | 2.74E+00 | 9.37E-01 | no | E230029C05Rik | downstream_gene_variant | MODIFIER |
| 7 | 90043381 | AX-170298301 | 0 | G | A | 0.132 | -1.66E-02 | 2.13E-01 | 2.66E+00 | 2.74E+00 | 9.37E-01 | no | E230029C05Rik | downstream_gene_variant | MODIFIER |
| 7 | 90045445 | AX-170295918 | 0 | T | C | 0.132 | -1.66E-02 | 2.13E-01 | 2.66E+00 | 2.74E+00 | 9.37E-01 | no | E230029C05Rik | intron_variant,non_coding_transcript_variant | MODIFIER |
| 7 | 90045876 | AX-170297966 | 0 | G | A | 0.132 | -1.66E-02 | 2.13E-01 | 2.66E+00 | 2.74E+00 | 9.37E-01 | no | E230029C05Rik | intron_variant,non_coding_transcript_variant | MODIFIER |
| 7 | 90052461 | AX-170187565 | 0 | T | C | 0.132 | -1.66E-02 | 2.13E-01 | 2.66E+00 | 2.74E+00 | 9.37E-01 | no | E230029C05Rik | downstream_gene_variant | MODIFIER |
| 7 | 90113793 | AX-170376198 | 0 | C | T | 0.132 | -1.66E-02 | 2.13E-01 | 2.66E+00 | 2.74E+00 | 9.37E-01 | no | | intergenic_variant | MODIFIER |
| 7 | 90125657 | AX-170029483 | 0 | A | G | 0.132 | -1.66E-02 | 2.13E-01 | 2.66E+00 | 2.74E+00 | 9.37E-01 | no | Picalm | upstream_gene_variant | MODIFIER |
| 7 | 90141888 | AX-170078028 | 0 | A | G | 0.132 | -1.66E-02 | 2.13E-01 | 2.66E+00 | 2.74E+00 | 9.37E-01 | no | Picalm | intron_variant | MODIFIER |
| 7 | 90142546 | AX-170078027 | 0 | T | C | 0.132 | -1.66E-02 | 2.13E-01 | 2.66E+00 | 2.74E+00 | 9.37E-01 | no | Picalm | intron_variant | MODIFIER |
| 7 | 90143389 | AX-170096019 | 0 | C | T | 0.132 | -1.66E-02 | 2.13E-01 | 2.66E+00 | 2.74E+00 | 9.37E-01 | no | Picalm | intron_variant | MODIFIER |
| 7 | 90214175 | AX-170164591 | 0 | A | G | 0.132 | -1.66E-02 | 2.13E-01 | 2.66E+00 | 2.74E+00 | 9.37E-01 | no | Picalm | downstream_gene_variant | MODIFIER |
| 7 | 90217274 | AX-170268085 | 0 | A | G | 0.132 | -1.66E-02 | 2.13E-01 | 2.66E+00 | 2.74E+00 | 9.37E-01 | no | Picalm | downstream_gene_variant | MODIFIER |
| 7 | 90217880 | AX-170357019 | 0 | C | T | 0.132 | -1.66E-02 | 2.13E-01 | 2.66E+00 | 2.74E+00 | 9.37E-01 | no | Picalm | downstream_gene_variant | MODIFIER |
| 7 | 90221068 | AX-170403857 | 0 | G | A | 0.132 | -1.66E-02 | 2.13E-01 | 2.66E+00 | 2.74E+00 | 9.37E-01 | no | Ccdc83 | downstream_gene_variant | MODIFIER |
| 7 | 90221115 | AX-170439511 | 0 | A | G | 0.132 | -1.66E-02 | 2.13E-01 | 2.66E+00 | 2.74E+00 | 9.37E-01 | no | Ccdc83 | downstream_gene_variant | MODIFIER |
| 7 | 90245501 | AX-170078029 | 0 | C | T | 0.132 | -1.66E-02 | 2.13E-01 | 2.66E+00 | 2.74E+00 | 9.37E-01 | no | Ccdc83 | intron_variant | MODIFIER |
| 7 | 90245583 | AX-170078030 | 0 | A | G | 0.132 | -1.66E-02 | 2.13E-01 | 2.66E+00 | 2.74E+00 | 9.37E-01 | no | Ccdc83 | intron_variant | MODIFIER |
| 7 | 90262200 | AX-170106436 | 0 | A | G | 0.132 | -1.66E-02 | 2.13E-01 | 2.66E+00 | 2.74E+00 | 9.37E-01 | no | Ccdc83 | intron_variant | MODIFIER |
| 7 | 90264504 | AX-170208842 | 0 | A | G | 0.132 | -1.66E-02 | 2.13E-01 | 2.66E+00 | 2.74E+00 | 9.37E-01 | no | Ccdc83 | intron_variant | MODIFIER |
| 7 | 90282495 | AX-170232867 | 0 | C | T | 0.132 | -1.66E-02 | 2.13E-01 | 2.66E+00 | 2.74E+00 | 9.37E-01 | no | | regulatory_region_variant | MODIFIER |
| 7 | 90319102 | AX-97855149 | 0 | A | G | 0.132 | -1.66E-02 | 2.13E-01 | 2.66E+00 | 2.74E+00 | 9.37E-01 | no | Sytl2 | intron_variant | MODIFIER |
| 7 | 90403660 | AX-97835432 | 0 | A | G | 0.132 | -1.66E-02 | 2.13E-01 | 2.66E+00 | 2.74E+00 | 9.37E-01 | no | Sytl2 | intron_variant,non_coding_transcript_variant | MODIFIER |
| 7 | 90409199 | AX-170274416 | 0 | G | T | 0.132 | -1.66E-02 | 2.13E-01 | 2.66E+00 | 2.74E+00 | 9.37E-01 | no | Sytl2 | intron_variant,non_coding_transcript_variant | MODIFIER |
| 7 | 90429990 | AX-170387541 | 0 | C | T | 0.132 | -1.66E-02 | 2.13E-01 | 2.66E+00 | 2.74E+00 | 9.37E-01 | no | Ccdc89 | downstream_gene_variant | MODIFIER |
| 7 | 90430538 | AX-97843736 | 0 | T | C | 0.132 | -1.66E-02 | 2.13E-01 | 2.66E+00 | 2.74E+00 | 9.37E-01 | no | Ccdc89 | downstream_gene_variant | MODIFIER |
| 7 | 90430770 | AX-170152270 | 0 | C | T | 0.132 | -1.66E-02 | 2.13E-01 | 2.66E+00 | 2.74E+00 | 9.37E-01 | no | Ccdc89 | downstream_gene_variant | MODIFIER |
| 7 | 90451777 | AX-97798518 | 0 | G | T | 0.132 | -1.66E-02 | 2.13E-01 | 2.66E+00 | 2.74E+00 | 9.37E-01 | no | Tmem126a | intron_variant | MODIFIER |
| 7 | 90473439 | AX-170362927 | 0 | C | G | 0.132 | -1.66E-02 | 2.13E-01 | 2.66E+00 | 2.74E+00 | 9.37E-01 | no | Tmem126b | intron_variant | MODIFIER |
| 7 | 90477577 | AX-97855841 | 0 | G | A | 0.132 | -1.66E-02 | 2.13E-01 | 2.66E+00 | 2.74E+00 | 9.37E-01 | no | Tmem126b | upstream_gene_variant | MODIFIER |
| 7 | 90489269 | AX-170078036 | 0 | T | C | 0.132 | -1.66E-02 | 2.13E-01 | 2.66E+00 | 2.74E+00 | 9.37E-01 | no | Dlg2 | intron_variant,non_coding_transcript_variant | MODIFIER |
| 7 | 90502488 | AX-170485728 | 0 | C | T | 0.132 | -1.66E-02 | 2.13E-01 | 2.66E+00 | 2.74E+00 | 9.37E-01 | no | Dlg2 | intron_variant,non_coding_transcript_variant | MODIFIER |
| 7 | 90539354 | AX-170123270 | 0 | G | A | 0.132 | -1.66E-02 | 2.13E-01 | 2.66E+00 | 2.74E+00 | 9.37E-01 | no | Dlg2 | intron_variant | MODIFIER |
| 7 | 90550915 | AX-170138881 | 0 | A | G | 0.132 | -1.66E-02 | 2.13E-01 | 2.66E+00 | 2.74E+00 | 9.37E-01 | no | Dlg2 | intron_variant | MODIFIER |
| 7 | 90555277 | AX-170197755 | 0 | C | T | 0.132 | -1.66E-02 | 2.13E-01 | 2.66E+00 | 2.74E+00 | 9.37E-01 | no | Dlg2 | intron_variant | MODIFIER |
| 7 | 90566589 | AX-170322420 | 0 | C | T | 0.132 | -1.66E-02 | 2.13E-01 | 2.66E+00 | 2.74E+00 | 9.37E-01 | no | Dlg2 | intron_variant | MODIFIER |
| 7 | 90566776 | AX-170477460 | 0 | T | C | 0.132 | -1.66E-02 | 2.13E-01 | 2.66E+00 | 2.74E+00 | 9.37E-01 | no | Dlg2 | intron_variant | MODIFIER |
| 7 | 90631544 | AX-170512641 | 0 | T | C | 0.132 | -1.66E-02 | 2.13E-01 | 2.66E+00 | 2.74E+00 | 9.37E-01 | no | Dlg2 | intron_variant | MODIFIER |
| 7 | 90901487 | AX-97846921 | 0 | A | G | 0.132 | -1.66E-02 | 2.13E-01 | 2.66E+00 | 2.74E+00 | 9.37E-01 | no | Gm45159 | intron_variant,non_coding_transcript_variant | MODIFIER |
| 7 | 91056928 | AX-170213485 | 0 | G | T | 0.132 | -1.66E-02 | 2.13E-01 | 2.66E+00 | 2.74E+00 | 9.37E-01 | no | Gm45179 | downstream_gene_variant | MODIFIER |
| 7 | 91062591 | AX-170097877 | 0 | G | C | 0.132 | -1.66E-02 | 2.13E-01 | 2.66E+00 | 2.74E+00 | 9.37E-01 | no | Dlg2 | intron_variant | MODIFIER |
| 7 | 91069906 | AX-170475204 | 0 | G | A | 0.132 | -1.66E-02 | 2.13E-01 | 2.66E+00 | 2.74E+00 | 9.37E-01 | no | Dlg2 | intron_variant | MODIFIER |
| 7 | 91069929 | AX-170509428 | 0 | G | A | 0.132 | -1.66E-02 | 2.13E-01 | 2.66E+00 | 2.74E+00 | 9.37E-01 | no | Dlg2 | intron_variant | MODIFIER |
| 7 | 91071927 | AX-170366752 | 0 | T | C | 0.132 | -1.66E-02 | 2.13E-01 | 2.66E+00 | 2.74E+00 | 9.37E-01 | no | Dlg2 | intron_variant | MODIFIER |
| 7 | 91074068 | AX-170141690 | 0 | G | T | 0.132 | -1.66E-02 | 2.13E-01 | 2.66E+00 | 2.74E+00 | 9.37E-01 | no | Dlg2 | intron_variant | MODIFIER |
| 7 | 91075638 | AX-97845096 | 0 | G | A | 0.132 | -1.66E-02 | 2.13E-01 | 2.66E+00 | 2.74E+00 | 9.37E-01 | no | Dlg2 | intron_variant | MODIFIER |
| 7 | 91094321 | AX-170078049 | 0 | T | A | 0.132 | -1.66E-02 | 2.13E-01 | 2.66E+00 | 2.74E+00 | 9.37E-01 | no | Dlg2 | intron_variant | MODIFIER |

| | | | | | | | | | | | | | | | | |
|---|---|---|---|---|---|---|---|---|---|---|---|---|---|---|---|---|
| 7 | 91100086 | AX-170317893 | 0 | C | T | 0.132 | -1.66E-02 | 2.13E-01 | 2.66E+00 | 2.74E+00 | 9.37E-01 | no | Dlg2 | | intron_variant | MODIFIER |
| 7 | 91100918 | AX-170471624 | 0 | G | A | 0.132 | -1.66E-02 | 2.13E-01 | 2.66E+00 | 2.74E+00 | 9.37E-01 | no | Dlg2 | | intron_variant | MODIFIER |
| 7 | 91332809 | AX-170488034 | 0 | A | T | 0.132 | -1.66E-02 | 2.13E-01 | 2.66E+00 | 2.74E+00 | 9.37E-01 | no | Dlg2 | | intron_variant | MODIFIER |
| 7 | 91371095 | AX-170405925 | 0 | C | T | 0.132 | -1.66E-02 | 2.13E-01 | 2.66E+00 | 2.74E+00 | 9.37E-01 | no | Dlg2 | | intron_variant | MODIFIER |
| 7 | 91382908 | AX-170117905 | 0 | T | C | 0.132 | -1.66E-02 | 2.13E-01 | 2.66E+00 | 2.74E+00 | 9.37E-01 | no | Dlg2 | | intron_variant | MODIFIER |
| 7 | 91386983 | AX-170456902 | 0 | T | C | 0.132 | -1.66E-02 | 2.13E-01 | 2.66E+00 | 2.74E+00 | 9.37E-01 | no | Dlg2 | | intron_variant | MODIFIER |
| 7 | 91400371 | AX-170101892 | 0 | T | C | 0.132 | -1.66E-02 | 2.13E-01 | 2.66E+00 | 2.74E+00 | 9.37E-01 | no | Dlg2 | | intron_variant | MODIFIER |
| 7 | 91400733 | AX-170359656 | 0 | C | T | 0.132 | -1.66E-02 | 2.13E-01 | 2.66E+00 | 2.74E+00 | 9.37E-01 | no | Dlg2 | | intron_variant | MODIFIER |
| 7 | 91415220 | AX-97846374 | 0 | C | T | 0.132 | -1.66E-02 | 2.13E-01 | 2.66E+00 | 2.74E+00 | 9.37E-01 | no | Dlg2 | | intron_variant | MODIFIER |
| 7 | 91424003 | AX-170430505 | 0 | G | C | 0.132 | -1.66E-02 | 2.13E-01 | 2.66E+00 | 2.74E+00 | 9.37E-01 | no | Dlg2 | | intron_variant | MODIFIER |
| 7 | 91455341 | AX-170078061 | 0 | T | C | 0.132 | -1.66E-02 | 2.13E-01 | 2.66E+00 | 2.74E+00 | 9.37E-01 | no | Dlg2 | | intron_variant | MODIFIER |
| 7 | 91457541 | AX-170078059 | 0 | T | A | 0.132 | -1.66E-02 | 2.13E-01 | 2.66E+00 | 2.74E+00 | 9.37E-01 | no | Dlg2 | | intron_variant | MODIFIER |
| 7 | 91460050 | AX-170472905 | 0 | T | C | 0.132 | -1.66E-02 | 2.13E-01 | 2.66E+00 | 2.74E+00 | 9.37E-01 | no | Dlg2 | | intron_variant | MODIFIER |
| 7 | 91466462 | AX-170148815 | 0 | G | A | 0.132 | -1.66E-02 | 2.13E-01 | 2.66E+00 | 2.74E+00 | 9.37E-01 | no | Dlg2 | | intron_variant | MODIFIER |
| 7 | 91468229 | AX-170520916 | 0 | T | C | 0.132 | -1.66E-02 | 2.13E-01 | 2.66E+00 | 2.74E+00 | 9.37E-01 | no | Dlg2 | | intron_variant | MODIFIER |
| 7 | 91471847 | AX-170411360 | 0 | C | T | 0.132 | -1.66E-02 | 2.13E-01 | 2.66E+00 | 2.74E+00 | 9.37E-01 | no | Dlg2 | | intron_variant | MODIFIER |
| 7 | 91478174 | AX-170265421 | 0 | T | C | 0.132 | -1.66E-02 | 2.13E-01 | 2.66E+00 | 2.74E+00 | 9.37E-01 | no | Dlg2 | | intron_variant | MODIFIER |
| 7 | 91506818 | AX-170228486 | 0 | A | G | 0.132 | -1.66E-02 | 2.13E-01 | 2.66E+00 | 2.74E+00 | 9.37E-01 | no | Dlg2 | | intron_variant | MODIFIER |
| 7 | 91508770 | AX-170446979 | 0 | C | T | 0.132 | -1.66E-02 | 2.13E-01 | 2.66E+00 | 2.74E+00 | 9.37E-01 | no | Dlg2 | | intron_variant | MODIFIER |
| 7 | 91556408 | AX-170246186 | 0 | T | A | 0.132 | -1.66E-02 | 2.13E-01 | 2.66E+00 | 2.74E+00 | 9.37E-01 | no | Dlg2 | | intron_variant | MODIFIER |
| 7 | 91559218 | AX-170532589 | 0 | C | A | 0.132 | -1.66E-02 | 2.13E-01 | 2.66E+00 | 2.74E+00 | 9.37E-01 | no | Dlg2 | | intron_variant | MODIFIER |
| 7 | 91559312 | AX-170163168 | 0 | G | T | 0.132 | -1.66E-02 | 2.13E-01 | 2.66E+00 | 2.74E+00 | 9.37E-01 | no | Dlg2 | | intron_variant | MODIFIER |
| 7 | 91575444 | AX-170339196 | 0 | T | A | 0.132 | -1.66E-02 | 2.13E-01 | 2.66E+00 | 2.74E+00 | 9.37E-01 | no | Dlg2 | | intron_variant | MODIFIER |
| 7 | 91600010 | AX-170078062 | 0 | C | T | 0.132 | -1.66E-02 | 2.13E-01 | 2.66E+00 | 2.74E+00 | 9.37E-01 | no | Dlg2 | | intron_variant | MODIFIER |
| 7 | 91600200 | AX-170128439 | 0 | G | A | 0.132 | -1.66E-02 | 2.13E-01 | 2.66E+00 | 2.74E+00 | 9.37E-01 | no | Dlg2 | | intron_variant | MODIFIER |
| 7 | 91602844 | AX-170487505 | 0 | G | T | 0.132 | -1.66E-02 | 2.13E-01 | 2.66E+00 | 2.74E+00 | 9.37E-01 | no | Dlg2 | | intron_variant | MODIFIER |
| 7 | 91603387 | AX-170394659 | 0 | G | A | 0.132 | -1.66E-02 | 2.13E-01 | 2.66E+00 | 2.74E+00 | 9.37E-01 | no | Dlg2 | | intron_variant | MODIFIER |
| 7 | 91603755 | AX-170122886 | 0 | G | A | 0.132 | -1.66E-02 | 2.13E-01 | 2.66E+00 | 2.74E+00 | 9.37E-01 | no | Dlg2 | | intron_variant | MODIFIER |
| 7 | 91635850 | AX-170376475 | 0 | T | C | 0.132 | -1.66E-02 | 2.13E-01 | 2.66E+00 | 2.74E+00 | 9.37E-01 | no | Dlg2 | | intron_variant | MODIFIER |
| 7 | 91654046 | AX-170522993 | 0 | G | A | 0.132 | -1.66E-02 | 2.13E-01 | 2.66E+00 | 2.74E+00 | 9.37E-01 | no | Dlg2 | | intron_variant | MODIFIER |
| 7 | 91664726 | AX-170326265 | 0 | G | C | 0.132 | -1.66E-02 | 2.13E-01 | 2.66E+00 | 2.74E+00 | 9.37E-01 | no | Dlg2 | | intron_variant | MODIFIER |
| 7 | 91669965 | AX-170029500 | 0 | A | G | 0.132 | -1.66E-02 | 2.13E-01 | 2.66E+00 | 2.74E+00 | 9.37E-01 | no | Dlg2 | | intron_variant | MODIFIER |
| 7 | 91701051 | AX-97820553 | 0 | T | C | 0.132 | -1.66E-02 | 2.13E-01 | 2.66E+00 | 2.74E+00 | 9.37E-01 | no | Dlg2 | | intron_variant | MODIFIER |
| 7 | 91703684 | AX-170029501 | 0 | G | A | 0.132 | -1.66E-02 | 2.13E-01 | 2.66E+00 | 2.74E+00 | 9.37E-01 | no | Dlg2 | | intron_variant | MODIFIER |
| 7 | 91762039 | AX-170029502 | 0 | T | C | 0.132 | -1.66E-02 | 2.13E-01 | 2.66E+00 | 2.74E+00 | 9.37E-01 | no | Dlg2 | | intron_variant | MODIFIER |
| 7 | 91790901 | AX-170442350 | 0 | T | G | 0.132 | -1.66E-02 | 2.13E-01 | 2.66E+00 | 2.74E+00 | 9.37E-01 | no | Dlg2 | | intron_variant | MODIFIER |
| 7 | 91815917 | AX-170382502 | 0 | A | G | 0.132 | -1.66E-02 | 2.13E-01 | 2.66E+00 | 2.74E+00 | 9.37E-01 | no | Dlg2 | | intron_variant | MODIFIER |
| 7 | 91974166 | AX-170417418 | 0 | C | T | 0.132 | -1.66E-02 | 2.13E-01 | 2.66E+00 | 2.74E+00 | 9.37E-01 | no | Dlg2 | | intron_variant | MODIFIER |
| 7 | 92034769 | AX-170438905 | 0 | C | T | 0.132 | -1.66E-02 | 2.13E-01 | 2.66E+00 | 2.74E+00 | 9.37E-01 | no | Dlg2 | | intron_variant | MODIFIER |
| 7 | 92038625 | AX-97824411 | 0 | A | G | 0.132 | -1.66E-02 | 2.13E-01 | 2.66E+00 | 2.74E+00 | 9.37E-01 | no | Dlg2 | | intron_variant | MODIFIER |
| 7 | 92051223 | AX-170528273 | 0 | G | A | 0.132 | -1.66E-02 | 2.13E-01 | 2.66E+00 | 2.74E+00 | 9.37E-01 | no | Dlg2 | | intron_variant | MODIFIER |
| 7 | 92098618 | AX-170361977 | 0 | G | A | 0.132 | -1.66E-02 | 2.13E-01 | 2.66E+00 | 2.74E+00 | 9.37E-01 | no | Dlg2 | | intron_variant | MODIFIER |
| 7 | 92121849 | AX-97822524 | 0 | A | G | 0.132 | -1.66E-02 | 2.13E-01 | 2.66E+00 | 2.74E+00 | 9.37E-01 | no | Dlg2 | | intron_variant | MODIFIER |
| 7 | 92125853 | AX-170445664 | 0 | T | C | 0.132 | -1.66E-02 | 2.13E-01 | 2.66E+00 | 2.74E+00 | 9.37E-01 | no | Dlg2 | | intron_variant | MODIFIER |
| 7 | 92154955 | AX-170169074 | 0 | C | A | 0.132 | -1.66E-02 | 2.13E-01 | 2.66E+00 | 2.74E+00 | 9.37E-01 | no | Dlg2 | | intron_variant | MODIFIER |
| 7 | 92163480 | AX-170508103 | 0 | A | G | 0.132 | -1.66E-02 | 2.13E-01 | 2.66E+00 | 2.74E+00 | 9.37E-01 | no | Dlg2 | | intron_variant | MODIFIER |
| 7 | 92163711 | AX-170029509 | 0 | G | A | 0.132 | -1.66E-02 | 2.13E-01 | 2.66E+00 | 2.74E+00 | 9.37E-01 | no | Dlg2 | | intron_variant | MODIFIER |
| 7 | 92165768 | AX-170465441 | 0 | T | C | 0.132 | -1.66E-02 | 2.13E-01 | 2.66E+00 | 2.74E+00 | 9.37E-01 | no | Dlg2 | | intron_variant | MODIFIER |
| 7 | 92170477 | AX-170353115 | 0 | T | C | 0.132 | -1.66E-02 | 2.13E-01 | 2.66E+00 | 2.74E+00 | 9.37E-01 | no | Dlg2 | | intron_variant | MODIFIER |
| 7 | 92197474 | AX-170377298 | 0 | T | C | 0.132 | -1.66E-02 | 2.13E-01 | 2.66E+00 | 2.74E+00 | 9.37E-01 | no | Dlg2 | | intron_variant | MODIFIER |
| 7 | 92198036 | AX-170029514 | 0 | C | T | 0.132 | -1.66E-02 | 2.13E-01 | 2.66E+00 | 2.74E+00 | 9.37E-01 | no | Dlg2 | | intron_variant | MODIFIER |
| 7 | 92302828 | AX-97831514 | 0 | G | A | 0.132 | -1.66E-02 | 2.13E-01 | 2.66E+00 | 2.74E+00 | 9.37E-01 | no | Dlg2 | | intron_variant | MODIFIER |
| 7 | 92359865 | AX-170157060 | 0 | A | C | 0.132 | -1.66E-02 | 2.13E-01 | 2.66E+00 | 2.74E+00 | 9.37E-01 | no | Dlg2 | | intron_variant | MODIFIER |
| 7 | 92414396 | AX-97802778 | 0 | A | G | 0.132 | -1.66E-02 | 2.13E-01 | 2.66E+00 | 2.74E+00 | 9.37E-01 | no | Dlg2 | | intron_variant | MODIFIER |

| | | | | | | | | | | | | | | | |
|---|---|---|---|---|---|---|---|---|---|---|---|---|---|---|---|
| 7 | 92414443 | AX-170371066 | 0 | A | G | 0.132 | -1.66E-02 | 2.13E-01 | 2.66E+00 | 2.74E+00 | 9.37E-01 | no | Dlg2 | intron_variant | MODIFIER |
| 7 | 92562251 | AX-170523131 | 0 | A | G | 0.132 | -1.66E-02 | 2.13E-01 | 2.66E+00 | 2.74E+00 | 9.37E-01 | no | Ccdc90b | intron_variant | MODIFIER |
| 7 | 92666795 | AX-170029526 | 0 | C | G | 0.132 | -1.66E-02 | 2.13E-01 | 2.66E+00 | 2.74E+00 | 9.37E-01 | no | Pcf11 | intron_variant | MODIFIER |
| 7 | 93020980 | AX-97787065 | 0 | A | T | 0.132 | -1.66E-02 | 2.13E-01 | 2.66E+00 | 2.74E+00 | 9.37E-01 | no | Gm26862 | intron_variant,non_coding_transcript_variant | MODIFIER |
| 7 | 93025342 | AX-170029524 | 0 | A | G | 0.132 | -1.66E-02 | 2.13E-01 | 2.66E+00 | 2.74E+00 | 9.37E-01 | no | Gm26862 | intron_variant,non_coding_transcript_variant | MODIFIER |
| 7 | 93052319 | AX-170340193 | 0 | G | A | 0.132 | -1.66E-02 | 2.13E-01 | 2.66E+00 | 2.74E+00 | 9.37E-01 | no | Gm9934 | non_coding_transcript_exon_variant | MODIFIER |
| 7 | 93064355 | AX-170192765 | 0 | A | G | 0.132 | -1.66E-02 | 2.13E-01 | 2.66E+00 | 2.74E+00 | 9.37E-01 | no | Gm9934 | intron_variant,non_coding_transcript_variant | MODIFIER |
| 7 | 93064427 | AX-170402419 | 0 | A | G | 0.132 | -1.66E-02 | 2.13E-01 | 2.66E+00 | 2.74E+00 | 9.37E-01 | no | Gm9934 | intron_variant,non_coding_transcript_variant | MODIFIER |
| 7 | 93064558 | AX-170491640 | 0 | A | G | 0.132 | -1.66E-02 | 2.13E-01 | 2.66E+00 | 2.74E+00 | 9.37E-01 | no | Gm9934 | intron_variant,non_coding_transcript_variant | MODIFIER |
| 7 | 93075230 | AX-97815543 | 0 | T | C | 0.132 | -1.66E-02 | 2.13E-01 | 2.66E+00 | 2.74E+00 | 9.37E-01 | no | Fam181b | upstream_gene_variant | MODIFIER |
| 7 | 93091859 | AX-170421892 | 0 | T | G | 0.132 | -1.66E-02 | 2.13E-01 | 2.66E+00 | 2.74E+00 | 9.37E-01 | no | Gm26862 | intron_variant,non_coding_transcript_variant | MODIFIER |
| 7 | 93109718 | AX-97812726 | 0 | C | T | 0.132 | -1.66E-02 | 2.13E-01 | 2.66E+00 | 2.74E+00 | 9.37E-01 | no | Gm26862 | downstream_gene_variant | MODIFIER |
| 7 | 93124484 | AX-170273455 | 0 | G | A | 0.132 | -1.66E-02 | 2.13E-01 | 2.66E+00 | 2.74E+00 | 9.37E-01 | no | | intergenic_variant | MODIFIER |
| 7 | 93133248 | AX-97787428 | 0 | T | C | 0.132 | -1.66E-02 | 2.13E-01 | 2.66E+00 | 2.74E+00 | 9.37E-01 | no | | intergenic_variant | MODIFIER |
| 7 | 93134504 | AX-170360432 | 0 | T | C | 0.132 | -1.66E-02 | 2.13E-01 | 2.66E+00 | 2.74E+00 | 9.37E-01 | no | | intergenic_variant | MODIFIER |
| 7 | 93248773 | AX-97806887 | 0 | C | T | 0.132 | -1.66E-02 | 2.13E-01 | 2.66E+00 | 2.74E+00 | 9.37E-01 | no | | intergenic_variant | MODIFIER |
| 7 | 93307832 | AX-97791943 | 0 | C | A | 0.132 | -1.66E-02 | 2.13E-01 | 2.66E+00 | 2.74E+00 | 9.37E-01 | no | Gm44947 | intron_variant,non_coding_transcript_variant | MODIFIER |
| 7 | 93342891 | AX-170089980 | 0 | T | C | 0.132 | -1.66E-02 | 2.13E-01 | 2.66E+00 | 2.74E+00 | 9.37E-01 | no | | intergenic_variant | MODIFIER |
| 7 | 93409353 | AX-170192816 | 0 | A | T | 0.132 | -1.66E-02 | 2.13E-01 | 2.66E+00 | 2.74E+00 | 9.37E-01 | no | | intergenic_variant | MODIFIER |
| 7 | 93612454 | AX-170301339 | 0 | A | G | 0.132 | -1.66E-02 | 2.13E-01 | 2.66E+00 | 2.74E+00 | 9.37E-01 | no | Gm45039 | upstream_gene_variant | MODIFIER |
| 7 | 93648454 | AX-170402872 | 0 | T | A | 0.132 | -1.66E-02 | 2.13E-01 | 2.66E+00 | 2.74E+00 | 9.37E-01 | no | | intergenic_variant | MODIFIER |
| 7 | 93652376 | AX-170384581 | 0 | T | G | 0.132 | -1.66E-02 | 2.13E-01 | 2.66E+00 | 2.74E+00 | 9.37E-01 | no | | intergenic_variant | MODIFIER |
| 7 | 93653306 | AX-170510958 | 0 | T | C | 0.132 | -1.66E-02 | 2.13E-01 | 2.66E+00 | 2.74E+00 | 9.37E-01 | no | | intergenic_variant | MODIFIER |
| 7 | 93657582 | AX-170310707 | 0 | G | A | 0.132 | -1.66E-02 | 2.13E-01 | 2.66E+00 | 2.74E+00 | 9.37E-01 | no | Bc1-ps1 | downstream_gene_variant | MODIFIER |
| 7 | 93693137 | AX-170423088 | 0 | C | T | 0.132 | -1.66E-02 | 2.13E-01 | 2.66E+00 | 2.74E+00 | 9.37E-01 | no | | intergenic_variant | MODIFIER |
| 7 | 93779154 | AX-170029541 | 0 | A | G | 0.132 | -1.66E-02 | 2.13E-01 | 2.66E+00 | 2.74E+00 | 9.37E-01 | no | | intergenic_variant | MODIFIER |
| 7 | 93807780 | AX-170029542 | 0 | C | A | 0.132 | -1.66E-02 | 2.13E-01 | 2.66E+00 | 2.74E+00 | 9.37E-01 | no | | intergenic_variant | MODIFIER |
| 7 | 93813194 | AX-170029537 | 0 | C | A | 0.132 | -1.66E-02 | 2.13E-01 | 2.66E+00 | 2.74E+00 | 9.37E-01 | no | | intergenic_variant | MODIFIER |
| 7 | 93960675 | AX-170335726 | 0 | T | C | 0.132 | -1.66E-02 | 2.13E-01 | 2.66E+00 | 2.74E+00 | 9.37E-01 | no | | intergenic_variant | MODIFIER |
| 7 | 93993550 | AX-170155349 | 0 | T | G | 0.132 | -1.66E-02 | 2.13E-01 | 2.66E+00 | 2.74E+00 | 9.37E-01 | no | | intergenic_variant | MODIFIER |
| 7 | 93993570 | AX-170528700 | 0 | A | G | 0.132 | -1.66E-02 | 2.13E-01 | 2.66E+00 | 2.74E+00 | 9.37E-01 | no | | intergenic_variant | MODIFIER |
| 7 | 94159750 | AX-170078106 | 0 | C | A | 0.132 | -1.66E-02 | 2.13E-01 | 2.66E+00 | 2.74E+00 | 9.37E-01 | no | Gm32647 | intron_variant,non_coding_transcript_variant | MODIFIER |
| 7 | 94167391 | AX-170242786 | 0 | T | C | 0.132 | -1.66E-02 | 2.13E-01 | 2.66E+00 | 2.74E+00 | 9.37E-01 | no | Gm32647 | intron_variant,non_coding_transcript_variant | MODIFIER |
| 7 | 94217585 | AX-170414706 | 0 | A | G | 0.132 | -1.66E-02 | 2.13E-01 | 2.66E+00 | 2.74E+00 | 9.37E-01 | no | Gm32647 | intron_variant,non_coding_transcript_variant | MODIFIER |
| 7 | 94237071 | AX-170125989 | 0 | A | C | 0.132 | -1.66E-02 | 2.13E-01 | 2.66E+00 | 2.74E+00 | 9.37E-01 | no | Gm32647 | intron_variant,non_coding_transcript_variant | MODIFIER |
| 7 | 94254014 | AX-170535058 | 0 | G | T | 0.132 | -1.66E-02 | 2.13E-01 | 2.66E+00 | 2.74E+00 | 9.37E-01 | no | Gm32647 | intron_variant,non_coding_transcript_variant | MODIFIER |
| 7 | 94304062 | AX-170078109 | 0 | A | T | 0.132 | -1.66E-02 | 2.13E-01 | 2.66E+00 | 2.74E+00 | 9.37E-01 | no | Gm32647 | intron_variant,non_coding_transcript_variant | MODIFIER |
| 7 | 94437838 | AX-170029544 | 0 | T | C | 0.132 | -1.66E-02 | 2.13E-01 | 2.66E+00 | 2.74E+00 | 9.37E-01 | no | Gm32647 | intron_variant,non_coding_transcript_variant | MODIFIER |
| 7 | 94438179 | AX-97782326 | 0 | A | G | 0.132 | -1.66E-02 | 2.13E-01 | 2.66E+00 | 2.74E+00 | 9.37E-01 | no | Gm32647 | intron_variant,non_coding_transcript_variant | MODIFIER |
| 7 | 94438427 | AX-170029546 | 0 | A | C | 0.132 | -1.66E-02 | 2.13E-01 | 2.66E+00 | 2.74E+00 | 9.37E-01 | no | Gm32647 | intron_variant,non_coding_transcript_variant | MODIFIER |
| 7 | 94508363 | AX-170467460 | 0 | G | A | 0.132 | -1.66E-02 | 2.13E-01 | 2.66E+00 | 2.74E+00 | 9.37E-01 | no | Gm32647 | intron_variant,non_coding_transcript_variant | MODIFIER |
| 7 | 94582352 | AX-170326055 | 0 | T | C | 0.132 | -1.66E-02 | 2.13E-01 | 2.66E+00 | 2.74E+00 | 9.37E-01 | no | Gm32647 | intron_variant,non_coding_transcript_variant | MODIFIER |
| 7 | 94604357 | AX-170410738 | 0 | T | G | 0.132 | -1.66E-02 | 2.13E-01 | 2.66E+00 | 2.74E+00 | 9.37E-01 | no | Gm19761 | upstream_gene_variant | MODIFIER |
| 7 | 94607819 | AX-170466014 | 0 | T | G | 0.132 | -1.66E-02 | 2.13E-01 | 2.66E+00 | 2.74E+00 | 9.37E-01 | no | Gm32647 | intron_variant,non_coding_transcript_variant | MODIFIER |
| 7 | 94614320 | AX-170428719 | 0 | G | T | 0.132 | -1.66E-02 | 2.13E-01 | 2.66E+00 | 2.74E+00 | 9.37E-01 | no | Gm32647 | intron_variant,non_coding_transcript_variant | MODIFIER |
| 7 | 94620224 | AX-170152434 | 0 | A | G | 0.132 | -1.66E-02 | 2.13E-01 | 2.66E+00 | 2.74E+00 | 9.37E-01 | no | Gm32647 | intron_variant,non_coding_transcript_variant | MODIFIER |
| 7 | 94833279 | AX-170485577 | 0 | C | T | 0.132 | -1.66E-02 | 2.13E-01 | 2.66E+00 | 2.74E+00 | 9.37E-01 | no | | intergenic_variant | MODIFIER |
| 7 | 94892151 | AX-170511561 | 0 | T | C | 0.132 | -1.66E-02 | 2.13E-01 | 2.66E+00 | 2.74E+00 | 9.37E-01 | no | | intergenic_variant | MODIFIER |
| 7 | 94895602 | AX-170029558 | 0 | G | A | 0.132 | -1.66E-02 | 2.13E-01 | 2.66E+00 | 2.74E+00 | 9.37E-01 | no | | intergenic_variant | MODIFIER |
| 7 | 94922091 | AX-170078118 | 0 | A | G | 0.132 | -1.66E-02 | 2.13E-01 | 2.66E+00 | 2.74E+00 | 9.37E-01 | no | | intergenic_variant | MODIFIER |
| 7 | 94929853 | AX-170168370 | 0 | T | G | 0.132 | -1.66E-02 | 2.13E-01 | 2.66E+00 | 2.74E+00 | 9.37E-01 | no | | intergenic_variant | MODIFIER |
| 7 | 94938065 | AX-170078119 | 0 | A | G | 0.132 | -1.66E-02 | 2.13E-01 | 2.66E+00 | 2.74E+00 | 9.37E-01 | no | | intergenic_variant | MODIFIER |
| 7 | 94943080 | AX-170029557 | 0 | G | T | 0.132 | -1.66E-02 | 2.13E-01 | 2.66E+00 | 2.74E+00 | 9.37E-01 | no | | intergenic_variant | MODIFIER |
| 7 | 94943172 | AX-170285922 | 0 | T | C | 0.132 | -1.66E-02 | 2.13E-01 | 2.66E+00 | 2.74E+00 | 9.37E-01 | no | | intergenic_variant | MODIFIER |

| | | | | | | | | | | | | | | | |
|---|---|---|---|---|---|---|---|---|---|---|---|---|---|---|---|
| 7 | 94943276 | AX-170534476 | 0 | G | A | 0.132 | -1.66E-02 | 2.13E-01 | 2.66E+00 | 2.74E+00 | 9.37E-01 | no | | intergenic_variant | MODIFIER |
| 7 | 94980636 | AX-170029560 | 0 | T | C | 0.132 | -1.66E-02 | 2.13E-01 | 2.66E+00 | 2.74E+00 | 9.37E-01 | no | | intergenic_variant | MODIFIER |
| 7 | 94989797 | AX-170240619 | 0 | T | C | 0.132 | -1.66E-02 | 2.13E-01 | 2.66E+00 | 2.74E+00 | 9.37E-01 | no | | intergenic_variant | MODIFIER |
| 7 | 94992290 | AX-170029553 | 0 | G | A | 0.132 | -1.66E-02 | 2.13E-01 | 2.66E+00 | 2.74E+00 | 9.37E-01 | no | | intergenic_variant | MODIFIER |
| 7 | 95022826 | AX-170425000 | 0 | T | C | 0.132 | -1.66E-02 | 2.13E-01 | 2.66E+00 | 2.74E+00 | 9.37E-01 | no | | intergenic_variant | MODIFIER |
| 7 | 95058468 | AX-97854860 | 0 | C | A | 0.132 | -1.66E-02 | 2.13E-01 | 2.66E+00 | 2.74E+00 | 9.37E-01 | no | | intergenic_variant | MODIFIER |
| 7 | 95362348 | AX-170530934 | 0 | T | C | 0.132 | -1.66E-02 | 2.13E-01 | 2.66E+00 | 2.74E+00 | 9.37E-01 | no | | intergenic_variant | MODIFIER |
| 7 | 95362572 | AX-170166213 | 0 | T | C | 0.132 | -1.66E-02 | 2.13E-01 | 2.66E+00 | 2.74E+00 | 9.37E-01 | no | | intergenic_variant | MODIFIER |
| 7 | 95363948 | AX-170267213 | 0 | C | T | 0.132 | -1.66E-02 | 2.13E-01 | 2.66E+00 | 2.74E+00 | 9.37E-01 | no | | intergenic_variant | MODIFIER |
| 7 | 95368322 | AX-97824691 | 0 | A | G | 0.132 | -1.66E-02 | 2.13E-01 | 2.66E+00 | 2.74E+00 | 9.37E-01 | no | | intergenic_variant | MODIFIER |
| 7 | 95401245 | AX-170268091 | 0 | A | C | 0.132 | -1.66E-02 | 2.13E-01 | 2.66E+00 | 2.74E+00 | 9.37E-01 | no | | intergenic_variant | MODIFIER |
| 7 | 95401605 | AX-97831585 | 0 | G | T | 0.132 | -1.66E-02 | 2.13E-01 | 2.66E+00 | 2.74E+00 | 9.37E-01 | no | | intergenic_variant | MODIFIER |
| 7 | 95418631 | AX-170029568 | 0 | G | T | 0.132 | -1.66E-02 | 2.13E-01 | 2.66E+00 | 2.74E+00 | 9.37E-01 | no | | intergenic_variant | MODIFIER |
| 7 | 95439207 | AX-170307105 | 0 | T | C | 0.132 | -1.66E-02 | 2.13E-01 | 2.66E+00 | 2.74E+00 | 9.37E-01 | no | Gm25684 | downstream_gene_variant | MODIFIER |
| 7 | 95440206 | AX-170406984 | 0 | G | A | 0.132 | -1.66E-02 | 2.13E-01 | 2.66E+00 | 2.74E+00 | 9.37E-01 | no | Gm25684 | downstream_gene_variant | MODIFIER |
| 7 | 95470001 | AX-170172033 | 0 | A | G | 0.132 | -1.66E-02 | 2.13E-01 | 2.66E+00 | 2.74E+00 | 9.37E-01 | no | Gm44603 | upstream_gene_variant | MODIFIER |
| 7 | 95470462 | AX-97839194 | 0 | A | G | 0.132 | -1.66E-02 | 2.13E-01 | 2.66E+00 | 2.74E+00 | 9.37E-01 | no | Gm44603 | upstream_gene_variant | MODIFIER |
| 7 | 95480513 | AX-170376147 | 0 | T | C | 0.132 | -1.66E-02 | 2.13E-01 | 2.66E+00 | 2.74E+00 | 9.37E-01 | no | | regulatory_region_variant | MODIFIER |
| 7 | 95510877 | AX-170270605 | 0 | G | A | 0.132 | -1.66E-02 | 2.13E-01 | 2.66E+00 | 2.74E+00 | 9.37E-01 | no | | intergenic_variant | MODIFIER |
| 7 | 95641089 | AX-170346839 | 0 | G | A | 0.132 | -1.66E-02 | 2.13E-01 | 2.66E+00 | 2.74E+00 | 9.37E-01 | no | | intergenic_variant | MODIFIER |
| 7 | 95659570 | AX-170332183 | 0 | G | C | 0.132 | -1.66E-02 | 2.13E-01 | 2.66E+00 | 2.74E+00 | 9.37E-01 | no | | intergenic_variant | MODIFIER |
| 7 | 95765412 | AX-170148297 | 0 | A | G | 0.132 | -1.66E-02 | 2.13E-01 | 2.66E+00 | 2.74E+00 | 9.37E-01 | no | | intergenic_variant | MODIFIER |
| 7 | 95770440 | AX-170431717 | 0 | C | G | 0.132 | -1.66E-02 | 2.13E-01 | 2.66E+00 | 2.74E+00 | 9.37E-01 | no | | intergenic_variant | MODIFIER |
| 7 | 95825798 | AX-170116262 | 0 | G | T | 0.132 | -1.66E-02 | 2.13E-01 | 2.66E+00 | 2.74E+00 | 9.37E-01 | no | | intergenic_variant | MODIFIER |
| 7 | 95826810 | AX-170449804 | 0 | G | A | 0.132 | -1.66E-02 | 2.13E-01 | 2.66E+00 | 2.74E+00 | 9.37E-01 | no | | intergenic_variant | MODIFIER |
| 7 | 95827595 | AX-170430824 | 0 | A | G | 0.132 | -1.66E-02 | 2.13E-01 | 2.66E+00 | 2.74E+00 | 9.37E-01 | no | | intergenic_variant | MODIFIER |
| 7 | 95855919 | AX-170411497 | 0 | G | A | 0.132 | -1.66E-02 | 2.13E-01 | 2.66E+00 | 2.74E+00 | 9.37E-01 | no | | intergenic_variant | MODIFIER |
| 7 | 95857776 | AX-170029569 | 0 | G | C | 0.132 | -1.66E-02 | 2.13E-01 | 2.66E+00 | 2.74E+00 | 9.37E-01 | no | | intergenic_variant | MODIFIER |
| 7 | 95894801 | AX-170029572 | 0 | C | A | 0.132 | -1.66E-02 | 2.13E-01 | 2.66E+00 | 2.74E+00 | 9.37E-01 | no | | intergenic_variant | MODIFIER |
| 7 | 95900126 | AX-97806925 | 0 | A | G | 0.132 | -1.66E-02 | 2.13E-01 | 2.66E+00 | 2.74E+00 | 9.37E-01 | no | | intergenic_variant | MODIFIER |
| 7 | 95904515 | AX-170029571 | 0 | A | G | 0.132 | -1.66E-02 | 2.13E-01 | 2.66E+00 | 2.74E+00 | 9.37E-01 | no | | intergenic_variant | MODIFIER |
| 7 | 95904818 | AX-170509254 | 0 | T | C | 0.132 | -1.66E-02 | 2.13E-01 | 2.66E+00 | 2.74E+00 | 9.37E-01 | no | | intergenic_variant | MODIFIER |
| 7 | 95915930 | AX-170326992 | 0 | A | G | 0.132 | -1.66E-02 | 2.13E-01 | 2.66E+00 | 2.74E+00 | 9.37E-01 | no | Gm5037 | upstream_gene_variant | MODIFIER |
| 7 | 95943585 | AX-170271576 | 0 | A | G | 0.132 | -1.66E-02 | 2.13E-01 | 2.66E+00 | 2.74E+00 | 9.37E-01 | no | | regulatory_region_variant | MODIFIER |
| 7 | 95955614 | AX-170334435 | 0 | G | A | 0.132 | -1.66E-02 | 2.13E-01 | 2.66E+00 | 2.74E+00 | 9.37E-01 | no | | intergenic_variant | MODIFIER |
| 7 | 95960254 | AX-170127646 | 0 | T | C | 0.132 | -1.66E-02 | 2.13E-01 | 2.66E+00 | 2.74E+00 | 9.37E-01 | no | | intergenic_variant | MODIFIER |
| 7 | 95964331 | AX-170123493 | 0 | G | A | 0.132 | -1.66E-02 | 2.13E-01 | 2.66E+00 | 2.74E+00 | 9.37E-01 | no | | intergenic_variant | MODIFIER |
| 7 | 95964359 | AX-170460266 | 0 | A | G | 0.132 | -1.66E-02 | 2.13E-01 | 2.66E+00 | 2.74E+00 | 9.37E-01 | no | | intergenic_variant | MODIFIER |
| 7 | 95966245 | AX-170189434 | 0 | T | G | 0.132 | -1.66E-02 | 2.13E-01 | 2.66E+00 | 2.74E+00 | 9.37E-01 | no | | intergenic_variant | MODIFIER |
| 7 | 95971674 | AX-170368899 | 0 | A | G | 0.132 | -1.66E-02 | 2.13E-01 | 2.66E+00 | 2.74E+00 | 9.37E-01 | no | | intergenic_variant | MODIFIER |
| 7 | 95971756 | AX-170384821 | 0 | A | G | 0.132 | -1.66E-02 | 2.13E-01 | 2.66E+00 | 2.74E+00 | 9.37E-01 | no | | intergenic_variant | MODIFIER |
| 7 | 96041488 | AX-170491929 | 0 | A | C | 0.132 | -1.66E-02 | 2.13E-01 | 2.66E+00 | 2.74E+00 | 9.37E-01 | no | | intergenic_variant | MODIFIER |
| 7 | 96042979 | AX-170223353 | 0 | G | A | 0.132 | -1.66E-02 | 2.13E-01 | 2.66E+00 | 2.74E+00 | 9.37E-01 | no | | intergenic_variant | MODIFIER |
| 7 | 96044285 | AX-170290762 | 0 | T | C | 0.132 | -1.66E-02 | 2.13E-01 | 2.66E+00 | 2.74E+00 | 9.37E-01 | no | | intergenic_variant | MODIFIER |
| 7 | 96156652 | AX-170407613 | 0 | A | G | 0.132 | -1.66E-02 | 2.13E-01 | 2.66E+00 | 2.74E+00 | 9.37E-01 | no | | intergenic_variant | MODIFIER |
| 7 | 96238153 | AX-97833532 | 0 | T | G | 0.132 | -1.66E-02 | 2.13E-01 | 2.66E+00 | 2.74E+00 | 9.37E-01 | no | Tenm4 | intron_variant | MODIFIER |
| 7 | 96283871 | AX-170247619 | 0 | T | C | 0.132 | -1.66E-02 | 2.13E-01 | 2.66E+00 | 2.74E+00 | 9.37E-01 | no | Tenm4 | intron_variant | MODIFIER |
| 7 | 96305405 | AX-170518481 | 0 | T | G | 0.132 | -1.66E-02 | 2.13E-01 | 2.66E+00 | 2.74E+00 | 9.37E-01 | no | Tenm4 | intron_variant | MODIFIER |
| 7 | 96312689 | AX-170334884 | 0 | G | A | 0.132 | -1.66E-02 | 2.13E-01 | 2.66E+00 | 2.74E+00 | 9.37E-01 | no | Tenm4 | intron_variant | MODIFIER |
| 7 | 96392737 | AX-170118891 | 0 | T | A | 0.132 | -1.66E-02 | 2.13E-01 | 2.66E+00 | 2.74E+00 | 9.37E-01 | no | Tenm4 | intron_variant | MODIFIER |
| 7 | 96450882 | AX-170314193 | 0 | T | C | 0.132 | -1.66E-02 | 2.13E-01 | 2.66E+00 | 2.74E+00 | 9.37E-01 | no | Tenm4 | intron_variant | MODIFIER |
| 7 | 96459178 | AX-97818195 | 0 | A | C | 0.132 | -1.66E-02 | 2.13E-01 | 2.66E+00 | 2.74E+00 | 9.37E-01 | no | Tenm4 | intron_variant | MODIFIER |
| 7 | 96585248 | AX-170416137 | 0 | A | T | 0.132 | -1.66E-02 | 2.13E-01 | 2.66E+00 | 2.74E+00 | 9.37E-01 | no | Tenm4 | intron_variant | MODIFIER |
| 7 | 96748598 | AX-170312229 | 0 | A | G | 0.132 | -1.66E-02 | 2.13E-01 | 2.66E+00 | 2.74E+00 | 9.37E-01 | no | Tenm4 | intron_variant | MODIFIER |

| | | | | | | | | | | | | | | | |
|---|---|---|---|---|---|---|---|---|---|---|---|---|---|---|---|
| 7 | 96749414 | AX-97802832 | 0 | C | A | 0.132 | -1.66E-02 | 2.13E-01 | 2.66E+00 | 2.74E+00 | 9.37E-01 | no | Tenm4 | intron_variant | MODIFIER |
| 7 | 96753842 | AX-170349507 | 0 | C | T | 0.132 | -1.66E-02 | 2.13E-01 | 2.66E+00 | 2.74E+00 | 9.37E-01 | no | Tenm4 | intron_variant | MODIFIER |
| 7 | 96798210 | AX-170222584 | 0 | A | T | 0.132 | -1.66E-02 | 2.13E-01 | 2.66E+00 | 2.74E+00 | 9.37E-01 | no | Tenm4 | intron_variant | MODIFIER |
| 7 | 96801181 | AX-170159071 | 0 | T | C | 0.132 | -1.66E-02 | 2.13E-01 | 2.66E+00 | 2.74E+00 | 9.37E-01 | no | Tenm4 | intron_variant | MODIFIER |
| 7 | 96914113 | AX-170291448 | 0 | T | A | 0.132 | -1.66E-02 | 2.13E-01 | 2.66E+00 | 2.74E+00 | 9.37E-01 | no | Tenm4 | downstream_gene_variant | MODIFIER |
| 7 | 96931456 | AX-170110997 | 0 | A | G | 0.132 | -1.66E-02 | 2.13E-01 | 2.66E+00 | 2.74E+00 | 9.37E-01 | no | C230038L03Rik | intron_variant,non_coding_transcript_variant | MODIFIER |
| 7 | 96943927 | AX-170397317 | 0 | T | C | 0.132 | -1.66E-02 | 2.13E-01 | 2.66E+00 | 2.74E+00 | 9.37E-01 | no | C230038L03Rik | downstream_gene_variant | MODIFIER |
| 7 | 96953990 | AX-170117216 | 0 | T | C | 0.132 | -1.66E-02 | 2.13E-01 | 2.66E+00 | 2.74E+00 | 9.37E-01 | no | Nars2 | intron_variant | MODIFIER |
| 7 | 97000590 | AX-170238383 | 0 | T | C | 0.132 | -1.66E-02 | 2.13E-01 | 2.66E+00 | 2.74E+00 | 9.37E-01 | no | Nars2 | intron_variant | MODIFIER |
| 7 | 97001231 | AX-170502410 | 0 | A | C | 0.132 | -1.66E-02 | 2.13E-01 | 2.66E+00 | 2.74E+00 | 9.37E-01 | no | Nars2 | intron_variant | MODIFIER |
| 7 | 97005781 | AX-170335762 | 0 | A | T | 0.132 | -1.66E-02 | 2.13E-01 | 2.66E+00 | 2.74E+00 | 9.37E-01 | no | Nars2 | intron_variant | MODIFIER |
| 7 | 97055999 | AX-170029577 | 0 | T | C | 0.132 | -1.66E-02 | 2.13E-01 | 2.66E+00 | 2.74E+00 | 9.37E-01 | no | Nars2 | intron_variant | MODIFIER |
| 7 | 97084165 | AX-170097614 | 0 | G | A | 0.132 | -1.66E-02 | 2.13E-01 | 2.66E+00 | 2.74E+00 | 9.37E-01 | no | Gab2 | intron_variant | MODIFIER |
| 7 | 97084193 | AX-170154207 | 0 | C | T | 0.132 | -1.66E-02 | 2.13E-01 | 2.66E+00 | 2.74E+00 | 9.37E-01 | no | Gab2 | intron_variant | MODIFIER |
| 7 | 97095356 | AX-97780585 | 0 | A | G | 0.132 | -1.66E-02 | 2.13E-01 | 2.66E+00 | 2.74E+00 | 9.37E-01 | no | Gab2 | intron_variant | MODIFIER |
| 7 | 97102003 | AX-97788248 | 0 | A | T | 0.132 | -1.66E-02 | 2.13E-01 | 2.66E+00 | 2.74E+00 | 9.37E-01 | no | Gab2 | intron_variant | MODIFIER |
| 7 | 97127857 | AX-170029581 | 0 | A | G | 0.132 | -1.66E-02 | 2.13E-01 | 2.66E+00 | 2.74E+00 | 9.37E-01 | no | Gab2 | intron_variant | MODIFIER |
| 7 | 97150284 | AX-170029580 | 0 | T | C | 0.132 | -1.66E-02 | 2.13E-01 | 2.66E+00 | 2.74E+00 | 9.37E-01 | no | Gab2 | intron_variant | MODIFIER |
| 7 | 97213623 | AX-170029582 | 0 | T | C | 0.132 | -1.66E-02 | 2.13E-01 | 2.66E+00 | 2.74E+00 | 9.37E-01 | no | Gab2 | intron_variant | MODIFIER |
| 7 | 97266662 | AX-170029583 | 0 | C | T | 0.132 | -1.66E-02 | 2.13E-01 | 2.66E+00 | 2.74E+00 | 9.37E-01 | no | Gab2 | intron_variant | MODIFIER |
| 7 | 97299811 | AX-170221669 | 0 | T | G | 0.132 | -1.66E-02 | 2.13E-01 | 2.66E+00 | 2.74E+00 | 9.37E-01 | no | Gab2 | intron_variant | MODIFIER |
| 7 | 97331083 | AX-97827483 | 0 | C | T | 0.132 | -1.66E-02 | 2.13E-01 | 2.66E+00 | 2.74E+00 | 9.37E-01 | no | Kctd21 | upstream_gene_variant | MODIFIER |
| 7 | 97351698 | AX-170423108 | 0 | G | A | 0.132 | -1.66E-02 | 2.13E-01 | 2.66E+00 | 2.74E+00 | 9.37E-01 | no | Kctd21 | downstream_gene_variant | MODIFIER |
| 7 | 97386479 | AX-170216588 | 0 | G | A | 0.132 | -1.66E-02 | 2.13E-01 | 2.66E+00 | 2.74E+00 | 9.37E-01 | no | Alg8 | intron_variant | MODIFIER |
| 7 | 97411715 | AX-170179917 | 0 | G | A | 0.132 | -1.66E-02 | 2.13E-01 | 2.66E+00 | 2.74E+00 | 9.37E-01 | no | Ndufc2 | downstream_gene_variant | MODIFIER |
| 7 | 97415500 | AX-170374843 | 0 | T | G | 0.132 | -1.66E-02 | 2.13E-01 | 2.66E+00 | 2.74E+00 | 9.37E-01 | no | Thrsp | intron_variant | MODIFIER |
| 7 | 97419374 | AX-170267544 | 0 | G | A | 0.132 | -1.66E-02 | 2.13E-01 | 2.66E+00 | 2.74E+00 | 9.37E-01 | no | Thrsp | upstream_gene_variant | MODIFIER |
| 7 | 97420955 | AX-170310190 | 0 | C | T | 0.132 | -1.66E-02 | 2.13E-01 | 2.66E+00 | 2.74E+00 | 9.37E-01 | no | Thrsp | upstream_gene_variant | MODIFIER |
| 7 | 97421077 | AX-170234441 | 0 | T | G | 0.132 | -1.66E-02 | 2.13E-01 | 2.66E+00 | 2.74E+00 | 9.37E-01 | no | Thrsp | upstream_gene_variant | MODIFIER |
| 7 | 97423892 | AX-170331461 | 0 | C | T | 0.132 | -1.66E-02 | 2.13E-01 | 2.66E+00 | 2.74E+00 | 9.37E-01 | no | | regulatory_region_variant | MODIFIER |
| 7 | 97441711 | AX-170369202 | 0 | A | G | 0.132 | -1.66E-02 | 2.13E-01 | 2.66E+00 | 2.74E+00 | 9.37E-01 | no | | intergenic_variant | MODIFIER |
| 7 | 97447452 | AX-170334688 | 0 | A | G | 0.132 | -1.66E-02 | 2.13E-01 | 2.66E+00 | 2.74E+00 | 9.37E-01 | no | Kctd14 | upstream_gene_variant | MODIFIER |
| 7 | 97457801 | AX-170153042 | 0 | A | G | 0.132 | -1.66E-02 | 2.13E-01 | 2.66E+00 | 2.74E+00 | 9.37E-01 | no | Kctd14 | missense_variant | MODERATE |
| 7 | 97457911 | AX-170205824 | 0 | G | A | 0.132 | -1.66E-02 | 2.13E-01 | 2.66E+00 | 2.74E+00 | 9.37E-01 | no | Kctd14 | synonymous_variant | LOW |
| 7 | 97457986 | AX-170517533 | 0 | C | G | 0.132 | -1.66E-02 | 2.13E-01 | 2.66E+00 | 2.74E+00 | 9.37E-01 | no | Kctd14 | synonymous_variant | LOW |
| 7 | 97458226 | AX-170291854 | 0 | C | T | 0.132 | -1.66E-02 | 2.13E-01 | 2.66E+00 | 2.74E+00 | 9.37E-01 | no | Kctd14 | synonymous_variant | LOW |
| 7 | 97464328 | AX-170334846 | 0 | T | C | 0.132 | -1.66E-02 | 2.13E-01 | 2.66E+00 | 2.74E+00 | 9.37E-01 | no | Kctd14 | downstream_gene_variant | MODIFIER |
| 7 | 97489666 | AX-97808862 | 0 | T | G | 0.132 | -1.66E-02 | 2.13E-01 | 2.66E+00 | 2.74E+00 | 9.37E-01 | no | Ints4 | intron_variant | MODIFIER |
| 7 | 97503151 | AX-170116650 | 0 | A | G | 0.132 | -1.66E-02 | 2.13E-01 | 2.66E+00 | 2.74E+00 | 9.37E-01 | no | Ints4 | intron_variant | MODIFIER |
| 7 | 97529431 | AX-170235980 | 0 | T | C | 0.132 | -1.66E-02 | 2.13E-01 | 2.66E+00 | 2.74E+00 | 9.37E-01 | no | Ints4 | intron_variant | MODIFIER |
| 7 | 97542979 | AX-170029589 | 0 | T | C | 0.132 | -1.66E-02 | 2.13E-01 | 2.66E+00 | 2.74E+00 | 9.37E-01 | no | Ints4 | downstream_gene_variant | MODIFIER |
| 7 | 97617548 | AX-170029590 | 0 | G | A | 0.132 | -1.66E-02 | 2.13E-01 | 2.66E+00 | 2.74E+00 | 9.37E-01 | no | Rsf1 | intron_variant | MODIFIER |
| 7 | 97629622 | AX-170029592 | 0 | C | A | 0.132 | -1.66E-02 | 2.13E-01 | 2.66E+00 | 2.74E+00 | 9.37E-01 | no | Rsf1 | intron_variant | MODIFIER |
| 7 | 97665353 | AX-170213661 | 0 | A | G | 0.132 | -1.66E-02 | 2.13E-01 | 2.66E+00 | 2.74E+00 | 9.37E-01 | no | Rsf1 | intron_variant | MODIFIER |
| 7 | 97678840 | AX-170524804 | 0 | G | A | 0.132 | -1.66E-02 | 2.13E-01 | 2.66E+00 | 2.74E+00 | 9.37E-01 | no | Rsf1 | intron_variant | MODIFIER |
| 7 | 97687361 | AX-170437933 | 0 | A | T | 0.132 | -1.66E-02 | 2.13E-01 | 2.66E+00 | 2.74E+00 | 9.37E-01 | no | Rsf1 | 3_prime_UTR_variant | MODIFIER |
| 7 | 97690555 | AX-170149459 | 0 | T | C | 0.132 | -1.66E-02 | 2.13E-01 | 2.66E+00 | 2.74E+00 | 9.37E-01 | no | Rsf1 | 3_prime_UTR_variant | MODIFIER |
| 7 | 97691276 | AX-170487238 | 0 | G | A | 0.132 | -1.66E-02 | 2.13E-01 | 2.66E+00 | 2.74E+00 | 9.37E-01 | no | Rsf1 | 3_prime_UTR_variant | MODIFIER |
| 7 | 97691706 | AX-97845118 | 0 | T | C | 0.132 | -1.66E-02 | 2.13E-01 | 2.66E+00 | 2.74E+00 | 9.37E-01 | no | Clns1a | upstream_gene_variant | MODIFIER |
| 7 | 97703997 | AX-170390182 | 0 | T | G | 0.132 | -1.66E-02 | 2.13E-01 | 2.66E+00 | 2.74E+00 | 9.37E-01 | no | Clns1a | intron_variant | MODIFIER |
| 7 | 97712733 | AX-170329731 | 0 | C | T | 0.132 | -1.66E-02 | 2.13E-01 | 2.66E+00 | 2.74E+00 | 9.37E-01 | no | Clns1a | splice_region_variant,synonymous_variant | LOW |
| 7 | 97765598 | AX-170294786 | 0 | T | C | 0.132 | -1.66E-02 | 2.13E-01 | 2.66E+00 | 2.74E+00 | 9.37E-01 | no | | regulatory_region_variant | MODIFIER |
| 7 | 97775567 | AX-170528982 | 0 | G | A | 0.132 | -1.66E-02 | 2.13E-01 | 2.66E+00 | 2.74E+00 | 9.37E-01 | no | | regulatory_region_variant | MODIFIER |
| 7 | 97798842 | AX-170304205 | 0 | C | T | 0.132 | -1.66E-02 | 2.13E-01 | 2.66E+00 | 2.74E+00 | 9.37E-01 | no | Pak1 | intron_variant,non_coding_transcript_variant | MODIFIER |

| | | | | | | | | | | | | | | | |
|---|---|---|---|---|---|---|---|---|---|---|---|---|---|---|---|
| 7 | 97841032 | AX-170145060 | 0 | A | G | 0.132 | -1.66E-02 | 2.13E-01 | 2.66E+00 | 2.74E+00 | 9.37E-01 | no | Pak1 | upstream_gene_variant | MODIFIER |
| 7 | 97875892 | AX-97803427 | 0 | T | C | 0.132 | -1.66E-02 | 2.13E-01 | 2.66E+00 | 2.74E+00 | 9.37E-01 | no | Pak1 | intron_variant | MODIFIER |
| 7 | 97903794 | AX-97836021 | 0 | G | T | 0.132 | -1.66E-02 | 2.13E-01 | 2.66E+00 | 2.74E+00 | 9.37E-01 | no | Pak1 | intron_variant | MODIFIER |
| 7 | 97910029 | AX-170208187 | 0 | T | C | 0.132 | -1.66E-02 | 2.13E-01 | 2.66E+00 | 2.74E+00 | 9.37E-01 | no | Pak1 | intron_variant | MODIFIER |
| 7 | 97910621 | AX-170422464 | 0 | G | A | 0.132 | -1.66E-02 | 2.13E-01 | 2.66E+00 | 2.74E+00 | 9.37E-01 | no | Pak1 | intron_variant | MODIFIER |
| 7 | 97911353 | AX-170452794 | 0 | T | A | 0.132 | -1.66E-02 | 2.13E-01 | 2.66E+00 | 2.74E+00 | 9.37E-01 | no | Pak1 | 3_prime_UTR_variant | MODIFIER |
| 7 | 97912158 | AX-170182831 | 0 | T | C | 0.132 | -1.66E-02 | 2.13E-01 | 2.66E+00 | 2.74E+00 | 9.37E-01 | no | Pak1 | 3_prime_UTR_variant | MODIFIER |
| 7 | 97956406 | AX-170202135 | 0 | T | C | 0.132 | -1.66E-02 | 2.13E-01 | 2.66E+00 | 2.74E+00 | 9.37E-01 | no | Gdpd4 | intron_variant | MODIFIER |
| 7 | 97965357 | AX-170430865 | 0 | G | A | 0.132 | -1.66E-02 | 2.13E-01 | 2.66E+00 | 2.74E+00 | 9.37E-01 | no | Gdpd4 | intron_variant | MODIFIER |
| 7 | 97981034 | AX-97789659 | 0 | A | G | 0.132 | -1.66E-02 | 2.13E-01 | 2.66E+00 | 2.74E+00 | 9.37E-01 | no | Gdpd4 | intron_variant | MODIFIER |
| 7 | 97982401 | AX-170227577 | 0 | T | G | 0.132 | -1.66E-02 | 2.13E-01 | 2.66E+00 | 2.74E+00 | 9.37E-01 | no | Gdpd4 | intron_variant | MODIFIER |
| 7 | 97998458 | AX-170386968 | 0 | C | T | 0.132 | -1.66E-02 | 2.13E-01 | 2.66E+00 | 2.74E+00 | 9.37E-01 | no | Gdpd4 | synonymous_variant | LOW |
| 7 | 98019482 | AX-170489652 | 0 | T | C | 0.132 | -1.66E-02 | 2.13E-01 | 2.66E+00 | 2.74E+00 | 9.37E-01 | no | Gdpd4 | intron_variant | MODIFIER |
| 7 | 98019809 | AX-170392509 | 0 | G | A | 0.132 | -1.66E-02 | 2.13E-01 | 2.66E+00 | 2.74E+00 | 9.37E-01 | no | Gdpd4 | intron_variant | MODIFIER |
| 7 | 98019967 | AX-170127674 | 0 | T | C | 0.132 | -1.66E-02 | 2.13E-01 | 2.66E+00 | 2.74E+00 | 9.37E-01 | no | Gdpd4 | intron_variant | MODIFIER |
| 7 | 98051866 | AX-170395645 | 0 | A | C | 0.132 | -1.66E-02 | 2.13E-01 | 2.66E+00 | 2.74E+00 | 9.37E-01 | no | Gdpd4 | downstream_gene_variant | MODIFIER |
| 7 | 98065035 | AX-97854152 | 0 | T | C | 0.132 | -1.66E-02 | 2.13E-01 | 2.66E+00 | 2.74E+00 | 9.37E-01 | no | Myo7a | intron_variant | MODIFIER |
| 7 | 98069423 | AX-170211140 | 0 | T | C | 0.132 | -1.66E-02 | 2.13E-01 | 2.66E+00 | 2.74E+00 | 9.37E-01 | no | Myo7a | synonymous_variant | LOW |
| 7 | 98070394 | AX-170171320 | 0 | A | G | 0.132 | -1.66E-02 | 2.13E-01 | 2.66E+00 | 2.74E+00 | 9.37E-01 | no | Myo7a | intron_variant | MODIFIER |
| 7 | 98071517 | AX-170432353 | 0 | G | A | 0.132 | -1.66E-02 | 2.13E-01 | 2.66E+00 | 2.74E+00 | 9.37E-01 | no | Myo7a | intron_variant | MODIFIER |
| 7 | 98085758 | AX-170283865 | 0 | A | G | 0.132 | -1.66E-02 | 2.13E-01 | 2.66E+00 | 2.74E+00 | 9.37E-01 | no | Myo7a | intron_variant | MODIFIER |
| 7 | 98087635 | AX-97835531 | 0 | G | A | 0.132 | -1.66E-02 | 2.13E-01 | 2.66E+00 | 2.74E+00 | 9.37E-01 | no | Myo7a | intron_variant | MODIFIER |
| 7 | 98092508 | AX-170029595 | 0 | G | A | 0.132 | -1.66E-02 | 2.13E-01 | 2.66E+00 | 2.74E+00 | 9.37E-01 | no | Myo7a | synonymous_variant | LOW |
| 7 | 98093406 | AX-170029597 | 0 | T | C | 0.132 | -1.66E-02 | 2.13E-01 | 2.66E+00 | 2.74E+00 | 9.37E-01 | no | Myo7a | intron_variant | MODIFIER |
| 7 | 98128885 | AX-170029594 | 0 | G | A | 0.132 | -1.66E-02 | 2.13E-01 | 2.66E+00 | 2.74E+00 | 9.37E-01 | no | Capn5 | intron_variant | MODIFIER |
| 7 | 98135534 | AX-170078180 | 0 | T | C | 0.132 | -1.66E-02 | 2.13E-01 | 2.66E+00 | 2.74E+00 | 9.37E-01 | no | Capn5 | intron_variant | MODIFIER |
| 7 | 98143556 | AX-170347133 | 0 | A | G | 0.132 | -1.66E-02 | 2.13E-01 | 2.66E+00 | 2.74E+00 | 9.37E-01 | no | Capn5 | intron_variant | MODIFIER |
| 7 | 98144239 | AX-170127820 | 0 | A | G | 0.132 | -1.66E-02 | 2.13E-01 | 2.66E+00 | 2.74E+00 | 9.37E-01 | no | Capn5 | intron_variant | MODIFIER |
| 7 | 98144334 | AX-170175046 | 0 | A | G | 0.132 | -1.66E-02 | 2.13E-01 | 2.66E+00 | 2.74E+00 | 9.37E-01 | no | Capn5 | intron_variant | MODIFIER |
| 7 | 98149333 | AX-170436085 | 0 | A | G | 0.132 | -1.66E-02 | 2.13E-01 | 2.66E+00 | 2.74E+00 | 9.37E-01 | no | Capn5 | intron_variant | MODIFIER |
| 7 | 98370565 | AX-170102841 | 0 | T | A | 0.132 | -1.66E-02 | 2.13E-01 | 2.66E+00 | 2.74E+00 | 9.37E-01 | no | | intergenic_variant | MODIFIER |
| 7 | 98377728 | AX-170366577 | 0 | T | C | 0.132 | -1.66E-02 | 2.13E-01 | 2.66E+00 | 2.74E+00 | 9.37E-01 | no | | regulatory_region_variant | MODIFIER |
| 7 | 98381340 | AX-170119597 | 0 | T | C | 0.132 | -1.66E-02 | 2.13E-01 | 2.66E+00 | 2.74E+00 | 9.37E-01 | no | | regulatory_region_variant | MODIFIER |
| 7 | 98384402 | AX-170439854 | 0 | A | G | 0.132 | -1.66E-02 | 2.13E-01 | 2.66E+00 | 2.74E+00 | 9.37E-01 | no | Gm44507 | downstream_gene_variant | MODIFIER |
| 7 | 98391621 | AX-170478858 | 0 | T | C | 0.132 | -1.66E-02 | 2.13E-01 | 2.66E+00 | 2.74E+00 | 9.37E-01 | no | Gm44507 | intron_variant,non_coding_transcript_variant | MODIFIER |
| 7 | 98391768 | AX-170312384 | 0 | C | T | 0.132 | -1.66E-02 | 2.13E-01 | 2.66E+00 | 2.74E+00 | 9.37E-01 | no | Gm44507 | intron_variant,non_coding_transcript_variant | MODIFIER |
| 7 | 98429493 | AX-170275022 | 0 | T | C | 0.132 | -1.66E-02 | 2.13E-01 | 2.66E+00 | 2.74E+00 | 9.37E-01 | no | | intergenic_variant | MODIFIER |
| 7 | 98433022 | AX-170136649 | 0 | A | G | 0.132 | -1.66E-02 | 2.13E-01 | 2.66E+00 | 2.74E+00 | 9.37E-01 | no | Gm44980 | downstream_gene_variant | MODIFIER |
| 7 | 98433784 | AX-170268386 | 0 | C | T | 0.132 | -1.66E-02 | 2.13E-01 | 2.66E+00 | 2.74E+00 | 9.37E-01 | no | Gm44980 | downstream_gene_variant | MODIFIER |
| 7 | 98452577 | AX-170211132 | 0 | T | C | 0.132 | -1.66E-02 | 2.13E-01 | 2.66E+00 | 2.74E+00 | 9.37E-01 | no | Gucy2d | intron_variant | MODIFIER |
| 7 | 98452942 | AX-170299134 | 0 | T | C | 0.132 | -1.66E-02 | 2.13E-01 | 2.66E+00 | 2.74E+00 | 9.37E-01 | no | Gucy2d | intron_variant | MODIFIER |
| 7 | 98487236 | AX-170393963 | 0 | C | T | 0.132 | -1.66E-02 | 2.13E-01 | 2.66E+00 | 2.74E+00 | 9.37E-01 | no | Lrrc32 | upstream_gene_variant | MODIFIER |
| 7 | 98521442 | AX-170299861 | 0 | C | A | 0.132 | -1.66E-02 | 2.13E-01 | 2.66E+00 | 2.74E+00 | 9.37E-01 | no | | intergenic_variant | MODIFIER |
| 7 | 98521559 | AX-170478769 | 0 | G | A | 0.132 | -1.66E-02 | 2.13E-01 | 2.66E+00 | 2.74E+00 | 9.37E-01 | no | | intergenic_variant | MODIFIER |
| 7 | 98523676 | AX-170461470 | 0 | A | T | 0.132 | -1.66E-02 | 2.13E-01 | 2.66E+00 | 2.74E+00 | 9.37E-01 | no | | regulatory_region_variant | MODIFIER |
| 7 | 98530013 | AX-170216700 | 0 | G | C | 0.132 | -1.66E-02 | 2.13E-01 | 2.66E+00 | 2.74E+00 | 9.37E-01 | no | | intergenic_variant | MODIFIER |
| 7 | 98563522 | AX-170485913 | 0 | T | C | 0.132 | -1.66E-02 | 2.13E-01 | 2.66E+00 | 2.74E+00 | 9.37E-01 | no | A630091E08Rik | upstream_gene_variant | MODIFIER |
| 7 | 98566747 | AX-97844650 | 0 | T | C | 0.132 | -1.66E-02 | 2.13E-01 | 2.66E+00 | 2.74E+00 | 9.37E-01 | no | A630091E08Rik | upstream_gene_variant | MODIFIER |
| 7 | 98593060 | AX-170411680 | 0 | A | G | 0.132 | -1.66E-02 | 2.13E-01 | 2.66E+00 | 2.74E+00 | 9.37E-01 | no | Emsy | intron_variant | MODIFIER |
| 7 | 98642918 | AX-170029604 | 0 | C | T | 0.132 | -1.66E-02 | 2.13E-01 | 2.66E+00 | 2.74E+00 | 9.37E-01 | no | Emsy | intron_variant | MODIFIER |
| 7 | 98647356 | AX-170168119 | 0 | C | A | 0.132 | -1.66E-02 | 2.13E-01 | 2.66E+00 | 2.74E+00 | 9.37E-01 | no | Emsy | intron_variant | MODIFIER |
| 7 | 98653431 | AX-170124585 | 0 | A | C | 0.132 | -1.66E-02 | 2.13E-01 | 2.66E+00 | 2.74E+00 | 9.37E-01 | no | Emsy | intron_variant | MODIFIER |
| 7 | 98668497 | AX-170199170 | 0 | T | C | 0.132 | -1.66E-02 | 2.13E-01 | 2.66E+00 | 2.74E+00 | 9.37E-01 | no | | regulatory_region_variant | MODIFIER |
| 7 | 98678141 | AX-170029603 | 0 | G | A | 0.132 | -1.66E-02 | 2.13E-01 | 2.66E+00 | 2.74E+00 | 9.37E-01 | no | Gm15506 | downstream_gene_variant | MODIFIER |

| | | | | | | | | | | | | | | | |
|---|---|---|---|---|---|---|---|---|---|---|---|---|---|---|---|
| 7 | 98679605 | AX-170029602 | 0 | T | C | 0.132 | -1.66E-02 | 2.13E-01 | 2.66E+00 | 2.74E+00 | 9.37E-01 | no | Gm15506 | downstream_gene_variant | MODIFIER |
| 1 | 148809666 | AX-170288921 | 1 | A | G | 0.196 | -9.15E-03 | 1.28E-01 | 2.66E+00 | 2.75E+00 | 9.37E-01 | no | | intergenic_variant | MODIFIER |
| 12 | 109862111 | AX-170344139 | 0 | T | C | 0.132 | -1.38E-02 | 1.84E-01 | 2.67E+00 | 2.74E+00 | 9.37E-01 | no | | intergenic_variant | MODIFIER |
| 12 | 109887478 | AX-170503331 | 0 | C | T | 0.132 | -1.38E-02 | 1.84E-01 | 2.67E+00 | 2.74E+00 | 9.37E-01 | no | | intergenic_variant | MODIFIER |
| 12 | 109901926 | AX-170406404 | 0 | C | G | 0.132 | -1.38E-02 | 1.84E-01 | 2.67E+00 | 2.74E+00 | 9.37E-01 | no | | intergenic_variant | MODIFIER |
| 12 | 109909377 | AX-170528515 | 0 | C | T | 0.132 | -1.38E-02 | 1.84E-01 | 2.67E+00 | 2.74E+00 | 9.37E-01 | no | 4930511J24Rik | upstream_gene_variant | MODIFIER |
| 12 | 109914638 | AX-170171293 | 0 | A | C | 0.132 | -1.38E-02 | 1.84E-01 | 2.67E+00 | 2.74E+00 | 9.37E-01 | no | 4930511J24Rik | downstream_gene_variant | MODIFIER |
| 12 | 109952181 | AX-170007517 | 0 | T | C | 0.132 | -1.38E-02 | 1.84E-01 | 2.67E+00 | 2.74E+00 | 9.37E-01 | no | | regulatory_region_variant | MODIFIER |
| 12 | 109980675 | AX-170282033 | 0 | G | C | 0.132 | -1.38E-02 | 1.84E-01 | 2.67E+00 | 2.74E+00 | 9.37E-01 | no | | intergenic_variant | MODIFIER |
| 12 | 113801866 | AX-170450041 | 0 | T | G | 0.132 | -1.38E-02 | 1.84E-01 | 2.67E+00 | 2.74E+00 | 9.37E-01 | no | Gm7003 | downstream_gene_variant | MODIFIER |
| 12 | 117694211 | AX-170007606 | 0 | T | C | 0.132 | -1.38E-02 | 1.84E-01 | 2.67E+00 | 2.74E+00 | 9.37E-01 | no | Rapgef5 | intron_variant | MODIFIER |
| 12 | 117717406 | AX-170104782 | 0 | C | T | 0.132 | -1.38E-02 | 1.84E-01 | 2.67E+00 | 2.74E+00 | 9.37E-01 | no | Rapgef5 | intron_variant | MODIFIER |
| 12 | 117719220 | AX-170318310 | 0 | T | C | 0.132 | -1.38E-02 | 1.84E-01 | 2.67E+00 | 2.74E+00 | 9.37E-01 | no | Rapgef5 | intron_variant | MODIFIER |
| 12 | 117744208 | AX-97824794 | 0 | A | G | 0.132 | -1.38E-02 | 1.84E-01 | 2.67E+00 | 2.74E+00 | 9.37E-01 | no | Rapgef5 | synonymous_variant | LOW |
| 12 | 117749772 | AX-170007608 | 0 | A | G | 0.132 | -1.38E-02 | 1.84E-01 | 2.67E+00 | 2.74E+00 | 9.37E-01 | no | Rapgef5 | intron_variant | MODIFIER |
| 12 | 118131408 | AX-170232242 | 0 | T | C | 0.132 | -1.38E-02 | 1.84E-01 | 2.67E+00 | 2.74E+00 | 9.37E-01 | no | Dnah11 | intron_variant | MODIFIER |
| 12 | 118197350 | AX-170092630 | 0 | A | G | 0.132 | -1.38E-02 | 1.84E-01 | 2.67E+00 | 2.74E+00 | 9.37E-01 | no | Dnah11 | intron_variant | MODIFIER |
| 12 | 118198480 | AX-170361169 | 0 | T | G | 0.132 | -1.38E-02 | 1.84E-01 | 2.67E+00 | 2.74E+00 | 9.37E-01 | no | Dnah11 | intron_variant | MODIFIER |
| 12 | 118756160 | AX-170007624 | 0 | C | A | 0.132 | -1.38E-02 | 1.84E-01 | 2.67E+00 | 2.74E+00 | 9.37E-01 | no | Gm22537 | downstream_gene_variant | MODIFIER |
| 1 | 7566639 | AX-170315373 | 0 | C | T | 0.421 | 6.33E-03 | 7.01E-02 | 2.65E+00 | 2.72E+00 | 9.37E-01 | no | | intergenic_variant | MODIFIER |
| 18 | 68531162 | AX-170373162 | 0 | G | A | 0.096 | 1.47E-02 | 1.70E-01 | 2.62E+00 | 2.73E+00 | 9.37E-01 | no | | intergenic_variant | MODIFIER |
| 1 | 9750259 | AX-169999918 | 1 | C | T | 0.527 | 6.70E-03 | 7.14E-02 | 2.64E+00 | 2.71E+00 | 9.37E-01 | no | Vcpip1 | upstream_gene_variant | MODIFIER |
| 7 | 102028399 | AX-170228459 | 0 | G | A | 0.456 | 8.19E-03 | 9.53E-02 | 2.66E+00 | 2.72E+00 | 9.37E-01 | no | Rnf121 | intron_variant | MODIFIER |
| 12 | 24551374 | AX-170048515 | 0 | G | T | 0.044 | -9.23E-03 | 1.31E-01 | 2.58E+00 | 2.74E+00 | 9.37E-01 | no | Taf1b | intron_variant | MODIFIER |
| 18 | 66111780 | AX-170526546 | 2 | A | G | 0.391 | 8.60E-03 | 8.35E-02 | 2.61E+00 | 2.71E+00 | 9.37E-01 | no | Ccbe1 | intron_variant,NMD_transcript_variant | MODIFIER |
| 18 | 58334564 | AX-170243835 | 1 | C | G | 0.188 | -9.47E-03 | 1.14E-01 | 2.65E+00 | 2.73E+00 | 9.37E-01 | no | | intergenic_variant | MODIFIER |
| 7 | 136820605 | AX-170030062 | 0 | G | A | 0.307 | -6.99E-03 | 7.96E-02 | 2.63E+00 | 2.73E+00 | 9.37E-01 | no | | intergenic_variant | MODIFIER |
| 9 | 86622220 | AX-170489475 | 2 | T | C | 0.136 | -1.12E-02 | 1.28E-01 | 2.64E+00 | 2.72E+00 | 9.37E-01 | no | Me1 | intron_variant | MODIFIER |
| 4 | 155905260 | AX-170388753 | 1 | G | A | 0.107 | -1.04E-02 | 1.43E-01 | 2.61E+00 | 2.74E+00 | 9.37E-01 | no | Acap3 | intron_variant | MODIFIER |
| 11 | 44855093 | AX-170223824 | 0 | G | C | 0.088 | -1.56E-02 | 1.83E-01 | 2.62E+00 | 2.73E+00 | 9.37E-01 | no | Ebf1 | intron_variant | MODIFIER |
| 16 | 92893217 | AX-170165436 | 1 | T | C | 0.482 | -6.01E-03 | 7.98E-02 | 2.66E+00 | 2.74E+00 | 9.37E-01 | no | | intergenic_variant | MODIFIER |
| 7 | 56114773 | AX-97819911 | 1 | C | T | 0.42 | 8.55E-03 | 9.89E-02 | 2.65E+00 | 2.72E+00 | 9.37E-01 | no | Herc2 | synonymous_variant | LOW |
| 8 | 55036653 | AX-170344348 | 0 | G | A | 0.158 | -1.20E-02 | 1.38E-01 | 2.65E+00 | 2.72E+00 | 9.37E-01 | no | Gpm6a | intron_variant | MODIFIER |
| 8 | 56861438 | AX-170376580 | 0 | G | A | 0.158 | -1.20E-02 | 1.38E-01 | 2.65E+00 | 2.72E+00 | 9.37E-01 | no | Gm33501 | intron_variant,non_coding_transcript_variant | MODIFIER |
| 8 | 57356510 | AX-97841816 | 0 | C | T | 0.158 | -1.20E-02 | 1.38E-01 | 2.65E+00 | 2.72E+00 | 9.37E-01 | no | 5033428I22Rik | downstream_gene_variant | MODIFIER |
| 8 | 57496608 | AX-97857368 | 0 | C | T | 0.158 | -1.20E-02 | 1.38E-01 | 2.65E+00 | 2.72E+00 | 9.37E-01 | no | 2500002B13Rik | intron_variant,non_coding_transcript_variant | MODIFIER |
| 11 | 18016815 | AX-170004398 | 1 | A | T | 0.062 | 1.55E-02 | 2.14E-01 | 2.65E+00 | 2.75E+00 | 9.38E-01 | no | | intergenic_variant | MODIFIER |
| 11 | 34506509 | AX-170432929 | 1 | T | A | 0.098 | -1.33E-02 | 1.88E-01 | 2.66E+00 | 2.75E+00 | 9.38E-01 | no | Dock2 | intron_variant | MODIFIER |
| 11 | 35196079 | AX-170251938 | 1 | C | T | 0.098 | -1.33E-02 | 1.88E-01 | 2.66E+00 | 2.75E+00 | 9.38E-01 | no | Slit3 | intron_variant | MODIFIER |
| 11 | 32982555 | AX-170399239 | 0 | G | A | 0.096 | -1.34E-02 | 1.88E-01 | 2.66E+00 | 2.75E+00 | 9.38E-01 | no | Gm26052 | downstream_gene_variant | MODIFIER |
| 11 | 33956570 | AX-97817932 | 0 | A | C | 0.096 | -1.34E-02 | 1.88E-01 | 2.66E+00 | 2.75E+00 | 9.38E-01 | no | Kcnip1 | intron_variant,non_coding_transcript_variant | MODIFIER |
| 11 | 33957712 | AX-170351506 | 0 | G | A | 0.096 | -1.34E-02 | 1.88E-01 | 2.66E+00 | 2.75E+00 | 9.38E-01 | no | Kcnip1 | intron_variant,non_coding_transcript_variant | MODIFIER |
| 11 | 33970591 | AX-170046745 | 0 | A | G | 0.096 | -1.34E-02 | 1.88E-01 | 2.66E+00 | 2.75E+00 | 9.38E-01 | no | Kcnmb1 | 3_prime_UTR_variant | MODIFIER |
| 11 | 33970643 | AX-170046748 | 0 | T | C | 0.096 | -1.34E-02 | 1.88E-01 | 2.66E+00 | 2.75E+00 | 9.38E-01 | no | Kcnmb1 | 3_prime_UTR_variant | MODIFIER |
| 11 | 33972900 | AX-170046746 | 0 | A | G | 0.096 | -1.34E-02 | 1.88E-01 | 2.66E+00 | 2.75E+00 | 9.38E-01 | no | Kcnmb1 | 3_prime_UTR_variant | MODIFIER |
| 11 | 33991153 | AX-170255814 | 0 | A | G | 0.096 | -1.34E-02 | 1.88E-01 | 2.66E+00 | 2.75E+00 | 9.38E-01 | no | Kcnip1 | intron_variant,non_coding_transcript_variant | MODIFIER |
| 11 | 33992450 | AX-170194427 | 0 | T | C | 0.096 | -1.34E-02 | 1.88E-01 | 2.66E+00 | 2.75E+00 | 9.38E-01 | no | Kcnip1 | intron_variant,non_coding_transcript_variant | MODIFIER |
| 11 | 33993746 | AX-170370195 | 0 | G | A | 0.096 | -1.34E-02 | 1.88E-01 | 2.66E+00 | 2.75E+00 | 9.38E-01 | no | Kcnip1 | upstream_gene_variant | MODIFIER |
| 11 | 33995148 | AX-170167709 | 0 | G | A | 0.096 | -1.34E-02 | 1.88E-01 | 2.66E+00 | 2.75E+00 | 9.38E-01 | no | Kcnip1 | upstream_gene_variant | MODIFIER |
| 11 | 33996940 | AX-97854364 | 0 | T | C | 0.096 | -1.34E-02 | 1.88E-01 | 2.66E+00 | 2.75E+00 | 9.38E-01 | no | Kcnip1 | upstream_gene_variant | MODIFIER |
| 11 | 34043992 | AX-97851506 | 0 | T | A | 0.096 | -1.34E-02 | 1.88E-01 | 2.66E+00 | 2.75E+00 | 9.38E-01 | no | Lcp2 | upstream_gene_variant | MODIFIER |
| 11 | 34067862 | AX-170092214 | 0 | A | G | 0.096 | -1.34E-02 | 1.88E-01 | 2.66E+00 | 2.75E+00 | 9.38E-01 | no | Lcp2 | intron_variant | MODIFIER |
| 11 | 34082185 | AX-170127123 | 0 | A | G | 0.096 | -1.34E-02 | 1.88E-01 | 2.66E+00 | 2.75E+00 | 9.38E-01 | no | Lcp2 | intron_variant | MODIFIER |
| 11 | 34084042 | AX-170513359 | 0 | G | T | 0.096 | -1.34E-02 | 1.88E-01 | 2.66E+00 | 2.75E+00 | 9.38E-01 | no | Lcp2 | intron_variant | MODIFIER |

| | | | | | | | | | | | | | | | |
|---|---|---|---|---|---|---|---|---|---|---|---|---|---|---|---|
| 11 | 34108032 | AX-97806348 | 0 | A | T | 0.096 | -1.34E-02 | 1.88E-01 | 2.66E+00 | 2.75E+00 | 9.38E-01 | no | | intergenic_variant | MODIFIER |
| 11 | 34138337 | AX-170527723 | 0 | C | T | 0.096 | -1.34E-02 | 1.88E-01 | 2.66E+00 | 2.75E+00 | 9.38E-01 | no | Gm12121 | upstream_gene_variant | MODIFIER |
| 11 | 34146793 | AX-97793784 | 0 | T | C | 0.096 | -1.34E-02 | 1.88E-01 | 2.66E+00 | 2.75E+00 | 9.38E-01 | no | Gm12121 | downstream_gene_variant | MODIFIER |
| 11 | 34147324 | AX-170411667 | 0 | C | T | 0.096 | -1.34E-02 | 1.88E-01 | 2.66E+00 | 2.75E+00 | 9.38E-01 | no | Gm12121 | downstream_gene_variant | MODIFIER |
| 11 | 34151057 | AX-170004671 | 0 | G | A | 0.096 | -1.34E-02 | 1.88E-01 | 2.66E+00 | 2.75E+00 | 9.38E-01 | no | | intergenic_variant | MODIFIER |
| 11 | 34151276 | AX-170457416 | 0 | G | A | 0.096 | -1.34E-02 | 1.88E-01 | 2.66E+00 | 2.75E+00 | 9.38E-01 | no | | intergenic_variant | MODIFIER |
| 11 | 34153615 | AX-170201070 | 0 | A | G | 0.096 | -1.34E-02 | 1.88E-01 | 2.66E+00 | 2.75E+00 | 9.38E-01 | no | | intergenic_variant | MODIFIER |
| 11 | 34187289 | AX-170254173 | 0 | G | A | 0.096 | -1.34E-02 | 1.88E-01 | 2.66E+00 | 2.75E+00 | 9.38E-01 | no | | intergenic_variant | MODIFIER |
| 11 | 34190822 | AX-170125495 | 0 | A | G | 0.096 | -1.34E-02 | 1.88E-01 | 2.66E+00 | 2.75E+00 | 9.38E-01 | no | | intergenic_variant | MODIFIER |
| 11 | 34192154 | AX-170046749 | 0 | A | C | 0.096 | -1.34E-02 | 1.88E-01 | 2.66E+00 | 2.75E+00 | 9.38E-01 | no | | intergenic_variant | MODIFIER |
| 11 | 34192457 | AX-170479235 | 0 | C | T | 0.096 | -1.34E-02 | 1.88E-01 | 2.66E+00 | 2.75E+00 | 9.38E-01 | no | | intergenic_variant | MODIFIER |
| 11 | 34192511 | AX-170164561 | 0 | G | A | 0.096 | -1.34E-02 | 1.88E-01 | 2.66E+00 | 2.75E+00 | 9.38E-01 | no | | intergenic_variant | MODIFIER |
| 11 | 34195104 | AX-170378407 | 0 | A | G | 0.096 | -1.34E-02 | 1.88E-01 | 2.66E+00 | 2.75E+00 | 9.38E-01 | no | | intergenic_variant | MODIFIER |
| 11 | 34234119 | AX-97833348 | 0 | G | A | 0.096 | -1.34E-02 | 1.88E-01 | 2.66E+00 | 2.75E+00 | 9.38E-01 | no | Dock2 | intron_variant | MODIFIER |
| 11 | 34236008 | AX-170212557 | 0 | A | C | 0.096 | -1.34E-02 | 1.88E-01 | 2.66E+00 | 2.75E+00 | 9.38E-01 | no | Dock2 | intron_variant | MODIFIER |
| 11 | 34254156 | AX-170429921 | 0 | A | G | 0.096 | -1.34E-02 | 1.88E-01 | 2.66E+00 | 2.75E+00 | 9.38E-01 | no | Dock2 | intron_variant | MODIFIER |
| 11 | 34258405 | AX-97783693 | 0 | C | T | 0.096 | -1.34E-02 | 1.88E-01 | 2.66E+00 | 2.75E+00 | 9.38E-01 | no | Dock2 | intron_variant | MODIFIER |
| 11 | 34259576 | AX-170330464 | 0 | T | C | 0.096 | -1.34E-02 | 1.88E-01 | 2.66E+00 | 2.75E+00 | 9.38E-01 | no | Dock2 | intron_variant | MODIFIER |
| 11 | 34286282 | AX-97818193 | 0 | T | G | 0.096 | -1.34E-02 | 1.88E-01 | 2.66E+00 | 2.75E+00 | 9.38E-01 | no | Dock2 | intron_variant | MODIFIER |
| 11 | 34293325 | AX-170465124 | 0 | A | G | 0.096 | -1.34E-02 | 1.88E-01 | 2.66E+00 | 2.75E+00 | 9.38E-01 | no | Dock2 | intron_variant | MODIFIER |
| 11 | 34296382 | AX-97820502 | 0 | C | T | 0.096 | -1.34E-02 | 1.88E-01 | 2.66E+00 | 2.75E+00 | 9.38E-01 | no | Dock2 | intron_variant | MODIFIER |
| 11 | 34297664 | AX-170276729 | 0 | C | T | 0.096 | -1.34E-02 | 1.88E-01 | 2.66E+00 | 2.75E+00 | 9.38E-01 | no | Dock2 | intron_variant | MODIFIER |
| 11 | 34298955 | AX-170151110 | 0 | A | G | 0.096 | -1.34E-02 | 1.88E-01 | 2.66E+00 | 2.75E+00 | 9.38E-01 | no | Dock2 | intron_variant | MODIFIER |
| 11 | 34299560 | AX-170194478 | 0 | G | T | 0.096 | -1.34E-02 | 1.88E-01 | 2.66E+00 | 2.75E+00 | 9.38E-01 | no | Dock2 | intron_variant | MODIFIER |
| 11 | 34299886 | AX-170365917 | 0 | T | G | 0.096 | -1.34E-02 | 1.88E-01 | 2.66E+00 | 2.75E+00 | 9.38E-01 | no | Dock2 | intron_variant | MODIFIER |
| 11 | 34300307 | AX-170319152 | 0 | G | A | 0.096 | -1.34E-02 | 1.88E-01 | 2.66E+00 | 2.75E+00 | 9.38E-01 | no | Dock2 | intron_variant | MODIFIER |
| 11 | 34301437 | AX-170528866 | 0 | C | T | 0.096 | -1.34E-02 | 1.88E-01 | 2.66E+00 | 2.75E+00 | 9.38E-01 | no | Dock2 | intron_variant | MODIFIER |
| 11 | 34301901 | AX-170379519 | 0 | C | T | 0.096 | -1.34E-02 | 1.88E-01 | 2.66E+00 | 2.75E+00 | 9.38E-01 | no | Dock2 | intron_variant | MODIFIER |
| 11 | 34302413 | AX-170295015 | 0 | C | T | 0.096 | -1.34E-02 | 1.88E-01 | 2.66E+00 | 2.75E+00 | 9.38E-01 | no | Dock2 | intron_variant | MODIFIER |
| 11 | 34302743 | AX-170361019 | 0 | A | C | 0.096 | -1.34E-02 | 1.88E-01 | 2.66E+00 | 2.75E+00 | 9.38E-01 | no | Dock2 | intron_variant | MODIFIER |
| 11 | 34312943 | AX-170425900 | 0 | T | C | 0.096 | -1.34E-02 | 1.88E-01 | 2.66E+00 | 2.75E+00 | 9.38E-01 | no | Fam196b | upstream_gene_variant | MODIFIER |
| 11 | 34320034 | AX-97824241 | 0 | A | G | 0.096 | -1.34E-02 | 1.88E-01 | 2.66E+00 | 2.75E+00 | 9.38E-01 | no | Fam196b | intron_variant | MODIFIER |
| 11 | 34353411 | AX-170470409 | 0 | G | A | 0.096 | -1.34E-02 | 1.88E-01 | 2.66E+00 | 2.75E+00 | 9.38E-01 | no | Fam196b | intron_variant | MODIFIER |
| 11 | 34354460 | AX-170409284 | 0 | T | C | 0.096 | -1.34E-02 | 1.88E-01 | 2.66E+00 | 2.75E+00 | 9.38E-01 | no | Fam196b | intron_variant | MODIFIER |
| 11 | 34360105 | AX-170489588 | 0 | T | G | 0.096 | -1.34E-02 | 1.88E-01 | 2.66E+00 | 2.75E+00 | 9.38E-01 | no | Fam196b | intron_variant | MODIFIER |
| 11 | 34360455 | AX-170046758 | 0 | A | G | 0.096 | -1.34E-02 | 1.88E-01 | 2.66E+00 | 2.75E+00 | 9.38E-01 | no | Fam196b | intron_variant | MODIFIER |
| 11 | 34377355 | AX-170392182 | 0 | C | T | 0.096 | -1.34E-02 | 1.88E-01 | 2.66E+00 | 2.75E+00 | 9.38E-01 | no | Fam196b | intron_variant | MODIFIER |
| 11 | 34387996 | AX-170250543 | 0 | T | G | 0.096 | -1.34E-02 | 1.88E-01 | 2.66E+00 | 2.75E+00 | 9.38E-01 | no | Fam196b | intron_variant | MODIFIER |
| 11 | 34389089 | AX-170438375 | 0 | T | C | 0.096 | -1.34E-02 | 1.88E-01 | 2.66E+00 | 2.75E+00 | 9.38E-01 | no | Fam196b | intron_variant | MODIFIER |
| 11 | 34389252 | AX-170423287 | 0 | C | A | 0.096 | -1.34E-02 | 1.88E-01 | 2.66E+00 | 2.75E+00 | 9.38E-01 | no | Fam196b | intron_variant | MODIFIER |
| 11 | 34389868 | AX-170319257 | 0 | T | C | 0.096 | -1.34E-02 | 1.88E-01 | 2.66E+00 | 2.75E+00 | 9.38E-01 | no | Fam196b | intron_variant | MODIFIER |
| 11 | 34394423 | AX-97794329 | 0 | T | C | 0.096 | -1.34E-02 | 1.88E-01 | 2.66E+00 | 2.75E+00 | 9.38E-01 | no | Fam196b | intron_variant | MODIFIER |
| 11 | 34401313 | AX-170523849 | 0 | G | A | 0.096 | -1.34E-02 | 1.88E-01 | 2.66E+00 | 2.75E+00 | 9.38E-01 | no | Fam196b | intron_variant | MODIFIER |
| 11 | 34401701 | AX-170432195 | 0 | C | T | 0.096 | -1.34E-02 | 1.88E-01 | 2.66E+00 | 2.75E+00 | 9.38E-01 | no | Fam196b | 5_prime_UTR_variant | MODIFIER |
| 11 | 34413058 | AX-170390077 | 0 | T | A | 0.096 | -1.34E-02 | 1.88E-01 | 2.66E+00 | 2.75E+00 | 9.38E-01 | no | Fam196b | intron_variant | MODIFIER |
| 11 | 34433642 | AX-97840260 | 0 | C | T | 0.096 | -1.34E-02 | 1.88E-01 | 2.66E+00 | 2.75E+00 | 9.38E-01 | no | Dock2 | intron_variant | MODIFIER |
| 11 | 34434367 | AX-170338368 | 0 | G | C | 0.096 | -1.34E-02 | 1.88E-01 | 2.66E+00 | 2.75E+00 | 9.38E-01 | no | Dock2 | intron_variant | MODIFIER |
| 11 | 34471344 | AX-170280595 | 0 | C | T | 0.096 | -1.34E-02 | 1.88E-01 | 2.66E+00 | 2.75E+00 | 9.38E-01 | no | Dock2 | intron_variant | MODIFIER |
| 11 | 34479292 | AX-170219702 | 0 | G | A | 0.096 | -1.34E-02 | 1.88E-01 | 2.66E+00 | 2.75E+00 | 9.38E-01 | no | Dock2 | intron_variant | MODIFIER |
| 11 | 34479852 | AX-170092215 | 0 | G | A | 0.096 | -1.34E-02 | 1.88E-01 | 2.66E+00 | 2.75E+00 | 9.38E-01 | no | Dock2 | intron_variant | MODIFIER |
| 11 | 34507599 | AX-170158109 | 0 | T | C | 0.096 | -1.34E-02 | 1.88E-01 | 2.66E+00 | 2.75E+00 | 9.38E-01 | no | Dock2 | intron_variant | MODIFIER |
| 11 | 34508346 | AX-170291839 | 0 | A | G | 0.096 | -1.34E-02 | 1.88E-01 | 2.66E+00 | 2.75E+00 | 9.38E-01 | no | Dock2 | intron_variant | MODIFIER |
| 11 | 34512175 | AX-170188475 | 0 | C | T | 0.096 | -1.34E-02 | 1.88E-01 | 2.66E+00 | 2.75E+00 | 9.38E-01 | no | Dock2 | intron_variant | MODIFIER |
| 11 | 34512917 | AX-170270814 | 0 | C | G | 0.096 | -1.34E-02 | 1.88E-01 | 2.66E+00 | 2.75E+00 | 9.38E-01 | no | Dock2 | intron_variant | MODIFIER |

| | | | | | | | | | | | | | | | |
|---|---|---|---|---|---|---|---|---|---|---|---|---|---|---|---|
| 11 | 34534819 | AX-170175958 | 0 | A | T | 0.096 | -1.34E-02 | 1.88E-01 | 2.66E+00 | 2.75E+00 | 9.38E-01 | no | Dock2 | intron_variant | MODIFIER |
| 11 | 34536033 | AX-170493478 | 0 | C | T | 0.096 | -1.34E-02 | 1.88E-01 | 2.66E+00 | 2.75E+00 | 9.38E-01 | no | Dock2 | intron_variant | MODIFIER |
| 11 | 34536161 | AX-170398964 | 0 | G | A | 0.096 | -1.34E-02 | 1.88E-01 | 2.66E+00 | 2.75E+00 | 9.38E-01 | no | Dock2 | intron_variant | MODIFIER |
| 11 | 34536962 | AX-170161100 | 0 | A | G | 0.096 | -1.34E-02 | 1.88E-01 | 2.66E+00 | 2.75E+00 | 9.38E-01 | no | Dock2 | intron_variant | MODIFIER |
| 11 | 34697567 | AX-170398197 | 0 | A | G | 0.096 | -1.34E-02 | 1.88E-01 | 2.66E+00 | 2.75E+00 | 9.38E-01 | no | Dock2 | intron_variant | MODIFIER |
| 11 | 34698880 | AX-170458811 | 0 | T | C | 0.096 | -1.34E-02 | 1.88E-01 | 2.66E+00 | 2.75E+00 | 9.38E-01 | no | Dock2 | intron_variant | MODIFIER |
| 11 | 34700335 | AX-170440119 | 0 | G | A | 0.096 | -1.34E-02 | 1.88E-01 | 2.66E+00 | 2.75E+00 | 9.38E-01 | no | Dock2 | intron_variant | MODIFIER |
| 11 | 34704180 | AX-170485910 | 0 | A | G | 0.096 | -1.34E-02 | 1.88E-01 | 2.66E+00 | 2.75E+00 | 9.38E-01 | no | Dock2 | intron_variant | MODIFIER |
| 11 | 34705846 | AX-170143163 | 0 | G | A | 0.096 | -1.34E-02 | 1.88E-01 | 2.66E+00 | 2.75E+00 | 9.38E-01 | no | Dock2 | synonymous_variant | LOW |
| 11 | 34708466 | AX-170369577 | 0 | G | A | 0.096 | -1.34E-02 | 1.88E-01 | 2.66E+00 | 2.75E+00 | 9.38E-01 | no | Dock2 | intron_variant | MODIFIER |
| 11 | 34709631 | AX-170482313 | 0 | G | A | 0.096 | -1.34E-02 | 1.88E-01 | 2.66E+00 | 2.75E+00 | 9.38E-01 | no | Dock2 | intron_variant | MODIFIER |
| 11 | 34712073 | AX-97847428 | 0 | G | A | 0.096 | -1.34E-02 | 1.88E-01 | 2.66E+00 | 2.75E+00 | 9.38E-01 | no | Dock2 | intron_variant | MODIFIER |
| 11 | 34712482 | AX-170165428 | 0 | C | A | 0.096 | -1.34E-02 | 1.88E-01 | 2.66E+00 | 2.75E+00 | 9.38E-01 | no | Dock2 | intron_variant | MODIFIER |
| 11 | 34713130 | AX-170116195 | 0 | A | G | 0.096 | -1.34E-02 | 1.88E-01 | 2.66E+00 | 2.75E+00 | 9.38E-01 | no | Dock2 | intron_variant | MODIFIER |
| 11 | 34717304 | AX-170225506 | 0 | C | G | 0.096 | -1.34E-02 | 1.88E-01 | 2.66E+00 | 2.75E+00 | 9.38E-01 | no | Dock2 | intron_variant | MODIFIER |
| 11 | 34722777 | AX-170099823 | 0 | C | A | 0.096 | -1.34E-02 | 1.88E-01 | 2.66E+00 | 2.75E+00 | 9.38E-01 | no | Dock2 | intron_variant | MODIFIER |
| 11 | 34729757 | AX-170111440 | 0 | C | T | 0.096 | -1.34E-02 | 1.88E-01 | 2.66E+00 | 2.75E+00 | 9.38E-01 | no | Dock2 | intron_variant | MODIFIER |
| 11 | 34730590 | AX-170101334 | 0 | T | C | 0.096 | -1.34E-02 | 1.88E-01 | 2.66E+00 | 2.75E+00 | 9.38E-01 | no | Dock2 | intron_variant | MODIFIER |
| 11 | 34740512 | AX-170308303 | 0 | C | T | 0.096 | -1.34E-02 | 1.88E-01 | 2.66E+00 | 2.75E+00 | 9.38E-01 | no | Dock2 | intron_variant | MODIFIER |
| 11 | 34756292 | AX-170046767 | 0 | A | G | 0.096 | -1.34E-02 | 1.88E-01 | 2.66E+00 | 2.75E+00 | 9.38E-01 | no | Dock2 | intron_variant | MODIFIER |
| 11 | 34760390 | AX-170046762 | 0 | G | A | 0.096 | -1.34E-02 | 1.88E-01 | 2.66E+00 | 2.75E+00 | 9.38E-01 | no | Dock2 | intron_variant | MODIFIER |
| 11 | 34769953 | AX-170259000 | 0 | G | T | 0.096 | -1.34E-02 | 1.88E-01 | 2.66E+00 | 2.75E+00 | 9.38E-01 | no | Dock2 | intron_variant | MODIFIER |
| 11 | 34770248 | AX-170471061 | 0 | G | A | 0.096 | -1.34E-02 | 1.88E-01 | 2.66E+00 | 2.75E+00 | 9.38E-01 | no | Dock2 | intron_variant | MODIFIER |
| 11 | 34774826 | AX-97819600 | 0 | A | G | 0.096 | -1.34E-02 | 1.88E-01 | 2.66E+00 | 2.75E+00 | 9.38E-01 | no | Dock2 | intron_variant | MODIFIER |
| 11 | 34817338 | AX-170454989 | 0 | A | C | 0.096 | -1.34E-02 | 1.88E-01 | 2.66E+00 | 2.75E+00 | 9.38E-01 | no | Spdl1 | intron_variant | MODIFIER |
| 11 | 34821964 | AX-97856071 | 0 | A | G | 0.096 | -1.34E-02 | 1.88E-01 | 2.66E+00 | 2.75E+00 | 9.38E-01 | no | Spdl1 | intron_variant | MODIFIER |
| 11 | 34823116 | AX-170004674 | 0 | C | G | 0.096 | -1.34E-02 | 1.88E-01 | 2.66E+00 | 2.75E+00 | 9.38E-01 | no | Spdl1 | intron_variant | MODIFIER |
| 11 | 34836673 | AX-170420067 | 0 | G | A | 0.096 | -1.34E-02 | 1.88E-01 | 2.66E+00 | 2.75E+00 | 9.38E-01 | no | Spdl1 | upstream_gene_variant | MODIFIER |
| 11 | 34837552 | AX-97785790 | 0 | C | T | 0.096 | -1.34E-02 | 1.88E-01 | 2.66E+00 | 2.75E+00 | 9.38E-01 | no | Spdl1 | upstream_gene_variant | MODIFIER |
| 11 | 34882714 | AX-97822720 | 0 | T | C | 0.096 | -1.34E-02 | 1.88E-01 | 2.66E+00 | 2.75E+00 | 9.38E-01 | no | Gm12711 | downstream_gene_variant | MODIFIER |
| 11 | 34894220 | AX-97847276 | 0 | C | T | 0.096 | -1.34E-02 | 1.88E-01 | 2.66E+00 | 2.75E+00 | 9.38E-01 | no | Gm25799 | upstream_gene_variant | MODIFIER |
| 11 | 34935707 | AX-170462936 | 0 | T | C | 0.096 | -1.34E-02 | 1.88E-01 | 2.66E+00 | 2.75E+00 | 9.38E-01 | no | | intergenic_variant | MODIFIER |
| 11 | 34946646 | AX-97817398 | 0 | T | A | 0.096 | -1.34E-02 | 1.88E-01 | 2.66E+00 | 2.75E+00 | 9.38E-01 | no | | intergenic_variant | MODIFIER |
| 11 | 34973268 | AX-170323329 | 0 | T | C | 0.096 | -1.34E-02 | 1.88E-01 | 2.66E+00 | 2.75E+00 | 9.38E-01 | no | | regulatory_region_variant | MODIFIER |
| 11 | 34973684 | AX-170355515 | 0 | G | A | 0.096 | -1.34E-02 | 1.88E-01 | 2.66E+00 | 2.75E+00 | 9.38E-01 | no | | regulatory_region_variant | MODIFIER |
| 11 | 34977079 | AX-97817990 | 0 | G | A | 0.096 | -1.34E-02 | 1.88E-01 | 2.66E+00 | 2.75E+00 | 9.38E-01 | no | | intergenic_variant | MODIFIER |
| 11 | 34995982 | AX-97853689 | 0 | G | A | 0.096 | -1.34E-02 | 1.88E-01 | 2.66E+00 | 2.75E+00 | 9.38E-01 | no | | intergenic_variant | MODIFIER |
| 11 | 35015460 | AX-170256825 | 0 | C | T | 0.096 | -1.34E-02 | 1.88E-01 | 2.66E+00 | 2.75E+00 | 9.38E-01 | no | | intergenic_variant | MODIFIER |
| 11 | 35017718 | AX-170154783 | 0 | G | A | 0.096 | -1.34E-02 | 1.88E-01 | 2.66E+00 | 2.75E+00 | 9.38E-01 | no | | intergenic_variant | MODIFIER |
| 11 | 35017840 | AX-170264568 | 0 | C | G | 0.096 | -1.34E-02 | 1.88E-01 | 2.66E+00 | 2.75E+00 | 9.38E-01 | no | | intergenic_variant | MODIFIER |
| 11 | 35018702 | AX-170160757 | 0 | C | G | 0.096 | -1.34E-02 | 1.88E-01 | 2.66E+00 | 2.75E+00 | 9.38E-01 | no | | intergenic_variant | MODIFIER |
| 11 | 35019057 | AX-170469673 | 0 | C | T | 0.096 | -1.34E-02 | 1.88E-01 | 2.66E+00 | 2.75E+00 | 9.38E-01 | no | | intergenic_variant | MODIFIER |
| 11 | 35019100 | AX-170251981 | 0 | G | A | 0.096 | -1.34E-02 | 1.88E-01 | 2.66E+00 | 2.75E+00 | 9.38E-01 | no | | intergenic_variant | MODIFIER |
| 11 | 35019488 | AX-170488556 | 0 | C | T | 0.096 | -1.34E-02 | 1.88E-01 | 2.66E+00 | 2.75E+00 | 9.38E-01 | no | | intergenic_variant | MODIFIER |
| 11 | 35023075 | AX-170114666 | 0 | G | A | 0.096 | -1.34E-02 | 1.88E-01 | 2.66E+00 | 2.75E+00 | 9.38E-01 | no | | intergenic_variant | MODIFIER |
| 11 | 35024976 | AX-170046771 | 0 | G | A | 0.096 | -1.34E-02 | 1.88E-01 | 2.66E+00 | 2.75E+00 | 9.38E-01 | no | | regulatory_region_variant | MODIFIER |
| 11 | 35027709 | AX-170046768 | 0 | G | C | 0.096 | -1.34E-02 | 1.88E-01 | 2.66E+00 | 2.75E+00 | 9.38E-01 | no | | intergenic_variant | MODIFIER |
| 11 | 35033817 | AX-170046770 | 0 | C | A | 0.096 | -1.34E-02 | 1.88E-01 | 2.66E+00 | 2.75E+00 | 9.38E-01 | no | | intergenic_variant | MODIFIER |
| 11 | 35034901 | AX-170462489 | 0 | A | G | 0.096 | -1.34E-02 | 1.88E-01 | 2.66E+00 | 2.75E+00 | 9.38E-01 | no | | intergenic_variant | MODIFIER |
| 11 | 35034976 | AX-170384232 | 0 | G | A | 0.096 | -1.34E-02 | 1.88E-01 | 2.66E+00 | 2.75E+00 | 9.38E-01 | no | | intergenic_variant | MODIFIER |
| 11 | 35036367 | AX-170342422 | 0 | T | C | 0.096 | -1.34E-02 | 1.88E-01 | 2.66E+00 | 2.75E+00 | 9.38E-01 | no | | intergenic_variant | MODIFIER |
| 11 | 35037192 | AX-170495468 | 0 | C | T | 0.096 | -1.34E-02 | 1.88E-01 | 2.66E+00 | 2.75E+00 | 9.38E-01 | no | | intergenic_variant | MODIFIER |
| 11 | 35058418 | AX-170311904 | 0 | G | A | 0.096 | -1.34E-02 | 1.88E-01 | 2.66E+00 | 2.75E+00 | 9.38E-01 | no | | intergenic_variant | MODIFIER |
| 11 | 35070352 | AX-170120931 | 0 | A | T | 0.096 | -1.34E-02 | 1.88E-01 | 2.66E+00 | 2.75E+00 | 9.38E-01 | no | | intergenic_variant | MODIFIER |

| | | | | | | | | | | | | | | | |
|---|---|---|---|---|---|---|---|---|---|---|---|---|---|---|---|
| 11 | 35093506 | AX-97811744 | 0 | T | C | 0.096 | -1.34E-02 | 1.88E-01 | 2.66E+00 | 2.75E+00 | 9.38E-01 | no | | intergenic_variant | MODIFIER |
| 11 | 35125519 | AX-97799511 | 0 | A | G | 0.096 | -1.34E-02 | 1.88E-01 | 2.66E+00 | 2.75E+00 | 9.38E-01 | no | Slit3 | intron_variant | MODIFIER |
| 11 | 35126638 | AX-170308230 | 0 | G | A | 0.096 | -1.34E-02 | 1.88E-01 | 2.66E+00 | 2.75E+00 | 9.38E-01 | no | Slit3 | intron_variant | MODIFIER |
| 11 | 35129902 | AX-170504384 | 0 | G | A | 0.096 | -1.34E-02 | 1.88E-01 | 2.66E+00 | 2.75E+00 | 9.38E-01 | no | Slit3 | intron_variant | MODIFIER |
| 11 | 35143733 | AX-170497983 | 0 | G | A | 0.096 | -1.34E-02 | 1.88E-01 | 2.66E+00 | 2.75E+00 | 9.38E-01 | no | Slit3 | intron_variant | MODIFIER |
| 11 | 35143916 | AX-170318925 | 0 | C | T | 0.096 | -1.34E-02 | 1.88E-01 | 2.66E+00 | 2.75E+00 | 9.38E-01 | no | Slit3 | intron_variant | MODIFIER |
| 11 | 35143972 | AX-170448130 | 0 | C | T | 0.096 | -1.34E-02 | 1.88E-01 | 2.66E+00 | 2.75E+00 | 9.38E-01 | no | Slit3 | intron_variant | MODIFIER |
| 11 | 35143988 | AX-170257767 | 0 | A | G | 0.096 | -1.34E-02 | 1.88E-01 | 2.66E+00 | 2.75E+00 | 9.38E-01 | no | Slit3 | intron_variant | MODIFIER |
| 11 | 35157621 | AX-170036469 | 0 | A | G | 0.096 | -1.34E-02 | 1.88E-01 | 2.66E+00 | 2.75E+00 | 9.38E-01 | no | Slit3 | intron_variant | MODIFIER |
| 11 | 35172969 | AX-97782706 | 0 | G | A | 0.096 | -1.34E-02 | 1.88E-01 | 2.66E+00 | 2.75E+00 | 9.38E-01 | no | Slit3 | intron_variant | MODIFIER |
| 11 | 35200038 | AX-170274797 | 0 | G | A | 0.096 | -1.34E-02 | 1.88E-01 | 2.66E+00 | 2.75E+00 | 9.38E-01 | no | Slit3 | intron_variant | MODIFIER |
| 11 | 35200411 | AX-170046774 | 0 | C | T | 0.096 | -1.34E-02 | 1.88E-01 | 2.66E+00 | 2.75E+00 | 9.38E-01 | no | Slit3 | intron_variant | MODIFIER |
| 11 | 35299383 | AX-170374603 | 0 | G | A | 0.096 | -1.34E-02 | 1.88E-01 | 2.66E+00 | 2.75E+00 | 9.38E-01 | no | Slit3 | intron_variant | MODIFIER |
| 11 | 35336607 | AX-170004678 | 0 | C | T | 0.096 | -1.34E-02 | 1.88E-01 | 2.66E+00 | 2.75E+00 | 9.38E-01 | no | Slit3 | intron_variant | MODIFIER |
| 11 | 35338353 | AX-170463139 | 0 | G | A | 0.096 | -1.34E-02 | 1.88E-01 | 2.66E+00 | 2.75E+00 | 9.38E-01 | no | Slit3 | intron_variant | MODIFIER |
| 11 | 35356671 | AX-97813012 | 0 | C | T | 0.096 | -1.34E-02 | 1.88E-01 | 2.66E+00 | 2.75E+00 | 9.38E-01 | no | Slit3 | intron_variant | MODIFIER |
| 11 | 35359700 | AX-170180945 | 0 | C | T | 0.096 | -1.34E-02 | 1.88E-01 | 2.66E+00 | 2.75E+00 | 9.38E-01 | no | Slit3 | intron_variant | MODIFIER |
| 11 | 35362157 | AX-170120538 | 0 | G | T | 0.096 | -1.34E-02 | 1.88E-01 | 2.66E+00 | 2.75E+00 | 9.38E-01 | no | Slit3 | intron_variant | MODIFIER |
| 11 | 35368053 | AX-170120611 | 0 | A | G | 0.096 | -1.34E-02 | 1.88E-01 | 2.66E+00 | 2.75E+00 | 9.38E-01 | no | Slit3 | intron_variant | MODIFIER |
| 11 | 35369706 | AX-170448252 | 0 | C | T | 0.096 | -1.34E-02 | 1.88E-01 | 2.66E+00 | 2.75E+00 | 9.38E-01 | no | Slit3 | intron_variant | MODIFIER |
| 11 | 35378248 | AX-170046777 | 0 | A | G | 0.096 | -1.34E-02 | 1.88E-01 | 2.66E+00 | 2.75E+00 | 9.38E-01 | no | Slit3 | intron_variant | MODIFIER |
| 11 | 35380801 | AX-170402027 | 0 | A | G | 0.096 | -1.34E-02 | 1.88E-01 | 2.66E+00 | 2.75E+00 | 9.38E-01 | no | Slit3 | intron_variant | MODIFIER |
| 11 | 35381052 | AX-170482814 | 0 | A | G | 0.096 | -1.34E-02 | 1.88E-01 | 2.66E+00 | 2.75E+00 | 9.38E-01 | no | Slit3 | intron_variant | MODIFIER |
| 11 | 35381394 | AX-170511458 | 0 | A | G | 0.096 | -1.34E-02 | 1.88E-01 | 2.66E+00 | 2.75E+00 | 9.38E-01 | no | Slit3 | intron_variant | MODIFIER |
| 5 | 99589884 | AX-170277416 | 1 | T | A | 0.42 | -7.51E-03 | 1.05E-01 | 2.69E+00 | 2.75E+00 | 9.38E-01 | no | Rasgef1b | intron_variant,non_coding_transcript_variant | MODIFIER |
| 9 | 108150453 | AX-170165983 | 0 | G | A | 0.132 | -1.01E-02 | 1.14E-01 | 2.65E+00 | 2.72E+00 | 9.38E-01 | no | Bsn | intron_variant | MODIFIER |
| 1 | 11670767 | AX-170210883 | 1 | G | T | 0.393 | -5.05E-03 | 7.30E-02 | 2.69E+00 | 2.76E+00 | 9.38E-01 | no | A830018L16Rik | intron_variant | MODIFIER |
| 4 | 140903948 | AX-170024121 | 0 | T | C | 0.096 | 1.19E-02 | 1.46E-01 | 2.63E+00 | 2.73E+00 | 9.38E-01 | no | Padi2 | upstream_gene_variant | MODIFIER |
| 18 | 18583440 | AX-170104945 | 1 | C | T | 0.054 | -1.45E-02 | 1.86E-01 | 2.64E+00 | 2.74E+00 | 9.38E-01 | no | | intergenic_variant | MODIFIER |
| 2 | 123821292 | AX-170153817 | 1 | T | A | 0.036 | -9.88E-03 | 1.29E-01 | 2.57E+00 | 2.74E+00 | 9.38E-01 | no | | intergenic_variant | MODIFIER |
| 1 | 72948486 | AX-170462626 | 0 | T | G | 0.053 | 1.44E-02 | 1.90E-01 | 2.65E+00 | 2.74E+00 | 9.38E-01 | no | | intergenic_variant | MODIFIER |
| 20 | 70051200 | AX-170090538 | 0 | C | T | 0.053 | 1.44E-02 | 1.90E-01 | 2.65E+00 | 2.74E+00 | 9.38E-01 | no | 1700111N16Rik | intron_variant,non_coding_transcript_variant | MODIFIER |
| 7 | 53002808 | AX-170525856 | 2 | A | T | 0.427 | -7.14E-03 | 1.01E-01 | 2.68E+00 | 2.75E+00 | 9.38E-01 | no | | intergenic_variant | MODIFIER |
| 10 | 49450668 | AX-97812097 | 0 | G | A | 0.026 | -1.36E-02 | 1.36E-01 | 2.53E+00 | 2.72E+00 | 9.38E-01 | no | Grik2 | intron_variant | MODIFIER |
| 9 | 80214858 | AX-170334732 | 2 | C | T | 0.209 | 9.45E-03 | 1.29E-01 | 2.70E+00 | 2.75E+00 | 9.38E-01 | no | Myo6 | intron_variant | MODIFIER |
| 7 | 107148456 | AX-97794182 | 1 | A | T | 0.446 | -6.32E-03 | 9.10E-02 | 2.68E+00 | 2.75E+00 | 9.38E-01 | no | Olfr716 | downstream_gene_variant | MODIFIER |
| 10 | 9627726 | AX-170044115 | 0 | G | T | 0.228 | 7.20E-03 | 1.02E-01 | 2.67E+00 | 2.75E+00 | 9.38E-01 | no | Samd5 | 3_prime_UTR_variant | MODIFIER |
| 10 | 9628180 | AX-170044117 | 0 | G | A | 0.228 | 7.20E-03 | 1.02E-01 | 2.67E+00 | 2.75E+00 | 9.38E-01 | no | Samd5 | 3_prime_UTR_variant | MODIFIER |
| 10 | 10046762 | AX-97823626 | 0 | C | T | 0.228 | 7.20E-03 | 1.02E-01 | 2.67E+00 | 2.75E+00 | 9.38E-01 | no | Gm28905 | intron_variant,non_coding_transcript_variant | MODIFIER |
| 10 | 10581678 | AX-97817296 | 0 | T | C | 0.228 | 7.20E-03 | 1.02E-01 | 2.67E+00 | 2.75E+00 | 9.38E-01 | no | | intergenic_variant | MODIFIER |
| 10 | 10608096 | AX-170250486 | 0 | T | C | 0.228 | 7.20E-03 | 1.02E-01 | 2.67E+00 | 2.75E+00 | 9.38E-01 | no | Gm5177 | upstream_gene_variant | MODIFIER |
| 10 | 10609012 | AX-170255778 | 0 | G | A | 0.228 | 7.20E-03 | 1.02E-01 | 2.67E+00 | 2.75E+00 | 9.38E-01 | no | Gm5177 | upstream_gene_variant | MODIFIER |
| 10 | 10619349 | AX-170417934 | 0 | A | G | 0.228 | 7.20E-03 | 1.02E-01 | 2.67E+00 | 2.75E+00 | 9.38E-01 | no | Gm5177 | downstream_gene_variant | MODIFIER |
| 10 | 10628907 | AX-170415347 | 0 | T | G | 0.228 | 7.20E-03 | 1.02E-01 | 2.67E+00 | 2.75E+00 | 9.38E-01 | no | | intergenic_variant | MODIFIER |
| 10 | 10648174 | AX-170126276 | 0 | T | C | 0.228 | 7.20E-03 | 1.02E-01 | 2.67E+00 | 2.75E+00 | 9.38E-01 | no | | intergenic_variant | MODIFIER |
| 10 | 10821511 | AX-170490178 | 0 | A | T | 0.228 | 7.20E-03 | 1.02E-01 | 2.67E+00 | 2.75E+00 | 9.38E-01 | no | Grm1 | intron_variant | MODIFIER |
| 10 | 10828092 | AX-170301452 | 0 | A | G | 0.228 | 7.20E-03 | 1.02E-01 | 2.67E+00 | 2.75E+00 | 9.38E-01 | no | Grm1 | intron_variant | MODIFIER |
| 10 | 11278102 | AX-97815462 | 0 | G | A | 0.228 | 7.20E-03 | 1.02E-01 | 2.67E+00 | 2.75E+00 | 9.38E-01 | no | Fbxo30 | upstream_gene_variant | MODIFIER |
| 10 | 11345820 | AX-97795362 | 0 | G | A | 0.228 | 7.20E-03 | 1.02E-01 | 2.67E+00 | 2.75E+00 | 9.38E-01 | no | Epm2a | intron_variant | MODIFIER |
| 4 | 94654852 | AX-170533275 | 0 | A | G | 0.333 | 8.73E-03 | 1.07E-01 | 2.66E+00 | 2.73E+00 | 9.38E-01 | no | Ift74 | synonymous_variant | LOW |
| 10 | 14372896 | AX-97799830 | 0 | C | T | 0.211 | -8.24E-03 | 9.72E-02 | 2.65E+00 | 2.73E+00 | 9.38E-01 | no | | intergenic_variant | MODIFIER |
| 2 | 163784863 | AX-170220731 | 2 | C | T | 0.136 | 1.13E-02 | 1.51E-01 | 2.63E+00 | 2.74E+00 | 9.38E-01 | no | | intergenic_variant | MODIFIER |
| 10 | 58514726 | AX-170487082 | 1 | T | C | 0.277 | 4.03E-03 | 8.13E-02 | 2.65E+00 | 2.78E+00 | 9.38E-01 | no | Ccdc138 | intron_variant | MODIFIER |
| 11 | 63451667 | AX-97846403 | 1 | C | T | 0.045 | -1.27E-02 | 1.45E-01 | 2.57E+00 | 2.73E+00 | 9.38E-01 | no | | intergenic_variant | MODIFIER |

| | | | | | | | | | | | | | | | |
|---|---|---|---|---|---|---|---|---|---|---|---|---|---|---|---|
| 1 | 168332862 | AX-170294366 | 1 | G | T | 0.223 | -9.17E-03 | 1.18E-01 | 2.64E+00 | 2.74E+00 | 9.38E-01 | no | Pbx1 | intron_variant | MODIFIER |
| 7 | 93552278 | AX-170390988 | 1 | G | A | 0.134 | -1.58E-02 | 2.11E-01 | 2.66E+00 | 2.74E+00 | 9.38E-01 | no | | intergenic_variant | MODIFIER |
| 7 | 97733300 | AX-170234268 | 1 | G | A | 0.134 | -1.58E-02 | 2.11E-01 | 2.66E+00 | 2.74E+00 | 9.38E-01 | no | Aqp11 | intron_variant | MODIFIER |
| 4 | 127377154 | AX-170213612 | 0 | A | G | 0.149 | 1.00E-02 | 1.04E-01 | 2.60E+00 | 2.72E+00 | 9.39E-01 | no | | intergenic_variant | MODIFIER |
| 10 | 56172135 | AX-170003282 | 1 | A | G | 0.286 | 4.15E-03 | 7.91E-02 | 2.66E+00 | 2.77E+00 | 9.39E-01 | no | Tbc1d32 | intron_variant | MODIFIER |
| 1 | 73249028 | AX-170292336 | 1 | G | C | 0.098 | 1.60E-02 | 2.06E-01 | 2.62E+00 | 2.74E+00 | 9.39E-01 | no | | intergenic_variant | MODIFIER |
| 5 | 127970182 | AX-170361785 | 1 | A | C | 0.42 | -5.76E-03 | 8.40E-02 | 2.64E+00 | 2.75E+00 | 9.39E-01 | no | Tmem132d | intron_variant | MODIFIER |
| 12 | 113918514 | AX-97783146 | 2 | G | A | 0.127 | -1.23E-02 | 1.61E-01 | 2.63E+00 | 2.74E+00 | 9.39E-01 | no | | intergenic_variant | MODIFIER |
| 11 | 34482837 | AX-170218410 | 1 | A | G | 0.098 | -1.29E-02 | 1.86E-01 | 2.66E+00 | 2.75E+00 | 9.39E-01 | no | Dock2 | intron_variant | MODIFIER |
| 11 | 35105886 | AX-170400086 | 1 | A | G | 0.098 | -1.29E-02 | 1.86E-01 | 2.66E+00 | 2.75E+00 | 9.39E-01 | no | | intergenic_variant | MODIFIER |
| 18 | 45684710 | AX-170501304 | 1 | G | A | 0.062 | 2.24E-02 | 3.01E-01 | 2.73E+00 | 2.75E+00 | 9.39E-01 | no | Kcnn2 | intron_variant | MODIFIER |
| 10 | 114408994 | AX-170323873 | 0 | G | A | 0.368 | 5.50E-03 | 7.46E-02 | 2.65E+00 | 2.74E+00 | 9.39E-01 | no | Trhde | intron_variant | MODIFIER |
| 1 | 95731373 | AX-97812398 | 0 | C | A | 0.158 | -9.57E-03 | 1.56E-01 | 2.62E+00 | 2.75E+00 | 9.39E-01 | no | | intergenic_variant | MODIFIER |
| 5 | 130472801 | AX-170302005 | 1 | T | C | 0.045 | 1.05E-02 | 1.23E-01 | 2.56E+00 | 2.73E+00 | 9.39E-01 | no | Caln1 | intron_variant | MODIFIER |
| 8 | 81711313 | AX-170031126 | 1 | A | G | 0.196 | -9.73E-03 | 1.32E-01 | 2.68E+00 | 2.74E+00 | 9.39E-01 | no | Inpp4b | upstream_gene_variant | MODIFIER |
| 8 | 47090835 | AX-97832547 | 1 | G | A | 0.152 | 1.06E-02 | 1.47E-01 | 2.69E+00 | 2.75E+00 | 9.39E-01 | no | Enpp6 | intron_variant | MODIFIER |
| 5 | 101039863 | AX-170452256 | 2 | C | T | 0.409 | -7.45E-03 | 1.05E-01 | 2.69E+00 | 2.75E+00 | 9.39E-01 | no | | intergenic_variant | MODIFIER |
| 2 | 93020440 | AX-170313825 | 0 | C | T | 0.061 | 1.75E-02 | 2.38E-01 | 2.67E+00 | 2.74E+00 | 9.39E-01 | no | Prdm11 | intron_variant | MODIFIER |
| 11 | 47322584 | AX-97833240 | 1 | C | T | 0.071 | -1.52E-02 | 1.92E-01 | 2.62E+00 | 2.73E+00 | 9.39E-01 | no | Sgcd | intron_variant | MODIFIER |
| 1 | 50120391 | AX-170000467 | 2 | A | G | 0.109 | 1.83E-02 | 2.40E-01 | 2.69E+00 | 2.74E+00 | 9.39E-01 | no | | intergenic_variant | MODIFIER |
| 10 | 9261488 | AX-170213645 | 0 | G | A | 0.202 | 6.85E-03 | 1.03E-01 | 2.67E+00 | 2.75E+00 | 9.39E-01 | no | | intergenic_variant | MODIFIER |
| 10 | 9266673 | AX-170504692 | 0 | A | G | 0.202 | 6.85E-03 | 1.03E-01 | 2.67E+00 | 2.75E+00 | 9.39E-01 | no | | intergenic_variant | MODIFIER |
| 10 | 9275549 | AX-97851322 | 0 | G | A | 0.202 | 6.85E-03 | 1.03E-01 | 2.67E+00 | 2.75E+00 | 9.39E-01 | no | | intergenic_variant | MODIFIER |
| 10 | 9280142 | AX-170228880 | 0 | G | A | 0.202 | 6.85E-03 | 1.03E-01 | 2.67E+00 | 2.75E+00 | 9.39E-01 | no | | intergenic_variant | MODIFIER |
| 10 | 9283707 | AX-170373265 | 0 | G | A | 0.202 | 6.85E-03 | 1.03E-01 | 2.67E+00 | 2.75E+00 | 9.39E-01 | no | | intergenic_variant | MODIFIER |
| 10 | 9289289 | AX-97830294 | 0 | C | T | 0.202 | 6.85E-03 | 1.03E-01 | 2.67E+00 | 2.75E+00 | 9.39E-01 | no | | intergenic_variant | MODIFIER |
| 10 | 9305921 | AX-170091856 | 0 | G | A | 0.202 | 6.85E-03 | 1.03E-01 | 2.67E+00 | 2.75E+00 | 9.39E-01 | no | | intergenic_variant | MODIFIER |
| 10 | 9314858 | AX-170515536 | 0 | T | A | 0.202 | 6.85E-03 | 1.03E-01 | 2.67E+00 | 2.75E+00 | 9.39E-01 | no | | intergenic_variant | MODIFIER |
| 10 | 9321281 | AX-170309327 | 0 | C | T | 0.202 | 6.85E-03 | 1.03E-01 | 2.67E+00 | 2.75E+00 | 9.39E-01 | no | | intergenic_variant | MODIFIER |
| 10 | 9325362 | AX-170002687 | 0 | G | A | 0.202 | 6.85E-03 | 1.03E-01 | 2.67E+00 | 2.75E+00 | 9.39E-01 | no | Gm48748 | downstream_gene_variant | MODIFIER |
| 10 | 9338881 | AX-170485005 | 0 | G | A | 0.202 | 6.85E-03 | 1.03E-01 | 2.67E+00 | 2.75E+00 | 9.39E-01 | no | Gm48749 | intron_variant,non_coding_transcript_variant | MODIFIER |
| 10 | 9338941 | AX-170342002 | 0 | T | C | 0.202 | 6.85E-03 | 1.03E-01 | 2.67E+00 | 2.75E+00 | 9.39E-01 | no | Gm48749 | intron_variant,non_coding_transcript_variant | MODIFIER |
| 10 | 9350696 | AX-170273998 | 0 | C | T | 0.202 | 6.85E-03 | 1.03E-01 | 2.67E+00 | 2.75E+00 | 9.39E-01 | no | Gm48749 | intron_variant,non_coding_transcript_variant | MODIFIER |
| 10 | 9370376 | AX-170002692 | 0 | A | G | 0.202 | 6.85E-03 | 1.03E-01 | 2.67E+00 | 2.75E+00 | 9.39E-01 | no | Gm48749 | intron_variant,non_coding_transcript_variant | MODIFIER |
| 10 | 9370616 | AX-97787716 | 0 | C | T | 0.202 | 6.85E-03 | 1.03E-01 | 2.67E+00 | 2.75E+00 | 9.39E-01 | no | Gm48749 | intron_variant,non_coding_transcript_variant | MODIFIER |
| 10 | 9398400 | AX-170330562 | 0 | G | A | 0.202 | 6.85E-03 | 1.03E-01 | 2.67E+00 | 2.75E+00 | 9.39E-01 | no | Gm48749 | downstream_gene_variant | MODIFIER |
| 11 | 47168671 | AX-170433382 | 0 | G | A | 0.07 | 1.25E-02 | 1.85E-01 | 2.63E+00 | 2.75E+00 | 9.39E-01 | no | Sgcd | intron_variant | MODIFIER |
| 7 | 94329148 | AX-97789497 | 1 | G | A | 0.116 | 1.66E-02 | 2.12E-01 | 2.65E+00 | 2.73E+00 | 9.39E-01 | no | Gm32647 | intron_variant,non_coding_transcript_variant | MODIFIER |
| 4 | 143094064 | AX-170024157 | 1 | A | T | 0.098 | 1.48E-02 | 1.94E-01 | 2.65E+00 | 2.74E+00 | 9.39E-01 | no | | intergenic_variant | MODIFIER |
| 20 | 76748609 | AX-170096813 | 2 | G | A | 0.236 | 5.03E-03 | 6.47E-02 | 2.63E+00 | 2.73E+00 | 9.39E-01 | no | | intergenic_variant | MODIFIER |
| 5 | 38619675 | AX-97802074 | 0 | A | G | 0.333 | 8.28E-03 | 1.03E-01 | 2.65E+00 | 2.73E+00 | 9.40E-01 | no | | regulatory_region_variant | MODIFIER |
| 11 | 104011458 | AX-170102015 | 0 | A | G | 0.053 | 1.39E-02 | 1.88E-01 | 2.62E+00 | 2.74E+00 | 9.40E-01 | no | | intergenic_variant | MODIFIER |
| 13 | 54874042 | AX-170247004 | 1 | T | A | 0.518 | 5.87E-03 | 6.44E-02 | 2.60E+00 | 2.72E+00 | 9.40E-01 | no | | regulatory_region_variant | MODIFIER |
| 1 | 74444552 | AX-170041680 | 0 | C | T | 0.149 | 1.55E-02 | 1.96E-01 | 2.63E+00 | 2.73E+00 | 9.40E-01 | no | Usp37 | intron_variant | MODIFIER |
| 16 | 92804308 | AX-170205089 | 1 | C | T | 0.473 | -5.81E-03 | 7.86E-02 | 2.66E+00 | 2.74E+00 | 9.40E-01 | no | Runx1 | intron_variant | MODIFIER |
| 7 | 138078361 | AX-170366459 | 1 | G | C | 0.304 | -6.28E-03 | 7.82E-02 | 2.63E+00 | 2.73E+00 | 9.40E-01 | no | | intergenic_variant | MODIFIER |
| 5 | 73727941 | AX-170307972 | 1 | C | T | 0.455 | 6.80E-03 | 9.16E-02 | 2.64E+00 | 2.74E+00 | 9.40E-01 | no | | intergenic_variant | MODIFIER |
| 4 | 55966009 | AX-170318164 | 1 | C | T | 0.366 | -7.54E-03 | 1.04E-01 | 2.66E+00 | 2.74E+00 | 9.40E-01 | no | Gm12519 | downstream_gene_variant | MODIFIER |
| 1 | 54766001 | AX-97811581 | 1 | G | A | 0.107 | 1.74E-02 | 2.33E-01 | 2.69E+00 | 2.74E+00 | 9.40E-01 | no | Ankrd44 | intron_variant | MODIFIER |
| 20 | 134184726 | AX-170035000 | 1 | A | G | 0.152 | 7.49E-03 | 9.10E-02 | 2.63E+00 | 2.73E+00 | 9.40E-01 | no | Arl13a | upstream_gene_variant | MODIFIER |
| 16 | 92912788 | AX-170508728 | 0 | G | A | 0.474 | -5.77E-03 | 7.87E-02 | 2.66E+00 | 2.74E+00 | 9.40E-01 | no | | intergenic_variant | MODIFIER |
| 9 | 55934502 | AX-170110543 | 2 | C | T | 0.2 | 8.31E-03 | 1.10E-01 | 2.65E+00 | 2.74E+00 | 9.40E-01 | no | Gm7435 | upstream_gene_variant | MODIFIER |
| 16 | 92816359 | AX-170111010 | 1 | G | C | 0.464 | -5.81E-03 | 7.81E-02 | 2.65E+00 | 2.74E+00 | 9.40E-01 | no | Runx1 | intron_variant | MODIFIER |
| 10 | 13876354 | AX-97796232 | 1 | C | T | 0.196 | -8.85E-03 | 1.10E-01 | 2.65E+00 | 2.73E+00 | 9.40E-01 | no | Gm32105 | downstream_gene_variant | MODIFIER |

| | | | | | | | | | | | | | | | |
|---|---|---|---|---|---|---|---|---|---|---|---|---|---|---|---|
| 10 | 13627244 | AX-170002773 | 0 | C | T | 0.193 | -8.89E-03 | 1.09E-01 | 2.65E+00 | 2.73E+00 | 9.40E-01 | no | Gm48773 | non_coding_transcript_exon_variant | MODIFIER |
| 10 | 13833549 | AX-170467705 | 0 | T | G | 0.193 | -8.89E-03 | 1.09E-01 | 2.65E+00 | 2.73E+00 | 9.40E-01 | no | Aig1 | intron_variant | MODIFIER |
| 10 | 13836598 | AX-170250664 | 0 | T | C | 0.193 | -8.89E-03 | 1.09E-01 | 2.65E+00 | 2.73E+00 | 9.40E-01 | no | Aig1 | intron_variant | MODIFIER |
| 10 | 13837083 | AX-170519259 | 0 | G | A | 0.193 | -8.89E-03 | 1.09E-01 | 2.65E+00 | 2.73E+00 | 9.40E-01 | no | Aig1 | intron_variant | MODIFIER |
| 10 | 13842934 | AX-170134358 | 0 | T | C | 0.193 | -8.89E-03 | 1.09E-01 | 2.65E+00 | 2.73E+00 | 9.40E-01 | no | Aig1 | intron_variant | MODIFIER |
| 10 | 13844405 | AX-170261657 | 0 | A | G | 0.193 | -8.89E-03 | 1.09E-01 | 2.65E+00 | 2.73E+00 | 9.40E-01 | no | Aig1 | intron_variant | MODIFIER |
| 10 | 13844801 | AX-170370978 | 0 | A | G | 0.193 | -8.89E-03 | 1.09E-01 | 2.65E+00 | 2.73E+00 | 9.40E-01 | no | Aig1 | intron_variant | MODIFIER |
| 10 | 13847139 | AX-170437963 | 0 | G | A | 0.193 | -8.89E-03 | 1.09E-01 | 2.65E+00 | 2.73E+00 | 9.40E-01 | no | Aig1 | intron_variant | MODIFIER |
| 10 | 13866564 | AX-170231670 | 0 | T | C | 0.193 | -8.89E-03 | 1.09E-01 | 2.65E+00 | 2.73E+00 | 9.40E-01 | no | Aig1 | intron_variant | MODIFIER |
| 10 | 13874651 | AX-170103615 | 0 | A | G | 0.193 | -8.89E-03 | 1.09E-01 | 2.65E+00 | 2.73E+00 | 9.40E-01 | no | Gm32105 | downstream_gene_variant | MODIFIER |
| 10 | 13939976 | AX-97853422 | 0 | C | T | 0.193 | -8.89E-03 | 1.09E-01 | 2.65E+00 | 2.73E+00 | 9.40E-01 | no | | intergenic_variant | MODIFIER |
| 10 | 13945488 | AX-170264573 | 0 | C | T | 0.193 | -8.89E-03 | 1.09E-01 | 2.65E+00 | 2.73E+00 | 9.40E-01 | no | | intergenic_variant | MODIFIER |
| 10 | 13975485 | AX-170521185 | 0 | C | T | 0.193 | -8.89E-03 | 1.09E-01 | 2.65E+00 | 2.73E+00 | 9.40E-01 | no | Hivep2 | intron_variant | MODIFIER |
| 10 | 13988164 | AX-97814825 | 0 | T | C | 0.193 | -8.89E-03 | 1.09E-01 | 2.65E+00 | 2.73E+00 | 9.40E-01 | no | Hivep2 | intron_variant | MODIFIER |
| 10 | 14380725 | AX-170526275 | 0 | C | T | 0.193 | -8.89E-03 | 1.09E-01 | 2.65E+00 | 2.73E+00 | 9.40E-01 | no | | intergenic_variant | MODIFIER |
| 10 | 14387038 | AX-170355287 | 0 | G | A | 0.193 | -8.89E-03 | 1.09E-01 | 2.65E+00 | 2.73E+00 | 9.40E-01 | no | | intergenic_variant | MODIFIER |
| 10 | 14387486 | AX-170121401 | 0 | G | A | 0.193 | -8.89E-03 | 1.09E-01 | 2.65E+00 | 2.73E+00 | 9.40E-01 | no | | intergenic_variant | MODIFIER |
| 10 | 14393681 | AX-170292920 | 0 | G | T | 0.193 | -8.89E-03 | 1.09E-01 | 2.65E+00 | 2.73E+00 | 9.40E-01 | no | | intergenic_variant | MODIFIER |
| 15 | 93677417 | AX-97839290 | 1 | G | A | 0.143 | 1.34E-02 | 1.67E-01 | 2.64E+00 | 2.73E+00 | 9.41E-01 | no | Zfp42-ps1 | downstream_gene_variant | MODIFIER |
| 9 | 52200126 | AX-170190839 | 0 | G | A | 0.219 | -8.65E-03 | 1.18E-01 | 2.68E+00 | 2.74E+00 | 9.41E-01 | no | | intergenic_variant | MODIFIER |
| 9 | 52449005 | AX-97783828 | 0 | T | G | 0.219 | -8.65E-03 | 1.18E-01 | 2.68E+00 | 2.74E+00 | 9.41E-01 | no | AC166902.1 | intron_variant,non_coding_transcript_variant | MODIFIER |
| 9 | 52457083 | AX-170528551 | 0 | T | C | 0.219 | -8.65E-03 | 1.18E-01 | 2.68E+00 | 2.74E+00 | 9.41E-01 | no | AC166902.1 | intron_variant,non_coding_transcript_variant | MODIFIER |
| 9 | 52458769 | AX-170236437 | 0 | A | G | 0.219 | -8.65E-03 | 1.18E-01 | 2.68E+00 | 2.74E+00 | 9.41E-01 | no | AC166902.1 | intron_variant,non_coding_transcript_variant | MODIFIER |
| 9 | 52462602 | AX-170271602 | 0 | G | A | 0.219 | -8.65E-03 | 1.18E-01 | 2.68E+00 | 2.74E+00 | 9.41E-01 | no | AC166902.1 | intron_variant,non_coding_transcript_variant | MODIFIER |
| 9 | 52473287 | AX-170228631 | 0 | G | T | 0.219 | -8.65E-03 | 1.18E-01 | 2.68E+00 | 2.74E+00 | 9.41E-01 | no | AC166902.1 | downstream_gene_variant | MODIFIER |
| 9 | 52473875 | AX-170171010 | 0 | T | C | 0.219 | -8.65E-03 | 1.18E-01 | 2.68E+00 | 2.74E+00 | 9.41E-01 | no | AC166902.1 | downstream_gene_variant | MODIFIER |
| 9 | 52480127 | AX-170082453 | 0 | T | C | 0.219 | -8.65E-03 | 1.18E-01 | 2.68E+00 | 2.74E+00 | 9.41E-01 | no | | intergenic_variant | MODIFIER |
| 9 | 52492708 | AX-170337723 | 0 | T | C | 0.219 | -8.65E-03 | 1.18E-01 | 2.68E+00 | 2.74E+00 | 9.41E-01 | no | | regulatory_region_variant | MODIFIER |
| 9 | 52511288 | AX-170115305 | 0 | C | A | 0.219 | -8.65E-03 | 1.18E-01 | 2.68E+00 | 2.74E+00 | 9.41E-01 | no | | intergenic_variant | MODIFIER |
| 9 | 52545228 | AX-170096548 | 0 | C | T | 0.219 | -8.65E-03 | 1.18E-01 | 2.68E+00 | 2.74E+00 | 9.41E-01 | no | | intergenic_variant | MODIFIER |
| 9 | 52563848 | AX-170488399 | 0 | G | A | 0.219 | -8.65E-03 | 1.18E-01 | 2.68E+00 | 2.74E+00 | 9.41E-01 | no | | intergenic_variant | MODIFIER |
| 9 | 52564945 | AX-170310368 | 0 | T | C | 0.219 | -8.65E-03 | 1.18E-01 | 2.68E+00 | 2.74E+00 | 9.41E-01 | no | | intergenic_variant | MODIFIER |
| 9 | 52570381 | AX-170435417 | 0 | C | T | 0.219 | -8.65E-03 | 1.18E-01 | 2.68E+00 | 2.74E+00 | 9.41E-01 | no | | intergenic_variant | MODIFIER |
| 9 | 52594471 | AX-170135801 | 0 | T | G | 0.219 | -8.65E-03 | 1.18E-01 | 2.68E+00 | 2.74E+00 | 9.41E-01 | no | | intergenic_variant | MODIFIER |
| 9 | 52618627 | AX-97832325 | 0 | G | A | 0.219 | -8.65E-03 | 1.18E-01 | 2.68E+00 | 2.74E+00 | 9.41E-01 | no | | intergenic_variant | MODIFIER |
| 9 | 52623060 | AX-170228584 | 0 | T | C | 0.219 | -8.65E-03 | 1.18E-01 | 2.68E+00 | 2.74E+00 | 9.41E-01 | no | | intergenic_variant | MODIFIER |
| 9 | 52628269 | AX-170486308 | 0 | C | T | 0.219 | -8.65E-03 | 1.18E-01 | 2.68E+00 | 2.74E+00 | 9.41E-01 | no | | intergenic_variant | MODIFIER |
| 9 | 52629546 | AX-170172282 | 0 | G | A | 0.219 | -8.65E-03 | 1.18E-01 | 2.68E+00 | 2.74E+00 | 9.41E-01 | no | | intergenic_variant | MODIFIER |
| 9 | 52637487 | AX-170474041 | 0 | G | C | 0.219 | -8.65E-03 | 1.18E-01 | 2.68E+00 | 2.74E+00 | 9.41E-01 | no | | intergenic_variant | MODIFIER |
| 9 | 52637814 | AX-170345315 | 0 | A | G | 0.219 | -8.65E-03 | 1.18E-01 | 2.68E+00 | 2.74E+00 | 9.41E-01 | no | | intergenic_variant | MODIFIER |
| 9 | 52653127 | AX-170210413 | 0 | G | C | 0.219 | -8.65E-03 | 1.18E-01 | 2.68E+00 | 2.74E+00 | 9.41E-01 | no | | intergenic_variant | MODIFIER |
| 9 | 52653619 | AX-170251428 | 0 | C | G | 0.219 | -8.65E-03 | 1.18E-01 | 2.68E+00 | 2.74E+00 | 9.41E-01 | no | | intergenic_variant | MODIFIER |
| 9 | 52656104 | AX-170309152 | 0 | G | A | 0.219 | -8.65E-03 | 1.18E-01 | 2.68E+00 | 2.74E+00 | 9.41E-01 | no | | intergenic_variant | MODIFIER |
| 9 | 52659575 | AX-170106845 | 0 | T | G | 0.219 | -8.65E-03 | 1.18E-01 | 2.68E+00 | 2.74E+00 | 9.41E-01 | no | | regulatory_region_variant | MODIFIER |
| 9 | 52662783 | AX-170082459 | 0 | T | C | 0.219 | -8.65E-03 | 1.18E-01 | 2.68E+00 | 2.74E+00 | 9.41E-01 | no | | intergenic_variant | MODIFIER |
| 9 | 52662816 | AX-170082460 | 0 | T | C | 0.219 | -8.65E-03 | 1.18E-01 | 2.68E+00 | 2.74E+00 | 9.41E-01 | no | | intergenic_variant | MODIFIER |
| 9 | 52666171 | AX-170491150 | 0 | A | G | 0.219 | -8.65E-03 | 1.18E-01 | 2.68E+00 | 2.74E+00 | 9.41E-01 | no | | intergenic_variant | MODIFIER |
| 9 | 52673205 | AX-170394637 | 0 | C | T | 0.219 | -8.65E-03 | 1.18E-01 | 2.68E+00 | 2.74E+00 | 9.41E-01 | no | AI593442 | 3_prime_UTR_variant | MODIFIER |
| 9 | 52690773 | AX-170195858 | 0 | A | G | 0.219 | -8.65E-03 | 1.18E-01 | 2.68E+00 | 2.74E+00 | 9.41E-01 | no | Gm1715 | intron_variant,non_coding_transcript_variant | MODIFIER |
| 9 | 52700733 | AX-170385319 | 0 | T | C | 0.219 | -8.65E-03 | 1.18E-01 | 2.68E+00 | 2.74E+00 | 9.41E-01 | no | Gm1715 | intron_variant,non_coding_transcript_variant | MODIFIER |
| 9 | 52701051 | AX-170404123 | 0 | A | G | 0.219 | -8.65E-03 | 1.18E-01 | 2.68E+00 | 2.74E+00 | 9.41E-01 | no | Gm1715 | intron_variant,non_coding_transcript_variant | MODIFIER |
| 9 | 52701597 | AX-170275402 | 0 | T | A | 0.219 | -8.65E-03 | 1.18E-01 | 2.68E+00 | 2.74E+00 | 9.41E-01 | no | Gm1715 | intron_variant,non_coding_transcript_variant | MODIFIER |
| 9 | 52718036 | AX-170255985 | 0 | T | C | 0.219 | -8.65E-03 | 1.18E-01 | 2.68E+00 | 2.74E+00 | 9.41E-01 | no | Gm1715 | intron_variant,non_coding_transcript_variant | MODIFIER |
| 9 | 52761292 | AX-97829802 | 0 | C | T | 0.219 | -8.65E-03 | 1.18E-01 | 2.68E+00 | 2.74E+00 | 9.41E-01 | no | Gm22242 | downstream_gene_variant | MODIFIER |

| | | | | | | | | | | | | | | | |
|---|---|---|---|---|---|---|---|---|---|---|---|---|---|---|---|
| 9 | 52772988 | AX-170032365 | 0 | T | A | 0.219 | -8.65E-03 | 1.18E-01 | 2.68E+00 | 2.74E+00 | 9.41E-01 | no | Gm1715 | intron_variant,non_coding_transcript_variant | MODIFIER |
| 9 | 52878272 | AX-170210275 | 0 | T | C | 0.219 | -8.65E-03 | 1.18E-01 | 2.68E+00 | 2.74E+00 | 9.41E-01 | no | | intergenic_variant | MODIFIER |
| 9 | 52913201 | AX-170032366 | 0 | A | G | 0.219 | -8.65E-03 | 1.18E-01 | 2.68E+00 | 2.74E+00 | 9.41E-01 | no | | intergenic_variant | MODIFIER |
| 9 | 52913595 | AX-170032361 | 0 | A | G | 0.219 | -8.65E-03 | 1.18E-01 | 2.68E+00 | 2.74E+00 | 9.41E-01 | no | | intergenic_variant | MODIFIER |
| 9 | 52913888 | AX-170310129 | 0 | A | C | 0.219 | -8.65E-03 | 1.18E-01 | 2.68E+00 | 2.74E+00 | 9.41E-01 | no | | intergenic_variant | MODIFIER |
| 9 | 52976964 | AX-170449739 | 0 | C | T | 0.219 | -8.65E-03 | 1.18E-01 | 2.68E+00 | 2.74E+00 | 9.41E-01 | no | | intergenic_variant | MODIFIER |
| 9 | 53004093 | AX-170082462 | 0 | T | C | 0.219 | -8.65E-03 | 1.18E-01 | 2.68E+00 | 2.74E+00 | 9.41E-01 | no | | intergenic_variant | MODIFIER |
| 9 | 53027965 | AX-97801658 | 0 | C | T | 0.219 | -8.65E-03 | 1.18E-01 | 2.68E+00 | 2.74E+00 | 9.41E-01 | no | | intergenic_variant | MODIFIER |
| 9 | 53072269 | AX-170112442 | 0 | C | G | 0.219 | -8.65E-03 | 1.18E-01 | 2.68E+00 | 2.74E+00 | 9.41E-01 | no | | regulatory_region_variant | MODIFIER |
| 9 | 53072589 | AX-97845954 | 0 | A | G | 0.219 | -8.65E-03 | 1.18E-01 | 2.68E+00 | 2.74E+00 | 9.41E-01 | no | | regulatory_region_variant | MODIFIER |
| 9 | 53074285 | AX-170478581 | 0 | G | A | 0.219 | -8.65E-03 | 1.18E-01 | 2.68E+00 | 2.74E+00 | 9.41E-01 | no | | regulatory_region_variant | MODIFIER |
| 9 | 53075132 | AX-170200699 | 0 | T | C | 0.219 | -8.65E-03 | 1.18E-01 | 2.68E+00 | 2.74E+00 | 9.41E-01 | no | | intergenic_variant | MODIFIER |
| 9 | 53089090 | AX-170144249 | 0 | A | G | 0.219 | -8.65E-03 | 1.18E-01 | 2.68E+00 | 2.74E+00 | 9.41E-01 | no | | intergenic_variant | MODIFIER |
| 9 | 53096237 | AX-97847319 | 0 | T | C | 0.219 | -8.65E-03 | 1.18E-01 | 2.68E+00 | 2.74E+00 | 9.41E-01 | no | Ddx10 | downstream_gene_variant | MODIFIER |
| 9 | 53460453 | AX-170256852 | 0 | G | A | 0.219 | -8.65E-03 | 1.18E-01 | 2.68E+00 | 2.74E+00 | 9.41E-01 | no | Atm | intron_variant | MODIFIER |
| 9 | 53467001 | AX-97830393 | 0 | C | A | 0.219 | -8.65E-03 | 1.18E-01 | 2.68E+00 | 2.74E+00 | 9.41E-01 | no | Atm | intron_variant | MODIFIER |
| 10 | 31474842 | AX-170221676 | 0 | G | A | 0.395 | -5.45E-03 | 7.28E-02 | 2.62E+00 | 2.74E+00 | 9.41E-01 | no | | intergenic_variant | MODIFIER |
| 7 | 51405690 | AX-170227559 | 1 | A | G | 0.42 | 7.64E-03 | 9.38E-02 | 2.63E+00 | 2.73E+00 | 9.41E-01 | no | | intergenic_variant | MODIFIER |
| 18 | 75928764 | AX-170321182 | 1 | T | A | 0.384 | -6.47E-03 | 8.24E-02 | 2.63E+00 | 2.73E+00 | 9.41E-01 | no | Zbtb7c | intron_variant | MODIFIER |
| 2 | 124524108 | AX-170390289 | 1 | C | T | 0.143 | 1.41E-02 | 1.73E-01 | 2.64E+00 | 2.73E+00 | 9.41E-01 | no | Sema6d | intron_variant | MODIFIER |
| 8 | 128482576 | AX-170081526 | 0 | C | T | 0.053 | -1.38E-02 | 1.96E-01 | 2.63E+00 | 2.74E+00 | 9.41E-01 | no | Nrp1 | intron_variant | MODIFIER |
| 7 | 106510434 | AX-170096056 | 2 | G | T | 0.482 | -5.92E-03 | 9.42E-02 | 2.68E+00 | 2.76E+00 | 9.41E-01 | no | Gm8995 | non_coding_transcript_exon_variant | MODIFIER |
| 12 | 116467398 | AX-170448602 | 1 | G | T | 0.125 | -1.37E-02 | 1.99E-01 | 2.68E+00 | 2.75E+00 | 9.42E-01 | no | Ncapg2 | downstream_gene_variant | MODIFIER |
| 9 | 85568641 | AX-170032872 | 2 | A | T | 0.2 | 8.42E-03 | 1.19E-01 | 2.66E+00 | 2.74E+00 | 9.42E-01 | no | | intergenic_variant | MODIFIER |
| 4 | 88967942 | AX-170323225 | 0 | C | A | 0.325 | -7.14E-03 | 1.08E-01 | 2.68E+00 | 2.75E+00 | 9.42E-01 | no | | intergenic_variant | MODIFIER |
| 1 | 25345543 | AX-170527894 | 1 | C | T | 0.42 | -5.42E-03 | 8.15E-02 | 2.68E+00 | 2.75E+00 | 9.42E-01 | no | Adgrb3 | intron_variant | MODIFIER |
| 7 | 68679716 | AX-170186716 | 0 | G | A | 0.412 | -6.04E-03 | 9.31E-02 | 2.66E+00 | 2.75E+00 | 9.42E-01 | no | | intergenic_variant | MODIFIER |
| 7 | 45022621 | AX-170262541 | 0 | G | T | 0.333 | 8.01E-03 | 9.13E-02 | 2.63E+00 | 2.72E+00 | 9.42E-01 | no | Rras | downstream_gene_variant | MODIFIER |
| 10 | 6388569 | AX-170533828 | 0 | C | T | 0.211 | 6.03E-03 | 9.49E-02 | 2.62E+00 | 2.75E+00 | 9.42E-01 | no | | intergenic_variant | MODIFIER |
| 13 | 40957892 | AX-170276634 | 0 | A | G | 0.518 | 5.48E-03 | 6.17E-02 | 2.61E+00 | 2.72E+00 | 9.42E-01 | no | Gcnt2 | intron_variant | MODIFIER |
| 13 | 40967428 | AX-97856406 | 0 | A | G | 0.518 | 5.48E-03 | 6.17E-02 | 2.61E+00 | 2.72E+00 | 9.42E-01 | no | A730081D07Rik | intron_variant,non_coding_transcript_variant | MODIFIER |
| 13 | 40984998 | AX-97817853 | 0 | T | C | 0.518 | 5.48E-03 | 6.17E-02 | 2.61E+00 | 2.72E+00 | 9.42E-01 | no | A730081D07Rik | intron_variant,non_coding_transcript_variant | MODIFIER |
| 13 | 40989325 | AX-170106160 | 0 | A | G | 0.518 | 5.48E-03 | 6.17E-02 | 2.61E+00 | 2.72E+00 | 9.42E-01 | no | A730081D07Rik | intron_variant,non_coding_transcript_variant | MODIFIER |
| 13 | 41008294 | AX-170395986 | 0 | T | C | 0.518 | 5.48E-03 | 6.17E-02 | 2.61E+00 | 2.72E+00 | 9.42E-01 | no | Pak1ip1 | intron_variant | MODIFIER |
| 13 | 41030456 | AX-170490082 | 0 | T | C | 0.518 | 5.48E-03 | 6.17E-02 | 2.61E+00 | 2.72E+00 | 9.42E-01 | no | Mak | intron_variant | MODIFIER |
| 13 | 41030825 | AX-97828460 | 0 | T | C | 0.518 | 5.48E-03 | 6.17E-02 | 2.61E+00 | 2.72E+00 | 9.42E-01 | no | Mak | intron_variant | MODIFIER |
| 13 | 41046551 | AX-97821398 | 0 | T | C | 0.518 | 5.48E-03 | 6.17E-02 | 2.61E+00 | 2.72E+00 | 9.42E-01 | no | Mak | intron_variant | MODIFIER |
| 13 | 41046851 | AX-170285695 | 0 | T | C | 0.518 | 5.48E-03 | 6.17E-02 | 2.61E+00 | 2.72E+00 | 9.42E-01 | no | Mak | intron_variant | MODIFIER |
| 13 | 41064667 | AX-170432945 | 0 | A | C | 0.518 | 5.48E-03 | 6.17E-02 | 2.61E+00 | 2.72E+00 | 9.42E-01 | no | Mak | intron_variant | MODIFIER |
| 13 | 41072909 | AX-170306858 | 0 | C | T | 0.518 | 5.48E-03 | 6.17E-02 | 2.61E+00 | 2.72E+00 | 9.42E-01 | no | Mak | upstream_gene_variant | MODIFIER |
| 13 | 41072934 | AX-97817852 | 0 | T | C | 0.518 | 5.48E-03 | 6.17E-02 | 2.61E+00 | 2.72E+00 | 9.42E-01 | no | Mak | upstream_gene_variant | MODIFIER |
| 13 | 41118982 | AX-170307574 | 0 | C | A | 0.518 | 5.48E-03 | 6.17E-02 | 2.61E+00 | 2.72E+00 | 9.42E-01 | no | Sycp2l | intron_variant | MODIFIER |
| 13 | 41129060 | AX-170188655 | 0 | G | C | 0.518 | 5.48E-03 | 6.17E-02 | 2.61E+00 | 2.72E+00 | 9.42E-01 | no | Sycp2l | intron_variant | MODIFIER |
| 13 | 41129083 | AX-170408850 | 0 | G | A | 0.518 | 5.48E-03 | 6.17E-02 | 2.61E+00 | 2.72E+00 | 9.42E-01 | no | Sycp2l | intron_variant | MODIFIER |
| 9 | 108824534 | AX-170124362 | 1 | G | C | 0.125 | 7.28E-03 | 1.18E-01 | 2.69E+00 | 2.76E+00 | 9.42E-01 | no | Celsr3 | upstream_gene_variant | MODIFIER |
| 5 | 28664822 | AX-170522975 | 0 | T | C | 0.36 | 7.13E-03 | 9.94E-02 | 2.67E+00 | 2.74E+00 | 9.42E-01 | no | 9530036O11Rik | intron_variant,non_coding_transcript_variant | MODIFIER |
| 10 | 7730085 | AX-170447000 | 1 | G | A | 0.214 | 6.98E-03 | 1.07E-01 | 2.67E+00 | 2.75E+00 | 9.42E-01 | no | Katna1 | intron_variant | MODIFIER |
| 4 | 80667126 | AX-170278794 | 1 | G | A | 0.232 | -5.90E-03 | 1.04E-01 | 2.68E+00 | 2.76E+00 | 9.42E-01 | no | | intergenic_variant | MODIFIER |
| 12 | 72842223 | AX-97797724 | 1 | G | A | 0.062 | -2.20E-02 | 3.01E-01 | 2.71E+00 | 2.73E+00 | 9.42E-01 | no | | intergenic_variant | MODIFIER |
| 12 | 101930620 | AX-170238420 | 0 | G | A | 0.07 | -1.16E-02 | 1.67E-01 | 2.66E+00 | 2.74E+00 | 9.42E-01 | no | Atxn3 | intron_variant | MODIFIER |
| 12 | 109472086 | AX-170007502 | 0 | C | T | 0.123 | -1.34E-02 | 1.97E-01 | 2.67E+00 | 2.74E+00 | 9.42E-01 | no | | intergenic_variant | MODIFIER |
| 12 | 109497275 | AX-170291941 | 0 | T | G | 0.246 | -6.68E-03 | 9.83E-02 | 2.67E+00 | 2.74E+00 | 9.42E-01 | no | Gm34081 | intron_variant,non_coding_transcript_variant | MODIFIER |
| 12 | 109512041 | AX-170108737 | 0 | A | G | 0.123 | -1.34E-02 | 1.97E-01 | 2.67E+00 | 2.74E+00 | 9.42E-01 | no | Gm34081 | intron_variant,non_coding_transcript_variant | MODIFIER |
| 12 | 109516838 | AX-170154891 | 0 | T | A | 0.123 | -1.34E-02 | 1.97E-01 | 2.67E+00 | 2.74E+00 | 9.42E-01 | no | Gm34081 | upstream_gene_variant | MODIFIER |

| | | | | | | | | | | | | | | | |
|---|---|---|---|---|---|---|---|---|---|---|---|---|---|---|---|
| 12 | 109523377 | AX-170485868 | 0 | C | T | 0.123 | -1.34E-02 | 1.97E-01 | 2.67E+00 | 2.74E+00 | 9.42E-01 | no | | regulatory_region_variant | MODIFIER |
| 12 | 109526192 | AX-170185290 | 0 | T | C | 0.123 | -1.34E-02 | 1.97E-01 | 2.67E+00 | 2.74E+00 | 9.42E-01 | no | Gm27528 | upstream_gene_variant | MODIFIER |
| 12 | 109557445 | AX-170358563 | 0 | C | G | 0.123 | -1.34E-02 | 1.97E-01 | 2.67E+00 | 2.74E+00 | 9.42E-01 | no | Meg3 | intron_variant,non_coding_transcript_variant | MODIFIER |
| 12 | 109575973 | AX-170529595 | 0 | T | C | 0.123 | -1.34E-02 | 1.97E-01 | 2.67E+00 | 2.74E+00 | 9.42E-01 | no | Mir673 | downstream_gene_variant | MODIFIER |
| 12 | 109576115 | AX-170463316 | 0 | T | C | 0.123 | -1.34E-02 | 1.97E-01 | 2.67E+00 | 2.74E+00 | 9.42E-01 | no | Mir673 | downstream_gene_variant | MODIFIER |
| 12 | 109583938 | AX-170150772 | 0 | T | C | 0.123 | -1.34E-02 | 1.97E-01 | 2.67E+00 | 2.74E+00 | 9.42E-01 | no | Mir337 | upstream_gene_variant | MODIFIER |
| 12 | 109584632 | AX-170296078 | 0 | T | C | 0.123 | -1.34E-02 | 1.97E-01 | 2.67E+00 | 2.74E+00 | 9.42E-01 | no | Mir337 | upstream_gene_variant | MODIFIER |
| 12 | 109622948 | AX-97822314 | 0 | A | G | 0.123 | -1.34E-02 | 1.97E-01 | 2.67E+00 | 2.74E+00 | 9.42E-01 | no | Mir370 | downstream_gene_variant | MODIFIER |
| 12 | 109627488 | AX-170167657 | 0 | G | A | 0.123 | -1.34E-02 | 1.97E-01 | 2.67E+00 | 2.74E+00 | 9.42E-01 | no | Gm24564 | upstream_gene_variant | MODIFIER |
| 12 | 109645790 | AX-170050164 | 0 | A | G | 0.123 | -1.34E-02 | 1.97E-01 | 2.67E+00 | 2.74E+00 | 9.42E-01 | no | DQ267100 | upstream_gene_variant | MODIFIER |
| 12 | 109649134 | AX-170105843 | 0 | T | C | 0.123 | -1.34E-02 | 1.97E-01 | 2.67E+00 | 2.74E+00 | 9.42E-01 | no | DQ267101 | upstream_gene_variant | MODIFIER |
| 12 | 109649296 | AX-170302514 | 0 | G | A | 0.123 | -1.34E-02 | 1.97E-01 | 2.67E+00 | 2.74E+00 | 9.42E-01 | no | DQ267101 | upstream_gene_variant | MODIFIER |
| 12 | 109663751 | AX-170501941 | 0 | T | G | 0.123 | -1.34E-02 | 1.97E-01 | 2.67E+00 | 2.74E+00 | 9.42E-01 | no | Rian | downstream_gene_variant | MODIFIER |
| 12 | 109665895 | AX-170100568 | 0 | A | C | 0.123 | -1.34E-02 | 1.97E-01 | 2.67E+00 | 2.74E+00 | 9.42E-01 | no | Rian | downstream_gene_variant | MODIFIER |
| 12 | 109667753 | AX-170426300 | 0 | C | T | 0.123 | -1.34E-02 | 1.97E-01 | 2.67E+00 | 2.74E+00 | 9.42E-01 | no | Gm37899 | upstream_gene_variant | MODIFIER |
| 12 | 109688310 | AX-170188128 | 0 | T | C | 0.123 | -1.34E-02 | 1.97E-01 | 2.67E+00 | 2.74E+00 | 9.42E-01 | no | Gm25854 | downstream_gene_variant | MODIFIER |
| 12 | 109689131 | AX-97840803 | 0 | A | G | 0.123 | -1.34E-02 | 1.97E-01 | 2.67E+00 | 2.74E+00 | 9.42E-01 | no | Gm25854 | downstream_gene_variant | MODIFIER |
| 12 | 109694272 | AX-170206256 | 0 | A | G | 0.123 | -1.34E-02 | 1.97E-01 | 2.67E+00 | 2.74E+00 | 9.42E-01 | no | Gm25224 | downstream_gene_variant | MODIFIER |
| 12 | 109696006 | AX-170157323 | 0 | C | T | 0.123 | -1.34E-02 | 1.97E-01 | 2.67E+00 | 2.74E+00 | 9.42E-01 | no | B830012L14Rik | non_coding_transcript_exon_variant | MODIFIER |
| 12 | 109720205 | AX-170401201 | 0 | T | C | 0.123 | -1.34E-02 | 1.97E-01 | 2.67E+00 | 2.74E+00 | 9.42E-01 | no | Mir300 | upstream_gene_variant | MODIFIER |
| 12 | 109720892 | AX-97784784 | 0 | C | T | 0.123 | -1.34E-02 | 1.97E-01 | 2.67E+00 | 2.74E+00 | 9.42E-01 | no | Mir300 | upstream_gene_variant | MODIFIER |
| 12 | 109721353 | AX-170232609 | 0 | T | C | 0.123 | -1.34E-02 | 1.97E-01 | 2.67E+00 | 2.74E+00 | 9.42E-01 | no | Mir300 | upstream_gene_variant | MODIFIER |
| 12 | 109721982 | AX-170297297 | 0 | A | G | 0.123 | -1.34E-02 | 1.97E-01 | 2.67E+00 | 2.74E+00 | 9.42E-01 | no | Mir300 | upstream_gene_variant | MODIFIER |
| 12 | 109730299 | AX-170280749 | 0 | T | C | 0.123 | -1.34E-02 | 1.97E-01 | 2.67E+00 | 2.74E+00 | 9.42E-01 | no | Mir134 | upstream_gene_variant | MODIFIER |
| 12 | 109756597 | AX-97843657 | 0 | A | G | 0.123 | -1.34E-02 | 1.97E-01 | 2.67E+00 | 2.74E+00 | 9.42E-01 | no | | intergenic_variant | MODIFIER |
| 12 | 109756751 | AX-170275479 | 0 | A | G | 0.123 | -1.34E-02 | 1.97E-01 | 2.67E+00 | 2.74E+00 | 9.42E-01 | no | | intergenic_variant | MODIFIER |
| 12 | 109764972 | AX-170323979 | 0 | G | T | 0.123 | -1.34E-02 | 1.97E-01 | 2.67E+00 | 2.74E+00 | 9.42E-01 | no | | intergenic_variant | MODIFIER |
| 12 | 109785590 | AX-97824994 | 0 | A | G | 0.123 | -1.34E-02 | 1.97E-01 | 2.67E+00 | 2.74E+00 | 9.42E-01 | no | | intergenic_variant | MODIFIER |
| 12 | 109808274 | AX-97842153 | 0 | A | C | 0.123 | -1.34E-02 | 1.97E-01 | 2.67E+00 | 2.74E+00 | 9.42E-01 | no | | intergenic_variant | MODIFIER |
| 12 | 109858443 | AX-170220453 | 0 | C | T | 0.123 | -1.34E-02 | 1.97E-01 | 2.67E+00 | 2.74E+00 | 9.42E-01 | no | | intergenic_variant | MODIFIER |
| 12 | 109899627 | AX-97807043 | 0 | A | G | 0.123 | -1.34E-02 | 1.97E-01 | 2.67E+00 | 2.74E+00 | 9.42E-01 | no | Gm3379 | upstream_gene_variant | MODIFIER |
| 12 | 109900197 | AX-170050173 | 0 | T | C | 0.123 | -1.34E-02 | 1.97E-01 | 2.67E+00 | 2.74E+00 | 9.42E-01 | no | | intergenic_variant | MODIFIER |
| 12 | 109921839 | AX-170211418 | 0 | A | G | 0.123 | -1.34E-02 | 1.97E-01 | 2.67E+00 | 2.74E+00 | 9.42E-01 | no | | intergenic_variant | MODIFIER |
| 12 | 109928838 | AX-170346778 | 0 | T | C | 0.123 | -1.34E-02 | 1.97E-01 | 2.67E+00 | 2.74E+00 | 9.42E-01 | no | | intergenic_variant | MODIFIER |
| 12 | 109935573 | AX-170143414 | 0 | A | G | 0.123 | -1.34E-02 | 1.97E-01 | 2.67E+00 | 2.74E+00 | 9.42E-01 | no | | intergenic_variant | MODIFIER |
| 12 | 109968213 | AX-97852414 | 0 | C | T | 0.123 | -1.34E-02 | 1.97E-01 | 2.67E+00 | 2.74E+00 | 9.42E-01 | no | | intergenic_variant | MODIFIER |
| 12 | 109973807 | AX-170351785 | 0 | T | C | 0.123 | -1.34E-02 | 1.97E-01 | 2.67E+00 | 2.74E+00 | 9.42E-01 | no | | intergenic_variant | MODIFIER |
| 12 | 109978217 | AX-170432809 | 0 | G | A | 0.123 | -1.34E-02 | 1.97E-01 | 2.67E+00 | 2.74E+00 | 9.42E-01 | no | | intergenic_variant | MODIFIER |
| 12 | 109980818 | AX-170149475 | 0 | A | G | 0.123 | -1.34E-02 | 1.97E-01 | 2.67E+00 | 2.74E+00 | 9.42E-01 | no | | intergenic_variant | MODIFIER |
| 12 | 109985006 | AX-170138849 | 0 | G | T | 0.123 | -1.34E-02 | 1.97E-01 | 2.67E+00 | 2.74E+00 | 9.42E-01 | no | | intergenic_variant | MODIFIER |
| 12 | 110797995 | AX-170494334 | 0 | A | G | 0.123 | -1.34E-02 | 1.97E-01 | 2.67E+00 | 2.74E+00 | 9.42E-01 | no | Wdr20 | downstream_gene_variant | MODIFIER |
| 12 | 110941321 | AX-97838586 | 0 | G | A | 0.123 | -1.34E-02 | 1.97E-01 | 2.67E+00 | 2.74E+00 | 9.42E-01 | no | Tecpr2 | intron_variant | MODIFIER |
| 12 | 111093078 | AX-170478988 | 0 | T | C | 0.123 | -1.34E-02 | 1.97E-01 | 2.67E+00 | 2.74E+00 | 9.42E-01 | no | Rcor1 | splice_region_variant,intron_variant | LOW |
| 12 | 111093606 | AX-170092615 | 0 | T | G | 0.123 | -1.34E-02 | 1.97E-01 | 2.67E+00 | 2.74E+00 | 9.42E-01 | no | Rcor1 | intron_variant | MODIFIER |
| 12 | 111093695 | AX-170244057 | 0 | A | G | 0.123 | -1.34E-02 | 1.97E-01 | 2.67E+00 | 2.74E+00 | 9.42E-01 | no | Rcor1 | intron_variant | MODIFIER |
| 12 | 111115042 | AX-170007525 | 0 | C | T | 0.123 | -1.34E-02 | 1.97E-01 | 2.67E+00 | 2.74E+00 | 9.42E-01 | no | Rcor1 | 3_prime_UTR_variant | MODIFIER |
| 12 | 111121560 | AX-97781823 | 0 | A | G | 0.123 | -1.34E-02 | 1.97E-01 | 2.67E+00 | 2.74E+00 | 9.42E-01 | no | | intergenic_variant | MODIFIER |
| 12 | 111134204 | AX-170050201 | 0 | T | C | 0.123 | -1.34E-02 | 1.97E-01 | 2.67E+00 | 2.74E+00 | 9.42E-01 | no | | regulatory_region_variant | MODIFIER |
| 12 | 111147452 | AX-170177929 | 0 | T | C | 0.123 | -1.34E-02 | 1.97E-01 | 2.67E+00 | 2.74E+00 | 9.42E-01 | no | | intergenic_variant | MODIFIER |
| 12 | 111154101 | AX-170224889 | 0 | T | C | 0.123 | -1.34E-02 | 1.97E-01 | 2.67E+00 | 2.74E+00 | 9.42E-01 | no | | regulatory_region_variant | MODIFIER |
| 12 | 111209098 | AX-170155782 | 0 | A | G | 0.123 | -1.34E-02 | 1.97E-01 | 2.67E+00 | 2.74E+00 | 9.42E-01 | no | Traf3 | intron_variant | MODIFIER |
| 12 | 111415089 | AX-170354079 | 0 | T | C | 0.123 | -1.34E-02 | 1.97E-01 | 2.67E+00 | 2.74E+00 | 9.42E-01 | no | Exoc3l4 | upstream_gene_variant | MODIFIER |
| 12 | 111502547 | AX-97826854 | 0 | A | G | 0.123 | -1.34E-02 | 1.97E-01 | 2.67E+00 | 2.74E+00 | 9.42E-01 | no | Gm40578 | upstream_gene_variant | MODIFIER |
| 12 | 111514301 | AX-170092616 | 0 | C | G | 0.123 | -1.34E-02 | 1.97E-01 | 2.67E+00 | 2.74E+00 | 9.42E-01 | no | | regulatory_region_variant | MODIFIER |

| | | | | | | | | | | | | | | | |
|---|---|---|---|---|---|---|---|---|---|---|---|---|---|---|---|
| 12 | 111518518 | AX-170374177 | 0 | T | C | 0.123 | -1.34E-02 | 1.97E-01 | 2.67E+00 | 2.74E+00 | 9.42E-01 | no | | regulatory_region_variant | MODIFIER |
| 12 | 111575254 | AX-97811766 | 0 | T | C | 0.123 | -1.34E-02 | 1.97E-01 | 2.67E+00 | 2.74E+00 | 9.42E-01 | no | Mark3 | intron_variant | MODIFIER |
| 12 | 111634174 | AX-170050215 | 0 | G | A | 0.123 | -1.34E-02 | 1.97E-01 | 2.67E+00 | 2.74E+00 | 9.42E-01 | no | Mark3 | intron_variant | MODIFIER |
| 12 | 111648962 | AX-170050213 | 0 | A | T | 0.123 | -1.34E-02 | 1.97E-01 | 2.67E+00 | 2.74E+00 | 9.42E-01 | no | Mark3 | intron_variant | MODIFIER |
| 12 | 111652515 | AX-170272328 | 0 | T | C | 0.123 | -1.34E-02 | 1.97E-01 | 2.67E+00 | 2.74E+00 | 9.42E-01 | no | Mark3 | intron_variant | MODIFIER |
| 12 | 111653072 | AX-170523587 | 0 | A | G | 0.123 | -1.34E-02 | 1.97E-01 | 2.67E+00 | 2.74E+00 | 9.42E-01 | no | Mark3 | intron_variant | MODIFIER |
| 12 | 111654942 | AX-170333882 | 0 | T | C | 0.123 | -1.34E-02 | 1.97E-01 | 2.67E+00 | 2.74E+00 | 9.42E-01 | no | Mark3 | intron_variant | MODIFIER |
| 12 | 111683711 | AX-170303916 | 0 | C | T | 0.123 | -1.34E-02 | 1.97E-01 | 2.67E+00 | 2.74E+00 | 9.42E-01 | no | Trmt61a | 3_prime_UTR_variant | MODIFIER |
| 12 | 111687974 | AX-170007531 | 0 | T | C | 0.123 | -1.34E-02 | 1.97E-01 | 2.67E+00 | 2.74E+00 | 9.42E-01 | no | Trmt61a | downstream_gene_variant | MODIFIER |
| 12 | 111711508 | AX-170154834 | 0 | T | C | 0.123 | -1.34E-02 | 1.97E-01 | 2.67E+00 | 2.74E+00 | 9.42E-01 | no | Apopt1 | upstream_gene_variant | MODIFIER |
| 12 | 111714444 | AX-97831737 | 0 | A | G | 0.123 | -1.34E-02 | 1.97E-01 | 2.67E+00 | 2.74E+00 | 9.42E-01 | no | Apopt1 | intron_variant | MODIFIER |
| 12 | 111767565 | AX-97826852 | 0 | T | C | 0.123 | -1.34E-02 | 1.97E-01 | 2.67E+00 | 2.74E+00 | 9.42E-01 | no | Klc1 | intron_variant | MODIFIER |
| 12 | 111831395 | AX-170250432 | 0 | A | C | 0.123 | -1.34E-02 | 1.97E-01 | 2.67E+00 | 2.74E+00 | 9.42E-01 | no | Zfyve21 | downstream_gene_variant | MODIFIER |
| 12 | 111843313 | AX-170359277 | 0 | T | C | 0.123 | -1.34E-02 | 1.97E-01 | 2.67E+00 | 2.74E+00 | 9.42E-01 | no | Ppp1r13b | intron_variant | MODIFIER |
| 12 | 111843953 | AX-170175328 | 0 | T | G | 0.123 | -1.34E-02 | 1.97E-01 | 2.67E+00 | 2.74E+00 | 9.42E-01 | no | Ppp1r13b | intron_variant | MODIFIER |
| 12 | 111884801 | AX-170357756 | 0 | C | T | 0.123 | -1.34E-02 | 1.97E-01 | 2.67E+00 | 2.74E+00 | 9.42E-01 | no | Ppp1r13b | intron_variant | MODIFIER |
| 12 | 111887341 | AX-170149044 | 0 | A | G | 0.123 | -1.34E-02 | 1.97E-01 | 2.67E+00 | 2.74E+00 | 9.42E-01 | no | Ppp1r13b | intron_variant | MODIFIER |
| 12 | 111887480 | AX-170421669 | 0 | A | G | 0.123 | -1.34E-02 | 1.97E-01 | 2.67E+00 | 2.74E+00 | 9.42E-01 | no | Ppp1r13b | intron_variant | MODIFIER |
| 12 | 111890210 | AX-170350505 | 0 | A | G | 0.123 | -1.34E-02 | 1.97E-01 | 2.67E+00 | 2.74E+00 | 9.42E-01 | no | Ppp1r13b | intron_variant | MODIFIER |
| 12 | 111949331 | AX-170377910 | 0 | T | A | 0.123 | -1.34E-02 | 1.97E-01 | 2.67E+00 | 2.74E+00 | 9.42E-01 | no | 5033406O09Rik | upstream_gene_variant | MODIFIER |
| 12 | 111951693 | AX-170365006 | 0 | A | G | 0.123 | -1.34E-02 | 1.97E-01 | 2.67E+00 | 2.74E+00 | 9.42E-01 | no | Gm36757 | non_coding_transcript_exon_variant | MODIFIER |
| 12 | 111952351 | AX-170462635 | 0 | G | A | 0.123 | -1.34E-02 | 1.97E-01 | 2.67E+00 | 2.74E+00 | 9.42E-01 | no | Gm36757 | non_coding_transcript_exon_variant | MODIFIER |
| 12 | 111955062 | AX-170338569 | 0 | C | T | 0.123 | -1.34E-02 | 1.97E-01 | 2.67E+00 | 2.74E+00 | 9.42E-01 | no | Gm36757 | downstream_gene_variant | MODIFIER |
| 12 | 111977497 | AX-170492970 | 0 | A | G | 0.123 | -1.34E-02 | 1.97E-01 | 2.67E+00 | 2.74E+00 | 9.42E-01 | no | Tdrd9 | intron_variant | MODIFIER |
| 12 | 112011751 | AX-170234368 | 0 | G | T | 0.123 | -1.34E-02 | 1.97E-01 | 2.67E+00 | 2.74E+00 | 9.42E-01 | no | Tdrd9 | intron_variant | MODIFIER |
| 12 | 112046718 | AX-97822433 | 0 | C | T | 0.123 | -1.34E-02 | 1.97E-01 | 2.67E+00 | 2.74E+00 | 9.42E-01 | no | Tdrd9 | intron_variant | MODIFIER |
| 12 | 112062926 | AX-170504816 | 0 | T | C | 0.123 | -1.34E-02 | 1.97E-01 | 2.67E+00 | 2.74E+00 | 9.42E-01 | no | Tdrd9 | intron_variant | MODIFIER |
| 12 | 112062981 | AX-170152206 | 0 | A | G | 0.123 | -1.34E-02 | 1.97E-01 | 2.67E+00 | 2.74E+00 | 9.42E-01 | no | Tdrd9 | intron_variant | MODIFIER |
| 12 | 112063027 | AX-170337697 | 0 | A | C | 0.123 | -1.34E-02 | 1.97E-01 | 2.67E+00 | 2.74E+00 | 9.42E-01 | no | Tdrd9 | intron_variant | MODIFIER |
| 12 | 112067773 | AX-170327946 | 0 | A | G | 0.123 | -1.34E-02 | 1.97E-01 | 2.67E+00 | 2.74E+00 | 9.42E-01 | no | Tdrd9 | intron_variant | MODIFIER |
| 12 | 112075585 | AX-170331845 | 0 | G | A | 0.123 | -1.34E-02 | 1.97E-01 | 2.67E+00 | 2.74E+00 | 9.42E-01 | no | | intergenic_variant | MODIFIER |
| 12 | 112114062 | AX-170371758 | 0 | A | G | 0.123 | -1.34E-02 | 1.97E-01 | 2.67E+00 | 2.74E+00 | 9.42E-01 | no | Aspg | intron_variant | MODIFIER |
| 12 | 112117663 | AX-170007540 | 0 | G | T | 0.123 | -1.34E-02 | 1.97E-01 | 2.67E+00 | 2.74E+00 | 9.42E-01 | no | Aspg | intron_variant | MODIFIER |
| 12 | 112174833 | AX-170356518 | 0 | T | C | 0.123 | -1.34E-02 | 1.97E-01 | 2.67E+00 | 2.74E+00 | 9.42E-01 | no | Kif26a | missense_variant | MODERATE |
| 12 | 112187611 | AX-170383513 | 0 | G | A | 0.123 | -1.34E-02 | 1.97E-01 | 2.67E+00 | 2.74E+00 | 9.42E-01 | no | | intergenic_variant | MODIFIER |
| 12 | 112201374 | AX-170318185 | 0 | T | C | 0.123 | -1.34E-02 | 1.97E-01 | 2.67E+00 | 2.74E+00 | 9.42E-01 | no | Gm20368 | downstream_gene_variant | MODIFIER |
| 12 | 112217200 | AX-170050226 | 0 | A | G | 0.123 | -1.34E-02 | 1.97E-01 | 2.67E+00 | 2.74E+00 | 9.42E-01 | no | | regulatory_region_variant | MODIFIER |
| 12 | 112221602 | AX-170470849 | 0 | C | T | 0.123 | -1.34E-02 | 1.97E-01 | 2.67E+00 | 2.74E+00 | 9.42E-01 | no | | regulatory_region_variant | MODIFIER |
| 12 | 112265780 | AX-170372712 | 0 | T | G | 0.123 | -1.34E-02 | 1.97E-01 | 2.67E+00 | 2.74E+00 | 9.42E-01 | no | | regulatory_region_variant | MODIFIER |
| 12 | 112276291 | AX-170007549 | 0 | T | G | 0.123 | -1.34E-02 | 1.97E-01 | 2.67E+00 | 2.74E+00 | 9.42E-01 | no | | regulatory_region_variant | MODIFIER |
| 12 | 112317453 | AX-170007545 | 0 | C | T | 0.123 | -1.34E-02 | 1.97E-01 | 2.67E+00 | 2.74E+00 | 9.42E-01 | no | | regulatory_region_variant | MODIFIER |
| 12 | 112317700 | AX-170007541 | 0 | G | C | 0.123 | -1.34E-02 | 1.97E-01 | 2.67E+00 | 2.74E+00 | 9.42E-01 | no | | intergenic_variant | MODIFIER |
| 12 | 112325540 | AX-170374174 | 0 | T | C | 0.123 | -1.34E-02 | 1.97E-01 | 2.67E+00 | 2.74E+00 | 9.42E-01 | no | | intergenic_variant | MODIFIER |
| 12 | 112371870 | AX-170007548 | 0 | A | G | 0.123 | -1.34E-02 | 1.97E-01 | 2.67E+00 | 2.74E+00 | 9.42E-01 | no | | intergenic_variant | MODIFIER |
| 12 | 112372955 | AX-170288864 | 0 | A | G | 0.123 | -1.34E-02 | 1.97E-01 | 2.67E+00 | 2.74E+00 | 9.42E-01 | no | | regulatory_region_variant | MODIFIER |
| 12 | 112401806 | AX-97788899 | 0 | A | C | 0.123 | -1.34E-02 | 1.97E-01 | 2.67E+00 | 2.74E+00 | 9.42E-01 | no | E330035G20Rik | intron_variant,non_coding_transcript_variant | MODIFIER |
| 12 | 112406352 | AX-170473009 | 0 | C | G | 0.123 | -1.34E-02 | 1.97E-01 | 2.67E+00 | 2.74E+00 | 9.42E-01 | no | E330035G20Rik | upstream_gene_variant | MODIFIER |
| 12 | 112409281 | AX-170188049 | 0 | G | A | 0.123 | -1.34E-02 | 1.97E-01 | 2.67E+00 | 2.74E+00 | 9.42E-01 | no | A730018C14Rik | downstream_gene_variant | MODIFIER |
| 12 | 112413158 | AX-170510878 | 0 | C | T | 0.123 | -1.34E-02 | 1.97E-01 | 2.67E+00 | 2.74E+00 | 9.42E-01 | no | A730018C14Rik | downstream_gene_variant | MODIFIER |
| 12 | 112415138 | AX-170108977 | 0 | A | G | 0.123 | -1.34E-02 | 1.97E-01 | 2.67E+00 | 2.74E+00 | 9.42E-01 | no | A730018C14Rik | non_coding_transcript_exon_variant | MODIFIER |
| 12 | 112538842 | AX-170476022 | 0 | C | T | 0.123 | -1.34E-02 | 1.97E-01 | 2.67E+00 | 2.74E+00 | 9.42E-01 | no | | intergenic_variant | MODIFIER |
| 12 | 112538864 | AX-170114894 | 0 | T | G | 0.123 | -1.34E-02 | 1.97E-01 | 2.67E+00 | 2.74E+00 | 9.42E-01 | no | | intergenic_variant | MODIFIER |
| 12 | 112553592 | AX-170458998 | 0 | T | C | 0.123 | -1.34E-02 | 1.97E-01 | 2.67E+00 | 2.74E+00 | 9.42E-01 | no | | intergenic_variant | MODIFIER |
| 12 | 112568091 | AX-170447899 | 0 | G | A | 0.123 | -1.34E-02 | 1.97E-01 | 2.67E+00 | 2.74E+00 | 9.42E-01 | no | | intergenic_variant | MODIFIER |

| | | | | | | | | | | | | | | | |
|---|---|---|---|---|---|---|---|---|---|---|---|---|---|---|---|
| 12 | 112574668 | AX-170459434 | 0 | T | C | 0.123 | -1.34E-02 | 1.97E-01 | 2.67E+00 | 2.74E+00 | 9.42E-01 | no | | intergenic_variant | MODIFIER |
| 12 | 112614770 | AX-170293564 | 0 | G | A | 0.123 | -1.34E-02 | 1.97E-01 | 2.67E+00 | 2.74E+00 | 9.42E-01 | no | Inf2 | intron_variant | MODIFIER |
| 12 | 113164303 | AX-170175209 | 0 | A | G | 0.123 | -1.34E-02 | 1.97E-01 | 2.67E+00 | 2.74E+00 | 9.42E-01 | no | Tedc1 | 3_prime_UTR_variant | MODIFIER |
| 12 | 113165039 | AX-170299216 | 0 | A | G | 0.123 | -1.34E-02 | 1.97E-01 | 2.67E+00 | 2.74E+00 | 9.42E-01 | no | Tedc1 | downstream_gene_variant | MODIFIER |
| 12 | 113165352 | AX-168389327 | 0 | C | A | 0.123 | -1.34E-02 | 1.97E-01 | 2.67E+00 | 2.74E+00 | 9.42E-01 | no | Tedc1 | downstream_gene_variant | MODIFIER |
| 12 | 113188554 | AX-170493445 | 0 | T | C | 0.123 | -1.34E-02 | 1.97E-01 | 2.67E+00 | 2.74E+00 | 9.42E-01 | no | Tmem121 | synonymous_variant | LOW |
| 12 | 113191095 | AX-97795973 | 0 | A | G | 0.123 | -1.34E-02 | 1.97E-01 | 2.67E+00 | 2.74E+00 | 9.42E-01 | no | Tmem121 | downstream_gene_variant | MODIFIER |
| 12 | 113192794 | AX-170007567 | 0 | A | G | 0.123 | -1.34E-02 | 1.97E-01 | 2.67E+00 | 2.74E+00 | 9.42E-01 | no | Tmem121 | downstream_gene_variant | MODIFIER |
| 12 | 113214607 | AX-170427382 | 0 | T | C | 0.246 | -6.68E-03 | 9.83E-02 | 2.67E+00 | 2.74E+00 | 9.42E-01 | no | Gm25622 | downstream_gene_variant | MODIFIER |
| 12 | 113221855 | AX-170398507 | 0 | T | C | 0.123 | -1.34E-02 | 1.97E-01 | 2.67E+00 | 2.74E+00 | 9.42E-01 | no | | regulatory_region_variant | MODIFIER |
| 12 | 113223771 | AX-170050251 | 0 | C | T | 0.246 | -6.68E-03 | 9.83E-02 | 2.67E+00 | 2.74E+00 | 9.42E-01 | no | | intergenic_variant | MODIFIER |
| 12 | 113223892 | AX-170050250 | 0 | G | T | 0.123 | -1.34E-02 | 1.97E-01 | 2.67E+00 | 2.74E+00 | 9.42E-01 | no | | intergenic_variant | MODIFIER |
| 12 | 113305777 | AX-97796863 | 0 | C | T | 0.123 | -1.34E-02 | 1.97E-01 | 2.67E+00 | 2.74E+00 | 9.42E-01 | no | Ighg2b | downstream_gene_variant | MODIFIER |
| 12 | 113306218 | AX-170239229 | 0 | T | C | 0.123 | -1.34E-02 | 1.97E-01 | 2.67E+00 | 2.74E+00 | 9.42E-01 | no | Ighg2b | downstream_gene_variant | MODIFIER |
| 12 | 113330061 | AX-170087338 | 0 | C | T | 0.123 | -1.34E-02 | 1.97E-01 | 2.67E+00 | 2.74E+00 | 9.42E-01 | no | Ighg1 | intron_variant | MODIFIER |
| 12 | 113491787 | AX-170369244 | 0 | T | A | 0.123 | -1.34E-02 | 1.97E-01 | 2.67E+00 | 2.74E+00 | 9.42E-01 | no | Adam6b | missense_variant | MODERATE |
| 12 | 113526683 | AX-97822383 | 0 | T | G | 0.123 | -1.34E-02 | 1.97E-01 | 2.67E+00 | 2.74E+00 | 9.42E-01 | no | Gm16968 | downstream_gene_variant | MODIFIER |
| 12 | 113588909 | AX-170481227 | 0 | G | C | 0.123 | -1.34E-02 | 1.97E-01 | 2.67E+00 | 2.74E+00 | 9.42E-01 | no | Ighv2-2 | upstream_gene_variant | MODIFIER |
| 12 | 113880988 | AX-170130398 | 0 | T | G | 0.123 | -1.34E-02 | 1.97E-01 | 2.67E+00 | 2.74E+00 | 9.42E-01 | no | Ighv2-9 | upstream_gene_variant | MODIFIER |
| 12 | 113897150 | AX-170435059 | 0 | T | C | 0.123 | -1.34E-02 | 1.97E-01 | 2.67E+00 | 2.74E+00 | 9.42E-01 | no | Ighv7-1 | upstream_gene_variant | MODIFIER |
| 12 | 113928386 | AX-170007572 | 0 | T | C | 0.123 | -1.34E-02 | 1.97E-01 | 2.67E+00 | 2.74E+00 | 9.42E-01 | no | Ighv14-1 | downstream_gene_variant | MODIFIER |
| 12 | 114175470 | AX-170265450 | 0 | G | C | 0.123 | -1.34E-02 | 1.97E-01 | 2.67E+00 | 2.74E+00 | 9.42E-01 | no | Ighv14-4 | downstream_gene_variant | MODIFIER |
| 12 | 114175991 | AX-170265979 | 0 | C | T | 0.123 | -1.34E-02 | 1.97E-01 | 2.67E+00 | 2.74E+00 | 9.42E-01 | no | Ighv14-4 | downstream_gene_variant | MODIFIER |
| 12 | 114209604 | AX-170349812 | 0 | A | G | 0.123 | -1.34E-02 | 1.97E-01 | 2.67E+00 | 2.74E+00 | 9.42E-01 | no | | intergenic_variant | MODIFIER |
| 12 | 114209639 | AX-170364510 | 0 | A | G | 0.123 | -1.34E-02 | 1.97E-01 | 2.67E+00 | 2.74E+00 | 9.42E-01 | no | | intergenic_variant | MODIFIER |
| 12 | 114283890 | AX-170523200 | 0 | T | G | 0.123 | -1.34E-02 | 1.97E-01 | 2.67E+00 | 2.74E+00 | 9.42E-01 | no | Ighv3-6 | downstream_gene_variant | MODIFIER |
| 12 | 114372092 | AX-97850614 | 0 | A | G | 0.123 | -1.34E-02 | 1.97E-01 | 2.67E+00 | 2.74E+00 | 9.42E-01 | no | | intergenic_variant | MODIFIER |
| 12 | 114433367 | AX-170314090 | 0 | T | A | 0.123 | -1.34E-02 | 1.97E-01 | 2.67E+00 | 2.74E+00 | 9.42E-01 | no | Ighv6-6 | downstream_gene_variant | MODIFIER |
| 12 | 114598004 | AX-170248257 | 0 | T | C | 0.123 | -1.34E-02 | 1.97E-01 | 2.67E+00 | 2.74E+00 | 9.42E-01 | no | Ighv1-10 | non_coding_transcript_exon_variant | MODIFIER |
| 12 | 114619001 | AX-97823672 | 0 | A | G | 0.123 | -1.34E-02 | 1.97E-01 | 2.67E+00 | 2.74E+00 | 9.42E-01 | no | Ighv1-12 | upstream_gene_variant | MODIFIER |
| 12 | 114642215 | AX-170410379 | 0 | C | T | 0.123 | -1.34E-02 | 1.97E-01 | 2.67E+00 | 2.74E+00 | 9.42E-01 | no | Ighv1-14 | downstream_gene_variant | MODIFIER |
| 12 | 114809461 | AX-170007576 | 0 | C | T | 0.123 | -1.34E-02 | 1.97E-01 | 2.67E+00 | 2.74E+00 | 9.42E-01 | no | Ighv1-28 | non_coding_transcript_exon_variant | MODIFIER |
| 12 | 114840369 | AX-170007575 | 0 | C | G | 0.123 | -1.34E-02 | 1.97E-01 | 2.67E+00 | 2.74E+00 | 9.42E-01 | no | Ighv1-33 | downstream_gene_variant | MODIFIER |
| 12 | 114925972 | AX-170219457 | 0 | T | C | 0.123 | -1.34E-02 | 1.97E-01 | 2.67E+00 | 2.74E+00 | 9.42E-01 | no | Ighv1-40 | upstream_gene_variant | MODIFIER |
| 12 | 114955828 | AX-170523388 | 0 | T | G | 0.123 | -1.34E-02 | 1.97E-01 | 2.67E+00 | 2.74E+00 | 9.42E-01 | no | | intergenic_variant | MODIFIER |
| 12 | 114955886 | AX-170157019 | 0 | A | C | 0.123 | -1.34E-02 | 1.97E-01 | 2.67E+00 | 2.74E+00 | 9.42E-01 | no | | intergenic_variant | MODIFIER |
| 12 | 115565410 | AX-170200388 | 0 | C | T | 0.123 | -1.34E-02 | 1.97E-01 | 2.67E+00 | 2.74E+00 | 9.42E-01 | no | Ighv8-11 | downstream_gene_variant | MODIFIER |
| 12 | 115614172 | AX-97826894 | 0 | T | C | 0.123 | -1.34E-02 | 1.97E-01 | 2.67E+00 | 2.74E+00 | 9.42E-01 | no | Gm5660 | upstream_gene_variant | MODIFIER |
| 12 | 116032027 | AX-170257303 | 0 | G | T | 0.123 | -1.34E-02 | 1.97E-01 | 2.67E+00 | 2.74E+00 | 9.42E-01 | no | | intergenic_variant | MODIFIER |
| 12 | 116099481 | AX-170494219 | 0 | A | T | 0.123 | -1.34E-02 | 1.97E-01 | 2.67E+00 | 2.74E+00 | 9.42E-01 | no | Vipr2 | intron_variant | MODIFIER |
| 12 | 116100543 | AX-170420471 | 0 | C | T | 0.123 | -1.34E-02 | 1.97E-01 | 2.67E+00 | 2.74E+00 | 9.42E-01 | no | Vipr2 | intron_variant | MODIFIER |
| 12 | 116100816 | AX-170503303 | 0 | T | C | 0.123 | -1.34E-02 | 1.97E-01 | 2.67E+00 | 2.74E+00 | 9.42E-01 | no | Vipr2 | intron_variant | MODIFIER |
| 12 | 116119835 | AX-170447862 | 0 | A | C | 0.123 | -1.34E-02 | 1.97E-01 | 2.67E+00 | 2.74E+00 | 9.42E-01 | no | Vipr2 | intron_variant | MODIFIER |
| 12 | 116119955 | AX-170175490 | 0 | G | C | 0.123 | -1.34E-02 | 1.97E-01 | 2.67E+00 | 2.74E+00 | 9.42E-01 | no | Vipr2 | intron_variant | MODIFIER |
| 12 | 116120012 | AX-170330368 | 0 | A | G | 0.123 | -1.34E-02 | 1.97E-01 | 2.67E+00 | 2.74E+00 | 9.42E-01 | no | Vipr2 | intron_variant | MODIFIER |
| 12 | 116120173 | AX-170190151 | 0 | T | C | 0.123 | -1.34E-02 | 1.97E-01 | 2.67E+00 | 2.74E+00 | 9.42E-01 | no | Vipr2 | intron_variant | MODIFIER |
| 12 | 116122108 | AX-170313792 | 0 | A | G | 0.123 | -1.34E-02 | 1.97E-01 | 2.67E+00 | 2.74E+00 | 9.42E-01 | no | Vipr2 | intron_variant | MODIFIER |
| 12 | 116123804 | AX-170456664 | 0 | A | G | 0.123 | -1.34E-02 | 1.97E-01 | 2.67E+00 | 2.74E+00 | 9.42E-01 | no | Vipr2 | intron_variant | MODIFIER |
| 12 | 116124007 | AX-170294183 | 0 | A | T | 0.123 | -1.34E-02 | 1.97E-01 | 2.67E+00 | 2.74E+00 | 9.42E-01 | no | Vipr2 | intron_variant | MODIFIER |
| 12 | 116124704 | AX-170246830 | 0 | A | G | 0.123 | -1.34E-02 | 1.97E-01 | 2.67E+00 | 2.74E+00 | 9.42E-01 | no | Vipr2 | intron_variant | MODIFIER |
| 12 | 116137857 | AX-170372206 | 0 | T | C | 0.123 | -1.34E-02 | 1.97E-01 | 2.67E+00 | 2.74E+00 | 9.42E-01 | no | Vipr2 | intron_variant | MODIFIER |
| 12 | 116140908 | AX-170139405 | 0 | G | A | 0.123 | -1.34E-02 | 1.97E-01 | 2.67E+00 | 2.74E+00 | 9.42E-01 | no | Vipr2 | intron_variant | MODIFIER |
| 12 | 116167373 | AX-170050314 | 0 | T | C | 0.123 | -1.34E-02 | 1.97E-01 | 2.67E+00 | 2.74E+00 | 9.42E-01 | no | Gm48620 | upstream_gene_variant | MODIFIER |
| 12 | 116468993 | AX-170484581 | 0 | T | G | 0.123 | -1.34E-02 | 1.97E-01 | 2.67E+00 | 2.74E+00 | 9.42E-01 | no | | intergenic_variant | MODIFIER |

| | | | | | | | | | | | | | | | |
|---|---|---|---|---|---|---|---|---|---|---|---|---|---|---|---|
| 12 | 116579174 | AX-170315926 | 0 | T | G | 0.123 | -1.34E-02 | 1.97E-01 | 2.67E+00 | 2.74E+00 | 9.42E-01 | no | Ptprn2 | intron_variant | MODIFIER |
| 12 | 116580706 | AX-170520576 | 0 | A | G | 0.123 | -1.34E-02 | 1.97E-01 | 2.67E+00 | 2.74E+00 | 9.42E-01 | no | Ptprn2 | intron_variant | MODIFIER |
| 12 | 116769455 | AX-170244328 | 0 | A | G | 0.123 | -1.34E-02 | 1.97E-01 | 2.67E+00 | 2.74E+00 | 9.42E-01 | no | Ptprn2 | intron_variant | MODIFIER |
| 12 | 116779111 | AX-170449179 | 0 | C | G | 0.123 | -1.34E-02 | 1.97E-01 | 2.67E+00 | 2.74E+00 | 9.42E-01 | no | Ptprn2 | intron_variant | MODIFIER |
| 12 | 116779691 | AX-170413886 | 0 | A | C | 0.123 | -1.34E-02 | 1.97E-01 | 2.67E+00 | 2.74E+00 | 9.42E-01 | no | Ptprn2 | intron_variant | MODIFIER |
| 12 | 116804704 | AX-170277038 | 0 | T | C | 0.123 | -1.34E-02 | 1.97E-01 | 2.67E+00 | 2.74E+00 | 9.42E-01 | no | Ptprn2 | intron_variant | MODIFIER |
| 12 | 116809231 | AX-170347307 | 0 | C | T | 0.123 | -1.34E-02 | 1.97E-01 | 2.67E+00 | 2.74E+00 | 9.42E-01 | no | Ptprn2 | intron_variant | MODIFIER |
| 12 | 116834785 | AX-170141764 | 0 | T | A | 0.123 | -1.34E-02 | 1.97E-01 | 2.67E+00 | 2.74E+00 | 9.42E-01 | no | Ptprn2 | intron_variant | MODIFIER |
| 12 | 116839373 | AX-170421690 | 0 | T | G | 0.123 | -1.34E-02 | 1.97E-01 | 2.67E+00 | 2.74E+00 | 9.42E-01 | no | Ptprn2 | intron_variant | MODIFIER |
| 12 | 116839858 | AX-170444888 | 0 | C | A | 0.123 | -1.34E-02 | 1.97E-01 | 2.67E+00 | 2.74E+00 | 9.42E-01 | no | Ptprn2 | intron_variant | MODIFIER |
| 12 | 116841510 | AX-170144456 | 0 | T | C | 0.123 | -1.34E-02 | 1.97E-01 | 2.67E+00 | 2.74E+00 | 9.42E-01 | no | Ptprn2 | intron_variant | MODIFIER |
| 12 | 116841544 | AX-170358730 | 0 | A | G | 0.123 | -1.34E-02 | 1.97E-01 | 2.67E+00 | 2.74E+00 | 9.42E-01 | no | Ptprn2 | intron_variant | MODIFIER |
| 12 | 116857787 | AX-170242228 | 0 | G | T | 0.123 | -1.34E-02 | 1.97E-01 | 2.67E+00 | 2.74E+00 | 9.42E-01 | no | Ptprn2 | intron_variant | MODIFIER |
| 12 | 116940226 | AX-170489945 | 0 | C | T | 0.123 | -1.34E-02 | 1.97E-01 | 2.67E+00 | 2.74E+00 | 9.42E-01 | no | Ptprn2 | intron_variant | MODIFIER |
| 12 | 116963122 | AX-170092624 | 0 | A | G | 0.123 | -1.34E-02 | 1.97E-01 | 2.67E+00 | 2.74E+00 | 9.42E-01 | no | Ptprn2 | intron_variant | MODIFIER |
| 12 | 117023061 | AX-170451197 | 0 | G | A | 0.123 | -1.34E-02 | 1.97E-01 | 2.67E+00 | 2.74E+00 | 9.42E-01 | no | Ptprn2 | intron_variant | MODIFIER |
| 12 | 117025234 | AX-170320790 | 0 | T | C | 0.123 | -1.34E-02 | 1.97E-01 | 2.67E+00 | 2.74E+00 | 9.42E-01 | no | Ptprn2 | intron_variant | MODIFIER |
| 12 | 117028002 | AX-170493552 | 0 | T | C | 0.123 | -1.34E-02 | 1.97E-01 | 2.67E+00 | 2.74E+00 | 9.42E-01 | no | Ptprn2 | intron_variant | MODIFIER |
| 12 | 117030342 | AX-170415239 | 0 | A | G | 0.123 | -1.34E-02 | 1.97E-01 | 2.67E+00 | 2.74E+00 | 9.42E-01 | no | Ptprn2 | intron_variant | MODIFIER |
| 12 | 117031240 | AX-170361852 | 0 | G | T | 0.123 | -1.34E-02 | 1.97E-01 | 2.67E+00 | 2.74E+00 | 9.42E-01 | no | Ptprn2 | intron_variant | MODIFIER |
| 12 | 117032364 | AX-170423222 | 0 | C | G | 0.123 | -1.34E-02 | 1.97E-01 | 2.67E+00 | 2.74E+00 | 9.42E-01 | no | Ptprn2 | intron_variant | MODIFIER |
| 12 | 117037885 | AX-170183536 | 0 | A | G | 0.123 | -1.34E-02 | 1.97E-01 | 2.67E+00 | 2.74E+00 | 9.42E-01 | no | Ptprn2 | intron_variant | MODIFIER |
| 12 | 117094100 | AX-97850771 | 0 | C | A | 0.123 | -1.34E-02 | 1.97E-01 | 2.67E+00 | 2.74E+00 | 9.42E-01 | no | Ptprn2 | intron_variant | MODIFIER |
| 12 | 117193039 | AX-170007594 | 0 | G | A | 0.123 | -1.34E-02 | 1.97E-01 | 2.67E+00 | 2.74E+00 | 9.42E-01 | no | Ptprn2 | intron_variant | MODIFIER |
| 12 | 117194323 | AX-97792751 | 0 | G | A | 0.123 | -1.34E-02 | 1.97E-01 | 2.67E+00 | 2.74E+00 | 9.42E-01 | no | Ptprn2 | intron_variant | MODIFIER |
| 12 | 117195045 | AX-170312772 | 0 | A | G | 0.123 | -1.34E-02 | 1.97E-01 | 2.67E+00 | 2.74E+00 | 9.42E-01 | no | Ptprn2 | intron_variant | MODIFIER |
| 12 | 117195507 | AX-170110079 | 0 | G | C | 0.123 | -1.34E-02 | 1.97E-01 | 2.67E+00 | 2.74E+00 | 9.42E-01 | no | Ptprn2 | intron_variant | MODIFIER |
| 12 | 117202963 | AX-170007595 | 0 | A | G | 0.123 | -1.34E-02 | 1.97E-01 | 2.67E+00 | 2.74E+00 | 9.42E-01 | no | Ptprn2 | intron_variant | MODIFIER |
| 12 | 117230856 | AX-170205286 | 0 | A | G | 0.123 | -1.34E-02 | 1.97E-01 | 2.67E+00 | 2.74E+00 | 9.42E-01 | no | Ptprn2 | intron_variant | MODIFIER |
| 12 | 117243091 | AX-97853526 | 0 | G | A | 0.123 | -1.34E-02 | 1.97E-01 | 2.67E+00 | 2.74E+00 | 9.42E-01 | no | Ptprn2 | intron_variant | MODIFIER |
| 12 | 117260344 | AX-170242479 | 0 | T | C | 0.123 | -1.34E-02 | 1.97E-01 | 2.67E+00 | 2.74E+00 | 9.42E-01 | no | Ptprn2 | intron_variant | MODIFIER |
| 12 | 117331481 | AX-170136201 | 0 | G | A | 0.123 | -1.34E-02 | 1.97E-01 | 2.67E+00 | 2.74E+00 | 9.42E-01 | no | Gm5441 | intron_variant,non_coding_transcript_variant | MODIFIER |
| 12 | 117332783 | AX-170143104 | 0 | T | A | 0.123 | -1.34E-02 | 1.97E-01 | 2.67E+00 | 2.74E+00 | 9.42E-01 | no | Gm5441 | intron_variant,non_coding_transcript_variant | MODIFIER |
| 12 | 117503380 | AX-97822548 | 0 | G | A | 0.123 | -1.34E-02 | 1.97E-01 | 2.67E+00 | 2.74E+00 | 9.42E-01 | no | | intergenic_variant | MODIFIER |
| 12 | 117504160 | AX-170449208 | 0 | T | G | 0.123 | -1.34E-02 | 1.97E-01 | 2.67E+00 | 2.74E+00 | 9.42E-01 | no | | intergenic_variant | MODIFIER |
| 12 | 117504197 | AX-170050345 | 0 | T | C | 0.123 | -1.34E-02 | 1.97E-01 | 2.67E+00 | 2.74E+00 | 9.42E-01 | no | | intergenic_variant | MODIFIER |
| 12 | 117505999 | AX-170422114 | 0 | C | T | 0.123 | -1.34E-02 | 1.97E-01 | 2.67E+00 | 2.74E+00 | 9.42E-01 | no | | intergenic_variant | MODIFIER |
| 12 | 117509016 | AX-97808882 | 0 | T | C | 0.123 | -1.34E-02 | 1.97E-01 | 2.67E+00 | 2.74E+00 | 9.42E-01 | no | | intergenic_variant | MODIFIER |
| 12 | 117542935 | AX-170380827 | 0 | G | C | 0.123 | -1.34E-02 | 1.97E-01 | 2.67E+00 | 2.74E+00 | 9.42E-01 | no | Rapgef5 | intron_variant | MODIFIER |
| 12 | 117543025 | AX-97840226 | 0 | T | C | 0.123 | -1.34E-02 | 1.97E-01 | 2.67E+00 | 2.74E+00 | 9.42E-01 | no | Rapgef5 | intron_variant | MODIFIER |
| 12 | 117560478 | AX-170263409 | 0 | T | C | 0.123 | -1.34E-02 | 1.97E-01 | 2.67E+00 | 2.74E+00 | 9.42E-01 | no | Rapgef5 | intron_variant | MODIFIER |
| 12 | 117562669 | AX-170246530 | 0 | C | A | 0.123 | -1.34E-02 | 1.97E-01 | 2.67E+00 | 2.74E+00 | 9.42E-01 | no | Rapgef5 | intron_variant | MODIFIER |
| 12 | 117565200 | AX-170188928 | 0 | A | G | 0.123 | -1.34E-02 | 1.97E-01 | 2.67E+00 | 2.74E+00 | 9.42E-01 | no | Rapgef5 | intron_variant | MODIFIER |
| 12 | 117659427 | AX-170526591 | 0 | G | C | 0.123 | -1.34E-02 | 1.97E-01 | 2.67E+00 | 2.74E+00 | 9.42E-01 | no | Rapgef5 | intron_variant | MODIFIER |
| 12 | 117659529 | AX-170007601 | 0 | A | C | 0.123 | -1.34E-02 | 1.97E-01 | 2.67E+00 | 2.74E+00 | 9.42E-01 | no | Rapgef5 | intron_variant | MODIFIER |
| 12 | 117665814 | AX-170007605 | 0 | C | T | 0.123 | -1.34E-02 | 1.97E-01 | 2.67E+00 | 2.74E+00 | 9.42E-01 | no | Rapgef5 | intron_variant | MODIFIER |
| 12 | 117665946 | AX-170050352 | 0 | C | T | 0.123 | -1.34E-02 | 1.97E-01 | 2.67E+00 | 2.74E+00 | 9.42E-01 | no | Rapgef5 | intron_variant | MODIFIER |
| 12 | 117667203 | AX-170482048 | 0 | A | G | 0.123 | -1.34E-02 | 1.97E-01 | 2.67E+00 | 2.74E+00 | 9.42E-01 | no | Rapgef5 | intron_variant | MODIFIER |
| 12 | 117667858 | AX-170050350 | 0 | T | C | 0.123 | -1.34E-02 | 1.97E-01 | 2.67E+00 | 2.74E+00 | 9.42E-01 | no | Rapgef5 | intron_variant | MODIFIER |
| 12 | 117672900 | AX-170481254 | 0 | C | T | 0.123 | -1.34E-02 | 1.97E-01 | 2.67E+00 | 2.74E+00 | 9.42E-01 | no | Rapgef5 | intron_variant | MODIFIER |
| 12 | 117693758 | AX-170519858 | 0 | G | A | 0.123 | -1.34E-02 | 1.97E-01 | 2.67E+00 | 2.74E+00 | 9.42E-01 | no | Rapgef5 | intron_variant | MODIFIER |
| 12 | 117694370 | AX-170215666 | 0 | A | G | 0.123 | -1.34E-02 | 1.97E-01 | 2.67E+00 | 2.74E+00 | 9.42E-01 | no | Rapgef5 | intron_variant | MODIFIER |
| 12 | 117746372 | AX-170183668 | 0 | C | T | 0.123 | -1.34E-02 | 1.97E-01 | 2.67E+00 | 2.74E+00 | 9.42E-01 | no | Rapgef5 | intron_variant | MODIFIER |
| 12 | 117891005 | AX-170345097 | 0 | A | T | 0.123 | -1.34E-02 | 1.97E-01 | 2.67E+00 | 2.74E+00 | 9.42E-01 | no | Dnah11 | intron_variant | MODIFIER |

| | | | | | | | | | | | | | | | |
|---|---|---|---|---|---|---|---|---|---|---|---|---|---|---|---|
| 12 | 117916789 | AX-170092629 | 0 | T | C | 0.123 | -1.34E-02 | 1.97E-01 | 2.67E+00 | 2.74E+00 | 9.42E-01 | no | Dnah11 | synonymous_variant | LOW |
| 12 | 117926376 | AX-170403519 | 0 | C | T | 0.123 | -1.34E-02 | 1.97E-01 | 2.67E+00 | 2.74E+00 | 9.42E-01 | no | Dnah11 | intron_variant | MODIFIER |
| 12 | 117929312 | AX-170348059 | 0 | A | G | 0.123 | -1.34E-02 | 1.97E-01 | 2.67E+00 | 2.74E+00 | 9.42E-01 | no | Dnah11 | intron_variant | MODIFIER |
| 12 | 117934064 | AX-170460324 | 0 | T | C | 0.123 | -1.34E-02 | 1.97E-01 | 2.67E+00 | 2.74E+00 | 9.42E-01 | no | Dnah11 | intron_variant | MODIFIER |
| 12 | 117935435 | AX-170437861 | 0 | A | T | 0.123 | -1.34E-02 | 1.97E-01 | 2.67E+00 | 2.74E+00 | 9.42E-01 | no | Dnah11 | intron_variant | MODIFIER |
| 12 | 117972509 | AX-168399458 | 0 | A | G | 0.123 | -1.34E-02 | 1.97E-01 | 2.67E+00 | 2.74E+00 | 9.42E-01 | no | Dnah11 | intron_variant | MODIFIER |
| 12 | 118367684 | AX-170229650 | 0 | T | G | 0.123 | -1.34E-02 | 1.97E-01 | 2.67E+00 | 2.74E+00 | 9.42E-01 | no | | intergenic_variant | MODIFIER |
| 12 | 118513103 | AX-170050369 | 0 | A | G | 0.123 | -1.34E-02 | 1.97E-01 | 2.67E+00 | 2.74E+00 | 9.42E-01 | no | | intergenic_variant | MODIFIER |
| 12 | 118694341 | AX-97855767 | 0 | G | A | 0.123 | -1.34E-02 | 1.97E-01 | 2.67E+00 | 2.74E+00 | 9.42E-01 | no | | intergenic_variant | MODIFIER |
| 12 | 118709019 | AX-97812154 | 0 | C | T | 0.123 | -1.34E-02 | 1.97E-01 | 2.67E+00 | 2.74E+00 | 9.42E-01 | no | | intergenic_variant | MODIFIER |
| 7 | 113026168 | AX-170134298 | 0 | C | G | 0.439 | 6.81E-03 | 9.38E-02 | 2.68E+00 | 2.74E+00 | 9.42E-01 | no | | intergenic_variant | MODIFIER |
| 13 | 54854933 | AX-170460008 | 0 | C | A | 0.526 | 5.76E-03 | 6.48E-02 | 2.61E+00 | 2.72E+00 | 9.42E-01 | no | | intergenic_variant | MODIFIER |
| 13 | 54857623 | AX-170484084 | 0 | T | C | 0.526 | 5.76E-03 | 6.48E-02 | 2.61E+00 | 2.72E+00 | 9.42E-01 | no | | intergenic_variant | MODIFIER |
| 13 | 54894200 | AX-170464243 | 0 | G | A | 0.526 | 5.76E-03 | 6.48E-02 | 2.61E+00 | 2.72E+00 | 9.42E-01 | no | | intergenic_variant | MODIFIER |
| 13 | 54895491 | AX-170008352 | 0 | T | C | 0.526 | 5.76E-03 | 6.48E-02 | 2.61E+00 | 2.72E+00 | 9.42E-01 | no | | intergenic_variant | MODIFIER |
| 13 | 54912035 | AX-170146535 | 0 | C | A | 0.526 | 5.76E-03 | 6.48E-02 | 2.61E+00 | 2.72E+00 | 9.42E-01 | no | 5330429C05Rik | upstream_gene_variant | MODIFIER |
| 13 | 54946934 | AX-97823656 | 0 | A | G | 0.526 | 5.76E-03 | 6.48E-02 | 2.61E+00 | 2.72E+00 | 9.42E-01 | no | Unc5a | upstream_gene_variant | MODIFIER |
| 13 | 54982696 | AX-97815309 | 0 | G | A | 0.526 | 5.76E-03 | 6.48E-02 | 2.61E+00 | 2.72E+00 | 9.42E-01 | no | Unc5a | intron_variant | MODIFIER |
| 13 | 54991657 | AX-170378309 | 0 | G | A | 0.526 | 5.76E-03 | 6.48E-02 | 2.61E+00 | 2.72E+00 | 9.42E-01 | no | Unc5a | intron_variant | MODIFIER |
| 13 | 55040169 | AX-170364037 | 0 | G | C | 0.526 | 5.76E-03 | 6.48E-02 | 2.61E+00 | 2.72E+00 | 9.42E-01 | no | Uimc1 | intron_variant | MODIFIER |
| 13 | 55040584 | AX-170008355 | 0 | C | T | 0.526 | 5.76E-03 | 6.48E-02 | 2.61E+00 | 2.72E+00 | 9.42E-01 | no | Uimc1 | missense_variant | MODERATE |
| 13 | 55060853 | AX-170193011 | 0 | T | C | 0.526 | 5.76E-03 | 6.48E-02 | 2.61E+00 | 2.72E+00 | 9.42E-01 | no | Uimc1 | intron_variant | MODIFIER |
| 13 | 55111065 | AX-170008351 | 0 | A | G | 0.526 | 5.76E-03 | 6.48E-02 | 2.61E+00 | 2.72E+00 | 9.42E-01 | no | Zfp346 | intron_variant | MODIFIER |
| 13 | 55123974 | AX-170143010 | 0 | A | C | 0.526 | 5.76E-03 | 6.48E-02 | 2.61E+00 | 2.72E+00 | 9.42E-01 | no | Zfp346 | intron_variant | MODIFIER |
| 13 | 55125070 | AX-170476667 | 0 | T | C | 0.526 | 5.76E-03 | 6.48E-02 | 2.61E+00 | 2.72E+00 | 9.42E-01 | no | Zfp346 | intron_variant | MODIFIER |
| 13 | 55129795 | AX-170141042 | 0 | T | C | 0.526 | 5.76E-03 | 6.48E-02 | 2.61E+00 | 2.72E+00 | 9.42E-01 | no | Zfp346 | intron_variant | MODIFIER |
| 13 | 55167123 | AX-97827254 | 0 | G | A | 0.526 | 5.76E-03 | 6.48E-02 | 2.61E+00 | 2.72E+00 | 9.42E-01 | no | Fgfr4 | synonymous_variant | LOW |
| 13 | 55172337 | AX-170051354 | 0 | T | G | 0.526 | 5.76E-03 | 6.48E-02 | 2.61E+00 | 2.72E+00 | 9.42E-01 | no | Fgfr4 | downstream_gene_variant | MODIFIER |
| 13 | 55174305 | AX-170306067 | 0 | G | A | 0.526 | 5.76E-03 | 6.48E-02 | 2.61E+00 | 2.72E+00 | 9.42E-01 | no | | intergenic_variant | MODIFIER |
| 13 | 55287549 | AX-170162947 | 0 | T | C | 0.526 | 5.76E-03 | 6.48E-02 | 2.61E+00 | 2.72E+00 | 9.42E-01 | no | Nsd1 | intron_variant | MODIFIER |
| 13 | 55292434 | AX-97797215 | 0 | G | A | 0.526 | 5.76E-03 | 6.48E-02 | 2.61E+00 | 2.72E+00 | 9.42E-01 | no | Nsd1 | intron_variant | MODIFIER |
| 5 | 123889677 | AX-97789173 | 1 | G | A | 0.455 | -7.58E-03 | 1.12E-01 | 2.70E+00 | 2.75E+00 | 9.42E-01 | no | | intergenic_variant | MODIFIER |
| 2 | 122056445 | AX-170018881 | 0 | G | A | 0.149 | -1.07E-02 | 1.78E-01 | 2.67E+00 | 2.76E+00 | 9.42E-01 | no | Eif3j1 | 3_prime_UTR_variant | MODIFIER |
| 1 | 67636344 | AX-170129617 | 0 | A | G | 0.114 | -1.61E-02 | 2.31E-01 | 2.69E+00 | 2.74E+00 | 9.42E-01 | no | | intergenic_variant | MODIFIER |
| 7 | 47155453 | AX-170035750 | 0 | T | C | 0.342 | -5.99E-03 | 9.44E-02 | 2.67E+00 | 2.75E+00 | 9.42E-01 | no | | regulatory_region_variant | MODIFIER |
| 3 | 25541605 | AX-170190675 | 0 | G | A | 0.474 | -2.42E-03 | 3.52E-02 | 2.58E+00 | 2.74E+00 | 9.43E-01 | no | Nlgn1 | intron_variant | MODIFIER |
| 7 | 119732937 | AX-97845237 | 0 | A | G | 0.237 | -4.84E-03 | 7.04E-02 | 2.58E+00 | 2.74E+00 | 9.43E-01 | no | Gm44866 | intron_variant,non_coding_transcript_variant | MODIFIER |
| 14 | 61181813 | AX-170532483 | 0 | T | C | 0.237 | -4.84E-03 | 7.04E-02 | 2.58E+00 | 2.74E+00 | 9.43E-01 | no | Sacs | intron_variant | MODIFIER |
| 14 | 61191513 | AX-170009934 | 0 | T | C | 0.237 | -4.84E-03 | 7.04E-02 | 2.58E+00 | 2.74E+00 | 9.43E-01 | no | Sacs | synonymous_variant | LOW |
| 5 | 71415139 | AX-170387037 | 1 | T | A | 0.446 | 7.53E-03 | 9.67E-02 | 2.65E+00 | 2.73E+00 | 9.43E-01 | no | | intergenic_variant | MODIFIER |
| 1 | 5070719 | AX-170153298 | 0 | G | A | 0.421 | -3.94E-03 | 6.84E-02 | 2.67E+00 | 2.76E+00 | 9.43E-01 | no | Rgs20 | upstream_gene_variant | MODIFIER |
| 13 | 38116284 | AX-170163089 | 0 | G | C | 0.175 | -9.11E-03 | 1.16E-01 | 2.65E+00 | 2.73E+00 | 9.43E-01 | no | | intergenic_variant | MODIFIER |
| 9 | 32806728 | AX-97843594 | 0 | C | T | 0.421 | 6.50E-03 | 7.87E-02 | 2.63E+00 | 2.72E+00 | 9.43E-01 | no | | intergenic_variant | MODIFIER |
| 13 | 37597304 | AX-170451107 | 1 | C | A | 0.196 | -9.24E-03 | 1.14E-01 | 2.65E+00 | 2.72E+00 | 9.43E-01 | no | | regulatory_region_variant | MODIFIER |
| 7 | 82838341 | AX-170250089 | 1 | G | A | 0.116 | 1.56E-02 | 2.13E-01 | 2.65E+00 | 2.73E+00 | 9.43E-01 | no | | intergenic_variant | MODIFIER |
| 6 | 139944348 | AX-170463978 | 1 | C | T | 0.08 | 1.49E-02 | 1.99E-01 | 2.63E+00 | 2.73E+00 | 9.43E-01 | no | Pik3c2g | intron_variant | MODIFIER |
| 5 | 101196607 | AX-170205530 | 0 | C | T | 0.421 | -6.44E-03 | 1.01E-01 | 2.68E+00 | 2.75E+00 | 9.43E-01 | no | | intergenic_variant | MODIFIER |
| 3 | 146636602 | AX-170035669 | 1 | A | G | 0.339 | 5.52E-03 | 7.34E-02 | 2.66E+00 | 2.73E+00 | 9.43E-01 | no | Uox | downstream_gene_variant | MODIFIER |
| 12 | 111881101 | AX-170508515 | 1 | A | T | 0.134 | -1.28E-02 | 1.80E-01 | 2.66E+00 | 2.74E+00 | 9.43E-01 | no | Ppp1r13b | intron_variant | MODIFIER |
| 12 | 112434970 | AX-170255195 | 1 | C | G | 0.134 | -1.28E-02 | 1.80E-01 | 2.66E+00 | 2.74E+00 | 9.43E-01 | no | B020018J22Rik | non_coding_transcript_exon_variant | MODIFIER |
| 12 | 112623421 | AX-170517194 | 1 | C | A | 0.134 | -1.28E-02 | 1.80E-01 | 2.66E+00 | 2.74E+00 | 9.43E-01 | no | Adssl1 | intron_variant | MODIFIER |
| 11 | 13082006 | AX-170086950 | 1 | A | T | 0.054 | -1.36E-02 | 2.02E-01 | 2.61E+00 | 2.74E+00 | 9.43E-01 | no | | intergenic_variant | MODIFIER |
| 16 | 96488580 | AX-97830152 | 1 | A | G | 0.455 | -6.61E-03 | 7.45E-02 | 2.61E+00 | 2.72E+00 | 9.43E-01 | no | Pcp4 | intron_variant | MODIFIER |
| 7 | 91752618 | AX-170270431 | 0 | T | C | 0.14 | -1.28E-02 | 1.94E-01 | 2.65E+00 | 2.74E+00 | 9.43E-01 | no | Dlg2 | intron_variant | MODIFIER |

| | | | | | | | | | | | | | | | |
|---|---|---|---|---|---|---|---|---|---|---|---|---|---|---|---|
| 8 | 75405298 | AX-170264582 | 2 | T | C | 0.182 | -9.60E-03 | 1.26E-01 | 2.64E+00 | 2.73E+00 | 9.43E-01 | no | Gm19885 | downstream_gene_variant | MODIFIER |
| 7 | 53778809 | AX-170282531 | 1 | T | C | 0.411 | -6.39E-03 | 9.61E-02 | 2.65E+00 | 2.74E+00 | 9.43E-01 | no | | intergenic_variant | MODIFIER |
| 18 | 69953314 | AX-170243373 | 1 | A | G | 0.259 | -7.39E-03 | 9.77E-02 | 2.66E+00 | 2.73E+00 | 9.43E-01 | no | Ccdc68 | intron_variant | MODIFIER |
| 5 | 37394164 | AX-170307651 | 0 | A | G | 0.325 | -7.13E-03 | 1.09E-01 | 2.68E+00 | 2.75E+00 | 9.43E-01 | no | Evc2 | intron_variant | MODIFIER |
| 1 | 170556903 | AX-170533529 | 0 | T | C | 0.211 | -8.39E-03 | 1.27E-01 | 2.69E+00 | 2.75E+00 | 9.43E-01 | no | Nos1ap | intron_variant,non_coding_transcript_variant | MODIFIER |
| 5 | 78954660 | AX-170095369 | 0 | C | T | 0.412 | -6.34E-03 | 9.72E-02 | 2.66E+00 | 2.75E+00 | 9.43E-01 | no | | intergenic_variant | MODIFIER |
| 1 | 10036420 | AX-170446797 | 0 | T | C | 0.421 | 5.93E-03 | 7.04E-02 | 2.64E+00 | 2.72E+00 | 9.43E-01 | no | Cops5 | intron_variant | MODIFIER |
| 6 | 102314230 | AX-170095730 | 1 | C | A | 0.071 | 1.93E-02 | 2.75E-01 | 2.70E+00 | 2.74E+00 | 9.43E-01 | no | Cntn3 | intron_variant | MODIFIER |
| 1 | 152534421 | AX-170390291 | 0 | G | C | 0.211 | 1.03E-02 | 1.25E-01 | 2.64E+00 | 2.72E+00 | 9.43E-01 | no | Rgl1 | intron_variant | MODIFIER |
| 12 | 111197028 | AX-170531896 | 1 | C | T | 0.125 | -1.30E-02 | 1.96E-01 | 2.67E+00 | 2.75E+00 | 9.43E-01 | no | Traf3 | intron_variant | MODIFIER |
| 9 | 61501450 | AX-170032491 | 1 | C | T | 0.205 | -8.95E-03 | 1.27E-01 | 2.69E+00 | 2.74E+00 | 9.43E-01 | no | | intergenic_variant | MODIFIER |
| 2 | 124888399 | AX-170334515 | 1 | C | T | 0.143 | 1.64E-02 | 2.27E-01 | 2.70E+00 | 2.73E+00 | 9.43E-01 | no | | regulatory_region_variant | MODIFIER |
| 2 | 128511391 | AX-170226712 | 1 | G | T | 0.143 | 1.64E-02 | 2.27E-01 | 2.70E+00 | 2.73E+00 | 9.43E-01 | no | | intergenic_variant | MODIFIER |
| 8 | 80995583 | AX-170414936 | 1 | C | T | 0.205 | -7.36E-03 | 1.12E-01 | 2.64E+00 | 2.74E+00 | 9.43E-01 | no | Usp38 | intron_variant | MODIFIER |
| 1 | 60629788 | AX-170511848 | 1 | C | T | 0.107 | 1.66E-02 | 2.35E-01 | 2.69E+00 | 2.74E+00 | 9.44E-01 | no | | intergenic_variant | MODIFIER |
| 1 | 12367881 | AX-170376185 | 0 | C | T | 0.386 | -4.63E-03 | 7.46E-02 | 2.69E+00 | 2.76E+00 | 9.44E-01 | no | | intergenic_variant | MODIFIER |
| 7 | 113825406 | AX-170400498 | 0 | C | A | 0.43 | 6.59E-03 | 9.28E-02 | 2.67E+00 | 2.74E+00 | 9.44E-01 | no | Spon1 | intron_variant | MODIFIER |
| 4 | 148337097 | AX-170347841 | 0 | T | C | 0.105 | -1.19E-02 | 1.83E-01 | 2.66E+00 | 2.74E+00 | 9.44E-01 | no | | intergenic_variant | MODIFIER |
| 9 | 53076805 | AX-170119111 | 1 | T | C | 0.223 | -7.96E-03 | 1.16E-01 | 2.68E+00 | 2.74E+00 | 9.44E-01 | no | | intergenic_variant | MODIFIER |
| 13 | 47657455 | AX-170214327 | 0 | T | C | 0.5 | 5.18E-03 | 6.61E-02 | 2.63E+00 | 2.73E+00 | 9.44E-01 | no | | intergenic_variant | MODIFIER |
| 10 | 43487623 | AX-170346538 | 1 | C | T | 0.375 | 4.80E-03 | 7.52E-02 | 2.64E+00 | 2.75E+00 | 9.44E-01 | no | Bend3 | intron_variant | MODIFIER |
| 10 | 43494416 | AX-170044716 | 1 | G | T | 0.375 | 4.80E-03 | 7.52E-02 | 2.64E+00 | 2.75E+00 | 9.44E-01 | no | Bend3 | intron_variant | MODIFIER |
| 12 | 63734198 | AX-97821431 | 0 | A | C | 0.061 | -2.14E-02 | 3.02E-01 | 2.71E+00 | 2.73E+00 | 9.44E-01 | no | | intergenic_variant | MODIFIER |
| 12 | 69477855 | AX-170006988 | 0 | C | T | 0.061 | -2.14E-02 | 3.02E-01 | 2.71E+00 | 2.73E+00 | 9.44E-01 | no | 5830428M24Rik | upstream_gene_variant | MODIFIER |
| 18 | 42009712 | AX-170015600 | 0 | G | A | 0.061 | -2.14E-02 | 3.02E-01 | 2.71E+00 | 2.73E+00 | 9.44E-01 | no | | regulatory_region_variant | MODIFIER |
| 1 | 36163528 | AX-170246937 | 0 | A | T | 0.456 | -5.54E-03 | 7.95E-02 | 2.65E+00 | 2.74E+00 | 9.44E-01 | no | Uggt1 | intron_variant | MODIFIER |
| 1 | 11381341 | AX-97836153 | 1 | A | G | 0.393 | -4.43E-03 | 7.46E-02 | 2.69E+00 | 2.76E+00 | 9.44E-01 | no | | intergenic_variant | MODIFIER |
| 2 | 124973742 | AX-170064811 | 0 | A | G | 0.14 | 1.55E-02 | 2.21E-01 | 2.70E+00 | 2.73E+00 | 9.44E-01 | no | | intergenic_variant | MODIFIER |
| 2 | 146844614 | AX-170447680 | 0 | C | T | 0.14 | 1.55E-02 | 2.21E-01 | 2.70E+00 | 2.73E+00 | 9.44E-01 | no | Gm14114 | upstream_gene_variant | MODIFIER |
| 7 | 102757499 | AX-170341712 | 1 | T | C | 0.464 | -5.70E-03 | 9.26E-02 | 2.68E+00 | 2.75E+00 | 9.44E-01 | no | Olfr78 | intron_variant | MODIFIER |
| 5 | 122684136 | AX-170283300 | 0 | G | A | 0.456 | 7.87E-03 | 1.05E-01 | 2.65E+00 | 2.73E+00 | 9.44E-01 | no | P2rx7 | intron_variant | MODIFIER |
| 19 | 29719767 | AX-168390215 | 0 | C | T | 0.053 | -1.17E-02 | 1.67E-01 | 2.62E+00 | 2.74E+00 | 9.44E-01 | no | 9930021J03Rik | intron_variant | MODIFIER |
| 8 | 73199042 | AX-170189708 | 0 | C | T | 0.193 | -9.33E-03 | 1.35E-01 | 2.68E+00 | 2.74E+00 | 9.44E-01 | no | Large1 | intron_variant | MODIFIER |
| 12 | 115294914 | AX-170422344 | 1 | C | A | 0.143 | -1.29E-02 | 1.92E-01 | 2.67E+00 | 2.74E+00 | 9.44E-01 | no | Ighv8-8 | upstream_gene_variant | MODIFIER |
| 1 | 118518545 | AX-170042445 | 0 | A | T | 0.158 | 1.03E-02 | 1.40E-01 | 2.59E+00 | 2.73E+00 | 9.45E-01 | no | Clasp1 | intron_variant | MODIFIER |
| 10 | 43405222 | AX-170109725 | 1 | G | A | 0.366 | -5.63E-03 | 7.58E-02 | 2.64E+00 | 2.73E+00 | 9.45E-01 | no | Pdss2 | intron_variant | MODIFIER |
| 10 | 6041927 | AX-170002634 | 1 | C | T | 0.214 | -7.69E-03 | 1.05E-01 | 2.64E+00 | 2.73E+00 | 9.45E-01 | no | | intergenic_variant | MODIFIER |
| 16 | 87294833 | AX-170013591 | 0 | A | T | 0.43 | 5.29E-03 | 7.90E-02 | 2.66E+00 | 2.74E+00 | 9.45E-01 | no | | intergenic_variant | MODIFIER |
| 8 | 81069193 | AX-97846586 | 1 | A | C | 0.196 | -8.65E-03 | 1.28E-01 | 2.67E+00 | 2.74E+00 | 9.45E-01 | no | Gm45538 | intron_variant,non_coding_transcript_variant | MODIFIER |
| 1 | 135843549 | AX-170176454 | 0 | T | C | 0.211 | -6.74E-03 | 1.17E-01 | 2.65E+00 | 2.75E+00 | 9.45E-01 | no | Tnnt2 | intron_variant | MODIFIER |
| 1 | 136001823 | AX-170217819 | 0 | T | C | 0.211 | -6.74E-03 | 1.17E-01 | 2.65E+00 | 2.75E+00 | 9.45E-01 | no | Igfn1 | intron_variant | MODIFIER |
| 1 | 136590111 | AX-170507018 | 0 | A | G | 0.211 | -6.74E-03 | 1.17E-01 | 2.65E+00 | 2.75E+00 | 9.45E-01 | no | | intergenic_variant | MODIFIER |
| 1 | 78724223 | AX-170284835 | 0 | G | T | 0.132 | -1.29E-02 | 1.82E-01 | 2.64E+00 | 2.73E+00 | 9.45E-01 | no | | regulatory_region_variant | MODIFIER |
| 12 | 60167175 | AX-170006839 | 1 | C | G | 0.062 | 2.00E-02 | 3.01E-01 | 2.73E+00 | 2.75E+00 | 9.45E-01 | no | | intergenic_variant | MODIFIER |
| 12 | 70668456 | AX-97815730 | 1 | C | T | 0.062 | 2.00E-02 | 3.01E-01 | 2.73E+00 | 2.75E+00 | 9.45E-01 | no | Gm24474 | intron_variant,non_coding_transcript_variant | MODIFIER |
| 18 | 34520242 | AX-97817753 | 1 | G | A | 0.062 | 2.00E-02 | 3.01E-01 | 2.73E+00 | 2.75E+00 | 9.45E-01 | no | | regulatory_region_variant | MODIFIER |
| 10 | 42988017 | AX-170446075 | 0 | C | T | 0.368 | 4.57E-03 | 7.52E-02 | 2.65E+00 | 2.75E+00 | 9.45E-01 | no | | regulatory_region_variant | MODIFIER |
| 10 | 43005585 | AX-170122769 | 0 | G | T | 0.368 | 4.57E-03 | 7.52E-02 | 2.65E+00 | 2.75E+00 | 9.45E-01 | no | Sobp | intron_variant | MODIFIER |
| 10 | 43053790 | AX-170044703 | 0 | C | T | 0.368 | 4.57E-03 | 7.52E-02 | 2.65E+00 | 2.75E+00 | 9.45E-01 | no | Sobp | intron_variant | MODIFIER |
| 10 | 43054582 | AX-170044707 | 0 | A | G | 0.368 | 4.57E-03 | 7.52E-02 | 2.65E+00 | 2.75E+00 | 9.45E-01 | no | Sobp | intron_variant | MODIFIER |
| 10 | 43138932 | AX-170209160 | 0 | A | C | 0.368 | 4.57E-03 | 7.52E-02 | 2.65E+00 | 2.75E+00 | 9.45E-01 | no | Sobp | intron_variant | MODIFIER |
| 10 | 43157284 | AX-170138064 | 0 | A | G | 0.368 | 4.57E-03 | 7.52E-02 | 2.65E+00 | 2.75E+00 | 9.45E-01 | no | Sobp | intron_variant | MODIFIER |
| 10 | 43160785 | AX-97845564 | 0 | C | T | 0.368 | 4.57E-03 | 7.52E-02 | 2.65E+00 | 2.75E+00 | 9.45E-01 | no | Sobp | synonymous_variant | LOW |
| 10 | 43340205 | AX-170523266 | 0 | G | T | 0.368 | 4.57E-03 | 7.52E-02 | 2.65E+00 | 2.75E+00 | 9.45E-01 | no | Pdss2 | intron_variant | MODIFIER |

| | | | | | | | | | | | | | | | |
|---|---|---|---|---|---|---|---|---|---|---|---|---|---|---|---|
| 10 | 43415116 | AX-170370852 | 0 | G | A | 0.368 | 4.57E-03 | 7.52E-02 | 2.65E+00 | 2.75E+00 | 9.45E-01 | no | Pdss2 | intron_variant | MODIFIER |
| 10 | 43420869 | AX-170474451 | 0 | A | C | 0.368 | 4.57E-03 | 7.52E-02 | 2.65E+00 | 2.75E+00 | 9.45E-01 | no | Pdss2 | intron_variant | MODIFIER |
| 10 | 43425105 | AX-97786824 | 0 | T | C | 0.368 | 4.57E-03 | 7.52E-02 | 2.65E+00 | 2.75E+00 | 9.45E-01 | no | Pdss2 | intron_variant | MODIFIER |
| 10 | 43425241 | AX-170227890 | 0 | G | A | 0.368 | 4.57E-03 | 7.52E-02 | 2.65E+00 | 2.75E+00 | 9.45E-01 | no | Pdss2 | intron_variant | MODIFIER |
| 10 | 43425738 | AX-170370431 | 0 | G | A | 0.368 | 4.57E-03 | 7.52E-02 | 2.65E+00 | 2.75E+00 | 9.45E-01 | no | Pdss2 | intron_variant | MODIFIER |
| 10 | 43490032 | AX-170003128 | 0 | T | C | 0.368 | 4.57E-03 | 7.52E-02 | 2.65E+00 | 2.75E+00 | 9.45E-01 | no | Bend3 | intron_variant | MODIFIER |
| 10 | 43492817 | AX-170044717 | 0 | C | T | 0.368 | 4.57E-03 | 7.52E-02 | 2.65E+00 | 2.75E+00 | 9.45E-01 | no | Bend3 | intron_variant | MODIFIER |
| 10 | 43493228 | AX-170283788 | 0 | A | G | 0.368 | 4.57E-03 | 7.52E-02 | 2.65E+00 | 2.75E+00 | 9.45E-01 | no | Bend3 | intron_variant | MODIFIER |
| 10 | 43494189 | AX-170044714 | 0 | G | C | 0.368 | 4.57E-03 | 7.52E-02 | 2.65E+00 | 2.75E+00 | 9.45E-01 | no | Bend3 | synonymous_variant | LOW |
| 10 | 43496112 | AX-170044715 | 0 | A | G | 0.368 | 4.57E-03 | 7.52E-02 | 2.65E+00 | 2.75E+00 | 9.45E-01 | no | Bend3 | intron_variant | MODIFIER |
| 10 | 43510224 | AX-170202621 | 0 | A | G | 0.368 | 4.57E-03 | 7.52E-02 | 2.65E+00 | 2.75E+00 | 9.45E-01 | no | Bend3 | synonymous_variant | LOW |
| 10 | 43513826 | AX-170003127 | 0 | T | A | 0.368 | 4.57E-03 | 7.52E-02 | 2.65E+00 | 2.75E+00 | 9.45E-01 | no | Bend3 | intron_variant | MODIFIER |
| 10 | 43533951 | AX-170171043 | 0 | C | A | 0.368 | 4.57E-03 | 7.52E-02 | 2.65E+00 | 2.75E+00 | 9.45E-01 | no | 1700021F05Rik | intron_variant,non_coding_transcript_variant | MODIFIER |
| 10 | 43619097 | AX-170186152 | 0 | G | A | 0.368 | 4.57E-03 | 7.52E-02 | 2.65E+00 | 2.75E+00 | 9.45E-01 | no | F930017D23Rik | intron_variant,non_coding_transcript_variant | MODIFIER |
| 10 | 43627147 | AX-170116559 | 0 | A | G | 0.368 | 4.57E-03 | 7.52E-02 | 2.65E+00 | 2.75E+00 | 9.45E-01 | no | F930017D23Rik | intron_variant,non_coding_transcript_variant | MODIFIER |
| 10 | 43635824 | AX-170507612 | 0 | T | G | 0.368 | 4.57E-03 | 7.52E-02 | 2.65E+00 | 2.75E+00 | 9.45E-01 | no | F930017D23Rik | downstream_gene_variant | MODIFIER |
| 10 | 43635982 | AX-170505736 | 0 | G | T | 0.368 | 4.57E-03 | 7.52E-02 | 2.65E+00 | 2.75E+00 | 9.45E-01 | no | Rpl26-ps1 | upstream_gene_variant | MODIFIER |
| 10 | 43640835 | AX-170291986 | 0 | T | C | 0.368 | 4.57E-03 | 7.52E-02 | 2.65E+00 | 2.75E+00 | 9.45E-01 | no | Rpl26-ps1 | downstream_gene_variant | MODIFIER |
| 10 | 43641109 | AX-170322647 | 0 | A | G | 0.368 | 4.57E-03 | 7.52E-02 | 2.65E+00 | 2.75E+00 | 9.45E-01 | no | Rpl26-ps1 | downstream_gene_variant | MODIFIER |
| 10 | 43644834 | AX-170530680 | 0 | T | C | 0.368 | 4.57E-03 | 7.52E-02 | 2.65E+00 | 2.75E+00 | 9.45E-01 | no | F930017D23Rik | intron_variant,non_coding_transcript_variant | MODIFIER |
| 10 | 43664960 | AX-97825840 | 0 | C | T | 0.368 | 4.57E-03 | 7.52E-02 | 2.65E+00 | 2.75E+00 | 9.45E-01 | no | F930017D23Rik | intron_variant,non_coding_transcript_variant | MODIFIER |
| 10 | 43676998 | AX-170044721 | 0 | A | G | 0.368 | 4.57E-03 | 7.52E-02 | 2.65E+00 | 2.75E+00 | 9.45E-01 | no | F930017D23Rik | intron_variant,non_coding_transcript_variant | MODIFIER |
| 10 | 43686248 | AX-170224275 | 0 | C | T | 0.368 | 4.57E-03 | 7.52E-02 | 2.65E+00 | 2.75E+00 | 9.45E-01 | no | F930017D23Rik | intron_variant,non_coding_transcript_variant | MODIFIER |
| 10 | 43985125 | AX-170003130 | 0 | G | A | 0.368 | 4.57E-03 | 7.52E-02 | 2.65E+00 | 2.75E+00 | 9.45E-01 | no | Crybg1 | intron_variant | MODIFIER |
| 10 | 44036261 | AX-170380585 | 0 | A | T | 0.368 | 4.57E-03 | 7.52E-02 | 2.65E+00 | 2.75E+00 | 9.45E-01 | no | Crybg1 | intron_variant | MODIFIER |
| 12 | 101500181 | AX-170163551 | 0 | G | A | 0.088 | -1.30E-02 | 1.95E-01 | 2.68E+00 | 2.74E+00 | 9.45E-01 | no | Catsperb | intron_variant | MODIFIER |
| 9 | 59301328 | AX-97839905 | 2 | G | C | 0.218 | -8.18E-03 | 1.23E-01 | 2.69E+00 | 2.74E+00 | 9.45E-01 | no | Adpgk | intron_variant | MODIFIER |
| 10 | 23285382 | AX-170483705 | 0 | G | C | 0.579 | -2.85E-03 | 7.04E-02 | 2.63E+00 | 2.76E+00 | 9.46E-01 | no | Eya4 | intron_variant | MODIFIER |
| 10 | 117554290 | AX-170103899 | 1 | A | C | 0.33 | 4.63E-03 | 7.68E-02 | 2.66E+00 | 2.75E+00 | 9.46E-01 | no | Gm32235 | upstream_gene_variant | MODIFIER |
| 9 | 53075079 | AX-170384996 | 0 | T | C | 0.228 | 7.58E-03 | 1.08E-01 | 2.66E+00 | 2.73E+00 | 9.46E-01 | no | | intergenic_variant | MODIFIER |
| 10 | 115966675 | AX-170498238 | 0 | C | T | 0.386 | -5.00E-03 | 7.77E-02 | 2.65E+00 | 2.74E+00 | 9.46E-01 | no | | intergenic_variant | MODIFIER |
| 3 | 25536759 | AX-170066129 | 1 | G | A | 0.482 | -2.27E-03 | 3.54E-02 | 2.58E+00 | 2.74E+00 | 9.46E-01 | no | Nlgn1 | intron_variant | MODIFIER |
| 1 | 21044944 | AX-170516224 | 0 | A | G | 0.404 | 6.65E-03 | 8.12E-02 | 2.65E+00 | 2.72E+00 | 9.46E-01 | no | Tram2 | intron_variant | MODIFIER |
| 4 | 143745679 | AX-97799093 | 2 | T | C | 0.073 | -1.46E-02 | 2.13E-01 | 2.63E+00 | 2.74E+00 | 9.46E-01 | no | Gm13081 | upstream_gene_variant | MODIFIER |
| 1 | 25426931 | AX-168407211 | 0 | G | T | 0.412 | -4.94E-03 | 8.07E-02 | 2.68E+00 | 2.75E+00 | 9.46E-01 | no | Adgrb3 | intron_variant | MODIFIER |
| 8 | 35244393 | AX-170096222 | 0 | A | G | 0.149 | -1.06E-02 | 1.44E-01 | 2.67E+00 | 2.73E+00 | 9.46E-01 | no | Gm34597 | intron_variant,non_coding_transcript_variant | MODIFIER |
| 5 | 27437716 | AX-97803798 | 0 | T | C | 0.061 | -1.43E-02 | 2.19E-01 | 2.68E+00 | 2.74E+00 | 9.46E-01 | no | Dpp6 | intron_variant | MODIFIER |
| 10 | 31054379 | AX-170298932 | 0 | A | C | 0.395 | -4.99E-03 | 7.11E-02 | 2.62E+00 | 2.74E+00 | 9.46E-01 | no | Gm30676 | intron_variant,non_coding_transcript_variant | MODIFIER |
| 8 | 32285299 | AX-170278349 | 0 | G | T | 0.149 | 7.67E-03 | 1.24E-01 | 2.67E+00 | 2.75E+00 | 9.46E-01 | no | Nrg1 | intron_variant | MODIFIER |
| 1 | 51792370 | AX-170000490 | 0 | T | C | 0.105 | -1.56E-02 | 2.34E-01 | 2.69E+00 | 2.74E+00 | 9.46E-01 | no | Myo1b | intron_variant | MODIFIER |
| 1 | 54773535 | AX-170335684 | 0 | G | A | 0.105 | -1.56E-02 | 2.34E-01 | 2.69E+00 | 2.74E+00 | 9.46E-01 | no | Ankrd44 | intron_variant | MODIFIER |
| 9 | 62374551 | AX-97838370 | 0 | G | A | 0.202 | -9.46E-03 | 1.26E-01 | 2.66E+00 | 2.72E+00 | 9.46E-01 | no | Anp32a | intron_variant | MODIFIER |
| 9 | 69372562 | AX-170297438 | 0 | G | A | 0.202 | -9.46E-03 | 1.26E-01 | 2.66E+00 | 2.72E+00 | 9.46E-01 | no | Rora | intron_variant | MODIFIER |
| 5 | 120281325 | AX-170073306 | 1 | G | T | 0.455 | 7.94E-03 | 1.09E-01 | 2.68E+00 | 2.73E+00 | 9.46E-01 | no | | regulatory_region_variant | MODIFIER |
| 5 | 121567876 | AX-170026031 | 1 | G | A | 0.455 | 7.94E-03 | 1.09E-01 | 2.68E+00 | 2.73E+00 | 9.46E-01 | no | Aldh2 | 3_prime_UTR_variant | MODIFIER |
| 15 | 93848510 | AX-97821974 | 0 | G | A | 0.149 | 1.29E-02 | 1.78E-01 | 2.66E+00 | 2.73E+00 | 9.46E-01 | no | Gm49445 | downstream_gene_variant | MODIFIER |
| 15 | 96953249 | AX-170168162 | 0 | A | G | 0.149 | 1.29E-02 | 1.78E-01 | 2.66E+00 | 2.73E+00 | 9.46E-01 | no | | regulatory_region_variant | MODIFIER |
| 3 | 148605466 | AX-170021888 | 1 | G | A | 0.304 | -4.58E-03 | 7.55E-02 | 2.65E+00 | 2.75E+00 | 9.46E-01 | no | | intergenic_variant | MODIFIER |
| 12 | 68968045 | AX-170186565 | 0 | C | G | 0.061 | -2.05E-02 | 3.00E-01 | 2.71E+00 | 2.73E+00 | 9.46E-01 | no | Gm47515 | intron_variant,non_coding_transcript_variant | MODIFIER |
| 9 | 52066469 | AX-170328898 | 0 | C | A | 0.211 | -7.86E-03 | 1.18E-01 | 2.68E+00 | 2.74E+00 | 9.46E-01 | no | Rdx | intron_variant | MODIFIER |
| 9 | 52066536 | AX-170530858 | 0 | C | T | 0.211 | -7.86E-03 | 1.18E-01 | 2.68E+00 | 2.74E+00 | 9.46E-01 | no | Rdx | intron_variant | MODIFIER |
| 9 | 52070691 | AX-170306639 | 0 | C | G | 0.211 | -7.86E-03 | 1.18E-01 | 2.68E+00 | 2.74E+00 | 9.46E-01 | no | Rdx | intron_variant | MODIFIER |
| 9 | 52079956 | AX-170229440 | 0 | A | G | 0.211 | -7.86E-03 | 1.18E-01 | 2.68E+00 | 2.74E+00 | 9.46E-01 | no | Rdx | intron_variant | MODIFIER |
| 9 | 52100631 | AX-170466238 | 0 | G | A | 0.211 | -7.86E-03 | 1.18E-01 | 2.68E+00 | 2.74E+00 | 9.46E-01 | no | Gm27686 | downstream_gene_variant | MODIFIER |

| | | | | | | | | | | | | | | | |
|---|---|---|---|---|---|---|---|---|---|---|---|---|---|---|---|
| 9 | 52592601 | AX-170382180 | 0 | T | C | 0.211 | -7.86E-03 | 1.18E-01 | 2.68E+00 | 2.74E+00 | 9.46E-01 | no | | intergenic_variant | MODIFIER |
| 1 | 53010883 | AX-170366251 | 2 | G | A | 0.1 | -1.38E-02 | 2.12E-01 | 2.65E+00 | 2.74E+00 | 9.46E-01 | no | 1700019D03Rik | intron_variant | MODIFIER |
| 10 | 27746714 | AX-170240688 | 0 | G | A | 0.167 | 5.46E-03 | 1.01E-01 | 2.66E+00 | 2.75E+00 | 9.46E-01 | no | Gm30228 | intron_variant,non_coding_transcript_variant | MODIFIER |
| 5 | 109832694 | AX-170025871 | 2 | G | A | 0.436 | -6.16E-03 | 1.07E-01 | 2.68E+00 | 2.75E+00 | 9.46E-01 | no | Gm10416 | downstream_gene_variant | MODIFIER |
| 7 | 101860057 | AX-97834939 | 0 | T | A | 0.465 | 6.75E-03 | 9.02E-02 | 2.66E+00 | 2.73E+00 | 9.46E-01 | no | Folr1 | intron_variant | MODIFIER |
| 6 | 114793990 | AX-170147737 | 1 | G | A | 0.08 | -1.25E-02 | 2.03E-01 | 2.63E+00 | 2.74E+00 | 9.47E-01 | no | Atg7 | intron_variant | MODIFIER |
| 4 | 145632616 | AX-170024192 | 1 | T | G | 0.08 | 1.44E-02 | 2.03E-01 | 2.64E+00 | 2.73E+00 | 9.47E-01 | no | Gm13234 | non_coding_transcript_exon_variant | MODIFIER |
| 9 | 86045392 | AX-170524197 | 1 | T | C | 0.125 | 7.02E-03 | 1.26E-01 | 2.66E+00 | 2.75E+00 | 9.47E-01 | no | | intergenic_variant | MODIFIER |
| 12 | 69583157 | AX-170262775 | 0 | T | A | 0.061 | -2.02E-02 | 3.00E-01 | 2.71E+00 | 2.73E+00 | 9.47E-01 | no | Sos2 | downstream_gene_variant | MODIFIER |
| 12 | 72379972 | AX-170511509 | 0 | C | T | 0.061 | -2.02E-02 | 3.00E-01 | 2.71E+00 | 2.73E+00 | 9.47E-01 | no | Rtn1 | intron_variant | MODIFIER |
| 11 | 8119411 | AX-97802666 | 0 | G | A | 0.053 | -1.25E-02 | 2.00E-01 | 2.61E+00 | 2.74E+00 | 9.47E-01 | no | | intergenic_variant | MODIFIER |
| 1 | 67659910 | AX-170205004 | 0 | C | T | 0.114 | -1.32E-02 | 1.80E-01 | 2.63E+00 | 2.73E+00 | 9.47E-01 | no | | intergenic_variant | MODIFIER |
| 9 | 79453598 | AX-170406897 | 0 | G | A | 0.211 | 7.36E-03 | 1.23E-01 | 2.68E+00 | 2.75E+00 | 9.47E-01 | no | Gm10635 | intron_variant,non_coding_transcript_variant | MODIFIER |
| 11 | 17743030 | AX-170254414 | 1 | G | A | 0.062 | 1.30E-02 | 2.14E-01 | 2.65E+00 | 2.74E+00 | 9.47E-01 | no | | intergenic_variant | MODIFIER |
| 2 | 123820067 | AX-170215296 | 0 | A | G | 0.14 | -1.19E-02 | 1.86E-01 | 2.67E+00 | 2.74E+00 | 9.47E-01 | no | | intergenic_variant | MODIFIER |
| 7 | 67522780 | AX-97853034 | 1 | A | T | 0.411 | -5.52E-03 | 9.36E-02 | 2.67E+00 | 2.75E+00 | 9.47E-01 | no | Lrrc28 | intron_variant | MODIFIER |
| 1 | 44533601 | AX-97811452 | 0 | C | T | 0.105 | -1.01E-02 | 1.80E-01 | 2.64E+00 | 2.75E+00 | 9.47E-01 | no | | intergenic_variant | MODIFIER |
| 13 | 25367191 | AX-170007952 | 0 | T | C | 0.175 | -1.08E-02 | 1.19E-01 | 2.62E+00 | 2.71E+00 | 9.47E-01 | no | | intergenic_variant | MODIFIER |
| 4 | 142960139 | AX-170323620 | 1 | G | A | 0.08 | 1.05E-02 | 1.68E-01 | 2.65E+00 | 2.74E+00 | 9.47E-01 | no | 6330411D24Rik | intron_variant,non_coding_transcript_variant | MODIFIER |
| 1 | 76468229 | AX-170250167 | 0 | G | T | 0.14 | 1.55E-02 | 2.35E-01 | 2.68E+00 | 2.74E+00 | 9.48E-01 | no | Gm29069 | intron_variant,non_coding_transcript_variant | MODIFIER |
| 3 | 147621085 | AX-170089103 | 1 | G | A | 0.348 | 5.10E-03 | 7.30E-02 | 2.66E+00 | 2.73E+00 | 9.48E-01 | no | Gm36888 | non_coding_transcript_exon_variant | MODIFIER |
| 8 | 84851581 | AX-170121121 | 0 | C | T | 0.202 | 8.40E-03 | 1.23E-01 | 2.65E+00 | 2.73E+00 | 9.48E-01 | no | Calr | upstream_gene_variant | MODIFIER |
| 12 | 65987001 | AX-170265625 | 0 | G | A | 0.018 | -1.90E-02 | 1.58E-01 | 2.51E+00 | 2.71E+00 | 9.48E-01 | no | | intergenic_variant | MODIFIER |
| 5 | 69550835 | AX-170318797 | 0 | T | C | 0.447 | 6.93E-03 | 9.74E-02 | 2.65E+00 | 2.73E+00 | 9.48E-01 | no | | intergenic_variant | MODIFIER |
| 5 | 69856289 | AX-170332203 | 0 | A | G | 0.447 | 6.93E-03 | 9.74E-02 | 2.65E+00 | 2.73E+00 | 9.48E-01 | no | | intergenic_variant | MODIFIER |
| 5 | 69939292 | AX-170203809 | 0 | A | C | 0.447 | 6.93E-03 | 9.74E-02 | 2.65E+00 | 2.73E+00 | 9.48E-01 | no | | intergenic_variant | MODIFIER |
| 5 | 69939486 | AX-170511697 | 0 | T | A | 0.447 | 6.93E-03 | 9.74E-02 | 2.65E+00 | 2.73E+00 | 9.48E-01 | no | | intergenic_variant | MODIFIER |
| 5 | 69939808 | AX-170249379 | 0 | C | T | 0.447 | 6.93E-03 | 9.74E-02 | 2.65E+00 | 2.73E+00 | 9.48E-01 | no | | intergenic_variant | MODIFIER |
| 5 | 69940345 | AX-97852911 | 0 | C | A | 0.447 | 6.93E-03 | 9.74E-02 | 2.65E+00 | 2.73E+00 | 9.48E-01 | no | | intergenic_variant | MODIFIER |
| 5 | 69944015 | AX-170072409 | 0 | G | A | 0.447 | 6.93E-03 | 9.74E-02 | 2.65E+00 | 2.73E+00 | 9.48E-01 | no | | intergenic_variant | MODIFIER |
| 5 | 69971436 | AX-97821488 | 0 | A | T | 0.447 | 6.93E-03 | 9.74E-02 | 2.65E+00 | 2.73E+00 | 9.48E-01 | no | | intergenic_variant | MODIFIER |
| 5 | 69991652 | AX-170025225 | 0 | C | A | 0.447 | 6.93E-03 | 9.74E-02 | 2.65E+00 | 2.73E+00 | 9.48E-01 | no | | intergenic_variant | MODIFIER |
| 5 | 69998033 | AX-170072410 | 0 | A | G | 0.447 | 6.93E-03 | 9.74E-02 | 2.65E+00 | 2.73E+00 | 9.48E-01 | no | | intergenic_variant | MODIFIER |
| 5 | 70001612 | AX-97783418 | 0 | C | G | 0.447 | 6.93E-03 | 9.74E-02 | 2.65E+00 | 2.73E+00 | 9.48E-01 | no | | intergenic_variant | MODIFIER |
| 5 | 70015964 | AX-170039029 | 0 | T | C | 0.447 | 6.93E-03 | 9.74E-02 | 2.65E+00 | 2.73E+00 | 9.48E-01 | no | | intergenic_variant | MODIFIER |
| 5 | 70027530 | AX-170089513 | 0 | C | T | 0.447 | 6.93E-03 | 9.74E-02 | 2.65E+00 | 2.73E+00 | 9.48E-01 | no | | intergenic_variant | MODIFIER |
| 5 | 70069621 | AX-170530370 | 0 | C | T | 0.447 | 6.93E-03 | 9.74E-02 | 2.65E+00 | 2.73E+00 | 9.48E-01 | no | | intergenic_variant | MODIFIER |
| 5 | 70069643 | AX-170425079 | 0 | A | G | 0.447 | 6.93E-03 | 9.74E-02 | 2.65E+00 | 2.73E+00 | 9.48E-01 | no | | intergenic_variant | MODIFIER |
| 5 | 70069699 | AX-170135748 | 0 | C | T | 0.447 | 6.93E-03 | 9.74E-02 | 2.65E+00 | 2.73E+00 | 9.48E-01 | no | | intergenic_variant | MODIFIER |
| 5 | 70079805 | AX-97849977 | 0 | A | G | 0.447 | 6.93E-03 | 9.74E-02 | 2.65E+00 | 2.73E+00 | 9.48E-01 | no | | intergenic_variant | MODIFIER |
| 5 | 70108590 | AX-97781957 | 0 | G | C | 0.447 | 6.93E-03 | 9.74E-02 | 2.65E+00 | 2.73E+00 | 9.48E-01 | no | | intergenic_variant | MODIFIER |
| 5 | 70148792 | AX-170025226 | 0 | G | T | 0.447 | 6.93E-03 | 9.74E-02 | 2.65E+00 | 2.73E+00 | 9.48E-01 | no | | intergenic_variant | MODIFIER |
| 5 | 70200152 | AX-170103746 | 0 | C | G | 0.447 | 6.93E-03 | 9.74E-02 | 2.65E+00 | 2.73E+00 | 9.48E-01 | no | | intergenic_variant | MODIFIER |
| 5 | 70299680 | AX-170504943 | 0 | T | G | 0.447 | 6.93E-03 | 9.74E-02 | 2.65E+00 | 2.73E+00 | 9.48E-01 | no | | intergenic_variant | MODIFIER |
| 5 | 70300718 | AX-170089515 | 0 | C | T | 0.447 | 6.93E-03 | 9.74E-02 | 2.65E+00 | 2.73E+00 | 9.48E-01 | no | | intergenic_variant | MODIFIER |
| 5 | 70336837 | AX-170494135 | 0 | C | T | 0.447 | 6.93E-03 | 9.74E-02 | 2.65E+00 | 2.73E+00 | 9.48E-01 | no | | intergenic_variant | MODIFIER |
| 5 | 70392163 | AX-170529860 | 0 | T | G | 0.447 | 6.93E-03 | 9.74E-02 | 2.65E+00 | 2.73E+00 | 9.48E-01 | no | | intergenic_variant | MODIFIER |
| 5 | 70401150 | AX-170365118 | 0 | T | C | 0.447 | 6.93E-03 | 9.74E-02 | 2.65E+00 | 2.73E+00 | 9.48E-01 | no | | intergenic_variant | MODIFIER |
| 5 | 70612927 | AX-170490615 | 0 | C | A | 0.447 | 6.93E-03 | 9.74E-02 | 2.65E+00 | 2.73E+00 | 9.48E-01 | no | | intergenic_variant | MODIFIER |
| 5 | 70665531 | AX-170535738 | 0 | T | G | 0.447 | 6.93E-03 | 9.74E-02 | 2.65E+00 | 2.73E+00 | 9.48E-01 | no | | intergenic_variant | MODIFIER |
| 5 | 70673532 | AX-170072421 | 0 | G | A | 0.447 | 6.93E-03 | 9.74E-02 | 2.65E+00 | 2.73E+00 | 9.48E-01 | no | | intergenic_variant | MODIFIER |
| 5 | 70674048 | AX-170072422 | 0 | T | G | 0.447 | 6.93E-03 | 9.74E-02 | 2.65E+00 | 2.73E+00 | 9.48E-01 | no | | intergenic_variant | MODIFIER |
| 5 | 70742061 | AX-170489607 | 0 | A | C | 0.447 | 6.93E-03 | 9.74E-02 | 2.65E+00 | 2.73E+00 | 9.48E-01 | no | | intergenic_variant | MODIFIER |
| 5 | 70823599 | AX-170429494 | 0 | C | T | 0.447 | 6.93E-03 | 9.74E-02 | 2.65E+00 | 2.73E+00 | 9.48E-01 | no | Gabrg1 | intron_variant | MODIFIER |

| | | | | | | | | | | | | | | | |
|---|---|---|---|---|---|---|---|---|---|---|---|---|---|---|---|
| 5 | 70918262 | AX-170126888 | 0 | A | T | 0.447 | 6.93E-03 | 9.74E-02 | 2.65E+00 | 2.73E+00 | 9.48E-01 | no | | intergenic_variant | MODIFIER |
| 5 | 71427144 | AX-170177366 | 0 | T | A | 0.447 | 6.93E-03 | 9.74E-02 | 2.65E+00 | 2.73E+00 | 9.48E-01 | no | | intergenic_variant | MODIFIER |
| 5 | 71609621 | AX-97850651 | 0 | T | C | 0.447 | 6.93E-03 | 9.74E-02 | 2.65E+00 | 2.73E+00 | 9.48E-01 | no | Gabra4 | intron_variant | MODIFIER |
| 5 | 71673198 | AX-170427119 | 0 | G | A | 0.447 | 6.93E-03 | 9.74E-02 | 2.65E+00 | 2.73E+00 | 9.48E-01 | no | Gabrb1 | intron_variant | MODIFIER |
| 5 | 71714621 | AX-97847660 | 0 | T | G | 0.447 | 6.93E-03 | 9.74E-02 | 2.65E+00 | 2.73E+00 | 9.48E-01 | no | Gabrb1 | intron_variant | MODIFIER |
| 5 | 72264471 | AX-170487339 | 0 | C | T | 0.447 | 6.93E-03 | 9.74E-02 | 2.65E+00 | 2.73E+00 | 9.48E-01 | no | Atp10d | stop_lost | HIGH |
| 5 | 72985885 | AX-170025272 | 0 | C | T | 0.447 | 6.93E-03 | 9.74E-02 | 2.65E+00 | 2.73E+00 | 9.48E-01 | no | | intergenic_variant | MODIFIER |
| 5 | 73022870 | AX-170025271 | 0 | C | G | 0.447 | 6.93E-03 | 9.74E-02 | 2.65E+00 | 2.73E+00 | 9.48E-01 | no | Fryl | intron_variant | MODIFIER |
| 5 | 73032881 | AX-97804021 | 0 | C | T | 0.447 | 6.93E-03 | 9.74E-02 | 2.65E+00 | 2.73E+00 | 9.48E-01 | no | Fryl | intron_variant | MODIFIER |
| 5 | 73032975 | AX-170025276 | 0 | G | C | 0.447 | 6.93E-03 | 9.74E-02 | 2.65E+00 | 2.73E+00 | 9.48E-01 | no | Fryl | intron_variant | MODIFIER |
| 5 | 73046296 | AX-170408650 | 0 | A | C | 0.447 | 6.93E-03 | 9.74E-02 | 2.65E+00 | 2.73E+00 | 9.48E-01 | no | Fryl | intron_variant | MODIFIER |
| 5 | 73133443 | AX-170072455 | 0 | A | G | 0.447 | 6.93E-03 | 9.74E-02 | 2.65E+00 | 2.73E+00 | 9.48E-01 | no | Fryl | intron_variant | MODIFIER |
| 5 | 73148891 | AX-170339063 | 0 | A | T | 0.447 | 6.93E-03 | 9.74E-02 | 2.65E+00 | 2.73E+00 | 9.48E-01 | no | Fryl | upstream_gene_variant | MODIFIER |
| 5 | 73161730 | AX-170095357 | 0 | C | A | 0.447 | 6.93E-03 | 9.74E-02 | 2.65E+00 | 2.73E+00 | 9.48E-01 | no | Fryl | intron_variant | MODIFIER |
| 5 | 73188070 | AX-170291051 | 0 | G | A | 0.447 | 6.93E-03 | 9.74E-02 | 2.65E+00 | 2.73E+00 | 9.48E-01 | no | Fryl | intron_variant | MODIFIER |
| 5 | 73194021 | AX-170401907 | 0 | T | C | 0.447 | 6.93E-03 | 9.74E-02 | 2.65E+00 | 2.73E+00 | 9.48E-01 | no | Fryl | intron_variant | MODIFIER |
| 5 | 73194077 | AX-170272078 | 0 | A | T | 0.447 | 6.93E-03 | 9.74E-02 | 2.65E+00 | 2.73E+00 | 9.48E-01 | no | Fryl | intron_variant | MODIFIER |
| 5 | 73203397 | AX-170179859 | 0 | A | G | 0.447 | 6.93E-03 | 9.74E-02 | 2.65E+00 | 2.73E+00 | 9.48E-01 | no | Fryl | intron_variant | MODIFIER |
| 5 | 73232448 | AX-170419895 | 0 | A | G | 0.447 | 6.93E-03 | 9.74E-02 | 2.65E+00 | 2.73E+00 | 9.48E-01 | no | Fryl | intron_variant | MODIFIER |
| 5 | 73242431 | AX-170369317 | 0 | G | A | 0.447 | 6.93E-03 | 9.74E-02 | 2.65E+00 | 2.73E+00 | 9.48E-01 | no | Fryl | intron_variant | MODIFIER |
| 5 | 73263405 | AX-170175827 | 0 | C | T | 0.447 | 6.93E-03 | 9.74E-02 | 2.65E+00 | 2.73E+00 | 9.48E-01 | no | Gm34411 | upstream_gene_variant | MODIFIER |
| 5 | 73313614 | AX-170249025 | 0 | G | A | 0.447 | 6.93E-03 | 9.74E-02 | 2.65E+00 | 2.73E+00 | 9.48E-01 | no | Ociad1 | 3_prime_UTR_variant | MODIFIER |
| 5 | 73314559 | AX-170220871 | 0 | T | C | 0.447 | 6.93E-03 | 9.74E-02 | 2.65E+00 | 2.73E+00 | 9.48E-01 | no | Ociad1 | downstream_gene_variant | MODIFIER |
| 5 | 73323871 | AX-97792393 | 0 | A | G | 0.447 | 6.93E-03 | 9.74E-02 | 2.65E+00 | 2.73E+00 | 9.48E-01 | no | Ociad2 | intron_variant | MODIFIER |
| 8 | 91055465 | AX-170031251 | 0 | T | C | 0.184 | -8.35E-03 | 1.18E-01 | 2.65E+00 | 2.73E+00 | 9.48E-01 | no | Chd9 | downstream_gene_variant | MODIFIER |
| 5 | 73143009 | AX-170167392 | 0 | G | A | 0.456 | 7.07E-03 | 9.63E-02 | 2.64E+00 | 2.73E+00 | 9.48E-01 | no | Fryl | intron_variant | MODIFIER |
| 8 | 84169095 | AX-170080622 | 0 | C | T | 0.193 | 8.61E-03 | 1.32E-01 | 2.68E+00 | 2.74E+00 | 9.48E-01 | no | Nanos3 | downstream_gene_variant | MODIFIER |
| 8 | 84170381 | AX-170080624 | 0 | C | T | 0.193 | 8.61E-03 | 1.32E-01 | 2.68E+00 | 2.74E+00 | 9.48E-01 | no | Nanos3 | downstream_gene_variant | MODIFIER |
| 8 | 84177336 | AX-170467208 | 0 | A | G | 0.193 | 8.61E-03 | 1.32E-01 | 2.68E+00 | 2.74E+00 | 9.48E-01 | no | Nanos3 | upstream_gene_variant | MODIFIER |
| 8 | 84179297 | AX-170150267 | 0 | A | T | 0.193 | 8.61E-03 | 1.32E-01 | 2.68E+00 | 2.74E+00 | 9.48E-01 | no | Nanos3 | upstream_gene_variant | MODIFIER |
| 8 | 84211829 | AX-170213276 | 0 | C | T | 0.193 | 8.61E-03 | 1.32E-01 | 2.68E+00 | 2.74E+00 | 9.48E-01 | no | Zswim4 | 3_prime_UTR_variant | MODIFIER |
| 8 | 84225265 | AX-170171547 | 0 | A | G | 0.193 | 8.61E-03 | 1.32E-01 | 2.68E+00 | 2.74E+00 | 9.48E-01 | no | Zswim4 | intron_variant | MODIFIER |
| 8 | 84229686 | AX-170266557 | 0 | T | C | 0.193 | 8.61E-03 | 1.32E-01 | 2.68E+00 | 2.74E+00 | 9.48E-01 | no | Zswim4 | intron_variant | MODIFIER |
| 8 | 84252297 | AX-97781421 | 0 | T | C | 0.193 | 8.61E-03 | 1.32E-01 | 2.68E+00 | 2.74E+00 | 9.48E-01 | no | Mri1 | downstream_gene_variant | MODIFIER |
| 8 | 84293008 | AX-170344596 | 0 | C | T | 0.193 | 8.61E-03 | 1.32E-01 | 2.68E+00 | 2.74E+00 | 9.48E-01 | no | | intergenic_variant | MODIFIER |
| 8 | 84296545 | AX-170288852 | 0 | T | C | 0.193 | 8.61E-03 | 1.32E-01 | 2.68E+00 | 2.74E+00 | 9.48E-01 | no | | intergenic_variant | MODIFIER |
| 8 | 84304359 | AX-170481414 | 0 | A | G | 0.193 | 8.61E-03 | 1.32E-01 | 2.68E+00 | 2.74E+00 | 9.48E-01 | no | Gm45608 | upstream_gene_variant | MODIFIER |
| 8 | 84317045 | AX-170371502 | 0 | A | G | 0.193 | 8.61E-03 | 1.32E-01 | 2.68E+00 | 2.74E+00 | 9.48E-01 | no | | intergenic_variant | MODIFIER |
| 8 | 84411304 | AX-170031152 | 0 | A | C | 0.193 | 8.61E-03 | 1.32E-01 | 2.68E+00 | 2.74E+00 | 9.48E-01 | no | Cacna1a | upstream_gene_variant | MODIFIER |
| 8 | 84424727 | AX-170389122 | 0 | G | T | 0.193 | 8.61E-03 | 1.32E-01 | 2.68E+00 | 2.74E+00 | 9.48E-01 | no | Cacna1a | intron_variant | MODIFIER |
| 8 | 84425073 | AX-170389495 | 0 | T | C | 0.193 | 8.61E-03 | 1.32E-01 | 2.68E+00 | 2.74E+00 | 9.48E-01 | no | Cacna1a | intron_variant | MODIFIER |
| 8 | 84468923 | AX-170136821 | 0 | A | G | 0.193 | 8.61E-03 | 1.32E-01 | 2.68E+00 | 2.74E+00 | 9.48E-01 | no | Cacna1a | intron_variant | MODIFIER |
| 8 | 84523664 | AX-170281317 | 0 | C | A | 0.193 | 8.61E-03 | 1.32E-01 | 2.68E+00 | 2.74E+00 | 9.48E-01 | no | Cacna1a | intron_variant | MODIFIER |
| 8 | 84527478 | AX-170487244 | 0 | G | A | 0.193 | 8.61E-03 | 1.32E-01 | 2.68E+00 | 2.74E+00 | 9.48E-01 | no | Cacna1a | intron_variant | MODIFIER |
| 8 | 84529364 | AX-170204341 | 0 | A | G | 0.193 | 8.61E-03 | 1.32E-01 | 2.68E+00 | 2.74E+00 | 9.48E-01 | no | Cacna1a | intron_variant | MODIFIER |
| 8 | 84569630 | AX-170080632 | 0 | C | T | 0.193 | 8.61E-03 | 1.32E-01 | 2.68E+00 | 2.74E+00 | 9.48E-01 | no | Cacna1a | intron_variant | MODIFIER |
| 8 | 84571496 | AX-170185965 | 0 | A | G | 0.193 | 8.61E-03 | 1.32E-01 | 2.68E+00 | 2.74E+00 | 9.48E-01 | no | Cacna1a | intron_variant | MODIFIER |
| 8 | 84572682 | AX-170393321 | 0 | T | C | 0.193 | 8.61E-03 | 1.32E-01 | 2.68E+00 | 2.74E+00 | 9.48E-01 | no | Cacna1a | intron_variant | MODIFIER |
| 8 | 84573383 | AX-170080628 | 0 | T | C | 0.193 | 8.61E-03 | 1.32E-01 | 2.68E+00 | 2.74E+00 | 9.48E-01 | no | Cacna1a | intron_variant | MODIFIER |
| 8 | 84577803 | AX-170325712 | 0 | A | G | 0.193 | 8.61E-03 | 1.32E-01 | 2.68E+00 | 2.74E+00 | 9.48E-01 | no | Cacna1a | intron_variant | MODIFIER |
| 8 | 84591828 | AX-170338133 | 0 | A | G | 0.193 | 8.61E-03 | 1.32E-01 | 2.68E+00 | 2.74E+00 | 9.48E-01 | no | Cacna1a | intron_variant | MODIFIER |
| 8 | 84614595 | AX-170280177 | 0 | G | A | 0.193 | 8.61E-03 | 1.32E-01 | 2.68E+00 | 2.74E+00 | 9.48E-01 | no | Cacna1a | intron_variant | MODIFIER |
| 8 | 84645314 | AX-170107955 | 0 | G | A | 0.193 | 8.61E-03 | 1.32E-01 | 2.68E+00 | 2.74E+00 | 9.48E-01 | no | | regulatory_region_variant | MODIFIER |
| 8 | 84697469 | AX-170267921 | 0 | T | C | 0.193 | 8.61E-03 | 1.32E-01 | 2.68E+00 | 2.74E+00 | 9.48E-01 | no | Trmt1 | synonymous_variant | LOW |

| | | | | | | | | | | | | | | | |
|---|---|---|---|---|---|---|---|---|---|---|---|---|---|---|---|
| 8 | 84725671 | AX-170439679 | 0 | G | A | 0.193 | 8.61E-03 | 1.32E-01 | 2.68E+00 | 2.74E+00 | 9.48E-01 | no | Nfix | intron_variant | MODIFIER |
| 8 | 84733450 | AX-170137094 | 0 | T | C | 0.193 | 8.61E-03 | 1.32E-01 | 2.68E+00 | 2.74E+00 | 9.48E-01 | no | Nfix | intron_variant | MODIFIER |
| 8 | 84733890 | AX-170489556 | 0 | T | C | 0.193 | 8.61E-03 | 1.32E-01 | 2.68E+00 | 2.74E+00 | 9.48E-01 | no | Nfix | intron_variant | MODIFIER |
| 8 | 84752666 | AX-170305700 | 0 | C | T | 0.193 | 8.61E-03 | 1.32E-01 | 2.68E+00 | 2.74E+00 | 9.48E-01 | no | Nfix | intron_variant | MODIFIER |
| 8 | 84754420 | AX-170117323 | 0 | A | G | 0.193 | 8.61E-03 | 1.32E-01 | 2.68E+00 | 2.74E+00 | 9.48E-01 | no | Nfix | intron_variant | MODIFIER |
| 8 | 84757024 | AX-170240167 | 0 | A | G | 0.193 | 8.61E-03 | 1.32E-01 | 2.68E+00 | 2.74E+00 | 9.48E-01 | no | Nfix | intron_variant | MODIFIER |
| 8 | 84766556 | AX-170304867 | 0 | C | T | 0.193 | 8.61E-03 | 1.32E-01 | 2.68E+00 | 2.74E+00 | 9.48E-01 | no | Nfix | intron_variant | MODIFIER |
| 8 | 84792871 | AX-170213020 | 0 | G | A | 0.193 | 8.61E-03 | 1.32E-01 | 2.68E+00 | 2.74E+00 | 9.48E-01 | no | Nfix | intron_variant | MODIFIER |
| 8 | 84792885 | AX-170245954 | 0 | A | G | 0.193 | 8.61E-03 | 1.32E-01 | 2.68E+00 | 2.74E+00 | 9.48E-01 | no | Nfix | intron_variant | MODIFIER |
| 8 | 84809048 | AX-170189231 | 0 | C | T | 0.193 | 8.61E-03 | 1.32E-01 | 2.68E+00 | 2.74E+00 | 9.48E-01 | no | | intergenic_variant | MODIFIER |
| 8 | 84845911 | AX-170435500 | 0 | A | G | 0.193 | 8.61E-03 | 1.32E-01 | 2.68E+00 | 2.74E+00 | 9.48E-01 | no | Calr | synonymous_variant | LOW |
| 8 | 84846112 | AX-170090165 | 0 | A | T | 0.193 | 8.61E-03 | 1.32E-01 | 2.68E+00 | 2.74E+00 | 9.48E-01 | no | Calr | intron_variant | MODIFIER |
| 8 | 84854427 | AX-170125364 | 0 | T | C | 0.193 | 8.61E-03 | 1.32E-01 | 2.68E+00 | 2.74E+00 | 9.48E-01 | no | Farsa | upstream_gene_variant | MODIFIER |
| 8 | 84881943 | AX-170344903 | 0 | C | T | 0.193 | 8.61E-03 | 1.32E-01 | 2.68E+00 | 2.74E+00 | 9.48E-01 | no | Gcdh | downstream_gene_variant | MODIFIER |
| 8 | 84887651 | AX-170528550 | 0 | C | T | 0.193 | 8.61E-03 | 1.32E-01 | 2.68E+00 | 2.74E+00 | 9.48E-01 | no | Gcdh | intron_variant | MODIFIER |
| 8 | 84919655 | AX-170149961 | 0 | C | T | 0.193 | 8.61E-03 | 1.32E-01 | 2.68E+00 | 2.74E+00 | 9.48E-01 | no | Mast1 | intron_variant | MODIFIER |
| 8 | 84947384 | AX-170422885 | 0 | T | C | 0.193 | 8.61E-03 | 1.32E-01 | 2.68E+00 | 2.74E+00 | 9.48E-01 | no | Rtbdn | intron_variant | MODIFIER |
| 8 | 84957636 | AX-170529744 | 0 | C | T | 0.193 | 8.61E-03 | 1.32E-01 | 2.68E+00 | 2.74E+00 | 9.48E-01 | no | Rnaseh2a | downstream_gene_variant | MODIFIER |
| 8 | 84974892 | AX-170535778 | 0 | C | T | 0.193 | 8.61E-03 | 1.32E-01 | 2.68E+00 | 2.74E+00 | 9.48E-01 | no | Prdx2 | downstream_gene_variant | MODIFIER |
| 8 | 85014874 | AX-170494169 | 0 | T | C | 0.193 | 8.61E-03 | 1.32E-01 | 2.68E+00 | 2.74E+00 | 9.48E-01 | no | Best2 | upstream_gene_variant | MODIFIER |
| 8 | 85054103 | AX-170311046 | 0 | A | T | 0.193 | 8.61E-03 | 1.32E-01 | 2.68E+00 | 2.74E+00 | 9.48E-01 | no | Tnpo2 | intron_variant | MODIFIER |
| 8 | 85055668 | AX-170376870 | 0 | G | C | 0.193 | 8.61E-03 | 1.32E-01 | 2.68E+00 | 2.74E+00 | 9.48E-01 | no | Tnpo2 | 3_prime_UTR_variant | MODIFIER |
| 8 | 85109422 | AX-170274059 | 0 | C | G | 0.193 | 8.61E-03 | 1.32E-01 | 2.68E+00 | 2.74E+00 | 9.48E-01 | no | Zfp791 | downstream_gene_variant | MODIFIER |
| 8 | 85135859 | AX-170480684 | 0 | G | A | 0.193 | 8.61E-03 | 1.32E-01 | 2.68E+00 | 2.74E+00 | 9.48E-01 | no | Gm42031 | upstream_gene_variant | MODIFIER |
| 8 | 85144364 | AX-170369368 | 0 | A | T | 0.193 | 8.61E-03 | 1.32E-01 | 2.68E+00 | 2.74E+00 | 9.48E-01 | no | | intergenic_variant | MODIFIER |
| 8 | 85157579 | AX-170410733 | 0 | A | G | 0.193 | 8.61E-03 | 1.32E-01 | 2.68E+00 | 2.74E+00 | 9.48E-01 | no | Cks1brt | intron_variant | MODIFIER |
| 8 | 85173649 | AX-170467076 | 0 | T | C | 0.193 | 8.61E-03 | 1.32E-01 | 2.68E+00 | 2.74E+00 | 9.48E-01 | no | Cks1brt | downstream_gene_variant | MODIFIER |
| 8 | 85223572 | AX-170201319 | 0 | T | C | 0.193 | 8.61E-03 | 1.32E-01 | 2.68E+00 | 2.74E+00 | 9.48E-01 | no | Olfr371 | upstream_gene_variant | MODIFIER |
| 8 | 85227397 | AX-170385158 | 0 | A | C | 0.193 | 8.61E-03 | 1.32E-01 | 2.68E+00 | 2.74E+00 | 9.48E-01 | no | Olfr371 | upstream_gene_variant | MODIFIER |
| 8 | 85284813 | AX-170509682 | 0 | T | G | 0.193 | 8.61E-03 | 1.32E-01 | 2.68E+00 | 2.74E+00 | 9.48E-01 | no | Vps35 | intron_variant | MODIFIER |
| 8 | 85328776 | AX-170158596 | 0 | G | A | 0.193 | 8.61E-03 | 1.32E-01 | 2.68E+00 | 2.74E+00 | 9.48E-01 | no | Mylk3 | intron_variant | MODIFIER |
| 8 | 85339622 | AX-97828116 | 0 | G | T | 0.193 | 8.61E-03 | 1.32E-01 | 2.68E+00 | 2.74E+00 | 9.48E-01 | no | Mylk3 | intron_variant | MODIFIER |
| 8 | 85360465 | AX-170487690 | 0 | G | A | 0.193 | 8.61E-03 | 1.32E-01 | 2.68E+00 | 2.74E+00 | 9.48E-01 | no | Mylk3 | intron_variant | MODIFIER |
| 8 | 85385745 | AX-170080638 | 0 | A | C | 0.193 | 8.61E-03 | 1.32E-01 | 2.68E+00 | 2.74E+00 | 9.48E-01 | no | Mylk3 | upstream_gene_variant | MODIFIER |
| 9 | 52473220 | AX-97804038 | 0 | G | A | 0.228 | -6.85E-03 | 1.10E-01 | 2.66E+00 | 2.74E+00 | 9.48E-01 | no | AC166902.1 | downstream_gene_variant | MODIFIER |
| 9 | 52573931 | AX-170258273 | 0 | G | A | 0.228 | -6.85E-03 | 1.10E-01 | 2.66E+00 | 2.74E+00 | 9.48E-01 | no | | intergenic_variant | MODIFIER |
| 12 | 114972852 | AX-170092620 | 0 | G | A | 0.149 | 1.13E-02 | 1.62E-01 | 2.63E+00 | 2.73E+00 | 9.48E-01 | no | Ighv1-45 | upstream_gene_variant | MODIFIER |
| 10 | 3908340 | AX-97814639 | 1 | G | C | 0.205 | 5.79E-03 | 9.47E-02 | 2.63E+00 | 2.74E+00 | 9.48E-01 | no | Plekhg1 | intron_variant | MODIFIER |
| 19 | 27672674 | AX-97835627 | 0 | C | A | 0.053 | 1.07E-02 | 1.58E-01 | 2.60E+00 | 2.73E+00 | 9.48E-01 | no | | intergenic_variant | MODIFIER |
| 7 | 81161677 | AX-170382467 | 1 | A | G | 0.098 | 1.43E-02 | 2.12E-01 | 2.63E+00 | 2.73E+00 | 9.48E-01 | no | Slc28a1 | intron_variant | MODIFIER |
| 1 | 176703609 | AX-170341742 | 0 | A | G | 0.219 | 8.55E-03 | 1.14E-01 | 2.64E+00 | 2.72E+00 | 9.48E-01 | no | Gm26104 | downstream_gene_variant | MODIFIER |
| 5 | 118759589 | AX-170073277 | 1 | T | C | 0.464 | 7.66E-03 | 1.11E-01 | 2.68E+00 | 2.73E+00 | 9.49E-01 | no | Med13l | intron_variant | MODIFIER |
| 5 | 122603003 | AX-97836504 | 1 | A | G | 0.464 | 7.66E-03 | 1.11E-01 | 2.68E+00 | 2.73E+00 | 9.49E-01 | no | Ift81 | intron_variant | MODIFIER |
| 5 | 125122389 | AX-170348716 | 1 | T | C | 0.464 | 7.66E-03 | 1.11E-01 | 2.68E+00 | 2.73E+00 | 9.49E-01 | no | Ncor2 | intron_variant | MODIFIER |
| 13 | 37982058 | AX-170486032 | 1 | T | A | 0.179 | -8.69E-03 | 1.16E-01 | 2.65E+00 | 2.72E+00 | 9.49E-01 | no | Ssr1 | intron_variant | MODIFIER |
| 6 | 138736882 | AX-97786025 | 1 | G | A | 0.071 | 1.75E-02 | 2.78E-01 | 2.70E+00 | 2.74E+00 | 9.49E-01 | no | | intergenic_variant | MODIFIER |
| 13 | 41129934 | AX-170152173 | 0 | C | T | 0.518 | 4.60E-03 | 6.01E-02 | 2.61E+00 | 2.73E+00 | 9.49E-01 | no | Sycp2l | intron_variant | MODIFIER |
| 8 | 85224311 | AX-170031162 | 1 | A | C | 0.188 | -8.05E-03 | 1.29E-01 | 2.67E+00 | 2.74E+00 | 9.49E-01 | no | Olfr371 | upstream_gene_variant | MODIFIER |
| 9 | 60097651 | AX-170345611 | 1 | G | A | 0.223 | 7.65E-03 | 1.21E-01 | 2.68E+00 | 2.74E+00 | 9.49E-01 | no | Thsd4 | intron_variant | MODIFIER |
| 8 | 78900840 | AX-170423969 | 2 | T | C | 0.209 | 7.89E-03 | 1.26E-01 | 2.67E+00 | 2.74E+00 | 9.49E-01 | no | | intergenic_variant | MODIFIER |
| 7 | 103338643 | AX-170361315 | 1 | A | T | 0.455 | -5.20E-03 | 9.42E-02 | 2.69E+00 | 2.75E+00 | 9.49E-01 | no | Olfr599 | synonymous_variant | LOW |
| 9 | 52096048 | AX-170096547 | 1 | C | T | 0.214 | -7.21E-03 | 1.17E-01 | 2.68E+00 | 2.74E+00 | 9.49E-01 | no | | regulatory_region_variant | MODIFIER |
| 5 | 34920746 | AX-170442865 | 0 | G | A | 0.333 | 6.94E-03 | 1.08E-01 | 2.68E+00 | 2.74E+00 | 9.49E-01 | no | Msantd1 | intron_variant | MODIFIER |
| 5 | 121963059 | AX-170266239 | 1 | T | C | 0.446 | 7.61E-03 | 1.15E-01 | 2.69E+00 | 2.73E+00 | 9.49E-01 | no | Cux2 | intron_variant | MODIFIER |

| | | | | | | | | | | | | | | | |
|---|---|---|---|---|---|---|---|---|---|---|---|---|---|---|---|
| 7 | 51887792 | AX-170370896 | 1 | T | C | 0.438 | 7.12E-03 | 1.01E-01 | 2.65E+00 | 2.73E+00 | 9.49E-01 | no | Gas2 | intron_variant | MODIFIER |
| 7 | 56152017 | AX-170029062 | 1 | G | A | 0.438 | 7.12E-03 | 1.01E-01 | 2.65E+00 | 2.73E+00 | 9.49E-01 | no | Herc2 | intron_variant | MODIFIER |
| 12 | 113847963 | AX-170221441 | 0 | G | C | 0.14 | -1.21E-02 | 1.92E-01 | 2.67E+00 | 2.74E+00 | 9.49E-01 | no | | intergenic_variant | MODIFIER |
| 7 | 93282483 | AX-97824966 | 0 | T | G | 0.123 | -1.25E-02 | 2.04E-01 | 2.66E+00 | 2.74E+00 | 9.50E-01 | no | | intergenic_variant | MODIFIER |
| 13 | 53282183 | AX-170008327 | 1 | C | T | 0.509 | 5.08E-03 | 6.53E-02 | 2.61E+00 | 2.72E+00 | 9.50E-01 | no | Ror2 | intron_variant | MODIFIER |
| 11 | 8889145 | AX-170400588 | 1 | G | A | 0.062 | 1.24E-02 | 2.16E-01 | 2.65E+00 | 2.74E+00 | 9.50E-01 | no | Gm11991 | upstream_gene_variant | MODIFIER |
| 1 | 28802659 | AX-170512775 | 1 | A | C | 0.393 | 5.10E-03 | 8.10E-02 | 2.65E+00 | 2.74E+00 | 9.50E-01 | no | Gm5697 | non_coding_transcript_exon_variant | MODIFIER |
| 1 | 99853097 | AX-170411007 | 0 | C | G | 0.14 | 9.67E-03 | 1.48E-01 | 2.62E+00 | 2.74E+00 | 9.50E-01 | no | Cntnap5b | intron_variant | MODIFIER |
| 13 | 47055172 | AX-170425388 | 0 | C | A | 0.474 | 4.51E-03 | 6.42E-02 | 2.63E+00 | 2.73E+00 | 9.50E-01 | no | Kdm1b | intron_variant | MODIFIER |
| 1 | 100395488 | AX-170472286 | 0 | A | C | 0.158 | -9.84E-03 | 1.76E-01 | 2.65E+00 | 2.75E+00 | 9.50E-01 | no | Cntnap5b | intron_variant | MODIFIER |
| 8 | 101414513 | AX-97813359 | 0 | T | G | 0.219 | 7.18E-03 | 1.10E-01 | 2.65E+00 | 2.73E+00 | 9.50E-01 | no | | intergenic_variant | MODIFIER |
| 12 | 111599751 | AX-170400556 | 1 | C | T | 0.143 | -1.15E-02 | 1.92E-01 | 2.67E+00 | 2.74E+00 | 9.50E-01 | no | Mark3 | intron_variant | MODIFIER |
| 16 | 19534110 | AX-170056836 | 0 | T | A | 0.132 | 5.10E-03 | 8.81E-02 | 2.60E+00 | 2.74E+00 | 9.50E-01 | no | Olfr168 | upstream_gene_variant | MODIFIER |
| 9 | 80763239 | AX-170169805 | 0 | T | C | 0.202 | 7.13E-03 | 1.29E-01 | 2.70E+00 | 2.75E+00 | 9.50E-01 | no | | intergenic_variant | MODIFIER |
| 8 | 90651545 | AX-170252362 | 0 | C | T | 0.211 | 6.02E-03 | 1.09E-01 | 2.64E+00 | 2.75E+00 | 9.50E-01 | no | | intergenic_variant | MODIFIER |
| 7 | 118149041 | AX-170420462 | 1 | A | G | 0.393 | -4.80E-03 | 8.79E-02 | 2.67E+00 | 2.75E+00 | 9.50E-01 | no | Smg1 | intron_variant | MODIFIER |
| 12 | 115453066 | AX-170108520 | 1 | T | C | 0.125 | 1.26E-02 | 1.96E-01 | 2.66E+00 | 2.73E+00 | 9.50E-01 | no | | intergenic_variant | MODIFIER |
| 13 | 54978476 | AX-170008347 | 1 | T | A | 0.518 | 4.96E-03 | 6.42E-02 | 2.61E+00 | 2.72E+00 | 9.50E-01 | no | Unc5a | intron_variant | MODIFIER |
| 9 | 85685772 | AX-170460780 | 1 | A | G | 0.152 | 6.58E-03 | 1.28E-01 | 2.68E+00 | 2.76E+00 | 9.50E-01 | no | Ibtk | downstream_gene_variant | MODIFIER |
| 3 | 36906597 | AX-170310755 | 1 | G | T | 0.188 | -5.53E-03 | 8.85E-02 | 2.63E+00 | 2.74E+00 | 9.51E-01 | no | 4932438A13Rik | intron_variant | MODIFIER |
| 12 | 117672823 | AX-170517041 | 1 | C | T | 0.107 | -1.18E-02 | 1.63E-01 | 2.62E+00 | 2.73E+00 | 9.51E-01 | no | Rapgef5 | intron_variant | MODIFIER |
| 8 | 81120345 | AX-170399083 | 0 | C | T | 0.202 | -8.04E-03 | 1.30E-01 | 2.67E+00 | 2.74E+00 | 9.51E-01 | no | | intergenic_variant | MODIFIER |
| 9 | 33087801 | AX-170313071 | 0 | A | G | 0.43 | 6.00E-03 | 8.60E-02 | 2.66E+00 | 2.73E+00 | 9.51E-01 | no | Gm27166 | intron_variant,non_coding_transcript_variant | MODIFIER |
| 9 | 86868147 | AX-170463261 | 0 | T | G | 0.14 | -8.31E-03 | 1.24E-01 | 2.64E+00 | 2.73E+00 | 9.51E-01 | no | Snap91 | intron_variant | MODIFIER |
| 3 | 147425366 | AX-170390957 | 1 | T | G | 0.33 | 4.87E-03 | 7.60E-02 | 2.65E+00 | 2.73E+00 | 9.51E-01 | no | Gm37041 | upstream_gene_variant | MODIFIER |
| 16 | 92682450 | AX-170037540 | 0 | A | G | 0.474 | 4.56E-03 | 7.73E-02 | 2.65E+00 | 2.74E+00 | 9.51E-01 | no | Runx1 | intron_variant | MODIFIER |
| 12 | 64742309 | AX-168374612 | 1 | G | A | 0.062 | -1.88E-02 | 3.01E-01 | 2.71E+00 | 2.73E+00 | 9.51E-01 | no | | intergenic_variant | MODIFIER |
| 18 | 43259047 | AX-170112861 | 1 | C | T | 0.062 | 1.78E-02 | 2.99E-01 | 2.72E+00 | 2.75E+00 | 9.51E-01 | no | Stk32a | intron_variant | MODIFIER |
| 7 | 94249714 | AX-170029543 | 0 | T | G | 0.123 | 1.03E-02 | 1.78E-01 | 2.64E+00 | 2.74E+00 | 9.51E-01 | no | Gm32647 | intron_variant,non_coding_transcript_variant | MODIFIER |
| 9 | 29622474 | AX-170221456 | 0 | C | T | 0.447 | -4.30E-03 | 8.37E-02 | 2.67E+00 | 2.75E+00 | 9.51E-01 | no | Ntm | intron_variant | MODIFIER |
| 1 | 39335968 | AX-170476003 | 1 | G | A | 0.402 | 3.99E-03 | 7.53E-02 | 2.67E+00 | 2.75E+00 | 9.51E-01 | no | Npas2 | intron_variant | MODIFIER |
| 4 | 144138479 | AX-97831700 | 0 | G | C | 0.105 | -1.20E-02 | 2.04E-01 | 2.68E+00 | 2.74E+00 | 9.51E-01 | no | | intergenic_variant | MODIFIER |
| 7 | 70569406 | AX-170113410 | 0 | A | C | 0.386 | 6.51E-03 | 1.03E-01 | 2.65E+00 | 2.74E+00 | 9.51E-01 | no | Gm44813 | upstream_gene_variant | MODIFIER |
| 15 | 87175819 | AX-170139791 | 0 | C | T | 0.14 | 1.06E-02 | 1.46E-01 | 2.64E+00 | 2.72E+00 | 9.51E-01 | no | Gm49470 | intron_variant,non_coding_transcript_variant | MODIFIER |
| 6 | 140008134 | AX-97839241 | 0 | A | T | 0.07 | -1.68E-02 | 2.77E-01 | 2.70E+00 | 2.74E+00 | 9.51E-01 | no | Plcz1 | intron_variant | MODIFIER |
| 13 | 46958575 | AX-170419188 | 0 | G | T | 0.465 | -3.31E-03 | 6.38E-02 | 2.64E+00 | 2.75E+00 | 9.51E-01 | no | | regulatory_region_variant | MODIFIER |
| 6 | 145045598 | AX-97791461 | 0 | A | G | 0.07 | -1.68E-02 | 2.77E-01 | 2.70E+00 | 2.74E+00 | 9.51E-01 | no | Bcat1 | intron_variant | MODIFIER |
| 11 | 13519231 | AX-170356893 | 1 | G | A | 0.054 | -1.43E-02 | 2.38E-01 | 2.67E+00 | 2.74E+00 | 9.51E-01 | no | | intergenic_variant | MODIFIER |
| 11 | 24488303 | AX-170210772 | 1 | A | T | 0.054 | -1.43E-02 | 2.38E-01 | 2.67E+00 | 2.74E+00 | 9.51E-01 | no | 4933430M04Rik | intron_variant,non_coding_transcript_variant | MODIFIER |
| 9 | 67621606 | AX-170267571 | 1 | T | A | 0.196 | 7.93E-03 | 1.35E-01 | 2.71E+00 | 2.74E+00 | 9.51E-01 | no | | regulatory_region_variant | MODIFIER |
| 9 | 68536453 | AX-170380670 | 1 | C | T | 0.196 | 7.93E-03 | 1.35E-01 | 2.71E+00 | 2.74E+00 | 9.51E-01 | no | | intergenic_variant | MODIFIER |
| 9 | 70312893 | AX-170533279 | 1 | G | A | 0.196 | 7.93E-03 | 1.35E-01 | 2.71E+00 | 2.74E+00 | 9.51E-01 | no | Myo1e | intron_variant | MODIFIER |
| 9 | 74916015 | AX-170437093 | 1 | C | T | 0.196 | 7.93E-03 | 1.35E-01 | 2.71E+00 | 2.74E+00 | 9.51E-01 | no | | regulatory_region_variant | MODIFIER |
| 2 | 137168787 | AX-170019152 | 0 | T | G | 0.14 | 1.37E-02 | 2.23E-01 | 2.71E+00 | 2.73E+00 | 9.51E-01 | no | | intergenic_variant | MODIFIER |
| 1 | 116356241 | AX-170229985 | 2 | A | G | 0.155 | 1.38E-02 | 2.02E-01 | 2.65E+00 | 2.73E+00 | 9.51E-01 | no | Cntnap5a | intron_variant | MODIFIER |
| 1 | 13779975 | AX-170364456 | 0 | T | C | 0.368 | 5.07E-03 | 7.50E-02 | 2.66E+00 | 2.73E+00 | 9.51E-01 | no | | intergenic_variant | MODIFIER |
| 11 | 5607639 | AX-170388539 | 2 | T | C | 0.055 | -1.41E-02 | 2.37E-01 | 2.67E+00 | 2.74E+00 | 9.52E-01 | no | Ankrd36 | intron_variant | MODIFIER |
| 1 | 8083777 | AX-170235725 | 0 | A | G | 0.088 | -9.43E-03 | 1.83E-01 | 2.63E+00 | 2.75E+00 | 9.52E-01 | no | | intergenic_variant | MODIFIER |
| 5 | 119998584 | AX-97822165 | 1 | T | C | 0.446 | -6.36E-03 | 1.13E-01 | 2.71E+00 | 2.75E+00 | 9.52E-01 | no | | intergenic_variant | MODIFIER |
| 9 | 66178654 | AX-170082679 | 1 | G | A | 0.196 | 8.18E-03 | 1.39E-01 | 2.72E+00 | 2.74E+00 | 9.52E-01 | no | Dapk2 | intron_variant | MODIFIER |
| 9 | 69922158 | AX-97848936 | 1 | G | T | 0.196 | 8.18E-03 | 1.39E-01 | 2.72E+00 | 2.74E+00 | 9.52E-01 | no | | intergenic_variant | MODIFIER |
| 9 | 78122586 | AX-170361636 | 1 | T | C | 0.196 | 8.18E-03 | 1.39E-01 | 2.72E+00 | 2.74E+00 | 9.52E-01 | no | Ick | intron_variant | MODIFIER |
| 7 | 79891124 | AX-170122240 | 1 | A | G | 0.107 | 1.23E-02 | 1.99E-01 | 2.66E+00 | 2.73E+00 | 9.52E-01 | no | Ap3s2 | intron_variant | MODIFIER |
| 1 | 69076295 | AX-170289268 | 0 | A | T | 0.14 | 1.42E-02 | 2.36E-01 | 2.68E+00 | 2.74E+00 | 9.52E-01 | no | Erbb4 | intron_variant | MODIFIER |

| | | | | | | | | | | | | | | | |
|---|---|---|---|---|---|---|---|---|---|---|---|---|---|---|---|
| 13 | 45279653 | AX-170162229 | 1 | T | C | 0.473 | -3.05E-03 | 6.18E-02 | 2.64E+00 | 2.75E+00 | 9.52E-01 | no | Gm47448 | intron_variant,non_coding_transcript_variant | MODIFIER |
| 7 | 95004442 | AX-170509727 | 1 | T | G | 0.134 | -1.30E-02 | 2.22E-01 | 2.67E+00 | 2.74E+00 | 9.52E-01 | no | | intergenic_variant | MODIFIER |
| 9 | 112978326 | AX-170457015 | 1 | A | G | 0.125 | -6.88E-03 | 1.07E-01 | 2.65E+00 | 2.73E+00 | 9.52E-01 | no | | intergenic_variant | MODIFIER |
| 5 | 88006694 | AX-97791986 | 0 | G | A | 0.43 | 6.36E-03 | 1.10E-01 | 2.69E+00 | 2.74E+00 | 9.52E-01 | no | Smr3a | intron_variant | MODIFIER |
| 18 | 60822951 | AX-170329988 | 1 | C | T | 0.42 | 3.93E-03 | 8.20E-02 | 2.68E+00 | 2.76E+00 | 9.52E-01 | no | Tcof1 | missense_variant,NMD_transcript_variant | MODERATE |
| 1 | 53866101 | AX-170155557 | 2 | G | T | 0.091 | 1.19E-02 | 1.95E-01 | 2.65E+00 | 2.74E+00 | 9.52E-01 | no | Hecw2 | intron_variant | MODIFIER |
| 9 | 61135119 | AX-170131487 | 0 | A | G | 0.193 | 8.03E-03 | 1.38E-01 | 2.72E+00 | 2.74E+00 | 9.52E-01 | no | I730028E13Rik | upstream_gene_variant | MODIFIER |
| 9 | 61154651 | AX-170177370 | 0 | C | T | 0.193 | 8.03E-03 | 1.38E-01 | 2.72E+00 | 2.74E+00 | 9.52E-01 | no | | intergenic_variant | MODIFIER |
| 9 | 61158585 | AX-170405739 | 0 | G | A | 0.193 | 8.03E-03 | 1.38E-01 | 2.72E+00 | 2.74E+00 | 9.52E-01 | no | | regulatory_region_variant | MODIFIER |
| 9 | 61158965 | AX-170366280 | 0 | G | A | 0.193 | 8.03E-03 | 1.38E-01 | 2.72E+00 | 2.74E+00 | 9.52E-01 | no | | regulatory_region_variant | MODIFIER |
| 9 | 61172885 | AX-170082592 | 0 | G | A | 0.193 | 8.03E-03 | 1.38E-01 | 2.72E+00 | 2.74E+00 | 9.52E-01 | no | | intergenic_variant | MODIFIER |
| 9 | 61193908 | AX-170439954 | 0 | T | C | 0.193 | 8.03E-03 | 1.38E-01 | 2.72E+00 | 2.74E+00 | 9.52E-01 | no | | intergenic_variant | MODIFIER |
| 9 | 61198855 | AX-170482539 | 0 | C | A | 0.193 | 8.03E-03 | 1.38E-01 | 2.72E+00 | 2.74E+00 | 9.52E-01 | no | | intergenic_variant | MODIFIER |
| 9 | 61212686 | AX-170472344 | 0 | G | A | 0.193 | 8.03E-03 | 1.38E-01 | 2.72E+00 | 2.74E+00 | 9.52E-01 | no | Gm34105 | intron_variant,non_coding_transcript_variant | MODIFIER |
| 9 | 61213723 | AX-170487974 | 0 | C | A | 0.193 | 8.03E-03 | 1.38E-01 | 2.72E+00 | 2.74E+00 | 9.52E-01 | no | Gm34105 | downstream_gene_variant | MODIFIER |
| 9 | 61214919 | AX-170501009 | 0 | C | T | 0.193 | 8.03E-03 | 1.38E-01 | 2.72E+00 | 2.74E+00 | 9.52E-01 | no | Gm34105 | downstream_gene_variant | MODIFIER |
| 9 | 61242192 | AX-170190448 | 0 | A | G | 0.193 | 8.03E-03 | 1.38E-01 | 2.72E+00 | 2.74E+00 | 9.52E-01 | no | | regulatory_region_variant | MODIFIER |
| 9 | 61286910 | AX-170114418 | 0 | A | G | 0.193 | 8.03E-03 | 1.38E-01 | 2.72E+00 | 2.74E+00 | 9.52E-01 | no | B930092H01Rik | intron_variant,non_coding_transcript_variant | MODIFIER |
| 9 | 61477160 | AX-170082594 | 0 | C | T | 0.193 | 8.03E-03 | 1.38E-01 | 2.72E+00 | 2.74E+00 | 9.52E-01 | no | Gm47240 | upstream_gene_variant | MODIFIER |
| 9 | 61477359 | AX-170082596 | 0 | C | T | 0.193 | 8.03E-03 | 1.38E-01 | 2.72E+00 | 2.74E+00 | 9.52E-01 | no | Gm47240 | upstream_gene_variant | MODIFIER |
| 9 | 61477928 | AX-170293456 | 0 | A | G | 0.193 | 8.03E-03 | 1.38E-01 | 2.72E+00 | 2.74E+00 | 9.52E-01 | no | Gm47240 | upstream_gene_variant | MODIFIER |
| 9 | 61478006 | AX-170352298 | 0 | C | T | 0.193 | 8.03E-03 | 1.38E-01 | 2.72E+00 | 2.74E+00 | 9.52E-01 | no | | intergenic_variant | MODIFIER |
| 9 | 61479942 | AX-170178896 | 0 | T | C | 0.193 | 8.03E-03 | 1.38E-01 | 2.72E+00 | 2.74E+00 | 9.52E-01 | no | | regulatory_region_variant | MODIFIER |
| 9 | 61486260 | AX-170246334 | 0 | T | G | 0.193 | 8.03E-03 | 1.38E-01 | 2.72E+00 | 2.74E+00 | 9.52E-01 | no | | intergenic_variant | MODIFIER |
| 9 | 61504960 | AX-170423962 | 0 | T | C | 0.193 | 8.03E-03 | 1.38E-01 | 2.72E+00 | 2.74E+00 | 9.52E-01 | no | Gm34424 | upstream_gene_variant | MODIFIER |
| 9 | 61509757 | AX-97805166 | 0 | T | C | 0.193 | 8.03E-03 | 1.38E-01 | 2.72E+00 | 2.74E+00 | 9.52E-01 | no | Gm34424 | non_coding_transcript_exon_variant | MODIFIER |
| 9 | 61509816 | AX-170511564 | 0 | T | C | 0.193 | 8.03E-03 | 1.38E-01 | 2.72E+00 | 2.74E+00 | 9.52E-01 | no | Gm34424 | non_coding_transcript_exon_variant | MODIFIER |
| 9 | 61510489 | AX-170032494 | 0 | A | G | 0.193 | 8.03E-03 | 1.38E-01 | 2.72E+00 | 2.74E+00 | 9.52E-01 | no | Gm34424 | intron_variant,non_coding_transcript_variant | MODIFIER |
| 9 | 61544337 | AX-170096559 | 0 | G | A | 0.193 | 8.03E-03 | 1.38E-01 | 2.72E+00 | 2.74E+00 | 9.52E-01 | no | | regulatory_region_variant | MODIFIER |
| 9 | 61575525 | AX-170138514 | 0 | T | C | 0.193 | 8.03E-03 | 1.38E-01 | 2.72E+00 | 2.74E+00 | 9.52E-01 | no | | intergenic_variant | MODIFIER |
| 9 | 61582149 | AX-170032500 | 0 | T | C | 0.193 | 8.03E-03 | 1.38E-01 | 2.72E+00 | 2.74E+00 | 9.52E-01 | no | | intergenic_variant | MODIFIER |
| 9 | 61582923 | AX-170339298 | 0 | G | A | 0.193 | 8.03E-03 | 1.38E-01 | 2.72E+00 | 2.74E+00 | 9.52E-01 | no | | regulatory_region_variant | MODIFIER |
| 9 | 61586398 | AX-170442406 | 0 | A | G | 0.193 | 8.03E-03 | 1.38E-01 | 2.72E+00 | 2.74E+00 | 9.52E-01 | no | | intergenic_variant | MODIFIER |
| 9 | 61586529 | AX-97836273 | 0 | A | G | 0.193 | 8.03E-03 | 1.38E-01 | 2.72E+00 | 2.74E+00 | 9.52E-01 | no | | intergenic_variant | MODIFIER |
| 9 | 61588086 | AX-170422285 | 0 | T | A | 0.193 | 8.03E-03 | 1.38E-01 | 2.72E+00 | 2.74E+00 | 9.52E-01 | no | | intergenic_variant | MODIFIER |
| 9 | 61590087 | AX-170487312 | 0 | C | T | 0.193 | 8.03E-03 | 1.38E-01 | 2.72E+00 | 2.74E+00 | 9.52E-01 | no | | intergenic_variant | MODIFIER |
| 9 | 61618782 | AX-97809100 | 0 | C | T | 0.193 | 8.03E-03 | 1.38E-01 | 2.72E+00 | 2.74E+00 | 9.52E-01 | no | | regulatory_region_variant | MODIFIER |
| 9 | 61622599 | AX-170234642 | 0 | T | C | 0.193 | 8.03E-03 | 1.38E-01 | 2.72E+00 | 2.74E+00 | 9.52E-01 | no | | regulatory_region_variant | MODIFIER |
| 9 | 61624259 | AX-170510351 | 0 | A | G | 0.193 | 8.03E-03 | 1.38E-01 | 2.72E+00 | 2.74E+00 | 9.52E-01 | no | | intergenic_variant | MODIFIER |
| 9 | 61630451 | AX-170145908 | 0 | C | T | 0.193 | 8.03E-03 | 1.38E-01 | 2.72E+00 | 2.74E+00 | 9.52E-01 | no | | intergenic_variant | MODIFIER |
| 9 | 61632855 | AX-170082598 | 0 | C | T | 0.193 | 8.03E-03 | 1.38E-01 | 2.72E+00 | 2.74E+00 | 9.52E-01 | no | | intergenic_variant | MODIFIER |
| 9 | 61668720 | AX-170286015 | 0 | A | T | 0.193 | 8.03E-03 | 1.38E-01 | 2.72E+00 | 2.74E+00 | 9.52E-01 | no | | regulatory_region_variant | MODIFIER |
| 9 | 61668732 | AX-170445363 | 0 | T | A | 0.193 | 8.03E-03 | 1.38E-01 | 2.72E+00 | 2.74E+00 | 9.52E-01 | no | | regulatory_region_variant | MODIFIER |
| 9 | 61669013 | AX-170405108 | 0 | T | C | 0.193 | 8.03E-03 | 1.38E-01 | 2.72E+00 | 2.74E+00 | 9.52E-01 | no | | regulatory_region_variant | MODIFIER |
| 9 | 61671177 | AX-97792378 | 0 | A | G | 0.193 | 8.03E-03 | 1.38E-01 | 2.72E+00 | 2.74E+00 | 9.52E-01 | no | | regulatory_region_variant | MODIFIER |
| 9 | 61673197 | AX-170484411 | 0 | C | A | 0.193 | 8.03E-03 | 1.38E-01 | 2.72E+00 | 2.74E+00 | 9.52E-01 | no | | regulatory_region_variant | MODIFIER |
| 9 | 61674745 | AX-170213902 | 0 | A | G | 0.193 | 8.03E-03 | 1.38E-01 | 2.72E+00 | 2.74E+00 | 9.52E-01 | no | | intergenic_variant | MODIFIER |
| 9 | 61676100 | AX-170379409 | 0 | C | T | 0.193 | 8.03E-03 | 1.38E-01 | 2.72E+00 | 2.74E+00 | 9.52E-01 | no | | intergenic_variant | MODIFIER |
| 9 | 61698599 | AX-97850533 | 0 | A | G | 0.193 | 8.03E-03 | 1.38E-01 | 2.72E+00 | 2.74E+00 | 9.52E-01 | no | | intergenic_variant | MODIFIER |
| 9 | 61742731 | AX-170364601 | 0 | A | G | 0.193 | 8.03E-03 | 1.38E-01 | 2.72E+00 | 2.74E+00 | 9.52E-01 | no | | intergenic_variant | MODIFIER |
| 9 | 61743177 | AX-170186824 | 0 | C | T | 0.193 | 8.03E-03 | 1.38E-01 | 2.72E+00 | 2.74E+00 | 9.52E-01 | no | | intergenic_variant | MODIFIER |
| 9 | 61743494 | AX-170323806 | 0 | C | T | 0.193 | 8.03E-03 | 1.38E-01 | 2.72E+00 | 2.74E+00 | 9.52E-01 | no | | intergenic_variant | MODIFIER |
| 9 | 61745894 | AX-170151772 | 0 | T | C | 0.193 | 8.03E-03 | 1.38E-01 | 2.72E+00 | 2.74E+00 | 9.52E-01 | no | | regulatory_region_variant | MODIFIER |
| 9 | 61746160 | AX-170245362 | 0 | A | G | 0.193 | 8.03E-03 | 1.38E-01 | 2.72E+00 | 2.74E+00 | 9.52E-01 | no | | regulatory_region_variant | MODIFIER |

| | | | | | | | | | | | | | | | |
|---|---|---|---|---|---|---|---|---|---|---|---|---|---|---|---|
| 9 | 61747228 | AX-97829679 | 0 | T | C | 0.193 | 8.03E-03 | 1.38E-01 | 2.72E+00 | 2.74E+00 | 9.52E-01 | no | | intergenic_variant | MODIFIER |
| 9 | 61751447 | AX-170176507 | 0 | C | T | 0.193 | 8.03E-03 | 1.38E-01 | 2.72E+00 | 2.74E+00 | 9.52E-01 | no | | intergenic_variant | MODIFIER |
| 9 | 61752549 | AX-170129306 | 0 | C | T | 0.193 | 8.03E-03 | 1.38E-01 | 2.72E+00 | 2.74E+00 | 9.52E-01 | no | | intergenic_variant | MODIFIER |
| 9 | 61763108 | AX-170082602 | 0 | G | A | 0.193 | 8.03E-03 | 1.38E-01 | 2.72E+00 | 2.74E+00 | 9.52E-01 | no | Gm47241 | non_coding_transcript_exon_variant | MODIFIER |
| 9 | 61785637 | AX-170439443 | 0 | A | C | 0.193 | 8.03E-03 | 1.38E-01 | 2.72E+00 | 2.74E+00 | 9.52E-01 | no | | intergenic_variant | MODIFIER |
| 9 | 61788280 | AX-170340078 | 0 | T | C | 0.193 | 8.03E-03 | 1.38E-01 | 2.72E+00 | 2.74E+00 | 9.52E-01 | no | | intergenic_variant | MODIFIER |
| 9 | 61794483 | AX-170454203 | 0 | G | C | 0.193 | 8.03E-03 | 1.38E-01 | 2.72E+00 | 2.74E+00 | 9.52E-01 | no | | intergenic_variant | MODIFIER |
| 9 | 61808789 | AX-170354296 | 0 | G | C | 0.193 | 8.03E-03 | 1.38E-01 | 2.72E+00 | 2.74E+00 | 9.52E-01 | no | | regulatory_region_variant | MODIFIER |
| 9 | 61809212 | AX-170152703 | 0 | G | A | 0.193 | 8.03E-03 | 1.38E-01 | 2.72E+00 | 2.74E+00 | 9.52E-01 | no | | regulatory_region_variant | MODIFIER |
| 9 | 61810410 | AX-170535782 | 0 | G | A | 0.193 | 8.03E-03 | 1.38E-01 | 2.72E+00 | 2.74E+00 | 9.52E-01 | no | | intergenic_variant | MODIFIER |
| 9 | 61821409 | AX-97833368 | 0 | T | C | 0.193 | 8.03E-03 | 1.38E-01 | 2.72E+00 | 2.74E+00 | 9.52E-01 | no | | intergenic_variant | MODIFIER |
| 9 | 61822052 | AX-170114902 | 0 | G | A | 0.193 | 8.03E-03 | 1.38E-01 | 2.72E+00 | 2.74E+00 | 9.52E-01 | no | | intergenic_variant | MODIFIER |
| 9 | 61824118 | AX-170148565 | 0 | A | G | 0.193 | 8.03E-03 | 1.38E-01 | 2.72E+00 | 2.74E+00 | 9.52E-01 | no | | intergenic_variant | MODIFIER |
| 9 | 61824791 | AX-170274456 | 0 | C | T | 0.193 | 8.03E-03 | 1.38E-01 | 2.72E+00 | 2.74E+00 | 9.52E-01 | no | | intergenic_variant | MODIFIER |
| 9 | 61855196 | AX-170335247 | 0 | A | T | 0.193 | 8.03E-03 | 1.38E-01 | 2.72E+00 | 2.74E+00 | 9.52E-01 | no | Gm19208 | downstream_gene_variant | MODIFIER |
| 9 | 61855859 | AX-170408144 | 0 | C | T | 0.193 | 8.03E-03 | 1.38E-01 | 2.72E+00 | 2.74E+00 | 9.52E-01 | no | Gm19208 | downstream_gene_variant | MODIFIER |
| 9 | 61858432 | AX-170237554 | 0 | G | A | 0.193 | 8.03E-03 | 1.38E-01 | 2.72E+00 | 2.74E+00 | 9.52E-01 | no | | intergenic_variant | MODIFIER |
| 9 | 61872834 | AX-97815466 | 0 | A | G | 0.193 | 8.03E-03 | 1.38E-01 | 2.72E+00 | 2.74E+00 | 9.52E-01 | no | | regulatory_region_variant | MODIFIER |
| 9 | 61887363 | AX-170082605 | 0 | T | G | 0.193 | 8.03E-03 | 1.38E-01 | 2.72E+00 | 2.74E+00 | 9.52E-01 | no | | intergenic_variant | MODIFIER |
| 9 | 61895507 | AX-170032504 | 0 | T | C | 0.193 | 8.03E-03 | 1.38E-01 | 2.72E+00 | 2.74E+00 | 9.52E-01 | no | | regulatory_region_variant | MODIFIER |
| 9 | 61895556 | AX-97841341 | 0 | G | A | 0.193 | 8.03E-03 | 1.38E-01 | 2.72E+00 | 2.74E+00 | 9.52E-01 | no | | regulatory_region_variant | MODIFIER |
| 9 | 61905624 | AX-170032501 | 0 | T | G | 0.193 | 8.03E-03 | 1.38E-01 | 2.72E+00 | 2.74E+00 | 9.52E-01 | no | | intergenic_variant | MODIFIER |
| 9 | 62174395 | AX-97810721 | 0 | A | G | 0.193 | 8.03E-03 | 1.38E-01 | 2.72E+00 | 2.74E+00 | 9.52E-01 | no | | intergenic_variant | MODIFIER |
| 9 | 62182405 | AX-170148505 | 0 | G | A | 0.193 | 8.03E-03 | 1.38E-01 | 2.72E+00 | 2.74E+00 | 9.52E-01 | no | | regulatory_region_variant | MODIFIER |
| 9 | 62185923 | AX-170082613 | 0 | A | G | 0.193 | 8.03E-03 | 1.38E-01 | 2.72E+00 | 2.74E+00 | 9.52E-01 | no | | intergenic_variant | MODIFIER |
| 9 | 62191605 | AX-170261246 | 0 | C | G | 0.193 | 8.03E-03 | 1.38E-01 | 2.72E+00 | 2.74E+00 | 9.52E-01 | no | | regulatory_region_variant | MODIFIER |
| 9 | 62264541 | AX-170379630 | 0 | C | A | 0.193 | 8.03E-03 | 1.38E-01 | 2.72E+00 | 2.74E+00 | 9.52E-01 | no | | regulatory_region_variant | MODIFIER |
| 9 | 62271334 | AX-170411438 | 0 | C | A | 0.193 | 8.03E-03 | 1.38E-01 | 2.72E+00 | 2.74E+00 | 9.52E-01 | no | Spesp1 | 3_prime_UTR_variant | MODIFIER |
| 9 | 62277122 | AX-170139540 | 0 | A | G | 0.193 | 8.03E-03 | 1.38E-01 | 2.72E+00 | 2.74E+00 | 9.52E-01 | no | Spesp1 | intron_variant | MODIFIER |
| 9 | 62282075 | AX-170345090 | 0 | G | C | 0.193 | 8.03E-03 | 1.38E-01 | 2.72E+00 | 2.74E+00 | 9.52E-01 | no | Spesp1 | 5_prime_UTR_variant | MODIFIER |
| 9 | 62335867 | AX-97853451 | 0 | A | G | 0.193 | 8.03E-03 | 1.38E-01 | 2.72E+00 | 2.74E+00 | 9.52E-01 | no | Gm35288 | non_coding_transcript_exon_variant | MODIFIER |
| 9 | 62338550 | AX-170126495 | 0 | C | A | 0.193 | 8.03E-03 | 1.38E-01 | 2.72E+00 | 2.74E+00 | 9.52E-01 | no | Anp32a | upstream_gene_variant | MODIFIER |
| 9 | 62338926 | AX-170032514 | 0 | C | G | 0.193 | 8.03E-03 | 1.38E-01 | 2.72E+00 | 2.74E+00 | 9.52E-01 | no | Anp32a | upstream_gene_variant | MODIFIER |
| 9 | 62372733 | AX-170527206 | 0 | T | C | 0.193 | 8.03E-03 | 1.38E-01 | 2.72E+00 | 2.74E+00 | 9.52E-01 | no | Anp32a | intron_variant | MODIFIER |
| 9 | 62382072 | AX-170405423 | 0 | A | G | 0.193 | 8.03E-03 | 1.38E-01 | 2.72E+00 | 2.74E+00 | 9.52E-01 | no | Anp32a | downstream_gene_variant | MODIFIER |
| 9 | 62411493 | AX-170032510 | 0 | A | G | 0.193 | 8.03E-03 | 1.38E-01 | 2.72E+00 | 2.74E+00 | 9.52E-01 | no | | intergenic_variant | MODIFIER |
| 9 | 62417817 | AX-170032513 | 0 | T | C | 0.193 | 8.03E-03 | 1.38E-01 | 2.72E+00 | 2.74E+00 | 9.52E-01 | no | Coro2b | downstream_gene_variant | MODIFIER |
| 9 | 62428746 | AX-170392652 | 0 | G | A | 0.193 | 8.03E-03 | 1.38E-01 | 2.72E+00 | 2.74E+00 | 9.52E-01 | no | Coro2b | intron_variant | MODIFIER |
| 9 | 62428999 | AX-170334177 | 0 | G | A | 0.193 | 8.03E-03 | 1.38E-01 | 2.72E+00 | 2.74E+00 | 9.52E-01 | no | Coro2b | synonymous_variant | LOW |
| 9 | 62432979 | AX-170459892 | 0 | T | C | 0.193 | 8.03E-03 | 1.38E-01 | 2.72E+00 | 2.74E+00 | 9.52E-01 | no | Coro2b | intron_variant | MODIFIER |
| 9 | 62498963 | AX-170096560 | 0 | T | C | 0.193 | 8.03E-03 | 1.38E-01 | 2.72E+00 | 2.74E+00 | 9.52E-01 | no | Coro2b | intron_variant | MODIFIER |
| 9 | 62505866 | AX-170343749 | 0 | A | G | 0.193 | 8.03E-03 | 1.38E-01 | 2.72E+00 | 2.74E+00 | 9.52E-01 | no | Coro2b | intron_variant | MODIFIER |
| 9 | 62507468 | AX-170313704 | 0 | G | A | 0.193 | 8.03E-03 | 1.38E-01 | 2.72E+00 | 2.74E+00 | 9.52E-01 | no | Coro2b | intron_variant | MODIFIER |
| 9 | 62514407 | AX-170171559 | 0 | C | T | 0.193 | 8.03E-03 | 1.38E-01 | 2.72E+00 | 2.74E+00 | 9.52E-01 | no | Coro2b | intron_variant | MODIFIER |
| 9 | 62521249 | AX-170353399 | 0 | C | G | 0.193 | 8.03E-03 | 1.38E-01 | 2.72E+00 | 2.74E+00 | 9.52E-01 | no | Coro2b | intron_variant | MODIFIER |
| 9 | 63916475 | AX-170527108 | 0 | G | A | 0.193 | 8.03E-03 | 1.38E-01 | 2.72E+00 | 2.74E+00 | 9.52E-01 | no | | regulatory_region_variant | MODIFIER |
| 9 | 63924848 | AX-170407685 | 0 | G | T | 0.193 | 8.03E-03 | 1.38E-01 | 2.72E+00 | 2.74E+00 | 9.52E-01 | no | Gm36033 | downstream_gene_variant | MODIFIER |
| 9 | 63926075 | AX-170361108 | 0 | T | C | 0.193 | 8.03E-03 | 1.38E-01 | 2.72E+00 | 2.74E+00 | 9.52E-01 | no | Gm36033 | downstream_gene_variant | MODIFIER |
| 9 | 63937726 | AX-170442054 | 0 | C | T | 0.193 | 8.03E-03 | 1.38E-01 | 2.72E+00 | 2.74E+00 | 9.52E-01 | no | Gm36033 | upstream_gene_variant | MODIFIER |
| 9 | 63954533 | AX-170082639 | 0 | T | C | 0.193 | 8.03E-03 | 1.38E-01 | 2.72E+00 | 2.74E+00 | 9.52E-01 | no | Smad6 | intron_variant | MODIFIER |
| 9 | 63976809 | AX-170032543 | 0 | T | C | 0.193 | 8.03E-03 | 1.38E-01 | 2.72E+00 | 2.74E+00 | 9.52E-01 | no | Smad6 | intron_variant | MODIFIER |
| 9 | 63991125 | AX-170114509 | 0 | T | C | 0.193 | 8.03E-03 | 1.38E-01 | 2.72E+00 | 2.74E+00 | 9.52E-01 | no | Smad6 | intron_variant | MODIFIER |
| 9 | 63999207 | AX-170082638 | 0 | T | C | 0.193 | 8.03E-03 | 1.38E-01 | 2.72E+00 | 2.74E+00 | 9.52E-01 | no | Smad6 | intron_variant | MODIFIER |
| 9 | 64026653 | AX-170032539 | 0 | G | A | 0.193 | 8.03E-03 | 1.38E-01 | 2.72E+00 | 2.74E+00 | 9.52E-01 | no | Smad6 | upstream_gene_variant | MODIFIER |

| | | | | | | | | | | | | | | | |
|---|---|---|---|---|---|---|---|---|---|---|---|---|---|---|---|
| 9 | 64032827 | AX-97845228 | 0 | T | C | 0.193 | 8.03E-03 | 1.38E-01 | 2.72E+00 | 2.74E+00 | 9.52E-01 | no | | intergenic_variant | MODIFIER |
| 9 | 64093866 | AX-170032538 | 0 | G | A | 0.193 | 8.03E-03 | 1.38E-01 | 2.72E+00 | 2.74E+00 | 9.52E-01 | no | | regulatory_region_variant | MODIFIER |
| 9 | 64193595 | AX-97804305 | 0 | T | C | 0.193 | 8.03E-03 | 1.38E-01 | 2.72E+00 | 2.74E+00 | 9.52E-01 | no | Map2k1 | intron_variant | MODIFIER |
| 9 | 64200704 | AX-170167266 | 0 | A | G | 0.193 | 8.03E-03 | 1.38E-01 | 2.72E+00 | 2.74E+00 | 9.52E-01 | no | Map2k1 | intron_variant | MODIFIER |
| 9 | 64203403 | AX-170494321 | 0 | C | T | 0.193 | 8.03E-03 | 1.38E-01 | 2.72E+00 | 2.74E+00 | 9.52E-01 | no | Map2k1 | intron_variant | MODIFIER |
| 9 | 64205272 | AX-170227594 | 0 | T | C | 0.193 | 8.03E-03 | 1.38E-01 | 2.72E+00 | 2.74E+00 | 9.52E-01 | no | Map2k1 | intron_variant | MODIFIER |
| 9 | 64208038 | AX-170456051 | 0 | T | C | 0.193 | 8.03E-03 | 1.38E-01 | 2.72E+00 | 2.74E+00 | 9.52E-01 | no | Map2k1 | intron_variant | MODIFIER |
| 9 | 64218595 | AX-170082645 | 0 | G | A | 0.193 | 8.03E-03 | 1.38E-01 | 2.72E+00 | 2.74E+00 | 9.52E-01 | no | Map2k1 | intron_variant | MODIFIER |
| 9 | 64220734 | AX-170082644 | 0 | G | A | 0.193 | 8.03E-03 | 1.38E-01 | 2.72E+00 | 2.74E+00 | 9.52E-01 | no | Map2k1 | intron_variant | MODIFIER |
| 9 | 64310654 | AX-170231278 | 0 | A | C | 0.193 | 8.03E-03 | 1.38E-01 | 2.72E+00 | 2.74E+00 | 9.52E-01 | no | Dis3l | intron_variant | MODIFIER |
| 9 | 64313555 | AX-97793585 | 0 | C | T | 0.193 | 8.03E-03 | 1.38E-01 | 2.72E+00 | 2.74E+00 | 9.52E-01 | no | Dis3l | intron_variant | MODIFIER |
| 9 | 64357669 | AX-170266059 | 0 | T | C | 0.193 | 8.03E-03 | 1.38E-01 | 2.72E+00 | 2.74E+00 | 9.52E-01 | no | | regulatory_region_variant | MODIFIER |
| 9 | 64376412 | AX-97786991 | 0 | C | T | 0.193 | 8.03E-03 | 1.38E-01 | 2.72E+00 | 2.74E+00 | 9.52E-01 | no | | intergenic_variant | MODIFIER |
| 9 | 64394432 | AX-170436215 | 0 | A | T | 0.193 | 8.03E-03 | 1.38E-01 | 2.72E+00 | 2.74E+00 | 9.52E-01 | no | Megf11 | intron_variant | MODIFIER |
| 9 | 64394467 | AX-97839537 | 0 | T | C | 0.193 | 8.03E-03 | 1.38E-01 | 2.72E+00 | 2.74E+00 | 9.52E-01 | no | Megf11 | intron_variant | MODIFIER |
| 9 | 64401421 | AX-170466575 | 0 | G | C | 0.193 | 8.03E-03 | 1.38E-01 | 2.72E+00 | 2.74E+00 | 9.52E-01 | no | Megf11 | intron_variant | MODIFIER |
| 9 | 64405890 | AX-170276824 | 0 | C | T | 0.193 | 8.03E-03 | 1.38E-01 | 2.72E+00 | 2.74E+00 | 9.52E-01 | no | Megf11 | intron_variant | MODIFIER |
| 9 | 64407119 | AX-170336261 | 0 | G | A | 0.193 | 8.03E-03 | 1.38E-01 | 2.72E+00 | 2.74E+00 | 9.52E-01 | no | Megf11 | intron_variant | MODIFIER |
| 9 | 64422937 | AX-97804259 | 0 | A | G | 0.193 | 8.03E-03 | 1.38E-01 | 2.72E+00 | 2.74E+00 | 9.52E-01 | no | Megf11 | intron_variant | MODIFIER |
| 9 | 64431952 | AX-170032550 | 0 | G | C | 0.193 | 8.03E-03 | 1.38E-01 | 2.72E+00 | 2.74E+00 | 9.52E-01 | no | Megf11 | intron_variant | MODIFIER |
| 9 | 64461492 | AX-97854985 | 0 | A | G | 0.193 | 8.03E-03 | 1.38E-01 | 2.72E+00 | 2.74E+00 | 9.52E-01 | no | Megf11 | intron_variant | MODIFIER |
| 9 | 64612393 | AX-170284934 | 0 | T | G | 0.193 | 8.03E-03 | 1.38E-01 | 2.72E+00 | 2.74E+00 | 9.52E-01 | no | Megf11 | intron_variant | MODIFIER |
| 9 | 64614203 | AX-170450262 | 0 | C | T | 0.193 | 8.03E-03 | 1.38E-01 | 2.72E+00 | 2.74E+00 | 9.52E-01 | no | Megf11 | intron_variant | MODIFIER |
| 9 | 64615240 | AX-170448022 | 0 | T | C | 0.193 | 8.03E-03 | 1.38E-01 | 2.72E+00 | 2.74E+00 | 9.52E-01 | no | Megf11 | intron_variant | MODIFIER |
| 9 | 64615704 | AX-170231687 | 0 | G | T | 0.193 | 8.03E-03 | 1.38E-01 | 2.72E+00 | 2.74E+00 | 9.52E-01 | no | Megf11 | intron_variant | MODIFIER |
| 9 | 64626694 | AX-97805022 | 0 | A | G | 0.193 | 8.03E-03 | 1.38E-01 | 2.72E+00 | 2.74E+00 | 9.52E-01 | no | Megf11 | intron_variant | MODIFIER |
| 9 | 64639704 | AX-170399063 | 0 | C | G | 0.193 | 8.03E-03 | 1.38E-01 | 2.72E+00 | 2.74E+00 | 9.52E-01 | no | Megf11 | intron_variant | MODIFIER |
| 9 | 64641254 | AX-170362304 | 0 | A | G | 0.193 | 8.03E-03 | 1.38E-01 | 2.72E+00 | 2.74E+00 | 9.52E-01 | no | Megf11 | intron_variant | MODIFIER |
| 9 | 64661999 | AX-170237631 | 0 | G | A | 0.193 | 8.03E-03 | 1.38E-01 | 2.72E+00 | 2.74E+00 | 9.52E-01 | no | Megf11 | intron_variant | MODIFIER |
| 9 | 64664381 | AX-170448636 | 0 | G | C | 0.193 | 8.03E-03 | 1.38E-01 | 2.72E+00 | 2.74E+00 | 9.52E-01 | no | Megf11 | intron_variant | MODIFIER |
| 9 | 64665029 | AX-170416467 | 0 | T | A | 0.193 | 8.03E-03 | 1.38E-01 | 2.72E+00 | 2.74E+00 | 9.52E-01 | no | Megf11 | intron_variant | MODIFIER |
| 9 | 64680669 | AX-170262144 | 0 | A | G | 0.193 | 8.03E-03 | 1.38E-01 | 2.72E+00 | 2.74E+00 | 9.52E-01 | no | Megf11 | intron_variant | MODIFIER |
| 9 | 64838124 | AX-170082655 | 0 | A | T | 0.193 | 8.03E-03 | 1.38E-01 | 2.72E+00 | 2.74E+00 | 9.52E-01 | no | Dennd4a | intron_variant | MODIFIER |
| 9 | 64845498 | AX-170296392 | 0 | G | T | 0.193 | 8.03E-03 | 1.38E-01 | 2.72E+00 | 2.74E+00 | 9.52E-01 | no | Dennd4a | intron_variant | MODIFIER |
| 9 | 64858404 | AX-170517638 | 0 | A | G | 0.193 | 8.03E-03 | 1.38E-01 | 2.72E+00 | 2.74E+00 | 9.52E-01 | no | Dennd4a | intron_variant | MODIFIER |
| 9 | 64892427 | AX-170262237 | 0 | T | A | 0.193 | 8.03E-03 | 1.38E-01 | 2.72E+00 | 2.74E+00 | 9.52E-01 | no | Dennd4a | intron_variant | MODIFIER |
| 9 | 64902472 | AX-170032559 | 0 | A | G | 0.193 | 8.03E-03 | 1.38E-01 | 2.72E+00 | 2.74E+00 | 9.52E-01 | no | Dennd4a | intron_variant | MODIFIER |
| 9 | 64904995 | AX-170223140 | 0 | C | T | 0.193 | 8.03E-03 | 1.38E-01 | 2.72E+00 | 2.74E+00 | 9.52E-01 | no | Dennd4a | intron_variant | MODIFIER |
| 9 | 64924228 | AX-170476757 | 0 | C | T | 0.193 | 8.03E-03 | 1.38E-01 | 2.72E+00 | 2.74E+00 | 9.52E-01 | no | Slc24a1 | 3_prime_UTR_variant | MODIFIER |
| 9 | 65027384 | AX-97857150 | 0 | T | C | 0.193 | 8.03E-03 | 1.38E-01 | 2.72E+00 | 2.74E+00 | 9.52E-01 | no | | intergenic_variant | MODIFIER |
| 9 | 65190407 | AX-170364116 | 0 | T | C | 0.193 | 8.03E-03 | 1.38E-01 | 2.72E+00 | 2.74E+00 | 9.52E-01 | no | Igdcc3 | downstream_gene_variant | MODIFIER |
| 9 | 65218654 | AX-97810425 | 0 | C | T | 0.193 | 8.03E-03 | 1.38E-01 | 2.72E+00 | 2.74E+00 | 9.52E-01 | no | Parp16 | intron_variant | MODIFIER |
| 9 | 65234789 | AX-170252957 | 0 | G | A | 0.193 | 8.03E-03 | 1.38E-01 | 2.72E+00 | 2.74E+00 | 9.52E-01 | no | Parp16 | intron_variant | MODIFIER |
| 9 | 65235598 | AX-170309533 | 0 | C | T | 0.193 | 8.03E-03 | 1.38E-01 | 2.72E+00 | 2.74E+00 | 9.52E-01 | no | Parp16 | intron_variant | MODIFIER |
| 9 | 65330426 | AX-170151277 | 0 | G | A | 0.193 | 8.03E-03 | 1.38E-01 | 2.72E+00 | 2.74E+00 | 9.52E-01 | no | Clpx | 3_prime_UTR_variant | MODIFIER |
| 9 | 65435024 | AX-170492556 | 0 | T | C | 0.193 | 8.03E-03 | 1.38E-01 | 2.72E+00 | 2.74E+00 | 9.52E-01 | no | Mtfmt | upstream_gene_variant | MODIFIER |
| 9 | 65435185 | AX-169999741 | 0 | G | C | 0.193 | 8.03E-03 | 1.38E-01 | 2.72E+00 | 2.74E+00 | 9.52E-01 | no | Mtfmt | upstream_gene_variant | MODIFIER |
| 9 | 65435210 | AX-97797535 | 0 | G | A | 0.193 | 8.03E-03 | 1.38E-01 | 2.72E+00 | 2.74E+00 | 9.52E-01 | no | Mtfmt | upstream_gene_variant | MODIFIER |
| 9 | 65464836 | AX-170174310 | 0 | G | C | 0.193 | 8.03E-03 | 1.38E-01 | 2.72E+00 | 2.74E+00 | 9.52E-01 | no | Spg21 | intron_variant | MODIFIER |
| 9 | 65465056 | AX-97805714 | 0 | C | T | 0.193 | 8.03E-03 | 1.38E-01 | 2.72E+00 | 2.74E+00 | 9.52E-01 | no | Spg21 | intron_variant | MODIFIER |
| 9 | 65465164 | AX-170032568 | 0 | T | C | 0.193 | 8.03E-03 | 1.38E-01 | 2.72E+00 | 2.74E+00 | 9.52E-01 | no | Spg21 | intron_variant | MODIFIER |
| 9 | 65478059 | AX-170298360 | 0 | A | G | 0.193 | 8.03E-03 | 1.38E-01 | 2.72E+00 | 2.74E+00 | 9.52E-01 | no | Spg21 | intron_variant | MODIFIER |
| 9 | 65480295 | AX-170378302 | 0 | C | T | 0.193 | 8.03E-03 | 1.38E-01 | 2.72E+00 | 2.74E+00 | 9.52E-01 | no | Spg21 | intron_variant | MODIFIER |
| 9 | 65484105 | AX-170359298 | 0 | G | A | 0.193 | 8.03E-03 | 1.38E-01 | 2.72E+00 | 2.74E+00 | 9.52E-01 | no | Spg21 | intron_variant | MODIFIER |

| | | | | | | | | | | | | | | | |
|---|---|---|---|---|---|---|---|---|---|---|---|---|---|---|---|
| 9 | 65489225 | AX-170366143 | 0 | T | C | 0.193 | 8.03E-03 | 1.38E-01 | 2.72E+00 | 2.74E+00 | 9.52E-01 | no | Spg21 | downstream_gene_variant | MODIFIER |
| 9 | 65489861 | AX-170296333 | 0 | C | A | 0.193 | 8.03E-03 | 1.38E-01 | 2.72E+00 | 2.74E+00 | 9.52E-01 | no | Spg21 | downstream_gene_variant | MODIFIER |
| 9 | 65490892 | AX-97842818 | 0 | C | T | 0.193 | 8.03E-03 | 1.38E-01 | 2.72E+00 | 2.74E+00 | 9.52E-01 | no | Spg21 | downstream_gene_variant | MODIFIER |
| 9 | 65495312 | AX-170142222 | 0 | G | A | 0.193 | 8.03E-03 | 1.38E-01 | 2.72E+00 | 2.74E+00 | 9.52E-01 | no | Ankdd1a | intron_variant,non_coding_transcript_variant | MODIFIER |
| 9 | 65501988 | AX-170190747 | 0 | C | T | 0.193 | 8.03E-03 | 1.38E-01 | 2.72E+00 | 2.74E+00 | 9.52E-01 | no | Ankdd1a | intron_variant | MODIFIER |
| 9 | 65508365 | AX-170032567 | 0 | A | G | 0.193 | 8.03E-03 | 1.38E-01 | 2.72E+00 | 2.74E+00 | 9.52E-01 | no | Ankdd1a | intron_variant | MODIFIER |
| 9 | 65515504 | AX-97829829 | 0 | C | A | 0.193 | 8.03E-03 | 1.38E-01 | 2.72E+00 | 2.74E+00 | 9.52E-01 | no | Ankdd1a | intron_variant | MODIFIER |
| 9 | 65522695 | AX-170354910 | 0 | A | G | 0.193 | 8.03E-03 | 1.38E-01 | 2.72E+00 | 2.74E+00 | 9.52E-01 | no | Gm17749 | intron_variant,non_coding_transcript_variant | MODIFIER |
| 9 | 65545829 | AX-170335339 | 0 | T | C | 0.193 | 8.03E-03 | 1.38E-01 | 2.72E+00 | 2.74E+00 | 9.52E-01 | no | | regulatory_region_variant | MODIFIER |
| 9 | 65547668 | AX-170258292 | 0 | C | T | 0.193 | 8.03E-03 | 1.38E-01 | 2.72E+00 | 2.74E+00 | 9.52E-01 | no | | regulatory_region_variant | MODIFIER |
| 9 | 65549150 | AX-170138894 | 0 | T | C | 0.193 | 8.03E-03 | 1.38E-01 | 2.72E+00 | 2.74E+00 | 9.52E-01 | no | Plekho2 | downstream_gene_variant | MODIFIER |
| 9 | 65558168 | AX-170257587 | 0 | A | G | 0.193 | 8.03E-03 | 1.38E-01 | 2.72E+00 | 2.74E+00 | 9.52E-01 | no | Plekho2 | intron_variant | MODIFIER |
| 9 | 65575044 | AX-97788958 | 0 | C | T | 0.193 | 8.03E-03 | 1.38E-01 | 2.72E+00 | 2.74E+00 | 9.52E-01 | no | Plekho2 | intron_variant | MODIFIER |
| 9 | 65613357 | AX-97821856 | 0 | A | C | 0.193 | 8.03E-03 | 1.38E-01 | 2.72E+00 | 2.74E+00 | 9.52E-01 | no | | regulatory_region_variant | MODIFIER |
| 9 | 65615890 | AX-170211768 | 0 | A | G | 0.193 | 8.03E-03 | 1.38E-01 | 2.72E+00 | 2.74E+00 | 9.52E-01 | no | | intergenic_variant | MODIFIER |
| 9 | 65634229 | AX-97855814 | 0 | C | A | 0.193 | 8.03E-03 | 1.38E-01 | 2.72E+00 | 2.74E+00 | 9.52E-01 | no | Rbpms2 | intron_variant | MODIFIER |
| 9 | 65634821 | AX-170520880 | 0 | A | G | 0.193 | 8.03E-03 | 1.38E-01 | 2.72E+00 | 2.74E+00 | 9.52E-01 | no | Rbpms2 | intron_variant | MODIFIER |
| 9 | 65836433 | AX-170285637 | 0 | A | G | 0.193 | 8.03E-03 | 1.38E-01 | 2.72E+00 | 2.74E+00 | 9.52E-01 | no | n-R5s85 | non_coding_transcript_exon_variant | MODIFIER |
| 9 | 65950570 | AX-170032575 | 0 | T | G | 0.193 | 8.03E-03 | 1.38E-01 | 2.72E+00 | 2.74E+00 | 9.52E-01 | no | Csnk1g1 | intron_variant | MODIFIER |
| 9 | 65963636 | AX-170135683 | 0 | T | A | 0.193 | 8.03E-03 | 1.38E-01 | 2.72E+00 | 2.74E+00 | 9.52E-01 | no | Csnk1g1 | intron_variant | MODIFIER |
| 9 | 65992278 | AX-170032574 | 0 | A | G | 0.193 | 8.03E-03 | 1.38E-01 | 2.72E+00 | 2.74E+00 | 9.52E-01 | no | Csnk1g1 | intron_variant | MODIFIER |
| 9 | 65992399 | AX-97837441 | 0 | C | T | 0.193 | 8.03E-03 | 1.38E-01 | 2.72E+00 | 2.74E+00 | 9.52E-01 | no | Csnk1g1 | intron_variant | MODIFIER |
| 9 | 66048380 | AX-170032580 | 0 | C | G | 0.193 | 8.03E-03 | 1.38E-01 | 2.72E+00 | 2.74E+00 | 9.52E-01 | no | Csnk1g1 | downstream_gene_variant | MODIFIER |
| 9 | 66060781 | AX-170232733 | 0 | G | T | 0.193 | 8.03E-03 | 1.38E-01 | 2.72E+00 | 2.74E+00 | 9.52E-01 | no | Ppib | intron_variant | MODIFIER |
| 9 | 66064643 | AX-170514047 | 0 | A | G | 0.193 | 8.03E-03 | 1.38E-01 | 2.72E+00 | 2.74E+00 | 9.52E-01 | no | Ppib | intron_variant | MODIFIER |
| 9 | 66078312 | AX-170111419 | 0 | G | C | 0.193 | 8.03E-03 | 1.38E-01 | 2.72E+00 | 2.74E+00 | 9.52E-01 | no | | regulatory_region_variant | MODIFIER |
| 9 | 66085410 | AX-170112063 | 0 | A | G | 0.193 | 8.03E-03 | 1.38E-01 | 2.72E+00 | 2.74E+00 | 9.52E-01 | no | Snx1 | downstream_gene_variant | MODIFIER |
| 9 | 66121026 | AX-170219751 | 0 | A | G | 0.193 | 8.03E-03 | 1.38E-01 | 2.72E+00 | 2.74E+00 | 9.52E-01 | no | Snx1 | intron_variant | MODIFIER |
| 9 | 66124830 | AX-170111265 | 0 | C | T | 0.193 | 8.03E-03 | 1.38E-01 | 2.72E+00 | 2.74E+00 | 9.52E-01 | no | Ciao2a | upstream_gene_variant | MODIFIER |
| 9 | 66169063 | AX-170371347 | 0 | C | T | 0.193 | 8.03E-03 | 1.38E-01 | 2.72E+00 | 2.74E+00 | 9.52E-01 | no | Dapk2 | intron_variant | MODIFIER |
| 9 | 66169085 | AX-170442237 | 0 | G | A | 0.193 | 8.03E-03 | 1.38E-01 | 2.72E+00 | 2.74E+00 | 9.52E-01 | no | Dapk2 | intron_variant | MODIFIER |
| 9 | 66173074 | AX-170157253 | 0 | C | A | 0.193 | 8.03E-03 | 1.38E-01 | 2.72E+00 | 2.74E+00 | 9.52E-01 | no | Dapk2 | intron_variant | MODIFIER |
| 9 | 66174757 | AX-97798501 | 0 | A | G | 0.193 | 8.03E-03 | 1.38E-01 | 2.72E+00 | 2.74E+00 | 9.52E-01 | no | Dapk2 | intron_variant | MODIFIER |
| 9 | 66177618 | AX-170352963 | 0 | G | A | 0.193 | 8.03E-03 | 1.38E-01 | 2.72E+00 | 2.74E+00 | 9.52E-01 | no | Dapk2 | intron_variant | MODIFIER |
| 9 | 66178180 | AX-170489643 | 0 | C | T | 0.193 | 8.03E-03 | 1.38E-01 | 2.72E+00 | 2.74E+00 | 9.52E-01 | no | Dapk2 | intron_variant | MODIFIER |
| 9 | 66181619 | AX-170456140 | 0 | T | C | 0.193 | 8.03E-03 | 1.38E-01 | 2.72E+00 | 2.74E+00 | 9.52E-01 | no | Dapk2 | intron_variant | MODIFIER |
| 9 | 66201958 | AX-170386832 | 0 | C | T | 0.193 | 8.03E-03 | 1.38E-01 | 2.72E+00 | 2.74E+00 | 9.52E-01 | no | Dapk2 | intron_variant | MODIFIER |
| 9 | 66202500 | AX-170352398 | 0 | A | C | 0.193 | 8.03E-03 | 1.38E-01 | 2.72E+00 | 2.74E+00 | 9.52E-01 | no | Dapk2 | intron_variant | MODIFIER |
| 9 | 66202555 | AX-170155776 | 0 | A | T | 0.193 | 8.03E-03 | 1.38E-01 | 2.72E+00 | 2.74E+00 | 9.52E-01 | no | Dapk2 | intron_variant | MODIFIER |
| 9 | 66206277 | AX-170394168 | 0 | A | G | 0.193 | 8.03E-03 | 1.38E-01 | 2.72E+00 | 2.74E+00 | 9.52E-01 | no | Dapk2 | intron_variant | MODIFIER |
| 9 | 66213703 | AX-170296471 | 0 | A | C | 0.193 | 8.03E-03 | 1.38E-01 | 2.72E+00 | 2.74E+00 | 9.52E-01 | no | Dapk2 | intron_variant | MODIFIER |
| 9 | 66217724 | AX-97838433 | 0 | A | G | 0.193 | 8.03E-03 | 1.38E-01 | 2.72E+00 | 2.74E+00 | 9.52E-01 | no | Dapk2 | intron_variant | MODIFIER |
| 9 | 66275014 | AX-170265873 | 0 | G | A | 0.193 | 8.03E-03 | 1.38E-01 | 2.72E+00 | 2.74E+00 | 9.52E-01 | no | Dapk2 | downstream_gene_variant | MODIFIER |
| 9 | 66300543 | AX-170277242 | 0 | T | C | 0.193 | 8.03E-03 | 1.38E-01 | 2.72E+00 | 2.74E+00 | 9.52E-01 | no | | intergenic_variant | MODIFIER |
| 9 | 66302153 | AX-170285588 | 0 | T | G | 0.193 | 8.03E-03 | 1.38E-01 | 2.72E+00 | 2.74E+00 | 9.52E-01 | no | | intergenic_variant | MODIFIER |
| 9 | 66303070 | AX-170168137 | 0 | T | C | 0.193 | 8.03E-03 | 1.38E-01 | 2.72E+00 | 2.74E+00 | 9.52E-01 | no | | intergenic_variant | MODIFIER |
| 9 | 66307731 | AX-170169825 | 0 | C | T | 0.193 | 8.03E-03 | 1.38E-01 | 2.72E+00 | 2.74E+00 | 9.52E-01 | no | | regulatory_region_variant | MODIFIER |
| 9 | 66307793 | AX-170316920 | 0 | G | T | 0.193 | 8.03E-03 | 1.38E-01 | 2.72E+00 | 2.74E+00 | 9.52E-01 | no | | regulatory_region_variant | MODIFIER |
| 9 | 66309240 | AX-170116678 | 0 | C | G | 0.193 | 8.03E-03 | 1.38E-01 | 2.72E+00 | 2.74E+00 | 9.52E-01 | no | | intergenic_variant | MODIFIER |
| 9 | 66657534 | AX-170534147 | 0 | T | C | 0.193 | 8.03E-03 | 1.38E-01 | 2.72E+00 | 2.74E+00 | 9.52E-01 | no | | intergenic_variant | MODIFIER |
| 9 | 66699931 | AX-170082687 | 0 | A | G | 0.193 | 8.03E-03 | 1.38E-01 | 2.72E+00 | 2.74E+00 | 9.52E-01 | no | | intergenic_variant | MODIFIER |
| 9 | 66736766 | AX-170291886 | 0 | T | C | 0.193 | 8.03E-03 | 1.38E-01 | 2.72E+00 | 2.74E+00 | 9.52E-01 | no | Car12 | intron_variant | MODIFIER |
| 9 | 66747932 | AX-170192391 | 0 | G | A | 0.193 | 8.03E-03 | 1.38E-01 | 2.72E+00 | 2.74E+00 | 9.52E-01 | no | Car12 | intron_variant | MODIFIER |
| 9 | 66749157 | AX-170486842 | 0 | T | G | 0.193 | 8.03E-03 | 1.38E-01 | 2.72E+00 | 2.74E+00 | 9.52E-01 | no | Car12 | intron_variant | MODIFIER |

| | | | | | | | | | | | | | | | |
|---|---|---|---|---|---|---|---|---|---|---|---|---|---|---|---|
| 9 | 66751922 | AX-170453664 | 0 | C | A | 0.193 | 8.03E-03 | 1.38E-01 | 2.72E+00 | 2.74E+00 | 9.52E-01 | no | Car12 | intron_variant | MODIFIER |
| 9 | 66755826 | AX-170529797 | 0 | T | C | 0.193 | 8.03E-03 | 1.38E-01 | 2.72E+00 | 2.74E+00 | 9.52E-01 | no | Car12 | intron_variant | MODIFIER |
| 9 | 66758869 | AX-170452737 | 0 | A | G | 0.193 | 8.03E-03 | 1.38E-01 | 2.72E+00 | 2.74E+00 | 9.52E-01 | no | Car12 | intron_variant | MODIFIER |
| 9 | 66768662 | AX-170428296 | 0 | A | G | 0.193 | 8.03E-03 | 1.38E-01 | 2.72E+00 | 2.74E+00 | 9.52E-01 | no | Car12 | downstream_gene_variant | MODIFIER |
| 9 | 66989778 | AX-170032590 | 0 | C | T | 0.193 | 8.03E-03 | 1.38E-01 | 2.72E+00 | 2.74E+00 | 9.52E-01 | no | | regulatory_region_variant | MODIFIER |
| 9 | 66990365 | AX-97835184 | 0 | A | G | 0.193 | 8.03E-03 | 1.38E-01 | 2.72E+00 | 2.74E+00 | 9.52E-01 | no | | regulatory_region_variant | MODIFIER |
| 9 | 67008596 | AX-170177089 | 0 | T | C | 0.193 | 8.03E-03 | 1.38E-01 | 2.72E+00 | 2.74E+00 | 9.52E-01 | no | | regulatory_region_variant | MODIFIER |
| 9 | 67010138 | AX-97827757 | 0 | G | T | 0.193 | 8.03E-03 | 1.38E-01 | 2.72E+00 | 2.74E+00 | 9.52E-01 | no | | intergenic_variant | MODIFIER |
| 9 | 67016000 | AX-170292097 | 0 | G | T | 0.193 | 8.03E-03 | 1.38E-01 | 2.72E+00 | 2.74E+00 | 9.52E-01 | no | | regulatory_region_variant | MODIFIER |
| 9 | 67023057 | AX-170032592 | 0 | A | T | 0.193 | 8.03E-03 | 1.38E-01 | 2.72E+00 | 2.74E+00 | 9.52E-01 | no | Tpm1 | 3_prime_UTR_variant | MODIFIER |
| 9 | 67053066 | AX-97806227 | 0 | A | C | 0.193 | 8.03E-03 | 1.38E-01 | 2.72E+00 | 2.74E+00 | 9.52E-01 | no | Tpm1 | upstream_gene_variant | MODIFIER |
| 9 | 67080370 | AX-170521634 | 0 | C | T | 0.193 | 8.03E-03 | 1.38E-01 | 2.72E+00 | 2.74E+00 | 9.52E-01 | no | | intergenic_variant | MODIFIER |
| 9 | 67100882 | AX-170176787 | 0 | A | C | 0.193 | 8.03E-03 | 1.38E-01 | 2.72E+00 | 2.74E+00 | 9.52E-01 | no | Gm22145 | upstream_gene_variant | MODIFIER |
| 9 | 67112537 | AX-170174631 | 0 | A | T | 0.193 | 8.03E-03 | 1.38E-01 | 2.72E+00 | 2.74E+00 | 9.52E-01 | no | Gm10646 | downstream_gene_variant | MODIFIER |
| 9 | 67136805 | AX-170245133 | 0 | C | T | 0.193 | 8.03E-03 | 1.38E-01 | 2.72E+00 | 2.74E+00 | 9.52E-01 | no | Gm19299 | intron_variant,non_coding_transcript_variant | MODIFIER |
| 9 | 67146778 | AX-97850730 | 0 | T | A | 0.193 | 8.03E-03 | 1.38E-01 | 2.72E+00 | 2.74E+00 | 9.52E-01 | no | Gm19299 | intron_variant,non_coding_transcript_variant | MODIFIER |
| 9 | 67148139 | AX-170421933 | 0 | A | G | 0.193 | 8.03E-03 | 1.38E-01 | 2.72E+00 | 2.74E+00 | 9.52E-01 | no | Gm19299 | intron_variant,non_coding_transcript_variant | MODIFIER |
| 9 | 67153016 | AX-170379176 | 0 | A | G | 0.193 | 8.03E-03 | 1.38E-01 | 2.72E+00 | 2.74E+00 | 9.52E-01 | no | Gm19299 | intron_variant,non_coding_transcript_variant | MODIFIER |
| 9 | 67153193 | AX-170237116 | 0 | C | T | 0.193 | 8.03E-03 | 1.38E-01 | 2.72E+00 | 2.74E+00 | 9.52E-01 | no | Gm19299 | intron_variant,non_coding_transcript_variant | MODIFIER |
| 9 | 67153428 | AX-170106311 | 0 | A | G | 0.193 | 8.03E-03 | 1.38E-01 | 2.72E+00 | 2.74E+00 | 9.52E-01 | no | Gm19299 | intron_variant,non_coding_transcript_variant | MODIFIER |
| 9 | 67158027 | AX-170396045 | 0 | A | G | 0.193 | 8.03E-03 | 1.38E-01 | 2.72E+00 | 2.74E+00 | 9.52E-01 | no | Gm19299 | intron_variant,non_coding_transcript_variant | MODIFIER |
| 9 | 67158414 | AX-170489888 | 0 | C | G | 0.193 | 8.03E-03 | 1.38E-01 | 2.72E+00 | 2.74E+00 | 9.52E-01 | no | Gm19299 | intron_variant,non_coding_transcript_variant | MODIFIER |
| 9 | 67178087 | AX-170392925 | 0 | A | G | 0.193 | 8.03E-03 | 1.38E-01 | 2.72E+00 | 2.74E+00 | 9.52E-01 | no | Gm19299 | intron_variant,non_coding_transcript_variant | MODIFIER |
| 9 | 67219793 | AX-170515072 | 0 | C | T | 0.193 | 8.03E-03 | 1.38E-01 | 2.72E+00 | 2.74E+00 | 9.52E-01 | no | Tln2 | downstream_gene_variant | MODIFIER |
| 9 | 67246219 | AX-170349115 | 0 | T | C | 0.193 | 8.03E-03 | 1.38E-01 | 2.72E+00 | 2.74E+00 | 9.52E-01 | no | Tln2 | intron_variant | MODIFIER |
| 9 | 67247812 | AX-170249151 | 0 | A | G | 0.193 | 8.03E-03 | 1.38E-01 | 2.72E+00 | 2.74E+00 | 9.52E-01 | no | Tln2 | intron_variant | MODIFIER |
| 9 | 67255136 | AX-170316797 | 0 | G | A | 0.193 | 8.03E-03 | 1.38E-01 | 2.72E+00 | 2.74E+00 | 9.52E-01 | no | Tln2 | intron_variant | MODIFIER |
| 9 | 67259235 | AX-170132853 | 0 | A | G | 0.193 | 8.03E-03 | 1.38E-01 | 2.72E+00 | 2.74E+00 | 9.52E-01 | no | Tln2 | intron_variant | MODIFIER |
| 9 | 67321288 | AX-170335007 | 0 | T | C | 0.193 | 8.03E-03 | 1.38E-01 | 2.72E+00 | 2.74E+00 | 9.52E-01 | no | Tln2 | intron_variant | MODIFIER |
| 9 | 67345380 | AX-170136354 | 0 | T | A | 0.193 | 8.03E-03 | 1.38E-01 | 2.72E+00 | 2.74E+00 | 9.52E-01 | no | Tln2 | intron_variant | MODIFIER |
| 9 | 67346220 | AX-170151975 | 0 | A | G | 0.193 | 8.03E-03 | 1.38E-01 | 2.72E+00 | 2.74E+00 | 9.52E-01 | no | Tln2 | intron_variant | MODIFIER |
| 9 | 67346504 | AX-170282096 | 0 | T | C | 0.193 | 8.03E-03 | 1.38E-01 | 2.72E+00 | 2.74E+00 | 9.52E-01 | no | Tln2 | missense_variant | MODERATE |
| 9 | 67361651 | AX-170124472 | 0 | G | A | 0.193 | 8.03E-03 | 1.38E-01 | 2.72E+00 | 2.74E+00 | 9.52E-01 | no | Tln2 | intron_variant | MODIFIER |
| 9 | 67365853 | AX-97832450 | 0 | C | T | 0.193 | 8.03E-03 | 1.38E-01 | 2.72E+00 | 2.74E+00 | 9.52E-01 | no | Tln2 | intron_variant | MODIFIER |
| 9 | 67388352 | AX-170501296 | 0 | T | C | 0.193 | 8.03E-03 | 1.38E-01 | 2.72E+00 | 2.74E+00 | 9.52E-01 | no | Tln2 | intron_variant | MODIFIER |
| 9 | 67399891 | AX-170457168 | 0 | T | C | 0.193 | 8.03E-03 | 1.38E-01 | 2.72E+00 | 2.74E+00 | 9.52E-01 | no | Tln2 | intron_variant | MODIFIER |
| 9 | 67400087 | AX-170292395 | 0 | C | T | 0.193 | 8.03E-03 | 1.38E-01 | 2.72E+00 | 2.74E+00 | 9.52E-01 | no | Tln2 | intron_variant | MODIFIER |
| 9 | 67401863 | AX-170082700 | 0 | G | A | 0.193 | 8.03E-03 | 1.38E-01 | 2.72E+00 | 2.74E+00 | 9.52E-01 | no | Tln2 | intron_variant | MODIFIER |
| 9 | 67401929 | AX-170082703 | 0 | G | A | 0.193 | 8.03E-03 | 1.38E-01 | 2.72E+00 | 2.74E+00 | 9.52E-01 | no | Tln2 | intron_variant | MODIFIER |
| 9 | 67402747 | AX-170082701 | 0 | A | C | 0.193 | 8.03E-03 | 1.38E-01 | 2.72E+00 | 2.74E+00 | 9.52E-01 | no | Tln2 | intron_variant | MODIFIER |
| 9 | 67403748 | AX-170082704 | 0 | C | T | 0.193 | 8.03E-03 | 1.38E-01 | 2.72E+00 | 2.74E+00 | 9.52E-01 | no | Tln2 | intron_variant | MODIFIER |
| 9 | 67404340 | AX-170082702 | 0 | G | T | 0.193 | 8.03E-03 | 1.38E-01 | 2.72E+00 | 2.74E+00 | 9.52E-01 | no | Tln2 | intron_variant | MODIFIER |
| 9 | 67412665 | AX-170442003 | 0 | G | T | 0.193 | 8.03E-03 | 1.38E-01 | 2.72E+00 | 2.74E+00 | 9.52E-01 | no | Tln2 | intron_variant | MODIFIER |
| 9 | 67432084 | AX-170248373 | 0 | G | A | 0.193 | 8.03E-03 | 1.38E-01 | 2.72E+00 | 2.74E+00 | 9.52E-01 | no | Tln2 | intron_variant | MODIFIER |
| 9 | 67549991 | AX-170032598 | 0 | T | G | 0.193 | 8.03E-03 | 1.38E-01 | 2.72E+00 | 2.74E+00 | 9.52E-01 | no | Tln2 | intron_variant | MODIFIER |
| 9 | 67568133 | AX-170398543 | 0 | T | G | 0.193 | 8.03E-03 | 1.38E-01 | 2.72E+00 | 2.74E+00 | 9.52E-01 | no | | intergenic_variant | MODIFIER |
| 9 | 67615404 | AX-170266714 | 0 | T | C | 0.193 | 8.03E-03 | 1.38E-01 | 2.72E+00 | 2.74E+00 | 9.52E-01 | no | | regulatory_region_variant | MODIFIER |
| 9 | 67825189 | AX-97844398 | 0 | G | A | 0.193 | 8.03E-03 | 1.38E-01 | 2.72E+00 | 2.74E+00 | 9.52E-01 | no | | intergenic_variant | MODIFIER |
| 9 | 67825268 | AX-170317435 | 0 | A | G | 0.193 | 8.03E-03 | 1.38E-01 | 2.72E+00 | 2.74E+00 | 9.52E-01 | no | | intergenic_variant | MODIFIER |
| 9 | 67847916 | AX-170305018 | 0 | A | T | 0.193 | 8.03E-03 | 1.38E-01 | 2.72E+00 | 2.74E+00 | 9.52E-01 | no | Vps13c | intron_variant | MODIFIER |
| 9 | 67848010 | AX-170458862 | 0 | C | T | 0.193 | 8.03E-03 | 1.38E-01 | 2.72E+00 | 2.74E+00 | 9.52E-01 | no | Vps13c | intron_variant | MODIFIER |
| 9 | 67850296 | AX-170307575 | 0 | A | G | 0.193 | 8.03E-03 | 1.38E-01 | 2.72E+00 | 2.74E+00 | 9.52E-01 | no | Vps13c | intron_variant | MODIFIER |
| 9 | 67865135 | AX-170209916 | 0 | A | G | 0.193 | 8.03E-03 | 1.38E-01 | 2.72E+00 | 2.74E+00 | 9.52E-01 | no | Vps13c | intron_variant | MODIFIER |
| 9 | 67865904 | AX-170169636 | 0 | A | G | 0.193 | 8.03E-03 | 1.38E-01 | 2.72E+00 | 2.74E+00 | 9.52E-01 | no | Vps13c | intron_variant | MODIFIER |

| | | | | | | | | | | | | | | | |
|---|---|---|---|---|---|---|---|---|---|---|---|---|---|---|---|
| 9 | 67866036 | AX-97784804 | 0 | A | C | 0.193 | 8.03E-03 | 1.38E-01 | 2.72E+00 | 2.74E+00 | 9.52E-01 | no | Vps13c | intron_variant | MODIFIER |
| 9 | 67875251 | AX-170340236 | 0 | T | G | 0.193 | 8.03E-03 | 1.38E-01 | 2.72E+00 | 2.74E+00 | 9.52E-01 | no | Vps13c | intron_variant | MODIFIER |
| 9 | 67890913 | AX-170200440 | 0 | A | G | 0.193 | 8.03E-03 | 1.38E-01 | 2.72E+00 | 2.74E+00 | 9.52E-01 | no | Vps13c | intron_variant | MODIFIER |
| 9 | 67890932 | AX-170132327 | 0 | C | T | 0.193 | 8.03E-03 | 1.38E-01 | 2.72E+00 | 2.74E+00 | 9.52E-01 | no | Vps13c | intron_variant | MODIFIER |
| 9 | 67897124 | AX-170229360 | 0 | A | G | 0.193 | 8.03E-03 | 1.38E-01 | 2.72E+00 | 2.74E+00 | 9.52E-01 | no | Vps13c | synonymous_variant | LOW |
| 9 | 67897331 | AX-170082714 | 0 | G | A | 0.193 | 8.03E-03 | 1.38E-01 | 2.72E+00 | 2.74E+00 | 9.52E-01 | no | Vps13c | intron_variant | MODIFIER |
| 9 | 67897363 | AX-170082712 | 0 | G | A | 0.193 | 8.03E-03 | 1.38E-01 | 2.72E+00 | 2.74E+00 | 9.52E-01 | no | Vps13c | intron_variant | MODIFIER |
| 9 | 67906678 | AX-170253564 | 0 | A | G | 0.193 | 8.03E-03 | 1.38E-01 | 2.72E+00 | 2.74E+00 | 9.52E-01 | no | Vps13c | intron_variant | MODIFIER |
| 9 | 67929030 | AX-170152887 | 0 | T | C | 0.193 | 8.03E-03 | 1.38E-01 | 2.72E+00 | 2.74E+00 | 9.52E-01 | no | Vps13c | intron_variant | MODIFIER |
| 9 | 67962914 | AX-97824021 | 0 | A | G | 0.193 | 8.03E-03 | 1.38E-01 | 2.72E+00 | 2.74E+00 | 9.52E-01 | no | Vps13c | intron_variant | MODIFIER |
| 9 | 67968396 | AX-170032604 | 0 | A | G | 0.193 | 8.03E-03 | 1.38E-01 | 2.72E+00 | 2.74E+00 | 9.52E-01 | no | Vps13c | intron_variant | MODIFIER |
| 9 | 67986817 | AX-170283645 | 0 | C | T | 0.193 | 8.03E-03 | 1.38E-01 | 2.72E+00 | 2.74E+00 | 9.52E-01 | no | Vps13c | intron_variant | MODIFIER |
| 9 | 67987086 | AX-170032605 | 0 | T | C | 0.193 | 8.03E-03 | 1.38E-01 | 2.72E+00 | 2.74E+00 | 9.52E-01 | no | Vps13c | intron_variant | MODIFIER |
| 9 | 68009384 | AX-170426395 | 0 | C | T | 0.193 | 8.03E-03 | 1.38E-01 | 2.72E+00 | 2.74E+00 | 9.52E-01 | no | | intergenic_variant | MODIFIER |
| 9 | 68011656 | AX-170032603 | 0 | T | C | 0.193 | 8.03E-03 | 1.38E-01 | 2.72E+00 | 2.74E+00 | 9.52E-01 | no | | intergenic_variant | MODIFIER |
| 9 | 68021025 | AX-97830880 | 0 | C | T | 0.193 | 8.03E-03 | 1.38E-01 | 2.72E+00 | 2.74E+00 | 9.52E-01 | no | Gm20745 | non_coding_transcript_exon_variant | MODIFIER |
| 9 | 68038807 | AX-97833987 | 0 | A | C | 0.193 | 8.03E-03 | 1.38E-01 | 2.72E+00 | 2.74E+00 | 9.52E-01 | no | | intergenic_variant | MODIFIER |
| 9 | 68058100 | AX-170198092 | 0 | C | T | 0.193 | 8.03E-03 | 1.38E-01 | 2.72E+00 | 2.74E+00 | 9.52E-01 | no | | intergenic_variant | MODIFIER |
| 9 | 68074061 | AX-170105582 | 0 | G | A | 0.193 | 8.03E-03 | 1.38E-01 | 2.72E+00 | 2.74E+00 | 9.52E-01 | no | | intergenic_variant | MODIFIER |
| 9 | 68093420 | AX-170360334 | 0 | A | G | 0.193 | 8.03E-03 | 1.38E-01 | 2.72E+00 | 2.74E+00 | 9.52E-01 | no | | intergenic_variant | MODIFIER |
| 9 | 68097682 | AX-170231730 | 0 | G | T | 0.193 | 8.03E-03 | 1.38E-01 | 2.72E+00 | 2.74E+00 | 9.52E-01 | no | | intergenic_variant | MODIFIER |
| 9 | 68112318 | AX-170082719 | 0 | C | T | 0.193 | 8.03E-03 | 1.38E-01 | 2.72E+00 | 2.74E+00 | 9.52E-01 | no | | intergenic_variant | MODIFIER |
| 9 | 68113767 | AX-170366678 | 0 | C | T | 0.193 | 8.03E-03 | 1.38E-01 | 2.72E+00 | 2.74E+00 | 9.52E-01 | no | | intergenic_variant | MODIFIER |
| 9 | 68123599 | AX-170228027 | 0 | T | C | 0.193 | 8.03E-03 | 1.38E-01 | 2.72E+00 | 2.74E+00 | 9.52E-01 | no | | intergenic_variant | MODIFIER |
| 9 | 68123862 | AX-170219170 | 0 | C | T | 0.193 | 8.03E-03 | 1.38E-01 | 2.72E+00 | 2.74E+00 | 9.52E-01 | no | | intergenic_variant | MODIFIER |
| 9 | 68123966 | AX-170526172 | 0 | T | G | 0.193 | 8.03E-03 | 1.38E-01 | 2.72E+00 | 2.74E+00 | 9.52E-01 | no | | intergenic_variant | MODIFIER |
| 9 | 68124491 | AX-170523887 | 0 | T | C | 0.193 | 8.03E-03 | 1.38E-01 | 2.72E+00 | 2.74E+00 | 9.52E-01 | no | | intergenic_variant | MODIFIER |
| 9 | 68125153 | AX-170459094 | 0 | T | C | 0.193 | 8.03E-03 | 1.38E-01 | 2.72E+00 | 2.74E+00 | 9.52E-01 | no | | intergenic_variant | MODIFIER |
| 9 | 68125955 | AX-170387466 | 0 | G | C | 0.193 | 8.03E-03 | 1.38E-01 | 2.72E+00 | 2.74E+00 | 9.52E-01 | no | | intergenic_variant | MODIFIER |
| 9 | 68126695 | AX-170096571 | 0 | G | A | 0.193 | 8.03E-03 | 1.38E-01 | 2.72E+00 | 2.74E+00 | 9.52E-01 | no | | intergenic_variant | MODIFIER |
| 9 | 68131553 | AX-170487734 | 0 | C | T | 0.193 | 8.03E-03 | 1.38E-01 | 2.72E+00 | 2.74E+00 | 9.52E-01 | no | | regulatory_region_variant | MODIFIER |
| 9 | 68190552 | AX-170390643 | 0 | G | C | 0.193 | 8.03E-03 | 1.38E-01 | 2.72E+00 | 2.74E+00 | 9.52E-01 | no | | intergenic_variant | MODIFIER |
| 9 | 68192155 | AX-170422458 | 0 | A | G | 0.193 | 8.03E-03 | 1.38E-01 | 2.72E+00 | 2.74E+00 | 9.52E-01 | no | | intergenic_variant | MODIFIER |
| 9 | 68217491 | AX-170481055 | 0 | G | A | 0.193 | 8.03E-03 | 1.38E-01 | 2.72E+00 | 2.74E+00 | 9.52E-01 | no | | intergenic_variant | MODIFIER |
| 9 | 68262457 | AX-170123994 | 0 | C | T | 0.193 | 8.03E-03 | 1.38E-01 | 2.72E+00 | 2.74E+00 | 9.52E-01 | no | | intergenic_variant | MODIFIER |
| 9 | 68265896 | AX-170170238 | 0 | G | A | 0.193 | 8.03E-03 | 1.38E-01 | 2.72E+00 | 2.74E+00 | 9.52E-01 | no | | intergenic_variant | MODIFIER |
| 9 | 68278955 | AX-170450786 | 0 | T | C | 0.193 | 8.03E-03 | 1.38E-01 | 2.72E+00 | 2.74E+00 | 9.52E-01 | no | Gm22382 | upstream_gene_variant | MODIFIER |
| 9 | 68288667 | AX-170339469 | 0 | G | A | 0.193 | 8.03E-03 | 1.38E-01 | 2.72E+00 | 2.74E+00 | 9.52E-01 | no | | intergenic_variant | MODIFIER |
| 9 | 68381775 | AX-170111528 | 0 | T | C | 0.193 | 8.03E-03 | 1.38E-01 | 2.72E+00 | 2.74E+00 | 9.52E-01 | no | | intergenic_variant | MODIFIER |
| 9 | 68403735 | AX-97802071 | 0 | G | A | 0.193 | 8.03E-03 | 1.38E-01 | 2.72E+00 | 2.74E+00 | 9.52E-01 | no | | intergenic_variant | MODIFIER |
| 9 | 68426034 | AX-170181675 | 0 | T | C | 0.193 | 8.03E-03 | 1.38E-01 | 2.72E+00 | 2.74E+00 | 9.52E-01 | no | Gm19987 | upstream_gene_variant | MODIFIER |
| 9 | 68448069 | AX-170032609 | 0 | C | T | 0.193 | 8.03E-03 | 1.38E-01 | 2.72E+00 | 2.74E+00 | 9.52E-01 | no | | intergenic_variant | MODIFIER |
| 9 | 68449491 | AX-170314200 | 0 | G | T | 0.193 | 8.03E-03 | 1.38E-01 | 2.72E+00 | 2.74E+00 | 9.52E-01 | no | | intergenic_variant | MODIFIER |
| 9 | 68449856 | AX-170032611 | 0 | T | C | 0.193 | 8.03E-03 | 1.38E-01 | 2.72E+00 | 2.74E+00 | 9.52E-01 | no | | intergenic_variant | MODIFIER |
| 9 | 68505424 | AX-170323933 | 0 | T | C | 0.193 | 8.03E-03 | 1.38E-01 | 2.72E+00 | 2.74E+00 | 9.52E-01 | no | Gm22962 | upstream_gene_variant | MODIFIER |
| 9 | 68571111 | AX-170245943 | 0 | G | A | 0.193 | 8.03E-03 | 1.38E-01 | 2.72E+00 | 2.74E+00 | 9.52E-01 | no | | regulatory_region_variant | MODIFIER |
| 9 | 68573486 | AX-170193173 | 0 | G | A | 0.193 | 8.03E-03 | 1.38E-01 | 2.72E+00 | 2.74E+00 | 9.52E-01 | no | | intergenic_variant | MODIFIER |
| 9 | 68573831 | AX-170481874 | 0 | A | G | 0.193 | 8.03E-03 | 1.38E-01 | 2.72E+00 | 2.74E+00 | 9.52E-01 | no | | intergenic_variant | MODIFIER |
| 9 | 68573890 | AX-170363107 | 0 | C | T | 0.193 | 8.03E-03 | 1.38E-01 | 2.72E+00 | 2.74E+00 | 9.52E-01 | no | | intergenic_variant | MODIFIER |
| 9 | 68584582 | AX-170173135 | 0 | C | T | 0.193 | 8.03E-03 | 1.38E-01 | 2.72E+00 | 2.74E+00 | 9.52E-01 | no | | intergenic_variant | MODIFIER |
| 9 | 68608451 | AX-170233918 | 0 | A | G | 0.193 | 8.03E-03 | 1.38E-01 | 2.72E+00 | 2.74E+00 | 9.52E-01 | no | | intergenic_variant | MODIFIER |
| 9 | 68620360 | AX-170262823 | 0 | T | C | 0.193 | 8.03E-03 | 1.38E-01 | 2.72E+00 | 2.74E+00 | 9.52E-01 | no | | regulatory_region_variant | MODIFIER |
| 9 | 68664040 | AX-170411768 | 0 | A | G | 0.193 | 8.03E-03 | 1.38E-01 | 2.72E+00 | 2.74E+00 | 9.52E-01 | no | Rora | intron_variant | MODIFIER |
| 9 | 68666882 | AX-170324347 | 0 | C | G | 0.193 | 8.03E-03 | 1.38E-01 | 2.72E+00 | 2.74E+00 | 9.52E-01 | no | Rora | intron_variant | MODIFIER |

| | | | | | | | | | | | | | | | |
|---|---|---|---|---|---|---|---|---|---|---|---|---|---|---|---|
| 9 | 68667492 | AX-170307891 | 0 | A | G | 0.193 | 8.03E-03 | 1.38E-01 | 2.72E+00 | 2.74E+00 | 9.52E-01 | no | Rora | intron_variant | MODIFIER |
| 9 | 68696785 | AX-170372909 | 0 | A | G | 0.193 | 8.03E-03 | 1.38E-01 | 2.72E+00 | 2.74E+00 | 9.52E-01 | no | Rora | intron_variant | MODIFIER |
| 9 | 68696844 | AX-170396884 | 0 | T | G | 0.193 | 8.03E-03 | 1.38E-01 | 2.72E+00 | 2.74E+00 | 9.52E-01 | no | Rora | intron_variant | MODIFIER |
| 9 | 68722786 | AX-97837667 | 0 | G | A | 0.193 | 8.03E-03 | 1.38E-01 | 2.72E+00 | 2.74E+00 | 9.52E-01 | no | Rora | intron_variant | MODIFIER |
| 9 | 68724513 | AX-170273168 | 0 | T | C | 0.193 | 8.03E-03 | 1.38E-01 | 2.72E+00 | 2.74E+00 | 9.52E-01 | no | Rora | intron_variant | MODIFIER |
| 9 | 68738905 | AX-97808774 | 0 | A | C | 0.193 | 8.03E-03 | 1.38E-01 | 2.72E+00 | 2.74E+00 | 9.52E-01 | no | Rora | intron_variant | MODIFIER |
| 9 | 68740201 | AX-170301159 | 0 | C | G | 0.193 | 8.03E-03 | 1.38E-01 | 2.72E+00 | 2.74E+00 | 9.52E-01 | no | Rora | intron_variant | MODIFIER |
| 9 | 68756252 | AX-170398551 | 0 | A | G | 0.193 | 8.03E-03 | 1.38E-01 | 2.72E+00 | 2.74E+00 | 9.52E-01 | no | Rora | intron_variant | MODIFIER |
| 9 | 68776594 | AX-170284254 | 0 | A | G | 0.193 | 8.03E-03 | 1.38E-01 | 2.72E+00 | 2.74E+00 | 9.52E-01 | no | Rora | intron_variant | MODIFIER |
| 9 | 68777360 | AX-97852715 | 0 | C | A | 0.193 | 8.03E-03 | 1.38E-01 | 2.72E+00 | 2.74E+00 | 9.52E-01 | no | Rora | intron_variant | MODIFIER |
| 9 | 68782922 | AX-170279422 | 0 | A | G | 0.193 | 8.03E-03 | 1.38E-01 | 2.72E+00 | 2.74E+00 | 9.52E-01 | no | Rora | intron_variant | MODIFIER |
| 9 | 68787363 | AX-170390618 | 0 | C | T | 0.193 | 8.03E-03 | 1.38E-01 | 2.72E+00 | 2.74E+00 | 9.52E-01 | no | Rora | intron_variant | MODIFIER |
| 9 | 68787645 | AX-170393941 | 0 | C | A | 0.193 | 8.03E-03 | 1.38E-01 | 2.72E+00 | 2.74E+00 | 9.52E-01 | no | Rora | intron_variant | MODIFIER |
| 9 | 68789023 | AX-170278250 | 0 | T | A | 0.193 | 8.03E-03 | 1.38E-01 | 2.72E+00 | 2.74E+00 | 9.52E-01 | no | Rora | intron_variant | MODIFIER |
| 9 | 68790217 | AX-170193632 | 0 | A | G | 0.193 | 8.03E-03 | 1.38E-01 | 2.72E+00 | 2.74E+00 | 9.52E-01 | no | Rora | intron_variant | MODIFIER |
| 9 | 68798646 | AX-170165117 | 0 | A | G | 0.193 | 8.03E-03 | 1.38E-01 | 2.72E+00 | 2.74E+00 | 9.52E-01 | no | Rora | intron_variant | MODIFIER |
| 9 | 68802971 | AX-97815931 | 0 | A | G | 0.193 | 8.03E-03 | 1.38E-01 | 2.72E+00 | 2.74E+00 | 9.52E-01 | no | Rora | intron_variant | MODIFIER |
| 9 | 68803034 | AX-170211176 | 0 | T | C | 0.193 | 8.03E-03 | 1.38E-01 | 2.72E+00 | 2.74E+00 | 9.52E-01 | no | Rora | intron_variant | MODIFIER |
| 9 | 68825270 | AX-97810920 | 0 | T | C | 0.193 | 8.03E-03 | 1.38E-01 | 2.72E+00 | 2.74E+00 | 9.52E-01 | no | Rora | intron_variant | MODIFIER |
| 9 | 68832884 | AX-168403606 | 0 | A | G | 0.193 | 8.03E-03 | 1.38E-01 | 2.72E+00 | 2.74E+00 | 9.52E-01 | no | Rora | intron_variant | MODIFIER |
| 9 | 68835219 | AX-170096574 | 0 | T | A | 0.193 | 8.03E-03 | 1.38E-01 | 2.72E+00 | 2.74E+00 | 9.52E-01 | no | Rora | intron_variant | MODIFIER |
| 9 | 68839035 | AX-170330725 | 0 | G | T | 0.193 | 8.03E-03 | 1.38E-01 | 2.72E+00 | 2.74E+00 | 9.52E-01 | no | Rora | intron_variant | MODIFIER |
| 9 | 68887903 | AX-170494963 | 0 | G | A | 0.193 | 8.03E-03 | 1.38E-01 | 2.72E+00 | 2.74E+00 | 9.52E-01 | no | Rora | intron_variant | MODIFIER |
| 9 | 68892009 | AX-170522210 | 0 | C | A | 0.193 | 8.03E-03 | 1.38E-01 | 2.72E+00 | 2.74E+00 | 9.52E-01 | no | Rora | intron_variant | MODIFIER |
| 9 | 68906118 | AX-170419705 | 0 | C | T | 0.193 | 8.03E-03 | 1.38E-01 | 2.72E+00 | 2.74E+00 | 9.52E-01 | no | Rora | intron_variant | MODIFIER |
| 9 | 68911815 | AX-170352594 | 0 | G | C | 0.193 | 8.03E-03 | 1.38E-01 | 2.72E+00 | 2.74E+00 | 9.52E-01 | no | Rora | intron_variant | MODIFIER |
| 9 | 68933773 | AX-170475839 | 0 | C | T | 0.193 | 8.03E-03 | 1.38E-01 | 2.72E+00 | 2.74E+00 | 9.52E-01 | no | Rora | intron_variant | MODIFIER |
| 9 | 68933995 | AX-170082739 | 0 | C | T | 0.193 | 8.03E-03 | 1.38E-01 | 2.72E+00 | 2.74E+00 | 9.52E-01 | no | Rora | intron_variant | MODIFIER |
| 9 | 68941692 | AX-170090323 | 0 | C | G | 0.193 | 8.03E-03 | 1.38E-01 | 2.72E+00 | 2.74E+00 | 9.52E-01 | no | Rora | intron_variant | MODIFIER |
| 9 | 68943292 | AX-170197471 | 0 | T | C | 0.193 | 8.03E-03 | 1.38E-01 | 2.72E+00 | 2.74E+00 | 9.52E-01 | no | Rora | intron_variant | MODIFIER |
| 9 | 68946074 | AX-170214526 | 0 | C | A | 0.193 | 8.03E-03 | 1.38E-01 | 2.72E+00 | 2.74E+00 | 9.52E-01 | no | Rora | intron_variant | MODIFIER |
| 9 | 69014324 | AX-170032617 | 0 | G | A | 0.193 | 8.03E-03 | 1.38E-01 | 2.72E+00 | 2.74E+00 | 9.52E-01 | no | Rora | intron_variant | MODIFIER |
| 9 | 69022630 | AX-170147597 | 0 | G | A | 0.193 | 8.03E-03 | 1.38E-01 | 2.72E+00 | 2.74E+00 | 9.52E-01 | no | Rora | intron_variant | MODIFIER |
| 9 | 69025369 | AX-170032614 | 0 | T | C | 0.193 | 8.03E-03 | 1.38E-01 | 2.72E+00 | 2.74E+00 | 9.52E-01 | no | Rora | intron_variant | MODIFIER |
| 9 | 69027028 | AX-170154054 | 0 | G | C | 0.193 | 8.03E-03 | 1.38E-01 | 2.72E+00 | 2.74E+00 | 9.52E-01 | no | Rora | intron_variant | MODIFIER |
| 9 | 69029190 | AX-170309745 | 0 | G | A | 0.193 | 8.03E-03 | 1.38E-01 | 2.72E+00 | 2.74E+00 | 9.52E-01 | no | Rora | intron_variant | MODIFIER |
| 9 | 69031906 | AX-170258441 | 0 | T | A | 0.193 | 8.03E-03 | 1.38E-01 | 2.72E+00 | 2.74E+00 | 9.52E-01 | no | Rora | intron_variant | MODIFIER |
| 9 | 69034469 | AX-170427674 | 0 | A | G | 0.193 | 8.03E-03 | 1.38E-01 | 2.72E+00 | 2.74E+00 | 9.52E-01 | no | Rora | intron_variant | MODIFIER |
| 9 | 69056873 | AX-170259356 | 0 | C | A | 0.193 | 8.03E-03 | 1.38E-01 | 2.72E+00 | 2.74E+00 | 9.52E-01 | no | Rora | intron_variant | MODIFIER |
| 9 | 69058472 | AX-170032618 | 0 | G | A | 0.193 | 8.03E-03 | 1.38E-01 | 2.72E+00 | 2.74E+00 | 9.52E-01 | no | Rora | intron_variant | MODIFIER |
| 9 | 69062617 | AX-170032615 | 0 | G | C | 0.193 | 8.03E-03 | 1.38E-01 | 2.72E+00 | 2.74E+00 | 9.52E-01 | no | Rora | intron_variant | MODIFIER |
| 9 | 69068589 | AX-170341015 | 0 | C | T | 0.193 | 8.03E-03 | 1.38E-01 | 2.72E+00 | 2.74E+00 | 9.52E-01 | no | Rora | intron_variant | MODIFIER |
| 9 | 69068953 | AX-170449303 | 0 | A | G | 0.193 | 8.03E-03 | 1.38E-01 | 2.72E+00 | 2.74E+00 | 9.52E-01 | no | Rora | intron_variant | MODIFIER |
| 9 | 69070866 | AX-170211145 | 0 | A | G | 0.193 | 8.03E-03 | 1.38E-01 | 2.72E+00 | 2.74E+00 | 9.52E-01 | no | Rora | intron_variant | MODIFIER |
| 9 | 69074140 | AX-170235786 | 0 | G | A | 0.193 | 8.03E-03 | 1.38E-01 | 2.72E+00 | 2.74E+00 | 9.52E-01 | no | Rora | intron_variant | MODIFIER |
| 9 | 69105174 | AX-170032620 | 0 | G | A | 0.193 | 8.03E-03 | 1.38E-01 | 2.72E+00 | 2.74E+00 | 9.52E-01 | no | Rora | intron_variant | MODIFIER |
| 9 | 69105857 | AX-97855646 | 0 | C | T | 0.193 | 8.03E-03 | 1.38E-01 | 2.72E+00 | 2.74E+00 | 9.52E-01 | no | Rora | intron_variant | MODIFIER |
| 9 | 69129261 | AX-97822123 | 0 | G | A | 0.193 | 8.03E-03 | 1.38E-01 | 2.72E+00 | 2.74E+00 | 9.52E-01 | no | Rora | intron_variant | MODIFIER |
| 9 | 69135679 | AX-170405578 | 0 | A | G | 0.193 | 8.03E-03 | 1.38E-01 | 2.72E+00 | 2.74E+00 | 9.52E-01 | no | Rora | intron_variant | MODIFIER |
| 9 | 69147292 | AX-170343164 | 0 | A | G | 0.193 | 8.03E-03 | 1.38E-01 | 2.72E+00 | 2.74E+00 | 9.52E-01 | no | Rora | intron_variant | MODIFIER |
| 9 | 69154770 | AX-170319515 | 0 | A | G | 0.193 | 8.03E-03 | 1.38E-01 | 2.72E+00 | 2.74E+00 | 9.52E-01 | no | Rora | intron_variant | MODIFIER |
| 9 | 69272245 | AX-170090326 | 0 | C | A | 0.193 | 8.03E-03 | 1.38E-01 | 2.72E+00 | 2.74E+00 | 9.52E-01 | no | Rora | intron_variant | MODIFIER |
| 9 | 69272438 | AX-170473243 | 0 | C | G | 0.193 | 8.03E-03 | 1.38E-01 | 2.72E+00 | 2.74E+00 | 9.52E-01 | no | Rora | intron_variant | MODIFIER |
| 9 | 69272463 | AX-170289862 | 0 | A | G | 0.193 | 8.03E-03 | 1.38E-01 | 2.72E+00 | 2.74E+00 | 9.52E-01 | no | Rora | intron_variant | MODIFIER |

| | | | | | | | | | | | | | | | |
|---|---|---|---|---|---|---|---|---|---|---|---|---|---|---|---|
| 9 | 69341176 | AX-170271380 | 0 | T | C | 0.193 | 8.03E-03 | 1.38E-01 | 2.72E+00 | 2.74E+00 | 9.52E-01 | no | Rora | intron_variant | MODIFIER |
| 9 | 69360999 | AX-170186348 | 0 | C | T | 0.193 | 8.03E-03 | 1.38E-01 | 2.72E+00 | 2.74E+00 | 9.52E-01 | no | Rora | intron_variant | MODIFIER |
| 9 | 69493454 | AX-170226380 | 0 | A | C | 0.193 | 8.03E-03 | 1.38E-01 | 2.72E+00 | 2.74E+00 | 9.52E-01 | no | Anxa2 | downstream_gene_variant | MODIFIER |
| 9 | 69496029 | AX-170130209 | 0 | T | A | 0.193 | 8.03E-03 | 1.38E-01 | 2.72E+00 | 2.74E+00 | 9.52E-01 | no | Anxa2 | downstream_gene_variant | MODIFIER |
| 9 | 69496697 | AX-170082750 | 0 | T | A | 0.193 | 8.03E-03 | 1.38E-01 | 2.72E+00 | 2.74E+00 | 9.52E-01 | no | Anxa2 | downstream_gene_variant | MODIFIER |
| 9 | 69501346 | AX-170096576 | 0 | T | C | 0.193 | 8.03E-03 | 1.38E-01 | 2.72E+00 | 2.74E+00 | 9.52E-01 | no | | intergenic_variant | MODIFIER |
| 9 | 69514615 | AX-170248407 | 0 | T | C | 0.193 | 8.03E-03 | 1.38E-01 | 2.72E+00 | 2.74E+00 | 9.52E-01 | no | | intergenic_variant | MODIFIER |
| 9 | 69522361 | AX-170364471 | 0 | A | G | 0.193 | 8.03E-03 | 1.38E-01 | 2.72E+00 | 2.74E+00 | 9.52E-01 | no | | intergenic_variant | MODIFIER |
| 9 | 69526800 | AX-170159862 | 0 | T | A | 0.193 | 8.03E-03 | 1.38E-01 | 2.72E+00 | 2.74E+00 | 9.52E-01 | no | | intergenic_variant | MODIFIER |
| 9 | 69537012 | AX-170032624 | 0 | G | A | 0.193 | 8.03E-03 | 1.38E-01 | 2.72E+00 | 2.74E+00 | 9.52E-01 | no | | intergenic_variant | MODIFIER |
| 9 | 69539298 | AX-170114729 | 0 | C | T | 0.193 | 8.03E-03 | 1.38E-01 | 2.72E+00 | 2.74E+00 | 9.52E-01 | no | | intergenic_variant | MODIFIER |
| 9 | 69565548 | AX-170286599 | 0 | G | A | 0.193 | 8.03E-03 | 1.38E-01 | 2.72E+00 | 2.74E+00 | 9.52E-01 | no | | intergenic_variant | MODIFIER |
| 9 | 69574982 | AX-170526893 | 0 | A | G | 0.193 | 8.03E-03 | 1.38E-01 | 2.72E+00 | 2.74E+00 | 9.52E-01 | no | | intergenic_variant | MODIFIER |
| 9 | 69577484 | AX-170405055 | 0 | A | G | 0.193 | 8.03E-03 | 1.38E-01 | 2.72E+00 | 2.74E+00 | 9.52E-01 | no | | intergenic_variant | MODIFIER |
| 9 | 69579436 | AX-170297192 | 0 | G | C | 0.193 | 8.03E-03 | 1.38E-01 | 2.72E+00 | 2.74E+00 | 9.52E-01 | no | | intergenic_variant | MODIFIER |
| 9 | 69581100 | AX-170203234 | 0 | G | A | 0.193 | 8.03E-03 | 1.38E-01 | 2.72E+00 | 2.74E+00 | 9.52E-01 | no | | intergenic_variant | MODIFIER |
| 9 | 69614717 | AX-97839392 | 0 | A | G | 0.193 | 8.03E-03 | 1.38E-01 | 2.72E+00 | 2.74E+00 | 9.52E-01 | no | | intergenic_variant | MODIFIER |
| 9 | 69614913 | AX-170039966 | 0 | A | G | 0.193 | 8.03E-03 | 1.38E-01 | 2.72E+00 | 2.74E+00 | 9.52E-01 | no | | intergenic_variant | MODIFIER |
| 9 | 69616557 | AX-170484310 | 0 | G | C | 0.193 | 8.03E-03 | 1.38E-01 | 2.72E+00 | 2.74E+00 | 9.52E-01 | no | | intergenic_variant | MODIFIER |
| 9 | 69630138 | AX-170463223 | 0 | T | C | 0.193 | 8.03E-03 | 1.38E-01 | 2.72E+00 | 2.74E+00 | 9.52E-01 | no | | intergenic_variant | MODIFIER |
| 9 | 69632365 | AX-170528891 | 0 | A | G | 0.193 | 8.03E-03 | 1.38E-01 | 2.72E+00 | 2.74E+00 | 9.52E-01 | no | | intergenic_variant | MODIFIER |
| 9 | 69637008 | AX-170326522 | 0 | T | C | 0.193 | 8.03E-03 | 1.38E-01 | 2.72E+00 | 2.74E+00 | 9.52E-01 | no | | intergenic_variant | MODIFIER |
| 9 | 69639212 | AX-170470441 | 0 | G | T | 0.193 | 8.03E-03 | 1.38E-01 | 2.72E+00 | 2.74E+00 | 9.52E-01 | no | | intergenic_variant | MODIFIER |
| 9 | 69650620 | AX-170096577 | 0 | C | T | 0.193 | 8.03E-03 | 1.38E-01 | 2.72E+00 | 2.74E+00 | 9.52E-01 | no | | intergenic_variant | MODIFIER |
| 9 | 69650665 | AX-170108893 | 0 | A | G | 0.193 | 8.03E-03 | 1.38E-01 | 2.72E+00 | 2.74E+00 | 9.52E-01 | no | | intergenic_variant | MODIFIER |
| 9 | 69653499 | AX-170396807 | 0 | C | T | 0.193 | 8.03E-03 | 1.38E-01 | 2.72E+00 | 2.74E+00 | 9.52E-01 | no | | intergenic_variant | MODIFIER |
| 9 | 69654970 | AX-170463946 | 0 | T | C | 0.193 | 8.03E-03 | 1.38E-01 | 2.72E+00 | 2.74E+00 | 9.52E-01 | no | | intergenic_variant | MODIFIER |
| 9 | 69666332 | AX-170116128 | 0 | C | T | 0.193 | 8.03E-03 | 1.38E-01 | 2.72E+00 | 2.74E+00 | 9.52E-01 | no | | intergenic_variant | MODIFIER |
| 9 | 69666371 | AX-170398096 | 0 | A | G | 0.193 | 8.03E-03 | 1.38E-01 | 2.72E+00 | 2.74E+00 | 9.52E-01 | no | | intergenic_variant | MODIFIER |
| 9 | 69677168 | AX-170317147 | 0 | C | T | 0.193 | 8.03E-03 | 1.38E-01 | 2.72E+00 | 2.74E+00 | 9.52E-01 | no | | intergenic_variant | MODIFIER |
| 9 | 69679153 | AX-170532662 | 0 | A | C | 0.193 | 8.03E-03 | 1.38E-01 | 2.72E+00 | 2.74E+00 | 9.52E-01 | no | | intergenic_variant | MODIFIER |
| 9 | 69697060 | AX-97792987 | 0 | G | T | 0.193 | 8.03E-03 | 1.38E-01 | 2.72E+00 | 2.74E+00 | 9.52E-01 | no | | intergenic_variant | MODIFIER |
| 9 | 69697246 | AX-170447983 | 0 | T | C | 0.193 | 8.03E-03 | 1.38E-01 | 2.72E+00 | 2.74E+00 | 9.52E-01 | no | | intergenic_variant | MODIFIER |
| 9 | 69697608 | AX-170118433 | 0 | T | C | 0.193 | 8.03E-03 | 1.38E-01 | 2.72E+00 | 2.74E+00 | 9.52E-01 | no | | intergenic_variant | MODIFIER |
| 9 | 69700792 | AX-170534822 | 0 | T | C | 0.193 | 8.03E-03 | 1.38E-01 | 2.72E+00 | 2.74E+00 | 9.52E-01 | no | | intergenic_variant | MODIFIER |
| 9 | 69704789 | AX-170278452 | 0 | A | G | 0.193 | 8.03E-03 | 1.38E-01 | 2.72E+00 | 2.74E+00 | 9.52E-01 | no | | intergenic_variant | MODIFIER |
| 9 | 69714962 | AX-170452113 | 0 | T | C | 0.193 | 8.03E-03 | 1.38E-01 | 2.72E+00 | 2.74E+00 | 9.52E-01 | no | | regulatory_region_variant | MODIFIER |
| 9 | 69733967 | AX-170082757 | 0 | A | G | 0.193 | 8.03E-03 | 1.38E-01 | 2.72E+00 | 2.74E+00 | 9.52E-01 | no | | intergenic_variant | MODIFIER |
| 9 | 69737553 | AX-97854909 | 0 | G | A | 0.193 | 8.03E-03 | 1.38E-01 | 2.72E+00 | 2.74E+00 | 9.52E-01 | no | | intergenic_variant | MODIFIER |
| 9 | 69740740 | AX-170531930 | 0 | C | T | 0.193 | 8.03E-03 | 1.38E-01 | 2.72E+00 | 2.74E+00 | 9.52E-01 | no | | intergenic_variant | MODIFIER |
| 9 | 69742549 | AX-170082753 | 0 | C | T | 0.193 | 8.03E-03 | 1.38E-01 | 2.72E+00 | 2.74E+00 | 9.52E-01 | no | | intergenic_variant | MODIFIER |
| 9 | 69780486 | AX-170517916 | 0 | C | A | 0.193 | 8.03E-03 | 1.38E-01 | 2.72E+00 | 2.74E+00 | 9.52E-01 | no | B230323A14Rik | intron_variant,non_coding_transcript_variant | MODIFIER |
| 9 | 69790450 | AX-170146901 | 0 | A | G | 0.193 | 8.03E-03 | 1.38E-01 | 2.72E+00 | 2.74E+00 | 9.52E-01 | no | B230323A14Rik | intron_variant,non_coding_transcript_variant | MODIFIER |
| 9 | 69814716 | AX-170349230 | 0 | T | G | 0.193 | 8.03E-03 | 1.38E-01 | 2.72E+00 | 2.74E+00 | 9.52E-01 | no | B230323A14Rik | intron_variant,non_coding_transcript_variant | MODIFIER |
| 9 | 69814973 | AX-170216354 | 0 | C | T | 0.193 | 8.03E-03 | 1.38E-01 | 2.72E+00 | 2.74E+00 | 9.52E-01 | no | B230323A14Rik | intron_variant,non_coding_transcript_variant | MODIFIER |
| 9 | 69822731 | AX-170469850 | 0 | C | T | 0.193 | 8.03E-03 | 1.38E-01 | 2.72E+00 | 2.74E+00 | 9.52E-01 | no | B230323A14Rik | intron_variant,non_coding_transcript_variant | MODIFIER |
| 9 | 69834345 | AX-170286030 | 0 | T | C | 0.193 | 8.03E-03 | 1.38E-01 | 2.72E+00 | 2.74E+00 | 9.52E-01 | no | B230323A14Rik | downstream_gene_variant | MODIFIER |
| 9 | 69848269 | AX-170522875 | 0 | A | T | 0.193 | 8.03E-03 | 1.38E-01 | 2.72E+00 | 2.74E+00 | 9.52E-01 | no | | intergenic_variant | MODIFIER |
| 9 | 69848321 | AX-170527184 | 0 | T | C | 0.193 | 8.03E-03 | 1.38E-01 | 2.72E+00 | 2.74E+00 | 9.52E-01 | no | | intergenic_variant | MODIFIER |
| 9 | 69910680 | AX-170168793 | 0 | C | T | 0.193 | 8.03E-03 | 1.38E-01 | 2.72E+00 | 2.74E+00 | 9.52E-01 | no | | regulatory_region_variant | MODIFIER |
| 9 | 69911332 | AX-170103214 | 0 | G | A | 0.193 | 8.03E-03 | 1.38E-01 | 2.72E+00 | 2.74E+00 | 9.52E-01 | no | | intergenic_variant | MODIFIER |
| 9 | 69917316 | AX-170305790 | 0 | C | T | 0.193 | 8.03E-03 | 1.38E-01 | 2.72E+00 | 2.74E+00 | 9.52E-01 | no | | intergenic_variant | MODIFIER |
| 9 | 69921808 | AX-170476818 | 0 | T | C | 0.193 | 8.03E-03 | 1.38E-01 | 2.72E+00 | 2.74E+00 | 9.52E-01 | no | | intergenic_variant | MODIFIER |
| 9 | 69942218 | AX-97841468 | 0 | A | G | 0.193 | 8.03E-03 | 1.38E-01 | 2.72E+00 | 2.74E+00 | 9.52E-01 | no | | intergenic_variant | MODIFIER |

| | | | | | | | | | | | | | | | |
|---|---|---|---|---|---|---|---|---|---|---|---|---|---|---|---|
| 9 | 69949346 | AX-170503401 | 0 | T | A | 0.193 | 8.03E-03 | 1.38E-01 | 2.72E+00 | 2.74E+00 | 9.52E-01 | no | | regulatory_region_variant | MODIFIER |
| 9 | 69962246 | AX-170154220 | 0 | G | A | 0.193 | 8.03E-03 | 1.38E-01 | 2.72E+00 | 2.74E+00 | 9.52E-01 | no | | intergenic_variant | MODIFIER |
| 9 | 69971761 | AX-97834887 | 0 | G | A | 0.193 | 8.03E-03 | 1.38E-01 | 2.72E+00 | 2.74E+00 | 9.52E-01 | no | | intergenic_variant | MODIFIER |
| 9 | 69986232 | AX-170373903 | 0 | C | G | 0.193 | 8.03E-03 | 1.38E-01 | 2.72E+00 | 2.74E+00 | 9.52E-01 | no | Bnip2 | upstream_gene_variant | MODIFIER |
| 9 | 69990940 | AX-97805768 | 0 | G | A | 0.193 | 8.03E-03 | 1.38E-01 | 2.72E+00 | 2.74E+00 | 9.52E-01 | no | Bnip2 | intron_variant | MODIFIER |
| 9 | 70123632 | AX-170032633 | 0 | G | A | 0.193 | 8.03E-03 | 1.38E-01 | 2.72E+00 | 2.74E+00 | 9.52E-01 | no | Fam81a | intron_variant | MODIFIER |
| 9 | 70130853 | AX-97820254 | 0 | G | A | 0.193 | 8.03E-03 | 1.38E-01 | 2.72E+00 | 2.74E+00 | 9.52E-01 | no | Fam81a | intron_variant | MODIFIER |
| 9 | 70135265 | AX-170032626 | 0 | C | T | 0.193 | 8.03E-03 | 1.38E-01 | 2.72E+00 | 2.74E+00 | 9.52E-01 | no | Fam81a | intron_variant | MODIFIER |
| 9 | 70137973 | AX-97849532 | 0 | G | A | 0.193 | 8.03E-03 | 1.38E-01 | 2.72E+00 | 2.74E+00 | 9.52E-01 | no | Fam81a | intron_variant | MODIFIER |
| 9 | 70149597 | AX-170266095 | 0 | A | G | 0.193 | 8.03E-03 | 1.38E-01 | 2.72E+00 | 2.74E+00 | 9.52E-01 | no | Gm24257 | upstream_gene_variant | MODIFIER |
| 9 | 70268150 | AX-170415130 | 0 | T | G | 0.193 | 8.03E-03 | 1.38E-01 | 2.72E+00 | 2.74E+00 | 9.52E-01 | no | Myo1e | intron_variant | MODIFIER |
| 9 | 70268256 | AX-170310892 | 0 | G | C | 0.193 | 8.03E-03 | 1.38E-01 | 2.72E+00 | 2.74E+00 | 9.52E-01 | no | Myo1e | intron_variant | MODIFIER |
| 9 | 70299950 | AX-170421119 | 0 | T | C | 0.193 | 8.03E-03 | 1.38E-01 | 2.72E+00 | 2.74E+00 | 9.52E-01 | no | Myo1e | intron_variant | MODIFIER |
| 9 | 70301181 | AX-170224966 | 0 | T | C | 0.193 | 8.03E-03 | 1.38E-01 | 2.72E+00 | 2.74E+00 | 9.52E-01 | no | Myo1e | intron_variant | MODIFIER |
| 9 | 70301731 | AX-97805677 | 0 | T | C | 0.193 | 8.03E-03 | 1.38E-01 | 2.72E+00 | 2.74E+00 | 9.52E-01 | no | Myo1e | synonymous_variant | LOW |
| 9 | 70302261 | AX-170210584 | 0 | G | T | 0.193 | 8.03E-03 | 1.38E-01 | 2.72E+00 | 2.74E+00 | 9.52E-01 | no | Myo1e | intron_variant | MODIFIER |
| 9 | 70331645 | AX-97813544 | 0 | A | C | 0.193 | 8.03E-03 | 1.38E-01 | 2.72E+00 | 2.74E+00 | 9.52E-01 | no | Myo1e | intron_variant | MODIFIER |
| 9 | 70331695 | AX-170420129 | 0 | T | C | 0.193 | 8.03E-03 | 1.38E-01 | 2.72E+00 | 2.74E+00 | 9.52E-01 | no | Myo1e | intron_variant | MODIFIER |
| 9 | 70332181 | AX-170497572 | 0 | A | G | 0.193 | 8.03E-03 | 1.38E-01 | 2.72E+00 | 2.74E+00 | 9.52E-01 | no | Myo1e | intron_variant | MODIFIER |
| 9 | 70335328 | AX-97818805 | 0 | G | A | 0.193 | 8.03E-03 | 1.38E-01 | 2.72E+00 | 2.74E+00 | 9.52E-01 | no | Myo1e | intron_variant | MODIFIER |
| 9 | 70371327 | AX-97794252 | 0 | T | C | 0.193 | 8.03E-03 | 1.38E-01 | 2.72E+00 | 2.74E+00 | 9.52E-01 | no | Myo1e | intron_variant | MODIFIER |
| 9 | 70378198 | AX-170450999 | 0 | A | G | 0.193 | 8.03E-03 | 1.38E-01 | 2.72E+00 | 2.74E+00 | 9.52E-01 | no | Myo1e | intron_variant | MODIFIER |
| 9 | 70386910 | AX-170524041 | 0 | A | G | 0.193 | 8.03E-03 | 1.38E-01 | 2.72E+00 | 2.74E+00 | 9.52E-01 | no | Myo1e | intron_variant | MODIFIER |
| 9 | 70406871 | AX-97804727 | 0 | A | G | 0.193 | 8.03E-03 | 1.38E-01 | 2.72E+00 | 2.74E+00 | 9.52E-01 | no | Ccnb2 | downstream_gene_variant | MODIFIER |
| 9 | 70459462 | AX-170157809 | 0 | T | C | 0.193 | 8.03E-03 | 1.38E-01 | 2.72E+00 | 2.74E+00 | 9.52E-01 | no | Rnf111 | intron_variant | MODIFIER |
| 9 | 70471782 | AX-97799707 | 0 | T | C | 0.193 | 8.03E-03 | 1.38E-01 | 2.72E+00 | 2.74E+00 | 9.52E-01 | no | Rnf111 | intron_variant | MODIFIER |
| 9 | 70508381 | AX-97823927 | 0 | T | C | 0.193 | 8.03E-03 | 1.38E-01 | 2.72E+00 | 2.74E+00 | 9.52E-01 | no | Rnf111 | upstream_gene_variant | MODIFIER |
| 9 | 70826986 | AX-97803644 | 0 | G | A | 0.193 | 8.03E-03 | 1.38E-01 | 2.72E+00 | 2.74E+00 | 9.52E-01 | no | Lipc | intron_variant | MODIFIER |
| 9 | 71404472 | AX-97788750 | 0 | C | T | 0.193 | 8.03E-03 | 1.38E-01 | 2.72E+00 | 2.74E+00 | 9.52E-01 | no | | intergenic_variant | MODIFIER |
| 9 | 71413025 | AX-170476697 | 0 | C | T | 0.193 | 8.03E-03 | 1.38E-01 | 2.72E+00 | 2.74E+00 | 9.52E-01 | no | | intergenic_variant | MODIFIER |
| 9 | 71414567 | AX-97792761 | 0 | A | G | 0.193 | 8.03E-03 | 1.38E-01 | 2.72E+00 | 2.74E+00 | 9.52E-01 | no | | intergenic_variant | MODIFIER |
| 9 | 71434833 | AX-170251135 | 0 | C | T | 0.193 | 8.03E-03 | 1.38E-01 | 2.72E+00 | 2.74E+00 | 9.52E-01 | no | | intergenic_variant | MODIFIER |
| 9 | 71435824 | AX-170469005 | 0 | A | G | 0.193 | 8.03E-03 | 1.38E-01 | 2.72E+00 | 2.74E+00 | 9.52E-01 | no | | intergenic_variant | MODIFIER |
| 9 | 71439947 | AX-97805061 | 0 | C | T | 0.193 | 8.03E-03 | 1.38E-01 | 2.72E+00 | 2.74E+00 | 9.52E-01 | no | | intergenic_variant | MODIFIER |
| 9 | 71478658 | AX-170111902 | 0 | G | A | 0.193 | 8.03E-03 | 1.38E-01 | 2.72E+00 | 2.74E+00 | 9.52E-01 | no | Polr2m | 3_prime_UTR_variant | MODIFIER |
| 9 | 71492852 | AX-170425029 | 0 | A | G | 0.193 | 8.03E-03 | 1.38E-01 | 2.72E+00 | 2.74E+00 | 9.52E-01 | no | | regulatory_region_variant | MODIFIER |
| 9 | 71504223 | AX-97799429 | 0 | T | C | 0.193 | 8.03E-03 | 1.38E-01 | 2.72E+00 | 2.74E+00 | 9.52E-01 | no | Myzap | downstream_gene_variant | MODIFIER |
| 9 | 71529646 | AX-170443363 | 0 | T | G | 0.193 | 8.03E-03 | 1.38E-01 | 2.72E+00 | 2.74E+00 | 9.52E-01 | no | Myzap | intron_variant | MODIFIER |
| 9 | 71543637 | AX-170303689 | 0 | C | T | 0.193 | 8.03E-03 | 1.38E-01 | 2.72E+00 | 2.74E+00 | 9.52E-01 | no | Myzap | intron_variant | MODIFIER |
| 9 | 71543716 | AX-170311210 | 0 | G | C | 0.193 | 8.03E-03 | 1.38E-01 | 2.72E+00 | 2.74E+00 | 9.52E-01 | no | Myzap | intron_variant | MODIFIER |
| 9 | 71550209 | AX-97840791 | 0 | T | C | 0.193 | 8.03E-03 | 1.38E-01 | 2.72E+00 | 2.74E+00 | 9.52E-01 | no | Myzap | synonymous_variant | LOW |
| 9 | 71552648 | AX-170326459 | 0 | T | C | 0.193 | 8.03E-03 | 1.38E-01 | 2.72E+00 | 2.74E+00 | 9.52E-01 | no | Myzap | intron_variant | MODIFIER |
| 9 | 71555818 | AX-170232803 | 0 | T | C | 0.193 | 8.03E-03 | 1.38E-01 | 2.72E+00 | 2.74E+00 | 9.52E-01 | no | Myzap | intron_variant | MODIFIER |
| 9 | 71582952 | AX-97804372 | 0 | T | C | 0.193 | 8.03E-03 | 1.38E-01 | 2.72E+00 | 2.74E+00 | 9.52E-01 | no | Myzap | intron_variant | MODIFIER |
| 9 | 71598800 | AX-97811937 | 0 | G | A | 0.193 | 8.03E-03 | 1.38E-01 | 2.72E+00 | 2.74E+00 | 9.52E-01 | no | Gm6018 | downstream_gene_variant | MODIFIER |
| 9 | 71606802 | AX-170283536 | 0 | T | G | 0.193 | 8.03E-03 | 1.38E-01 | 2.72E+00 | 2.74E+00 | 9.52E-01 | no | | regulatory_region_variant | MODIFIER |
| 9 | 71609176 | AX-170032665 | 0 | A | G | 0.193 | 8.03E-03 | 1.38E-01 | 2.72E+00 | 2.74E+00 | 9.52E-01 | no | | regulatory_region_variant | MODIFIER |
| 9 | 71611436 | AX-170453205 | 0 | C | T | 0.193 | 8.03E-03 | 1.38E-01 | 2.72E+00 | 2.74E+00 | 9.52E-01 | no | | regulatory_region_variant | MODIFIER |
| 9 | 71624484 | AX-170032657 | 0 | T | A | 0.193 | 8.03E-03 | 1.38E-01 | 2.72E+00 | 2.74E+00 | 9.52E-01 | no | Cgnl1 | downstream_gene_variant | MODIFIER |
| 9 | 71627435 | AX-170245284 | 0 | C | T | 0.193 | 8.03E-03 | 1.38E-01 | 2.72E+00 | 2.74E+00 | 9.52E-01 | no | Cgnl1 | 3_prime_UTR_variant | MODIFIER |
| 9 | 71633552 | AX-97824688 | 0 | A | G | 0.193 | 8.03E-03 | 1.38E-01 | 2.72E+00 | 2.74E+00 | 9.52E-01 | no | Cgnl1 | intron_variant | MODIFIER |
| 9 | 71656055 | AX-170032660 | 0 | T | C | 0.193 | 8.03E-03 | 1.38E-01 | 2.72E+00 | 2.74E+00 | 9.52E-01 | no | Cgnl1 | synonymous_variant | LOW |
| 9 | 71797707 | AX-97784694 | 0 | G | A | 0.193 | 8.03E-03 | 1.38E-01 | 2.72E+00 | 2.74E+00 | 9.52E-01 | no | | intergenic_variant | MODIFIER |
| 9 | 71871847 | AX-170082786 | 0 | T | G | 0.193 | 8.03E-03 | 1.38E-01 | 2.72E+00 | 2.74E+00 | 9.52E-01 | no | Tcf12 | intron_variant | MODIFIER |

| | | | | | | | | | | | | | | | |
|---|---|---|---|---|---|---|---|---|---|---|---|---|---|---|---|
| 9 | 71900799 | AX-170098650 | 0 | T | C | 0.193 | 8.03E-03 | 1.38E-01 | 2.72E+00 | 2.74E+00 | 9.52E-01 | no | Tcf12 | intron_variant | MODIFIER |
| 9 | 71912969 | AX-170123329 | 0 | G | C | 0.193 | 8.03E-03 | 1.38E-01 | 2.72E+00 | 2.74E+00 | 9.52E-01 | no | Tcf12 | intron_variant | MODIFIER |
| 9 | 71938428 | AX-170409904 | 0 | C | G | 0.193 | 8.03E-03 | 1.38E-01 | 2.72E+00 | 2.74E+00 | 9.52E-01 | no | Tcf12 | intron_variant | MODIFIER |
| 9 | 72020615 | AX-97799342 | 0 | A | C | 0.193 | 8.03E-03 | 1.38E-01 | 2.72E+00 | 2.74E+00 | 9.52E-01 | no | Tcf12 | intron_variant | MODIFIER |
| 9 | 72027534 | AX-170197700 | 0 | G | T | 0.193 | 8.03E-03 | 1.38E-01 | 2.72E+00 | 2.74E+00 | 9.52E-01 | no | Tcf12 | intron_variant | MODIFIER |
| 9 | 72039541 | AX-170517723 | 0 | T | C | 0.193 | 8.03E-03 | 1.38E-01 | 2.72E+00 | 2.74E+00 | 9.52E-01 | no | Tcf12 | intron_variant | MODIFIER |
| 9 | 72073523 | AX-170090330 | 0 | G | A | 0.193 | 8.03E-03 | 1.38E-01 | 2.72E+00 | 2.74E+00 | 9.52E-01 | no | Tcf12 | intron_variant | MODIFIER |
| 9 | 72079462 | AX-170471820 | 0 | T | C | 0.193 | 8.03E-03 | 1.38E-01 | 2.72E+00 | 2.74E+00 | 9.52E-01 | no | Tcf12 | intron_variant | MODIFIER |
| 9 | 72405773 | AX-170402254 | 0 | A | G | 0.193 | 8.03E-03 | 1.38E-01 | 2.72E+00 | 2.74E+00 | 9.52E-01 | no | BC065403 | intron_variant,non_coding_transcript_variant | MODIFIER |
| 9 | 72468495 | AX-170522433 | 0 | G | A | 0.193 | 8.03E-03 | 1.38E-01 | 2.72E+00 | 2.74E+00 | 9.52E-01 | no | Tex9 | intron_variant | MODIFIER |
| 9 | 72468524 | AX-170143144 | 0 | T | C | 0.193 | 8.03E-03 | 1.38E-01 | 2.72E+00 | 2.74E+00 | 9.52E-01 | no | Tex9 | intron_variant | MODIFIER |
| 9 | 72468830 | AX-170439569 | 0 | C | T | 0.193 | 8.03E-03 | 1.38E-01 | 2.72E+00 | 2.74E+00 | 9.52E-01 | no | Tex9 | intron_variant | MODIFIER |
| 9 | 72598502 | AX-170370393 | 0 | T | C | 0.193 | 8.03E-03 | 1.38E-01 | 2.72E+00 | 2.74E+00 | 9.52E-01 | no | Rfx7 | intron_variant | MODIFIER |
| 9 | 72653596 | AX-170154487 | 0 | T | C | 0.193 | 8.03E-03 | 1.38E-01 | 2.72E+00 | 2.74E+00 | 9.52E-01 | no | | intergenic_variant | MODIFIER |
| 9 | 72942717 | AX-170380668 | 0 | G | C | 0.193 | 8.03E-03 | 1.38E-01 | 2.72E+00 | 2.74E+00 | 9.52E-01 | no | Pygo1 | intron_variant | MODIFIER |
| 9 | 73029705 | AX-170232479 | 0 | C | T | 0.193 | 8.03E-03 | 1.38E-01 | 2.72E+00 | 2.74E+00 | 9.52E-01 | no | Pigb | missense_variant,splice_region_variant | MODERATE |
| 9 | 73053808 | AX-170529533 | 0 | G | A | 0.193 | 8.03E-03 | 1.38E-01 | 2.72E+00 | 2.74E+00 | 9.52E-01 | no | Rab27a | intron_variant | MODIFIER |
| 9 | 73104552 | AX-170032694 | 0 | T | C | 0.193 | 8.03E-03 | 1.38E-01 | 2.72E+00 | 2.74E+00 | 9.52E-01 | no | Khdc3 | downstream_gene_variant | MODIFIER |
| 9 | 73493919 | AX-97784188 | 0 | C | T | 0.193 | 8.03E-03 | 1.38E-01 | 2.72E+00 | 2.74E+00 | 9.52E-01 | no | Unc13c | intron_variant | MODIFIER |
| 9 | 73807237 | AX-170032704 | 0 | T | A | 0.193 | 8.03E-03 | 1.38E-01 | 2.72E+00 | 2.74E+00 | 9.52E-01 | no | Unc13c | intron_variant | MODIFIER |
| 9 | 73884420 | AX-170271558 | 0 | T | C | 0.193 | 8.03E-03 | 1.38E-01 | 2.72E+00 | 2.74E+00 | 9.52E-01 | no | Unc13c | intron_variant | MODIFIER |
| 9 | 74106817 | AX-97797450 | 0 | C | G | 0.193 | 8.03E-03 | 1.38E-01 | 2.72E+00 | 2.74E+00 | 9.52E-01 | no | Wdr72 | upstream_gene_variant | MODIFIER |
| 9 | 74132644 | AX-170304876 | 0 | T | C | 0.193 | 8.03E-03 | 1.38E-01 | 2.72E+00 | 2.74E+00 | 9.52E-01 | no | Wdr72 | intron_variant | MODIFIER |
| 9 | 74136735 | AX-170371159 | 0 | C | T | 0.193 | 8.03E-03 | 1.38E-01 | 2.72E+00 | 2.74E+00 | 9.52E-01 | no | Wdr72 | intron_variant | MODIFIER |
| 9 | 74141014 | AX-170185170 | 0 | C | T | 0.193 | 8.03E-03 | 1.38E-01 | 2.72E+00 | 2.74E+00 | 9.52E-01 | no | Wdr72 | intron_variant | MODIFIER |
| 9 | 74254111 | AX-170275460 | 0 | A | G | 0.193 | 8.03E-03 | 1.38E-01 | 2.72E+00 | 2.74E+00 | 9.52E-01 | no | Wdr72 | intron_variant | MODIFIER |
| 9 | 74255266 | AX-170428306 | 0 | T | C | 0.193 | 8.03E-03 | 1.38E-01 | 2.72E+00 | 2.74E+00 | 9.52E-01 | no | Wdr72 | intron_variant | MODIFIER |
| 9 | 74256001 | AX-170159818 | 0 | T | C | 0.193 | 8.03E-03 | 1.38E-01 | 2.72E+00 | 2.74E+00 | 9.52E-01 | no | Wdr72 | intron_variant | MODIFIER |
| 9 | 74305218 | AX-170299583 | 0 | A | G | 0.193 | 8.03E-03 | 1.38E-01 | 2.72E+00 | 2.74E+00 | 9.52E-01 | no | | intergenic_variant | MODIFIER |
| 9 | 74329104 | AX-170032711 | 0 | A | C | 0.193 | 8.03E-03 | 1.38E-01 | 2.72E+00 | 2.74E+00 | 9.52E-01 | no | | regulatory_region_variant | MODIFIER |
| 9 | 74438242 | AX-170293385 | 0 | C | T | 0.193 | 8.03E-03 | 1.38E-01 | 2.72E+00 | 2.74E+00 | 9.52E-01 | no | | intergenic_variant | MODIFIER |
| 9 | 74542365 | AX-170517472 | 0 | T | C | 0.193 | 8.03E-03 | 1.38E-01 | 2.72E+00 | 2.74E+00 | 9.52E-01 | no | | intergenic_variant | MODIFIER |
| 9 | 74573171 | AX-170113156 | 0 | A | G | 0.193 | 8.03E-03 | 1.38E-01 | 2.72E+00 | 2.74E+00 | 9.52E-01 | no | Gm27189 | downstream_gene_variant | MODIFIER |
| 9 | 74573390 | AX-170349923 | 0 | A | G | 0.193 | 8.03E-03 | 1.38E-01 | 2.72E+00 | 2.74E+00 | 9.52E-01 | no | Gm27189 | downstream_gene_variant | MODIFIER |
| 9 | 74573406 | AX-170465388 | 0 | A | G | 0.193 | 8.03E-03 | 1.38E-01 | 2.72E+00 | 2.74E+00 | 9.52E-01 | no | Gm27189 | downstream_gene_variant | MODIFIER |
| 9 | 74674430 | AX-170489751 | 0 | T | C | 0.193 | 8.03E-03 | 1.38E-01 | 2.72E+00 | 2.74E+00 | 9.52E-01 | no | | intergenic_variant | MODIFIER |
| 9 | 74698120 | AX-170032716 | 0 | T | C | 0.193 | 8.03E-03 | 1.38E-01 | 2.72E+00 | 2.74E+00 | 9.52E-01 | no | | intergenic_variant | MODIFIER |
| 9 | 74759006 | AX-170082822 | 0 | C | T | 0.193 | 8.03E-03 | 1.38E-01 | 2.72E+00 | 2.74E+00 | 9.52E-01 | no | | intergenic_variant | MODIFIER |
| 9 | 74794051 | AX-170340200 | 0 | T | C | 0.193 | 8.03E-03 | 1.38E-01 | 2.72E+00 | 2.74E+00 | 9.52E-01 | no | | intergenic_variant | MODIFIER |
| 9 | 74799716 | AX-97796402 | 0 | G | A | 0.193 | 8.03E-03 | 1.38E-01 | 2.72E+00 | 2.74E+00 | 9.52E-01 | no | | intergenic_variant | MODIFIER |
| 9 | 74804048 | AX-170032722 | 0 | T | C | 0.193 | 8.03E-03 | 1.38E-01 | 2.72E+00 | 2.74E+00 | 9.52E-01 | no | | intergenic_variant | MODIFIER |
| 9 | 74878003 | AX-170249288 | 0 | T | G | 0.193 | 8.03E-03 | 1.38E-01 | 2.72E+00 | 2.74E+00 | 9.52E-01 | no | Onecut1 | intron_variant | MODIFIER |
| 9 | 74885661 | AX-170134236 | 0 | A | G | 0.193 | 8.03E-03 | 1.38E-01 | 2.72E+00 | 2.74E+00 | 9.52E-01 | no | Onecut1 | intron_variant | MODIFIER |
| 9 | 74887469 | AX-170251939 | 0 | G | T | 0.193 | 8.03E-03 | 1.38E-01 | 2.72E+00 | 2.74E+00 | 9.52E-01 | no | Onecut1 | intron_variant | MODIFIER |
| 9 | 74887497 | AX-170383424 | 0 | A | G | 0.193 | 8.03E-03 | 1.38E-01 | 2.72E+00 | 2.74E+00 | 9.52E-01 | no | Onecut1 | intron_variant | MODIFIER |
| 9 | 74887588 | AX-170354372 | 0 | T | C | 0.193 | 8.03E-03 | 1.38E-01 | 2.72E+00 | 2.74E+00 | 9.52E-01 | no | Onecut1 | intron_variant | MODIFIER |
| 9 | 74905052 | AX-170356683 | 0 | C | A | 0.193 | 8.03E-03 | 1.38E-01 | 2.72E+00 | 2.74E+00 | 9.52E-01 | no | | regulatory_region_variant | MODIFIER |
| 9 | 74906102 | AX-97805086 | 0 | G | C | 0.193 | 8.03E-03 | 1.38E-01 | 2.72E+00 | 2.74E+00 | 9.52E-01 | no | | regulatory_region_variant | MODIFIER |
| 9 | 74909503 | AX-170377573 | 0 | C | T | 0.193 | 8.03E-03 | 1.38E-01 | 2.72E+00 | 2.74E+00 | 9.52E-01 | no | | regulatory_region_variant | MODIFIER |
| 9 | 74913204 | AX-170466283 | 0 | T | C | 0.193 | 8.03E-03 | 1.38E-01 | 2.72E+00 | 2.74E+00 | 9.52E-01 | no | | regulatory_region_variant | MODIFIER |
| 9 | 74939426 | AX-97794913 | 0 | A | G | 0.193 | 8.03E-03 | 1.38E-01 | 2.72E+00 | 2.74E+00 | 9.52E-01 | no | | regulatory_region_variant | MODIFIER |
| 9 | 74946894 | AX-170205391 | 0 | A | G | 0.193 | 8.03E-03 | 1.38E-01 | 2.72E+00 | 2.74E+00 | 9.52E-01 | no | | regulatory_region_variant | MODIFIER |
| 9 | 74965483 | AX-170090335 | 0 | T | A | 0.193 | 8.03E-03 | 1.38E-01 | 2.72E+00 | 2.74E+00 | 9.52E-01 | no | Fam214a | intron_variant | MODIFIER |
| 9 | 74988380 | AX-97806024 | 0 | C | T | 0.193 | 8.03E-03 | 1.38E-01 | 2.72E+00 | 2.74E+00 | 9.52E-01 | no | Fam214a | intron_variant | MODIFIER |

| | | | | | | | | | | | | | | | |
|---|---|---|---|---|---|---|---|---|---|---|---|---|---|---|---|
| 9 | 74994443 | AX-170431817 | 0 | T | C | 0.193 | 8.03E-03 | 1.38E-01 | 2.72E+00 | 2.74E+00 | 9.52E-01 | no | Fam214a | intron_variant | MODIFIER |
| 9 | 75024316 | AX-97805830 | 0 | T | C | 0.193 | 8.03E-03 | 1.38E-01 | 2.72E+00 | 2.74E+00 | 9.52E-01 | no | Fam214a | intron_variant | MODIFIER |
| 9 | 75031362 | AX-170373053 | 0 | G | A | 0.193 | 8.03E-03 | 1.38E-01 | 2.72E+00 | 2.74E+00 | 9.52E-01 | no | Fam214a | intron_variant | MODIFIER |
| 9 | 75033717 | AX-170448911 | 0 | G | C | 0.193 | 8.03E-03 | 1.38E-01 | 2.72E+00 | 2.74E+00 | 9.52E-01 | no | Arpp19 | upstream_gene_variant | MODIFIER |
| 9 | 75056053 | AX-170217455 | 0 | T | G | 0.193 | 8.03E-03 | 1.38E-01 | 2.72E+00 | 2.74E+00 | 9.52E-01 | no | Arpp19 | intron_variant | MODIFIER |
| 9 | 75056223 | AX-97805150 | 0 | C | T | 0.193 | 8.03E-03 | 1.38E-01 | 2.72E+00 | 2.74E+00 | 9.52E-01 | no | Arpp19 | intron_variant | MODIFIER |
| 9 | 75283090 | AX-170359574 | 0 | G | A | 0.193 | 8.03E-03 | 1.38E-01 | 2.72E+00 | 2.74E+00 | 9.52E-01 | no | Myo5c | intron_variant | MODIFIER |
| 9 | 75290643 | AX-170481063 | 0 | G | A | 0.193 | 8.03E-03 | 1.38E-01 | 2.72E+00 | 2.74E+00 | 9.52E-01 | no | Myo5c | intron_variant | MODIFIER |
| 9 | 75294839 | AX-170521044 | 0 | G | T | 0.193 | 8.03E-03 | 1.38E-01 | 2.72E+00 | 2.74E+00 | 9.52E-01 | no | Myo5c | intron_variant | MODIFIER |
| 9 | 75363759 | AX-170177626 | 0 | C | T | 0.193 | 8.03E-03 | 1.38E-01 | 2.72E+00 | 2.74E+00 | 9.52E-01 | no | | intergenic_variant | MODIFIER |
| 9 | 75447893 | AX-170082833 | 0 | A | C | 0.193 | 8.03E-03 | 1.38E-01 | 2.72E+00 | 2.74E+00 | 9.52E-01 | no | Leo1 | intron_variant | MODIFIER |
| 9 | 75448728 | AX-97830620 | 0 | A | G | 0.193 | 8.03E-03 | 1.38E-01 | 2.72E+00 | 2.74E+00 | 9.52E-01 | no | Leo1 | intron_variant | MODIFIER |
| 9 | 75455620 | AX-170082837 | 0 | T | C | 0.193 | 8.03E-03 | 1.38E-01 | 2.72E+00 | 2.74E+00 | 9.52E-01 | no | Leo1 | intron_variant | MODIFIER |
| 9 | 75461046 | AX-170281308 | 0 | C | A | 0.193 | 8.03E-03 | 1.38E-01 | 2.72E+00 | 2.74E+00 | 9.52E-01 | no | Leo1 | synonymous_variant | LOW |
| 9 | 75463911 | AX-170390012 | 0 | C | G | 0.193 | 8.03E-03 | 1.38E-01 | 2.72E+00 | 2.74E+00 | 9.52E-01 | no | Leo1 | intron_variant | MODIFIER |
| 9 | 75479667 | AX-170535328 | 0 | A | G | 0.193 | 8.03E-03 | 1.38E-01 | 2.72E+00 | 2.74E+00 | 9.52E-01 | no | Gm7972 | splice_region_variant,intron_variant,non_coding_transcript_variant | LOW |
| 9 | 75481056 | AX-170267236 | 0 | T | C | 0.193 | 8.03E-03 | 1.38E-01 | 2.72E+00 | 2.74E+00 | 9.52E-01 | no | Gm7972 | intron_variant,non_coding_transcript_variant | MODIFIER |
| 9 | 75494501 | AX-170444527 | 0 | C | A | 0.193 | 8.03E-03 | 1.38E-01 | 2.72E+00 | 2.74E+00 | 9.52E-01 | no | Tmod3 | downstream_gene_variant | MODIFIER |
| 9 | 75527758 | AX-97823041 | 0 | T | G | 0.193 | 8.03E-03 | 1.38E-01 | 2.72E+00 | 2.74E+00 | 9.52E-01 | no | Tmod3 | intron_variant | MODIFIER |
| 9 | 75554058 | AX-170242356 | 0 | T | C | 0.193 | 8.03E-03 | 1.38E-01 | 2.72E+00 | 2.74E+00 | 9.52E-01 | no | Tmod3 | intron_variant | MODIFIER |
| 9 | 75672453 | AX-170302705 | 0 | T | C | 0.193 | 8.03E-03 | 1.38E-01 | 2.72E+00 | 2.74E+00 | 9.52E-01 | no | Scg3 | intron_variant | MODIFIER |
| 9 | 75673913 | AX-170032731 | 0 | G | A | 0.193 | 8.03E-03 | 1.38E-01 | 2.72E+00 | 2.74E+00 | 9.52E-01 | no | Scg3 | intron_variant | MODIFIER |
| 9 | 76009745 | AX-170317417 | 0 | A | C | 0.193 | 8.03E-03 | 1.38E-01 | 2.72E+00 | 2.74E+00 | 9.52E-01 | no | Gm26903 | intron_variant,non_coding_transcript_variant | MODIFIER |
| 9 | 76013245 | AX-170331513 | 0 | T | C | 0.193 | 8.03E-03 | 1.38E-01 | 2.72E+00 | 2.74E+00 | 9.52E-01 | no | Hmgcll1 | upstream_gene_variant | MODIFIER |
| 9 | 76023178 | AX-170482162 | 0 | G | A | 0.193 | 8.03E-03 | 1.38E-01 | 2.72E+00 | 2.74E+00 | 9.52E-01 | no | Hmgcll1 | intron_variant | MODIFIER |
| 9 | 76254573 | AX-170138166 | 0 | G | A | 0.193 | 8.03E-03 | 1.38E-01 | 2.72E+00 | 2.74E+00 | 9.52E-01 | no | Hcrtr2 | synonymous_variant | LOW |
| 9 | 76254727 | AX-170032735 | 0 | C | T | 0.193 | 8.03E-03 | 1.38E-01 | 2.72E+00 | 2.74E+00 | 9.52E-01 | no | Hcrtr2 | intron_variant | MODIFIER |
| 9 | 76467043 | AX-170372487 | 0 | A | G | 0.193 | 8.03E-03 | 1.38E-01 | 2.72E+00 | 2.74E+00 | 9.52E-01 | no | | intergenic_variant | MODIFIER |
| 9 | 76681943 | AX-170250364 | 0 | A | G | 0.193 | 8.03E-03 | 1.38E-01 | 2.72E+00 | 2.74E+00 | 9.52E-01 | no | | intergenic_variant | MODIFIER |
| 9 | 76691194 | AX-170414652 | 0 | T | A | 0.193 | 8.03E-03 | 1.38E-01 | 2.72E+00 | 2.74E+00 | 9.52E-01 | no | | intergenic_variant | MODIFIER |
| 9 | 76696719 | AX-170419846 | 0 | A | G | 0.193 | 8.03E-03 | 1.38E-01 | 2.72E+00 | 2.74E+00 | 9.52E-01 | no | | intergenic_variant | MODIFIER |
| 9 | 76697232 | AX-170441000 | 0 | T | C | 0.193 | 8.03E-03 | 1.38E-01 | 2.72E+00 | 2.74E+00 | 9.52E-01 | no | | intergenic_variant | MODIFIER |
| 9 | 76732937 | AX-170172888 | 0 | T | C | 0.193 | 8.03E-03 | 1.38E-01 | 2.72E+00 | 2.74E+00 | 9.52E-01 | no | | intergenic_variant | MODIFIER |
| 9 | 76733026 | AX-170032744 | 0 | T | A | 0.193 | 8.03E-03 | 1.38E-01 | 2.72E+00 | 2.74E+00 | 9.52E-01 | no | | intergenic_variant | MODIFIER |
| 9 | 76756585 | AX-170032747 | 0 | A | G | 0.193 | 8.03E-03 | 1.38E-01 | 2.72E+00 | 2.74E+00 | 9.52E-01 | no | Gm27155 | downstream_gene_variant | MODIFIER |
| 9 | 76787644 | AX-170515758 | 0 | A | C | 0.193 | 8.03E-03 | 1.38E-01 | 2.72E+00 | 2.74E+00 | 9.52E-01 | no | Gm34158 | downstream_gene_variant | MODIFIER |
| 9 | 76787825 | AX-170119133 | 0 | A | C | 0.193 | 8.03E-03 | 1.38E-01 | 2.72E+00 | 2.74E+00 | 9.52E-01 | no | Gm34158 | downstream_gene_variant | MODIFIER |
| 9 | 76790261 | AX-170082862 | 0 | A | G | 0.193 | 8.03E-03 | 1.38E-01 | 2.72E+00 | 2.74E+00 | 9.52E-01 | no | Gm34158 | downstream_gene_variant | MODIFIER |
| 9 | 76790551 | AX-170082863 | 0 | A | G | 0.193 | 8.03E-03 | 1.38E-01 | 2.72E+00 | 2.74E+00 | 9.52E-01 | no | Gm34158 | downstream_gene_variant | MODIFIER |
| 9 | 76801459 | AX-170032746 | 0 | A | C | 0.193 | 8.03E-03 | 1.38E-01 | 2.72E+00 | 2.74E+00 | 9.52E-01 | no | Gm47800 | downstream_gene_variant | MODIFIER |
| 9 | 76843214 | AX-170032743 | 0 | C | A | 0.193 | 8.03E-03 | 1.38E-01 | 2.72E+00 | 2.74E+00 | 9.52E-01 | no | | intergenic_variant | MODIFIER |
| 9 | 76851458 | AX-170138186 | 0 | T | C | 0.193 | 8.03E-03 | 1.38E-01 | 2.72E+00 | 2.74E+00 | 9.52E-01 | no | | intergenic_variant | MODIFIER |
| 9 | 76884430 | AX-170247007 | 0 | C | G | 0.193 | 8.03E-03 | 1.38E-01 | 2.72E+00 | 2.74E+00 | 9.52E-01 | no | | intergenic_variant | MODIFIER |
| 9 | 76892677 | AX-170435776 | 0 | T | G | 0.193 | 8.03E-03 | 1.38E-01 | 2.72E+00 | 2.74E+00 | 9.52E-01 | no | | intergenic_variant | MODIFIER |
| 9 | 76894486 | AX-170374009 | 0 | A | T | 0.193 | 8.03E-03 | 1.38E-01 | 2.72E+00 | 2.74E+00 | 9.52E-01 | no | | intergenic_variant | MODIFIER |
| 9 | 77018252 | AX-170516353 | 0 | A | G | 0.193 | 8.03E-03 | 1.38E-01 | 2.72E+00 | 2.74E+00 | 9.52E-01 | no | Tinag | intron_variant | MODIFIER |
| 9 | 77071673 | AX-170380118 | 0 | A | G | 0.193 | 8.03E-03 | 1.38E-01 | 2.72E+00 | 2.74E+00 | 9.52E-01 | no | | intergenic_variant | MODIFIER |
| 9 | 77071883 | AX-170460167 | 0 | C | T | 0.193 | 8.03E-03 | 1.38E-01 | 2.72E+00 | 2.74E+00 | 9.52E-01 | no | | intergenic_variant | MODIFIER |
| 9 | 77102178 | AX-170361726 | 0 | C | T | 0.193 | 8.03E-03 | 1.38E-01 | 2.72E+00 | 2.74E+00 | 9.52E-01 | no | Mlip | 3_prime_UTR_variant | MODIFIER |
| 9 | 77114575 | AX-170488697 | 0 | A | T | 0.193 | 8.03E-03 | 1.38E-01 | 2.72E+00 | 2.74E+00 | 9.52E-01 | no | Mlip | intron_variant | MODIFIER |
| 9 | 77265556 | AX-170206065 | 0 | C | T | 0.193 | 8.03E-03 | 1.38E-01 | 2.72E+00 | 2.74E+00 | 9.52E-01 | no | Mlip | intron_variant | MODIFIER |
| 9 | 77305686 | AX-170426228 | 0 | C | T | 0.193 | 8.03E-03 | 1.38E-01 | 2.72E+00 | 2.74E+00 | 9.52E-01 | no | Mlip | intron_variant | MODIFIER |
| 9 | 77308191 | AX-170328339 | 0 | A | G | 0.193 | 8.03E-03 | 1.38E-01 | 2.72E+00 | 2.74E+00 | 9.52E-01 | no | Mlip | intron_variant | MODIFIER |
| 9 | 77354397 | AX-170032748 | 0 | T | A | 0.193 | 8.03E-03 | 1.38E-01 | 2.72E+00 | 2.74E+00 | 9.52E-01 | no | Mlip | upstream_gene_variant | MODIFIER |

| | | | | | | | | | | | | | | | |
|---|---|---|---|---|---|---|---|---|---|---|---|---|---|---|---|
| 9 | 77355081 | AX-170032756 | 0 | G | A | 0.193 | 8.03E-03 | 1.38E-01 | 2.72E+00 | 2.74E+00 | 9.52E-01 | no | Mlip | upstream_gene_variant | MODIFIER |
| 9 | 77369478 | AX-170086153 | 0 | A | G | 0.193 | 8.03E-03 | 1.38E-01 | 2.72E+00 | 2.74E+00 | 9.52E-01 | no | 5730403I07Rik | intron_variant,non_coding_transcript_variant | MODIFIER |
| 9 | 77433750 | AX-97805181 | 0 | T | G | 0.193 | 8.03E-03 | 1.38E-01 | 2.72E+00 | 2.74E+00 | 9.52E-01 | no | Lrrc1 | intron_variant | MODIFIER |
| 9 | 77441630 | AX-170271969 | 0 | A | G | 0.193 | 8.03E-03 | 1.38E-01 | 2.72E+00 | 2.74E+00 | 9.52E-01 | no | Lrrc1 | intron_variant | MODIFIER |
| 9 | 77447337 | AX-170319437 | 0 | T | C | 0.193 | 8.03E-03 | 1.38E-01 | 2.72E+00 | 2.74E+00 | 9.52E-01 | no | Lrrc1 | intron_variant | MODIFIER |
| 9 | 77447458 | AX-170533473 | 0 | C | T | 0.193 | 8.03E-03 | 1.38E-01 | 2.72E+00 | 2.74E+00 | 9.52E-01 | no | Lrrc1 | intron_variant | MODIFIER |
| 9 | 77453426 | AX-170341014 | 0 | G | A | 0.193 | 8.03E-03 | 1.38E-01 | 2.72E+00 | 2.74E+00 | 9.52E-01 | no | Lrrc1 | intron_variant | MODIFIER |
| 9 | 77453475 | AX-170255527 | 0 | C | T | 0.193 | 8.03E-03 | 1.38E-01 | 2.72E+00 | 2.74E+00 | 9.52E-01 | no | Lrrc1 | intron_variant | MODIFIER |
| 9 | 77631978 | AX-170143084 | 0 | C | T | 0.193 | 8.03E-03 | 1.38E-01 | 2.72E+00 | 2.74E+00 | 9.52E-01 | no | Klhl31 | upstream_gene_variant | MODIFIER |
| 9 | 77637970 | AX-170378539 | 0 | A | G | 0.193 | 8.03E-03 | 1.38E-01 | 2.72E+00 | 2.74E+00 | 9.52E-01 | no | Klhl31 | intron_variant | MODIFIER |
| 9 | 77664844 | AX-170106917 | 0 | G | A | 0.193 | 8.03E-03 | 1.38E-01 | 2.72E+00 | 2.74E+00 | 9.52E-01 | no | Klhl31 | downstream_gene_variant | MODIFIER |
| 9 | 77673522 | AX-97829769 | 0 | C | T | 0.193 | 8.03E-03 | 1.38E-01 | 2.72E+00 | 2.74E+00 | 9.52E-01 | no | | intergenic_variant | MODIFIER |
| 9 | 77698281 | AX-170082873 | 0 | T | C | 0.193 | 8.03E-03 | 1.38E-01 | 2.72E+00 | 2.74E+00 | 9.52E-01 | no | Gm47790 | intron_variant,non_coding_transcript_variant | MODIFIER |
| 9 | 77705855 | AX-170420949 | 0 | G | C | 0.193 | 8.03E-03 | 1.38E-01 | 2.72E+00 | 2.74E+00 | 9.52E-01 | no | Gm47789 | downstream_gene_variant | MODIFIER |
| 9 | 77706081 | AX-170082877 | 0 | T | C | 0.193 | 8.03E-03 | 1.38E-01 | 2.72E+00 | 2.74E+00 | 9.52E-01 | no | Gm47789 | downstream_gene_variant | MODIFIER |
| 9 | 77711879 | AX-170032765 | 0 | T | C | 0.193 | 8.03E-03 | 1.38E-01 | 2.72E+00 | 2.74E+00 | 9.52E-01 | no | Gm47789 | upstream_gene_variant | MODIFIER |
| 9 | 77713360 | AX-170032757 | 0 | A | G | 0.193 | 8.03E-03 | 1.38E-01 | 2.72E+00 | 2.74E+00 | 9.52E-01 | no | Gm47790 | intron_variant,non_coding_transcript_variant | MODIFIER |
| 9 | 77719288 | AX-170121649 | 0 | C | T | 0.193 | 8.03E-03 | 1.38E-01 | 2.72E+00 | 2.74E+00 | 9.52E-01 | no | Gm47790 | intron_variant,non_coding_transcript_variant | MODIFIER |
| 9 | 77720698 | AX-170096594 | 0 | A | G | 0.193 | 8.03E-03 | 1.38E-01 | 2.72E+00 | 2.74E+00 | 9.52E-01 | no | Gm47790 | intron_variant,non_coding_transcript_variant | MODIFIER |
| 9 | 77725167 | AX-170255526 | 0 | G | T | 0.193 | 8.03E-03 | 1.38E-01 | 2.72E+00 | 2.74E+00 | 9.52E-01 | no | Gm47790 | intron_variant,non_coding_transcript_variant | MODIFIER |
| 9 | 77766255 | AX-170032758 | 0 | G | A | 0.193 | 8.03E-03 | 1.38E-01 | 2.72E+00 | 2.74E+00 | 9.52E-01 | no | Gclc | intron_variant | MODIFIER |
| 9 | 77782880 | AX-170357208 | 0 | T | C | 0.193 | 8.03E-03 | 1.38E-01 | 2.72E+00 | 2.74E+00 | 9.52E-01 | no | Gclc | intron_variant | MODIFIER |
| 9 | 77783047 | AX-170416270 | 0 | A | C | 0.193 | 8.03E-03 | 1.38E-01 | 2.72E+00 | 2.74E+00 | 9.52E-01 | no | Gclc | intron_variant | MODIFIER |
| 9 | 77783235 | AX-170106755 | 0 | G | A | 0.193 | 8.03E-03 | 1.38E-01 | 2.72E+00 | 2.74E+00 | 9.52E-01 | no | Gclc | intron_variant | MODIFIER |
| 9 | 77784124 | AX-170032760 | 0 | A | G | 0.193 | 8.03E-03 | 1.38E-01 | 2.72E+00 | 2.74E+00 | 9.52E-01 | no | Gclc | intron_variant | MODIFIER |
| 9 | 77799810 | AX-97856222 | 0 | C | T | 0.193 | 8.03E-03 | 1.38E-01 | 2.72E+00 | 2.74E+00 | 9.52E-01 | no | | intergenic_variant | MODIFIER |
| 9 | 77805100 | AX-170032764 | 0 | T | C | 0.193 | 8.03E-03 | 1.38E-01 | 2.72E+00 | 2.74E+00 | 9.52E-01 | no | Gm34654 | downstream_gene_variant | MODIFIER |
| 9 | 77845586 | AX-170090341 | 0 | G | C | 0.193 | 8.03E-03 | 1.38E-01 | 2.72E+00 | 2.74E+00 | 9.52E-01 | no | Gm47818 | intron_variant,non_coding_transcript_variant | MODIFIER |
| 9 | 77846189 | AX-170032762 | 0 | G | A | 0.193 | 8.03E-03 | 1.38E-01 | 2.72E+00 | 2.74E+00 | 9.52E-01 | no | Gm47818 | non_coding_transcript_exon_variant | MODIFIER |
| 9 | 77884716 | AX-170032759 | 0 | A | T | 0.193 | 8.03E-03 | 1.38E-01 | 2.72E+00 | 2.74E+00 | 9.52E-01 | no | | intergenic_variant | MODIFIER |
| 9 | 77957030 | AX-170380933 | 0 | C | T | 0.193 | 8.03E-03 | 1.38E-01 | 2.72E+00 | 2.74E+00 | 9.52E-01 | no | Elovl5 | intron_variant | MODIFIER |
| 9 | 78039622 | AX-170479472 | 0 | T | C | 0.193 | 8.03E-03 | 1.38E-01 | 2.72E+00 | 2.74E+00 | 9.52E-01 | no | | regulatory_region_variant | MODIFIER |
| 9 | 78039730 | AX-170499540 | 0 | G | A | 0.193 | 8.03E-03 | 1.38E-01 | 2.72E+00 | 2.74E+00 | 9.52E-01 | no | | regulatory_region_variant | MODIFIER |
| 9 | 78039867 | AX-170423243 | 0 | G | A | 0.193 | 8.03E-03 | 1.38E-01 | 2.72E+00 | 2.74E+00 | 9.52E-01 | no | | intergenic_variant | MODIFIER |
| 9 | 78039902 | AX-170361983 | 0 | G | A | 0.193 | 8.03E-03 | 1.38E-01 | 2.72E+00 | 2.74E+00 | 9.52E-01 | no | | intergenic_variant | MODIFIER |
| 9 | 78039994 | AX-170511270 | 0 | A | G | 0.193 | 8.03E-03 | 1.38E-01 | 2.72E+00 | 2.74E+00 | 9.52E-01 | no | | intergenic_variant | MODIFIER |
| 9 | 78054660 | AX-97792618 | 0 | A | G | 0.193 | 8.03E-03 | 1.38E-01 | 2.72E+00 | 2.74E+00 | 9.52E-01 | no | Gcm1 | intron_variant | MODIFIER |
| 9 | 78084759 | AX-97795769 | 0 | C | T | 0.193 | 8.03E-03 | 1.38E-01 | 2.72E+00 | 2.74E+00 | 9.52E-01 | no | Fbxo9 | intron_variant | MODIFIER |
| 9 | 78148875 | AX-97838556 | 0 | A | G | 0.193 | 8.03E-03 | 1.38E-01 | 2.72E+00 | 2.74E+00 | 9.52E-01 | no | Ick | intron_variant | MODIFIER |
| 9 | 78169489 | AX-97808499 | 0 | G | A | 0.193 | 8.03E-03 | 1.38E-01 | 2.72E+00 | 2.74E+00 | 9.52E-01 | no | Ick | 3_prime_UTR_variant | MODIFIER |
| 9 | 78169746 | AX-170361406 | 0 | A | G | 0.193 | 8.03E-03 | 1.38E-01 | 2.72E+00 | 2.74E+00 | 9.52E-01 | no | Ick | 3_prime_UTR_variant | MODIFIER |
| 9 | 78177837 | AX-170315955 | 0 | G | A | 0.193 | 8.03E-03 | 1.38E-01 | 2.72E+00 | 2.74E+00 | 9.52E-01 | no | Rn7sk | upstream_gene_variant | MODIFIER |
| 9 | 78183922 | AX-170494977 | 0 | T | C | 0.193 | 8.03E-03 | 1.38E-01 | 2.72E+00 | 2.74E+00 | 9.52E-01 | no | Gsta4 | intron_variant | MODIFIER |
| 9 | 78184011 | AX-170292446 | 0 | A | C | 0.193 | 8.03E-03 | 1.38E-01 | 2.72E+00 | 2.74E+00 | 9.52E-01 | no | Gsta4 | intron_variant | MODIFIER |
| 9 | 78201795 | AX-170203530 | 0 | G | A | 0.193 | 8.03E-03 | 1.38E-01 | 2.72E+00 | 2.74E+00 | 9.52E-01 | no | Gsta4 | intron_variant | MODIFIER |
| 9 | 78201926 | AX-170151565 | 0 | A | T | 0.193 | 8.03E-03 | 1.38E-01 | 2.72E+00 | 2.74E+00 | 9.52E-01 | no | Gsta4 | intron_variant | MODIFIER |
| 9 | 78204567 | AX-170187170 | 0 | G | A | 0.193 | 8.03E-03 | 1.38E-01 | 2.72E+00 | 2.74E+00 | 9.52E-01 | no | Gsta4 | intron_variant | MODIFIER |
| 9 | 78210233 | AX-170283921 | 0 | A | G | 0.193 | 8.03E-03 | 1.38E-01 | 2.72E+00 | 2.74E+00 | 9.52E-01 | no | Gsta4 | downstream_gene_variant | MODIFIER |
| 9 | 78217038 | AX-170352169 | 0 | T | C | 0.193 | 8.03E-03 | 1.38E-01 | 2.72E+00 | 2.74E+00 | 9.52E-01 | no | Gm3126 | upstream_gene_variant | MODIFIER |
| 9 | 78221273 | AX-170410993 | 0 | C | A | 0.193 | 8.03E-03 | 1.38E-01 | 2.72E+00 | 2.74E+00 | 9.52E-01 | no | | intergenic_variant | MODIFIER |
| 9 | 78315011 | AX-170032771 | 0 | G | C | 0.193 | 8.03E-03 | 1.38E-01 | 2.72E+00 | 2.74E+00 | 9.52E-01 | no | Omt2a | upstream_gene_variant | MODIFIER |
| 9 | 78618066 | AX-170209092 | 0 | T | A | 0.193 | 8.03E-03 | 1.38E-01 | 2.72E+00 | 2.74E+00 | 9.52E-01 | no | Cd109 | intron_variant | MODIFIER |
| 9 | 78618785 | AX-170211537 | 0 | T | C | 0.193 | 8.03E-03 | 1.38E-01 | 2.72E+00 | 2.74E+00 | 9.52E-01 | no | Cd109 | intron_variant | MODIFIER |
| 9 | 78638545 | AX-170102919 | 0 | T | C | 0.193 | 8.03E-03 | 1.38E-01 | 2.72E+00 | 2.74E+00 | 9.52E-01 | no | Cd109 | intron_variant | MODIFIER |

| | | | | | | | | | | | | | | | |
|---|---|---|---|---|---|---|---|---|---|---|---|---|---|---|---|
| 9 | 78662875 | AX-170328873 | 0 | T | C | 0.193 | 8.03E-03 | 1.38E-01 | 2.72E+00 | 2.74E+00 | 9.52E-01 | no | Cd109 | intron_variant | MODIFIER |
| 9 | 78674277 | AX-97825418 | 0 | C | G | 0.193 | 8.03E-03 | 1.38E-01 | 2.72E+00 | 2.74E+00 | 9.52E-01 | no | Cd109 | intron_variant | MODIFIER |
| 9 | 78676171 | AX-170377413 | 0 | A | G | 0.193 | 8.03E-03 | 1.38E-01 | 2.72E+00 | 2.74E+00 | 9.52E-01 | no | Cd109 | intron_variant | MODIFIER |
| 9 | 78708979 | AX-170456755 | 0 | A | G | 0.193 | 8.03E-03 | 1.38E-01 | 2.72E+00 | 2.74E+00 | 9.52E-01 | no | Cd109 | intron_variant | MODIFIER |
| 9 | 78710022 | AX-170359046 | 0 | A | T | 0.193 | 8.03E-03 | 1.38E-01 | 2.72E+00 | 2.74E+00 | 9.52E-01 | no | Cd109 | intron_variant | MODIFIER |
| 9 | 78711692 | AX-170039989 | 0 | G | C | 0.193 | 8.03E-03 | 1.38E-01 | 2.72E+00 | 2.74E+00 | 9.52E-01 | no | Cd109 | intron_variant | MODIFIER |
| 9 | 78788106 | AX-170032773 | 0 | T | G | 0.193 | 8.03E-03 | 1.38E-01 | 2.72E+00 | 2.74E+00 | 9.52E-01 | no | | regulatory_region_variant | MODIFIER |
| 16 | 77178380 | AX-170013422 | 0 | A | T | 0.456 | -3.38E-03 | 7.59E-02 | 2.67E+00 | 2.76E+00 | 9.52E-01 | no | | intergenic_variant | MODIFIER |
| 16 | 89222214 | AX-170424417 | 0 | C | T | 0.474 | -4.36E-03 | 7.79E-02 | 2.66E+00 | 2.74E+00 | 9.52E-01 | no | | intergenic_variant | MODIFIER |
| 7 | 103841592 | AX-170514023 | 0 | G | T | 0.456 | -4.82E-03 | 9.23E-02 | 2.68E+00 | 2.75E+00 | 9.52E-01 | no | Hbb-bh1 | downstream_gene_variant | MODIFIER |
| 9 | 29748093 | AX-170345242 | 0 | A | G | 0.439 | -4.40E-03 | 8.46E-02 | 2.68E+00 | 2.75E+00 | 9.52E-01 | no | Ntm | intron_variant | MODIFIER |
| 9 | 34170306 | AX-170128375 | 0 | C | T | 0.439 | -4.40E-03 | 8.46E-02 | 2.68E+00 | 2.75E+00 | 9.52E-01 | no | | intergenic_variant | MODIFIER |
| 7 | 57279966 | AX-170125728 | 0 | G | A | 0.43 | 5.96E-03 | 9.38E-02 | 2.64E+00 | 2.73E+00 | 9.53E-01 | no | Gabrg3 | intron_variant | MODIFIER |
| 7 | 87688681 | AX-170324863 | 0 | A | G | 0.114 | -1.18E-02 | 2.11E-01 | 2.66E+00 | 2.74E+00 | 9.53E-01 | no | Grm5 | intron_variant | MODIFIER |
| 1 | 47093211 | AX-170457750 | 1 | G | A | 0.107 | 1.40E-02 | 2.37E-01 | 2.69E+00 | 2.74E+00 | 9.53E-01 | no | | intergenic_variant | MODIFIER |
| 1 | 48132060 | AX-170187085 | 1 | G | A | 0.107 | 1.40E-02 | 2.37E-01 | 2.69E+00 | 2.74E+00 | 9.53E-01 | no | | intergenic_variant | MODIFIER |
| 1 | 53558557 | AX-170332675 | 1 | G | T | 0.107 | 1.40E-02 | 2.37E-01 | 2.69E+00 | 2.74E+00 | 9.53E-01 | no | Dnah7a | intron_variant | MODIFIER |
| 1 | 53846707 | AX-170205448 | 1 | A | T | 0.107 | 1.40E-02 | 2.37E-01 | 2.69E+00 | 2.74E+00 | 9.53E-01 | no | Hecw2 | intron_variant | MODIFIER |
| 9 | 33259281 | AX-170254505 | 2 | A | G | 0.418 | -4.17E-03 | 8.49E-02 | 2.67E+00 | 2.75E+00 | 9.53E-01 | no | Gm32171 | intron_variant,non_coding_transcript_variant | MODIFIER |
| 5 | 72196370 | AX-170535302 | 2 | G | A | 0.455 | -4.83E-03 | 9.72E-02 | 2.66E+00 | 2.75E+00 | 9.53E-01 | no | | intergenic_variant | MODIFIER |
| 2 | 155287523 | AX-170019432 | 0 | A | G | 0.132 | 1.27E-02 | 2.06E-01 | 2.69E+00 | 2.73E+00 | 9.53E-01 | no | Pigu | intron_variant | MODIFIER |
| 9 | 74947009 | AX-170277326 | 0 | T | C | 0.193 | 7.71E-03 | 1.35E-01 | 2.71E+00 | 2.74E+00 | 9.53E-01 | no | | regulatory_region_variant | MODIFIER |
| 9 | 61548914 | AX-170221247 | 0 | C | T | 0.202 | 7.12E-03 | 1.25E-01 | 2.68E+00 | 2.74E+00 | 9.53E-01 | no | | intergenic_variant | MODIFIER |
| 9 | 65565418 | AX-170032572 | 0 | C | T | 0.202 | 7.12E-03 | 1.25E-01 | 2.68E+00 | 2.74E+00 | 9.53E-01 | no | Plekho2 | intron_variant | MODIFIER |
| 9 | 66088936 | AX-170464735 | 0 | G | A | 0.202 | 7.12E-03 | 1.25E-01 | 2.68E+00 | 2.74E+00 | 9.53E-01 | no | Snx1 | intron_variant | MODIFIER |
| 9 | 67089367 | AX-170513401 | 0 | C | T | 0.202 | 7.12E-03 | 1.25E-01 | 2.68E+00 | 2.74E+00 | 9.53E-01 | no | | regulatory_region_variant | MODIFIER |
| 9 | 67932282 | AX-170483843 | 0 | C | T | 0.202 | 7.12E-03 | 1.25E-01 | 2.68E+00 | 2.74E+00 | 9.53E-01 | no | Vps13c | intron_variant | MODIFIER |
| 9 | 68968816 | AX-170160237 | 0 | C | T | 0.202 | 7.12E-03 | 1.25E-01 | 2.68E+00 | 2.74E+00 | 9.53E-01 | no | Rora | intron_variant | MODIFIER |
| 9 | 74998170 | AX-170449227 | 0 | T | A | 0.202 | 7.12E-03 | 1.25E-01 | 2.68E+00 | 2.74E+00 | 9.53E-01 | no | Fam214a | intron_variant | MODIFIER |
| 9 | 77459118 | AX-170387614 | 0 | C | T | 0.202 | 7.12E-03 | 1.25E-01 | 2.68E+00 | 2.74E+00 | 9.53E-01 | no | Lrrc1 | intron_variant | MODIFIER |
| 9 | 105897710 | AX-170033211 | 0 | T | A | 0.132 | -8.10E-03 | 1.13E-01 | 2.65E+00 | 2.72E+00 | 9.53E-01 | no | Col6a5 | intron_variant | MODIFIER |
| 7 | 50800990 | AX-170090791 | 0 | T | G | 0.421 | -5.15E-03 | 1.02E-01 | 2.67E+00 | 2.75E+00 | 9.53E-01 | no | Nell1 | intron_variant | MODIFIER |
| 7 | 51583633 | AX-170089916 | 0 | A | C | 0.421 | -5.15E-03 | 1.02E-01 | 2.67E+00 | 2.75E+00 | 9.53E-01 | no | Ano5 | intron_variant | MODIFIER |
| 7 | 51997559 | AX-97790083 | 0 | A | G | 0.421 | -5.15E-03 | 1.02E-01 | 2.67E+00 | 2.75E+00 | 9.53E-01 | no | Gas2 | downstream_gene_variant | MODIFIER |
| 7 | 52407757 | AX-170077350 | 0 | A | G | 0.421 | -5.15E-03 | 1.02E-01 | 2.67E+00 | 2.75E+00 | 9.53E-01 | no | Gm20074 | upstream_gene_variant | MODIFIER |
| 7 | 57247377 | AX-170077437 | 0 | A | G | 0.421 | -5.15E-03 | 1.02E-01 | 2.67E+00 | 2.75E+00 | 9.53E-01 | no | Gabrg3 | intron_variant | MODIFIER |
| 7 | 57510736 | AX-170231487 | 0 | C | T | 0.421 | -5.15E-03 | 1.02E-01 | 2.67E+00 | 2.75E+00 | 9.53E-01 | no | Gabra5 | upstream_gene_variant | MODIFIER |
| 8 | 47799715 | AX-97843029 | 0 | G | A | 0.158 | 6.91E-03 | 1.34E-01 | 2.68E+00 | 2.75E+00 | 9.53E-01 | no | | regulatory_region_variant | MODIFIER |
| 18 | 43134397 | AX-97785627 | 0 | G | A | 0.061 | -1.80E-02 | 3.00E-01 | 2.71E+00 | 2.73E+00 | 9.53E-01 | no | AC099605.1 | intron_variant,non_coding_transcript_variant | MODIFIER |
| 5 | 101482194 | AX-170025742 | 1 | C | G | 0.438 | -5.47E-03 | 1.01E-01 | 2.67E+00 | 2.74E+00 | 9.53E-01 | no | | intergenic_variant | MODIFIER |
| 20 | 108824687 | AX-97806662 | 0 | A | G | 0.298 | -3.10E-03 | 5.35E-02 | 2.62E+00 | 2.74E+00 | 9.53E-01 | no | Brwd3 | intron_variant,NMD_transcript_variant | MODIFIER |
| 20 | 108968266 | AX-170518926 | 0 | C | G | 0.298 | -3.10E-03 | 5.35E-02 | 2.62E+00 | 2.74E+00 | 9.53E-01 | no | 2810403D21Rik | intron_variant,non_coding_transcript_variant | MODIFIER |
| 20 | 108968353 | AX-170513725 | 0 | G | C | 0.298 | -3.10E-03 | 5.35E-02 | 2.62E+00 | 2.74E+00 | 9.53E-01 | no | 2810403D21Rik | intron_variant,non_coding_transcript_variant | MODIFIER |
| 20 | 108973501 | AX-97840191 | 0 | T | G | 0.298 | -3.10E-03 | 5.35E-02 | 2.62E+00 | 2.74E+00 | 9.53E-01 | no | 2810403D21Rik | intron_variant,non_coding_transcript_variant | MODIFIER |
| 20 | 108980044 | AX-170445822 | 0 | T | A | 0.298 | -3.10E-03 | 5.35E-02 | 2.62E+00 | 2.74E+00 | 9.53E-01 | no | | intergenic_variant | MODIFIER |
| 20 | 109003457 | AX-97797309 | 0 | C | T | 0.298 | -3.10E-03 | 5.35E-02 | 2.62E+00 | 2.74E+00 | 9.53E-01 | no | Hmgn5 | downstream_gene_variant | MODIFIER |
| 20 | 109024014 | AX-97848894 | 0 | A | G | 0.298 | -3.10E-03 | 5.35E-02 | 2.62E+00 | 2.74E+00 | 9.53E-01 | no | Gm25797 | upstream_gene_variant | MODIFIER |
| 20 | 109067875 | AX-97783009 | 0 | G | T | 0.298 | -3.10E-03 | 5.35E-02 | 2.62E+00 | 2.74E+00 | 9.53E-01 | no | | intergenic_variant | MODIFIER |
| 20 | 109087480 | AX-97791827 | 0 | C | T | 0.298 | -3.10E-03 | 5.35E-02 | 2.62E+00 | 2.74E+00 | 9.53E-01 | no | | intergenic_variant | MODIFIER |
| 20 | 109110800 | AX-97798244 | 0 | G | A | 0.298 | -3.10E-03 | 5.35E-02 | 2.62E+00 | 2.74E+00 | 9.53E-01 | no | Sh3bgrl | intron_variant | MODIFIER |
| 20 | 109120056 | AX-170034680 | 0 | T | C | 0.298 | -3.10E-03 | 5.35E-02 | 2.62E+00 | 2.74E+00 | 9.53E-01 | no | Sh3bgrl | intron_variant | MODIFIER |
| 20 | 109786752 | AX-97792596 | 0 | A | C | 0.298 | -3.10E-03 | 5.35E-02 | 2.62E+00 | 2.74E+00 | 9.53E-01 | no | | intergenic_variant | MODIFIER |
| 20 | 110122079 | AX-97802502 | 0 | A | C | 0.298 | -3.10E-03 | 5.35E-02 | 2.62E+00 | 2.74E+00 | 9.53E-01 | no | | intergenic_variant | MODIFIER |
| 20 | 110667777 | AX-97844288 | 0 | T | C | 0.298 | -3.10E-03 | 5.35E-02 | 2.62E+00 | 2.74E+00 | 9.53E-01 | no | Gm44593 | intron_variant,non_coding_transcript_variant | MODIFIER |

| | | | | | | | | | | | | | | | |
|---|---|---|---|---|---|---|---|---|---|---|---|---|---|---|---|
| 20 | 111222721 | AX-170360131 | 0 | G | T | 0.298 | -3.10E-03 | 5.35E-02 | 2.62E+00 | 2.74E+00 | 9.53E-01 | no | | intergenic_variant | MODIFIER |
| 20 | 111375129 | AX-97823555 | 0 | C | T | 0.298 | -3.10E-03 | 5.35E-02 | 2.62E+00 | 2.74E+00 | 9.53E-01 | no | | intergenic_variant | MODIFIER |
| 20 | 111392334 | AX-168414086 | 0 | G | A | 0.298 | -3.10E-03 | 5.35E-02 | 2.62E+00 | 2.74E+00 | 9.53E-01 | no | Rps6ka6 | intron_variant | MODIFIER |
| 20 | 111395792 | AX-170276217 | 0 | A | G | 0.298 | -3.10E-03 | 5.35E-02 | 2.62E+00 | 2.74E+00 | 9.53E-01 | no | Rps6ka6 | intron_variant | MODIFIER |
| 20 | 111403230 | AX-97807216 | 0 | A | G | 0.298 | -3.10E-03 | 5.35E-02 | 2.62E+00 | 2.74E+00 | 9.53E-01 | no | Rps6ka6 | intron_variant | MODIFIER |
| 20 | 111427487 | AX-170034729 | 0 | G | A | 0.298 | -3.10E-03 | 5.35E-02 | 2.62E+00 | 2.74E+00 | 9.53E-01 | no | Rps6ka6 | intron_variant | MODIFIER |
| 20 | 111494241 | AX-97852941 | 0 | C | T | 0.298 | -3.10E-03 | 5.35E-02 | 2.62E+00 | 2.74E+00 | 9.53E-01 | no | Rps6ka6 | intron_variant | MODIFIER |
| 20 | 111517912 | AX-170034728 | 0 | C | G | 0.298 | -3.10E-03 | 5.35E-02 | 2.62E+00 | 2.74E+00 | 9.53E-01 | no | Rps6ka6 | intron_variant | MODIFIER |
| 20 | 111519187 | AX-170034726 | 0 | T | G | 0.298 | -3.10E-03 | 5.35E-02 | 2.62E+00 | 2.74E+00 | 9.53E-01 | no | Rps6ka6 | intron_variant | MODIFIER |
| 20 | 111520871 | AX-97843530 | 0 | G | A | 0.298 | -3.10E-03 | 5.35E-02 | 2.62E+00 | 2.74E+00 | 9.53E-01 | no | Rps6ka6 | intron_variant | MODIFIER |
| 20 | 111528982 | AX-170034725 | 0 | C | T | 0.298 | -3.10E-03 | 5.35E-02 | 2.62E+00 | 2.74E+00 | 9.53E-01 | no | Rps6ka6 | intron_variant | MODIFIER |
| 20 | 111532914 | AX-170034732 | 0 | C | G | 0.298 | -3.10E-03 | 5.35E-02 | 2.62E+00 | 2.74E+00 | 9.53E-01 | no | Rps6ka6 | intron_variant | MODIFIER |
| 18 | 65294018 | AX-170015869 | 1 | A | T | 0.393 | -5.80E-03 | 7.85E-02 | 2.64E+00 | 2.72E+00 | 9.53E-01 | no | Alpk2 | intron_variant | MODIFIER |
| 1 | 9768162 | AX-169999913 | 0 | T | C | 0.509 | 5.12E-03 | 7.20E-02 | 2.65E+00 | 2.72E+00 | 9.53E-01 | no | 1700034P13Rik | intron_variant,non_coding_transcript_variant | MODIFIER |
| 13 | 52761773 | AX-170205795 | 0 | C | T | 0.518 | 4.82E-03 | 6.59E-02 | 2.62E+00 | 2.73E+00 | 9.53E-01 | no | BB123696 | upstream_gene_variant | MODIFIER |
| 13 | 52763504 | AX-170372216 | 0 | G | A | 0.518 | 4.82E-03 | 6.59E-02 | 2.62E+00 | 2.73E+00 | 9.53E-01 | no | | intergenic_variant | MODIFIER |
| 13 | 52834256 | AX-97823502 | 0 | C | T | 0.518 | 4.82E-03 | 6.59E-02 | 2.62E+00 | 2.73E+00 | 9.53E-01 | no | Auh | downstream_gene_variant | MODIFIER |
| 13 | 53068432 | AX-170463704 | 0 | T | C | 0.518 | 4.82E-03 | 6.59E-02 | 2.62E+00 | 2.73E+00 | 9.53E-01 | no | | regulatory_region_variant | MODIFIER |
| 13 | 53100009 | AX-170255771 | 0 | G | C | 0.518 | 4.82E-03 | 6.59E-02 | 2.62E+00 | 2.73E+00 | 9.53E-01 | no | | regulatory_region_variant | MODIFIER |
| 13 | 53163845 | AX-170266302 | 0 | G | A | 0.518 | 4.82E-03 | 6.59E-02 | 2.62E+00 | 2.73E+00 | 9.53E-01 | no | Ror2 | intron_variant | MODIFIER |
| 13 | 53164218 | AX-170279463 | 0 | T | G | 0.518 | 4.82E-03 | 6.59E-02 | 2.62E+00 | 2.73E+00 | 9.53E-01 | no | Ror2 | intron_variant | MODIFIER |
| 13 | 53171313 | AX-170211248 | 0 | C | T | 0.518 | 4.82E-03 | 6.59E-02 | 2.62E+00 | 2.73E+00 | 9.53E-01 | no | Ror2 | intron_variant | MODIFIER |
| 13 | 53171862 | AX-170159091 | 0 | T | C | 0.518 | 4.82E-03 | 6.59E-02 | 2.62E+00 | 2.73E+00 | 9.53E-01 | no | Ror2 | intron_variant | MODIFIER |
| 13 | 53302075 | AX-170203347 | 0 | A | G | 0.518 | 4.82E-03 | 6.59E-02 | 2.62E+00 | 2.73E+00 | 9.53E-01 | no | | regulatory_region_variant | MODIFIER |
| 13 | 53304582 | AX-170206157 | 0 | T | C | 0.518 | 4.82E-03 | 6.59E-02 | 2.62E+00 | 2.73E+00 | 9.53E-01 | no | | intergenic_variant | MODIFIER |
| 13 | 53305311 | AX-170358931 | 0 | C | T | 0.518 | 4.82E-03 | 6.59E-02 | 2.62E+00 | 2.73E+00 | 9.53E-01 | no | | intergenic_variant | MODIFIER |
| 13 | 53330792 | AX-170252119 | 0 | G | A | 0.518 | 4.82E-03 | 6.59E-02 | 2.62E+00 | 2.73E+00 | 9.53E-01 | no | Sptlc1 | downstream_gene_variant | MODIFIER |
| 13 | 53350261 | AX-170490165 | 0 | C | A | 0.518 | 4.82E-03 | 6.59E-02 | 2.62E+00 | 2.73E+00 | 9.53E-01 | no | Sptlc1 | intron_variant | MODIFIER |
| 13 | 53352064 | AX-170482091 | 0 | C | T | 0.518 | 4.82E-03 | 6.59E-02 | 2.62E+00 | 2.73E+00 | 9.53E-01 | no | Sptlc1 | intron_variant | MODIFIER |
| 13 | 53372762 | AX-170185446 | 0 | A | G | 0.518 | 4.82E-03 | 6.59E-02 | 2.62E+00 | 2.73E+00 | 9.53E-01 | no | Sptlc1 | intron_variant | MODIFIER |
| 13 | 53373713 | AX-170051313 | 0 | C | T | 0.518 | 4.82E-03 | 6.59E-02 | 2.62E+00 | 2.73E+00 | 9.53E-01 | no | Sptlc1 | intron_variant | MODIFIER |
| 13 | 53414132 | AX-170485703 | 0 | G | A | 0.518 | 4.82E-03 | 6.59E-02 | 2.62E+00 | 2.73E+00 | 9.53E-01 | no | Gm2762 | intron_variant,non_coding_transcript_variant | MODIFIER |
| 13 | 53414387 | AX-170361281 | 0 | A | G | 0.518 | 4.82E-03 | 6.59E-02 | 2.62E+00 | 2.73E+00 | 9.53E-01 | no | Gm2762 | intron_variant,non_coding_transcript_variant | MODIFIER |
| 13 | 53433384 | AX-170179109 | 0 | G | A | 0.518 | 4.82E-03 | 6.59E-02 | 2.62E+00 | 2.73E+00 | 9.53E-01 | no | Gm2762 | intron_variant,non_coding_transcript_variant | MODIFIER |
| 13 | 53442568 | AX-170284432 | 0 | A | G | 0.518 | 4.82E-03 | 6.59E-02 | 2.62E+00 | 2.73E+00 | 9.53E-01 | no | Gm2762 | intron_variant,non_coding_transcript_variant | MODIFIER |
| 13 | 53450389 | AX-170390762 | 0 | C | T | 0.518 | 4.82E-03 | 6.59E-02 | 2.62E+00 | 2.73E+00 | 9.53E-01 | no | Gm2762 | intron_variant,non_coding_transcript_variant | MODIFIER |
| 13 | 53453592 | AX-170285737 | 0 | A | G | 0.518 | 4.82E-03 | 6.59E-02 | 2.62E+00 | 2.73E+00 | 9.53E-01 | no | Gm2762 | intron_variant,non_coding_transcript_variant | MODIFIER |
| 13 | 53476849 | AX-170322255 | 0 | T | C | 0.518 | 4.82E-03 | 6.59E-02 | 2.62E+00 | 2.73E+00 | 9.53E-01 | no | Msx2 | upstream_gene_variant | MODIFIER |
| 8 | 81134589 | AX-170080565 | 2 | G | A | 0.2 | 6.76E-03 | 1.11E-01 | 2.63E+00 | 2.74E+00 | 9.53E-01 | no | | intergenic_variant | MODIFIER |
| 1 | 47639992 | AX-170000424 | 1 | G | A | 0.107 | 1.39E-02 | 2.36E-01 | 2.69E+00 | 2.74E+00 | 9.53E-01 | no | | intergenic_variant | MODIFIER |
| 1 | 51203048 | AX-97811531 | 1 | C | T | 0.107 | 1.39E-02 | 2.36E-01 | 2.69E+00 | 2.74E+00 | 9.53E-01 | no | | intergenic_variant | MODIFIER |
| 1 | 52915151 | AX-170035923 | 1 | T | A | 0.107 | 1.39E-02 | 2.36E-01 | 2.69E+00 | 2.74E+00 | 9.53E-01 | no | Hibch | intron_variant | MODIFIER |
| 16 | 51503007 | AX-170013035 | 1 | G | A | 0.491 | -5.48E-03 | 7.74E-02 | 2.65E+00 | 2.72E+00 | 9.53E-01 | no | | intergenic_variant | MODIFIER |
| 8 | 83048310 | AX-170257774 | 0 | C | A | 0.202 | -7.77E-03 | 1.24E-01 | 2.65E+00 | 2.73E+00 | 9.53E-01 | no | Rnf150 | intron_variant | MODIFIER |
| 10 | 117459963 | AX-170092126 | 1 | T | C | 0.321 | -4.93E-03 | 7.46E-02 | 2.63E+00 | 2.73E+00 | 9.53E-01 | no | | regulatory_region_variant | MODIFIER |
| 9 | 68665945 | AX-170417338 | 0 | A | G | 0.202 | 7.62E-03 | 1.34E-01 | 2.71E+00 | 2.74E+00 | 9.53E-01 | no | Rora | intron_variant | MODIFIER |
| 9 | 72428932 | AX-170415690 | 0 | G | C | 0.202 | 7.62E-03 | 1.34E-01 | 2.71E+00 | 2.74E+00 | 9.53E-01 | no | Gm27254 | downstream_gene_variant | MODIFIER |
| 7 | 58991472 | AX-97812642 | 2 | G | A | 0.436 | 6.07E-03 | 9.82E-02 | 2.66E+00 | 2.73E+00 | 9.53E-01 | no | | intergenic_variant | MODIFIER |
| 1 | 10622427 | AX-170321046 | 0 | G | T | 0.395 | -3.60E-03 | 7.51E-02 | 2.68E+00 | 2.75E+00 | 9.53E-01 | no | Cpa6 | intron_variant | MODIFIER |
| 10 | 19083739 | AX-170292211 | 0 | T | C | 0.193 | 4.39E-03 | 9.32E-02 | 2.64E+00 | 2.75E+00 | 9.54E-01 | no | 1700124M09Rik | downstream_gene_variant | MODIFIER |
| 1 | 54707909 | AX-170418434 | 0 | T | C | 0.096 | 1.19E-02 | 2.01E-01 | 2.64E+00 | 2.74E+00 | 9.54E-01 | no | Ankrd44 | intron_variant | MODIFIER |
| 1 | 60364033 | AX-97811677 | 0 | G | A | 0.096 | 1.19E-02 | 2.01E-01 | 2.64E+00 | 2.74E+00 | 9.54E-01 | no | Cyp20a1 | intron_variant | MODIFIER |
| 9 | 55932058 | AX-170533589 | 1 | C | A | 0.205 | -6.92E-03 | 1.21E-01 | 2.68E+00 | 2.74E+00 | 9.54E-01 | no | Scaper | intron_variant | MODIFIER |
| 8 | 81484341 | AX-170173022 | 1 | G | A | 0.205 | 7.63E-03 | 1.29E-01 | 2.67E+00 | 2.74E+00 | 9.54E-01 | no | Inpp4b | intron_variant,NMD_transcript_variant | MODIFIER |

| | | | | | | | | | | | | | | | |
|---|---|---|---|---|---|---|---|---|---|---|---|---|---|---|---|
| 8 | 85616992 | AX-170031175 | 1 | T | C | 0.205 | 7.63E-03 | 1.29E-01 | 2.67E+00 | 2.74E+00 | 9.54E-01 | no | | intergenic_variant | MODIFIER |
| 16 | 93435013 | AX-170521802 | 0 | C | T | 0.412 | -5.72E-03 | 8.25E-02 | 2.63E+00 | 2.73E+00 | 9.54E-01 | no | 1700029J03Rik | intron_variant,non_coding_transcript_variant | MODIFIER |
| 16 | 93446416 | AX-170405599 | 0 | C | T | 0.412 | -5.72E-03 | 8.25E-02 | 2.63E+00 | 2.73E+00 | 9.54E-01 | no | 1700029J03Rik | intron_variant,non_coding_transcript_variant | MODIFIER |
| 16 | 93448014 | AX-170294335 | 0 | T | C | 0.412 | -5.72E-03 | 8.25E-02 | 2.63E+00 | 2.73E+00 | 9.54E-01 | no | 1700029J03Rik | intron_variant,non_coding_transcript_variant | MODIFIER |
| 16 | 93487325 | AX-170189513 | 0 | T | C | 0.412 | -5.72E-03 | 8.25E-02 | 2.63E+00 | 2.73E+00 | 9.54E-01 | no | Gm30695 | upstream_gene_variant | MODIFIER |
| 16 | 93487402 | AX-170266640 | 0 | C | T | 0.412 | -5.72E-03 | 8.25E-02 | 2.63E+00 | 2.73E+00 | 9.54E-01 | no | Gm30695 | upstream_gene_variant | MODIFIER |
| 16 | 93494717 | AX-170442378 | 0 | G | A | 0.412 | -5.72E-03 | 8.25E-02 | 2.63E+00 | 2.73E+00 | 9.54E-01 | no | | intergenic_variant | MODIFIER |
| 16 | 93495169 | AX-97794690 | 0 | G | A | 0.412 | -5.72E-03 | 8.25E-02 | 2.63E+00 | 2.73E+00 | 9.54E-01 | no | | intergenic_variant | MODIFIER |
| 16 | 93495213 | AX-170162336 | 0 | A | G | 0.412 | -5.72E-03 | 8.25E-02 | 2.63E+00 | 2.73E+00 | 9.54E-01 | no | | intergenic_variant | MODIFIER |
| 16 | 93495447 | AX-170413686 | 0 | G | C | 0.412 | -5.72E-03 | 8.25E-02 | 2.63E+00 | 2.73E+00 | 9.54E-01 | no | | intergenic_variant | MODIFIER |
| 16 | 93504391 | AX-170336214 | 0 | A | C | 0.412 | -5.72E-03 | 8.25E-02 | 2.63E+00 | 2.73E+00 | 9.54E-01 | no | | intergenic_variant | MODIFIER |
| 16 | 93513459 | AX-170360202 | 0 | A | G | 0.412 | -5.72E-03 | 8.25E-02 | 2.63E+00 | 2.73E+00 | 9.54E-01 | no | | intergenic_variant | MODIFIER |
| 16 | 93518540 | AX-170279886 | 0 | A | G | 0.412 | -5.72E-03 | 8.25E-02 | 2.63E+00 | 2.73E+00 | 9.54E-01 | no | | intergenic_variant | MODIFIER |
| 16 | 93519491 | AX-170467007 | 0 | G | T | 0.412 | -5.72E-03 | 8.25E-02 | 2.63E+00 | 2.73E+00 | 9.54E-01 | no | | intergenic_variant | MODIFIER |
| 16 | 93519744 | AX-170090766 | 0 | C | T | 0.412 | -5.72E-03 | 8.25E-02 | 2.63E+00 | 2.73E+00 | 9.54E-01 | no | | intergenic_variant | MODIFIER |
| 16 | 93520975 | AX-170374663 | 0 | G | A | 0.412 | -5.72E-03 | 8.25E-02 | 2.63E+00 | 2.73E+00 | 9.54E-01 | no | | intergenic_variant | MODIFIER |
| 16 | 93522332 | AX-170142650 | 0 | C | A | 0.412 | -5.72E-03 | 8.25E-02 | 2.63E+00 | 2.73E+00 | 9.54E-01 | no | | intergenic_variant | MODIFIER |
| 16 | 93522499 | AX-170414259 | 0 | T | C | 0.412 | -5.72E-03 | 8.25E-02 | 2.63E+00 | 2.73E+00 | 9.54E-01 | no | | regulatory_region_variant | MODIFIER |
| 16 | 93526871 | AX-170345694 | 0 | A | G | 0.412 | -5.72E-03 | 8.25E-02 | 2.63E+00 | 2.73E+00 | 9.54E-01 | no | | intergenic_variant | MODIFIER |
| 16 | 93529497 | AX-170058143 | 0 | A | G | 0.412 | -5.72E-03 | 8.25E-02 | 2.63E+00 | 2.73E+00 | 9.54E-01 | no | Gm49725 | upstream_gene_variant | MODIFIER |
| 16 | 93544070 | AX-170093568 | 0 | G | T | 0.412 | -5.72E-03 | 8.25E-02 | 2.63E+00 | 2.73E+00 | 9.54E-01 | no | | regulatory_region_variant | MODIFIER |
| 16 | 93544552 | AX-170331767 | 0 | T | C | 0.412 | -5.72E-03 | 8.25E-02 | 2.63E+00 | 2.73E+00 | 9.54E-01 | no | | intergenic_variant | MODIFIER |
| 16 | 93596307 | AX-170290880 | 0 | T | C | 0.412 | -5.72E-03 | 8.25E-02 | 2.63E+00 | 2.73E+00 | 9.54E-01 | no | Setd4 | synonymous_variant | LOW |
| 16 | 93597056 | AX-170218368 | 0 | T | C | 0.412 | -5.72E-03 | 8.25E-02 | 2.63E+00 | 2.73E+00 | 9.54E-01 | no | Setd4 | intron_variant | MODIFIER |
| 16 | 93606260 | AX-97853600 | 0 | A | G | 0.412 | -5.72E-03 | 8.25E-02 | 2.63E+00 | 2.73E+00 | 9.54E-01 | no | Setd4 | upstream_gene_variant | MODIFIER |
| 16 | 93612703 | AX-170147197 | 0 | T | G | 0.412 | -5.72E-03 | 8.25E-02 | 2.63E+00 | 2.73E+00 | 9.54E-01 | no | Cbr1 | downstream_gene_variant | MODIFIER |
| 16 | 93643402 | AX-97832176 | 0 | A | C | 0.412 | -5.72E-03 | 8.25E-02 | 2.63E+00 | 2.73E+00 | 9.54E-01 | no | | intergenic_variant | MODIFIER |
| 16 | 93660712 | AX-97803814 | 0 | A | G | 0.412 | -5.72E-03 | 8.25E-02 | 2.63E+00 | 2.73E+00 | 9.54E-01 | no | | intergenic_variant | MODIFIER |
| 16 | 93667702 | AX-170521016 | 0 | C | T | 0.412 | -5.72E-03 | 8.25E-02 | 2.63E+00 | 2.73E+00 | 9.54E-01 | no | | intergenic_variant | MODIFIER |
| 16 | 93670025 | AX-170525076 | 0 | G | A | 0.412 | -5.72E-03 | 8.25E-02 | 2.63E+00 | 2.73E+00 | 9.54E-01 | no | | intergenic_variant | MODIFIER |
| 16 | 93672898 | AX-170137749 | 0 | C | T | 0.412 | -5.72E-03 | 8.25E-02 | 2.63E+00 | 2.73E+00 | 9.54E-01 | no | | intergenic_variant | MODIFIER |
| 16 | 93691371 | AX-170320431 | 0 | G | A | 0.412 | -5.72E-03 | 8.25E-02 | 2.63E+00 | 2.73E+00 | 9.54E-01 | no | Cbr3 | downstream_gene_variant | MODIFIER |
| 16 | 93691406 | AX-170145616 | 0 | A | G | 0.412 | -5.72E-03 | 8.25E-02 | 2.63E+00 | 2.73E+00 | 9.54E-01 | no | Cbr3 | downstream_gene_variant | MODIFIER |
| 13 | 36959619 | AX-170008123 | 1 | A | G | 0.188 | -7.79E-03 | 1.15E-01 | 2.64E+00 | 2.73E+00 | 9.54E-01 | no | F13a1 | intron_variant | MODIFIER |
| 4 | 144746543 | AX-170399021 | 1 | A | G | 0.098 | 1.10E-02 | 2.06E-01 | 2.66E+00 | 2.74E+00 | 9.54E-01 | no | | intergenic_variant | MODIFIER |
| 4 | 144955099 | AX-170337435 | 1 | G | A | 0.098 | 1.10E-02 | 2.06E-01 | 2.66E+00 | 2.74E+00 | 9.54E-01 | no | Gm38074 | downstream_gene_variant | MODIFIER |
| 4 | 73033138 | AX-170069675 | 0 | G | A | 0.237 | -4.70E-03 | 1.05E-01 | 2.67E+00 | 2.76E+00 | 9.54E-01 | no | Gm11251 | non_coding_transcript_exon_variant | MODIFIER |
| 8 | 78840120 | AX-97853777 | 1 | T | C | 0.196 | -7.92E-03 | 1.34E-01 | 2.68E+00 | 2.74E+00 | 9.54E-01 | no | | intergenic_variant | MODIFIER |
| 7 | 69935742 | AX-97801041 | 0 | C | A | 0.404 | 5.34E-03 | 8.76E-02 | 2.64E+00 | 2.73E+00 | 9.54E-01 | no | | intergenic_variant | MODIFIER |
| 7 | 102676496 | AX-170177796 | 1 | T | C | 0.464 | -4.81E-03 | 9.39E-02 | 2.68E+00 | 2.75E+00 | 9.54E-01 | no | Trim68 | downstream_gene_variant | MODIFIER |
| 6 | 102799981 | AX-97822983 | 0 | G | A | 0.053 | 1.75E-02 | 2.96E-01 | 2.67E+00 | 2.74E+00 | 9.54E-01 | no | | intergenic_variant | MODIFIER |
| 8 | 47519512 | AX-170168510 | 1 | G | T | 0.152 | -9.07E-03 | 1.48E-01 | 2.68E+00 | 2.73E+00 | 9.54E-01 | no | Trappc11 | intron_variant | MODIFIER |
| 5 | 120043747 | AX-170146195 | 1 | C | T | 0.455 | -5.06E-03 | 1.05E-01 | 2.69E+00 | 2.75E+00 | 9.54E-01 | no | | intergenic_variant | MODIFIER |
| 9 | 35516151 | AX-170313353 | 1 | C | G | 0.438 | -4.44E-03 | 8.75E-02 | 2.68E+00 | 2.75E+00 | 9.54E-01 | no | Gm1113 | intron_variant | MODIFIER |
| 9 | 61823771 | AX-170489237 | 1 | T | C | 0.196 | -7.76E-03 | 1.32E-01 | 2.69E+00 | 2.73E+00 | 9.54E-01 | no | | intergenic_variant | MODIFIER |
| 1 | 40405267 | AX-170143819 | 1 | T | C | 0.393 | -4.82E-03 | 7.64E-02 | 2.65E+00 | 2.73E+00 | 9.54E-01 | no | | intergenic_variant | MODIFIER |
| 7 | 57124463 | AX-170306739 | 0 | T | C | 0.421 | -5.11E-03 | 9.95E-02 | 2.67E+00 | 2.75E+00 | 9.54E-01 | no | Gabrg3 | intron_variant | MODIFIER |
| 13 | 53476831 | AX-170448629 | 2 | A | G | 0.509 | 4.68E-03 | 6.53E-02 | 2.62E+00 | 2.73E+00 | 9.54E-01 | no | Msx2 | upstream_gene_variant | MODIFIER |
| 10 | 5868376 | AX-170412828 | 1 | C | T | 0.188 | -6.66E-03 | 1.07E-01 | 2.64E+00 | 2.73E+00 | 9.54E-01 | no | Rgs17 | intron_variant | MODIFIER |
| 11 | 34195119 | AX-170172262 | 1 | T | C | 0.071 | -8.17E-03 | 1.81E-01 | 2.62E+00 | 2.75E+00 | 9.54E-01 | no | | intergenic_variant | MODIFIER |
| 10 | 40946414 | AX-97782752 | 0 | C | T | 0.386 | 3.59E-03 | 7.55E-02 | 2.64E+00 | 2.75E+00 | 9.54E-01 | no | | intergenic_variant | MODIFIER |
| 8 | 83000153 | AX-170498968 | 0 | G | A | 0.211 | 7.28E-03 | 1.22E-01 | 2.66E+00 | 2.73E+00 | 9.54E-01 | no | Rnf150 | intron_variant | MODIFIER |
| 1 | 9160752 | AX-170163660 | 1 | A | G | 0.509 | 5.00E-03 | 7.20E-02 | 2.65E+00 | 2.72E+00 | 9.54E-01 | no | Sntg1 | intron_variant,non_coding_transcript_variant | MODIFIER |
| 2 | 113704619 | AX-97820734 | 0 | T | G | 0.14 | -1.21E-02 | 2.21E-01 | 2.72E+00 | 2.75E+00 | 9.54E-01 | no | Fmn1 | intron_variant | MODIFIER |

| | | | | | | | | | | | | | | |
|---|---|---|---|---|---|---|---|---|---|---|---|---|---|---|
| 2 | 132540430 | AX-170147929 | 0 | G | T | 0.14 | -1.21E-02 | 2.21E-01 | 2.72E+00 | 2.75E+00 | 9.54E-01 | no | Gpcpd1 | intron_variant | MODIFIER |
| 2 | 142120636 | AX-170481695 | 0 | C | T | 0.14 | -1.21E-02 | 2.21E-01 | 2.72E+00 | 2.75E+00 | 9.54E-01 | no | Macrod2 | intron_variant | MODIFIER |
| 2 | 157465910 | AX-97806865 | 0 | G | A | 0.14 | -1.21E-02 | 2.21E-01 | 2.72E+00 | 2.75E+00 | 9.54E-01 | no | Src | synonymous_variant | LOW |
| 1 | 6823604 | AX-170294527 | 0 | C | T | 0.509 | 4.99E-03 | 7.20E-02 | 2.65E+00 | 2.72E+00 | 9.54E-01 | no | St18 | intron_variant | MODIFIER |
| 1 | 6871585 | AX-169999880 | 0 | C | T | 0.509 | 4.99E-03 | 7.20E-02 | 2.65E+00 | 2.72E+00 | 9.54E-01 | no | | intergenic_variant | MODIFIER |
| 1 | 6882384 | AX-170387625 | 0 | A | G | 0.509 | 4.99E-03 | 7.20E-02 | 2.65E+00 | 2.72E+00 | 9.54E-01 | no | | intergenic_variant | MODIFIER |
| 1 | 6890939 | AX-97810622 | 0 | C | A | 0.509 | 4.99E-03 | 7.20E-02 | 2.65E+00 | 2.72E+00 | 9.54E-01 | no | | intergenic_variant | MODIFIER |
| 1 | 6890959 | AX-169999879 | 0 | G | T | 0.509 | 4.99E-03 | 7.20E-02 | 2.65E+00 | 2.72E+00 | 9.54E-01 | no | | intergenic_variant | MODIFIER |
| 1 | 6907979 | AX-170472164 | 0 | C | T | 0.509 | 4.99E-03 | 7.20E-02 | 2.65E+00 | 2.72E+00 | 9.54E-01 | no | Gm7449 | upstream_gene_variant | MODIFIER |
| 1 | 6928937 | AX-170040301 | 0 | G | T | 0.509 | 4.99E-03 | 7.20E-02 | 2.65E+00 | 2.72E+00 | 9.54E-01 | no | Gm37275 | upstream_gene_variant | MODIFIER |
| 1 | 6929299 | AX-170040300 | 0 | A | C | 0.509 | 4.99E-03 | 7.20E-02 | 2.65E+00 | 2.72E+00 | 9.54E-01 | no | Gm37275 | upstream_gene_variant | MODIFIER |
| 1 | 6935240 | AX-169999876 | 0 | T | G | 0.509 | 4.99E-03 | 7.20E-02 | 2.65E+00 | 2.72E+00 | 9.54E-01 | no | | intergenic_variant | MODIFIER |
| 1 | 7469098 | AX-170532956 | 0 | C | G | 0.509 | 4.99E-03 | 7.20E-02 | 2.65E+00 | 2.72E+00 | 9.54E-01 | no | | intergenic_variant | MODIFIER |
| 1 | 8008753 | AX-169999888 | 0 | A | G | 0.509 | 4.99E-03 | 7.20E-02 | 2.65E+00 | 2.72E+00 | 9.54E-01 | no | | intergenic_variant | MODIFIER |
| 1 | 8011731 | AX-170505328 | 0 | G | A | 0.509 | 4.99E-03 | 7.20E-02 | 2.65E+00 | 2.72E+00 | 9.54E-01 | no | | intergenic_variant | MODIFIER |
| 1 | 8035287 | AX-170340964 | 0 | A | G | 0.509 | 4.99E-03 | 7.20E-02 | 2.65E+00 | 2.72E+00 | 9.54E-01 | no | | intergenic_variant | MODIFIER |
| 1 | 8263888 | AX-97798591 | 0 | T | C | 0.509 | 4.99E-03 | 7.20E-02 | 2.65E+00 | 2.72E+00 | 9.54E-01 | no | | intergenic_variant | MODIFIER |
| 1 | 8263913 | AX-170040325 | 0 | C | T | 0.509 | 4.99E-03 | 7.20E-02 | 2.65E+00 | 2.72E+00 | 9.54E-01 | no | | intergenic_variant | MODIFIER |
| 1 | 8265198 | AX-170100114 | 0 | T | A | 0.509 | 4.99E-03 | 7.20E-02 | 2.65E+00 | 2.72E+00 | 9.54E-01 | no | | intergenic_variant | MODIFIER |
| 1 | 8266073 | AX-170487334 | 0 | C | G | 0.509 | 4.99E-03 | 7.20E-02 | 2.65E+00 | 2.72E+00 | 9.54E-01 | no | | intergenic_variant | MODIFIER |
| 1 | 8289124 | AX-170146352 | 0 | C | T | 0.509 | 4.99E-03 | 7.20E-02 | 2.65E+00 | 2.72E+00 | 9.54E-01 | no | | intergenic_variant | MODIFIER |
| 1 | 8360977 | AX-170145779 | 0 | C | G | 0.509 | 4.99E-03 | 7.20E-02 | 2.65E+00 | 2.72E+00 | 9.54E-01 | no | Sntg1 | downstream_gene_variant | MODIFIER |
| 1 | 8361104 | AX-170290178 | 0 | T | C | 0.509 | 4.99E-03 | 7.20E-02 | 2.65E+00 | 2.72E+00 | 9.54E-01 | no | Sntg1 | downstream_gene_variant | MODIFIER |
| 1 | 8448543 | AX-170523321 | 0 | T | C | 0.509 | 4.99E-03 | 7.20E-02 | 2.65E+00 | 2.72E+00 | 9.54E-01 | no | Sntg1 | intron_variant,non_coding_transcript_variant | MODIFIER |
| 1 | 9035809 | AX-97810601 | 0 | C | T | 0.509 | 4.99E-03 | 7.20E-02 | 2.65E+00 | 2.72E+00 | 9.54E-01 | no | Sntg1 | intron_variant,non_coding_transcript_variant | MODIFIER |
| 1 | 9037356 | AX-170246917 | 0 | A | G | 0.509 | 4.99E-03 | 7.20E-02 | 2.65E+00 | 2.72E+00 | 9.54E-01 | no | Sntg1 | intron_variant,non_coding_transcript_variant | MODIFIER |
| 1 | 9051669 | AX-170213026 | 0 | A | G | 0.509 | 4.99E-03 | 7.20E-02 | 2.65E+00 | 2.72E+00 | 9.54E-01 | no | Sntg1 | intron_variant,non_coding_transcript_variant | MODIFIER |
| 1 | 9088647 | AX-170195081 | 0 | T | C | 0.509 | 4.99E-03 | 7.20E-02 | 2.65E+00 | 2.72E+00 | 9.54E-01 | no | Sntg1 | intron_variant,non_coding_transcript_variant | MODIFIER |
| 1 | 9102112 | AX-170040341 | 0 | T | C | 0.509 | 4.99E-03 | 7.20E-02 | 2.65E+00 | 2.72E+00 | 9.54E-01 | no | Sntg1 | intron_variant,non_coding_transcript_variant | MODIFIER |
| 1 | 9108247 | AX-97810599 | 0 | T | G | 0.509 | 4.99E-03 | 7.20E-02 | 2.65E+00 | 2.72E+00 | 9.54E-01 | no | Sntg1 | intron_variant,non_coding_transcript_variant | MODIFIER |
| 1 | 9108300 | AX-170205002 | 0 | C | T | 0.509 | 4.99E-03 | 7.20E-02 | 2.65E+00 | 2.72E+00 | 9.54E-01 | no | Sntg1 | intron_variant,non_coding_transcript_variant | MODIFIER |
| 1 | 9125385 | AX-170316676 | 0 | C | T | 0.509 | 4.99E-03 | 7.20E-02 | 2.65E+00 | 2.72E+00 | 9.54E-01 | no | Sntg1 | intron_variant,non_coding_transcript_variant | MODIFIER |
| 1 | 9151802 | AX-169999907 | 0 | G | C | 0.509 | 4.99E-03 | 7.20E-02 | 2.65E+00 | 2.72E+00 | 9.54E-01 | no | Sntg1 | intron_variant,non_coding_transcript_variant | MODIFIER |
| 1 | 9170202 | AX-169999908 | 0 | T | C | 0.509 | 4.99E-03 | 7.20E-02 | 2.65E+00 | 2.72E+00 | 9.54E-01 | no | Sntg1 | intron_variant,non_coding_transcript_variant | MODIFIER |
| 1 | 9170299 | AX-169999905 | 0 | G | A | 0.509 | 4.99E-03 | 7.20E-02 | 2.65E+00 | 2.72E+00 | 9.54E-01 | no | Sntg1 | intron_variant,non_coding_transcript_variant | MODIFIER |
| 1 | 9190499 | AX-170408275 | 0 | T | A | 0.509 | 4.99E-03 | 7.20E-02 | 2.65E+00 | 2.72E+00 | 9.54E-01 | no | Sntg1 | intron_variant,non_coding_transcript_variant | MODIFIER |
| 1 | 9208170 | AX-97797887 | 0 | C | T | 0.509 | 4.99E-03 | 7.20E-02 | 2.65E+00 | 2.72E+00 | 9.54E-01 | no | Sntg1 | intron_variant,non_coding_transcript_variant | MODIFIER |
| 1 | 9214736 | AX-170126085 | 0 | C | T | 0.509 | 4.99E-03 | 7.20E-02 | 2.65E+00 | 2.72E+00 | 9.54E-01 | no | Sntg1 | intron_variant,non_coding_transcript_variant | MODIFIER |
| 1 | 9225076 | AX-97821446 | 0 | A | G | 0.509 | 4.99E-03 | 7.20E-02 | 2.65E+00 | 2.72E+00 | 9.54E-01 | no | Sntg1 | intron_variant,non_coding_transcript_variant | MODIFIER |
| 1 | 9249385 | AX-170426366 | 0 | T | G | 0.509 | 4.99E-03 | 7.20E-02 | 2.65E+00 | 2.72E+00 | 9.54E-01 | no | Sntg1 | non_coding_transcript_exon_variant | MODIFIER |
| 1 | 9279470 | AX-170120163 | 0 | G | A | 0.509 | 4.99E-03 | 7.20E-02 | 2.65E+00 | 2.72E+00 | 9.54E-01 | no | Sntg1 | intron_variant,non_coding_transcript_variant | MODIFIER |
| 1 | 9279560 | AX-170148046 | 0 | G | C | 0.509 | 4.99E-03 | 7.20E-02 | 2.65E+00 | 2.72E+00 | 9.54E-01 | no | Sntg1 | intron_variant,non_coding_transcript_variant | MODIFIER |
| 1 | 9294996 | AX-170261960 | 0 | G | A | 0.509 | 4.99E-03 | 7.20E-02 | 2.65E+00 | 2.72E+00 | 9.54E-01 | no | Sntg1 | intron_variant,non_coding_transcript_variant | MODIFIER |
| 1 | 9296275 | AX-170413933 | 0 | C | T | 0.509 | 4.99E-03 | 7.20E-02 | 2.65E+00 | 2.72E+00 | 9.54E-01 | no | Sntg1 | intron_variant,non_coding_transcript_variant | MODIFIER |
| 1 | 9320991 | AX-170040344 | 0 | C | T | 0.509 | 4.99E-03 | 7.20E-02 | 2.65E+00 | 2.72E+00 | 9.54E-01 | no | | intergenic_variant | MODIFIER |
| 1 | 9777083 | AX-170040351 | 0 | C | G | 0.509 | 4.99E-03 | 7.20E-02 | 2.65E+00 | 2.72E+00 | 9.54E-01 | no | 1700034P13Rik | intron_variant,non_coding_transcript_variant | MODIFIER |
| 1 | 9802627 | AX-169999916 | 0 | G | C | 0.509 | 4.99E-03 | 7.20E-02 | 2.65E+00 | 2.72E+00 | 9.54E-01 | no | Gm6195 | upstream_gene_variant | MODIFIER |
| 1 | 9802793 | AX-97808573 | 0 | C | G | 0.509 | 4.99E-03 | 7.20E-02 | 2.65E+00 | 2.72E+00 | 9.54E-01 | no | Gm6195 | upstream_gene_variant | MODIFIER |
| 1 | 9874951 | AX-169999921 | 0 | C | T | 0.509 | 4.99E-03 | 7.20E-02 | 2.65E+00 | 2.72E+00 | 9.54E-01 | no | Sgk3 | intron_variant | MODIFIER |
| 1 | 9929081 | AX-170210850 | 0 | A | G | 0.509 | 4.99E-03 | 7.20E-02 | 2.65E+00 | 2.72E+00 | 9.54E-01 | no | Mcmdc2 | intron_variant | MODIFIER |
| 1 | 9936854 | AX-170035607 | 0 | T | G | 0.509 | 4.99E-03 | 7.20E-02 | 2.65E+00 | 2.72E+00 | 9.54E-01 | no | Mcmdc2 | intron_variant | MODIFIER |
| 1 | 10020165 | AX-170187322 | 0 | C | T | 0.509 | 4.99E-03 | 7.20E-02 | 2.65E+00 | 2.72E+00 | 9.54E-01 | no | Cops5 | downstream_gene_variant | MODIFIER |
| 1 | 10020904 | AX-170303822 | 0 | T | C | 0.509 | 4.99E-03 | 7.20E-02 | 2.65E+00 | 2.72E+00 | 9.54E-01 | no | Cops5 | downstream_gene_variant | MODIFIER |
| 1 | 10037549 | AX-170287312 | 0 | T | A | 0.509 | 4.99E-03 | 7.20E-02 | 2.65E+00 | 2.72E+00 | 9.54E-01 | no | Cops5 | intron_variant | MODIFIER |

| | | | | | | | | | | | | | | | |
|---|---|---|---|---|---|---|---|---|---|---|---|---|---|---|---|
| 10 | 120077704 | AX-170532557 | 1 | G | A | 0.33 | -3.77E-03 | 6.90E-02 | 2.64E+00 | 2.74E+00 | 9.54E-01 | no | Grip1 | downstream_gene_variant | MODIFIER |
| 1 | 36899458 | AX-170193316 | 0 | G | A | 0.465 | -4.13E-03 | 7.25E-02 | 2.63E+00 | 2.74E+00 | 9.54E-01 | no | Tmem131 | intron_variant | MODIFIER |
| 1 | 39949598 | AX-170155440 | 0 | G | A | 0.395 | -4.75E-03 | 7.65E-02 | 2.66E+00 | 2.73E+00 | 9.54E-01 | no | Map4k4 | intron_variant | MODIFIER |
| 1 | 39959750 | AX-97811303 | 0 | G | A | 0.395 | -4.75E-03 | 7.65E-02 | 2.66E+00 | 2.73E+00 | 9.54E-01 | no | Map4k4 | intron_variant | MODIFIER |
| 1 | 40095023 | AX-170372310 | 0 | C | T | 0.395 | -4.75E-03 | 7.65E-02 | 2.66E+00 | 2.73E+00 | 9.54E-01 | no | Il1r2 | intron_variant | MODIFIER |
| 1 | 40432949 | AX-170431056 | 0 | T | C | 0.395 | -4.75E-03 | 7.65E-02 | 2.66E+00 | 2.73E+00 | 9.54E-01 | no | Il1rl1 | upstream_gene_variant | MODIFIER |
| 1 | 40439205 | AX-170221924 | 0 | C | A | 0.395 | -4.75E-03 | 7.65E-02 | 2.66E+00 | 2.73E+00 | 9.54E-01 | no | Il1rl1 | upstream_gene_variant | MODIFIER |
| 1 | 40439384 | AX-170456969 | 0 | G | A | 0.395 | -4.75E-03 | 7.65E-02 | 2.66E+00 | 2.73E+00 | 9.54E-01 | no | Il1rl1 | upstream_gene_variant | MODIFIER |
| 1 | 40484666 | AX-170520573 | 0 | G | C | 0.395 | -4.75E-03 | 7.65E-02 | 2.66E+00 | 2.73E+00 | 9.54E-01 | no | Il18r1 | intron_variant | MODIFIER |
| 7 | 98719761 | AX-170348883 | 0 | G | A | 0.14 | -8.71E-03 | 1.72E-01 | 2.63E+00 | 2.74E+00 | 9.54E-01 | no | Thap12 | downstream_gene_variant | MODIFIER |
| 1 | 71128475 | AX-170086493 | 0 | G | T | 0.14 | 1.30E-02 | 2.34E-01 | 2.68E+00 | 2.74E+00 | 9.55E-01 | no | | intergenic_variant | MODIFIER |
| 6 | 110686799 | AX-170443608 | 0 | G | A | 0.07 | -1.55E-02 | 2.76E-01 | 2.70E+00 | 2.74E+00 | 9.55E-01 | no | Grm7 | intron_variant | MODIFIER |
| 7 | 69063558 | AX-97805418 | 1 | T | C | 0.402 | -4.78E-03 | 9.71E-02 | 2.67E+00 | 2.75E+00 | 9.55E-01 | no | | intergenic_variant | MODIFIER |
| 11 | 6643926 | AX-170390062 | 1 | C | T | 0.062 | 1.10E-02 | 2.14E-01 | 2.65E+00 | 2.74E+00 | 9.55E-01 | no | | intergenic_variant | MODIFIER |
| 9 | 104364685 | AX-170160195 | 1 | A | T | 0.125 | 5.27E-03 | 1.17E-01 | 2.69E+00 | 2.76E+00 | 9.55E-01 | no | Gm28548 | intron_variant,non_coding_transcript_variant | MODIFIER |
| 7 | 112199625 | AX-170149705 | 0 | T | C | 0.456 | 5.63E-03 | 9.71E-02 | 2.68E+00 | 2.73E+00 | 9.55E-01 | no | | intergenic_variant | MODIFIER |
| 7 | 112234522 | AX-170532035 | 0 | C | G | 0.456 | 5.63E-03 | 9.71E-02 | 2.68E+00 | 2.73E+00 | 9.55E-01 | no | Mical2 | intron_variant | MODIFIER |
| 7 | 112236474 | AX-170508867 | 0 | T | A | 0.456 | 5.63E-03 | 9.71E-02 | 2.68E+00 | 2.73E+00 | 9.55E-01 | no | Mical2 | intron_variant | MODIFIER |
| 7 | 112239540 | AX-170322041 | 0 | A | G | 0.456 | 5.63E-03 | 9.71E-02 | 2.68E+00 | 2.73E+00 | 9.55E-01 | no | Mical2 | intron_variant | MODIFIER |
| 7 | 112240126 | AX-97813821 | 0 | A | G | 0.456 | 5.63E-03 | 9.71E-02 | 2.68E+00 | 2.73E+00 | 9.55E-01 | no | Mical2 | intron_variant | MODIFIER |
| 7 | 112246044 | AX-170202970 | 0 | G | T | 0.456 | 5.63E-03 | 9.71E-02 | 2.68E+00 | 2.73E+00 | 9.55E-01 | no | Mical2 | intron_variant | MODIFIER |
| 7 | 112246100 | AX-170480004 | 0 | G | T | 0.456 | 5.63E-03 | 9.71E-02 | 2.68E+00 | 2.73E+00 | 9.55E-01 | no | Mical2 | intron_variant | MODIFIER |
| 7 | 112251403 | AX-170176881 | 0 | A | G | 0.456 | 5.63E-03 | 9.71E-02 | 2.68E+00 | 2.73E+00 | 9.55E-01 | no | Mical2 | intron_variant | MODIFIER |
| 7 | 112253848 | AX-97818609 | 0 | G | A | 0.456 | 5.63E-03 | 9.71E-02 | 2.68E+00 | 2.73E+00 | 9.55E-01 | no | Mical2 | intron_variant | MODIFIER |
| 7 | 112261785 | AX-170120916 | 0 | T | C | 0.456 | 5.63E-03 | 9.71E-02 | 2.68E+00 | 2.73E+00 | 9.55E-01 | no | Mical2 | intron_variant | MODIFIER |
| 7 | 112271149 | AX-170423413 | 0 | G | A | 0.456 | 5.63E-03 | 9.71E-02 | 2.68E+00 | 2.73E+00 | 9.55E-01 | no | Mical2 | upstream_gene_variant | MODIFIER |
| 7 | 112277611 | AX-170237168 | 0 | A | C | 0.456 | 5.63E-03 | 9.71E-02 | 2.68E+00 | 2.73E+00 | 9.55E-01 | no | Mical2 | intron_variant | MODIFIER |
| 7 | 112290690 | AX-170328444 | 0 | C | T | 0.456 | 5.63E-03 | 9.71E-02 | 2.68E+00 | 2.73E+00 | 9.55E-01 | no | Mical2 | intron_variant | MODIFIER |
| 7 | 112306376 | AX-170145676 | 0 | G | C | 0.456 | 5.63E-03 | 9.71E-02 | 2.68E+00 | 2.73E+00 | 9.55E-01 | no | Mical2 | intron_variant | MODIFIER |
| 7 | 112307352 | AX-170503639 | 0 | T | C | 0.456 | 5.63E-03 | 9.71E-02 | 2.68E+00 | 2.73E+00 | 9.55E-01 | no | Mical2 | intron_variant | MODIFIER |
| 7 | 112330645 | AX-170172320 | 0 | T | G | 0.456 | 5.63E-03 | 9.71E-02 | 2.68E+00 | 2.73E+00 | 9.55E-01 | no | Mical2 | intron_variant | MODIFIER |
| 7 | 112340521 | AX-170432768 | 0 | G | T | 0.456 | 5.63E-03 | 9.71E-02 | 2.68E+00 | 2.73E+00 | 9.55E-01 | no | Mical2 | intron_variant | MODIFIER |
| 7 | 112346901 | AX-170351703 | 0 | T | C | 0.456 | 5.63E-03 | 9.71E-02 | 2.68E+00 | 2.73E+00 | 9.55E-01 | no | Mical2 | synonymous_variant | LOW |
| 7 | 112372681 | AX-170113363 | 0 | C | T | 0.456 | 5.63E-03 | 9.71E-02 | 2.68E+00 | 2.73E+00 | 9.55E-01 | no | Mical2 | upstream_gene_variant | MODIFIER |
| 7 | 112388735 | AX-170029785 | 0 | G | A | 0.456 | 5.63E-03 | 9.71E-02 | 2.68E+00 | 2.73E+00 | 9.55E-01 | no | Mical2 | intron_variant | MODIFIER |
| 7 | 112389727 | AX-170507632 | 0 | G | A | 0.456 | 5.63E-03 | 9.71E-02 | 2.68E+00 | 2.73E+00 | 9.55E-01 | no | Mical2 | intron_variant | MODIFIER |
| 7 | 112407017 | AX-170368998 | 0 | T | C | 0.456 | 5.63E-03 | 9.71E-02 | 2.68E+00 | 2.73E+00 | 9.55E-01 | no | Mical2 | intron_variant | MODIFIER |
| 7 | 112413110 | AX-170029784 | 0 | G | T | 0.456 | 5.63E-03 | 9.71E-02 | 2.68E+00 | 2.73E+00 | 9.55E-01 | no | Mical2 | downstream_gene_variant | MODIFIER |
| 7 | 112447391 | AX-170218897 | 0 | A | G | 0.456 | 5.63E-03 | 9.71E-02 | 2.68E+00 | 2.73E+00 | 9.55E-01 | no | Parva | intron_variant | MODIFIER |
| 7 | 112453066 | AX-170478998 | 0 | A | T | 0.456 | 5.63E-03 | 9.71E-02 | 2.68E+00 | 2.73E+00 | 9.55E-01 | no | Parva | intron_variant | MODIFIER |
| 7 | 112454919 | AX-170078448 | 0 | G | A | 0.456 | 5.63E-03 | 9.71E-02 | 2.68E+00 | 2.73E+00 | 9.55E-01 | no | Parva | intron_variant | MODIFIER |
| 7 | 112467695 | AX-170176846 | 0 | G | T | 0.456 | 5.63E-03 | 9.71E-02 | 2.68E+00 | 2.73E+00 | 9.55E-01 | no | Parva | intron_variant | MODIFIER |
| 7 | 112468546 | AX-170202854 | 0 | G | A | 0.456 | 5.63E-03 | 9.71E-02 | 2.68E+00 | 2.73E+00 | 9.55E-01 | no | Parva | intron_variant | MODIFIER |
| 7 | 112469355 | AX-170284477 | 0 | A | G | 0.456 | 5.63E-03 | 9.71E-02 | 2.68E+00 | 2.73E+00 | 9.55E-01 | no | Parva | intron_variant | MODIFIER |
| 7 | 112469487 | AX-170512806 | 0 | C | T | 0.456 | 5.63E-03 | 9.71E-02 | 2.68E+00 | 2.73E+00 | 9.55E-01 | no | Parva | intron_variant | MODIFIER |
| 7 | 112471312 | AX-170242207 | 0 | G | A | 0.456 | 5.63E-03 | 9.71E-02 | 2.68E+00 | 2.73E+00 | 9.55E-01 | no | Parva | intron_variant | MODIFIER |
| 7 | 112502376 | AX-170473963 | 0 | A | G | 0.456 | 5.63E-03 | 9.71E-02 | 2.68E+00 | 2.73E+00 | 9.55E-01 | no | Parva | intron_variant | MODIFIER |
| 7 | 112520124 | AX-170377924 | 0 | T | C | 0.456 | 5.63E-03 | 9.71E-02 | 2.68E+00 | 2.73E+00 | 9.55E-01 | no | Parva | intron_variant | MODIFIER |
| 7 | 112542779 | AX-170251094 | 0 | T | C | 0.456 | 5.63E-03 | 9.71E-02 | 2.68E+00 | 2.73E+00 | 9.55E-01 | no | Parva | intron_variant | MODIFIER |
| 7 | 112549830 | AX-170256137 | 0 | C | G | 0.456 | 5.63E-03 | 9.71E-02 | 2.68E+00 | 2.73E+00 | 9.55E-01 | no | Parva | intron_variant | MODIFIER |
| 7 | 112550040 | AX-170158881 | 0 | C | T | 0.456 | 5.63E-03 | 9.71E-02 | 2.68E+00 | 2.73E+00 | 9.55E-01 | no | Parva | intron_variant | MODIFIER |
| 7 | 112555491 | AX-170269119 | 0 | T | C | 0.456 | 5.63E-03 | 9.71E-02 | 2.68E+00 | 2.73E+00 | 9.55E-01 | no | Parva | intron_variant | MODIFIER |
| 7 | 112565527 | AX-170229539 | 0 | A | G | 0.456 | 5.63E-03 | 9.71E-02 | 2.68E+00 | 2.73E+00 | 9.55E-01 | no | Parva | intron_variant | MODIFIER |
| 7 | 112570834 | AX-170078454 | 0 | G | A | 0.456 | 5.63E-03 | 9.71E-02 | 2.68E+00 | 2.73E+00 | 9.55E-01 | no | Parva | intron_variant | MODIFIER |

| | | | | | | | | | | | | | | | |
|---|---|---|---|---|---|---|---|---|---|---|---|---|---|---|---|
| 7 | 75612728 | AX-170307900 | 1 | G | A | 0.393 | 5.51E-03 | 9.67E-02 | 2.63E+00 | 2.74E+00 | 9.55E-01 | no | Akap13 | intron_variant | MODIFIER |
| 7 | 40370186 | AX-170102489 | 0 | G | A | 0.325 | 6.10E-03 | 9.41E-02 | 2.65E+00 | 2.73E+00 | 9.55E-01 | no | Gm28807 | downstream_gene_variant | MODIFIER |
| 7 | 95262370 | AX-170159030 | 0 | T | C | 0.105 | 1.06E-02 | 1.77E-01 | 2.64E+00 | 2.73E+00 | 9.55E-01 | no | | intergenic_variant | MODIFIER |
| 7 | 32434437 | AX-97802260 | 0 | C | A | 0.491 | 2.66E-02 | 4.67E-01 | 2.65E+00 | 2.74E+00 | 9.55E-01 | no | | intergenic_variant | MODIFIER |
| 11 | 74214619 | AX-170504446 | 0 | G | A | 0.263 | 3.66E-03 | 7.12E-02 | 2.60E+00 | 2.74E+00 | 9.55E-01 | no | Olfr43 | upstream_gene_variant | MODIFIER |
| 8 | 82505883 | AX-170463716 | 2 | T | A | 0.209 | 7.27E-03 | 1.29E-01 | 2.68E+00 | 2.74E+00 | 9.55E-01 | no | | intergenic_variant | MODIFIER |
| 19 | 37670911 | AX-170291082 | 1 | C | T | 0.045 | -1.04E-02 | 1.67E-01 | 2.61E+00 | 2.73E+00 | 9.55E-01 | no | Exoc6 | intron_variant | MODIFIER |
| 8 | 79033219 | AX-170381165 | 0 | T | C | 0.202 | 7.36E-03 | 1.31E-01 | 2.68E+00 | 2.74E+00 | 9.55E-01 | no | Zfp827 | intron_variant | MODIFIER |
| 8 | 79084569 | AX-97783322 | 0 | G | A | 0.202 | 7.36E-03 | 1.31E-01 | 2.68E+00 | 2.74E+00 | 9.55E-01 | no | Zfp827 | intron_variant | MODIFIER |
| 8 | 80883754 | AX-170285051 | 0 | A | G | 0.202 | 7.36E-03 | 1.31E-01 | 2.68E+00 | 2.74E+00 | 9.55E-01 | no | Gab1 | upstream_gene_variant | MODIFIER |
| 8 | 80884212 | AX-170227915 | 0 | C | T | 0.202 | 7.36E-03 | 1.31E-01 | 2.68E+00 | 2.74E+00 | 9.55E-01 | no | Gab1 | upstream_gene_variant | MODIFIER |
| 8 | 80892314 | AX-170444519 | 0 | A | G | 0.202 | 7.36E-03 | 1.31E-01 | 2.68E+00 | 2.74E+00 | 9.55E-01 | no | | intergenic_variant | MODIFIER |
| 8 | 80903849 | AX-170511521 | 0 | G | T | 0.202 | 7.36E-03 | 1.31E-01 | 2.68E+00 | 2.74E+00 | 9.55E-01 | no | | intergenic_variant | MODIFIER |
| 8 | 80905105 | AX-170131655 | 0 | C | G | 0.202 | 7.36E-03 | 1.31E-01 | 2.68E+00 | 2.74E+00 | 9.55E-01 | no | | intergenic_variant | MODIFIER |
| 8 | 80905926 | AX-170272870 | 0 | A | G | 0.202 | 7.36E-03 | 1.31E-01 | 2.68E+00 | 2.74E+00 | 9.55E-01 | no | | intergenic_variant | MODIFIER |
| 8 | 80906344 | AX-170524684 | 0 | T | C | 0.202 | 7.36E-03 | 1.31E-01 | 2.68E+00 | 2.74E+00 | 9.55E-01 | no | | intergenic_variant | MODIFIER |
| 8 | 80936785 | AX-170395574 | 0 | C | T | 0.202 | 7.36E-03 | 1.31E-01 | 2.68E+00 | 2.74E+00 | 9.55E-01 | no | Gm4891 | downstream_gene_variant | MODIFIER |
| 8 | 80951813 | AX-170104648 | 0 | C | T | 0.202 | 7.36E-03 | 1.31E-01 | 2.68E+00 | 2.74E+00 | 9.55E-01 | no | | regulatory_region_variant | MODIFIER |
| 8 | 80962284 | AX-170080564 | 0 | T | C | 0.202 | 7.36E-03 | 1.31E-01 | 2.68E+00 | 2.74E+00 | 9.55E-01 | no | | regulatory_region_variant | MODIFIER |
| 8 | 80970467 | AX-170527800 | 0 | G | A | 0.202 | 7.36E-03 | 1.31E-01 | 2.68E+00 | 2.74E+00 | 9.55E-01 | no | | intergenic_variant | MODIFIER |
| 8 | 80976238 | AX-97807885 | 0 | A | G | 0.202 | 7.36E-03 | 1.31E-01 | 2.68E+00 | 2.74E+00 | 9.55E-01 | no | Usp38 | downstream_gene_variant | MODIFIER |
| 8 | 80979430 | AX-170216985 | 0 | G | A | 0.202 | 7.36E-03 | 1.31E-01 | 2.68E+00 | 2.74E+00 | 9.55E-01 | no | Usp38 | downstream_gene_variant | MODIFIER |
| 8 | 80987953 | AX-97845458 | 0 | G | A | 0.202 | 7.36E-03 | 1.31E-01 | 2.68E+00 | 2.74E+00 | 9.55E-01 | no | Usp38 | intron_variant | MODIFIER |
| 8 | 80995675 | AX-170444171 | 0 | A | G | 0.202 | 7.36E-03 | 1.31E-01 | 2.68E+00 | 2.74E+00 | 9.55E-01 | no | Usp38 | splice_region_variant,intron_variant | LOW |
| 8 | 80996855 | AX-170503790 | 0 | T | A | 0.202 | 7.36E-03 | 1.31E-01 | 2.68E+00 | 2.74E+00 | 9.55E-01 | no | Usp38 | intron_variant | MODIFIER |
| 8 | 80997909 | AX-170314771 | 0 | A | T | 0.202 | 7.36E-03 | 1.31E-01 | 2.68E+00 | 2.74E+00 | 9.55E-01 | no | Usp38 | intron_variant | MODIFIER |
| 8 | 81001068 | AX-170166914 | 0 | A | G | 0.202 | 7.36E-03 | 1.31E-01 | 2.68E+00 | 2.74E+00 | 9.55E-01 | no | Usp38 | synonymous_variant | LOW |
| 8 | 81001671 | AX-170401516 | 0 | C | T | 0.202 | 7.36E-03 | 1.31E-01 | 2.68E+00 | 2.74E+00 | 9.55E-01 | no | Usp38 | intron_variant | MODIFIER |
| 8 | 81006049 | AX-170200847 | 0 | C | G | 0.202 | 7.36E-03 | 1.31E-01 | 2.68E+00 | 2.74E+00 | 9.55E-01 | no | Usp38 | intron_variant | MODIFIER |
| 8 | 81035108 | AX-170341955 | 0 | C | T | 0.202 | 7.36E-03 | 1.31E-01 | 2.68E+00 | 2.74E+00 | 9.55E-01 | no | Gm9725 | intron_variant,non_coding_transcript_variant | MODIFIER |
| 8 | 81044458 | AX-97854791 | 0 | C | T | 0.202 | 7.36E-03 | 1.31E-01 | 2.68E+00 | 2.74E+00 | 9.55E-01 | no | Gm9725 | intron_variant,non_coding_transcript_variant | MODIFIER |
| 8 | 81053046 | AX-170435753 | 0 | A | G | 0.202 | 7.36E-03 | 1.31E-01 | 2.68E+00 | 2.74E+00 | 9.55E-01 | no | Gm9725 | intron_variant,non_coding_transcript_variant | MODIFIER |
| 8 | 81093807 | AX-97803964 | 0 | A | G | 0.202 | 7.36E-03 | 1.31E-01 | 2.68E+00 | 2.74E+00 | 9.55E-01 | no | | intergenic_variant | MODIFIER |
| 8 | 81101691 | AX-170416479 | 0 | G | A | 0.202 | 7.36E-03 | 1.31E-01 | 2.68E+00 | 2.74E+00 | 9.55E-01 | no | | intergenic_variant | MODIFIER |
| 8 | 81120319 | AX-170423374 | 0 | T | C | 0.202 | 7.36E-03 | 1.31E-01 | 2.68E+00 | 2.74E+00 | 9.55E-01 | no | | intergenic_variant | MODIFIER |
| 8 | 81120690 | AX-170295596 | 0 | A | G | 0.202 | 7.36E-03 | 1.31E-01 | 2.68E+00 | 2.74E+00 | 9.55E-01 | no | | intergenic_variant | MODIFIER |
| 8 | 81121622 | AX-97803965 | 0 | G | T | 0.202 | 7.36E-03 | 1.31E-01 | 2.68E+00 | 2.74E+00 | 9.55E-01 | no | | intergenic_variant | MODIFIER |
| 8 | 81126271 | AX-170447742 | 0 | T | C | 0.202 | 7.36E-03 | 1.31E-01 | 2.68E+00 | 2.74E+00 | 9.55E-01 | no | | intergenic_variant | MODIFIER |
| 8 | 81126610 | AX-170403052 | 0 | G | T | 0.202 | 7.36E-03 | 1.31E-01 | 2.68E+00 | 2.74E+00 | 9.55E-01 | no | | intergenic_variant | MODIFIER |
| 8 | 81146764 | AX-170330114 | 0 | T | C | 0.202 | 7.36E-03 | 1.31E-01 | 2.68E+00 | 2.74E+00 | 9.55E-01 | no | | intergenic_variant | MODIFIER |
| 8 | 81148459 | AX-97844194 | 0 | G | A | 0.202 | 7.36E-03 | 1.31E-01 | 2.68E+00 | 2.74E+00 | 9.55E-01 | no | | intergenic_variant | MODIFIER |
| 8 | 81227076 | AX-170316661 | 0 | T | C | 0.202 | 7.36E-03 | 1.31E-01 | 2.68E+00 | 2.74E+00 | 9.55E-01 | no | | intergenic_variant | MODIFIER |
| 8 | 81227711 | AX-170209894 | 0 | C | T | 0.202 | 7.36E-03 | 1.31E-01 | 2.68E+00 | 2.74E+00 | 9.55E-01 | no | | intergenic_variant | MODIFIER |
| 8 | 81243307 | AX-170031118 | 0 | C | T | 0.202 | 7.36E-03 | 1.31E-01 | 2.68E+00 | 2.74E+00 | 9.55E-01 | no | | intergenic_variant | MODIFIER |
| 8 | 81244617 | AX-170432460 | 0 | G | A | 0.202 | 7.36E-03 | 1.31E-01 | 2.68E+00 | 2.74E+00 | 9.55E-01 | no | | intergenic_variant | MODIFIER |
| 8 | 81244804 | AX-170124228 | 0 | C | G | 0.202 | 7.36E-03 | 1.31E-01 | 2.68E+00 | 2.74E+00 | 9.55E-01 | no | | intergenic_variant | MODIFIER |
| 8 | 81266176 | AX-170143304 | 0 | G | A | 0.202 | 7.36E-03 | 1.31E-01 | 2.68E+00 | 2.74E+00 | 9.55E-01 | no | | intergenic_variant | MODIFIER |
| 8 | 81286320 | AX-170121665 | 0 | G | C | 0.202 | 7.36E-03 | 1.31E-01 | 2.68E+00 | 2.74E+00 | 9.55E-01 | no | | intergenic_variant | MODIFIER |
| 8 | 81287474 | AX-170124582 | 0 | G | A | 0.202 | 7.36E-03 | 1.31E-01 | 2.68E+00 | 2.74E+00 | 9.55E-01 | no | | intergenic_variant | MODIFIER |
| 8 | 81287502 | AX-170137132 | 0 | G | A | 0.202 | 7.36E-03 | 1.31E-01 | 2.68E+00 | 2.74E+00 | 9.55E-01 | no | | intergenic_variant | MODIFIER |
| 8 | 81287687 | AX-170080567 | 0 | T | C | 0.202 | 7.36E-03 | 1.31E-01 | 2.68E+00 | 2.74E+00 | 9.55E-01 | no | | intergenic_variant | MODIFIER |
| 8 | 81295594 | AX-170320804 | 0 | A | G | 0.202 | 7.36E-03 | 1.31E-01 | 2.68E+00 | 2.74E+00 | 9.55E-01 | no | Gm45539 | upstream_gene_variant | MODIFIER |
| 8 | 81324460 | AX-170193125 | 0 | T | G | 0.202 | 7.36E-03 | 1.31E-01 | 2.68E+00 | 2.74E+00 | 9.55E-01 | no | | intergenic_variant | MODIFIER |
| 8 | 81325793 | AX-170374386 | 0 | T | C | 0.202 | 7.36E-03 | 1.31E-01 | 2.68E+00 | 2.74E+00 | 9.55E-01 | no | | intergenic_variant | MODIFIER |

| | | | | | | | | | | | | | | | |
|---|---|---|---|---|---|---|---|---|---|---|---|---|---|---|---|
| 8 | 81328392 | AX-170031122 | 0 | A | C | 0.202 | 7.36E-03 | 1.31E-01 | 2.68E+00 | 2.74E+00 | 9.55E-01 | no | | intergenic_variant | MODIFIER |
| 8 | 81328434 | AX-170031117 | 0 | C | T | 0.202 | 7.36E-03 | 1.31E-01 | 2.68E+00 | 2.74E+00 | 9.55E-01 | no | | intergenic_variant | MODIFIER |
| 8 | 81328456 | AX-170031123 | 0 | G | A | 0.202 | 7.36E-03 | 1.31E-01 | 2.68E+00 | 2.74E+00 | 9.55E-01 | no | | intergenic_variant | MODIFIER |
| 8 | 81352373 | AX-170454178 | 0 | G | A | 0.202 | 7.36E-03 | 1.31E-01 | 2.68E+00 | 2.74E+00 | 9.55E-01 | no | Inpp4b | intron_variant,NMD_transcript_variant | MODIFIER |
| 8 | 81361091 | AX-170031116 | 0 | T | A | 0.202 | 7.36E-03 | 1.31E-01 | 2.68E+00 | 2.74E+00 | 9.55E-01 | no | Inpp4b | intron_variant,NMD_transcript_variant | MODIFIER |
| 8 | 81361582 | AX-170157402 | 0 | C | A | 0.202 | 7.36E-03 | 1.31E-01 | 2.68E+00 | 2.74E+00 | 9.55E-01 | no | Inpp4b | intron_variant,NMD_transcript_variant | MODIFIER |
| 8 | 81363429 | AX-97795038 | 0 | C | T | 0.202 | 7.36E-03 | 1.31E-01 | 2.68E+00 | 2.74E+00 | 9.55E-01 | no | Inpp4b | intron_variant,NMD_transcript_variant | MODIFIER |
| 8 | 81393982 | AX-170495645 | 0 | A | G | 0.202 | 7.36E-03 | 1.31E-01 | 2.68E+00 | 2.74E+00 | 9.55E-01 | no | Inpp4b | intron_variant,NMD_transcript_variant | MODIFIER |
| 8 | 81408191 | AX-170466797 | 0 | C | T | 0.202 | 7.36E-03 | 1.31E-01 | 2.68E+00 | 2.74E+00 | 9.55E-01 | no | Inpp4b | intron_variant,NMD_transcript_variant | MODIFIER |
| 8 | 81408813 | AX-170428676 | 0 | T | C | 0.202 | 7.36E-03 | 1.31E-01 | 2.68E+00 | 2.74E+00 | 9.55E-01 | no | Inpp4b | intron_variant,NMD_transcript_variant | MODIFIER |
| 8 | 81409980 | AX-170256117 | 0 | A | C | 0.202 | 7.36E-03 | 1.31E-01 | 2.68E+00 | 2.74E+00 | 9.55E-01 | no | Inpp4b | intron_variant,NMD_transcript_variant | MODIFIER |
| 8 | 81411765 | AX-170467138 | 0 | A | C | 0.202 | 7.36E-03 | 1.31E-01 | 2.68E+00 | 2.74E+00 | 9.55E-01 | no | Inpp4b | intron_variant,NMD_transcript_variant | MODIFIER |
| 8 | 81420974 | AX-170115655 | 0 | G | A | 0.202 | 7.36E-03 | 1.31E-01 | 2.68E+00 | 2.74E+00 | 9.55E-01 | no | Inpp4b | intron_variant,NMD_transcript_variant | MODIFIER |
| 8 | 81428331 | AX-170534811 | 0 | A | G | 0.202 | 7.36E-03 | 1.31E-01 | 2.68E+00 | 2.74E+00 | 9.55E-01 | no | Inpp4b | intron_variant,NMD_transcript_variant | MODIFIER |
| 8 | 81444833 | AX-170441227 | 0 | C | A | 0.202 | 7.36E-03 | 1.31E-01 | 2.68E+00 | 2.74E+00 | 9.55E-01 | no | Inpp4b | intron_variant,NMD_transcript_variant | MODIFIER |
| 8 | 81482764 | AX-170428392 | 0 | T | A | 0.202 | 7.36E-03 | 1.31E-01 | 2.68E+00 | 2.74E+00 | 9.55E-01 | no | Inpp4b | intron_variant,NMD_transcript_variant | MODIFIER |
| 8 | 81505604 | AX-170189755 | 0 | A | C | 0.202 | 7.36E-03 | 1.31E-01 | 2.68E+00 | 2.74E+00 | 9.55E-01 | no | Inpp4b | intron_variant,NMD_transcript_variant | MODIFIER |
| 8 | 81512087 | AX-170487558 | 0 | A | G | 0.202 | 7.36E-03 | 1.31E-01 | 2.68E+00 | 2.74E+00 | 9.55E-01 | no | Inpp4b | intron_variant,NMD_transcript_variant | MODIFIER |
| 8 | 81512235 | AX-170158001 | 0 | G | A | 0.202 | 7.36E-03 | 1.31E-01 | 2.68E+00 | 2.74E+00 | 9.55E-01 | no | Inpp4b | intron_variant,NMD_transcript_variant | MODIFIER |
| 8 | 81512786 | AX-97843859 | 0 | T | C | 0.202 | 7.36E-03 | 1.31E-01 | 2.68E+00 | 2.74E+00 | 9.55E-01 | no | Inpp4b | intron_variant,NMD_transcript_variant | MODIFIER |
| 8 | 81512968 | AX-170080571 | 0 | G | A | 0.202 | 7.36E-03 | 1.31E-01 | 2.68E+00 | 2.74E+00 | 9.55E-01 | no | Inpp4b | intron_variant,NMD_transcript_variant | MODIFIER |
| 8 | 81514134 | AX-170418731 | 0 | G | A | 0.202 | 7.36E-03 | 1.31E-01 | 2.68E+00 | 2.74E+00 | 9.55E-01 | no | Inpp4b | intron_variant,NMD_transcript_variant | MODIFIER |
| 8 | 81521931 | AX-170227164 | 0 | A | T | 0.202 | 7.36E-03 | 1.31E-01 | 2.68E+00 | 2.74E+00 | 9.55E-01 | no | Inpp4b | intron_variant,NMD_transcript_variant | MODIFIER |
| 8 | 81563305 | AX-170317398 | 0 | G | A | 0.202 | 7.36E-03 | 1.31E-01 | 2.68E+00 | 2.74E+00 | 9.55E-01 | no | Inpp4b | intron_variant,NMD_transcript_variant | MODIFIER |
| 8 | 81590486 | AX-170197493 | 0 | A | C | 0.202 | 7.36E-03 | 1.31E-01 | 2.68E+00 | 2.74E+00 | 9.55E-01 | no | Inpp4b | intron_variant,NMD_transcript_variant | MODIFIER |
| 8 | 81590717 | AX-97822157 | 0 | G | A | 0.202 | 7.36E-03 | 1.31E-01 | 2.68E+00 | 2.74E+00 | 9.55E-01 | no | Inpp4b | intron_variant,NMD_transcript_variant | MODIFIER |
| 8 | 81612837 | AX-170440448 | 0 | C | T | 0.202 | 7.36E-03 | 1.31E-01 | 2.68E+00 | 2.74E+00 | 9.55E-01 | no | Inpp4b | intron_variant,NMD_transcript_variant | MODIFIER |
| 8 | 81616447 | AX-170107620 | 0 | T | A | 0.202 | 7.36E-03 | 1.31E-01 | 2.68E+00 | 2.74E+00 | 9.55E-01 | no | Inpp4b | intron_variant,NMD_transcript_variant | MODIFIER |
| 8 | 81619449 | AX-170266233 | 0 | A | T | 0.202 | 7.36E-03 | 1.31E-01 | 2.68E+00 | 2.74E+00 | 9.55E-01 | no | Inpp4b | intron_variant,NMD_transcript_variant | MODIFIER |
| 8 | 81629913 | AX-170487111 | 0 | T | C | 0.202 | 7.36E-03 | 1.31E-01 | 2.68E+00 | 2.74E+00 | 9.55E-01 | no | Inpp4b | intron_variant,NMD_transcript_variant | MODIFIER |
| 8 | 81656130 | AX-170464616 | 0 | G | C | 0.202 | 7.36E-03 | 1.31E-01 | 2.68E+00 | 2.74E+00 | 9.55E-01 | no | Inpp4b | intron_variant,NMD_transcript_variant | MODIFIER |
| 8 | 81656895 | AX-97819461 | 0 | G | A | 0.202 | 7.36E-03 | 1.31E-01 | 2.68E+00 | 2.74E+00 | 9.55E-01 | no | Inpp4b | intron_variant,NMD_transcript_variant | MODIFIER |
| 8 | 81690109 | AX-170031129 | 0 | G | A | 0.202 | 7.36E-03 | 1.31E-01 | 2.68E+00 | 2.74E+00 | 9.55E-01 | no | Inpp4b | intron_variant,NMD_transcript_variant | MODIFIER |
| 8 | 81690160 | AX-170263890 | 0 | G | A | 0.202 | 7.36E-03 | 1.31E-01 | 2.68E+00 | 2.74E+00 | 9.55E-01 | no | Inpp4b | intron_variant,NMD_transcript_variant | MODIFIER |
| 8 | 81692580 | AX-170234786 | 0 | G | A | 0.202 | 7.36E-03 | 1.31E-01 | 2.68E+00 | 2.74E+00 | 9.55E-01 | no | Inpp4b | intron_variant,NMD_transcript_variant | MODIFIER |
| 8 | 81692637 | AX-170179546 | 0 | T | C | 0.202 | 7.36E-03 | 1.31E-01 | 2.68E+00 | 2.74E+00 | 9.55E-01 | no | Inpp4b | intron_variant,NMD_transcript_variant | MODIFIER |
| 8 | 81695215 | AX-97799499 | 0 | T | C | 0.202 | 7.36E-03 | 1.31E-01 | 2.68E+00 | 2.74E+00 | 9.55E-01 | no | Inpp4b | intron_variant,NMD_transcript_variant | MODIFIER |
| 8 | 81711171 | AX-97786087 | 0 | T | G | 0.202 | 7.36E-03 | 1.31E-01 | 2.68E+00 | 2.74E+00 | 9.55E-01 | no | Inpp4b | upstream_gene_variant | MODIFIER |
| 8 | 81711229 | AX-170208495 | 0 | A | G | 0.202 | 7.36E-03 | 1.31E-01 | 2.68E+00 | 2.74E+00 | 9.55E-01 | no | Inpp4b | upstream_gene_variant | MODIFIER |
| 8 | 81717922 | AX-170108488 | 0 | T | C | 0.202 | 7.36E-03 | 1.31E-01 | 2.68E+00 | 2.74E+00 | 9.55E-01 | no | Inpp4b | intron_variant | MODIFIER |
| 8 | 81718838 | AX-170031127 | 0 | C | T | 0.202 | 7.36E-03 | 1.31E-01 | 2.68E+00 | 2.74E+00 | 9.55E-01 | no | Inpp4b | intron_variant | MODIFIER |
| 8 | 81730703 | AX-170274173 | 0 | G | T | 0.202 | 7.36E-03 | 1.31E-01 | 2.68E+00 | 2.74E+00 | 9.55E-01 | no | Inpp4b | intron_variant | MODIFIER |
| 8 | 81759678 | AX-170190548 | 0 | A | G | 0.202 | 7.36E-03 | 1.31E-01 | 2.68E+00 | 2.74E+00 | 9.55E-01 | no | Inpp4b | intron_variant | MODIFIER |
| 8 | 81759708 | AX-170321919 | 0 | C | G | 0.202 | 7.36E-03 | 1.31E-01 | 2.68E+00 | 2.74E+00 | 9.55E-01 | no | Inpp4b | intron_variant | MODIFIER |
| 8 | 81773026 | AX-97789009 | 0 | G | A | 0.202 | 7.36E-03 | 1.31E-01 | 2.68E+00 | 2.74E+00 | 9.55E-01 | no | Inpp4b | intron_variant | MODIFIER |
| 8 | 81792805 | AX-170483670 | 0 | C | T | 0.202 | 7.36E-03 | 1.31E-01 | 2.68E+00 | 2.74E+00 | 9.55E-01 | no | Inpp4b | intron_variant | MODIFIER |
| 8 | 81825188 | AX-170080575 | 0 | A | C | 0.202 | 7.36E-03 | 1.31E-01 | 2.68E+00 | 2.74E+00 | 9.55E-01 | no | Inpp4b | intron_variant | MODIFIER |
| 8 | 81854092 | AX-170208624 | 0 | T | C | 0.202 | 7.36E-03 | 1.31E-01 | 2.68E+00 | 2.74E+00 | 9.55E-01 | no | Inpp4b | intron_variant | MODIFIER |
| 8 | 81854397 | AX-97849236 | 0 | G | T | 0.202 | 7.36E-03 | 1.31E-01 | 2.68E+00 | 2.74E+00 | 9.55E-01 | no | Inpp4b | intron_variant | MODIFIER |
| 8 | 81854661 | AX-170031128 | 0 | G | T | 0.202 | 7.36E-03 | 1.31E-01 | 2.68E+00 | 2.74E+00 | 9.55E-01 | no | Inpp4b | intron_variant | MODIFIER |
| 8 | 81858556 | AX-170531014 | 0 | G | A | 0.202 | 7.36E-03 | 1.31E-01 | 2.68E+00 | 2.74E+00 | 9.55E-01 | no | Inpp4b | intron_variant | MODIFIER |
| 8 | 81859434 | AX-170245370 | 0 | G | A | 0.202 | 7.36E-03 | 1.31E-01 | 2.68E+00 | 2.74E+00 | 9.55E-01 | no | Inpp4b | intron_variant | MODIFIER |
| 8 | 81871158 | AX-97835148 | 0 | C | T | 0.202 | 7.36E-03 | 1.31E-01 | 2.68E+00 | 2.74E+00 | 9.55E-01 | no | Inpp4b | intron_variant | MODIFIER |
| 8 | 81878820 | AX-170491576 | 0 | G | T | 0.202 | 7.36E-03 | 1.31E-01 | 2.68E+00 | 2.74E+00 | 9.55E-01 | no | Inpp4b | intron_variant | MODIFIER |

| | | | | | | | | | | | | | | | |
|---|---|---|---|---|---|---|---|---|---|---|---|---|---|---|---|
| 8 | 81886483 | AX-97838136 | 0 | G | T | 0.202 | 7.36E-03 | 1.31E-01 | 2.68E+00 | 2.74E+00 | 9.55E-01 | no | Inpp4b | intron_variant | MODIFIER |
| 8 | 81888621 | AX-170131586 | 0 | G | C | 0.202 | 7.36E-03 | 1.31E-01 | 2.68E+00 | 2.74E+00 | 9.55E-01 | no | Inpp4b | intron_variant | MODIFIER |
| 8 | 81907202 | AX-170407869 | 0 | G | A | 0.202 | 7.36E-03 | 1.31E-01 | 2.68E+00 | 2.74E+00 | 9.55E-01 | no | Inpp4b | intron_variant | MODIFIER |
| 8 | 81926847 | AX-170125232 | 0 | G | T | 0.202 | 7.36E-03 | 1.31E-01 | 2.68E+00 | 2.74E+00 | 9.55E-01 | no | Inpp4b | intron_variant | MODIFIER |
| 8 | 81942731 | AX-97783766 | 0 | C | T | 0.202 | 7.36E-03 | 1.31E-01 | 2.68E+00 | 2.74E+00 | 9.55E-01 | no | Inpp4b | intron_variant | MODIFIER |
| 8 | 81942759 | AX-170392146 | 0 | G | C | 0.202 | 7.36E-03 | 1.31E-01 | 2.68E+00 | 2.74E+00 | 9.55E-01 | no | Inpp4b | intron_variant | MODIFIER |
| 8 | 81951757 | AX-170482862 | 0 | T | C | 0.202 | 7.36E-03 | 1.31E-01 | 2.68E+00 | 2.74E+00 | 9.55E-01 | no | Inpp4b | intron_variant | MODIFIER |
| 8 | 81953162 | AX-170511944 | 0 | T | C | 0.202 | 7.36E-03 | 1.31E-01 | 2.68E+00 | 2.74E+00 | 9.55E-01 | no | Inpp4b | intron_variant | MODIFIER |
| 8 | 82004676 | AX-170412285 | 0 | A | G | 0.202 | 7.36E-03 | 1.31E-01 | 2.68E+00 | 2.74E+00 | 9.55E-01 | no | Inpp4b | intron_variant | MODIFIER |
| 8 | 82017253 | AX-97847795 | 0 | T | C | 0.202 | 7.36E-03 | 1.31E-01 | 2.68E+00 | 2.74E+00 | 9.55E-01 | no | Inpp4b | intron_variant | MODIFIER |
| 8 | 82042308 | AX-170347979 | 0 | G | T | 0.202 | 7.36E-03 | 1.31E-01 | 2.68E+00 | 2.74E+00 | 9.55E-01 | no | Inpp4b | intron_variant | MODIFIER |
| 8 | 82042750 | AX-170090162 | 0 | C | A | 0.202 | 7.36E-03 | 1.31E-01 | 2.68E+00 | 2.74E+00 | 9.55E-01 | no | Inpp4b | intron_variant | MODIFIER |
| 8 | 82048124 | AX-170478070 | 0 | A | G | 0.202 | 7.36E-03 | 1.31E-01 | 2.68E+00 | 2.74E+00 | 9.55E-01 | no | Inpp4b | intron_variant | MODIFIER |
| 8 | 82065423 | AX-170495189 | 0 | A | C | 0.202 | 7.36E-03 | 1.31E-01 | 2.68E+00 | 2.74E+00 | 9.55E-01 | no | Inpp4b | intron_variant | MODIFIER |
| 8 | 82079715 | AX-170302160 | 0 | G | A | 0.202 | 7.36E-03 | 1.31E-01 | 2.68E+00 | 2.74E+00 | 9.55E-01 | no | Inpp4b | intron_variant | MODIFIER |
| 8 | 82123489 | AX-170302378 | 0 | A | G | 0.202 | 7.36E-03 | 1.31E-01 | 2.68E+00 | 2.74E+00 | 9.55E-01 | no | Inpp4b | downstream_gene_variant | MODIFIER |
| 8 | 82128273 | AX-97791821 | 0 | A | T | 0.202 | 7.36E-03 | 1.31E-01 | 2.68E+00 | 2.74E+00 | 9.55E-01 | no | Inpp4b | downstream_gene_variant | MODIFIER |
| 8 | 82140805 | AX-170100886 | 0 | T | C | 0.202 | 7.36E-03 | 1.31E-01 | 2.68E+00 | 2.74E+00 | 9.55E-01 | no | | intergenic_variant | MODIFIER |
| 8 | 82159251 | AX-97790248 | 0 | C | T | 0.202 | 7.36E-03 | 1.31E-01 | 2.68E+00 | 2.74E+00 | 9.55E-01 | no | Gm9520 | upstream_gene_variant | MODIFIER |
| 8 | 82166562 | AX-170120952 | 0 | G | A | 0.202 | 7.36E-03 | 1.31E-01 | 2.68E+00 | 2.74E+00 | 9.55E-01 | no | Gm9520 | downstream_gene_variant | MODIFIER |
| 8 | 82168443 | AX-170409371 | 0 | T | C | 0.202 | 7.36E-03 | 1.31E-01 | 2.68E+00 | 2.74E+00 | 9.55E-01 | no | Gm9520 | downstream_gene_variant | MODIFIER |
| 8 | 82198727 | AX-97840239 | 0 | G | A | 0.202 | 7.36E-03 | 1.31E-01 | 2.68E+00 | 2.74E+00 | 9.55E-01 | no | Gm38590 | intron_variant,non_coding_transcript_variant | MODIFIER |
| 8 | 82198840 | AX-170391894 | 0 | G | A | 0.202 | 7.36E-03 | 1.31E-01 | 2.68E+00 | 2.74E+00 | 9.55E-01 | no | Gm38590 | intron_variant,non_coding_transcript_variant | MODIFIER |
| 8 | 82207671 | AX-170437621 | 0 | A | G | 0.202 | 7.36E-03 | 1.31E-01 | 2.68E+00 | 2.74E+00 | 9.55E-01 | no | Gm38590 | upstream_gene_variant | MODIFIER |
| 8 | 82232543 | AX-170125694 | 0 | G | T | 0.202 | 7.36E-03 | 1.31E-01 | 2.68E+00 | 2.74E+00 | 9.55E-01 | no | | intergenic_variant | MODIFIER |
| 8 | 82234063 | AX-97847228 | 0 | T | A | 0.202 | 7.36E-03 | 1.31E-01 | 2.68E+00 | 2.74E+00 | 9.55E-01 | no | | intergenic_variant | MODIFIER |
| 8 | 82234389 | AX-170080586 | 0 | T | C | 0.202 | 7.36E-03 | 1.31E-01 | 2.68E+00 | 2.74E+00 | 9.55E-01 | no | | intergenic_variant | MODIFIER |
| 8 | 82251697 | AX-170472231 | 0 | G | A | 0.202 | 7.36E-03 | 1.31E-01 | 2.68E+00 | 2.74E+00 | 9.55E-01 | no | | intergenic_variant | MODIFIER |
| 8 | 82253995 | AX-170168897 | 0 | T | C | 0.202 | 7.36E-03 | 1.31E-01 | 2.68E+00 | 2.74E+00 | 9.55E-01 | no | | intergenic_variant | MODIFIER |
| 8 | 82266178 | AX-170080584 | 0 | C | T | 0.202 | 7.36E-03 | 1.31E-01 | 2.68E+00 | 2.74E+00 | 9.55E-01 | no | | intergenic_variant | MODIFIER |
| 8 | 82266451 | AX-170441932 | 0 | C | T | 0.202 | 7.36E-03 | 1.31E-01 | 2.68E+00 | 2.74E+00 | 9.55E-01 | no | | intergenic_variant | MODIFIER |
| 8 | 82278780 | AX-170031133 | 0 | A | T | 0.202 | 7.36E-03 | 1.31E-01 | 2.68E+00 | 2.74E+00 | 9.55E-01 | no | | intergenic_variant | MODIFIER |
| 8 | 82278816 | AX-170425209 | 0 | A | C | 0.202 | 7.36E-03 | 1.31E-01 | 2.68E+00 | 2.74E+00 | 9.55E-01 | no | | intergenic_variant | MODIFIER |
| 8 | 82285558 | AX-170031134 | 0 | A | G | 0.202 | 7.36E-03 | 1.31E-01 | 2.68E+00 | 2.74E+00 | 9.55E-01 | no | | intergenic_variant | MODIFIER |
| 8 | 82308885 | AX-97849453 | 0 | G | A | 0.202 | 7.36E-03 | 1.31E-01 | 2.68E+00 | 2.74E+00 | 9.55E-01 | no | | intergenic_variant | MODIFIER |
| 8 | 82329542 | AX-170369277 | 0 | C | T | 0.202 | 7.36E-03 | 1.31E-01 | 2.68E+00 | 2.74E+00 | 9.55E-01 | no | Il15 | downstream_gene_variant | MODIFIER |
| 8 | 82368232 | AX-170152868 | 0 | C | T | 0.202 | 7.36E-03 | 1.31E-01 | 2.68E+00 | 2.74E+00 | 9.55E-01 | no | Il15 | intron_variant | MODIFIER |
| 8 | 82372714 | AX-170264356 | 0 | G | A | 0.202 | 7.36E-03 | 1.31E-01 | 2.68E+00 | 2.74E+00 | 9.55E-01 | no | Il15 | intron_variant | MODIFIER |
| 8 | 82377882 | AX-97784027 | 0 | A | C | 0.202 | 7.36E-03 | 1.31E-01 | 2.68E+00 | 2.74E+00 | 9.55E-01 | no | Il15 | intron_variant | MODIFIER |
| 8 | 82378530 | AX-170355342 | 0 | G | A | 0.202 | 7.36E-03 | 1.31E-01 | 2.68E+00 | 2.74E+00 | 9.55E-01 | no | Il15 | intron_variant | MODIFIER |
| 8 | 82379912 | AX-170138236 | 0 | C | G | 0.202 | 7.36E-03 | 1.31E-01 | 2.68E+00 | 2.74E+00 | 9.55E-01 | no | Il15 | intron_variant | MODIFIER |
| 8 | 82413240 | AX-97849999 | 0 | A | G | 0.202 | 7.36E-03 | 1.31E-01 | 2.68E+00 | 2.74E+00 | 9.55E-01 | no | Gm25138 | downstream_gene_variant | MODIFIER |
| 8 | 82443417 | AX-170291395 | 0 | A | G | 0.202 | 7.36E-03 | 1.31E-01 | 2.68E+00 | 2.74E+00 | 9.55E-01 | no | | intergenic_variant | MODIFIER |
| 8 | 82460408 | AX-170295661 | 0 | T | G | 0.202 | 7.36E-03 | 1.31E-01 | 2.68E+00 | 2.74E+00 | 9.55E-01 | no | | intergenic_variant | MODIFIER |
| 8 | 82502271 | AX-170354301 | 0 | G | C | 0.202 | 7.36E-03 | 1.31E-01 | 2.68E+00 | 2.74E+00 | 9.55E-01 | no | | intergenic_variant | MODIFIER |
| 8 | 82527509 | AX-170412919 | 0 | A | G | 0.202 | 7.36E-03 | 1.31E-01 | 2.68E+00 | 2.74E+00 | 9.55E-01 | no | | intergenic_variant | MODIFIER |
| 8 | 82528166 | AX-170362396 | 0 | G | A | 0.202 | 7.36E-03 | 1.31E-01 | 2.68E+00 | 2.74E+00 | 9.55E-01 | no | | intergenic_variant | MODIFIER |
| 8 | 82529515 | AX-170219217 | 0 | G | T | 0.202 | 7.36E-03 | 1.31E-01 | 2.68E+00 | 2.74E+00 | 9.55E-01 | no | | regulatory_region_variant | MODIFIER |
| 8 | 82530017 | AX-170227748 | 0 | G | A | 0.202 | 7.36E-03 | 1.31E-01 | 2.68E+00 | 2.74E+00 | 9.55E-01 | no | | intergenic_variant | MODIFIER |
| 8 | 82531121 | AX-97857421 | 0 | A | G | 0.202 | 7.36E-03 | 1.31E-01 | 2.68E+00 | 2.74E+00 | 9.55E-01 | no | | intergenic_variant | MODIFIER |
| 8 | 82537985 | AX-170080588 | 0 | T | C | 0.202 | 7.36E-03 | 1.31E-01 | 2.68E+00 | 2.74E+00 | 9.55E-01 | no | | intergenic_variant | MODIFIER |
| 8 | 82557549 | AX-97793908 | 0 | T | C | 0.202 | 7.36E-03 | 1.31E-01 | 2.68E+00 | 2.74E+00 | 9.55E-01 | no | | intergenic_variant | MODIFIER |
| 8 | 82568200 | AX-170375748 | 0 | T | G | 0.202 | 7.36E-03 | 1.31E-01 | 2.68E+00 | 2.74E+00 | 9.55E-01 | no | | intergenic_variant | MODIFIER |
| 8 | 82572817 | AX-170458143 | 0 | G | C | 0.202 | 7.36E-03 | 1.31E-01 | 2.68E+00 | 2.74E+00 | 9.55E-01 | no | Gm8167 | upstream_gene_variant | MODIFIER |

| | | | | | | | | | | | | | | | |
|---|---|---|---|---|---|---|---|---|---|---|---|---|---|---|---|
| 8 | 82591707 | AX-170492963 | 0 | G | A | 0.202 | 7.36E-03 | 1.31E-01 | 2.68E+00 | 2.74E+00 | 9.55E-01 | no | | intergenic_variant | MODIFIER |
| 8 | 82608943 | AX-97788399 | 0 | C | T | 0.202 | 7.36E-03 | 1.31E-01 | 2.68E+00 | 2.74E+00 | 9.55E-01 | no | | intergenic_variant | MODIFIER |
| 8 | 82629463 | AX-170419293 | 0 | A | C | 0.202 | 7.36E-03 | 1.31E-01 | 2.68E+00 | 2.74E+00 | 9.55E-01 | no | | intergenic_variant | MODIFIER |
| 8 | 82629821 | AX-170440327 | 0 | A | G | 0.202 | 7.36E-03 | 1.31E-01 | 2.68E+00 | 2.74E+00 | 9.55E-01 | no | | intergenic_variant | MODIFIER |
| 8 | 82630499 | AX-170150464 | 0 | G | A | 0.202 | 7.36E-03 | 1.31E-01 | 2.68E+00 | 2.74E+00 | 9.55E-01 | no | | intergenic_variant | MODIFIER |
| 8 | 82671371 | AX-170285615 | 0 | G | A | 0.202 | 7.36E-03 | 1.31E-01 | 2.68E+00 | 2.74E+00 | 9.55E-01 | no | | intergenic_variant | MODIFIER |
| 8 | 82692541 | AX-170497263 | 0 | A | G | 0.202 | 7.36E-03 | 1.31E-01 | 2.68E+00 | 2.74E+00 | 9.55E-01 | no | | intergenic_variant | MODIFIER |
| 8 | 82694786 | AX-170185845 | 0 | A | G | 0.202 | 7.36E-03 | 1.31E-01 | 2.68E+00 | 2.74E+00 | 9.55E-01 | no | | intergenic_variant | MODIFIER |
| 8 | 82696217 | AX-170183264 | 0 | C | T | 0.202 | 7.36E-03 | 1.31E-01 | 2.68E+00 | 2.74E+00 | 9.55E-01 | no | | intergenic_variant | MODIFIER |
| 8 | 82701080 | AX-170502648 | 0 | G | A | 0.202 | 7.36E-03 | 1.31E-01 | 2.68E+00 | 2.74E+00 | 9.55E-01 | no | | intergenic_variant | MODIFIER |
| 8 | 82728196 | AX-97781151 | 0 | C | T | 0.202 | 7.36E-03 | 1.31E-01 | 2.68E+00 | 2.74E+00 | 9.55E-01 | no | | intergenic_variant | MODIFIER |
| 8 | 82765534 | AX-170162507 | 0 | G | C | 0.202 | 7.36E-03 | 1.31E-01 | 2.68E+00 | 2.74E+00 | 9.55E-01 | no | Zfp330 | intron_variant | MODIFIER |
| 8 | 82766863 | AX-170101065 | 0 | T | A | 0.202 | 7.36E-03 | 1.31E-01 | 2.68E+00 | 2.74E+00 | 9.55E-01 | no | Zfp330 | intron_variant | MODIFIER |
| 8 | 82770165 | AX-170233935 | 0 | G | A | 0.202 | 7.36E-03 | 1.31E-01 | 2.68E+00 | 2.74E+00 | 9.55E-01 | no | Zfp330 | intron_variant | MODIFIER |
| 8 | 82829536 | AX-170349709 | 0 | A | C | 0.202 | 7.36E-03 | 1.31E-01 | 2.68E+00 | 2.74E+00 | 9.55E-01 | no | | intergenic_variant | MODIFIER |
| 8 | 82834430 | AX-170224426 | 0 | A | G | 0.202 | 7.36E-03 | 1.31E-01 | 2.68E+00 | 2.74E+00 | 9.55E-01 | no | | intergenic_variant | MODIFIER |
| 8 | 82855168 | AX-170031140 | 0 | G | A | 0.202 | 7.36E-03 | 1.31E-01 | 2.68E+00 | 2.74E+00 | 9.55E-01 | no | | intergenic_variant | MODIFIER |
| 8 | 82881176 | AX-170244578 | 0 | T | C | 0.202 | 7.36E-03 | 1.31E-01 | 2.68E+00 | 2.74E+00 | 9.55E-01 | no | Rnf150 | intron_variant | MODIFIER |
| 8 | 82892460 | AX-97798101 | 0 | C | G | 0.202 | 7.36E-03 | 1.31E-01 | 2.68E+00 | 2.74E+00 | 9.55E-01 | no | Rnf150 | intron_variant | MODIFIER |
| 8 | 82893086 | AX-170080595 | 0 | T | C | 0.202 | 7.36E-03 | 1.31E-01 | 2.68E+00 | 2.74E+00 | 9.55E-01 | no | Rnf150 | intron_variant | MODIFIER |
| 8 | 82893631 | AX-170080592 | 0 | G | A | 0.202 | 7.36E-03 | 1.31E-01 | 2.68E+00 | 2.74E+00 | 9.55E-01 | no | Rnf150 | intron_variant | MODIFIER |
| 8 | 82920552 | AX-97786174 | 0 | A | G | 0.202 | 7.36E-03 | 1.31E-01 | 2.68E+00 | 2.74E+00 | 9.55E-01 | no | Rnf150 | intron_variant | MODIFIER |
| 8 | 82923549 | AX-170080593 | 0 | C | T | 0.202 | 7.36E-03 | 1.31E-01 | 2.68E+00 | 2.74E+00 | 9.55E-01 | no | Rnf150 | intron_variant | MODIFIER |
| 8 | 82927385 | AX-170430343 | 0 | T | C | 0.202 | 7.36E-03 | 1.31E-01 | 2.68E+00 | 2.74E+00 | 9.55E-01 | no | Rnf150 | intron_variant | MODIFIER |
| 8 | 82929477 | AX-170132472 | 0 | T | G | 0.202 | 7.36E-03 | 1.31E-01 | 2.68E+00 | 2.74E+00 | 9.55E-01 | no | Rnf150 | intron_variant | MODIFIER |
| 8 | 82936158 | AX-170031141 | 0 | C | T | 0.202 | 7.36E-03 | 1.31E-01 | 2.68E+00 | 2.74E+00 | 9.55E-01 | no | Rnf150 | intron_variant | MODIFIER |
| 8 | 82939602 | AX-97840930 | 0 | G | T | 0.202 | 7.36E-03 | 1.31E-01 | 2.68E+00 | 2.74E+00 | 9.55E-01 | no | Rnf150 | intron_variant | MODIFIER |
| 8 | 82960916 | AX-170416135 | 0 | A | G | 0.202 | 7.36E-03 | 1.31E-01 | 2.68E+00 | 2.74E+00 | 9.55E-01 | no | Rnf150 | intron_variant | MODIFIER |
| 8 | 82985924 | AX-170429319 | 0 | T | G | 0.202 | 7.36E-03 | 1.31E-01 | 2.68E+00 | 2.74E+00 | 9.55E-01 | no | Rnf150 | intron_variant | MODIFIER |
| 8 | 83000494 | AX-170239288 | 0 | A | T | 0.202 | 7.36E-03 | 1.31E-01 | 2.68E+00 | 2.74E+00 | 9.55E-01 | no | Rnf150 | intron_variant | MODIFIER |
| 8 | 83005410 | AX-170296754 | 0 | T | C | 0.202 | 7.36E-03 | 1.31E-01 | 2.68E+00 | 2.74E+00 | 9.55E-01 | no | Rnf150 | intron_variant | MODIFIER |
| 8 | 83006485 | AX-170241216 | 0 | A | G | 0.202 | 7.36E-03 | 1.31E-01 | 2.68E+00 | 2.74E+00 | 9.55E-01 | no | Rnf150 | intron_variant | MODIFIER |
| 8 | 83045228 | AX-170343182 | 0 | T | C | 0.202 | 7.36E-03 | 1.31E-01 | 2.68E+00 | 2.74E+00 | 9.55E-01 | no | Rnf150 | intron_variant | MODIFIER |
| 8 | 83046805 | AX-170532099 | 0 | T | C | 0.202 | 7.36E-03 | 1.31E-01 | 2.68E+00 | 2.74E+00 | 9.55E-01 | no | Rnf150 | intron_variant | MODIFIER |
| 8 | 83054967 | AX-170300556 | 0 | C | A | 0.202 | 7.36E-03 | 1.31E-01 | 2.68E+00 | 2.74E+00 | 9.55E-01 | no | Rnf150 | intron_variant | MODIFIER |
| 8 | 83101177 | AX-170303806 | 0 | C | A | 0.202 | 7.36E-03 | 1.31E-01 | 2.68E+00 | 2.74E+00 | 9.55E-01 | no | | intergenic_variant | MODIFIER |
| 8 | 83108615 | AX-170277405 | 0 | G | A | 0.202 | 7.36E-03 | 1.31E-01 | 2.68E+00 | 2.74E+00 | 9.55E-01 | no | | intergenic_variant | MODIFIER |
| 8 | 83110291 | AX-170481101 | 0 | G | T | 0.202 | 7.36E-03 | 1.31E-01 | 2.68E+00 | 2.74E+00 | 9.55E-01 | no | | regulatory_region_variant | MODIFIER |
| 8 | 83110725 | AX-170186778 | 0 | C | T | 0.202 | 7.36E-03 | 1.31E-01 | 2.68E+00 | 2.74E+00 | 9.55E-01 | no | | regulatory_region_variant | MODIFIER |
| 8 | 83111364 | AX-170113295 | 0 | G | A | 0.202 | 7.36E-03 | 1.31E-01 | 2.68E+00 | 2.74E+00 | 9.55E-01 | no | | intergenic_variant | MODIFIER |
| 8 | 83111462 | AX-170430517 | 0 | T | C | 0.202 | 7.36E-03 | 1.31E-01 | 2.68E+00 | 2.74E+00 | 9.55E-01 | no | | intergenic_variant | MODIFIER |
| 8 | 83115107 | AX-170080597 | 0 | A | G | 0.202 | 7.36E-03 | 1.31E-01 | 2.68E+00 | 2.74E+00 | 9.55E-01 | no | | intergenic_variant | MODIFIER |
| 8 | 83149829 | AX-97784507 | 0 | T | C | 0.202 | 7.36E-03 | 1.31E-01 | 2.68E+00 | 2.74E+00 | 9.55E-01 | no | | intergenic_variant | MODIFIER |
| 8 | 83149869 | AX-170080600 | 0 | A | G | 0.202 | 7.36E-03 | 1.31E-01 | 2.68E+00 | 2.74E+00 | 9.55E-01 | no | | intergenic_variant | MODIFIER |
| 8 | 83150755 | AX-170080599 | 0 | T | C | 0.202 | 7.36E-03 | 1.31E-01 | 2.68E+00 | 2.74E+00 | 9.55E-01 | no | | intergenic_variant | MODIFIER |
| 8 | 83150972 | AX-170406739 | 0 | A | G | 0.202 | 7.36E-03 | 1.31E-01 | 2.68E+00 | 2.74E+00 | 9.55E-01 | no | | intergenic_variant | MODIFIER |
| 8 | 83158422 | AX-170111517 | 0 | C | T | 0.202 | 7.36E-03 | 1.31E-01 | 2.68E+00 | 2.74E+00 | 9.55E-01 | no | | intergenic_variant | MODIFIER |
| 8 | 83161196 | AX-170080601 | 0 | T | C | 0.202 | 7.36E-03 | 1.31E-01 | 2.68E+00 | 2.74E+00 | 9.55E-01 | no | Tbc1d9 | upstream_gene_variant | MODIFIER |
| 8 | 83196551 | AX-170080602 | 0 | G | C | 0.202 | 7.36E-03 | 1.31E-01 | 2.68E+00 | 2.74E+00 | 9.55E-01 | no | Tbc1d9 | intron_variant | MODIFIER |
| 8 | 83196658 | AX-170123326 | 0 | G | A | 0.202 | 7.36E-03 | 1.31E-01 | 2.68E+00 | 2.74E+00 | 9.55E-01 | no | Tbc1d9 | intron_variant | MODIFIER |
| 8 | 83204586 | AX-170438958 | 0 | C | T | 0.202 | 7.36E-03 | 1.31E-01 | 2.68E+00 | 2.74E+00 | 9.55E-01 | no | Tbc1d9 | intron_variant | MODIFIER |
| 8 | 83204722 | AX-170213348 | 0 | C | G | 0.202 | 7.36E-03 | 1.31E-01 | 2.68E+00 | 2.74E+00 | 9.55E-01 | no | Tbc1d9 | intron_variant | MODIFIER |
| 8 | 83223522 | AX-170495007 | 0 | G | C | 0.202 | 7.36E-03 | 1.31E-01 | 2.68E+00 | 2.74E+00 | 9.55E-01 | no | Tbc1d9 | intron_variant | MODIFIER |
| 8 | 83227754 | AX-170255652 | 0 | C | G | 0.202 | 7.36E-03 | 1.31E-01 | 2.68E+00 | 2.74E+00 | 9.55E-01 | no | Tbc1d9 | intron_variant | MODIFIER |

| | | | | | | | | | | | | | | | |
|---|---|---|---|---|---|---|---|---|---|---|---|---|---|---|---|
| 8 | 83250007 | AX-170126958 | 0 | C | T | 0.202 | 7.36E-03 | 1.31E-01 | 2.68E+00 | 2.74E+00 | 9.55E-01 | no | Tbc1d9 | intron_variant | MODIFIER |
| 8 | 83250282 | AX-170150066 | 0 | A | G | 0.202 | 7.36E-03 | 1.31E-01 | 2.68E+00 | 2.74E+00 | 9.55E-01 | no | Tbc1d9 | intron_variant | MODIFIER |
| 8 | 83254347 | AX-170347941 | 0 | C | G | 0.202 | 7.36E-03 | 1.31E-01 | 2.68E+00 | 2.74E+00 | 9.55E-01 | no | Tbc1d9 | intron_variant | MODIFIER |
| 8 | 83280916 | AX-170351660 | 0 | A | C | 0.202 | 7.36E-03 | 1.31E-01 | 2.68E+00 | 2.74E+00 | 9.55E-01 | no | | intergenic_variant | MODIFIER |
| 8 | 83289026 | AX-97857344 | 0 | C | T | 0.202 | 7.36E-03 | 1.31E-01 | 2.68E+00 | 2.74E+00 | 9.55E-01 | no | Ucp1 | upstream_gene_variant | MODIFIER |
| 8 | 83301308 | AX-170442828 | 0 | A | G | 0.202 | 7.36E-03 | 1.31E-01 | 2.68E+00 | 2.74E+00 | 9.55E-01 | no | Ucp1 | downstream_gene_variant | MODIFIER |
| 8 | 83319822 | AX-97795759 | 0 | G | A | 0.202 | 7.36E-03 | 1.31E-01 | 2.68E+00 | 2.74E+00 | 9.55E-01 | no | Elmod2 | intron_variant | MODIFIER |
| 8 | 83328645 | AX-170140976 | 0 | C | G | 0.202 | 7.36E-03 | 1.31E-01 | 2.68E+00 | 2.74E+00 | 9.55E-01 | no | Elmod2 | intron_variant | MODIFIER |
| 8 | 83339428 | AX-170147959 | 0 | T | C | 0.202 | 7.36E-03 | 1.31E-01 | 2.68E+00 | 2.74E+00 | 9.55E-01 | no | | intergenic_variant | MODIFIER |
| 8 | 83349618 | AX-170344782 | 0 | T | C | 0.202 | 7.36E-03 | 1.31E-01 | 2.68E+00 | 2.74E+00 | 9.55E-01 | no | Mgat4d | intron_variant | MODIFIER |
| 8 | 83349740 | AX-170383108 | 0 | G | A | 0.202 | 7.36E-03 | 1.31E-01 | 2.68E+00 | 2.74E+00 | 9.55E-01 | no | Mgat4d | intron_variant | MODIFIER |
| 8 | 83365942 | AX-170311997 | 0 | A | G | 0.202 | 7.36E-03 | 1.31E-01 | 2.68E+00 | 2.74E+00 | 9.55E-01 | no | Mgat4d | intron_variant | MODIFIER |
| 8 | 83366151 | AX-170031144 | 0 | A | G | 0.202 | 7.36E-03 | 1.31E-01 | 2.68E+00 | 2.74E+00 | 9.55E-01 | no | Mgat4d | intron_variant | MODIFIER |
| 8 | 83397928 | AX-170080607 | 0 | C | T | 0.202 | 7.36E-03 | 1.31E-01 | 2.68E+00 | 2.74E+00 | 9.55E-01 | no | Clgn | intron_variant | MODIFIER |
| 8 | 83398285 | AX-170080606 | 0 | T | C | 0.202 | 7.36E-03 | 1.31E-01 | 2.68E+00 | 2.74E+00 | 9.55E-01 | no | Clgn | intron_variant | MODIFIER |
| 8 | 83398715 | AX-170031142 | 0 | C | T | 0.202 | 7.36E-03 | 1.31E-01 | 2.68E+00 | 2.74E+00 | 9.55E-01 | no | Clgn | intron_variant | MODIFIER |
| 8 | 83432039 | AX-170405597 | 0 | G | C | 0.202 | 7.36E-03 | 1.31E-01 | 2.68E+00 | 2.74E+00 | 9.55E-01 | no | Clgn | downstream_gene_variant | MODIFIER |
| 8 | 83432136 | AX-97796851 | 0 | T | C | 0.202 | 7.36E-03 | 1.31E-01 | 2.68E+00 | 2.74E+00 | 9.55E-01 | no | Clgn | downstream_gene_variant | MODIFIER |
| 8 | 83437854 | AX-170234176 | 0 | C | A | 0.202 | 7.36E-03 | 1.31E-01 | 2.68E+00 | 2.74E+00 | 9.55E-01 | no | Scoc | intron_variant | MODIFIER |
| 8 | 83438559 | AX-170143722 | 0 | A | G | 0.202 | 7.36E-03 | 1.31E-01 | 2.68E+00 | 2.74E+00 | 9.55E-01 | no | Scoc | intron_variant | MODIFIER |
| 8 | 83449714 | AX-170476902 | 0 | T | C | 0.202 | 7.36E-03 | 1.31E-01 | 2.68E+00 | 2.74E+00 | 9.55E-01 | no | Scoc | intron_variant | MODIFIER |
| 8 | 83490220 | AX-170109922 | 0 | C | G | 0.202 | 7.36E-03 | 1.31E-01 | 2.68E+00 | 2.74E+00 | 9.55E-01 | no | Gm33242 | non_coding_transcript_exon_variant | MODIFIER |
| 8 | 83499247 | AX-170090164 | 0 | G | A | 0.202 | 7.36E-03 | 1.31E-01 | 2.68E+00 | 2.74E+00 | 9.55E-01 | no | | intergenic_variant | MODIFIER |
| 8 | 83499340 | AX-170251278 | 0 | C | G | 0.202 | 7.36E-03 | 1.31E-01 | 2.68E+00 | 2.74E+00 | 9.55E-01 | no | | intergenic_variant | MODIFIER |
| 8 | 83535328 | AX-170166053 | 0 | G | T | 0.202 | 7.36E-03 | 1.31E-01 | 2.68E+00 | 2.74E+00 | 9.55E-01 | no | Olfr370 | upstream_gene_variant | MODIFIER |
| 8 | 83542087 | AX-97793997 | 0 | C | T | 0.202 | 7.36E-03 | 1.31E-01 | 2.68E+00 | 2.74E+00 | 9.55E-01 | no | Olfr370 | synonymous_variant | LOW |
| 8 | 83544297 | AX-170116301 | 0 | C | A | 0.202 | 7.36E-03 | 1.31E-01 | 2.68E+00 | 2.74E+00 | 9.55E-01 | no | Olfr370 | downstream_gene_variant | MODIFIER |
| 8 | 83574368 | AX-170114457 | 0 | T | C | 0.202 | 7.36E-03 | 1.31E-01 | 2.68E+00 | 2.74E+00 | 9.55E-01 | no | Tecr | intron_variant | MODIFIER |
| 8 | 83578418 | AX-170346397 | 0 | G | A | 0.202 | 7.36E-03 | 1.31E-01 | 2.68E+00 | 2.74E+00 | 9.55E-01 | no | Tecr | intron_variant | MODIFIER |
| 8 | 83582589 | AX-170209448 | 0 | G | T | 0.202 | 7.36E-03 | 1.31E-01 | 2.68E+00 | 2.74E+00 | 9.55E-01 | no | Tecr | intron_variant | MODIFIER |
| 8 | 83584044 | AX-170080609 | 0 | G | A | 0.202 | 7.36E-03 | 1.31E-01 | 2.68E+00 | 2.74E+00 | 9.55E-01 | no | Tecr | intron_variant | MODIFIER |
| 8 | 83588086 | AX-170459689 | 0 | G | A | 0.202 | 7.36E-03 | 1.31E-01 | 2.68E+00 | 2.74E+00 | 9.55E-01 | no | Tecr | intron_variant | MODIFIER |
| 8 | 83588121 | AX-170346551 | 0 | A | G | 0.202 | 7.36E-03 | 1.31E-01 | 2.68E+00 | 2.74E+00 | 9.55E-01 | no | Tecr | intron_variant | MODIFIER |
| 8 | 83620091 | AX-170080610 | 0 | G | A | 0.202 | 7.36E-03 | 1.31E-01 | 2.68E+00 | 2.74E+00 | 9.55E-01 | no | | intergenic_variant | MODIFIER |
| 8 | 83624079 | AX-170298541 | 0 | T | A | 0.202 | 7.36E-03 | 1.31E-01 | 2.68E+00 | 2.74E+00 | 9.55E-01 | no | | regulatory_region_variant | MODIFIER |
| 8 | 83655261 | AX-170274391 | 0 | A | G | 0.202 | 7.36E-03 | 1.31E-01 | 2.68E+00 | 2.74E+00 | 9.55E-01 | no | Gipc1 | intron_variant | MODIFIER |
| 8 | 83669953 | AX-170266418 | 0 | T | C | 0.202 | 7.36E-03 | 1.31E-01 | 2.68E+00 | 2.74E+00 | 9.55E-01 | no | Pkn1 | 3_prime_UTR_variant | MODIFIER |
| 8 | 83673180 | AX-170368500 | 0 | T | C | 0.202 | 7.36E-03 | 1.31E-01 | 2.68E+00 | 2.74E+00 | 9.55E-01 | no | Pkn1 | intron_variant | MODIFIER |
| 8 | 83674087 | AX-170434269 | 0 | G | A | 0.202 | 7.36E-03 | 1.31E-01 | 2.68E+00 | 2.74E+00 | 9.55E-01 | no | Pkn1 | intron_variant | MODIFIER |
| 8 | 83677563 | AX-170301309 | 0 | T | C | 0.202 | 7.36E-03 | 1.31E-01 | 2.68E+00 | 2.74E+00 | 9.55E-01 | no | Pkn1 | synonymous_variant | LOW |
| 8 | 83691679 | AX-170138494 | 0 | A | G | 0.202 | 7.36E-03 | 1.31E-01 | 2.68E+00 | 2.74E+00 | 9.55E-01 | no | Pkn1 | intron_variant | MODIFIER |
| 8 | 83720953 | AX-97809293 | 0 | A | G | 0.202 | 7.36E-03 | 1.31E-01 | 2.68E+00 | 2.74E+00 | 9.55E-01 | no | Adgre5 | downstream_gene_variant | MODIFIER |
| 8 | 83722733 | AX-170357334 | 0 | G | A | 0.202 | 7.36E-03 | 1.31E-01 | 2.68E+00 | 2.74E+00 | 9.55E-01 | no | Adgre5 | downstream_gene_variant | MODIFIER |
| 8 | 83725032 | AX-170150665 | 0 | C | G | 0.202 | 7.36E-03 | 1.31E-01 | 2.68E+00 | 2.74E+00 | 9.55E-01 | no | Adgre5 | intron_variant | MODIFIER |
| 8 | 83729093 | AX-170482029 | 0 | A | T | 0.202 | 7.36E-03 | 1.31E-01 | 2.68E+00 | 2.74E+00 | 9.55E-01 | no | Adgre5 | synonymous_variant | LOW |
| 8 | 83731564 | AX-170533662 | 0 | A | C | 0.202 | 7.36E-03 | 1.31E-01 | 2.68E+00 | 2.74E+00 | 9.55E-01 | no | Adgre5 | intron_variant | MODIFIER |
| 8 | 83739033 | AX-97837115 | 0 | C | T | 0.202 | 7.36E-03 | 1.31E-01 | 2.68E+00 | 2.74E+00 | 9.55E-01 | no | Adgre5 | intron_variant | MODIFIER |
| 8 | 83768924 | AX-170419720 | 0 | G | C | 0.202 | 7.36E-03 | 1.31E-01 | 2.68E+00 | 2.74E+00 | 9.55E-01 | no | | regulatory_region_variant | MODIFIER |
| 8 | 83785356 | AX-170080615 | 0 | G | C | 0.202 | 7.36E-03 | 1.31E-01 | 2.68E+00 | 2.74E+00 | 9.55E-01 | no | | regulatory_region_variant | MODIFIER |
| 8 | 83787992 | AX-170080619 | 0 | T | C | 0.202 | 7.36E-03 | 1.31E-01 | 2.68E+00 | 2.74E+00 | 9.55E-01 | no | | intergenic_variant | MODIFIER |
| 8 | 83795290 | AX-170080614 | 0 | A | G | 0.202 | 7.36E-03 | 1.31E-01 | 2.68E+00 | 2.74E+00 | 9.55E-01 | no | | intergenic_variant | MODIFIER |
| 8 | 83799820 | AX-170326895 | 0 | G | A | 0.202 | 7.36E-03 | 1.31E-01 | 2.68E+00 | 2.74E+00 | 9.55E-01 | no | | regulatory_region_variant | MODIFIER |
| 8 | 83808170 | AX-170374085 | 0 | T | C | 0.202 | 7.36E-03 | 1.31E-01 | 2.68E+00 | 2.74E+00 | 9.55E-01 | no | | intergenic_variant | MODIFIER |
| 8 | 83840051 | AX-170490415 | 0 | A | T | 0.202 | 7.36E-03 | 1.31E-01 | 2.68E+00 | 2.74E+00 | 9.55E-01 | no | | intergenic_variant | MODIFIER |

| | | | | | | | | | | | | | | | |
|---|---|---|---|---|---|---|---|---|---|---|---|---|---|---|---|
| 8 | 83871742 | AX-97793962 | 0 | A | T | 0.202 | 7.36E-03 | 1.31E-01 | 2.68E+00 | 2.74E+00 | 9.55E-01 | no | | intergenic_variant | MODIFIER |
| 8 | 83889821 | AX-170139822 | 0 | C | T | 0.202 | 7.36E-03 | 1.31E-01 | 2.68E+00 | 2.74E+00 | 9.55E-01 | no | Gm26721 | intron_variant,non_coding_transcript_variant | MODIFIER |
| 8 | 83892553 | AX-97813334 | 0 | C | A | 0.202 | 7.36E-03 | 1.31E-01 | 2.68E+00 | 2.74E+00 | 9.55E-01 | no | Gm26721 | intron_variant,non_coding_transcript_variant | MODIFIER |
| 8 | 83907733 | AX-170451677 | 0 | G | A | 0.202 | 7.36E-03 | 1.31E-01 | 2.68E+00 | 2.74E+00 | 9.55E-01 | no | Adgrl1 | intron_variant | MODIFIER |
| 8 | 83911103 | AX-170339507 | 0 | T | C | 0.202 | 7.36E-03 | 1.31E-01 | 2.68E+00 | 2.74E+00 | 9.55E-01 | no | Adgrl1 | intron_variant | MODIFIER |
| 8 | 83915913 | AX-170276981 | 0 | A | G | 0.202 | 7.36E-03 | 1.31E-01 | 2.68E+00 | 2.74E+00 | 9.55E-01 | no | Adgrl1 | intron_variant | MODIFIER |
| 8 | 83926528 | AX-170031149 | 0 | T | G | 0.202 | 7.36E-03 | 1.31E-01 | 2.68E+00 | 2.74E+00 | 9.55E-01 | no | Adgrl1 | intron_variant | MODIFIER |
| 8 | 83931363 | AX-170031147 | 0 | C | T | 0.202 | 7.36E-03 | 1.31E-01 | 2.68E+00 | 2.74E+00 | 9.55E-01 | no | Adgrl1 | intron_variant | MODIFIER |
| 8 | 83986728 | AX-170145000 | 0 | G | C | 0.202 | 7.36E-03 | 1.31E-01 | 2.68E+00 | 2.74E+00 | 9.55E-01 | no | Prkaca | intron_variant | MODIFIER |
| 8 | 83991814 | AX-170443475 | 0 | C | T | 0.202 | 7.36E-03 | 1.31E-01 | 2.68E+00 | 2.74E+00 | 9.55E-01 | no | Prkaca | intron_variant | MODIFIER |
| 8 | 83991913 | AX-170242828 | 0 | G | A | 0.202 | 7.36E-03 | 1.31E-01 | 2.68E+00 | 2.74E+00 | 9.55E-01 | no | Prkaca | intron_variant | MODIFIER |
| 8 | 84008536 | AX-170080621 | 0 | C | T | 0.202 | 7.36E-03 | 1.31E-01 | 2.68E+00 | 2.74E+00 | 9.55E-01 | no | 2210011C24Rik | downstream_gene_variant | MODIFIER |
| 8 | 84019758 | AX-97834431 | 0 | G | A | 0.202 | 7.36E-03 | 1.31E-01 | 2.68E+00 | 2.74E+00 | 9.55E-01 | no | Palm3 | upstream_gene_variant | MODIFIER |
| 8 | 84036670 | AX-170173999 | 0 | C | T | 0.202 | 7.36E-03 | 1.31E-01 | 2.68E+00 | 2.74E+00 | 9.55E-01 | no | Il27ra | intron_variant | MODIFIER |
| 8 | 84069352 | AX-170143711 | 0 | C | T | 0.202 | 7.36E-03 | 1.31E-01 | 2.68E+00 | 2.74E+00 | 9.55E-01 | no | Rfx1 | intron_variant | MODIFIER |
| 8 | 84070968 | AX-170439594 | 0 | G | A | 0.202 | 7.36E-03 | 1.31E-01 | 2.68E+00 | 2.74E+00 | 9.55E-01 | no | Rfx1 | intron_variant | MODIFIER |
| 8 | 84072089 | AX-97793746 | 0 | C | T | 0.202 | 7.36E-03 | 1.31E-01 | 2.68E+00 | 2.74E+00 | 9.55E-01 | no | Rfx1 | intron_variant | MODIFIER |
| 8 | 84073148 | AX-170135284 | 0 | A | G | 0.202 | 7.36E-03 | 1.31E-01 | 2.68E+00 | 2.74E+00 | 9.55E-01 | no | Rfx1 | intron_variant | MODIFIER |
| 8 | 84080968 | AX-170380719 | 0 | T | C | 0.202 | 7.36E-03 | 1.31E-01 | 2.68E+00 | 2.74E+00 | 9.55E-01 | no | Rfx1 | intron_variant | MODIFIER |
| 8 | 84089452 | AX-170262658 | 0 | T | C | 0.202 | 7.36E-03 | 1.31E-01 | 2.68E+00 | 2.74E+00 | 9.55E-01 | no | Rfx1 | intron_variant | MODIFIER |
| 8 | 84092144 | AX-170501262 | 0 | G | T | 0.202 | 7.36E-03 | 1.31E-01 | 2.68E+00 | 2.74E+00 | 9.55E-01 | no | Rfx1 | intron_variant | MODIFIER |
| 8 | 84092317 | AX-170376910 | 0 | G | A | 0.202 | 7.36E-03 | 1.31E-01 | 2.68E+00 | 2.74E+00 | 9.55E-01 | no | Rfx1 | intron_variant | MODIFIER |
| 8 | 84092549 | AX-170445082 | 0 | T | C | 0.202 | 7.36E-03 | 1.31E-01 | 2.68E+00 | 2.74E+00 | 9.55E-01 | no | Rfx1 | synonymous_variant | LOW |
| 8 | 84094855 | AX-170346280 | 0 | G | C | 0.202 | 7.36E-03 | 1.31E-01 | 2.68E+00 | 2.74E+00 | 9.55E-01 | no | Rfx1 | synonymous_variant | LOW |
| 8 | 84096477 | AX-170494098 | 0 | T | C | 0.202 | 7.36E-03 | 1.31E-01 | 2.68E+00 | 2.74E+00 | 9.55E-01 | no | Rfx1 | 3_prime_UTR_variant | MODIFIER |
| 8 | 84099499 | AX-170209719 | 0 | C | T | 0.202 | 7.36E-03 | 1.31E-01 | 2.68E+00 | 2.74E+00 | 9.55E-01 | no | Rfx1 | downstream_gene_variant | MODIFIER |
| 8 | 84103291 | AX-97810087 | 0 | C | A | 0.202 | 7.36E-03 | 1.31E-01 | 2.68E+00 | 2.74E+00 | 9.55E-01 | no | Dcaf15 | intron_variant | MODIFIER |
| 8 | 84135704 | AX-170368583 | 0 | C | A | 0.202 | 7.36E-03 | 1.31E-01 | 2.68E+00 | 2.74E+00 | 9.55E-01 | no | Cc2d1a | intron_variant | MODIFIER |
| 8 | 84143469 | AX-170209236 | 0 | T | C | 0.202 | 7.36E-03 | 1.31E-01 | 2.68E+00 | 2.74E+00 | 9.55E-01 | no | Cc2d1a | missense_variant | MODERATE |
| 8 | 84143936 | AX-170319282 | 0 | G | A | 0.202 | 7.36E-03 | 1.31E-01 | 2.68E+00 | 2.74E+00 | 9.55E-01 | no | Cc2d1a | intron_variant | MODIFIER |
| 8 | 84144753 | AX-170413599 | 0 | A | G | 0.202 | 7.36E-03 | 1.31E-01 | 2.68E+00 | 2.74E+00 | 9.55E-01 | no | Cc2d1a | intron_variant | MODIFIER |
| 8 | 84171072 | AX-97809794 | 0 | A | G | 0.202 | 7.36E-03 | 1.31E-01 | 2.68E+00 | 2.74E+00 | 9.55E-01 | no | Nanos3 | downstream_gene_variant | MODIFIER |
| 8 | 84171908 | AX-170143096 | 0 | G | C | 0.202 | 7.36E-03 | 1.31E-01 | 2.68E+00 | 2.74E+00 | 9.55E-01 | no | Nanos3 | downstream_gene_variant | MODIFIER |
| 8 | 84173382 | AX-170276104 | 0 | G | A | 0.202 | 7.36E-03 | 1.31E-01 | 2.68E+00 | 2.74E+00 | 9.55E-01 | no | Nanos3 | downstream_gene_variant | MODIFIER |
| 8 | 84174284 | AX-170182203 | 0 | T | C | 0.202 | 7.36E-03 | 1.31E-01 | 2.68E+00 | 2.74E+00 | 9.55E-01 | no | Nanos3 | intron_variant | MODIFIER |
| 8 | 84174324 | AX-170457693 | 0 | G | A | 0.202 | 7.36E-03 | 1.31E-01 | 2.68E+00 | 2.74E+00 | 9.55E-01 | no | Nanos3 | intron_variant | MODIFIER |
| 8 | 84174374 | AX-170281364 | 0 | A | G | 0.202 | 7.36E-03 | 1.31E-01 | 2.68E+00 | 2.74E+00 | 9.55E-01 | no | Nanos3 | intron_variant | MODIFIER |
| 8 | 84175110 | AX-170398266 | 0 | C | T | 0.202 | 7.36E-03 | 1.31E-01 | 2.68E+00 | 2.74E+00 | 9.55E-01 | no | Nanos3 | intron_variant | MODIFIER |
| 8 | 84331433 | AX-170307546 | 0 | T | C | 0.202 | 7.36E-03 | 1.31E-01 | 2.68E+00 | 2.74E+00 | 9.55E-01 | no | | intergenic_variant | MODIFIER |
| 8 | 84332837 | AX-170265909 | 0 | A | G | 0.202 | 7.36E-03 | 1.31E-01 | 2.68E+00 | 2.74E+00 | 9.55E-01 | no | | intergenic_variant | MODIFIER |
| 8 | 84335884 | AX-170345770 | 0 | G | C | 0.202 | 7.36E-03 | 1.31E-01 | 2.68E+00 | 2.74E+00 | 9.55E-01 | no | | intergenic_variant | MODIFIER |
| 8 | 84365529 | AX-170031155 | 0 | A | G | 0.202 | 7.36E-03 | 1.31E-01 | 2.68E+00 | 2.74E+00 | 9.55E-01 | no | | intergenic_variant | MODIFIER |
| 8 | 84369484 | AX-170031158 | 0 | G | A | 0.202 | 7.36E-03 | 1.31E-01 | 2.68E+00 | 2.74E+00 | 9.55E-01 | no | | intergenic_variant | MODIFIER |
| 8 | 84467684 | AX-170442782 | 0 | G | T | 0.202 | 7.36E-03 | 1.31E-01 | 2.68E+00 | 2.74E+00 | 9.55E-01 | no | Cacna1a | intron_variant | MODIFIER |
| 8 | 84483222 | AX-170212764 | 0 | A | G | 0.202 | 7.36E-03 | 1.31E-01 | 2.68E+00 | 2.74E+00 | 9.55E-01 | no | Cacna1a | intron_variant | MODIFIER |
| 8 | 84486707 | AX-170031156 | 0 | A | G | 0.202 | 7.36E-03 | 1.31E-01 | 2.68E+00 | 2.74E+00 | 9.55E-01 | no | Cacna1a | intron_variant | MODIFIER |
| 8 | 84491625 | AX-97784574 | 0 | G | A | 0.202 | 7.36E-03 | 1.31E-01 | 2.68E+00 | 2.74E+00 | 9.55E-01 | no | Cacna1a | intron_variant | MODIFIER |
| 8 | 84540748 | AX-170191684 | 0 | C | A | 0.202 | 7.36E-03 | 1.31E-01 | 2.68E+00 | 2.74E+00 | 9.55E-01 | no | Cacna1a | intron_variant | MODIFIER |
| 8 | 84542010 | AX-170395668 | 0 | G | A | 0.202 | 7.36E-03 | 1.31E-01 | 2.68E+00 | 2.74E+00 | 9.55E-01 | no | Cacna1a | intron_variant | MODIFIER |
| 8 | 84545405 | AX-97785036 | 0 | G | A | 0.202 | 7.36E-03 | 1.31E-01 | 2.68E+00 | 2.74E+00 | 9.55E-01 | no | Cacna1a | intron_variant | MODIFIER |
| 8 | 84867550 | AX-97807936 | 0 | T | C | 0.202 | 7.36E-03 | 1.31E-01 | 2.68E+00 | 2.74E+00 | 9.55E-01 | no | Farsa | missense_variant,splice_region_variant | MODERATE |
| 8 | 84893047 | AX-97807749 | 0 | C | T | 0.202 | 7.36E-03 | 1.31E-01 | 2.68E+00 | 2.74E+00 | 9.55E-01 | no | Gcdh | missense_variant | MODERATE |
| 8 | 84910437 | AX-97815850 | 0 | C | T | 0.202 | 7.36E-03 | 1.31E-01 | 2.68E+00 | 2.74E+00 | 9.55E-01 | no | Dnase2a | intron_variant | MODIFIER |
| 8 | 84946560 | AX-97786231 | 0 | C | T | 0.202 | 7.36E-03 | 1.31E-01 | 2.68E+00 | 2.74E+00 | 9.55E-01 | no | Rtbdn | upstream_gene_variant | MODIFIER |

| | | | | | | | | | | | | | | | |
|---|---|---|---|---|---|---|---|---|---|---|---|---|---|---|---|
| 8 | 84987230 | AX-170195838 | 0 | C | T | 0.202 | 7.36E-03 | 1.31E-01 | 2.68E+00 | 2.74E+00 | 9.55E-01 | no | Hook2 | upstream_gene_variant | MODIFIER |
| 8 | 84990242 | AX-97790237 | 0 | C | T | 0.202 | 7.36E-03 | 1.31E-01 | 2.68E+00 | 2.74E+00 | 9.55E-01 | no | Hook2 | upstream_gene_variant | MODIFIER |
| 8 | 85022849 | AX-170031168 | 0 | C | T | 0.202 | 7.36E-03 | 1.31E-01 | 2.68E+00 | 2.74E+00 | 9.55E-01 | no | Trir | upstream_gene_variant | MODIFIER |
| 8 | 85033201 | AX-170031160 | 0 | A | T | 0.202 | 7.36E-03 | 1.31E-01 | 2.68E+00 | 2.74E+00 | 9.55E-01 | no | Trir | downstream_gene_variant | MODIFIER |
| 8 | 85070850 | AX-170204037 | 0 | C | A | 0.202 | 7.36E-03 | 1.31E-01 | 2.68E+00 | 2.74E+00 | 9.55E-01 | no | Dhps | upstream_gene_variant | MODIFIER |
| 8 | 85092100 | AX-170090167 | 0 | G | A | 0.202 | 7.36E-03 | 1.31E-01 | 2.68E+00 | 2.74E+00 | 9.55E-01 | no | Man2b1 | synonymous_variant | LOW |
| 8 | 85297097 | AX-170460247 | 0 | G | A | 0.202 | 7.36E-03 | 1.31E-01 | 2.68E+00 | 2.74E+00 | 9.55E-01 | no | Vps35 | intron_variant | MODIFIER |
| 8 | 85305403 | AX-170333201 | 0 | G | A | 0.202 | 7.36E-03 | 1.31E-01 | 2.68E+00 | 2.74E+00 | 9.55E-01 | no | Orc6 | intron_variant | MODIFIER |
| 8 | 85397517 | AX-97839025 | 0 | C | T | 0.202 | 7.36E-03 | 1.31E-01 | 2.68E+00 | 2.74E+00 | 9.55E-01 | no | | regulatory_region_variant | MODIFIER |
| 8 | 85519136 | AX-97822942 | 0 | T | G | 0.202 | 7.36E-03 | 1.31E-01 | 2.68E+00 | 2.74E+00 | 9.55E-01 | no | Gpt2 | intron_variant | MODIFIER |
| 8 | 85541961 | AX-170203092 | 0 | A | G | 0.202 | 7.36E-03 | 1.31E-01 | 2.68E+00 | 2.74E+00 | 9.55E-01 | no | Dnaja2 | intron_variant | MODIFIER |
| 8 | 85546910 | AX-97855192 | 0 | T | C | 0.202 | 7.36E-03 | 1.31E-01 | 2.68E+00 | 2.74E+00 | 9.55E-01 | no | Dnaja2 | intron_variant | MODIFIER |
| 8 | 85547561 | AX-170509144 | 0 | T | C | 0.202 | 7.36E-03 | 1.31E-01 | 2.68E+00 | 2.74E+00 | 9.55E-01 | no | Dnaja2 | intron_variant | MODIFIER |
| 8 | 85550927 | AX-170031179 | 0 | C | T | 0.202 | 7.36E-03 | 1.31E-01 | 2.68E+00 | 2.74E+00 | 9.55E-01 | no | Dnaja2 | intron_variant | MODIFIER |
| 8 | 85579877 | AX-97853877 | 0 | T | C | 0.202 | 7.36E-03 | 1.31E-01 | 2.68E+00 | 2.74E+00 | 9.55E-01 | no | | intergenic_variant | MODIFIER |
| 8 | 85607034 | AX-170031173 | 0 | C | T | 0.202 | 7.36E-03 | 1.31E-01 | 2.68E+00 | 2.74E+00 | 9.55E-01 | no | Gm45424 | downstream_gene_variant | MODIFIER |
| 8 | 85609576 | AX-170420425 | 0 | A | T | 0.202 | 7.36E-03 | 1.31E-01 | 2.68E+00 | 2.74E+00 | 9.55E-01 | no | Gm45425 | intron_variant,non_coding_transcript_variant | MODIFIER |
| 8 | 85680938 | AX-97806824 | 0 | A | G | 0.202 | 7.36E-03 | 1.31E-01 | 2.68E+00 | 2.74E+00 | 9.55E-01 | no | Neto2 | intron_variant | MODIFIER |
| 7 | 111866391 | AX-170029779 | 1 | G | A | 0.455 | 5.52E-03 | 9.65E-02 | 2.68E+00 | 2.74E+00 | 9.55E-01 | no | | intergenic_variant | MODIFIER |
| 7 | 111680092 | AX-170520040 | 0 | G | A | 0.465 | 5.60E-03 | 9.46E-02 | 2.68E+00 | 2.73E+00 | 9.55E-01 | no | Galnt18 | intron_variant | MODIFIER |
| 7 | 111683462 | AX-170476834 | 0 | G | T | 0.465 | 5.60E-03 | 9.46E-02 | 2.68E+00 | 2.73E+00 | 9.55E-01 | no | Galnt18 | intron_variant | MODIFIER |
| 7 | 111707135 | AX-170188491 | 0 | T | C | 0.465 | 5.60E-03 | 9.46E-02 | 2.68E+00 | 2.73E+00 | 9.55E-01 | no | Galnt18 | intron_variant | MODIFIER |
| 7 | 111715210 | AX-170442720 | 0 | G | A | 0.465 | 5.60E-03 | 9.46E-02 | 2.68E+00 | 2.73E+00 | 9.55E-01 | no | Galnt18 | intron_variant | MODIFIER |
| 7 | 111722083 | AX-97836763 | 0 | T | C | 0.465 | 5.60E-03 | 9.46E-02 | 2.68E+00 | 2.73E+00 | 9.55E-01 | no | Galnt18 | intron_variant | MODIFIER |
| 7 | 111723217 | AX-170504281 | 0 | C | T | 0.465 | 5.60E-03 | 9.46E-02 | 2.68E+00 | 2.73E+00 | 9.55E-01 | no | Galnt18 | intron_variant | MODIFIER |
| 7 | 111725177 | AX-97849770 | 0 | A | T | 0.465 | 5.60E-03 | 9.46E-02 | 2.68E+00 | 2.73E+00 | 9.55E-01 | no | Galnt18 | intron_variant | MODIFIER |
| 7 | 111725244 | AX-170340350 | 0 | T | C | 0.465 | 5.60E-03 | 9.46E-02 | 2.68E+00 | 2.73E+00 | 9.55E-01 | no | Galnt18 | intron_variant | MODIFIER |
| 7 | 111727095 | AX-170182384 | 0 | C | A | 0.465 | 5.60E-03 | 9.46E-02 | 2.68E+00 | 2.73E+00 | 9.55E-01 | no | Galnt18 | intron_variant | MODIFIER |
| 7 | 111734188 | AX-170097936 | 0 | T | G | 0.465 | 5.60E-03 | 9.46E-02 | 2.68E+00 | 2.73E+00 | 9.55E-01 | no | Galnt18 | intron_variant | MODIFIER |
| 7 | 111734296 | AX-170078432 | 0 | G | C | 0.465 | 5.60E-03 | 9.46E-02 | 2.68E+00 | 2.73E+00 | 9.55E-01 | no | Galnt18 | intron_variant | MODIFIER |
| 7 | 111734395 | AX-170078431 | 0 | A | G | 0.465 | 5.60E-03 | 9.46E-02 | 2.68E+00 | 2.73E+00 | 9.55E-01 | no | Galnt18 | intron_variant | MODIFIER |
| 7 | 111736368 | AX-170353033 | 0 | T | C | 0.465 | 5.60E-03 | 9.46E-02 | 2.68E+00 | 2.73E+00 | 9.55E-01 | no | Galnt18 | intron_variant | MODIFIER |
| 7 | 111789692 | AX-170029782 | 0 | T | A | 0.465 | 5.60E-03 | 9.46E-02 | 2.68E+00 | 2.73E+00 | 9.55E-01 | no | Gm25787 | downstream_gene_variant | MODIFIER |
| 7 | 111791040 | AX-170029781 | 0 | G | A | 0.465 | 5.60E-03 | 9.46E-02 | 2.68E+00 | 2.73E+00 | 9.55E-01 | no | Gm25787 | downstream_gene_variant | MODIFIER |
| 7 | 111791515 | AX-97786456 | 0 | C | T | 0.465 | 5.60E-03 | 9.46E-02 | 2.68E+00 | 2.73E+00 | 9.55E-01 | no | Gm25787 | downstream_gene_variant | MODIFIER |
| 7 | 111827743 | AX-170035766 | 0 | G | A | 0.465 | 5.60E-03 | 9.46E-02 | 2.68E+00 | 2.73E+00 | 9.55E-01 | no | | intergenic_variant | MODIFIER |
| 7 | 111831692 | AX-170029783 | 0 | G | A | 0.465 | 5.60E-03 | 9.46E-02 | 2.68E+00 | 2.73E+00 | 9.55E-01 | no | | intergenic_variant | MODIFIER |
| 7 | 111878885 | AX-170078435 | 0 | T | C | 0.465 | 5.60E-03 | 9.46E-02 | 2.68E+00 | 2.73E+00 | 9.55E-01 | no | | intergenic_variant | MODIFIER |
| 7 | 111880758 | AX-170424469 | 0 | C | T | 0.465 | 5.60E-03 | 9.46E-02 | 2.68E+00 | 2.73E+00 | 9.55E-01 | no | | intergenic_variant | MODIFIER |
| 7 | 111882752 | AX-170305116 | 0 | A | C | 0.465 | 5.60E-03 | 9.46E-02 | 2.68E+00 | 2.73E+00 | 9.55E-01 | no | | intergenic_variant | MODIFIER |
| 7 | 111902685 | AX-170132233 | 0 | A | G | 0.465 | 5.60E-03 | 9.46E-02 | 2.68E+00 | 2.73E+00 | 9.55E-01 | no | | intergenic_variant | MODIFIER |
| 7 | 111919057 | AX-170506595 | 0 | A | G | 0.465 | 5.60E-03 | 9.46E-02 | 2.68E+00 | 2.73E+00 | 9.55E-01 | no | | intergenic_variant | MODIFIER |
| 7 | 111940894 | AX-170125842 | 0 | T | C | 0.465 | 5.60E-03 | 9.46E-02 | 2.68E+00 | 2.73E+00 | 9.55E-01 | no | | intergenic_variant | MODIFIER |
| 7 | 111945623 | AX-170137448 | 0 | G | A | 0.465 | 5.60E-03 | 9.46E-02 | 2.68E+00 | 2.73E+00 | 9.55E-01 | no | Gm23345 | downstream_gene_variant | MODIFIER |
| 7 | 111946604 | AX-170209319 | 0 | A | G | 0.465 | 5.60E-03 | 9.46E-02 | 2.68E+00 | 2.73E+00 | 9.55E-01 | no | Gm23345 | downstream_gene_variant | MODIFIER |
| 7 | 111974173 | AX-97850269 | 0 | T | C | 0.465 | 5.60E-03 | 9.46E-02 | 2.68E+00 | 2.73E+00 | 9.55E-01 | no | | intergenic_variant | MODIFIER |
| 7 | 111977118 | AX-170491356 | 0 | C | T | 0.465 | 5.60E-03 | 9.46E-02 | 2.68E+00 | 2.73E+00 | 9.55E-01 | no | | regulatory_region_variant | MODIFIER |
| 7 | 111978634 | AX-170386430 | 0 | A | G | 0.465 | 5.60E-03 | 9.46E-02 | 2.68E+00 | 2.73E+00 | 9.55E-01 | no | | intergenic_variant | MODIFIER |
| 7 | 111982229 | AX-170493234 | 0 | T | C | 0.465 | 5.60E-03 | 9.46E-02 | 2.68E+00 | 2.73E+00 | 9.55E-01 | no | | intergenic_variant | MODIFIER |
| 7 | 111982307 | AX-170352469 | 0 | T | C | 0.465 | 5.60E-03 | 9.46E-02 | 2.68E+00 | 2.73E+00 | 9.55E-01 | no | | intergenic_variant | MODIFIER |
| 7 | 111999550 | AX-170213232 | 0 | A | G | 0.465 | 5.60E-03 | 9.46E-02 | 2.68E+00 | 2.73E+00 | 9.55E-01 | no | | intergenic_variant | MODIFIER |
| 7 | 112002315 | AX-170078438 | 0 | G | A | 0.465 | 5.60E-03 | 9.46E-02 | 2.68E+00 | 2.73E+00 | 9.55E-01 | no | | intergenic_variant | MODIFIER |
| 7 | 112002881 | AX-97834780 | 0 | C | T | 0.465 | 5.60E-03 | 9.46E-02 | 2.68E+00 | 2.73E+00 | 9.55E-01 | no | | intergenic_variant | MODIFIER |
| 7 | 112055823 | AX-97792760 | 0 | C | T | 0.465 | 5.60E-03 | 9.46E-02 | 2.68E+00 | 2.73E+00 | 9.55E-01 | no | Usp47 | intron_variant | MODIFIER |

| | | | | | | | | | | | | | | | |
|---|---|---|---|---|---|---|---|---|---|---|---|---|---|---|---|
| 7 | 112188636 | AX-170285435 | 0 | T | C | 0.465 | 5.60E-03 | 9.46E-02 | 2.68E+00 | 2.73E+00 | 9.55E-01 | no | | regulatory_region_variant | MODIFIER |
| 5 | 100190537 | AX-97833042 | 1 | G | A | 0.438 | 6.17E-03 | 1.05E-01 | 2.67E+00 | 2.73E+00 | 9.55E-01 | no | | intergenic_variant | MODIFIER |
| 5 | 100905910 | AX-170072887 | 1 | G | C | 0.438 | 6.17E-03 | 1.05E-01 | 2.67E+00 | 2.73E+00 | 9.55E-01 | no | Gm36793 | downstream_gene_variant | MODIFIER |
| 8 | 21906576 | AX-170477455 | 1 | A | C | 0.17 | -6.18E-03 | 1.11E-01 | 2.65E+00 | 2.74E+00 | 9.55E-01 | no | Defb11 | upstream_gene_variant | MODIFIER |
| 4 | 63671033 | AX-170253771 | 1 | C | T | 0.348 | -5.11E-03 | 1.08E-01 | 2.67E+00 | 2.75E+00 | 9.55E-01 | no | | intergenic_variant | MODIFIER |
| 10 | 17793092 | AX-170002841 | 0 | T | G | 0.219 | 4.03E-03 | 9.03E-02 | 2.63E+00 | 2.75E+00 | 9.56E-01 | no | Txlnb | upstream_gene_variant | MODIFIER |
| 20 | 59391587 | AX-97843149 | 0 | C | A | 0.246 | 3.60E-03 | 6.26E-02 | 2.64E+00 | 2.74E+00 | 9.56E-01 | no | Fgf13 | intron_variant | MODIFIER |
| 18 | 41827479 | AX-170353808 | 1 | G | A | 0.062 | 1.62E-02 | 3.02E-01 | 2.72E+00 | 2.75E+00 | 9.56E-01 | no | AC154172.2 | intron_variant,non_coding_transcript_variant | MODIFIER |
| 8 | 85518168 | AX-170458092 | 1 | A | G | 0.205 | 7.26E-03 | 1.29E-01 | 2.67E+00 | 2.74E+00 | 9.56E-01 | no | Gpt2 | intron_variant | MODIFIER |
| 18 | 57725815 | AX-97857179 | 1 | G | A | 0.393 | 7.49E-03 | 9.44E-02 | 2.63E+00 | 2.71E+00 | 9.56E-01 | no | Ccdc192 | intron_variant | MODIFIER |
| 1 | 9397195 | AX-170430526 | 1 | T | C | 0.5 | 4.73E-03 | 7.04E-02 | 2.65E+00 | 2.72E+00 | 9.56E-01 | no | Gm6187 | non_coding_transcript_exon_variant | MODIFIER |
| 1 | 6075485 | AX-170040286 | 0 | C | A | 0.088 | -8.52E-03 | 1.66E-01 | 2.63E+00 | 2.74E+00 | 9.56E-01 | no | | intergenic_variant | MODIFIER |
| 13 | 54968023 | AX-170166956 | 1 | G | A | 0.527 | 4.68E-03 | 6.49E-02 | 2.61E+00 | 2.72E+00 | 9.56E-01 | no | Unc5a | intron_variant | MODIFIER |
| 8 | 93392348 | AX-170514786 | 1 | G | C | 0.214 | 6.07E-03 | 1.17E-01 | 2.68E+00 | 2.74E+00 | 9.56E-01 | no | Gm45765 | intron_variant,non_coding_transcript_variant | MODIFIER |
| 13 | 40895164 | AX-170487589 | 0 | G | A | 0.175 | 3.14E-03 | 1.05E-01 | 2.65E+00 | 2.76E+00 | 9.56E-01 | no | Gcnt2 | intron_variant | MODIFIER |
| 2 | 119104989 | AX-170123311 | 0 | C | T | 0.14 | 1.24E-02 | 2.20E-01 | 2.70E+00 | 2.73E+00 | 9.56E-01 | no | Knl1 | 3_prime_UTR_variant | MODIFIER |
| 2 | 132639373 | AX-168419979 | 0 | C | T | 0.14 | 1.24E-02 | 2.20E-01 | 2.70E+00 | 2.73E+00 | 9.56E-01 | no | AU019990 | intron_variant,non_coding_transcript_variant | MODIFIER |
| 2 | 148750491 | AX-170485951 | 0 | G | A | 0.14 | 1.24E-02 | 2.20E-01 | 2.70E+00 | 2.73E+00 | 9.56E-01 | no | Cstl1 | intron_variant | MODIFIER |
| 9 | 75168451 | AX-170419423 | 1 | T | A | 0.196 | 7.26E-03 | 1.36E-01 | 2.71E+00 | 2.74E+00 | 9.56E-01 | no | Myo5a | intron_variant | MODIFIER |
| 2 | 157862428 | AX-97795966 | 2 | G | A | 0.136 | 1.26E-02 | 2.15E-01 | 2.68E+00 | 2.73E+00 | 9.56E-01 | no | Ctnnbl1 | intron_variant | MODIFIER |
| 2 | 147392559 | AX-170019317 | 0 | G | A | 0.14 | -1.17E-02 | 2.21E-01 | 2.71E+00 | 2.74E+00 | 9.56E-01 | no | Pax1 | non_coding_transcript_exon_variant | MODIFIER |
| 11 | 5415430 | AX-170427553 | 1 | C | T | 0.062 | 1.08E-02 | 2.14E-01 | 2.65E+00 | 2.74E+00 | 9.56E-01 | no | Znrf3 | intron_variant | MODIFIER |
| 20 | 87601947 | AX-170222031 | 1 | A | G | 0.277 | 3.68E-03 | 6.06E-02 | 2.64E+00 | 2.73E+00 | 9.56E-01 | no | Il1rapl1 | intron_variant | MODIFIER |
| 8 | 26690117 | AX-170090068 | 1 | G | A | 0.17 | -6.36E-03 | 1.12E-01 | 2.65E+00 | 2.74E+00 | 9.56E-01 | no | | regulatory_region_variant | MODIFIER |
| 20 | 56958456 | AX-170154275 | 1 | A | T | 0.25 | 3.59E-03 | 6.25E-02 | 2.64E+00 | 2.74E+00 | 9.56E-01 | no | Adgrg4 | upstream_gene_variant | MODIFIER |
| 20 | 87580283 | AX-170177720 | 0 | T | C | 0.272 | 3.61E-03 | 6.06E-02 | 2.64E+00 | 2.73E+00 | 9.56E-01 | no | Il1rapl1 | intron_variant | MODIFIER |
| 20 | 87585991 | AX-170223035 | 0 | G | C | 0.272 | 3.61E-03 | 6.06E-02 | 2.64E+00 | 2.73E+00 | 9.56E-01 | no | Il1rapl1 | intron_variant | MODIFIER |
| 20 | 87607849 | AX-97783482 | 0 | C | T | 0.272 | 3.61E-03 | 6.06E-02 | 2.64E+00 | 2.73E+00 | 9.56E-01 | no | Il1rapl1 | intron_variant | MODIFIER |
| 20 | 87887850 | AX-97852937 | 0 | A | G | 0.272 | 3.61E-03 | 6.06E-02 | 2.64E+00 | 2.73E+00 | 9.56E-01 | no | Il1rapl1 | intron_variant | MODIFIER |
| 20 | 87896767 | AX-170034354 | 0 | A | G | 0.272 | 3.61E-03 | 6.06E-02 | 2.64E+00 | 2.73E+00 | 9.56E-01 | no | Il1rapl1 | intron_variant | MODIFIER |
| 20 | 88224655 | AX-97789797 | 0 | A | G | 0.272 | 3.61E-03 | 6.06E-02 | 2.64E+00 | 2.73E+00 | 9.56E-01 | no | | intergenic_variant | MODIFIER |
| 20 | 88597919 | AX-170090579 | 0 | T | C | 0.272 | 3.61E-03 | 6.06E-02 | 2.64E+00 | 2.73E+00 | 9.56E-01 | no | | intergenic_variant | MODIFIER |
| 20 | 88827979 | AX-170148282 | 0 | A | T | 0.272 | 3.61E-03 | 6.06E-02 | 2.64E+00 | 2.73E+00 | 9.56E-01 | no | | intergenic_variant | MODIFIER |
| 20 | 88834661 | AX-170248012 | 0 | T | C | 0.272 | 3.61E-03 | 6.06E-02 | 2.64E+00 | 2.73E+00 | 9.56E-01 | no | | intergenic_variant | MODIFIER |
| 20 | 88866735 | AX-170260354 | 0 | T | C | 0.272 | 3.61E-03 | 6.06E-02 | 2.64E+00 | 2.73E+00 | 9.56E-01 | no | | intergenic_variant | MODIFIER |
| 20 | 88938307 | AX-170521806 | 0 | G | T | 0.272 | 3.61E-03 | 6.06E-02 | 2.64E+00 | 2.73E+00 | 9.56E-01 | no | | intergenic_variant | MODIFIER |
| 20 | 88941079 | AX-170211033 | 0 | T | C | 0.272 | 3.61E-03 | 6.06E-02 | 2.64E+00 | 2.73E+00 | 9.56E-01 | no | | intergenic_variant | MODIFIER |
| 20 | 89131766 | AX-170084736 | 0 | A | G | 0.272 | 3.61E-03 | 6.06E-02 | 2.64E+00 | 2.73E+00 | 9.56E-01 | no | | intergenic_variant | MODIFIER |
| 20 | 89179057 | AX-170034384 | 0 | G | A | 0.272 | 3.61E-03 | 6.06E-02 | 2.64E+00 | 2.73E+00 | 9.56E-01 | no | | intergenic_variant | MODIFIER |
| 20 | 90177801 | AX-97807869 | 0 | T | C | 0.272 | 3.61E-03 | 6.06E-02 | 2.64E+00 | 2.73E+00 | 9.56E-01 | no | | intergenic_variant | MODIFIER |
| 20 | 90734981 | AX-170084760 | 0 | T | C | 0.272 | 3.61E-03 | 6.06E-02 | 2.64E+00 | 2.73E+00 | 9.56E-01 | no | | intergenic_variant | MODIFIER |
| 20 | 92685669 | AX-170034432 | 0 | C | G | 0.272 | 3.61E-03 | 6.06E-02 | 2.64E+00 | 2.73E+00 | 9.56E-01 | no | | intergenic_variant | MODIFIER |
| 20 | 92697004 | AX-97829182 | 0 | C | T | 0.272 | 3.61E-03 | 6.06E-02 | 2.64E+00 | 2.73E+00 | 9.56E-01 | no | | intergenic_variant | MODIFIER |
| 20 | 92735294 | AX-168407092 | 0 | C | T | 0.272 | 3.61E-03 | 6.06E-02 | 2.64E+00 | 2.73E+00 | 9.56E-01 | no | | intergenic_variant | MODIFIER |
| 20 | 92858648 | AX-170164576 | 0 | A | G | 0.272 | 3.61E-03 | 6.06E-02 | 2.64E+00 | 2.73E+00 | 9.56E-01 | no | | intergenic_variant | MODIFIER |
| 20 | 92858897 | AX-97829747 | 0 | T | A | 0.272 | 3.61E-03 | 6.06E-02 | 2.64E+00 | 2.73E+00 | 9.56E-01 | no | | intergenic_variant | MODIFIER |
| 20 | 93278461 | AX-97818175 | 0 | G | A | 0.272 | 3.61E-03 | 6.06E-02 | 2.64E+00 | 2.73E+00 | 9.56E-01 | no | | intergenic_variant | MODIFIER |
| 20 | 93465119 | AX-168376835 | 0 | G | A | 0.272 | 3.61E-03 | 6.06E-02 | 2.64E+00 | 2.73E+00 | 9.56E-01 | no | Pola1 | intron_variant | MODIFIER |
| 20 | 93486170 | AX-170523456 | 0 | G | C | 0.272 | 3.61E-03 | 6.06E-02 | 2.64E+00 | 2.73E+00 | 9.56E-01 | no | Pola1 | intron_variant | MODIFIER |
| 20 | 93699907 | AX-170034446 | 0 | G | A | 0.272 | 3.61E-03 | 6.06E-02 | 2.64E+00 | 2.73E+00 | 9.56E-01 | no | Pcyt1b | intron_variant | MODIFIER |
| 20 | 95069663 | AX-170449024 | 0 | G | T | 0.272 | 3.61E-03 | 6.06E-02 | 2.64E+00 | 2.73E+00 | 9.56E-01 | no | Arhgef9 | intron_variant | MODIFIER |
| 20 | 95092461 | AX-97801232 | 0 | C | T | 0.272 | 3.61E-03 | 6.06E-02 | 2.64E+00 | 2.73E+00 | 9.56E-01 | no | Arhgef9 | intron_variant | MODIFIER |
| 20 | 95426966 | AX-97849376 | 0 | T | C | 0.272 | 3.61E-03 | 6.06E-02 | 2.64E+00 | 2.73E+00 | 9.56E-01 | no | Amer1 | synonymous_variant | LOW |
| 12 | 117260616 | AX-97822519 | 0 | G | T | 0.132 | -7.59E-03 | 1.67E-01 | 2.64E+00 | 2.75E+00 | 9.56E-01 | no | Ptprn2 | intron_variant | MODIFIER |

| | | | | | | | | | | | | | | | |
|---|---|---|---|---|---|---|---|---|---|---|---|---|---|---|---|
| 7 | 80219888 | AX-170387526 | 2 | C | A | 0.118 | 1.24E-02 | 2.14E-01 | 2.65E+00 | 2.73E+00 | 9.56E-01 | no | Sema4b | intron_variant | MODIFIER |
| 8 | 41252175 | AX-170266833 | 0 | G | A | 0.158 | -8.21E-03 | 1.30E-01 | 2.64E+00 | 2.73E+00 | 9.56E-01 | no | Pcm1 | intron_variant | MODIFIER |
| 1 | 190012126 | AX-170323221 | 1 | A | G | 0.241 | 8.03E-03 | 9.67E-02 | 2.61E+00 | 2.71E+00 | 9.56E-01 | no | | intergenic_variant | MODIFIER |
| 9 | 38313486 | AX-170327656 | 0 | T | G | 0.5 | -3.89E-03 | 7.54E-02 | 2.65E+00 | 2.74E+00 | 9.57E-01 | no | Olfr897-ps1 | downstream_gene_variant | MODIFIER |
| 1 | 38560374 | AX-170136622 | 1 | A | G | 0.455 | -4.48E-03 | 7.82E-02 | 2.66E+00 | 2.73E+00 | 9.57E-01 | no | Aff3 | intron_variant,non_coding_transcript_variant | MODIFIER |
| 4 | 10213838 | AX-170134758 | 0 | C | A | 0.07 | 6.17E-03 | 1.15E-01 | 2.59E+00 | 2.74E+00 | 9.57E-01 | no | | intergenic_variant | MODIFIER |
| 12 | 115804429 | AX-170395995 | 0 | A | G | 0.114 | -8.93E-03 | 1.79E-01 | 2.67E+00 | 2.74E+00 | 9.57E-01 | no | Ighv1-74 | upstream_gene_variant | MODIFIER |
| 8 | 37103244 | AX-170388365 | 2 | C | T | 0.136 | -8.91E-03 | 1.42E-01 | 2.66E+00 | 2.73E+00 | 9.57E-01 | no | Gm3433 | upstream_gene_variant | MODIFIER |
| 20 | 93441811 | AX-97808186 | 0 | G | C | 0.272 | 3.52E-03 | 6.06E-02 | 2.64E+00 | 2.73E+00 | 9.57E-01 | no | Pola1 | intron_variant | MODIFIER |
| 10 | 43408172 | AX-170144765 | 1 | G | A | 0.366 | -4.48E-03 | 7.52E-02 | 2.63E+00 | 2.73E+00 | 9.57E-01 | no | Pdss2 | intron_variant | MODIFIER |
| 7 | 80517473 | AX-170099514 | 0 | T | C | 0.114 | -1.07E-02 | 2.12E-01 | 2.66E+00 | 2.74E+00 | 9.57E-01 | no | Blm | intron_variant | MODIFIER |
| 7 | 81097193 | AX-170077859 | 1 | T | A | 0.116 | 1.20E-02 | 2.12E-01 | 2.65E+00 | 2.73E+00 | 9.57E-01 | no | Alpk3 | intron_variant | MODIFIER |
| 7 | 94001708 | AX-170341830 | 1 | G | A | 0.116 | 1.20E-02 | 2.12E-01 | 2.65E+00 | 2.73E+00 | 9.57E-01 | no | | intergenic_variant | MODIFIER |
| 7 | 94683289 | AX-170205874 | 1 | G | A | 0.116 | 1.20E-02 | 2.12E-01 | 2.65E+00 | 2.73E+00 | 9.57E-01 | no | Gm32647 | intron_variant,non_coding_transcript_variant | MODIFIER |
| 8 | 48603188 | AX-170299958 | 1 | G | C | 0.152 | 6.67E-03 | 1.48E-01 | 2.69E+00 | 2.75E+00 | 9.57E-01 | no | Tenm3 | intron_variant | MODIFIER |
| 8 | 49016565 | AX-170030637 | 1 | G | T | 0.152 | 6.67E-03 | 1.48E-01 | 2.69E+00 | 2.75E+00 | 9.57E-01 | no | | intergenic_variant | MODIFIER |
| 9 | 77448351 | AX-170360799 | 0 | T | C | 0.211 | 6.06E-03 | 1.14E-01 | 2.65E+00 | 2.74E+00 | 9.57E-01 | no | Lrrc1 | intron_variant | MODIFIER |
| 5 | 27487387 | AX-170113473 | 0 | C | T | 0.061 | 1.24E-02 | 2.13E-01 | 2.63E+00 | 2.73E+00 | 9.57E-01 | no | Dpp6 | intron_variant | MODIFIER |
| 7 | 91690768 | AX-97841215 | 1 | G | A | 0.134 | -1.07E-02 | 2.10E-01 | 2.66E+00 | 2.74E+00 | 9.57E-01 | no | Dlg2 | intron_variant | MODIFIER |
| 7 | 102805933 | AX-170029659 | 0 | C | A | 0.456 | 5.83E-03 | 9.45E-02 | 2.66E+00 | 2.73E+00 | 9.57E-01 | no | Olfr564 | downstream_gene_variant | MODIFIER |
| 7 | 27237421 | AX-170400852 | 0 | C | T | 0.395 | 5.14E-03 | 8.67E-02 | 2.62E+00 | 2.73E+00 | 9.57E-01 | no | Coq8b | synonymous_variant | LOW |
| 1 | 53549413 | AX-170086469 | 1 | G | T | 0.393 | 1.22E-02 | 2.35E-01 | 2.69E+00 | 2.74E+00 | 9.57E-01 | no | Dnah7a | intron_variant | MODIFIER |
| 7 | 55448515 | AX-97809966 | 0 | A | G | 0.421 | 6.27E-03 | 1.02E-01 | 2.65E+00 | 2.73E+00 | 9.57E-01 | no | | intergenic_variant | MODIFIER |
| 13 | 56224663 | AX-170333224 | 0 | C | T | 0.509 | 4.34E-03 | 6.49E-02 | 2.61E+00 | 2.73E+00 | 9.57E-01 | no | 4930550C17Rik | intron_variant,non_coding_transcript_variant | MODIFIER |
| 7 | 53034568 | AX-170208109 | 0 | T | C | 0.421 | 6.23E-03 | 1.02E-01 | 2.65E+00 | 2.73E+00 | 9.57E-01 | no | | intergenic_variant | MODIFIER |
| 11 | 33334891 | AX-170405093 | 0 | A | T | 0.114 | 7.69E-03 | 1.46E-01 | 2.62E+00 | 2.74E+00 | 9.57E-01 | no | Ranbp17 | intron_variant | MODIFIER |
| 15 | 92190779 | AX-170375620 | 1 | T | G | 0.125 | 9.07E-03 | 1.55E-01 | 2.66E+00 | 2.73E+00 | 9.57E-01 | no | Cntn1 | intron_variant | MODIFIER |
| 1 | 8266045 | AX-170406222 | 1 | A | G | 0.509 | 4.72E-03 | 7.20E-02 | 2.65E+00 | 2.72E+00 | 9.57E-01 | no | | intergenic_variant | MODIFIER |
| 12 | 107169356 | AX-170100059 | 0 | A | G | 0.123 | 9.04E-03 | 1.62E-01 | 2.62E+00 | 2.74E+00 | 9.57E-01 | no | | intergenic_variant | MODIFIER |
| 18 | 53372500 | AX-97854338 | 2 | G | T | 0.045 | -1.27E-02 | 2.26E-01 | 2.61E+00 | 2.74E+00 | 9.57E-01 | no | Snx24 | intron_variant | MODIFIER |
| 8 | 49808469 | AX-170136750 | 2 | G | A | 0.155 | 6.81E-03 | 1.49E-01 | 2.70E+00 | 2.75E+00 | 9.58E-01 | no | | intergenic_variant | MODIFIER |
| 9 | 64674411 | AX-170169576 | 2 | T | C | 0.191 | -7.34E-03 | 1.36E-01 | 2.71E+00 | 2.73E+00 | 9.58E-01 | no | Megf11 | intron_variant | MODIFIER |
| 20 | 110831953 | AX-170034720 | 0 | G | A | 0.281 | -2.81E-03 | 5.45E-02 | 2.62E+00 | 2.74E+00 | 9.58E-01 | no | | intergenic_variant | MODIFIER |
| 11 | 33648895 | AX-97851937 | 0 | G | A | 0.105 | -8.86E-03 | 1.88E-01 | 2.67E+00 | 2.74E+00 | 9.58E-01 | no | Kcnip1 | intron_variant | MODIFIER |
| 1 | 191296530 | AX-170265541 | 0 | T | C | 0.254 | 8.84E-03 | 9.23E-02 | 2.58E+00 | 2.71E+00 | 9.58E-01 | no | | regulatory_region_variant | MODIFIER |
| 9 | 71437644 | AX-170398060 | 0 | T | C | 0.193 | 6.05E-03 | 1.25E-01 | 2.70E+00 | 2.75E+00 | 9.58E-01 | no | | intergenic_variant | MODIFIER |
| 8 | 21894052 | AX-170168993 | 0 | C | T | 0.175 | -5.65E-03 | 1.10E-01 | 2.66E+00 | 2.74E+00 | 9.58E-01 | no | | intergenic_variant | MODIFIER |
| 8 | 21914979 | AX-170473191 | 0 | A | G | 0.175 | -5.65E-03 | 1.10E-01 | 2.66E+00 | 2.74E+00 | 9.58E-01 | no | Gm36879 | intron_variant,non_coding_transcript_variant | MODIFIER |
| 8 | 21924100 | AX-170096194 | 0 | A | G | 0.175 | -5.65E-03 | 1.10E-01 | 2.66E+00 | 2.74E+00 | 9.58E-01 | no | Gm36879 | intron_variant,non_coding_transcript_variant | MODIFIER |
| 8 | 21928995 | AX-170292842 | 0 | G | A | 0.175 | -5.65E-03 | 1.10E-01 | 2.66E+00 | 2.74E+00 | 9.58E-01 | no | Defb15 | downstream_gene_variant | MODIFIER |
| 8 | 21942439 | AX-97810961 | 0 | C | T | 0.175 | -5.65E-03 | 1.10E-01 | 2.66E+00 | 2.74E+00 | 9.58E-01 | no | Defb13 | upstream_gene_variant | MODIFIER |
| 8 | 21960955 | AX-170452801 | 0 | C | T | 0.175 | -5.65E-03 | 1.10E-01 | 2.66E+00 | 2.74E+00 | 9.58E-01 | no | Fam90a1a | intron_variant | MODIFIER |
| 8 | 21965206 | AX-170496936 | 0 | T | C | 0.175 | -5.65E-03 | 1.10E-01 | 2.66E+00 | 2.74E+00 | 9.58E-01 | no | Ccdc70 | upstream_gene_variant | MODIFIER |
| 8 | 21965231 | AX-170231397 | 0 | A | T | 0.175 | -5.65E-03 | 1.10E-01 | 2.66E+00 | 2.74E+00 | 9.58E-01 | no | Ccdc70 | upstream_gene_variant | MODIFIER |
| 8 | 21976101 | AX-170242779 | 0 | A | G | 0.175 | -5.65E-03 | 1.10E-01 | 2.66E+00 | 2.74E+00 | 9.58E-01 | no | Ccdc70 | downstream_gene_variant | MODIFIER |
| 8 | 21977228 | AX-170096195 | 0 | G | A | 0.175 | -5.65E-03 | 1.10E-01 | 2.66E+00 | 2.74E+00 | 9.58E-01 | no | Ccdc70 | downstream_gene_variant | MODIFIER |
| 8 | 21991820 | AX-170492352 | 0 | A | G | 0.175 | -5.65E-03 | 1.10E-01 | 2.66E+00 | 2.74E+00 | 9.58E-01 | no | Atp7b | downstream_gene_variant | MODIFIER |
| 8 | 22016962 | AX-170202197 | 0 | G | A | 0.175 | -5.65E-03 | 1.10E-01 | 2.66E+00 | 2.74E+00 | 9.58E-01 | no | Atp7b | intron_variant | MODIFIER |
| 8 | 22017268 | AX-170121599 | 0 | T | C | 0.175 | -5.65E-03 | 1.10E-01 | 2.66E+00 | 2.74E+00 | 9.58E-01 | no | Atp7b | intron_variant | MODIFIER |
| 8 | 22027217 | AX-170444307 | 0 | A | C | 0.175 | -5.65E-03 | 1.10E-01 | 2.66E+00 | 2.74E+00 | 9.58E-01 | no | Atp7b | intron_variant | MODIFIER |
| 8 | 22027521 | AX-170119257 | 0 | A | G | 0.175 | -5.65E-03 | 1.10E-01 | 2.66E+00 | 2.74E+00 | 9.58E-01 | no | Atp7b | intron_variant | MODIFIER |
| 8 | 22061152 | AX-170436304 | 0 | C | T | 0.175 | -5.65E-03 | 1.10E-01 | 2.66E+00 | 2.74E+00 | 9.58E-01 | no | Atp7b | upstream_gene_variant | MODIFIER |
| 8 | 22065066 | AX-170383817 | 0 | T | C | 0.175 | -5.65E-03 | 1.10E-01 | 2.66E+00 | 2.74E+00 | 9.58E-01 | no | Atp7b | upstream_gene_variant | MODIFIER |
| 8 | 22066545 | AX-170300989 | 0 | A | G | 0.175 | -5.65E-03 | 1.10E-01 | 2.66E+00 | 2.74E+00 | 9.58E-01 | no | Alg11 | intron_variant | MODIFIER |

| | | | | | | | | | | | | | | | |
|---|---|---|---|---|---|---|---|---|---|---|---|---|---|---|---|
| 8 | 22067888 | AX-170079495 | 0 | C | T | 0.175 | -5.65E-03 | 1.10E-01 | 2.66E+00 | 2.74E+00 | 9.58E-01 | no | Alg11 | synonymous_variant | LOW |
| 8 | 22074148 | AX-97780595 | 0 | G | A | 0.175 | -5.65E-03 | 1.10E-01 | 2.66E+00 | 2.74E+00 | 9.58E-01 | no | Alg11 | downstream_gene_variant | MODIFIER |
| 8 | 22119597 | AX-170415673 | 0 | A | C | 0.175 | -5.65E-03 | 1.10E-01 | 2.66E+00 | 2.74E+00 | 9.58E-01 | no | Nek5 | intron_variant | MODIFIER |
| 8 | 22120577 | AX-170146786 | 0 | A | C | 0.175 | -5.65E-03 | 1.10E-01 | 2.66E+00 | 2.74E+00 | 9.58E-01 | no | Nek5 | intron_variant | MODIFIER |
| 8 | 22120753 | AX-170520100 | 0 | A | C | 0.175 | -5.65E-03 | 1.10E-01 | 2.66E+00 | 2.74E+00 | 9.58E-01 | no | Nek5 | intron_variant | MODIFIER |
| 8 | 22127734 | AX-170342473 | 0 | A | G | 0.175 | -5.65E-03 | 1.10E-01 | 2.66E+00 | 2.74E+00 | 9.58E-01 | no | Nek3 | downstream_gene_variant | MODIFIER |
| 8 | 22141712 | AX-170516132 | 0 | C | T | 0.175 | -5.65E-03 | 1.10E-01 | 2.66E+00 | 2.74E+00 | 9.58E-01 | no | Nek3 | intron_variant | MODIFIER |
| 8 | 22157163 | AX-170146124 | 0 | G | A | 0.175 | -5.65E-03 | 1.10E-01 | 2.66E+00 | 2.74E+00 | 9.58E-01 | no | Nek3 | missense_variant | MODERATE |
| 8 | 22196835 | AX-170260821 | 0 | G | T | 0.175 | -5.65E-03 | 1.10E-01 | 2.66E+00 | 2.74E+00 | 9.58E-01 | no | Vps36 | intron_variant | MODIFIER |
| 8 | 22197390 | AX-170208496 | 0 | A | G | 0.175 | -5.65E-03 | 1.10E-01 | 2.66E+00 | 2.74E+00 | 9.58E-01 | no | Vps36 | intron_variant | MODIFIER |
| 8 | 22215407 | AX-170381528 | 0 | A | G | 0.175 | -5.65E-03 | 1.10E-01 | 2.66E+00 | 2.74E+00 | 9.58E-01 | no | Vps36 | intron_variant | MODIFIER |
| 8 | 22219092 | AX-170332991 | 0 | G | C | 0.175 | -5.65E-03 | 1.10E-01 | 2.66E+00 | 2.74E+00 | 9.58E-01 | no | Vps36 | 3_prime_UTR_variant | MODIFIER |
| 8 | 22228463 | AX-170398166 | 0 | C | G | 0.175 | -5.65E-03 | 1.10E-01 | 2.66E+00 | 2.74E+00 | 9.58E-01 | no | Thsd1 | intron_variant | MODIFIER |
| 8 | 22228648 | AX-170260385 | 0 | A | T | 0.175 | -5.65E-03 | 1.10E-01 | 2.66E+00 | 2.74E+00 | 9.58E-01 | no | Thsd1 | intron_variant | MODIFIER |
| 8 | 22228845 | AX-170365538 | 0 | A | T | 0.175 | -5.65E-03 | 1.10E-01 | 2.66E+00 | 2.74E+00 | 9.58E-01 | no | Thsd1 | intron_variant | MODIFIER |
| 8 | 22251315 | AX-170273934 | 0 | C | A | 0.175 | -5.65E-03 | 1.10E-01 | 2.66E+00 | 2.74E+00 | 9.58E-01 | no | Thsd1 | intron_variant | MODIFIER |
| 8 | 22255865 | AX-170357620 | 0 | G | A | 0.175 | -5.65E-03 | 1.10E-01 | 2.66E+00 | 2.74E+00 | 9.58E-01 | no | Thsd1 | intron_variant | MODIFIER |
| 8 | 22257970 | AX-170079497 | 0 | C | T | 0.175 | -5.65E-03 | 1.10E-01 | 2.66E+00 | 2.74E+00 | 9.58E-01 | no | Thsd1 | intron_variant | MODIFIER |
| 8 | 22270209 | AX-170380016 | 0 | T | C | 0.175 | -5.65E-03 | 1.10E-01 | 2.66E+00 | 2.74E+00 | 9.58E-01 | no | | intergenic_variant | MODIFIER |
| 8 | 22271020 | AX-170459568 | 0 | A | G | 0.175 | -5.65E-03 | 1.10E-01 | 2.66E+00 | 2.74E+00 | 9.58E-01 | no | | intergenic_variant | MODIFIER |
| 8 | 22322500 | AX-170124333 | 0 | C | T | 0.175 | -5.65E-03 | 1.10E-01 | 2.66E+00 | 2.74E+00 | 9.58E-01 | no | Tpte | intron_variant | MODIFIER |
| 8 | 22325138 | AX-170251906 | 0 | A | C | 0.175 | -5.65E-03 | 1.10E-01 | 2.66E+00 | 2.74E+00 | 9.58E-01 | no | Tpte | intron_variant | MODIFIER |
| 8 | 22328333 | AX-170509160 | 0 | A | G | 0.175 | -5.65E-03 | 1.10E-01 | 2.66E+00 | 2.74E+00 | 9.58E-01 | no | Tpte | intron_variant | MODIFIER |
| 8 | 22364986 | AX-170385886 | 0 | T | C | 0.175 | -5.65E-03 | 1.10E-01 | 2.66E+00 | 2.74E+00 | 9.58E-01 | no | Tpte | intron_variant | MODIFIER |
| 8 | 22365516 | AX-170334746 | 0 | C | T | 0.175 | -5.65E-03 | 1.10E-01 | 2.66E+00 | 2.74E+00 | 9.58E-01 | no | Tpte | intron_variant | MODIFIER |
| 8 | 22372361 | AX-97794199 | 0 | T | G | 0.175 | -5.65E-03 | 1.10E-01 | 2.66E+00 | 2.74E+00 | 9.58E-01 | no | Slc25a15 | downstream_gene_variant | MODIFIER |
| 8 | 22378080 | AX-170138763 | 0 | C | T | 0.175 | -5.65E-03 | 1.10E-01 | 2.66E+00 | 2.74E+00 | 9.58E-01 | no | Slc25a15 | intron_variant | MODIFIER |
| 8 | 22387065 | AX-170263484 | 0 | T | C | 0.175 | -5.65E-03 | 1.10E-01 | 2.66E+00 | 2.74E+00 | 9.58E-01 | no | Slc25a15 | intron_variant | MODIFIER |
| 8 | 22393010 | AX-170344898 | 0 | T | C | 0.175 | -5.65E-03 | 1.10E-01 | 2.66E+00 | 2.74E+00 | 9.58E-01 | no | Slc25a15 | intron_variant | MODIFIER |
| 8 | 22420265 | AX-170426813 | 0 | A | G | 0.175 | -5.65E-03 | 1.10E-01 | 2.66E+00 | 2.74E+00 | 9.58E-01 | no | Mrps31 | intron_variant | MODIFIER |
| 8 | 22423258 | AX-170109297 | 0 | T | C | 0.175 | -5.65E-03 | 1.10E-01 | 2.66E+00 | 2.74E+00 | 9.58E-01 | no | Mrps31 | intron_variant | MODIFIER |
| 8 | 22434759 | AX-170030339 | 0 | A | G | 0.175 | -5.65E-03 | 1.10E-01 | 2.66E+00 | 2.74E+00 | 9.58E-01 | no | | regulatory_region_variant | MODIFIER |
| 7 | 78600074 | AX-170077812 | 0 | T | C | 0.114 | 1.16E-02 | 2.12E-01 | 2.66E+00 | 2.74E+00 | 9.58E-01 | no | Ntrk3 | intron_variant,non_coding_transcript_variant | MODIFIER |
| 7 | 78829870 | AX-97803013 | 0 | A | G | 0.114 | 1.16E-02 | 2.12E-01 | 2.66E+00 | 2.74E+00 | 9.58E-01 | no | Det1 | intron_variant | MODIFIER |
| 7 | 78851934 | AX-170345022 | 0 | G | A | 0.114 | 1.16E-02 | 2.12E-01 | 2.66E+00 | 2.74E+00 | 9.58E-01 | no | Det1 | upstream_gene_variant | MODIFIER |
| 7 | 78857680 | AX-170344719 | 0 | C | T | 0.114 | 1.16E-02 | 2.12E-01 | 2.66E+00 | 2.74E+00 | 9.58E-01 | no | Gm44730 | intron_variant,non_coding_transcript_variant | MODIFIER |
| 7 | 78862351 | AX-170343065 | 0 | C | T | 0.114 | 1.16E-02 | 2.12E-01 | 2.66E+00 | 2.74E+00 | 9.58E-01 | no | Gm44730 | upstream_gene_variant | MODIFIER |
| 7 | 78864405 | AX-170267146 | 0 | T | C | 0.114 | 1.16E-02 | 2.12E-01 | 2.66E+00 | 2.74E+00 | 9.58E-01 | no | | regulatory_region_variant | MODIFIER |
| 7 | 78869590 | AX-170212607 | 0 | A | G | 0.114 | 1.16E-02 | 2.12E-01 | 2.66E+00 | 2.74E+00 | 9.58E-01 | no | | intergenic_variant | MODIFIER |
| 7 | 78878070 | AX-170355657 | 0 | C | T | 0.114 | 1.16E-02 | 2.12E-01 | 2.66E+00 | 2.74E+00 | 9.58E-01 | no | | intergenic_variant | MODIFIER |
| 7 | 78886014 | AX-97809371 | 0 | G | A | 0.114 | 1.16E-02 | 2.12E-01 | 2.66E+00 | 2.74E+00 | 9.58E-01 | no | Mir7-2 | upstream_gene_variant | MODIFIER |
| 7 | 78902501 | AX-170395371 | 0 | T | G | 0.114 | 1.16E-02 | 2.12E-01 | 2.66E+00 | 2.74E+00 | 9.58E-01 | no | Aen | missense_variant | MODERATE |
| 7 | 78919452 | AX-97834066 | 0 | G | T | 0.114 | 1.16E-02 | 2.12E-01 | 2.66E+00 | 2.74E+00 | 9.58E-01 | no | Isg20 | intron_variant | MODIFIER |
| 7 | 79094004 | AX-170297336 | 0 | T | C | 0.114 | 1.16E-02 | 2.12E-01 | 2.66E+00 | 2.74E+00 | 9.58E-01 | no | Acan | synonymous_variant | LOW |
| 7 | 79095130 | AX-170472678 | 0 | A | G | 0.114 | 1.16E-02 | 2.12E-01 | 2.66E+00 | 2.74E+00 | 9.58E-01 | no | Acan | intron_variant | MODIFIER |
| 7 | 79108346 | AX-170188369 | 0 | T | C | 0.114 | 1.16E-02 | 2.12E-01 | 2.66E+00 | 2.74E+00 | 9.58E-01 | no | Acan | intron_variant | MODIFIER |
| 7 | 79108560 | AX-170146565 | 0 | G | A | 0.114 | 1.16E-02 | 2.12E-01 | 2.66E+00 | 2.74E+00 | 9.58E-01 | no | Acan | intron_variant | MODIFIER |
| 7 | 79184421 | AX-170276740 | 0 | A | G | 0.114 | 1.16E-02 | 2.12E-01 | 2.66E+00 | 2.74E+00 | 9.58E-01 | no | | intergenic_variant | MODIFIER |
| 7 | 79202848 | AX-170360738 | 0 | A | G | 0.114 | 1.16E-02 | 2.12E-01 | 2.66E+00 | 2.74E+00 | 9.58E-01 | no | | regulatory_region_variant | MODIFIER |
| 7 | 79251520 | AX-170175180 | 0 | G | A | 0.114 | 1.16E-02 | 2.12E-01 | 2.66E+00 | 2.74E+00 | 9.58E-01 | no | | intergenic_variant | MODIFIER |
| 7 | 79253450 | AX-170211519 | 0 | C | A | 0.114 | 1.16E-02 | 2.12E-01 | 2.66E+00 | 2.74E+00 | 9.58E-01 | no | | intergenic_variant | MODIFIER |
| 7 | 79263125 | AX-170226592 | 0 | G | A | 0.114 | 1.16E-02 | 2.12E-01 | 2.66E+00 | 2.74E+00 | 9.58E-01 | no | | regulatory_region_variant | MODIFIER |
| 7 | 79275561 | AX-170469678 | 0 | C | T | 0.114 | 1.16E-02 | 2.12E-01 | 2.66E+00 | 2.74E+00 | 9.58E-01 | no | Abhd2 | intron_variant | MODIFIER |
| 7 | 79277284 | AX-170150673 | 0 | G | C | 0.114 | 1.16E-02 | 2.12E-01 | 2.66E+00 | 2.74E+00 | 9.58E-01 | no | Abhd2 | intron_variant | MODIFIER |

| | | | | | | | | | | | | | | | |
|---|---|---|---|---|---|---|---|---|---|---|---|---|---|---|---|
| 7 | 79279513 | AX-170077824 | 0 | C | T | 0.114 | 1.16E-02 | 2.12E-01 | 2.66E+00 | 2.74E+00 | 9.58E-01 | no | Abhd2 | intron_variant | MODIFIER |
| 7 | 79285185 | AX-97783170 | 0 | A | G | 0.114 | 1.16E-02 | 2.12E-01 | 2.66E+00 | 2.74E+00 | 9.58E-01 | no | Abhd2 | intron_variant | MODIFIER |
| 7 | 79307719 | AX-170452564 | 0 | T | C | 0.114 | 1.16E-02 | 2.12E-01 | 2.66E+00 | 2.74E+00 | 9.58E-01 | no | Abhd2 | intron_variant | MODIFIER |
| 7 | 79311634 | AX-97802703 | 0 | C | T | 0.114 | 1.16E-02 | 2.12E-01 | 2.66E+00 | 2.74E+00 | 9.58E-01 | no | Abhd2 | intron_variant | MODIFIER |
| 7 | 79331234 | AX-97801648 | 0 | G | A | 0.114 | 1.16E-02 | 2.12E-01 | 2.66E+00 | 2.74E+00 | 9.58E-01 | no | Abhd2 | intron_variant | MODIFIER |
| 7 | 79343795 | AX-170503371 | 0 | G | A | 0.114 | 1.16E-02 | 2.12E-01 | 2.66E+00 | 2.74E+00 | 9.58E-01 | no | Abhd2 | intron_variant | MODIFIER |
| 7 | 79344065 | AX-170404512 | 0 | C | T | 0.114 | 1.16E-02 | 2.12E-01 | 2.66E+00 | 2.74E+00 | 9.58E-01 | no | Abhd2 | intron_variant | MODIFIER |
| 7 | 79347797 | AX-170521423 | 0 | T | C | 0.114 | 1.16E-02 | 2.12E-01 | 2.66E+00 | 2.74E+00 | 9.58E-01 | no | Abhd2 | intron_variant | MODIFIER |
| 7 | 79348757 | AX-170182218 | 0 | A | C | 0.114 | 1.16E-02 | 2.12E-01 | 2.66E+00 | 2.74E+00 | 9.58E-01 | no | Abhd2 | intron_variant | MODIFIER |
| 7 | 79348888 | AX-97854664 | 0 | G | A | 0.114 | 1.16E-02 | 2.12E-01 | 2.66E+00 | 2.74E+00 | 9.58E-01 | no | Abhd2 | intron_variant | MODIFIER |
| 7 | 79359222 | AX-170412664 | 0 | A | G | 0.114 | 1.16E-02 | 2.12E-01 | 2.66E+00 | 2.74E+00 | 9.58E-01 | no | Abhd2 | intron_variant | MODIFIER |
| 7 | 79381643 | AX-170288141 | 0 | T | C | 0.114 | 1.16E-02 | 2.12E-01 | 2.66E+00 | 2.74E+00 | 9.58E-01 | no | Rlbp1 | synonymous_variant | LOW |
| 7 | 79392683 | AX-170265583 | 0 | T | G | 0.114 | 1.16E-02 | 2.12E-01 | 2.66E+00 | 2.74E+00 | 9.58E-01 | no | Fanci | intron_variant | MODIFIER |
| 7 | 79420906 | AX-170381934 | 0 | G | A | 0.114 | 1.16E-02 | 2.12E-01 | 2.66E+00 | 2.74E+00 | 9.58E-01 | no | Fanci | intron_variant | MODIFIER |
| 7 | 79429127 | AX-170395900 | 0 | C | T | 0.114 | 1.16E-02 | 2.12E-01 | 2.66E+00 | 2.74E+00 | 9.58E-01 | no | Fanci | intron_variant | MODIFIER |
| 7 | 79432337 | AX-97809151 | 0 | T | C | 0.114 | 1.16E-02 | 2.12E-01 | 2.66E+00 | 2.74E+00 | 9.58E-01 | no | Fanci | intron_variant | MODIFIER |
| 7 | 79447597 | AX-170099261 | 0 | C | T | 0.114 | 1.16E-02 | 2.12E-01 | 2.66E+00 | 2.74E+00 | 9.58E-01 | no | Fanci | intron_variant | MODIFIER |
| 7 | 79448743 | AX-170201403 | 0 | A | G | 0.114 | 1.16E-02 | 2.12E-01 | 2.66E+00 | 2.74E+00 | 9.58E-01 | no | Fanci | intron_variant | MODIFIER |
| 7 | 79456727 | AX-170241454 | 0 | T | C | 0.114 | 1.16E-02 | 2.12E-01 | 2.66E+00 | 2.74E+00 | 9.58E-01 | no | Polg | missense_variant | MODERATE |
| 7 | 79471304 | AX-97814903 | 0 | A | G | 0.114 | 1.16E-02 | 2.12E-01 | 2.66E+00 | 2.74E+00 | 9.58E-01 | no | Polg | upstream_gene_variant | MODIFIER |
| 7 | 79471422 | AX-170318251 | 0 | T | C | 0.114 | 1.16E-02 | 2.12E-01 | 2.66E+00 | 2.74E+00 | 9.58E-01 | no | Gm10616 | downstream_gene_variant | MODIFIER |
| 7 | 79486248 | AX-97802667 | 0 | T | C | 0.114 | 1.16E-02 | 2.12E-01 | 2.66E+00 | 2.74E+00 | 9.58E-01 | no | | intergenic_variant | MODIFIER |
| 7 | 79487590 | AX-170077827 | 0 | A | C | 0.114 | 1.16E-02 | 2.12E-01 | 2.66E+00 | 2.74E+00 | 9.58E-01 | no | | intergenic_variant | MODIFIER |
| 7 | 79494328 | AX-170468525 | 0 | C | T | 0.114 | 1.16E-02 | 2.12E-01 | 2.66E+00 | 2.74E+00 | 9.58E-01 | no | | regulatory_region_variant | MODIFIER |
| 7 | 79494521 | AX-170316854 | 0 | C | T | 0.114 | 1.16E-02 | 2.12E-01 | 2.66E+00 | 2.74E+00 | 9.58E-01 | no | | intergenic_variant | MODIFIER |
| 7 | 79502462 | AX-170514795 | 0 | G | A | 0.114 | 1.16E-02 | 2.12E-01 | 2.66E+00 | 2.74E+00 | 9.58E-01 | no | Mir9-3 | upstream_gene_variant | MODIFIER |
| 7 | 79537975 | AX-170293709 | 0 | G | A | 0.114 | 1.16E-02 | 2.12E-01 | 2.66E+00 | 2.74E+00 | 9.58E-01 | no | Mir9-3hg | downstream_gene_variant | MODIFIER |
| 7 | 79559353 | AX-170304582 | 0 | G | A | 0.114 | 1.16E-02 | 2.12E-01 | 2.66E+00 | 2.74E+00 | 9.58E-01 | no | Gm35040 | non_coding_transcript_exon_variant | MODIFIER |
| 7 | 79575676 | AX-170101109 | 0 | G | A | 0.114 | 1.16E-02 | 2.12E-01 | 2.66E+00 | 2.74E+00 | 9.58E-01 | no | Gm45168 | downstream_gene_variant | MODIFIER |
| 7 | 79578195 | AX-170281138 | 0 | T | C | 0.114 | 1.16E-02 | 2.12E-01 | 2.66E+00 | 2.74E+00 | 9.58E-01 | no | Gm45168 | splice_region_variant,non_coding_transcript_exon_variant | LOW |
| 7 | 79581698 | AX-170376138 | 0 | G | A | 0.114 | 1.16E-02 | 2.12E-01 | 2.66E+00 | 2.74E+00 | 9.58E-01 | no | Gm45168 | upstream_gene_variant | MODIFIER |
| 7 | 79667697 | AX-170029384 | 0 | A | G | 0.114 | 1.16E-02 | 2.12E-01 | 2.66E+00 | 2.74E+00 | 9.58E-01 | no | Ticrr | synonymous_variant | LOW |
| 7 | 80029789 | AX-97798999 | 0 | T | C | 0.114 | 1.16E-02 | 2.12E-01 | 2.66E+00 | 2.74E+00 | 9.58E-01 | no | Zfp710 | intron_variant | MODIFIER |
| 7 | 80045376 | AX-170077841 | 0 | A | G | 0.114 | 1.16E-02 | 2.12E-01 | 2.66E+00 | 2.74E+00 | 9.58E-01 | no | Zfp710 | intron_variant | MODIFIER |
| 7 | 80046301 | AX-170281385 | 0 | A | G | 0.114 | 1.16E-02 | 2.12E-01 | 2.66E+00 | 2.74E+00 | 9.58E-01 | no | Zfp710 | intron_variant | MODIFIER |
| 7 | 80047066 | AX-170270648 | 0 | C | G | 0.114 | 1.16E-02 | 2.12E-01 | 2.66E+00 | 2.74E+00 | 9.58E-01 | no | Zfp710 | intron_variant | MODIFIER |
| 7 | 80049580 | AX-170295295 | 0 | C | G | 0.114 | 1.16E-02 | 2.12E-01 | 2.66E+00 | 2.74E+00 | 9.58E-01 | no | Zfp710 | intron_variant | MODIFIER |
| 7 | 80052256 | AX-170199965 | 0 | T | G | 0.114 | 1.16E-02 | 2.12E-01 | 2.66E+00 | 2.74E+00 | 9.58E-01 | no | Zfp710 | intron_variant | MODIFIER |
| 7 | 80071819 | AX-170282289 | 0 | A | G | 0.114 | 1.16E-02 | 2.12E-01 | 2.66E+00 | 2.74E+00 | 9.58E-01 | no | Zfp710 | intron_variant | MODIFIER |
| 7 | 80071842 | AX-170284693 | 0 | T | C | 0.114 | 1.16E-02 | 2.12E-01 | 2.66E+00 | 2.74E+00 | 9.58E-01 | no | Zfp710 | intron_variant | MODIFIER |
| 7 | 80099754 | AX-170248921 | 0 | T | C | 0.114 | 1.16E-02 | 2.12E-01 | 2.66E+00 | 2.74E+00 | 9.58E-01 | no | Idh2 | intron_variant | MODIFIER |
| 7 | 80159054 | AX-170029394 | 0 | A | G | 0.114 | 1.16E-02 | 2.12E-01 | 2.66E+00 | 2.74E+00 | 9.58E-01 | no | Gm45202 | downstream_gene_variant | MODIFIER |
| 7 | 80159556 | AX-170029389 | 0 | G | A | 0.114 | 1.16E-02 | 2.12E-01 | 2.66E+00 | 2.74E+00 | 9.58E-01 | no | Gm45202 | downstream_gene_variant | MODIFIER |
| 7 | 80169003 | AX-170318316 | 0 | A | T | 0.114 | 1.16E-02 | 2.12E-01 | 2.66E+00 | 2.74E+00 | 9.58E-01 | no | | intergenic_variant | MODIFIER |
| 7 | 80191026 | AX-170029391 | 0 | A | G | 0.114 | 1.16E-02 | 2.12E-01 | 2.66E+00 | 2.74E+00 | 9.58E-01 | no | Sema4b | intron_variant | MODIFIER |
| 7 | 80192928 | AX-170089963 | 0 | G | A | 0.114 | 1.16E-02 | 2.12E-01 | 2.66E+00 | 2.74E+00 | 9.58E-01 | no | Sema4b | intron_variant | MODIFIER |
| 7 | 80213588 | AX-97835307 | 0 | T | C | 0.114 | 1.16E-02 | 2.12E-01 | 2.66E+00 | 2.74E+00 | 9.58E-01 | no | Sema4b | intron_variant | MODIFIER |
| 7 | 80228388 | AX-170029392 | 0 | A | C | 0.114 | 1.16E-02 | 2.12E-01 | 2.66E+00 | 2.74E+00 | 9.58E-01 | no | Sema4b | downstream_gene_variant | MODIFIER |
| 7 | 80228887 | AX-170029395 | 0 | A | G | 0.114 | 1.16E-02 | 2.12E-01 | 2.66E+00 | 2.74E+00 | 9.58E-01 | no | Sema4b | downstream_gene_variant | MODIFIER |
| 7 | 80236501 | AX-170077843 | 0 | G | A | 0.114 | 1.16E-02 | 2.12E-01 | 2.66E+00 | 2.74E+00 | 9.58E-01 | no | Gdpgp1 | intron_variant | MODIFIER |
| 7 | 80266065 | AX-170384389 | 0 | G | T | 0.114 | 1.16E-02 | 2.12E-01 | 2.66E+00 | 2.74E+00 | 9.58E-01 | no | Vps33b | upstream_gene_variant | MODIFIER |
| 7 | 80266101 | AX-170413615 | 0 | C | G | 0.114 | 1.16E-02 | 2.12E-01 | 2.66E+00 | 2.74E+00 | 9.58E-01 | no | Vps33b | upstream_gene_variant | MODIFIER |
| 7 | 80270349 | AX-170303144 | 0 | T | C | 0.114 | 1.16E-02 | 2.12E-01 | 2.66E+00 | 2.74E+00 | 9.58E-01 | no | Vps33b | intron_variant | MODIFIER |
| 7 | 80272472 | AX-97829418 | 0 | C | T | 0.114 | 1.16E-02 | 2.12E-01 | 2.66E+00 | 2.74E+00 | 9.58E-01 | no | Vps33b | intron_variant | MODIFIER |

| | | | | | | | | | | | | | | | |
|---|---|---|---|---|---|---|---|---|---|---|---|---|---|---|---|
| 7 | 80286466 | AX-170120748 | 0 | A | G | 0.114 | 1.16E-02 | 2.12E-01 | 2.66E+00 | 2.74E+00 | 9.58E-01 | no | Vps33b | intron_variant | MODIFIER |
| 7 | 80297876 | AX-170336917 | 0 | G | A | 0.114 | 1.16E-02 | 2.12E-01 | 2.66E+00 | 2.74E+00 | 9.58E-01 | no | Prc1 | intron_variant | MODIFIER |
| 7 | 80301872 | AX-170029396 | 0 | G | A | 0.114 | 1.16E-02 | 2.12E-01 | 2.66E+00 | 2.74E+00 | 9.58E-01 | no | Prc1 | intron_variant | MODIFIER |
| 7 | 80304779 | AX-97820002 | 0 | A | G | 0.114 | 1.16E-02 | 2.12E-01 | 2.66E+00 | 2.74E+00 | 9.58E-01 | no | Prc1 | synonymous_variant | LOW |
| 7 | 80329678 | AX-170351720 | 0 | G | A | 0.114 | 1.16E-02 | 2.12E-01 | 2.66E+00 | 2.74E+00 | 9.58E-01 | no | Unc45a | intron_variant | MODIFIER |
| 7 | 80331297 | AX-170161478 | 0 | T | C | 0.114 | 1.16E-02 | 2.12E-01 | 2.66E+00 | 2.74E+00 | 9.58E-01 | no | Unc45a | intron_variant | MODIFIER |
| 7 | 80357462 | AX-170242990 | 0 | C | T | 0.114 | 1.16E-02 | 2.12E-01 | 2.66E+00 | 2.74E+00 | 9.58E-01 | no | Man2a2 | intron_variant | MODIFIER |
| 7 | 80362702 | AX-170374979 | 0 | A | G | 0.114 | 1.16E-02 | 2.12E-01 | 2.66E+00 | 2.74E+00 | 9.58E-01 | no | Man2a2 | intron_variant | MODIFIER |
| 7 | 80433516 | AX-170439564 | 0 | G | A | 0.114 | 1.16E-02 | 2.12E-01 | 2.66E+00 | 2.74E+00 | 9.58E-01 | no | Gm15544 | downstream_gene_variant | MODIFIER |
| 7 | 80435900 | AX-170397337 | 0 | A | G | 0.114 | 1.16E-02 | 2.12E-01 | 2.66E+00 | 2.74E+00 | 9.58E-01 | no | | regulatory_region_variant | MODIFIER |
| 7 | 80454932 | AX-170434016 | 0 | T | C | 0.114 | 1.16E-02 | 2.12E-01 | 2.66E+00 | 2.74E+00 | 9.58E-01 | no | Blm | downstream_gene_variant | MODIFIER |
| 7 | 80457935 | AX-170530877 | 0 | A | G | 0.114 | 1.16E-02 | 2.12E-01 | 2.66E+00 | 2.74E+00 | 9.58E-01 | no | Blm | intron_variant | MODIFIER |
| 7 | 80461004 | AX-97835306 | 0 | T | C | 0.114 | 1.16E-02 | 2.12E-01 | 2.66E+00 | 2.74E+00 | 9.58E-01 | no | Blm | intron_variant | MODIFIER |
| 7 | 80473638 | AX-170174368 | 0 | C | G | 0.114 | 1.16E-02 | 2.12E-01 | 2.66E+00 | 2.74E+00 | 9.58E-01 | no | Blm | intron_variant | MODIFIER |
| 7 | 80499603 | AX-170466192 | 0 | T | G | 0.114 | 1.16E-02 | 2.12E-01 | 2.66E+00 | 2.74E+00 | 9.58E-01 | no | Blm | intron_variant | MODIFIER |
| 7 | 80580703 | AX-170476769 | 0 | C | T | 0.114 | 1.16E-02 | 2.12E-01 | 2.66E+00 | 2.74E+00 | 9.58E-01 | no | | regulatory_region_variant | MODIFIER |
| 7 | 80585214 | AX-170266653 | 0 | A | G | 0.114 | 1.16E-02 | 2.12E-01 | 2.66E+00 | 2.74E+00 | 9.58E-01 | no | Crtc3 | downstream_gene_variant | MODIFIER |
| 7 | 80600429 | AX-170256266 | 0 | A | G | 0.114 | 1.16E-02 | 2.12E-01 | 2.66E+00 | 2.74E+00 | 9.58E-01 | no | Crtc3 | intron_variant | MODIFIER |
| 7 | 80609910 | AX-170266568 | 0 | G | A | 0.114 | 1.16E-02 | 2.12E-01 | 2.66E+00 | 2.74E+00 | 9.58E-01 | no | Crtc3 | synonymous_variant | LOW |
| 7 | 80631582 | AX-170488729 | 0 | T | C | 0.114 | 1.16E-02 | 2.12E-01 | 2.66E+00 | 2.74E+00 | 9.58E-01 | no | Crtc3 | intron_variant | MODIFIER |
| 7 | 80632588 | AX-170323207 | 0 | A | G | 0.114 | 1.16E-02 | 2.12E-01 | 2.66E+00 | 2.74E+00 | 9.58E-01 | no | Crtc3 | intron_variant | MODIFIER |
| 7 | 80670468 | AX-170148589 | 0 | G | A | 0.114 | 1.16E-02 | 2.12E-01 | 2.66E+00 | 2.74E+00 | 9.58E-01 | no | Crtc3 | intron_variant | MODIFIER |
| 7 | 80706797 | AX-97843142 | 0 | T | C | 0.114 | 1.16E-02 | 2.12E-01 | 2.66E+00 | 2.74E+00 | 9.58E-01 | no | Iqgap1 | downstream_gene_variant | MODIFIER |
| 7 | 80752238 | AX-170478568 | 0 | G | A | 0.114 | 1.16E-02 | 2.12E-01 | 2.66E+00 | 2.74E+00 | 9.58E-01 | no | Iqgap1 | synonymous_variant | LOW |
| 7 | 80757855 | AX-170330872 | 0 | T | C | 0.114 | 1.16E-02 | 2.12E-01 | 2.66E+00 | 2.74E+00 | 9.58E-01 | no | Iqgap1 | intron_variant | MODIFIER |
| 7 | 80819066 | AX-170475558 | 0 | G | A | 0.114 | 1.16E-02 | 2.12E-01 | 2.66E+00 | 2.74E+00 | 9.58E-01 | no | Iqgap1 | intron_variant | MODIFIER |
| 7 | 80819550 | AX-170306468 | 0 | G | A | 0.114 | 1.16E-02 | 2.12E-01 | 2.66E+00 | 2.74E+00 | 9.58E-01 | no | Iqgap1 | intron_variant | MODIFIER |
| 7 | 81029509 | AX-170475941 | 0 | G | A | 0.114 | 1.16E-02 | 2.12E-01 | 2.66E+00 | 2.74E+00 | 9.58E-01 | no | Zfp592 | missense_variant | MODERATE |
| 7 | 81029641 | AX-170209400 | 0 | G | C | 0.114 | 1.16E-02 | 2.12E-01 | 2.66E+00 | 2.74E+00 | 9.58E-01 | no | Zfp592 | intron_variant | MODIFIER |
| 7 | 81031139 | AX-170188664 | 0 | T | C | 0.114 | 1.16E-02 | 2.12E-01 | 2.66E+00 | 2.74E+00 | 9.58E-01 | no | Zfp592 | intron_variant | MODIFIER |
| 7 | 81033161 | AX-170396916 | 0 | C | T | 0.114 | 1.16E-02 | 2.12E-01 | 2.66E+00 | 2.74E+00 | 9.58E-01 | no | Zfp592 | intron_variant | MODIFIER |
| 7 | 81040449 | AX-170469224 | 0 | C | G | 0.114 | 1.16E-02 | 2.12E-01 | 2.66E+00 | 2.74E+00 | 9.58E-01 | no | Zfp592 | intron_variant | MODIFIER |
| 7 | 81045479 | AX-170163030 | 0 | T | C | 0.114 | 1.16E-02 | 2.12E-01 | 2.66E+00 | 2.74E+00 | 9.58E-01 | no | Zfp592 | downstream_gene_variant | MODIFIER |
| 7 | 81055007 | AX-170484241 | 0 | A | G | 0.114 | 1.16E-02 | 2.12E-01 | 2.66E+00 | 2.74E+00 | 9.58E-01 | no | Alpk3 | upstream_gene_variant | MODIFIER |
| 7 | 81057153 | AX-170365053 | 0 | G | A | 0.114 | 1.16E-02 | 2.12E-01 | 2.66E+00 | 2.74E+00 | 9.58E-01 | no | Alpk3 | upstream_gene_variant | MODIFIER |
| 7 | 81062319 | AX-170398980 | 0 | A | G | 0.114 | 1.16E-02 | 2.12E-01 | 2.66E+00 | 2.74E+00 | 9.58E-01 | no | Alpk3 | intron_variant | MODIFIER |
| 7 | 81065490 | AX-170315253 | 0 | T | A | 0.114 | 1.16E-02 | 2.12E-01 | 2.66E+00 | 2.74E+00 | 9.58E-01 | no | Alpk3 | intron_variant | MODIFIER |
| 7 | 81095134 | AX-97844224 | 0 | A | C | 0.114 | 1.16E-02 | 2.12E-01 | 2.66E+00 | 2.74E+00 | 9.58E-01 | no | Alpk3 | intron_variant | MODIFIER |
| 7 | 81098720 | AX-170289844 | 0 | G | T | 0.114 | 1.16E-02 | 2.12E-01 | 2.66E+00 | 2.74E+00 | 9.58E-01 | no | Alpk3 | intron_variant | MODIFIER |
| 7 | 81111584 | AX-170295867 | 0 | T | C | 0.114 | 1.16E-02 | 2.12E-01 | 2.66E+00 | 2.74E+00 | 9.58E-01 | no | Slc28a1 | upstream_gene_variant | MODIFIER |
| 7 | 81112132 | AX-170383776 | 0 | T | C | 0.114 | 1.16E-02 | 2.12E-01 | 2.66E+00 | 2.74E+00 | 9.58E-01 | no | Slc28a1 | upstream_gene_variant | MODIFIER |
| 7 | 81112736 | AX-170453685 | 0 | G | T | 0.114 | 1.16E-02 | 2.12E-01 | 2.66E+00 | 2.74E+00 | 9.58E-01 | no | Slc28a1 | upstream_gene_variant | MODIFIER |
| 7 | 81372314 | AX-170077864 | 0 | C | T | 0.114 | 1.16E-02 | 2.12E-01 | 2.66E+00 | 2.74E+00 | 9.58E-01 | no | Cpeb1 | intron_variant | MODIFIER |
| 7 | 81518193 | AX-170199832 | 0 | A | G | 0.114 | 1.16E-02 | 2.12E-01 | 2.66E+00 | 2.74E+00 | 9.58E-01 | no | | intergenic_variant | MODIFIER |
| 7 | 81519056 | AX-170118322 | 0 | A | C | 0.114 | 1.16E-02 | 2.12E-01 | 2.66E+00 | 2.74E+00 | 9.58E-01 | no | 2900076A07Rik | upstream_gene_variant | MODIFIER |
| 7 | 81523675 | AX-170519646 | 0 | C | A | 0.114 | 1.16E-02 | 2.12E-01 | 2.66E+00 | 2.74E+00 | 9.58E-01 | no | 2900076A07Rik | splice_region_variant,non_coding_transcript_exon_variant | LOW |
| 7 | 81524486 | AX-170424973 | 0 | T | C | 0.114 | 1.16E-02 | 2.12E-01 | 2.66E+00 | 2.74E+00 | 9.58E-01 | no | 2900076A07Rik | intron_variant,non_coding_transcript_variant | MODIFIER |
| 7 | 81538902 | AX-170461677 | 0 | G | A | 0.114 | 1.16E-02 | 2.12E-01 | 2.66E+00 | 2.74E+00 | 9.58E-01 | no | Fsd2 | intron_variant | MODIFIER |
| 7 | 81539761 | AX-170193501 | 0 | G | A | 0.114 | 1.16E-02 | 2.12E-01 | 2.66E+00 | 2.74E+00 | 9.58E-01 | no | Fsd2 | intron_variant | MODIFIER |
| 7 | 81540434 | AX-170392523 | 0 | A | G | 0.114 | 1.16E-02 | 2.12E-01 | 2.66E+00 | 2.74E+00 | 9.58E-01 | no | Fsd2 | synonymous_variant | LOW |
| 7 | 81540506 | AX-170268116 | 0 | C | T | 0.114 | 1.16E-02 | 2.12E-01 | 2.66E+00 | 2.74E+00 | 9.58E-01 | no | Fsd2 | intron_variant | MODIFIER |
| 7 | 81574301 | AX-170202134 | 0 | A | G | 0.114 | 1.16E-02 | 2.12E-01 | 2.66E+00 | 2.74E+00 | 9.58E-01 | no | Whamm | intron_variant | MODIFIER |
| 7 | 81612063 | AX-170287326 | 0 | A | G | 0.114 | 1.16E-02 | 2.12E-01 | 2.66E+00 | 2.74E+00 | 9.58E-01 | no | Homer2 | intron_variant | MODIFIER |
| 7 | 81617149 | AX-170236536 | 0 | T | C | 0.114 | 1.16E-02 | 2.12E-01 | 2.66E+00 | 2.74E+00 | 9.58E-01 | no | Homer2 | intron_variant | MODIFIER |

| | | | | | | | | | | | | | | | |
|---|---|---|---|---|---|---|---|---|---|---|---|---|---|---|---|
| 7 | 81619012 | AX-170104016 | 0 | G | A | 0.114 | 1.16E-02 | 2.12E-01 | 2.66E+00 | 2.74E+00 | 9.58E-01 | no | Homer2 | intron_variant | MODIFIER |
| 7 | 81621553 | AX-97844630 | 0 | A | G | 0.114 | 1.16E-02 | 2.12E-01 | 2.66E+00 | 2.74E+00 | 9.58E-01 | no | Homer2 | intron_variant | MODIFIER |
| 7 | 81644900 | AX-97787160 | 0 | G | T | 0.114 | 1.16E-02 | 2.12E-01 | 2.66E+00 | 2.74E+00 | 9.58E-01 | no | Homer2 | intron_variant | MODIFIER |
| 7 | 81652311 | AX-170207431 | 0 | C | T | 0.114 | 1.16E-02 | 2.12E-01 | 2.66E+00 | 2.74E+00 | 9.58E-01 | no | Homer2 | intron_variant | MODIFIER |
| 7 | 81679615 | AX-170368293 | 0 | A | G | 0.114 | 1.16E-02 | 2.12E-01 | 2.66E+00 | 2.74E+00 | 9.58E-01 | no | Homer2 | intron_variant | MODIFIER |
| 7 | 81700901 | AX-170518813 | 0 | A | G | 0.114 | 1.16E-02 | 2.12E-01 | 2.66E+00 | 2.74E+00 | 9.58E-01 | no | Homer2 | intron_variant | MODIFIER |
| 7 | 81736110 | AX-170108007 | 0 | T | C | 0.114 | 1.16E-02 | 2.12E-01 | 2.66E+00 | 2.74E+00 | 9.58E-01 | no | | intergenic_variant | MODIFIER |
| 7 | 81845571 | AX-170029417 | 0 | A | G | 0.114 | 1.16E-02 | 2.12E-01 | 2.66E+00 | 2.74E+00 | 9.58E-01 | no | | intergenic_variant | MODIFIER |
| 7 | 81858165 | AX-97824723 | 0 | G | A | 0.114 | 1.16E-02 | 2.12E-01 | 2.66E+00 | 2.74E+00 | 9.58E-01 | no | Tm6sf1 | upstream_gene_variant | MODIFIER |
| 7 | 81902543 | AX-170029413 | 0 | A | G | 0.114 | 1.16E-02 | 2.12E-01 | 2.66E+00 | 2.74E+00 | 9.58E-01 | no | Hdgfl3 | intron_variant | MODIFIER |
| 7 | 81923457 | AX-170255411 | 0 | A | G | 0.114 | 1.16E-02 | 2.12E-01 | 2.66E+00 | 2.74E+00 | 9.58E-01 | no | Hdgfl3 | intron_variant | MODIFIER |
| 7 | 81927058 | AX-170156035 | 0 | T | A | 0.114 | 1.16E-02 | 2.12E-01 | 2.66E+00 | 2.74E+00 | 9.58E-01 | no | Hdgfl3 | intron_variant | MODIFIER |
| 7 | 81954612 | AX-170370021 | 0 | G | T | 0.114 | 1.16E-02 | 2.12E-01 | 2.66E+00 | 2.74E+00 | 9.58E-01 | no | | regulatory_region_variant | MODIFIER |
| 7 | 81956234 | AX-170304531 | 0 | G | T | 0.114 | 1.16E-02 | 2.12E-01 | 2.66E+00 | 2.74E+00 | 9.58E-01 | no | | intergenic_variant | MODIFIER |
| 7 | 81956858 | AX-170325352 | 0 | T | G | 0.114 | 1.16E-02 | 2.12E-01 | 2.66E+00 | 2.74E+00 | 9.58E-01 | no | | intergenic_variant | MODIFIER |
| 7 | 81959795 | AX-170182016 | 0 | C | T | 0.114 | 1.16E-02 | 2.12E-01 | 2.66E+00 | 2.74E+00 | 9.58E-01 | no | | regulatory_region_variant | MODIFIER |
| 7 | 81960587 | AX-170125941 | 0 | A | G | 0.114 | 1.16E-02 | 2.12E-01 | 2.66E+00 | 2.74E+00 | 9.58E-01 | no | | intergenic_variant | MODIFIER |
| 7 | 81962767 | AX-170326387 | 0 | A | G | 0.114 | 1.16E-02 | 2.12E-01 | 2.66E+00 | 2.74E+00 | 9.58E-01 | no | Bnc1 | downstream_gene_variant | MODIFIER |
| 7 | 81963941 | AX-170132956 | 0 | G | T | 0.114 | 1.16E-02 | 2.12E-01 | 2.66E+00 | 2.74E+00 | 9.58E-01 | no | Bnc1 | downstream_gene_variant | MODIFIER |
| 7 | 81982501 | AX-170530759 | 0 | A | G | 0.114 | 1.16E-02 | 2.12E-01 | 2.66E+00 | 2.74E+00 | 9.58E-01 | no | Bnc1 | intron_variant | MODIFIER |
| 7 | 81986573 | AX-170353126 | 0 | T | A | 0.114 | 1.16E-02 | 2.12E-01 | 2.66E+00 | 2.74E+00 | 9.58E-01 | no | Bnc1 | intron_variant | MODIFIER |
| 7 | 81986851 | AX-170416441 | 0 | C | T | 0.114 | 1.16E-02 | 2.12E-01 | 2.66E+00 | 2.74E+00 | 9.58E-01 | no | Bnc1 | intron_variant | MODIFIER |
| 7 | 82005578 | AX-170239838 | 0 | C | T | 0.114 | 1.16E-02 | 2.12E-01 | 2.66E+00 | 2.74E+00 | 9.58E-01 | no | | intergenic_variant | MODIFIER |
| 7 | 82027792 | AX-170104198 | 0 | C | T | 0.114 | 1.16E-02 | 2.12E-01 | 2.66E+00 | 2.74E+00 | 9.58E-01 | no | | intergenic_variant | MODIFIER |
| 7 | 82038691 | AX-170077878 | 0 | T | C | 0.114 | 1.16E-02 | 2.12E-01 | 2.66E+00 | 2.74E+00 | 9.58E-01 | no | | intergenic_variant | MODIFIER |
| 7 | 82062866 | AX-170402662 | 0 | C | T | 0.114 | 1.16E-02 | 2.12E-01 | 2.66E+00 | 2.74E+00 | 9.58E-01 | no | | intergenic_variant | MODIFIER |
| 7 | 82067326 | AX-170493322 | 0 | C | T | 0.114 | 1.16E-02 | 2.12E-01 | 2.66E+00 | 2.74E+00 | 9.58E-01 | no | Gm20744 | upstream_gene_variant | MODIFIER |
| 7 | 82067671 | AX-170226966 | 0 | C | A | 0.114 | 1.16E-02 | 2.12E-01 | 2.66E+00 | 2.74E+00 | 9.58E-01 | no | Gm20744 | upstream_gene_variant | MODIFIER |
| 7 | 82068034 | AX-170242535 | 0 | G | T | 0.114 | 1.16E-02 | 2.12E-01 | 2.66E+00 | 2.74E+00 | 9.58E-01 | no | Gm20744 | upstream_gene_variant | MODIFIER |
| 7 | 82070519 | AX-170433792 | 0 | C | T | 0.114 | 1.16E-02 | 2.12E-01 | 2.66E+00 | 2.74E+00 | 9.58E-01 | no | Gm20744 | intron_variant,non_coding_transcript_variant | MODIFIER |
| 7 | 82113163 | AX-170352343 | 0 | C | T | 0.114 | 1.16E-02 | 2.12E-01 | 2.66E+00 | 2.74E+00 | 9.58E-01 | no | Gm45056 | downstream_gene_variant | MODIFIER |
| 7 | 82244964 | AX-170361392 | 0 | G | A | 0.114 | 1.16E-02 | 2.12E-01 | 2.66E+00 | 2.74E+00 | 9.58E-01 | no | Sh3gl3 | intron_variant | MODIFIER |
| 7 | 82245934 | AX-170415960 | 0 | T | C | 0.114 | 1.16E-02 | 2.12E-01 | 2.66E+00 | 2.74E+00 | 9.58E-01 | no | Sh3gl3 | intron_variant | MODIFIER |
| 7 | 82254532 | AX-170163052 | 0 | A | C | 0.114 | 1.16E-02 | 2.12E-01 | 2.66E+00 | 2.74E+00 | 9.58E-01 | no | Sh3gl3 | intron_variant | MODIFIER |
| 7 | 82254904 | AX-170231386 | 0 | C | T | 0.114 | 1.16E-02 | 2.12E-01 | 2.66E+00 | 2.74E+00 | 9.58E-01 | no | Sh3gl3 | intron_variant | MODIFIER |
| 7 | 82259766 | AX-170256547 | 0 | T | C | 0.114 | 1.16E-02 | 2.12E-01 | 2.66E+00 | 2.74E+00 | 9.58E-01 | no | Sh3gl3 | intron_variant | MODIFIER |
| 7 | 82277960 | AX-170295434 | 0 | A | T | 0.114 | 1.16E-02 | 2.12E-01 | 2.66E+00 | 2.74E+00 | 9.58E-01 | no | Sh3gl3 | intron_variant | MODIFIER |
| 7 | 82299981 | AX-170207565 | 0 | A | G | 0.114 | 1.16E-02 | 2.12E-01 | 2.66E+00 | 2.74E+00 | 9.58E-01 | no | Sh3gl3 | intron_variant | MODIFIER |
| 7 | 82324940 | AX-170256275 | 0 | C | T | 0.114 | 1.16E-02 | 2.12E-01 | 2.66E+00 | 2.74E+00 | 9.58E-01 | no | | intergenic_variant | MODIFIER |
| 7 | 82328037 | AX-170447384 | 0 | C | T | 0.114 | 1.16E-02 | 2.12E-01 | 2.66E+00 | 2.74E+00 | 9.58E-01 | no | | intergenic_variant | MODIFIER |
| 7 | 82336525 | AX-170271057 | 0 | T | C | 0.114 | 1.16E-02 | 2.12E-01 | 2.66E+00 | 2.74E+00 | 9.58E-01 | no | Adamtsl3 | intron_variant,non_coding_transcript_variant | MODIFIER |
| 7 | 82342253 | AX-170029419 | 0 | C | T | 0.114 | 1.16E-02 | 2.12E-01 | 2.66E+00 | 2.74E+00 | 9.58E-01 | no | Adamtsl3 | downstream_gene_variant | MODIFIER |
| 7 | 82361487 | AX-170029420 | 0 | C | T | 0.114 | 1.16E-02 | 2.12E-01 | 2.66E+00 | 2.74E+00 | 9.58E-01 | no | Adamtsl3 | intron_variant | MODIFIER |
| 7 | 82383875 | AX-170077884 | 0 | T | C | 0.114 | 1.16E-02 | 2.12E-01 | 2.66E+00 | 2.74E+00 | 9.58E-01 | no | Adamtsl3 | intron_variant | MODIFIER |
| 7 | 82387482 | AX-170077883 | 0 | T | C | 0.114 | 1.16E-02 | 2.12E-01 | 2.66E+00 | 2.74E+00 | 9.58E-01 | no | Adamtsl3 | intron_variant | MODIFIER |
| 7 | 82411386 | AX-170311648 | 0 | T | C | 0.114 | 1.16E-02 | 2.12E-01 | 2.66E+00 | 2.74E+00 | 9.58E-01 | no | Adamtsl3 | intron_variant | MODIFIER |
| 7 | 82528828 | AX-97808032 | 0 | C | T | 0.114 | 1.16E-02 | 2.12E-01 | 2.66E+00 | 2.74E+00 | 9.58E-01 | no | Adamtsl3 | splice_region_variant,intron_variant | LOW |
| 7 | 82548805 | AX-170227234 | 0 | A | G | 0.114 | 1.16E-02 | 2.12E-01 | 2.66E+00 | 2.74E+00 | 9.58E-01 | no | Adamtsl3 | intron_variant | MODIFIER |
| 7 | 82588610 | AX-170077887 | 0 | A | G | 0.114 | 1.16E-02 | 2.12E-01 | 2.66E+00 | 2.74E+00 | 9.58E-01 | no | Adamtsl3 | intron_variant | MODIFIER |
| 7 | 82598860 | AX-170408728 | 0 | G | T | 0.114 | 1.16E-02 | 2.12E-01 | 2.66E+00 | 2.74E+00 | 9.58E-01 | no | Adamtsl3 | intron_variant | MODIFIER |
| 7 | 82603325 | AX-170077891 | 0 | C | T | 0.114 | 1.16E-02 | 2.12E-01 | 2.66E+00 | 2.74E+00 | 9.58E-01 | no | Adamtsl3 | intron_variant | MODIFIER |
| 7 | 82611924 | AX-170183961 | 0 | T | C | 0.114 | 1.16E-02 | 2.12E-01 | 2.66E+00 | 2.74E+00 | 9.58E-01 | no | Adamtsl3 | intron_variant | MODIFIER |
| 7 | 82611986 | AX-170182876 | 0 | T | A | 0.114 | 1.16E-02 | 2.12E-01 | 2.66E+00 | 2.74E+00 | 9.58E-01 | no | Adamtsl3 | intron_variant | MODIFIER |
| 7 | 82634205 | AX-170224948 | 0 | C | T | 0.114 | 1.16E-02 | 2.12E-01 | 2.66E+00 | 2.74E+00 | 9.58E-01 | no | Saxo2 | 3_prime_UTR_variant | MODIFIER |

| | | | | | | | | | | | | | | | |
|---|---|---|---|---|---|---|---|---|---|---|---|---|---|---|---|
| 7 | 82665738 | AX-170294432 | 0 | A | C | 0.114 | 1.16E-02 | 2.12E-01 | 2.66E+00 | 2.74E+00 | 9.58E-01 | no | Efl1 | intron_variant | MODIFIER |
| 7 | 82670492 | AX-170403298 | 0 | G | A | 0.114 | 1.16E-02 | 2.12E-01 | 2.66E+00 | 2.74E+00 | 9.58E-01 | no | Efl1 | intron_variant | MODIFIER |
| 7 | 82702827 | AX-170509125 | 0 | G | A | 0.114 | 1.16E-02 | 2.12E-01 | 2.66E+00 | 2.74E+00 | 9.58E-01 | no | Efl1 | intron_variant | MODIFIER |
| 7 | 82720705 | AX-170263440 | 0 | T | A | 0.114 | 1.16E-02 | 2.12E-01 | 2.66E+00 | 2.74E+00 | 9.58E-01 | no | Efl1 | intron_variant | MODIFIER |
| 7 | 82742457 | AX-170498269 | 0 | T | C | 0.114 | 1.16E-02 | 2.12E-01 | 2.66E+00 | 2.74E+00 | 9.58E-01 | no | Efl1 | intron_variant | MODIFIER |
| 7 | 82766402 | AX-170398742 | 0 | T | C | 0.114 | 1.16E-02 | 2.12E-01 | 2.66E+00 | 2.74E+00 | 9.58E-01 | no | Efl1 | intron_variant | MODIFIER |
| 7 | 82788064 | AX-170442386 | 0 | A | G | 0.114 | 1.16E-02 | 2.12E-01 | 2.66E+00 | 2.74E+00 | 9.58E-01 | no | | intergenic_variant | MODIFIER |
| 7 | 82810161 | AX-170077895 | 0 | C | T | 0.114 | 1.16E-02 | 2.12E-01 | 2.66E+00 | 2.74E+00 | 9.58E-01 | no | | intergenic_variant | MODIFIER |
| 7 | 82833967 | AX-170296067 | 0 | A | G | 0.114 | 1.16E-02 | 2.12E-01 | 2.66E+00 | 2.74E+00 | 9.58E-01 | no | | intergenic_variant | MODIFIER |
| 7 | 82847408 | AX-170492597 | 0 | G | A | 0.114 | 1.16E-02 | 2.12E-01 | 2.66E+00 | 2.74E+00 | 9.58E-01 | no | 1700010L04Rik | upstream_gene_variant | MODIFIER |
| 7 | 82851247 | AX-97812316 | 0 | A | C | 0.114 | 1.16E-02 | 2.12E-01 | 2.66E+00 | 2.74E+00 | 9.58E-01 | no | 1700010L04Rik | upstream_gene_variant | MODIFIER |
| 7 | 82936666 | AX-170029427 | 0 | C | T | 0.114 | 1.16E-02 | 2.12E-01 | 2.66E+00 | 2.74E+00 | 9.58E-01 | no | 4933430H16Rik | downstream_gene_variant | MODIFIER |
| 7 | 87429505 | AX-170317945 | 0 | C | T | 0.114 | 1.16E-02 | 2.12E-01 | 2.66E+00 | 2.74E+00 | 9.58E-01 | no | Tyr | intron_variant | MODIFIER |
| 7 | 87432027 | AX-170125693 | 0 | G | T | 0.114 | 1.16E-02 | 2.12E-01 | 2.66E+00 | 2.74E+00 | 9.58E-01 | no | Tyr | intron_variant | MODIFIER |
| 7 | 87435149 | AX-170184917 | 0 | T | G | 0.114 | 1.16E-02 | 2.12E-01 | 2.66E+00 | 2.74E+00 | 9.58E-01 | no | Tyr | intron_variant | MODIFIER |
| 7 | 87457848 | AX-170504697 | 0 | A | C | 0.114 | 1.16E-02 | 2.12E-01 | 2.66E+00 | 2.74E+00 | 9.58E-01 | no | Tyr | intron_variant | MODIFIER |
| 7 | 87508115 | AX-170479829 | 0 | T | A | 0.114 | 1.16E-02 | 2.12E-01 | 2.66E+00 | 2.74E+00 | 9.58E-01 | no | | intergenic_variant | MODIFIER |
| 7 | 87610374 | AX-170185327 | 0 | G | T | 0.114 | 1.16E-02 | 2.12E-01 | 2.66E+00 | 2.74E+00 | 9.58E-01 | no | Grm5 | intron_variant | MODIFIER |
| 7 | 87650685 | AX-170077980 | 0 | G | C | 0.114 | 1.16E-02 | 2.12E-01 | 2.66E+00 | 2.74E+00 | 9.58E-01 | no | Grm5 | intron_variant | MODIFIER |
| 7 | 87663369 | AX-170528196 | 0 | A | G | 0.114 | 1.16E-02 | 2.12E-01 | 2.66E+00 | 2.74E+00 | 9.58E-01 | no | Grm5 | intron_variant | MODIFIER |
| 7 | 87688795 | AX-170258020 | 0 | C | T | 0.114 | 1.16E-02 | 2.12E-01 | 2.66E+00 | 2.74E+00 | 9.58E-01 | no | Grm5 | intron_variant | MODIFIER |
| 7 | 87689885 | AX-170210515 | 0 | C | T | 0.114 | 1.16E-02 | 2.12E-01 | 2.66E+00 | 2.74E+00 | 9.58E-01 | no | Grm5 | intron_variant | MODIFIER |
| 7 | 87715124 | AX-170168263 | 0 | A | G | 0.114 | 1.16E-02 | 2.12E-01 | 2.66E+00 | 2.74E+00 | 9.58E-01 | no | Grm5 | intron_variant | MODIFIER |
| 7 | 87727054 | AX-170465084 | 0 | A | C | 0.114 | 1.16E-02 | 2.12E-01 | 2.66E+00 | 2.74E+00 | 9.58E-01 | no | Grm5 | intron_variant | MODIFIER |
| 7 | 87728606 | AX-170388412 | 0 | A | G | 0.114 | 1.16E-02 | 2.12E-01 | 2.66E+00 | 2.74E+00 | 9.58E-01 | no | Grm5 | intron_variant | MODIFIER |
| 7 | 87731929 | AX-170128660 | 0 | C | T | 0.114 | 1.16E-02 | 2.12E-01 | 2.66E+00 | 2.74E+00 | 9.58E-01 | no | Grm5 | intron_variant | MODIFIER |
| 7 | 87745587 | AX-170408878 | 0 | G | T | 0.114 | 1.16E-02 | 2.12E-01 | 2.66E+00 | 2.74E+00 | 9.58E-01 | no | Grm5 | intron_variant | MODIFIER |
| 7 | 87787604 | AX-170295347 | 0 | T | C | 0.114 | 1.16E-02 | 2.12E-01 | 2.66E+00 | 2.74E+00 | 9.58E-01 | no | Grm5 | intron_variant | MODIFIER |
| 7 | 87819293 | AX-170247631 | 0 | T | G | 0.114 | 1.16E-02 | 2.12E-01 | 2.66E+00 | 2.74E+00 | 9.58E-01 | no | Grm5 | intron_variant | MODIFIER |
| 7 | 91515708 | AX-97809897 | 0 | G | T | 0.114 | 1.16E-02 | 2.12E-01 | 2.66E+00 | 2.74E+00 | 9.58E-01 | no | Dlg2 | intron_variant | MODIFIER |
| 7 | 92090940 | AX-97812207 | 0 | A | G | 0.114 | 1.16E-02 | 2.12E-01 | 2.66E+00 | 2.74E+00 | 9.58E-01 | no | Dlg2 | intron_variant | MODIFIER |
| 7 | 92158542 | AX-97834879 | 0 | A | G | 0.114 | 1.16E-02 | 2.12E-01 | 2.66E+00 | 2.74E+00 | 9.58E-01 | no | Dlg2 | intron_variant | MODIFIER |
| 7 | 92194007 | AX-97802777 | 0 | T | C | 0.114 | 1.16E-02 | 2.12E-01 | 2.66E+00 | 2.74E+00 | 9.58E-01 | no | Dlg2 | intron_variant | MODIFIER |
| 7 | 92210041 | AX-97825750 | 0 | C | A | 0.114 | 1.16E-02 | 2.12E-01 | 2.66E+00 | 2.74E+00 | 9.58E-01 | no | Dlg2 | intron_variant | MODIFIER |
| 7 | 92215465 | AX-170512026 | 0 | T | G | 0.114 | 1.16E-02 | 2.12E-01 | 2.66E+00 | 2.74E+00 | 9.58E-01 | no | Dlg2 | intron_variant | MODIFIER |
| 7 | 92215926 | AX-170078075 | 0 | A | T | 0.114 | 1.16E-02 | 2.12E-01 | 2.66E+00 | 2.74E+00 | 9.58E-01 | no | Dlg2 | intron_variant | MODIFIER |
| 7 | 92283715 | AX-170029510 | 0 | T | A | 0.114 | 1.16E-02 | 2.12E-01 | 2.66E+00 | 2.74E+00 | 9.58E-01 | no | Dlg2 | intron_variant | MODIFIER |
| 7 | 92345354 | AX-97802675 | 0 | T | A | 0.114 | 1.16E-02 | 2.12E-01 | 2.66E+00 | 2.74E+00 | 9.58E-01 | no | Dlg2 | intron_variant | MODIFIER |
| 7 | 92381623 | AX-170477828 | 0 | A | G | 0.114 | 1.16E-02 | 2.12E-01 | 2.66E+00 | 2.74E+00 | 9.58E-01 | no | Dlg2 | intron_variant | MODIFIER |
| 7 | 92393316 | AX-170198079 | 0 | T | C | 0.114 | 1.16E-02 | 2.12E-01 | 2.66E+00 | 2.74E+00 | 9.58E-01 | no | Dlg2 | intron_variant | MODIFIER |
| 7 | 92395561 | AX-97785259 | 0 | C | T | 0.114 | 1.16E-02 | 2.12E-01 | 2.66E+00 | 2.74E+00 | 9.58E-01 | no | Dlg2 | intron_variant | MODIFIER |
| 7 | 92399689 | AX-170078077 | 0 | T | C | 0.114 | 1.16E-02 | 2.12E-01 | 2.66E+00 | 2.74E+00 | 9.58E-01 | no | Dlg2 | intron_variant | MODIFIER |
| 7 | 92417739 | AX-170388604 | 0 | A | G | 0.114 | 1.16E-02 | 2.12E-01 | 2.66E+00 | 2.74E+00 | 9.58E-01 | no | Dlg2 | intron_variant | MODIFIER |
| 7 | 92439662 | AX-170214093 | 0 | G | C | 0.114 | 1.16E-02 | 2.12E-01 | 2.66E+00 | 2.74E+00 | 9.58E-01 | no | Dlg2 | intron_variant | MODIFIER |
| 7 | 92440190 | AX-170237417 | 0 | A | T | 0.114 | 1.16E-02 | 2.12E-01 | 2.66E+00 | 2.74E+00 | 9.58E-01 | no | Dlg2 | intron_variant | MODIFIER |
| 7 | 92444271 | AX-170133193 | 0 | T | C | 0.114 | 1.16E-02 | 2.12E-01 | 2.66E+00 | 2.74E+00 | 9.58E-01 | no | Dlg2 | intron_variant | MODIFIER |
| 7 | 92460764 | AX-170089978 | 0 | T | G | 0.114 | 1.16E-02 | 2.12E-01 | 2.66E+00 | 2.74E+00 | 9.58E-01 | no | | intergenic_variant | MODIFIER |
| 7 | 92475805 | AX-170089979 | 0 | T | G | 0.114 | 1.16E-02 | 2.12E-01 | 2.66E+00 | 2.74E+00 | 9.58E-01 | no | | intergenic_variant | MODIFIER |
| 7 | 92476053 | AX-170339988 | 0 | T | C | 0.114 | 1.16E-02 | 2.12E-01 | 2.66E+00 | 2.74E+00 | 9.58E-01 | no | | intergenic_variant | MODIFIER |
| 7 | 92507548 | AX-170297616 | 0 | C | T | 0.114 | 1.16E-02 | 2.12E-01 | 2.66E+00 | 2.74E+00 | 9.58E-01 | no | | intergenic_variant | MODIFIER |
| 7 | 92516061 | AX-97851006 | 0 | A | G | 0.114 | 1.16E-02 | 2.12E-01 | 2.66E+00 | 2.74E+00 | 9.58E-01 | no | | intergenic_variant | MODIFIER |
| 7 | 92518809 | AX-170157477 | 0 | A | C | 0.114 | 1.16E-02 | 2.12E-01 | 2.66E+00 | 2.74E+00 | 9.58E-01 | no | | intergenic_variant | MODIFIER |
| 7 | 92699899 | AX-170180764 | 0 | A | G | 0.114 | 1.16E-02 | 2.12E-01 | 2.66E+00 | 2.74E+00 | 9.58E-01 | no | | intergenic_variant | MODIFIER |
| 7 | 92701780 | AX-97789502 | 0 | A | G | 0.114 | 1.16E-02 | 2.12E-01 | 2.66E+00 | 2.74E+00 | 9.58E-01 | no | Gm45199 | upstream_gene_variant | MODIFIER |

| | | | | | | | | | | | | | | | |
|---|---|---|---|---|---|---|---|---|---|---|---|---|---|---|---|
| 7 | 93547525 | AX-97835997 | 0 | A | G | 0.114 | 1.16E-02 | 2.12E-01 | 2.66E+00 | 2.74E+00 | 9.58E-01 | no | | intergenic_variant | MODIFIER |
| 7 | 93575577 | AX-170187571 | 0 | T | C | 0.114 | 1.16E-02 | 2.12E-01 | 2.66E+00 | 2.74E+00 | 9.58E-01 | no | | intergenic_variant | MODIFIER |
| 7 | 93647231 | AX-170230028 | 0 | C | T | 0.114 | 1.16E-02 | 2.12E-01 | 2.66E+00 | 2.74E+00 | 9.58E-01 | no | | intergenic_variant | MODIFIER |
| 7 | 93648205 | AX-170212288 | 0 | A | C | 0.114 | 1.16E-02 | 2.12E-01 | 2.66E+00 | 2.74E+00 | 9.58E-01 | no | | intergenic_variant | MODIFIER |
| 7 | 93676845 | AX-170193426 | 0 | A | T | 0.114 | 1.16E-02 | 2.12E-01 | 2.66E+00 | 2.74E+00 | 9.58E-01 | no | | intergenic_variant | MODIFIER |
| 7 | 93719609 | AX-97818948 | 0 | A | G | 0.114 | 1.16E-02 | 2.12E-01 | 2.66E+00 | 2.74E+00 | 9.58E-01 | no | | intergenic_variant | MODIFIER |
| 7 | 93735344 | AX-170497073 | 0 | G | T | 0.114 | 1.16E-02 | 2.12E-01 | 2.66E+00 | 2.74E+00 | 9.58E-01 | no | | intergenic_variant | MODIFIER |
| 7 | 93740236 | AX-170493694 | 0 | A | C | 0.114 | 1.16E-02 | 2.12E-01 | 2.66E+00 | 2.74E+00 | 9.58E-01 | no | | intergenic_variant | MODIFIER |
| 7 | 93820578 | AX-170490101 | 0 | T | A | 0.114 | 1.16E-02 | 2.12E-01 | 2.66E+00 | 2.74E+00 | 9.58E-01 | no | | intergenic_variant | MODIFIER |
| 7 | 93868775 | AX-170089981 | 0 | C | T | 0.114 | 1.16E-02 | 2.12E-01 | 2.66E+00 | 2.74E+00 | 9.58E-01 | no | | intergenic_variant | MODIFIER |
| 7 | 93901561 | AX-170078098 | 0 | A | T | 0.114 | 1.16E-02 | 2.12E-01 | 2.66E+00 | 2.74E+00 | 9.58E-01 | no | | intergenic_variant | MODIFIER |
| 7 | 94002758 | AX-97799478 | 0 | G | A | 0.114 | 1.16E-02 | 2.12E-01 | 2.66E+00 | 2.74E+00 | 9.58E-01 | no | | intergenic_variant | MODIFIER |
| 7 | 94013145 | AX-170197122 | 0 | G | C | 0.114 | 1.16E-02 | 2.12E-01 | 2.66E+00 | 2.74E+00 | 9.58E-01 | no | | intergenic_variant | MODIFIER |
| 7 | 94043710 | AX-97838695 | 0 | T | C | 0.114 | 1.16E-02 | 2.12E-01 | 2.66E+00 | 2.74E+00 | 9.58E-01 | no | Gm32647 | intron_variant,non_coding_transcript_variant | MODIFIER |
| 7 | 94089291 | AX-170349124 | 0 | A | G | 0.114 | 1.16E-02 | 2.12E-01 | 2.66E+00 | 2.74E+00 | 9.58E-01 | no | Gm32647 | intron_variant,non_coding_transcript_variant | MODIFIER |
| 7 | 94089833 | AX-170159503 | 0 | T | A | 0.114 | 1.16E-02 | 2.12E-01 | 2.66E+00 | 2.74E+00 | 9.58E-01 | no | Gm32647 | intron_variant,non_coding_transcript_variant | MODIFIER |
| 7 | 94090432 | AX-170111773 | 0 | T | C | 0.114 | 1.16E-02 | 2.12E-01 | 2.66E+00 | 2.74E+00 | 9.58E-01 | no | Gm32647 | intron_variant,non_coding_transcript_variant | MODIFIER |
| 7 | 94163187 | AX-170111803 | 0 | G | A | 0.114 | 1.16E-02 | 2.12E-01 | 2.66E+00 | 2.74E+00 | 9.58E-01 | no | Gm32647 | intron_variant,non_coding_transcript_variant | MODIFIER |
| 7 | 94165414 | AX-170193625 | 0 | T | C | 0.114 | 1.16E-02 | 2.12E-01 | 2.66E+00 | 2.74E+00 | 9.58E-01 | no | Gm32647 | intron_variant,non_coding_transcript_variant | MODIFIER |
| 7 | 94168432 | AX-170399342 | 0 | G | A | 0.114 | 1.16E-02 | 2.12E-01 | 2.66E+00 | 2.74E+00 | 9.58E-01 | no | Gm32647 | intron_variant,non_coding_transcript_variant | MODIFIER |
| 7 | 94278567 | AX-97806979 | 0 | C | A | 0.114 | 1.16E-02 | 2.12E-01 | 2.66E+00 | 2.74E+00 | 9.58E-01 | no | Gm32647 | intron_variant,non_coding_transcript_variant | MODIFIER |
| 7 | 94320190 | AX-170235527 | 0 | A | G | 0.114 | 1.16E-02 | 2.12E-01 | 2.66E+00 | 2.74E+00 | 9.58E-01 | no | Gm32647 | intron_variant,non_coding_transcript_variant | MODIFIER |
| 7 | 94322127 | AX-170231609 | 0 | T | C | 0.114 | 1.16E-02 | 2.12E-01 | 2.66E+00 | 2.74E+00 | 9.58E-01 | no | Gm32647 | intron_variant,non_coding_transcript_variant | MODIFIER |
| 7 | 94332503 | AX-170166379 | 0 | G | A | 0.114 | 1.16E-02 | 2.12E-01 | 2.66E+00 | 2.74E+00 | 9.58E-01 | no | Gm32647 | intron_variant,non_coding_transcript_variant | MODIFIER |
| 7 | 94333481 | AX-170029549 | 0 | G | A | 0.114 | 1.16E-02 | 2.12E-01 | 2.66E+00 | 2.74E+00 | 9.58E-01 | no | Gm32647 | intron_variant,non_coding_transcript_variant | MODIFIER |
| 7 | 94541505 | AX-170096030 | 0 | C | A | 0.114 | 1.16E-02 | 2.12E-01 | 2.66E+00 | 2.74E+00 | 9.58E-01 | no | Gm32647 | intron_variant,non_coding_transcript_variant | MODIFIER |
| 7 | 94543162 | AX-170273862 | 0 | T | C | 0.114 | 1.16E-02 | 2.12E-01 | 2.66E+00 | 2.74E+00 | 9.58E-01 | no | Gm32647 | intron_variant,non_coding_transcript_variant | MODIFIER |
| 7 | 94545640 | AX-170272056 | 0 | G | A | 0.114 | 1.16E-02 | 2.12E-01 | 2.66E+00 | 2.74E+00 | 9.58E-01 | no | Gm32647 | intron_variant,non_coding_transcript_variant | MODIFIER |
| 7 | 94546008 | AX-170352415 | 0 | A | G | 0.114 | 1.16E-02 | 2.12E-01 | 2.66E+00 | 2.74E+00 | 9.58E-01 | no | Gm32647 | intron_variant,non_coding_transcript_variant | MODIFIER |
| 7 | 94547627 | AX-170237914 | 0 | A | G | 0.114 | 1.16E-02 | 2.12E-01 | 2.66E+00 | 2.74E+00 | 9.58E-01 | no | Gm32647 | intron_variant,non_coding_transcript_variant | MODIFIER |
| 7 | 94551278 | AX-170193712 | 0 | A | T | 0.114 | 1.16E-02 | 2.12E-01 | 2.66E+00 | 2.74E+00 | 9.58E-01 | no | Gm32647 | intron_variant,non_coding_transcript_variant | MODIFIER |
| 7 | 94558605 | AX-97856548 | 0 | A | G | 0.114 | 1.16E-02 | 2.12E-01 | 2.66E+00 | 2.74E+00 | 9.58E-01 | no | Gm32647 | intron_variant,non_coding_transcript_variant | MODIFIER |
| 7 | 94577573 | AX-170166020 | 0 | G | C | 0.114 | 1.16E-02 | 2.12E-01 | 2.66E+00 | 2.74E+00 | 9.58E-01 | no | Gm32647 | intron_variant,non_coding_transcript_variant | MODIFIER |
| 7 | 94587385 | AX-97812503 | 0 | T | C | 0.114 | 1.16E-02 | 2.12E-01 | 2.66E+00 | 2.74E+00 | 9.58E-01 | no | Gm32647 | intron_variant,non_coding_transcript_variant | MODIFIER |
| 7 | 94682507 | AX-170244160 | 0 | C | T | 0.114 | 1.16E-02 | 2.12E-01 | 2.66E+00 | 2.74E+00 | 9.58E-01 | no | Gm32647 | intron_variant,non_coding_transcript_variant | MODIFIER |
| 7 | 94692237 | AX-97824023 | 0 | G | T | 0.114 | 1.16E-02 | 2.12E-01 | 2.66E+00 | 2.74E+00 | 9.58E-01 | no | Gm32647 | intron_variant,non_coding_transcript_variant | MODIFIER |
| 7 | 94751585 | AX-170301317 | 0 | T | C | 0.114 | 1.16E-02 | 2.12E-01 | 2.66E+00 | 2.74E+00 | 9.58E-01 | no | | intergenic_variant | MODIFIER |
| 7 | 94761730 | AX-170456310 | 0 | A | G | 0.114 | 1.16E-02 | 2.12E-01 | 2.66E+00 | 2.74E+00 | 9.58E-01 | no | | intergenic_variant | MODIFIER |
| 7 | 94765371 | AX-170399285 | 0 | C | T | 0.114 | 1.16E-02 | 2.12E-01 | 2.66E+00 | 2.74E+00 | 9.58E-01 | no | | intergenic_variant | MODIFIER |
| 7 | 94767003 | AX-170415484 | 0 | G | C | 0.114 | 1.16E-02 | 2.12E-01 | 2.66E+00 | 2.74E+00 | 9.58E-01 | no | | intergenic_variant | MODIFIER |
| 7 | 94774750 | AX-170499137 | 0 | C | T | 0.114 | 1.16E-02 | 2.12E-01 | 2.66E+00 | 2.74E+00 | 9.58E-01 | no | | intergenic_variant | MODIFIER |
| 7 | 94798147 | AX-170078114 | 0 | A | G | 0.114 | 1.16E-02 | 2.12E-01 | 2.66E+00 | 2.74E+00 | 9.58E-01 | no | | regulatory_region_variant | MODIFIER |
| 7 | 94798199 | AX-170306474 | 0 | A | T | 0.114 | 1.16E-02 | 2.12E-01 | 2.66E+00 | 2.74E+00 | 9.58E-01 | no | | regulatory_region_variant | MODIFIER |
| 7 | 94849677 | AX-170446573 | 0 | C | T | 0.114 | 1.16E-02 | 2.12E-01 | 2.66E+00 | 2.74E+00 | 9.58E-01 | no | | intergenic_variant | MODIFIER |
| 7 | 94861578 | AX-170029554 | 0 | T | C | 0.114 | 1.16E-02 | 2.12E-01 | 2.66E+00 | 2.74E+00 | 9.58E-01 | no | | intergenic_variant | MODIFIER |
| 7 | 94879498 | AX-170429316 | 0 | G | A | 0.114 | 1.16E-02 | 2.12E-01 | 2.66E+00 | 2.74E+00 | 9.58E-01 | no | | intergenic_variant | MODIFIER |
| 7 | 94880944 | AX-170162621 | 0 | T | A | 0.114 | 1.16E-02 | 2.12E-01 | 2.66E+00 | 2.74E+00 | 9.58E-01 | no | | intergenic_variant | MODIFIER |
| 7 | 94881691 | AX-170029556 | 0 | G | T | 0.114 | 1.16E-02 | 2.12E-01 | 2.66E+00 | 2.74E+00 | 9.58E-01 | no | | intergenic_variant | MODIFIER |
| 7 | 94967999 | AX-97802831 | 0 | G | T | 0.114 | 1.16E-02 | 2.12E-01 | 2.66E+00 | 2.74E+00 | 9.58E-01 | no | | intergenic_variant | MODIFIER |
| 7 | 95123864 | AX-170533917 | 0 | T | C | 0.114 | 1.16E-02 | 2.12E-01 | 2.66E+00 | 2.74E+00 | 9.58E-01 | no | | intergenic_variant | MODIFIER |
| 7 | 95230723 | AX-170445558 | 0 | T | C | 0.114 | 1.16E-02 | 2.12E-01 | 2.66E+00 | 2.74E+00 | 9.58E-01 | no | | intergenic_variant | MODIFIER |
| 7 | 95354899 | AX-170469906 | 0 | T | C | 0.114 | 1.16E-02 | 2.12E-01 | 2.66E+00 | 2.74E+00 | 9.58E-01 | no | | intergenic_variant | MODIFIER |
| 7 | 95401054 | AX-170029563 | 0 | A | C | 0.114 | 1.16E-02 | 2.12E-01 | 2.66E+00 | 2.74E+00 | 9.58E-01 | no | | regulatory_region_variant | MODIFIER |
| 7 | 95547673 | AX-170327651 | 0 | A | C | 0.114 | 1.16E-02 | 2.12E-01 | 2.66E+00 | 2.74E+00 | 9.58E-01 | no | | intergenic_variant | MODIFIER |

| | | | | | | | | | | | | | | | |
|---|---|---|---|---|---|---|---|---|---|---|---|---|---|---|---|
| 7 | 95549925 | AX-170493635 | 0 | T | C | 0.114 | 1.16E-02 | 2.12E-01 | 2.66E+00 | 2.74E+00 | 9.58E-01 | no | | intergenic_variant | MODIFIER |
| 7 | 95570016 | AX-170362820 | 0 | T | C | 0.114 | 1.16E-02 | 2.12E-01 | 2.66E+00 | 2.74E+00 | 9.58E-01 | no | | intergenic_variant | MODIFIER |
| 7 | 95613996 | AX-170249135 | 0 | A | G | 0.114 | 1.16E-02 | 2.12E-01 | 2.66E+00 | 2.74E+00 | 9.58E-01 | no | | intergenic_variant | MODIFIER |
| 7 | 95660870 | AX-97836601 | 0 | A | G | 0.114 | 1.16E-02 | 2.12E-01 | 2.66E+00 | 2.74E+00 | 9.58E-01 | no | | intergenic_variant | MODIFIER |
| 7 | 95680408 | AX-170201146 | 0 | C | T | 0.114 | 1.16E-02 | 2.12E-01 | 2.66E+00 | 2.74E+00 | 9.58E-01 | no | | intergenic_variant | MODIFIER |
| 7 | 95681036 | AX-97848444 | 0 | C | T | 0.114 | 1.16E-02 | 2.12E-01 | 2.66E+00 | 2.74E+00 | 9.58E-01 | no | | intergenic_variant | MODIFIER |
| 7 | 95705509 | AX-170495597 | 0 | C | T | 0.114 | 1.16E-02 | 2.12E-01 | 2.66E+00 | 2.74E+00 | 9.58E-01 | no | | intergenic_variant | MODIFIER |
| 7 | 95733936 | AX-97836594 | 0 | G | A | 0.114 | 1.16E-02 | 2.12E-01 | 2.66E+00 | 2.74E+00 | 9.58E-01 | no | | intergenic_variant | MODIFIER |
| 7 | 95775226 | AX-170507144 | 0 | T | C | 0.114 | 1.16E-02 | 2.12E-01 | 2.66E+00 | 2.74E+00 | 9.58E-01 | no | | intergenic_variant | MODIFIER |
| 7 | 95801713 | AX-170362019 | 0 | C | T | 0.114 | 1.16E-02 | 2.12E-01 | 2.66E+00 | 2.74E+00 | 9.58E-01 | no | | intergenic_variant | MODIFIER |
| 7 | 95875409 | AX-170438302 | 0 | A | T | 0.114 | 1.16E-02 | 2.12E-01 | 2.66E+00 | 2.74E+00 | 9.58E-01 | no | | intergenic_variant | MODIFIER |
| 7 | 95995347 | AX-170317731 | 0 | A | G | 0.114 | 1.16E-02 | 2.12E-01 | 2.66E+00 | 2.74E+00 | 9.58E-01 | no | | intergenic_variant | MODIFIER |
| 7 | 96007244 | AX-170396940 | 0 | C | T | 0.114 | 1.16E-02 | 2.12E-01 | 2.66E+00 | 2.74E+00 | 9.58E-01 | no | | intergenic_variant | MODIFIER |
| 7 | 96037457 | AX-170430314 | 0 | A | G | 0.114 | 1.16E-02 | 2.12E-01 | 2.66E+00 | 2.74E+00 | 9.58E-01 | no | | intergenic_variant | MODIFIER |
| 7 | 96037514 | AX-170504841 | 0 | T | C | 0.114 | 1.16E-02 | 2.12E-01 | 2.66E+00 | 2.74E+00 | 9.58E-01 | no | | intergenic_variant | MODIFIER |
| 7 | 96078881 | AX-170143275 | 0 | T | G | 0.114 | 1.16E-02 | 2.12E-01 | 2.66E+00 | 2.74E+00 | 9.58E-01 | no | | intergenic_variant | MODIFIER |
| 7 | 96242281 | AX-170381970 | 0 | A | C | 0.114 | 1.16E-02 | 2.12E-01 | 2.66E+00 | 2.74E+00 | 9.58E-01 | no | Tenm4 | intron_variant | MODIFIER |
| 7 | 96253608 | AX-170214348 | 0 | C | G | 0.114 | 1.16E-02 | 2.12E-01 | 2.66E+00 | 2.74E+00 | 9.58E-01 | no | Mir708 | downstream_gene_variant | MODIFIER |
| 7 | 96295403 | AX-170265378 | 0 | T | C | 0.114 | 1.16E-02 | 2.12E-01 | 2.66E+00 | 2.74E+00 | 9.58E-01 | no | Tenm4 | intron_variant | MODIFIER |
| 7 | 96327577 | AX-170251918 | 0 | A | C | 0.114 | 1.16E-02 | 2.12E-01 | 2.66E+00 | 2.74E+00 | 9.58E-01 | no | Tenm4 | intron_variant | MODIFIER |
| 7 | 96334908 | AX-170362614 | 0 | A | G | 0.114 | 1.16E-02 | 2.12E-01 | 2.66E+00 | 2.74E+00 | 9.58E-01 | no | Tenm4 | intron_variant | MODIFIER |
| 7 | 96376908 | AX-170333383 | 0 | T | C | 0.114 | 1.16E-02 | 2.12E-01 | 2.66E+00 | 2.74E+00 | 9.58E-01 | no | Tenm4 | intron_variant | MODIFIER |
| 7 | 96409159 | AX-170434010 | 0 | T | G | 0.114 | 1.16E-02 | 2.12E-01 | 2.66E+00 | 2.74E+00 | 9.58E-01 | no | Tenm4 | intron_variant | MODIFIER |
| 7 | 96411103 | AX-170437396 | 0 | C | T | 0.114 | 1.16E-02 | 2.12E-01 | 2.66E+00 | 2.74E+00 | 9.58E-01 | no | Tenm4 | intron_variant | MODIFIER |
| 7 | 96411625 | AX-170096034 | 0 | G | C | 0.114 | 1.16E-02 | 2.12E-01 | 2.66E+00 | 2.74E+00 | 9.58E-01 | no | Tenm4 | intron_variant | MODIFIER |
| 7 | 96426219 | AX-170114573 | 0 | T | G | 0.114 | 1.16E-02 | 2.12E-01 | 2.66E+00 | 2.74E+00 | 9.58E-01 | no | Tenm4 | intron_variant | MODIFIER |
| 7 | 96457111 | AX-170281051 | 0 | T | G | 0.114 | 1.16E-02 | 2.12E-01 | 2.66E+00 | 2.74E+00 | 9.58E-01 | no | Tenm4 | intron_variant | MODIFIER |
| 7 | 96534729 | AX-97808030 | 0 | C | T | 0.114 | 1.16E-02 | 2.12E-01 | 2.66E+00 | 2.74E+00 | 9.58E-01 | no | Tenm4 | intron_variant | MODIFIER |
| 7 | 96554827 | AX-170303024 | 0 | A | C | 0.114 | 1.16E-02 | 2.12E-01 | 2.66E+00 | 2.74E+00 | 9.58E-01 | no | Tenm4 | intron_variant | MODIFIER |
| 7 | 96561182 | AX-170488633 | 0 | A | C | 0.114 | 1.16E-02 | 2.12E-01 | 2.66E+00 | 2.74E+00 | 9.58E-01 | no | Tenm4 | intron_variant | MODIFIER |
| 7 | 96575014 | AX-170158558 | 0 | A | G | 0.114 | 1.16E-02 | 2.12E-01 | 2.66E+00 | 2.74E+00 | 9.58E-01 | no | Tenm4 | intron_variant | MODIFIER |
| 7 | 96575688 | AX-170366874 | 0 | T | C | 0.114 | 1.16E-02 | 2.12E-01 | 2.66E+00 | 2.74E+00 | 9.58E-01 | no | Tenm4 | intron_variant | MODIFIER |
| 7 | 96579575 | AX-170530711 | 0 | G | A | 0.114 | 1.16E-02 | 2.12E-01 | 2.66E+00 | 2.74E+00 | 9.58E-01 | no | Tenm4 | intron_variant | MODIFIER |
| 7 | 96589160 | AX-170320352 | 0 | G | A | 0.114 | 1.16E-02 | 2.12E-01 | 2.66E+00 | 2.74E+00 | 9.58E-01 | no | Tenm4 | intron_variant | MODIFIER |
| 7 | 96607777 | AX-170078157 | 0 | G | A | 0.114 | 1.16E-02 | 2.12E-01 | 2.66E+00 | 2.74E+00 | 9.58E-01 | no | Tenm4 | intron_variant | MODIFIER |
| 7 | 96660783 | AX-170106841 | 0 | G | C | 0.114 | 1.16E-02 | 2.12E-01 | 2.66E+00 | 2.74E+00 | 9.58E-01 | no | Tenm4 | intron_variant | MODIFIER |
| 7 | 96660822 | AX-170286666 | 0 | C | T | 0.114 | 1.16E-02 | 2.12E-01 | 2.66E+00 | 2.74E+00 | 9.58E-01 | no | Tenm4 | intron_variant | MODIFIER |
| 7 | 96660896 | AX-170441165 | 0 | A | G | 0.114 | 1.16E-02 | 2.12E-01 | 2.66E+00 | 2.74E+00 | 9.58E-01 | no | Tenm4 | intron_variant | MODIFIER |
| 7 | 96702983 | AX-170416913 | 0 | T | C | 0.114 | 1.16E-02 | 2.12E-01 | 2.66E+00 | 2.74E+00 | 9.58E-01 | no | Tenm4 | intron_variant | MODIFIER |
| 7 | 96703471 | AX-170265684 | 0 | T | C | 0.114 | 1.16E-02 | 2.12E-01 | 2.66E+00 | 2.74E+00 | 9.58E-01 | no | Tenm4 | intron_variant | MODIFIER |
| 7 | 96721239 | AX-170096035 | 0 | A | T | 0.114 | 1.16E-02 | 2.12E-01 | 2.66E+00 | 2.74E+00 | 9.58E-01 | no | Tenm4 | intron_variant | MODIFIER |
| 7 | 96721731 | AX-170286747 | 0 | T | C | 0.114 | 1.16E-02 | 2.12E-01 | 2.66E+00 | 2.74E+00 | 9.58E-01 | no | Tenm4 | intron_variant | MODIFIER |
| 7 | 96723554 | AX-170314860 | 0 | A | T | 0.114 | 1.16E-02 | 2.12E-01 | 2.66E+00 | 2.74E+00 | 9.58E-01 | no | Tenm4 | intron_variant | MODIFIER |
| 7 | 96738324 | AX-170511313 | 0 | A | G | 0.114 | 1.16E-02 | 2.12E-01 | 2.66E+00 | 2.74E+00 | 9.58E-01 | no | Tenm4 | intron_variant | MODIFIER |
| 7 | 96815296 | AX-170455934 | 0 | A | G | 0.114 | 1.16E-02 | 2.12E-01 | 2.66E+00 | 2.74E+00 | 9.58E-01 | no | Tenm4 | intron_variant | MODIFIER |
| 7 | 96820877 | AX-97831942 | 0 | A | G | 0.114 | 1.16E-02 | 2.12E-01 | 2.66E+00 | 2.74E+00 | 9.58E-01 | no | Tenm4 | intron_variant | MODIFIER |
| 8 | 95972161 | AX-170080889 | 0 | G | T | 0.228 | 6.05E-03 | 1.17E-01 | 2.68E+00 | 2.74E+00 | 9.58E-01 | no | | intergenic_variant | MODIFIER |
| 13 | 45200458 | AX-170008237 | 0 | G | T | 0.474 | -2.55E-03 | 6.16E-02 | 2.64E+00 | 2.75E+00 | 9.58E-01 | no | | intergenic_variant | MODIFIER |
| 13 | 45223438 | AX-97824246 | 0 | C | T | 0.474 | -2.55E-03 | 6.16E-02 | 2.64E+00 | 2.75E+00 | 9.58E-01 | no | | intergenic_variant | MODIFIER |
| 13 | 45225023 | AX-170453134 | 0 | G | A | 0.474 | -2.55E-03 | 6.16E-02 | 2.64E+00 | 2.75E+00 | 9.58E-01 | no | | intergenic_variant | MODIFIER |
| 13 | 45231592 | AX-170153377 | 0 | G | A | 0.474 | -2.55E-03 | 6.16E-02 | 2.64E+00 | 2.75E+00 | 9.58E-01 | no | Gm47448 | upstream_gene_variant | MODIFIER |
| 13 | 45245803 | AX-170371641 | 0 | T | G | 0.474 | -2.55E-03 | 6.16E-02 | 2.64E+00 | 2.75E+00 | 9.58E-01 | no | Gm47448 | intron_variant,non_coding_transcript_variant | MODIFIER |
| 13 | 45250710 | AX-170288507 | 0 | T | G | 0.474 | -2.55E-03 | 6.16E-02 | 2.64E+00 | 2.75E+00 | 9.58E-01 | no | Gm47448 | intron_variant,non_coding_transcript_variant | MODIFIER |
| 13 | 45252102 | AX-97829693 | 0 | A | G | 0.474 | -2.55E-03 | 6.16E-02 | 2.64E+00 | 2.75E+00 | 9.58E-01 | no | Gm47448 | intron_variant,non_coding_transcript_variant | MODIFIER |

| | | | | | | | | | | | | | | | |
|---|---|---|---|---|---|---|---|---|---|---|---|---|---|---|---|
| 13 | 45269790 | AX-97817893 | 0 | A | G | 0.474 | -2.55E-03 | 6.16E-02 | 2.64E+00 | 2.75E+00 | 9.58E-01 | no | Gm47448 | intron_variant,non_coding_transcript_variant | MODIFIER |
| 13 | 45270059 | AX-170492610 | 0 | T | G | 0.474 | -2.55E-03 | 6.16E-02 | 2.64E+00 | 2.75E+00 | 9.58E-01 | no | Gm47448 | intron_variant,non_coding_transcript_variant | MODIFIER |
| 13 | 45275528 | AX-170141248 | 0 | T | G | 0.474 | -2.55E-03 | 6.16E-02 | 2.64E+00 | 2.75E+00 | 9.58E-01 | no | Gm47448 | intron_variant,non_coding_transcript_variant | MODIFIER |
| 13 | 45278350 | AX-170309135 | 0 | G | A | 0.474 | -2.55E-03 | 6.16E-02 | 2.64E+00 | 2.75E+00 | 9.58E-01 | no | Gm47448 | intron_variant,non_coding_transcript_variant | MODIFIER |
| 13 | 45281420 | AX-170530924 | 0 | C | T | 0.474 | -2.55E-03 | 6.16E-02 | 2.64E+00 | 2.75E+00 | 9.58E-01 | no | Gm47448 | intron_variant,non_coding_transcript_variant | MODIFIER |
| 13 | 45290886 | AX-170189048 | 0 | G | A | 0.474 | -2.55E-03 | 6.16E-02 | 2.64E+00 | 2.75E+00 | 9.58E-01 | no | Gm23755 | downstream_gene_variant | MODIFIER |
| 13 | 45298132 | AX-170482251 | 0 | C | T | 0.474 | -2.55E-03 | 6.16E-02 | 2.64E+00 | 2.75E+00 | 9.58E-01 | no | Gm23755 | upstream_gene_variant | MODIFIER |
| 13 | 45338700 | AX-170220681 | 0 | G | A | 0.474 | -2.55E-03 | 6.16E-02 | 2.64E+00 | 2.75E+00 | 9.58E-01 | no | Gm47448 | downstream_gene_variant | MODIFIER |
| 13 | 45349804 | AX-170427548 | 0 | A | G | 0.474 | -2.55E-03 | 6.16E-02 | 2.64E+00 | 2.75E+00 | 9.58E-01 | no | | intergenic_variant | MODIFIER |
| 13 | 45350053 | AX-170404438 | 0 | A | G | 0.474 | -2.55E-03 | 6.16E-02 | 2.64E+00 | 2.75E+00 | 9.58E-01 | no | | intergenic_variant | MODIFIER |
| 13 | 45351803 | AX-170482912 | 0 | T | A | 0.474 | -2.55E-03 | 6.16E-02 | 2.64E+00 | 2.75E+00 | 9.58E-01 | no | | intergenic_variant | MODIFIER |
| 13 | 45363333 | AX-170472774 | 0 | G | A | 0.474 | -2.55E-03 | 6.16E-02 | 2.64E+00 | 2.75E+00 | 9.58E-01 | no | | intergenic_variant | MODIFIER |
| 13 | 45366396 | AX-170294678 | 0 | G | A | 0.474 | -2.55E-03 | 6.16E-02 | 2.64E+00 | 2.75E+00 | 9.58E-01 | no | | intergenic_variant | MODIFIER |
| 13 | 45372511 | AX-170529467 | 0 | T | G | 0.474 | -2.55E-03 | 6.16E-02 | 2.64E+00 | 2.75E+00 | 9.58E-01 | no | | intergenic_variant | MODIFIER |
| 13 | 45424936 | AX-170420635 | 0 | G | A | 0.474 | -2.55E-03 | 6.16E-02 | 2.64E+00 | 2.75E+00 | 9.58E-01 | no | | intergenic_variant | MODIFIER |
| 11 | 46023324 | AX-170381486 | 0 | G | A | 0.088 | -1.09E-02 | 1.82E-01 | 2.63E+00 | 2.73E+00 | 9.58E-01 | no | | intergenic_variant | MODIFIER |
| 9 | 29747785 | AX-170370167 | 0 | G | A | 0.439 | 4.97E-03 | 8.52E-02 | 2.66E+00 | 2.73E+00 | 9.58E-01 | no | Ntm | intron_variant | MODIFIER |
| 5 | 70714881 | AX-170422998 | 1 | A | T | 0.446 | 5.66E-03 | 9.70E-02 | 2.65E+00 | 2.73E+00 | 9.58E-01 | no | | intergenic_variant | MODIFIER |
| 8 | 34939742 | AX-170030493 | 0 | G | A | 0.158 | -8.50E-03 | 1.39E-01 | 2.66E+00 | 2.73E+00 | 9.58E-01 | no | Tnks | intron_variant | MODIFIER |
| 8 | 19168378 | AX-170513674 | 1 | C | G | 0.179 | -5.44E-03 | 1.09E-01 | 2.66E+00 | 2.74E+00 | 9.58E-01 | no | Gm39139 | upstream_gene_variant | MODIFIER |
| 9 | 106825940 | AX-170265051 | 0 | T | A | 0.14 | -6.38E-03 | 1.01E-01 | 2.64E+00 | 2.73E+00 | 9.58E-01 | no | Dcaf1 | upstream_gene_variant | MODIFIER |
| 1 | 39572889 | AX-97841942 | 0 | C | T | 0.412 | 3.51E-03 | 7.61E-02 | 2.67E+00 | 2.74E+00 | 9.59E-01 | no | Rnf149 | intron_variant | MODIFIER |
| 1 | 53715656 | AX-170000515 | 1 | A | G | 0.098 | 1.11E-02 | 2.10E-01 | 2.65E+00 | 2.74E+00 | 9.59E-01 | no | | regulatory_region_variant | MODIFIER |
| 1 | 55227912 | AX-170035925 | 1 | A | G | 0.098 | 1.11E-02 | 2.10E-01 | 2.65E+00 | 2.74E+00 | 9.59E-01 | no | Rftn2 | upstream_gene_variant | MODIFIER |
| 1 | 55400349 | AX-170417554 | 1 | C | T | 0.098 | 1.11E-02 | 2.10E-01 | 2.65E+00 | 2.74E+00 | 9.59E-01 | no | | intergenic_variant | MODIFIER |
| 18 | 58293919 | AX-170088349 | 1 | C | T | 0.393 | 7.40E-03 | 9.30E-02 | 2.62E+00 | 2.71E+00 | 9.59E-01 | no | | intergenic_variant | MODIFIER |
| 8 | 97662869 | AX-170175877 | 0 | T | C | 0.219 | 5.47E-03 | 1.08E-01 | 2.66E+00 | 2.74E+00 | 9.59E-01 | no | | intergenic_variant | MODIFIER |
| 10 | 15713165 | AX-170002805 | 0 | T | C | 0.184 | 4.47E-03 | 1.08E-01 | 2.67E+00 | 2.75E+00 | 9.59E-01 | no | Gm40604 | intron_variant,non_coding_transcript_variant | MODIFIER |
| 10 | 15765155 | AX-170002807 | 0 | A | C | 0.184 | 4.47E-03 | 1.08E-01 | 2.67E+00 | 2.75E+00 | 9.59E-01 | no | Gm47712 | intron_variant,non_coding_transcript_variant | MODIFIER |
| 1 | 69844871 | AX-170000759 | 1 | G | A | 0.107 | -9.52E-03 | 1.94E-01 | 2.64E+00 | 2.74E+00 | 9.59E-01 | no | Spag16 | intron_variant | MODIFIER |
| 1 | 11141426 | AX-170189552 | 0 | A | G | 0.395 | -3.10E-03 | 7.29E-02 | 2.69E+00 | 2.75E+00 | 9.59E-01 | no | Prex2 | intron_variant | MODIFIER |
| 1 | 11180533 | AX-170040373 | 0 | T | C | 0.395 | -3.10E-03 | 7.29E-02 | 2.69E+00 | 2.75E+00 | 9.59E-01 | no | Prex2 | intron_variant | MODIFIER |
| 1 | 11186773 | AX-170040372 | 0 | A | C | 0.395 | -3.10E-03 | 7.29E-02 | 2.69E+00 | 2.75E+00 | 9.59E-01 | no | Prex2 | splice_region_variant,intron_variant | LOW |
| 1 | 11186867 | AX-170105714 | 0 | C | G | 0.395 | -3.10E-03 | 7.29E-02 | 2.69E+00 | 2.75E+00 | 9.59E-01 | no | Prex2 | intron_variant | MODIFIER |
| 1 | 11197722 | AX-170270303 | 0 | T | C | 0.395 | -3.10E-03 | 7.29E-02 | 2.69E+00 | 2.75E+00 | 9.59E-01 | no | Prex2 | intron_variant | MODIFIER |
| 1 | 11199663 | AX-170154789 | 0 | G | T | 0.395 | -3.10E-03 | 7.29E-02 | 2.69E+00 | 2.75E+00 | 9.59E-01 | no | Prex2 | intron_variant | MODIFIER |
| 1 | 11202757 | AX-170207319 | 0 | C | T | 0.395 | -3.10E-03 | 7.29E-02 | 2.69E+00 | 2.75E+00 | 9.59E-01 | no | Prex2 | intron_variant | MODIFIER |
| 1 | 11288572 | AX-170227930 | 0 | T | C | 0.395 | -3.10E-03 | 7.29E-02 | 2.69E+00 | 2.75E+00 | 9.59E-01 | no | Prex2 | intron_variant | MODIFIER |
| 1 | 11305877 | AX-170245345 | 0 | C | G | 0.395 | -3.10E-03 | 7.29E-02 | 2.69E+00 | 2.75E+00 | 9.59E-01 | no | Prex2 | downstream_gene_variant | MODIFIER |
| 1 | 11357180 | AX-170418664 | 0 | T | C | 0.395 | -3.10E-03 | 7.29E-02 | 2.69E+00 | 2.75E+00 | 9.59E-01 | no | | intergenic_variant | MODIFIER |
| 1 | 11380074 | AX-170390272 | 0 | G | T | 0.395 | -3.10E-03 | 7.29E-02 | 2.69E+00 | 2.75E+00 | 9.59E-01 | no | | intergenic_variant | MODIFIER |
| 1 | 11421985 | AX-170451528 | 0 | C | T | 0.395 | -3.10E-03 | 7.29E-02 | 2.69E+00 | 2.75E+00 | 9.59E-01 | no | A830018L16Rik | intron_variant | MODIFIER |
| 1 | 11423254 | AX-170256465 | 0 | T | C | 0.395 | -3.10E-03 | 7.29E-02 | 2.69E+00 | 2.75E+00 | 9.59E-01 | no | A830018L16Rik | intron_variant | MODIFIER |
| 1 | 11435898 | AX-97829775 | 0 | G | A | 0.395 | -3.10E-03 | 7.29E-02 | 2.69E+00 | 2.75E+00 | 9.59E-01 | no | A830018L16Rik | intron_variant | MODIFIER |
| 1 | 11452131 | AX-170103230 | 0 | T | C | 0.395 | -3.10E-03 | 7.29E-02 | 2.69E+00 | 2.75E+00 | 9.59E-01 | no | A830018L16Rik | intron_variant | MODIFIER |
| 1 | 11464962 | AX-170177035 | 0 | A | G | 0.395 | -3.10E-03 | 7.29E-02 | 2.69E+00 | 2.75E+00 | 9.59E-01 | no | A830018L16Rik | intron_variant | MODIFIER |
| 1 | 11504847 | AX-97795868 | 0 | A | G | 0.395 | -3.10E-03 | 7.29E-02 | 2.69E+00 | 2.75E+00 | 9.59E-01 | no | A830018L16Rik | intron_variant | MODIFIER |
| 1 | 11515339 | AX-170311474 | 0 | C | A | 0.395 | -3.10E-03 | 7.29E-02 | 2.69E+00 | 2.75E+00 | 9.59E-01 | no | A830018L16Rik | intron_variant | MODIFIER |
| 1 | 11516744 | AX-170280952 | 0 | A | G | 0.395 | -3.10E-03 | 7.29E-02 | 2.69E+00 | 2.75E+00 | 9.59E-01 | no | A830018L16Rik | intron_variant | MODIFIER |
| 1 | 11516776 | AX-170420147 | 0 | T | C | 0.395 | -3.10E-03 | 7.29E-02 | 2.69E+00 | 2.75E+00 | 9.59E-01 | no | A830018L16Rik | intron_variant | MODIFIER |
| 1 | 11516996 | AX-170040379 | 0 | C | T | 0.395 | -3.10E-03 | 7.29E-02 | 2.69E+00 | 2.75E+00 | 9.59E-01 | no | A830018L16Rik | intron_variant | MODIFIER |
| 1 | 11552923 | AX-170450967 | 0 | A | G | 0.395 | -3.10E-03 | 7.29E-02 | 2.69E+00 | 2.75E+00 | 9.59E-01 | no | A830018L16Rik | intron_variant | MODIFIER |
| 1 | 11555709 | AX-170337649 | 0 | T | C | 0.395 | -3.10E-03 | 7.29E-02 | 2.69E+00 | 2.75E+00 | 9.59E-01 | no | A830018L16Rik | intron_variant | MODIFIER |
| 1 | 11555973 | AX-97821935 | 0 | G | A | 0.395 | -3.10E-03 | 7.29E-02 | 2.69E+00 | 2.75E+00 | 9.59E-01 | no | A830018L16Rik | intron_variant | MODIFIER |

| | | | | | | | | | | | | | | | |
|---|---|---|---|---|---|---|---|---|---|---|---|---|---|---|---|
| 1 | 11596048 | AX-170431432 | 0 | T | G | 0.395 | -3.10E-03 | 7.29E-02 | 2.69E+00 | 2.75E+00 | 9.59E-01 | no | A830018L16Rik | intron_variant | MODIFIER |
| 1 | 11606500 | AX-97848739 | 0 | G | T | 0.395 | -3.10E-03 | 7.29E-02 | 2.69E+00 | 2.75E+00 | 9.59E-01 | no | A830018L16Rik | intron_variant | MODIFIER |
| 1 | 11633153 | AX-170171245 | 0 | T | C | 0.395 | -3.10E-03 | 7.29E-02 | 2.69E+00 | 2.75E+00 | 9.59E-01 | no | A830018L16Rik | intron_variant | MODIFIER |
| 1 | 11633594 | AX-170383838 | 0 | A | G | 0.395 | -3.10E-03 | 7.29E-02 | 2.69E+00 | 2.75E+00 | 9.59E-01 | no | A830018L16Rik | intron_variant | MODIFIER |
| 1 | 11653006 | AX-170507478 | 0 | G | A | 0.395 | -3.10E-03 | 7.29E-02 | 2.69E+00 | 2.75E+00 | 9.59E-01 | no | A830018L16Rik | intron_variant | MODIFIER |
| 1 | 11696282 | AX-170178549 | 0 | A | G | 0.395 | -3.10E-03 | 7.29E-02 | 2.69E+00 | 2.75E+00 | 9.59E-01 | no | A830018L16Rik | intron_variant | MODIFIER |
| 1 | 7940508 | AX-170482892 | 1 | G | A | 0.42 | -2.86E-03 | 7.13E-02 | 2.68E+00 | 2.75E+00 | 9.59E-01 | no | | intergenic_variant | MODIFIER |
| 8 | 32371189 | AX-170354364 | 0 | T | C | 0.149 | 5.62E-03 | 1.18E-01 | 2.64E+00 | 2.74E+00 | 9.59E-01 | no | Nrg1 | intron_variant | MODIFIER |
| 8 | 55726159 | AX-170030762 | 1 | T | C | 0.152 | 6.36E-03 | 1.51E-01 | 2.70E+00 | 2.75E+00 | 9.59E-01 | no | | intergenic_variant | MODIFIER |
| 11 | 34861405 | AX-170004673 | 1 | G | C | 0.098 | -8.44E-03 | 1.89E-01 | 2.66E+00 | 2.74E+00 | 9.59E-01 | no | | intergenic_variant | MODIFIER |
| 1 | 187684747 | AX-170182631 | 0 | A | C | 0.246 | -1.57E-03 | 9.40E-02 | 2.62E+00 | 2.76E+00 | 9.59E-01 | no | Esrrg | intron_variant | MODIFIER |
| 7 | 87672904 | AX-97796465 | 1 | C | T | 0.116 | 1.12E-02 | 2.12E-01 | 2.65E+00 | 2.74E+00 | 9.59E-01 | no | Grm5 | intron_variant | MODIFIER |
| 12 | 34878343 | AX-170325453 | 1 | A | G | 0.036 | 8.64E-03 | 1.30E-01 | 2.55E+00 | 2.73E+00 | 9.60E-01 | no | 4921508M14Rik | upstream_gene_variant | MODIFIER |
| 8 | 29928070 | AX-170030432 | 0 | A | G | 0.167 | -6.04E-03 | 1.10E-01 | 2.65E+00 | 2.73E+00 | 9.60E-01 | no | | intergenic_variant | MODIFIER |
| 7 | 87437537 | AX-170029460 | 0 | G | A | 0.123 | 1.02E-02 | 1.74E-01 | 2.61E+00 | 2.73E+00 | 9.60E-01 | no | Tyr | intron_variant | MODIFIER |
| 10 | 32415917 | AX-170003012 | 0 | A | G | 0.395 | 3.49E-03 | 7.38E-02 | 2.62E+00 | 2.74E+00 | 9.60E-01 | no | Nkain2 | intron_variant | MODIFIER |
| 9 | 61200246 | AX-170106151 | 1 | G | A | 0.205 | 6.59E-03 | 1.36E-01 | 2.71E+00 | 2.74E+00 | 9.60E-01 | no | | regulatory_region_variant | MODIFIER |
| 9 | 62643078 | AX-97833631 | 1 | G | A | 0.205 | 6.59E-03 | 1.36E-01 | 2.71E+00 | 2.74E+00 | 9.60E-01 | no | | intergenic_variant | MODIFIER |
| 3 | 147742667 | AX-170171520 | 0 | G | A | 0.36 | -3.06E-03 | 7.13E-02 | 2.66E+00 | 2.75E+00 | 9.60E-01 | no | | intergenic_variant | MODIFIER |
| 16 | 18864939 | AX-97781500 | 0 | T | G | 0.132 | -4.27E-03 | 8.78E-02 | 2.59E+00 | 2.74E+00 | 9.60E-01 | no | Gm28539 | intron_variant,NMD_transcript_variant | MODIFIER |
| 9 | 77664863 | AX-170160914 | 1 | G | C | 0.188 | 6.55E-03 | 1.37E-01 | 2.72E+00 | 2.74E+00 | 9.60E-01 | no | Klhl31 | downstream_gene_variant | MODIFIER |
| 8 | 35517639 | AX-170079738 | 0 | T | C | 0.158 | -7.76E-03 | 1.34E-01 | 2.65E+00 | 2.73E+00 | 9.60E-01 | no | | regulatory_region_variant | MODIFIER |
| 1 | 70073706 | AX-170041576 | 1 | A | G | 0.143 | 1.19E-02 | 2.36E-01 | 2.68E+00 | 2.74E+00 | 9.60E-01 | no | Spag16 | intron_variant | MODIFIER |
| 9 | 53956762 | AX-170286375 | 1 | G | T | 0.205 | 5.67E-03 | 1.16E-01 | 2.67E+00 | 2.74E+00 | 9.60E-01 | no | Elmod1 | intron_variant | MODIFIER |
| 1 | 41424789 | AX-170413838 | 1 | C | T | 0.429 | -3.98E-03 | 7.50E-02 | 2.65E+00 | 2.73E+00 | 9.60E-01 | no | | intergenic_variant | MODIFIER |
| 18 | 67005380 | AX-170463568 | 1 | G | A | 0.286 | 4.52E-03 | 9.41E-02 | 2.66E+00 | 2.74E+00 | 9.60E-01 | no | | intergenic_variant | MODIFIER |
| 1 | 41706086 | AX-97810095 | 0 | T | C | 0.421 | 3.55E-03 | 7.58E-02 | 2.65E+00 | 2.74E+00 | 9.60E-01 | no | | intergenic_variant | MODIFIER |
| 9 | 85977708 | AX-170371082 | 1 | C | T | 0.143 | 4.81E-03 | 1.25E-01 | 2.67E+00 | 2.75E+00 | 9.60E-01 | no | | intergenic_variant | MODIFIER |
| 9 | 60171780 | AX-170395810 | 0 | T | C | 0.202 | 6.41E-03 | 1.35E-01 | 2.71E+00 | 2.74E+00 | 9.60E-01 | no | Thsd4 | intron_variant | MODIFIER |
| 9 | 60180787 | AX-97804077 | 0 | A | G | 0.202 | 6.41E-03 | 1.35E-01 | 2.71E+00 | 2.74E+00 | 9.60E-01 | no | Thsd4 | intron_variant | MODIFIER |
| 9 | 60197308 | AX-170128006 | 0 | G | C | 0.202 | 6.41E-03 | 1.35E-01 | 2.71E+00 | 2.74E+00 | 9.60E-01 | no | Thsd4 | intron_variant | MODIFIER |
| 9 | 60197580 | AX-170372328 | 0 | A | C | 0.202 | 6.41E-03 | 1.35E-01 | 2.71E+00 | 2.74E+00 | 9.60E-01 | no | Thsd4 | intron_variant | MODIFIER |
| 9 | 60197745 | AX-97857310 | 0 | G | A | 0.202 | 6.41E-03 | 1.35E-01 | 2.71E+00 | 2.74E+00 | 9.60E-01 | no | Thsd4 | intron_variant | MODIFIER |
| 9 | 60198185 | AX-170235681 | 0 | C | T | 0.202 | 6.41E-03 | 1.35E-01 | 2.71E+00 | 2.74E+00 | 9.60E-01 | no | Thsd4 | intron_variant | MODIFIER |
| 9 | 60198270 | AX-170131322 | 0 | A | G | 0.202 | 6.41E-03 | 1.35E-01 | 2.71E+00 | 2.74E+00 | 9.60E-01 | no | Thsd4 | intron_variant | MODIFIER |
| 9 | 60198577 | AX-170335423 | 0 | A | G | 0.202 | 6.41E-03 | 1.35E-01 | 2.71E+00 | 2.74E+00 | 9.60E-01 | no | Thsd4 | intron_variant | MODIFIER |
| 9 | 60209141 | AX-170297567 | 0 | A | G | 0.202 | 6.41E-03 | 1.35E-01 | 2.71E+00 | 2.74E+00 | 9.60E-01 | no | Thsd4 | intron_variant | MODIFIER |
| 9 | 60221308 | AX-97844613 | 0 | A | G | 0.202 | 6.41E-03 | 1.35E-01 | 2.71E+00 | 2.74E+00 | 9.60E-01 | no | Thsd4 | intron_variant | MODIFIER |
| 9 | 60224301 | AX-170312269 | 0 | A | G | 0.202 | 6.41E-03 | 1.35E-01 | 2.71E+00 | 2.74E+00 | 9.60E-01 | no | Thsd4 | intron_variant | MODIFIER |
| 9 | 60251079 | AX-170385667 | 0 | C | A | 0.202 | 6.41E-03 | 1.35E-01 | 2.71E+00 | 2.74E+00 | 9.60E-01 | no | Thsd4 | intron_variant | MODIFIER |
| 9 | 60251742 | AX-97832426 | 0 | G | T | 0.202 | 6.41E-03 | 1.35E-01 | 2.71E+00 | 2.74E+00 | 9.60E-01 | no | Thsd4 | intron_variant | MODIFIER |
| 9 | 60269846 | AX-170450249 | 0 | A | T | 0.202 | 6.41E-03 | 1.35E-01 | 2.71E+00 | 2.74E+00 | 9.60E-01 | no | Thsd4 | intron_variant | MODIFIER |
| 9 | 60273272 | AX-170146364 | 0 | C | A | 0.202 | 6.41E-03 | 1.35E-01 | 2.71E+00 | 2.74E+00 | 9.60E-01 | no | Thsd4 | intron_variant | MODIFIER |
| 9 | 60280519 | AX-170229928 | 0 | A | G | 0.202 | 6.41E-03 | 1.35E-01 | 2.71E+00 | 2.74E+00 | 9.60E-01 | no | Thsd4 | intron_variant | MODIFIER |
| 9 | 60282601 | AX-170219849 | 0 | G | A | 0.202 | 6.41E-03 | 1.35E-01 | 2.71E+00 | 2.74E+00 | 9.60E-01 | no | Thsd4 | intron_variant | MODIFIER |
| 9 | 60290421 | AX-97805659 | 0 | A | T | 0.202 | 6.41E-03 | 1.35E-01 | 2.71E+00 | 2.74E+00 | 9.60E-01 | no | Thsd4 | intron_variant | MODIFIER |
| 9 | 60321248 | AX-170438016 | 0 | A | G | 0.202 | 6.41E-03 | 1.35E-01 | 2.71E+00 | 2.74E+00 | 9.60E-01 | no | Thsd4 | intron_variant | MODIFIER |
| 9 | 60321618 | AX-170515780 | 0 | G | A | 0.202 | 6.41E-03 | 1.35E-01 | 2.71E+00 | 2.74E+00 | 9.60E-01 | no | Thsd4 | intron_variant | MODIFIER |
| 9 | 60327328 | AX-170082573 | 0 | G | C | 0.202 | 6.41E-03 | 1.35E-01 | 2.71E+00 | 2.74E+00 | 9.60E-01 | no | Thsd4 | intron_variant | MODIFIER |
| 9 | 60327487 | AX-170082575 | 0 | A | G | 0.202 | 6.41E-03 | 1.35E-01 | 2.71E+00 | 2.74E+00 | 9.60E-01 | no | Thsd4 | intron_variant | MODIFIER |
| 9 | 60330457 | AX-97853582 | 0 | G | A | 0.202 | 6.41E-03 | 1.35E-01 | 2.71E+00 | 2.74E+00 | 9.60E-01 | no | Thsd4 | intron_variant | MODIFIER |
| 9 | 60331005 | AX-170082574 | 0 | A | G | 0.202 | 6.41E-03 | 1.35E-01 | 2.71E+00 | 2.74E+00 | 9.60E-01 | no | Thsd4 | intron_variant | MODIFIER |
| 9 | 60331545 | AX-170501917 | 0 | A | C | 0.202 | 6.41E-03 | 1.35E-01 | 2.71E+00 | 2.74E+00 | 9.60E-01 | no | Thsd4 | intron_variant | MODIFIER |
| 9 | 60331700 | AX-170288822 | 0 | G | T | 0.202 | 6.41E-03 | 1.35E-01 | 2.71E+00 | 2.74E+00 | 9.60E-01 | no | Thsd4 | intron_variant | MODIFIER |

| | | | | | | | | | | | | | | | |
|---|---|---|---|---|---|---|---|---|---|---|---|---|---|---|---|
| 9 | 60332498 | AX-170032479 | 0 | G | T | 0.202 | 6.41E-03 | 1.35E-01 | 2.71E+00 | 2.74E+00 | 9.60E-01 | no | Thsd4 | intron_variant | MODIFIER |
| 9 | 60332812 | AX-170032482 | 0 | A | G | 0.202 | 6.41E-03 | 1.35E-01 | 2.71E+00 | 2.74E+00 | 9.60E-01 | no | Thsd4 | intron_variant | MODIFIER |
| 9 | 60349213 | AX-170493933 | 0 | G | T | 0.202 | 6.41E-03 | 1.35E-01 | 2.71E+00 | 2.74E+00 | 9.60E-01 | no | Thsd4 | intron_variant | MODIFIER |
| 9 | 60351981 | AX-170032477 | 0 | T | C | 0.202 | 6.41E-03 | 1.35E-01 | 2.71E+00 | 2.74E+00 | 9.60E-01 | no | Thsd4 | intron_variant | MODIFIER |
| 9 | 60354049 | AX-97793609 | 0 | T | C | 0.202 | 6.41E-03 | 1.35E-01 | 2.71E+00 | 2.74E+00 | 9.60E-01 | no | Thsd4 | intron_variant | MODIFIER |
| 9 | 60369746 | AX-170267001 | 0 | G | C | 0.202 | 6.41E-03 | 1.35E-01 | 2.71E+00 | 2.74E+00 | 9.60E-01 | no | Thsd4 | intron_variant | MODIFIER |
| 9 | 60372103 | AX-97849341 | 0 | T | G | 0.202 | 6.41E-03 | 1.35E-01 | 2.71E+00 | 2.74E+00 | 9.60E-01 | no | Thsd4 | intron_variant | MODIFIER |
| 9 | 60383334 | AX-170032478 | 0 | A | G | 0.202 | 6.41E-03 | 1.35E-01 | 2.71E+00 | 2.74E+00 | 9.60E-01 | no | Thsd4 | intron_variant | MODIFIER |
| 9 | 60386093 | AX-170032480 | 0 | A | G | 0.202 | 6.41E-03 | 1.35E-01 | 2.71E+00 | 2.74E+00 | 9.60E-01 | no | Thsd4 | intron_variant | MODIFIER |
| 9 | 60395815 | AX-170298291 | 0 | A | C | 0.202 | 6.41E-03 | 1.35E-01 | 2.71E+00 | 2.74E+00 | 9.60E-01 | no | Thsd4 | intron_variant | MODIFIER |
| 9 | 60401884 | AX-170419466 | 0 | C | T | 0.202 | 6.41E-03 | 1.35E-01 | 2.71E+00 | 2.74E+00 | 9.60E-01 | no | Thsd4 | intron_variant | MODIFIER |
| 9 | 60411314 | AX-170279821 | 0 | G | A | 0.202 | 6.41E-03 | 1.35E-01 | 2.71E+00 | 2.74E+00 | 9.60E-01 | no | Thsd4 | intron_variant | MODIFIER |
| 9 | 60412421 | AX-170327709 | 0 | G | A | 0.202 | 6.41E-03 | 1.35E-01 | 2.71E+00 | 2.74E+00 | 9.60E-01 | no | Thsd4 | intron_variant | MODIFIER |
| 9 | 60413635 | AX-97834188 | 0 | T | C | 0.202 | 6.41E-03 | 1.35E-01 | 2.71E+00 | 2.74E+00 | 9.60E-01 | no | Thsd4 | intron_variant | MODIFIER |
| 9 | 60423104 | AX-170291136 | 0 | A | G | 0.202 | 6.41E-03 | 1.35E-01 | 2.71E+00 | 2.74E+00 | 9.60E-01 | no | Thsd4 | intron_variant | MODIFIER |
| 9 | 60439915 | AX-170082578 | 0 | G | C | 0.202 | 6.41E-03 | 1.35E-01 | 2.71E+00 | 2.74E+00 | 9.60E-01 | no | Thsd4 | intron_variant | MODIFIER |
| 9 | 60439961 | AX-97851597 | 0 | G | A | 0.202 | 6.41E-03 | 1.35E-01 | 2.71E+00 | 2.74E+00 | 9.60E-01 | no | Thsd4 | intron_variant | MODIFIER |
| 9 | 60460257 | AX-170500530 | 0 | C | A | 0.202 | 6.41E-03 | 1.35E-01 | 2.71E+00 | 2.74E+00 | 9.60E-01 | no | Thsd4 | intron_variant | MODIFIER |
| 9 | 60466788 | AX-170129869 | 0 | C | T | 0.202 | 6.41E-03 | 1.35E-01 | 2.71E+00 | 2.74E+00 | 9.60E-01 | no | Thsd4 | intron_variant | MODIFIER |
| 9 | 60494703 | AX-97826055 | 0 | A | G | 0.202 | 6.41E-03 | 1.35E-01 | 2.71E+00 | 2.74E+00 | 9.60E-01 | no | Thsd4 | intron_variant | MODIFIER |
| 9 | 60505249 | AX-170105858 | 0 | T | C | 0.202 | 6.41E-03 | 1.35E-01 | 2.71E+00 | 2.74E+00 | 9.60E-01 | no | Thsd4 | intron_variant | MODIFIER |
| 9 | 60505742 | AX-170397681 | 0 | G | A | 0.202 | 6.41E-03 | 1.35E-01 | 2.71E+00 | 2.74E+00 | 9.60E-01 | no | Thsd4 | intron_variant | MODIFIER |
| 9 | 60508129 | AX-170407010 | 0 | G | A | 0.202 | 6.41E-03 | 1.35E-01 | 2.71E+00 | 2.74E+00 | 9.60E-01 | no | Thsd4 | intron_variant | MODIFIER |
| 9 | 60585873 | AX-170300876 | 0 | G | C | 0.202 | 6.41E-03 | 1.35E-01 | 2.71E+00 | 2.74E+00 | 9.60E-01 | no | Lrrc49 | intron_variant | MODIFIER |
| 9 | 60586721 | AX-170355331 | 0 | C | A | 0.202 | 6.41E-03 | 1.35E-01 | 2.71E+00 | 2.74E+00 | 9.60E-01 | no | Lrrc49 | intron_variant | MODIFIER |
| 9 | 60588345 | AX-97797182 | 0 | T | C | 0.202 | 6.41E-03 | 1.35E-01 | 2.71E+00 | 2.74E+00 | 9.60E-01 | no | Lrrc49 | intron_variant | MODIFIER |
| 9 | 60605798 | AX-170393213 | 0 | G | A | 0.202 | 6.41E-03 | 1.35E-01 | 2.71E+00 | 2.74E+00 | 9.60E-01 | no | Lrrc49 | intron_variant | MODIFIER |
| 9 | 60801670 | AX-97821628 | 0 | C | T | 0.202 | 6.41E-03 | 1.35E-01 | 2.71E+00 | 2.74E+00 | 9.60E-01 | no | Uaca | intron_variant | MODIFIER |
| 9 | 60921922 | AX-170032489 | 0 | A | G | 0.202 | 6.41E-03 | 1.35E-01 | 2.71E+00 | 2.74E+00 | 9.60E-01 | no | Gm47923 | intron_variant,non_coding_transcript_variant | MODIFIER |
| 9 | 60930265 | AX-170465947 | 0 | A | G | 0.202 | 6.41E-03 | 1.35E-01 | 2.71E+00 | 2.74E+00 | 9.60E-01 | no | Gm47923 | intron_variant,non_coding_transcript_variant | MODIFIER |
| 9 | 60930969 | AX-170274635 | 0 | T | C | 0.202 | 6.41E-03 | 1.35E-01 | 2.71E+00 | 2.74E+00 | 9.60E-01 | no | Gm47923 | intron_variant,non_coding_transcript_variant | MODIFIER |
| 9 | 60931726 | AX-170271539 | 0 | C | A | 0.202 | 6.41E-03 | 1.35E-01 | 2.71E+00 | 2.74E+00 | 9.60E-01 | no | Gm47923 | intron_variant,non_coding_transcript_variant | MODIFIER |
| 9 | 60935702 | AX-170032486 | 0 | C | G | 0.202 | 6.41E-03 | 1.35E-01 | 2.71E+00 | 2.74E+00 | 9.60E-01 | no | Gm47923 | intron_variant,non_coding_transcript_variant | MODIFIER |
| 9 | 60942148 | AX-170032488 | 0 | G | T | 0.202 | 6.41E-03 | 1.35E-01 | 2.71E+00 | 2.74E+00 | 9.60E-01 | no | Gm47923 | intron_variant,non_coding_transcript_variant | MODIFIER |
| 9 | 60942399 | AX-170462470 | 0 | C | T | 0.202 | 6.41E-03 | 1.35E-01 | 2.71E+00 | 2.74E+00 | 9.60E-01 | no | Gm47923 | intron_variant,non_coding_transcript_variant | MODIFIER |
| 9 | 61191311 | AX-170179029 | 0 | T | C | 0.202 | 6.41E-03 | 1.35E-01 | 2.71E+00 | 2.74E+00 | 9.60E-01 | no | | intergenic_variant | MODIFIER |
| 9 | 61292316 | AX-170449935 | 0 | G | A | 0.202 | 6.41E-03 | 1.35E-01 | 2.71E+00 | 2.74E+00 | 9.60E-01 | no | B930092H01Rik | intron_variant,non_coding_transcript_variant | MODIFIER |
| 9 | 61892874 | AX-170082604 | 0 | G | A | 0.202 | 6.41E-03 | 1.35E-01 | 2.71E+00 | 2.74E+00 | 9.60E-01 | no | | intergenic_variant | MODIFIER |
| 9 | 61930349 | AX-97838378 | 0 | T | C | 0.202 | 6.41E-03 | 1.35E-01 | 2.71E+00 | 2.74E+00 | 9.60E-01 | no | Kif23 | intron_variant | MODIFIER |
| 9 | 62150330 | AX-97838801 | 0 | A | G | 0.202 | 6.41E-03 | 1.35E-01 | 2.71E+00 | 2.74E+00 | 9.60E-01 | no | | intergenic_variant | MODIFIER |
| 9 | 62154886 | AX-170347865 | 0 | T | C | 0.202 | 6.41E-03 | 1.35E-01 | 2.71E+00 | 2.74E+00 | 9.60E-01 | no | | regulatory_region_variant | MODIFIER |
| 9 | 62175805 | AX-170345801 | 0 | C | T | 0.202 | 6.41E-03 | 1.35E-01 | 2.71E+00 | 2.74E+00 | 9.60E-01 | no | | intergenic_variant | MODIFIER |
| 9 | 62182120 | AX-168408544 | 0 | C | T | 0.202 | 6.41E-03 | 1.35E-01 | 2.71E+00 | 2.74E+00 | 9.60E-01 | no | | regulatory_region_variant | MODIFIER |
| 9 | 62193456 | AX-170136067 | 0 | T | A | 0.202 | 6.41E-03 | 1.35E-01 | 2.71E+00 | 2.74E+00 | 9.60E-01 | no | | regulatory_region_variant | MODIFIER |
| 9 | 62220574 | AX-97809291 | 0 | G | A | 0.202 | 6.41E-03 | 1.35E-01 | 2.71E+00 | 2.74E+00 | 9.60E-01 | no | | regulatory_region_variant | MODIFIER |
| 9 | 62283163 | AX-170395907 | 0 | T | C | 0.202 | 6.41E-03 | 1.35E-01 | 2.71E+00 | 2.74E+00 | 9.60E-01 | no | Spesp1 | upstream_gene_variant | MODIFIER |
| 9 | 62289370 | AX-170226344 | 0 | A | G | 0.202 | 6.41E-03 | 1.35E-01 | 2.71E+00 | 2.74E+00 | 9.60E-01 | no | | intergenic_variant | MODIFIER |
| 9 | 62290553 | AX-170368656 | 0 | T | G | 0.202 | 6.41E-03 | 1.35E-01 | 2.71E+00 | 2.74E+00 | 9.60E-01 | no | | intergenic_variant | MODIFIER |
| 9 | 62290725 | AX-170175190 | 0 | G | A | 0.202 | 6.41E-03 | 1.35E-01 | 2.71E+00 | 2.74E+00 | 9.60E-01 | no | | intergenic_variant | MODIFIER |
| 9 | 62312549 | AX-170353897 | 0 | A | G | 0.202 | 6.41E-03 | 1.35E-01 | 2.71E+00 | 2.74E+00 | 9.60E-01 | no | Gm35288 | intron_variant,non_coding_transcript_variant | MODIFIER |
| 9 | 62380250 | AX-170032515 | 0 | G | A | 0.202 | 6.41E-03 | 1.35E-01 | 2.71E+00 | 2.74E+00 | 9.60E-01 | no | Anp32a | downstream_gene_variant | MODIFIER |
| 9 | 62431320 | AX-97839507 | 0 | T | G | 0.202 | 6.41E-03 | 1.35E-01 | 2.71E+00 | 2.74E+00 | 9.60E-01 | no | Coro2b | synonymous_variant | LOW |
| 9 | 62469789 | AX-170332456 | 0 | T | C | 0.202 | 6.41E-03 | 1.35E-01 | 2.71E+00 | 2.74E+00 | 9.60E-01 | no | Coro2b | intron_variant | MODIFIER |
| 9 | 62484473 | AX-170173083 | 0 | C | T | 0.202 | 6.41E-03 | 1.35E-01 | 2.71E+00 | 2.74E+00 | 9.60E-01 | no | Coro2b | intron_variant | MODIFIER |

| | | | | | | | | | | | | | | | |
|---|---|---|---|---|---|---|---|---|---|---|---|---|---|---|---|
| 9 | 62505489 | AX-170426790 | 0 | A | G | 0.202 | 6.41E-03 | 1.35E-01 | 2.71E+00 | 2.74E+00 | 9.60E-01 | no | Coro2b | intron_variant | MODIFIER |
| 9 | 62516190 | AX-170307831 | 0 | A | G | 0.202 | 6.41E-03 | 1.35E-01 | 2.71E+00 | 2.74E+00 | 9.60E-01 | no | Coro2b | intron_variant | MODIFIER |
| 9 | 62524404 | AX-97826738 | 0 | C | T | 0.202 | 6.41E-03 | 1.35E-01 | 2.71E+00 | 2.74E+00 | 9.60E-01 | no | Coro2b | intron_variant | MODIFIER |
| 9 | 62527719 | AX-170161009 | 0 | G | A | 0.202 | 6.41E-03 | 1.35E-01 | 2.71E+00 | 2.74E+00 | 9.60E-01 | no | Coro2b | intron_variant | MODIFIER |
| 9 | 62538564 | AX-170204600 | 0 | G | A | 0.202 | 6.41E-03 | 1.35E-01 | 2.71E+00 | 2.74E+00 | 9.60E-01 | no | Coro2b | upstream_gene_variant | MODIFIER |
| 9 | 62542768 | AX-97790582 | 0 | C | T | 0.202 | 6.41E-03 | 1.35E-01 | 2.71E+00 | 2.74E+00 | 9.60E-01 | no | | regulatory_region_variant | MODIFIER |
| 9 | 62628712 | AX-170243878 | 0 | C | T | 0.202 | 6.41E-03 | 1.35E-01 | 2.71E+00 | 2.74E+00 | 9.60E-01 | no | | intergenic_variant | MODIFIER |
| 9 | 62665990 | AX-170165348 | 0 | G | C | 0.202 | 6.41E-03 | 1.35E-01 | 2.71E+00 | 2.74E+00 | 9.60E-01 | no | | regulatory_region_variant | MODIFIER |
| 9 | 62670171 | AX-97853613 | 0 | G | A | 0.202 | 6.41E-03 | 1.35E-01 | 2.71E+00 | 2.74E+00 | 9.60E-01 | no | | intergenic_variant | MODIFIER |
| 9 | 62674340 | AX-170397163 | 0 | T | G | 0.202 | 6.41E-03 | 1.35E-01 | 2.71E+00 | 2.74E+00 | 9.60E-01 | no | Itga11 | upstream_gene_variant | MODIFIER |
| 9 | 62684655 | AX-170134928 | 0 | T | C | 0.202 | 6.41E-03 | 1.35E-01 | 2.71E+00 | 2.74E+00 | 9.60E-01 | no | Itga11 | intron_variant | MODIFIER |
| 9 | 62702352 | AX-170231422 | 0 | T | C | 0.202 | 6.41E-03 | 1.35E-01 | 2.71E+00 | 2.74E+00 | 9.60E-01 | no | Itga11 | intron_variant | MODIFIER |
| 9 | 62719371 | AX-97803894 | 0 | A | C | 0.202 | 6.41E-03 | 1.35E-01 | 2.71E+00 | 2.74E+00 | 9.60E-01 | no | Itga11 | intron_variant | MODIFIER |
| 9 | 62727748 | AX-170113759 | 0 | A | G | 0.202 | 6.41E-03 | 1.35E-01 | 2.71E+00 | 2.74E+00 | 9.60E-01 | no | Itga11 | intron_variant | MODIFIER |
| 9 | 62730443 | AX-170236231 | 0 | A | G | 0.202 | 6.41E-03 | 1.35E-01 | 2.71E+00 | 2.74E+00 | 9.60E-01 | no | Itga11 | intron_variant | MODIFIER |
| 9 | 62748616 | AX-170310713 | 0 | C | A | 0.202 | 6.41E-03 | 1.35E-01 | 2.71E+00 | 2.74E+00 | 9.60E-01 | no | Itga11 | intron_variant | MODIFIER |
| 9 | 62749165 | AX-170411557 | 0 | C | T | 0.202 | 6.41E-03 | 1.35E-01 | 2.71E+00 | 2.74E+00 | 9.60E-01 | no | Itga11 | intron_variant | MODIFIER |
| 9 | 62764652 | AX-170082622 | 0 | G | A | 0.202 | 6.41E-03 | 1.35E-01 | 2.71E+00 | 2.74E+00 | 9.60E-01 | no | Itga11 | intron_variant | MODIFIER |
| 9 | 62764860 | AX-97805692 | 0 | C | T | 0.202 | 6.41E-03 | 1.35E-01 | 2.71E+00 | 2.74E+00 | 9.60E-01 | no | Itga11 | intron_variant | MODIFIER |
| 9 | 62765722 | AX-170361059 | 0 | G | A | 0.202 | 6.41E-03 | 1.35E-01 | 2.71E+00 | 2.74E+00 | 9.60E-01 | no | Itga11 | intron_variant | MODIFIER |
| 9 | 62767410 | AX-170356479 | 0 | A | C | 0.202 | 6.41E-03 | 1.35E-01 | 2.71E+00 | 2.74E+00 | 9.60E-01 | no | Itga11 | intron_variant | MODIFIER |
| 9 | 62783317 | AX-170109024 | 0 | T | C | 0.202 | 6.41E-03 | 1.35E-01 | 2.71E+00 | 2.74E+00 | 9.60E-01 | no | Itga11 | 3_prime_UTR_variant | MODIFIER |
| 9 | 63797762 | AX-170267951 | 0 | G | A | 0.202 | 6.41E-03 | 1.35E-01 | 2.71E+00 | 2.74E+00 | 9.60E-01 | no | Gm2553 | downstream_gene_variant | MODIFIER |
| 9 | 63811947 | AX-170262335 | 0 | A | G | 0.202 | 6.41E-03 | 1.35E-01 | 2.71E+00 | 2.74E+00 | 9.60E-01 | no | | regulatory_region_variant | MODIFIER |
| 9 | 63819519 | AX-170090317 | 0 | C | T | 0.202 | 6.41E-03 | 1.35E-01 | 2.71E+00 | 2.74E+00 | 9.60E-01 | no | | intergenic_variant | MODIFIER |
| 9 | 63824367 | AX-170464125 | 0 | T | C | 0.202 | 6.41E-03 | 1.35E-01 | 2.71E+00 | 2.74E+00 | 9.60E-01 | no | | intergenic_variant | MODIFIER |
| 9 | 63836496 | AX-170136587 | 0 | T | C | 0.202 | 6.41E-03 | 1.35E-01 | 2.71E+00 | 2.74E+00 | 9.60E-01 | no | | regulatory_region_variant | MODIFIER |
| 9 | 63881813 | AX-170493249 | 0 | G | A | 0.202 | 6.41E-03 | 1.35E-01 | 2.71E+00 | 2.74E+00 | 9.60E-01 | no | | regulatory_region_variant | MODIFIER |
| 9 | 63886585 | AX-170449934 | 0 | A | G | 0.202 | 6.41E-03 | 1.35E-01 | 2.71E+00 | 2.74E+00 | 9.60E-01 | no | | intergenic_variant | MODIFIER |
| 9 | 64053442 | AX-97832039 | 0 | T | C | 0.202 | 6.41E-03 | 1.35E-01 | 2.71E+00 | 2.74E+00 | 9.60E-01 | no | Gm25606 | downstream_gene_variant | MODIFIER |
| 9 | 64133111 | AX-97839535 | 0 | C | T | 0.202 | 6.41E-03 | 1.35E-01 | 2.71E+00 | 2.74E+00 | 9.60E-01 | no | Lctl | synonymous_variant | LOW |
| 9 | 64134271 | AX-170448900 | 0 | T | C | 0.202 | 6.41E-03 | 1.35E-01 | 2.71E+00 | 2.74E+00 | 9.60E-01 | no | Lctl | intron_variant | MODIFIER |
| 9 | 64144157 | AX-97849254 | 0 | T | C | 0.202 | 6.41E-03 | 1.35E-01 | 2.71E+00 | 2.74E+00 | 9.60E-01 | no | Zwilch | missense_variant | MODERATE |
| 9 | 64170566 | AX-170318875 | 0 | T | G | 0.202 | 6.41E-03 | 1.35E-01 | 2.71E+00 | 2.74E+00 | 9.60E-01 | no | Rpl4 | upstream_gene_variant | MODIFIER |
| 9 | 64192155 | AX-170145436 | 0 | G | C | 0.202 | 6.41E-03 | 1.35E-01 | 2.71E+00 | 2.74E+00 | 9.60E-01 | no | Map2k1 | intron_variant | MODIFIER |
| 9 | 64230367 | AX-170512952 | 0 | T | C | 0.202 | 6.41E-03 | 1.35E-01 | 2.71E+00 | 2.74E+00 | 9.60E-01 | no | Map2k1 | intron_variant | MODIFIER |
| 9 | 64231516 | AX-170403573 | 0 | G | A | 0.202 | 6.41E-03 | 1.35E-01 | 2.71E+00 | 2.74E+00 | 9.60E-01 | no | Map2k1 | intron_variant | MODIFIER |
| 9 | 64349571 | AX-170318478 | 0 | C | A | 0.202 | 6.41E-03 | 1.35E-01 | 2.71E+00 | 2.74E+00 | 9.60E-01 | no | | intergenic_variant | MODIFIER |
| 9 | 64379304 | AX-170450985 | 0 | T | C | 0.202 | 6.41E-03 | 1.35E-01 | 2.71E+00 | 2.74E+00 | 9.60E-01 | no | | intergenic_variant | MODIFIER |
| 9 | 64503314 | AX-170032553 | 0 | T | C | 0.202 | 6.41E-03 | 1.35E-01 | 2.71E+00 | 2.74E+00 | 9.60E-01 | no | Megf11 | intron_variant | MODIFIER |
| 9 | 69445005 | AX-170242112 | 0 | A | G | 0.202 | 6.41E-03 | 1.35E-01 | 2.71E+00 | 2.74E+00 | 9.60E-01 | no | Gm18587 | non_coding_transcript_exon_variant | MODIFIER |
| 9 | 69445872 | AX-170270732 | 0 | C | T | 0.202 | 6.41E-03 | 1.35E-01 | 2.71E+00 | 2.74E+00 | 9.60E-01 | no | Gm4978 | downstream_gene_variant | MODIFIER |
| 9 | 74583181 | AX-97795248 | 0 | A | G | 0.202 | 6.41E-03 | 1.35E-01 | 2.71E+00 | 2.74E+00 | 9.60E-01 | no | | intergenic_variant | MODIFIER |
| 9 | 74602811 | AX-97830988 | 0 | T | C | 0.202 | 6.41E-03 | 1.35E-01 | 2.71E+00 | 2.74E+00 | 9.60E-01 | no | | intergenic_variant | MODIFIER |
| 9 | 74609386 | AX-170210293 | 0 | G | A | 0.202 | 6.41E-03 | 1.35E-01 | 2.71E+00 | 2.74E+00 | 9.60E-01 | no | | intergenic_variant | MODIFIER |
| 9 | 74622787 | AX-170224917 | 0 | T | A | 0.202 | 6.41E-03 | 1.35E-01 | 2.71E+00 | 2.74E+00 | 9.60E-01 | no | | intergenic_variant | MODIFIER |
| 9 | 74623547 | AX-97792876 | 0 | C | A | 0.202 | 6.41E-03 | 1.35E-01 | 2.71E+00 | 2.74E+00 | 9.60E-01 | no | | intergenic_variant | MODIFIER |
| 9 | 74660741 | AX-97797372 | 0 | A | G | 0.202 | 6.41E-03 | 1.35E-01 | 2.71E+00 | 2.74E+00 | 9.60E-01 | no | | intergenic_variant | MODIFIER |
| 9 | 74890120 | AX-170283179 | 0 | G | A | 0.202 | 6.41E-03 | 1.35E-01 | 2.71E+00 | 2.74E+00 | 9.60E-01 | no | Onecut1 | 3_prime_UTR_variant | MODIFIER |
| 9 | 75168827 | AX-170032724 | 0 | A | G | 0.202 | 6.41E-03 | 1.35E-01 | 2.71E+00 | 2.74E+00 | 9.60E-01 | no | Myo5a | intron_variant | MODIFIER |
| 9 | 75327992 | AX-170499582 | 0 | T | C | 0.202 | 6.41E-03 | 1.35E-01 | 2.71E+00 | 2.74E+00 | 9.60E-01 | no | Gnb5 | intron_variant | MODIFIER |
| 9 | 75359345 | AX-97809996 | 0 | A | G | 0.202 | 6.41E-03 | 1.35E-01 | 2.71E+00 | 2.74E+00 | 9.60E-01 | no | | intergenic_variant | MODIFIER |
| 9 | 75362979 | AX-170383451 | 0 | G | A | 0.202 | 6.41E-03 | 1.35E-01 | 2.71E+00 | 2.74E+00 | 9.60E-01 | no | | intergenic_variant | MODIFIER |
| 9 | 75384883 | AX-170348610 | 0 | G | C | 0.202 | 6.41E-03 | 1.35E-01 | 2.71E+00 | 2.74E+00 | 9.60E-01 | no | Mapk6 | downstream_gene_variant | MODIFIER |

| | | | | | | | | | | | | | | | |
|---|---|---|---|---|---|---|---|---|---|---|---|---|---|---|---|
| 9 | 75401443 | AX-97801278 | 0 | C | A | 0.202 | 6.41E-03 | 1.35E-01 | 2.71E+00 | 2.74E+00 | 9.60E-01 | no | Mapk6 | intron_variant | MODIFIER |
| 9 | 75659801 | AX-170082841 | 0 | G | A | 0.202 | 6.41E-03 | 1.35E-01 | 2.71E+00 | 2.74E+00 | 9.60E-01 | no | Scg3 | intron_variant | MODIFIER |
| 9 | 75660134 | AX-170082840 | 0 | T | C | 0.202 | 6.41E-03 | 1.35E-01 | 2.71E+00 | 2.74E+00 | 9.60E-01 | no | Scg3 | intron_variant | MODIFIER |
| 9 | 75702438 | AX-170096588 | 0 | C | T | 0.202 | 6.41E-03 | 1.35E-01 | 2.71E+00 | 2.74E+00 | 9.60E-01 | no | | intergenic_variant | MODIFIER |
| 9 | 75720190 | AX-170486750 | 0 | G | C | 0.202 | 6.41E-03 | 1.35E-01 | 2.71E+00 | 2.74E+00 | 9.60E-01 | no | | regulatory_region_variant | MODIFIER |
| 9 | 75791810 | AX-97805829 | 0 | C | T | 0.202 | 6.41E-03 | 1.35E-01 | 2.71E+00 | 2.74E+00 | 9.60E-01 | no | Bmp5 | intron_variant | MODIFIER |
| 9 | 75796741 | AX-170032733 | 0 | A | G | 0.202 | 6.41E-03 | 1.35E-01 | 2.71E+00 | 2.74E+00 | 9.60E-01 | no | Bmp5 | intron_variant | MODIFIER |
| 9 | 75825038 | AX-170102578 | 0 | A | G | 0.202 | 6.41E-03 | 1.35E-01 | 2.71E+00 | 2.74E+00 | 9.60E-01 | no | Bmp5 | intron_variant | MODIFIER |
| 9 | 75827828 | AX-170262696 | 0 | A | G | 0.202 | 6.41E-03 | 1.35E-01 | 2.71E+00 | 2.74E+00 | 9.60E-01 | no | Bmp5 | intron_variant | MODIFIER |
| 9 | 75827981 | AX-170367536 | 0 | A | C | 0.202 | 6.41E-03 | 1.35E-01 | 2.71E+00 | 2.74E+00 | 9.60E-01 | no | Bmp5 | intron_variant | MODIFIER |
| 9 | 76241440 | AX-170032737 | 0 | G | C | 0.202 | 6.41E-03 | 1.35E-01 | 2.71E+00 | 2.74E+00 | 9.60E-01 | no | Hcrtr2 | intron_variant | MODIFIER |
| 9 | 76309719 | AX-97813032 | 0 | T | C | 0.202 | 6.41E-03 | 1.35E-01 | 2.71E+00 | 2.74E+00 | 9.60E-01 | no | Hcrtr2 | intron_variant | MODIFIER |
| 7 | 102457035 | AX-170029654 | 1 | G | A | 0.455 | 5.35E-03 | 9.44E-02 | 2.67E+00 | 2.73E+00 | 9.60E-01 | no | Rrm1 | intron_variant | MODIFIER |
| 5 | 70085363 | AX-170025230 | 1 | G | A | 0.446 | 5.38E-03 | 9.77E-02 | 2.65E+00 | 2.73E+00 | 9.60E-01 | no | | intergenic_variant | MODIFIER |
| 1 | 63622033 | AX-170349022 | 0 | T | C | 0.132 | -1.17E-02 | 2.38E-01 | 2.69E+00 | 2.74E+00 | 9.60E-01 | no | Dytn | downstream_gene_variant | MODIFIER |
| 9 | 87385771 | AX-170317733 | 1 | G | A | 0.134 | -7.67E-03 | 1.33E-01 | 2.66E+00 | 2.73E+00 | 9.60E-01 | no | | intergenic_variant | MODIFIER |
| 12 | 116833611 | AX-170183809 | 1 | C | T | 0.125 | -8.85E-03 | 1.98E-01 | 2.67E+00 | 2.74E+00 | 9.60E-01 | no | Ptprn2 | intron_variant | MODIFIER |
| 7 | 117758647 | AX-170467994 | 0 | T | C | 0.395 | 4.51E-03 | 8.83E-02 | 2.68E+00 | 2.74E+00 | 9.60E-01 | no | | intergenic_variant | MODIFIER |
| 11 | 35032944 | AX-97789248 | 0 | C | T | 0.096 | 1.02E-02 | 1.87E-01 | 2.65E+00 | 2.73E+00 | 9.60E-01 | no | | intergenic_variant | MODIFIER |
| 13 | 52906604 | AX-97820212 | 1 | A | G | 0.5 | 4.34E-03 | 6.63E-02 | 2.62E+00 | 2.73E+00 | 9.60E-01 | no | Auh | intron_variant | MODIFIER |
| 5 | 123719713 | AX-170026071 | 0 | C | A | 0.456 | 5.90E-03 | 1.11E-01 | 2.68E+00 | 2.73E+00 | 9.60E-01 | no | Zcchc8 | intron_variant | MODIFIER |
| 4 | 141036860 | AX-170211311 | 0 | G | C | 0.079 | -7.18E-03 | 1.60E-01 | 2.64E+00 | 2.74E+00 | 9.61E-01 | no | Crocc | intron_variant | MODIFIER |
| 1 | 67032441 | AX-170239211 | 0 | C | A | 0.114 | -9.30E-03 | 1.85E-01 | 2.64E+00 | 2.74E+00 | 9.61E-01 | no | Lancl1 | intron_variant | MODIFIER |
| 8 | 23819703 | AX-97807173 | 2 | G | A | 0.173 | 5.02E-03 | 1.12E-01 | 2.66E+00 | 2.74E+00 | 9.61E-01 | no | Zmat4 | intron_variant | MODIFIER |
| 9 | 66990196 | AX-170032589 | 0 | C | T | 0.193 | -6.90E-03 | 1.32E-01 | 2.69E+00 | 2.73E+00 | 9.61E-01 | no | | regulatory_region_variant | MODIFIER |
| 7 | 93905508 | AX-170128991 | 1 | A | T | 0.125 | 9.55E-03 | 2.00E-01 | 2.64E+00 | 2.74E+00 | 9.61E-01 | no | | intergenic_variant | MODIFIER |
| 1 | 173646649 | AX-170043629 | 1 | G | A | 0.232 | 8.19E-03 | 1.26E-01 | 2.65E+00 | 2.72E+00 | 9.61E-01 | no | Ifi209 | 3_prime_UTR_variant | MODIFIER |
| 8 | 78164978 | AX-170477534 | 0 | C | A | 0.202 | -6.39E-03 | 1.31E-01 | 2.68E+00 | 2.74E+00 | 9.61E-01 | no | | intergenic_variant | MODIFIER |
| 8 | 78212150 | AX-170283074 | 0 | A | C | 0.202 | -6.39E-03 | 1.31E-01 | 2.68E+00 | 2.74E+00 | 9.61E-01 | no | Ttc29 | upstream_gene_variant | MODIFIER |
| 8 | 78216877 | AX-170339161 | 0 | C | T | 0.202 | -6.39E-03 | 1.31E-01 | 2.68E+00 | 2.74E+00 | 9.61E-01 | no | Ttc29 | intron_variant | MODIFIER |
| 8 | 78218825 | AX-170266325 | 0 | A | G | 0.202 | -6.39E-03 | 1.31E-01 | 2.68E+00 | 2.74E+00 | 9.61E-01 | no | Ttc29 | missense_variant | MODERATE |
| 8 | 78230496 | AX-97841569 | 0 | T | C | 0.202 | -6.39E-03 | 1.31E-01 | 2.68E+00 | 2.74E+00 | 9.61E-01 | no | Ttc29 | intron_variant | MODIFIER |
| 8 | 78231077 | AX-170207531 | 0 | T | G | 0.202 | -6.39E-03 | 1.31E-01 | 2.68E+00 | 2.74E+00 | 9.61E-01 | no | Ttc29 | intron_variant | MODIFIER |
| 8 | 78234671 | AX-170321762 | 0 | A | G | 0.202 | -6.39E-03 | 1.31E-01 | 2.68E+00 | 2.74E+00 | 9.61E-01 | no | Ttc29 | intron_variant | MODIFIER |
| 8 | 78260892 | AX-170481820 | 0 | G | C | 0.202 | -6.39E-03 | 1.31E-01 | 2.68E+00 | 2.74E+00 | 9.61E-01 | no | Ttc29 | intron_variant | MODIFIER |
| 8 | 78260927 | AX-170387498 | 0 | T | A | 0.202 | -6.39E-03 | 1.31E-01 | 2.68E+00 | 2.74E+00 | 9.61E-01 | no | Ttc29 | intron_variant | MODIFIER |
| 8 | 78261269 | AX-170255629 | 0 | C | T | 0.202 | -6.39E-03 | 1.31E-01 | 2.68E+00 | 2.74E+00 | 9.61E-01 | no | Ttc29 | intron_variant | MODIFIER |
| 8 | 78264244 | AX-170147846 | 0 | T | G | 0.202 | -6.39E-03 | 1.31E-01 | 2.68E+00 | 2.74E+00 | 9.61E-01 | no | Ttc29 | intron_variant | MODIFIER |
| 8 | 78270834 | AX-170259674 | 0 | C | G | 0.202 | -6.39E-03 | 1.31E-01 | 2.68E+00 | 2.74E+00 | 9.61E-01 | no | Ttc29 | intron_variant | MODIFIER |
| 8 | 78282803 | AX-170406929 | 0 | G | A | 0.202 | -6.39E-03 | 1.31E-01 | 2.68E+00 | 2.74E+00 | 9.61E-01 | no | Ttc29 | intron_variant | MODIFIER |
| 8 | 78284521 | AX-170465289 | 0 | A | C | 0.202 | -6.39E-03 | 1.31E-01 | 2.68E+00 | 2.74E+00 | 9.61E-01 | no | Ttc29 | intron_variant | MODIFIER |
| 8 | 78287526 | AX-170502520 | 0 | T | A | 0.202 | -6.39E-03 | 1.31E-01 | 2.68E+00 | 2.74E+00 | 9.61E-01 | no | Ttc29 | intron_variant | MODIFIER |
| 8 | 78287733 | AX-97812291 | 0 | A | G | 0.202 | -6.39E-03 | 1.31E-01 | 2.68E+00 | 2.74E+00 | 9.61E-01 | no | Ttc29 | intron_variant | MODIFIER |
| 8 | 78310300 | AX-170167361 | 0 | G | A | 0.202 | -6.39E-03 | 1.31E-01 | 2.68E+00 | 2.74E+00 | 9.61E-01 | no | Ttc29 | intron_variant | MODIFIER |
| 8 | 78310523 | AX-170192491 | 0 | T | C | 0.202 | -6.39E-03 | 1.31E-01 | 2.68E+00 | 2.74E+00 | 9.61E-01 | no | Ttc29 | intron_variant | MODIFIER |
| 8 | 78311286 | AX-170080509 | 0 | A | G | 0.202 | -6.39E-03 | 1.31E-01 | 2.68E+00 | 2.74E+00 | 9.61E-01 | no | Ttc29 | intron_variant | MODIFIER |
| 8 | 78327236 | AX-170400540 | 0 | C | T | 0.202 | -6.39E-03 | 1.31E-01 | 2.68E+00 | 2.74E+00 | 9.61E-01 | no | Ttc29 | intron_variant | MODIFIER |
| 8 | 78328333 | AX-170397508 | 0 | A | G | 0.202 | -6.39E-03 | 1.31E-01 | 2.68E+00 | 2.74E+00 | 9.61E-01 | no | Ttc29 | intron_variant | MODIFIER |
| 8 | 78333044 | AX-170126440 | 0 | T | C | 0.202 | -6.39E-03 | 1.31E-01 | 2.68E+00 | 2.74E+00 | 9.61E-01 | no | Ttc29 | intron_variant | MODIFIER |
| 8 | 78350735 | AX-170485710 | 0 | T | C | 0.202 | -6.39E-03 | 1.31E-01 | 2.68E+00 | 2.74E+00 | 9.61E-01 | no | Ttc29 | intron_variant | MODIFIER |
| 8 | 78364532 | AX-170521049 | 0 | C | T | 0.202 | -6.39E-03 | 1.31E-01 | 2.68E+00 | 2.74E+00 | 9.61E-01 | no | Ttc29 | intron_variant | MODIFIER |
| 8 | 78368343 | AX-97824653 | 0 | G | A | 0.202 | -6.39E-03 | 1.31E-01 | 2.68E+00 | 2.74E+00 | 9.61E-01 | no | Ttc29 | intron_variant | MODIFIER |
| 8 | 78368475 | AX-170492453 | 0 | G | A | 0.202 | -6.39E-03 | 1.31E-01 | 2.68E+00 | 2.74E+00 | 9.61E-01 | no | Ttc29 | intron_variant | MODIFIER |
| 8 | 78374765 | AX-170516459 | 0 | G | T | 0.202 | -6.39E-03 | 1.31E-01 | 2.68E+00 | 2.74E+00 | 9.61E-01 | no | Ttc29 | intron_variant | MODIFIER |

| | | | | | | | | | | | | | | | |
|---|---|---|---|---|---|---|---|---|---|---|---|---|---|---|---|
| 8 | 78382092 | AX-170098185 | 0 | C | T | 0.202 | -6.39E-03 | 1.31E-01 | 2.68E+00 | 2.74E+00 | 9.61E-01 | no | Ttc29 | intron_variant | MODIFIER |
| 8 | 78542254 | AX-170031083 | 0 | A | G | 0.202 | -6.39E-03 | 1.31E-01 | 2.68E+00 | 2.74E+00 | 9.61E-01 | no | Slc10a7 | intron_variant | MODIFIER |
| 8 | 78567198 | AX-97855295 | 0 | A | G | 0.202 | -6.39E-03 | 1.31E-01 | 2.68E+00 | 2.74E+00 | 9.61E-01 | no | Slc10a7 | intron_variant | MODIFIER |
| 8 | 78641414 | AX-170363628 | 0 | C | T | 0.202 | -6.39E-03 | 1.31E-01 | 2.68E+00 | 2.74E+00 | 9.61E-01 | no | Slc10a7 | intron_variant | MODIFIER |
| 8 | 78653762 | AX-170310928 | 0 | C | T | 0.202 | -6.39E-03 | 1.31E-01 | 2.68E+00 | 2.74E+00 | 9.61E-01 | no | Slc10a7 | intron_variant | MODIFIER |
| 8 | 78659607 | AX-170243249 | 0 | C | G | 0.202 | -6.39E-03 | 1.31E-01 | 2.68E+00 | 2.74E+00 | 9.61E-01 | no | Slc10a7 | intron_variant | MODIFIER |
| 8 | 78672708 | AX-170248845 | 0 | G | T | 0.202 | -6.39E-03 | 1.31E-01 | 2.68E+00 | 2.74E+00 | 9.61E-01 | no | Slc10a7 | intron_variant | MODIFIER |
| 8 | 78681178 | AX-170311256 | 0 | C | T | 0.202 | -6.39E-03 | 1.31E-01 | 2.68E+00 | 2.74E+00 | 9.61E-01 | no | Slc10a7 | intron_variant | MODIFIER |
| 8 | 78685545 | AX-170336422 | 0 | G | C | 0.202 | -6.39E-03 | 1.31E-01 | 2.68E+00 | 2.74E+00 | 9.61E-01 | no | Slc10a7 | intron_variant | MODIFIER |
| 8 | 78685639 | AX-170120141 | 0 | G | A | 0.202 | -6.39E-03 | 1.31E-01 | 2.68E+00 | 2.74E+00 | 9.61E-01 | no | Slc10a7 | intron_variant | MODIFIER |
| 8 | 78760692 | AX-170133332 | 0 | A | G | 0.202 | -6.39E-03 | 1.31E-01 | 2.68E+00 | 2.74E+00 | 9.61E-01 | no | Gm47209 | intron_variant,non_coding_transcript_variant | MODIFIER |
| 8 | 78765255 | AX-170149966 | 0 | C | T | 0.202 | -6.39E-03 | 1.31E-01 | 2.68E+00 | 2.74E+00 | 9.61E-01 | no | Gm47209 | non_coding_transcript_exon_variant | MODIFIER |
| 8 | 78774683 | AX-97806540 | 0 | T | C | 0.202 | -6.39E-03 | 1.31E-01 | 2.68E+00 | 2.74E+00 | 9.61E-01 | no | Gm45446 | downstream_gene_variant | MODIFIER |
| 8 | 78780870 | AX-170165668 | 0 | A | C | 0.202 | -6.39E-03 | 1.31E-01 | 2.68E+00 | 2.74E+00 | 9.61E-01 | no | Gm45446 | upstream_gene_variant | MODIFIER |
| 8 | 78789685 | AX-170104083 | 0 | T | C | 0.202 | -6.39E-03 | 1.31E-01 | 2.68E+00 | 2.74E+00 | 9.61E-01 | no | | intergenic_variant | MODIFIER |
| 8 | 78805516 | AX-170158776 | 0 | T | C | 0.202 | -6.39E-03 | 1.31E-01 | 2.68E+00 | 2.74E+00 | 9.61E-01 | no | Lsm6 | downstream_gene_variant | MODIFIER |
| 8 | 78839536 | AX-168392060 | 0 | G | A | 0.202 | -6.39E-03 | 1.31E-01 | 2.68E+00 | 2.74E+00 | 9.61E-01 | no | | intergenic_variant | MODIFIER |
| 8 | 78860434 | AX-170256600 | 0 | T | C | 0.202 | -6.39E-03 | 1.31E-01 | 2.68E+00 | 2.74E+00 | 9.61E-01 | no | | intergenic_variant | MODIFIER |
| 8 | 78860465 | AX-170425177 | 0 | G | T | 0.202 | -6.39E-03 | 1.31E-01 | 2.68E+00 | 2.74E+00 | 9.61E-01 | no | | intergenic_variant | MODIFIER |
| 8 | 78862765 | AX-170249425 | 0 | A | G | 0.202 | -6.39E-03 | 1.31E-01 | 2.68E+00 | 2.74E+00 | 9.61E-01 | no | | intergenic_variant | MODIFIER |
| 8 | 78862811 | AX-97848343 | 0 | G | T | 0.202 | -6.39E-03 | 1.31E-01 | 2.68E+00 | 2.74E+00 | 9.61E-01 | no | | intergenic_variant | MODIFIER |
| 8 | 78867326 | AX-170471974 | 0 | C | T | 0.202 | -6.39E-03 | 1.31E-01 | 2.68E+00 | 2.74E+00 | 9.61E-01 | no | Gm8054 | downstream_gene_variant | MODIFIER |
| 8 | 78868095 | AX-170171751 | 0 | C | T | 0.202 | -6.39E-03 | 1.31E-01 | 2.68E+00 | 2.74E+00 | 9.61E-01 | no | Gm8054 | downstream_gene_variant | MODIFIER |
| 8 | 78900255 | AX-97828603 | 0 | A | G | 0.202 | -6.39E-03 | 1.31E-01 | 2.68E+00 | 2.74E+00 | 9.61E-01 | no | | regulatory_region_variant | MODIFIER |
| 8 | 78906350 | AX-170409029 | 0 | C | T | 0.202 | -6.39E-03 | 1.31E-01 | 2.68E+00 | 2.74E+00 | 9.61E-01 | no | | intergenic_variant | MODIFIER |
| 8 | 81950191 | AX-170170374 | 0 | C | T | 0.202 | -6.39E-03 | 1.31E-01 | 2.68E+00 | 2.74E+00 | 9.61E-01 | no | Inpp4b | intron_variant | MODIFIER |
| 1 | 73287124 | AX-170101764 | 0 | G | A | 0.114 | 1.13E-02 | 2.33E-01 | 2.69E+00 | 2.74E+00 | 9.61E-01 | no | Gm29185 | intron_variant,non_coding_transcript_variant | MODIFIER |
| 18 | 65133129 | AX-170061123 | 0 | T | C | 0.404 | -4.84E-03 | 8.10E-02 | 2.66E+00 | 2.72E+00 | 9.62E-01 | no | Nedd4l | intron_variant | MODIFIER |
| 18 | 65133850 | AX-170211628 | 0 | A | G | 0.404 | -4.84E-03 | 8.10E-02 | 2.66E+00 | 2.72E+00 | 9.62E-01 | no | Nedd4l | intron_variant | MODIFIER |
| 18 | 65170631 | AX-170179022 | 0 | C | T | 0.404 | -4.84E-03 | 8.10E-02 | 2.66E+00 | 2.72E+00 | 9.62E-01 | no | Nedd4l | intron_variant | MODIFIER |
| 18 | 65243135 | AX-170154659 | 0 | A | G | 0.404 | -4.84E-03 | 8.10E-02 | 2.66E+00 | 2.72E+00 | 9.62E-01 | no | AC157910.3 | upstream_gene_variant | MODIFIER |
| 18 | 65243409 | AX-170247966 | 0 | C | T | 0.404 | -4.84E-03 | 8.10E-02 | 2.66E+00 | 2.72E+00 | 9.62E-01 | no | AC157910.3 | upstream_gene_variant | MODIFIER |
| 18 | 65244521 | AX-170349298 | 0 | C | T | 0.404 | -4.84E-03 | 8.10E-02 | 2.66E+00 | 2.72E+00 | 9.62E-01 | no | Mir122 | upstream_gene_variant | MODIFIER |
| 18 | 65267275 | AX-170391272 | 0 | G | A | 0.404 | -4.84E-03 | 8.10E-02 | 2.66E+00 | 2.72E+00 | 9.62E-01 | no | Alpk2 | intron_variant | MODIFIER |
| 18 | 65289973 | AX-170149094 | 0 | A | C | 0.404 | -4.84E-03 | 8.10E-02 | 2.66E+00 | 2.72E+00 | 9.62E-01 | no | Alpk2 | intron_variant | MODIFIER |
| 18 | 65294997 | AX-170416046 | 0 | T | A | 0.404 | -4.84E-03 | 8.10E-02 | 2.66E+00 | 2.72E+00 | 9.62E-01 | no | Alpk2 | intron_variant | MODIFIER |
| 18 | 65397881 | AX-170015867 | 0 | A | G | 0.404 | -4.84E-03 | 8.10E-02 | 2.66E+00 | 2.72E+00 | 9.62E-01 | no | Alpk2 | upstream_gene_variant | MODIFIER |
| 18 | 65481216 | AX-97818248 | 0 | T | A | 0.404 | -4.84E-03 | 8.10E-02 | 2.66E+00 | 2.72E+00 | 9.62E-01 | no | Malt1 | downstream_gene_variant | MODIFIER |
| 18 | 65674943 | AX-170061135 | 0 | A | G | 0.404 | -4.84E-03 | 8.10E-02 | 2.66E+00 | 2.72E+00 | 9.62E-01 | no | Zfp532 | intron_variant | MODIFIER |
| 18 | 65703627 | AX-170139414 | 0 | T | C | 0.404 | -4.84E-03 | 8.10E-02 | 2.66E+00 | 2.72E+00 | 9.62E-01 | no | Oacyl | intron_variant | MODIFIER |
| 18 | 65711037 | AX-97812142 | 0 | A | G | 0.404 | -4.84E-03 | 8.10E-02 | 2.66E+00 | 2.72E+00 | 9.62E-01 | no | Oacyl | intron_variant | MODIFIER |
| 18 | 65711149 | AX-170417207 | 0 | G | A | 0.404 | -4.84E-03 | 8.10E-02 | 2.66E+00 | 2.72E+00 | 9.62E-01 | no | Oacyl | intron_variant | MODIFIER |
| 18 | 65735800 | AX-97835676 | 0 | G | A | 0.404 | -4.84E-03 | 8.10E-02 | 2.66E+00 | 2.72E+00 | 9.62E-01 | no | Oacyl | intron_variant | MODIFIER |
| 18 | 65741129 | AX-97835672 | 0 | G | A | 0.404 | -4.84E-03 | 8.10E-02 | 2.66E+00 | 2.72E+00 | 9.62E-01 | no | Oacyl | intron_variant | MODIFIER |
| 18 | 65749328 | AX-170374029 | 0 | A | T | 0.404 | -4.84E-03 | 8.10E-02 | 2.66E+00 | 2.72E+00 | 9.62E-01 | no | Oacyl | intron_variant | MODIFIER |
| 18 | 66113410 | AX-170469792 | 0 | T | C | 0.404 | -4.84E-03 | 8.10E-02 | 2.66E+00 | 2.72E+00 | 9.62E-01 | no | Ccbe1 | intron_variant,NMD_transcript_variant | MODIFIER |
| 12 | 117672853 | AX-170220289 | 1 | T | C | 0.125 | -8.45E-03 | 1.94E-01 | 2.67E+00 | 2.74E+00 | 9.62E-01 | no | Rapgef5 | intron_variant | MODIFIER |
| 9 | 69556880 | AX-170379839 | 0 | T | A | 0.202 | 5.86E-03 | 1.28E-01 | 2.69E+00 | 2.74E+00 | 9.62E-01 | no | | intergenic_variant | MODIFIER |
| 9 | 78775561 | AX-170269246 | 0 | T | C | 0.202 | 5.86E-03 | 1.28E-01 | 2.69E+00 | 2.74E+00 | 9.62E-01 | no | | regulatory_region_variant | MODIFIER |
| 9 | 111857345 | AX-170321224 | 1 | T | C | 0.107 | 4.57E-03 | 1.16E-01 | 2.67E+00 | 2.75E+00 | 9.62E-01 | no | | intergenic_variant | MODIFIER |
| 2 | 122333166 | AX-170268436 | 0 | G | A | 0.132 | 1.06E-02 | 2.21E-01 | 2.68E+00 | 2.74E+00 | 9.62E-01 | no | Duox1 | synonymous_variant | LOW |
| 7 | 51454882 | AX-170028998 | 2 | T | A | 0.427 | -4.02E-03 | 1.00E-01 | 2.67E+00 | 2.75E+00 | 9.62E-01 | no | Gm9354 | upstream_gene_variant | MODIFIER |
| 16 | 92999666 | AX-170118085 | 2 | G | A | 0.482 | -3.70E-03 | 7.86E-02 | 2.65E+00 | 2.74E+00 | 9.62E-01 | no | | intergenic_variant | MODIFIER |
| 7 | 118767824 | AX-170224135 | 1 | A | C | 0.384 | -3.62E-03 | 8.79E-02 | 2.67E+00 | 2.75E+00 | 9.62E-01 | no | Vps35l | intron_variant | MODIFIER |

| | | | | | | | | | | | | | | | |
|---|---|---|---|---|---|---|---|---|---|---|---|---|---|---|---|
| 7 | 118843719 | AX-170078597 | 1 | T | C | 0.384 | -3.62E-03 | 8.79E-02 | 2.67E+00 | 2.75E+00 | 9.62E-01 | no | Knop1 | downstream_gene_variant | MODIFIER |
| 6 | 140163108 | AX-170296238 | 1 | A | C | 0.071 | 1.30E-02 | 2.77E-01 | 2.70E+00 | 2.74E+00 | 9.62E-01 | no | Gm24174 | downstream_gene_variant | MODIFIER |
| 12 | 109977504 | AX-170425890 | 0 | C | T | 0.132 | 9.87E-03 | 1.65E-01 | 2.62E+00 | 2.73E+00 | 9.62E-01 | no | | intergenic_variant | MODIFIER |
| 1 | 70190939 | AX-97785504 | 0 | G | T | 0.114 | -1.05E-02 | 2.32E-01 | 2.69E+00 | 2.74E+00 | 9.62E-01 | no | Spag16 | intron_variant | MODIFIER |
| 13 | 45374267 | AX-170110419 | 1 | G | C | 0.464 | -2.24E-03 | 6.07E-02 | 2.63E+00 | 2.75E+00 | 9.62E-01 | no | | intergenic_variant | MODIFIER |
| 9 | 105920768 | AX-170456443 | 1 | C | T | 0.125 | -6.94E-03 | 1.19E-01 | 2.66E+00 | 2.73E+00 | 9.62E-01 | no | Col6a5 | intron_variant | MODIFIER |
| 1 | 54403697 | AX-97839946 | 1 | C | A | 0.107 | -1.08E-02 | 2.36E-01 | 2.69E+00 | 2.74E+00 | 9.62E-01 | no | Gtf3c3 | intron_variant | MODIFIER |
| 1 | 55791987 | AX-170090817 | 1 | G | T | 0.107 | -1.08E-02 | 2.36E-01 | 2.69E+00 | 2.74E+00 | 9.62E-01 | no | | intergenic_variant | MODIFIER |
| 15 | 100452626 | AX-170113674 | 1 | A | T | 0.089 | 9.95E-03 | 1.79E-01 | 2.65E+00 | 2.73E+00 | 9.62E-01 | no | n-R5s43 | downstream_gene_variant | MODIFIER |
| 4 | 134956493 | AX-97809250 | 1 | C | T | 0.17 | -3.33E-03 | 1.07E-01 | 2.68E+00 | 2.76E+00 | 9.62E-01 | no | | regulatory_region_variant | MODIFIER |
| 7 | 109498561 | AX-170078389 | 0 | G | A | 0.465 | -3.79E-03 | 9.31E-02 | 2.67E+00 | 2.75E+00 | 9.62E-01 | no | Trim66 | intron_variant | MODIFIER |
| 18 | 31980664 | AX-170352073 | 1 | C | T | 0.062 | 1.37E-02 | 3.01E-01 | 2.72E+00 | 2.74E+00 | 9.62E-01 | no | Myo7b | intron_variant | MODIFIER |
| 18 | 35149013 | AX-170504884 | 1 | C | T | 0.062 | 1.37E-02 | 3.01E-01 | 2.72E+00 | 2.74E+00 | 9.62E-01 | no | Ctnna1 | intron_variant | MODIFIER |
| 11 | 34697222 | AX-170404423 | 1 | G | A | 0.098 | 9.04E-03 | 1.82E-01 | 2.63E+00 | 2.74E+00 | 9.62E-01 | no | Dock2 | intron_variant | MODIFIER |
| 5 | 127120407 | AX-170026122 | 0 | G | A | 0.421 | 4.65E-03 | 8.86E-02 | 2.64E+00 | 2.73E+00 | 9.63E-01 | no | | intergenic_variant | MODIFIER |
| 3 | 147746900 | AX-97832349 | 0 | C | T | 0.351 | 3.82E-03 | 7.32E-02 | 2.66E+00 | 2.73E+00 | 9.63E-01 | no | | intergenic_variant | MODIFIER |
| 3 | 147747201 | AX-170190900 | 0 | C | T | 0.351 | 3.82E-03 | 7.32E-02 | 2.66E+00 | 2.73E+00 | 9.63E-01 | no | | intergenic_variant | MODIFIER |
| 12 | 78928077 | AX-170211663 | 0 | G | A | 0.053 | 1.03E-02 | 1.97E-01 | 2.62E+00 | 2.73E+00 | 9.63E-01 | no | | intergenic_variant | MODIFIER |
| 10 | 120336113 | AX-170004030 | 0 | A | G | 0.342 | -3.27E-03 | 7.06E-02 | 2.65E+00 | 2.74E+00 | 9.63E-01 | no | | intergenic_variant | MODIFIER |
| 9 | 84577478 | AX-97809207 | 0 | G | T | 0.211 | 4.47E-03 | 1.21E-01 | 2.67E+00 | 2.75E+00 | 9.63E-01 | no | Gm39388 | upstream_gene_variant | MODIFIER |
| 7 | 94930032 | AX-170078116 | 2 | C | A | 0.136 | -9.54E-03 | 2.14E-01 | 2.66E+00 | 2.74E+00 | 9.63E-01 | no | | intergenic_variant | MODIFIER |
| 7 | 52135854 | AX-170526499 | 0 | C | T | 0.43 | 5.10E-03 | 9.67E-02 | 2.65E+00 | 2.73E+00 | 9.63E-01 | no | | intergenic_variant | MODIFIER |
| 5 | 70016135 | AX-170211553 | 1 | C | G | 0.446 | -3.81E-03 | 9.79E-02 | 2.66E+00 | 2.75E+00 | 9.63E-01 | no | | intergenic_variant | MODIFIER |
| 10 | 4959876 | AX-170468444 | 1 | C | A | 0.196 | 4.43E-03 | 9.87E-02 | 2.64E+00 | 2.74E+00 | 9.63E-01 | no | Esr1 | intron_variant | MODIFIER |
| 13 | 42453906 | AX-170176576 | 0 | C | T | 0.491 | -2.03E-03 | 6.17E-02 | 2.64E+00 | 2.75E+00 | 9.63E-01 | no | | intergenic_variant | MODIFIER |
| 13 | 42455340 | AX-170463231 | 0 | G | T | 0.491 | -2.03E-03 | 6.17E-02 | 2.64E+00 | 2.75E+00 | 9.63E-01 | no | | intergenic_variant | MODIFIER |
| 13 | 42455738 | AX-170008185 | 0 | G | A | 0.491 | -2.03E-03 | 6.17E-02 | 2.64E+00 | 2.75E+00 | 9.63E-01 | no | | intergenic_variant | MODIFIER |
| 13 | 42479548 | AX-170337488 | 0 | T | A | 0.491 | -2.03E-03 | 6.17E-02 | 2.64E+00 | 2.75E+00 | 9.63E-01 | no | | intergenic_variant | MODIFIER |
| 13 | 42479609 | AX-170087412 | 0 | A | G | 0.491 | -2.03E-03 | 6.17E-02 | 2.64E+00 | 2.75E+00 | 9.63E-01 | no | | intergenic_variant | MODIFIER |
| 13 | 42482840 | AX-97821128 | 0 | T | C | 0.491 | -2.03E-03 | 6.17E-02 | 2.64E+00 | 2.75E+00 | 9.63E-01 | no | | intergenic_variant | MODIFIER |
| 13 | 42515468 | AX-168396522 | 0 | G | A | 0.491 | -2.03E-03 | 6.17E-02 | 2.64E+00 | 2.75E+00 | 9.63E-01 | no | | intergenic_variant | MODIFIER |
| 13 | 42530700 | AX-170008186 | 0 | G | A | 0.491 | -2.03E-03 | 6.17E-02 | 2.64E+00 | 2.75E+00 | 9.63E-01 | no | | intergenic_variant | MODIFIER |
| 13 | 42531485 | AX-168385272 | 0 | T | C | 0.491 | -2.03E-03 | 6.17E-02 | 2.64E+00 | 2.75E+00 | 9.63E-01 | no | | intergenic_variant | MODIFIER |
| 13 | 42544730 | AX-170051103 | 0 | C | G | 0.491 | -2.03E-03 | 6.17E-02 | 2.64E+00 | 2.75E+00 | 9.63E-01 | no | | regulatory_region_variant | MODIFIER |
| 13 | 42547214 | AX-97833781 | 0 | C | T | 0.491 | -2.03E-03 | 6.17E-02 | 2.64E+00 | 2.75E+00 | 9.63E-01 | no | | intergenic_variant | MODIFIER |
| 13 | 42603608 | AX-170087413 | 0 | C | T | 0.491 | -2.03E-03 | 6.17E-02 | 2.64E+00 | 2.75E+00 | 9.63E-01 | no | | intergenic_variant | MODIFIER |
| 13 | 42610343 | AX-170339156 | 0 | A | G | 0.491 | -2.03E-03 | 6.17E-02 | 2.64E+00 | 2.75E+00 | 9.63E-01 | no | | intergenic_variant | MODIFIER |
| 13 | 42610526 | AX-170171161 | 0 | G | C | 0.491 | -2.03E-03 | 6.17E-02 | 2.64E+00 | 2.75E+00 | 9.63E-01 | no | | intergenic_variant | MODIFIER |
| 13 | 42628474 | AX-97781277 | 0 | A | G | 0.491 | -2.03E-03 | 6.17E-02 | 2.64E+00 | 2.75E+00 | 9.63E-01 | no | Gm23086 | downstream_gene_variant | MODIFIER |
| 13 | 42646234 | AX-97856216 | 0 | T | C | 0.491 | -2.03E-03 | 6.17E-02 | 2.64E+00 | 2.75E+00 | 9.63E-01 | no | | intergenic_variant | MODIFIER |
| 13 | 42666133 | AX-97784394 | 0 | G | A | 0.491 | -2.03E-03 | 6.17E-02 | 2.64E+00 | 2.75E+00 | 9.63E-01 | no | | intergenic_variant | MODIFIER |
| 13 | 42726555 | AX-97790021 | 0 | A | G | 0.491 | -2.03E-03 | 6.17E-02 | 2.64E+00 | 2.75E+00 | 9.63E-01 | no | Phactr1 | intron_variant | MODIFIER |
| 13 | 42727117 | AX-170336276 | 0 | A | T | 0.491 | -2.03E-03 | 6.17E-02 | 2.64E+00 | 2.75E+00 | 9.63E-01 | no | Phactr1 | intron_variant | MODIFIER |
| 13 | 42736234 | AX-170051108 | 0 | T | C | 0.491 | -2.03E-03 | 6.17E-02 | 2.64E+00 | 2.75E+00 | 9.63E-01 | no | Phactr1 | intron_variant | MODIFIER |
| 13 | 42736706 | AX-170051109 | 0 | C | T | 0.491 | -2.03E-03 | 6.17E-02 | 2.64E+00 | 2.75E+00 | 9.63E-01 | no | Phactr1 | intron_variant | MODIFIER |
| 13 | 42738939 | AX-170373658 | 0 | C | T | 0.491 | -2.03E-03 | 6.17E-02 | 2.64E+00 | 2.75E+00 | 9.63E-01 | no | Phactr1 | intron_variant | MODIFIER |
| 6 | 139061747 | AX-170028366 | 1 | C | T | 0.071 | 1.26E-02 | 2.77E-01 | 2.70E+00 | 2.74E+00 | 9.63E-01 | no | | intergenic_variant | MODIFIER |
| 11 | 48097181 | AX-170455747 | 1 | G | T | 0.089 | -9.55E-03 | 1.83E-01 | 2.63E+00 | 2.73E+00 | 9.63E-01 | no | | intergenic_variant | MODIFIER |
| 8 | 77829473 | AX-97824178 | 0 | T | C | 0.193 | 5.57E-03 | 1.30E-01 | 2.67E+00 | 2.74E+00 | 9.63E-01 | no | | regulatory_region_variant | MODIFIER |
| 18 | 64846663 | AX-97855586 | 0 | A | C | 0.421 | 2.58E-03 | 8.04E-02 | 2.68E+00 | 2.75E+00 | 9.63E-01 | no | | intergenic_variant | MODIFIER |
| 8 | 75531492 | AX-170245026 | 0 | C | T | 0.193 | 6.18E-03 | 1.39E-01 | 2.69E+00 | 2.74E+00 | 9.63E-01 | no | Iqcm | intron_variant | MODIFIER |
| 4 | 57054056 | AX-170283906 | 0 | A | G | 0.368 | -4.73E-03 | 1.07E-01 | 2.66E+00 | 2.74E+00 | 9.63E-01 | no | Epb41l4b | intron_variant | MODIFIER |
| 8 | 76376670 | AX-170346750 | 0 | C | T | 0.193 | -6.19E-03 | 1.33E-01 | 2.67E+00 | 2.74E+00 | 9.63E-01 | no | Gm45740 | intron_variant,non_coding_transcript_variant | MODIFIER |
| 12 | 72907364 | AX-97827999 | 0 | A | G | 0.061 | 8.95E-03 | 2.18E-01 | 2.62E+00 | 2.74E+00 | 9.63E-01 | no | 4930447C04Rik | intron_variant | MODIFIER |

| | | | | | | | | | | | | | | | |
|---|---|---|---|---|---|---|---|---|---|---|---|---|---|---|---|
| 12 | 116355588 | AX-170433378 | 2 | T | C | 0.136 | -5.43E-03 | 1.38E-01 | 2.65E+00 | 2.74E+00 | 9.63E-01 | no | Esyt2 | intron_variant | MODIFIER |
| 10 | 43452645 | AX-170003126 | 1 | G | C | 0.375 | -3.70E-03 | 7.44E-02 | 2.63E+00 | 2.73E+00 | 9.64E-01 | no | Pdss2 | intron_variant | MODIFIER |
| 7 | 102585149 | AX-170516452 | 1 | T | A | 0.473 | -3.69E-03 | 9.59E-02 | 2.68E+00 | 2.75E+00 | 9.64E-01 | no | Olfr551 | downstream_gene_variant | MODIFIER |
| 1 | 39232378 | AX-170307225 | 0 | C | T | 0.386 | 2.91E-03 | 7.56E-02 | 2.67E+00 | 2.75E+00 | 9.64E-01 | no | Npas2 | intron_variant | MODIFIER |
| 1 | 39243848 | AX-170520455 | 0 | G | T | 0.386 | 2.91E-03 | 7.56E-02 | 2.67E+00 | 2.75E+00 | 9.64E-01 | no | Npas2 | intron_variant | MODIFIER |
| 1 | 39244101 | AX-170263028 | 0 | A | G | 0.386 | 2.91E-03 | 7.56E-02 | 2.67E+00 | 2.75E+00 | 9.64E-01 | no | Npas2 | intron_variant | MODIFIER |
| 1 | 39252351 | AX-97794025 | 0 | T | C | 0.386 | 2.91E-03 | 7.56E-02 | 2.67E+00 | 2.75E+00 | 9.64E-01 | no | Npas2 | intron_variant | MODIFIER |
| 1 | 39306594 | AX-170352638 | 0 | A | C | 0.386 | 2.91E-03 | 7.56E-02 | 2.67E+00 | 2.75E+00 | 9.64E-01 | no | Npas2 | intron_variant | MODIFIER |
| 1 | 39306969 | AX-170290157 | 0 | A | G | 0.386 | 2.91E-03 | 7.56E-02 | 2.67E+00 | 2.75E+00 | 9.64E-01 | no | Npas2 | intron_variant | MODIFIER |
| 1 | 39307757 | AX-170275807 | 0 | G | A | 0.386 | 2.91E-03 | 7.56E-02 | 2.67E+00 | 2.75E+00 | 9.64E-01 | no | Npas2 | intron_variant | MODIFIER |
| 1 | 39308547 | AX-170210037 | 0 | T | C | 0.386 | 2.91E-03 | 7.56E-02 | 2.67E+00 | 2.75E+00 | 9.64E-01 | no | Npas2 | intron_variant | MODIFIER |
| 1 | 39324204 | AX-170040958 | 0 | C | T | 0.386 | 2.91E-03 | 7.56E-02 | 2.67E+00 | 2.75E+00 | 9.64E-01 | no | Npas2 | intron_variant | MODIFIER |
| 1 | 39347535 | AX-170259304 | 0 | A | C | 0.386 | 2.91E-03 | 7.56E-02 | 2.67E+00 | 2.75E+00 | 9.64E-01 | no | Npas2 | synonymous_variant | LOW |
| 1 | 39348863 | AX-170241751 | 0 | G | A | 0.386 | 2.91E-03 | 7.56E-02 | 2.67E+00 | 2.75E+00 | 9.64E-01 | no | Npas2 | intron_variant | MODIFIER |
| 1 | 39349627 | AX-170369570 | 0 | C | T | 0.386 | 2.91E-03 | 7.56E-02 | 2.67E+00 | 2.75E+00 | 9.64E-01 | no | Npas2 | intron_variant | MODIFIER |
| 1 | 39633497 | AX-170000331 | 0 | C | T | 0.386 | 2.91E-03 | 7.56E-02 | 2.67E+00 | 2.75E+00 | 9.64E-01 | no | Creg2 | intron_variant | MODIFIER |
| 1 | 39633779 | AX-170132334 | 0 | C | T | 0.386 | 2.91E-03 | 7.56E-02 | 2.67E+00 | 2.75E+00 | 9.64E-01 | no | Creg2 | intron_variant | MODIFIER |
| 1 | 39637388 | AX-170437596 | 0 | G | T | 0.386 | 2.91E-03 | 7.56E-02 | 2.67E+00 | 2.75E+00 | 9.64E-01 | no | Creg2 | intron_variant | MODIFIER |
| 1 | 39638316 | AX-170000330 | 0 | A | G | 0.386 | 2.91E-03 | 7.56E-02 | 2.67E+00 | 2.75E+00 | 9.64E-01 | no | Creg2 | intron_variant | MODIFIER |
| 1 | 39666809 | AX-170277629 | 0 | C | T | 0.386 | 2.91E-03 | 7.56E-02 | 2.67E+00 | 2.75E+00 | 9.64E-01 | no | Rfx8 | intron_variant | MODIFIER |
| 1 | 39712542 | AX-170334834 | 0 | A | G | 0.386 | 2.91E-03 | 7.56E-02 | 2.67E+00 | 2.75E+00 | 9.64E-01 | no | Rfx8 | intron_variant | MODIFIER |
| 1 | 39714907 | AX-170374249 | 0 | C | A | 0.386 | 2.91E-03 | 7.56E-02 | 2.67E+00 | 2.75E+00 | 9.64E-01 | no | Rfx8 | intron_variant | MODIFIER |
| 1 | 39719384 | AX-170040966 | 0 | T | G | 0.386 | 2.91E-03 | 7.56E-02 | 2.67E+00 | 2.75E+00 | 9.64E-01 | no | Rfx8 | intron_variant | MODIFIER |
| 1 | 39732405 | AX-170279295 | 0 | G | A | 0.386 | 2.91E-03 | 7.56E-02 | 2.67E+00 | 2.75E+00 | 9.64E-01 | no | | regulatory_region_variant | MODIFIER |
| 1 | 39732957 | AX-170249497 | 0 | A | G | 0.386 | 2.91E-03 | 7.56E-02 | 2.67E+00 | 2.75E+00 | 9.64E-01 | no | | regulatory_region_variant | MODIFIER |
| 1 | 39749631 | AX-170371382 | 0 | A | G | 0.386 | 2.91E-03 | 7.56E-02 | 2.67E+00 | 2.75E+00 | 9.64E-01 | no | | regulatory_region_variant | MODIFIER |
| 1 | 39762541 | AX-170218150 | 0 | T | C | 0.386 | 2.91E-03 | 7.56E-02 | 2.67E+00 | 2.75E+00 | 9.64E-01 | no | | regulatory_region_variant | MODIFIER |
| 1 | 39766139 | AX-170166981 | 0 | G | A | 0.386 | 2.91E-03 | 7.56E-02 | 2.67E+00 | 2.75E+00 | 9.64E-01 | no | | intergenic_variant | MODIFIER |
| 1 | 39773809 | AX-97857176 | 0 | C | T | 0.386 | 2.91E-03 | 7.56E-02 | 2.67E+00 | 2.75E+00 | 9.64E-01 | no | | regulatory_region_variant | MODIFIER |
| 1 | 39785776 | AX-170518857 | 0 | T | G | 0.386 | 2.91E-03 | 7.56E-02 | 2.67E+00 | 2.75E+00 | 9.64E-01 | no | | intergenic_variant | MODIFIER |
| 1 | 39882244 | AX-170160901 | 0 | C | T | 0.386 | 2.91E-03 | 7.56E-02 | 2.67E+00 | 2.75E+00 | 9.64E-01 | no | | intergenic_variant | MODIFIER |
| 1 | 39889105 | AX-170108279 | 0 | T | A | 0.386 | 2.91E-03 | 7.56E-02 | 2.67E+00 | 2.75E+00 | 9.64E-01 | no | | intergenic_variant | MODIFIER |
| 1 | 39903525 | AX-170191144 | 0 | T | C | 0.386 | 2.91E-03 | 7.56E-02 | 2.67E+00 | 2.75E+00 | 9.64E-01 | no | Map4k4 | intron_variant | MODIFIER |
| 1 | 39909353 | AX-170086451 | 0 | G | A | 0.386 | 2.91E-03 | 7.56E-02 | 2.67E+00 | 2.75E+00 | 9.64E-01 | no | Map4k4 | intron_variant | MODIFIER |
| 1 | 39912024 | AX-170085968 | 0 | G | A | 0.386 | 2.91E-03 | 7.56E-02 | 2.67E+00 | 2.75E+00 | 9.64E-01 | no | Map4k4 | intron_variant | MODIFIER |
| 1 | 39912687 | AX-170212216 | 0 | T | C | 0.386 | 2.91E-03 | 7.56E-02 | 2.67E+00 | 2.75E+00 | 9.64E-01 | no | Map4k4 | intron_variant | MODIFIER |
| 11 | 74290222 | AX-170479872 | 1 | A | G | 0.268 | 2.92E-03 | 7.29E-02 | 2.61E+00 | 2.74E+00 | 9.64E-01 | no | Olfr59 | 3_prime_UTR_variant | MODIFIER |
| 12 | 107141124 | AX-170246492 | 0 | G | A | 0.114 | -8.23E-03 | 1.97E-01 | 2.67E+00 | 2.74E+00 | 9.64E-01 | no | | intergenic_variant | MODIFIER |
| 18 | 29836989 | AX-170015441 | 0 | A | G | 0.044 | -1.24E-02 | 2.28E-01 | 2.61E+00 | 2.73E+00 | 9.64E-01 | no | | intergenic_variant | MODIFIER |
| 4 | 58541677 | AX-170160171 | 1 | G | A | 0.411 | 5.03E-03 | 9.57E-02 | 2.61E+00 | 2.73E+00 | 9.64E-01 | no | Lpar1 | intron_variant | MODIFIER |
| 13 | 53400329 | AX-170230784 | 1 | C | T | 0.518 | 3.91E-03 | 6.59E-02 | 2.62E+00 | 2.73E+00 | 9.64E-01 | no | Gm2762 | intron_variant,non_coding_transcript_variant | MODIFIER |
| 10 | 32468775 | AX-97800989 | 1 | G | A | 0.393 | -3.75E-03 | 7.46E-02 | 2.62E+00 | 2.73E+00 | 9.64E-01 | no | Nkain2 | intron_variant | MODIFIER |
| 7 | 102003319 | AX-170485434 | 0 | C | T | 0.474 | -3.57E-03 | 9.56E-02 | 2.68E+00 | 2.75E+00 | 9.64E-01 | no | Numa1 | intron_variant | MODIFIER |
| 7 | 102341971 | AX-170029648 | 0 | C | T | 0.474 | -3.57E-03 | 9.56E-02 | 2.68E+00 | 2.75E+00 | 9.64E-01 | no | Stim1 | intron_variant | MODIFIER |
| 7 | 102363079 | AX-170136153 | 0 | T | A | 0.474 | -3.57E-03 | 9.56E-02 | 2.68E+00 | 2.75E+00 | 9.64E-01 | no | Stim1 | intron_variant | MODIFIER |
| 7 | 102390746 | AX-170089991 | 0 | G | T | 0.474 | -3.57E-03 | 9.56E-02 | 2.68E+00 | 2.75E+00 | 9.64E-01 | no | Stim1 | intron_variant | MODIFIER |
| 7 | 102455513 | AX-170527446 | 0 | A | G | 0.474 | -3.57E-03 | 9.56E-02 | 2.68E+00 | 2.75E+00 | 9.64E-01 | no | Rrm1 | intron_variant | MODIFIER |
| 7 | 102501975 | AX-170245178 | 0 | C | T | 0.474 | -3.57E-03 | 9.56E-02 | 2.68E+00 | 2.75E+00 | 9.64E-01 | no | Olfr545 | upstream_gene_variant | MODIFIER |
| 7 | 102541624 | AX-170397423 | 0 | G | A | 0.474 | -3.57E-03 | 9.56E-02 | 2.68E+00 | 2.75E+00 | 9.64E-01 | no | Olfr548-ps1 | upstream_gene_variant | MODIFIER |
| 7 | 102552358 | AX-170207608 | 0 | G | A | 0.474 | -3.57E-03 | 9.56E-02 | 2.68E+00 | 2.75E+00 | 9.64E-01 | no | Olfr549 | upstream_gene_variant | MODIFIER |
| 7 | 102606723 | AX-170111790 | 0 | T | A | 0.474 | -3.57E-03 | 9.56E-02 | 2.68E+00 | 2.75E+00 | 9.64E-01 | no | Olfr552 | downstream_gene_variant | MODIFIER |
| 7 | 102670048 | AX-170482919 | 0 | C | G | 0.474 | -3.57E-03 | 9.56E-02 | 2.68E+00 | 2.75E+00 | 9.64E-01 | no | Olfr556 | missense_variant | MODERATE |
| 7 | 102707621 | AX-170307078 | 0 | A | T | 0.474 | -3.57E-03 | 9.56E-02 | 2.68E+00 | 2.75E+00 | 9.64E-01 | no | Olfr558 | intron_variant | MODIFIER |
| 7 | 102817987 | AX-170029663 | 0 | C | A | 0.474 | -3.57E-03 | 9.56E-02 | 2.68E+00 | 2.75E+00 | 9.64E-01 | no | Olfr565-ps1 | non_coding_transcript_exon_variant | MODIFIER |

| | | | | | | | | | | | | | | | |
|---|---|---|---|---|---|---|---|---|---|---|---|---|---|---|---|
| 7 | 102864898 | AX-97803482 | 0 | C | A | 0.474 | -3.57E-03 | 9.56E-02 | 2.68E+00 | 2.75E+00 | 9.64E-01 | no | Olfr567-ps1 | upstream_gene_variant | MODIFIER |
| 7 | 102879933 | AX-170029664 | 0 | A | C | 0.474 | -3.57E-03 | 9.56E-02 | 2.68E+00 | 2.75E+00 | 9.64E-01 | no | Olfr568 | downstream_gene_variant | MODIFIER |
| 7 | 102880819 | AX-97799857 | 0 | G | A | 0.474 | -3.57E-03 | 9.56E-02 | 2.68E+00 | 2.75E+00 | 9.64E-01 | no | Olfr568 | downstream_gene_variant | MODIFIER |
| 7 | 102909194 | AX-170078287 | 0 | G | A | 0.474 | -3.57E-03 | 9.56E-02 | 2.68E+00 | 2.75E+00 | 9.64E-01 | no | Olfr571 | missense_variant | MODERATE |
| 7 | 102909944 | AX-170078286 | 0 | C | T | 0.474 | -3.57E-03 | 9.56E-02 | 2.68E+00 | 2.75E+00 | 9.64E-01 | no | Olfr571 | upstream_gene_variant | MODIFIER |
| 7 | 102913314 | AX-170452892 | 0 | G | C | 0.474 | -3.57E-03 | 9.56E-02 | 2.68E+00 | 2.75E+00 | 9.64E-01 | no | Olfr571 | upstream_gene_variant | MODIFIER |
| 7 | 102916460 | AX-170078285 | 0 | C | T | 0.474 | -3.57E-03 | 9.56E-02 | 2.68E+00 | 2.75E+00 | 9.64E-01 | no | Olfr571 | intron_variant | MODIFIER |
| 7 | 102949065 | AX-170264400 | 0 | T | A | 0.474 | -3.57E-03 | 9.56E-02 | 2.68E+00 | 2.75E+00 | 9.64E-01 | no | Olfr574 | missense_variant | MODERATE |
| 7 | 103581673 | AX-170190129 | 0 | A | T | 0.474 | -3.57E-03 | 9.56E-02 | 2.68E+00 | 2.75E+00 | 9.64E-01 | no | Olfr617 | upstream_gene_variant | MODIFIER |
| 7 | 103590270 | AX-170098114 | 0 | G | A | 0.474 | -3.57E-03 | 9.56E-02 | 2.68E+00 | 2.75E+00 | 9.64E-01 | no | Olfr618 | intron_variant | MODIFIER |
| 7 | 103628703 | AX-170308719 | 0 | C | G | 0.474 | -3.57E-03 | 9.56E-02 | 2.68E+00 | 2.75E+00 | 9.64E-01 | no | Olfr621-ps1 | downstream_gene_variant | MODIFIER |
| 7 | 103628943 | AX-170499913 | 0 | A | G | 0.474 | -3.57E-03 | 9.56E-02 | 2.68E+00 | 2.75E+00 | 9.64E-01 | no | Olfr621-ps1 | downstream_gene_variant | MODIFIER |
| 7 | 103629023 | AX-170524705 | 0 | A | G | 0.474 | -3.57E-03 | 9.56E-02 | 2.68E+00 | 2.75E+00 | 9.64E-01 | no | Olfr621-ps1 | non_coding_transcript_exon_variant | MODIFIER |
| 7 | 103637497 | AX-170097789 | 0 | T | C | 0.474 | -3.57E-03 | 9.56E-02 | 2.68E+00 | 2.75E+00 | 9.64E-01 | no | Olfr622 | downstream_gene_variant | MODIFIER |
| 7 | 103642975 | AX-170078299 | 0 | G | A | 0.474 | -3.57E-03 | 9.56E-02 | 2.68E+00 | 2.75E+00 | 9.64E-01 | no | Olfr622 | upstream_gene_variant | MODIFIER |
| 7 | 103659355 | AX-97822152 | 0 | T | C | 0.474 | -3.57E-03 | 9.56E-02 | 2.68E+00 | 2.75E+00 | 9.64E-01 | no | Olfr623 | downstream_gene_variant | MODIFIER |
| 7 | 103769126 | AX-170317130 | 0 | C | T | 0.474 | -3.57E-03 | 9.56E-02 | 2.68E+00 | 2.75E+00 | 9.64E-01 | no | Olfr69 | intron_variant | MODIFIER |
| 7 | 104230845 | AX-97802916 | 0 | A | G | 0.474 | -3.57E-03 | 9.56E-02 | 2.68E+00 | 2.75E+00 | 9.64E-01 | no | Trim6 | intron_variant | MODIFIER |
| 7 | 104239041 | AX-170029676 | 0 | A | G | 0.474 | -3.57E-03 | 9.56E-02 | 2.68E+00 | 2.75E+00 | 9.64E-01 | no | Trim6 | downstream_gene_variant | MODIFIER |
| 7 | 104296664 | AX-97824967 | 0 | C | A | 0.474 | -3.57E-03 | 9.56E-02 | 2.68E+00 | 2.75E+00 | 9.64E-01 | no | Trim12a | downstream_gene_variant | MODIFIER |
| 7 | 104364850 | AX-170199235 | 0 | A | G | 0.474 | -3.57E-03 | 9.56E-02 | 2.68E+00 | 2.75E+00 | 9.64E-01 | no | Trim30b | intron_variant | MODIFIER |
| 7 | 104479672 | AX-170472824 | 0 | C | G | 0.474 | -3.57E-03 | 9.56E-02 | 2.68E+00 | 2.75E+00 | 9.64E-01 | no | Trim30d | intron_variant | MODIFIER |
| 7 | 104486669 | AX-170113924 | 0 | A | G | 0.474 | -3.57E-03 | 9.56E-02 | 2.68E+00 | 2.75E+00 | 9.64E-01 | no | Trim30d | intron_variant | MODIFIER |
| 7 | 104551017 | AX-170163699 | 0 | A | G | 0.474 | -3.57E-03 | 9.56E-02 | 2.68E+00 | 2.75E+00 | 9.64E-01 | no | Olfr651 | upstream_gene_variant | MODIFIER |
| 7 | 104574845 | AX-170167504 | 0 | G | A | 0.474 | -3.57E-03 | 9.56E-02 | 2.68E+00 | 2.75E+00 | 9.64E-01 | no | Olfr653 | upstream_gene_variant | MODIFIER |
| 7 | 104598267 | AX-170199191 | 0 | G | C | 0.474 | -3.57E-03 | 9.56E-02 | 2.68E+00 | 2.75E+00 | 9.64E-01 | no | Olfr655 | upstream_gene_variant | MODIFIER |
| 7 | 105609899 | AX-170519158 | 0 | C | A | 0.474 | -3.57E-03 | 9.56E-02 | 2.68E+00 | 2.75E+00 | 9.64E-01 | no | Trim3 | downstream_gene_variant | MODIFIER |
| 7 | 105613778 | AX-170516613 | 0 | G | A | 0.474 | -3.57E-03 | 9.56E-02 | 2.68E+00 | 2.75E+00 | 9.64E-01 | no | Trim3 | intron_variant | MODIFIER |
| 7 | 105701891 | AX-97811983 | 0 | T | C | 0.474 | -3.57E-03 | 9.56E-02 | 2.68E+00 | 2.75E+00 | 9.64E-01 | no | Dnhd1 | intron_variant | MODIFIER |
| 7 | 105762942 | AX-97829652 | 0 | C | T | 0.474 | -3.57E-03 | 9.56E-02 | 2.68E+00 | 2.75E+00 | 9.64E-01 | no | Dchs1 | synonymous_variant | LOW |
| 7 | 106444512 | AX-97853614 | 0 | T | C | 0.474 | -3.57E-03 | 9.56E-02 | 2.68E+00 | 2.75E+00 | 9.64E-01 | no | | intergenic_variant | MODIFIER |
| 7 | 106448335 | AX-170120608 | 0 | A | G | 0.474 | -3.57E-03 | 9.56E-02 | 2.68E+00 | 2.75E+00 | 9.64E-01 | no | | intergenic_variant | MODIFIER |
| 7 | 106505327 | AX-170135173 | 0 | G | T | 0.474 | -3.57E-03 | 9.56E-02 | 2.68E+00 | 2.75E+00 | 9.64E-01 | no | Gm8995 | non_coding_transcript_exon_variant | MODIFIER |
| 7 | 106510982 | AX-170113179 | 0 | T | C | 0.474 | -3.57E-03 | 9.56E-02 | 2.68E+00 | 2.75E+00 | 9.64E-01 | no | Gm8995 | non_coding_transcript_exon_variant | MODIFIER |
| 7 | 106596816 | AX-170096057 | 0 | G | A | 0.474 | -3.57E-03 | 9.56E-02 | 2.68E+00 | 2.75E+00 | 9.64E-01 | no | Gm1966 | non_coding_transcript_exon_variant | MODIFIER |
| 7 | 106596963 | AX-170294735 | 0 | T | C | 0.474 | -3.57E-03 | 9.56E-02 | 2.68E+00 | 2.75E+00 | 9.64E-01 | no | Gm1966 | non_coding_transcript_exon_variant | MODIFIER |
| 7 | 106605536 | AX-97855846 | 0 | G | A | 0.474 | -3.57E-03 | 9.56E-02 | 2.68E+00 | 2.75E+00 | 9.64E-01 | no | Gm1966 | upstream_gene_variant | MODIFIER |
| 7 | 106640987 | AX-170178028 | 0 | T | A | 0.474 | -3.57E-03 | 9.56E-02 | 2.68E+00 | 2.75E+00 | 9.64E-01 | no | | intergenic_variant | MODIFIER |
| 7 | 106650507 | AX-170218006 | 0 | C | T | 0.474 | -3.57E-03 | 9.56E-02 | 2.68E+00 | 2.75E+00 | 9.64E-01 | no | | regulatory_region_variant | MODIFIER |
| 7 | 106650624 | AX-97842197 | 0 | T | C | 0.474 | -3.57E-03 | 9.56E-02 | 2.68E+00 | 2.75E+00 | 9.64E-01 | no | | regulatory_region_variant | MODIFIER |
| 7 | 106862487 | AX-170311123 | 0 | T | A | 0.474 | -3.57E-03 | 9.56E-02 | 2.68E+00 | 2.75E+00 | 9.64E-01 | no | Olfr704 | upstream_gene_variant | MODIFIER |
| 7 | 106864731 | AX-97852756 | 0 | A | G | 0.474 | -3.57E-03 | 9.56E-02 | 2.68E+00 | 2.75E+00 | 9.64E-01 | no | Olfr704 | intron_variant | MODIFIER |
| 7 | 106867290 | AX-170029721 | 0 | T | G | 0.474 | -3.57E-03 | 9.56E-02 | 2.68E+00 | 2.75E+00 | 9.64E-01 | no | Olfr704 | downstream_gene_variant | MODIFIER |
| 7 | 106899426 | AX-170396825 | 0 | A | G | 0.474 | -3.57E-03 | 9.56E-02 | 2.68E+00 | 2.75E+00 | 9.64E-01 | no | Olfr707 | upstream_gene_variant | MODIFIER |
| 7 | 106909896 | AX-170435978 | 0 | C | T | 0.474 | -3.57E-03 | 9.56E-02 | 2.68E+00 | 2.75E+00 | 9.64E-01 | no | Olfr1532-ps1 | upstream_gene_variant | MODIFIER |
| 7 | 106916204 | AX-170134409 | 0 | T | C | 0.474 | -3.57E-03 | 9.56E-02 | 2.68E+00 | 2.75E+00 | 9.64E-01 | no | Olfr1532-ps1 | downstream_gene_variant | MODIFIER |
| 7 | 106916390 | AX-170241363 | 0 | A | G | 0.474 | -3.57E-03 | 9.56E-02 | 2.68E+00 | 2.75E+00 | 9.64E-01 | no | Olfr1532-ps1 | downstream_gene_variant | MODIFIER |
| 7 | 106916578 | AX-170337042 | 0 | A | G | 0.474 | -3.57E-03 | 9.56E-02 | 2.68E+00 | 2.75E+00 | 9.64E-01 | no | Olfr1532-ps1 | downstream_gene_variant | MODIFIER |
| 7 | 106928532 | AX-170251768 | 0 | C | T | 0.474 | -3.57E-03 | 9.56E-02 | 2.68E+00 | 2.75E+00 | 9.64E-01 | no | Olfr709-ps1 | upstream_gene_variant | MODIFIER |
| 7 | 106932033 | AX-97821117 | 0 | T | C | 0.474 | -3.57E-03 | 9.56E-02 | 2.68E+00 | 2.75E+00 | 9.64E-01 | no | Olfr709-ps1 | upstream_gene_variant | MODIFIER |
| 7 | 106934210 | AX-170124354 | 0 | G | T | 0.474 | -3.57E-03 | 9.56E-02 | 2.68E+00 | 2.75E+00 | 9.64E-01 | no | Olfr709-ps1 | upstream_gene_variant | MODIFIER |
| 7 | 106956377 | AX-170029717 | 0 | G | A | 0.474 | -3.57E-03 | 9.56E-02 | 2.68E+00 | 2.75E+00 | 9.64E-01 | no | Olfr6 | synonymous_variant | LOW |
| 7 | 106956945 | AX-170237949 | 0 | C | G | 0.474 | -3.57E-03 | 9.56E-02 | 2.68E+00 | 2.75E+00 | 9.64E-01 | no | Olfr6 | 5_prime_UTR_variant | MODIFIER |
| 7 | 106970816 | AX-97848232 | 0 | C | T | 0.474 | -3.57E-03 | 9.56E-02 | 2.68E+00 | 2.75E+00 | 9.64E-01 | no | Olfr711 | 3_prime_UTR_variant | MODIFIER |

| | | | | | | | | | | | | | | | |
|---|---|---|---|---|---|---|---|---|---|---|---|---|---|---|---|
| 7 | 106971872 | AX-170239280 | 0 | G | A | 0.474 | -3.57E-03 | 9.56E-02 | 2.68E+00 | 2.75E+00 | 9.64E-01 | no | Olfr711 | synonymous_variant | LOW |
| 7 | 106975357 | AX-97803511 | 0 | A | G | 0.474 | -3.57E-03 | 9.56E-02 | 2.68E+00 | 2.75E+00 | 9.64E-01 | no | Olfr711 | 5_prime_UTR_variant | MODIFIER |
| 7 | 107001966 | AX-97800424 | 0 | A | G | 0.474 | -3.57E-03 | 9.56E-02 | 2.68E+00 | 2.75E+00 | 9.64E-01 | no | Olfr2 | 5_prime_UTR_variant | MODIFIER |
| 7 | 107027384 | AX-170147678 | 0 | T | C | 0.474 | -3.57E-03 | 9.56E-02 | 2.68E+00 | 2.75E+00 | 9.64E-01 | no | Olfr713 | upstream_gene_variant | MODIFIER |
| 7 | 107027665 | AX-97847961 | 0 | G | A | 0.474 | -3.57E-03 | 9.56E-02 | 2.68E+00 | 2.75E+00 | 9.64E-01 | no | Olfr713 | upstream_gene_variant | MODIFIER |
| 7 | 107030601 | AX-170199304 | 0 | C | T | 0.474 | -3.57E-03 | 9.56E-02 | 2.68E+00 | 2.75E+00 | 9.64E-01 | no | Olfr713 | upstream_gene_variant | MODIFIER |
| 7 | 109241501 | AX-170397260 | 0 | A | T | 0.474 | -3.57E-03 | 9.56E-02 | 2.68E+00 | 2.75E+00 | 9.64E-01 | no | | regulatory_region_variant | MODIFIER |
| 7 | 109246106 | AX-170432382 | 0 | T | C | 0.474 | -3.57E-03 | 9.56E-02 | 2.68E+00 | 2.75E+00 | 9.64E-01 | no | | intergenic_variant | MODIFIER |
| 7 | 109274161 | AX-170268334 | 0 | G | A | 0.474 | -3.57E-03 | 9.56E-02 | 2.68E+00 | 2.75E+00 | 9.64E-01 | no | | intergenic_variant | MODIFIER |
| 7 | 109274408 | AX-170111996 | 0 | C | T | 0.474 | -3.57E-03 | 9.56E-02 | 2.68E+00 | 2.75E+00 | 9.64E-01 | no | Stk33 | downstream_gene_variant | MODIFIER |
| 7 | 109276164 | AX-170029746 | 0 | T | C | 0.474 | -3.57E-03 | 9.56E-02 | 2.68E+00 | 2.75E+00 | 9.64E-01 | no | Stk33 | downstream_gene_variant | MODIFIER |
| 7 | 109281300 | AX-170029747 | 0 | C | T | 0.474 | -3.57E-03 | 9.56E-02 | 2.68E+00 | 2.75E+00 | 9.64E-01 | no | Stk33 | intron_variant | MODIFIER |
| 7 | 109292708 | AX-170096065 | 0 | T | C | 0.474 | -3.57E-03 | 9.56E-02 | 2.68E+00 | 2.75E+00 | 9.64E-01 | no | Stk33 | intron_variant | MODIFIER |
| 7 | 109309794 | AX-170029752 | 0 | A | C | 0.474 | -3.57E-03 | 9.56E-02 | 2.68E+00 | 2.75E+00 | 9.64E-01 | no | Stk33 | intron_variant | MODIFIER |
| 7 | 109322369 | AX-170146341 | 0 | A | C | 0.474 | -3.57E-03 | 9.56E-02 | 2.68E+00 | 2.75E+00 | 9.64E-01 | no | Stk33 | intron_variant | MODIFIER |
| 7 | 109391023 | AX-170096066 | 0 | A | C | 0.474 | -3.57E-03 | 9.56E-02 | 2.68E+00 | 2.75E+00 | 9.64E-01 | no | Stk33 | intron_variant | MODIFIER |
| 7 | 109392389 | AX-170104941 | 0 | G | T | 0.474 | -3.57E-03 | 9.56E-02 | 2.68E+00 | 2.75E+00 | 9.64E-01 | no | Stk33 | intron_variant | MODIFIER |
| 7 | 109409411 | AX-97850635 | 0 | A | C | 0.474 | -3.57E-03 | 9.56E-02 | 2.68E+00 | 2.75E+00 | 9.64E-01 | no | Stk33 | intron_variant | MODIFIER |
| 7 | 109438366 | AX-170284092 | 0 | C | T | 0.474 | -3.57E-03 | 9.56E-02 | 2.68E+00 | 2.75E+00 | 9.64E-01 | no | Stk33 | intron_variant | MODIFIER |
| 7 | 109441030 | AX-170096067 | 0 | T | G | 0.474 | -3.57E-03 | 9.56E-02 | 2.68E+00 | 2.75E+00 | 9.64E-01 | no | Stk33 | upstream_gene_variant | MODIFIER |
| 7 | 109479170 | AX-170382669 | 0 | G | A | 0.474 | -3.57E-03 | 9.56E-02 | 2.68E+00 | 2.75E+00 | 9.64E-01 | no | Trim66 | intron_variant | MODIFIER |
| 7 | 109479283 | AX-170362354 | 0 | C | T | 0.474 | -3.57E-03 | 9.56E-02 | 2.68E+00 | 2.75E+00 | 9.64E-01 | no | Trim66 | intron_variant | MODIFIER |
| 7 | 109496620 | AX-170414560 | 0 | T | G | 0.474 | -3.57E-03 | 9.56E-02 | 2.68E+00 | 2.75E+00 | 9.64E-01 | no | Trim66 | intron_variant | MODIFIER |
| 7 | 109500256 | AX-170078391 | 0 | A | G | 0.474 | -3.57E-03 | 9.56E-02 | 2.68E+00 | 2.75E+00 | 9.64E-01 | no | Trim66 | intron_variant | MODIFIER |
| 7 | 109501693 | AX-170078388 | 0 | T | A | 0.474 | -3.57E-03 | 9.56E-02 | 2.68E+00 | 2.75E+00 | 9.64E-01 | no | Trim66 | intron_variant | MODIFIER |
| 7 | 109533144 | AX-97802976 | 0 | C | T | 0.474 | -3.57E-03 | 9.56E-02 | 2.68E+00 | 2.75E+00 | 9.64E-01 | no | St5 | intron_variant | MODIFIER |
| 7 | 109545918 | AX-170316645 | 0 | A | G | 0.474 | -3.57E-03 | 9.56E-02 | 2.68E+00 | 2.75E+00 | 9.64E-01 | no | St5 | intron_variant | MODIFIER |
| 7 | 109547658 | AX-170281621 | 0 | G | A | 0.474 | -3.57E-03 | 9.56E-02 | 2.68E+00 | 2.75E+00 | 9.64E-01 | no | St5 | intron_variant | MODIFIER |
| 7 | 109554098 | AX-170099212 | 0 | T | C | 0.474 | -3.57E-03 | 9.56E-02 | 2.68E+00 | 2.75E+00 | 9.64E-01 | no | St5 | intron_variant | MODIFIER |
| 7 | 109569518 | AX-97851234 | 0 | T | C | 0.474 | -3.57E-03 | 9.56E-02 | 2.68E+00 | 2.75E+00 | 9.64E-01 | no | St5 | intron_variant | MODIFIER |
| 7 | 109580109 | AX-170221530 | 0 | A | G | 0.474 | -3.57E-03 | 9.56E-02 | 2.68E+00 | 2.75E+00 | 9.64E-01 | no | St5 | intron_variant | MODIFIER |
| 7 | 109585024 | AX-170244561 | 0 | G | A | 0.474 | -3.57E-03 | 9.56E-02 | 2.68E+00 | 2.75E+00 | 9.64E-01 | no | St5 | intron_variant | MODIFIER |
| 7 | 109586196 | AX-170096068 | 0 | C | T | 0.474 | -3.57E-03 | 9.56E-02 | 2.68E+00 | 2.75E+00 | 9.64E-01 | no | St5 | intron_variant | MODIFIER |
| 7 | 109586379 | AX-170236534 | 0 | C | T | 0.474 | -3.57E-03 | 9.56E-02 | 2.68E+00 | 2.75E+00 | 9.64E-01 | no | St5 | intron_variant | MODIFIER |
| 7 | 109706852 | AX-170393533 | 0 | G | A | 0.474 | -3.57E-03 | 9.56E-02 | 2.68E+00 | 2.75E+00 | 9.64E-01 | no | Akip1 | intron_variant | MODIFIER |
| 7 | 109709565 | AX-97823330 | 0 | C | T | 0.474 | -3.57E-03 | 9.56E-02 | 2.68E+00 | 2.75E+00 | 9.64E-01 | no | BC051019 | downstream_gene_variant | MODIFIER |
| 7 | 109713183 | AX-170078394 | 0 | A | G | 0.474 | -3.57E-03 | 9.56E-02 | 2.68E+00 | 2.75E+00 | 9.64E-01 | no | BC051019 | intron_variant | MODIFIER |
| 9 | 89812516 | AX-170119393 | 1 | A | T | 0.134 | -7.11E-03 | 1.30E-01 | 2.65E+00 | 2.73E+00 | 9.64E-01 | no | | intergenic_variant | MODIFIER |
| 7 | 98531105 | AX-170322812 | 1 | G | C | 0.116 | -7.45E-03 | 1.75E-01 | 2.66E+00 | 2.74E+00 | 9.64E-01 | no | | intergenic_variant | MODIFIER |
| 5 | 99477192 | AX-170025708 | 0 | A | G | 0.43 | 5.59E-03 | 1.10E-01 | 2.67E+00 | 2.73E+00 | 9.64E-01 | no | Rasgef1b | intron_variant | MODIFIER |
| 11 | 60787117 | AX-170506707 | 2 | G | C | 0.055 | -7.52E-03 | 1.52E-01 | 2.62E+00 | 2.73E+00 | 9.64E-01 | no | Shmt1 | downstream_gene_variant | MODIFIER |
| 13 | 53651181 | AX-170051324 | 0 | A | T | 0.509 | -2.15E-03 | 6.87E-02 | 2.64E+00 | 2.75E+00 | 9.64E-01 | no | 4930555G21Rik | downstream_gene_variant | MODIFIER |
| 5 | 127384037 | AX-170192349 | 0 | A | G | 0.421 | -3.04E-03 | 8.40E-02 | 2.64E+00 | 2.74E+00 | 9.64E-01 | no | Tmem132c | intron_variant | MODIFIER |
| 5 | 127389248 | AX-170035720 | 0 | A | T | 0.421 | -3.04E-03 | 8.40E-02 | 2.64E+00 | 2.74E+00 | 9.64E-01 | no | Tmem132c | intron_variant | MODIFIER |
| 5 | 127433263 | AX-97813468 | 0 | A | G | 0.421 | -3.04E-03 | 8.40E-02 | 2.64E+00 | 2.74E+00 | 9.64E-01 | no | Tmem132c | intron_variant | MODIFIER |
| 5 | 127454230 | AX-170470234 | 0 | G | A | 0.421 | -3.04E-03 | 8.40E-02 | 2.64E+00 | 2.74E+00 | 9.64E-01 | no | Tmem132c | intron_variant | MODIFIER |
| 5 | 127454552 | AX-170483629 | 0 | G | A | 0.421 | -3.04E-03 | 8.40E-02 | 2.64E+00 | 2.74E+00 | 9.64E-01 | no | Tmem132c | intron_variant | MODIFIER |
| 5 | 127463134 | AX-97833364 | 0 | G | A | 0.421 | -3.04E-03 | 8.40E-02 | 2.64E+00 | 2.74E+00 | 9.64E-01 | no | Tmem132c | intron_variant | MODIFIER |
| 5 | 127484104 | AX-170159537 | 0 | C | T | 0.421 | -3.04E-03 | 8.40E-02 | 2.64E+00 | 2.74E+00 | 9.64E-01 | no | Tmem132c | intron_variant | MODIFIER |
| 5 | 127500781 | AX-170491744 | 0 | C | T | 0.421 | -3.04E-03 | 8.40E-02 | 2.64E+00 | 2.74E+00 | 9.64E-01 | no | Tmem132c | intron_variant | MODIFIER |
| 5 | 127516901 | AX-170487451 | 0 | T | G | 0.421 | -3.04E-03 | 8.40E-02 | 2.64E+00 | 2.74E+00 | 9.64E-01 | no | Tmem132c | intron_variant | MODIFIER |
| 5 | 127646893 | AX-170311864 | 0 | G | A | 0.421 | -3.04E-03 | 8.40E-02 | 2.64E+00 | 2.74E+00 | 9.64E-01 | no | Glt1d1 | intron_variant | MODIFIER |
| 5 | 127657039 | AX-170292692 | 0 | G | T | 0.421 | -3.04E-03 | 8.40E-02 | 2.64E+00 | 2.74E+00 | 9.64E-01 | no | Glt1d1 | intron_variant | MODIFIER |
| 5 | 127660678 | AX-97794973 | 0 | C | T | 0.421 | -3.04E-03 | 8.40E-02 | 2.64E+00 | 2.74E+00 | 9.64E-01 | no | Glt1d1 | intron_variant | MODIFIER |

| | | | | | | | | | | | | | | | |
|---|---|---|---|---|---|---|---|---|---|---|---|---|---|---|---|
| 5 | 127661562 | AX-170460042 | 0 | G | A | 0.421 | -3.04E-03 | 8.40E-02 | 2.64E+00 | 2.74E+00 | 9.64E-01 | no | Glt1d1 | intron_variant | MODIFIER |
| 5 | 127692712 | AX-170026129 | 0 | G | A | 0.421 | -3.04E-03 | 8.40E-02 | 2.64E+00 | 2.74E+00 | 9.64E-01 | no | Glt1d1 | intron_variant | MODIFIER |
| 5 | 127812597 | AX-97854708 | 0 | A | G | 0.421 | -3.04E-03 | 8.40E-02 | 2.64E+00 | 2.74E+00 | 9.64E-01 | no | Tmem132d | intron_variant | MODIFIER |
| 5 | 127812797 | AX-170522090 | 0 | C | T | 0.421 | -3.04E-03 | 8.40E-02 | 2.64E+00 | 2.74E+00 | 9.64E-01 | no | Tmem132d | intron_variant | MODIFIER |
| 5 | 127815388 | AX-170376087 | 0 | C | G | 0.421 | -3.04E-03 | 8.40E-02 | 2.64E+00 | 2.74E+00 | 9.64E-01 | no | Tmem132d | intron_variant | MODIFIER |
| 5 | 127817882 | AX-170285449 | 0 | C | T | 0.421 | -3.04E-03 | 8.40E-02 | 2.64E+00 | 2.74E+00 | 9.64E-01 | no | Tmem132d | intron_variant | MODIFIER |
| 5 | 127826569 | AX-170448648 | 0 | G | A | 0.421 | -3.04E-03 | 8.40E-02 | 2.64E+00 | 2.74E+00 | 9.64E-01 | no | Tmem132d | intron_variant | MODIFIER |
| 5 | 127840064 | AX-170284448 | 0 | C | T | 0.421 | -3.04E-03 | 8.40E-02 | 2.64E+00 | 2.74E+00 | 9.64E-01 | no | Tmem132d | intron_variant | MODIFIER |
| 5 | 127841691 | AX-170354763 | 0 | A | C | 0.421 | -3.04E-03 | 8.40E-02 | 2.64E+00 | 2.74E+00 | 9.64E-01 | no | Tmem132d | intron_variant | MODIFIER |
| 5 | 127841986 | AX-170306864 | 0 | A | G | 0.421 | -3.04E-03 | 8.40E-02 | 2.64E+00 | 2.74E+00 | 9.64E-01 | no | Tmem132d | intron_variant | MODIFIER |
| 5 | 127945529 | AX-170151452 | 0 | G | T | 0.421 | -3.04E-03 | 8.40E-02 | 2.64E+00 | 2.74E+00 | 9.64E-01 | no | Tmem132d | intron_variant | MODIFIER |
| 5 | 127947617 | AX-170294670 | 0 | T | C | 0.421 | -3.04E-03 | 8.40E-02 | 2.64E+00 | 2.74E+00 | 9.64E-01 | no | Tmem132d | intron_variant | MODIFIER |
| 5 | 127952924 | AX-170527718 | 0 | C | T | 0.421 | -3.04E-03 | 8.40E-02 | 2.64E+00 | 2.74E+00 | 9.64E-01 | no | Tmem132d | intron_variant | MODIFIER |
| 5 | 127959393 | AX-170095461 | 0 | T | G | 0.421 | -3.04E-03 | 8.40E-02 | 2.64E+00 | 2.74E+00 | 9.64E-01 | no | Tmem132d | intron_variant | MODIFIER |
| 5 | 127967998 | AX-170121670 | 0 | T | G | 0.421 | -3.04E-03 | 8.40E-02 | 2.64E+00 | 2.74E+00 | 9.64E-01 | no | Tmem132d | intron_variant | MODIFIER |
| 5 | 127970726 | AX-170233163 | 0 | C | T | 0.421 | -3.04E-03 | 8.40E-02 | 2.64E+00 | 2.74E+00 | 9.64E-01 | no | Tmem132d | intron_variant | MODIFIER |
| 10 | 31474282 | AX-170523678 | 1 | G | C | 0.384 | 2.76E-03 | 7.44E-02 | 2.63E+00 | 2.74E+00 | 9.65E-01 | no | | intergenic_variant | MODIFIER |
| 2 | 120133905 | AX-170195183 | 0 | T | C | 0.14 | -8.43E-03 | 1.78E-01 | 2.66E+00 | 2.73E+00 | 9.65E-01 | no | Ehd4 | intron_variant | MODIFIER |
| 2 | 121684433 | AX-170514118 | 0 | G | A | 0.14 | -8.43E-03 | 1.78E-01 | 2.66E+00 | 2.73E+00 | 9.65E-01 | no | Frmd5 | intron_variant | MODIFIER |
| 2 | 124704866 | AX-170386252 | 0 | A | G | 0.14 | -8.43E-03 | 1.78E-01 | 2.66E+00 | 2.73E+00 | 9.65E-01 | no | Gm13994 | upstream_gene_variant | MODIFIER |
| 2 | 125176253 | AX-170018936 | 0 | A | G | 0.14 | -8.43E-03 | 1.78E-01 | 2.66E+00 | 2.73E+00 | 9.65E-01 | no | Slc12a1 | intron_variant | MODIFIER |
| 2 | 127408600 | AX-97817818 | 0 | G | A | 0.14 | -8.43E-03 | 1.78E-01 | 2.66E+00 | 2.73E+00 | 9.65E-01 | no | | intergenic_variant | MODIFIER |
| 6 | 133763637 | AX-97801342 | 0 | C | T | 0.561 | 4.65E-03 | 8.17E-02 | 2.65E+00 | 2.73E+00 | 9.65E-01 | no | Gm23375 | upstream_gene_variant | MODIFIER |
| 12 | 70833980 | AX-170219793 | 1 | G | A | 0.062 | 1.26E-02 | 3.00E-01 | 2.72E+00 | 2.74E+00 | 9.65E-01 | no | Frmd6 | intron_variant | MODIFIER |
| 12 | 71023818 | AX-97846838 | 1 | G | A | 0.062 | 1.26E-02 | 3.00E-01 | 2.72E+00 | 2.74E+00 | 9.65E-01 | no | Arid4a | intron_variant | MODIFIER |
| 10 | 7798587 | AX-170002663 | 1 | G | A | 0.214 | -5.20E-03 | 1.05E-01 | 2.65E+00 | 2.73E+00 | 9.65E-01 | no | Ppil4 | intron_variant | MODIFIER |
| 8 | 110077507 | AX-170416246 | 2 | A | G | 0.218 | 4.10E-03 | 1.11E-01 | 2.65E+00 | 2.74E+00 | 9.65E-01 | no | Zfp612 | upstream_gene_variant | MODIFIER |
| 1 | 53591212 | AX-170041245 | 1 | A | G | 0.107 | -9.78E-03 | 2.34E-01 | 2.69E+00 | 2.74E+00 | 9.65E-01 | no | Dnah7a | intron_variant | MODIFIER |
| 6 | 144633360 | AX-170468971 | 1 | A | T | 0.054 | 8.39E-03 | 1.92E-01 | 2.65E+00 | 2.74E+00 | 9.65E-01 | no | Sox5 | intron_variant,non_coding_transcript_variant | MODIFIER |
| 9 | 59500677 | AX-97822637 | 1 | C | T | 0.214 | 5.36E-03 | 1.23E-01 | 2.68E+00 | 2.74E+00 | 9.65E-01 | no | Gm33180 | intron_variant,non_coding_transcript_variant | MODIFIER |
| 15 | 88088899 | AX-170187509 | 0 | T | G | 0.132 | 9.52E-03 | 1.82E-01 | 2.64E+00 | 2.73E+00 | 9.65E-01 | no | | intergenic_variant | MODIFIER |
| 2 | 145569113 | AX-170331119 | 1 | G | T | 0.134 | 9.97E-03 | 2.29E-01 | 2.70E+00 | 2.74E+00 | 9.65E-01 | no | Slc24a3 | intron_variant | MODIFIER |
| 9 | 40425851 | AX-170467032 | 1 | C | T | 0.438 | 2.92E-03 | 7.34E-02 | 2.64E+00 | 2.74E+00 | 9.65E-01 | no | Gramd1b | intron_variant | MODIFIER |
| 10 | 16100034 | AX-97853989 | 0 | T | C | 0.193 | 3.44E-03 | 1.04E-01 | 2.67E+00 | 2.75E+00 | 9.65E-01 | no | | intergenic_variant | MODIFIER |
| 10 | 16120090 | AX-170002816 | 0 | G | A | 0.193 | 3.44E-03 | 1.04E-01 | 2.67E+00 | 2.75E+00 | 9.65E-01 | no | | intergenic_variant | MODIFIER |
| 10 | 16164158 | AX-97814901 | 0 | T | C | 0.193 | 3.44E-03 | 1.04E-01 | 2.67E+00 | 2.75E+00 | 9.65E-01 | no | | intergenic_variant | MODIFIER |
| 10 | 16175865 | AX-170002818 | 0 | C | A | 0.193 | 3.44E-03 | 1.04E-01 | 2.67E+00 | 2.75E+00 | 9.65E-01 | no | | intergenic_variant | MODIFIER |
| 10 | 16202664 | AX-97847455 | 0 | T | C | 0.193 | 3.44E-03 | 1.04E-01 | 2.67E+00 | 2.75E+00 | 9.65E-01 | no | | intergenic_variant | MODIFIER |
| 1 | 168814779 | AX-170119774 | 1 | T | A | 0.223 | 7.02E-03 | 1.19E-01 | 2.62E+00 | 2.73E+00 | 9.66E-01 | no | | intergenic_variant | MODIFIER |
| 8 | 55021344 | AX-97801237 | 1 | C | G | 0.152 | -7.61E-03 | 1.53E-01 | 2.68E+00 | 2.73E+00 | 9.66E-01 | no | Gpm6a | intron_variant | MODIFIER |
| 2 | 142899466 | AX-170158906 | 0 | G | C | 0.14 | 9.84E-03 | 2.22E-01 | 2.71E+00 | 2.73E+00 | 9.66E-01 | no | Kif16b | intron_variant | MODIFIER |
| 2 | 157481689 | AX-170509440 | 1 | T | A | 0.134 | -7.62E-03 | 1.73E-01 | 2.66E+00 | 2.74E+00 | 9.66E-01 | no | | regulatory_region_variant | MODIFIER |
| 11 | 11128651 | AX-170382832 | 1 | T | C | 0.062 | 8.16E-03 | 2.13E-01 | 2.65E+00 | 2.74E+00 | 9.66E-01 | no | Vwc2 | intron_variant | MODIFIER |
| 5 | 123476737 | AX-170026061 | 0 | C | G | 0.465 | 5.23E-03 | 1.12E-01 | 2.68E+00 | 2.73E+00 | 9.66E-01 | no | Il31 | downstream_gene_variant | MODIFIER |
| 2 | 95102747 | AX-170286226 | 0 | G | T | 0.061 | 9.40E-03 | 2.26E-01 | 2.65E+00 | 2.74E+00 | 9.66E-01 | no | | intergenic_variant | MODIFIER |
| 7 | 58939941 | AX-97786420 | 0 | T | C | 0.421 | -3.49E-03 | 9.41E-02 | 2.66E+00 | 2.74E+00 | 9.66E-01 | no | | intergenic_variant | MODIFIER |
| 13 | 37278848 | AX-170182415 | 1 | G | A | 0.179 | 3.96E-03 | 1.21E-01 | 2.68E+00 | 2.75E+00 | 9.66E-01 | no | | intergenic_variant | MODIFIER |
| 11 | 3334213 | AX-170389817 | 0 | T | C | 0.061 | 8.02E-03 | 2.14E-01 | 2.65E+00 | 2.74E+00 | 9.66E-01 | no | Pik3ip1 | intron_variant | MODIFIER |
| 11 | 3350241 | AX-170394489 | 0 | T | G | 0.061 | 8.02E-03 | 2.14E-01 | 2.65E+00 | 2.74E+00 | 9.66E-01 | no | Limk2 | intron_variant | MODIFIER |
| 11 | 4139872 | AX-170004172 | 0 | T | G | 0.061 | 8.02E-03 | 2.14E-01 | 2.65E+00 | 2.74E+00 | 9.66E-01 | no | Rnf215 | intron_variant | MODIFIER |
| 11 | 4355625 | AX-170331921 | 0 | T | C | 0.061 | 8.02E-03 | 2.14E-01 | 2.65E+00 | 2.74E+00 | 9.66E-01 | no | Hormad2 | intron_variant,NMD_transcript_variant | MODIFIER |
| 11 | 4400324 | AX-170304989 | 0 | T | C | 0.061 | 8.02E-03 | 2.14E-01 | 2.65E+00 | 2.74E+00 | 9.66E-01 | no | Hormad2 | intron_variant,NMD_transcript_variant | MODIFIER |
| 11 | 4718725 | AX-170288219 | 0 | G | T | 0.061 | 8.02E-03 | 2.14E-01 | 2.65E+00 | 2.74E+00 | 9.66E-01 | no | Zmat5 | intron_variant | MODIFIER |
| 11 | 4751158 | AX-170186719 | 0 | T | C | 0.061 | 8.02E-03 | 2.14E-01 | 2.65E+00 | 2.74E+00 | 9.66E-01 | no | Cabp7 | upstream_gene_variant | MODIFIER |

| | | | | | | | | | | | | | | | |
|---|---|---|---|---|---|---|---|---|---|---|---|---|---|---|---|
| 11 | 4751344 | AX-170245694 | 0 | C | T | 0.061 | 8.02E-03 | 2.14E-01 | 2.65E+00 | 2.74E+00 | 9.66E-01 | no | Cabp7 | upstream_gene_variant | MODIFIER |
| 11 | 4751425 | AX-170139170 | 0 | A | T | 0.061 | 8.02E-03 | 2.14E-01 | 2.65E+00 | 2.74E+00 | 9.66E-01 | no | Cabp7 | upstream_gene_variant | MODIFIER |
| 11 | 4752447 | AX-97821963 | 0 | T | C | 0.061 | 8.02E-03 | 2.14E-01 | 2.65E+00 | 2.74E+00 | 9.66E-01 | no | | intergenic_variant | MODIFIER |
| 11 | 4752502 | AX-170182260 | 0 | T | C | 0.061 | 8.02E-03 | 2.14E-01 | 2.65E+00 | 2.74E+00 | 9.66E-01 | no | | intergenic_variant | MODIFIER |
| 11 | 4761577 | AX-170238291 | 0 | A | G | 0.061 | 8.02E-03 | 2.14E-01 | 2.65E+00 | 2.74E+00 | 9.66E-01 | no | Nf2 | downstream_gene_variant | MODIFIER |
| 11 | 4766933 | AX-170294479 | 0 | G | T | 0.061 | 8.02E-03 | 2.14E-01 | 2.65E+00 | 2.74E+00 | 9.66E-01 | no | Nf2 | downstream_gene_variant | MODIFIER |
| 11 | 4772180 | AX-170500414 | 0 | T | C | 0.061 | 8.02E-03 | 2.14E-01 | 2.65E+00 | 2.74E+00 | 9.66E-01 | no | Nf2 | downstream_gene_variant | MODIFIER |
| 11 | 4779801 | AX-170381582 | 0 | C | T | 0.061 | 8.02E-03 | 2.14E-01 | 2.65E+00 | 2.74E+00 | 9.66E-01 | no | Nf2 | intron_variant | MODIFIER |
| 11 | 4787389 | AX-170306546 | 0 | T | C | 0.061 | 8.02E-03 | 2.14E-01 | 2.65E+00 | 2.74E+00 | 9.66E-01 | no | Nf2 | intron_variant | MODIFIER |
| 11 | 4789910 | AX-170046299 | 0 | T | C | 0.061 | 8.02E-03 | 2.14E-01 | 2.65E+00 | 2.74E+00 | 9.66E-01 | no | Nf2 | intron_variant | MODIFIER |
| 11 | 4805906 | AX-97815704 | 0 | T | G | 0.061 | 8.02E-03 | 2.14E-01 | 2.65E+00 | 2.74E+00 | 9.66E-01 | no | Nf2 | intron_variant | MODIFIER |
| 11 | 4834106 | AX-170302859 | 0 | G | A | 0.061 | 8.02E-03 | 2.14E-01 | 2.65E+00 | 2.74E+00 | 9.66E-01 | no | Nf2 | intron_variant | MODIFIER |
| 11 | 4836200 | AX-170282000 | 0 | G | A | 0.061 | 8.02E-03 | 2.14E-01 | 2.65E+00 | 2.74E+00 | 9.66E-01 | no | Nf2 | intron_variant | MODIFIER |
| 11 | 4836843 | AX-97801919 | 0 | T | G | 0.061 | 8.02E-03 | 2.14E-01 | 2.65E+00 | 2.74E+00 | 9.66E-01 | no | Nf2 | intron_variant | MODIFIER |
| 11 | 4923635 | AX-170421413 | 0 | C | T | 0.061 | 8.02E-03 | 2.14E-01 | 2.65E+00 | 2.74E+00 | 9.66E-01 | no | Thoc5 | intron_variant | MODIFIER |
| 11 | 4923873 | AX-170449308 | 0 | A | G | 0.061 | 8.02E-03 | 2.14E-01 | 2.65E+00 | 2.74E+00 | 9.66E-01 | no | Thoc5 | intron_variant | MODIFIER |
| 11 | 4931346 | AX-170507960 | 0 | G | A | 0.061 | 8.02E-03 | 2.14E-01 | 2.65E+00 | 2.74E+00 | 9.66E-01 | no | Thoc5 | downstream_gene_variant | MODIFIER |
| 11 | 4986526 | AX-97790682 | 0 | G | C | 0.061 | 8.02E-03 | 2.14E-01 | 2.65E+00 | 2.74E+00 | 9.66E-01 | no | Ap1b1 | upstream_gene_variant | MODIFIER |
| 11 | 4986590 | AX-170262170 | 0 | T | C | 0.061 | 8.02E-03 | 2.14E-01 | 2.65E+00 | 2.74E+00 | 9.66E-01 | no | Ap1b1 | upstream_gene_variant | MODIFIER |
| 11 | 4996607 | AX-170086933 | 0 | G | C | 0.061 | 8.02E-03 | 2.14E-01 | 2.65E+00 | 2.74E+00 | 9.66E-01 | no | Ap1b1 | intron_variant | MODIFIER |
| 11 | 5030237 | AX-170340148 | 0 | G | A | 0.061 | 8.02E-03 | 2.14E-01 | 2.65E+00 | 2.74E+00 | 9.66E-01 | no | Ap1b1 | intron_variant | MODIFIER |
| 11 | 5030301 | AX-170046304 | 0 | A | G | 0.061 | 8.02E-03 | 2.14E-01 | 2.65E+00 | 2.74E+00 | 9.66E-01 | no | Ap1b1 | intron_variant | MODIFIER |
| 11 | 5037888 | AX-97832301 | 0 | A | G | 0.061 | 8.02E-03 | 2.14E-01 | 2.65E+00 | 2.74E+00 | 9.66E-01 | no | Ap1b1 | intron_variant | MODIFIER |
| 11 | 5071257 | AX-97789515 | 0 | G | T | 0.061 | 8.02E-03 | 2.14E-01 | 2.65E+00 | 2.74E+00 | 9.66E-01 | no | Ewsr1 | intron_variant | MODIFIER |
| 11 | 5082516 | AX-170209514 | 0 | C | T | 0.061 | 8.02E-03 | 2.14E-01 | 2.65E+00 | 2.74E+00 | 9.66E-01 | no | Ewsr1 | intron_variant | MODIFIER |
| 11 | 5088235 | AX-170092163 | 0 | G | T | 0.061 | 8.02E-03 | 2.14E-01 | 2.65E+00 | 2.74E+00 | 9.66E-01 | no | Ewsr1 | downstream_gene_variant | MODIFIER |
| 11 | 5103915 | AX-170385847 | 0 | A | G | 0.061 | 8.02E-03 | 2.14E-01 | 2.65E+00 | 2.74E+00 | 9.66E-01 | no | Rhbdd3 | intron_variant | MODIFIER |
| 11 | 5126366 | AX-170459675 | 0 | G | A | 0.061 | 8.02E-03 | 2.14E-01 | 2.65E+00 | 2.74E+00 | 9.66E-01 | no | Emid1 | intron_variant | MODIFIER |
| 11 | 5192132 | AX-170004183 | 0 | A | G | 0.061 | 8.02E-03 | 2.14E-01 | 2.65E+00 | 2.74E+00 | 9.66E-01 | no | Kremen1 | 3_prime_UTR_variant | MODIFIER |
| 11 | 5192290 | AX-170270751 | 0 | T | C | 0.061 | 8.02E-03 | 2.14E-01 | 2.65E+00 | 2.74E+00 | 9.66E-01 | no | Kremen1 | 3_prime_UTR_variant | MODIFIER |
| 11 | 5194448 | AX-170382897 | 0 | C | T | 0.061 | 8.02E-03 | 2.14E-01 | 2.65E+00 | 2.74E+00 | 9.66E-01 | no | Kremen1 | 3_prime_UTR_variant | MODIFIER |
| 11 | 5198436 | AX-170004186 | 0 | C | T | 0.061 | 8.02E-03 | 2.14E-01 | 2.65E+00 | 2.74E+00 | 9.66E-01 | no | Kremen1 | intron_variant | MODIFIER |
| 11 | 5198559 | AX-170240880 | 0 | A | T | 0.061 | 8.02E-03 | 2.14E-01 | 2.65E+00 | 2.74E+00 | 9.66E-01 | no | Kremen1 | intron_variant | MODIFIER |
| 11 | 5201869 | AX-170426074 | 0 | T | C | 0.061 | 8.02E-03 | 2.14E-01 | 2.65E+00 | 2.74E+00 | 9.66E-01 | no | Kremen1 | synonymous_variant | LOW |
| 11 | 5204322 | AX-170004184 | 0 | C | G | 0.061 | 8.02E-03 | 2.14E-01 | 2.65E+00 | 2.74E+00 | 9.66E-01 | no | Kremen1 | intron_variant | MODIFIER |
| 11 | 5208798 | AX-170046308 | 0 | G | A | 0.061 | 8.02E-03 | 2.14E-01 | 2.65E+00 | 2.74E+00 | 9.66E-01 | no | Kremen1 | intron_variant | MODIFIER |
| 11 | 5208823 | AX-97841074 | 0 | C | T | 0.061 | 8.02E-03 | 2.14E-01 | 2.65E+00 | 2.74E+00 | 9.66E-01 | no | Kremen1 | intron_variant | MODIFIER |
| 11 | 5258270 | AX-97834972 | 0 | A | G | 0.061 | 8.02E-03 | 2.14E-01 | 2.65E+00 | 2.74E+00 | 9.66E-01 | no | Kremen1 | intron_variant | MODIFIER |
| 11 | 5278091 | AX-97852444 | 0 | A | G | 0.061 | 8.02E-03 | 2.14E-01 | 2.65E+00 | 2.74E+00 | 9.66E-01 | no | Znrf3 | 3_prime_UTR_variant | MODIFIER |
| 11 | 5288239 | AX-170440214 | 0 | C | G | 0.061 | 8.02E-03 | 2.14E-01 | 2.65E+00 | 2.74E+00 | 9.66E-01 | no | Znrf3 | intron_variant | MODIFIER |
| 11 | 5288548 | AX-170156558 | 0 | G | A | 0.061 | 8.02E-03 | 2.14E-01 | 2.65E+00 | 2.74E+00 | 9.66E-01 | no | Znrf3 | intron_variant | MODIFIER |
| 11 | 5310974 | AX-170280134 | 0 | A | G | 0.061 | 8.02E-03 | 2.14E-01 | 2.65E+00 | 2.74E+00 | 9.66E-01 | no | Znrf3 | intron_variant | MODIFIER |
| 11 | 5314361 | AX-97825608 | 0 | A | G | 0.061 | 8.02E-03 | 2.14E-01 | 2.65E+00 | 2.74E+00 | 9.66E-01 | no | Znrf3 | intron_variant | MODIFIER |
| 11 | 5320025 | AX-170399006 | 0 | C | G | 0.061 | 8.02E-03 | 2.14E-01 | 2.65E+00 | 2.74E+00 | 9.66E-01 | no | Znrf3 | intron_variant | MODIFIER |
| 11 | 5341601 | AX-97786925 | 0 | C | T | 0.061 | 8.02E-03 | 2.14E-01 | 2.65E+00 | 2.74E+00 | 9.66E-01 | no | Znrf3 | intron_variant | MODIFIER |
| 11 | 5360959 | AX-170412175 | 0 | A | C | 0.061 | 8.02E-03 | 2.14E-01 | 2.65E+00 | 2.74E+00 | 9.66E-01 | no | Znrf3 | intron_variant | MODIFIER |
| 11 | 5372351 | AX-170227978 | 0 | G | A | 0.061 | 8.02E-03 | 2.14E-01 | 2.65E+00 | 2.74E+00 | 9.66E-01 | no | Znrf3 | intron_variant | MODIFIER |
| 11 | 5380351 | AX-97814475 | 0 | A | G | 0.061 | 8.02E-03 | 2.14E-01 | 2.65E+00 | 2.74E+00 | 9.66E-01 | no | Znrf3 | intron_variant | MODIFIER |
| 11 | 5391122 | AX-170486600 | 0 | A | G | 0.061 | 8.02E-03 | 2.14E-01 | 2.65E+00 | 2.74E+00 | 9.66E-01 | no | Znrf3 | intron_variant | MODIFIER |
| 11 | 5400374 | AX-170310323 | 0 | T | C | 0.061 | 8.02E-03 | 2.14E-01 | 2.65E+00 | 2.74E+00 | 9.66E-01 | no | Znrf3 | intron_variant | MODIFIER |
| 11 | 5405877 | AX-170183938 | 0 | A | C | 0.061 | 8.02E-03 | 2.14E-01 | 2.65E+00 | 2.74E+00 | 9.66E-01 | no | Znrf3 | intron_variant | MODIFIER |
| 11 | 5422346 | AX-170097341 | 0 | G | A | 0.061 | 8.02E-03 | 2.14E-01 | 2.65E+00 | 2.74E+00 | 9.66E-01 | no | Znrf3 | intron_variant | MODIFIER |
| 11 | 5433273 | AX-170362246 | 0 | A | G | 0.061 | 8.02E-03 | 2.14E-01 | 2.65E+00 | 2.74E+00 | 9.66E-01 | no | Znrf3 | intron_variant | MODIFIER |
| 11 | 5442866 | AX-170126790 | 0 | G | A | 0.061 | 8.02E-03 | 2.14E-01 | 2.65E+00 | 2.74E+00 | 9.66E-01 | no | Znrf3 | intron_variant | MODIFIER |

| | | | | | | | | | | | | | | | |
|---|---|---|---|---|---|---|---|---|---|---|---|---|---|---|---|
| 11 | 5567122 | AX-170046310 | 0 | G | A | 0.061 | 8.02E-03 | 2.14E-01 | 2.65E+00 | 2.74E+00 | 9.66E-01 | no | Ankrd36 | upstream_gene_variant | MODIFIER |
| 11 | 5569790 | AX-170343416 | 0 | A | G | 0.061 | 8.02E-03 | 2.14E-01 | 2.65E+00 | 2.74E+00 | 9.66E-01 | no | Ankrd36 | 5_prime_UTR_variant | MODIFIER |
| 11 | 5577028 | AX-170471425 | 0 | A | G | 0.061 | 8.02E-03 | 2.14E-01 | 2.65E+00 | 2.74E+00 | 9.66E-01 | no | Ankrd36 | intron_variant | MODIFIER |
| 11 | 5577437 | AX-170427771 | 0 | A | G | 0.061 | 8.02E-03 | 2.14E-01 | 2.65E+00 | 2.74E+00 | 9.66E-01 | no | Ankrd36 | intron_variant | MODIFIER |
| 11 | 5589518 | AX-170389530 | 0 | C | T | 0.061 | 8.02E-03 | 2.14E-01 | 2.65E+00 | 2.74E+00 | 9.66E-01 | no | Ankrd36 | intron_variant | MODIFIER |
| 11 | 5589551 | AX-170458440 | 0 | A | G | 0.061 | 8.02E-03 | 2.14E-01 | 2.65E+00 | 2.74E+00 | 9.66E-01 | no | Ankrd36 | intron_variant | MODIFIER |
| 11 | 5589617 | AX-97820745 | 0 | T | C | 0.061 | 8.02E-03 | 2.14E-01 | 2.65E+00 | 2.74E+00 | 9.66E-01 | no | Ankrd36 | intron_variant | MODIFIER |
| 11 | 5590472 | AX-170296850 | 0 | C | A | 0.061 | 8.02E-03 | 2.14E-01 | 2.65E+00 | 2.74E+00 | 9.66E-01 | no | Ankrd36 | intron_variant | MODIFIER |
| 11 | 5605438 | AX-170340944 | 0 | G | A | 0.061 | 8.02E-03 | 2.14E-01 | 2.65E+00 | 2.74E+00 | 9.66E-01 | no | Ankrd36 | intron_variant | MODIFIER |
| 11 | 5621205 | AX-97826944 | 0 | T | G | 0.061 | 8.02E-03 | 2.14E-01 | 2.65E+00 | 2.74E+00 | 9.66E-01 | no | Ankrd36 | intron_variant | MODIFIER |
| 11 | 5650312 | AX-170402011 | 0 | T | G | 0.061 | 8.02E-03 | 2.14E-01 | 2.65E+00 | 2.74E+00 | 9.66E-01 | no | Ankrd36 | intron_variant | MODIFIER |
| 11 | 5684997 | AX-170004196 | 0 | C | T | 0.061 | 8.02E-03 | 2.14E-01 | 2.65E+00 | 2.74E+00 | 9.66E-01 | no | Ankrd36 | intron_variant | MODIFIER |
| 11 | 5721466 | AX-170004191 | 0 | T | G | 0.061 | 8.02E-03 | 2.14E-01 | 2.65E+00 | 2.74E+00 | 9.66E-01 | no | Urgcp | intron_variant | MODIFIER |
| 11 | 5728821 | AX-170443514 | 0 | G | C | 0.061 | 8.02E-03 | 2.14E-01 | 2.65E+00 | 2.74E+00 | 9.66E-01 | no | Urgcp | intron_variant | MODIFIER |
| 11 | 5730107 | AX-170461904 | 0 | G | A | 0.061 | 8.02E-03 | 2.14E-01 | 2.65E+00 | 2.74E+00 | 9.66E-01 | no | Urgcp | intron_variant | MODIFIER |
| 11 | 5730147 | AX-170177856 | 0 | G | A | 0.061 | 8.02E-03 | 2.14E-01 | 2.65E+00 | 2.74E+00 | 9.66E-01 | no | Urgcp | intron_variant | MODIFIER |
| 11 | 5730315 | AX-170348374 | 0 | T | A | 0.061 | 8.02E-03 | 2.14E-01 | 2.65E+00 | 2.74E+00 | 9.66E-01 | no | Urgcp | intron_variant | MODIFIER |
| 11 | 5756759 | AX-97817256 | 0 | T | C | 0.061 | 8.02E-03 | 2.14E-01 | 2.65E+00 | 2.74E+00 | 9.66E-01 | no | Urgcp | intron_variant | MODIFIER |
| 11 | 5756788 | AX-170233890 | 0 | C | T | 0.061 | 8.02E-03 | 2.14E-01 | 2.65E+00 | 2.74E+00 | 9.66E-01 | no | Urgcp | intron_variant | MODIFIER |
| 11 | 5757730 | AX-170004194 | 0 | G | A | 0.061 | 8.02E-03 | 2.14E-01 | 2.65E+00 | 2.74E+00 | 9.66E-01 | no | Urgcp | intron_variant | MODIFIER |
| 11 | 5805985 | AX-170004188 | 0 | C | T | 0.061 | 8.02E-03 | 2.14E-01 | 2.65E+00 | 2.74E+00 | 9.66E-01 | no | Pgam2 | upstream_gene_variant | MODIFIER |
| 11 | 5806941 | AX-170329024 | 0 | G | A | 0.061 | 8.02E-03 | 2.14E-01 | 2.65E+00 | 2.74E+00 | 9.66E-01 | no | Pgam2 | upstream_gene_variant | MODIFIER |
| 11 | 5817032 | AX-170481304 | 0 | T | C | 0.061 | 8.02E-03 | 2.14E-01 | 2.65E+00 | 2.74E+00 | 9.66E-01 | no | Gm11966 | downstream_gene_variant | MODIFIER |
| 11 | 5890670 | AX-170227792 | 0 | C | T | 0.061 | 8.02E-03 | 2.14E-01 | 2.65E+00 | 2.74E+00 | 9.66E-01 | no | | intergenic_variant | MODIFIER |
| 11 | 5978181 | AX-170046319 | 0 | G | A | 0.061 | 8.02E-03 | 2.14E-01 | 2.65E+00 | 2.74E+00 | 9.66E-01 | no | Camk2b | intron_variant | MODIFIER |
| 11 | 6000875 | AX-170046317 | 0 | G | A | 0.061 | 8.02E-03 | 2.14E-01 | 2.65E+00 | 2.74E+00 | 9.66E-01 | no | Camk2b | intron_variant | MODIFIER |
| 11 | 6015438 | AX-170348562 | 0 | C | T | 0.061 | 8.02E-03 | 2.14E-01 | 2.65E+00 | 2.74E+00 | 9.66E-01 | no | Camk2b | intron_variant | MODIFIER |
| 11 | 6015487 | AX-170214646 | 0 | C | T | 0.061 | 8.02E-03 | 2.14E-01 | 2.65E+00 | 2.74E+00 | 9.66E-01 | no | Camk2b | synonymous_variant | LOW |
| 11 | 6267403 | AX-97800495 | 0 | A | C | 0.061 | 8.02E-03 | 2.14E-01 | 2.65E+00 | 2.74E+00 | 9.66E-01 | no | Ddx56 | intron_variant | MODIFIER |
| 11 | 6268179 | AX-170004206 | 0 | T | C | 0.061 | 8.02E-03 | 2.14E-01 | 2.65E+00 | 2.74E+00 | 9.66E-01 | no | Ddx56 | upstream_gene_variant | MODIFIER |
| 11 | 6274186 | AX-170465900 | 0 | G | A | 0.061 | 8.02E-03 | 2.14E-01 | 2.65E+00 | 2.74E+00 | 9.66E-01 | no | Tmed4 | synonymous_variant | LOW |
| 11 | 6321720 | AX-170004203 | 0 | A | G | 0.061 | 8.02E-03 | 2.14E-01 | 2.65E+00 | 2.74E+00 | 9.66E-01 | no | Ogdh | intron_variant | MODIFIER |
| 11 | 6322093 | AX-170507975 | 0 | T | G | 0.061 | 8.02E-03 | 2.14E-01 | 2.65E+00 | 2.74E+00 | 9.66E-01 | no | Ogdh | intron_variant | MODIFIER |
| 11 | 6406812 | AX-170004200 | 0 | A | G | 0.061 | 8.02E-03 | 2.14E-01 | 2.65E+00 | 2.74E+00 | 9.66E-01 | no | Zmiz2 | downstream_gene_variant | MODIFIER |
| 11 | 6466657 | AX-170004201 | 0 | T | C | 0.061 | 8.02E-03 | 2.14E-01 | 2.65E+00 | 2.74E+00 | 9.66E-01 | no | Purb | downstream_gene_variant | MODIFIER |
| 11 | 6470622 | AX-170410764 | 0 | T | C | 0.061 | 8.02E-03 | 2.14E-01 | 2.65E+00 | 2.74E+00 | 9.66E-01 | no | Gm11973 | upstream_gene_variant | MODIFIER |
| 11 | 6478565 | AX-170046326 | 0 | G | A | 0.061 | 8.02E-03 | 2.14E-01 | 2.65E+00 | 2.74E+00 | 9.66E-01 | no | Gm11973 | intron_variant,non_coding_transcript_variant | MODIFIER |
| 11 | 6479169 | AX-97806717 | 0 | G | A | 0.061 | 8.02E-03 | 2.14E-01 | 2.65E+00 | 2.74E+00 | 9.66E-01 | no | Gm11973 | intron_variant,non_coding_transcript_variant | MODIFIER |
| 11 | 6481325 | AX-170330980 | 0 | G | A | 0.061 | 8.02E-03 | 2.14E-01 | 2.65E+00 | 2.74E+00 | 9.66E-01 | no | Gm11973 | intron_variant,non_coding_transcript_variant | MODIFIER |
| 11 | 6506472 | AX-170392239 | 0 | C | T | 0.061 | 8.02E-03 | 2.14E-01 | 2.65E+00 | 2.74E+00 | 9.66E-01 | no | Myo1g | downstream_gene_variant | MODIFIER |
| 11 | 6516557 | AX-170191791 | 0 | G | A | 0.061 | 8.02E-03 | 2.14E-01 | 2.65E+00 | 2.74E+00 | 9.66E-01 | no | Myo1g | intron_variant | MODIFIER |
| 11 | 6702035 | AX-170509034 | 0 | T | G | 0.061 | 8.02E-03 | 2.14E-01 | 2.65E+00 | 2.74E+00 | 9.66E-01 | no | | intergenic_variant | MODIFIER |
| 11 | 7130166 | AX-170241409 | 0 | A | G | 0.061 | 8.02E-03 | 2.14E-01 | 2.65E+00 | 2.74E+00 | 9.66E-01 | no | Adcy1 | intron_variant | MODIFIER |
| 11 | 8497583 | AX-170296956 | 0 | G | A | 0.061 | 8.02E-03 | 2.14E-01 | 2.65E+00 | 2.74E+00 | 9.66E-01 | no | Tns3 | intron_variant | MODIFIER |
| 11 | 8502194 | AX-170164739 | 0 | G | T | 0.061 | 8.02E-03 | 2.14E-01 | 2.65E+00 | 2.74E+00 | 9.66E-01 | no | Tns3 | intron_variant | MODIFIER |
| 11 | 8535560 | AX-170185120 | 0 | G | A | 0.061 | 8.02E-03 | 2.14E-01 | 2.65E+00 | 2.74E+00 | 9.66E-01 | no | Tns3 | intron_variant | MODIFIER |
| 11 | 8576483 | AX-97808071 | 0 | C | T | 0.061 | 8.02E-03 | 2.14E-01 | 2.65E+00 | 2.74E+00 | 9.66E-01 | no | Tns3 | intron_variant | MODIFIER |
| 11 | 8576781 | AX-170281897 | 0 | A | C | 0.061 | 8.02E-03 | 2.14E-01 | 2.65E+00 | 2.74E+00 | 9.66E-01 | no | Tns3 | intron_variant | MODIFIER |
| 11 | 8579596 | AX-170298182 | 0 | C | T | 0.061 | 8.02E-03 | 2.14E-01 | 2.65E+00 | 2.74E+00 | 9.66E-01 | no | Tns3 | intron_variant | MODIFIER |
| 11 | 8827156 | AX-170158450 | 0 | A | T | 0.061 | 8.02E-03 | 2.14E-01 | 2.65E+00 | 2.74E+00 | 9.66E-01 | no | Pkd1l1 | intron_variant | MODIFIER |
| 11 | 8886286 | AX-170121334 | 0 | T | C | 0.061 | 8.02E-03 | 2.14E-01 | 2.65E+00 | 2.74E+00 | 9.66E-01 | no | Gm11991 | upstream_gene_variant | MODIFIER |
| 11 | 8889410 | AX-170238692 | 0 | T | C | 0.061 | 8.02E-03 | 2.14E-01 | 2.65E+00 | 2.74E+00 | 9.66E-01 | no | Gm11991 | upstream_gene_variant | MODIFIER |
| 11 | 8976078 | AX-170046374 | 0 | C | T | 0.061 | 8.02E-03 | 2.14E-01 | 2.65E+00 | 2.74E+00 | 9.66E-01 | no | Pkd1l1 | upstream_gene_variant | MODIFIER |
| 11 | 8976826 | AX-170046375 | 0 | C | A | 0.061 | 8.02E-03 | 2.14E-01 | 2.65E+00 | 2.74E+00 | 9.66E-01 | no | Pkd1l1 | upstream_gene_variant | MODIFIER |

| | | | | | | | | | | | | | | | |
|---|---|---|---|---|---|---|---|---|---|---|---|---|---|---|---|
| 11 | 8996416 | AX-97789672 | 0 | T | C | 0.061 | 8.02E-03 | 2.14E-01 | 2.65E+00 | 2.74E+00 | 9.66E-01 | no | Hus1 | 3_prime_UTR_variant | MODIFIER |
| 11 | 8997046 | AX-170300417 | 0 | C | G | 0.061 | 8.02E-03 | 2.14E-01 | 2.65E+00 | 2.74E+00 | 9.66E-01 | no | Hus1 | intron_variant | MODIFIER |
| 11 | 9018317 | AX-170004242 | 0 | C | A | 0.061 | 8.02E-03 | 2.14E-01 | 2.65E+00 | 2.74E+00 | 9.66E-01 | no | Sun3 | intron_variant | MODIFIER |
| 11 | 9034179 | AX-170004237 | 0 | A | G | 0.061 | 8.02E-03 | 2.14E-01 | 2.65E+00 | 2.74E+00 | 9.66E-01 | no | Sun3 | intron_variant | MODIFIER |
| 11 | 9034316 | AX-97828476 | 0 | C | G | 0.061 | 8.02E-03 | 2.14E-01 | 2.65E+00 | 2.74E+00 | 9.66E-01 | no | Sun3 | intron_variant | MODIFIER |
| 11 | 9165457 | AX-170217784 | 0 | G | A | 0.061 | 8.02E-03 | 2.14E-01 | 2.65E+00 | 2.74E+00 | 9.66E-01 | no | | intergenic_variant | MODIFIER |
| 11 | 9165595 | AX-170268122 | 0 | A | G | 0.061 | 8.02E-03 | 2.14E-01 | 2.65E+00 | 2.74E+00 | 9.66E-01 | no | | intergenic_variant | MODIFIER |
| 11 | 9180058 | AX-170431751 | 0 | C | A | 0.061 | 8.02E-03 | 2.14E-01 | 2.65E+00 | 2.74E+00 | 9.66E-01 | no | | intergenic_variant | MODIFIER |
| 11 | 9181917 | AX-170394564 | 0 | G | A | 0.061 | 8.02E-03 | 2.14E-01 | 2.65E+00 | 2.74E+00 | 9.66E-01 | no | | intergenic_variant | MODIFIER |
| 11 | 9183609 | AX-170161182 | 0 | C | T | 0.061 | 8.02E-03 | 2.14E-01 | 2.65E+00 | 2.74E+00 | 9.66E-01 | no | | intergenic_variant | MODIFIER |
| 11 | 9193431 | AX-170194166 | 0 | A | G | 0.061 | 8.02E-03 | 2.14E-01 | 2.65E+00 | 2.74E+00 | 9.66E-01 | no | Abca13 | intron_variant | MODIFIER |
| 11 | 9215564 | AX-170046376 | 0 | C | T | 0.061 | 8.02E-03 | 2.14E-01 | 2.65E+00 | 2.74E+00 | 9.66E-01 | no | Abca13 | intron_variant | MODIFIER |
| 11 | 9215641 | AX-170046377 | 0 | C | T | 0.061 | 8.02E-03 | 2.14E-01 | 2.65E+00 | 2.74E+00 | 9.66E-01 | no | Abca13 | intron_variant | MODIFIER |
| 11 | 9229436 | AX-97846451 | 0 | T | C | 0.061 | 8.02E-03 | 2.14E-01 | 2.65E+00 | 2.74E+00 | 9.66E-01 | no | Abca13 | intron_variant | MODIFIER |
| 11 | 9293617 | AX-170239728 | 0 | T | G | 0.061 | 8.02E-03 | 2.14E-01 | 2.65E+00 | 2.74E+00 | 9.66E-01 | no | Abca13 | missense_variant | MODERATE |
| 11 | 9706684 | AX-170239696 | 0 | A | G | 0.061 | 8.02E-03 | 2.14E-01 | 2.65E+00 | 2.74E+00 | 9.66E-01 | no | Gm36954 | upstream_gene_variant | MODIFIER |
| 11 | 9717475 | AX-170374626 | 0 | C | T | 0.061 | 8.02E-03 | 2.14E-01 | 2.65E+00 | 2.74E+00 | 9.66E-01 | no | | intergenic_variant | MODIFIER |
| 11 | 9720812 | AX-170201018 | 0 | T | C | 0.061 | 8.02E-03 | 2.14E-01 | 2.65E+00 | 2.74E+00 | 9.66E-01 | no | | intergenic_variant | MODIFIER |
| 11 | 9722491 | AX-170325723 | 0 | G | A | 0.061 | 8.02E-03 | 2.14E-01 | 2.65E+00 | 2.74E+00 | 9.66E-01 | no | | intergenic_variant | MODIFIER |
| 11 | 9751095 | AX-170086946 | 0 | T | C | 0.061 | 8.02E-03 | 2.14E-01 | 2.65E+00 | 2.74E+00 | 9.66E-01 | no | | intergenic_variant | MODIFIER |
| 11 | 10038698 | AX-170456101 | 0 | T | C | 0.061 | 8.02E-03 | 2.14E-01 | 2.65E+00 | 2.74E+00 | 9.66E-01 | no | | intergenic_variant | MODIFIER |
| 11 | 10546053 | AX-97853146 | 0 | C | T | 0.061 | 8.02E-03 | 2.14E-01 | 2.65E+00 | 2.74E+00 | 9.66E-01 | no | | intergenic_variant | MODIFIER |
| 11 | 10552720 | AX-170425846 | 0 | T | G | 0.061 | 8.02E-03 | 2.14E-01 | 2.65E+00 | 2.74E+00 | 9.66E-01 | no | | intergenic_variant | MODIFIER |
| 11 | 10556023 | AX-170004265 | 0 | G | A | 0.061 | 8.02E-03 | 2.14E-01 | 2.65E+00 | 2.74E+00 | 9.66E-01 | no | | intergenic_variant | MODIFIER |
| 11 | 10556352 | AX-170004270 | 0 | A | G | 0.061 | 8.02E-03 | 2.14E-01 | 2.65E+00 | 2.74E+00 | 9.66E-01 | no | | intergenic_variant | MODIFIER |
| 11 | 10556995 | AX-170375261 | 0 | G | A | 0.061 | 8.02E-03 | 2.14E-01 | 2.65E+00 | 2.74E+00 | 9.66E-01 | no | | intergenic_variant | MODIFIER |
| 11 | 10927955 | AX-97857180 | 0 | T | C | 0.061 | 8.02E-03 | 2.14E-01 | 2.65E+00 | 2.74E+00 | 9.66E-01 | no | | intergenic_variant | MODIFIER |
| 11 | 10948737 | AX-97835226 | 0 | C | G | 0.061 | 8.02E-03 | 2.14E-01 | 2.65E+00 | 2.74E+00 | 9.66E-01 | no | | intergenic_variant | MODIFIER |
| 11 | 10956062 | AX-170004279 | 0 | A | G | 0.061 | 8.02E-03 | 2.14E-01 | 2.65E+00 | 2.74E+00 | 9.66E-01 | no | | intergenic_variant | MODIFIER |
| 11 | 11051128 | AX-170139057 | 0 | G | A | 0.061 | 8.02E-03 | 2.14E-01 | 2.65E+00 | 2.74E+00 | 9.66E-01 | no | | intergenic_variant | MODIFIER |
| 11 | 11052398 | AX-97796318 | 0 | T | G | 0.061 | 8.02E-03 | 2.14E-01 | 2.65E+00 | 2.74E+00 | 9.66E-01 | no | | intergenic_variant | MODIFIER |
| 11 | 11080577 | AX-97788749 | 0 | A | C | 0.061 | 8.02E-03 | 2.14E-01 | 2.65E+00 | 2.74E+00 | 9.66E-01 | no | | intergenic_variant | MODIFIER |
| 11 | 11089710 | AX-97799770 | 0 | A | G | 0.061 | 8.02E-03 | 2.14E-01 | 2.65E+00 | 2.74E+00 | 9.66E-01 | no | | intergenic_variant | MODIFIER |
| 11 | 11140106 | AX-97801825 | 0 | T | C | 0.061 | 8.02E-03 | 2.14E-01 | 2.65E+00 | 2.74E+00 | 9.66E-01 | no | Vwc2 | intron_variant | MODIFIER |
| 11 | 11155056 | AX-170004275 | 0 | G | A | 0.061 | 8.02E-03 | 2.14E-01 | 2.65E+00 | 2.74E+00 | 9.66E-01 | no | Vwc2 | intron_variant | MODIFIER |
| 11 | 11171518 | AX-170381830 | 0 | A | G | 0.061 | 8.02E-03 | 2.14E-01 | 2.65E+00 | 2.74E+00 | 9.66E-01 | no | Vwc2 | intron_variant | MODIFIER |
| 11 | 11171618 | AX-170133979 | 0 | C | G | 0.061 | 8.02E-03 | 2.14E-01 | 2.65E+00 | 2.74E+00 | 9.66E-01 | no | Vwc2 | intron_variant | MODIFIER |
| 11 | 11195444 | AX-170004277 | 0 | A | T | 0.061 | 8.02E-03 | 2.14E-01 | 2.65E+00 | 2.74E+00 | 9.66E-01 | no | Vwc2 | intron_variant | MODIFIER |
| 11 | 11225120 | AX-170455533 | 0 | T | C | 0.061 | 8.02E-03 | 2.14E-01 | 2.65E+00 | 2.74E+00 | 9.66E-01 | no | Vwc2 | intron_variant | MODIFIER |
| 11 | 11268829 | AX-170354403 | 0 | C | T | 0.061 | 8.02E-03 | 2.14E-01 | 2.65E+00 | 2.74E+00 | 9.66E-01 | no | Vwc2 | downstream_gene_variant | MODIFIER |
| 11 | 12243960 | AX-97854452 | 0 | A | G | 0.061 | 8.02E-03 | 2.14E-01 | 2.65E+00 | 2.74E+00 | 9.66E-01 | no | Cobl | intron_variant | MODIFIER |
| 11 | 12248827 | AX-170046417 | 0 | A | G | 0.061 | 8.02E-03 | 2.14E-01 | 2.65E+00 | 2.74E+00 | 9.66E-01 | no | Cobl | intron_variant | MODIFIER |
| 11 | 12252265 | AX-170242595 | 0 | C | T | 0.061 | 8.02E-03 | 2.14E-01 | 2.65E+00 | 2.74E+00 | 9.66E-01 | no | Cobl | intron_variant | MODIFIER |
| 11 | 12271162 | AX-170183772 | 0 | G | C | 0.061 | 8.02E-03 | 2.14E-01 | 2.65E+00 | 2.74E+00 | 9.66E-01 | no | Cobl | intron_variant | MODIFIER |
| 11 | 12271301 | AX-170178933 | 0 | G | A | 0.061 | 8.02E-03 | 2.14E-01 | 2.65E+00 | 2.74E+00 | 9.66E-01 | no | Cobl | intron_variant | MODIFIER |
| 11 | 12271595 | AX-170308281 | 0 | T | C | 0.061 | 8.02E-03 | 2.14E-01 | 2.65E+00 | 2.74E+00 | 9.66E-01 | no | Cobl | intron_variant | MODIFIER |
| 11 | 12274483 | AX-97830692 | 0 | T | C | 0.061 | 8.02E-03 | 2.14E-01 | 2.65E+00 | 2.74E+00 | 9.66E-01 | no | Cobl | intron_variant | MODIFIER |
| 11 | 12274709 | AX-170440040 | 0 | A | T | 0.061 | 8.02E-03 | 2.14E-01 | 2.65E+00 | 2.74E+00 | 9.66E-01 | no | Cobl | intron_variant | MODIFIER |
| 11 | 12286961 | AX-97783929 | 0 | A | G | 0.061 | 8.02E-03 | 2.14E-01 | 2.65E+00 | 2.74E+00 | 9.66E-01 | no | Cobl | intron_variant | MODIFIER |
| 11 | 12764324 | AX-170202120 | 0 | A | G | 0.061 | 8.02E-03 | 2.14E-01 | 2.65E+00 | 2.74E+00 | 9.66E-01 | no | | intergenic_variant | MODIFIER |
| 11 | 12838559 | AX-170479385 | 0 | G | A | 0.061 | 8.02E-03 | 2.14E-01 | 2.65E+00 | 2.74E+00 | 9.66E-01 | no | | intergenic_variant | MODIFIER |
| 11 | 12873556 | AX-170514798 | 0 | A | T | 0.061 | 8.02E-03 | 2.14E-01 | 2.65E+00 | 2.74E+00 | 9.66E-01 | no | | intergenic_variant | MODIFIER |
| 11 | 12919617 | AX-170119352 | 0 | T | G | 0.061 | 8.02E-03 | 2.14E-01 | 2.65E+00 | 2.74E+00 | 9.66E-01 | no | | intergenic_variant | MODIFIER |
| 11 | 12986448 | AX-170223198 | 0 | T | C | 0.061 | 8.02E-03 | 2.14E-01 | 2.65E+00 | 2.74E+00 | 9.66E-01 | no | | intergenic_variant | MODIFIER |

| | | | | | | | | | | | | | | | |
|---|---|---|---|---|---|---|---|---|---|---|---|---|---|---|---|
| 11 | 13043202 | AX-170393161 | 0 | G | A | 0.061 | 8.02E-03 | 2.14E-01 | 2.65E+00 | 2.74E+00 | 9.66E-01 | no | | intergenic_variant | MODIFIER |
| 11 | 13056513 | AX-168373898 | 0 | A | C | 0.061 | 8.02E-03 | 2.14E-01 | 2.65E+00 | 2.74E+00 | 9.66E-01 | no | | intergenic_variant | MODIFIER |
| 11 | 13100198 | AX-170046432 | 0 | C | T | 0.061 | 8.02E-03 | 2.14E-01 | 2.65E+00 | 2.74E+00 | 9.66E-01 | no | | intergenic_variant | MODIFIER |
| 11 | 13107029 | AX-170046434 | 0 | G | A | 0.061 | 8.02E-03 | 2.14E-01 | 2.65E+00 | 2.74E+00 | 9.66E-01 | no | | intergenic_variant | MODIFIER |
| 11 | 13113749 | AX-170145548 | 0 | C | A | 0.061 | 8.02E-03 | 2.14E-01 | 2.65E+00 | 2.74E+00 | 9.66E-01 | no | | intergenic_variant | MODIFIER |
| 11 | 13125371 | AX-170508549 | 0 | C | T | 0.061 | 8.02E-03 | 2.14E-01 | 2.65E+00 | 2.74E+00 | 9.66E-01 | no | | intergenic_variant | MODIFIER |
| 11 | 13434401 | AX-170171467 | 0 | G | A | 0.061 | 8.02E-03 | 2.14E-01 | 2.65E+00 | 2.74E+00 | 9.66E-01 | no | | intergenic_variant | MODIFIER |
| 11 | 13454445 | AX-170507040 | 0 | T | C | 0.061 | 8.02E-03 | 2.14E-01 | 2.65E+00 | 2.74E+00 | 9.66E-01 | no | | intergenic_variant | MODIFIER |
| 11 | 13460357 | AX-170422072 | 0 | G | A | 0.061 | 8.02E-03 | 2.14E-01 | 2.65E+00 | 2.74E+00 | 9.66E-01 | no | | intergenic_variant | MODIFIER |
| 11 | 13610477 | AX-170518074 | 0 | T | G | 0.061 | 8.02E-03 | 2.14E-01 | 2.65E+00 | 2.74E+00 | 9.66E-01 | no | | intergenic_variant | MODIFIER |
| 11 | 13611314 | AX-170295388 | 0 | G | T | 0.061 | 8.02E-03 | 2.14E-01 | 2.65E+00 | 2.74E+00 | 9.66E-01 | no | | intergenic_variant | MODIFIER |
| 11 | 13613209 | AX-170004321 | 0 | C | T | 0.061 | 8.02E-03 | 2.14E-01 | 2.65E+00 | 2.74E+00 | 9.66E-01 | no | | intergenic_variant | MODIFIER |
| 11 | 13623821 | AX-170319537 | 0 | G | A | 0.061 | 8.02E-03 | 2.14E-01 | 2.65E+00 | 2.74E+00 | 9.66E-01 | no | | intergenic_variant | MODIFIER |
| 11 | 13623901 | AX-170004322 | 0 | A | G | 0.061 | 8.02E-03 | 2.14E-01 | 2.65E+00 | 2.74E+00 | 9.66E-01 | no | | intergenic_variant | MODIFIER |
| 11 | 13624226 | AX-170004323 | 0 | G | A | 0.061 | 8.02E-03 | 2.14E-01 | 2.65E+00 | 2.74E+00 | 9.66E-01 | no | | intergenic_variant | MODIFIER |
| 11 | 13672870 | AX-170004320 | 0 | G | A | 0.061 | 8.02E-03 | 2.14E-01 | 2.65E+00 | 2.74E+00 | 9.66E-01 | no | | intergenic_variant | MODIFIER |
| 11 | 13710419 | AX-170367974 | 0 | T | C | 0.061 | 8.02E-03 | 2.14E-01 | 2.65E+00 | 2.74E+00 | 9.66E-01 | no | | intergenic_variant | MODIFIER |
| 11 | 13787288 | AX-170246739 | 0 | C | T | 0.061 | 8.02E-03 | 2.14E-01 | 2.65E+00 | 2.74E+00 | 9.66E-01 | no | | intergenic_variant | MODIFIER |
| 11 | 13806824 | AX-170377760 | 0 | C | T | 0.061 | 8.02E-03 | 2.14E-01 | 2.65E+00 | 2.74E+00 | 9.66E-01 | no | | intergenic_variant | MODIFIER |
| 11 | 13855547 | AX-170382188 | 0 | A | G | 0.061 | 8.02E-03 | 2.14E-01 | 2.65E+00 | 2.74E+00 | 9.66E-01 | no | | intergenic_variant | MODIFIER |
| 11 | 13881537 | AX-170129107 | 0 | C | T | 0.061 | 8.02E-03 | 2.14E-01 | 2.65E+00 | 2.74E+00 | 9.66E-01 | no | | intergenic_variant | MODIFIER |
| 11 | 14024096 | AX-170493571 | 0 | G | A | 0.061 | 8.02E-03 | 2.14E-01 | 2.65E+00 | 2.74E+00 | 9.66E-01 | no | | intergenic_variant | MODIFIER |
| 11 | 14025054 | AX-170254844 | 0 | G | C | 0.061 | 8.02E-03 | 2.14E-01 | 2.65E+00 | 2.74E+00 | 9.66E-01 | no | | intergenic_variant | MODIFIER |
| 11 | 14028598 | AX-170410696 | 0 | C | A | 0.061 | 8.02E-03 | 2.14E-01 | 2.65E+00 | 2.74E+00 | 9.66E-01 | no | | intergenic_variant | MODIFIER |
| 11 | 14225071 | AX-97806858 | 0 | T | A | 0.061 | 8.02E-03 | 2.14E-01 | 2.65E+00 | 2.74E+00 | 9.66E-01 | no | | intergenic_variant | MODIFIER |
| 11 | 14251022 | AX-97817432 | 0 | T | A | 0.061 | 8.02E-03 | 2.14E-01 | 2.65E+00 | 2.74E+00 | 9.66E-01 | no | | intergenic_variant | MODIFIER |
| 11 | 14271068 | AX-170173688 | 0 | A | T | 0.061 | 8.02E-03 | 2.14E-01 | 2.65E+00 | 2.74E+00 | 9.66E-01 | no | | intergenic_variant | MODIFIER |
| 11 | 15038809 | AX-170417814 | 0 | A | G | 0.061 | 8.02E-03 | 2.14E-01 | 2.65E+00 | 2.74E+00 | 9.66E-01 | no | | intergenic_variant | MODIFIER |
| 11 | 16906827 | AX-170046485 | 0 | A | G | 0.061 | 8.02E-03 | 2.14E-01 | 2.65E+00 | 2.74E+00 | 9.66E-01 | no | Egfr | synonymous_variant | LOW |
| 11 | 17499083 | AX-170380794 | 0 | C | T | 0.061 | 8.02E-03 | 2.14E-01 | 2.65E+00 | 2.74E+00 | 9.66E-01 | no | | intergenic_variant | MODIFIER |
| 11 | 17599492 | AX-97823993 | 0 | G | A | 0.061 | 8.02E-03 | 2.14E-01 | 2.65E+00 | 2.74E+00 | 9.66E-01 | no | | intergenic_variant | MODIFIER |
| 11 | 17629954 | AX-97793360 | 0 | G | A | 0.061 | 8.02E-03 | 2.14E-01 | 2.65E+00 | 2.74E+00 | 9.66E-01 | no | | regulatory_region_variant | MODIFIER |
| 11 | 17630607 | AX-170436358 | 0 | A | C | 0.061 | 8.02E-03 | 2.14E-01 | 2.65E+00 | 2.74E+00 | 9.66E-01 | no | | intergenic_variant | MODIFIER |
| 11 | 17632414 | AX-170328155 | 0 | T | C | 0.061 | 8.02E-03 | 2.14E-01 | 2.65E+00 | 2.74E+00 | 9.66E-01 | no | | intergenic_variant | MODIFIER |
| 11 | 17664300 | AX-170491026 | 0 | G | A | 0.061 | 8.02E-03 | 2.14E-01 | 2.65E+00 | 2.74E+00 | 9.66E-01 | no | | regulatory_region_variant | MODIFIER |
| 11 | 17681257 | AX-170298771 | 0 | C | T | 0.061 | 8.02E-03 | 2.14E-01 | 2.65E+00 | 2.74E+00 | 9.66E-01 | no | | regulatory_region_variant | MODIFIER |
| 11 | 17710694 | AX-170450929 | 0 | C | T | 0.061 | 8.02E-03 | 2.14E-01 | 2.65E+00 | 2.74E+00 | 9.66E-01 | no | | intergenic_variant | MODIFIER |
| 11 | 17710884 | AX-170521353 | 0 | G | A | 0.061 | 8.02E-03 | 2.14E-01 | 2.65E+00 | 2.74E+00 | 9.66E-01 | no | | intergenic_variant | MODIFIER |
| 11 | 17713806 | AX-170195406 | 0 | A | G | 0.061 | 8.02E-03 | 2.14E-01 | 2.65E+00 | 2.74E+00 | 9.66E-01 | no | | intergenic_variant | MODIFIER |
| 11 | 17797045 | AX-170099813 | 0 | G | A | 0.061 | 8.02E-03 | 2.14E-01 | 2.65E+00 | 2.74E+00 | 9.66E-01 | no | | intergenic_variant | MODIFIER |
| 11 | 17803850 | AX-170365784 | 0 | A | G | 0.061 | 8.02E-03 | 2.14E-01 | 2.65E+00 | 2.74E+00 | 9.66E-01 | no | | intergenic_variant | MODIFIER |
| 11 | 17860799 | AX-170086956 | 0 | T | G | 0.061 | 8.02E-03 | 2.14E-01 | 2.65E+00 | 2.74E+00 | 9.66E-01 | no | | intergenic_variant | MODIFIER |
| 11 | 17860998 | AX-170004399 | 0 | G | A | 0.061 | 8.02E-03 | 2.14E-01 | 2.65E+00 | 2.74E+00 | 9.66E-01 | no | | intergenic_variant | MODIFIER |
| 11 | 17902316 | AX-170529359 | 0 | C | T | 0.061 | 8.02E-03 | 2.14E-01 | 2.65E+00 | 2.74E+00 | 9.66E-01 | no | | intergenic_variant | MODIFIER |
| 11 | 17905352 | AX-170344019 | 0 | G | A | 0.061 | 8.02E-03 | 2.14E-01 | 2.65E+00 | 2.74E+00 | 9.66E-01 | no | | intergenic_variant | MODIFIER |
| 11 | 17948878 | AX-170004407 | 0 | T | A | 0.061 | 8.02E-03 | 2.14E-01 | 2.65E+00 | 2.74E+00 | 9.66E-01 | no | Etaa1 | intron_variant | MODIFIER |
| 11 | 17951112 | AX-170004402 | 0 | A | G | 0.061 | 8.02E-03 | 2.14E-01 | 2.65E+00 | 2.74E+00 | 9.66E-01 | no | Etaa1 | intron_variant | MODIFIER |
| 11 | 17989907 | AX-170532093 | 0 | G | C | 0.061 | 8.02E-03 | 2.14E-01 | 2.65E+00 | 2.74E+00 | 9.66E-01 | no | | intergenic_variant | MODIFIER |
| 11 | 18076638 | AX-170505901 | 0 | C | A | 0.061 | 8.02E-03 | 2.14E-01 | 2.65E+00 | 2.74E+00 | 9.66E-01 | no | | intergenic_variant | MODIFIER |
| 11 | 18085601 | AX-170413168 | 0 | G | C | 0.061 | 8.02E-03 | 2.14E-01 | 2.65E+00 | 2.74E+00 | 9.66E-01 | no | | intergenic_variant | MODIFIER |
| 11 | 18096223 | AX-170332823 | 0 | A | T | 0.061 | 8.02E-03 | 2.14E-01 | 2.65E+00 | 2.74E+00 | 9.66E-01 | no | Gm12018 | non_coding_transcript_exon_variant | MODIFIER |
| 11 | 18122441 | AX-170004396 | 0 | C | T | 0.061 | 8.02E-03 | 2.14E-01 | 2.65E+00 | 2.74E+00 | 9.66E-01 | no | | intergenic_variant | MODIFIER |
| 11 | 18122494 | AX-170004408 | 0 | A | G | 0.061 | 8.02E-03 | 2.14E-01 | 2.65E+00 | 2.74E+00 | 9.66E-01 | no | | intergenic_variant | MODIFIER |
| 11 | 18207451 | AX-170086957 | 0 | C | G | 0.061 | 8.02E-03 | 2.14E-01 | 2.65E+00 | 2.74E+00 | 9.66E-01 | no | Gm12019 | upstream_gene_variant | MODIFIER |

| | | | | | | | | | | | | | | | |
|---|---|---|---|---|---|---|---|---|---|---|---|---|---|---|---|
| 11 | 18207634 | AX-170315642 | 0 | A | C | 0.061 | 8.02E-03 | 2.14E-01 | 2.65E+00 | 2.74E+00 | 9.66E-01 | no | Gm12019 | upstream_gene_variant | MODIFIER |
| 11 | 18273132 | AX-170484995 | 0 | T | C | 0.061 | 8.02E-03 | 2.14E-01 | 2.65E+00 | 2.74E+00 | 9.66E-01 | no | | intergenic_variant | MODIFIER |
| 11 | 18277271 | AX-97802995 | 0 | T | C | 0.061 | 8.02E-03 | 2.14E-01 | 2.65E+00 | 2.74E+00 | 9.66E-01 | no | | intergenic_variant | MODIFIER |
| 11 | 18277387 | AX-170268099 | 0 | A | C | 0.061 | 8.02E-03 | 2.14E-01 | 2.65E+00 | 2.74E+00 | 9.66E-01 | no | | intergenic_variant | MODIFIER |
| 11 | 18283704 | AX-170223485 | 0 | C | T | 0.061 | 8.02E-03 | 2.14E-01 | 2.65E+00 | 2.74E+00 | 9.66E-01 | no | | intergenic_variant | MODIFIER |
| 11 | 18364090 | AX-97846273 | 0 | T | G | 0.061 | 8.02E-03 | 2.14E-01 | 2.65E+00 | 2.74E+00 | 9.66E-01 | no | | intergenic_variant | MODIFIER |
| 11 | 18371030 | AX-170190795 | 0 | T | C | 0.061 | 8.02E-03 | 2.14E-01 | 2.65E+00 | 2.74E+00 | 9.66E-01 | no | | regulatory_region_variant | MODIFIER |
| 11 | 18399666 | AX-97817430 | 0 | C | T | 0.061 | 8.02E-03 | 2.14E-01 | 2.65E+00 | 2.74E+00 | 9.66E-01 | no | | regulatory_region_variant | MODIFIER |
| 11 | 18407934 | AX-170004412 | 0 | T | A | 0.061 | 8.02E-03 | 2.14E-01 | 2.65E+00 | 2.74E+00 | 9.66E-01 | no | | intergenic_variant | MODIFIER |
| 11 | 18414079 | AX-170528040 | 0 | T | C | 0.061 | 8.02E-03 | 2.14E-01 | 2.65E+00 | 2.74E+00 | 9.66E-01 | no | | intergenic_variant | MODIFIER |
| 11 | 18414557 | AX-170120496 | 0 | C | T | 0.061 | 8.02E-03 | 2.14E-01 | 2.65E+00 | 2.74E+00 | 9.66E-01 | no | | intergenic_variant | MODIFIER |
| 11 | 18419282 | AX-170046505 | 0 | T | C | 0.061 | 8.02E-03 | 2.14E-01 | 2.65E+00 | 2.74E+00 | 9.66E-01 | no | | intergenic_variant | MODIFIER |
| 11 | 18421690 | AX-170046504 | 0 | C | A | 0.061 | 8.02E-03 | 2.14E-01 | 2.65E+00 | 2.74E+00 | 9.66E-01 | no | | intergenic_variant | MODIFIER |
| 11 | 18531319 | AX-170421188 | 0 | A | G | 0.061 | 8.02E-03 | 2.14E-01 | 2.65E+00 | 2.74E+00 | 9.66E-01 | no | | regulatory_region_variant | MODIFIER |
| 11 | 18538127 | AX-97853592 | 0 | G | A | 0.061 | 8.02E-03 | 2.14E-01 | 2.65E+00 | 2.74E+00 | 9.66E-01 | no | | intergenic_variant | MODIFIER |
| 11 | 18538430 | AX-170385325 | 0 | G | A | 0.061 | 8.02E-03 | 2.14E-01 | 2.65E+00 | 2.74E+00 | 9.66E-01 | no | | intergenic_variant | MODIFIER |
| 11 | 18546805 | AX-170004418 | 0 | C | G | 0.061 | 8.02E-03 | 2.14E-01 | 2.65E+00 | 2.74E+00 | 9.66E-01 | no | | intergenic_variant | MODIFIER |
| 11 | 18547029 | AX-170004410 | 0 | G | A | 0.061 | 8.02E-03 | 2.14E-01 | 2.65E+00 | 2.74E+00 | 9.66E-01 | no | | intergenic_variant | MODIFIER |
| 11 | 18559188 | AX-170004409 | 0 | C | T | 0.061 | 8.02E-03 | 2.14E-01 | 2.65E+00 | 2.74E+00 | 9.66E-01 | no | | intergenic_variant | MODIFIER |
| 11 | 18585284 | AX-170237037 | 0 | A | T | 0.061 | 8.02E-03 | 2.14E-01 | 2.65E+00 | 2.74E+00 | 9.66E-01 | no | | intergenic_variant | MODIFIER |
| 11 | 18586307 | AX-170350029 | 0 | A | G | 0.061 | 8.02E-03 | 2.14E-01 | 2.65E+00 | 2.74E+00 | 9.66E-01 | no | | intergenic_variant | MODIFIER |
| 11 | 18590814 | AX-170432646 | 0 | G | A | 0.061 | 8.02E-03 | 2.14E-01 | 2.65E+00 | 2.74E+00 | 9.66E-01 | no | Gm12023 | downstream_gene_variant | MODIFIER |
| 11 | 18657459 | AX-170396132 | 0 | G | A | 0.061 | 8.02E-03 | 2.14E-01 | 2.65E+00 | 2.74E+00 | 9.66E-01 | no | Gm12023 | intron_variant,non_coding_transcript_variant | MODIFIER |
| 11 | 18657599 | AX-170086958 | 0 | G | A | 0.061 | 8.02E-03 | 2.14E-01 | 2.65E+00 | 2.74E+00 | 9.66E-01 | no | Gm12023 | intron_variant,non_coding_transcript_variant | MODIFIER |
| 11 | 18668252 | AX-170513383 | 0 | T | G | 0.061 | 8.02E-03 | 2.14E-01 | 2.65E+00 | 2.74E+00 | 9.66E-01 | no | Gm12023 | intron_variant,non_coding_transcript_variant | MODIFIER |
| 11 | 18668405 | AX-170128018 | 0 | T | C | 0.061 | 8.02E-03 | 2.14E-01 | 2.65E+00 | 2.74E+00 | 9.66E-01 | no | Gm12023 | intron_variant,non_coding_transcript_variant | MODIFIER |
| 11 | 18671000 | AX-170268552 | 0 | C | T | 0.061 | 8.02E-03 | 2.14E-01 | 2.65E+00 | 2.74E+00 | 9.66E-01 | no | Gm12023 | intron_variant,non_coding_transcript_variant | MODIFIER |
| 11 | 18675911 | AX-170092188 | 0 | T | C | 0.061 | 8.02E-03 | 2.14E-01 | 2.65E+00 | 2.74E+00 | 9.66E-01 | no | Gm12023 | intron_variant,non_coding_transcript_variant | MODIFIER |
| 11 | 18683489 | AX-170509939 | 0 | C | T | 0.061 | 8.02E-03 | 2.14E-01 | 2.65E+00 | 2.74E+00 | 9.66E-01 | no | Gm12023 | intron_variant,non_coding_transcript_variant | MODIFIER |
| 11 | 18690231 | AX-170046507 | 0 | T | C | 0.061 | 8.02E-03 | 2.14E-01 | 2.65E+00 | 2.74E+00 | 9.66E-01 | no | Gm12023 | intron_variant,non_coding_transcript_variant | MODIFIER |
| 11 | 18715592 | AX-170149863 | 0 | C | T | 0.061 | 8.02E-03 | 2.14E-01 | 2.65E+00 | 2.74E+00 | 9.66E-01 | no | | intergenic_variant | MODIFIER |
| 11 | 18716846 | AX-170362999 | 0 | T | C | 0.061 | 8.02E-03 | 2.14E-01 | 2.65E+00 | 2.74E+00 | 9.66E-01 | no | | intergenic_variant | MODIFIER |
| 11 | 18724620 | AX-170216269 | 0 | G | A | 0.061 | 8.02E-03 | 2.14E-01 | 2.65E+00 | 2.74E+00 | 9.66E-01 | no | | intergenic_variant | MODIFIER |
| 11 | 18744715 | AX-170347914 | 0 | G | A | 0.061 | 8.02E-03 | 2.14E-01 | 2.65E+00 | 2.74E+00 | 9.66E-01 | no | | intergenic_variant | MODIFIER |
| 11 | 18746223 | AX-170118377 | 0 | C | T | 0.061 | 8.02E-03 | 2.14E-01 | 2.65E+00 | 2.74E+00 | 9.66E-01 | no | | regulatory_region_variant | MODIFIER |
| 11 | 18759637 | AX-170099724 | 0 | T | C | 0.061 | 8.02E-03 | 2.14E-01 | 2.65E+00 | 2.74E+00 | 9.66E-01 | no | Gm28401 | upstream_gene_variant | MODIFIER |
| 11 | 18766005 | AX-170380431 | 0 | G | A | 0.061 | 8.02E-03 | 2.14E-01 | 2.65E+00 | 2.74E+00 | 9.66E-01 | no | | intergenic_variant | MODIFIER |
| 11 | 18829441 | AX-170103897 | 0 | G | A | 0.061 | 8.02E-03 | 2.14E-01 | 2.65E+00 | 2.74E+00 | 9.66E-01 | no | | intergenic_variant | MODIFIER |
| 11 | 18829458 | AX-170351659 | 0 | C | G | 0.061 | 8.02E-03 | 2.14E-01 | 2.65E+00 | 2.74E+00 | 9.66E-01 | no | | intergenic_variant | MODIFIER |
| 11 | 18829916 | AX-170092189 | 0 | C | T | 0.061 | 8.02E-03 | 2.14E-01 | 2.65E+00 | 2.74E+00 | 9.66E-01 | no | | intergenic_variant | MODIFIER |
| 11 | 18830114 | AX-170240589 | 0 | T | C | 0.061 | 8.02E-03 | 2.14E-01 | 2.65E+00 | 2.74E+00 | 9.66E-01 | no | | intergenic_variant | MODIFIER |
| 11 | 18830178 | AX-170153470 | 0 | T | C | 0.061 | 8.02E-03 | 2.14E-01 | 2.65E+00 | 2.74E+00 | 9.66E-01 | no | | intergenic_variant | MODIFIER |
| 11 | 18830521 | AX-170324518 | 0 | G | A | 0.061 | 8.02E-03 | 2.14E-01 | 2.65E+00 | 2.74E+00 | 9.66E-01 | no | | intergenic_variant | MODIFIER |
| 11 | 18831037 | AX-170444953 | 0 | A | C | 0.061 | 8.02E-03 | 2.14E-01 | 2.65E+00 | 2.74E+00 | 9.66E-01 | no | | intergenic_variant | MODIFIER |
| 11 | 18837116 | AX-170460206 | 0 | G | T | 0.061 | 8.02E-03 | 2.14E-01 | 2.65E+00 | 2.74E+00 | 9.66E-01 | no | | intergenic_variant | MODIFIER |
| 11 | 18850087 | AX-170046512 | 0 | G | C | 0.061 | 8.02E-03 | 2.14E-01 | 2.65E+00 | 2.74E+00 | 9.66E-01 | no | 8430419K02Rik | intron_variant,non_coding_transcript_variant | MODIFIER |
| 11 | 18861116 | AX-170046514 | 0 | C | T | 0.061 | 8.02E-03 | 2.14E-01 | 2.65E+00 | 2.74E+00 | 9.66E-01 | no | 8430419K02Rik | intron_variant,non_coding_transcript_variant | MODIFIER |
| 11 | 19199516 | AX-97817635 | 0 | T | G | 0.061 | 8.02E-03 | 2.14E-01 | 2.65E+00 | 2.74E+00 | 9.66E-01 | no | | intergenic_variant | MODIFIER |
| 11 | 19199593 | AX-170474358 | 0 | G | T | 0.061 | 8.02E-03 | 2.14E-01 | 2.65E+00 | 2.74E+00 | 9.66E-01 | no | | intergenic_variant | MODIFIER |
| 11 | 19228223 | AX-97784749 | 0 | A | G | 0.061 | 8.02E-03 | 2.14E-01 | 2.65E+00 | 2.74E+00 | 9.66E-01 | no | | intergenic_variant | MODIFIER |
| 11 | 19234958 | AX-170486521 | 0 | T | A | 0.061 | 8.02E-03 | 2.14E-01 | 2.65E+00 | 2.74E+00 | 9.66E-01 | no | | intergenic_variant | MODIFIER |
| 11 | 19235018 | AX-170168914 | 0 | A | G | 0.061 | 8.02E-03 | 2.14E-01 | 2.65E+00 | 2.74E+00 | 9.66E-01 | no | | intergenic_variant | MODIFIER |
| 11 | 19256177 | AX-170454945 | 0 | A | G | 0.061 | 8.02E-03 | 2.14E-01 | 2.65E+00 | 2.74E+00 | 9.66E-01 | no | | intergenic_variant | MODIFIER |
| 11 | 20647221 | AX-170004441 | 0 | A | G | 0.061 | 8.02E-03 | 2.14E-01 | 2.65E+00 | 2.74E+00 | 9.66E-01 | no | Sertad2 | intron_variant | MODIFIER |

| | | | | | | | | | | | | | | | |
|---|---|---|---|---|---|---|---|---|---|---|---|---|---|---|---|
| 11 | 21140133 | AX-170035495 | 0 | G | A | 0.061 | 8.02E-03 | 2.14E-01 | 2.65E+00 | 2.74E+00 | 9.66E-01 | no | Peli1 | intron_variant | MODIFIER |
| 11 | 23005784 | AX-170004489 | 0 | C | T | 0.061 | 8.02E-03 | 2.14E-01 | 2.65E+00 | 2.74E+00 | 9.66E-01 | no | Cct4 | downstream_gene_variant | MODIFIER |
| 11 | 23488087 | AX-97817693 | 0 | A | G | 0.061 | 8.02E-03 | 2.14E-01 | 2.65E+00 | 2.74E+00 | 9.66E-01 | no | Ahsa2 | downstream_gene_variant | MODIFIER |
| 11 | 24147253 | AX-97818689 | 0 | A | G | 0.061 | 8.02E-03 | 2.14E-01 | 2.65E+00 | 2.74E+00 | 9.66E-01 | no | Bcl11a | intron_variant | MODIFIER |
| 11 | 24478976 | AX-170237795 | 0 | C | T | 0.061 | 8.02E-03 | 2.14E-01 | 2.65E+00 | 2.74E+00 | 9.66E-01 | no | 4933430M04Rik | upstream_gene_variant | MODIFIER |
| 11 | 24481395 | AX-170213341 | 0 | G | A | 0.061 | 8.02E-03 | 2.14E-01 | 2.65E+00 | 2.74E+00 | 9.66E-01 | no | 4933430M04Rik | upstream_gene_variant | MODIFIER |
| 11 | 24481797 | AX-170179879 | 0 | T | C | 0.061 | 8.02E-03 | 2.14E-01 | 2.65E+00 | 2.74E+00 | 9.66E-01 | no | 4933430M04Rik | upstream_gene_variant | MODIFIER |
| 11 | 24486320 | AX-170004507 | 0 | T | C | 0.061 | 8.02E-03 | 2.14E-01 | 2.65E+00 | 2.74E+00 | 9.66E-01 | no | 4933430M04Rik | intron_variant,non_coding_transcript_variant | MODIFIER |
| 11 | 24486409 | AX-170004508 | 0 | T | C | 0.061 | 8.02E-03 | 2.14E-01 | 2.65E+00 | 2.74E+00 | 9.66E-01 | no | 4933430M04Rik | intron_variant,non_coding_transcript_variant | MODIFIER |
| 11 | 24486584 | AX-170287252 | 0 | C | G | 0.061 | 8.02E-03 | 2.14E-01 | 2.65E+00 | 2.74E+00 | 9.66E-01 | no | 4933430M04Rik | intron_variant,non_coding_transcript_variant | MODIFIER |
| 11 | 24513986 | AX-170446311 | 0 | G | A | 0.061 | 8.02E-03 | 2.14E-01 | 2.65E+00 | 2.74E+00 | 9.66E-01 | no | 4933430M04Rik | intron_variant,non_coding_transcript_variant | MODIFIER |
| 11 | 24515229 | AX-170335129 | 0 | T | C | 0.061 | 8.02E-03 | 2.14E-01 | 2.65E+00 | 2.74E+00 | 9.66E-01 | no | 4933430M04Rik | intron_variant,non_coding_transcript_variant | MODIFIER |
| 11 | 24516834 | AX-170046596 | 0 | T | G | 0.061 | 8.02E-03 | 2.14E-01 | 2.65E+00 | 2.74E+00 | 9.66E-01 | no | 4933430M04Rik | intron_variant,non_coding_transcript_variant | MODIFIER |
| 11 | 24516946 | AX-170046597 | 0 | A | G | 0.061 | 8.02E-03 | 2.14E-01 | 2.65E+00 | 2.74E+00 | 9.66E-01 | no | 4933430M04Rik | intron_variant,non_coding_transcript_variant | MODIFIER |
| 11 | 24547865 | AX-170466651 | 0 | A | C | 0.061 | 8.02E-03 | 2.14E-01 | 2.65E+00 | 2.74E+00 | 9.66E-01 | no | Gm12067 | downstream_gene_variant | MODIFIER |
| 11 | 24569170 | AX-170046595 | 0 | G | A | 0.061 | 8.02E-03 | 2.14E-01 | 2.65E+00 | 2.74E+00 | 9.66E-01 | no | 4933430M04Rik | intron_variant,non_coding_transcript_variant | MODIFIER |
| 11 | 24570104 | AX-170516657 | 0 | T | C | 0.061 | 8.02E-03 | 2.14E-01 | 2.65E+00 | 2.74E+00 | 9.66E-01 | no | 4933430M04Rik | downstream_gene_variant | MODIFIER |
| 11 | 24639185 | AX-170351975 | 0 | T | C | 0.061 | 8.02E-03 | 2.14E-01 | 2.65E+00 | 2.74E+00 | 9.66E-01 | no | Gm12068 | intron_variant,non_coding_transcript_variant | MODIFIER |
| 3 | 57651654 | AX-170478114 | 2 | C | T | 0.273 | 2.93E-03 | 7.17E-02 | 2.58E+00 | 2.74E+00 | 9.66E-01 | no | Commd2 | 5_prime_UTR_variant,NMD_transcript_variant | MODIFIER |
| 1 | 33612796 | AX-170453790 | 0 | C | T | 0.421 | -3.33E-03 | 8.16E-02 | 2.66E+00 | 2.74E+00 | 9.66E-01 | no | Prim2 | intron_variant | MODIFIER |
| 7 | 117458266 | AX-170138520 | 0 | G | A | 0.386 | -3.51E-03 | 8.91E-02 | 2.68E+00 | 2.74E+00 | 9.66E-01 | no | Xylt1 | intron_variant | MODIFIER |
| 5 | 127052225 | AX-170304180 | 1 | T | C | 0.42 | -3.24E-03 | 8.83E-02 | 2.64E+00 | 2.74E+00 | 9.66E-01 | no | 4933438B17Rik | intron_variant,non_coding_transcript_variant | MODIFIER |
| 9 | 108724071 | AX-170452180 | 1 | C | T | 0.125 | -6.16E-03 | 1.18E-01 | 2.66E+00 | 2.73E+00 | 9.66E-01 | no | Prkar2a | intron_variant | MODIFIER |
| 5 | 122997634 | AX-170263016 | 1 | C | T | 0.455 | -4.18E-03 | 1.12E-01 | 2.70E+00 | 2.75E+00 | 9.66E-01 | no | A930024E05Rik | non_coding_transcript_exon_variant | MODIFIER |
| 4 | 145066504 | AX-97845000 | 2 | A | G | 0.082 | 9.64E-03 | 2.04E-01 | 2.64E+00 | 2.73E+00 | 9.66E-01 | no | Vps13d | intron_variant | MODIFIER |
| 6 | 147373575 | AX-170312834 | 0 | G | A | 0.061 | 1.03E-02 | 2.50E-01 | 2.67E+00 | 2.74E+00 | 9.66E-01 | no | | intergenic_variant | MODIFIER |
| 7 | 98429441 | AX-170343717 | 1 | T | C | 0.125 | 7.26E-03 | 1.86E-01 | 2.65E+00 | 2.74E+00 | 9.66E-01 | no | | intergenic_variant | MODIFIER |
| 10 | 12493997 | AX-170534925 | 1 | C | T | 0.214 | -4.95E-03 | 1.05E-01 | 2.65E+00 | 2.73E+00 | 9.66E-01 | no | Utrn | intron_variant | MODIFIER |
| 12 | 111885209 | AX-170345318 | 1 | C | T | 0.125 | 8.85E-03 | 1.96E-01 | 2.66E+00 | 2.73E+00 | 9.67E-01 | no | Ppp1r13b | intron_variant | MODIFIER |
| 9 | 61744018 | AX-170394686 | 1 | G | A | 0.196 | 4.69E-03 | 1.19E-01 | 2.67E+00 | 2.74E+00 | 9.67E-01 | no | | intergenic_variant | MODIFIER |
| 20 | 145005453 | AX-97813120 | 0 | C | T | 0.149 | 6.11E-03 | 1.35E-01 | 2.64E+00 | 2.74E+00 | 9.67E-01 | no | Rtl4 | intron_variant | MODIFIER |
| 11 | 4767290 | AX-170428946 | 1 | C | T | 0.062 | 7.86E-03 | 2.14E-01 | 2.65E+00 | 2.74E+00 | 9.67E-01 | no | Nf2 | downstream_gene_variant | MODIFIER |
| 8 | 35466200 | AX-170154534 | 0 | T | C | 0.149 | 5.68E-03 | 1.53E-01 | 2.71E+00 | 2.75E+00 | 9.67E-01 | no | Eri1 | 3_prime_UTR_variant | MODIFIER |
| 7 | 89631477 | AX-170348519 | 1 | C | A | 0.134 | 8.55E-03 | 2.11E-01 | 2.66E+00 | 2.74E+00 | 9.67E-01 | no | Me3 | upstream_gene_variant | MODIFIER |
| 2 | 124337514 | AX-170152504 | 0 | G | A | 0.14 | 9.32E-03 | 2.20E-01 | 2.70E+00 | 2.73E+00 | 9.67E-01 | no | Sema6d | intron_variant | MODIFIER |
| 2 | 130540735 | AX-170392549 | 0 | A | G | 0.14 | 9.32E-03 | 2.20E-01 | 2.70E+00 | 2.73E+00 | 9.67E-01 | no | Ptpra | intron_variant | MODIFIER |
| 4 | 143208777 | AX-170108591 | 0 | A | G | 0.114 | 7.90E-03 | 1.70E-01 | 2.63E+00 | 2.73E+00 | 9.67E-01 | no | Prdm2 | intron_variant | MODIFIER |
| 1 | 66354984 | AX-97845803 | 0 | G | T | 0.123 | -6.08E-03 | 1.85E-01 | 2.64E+00 | 2.74E+00 | 9.67E-01 | no | Map2 | intron_variant | MODIFIER |
| 4 | 141763100 | AX-170465933 | 1 | T | C | 0.089 | -7.18E-03 | 1.93E-01 | 2.65E+00 | 2.74E+00 | 9.67E-01 | no | Dnajc16 | 3_prime_UTR_variant | MODIFIER |
| 4 | 150016207 | AX-170512949 | 1 | C | T | 0.089 | -7.18E-03 | 1.93E-01 | 2.65E+00 | 2.74E+00 | 9.67E-01 | no | | intergenic_variant | MODIFIER |
| 1 | 155753814 | AX-170443317 | 1 | A | T | 0.214 | -4.77E-03 | 1.39E-01 | 2.70E+00 | 2.75E+00 | 9.67E-01 | no | Lhx4 | upstream_gene_variant | MODIFIER |
| 7 | 40288949 | AX-97834231 | 0 | G | A | 0.325 | 4.86E-03 | 9.63E-02 | 2.65E+00 | 2.73E+00 | 9.67E-01 | no | | intergenic_variant | MODIFIER |
| 1 | 37620925 | AX-170328615 | 0 | G | A | 0.456 | -3.43E-03 | 7.82E-02 | 2.66E+00 | 2.74E+00 | 9.67E-01 | no | 2010300C02Rik | intron_variant | MODIFIER |
| 1 | 37624592 | AX-170356526 | 0 | T | C | 0.456 | -3.43E-03 | 7.82E-02 | 2.66E+00 | 2.74E+00 | 9.67E-01 | no | 2010300C02Rik | missense_variant | MODERATE |
| 1 | 37626267 | AX-170209917 | 0 | A | C | 0.456 | -3.43E-03 | 7.82E-02 | 2.66E+00 | 2.74E+00 | 9.67E-01 | no | 2010300C02Rik | upstream_gene_variant | MODIFIER |
| 1 | 37632448 | AX-170475091 | 0 | C | A | 0.456 | -3.43E-03 | 7.82E-02 | 2.66E+00 | 2.74E+00 | 9.67E-01 | no | 2010300C02Rik | intron_variant | MODIFIER |
| 1 | 37632490 | AX-170306489 | 0 | G | A | 0.456 | -3.43E-03 | 7.82E-02 | 2.66E+00 | 2.74E+00 | 9.67E-01 | no | 2010300C02Rik | intron_variant | MODIFIER |
| 1 | 37643578 | AX-170223103 | 0 | C | T | 0.456 | -3.43E-03 | 7.82E-02 | 2.66E+00 | 2.74E+00 | 9.67E-01 | no | 2010300C02Rik | intron_variant | MODIFIER |
| 1 | 37644130 | AX-170180896 | 0 | C | T | 0.456 | -3.43E-03 | 7.82E-02 | 2.66E+00 | 2.74E+00 | 9.67E-01 | no | 2010300C02Rik | intron_variant | MODIFIER |
| 1 | 37647425 | AX-97835292 | 0 | T | C | 0.456 | -3.43E-03 | 7.82E-02 | 2.66E+00 | 2.74E+00 | 9.67E-01 | no | 2010300C02Rik | intron_variant | MODIFIER |
| 1 | 37706705 | AX-170040921 | 0 | A | G | 0.456 | -3.43E-03 | 7.82E-02 | 2.66E+00 | 2.74E+00 | 9.67E-01 | no | 2010300C02Rik | intron_variant | MODIFIER |
| 1 | 37710746 | AX-97836535 | 0 | C | T | 0.456 | -3.43E-03 | 7.82E-02 | 2.66E+00 | 2.74E+00 | 9.67E-01 | no | 2010300C02Rik | intron_variant | MODIFIER |
| 1 | 37714874 | AX-170243378 | 0 | A | T | 0.456 | -3.43E-03 | 7.82E-02 | 2.66E+00 | 2.74E+00 | 9.67E-01 | no | 2010300C02Rik | intron_variant | MODIFIER |
| 1 | 37889207 | AX-97809442 | 0 | G | A | 0.456 | -3.43E-03 | 7.82E-02 | 2.66E+00 | 2.74E+00 | 9.67E-01 | no | Mrpl30 | upstream_gene_variant | MODIFIER |

| | | | | | | | | | | | | | | | |
|---|---|---|---|---|---|---|---|---|---|---|---|---|---|---|---|
| 1 | 37907323 | AX-170000309 | 0 | T | C | 0.456 | -3.43E-03 | 7.82E-02 | 2.66E+00 | 2.74E+00 | 9.67E-01 | no | Lyg2 | 3_prime_UTR_variant | MODIFIER |
| 1 | 37907483 | AX-170000306 | 0 | G | A | 0.456 | -3.43E-03 | 7.82E-02 | 2.66E+00 | 2.74E+00 | 9.67E-01 | no | Lyg2 | 3_prime_UTR_variant | MODIFIER |
| 1 | 38048046 | AX-97842680 | 0 | G | T | 0.456 | -3.43E-03 | 7.82E-02 | 2.66E+00 | 2.74E+00 | 9.67E-01 | no | Rev1 | downstream_gene_variant | MODIFIER |
| 1 | 38157244 | AX-170534197 | 0 | T | C | 0.456 | -3.43E-03 | 7.82E-02 | 2.66E+00 | 2.74E+00 | 9.67E-01 | no | Gm5099 | intron_variant,non_coding_transcript_variant | MODIFIER |
| 1 | 38167249 | AX-170187737 | 0 | T | C | 0.456 | -3.43E-03 | 7.82E-02 | 2.66E+00 | 2.74E+00 | 9.67E-01 | no | | intergenic_variant | MODIFIER |
| 1 | 38179795 | AX-170307731 | 0 | A | G | 0.456 | -3.43E-03 | 7.82E-02 | 2.66E+00 | 2.74E+00 | 9.67E-01 | no | Aff3 | intron_variant | MODIFIER |
| 1 | 38181889 | AX-170091473 | 0 | C | T | 0.456 | -3.43E-03 | 7.82E-02 | 2.66E+00 | 2.74E+00 | 9.67E-01 | no | Aff3 | intron_variant | MODIFIER |
| 1 | 38213227 | AX-97816790 | 0 | T | C | 0.456 | -3.43E-03 | 7.82E-02 | 2.66E+00 | 2.74E+00 | 9.67E-01 | no | Aff3 | intron_variant | MODIFIER |
| 1 | 38215905 | AX-170113090 | 0 | C | T | 0.456 | -3.43E-03 | 7.82E-02 | 2.66E+00 | 2.74E+00 | 9.67E-01 | no | Aff3 | intron_variant | MODIFIER |
| 1 | 38216852 | AX-170366301 | 0 | A | G | 0.456 | -3.43E-03 | 7.82E-02 | 2.66E+00 | 2.74E+00 | 9.67E-01 | no | Aff3 | intron_variant | MODIFIER |
| 1 | 38262680 | AX-170234397 | 0 | G | A | 0.456 | -3.43E-03 | 7.82E-02 | 2.66E+00 | 2.74E+00 | 9.67E-01 | no | Aff3 | intron_variant | MODIFIER |
| 1 | 38394101 | AX-170139229 | 0 | G | A | 0.456 | -3.43E-03 | 7.82E-02 | 2.66E+00 | 2.74E+00 | 9.67E-01 | no | Aff3 | intron_variant,non_coding_transcript_variant | MODIFIER |
| 1 | 38476249 | AX-170451353 | 0 | C | T | 0.456 | -3.43E-03 | 7.82E-02 | 2.66E+00 | 2.74E+00 | 9.67E-01 | no | Aff3 | intron_variant,non_coding_transcript_variant | MODIFIER |
| 1 | 38477592 | AX-170000316 | 0 | T | G | 0.456 | -3.43E-03 | 7.82E-02 | 2.66E+00 | 2.74E+00 | 9.67E-01 | no | Aff3 | intron_variant,non_coding_transcript_variant | MODIFIER |
| 1 | 38477617 | AX-170000312 | 0 | C | T | 0.456 | -3.43E-03 | 7.82E-02 | 2.66E+00 | 2.74E+00 | 9.67E-01 | no | Aff3 | intron_variant,non_coding_transcript_variant | MODIFIER |
| 1 | 38490733 | AX-170000313 | 0 | T | C | 0.456 | -3.43E-03 | 7.82E-02 | 2.66E+00 | 2.74E+00 | 9.67E-01 | no | Aff3 | intron_variant,non_coding_transcript_variant | MODIFIER |
| 1 | 38491323 | AX-170490979 | 0 | G | A | 0.456 | -3.43E-03 | 7.82E-02 | 2.66E+00 | 2.74E+00 | 9.67E-01 | no | Aff3 | intron_variant,non_coding_transcript_variant | MODIFIER |
| 1 | 38507478 | AX-170000315 | 0 | C | T | 0.456 | -3.43E-03 | 7.82E-02 | 2.66E+00 | 2.74E+00 | 9.67E-01 | no | Aff3 | intron_variant,non_coding_transcript_variant | MODIFIER |
| 1 | 38516541 | AX-97827314 | 0 | G | T | 0.456 | -3.43E-03 | 7.82E-02 | 2.66E+00 | 2.74E+00 | 9.67E-01 | no | Aff3 | intron_variant,non_coding_transcript_variant | MODIFIER |
| 1 | 38517287 | AX-170000314 | 0 | A | G | 0.456 | -3.43E-03 | 7.82E-02 | 2.66E+00 | 2.74E+00 | 9.67E-01 | no | Aff3 | intron_variant,non_coding_transcript_variant | MODIFIER |
| 1 | 38521631 | AX-170171942 | 0 | T | G | 0.456 | -3.43E-03 | 7.82E-02 | 2.66E+00 | 2.74E+00 | 9.67E-01 | no | Aff3 | intron_variant,non_coding_transcript_variant | MODIFIER |
| 1 | 38526321 | AX-170194116 | 0 | G | A | 0.456 | -3.43E-03 | 7.82E-02 | 2.66E+00 | 2.74E+00 | 9.67E-01 | no | Aff3 | intron_variant,non_coding_transcript_variant | MODIFIER |
| 1 | 38539041 | AX-170117699 | 0 | A | G | 0.456 | -3.43E-03 | 7.82E-02 | 2.66E+00 | 2.74E+00 | 9.67E-01 | no | Aff3 | intron_variant,non_coding_transcript_variant | MODIFIER |
| 1 | 38546632 | AX-170166571 | 0 | C | T | 0.456 | -3.43E-03 | 7.82E-02 | 2.66E+00 | 2.74E+00 | 9.67E-01 | no | Aff3 | intron_variant,non_coding_transcript_variant | MODIFIER |
| 1 | 38550799 | AX-170272590 | 0 | G | A | 0.456 | -3.43E-03 | 7.82E-02 | 2.66E+00 | 2.74E+00 | 9.67E-01 | no | Aff3 | intron_variant,non_coding_transcript_variant | MODIFIER |
| 1 | 38563883 | AX-170245322 | 0 | G | A | 0.456 | -3.43E-03 | 7.82E-02 | 2.66E+00 | 2.74E+00 | 9.67E-01 | no | Aff3 | intron_variant,non_coding_transcript_variant | MODIFIER |
| 1 | 38584963 | AX-170408087 | 0 | T | G | 0.456 | -3.43E-03 | 7.82E-02 | 2.66E+00 | 2.74E+00 | 9.67E-01 | no | Aff3 | intron_variant,non_coding_transcript_variant | MODIFIER |
| 1 | 38586216 | AX-170040942 | 0 | G | A | 0.456 | -3.43E-03 | 7.82E-02 | 2.66E+00 | 2.74E+00 | 9.67E-01 | no | Aff3 | intron_variant,non_coding_transcript_variant | MODIFIER |
| 1 | 38587545 | AX-170040943 | 0 | A | G | 0.456 | -3.43E-03 | 7.82E-02 | 2.66E+00 | 2.74E+00 | 9.67E-01 | no | Aff3 | intron_variant,non_coding_transcript_variant | MODIFIER |
| 1 | 38587802 | AX-170323021 | 0 | A | G | 0.456 | -3.43E-03 | 7.82E-02 | 2.66E+00 | 2.74E+00 | 9.67E-01 | no | Aff3 | intron_variant,non_coding_transcript_variant | MODIFIER |
| 1 | 38682230 | AX-97811330 | 0 | G | A | 0.456 | -3.43E-03 | 7.82E-02 | 2.66E+00 | 2.74E+00 | 9.67E-01 | no | | intergenic_variant | MODIFIER |
| 1 | 38686463 | AX-170293691 | 0 | G | T | 0.456 | -3.43E-03 | 7.82E-02 | 2.66E+00 | 2.74E+00 | 9.67E-01 | no | | intergenic_variant | MODIFIER |
| 1 | 38686885 | AX-170370524 | 0 | A | G | 0.456 | -3.43E-03 | 7.82E-02 | 2.66E+00 | 2.74E+00 | 9.67E-01 | no | | intergenic_variant | MODIFIER |
| 1 | 38690999 | AX-170172625 | 0 | T | C | 0.456 | -3.43E-03 | 7.82E-02 | 2.66E+00 | 2.74E+00 | 9.67E-01 | no | | intergenic_variant | MODIFIER |
| 1 | 38701778 | AX-170450449 | 0 | C | T | 0.456 | -3.43E-03 | 7.82E-02 | 2.66E+00 | 2.74E+00 | 9.67E-01 | no | | intergenic_variant | MODIFIER |
| 1 | 38709967 | AX-170098125 | 0 | A | C | 0.456 | -3.43E-03 | 7.82E-02 | 2.66E+00 | 2.74E+00 | 9.67E-01 | no | | intergenic_variant | MODIFIER |
| 1 | 38712107 | AX-170086448 | 0 | T | C | 0.456 | -3.43E-03 | 7.82E-02 | 2.66E+00 | 2.74E+00 | 9.67E-01 | no | | intergenic_variant | MODIFIER |
| 1 | 38726592 | AX-170040945 | 0 | A | G | 0.456 | -3.43E-03 | 7.82E-02 | 2.66E+00 | 2.74E+00 | 9.67E-01 | no | Gm38135 | downstream_gene_variant | MODIFIER |
| 1 | 38739662 | AX-170513658 | 0 | T | C | 0.456 | -3.43E-03 | 7.82E-02 | 2.66E+00 | 2.74E+00 | 9.67E-01 | no | | intergenic_variant | MODIFIER |
| 1 | 38741496 | AX-170124507 | 0 | G | A | 0.456 | -3.43E-03 | 7.82E-02 | 2.66E+00 | 2.74E+00 | 9.67E-01 | no | | intergenic_variant | MODIFIER |
| 1 | 38742970 | AX-170125189 | 0 | C | T | 0.456 | -3.43E-03 | 7.82E-02 | 2.66E+00 | 2.74E+00 | 9.67E-01 | no | | intergenic_variant | MODIFIER |
| 1 | 38743128 | AX-97834235 | 0 | T | C | 0.456 | -3.43E-03 | 7.82E-02 | 2.66E+00 | 2.74E+00 | 9.67E-01 | no | | intergenic_variant | MODIFIER |
| 1 | 38743184 | AX-170460669 | 0 | C | A | 0.456 | -3.43E-03 | 7.82E-02 | 2.66E+00 | 2.74E+00 | 9.67E-01 | no | | intergenic_variant | MODIFIER |
| 1 | 38760847 | AX-170276256 | 0 | A | G | 0.456 | -3.43E-03 | 7.82E-02 | 2.66E+00 | 2.74E+00 | 9.67E-01 | no | | regulatory_region_variant | MODIFIER |
| 1 | 38779297 | AX-170444864 | 0 | G | T | 0.456 | -3.43E-03 | 7.82E-02 | 2.66E+00 | 2.74E+00 | 9.67E-01 | no | | intergenic_variant | MODIFIER |
| 1 | 38779731 | AX-170249251 | 0 | T | C | 0.456 | -3.43E-03 | 7.82E-02 | 2.66E+00 | 2.74E+00 | 9.67E-01 | no | | intergenic_variant | MODIFIER |
| 1 | 38785789 | AX-97791312 | 0 | G | T | 0.456 | -3.43E-03 | 7.82E-02 | 2.66E+00 | 2.74E+00 | 9.67E-01 | no | | intergenic_variant | MODIFIER |
| 1 | 38788921 | AX-170403969 | 0 | A | C | 0.456 | -3.43E-03 | 7.82E-02 | 2.66E+00 | 2.74E+00 | 9.67E-01 | no | Lonrf2 | downstream_gene_variant | MODIFIER |
| 1 | 38792079 | AX-170527647 | 0 | C | T | 0.456 | -3.43E-03 | 7.82E-02 | 2.66E+00 | 2.74E+00 | 9.67E-01 | no | Lonrf2 | downstream_gene_variant | MODIFIER |
| 1 | 38792949 | AX-170484041 | 0 | G | A | 0.456 | -3.43E-03 | 7.82E-02 | 2.66E+00 | 2.74E+00 | 9.67E-01 | no | Lonrf2 | downstream_gene_variant | MODIFIER |
| 1 | 38869490 | AX-170358952 | 0 | A | C | 0.456 | -3.43E-03 | 7.82E-02 | 2.66E+00 | 2.74E+00 | 9.67E-01 | no | Chst10 | intron_variant | MODIFIER |
| 1 | 38870584 | AX-170458264 | 0 | T | C | 0.456 | -3.43E-03 | 7.82E-02 | 2.66E+00 | 2.74E+00 | 9.67E-01 | no | Chst10 | intron_variant | MODIFIER |
| 1 | 38871204 | AX-170427645 | 0 | T | C | 0.456 | -3.43E-03 | 7.82E-02 | 2.66E+00 | 2.74E+00 | 9.67E-01 | no | Chst10 | intron_variant | MODIFIER |
| 1 | 38874601 | AX-170159903 | 0 | C | T | 0.456 | -3.43E-03 | 7.82E-02 | 2.66E+00 | 2.74E+00 | 9.67E-01 | no | Chst10 | intron_variant | MODIFIER |

| | | | | | | | | | | | | | | | |
|---|---|---|---|---|---|---|---|---|---|---|---|---|---|---|---|
| 1 | 38890945 | AX-170204791 | 0 | C | T | 0.456 | -3.43E-03 | 7.82E-02 | 2.66E+00 | 2.74E+00 | 9.67E-01 | no | Chst10 | intron_variant | MODIFIER |
| 1 | 38891090 | AX-170206369 | 0 | C | G | 0.456 | -3.43E-03 | 7.82E-02 | 2.66E+00 | 2.74E+00 | 9.67E-01 | no | Chst10 | intron_variant | MODIFIER |
| 1 | 38891578 | AX-170179274 | 0 | A | G | 0.456 | -3.43E-03 | 7.82E-02 | 2.66E+00 | 2.74E+00 | 9.67E-01 | no | Chst10 | intron_variant | MODIFIER |
| 1 | 38894687 | AX-170266340 | 0 | A | G | 0.456 | -3.43E-03 | 7.82E-02 | 2.66E+00 | 2.74E+00 | 9.67E-01 | no | Chst10 | intron_variant | MODIFIER |
| 1 | 38968470 | AX-170260955 | 0 | G | A | 0.456 | -3.43E-03 | 7.82E-02 | 2.66E+00 | 2.74E+00 | 9.67E-01 | no | | intergenic_variant | MODIFIER |
| 1 | 38969077 | AX-170395530 | 0 | C | G | 0.456 | -3.43E-03 | 7.82E-02 | 2.66E+00 | 2.74E+00 | 9.67E-01 | no | | intergenic_variant | MODIFIER |
| 1 | 38969126 | AX-97806486 | 0 | G | A | 0.456 | -3.43E-03 | 7.82E-02 | 2.66E+00 | 2.74E+00 | 9.67E-01 | no | | intergenic_variant | MODIFIER |
| 1 | 39005858 | AX-170518015 | 0 | G | T | 0.456 | -3.43E-03 | 7.82E-02 | 2.66E+00 | 2.74E+00 | 9.67E-01 | no | | regulatory_region_variant | MODIFIER |
| 1 | 39014729 | AX-97786855 | 0 | G | A | 0.456 | -3.43E-03 | 7.82E-02 | 2.66E+00 | 2.74E+00 | 9.67E-01 | no | | intergenic_variant | MODIFIER |
| 1 | 39014809 | AX-170000323 | 0 | A | C | 0.456 | -3.43E-03 | 7.82E-02 | 2.66E+00 | 2.74E+00 | 9.67E-01 | no | | intergenic_variant | MODIFIER |
| 1 | 39015261 | AX-170375776 | 0 | A | C | 0.456 | -3.43E-03 | 7.82E-02 | 2.66E+00 | 2.74E+00 | 9.67E-01 | no | | intergenic_variant | MODIFIER |
| 1 | 39016071 | AX-170322166 | 0 | C | T | 0.456 | -3.43E-03 | 7.82E-02 | 2.66E+00 | 2.74E+00 | 9.67E-01 | no | | intergenic_variant | MODIFIER |
| 1 | 39020746 | AX-170460358 | 0 | G | A | 0.456 | -3.43E-03 | 7.82E-02 | 2.66E+00 | 2.74E+00 | 9.67E-01 | no | | intergenic_variant | MODIFIER |
| 1 | 39100624 | AX-170116034 | 0 | C | A | 0.456 | -3.43E-03 | 7.82E-02 | 2.66E+00 | 2.74E+00 | 9.67E-01 | no | | regulatory_region_variant | MODIFIER |
| 1 | 39101952 | AX-170198501 | 0 | T | G | 0.456 | -3.43E-03 | 7.82E-02 | 2.66E+00 | 2.74E+00 | 9.67E-01 | no | | intergenic_variant | MODIFIER |
| 1 | 39102123 | AX-170527657 | 0 | A | G | 0.456 | -3.43E-03 | 7.82E-02 | 2.66E+00 | 2.74E+00 | 9.67E-01 | no | | intergenic_variant | MODIFIER |
| 1 | 39102672 | AX-170212836 | 0 | C | T | 0.456 | -3.43E-03 | 7.82E-02 | 2.66E+00 | 2.74E+00 | 9.67E-01 | no | | regulatory_region_variant | MODIFIER |
| 1 | 39104250 | AX-170040953 | 0 | C | A | 0.456 | -3.43E-03 | 7.82E-02 | 2.66E+00 | 2.74E+00 | 9.67E-01 | no | | regulatory_region_variant | MODIFIER |
| 1 | 39108689 | AX-170040952 | 0 | C | T | 0.456 | -3.43E-03 | 7.82E-02 | 2.66E+00 | 2.74E+00 | 9.67E-01 | no | | intergenic_variant | MODIFIER |
| 1 | 39108930 | AX-170102418 | 0 | T | C | 0.456 | -3.43E-03 | 7.82E-02 | 2.66E+00 | 2.74E+00 | 9.67E-01 | no | | intergenic_variant | MODIFIER |
| 1 | 39140420 | AX-170417476 | 0 | G | A | 0.456 | -3.43E-03 | 7.82E-02 | 2.66E+00 | 2.74E+00 | 9.67E-01 | no | | intergenic_variant | MODIFIER |
| 1 | 39190394 | AX-170000324 | 0 | A | G | 0.456 | -3.43E-03 | 7.82E-02 | 2.66E+00 | 2.74E+00 | 9.67E-01 | no | Npas2 | upstream_gene_variant | MODIFIER |
| 1 | 39196915 | AX-170000322 | 0 | T | G | 0.456 | -3.43E-03 | 7.82E-02 | 2.66E+00 | 2.74E+00 | 9.67E-01 | no | Npas2 | intron_variant | MODIFIER |
| 9 | 55815298 | AX-170434004 | 0 | G | A | 0.202 | -4.09E-03 | 1.00E-01 | 2.62E+00 | 2.74E+00 | 9.67E-01 | no | Scaper | intron_variant | MODIFIER |
| 1 | 66135303 | AX-170263700 | 0 | C | T | 0.14 | -9.11E-03 | 2.36E-01 | 2.69E+00 | 2.74E+00 | 9.68E-01 | no | | intergenic_variant | MODIFIER |
| 12 | 116838259 | AX-170387207 | 0 | G | A | 0.132 | -6.23E-03 | 1.63E-01 | 2.63E+00 | 2.74E+00 | 9.68E-01 | no | Ptprn2 | intron_variant | MODIFIER |
| 1 | 52056763 | AX-97818718 | 0 | G | T | 0.132 | 9.74E-03 | 2.42E-01 | 2.69E+00 | 2.74E+00 | 9.68E-01 | no | Stat4 | intron_variant | MODIFIER |
| 1 | 52057057 | AX-170518982 | 0 | T | C | 0.132 | 9.74E-03 | 2.42E-01 | 2.69E+00 | 2.74E+00 | 9.68E-01 | no | Stat4 | intron_variant | MODIFIER |
| 1 | 52057862 | AX-170228243 | 0 | A | G | 0.132 | 9.74E-03 | 2.42E-01 | 2.69E+00 | 2.74E+00 | 9.68E-01 | no | Stat4 | intron_variant | MODIFIER |
| 1 | 52067812 | AX-170123047 | 0 | A | G | 0.132 | 9.74E-03 | 2.42E-01 | 2.69E+00 | 2.74E+00 | 9.68E-01 | no | Stat4 | intron_variant | MODIFIER |
| 1 | 52071825 | AX-170490289 | 0 | A | G | 0.132 | 9.74E-03 | 2.42E-01 | 2.69E+00 | 2.74E+00 | 9.68E-01 | no | Stat4 | intron_variant | MODIFIER |
| 1 | 52087359 | AX-97849063 | 0 | G | A | 0.132 | 9.74E-03 | 2.42E-01 | 2.69E+00 | 2.74E+00 | 9.68E-01 | no | Stat4 | intron_variant | MODIFIER |
| 1 | 52093306 | AX-170147810 | 0 | A | G | 0.132 | 9.74E-03 | 2.42E-01 | 2.69E+00 | 2.74E+00 | 9.68E-01 | no | Stat4 | intron_variant | MODIFIER |
| 1 | 52100009 | AX-170463429 | 0 | G | A | 0.132 | 9.74E-03 | 2.42E-01 | 2.69E+00 | 2.74E+00 | 9.68E-01 | no | Stat4 | intron_variant | MODIFIER |
| 1 | 52102109 | AX-170295065 | 0 | C | A | 0.132 | 9.74E-03 | 2.42E-01 | 2.69E+00 | 2.74E+00 | 9.68E-01 | no | Stat4 | splice_region_variant,intron_variant | LOW |
| 1 | 52105347 | AX-170219574 | 0 | G | A | 0.132 | 9.74E-03 | 2.42E-01 | 2.69E+00 | 2.74E+00 | 9.68E-01 | no | Stat4 | intron_variant | MODIFIER |
| 1 | 52107410 | AX-170202076 | 0 | A | G | 0.132 | 9.74E-03 | 2.42E-01 | 2.69E+00 | 2.74E+00 | 9.68E-01 | no | Stat4 | downstream_gene_variant | MODIFIER |
| 1 | 52107661 | AX-170000498 | 0 | T | C | 0.132 | 9.74E-03 | 2.42E-01 | 2.69E+00 | 2.74E+00 | 9.68E-01 | no | Stat4 | downstream_gene_variant | MODIFIER |
| 1 | 52132720 | AX-97783870 | 0 | A | G | 0.132 | 9.74E-03 | 2.42E-01 | 2.69E+00 | 2.74E+00 | 9.68E-01 | no | Stat1 | intron_variant | MODIFIER |
| 1 | 52136979 | AX-170041221 | 0 | T | C | 0.132 | 9.74E-03 | 2.42E-01 | 2.69E+00 | 2.74E+00 | 9.68E-01 | no | Stat1 | intron_variant | MODIFIER |
| 1 | 52137380 | AX-170329264 | 0 | T | C | 0.132 | 9.74E-03 | 2.42E-01 | 2.69E+00 | 2.74E+00 | 9.68E-01 | no | Stat1 | synonymous_variant | LOW |
| 1 | 52137560 | AX-170474071 | 0 | G | A | 0.132 | 9.74E-03 | 2.42E-01 | 2.69E+00 | 2.74E+00 | 9.68E-01 | no | Stat1 | intron_variant | MODIFIER |
| 1 | 52138041 | AX-170423338 | 0 | G | T | 0.132 | 9.74E-03 | 2.42E-01 | 2.69E+00 | 2.74E+00 | 9.68E-01 | no | Stat1 | intron_variant | MODIFIER |
| 1 | 52156267 | AX-170000502 | 0 | C | G | 0.132 | 9.74E-03 | 2.42E-01 | 2.69E+00 | 2.74E+00 | 9.68E-01 | no | Stat1 | intron_variant | MODIFIER |
| 1 | 52162597 | AX-170000497 | 0 | G | A | 0.132 | 9.74E-03 | 2.42E-01 | 2.69E+00 | 2.74E+00 | 9.68E-01 | no | Stat1 | downstream_gene_variant | MODIFIER |
| 1 | 52166759 | AX-170444989 | 0 | T | C | 0.132 | 9.74E-03 | 2.42E-01 | 2.69E+00 | 2.74E+00 | 9.68E-01 | no | Stat1 | downstream_gene_variant | MODIFIER |
| 1 | 52168064 | AX-170257945 | 0 | C | G | 0.132 | 9.74E-03 | 2.42E-01 | 2.69E+00 | 2.74E+00 | 9.68E-01 | no | Gls | intron_variant | MODIFIER |
| 1 | 52168569 | AX-170293627 | 0 | A | G | 0.132 | 9.74E-03 | 2.42E-01 | 2.69E+00 | 2.74E+00 | 9.68E-01 | no | Gls | splice_region_variant,intron_variant | LOW |
| 1 | 52172827 | AX-97811476 | 0 | T | G | 0.132 | 9.74E-03 | 2.42E-01 | 2.69E+00 | 2.74E+00 | 9.68E-01 | no | Gls | intron_variant | MODIFIER |
| 1 | 52197577 | AX-170000500 | 0 | A | G | 0.132 | 9.74E-03 | 2.42E-01 | 2.69E+00 | 2.74E+00 | 9.68E-01 | no | Gls | intron_variant,non_coding_transcript_variant | MODIFIER |
| 1 | 52197611 | AX-170000499 | 0 | T | A | 0.132 | 9.74E-03 | 2.42E-01 | 2.69E+00 | 2.74E+00 | 9.68E-01 | no | Gls | intron_variant,non_coding_transcript_variant | MODIFIER |
| 1 | 52208807 | AX-170000496 | 0 | C | T | 0.132 | 9.74E-03 | 2.42E-01 | 2.69E+00 | 2.74E+00 | 9.68E-01 | no | Gls | intron_variant | MODIFIER |
| 1 | 52228117 | AX-170246267 | 0 | C | T | 0.132 | 9.74E-03 | 2.42E-01 | 2.69E+00 | 2.74E+00 | 9.68E-01 | no | Gls | intron_variant | MODIFIER |
| 1 | 52229525 | AX-170000501 | 0 | T | C | 0.132 | 9.74E-03 | 2.42E-01 | 2.69E+00 | 2.74E+00 | 9.68E-01 | no | Gls | intron_variant | MODIFIER |

| | | | | | | | | | | | | | | | |
|---|---|---|---|---|---|---|---|---|---|---|---|---|---|---|---|
| 1 | 52230586 | AX-170407739 | 0 | C | T | 0.132 | 9.74E-03 | 2.42E-01 | 2.69E+00 | 2.74E+00 | 9.68E-01 | no | Gls | intron_variant | MODIFIER |
| 1 | 52246821 | AX-170449727 | 0 | A | G | 0.132 | 9.74E-03 | 2.42E-01 | 2.69E+00 | 2.74E+00 | 9.68E-01 | no | | intergenic_variant | MODIFIER |
| 1 | 52247476 | AX-97783037 | 0 | A | T | 0.132 | 9.74E-03 | 2.42E-01 | 2.69E+00 | 2.74E+00 | 9.68E-01 | no | | intergenic_variant | MODIFIER |
| 1 | 52251171 | AX-170041222 | 0 | A | G | 0.132 | 9.74E-03 | 2.42E-01 | 2.69E+00 | 2.74E+00 | 9.68E-01 | no | | regulatory_region_variant | MODIFIER |
| 1 | 52270438 | AX-170349921 | 0 | T | G | 0.132 | 9.74E-03 | 2.42E-01 | 2.69E+00 | 2.74E+00 | 9.68E-01 | no | | regulatory_region_variant | MODIFIER |
| 1 | 52271830 | AX-170322748 | 0 | A | G | 0.132 | 9.74E-03 | 2.42E-01 | 2.69E+00 | 2.74E+00 | 9.68E-01 | no | | intergenic_variant | MODIFIER |
| 1 | 52276221 | AX-97832424 | 0 | G | A | 0.132 | 9.74E-03 | 2.42E-01 | 2.69E+00 | 2.74E+00 | 9.68E-01 | no | | intergenic_variant | MODIFIER |
| 1 | 52291393 | AX-170102006 | 0 | T | C | 0.132 | 9.74E-03 | 2.42E-01 | 2.69E+00 | 2.74E+00 | 9.68E-01 | no | Gm5975 | upstream_gene_variant | MODIFIER |
| 1 | 52300917 | AX-170144871 | 0 | T | A | 0.132 | 9.74E-03 | 2.42E-01 | 2.69E+00 | 2.74E+00 | 9.68E-01 | no | | intergenic_variant | MODIFIER |
| 1 | 52303885 | AX-170442005 | 0 | A | T | 0.132 | 9.74E-03 | 2.42E-01 | 2.69E+00 | 2.74E+00 | 9.68E-01 | no | | intergenic_variant | MODIFIER |
| 1 | 52307009 | AX-170464832 | 0 | A | G | 0.132 | 9.74E-03 | 2.42E-01 | 2.69E+00 | 2.74E+00 | 9.68E-01 | no | | regulatory_region_variant | MODIFIER |
| 1 | 52307044 | AX-97834093 | 0 | G | A | 0.132 | 9.74E-03 | 2.42E-01 | 2.69E+00 | 2.74E+00 | 9.68E-01 | no | | regulatory_region_variant | MODIFIER |
| 1 | 52307113 | AX-170229709 | 0 | A | T | 0.132 | 9.74E-03 | 2.42E-01 | 2.69E+00 | 2.74E+00 | 9.68E-01 | no | | regulatory_region_variant | MODIFIER |
| 1 | 52307168 | AX-170419608 | 0 | T | A | 0.132 | 9.74E-03 | 2.42E-01 | 2.69E+00 | 2.74E+00 | 9.68E-01 | no | | regulatory_region_variant | MODIFIER |
| 1 | 52307262 | AX-170327423 | 0 | G | T | 0.132 | 9.74E-03 | 2.42E-01 | 2.69E+00 | 2.74E+00 | 9.68E-01 | no | | regulatory_region_variant | MODIFIER |
| 1 | 52321313 | AX-170180860 | 0 | C | T | 0.132 | 9.74E-03 | 2.42E-01 | 2.69E+00 | 2.74E+00 | 9.68E-01 | no | | intergenic_variant | MODIFIER |
| 1 | 52336569 | AX-170370142 | 0 | T | C | 0.132 | 9.74E-03 | 2.42E-01 | 2.69E+00 | 2.74E+00 | 9.68E-01 | no | | intergenic_variant | MODIFIER |
| 1 | 52338049 | AX-170303601 | 0 | A | G | 0.132 | 9.74E-03 | 2.42E-01 | 2.69E+00 | 2.74E+00 | 9.68E-01 | no | | intergenic_variant | MODIFIER |
| 1 | 52338282 | AX-97817886 | 0 | A | G | 0.132 | 9.74E-03 | 2.42E-01 | 2.69E+00 | 2.74E+00 | 9.68E-01 | no | | intergenic_variant | MODIFIER |
| 1 | 52338850 | AX-170203625 | 0 | A | G | 0.132 | 9.74E-03 | 2.42E-01 | 2.69E+00 | 2.74E+00 | 9.68E-01 | no | | intergenic_variant | MODIFIER |
| 1 | 52339573 | AX-170097596 | 0 | C | T | 0.132 | 9.74E-03 | 2.42E-01 | 2.69E+00 | 2.74E+00 | 9.68E-01 | no | | intergenic_variant | MODIFIER |
| 1 | 52353785 | AX-97816488 | 0 | A | G | 0.132 | 9.74E-03 | 2.42E-01 | 2.69E+00 | 2.74E+00 | 9.68E-01 | no | | intergenic_variant | MODIFIER |
| 1 | 52357916 | AX-170373278 | 0 | A | G | 0.132 | 9.74E-03 | 2.42E-01 | 2.69E+00 | 2.74E+00 | 9.68E-01 | no | | intergenic_variant | MODIFIER |
| 1 | 52369445 | AX-170041223 | 0 | T | G | 0.132 | 9.74E-03 | 2.42E-01 | 2.69E+00 | 2.74E+00 | 9.68E-01 | no | | intergenic_variant | MODIFIER |
| 1 | 52390033 | AX-170150607 | 0 | T | C | 0.132 | 9.74E-03 | 2.42E-01 | 2.69E+00 | 2.74E+00 | 9.68E-01 | no | | intergenic_variant | MODIFIER |
| 1 | 52392981 | AX-97810843 | 0 | C | G | 0.132 | 9.74E-03 | 2.42E-01 | 2.69E+00 | 2.74E+00 | 9.68E-01 | no | | regulatory_region_variant | MODIFIER |
| 1 | 52423619 | AX-170484634 | 0 | A | C | 0.132 | 9.74E-03 | 2.42E-01 | 2.69E+00 | 2.74E+00 | 9.68E-01 | no | | intergenic_variant | MODIFIER |
| 1 | 52424902 | AX-170140383 | 0 | A | G | 0.132 | 9.74E-03 | 2.42E-01 | 2.69E+00 | 2.74E+00 | 9.68E-01 | no | | intergenic_variant | MODIFIER |
| 1 | 52429887 | AX-170411327 | 0 | A | G | 0.132 | 9.74E-03 | 2.42E-01 | 2.69E+00 | 2.74E+00 | 9.68E-01 | no | | intergenic_variant | MODIFIER |
| 1 | 52435848 | AX-170269885 | 0 | A | G | 0.132 | 9.74E-03 | 2.42E-01 | 2.69E+00 | 2.74E+00 | 9.68E-01 | no | | regulatory_region_variant | MODIFIER |
| 1 | 52440571 | AX-170137943 | 0 | C | G | 0.132 | 9.74E-03 | 2.42E-01 | 2.69E+00 | 2.74E+00 | 9.68E-01 | no | Gm23460 | downstream_gene_variant | MODIFIER |
| 1 | 52444933 | AX-170197896 | 0 | C | T | 0.132 | 9.74E-03 | 2.42E-01 | 2.69E+00 | 2.74E+00 | 9.68E-01 | no | Gm23460 | upstream_gene_variant | MODIFIER |
| 1 | 52466036 | AX-170378110 | 0 | A | G | 0.132 | 9.74E-03 | 2.42E-01 | 2.69E+00 | 2.74E+00 | 9.68E-01 | no | Nab1 | intron_variant | MODIFIER |
| 1 | 52468944 | AX-170409868 | 0 | T | C | 0.132 | 9.74E-03 | 2.42E-01 | 2.69E+00 | 2.74E+00 | 9.68E-01 | no | Nab1 | intron_variant | MODIFIER |
| 1 | 52470794 | AX-170454374 | 0 | G | T | 0.132 | 9.74E-03 | 2.42E-01 | 2.69E+00 | 2.74E+00 | 9.68E-01 | no | Nab1 | intron_variant | MODIFIER |
| 1 | 52471850 | AX-170105877 | 0 | C | T | 0.132 | 9.74E-03 | 2.42E-01 | 2.69E+00 | 2.74E+00 | 9.68E-01 | no | Nab1 | intron_variant | MODIFIER |
| 1 | 52472062 | AX-170276610 | 0 | G | A | 0.132 | 9.74E-03 | 2.42E-01 | 2.69E+00 | 2.74E+00 | 9.68E-01 | no | Nab1 | intron_variant | MODIFIER |
| 1 | 52478536 | AX-170105012 | 0 | A | G | 0.132 | 9.74E-03 | 2.42E-01 | 2.69E+00 | 2.74E+00 | 9.68E-01 | no | Nab1 | intron_variant | MODIFIER |
| 1 | 52530501 | AX-170230798 | 0 | A | C | 0.132 | 9.74E-03 | 2.42E-01 | 2.69E+00 | 2.74E+00 | 9.68E-01 | no | Gm553 | intron_variant,non_coding_transcript_variant | MODIFIER |
| 1 | 52530532 | AX-170329295 | 0 | A | G | 0.132 | 9.74E-03 | 2.42E-01 | 2.69E+00 | 2.74E+00 | 9.68E-01 | no | Gm553 | intron_variant,non_coding_transcript_variant | MODIFIER |
| 1 | 52535650 | AX-170519470 | 0 | T | C | 0.132 | 9.74E-03 | 2.42E-01 | 2.69E+00 | 2.74E+00 | 9.68E-01 | no | Gm553 | intron_variant,non_coding_transcript_variant | MODIFIER |
| 1 | 52536353 | AX-170175715 | 0 | A | T | 0.132 | 9.74E-03 | 2.42E-01 | 2.69E+00 | 2.74E+00 | 9.68E-01 | no | Gm553 | intron_variant,non_coding_transcript_variant | MODIFIER |
| 1 | 52537797 | AX-170141602 | 0 | A | G | 0.132 | 9.74E-03 | 2.42E-01 | 2.69E+00 | 2.74E+00 | 9.68E-01 | no | Gm553 | intron_variant,non_coding_transcript_variant | MODIFIER |
| 1 | 52539724 | AX-170285748 | 0 | T | G | 0.132 | 9.74E-03 | 2.42E-01 | 2.69E+00 | 2.74E+00 | 9.68E-01 | no | Gm553 | intron_variant,non_coding_transcript_variant | MODIFIER |
| 1 | 52539771 | AX-170314406 | 0 | A | C | 0.132 | 9.74E-03 | 2.42E-01 | 2.69E+00 | 2.74E+00 | 9.68E-01 | no | Gm553 | intron_variant,non_coding_transcript_variant | MODIFIER |
| 1 | 52543140 | AX-170201195 | 0 | T | G | 0.132 | 9.74E-03 | 2.42E-01 | 2.69E+00 | 2.74E+00 | 9.68E-01 | no | Gm553 | intron_variant,non_coding_transcript_variant | MODIFIER |
| 1 | 52543169 | AX-170163772 | 0 | A | G | 0.132 | 9.74E-03 | 2.42E-01 | 2.69E+00 | 2.74E+00 | 9.68E-01 | no | Gm553 | intron_variant,non_coding_transcript_variant | MODIFIER |
| 1 | 52543185 | AX-170403339 | 0 | G | A | 0.132 | 9.74E-03 | 2.42E-01 | 2.69E+00 | 2.74E+00 | 9.68E-01 | no | Gm553 | intron_variant,non_coding_transcript_variant | MODIFIER |
| 1 | 52554850 | AX-170233785 | 0 | C | T | 0.132 | 9.74E-03 | 2.42E-01 | 2.69E+00 | 2.74E+00 | 9.68E-01 | no | Gm553 | intron_variant,non_coding_transcript_variant | MODIFIER |
| 1 | 52562929 | AX-170188209 | 0 | T | C | 0.132 | 9.74E-03 | 2.42E-01 | 2.69E+00 | 2.74E+00 | 9.68E-01 | no | Gm553 | intron_variant,non_coding_transcript_variant | MODIFIER |
| 1 | 53011522 | AX-170273905 | 0 | C | T | 0.132 | 9.74E-03 | 2.42E-01 | 2.69E+00 | 2.74E+00 | 9.68E-01 | no | 1700019D03Rik | intron_variant | MODIFIER |
| 1 | 60716947 | AX-170362577 | 0 | C | G | 0.132 | 9.74E-03 | 2.42E-01 | 2.69E+00 | 2.74E+00 | 9.68E-01 | no | Cd28 | non_coding_transcript_exon_variant | MODIFIER |
| 1 | 62607724 | AX-170246228 | 0 | G | A | 0.132 | 9.74E-03 | 2.42E-01 | 2.69E+00 | 2.74E+00 | 9.68E-01 | no | Pard3b | intron_variant | MODIFIER |
| 1 | 62607859 | AX-170219227 | 0 | G | A | 0.132 | 9.74E-03 | 2.42E-01 | 2.69E+00 | 2.74E+00 | 9.68E-01 | no | Pard3b | intron_variant | MODIFIER |

| | | | | | | | | | | | | | | | |
|---|---|---|---|---|---|---|---|---|---|---|---|---|---|---|---|
| 1 | 62614851 | AX-170202167 | 0 | A | T | 0.132 | 9.74E-03 | 2.42E-01 | 2.69E+00 | 2.74E+00 | 9.68E-01 | no | Pard3b | intron_variant | MODIFIER |
| 1 | 62636451 | AX-170035612 | 0 | A | G | 0.132 | 9.74E-03 | 2.42E-01 | 2.69E+00 | 2.74E+00 | 9.68E-01 | no | Pard3b | intron_variant | MODIFIER |
| 1 | 62653218 | AX-170041428 | 0 | G | A | 0.132 | 9.74E-03 | 2.42E-01 | 2.69E+00 | 2.74E+00 | 9.68E-01 | no | Gm11600 | downstream_gene_variant | MODIFIER |
| 1 | 62687260 | AX-170460253 | 0 | T | C | 0.132 | 9.74E-03 | 2.42E-01 | 2.69E+00 | 2.74E+00 | 9.68E-01 | no | | intergenic_variant | MODIFIER |
| 1 | 62698621 | AX-170000643 | 0 | C | A | 0.132 | 9.74E-03 | 2.42E-01 | 2.69E+00 | 2.74E+00 | 9.68E-01 | no | Nrp2 | upstream_gene_variant | MODIFIER |
| 1 | 62791587 | AX-97849823 | 0 | T | C | 0.132 | 9.74E-03 | 2.42E-01 | 2.69E+00 | 2.74E+00 | 9.68E-01 | no | Nrp2 | intron_variant | MODIFIER |
| 1 | 62807358 | AX-97811761 | 0 | T | C | 0.132 | 9.74E-03 | 2.42E-01 | 2.69E+00 | 2.74E+00 | 9.68E-01 | no | Nrp2 | intron_variant | MODIFIER |
| 1 | 62838936 | AX-170517754 | 0 | G | A | 0.132 | 9.74E-03 | 2.42E-01 | 2.69E+00 | 2.74E+00 | 9.68E-01 | no | | intergenic_variant | MODIFIER |
| 1 | 62842218 | AX-170253879 | 0 | G | A | 0.132 | 9.74E-03 | 2.42E-01 | 2.69E+00 | 2.74E+00 | 9.68E-01 | no | | intergenic_variant | MODIFIER |
| 1 | 62844437 | AX-170324379 | 0 | G | C | 0.132 | 9.74E-03 | 2.42E-01 | 2.69E+00 | 2.74E+00 | 9.68E-01 | no | | intergenic_variant | MODIFIER |
| 1 | 62849178 | AX-170131934 | 0 | T | C | 0.132 | 9.74E-03 | 2.42E-01 | 2.69E+00 | 2.74E+00 | 9.68E-01 | no | | intergenic_variant | MODIFIER |
| 1 | 62854123 | AX-170251824 | 0 | T | C | 0.132 | 9.74E-03 | 2.42E-01 | 2.69E+00 | 2.74E+00 | 9.68E-01 | no | | intergenic_variant | MODIFIER |
| 1 | 62882460 | AX-97798221 | 0 | A | G | 0.132 | 9.74E-03 | 2.42E-01 | 2.69E+00 | 2.74E+00 | 9.68E-01 | no | | intergenic_variant | MODIFIER |
| 1 | 62882509 | AX-170041433 | 0 | C | G | 0.132 | 9.74E-03 | 2.42E-01 | 2.69E+00 | 2.74E+00 | 9.68E-01 | no | | intergenic_variant | MODIFIER |
| 1 | 62884934 | AX-170149280 | 0 | G | A | 0.132 | 9.74E-03 | 2.42E-01 | 2.69E+00 | 2.74E+00 | 9.68E-01 | no | | regulatory_region_variant | MODIFIER |
| 1 | 62900203 | AX-97846549 | 0 | C | A | 0.132 | 9.74E-03 | 2.42E-01 | 2.69E+00 | 2.74E+00 | 9.68E-01 | no | | regulatory_region_variant | MODIFIER |
| 1 | 62908077 | AX-170105092 | 0 | A | G | 0.132 | 9.74E-03 | 2.42E-01 | 2.69E+00 | 2.74E+00 | 9.68E-01 | no | | intergenic_variant | MODIFIER |
| 1 | 62908559 | AX-170373965 | 0 | G | A | 0.132 | 9.74E-03 | 2.42E-01 | 2.69E+00 | 2.74E+00 | 9.68E-01 | no | | regulatory_region_variant | MODIFIER |
| 1 | 62910393 | AX-170168043 | 0 | C | G | 0.132 | 9.74E-03 | 2.42E-01 | 2.69E+00 | 2.74E+00 | 9.68E-01 | no | | regulatory_region_variant | MODIFIER |
| 1 | 62910546 | AX-170502980 | 0 | C | T | 0.132 | 9.74E-03 | 2.42E-01 | 2.69E+00 | 2.74E+00 | 9.68E-01 | no | | intergenic_variant | MODIFIER |
| 1 | 63005312 | AX-97797695 | 0 | C | T | 0.132 | 9.74E-03 | 2.42E-01 | 2.69E+00 | 2.74E+00 | 9.68E-01 | no | Ino80d | intron_variant,NMD_transcript_variant | MODIFIER |
| 1 | 63263837 | AX-170321259 | 0 | C | T | 0.132 | 9.74E-03 | 2.42E-01 | 2.69E+00 | 2.74E+00 | 9.68E-01 | no | Gm11608 | downstream_gene_variant | MODIFIER |
| 1 | 63265798 | AX-170174801 | 0 | A | C | 0.132 | 9.74E-03 | 2.42E-01 | 2.69E+00 | 2.74E+00 | 9.68E-01 | no | Gm11608 | downstream_gene_variant | MODIFIER |
| 1 | 63271204 | AX-170000656 | 0 | T | C | 0.132 | 9.74E-03 | 2.42E-01 | 2.69E+00 | 2.74E+00 | 9.68E-01 | no | Zdbf2 | upstream_gene_variant | MODIFIER |
| 1 | 63293552 | AX-170379337 | 0 | G | A | 0.132 | 9.74E-03 | 2.42E-01 | 2.69E+00 | 2.74E+00 | 9.68E-01 | no | Zdbf2 | intron_variant | MODIFIER |
| 1 | 63361773 | AX-97802210 | 0 | A | G | 0.132 | 9.74E-03 | 2.42E-01 | 2.69E+00 | 2.74E+00 | 9.68E-01 | no | Gm11605 | downstream_gene_variant | MODIFIER |
| 1 | 63397194 | AX-97829658 | 0 | T | G | 0.132 | 9.74E-03 | 2.42E-01 | 2.69E+00 | 2.74E+00 | 9.68E-01 | no | | intergenic_variant | MODIFIER |
| 1 | 63410830 | AX-170309089 | 0 | C | T | 0.132 | 9.74E-03 | 2.42E-01 | 2.69E+00 | 2.74E+00 | 9.68E-01 | no | | intergenic_variant | MODIFIER |
| 1 | 63414925 | AX-170104905 | 0 | T | C | 0.132 | 9.74E-03 | 2.42E-01 | 2.69E+00 | 2.74E+00 | 9.68E-01 | no | | intergenic_variant | MODIFIER |
| 1 | 63415114 | AX-170231975 | 0 | T | C | 0.132 | 9.74E-03 | 2.42E-01 | 2.69E+00 | 2.74E+00 | 9.68E-01 | no | | intergenic_variant | MODIFIER |
| 1 | 63428968 | AX-170501093 | 0 | T | A | 0.132 | 9.74E-03 | 2.42E-01 | 2.69E+00 | 2.74E+00 | 9.68E-01 | no | | intergenic_variant | MODIFIER |
| 1 | 63430636 | AX-170481208 | 0 | G | A | 0.132 | 9.74E-03 | 2.42E-01 | 2.69E+00 | 2.74E+00 | 9.68E-01 | no | | regulatory_region_variant | MODIFIER |
| 1 | 63450140 | AX-170163680 | 0 | T | C | 0.132 | 9.74E-03 | 2.42E-01 | 2.69E+00 | 2.74E+00 | 9.68E-01 | no | Adam23 | intron_variant | MODIFIER |
| 1 | 63453031 | AX-97822128 | 0 | A | G | 0.132 | 9.74E-03 | 2.42E-01 | 2.69E+00 | 2.74E+00 | 9.68E-01 | no | Adam23 | intron_variant | MODIFIER |
| 1 | 63453454 | AX-170035614 | 0 | C | T | 0.132 | 9.74E-03 | 2.42E-01 | 2.69E+00 | 2.74E+00 | 9.68E-01 | no | Adam23 | intron_variant | MODIFIER |
| 1 | 63471551 | AX-170534193 | 0 | T | C | 0.132 | 9.74E-03 | 2.42E-01 | 2.69E+00 | 2.74E+00 | 9.68E-01 | no | Adam23 | intron_variant | MODIFIER |
| 1 | 63473071 | AX-170349431 | 0 | T | C | 0.132 | 9.74E-03 | 2.42E-01 | 2.69E+00 | 2.74E+00 | 9.68E-01 | no | Adam23 | intron_variant | MODIFIER |
| 1 | 63487040 | AX-97811728 | 0 | A | G | 0.132 | 9.74E-03 | 2.42E-01 | 2.69E+00 | 2.74E+00 | 9.68E-01 | no | Adam23 | intron_variant | MODIFIER |
| 1 | 63509472 | AX-97841772 | 0 | A | G | 0.132 | 9.74E-03 | 2.42E-01 | 2.69E+00 | 2.74E+00 | 9.68E-01 | no | Adam23 | intron_variant | MODIFIER |
| 1 | 63524685 | AX-170470426 | 0 | G | A | 0.132 | 9.74E-03 | 2.42E-01 | 2.69E+00 | 2.74E+00 | 9.68E-01 | no | Adam23 | intron_variant | MODIFIER |
| 1 | 63530277 | AX-170529586 | 0 | T | G | 0.132 | 9.74E-03 | 2.42E-01 | 2.69E+00 | 2.74E+00 | 9.68E-01 | no | Adam23 | synonymous_variant | LOW |
| 1 | 63550283 | AX-97785309 | 0 | C | T | 0.132 | 9.74E-03 | 2.42E-01 | 2.69E+00 | 2.74E+00 | 9.68E-01 | no | Adam23 | intron_variant | MODIFIER |
| 1 | 63573808 | AX-170490770 | 0 | T | A | 0.132 | 9.74E-03 | 2.42E-01 | 2.69E+00 | 2.74E+00 | 9.68E-01 | no | Adam23 | intron_variant | MODIFIER |
| 1 | 63579350 | AX-170319039 | 0 | T | G | 0.132 | 9.74E-03 | 2.42E-01 | 2.69E+00 | 2.74E+00 | 9.68E-01 | no | Adam23 | intron_variant | MODIFIER |
| 1 | 63579559 | AX-170509264 | 0 | A | G | 0.132 | 9.74E-03 | 2.42E-01 | 2.69E+00 | 2.74E+00 | 9.68E-01 | no | Adam23 | intron_variant | MODIFIER |
| 1 | 63583945 | AX-170431811 | 0 | A | G | 0.132 | 9.74E-03 | 2.42E-01 | 2.69E+00 | 2.74E+00 | 9.68E-01 | no | Adam23 | intron_variant | MODIFIER |
| 1 | 63586829 | AX-97853390 | 0 | C | T | 0.132 | 9.74E-03 | 2.42E-01 | 2.69E+00 | 2.74E+00 | 9.68E-01 | no | Adam23 | intron_variant | MODIFIER |
| 1 | 63620208 | AX-170041445 | 0 | G | A | 0.132 | 9.74E-03 | 2.42E-01 | 2.69E+00 | 2.74E+00 | 9.68E-01 | no | Dytn | downstream_gene_variant | MODIFIER |
| 1 | 63644792 | AX-170149648 | 0 | T | C | 0.132 | 9.74E-03 | 2.42E-01 | 2.69E+00 | 2.74E+00 | 9.68E-01 | no | Dytn | intron_variant | MODIFIER |
| 1 | 63656189 | AX-170133243 | 0 | T | G | 0.132 | 9.74E-03 | 2.42E-01 | 2.69E+00 | 2.74E+00 | 9.68E-01 | no | Dytn | intron_variant | MODIFIER |
| 1 | 63685144 | AX-170000659 | 0 | A | C | 0.132 | 9.74E-03 | 2.42E-01 | 2.69E+00 | 2.74E+00 | 9.68E-01 | no | Dytn | 5_prime_UTR_variant | MODIFIER |
| 1 | 63730569 | AX-97856576 | 0 | A | G | 0.132 | 9.74E-03 | 2.42E-01 | 2.69E+00 | 2.74E+00 | 9.68E-01 | no | Fastkd2 | upstream_gene_variant | MODIFIER |
| 1 | 63786290 | AX-97785180 | 0 | A | G | 0.132 | 9.74E-03 | 2.42E-01 | 2.69E+00 | 2.74E+00 | 9.68E-01 | no | | intergenic_variant | MODIFIER |
| 1 | 63843059 | AX-170171249 | 0 | C | G | 0.132 | 9.74E-03 | 2.42E-01 | 2.69E+00 | 2.74E+00 | 9.68E-01 | no | Gm13751 | downstream_gene_variant | MODIFIER |

| | | | | | | | | | | | | | | | |
|---|---|---|---|---|---|---|---|---|---|---|---|---|---|---|---|
| 1 | 63857837 | AX-170325883 | 0 | G | A | 0.132 | 9.74E-03 | 2.42E-01 | 2.69E+00 | 2.74E+00 | 9.68E-01 | no | | regulatory_region_variant | MODIFIER |
| 1 | 63901004 | AX-170220539 | 0 | C | G | 0.132 | 9.74E-03 | 2.42E-01 | 2.69E+00 | 2.74E+00 | 9.68E-01 | no | | intergenic_variant | MODIFIER |
| 1 | 63903509 | AX-170178830 | 0 | C | G | 0.132 | 9.74E-03 | 2.42E-01 | 2.69E+00 | 2.74E+00 | 9.68E-01 | no | | regulatory_region_variant | MODIFIER |
| 1 | 63937394 | AX-170151749 | 0 | T | G | 0.132 | 9.74E-03 | 2.42E-01 | 2.69E+00 | 2.74E+00 | 9.68E-01 | no | Gm13754 | downstream_gene_variant | MODIFIER |
| 1 | 63937970 | AX-170161826 | 0 | A | C | 0.132 | 9.74E-03 | 2.42E-01 | 2.69E+00 | 2.74E+00 | 9.68E-01 | no | Gm13754 | downstream_gene_variant | MODIFIER |
| 1 | 63967585 | AX-170352078 | 0 | C | T | 0.132 | 9.74E-03 | 2.42E-01 | 2.69E+00 | 2.74E+00 | 9.68E-01 | no | Gm13749 | intron_variant,non_coding_transcript_variant | MODIFIER |
| 1 | 63967604 | AX-170288837 | 0 | G | T | 0.132 | 9.74E-03 | 2.42E-01 | 2.69E+00 | 2.74E+00 | 9.68E-01 | no | Gm13749 | intron_variant,non_coding_transcript_variant | MODIFIER |
| 1 | 63967678 | AX-97807327 | 0 | C | G | 0.132 | 9.74E-03 | 2.42E-01 | 2.69E+00 | 2.74E+00 | 9.68E-01 | no | Gm13749 | non_coding_transcript_exon_variant | MODIFIER |
| 1 | 63967706 | AX-170210375 | 0 | A | C | 0.132 | 9.74E-03 | 2.42E-01 | 2.69E+00 | 2.74E+00 | 9.68E-01 | no | Gm13749 | non_coding_transcript_exon_variant | MODIFIER |
| 1 | 63969227 | AX-170101564 | 0 | C | G | 0.132 | 9.74E-03 | 2.42E-01 | 2.69E+00 | 2.74E+00 | 9.68E-01 | no | Gm13749 | downstream_gene_variant | MODIFIER |
| 1 | 64017720 | AX-97783499 | 0 | C | T | 0.132 | 9.74E-03 | 2.42E-01 | 2.69E+00 | 2.74E+00 | 9.68E-01 | no | | intergenic_variant | MODIFIER |
| 1 | 64023599 | AX-97846515 | 0 | A | G | 0.132 | 9.74E-03 | 2.42E-01 | 2.69E+00 | 2.74E+00 | 9.68E-01 | no | | intergenic_variant | MODIFIER |
| 1 | 64043098 | AX-170480993 | 0 | G | A | 0.132 | 9.74E-03 | 2.42E-01 | 2.69E+00 | 2.74E+00 | 9.68E-01 | no | Klf7 | intron_variant | MODIFIER |
| 1 | 64048457 | AX-170320756 | 0 | T | G | 0.132 | 9.74E-03 | 2.42E-01 | 2.69E+00 | 2.74E+00 | 9.68E-01 | no | Klf7 | intron_variant | MODIFIER |
| 1 | 64048508 | AX-170458379 | 0 | A | G | 0.132 | 9.74E-03 | 2.42E-01 | 2.69E+00 | 2.74E+00 | 9.68E-01 | no | Klf7 | intron_variant | MODIFIER |
| 1 | 64050451 | AX-97824560 | 0 | T | C | 0.132 | 9.74E-03 | 2.42E-01 | 2.69E+00 | 2.74E+00 | 9.68E-01 | no | Klf7 | intron_variant | MODIFIER |
| 1 | 64054258 | AX-170441856 | 0 | G | C | 0.132 | 9.74E-03 | 2.42E-01 | 2.69E+00 | 2.74E+00 | 9.68E-01 | no | Klf7 | intron_variant | MODIFIER |
| 1 | 64084795 | AX-170188464 | 0 | C | T | 0.132 | 9.74E-03 | 2.42E-01 | 2.69E+00 | 2.74E+00 | 9.68E-01 | no | Klf7 | intron_variant | MODIFIER |
| 1 | 64085097 | AX-170207606 | 0 | T | A | 0.132 | 9.74E-03 | 2.42E-01 | 2.69E+00 | 2.74E+00 | 9.68E-01 | no | Klf7 | intron_variant | MODIFIER |
| 1 | 64085286 | AX-97783786 | 0 | T | C | 0.132 | 9.74E-03 | 2.42E-01 | 2.69E+00 | 2.74E+00 | 9.68E-01 | no | Klf7 | intron_variant | MODIFIER |
| 1 | 64103457 | AX-170235777 | 0 | T | C | 0.132 | 9.74E-03 | 2.42E-01 | 2.69E+00 | 2.74E+00 | 9.68E-01 | no | Klf7 | intron_variant | MODIFIER |
| 1 | 64105141 | AX-170312608 | 0 | T | C | 0.132 | 9.74E-03 | 2.42E-01 | 2.69E+00 | 2.74E+00 | 9.68E-01 | no | Klf7 | intron_variant | MODIFIER |
| 1 | 64106177 | AX-170391228 | 0 | C | G | 0.132 | 9.74E-03 | 2.42E-01 | 2.69E+00 | 2.74E+00 | 9.68E-01 | no | Klf7 | intron_variant | MODIFIER |
| 1 | 64106217 | AX-170491755 | 0 | G | T | 0.132 | 9.74E-03 | 2.42E-01 | 2.69E+00 | 2.74E+00 | 9.68E-01 | no | Klf7 | intron_variant | MODIFIER |
| 1 | 64107713 | AX-170122504 | 0 | C | T | 0.132 | 9.74E-03 | 2.42E-01 | 2.69E+00 | 2.74E+00 | 9.68E-01 | no | Klf7 | intron_variant | MODIFIER |
| 1 | 64109910 | AX-97835230 | 0 | A | G | 0.132 | 9.74E-03 | 2.42E-01 | 2.69E+00 | 2.74E+00 | 9.68E-01 | no | Klf7 | intron_variant | MODIFIER |
| 1 | 64110163 | AX-170482339 | 0 | T | C | 0.132 | 9.74E-03 | 2.42E-01 | 2.69E+00 | 2.74E+00 | 9.68E-01 | no | Klf7 | intron_variant | MODIFIER |
| 1 | 64111431 | AX-170218948 | 0 | G | A | 0.132 | 9.74E-03 | 2.42E-01 | 2.69E+00 | 2.74E+00 | 9.68E-01 | no | Klf7 | intron_variant | MODIFIER |
| 1 | 64111929 | AX-170101271 | 0 | A | G | 0.132 | 9.74E-03 | 2.42E-01 | 2.69E+00 | 2.74E+00 | 9.68E-01 | no | Klf7 | intron_variant | MODIFIER |
| 1 | 64149720 | AX-170041458 | 0 | G | A | 0.132 | 9.74E-03 | 2.42E-01 | 2.69E+00 | 2.74E+00 | 9.68E-01 | no | Gm13750 | upstream_gene_variant | MODIFIER |
| 1 | 64153313 | AX-170041454 | 0 | G | T | 0.132 | 9.74E-03 | 2.42E-01 | 2.69E+00 | 2.74E+00 | 9.68E-01 | no | Gm13750 | downstream_gene_variant | MODIFIER |
| 1 | 64153349 | AX-170041455 | 0 | C | T | 0.132 | 9.74E-03 | 2.42E-01 | 2.69E+00 | 2.74E+00 | 9.68E-01 | no | Gm13750 | downstream_gene_variant | MODIFIER |
| 1 | 64153597 | AX-170041457 | 0 | C | G | 0.132 | 9.74E-03 | 2.42E-01 | 2.69E+00 | 2.74E+00 | 9.68E-01 | no | Gm13750 | downstream_gene_variant | MODIFIER |
| 1 | 64153998 | AX-170354683 | 0 | A | G | 0.132 | 9.74E-03 | 2.42E-01 | 2.69E+00 | 2.74E+00 | 9.68E-01 | no | Gm13750 | downstream_gene_variant | MODIFIER |
| 1 | 64166862 | AX-170000666 | 0 | T | G | 0.132 | 9.74E-03 | 2.42E-01 | 2.69E+00 | 2.74E+00 | 9.68E-01 | no | | intergenic_variant | MODIFIER |
| 1 | 64201441 | AX-170000669 | 0 | G | A | 0.132 | 9.74E-03 | 2.42E-01 | 2.69E+00 | 2.74E+00 | 9.68E-01 | no | Gm28981 | intron_variant,non_coding_transcript_variant | MODIFIER |
| 1 | 64238581 | AX-97782325 | 0 | T | C | 0.132 | 9.74E-03 | 2.42E-01 | 2.69E+00 | 2.74E+00 | 9.68E-01 | no | Gm28981 | intron_variant,non_coding_transcript_variant | MODIFIER |
| 9 | 29872548 | AX-170163796 | 0 | T | A | 0.43 | -2.77E-03 | 8.63E-02 | 2.68E+00 | 2.75E+00 | 9.68E-01 | no | Ntm | intron_variant | MODIFIER |
| 8 | 75681677 | AX-170531784 | 0 | A | T | 0.193 | -5.38E-03 | 1.32E-01 | 2.67E+00 | 2.74E+00 | 9.68E-01 | no | Iqcm | intron_variant | MODIFIER |
| 1 | 103473535 | AX-170001229 | 0 | C | T | 0.158 | -5.30E-03 | 1.55E-01 | 2.62E+00 | 2.74E+00 | 9.68E-01 | no | | intergenic_variant | MODIFIER |
| 4 | 55672927 | AX-170097759 | 1 | G | A | 0.375 | 4.25E-03 | 1.06E-01 | 2.66E+00 | 2.74E+00 | 9.68E-01 | no | | intergenic_variant | MODIFIER |
| 18 | 67427680 | AX-170163182 | 2 | T | C | 0.282 | -4.45E-03 | 9.94E-02 | 2.65E+00 | 2.73E+00 | 9.68E-01 | no | Afg3l2 | intron_variant | MODIFIER |
| 11 | 4752187 | AX-170348130 | 0 | C | T | 0.07 | 6.78E-03 | 1.68E-01 | 2.61E+00 | 2.74E+00 | 9.68E-01 | no | | intergenic_variant | MODIFIER |
| 11 | 8490254 | AX-170318518 | 0 | C | T | 0.07 | 6.78E-03 | 1.68E-01 | 2.61E+00 | 2.74E+00 | 9.68E-01 | no | Tns3 | intron_variant | MODIFIER |
| 11 | 11171143 | AX-170524610 | 0 | G | T | 0.07 | 6.78E-03 | 1.68E-01 | 2.61E+00 | 2.74E+00 | 9.68E-01 | no | Vwc2 | intron_variant | MODIFIER |
| 18 | 41969880 | AX-170015597 | 0 | A | T | 0.061 | -1.24E-02 | 3.01E-01 | 2.71E+00 | 2.73E+00 | 9.68E-01 | no | | intergenic_variant | MODIFIER |
| 1 | 37647267 | AX-170151776 | 0 | A | G | 0.456 | -3.29E-03 | 7.86E-02 | 2.65E+00 | 2.74E+00 | 9.68E-01 | no | 2010300C02Rik | intron_variant | MODIFIER |
| 9 | 105530538 | AX-170398267 | 0 | T | C | 0.123 | 3.44E-03 | 1.15E-01 | 2.67E+00 | 2.75E+00 | 9.68E-01 | no | Atp2c1 | upstream_gene_variant | MODIFIER |
| 1 | 106463331 | AX-170515943 | 0 | G | A | 0.158 | 9.72E-03 | 1.88E-01 | 2.64E+00 | 2.73E+00 | 9.68E-01 | no | | intergenic_variant | MODIFIER |
| 1 | 118602327 | AX-170126004 | 0 | C | A | 0.158 | 9.72E-03 | 1.88E-01 | 2.64E+00 | 2.73E+00 | 9.68E-01 | no | Clasp1 | intron_variant | MODIFIER |
| 1 | 119150332 | AX-97824709 | 0 | G | A | 0.158 | 9.72E-03 | 1.88E-01 | 2.64E+00 | 2.73E+00 | 9.68E-01 | no | | intergenic_variant | MODIFIER |
| 12 | 68982379 | AX-170087275 | 1 | C | A | 0.062 | -1.23E-02 | 3.01E-01 | 2.71E+00 | 2.73E+00 | 9.68E-01 | no | Gm47515 | intron_variant,non_coding_transcript_variant | MODIFIER |
| 8 | 29883095 | AX-170030428 | 1 | T | C | 0.179 | -4.05E-03 | 1.01E-01 | 2.63E+00 | 2.74E+00 | 9.68E-01 | no | | intergenic_variant | MODIFIER |
| 1 | 155726239 | AX-170461258 | 0 | T | C | 0.211 | 6.72E-03 | 1.46E-01 | 2.68E+00 | 2.73E+00 | 9.68E-01 | no | Lhx4 | intron_variant | MODIFIER |

| | | | | | | | | | | | | | | | |
|---|---|---|---|---|---|---|---|---|---|---|---|---|---|---|---|
| 1 | 52428070 | AX-170103583 | 0 | C | T | 0.132 | 9.34E-03 | 2.39E-01 | 2.69E+00 | 2.74E+00 | 9.68E-01 | no | | regulatory_region_variant | MODIFIER |
| 1 | 52537048 | AX-170322433 | 0 | T | A | 0.14 | 8.37E-03 | 1.88E-01 | 2.63E+00 | 2.73E+00 | 9.69E-01 | no | Gm553 | intron_variant,non_coding_transcript_variant | MODIFIER |
| 1 | 63257134 | AX-170000649 | 0 | A | T | 0.14 | 8.37E-03 | 1.88E-01 | 2.63E+00 | 2.73E+00 | 9.69E-01 | no | | intergenic_variant | MODIFIER |
| 9 | 65950232 | AX-170032579 | 0 | C | G | 0.202 | -5.29E-03 | 1.28E-01 | 2.69E+00 | 2.74E+00 | 9.69E-01 | no | Csnk1g1 | intron_variant | MODIFIER |
| 9 | 78620349 | AX-97789027 | 0 | A | G | 0.202 | -5.29E-03 | 1.28E-01 | 2.69E+00 | 2.74E+00 | 9.69E-01 | no | Cd109 | intron_variant | MODIFIER |
| 18 | 69322163 | AX-170374402 | 0 | T | C | 0.096 | 7.69E-03 | 1.71E-01 | 2.63E+00 | 2.73E+00 | 9.69E-01 | no | | intergenic_variant | MODIFIER |
| 1 | 70879767 | AX-170165489 | 0 | G | T | 0.14 | -8.95E-03 | 2.38E-01 | 2.69E+00 | 2.74E+00 | 9.69E-01 | no | Vwc2l | intron_variant | MODIFIER |
| 1 | 78593897 | AX-170113354 | 0 | G | T | 0.14 | -8.95E-03 | 2.38E-01 | 2.69E+00 | 2.74E+00 | 9.69E-01 | no | 5730419F03Rik | downstream_gene_variant | MODIFIER |
| 5 | 127340302 | AX-170484961 | 0 | T | C | 0.412 | -2.55E-03 | 8.36E-02 | 2.64E+00 | 2.74E+00 | 9.69E-01 | no | Tmem132c | intron_variant | MODIFIER |
| 5 | 127417165 | AX-97823803 | 0 | C | A | 0.412 | -2.55E-03 | 8.36E-02 | 2.64E+00 | 2.74E+00 | 9.69E-01 | no | Tmem132c | intron_variant | MODIFIER |
| 5 | 127427198 | AX-170458248 | 0 | C | T | 0.412 | -2.55E-03 | 8.36E-02 | 2.64E+00 | 2.74E+00 | 9.69E-01 | no | Tmem132c | intron_variant | MODIFIER |
| 5 | 127433350 | AX-170382655 | 0 | G | A | 0.412 | -2.55E-03 | 8.36E-02 | 2.64E+00 | 2.74E+00 | 9.69E-01 | no | Tmem132c | intron_variant | MODIFIER |
| 5 | 127465978 | AX-170415946 | 0 | G | A | 0.412 | -2.55E-03 | 8.36E-02 | 2.64E+00 | 2.74E+00 | 9.69E-01 | no | Tmem132c | intron_variant | MODIFIER |
| 5 | 127488642 | AX-170423254 | 0 | G | A | 0.412 | -2.55E-03 | 8.36E-02 | 2.64E+00 | 2.74E+00 | 9.69E-01 | no | Tmem132c | intron_variant | MODIFIER |
| 5 | 127498801 | AX-170143809 | 0 | A | G | 0.412 | -2.55E-03 | 8.36E-02 | 2.64E+00 | 2.74E+00 | 9.69E-01 | no | Tmem132c | intron_variant | MODIFIER |
| 5 | 127507333 | AX-170095460 | 0 | A | G | 0.412 | -2.55E-03 | 8.36E-02 | 2.64E+00 | 2.74E+00 | 9.69E-01 | no | Tmem132c | intron_variant | MODIFIER |
| 5 | 127516703 | AX-170250887 | 0 | A | G | 0.412 | -2.55E-03 | 8.36E-02 | 2.64E+00 | 2.74E+00 | 9.69E-01 | no | Tmem132c | intron_variant | MODIFIER |
| 5 | 127526181 | AX-97807139 | 0 | T | C | 0.412 | -2.55E-03 | 8.36E-02 | 2.64E+00 | 2.74E+00 | 9.69E-01 | no | Tmem132c | intron_variant | MODIFIER |
| 5 | 127674747 | AX-170308647 | 0 | A | G | 0.412 | -2.55E-03 | 8.36E-02 | 2.64E+00 | 2.74E+00 | 9.69E-01 | no | Glt1d1 | intron_variant | MODIFIER |
| 5 | 127813778 | AX-170341325 | 0 | C | G | 0.412 | -2.55E-03 | 8.36E-02 | 2.64E+00 | 2.74E+00 | 9.69E-01 | no | Tmem132d | intron_variant | MODIFIER |
| 5 | 127814635 | AX-170460914 | 0 | G | A | 0.412 | -2.55E-03 | 8.36E-02 | 2.64E+00 | 2.74E+00 | 9.69E-01 | no | Tmem132d | intron_variant | MODIFIER |
| 5 | 127842027 | AX-170228396 | 0 | G | A | 0.412 | -2.55E-03 | 8.36E-02 | 2.64E+00 | 2.74E+00 | 9.69E-01 | no | Tmem132d | intron_variant | MODIFIER |
| 5 | 127842092 | AX-170443576 | 0 | A | G | 0.412 | -2.55E-03 | 8.36E-02 | 2.64E+00 | 2.74E+00 | 9.69E-01 | no | Tmem132d | intron_variant | MODIFIER |
| 5 | 127842715 | AX-170316595 | 0 | G | A | 0.412 | -2.55E-03 | 8.36E-02 | 2.64E+00 | 2.74E+00 | 9.69E-01 | no | Tmem132d | intron_variant | MODIFIER |
| 5 | 128004526 | AX-170297384 | 0 | C | T | 0.412 | -2.55E-03 | 8.36E-02 | 2.64E+00 | 2.74E+00 | 9.69E-01 | no | Tmem132d | intron_variant | MODIFIER |
| 5 | 128011106 | AX-170344332 | 0 | G | A | 0.412 | -2.55E-03 | 8.36E-02 | 2.64E+00 | 2.74E+00 | 9.69E-01 | no | Tmem132d | intron_variant | MODIFIER |
| 5 | 128011387 | AX-170260667 | 0 | A | G | 0.412 | -2.55E-03 | 8.36E-02 | 2.64E+00 | 2.74E+00 | 9.69E-01 | no | Tmem132d | intron_variant | MODIFIER |
| 5 | 128011773 | AX-170343357 | 0 | G | A | 0.412 | -2.55E-03 | 8.36E-02 | 2.64E+00 | 2.74E+00 | 9.69E-01 | no | Tmem132d | intron_variant | MODIFIER |
| 5 | 128025859 | AX-170302042 | 0 | A | G | 0.412 | -2.55E-03 | 8.36E-02 | 2.64E+00 | 2.74E+00 | 9.69E-01 | no | Tmem132d | intron_variant | MODIFIER |
| 5 | 128027823 | AX-170213482 | 0 | T | C | 0.412 | -2.55E-03 | 8.36E-02 | 2.64E+00 | 2.74E+00 | 9.69E-01 | no | Tmem132d | intron_variant | MODIFIER |
| 5 | 128030312 | AX-170073459 | 0 | A | T | 0.412 | -2.55E-03 | 8.36E-02 | 2.64E+00 | 2.74E+00 | 9.69E-01 | no | Tmem132d | intron_variant | MODIFIER |
| 5 | 128030391 | AX-97782173 | 0 | T | C | 0.412 | -2.55E-03 | 8.36E-02 | 2.64E+00 | 2.74E+00 | 9.69E-01 | no | Tmem132d | intron_variant | MODIFIER |
| 5 | 128036029 | AX-170073463 | 0 | G | A | 0.412 | -2.55E-03 | 8.36E-02 | 2.64E+00 | 2.74E+00 | 9.69E-01 | no | Tmem132d | intron_variant | MODIFIER |
| 5 | 128040844 | AX-170035719 | 0 | G | A | 0.412 | -2.55E-03 | 8.36E-02 | 2.64E+00 | 2.74E+00 | 9.69E-01 | no | Tmem132d | intron_variant | MODIFIER |
| 5 | 128042764 | AX-170186310 | 0 | G | A | 0.412 | -2.55E-03 | 8.36E-02 | 2.64E+00 | 2.74E+00 | 9.69E-01 | no | Tmem132d | intron_variant | MODIFIER |
| 5 | 128073470 | AX-170211521 | 0 | T | C | 0.412 | -2.55E-03 | 8.36E-02 | 2.64E+00 | 2.74E+00 | 9.69E-01 | no | Tmem132d | intron_variant | MODIFIER |
| 5 | 128081795 | AX-170105701 | 0 | G | A | 0.412 | -2.55E-03 | 8.36E-02 | 2.64E+00 | 2.74E+00 | 9.69E-01 | no | Tmem132d | intron_variant | MODIFIER |
| 5 | 128092134 | AX-170481425 | 0 | T | C | 0.412 | -2.55E-03 | 8.36E-02 | 2.64E+00 | 2.74E+00 | 9.69E-01 | no | Tmem132d | intron_variant | MODIFIER |
| 5 | 128092374 | AX-170208763 | 0 | T | C | 0.412 | -2.55E-03 | 8.36E-02 | 2.64E+00 | 2.74E+00 | 9.69E-01 | no | Tmem132d | intron_variant | MODIFIER |
| 12 | 101968170 | AX-97822212 | 0 | G | T | 0.07 | -7.04E-03 | 1.78E-01 | 2.63E+00 | 2.74E+00 | 9.69E-01 | no | Ndufb1-ps | non_coding_transcript_exon_variant | MODIFIER |
| 1 | 53530963 | AX-170326839 | 0 | T | C | 0.105 | -8.98E-03 | 2.34E-01 | 2.69E+00 | 2.74E+00 | 9.69E-01 | no | Dnah7a | intron_variant | MODIFIER |
| 1 | 23287063 | AX-170287526 | 1 | A | T | 0.429 | 3.85E-03 | 8.27E-02 | 2.66E+00 | 2.73E+00 | 9.69E-01 | no | Mir30c-2 | upstream_gene_variant | MODIFIER |
| 10 | 14402686 | AX-170442431 | 0 | T | C | 0.193 | 2.61E-03 | 1.05E-01 | 2.66E+00 | 2.75E+00 | 9.69E-01 | no | Adgrg6 | 3_prime_UTR_variant | MODIFIER |
| 12 | 101654507 | AX-170007401 | 1 | C | T | 0.089 | 7.47E-03 | 1.94E-01 | 2.68E+00 | 2.74E+00 | 9.69E-01 | no | Tc2n | intron_variant,NMD_transcript_variant | MODIFIER |
| 1 | 25410156 | AX-170040650 | 0 | G | A | 0.421 | 3.47E-03 | 7.82E-02 | 2.65E+00 | 2.73E+00 | 9.69E-01 | no | Adgrb3 | intron_variant | MODIFIER |
| 15 | 92314285 | AX-170187735 | 0 | T | A | 0.149 | 7.86E-03 | 1.79E-01 | 2.67E+00 | 2.73E+00 | 9.69E-01 | no | Cntn1 | intron_variant | MODIFIER |
| 15 | 92945723 | AX-170144540 | 0 | G | C | 0.149 | 7.86E-03 | 1.79E-01 | 2.67E+00 | 2.73E+00 | 9.69E-01 | no | Gm49418 | intron_variant,non_coding_transcript_variant | MODIFIER |
| 15 | 94223774 | AX-170283386 | 0 | G | T | 0.149 | 7.89E-03 | 1.79E-01 | 2.67E+00 | 2.73E+00 | 9.69E-01 | no | | intergenic_variant | MODIFIER |
| 6 | 114276769 | AX-170028024 | 0 | T | C | 0.053 | 1.17E-02 | 2.95E-01 | 2.67E+00 | 2.74E+00 | 9.69E-01 | no | | intergenic_variant | MODIFIER |
| 1 | 146689005 | AX-170086639 | 1 | A | C | 0.188 | 5.81E-03 | 1.30E-01 | 2.66E+00 | 2.73E+00 | 9.69E-01 | no | Brinp3 | intron_variant | MODIFIER |
| 1 | 44598651 | AX-97791318 | 0 | T | A | 0.105 | 6.42E-03 | 1.74E-01 | 2.62E+00 | 2.74E+00 | 9.69E-01 | no | Gulp1 | intron_variant | MODIFIER |
| 7 | 51261411 | AX-97833183 | 1 | T | C | 0.411 | -3.31E-03 | 1.04E-01 | 2.66E+00 | 2.74E+00 | 9.70E-01 | no | | intergenic_variant | MODIFIER |
| 1 | 48128596 | AX-170179188 | 1 | C | T | 0.107 | 9.10E-03 | 2.36E-01 | 2.69E+00 | 2.74E+00 | 9.70E-01 | no | | intergenic_variant | MODIFIER |
| 6 | 118072426 | AX-170028070 | 0 | C | T | 0.07 | -1.06E-02 | 2.78E-01 | 2.70E+00 | 2.74E+00 | 9.70E-01 | no | Rasgef1a | intron_variant | MODIFIER |

| | | | | | | | | | | | | | | | |
|---|---|---|---|---|---|---|---|---|---|---|---|---|---|---|---|
| 10 | 30859109 | AX-97834062 | 1 | G | T | 0.179 | -4.13E-03 | 9.39E-02 | 2.62E+00 | 2.73E+00 | 9.70E-01 | no | | intergenic_variant | MODIFIER |
| 13 | 36368495 | AX-97823195 | 0 | C | T | 0.175 | 3.19E-03 | 1.20E-01 | 2.68E+00 | 2.75E+00 | 9.70E-01 | no | Fars2 | intron_variant | MODIFIER |
| 9 | 105527719 | AX-97789156 | 1 | C | T | 0.125 | 3.30E-03 | 1.18E-01 | 2.68E+00 | 2.75E+00 | 9.70E-01 | no | Atp2c1 | upstream_gene_variant | MODIFIER |
| 1 | 38550403 | AX-170199786 | 1 | A | G | 0.455 | -3.19E-03 | 7.81E-02 | 2.66E+00 | 2.74E+00 | 9.70E-01 | no | Aff3 | intron_variant,non_coding_transcript_variant | MODIFIER |
| 10 | 16226008 | AX-170189102 | 1 | G | T | 0.205 | 2.92E-03 | 1.07E-01 | 2.67E+00 | 2.75E+00 | 9.70E-01 | no | | intergenic_variant | MODIFIER |
| 12 | 116132899 | AX-170489226 | 1 | G | T | 0.134 | 6.63E-03 | 1.73E-01 | 2.64E+00 | 2.74E+00 | 9.70E-01 | no | Vipr2 | intron_variant | MODIFIER |
| 7 | 118453817 | AX-170097613 | 0 | C | T | 0.395 | -3.03E-03 | 9.10E-02 | 2.69E+00 | 2.74E+00 | 9.70E-01 | no | | intergenic_variant | MODIFIER |
| 15 | 94797195 | AX-97803294 | 1 | C | T | 0.152 | 7.86E-03 | 1.79E-01 | 2.67E+00 | 2.73E+00 | 9.70E-01 | no | Tmem117 | intron_variant | MODIFIER |
| 1 | 39432156 | AX-170282895 | 1 | A | T | 0.402 | -3.35E-03 | 7.57E-02 | 2.65E+00 | 2.73E+00 | 9.70E-01 | no | Tbc1d8 | intron_variant | MODIFIER |
| 2 | 129616516 | AX-170173470 | 0 | C | T | 0.14 | 8.49E-03 | 2.20E-01 | 2.70E+00 | 2.73E+00 | 9.70E-01 | no | Sirpa | missense_variant | MODERATE |
| 2 | 132813458 | AX-170301050 | 0 | A | G | 0.14 | 8.49E-03 | 2.20E-01 | 2.70E+00 | 2.73E+00 | 9.70E-01 | no | Mcm8 | upstream_gene_variant | MODIFIER |
| 2 | 138975324 | AX-170172206 | 0 | G | T | 0.14 | 8.49E-03 | 2.20E-01 | 2.70E+00 | 2.73E+00 | 9.70E-01 | no | | regulatory_region_variant | MODIFIER |
| 2 | 155879246 | AX-170361778 | 0 | C | T | 0.14 | 8.49E-03 | 2.20E-01 | 2.70E+00 | 2.73E+00 | 9.70E-01 | no | Uqcc1 | intron_variant | MODIFIER |
| 2 | 114039258 | AX-170280297 | 1 | C | A | 0.134 | 8.05E-03 | 2.03E-01 | 2.67E+00 | 2.74E+00 | 9.70E-01 | no | A530058N18Rik | intron_variant,non_coding_transcript_variant | MODIFIER |
| 8 | 36795694 | AX-170079776 | 0 | C | T | 0.149 | 4.88E-03 | 1.49E-01 | 2.69E+00 | 2.74E+00 | 9.70E-01 | no | Dlc1 | intron_variant | MODIFIER |
| 8 | 38121727 | AX-170250560 | 0 | A | T | 0.149 | 4.88E-03 | 1.49E-01 | 2.69E+00 | 2.74E+00 | 9.70E-01 | no | Sgcz | intron_variant | MODIFIER |
| 11 | 7416057 | AX-170123757 | 0 | A | G | 0.044 | -8.68E-03 | 1.98E-01 | 2.61E+00 | 2.73E+00 | 9.70E-01 | no | | intergenic_variant | MODIFIER |
| 1 | 68926953 | AX-97816710 | 0 | T | C | 0.114 | 8.84E-03 | 2.31E-01 | 2.69E+00 | 2.74E+00 | 9.70E-01 | no | Erbb4 | intron_variant | MODIFIER |
| 7 | 53940124 | AX-170029026 | 1 | G | T | 0.411 | 4.16E-03 | 9.70E-02 | 2.65E+00 | 2.73E+00 | 9.70E-01 | no | | intergenic_variant | MODIFIER |
| 10 | 5070456 | AX-170002647 | 0 | C | T | 0.211 | 3.21E-03 | 9.88E-02 | 2.64E+00 | 2.74E+00 | 9.70E-01 | no | Syne1 | upstream_gene_variant | MODIFIER |
| 10 | 5071448 | AX-170282042 | 0 | G | A | 0.211 | 3.21E-03 | 9.88E-02 | 2.64E+00 | 2.74E+00 | 9.70E-01 | no | Syne1 | upstream_gene_variant | MODIFIER |
| 10 | 5240421 | AX-170214602 | 0 | G | A | 0.211 | 3.21E-03 | 9.88E-02 | 2.64E+00 | 2.74E+00 | 9.70E-01 | no | Syne1 | intron_variant | MODIFIER |
| 10 | 5298460 | AX-97813901 | 0 | G | C | 0.211 | 3.21E-03 | 9.88E-02 | 2.64E+00 | 2.74E+00 | 9.70E-01 | no | Syne1 | intron_variant | MODIFIER |
| 10 | 5304167 | AX-170341908 | 0 | T | G | 0.211 | 3.21E-03 | 9.88E-02 | 2.64E+00 | 2.74E+00 | 9.70E-01 | no | Syne1 | intron_variant | MODIFIER |
| 10 | 5343832 | AX-170102706 | 0 | A | G | 0.211 | 3.21E-03 | 9.88E-02 | 2.64E+00 | 2.74E+00 | 9.70E-01 | no | Syne1 | intron_variant | MODIFIER |
| 10 | 5344960 | AX-170465690 | 0 | G | A | 0.211 | 3.21E-03 | 9.88E-02 | 2.64E+00 | 2.74E+00 | 9.70E-01 | no | Syne1 | splice_region_variant,synonymous_variant | LOW |
| 10 | 5354042 | AX-170134350 | 0 | C | T | 0.211 | 3.21E-03 | 9.88E-02 | 2.64E+00 | 2.74E+00 | 9.70E-01 | no | Syne1 | intron_variant | MODIFIER |
| 10 | 5354447 | AX-170335210 | 0 | G | A | 0.211 | 3.21E-03 | 9.88E-02 | 2.64E+00 | 2.74E+00 | 9.70E-01 | no | Syne1 | intron_variant | MODIFIER |
| 10 | 5367811 | AX-170461890 | 0 | T | C | 0.211 | 3.21E-03 | 9.88E-02 | 2.64E+00 | 2.74E+00 | 9.70E-01 | no | Syne1 | intron_variant | MODIFIER |
| 10 | 5368887 | AX-170147292 | 0 | A | G | 0.211 | 3.21E-03 | 9.88E-02 | 2.64E+00 | 2.74E+00 | 9.70E-01 | no | Syne1 | intron_variant | MODIFIER |
| 10 | 5369745 | AX-170369881 | 0 | C | T | 0.211 | 3.21E-03 | 9.88E-02 | 2.64E+00 | 2.74E+00 | 9.70E-01 | no | Syne1 | intron_variant | MODIFIER |
| 10 | 5405426 | AX-170465904 | 0 | C | T | 0.211 | 3.21E-03 | 9.88E-02 | 2.64E+00 | 2.74E+00 | 9.70E-01 | no | Syne1 | missense_variant,splice_region_variant | MODERATE |
| 10 | 5455943 | AX-170302090 | 0 | G | A | 0.211 | 3.21E-03 | 9.88E-02 | 2.64E+00 | 2.74E+00 | 9.70E-01 | no | Syne1 | intron_variant | MODIFIER |
| 10 | 5456327 | AX-170255186 | 0 | A | G | 0.211 | 3.21E-03 | 9.88E-02 | 2.64E+00 | 2.74E+00 | 9.70E-01 | no | Syne1 | intron_variant | MODIFIER |
| 10 | 5481758 | AX-170148302 | 0 | C | A | 0.211 | 3.21E-03 | 9.88E-02 | 2.64E+00 | 2.74E+00 | 9.70E-01 | no | Syne1 | intron_variant | MODIFIER |
| 10 | 5482030 | AX-170329512 | 0 | C | T | 0.211 | 3.21E-03 | 9.88E-02 | 2.64E+00 | 2.74E+00 | 9.70E-01 | no | Syne1 | intron_variant | MODIFIER |
| 10 | 5522967 | AX-170181771 | 0 | T | C | 0.211 | 3.21E-03 | 9.88E-02 | 2.64E+00 | 2.74E+00 | 9.70E-01 | no | Syne1 | intron_variant | MODIFIER |
| 10 | 5523171 | AX-97840908 | 0 | C | G | 0.211 | 3.21E-03 | 9.88E-02 | 2.64E+00 | 2.74E+00 | 9.70E-01 | no | Syne1 | intron_variant | MODIFIER |
| 10 | 5531105 | AX-170440564 | 0 | A | G | 0.211 | 3.21E-03 | 9.88E-02 | 2.64E+00 | 2.74E+00 | 9.70E-01 | no | Syne1 | intron_variant | MODIFIER |
| 10 | 5534236 | AX-170320104 | 0 | A | G | 0.211 | 3.21E-03 | 9.88E-02 | 2.64E+00 | 2.74E+00 | 9.70E-01 | no | Syne1 | intron_variant | MODIFIER |
| 10 | 5542724 | AX-170301728 | 0 | T | C | 0.211 | 3.21E-03 | 9.88E-02 | 2.64E+00 | 2.74E+00 | 9.70E-01 | no | Syne1 | intron_variant | MODIFIER |
| 10 | 5554356 | AX-170044063 | 0 | G | A | 0.211 | 3.21E-03 | 9.88E-02 | 2.64E+00 | 2.74E+00 | 9.70E-01 | no | Syne1 | upstream_gene_variant | MODIFIER |
| 10 | 5557681 | AX-97849382 | 0 | A | T | 0.211 | 3.21E-03 | 9.88E-02 | 2.64E+00 | 2.74E+00 | 9.70E-01 | no | | intergenic_variant | MODIFIER |
| 10 | 5560639 | AX-170433890 | 0 | C | A | 0.211 | 3.21E-03 | 9.88E-02 | 2.64E+00 | 2.74E+00 | 9.70E-01 | no | | intergenic_variant | MODIFIER |
| 10 | 5606075 | AX-170002636 | 0 | C | A | 0.211 | 3.21E-03 | 9.88E-02 | 2.64E+00 | 2.74E+00 | 9.70E-01 | no | Myct1 | 3_prime_UTR_variant | MODIFIER |
| 1 | 169448487 | AX-170043513 | 0 | A | G | 0.211 | -3.57E-03 | 1.31E-01 | 2.67E+00 | 2.75E+00 | 9.70E-01 | no | | intergenic_variant | MODIFIER |
| 13 | 47453602 | AX-170100971 | 0 | A | G | 0.465 | 2.97E-03 | 6.60E-02 | 2.63E+00 | 2.73E+00 | 9.70E-01 | no | | intergenic_variant | MODIFIER |
| 8 | 94216237 | AX-170080856 | 1 | G | A | 0.214 | -4.21E-03 | 1.17E-01 | 2.67E+00 | 2.74E+00 | 9.70E-01 | no | Nup93 | intron_variant | MODIFIER |
| 11 | 18586059 | AX-170035494 | 1 | C | T | 0.062 | 6.87E-03 | 2.15E-01 | 2.65E+00 | 2.74E+00 | 9.71E-01 | no | | intergenic_variant | MODIFIER |
| 11 | 13599235 | AX-170004324 | 0 | A | G | 0.07 | 6.59E-03 | 1.82E-01 | 2.63E+00 | 2.74E+00 | 9.71E-01 | no | | intergenic_variant | MODIFIER |
| 5 | 110590709 | AX-170444160 | 1 | T | G | 0.446 | 4.52E-03 | 1.12E-01 | 2.67E+00 | 2.73E+00 | 9.71E-01 | no | Galnt9 | intron_variant | MODIFIER |
| 9 | 86453803 | AX-97784064 | 1 | A | T | 0.134 | 2.69E-03 | 1.16E-01 | 2.65E+00 | 2.75E+00 | 9.71E-01 | no | Ube2cbp | intron_variant | MODIFIER |
| 8 | 83114342 | AX-170502523 | 0 | A | G | 0.211 | -4.25E-03 | 1.26E-01 | 2.67E+00 | 2.74E+00 | 9.71E-01 | no | | intergenic_variant | MODIFIER |
| 1 | 167518810 | AX-170448222 | 0 | A | G | 0.228 | -2.02E-03 | 1.10E-01 | 2.63E+00 | 2.75E+00 | 9.71E-01 | no | | intergenic_variant | MODIFIER |

| | | | | | | | | | | | | | | | |
|---|---|---|---|---|---|---|---|---|---|---|---|---|---|---|---|
| 1 | 19962830 | AX-170285339 | 0 | T | C | 0.404 | -2.19E-03 | 8.10E-02 | 2.69E+00 | 2.75E+00 | 9.71E-01 | no | | intergenic_variant | MODIFIER |
| 15 | 96931676 | AX-170254685 | 0 | G | A | 0.149 | -5.53E-03 | 1.80E-01 | 2.68E+00 | 2.74E+00 | 9.71E-01 | no | | intergenic_variant | MODIFIER |
| 4 | 149868293 | AX-170175581 | 0 | T | C | 0.096 | 7.53E-03 | 2.00E-01 | 2.67E+00 | 2.74E+00 | 9.71E-01 | no | | regulatory_region_variant | MODIFIER |
| 6 | 113269978 | AX-170389510 | 0 | C | T | 0.053 | -1.01E-02 | 2.08E-01 | 2.61E+00 | 2.73E+00 | 9.71E-01 | no | Mtmr14 | intron_variant | MODIFIER |
| 3 | 147685925 | AX-170461441 | 0 | A | G | 0.342 | -2.35E-03 | 7.43E-02 | 2.67E+00 | 2.74E+00 | 9.71E-01 | no | | intergenic_variant | MODIFIER |
| 13 | 53628302 | AX-170108512 | 1 | C | T | 0.5 | 3.61E-03 | 6.70E-02 | 2.62E+00 | 2.73E+00 | 9.71E-01 | no | | intergenic_variant | MODIFIER |
| 9 | 33693271 | AX-170370464 | 2 | G | A | 0.445 | 3.91E-03 | 8.77E-02 | 2.66E+00 | 2.73E+00 | 9.71E-01 | no | | intergenic_variant | MODIFIER |
| 5 | 127660973 | AX-170317133 | 1 | A | G | 0.42 | -2.15E-03 | 8.31E-02 | 2.64E+00 | 2.74E+00 | 9.71E-01 | no | Glt1d1 | intron_variant | MODIFIER |
| 20 | 140080162 | AX-97819885 | 0 | G | T | 0.149 | -4.29E-03 | 1.23E-01 | 2.64E+00 | 2.74E+00 | 9.71E-01 | no | Pih1h3b | intron_variant | MODIFIER |
| 7 | 106807711 | AX-97820039 | 1 | G | T | 0.464 | 3.81E-03 | 9.11E-02 | 2.66E+00 | 2.73E+00 | 9.71E-01 | no | Olfr700 | upstream_gene_variant | MODIFIER |
| 10 | 8998830 | AX-170262521 | 1 | A | T | 0.214 | -3.75E-03 | 9.47E-02 | 2.63E+00 | 2.74E+00 | 9.71E-01 | no | | regulatory_region_variant | MODIFIER |
| 1 | 13072211 | AX-170510193 | 1 | T | C | 0.366 | 3.30E-03 | 7.62E-02 | 2.66E+00 | 2.73E+00 | 9.71E-01 | no | Gm29283 | intron_variant,non_coding_transcript_variant | MODIFIER |
| 1 | 13278037 | AX-170419194 | 1 | T | C | 0.366 | 3.30E-03 | 7.62E-02 | 2.66E+00 | 2.73E+00 | 9.71E-01 | no | Ncoa2 | intron_variant | MODIFIER |
| 9 | 108968712 | AX-170453963 | 0 | G | A | 0.123 | -5.95E-03 | 1.18E-01 | 2.65E+00 | 2.73E+00 | 9.71E-01 | no | Col7a1 | intron_variant | MODIFIER |
| 7 | 103632687 | AX-170332659 | 0 | C | T | 0.491 | -2.21E-03 | 8.90E-02 | 2.65E+00 | 2.75E+00 | 9.71E-01 | no | Olfr621-ps1 | upstream_gene_variant | MODIFIER |
| 3 | 147546726 | AX-170021866 | 0 | C | G | 0.342 | -2.27E-03 | 7.34E-02 | 2.66E+00 | 2.74E+00 | 9.71E-01 | no | | intergenic_variant | MODIFIER |
| 4 | 56028381 | AX-170230687 | 0 | T | A | 0.368 | 3.67E-03 | 1.05E-01 | 2.66E+00 | 2.74E+00 | 9.71E-01 | no | | intergenic_variant | MODIFIER |
| 4 | 132429036 | AX-170123788 | 0 | A | T | 0.158 | -1.61E-03 | 1.03E-01 | 2.64E+00 | 2.75E+00 | 9.71E-01 | no | | regulatory_region_variant | MODIFIER |
| 4 | 136493989 | AX-170024031 | 0 | A | G | 0.158 | -1.61E-03 | 1.03E-01 | 2.64E+00 | 2.75E+00 | 9.71E-01 | no | Luzp1 | intron_variant | MODIFIER |
| 4 | 132365206 | AX-170210211 | 1 | G | A | 0.179 | 4.97E-03 | 1.06E-01 | 2.66E+00 | 2.73E+00 | 9.71E-01 | no | Phactr4 | intron_variant | MODIFIER |
| 1 | 25459510 | AX-97846146 | 0 | A | G | 0.412 | 3.62E-03 | 7.97E-02 | 2.64E+00 | 2.73E+00 | 9.72E-01 | no | Adgrb3 | intron_variant | MODIFIER |
| 7 | 94196551 | AX-170341458 | 0 | A | C | 0.123 | 7.13E-03 | 1.93E-01 | 2.65E+00 | 2.74E+00 | 9.72E-01 | no | Gm5899 | upstream_gene_variant | MODIFIER |
| 7 | 94943578 | AX-170029555 | 0 | T | G | 0.123 | 7.13E-03 | 1.93E-01 | 2.65E+00 | 2.74E+00 | 9.72E-01 | no | | intergenic_variant | MODIFIER |
| 1 | 39632984 | AX-170403365 | 1 | C | T | 0.393 | -3.26E-03 | 7.47E-02 | 2.65E+00 | 2.73E+00 | 9.72E-01 | no | Creg2 | intron_variant | MODIFIER |
| 1 | 59517034 | AX-170000595 | 0 | C | T | 0.105 | -8.10E-03 | 2.34E-01 | 2.69E+00 | 2.74E+00 | 9.72E-01 | no | Gm973 | intron_variant | MODIFIER |
| 16 | 27932607 | AX-170118884 | 0 | G | A | 0.246 | -2.60E-03 | 8.43E-02 | 2.61E+00 | 2.74E+00 | 9.72E-01 | no | | intergenic_variant | MODIFIER |
| 8 | 82319617 | AX-170395388 | 0 | A | T | 0.211 | -3.79E-03 | 1.13E-01 | 2.66E+00 | 2.74E+00 | 9.72E-01 | no | | intergenic_variant | MODIFIER |
| 9 | 105260590 | AX-170409197 | 1 | C | A | 0.125 | 2.88E-03 | 1.16E-01 | 2.67E+00 | 2.75E+00 | 9.72E-01 | no | Nek11 | intron_variant | MODIFIER |
| 12 | 72797135 | AX-97849897 | 1 | C | T | 0.062 | -1.09E-02 | 3.01E-01 | 2.71E+00 | 2.73E+00 | 9.72E-01 | no | Ppm1a | 3_prime_UTR_variant | MODIFIER |
| 15 | 86148114 | AX-170056191 | 0 | G | A | 0.518 | -1.10E-03 | 7.23E-02 | 2.64E+00 | 2.75E+00 | 9.72E-01 | no | Cerk | intron_variant | MODIFIER |
| 1 | 149158666 | AX-97844632 | 0 | C | T | 0.202 | -3.38E-03 | 1.30E-01 | 2.67E+00 | 2.75E+00 | 9.72E-01 | no | | intergenic_variant | MODIFIER |
| 1 | 150402686 | AX-170127252 | 0 | A | C | 0.202 | -3.38E-03 | 1.30E-01 | 2.67E+00 | 2.75E+00 | 9.72E-01 | no | Tpr | intron_variant | MODIFIER |
| 7 | 57155640 | AX-97781677 | 0 | C | T | 0.43 | -2.84E-03 | 9.89E-02 | 2.67E+00 | 2.74E+00 | 9.72E-01 | no | Gabrg3 | intron_variant | MODIFIER |
| 7 | 57630411 | AX-170422973 | 0 | T | C | 0.43 | -2.84E-03 | 9.89E-02 | 2.67E+00 | 2.74E+00 | 9.72E-01 | no | Gabrb3 | intron_variant | MODIFIER |
| 8 | 83381985 | AX-170080608 | 0 | G | A | 0.193 | -4.59E-03 | 1.26E-01 | 2.67E+00 | 2.74E+00 | 9.72E-01 | no | Mgat4d | downstream_gene_variant | MODIFIER |
| 6 | 118985646 | AX-170432234 | 0 | G | A | 0.07 | -9.51E-03 | 2.77E-01 | 2.70E+00 | 2.74E+00 | 9.72E-01 | no | Cacna1c | intron_variant | MODIFIER |
| 9 | 69921019 | AX-170191872 | 2 | T | C | 0.182 | -5.02E-03 | 1.37E-01 | 2.70E+00 | 2.73E+00 | 9.72E-01 | no | | intergenic_variant | MODIFIER |
| 7 | 41253994 | AX-170182083 | 2 | G | A | 0.327 | 4.11E-03 | 9.53E-02 | 2.65E+00 | 2.73E+00 | 9.72E-01 | no | Gm9244 | downstream_gene_variant | MODIFIER |
| 8 | 82544806 | AX-170080589 | 1 | G | A | 0.214 | -4.06E-03 | 1.25E-01 | 2.67E+00 | 2.74E+00 | 9.73E-01 | no | | intergenic_variant | MODIFIER |
| 7 | 10277476 | AX-170363766 | 0 | T | C | 0.167 | 1.12E-03 | 7.89E-02 | 2.60E+00 | 2.75E+00 | 9.73E-01 | no | Vmn1r66 | upstream_gene_variant | MODIFIER |
| 7 | 11194649 | AX-170432357 | 0 | A | G | 0.167 | 1.12E-03 | 7.89E-02 | 2.60E+00 | 2.75E+00 | 9.73E-01 | no | | intergenic_variant | MODIFIER |
| 1 | 53005920 | AX-170035922 | 0 | G | A | 0.096 | -7.49E-03 | 2.12E-01 | 2.67E+00 | 2.74E+00 | 9.73E-01 | no | 1700019D03Rik | intron_variant | MODIFIER |
| 7 | 111788328 | AX-170178725 | 2 | A | C | 0.473 | -2.72E-03 | 9.42E-02 | 2.68E+00 | 2.74E+00 | 9.73E-01 | no | Gm25787 | downstream_gene_variant | MODIFIER |
| 8 | 23321628 | AX-170374829 | 1 | A | G | 0.161 | -4.14E-03 | 1.10E-01 | 2.65E+00 | 2.74E+00 | 9.73E-01 | no | | regulatory_region_variant | MODIFIER |
| 13 | 25451095 | AX-170007947 | 1 | C | T | 0.17 | -6.87E-03 | 1.18E-01 | 2.63E+00 | 2.72E+00 | 9.73E-01 | no | Tpi-rs8 | upstream_gene_variant | MODIFIER |
| 8 | 81276906 | AX-170434530 | 1 | T | C | 0.205 | 4.44E-03 | 1.30E-01 | 2.68E+00 | 2.74E+00 | 9.73E-01 | no | | intergenic_variant | MODIFIER |
| 8 | 81527356 | AX-170424169 | 1 | C | A | 0.205 | 4.44E-03 | 1.30E-01 | 2.68E+00 | 2.74E+00 | 9.73E-01 | no | Inpp4b | intron_variant,NMD_transcript_variant | MODIFIER |
| 2 | 121121831 | AX-170361912 | 0 | A | G | 0.14 | 5.96E-03 | 1.84E-01 | 2.67E+00 | 2.74E+00 | 9.73E-01 | no | | intergenic_variant | MODIFIER |
| 4 | 137464207 | AX-170226115 | 0 | T | C | 0.167 | -1.64E-03 | 1.04E-01 | 2.65E+00 | 2.75E+00 | 9.73E-01 | no | Hspg2 | upstream_gene_variant | MODIFIER |
| 10 | 4699796 | AX-170369259 | 0 | A | G | 0.193 | -3.48E-03 | 9.44E-02 | 2.63E+00 | 2.74E+00 | 9.73E-01 | no | Esr1 | intron_variant | MODIFIER |
| 1 | 63397754 | AX-170448547 | 0 | C | T | 0.14 | 7.43E-03 | 1.96E-01 | 2.64E+00 | 2.74E+00 | 9.73E-01 | no | | intergenic_variant | MODIFIER |
| 6 | 145039288 | AX-170076536 | 1 | A | G | 0.062 | -8.16E-03 | 2.25E-01 | 2.65E+00 | 2.74E+00 | 9.73E-01 | no | Bcat1 | intron_variant | MODIFIER |
| 13 | 38500515 | AX-97826376 | 2 | C | T | 0.2 | -5.88E-03 | 1.09E-01 | 2.63E+00 | 2.73E+00 | 9.73E-01 | no | Txndc5 | 3_prime_UTR_variant | MODIFIER |
| 4 | 143706533 | AX-170070932 | 2 | A | T | 0.064 | -6.03E-03 | 1.91E-01 | 2.61E+00 | 2.74E+00 | 9.73E-01 | no | Gm13089 | upstream_gene_variant | MODIFIER |

| | | | | | | | | | | | | | | | |
|---|---|---|---|---|---|---|---|---|---|---|---|---|---|---|---|
| 3 | 148736093 | AX-170086000 | 0 | A | T | 0.316 | -2.00E-03 | 7.78E-02 | 2.66E+00 | 2.74E+00 | 9.73E-01 | no | | intergenic_variant | MODIFIER |
| 5 | 37519437 | AX-170218340 | 0 | G | A | 0.325 | -2.56E-03 | 1.03E-01 | 2.65E+00 | 2.74E+00 | 9.73E-01 | no | Stk32b | intron_variant | MODIFIER |
| 7 | 52979158 | AX-170077362 | 1 | C | T | 0.446 | -2.45E-03 | 9.69E-02 | 2.65E+00 | 2.74E+00 | 9.73E-01 | no | | intergenic_variant | MODIFIER |
| 11 | 49486010 | AX-97818361 | 0 | G | A | 0.088 | -7.51E-03 | 1.82E-01 | 2.63E+00 | 2.73E+00 | 9.74E-01 | no | Olfr1385 | upstream_gene_variant | MODIFIER |
| 8 | 93349082 | AX-170242761 | 1 | G | A | 0.214 | -4.00E-03 | 1.17E-01 | 2.67E+00 | 2.74E+00 | 9.74E-01 | no | Ces1h | downstream_gene_variant | MODIFIER |
| 7 | 69542362 | AX-97790229 | 0 | G | T | 0.421 | 4.19E-03 | 1.01E-01 | 2.66E+00 | 2.73E+00 | 9.74E-01 | no | 4930402F11Rik | intron_variant,non_coding_transcript_variant | MODIFIER |
| 7 | 69542954 | AX-170343414 | 0 | C | T | 0.421 | 4.19E-03 | 1.01E-01 | 2.66E+00 | 2.73E+00 | 9.74E-01 | no | 4930402F11Rik | intron_variant,non_coding_transcript_variant | MODIFIER |
| 7 | 69551968 | AX-170145067 | 0 | C | T | 0.421 | 4.19E-03 | 1.01E-01 | 2.66E+00 | 2.73E+00 | 9.74E-01 | no | 4930402F11Rik | downstream_gene_variant | MODIFIER |
| 7 | 69552684 | AX-170077650 | 0 | T | C | 0.421 | 4.19E-03 | 1.01E-01 | 2.66E+00 | 2.73E+00 | 9.74E-01 | no | 4930402F11Rik | downstream_gene_variant | MODIFIER |
| 7 | 69554483 | AX-170077652 | 0 | A | C | 0.421 | 4.19E-03 | 1.01E-01 | 2.66E+00 | 2.73E+00 | 9.74E-01 | no | 4930402F11Rik | downstream_gene_variant | MODIFIER |
| 7 | 69555407 | AX-170254606 | 0 | C | A | 0.421 | 4.19E-03 | 1.01E-01 | 2.66E+00 | 2.73E+00 | 9.74E-01 | no | 4930402F11Rik | downstream_gene_variant | MODIFIER |
| 7 | 69555443 | AX-170456095 | 0 | A | G | 0.421 | 4.19E-03 | 1.01E-01 | 2.66E+00 | 2.73E+00 | 9.74E-01 | no | 4930402F11Rik | downstream_gene_variant | MODIFIER |
| 7 | 69593034 | AX-170226929 | 0 | G | A | 0.421 | 4.19E-03 | 1.01E-01 | 2.66E+00 | 2.73E+00 | 9.74E-01 | no | | intergenic_variant | MODIFIER |
| 7 | 69766684 | AX-170035757 | 0 | C | T | 0.421 | 4.19E-03 | 1.01E-01 | 2.66E+00 | 2.73E+00 | 9.74E-01 | no | | regulatory_region_variant | MODIFIER |
| 7 | 69964133 | AX-170490640 | 0 | C | A | 0.421 | 4.19E-03 | 1.01E-01 | 2.66E+00 | 2.73E+00 | 9.74E-01 | no | | intergenic_variant | MODIFIER |
| 7 | 69983851 | AX-97837231 | 0 | T | C | 0.421 | 4.19E-03 | 1.01E-01 | 2.66E+00 | 2.73E+00 | 9.74E-01 | no | | intergenic_variant | MODIFIER |
| 7 | 70030120 | AX-170213423 | 0 | G | T | 0.421 | 4.19E-03 | 1.01E-01 | 2.66E+00 | 2.73E+00 | 9.74E-01 | no | | intergenic_variant | MODIFIER |
| 7 | 70030205 | AX-170190003 | 0 | A | C | 0.421 | 4.19E-03 | 1.01E-01 | 2.66E+00 | 2.73E+00 | 9.74E-01 | no | | intergenic_variant | MODIFIER |
| 10 | 3180584 | AX-170274385 | 0 | T | C | 0.202 | -3.27E-03 | 9.57E-02 | 2.63E+00 | 2.74E+00 | 9.74E-01 | no | Gm48097 | downstream_gene_variant | MODIFIER |
| 10 | 3195232 | AX-170292848 | 0 | C | T | 0.202 | -3.27E-03 | 9.57E-02 | 2.63E+00 | 2.74E+00 | 9.74E-01 | no | | intergenic_variant | MODIFIER |
| 10 | 3377740 | AX-170201946 | 0 | T | G | 0.202 | -3.27E-03 | 9.57E-02 | 2.63E+00 | 2.74E+00 | 9.74E-01 | no | Ppp1r14c | intron_variant | MODIFIER |
| 10 | 3381827 | AX-170044079 | 0 | T | G | 0.202 | -3.27E-03 | 9.57E-02 | 2.63E+00 | 2.74E+00 | 9.74E-01 | no | Ppp1r14c | intron_variant | MODIFIER |
| 10 | 3417409 | AX-170455208 | 0 | A | G | 0.202 | -3.27E-03 | 9.57E-02 | 2.63E+00 | 2.74E+00 | 9.74E-01 | no | Ppp1r14c | intron_variant | MODIFIER |
| 10 | 3419681 | AX-170209331 | 0 | C | T | 0.202 | -3.27E-03 | 9.57E-02 | 2.63E+00 | 2.74E+00 | 9.74E-01 | no | Ppp1r14c | intron_variant | MODIFIER |
| 10 | 3420310 | AX-170002652 | 0 | G | A | 0.202 | -3.27E-03 | 9.57E-02 | 2.63E+00 | 2.74E+00 | 9.74E-01 | no | Ppp1r14c | intron_variant | MODIFIER |
| 10 | 3444691 | AX-170002654 | 0 | C | T | 0.202 | -3.27E-03 | 9.57E-02 | 2.63E+00 | 2.74E+00 | 9.74E-01 | no | Ppp1r14c | intron_variant | MODIFIER |
| 10 | 3450230 | AX-170347490 | 0 | A | G | 0.202 | -3.27E-03 | 9.57E-02 | 2.63E+00 | 2.74E+00 | 9.74E-01 | no | Ppp1r14c | intron_variant | MODIFIER |
| 10 | 3454685 | AX-170523408 | 0 | C | A | 0.202 | -3.27E-03 | 9.57E-02 | 2.63E+00 | 2.74E+00 | 9.74E-01 | no | Ppp1r14c | intron_variant | MODIFIER |
| 10 | 3490545 | AX-170460833 | 0 | C | T | 0.202 | -3.27E-03 | 9.57E-02 | 2.63E+00 | 2.74E+00 | 9.74E-01 | no | | intergenic_variant | MODIFIER |
| 10 | 3514188 | AX-170507201 | 0 | G | A | 0.202 | -3.27E-03 | 9.57E-02 | 2.63E+00 | 2.74E+00 | 9.74E-01 | no | | intergenic_variant | MODIFIER |
| 10 | 3856986 | AX-170249693 | 0 | C | T | 0.202 | -3.27E-03 | 9.57E-02 | 2.63E+00 | 2.74E+00 | 9.74E-01 | no | Plekhg1 | intron_variant | MODIFIER |
| 10 | 3901930 | AX-170414170 | 0 | C | T | 0.202 | -3.27E-03 | 9.57E-02 | 2.63E+00 | 2.74E+00 | 9.74E-01 | no | Plekhg1 | intron_variant | MODIFIER |
| 10 | 3901982 | AX-170167370 | 0 | G | A | 0.202 | -3.27E-03 | 9.57E-02 | 2.63E+00 | 2.74E+00 | 9.74E-01 | no | Plekhg1 | intron_variant | MODIFIER |
| 10 | 3908415 | AX-170044077 | 0 | T | C | 0.202 | -3.27E-03 | 9.57E-02 | 2.63E+00 | 2.74E+00 | 9.74E-01 | no | Plekhg1 | intron_variant | MODIFIER |
| 10 | 3950733 | AX-170468349 | 0 | T | G | 0.202 | -3.27E-03 | 9.57E-02 | 2.63E+00 | 2.74E+00 | 9.74E-01 | no | Plekhg1 | intron_variant | MODIFIER |
| 10 | 4080358 | AX-170533724 | 0 | T | C | 0.202 | -3.27E-03 | 9.57E-02 | 2.63E+00 | 2.74E+00 | 9.74E-01 | no | Mthfd1l | intron_variant | MODIFIER |
| 10 | 4080404 | AX-170251755 | 0 | A | G | 0.202 | -3.27E-03 | 9.57E-02 | 2.63E+00 | 2.74E+00 | 9.74E-01 | no | Mthfd1l | intron_variant | MODIFIER |
| 10 | 4097937 | AX-170453717 | 0 | G | A | 0.202 | -3.27E-03 | 9.57E-02 | 2.63E+00 | 2.74E+00 | 9.74E-01 | no | Mthfd1l | intron_variant | MODIFIER |
| 12 | 61232407 | AX-170049198 | 1 | G | C | 0.062 | -1.01E-02 | 3.03E-01 | 2.72E+00 | 2.74E+00 | 9.74E-01 | no | Gm48301 | intron_variant,non_coding_transcript_variant | MODIFIER |
| 7 | 94506625 | AX-170437706 | 2 | C | T | 0.136 | -6.42E-03 | 2.13E-01 | 2.66E+00 | 2.74E+00 | 9.74E-01 | no | Gm32647 | intron_variant,non_coding_transcript_variant | MODIFIER |
| 18 | 60975856 | AX-170459190 | 0 | T | C | 0.412 | 1.86E-03 | 8.31E-02 | 2.68E+00 | 2.75E+00 | 9.74E-01 | no | Camk2a | intron_variant | MODIFIER |
| 1 | 25451498 | AX-170339719 | 1 | A | G | 0.393 | 2.98E-03 | 8.26E-02 | 2.66E+00 | 2.74E+00 | 9.74E-01 | no | Adgrb3 | intron_variant | MODIFIER |
| 8 | 101626823 | AX-170506090 | 2 | G | A | 0.227 | -2.89E-03 | 1.03E-01 | 2.65E+00 | 2.74E+00 | 9.74E-01 | no | | intergenic_variant | MODIFIER |
| 9 | 52429452 | AX-170225839 | 1 | A | G | 0.214 | -3.83E-03 | 1.16E-01 | 2.67E+00 | 2.74E+00 | 9.74E-01 | no | AC166902.1 | intron_variant,non_coding_transcript_variant | MODIFIER |
| 11 | 74754966 | AX-170461624 | 0 | A | G | 0.237 | -2.23E-03 | 7.14E-02 | 2.60E+00 | 2.74E+00 | 9.74E-01 | no | | intergenic_variant | MODIFIER |
| 1 | 9883920 | AX-170255346 | 1 | T | G | 0.402 | 2.95E-03 | 7.07E-02 | 2.66E+00 | 2.73E+00 | 9.74E-01 | no | Sgk3 | intron_variant | MODIFIER |
| 9 | 87302854 | AX-170483039 | 1 | A | G | 0.143 | 2.78E-03 | 1.25E-01 | 2.66E+00 | 2.75E+00 | 9.74E-01 | no | | intergenic_variant | MODIFIER |
| 20 | 92685999 | AX-170034422 | 1 | C | T | 0.277 | 2.27E-03 | 5.97E-02 | 2.63E+00 | 2.74E+00 | 9.74E-01 | no | | intergenic_variant | MODIFIER |
| 8 | 87125714 | AX-170101100 | 0 | A | G | 0.211 | -4.49E-03 | 1.16E-01 | 2.63E+00 | 2.73E+00 | 9.74E-01 | no | Gm27246 | intron_variant,non_coding_transcript_variant | MODIFIER |
| 7 | 88331377 | AX-170469648 | 1 | C | G | 0.125 | 6.01E-03 | 1.84E-01 | 2.63E+00 | 2.74E+00 | 9.74E-01 | no | | intergenic_variant | MODIFIER |
| 8 | 45072521 | AX-170417793 | 0 | A | T | 0.14 | 3.19E-03 | 1.38E-01 | 2.68E+00 | 2.75E+00 | 9.74E-01 | no | Mtnr1a | intron_variant | MODIFIER |
| 10 | 6554162 | AX-170514237 | 0 | T | C | 0.202 | -3.53E-03 | 1.03E-01 | 2.64E+00 | 2.74E+00 | 9.74E-01 | no | | intergenic_variant | MODIFIER |
| 1 | 54486428 | AX-170244845 | 1 | C | T | 0.098 | -7.15E-03 | 2.20E-01 | 2.68E+00 | 2.74E+00 | 9.74E-01 | no | Pgap1 | intron_variant | MODIFIER |
| 1 | 60609441 | AX-170041390 | 1 | T | A | 0.098 | -7.15E-03 | 2.20E-01 | 2.68E+00 | 2.74E+00 | 9.74E-01 | no | Gm23762 | downstream_gene_variant | MODIFIER |

| | | | | | | | | | | | | | | | |
|---|---|---|---|---|---|---|---|---|---|---|---|---|---|---|---|
| 16 | 92570970 | AX-170522786 | 1 | C | G | 0.464 | -2.35E-03 | 7.77E-02 | 2.66E+00 | 2.74E+00 | 9.75E-01 | no | | intergenic_variant | MODIFIER |
| 10 | 5578938 | AX-170238549 | 1 | C | T | 0.214 | -3.41E-03 | 9.76E-02 | 2.64E+00 | 2.74E+00 | 9.75E-01 | no | | intergenic_variant | MODIFIER |
| 4 | 143989482 | AX-170024167 | 0 | G | C | 0.061 | 5.07E-03 | 1.88E-01 | 2.62E+00 | 2.74E+00 | 9.75E-01 | no | Pramef17 | downstream_gene_variant | MODIFIER |
| 15 | 78288377 | AX-170056056 | 0 | C | A | 0.553 | -1.76E-03 | 8.21E-02 | 2.67E+00 | 2.75E+00 | 9.75E-01 | no | Csf2rb2 | intron_variant | MODIFIER |
| 13 | 53545895 | AX-170179518 | 0 | G | A | 0.526 | -1.09E-03 | 6.81E-02 | 2.64E+00 | 2.75E+00 | 9.75E-01 | no | | intergenic_variant | MODIFIER |
| 13 | 53721608 | AX-170242169 | 0 | C | T | 0.526 | -1.09E-03 | 6.81E-02 | 2.64E+00 | 2.75E+00 | 9.75E-01 | no | | intergenic_variant | MODIFIER |
| 13 | 53730436 | AX-97853331 | 0 | A | G | 0.526 | -1.09E-03 | 6.81E-02 | 2.64E+00 | 2.75E+00 | 9.75E-01 | no | | intergenic_variant | MODIFIER |
| 13 | 53736687 | AX-170180663 | 0 | G | A | 0.526 | -1.09E-03 | 6.81E-02 | 2.64E+00 | 2.75E+00 | 9.75E-01 | no | | intergenic_variant | MODIFIER |
| 13 | 53748413 | AX-170277288 | 0 | G | C | 0.526 | -1.09E-03 | 6.81E-02 | 2.64E+00 | 2.75E+00 | 9.75E-01 | no | | intergenic_variant | MODIFIER |
| 13 | 53748483 | AX-170092755 | 0 | A | G | 0.526 | -1.09E-03 | 6.81E-02 | 2.64E+00 | 2.75E+00 | 9.75E-01 | no | | intergenic_variant | MODIFIER |
| 13 | 53759256 | AX-170198514 | 0 | C | T | 0.526 | -1.09E-03 | 6.81E-02 | 2.64E+00 | 2.75E+00 | 9.75E-01 | no | | intergenic_variant | MODIFIER |
| 13 | 53759931 | AX-170165995 | 0 | T | C | 0.526 | -1.09E-03 | 6.81E-02 | 2.64E+00 | 2.75E+00 | 9.75E-01 | no | | intergenic_variant | MODIFIER |
| 13 | 53761064 | AX-97810514 | 0 | C | T | 0.526 | -1.09E-03 | 6.81E-02 | 2.64E+00 | 2.75E+00 | 9.75E-01 | no | | intergenic_variant | MODIFIER |
| 13 | 53763952 | AX-170278093 | 0 | C | T | 0.526 | -1.09E-03 | 6.81E-02 | 2.64E+00 | 2.75E+00 | 9.75E-01 | no | | intergenic_variant | MODIFIER |
| 13 | 53767440 | AX-170253572 | 0 | A | G | 0.526 | -1.09E-03 | 6.81E-02 | 2.64E+00 | 2.75E+00 | 9.75E-01 | no | | intergenic_variant | MODIFIER |
| 13 | 53782415 | AX-170360496 | 0 | G | A | 0.526 | -1.09E-03 | 6.81E-02 | 2.64E+00 | 2.75E+00 | 9.75E-01 | no | Gm48548 | downstream_gene_variant | MODIFIER |
| 13 | 53904112 | AX-170457840 | 0 | G | A | 0.526 | -1.09E-03 | 6.81E-02 | 2.64E+00 | 2.75E+00 | 9.75E-01 | no | | regulatory_region_variant | MODIFIER |
| 13 | 54251183 | AX-97815981 | 0 | A | C | 0.526 | -1.09E-03 | 6.81E-02 | 2.64E+00 | 2.75E+00 | 9.75E-01 | no | | intergenic_variant | MODIFIER |
| 13 | 54329182 | AX-170008344 | 0 | A | C | 0.526 | -1.09E-03 | 6.81E-02 | 2.64E+00 | 2.75E+00 | 9.75E-01 | no | Gm48622 | upstream_gene_variant | MODIFIER |
| 13 | 54344085 | AX-170244550 | 0 | G | A | 0.526 | -1.09E-03 | 6.81E-02 | 2.64E+00 | 2.75E+00 | 9.75E-01 | no | | intergenic_variant | MODIFIER |
| 13 | 54344364 | AX-97835135 | 0 | C | T | 0.526 | -1.09E-03 | 6.81E-02 | 2.64E+00 | 2.75E+00 | 9.75E-01 | no | | intergenic_variant | MODIFIER |
| 13 | 54345689 | AX-170485826 | 0 | G | A | 0.526 | -1.09E-03 | 6.81E-02 | 2.64E+00 | 2.75E+00 | 9.75E-01 | no | | intergenic_variant | MODIFIER |
| 13 | 54345830 | AX-170051337 | 0 | G | C | 0.526 | -1.09E-03 | 6.81E-02 | 2.64E+00 | 2.75E+00 | 9.75E-01 | no | | regulatory_region_variant | MODIFIER |
| 8 | 90417312 | AX-170345539 | 1 | G | A | 0.196 | 2.74E-03 | 1.16E-01 | 2.66E+00 | 2.74E+00 | 9.75E-01 | no | | intergenic_variant | MODIFIER |
| 9 | 105485685 | AX-170355191 | 1 | T | A | 0.125 | 2.43E-03 | 1.17E-01 | 2.68E+00 | 2.75E+00 | 9.75E-01 | no | Atp2c1 | intron_variant | MODIFIER |
| 5 | 73186660 | AX-97841521 | 0 | G | A | 0.447 | 4.24E-03 | 9.39E-02 | 2.63E+00 | 2.73E+00 | 9.75E-01 | no | Fryl | intron_variant | MODIFIER |
| 13 | 42472941 | AX-170459780 | 0 | T | A | 0.5 | 2.87E-03 | 5.97E-02 | 2.62E+00 | 2.73E+00 | 9.75E-01 | no | Gm6049 | upstream_gene_variant | MODIFIER |
| 10 | 114788365 | AX-170100619 | 0 | C | A | 0.36 | -2.63E-03 | 7.72E-02 | 2.65E+00 | 2.74E+00 | 9.75E-01 | no | Trhde | intron_variant | MODIFIER |
| 8 | 102616366 | AX-170090198 | 2 | A | G | 0.209 | -3.88E-03 | 1.19E-01 | 2.68E+00 | 2.74E+00 | 9.75E-01 | no | | intergenic_variant | MODIFIER |
| 2 | 155107749 | AX-97795922 | 1 | G | A | 0.143 | 6.82E-03 | 1.70E-01 | 2.64E+00 | 2.73E+00 | 9.75E-01 | no | | regulatory_region_variant | MODIFIER |
| 18 | 36759191 | AX-97787338 | 0 | A | G | 0.061 | -9.73E-03 | 2.98E-01 | 2.71E+00 | 2.73E+00 | 9.75E-01 | no | Ik | downstream_gene_variant | MODIFIER |
| 7 | 93388652 | AX-170029534 | 1 | T | C | 0.134 | -6.07E-03 | 2.11E-01 | 2.66E+00 | 2.74E+00 | 9.75E-01 | no | | intergenic_variant | MODIFIER |
| 8 | 73058746 | AX-170337192 | 1 | C | A | 0.196 | -4.36E-03 | 1.33E-01 | 2.67E+00 | 2.74E+00 | 9.75E-01 | no | Large1 | intron_variant | MODIFIER |
| 12 | 101159301 | AX-170146934 | 1 | C | T | 0.098 | -3.67E-03 | 1.65E-01 | 2.65E+00 | 2.74E+00 | 9.75E-01 | no | | intergenic_variant | MODIFIER |
| 5 | 73252271 | AX-97814692 | 1 | C | T | 0.438 | 3.74E-03 | 9.39E-02 | 2.64E+00 | 2.73E+00 | 9.75E-01 | no | Fryl | intron_variant | MODIFIER |
| 8 | 47372489 | AX-170212463 | 1 | G | A | 0.152 | 3.95E-03 | 1.50E-01 | 2.70E+00 | 2.74E+00 | 9.75E-01 | no | Stox2 | intron_variant,non_coding_transcript_variant | MODIFIER |
| 8 | 23329713 | AX-170289446 | 1 | G | A | 0.188 | 2.70E-03 | 9.96E-02 | 2.64E+00 | 2.74E+00 | 9.75E-01 | no | | intergenic_variant | MODIFIER |
| 2 | 117401542 | AX-170340415 | 1 | G | A | 0.125 | 6.38E-03 | 1.95E-01 | 2.67E+00 | 2.74E+00 | 9.75E-01 | no | | intergenic_variant | MODIFIER |
| 10 | 127246085 | AX-170427726 | 0 | C | T | 0.325 | 1.77E-03 | 7.25E-02 | 2.64E+00 | 2.74E+00 | 9.75E-01 | no | Kif5a | intron_variant | MODIFIER |
| 13 | 40134240 | AX-170313320 | 0 | A | C | 0.184 | 6.85E-04 | 9.93E-02 | 2.64E+00 | 2.75E+00 | 9.75E-01 | no | Ofcc1 | intron_variant | MODIFIER |
| 7 | 137860238 | AX-168415453 | 0 | A | G | 0.307 | -2.88E-03 | 7.74E-02 | 2.63E+00 | 2.73E+00 | 9.75E-01 | no | | intergenic_variant | MODIFIER |
| 7 | 138732565 | AX-170183797 | 0 | T | A | 0.307 | -2.88E-03 | 7.74E-02 | 2.63E+00 | 2.73E+00 | 9.75E-01 | no | | intergenic_variant | MODIFIER |
| 7 | 138742232 | AX-170079024 | 0 | T | G | 0.307 | -2.88E-03 | 7.74E-02 | 2.63E+00 | 2.73E+00 | 9.75E-01 | no | | regulatory_region_variant | MODIFIER |
| 1 | 59610624 | AX-170257864 | 1 | A | G | 0.107 | 6.08E-03 | 1.71E-01 | 2.61E+00 | 2.74E+00 | 9.76E-01 | no | Gm973 | intron_variant | MODIFIER |
| 9 | 78651173 | AX-170406722 | 1 | T | C | 0.196 | 3.95E-03 | 1.39E-01 | 2.72E+00 | 2.74E+00 | 9.76E-01 | no | Cd109 | intron_variant | MODIFIER |
| 15 | 77912558 | AX-170354803 | 0 | C | T | 0.553 | 3.47E-03 | 8.28E-02 | 2.66E+00 | 2.73E+00 | 9.76E-01 | no | Txn2 | downstream_gene_variant | MODIFIER |
| 15 | 77972006 | AX-170087916 | 0 | C | T | 0.553 | 3.47E-03 | 8.28E-02 | 2.66E+00 | 2.73E+00 | 9.76E-01 | no | Eif3d | upstream_gene_variant | MODIFIER |
| 15 | 78012444 | AX-97829788 | 0 | C | T | 0.553 | 3.47E-03 | 8.28E-02 | 2.66E+00 | 2.73E+00 | 9.76E-01 | no | Cacng2 | intron_variant | MODIFIER |
| 15 | 78065630 | AX-170374175 | 0 | C | A | 0.553 | 3.47E-03 | 8.28E-02 | 2.66E+00 | 2.73E+00 | 9.76E-01 | no | Cacng2 | intron_variant | MODIFIER |
| 15 | 78069369 | AX-97828322 | 0 | G | A | 0.553 | 3.47E-03 | 8.28E-02 | 2.66E+00 | 2.73E+00 | 9.76E-01 | no | Cacng2 | intron_variant | MODIFIER |
| 15 | 78102542 | AX-170012040 | 0 | C | A | 0.553 | 3.47E-03 | 8.28E-02 | 2.66E+00 | 2.73E+00 | 9.76E-01 | no | Cacng2 | intron_variant | MODIFIER |
| 15 | 78111142 | AX-97839662 | 0 | C | G | 0.553 | 3.47E-03 | 8.28E-02 | 2.66E+00 | 2.73E+00 | 9.76E-01 | no | Cacng2 | intron_variant | MODIFIER |
| 15 | 78153058 | AX-168410390 | 0 | G | A | 0.553 | 3.47E-03 | 8.28E-02 | 2.66E+00 | 2.73E+00 | 9.76E-01 | no | | regulatory_region_variant | MODIFIER |
| 15 | 78186940 | AX-170056054 | 0 | T | C | 0.553 | 3.47E-03 | 8.28E-02 | 2.66E+00 | 2.73E+00 | 9.76E-01 | no | Pvalb | downstream_gene_variant | MODIFIER |

| | | | | | | | | | | | | | | | |
|---|---|---|---|---|---|---|---|---|---|---|---|---|---|---|---|
| 15 | 78190293 | AX-170012045 | 0 | G | A | 0.553 | 3.47E-03 | 8.28E-02 | 2.66E+00 | 2.73E+00 | 9.76E-01 | no | Pvalb | downstream_gene_variant | MODIFIER |
| 15 | 78190745 | AX-170121590 | 0 | G | A | 0.553 | 3.47E-03 | 8.28E-02 | 2.66E+00 | 2.73E+00 | 9.76E-01 | no | Pvalb | downstream_gene_variant | MODIFIER |
| 15 | 78224968 | AX-170121805 | 0 | T | C | 0.553 | 3.47E-03 | 8.28E-02 | 2.66E+00 | 2.73E+00 | 9.76E-01 | no | | intergenic_variant | MODIFIER |
| 15 | 78234670 | AX-170304061 | 0 | T | A | 0.553 | 3.47E-03 | 8.28E-02 | 2.66E+00 | 2.73E+00 | 9.76E-01 | no | | regulatory_region_variant | MODIFIER |
| 15 | 78237257 | AX-170492598 | 0 | T | G | 0.553 | 3.47E-03 | 8.28E-02 | 2.66E+00 | 2.73E+00 | 9.76E-01 | no | | intergenic_variant | MODIFIER |
| 15 | 78237280 | AX-170391284 | 0 | T | C | 0.553 | 3.47E-03 | 8.28E-02 | 2.66E+00 | 2.73E+00 | 9.76E-01 | no | | intergenic_variant | MODIFIER |
| 15 | 78245697 | AX-170336102 | 0 | A | C | 0.553 | 3.47E-03 | 8.28E-02 | 2.66E+00 | 2.73E+00 | 9.76E-01 | no | Ncf4 | intron_variant | MODIFIER |
| 15 | 78245965 | AX-170492419 | 0 | T | C | 0.553 | 3.47E-03 | 8.28E-02 | 2.66E+00 | 2.73E+00 | 9.76E-01 | no | Ncf4 | intron_variant | MODIFIER |
| 15 | 78262111 | AX-170264627 | 0 | A | G | 0.553 | 3.47E-03 | 8.28E-02 | 2.66E+00 | 2.73E+00 | 9.76E-01 | no | Ncf4 | intron_variant | MODIFIER |
| 15 | 78264038 | AX-170281520 | 0 | A | G | 0.553 | 3.47E-03 | 8.28E-02 | 2.66E+00 | 2.73E+00 | 9.76E-01 | no | Ncf4 | downstream_gene_variant | MODIFIER |
| 15 | 78279750 | AX-97829724 | 0 | C | T | 0.553 | 3.47E-03 | 8.28E-02 | 2.66E+00 | 2.73E+00 | 9.76E-01 | no | Csf2rb2 | downstream_gene_variant | MODIFIER |
| 15 | 78281277 | AX-170279280 | 0 | C | G | 0.553 | 3.47E-03 | 8.28E-02 | 2.66E+00 | 2.73E+00 | 9.76E-01 | no | Csf2rb2 | downstream_gene_variant | MODIFIER |
| 15 | 78292885 | AX-170368809 | 0 | G | A | 0.553 | 3.47E-03 | 8.28E-02 | 2.66E+00 | 2.73E+00 | 9.76E-01 | no | Csf2rb2 | intron_variant | MODIFIER |
| 15 | 78300241 | AX-170181109 | 0 | C | A | 0.553 | 3.47E-03 | 8.28E-02 | 2.66E+00 | 2.73E+00 | 9.76E-01 | no | Csf2rb2 | intron_variant | MODIFIER |
| 15 | 78320285 | AX-170521235 | 0 | G | A | 0.553 | 3.47E-03 | 8.28E-02 | 2.66E+00 | 2.73E+00 | 9.76E-01 | no | | regulatory_region_variant | MODIFIER |
| 15 | 78332041 | AX-97812604 | 0 | A | G | 0.553 | 3.47E-03 | 8.28E-02 | 2.66E+00 | 2.73E+00 | 9.76E-01 | no | Csf2rb | intron_variant | MODIFIER |
| 15 | 78332283 | AX-170458277 | 0 | T | A | 0.553 | 3.47E-03 | 8.28E-02 | 2.66E+00 | 2.73E+00 | 9.76E-01 | no | Csf2rb | intron_variant | MODIFIER |
| 15 | 78338293 | AX-170282639 | 0 | T | C | 0.553 | 3.47E-03 | 8.28E-02 | 2.66E+00 | 2.73E+00 | 9.76E-01 | no | Csf2rb | intron_variant | MODIFIER |
| 15 | 78340721 | AX-170419821 | 0 | C | A | 0.553 | 3.47E-03 | 8.28E-02 | 2.66E+00 | 2.73E+00 | 9.76E-01 | no | Csf2rb | missense_variant | MODERATE |
| 15 | 78351738 | AX-97826098 | 0 | C | T | 0.553 | 3.47E-03 | 8.28E-02 | 2.66E+00 | 2.73E+00 | 9.76E-01 | no | Csf2rb | downstream_gene_variant | MODIFIER |
| 15 | 78381619 | AX-170161413 | 0 | A | G | 0.553 | 3.47E-03 | 8.28E-02 | 2.66E+00 | 2.73E+00 | 9.76E-01 | no | Tex33 | intron_variant | MODIFIER |
| 15 | 78516240 | AX-170489419 | 0 | A | G | 0.553 | 3.47E-03 | 8.28E-02 | 2.66E+00 | 2.73E+00 | 9.76E-01 | no | | regulatory_region_variant | MODIFIER |
| 15 | 78516336 | AX-170329965 | 0 | T | C | 0.553 | 3.47E-03 | 8.28E-02 | 2.66E+00 | 2.73E+00 | 9.76E-01 | no | | regulatory_region_variant | MODIFIER |
| 15 | 78520336 | AX-97830477 | 0 | C | T | 0.553 | 3.47E-03 | 8.28E-02 | 2.66E+00 | 2.73E+00 | 9.76E-01 | no | C1qtnf6 | downstream_gene_variant | MODIFIER |
| 15 | 78535845 | AX-170149101 | 0 | G | A | 0.553 | 3.47E-03 | 8.28E-02 | 2.66E+00 | 2.73E+00 | 9.76E-01 | no | Sstr3 | downstream_gene_variant | MODIFIER |
| 15 | 78539253 | AX-97817878 | 0 | C | A | 0.553 | 3.47E-03 | 8.28E-02 | 2.66E+00 | 2.73E+00 | 9.76E-01 | no | Sstr3 | 3_prime_UTR_variant | MODIFIER |
| 15 | 78539610 | AX-170121521 | 0 | A | G | 0.553 | 3.47E-03 | 8.28E-02 | 2.66E+00 | 2.73E+00 | 9.76E-01 | no | Sstr3 | synonymous_variant | LOW |
| 15 | 78539925 | AX-170372931 | 0 | A | G | 0.553 | 3.47E-03 | 8.28E-02 | 2.66E+00 | 2.73E+00 | 9.76E-01 | no | Sstr3 | synonymous_variant | LOW |
| 15 | 78541621 | AX-170301701 | 0 | A | G | 0.553 | 3.47E-03 | 8.28E-02 | 2.66E+00 | 2.73E+00 | 9.76E-01 | no | Sstr3 | intron_variant | MODIFIER |
| 15 | 78587072 | AX-170148966 | 0 | T | C | 0.553 | 3.47E-03 | 8.28E-02 | 2.66E+00 | 2.73E+00 | 9.76E-01 | no | | regulatory_region_variant | MODIFIER |
| 15 | 78587598 | AX-170012053 | 0 | C | T | 0.553 | 3.47E-03 | 8.28E-02 | 2.66E+00 | 2.73E+00 | 9.76E-01 | no | | regulatory_region_variant | MODIFIER |
| 15 | 78602006 | AX-168419953 | 0 | A | G | 0.553 | 3.47E-03 | 8.28E-02 | 2.66E+00 | 2.73E+00 | 9.76E-01 | no | Cyth4 | intron_variant | MODIFIER |
| 15 | 78604692 | AX-170056063 | 0 | T | C | 0.553 | 3.47E-03 | 8.28E-02 | 2.66E+00 | 2.73E+00 | 9.76E-01 | no | Cyth4 | synonymous_variant | LOW |
| 15 | 78608532 | AX-170012057 | 0 | C | T | 0.553 | 3.47E-03 | 8.28E-02 | 2.66E+00 | 2.73E+00 | 9.76E-01 | no | Cyth4 | intron_variant | MODIFIER |
| 15 | 78640384 | AX-170012056 | 0 | C | T | 0.553 | 3.47E-03 | 8.28E-02 | 2.66E+00 | 2.73E+00 | 9.76E-01 | no | | intergenic_variant | MODIFIER |
| 15 | 78642860 | AX-170012052 | 0 | A | G | 0.553 | 3.47E-03 | 8.28E-02 | 2.66E+00 | 2.73E+00 | 9.76E-01 | no | | intergenic_variant | MODIFIER |
| 15 | 78643117 | AX-170012058 | 0 | T | G | 0.553 | 3.47E-03 | 8.28E-02 | 2.66E+00 | 2.73E+00 | 9.76E-01 | no | | intergenic_variant | MODIFIER |
| 15 | 78643148 | AX-170119747 | 0 | C | T | 0.553 | 3.47E-03 | 8.28E-02 | 2.66E+00 | 2.73E+00 | 9.76E-01 | no | | intergenic_variant | MODIFIER |
| 15 | 78692169 | AX-170012051 | 0 | T | C | 0.553 | 3.47E-03 | 8.28E-02 | 2.66E+00 | 2.73E+00 | 9.76E-01 | no | Elfn2 | intron_variant | MODIFIER |
| 15 | 78694490 | AX-97829818 | 0 | G | A | 0.553 | 3.47E-03 | 8.28E-02 | 2.66E+00 | 2.73E+00 | 9.76E-01 | no | Elfn2 | intron_variant | MODIFIER |
| 15 | 78741359 | AX-170379152 | 0 | C | A | 0.553 | 3.47E-03 | 8.28E-02 | 2.66E+00 | 2.73E+00 | 9.76E-01 | no | | intergenic_variant | MODIFIER |
| 15 | 78741725 | AX-170109979 | 0 | T | C | 0.553 | 3.47E-03 | 8.28E-02 | 2.66E+00 | 2.73E+00 | 9.76E-01 | no | | intergenic_variant | MODIFIER |
| 15 | 78744075 | AX-170270597 | 0 | T | C | 0.553 | 3.47E-03 | 8.28E-02 | 2.66E+00 | 2.73E+00 | 9.76E-01 | no | | regulatory_region_variant | MODIFIER |
| 15 | 78745691 | AX-97806433 | 0 | T | G | 0.553 | 3.47E-03 | 8.28E-02 | 2.66E+00 | 2.73E+00 | 9.76E-01 | no | | intergenic_variant | MODIFIER |
| 12 | 71085750 | AX-170007011 | 1 | T | C | 0.062 | 8.67E-03 | 3.02E-01 | 2.72E+00 | 2.74E+00 | 9.76E-01 | no | Arid4a | intron_variant | MODIFIER |
| 20 | 76540395 | AX-170129223 | 0 | A | G | 0.254 | -1.52E-03 | 6.06E-02 | 2.64E+00 | 2.74E+00 | 9.76E-01 | no | Gm5793 | upstream_gene_variant | MODIFIER |
| 1 | 52964897 | AX-170000510 | 1 | G | A | 0.107 | -6.80E-03 | 2.35E-01 | 2.69E+00 | 2.74E+00 | 9.76E-01 | no | 1700019D03Rik | upstream_gene_variant | MODIFIER |
| 9 | 64834935 | AX-170145722 | 1 | G | A | 0.196 | -4.29E-03 | 1.37E-01 | 2.71E+00 | 2.74E+00 | 9.76E-01 | no | Dennd4a | intron_variant | MODIFIER |
| 9 | 66204953 | AX-170300028 | 1 | C | T | 0.196 | -4.29E-03 | 1.37E-01 | 2.71E+00 | 2.74E+00 | 9.76E-01 | no | Dapk2 | intron_variant | MODIFIER |
| 1 | 11083825 | AX-97810738 | 0 | C | A | 0.096 | 6.01E-03 | 1.71E-01 | 2.63E+00 | 2.74E+00 | 9.76E-01 | no | Prex2 | intron_variant | MODIFIER |
| 7 | 81069494 | AX-170316262 | 0 | A | T | 0.114 | -5.82E-03 | 2.10E-01 | 2.66E+00 | 2.74E+00 | 9.76E-01 | no | Alpk3 | intron_variant | MODIFIER |
| 15 | 102522602 | AX-97833784 | 0 | G | A | 0.132 | 5.21E-03 | 1.61E-01 | 2.67E+00 | 2.74E+00 | 9.76E-01 | no | Tarbp2 | intron_variant | MODIFIER |
| 11 | 50200290 | AX-170047013 | 1 | T | A | 0.089 | 4.01E-03 | 1.82E-01 | 2.64E+00 | 2.74E+00 | 9.76E-01 | no | Sqstm1 | 3_prime_UTR_variant | MODIFIER |
| 7 | 112503279 | AX-97827699 | 1 | A | G | 0.455 | 3.10E-03 | 9.66E-02 | 2.68E+00 | 2.74E+00 | 9.76E-01 | no | Parva | intron_variant | MODIFIER |

| | | | | | | | | | | | | | | | |
|---|---|---|---|---|---|---|---|---|---|---|---|---|---|---|---|
| 14 | 64105424 | AX-170320974 | 0 | T | C | 0.044 | -9.10E-03 | 2.36E-01 | 2.66E+00 | 2.73E+00 | 9.76E-01 | no | Prss52 | intron_variant | MODIFIER |
| 14 | 64530061 | AX-170173262 | 0 | T | C | 0.044 | -9.10E-03 | 2.36E-01 | 2.66E+00 | 2.73E+00 | 9.76E-01 | no | Gm47202 | downstream_gene_variant | MODIFIER |
| 14 | 65346493 | AX-170009998 | 0 | A | C | 0.044 | -9.10E-03 | 2.36E-01 | 2.66E+00 | 2.73E+00 | 9.76E-01 | no | Gm48449 | downstream_gene_variant | MODIFIER |
| 4 | 90748630 | AX-170284122 | 2 | C | T | 0.327 | -2.31E-03 | 1.06E-01 | 2.66E+00 | 2.74E+00 | 9.76E-01 | no | | intergenic_variant | MODIFIER |
| 9 | 80916690 | AX-170212554 | 1 | T | C | 0.205 | -4.27E-03 | 1.29E-01 | 2.69E+00 | 2.73E+00 | 9.76E-01 | no | | intergenic_variant | MODIFIER |
| 6 | 101422669 | AX-170250045 | 1 | C | T | 0.071 | 8.10E-03 | 2.76E-01 | 2.70E+00 | 2.74E+00 | 9.76E-01 | no | | intergenic_variant | MODIFIER |
| 6 | 118733338 | AX-170353297 | 1 | C | T | 0.071 | 8.10E-03 | 2.76E-01 | 2.70E+00 | 2.74E+00 | 9.76E-01 | no | Cacna1c | intron_variant | MODIFIER |
| 6 | 140097896 | AX-170318113 | 1 | A | G | 0.071 | 8.10E-03 | 2.76E-01 | 2.70E+00 | 2.74E+00 | 9.76E-01 | no | | intergenic_variant | MODIFIER |
| 14 | 67473749 | AX-170010027 | 1 | C | T | 0.036 | -9.37E-03 | 2.32E-01 | 2.63E+00 | 2.73E+00 | 9.76E-01 | no | Gm38409 | upstream_gene_variant | MODIFIER |
| 4 | 138749991 | AX-170372243 | 0 | T | C | 0.158 | -1.19E-03 | 1.02E-01 | 2.64E+00 | 2.75E+00 | 9.76E-01 | no | Pla2g2f | downstream_gene_variant | MODIFIER |
| 10 | 8138038 | AX-170315298 | 1 | C | A | 0.214 | -3.76E-03 | 1.04E-01 | 2.65E+00 | 2.73E+00 | 9.76E-01 | no | | intergenic_variant | MODIFIER |
| 4 | 71128955 | AX-170403122 | 0 | G | T | 0.228 | -1.61E-03 | 1.02E-01 | 2.65E+00 | 2.75E+00 | 9.76E-01 | no | Gm11228 | intron_variant,non_coding_transcript_variant | MODIFIER |
| 4 | 149700171 | AX-170121266 | 2 | G | A | 0.1 | -4.75E-03 | 1.67E-01 | 2.61E+00 | 2.74E+00 | 9.76E-01 | no | Pik3cd | upstream_gene_variant | MODIFIER |
| 1 | 10616456 | AX-97841522 | 0 | T | C | 0.491 | 3.10E-03 | 7.17E-02 | 2.66E+00 | 2.73E+00 | 9.76E-01 | no | Cpa6 | intron_variant | MODIFIER |
| 13 | 42450014 | AX-170008191 | 0 | C | A | 0.491 | -9.55E-04 | 6.19E-02 | 2.63E+00 | 2.74E+00 | 9.77E-01 | no | | intergenic_variant | MODIFIER |
| 4 | 141407134 | AX-170235324 | 0 | T | C | 0.114 | 5.53E-03 | 1.70E-01 | 2.66E+00 | 2.74E+00 | 9.77E-01 | no | Clcnkb | missense_variant | MODERATE |
| 8 | 94477421 | AX-170183502 | 0 | G | A | 0.211 | 2.46E-03 | 1.15E-01 | 2.66E+00 | 2.74E+00 | 9.77E-01 | no | Nlrc5 | intron_variant | MODIFIER |
| 6 | 118631443 | AX-170190920 | 0 | T | A | 0.079 | 6.60E-03 | 1.99E-01 | 2.63E+00 | 2.74E+00 | 9.77E-01 | no | Cacna1c | intron_variant | MODIFIER |
| 6 | 139803794 | AX-170445033 | 0 | T | A | 0.079 | 6.60E-03 | 1.99E-01 | 2.63E+00 | 2.74E+00 | 9.77E-01 | no | Pik3c2g | intron_variant | MODIFIER |
| 6 | 100799250 | AX-170182292 | 0 | C | T | 0.07 | 7.84E-03 | 2.78E-01 | 2.70E+00 | 2.74E+00 | 9.77E-01 | no | Gxylt2 | intron_variant | MODIFIER |
| 6 | 100803522 | AX-170299479 | 0 | A | C | 0.07 | 7.84E-03 | 2.78E-01 | 2.70E+00 | 2.74E+00 | 9.77E-01 | no | Gxylt2 | intron_variant | MODIFIER |
| 6 | 100803871 | AX-97834975 | 0 | A | G | 0.07 | 7.84E-03 | 2.78E-01 | 2.70E+00 | 2.74E+00 | 9.77E-01 | no | Gxylt2 | intron_variant | MODIFIER |
| 6 | 100805658 | AX-170027828 | 0 | G | T | 0.07 | 7.84E-03 | 2.78E-01 | 2.70E+00 | 2.74E+00 | 9.77E-01 | no | Gxylt2 | 3_prime_UTR_variant | MODIFIER |
| 6 | 100810164 | AX-170027826 | 0 | G | A | 0.07 | 7.84E-03 | 2.78E-01 | 2.70E+00 | 2.74E+00 | 9.77E-01 | no | Gxylt2 | 3_prime_UTR_variant | MODIFIER |
| 6 | 100830300 | AX-170027824 | 0 | A | T | 0.07 | 7.84E-03 | 2.78E-01 | 2.70E+00 | 2.74E+00 | 9.77E-01 | no | Ppp4r2 | upstream_gene_variant | MODIFIER |
| 6 | 100843278 | AX-170525255 | 0 | C | G | 0.07 | 7.84E-03 | 2.78E-01 | 2.70E+00 | 2.74E+00 | 9.77E-01 | no | Ppp4r2 | intron_variant | MODIFIER |
| 6 | 100843430 | AX-97802004 | 0 | A | C | 0.07 | 7.84E-03 | 2.78E-01 | 2.70E+00 | 2.74E+00 | 9.77E-01 | no | Ppp4r2 | intron_variant | MODIFIER |
| 6 | 100857215 | AX-170027829 | 0 | A | T | 0.07 | 7.84E-03 | 2.78E-01 | 2.70E+00 | 2.74E+00 | 9.77E-01 | no | Ppp4r2 | intron_variant | MODIFIER |
| 6 | 100869556 | AX-170174287 | 0 | T | G | 0.07 | 7.84E-03 | 2.78E-01 | 2.70E+00 | 2.74E+00 | 9.77E-01 | no | Ppp4r2 | 3_prime_UTR_variant | MODIFIER |
| 6 | 100872395 | AX-97790480 | 0 | T | C | 0.07 | 7.84E-03 | 2.78E-01 | 2.70E+00 | 2.74E+00 | 9.77E-01 | no | Ppp4r2 | downstream_gene_variant | MODIFIER |
| 6 | 100990289 | AX-170471558 | 0 | C | T | 0.07 | 7.84E-03 | 2.78E-01 | 2.70E+00 | 2.74E+00 | 9.77E-01 | no | | intergenic_variant | MODIFIER |
| 6 | 100991654 | AX-170126747 | 0 | T | C | 0.07 | 7.84E-03 | 2.78E-01 | 2.70E+00 | 2.74E+00 | 9.77E-01 | no | | intergenic_variant | MODIFIER |
| 6 | 100991790 | AX-170106802 | 0 | G | A | 0.07 | 7.84E-03 | 2.78E-01 | 2.70E+00 | 2.74E+00 | 9.77E-01 | no | | intergenic_variant | MODIFIER |
| 6 | 101016396 | AX-170141359 | 0 | T | C | 0.07 | 7.84E-03 | 2.78E-01 | 2.70E+00 | 2.74E+00 | 9.77E-01 | no | | intergenic_variant | MODIFIER |
| 6 | 101022710 | AX-170212912 | 0 | A | C | 0.07 | 7.84E-03 | 2.78E-01 | 2.70E+00 | 2.74E+00 | 9.77E-01 | no | | intergenic_variant | MODIFIER |
| 6 | 101026316 | AX-170411590 | 0 | T | C | 0.07 | 7.84E-03 | 2.78E-01 | 2.70E+00 | 2.74E+00 | 9.77E-01 | no | Gm43950 | upstream_gene_variant | MODIFIER |
| 6 | 101027969 | AX-170239332 | 0 | A | G | 0.07 | 7.84E-03 | 2.78E-01 | 2.70E+00 | 2.74E+00 | 9.77E-01 | no | Gm43950 | upstream_gene_variant | MODIFIER |
| 6 | 101028302 | AX-97811672 | 0 | T | C | 0.07 | 7.84E-03 | 2.78E-01 | 2.70E+00 | 2.74E+00 | 9.77E-01 | no | Gm43950 | non_coding_transcript_exon_variant | MODIFIER |
| 6 | 101028549 | AX-170374628 | 0 | C | T | 0.07 | 7.84E-03 | 2.78E-01 | 2.70E+00 | 2.74E+00 | 9.77E-01 | no | Gm43950 | non_coding_transcript_exon_variant | MODIFIER |
| 6 | 101028770 | AX-170359504 | 0 | C | G | 0.07 | 7.84E-03 | 2.78E-01 | 2.70E+00 | 2.74E+00 | 9.77E-01 | no | Gm43950 | intron_variant,non_coding_transcript_variant | MODIFIER |
| 6 | 101030624 | AX-170356454 | 0 | C | T | 0.07 | 7.84E-03 | 2.78E-01 | 2.70E+00 | 2.74E+00 | 9.77E-01 | no | Gm43950 | intron_variant,non_coding_transcript_variant | MODIFIER |
| 6 | 101060236 | AX-170504790 | 0 | A | G | 0.07 | 7.84E-03 | 2.78E-01 | 2.70E+00 | 2.74E+00 | 9.77E-01 | no | Gm43948 | intron_variant,non_coding_transcript_variant | MODIFIER |
| 6 | 101065300 | AX-97853352 | 0 | T | C | 0.07 | 7.84E-03 | 2.78E-01 | 2.70E+00 | 2.74E+00 | 9.77E-01 | no | Gm43948 | intron_variant,non_coding_transcript_variant | MODIFIER |
| 6 | 101081857 | AX-97833970 | 0 | T | C | 0.07 | 7.84E-03 | 2.78E-01 | 2.70E+00 | 2.74E+00 | 9.77E-01 | no | Gm43948 | intron_variant,non_coding_transcript_variant | MODIFIER |
| 6 | 101087898 | AX-170409023 | 0 | A | T | 0.07 | 7.84E-03 | 2.78E-01 | 2.70E+00 | 2.74E+00 | 9.77E-01 | no | Gm43948 | intron_variant,non_coding_transcript_variant | MODIFIER |
| 6 | 101088924 | AX-170306278 | 0 | C | T | 0.07 | 7.84E-03 | 2.78E-01 | 2.70E+00 | 2.74E+00 | 9.77E-01 | no | Gm43948 | intron_variant,non_coding_transcript_variant | MODIFIER |
| 6 | 101094599 | AX-170479055 | 0 | T | C | 0.07 | 7.84E-03 | 2.78E-01 | 2.70E+00 | 2.74E+00 | 9.77E-01 | no | Gm43948 | intron_variant,non_coding_transcript_variant | MODIFIER |
| 6 | 101094955 | AX-170534824 | 0 | G | C | 0.07 | 7.84E-03 | 2.78E-01 | 2.70E+00 | 2.74E+00 | 9.77E-01 | no | Gm43948 | intron_variant,non_coding_transcript_variant | MODIFIER |
| 6 | 101095508 | AX-170417284 | 0 | T | C | 0.07 | 7.84E-03 | 2.78E-01 | 2.70E+00 | 2.74E+00 | 9.77E-01 | no | Gm43948 | intron_variant,non_coding_transcript_variant | MODIFIER |
| 6 | 101103892 | AX-97840756 | 0 | T | C | 0.07 | 7.84E-03 | 2.78E-01 | 2.70E+00 | 2.74E+00 | 9.77E-01 | no | Gm43948 | intron_variant,non_coding_transcript_variant | MODIFIER |
| 6 | 101106427 | AX-170340742 | 0 | A | G | 0.07 | 7.84E-03 | 2.78E-01 | 2.70E+00 | 2.74E+00 | 9.77E-01 | no | Gm43948 | intron_variant,non_coding_transcript_variant | MODIFIER |
| 6 | 101122513 | AX-170337548 | 0 | C | T | 0.07 | 7.84E-03 | 2.78E-01 | 2.70E+00 | 2.74E+00 | 9.77E-01 | no | Gm43948 | intron_variant,non_coding_transcript_variant | MODIFIER |
| 6 | 101122616 | AX-170403454 | 0 | T | C | 0.07 | 7.84E-03 | 2.78E-01 | 2.70E+00 | 2.74E+00 | 9.77E-01 | no | Gm43948 | intron_variant,non_coding_transcript_variant | MODIFIER |
| 6 | 101161360 | AX-170183841 | 0 | T | C | 0.07 | 7.84E-03 | 2.78E-01 | 2.70E+00 | 2.74E+00 | 9.77E-01 | no | Pdzrn3 | intron_variant | MODIFIER |

| | | | | | | | | | | | | | | | |
|---|---|---|---|---|---|---|---|---|---|---|---|---|---|---|---|
| 6 | 101173218 | AX-97834211 | 0 | A | G | 0.07 | 7.84E-03 | 2.78E-01 | 2.70E+00 | 2.74E+00 | 9.77E-01 | no | Pdzrn3 | intron_variant | MODIFIER |
| 6 | 101271099 | AX-97785966 | 0 | A | G | 0.07 | 7.84E-03 | 2.78E-01 | 2.70E+00 | 2.74E+00 | 9.77E-01 | no | Pdzrn3 | intron_variant | MODIFIER |
| 6 | 101419893 | AX-170027838 | 0 | G | A | 0.07 | 7.84E-03 | 2.78E-01 | 2.70E+00 | 2.74E+00 | 9.77E-01 | no | | intergenic_variant | MODIFIER |
| 6 | 101421018 | AX-170339607 | 0 | T | C | 0.07 | 7.84E-03 | 2.78E-01 | 2.70E+00 | 2.74E+00 | 9.77E-01 | no | | intergenic_variant | MODIFIER |
| 6 | 101426954 | AX-170153790 | 0 | G | A | 0.07 | 7.84E-03 | 2.78E-01 | 2.70E+00 | 2.74E+00 | 9.77E-01 | no | | intergenic_variant | MODIFIER |
| 6 | 101462246 | AX-97832376 | 0 | C | T | 0.07 | 7.84E-03 | 2.78E-01 | 2.70E+00 | 2.74E+00 | 9.77E-01 | no | | intergenic_variant | MODIFIER |
| 6 | 101476308 | AX-170117759 | 0 | G | A | 0.07 | 7.84E-03 | 2.78E-01 | 2.70E+00 | 2.74E+00 | 9.77E-01 | no | | intergenic_variant | MODIFIER |
| 6 | 101481286 | AX-170098938 | 0 | T | C | 0.07 | 7.84E-03 | 2.78E-01 | 2.70E+00 | 2.74E+00 | 9.77E-01 | no | | intergenic_variant | MODIFIER |
| 6 | 101482940 | AX-170226801 | 0 | T | C | 0.07 | 7.84E-03 | 2.78E-01 | 2.70E+00 | 2.74E+00 | 9.77E-01 | no | | regulatory_region_variant | MODIFIER |
| 6 | 101484515 | AX-170354547 | 0 | T | C | 0.07 | 7.84E-03 | 2.78E-01 | 2.70E+00 | 2.74E+00 | 9.77E-01 | no | | regulatory_region_variant | MODIFIER |
| 6 | 101491043 | AX-97844799 | 0 | A | T | 0.07 | 7.84E-03 | 2.78E-01 | 2.70E+00 | 2.74E+00 | 9.77E-01 | no | | intergenic_variant | MODIFIER |
| 6 | 101655778 | AX-170202462 | 0 | C | A | 0.07 | 7.84E-03 | 2.78E-01 | 2.70E+00 | 2.74E+00 | 9.77E-01 | no | Gm32453 | downstream_gene_variant | MODIFIER |
| 6 | 102245965 | AX-170323368 | 0 | C | T | 0.07 | 7.84E-03 | 2.78E-01 | 2.70E+00 | 2.74E+00 | 9.77E-01 | no | Cntn3 | intron_variant | MODIFIER |
| 6 | 102277810 | AX-170362337 | 0 | G | T | 0.07 | 7.84E-03 | 2.78E-01 | 2.70E+00 | 2.74E+00 | 9.77E-01 | no | Cntn3 | intron_variant | MODIFIER |
| 6 | 102278121 | AX-170512819 | 0 | T | A | 0.07 | 7.84E-03 | 2.78E-01 | 2.70E+00 | 2.74E+00 | 9.77E-01 | no | Cntn3 | intron_variant | MODIFIER |
| 6 | 102282626 | AX-170417024 | 0 | C | T | 0.07 | 7.84E-03 | 2.78E-01 | 2.70E+00 | 2.74E+00 | 9.77E-01 | no | Cntn3 | intron_variant | MODIFIER |
| 6 | 102289288 | AX-97811150 | 0 | G | A | 0.07 | 7.84E-03 | 2.78E-01 | 2.70E+00 | 2.74E+00 | 9.77E-01 | no | Cntn3 | intron_variant | MODIFIER |
| 6 | 102488340 | AX-170027853 | 0 | C | T | 0.07 | 7.84E-03 | 2.78E-01 | 2.70E+00 | 2.74E+00 | 9.77E-01 | no | Cntn3 | intron_variant | MODIFIER |
| 6 | 102489070 | AX-170326981 | 0 | C | T | 0.07 | 7.84E-03 | 2.78E-01 | 2.70E+00 | 2.74E+00 | 9.77E-01 | no | Cntn3 | intron_variant | MODIFIER |
| 6 | 102501719 | AX-170027849 | 0 | A | G | 0.07 | 7.84E-03 | 2.78E-01 | 2.70E+00 | 2.74E+00 | 9.77E-01 | no | Cntn3 | intron_variant | MODIFIER |
| 6 | 103224797 | AX-97781621 | 0 | C | T | 0.07 | 7.84E-03 | 2.78E-01 | 2.70E+00 | 2.74E+00 | 9.77E-01 | no | | intergenic_variant | MODIFIER |
| 6 | 103290895 | AX-170317113 | 0 | A | G | 0.07 | 7.84E-03 | 2.78E-01 | 2.70E+00 | 2.74E+00 | 9.77E-01 | no | | intergenic_variant | MODIFIER |
| 6 | 103398840 | AX-168419016 | 0 | A | T | 0.07 | 7.84E-03 | 2.78E-01 | 2.70E+00 | 2.74E+00 | 9.77E-01 | no | | intergenic_variant | MODIFIER |
| 6 | 104205562 | AX-170027864 | 0 | A | G | 0.07 | 7.84E-03 | 2.78E-01 | 2.70E+00 | 2.74E+00 | 9.77E-01 | no | | intergenic_variant | MODIFIER |
| 6 | 104305719 | AX-170534423 | 0 | G | T | 0.07 | 7.84E-03 | 2.78E-01 | 2.70E+00 | 2.74E+00 | 9.77E-01 | no | | intergenic_variant | MODIFIER |
| 6 | 104317278 | AX-170194287 | 0 | A | G | 0.07 | 7.84E-03 | 2.78E-01 | 2.70E+00 | 2.74E+00 | 9.77E-01 | no | | intergenic_variant | MODIFIER |
| 6 | 104319481 | AX-170117520 | 0 | C | T | 0.07 | 7.84E-03 | 2.78E-01 | 2.70E+00 | 2.74E+00 | 9.77E-01 | no | | intergenic_variant | MODIFIER |
| 6 | 104454547 | AX-170192513 | 0 | T | C | 0.07 | 7.84E-03 | 2.78E-01 | 2.70E+00 | 2.74E+00 | 9.77E-01 | no | | intergenic_variant | MODIFIER |
| 6 | 104460634 | AX-170206839 | 0 | G | A | 0.07 | 7.84E-03 | 2.78E-01 | 2.70E+00 | 2.74E+00 | 9.77E-01 | no | | intergenic_variant | MODIFIER |
| 6 | 104469675 | AX-170129427 | 0 | T | C | 0.07 | 7.84E-03 | 2.78E-01 | 2.70E+00 | 2.74E+00 | 9.77E-01 | no | | intergenic_variant | MODIFIER |
| 6 | 104479335 | AX-170106456 | 0 | C | T | 0.07 | 7.84E-03 | 2.78E-01 | 2.70E+00 | 2.74E+00 | 9.77E-01 | no | | regulatory_region_variant | MODIFIER |
| 6 | 104538011 | AX-170227429 | 0 | T | C | 0.07 | 7.84E-03 | 2.78E-01 | 2.70E+00 | 2.74E+00 | 9.77E-01 | no | Cntn6 | intron_variant | MODIFIER |
| 6 | 104544121 | AX-97855977 | 0 | T | C | 0.07 | 7.84E-03 | 2.78E-01 | 2.70E+00 | 2.74E+00 | 9.77E-01 | no | Cntn6 | intron_variant | MODIFIER |
| 6 | 104568100 | AX-170075755 | 0 | A | G | 0.07 | 7.84E-03 | 2.78E-01 | 2.70E+00 | 2.74E+00 | 9.77E-01 | no | Cntn6 | intron_variant | MODIFIER |
| 6 | 104574387 | AX-170390649 | 0 | C | A | 0.07 | 7.84E-03 | 2.78E-01 | 2.70E+00 | 2.74E+00 | 9.77E-01 | no | Cntn6 | intron_variant | MODIFIER |
| 6 | 104607680 | AX-170466546 | 0 | T | C | 0.07 | 7.84E-03 | 2.78E-01 | 2.70E+00 | 2.74E+00 | 9.77E-01 | no | Cntn6 | intron_variant | MODIFIER |
| 6 | 104626086 | AX-170111235 | 0 | G | A | 0.07 | 7.84E-03 | 2.78E-01 | 2.70E+00 | 2.74E+00 | 9.77E-01 | no | Cntn6 | intron_variant | MODIFIER |
| 6 | 104644776 | AX-170089800 | 0 | A | G | 0.07 | 7.84E-03 | 2.78E-01 | 2.70E+00 | 2.74E+00 | 9.77E-01 | no | Cntn6 | intron_variant | MODIFIER |
| 6 | 104675080 | AX-170479839 | 0 | G | T | 0.07 | 7.84E-03 | 2.78E-01 | 2.70E+00 | 2.74E+00 | 9.77E-01 | no | Cntn6 | intron_variant | MODIFIER |
| 6 | 104733499 | AX-170463010 | 0 | A | C | 0.07 | 7.84E-03 | 2.78E-01 | 2.70E+00 | 2.74E+00 | 9.77E-01 | no | Cntn6 | intron_variant | MODIFIER |
| 6 | 104733933 | AX-170027876 | 0 | T | C | 0.07 | 7.84E-03 | 2.78E-01 | 2.70E+00 | 2.74E+00 | 9.77E-01 | no | Cntn6 | intron_variant | MODIFIER |
| 6 | 104770983 | AX-170027873 | 0 | G | A | 0.07 | 7.84E-03 | 2.78E-01 | 2.70E+00 | 2.74E+00 | 9.77E-01 | no | Cntn6 | intron_variant | MODIFIER |
| 6 | 104771159 | AX-170413266 | 0 | G | A | 0.07 | 7.84E-03 | 2.78E-01 | 2.70E+00 | 2.74E+00 | 9.77E-01 | no | Cntn6 | intron_variant | MODIFIER |
| 6 | 104773278 | AX-170135479 | 0 | T | C | 0.07 | 7.84E-03 | 2.78E-01 | 2.70E+00 | 2.74E+00 | 9.77E-01 | no | Cntn6 | intron_variant | MODIFIER |
| 6 | 104787923 | AX-170312664 | 0 | C | T | 0.07 | 7.84E-03 | 2.78E-01 | 2.70E+00 | 2.74E+00 | 9.77E-01 | no | Cntn6 | intron_variant | MODIFIER |
| 6 | 104856925 | AX-170405037 | 0 | C | T | 0.07 | 7.84E-03 | 2.78E-01 | 2.70E+00 | 2.74E+00 | 9.77E-01 | no | Cntn6 | intron_variant | MODIFIER |
| 6 | 104862718 | AX-170455673 | 0 | G | A | 0.07 | 7.84E-03 | 2.78E-01 | 2.70E+00 | 2.74E+00 | 9.77E-01 | no | Cntn6 | intron_variant | MODIFIER |
| 6 | 104938694 | AX-170300362 | 0 | G | C | 0.07 | 7.84E-03 | 2.78E-01 | 2.70E+00 | 2.74E+00 | 9.77E-01 | no | | intergenic_variant | MODIFIER |
| 6 | 105224953 | AX-97802036 | 0 | T | C | 0.07 | 7.84E-03 | 2.78E-01 | 2.70E+00 | 2.74E+00 | 9.77E-01 | no | | intergenic_variant | MODIFIER |
| 6 | 105225252 | AX-170027883 | 0 | A | G | 0.07 | 7.84E-03 | 2.78E-01 | 2.70E+00 | 2.74E+00 | 9.77E-01 | no | | intergenic_variant | MODIFIER |
| 6 | 105259436 | AX-170141624 | 0 | G | A | 0.07 | 7.84E-03 | 2.78E-01 | 2.70E+00 | 2.74E+00 | 9.77E-01 | no | | intergenic_variant | MODIFIER |
| 6 | 105284512 | AX-170027878 | 0 | A | C | 0.07 | 7.84E-03 | 2.78E-01 | 2.70E+00 | 2.74E+00 | 9.77E-01 | no | | intergenic_variant | MODIFIER |
| 6 | 105352995 | AX-170139950 | 0 | T | C | 0.07 | 7.84E-03 | 2.78E-01 | 2.70E+00 | 2.74E+00 | 9.77E-01 | no | | intergenic_variant | MODIFIER |
| 6 | 105396702 | AX-97795749 | 0 | C | T | 0.07 | 7.84E-03 | 2.78E-01 | 2.70E+00 | 2.74E+00 | 9.77E-01 | no | Gm25570 | upstream_gene_variant | MODIFIER |

| | | | | | | | | | | | | | | | |
|---|---|---|---|---|---|---|---|---|---|---|---|---|---|---|---|
| 6 | 105428325 | AX-97825610 | 0 | G | T | 0.07 | 7.84E-03 | 2.78E-01 | 2.70E+00 | 2.74E+00 | 9.77E-01 | no | | intergenic_variant | MODIFIER |
| 6 | 105428953 | AX-170157882 | 0 | C | G | 0.07 | 7.84E-03 | 2.78E-01 | 2.70E+00 | 2.74E+00 | 9.77E-01 | no | | intergenic_variant | MODIFIER |
| 6 | 105433884 | AX-170277959 | 0 | C | G | 0.07 | 7.84E-03 | 2.78E-01 | 2.70E+00 | 2.74E+00 | 9.77E-01 | no | | intergenic_variant | MODIFIER |
| 6 | 105574388 | AX-170525440 | 0 | G | C | 0.07 | 7.84E-03 | 2.78E-01 | 2.70E+00 | 2.74E+00 | 9.77E-01 | no | | intergenic_variant | MODIFIER |
| 6 | 105577809 | AX-170134063 | 0 | T | A | 0.07 | 7.84E-03 | 2.78E-01 | 2.70E+00 | 2.74E+00 | 9.77E-01 | no | | intergenic_variant | MODIFIER |
| 6 | 105609523 | AX-170506456 | 0 | G | T | 0.07 | 7.84E-03 | 2.78E-01 | 2.70E+00 | 2.74E+00 | 9.77E-01 | no | | intergenic_variant | MODIFIER |
| 6 | 105708381 | AX-170196834 | 0 | A | G | 0.07 | 7.84E-03 | 2.78E-01 | 2.70E+00 | 2.74E+00 | 9.77E-01 | no | Cntn4 | intron_variant | MODIFIER |
| 6 | 105708523 | AX-170497233 | 0 | A | G | 0.07 | 7.84E-03 | 2.78E-01 | 2.70E+00 | 2.74E+00 | 9.77E-01 | no | Cntn4 | intron_variant | MODIFIER |
| 6 | 105709961 | AX-170280719 | 0 | C | T | 0.07 | 7.84E-03 | 2.78E-01 | 2.70E+00 | 2.74E+00 | 9.77E-01 | no | Cntn4 | intron_variant | MODIFIER |
| 6 | 105716663 | AX-170223734 | 0 | G | T | 0.07 | 7.84E-03 | 2.78E-01 | 2.70E+00 | 2.74E+00 | 9.77E-01 | no | Cntn4 | intron_variant | MODIFIER |
| 6 | 105717408 | AX-170027885 | 0 | C | T | 0.07 | 7.84E-03 | 2.78E-01 | 2.70E+00 | 2.74E+00 | 9.77E-01 | no | Cntn4 | intron_variant | MODIFIER |
| 6 | 105731495 | AX-170166515 | 0 | C | T | 0.07 | 7.84E-03 | 2.78E-01 | 2.70E+00 | 2.74E+00 | 9.77E-01 | no | Cntn4 | intron_variant | MODIFIER |
| 6 | 105762058 | AX-170361862 | 0 | G | T | 0.07 | 7.84E-03 | 2.78E-01 | 2.70E+00 | 2.74E+00 | 9.77E-01 | no | Cntn4 | intron_variant | MODIFIER |
| 6 | 105779883 | AX-170169777 | 0 | A | T | 0.07 | 7.84E-03 | 2.78E-01 | 2.70E+00 | 2.74E+00 | 9.77E-01 | no | Cntn4 | intron_variant | MODIFIER |
| 6 | 105816559 | AX-170089803 | 0 | T | C | 0.07 | 7.84E-03 | 2.78E-01 | 2.70E+00 | 2.74E+00 | 9.77E-01 | no | Cntn4 | intron_variant | MODIFIER |
| 6 | 105861975 | AX-170027890 | 0 | T | A | 0.07 | 7.84E-03 | 2.78E-01 | 2.70E+00 | 2.74E+00 | 9.77E-01 | no | Cntn4 | intron_variant | MODIFIER |
| 6 | 105893901 | AX-170027889 | 0 | G | A | 0.07 | 7.84E-03 | 2.78E-01 | 2.70E+00 | 2.74E+00 | 9.77E-01 | no | Cntn4 | intron_variant | MODIFIER |
| 6 | 105894493 | AX-170241688 | 0 | G | A | 0.07 | 7.84E-03 | 2.78E-01 | 2.70E+00 | 2.74E+00 | 9.77E-01 | no | Cntn4 | intron_variant | MODIFIER |
| 6 | 105923230 | AX-97802034 | 0 | C | T | 0.07 | 7.84E-03 | 2.78E-01 | 2.70E+00 | 2.74E+00 | 9.77E-01 | no | Cntn4 | intron_variant | MODIFIER |
| 6 | 105949724 | AX-97802037 | 0 | G | A | 0.07 | 7.84E-03 | 2.78E-01 | 2.70E+00 | 2.74E+00 | 9.77E-01 | no | Cntn4 | intron_variant | MODIFIER |
| 6 | 105979981 | AX-170522635 | 0 | C | A | 0.07 | 7.84E-03 | 2.78E-01 | 2.70E+00 | 2.74E+00 | 9.77E-01 | no | Cntn4 | intron_variant | MODIFIER |
| 6 | 106012113 | AX-170466994 | 0 | A | T | 0.07 | 7.84E-03 | 2.78E-01 | 2.70E+00 | 2.74E+00 | 9.77E-01 | no | Cntn4 | intron_variant | MODIFIER |
| 6 | 106020615 | AX-170098545 | 0 | G | A | 0.07 | 7.84E-03 | 2.78E-01 | 2.70E+00 | 2.74E+00 | 9.77E-01 | no | Cntn4 | intron_variant | MODIFIER |
| 6 | 110485932 | AX-170446952 | 0 | T | G | 0.07 | 7.84E-03 | 2.78E-01 | 2.70E+00 | 2.74E+00 | 9.77E-01 | no | | intergenic_variant | MODIFIER |
| 6 | 110488612 | AX-170130794 | 0 | A | C | 0.07 | 7.84E-03 | 2.78E-01 | 2.70E+00 | 2.74E+00 | 9.77E-01 | no | | intergenic_variant | MODIFIER |
| 6 | 110574856 | AX-170445594 | 0 | T | C | 0.07 | 7.84E-03 | 2.78E-01 | 2.70E+00 | 2.74E+00 | 9.77E-01 | no | | intergenic_variant | MODIFIER |
| 6 | 110580995 | AX-170394229 | 0 | C | T | 0.07 | 7.84E-03 | 2.78E-01 | 2.70E+00 | 2.74E+00 | 9.77E-01 | no | | intergenic_variant | MODIFIER |
| 6 | 110592540 | AX-170239818 | 0 | G | C | 0.07 | 7.84E-03 | 2.78E-01 | 2.70E+00 | 2.74E+00 | 9.77E-01 | no | | intergenic_variant | MODIFIER |
| 6 | 110612773 | AX-170527413 | 0 | C | T | 0.07 | 7.84E-03 | 2.78E-01 | 2.70E+00 | 2.74E+00 | 9.77E-01 | no | | intergenic_variant | MODIFIER |
| 6 | 110644747 | AX-170392122 | 0 | G | A | 0.07 | 7.84E-03 | 2.78E-01 | 2.70E+00 | 2.74E+00 | 9.77E-01 | no | Grm7 | upstream_gene_variant | MODIFIER |
| 6 | 110710478 | AX-97807384 | 0 | T | C | 0.07 | 7.84E-03 | 2.78E-01 | 2.70E+00 | 2.74E+00 | 9.77E-01 | no | Grm7 | intron_variant | MODIFIER |
| 6 | 110737918 | AX-97793082 | 0 | A | G | 0.07 | 7.84E-03 | 2.78E-01 | 2.70E+00 | 2.74E+00 | 9.77E-01 | no | Grm7 | intron_variant | MODIFIER |
| 6 | 110766402 | AX-170397894 | 0 | G | C | 0.07 | 7.84E-03 | 2.78E-01 | 2.70E+00 | 2.74E+00 | 9.77E-01 | no | Grm7 | intron_variant | MODIFIER |
| 6 | 110775625 | AX-170405416 | 0 | A | C | 0.07 | 7.84E-03 | 2.78E-01 | 2.70E+00 | 2.74E+00 | 9.77E-01 | no | Grm7 | intron_variant | MODIFIER |
| 6 | 110775692 | AX-97802093 | 0 | A | C | 0.07 | 7.84E-03 | 2.78E-01 | 2.70E+00 | 2.74E+00 | 9.77E-01 | no | Grm7 | intron_variant | MODIFIER |
| 6 | 110824289 | AX-97827142 | 0 | T | C | 0.07 | 7.84E-03 | 2.78E-01 | 2.70E+00 | 2.74E+00 | 9.77E-01 | no | Grm7 | intron_variant | MODIFIER |
| 6 | 110928139 | AX-97836226 | 0 | A | C | 0.07 | 7.84E-03 | 2.78E-01 | 2.70E+00 | 2.74E+00 | 9.77E-01 | no | Grm7 | intron_variant | MODIFIER |
| 6 | 110962179 | AX-170502526 | 0 | A | T | 0.07 | 7.84E-03 | 2.78E-01 | 2.70E+00 | 2.74E+00 | 9.77E-01 | no | Grm7 | intron_variant | MODIFIER |
| 6 | 110964522 | AX-170346435 | 0 | C | T | 0.07 | 7.84E-03 | 2.78E-01 | 2.70E+00 | 2.74E+00 | 9.77E-01 | no | Grm7 | intron_variant | MODIFIER |
| 6 | 110968546 | AX-170027980 | 0 | G | A | 0.07 | 7.84E-03 | 2.78E-01 | 2.70E+00 | 2.74E+00 | 9.77E-01 | no | Grm7 | intron_variant | MODIFIER |
| 6 | 110982568 | AX-170027974 | 0 | G | A | 0.07 | 7.84E-03 | 2.78E-01 | 2.70E+00 | 2.74E+00 | 9.77E-01 | no | Grm7 | intron_variant | MODIFIER |
| 6 | 111100480 | AX-170027981 | 0 | T | C | 0.07 | 7.84E-03 | 2.78E-01 | 2.70E+00 | 2.74E+00 | 9.77E-01 | no | Grm7 | intron_variant | MODIFIER |
| 6 | 111210065 | AX-97829503 | 0 | G | A | 0.07 | 7.84E-03 | 2.78E-01 | 2.70E+00 | 2.74E+00 | 9.77E-01 | no | Grm7 | intron_variant | MODIFIER |
| 6 | 111233725 | AX-170403391 | 0 | C | T | 0.07 | 7.84E-03 | 2.78E-01 | 2.70E+00 | 2.74E+00 | 9.77E-01 | no | Grm7 | intron_variant | MODIFIER |
| 6 | 111234823 | AX-170451670 | 0 | T | C | 0.07 | 7.84E-03 | 2.78E-01 | 2.70E+00 | 2.74E+00 | 9.77E-01 | no | Grm7 | intron_variant | MODIFIER |
| 6 | 111312826 | AX-97826438 | 0 | C | A | 0.07 | 7.84E-03 | 2.78E-01 | 2.70E+00 | 2.74E+00 | 9.77E-01 | no | Grm7 | intron_variant | MODIFIER |
| 6 | 111323498 | AX-170404077 | 0 | G | T | 0.07 | 7.84E-03 | 2.78E-01 | 2.70E+00 | 2.74E+00 | 9.77E-01 | no | Grm7 | intron_variant | MODIFIER |
| 6 | 111329560 | AX-170128027 | 0 | G | T | 0.07 | 7.84E-03 | 2.78E-01 | 2.70E+00 | 2.74E+00 | 9.77E-01 | no | Grm7 | intron_variant | MODIFIER |
| 6 | 111331737 | AX-170192082 | 0 | C | T | 0.07 | 7.84E-03 | 2.78E-01 | 2.70E+00 | 2.74E+00 | 9.77E-01 | no | Grm7 | intron_variant | MODIFIER |
| 6 | 111336905 | AX-170408085 | 0 | G | A | 0.07 | 7.84E-03 | 2.78E-01 | 2.70E+00 | 2.74E+00 | 9.77E-01 | no | Grm7 | intron_variant | MODIFIER |
| 6 | 111341006 | AX-170301714 | 0 | T | C | 0.07 | 7.84E-03 | 2.78E-01 | 2.70E+00 | 2.74E+00 | 9.77E-01 | no | Grm7 | intron_variant | MODIFIER |
| 6 | 111343001 | AX-170418164 | 0 | T | C | 0.07 | 7.84E-03 | 2.78E-01 | 2.70E+00 | 2.74E+00 | 9.77E-01 | no | Grm7 | intron_variant | MODIFIER |
| 6 | 111358831 | AX-170406679 | 0 | A | G | 0.07 | 7.84E-03 | 2.78E-01 | 2.70E+00 | 2.74E+00 | 9.77E-01 | no | Grm7 | synonymous_variant | LOW |
| 6 | 111401594 | AX-170150327 | 0 | G | T | 0.07 | 7.84E-03 | 2.78E-01 | 2.70E+00 | 2.74E+00 | 9.77E-01 | no | Grm7 | intron_variant | MODIFIER |

| | | | | | | | | | | | | | | | |
|---|---|---|---|---|---|---|---|---|---|---|---|---|---|---|---|
| 6 | 111423571 | AX-170106060 | 0 | C | G | 0.07 | 7.84E-03 | 2.78E-01 | 2.70E+00 | 2.74E+00 | 9.77E-01 | no | Grm7 | intron_variant | MODIFIER |
| 6 | 111446984 | AX-170204054 | 0 | A | G | 0.07 | 7.84E-03 | 2.78E-01 | 2.70E+00 | 2.74E+00 | 9.77E-01 | no | Grm7 | intron_variant | MODIFIER |
| 6 | 111448722 | AX-170313583 | 0 | T | C | 0.07 | 7.84E-03 | 2.78E-01 | 2.70E+00 | 2.74E+00 | 9.77E-01 | no | Grm7 | intron_variant | MODIFIER |
| 6 | 111450507 | AX-170265349 | 0 | G | A | 0.07 | 7.84E-03 | 2.78E-01 | 2.70E+00 | 2.74E+00 | 9.77E-01 | no | Grm7 | intron_variant | MODIFIER |
| 6 | 111456873 | AX-170147611 | 0 | C | T | 0.07 | 7.84E-03 | 2.78E-01 | 2.70E+00 | 2.74E+00 | 9.77E-01 | no | Grm7 | intron_variant | MODIFIER |
| 6 | 111483352 | AX-97809394 | 0 | T | C | 0.07 | 7.84E-03 | 2.78E-01 | 2.70E+00 | 2.74E+00 | 9.77E-01 | no | Grm7 | intron_variant | MODIFIER |
| 6 | 111492764 | AX-97813569 | 0 | A | G | 0.07 | 7.84E-03 | 2.78E-01 | 2.70E+00 | 2.74E+00 | 9.77E-01 | no | Grm7 | intron_variant | MODIFIER |
| 6 | 111493294 | AX-170075876 | 0 | T | C | 0.07 | 7.84E-03 | 2.78E-01 | 2.70E+00 | 2.74E+00 | 9.77E-01 | no | Grm7 | intron_variant | MODIFIER |
| 6 | 112821522 | AX-97784988 | 0 | C | T | 0.07 | 7.84E-03 | 2.78E-01 | 2.70E+00 | 2.74E+00 | 9.77E-01 | no | Srgap3 | intron_variant,non_coding_transcript_variant | MODIFIER |
| 6 | 112860674 | AX-97830437 | 0 | C | T | 0.07 | 7.84E-03 | 2.78E-01 | 2.70E+00 | 2.74E+00 | 9.77E-01 | no | Srgap3 | intron_variant,non_coding_transcript_variant | MODIFIER |
| 6 | 112901793 | AX-170420902 | 0 | G | A | 0.07 | 7.84E-03 | 2.78E-01 | 2.70E+00 | 2.74E+00 | 9.77E-01 | no | Srgap3 | intron_variant,non_coding_transcript_variant | MODIFIER |
| 6 | 112912580 | AX-170373544 | 0 | G | T | 0.07 | 7.84E-03 | 2.78E-01 | 2.70E+00 | 2.74E+00 | 9.77E-01 | no | Srgap3 | intron_variant,non_coding_transcript_variant | MODIFIER |
| 6 | 113001941 | AX-170229100 | 0 | C | G | 0.07 | 7.84E-03 | 2.78E-01 | 2.70E+00 | 2.74E+00 | 9.77E-01 | no | | regulatory_region_variant | MODIFIER |
| 6 | 113031256 | AX-97812166 | 0 | A | C | 0.07 | 7.84E-03 | 2.78E-01 | 2.70E+00 | 2.74E+00 | 9.77E-01 | no | Gm23244 | downstream_gene_variant | MODIFIER |
| 6 | 113033330 | AX-170197374 | 0 | T | C | 0.07 | 7.84E-03 | 2.78E-01 | 2.70E+00 | 2.74E+00 | 9.77E-01 | no | Gm23244 | downstream_gene_variant | MODIFIER |
| 6 | 113093142 | AX-97844322 | 0 | C | T | 0.07 | 7.84E-03 | 2.78E-01 | 2.70E+00 | 2.74E+00 | 9.77E-01 | no | Setd5 | intron_variant | MODIFIER |
| 6 | 113096033 | AX-170028006 | 0 | G | A | 0.07 | 7.84E-03 | 2.78E-01 | 2.70E+00 | 2.74E+00 | 9.77E-01 | no | Setd5 | intron_variant | MODIFIER |
| 6 | 113134882 | AX-97837513 | 0 | A | C | 0.07 | 7.84E-03 | 2.78E-01 | 2.70E+00 | 2.74E+00 | 9.77E-01 | no | Setd5 | intron_variant | MODIFIER |
| 6 | 113135716 | AX-170028005 | 0 | G | A | 0.07 | 7.84E-03 | 2.78E-01 | 2.70E+00 | 2.74E+00 | 9.77E-01 | no | Setd5 | intron_variant | MODIFIER |
| 6 | 113167696 | AX-170306150 | 0 | T | C | 0.07 | 7.84E-03 | 2.78E-01 | 2.70E+00 | 2.74E+00 | 9.77E-01 | no | Lhfpl4 | downstream_gene_variant | MODIFIER |
| 6 | 114427813 | AX-170075935 | 0 | C | T | 0.07 | 7.84E-03 | 2.78E-01 | 2.70E+00 | 2.74E+00 | 9.77E-01 | no | Hrh1 | intron_variant | MODIFIER |
| 6 | 114427917 | AX-170416097 | 0 | A | G | 0.07 | 7.84E-03 | 2.78E-01 | 2.70E+00 | 2.74E+00 | 9.77E-01 | no | Hrh1 | intron_variant | MODIFIER |
| 6 | 114440522 | AX-170395790 | 0 | G | A | 0.07 | 7.84E-03 | 2.78E-01 | 2.70E+00 | 2.74E+00 | 9.77E-01 | no | Hrh1 | intron_variant | MODIFIER |
| 6 | 114445211 | AX-170403911 | 0 | C | T | 0.07 | 7.84E-03 | 2.78E-01 | 2.70E+00 | 2.74E+00 | 9.77E-01 | no | Hrh1 | intron_variant | MODIFIER |
| 6 | 114451211 | AX-170200243 | 0 | G | A | 0.07 | 7.84E-03 | 2.78E-01 | 2.70E+00 | 2.74E+00 | 9.77E-01 | no | Hrh1 | intron_variant | MODIFIER |
| 6 | 114451473 | AX-170089817 | 0 | T | C | 0.07 | 7.84E-03 | 2.78E-01 | 2.70E+00 | 2.74E+00 | 9.77E-01 | no | Hrh1 | intron_variant | MODIFIER |
| 6 | 114479303 | AX-97835117 | 0 | A | C | 0.07 | 7.84E-03 | 2.78E-01 | 2.70E+00 | 2.74E+00 | 9.77E-01 | no | Hrh1 | intron_variant | MODIFIER |
| 6 | 114481980 | AX-170228343 | 0 | G | A | 0.07 | 7.84E-03 | 2.78E-01 | 2.70E+00 | 2.74E+00 | 9.77E-01 | no | Hrh1 | 3_prime_UTR_variant | MODIFIER |
| 6 | 114482860 | AX-170441767 | 0 | A | G | 0.07 | 7.84E-03 | 2.78E-01 | 2.70E+00 | 2.74E+00 | 9.77E-01 | no | Hrh1 | 3_prime_UTR_variant | MODIFIER |
| 6 | 114643357 | AX-170256050 | 0 | G | C | 0.07 | 7.84E-03 | 2.78E-01 | 2.70E+00 | 2.74E+00 | 9.77E-01 | no | Atg7 | intron_variant | MODIFIER |
| 6 | 114649736 | AX-170327643 | 0 | C | A | 0.07 | 7.84E-03 | 2.78E-01 | 2.70E+00 | 2.74E+00 | 9.77E-01 | no | Atg7 | intron_variant | MODIFIER |
| 6 | 114690790 | AX-97855861 | 0 | G | A | 0.07 | 7.84E-03 | 2.78E-01 | 2.70E+00 | 2.74E+00 | 9.77E-01 | no | Atg7 | intron_variant | MODIFIER |
| 6 | 114713963 | AX-170443322 | 0 | A | T | 0.07 | 7.84E-03 | 2.78E-01 | 2.70E+00 | 2.74E+00 | 9.77E-01 | no | Atg7 | intron_variant | MODIFIER |
| 6 | 114726749 | AX-97844053 | 0 | C | T | 0.07 | 7.84E-03 | 2.78E-01 | 2.70E+00 | 2.74E+00 | 9.77E-01 | no | Atg7 | intron_variant | MODIFIER |
| 6 | 114781041 | AX-97781271 | 0 | C | T | 0.07 | 7.84E-03 | 2.78E-01 | 2.70E+00 | 2.74E+00 | 9.77E-01 | no | Atg7 | intron_variant | MODIFIER |
| 6 | 114799455 | AX-170427671 | 0 | G | A | 0.07 | 7.84E-03 | 2.78E-01 | 2.70E+00 | 2.74E+00 | 9.77E-01 | no | Atg7 | intron_variant | MODIFIER |
| 6 | 114803830 | AX-170378450 | 0 | G | A | 0.07 | 7.84E-03 | 2.78E-01 | 2.70E+00 | 2.74E+00 | 9.77E-01 | no | Atg7 | intron_variant | MODIFIER |
| 6 | 114986891 | AX-97804987 | 0 | C | T | 0.07 | 7.84E-03 | 2.78E-01 | 2.70E+00 | 2.74E+00 | 9.77E-01 | no | | intergenic_variant | MODIFIER |
| 6 | 116943004 | AX-170028061 | 0 | C | T | 0.07 | 7.84E-03 | 2.78E-01 | 2.70E+00 | 2.74E+00 | 9.77E-01 | no | | intergenic_variant | MODIFIER |
| 6 | 117470880 | AX-170167134 | 0 | G | C | 0.07 | 7.84E-03 | 2.78E-01 | 2.70E+00 | 2.74E+00 | 9.77E-01 | no | Gm6227 | downstream_gene_variant | MODIFIER |
| 6 | 118079442 | AX-97839856 | 0 | C | T | 0.07 | 7.84E-03 | 2.78E-01 | 2.70E+00 | 2.74E+00 | 9.77E-01 | no | Rasgef1a | intron_variant | MODIFIER |
| 6 | 118173020 | AX-170266786 | 0 | G | A | 0.07 | 7.84E-03 | 2.78E-01 | 2.70E+00 | 2.74E+00 | 9.77E-01 | no | Ret | intron_variant | MODIFIER |
| 6 | 118280071 | AX-97792525 | 0 | G | T | 0.07 | 7.84E-03 | 2.78E-01 | 2.70E+00 | 2.74E+00 | 9.77E-01 | no | | intergenic_variant | MODIFIER |
| 6 | 118295567 | AX-170204947 | 0 | G | A | 0.07 | 7.84E-03 | 2.78E-01 | 2.70E+00 | 2.74E+00 | 9.77E-01 | no | Gm30498 | downstream_gene_variant | MODIFIER |
| 6 | 118299640 | AX-170285962 | 0 | A | T | 0.07 | 7.84E-03 | 2.78E-01 | 2.70E+00 | 2.74E+00 | 9.77E-01 | no | Gm30498 | downstream_gene_variant | MODIFIER |
| 6 | 118336700 | AX-170376141 | 0 | C | T | 0.07 | 7.84E-03 | 2.78E-01 | 2.70E+00 | 2.74E+00 | 9.77E-01 | no | | regulatory_region_variant | MODIFIER |
| 6 | 118499853 | AX-170192021 | 0 | C | T | 0.07 | 7.84E-03 | 2.78E-01 | 2.70E+00 | 2.74E+00 | 9.77E-01 | no | Ankrd26 | downstream_gene_variant | MODIFIER |
| 6 | 118500448 | AX-170233604 | 0 | T | C | 0.07 | 7.84E-03 | 2.78E-01 | 2.70E+00 | 2.74E+00 | 9.77E-01 | no | Ankrd26 | downstream_gene_variant | MODIFIER |
| 6 | 118543090 | AX-168374496 | 0 | C | T | 0.07 | 7.84E-03 | 2.78E-01 | 2.70E+00 | 2.74E+00 | 9.77E-01 | no | Ankrd26 | intron_variant | MODIFIER |
| 6 | 118547534 | AX-170152168 | 0 | C | T | 0.07 | 7.84E-03 | 2.78E-01 | 2.70E+00 | 2.74E+00 | 9.77E-01 | no | Ankrd26 | intron_variant | MODIFIER |
| 6 | 118564325 | AX-170028078 | 0 | G | A | 0.07 | 7.84E-03 | 2.78E-01 | 2.70E+00 | 2.74E+00 | 9.77E-01 | no | Ankrd26 | upstream_gene_variant | MODIFIER |
| 6 | 118574829 | AX-170028075 | 0 | A | T | 0.07 | 7.84E-03 | 2.78E-01 | 2.70E+00 | 2.74E+00 | 9.77E-01 | no | | intergenic_variant | MODIFIER |
| 6 | 118625115 | AX-97823717 | 0 | A | G | 0.07 | 7.84E-03 | 2.78E-01 | 2.70E+00 | 2.74E+00 | 9.77E-01 | no | Cacna1c | intron_variant | MODIFIER |
| 6 | 118630643 | AX-170486007 | 0 | G | A | 0.07 | 7.84E-03 | 2.78E-01 | 2.70E+00 | 2.74E+00 | 9.77E-01 | no | Cacna1c | intron_variant | MODIFIER |

| | | | | | | | | | | | | | | | |
|---|---|---|---|---|---|---|---|---|---|---|---|---|---|---|---|
| 6 | 118632510 | AX-170151039 | 0 | T | C | 0.07 | 7.84E-03 | 2.78E-01 | 2.70E+00 | 2.74E+00 | 9.77E-01 | no | Cacna1c | intron_variant | MODIFIER |
| 6 | 118652400 | AX-170324945 | 0 | T | C | 0.07 | 7.84E-03 | 2.78E-01 | 2.70E+00 | 2.74E+00 | 9.77E-01 | no | Cacna1c | intron_variant | MODIFIER |
| 6 | 118653251 | AX-170218726 | 0 | A | T | 0.07 | 7.84E-03 | 2.78E-01 | 2.70E+00 | 2.74E+00 | 9.77E-01 | no | Cacna1c | intron_variant | MODIFIER |
| 6 | 118653973 | AX-170528820 | 0 | G | A | 0.07 | 7.84E-03 | 2.78E-01 | 2.70E+00 | 2.74E+00 | 9.77E-01 | no | Cacna1c | intron_variant | MODIFIER |
| 6 | 118695900 | AX-170486694 | 0 | C | G | 0.07 | 7.84E-03 | 2.78E-01 | 2.70E+00 | 2.74E+00 | 9.77E-01 | no | Cacna1c | intron_variant | MODIFIER |
| 6 | 118733361 | AX-170193048 | 0 | C | T | 0.07 | 7.84E-03 | 2.78E-01 | 2.70E+00 | 2.74E+00 | 9.77E-01 | no | Cacna1c | intron_variant | MODIFIER |
| 6 | 118733472 | AX-97843609 | 0 | C | T | 0.07 | 7.84E-03 | 2.78E-01 | 2.70E+00 | 2.74E+00 | 9.77E-01 | no | Cacna1c | intron_variant | MODIFIER |
| 6 | 118975237 | AX-170175021 | 0 | G | A | 0.07 | 7.84E-03 | 2.78E-01 | 2.70E+00 | 2.74E+00 | 9.77E-01 | no | Cacna1c | intron_variant | MODIFIER |
| 6 | 118977487 | AX-170342182 | 0 | G | A | 0.07 | 7.84E-03 | 2.78E-01 | 2.70E+00 | 2.74E+00 | 9.77E-01 | no | Cacna1c | intron_variant | MODIFIER |
| 6 | 118978012 | AX-170127206 | 0 | G | A | 0.07 | 7.84E-03 | 2.78E-01 | 2.70E+00 | 2.74E+00 | 9.77E-01 | no | Cacna1c | intron_variant | MODIFIER |
| 6 | 118978625 | AX-170116457 | 0 | A | G | 0.07 | 7.84E-03 | 2.78E-01 | 2.70E+00 | 2.74E+00 | 9.77E-01 | no | Cacna1c | intron_variant | MODIFIER |
| 6 | 118994301 | AX-170499760 | 0 | A | G | 0.07 | 7.84E-03 | 2.78E-01 | 2.70E+00 | 2.74E+00 | 9.77E-01 | no | Cacna1c | intron_variant | MODIFIER |
| 6 | 119000186 | AX-170396990 | 0 | A | G | 0.07 | 7.84E-03 | 2.78E-01 | 2.70E+00 | 2.74E+00 | 9.77E-01 | no | Cacna1c | intron_variant | MODIFIER |
| 6 | 119003920 | AX-170260419 | 0 | C | A | 0.07 | 7.84E-03 | 2.78E-01 | 2.70E+00 | 2.74E+00 | 9.77E-01 | no | Cacna1c | intron_variant | MODIFIER |
| 6 | 119004921 | AX-170513300 | 0 | G | A | 0.07 | 7.84E-03 | 2.78E-01 | 2.70E+00 | 2.74E+00 | 9.77E-01 | no | Cacna1c | intron_variant | MODIFIER |
| 6 | 119066647 | AX-170299623 | 0 | T | C | 0.07 | 7.84E-03 | 2.78E-01 | 2.70E+00 | 2.74E+00 | 9.77E-01 | no | Cacna1c | intron_variant | MODIFIER |
| 6 | 119066914 | AX-170028085 | 0 | A | G | 0.07 | 7.84E-03 | 2.78E-01 | 2.70E+00 | 2.74E+00 | 9.77E-01 | no | Cacna1c | intron_variant | MODIFIER |
| 6 | 119072418 | AX-170320331 | 0 | G | A | 0.07 | 7.84E-03 | 2.78E-01 | 2.70E+00 | 2.74E+00 | 9.77E-01 | no | Cacna1c | intron_variant | MODIFIER |
| 6 | 119075758 | AX-170112453 | 0 | C | A | 0.07 | 7.84E-03 | 2.78E-01 | 2.70E+00 | 2.74E+00 | 9.77E-01 | no | Cacna1c | intron_variant | MODIFIER |
| 6 | 119081262 | AX-170534356 | 0 | T | C | 0.07 | 7.84E-03 | 2.78E-01 | 2.70E+00 | 2.74E+00 | 9.77E-01 | no | Cacna1c | intron_variant | MODIFIER |
| 6 | 119084512 | AX-170028084 | 0 | G | A | 0.07 | 7.84E-03 | 2.78E-01 | 2.70E+00 | 2.74E+00 | 9.77E-01 | no | Cacna1c | intron_variant | MODIFIER |
| 6 | 119093964 | AX-170155873 | 0 | C | A | 0.07 | 7.84E-03 | 2.78E-01 | 2.70E+00 | 2.74E+00 | 9.77E-01 | no | Cacna1c | intron_variant | MODIFIER |
| 6 | 119098230 | AX-170228822 | 0 | A | G | 0.07 | 7.84E-03 | 2.78E-01 | 2.70E+00 | 2.74E+00 | 9.77E-01 | no | Cacna1c | intron_variant | MODIFIER |
| 6 | 119100387 | AX-170197554 | 0 | T | C | 0.07 | 7.84E-03 | 2.78E-01 | 2.70E+00 | 2.74E+00 | 9.77E-01 | no | Cacna1c | intron_variant | MODIFIER |
| 6 | 119138960 | AX-170028082 | 0 | G | C | 0.07 | 7.84E-03 | 2.78E-01 | 2.70E+00 | 2.74E+00 | 9.77E-01 | no | Cacna1c | intron_variant | MODIFIER |
| 6 | 119152771 | AX-97780407 | 0 | T | C | 0.07 | 7.84E-03 | 2.78E-01 | 2.70E+00 | 2.74E+00 | 9.77E-01 | no | Cacna1c | intron_variant | MODIFIER |
| 6 | 119176904 | AX-97789627 | 0 | T | C | 0.07 | 7.84E-03 | 2.78E-01 | 2.70E+00 | 2.74E+00 | 9.77E-01 | no | Dcp1b | intron_variant | MODIFIER |
| 6 | 119199924 | AX-170145958 | 0 | C | T | 0.07 | 7.84E-03 | 2.78E-01 | 2.70E+00 | 2.74E+00 | 9.77E-01 | no | Dcp1b | intron_variant | MODIFIER |
| 6 | 119215850 | AX-170488101 | 0 | G | A | 0.07 | 7.84E-03 | 2.78E-01 | 2.70E+00 | 2.74E+00 | 9.77E-01 | no | Dcp1b | intron_variant | MODIFIER |
| 6 | 119218245 | AX-170148830 | 0 | C | T | 0.07 | 7.84E-03 | 2.78E-01 | 2.70E+00 | 2.74E+00 | 9.77E-01 | no | Dcp1b | intron_variant | MODIFIER |
| 6 | 119223409 | AX-97826677 | 0 | C | T | 0.07 | 7.84E-03 | 2.78E-01 | 2.70E+00 | 2.74E+00 | 9.77E-01 | no | Dcp1b | downstream_gene_variant | MODIFIER |
| 6 | 119503340 | AX-170300216 | 0 | T | A | 0.07 | 7.84E-03 | 2.78E-01 | 2.70E+00 | 2.74E+00 | 9.77E-01 | no | Wnt5b | intron_variant | MODIFIER |
| 6 | 119525896 | AX-170334734 | 0 | C | T | 0.07 | 7.84E-03 | 2.78E-01 | 2.70E+00 | 2.74E+00 | 9.77E-01 | no | Wnt5b | intron_variant | MODIFIER |
| 6 | 119534633 | AX-170488672 | 0 | A | G | 0.07 | 7.84E-03 | 2.78E-01 | 2.70E+00 | 2.74E+00 | 9.77E-01 | no | Wnt5b | intron_variant | MODIFIER |
| 6 | 119534661 | AX-170028093 | 0 | C | T | 0.07 | 7.84E-03 | 2.78E-01 | 2.70E+00 | 2.74E+00 | 9.77E-01 | no | Wnt5b | intron_variant | MODIFIER |
| 6 | 119573641 | AX-97820682 | 0 | A | G | 0.07 | 7.84E-03 | 2.78E-01 | 2.70E+00 | 2.74E+00 | 9.77E-01 | no | Erc1 | 3_prime_UTR_variant | MODIFIER |
| 6 | 119574492 | AX-170185548 | 0 | C | T | 0.07 | 7.84E-03 | 2.78E-01 | 2.70E+00 | 2.74E+00 | 9.77E-01 | no | Erc1 | 3_prime_UTR_variant | MODIFIER |
| 6 | 119595777 | AX-170076052 | 0 | G | T | 0.07 | 7.84E-03 | 2.78E-01 | 2.70E+00 | 2.74E+00 | 9.77E-01 | no | Erc1 | intron_variant | MODIFIER |
| 6 | 119597005 | AX-170089824 | 0 | G | A | 0.07 | 7.84E-03 | 2.78E-01 | 2.70E+00 | 2.74E+00 | 9.77E-01 | no | Erc1 | intron_variant | MODIFIER |
| 6 | 119599676 | AX-170028096 | 0 | T | C | 0.07 | 7.84E-03 | 2.78E-01 | 2.70E+00 | 2.74E+00 | 9.77E-01 | no | Erc1 | intron_variant | MODIFIER |
| 6 | 119648106 | AX-97811770 | 0 | G | A | 0.07 | 7.84E-03 | 2.78E-01 | 2.70E+00 | 2.74E+00 | 9.77E-01 | no | Erc1 | intron_variant | MODIFIER |
| 6 | 119656766 | AX-170028097 | 0 | C | T | 0.07 | 7.84E-03 | 2.78E-01 | 2.70E+00 | 2.74E+00 | 9.77E-01 | no | Erc1 | intron_variant | MODIFIER |
| 6 | 119698879 | AX-170028098 | 0 | C | T | 0.07 | 7.84E-03 | 2.78E-01 | 2.70E+00 | 2.74E+00 | 9.77E-01 | no | Erc1 | intron_variant | MODIFIER |
| 6 | 119804799 | AX-170460601 | 0 | C | A | 0.07 | 7.84E-03 | 2.78E-01 | 2.70E+00 | 2.74E+00 | 9.77E-01 | no | Erc1 | intron_variant | MODIFIER |
| 6 | 119826601 | AX-170125243 | 0 | T | C | 0.07 | 7.84E-03 | 2.78E-01 | 2.70E+00 | 2.74E+00 | 9.77E-01 | no | Erc1 | upstream_gene_variant | MODIFIER |
| 6 | 119928828 | AX-97822131 | 0 | A | G | 0.07 | 7.84E-03 | 2.78E-01 | 2.70E+00 | 2.74E+00 | 9.77E-01 | no | Wnk1 | intron_variant | MODIFIER |
| 6 | 119931314 | AX-170300536 | 0 | G | T | 0.07 | 7.84E-03 | 2.78E-01 | 2.70E+00 | 2.74E+00 | 9.77E-01 | no | Wnk1 | intron_variant | MODIFIER |
| 6 | 119986173 | AX-170425020 | 0 | G | A | 0.07 | 7.84E-03 | 2.78E-01 | 2.70E+00 | 2.74E+00 | 9.77E-01 | no | Wnk1 | upstream_gene_variant | MODIFIER |
| 6 | 119995593 | AX-170137893 | 0 | G | A | 0.07 | 7.84E-03 | 2.78E-01 | 2.70E+00 | 2.74E+00 | 9.77E-01 | no | Wnk1 | intron_variant | MODIFIER |
| 6 | 120004619 | AX-97808889 | 0 | C | T | 0.07 | 7.84E-03 | 2.78E-01 | 2.70E+00 | 2.74E+00 | 9.77E-01 | no | Wnk1 | intron_variant | MODIFIER |
| 6 | 120016593 | AX-97837544 | 0 | T | C | 0.07 | 7.84E-03 | 2.78E-01 | 2.70E+00 | 2.74E+00 | 9.77E-01 | no | Wnk1 | intron_variant | MODIFIER |
| 6 | 120083565 | AX-170150970 | 0 | G | A | 0.07 | 7.84E-03 | 2.78E-01 | 2.70E+00 | 2.74E+00 | 9.77E-01 | no | | regulatory_region_variant | MODIFIER |
| 6 | 120085124 | AX-97851629 | 0 | T | C | 0.07 | 7.84E-03 | 2.78E-01 | 2.70E+00 | 2.74E+00 | 9.77E-01 | no | | regulatory_region_variant | MODIFIER |
| 6 | 120120680 | AX-97815963 | 0 | G | A | 0.07 | 7.84E-03 | 2.78E-01 | 2.70E+00 | 2.74E+00 | 9.77E-01 | no | Ninj2 | intron_variant | MODIFIER |

| | | | | | | | | | | | | | | | |
|---|---|---|---|---|---|---|---|---|---|---|---|---|---|---|---|
| 6 | 120152134 | AX-97790917 | 0 | T | C | 0.07 | 7.84E-03 | 2.78E-01 | 2.70E+00 | 2.74E+00 | 9.77E-01 | no | Ninj2 | intron_variant | MODIFIER |
| 6 | 120231818 | AX-170434178 | 0 | T | G | 0.07 | 7.84E-03 | 2.78E-01 | 2.70E+00 | 2.74E+00 | 9.77E-01 | no | B4galnt3 | intron_variant | MODIFIER |
| 6 | 120477863 | AX-97809126 | 0 | G | A | 0.07 | 7.84E-03 | 2.78E-01 | 2.70E+00 | 2.74E+00 | 9.77E-01 | no | Il17ra | intron_variant | MODIFIER |
| 6 | 120510375 | AX-97857627 | 0 | G | A | 0.07 | 7.84E-03 | 2.78E-01 | 2.70E+00 | 2.74E+00 | 9.77E-01 | no | Hdhd5 | synonymous_variant | LOW |
| 6 | 120527875 | AX-170117279 | 0 | A | C | 0.07 | 7.84E-03 | 2.78E-01 | 2.70E+00 | 2.74E+00 | 9.77E-01 | no | Hdhd5 | intron_variant | MODIFIER |
| 6 | 120600323 | AX-97852491 | 0 | G | A | 0.07 | 7.84E-03 | 2.78E-01 | 2.70E+00 | 2.74E+00 | 9.77E-01 | no | | regulatory_region_variant | MODIFIER |
| 6 | 120612668 | AX-170028118 | 0 | C | T | 0.07 | 7.84E-03 | 2.78E-01 | 2.70E+00 | 2.74E+00 | 9.77E-01 | no | | regulatory_region_variant | MODIFIER |
| 6 | 120622083 | AX-97855817 | 0 | A | G | 0.07 | 7.84E-03 | 2.78E-01 | 2.70E+00 | 2.74E+00 | 9.77E-01 | no | | intergenic_variant | MODIFIER |
| 6 | 120637738 | AX-170028110 | 0 | T | G | 0.07 | 7.84E-03 | 2.78E-01 | 2.70E+00 | 2.74E+00 | 9.77E-01 | no | | intergenic_variant | MODIFIER |
| 6 | 120643093 | AX-97801770 | 0 | A | G | 0.07 | 7.84E-03 | 2.78E-01 | 2.70E+00 | 2.74E+00 | 9.77E-01 | no | | regulatory_region_variant | MODIFIER |
| 6 | 120692057 | AX-170028112 | 0 | C | T | 0.07 | 7.84E-03 | 2.78E-01 | 2.70E+00 | 2.74E+00 | 9.77E-01 | no | Cecr2 | intron_variant | MODIFIER |
| 6 | 120694036 | AX-170028120 | 0 | T | C | 0.07 | 7.84E-03 | 2.78E-01 | 2.70E+00 | 2.74E+00 | 9.77E-01 | no | Cecr2 | intron_variant | MODIFIER |
| 6 | 120724484 | AX-170028117 | 0 | A | C | 0.07 | 7.84E-03 | 2.78E-01 | 2.70E+00 | 2.74E+00 | 9.77E-01 | no | Cecr2 | intron_variant | MODIFIER |
| 6 | 120777941 | AX-170028115 | 0 | A | G | 0.07 | 7.84E-03 | 2.78E-01 | 2.70E+00 | 2.74E+00 | 9.77E-01 | no | Slc25a18 | intron_variant | MODIFIER |
| 6 | 120892242 | AX-97832449 | 0 | G | A | 0.07 | 7.84E-03 | 2.78E-01 | 2.70E+00 | 2.74E+00 | 9.77E-01 | no | Bid | downstream_gene_variant | MODIFIER |
| 6 | 120938702 | AX-170484446 | 0 | G | A | 0.07 | 7.84E-03 | 2.78E-01 | 2.70E+00 | 2.74E+00 | 9.77E-01 | no | Mical3 | intron_variant,non_coding_transcript_variant | MODIFIER |
| 6 | 120959397 | AX-97837689 | 0 | C | T | 0.07 | 7.84E-03 | 2.78E-01 | 2.70E+00 | 2.74E+00 | 9.77E-01 | no | Mical3 | non_coding_transcript_exon_variant | MODIFIER |
| 6 | 120978365 | AX-170135184 | 0 | T | C | 0.07 | 7.84E-03 | 2.78E-01 | 2.70E+00 | 2.74E+00 | 9.77E-01 | no | Mical3 | intron_variant,non_coding_transcript_variant | MODIFIER |
| 6 | 120980606 | AX-170348617 | 0 | C | T | 0.07 | 7.84E-03 | 2.78E-01 | 2.70E+00 | 2.74E+00 | 9.77E-01 | no | Mical3 | intron_variant,non_coding_transcript_variant | MODIFIER |
| 6 | 121024482 | AX-170181499 | 0 | C | T | 0.07 | 7.84E-03 | 2.78E-01 | 2.70E+00 | 2.74E+00 | 9.77E-01 | no | Mical3 | intron_variant | MODIFIER |
| 6 | 121040826 | AX-170272348 | 0 | C | A | 0.07 | 7.84E-03 | 2.78E-01 | 2.70E+00 | 2.74E+00 | 9.77E-01 | no | Mical3 | intron_variant | MODIFIER |
| 6 | 121048415 | AX-170358360 | 0 | G | A | 0.07 | 7.84E-03 | 2.78E-01 | 2.70E+00 | 2.74E+00 | 9.77E-01 | no | Mical3 | intron_variant | MODIFIER |
| 6 | 121052508 | AX-170196740 | 0 | A | G | 0.07 | 7.84E-03 | 2.78E-01 | 2.70E+00 | 2.74E+00 | 9.77E-01 | no | Mical3 | intron_variant | MODIFIER |
| 6 | 121117599 | AX-170355976 | 0 | T | C | 0.07 | 7.84E-03 | 2.78E-01 | 2.70E+00 | 2.74E+00 | 9.77E-01 | no | Minpp1-ps | non_coding_transcript_exon_variant | MODIFIER |
| 6 | 121129328 | AX-170265990 | 0 | C | T | 0.07 | 7.84E-03 | 2.78E-01 | 2.70E+00 | 2.74E+00 | 9.77E-01 | no | Mical3 | intron_variant | MODIFIER |
| 6 | 121176339 | AX-97838117 | 0 | A | G | 0.07 | 7.84E-03 | 2.78E-01 | 2.70E+00 | 2.74E+00 | 9.77E-01 | no | | regulatory_region_variant | MODIFIER |
| 6 | 121243453 | AX-170483980 | 0 | A | T | 0.07 | 7.84E-03 | 2.78E-01 | 2.70E+00 | 2.74E+00 | 9.77E-01 | no | Usp18 | upstream_gene_variant | MODIFIER |
| 6 | 121326400 | AX-170076079 | 0 | C | T | 0.07 | 7.84E-03 | 2.78E-01 | 2.70E+00 | 2.74E+00 | 9.77E-01 | no | Slc6a13 | intron_variant | MODIFIER |
| 6 | 121328292 | AX-97808802 | 0 | G | T | 0.07 | 7.84E-03 | 2.78E-01 | 2.70E+00 | 2.74E+00 | 9.77E-01 | no | Slc6a13 | intron_variant | MODIFIER |
| 6 | 121334907 | AX-170399944 | 0 | C | T | 0.07 | 7.84E-03 | 2.78E-01 | 2.70E+00 | 2.74E+00 | 9.77E-01 | no | Slc6a13 | synonymous_variant | LOW |
| 6 | 121364141 | AX-170138921 | 0 | C | T | 0.07 | 7.84E-03 | 2.78E-01 | 2.70E+00 | 2.74E+00 | 9.77E-01 | no | Slc6a12 | intron_variant | MODIFIER |
| 6 | 121497253 | AX-170194020 | 0 | T | C | 0.07 | 7.84E-03 | 2.78E-01 | 2.70E+00 | 2.74E+00 | 9.77E-01 | no | | intergenic_variant | MODIFIER |
| 6 | 121500587 | AX-170119263 | 0 | C | A | 0.07 | 7.84E-03 | 2.78E-01 | 2.70E+00 | 2.74E+00 | 9.77E-01 | no | | intergenic_variant | MODIFIER |
| 6 | 121524116 | AX-170353517 | 0 | T | C | 0.07 | 7.84E-03 | 2.78E-01 | 2.70E+00 | 2.74E+00 | 9.77E-01 | no | | regulatory_region_variant | MODIFIER |
| 6 | 121529887 | AX-170241466 | 0 | A | G | 0.07 | 7.84E-03 | 2.78E-01 | 2.70E+00 | 2.74E+00 | 9.77E-01 | no | | intergenic_variant | MODIFIER |
| 6 | 121530406 | AX-170338125 | 0 | G | A | 0.07 | 7.84E-03 | 2.78E-01 | 2.70E+00 | 2.74E+00 | 9.77E-01 | no | | intergenic_variant | MODIFIER |
| 6 | 121530758 | AX-170222957 | 0 | A | T | 0.07 | 7.84E-03 | 2.78E-01 | 2.70E+00 | 2.74E+00 | 9.77E-01 | no | | intergenic_variant | MODIFIER |
| 6 | 121530908 | AX-170491342 | 0 | G | A | 0.07 | 7.84E-03 | 2.78E-01 | 2.70E+00 | 2.74E+00 | 9.77E-01 | no | | intergenic_variant | MODIFIER |
| 6 | 121580307 | AX-170203537 | 0 | T | C | 0.07 | 7.84E-03 | 2.78E-01 | 2.70E+00 | 2.74E+00 | 9.77E-01 | no | | intergenic_variant | MODIFIER |
| 6 | 121611864 | AX-170374388 | 0 | G | A | 0.07 | 7.84E-03 | 2.78E-01 | 2.70E+00 | 2.74E+00 | 9.77E-01 | no | | regulatory_region_variant | MODIFIER |
| 6 | 121665255 | AX-170152410 | 0 | C | T | 0.07 | 7.84E-03 | 2.78E-01 | 2.70E+00 | 2.74E+00 | 9.77E-01 | no | A2m | intron_variant | MODIFIER |
| 6 | 121665273 | AX-170184557 | 0 | T | C | 0.07 | 7.84E-03 | 2.78E-01 | 2.70E+00 | 2.74E+00 | 9.77E-01 | no | A2m | intron_variant | MODIFIER |
| 6 | 121669253 | AX-170097497 | 0 | T | C | 0.07 | 7.84E-03 | 2.78E-01 | 2.70E+00 | 2.74E+00 | 9.77E-01 | no | A2m | intron_variant | MODIFIER |
| 6 | 121677995 | AX-170524939 | 0 | C | T | 0.07 | 7.84E-03 | 2.78E-01 | 2.70E+00 | 2.74E+00 | 9.77E-01 | no | A2m | missense_variant | MODERATE |
| 6 | 121724231 | AX-170076088 | 0 | A | G | 0.07 | 7.84E-03 | 2.78E-01 | 2.70E+00 | 2.74E+00 | 9.77E-01 | no | | intergenic_variant | MODIFIER |
| 6 | 121728413 | AX-170028132 | 0 | C | T | 0.07 | 7.84E-03 | 2.78E-01 | 2.70E+00 | 2.74E+00 | 9.77E-01 | no | Gm7298 | upstream_gene_variant | MODIFIER |
| 6 | 121862124 | AX-97835347 | 0 | G | A | 0.07 | 7.84E-03 | 2.78E-01 | 2.70E+00 | 2.74E+00 | 9.77E-01 | no | Mug1 | intron_variant | MODIFIER |
| 6 | 121964432 | AX-170028130 | 0 | G | A | 0.07 | 7.84E-03 | 2.78E-01 | 2.70E+00 | 2.74E+00 | 9.77E-01 | no | | intergenic_variant | MODIFIER |
| 6 | 121965144 | AX-170028133 | 0 | C | T | 0.07 | 7.84E-03 | 2.78E-01 | 2.70E+00 | 2.74E+00 | 9.77E-01 | no | | intergenic_variant | MODIFIER |
| 6 | 122040103 | AX-170028136 | 0 | A | T | 0.07 | 7.84E-03 | 2.78E-01 | 2.70E+00 | 2.74E+00 | 9.77E-01 | no | Mug2 | intron_variant | MODIFIER |
| 6 | 122293797 | AX-97797171 | 0 | A | G | 0.07 | 7.84E-03 | 2.78E-01 | 2.70E+00 | 2.74E+00 | 9.77E-01 | no | Gm6637 | upstream_gene_variant | MODIFIER |
| 6 | 122311211 | AX-170414408 | 0 | T | A | 0.07 | 7.84E-03 | 2.78E-01 | 2.70E+00 | 2.74E+00 | 9.77E-01 | no | M6pr | intron_variant | MODIFIER |
| 6 | 122319474 | AX-170168863 | 0 | C | T | 0.07 | 7.84E-03 | 2.78E-01 | 2.70E+00 | 2.74E+00 | 9.77E-01 | no | M6pr | downstream_gene_variant | MODIFIER |
| 6 | 138732732 | AX-170106793 | 0 | A | G | 0.07 | 7.84E-03 | 2.78E-01 | 2.70E+00 | 2.74E+00 | 9.77E-01 | no | | intergenic_variant | MODIFIER |

| | | | | | | | | | | | | | | | |
|---|---|---|---|---|---|---|---|---|---|---|---|---|---|---|---|
| 6 | 138736688 | AX-170222215 | 0 | G | C | 0.07 | 7.84E-03 | 2.78E-01 | 2.70E+00 | 2.74E+00 | 9.77E-01 | no | | intergenic_variant | MODIFIER |
| 6 | 138737086 | AX-170445865 | 0 | C | A | 0.07 | 7.84E-03 | 2.78E-01 | 2.70E+00 | 2.74E+00 | 9.77E-01 | no | | intergenic_variant | MODIFIER |
| 6 | 138772546 | AX-97814330 | 0 | A | G | 0.07 | 7.84E-03 | 2.78E-01 | 2.70E+00 | 2.74E+00 | 9.77E-01 | no | | intergenic_variant | MODIFIER |
| 6 | 138784664 | AX-170076395 | 0 | G | C | 0.07 | 7.84E-03 | 2.78E-01 | 2.70E+00 | 2.74E+00 | 9.77E-01 | no | | intergenic_variant | MODIFIER |
| 6 | 138791429 | AX-170390090 | 0 | T | C | 0.07 | 7.84E-03 | 2.78E-01 | 2.70E+00 | 2.74E+00 | 9.77E-01 | no | | intergenic_variant | MODIFIER |
| 6 | 138794940 | AX-170108208 | 0 | T | A | 0.07 | 7.84E-03 | 2.78E-01 | 2.70E+00 | 2.74E+00 | 9.77E-01 | no | | intergenic_variant | MODIFIER |
| 6 | 138795932 | AX-170530348 | 0 | T | G | 0.07 | 7.84E-03 | 2.78E-01 | 2.70E+00 | 2.74E+00 | 9.77E-01 | no | | intergenic_variant | MODIFIER |
| 6 | 138798251 | AX-170175949 | 0 | C | G | 0.07 | 7.84E-03 | 2.78E-01 | 2.70E+00 | 2.74E+00 | 9.77E-01 | no | | intergenic_variant | MODIFIER |
| 6 | 138798428 | AX-97784233 | 0 | T | C | 0.07 | 7.84E-03 | 2.78E-01 | 2.70E+00 | 2.74E+00 | 9.77E-01 | no | | intergenic_variant | MODIFIER |
| 6 | 138807164 | AX-170273987 | 0 | C | T | 0.07 | 7.84E-03 | 2.78E-01 | 2.70E+00 | 2.74E+00 | 9.77E-01 | no | | intergenic_variant | MODIFIER |
| 6 | 138813549 | AX-170158643 | 0 | C | G | 0.07 | 7.84E-03 | 2.78E-01 | 2.70E+00 | 2.74E+00 | 9.77E-01 | no | | intergenic_variant | MODIFIER |
| 6 | 138824682 | AX-170503098 | 0 | T | G | 0.07 | 7.84E-03 | 2.78E-01 | 2.70E+00 | 2.74E+00 | 9.77E-01 | no | | intergenic_variant | MODIFIER |
| 6 | 138832263 | AX-170506247 | 0 | A | C | 0.07 | 7.84E-03 | 2.78E-01 | 2.70E+00 | 2.74E+00 | 9.77E-01 | no | | intergenic_variant | MODIFIER |
| 6 | 138843115 | AX-170532232 | 0 | T | C | 0.07 | 7.84E-03 | 2.78E-01 | 2.70E+00 | 2.74E+00 | 9.77E-01 | no | | intergenic_variant | MODIFIER |
| 6 | 138854508 | AX-97794774 | 0 | A | G | 0.07 | 7.84E-03 | 2.78E-01 | 2.70E+00 | 2.74E+00 | 9.77E-01 | no | | intergenic_variant | MODIFIER |
| 6 | 138858497 | AX-170271677 | 0 | A | G | 0.07 | 7.84E-03 | 2.78E-01 | 2.70E+00 | 2.74E+00 | 9.77E-01 | no | Gm9038 | downstream_gene_variant | MODIFIER |
| 6 | 138860056 | AX-170316821 | 0 | G | T | 0.07 | 7.84E-03 | 2.78E-01 | 2.70E+00 | 2.74E+00 | 9.77E-01 | no | Gm9038 | non_coding_transcript_exon_variant | MODIFIER |
| 6 | 138889286 | AX-170468161 | 0 | A | G | 0.07 | 7.84E-03 | 2.78E-01 | 2.70E+00 | 2.74E+00 | 9.77E-01 | no | | intergenic_variant | MODIFIER |
| 6 | 138889662 | AX-170089854 | 0 | A | C | 0.07 | 7.84E-03 | 2.78E-01 | 2.70E+00 | 2.74E+00 | 9.77E-01 | no | | intergenic_variant | MODIFIER |
| 6 | 138906733 | AX-97854829 | 0 | C | T | 0.07 | 7.84E-03 | 2.78E-01 | 2.70E+00 | 2.74E+00 | 9.77E-01 | no | | intergenic_variant | MODIFIER |
| 6 | 138926452 | AX-170159480 | 0 | T | G | 0.07 | 7.84E-03 | 2.78E-01 | 2.70E+00 | 2.74E+00 | 9.77E-01 | no | | intergenic_variant | MODIFIER |
| 6 | 138926592 | AX-170493315 | 0 | T | G | 0.07 | 7.84E-03 | 2.78E-01 | 2.70E+00 | 2.74E+00 | 9.77E-01 | no | | intergenic_variant | MODIFIER |
| 6 | 138927057 | AX-97807230 | 0 | T | C | 0.07 | 7.84E-03 | 2.78E-01 | 2.70E+00 | 2.74E+00 | 9.77E-01 | no | | intergenic_variant | MODIFIER |
| 6 | 138933420 | AX-170473609 | 0 | T | G | 0.07 | 7.84E-03 | 2.78E-01 | 2.70E+00 | 2.74E+00 | 9.77E-01 | no | | intergenic_variant | MODIFIER |
| 6 | 138933489 | AX-170169575 | 0 | A | T | 0.07 | 7.84E-03 | 2.78E-01 | 2.70E+00 | 2.74E+00 | 9.77E-01 | no | | intergenic_variant | MODIFIER |
| 6 | 138934949 | AX-170198377 | 0 | A | G | 0.07 | 7.84E-03 | 2.78E-01 | 2.70E+00 | 2.74E+00 | 9.77E-01 | no | | intergenic_variant | MODIFIER |
| 6 | 138959759 | AX-97785222 | 0 | G | A | 0.07 | 7.84E-03 | 2.78E-01 | 2.70E+00 | 2.74E+00 | 9.77E-01 | no | | intergenic_variant | MODIFIER |
| 6 | 138960053 | AX-170196731 | 0 | G | A | 0.07 | 7.84E-03 | 2.78E-01 | 2.70E+00 | 2.74E+00 | 9.77E-01 | no | | intergenic_variant | MODIFIER |
| 6 | 138967113 | AX-97798740 | 0 | T | G | 0.07 | 7.84E-03 | 2.78E-01 | 2.70E+00 | 2.74E+00 | 9.77E-01 | no | | intergenic_variant | MODIFIER |
| 6 | 138967498 | AX-170211070 | 0 | G | C | 0.07 | 7.84E-03 | 2.78E-01 | 2.70E+00 | 2.74E+00 | 9.77E-01 | no | | intergenic_variant | MODIFIER |
| 6 | 139000820 | AX-170378081 | 0 | T | G | 0.07 | 7.84E-03 | 2.78E-01 | 2.70E+00 | 2.74E+00 | 9.77E-01 | no | | intergenic_variant | MODIFIER |
| 6 | 139003342 | AX-97783765 | 0 | T | G | 0.07 | 7.84E-03 | 2.78E-01 | 2.70E+00 | 2.74E+00 | 9.77E-01 | no | | intergenic_variant | MODIFIER |
| 6 | 139065539 | AX-170028370 | 0 | T | G | 0.07 | 7.84E-03 | 2.78E-01 | 2.70E+00 | 2.74E+00 | 9.77E-01 | no | | intergenic_variant | MODIFIER |
| 6 | 139066703 | AX-170438416 | 0 | G | T | 0.07 | 7.84E-03 | 2.78E-01 | 2.70E+00 | 2.74E+00 | 9.77E-01 | no | | intergenic_variant | MODIFIER |
| 6 | 139067976 | AX-170028369 | 0 | C | T | 0.07 | 7.84E-03 | 2.78E-01 | 2.70E+00 | 2.74E+00 | 9.77E-01 | no | | intergenic_variant | MODIFIER |
| 6 | 139071903 | AX-97841165 | 0 | G | A | 0.07 | 7.84E-03 | 2.78E-01 | 2.70E+00 | 2.74E+00 | 9.77E-01 | no | | intergenic_variant | MODIFIER |
| 6 | 139087381 | AX-170211951 | 0 | A | T | 0.07 | 7.84E-03 | 2.78E-01 | 2.70E+00 | 2.74E+00 | 9.77E-01 | no | | intergenic_variant | MODIFIER |
| 6 | 139089176 | AX-170028367 | 0 | C | T | 0.07 | 7.84E-03 | 2.78E-01 | 2.70E+00 | 2.74E+00 | 9.77E-01 | no | | intergenic_variant | MODIFIER |
| 6 | 139103104 | AX-97804408 | 0 | A | G | 0.07 | 7.84E-03 | 2.78E-01 | 2.70E+00 | 2.74E+00 | 9.77E-01 | no | | intergenic_variant | MODIFIER |
| 6 | 139108399 | AX-170495726 | 0 | G | T | 0.07 | 7.84E-03 | 2.78E-01 | 2.70E+00 | 2.74E+00 | 9.77E-01 | no | | intergenic_variant | MODIFIER |
| 6 | 139119306 | AX-170317403 | 0 | A | G | 0.07 | 7.84E-03 | 2.78E-01 | 2.70E+00 | 2.74E+00 | 9.77E-01 | no | | intergenic_variant | MODIFIER |
| 6 | 139134862 | AX-170076401 | 0 | A | T | 0.07 | 7.84E-03 | 2.78E-01 | 2.70E+00 | 2.74E+00 | 9.77E-01 | no | Gm44267 | upstream_gene_variant | MODIFIER |
| 6 | 139237208 | AX-170271087 | 0 | C | A | 0.07 | 7.84E-03 | 2.78E-01 | 2.70E+00 | 2.74E+00 | 9.77E-01 | no | | intergenic_variant | MODIFIER |
| 6 | 139285730 | AX-170285734 | 0 | C | G | 0.07 | 7.84E-03 | 2.78E-01 | 2.70E+00 | 2.74E+00 | 9.77E-01 | no | | intergenic_variant | MODIFIER |
| 6 | 139494033 | AX-170035743 | 0 | G | A | 0.07 | 7.84E-03 | 2.78E-01 | 2.70E+00 | 2.74E+00 | 9.77E-01 | no | Rergl | intron_variant | MODIFIER |
| 6 | 139716080 | AX-170452374 | 0 | A | T | 0.07 | 7.84E-03 | 2.78E-01 | 2.70E+00 | 2.74E+00 | 9.77E-01 | no | Pik3c2g | intron_variant | MODIFIER |
| 6 | 139718135 | AX-170465337 | 0 | G | A | 0.07 | 7.84E-03 | 2.78E-01 | 2.70E+00 | 2.74E+00 | 9.77E-01 | no | Pik3c2g | intron_variant | MODIFIER |
| 6 | 139719354 | AX-170453008 | 0 | T | C | 0.07 | 7.84E-03 | 2.78E-01 | 2.70E+00 | 2.74E+00 | 9.77E-01 | no | Pik3c2g | intron_variant | MODIFIER |
| 6 | 139760010 | AX-170103726 | 0 | T | A | 0.07 | 7.84E-03 | 2.78E-01 | 2.70E+00 | 2.74E+00 | 9.77E-01 | no | Pik3c2g | intron_variant | MODIFIER |
| 6 | 139767439 | AX-170229641 | 0 | A | G | 0.07 | 7.84E-03 | 2.78E-01 | 2.70E+00 | 2.74E+00 | 9.77E-01 | no | Pik3c2g | intron_variant | MODIFIER |
| 6 | 139800401 | AX-170318852 | 0 | A | C | 0.07 | 7.84E-03 | 2.78E-01 | 2.70E+00 | 2.74E+00 | 9.77E-01 | no | Pik3c2g | intron_variant | MODIFIER |
| 6 | 139803912 | AX-170188613 | 0 | C | T | 0.07 | 7.84E-03 | 2.78E-01 | 2.70E+00 | 2.74E+00 | 9.77E-01 | no | Pik3c2g | intron_variant | MODIFIER |
| 6 | 139804044 | AX-170360260 | 0 | T | C | 0.07 | 7.84E-03 | 2.78E-01 | 2.70E+00 | 2.74E+00 | 9.77E-01 | no | Pik3c2g | intron_variant | MODIFIER |
| 6 | 139833813 | AX-170450908 | 0 | G | A | 0.07 | 7.84E-03 | 2.78E-01 | 2.70E+00 | 2.74E+00 | 9.77E-01 | no | Pik3c2g | intron_variant | MODIFIER |

| | | | | | | | | | | | | | | | |
|---|---|---|---|---|---|---|---|---|---|---|---|---|---|---|---|
| 6 | 139833920 | AX-170339300 | 0 | A | G | 0.07 | 7.84E-03 | 2.78E-01 | 2.70E+00 | 2.74E+00 | 9.77E-01 | no | Pik3c2g | intron_variant | MODIFIER |
| 6 | 139834072 | AX-170263093 | 0 | A | G | 0.07 | 7.84E-03 | 2.78E-01 | 2.70E+00 | 2.74E+00 | 9.77E-01 | no | Pik3c2g | intron_variant | MODIFIER |
| 6 | 139834392 | AX-170366053 | 0 | A | G | 0.07 | 7.84E-03 | 2.78E-01 | 2.70E+00 | 2.74E+00 | 9.77E-01 | no | Pik3c2g | intron_variant | MODIFIER |
| 6 | 139840842 | AX-170154251 | 0 | C | G | 0.07 | 7.84E-03 | 2.78E-01 | 2.70E+00 | 2.74E+00 | 9.77E-01 | no | Pik3c2g | upstream_gene_variant | MODIFIER |
| 6 | 139841368 | AX-170115301 | 0 | T | G | 0.07 | 7.84E-03 | 2.78E-01 | 2.70E+00 | 2.74E+00 | 9.77E-01 | no | Pik3c2g | upstream_gene_variant | MODIFIER |
| 6 | 139893335 | AX-170534732 | 0 | A | G | 0.07 | 7.84E-03 | 2.78E-01 | 2.70E+00 | 2.74E+00 | 9.77E-01 | no | Pik3c2g | intron_variant | MODIFIER |
| 6 | 139942295 | AX-170028391 | 0 | C | T | 0.07 | 7.84E-03 | 2.78E-01 | 2.70E+00 | 2.74E+00 | 9.77E-01 | no | Pik3c2g | intron_variant | MODIFIER |
| 6 | 139943967 | AX-97798215 | 0 | C | A | 0.07 | 7.84E-03 | 2.78E-01 | 2.70E+00 | 2.74E+00 | 9.77E-01 | no | Pik3c2g | intron_variant | MODIFIER |
| 6 | 139977915 | AX-170223210 | 0 | A | G | 0.07 | 7.84E-03 | 2.78E-01 | 2.70E+00 | 2.74E+00 | 9.77E-01 | no | | intergenic_variant | MODIFIER |
| 6 | 140017437 | AX-170028389 | 0 | A | G | 0.07 | 7.84E-03 | 2.78E-01 | 2.70E+00 | 2.74E+00 | 9.77E-01 | no | Plcz1 | intron_variant | MODIFIER |
| 6 | 140022173 | AX-170466368 | 0 | T | G | 0.07 | 7.84E-03 | 2.78E-01 | 2.70E+00 | 2.74E+00 | 9.77E-01 | no | Plcz1 | intron_variant | MODIFIER |
| 6 | 140027447 | AX-170380351 | 0 | C | T | 0.07 | 7.84E-03 | 2.78E-01 | 2.70E+00 | 2.74E+00 | 9.77E-01 | no | Plcz1 | intron_variant | MODIFIER |
| 6 | 140028667 | AX-170333491 | 0 | G | A | 0.07 | 7.84E-03 | 2.78E-01 | 2.70E+00 | 2.74E+00 | 9.77E-01 | no | Plcz1 | intron_variant | MODIFIER |
| 6 | 140028764 | AX-170106482 | 0 | C | T | 0.07 | 7.84E-03 | 2.78E-01 | 2.70E+00 | 2.74E+00 | 9.77E-01 | no | Plcz1 | intron_variant | MODIFIER |
| 6 | 140065547 | AX-170130426 | 0 | C | G | 0.07 | 7.84E-03 | 2.78E-01 | 2.70E+00 | 2.74E+00 | 9.77E-01 | no | Gm44158 | downstream_gene_variant | MODIFIER |
| 6 | 140071257 | AX-170473518 | 0 | C | T | 0.07 | 7.84E-03 | 2.78E-01 | 2.70E+00 | 2.74E+00 | 9.77E-01 | no | Gm44158 | upstream_gene_variant | MODIFIER |
| 6 | 140077953 | AX-170076420 | 0 | T | C | 0.07 | 7.84E-03 | 2.78E-01 | 2.70E+00 | 2.74E+00 | 9.77E-01 | no | | intergenic_variant | MODIFIER |
| 6 | 140078348 | AX-170076418 | 0 | G | T | 0.07 | 7.84E-03 | 2.78E-01 | 2.70E+00 | 2.74E+00 | 9.77E-01 | no | | intergenic_variant | MODIFIER |
| 6 | 140089042 | AX-170172091 | 0 | T | C | 0.07 | 7.84E-03 | 2.78E-01 | 2.70E+00 | 2.74E+00 | 9.77E-01 | no | | intergenic_variant | MODIFIER |
| 6 | 140089245 | AX-170138652 | 0 | C | A | 0.07 | 7.84E-03 | 2.78E-01 | 2.70E+00 | 2.74E+00 | 9.77E-01 | no | | intergenic_variant | MODIFIER |
| 6 | 140091504 | AX-170459493 | 0 | T | C | 0.07 | 7.84E-03 | 2.78E-01 | 2.70E+00 | 2.74E+00 | 9.77E-01 | no | | intergenic_variant | MODIFIER |
| 6 | 140095136 | AX-170523794 | 0 | C | A | 0.07 | 7.84E-03 | 2.78E-01 | 2.70E+00 | 2.74E+00 | 9.77E-01 | no | | intergenic_variant | MODIFIER |
| 6 | 140105131 | AX-170451190 | 0 | T | C | 0.07 | 7.84E-03 | 2.78E-01 | 2.70E+00 | 2.74E+00 | 9.77E-01 | no | | intergenic_variant | MODIFIER |
| 6 | 140109238 | AX-97785911 | 0 | T | C | 0.07 | 7.84E-03 | 2.78E-01 | 2.70E+00 | 2.74E+00 | 9.77E-01 | no | | intergenic_variant | MODIFIER |
| 6 | 140118062 | AX-170214576 | 0 | A | G | 0.07 | 7.84E-03 | 2.78E-01 | 2.70E+00 | 2.74E+00 | 9.77E-01 | no | | intergenic_variant | MODIFIER |
| 6 | 140118090 | AX-97802413 | 0 | A | G | 0.07 | 7.84E-03 | 2.78E-01 | 2.70E+00 | 2.74E+00 | 9.77E-01 | no | | intergenic_variant | MODIFIER |
| 6 | 140143037 | AX-97802382 | 0 | T | C | 0.07 | 7.84E-03 | 2.78E-01 | 2.70E+00 | 2.74E+00 | 9.77E-01 | no | | intergenic_variant | MODIFIER |
| 6 | 140170992 | AX-170089856 | 0 | G | C | 0.07 | 7.84E-03 | 2.78E-01 | 2.70E+00 | 2.74E+00 | 9.77E-01 | no | Gm24174 | upstream_gene_variant | MODIFIER |
| 6 | 144100198 | AX-170532895 | 0 | A | G | 0.07 | 7.84E-03 | 2.78E-01 | 2.70E+00 | 2.74E+00 | 9.77E-01 | no | Sox5 | intron_variant | MODIFIER |
| 6 | 144100939 | AX-170465946 | 0 | G | A | 0.07 | 7.84E-03 | 2.78E-01 | 2.70E+00 | 2.74E+00 | 9.77E-01 | no | Sox5 | intron_variant | MODIFIER |
| 6 | 144118204 | AX-170408740 | 0 | G | A | 0.07 | 7.84E-03 | 2.78E-01 | 2.70E+00 | 2.74E+00 | 9.77E-01 | no | Sox5 | intron_variant | MODIFIER |
| 6 | 144119260 | AX-170449720 | 0 | C | T | 0.07 | 7.84E-03 | 2.78E-01 | 2.70E+00 | 2.74E+00 | 9.77E-01 | no | Sox5 | intron_variant | MODIFIER |
| 6 | 144121263 | AX-170419982 | 0 | C | G | 0.07 | 7.84E-03 | 2.78E-01 | 2.70E+00 | 2.74E+00 | 9.77E-01 | no | Sox5 | intron_variant | MODIFIER |
| 6 | 144263031 | AX-97783219 | 0 | T | C | 0.07 | 7.84E-03 | 2.78E-01 | 2.70E+00 | 2.74E+00 | 9.77E-01 | no | Sox5 | intron_variant,non_coding_transcript_variant | MODIFIER |
| 6 | 144335646 | AX-170375657 | 0 | G | A | 0.07 | 7.84E-03 | 2.78E-01 | 2.70E+00 | 2.74E+00 | 9.77E-01 | no | Sox5 | intron_variant,non_coding_transcript_variant | MODIFIER |
| 6 | 144360384 | AX-170453136 | 0 | G | A | 0.07 | 7.84E-03 | 2.78E-01 | 2.70E+00 | 2.74E+00 | 9.77E-01 | no | Sox5 | intron_variant,non_coding_transcript_variant | MODIFIER |
| 6 | 144380354 | AX-170306473 | 0 | G | A | 0.07 | 7.84E-03 | 2.78E-01 | 2.70E+00 | 2.74E+00 | 9.77E-01 | no | Sox5 | intron_variant,non_coding_transcript_variant | MODIFIER |
| 6 | 144380422 | AX-170291932 | 0 | T | C | 0.07 | 7.84E-03 | 2.78E-01 | 2.70E+00 | 2.74E+00 | 9.77E-01 | no | Sox5 | intron_variant,non_coding_transcript_variant | MODIFIER |
| 6 | 144381220 | AX-170281154 | 0 | G | A | 0.07 | 7.84E-03 | 2.78E-01 | 2.70E+00 | 2.74E+00 | 9.77E-01 | no | Sox5 | intron_variant,non_coding_transcript_variant | MODIFIER |
| 6 | 144405208 | AX-170144336 | 0 | T | C | 0.07 | 7.84E-03 | 2.78E-01 | 2.70E+00 | 2.74E+00 | 9.77E-01 | no | Sox5 | intron_variant,non_coding_transcript_variant | MODIFIER |
| 6 | 144405470 | AX-170230983 | 0 | G | T | 0.07 | 7.84E-03 | 2.78E-01 | 2.70E+00 | 2.74E+00 | 9.77E-01 | no | Sox5 | intron_variant,non_coding_transcript_variant | MODIFIER |
| 6 | 144405498 | AX-170336568 | 0 | A | G | 0.07 | 7.84E-03 | 2.78E-01 | 2.70E+00 | 2.74E+00 | 9.77E-01 | no | Sox5 | intron_variant,non_coding_transcript_variant | MODIFIER |
| 6 | 144457109 | AX-97823713 | 0 | G | T | 0.07 | 7.84E-03 | 2.78E-01 | 2.70E+00 | 2.74E+00 | 9.77E-01 | no | Sox5 | intron_variant,non_coding_transcript_variant | MODIFIER |
| 6 | 144458297 | AX-170406477 | 0 | A | G | 0.07 | 7.84E-03 | 2.78E-01 | 2.70E+00 | 2.74E+00 | 9.77E-01 | no | Sox5 | intron_variant,non_coding_transcript_variant | MODIFIER |
| 6 | 144463250 | AX-170454739 | 0 | C | T | 0.07 | 7.84E-03 | 2.78E-01 | 2.70E+00 | 2.74E+00 | 9.77E-01 | no | Sox5 | intron_variant,non_coding_transcript_variant | MODIFIER |
| 6 | 144463505 | AX-170156202 | 0 | A | G | 0.07 | 7.84E-03 | 2.78E-01 | 2.70E+00 | 2.74E+00 | 9.77E-01 | no | Sox5 | intron_variant,non_coding_transcript_variant | MODIFIER |
| 6 | 144497028 | AX-170204790 | 0 | C | T | 0.07 | 7.84E-03 | 2.78E-01 | 2.70E+00 | 2.74E+00 | 9.77E-01 | no | Sox5 | intron_variant,non_coding_transcript_variant | MODIFIER |
| 6 | 144631787 | AX-97834443 | 0 | T | C | 0.07 | 7.84E-03 | 2.78E-01 | 2.70E+00 | 2.74E+00 | 9.77E-01 | no | Sox5 | intron_variant,non_coding_transcript_variant | MODIFIER |
| 6 | 144631958 | AX-170449765 | 0 | G | T | 0.07 | 7.84E-03 | 2.78E-01 | 2.70E+00 | 2.74E+00 | 9.77E-01 | no | Sox5 | intron_variant,non_coding_transcript_variant | MODIFIER |
| 6 | 144632072 | AX-170357379 | 0 | A | G | 0.07 | 7.84E-03 | 2.78E-01 | 2.70E+00 | 2.74E+00 | 9.77E-01 | no | Sox5 | intron_variant,non_coding_transcript_variant | MODIFIER |
| 6 | 144634178 | AX-170396428 | 0 | G | A | 0.07 | 7.84E-03 | 2.78E-01 | 2.70E+00 | 2.74E+00 | 9.77E-01 | no | Sox5 | intron_variant,non_coding_transcript_variant | MODIFIER |
| 6 | 144634769 | AX-170254519 | 0 | A | G | 0.07 | 7.84E-03 | 2.78E-01 | 2.70E+00 | 2.74E+00 | 9.77E-01 | no | Sox5 | intron_variant,non_coding_transcript_variant | MODIFIER |
| 6 | 144682906 | AX-170412628 | 0 | A | G | 0.07 | 7.84E-03 | 2.78E-01 | 2.70E+00 | 2.74E+00 | 9.77E-01 | no | Sox5os3 | intron_variant,non_coding_transcript_variant | MODIFIER |
| 6 | 144742407 | AX-170421130 | 0 | T | C | 0.07 | 7.84E-03 | 2.78E-01 | 2.70E+00 | 2.74E+00 | 9.77E-01 | no | Sox5 | intron_variant,non_coding_transcript_variant | MODIFIER |

| | | | | | | | | | | | | | | | |
|---|---|---|---|---|---|---|---|---|---|---|---|---|---|---|---|
| 6 | 144744290 | AX-170504118 | 0 | G | C | 0.07 | 7.84E-03 | 2.78E-01 | 2.70E+00 | 2.74E+00 | 9.77E-01 | no | Sox5 | intron_variant,non_coding_transcript_variant | MODIFIER |
| 6 | 144745209 | AX-97834441 | 0 | T | C | 0.07 | 7.84E-03 | 2.78E-01 | 2.70E+00 | 2.74E+00 | 9.77E-01 | no | Sox5 | intron_variant,non_coding_transcript_variant | MODIFIER |
| 6 | 144804809 | AX-170299050 | 0 | G | C | 0.07 | 7.84E-03 | 2.78E-01 | 2.70E+00 | 2.74E+00 | 9.77E-01 | no | Gm22792 | upstream_gene_variant | MODIFIER |
| 6 | 144805088 | AX-170488943 | 0 | C | T | 0.07 | 7.84E-03 | 2.78E-01 | 2.70E+00 | 2.74E+00 | 9.77E-01 | no | Gm22792 | upstream_gene_variant | MODIFIER |
| 6 | 144809836 | AX-170392014 | 0 | A | G | 0.07 | 7.84E-03 | 2.78E-01 | 2.70E+00 | 2.74E+00 | 9.77E-01 | no | | intergenic_variant | MODIFIER |
| 6 | 144958793 | AX-170483529 | 0 | C | T | 0.07 | 7.84E-03 | 2.78E-01 | 2.70E+00 | 2.74E+00 | 9.77E-01 | no | | intergenic_variant | MODIFIER |
| 6 | 144989181 | AX-97797620 | 0 | A | C | 0.07 | 7.84E-03 | 2.78E-01 | 2.70E+00 | 2.74E+00 | 9.77E-01 | no | Bcat1 | downstream_gene_variant | MODIFIER |
| 6 | 144989378 | AX-170028452 | 0 | C | G | 0.07 | 7.84E-03 | 2.78E-01 | 2.70E+00 | 2.74E+00 | 9.77E-01 | no | Bcat1 | downstream_gene_variant | MODIFIER |
| 6 | 144991107 | AX-97826662 | 0 | T | G | 0.07 | 7.84E-03 | 2.78E-01 | 2.70E+00 | 2.74E+00 | 9.77E-01 | no | Bcat1 | downstream_gene_variant | MODIFIER |
| 6 | 144993837 | AX-170114930 | 0 | T | C | 0.07 | 7.84E-03 | 2.78E-01 | 2.70E+00 | 2.74E+00 | 9.77E-01 | no | Bcat1 | 3_prime_UTR_variant | MODIFIER |
| 6 | 144998536 | AX-170293304 | 0 | C | T | 0.07 | 7.84E-03 | 2.78E-01 | 2.70E+00 | 2.74E+00 | 9.77E-01 | no | Bcat1 | 3_prime_UTR_variant | MODIFIER |
| 6 | 145035353 | AX-170366382 | 0 | G | A | 0.07 | 7.84E-03 | 2.78E-01 | 2.70E+00 | 2.74E+00 | 9.77E-01 | no | Bcat1 | intron_variant | MODIFIER |
| 6 | 145035384 | AX-97784486 | 0 | A | G | 0.07 | 7.84E-03 | 2.78E-01 | 2.70E+00 | 2.74E+00 | 9.77E-01 | no | Bcat1 | intron_variant | MODIFIER |
| 6 | 145042165 | AX-170076539 | 0 | G | A | 0.07 | 7.84E-03 | 2.78E-01 | 2.70E+00 | 2.74E+00 | 9.77E-01 | no | Bcat1 | intron_variant | MODIFIER |
| 6 | 145042421 | AX-170028451 | 0 | T | C | 0.07 | 7.84E-03 | 2.78E-01 | 2.70E+00 | 2.74E+00 | 9.77E-01 | no | Bcat1 | intron_variant | MODIFIER |
| 6 | 145045127 | AX-170151706 | 0 | T | G | 0.07 | 7.84E-03 | 2.78E-01 | 2.70E+00 | 2.74E+00 | 9.77E-01 | no | Bcat1 | intron_variant | MODIFIER |
| 6 | 145063301 | AX-170263750 | 0 | A | G | 0.07 | 7.84E-03 | 2.78E-01 | 2.70E+00 | 2.74E+00 | 9.77E-01 | no | Bcat1 | intron_variant | MODIFIER |
| 6 | 145063732 | AX-170398355 | 0 | A | G | 0.07 | 7.84E-03 | 2.78E-01 | 2.70E+00 | 2.74E+00 | 9.77E-01 | no | Bcat1 | intron_variant | MODIFIER |
| 6 | 145074578 | AX-170097526 | 0 | C | A | 0.07 | 7.84E-03 | 2.78E-01 | 2.70E+00 | 2.74E+00 | 9.77E-01 | no | Bcat1 | intron_variant | MODIFIER |
| 6 | 145075261 | AX-170269520 | 0 | C | G | 0.07 | 7.84E-03 | 2.78E-01 | 2.70E+00 | 2.74E+00 | 9.77E-01 | no | Bcat1 | intron_variant | MODIFIER |
| 6 | 145077916 | AX-170267159 | 0 | G | A | 0.07 | 7.84E-03 | 2.78E-01 | 2.70E+00 | 2.74E+00 | 9.77E-01 | no | Bcat1 | upstream_gene_variant | MODIFIER |
| 6 | 145114208 | AX-97823952 | 0 | T | G | 0.07 | 7.84E-03 | 2.78E-01 | 2.70E+00 | 2.74E+00 | 9.77E-01 | no | Lrmp | upstream_gene_variant | MODIFIER |
| 6 | 145114379 | AX-170409858 | 0 | T | C | 0.07 | 7.84E-03 | 2.78E-01 | 2.70E+00 | 2.74E+00 | 9.77E-01 | no | Lrmp | upstream_gene_variant | MODIFIER |
| 6 | 145116047 | AX-170481466 | 0 | A | G | 0.07 | 7.84E-03 | 2.78E-01 | 2.70E+00 | 2.74E+00 | 9.77E-01 | no | Lrmp | intron_variant,NMD_transcript_variant | MODIFIER |
| 6 | 145116786 | AX-170382744 | 0 | T | C | 0.07 | 7.84E-03 | 2.78E-01 | 2.70E+00 | 2.74E+00 | 9.77E-01 | no | Lrmp | upstream_gene_variant | MODIFIER |
| 6 | 145137191 | AX-170459832 | 0 | T | C | 0.07 | 7.84E-03 | 2.78E-01 | 2.70E+00 | 2.74E+00 | 9.77E-01 | no | Lrmp | intron_variant | MODIFIER |
| 6 | 145138850 | AX-170178528 | 0 | G | A | 0.07 | 7.84E-03 | 2.78E-01 | 2.70E+00 | 2.74E+00 | 9.77E-01 | no | Lrmp | intron_variant | MODIFIER |
| 6 | 145140565 | AX-170394836 | 0 | C | T | 0.07 | 7.84E-03 | 2.78E-01 | 2.70E+00 | 2.74E+00 | 9.77E-01 | no | Lrmp | intron_variant | MODIFIER |
| 6 | 145174235 | AX-170076541 | 0 | T | C | 0.07 | 7.84E-03 | 2.78E-01 | 2.70E+00 | 2.74E+00 | 9.77E-01 | no | Lrmp | intron_variant | MODIFIER |
| 6 | 145174670 | AX-170211158 | 0 | C | T | 0.07 | 7.84E-03 | 2.78E-01 | 2.70E+00 | 2.74E+00 | 9.77E-01 | no | Lrmp | missense_variant | MODERATE |
| 6 | 145179183 | AX-97854973 | 0 | C | T | 0.07 | 7.84E-03 | 2.78E-01 | 2.70E+00 | 2.74E+00 | 9.77E-01 | no | Lrmp | downstream_gene_variant | MODIFIER |
| 6 | 145206182 | AX-97791440 | 0 | C | T | 0.07 | 7.84E-03 | 2.78E-01 | 2.70E+00 | 2.74E+00 | 9.77E-01 | no | Etfrf1 | upstream_gene_variant | MODIFIER |
| 6 | 145219190 | AX-170258673 | 0 | T | C | 0.07 | 7.84E-03 | 2.78E-01 | 2.70E+00 | 2.74E+00 | 9.77E-01 | no | Kras | 3_prime_UTR_variant | MODIFIER |
| 6 | 145222558 | AX-170388685 | 0 | C | T | 0.07 | 7.84E-03 | 2.78E-01 | 2.70E+00 | 2.74E+00 | 9.77E-01 | no | Kras | intron_variant | MODIFIER |
| 6 | 145223273 | AX-170523807 | 0 | A | G | 0.07 | 7.84E-03 | 2.78E-01 | 2.70E+00 | 2.74E+00 | 9.77E-01 | no | Kras | intron_variant | MODIFIER |
| 6 | 145255978 | AX-170089864 | 0 | C | T | 0.07 | 7.84E-03 | 2.78E-01 | 2.70E+00 | 2.74E+00 | 9.77E-01 | no | Gm15706 | downstream_gene_variant | MODIFIER |
| 6 | 145261256 | AX-170110479 | 0 | C | T | 0.07 | 7.84E-03 | 2.78E-01 | 2.70E+00 | 2.74E+00 | 9.77E-01 | no | | intergenic_variant | MODIFIER |
| 6 | 145262833 | AX-170224494 | 0 | T | C | 0.07 | 7.84E-03 | 2.78E-01 | 2.70E+00 | 2.74E+00 | 9.77E-01 | no | Rps25-ps1 | upstream_gene_variant | MODIFIER |
| 6 | 145262976 | AX-170228445 | 0 | G | A | 0.07 | 7.84E-03 | 2.78E-01 | 2.70E+00 | 2.74E+00 | 9.77E-01 | no | Rps25-ps1 | upstream_gene_variant | MODIFIER |
| 6 | 145263498 | AX-170156907 | 0 | T | C | 0.07 | 7.84E-03 | 2.78E-01 | 2.70E+00 | 2.74E+00 | 9.77E-01 | no | Rps25-ps1 | upstream_gene_variant | MODIFIER |
| 6 | 145267295 | AX-170533573 | 0 | T | C | 0.07 | 7.84E-03 | 2.78E-01 | 2.70E+00 | 2.74E+00 | 9.77E-01 | no | Rps25-ps1 | downstream_gene_variant | MODIFIER |
| 6 | 145267766 | AX-170325949 | 0 | T | C | 0.07 | 7.84E-03 | 2.78E-01 | 2.70E+00 | 2.74E+00 | 9.77E-01 | no | Rps25-ps1 | downstream_gene_variant | MODIFIER |
| 6 | 145271113 | AX-170352377 | 0 | T | C | 0.07 | 7.84E-03 | 2.78E-01 | 2.70E+00 | 2.74E+00 | 9.77E-01 | no | Rps25-ps1 | downstream_gene_variant | MODIFIER |
| 6 | 145276623 | AX-170134527 | 0 | C | T | 0.07 | 7.84E-03 | 2.78E-01 | 2.70E+00 | 2.74E+00 | 9.77E-01 | no | | regulatory_region_variant | MODIFIER |
| 6 | 145276961 | AX-97827910 | 0 | T | G | 0.07 | 7.84E-03 | 2.78E-01 | 2.70E+00 | 2.74E+00 | 9.77E-01 | no | | regulatory_region_variant | MODIFIER |
| 6 | 145289340 | AX-170267694 | 0 | C | G | 0.07 | 7.84E-03 | 2.78E-01 | 2.70E+00 | 2.74E+00 | 9.77E-01 | no | Gm22793 | upstream_gene_variant | MODIFIER |
| 6 | 145294624 | AX-170427085 | 0 | G | C | 0.07 | 7.84E-03 | 2.78E-01 | 2.70E+00 | 2.74E+00 | 9.77E-01 | no | | intergenic_variant | MODIFIER |
| 6 | 145299032 | AX-97839721 | 0 | A | G | 0.07 | 7.84E-03 | 2.78E-01 | 2.70E+00 | 2.74E+00 | 9.77E-01 | no | 2010013B24Rik | upstream_gene_variant | MODIFIER |
| 6 | 145308627 | AX-170142207 | 0 | T | C | 0.07 | 7.84E-03 | 2.78E-01 | 2.70E+00 | 2.74E+00 | 9.77E-01 | no | 2010013B24Rik | non_coding_transcript_exon_variant | MODIFIER |
| 6 | 145309110 | AX-170172028 | 0 | T | C | 0.07 | 7.84E-03 | 2.78E-01 | 2.70E+00 | 2.74E+00 | 9.77E-01 | no | 2010013B24Rik | downstream_gene_variant | MODIFIER |
| 6 | 145344740 | AX-170132066 | 0 | C | T | 0.07 | 7.84E-03 | 2.78E-01 | 2.70E+00 | 2.74E+00 | 9.77E-01 | no | | regulatory_region_variant | MODIFIER |
| 6 | 145360638 | AX-170076543 | 0 | G | T | 0.07 | 7.84E-03 | 2.78E-01 | 2.70E+00 | 2.74E+00 | 9.77E-01 | no | Lmntd1 | downstream_gene_variant | MODIFIER |
| 6 | 145368993 | AX-170299274 | 0 | A | G | 0.07 | 7.84E-03 | 2.78E-01 | 2.70E+00 | 2.74E+00 | 9.77E-01 | no | Lmntd1 | intron_variant,NMD_transcript_variant | MODIFIER |
| 6 | 145377356 | AX-170377147 | 0 | C | T | 0.07 | 7.84E-03 | 2.78E-01 | 2.70E+00 | 2.74E+00 | 9.77E-01 | no | Lmntd1 | intron_variant,NMD_transcript_variant | MODIFIER |

| | | | | | | | | | | | | | | | |
|---|---|---|---|---|---|---|---|---|---|---|---|---|---|---|---|
| 6 | 145377411 | AX-97838203 | 0 | A | G | 0.07 | 7.84E-03 | 2.78E-01 | 2.70E+00 | 2.74E+00 | 9.77E-01 | no | Lmntd1 | intron_variant,NMD_transcript_variant | MODIFIER |
| 6 | 145378542 | AX-170123375 | 0 | T | C | 0.07 | 7.84E-03 | 2.78E-01 | 2.70E+00 | 2.74E+00 | 9.77E-01 | no | Lmntd1 | intron_variant,NMD_transcript_variant | MODIFIER |
| 6 | 145396428 | AX-170028462 | 0 | C | T | 0.07 | 7.84E-03 | 2.78E-01 | 2.70E+00 | 2.74E+00 | 9.77E-01 | no | Lmntd1 | intron_variant | MODIFIER |
| 6 | 145397880 | AX-170523436 | 0 | T | C | 0.07 | 7.84E-03 | 2.78E-01 | 2.70E+00 | 2.74E+00 | 9.77E-01 | no | Lmntd1 | intron_variant | MODIFIER |
| 6 | 145400014 | AX-97808221 | 0 | C | T | 0.07 | 7.84E-03 | 2.78E-01 | 2.70E+00 | 2.74E+00 | 9.77E-01 | no | Lmntd1 | intron_variant | MODIFIER |
| 6 | 145422819 | AX-170028464 | 0 | T | C | 0.07 | 7.84E-03 | 2.78E-01 | 2.70E+00 | 2.74E+00 | 9.77E-01 | no | Lmntd1 | intron_variant | MODIFIER |
| 6 | 145424032 | AX-170028463 | 0 | A | G | 0.07 | 7.84E-03 | 2.78E-01 | 2.70E+00 | 2.74E+00 | 9.77E-01 | no | Lmntd1 | intron_variant | MODIFIER |
| 6 | 145440984 | AX-170458849 | 0 | A | G | 0.07 | 7.84E-03 | 2.78E-01 | 2.70E+00 | 2.74E+00 | 9.77E-01 | no | Lmntd1 | intron_variant | MODIFIER |
| 6 | 145446541 | AX-170076548 | 0 | G | T | 0.07 | 7.84E-03 | 2.78E-01 | 2.70E+00 | 2.74E+00 | 9.77E-01 | no | Lmntd1 | intron_variant | MODIFIER |
| 6 | 145446630 | AX-170076547 | 0 | C | T | 0.07 | 7.84E-03 | 2.78E-01 | 2.70E+00 | 2.74E+00 | 9.77E-01 | no | Lmntd1 | intron_variant | MODIFIER |
| 6 | 145447078 | AX-170453770 | 0 | T | C | 0.07 | 7.84E-03 | 2.78E-01 | 2.70E+00 | 2.74E+00 | 9.77E-01 | no | Lmntd1 | intron_variant | MODIFIER |
| 6 | 145448676 | AX-170394493 | 0 | T | C | 0.07 | 7.84E-03 | 2.78E-01 | 2.70E+00 | 2.74E+00 | 9.77E-01 | no | Lmntd1 | intron_variant | MODIFIER |
| 6 | 145449254 | AX-170233473 | 0 | C | T | 0.07 | 7.84E-03 | 2.78E-01 | 2.70E+00 | 2.74E+00 | 9.77E-01 | no | Lmntd1 | intron_variant | MODIFIER |
| 6 | 145449279 | AX-97840411 | 0 | C | T | 0.07 | 7.84E-03 | 2.78E-01 | 2.70E+00 | 2.74E+00 | 9.77E-01 | no | Lmntd1 | intron_variant | MODIFIER |
| 6 | 146427668 | AX-170168398 | 0 | A | G | 0.07 | 7.84E-03 | 2.78E-01 | 2.70E+00 | 2.74E+00 | 9.77E-01 | no | Itpr2 | intron_variant | MODIFIER |
| 6 | 146431520 | AX-170533222 | 0 | C | T | 0.07 | 7.84E-03 | 2.78E-01 | 2.70E+00 | 2.74E+00 | 9.77E-01 | no | Itpr2 | intron_variant | MODIFIER |
| 6 | 146437022 | AX-97818657 | 0 | C | T | 0.07 | 7.84E-03 | 2.78E-01 | 2.70E+00 | 2.74E+00 | 9.77E-01 | no | Itpr2 | intron_variant | MODIFIER |
| 6 | 146446487 | AX-170102475 | 0 | T | C | 0.07 | 7.84E-03 | 2.78E-01 | 2.70E+00 | 2.74E+00 | 9.77E-01 | no | Itpr2 | intron_variant | MODIFIER |
| 6 | 146455705 | AX-97839148 | 0 | C | T | 0.07 | 7.84E-03 | 2.78E-01 | 2.70E+00 | 2.74E+00 | 9.77E-01 | no | Itpr2 | intron_variant | MODIFIER |
| 6 | 146456342 | AX-170532311 | 0 | G | A | 0.07 | 7.84E-03 | 2.78E-01 | 2.70E+00 | 2.74E+00 | 9.77E-01 | no | Itpr2 | intron_variant | MODIFIER |
| 6 | 146456376 | AX-170432895 | 0 | C | T | 0.07 | 7.84E-03 | 2.78E-01 | 2.70E+00 | 2.74E+00 | 9.77E-01 | no | Itpr2 | intron_variant | MODIFIER |
| 6 | 146461739 | AX-170165745 | 0 | A | C | 0.07 | 7.84E-03 | 2.78E-01 | 2.70E+00 | 2.74E+00 | 9.77E-01 | no | Itpr2 | intron_variant | MODIFIER |
| 6 | 146466621 | AX-170157652 | 0 | T | C | 0.07 | 7.84E-03 | 2.78E-01 | 2.70E+00 | 2.74E+00 | 9.77E-01 | no | Itpr2 | intron_variant | MODIFIER |
| 6 | 146476797 | AX-170181758 | 0 | C | T | 0.07 | 7.84E-03 | 2.78E-01 | 2.70E+00 | 2.74E+00 | 9.77E-01 | no | Itpr2 | intron_variant | MODIFIER |
| 6 | 146480195 | AX-97845777 | 0 | C | T | 0.07 | 7.84E-03 | 2.78E-01 | 2.70E+00 | 2.74E+00 | 9.77E-01 | no | Itpr2 | intron_variant | MODIFIER |
| 6 | 146498209 | AX-97821311 | 0 | A | G | 0.07 | 7.84E-03 | 2.78E-01 | 2.70E+00 | 2.74E+00 | 9.77E-01 | no | Itpr2 | intron_variant | MODIFIER |
| 6 | 146516299 | AX-97786159 | 0 | T | C | 0.07 | 7.84E-03 | 2.78E-01 | 2.70E+00 | 2.74E+00 | 9.77E-01 | no | | regulatory_region_variant | MODIFIER |
| 6 | 146523818 | AX-170452030 | 0 | T | G | 0.07 | 7.84E-03 | 2.78E-01 | 2.70E+00 | 2.74E+00 | 9.77E-01 | no | | intergenic_variant | MODIFIER |
| 6 | 146553554 | AX-170196203 | 0 | A | T | 0.07 | 7.84E-03 | 2.78E-01 | 2.70E+00 | 2.74E+00 | 9.77E-01 | no | Ints13 | intron_variant | MODIFIER |
| 6 | 146556391 | AX-170200332 | 0 | A | T | 0.07 | 7.84E-03 | 2.78E-01 | 2.70E+00 | 2.74E+00 | 9.77E-01 | no | Ints13 | intron_variant | MODIFIER |
| 6 | 146556662 | AX-170028487 | 0 | C | T | 0.07 | 7.84E-03 | 2.78E-01 | 2.70E+00 | 2.74E+00 | 9.77E-01 | no | Ints13 | intron_variant | MODIFIER |
| 6 | 146576859 | AX-170028484 | 0 | T | G | 0.07 | 7.84E-03 | 2.78E-01 | 2.70E+00 | 2.74E+00 | 9.77E-01 | no | Ints13 | intron_variant | MODIFIER |
| 6 | 146588700 | AX-97849706 | 0 | G | T | 0.07 | 7.84E-03 | 2.78E-01 | 2.70E+00 | 2.74E+00 | 9.77E-01 | no | Fgfr1op2 | intron_variant | MODIFIER |
| 6 | 146602787 | AX-170329499 | 0 | A | G | 0.07 | 7.84E-03 | 2.78E-01 | 2.70E+00 | 2.74E+00 | 9.77E-01 | no | Tm7sf3 | 3_prime_UTR_variant | MODIFIER |
| 6 | 146605393 | AX-170076571 | 0 | A | G | 0.07 | 7.84E-03 | 2.78E-01 | 2.70E+00 | 2.74E+00 | 9.77E-01 | no | Tm7sf3 | intron_variant | MODIFIER |
| 6 | 146605900 | AX-97849642 | 0 | T | C | 0.07 | 7.84E-03 | 2.78E-01 | 2.70E+00 | 2.74E+00 | 9.77E-01 | no | Tm7sf3 | intron_variant | MODIFIER |
| 6 | 146613807 | AX-170344746 | 0 | A | G | 0.07 | 7.84E-03 | 2.78E-01 | 2.70E+00 | 2.74E+00 | 9.77E-01 | no | Tm7sf3 | intron_variant | MODIFIER |
| 6 | 146625521 | AX-97820127 | 0 | C | T | 0.07 | 7.84E-03 | 2.78E-01 | 2.70E+00 | 2.74E+00 | 9.77E-01 | no | Tm7sf3 | intron_variant | MODIFIER |
| 6 | 146629057 | AX-170028480 | 0 | T | C | 0.07 | 7.84E-03 | 2.78E-01 | 2.70E+00 | 2.74E+00 | 9.77E-01 | no | Tm7sf3 | intron_variant | MODIFIER |
| 6 | 146630051 | AX-170100305 | 0 | G | A | 0.07 | 7.84E-03 | 2.78E-01 | 2.70E+00 | 2.74E+00 | 9.77E-01 | no | Tm7sf3 | intron_variant | MODIFIER |
| 6 | 146657570 | AX-97850025 | 0 | A | G | 0.07 | 7.84E-03 | 2.78E-01 | 2.70E+00 | 2.74E+00 | 9.77E-01 | no | 4930479D17Rik | intron_variant,non_coding_transcript_variant | MODIFIER |
| 6 | 146661276 | AX-170149298 | 0 | T | C | 0.07 | 7.84E-03 | 2.78E-01 | 2.70E+00 | 2.74E+00 | 9.77E-01 | no | 4930479D17Rik | intron_variant,non_coding_transcript_variant | MODIFIER |
| 6 | 146661433 | AX-170125690 | 0 | A | G | 0.07 | 7.84E-03 | 2.78E-01 | 2.70E+00 | 2.74E+00 | 9.77E-01 | no | 4930479D17Rik | intron_variant,non_coding_transcript_variant | MODIFIER |
| 6 | 146661879 | AX-170400848 | 0 | G | A | 0.07 | 7.84E-03 | 2.78E-01 | 2.70E+00 | 2.74E+00 | 9.77E-01 | no | 4930479D17Rik | intron_variant,non_coding_transcript_variant | MODIFIER |
| 6 | 146679074 | AX-170028485 | 0 | T | C | 0.07 | 7.84E-03 | 2.78E-01 | 2.70E+00 | 2.74E+00 | 9.77E-01 | no | | regulatory_region_variant | MODIFIER |
| 6 | 146683304 | AX-97805855 | 0 | T | C | 0.07 | 7.84E-03 | 2.78E-01 | 2.70E+00 | 2.74E+00 | 9.77E-01 | no | | regulatory_region_variant | MODIFIER |
| 6 | 146683599 | AX-170461833 | 0 | A | C | 0.07 | 7.84E-03 | 2.78E-01 | 2.70E+00 | 2.74E+00 | 9.77E-01 | no | | regulatory_region_variant | MODIFIER |
| 6 | 146685057 | AX-170028478 | 0 | A | C | 0.07 | 7.84E-03 | 2.78E-01 | 2.70E+00 | 2.74E+00 | 9.77E-01 | no | | regulatory_region_variant | MODIFIER |
| 6 | 146685239 | AX-170028489 | 0 | A | G | 0.07 | 7.84E-03 | 2.78E-01 | 2.70E+00 | 2.74E+00 | 9.77E-01 | no | | regulatory_region_variant | MODIFIER |
| 6 | 146685471 | AX-170187609 | 0 | T | C | 0.07 | 7.84E-03 | 2.78E-01 | 2.70E+00 | 2.74E+00 | 9.77E-01 | no | | regulatory_region_variant | MODIFIER |
| 6 | 146687902 | AX-170455969 | 0 | T | G | 0.07 | 7.84E-03 | 2.78E-01 | 2.70E+00 | 2.74E+00 | 9.77E-01 | no | | regulatory_region_variant | MODIFIER |
| 6 | 146700556 | AX-97837943 | 0 | T | C | 0.07 | 7.84E-03 | 2.78E-01 | 2.70E+00 | 2.74E+00 | 9.77E-01 | no | | intergenic_variant | MODIFIER |
| 6 | 146727679 | AX-97824210 | 0 | C | T | 0.07 | 7.84E-03 | 2.78E-01 | 2.70E+00 | 2.74E+00 | 9.77E-01 | no | Stk38l | intron_variant | MODIFIER |
| 6 | 146741667 | AX-170365435 | 0 | G | T | 0.07 | 7.84E-03 | 2.78E-01 | 2.70E+00 | 2.74E+00 | 9.77E-01 | no | Stk38l | intron_variant | MODIFIER |

| | | | | | | | | | | | | | | | |
|---|---|---|---|---|---|---|---|---|---|---|---|---|---|---|---|
| 6 | 146753421 | AX-170149761 | 0 | T | G | 0.07 | 7.84E-03 | 2.78E-01 | 2.70E+00 | 2.74E+00 | 9.77E-01 | no | Stk38l | intron_variant | MODIFIER |
| 6 | 146754616 | AX-97823572 | 0 | C | T | 0.07 | 7.84E-03 | 2.78E-01 | 2.70E+00 | 2.74E+00 | 9.77E-01 | no | Stk38l | intron_variant | MODIFIER |
| 6 | 146765330 | AX-170529322 | 0 | T | C | 0.07 | 7.84E-03 | 2.78E-01 | 2.70E+00 | 2.74E+00 | 9.77E-01 | no | Stk38l | intron_variant | MODIFIER |
| 6 | 146766353 | AX-170028481 | 0 | A | C | 0.07 | 7.84E-03 | 2.78E-01 | 2.70E+00 | 2.74E+00 | 9.77E-01 | no | Stk38l | intron_variant | MODIFIER |
| 6 | 146767202 | AX-97787406 | 0 | C | A | 0.07 | 7.84E-03 | 2.78E-01 | 2.70E+00 | 2.74E+00 | 9.77E-01 | no | Stk38l | intron_variant | MODIFIER |
| 6 | 146771404 | AX-170324010 | 0 | T | C | 0.07 | 7.84E-03 | 2.78E-01 | 2.70E+00 | 2.74E+00 | 9.77E-01 | no | Stk38l | intron_variant | MODIFIER |
| 6 | 146797630 | AX-170269424 | 0 | C | T | 0.07 | 7.84E-03 | 2.78E-01 | 2.70E+00 | 2.74E+00 | 9.77E-01 | no | Arntl2 | intron_variant | MODIFIER |
| 6 | 146805714 | AX-170272238 | 0 | C | G | 0.07 | 7.84E-03 | 2.78E-01 | 2.70E+00 | 2.74E+00 | 9.77E-01 | no | Arntl2 | intron_variant | MODIFIER |
| 6 | 146806513 | AX-170089865 | 0 | G | A | 0.07 | 7.84E-03 | 2.78E-01 | 2.70E+00 | 2.74E+00 | 9.77E-01 | no | Arntl2 | synonymous_variant | LOW |
| 6 | 146809710 | AX-170394556 | 0 | A | C | 0.07 | 7.84E-03 | 2.78E-01 | 2.70E+00 | 2.74E+00 | 9.77E-01 | no | Arntl2 | missense_variant | MODERATE |
| 6 | 146837649 | AX-97852103 | 0 | C | T | 0.07 | 7.84E-03 | 2.78E-01 | 2.70E+00 | 2.74E+00 | 9.77E-01 | no | Arntl2 | downstream_gene_variant | MODIFIER |
| 6 | 146847336 | AX-170189872 | 0 | G | T | 0.07 | 7.84E-03 | 2.78E-01 | 2.70E+00 | 2.74E+00 | 9.77E-01 | no | Smco2 | upstream_gene_variant | MODIFIER |
| 6 | 146849272 | AX-170509894 | 0 | G | A | 0.07 | 7.84E-03 | 2.78E-01 | 2.70E+00 | 2.74E+00 | 9.77E-01 | no | Smco2 | upstream_gene_variant | MODIFIER |
| 6 | 146852072 | AX-170432208 | 0 | G | A | 0.07 | 7.84E-03 | 2.78E-01 | 2.70E+00 | 2.74E+00 | 9.77E-01 | no | Smco2 | intron_variant | MODIFIER |
| 6 | 146852102 | AX-170423706 | 0 | C | T | 0.07 | 7.84E-03 | 2.78E-01 | 2.70E+00 | 2.74E+00 | 9.77E-01 | no | Smco2 | intron_variant | MODIFIER |
| 6 | 146852788 | AX-97835497 | 0 | T | G | 0.07 | 7.84E-03 | 2.78E-01 | 2.70E+00 | 2.74E+00 | 9.77E-01 | no | Smco2 | intron_variant | MODIFIER |
| 6 | 146865664 | AX-170451399 | 0 | A | G | 0.07 | 7.84E-03 | 2.78E-01 | 2.70E+00 | 2.74E+00 | 9.77E-01 | no | Smco2 | intron_variant | MODIFIER |
| 6 | 146866700 | AX-170207566 | 0 | C | T | 0.07 | 7.84E-03 | 2.78E-01 | 2.70E+00 | 2.74E+00 | 9.77E-01 | no | Smco2 | intron_variant | MODIFIER |
| 6 | 146883128 | AX-97820685 | 0 | A | G | 0.07 | 7.84E-03 | 2.78E-01 | 2.70E+00 | 2.74E+00 | 9.77E-01 | no | | regulatory_region_variant | MODIFIER |
| 6 | 146916583 | AX-170257304 | 0 | A | G | 0.07 | 7.84E-03 | 2.78E-01 | 2.70E+00 | 2.74E+00 | 9.77E-01 | no | Ppfibp1 | intron_variant | MODIFIER |
| 6 | 146928261 | AX-170392136 | 0 | G | A | 0.07 | 7.84E-03 | 2.78E-01 | 2.70E+00 | 2.74E+00 | 9.77E-01 | no | Ppfibp1 | intron_variant | MODIFIER |
| 6 | 146928541 | AX-170182303 | 0 | C | T | 0.07 | 7.84E-03 | 2.78E-01 | 2.70E+00 | 2.74E+00 | 9.77E-01 | no | Ppfibp1 | intron_variant | MODIFIER |
| 6 | 146938910 | AX-170095844 | 0 | G | A | 0.07 | 7.84E-03 | 2.78E-01 | 2.70E+00 | 2.74E+00 | 9.77E-01 | no | Ppfibp1 | 5_prime_UTR_variant | MODIFIER |
| 6 | 146939478 | AX-170441915 | 0 | T | C | 0.07 | 7.84E-03 | 2.78E-01 | 2.70E+00 | 2.74E+00 | 9.77E-01 | no | Ppfibp1 | intron_variant | MODIFIER |
| 6 | 146941240 | AX-170254735 | 0 | G | A | 0.07 | 7.84E-03 | 2.78E-01 | 2.70E+00 | 2.74E+00 | 9.77E-01 | no | Ppfibp1 | intron_variant | MODIFIER |
| 6 | 146945122 | AX-170371545 | 0 | G | A | 0.07 | 7.84E-03 | 2.78E-01 | 2.70E+00 | 2.74E+00 | 9.77E-01 | no | Ppfibp1 | intron_variant | MODIFIER |
| 6 | 146946347 | AX-170306725 | 0 | A | T | 0.07 | 7.84E-03 | 2.78E-01 | 2.70E+00 | 2.74E+00 | 9.77E-01 | no | Ppfibp1 | intron_variant | MODIFIER |
| 6 | 146948008 | AX-170434499 | 0 | G | A | 0.07 | 7.84E-03 | 2.78E-01 | 2.70E+00 | 2.74E+00 | 9.77E-01 | no | Ppfibp1 | intron_variant | MODIFIER |
| 6 | 146954996 | AX-97787193 | 0 | C | T | 0.07 | 7.84E-03 | 2.78E-01 | 2.70E+00 | 2.74E+00 | 9.77E-01 | no | Ppfibp1 | intron_variant | MODIFIER |
| 6 | 146958335 | AX-170323305 | 0 | A | T | 0.07 | 7.84E-03 | 2.78E-01 | 2.70E+00 | 2.74E+00 | 9.77E-01 | no | Ppfibp1 | intron_variant | MODIFIER |
| 6 | 146964997 | AX-170354661 | 0 | G | A | 0.07 | 7.84E-03 | 2.78E-01 | 2.70E+00 | 2.74E+00 | 9.77E-01 | no | Ppfibp1 | intron_variant | MODIFIER |
| 6 | 146969299 | AX-170376429 | 0 | T | G | 0.07 | 7.84E-03 | 2.78E-01 | 2.70E+00 | 2.74E+00 | 9.77E-01 | no | Ppfibp1 | intron_variant | MODIFIER |
| 6 | 146969544 | AX-170177257 | 0 | G | A | 0.07 | 7.84E-03 | 2.78E-01 | 2.70E+00 | 2.74E+00 | 9.77E-01 | no | Ppfibp1 | intron_variant | MODIFIER |
| 6 | 146977943 | AX-97837995 | 0 | C | T | 0.07 | 7.84E-03 | 2.78E-01 | 2.70E+00 | 2.74E+00 | 9.77E-01 | no | Ppfibp1 | intron_variant | MODIFIER |
| 6 | 146985643 | AX-170076574 | 0 | G | T | 0.07 | 7.84E-03 | 2.78E-01 | 2.70E+00 | 2.74E+00 | 9.77E-01 | no | Ppfibp1 | intron_variant | MODIFIER |
| 6 | 146996724 | AX-170244547 | 0 | C | T | 0.07 | 7.84E-03 | 2.78E-01 | 2.70E+00 | 2.74E+00 | 9.77E-01 | no | Ppfibp1 | intron_variant | MODIFIER |
| 6 | 147002046 | AX-170412064 | 0 | G | C | 0.07 | 7.84E-03 | 2.78E-01 | 2.70E+00 | 2.74E+00 | 9.77E-01 | no | Ppfibp1 | intron_variant | MODIFIER |
| 6 | 147010124 | AX-97847121 | 0 | T | C | 0.07 | 7.84E-03 | 2.78E-01 | 2.70E+00 | 2.74E+00 | 9.77E-01 | no | Ppfibp1 | intron_variant | MODIFIER |
| 6 | 147020331 | AX-170334694 | 0 | T | C | 0.07 | 7.84E-03 | 2.78E-01 | 2.70E+00 | 2.74E+00 | 9.77E-01 | no | Ppfibp1 | intron_variant | MODIFIER |
| 6 | 147023505 | AX-170277057 | 0 | T | C | 0.07 | 7.84E-03 | 2.78E-01 | 2.70E+00 | 2.74E+00 | 9.77E-01 | no | Ppfibp1 | intron_variant | MODIFIER |
| 6 | 147025719 | AX-170028497 | 0 | C | G | 0.07 | 7.84E-03 | 2.78E-01 | 2.70E+00 | 2.74E+00 | 9.77E-01 | no | Ppfibp1 | intron_variant | MODIFIER |
| 6 | 147029353 | AX-170225600 | 0 | C | G | 0.07 | 7.84E-03 | 2.78E-01 | 2.70E+00 | 2.74E+00 | 9.77E-01 | no | Ppfibp1 | intron_variant | MODIFIER |
| 6 | 147032754 | AX-170352777 | 0 | C | T | 0.07 | 7.84E-03 | 2.78E-01 | 2.70E+00 | 2.74E+00 | 9.77E-01 | no | Ppfibp1 | downstream_gene_variant | MODIFIER |
| 6 | 147045193 | AX-170386451 | 0 | T | G | 0.07 | 7.84E-03 | 2.78E-01 | 2.70E+00 | 2.74E+00 | 9.77E-01 | no | Mrps35 | intron_variant | MODIFIER |
| 6 | 147048892 | AX-170028493 | 0 | A | C | 0.07 | 7.84E-03 | 2.78E-01 | 2.70E+00 | 2.74E+00 | 9.77E-01 | no | Mrps35 | intron_variant | MODIFIER |
| 6 | 147054858 | AX-170280960 | 0 | C | T | 0.07 | 7.84E-03 | 2.78E-01 | 2.70E+00 | 2.74E+00 | 9.77E-01 | no | Mrps35 | intron_variant | MODIFIER |
| 6 | 147060475 | AX-170095845 | 0 | T | C | 0.07 | 7.84E-03 | 2.78E-01 | 2.70E+00 | 2.74E+00 | 9.77E-01 | no | Mrps35 | intron_variant | MODIFIER |
| 6 | 147061377 | AX-170165640 | 0 | T | A | 0.07 | 7.84E-03 | 2.78E-01 | 2.70E+00 | 2.74E+00 | 9.77E-01 | no | Mrps35 | intron_variant | MODIFIER |
| 6 | 147061446 | AX-170028498 | 0 | A | G | 0.07 | 7.84E-03 | 2.78E-01 | 2.70E+00 | 2.74E+00 | 9.77E-01 | no | Mrps35 | missense_variant | MODERATE |
| 6 | 147067119 | AX-97853664 | 0 | A | G | 0.07 | 7.84E-03 | 2.78E-01 | 2.70E+00 | 2.74E+00 | 9.77E-01 | no | Mrps35 | intron_variant | MODIFIER |
| 6 | 147094795 | AX-170028495 | 0 | A | G | 0.07 | 7.84E-03 | 2.78E-01 | 2.70E+00 | 2.74E+00 | 9.77E-01 | no | Klhl42 | intron_variant | MODIFIER |
| 6 | 147114253 | AX-170331287 | 0 | T | C | 0.07 | 7.84E-03 | 2.78E-01 | 2.70E+00 | 2.74E+00 | 9.77E-01 | no | Klhl42 | downstream_gene_variant | MODIFIER |
| 6 | 147136848 | AX-170028492 | 0 | A | G | 0.07 | 7.84E-03 | 2.78E-01 | 2.70E+00 | 2.74E+00 | 9.77E-01 | no | | intergenic_variant | MODIFIER |
| 6 | 147137022 | AX-170470468 | 0 | T | C | 0.07 | 7.84E-03 | 2.78E-01 | 2.70E+00 | 2.74E+00 | 9.77E-01 | no | | intergenic_variant | MODIFIER |

| | | | | | | | | | | | | | | | |
|---|---|---|---|---|---|---|---|---|---|---|---|---|---|---|---|
| 6 | 147143892 | AX-170076580 | 0 | A | G | 0.07 | 7.84E-03 | 2.78E-01 | 2.70E+00 | 2.74E+00 | 9.77E-01 | no | | intergenic_variant | MODIFIER |
| 6 | 147146153 | AX-170239008 | 0 | A | T | 0.07 | 7.84E-03 | 2.78E-01 | 2.70E+00 | 2.74E+00 | 9.77E-01 | no | | regulatory_region_variant | MODIFIER |
| 6 | 147149013 | AX-170180877 | 0 | C | T | 0.07 | 7.84E-03 | 2.78E-01 | 2.70E+00 | 2.74E+00 | 9.77E-01 | no | | intergenic_variant | MODIFIER |
| 6 | 147181824 | AX-170089866 | 0 | A | G | 0.07 | 7.84E-03 | 2.78E-01 | 2.70E+00 | 2.74E+00 | 9.77E-01 | no | E330012B07Rik | intron_variant,non_coding_transcript_variant | MODIFIER |
| 6 | 147181860 | AX-170438465 | 0 | C | T | 0.07 | 7.84E-03 | 2.78E-01 | 2.70E+00 | 2.74E+00 | 9.77E-01 | no | E330012B07Rik | intron_variant,non_coding_transcript_variant | MODIFIER |
| 6 | 147195806 | AX-97845817 | 0 | A | G | 0.07 | 7.84E-03 | 2.78E-01 | 2.70E+00 | 2.74E+00 | 9.77E-01 | no | E330012B07Rik | intron_variant,non_coding_transcript_variant | MODIFIER |
| 6 | 147210803 | AX-170501667 | 0 | T | A | 0.07 | 7.84E-03 | 2.78E-01 | 2.70E+00 | 2.74E+00 | 9.77E-01 | no | E330012B07Rik | downstream_gene_variant | MODIFIER |
| 6 | 147219570 | AX-97824610 | 0 | C | T | 0.07 | 7.84E-03 | 2.78E-01 | 2.70E+00 | 2.74E+00 | 9.77E-01 | no | | regulatory_region_variant | MODIFIER |
| 6 | 147224996 | AX-170476331 | 0 | C | T | 0.07 | 7.84E-03 | 2.78E-01 | 2.70E+00 | 2.74E+00 | 9.77E-01 | no | | intergenic_variant | MODIFIER |
| 6 | 147228098 | AX-170466612 | 0 | T | G | 0.07 | 7.84E-03 | 2.78E-01 | 2.70E+00 | 2.74E+00 | 9.77E-01 | no | | intergenic_variant | MODIFIER |
| 6 | 147229066 | AX-170386025 | 0 | C | T | 0.07 | 7.84E-03 | 2.78E-01 | 2.70E+00 | 2.74E+00 | 9.77E-01 | no | | intergenic_variant | MODIFIER |
| 6 | 147230089 | AX-170253261 | 0 | T | C | 0.07 | 7.84E-03 | 2.78E-01 | 2.70E+00 | 2.74E+00 | 9.77E-01 | no | | intergenic_variant | MODIFIER |
| 6 | 147234746 | AX-170303291 | 0 | T | A | 0.07 | 7.84E-03 | 2.78E-01 | 2.70E+00 | 2.74E+00 | 9.77E-01 | no | | intergenic_variant | MODIFIER |
| 6 | 147247836 | AX-170208095 | 0 | T | A | 0.07 | 7.84E-03 | 2.78E-01 | 2.70E+00 | 2.74E+00 | 9.77E-01 | no | Pthlh | downstream_gene_variant | MODIFIER |
| 6 | 147254407 | AX-170409745 | 0 | C | T | 0.07 | 7.84E-03 | 2.78E-01 | 2.70E+00 | 2.74E+00 | 9.77E-01 | no | Pthlh | intron_variant | MODIFIER |
| 6 | 147254445 | AX-97783053 | 0 | C | T | 0.07 | 7.84E-03 | 2.78E-01 | 2.70E+00 | 2.74E+00 | 9.77E-01 | no | Pthlh | intron_variant | MODIFIER |
| 6 | 147261387 | AX-170481311 | 0 | C | T | 0.07 | 7.84E-03 | 2.78E-01 | 2.70E+00 | 2.74E+00 | 9.77E-01 | no | Pthlh | intron_variant | MODIFIER |
| 6 | 147270243 | AX-170329807 | 0 | C | T | 0.07 | 7.84E-03 | 2.78E-01 | 2.70E+00 | 2.74E+00 | 9.77E-01 | no | Gm15766 | upstream_gene_variant | MODIFIER |
| 6 | 147273457 | AX-170401346 | 0 | G | T | 0.07 | 7.84E-03 | 2.78E-01 | 2.70E+00 | 2.74E+00 | 9.77E-01 | no | Gm15766 | downstream_gene_variant | MODIFIER |
| 6 | 147278033 | AX-97837991 | 0 | T | G | 0.07 | 7.84E-03 | 2.78E-01 | 2.70E+00 | 2.74E+00 | 9.77E-01 | no | | intergenic_variant | MODIFIER |
| 6 | 147279465 | AX-170517357 | 0 | T | C | 0.07 | 7.84E-03 | 2.78E-01 | 2.70E+00 | 2.74E+00 | 9.77E-01 | no | | intergenic_variant | MODIFIER |
| 6 | 147289323 | AX-170491877 | 0 | C | T | 0.07 | 7.84E-03 | 2.78E-01 | 2.70E+00 | 2.74E+00 | 9.77E-01 | no | | intergenic_variant | MODIFIER |
| 6 | 147299112 | AX-170133856 | 0 | C | T | 0.07 | 7.84E-03 | 2.78E-01 | 2.70E+00 | 2.74E+00 | 9.77E-01 | no | | intergenic_variant | MODIFIER |
| 6 | 147310820 | AX-170198101 | 0 | T | G | 0.07 | 7.84E-03 | 2.78E-01 | 2.70E+00 | 2.74E+00 | 9.77E-01 | no | | intergenic_variant | MODIFIER |
| 6 | 147322519 | AX-170076584 | 0 | C | A | 0.07 | 7.84E-03 | 2.78E-01 | 2.70E+00 | 2.74E+00 | 9.77E-01 | no | | intergenic_variant | MODIFIER |
| 6 | 147325953 | AX-170095846 | 0 | A | C | 0.07 | 7.84E-03 | 2.78E-01 | 2.70E+00 | 2.74E+00 | 9.77E-01 | no | | intergenic_variant | MODIFIER |
| 6 | 147334288 | AX-170101751 | 0 | T | C | 0.07 | 7.84E-03 | 2.78E-01 | 2.70E+00 | 2.74E+00 | 9.77E-01 | no | | intergenic_variant | MODIFIER |
| 6 | 147336576 | AX-170163570 | 0 | A | G | 0.07 | 7.84E-03 | 2.78E-01 | 2.70E+00 | 2.74E+00 | 9.77E-01 | no | | intergenic_variant | MODIFIER |
| 6 | 147336722 | AX-170250799 | 0 | T | C | 0.07 | 7.84E-03 | 2.78E-01 | 2.70E+00 | 2.74E+00 | 9.77E-01 | no | | intergenic_variant | MODIFIER |
| 6 | 147341192 | AX-170496751 | 0 | C | T | 0.07 | 7.84E-03 | 2.78E-01 | 2.70E+00 | 2.74E+00 | 9.77E-01 | no | | intergenic_variant | MODIFIER |
| 6 | 147348061 | AX-170213399 | 0 | T | C | 0.07 | 7.84E-03 | 2.78E-01 | 2.70E+00 | 2.74E+00 | 9.77E-01 | no | | intergenic_variant | MODIFIER |
| 6 | 147348619 | AX-170359627 | 0 | C | T | 0.07 | 7.84E-03 | 2.78E-01 | 2.70E+00 | 2.74E+00 | 9.77E-01 | no | | intergenic_variant | MODIFIER |
| 6 | 147357428 | AX-170453350 | 0 | A | G | 0.07 | 7.84E-03 | 2.78E-01 | 2.70E+00 | 2.74E+00 | 9.77E-01 | no | | regulatory_region_variant | MODIFIER |
| 6 | 147359156 | AX-170465444 | 0 | T | C | 0.07 | 7.84E-03 | 2.78E-01 | 2.70E+00 | 2.74E+00 | 9.77E-01 | no | | intergenic_variant | MODIFIER |
| 6 | 147359345 | AX-170432711 | 0 | T | C | 0.07 | 7.84E-03 | 2.78E-01 | 2.70E+00 | 2.74E+00 | 9.77E-01 | no | | intergenic_variant | MODIFIER |
| 6 | 147360768 | AX-97853957 | 0 | G | T | 0.07 | 7.84E-03 | 2.78E-01 | 2.70E+00 | 2.74E+00 | 9.77E-01 | no | | intergenic_variant | MODIFIER |
| 6 | 147368405 | AX-170463746 | 0 | C | T | 0.07 | 7.84E-03 | 2.78E-01 | 2.70E+00 | 2.74E+00 | 9.77E-01 | no | | intergenic_variant | MODIFIER |
| 6 | 147368562 | AX-170361953 | 0 | G | A | 0.07 | 7.84E-03 | 2.78E-01 | 2.70E+00 | 2.74E+00 | 9.77E-01 | no | | intergenic_variant | MODIFIER |
| 6 | 147377340 | AX-170192782 | 0 | C | T | 0.07 | 7.84E-03 | 2.78E-01 | 2.70E+00 | 2.74E+00 | 9.77E-01 | no | | intergenic_variant | MODIFIER |
| 6 | 147382377 | AX-170076585 | 0 | C | T | 0.07 | 7.84E-03 | 2.78E-01 | 2.70E+00 | 2.74E+00 | 9.77E-01 | no | | intergenic_variant | MODIFIER |
| 9 | 58924060 | AX-170096555 | 0 | C | T | 0.219 | -2.79E-03 | 1.18E-01 | 2.67E+00 | 2.74E+00 | 9.77E-01 | no | Neo1 | intron_variant | MODIFIER |
| 6 | 107021551 | AX-170098418 | 1 | A | G | 0.062 | -5.76E-03 | 2.32E-01 | 2.62E+00 | 2.74E+00 | 9.77E-01 | no | | intergenic_variant | MODIFIER |
| 18 | 38735976 | AX-170119318 | 0 | C | T | 0.061 | -9.13E-03 | 3.01E-01 | 2.71E+00 | 2.73E+00 | 9.77E-01 | no | AC133601.1 | intron_variant,non_coding_transcript_variant | MODIFIER |
| 17 | 81578579 | AX-170145904 | 0 | C | T | 0.044 | 3.56E-03 | 1.14E-01 | 2.55E+00 | 2.74E+00 | 9.77E-01 | no | Slc8a1 | intron_variant | MODIFIER |
| 18 | 34365272 | AX-170396814 | 1 | C | T | 0.062 | 8.21E-03 | 3.01E-01 | 2.72E+00 | 2.74E+00 | 9.77E-01 | no | Reep5 | intron_variant | MODIFIER |
| 6 | 121590228 | AX-170483438 | 0 | G | T | 0.07 | 7.70E-03 | 2.78E-01 | 2.70E+00 | 2.74E+00 | 9.77E-01 | no | | intergenic_variant | MODIFIER |
| 11 | 35150508 | AX-97831259 | 1 | A | G | 0.098 | -4.15E-03 | 1.88E-01 | 2.66E+00 | 2.74E+00 | 9.77E-01 | no | Slit3 | intron_variant | MODIFIER |
| 11 | 34771212 | AX-170046766 | 1 | A | G | 0.098 | -4.25E-03 | 1.88E-01 | 2.66E+00 | 2.74E+00 | 9.77E-01 | no | Dock2 | intron_variant | MODIFIER |
| 15 | 78285523 | AX-170056057 | 0 | G | C | 0.544 | 3.05E-03 | 7.89E-02 | 2.66E+00 | 2.73E+00 | 9.77E-01 | no | Csf2rb2 | intron_variant | MODIFIER |
| 5 | 78608061 | AX-97794226 | 0 | C | A | 0.421 | 3.24E-03 | 1.02E-01 | 2.67E+00 | 2.74E+00 | 9.77E-01 | no | | intergenic_variant | MODIFIER |
| 5 | 79872209 | AX-170522849 | 0 | A | G | 0.421 | 3.24E-03 | 1.02E-01 | 2.67E+00 | 2.74E+00 | 9.77E-01 | no | | intergenic_variant | MODIFIER |
| 12 | 57418450 | AX-170314670 | 2 | C | T | 0.064 | 8.07E-03 | 3.00E-01 | 2.72E+00 | 2.74E+00 | 9.77E-01 | no | Mipol1 | intron_variant | MODIFIER |
| 8 | 84530018 | AX-170185513 | 0 | C | T | 0.202 | -2.97E-03 | 1.12E-01 | 2.64E+00 | 2.74E+00 | 9.77E-01 | no | Cacna1a | intron_variant | MODIFIER |
| 5 | 125138824 | AX-170356169 | 0 | C | T | 0.465 | 3.66E-03 | 1.12E-01 | 2.68E+00 | 2.73E+00 | 9.77E-01 | no | Ncor2 | intron_variant | MODIFIER |

| | | | | | | | | | | | | | | | |
|---|---|---|---|---|---|---|---|---|---|---|---|---|---|---|---|
| 4 | 139683386 | AX-170386429 | 0 | T | C | 0.175 | -1.27E-03 | 9.62E-02 | 2.64E+00 | 2.75E+00 | 9.77E-01 | no | | intergenic_variant | MODIFIER |
| 1 | 38725231 | AX-170509984 | 0 | T | G | 0.456 | -2.47E-03 | 7.84E-02 | 2.66E+00 | 2.74E+00 | 9.77E-01 | no | Gm38135 | downstream_gene_variant | MODIFIER |
| 2 | 124915271 | AX-170277693 | 2 | T | C | 0.136 | -3.71E-03 | 1.50E-01 | 2.61E+00 | 2.74E+00 | 9.77E-01 | no | | intergenic_variant | MODIFIER |
| 11 | 9183469 | AX-170436235 | 0 | G | A | 0.061 | 4.92E-03 | 2.14E-01 | 2.65E+00 | 2.74E+00 | 9.77E-01 | no | | intergenic_variant | MODIFIER |
| 7 | 96555353 | AX-170113269 | 2 | C | T | 0.109 | 5.43E-03 | 1.91E-01 | 2.64E+00 | 2.74E+00 | 9.78E-01 | no | Tenm4 | intron_variant | MODIFIER |
| 1 | 3481133 | AX-170103557 | 1 | T | C | 0.509 | -6.52E-04 | 6.71E-02 | 2.65E+00 | 2.75E+00 | 9.78E-01 | no | Xkr4 | intron_variant | MODIFIER |
| 1 | 40484126 | AX-97840964 | 0 | A | G | 0.395 | 1.72E-03 | 7.69E-02 | 2.67E+00 | 2.74E+00 | 9.78E-01 | no | Il18r1 | intron_variant | MODIFIER |
| 9 | 105597133 | AX-97834827 | 1 | T | G | 0.125 | -4.60E-03 | 1.17E-01 | 2.66E+00 | 2.73E+00 | 9.78E-01 | no | | intergenic_variant | MODIFIER |
| 9 | 107405111 | AX-170355589 | 1 | C | A | 0.125 | -4.60E-03 | 1.17E-01 | 2.66E+00 | 2.73E+00 | 9.78E-01 | no | Cacna2d2 | intron_variant | MODIFIER |
| 13 | 45212729 | AX-170216639 | 1 | G | A | 0.464 | -9.53E-04 | 6.00E-02 | 2.63E+00 | 2.74E+00 | 9.78E-01 | no | | intergenic_variant | MODIFIER |
| 2 | 122986567 | AX-170064777 | 2 | C | T | 0.127 | 6.11E-03 | 1.96E-01 | 2.68E+00 | 2.74E+00 | 9.78E-01 | no | | intergenic_variant | MODIFIER |
| 18 | 60943019 | AX-170426988 | 0 | G | A | 0.421 | -3.25E-03 | 8.27E-02 | 2.66E+00 | 2.73E+00 | 9.78E-01 | no | Camk2a | intron_variant | MODIFIER |
| 4 | 149729033 | AX-170368467 | 0 | G | C | 0.096 | -4.99E-03 | 1.99E-01 | 2.68E+00 | 2.74E+00 | 9.78E-01 | no | Tmem201 | intron_variant | MODIFIER |
| 8 | 84458620 | AX-170312255 | 1 | G | A | 0.196 | 3.58E-03 | 1.29E-01 | 2.67E+00 | 2.74E+00 | 9.78E-01 | no | Cacna1a | intron_variant | MODIFIER |
| 6 | 147316826 | AX-170147302 | 1 | C | G | 0.071 | 7.44E-03 | 2.77E-01 | 2.70E+00 | 2.74E+00 | 9.78E-01 | no | | regulatory_region_variant | MODIFIER |
| 1 | 11099540 | AX-170143222 | 0 | C | T | 0.088 | 6.16E-03 | 1.82E-01 | 2.62E+00 | 2.74E+00 | 9.78E-01 | no | Prex2 | intron_variant | MODIFIER |
| 12 | 111507160 | AX-170523969 | 1 | A | T | 0.116 | -3.82E-03 | 1.57E-01 | 2.62E+00 | 2.74E+00 | 9.78E-01 | no | | regulatory_region_variant | MODIFIER |
| 7 | 82837401 | AX-170467231 | 1 | T | G | 0.107 | 5.10E-03 | 1.92E-01 | 2.64E+00 | 2.74E+00 | 9.78E-01 | no | | intergenic_variant | MODIFIER |
| 9 | 53076354 | AX-170265587 | 0 | A | G | 0.219 | 3.26E-03 | 1.17E-01 | 2.67E+00 | 2.74E+00 | 9.78E-01 | no | | regulatory_region_variant | MODIFIER |
| 12 | 115520923 | AX-170244793 | 2 | A | G | 0.109 | 5.60E-03 | 1.78E-01 | 2.67E+00 | 2.74E+00 | 9.78E-01 | no | Ighv8-10 | upstream_gene_variant | MODIFIER |
| 1 | 113755296 | AX-170467475 | 1 | T | A | 0.152 | 4.70E-03 | 1.32E-01 | 2.62E+00 | 2.73E+00 | 9.78E-01 | no | | intergenic_variant | MODIFIER |
| 9 | 57504392 | AX-170122901 | 1 | C | T | 0.214 | -3.11E-03 | 1.22E-01 | 2.68E+00 | 2.74E+00 | 9.79E-01 | no | Rpp25 | synonymous_variant | LOW |
| 7 | 117660327 | AX-170289208 | 0 | A | G | 0.386 | 2.87E-03 | 9.00E-02 | 2.67E+00 | 2.73E+00 | 9.79E-01 | no | Xylt1 | intron_variant | MODIFIER |
| 7 | 118469979 | AX-170113164 | 0 | T | C | 0.386 | 2.87E-03 | 9.00E-02 | 2.67E+00 | 2.73E+00 | 9.79E-01 | no | | regulatory_region_variant | MODIFIER |
| 14 | 112387702 | AX-170130751 | 0 | C | A | 0.421 | -1.80E-03 | 4.21E-02 | 2.61E+00 | 2.73E+00 | 9.79E-01 | no | | intergenic_variant | MODIFIER |
| 9 | 71522346 | AX-170492676 | 0 | A | C | 0.211 | -3.97E-03 | 1.41E-01 | 2.71E+00 | 2.73E+00 | 9.79E-01 | no | Myzap | intron_variant | MODIFIER |
| 9 | 71779514 | AX-170032656 | 0 | C | T | 0.211 | -3.97E-03 | 1.41E-01 | 2.71E+00 | 2.73E+00 | 9.79E-01 | no | | intergenic_variant | MODIFIER |
| 9 | 71779713 | AX-170237768 | 0 | C | T | 0.211 | -3.97E-03 | 1.41E-01 | 2.71E+00 | 2.73E+00 | 9.79E-01 | no | | intergenic_variant | MODIFIER |
| 9 | 71842001 | AX-170470312 | 0 | A | G | 0.211 | -3.97E-03 | 1.41E-01 | 2.71E+00 | 2.73E+00 | 9.79E-01 | no | Tcf12 | downstream_gene_variant | MODIFIER |
| 9 | 71871698 | AX-170082785 | 0 | T | C | 0.211 | -3.97E-03 | 1.41E-01 | 2.71E+00 | 2.73E+00 | 9.79E-01 | no | Tcf12 | intron_variant | MODIFIER |
| 9 | 71956807 | AX-170346968 | 0 | C | G | 0.211 | -3.97E-03 | 1.41E-01 | 2.71E+00 | 2.73E+00 | 9.79E-01 | no | Tcf12 | intron_variant | MODIFIER |
| 9 | 71973081 | AX-170188545 | 0 | G | A | 0.211 | -3.97E-03 | 1.41E-01 | 2.71E+00 | 2.73E+00 | 9.79E-01 | no | Tcf12 | intron_variant | MODIFIER |
| 9 | 72064393 | AX-170494929 | 0 | T | C | 0.211 | -3.97E-03 | 1.41E-01 | 2.71E+00 | 2.73E+00 | 9.79E-01 | no | Tcf12 | intron_variant | MODIFIER |
| 9 | 72121219 | AX-97812962 | 0 | A | G | 0.211 | -3.97E-03 | 1.41E-01 | 2.71E+00 | 2.73E+00 | 9.79E-01 | no | Gm25045 | downstream_gene_variant | MODIFIER |
| 9 | 72123051 | AX-170503144 | 0 | C | A | 0.211 | -3.97E-03 | 1.41E-01 | 2.71E+00 | 2.73E+00 | 9.79E-01 | no | Gm25045 | upstream_gene_variant | MODIFIER |
| 9 | 72149382 | AX-170032673 | 0 | G | C | 0.211 | -3.97E-03 | 1.41E-01 | 2.71E+00 | 2.73E+00 | 9.79E-01 | no | | intergenic_variant | MODIFIER |
| 9 | 72409660 | AX-97784367 | 0 | G | A | 0.211 | -3.97E-03 | 1.41E-01 | 2.71E+00 | 2.73E+00 | 9.79E-01 | no | BC065403 | intron_variant,non_coding_transcript_variant | MODIFIER |
| 9 | 72459797 | AX-170032678 | 0 | C | G | 0.211 | -3.97E-03 | 1.41E-01 | 2.71E+00 | 2.73E+00 | 9.79E-01 | no | Mns1 | 3_prime_UTR_variant | MODIFIER |
| 9 | 72464366 | AX-170479483 | 0 | G | C | 0.211 | -3.97E-03 | 1.41E-01 | 2.71E+00 | 2.73E+00 | 9.79E-01 | no | Mns1 | downstream_gene_variant | MODIFIER |
| 9 | 72471709 | AX-170458063 | 0 | C | T | 0.211 | -3.97E-03 | 1.41E-01 | 2.71E+00 | 2.73E+00 | 9.79E-01 | no | Tex9 | intron_variant | MODIFIER |
| 9 | 72481245 | AX-170184044 | 0 | C | T | 0.211 | -3.97E-03 | 1.41E-01 | 2.71E+00 | 2.73E+00 | 9.79E-01 | no | Tex9 | intron_variant | MODIFIER |
| 9 | 72548941 | AX-170139554 | 0 | T | G | 0.211 | -3.97E-03 | 1.41E-01 | 2.71E+00 | 2.73E+00 | 9.79E-01 | no | Rfx7 | intron_variant | MODIFIER |
| 9 | 72555240 | AX-97852897 | 0 | C | T | 0.211 | -3.97E-03 | 1.41E-01 | 2.71E+00 | 2.73E+00 | 9.79E-01 | no | Rfx7 | intron_variant | MODIFIER |
| 9 | 72577097 | AX-97789870 | 0 | C | A | 0.211 | -3.97E-03 | 1.41E-01 | 2.71E+00 | 2.73E+00 | 9.79E-01 | no | Rfx7 | intron_variant | MODIFIER |
| 9 | 72584031 | AX-170315474 | 0 | T | A | 0.211 | -3.97E-03 | 1.41E-01 | 2.71E+00 | 2.73E+00 | 9.79E-01 | no | Rfx7 | intron_variant | MODIFIER |
| 9 | 72641493 | AX-97833649 | 0 | T | G | 0.211 | -3.97E-03 | 1.41E-01 | 2.71E+00 | 2.73E+00 | 9.79E-01 | no | | regulatory_region_variant | MODIFIER |
| 9 | 72656256 | AX-170082792 | 0 | A | T | 0.211 | -3.97E-03 | 1.41E-01 | 2.71E+00 | 2.73E+00 | 9.79E-01 | no | | intergenic_variant | MODIFIER |
| 9 | 72947699 | AX-97842035 | 0 | C | G | 0.211 | -3.97E-03 | 1.41E-01 | 2.71E+00 | 2.73E+00 | 9.79E-01 | no | Pygo1 | 3_prime_UTR_variant | MODIFIER |
| 9 | 72948039 | AX-170097667 | 0 | C | G | 0.211 | -3.97E-03 | 1.41E-01 | 2.71E+00 | 2.73E+00 | 9.79E-01 | no | Pygo1 | 3_prime_UTR_variant | MODIFIER |
| 9 | 73218794 | AX-170082799 | 0 | A | G | 0.211 | -3.97E-03 | 1.41E-01 | 2.71E+00 | 2.73E+00 | 9.79E-01 | no | Gm27211 | intron_variant,non_coding_transcript_variant | MODIFIER |
| 9 | 73222403 | AX-97814867 | 0 | T | C | 0.211 | -3.97E-03 | 1.41E-01 | 2.71E+00 | 2.73E+00 | 9.79E-01 | no | Gm27211 | intron_variant,non_coding_transcript_variant | MODIFIER |
| 9 | 73305725 | AX-170410252 | 0 | A | T | 0.211 | -3.97E-03 | 1.41E-01 | 2.71E+00 | 2.73E+00 | 9.79E-01 | no | | intergenic_variant | MODIFIER |
| 9 | 73433381 | AX-170342869 | 0 | C | T | 0.211 | -3.97E-03 | 1.41E-01 | 2.71E+00 | 2.73E+00 | 9.79E-01 | no | | intergenic_variant | MODIFIER |
| 9 | 73462066 | AX-170451172 | 0 | T | C | 0.211 | -3.97E-03 | 1.41E-01 | 2.71E+00 | 2.73E+00 | 9.79E-01 | no | | intergenic_variant | MODIFIER |

| | | | | | | | | | | | | | | | |
|---|---|---|---|---|---|---|---|---|---|---|---|---|---|---|---|
| 9 | 73483742 | AX-112893139 | 0 | C | T | 0.211 | -3.97E-03 | 1.41E-01 | 2.71E+00 | 2.73E+00 | 9.79E-01 | no | Unc13c | missense_variant | MODERATE |
| 9 | 73699927 | AX-170032702 | 0 | G | A | 0.211 | -3.97E-03 | 1.41E-01 | 2.71E+00 | 2.73E+00 | 9.79E-01 | no | Unc13c | intron_variant | MODIFIER |
| 9 | 73751705 | AX-97783149 | 0 | A | G | 0.211 | -3.97E-03 | 1.41E-01 | 2.71E+00 | 2.73E+00 | 9.79E-01 | no | Unc13c | intron_variant | MODIFIER |
| 9 | 73758470 | AX-170443764 | 0 | T | G | 0.211 | -3.97E-03 | 1.41E-01 | 2.71E+00 | 2.73E+00 | 9.79E-01 | no | Unc13c | intron_variant | MODIFIER |
| 9 | 73758742 | AX-170142904 | 0 | T | C | 0.211 | -3.97E-03 | 1.41E-01 | 2.71E+00 | 2.73E+00 | 9.79E-01 | no | Unc13c | intron_variant | MODIFIER |
| 9 | 73760429 | AX-170119745 | 0 | C | T | 0.211 | -3.97E-03 | 1.41E-01 | 2.71E+00 | 2.73E+00 | 9.79E-01 | no | Unc13c | intron_variant | MODIFIER |
| 9 | 73841429 | AX-170158376 | 0 | G | C | 0.211 | -3.97E-03 | 1.41E-01 | 2.71E+00 | 2.73E+00 | 9.79E-01 | no | Unc13c | intron_variant | MODIFIER |
| 9 | 73930214 | AX-170285717 | 0 | C | A | 0.211 | -3.97E-03 | 1.41E-01 | 2.71E+00 | 2.73E+00 | 9.79E-01 | no | Unc13c | intron_variant | MODIFIER |
| 9 | 73934440 | AX-97826737 | 0 | T | C | 0.211 | -3.97E-03 | 1.41E-01 | 2.71E+00 | 2.73E+00 | 9.79E-01 | no | Unc13c | upstream_gene_variant | MODIFIER |
| 9 | 73951346 | AX-170147040 | 0 | T | G | 0.211 | -3.97E-03 | 1.41E-01 | 2.71E+00 | 2.73E+00 | 9.79E-01 | no | Unc13c | intron_variant | MODIFIER |
| 9 | 73966743 | AX-170364921 | 0 | A | G | 0.211 | -3.97E-03 | 1.41E-01 | 2.71E+00 | 2.73E+00 | 9.79E-01 | no | Unc13c | intron_variant | MODIFIER |
| 9 | 74010277 | AX-170294521 | 0 | T | G | 0.211 | -3.97E-03 | 1.41E-01 | 2.71E+00 | 2.73E+00 | 9.79E-01 | no | | intergenic_variant | MODIFIER |
| 9 | 74021006 | AX-170527327 | 0 | T | C | 0.211 | -3.97E-03 | 1.41E-01 | 2.71E+00 | 2.73E+00 | 9.79E-01 | no | | intergenic_variant | MODIFIER |
| 9 | 74022676 | AX-170202347 | 0 | T | C | 0.211 | -3.97E-03 | 1.41E-01 | 2.71E+00 | 2.73E+00 | 9.79E-01 | no | | intergenic_variant | MODIFIER |
| 9 | 74022784 | AX-170082807 | 0 | G | A | 0.211 | -3.97E-03 | 1.41E-01 | 2.71E+00 | 2.73E+00 | 9.79E-01 | no | | intergenic_variant | MODIFIER |
| 9 | 74022801 | AX-170082809 | 0 | A | G | 0.211 | -3.97E-03 | 1.41E-01 | 2.71E+00 | 2.73E+00 | 9.79E-01 | no | | intergenic_variant | MODIFIER |
| 9 | 74034250 | AX-170271757 | 0 | A | G | 0.211 | -3.97E-03 | 1.41E-01 | 2.71E+00 | 2.73E+00 | 9.79E-01 | no | | intergenic_variant | MODIFIER |
| 9 | 74067251 | AX-170236140 | 0 | G | C | 0.211 | -3.97E-03 | 1.41E-01 | 2.71E+00 | 2.73E+00 | 9.79E-01 | no | | regulatory_region_variant | MODIFIER |
| 9 | 74067279 | AX-170394494 | 0 | G | A | 0.211 | -3.97E-03 | 1.41E-01 | 2.71E+00 | 2.73E+00 | 9.79E-01 | no | | intergenic_variant | MODIFIER |
| 9 | 74092532 | AX-170174302 | 0 | T | C | 0.211 | -3.97E-03 | 1.41E-01 | 2.71E+00 | 2.73E+00 | 9.79E-01 | no | | intergenic_variant | MODIFIER |
| 9 | 74109574 | AX-170303041 | 0 | A | T | 0.211 | -3.97E-03 | 1.41E-01 | 2.71E+00 | 2.73E+00 | 9.79E-01 | no | Wdr72 | upstream_gene_variant | MODIFIER |
| 9 | 74122123 | AX-170096585 | 0 | G | A | 0.211 | -3.97E-03 | 1.41E-01 | 2.71E+00 | 2.73E+00 | 9.79E-01 | no | Wdr72 | intron_variant | MODIFIER |
| 9 | 74141207 | AX-170389310 | 0 | A | G | 0.211 | -3.97E-03 | 1.41E-01 | 2.71E+00 | 2.73E+00 | 9.79E-01 | no | Wdr72 | intron_variant | MODIFIER |
| 9 | 74142729 | AX-170096586 | 0 | T | C | 0.211 | -3.97E-03 | 1.41E-01 | 2.71E+00 | 2.73E+00 | 9.79E-01 | no | Wdr72 | intron_variant | MODIFIER |
| 9 | 74158941 | AX-170357733 | 0 | T | C | 0.211 | -3.97E-03 | 1.41E-01 | 2.71E+00 | 2.73E+00 | 9.79E-01 | no | Wdr72 | intron_variant | MODIFIER |
| 9 | 74192922 | AX-170404465 | 0 | T | G | 0.211 | -3.97E-03 | 1.41E-01 | 2.71E+00 | 2.73E+00 | 9.79E-01 | no | Wdr72 | intron_variant | MODIFIER |
| 9 | 74259537 | AX-170262615 | 0 | T | C | 0.211 | -3.97E-03 | 1.41E-01 | 2.71E+00 | 2.73E+00 | 9.79E-01 | no | Wdr72 | intron_variant | MODIFIER |
| 9 | 74260591 | AX-170461014 | 0 | T | A | 0.211 | -3.97E-03 | 1.41E-01 | 2.71E+00 | 2.73E+00 | 9.79E-01 | no | Wdr72 | intron_variant | MODIFIER |
| 9 | 74377989 | AX-170162318 | 0 | T | A | 0.211 | -3.97E-03 | 1.41E-01 | 2.71E+00 | 2.73E+00 | 9.79E-01 | no | | intergenic_variant | MODIFIER |
| 9 | 74390027 | AX-170324614 | 0 | T | C | 0.211 | -3.97E-03 | 1.41E-01 | 2.71E+00 | 2.73E+00 | 9.79E-01 | no | | intergenic_variant | MODIFIER |
| 9 | 74431970 | AX-170456817 | 0 | T | G | 0.211 | -3.97E-03 | 1.41E-01 | 2.71E+00 | 2.73E+00 | 9.79E-01 | no | | intergenic_variant | MODIFIER |
| 9 | 74436427 | AX-170117616 | 0 | T | C | 0.211 | -3.97E-03 | 1.41E-01 | 2.71E+00 | 2.73E+00 | 9.79E-01 | no | | intergenic_variant | MODIFIER |
| 9 | 74513124 | AX-170384076 | 0 | A | T | 0.211 | -3.97E-03 | 1.41E-01 | 2.71E+00 | 2.73E+00 | 9.79E-01 | no | | intergenic_variant | MODIFIER |
| 9 | 74517326 | AX-170321082 | 0 | G | A | 0.211 | -3.97E-03 | 1.41E-01 | 2.71E+00 | 2.73E+00 | 9.79E-01 | no | | intergenic_variant | MODIFIER |
| 2 | 127033403 | AX-97839324 | 0 | G | A | 0.14 | -4.69E-03 | 1.81E-01 | 2.66E+00 | 2.74E+00 | 9.79E-01 | no | Ap4e1 | intron_variant | MODIFIER |
| 2 | 145525941 | AX-170286099 | 0 | G | A | 0.14 | -4.69E-03 | 1.81E-01 | 2.66E+00 | 2.74E+00 | 9.79E-01 | no | Slc24a3 | intron_variant | MODIFIER |
| 2 | 149247184 | AX-170338140 | 0 | G | A | 0.14 | -4.69E-03 | 1.81E-01 | 2.66E+00 | 2.74E+00 | 9.79E-01 | no | | intergenic_variant | MODIFIER |
| 2 | 150010138 | AX-97825991 | 0 | G | A | 0.14 | -4.69E-03 | 1.81E-01 | 2.66E+00 | 2.74E+00 | 9.79E-01 | no | | intergenic_variant | MODIFIER |
| 2 | 157963674 | AX-170019493 | 0 | C | T | 0.14 | -4.69E-03 | 1.81E-01 | 2.66E+00 | 2.74E+00 | 9.79E-01 | no | | intergenic_variant | MODIFIER |
| 6 | 125126875 | AX-170500901 | 0 | C | T | 0.202 | 2.22E-03 | 8.17E-02 | 2.61E+00 | 2.74E+00 | 9.79E-01 | no | Chd4 | intron_variant | MODIFIER |
| 4 | 149854892 | AX-170024227 | 0 | G | A | 0.096 | 4.08E-03 | 1.70E-01 | 2.65E+00 | 2.74E+00 | 9.79E-01 | no | | regulatory_region_variant | MODIFIER |
| 9 | 34152694 | AX-170291628 | 0 | T | C | 0.43 | 2.88E-03 | 8.64E-02 | 2.66E+00 | 2.73E+00 | 9.79E-01 | no | | intergenic_variant | MODIFIER |
| 7 | 78350013 | AX-97842352 | 0 | G | A | 0.114 | 5.88E-03 | 2.11E-01 | 2.66E+00 | 2.74E+00 | 9.79E-01 | no | Ntrk3 | intron_variant | MODIFIER |
| 7 | 80577056 | AX-97802704 | 0 | G | A | 0.114 | 5.88E-03 | 2.11E-01 | 2.66E+00 | 2.74E+00 | 9.79E-01 | no | | regulatory_region_variant | MODIFIER |
| 7 | 94769305 | AX-170107893 | 0 | T | G | 0.114 | 5.88E-03 | 2.11E-01 | 2.66E+00 | 2.74E+00 | 9.79E-01 | no | | intergenic_variant | MODIFIER |
| 5 | 127413473 | AX-170493223 | 0 | T | A | 0.421 | -1.42E-03 | 8.36E-02 | 2.63E+00 | 2.74E+00 | 9.79E-01 | no | Tmem132c | intron_variant | MODIFIER |
| 5 | 90280241 | AX-97807418 | 1 | G | A | 0.429 | 2.73E-03 | 1.11E-01 | 2.69E+00 | 2.74E+00 | 9.79E-01 | no | Ankrd17 | intron_variant | MODIFIER |
| 12 | 63369282 | AX-170213040 | 0 | G | A | 0.061 | -8.28E-03 | 3.01E-01 | 2.71E+00 | 2.74E+00 | 9.79E-01 | no | | intergenic_variant | MODIFIER |
| 3 | 145768323 | AX-170362430 | 0 | G | A | 0.36 | -1.43E-03 | 7.30E-02 | 2.66E+00 | 2.74E+00 | 9.79E-01 | no | Ddah1 | intron_variant | MODIFIER |
| 7 | 137863701 | AX-97850951 | 0 | C | T | 0.298 | -2.53E-03 | 7.81E-02 | 2.63E+00 | 2.74E+00 | 9.79E-01 | no | | intergenic_variant | MODIFIER |
| 7 | 137867490 | AX-170030079 | 0 | A | G | 0.298 | -2.53E-03 | 7.81E-02 | 2.63E+00 | 2.74E+00 | 9.79E-01 | no | | intergenic_variant | MODIFIER |
| 7 | 137867812 | AX-170121621 | 0 | C | T | 0.298 | -2.53E-03 | 7.81E-02 | 2.63E+00 | 2.74E+00 | 9.79E-01 | no | | intergenic_variant | MODIFIER |
| 7 | 137871389 | AX-170382359 | 0 | A | G | 0.298 | -2.53E-03 | 7.81E-02 | 2.63E+00 | 2.74E+00 | 9.79E-01 | no | | intergenic_variant | MODIFIER |
| 7 | 137871448 | AX-170486726 | 0 | C | A | 0.298 | -2.53E-03 | 7.81E-02 | 2.63E+00 | 2.74E+00 | 9.79E-01 | no | | intergenic_variant | MODIFIER |

| | | | | | | | | | | | | | | | |
|---|---|---|---|---|---|---|---|---|---|---|---|---|---|---|---|
| 7 | 137873424 | AX-170179797 | 0 | G | C | 0.298 | -2.53E-03 | 7.81E-02 | 2.63E+00 | 2.74E+00 | 9.79E-01 | no | | intergenic_variant | MODIFIER |
| 7 | 137878307 | AX-170367086 | 0 | G | A | 0.298 | -2.53E-03 | 7.81E-02 | 2.63E+00 | 2.74E+00 | 9.79E-01 | no | | intergenic_variant | MODIFIER |
| 7 | 137882931 | AX-170184461 | 0 | T | C | 0.298 | -2.53E-03 | 7.81E-02 | 2.63E+00 | 2.74E+00 | 9.79E-01 | no | | regulatory_region_variant | MODIFIER |
| 7 | 137883020 | AX-170304258 | 0 | G | T | 0.298 | -2.53E-03 | 7.81E-02 | 2.63E+00 | 2.74E+00 | 9.79E-01 | no | | regulatory_region_variant | MODIFIER |
| 7 | 137883441 | AX-170096139 | 0 | T | G | 0.298 | -2.53E-03 | 7.81E-02 | 2.63E+00 | 2.74E+00 | 9.79E-01 | no | | intergenic_variant | MODIFIER |
| 7 | 137889900 | AX-170139499 | 0 | A | G | 0.298 | -2.53E-03 | 7.81E-02 | 2.63E+00 | 2.74E+00 | 9.79E-01 | no | | intergenic_variant | MODIFIER |
| 7 | 137906730 | AX-97826942 | 0 | A | C | 0.298 | -2.53E-03 | 7.81E-02 | 2.63E+00 | 2.74E+00 | 9.79E-01 | no | | regulatory_region_variant | MODIFIER |
| 7 | 137929102 | AX-97798984 | 0 | G | A | 0.298 | -2.53E-03 | 7.81E-02 | 2.63E+00 | 2.74E+00 | 9.79E-01 | no | | intergenic_variant | MODIFIER |
| 7 | 137952829 | AX-170488057 | 0 | T | A | 0.298 | -2.53E-03 | 7.81E-02 | 2.63E+00 | 2.74E+00 | 9.79E-01 | no | | regulatory_region_variant | MODIFIER |
| 7 | 137967075 | AX-170127973 | 0 | T | A | 0.298 | -2.53E-03 | 7.81E-02 | 2.63E+00 | 2.74E+00 | 9.79E-01 | no | | intergenic_variant | MODIFIER |
| 7 | 137970966 | AX-97851802 | 0 | C | T | 0.298 | -2.53E-03 | 7.81E-02 | 2.63E+00 | 2.74E+00 | 9.79E-01 | no | | intergenic_variant | MODIFIER |
| 7 | 137996389 | AX-97853332 | 0 | T | C | 0.298 | -2.53E-03 | 7.81E-02 | 2.63E+00 | 2.74E+00 | 9.79E-01 | no | | intergenic_variant | MODIFIER |
| 7 | 138010646 | AX-97848674 | 0 | A | G | 0.298 | -2.53E-03 | 7.81E-02 | 2.63E+00 | 2.74E+00 | 9.79E-01 | no | | intergenic_variant | MODIFIER |
| 7 | 138036751 | AX-97788056 | 0 | C | A | 0.298 | -2.53E-03 | 7.81E-02 | 2.63E+00 | 2.74E+00 | 9.79E-01 | no | | intergenic_variant | MODIFIER |
| 7 | 138076567 | AX-170406636 | 0 | G | A | 0.298 | -2.53E-03 | 7.81E-02 | 2.63E+00 | 2.74E+00 | 9.79E-01 | no | | intergenic_variant | MODIFIER |
| 7 | 138077416 | AX-170315727 | 0 | A | G | 0.298 | -2.53E-03 | 7.81E-02 | 2.63E+00 | 2.74E+00 | 9.79E-01 | no | | intergenic_variant | MODIFIER |
| 7 | 138732624 | AX-97781744 | 0 | G | A | 0.298 | -2.53E-03 | 7.81E-02 | 2.63E+00 | 2.74E+00 | 9.79E-01 | no | | intergenic_variant | MODIFIER |
| 7 | 138922940 | AX-170030094 | 0 | G | A | 0.298 | -2.53E-03 | 7.81E-02 | 2.63E+00 | 2.74E+00 | 9.79E-01 | no | | regulatory_region_variant | MODIFIER |
| 7 | 138984427 | AX-97783583 | 0 | C | T | 0.298 | -2.53E-03 | 7.81E-02 | 2.63E+00 | 2.74E+00 | 9.79E-01 | no | Jakmip3 | intron_variant | MODIFIER |
| 7 | 139068052 | AX-97803757 | 0 | G | A | 0.298 | -2.53E-03 | 7.81E-02 | 2.63E+00 | 2.74E+00 | 9.79E-01 | no | Jakmip3 | intron_variant | MODIFIER |
| 7 | 139076251 | AX-170478985 | 0 | A | G | 0.298 | -2.53E-03 | 7.81E-02 | 2.63E+00 | 2.74E+00 | 9.79E-01 | no | Jakmip3 | intron_variant | MODIFIER |
| 7 | 139077708 | AX-97816257 | 0 | G | A | 0.298 | -2.53E-03 | 7.81E-02 | 2.63E+00 | 2.74E+00 | 9.79E-01 | no | Jakmip3 | intron_variant | MODIFIER |
| 7 | 139079138 | AX-170360091 | 0 | T | C | 0.298 | -2.53E-03 | 7.81E-02 | 2.63E+00 | 2.74E+00 | 9.79E-01 | no | Jakmip3 | intron_variant | MODIFIER |
| 7 | 139095291 | AX-170366816 | 0 | A | G | 0.298 | -2.53E-03 | 7.81E-02 | 2.63E+00 | 2.74E+00 | 9.79E-01 | no | Dpysl4 | intron_variant | MODIFIER |
| 7 | 139127313 | AX-170370845 | 0 | A | G | 0.298 | -2.53E-03 | 7.81E-02 | 2.63E+00 | 2.74E+00 | 9.79E-01 | no | Stk32c | intron_variant | MODIFIER |
| 7 | 60926598 | AX-97797155 | 1 | A | G | 0.42 | -1.87E-03 | 9.22E-02 | 2.66E+00 | 2.74E+00 | 9.79E-01 | no | Gm44644 | upstream_gene_variant | MODIFIER |
| 7 | 91378006 | AX-170488349 | 2 | C | T | 0.136 | -5.07E-03 | 2.13E-01 | 2.66E+00 | 2.74E+00 | 9.79E-01 | no | Dlg2 | intron_variant | MODIFIER |
| 8 | 20922194 | AX-170098175 | 0 | C | T | 0.184 | 3.08E-03 | 9.89E-02 | 2.64E+00 | 2.74E+00 | 9.79E-01 | no | Gm6040 | upstream_gene_variant | MODIFIER |
| 7 | 102280567 | AX-97857621 | 1 | T | C | 0.455 | 3.14E-03 | 9.41E-02 | 2.67E+00 | 2.73E+00 | 9.79E-01 | no | Stim1 | intron_variant | MODIFIER |
| 10 | 6794285 | AX-170149439 | 1 | G | A | 0.205 | 1.94E-03 | 1.03E-01 | 2.65E+00 | 2.74E+00 | 9.80E-01 | no | Oprm1 | intron_variant | MODIFIER |
| 6 | 105110865 | AX-170338132 | 1 | C | T | 0.08 | -1.94E-03 | 1.75E-01 | 2.61E+00 | 2.74E+00 | 9.80E-01 | no | | intergenic_variant | MODIFIER |
| 1 | 54145663 | AX-170486196 | 1 | G | A | 0.08 | -5.48E-03 | 1.90E-01 | 2.61E+00 | 2.74E+00 | 9.80E-01 | no | Hecw2 | intron_variant | MODIFIER |
| 12 | 112102721 | AX-170160894 | 1 | C | G | 0.125 | 5.39E-03 | 1.96E-01 | 2.67E+00 | 2.74E+00 | 9.80E-01 | no | Aspg | upstream_gene_variant | MODIFIER |
| 12 | 116595185 | AX-170436570 | 1 | C | G | 0.125 | 5.39E-03 | 1.96E-01 | 2.67E+00 | 2.74E+00 | 9.80E-01 | no | Ptprn2 | intron_variant | MODIFIER |
| 12 | 56927044 | AX-170006803 | 1 | C | A | 0.062 | -7.90E-03 | 3.03E-01 | 2.71E+00 | 2.74E+00 | 9.80E-01 | no | Slc25a21 | intron_variant | MODIFIER |
| 4 | 148052081 | AX-170365693 | 0 | C | T | 0.114 | 4.33E-03 | 1.64E-01 | 2.64E+00 | 2.74E+00 | 9.80E-01 | no | Mthfr | intron_variant | MODIFIER |
| 10 | 31604926 | AX-97826254 | 0 | T | C | 0.377 | -2.27E-03 | 7.39E-02 | 2.62E+00 | 2.74E+00 | 9.80E-01 | no | Rnf217 | intron_variant | MODIFIER |
| 11 | 33419686 | AX-170512557 | 0 | A | G | 0.105 | 5.43E-03 | 1.89E-01 | 2.66E+00 | 2.74E+00 | 9.80E-01 | no | Ranbp17 | intron_variant | MODIFIER |
| 11 | 13081869 | AX-170004310 | 0 | A | G | 0.07 | -5.75E-03 | 1.91E-01 | 2.62E+00 | 2.74E+00 | 9.80E-01 | no | | intergenic_variant | MODIFIER |
| 11 | 23005494 | AX-170092199 | 0 | G | A | 0.07 | -5.75E-03 | 1.91E-01 | 2.62E+00 | 2.74E+00 | 9.80E-01 | no | Cct4 | downstream_gene_variant | MODIFIER |
| 2 | 88154192 | AX-97786339 | 1 | C | T | 0.054 | 4.77E-03 | 2.27E-01 | 2.65E+00 | 2.74E+00 | 9.80E-01 | no | Olfr1167 | upstream_gene_variant | MODIFIER |
| 9 | 35488174 | AX-170228049 | 1 | C | T | 0.438 | -1.38E-03 | 8.74E-02 | 2.67E+00 | 2.74E+00 | 9.80E-01 | no | Cdon | intron_variant | MODIFIER |
| 9 | 111431495 | AX-170504505 | 0 | G | A | 0.105 | 1.84E-03 | 1.16E-01 | 2.67E+00 | 2.74E+00 | 9.81E-01 | no | | intergenic_variant | MODIFIER |
| 9 | 111455945 | AX-97815722 | 0 | G | A | 0.105 | 1.84E-03 | 1.16E-01 | 2.67E+00 | 2.74E+00 | 9.81E-01 | no | Dclk3 | intron_variant | MODIFIER |
| 9 | 111463773 | AX-170324362 | 0 | G | T | 0.105 | 1.84E-03 | 1.16E-01 | 2.67E+00 | 2.74E+00 | 9.81E-01 | no | Dclk3 | intron_variant | MODIFIER |
| 9 | 111464310 | AX-170374970 | 0 | T | C | 0.105 | 1.84E-03 | 1.16E-01 | 2.67E+00 | 2.74E+00 | 9.81E-01 | no | Dclk3 | intron_variant | MODIFIER |
| 9 | 111464378 | AX-170404738 | 0 | A | G | 0.105 | 1.84E-03 | 1.16E-01 | 2.67E+00 | 2.74E+00 | 9.81E-01 | no | Dclk3 | intron_variant | MODIFIER |
| 9 | 111464735 | AX-170473379 | 0 | C | T | 0.105 | 1.84E-03 | 1.16E-01 | 2.67E+00 | 2.74E+00 | 9.81E-01 | no | Dclk3 | intron_variant | MODIFIER |
| 9 | 111465067 | AX-170158851 | 0 | T | C | 0.105 | 1.84E-03 | 1.16E-01 | 2.67E+00 | 2.74E+00 | 9.81E-01 | no | Dclk3 | intron_variant | MODIFIER |
| 9 | 111466508 | AX-170414290 | 0 | C | T | 0.105 | 1.84E-03 | 1.16E-01 | 2.67E+00 | 2.74E+00 | 9.81E-01 | no | Dclk3 | intron_variant | MODIFIER |
| 9 | 111474901 | AX-170325346 | 0 | T | A | 0.105 | 1.84E-03 | 1.16E-01 | 2.67E+00 | 2.74E+00 | 9.81E-01 | no | Dclk3 | intron_variant | MODIFIER |
| 9 | 111493423 | AX-97800192 | 0 | A | G | 0.105 | 1.84E-03 | 1.16E-01 | 2.67E+00 | 2.74E+00 | 9.81E-01 | no | Dclk3 | downstream_gene_variant | MODIFIER |
| 9 | 111501814 | AX-170465732 | 0 | C | T | 0.105 | 1.84E-03 | 1.16E-01 | 2.67E+00 | 2.74E+00 | 9.81E-01 | no | | intergenic_variant | MODIFIER |
| 9 | 111513332 | AX-170297671 | 0 | A | G | 0.105 | 1.84E-03 | 1.16E-01 | 2.67E+00 | 2.74E+00 | 9.81E-01 | no | | intergenic_variant | MODIFIER |

| | | | | | | | | | | | | | | | |
|---|---|---|---|---|---|---|---|---|---|---|---|---|---|---|---|
| 9 | 111627184 | AX-170481373 | 0 | T | A | 0.105 | 1.84E-03 | 1.16E-01 | 2.67E+00 | 2.74E+00 | 9.81E-01 | no | Stac | intron_variant | MODIFIER |
| 9 | 111627659 | AX-170243221 | 0 | C | T | 0.105 | 1.84E-03 | 1.16E-01 | 2.67E+00 | 2.74E+00 | 9.81E-01 | no | Stac | intron_variant | MODIFIER |
| 9 | 111628602 | AX-170033282 | 0 | G | A | 0.105 | 1.84E-03 | 1.16E-01 | 2.67E+00 | 2.74E+00 | 9.81E-01 | no | Stac | intron_variant | MODIFIER |
| 9 | 111633537 | AX-170396514 | 0 | C | G | 0.105 | 1.84E-03 | 1.16E-01 | 2.67E+00 | 2.74E+00 | 9.81E-01 | no | Stac | intron_variant | MODIFIER |
| 9 | 111633983 | AX-170215002 | 0 | T | C | 0.105 | 1.84E-03 | 1.16E-01 | 2.67E+00 | 2.74E+00 | 9.81E-01 | no | Stac | intron_variant | MODIFIER |
| 9 | 111634028 | AX-170335919 | 0 | G | C | 0.105 | 1.84E-03 | 1.16E-01 | 2.67E+00 | 2.74E+00 | 9.81E-01 | no | Stac | intron_variant | MODIFIER |
| 9 | 111657810 | AX-170033286 | 0 | A | C | 0.105 | 1.84E-03 | 1.16E-01 | 2.67E+00 | 2.74E+00 | 9.81E-01 | no | Stac | intron_variant | MODIFIER |
| 9 | 111672064 | AX-170033283 | 0 | C | G | 0.105 | 1.84E-03 | 1.16E-01 | 2.67E+00 | 2.74E+00 | 9.81E-01 | no | Stac | intron_variant | MODIFIER |
| 9 | 111676306 | AX-170417385 | 0 | T | G | 0.105 | 1.84E-03 | 1.16E-01 | 2.67E+00 | 2.74E+00 | 9.81E-01 | no | Stac | intron_variant | MODIFIER |
| 9 | 111692208 | AX-170033284 | 0 | T | C | 0.105 | 1.84E-03 | 1.16E-01 | 2.67E+00 | 2.74E+00 | 9.81E-01 | no | Stac | upstream_gene_variant | MODIFIER |
| 9 | 111705323 | AX-170339891 | 0 | T | A | 0.105 | 1.84E-03 | 1.16E-01 | 2.67E+00 | 2.74E+00 | 9.81E-01 | no | | intergenic_variant | MODIFIER |
| 9 | 111713630 | AX-170450510 | 0 | G | C | 0.105 | 1.84E-03 | 1.16E-01 | 2.67E+00 | 2.74E+00 | 9.81E-01 | no | | intergenic_variant | MODIFIER |
| 9 | 111715111 | AX-170090417 | 0 | G | A | 0.105 | 1.84E-03 | 1.16E-01 | 2.67E+00 | 2.74E+00 | 9.81E-01 | no | | intergenic_variant | MODIFIER |
| 9 | 111717282 | AX-170409156 | 0 | A | G | 0.105 | 1.84E-03 | 1.16E-01 | 2.67E+00 | 2.74E+00 | 9.81E-01 | no | | intergenic_variant | MODIFIER |
| 9 | 111760359 | AX-170206416 | 0 | A | G | 0.105 | 1.84E-03 | 1.16E-01 | 2.67E+00 | 2.74E+00 | 9.81E-01 | no | | intergenic_variant | MODIFIER |
| 9 | 111768996 | AX-170083470 | 0 | A | G | 0.105 | 1.84E-03 | 1.16E-01 | 2.67E+00 | 2.74E+00 | 9.81E-01 | no | | intergenic_variant | MODIFIER |
| 9 | 111769390 | AX-170245860 | 0 | A | T | 0.105 | 1.84E-03 | 1.16E-01 | 2.67E+00 | 2.74E+00 | 9.81E-01 | no | | intergenic_variant | MODIFIER |
| 9 | 111769651 | AX-170347821 | 0 | A | G | 0.105 | 1.84E-03 | 1.16E-01 | 2.67E+00 | 2.74E+00 | 9.81E-01 | no | | intergenic_variant | MODIFIER |
| 9 | 111769888 | AX-170349258 | 0 | A | G | 0.105 | 1.84E-03 | 1.16E-01 | 2.67E+00 | 2.74E+00 | 9.81E-01 | no | | intergenic_variant | MODIFIER |
| 9 | 111771099 | AX-97823852 | 0 | T | C | 0.105 | 1.84E-03 | 1.16E-01 | 2.67E+00 | 2.74E+00 | 9.81E-01 | no | | intergenic_variant | MODIFIER |
| 9 | 111772377 | AX-170277619 | 0 | C | T | 0.105 | 1.84E-03 | 1.16E-01 | 2.67E+00 | 2.74E+00 | 9.81E-01 | no | | intergenic_variant | MODIFIER |
| 9 | 111813114 | AX-170256154 | 0 | T | C | 0.105 | 1.84E-03 | 1.16E-01 | 2.67E+00 | 2.74E+00 | 9.81E-01 | no | | intergenic_variant | MODIFIER |
| 9 | 111813606 | AX-170401582 | 0 | C | T | 0.105 | 1.84E-03 | 1.16E-01 | 2.67E+00 | 2.74E+00 | 9.81E-01 | no | | intergenic_variant | MODIFIER |
| 9 | 111814784 | AX-170242270 | 0 | G | C | 0.105 | 1.84E-03 | 1.16E-01 | 2.67E+00 | 2.74E+00 | 9.81E-01 | no | | intergenic_variant | MODIFIER |
| 9 | 111817953 | AX-97832714 | 0 | G | A | 0.105 | 1.84E-03 | 1.16E-01 | 2.67E+00 | 2.74E+00 | 9.81E-01 | no | | intergenic_variant | MODIFIER |
| 9 | 111818059 | AX-170214118 | 0 | T | C | 0.105 | 1.84E-03 | 1.16E-01 | 2.67E+00 | 2.74E+00 | 9.81E-01 | no | | intergenic_variant | MODIFIER |
| 9 | 111855074 | AX-170396820 | 0 | G | A | 0.105 | 1.84E-03 | 1.16E-01 | 2.67E+00 | 2.74E+00 | 9.81E-01 | no | | intergenic_variant | MODIFIER |
| 9 | 111857784 | AX-170202824 | 0 | A | G | 0.105 | 1.84E-03 | 1.16E-01 | 2.67E+00 | 2.74E+00 | 9.81E-01 | no | | intergenic_variant | MODIFIER |
| 9 | 111858398 | AX-170173716 | 0 | C | T | 0.105 | 1.84E-03 | 1.16E-01 | 2.67E+00 | 2.74E+00 | 9.81E-01 | no | | intergenic_variant | MODIFIER |
| 9 | 111884848 | AX-170205457 | 0 | C | G | 0.105 | 1.84E-03 | 1.16E-01 | 2.67E+00 | 2.74E+00 | 9.81E-01 | no | | intergenic_variant | MODIFIER |
| 9 | 111885098 | AX-170249642 | 0 | T | C | 0.105 | 1.84E-03 | 1.16E-01 | 2.67E+00 | 2.74E+00 | 9.81E-01 | no | | regulatory_region_variant | MODIFIER |
| 9 | 111939922 | AX-170388671 | 0 | T | C | 0.105 | 1.84E-03 | 1.16E-01 | 2.67E+00 | 2.74E+00 | 9.81E-01 | no | | regulatory_region_variant | MODIFIER |
| 9 | 111940179 | AX-170140712 | 0 | T | C | 0.105 | 1.84E-03 | 1.16E-01 | 2.67E+00 | 2.74E+00 | 9.81E-01 | no | | intergenic_variant | MODIFIER |
| 9 | 111942411 | AX-170408954 | 0 | C | T | 0.105 | 1.84E-03 | 1.16E-01 | 2.67E+00 | 2.74E+00 | 9.81E-01 | no | | intergenic_variant | MODIFIER |
| 9 | 111943490 | AX-170120108 | 0 | G | C | 0.105 | 1.84E-03 | 1.16E-01 | 2.67E+00 | 2.74E+00 | 9.81E-01 | no | | intergenic_variant | MODIFIER |
| 9 | 112008605 | AX-170530225 | 0 | A | C | 0.105 | 1.84E-03 | 1.16E-01 | 2.67E+00 | 2.74E+00 | 9.81E-01 | no | | regulatory_region_variant | MODIFIER |
| 9 | 112037606 | AX-170463171 | 0 | A | G | 0.105 | 1.84E-03 | 1.16E-01 | 2.67E+00 | 2.74E+00 | 9.81E-01 | no | | regulatory_region_variant | MODIFIER |
| 9 | 112079279 | AX-170185006 | 0 | A | G | 0.105 | 1.84E-03 | 1.16E-01 | 2.67E+00 | 2.74E+00 | 9.81E-01 | no | Arpp21 | intron_variant | MODIFIER |
| 9 | 112145232 | AX-97846687 | 0 | A | G | 0.105 | 1.84E-03 | 1.16E-01 | 2.67E+00 | 2.74E+00 | 9.81E-01 | no | Arpp21 | intron_variant | MODIFIER |
| 9 | 112149361 | AX-170033293 | 0 | T | G | 0.105 | 1.84E-03 | 1.16E-01 | 2.67E+00 | 2.74E+00 | 9.81E-01 | no | Arpp21 | synonymous_variant | LOW |
| 1 | 38585794 | AX-170202843 | 1 | A | T | 0.446 | 1.51E-03 | 7.70E-02 | 2.65E+00 | 2.74E+00 | 9.81E-01 | no | Aff3 | intron_variant,non_coding_transcript_variant | MODIFIER |
| 7 | 98881366 | AX-170126773 | 0 | T | C | 0.123 | 3.33E-03 | 1.63E-01 | 2.62E+00 | 2.74E+00 | 9.81E-01 | no | Uvrag | downstream_gene_variant | MODIFIER |
| 8 | 40550379 | AX-170393031 | 1 | G | A | 0.143 | -4.10E-03 | 1.45E-01 | 2.68E+00 | 2.73E+00 | 9.81E-01 | no | Mtmr7 | downstream_gene_variant | MODIFIER |
| 1 | 39714970 | AX-170360284 | 1 | G | A | 0.393 | -2.20E-03 | 7.45E-02 | 2.65E+00 | 2.74E+00 | 9.81E-01 | no | Rfx8 | intron_variant | MODIFIER |
| 15 | 94673028 | AX-170012305 | 2 | T | A | 0.127 | 4.97E-03 | 1.57E-01 | 2.63E+00 | 2.74E+00 | 9.81E-01 | no | Tmem117 | intron_variant | MODIFIER |
| 5 | 77057707 | AX-97809346 | 1 | T | C | 0.42 | -1.40E-03 | 9.63E-02 | 2.66E+00 | 2.74E+00 | 9.81E-01 | no | Thegl | intron_variant | MODIFIER |
| 7 | 102428725 | AX-170150975 | 1 | T | C | 0.455 | -1.51E-03 | 9.46E-02 | 2.68E+00 | 2.74E+00 | 9.81E-01 | no | Stim1 | intron_variant | MODIFIER |
| 16 | 19628441 | AX-97823915 | 1 | G | C | 0.125 | 1.91E-03 | 9.58E-02 | 2.61E+00 | 2.74E+00 | 9.81E-01 | no | Olfr171 | upstream_gene_variant | MODIFIER |
| 1 | 6024654 | AX-170178342 | 0 | T | G | 0.105 | 3.92E-03 | 1.42E-01 | 2.61E+00 | 2.74E+00 | 9.81E-01 | no | | intergenic_variant | MODIFIER |
| 1 | 25345334 | AX-97845190 | 0 | A | G | 0.404 | -1.34E-03 | 8.26E-02 | 2.67E+00 | 2.74E+00 | 9.81E-01 | no | Adgrb3 | intron_variant | MODIFIER |
| 1 | 25368841 | AX-170040652 | 0 | T | C | 0.404 | -1.34E-03 | 8.26E-02 | 2.67E+00 | 2.74E+00 | 9.81E-01 | no | Adgrb3 | intron_variant | MODIFIER |
| 1 | 25440501 | AX-170491652 | 0 | A | T | 0.404 | -1.34E-03 | 8.26E-02 | 2.67E+00 | 2.74E+00 | 9.81E-01 | no | Adgrb3 | intron_variant | MODIFIER |
| 1 | 25442443 | AX-170136245 | 0 | A | G | 0.404 | -1.34E-03 | 8.26E-02 | 2.67E+00 | 2.74E+00 | 9.81E-01 | no | Adgrb3 | intron_variant | MODIFIER |
| 1 | 25444091 | AX-170515618 | 0 | T | C | 0.404 | -1.34E-03 | 8.26E-02 | 2.67E+00 | 2.74E+00 | 9.81E-01 | no | Adgrb3 | intron_variant | MODIFIER |

| | | | | | | | | | | | | | | | |
|---|---|---|---|---|---|---|---|---|---|---|---|---|---|---|---|
| 1 | 25444461 | AX-170231233 | 0 | A | C | 0.404 | -1.34E-03 | 8.26E-02 | 2.67E+00 | 2.74E+00 | 9.81E-01 | no | Adgrb3 | intron_variant | MODIFIER |
| 1 | 25473333 | AX-170277970 | 0 | T | G | 0.404 | -1.34E-03 | 8.26E-02 | 2.67E+00 | 2.74E+00 | 9.81E-01 | no | Adgrb3 | intron_variant | MODIFIER |
| 1 | 25474349 | AX-170386958 | 0 | A | G | 0.404 | -1.34E-03 | 8.26E-02 | 2.67E+00 | 2.74E+00 | 9.81E-01 | no | Adgrb3 | intron_variant | MODIFIER |
| 1 | 25489787 | AX-97783945 | 0 | G | A | 0.404 | -1.34E-03 | 8.26E-02 | 2.67E+00 | 2.74E+00 | 9.81E-01 | no | Adgrb3 | intron_variant | MODIFIER |
| 1 | 25490270 | AX-170523770 | 0 | G | T | 0.404 | -1.34E-03 | 8.26E-02 | 2.67E+00 | 2.74E+00 | 9.81E-01 | no | Adgrb3 | intron_variant | MODIFIER |
| 1 | 25507121 | AX-170466773 | 0 | T | G | 0.404 | -1.34E-03 | 8.26E-02 | 2.67E+00 | 2.74E+00 | 9.81E-01 | no | Adgrb3 | intron_variant | MODIFIER |
| 1 | 73982445 | AX-97819512 | 0 | A | G | 0.044 | -3.55E-03 | 1.33E-01 | 2.62E+00 | 2.74E+00 | 9.81E-01 | no | Tns1 | intron_variant | MODIFIER |
| 11 | 44862531 | AX-170437376 | 0 | A | G | 0.088 | -5.57E-03 | 1.83E-01 | 2.63E+00 | 2.74E+00 | 9.82E-01 | no | Ebf1 | intron_variant | MODIFIER |
| 11 | 44895211 | AX-170400527 | 0 | C | T | 0.088 | -5.57E-03 | 1.83E-01 | 2.63E+00 | 2.74E+00 | 9.82E-01 | no | Ebf1 | intron_variant | MODIFIER |
| 11 | 44895483 | AX-170271336 | 0 | T | C | 0.088 | -5.57E-03 | 1.83E-01 | 2.63E+00 | 2.74E+00 | 9.82E-01 | no | Ebf1 | intron_variant | MODIFIER |
| 11 | 44897363 | AX-170123607 | 0 | A | C | 0.088 | -5.57E-03 | 1.83E-01 | 2.63E+00 | 2.74E+00 | 9.82E-01 | no | Ebf1 | intron_variant | MODIFIER |
| 11 | 44902691 | AX-97819445 | 0 | T | C | 0.088 | -5.57E-03 | 1.83E-01 | 2.63E+00 | 2.74E+00 | 9.82E-01 | no | Ebf1 | intron_variant | MODIFIER |
| 11 | 44913951 | AX-170141313 | 0 | T | C | 0.088 | -5.57E-03 | 1.83E-01 | 2.63E+00 | 2.74E+00 | 9.82E-01 | no | Ebf1 | intron_variant | MODIFIER |
| 11 | 44914111 | AX-170046933 | 0 | G | A | 0.088 | -5.57E-03 | 1.83E-01 | 2.63E+00 | 2.74E+00 | 9.82E-01 | no | Ebf1 | intron_variant | MODIFIER |
| 11 | 44918276 | AX-170317603 | 0 | G | C | 0.088 | -5.57E-03 | 1.83E-01 | 2.63E+00 | 2.74E+00 | 9.82E-01 | no | Ebf1 | intron_variant | MODIFIER |
| 11 | 44920842 | AX-170181178 | 0 | C | T | 0.088 | -5.57E-03 | 1.83E-01 | 2.63E+00 | 2.74E+00 | 9.82E-01 | no | Ebf1 | intron_variant | MODIFIER |
| 11 | 44921953 | AX-170186505 | 0 | A | G | 0.088 | -5.57E-03 | 1.83E-01 | 2.63E+00 | 2.74E+00 | 9.82E-01 | no | Ebf1 | intron_variant | MODIFIER |
| 11 | 44923951 | AX-97824800 | 0 | C | T | 0.088 | -5.57E-03 | 1.83E-01 | 2.63E+00 | 2.74E+00 | 9.82E-01 | no | Ebf1 | intron_variant | MODIFIER |
| 11 | 44927048 | AX-170507875 | 0 | T | G | 0.088 | -5.57E-03 | 1.83E-01 | 2.63E+00 | 2.74E+00 | 9.82E-01 | no | Ebf1 | intron_variant | MODIFIER |
| 11 | 45023447 | AX-170269060 | 0 | A | G | 0.088 | -5.57E-03 | 1.83E-01 | 2.63E+00 | 2.74E+00 | 9.82E-01 | no | | intergenic_variant | MODIFIER |
| 11 | 45029566 | AX-97804275 | 0 | T | C | 0.088 | -5.57E-03 | 1.83E-01 | 2.63E+00 | 2.74E+00 | 9.82E-01 | no | | intergenic_variant | MODIFIER |
| 11 | 45036382 | AX-170004844 | 0 | C | T | 0.088 | -5.57E-03 | 1.83E-01 | 2.63E+00 | 2.74E+00 | 9.82E-01 | no | | intergenic_variant | MODIFIER |
| 11 | 45068947 | AX-97798719 | 0 | T | C | 0.088 | -5.57E-03 | 1.83E-01 | 2.63E+00 | 2.74E+00 | 9.82E-01 | no | | regulatory_region_variant | MODIFIER |
| 11 | 45082183 | AX-170004848 | 0 | A | G | 0.088 | -5.57E-03 | 1.83E-01 | 2.63E+00 | 2.74E+00 | 9.82E-01 | no | | intergenic_variant | MODIFIER |
| 11 | 45121662 | AX-97851072 | 0 | A | G | 0.088 | -5.57E-03 | 1.83E-01 | 2.63E+00 | 2.74E+00 | 9.82E-01 | no | Gm12159 | upstream_gene_variant | MODIFIER |
| 11 | 45144001 | AX-170265999 | 0 | G | A | 0.088 | -5.57E-03 | 1.83E-01 | 2.63E+00 | 2.74E+00 | 9.82E-01 | no | | intergenic_variant | MODIFIER |
| 11 | 45153582 | AX-170092232 | 0 | A | G | 0.088 | -5.57E-03 | 1.83E-01 | 2.63E+00 | 2.74E+00 | 9.82E-01 | no | | intergenic_variant | MODIFIER |
| 11 | 45154726 | AX-170525684 | 0 | T | C | 0.088 | -5.57E-03 | 1.83E-01 | 2.63E+00 | 2.74E+00 | 9.82E-01 | no | | regulatory_region_variant | MODIFIER |
| 11 | 45164421 | AX-170309458 | 0 | C | T | 0.088 | -5.57E-03 | 1.83E-01 | 2.63E+00 | 2.74E+00 | 9.82E-01 | no | Gm12160 | intron_variant,non_coding_transcript_variant | MODIFIER |
| 11 | 45171493 | AX-170426225 | 0 | C | T | 0.088 | -5.57E-03 | 1.83E-01 | 2.63E+00 | 2.74E+00 | 9.82E-01 | no | Gm12160 | intron_variant,non_coding_transcript_variant | MODIFIER |
| 11 | 45176264 | AX-170495088 | 0 | G | A | 0.088 | -5.57E-03 | 1.83E-01 | 2.63E+00 | 2.74E+00 | 9.82E-01 | no | Gm12160 | intron_variant,non_coding_transcript_variant | MODIFIER |
| 11 | 45176462 | AX-170446000 | 0 | G | A | 0.088 | -5.57E-03 | 1.83E-01 | 2.63E+00 | 2.74E+00 | 9.82E-01 | no | Gm12160 | intron_variant,non_coding_transcript_variant | MODIFIER |
| 11 | 45178856 | AX-170427338 | 0 | T | C | 0.088 | -5.57E-03 | 1.83E-01 | 2.63E+00 | 2.74E+00 | 9.82E-01 | no | Gm12160 | intron_variant,non_coding_transcript_variant | MODIFIER |
| 11 | 45185677 | AX-170296045 | 0 | C | T | 0.088 | -5.57E-03 | 1.83E-01 | 2.63E+00 | 2.74E+00 | 9.82E-01 | no | Gm12160 | intron_variant,non_coding_transcript_variant | MODIFIER |
| 11 | 45196208 | AX-170490582 | 0 | T | A | 0.088 | -5.57E-03 | 1.83E-01 | 2.63E+00 | 2.74E+00 | 9.82E-01 | no | Gm12160 | intron_variant,non_coding_transcript_variant | MODIFIER |
| 11 | 45232233 | AX-170498666 | 0 | G | C | 0.088 | -5.57E-03 | 1.83E-01 | 2.63E+00 | 2.74E+00 | 9.82E-01 | no | | intergenic_variant | MODIFIER |
| 11 | 45233399 | AX-170300109 | 0 | G | A | 0.088 | -5.57E-03 | 1.83E-01 | 2.63E+00 | 2.74E+00 | 9.82E-01 | no | | intergenic_variant | MODIFIER |
| 11 | 45237015 | AX-170238615 | 0 | A | G | 0.088 | -5.57E-03 | 1.83E-01 | 2.63E+00 | 2.74E+00 | 9.82E-01 | no | | intergenic_variant | MODIFIER |
| 11 | 45237454 | AX-97836406 | 0 | T | C | 0.088 | -5.57E-03 | 1.83E-01 | 2.63E+00 | 2.74E+00 | 9.82E-01 | no | | regulatory_region_variant | MODIFIER |
| 11 | 45240857 | AX-170276469 | 0 | G | A | 0.088 | -5.57E-03 | 1.83E-01 | 2.63E+00 | 2.74E+00 | 9.82E-01 | no | | regulatory_region_variant | MODIFIER |
| 11 | 45241485 | AX-170516522 | 0 | C | T | 0.088 | -5.57E-03 | 1.83E-01 | 2.63E+00 | 2.74E+00 | 9.82E-01 | no | | intergenic_variant | MODIFIER |
| 11 | 45242981 | AX-170235807 | 0 | T | A | 0.088 | -5.57E-03 | 1.83E-01 | 2.63E+00 | 2.74E+00 | 9.82E-01 | no | | intergenic_variant | MODIFIER |
| 11 | 45246496 | AX-170184940 | 0 | C | G | 0.088 | -5.57E-03 | 1.83E-01 | 2.63E+00 | 2.74E+00 | 9.82E-01 | no | | intergenic_variant | MODIFIER |
| 11 | 45273073 | AX-170438534 | 0 | C | T | 0.088 | -5.57E-03 | 1.83E-01 | 2.63E+00 | 2.74E+00 | 9.82E-01 | no | Gm37312 | downstream_gene_variant | MODIFIER |
| 11 | 45276906 | AX-170240943 | 0 | C | T | 0.088 | -5.57E-03 | 1.83E-01 | 2.63E+00 | 2.74E+00 | 9.82E-01 | no | Gm12161 | downstream_gene_variant | MODIFIER |
| 11 | 45330865 | AX-170412627 | 0 | A | G | 0.088 | -5.57E-03 | 1.83E-01 | 2.63E+00 | 2.74E+00 | 9.82E-01 | no | | intergenic_variant | MODIFIER |
| 11 | 45525510 | AX-170166696 | 0 | G | A | 0.088 | -5.57E-03 | 1.83E-01 | 2.63E+00 | 2.74E+00 | 9.82E-01 | no | | intergenic_variant | MODIFIER |
| 11 | 45526303 | AX-170177556 | 0 | C | G | 0.088 | -5.57E-03 | 1.83E-01 | 2.63E+00 | 2.74E+00 | 9.82E-01 | no | | intergenic_variant | MODIFIER |
| 11 | 45526439 | AX-97787931 | 0 | A | G | 0.088 | -5.57E-03 | 1.83E-01 | 2.63E+00 | 2.74E+00 | 9.82E-01 | no | | intergenic_variant | MODIFIER |
| 11 | 45597061 | AX-97801879 | 0 | A | G | 0.088 | -5.57E-03 | 1.83E-01 | 2.63E+00 | 2.74E+00 | 9.82E-01 | no | | intergenic_variant | MODIFIER |
| 11 | 45600427 | AX-170197331 | 0 | G | A | 0.088 | -5.57E-03 | 1.83E-01 | 2.63E+00 | 2.74E+00 | 9.82E-01 | no | | intergenic_variant | MODIFIER |
| 11 | 45626347 | AX-97853149 | 0 | A | G | 0.088 | -5.57E-03 | 1.83E-01 | 2.63E+00 | 2.74E+00 | 9.82E-01 | no | | intergenic_variant | MODIFIER |
| 11 | 45642820 | AX-170322825 | 0 | A | C | 0.088 | -5.57E-03 | 1.83E-01 | 2.63E+00 | 2.74E+00 | 9.82E-01 | no | Gm22284 | downstream_gene_variant | MODIFIER |
| 11 | 45647518 | AX-97810168 | 0 | C | T | 0.088 | -5.57E-03 | 1.83E-01 | 2.63E+00 | 2.74E+00 | 9.82E-01 | no | | intergenic_variant | MODIFIER |

| | | | | | | | | | | | | | | | |
|---|---|---|---|---|---|---|---|---|---|---|---|---|---|---|---|
| 11 | 45701298 | AX-97785552 | 0 | C | T | 0.088 | -5.57E-03 | 1.83E-01 | 2.63E+00 | 2.74E+00 | 9.82E-01 | no | | intergenic_variant | MODIFIER |
| 11 | 46033309 | AX-170174981 | 0 | G | A | 0.088 | -5.57E-03 | 1.83E-01 | 2.63E+00 | 2.74E+00 | 9.82E-01 | no | Gm12165 | upstream_gene_variant | MODIFIER |
| 11 | 46231677 | AX-170326191 | 0 | T | C | 0.088 | -5.57E-03 | 1.83E-01 | 2.63E+00 | 2.74E+00 | 9.82E-01 | no | Fndc9 | upstream_gene_variant | MODIFIER |
| 11 | 46283411 | AX-97818288 | 0 | C | T | 0.088 | -5.57E-03 | 1.83E-01 | 2.63E+00 | 2.74E+00 | 9.82E-01 | no | Cyfip2 | intron_variant | MODIFIER |
| 11 | 46290299 | AX-170111858 | 0 | C | T | 0.088 | -5.57E-03 | 1.83E-01 | 2.63E+00 | 2.74E+00 | 9.82E-01 | no | Cyfip2 | intron_variant | MODIFIER |
| 11 | 46317084 | AX-170477896 | 0 | G | A | 0.088 | -5.57E-03 | 1.83E-01 | 2.63E+00 | 2.74E+00 | 9.82E-01 | no | Cyfip2 | upstream_gene_variant | MODIFIER |
| 11 | 46325930 | AX-170244691 | 0 | A | T | 0.088 | -5.57E-03 | 1.83E-01 | 2.63E+00 | 2.74E+00 | 9.82E-01 | no | Itk | 3_prime_UTR_variant | MODIFIER |
| 11 | 46329911 | AX-170226654 | 0 | C | G | 0.088 | -5.57E-03 | 1.83E-01 | 2.63E+00 | 2.74E+00 | 9.82E-01 | no | Itk | intron_variant | MODIFIER |
| 11 | 46331034 | AX-170418707 | 0 | C | T | 0.088 | -5.57E-03 | 1.83E-01 | 2.63E+00 | 2.74E+00 | 9.82E-01 | no | Itk | intron_variant | MODIFIER |
| 11 | 46332133 | AX-97808026 | 0 | C | A | 0.088 | -5.57E-03 | 1.83E-01 | 2.63E+00 | 2.74E+00 | 9.82E-01 | no | Itk | intron_variant | MODIFIER |
| 11 | 46333563 | AX-170433781 | 0 | A | G | 0.088 | -5.57E-03 | 1.83E-01 | 2.63E+00 | 2.74E+00 | 9.82E-01 | no | Itk | intron_variant | MODIFIER |
| 11 | 46333867 | AX-170304107 | 0 | A | G | 0.088 | -5.57E-03 | 1.83E-01 | 2.63E+00 | 2.74E+00 | 9.82E-01 | no | Itk | intron_variant | MODIFIER |
| 11 | 46364769 | AX-170277799 | 0 | T | C | 0.088 | -5.57E-03 | 1.83E-01 | 2.63E+00 | 2.74E+00 | 9.82E-01 | no | Itk | intron_variant | MODIFIER |
| 11 | 46368159 | AX-170509856 | 0 | T | C | 0.088 | -5.57E-03 | 1.83E-01 | 2.63E+00 | 2.74E+00 | 9.82E-01 | no | Itk | intron_variant | MODIFIER |
| 11 | 46379659 | AX-170230448 | 0 | C | A | 0.088 | -5.57E-03 | 1.83E-01 | 2.63E+00 | 2.74E+00 | 9.82E-01 | no | Itk | intron_variant | MODIFIER |
| 11 | 46385126 | AX-170533178 | 0 | G | A | 0.088 | -5.57E-03 | 1.83E-01 | 2.63E+00 | 2.74E+00 | 9.82E-01 | no | Itk | intron_variant | MODIFIER |
| 11 | 46396694 | AX-170046964 | 0 | A | G | 0.088 | -5.57E-03 | 1.83E-01 | 2.63E+00 | 2.74E+00 | 9.82E-01 | no | Fam71b | non_coding_transcript_exon_variant | MODIFIER |
| 11 | 46397587 | AX-170248040 | 0 | A | G | 0.088 | -5.57E-03 | 1.83E-01 | 2.63E+00 | 2.74E+00 | 9.82E-01 | no | Fam71b | intron_variant,non_coding_transcript_variant | MODIFIER |
| 11 | 46399193 | AX-170345990 | 0 | T | C | 0.088 | -5.57E-03 | 1.83E-01 | 2.63E+00 | 2.74E+00 | 9.82E-01 | no | Fam71b | intron_variant,non_coding_transcript_variant | MODIFIER |
| 11 | 46421368 | AX-170147430 | 0 | A | G | 0.088 | -5.57E-03 | 1.83E-01 | 2.63E+00 | 2.74E+00 | 9.82E-01 | no | | regulatory_region_variant | MODIFIER |
| 11 | 46421615 | AX-170372386 | 0 | G | C | 0.088 | -5.57E-03 | 1.83E-01 | 2.63E+00 | 2.74E+00 | 9.82E-01 | no | | regulatory_region_variant | MODIFIER |
| 11 | 46421848 | AX-170183077 | 0 | A | G | 0.088 | -5.57E-03 | 1.83E-01 | 2.63E+00 | 2.74E+00 | 9.82E-01 | no | | regulatory_region_variant | MODIFIER |
| 11 | 46424204 | AX-170162680 | 0 | A | G | 0.088 | -5.57E-03 | 1.83E-01 | 2.63E+00 | 2.74E+00 | 9.82E-01 | no | | intergenic_variant | MODIFIER |
| 11 | 46424224 | AX-170142048 | 0 | C | T | 0.088 | -5.57E-03 | 1.83E-01 | 2.63E+00 | 2.74E+00 | 9.82E-01 | no | | intergenic_variant | MODIFIER |
| 11 | 46478901 | AX-170316817 | 0 | A | G | 0.088 | -5.57E-03 | 1.83E-01 | 2.63E+00 | 2.74E+00 | 9.82E-01 | no | Havcr2 | intron_variant | MODIFIER |
| 11 | 46479834 | AX-170183917 | 0 | C | A | 0.088 | -5.57E-03 | 1.83E-01 | 2.63E+00 | 2.74E+00 | 9.82E-01 | no | Havcr2 | 3_prime_UTR_variant | MODIFIER |
| 11 | 46489067 | AX-170296458 | 0 | T | C | 0.088 | -5.57E-03 | 1.83E-01 | 2.63E+00 | 2.74E+00 | 9.82E-01 | no | | intergenic_variant | MODIFIER |
| 11 | 46509767 | AX-170177109 | 0 | A | C | 0.088 | -5.57E-03 | 1.83E-01 | 2.63E+00 | 2.74E+00 | 9.82E-01 | no | | intergenic_variant | MODIFIER |
| 11 | 46511864 | AX-170427543 | 0 | G | A | 0.088 | -5.57E-03 | 1.83E-01 | 2.63E+00 | 2.74E+00 | 9.82E-01 | no | | intergenic_variant | MODIFIER |
| 11 | 46520801 | AX-97818313 | 0 | T | C | 0.088 | -5.57E-03 | 1.83E-01 | 2.63E+00 | 2.74E+00 | 9.82E-01 | no | Gm12169 | upstream_gene_variant | MODIFIER |
| 11 | 46520891 | AX-170255839 | 0 | T | G | 0.088 | -5.57E-03 | 1.83E-01 | 2.63E+00 | 2.74E+00 | 9.82E-01 | no | Gm12169 | upstream_gene_variant | MODIFIER |
| 11 | 46522106 | AX-170004868 | 0 | T | C | 0.088 | -5.57E-03 | 1.83E-01 | 2.63E+00 | 2.74E+00 | 9.82E-01 | no | Gm12169 | upstream_gene_variant | MODIFIER |
| 11 | 46554227 | AX-170472647 | 0 | C | G | 0.088 | -5.57E-03 | 1.83E-01 | 2.63E+00 | 2.74E+00 | 9.82E-01 | no | Gm12171 | intron_variant,non_coding_transcript_variant | MODIFIER |
| 11 | 46565562 | AX-170004863 | 0 | T | C | 0.088 | -5.57E-03 | 1.83E-01 | 2.63E+00 | 2.74E+00 | 9.82E-01 | no | Rpl9-ps2 | downstream_gene_variant | MODIFIER |
| 11 | 46575430 | AX-170492616 | 0 | T | C | 0.088 | -5.57E-03 | 1.83E-01 | 2.63E+00 | 2.74E+00 | 9.82E-01 | no | BC053393 | intron_variant | MODIFIER |
| 11 | 46580912 | AX-170220851 | 0 | A | G | 0.088 | -5.57E-03 | 1.83E-01 | 2.63E+00 | 2.74E+00 | 9.82E-01 | no | BC053393 | intron_variant | MODIFIER |
| 11 | 46581114 | AX-170444455 | 0 | G | T | 0.088 | -5.57E-03 | 1.83E-01 | 2.63E+00 | 2.74E+00 | 9.82E-01 | no | BC053393 | intron_variant | MODIFIER |
| 11 | 46583493 | AX-170151453 | 0 | T | C | 0.088 | -5.57E-03 | 1.83E-01 | 2.63E+00 | 2.74E+00 | 9.82E-01 | no | BC053393 | intron_variant | MODIFIER |
| 11 | 46609784 | AX-170046966 | 0 | G | A | 0.088 | -5.57E-03 | 1.83E-01 | 2.63E+00 | 2.74E+00 | 9.82E-01 | no | Dppa1 | intron_variant | MODIFIER |
| 11 | 46612556 | AX-170046967 | 0 | C | T | 0.088 | -5.57E-03 | 1.83E-01 | 2.63E+00 | 2.74E+00 | 9.82E-01 | no | Dppa1 | intron_variant | MODIFIER |
| 11 | 46630271 | AX-97854675 | 0 | G | A | 0.088 | -5.57E-03 | 1.83E-01 | 2.63E+00 | 2.74E+00 | 9.82E-01 | no | Dppa1 | upstream_gene_variant | MODIFIER |
| 11 | 46648969 | AX-170508099 | 0 | C | G | 0.088 | -5.57E-03 | 1.83E-01 | 2.63E+00 | 2.74E+00 | 9.82E-01 | no | Gm12173 | upstream_gene_variant | MODIFIER |
| 11 | 46679927 | AX-170359890 | 0 | A | C | 0.088 | -5.57E-03 | 1.83E-01 | 2.63E+00 | 2.74E+00 | 9.82E-01 | no | Timd2 | intron_variant | MODIFIER |
| 11 | 46680979 | AX-170092237 | 0 | C | T | 0.088 | -5.57E-03 | 1.83E-01 | 2.63E+00 | 2.74E+00 | 9.82E-01 | no | Timd2 | intron_variant | MODIFIER |
| 11 | 46755717 | AX-170504640 | 0 | A | G | 0.088 | -5.57E-03 | 1.83E-01 | 2.63E+00 | 2.74E+00 | 9.82E-01 | no | Havcr1 | intron_variant | MODIFIER |
| 11 | 46773145 | AX-97818365 | 0 | A | G | 0.088 | -5.57E-03 | 1.83E-01 | 2.63E+00 | 2.74E+00 | 9.82E-01 | no | Havcr1 | intron_variant | MODIFIER |
| 11 | 46809188 | AX-170351631 | 0 | T | A | 0.088 | -5.57E-03 | 1.83E-01 | 2.63E+00 | 2.74E+00 | 9.82E-01 | no | Timd4 | upstream_gene_variant | MODIFIER |
| 11 | 46836963 | AX-170046969 | 0 | A | G | 0.088 | -5.57E-03 | 1.83E-01 | 2.63E+00 | 2.74E+00 | 9.82E-01 | no | Timd4 | intron_variant | MODIFIER |
| 11 | 46853362 | AX-170355776 | 0 | A | C | 0.088 | -5.57E-03 | 1.83E-01 | 2.63E+00 | 2.74E+00 | 9.82E-01 | no | | intergenic_variant | MODIFIER |
| 11 | 46893807 | AX-170464493 | 0 | C | T | 0.088 | -5.57E-03 | 1.83E-01 | 2.63E+00 | 2.74E+00 | 9.82E-01 | no | Sgcd | downstream_gene_variant | MODIFIER |
| 11 | 46977298 | AX-170393363 | 0 | T | A | 0.088 | -5.57E-03 | 1.83E-01 | 2.63E+00 | 2.74E+00 | 9.82E-01 | no | Sgcd | downstream_gene_variant | MODIFIER |
| 11 | 46977475 | AX-170314016 | 0 | C | A | 0.088 | -5.57E-03 | 1.83E-01 | 2.63E+00 | 2.74E+00 | 9.82E-01 | no | Sgcd | downstream_gene_variant | MODIFIER |
| 11 | 46982744 | AX-170155617 | 0 | G | A | 0.088 | -5.57E-03 | 1.83E-01 | 2.63E+00 | 2.74E+00 | 9.82E-01 | no | Sgcd | intron_variant | MODIFIER |
| 11 | 46985510 | AX-170335736 | 0 | G | A | 0.088 | -5.57E-03 | 1.83E-01 | 2.63E+00 | 2.74E+00 | 9.82E-01 | no | Sgcd | intron_variant | MODIFIER |

| | | | | | | | | | | | | | | | |
|---|---|---|---|---|---|---|---|---|---|---|---|---|---|---|---|
| 11 | 47009247 | AX-170303135 | 0 | A | G | 0.088 | -5.57E-03 | 1.83E-01 | 2.63E+00 | 2.74E+00 | 9.82E-01 | no | Sgcd | intron_variant | MODIFIER |
| 11 | 47057236 | AX-170112879 | 0 | C | T | 0.088 | -5.57E-03 | 1.83E-01 | 2.63E+00 | 2.74E+00 | 9.82E-01 | no | Sgcd | intron_variant | MODIFIER |
| 11 | 47057990 | AX-170249688 | 0 | G | A | 0.088 | -5.57E-03 | 1.83E-01 | 2.63E+00 | 2.74E+00 | 9.82E-01 | no | Sgcd | intron_variant | MODIFIER |
| 11 | 47080348 | AX-170183252 | 0 | T | C | 0.088 | -5.57E-03 | 1.83E-01 | 2.63E+00 | 2.74E+00 | 9.82E-01 | no | Sgcd | intron_variant | MODIFIER |
| 11 | 47084123 | AX-97818363 | 0 | T | G | 0.088 | -5.57E-03 | 1.83E-01 | 2.63E+00 | 2.74E+00 | 9.82E-01 | no | Sgcd | intron_variant | MODIFIER |
| 11 | 47100770 | AX-97807668 | 0 | C | T | 0.088 | -5.57E-03 | 1.83E-01 | 2.63E+00 | 2.74E+00 | 9.82E-01 | no | Sgcd | intron_variant | MODIFIER |
| 11 | 47106774 | AX-170189087 | 0 | T | A | 0.088 | -5.57E-03 | 1.83E-01 | 2.63E+00 | 2.74E+00 | 9.82E-01 | no | Sgcd | intron_variant | MODIFIER |
| 11 | 47133730 | AX-170004872 | 0 | G | A | 0.088 | -5.57E-03 | 1.83E-01 | 2.63E+00 | 2.74E+00 | 9.82E-01 | no | Sgcd | intron_variant | MODIFIER |
| 11 | 47164882 | AX-170004876 | 0 | A | T | 0.088 | -5.57E-03 | 1.83E-01 | 2.63E+00 | 2.74E+00 | 9.82E-01 | no | Sgcd | intron_variant | MODIFIER |
| 11 | 47169596 | AX-170004873 | 0 | T | G | 0.088 | -5.57E-03 | 1.83E-01 | 2.63E+00 | 2.74E+00 | 9.82E-01 | no | Sgcd | intron_variant | MODIFIER |
| 11 | 47188959 | AX-170473610 | 0 | C | T | 0.088 | -5.57E-03 | 1.83E-01 | 2.63E+00 | 2.74E+00 | 9.82E-01 | no | Sgcd | intron_variant | MODIFIER |
| 11 | 47189043 | AX-170151281 | 0 | T | C | 0.088 | -5.57E-03 | 1.83E-01 | 2.63E+00 | 2.74E+00 | 9.82E-01 | no | Sgcd | intron_variant | MODIFIER |
| 11 | 47189367 | AX-170192861 | 0 | C | T | 0.088 | -5.57E-03 | 1.83E-01 | 2.63E+00 | 2.74E+00 | 9.82E-01 | no | Sgcd | intron_variant | MODIFIER |
| 11 | 47190672 | AX-170143378 | 0 | G | A | 0.088 | -5.57E-03 | 1.83E-01 | 2.63E+00 | 2.74E+00 | 9.82E-01 | no | Sgcd | intron_variant | MODIFIER |
| 11 | 47193565 | AX-170004871 | 0 | T | C | 0.088 | -5.57E-03 | 1.83E-01 | 2.63E+00 | 2.74E+00 | 9.82E-01 | no | Sgcd | intron_variant | MODIFIER |
| 11 | 47325055 | AX-170308298 | 0 | T | C | 0.088 | -5.57E-03 | 1.83E-01 | 2.63E+00 | 2.74E+00 | 9.82E-01 | no | Sgcd | intron_variant | MODIFIER |
| 11 | 47540941 | AX-170142872 | 0 | A | C | 0.088 | -5.57E-03 | 1.83E-01 | 2.63E+00 | 2.74E+00 | 9.82E-01 | no | Sgcd | intron_variant | MODIFIER |
| 11 | 47553207 | AX-170405035 | 0 | G | A | 0.088 | -5.57E-03 | 1.83E-01 | 2.63E+00 | 2.74E+00 | 9.82E-01 | no | Sgcd | intron_variant | MODIFIER |
| 11 | 47573124 | AX-97793523 | 0 | C | T | 0.088 | -5.57E-03 | 1.83E-01 | 2.63E+00 | 2.74E+00 | 9.82E-01 | no | Sgcd | intron_variant | MODIFIER |
| 11 | 47641862 | AX-170087015 | 0 | A | G | 0.088 | -5.57E-03 | 1.83E-01 | 2.63E+00 | 2.74E+00 | 9.82E-01 | no | Sgcd | intron_variant,non_coding_transcript_variant | MODIFIER |
| 11 | 47701250 | AX-97821936 | 0 | C | A | 0.088 | -5.57E-03 | 1.83E-01 | 2.63E+00 | 2.74E+00 | 9.82E-01 | no | Sgcd | intron_variant,non_coding_transcript_variant | MODIFIER |
| 11 | 47924940 | AX-170136598 | 0 | T | C | 0.088 | -5.57E-03 | 1.83E-01 | 2.63E+00 | 2.74E+00 | 9.82E-01 | no | Sgcd | intron_variant,non_coding_transcript_variant | MODIFIER |
| 11 | 47932673 | AX-170420587 | 0 | T | C | 0.088 | -5.57E-03 | 1.83E-01 | 2.63E+00 | 2.74E+00 | 9.82E-01 | no | Sgcd | intron_variant,non_coding_transcript_variant | MODIFIER |
| 11 | 47982190 | AX-97794153 | 0 | T | C | 0.088 | -5.57E-03 | 1.83E-01 | 2.63E+00 | 2.74E+00 | 9.82E-01 | no | Sgcd | intron_variant,non_coding_transcript_variant | MODIFIER |
| 11 | 47982222 | AX-170457061 | 0 | A | G | 0.088 | -5.57E-03 | 1.83E-01 | 2.63E+00 | 2.74E+00 | 9.82E-01 | no | Sgcd | intron_variant,non_coding_transcript_variant | MODIFIER |
| 11 | 47983430 | AX-170409629 | 0 | T | C | 0.088 | -5.57E-03 | 1.83E-01 | 2.63E+00 | 2.74E+00 | 9.82E-01 | no | Sgcd | intron_variant,non_coding_transcript_variant | MODIFIER |
| 11 | 47984221 | AX-170437528 | 0 | T | C | 0.088 | -5.57E-03 | 1.83E-01 | 2.63E+00 | 2.74E+00 | 9.82E-01 | no | Sgcd | intron_variant,non_coding_transcript_variant | MODIFIER |
| 11 | 47996536 | AX-170404202 | 0 | G | A | 0.088 | -5.57E-03 | 1.83E-01 | 2.63E+00 | 2.74E+00 | 9.82E-01 | no | | intergenic_variant | MODIFIER |
| 11 | 48002926 | AX-97818596 | 0 | G | A | 0.088 | -5.57E-03 | 1.83E-01 | 2.63E+00 | 2.74E+00 | 9.82E-01 | no | | intergenic_variant | MODIFIER |
| 11 | 48039045 | AX-170374058 | 0 | G | A | 0.088 | -5.57E-03 | 1.83E-01 | 2.63E+00 | 2.74E+00 | 9.82E-01 | no | | intergenic_variant | MODIFIER |
| 11 | 48053990 | AX-170495770 | 0 | C | G | 0.088 | -5.57E-03 | 1.83E-01 | 2.63E+00 | 2.74E+00 | 9.82E-01 | no | | intergenic_variant | MODIFIER |
| 11 | 48054759 | AX-170235296 | 0 | A | G | 0.088 | -5.57E-03 | 1.83E-01 | 2.63E+00 | 2.74E+00 | 9.82E-01 | no | | intergenic_variant | MODIFIER |
| 11 | 48055048 | AX-170397327 | 0 | T | C | 0.088 | -5.57E-03 | 1.83E-01 | 2.63E+00 | 2.74E+00 | 9.82E-01 | no | | intergenic_variant | MODIFIER |
| 11 | 48090874 | AX-170117533 | 0 | C | T | 0.088 | -5.57E-03 | 1.83E-01 | 2.63E+00 | 2.74E+00 | 9.82E-01 | no | | intergenic_variant | MODIFIER |
| 11 | 48091415 | AX-170448508 | 0 | A | G | 0.088 | -5.57E-03 | 1.83E-01 | 2.63E+00 | 2.74E+00 | 9.82E-01 | no | | intergenic_variant | MODIFIER |
| 11 | 48097937 | AX-170273774 | 0 | C | T | 0.088 | -5.57E-03 | 1.83E-01 | 2.63E+00 | 2.74E+00 | 9.82E-01 | no | | intergenic_variant | MODIFIER |
| 11 | 48110554 | AX-170269966 | 0 | A | G | 0.088 | -5.57E-03 | 1.83E-01 | 2.63E+00 | 2.74E+00 | 9.82E-01 | no | | intergenic_variant | MODIFIER |
| 11 | 48116911 | AX-170497576 | 0 | C | T | 0.088 | -5.57E-03 | 1.83E-01 | 2.63E+00 | 2.74E+00 | 9.82E-01 | no | | intergenic_variant | MODIFIER |
| 11 | 48117382 | AX-170410130 | 0 | C | T | 0.088 | -5.57E-03 | 1.83E-01 | 2.63E+00 | 2.74E+00 | 9.82E-01 | no | | intergenic_variant | MODIFIER |
| 11 | 48130971 | AX-170119751 | 0 | G | A | 0.088 | -5.57E-03 | 1.83E-01 | 2.63E+00 | 2.74E+00 | 9.82E-01 | no | | intergenic_variant | MODIFIER |
| 11 | 48131527 | AX-170432073 | 0 | C | T | 0.088 | -5.57E-03 | 1.83E-01 | 2.63E+00 | 2.74E+00 | 9.82E-01 | no | | intergenic_variant | MODIFIER |
| 11 | 48137666 | AX-97815460 | 0 | C | T | 0.088 | -5.57E-03 | 1.83E-01 | 2.63E+00 | 2.74E+00 | 9.82E-01 | no | | intergenic_variant | MODIFIER |
| 11 | 48142686 | AX-170251189 | 0 | T | C | 0.088 | -5.57E-03 | 1.83E-01 | 2.63E+00 | 2.74E+00 | 9.82E-01 | no | | intergenic_variant | MODIFIER |
| 11 | 48166918 | AX-170319062 | 0 | C | T | 0.088 | -5.57E-03 | 1.83E-01 | 2.63E+00 | 2.74E+00 | 9.82E-01 | no | | intergenic_variant | MODIFIER |
| 11 | 48170340 | AX-170436628 | 0 | A | C | 0.088 | -5.57E-03 | 1.83E-01 | 2.63E+00 | 2.74E+00 | 9.82E-01 | no | | intergenic_variant | MODIFIER |
| 11 | 48171638 | AX-170004888 | 0 | G | T | 0.088 | -5.57E-03 | 1.83E-01 | 2.63E+00 | 2.74E+00 | 9.82E-01 | no | | intergenic_variant | MODIFIER |
| 11 | 48171784 | AX-97818594 | 0 | A | G | 0.088 | -5.57E-03 | 1.83E-01 | 2.63E+00 | 2.74E+00 | 9.82E-01 | no | | intergenic_variant | MODIFIER |
| 11 | 48178537 | AX-170004886 | 0 | T | C | 0.088 | -5.57E-03 | 1.83E-01 | 2.63E+00 | 2.74E+00 | 9.82E-01 | no | | regulatory_region_variant | MODIFIER |
| 11 | 48215066 | AX-170004885 | 0 | A | T | 0.088 | -5.57E-03 | 1.83E-01 | 2.63E+00 | 2.74E+00 | 9.82E-01 | no | | intergenic_variant | MODIFIER |
| 11 | 48218739 | AX-97848934 | 0 | T | C | 0.088 | -5.57E-03 | 1.83E-01 | 2.63E+00 | 2.74E+00 | 9.82E-01 | no | | intergenic_variant | MODIFIER |
| 11 | 48245994 | AX-170312763 | 0 | A | C | 0.088 | -5.57E-03 | 1.83E-01 | 2.63E+00 | 2.74E+00 | 9.82E-01 | no | | intergenic_variant | MODIFIER |
| 11 | 48246716 | AX-170004889 | 0 | A | G | 0.088 | -5.57E-03 | 1.83E-01 | 2.63E+00 | 2.74E+00 | 9.82E-01 | no | | intergenic_variant | MODIFIER |
| 11 | 48247647 | AX-170326216 | 0 | T | C | 0.088 | -5.57E-03 | 1.83E-01 | 2.63E+00 | 2.74E+00 | 9.82E-01 | no | | intergenic_variant | MODIFIER |
| 11 | 48248787 | AX-97851818 | 0 | T | C | 0.088 | -5.57E-03 | 1.83E-01 | 2.63E+00 | 2.74E+00 | 9.82E-01 | no | | intergenic_variant | MODIFIER |

| | | | | | | | | | | | | | | | |
|---|---|---|---|---|---|---|---|---|---|---|---|---|---|---|---|
| 11 | 48252297 | AX-170445261 | 0 | C | T | 0.088 | -5.57E-03 | 1.83E-01 | 2.63E+00 | 2.74E+00 | 9.82E-01 | no | | intergenic_variant | MODIFIER |
| 11 | 48290847 | AX-97818591 | 0 | G | A | 0.088 | -5.57E-03 | 1.83E-01 | 2.63E+00 | 2.74E+00 | 9.82E-01 | no | Gm12178 | upstream_gene_variant | MODIFIER |
| 11 | 48308629 | AX-170389413 | 0 | G | A | 0.088 | -5.57E-03 | 1.83E-01 | 2.63E+00 | 2.74E+00 | 9.82E-01 | no | | intergenic_variant | MODIFIER |
| 11 | 48358040 | AX-170467276 | 0 | A | C | 0.088 | -5.57E-03 | 1.83E-01 | 2.63E+00 | 2.74E+00 | 9.82E-01 | no | | intergenic_variant | MODIFIER |
| 11 | 48362313 | AX-170289470 | 0 | A | C | 0.088 | -5.57E-03 | 1.83E-01 | 2.63E+00 | 2.74E+00 | 9.82E-01 | no | | intergenic_variant | MODIFIER |
| 11 | 48696355 | AX-170180040 | 0 | A | G | 0.088 | -5.57E-03 | 1.83E-01 | 2.63E+00 | 2.74E+00 | 9.82E-01 | no | | regulatory_region_variant | MODIFIER |
| 11 | 48803098 | AX-97855750 | 0 | G | T | 0.088 | -5.57E-03 | 1.83E-01 | 2.63E+00 | 2.74E+00 | 9.82E-01 | no | Rack1 | intron_variant | MODIFIER |
| 11 | 48803629 | AX-170113289 | 0 | T | G | 0.088 | -5.57E-03 | 1.83E-01 | 2.63E+00 | 2.74E+00 | 9.82E-01 | no | Rack1 | intron_variant | MODIFIER |
| 11 | 48812706 | AX-97810704 | 0 | A | T | 0.088 | -5.57E-03 | 1.83E-01 | 2.63E+00 | 2.74E+00 | 9.82E-01 | no | Trim41 | intron_variant | MODIFIER |
| 11 | 49095589 | AX-170174055 | 0 | G | T | 0.088 | -5.57E-03 | 1.83E-01 | 2.63E+00 | 2.74E+00 | 9.82E-01 | no | Ifi47 | missense_variant | MODERATE |
| 11 | 49096052 | AX-170363153 | 0 | T | A | 0.088 | -5.57E-03 | 1.83E-01 | 2.63E+00 | 2.74E+00 | 9.82E-01 | no | Ifi47 | synonymous_variant | LOW |
| 11 | 49096098 | AX-170468279 | 0 | C | T | 0.088 | -5.57E-03 | 1.83E-01 | 2.63E+00 | 2.74E+00 | 9.82E-01 | no | Ifi47 | missense_variant | MODERATE |
| 11 | 49096505 | AX-170513860 | 0 | A | G | 0.088 | -5.57E-03 | 1.83E-01 | 2.63E+00 | 2.74E+00 | 9.82E-01 | no | Ifi47 | synonymous_variant | LOW |
| 11 | 49110660 | AX-170218445 | 0 | G | A | 0.088 | -5.57E-03 | 1.83E-01 | 2.63E+00 | 2.74E+00 | 9.82E-01 | no | Olfr56 | intron_variant | MODIFIER |
| 11 | 49110922 | AX-170124165 | 0 | G | A | 0.088 | -5.57E-03 | 1.83E-01 | 2.63E+00 | 2.74E+00 | 9.82E-01 | no | Olfr56 | intron_variant | MODIFIER |
| 11 | 49115140 | AX-170046998 | 0 | T | C | 0.088 | -5.57E-03 | 1.83E-01 | 2.63E+00 | 2.74E+00 | 9.82E-01 | no | Olfr56 | intron_variant | MODIFIER |
| 11 | 49115207 | AX-170046996 | 0 | T | C | 0.088 | -5.57E-03 | 1.83E-01 | 2.63E+00 | 2.74E+00 | 9.82E-01 | no | Olfr56 | intron_variant | MODIFIER |
| 11 | 49165632 | AX-97789261 | 0 | C | T | 0.088 | -5.57E-03 | 1.83E-01 | 2.63E+00 | 2.74E+00 | 9.82E-01 | no | Btnl9 | 3_prime_UTR_variant | MODIFIER |
| 11 | 49183924 | AX-170426508 | 0 | T | A | 0.088 | -5.57E-03 | 1.83E-01 | 2.63E+00 | 2.74E+00 | 9.82E-01 | no | Btnl9 | intron_variant | MODIFIER |
| 11 | 49184362 | AX-170433227 | 0 | C | T | 0.088 | -5.57E-03 | 1.83E-01 | 2.63E+00 | 2.74E+00 | 9.82E-01 | no | Btnl9 | intron_variant | MODIFIER |
| 11 | 49184412 | AX-170247185 | 0 | T | C | 0.088 | -5.57E-03 | 1.83E-01 | 2.63E+00 | 2.74E+00 | 9.82E-01 | no | Btnl9 | intron_variant | MODIFIER |
| 11 | 49187376 | AX-97812108 | 0 | C | G | 0.088 | -5.57E-03 | 1.83E-01 | 2.63E+00 | 2.74E+00 | 9.82E-01 | no | Btnl9 | upstream_gene_variant | MODIFIER |
| 11 | 49206919 | AX-170198082 | 0 | C | G | 0.088 | -5.57E-03 | 1.83E-01 | 2.63E+00 | 2.74E+00 | 9.82E-01 | no | Zfp62 | intron_variant | MODIFIER |
| 11 | 49213917 | AX-170102700 | 0 | G | A | 0.088 | -5.57E-03 | 1.83E-01 | 2.63E+00 | 2.74E+00 | 9.82E-01 | no | Zfp62 | upstream_gene_variant | MODIFIER |
| 11 | 49218562 | AX-170324864 | 0 | A | T | 0.088 | -5.57E-03 | 1.83E-01 | 2.63E+00 | 2.74E+00 | 9.82E-01 | no | Zfp62 | 3_prime_UTR_variant | MODIFIER |
| 11 | 49237711 | AX-170397978 | 0 | A | G | 0.088 | -5.57E-03 | 1.83E-01 | 2.63E+00 | 2.74E+00 | 9.82E-01 | no | | regulatory_region_variant | MODIFIER |
| 11 | 49254423 | AX-97805665 | 0 | T | C | 0.088 | -5.57E-03 | 1.83E-01 | 2.63E+00 | 2.74E+00 | 9.82E-01 | no | Mgat1 | intron_variant | MODIFIER |
| 11 | 49255752 | AX-170099510 | 0 | G | A | 0.088 | -5.57E-03 | 1.83E-01 | 2.63E+00 | 2.74E+00 | 9.82E-01 | no | Mgat1 | intron_variant | MODIFIER |
| 11 | 49267397 | AX-170266874 | 0 | C | T | 0.088 | -5.57E-03 | 1.83E-01 | 2.63E+00 | 2.74E+00 | 9.82E-01 | no | Mgat1 | downstream_gene_variant | MODIFIER |
| 11 | 49267517 | AX-170168925 | 0 | G | T | 0.088 | -5.57E-03 | 1.83E-01 | 2.63E+00 | 2.74E+00 | 9.82E-01 | no | Mgat1 | downstream_gene_variant | MODIFIER |
| 11 | 49267646 | AX-97804644 | 0 | T | C | 0.088 | -5.57E-03 | 1.83E-01 | 2.63E+00 | 2.74E+00 | 9.82E-01 | no | Mgat1 | downstream_gene_variant | MODIFIER |
| 11 | 49282085 | AX-170193984 | 0 | A | C | 0.088 | -5.57E-03 | 1.83E-01 | 2.63E+00 | 2.74E+00 | 9.82E-01 | no | Olfr1393 | downstream_gene_variant | MODIFIER |
| 11 | 49335912 | AX-97843472 | 0 | C | T | 0.088 | -5.57E-03 | 1.83E-01 | 2.63E+00 | 2.74E+00 | 9.82E-01 | no | Olfr1390 | upstream_gene_variant | MODIFIER |
| 11 | 49336430 | AX-170004892 | 0 | G | C | 0.088 | -5.57E-03 | 1.83E-01 | 2.63E+00 | 2.74E+00 | 9.82E-01 | no | Olfr1390 | upstream_gene_variant | MODIFIER |
| 11 | 49336643 | AX-170004893 | 0 | C | A | 0.088 | -5.57E-03 | 1.83E-01 | 2.63E+00 | 2.74E+00 | 9.82E-01 | no | Olfr1390 | upstream_gene_variant | MODIFIER |
| 11 | 49338276 | AX-97826490 | 0 | A | G | 0.088 | -5.57E-03 | 1.83E-01 | 2.63E+00 | 2.74E+00 | 9.82E-01 | no | Olfr1390 | upstream_gene_variant | MODIFIER |
| 11 | 49393762 | AX-170004897 | 0 | G | A | 0.088 | -5.57E-03 | 1.83E-01 | 2.63E+00 | 2.74E+00 | 9.82E-01 | no | | intergenic_variant | MODIFIER |
| 11 | 49423271 | AX-170087019 | 0 | C | T | 0.088 | -5.57E-03 | 1.83E-01 | 2.63E+00 | 2.74E+00 | 9.82E-01 | no | Olfr1389 | intron_variant | MODIFIER |
| 11 | 49462914 | AX-170004894 | 0 | A | G | 0.088 | -5.57E-03 | 1.83E-01 | 2.63E+00 | 2.74E+00 | 9.82E-01 | no | Olfr1387 | downstream_gene_variant | MODIFIER |
| 11 | 49557498 | AX-170381454 | 0 | G | C | 0.088 | -5.57E-03 | 1.83E-01 | 2.63E+00 | 2.74E+00 | 9.82E-01 | no | Olfr1381 | downstream_gene_variant | MODIFIER |
| 11 | 49776384 | AX-170322200 | 0 | T | A | 0.088 | -5.57E-03 | 1.83E-01 | 2.63E+00 | 2.74E+00 | 9.82E-01 | no | Gm12193 | upstream_gene_variant | MODIFIER |
| 11 | 49778191 | AX-97849224 | 0 | C | T | 0.088 | -5.57E-03 | 1.83E-01 | 2.63E+00 | 2.74E+00 | 9.82E-01 | no | Gm12193 | downstream_gene_variant | MODIFIER |
| 11 | 49800473 | AX-97793090 | 0 | G | A | 0.088 | -5.57E-03 | 1.83E-01 | 2.63E+00 | 2.74E+00 | 9.82E-01 | no | Gfpt2 | intron_variant | MODIFIER |
| 11 | 49834420 | AX-97806241 | 0 | T | C | 0.088 | -5.57E-03 | 1.83E-01 | 2.63E+00 | 2.74E+00 | 9.82E-01 | no | Gfpt2 | intron_variant | MODIFIER |
| 11 | 49857748 | AX-97830499 | 0 | C | T | 0.088 | -5.57E-03 | 1.83E-01 | 2.63E+00 | 2.74E+00 | 9.82E-01 | no | Mapk9 | intron_variant | MODIFIER |
| 11 | 49976548 | AX-170004899 | 0 | A | G | 0.088 | -5.57E-03 | 1.83E-01 | 2.63E+00 | 2.74E+00 | 9.82E-01 | no | Rasgef1c | intron_variant | MODIFIER |
| 11 | 50019847 | AX-97805890 | 0 | G | A | 0.088 | -5.57E-03 | 1.83E-01 | 2.63E+00 | 2.74E+00 | 9.82E-01 | no | | intergenic_variant | MODIFIER |
| 11 | 50020403 | AX-170125995 | 0 | A | G | 0.088 | -5.57E-03 | 1.83E-01 | 2.63E+00 | 2.74E+00 | 9.82E-01 | no | Rnf130 | upstream_gene_variant | MODIFIER |
| 11 | 50057758 | AX-170004900 | 0 | T | C | 0.088 | -5.57E-03 | 1.83E-01 | 2.63E+00 | 2.74E+00 | 9.82E-01 | no | Rnf130 | intron_variant | MODIFIER |
| 11 | 50058387 | AX-97818364 | 0 | T | C | 0.088 | -5.57E-03 | 1.83E-01 | 2.63E+00 | 2.74E+00 | 9.82E-01 | no | Rnf130 | intron_variant | MODIFIER |
| 11 | 50058453 | AX-170004907 | 0 | C | T | 0.088 | -5.57E-03 | 1.83E-01 | 2.63E+00 | 2.74E+00 | 9.82E-01 | no | Rnf130 | intron_variant | MODIFIER |
| 11 | 50098572 | AX-97818593 | 0 | C | T | 0.088 | -5.57E-03 | 1.83E-01 | 2.63E+00 | 2.74E+00 | 9.82E-01 | no | Rnf130 | intron_variant | MODIFIER |
| 11 | 50177506 | AX-97818618 | 0 | G | A | 0.088 | -5.57E-03 | 1.83E-01 | 2.63E+00 | 2.74E+00 | 9.82E-01 | no | Mrnip | intron_variant | MODIFIER |
| 11 | 50180953 | AX-170347823 | 0 | A | C | 0.088 | -5.57E-03 | 1.83E-01 | 2.63E+00 | 2.74E+00 | 9.82E-01 | no | Mrnip | intron_variant | MODIFIER |

| | | | | | | | | | | | | | | | |
|---|---|---|---|---|---|---|---|---|---|---|---|---|---|---|---|
| 11 | 50181208 | AX-170270209 | 0 | C | T | 0.088 | -5.57E-03 | 1.83E-01 | 2.63E+00 | 2.74E+00 | 9.82E-01 | no | Mrnip | intron_variant | MODIFIER |
| 11 | 50182997 | AX-170047014 | 0 | A | G | 0.088 | -5.57E-03 | 1.83E-01 | 2.63E+00 | 2.74E+00 | 9.82E-01 | no | Mrnip | intron_variant | MODIFIER |
| 11 | 50187548 | AX-97835819 | 0 | T | A | 0.088 | -5.57E-03 | 1.83E-01 | 2.63E+00 | 2.74E+00 | 9.82E-01 | no | Mrnip | intron_variant | MODIFIER |
| 9 | 34948067 | AX-170300163 | 0 | T | C | 0.43 | 2.65E-03 | 8.49E-02 | 2.66E+00 | 2.73E+00 | 9.82E-01 | no | Kirrel3 | intron_variant | MODIFIER |
| 16 | 92726919 | AX-170272119 | 0 | G | A | 0.465 | -1.77E-03 | 7.62E-02 | 2.66E+00 | 2.74E+00 | 9.82E-01 | no | Runx1 | intron_variant | MODIFIER |
| 10 | 31474769 | AX-170091909 | 0 | C | T | 0.386 | 1.08E-03 | 7.52E-02 | 2.63E+00 | 2.74E+00 | 9.82E-01 | no | | intergenic_variant | MODIFIER |
| 16 | 92625814 | AX-170515769 | 1 | C | T | 0.482 | -1.75E-03 | 7.84E-02 | 2.65E+00 | 2.74E+00 | 9.82E-01 | no | Runx1 | intron_variant | MODIFIER |
| 8 | 23745600 | AX-97810102 | 1 | C | T | 0.17 | -2.99E-03 | 1.14E-01 | 2.66E+00 | 2.74E+00 | 9.82E-01 | no | Zmat4 | intron_variant | MODIFIER |
| 8 | 23817758 | AX-170445028 | 1 | G | A | 0.17 | -2.99E-03 | 1.14E-01 | 2.66E+00 | 2.74E+00 | 9.82E-01 | no | Zmat4 | intron_variant | MODIFIER |
| 8 | 26643511 | AX-170428546 | 1 | G | A | 0.17 | -2.99E-03 | 1.14E-01 | 2.66E+00 | 2.74E+00 | 9.82E-01 | no | Gm45584 | downstream_gene_variant | MODIFIER |
| 11 | 48132964 | AX-170170533 | 1 | C | T | 0.089 | -5.38E-03 | 1.82E-01 | 2.63E+00 | 2.74E+00 | 9.82E-01 | no | | intergenic_variant | MODIFIER |
| 13 | 48607862 | AX-170476228 | 0 | A | G | 0.509 | -8.65E-04 | 6.74E-02 | 2.63E+00 | 2.74E+00 | 9.82E-01 | no | Ptpdc1 | intron_variant | MODIFIER |
| 9 | 71512563 | AX-170297034 | 2 | A | G | 0.191 | 2.83E-03 | 1.34E-01 | 2.70E+00 | 2.74E+00 | 9.82E-01 | no | Myzap | intron_variant | MODIFIER |
| 1 | 167503445 | AX-170002243 | 0 | T | C | 0.219 | 4.18E-03 | 1.19E-01 | 2.63E+00 | 2.73E+00 | 9.82E-01 | no | | intergenic_variant | MODIFIER |
| 9 | 77471555 | AX-170198724 | 0 | T | A | 0.193 | 2.71E-03 | 1.36E-01 | 2.71E+00 | 2.74E+00 | 9.82E-01 | no | Lrrc1 | intron_variant | MODIFIER |
| 1 | 169148853 | AX-170389306 | 0 | T | G | 0.211 | 3.36E-03 | 1.28E-01 | 2.68E+00 | 2.74E+00 | 9.82E-01 | no | | intergenic_variant | MODIFIER |
| 2 | 50815887 | AX-170017662 | 1 | G | A | 0.054 | -4.67E-03 | 1.35E-01 | 2.59E+00 | 2.74E+00 | 9.82E-01 | no | | regulatory_region_variant | MODIFIER |
| 12 | 59781524 | AX-170049168 | 1 | G | A | 0.062 | -7.08E-03 | 3.01E-01 | 2.71E+00 | 2.74E+00 | 9.82E-01 | no | | intergenic_variant | MODIFIER |
| 12 | 61444182 | AX-170006860 | 1 | C | T | 0.062 | -7.08E-03 | 3.01E-01 | 2.71E+00 | 2.74E+00 | 9.82E-01 | no | | intergenic_variant | MODIFIER |
| 9 | 61474583 | AX-97839016 | 1 | T | C | 0.196 | 2.86E-03 | 1.39E-01 | 2.72E+00 | 2.74E+00 | 9.82E-01 | no | Gm47240 | upstream_gene_variant | MODIFIER |
| 9 | 77780487 | AX-170130258 | 1 | C | T | 0.196 | 2.86E-03 | 1.39E-01 | 2.72E+00 | 2.74E+00 | 9.82E-01 | no | Gclc | intron_variant | MODIFIER |
| 11 | 46951217 | AX-170445438 | 0 | C | T | 0.088 | -5.49E-03 | 1.60E-01 | 2.61E+00 | 2.73E+00 | 9.82E-01 | no | Sgcd | intron_variant | MODIFIER |
| 11 | 45391966 | AX-170046941 | 1 | G | A | 0.089 | -5.23E-03 | 1.82E-01 | 2.63E+00 | 2.74E+00 | 9.82E-01 | no | | intergenic_variant | MODIFIER |
| 11 | 46315431 | AX-170242719 | 1 | T | A | 0.089 | -5.23E-03 | 1.82E-01 | 2.63E+00 | 2.74E+00 | 9.82E-01 | no | Cyfip2 | upstream_gene_variant | MODIFIER |
| 11 | 49154139 | AX-170240264 | 1 | G | A | 0.089 | -5.23E-03 | 1.82E-01 | 2.63E+00 | 2.74E+00 | 9.82E-01 | no | Olfr1395 | downstream_gene_variant | MODIFIER |
| 8 | 90332815 | AX-170504742 | 1 | T | C | 0.196 | -3.78E-03 | 1.15E-01 | 2.65E+00 | 2.73E+00 | 9.82E-01 | no | Tox3 | intron_variant | MODIFIER |
| 1 | 70927503 | AX-170198948 | 1 | C | G | 0.143 | -4.43E-03 | 2.34E-01 | 2.68E+00 | 2.74E+00 | 9.83E-01 | no | | intergenic_variant | MODIFIER |
| 1 | 176906543 | AX-170116977 | 0 | C | T | 0.219 | 4.15E-03 | 1.23E-01 | 2.66E+00 | 2.73E+00 | 9.83E-01 | no | Sdccag8 | intron_variant | MODIFIER |
| 11 | 90415030 | AX-170335037 | 0 | G | A | 0.053 | 3.50E-03 | 1.74E-01 | 2.63E+00 | 2.74E+00 | 9.83E-01 | no | | intergenic_variant | MODIFIER |
| 8 | 115918525 | AX-170459324 | 0 | A | G | 0.263 | -2.32E-03 | 9.38E-02 | 2.62E+00 | 2.74E+00 | 9.83E-01 | no | | intergenic_variant | MODIFIER |
| 10 | 7816589 | AX-170438511 | 1 | G | T | 0.214 | -2.85E-03 | 1.04E-01 | 2.65E+00 | 2.74E+00 | 9.83E-01 | no | Ppil4 | intron_variant | MODIFIER |
| 10 | 43997311 | AX-170531365 | 0 | T | C | 0.386 | 9.99E-04 | 7.47E-02 | 2.64E+00 | 2.74E+00 | 9.83E-01 | no | Crybg1 | missense_variant | MODERATE |
| 10 | 45812831 | AX-97845020 | 0 | C | T | 0.386 | 9.99E-04 | 7.47E-02 | 2.64E+00 | 2.74E+00 | 9.83E-01 | no | Gm27410 | downstream_gene_variant | MODIFIER |
| 1 | 66984285 | AX-97855579 | 0 | A | G | 0.123 | -2.60E-03 | 1.85E-01 | 2.64E+00 | 2.74E+00 | 9.83E-01 | no | | intergenic_variant | MODIFIER |
| 4 | 141152327 | AX-170097455 | 0 | G | A | 0.096 | -2.57E-03 | 1.60E-01 | 2.66E+00 | 2.74E+00 | 9.83E-01 | no | Fbxo42 | intron_variant | MODIFIER |
| 1 | 39948194 | AX-170308392 | 1 | T | C | 0.402 | -2.07E-03 | 7.67E-02 | 2.66E+00 | 2.74E+00 | 9.83E-01 | no | Map4k4 | intron_variant | MODIFIER |
| 20 | 144508820 | AX-97819426 | 1 | A | T | 0.143 | -2.12E-03 | 1.29E-01 | 2.64E+00 | 2.74E+00 | 9.83E-01 | no | Trpc5 | intron_variant | MODIFIER |
| 11 | 22238945 | AX-170363581 | 2 | A | G | 0.055 | -3.27E-03 | 1.61E-01 | 2.58E+00 | 2.74E+00 | 9.83E-01 | no | Ehbp1 | intron_variant | MODIFIER |
| 8 | 80937346 | AX-170110624 | 1 | G | T | 0.205 | -2.55E-03 | 1.32E-01 | 2.68E+00 | 2.74E+00 | 9.83E-01 | no | Gm4891 | downstream_gene_variant | MODIFIER |
| 8 | 82373432 | AX-170401604 | 1 | T | C | 0.205 | -2.55E-03 | 1.32E-01 | 2.68E+00 | 2.74E+00 | 9.83E-01 | no | Il15 | intron_variant | MODIFIER |
| 7 | 117998686 | AX-170261430 | 1 | C | T | 0.402 | -1.39E-03 | 8.79E-02 | 2.67E+00 | 2.74E+00 | 9.83E-01 | no | | intergenic_variant | MODIFIER |
| 7 | 67711820 | AX-97812362 | 0 | A | C | 0.404 | -1.37E-03 | 9.86E-02 | 2.67E+00 | 2.74E+00 | 9.83E-01 | no | Ttc23 | intron_variant | MODIFIER |
| 7 | 70057002 | AX-170143493 | 0 | G | A | 0.404 | -1.37E-03 | 9.86E-02 | 2.67E+00 | 2.74E+00 | 9.83E-01 | no | | intergenic_variant | MODIFIER |
| 9 | 59471656 | AX-170475470 | 0 | T | C | 0.219 | -2.11E-03 | 1.18E-01 | 2.67E+00 | 2.74E+00 | 9.83E-01 | no | Arih1 | intron_variant,NMD_transcript_variant | MODIFIER |
| 7 | 97958029 | AX-97815912 | 1 | T | C | 0.134 | 4.20E-03 | 2.10E-01 | 2.66E+00 | 2.74E+00 | 9.83E-01 | no | Gdpd4 | intron_variant | MODIFIER |
| 9 | 87537555 | AX-170111834 | 1 | C | T | 0.143 | 1.15E-03 | 1.25E-01 | 2.66E+00 | 2.74E+00 | 9.83E-01 | no | | intergenic_variant | MODIFIER |
| 11 | 48055635 | AX-170391224 | 1 | A | C | 0.089 | 2.28E-03 | 1.82E-01 | 2.64E+00 | 2.74E+00 | 9.83E-01 | no | | intergenic_variant | MODIFIER |
| 11 | 48359029 | AX-170278733 | 1 | C | G | 0.089 | 2.36E-03 | 1.82E-01 | 2.64E+00 | 2.74E+00 | 9.83E-01 | no | | intergenic_variant | MODIFIER |
| 12 | 70302440 | AX-170323013 | 1 | C | T | 0.062 | -6.50E-03 | 3.02E-01 | 2.71E+00 | 2.74E+00 | 9.83E-01 | no | Trim9 | intron_variant | MODIFIER |
| 1 | 78414880 | AX-170235425 | 0 | G | T | 0.14 | 5.02E-03 | 2.35E-01 | 2.68E+00 | 2.74E+00 | 9.84E-01 | no | Sgpp2 | intron_variant | MODIFIER |
| 1 | 78795782 | AX-170000874 | 0 | C | T | 0.14 | 5.06E-03 | 2.35E-01 | 2.68E+00 | 2.74E+00 | 9.84E-01 | no | Gm29536 | intron_variant,non_coding_transcript_variant | MODIFIER |
| 12 | 100980188 | AX-170353748 | 0 | C | T | 0.088 | -3.92E-03 | 1.94E-01 | 2.68E+00 | 2.74E+00 | 9.84E-01 | no | Ccdc88c | intron_variant | MODIFIER |
| 1 | 7564522 | AX-170430087 | 0 | T | A | 0.412 | 2.18E-03 | 7.11E-02 | 2.66E+00 | 2.73E+00 | 9.84E-01 | no | | intergenic_variant | MODIFIER |
| 1 | 7594314 | AX-97783484 | 0 | A | G | 0.412 | 2.18E-03 | 7.11E-02 | 2.66E+00 | 2.73E+00 | 9.84E-01 | no | Gm7470 | upstream_gene_variant | MODIFIER |

| | | | | | | | | | | | | | | | |
|---|---|---|---|---|---|---|---|---|---|---|---|---|---|---|---|
| 1 | 7599821 | AX-97853082 | 0 | T | C | 0.412 | 2.18E-03 | 7.11E-02 | 2.66E+00 | 2.73E+00 | 9.84E-01 | no | Gm7470 | downstream_gene_variant | MODIFIER |
| 1 | 7614022 | AX-170140462 | 0 | A | G | 0.412 | 2.18E-03 | 7.11E-02 | 2.66E+00 | 2.73E+00 | 9.84E-01 | no | | intergenic_variant | MODIFIER |
| 1 | 7669185 | AX-170040308 | 0 | G | A | 0.412 | 2.18E-03 | 7.11E-02 | 2.66E+00 | 2.73E+00 | 9.84E-01 | no | Rps2-ps2 | downstream_gene_variant | MODIFIER |
| 1 | 7669308 | AX-97848243 | 0 | A | G | 0.412 | 2.18E-03 | 7.11E-02 | 2.66E+00 | 2.73E+00 | 9.84E-01 | no | Rps2-ps2 | downstream_gene_variant | MODIFIER |
| 1 | 7669592 | AX-170494697 | 0 | T | C | 0.412 | 2.18E-03 | 7.11E-02 | 2.66E+00 | 2.73E+00 | 9.84E-01 | no | Rps2-ps2 | downstream_gene_variant | MODIFIER |
| 1 | 7677513 | AX-170040311 | 0 | T | C | 0.412 | 2.18E-03 | 7.11E-02 | 2.66E+00 | 2.73E+00 | 9.84E-01 | no | | intergenic_variant | MODIFIER |
| 1 | 7698048 | AX-97846057 | 0 | G | A | 0.412 | 2.18E-03 | 7.11E-02 | 2.66E+00 | 2.73E+00 | 9.84E-01 | no | | intergenic_variant | MODIFIER |
| 1 | 7782007 | AX-170342698 | 0 | T | C | 0.412 | 2.18E-03 | 7.11E-02 | 2.66E+00 | 2.73E+00 | 9.84E-01 | no | | intergenic_variant | MODIFIER |
| 1 | 7818432 | AX-170137415 | 0 | C | A | 0.412 | 2.18E-03 | 7.11E-02 | 2.66E+00 | 2.73E+00 | 9.84E-01 | no | | intergenic_variant | MODIFIER |
| 1 | 7823013 | AX-170253688 | 0 | G | A | 0.412 | 2.18E-03 | 7.11E-02 | 2.66E+00 | 2.73E+00 | 9.84E-01 | no | | intergenic_variant | MODIFIER |
| 1 | 7826771 | AX-169999889 | 0 | T | C | 0.412 | 2.18E-03 | 7.11E-02 | 2.66E+00 | 2.73E+00 | 9.84E-01 | no | | intergenic_variant | MODIFIER |
| 1 | 7871667 | AX-169999892 | 0 | G | A | 0.412 | 2.18E-03 | 7.11E-02 | 2.66E+00 | 2.73E+00 | 9.84E-01 | no | | intergenic_variant | MODIFIER |
| 1 | 7904705 | AX-170086403 | 0 | A | G | 0.412 | 2.18E-03 | 7.11E-02 | 2.66E+00 | 2.73E+00 | 9.84E-01 | no | | intergenic_variant | MODIFIER |
| 1 | 7966419 | AX-169999891 | 0 | G | T | 0.412 | 2.18E-03 | 7.11E-02 | 2.66E+00 | 2.73E+00 | 9.84E-01 | no | | intergenic_variant | MODIFIER |
| 1 | 7976936 | AX-170086404 | 0 | C | A | 0.412 | 2.18E-03 | 7.11E-02 | 2.66E+00 | 2.73E+00 | 9.84E-01 | no | | intergenic_variant | MODIFIER |
| 1 | 8026452 | AX-170040315 | 0 | A | G | 0.412 | 2.18E-03 | 7.11E-02 | 2.66E+00 | 2.73E+00 | 9.84E-01 | no | | intergenic_variant | MODIFIER |
| 1 | 8082792 | AX-170175380 | 0 | A | G | 0.412 | 2.18E-03 | 7.11E-02 | 2.66E+00 | 2.73E+00 | 9.84E-01 | no | | intergenic_variant | MODIFIER |
| 1 | 8083854 | AX-170240380 | 0 | T | C | 0.412 | 2.18E-03 | 7.11E-02 | 2.66E+00 | 2.73E+00 | 9.84E-01 | no | | intergenic_variant | MODIFIER |
| 1 | 8246868 | AX-170040320 | 0 | G | C | 0.412 | 2.18E-03 | 7.11E-02 | 2.66E+00 | 2.73E+00 | 9.84E-01 | no | | intergenic_variant | MODIFIER |
| 1 | 8259804 | AX-170040324 | 0 | A | T | 0.412 | 2.18E-03 | 7.11E-02 | 2.66E+00 | 2.73E+00 | 9.84E-01 | no | | intergenic_variant | MODIFIER |
| 1 | 9897624 | AX-170239521 | 0 | G | A | 0.412 | 2.18E-03 | 7.11E-02 | 2.66E+00 | 2.73E+00 | 9.84E-01 | no | Sgk3 | intron_variant | MODIFIER |
| 1 | 9900096 | AX-170396694 | 0 | A | G | 0.412 | 2.18E-03 | 7.11E-02 | 2.66E+00 | 2.73E+00 | 9.84E-01 | no | Sgk3 | downstream_gene_variant | MODIFIER |
| 1 | 9900222 | AX-170376256 | 0 | G | A | 0.412 | 2.18E-03 | 7.11E-02 | 2.66E+00 | 2.73E+00 | 9.84E-01 | no | Sgk3 | downstream_gene_variant | MODIFIER |
| 1 | 9903974 | AX-169999915 | 0 | A | G | 0.412 | 2.18E-03 | 7.11E-02 | 2.66E+00 | 2.73E+00 | 9.84E-01 | no | Mcmdc2 | upstream_gene_variant | MODIFIER |
| 1 | 10018158 | AX-170164991 | 0 | G | A | 0.412 | 2.18E-03 | 7.11E-02 | 2.66E+00 | 2.73E+00 | 9.84E-01 | no | | intergenic_variant | MODIFIER |
| 1 | 10027837 | AX-170533548 | 0 | T | C | 0.412 | 2.18E-03 | 7.11E-02 | 2.66E+00 | 2.73E+00 | 9.84E-01 | no | Cops5 | intron_variant | MODIFIER |
| 1 | 10033421 | AX-97810679 | 0 | A | G | 0.412 | 2.18E-03 | 7.11E-02 | 2.66E+00 | 2.73E+00 | 9.84E-01 | no | Cops5 | intron_variant | MODIFIER |
| 1 | 10179462 | AX-169999929 | 0 | A | G | 0.412 | 2.18E-03 | 7.11E-02 | 2.66E+00 | 2.73E+00 | 9.84E-01 | no | Arfgef1 | intron_variant | MODIFIER |
| 2 | 129717072 | AX-97789176 | 2 | C | A | 0.145 | 5.11E-03 | 2.20E-01 | 2.70E+00 | 2.74E+00 | 9.84E-01 | no | | intergenic_variant | MODIFIER |
| 11 | 8095548 | AX-170390557 | 0 | C | T | 0.044 | -5.91E-03 | 1.92E-01 | 2.60E+00 | 2.74E+00 | 9.84E-01 | no | | intergenic_variant | MODIFIER |
| 1 | 52538617 | AX-170194937 | 0 | C | T | 0.132 | 4.78E-03 | 2.42E-01 | 2.69E+00 | 2.74E+00 | 9.84E-01 | no | Gm553 | intron_variant,non_coding_transcript_variant | MODIFIER |
| 18 | 69725347 | AX-170380355 | 0 | G | A | 0.272 | -2.66E-03 | 9.67E-02 | 2.66E+00 | 2.73E+00 | 9.84E-01 | no | | intergenic_variant | MODIFIER |
| 9 | 73158300 | AX-170313003 | 1 | G | A | 0.214 | 2.60E-03 | 1.39E-01 | 2.71E+00 | 2.74E+00 | 9.84E-01 | no | | regulatory_region_variant | MODIFIER |
| 9 | 39643575 | AX-170380106 | 0 | G | A | 0.43 | -1.65E-03 | 7.39E-02 | 2.65E+00 | 2.74E+00 | 9.84E-01 | no | Olfr961 | upstream_gene_variant | MODIFIER |
| 2 | 121248143 | AX-170217824 | 1 | G | A | 0.143 | 4.97E-03 | 2.21E-01 | 2.71E+00 | 2.74E+00 | 9.84E-01 | no | Trp53bp1 | intron_variant | MODIFIER |
| 2 | 143496518 | AX-170219780 | 1 | A | G | 0.143 | 4.97E-03 | 2.21E-01 | 2.71E+00 | 2.74E+00 | 9.84E-01 | no | | intergenic_variant | MODIFIER |
| 8 | 30904926 | AX-170517297 | 1 | C | A | 0.152 | 1.75E-03 | 1.15E-01 | 2.65E+00 | 2.74E+00 | 9.84E-01 | no | | intergenic_variant | MODIFIER |
| 16 | 98029840 | AX-170318435 | 1 | T | C | 0.455 | -2.52E-03 | 8.16E-02 | 2.64E+00 | 2.73E+00 | 9.84E-01 | no | B230307C23Rik | intron_variant | MODIFIER |
| 8 | 42981287 | AX-170508133 | 1 | C | T | 0.152 | 1.62E-03 | 1.26E-01 | 2.65E+00 | 2.74E+00 | 9.84E-01 | no | | intergenic_variant | MODIFIER |
| 9 | 58457114 | AX-170432088 | 0 | G | C | 0.228 | -2.08E-03 | 1.10E-01 | 2.67E+00 | 2.74E+00 | 9.84E-01 | no | 4930461G14Rik | intron_variant,non_coding_transcript_variant | MODIFIER |
| 10 | 14280818 | AX-170086750 | 1 | C | T | 0.205 | 1.11E-03 | 1.07E-01 | 2.67E+00 | 2.74E+00 | 9.84E-01 | no | | intergenic_variant | MODIFIER |
| 7 | 53044587 | AX-170325924 | 1 | A | T | 0.429 | -1.25E-03 | 1.00E-01 | 2.67E+00 | 2.74E+00 | 9.85E-01 | no | | intergenic_variant | MODIFIER |
| 11 | 45821308 | AX-170405969 | 1 | G | A | 0.098 | -3.37E-03 | 1.52E-01 | 2.60E+00 | 2.74E+00 | 9.85E-01 | no | F630206G17Rik | intron_variant,non_coding_transcript_variant | MODIFIER |
| 12 | 113188295 | AX-170113877 | 1 | C | T | 0.125 | -2.98E-03 | 1.96E-01 | 2.67E+00 | 2.74E+00 | 9.85E-01 | no | Tmem121 | synonymous_variant | LOW |
| 1 | 169416012 | AX-170346166 | 0 | C | T | 0.237 | -1.85E-03 | 1.28E-01 | 2.69E+00 | 2.74E+00 | 9.85E-01 | no | | intergenic_variant | MODIFIER |
| 5 | 121100986 | AX-170440101 | 1 | C | G | 0.455 | 2.57E-03 | 1.12E-01 | 2.69E+00 | 2.74E+00 | 9.85E-01 | no | | intergenic_variant | MODIFIER |
| 9 | 54184874 | AX-170308926 | 0 | G | T | 0.211 | 2.48E-03 | 1.22E-01 | 2.68E+00 | 2.74E+00 | 9.85E-01 | no | Cyp19a1 | intron_variant | MODIFIER |
| 1 | 70207338 | AX-170297389 | 0 | C | A | 0.132 | -3.98E-03 | 2.15E-01 | 2.69E+00 | 2.74E+00 | 9.85E-01 | no | Spag16 | intron_variant | MODIFIER |
| 8 | 85612289 | AX-97810319 | 2 | C | T | 0.209 | 2.86E-03 | 1.25E-01 | 2.66E+00 | 2.74E+00 | 9.85E-01 | no | Gm45425 | downstream_gene_variant | MODIFIER |
| 8 | 82079229 | AX-97826899 | 1 | A | T | 0.205 | 2.49E-03 | 1.32E-01 | 2.68E+00 | 2.74E+00 | 9.85E-01 | no | Inpp4b | intron_variant | MODIFIER |
| 12 | 72956704 | AX-97853814 | 1 | C | T | 0.071 | 4.09E-03 | 2.08E-01 | 2.64E+00 | 2.74E+00 | 9.85E-01 | no | Gm26709 | intron_variant,non_coding_transcript_variant | MODIFIER |
| 8 | 84228778 | AX-170389187 | 0 | A | G | 0.202 | -1.71E-03 | 1.19E-01 | 2.65E+00 | 2.74E+00 | 9.85E-01 | no | Zswim4 | intron_variant | MODIFIER |
| 8 | 16654701 | AX-170467753 | 1 | G | A | 0.232 | 4.35E-04 | 7.38E-02 | 2.63E+00 | 2.74E+00 | 9.85E-01 | no | Csmd1 | intron_variant | MODIFIER |
| 1 | 36145976 | AX-170168818 | 0 | C | T | 0.456 | 1.22E-03 | 7.92E-02 | 2.66E+00 | 2.74E+00 | 9.85E-01 | no | Uggt1 | intron_variant | MODIFIER |

| | | | | | | | | | | | | | | | |
|---|---|---|---|---|---|---|---|---|---|---|---|---|---|---|---|
| 11 | 23028398 | AX-170004480 | 1 | T | A | 0.062 | 3.02E-03 | 2.14E-01 | 2.65E+00 | 2.74E+00 | 9.85E-01 | no | Fam161a | downstream_gene_variant | MODIFIER |
| 9 | 39726409 | AX-170289321 | 0 | G | A | 0.43 | -1.38E-03 | 7.22E-02 | 2.64E+00 | 2.74E+00 | 9.85E-01 | no | Olfr966-ps1 | downstream_gene_variant | MODIFIER |
| 14 | 61180567 | AX-97834839 | 1 | G | A | 0.241 | 1.35E-03 | 7.03E-02 | 2.58E+00 | 2.74E+00 | 9.85E-01 | no | Sacs | intron_variant | MODIFIER |
| 15 | 91482436 | AX-170107190 | 0 | C | T | 0.132 | 2.56E-03 | 1.38E-01 | 2.65E+00 | 2.74E+00 | 9.85E-01 | no | Slc2a13 | intron_variant | MODIFIER |
| 15 | 92004879 | AX-170346112 | 0 | G | A | 0.132 | 2.56E-03 | 1.38E-01 | 2.65E+00 | 2.74E+00 | 9.85E-01 | no | | intergenic_variant | MODIFIER |
| 13 | 46331406 | AX-170008253 | 0 | C | T | 0.5 | -5.75E-04 | 6.20E-02 | 2.63E+00 | 2.74E+00 | 9.85E-01 | no | | intergenic_variant | MODIFIER |
| 7 | 81845642 | AX-170029416 | 1 | C | T | 0.116 | 4.58E-03 | 2.12E-01 | 2.66E+00 | 2.74E+00 | 9.86E-01 | no | | intergenic_variant | MODIFIER |
| 1 | 78223566 | AX-170503870 | 0 | G | A | 0.14 | 4.32E-03 | 2.34E-01 | 2.68E+00 | 2.74E+00 | 9.86E-01 | no | | intergenic_variant | MODIFIER |
| 9 | 64185363 | AX-170428347 | 0 | C | A | 0.184 | -2.68E-03 | 1.30E-01 | 2.69E+00 | 2.74E+00 | 9.86E-01 | no | Map2k1 | downstream_gene_variant | MODIFIER |
| 10 | 9409922 | AX-170044114 | 0 | A | C | 0.211 | 8.73E-04 | 1.05E-01 | 2.66E+00 | 2.74E+00 | 9.86E-01 | no | | intergenic_variant | MODIFIER |
| 10 | 9411646 | AX-170002693 | 0 | G | A | 0.211 | 8.73E-04 | 1.05E-01 | 2.66E+00 | 2.74E+00 | 9.86E-01 | no | | intergenic_variant | MODIFIER |
| 10 | 9443969 | AX-170086738 | 0 | G | A | 0.211 | 8.73E-04 | 1.05E-01 | 2.66E+00 | 2.74E+00 | 9.86E-01 | no | | intergenic_variant | MODIFIER |
| 9 | 52446735 | AX-170381080 | 1 | G | A | 0.232 | -2.29E-03 | 1.22E-01 | 2.68E+00 | 2.74E+00 | 9.86E-01 | no | AC166902.1 | intron_variant,non_coding_transcript_variant | MODIFIER |
| 10 | 12283865 | AX-97821832 | 0 | A | G | 0.211 | 8.47E-04 | 9.99E-02 | 2.64E+00 | 2.74E+00 | 9.86E-01 | no | | intergenic_variant | MODIFIER |
| 1 | 55752313 | AX-170173361 | 0 | T | C | 0.114 | -2.44E-03 | 1.88E-01 | 2.64E+00 | 2.74E+00 | 9.86E-01 | no | Plcl1 | 3_prime_UTR_variant | MODIFIER |
| 6 | 138856093 | AX-170508743 | 0 | A | T | 0.07 | -5.05E-03 | 2.77E-01 | 2.70E+00 | 2.74E+00 | 9.86E-01 | no | Gm9038 | downstream_gene_variant | MODIFIER |
| 12 | 113588844 | AX-97781771 | 2 | C | T | 0.1 | 3.12E-03 | 1.81E-01 | 2.64E+00 | 2.74E+00 | 9.86E-01 | no | Ighv2-2 | upstream_gene_variant | MODIFIER |
| 3 | 147238292 | AX-170127560 | 1 | T | A | 0.339 | -9.03E-04 | 7.38E-02 | 2.67E+00 | 2.74E+00 | 9.86E-01 | no | | intergenic_variant | MODIFIER |
| 1 | 4642252 | AX-170108568 | 0 | A | G | 0.509 | 8.60E-05 | 6.72E-02 | 2.65E+00 | 2.74E+00 | 9.86E-01 | no | | intergenic_variant | MODIFIER |
| 2 | 146869738 | AX-97804512 | 1 | G | A | 0.134 | 3.98E-03 | 1.92E-01 | 2.66E+00 | 2.74E+00 | 9.86E-01 | no | Kiz | intron_variant | MODIFIER |
| 8 | 19233305 | AX-170468025 | 0 | T | C | 0.184 | 1.94E-03 | 9.95E-02 | 2.63E+00 | 2.74E+00 | 9.86E-01 | no | | intergenic_variant | MODIFIER |
| 9 | 77252090 | AX-170376861 | 0 | T | C | 0.202 | -2.54E-03 | 1.35E-01 | 2.71E+00 | 2.74E+00 | 9.86E-01 | no | Mlip | intron_variant | MODIFIER |
| 9 | 77265180 | AX-97786983 | 0 | A | G | 0.202 | -2.54E-03 | 1.35E-01 | 2.71E+00 | 2.74E+00 | 9.86E-01 | no | Mlip | intron_variant | MODIFIER |
| 9 | 77301573 | AX-170090339 | 0 | G | A | 0.202 | -2.54E-03 | 1.35E-01 | 2.71E+00 | 2.74E+00 | 9.86E-01 | no | Mlip | intron_variant | MODIFIER |
| 9 | 77450028 | AX-97783780 | 0 | A | T | 0.202 | -2.54E-03 | 1.35E-01 | 2.71E+00 | 2.74E+00 | 9.86E-01 | no | Lrrc1 | intron_variant | MODIFIER |
| 9 | 77468442 | AX-170333776 | 0 | G | A | 0.202 | -2.54E-03 | 1.35E-01 | 2.71E+00 | 2.74E+00 | 9.86E-01 | no | Lrrc1 | intron_variant | MODIFIER |
| 9 | 77473588 | AX-170086240 | 0 | T | C | 0.202 | -2.54E-03 | 1.35E-01 | 2.71E+00 | 2.74E+00 | 9.86E-01 | no | Lrrc1 | intron_variant | MODIFIER |
| 9 | 77474148 | AX-170340072 | 0 | C | G | 0.202 | -2.54E-03 | 1.35E-01 | 2.71E+00 | 2.74E+00 | 9.86E-01 | no | Lrrc1 | intron_variant | MODIFIER |
| 9 | 77486976 | AX-170291349 | 0 | A | T | 0.202 | -2.54E-03 | 1.35E-01 | 2.71E+00 | 2.74E+00 | 9.86E-01 | no | Lrrc1 | intron_variant | MODIFIER |
| 9 | 77487601 | AX-170187040 | 0 | T | G | 0.202 | -2.54E-03 | 1.35E-01 | 2.71E+00 | 2.74E+00 | 9.86E-01 | no | Lrrc1 | intron_variant | MODIFIER |
| 9 | 77503567 | AX-170096593 | 0 | A | C | 0.202 | -2.54E-03 | 1.35E-01 | 2.71E+00 | 2.74E+00 | 9.86E-01 | no | Lrrc1 | intron_variant | MODIFIER |
| 9 | 77509518 | AX-170099117 | 0 | C | A | 0.202 | -2.54E-03 | 1.35E-01 | 2.71E+00 | 2.74E+00 | 9.86E-01 | no | Lrrc1 | intron_variant | MODIFIER |
| 9 | 77509763 | AX-170082867 | 0 | G | A | 0.202 | -2.54E-03 | 1.35E-01 | 2.71E+00 | 2.74E+00 | 9.86E-01 | no | Lrrc1 | intron_variant | MODIFIER |
| 9 | 77972296 | AX-170245681 | 0 | C | T | 0.202 | -2.54E-03 | 1.35E-01 | 2.71E+00 | 2.74E+00 | 9.86E-01 | no | Elovl5 | intron_variant | MODIFIER |
| 8 | 85299399 | AX-170354791 | 1 | A | G | 0.205 | 2.48E-03 | 1.27E-01 | 2.67E+00 | 2.74E+00 | 9.87E-01 | no | Vps35 | intron_variant | MODIFIER |
| 15 | 82692525 | AX-170234495 | 0 | T | C | 0.544 | 2.32E-03 | 7.52E-02 | 2.64E+00 | 2.73E+00 | 9.87E-01 | no | Cyp2d37-ps | intron_variant,non_coding_transcript_variant | MODIFIER |
| 8 | 54154317 | AX-170236826 | 0 | T | C | 0.149 | -3.48E-03 | 1.51E-01 | 2.68E+00 | 2.73E+00 | 9.87E-01 | no | Vegfc | intron_variant | MODIFIER |
| 7 | 116458132 | AX-170419704 | 0 | T | C | 0.412 | -1.09E-03 | 8.81E-02 | 2.68E+00 | 2.74E+00 | 9.87E-01 | no | | regulatory_region_variant | MODIFIER |
| 20 | 87716492 | AX-97843641 | 1 | G | A | 0.277 | -5.82E-04 | 6.06E-02 | 2.64E+00 | 2.74E+00 | 9.87E-01 | no | Il1rapl1 | intron_variant | MODIFIER |
| 8 | 84351603 | AX-170486782 | 0 | C | G | 0.211 | -2.10E-03 | 1.29E-01 | 2.68E+00 | 2.74E+00 | 9.87E-01 | no | | intergenic_variant | MODIFIER |
| 1 | 70112610 | AX-170041578 | 1 | T | A | 0.143 | -3.25E-03 | 2.31E-01 | 2.67E+00 | 2.74E+00 | 9.87E-01 | no | Spag16 | intron_variant | MODIFIER |
| 7 | 102522064 | AX-170238428 | 1 | G | T | 0.455 | 2.26E-03 | 9.47E-02 | 2.67E+00 | 2.73E+00 | 9.87E-01 | no | Olfr546-ps1 | downstream_gene_variant | MODIFIER |
| 15 | 78047418 | AX-97829733 | 1 | C | T | 0.554 | 2.30E-03 | 8.28E-02 | 2.66E+00 | 2.73E+00 | 9.87E-01 | no | Cacng2 | intron_variant | MODIFIER |
| 15 | 78563914 | AX-170087917 | 1 | A | G | 0.554 | 2.29E-03 | 8.27E-02 | 2.66E+00 | 2.73E+00 | 9.87E-01 | no | Rac2 | intron_variant | MODIFIER |
| 15 | 92197718 | AX-170525746 | 0 | A | C | 0.14 | -1.05E-03 | 1.34E-01 | 2.67E+00 | 2.74E+00 | 9.87E-01 | no | Cntn1 | intron_variant | MODIFIER |
| 6 | 139586392 | AX-170500055 | 0 | C | T | 0.061 | 5.23E-03 | 2.08E-01 | 2.61E+00 | 2.74E+00 | 9.87E-01 | no | Pik3c2g | upstream_gene_variant | MODIFIER |
| 7 | 44851401 | AX-170152553 | 1 | G | C | 0.33 | -6.81E-04 | 9.68E-02 | 2.67E+00 | 2.74E+00 | 9.87E-01 | no | Tbc1d17 | upstream_gene_variant | MODIFIER |
| 7 | 53755665 | AX-97818084 | 1 | T | C | 0.411 | 2.33E-03 | 9.52E-02 | 2.65E+00 | 2.74E+00 | 9.87E-01 | no | | intergenic_variant | MODIFIER |
| 10 | 15818171 | AX-170286260 | 0 | G | C | 0.202 | -2.78E-03 | 1.05E-01 | 2.66E+00 | 2.73E+00 | 9.87E-01 | no | | intergenic_variant | MODIFIER |
| 10 | 16052723 | AX-170270223 | 0 | G | A | 0.202 | -2.78E-03 | 1.05E-01 | 2.66E+00 | 2.73E+00 | 9.87E-01 | no | | regulatory_region_variant | MODIFIER |
| 10 | 16099950 | AX-170405817 | 0 | T | C | 0.202 | -2.78E-03 | 1.05E-01 | 2.66E+00 | 2.73E+00 | 9.87E-01 | no | | intergenic_variant | MODIFIER |
| 10 | 16138293 | AX-170002815 | 0 | G | A | 0.202 | -2.78E-03 | 1.05E-01 | 2.66E+00 | 2.73E+00 | 9.87E-01 | no | | intergenic_variant | MODIFIER |
| 10 | 16203453 | AX-170002819 | 0 | T | G | 0.202 | -2.78E-03 | 1.05E-01 | 2.66E+00 | 2.73E+00 | 9.87E-01 | no | | intergenic_variant | MODIFIER |
| 4 | 147108645 | AX-170259020 | 0 | G | A | 0.105 | 2.23E-03 | 1.72E-01 | 2.65E+00 | 2.74E+00 | 9.87E-01 | no | Gm13142 | downstream_gene_variant | MODIFIER |

| | | | | | | | | | | | | | | | |
|---|---|---|---|---|---|---|---|---|---|---|---|---|---|---|---|
| 5 | 126425272 | AX-170415164 | 1 | C | T | 0.473 | -1.09E-03 | 9.13E-02 | 2.64E+00 | 2.74E+00 | 9.87E-01 | no | | intergenic_variant | MODIFIER |
| 2 | 125068536 | AX-170182708 | 1 | A | T | 0.143 | -2.95E-03 | 2.22E-01 | 2.71E+00 | 2.74E+00 | 9.87E-01 | no | Slc24a5 | intron_variant | MODIFIER |
| 13 | 53906341 | AX-170191909 | 1 | A | G | 0.5 | 2.16E-03 | 6.76E-02 | 2.62E+00 | 2.73E+00 | 9.88E-01 | no | | intergenic_variant | MODIFIER |
| 11 | 5721272 | AX-170004189 | 0 | T | C | 0.061 | -4.22E-03 | 2.13E-01 | 2.64E+00 | 2.74E+00 | 9.88E-01 | no | Urgcp | intron_variant | MODIFIER |
| 5 | 126245556 | AX-168406744 | 0 | G | A | 0.456 | -1.11E-03 | 1.06E-01 | 2.67E+00 | 2.74E+00 | 9.88E-01 | no | | intergenic_variant | MODIFIER |
| 18 | 57853488 | AX-170223826 | 1 | A | G | 0.42 | -2.14E-03 | 8.58E-02 | 2.67E+00 | 2.73E+00 | 9.88E-01 | no | | intergenic_variant | MODIFIER |
| 6 | 139748470 | AX-170144883 | 0 | C | T | 0.053 | -4.11E-03 | 2.12E-01 | 2.62E+00 | 2.74E+00 | 9.88E-01 | no | Pik3c2g | intron_variant | MODIFIER |
| 2 | 157778050 | AX-170019491 | 0 | T | C | 0.149 | -1.89E-03 | 1.75E-01 | 2.66E+00 | 2.74E+00 | 9.88E-01 | no | Ctnnbl1 | intron_variant | MODIFIER |
| 7 | 101665909 | AX-170096048 | 0 | C | T | 0.456 | 2.15E-03 | 9.47E-02 | 2.67E+00 | 2.74E+00 | 9.88E-01 | no | Clpb | intron_variant | MODIFIER |
| 7 | 101666027 | AX-170231420 | 0 | C | A | 0.456 | 2.15E-03 | 9.47E-02 | 2.67E+00 | 2.74E+00 | 9.88E-01 | no | Clpb | intron_variant | MODIFIER |
| 7 | 101702748 | AX-170119998 | 0 | G | T | 0.456 | 2.15E-03 | 9.47E-02 | 2.67E+00 | 2.74E+00 | 9.88E-01 | no | Clpb | intron_variant | MODIFIER |
| 7 | 101703226 | AX-170078260 | 0 | T | C | 0.456 | 2.15E-03 | 9.47E-02 | 2.67E+00 | 2.74E+00 | 9.88E-01 | no | Clpb | intron_variant | MODIFIER |
| 7 | 101706637 | AX-170269324 | 0 | A | C | 0.456 | 2.15E-03 | 9.47E-02 | 2.67E+00 | 2.74E+00 | 9.88E-01 | no | Clpb | intron_variant | MODIFIER |
| 7 | 101710478 | AX-170193386 | 0 | G | T | 0.456 | 2.15E-03 | 9.47E-02 | 2.67E+00 | 2.74E+00 | 9.88E-01 | no | Clpb | intron_variant | MODIFIER |
| 7 | 101733331 | AX-97791499 | 0 | T | C | 0.456 | 2.15E-03 | 9.47E-02 | 2.67E+00 | 2.74E+00 | 9.88E-01 | no | Clpb | intron_variant | MODIFIER |
| 7 | 101735918 | AX-170263688 | 0 | A | G | 0.456 | 2.15E-03 | 9.47E-02 | 2.67E+00 | 2.74E+00 | 9.88E-01 | no | Clpb | intron_variant | MODIFIER |
| 7 | 101738347 | AX-170372904 | 0 | G | T | 0.456 | 2.15E-03 | 9.47E-02 | 2.67E+00 | 2.74E+00 | 9.88E-01 | no | Clpb | intron_variant | MODIFIER |
| 7 | 101752146 | AX-170476515 | 0 | T | C | 0.456 | 2.15E-03 | 9.47E-02 | 2.67E+00 | 2.74E+00 | 9.88E-01 | no | Clpb | intron_variant | MODIFIER |
| 7 | 101752485 | AX-170287699 | 0 | A | G | 0.456 | 2.15E-03 | 9.47E-02 | 2.67E+00 | 2.74E+00 | 9.88E-01 | no | Clpb | intron_variant | MODIFIER |
| 7 | 101753799 | AX-170432029 | 0 | A | G | 0.456 | 2.15E-03 | 9.47E-02 | 2.67E+00 | 2.74E+00 | 9.88E-01 | no | Clpb | intron_variant | MODIFIER |
| 7 | 101755925 | AX-170309406 | 0 | T | C | 0.456 | 2.15E-03 | 9.47E-02 | 2.67E+00 | 2.74E+00 | 9.88E-01 | no | Clpb | intron_variant | MODIFIER |
| 7 | 101767435 | AX-170297077 | 0 | G | A | 0.456 | 2.15E-03 | 9.47E-02 | 2.67E+00 | 2.74E+00 | 9.88E-01 | no | Clpb | intron_variant | MODIFIER |
| 7 | 101769676 | AX-97793637 | 0 | G | T | 0.456 | 2.15E-03 | 9.47E-02 | 2.67E+00 | 2.74E+00 | 9.88E-01 | no | Clpb | intron_variant | MODIFIER |
| 7 | 101787964 | AX-97817564 | 0 | A | G | 0.456 | 2.15E-03 | 9.47E-02 | 2.67E+00 | 2.74E+00 | 9.88E-01 | no | Clpb | 3_prime_UTR_variant | MODIFIER |
| 7 | 101991515 | AX-97792505 | 0 | G | A | 0.456 | 2.15E-03 | 9.47E-02 | 2.67E+00 | 2.74E+00 | 9.88E-01 | no | Numa1 | intron_variant | MODIFIER |
| 7 | 102016307 | AX-97811032 | 0 | A | G | 0.456 | 2.15E-03 | 9.47E-02 | 2.67E+00 | 2.74E+00 | 9.88E-01 | no | Numa1 | downstream_gene_variant | MODIFIER |
| 7 | 102029616 | AX-170157263 | 0 | A | G | 0.456 | 2.15E-03 | 9.47E-02 | 2.67E+00 | 2.74E+00 | 9.88E-01 | no | Rnf121 | intron_variant | MODIFIER |
| 7 | 102035551 | AX-170261591 | 0 | T | C | 0.456 | 2.15E-03 | 9.47E-02 | 2.67E+00 | 2.74E+00 | 9.88E-01 | no | Rnf121 | intron_variant | MODIFIER |
| 7 | 102120031 | AX-170470448 | 0 | A | T | 0.456 | 2.15E-03 | 9.47E-02 | 2.67E+00 | 2.74E+00 | 9.88E-01 | no | Chrna10 | upstream_gene_variant | MODIFIER |
| 7 | 102121287 | AX-97797384 | 0 | G | A | 0.456 | 2.15E-03 | 9.47E-02 | 2.67E+00 | 2.74E+00 | 9.88E-01 | no | Chrna10 | upstream_gene_variant | MODIFIER |
| 7 | 102141075 | AX-170169539 | 0 | G | C | 0.456 | 2.15E-03 | 9.47E-02 | 2.67E+00 | 2.74E+00 | 9.88E-01 | no | Nup98 | intron_variant | MODIFIER |
| 7 | 102142368 | AX-97853165 | 0 | C | T | 0.456 | 2.15E-03 | 9.47E-02 | 2.67E+00 | 2.74E+00 | 9.88E-01 | no | Nup98 | intron_variant | MODIFIER |
| 7 | 102142844 | AX-170380826 | 0 | C | T | 0.456 | 2.15E-03 | 9.47E-02 | 2.67E+00 | 2.74E+00 | 9.88E-01 | no | Nup98 | intron_variant | MODIFIER |
| 7 | 102142901 | AX-170242058 | 0 | T | A | 0.456 | 2.15E-03 | 9.47E-02 | 2.67E+00 | 2.74E+00 | 9.88E-01 | no | Nup98 | intron_variant | MODIFIER |
| 7 | 102152386 | AX-168419901 | 0 | A | G | 0.456 | 2.15E-03 | 9.47E-02 | 2.67E+00 | 2.74E+00 | 9.88E-01 | no | Nup98 | synonymous_variant | LOW |
| 7 | 102177370 | AX-97806968 | 0 | T | G | 0.456 | 2.15E-03 | 9.47E-02 | 2.67E+00 | 2.74E+00 | 9.88E-01 | no | Nup98 | intron_variant | MODIFIER |
| 7 | 102226175 | AX-170301413 | 0 | A | G | 0.456 | 2.15E-03 | 9.47E-02 | 2.67E+00 | 2.74E+00 | 9.88E-01 | no | Pgap2 | intron_variant | MODIFIER |
| 7 | 102228162 | AX-170518160 | 0 | G | T | 0.456 | 2.15E-03 | 9.47E-02 | 2.67E+00 | 2.74E+00 | 9.88E-01 | no | Pgap2 | intron_variant | MODIFIER |
| 7 | 102233179 | AX-170338161 | 0 | C | T | 0.456 | 2.15E-03 | 9.47E-02 | 2.67E+00 | 2.74E+00 | 9.88E-01 | no | Pgap2 | intron_variant | MODIFIER |
| 7 | 102236653 | AX-97857538 | 0 | A | G | 0.456 | 2.15E-03 | 9.47E-02 | 2.67E+00 | 2.74E+00 | 9.88E-01 | no | Pgap2 | intron_variant | MODIFIER |
| 7 | 102335395 | AX-97830738 | 0 | C | T | 0.456 | 2.15E-03 | 9.47E-02 | 2.67E+00 | 2.74E+00 | 9.88E-01 | no | Stim1 | intron_variant | MODIFIER |
| 7 | 102384360 | AX-170029650 | 0 | C | T | 0.456 | 2.15E-03 | 9.47E-02 | 2.67E+00 | 2.74E+00 | 9.88E-01 | no | Stim1 | intron_variant | MODIFIER |
| 7 | 102434839 | AX-97845253 | 0 | T | C | 0.456 | 2.15E-03 | 9.47E-02 | 2.67E+00 | 2.74E+00 | 9.88E-01 | no | Stim1 | intron_variant | MODIFIER |
| 7 | 102441185 | AX-170029655 | 0 | C | T | 0.456 | 2.15E-03 | 9.47E-02 | 2.67E+00 | 2.74E+00 | 9.88E-01 | no | Rrm1 | upstream_gene_variant | MODIFIER |
| 7 | 102455463 | AX-170480060 | 0 | G | T | 0.456 | 2.15E-03 | 9.47E-02 | 2.67E+00 | 2.74E+00 | 9.88E-01 | no | Rrm1 | intron_variant | MODIFIER |
| 7 | 102456608 | AX-170029649 | 0 | T | C | 0.456 | 2.15E-03 | 9.47E-02 | 2.67E+00 | 2.74E+00 | 9.88E-01 | no | Rrm1 | intron_variant | MODIFIER |
| 7 | 102479460 | AX-170319886 | 0 | A | G | 0.456 | 2.15E-03 | 9.47E-02 | 2.67E+00 | 2.74E+00 | 9.88E-01 | no | Olfr544 | downstream_gene_variant | MODIFIER |
| 7 | 102493485 | AX-170218166 | 0 | G | C | 0.456 | 2.15E-03 | 9.47E-02 | 2.67E+00 | 2.74E+00 | 9.88E-01 | no | Olfr545 | downstream_gene_variant | MODIFIER |
| 7 | 102535510 | AX-170477247 | 0 | C | T | 0.456 | 2.15E-03 | 9.47E-02 | 2.67E+00 | 2.74E+00 | 9.88E-01 | no | Olfr547 | synonymous_variant | LOW |
| 7 | 102535834 | AX-97803481 | 0 | C | T | 0.456 | 2.15E-03 | 9.47E-02 | 2.67E+00 | 2.74E+00 | 9.88E-01 | no | Olfr547 | downstream_gene_variant | MODIFIER |
| 7 | 102560975 | AX-170240692 | 0 | T | C | 0.456 | 2.15E-03 | 9.47E-02 | 2.67E+00 | 2.74E+00 | 9.88E-01 | no | Trim21 | intron_variant | MODIFIER |
| 7 | 102573731 | AX-170407468 | 0 | T | C | 0.456 | 2.15E-03 | 9.47E-02 | 2.67E+00 | 2.74E+00 | 9.88E-01 | no | Olfr550 | upstream_gene_variant | MODIFIER |
| 7 | 102581418 | AX-170409341 | 0 | A | T | 0.456 | 2.15E-03 | 9.47E-02 | 2.67E+00 | 2.74E+00 | 9.88E-01 | no | Olfr550 | downstream_gene_variant | MODIFIER |
| 7 | 102588640 | AX-97827617 | 0 | A | G | 0.456 | 2.15E-03 | 9.47E-02 | 2.67E+00 | 2.74E+00 | 9.88E-01 | no | Olfr551 | synonymous_variant | LOW |

| | | | | | | | | | | | | | | | |
|---|---|---|---|---|---|---|---|---|---|---|---|---|---|---|---|
| 7 | 102595839 | AX-170388166 | 0 | G | T | 0.456 | 2.15E-03 | 9.47E-02 | 2.67E+00 | 2.74E+00 | 9.88E-01 | no | Olfr551 | upstream_gene_variant | MODIFIER |
| 7 | 102598504 | AX-170302539 | 0 | A | G | 0.456 | 2.15E-03 | 9.47E-02 | 2.67E+00 | 2.74E+00 | 9.88E-01 | no | Olfr552 | intron_variant | MODIFIER |
| 7 | 102695415 | AX-170096050 | 0 | G | T | 0.456 | 2.15E-03 | 9.47E-02 | 2.67E+00 | 2.74E+00 | 9.88E-01 | no | Olfr557 | upstream_gene_variant | MODIFIER |
| 7 | 102737690 | AX-97841677 | 0 | A | C | 0.456 | 2.15E-03 | 9.47E-02 | 2.67E+00 | 2.74E+00 | 9.88E-01 | no | Olfr78 | downstream_gene_variant | MODIFIER |
| 7 | 102754527 | AX-170089992 | 0 | A | G | 0.456 | 2.15E-03 | 9.47E-02 | 2.67E+00 | 2.74E+00 | 9.88E-01 | no | Olfr78 | intron_variant | MODIFIER |
| 7 | 102848936 | AX-170158502 | 0 | T | G | 0.456 | 2.15E-03 | 9.47E-02 | 2.67E+00 | 2.74E+00 | 9.88E-01 | no | | intergenic_variant | MODIFIER |
| 7 | 102900506 | AX-170029658 | 0 | T | C | 0.456 | 2.15E-03 | 9.47E-02 | 2.67E+00 | 2.74E+00 | 9.88E-01 | no | Olfr570 | synonymous_variant | LOW |
| 7 | 102909585 | AX-170078284 | 0 | G | A | 0.456 | 2.15E-03 | 9.47E-02 | 2.67E+00 | 2.74E+00 | 9.88E-01 | no | Olfr571 | synonymous_variant | LOW |
| 7 | 102909705 | AX-97824744 | 0 | A | G | 0.456 | 2.15E-03 | 9.47E-02 | 2.67E+00 | 2.74E+00 | 9.88E-01 | no | Olfr571 | missense_variant | MODERATE |
| 7 | 102913452 | AX-170029662 | 0 | A | G | 0.456 | 2.15E-03 | 9.47E-02 | 2.67E+00 | 2.74E+00 | 9.88E-01 | no | Olfr571 | upstream_gene_variant | MODIFIER |
| 7 | 102915660 | AX-168396267 | 0 | C | T | 0.456 | 2.15E-03 | 9.47E-02 | 2.67E+00 | 2.74E+00 | 9.88E-01 | no | Olfr571 | intron_variant | MODIFIER |
| 7 | 102941674 | AX-170516203 | 0 | T | C | 0.456 | 2.15E-03 | 9.47E-02 | 2.67E+00 | 2.74E+00 | 9.88E-01 | no | Olfr573-ps1 | downstream_gene_variant | MODIFIER |
| 7 | 102942231 | AX-170474680 | 0 | A | G | 0.456 | 2.15E-03 | 9.47E-02 | 2.67E+00 | 2.74E+00 | 9.88E-01 | no | Olfr573-ps1 | synonymous_variant | LOW |
| 7 | 102949452 | AX-170262577 | 0 | T | G | 0.456 | 2.15E-03 | 9.47E-02 | 2.67E+00 | 2.74E+00 | 9.88E-01 | no | Olfr574 | missense_variant | MODERATE |
| 7 | 102949492 | AX-170179062 | 0 | G | A | 0.456 | 2.15E-03 | 9.47E-02 | 2.67E+00 | 2.74E+00 | 9.88E-01 | no | Olfr574 | synonymous_variant | LOW |
| 7 | 103347156 | AX-170343883 | 0 | G | C | 0.456 | 2.15E-03 | 9.47E-02 | 2.67E+00 | 2.74E+00 | 9.88E-01 | no | Olfr600 | upstream_gene_variant | MODIFIER |
| 7 | 103468432 | AX-170029666 | 0 | A | G | 0.456 | 2.15E-03 | 9.47E-02 | 2.67E+00 | 2.74E+00 | 9.88E-01 | no | Olfr608 | upstream_gene_variant | MODIFIER |
| 7 | 103504391 | AX-170029667 | 0 | C | T | 0.456 | 2.15E-03 | 9.47E-02 | 2.67E+00 | 2.74E+00 | 9.88E-01 | no | Olfr610 | downstream_gene_variant | MODIFIER |
| 7 | 103506639 | AX-97822208 | 0 | C | T | 0.456 | 2.15E-03 | 9.47E-02 | 2.67E+00 | 2.74E+00 | 9.88E-01 | no | Olfr610 | synonymous_variant | LOW |
| 7 | 103554214 | AX-170029673 | 0 | T | A | 0.456 | 2.15E-03 | 9.47E-02 | 2.67E+00 | 2.74E+00 | 9.88E-01 | no | Olfr613 | 3_prime_UTR_variant | MODIFIER |
| 7 | 103567205 | AX-97796916 | 0 | A | G | 0.456 | 2.15E-03 | 9.47E-02 | 2.67E+00 | 2.74E+00 | 9.88E-01 | no | Olfr616 | upstream_gene_variant | MODIFIER |
| 7 | 103713859 | AX-97786074 | 0 | G | A | 0.456 | 2.15E-03 | 9.47E-02 | 2.67E+00 | 2.74E+00 | 9.88E-01 | no | Olfr243 | upstream_gene_variant | MODIFIER |
| 7 | 103766911 | AX-170096053 | 0 | G | T | 0.456 | 2.15E-03 | 9.47E-02 | 2.67E+00 | 2.74E+00 | 9.88E-01 | no | Olfr69 | 3_prime_UTR_variant | MODIFIER |
| 7 | 103768406 | AX-170165062 | 0 | C | T | 0.456 | 2.15E-03 | 9.47E-02 | 2.67E+00 | 2.74E+00 | 9.88E-01 | no | Olfr69 | 5_prime_UTR_variant | MODIFIER |
| 7 | 103811262 | AX-97799905 | 0 | G | C | 0.456 | 2.15E-03 | 9.47E-02 | 2.67E+00 | 2.74E+00 | 9.88E-01 | no | Hbb-bt | downstream_gene_variant | MODIFIER |
| 7 | 103828588 | AX-170488857 | 0 | G | A | 0.456 | 2.15E-03 | 9.47E-02 | 2.67E+00 | 2.74E+00 | 9.88E-01 | no | Hbb-bs | upstream_gene_variant | MODIFIER |
| 7 | 103841449 | AX-97815657 | 0 | G | T | 0.456 | 2.15E-03 | 9.47E-02 | 2.67E+00 | 2.74E+00 | 9.88E-01 | no | Hbb-bh1 | downstream_gene_variant | MODIFIER |
| 7 | 103866840 | AX-97781388 | 0 | T | G | 0.456 | 2.15E-03 | 9.47E-02 | 2.67E+00 | 2.74E+00 | 9.88E-01 | no | | regulatory_region_variant | MODIFIER |
| 7 | 103867162 | AX-170401669 | 0 | A | G | 0.456 | 2.15E-03 | 9.47E-02 | 2.67E+00 | 2.74E+00 | 9.88E-01 | no | | regulatory_region_variant | MODIFIER |
| 7 | 103885020 | AX-97796095 | 0 | G | A | 0.456 | 2.15E-03 | 9.47E-02 | 2.67E+00 | 2.74E+00 | 9.88E-01 | no | Olfr66 | upstream_gene_variant | MODIFIER |
| 7 | 103893539 | AX-170372460 | 0 | T | C | 0.456 | 2.15E-03 | 9.47E-02 | 2.67E+00 | 2.74E+00 | 9.88E-01 | no | Olfr64 | synonymous_variant | LOW |
| 7 | 103895820 | AX-170445708 | 0 | G | T | 0.456 | 2.15E-03 | 9.47E-02 | 2.67E+00 | 2.74E+00 | 9.88E-01 | no | Olfr64 | upstream_gene_variant | MODIFIER |
| 7 | 103904393 | AX-170264549 | 0 | A | G | 0.456 | 2.15E-03 | 9.47E-02 | 2.67E+00 | 2.74E+00 | 9.88E-01 | no | Olfr65 | 5_prime_UTR_variant | MODIFIER |
| 7 | 103904436 | AX-170387710 | 0 | G | C | 0.456 | 2.15E-03 | 9.47E-02 | 2.67E+00 | 2.74E+00 | 9.88E-01 | no | Olfr65 | intron_variant | MODIFIER |
| 7 | 103915789 | AX-170426406 | 0 | T | G | 0.456 | 2.15E-03 | 9.47E-02 | 2.67E+00 | 2.74E+00 | 9.88E-01 | no | | regulatory_region_variant | MODIFIER |
| 7 | 103919951 | AX-170407858 | 0 | T | C | 0.456 | 2.15E-03 | 9.47E-02 | 2.67E+00 | 2.74E+00 | 9.88E-01 | no | | intergenic_variant | MODIFIER |
| 7 | 104362662 | AX-97820354 | 0 | T | C | 0.456 | 2.15E-03 | 9.47E-02 | 2.67E+00 | 2.74E+00 | 9.88E-01 | no | Trim30b | intron_variant | MODIFIER |
| 7 | 104387147 | AX-170213861 | 0 | C | A | 0.456 | 2.15E-03 | 9.47E-02 | 2.67E+00 | 2.74E+00 | 9.88E-01 | no | Trim30c | intron_variant | MODIFIER |
| 7 | 104400779 | AX-97800774 | 0 | T | C | 0.456 | 2.15E-03 | 9.47E-02 | 2.67E+00 | 2.74E+00 | 9.88E-01 | no | Trim30c | 5_prime_UTR_variant | MODIFIER |
| 7 | 104408283 | AX-170265226 | 0 | G | C | 0.456 | 2.15E-03 | 9.47E-02 | 2.67E+00 | 2.74E+00 | 9.88E-01 | no | Trim30a | downstream_gene_variant | MODIFIER |
| 7 | 104487473 | AX-97791696 | 0 | G | A | 0.456 | 2.15E-03 | 9.47E-02 | 2.67E+00 | 2.74E+00 | 9.88E-01 | no | Trim30d | intron_variant | MODIFIER |
| 7 | 104550201 | AX-170129764 | 0 | G | A | 0.456 | 2.15E-03 | 9.47E-02 | 2.67E+00 | 2.74E+00 | 9.88E-01 | no | Olfr651 | upstream_gene_variant | MODIFIER |
| 7 | 104560818 | AX-170096054 | 0 | A | G | 0.456 | 2.15E-03 | 9.47E-02 | 2.67E+00 | 2.74E+00 | 9.88E-01 | no | Olfr652 | upstream_gene_variant | MODIFIER |
| 7 | 104571532 | AX-170221134 | 0 | G | T | 0.456 | 2.15E-03 | 9.47E-02 | 2.67E+00 | 2.74E+00 | 9.88E-01 | no | Olfr652 | downstream_gene_variant | MODIFIER |
| 7 | 104584688 | AX-170232413 | 0 | A | G | 0.456 | 2.15E-03 | 9.47E-02 | 2.67E+00 | 2.74E+00 | 9.88E-01 | no | Olfr654 | upstream_gene_variant | MODIFIER |
| 7 | 104585687 | AX-170029693 | 0 | A | G | 0.456 | 2.15E-03 | 9.47E-02 | 2.67E+00 | 2.74E+00 | 9.88E-01 | no | Olfr654 | upstream_gene_variant | MODIFIER |
| 7 | 104618852 | AX-170089993 | 0 | C | T | 0.456 | 2.15E-03 | 9.47E-02 | 2.67E+00 | 2.74E+00 | 9.88E-01 | no | Olfr656 | downstream_gene_variant | MODIFIER |
| 7 | 105438136 | AX-168419916 | 0 | T | C | 0.456 | 2.15E-03 | 9.47E-02 | 2.67E+00 | 2.74E+00 | 9.88E-01 | no | Cckbr | downstream_gene_variant | MODIFIER |
| 7 | 105612828 | AX-97830775 | 0 | A | G | 0.456 | 2.15E-03 | 9.47E-02 | 2.67E+00 | 2.74E+00 | 9.88E-01 | no | Trim3 | intron_variant | MODIFIER |
| 7 | 105615949 | AX-170172012 | 0 | C | G | 0.456 | 2.15E-03 | 9.47E-02 | 2.67E+00 | 2.74E+00 | 9.88E-01 | no | Trim3 | intron_variant | MODIFIER |
| 7 | 105622423 | AX-170296029 | 0 | T | C | 0.456 | 2.15E-03 | 9.47E-02 | 2.67E+00 | 2.74E+00 | 9.88E-01 | no | Trim3 | intron_variant | MODIFIER |
| 7 | 107030769 | AX-170349325 | 0 | G | T | 0.456 | 2.15E-03 | 9.47E-02 | 2.67E+00 | 2.74E+00 | 9.88E-01 | no | Olfr713 | upstream_gene_variant | MODIFIER |
| 7 | 107035824 | AX-170202509 | 0 | C | A | 0.456 | 2.15E-03 | 9.47E-02 | 2.67E+00 | 2.74E+00 | 9.88E-01 | no | Olfr713 | upstream_gene_variant | MODIFIER |
| 7 | 107067808 | AX-97821983 | 0 | T | G | 0.456 | 2.15E-03 | 9.47E-02 | 2.67E+00 | 2.74E+00 | 9.88E-01 | no | Gm17991 | upstream_gene_variant | MODIFIER |

| | | | | | | | | | | | | | | | |
|---|---|---|---|---|---|---|---|---|---|---|---|---|---|---|---|
| 7 | 107076436 | AX-170089999 | 0 | C | T | 0.456 | 2.15E-03 | 9.47E-02 | 2.67E+00 | 2.74E+00 | 9.88E-01 | no | Olfr714 | downstream_gene_variant | MODIFIER |
| 7 | 107096735 | AX-170310924 | 0 | A | G | 0.456 | 2.15E-03 | 9.47E-02 | 2.67E+00 | 2.74E+00 | 9.88E-01 | no | Olfr17 | upstream_gene_variant | MODIFIER |
| 7 | 107099808 | AX-170078348 | 0 | G | C | 0.456 | 2.15E-03 | 9.47E-02 | 2.67E+00 | 2.74E+00 | 9.88E-01 | no | Olfr17 | downstream_gene_variant | MODIFIER |
| 7 | 107100133 | AX-170511116 | 0 | G | A | 0.456 | 2.15E-03 | 9.47E-02 | 2.67E+00 | 2.74E+00 | 9.88E-01 | no | Olfr17 | downstream_gene_variant | MODIFIER |
| 7 | 107100687 | AX-170134118 | 0 | C | T | 0.456 | 2.15E-03 | 9.47E-02 | 2.67E+00 | 2.74E+00 | 9.88E-01 | no | Olfr17 | downstream_gene_variant | MODIFIER |
| 7 | 107100725 | AX-97824871 | 0 | T | C | 0.456 | 2.15E-03 | 9.47E-02 | 2.67E+00 | 2.74E+00 | 9.88E-01 | no | Olfr17 | downstream_gene_variant | MODIFIER |
| 7 | 107183371 | AX-170168397 | 0 | A | T | 0.456 | 2.15E-03 | 9.47E-02 | 2.67E+00 | 2.74E+00 | 9.88E-01 | no | Nlrp14 | intron_variant | MODIFIER |
| 7 | 107211007 | AX-170096059 | 0 | G | A | 0.456 | 2.15E-03 | 9.47E-02 | 2.67E+00 | 2.74E+00 | 9.88E-01 | no | Rbmxl2 | downstream_gene_variant | MODIFIER |
| 7 | 107229236 | AX-170206747 | 0 | A | G | 0.456 | 2.15E-03 | 9.47E-02 | 2.67E+00 | 2.74E+00 | 9.88E-01 | no | | intergenic_variant | MODIFIER |
| 7 | 107229318 | AX-170533592 | 0 | C | T | 0.456 | 2.15E-03 | 9.47E-02 | 2.67E+00 | 2.74E+00 | 9.88E-01 | no | | intergenic_variant | MODIFIER |
| 7 | 107235532 | AX-97834179 | 0 | T | C | 0.456 | 2.15E-03 | 9.47E-02 | 2.67E+00 | 2.74E+00 | 9.88E-01 | no | | intergenic_variant | MODIFIER |
| 7 | 107235952 | AX-170316397 | 0 | G | A | 0.456 | 2.15E-03 | 9.47E-02 | 2.67E+00 | 2.74E+00 | 9.88E-01 | no | | intergenic_variant | MODIFIER |
| 7 | 107258340 | AX-170524471 | 0 | G | A | 0.456 | 2.15E-03 | 9.47E-02 | 2.67E+00 | 2.74E+00 | 9.88E-01 | no | Platr28 | intron_variant,non_coding_transcript_variant | MODIFIER |
| 7 | 107263632 | AX-170390982 | 0 | C | A | 0.456 | 2.15E-03 | 9.47E-02 | 2.67E+00 | 2.74E+00 | 9.88E-01 | no | Platr28 | intron_variant,non_coding_transcript_variant | MODIFIER |
| 7 | 107273270 | AX-170443385 | 0 | A | C | 0.456 | 2.15E-03 | 9.47E-02 | 2.67E+00 | 2.74E+00 | 9.88E-01 | no | Platr28 | intron_variant,non_coding_transcript_variant | MODIFIER |
| 7 | 107273305 | AX-170260193 | 0 | T | C | 0.456 | 2.15E-03 | 9.47E-02 | 2.67E+00 | 2.74E+00 | 9.88E-01 | no | Platr28 | intron_variant,non_coding_transcript_variant | MODIFIER |
| 7 | 107275003 | AX-170108415 | 0 | C | T | 0.456 | 2.15E-03 | 9.47E-02 | 2.67E+00 | 2.74E+00 | 9.88E-01 | no | Platr28 | intron_variant,non_coding_transcript_variant | MODIFIER |
| 7 | 107294979 | AX-170335080 | 0 | A | C | 0.456 | 2.15E-03 | 9.47E-02 | 2.67E+00 | 2.74E+00 | 9.88E-01 | no | Platr28 | intron_variant,non_coding_transcript_variant | MODIFIER |
| 7 | 107298856 | AX-170035764 | 0 | C | T | 0.456 | 2.15E-03 | 9.47E-02 | 2.67E+00 | 2.74E+00 | 9.88E-01 | no | Platr28 | intron_variant,non_coding_transcript_variant | MODIFIER |
| 7 | 107300747 | AX-170178653 | 0 | C | T | 0.456 | 2.15E-03 | 9.47E-02 | 2.67E+00 | 2.74E+00 | 9.88E-01 | no | Platr28 | downstream_gene_variant | MODIFIER |
| 7 | 107300818 | AX-170328648 | 0 | T | C | 0.456 | 2.15E-03 | 9.47E-02 | 2.67E+00 | 2.74E+00 | 9.88E-01 | no | Platr28 | downstream_gene_variant | MODIFIER |
| 7 | 107312889 | AX-170232567 | 0 | G | A | 0.456 | 2.15E-03 | 9.47E-02 | 2.67E+00 | 2.74E+00 | 9.88E-01 | no | | intergenic_variant | MODIFIER |
| 7 | 107313081 | AX-170122610 | 0 | A | G | 0.456 | 2.15E-03 | 9.47E-02 | 2.67E+00 | 2.74E+00 | 9.88E-01 | no | | intergenic_variant | MODIFIER |
| 7 | 107313542 | AX-170214183 | 0 | G | A | 0.456 | 2.15E-03 | 9.47E-02 | 2.67E+00 | 2.74E+00 | 9.88E-01 | no | | intergenic_variant | MODIFIER |
| 7 | 107316156 | AX-170299886 | 0 | C | A | 0.456 | 2.15E-03 | 9.47E-02 | 2.67E+00 | 2.74E+00 | 9.88E-01 | no | Gm24932 | downstream_gene_variant | MODIFIER |
| 7 | 107323030 | AX-170029725 | 0 | C | A | 0.456 | 2.15E-03 | 9.47E-02 | 2.67E+00 | 2.74E+00 | 9.88E-01 | no | Gm24932 | upstream_gene_variant | MODIFIER |
| 7 | 107335375 | AX-170029731 | 0 | A | G | 0.456 | 2.15E-03 | 9.47E-02 | 2.67E+00 | 2.74E+00 | 9.88E-01 | no | | intergenic_variant | MODIFIER |
| 7 | 109127274 | AX-170526904 | 0 | T | C | 0.456 | 2.15E-03 | 9.47E-02 | 2.67E+00 | 2.74E+00 | 9.88E-01 | no | | intergenic_variant | MODIFIER |
| 7 | 109143273 | AX-97796252 | 0 | T | C | 0.456 | 2.15E-03 | 9.47E-02 | 2.67E+00 | 2.74E+00 | 9.88E-01 | no | Lmo1 | intron_variant | MODIFIER |
| 7 | 109154544 | AX-170105455 | 0 | T | C | 0.456 | 2.15E-03 | 9.47E-02 | 2.67E+00 | 2.74E+00 | 9.88E-01 | no | Lmo1 | intron_variant | MODIFIER |
| 7 | 109164266 | AX-97805186 | 0 | C | T | 0.456 | 2.15E-03 | 9.47E-02 | 2.67E+00 | 2.74E+00 | 9.88E-01 | no | Lmo1 | intron_variant | MODIFIER |
| 7 | 109188637 | AX-97848958 | 0 | C | T | 0.456 | 2.15E-03 | 9.47E-02 | 2.67E+00 | 2.74E+00 | 9.88E-01 | no | | intergenic_variant | MODIFIER |
| 7 | 109188887 | AX-170370449 | 0 | C | T | 0.456 | 2.15E-03 | 9.47E-02 | 2.67E+00 | 2.74E+00 | 9.88E-01 | no | | intergenic_variant | MODIFIER |
| 7 | 109189345 | AX-170172095 | 0 | A | G | 0.456 | 2.15E-03 | 9.47E-02 | 2.67E+00 | 2.74E+00 | 9.88E-01 | no | | intergenic_variant | MODIFIER |
| 7 | 109198572 | AX-170029753 | 0 | A | G | 0.456 | 2.15E-03 | 9.47E-02 | 2.67E+00 | 2.74E+00 | 9.88E-01 | no | | intergenic_variant | MODIFIER |
| 7 | 109207382 | AX-170039591 | 0 | G | A | 0.456 | 2.15E-03 | 9.47E-02 | 2.67E+00 | 2.74E+00 | 9.88E-01 | no | Gm45024 | downstream_gene_variant | MODIFIER |
| 7 | 109222875 | AX-170029748 | 0 | T | A | 0.456 | 2.15E-03 | 9.47E-02 | 2.67E+00 | 2.74E+00 | 9.88E-01 | no | Gm45024 | intron_variant,non_coding_transcript_variant | MODIFIER |
| 7 | 109223019 | AX-97824814 | 0 | C | T | 0.456 | 2.15E-03 | 9.47E-02 | 2.67E+00 | 2.74E+00 | 9.88E-01 | no | Gm45024 | intron_variant,non_coding_transcript_variant | MODIFIER |
| 7 | 109223198 | AX-170029754 | 0 | G | A | 0.456 | 2.15E-03 | 9.47E-02 | 2.67E+00 | 2.74E+00 | 9.88E-01 | no | Gm45024 | intron_variant,non_coding_transcript_variant | MODIFIER |
| 1 | 48391020 | AX-97790747 | 1 | C | G | 0.107 | 3.88E-03 | 2.40E-01 | 2.70E+00 | 2.74E+00 | 9.88E-01 | no | | intergenic_variant | MODIFIER |
| 1 | 53799087 | AX-170511232 | 1 | C | T | 0.107 | 3.88E-03 | 2.40E-01 | 2.70E+00 | 2.74E+00 | 9.88E-01 | no | | intergenic_variant | MODIFIER |
| 1 | 54435788 | AX-97811606 | 1 | G | A | 0.107 | 3.88E-03 | 2.40E-01 | 2.70E+00 | 2.74E+00 | 9.88E-01 | no | Gtf3c3 | intron_variant | MODIFIER |
| 1 | 54666372 | AX-170529892 | 1 | G | A | 0.107 | 3.88E-03 | 2.40E-01 | 2.70E+00 | 2.74E+00 | 9.88E-01 | no | Ankrd44 | intron_variant | MODIFIER |
| 2 | 121453976 | AX-170018872 | 0 | G | A | 0.132 | -2.55E-03 | 1.92E-01 | 2.66E+00 | 2.74E+00 | 9.88E-01 | no | Hypk | upstream_gene_variant | MODIFIER |
| 15 | 90939620 | AX-170431882 | 1 | G | A | 0.125 | -4.37E-04 | 1.26E-01 | 2.63E+00 | 2.74E+00 | 9.88E-01 | no | Kif21a | intron_variant | MODIFIER |
| 1 | 48129420 | AX-170451576 | 1 | C | T | 0.098 | -2.72E-03 | 2.09E-01 | 2.65E+00 | 2.74E+00 | 9.88E-01 | no | | intergenic_variant | MODIFIER |
| 8 | 90861965 | AX-170381470 | 0 | G | A | 0.202 | 8.66E-04 | 1.15E-01 | 2.66E+00 | 2.74E+00 | 9.88E-01 | no | Chd9 | intron_variant | MODIFIER |
| 7 | 93928158 | AX-97848452 | 1 | T | C | 0.116 | 3.82E-03 | 2.13E-01 | 2.66E+00 | 2.74E+00 | 9.88E-01 | no | | intergenic_variant | MODIFIER |
| 12 | 105548905 | AX-170007457 | 0 | C | T | 0.132 | -1.11E-03 | 1.56E-01 | 2.62E+00 | 2.74E+00 | 9.88E-01 | no | | regulatory_region_variant | MODIFIER |
| 5 | 108708808 | AX-168379929 | 0 | T | C | 0.447 | 2.13E-03 | 1.16E-01 | 2.70E+00 | 2.74E+00 | 9.88E-01 | no | Fgfrl1 | downstream_gene_variant | MODIFIER |
| 1 | 29363667 | AX-170000193 | 0 | C | A | 0.404 | 1.35E-03 | 8.28E-02 | 2.66E+00 | 2.74E+00 | 9.88E-01 | no | Gm23785 | upstream_gene_variant | MODIFIER |
| 1 | 33058410 | AX-170497552 | 0 | G | A | 0.404 | 1.35E-03 | 8.28E-02 | 2.66E+00 | 2.74E+00 | 9.88E-01 | no | | intergenic_variant | MODIFIER |
| 4 | 63556202 | AX-170228474 | 1 | C | T | 0.348 | -1.27E-03 | 1.05E-01 | 2.65E+00 | 2.74E+00 | 9.88E-01 | no | Tmem268 | upstream_gene_variant | MODIFIER |
| 3 | 144675001 | AX-170068336 | 1 | A | G | 0.366 | -7.02E-04 | 7.05E-02 | 2.66E+00 | 2.74E+00 | 9.88E-01 | no | | intergenic_variant | MODIFIER |

| | | | | | | | | | | | | | | | |
|---|---|---|---|---|---|---|---|---|---|---|---|---|---|---|---|
| 8 | 57457630 | AX-170498810 | 0 | G | A | 0.158 | 1.26E-03 | 1.46E-01 | 2.69E+00 | 2.74E+00 | 9.88E-01 | no | Scrg1 | intron_variant | MODIFIER |
| 12 | 100581996 | AX-170007385 | 0 | T | G | 0.07 | 2.71E-03 | 1.67E-01 | 2.65E+00 | 2.74E+00 | 9.88E-01 | no | Rps6ka5 | intron_variant | MODIFIER |
| 12 | 101780082 | AX-97844075 | 0 | T | C | 0.07 | 2.71E-03 | 1.67E-01 | 2.65E+00 | 2.74E+00 | 9.88E-01 | no | Fbln5 | intron_variant | MODIFIER |
| 12 | 101781706 | AX-168394723 | 0 | G | T | 0.07 | 2.71E-03 | 1.67E-01 | 2.65E+00 | 2.74E+00 | 9.88E-01 | no | Fbln5 | intron_variant | MODIFIER |
| 12 | 101785492 | AX-170508969 | 0 | G | C | 0.07 | 2.71E-03 | 1.67E-01 | 2.65E+00 | 2.74E+00 | 9.88E-01 | no | Fbln5 | intron_variant | MODIFIER |
| 12 | 101846828 | AX-170356377 | 0 | A | G | 0.07 | 2.71E-03 | 1.67E-01 | 2.65E+00 | 2.74E+00 | 9.88E-01 | no | Trip11 | intron_variant | MODIFIER |
| 12 | 101859457 | AX-168397379 | 0 | G | A | 0.07 | 2.71E-03 | 1.67E-01 | 2.65E+00 | 2.74E+00 | 9.88E-01 | no | Trip11 | intron_variant | MODIFIER |
| 12 | 101860090 | AX-170419721 | 0 | G | A | 0.07 | 2.71E-03 | 1.67E-01 | 2.65E+00 | 2.74E+00 | 9.88E-01 | no | Trip11 | intron_variant | MODIFIER |
| 12 | 101884363 | AX-170110363 | 0 | G | A | 0.07 | 2.71E-03 | 1.67E-01 | 2.65E+00 | 2.74E+00 | 9.88E-01 | no | Trip11 | synonymous_variant | LOW |
| 12 | 101893404 | AX-97818323 | 0 | T | C | 0.07 | 2.71E-03 | 1.67E-01 | 2.65E+00 | 2.74E+00 | 9.88E-01 | no | Trip11 | intron_variant | MODIFIER |
| 12 | 101905459 | AX-170120452 | 0 | A | G | 0.07 | 2.71E-03 | 1.67E-01 | 2.65E+00 | 2.74E+00 | 9.88E-01 | no | Trip11 | intron_variant | MODIFIER |
| 12 | 101922315 | AX-170417162 | 0 | A | T | 0.07 | 2.71E-03 | 1.67E-01 | 2.65E+00 | 2.74E+00 | 9.88E-01 | no | Atxn3 | 3_prime_UTR_variant | MODIFIER |
| 12 | 101924472 | AX-170490639 | 0 | G | A | 0.07 | 2.71E-03 | 1.67E-01 | 2.65E+00 | 2.74E+00 | 9.88E-01 | no | Atxn3 | intron_variant | MODIFIER |
| 12 | 101930596 | AX-170437236 | 0 | T | C | 0.07 | 2.71E-03 | 1.67E-01 | 2.65E+00 | 2.74E+00 | 9.88E-01 | no | Atxn3 | intron_variant | MODIFIER |
| 12 | 101940829 | AX-170226310 | 0 | T | G | 0.07 | 2.71E-03 | 1.67E-01 | 2.65E+00 | 2.74E+00 | 9.88E-01 | no | Atxn3 | intron_variant | MODIFIER |
| 12 | 102170077 | AX-170231655 | 0 | A | G | 0.07 | 2.71E-03 | 1.67E-01 | 2.65E+00 | 2.74E+00 | 9.88E-01 | no | Slc24a4 | intron_variant | MODIFIER |
| 12 | 102170342 | AX-170309952 | 0 | T | C | 0.07 | 2.71E-03 | 1.67E-01 | 2.65E+00 | 2.74E+00 | 9.88E-01 | no | Slc24a4 | intron_variant | MODIFIER |
| 12 | 102170738 | AX-170390534 | 0 | A | G | 0.07 | 2.71E-03 | 1.67E-01 | 2.65E+00 | 2.74E+00 | 9.88E-01 | no | Slc24a4 | intron_variant | MODIFIER |
| 12 | 102171376 | AX-97826702 | 0 | C | T | 0.07 | 2.71E-03 | 1.67E-01 | 2.65E+00 | 2.74E+00 | 9.88E-01 | no | Slc24a4 | intron_variant | MODIFIER |
| 12 | 102174991 | AX-170531614 | 0 | C | T | 0.07 | 2.71E-03 | 1.67E-01 | 2.65E+00 | 2.74E+00 | 9.88E-01 | no | Slc24a4 | intron_variant | MODIFIER |
| 12 | 102182441 | AX-170145089 | 0 | G | A | 0.07 | 2.71E-03 | 1.67E-01 | 2.65E+00 | 2.74E+00 | 9.88E-01 | no | Slc24a4 | intron_variant | MODIFIER |
| 12 | 102193816 | AX-170007415 | 0 | G | A | 0.07 | 2.71E-03 | 1.67E-01 | 2.65E+00 | 2.74E+00 | 9.88E-01 | no | Slc24a4 | intron_variant | MODIFIER |
| 12 | 102199428 | AX-170480257 | 0 | A | G | 0.07 | 2.71E-03 | 1.67E-01 | 2.65E+00 | 2.74E+00 | 9.88E-01 | no | Slc24a4 | intron_variant | MODIFIER |
| 12 | 102199492 | AX-170092595 | 0 | A | G | 0.07 | 2.71E-03 | 1.67E-01 | 2.65E+00 | 2.74E+00 | 9.88E-01 | no | Slc24a4 | intron_variant | MODIFIER |
| 12 | 102201122 | AX-170007413 | 0 | C | G | 0.07 | 2.71E-03 | 1.67E-01 | 2.65E+00 | 2.74E+00 | 9.88E-01 | no | Slc24a4 | intron_variant | MODIFIER |
| 12 | 102201363 | AX-170007408 | 0 | A | G | 0.07 | 2.71E-03 | 1.67E-01 | 2.65E+00 | 2.74E+00 | 9.88E-01 | no | Slc24a4 | intron_variant | MODIFIER |
| 12 | 102203963 | AX-170275290 | 0 | G | A | 0.07 | 2.71E-03 | 1.67E-01 | 2.65E+00 | 2.74E+00 | 9.88E-01 | no | Slc24a4 | intron_variant | MODIFIER |
| 12 | 102204214 | AX-170289413 | 0 | T | C | 0.07 | 2.71E-03 | 1.67E-01 | 2.65E+00 | 2.74E+00 | 9.88E-01 | no | Slc24a4 | intron_variant | MODIFIER |
| 12 | 102205180 | AX-170289785 | 0 | G | A | 0.07 | 2.71E-03 | 1.67E-01 | 2.65E+00 | 2.74E+00 | 9.88E-01 | no | Slc24a4 | intron_variant | MODIFIER |
| 12 | 102393261 | AX-97812951 | 0 | T | C | 0.07 | 2.71E-03 | 1.67E-01 | 2.65E+00 | 2.74E+00 | 9.88E-01 | no | Lgmn | downstream_gene_variant | MODIFIER |
| 12 | 102415807 | AX-170126377 | 0 | G | A | 0.07 | 2.71E-03 | 1.67E-01 | 2.65E+00 | 2.74E+00 | 9.88E-01 | no | Lgmn | intron_variant | MODIFIER |
| 12 | 102417914 | AX-170205326 | 0 | T | G | 0.07 | 2.71E-03 | 1.67E-01 | 2.65E+00 | 2.74E+00 | 9.88E-01 | no | Lgmn | intron_variant | MODIFIER |
| 12 | 102420293 | AX-170356363 | 0 | A | G | 0.07 | 2.71E-03 | 1.67E-01 | 2.65E+00 | 2.74E+00 | 9.88E-01 | no | Lgmn | intron_variant | MODIFIER |
| 12 | 102420410 | AX-97803518 | 0 | T | C | 0.07 | 2.71E-03 | 1.67E-01 | 2.65E+00 | 2.74E+00 | 9.88E-01 | no | Lgmn | intron_variant | MODIFIER |
| 12 | 102420571 | AX-170230728 | 0 | G | A | 0.07 | 2.71E-03 | 1.67E-01 | 2.65E+00 | 2.74E+00 | 9.88E-01 | no | Lgmn | intron_variant | MODIFIER |
| 12 | 102422230 | AX-170200341 | 0 | T | C | 0.07 | 2.71E-03 | 1.67E-01 | 2.65E+00 | 2.74E+00 | 9.88E-01 | no | Lgmn | intron_variant | MODIFIER |
| 9 | 110986921 | AX-170246411 | 1 | C | T | 0.107 | -2.63E-03 | 1.07E-01 | 2.65E+00 | 2.74E+00 | 9.89E-01 | no | Lrrc2 | downstream_gene_variant | MODIFIER |
| 5 | 89056277 | AX-170427181 | 2 | C | G | 0.436 | -1.56E-03 | 1.05E-01 | 2.68E+00 | 2.74E+00 | 9.89E-01 | no | Slc4a4 | intron_variant | MODIFIER |
| 2 | 130046935 | AX-170133845 | 0 | A | G | 0.149 | 4.70E-03 | 1.84E-01 | 2.65E+00 | 2.73E+00 | 9.89E-01 | no | Tgm3 | intron_variant | MODIFIER |
| 7 | 10125090 | AX-170411210 | 0 | A | G | 0.167 | -2.61E-05 | 7.88E-02 | 2.60E+00 | 2.74E+00 | 9.89E-01 | no | | intergenic_variant | MODIFIER |
| 16 | 44430516 | AX-170012935 | 0 | T | C | 0.474 | -2.23E-03 | 7.98E-02 | 2.67E+00 | 2.73E+00 | 9.89E-01 | no | Cfap44 | intron_variant | MODIFIER |
| 9 | 54668096 | AX-170166636 | 1 | G | C | 0.214 | -1.52E-03 | 1.18E-01 | 2.68E+00 | 2.74E+00 | 9.89E-01 | no | | intergenic_variant | MODIFIER |
| 8 | 26532039 | AX-170167556 | 1 | T | C | 0.17 | 1.05E-03 | 1.12E-01 | 2.66E+00 | 2.74E+00 | 9.89E-01 | no | | intergenic_variant | MODIFIER |
| 6 | 121524240 | AX-170384174 | 1 | G | A | 0.071 | 3.79E-03 | 2.77E-01 | 2.70E+00 | 2.74E+00 | 9.89E-01 | no | | regulatory_region_variant | MODIFIER |
| 3 | 120489018 | AX-97824371 | 0 | C | T | 0.044 | 1.69E-03 | 1.68E-01 | 2.62E+00 | 2.74E+00 | 9.89E-01 | no | | intergenic_variant | MODIFIER |
| 12 | 64350326 | AX-170199514 | 1 | G | A | 0.062 | 3.57E-03 | 3.01E-01 | 2.72E+00 | 2.74E+00 | 9.89E-01 | no | | intergenic_variant | MODIFIER |
| 18 | 58831091 | AX-170214260 | 1 | A | G | 0.411 | -2.28E-03 | 8.56E-02 | 2.66E+00 | 2.73E+00 | 9.89E-01 | no | | regulatory_region_variant | MODIFIER |
| 7 | 112374458 | AX-170356679 | 0 | C | A | 0.456 | 1.19E-03 | 9.39E-02 | 2.67E+00 | 2.74E+00 | 9.90E-01 | no | Mical2 | 5_prime_UTR_variant | MODIFIER |
| 8 | 75056533 | AX-170317278 | 2 | G | A | 0.209 | -1.80E-03 | 1.19E-01 | 2.65E+00 | 2.74E+00 | 9.90E-01 | no | Tom1 | intron_variant | MODIFIER |
| 11 | 46679488 | AX-170406262 | 0 | G | C | 0.096 | -2.34E-03 | 1.51E-01 | 2.60E+00 | 2.74E+00 | 9.90E-01 | no | Timd2 | intron_variant | MODIFIER |
| 8 | 93634646 | AX-170522280 | 0 | T | C | 0.211 | 1.28E-03 | 1.17E-01 | 2.67E+00 | 2.74E+00 | 9.90E-01 | no | | intergenic_variant | MODIFIER |
| 8 | 42264607 | AX-170079893 | 0 | G | C | 0.149 | -2.89E-03 | 1.39E-01 | 2.67E+00 | 2.74E+00 | 9.90E-01 | no | | intergenic_variant | MODIFIER |
| 9 | 60183851 | AX-170292795 | 1 | C | T | 0.205 | 1.50E-03 | 1.40E-01 | 2.72E+00 | 2.74E+00 | 9.90E-01 | no | Thsd4 | intron_variant | MODIFIER |
| 16 | 19493705 | AX-97818684 | 1 | C | T | 0.134 | -8.36E-04 | 8.85E-02 | 2.59E+00 | 2.74E+00 | 9.90E-01 | no | Olfr166 | downstream_gene_variant | MODIFIER |

| | | | | | | | | | | | | | | | |
|---|---|---|---|---|---|---|---|---|---|---|---|---|---|---|---|
| 1 | 52513755 | AX-170395814 | 1 | C | T | 0.134 | 3.14E-03 | 2.39E-01 | 2.69E+00 | 2.74E+00 | 9.90E-01 | no | | intergenic_variant | MODIFIER |
| 18 | 52186651 | AX-170168681 | 0 | T | C | 0.184 | -2.20E-03 | 9.45E-02 | 2.61E+00 | 2.74E+00 | 9.90E-01 | no | | intergenic_variant | MODIFIER |
| 5 | 101705142 | AX-97853961 | 0 | G | A | 0.412 | -1.32E-03 | 1.06E-01 | 2.67E+00 | 2.74E+00 | 9.90E-01 | no | Gm42133 | upstream_gene_variant | MODIFIER |
| 18 | 68469945 | AX-170241903 | 2 | T | C | 0.073 | 1.65E-03 | 1.44E-01 | 2.59E+00 | 2.74E+00 | 9.90E-01 | no | | intergenic_variant | MODIFIER |
| 12 | 56509556 | AX-97834045 | 0 | G | T | 0.053 | 2.50E-03 | 2.75E-01 | 2.68E+00 | 2.74E+00 | 9.90E-01 | no | Sfta3-ps | intron_variant,non_coding_transcript_variant | MODIFIER |
| 12 | 56509982 | AX-170299518 | 0 | A | G | 0.053 | 2.50E-03 | 2.75E-01 | 2.68E+00 | 2.74E+00 | 9.90E-01 | no | Sfta3-ps | intron_variant,non_coding_transcript_variant | MODIFIER |
| 12 | 56537361 | AX-170478202 | 0 | T | A | 0.053 | 2.50E-03 | 2.75E-01 | 2.68E+00 | 2.74E+00 | 9.90E-01 | no | Nkx2-1 | upstream_gene_variant | MODIFIER |
| 12 | 56542940 | AX-97821275 | 0 | T | C | 0.053 | 2.50E-03 | 2.75E-01 | 2.68E+00 | 2.74E+00 | 9.90E-01 | no | Gm26973 | intron_variant,non_coding_transcript_variant | MODIFIER |
| 12 | 56820272 | AX-170349748 | 0 | G | T | 0.053 | 2.50E-03 | 2.75E-01 | 2.68E+00 | 2.74E+00 | 9.90E-01 | no | Slc25a21 | intron_variant | MODIFIER |
| 12 | 56861066 | AX-170276941 | 0 | G | T | 0.053 | 2.50E-03 | 2.75E-01 | 2.68E+00 | 2.74E+00 | 9.90E-01 | no | Slc25a21 | intron_variant | MODIFIER |
| 12 | 56922763 | AX-170453709 | 0 | A | T | 0.053 | 2.50E-03 | 2.75E-01 | 2.68E+00 | 2.74E+00 | 9.90E-01 | no | Slc25a21 | intron_variant | MODIFIER |
| 12 | 56929936 | AX-170006806 | 0 | A | G | 0.053 | 2.50E-03 | 2.75E-01 | 2.68E+00 | 2.74E+00 | 9.90E-01 | no | Slc25a21 | intron_variant | MODIFIER |
| 12 | 61455701 | AX-170006864 | 0 | A | G | 0.053 | 2.50E-03 | 2.75E-01 | 2.68E+00 | 2.74E+00 | 9.90E-01 | no | Nanog-ps2 | upstream_gene_variant | MODIFIER |
| 12 | 72837796 | AX-170366435 | 0 | T | C | 0.053 | 2.50E-03 | 2.75E-01 | 2.68E+00 | 2.74E+00 | 9.90E-01 | no | | intergenic_variant | MODIFIER |
| 18 | 30757183 | AX-170163462 | 0 | C | T | 0.053 | 2.50E-03 | 2.75E-01 | 2.68E+00 | 2.74E+00 | 9.90E-01 | no | | intergenic_variant | MODIFIER |
| 18 | 39258725 | AX-170417985 | 0 | T | C | 0.053 | 2.50E-03 | 2.75E-01 | 2.68E+00 | 2.74E+00 | 9.90E-01 | no | Arhgap26 | intron_variant | MODIFIER |
| 2 | 129865240 | AX-170019018 | 2 | T | A | 0.145 | 3.40E-03 | 2.18E-01 | 2.70E+00 | 2.74E+00 | 9.90E-01 | no | | intergenic_variant | MODIFIER |
| 7 | 104387047 | AX-97820993 | 1 | T | C | 0.455 | -4.19E-04 | 9.50E-02 | 2.68E+00 | 2.74E+00 | 9.91E-01 | no | Trim30c | intron_variant | MODIFIER |
| 7 | 69549184 | AX-170468476 | 2 | A | T | 0.409 | 1.94E-03 | 1.00E-01 | 2.66E+00 | 2.74E+00 | 9.91E-01 | no | 4930402F11Rik | intron_variant,non_coding_transcript_variant | MODIFIER |
| 9 | 33631164 | AX-170032090 | 1 | C | T | 0.429 | 1.75E-03 | 8.46E-02 | 2.66E+00 | 2.74E+00 | 9.91E-01 | no | | intergenic_variant | MODIFIER |
| 7 | 136963277 | AX-170401296 | 0 | T | C | 0.289 | -1.70E-03 | 7.91E-02 | 2.63E+00 | 2.74E+00 | 9.91E-01 | no | Mgmt | intron_variant | MODIFIER |
| 12 | 109490693 | AX-97817801 | 0 | A | G | 0.114 | -2.14E-03 | 1.90E-01 | 2.64E+00 | 2.74E+00 | 9.91E-01 | no | Gm34081 | downstream_gene_variant | MODIFIER |
| 10 | 31769764 | AX-170385058 | 2 | T | C | 0.382 | -1.16E-03 | 7.36E-02 | 2.62E+00 | 2.74E+00 | 9.91E-01 | no | Nkain2 | intron_variant | MODIFIER |
| 12 | 58697616 | AX-170298314 | 1 | G | A | 0.062 | 3.06E-03 | 3.01E-01 | 2.72E+00 | 2.74E+00 | 9.91E-01 | no | | intergenic_variant | MODIFIER |
| 7 | 102943167 | AX-170098375 | 0 | C | T | 0.474 | 1.78E-03 | 9.61E-02 | 2.67E+00 | 2.74E+00 | 9.91E-01 | no | Olfr573-ps1 | upstream_gene_variant | MODIFIER |
| 7 | 103594092 | AX-170395919 | 0 | A | G | 0.474 | 1.78E-03 | 9.61E-02 | 2.67E+00 | 2.74E+00 | 9.91E-01 | no | Olfr618 | upstream_gene_variant | MODIFIER |
| 11 | 32991124 | AX-170347722 | 0 | C | T | 0.114 | 3.27E-03 | 1.65E-01 | 2.64E+00 | 2.74E+00 | 9.91E-01 | no | | intergenic_variant | MODIFIER |
| 13 | 53548948 | AX-170402105 | 0 | G | T | 0.509 | 1.94E-03 | 6.85E-02 | 2.63E+00 | 2.74E+00 | 9.91E-01 | no | | intergenic_variant | MODIFIER |
| 13 | 53621646 | AX-170467993 | 0 | A | G | 0.509 | 1.94E-03 | 6.85E-02 | 2.63E+00 | 2.74E+00 | 9.91E-01 | no | | intergenic_variant | MODIFIER |
| 13 | 53629083 | AX-170222372 | 0 | C | T | 0.509 | 1.94E-03 | 6.85E-02 | 2.63E+00 | 2.74E+00 | 9.91E-01 | no | | intergenic_variant | MODIFIER |
| 13 | 53632658 | AX-170222485 | 0 | A | C | 0.509 | 1.94E-03 | 6.85E-02 | 2.63E+00 | 2.74E+00 | 9.91E-01 | no | 9530014B07Rik | downstream_gene_variant | MODIFIER |
| 13 | 53632684 | AX-170329797 | 0 | C | T | 0.509 | 1.94E-03 | 6.85E-02 | 2.63E+00 | 2.74E+00 | 9.91E-01 | no | 9530014B07Rik | downstream_gene_variant | MODIFIER |
| 13 | 53673038 | AX-170483994 | 0 | G | A | 0.509 | 1.94E-03 | 6.85E-02 | 2.63E+00 | 2.74E+00 | 9.91E-01 | no | | intergenic_variant | MODIFIER |
| 13 | 53674093 | AX-170368335 | 0 | G | A | 0.509 | 1.94E-03 | 6.85E-02 | 2.63E+00 | 2.74E+00 | 9.91E-01 | no | | intergenic_variant | MODIFIER |
| 13 | 53816807 | AX-170188299 | 0 | A | G | 0.509 | 1.94E-03 | 6.85E-02 | 2.63E+00 | 2.74E+00 | 9.91E-01 | no | | intergenic_variant | MODIFIER |
| 13 | 53818439 | AX-170474273 | 0 | C | T | 0.509 | 1.94E-03 | 6.85E-02 | 2.63E+00 | 2.74E+00 | 9.91E-01 | no | | intergenic_variant | MODIFIER |
| 13 | 53819234 | AX-170390466 | 0 | A | G | 0.509 | 1.94E-03 | 6.85E-02 | 2.63E+00 | 2.74E+00 | 9.91E-01 | no | | intergenic_variant | MODIFIER |
| 13 | 53821268 | AX-170316106 | 0 | G | A | 0.509 | 1.94E-03 | 6.85E-02 | 2.63E+00 | 2.74E+00 | 9.91E-01 | no | | intergenic_variant | MODIFIER |
| 13 | 53822383 | AX-170324028 | 0 | T | C | 0.509 | 1.94E-03 | 6.85E-02 | 2.63E+00 | 2.74E+00 | 9.91E-01 | no | | regulatory_region_variant | MODIFIER |
| 13 | 53854164 | AX-170087428 | 0 | C | T | 0.509 | 1.94E-03 | 6.85E-02 | 2.63E+00 | 2.74E+00 | 9.91E-01 | no | | intergenic_variant | MODIFIER |
| 13 | 53858861 | AX-170402438 | 0 | A | G | 0.509 | 1.94E-03 | 6.85E-02 | 2.63E+00 | 2.74E+00 | 9.91E-01 | no | | intergenic_variant | MODIFIER |
| 13 | 53886051 | AX-97850780 | 0 | C | A | 0.509 | 1.94E-03 | 6.85E-02 | 2.63E+00 | 2.74E+00 | 9.91E-01 | no | Gm18974 | upstream_gene_variant | MODIFIER |
| 13 | 53898084 | AX-170343723 | 0 | A | G | 0.509 | 1.94E-03 | 6.85E-02 | 2.63E+00 | 2.74E+00 | 9.91E-01 | no | | intergenic_variant | MODIFIER |
| 13 | 53900354 | AX-170333134 | 0 | G | A | 0.509 | 1.94E-03 | 6.85E-02 | 2.63E+00 | 2.74E+00 | 9.91E-01 | no | | regulatory_region_variant | MODIFIER |
| 13 | 53908579 | AX-170310270 | 0 | A | T | 0.509 | 1.94E-03 | 6.85E-02 | 2.63E+00 | 2.74E+00 | 9.91E-01 | no | E130119H09Rik | downstream_gene_variant | MODIFIER |
| 9 | 106033740 | AX-170210393 | 1 | C | G | 0.125 | 8.38E-05 | 1.17E-01 | 2.67E+00 | 2.74E+00 | 9.91E-01 | no | Col6a4 | intron_variant | MODIFIER |
| 10 | 114217971 | AX-170173057 | 0 | T | C | 0.351 | -1.06E-03 | 7.97E-02 | 2.66E+00 | 2.74E+00 | 9.91E-01 | no | | intergenic_variant | MODIFIER |
| 15 | 94106002 | AX-170012299 | 0 | C | A | 0.149 | -8.95E-04 | 1.79E-01 | 2.68E+00 | 2.74E+00 | 9.91E-01 | no | | intergenic_variant | MODIFIER |
| 15 | 75228154 | AX-97829701 | 0 | G | A | 0.553 | 2.35E-03 | 8.21E-02 | 2.65E+00 | 2.73E+00 | 9.91E-01 | no | | intergenic_variant | MODIFIER |
| 1 | 78826798 | AX-170401324 | 0 | G | C | 0.14 | 2.46E-03 | 1.76E-01 | 2.64E+00 | 2.74E+00 | 9.91E-01 | no | Gm29187 | downstream_gene_variant | MODIFIER |
| 8 | 84614102 | AX-170502586 | 1 | A | G | 0.196 | -1.19E-03 | 1.32E-01 | 2.68E+00 | 2.74E+00 | 9.91E-01 | no | Cacna1a | intron_variant | MODIFIER |
| 4 | 56952883 | AX-170205250 | 1 | G | A | 0.375 | -1.16E-03 | 1.06E-01 | 2.66E+00 | 2.74E+00 | 9.91E-01 | no | Frrs1l | downstream_gene_variant | MODIFIER |
| 11 | 48003302 | AX-170160444 | 0 | T | C | 0.079 | -3.15E-03 | 1.76E-01 | 2.63E+00 | 2.74E+00 | 9.91E-01 | no | | intergenic_variant | MODIFIER |
| 11 | 50180428 | AX-170120747 | 0 | C | G | 0.079 | -3.15E-03 | 1.76E-01 | 2.63E+00 | 2.74E+00 | 9.91E-01 | no | Mrnip | intron_variant | MODIFIER |

| | | | | | | | | | | | | | | | |
|---|---|---|---|---|---|---|---|---|---|---|---|---|---|---|---|
| 8 | 41397857 | AX-170436807 | 0 | C | T | 0.158 | -2.06E-03 | 1.37E-01 | 2.67E+00 | 2.74E+00 | 9.91E-01 | no | Frg1 | intron_variant | MODIFIER |
| 1 | 66681090 | AX-170350141 | 1 | C | G | 0.107 | 2.78E-03 | 2.44E-01 | 2.70E+00 | 2.74E+00 | 9.91E-01 | no | Unc80 | intron_variant | MODIFIER |
| 1 | 7771532 | AX-170429665 | 1 | G | A | 0.429 | 1.28E-04 | 7.01E-02 | 2.66E+00 | 2.74E+00 | 9.91E-01 | no | | intergenic_variant | MODIFIER |
| 11 | 13524100 | AX-170420399 | 0 | A | G | 0.061 | 1.17E-03 | 2.13E-01 | 2.65E+00 | 2.74E+00 | 9.91E-01 | no | | intergenic_variant | MODIFIER |
| 11 | 17750062 | AX-170097730 | 0 | T | C | 0.061 | 1.17E-03 | 2.13E-01 | 2.65E+00 | 2.74E+00 | 9.91E-01 | no | | intergenic_variant | MODIFIER |
| 4 | 145369141 | AX-170436101 | 1 | A | T | 0.098 | -1.61E-03 | 1.98E-01 | 2.67E+00 | 2.74E+00 | 9.91E-01 | no | | intergenic_variant | MODIFIER |
| 3 | 28797325 | AX-168394528 | 0 | G | A | 0.184 | 5.65E-04 | 9.65E-02 | 2.66E+00 | 2.74E+00 | 9.91E-01 | no | Eif5a2 | 3_prime_UTR_variant | MODIFIER |
| 7 | 101702779 | AX-170078259 | 1 | C | T | 0.455 | -2.56E-04 | 9.42E-02 | 2.68E+00 | 2.74E+00 | 9.91E-01 | no | Clpb | intron_variant | MODIFIER |
| 18 | 70943288 | AX-170035589 | 1 | G | A | 0.17 | 1.67E-05 | 1.12E-01 | 2.65E+00 | 2.74E+00 | 9.92E-01 | no | | intergenic_variant | MODIFIER |
| 7 | 37103086 | AX-170305413 | 0 | T | C | 0.342 | 2.26E-03 | 9.04E-02 | 2.65E+00 | 2.74E+00 | 9.92E-01 | no | | intergenic_variant | MODIFIER |
| 1 | 11230419 | AX-170322069 | 0 | A | T | 0.386 | -6.33E-05 | 7.46E-02 | 2.68E+00 | 2.74E+00 | 9.92E-01 | no | Prex2 | intron_variant | MODIFIER |
| 1 | 11230703 | AX-170210360 | 0 | C | T | 0.386 | -6.33E-05 | 7.46E-02 | 2.68E+00 | 2.74E+00 | 9.92E-01 | no | Prex2 | intron_variant | MODIFIER |
| 1 | 11242221 | AX-170370531 | 0 | G | T | 0.386 | -6.33E-05 | 7.46E-02 | 2.68E+00 | 2.74E+00 | 9.92E-01 | no | Prex2 | intron_variant | MODIFIER |
| 1 | 11253148 | AX-169999942 | 0 | A | G | 0.386 | -6.33E-05 | 7.46E-02 | 2.68E+00 | 2.74E+00 | 9.92E-01 | no | Prex2 | intron_variant | MODIFIER |
| 1 | 11274151 | AX-170489581 | 0 | G | A | 0.386 | -6.33E-05 | 7.46E-02 | 2.68E+00 | 2.74E+00 | 9.92E-01 | no | Prex2 | intron_variant | MODIFIER |
| 1 | 11304442 | AX-169999948 | 0 | A | G | 0.386 | -6.33E-05 | 7.46E-02 | 2.68E+00 | 2.74E+00 | 9.92E-01 | no | Prex2 | downstream_gene_variant | MODIFIER |
| 1 | 11345094 | AX-97822858 | 0 | T | G | 0.386 | -6.33E-05 | 7.46E-02 | 2.68E+00 | 2.74E+00 | 9.92E-01 | no | | intergenic_variant | MODIFIER |
| 1 | 11361445 | AX-170434291 | 0 | C | T | 0.386 | -6.33E-05 | 7.46E-02 | 2.68E+00 | 2.74E+00 | 9.92E-01 | no | Gm37410 | downstream_gene_variant | MODIFIER |
| 1 | 11380016 | AX-170497723 | 0 | T | C | 0.386 | -6.33E-05 | 7.46E-02 | 2.68E+00 | 2.74E+00 | 9.92E-01 | no | | intergenic_variant | MODIFIER |
| 1 | 11380312 | AX-170341187 | 0 | C | T | 0.386 | -6.33E-05 | 7.46E-02 | 2.68E+00 | 2.74E+00 | 9.92E-01 | no | | intergenic_variant | MODIFIER |
| 1 | 11412172 | AX-97810736 | 0 | T | C | 0.386 | -6.33E-05 | 7.46E-02 | 2.68E+00 | 2.74E+00 | 9.92E-01 | no | A830018L16Rik | upstream_gene_variant | MODIFIER |
| 1 | 11436159 | AX-169999944 | 0 | T | C | 0.386 | -6.33E-05 | 7.46E-02 | 2.68E+00 | 2.74E+00 | 9.92E-01 | no | A830018L16Rik | intron_variant | MODIFIER |
| 1 | 12368506 | AX-170365079 | 0 | T | A | 0.386 | -6.33E-05 | 7.46E-02 | 2.68E+00 | 2.74E+00 | 9.92E-01 | no | | intergenic_variant | MODIFIER |
| 1 | 12797959 | AX-170169702 | 0 | A | G | 0.386 | -6.33E-05 | 7.46E-02 | 2.68E+00 | 2.74E+00 | 9.92E-01 | no | Sulf1 | intron_variant | MODIFIER |
| 1 | 12806739 | AX-170285066 | 0 | T | C | 0.386 | -6.33E-05 | 7.46E-02 | 2.68E+00 | 2.74E+00 | 9.92E-01 | no | Sulf1 | intron_variant | MODIFIER |
| 1 | 12888028 | AX-169999969 | 0 | T | C | 0.386 | -6.33E-05 | 7.46E-02 | 2.68E+00 | 2.74E+00 | 9.92E-01 | no | Slco5a1 | intron_variant | MODIFIER |
| 1 | 12890213 | AX-169999968 | 0 | G | A | 0.386 | -6.33E-05 | 7.46E-02 | 2.68E+00 | 2.74E+00 | 9.92E-01 | no | Slco5a1 | intron_variant | MODIFIER |
| 1 | 12894720 | AX-97852115 | 0 | C | G | 0.386 | -6.33E-05 | 7.46E-02 | 2.68E+00 | 2.74E+00 | 9.92E-01 | no | Slco5a1 | intron_variant | MODIFIER |
| 1 | 12920189 | AX-169999967 | 0 | A | T | 0.386 | -6.33E-05 | 7.46E-02 | 2.68E+00 | 2.74E+00 | 9.92E-01 | no | Slco5a1 | intron_variant | MODIFIER |
| 1 | 12925008 | AX-170170611 | 0 | T | A | 0.386 | -6.33E-05 | 7.46E-02 | 2.68E+00 | 2.74E+00 | 9.92E-01 | no | Slco5a1 | intron_variant | MODIFIER |
| 1 | 12925028 | AX-170355118 | 0 | G | A | 0.386 | -6.33E-05 | 7.46E-02 | 2.68E+00 | 2.74E+00 | 9.92E-01 | no | Slco5a1 | intron_variant | MODIFIER |
| 1 | 12929645 | AX-168399818 | 0 | A | C | 0.386 | -6.33E-05 | 7.46E-02 | 2.68E+00 | 2.74E+00 | 9.92E-01 | no | Slco5a1 | intron_variant | MODIFIER |
| 1 | 12930271 | AX-170509860 | 0 | T | G | 0.386 | -6.33E-05 | 7.46E-02 | 2.68E+00 | 2.74E+00 | 9.92E-01 | no | Slco5a1 | intron_variant | MODIFIER |
| 1 | 12932239 | AX-170280729 | 0 | A | G | 0.386 | -6.33E-05 | 7.46E-02 | 2.68E+00 | 2.74E+00 | 9.92E-01 | no | Slco5a1 | intron_variant | MODIFIER |
| 1 | 12934887 | AX-170127767 | 0 | G | A | 0.386 | -6.33E-05 | 7.46E-02 | 2.68E+00 | 2.74E+00 | 9.92E-01 | no | Slco5a1 | intron_variant | MODIFIER |
| 1 | 12953951 | AX-170394016 | 0 | A | G | 0.386 | -6.33E-05 | 7.46E-02 | 2.68E+00 | 2.74E+00 | 9.92E-01 | no | Slco5a1 | intron_variant | MODIFIER |
| 1 | 12954705 | AX-170528058 | 0 | T | A | 0.386 | -6.33E-05 | 7.46E-02 | 2.68E+00 | 2.74E+00 | 9.92E-01 | no | Slco5a1 | intron_variant | MODIFIER |
| 1 | 12966263 | AX-170358220 | 0 | A | G | 0.386 | -6.33E-05 | 7.46E-02 | 2.68E+00 | 2.74E+00 | 9.92E-01 | no | Slco5a1 | intron_variant | MODIFIER |
| 1 | 12966532 | AX-170350709 | 0 | A | G | 0.386 | -6.33E-05 | 7.46E-02 | 2.68E+00 | 2.74E+00 | 9.92E-01 | no | Slco5a1 | intron_variant | MODIFIER |
| 1 | 12977461 | AX-170035606 | 0 | G | T | 0.386 | -6.33E-05 | 7.46E-02 | 2.68E+00 | 2.74E+00 | 9.92E-01 | no | Slco5a1 | intron_variant | MODIFIER |
| 1 | 12989086 | AX-170518816 | 0 | A | G | 0.386 | -6.33E-05 | 7.46E-02 | 2.68E+00 | 2.74E+00 | 9.92E-01 | no | Slco5a1 | intron_variant | MODIFIER |
| 1 | 12995223 | AX-170321220 | 0 | A | G | 0.386 | -6.33E-05 | 7.46E-02 | 2.68E+00 | 2.74E+00 | 9.92E-01 | no | Slco5a1 | upstream_gene_variant | MODIFIER |
| 1 | 12995266 | AX-170288814 | 0 | A | C | 0.386 | -6.33E-05 | 7.46E-02 | 2.68E+00 | 2.74E+00 | 9.92E-01 | no | Slco5a1 | upstream_gene_variant | MODIFIER |
| 1 | 12995339 | AX-97850885 | 0 | C | T | 0.386 | -6.33E-05 | 7.46E-02 | 2.68E+00 | 2.74E+00 | 9.92E-01 | no | Slco5a1 | upstream_gene_variant | MODIFIER |
| 1 | 13038248 | AX-170459011 | 0 | A | G | 0.386 | -6.33E-05 | 7.46E-02 | 2.68E+00 | 2.74E+00 | 9.92E-01 | no | | intergenic_variant | MODIFIER |
| 1 | 13038807 | AX-170347085 | 0 | T | A | 0.386 | -6.33E-05 | 7.46E-02 | 2.68E+00 | 2.74E+00 | 9.92E-01 | no | | intergenic_variant | MODIFIER |
| 12 | 68792248 | AX-170129468 | 1 | G | A | 0.08 | 1.10E-03 | 1.52E-01 | 2.59E+00 | 2.74E+00 | 9.92E-01 | no | | intergenic_variant | MODIFIER |
| 10 | 32101368 | AX-170463919 | 0 | A | T | 0.386 | -1.11E-03 | 7.40E-02 | 2.62E+00 | 2.74E+00 | 9.92E-01 | no | Nkain2 | intron_variant | MODIFIER |
| 10 | 32559617 | AX-97843336 | 0 | T | A | 0.386 | -1.11E-03 | 7.40E-02 | 2.62E+00 | 2.74E+00 | 9.92E-01 | no | Nkain2 | intron_variant | MODIFIER |
| 7 | 103746679 | AX-170472000 | 2 | A | G | 0.455 | 1.93E-03 | 9.25E-02 | 2.67E+00 | 2.74E+00 | 9.92E-01 | no | Olfr629 | upstream_gene_variant | MODIFIER |
| 18 | 41203011 | AX-170252843 | 2 | C | T | 0.073 | -3.69E-03 | 2.27E-01 | 2.63E+00 | 2.74E+00 | 9.92E-01 | no | | intergenic_variant | MODIFIER |
| 9 | 37726640 | AX-97780393 | 0 | A | G | 0.5 | 1.13E-03 | 7.55E-02 | 2.64E+00 | 2.74E+00 | 9.92E-01 | no | Olfr151 | downstream_gene_variant | MODIFIER |
| 10 | 120528396 | AX-97793580 | 1 | A | G | 0.339 | 5.95E-04 | 6.83E-02 | 2.64E+00 | 2.74E+00 | 9.92E-01 | no | | intergenic_variant | MODIFIER |
| 9 | 105127003 | AX-170511972 | 0 | C | T | 0.132 | -9.86E-05 | 1.09E-01 | 2.66E+00 | 2.74E+00 | 9.92E-01 | no | Nudt16 | downstream_gene_variant | MODIFIER |

| | | | | | | | | | | | | | | | |
|---|---|---|---|---|---|---|---|---|---|---|---|---|---|---|---|
| 9 | 105762873 | AX-170439104 | 0 | G | A | 0.132 | -9.86E-05 | 1.09E-01 | 2.66E+00 | 2.74E+00 | 9.92E-01 | no | Col6a6 | synonymous_variant | LOW |
| 9 | 105944676 | AX-170033209 | 0 | G | A | 0.132 | -9.86E-05 | 1.09E-01 | 2.66E+00 | 2.74E+00 | 9.92E-01 | no | Col6a5 | intron_variant | MODIFIER |
| 13 | 117082348 | AX-170427881 | 2 | C | T | 0.055 | 1.50E-03 | 1.08E-01 | 2.57E+00 | 2.74E+00 | 9.92E-01 | no | Gm48026 | downstream_gene_variant | MODIFIER |
| 7 | 104247856 | AX-170029683 | 0 | C | G | 0.404 | -9.58E-05 | 1.28E-01 | 2.59E+00 | 2.74E+00 | 9.92E-01 | no | Trim34a | missense_variant | MODERATE |
| 1 | 38193522 | AX-170353880 | 1 | T | A | 0.455 | 2.41E-04 | 7.47E-02 | 2.65E+00 | 2.74E+00 | 9.92E-01 | no | Aff3 | intron_variant | MODIFIER |
| 15 | 97511208 | AX-170319151 | 0 | C | T | 0.149 | -6.96E-04 | 1.79E-01 | 2.68E+00 | 2.74E+00 | 9.92E-01 | no | | intergenic_variant | MODIFIER |
| 1 | 40925736 | AX-170128542 | 0 | T | G | 0.412 | 5.00E-04 | 7.52E-02 | 2.65E+00 | 2.74E+00 | 9.92E-01 | no | | intergenic_variant | MODIFIER |
| 12 | 118199007 | AX-170050365 | 1 | T | G | 0.134 | -1.04E-03 | 1.84E-01 | 2.66E+00 | 2.74E+00 | 9.92E-01 | no | Dnah11 | 5_prime_UTR_variant | MODIFIER |
| 5 | 96871067 | AX-170072822 | 0 | A | C | 0.439 | -8.12E-04 | 1.08E-01 | 2.68E+00 | 2.74E+00 | 9.93E-01 | no | | regulatory_region_variant | MODIFIER |
| 8 | 78164232 | AX-170276759 | 1 | T | C | 0.205 | 6.53E-04 | 1.29E-01 | 2.67E+00 | 2.74E+00 | 9.93E-01 | no | | intergenic_variant | MODIFIER |
| 5 | 108729704 | AX-97836209 | 1 | C | A | 0.455 | 1.65E-03 | 1.11E-01 | 2.69E+00 | 2.74E+00 | 9.93E-01 | no | Rnf212 | downstream_gene_variant | MODIFIER |
| 18 | 57602056 | AX-170117727 | 0 | G | A | 0.412 | -3.40E-04 | 8.61E-02 | 2.67E+00 | 2.74E+00 | 9.93E-01 | no | Ccdc192 | intron_variant | MODIFIER |
| 18 | 39483786 | AX-97834522 | 1 | C | T | 0.062 | -3.29E-03 | 3.00E-01 | 2.71E+00 | 2.74E+00 | 9.93E-01 | no | Nr3c1 | intron_variant | MODIFIER |
| 2 | 71799576 | AX-170272100 | 0 | G | C | 0.053 | 9.19E-04 | 1.94E-01 | 2.64E+00 | 2.74E+00 | 9.93E-01 | no | Itga6 | intron_variant | MODIFIER |
| 2 | 72307007 | AX-170063968 | 0 | T | C | 0.053 | 9.19E-04 | 1.94E-01 | 2.64E+00 | 2.74E+00 | 9.93E-01 | no | Map3k20 | intron_variant | MODIFIER |
| 18 | 44037864 | AX-170130088 | 1 | G | A | 0.062 | 2.10E-03 | 2.98E-01 | 2.71E+00 | 2.74E+00 | 9.93E-01 | no | | intergenic_variant | MODIFIER |
| 20 | 168735936 | AX-170443910 | 1 | A | T | 0.17 | 1.93E-03 | 1.18E-01 | 2.64E+00 | 2.74E+00 | 9.93E-01 | no | | intergenic_variant | MODIFIER |
| 7 | 102320165 | AX-97790119 | 0 | G | T | 0.465 | 1.53E-03 | 9.26E-02 | 2.67E+00 | 2.74E+00 | 9.93E-01 | no | Stim1 | intron_variant | MODIFIER |
| 7 | 102534278 | AX-170078278 | 0 | A | G | 0.465 | 1.53E-03 | 9.26E-02 | 2.67E+00 | 2.74E+00 | 9.93E-01 | no | Olfr547 | upstream_gene_variant | MODIFIER |
| 7 | 102896544 | AX-170029660 | 0 | A | G | 0.465 | 1.53E-03 | 9.26E-02 | 2.67E+00 | 2.74E+00 | 9.93E-01 | no | Olfr570 | upstream_gene_variant | MODIFIER |
| 1 | 43694508 | AX-170000384 | 0 | G | A | 0.105 | -1.87E-03 | 2.36E-01 | 2.69E+00 | 2.74E+00 | 9.93E-01 | no | | regulatory_region_variant | MODIFIER |
| 1 | 46531184 | AX-170395314 | 0 | C | T | 0.105 | -1.87E-03 | 2.36E-01 | 2.69E+00 | 2.74E+00 | 9.93E-01 | no | Dnah7c | intron_variant | MODIFIER |
| 1 | 46926006 | AX-170147080 | 0 | G | A | 0.105 | -1.87E-03 | 2.36E-01 | 2.69E+00 | 2.74E+00 | 9.93E-01 | no | | intergenic_variant | MODIFIER |
| 1 | 46926036 | AX-170285963 | 0 | A | G | 0.105 | -1.87E-03 | 2.36E-01 | 2.69E+00 | 2.74E+00 | 9.93E-01 | no | | intergenic_variant | MODIFIER |
| 1 | 46926519 | AX-170114079 | 0 | A | G | 0.105 | -1.87E-03 | 2.36E-01 | 2.69E+00 | 2.74E+00 | 9.93E-01 | no | | intergenic_variant | MODIFIER |
| 1 | 47059814 | AX-170414944 | 0 | C | T | 0.105 | -1.87E-03 | 2.36E-01 | 2.69E+00 | 2.74E+00 | 9.93E-01 | no | | intergenic_variant | MODIFIER |
| 1 | 47079733 | AX-170439539 | 0 | T | C | 0.105 | -1.87E-03 | 2.36E-01 | 2.69E+00 | 2.74E+00 | 9.93E-01 | no | | intergenic_variant | MODIFIER |
| 1 | 47079745 | AX-170344966 | 0 | A | G | 0.105 | -1.87E-03 | 2.36E-01 | 2.69E+00 | 2.74E+00 | 9.93E-01 | no | | intergenic_variant | MODIFIER |
| 1 | 47101802 | AX-170106093 | 0 | C | T | 0.105 | -1.87E-03 | 2.36E-01 | 2.69E+00 | 2.74E+00 | 9.93E-01 | no | | intergenic_variant | MODIFIER |
| 1 | 47104131 | AX-170532819 | 0 | A | G | 0.105 | -1.87E-03 | 2.36E-01 | 2.69E+00 | 2.74E+00 | 9.93E-01 | no | | regulatory_region_variant | MODIFIER |
| 1 | 47164523 | AX-170000419 | 0 | A | G | 0.105 | -1.87E-03 | 2.36E-01 | 2.69E+00 | 2.74E+00 | 9.93E-01 | no | Gm28826 | intron_variant,non_coding_transcript_variant | MODIFIER |
| 1 | 47173384 | AX-170000418 | 0 | T | C | 0.105 | -1.87E-03 | 2.36E-01 | 2.69E+00 | 2.74E+00 | 9.93E-01 | no | Gm28826 | intron_variant,non_coding_transcript_variant | MODIFIER |
| 1 | 47204083 | AX-170220189 | 0 | A | G | 0.105 | -1.87E-03 | 2.36E-01 | 2.69E+00 | 2.74E+00 | 9.93E-01 | no | | intergenic_variant | MODIFIER |
| 1 | 47204173 | AX-170175343 | 0 | T | C | 0.105 | -1.87E-03 | 2.36E-01 | 2.69E+00 | 2.74E+00 | 9.93E-01 | no | | intergenic_variant | MODIFIER |
| 1 | 47216411 | AX-170186171 | 0 | G | A | 0.105 | -1.87E-03 | 2.36E-01 | 2.69E+00 | 2.74E+00 | 9.93E-01 | no | | intergenic_variant | MODIFIER |
| 1 | 47219885 | AX-170349328 | 0 | A | C | 0.105 | -1.87E-03 | 2.36E-01 | 2.69E+00 | 2.74E+00 | 9.93E-01 | no | | intergenic_variant | MODIFIER |
| 1 | 47219999 | AX-170144451 | 0 | G | A | 0.105 | -1.87E-03 | 2.36E-01 | 2.69E+00 | 2.74E+00 | 9.93E-01 | no | | intergenic_variant | MODIFIER |
| 1 | 47229371 | AX-170532430 | 0 | T | C | 0.105 | -1.87E-03 | 2.36E-01 | 2.69E+00 | 2.74E+00 | 9.93E-01 | no | | intergenic_variant | MODIFIER |
| 1 | 47264045 | AX-170180337 | 0 | G | C | 0.105 | -1.87E-03 | 2.36E-01 | 2.69E+00 | 2.74E+00 | 9.93E-01 | no | | intergenic_variant | MODIFIER |
| 1 | 47264380 | AX-170411078 | 0 | A | T | 0.105 | -1.87E-03 | 2.36E-01 | 2.69E+00 | 2.74E+00 | 9.93E-01 | no | | intergenic_variant | MODIFIER |
| 1 | 47268609 | AX-170386150 | 0 | T | C | 0.105 | -1.87E-03 | 2.36E-01 | 2.69E+00 | 2.74E+00 | 9.93E-01 | no | | intergenic_variant | MODIFIER |
| 1 | 47498087 | AX-170041139 | 0 | T | G | 0.105 | -1.87E-03 | 2.36E-01 | 2.69E+00 | 2.74E+00 | 9.93E-01 | no | | intergenic_variant | MODIFIER |
| 1 | 47606095 | AX-170283651 | 0 | C | T | 0.105 | -1.87E-03 | 2.36E-01 | 2.69E+00 | 2.74E+00 | 9.93E-01 | no | Gm37196 | non_coding_transcript_exon_variant | MODIFIER |
| 1 | 47606121 | AX-170104743 | 0 | T | G | 0.105 | -1.87E-03 | 2.36E-01 | 2.69E+00 | 2.74E+00 | 9.93E-01 | no | Gm37196 | non_coding_transcript_exon_variant | MODIFIER |
| 1 | 47606318 | AX-170507016 | 0 | T | G | 0.105 | -1.87E-03 | 2.36E-01 | 2.69E+00 | 2.74E+00 | 9.93E-01 | no | Gm37196 | non_coding_transcript_exon_variant | MODIFIER |
| 1 | 47609952 | AX-170000423 | 0 | T | C | 0.105 | -1.87E-03 | 2.36E-01 | 2.69E+00 | 2.74E+00 | 9.93E-01 | no | Gm37196 | downstream_gene_variant | MODIFIER |
| 1 | 47609985 | AX-170000422 | 0 | T | C | 0.105 | -1.87E-03 | 2.36E-01 | 2.69E+00 | 2.74E+00 | 9.93E-01 | no | Gm37196 | downstream_gene_variant | MODIFIER |
| 1 | 47621676 | AX-170041144 | 0 | G | C | 0.105 | -1.87E-03 | 2.36E-01 | 2.69E+00 | 2.74E+00 | 9.93E-01 | no | | intergenic_variant | MODIFIER |
| 1 | 47621850 | AX-170041143 | 0 | C | T | 0.105 | -1.87E-03 | 2.36E-01 | 2.69E+00 | 2.74E+00 | 9.93E-01 | no | | intergenic_variant | MODIFIER |
| 1 | 47621964 | AX-170041146 | 0 | A | C | 0.105 | -1.87E-03 | 2.36E-01 | 2.69E+00 | 2.74E+00 | 9.93E-01 | no | | intergenic_variant | MODIFIER |
| 1 | 47644771 | AX-170000421 | 0 | C | G | 0.105 | -1.87E-03 | 2.36E-01 | 2.69E+00 | 2.74E+00 | 9.93E-01 | no | | intergenic_variant | MODIFIER |
| 1 | 47829101 | AX-170129452 | 0 | T | C | 0.105 | -1.87E-03 | 2.36E-01 | 2.69E+00 | 2.74E+00 | 9.93E-01 | no | | intergenic_variant | MODIFIER |
| 1 | 47865227 | AX-170135892 | 0 | A | C | 0.105 | -1.87E-03 | 2.36E-01 | 2.69E+00 | 2.74E+00 | 9.93E-01 | no | | intergenic_variant | MODIFIER |
| 1 | 47985153 | AX-170189029 | 0 | C | T | 0.105 | -1.87E-03 | 2.36E-01 | 2.69E+00 | 2.74E+00 | 9.93E-01 | no | | intergenic_variant | MODIFIER |

| | | | | | | | | | | | | | | | |
|---|---|---|---|---|---|---|---|---|---|---|---|---|---|---|---|
| 1 | 48105209 | AX-170130363 | 0 | C | A | 0.105 | -1.87E-03 | 2.36E-01 | 2.69E+00 | 2.74E+00 | 9.93E-01 | no | | intergenic_variant | MODIFIER |
| 1 | 48108134 | AX-97804450 | 0 | C | A | 0.105 | -1.87E-03 | 2.36E-01 | 2.69E+00 | 2.74E+00 | 9.93E-01 | no | | intergenic_variant | MODIFIER |
| 1 | 48129245 | AX-170464278 | 0 | C | T | 0.105 | -1.87E-03 | 2.36E-01 | 2.69E+00 | 2.74E+00 | 9.93E-01 | no | | intergenic_variant | MODIFIER |
| 1 | 48133190 | AX-170520669 | 0 | A | G | 0.105 | -1.87E-03 | 2.36E-01 | 2.69E+00 | 2.74E+00 | 9.93E-01 | no | | intergenic_variant | MODIFIER |
| 1 | 48142653 | AX-170106435 | 0 | G | A | 0.105 | -1.87E-03 | 2.36E-01 | 2.69E+00 | 2.74E+00 | 9.93E-01 | no | | intergenic_variant | MODIFIER |
| 1 | 48205329 | AX-97787522 | 0 | T | C | 0.105 | -1.87E-03 | 2.36E-01 | 2.69E+00 | 2.74E+00 | 9.93E-01 | no | | intergenic_variant | MODIFIER |
| 1 | 48207945 | AX-170300251 | 0 | G | T | 0.105 | -1.87E-03 | 2.36E-01 | 2.69E+00 | 2.74E+00 | 9.93E-01 | no | | intergenic_variant | MODIFIER |
| 1 | 48208488 | AX-170000433 | 0 | C | G | 0.105 | -1.87E-03 | 2.36E-01 | 2.69E+00 | 2.74E+00 | 9.93E-01 | no | | intergenic_variant | MODIFIER |
| 1 | 48215994 | AX-170000432 | 0 | C | T | 0.105 | -1.87E-03 | 2.36E-01 | 2.69E+00 | 2.74E+00 | 9.93E-01 | no | | intergenic_variant | MODIFIER |
| 1 | 48240297 | AX-170000430 | 0 | T | G | 0.105 | -1.87E-03 | 2.36E-01 | 2.69E+00 | 2.74E+00 | 9.93E-01 | no | | intergenic_variant | MODIFIER |
| 1 | 48262558 | AX-170000434 | 0 | G | C | 0.105 | -1.87E-03 | 2.36E-01 | 2.69E+00 | 2.74E+00 | 9.93E-01 | no | | intergenic_variant | MODIFIER |
| 1 | 48269646 | AX-170000427 | 0 | G | A | 0.105 | -1.87E-03 | 2.36E-01 | 2.69E+00 | 2.74E+00 | 9.93E-01 | no | | intergenic_variant | MODIFIER |
| 1 | 48329900 | AX-170120424 | 0 | A | T | 0.105 | -1.87E-03 | 2.36E-01 | 2.69E+00 | 2.74E+00 | 9.93E-01 | no | | intergenic_variant | MODIFIER |
| 1 | 48390798 | AX-170160780 | 0 | G | C | 0.105 | -1.87E-03 | 2.36E-01 | 2.69E+00 | 2.74E+00 | 9.93E-01 | no | | intergenic_variant | MODIFIER |
| 1 | 48405811 | AX-170470564 | 0 | T | C | 0.105 | -1.87E-03 | 2.36E-01 | 2.69E+00 | 2.74E+00 | 9.93E-01 | no | | intergenic_variant | MODIFIER |
| 1 | 48426410 | AX-170369664 | 0 | G | A | 0.105 | -1.87E-03 | 2.36E-01 | 2.69E+00 | 2.74E+00 | 9.93E-01 | no | | intergenic_variant | MODIFIER |
| 1 | 48477262 | AX-97841956 | 0 | A | G | 0.105 | -1.87E-03 | 2.36E-01 | 2.69E+00 | 2.74E+00 | 9.93E-01 | no | | intergenic_variant | MODIFIER |
| 1 | 48515947 | AX-170132874 | 0 | G | T | 0.105 | -1.87E-03 | 2.36E-01 | 2.69E+00 | 2.74E+00 | 9.93E-01 | no | | intergenic_variant | MODIFIER |
| 1 | 48516038 | AX-170218053 | 0 | T | C | 0.105 | -1.87E-03 | 2.36E-01 | 2.69E+00 | 2.74E+00 | 9.93E-01 | no | | intergenic_variant | MODIFIER |
| 1 | 48516127 | AX-170457617 | 0 | A | G | 0.105 | -1.87E-03 | 2.36E-01 | 2.69E+00 | 2.74E+00 | 9.93E-01 | no | | intergenic_variant | MODIFIER |
| 1 | 48542969 | AX-97782067 | 0 | C | A | 0.105 | -1.87E-03 | 2.36E-01 | 2.69E+00 | 2.74E+00 | 9.93E-01 | no | | intergenic_variant | MODIFIER |
| 1 | 48605866 | AX-97785061 | 0 | T | A | 0.105 | -1.87E-03 | 2.36E-01 | 2.69E+00 | 2.74E+00 | 9.93E-01 | no | | intergenic_variant | MODIFIER |
| 1 | 48644672 | AX-170086460 | 0 | G | A | 0.105 | -1.87E-03 | 2.36E-01 | 2.69E+00 | 2.74E+00 | 9.93E-01 | no | | intergenic_variant | MODIFIER |
| 1 | 48646667 | AX-97801875 | 0 | C | T | 0.105 | -1.87E-03 | 2.36E-01 | 2.69E+00 | 2.74E+00 | 9.93E-01 | no | | intergenic_variant | MODIFIER |
| 1 | 48647405 | AX-170477682 | 0 | A | G | 0.105 | -1.87E-03 | 2.36E-01 | 2.69E+00 | 2.74E+00 | 9.93E-01 | no | | intergenic_variant | MODIFIER |
| 1 | 48678817 | AX-170000438 | 0 | C | T | 0.105 | -1.87E-03 | 2.36E-01 | 2.69E+00 | 2.74E+00 | 9.93E-01 | no | | intergenic_variant | MODIFIER |
| 1 | 48729521 | AX-170000441 | 0 | A | G | 0.105 | -1.87E-03 | 2.36E-01 | 2.69E+00 | 2.74E+00 | 9.93E-01 | no | Gm8354 | downstream_gene_variant | MODIFIER |
| 1 | 48840911 | AX-97794422 | 0 | A | G | 0.105 | -1.87E-03 | 2.36E-01 | 2.69E+00 | 2.74E+00 | 9.93E-01 | no | | intergenic_variant | MODIFIER |
| 1 | 48841144 | AX-170000435 | 0 | C | T | 0.105 | -1.87E-03 | 2.36E-01 | 2.69E+00 | 2.74E+00 | 9.93E-01 | no | | intergenic_variant | MODIFIER |
| 1 | 48853184 | AX-170000443 | 0 | T | A | 0.105 | -1.87E-03 | 2.36E-01 | 2.69E+00 | 2.74E+00 | 9.93E-01 | no | | intergenic_variant | MODIFIER |
| 1 | 48951679 | AX-170329534 | 0 | C | G | 0.105 | -1.87E-03 | 2.36E-01 | 2.69E+00 | 2.74E+00 | 9.93E-01 | no | | intergenic_variant | MODIFIER |
| 1 | 49007812 | AX-97852871 | 0 | A | G | 0.105 | -1.87E-03 | 2.36E-01 | 2.69E+00 | 2.74E+00 | 9.93E-01 | no | | intergenic_variant | MODIFIER |
| 1 | 49198846 | AX-97847617 | 0 | C | T | 0.105 | -1.87E-03 | 2.36E-01 | 2.69E+00 | 2.74E+00 | 9.93E-01 | no | | intergenic_variant | MODIFIER |
| 1 | 49244590 | AX-170157059 | 0 | T | C | 0.105 | -1.87E-03 | 2.36E-01 | 2.69E+00 | 2.74E+00 | 9.93E-01 | no | C230029F24Rik | upstream_gene_variant | MODIFIER |
| 1 | 49278026 | AX-170446900 | 0 | G | A | 0.105 | -1.87E-03 | 2.36E-01 | 2.69E+00 | 2.74E+00 | 9.93E-01 | no | C230029F24Rik | intron_variant,non_coding_transcript_variant | MODIFIER |
| 1 | 49280676 | AX-97811062 | 0 | G | A | 0.105 | -1.87E-03 | 2.36E-01 | 2.69E+00 | 2.74E+00 | 9.93E-01 | no | C230029F24Rik | intron_variant,non_coding_transcript_variant | MODIFIER |
| 1 | 49369126 | AX-97826613 | 0 | T | C | 0.105 | -1.87E-03 | 2.36E-01 | 2.69E+00 | 2.74E+00 | 9.93E-01 | no | | intergenic_variant | MODIFIER |
| 1 | 49376356 | AX-97811501 | 0 | T | C | 0.105 | -1.87E-03 | 2.36E-01 | 2.69E+00 | 2.74E+00 | 9.93E-01 | no | | intergenic_variant | MODIFIER |
| 1 | 49623477 | AX-97805834 | 0 | T | C | 0.105 | -1.87E-03 | 2.36E-01 | 2.69E+00 | 2.74E+00 | 9.93E-01 | no | | intergenic_variant | MODIFIER |
| 1 | 50090028 | AX-97843188 | 0 | T | C | 0.105 | -1.87E-03 | 2.36E-01 | 2.69E+00 | 2.74E+00 | 9.93E-01 | no | | intergenic_variant | MODIFIER |
| 1 | 50090640 | AX-170243029 | 0 | C | T | 0.105 | -1.87E-03 | 2.36E-01 | 2.69E+00 | 2.74E+00 | 9.93E-01 | no | | intergenic_variant | MODIFIER |
| 1 | 50115209 | AX-97807326 | 0 | T | C | 0.105 | -1.87E-03 | 2.36E-01 | 2.69E+00 | 2.74E+00 | 9.93E-01 | no | | intergenic_variant | MODIFIER |
| 1 | 50183577 | AX-170086465 | 0 | T | C | 0.105 | -1.87E-03 | 2.36E-01 | 2.69E+00 | 2.74E+00 | 9.93E-01 | no | | intergenic_variant | MODIFIER |
| 1 | 50223125 | AX-170109998 | 0 | C | T | 0.105 | -1.87E-03 | 2.36E-01 | 2.69E+00 | 2.74E+00 | 9.93E-01 | no | Gm27330 | upstream_gene_variant | MODIFIER |
| 1 | 50238354 | AX-170252352 | 0 | A | T | 0.105 | -1.87E-03 | 2.36E-01 | 2.69E+00 | 2.74E+00 | 9.93E-01 | no | | intergenic_variant | MODIFIER |
| 1 | 50238865 | AX-170191779 | 0 | A | G | 0.105 | -1.87E-03 | 2.36E-01 | 2.69E+00 | 2.74E+00 | 9.93E-01 | no | | intergenic_variant | MODIFIER |
| 1 | 50238942 | AX-170207585 | 0 | T | A | 0.105 | -1.87E-03 | 2.36E-01 | 2.69E+00 | 2.74E+00 | 9.93E-01 | no | | intergenic_variant | MODIFIER |
| 1 | 50239066 | AX-170215159 | 0 | T | G | 0.105 | -1.87E-03 | 2.36E-01 | 2.69E+00 | 2.74E+00 | 9.93E-01 | no | | intergenic_variant | MODIFIER |
| 1 | 50240045 | AX-170000464 | 0 | T | G | 0.105 | -1.87E-03 | 2.36E-01 | 2.69E+00 | 2.74E+00 | 9.93E-01 | no | | intergenic_variant | MODIFIER |
| 1 | 50240137 | AX-170000456 | 0 | T | C | 0.105 | -1.87E-03 | 2.36E-01 | 2.69E+00 | 2.74E+00 | 9.93E-01 | no | | intergenic_variant | MODIFIER |
| 1 | 50350041 | AX-170300592 | 0 | A | G | 0.105 | -1.87E-03 | 2.36E-01 | 2.69E+00 | 2.74E+00 | 9.93E-01 | no | | intergenic_variant | MODIFIER |
| 1 | 50392859 | AX-170469727 | 0 | T | G | 0.105 | -1.87E-03 | 2.36E-01 | 2.69E+00 | 2.74E+00 | 9.93E-01 | no | Gm29665 | downstream_gene_variant | MODIFIER |
| 1 | 50955680 | AX-170412058 | 0 | G | A | 0.105 | -1.87E-03 | 2.36E-01 | 2.69E+00 | 2.74E+00 | 9.93E-01 | no | Tmeff2 | intron_variant | MODIFIER |
| 1 | 51065798 | AX-170167938 | 0 | A | T | 0.105 | -1.87E-03 | 2.36E-01 | 2.69E+00 | 2.74E+00 | 9.93E-01 | no | Tmeff2 | intron_variant | MODIFIER |

| | | | | | | | | | | | | | | | |
|---|---|---|---|---|---|---|---|---|---|---|---|---|---|---|---|
| 1 | 51071575 | AX-170398154 | 0 | G | A | 0.105 | -1.87E-03 | 2.36E-01 | 2.69E+00 | 2.74E+00 | 9.93E-01 | no | Tmeff2 | intron_variant | MODIFIER |
| 1 | 51074400 | AX-170490962 | 0 | T | C | 0.105 | -1.87E-03 | 2.36E-01 | 2.69E+00 | 2.74E+00 | 9.93E-01 | no | Tmeff2 | intron_variant | MODIFIER |
| 1 | 51076703 | AX-97811532 | 0 | A | G | 0.105 | -1.87E-03 | 2.36E-01 | 2.69E+00 | 2.74E+00 | 9.93E-01 | no | Tmeff2 | intron_variant | MODIFIER |
| 1 | 51077882 | AX-170000483 | 0 | T | G | 0.105 | -1.87E-03 | 2.36E-01 | 2.69E+00 | 2.74E+00 | 9.93E-01 | no | Tmeff2 | intron_variant | MODIFIER |
| 1 | 51249392 | AX-170527912 | 0 | T | A | 0.105 | -1.87E-03 | 2.36E-01 | 2.69E+00 | 2.74E+00 | 9.93E-01 | no | 9330175M20Rik | intron_variant,non_coding_transcript_variant | MODIFIER |
| 1 | 51261579 | AX-170107760 | 0 | T | C | 0.105 | -1.87E-03 | 2.36E-01 | 2.69E+00 | 2.74E+00 | 9.93E-01 | no | 9330175M20Rik | intron_variant,non_coding_transcript_variant | MODIFIER |
| 1 | 51261712 | AX-170379257 | 0 | A | G | 0.105 | -1.87E-03 | 2.36E-01 | 2.69E+00 | 2.74E+00 | 9.93E-01 | no | 9330175M20Rik | intron_variant,non_coding_transcript_variant | MODIFIER |
| 1 | 51268033 | AX-170346754 | 0 | C | T | 0.105 | -1.87E-03 | 2.36E-01 | 2.69E+00 | 2.74E+00 | 9.93E-01 | no | 9330175M20Rik | intron_variant,non_coding_transcript_variant | MODIFIER |
| 1 | 51354680 | AX-97811528 | 0 | T | G | 0.105 | -1.87E-03 | 2.36E-01 | 2.69E+00 | 2.74E+00 | 9.93E-01 | no | | intergenic_variant | MODIFIER |
| 1 | 51369255 | AX-168397630 | 0 | C | G | 0.105 | -1.87E-03 | 2.36E-01 | 2.69E+00 | 2.74E+00 | 9.93E-01 | no | | intergenic_variant | MODIFIER |
| 1 | 51407207 | AX-170274830 | 0 | T | C | 0.105 | -1.87E-03 | 2.36E-01 | 2.69E+00 | 2.74E+00 | 9.93E-01 | no | | intergenic_variant | MODIFIER |
| 1 | 51407831 | AX-170187447 | 0 | T | C | 0.105 | -1.87E-03 | 2.36E-01 | 2.69E+00 | 2.74E+00 | 9.93E-01 | no | | intergenic_variant | MODIFIER |
| 1 | 51407929 | AX-97789094 | 0 | A | G | 0.105 | -1.87E-03 | 2.36E-01 | 2.69E+00 | 2.74E+00 | 9.93E-01 | no | | intergenic_variant | MODIFIER |
| 1 | 51426954 | AX-97842184 | 0 | T | G | 0.105 | -1.87E-03 | 2.36E-01 | 2.69E+00 | 2.74E+00 | 9.93E-01 | no | | intergenic_variant | MODIFIER |
| 1 | 51488037 | AX-97789093 | 0 | T | C | 0.105 | -1.87E-03 | 2.36E-01 | 2.69E+00 | 2.74E+00 | 9.93E-01 | no | | intergenic_variant | MODIFIER |
| 1 | 51507962 | AX-170345081 | 0 | A | G | 0.105 | -1.87E-03 | 2.36E-01 | 2.69E+00 | 2.74E+00 | 9.93E-01 | no | Gm17767 | downstream_gene_variant | MODIFIER |
| 1 | 51533756 | AX-97816184 | 0 | T | C | 0.105 | -1.87E-03 | 2.36E-01 | 2.69E+00 | 2.74E+00 | 9.93E-01 | no | Gm17767 | intron_variant,non_coding_transcript_variant | MODIFIER |
| 1 | 51539355 | AX-170269053 | 0 | T | C | 0.105 | -1.87E-03 | 2.36E-01 | 2.69E+00 | 2.74E+00 | 9.93E-01 | no | Gm17767 | intron_variant,non_coding_transcript_variant | MODIFIER |
| 1 | 51553699 | AX-97810841 | 0 | C | G | 0.105 | -1.87E-03 | 2.36E-01 | 2.69E+00 | 2.74E+00 | 9.93E-01 | no | Gm17767 | intron_variant,non_coding_transcript_variant | MODIFIER |
| 1 | 51595532 | AX-97816275 | 0 | T | C | 0.105 | -1.87E-03 | 2.36E-01 | 2.69E+00 | 2.74E+00 | 9.93E-01 | no | Gm17767 | intron_variant,non_coding_transcript_variant | MODIFIER |
| 1 | 51754523 | AX-97834918 | 0 | T | C | 0.105 | -1.87E-03 | 2.36E-01 | 2.69E+00 | 2.74E+00 | 9.93E-01 | no | Myo1b | intron_variant | MODIFIER |
| 1 | 51760020 | AX-170472800 | 0 | A | G | 0.105 | -1.87E-03 | 2.36E-01 | 2.69E+00 | 2.74E+00 | 9.93E-01 | no | Myo1b | intron_variant | MODIFIER |
| 1 | 51768193 | AX-170000493 | 0 | C | A | 0.105 | -1.87E-03 | 2.36E-01 | 2.69E+00 | 2.74E+00 | 9.93E-01 | no | Myo1b | intron_variant | MODIFIER |
| 1 | 51769109 | AX-170281856 | 0 | C | G | 0.105 | -1.87E-03 | 2.36E-01 | 2.69E+00 | 2.74E+00 | 9.93E-01 | no | Myo1b | intron_variant | MODIFIER |
| 1 | 51769463 | AX-170165103 | 0 | A | C | 0.105 | -1.87E-03 | 2.36E-01 | 2.69E+00 | 2.74E+00 | 9.93E-01 | no | Myo1b | intron_variant | MODIFIER |
| 1 | 51772703 | AX-170136478 | 0 | A | G | 0.105 | -1.87E-03 | 2.36E-01 | 2.69E+00 | 2.74E+00 | 9.93E-01 | no | Myo1b | intron_variant | MODIFIER |
| 1 | 51773878 | AX-170335001 | 0 | T | G | 0.105 | -1.87E-03 | 2.36E-01 | 2.69E+00 | 2.74E+00 | 9.93E-01 | no | Myo1b | intron_variant | MODIFIER |
| 1 | 51774238 | AX-170359055 | 0 | C | T | 0.105 | -1.87E-03 | 2.36E-01 | 2.69E+00 | 2.74E+00 | 9.93E-01 | no | Myo1b | intron_variant | MODIFIER |
| 1 | 51786962 | AX-170350518 | 0 | A | G | 0.105 | -1.87E-03 | 2.36E-01 | 2.69E+00 | 2.74E+00 | 9.93E-01 | no | Myo1b | intron_variant | MODIFIER |
| 1 | 51792230 | AX-170314714 | 0 | A | G | 0.105 | -1.87E-03 | 2.36E-01 | 2.69E+00 | 2.74E+00 | 9.93E-01 | no | Myo1b | intron_variant | MODIFIER |
| 1 | 51792387 | AX-170000495 | 0 | G | A | 0.105 | -1.87E-03 | 2.36E-01 | 2.69E+00 | 2.74E+00 | 9.93E-01 | no | Myo1b | intron_variant | MODIFIER |
| 1 | 51996595 | AX-170225138 | 0 | C | T | 0.105 | -1.87E-03 | 2.36E-01 | 2.69E+00 | 2.74E+00 | 9.93E-01 | no | Stat4 | intron_variant | MODIFIER |
| 1 | 51997024 | AX-170000492 | 0 | A | G | 0.105 | -1.87E-03 | 2.36E-01 | 2.69E+00 | 2.74E+00 | 9.93E-01 | no | Stat4 | intron_variant | MODIFIER |
| 1 | 51997261 | AX-170041216 | 0 | A | C | 0.105 | -1.87E-03 | 2.36E-01 | 2.69E+00 | 2.74E+00 | 9.93E-01 | no | Stat4 | intron_variant | MODIFIER |
| 1 | 52008386 | AX-97841444 | 0 | A | G | 0.105 | -1.87E-03 | 2.36E-01 | 2.69E+00 | 2.74E+00 | 9.93E-01 | no | Stat4 | 5_prime_UTR_variant | MODIFIER |
| 1 | 52010009 | AX-170522213 | 0 | A | G | 0.105 | -1.87E-03 | 2.36E-01 | 2.69E+00 | 2.74E+00 | 9.93E-01 | no | Stat4 | intron_variant | MODIFIER |
| 1 | 52011530 | AX-170370340 | 0 | T | C | 0.105 | -1.87E-03 | 2.36E-01 | 2.69E+00 | 2.74E+00 | 9.93E-01 | no | Stat4 | intron_variant | MODIFIER |
| 1 | 52011702 | AX-170269181 | 0 | A | G | 0.105 | -1.87E-03 | 2.36E-01 | 2.69E+00 | 2.74E+00 | 9.93E-01 | no | Stat4 | intron_variant | MODIFIER |
| 1 | 52017775 | AX-170114011 | 0 | G | T | 0.105 | -1.87E-03 | 2.36E-01 | 2.69E+00 | 2.74E+00 | 9.93E-01 | no | Stat4 | intron_variant | MODIFIER |
| 1 | 52565009 | AX-97784184 | 0 | G | A | 0.105 | -1.87E-03 | 2.36E-01 | 2.69E+00 | 2.74E+00 | 9.93E-01 | no | Gm553 | intron_variant,non_coding_transcript_variant | MODIFIER |
| 1 | 52565259 | AX-170230005 | 0 | G | T | 0.105 | -1.87E-03 | 2.36E-01 | 2.69E+00 | 2.74E+00 | 9.93E-01 | no | Gm28178 | downstream_gene_variant | MODIFIER |
| 1 | 52586270 | AX-170178850 | 0 | A | G | 0.105 | -1.87E-03 | 2.36E-01 | 2.69E+00 | 2.74E+00 | 9.93E-01 | no | Gm5527 | downstream_gene_variant | MODIFIER |
| 1 | 52594475 | AX-97824480 | 0 | G | A | 0.105 | -1.87E-03 | 2.36E-01 | 2.69E+00 | 2.74E+00 | 9.93E-01 | no | | intergenic_variant | MODIFIER |
| 1 | 52604111 | AX-170443123 | 0 | T | C | 0.105 | -1.87E-03 | 2.36E-01 | 2.69E+00 | 2.74E+00 | 9.93E-01 | no | | intergenic_variant | MODIFIER |
| 1 | 52604512 | AX-97793545 | 0 | G | A | 0.105 | -1.87E-03 | 2.36E-01 | 2.69E+00 | 2.74E+00 | 9.93E-01 | no | | intergenic_variant | MODIFIER |
| 1 | 52633040 | AX-170041228 | 0 | T | A | 0.105 | -1.87E-03 | 2.36E-01 | 2.69E+00 | 2.74E+00 | 9.93E-01 | no | Nemp2 | intron_variant | MODIFIER |
| 1 | 52641052 | AX-170496957 | 0 | A | G | 0.105 | -1.87E-03 | 2.36E-01 | 2.69E+00 | 2.74E+00 | 9.93E-01 | no | Nemp2 | missense_variant | MODERATE |
| 1 | 52651656 | AX-170186161 | 0 | G | A | 0.105 | -1.87E-03 | 2.36E-01 | 2.69E+00 | 2.74E+00 | 9.93E-01 | no | Mfsd6 | downstream_gene_variant | MODIFIER |
| 1 | 52651813 | AX-170284050 | 0 | G | A | 0.105 | -1.87E-03 | 2.36E-01 | 2.69E+00 | 2.74E+00 | 9.93E-01 | no | Mfsd6 | downstream_gene_variant | MODIFIER |
| 1 | 52652048 | AX-170463932 | 0 | G | A | 0.105 | -1.87E-03 | 2.36E-01 | 2.69E+00 | 2.74E+00 | 9.93E-01 | no | Mfsd6 | downstream_gene_variant | MODIFIER |
| 1 | 52660466 | AX-97792597 | 0 | T | C | 0.105 | -1.87E-03 | 2.36E-01 | 2.69E+00 | 2.74E+00 | 9.93E-01 | no | Mfsd6 | 3_prime_UTR_variant | MODIFIER |
| 1 | 52684578 | AX-170178315 | 0 | C | G | 0.105 | -1.87E-03 | 2.36E-01 | 2.69E+00 | 2.74E+00 | 9.93E-01 | no | Mfsd6 | intron_variant | MODIFIER |
| 1 | 52685485 | AX-170456324 | 0 | T | C | 0.105 | -1.87E-03 | 2.36E-01 | 2.69E+00 | 2.74E+00 | 9.93E-01 | no | Mfsd6 | intron_variant | MODIFIER |
| 1 | 52686120 | AX-170403385 | 0 | G | A | 0.105 | -1.87E-03 | 2.36E-01 | 2.69E+00 | 2.74E+00 | 9.93E-01 | no | Mfsd6 | intron_variant | MODIFIER |

| | | | | | | | | | | | | | | | |
|---|---|---|---|---|---|---|---|---|---|---|---|---|---|---|---|
| 1 | 52686188 | AX-170168891 | 0 | A | G | 0.105 | -1.87E-03 | 2.36E-01 | 2.69E+00 | 2.74E+00 | 9.93E-01 | no | Mfsd6 | intron_variant | MODIFIER |
| 1 | 52688213 | AX-170231885 | 0 | C | T | 0.105 | -1.87E-03 | 2.36E-01 | 2.69E+00 | 2.74E+00 | 9.93E-01 | no | Mfsd6 | intron_variant | MODIFIER |
| 1 | 52692855 | AX-170447814 | 0 | A | T | 0.105 | -1.87E-03 | 2.36E-01 | 2.69E+00 | 2.74E+00 | 9.93E-01 | no | Mfsd6 | intron_variant | MODIFIER |
| 1 | 52693507 | AX-170294993 | 0 | T | C | 0.105 | -1.87E-03 | 2.36E-01 | 2.69E+00 | 2.74E+00 | 9.93E-01 | no | Mfsd6 | intron_variant | MODIFIER |
| 1 | 52705338 | AX-170041231 | 0 | A | G | 0.105 | -1.87E-03 | 2.36E-01 | 2.69E+00 | 2.74E+00 | 9.93E-01 | no | Mfsd6 | intron_variant | MODIFIER |
| 1 | 52707428 | AX-170389001 | 0 | C | T | 0.105 | -1.87E-03 | 2.36E-01 | 2.69E+00 | 2.74E+00 | 9.93E-01 | no | Mfsd6 | intron_variant | MODIFIER |
| 1 | 52717673 | AX-170365050 | 0 | A | G | 0.105 | -1.87E-03 | 2.36E-01 | 2.69E+00 | 2.74E+00 | 9.93E-01 | no | Mfsd6 | intron_variant | MODIFIER |
| 1 | 52717743 | AX-170110924 | 0 | A | T | 0.105 | -1.87E-03 | 2.36E-01 | 2.69E+00 | 2.74E+00 | 9.93E-01 | no | Mfsd6 | intron_variant | MODIFIER |
| 1 | 52718363 | AX-170358312 | 0 | T | C | 0.105 | -1.87E-03 | 2.36E-01 | 2.69E+00 | 2.74E+00 | 9.93E-01 | no | Mfsd6 | intron_variant | MODIFIER |
| 1 | 52718540 | AX-170208925 | 0 | T | C | 0.105 | -1.87E-03 | 2.36E-01 | 2.69E+00 | 2.74E+00 | 9.93E-01 | no | Mfsd6 | intron_variant | MODIFIER |
| 1 | 52791711 | AX-170000511 | 0 | T | C | 0.105 | -1.87E-03 | 2.36E-01 | 2.69E+00 | 2.74E+00 | 9.93E-01 | no | Inpp1 | synonymous_variant | LOW |
| 1 | 52793371 | AX-170000505 | 0 | G | A | 0.105 | -1.87E-03 | 2.36E-01 | 2.69E+00 | 2.74E+00 | 9.93E-01 | no | Inpp1 | intron_variant | MODIFIER |
| 1 | 52831128 | AX-170311475 | 0 | G | A | 0.105 | -1.87E-03 | 2.36E-01 | 2.69E+00 | 2.74E+00 | 9.93E-01 | no | | intergenic_variant | MODIFIER |
| 1 | 52833121 | AX-170473923 | 0 | G | C | 0.105 | -1.87E-03 | 2.36E-01 | 2.69E+00 | 2.74E+00 | 9.93E-01 | no | | intergenic_variant | MODIFIER |
| 1 | 52860015 | AX-97816816 | 0 | T | C | 0.105 | -1.87E-03 | 2.36E-01 | 2.69E+00 | 2.74E+00 | 9.93E-01 | no | Hibch | intron_variant | MODIFIER |
| 1 | 52871271 | AX-170214853 | 0 | T | G | 0.105 | -1.87E-03 | 2.36E-01 | 2.69E+00 | 2.74E+00 | 9.93E-01 | no | Hibch | intron_variant | MODIFIER |
| 1 | 52876048 | AX-170313570 | 0 | T | C | 0.105 | -1.87E-03 | 2.36E-01 | 2.69E+00 | 2.74E+00 | 9.93E-01 | no | Hibch | intron_variant | MODIFIER |
| 1 | 52884930 | AX-170174340 | 0 | G | C | 0.105 | -1.87E-03 | 2.36E-01 | 2.69E+00 | 2.74E+00 | 9.93E-01 | no | Hibch | intron_variant | MODIFIER |
| 1 | 52885455 | AX-170492541 | 0 | T | C | 0.105 | -1.87E-03 | 2.36E-01 | 2.69E+00 | 2.74E+00 | 9.93E-01 | no | Hibch | intron_variant | MODIFIER |
| 1 | 52911635 | AX-170041235 | 0 | G | A | 0.105 | -1.87E-03 | 2.36E-01 | 2.69E+00 | 2.74E+00 | 9.93E-01 | no | Hibch | intron_variant | MODIFIER |
| 1 | 52918684 | AX-170107666 | 0 | C | T | 0.105 | -1.87E-03 | 2.36E-01 | 2.69E+00 | 2.74E+00 | 9.93E-01 | no | Hibch | intron_variant | MODIFIER |
| 1 | 52919199 | AX-170197206 | 0 | G | A | 0.105 | -1.87E-03 | 2.36E-01 | 2.69E+00 | 2.74E+00 | 9.93E-01 | no | Hibch | intron_variant | MODIFIER |
| 1 | 52934076 | AX-170107548 | 0 | C | T | 0.105 | -1.87E-03 | 2.36E-01 | 2.69E+00 | 2.74E+00 | 9.93E-01 | no | 1700019D03Rik | intron_variant | MODIFIER |
| 1 | 52958714 | AX-170000509 | 0 | G | C | 0.105 | -1.87E-03 | 2.36E-01 | 2.69E+00 | 2.74E+00 | 9.93E-01 | no | 1700019D03Rik | intron_variant | MODIFIER |
| 1 | 52972142 | AX-170000507 | 0 | T | C | 0.105 | -1.87E-03 | 2.36E-01 | 2.69E+00 | 2.74E+00 | 9.93E-01 | no | 1700019D03Rik | upstream_gene_variant | MODIFIER |
| 1 | 52973073 | AX-170313577 | 0 | A | G | 0.105 | -1.87E-03 | 2.36E-01 | 2.69E+00 | 2.74E+00 | 9.93E-01 | no | 1700019D03Rik | upstream_gene_variant | MODIFIER |
| 1 | 52974274 | AX-170433968 | 0 | G | A | 0.105 | -1.87E-03 | 2.36E-01 | 2.69E+00 | 2.74E+00 | 9.93E-01 | no | 1700019D03Rik | upstream_gene_variant | MODIFIER |
| 1 | 52974440 | AX-170511312 | 0 | A | T | 0.105 | -1.87E-03 | 2.36E-01 | 2.69E+00 | 2.74E+00 | 9.93E-01 | no | 1700019D03Rik | upstream_gene_variant | MODIFIER |
| 1 | 53011348 | AX-170390285 | 0 | A | G | 0.105 | -1.87E-03 | 2.36E-01 | 2.69E+00 | 2.74E+00 | 9.93E-01 | no | 1700019D03Rik | intron_variant | MODIFIER |
| 1 | 53011400 | AX-170534253 | 0 | C | A | 0.105 | -1.87E-03 | 2.36E-01 | 2.69E+00 | 2.74E+00 | 9.93E-01 | no | 1700019D03Rik | intron_variant | MODIFIER |
| 1 | 53012591 | AX-170530634 | 0 | C | T | 0.105 | -1.87E-03 | 2.36E-01 | 2.69E+00 | 2.74E+00 | 9.93E-01 | no | 1700019D03Rik | intron_variant | MODIFIER |
| 1 | 53019140 | AX-170529321 | 0 | T | A | 0.105 | -1.87E-03 | 2.36E-01 | 2.69E+00 | 2.74E+00 | 9.93E-01 | no | 1700019D03Rik | intron_variant | MODIFIER |
| 1 | 53023598 | AX-170160207 | 0 | G | A | 0.105 | -1.87E-03 | 2.36E-01 | 2.69E+00 | 2.74E+00 | 9.93E-01 | no | 1700019D03Rik | upstream_gene_variant | MODIFIER |
| 1 | 53034548 | AX-97811608 | 0 | T | G | 0.105 | -1.87E-03 | 2.36E-01 | 2.69E+00 | 2.74E+00 | 9.93E-01 | no | | intergenic_variant | MODIFIER |
| 1 | 53034895 | AX-170504011 | 0 | C | A | 0.105 | -1.87E-03 | 2.36E-01 | 2.69E+00 | 2.74E+00 | 9.93E-01 | no | | intergenic_variant | MODIFIER |
| 1 | 53055145 | AX-170346072 | 0 | A | G | 0.105 | -1.87E-03 | 2.36E-01 | 2.69E+00 | 2.74E+00 | 9.93E-01 | no | | intergenic_variant | MODIFIER |
| 1 | 53073728 | AX-170222159 | 0 | A | G | 0.105 | -1.87E-03 | 2.36E-01 | 2.69E+00 | 2.74E+00 | 9.93E-01 | no | | intergenic_variant | MODIFIER |
| 1 | 53109601 | AX-170155159 | 0 | G | A | 0.105 | -1.87E-03 | 2.36E-01 | 2.69E+00 | 2.74E+00 | 9.93E-01 | no | | intergenic_variant | MODIFIER |
| 1 | 53110092 | AX-170315331 | 0 | G | A | 0.105 | -1.87E-03 | 2.36E-01 | 2.69E+00 | 2.74E+00 | 9.93E-01 | no | | intergenic_variant | MODIFIER |
| 1 | 53112617 | AX-170418696 | 0 | T | G | 0.105 | -1.87E-03 | 2.36E-01 | 2.69E+00 | 2.74E+00 | 9.93E-01 | no | | intergenic_variant | MODIFIER |
| 1 | 53112755 | AX-170400655 | 0 | T | G | 0.211 | -9.34E-04 | 1.18E-01 | 2.69E+00 | 2.74E+00 | 9.93E-01 | no | | intergenic_variant | MODIFIER |
| 1 | 53112935 | AX-170186701 | 0 | G | C | 0.105 | -1.87E-03 | 2.36E-01 | 2.69E+00 | 2.74E+00 | 9.93E-01 | no | | intergenic_variant | MODIFIER |
| 1 | 53123605 | AX-170531574 | 0 | A | G | 0.105 | -1.87E-03 | 2.36E-01 | 2.69E+00 | 2.74E+00 | 9.93E-01 | no | | intergenic_variant | MODIFIER |
| 1 | 53138771 | AX-170383804 | 0 | A | T | 0.105 | -1.87E-03 | 2.36E-01 | 2.69E+00 | 2.74E+00 | 9.93E-01 | no | | intergenic_variant | MODIFIER |
| 1 | 53139035 | AX-170086468 | 0 | C | A | 0.105 | -1.87E-03 | 2.36E-01 | 2.69E+00 | 2.74E+00 | 9.93E-01 | no | | intergenic_variant | MODIFIER |
| 1 | 53145986 | AX-170500968 | 0 | T | C | 0.105 | -1.87E-03 | 2.36E-01 | 2.69E+00 | 2.74E+00 | 9.93E-01 | no | | intergenic_variant | MODIFIER |
| 1 | 53149466 | AX-170269474 | 0 | A | C | 0.105 | -1.87E-03 | 2.36E-01 | 2.69E+00 | 2.74E+00 | 9.93E-01 | no | | intergenic_variant | MODIFIER |
| 1 | 53184540 | AX-170398380 | 0 | C | T | 0.105 | -1.87E-03 | 2.36E-01 | 2.69E+00 | 2.74E+00 | 9.93E-01 | no | Pms1 | downstream_gene_variant | MODIFIER |
| 1 | 53188738 | AX-97819102 | 0 | G | A | 0.105 | -1.87E-03 | 2.36E-01 | 2.69E+00 | 2.74E+00 | 9.93E-01 | no | Pms1 | downstream_gene_variant | MODIFIER |
| 1 | 53194537 | AX-170118701 | 0 | G | C | 0.105 | -1.87E-03 | 2.36E-01 | 2.69E+00 | 2.74E+00 | 9.93E-01 | no | Pms1 | intron_variant | MODIFIER |
| 1 | 53209004 | AX-97817476 | 0 | C | A | 0.105 | -1.87E-03 | 2.36E-01 | 2.69E+00 | 2.74E+00 | 9.93E-01 | no | Pms1 | intron_variant | MODIFIER |
| 1 | 53215644 | AX-170128583 | 0 | C | T | 0.105 | -1.87E-03 | 2.36E-01 | 2.69E+00 | 2.74E+00 | 9.93E-01 | no | Pms1 | intron_variant | MODIFIER |
| 1 | 53216029 | AX-170199885 | 0 | T | G | 0.105 | -1.87E-03 | 2.36E-01 | 2.69E+00 | 2.74E+00 | 9.93E-01 | no | Pms1 | intron_variant | MODIFIER |
| 1 | 53216471 | AX-170279566 | 0 | T | C | 0.105 | -1.87E-03 | 2.36E-01 | 2.69E+00 | 2.74E+00 | 9.93E-01 | no | Pms1 | intron_variant | MODIFIER |

| | | | | | | | | | | | | | | | |
|---|---|---|---|---|---|---|---|---|---|---|---|---|---|---|---|
| 1 | 53230619 | AX-170140301 | 0 | A | G | 0.105 | -1.87E-03 | 2.36E-01 | 2.69E+00 | 2.74E+00 | 9.93E-01 | no | Pms1 | intron_variant | MODIFIER |
| 1 | 53262849 | AX-170310871 | 0 | A | G | 0.105 | -1.87E-03 | 2.36E-01 | 2.69E+00 | 2.74E+00 | 9.93E-01 | no | Pms1 | intron_variant | MODIFIER |
| 1 | 53294205 | AX-170358781 | 0 | T | C | 0.105 | -1.87E-03 | 2.36E-01 | 2.69E+00 | 2.74E+00 | 9.93E-01 | no | Ormdl1 | upstream_gene_variant | MODIFIER |
| 1 | 53323119 | AX-170225037 | 0 | G | A | 0.105 | -1.87E-03 | 2.36E-01 | 2.69E+00 | 2.74E+00 | 9.93E-01 | no | Osgepl1 | intron_variant | MODIFIER |
| 1 | 53323253 | AX-170186034 | 0 | T | C | 0.105 | -1.87E-03 | 2.36E-01 | 2.69E+00 | 2.74E+00 | 9.93E-01 | no | Osgepl1 | 3_prime_UTR_variant | MODIFIER |
| 1 | 53330852 | AX-170494881 | 0 | G | A | 0.105 | -1.87E-03 | 2.36E-01 | 2.69E+00 | 2.74E+00 | 9.93E-01 | no | Osgepl1 | downstream_gene_variant | MODIFIER |
| 1 | 53331015 | AX-170175447 | 0 | G | T | 0.105 | -1.87E-03 | 2.36E-01 | 2.69E+00 | 2.74E+00 | 9.93E-01 | no | Osgepl1 | downstream_gene_variant | MODIFIER |
| 1 | 53350039 | AX-170223598 | 0 | A | G | 0.105 | -1.87E-03 | 2.36E-01 | 2.69E+00 | 2.74E+00 | 9.93E-01 | no | Asnsd1 | 5_prime_UTR_variant | MODIFIER |
| 1 | 53429115 | AX-170323727 | 0 | C | T | 0.105 | -1.87E-03 | 2.36E-01 | 2.69E+00 | 2.74E+00 | 9.93E-01 | no | Dnah7a | intron_variant | MODIFIER |
| 1 | 53442860 | AX-97849179 | 0 | C | T | 0.105 | -1.87E-03 | 2.36E-01 | 2.69E+00 | 2.74E+00 | 9.93E-01 | no | Dnah7a | intron_variant | MODIFIER |
| 1 | 53453975 | AX-97784588 | 0 | C | T | 0.105 | -1.87E-03 | 2.36E-01 | 2.69E+00 | 2.74E+00 | 9.93E-01 | no | Dnah7a | intron_variant | MODIFIER |
| 1 | 53472201 | AX-170514836 | 0 | T | C | 0.105 | -1.87E-03 | 2.36E-01 | 2.69E+00 | 2.74E+00 | 9.93E-01 | no | Dnah7a | intron_variant | MODIFIER |
| 1 | 53472838 | AX-170369271 | 0 | A | G | 0.105 | -1.87E-03 | 2.36E-01 | 2.69E+00 | 2.74E+00 | 9.93E-01 | no | Dnah7a | synonymous_variant | LOW |
| 1 | 53476735 | AX-170467097 | 0 | A | G | 0.105 | -1.87E-03 | 2.36E-01 | 2.69E+00 | 2.74E+00 | 9.93E-01 | no | Dnah7a | intron_variant | MODIFIER |
| 1 | 53518067 | AX-170000516 | 0 | T | C | 0.105 | -1.87E-03 | 2.36E-01 | 2.69E+00 | 2.74E+00 | 9.93E-01 | no | Dnah7a | intron_variant | MODIFIER |
| 1 | 53557027 | AX-170000518 | 0 | G | C | 0.105 | -1.87E-03 | 2.36E-01 | 2.69E+00 | 2.74E+00 | 9.93E-01 | no | Dnah7a | intron_variant | MODIFIER |
| 1 | 53635044 | AX-170000520 | 0 | G | A | 0.105 | -1.87E-03 | 2.36E-01 | 2.69E+00 | 2.74E+00 | 9.93E-01 | no | Dnah7a | intron_variant | MODIFIER |
| 1 | 53646551 | AX-170230156 | 0 | A | G | 0.105 | -1.87E-03 | 2.36E-01 | 2.69E+00 | 2.74E+00 | 9.93E-01 | no | Dnah7a | intron_variant | MODIFIER |
| 1 | 53711039 | AX-170260813 | 0 | G | C | 0.105 | -1.87E-03 | 2.36E-01 | 2.69E+00 | 2.74E+00 | 9.93E-01 | no | Dnah7a | upstream_gene_variant | MODIFIER |
| 1 | 53719190 | AX-97855267 | 0 | C | T | 0.105 | -1.87E-03 | 2.36E-01 | 2.69E+00 | 2.74E+00 | 9.93E-01 | no | | intergenic_variant | MODIFIER |
| 1 | 53726527 | AX-170281045 | 0 | T | C | 0.105 | -1.87E-03 | 2.36E-01 | 2.69E+00 | 2.74E+00 | 9.93E-01 | no | | intergenic_variant | MODIFIER |
| 1 | 53745456 | AX-170134523 | 0 | A | T | 0.105 | -1.87E-03 | 2.36E-01 | 2.69E+00 | 2.74E+00 | 9.93E-01 | no | | intergenic_variant | MODIFIER |
| 1 | 53745850 | AX-170177950 | 0 | G | C | 0.105 | -1.87E-03 | 2.36E-01 | 2.69E+00 | 2.74E+00 | 9.93E-01 | no | | intergenic_variant | MODIFIER |
| 1 | 53756383 | AX-170261293 | 0 | G | A | 0.105 | -1.87E-03 | 2.36E-01 | 2.69E+00 | 2.74E+00 | 9.93E-01 | no | Stk17b | 3_prime_UTR_variant | MODIFIER |
| 1 | 53789116 | AX-170041252 | 0 | T | G | 0.105 | -1.87E-03 | 2.36E-01 | 2.69E+00 | 2.74E+00 | 9.93E-01 | no | Stk17b | upstream_gene_variant | MODIFIER |
| 1 | 53797528 | AX-170444487 | 0 | G | A | 0.105 | -1.87E-03 | 2.36E-01 | 2.69E+00 | 2.74E+00 | 9.93E-01 | no | Gm37863 | downstream_gene_variant | MODIFIER |
| 1 | 53797590 | AX-170167785 | 0 | C | T | 0.105 | -1.87E-03 | 2.36E-01 | 2.69E+00 | 2.74E+00 | 9.93E-01 | no | Gm37863 | downstream_gene_variant | MODIFIER |
| 1 | 53797748 | AX-170493277 | 0 | C | T | 0.105 | -1.87E-03 | 2.36E-01 | 2.69E+00 | 2.74E+00 | 9.93E-01 | no | | regulatory_region_variant | MODIFIER |
| 1 | 53799250 | AX-170370159 | 0 | A | G | 0.105 | -1.87E-03 | 2.36E-01 | 2.69E+00 | 2.74E+00 | 9.93E-01 | no | | intergenic_variant | MODIFIER |
| 1 | 53808283 | AX-170535557 | 0 | A | G | 0.105 | -1.87E-03 | 2.36E-01 | 2.69E+00 | 2.74E+00 | 9.93E-01 | no | Hecw2 | 3_prime_UTR_variant | MODIFIER |
| 1 | 53831752 | AX-97850310 | 0 | C | T | 0.105 | -1.87E-03 | 2.36E-01 | 2.69E+00 | 2.74E+00 | 9.93E-01 | no | Hecw2 | intron_variant | MODIFIER |
| 1 | 53832813 | AX-170142411 | 0 | A | G | 0.105 | -1.87E-03 | 2.36E-01 | 2.69E+00 | 2.74E+00 | 9.93E-01 | no | Hecw2 | intron_variant | MODIFIER |
| 1 | 53863060 | AX-170463684 | 0 | A | G | 0.105 | -1.87E-03 | 2.36E-01 | 2.69E+00 | 2.74E+00 | 9.93E-01 | no | Hecw2 | intron_variant | MODIFIER |
| 1 | 53870497 | AX-170249761 | 0 | G | A | 0.105 | -1.87E-03 | 2.36E-01 | 2.69E+00 | 2.74E+00 | 9.93E-01 | no | Hecw2 | intron_variant | MODIFIER |
| 1 | 53890943 | AX-170258406 | 0 | T | C | 0.105 | -1.87E-03 | 2.36E-01 | 2.69E+00 | 2.74E+00 | 9.93E-01 | no | Hecw2 | intron_variant | MODIFIER |
| 1 | 53892324 | AX-170441104 | 0 | T | C | 0.105 | -1.87E-03 | 2.36E-01 | 2.69E+00 | 2.74E+00 | 9.93E-01 | no | Hecw2 | intron_variant | MODIFIER |
| 1 | 53893527 | AX-170466862 | 0 | T | A | 0.105 | -1.87E-03 | 2.36E-01 | 2.69E+00 | 2.74E+00 | 9.93E-01 | no | Hecw2 | intron_variant | MODIFIER |
| 1 | 53931514 | AX-170180401 | 0 | C | T | 0.105 | -1.87E-03 | 2.36E-01 | 2.69E+00 | 2.74E+00 | 9.93E-01 | no | Hecw2 | intron_variant | MODIFIER |
| 1 | 53952219 | AX-170250849 | 0 | G | A | 0.105 | -1.87E-03 | 2.36E-01 | 2.69E+00 | 2.74E+00 | 9.93E-01 | no | Hecw2 | intron_variant | MODIFIER |
| 1 | 54004316 | AX-170501144 | 0 | G | A | 0.105 | -1.87E-03 | 2.36E-01 | 2.69E+00 | 2.74E+00 | 9.93E-01 | no | Hecw2 | intron_variant | MODIFIER |
| 1 | 54009856 | AX-170214008 | 0 | G | A | 0.105 | -1.87E-03 | 2.36E-01 | 2.69E+00 | 2.74E+00 | 9.93E-01 | no | Hecw2 | intron_variant | MODIFIER |
| 1 | 54013571 | AX-170200717 | 0 | A | G | 0.105 | -1.87E-03 | 2.36E-01 | 2.69E+00 | 2.74E+00 | 9.93E-01 | no | Hecw2 | intron_variant | MODIFIER |
| 1 | 54024190 | AX-170401427 | 0 | G | A | 0.105 | -1.87E-03 | 2.36E-01 | 2.69E+00 | 2.74E+00 | 9.93E-01 | no | Hecw2 | intron_variant | MODIFIER |
| 1 | 54024436 | AX-170203762 | 0 | G | A | 0.105 | -1.87E-03 | 2.36E-01 | 2.69E+00 | 2.74E+00 | 9.93E-01 | no | Hecw2 | intron_variant | MODIFIER |
| 1 | 54040089 | AX-170364497 | 0 | C | T | 0.105 | -1.87E-03 | 2.36E-01 | 2.69E+00 | 2.74E+00 | 9.93E-01 | no | Hecw2 | intron_variant | MODIFIER |
| 1 | 54041666 | AX-170142467 | 0 | A | C | 0.105 | -1.87E-03 | 2.36E-01 | 2.69E+00 | 2.74E+00 | 9.93E-01 | no | Hecw2 | intron_variant | MODIFIER |
| 1 | 54058379 | AX-170492292 | 0 | T | C | 0.105 | -1.87E-03 | 2.36E-01 | 2.69E+00 | 2.74E+00 | 9.93E-01 | no | Hecw2 | intron_variant | MODIFIER |
| 1 | 54079130 | AX-170260338 | 0 | C | T | 0.105 | -1.87E-03 | 2.36E-01 | 2.69E+00 | 2.74E+00 | 9.93E-01 | no | Hecw2 | intron_variant | MODIFIER |
| 1 | 54079171 | AX-170404262 | 0 | T | C | 0.105 | -1.87E-03 | 2.36E-01 | 2.69E+00 | 2.74E+00 | 9.93E-01 | no | Hecw2 | intron_variant | MODIFIER |
| 1 | 54105413 | AX-170168497 | 0 | C | T | 0.105 | -1.87E-03 | 2.36E-01 | 2.69E+00 | 2.74E+00 | 9.93E-01 | no | Hecw2 | intron_variant | MODIFIER |
| 1 | 54105627 | AX-170256447 | 0 | G | C | 0.105 | -1.87E-03 | 2.36E-01 | 2.69E+00 | 2.74E+00 | 9.93E-01 | no | Hecw2 | intron_variant | MODIFIER |
| 1 | 54124483 | AX-170041258 | 0 | C | A | 0.105 | -1.87E-03 | 2.36E-01 | 2.69E+00 | 2.74E+00 | 9.93E-01 | no | Hecw2 | intron_variant | MODIFIER |
| 1 | 54132171 | AX-170453798 | 0 | G | A | 0.105 | -1.87E-03 | 2.36E-01 | 2.69E+00 | 2.74E+00 | 9.93E-01 | no | Hecw2 | intron_variant | MODIFIER |
| 1 | 54150762 | AX-170470717 | 0 | G | A | 0.105 | -1.87E-03 | 2.36E-01 | 2.69E+00 | 2.74E+00 | 9.93E-01 | no | Hecw2 | intron_variant | MODIFIER |

| | | | | | | | | | | | | | | | |
|---|---|---|---|---|---|---|---|---|---|---|---|---|---|---|---|
| 1 | 54153191 | AX-170326912 | 0 | T | G | 0.105 | -1.87E-03 | 2.36E-01 | 2.69E+00 | 2.74E+00 | 9.93E-01 | no | Hecw2 | intron_variant | MODIFIER |
| 1 | 54161302 | AX-170340159 | 0 | A | G | 0.105 | -1.87E-03 | 2.36E-01 | 2.69E+00 | 2.74E+00 | 9.93E-01 | no | Hecw2 | intron_variant | MODIFIER |
| 1 | 54162763 | AX-170000523 | 0 | G | A | 0.105 | -1.87E-03 | 2.36E-01 | 2.69E+00 | 2.74E+00 | 9.93E-01 | no | Hecw2 | intron_variant | MODIFIER |
| 1 | 54163054 | AX-170438256 | 0 | T | C | 0.105 | -1.87E-03 | 2.36E-01 | 2.69E+00 | 2.74E+00 | 9.93E-01 | no | Hecw2 | intron_variant | MODIFIER |
| 1 | 54173315 | AX-170183916 | 0 | T | C | 0.105 | -1.87E-03 | 2.36E-01 | 2.69E+00 | 2.74E+00 | 9.93E-01 | no | Hecw2 | intron_variant | MODIFIER |
| 1 | 54173451 | AX-170264718 | 0 | A | G | 0.105 | -1.87E-03 | 2.36E-01 | 2.69E+00 | 2.74E+00 | 9.93E-01 | no | Hecw2 | intron_variant | MODIFIER |
| 1 | 54192100 | AX-97801826 | 0 | G | T | 0.105 | -1.87E-03 | 2.36E-01 | 2.69E+00 | 2.74E+00 | 9.93E-01 | no | Hecw2 | intron_variant | MODIFIER |
| 1 | 54225272 | AX-170414619 | 0 | C | A | 0.105 | -1.87E-03 | 2.36E-01 | 2.69E+00 | 2.74E+00 | 9.93E-01 | no | | intergenic_variant | MODIFIER |
| 1 | 54251380 | AX-170374155 | 0 | T | C | 0.105 | -1.87E-03 | 2.36E-01 | 2.69E+00 | 2.74E+00 | 9.93E-01 | no | Ccdc150 | intron_variant | MODIFIER |
| 1 | 54260964 | AX-170344309 | 0 | T | G | 0.105 | -1.87E-03 | 2.36E-01 | 2.69E+00 | 2.74E+00 | 9.93E-01 | no | Ccdc150 | intron_variant | MODIFIER |
| 1 | 54264478 | AX-97800074 | 0 | T | C | 0.105 | -1.87E-03 | 2.36E-01 | 2.69E+00 | 2.74E+00 | 9.93E-01 | no | Ccdc150 | intron_variant | MODIFIER |
| 1 | 54272427 | AX-97811585 | 0 | G | A | 0.105 | -1.87E-03 | 2.36E-01 | 2.69E+00 | 2.74E+00 | 9.93E-01 | no | Ccdc150 | intron_variant | MODIFIER |
| 1 | 54272826 | AX-170320881 | 0 | A | T | 0.105 | -1.87E-03 | 2.36E-01 | 2.69E+00 | 2.74E+00 | 9.93E-01 | no | Ccdc150 | intron_variant | MODIFIER |
| 1 | 54279621 | AX-170403434 | 0 | C | T | 0.105 | -1.87E-03 | 2.36E-01 | 2.69E+00 | 2.74E+00 | 9.93E-01 | no | Ccdc150 | intron_variant | MODIFIER |
| 1 | 54305042 | AX-97826049 | 0 | G | A | 0.105 | -1.87E-03 | 2.36E-01 | 2.69E+00 | 2.74E+00 | 9.93E-01 | no | Ccdc150 | intron_variant | MODIFIER |
| 1 | 54314429 | AX-170000524 | 0 | G | A | 0.105 | -1.87E-03 | 2.36E-01 | 2.69E+00 | 2.74E+00 | 9.93E-01 | no | Ccdc150 | intron_variant | MODIFIER |
| 1 | 54327797 | AX-170199178 | 0 | C | T | 0.105 | -1.87E-03 | 2.36E-01 | 2.69E+00 | 2.74E+00 | 9.93E-01 | no | Ccdc150 | intron_variant | MODIFIER |
| 1 | 54357627 | AX-170179404 | 0 | G | A | 0.105 | -1.87E-03 | 2.36E-01 | 2.69E+00 | 2.74E+00 | 9.93E-01 | no | Ccdc150 | intron_variant | MODIFIER |
| 1 | 54360504 | AX-97825312 | 0 | C | T | 0.105 | -1.87E-03 | 2.36E-01 | 2.69E+00 | 2.74E+00 | 9.93E-01 | no | Ccdc150 | intron_variant | MODIFIER |
| 1 | 54370492 | AX-170113459 | 0 | A | G | 0.105 | -1.87E-03 | 2.36E-01 | 2.69E+00 | 2.74E+00 | 9.93E-01 | no | Ccdc150 | downstream_gene_variant | MODIFIER |
| 1 | 54380723 | AX-170041262 | 0 | G | A | 0.105 | -1.87E-03 | 2.36E-01 | 2.69E+00 | 2.74E+00 | 9.93E-01 | no | | intergenic_variant | MODIFIER |
| 1 | 54380797 | AX-97811582 | 0 | A | C | 0.105 | -1.87E-03 | 2.36E-01 | 2.69E+00 | 2.74E+00 | 9.93E-01 | no | | intergenic_variant | MODIFIER |
| 1 | 54398285 | AX-170470286 | 0 | A | C | 0.105 | -1.87E-03 | 2.36E-01 | 2.69E+00 | 2.74E+00 | 9.93E-01 | no | Gtf3c3 | intron_variant | MODIFIER |
| 1 | 54422558 | AX-170041263 | 0 | A | C | 0.105 | -1.87E-03 | 2.36E-01 | 2.69E+00 | 2.74E+00 | 9.93E-01 | no | Gtf3c3 | intron_variant | MODIFIER |
| 1 | 54424628 | AX-170041261 | 0 | A | G | 0.105 | -1.87E-03 | 2.36E-01 | 2.69E+00 | 2.74E+00 | 9.93E-01 | no | Gtf3c3 | intron_variant | MODIFIER |
| 1 | 54448258 | AX-170377444 | 0 | C | A | 0.105 | -1.87E-03 | 2.36E-01 | 2.69E+00 | 2.74E+00 | 9.93E-01 | no | Gm34066 | intron_variant,non_coding_transcript_variant | MODIFIER |
| 1 | 54449296 | AX-170241151 | 0 | T | C | 0.105 | -1.87E-03 | 2.36E-01 | 2.69E+00 | 2.74E+00 | 9.93E-01 | no | Gm34066 | intron_variant,non_coding_transcript_variant | MODIFIER |
| 1 | 54475451 | AX-170410891 | 0 | C | T | 0.105 | -1.87E-03 | 2.36E-01 | 2.69E+00 | 2.74E+00 | 9.93E-01 | no | Pgap1 | 3_prime_UTR_variant | MODIFIER |
| 1 | 54477043 | AX-170255786 | 0 | T | C | 0.105 | -1.87E-03 | 2.36E-01 | 2.69E+00 | 2.74E+00 | 9.93E-01 | no | Pgap1 | 3_prime_UTR_variant | MODIFIER |
| 1 | 54492979 | AX-170381668 | 0 | C | T | 0.105 | -1.87E-03 | 2.36E-01 | 2.69E+00 | 2.74E+00 | 9.93E-01 | no | Pgap1 | intron_variant | MODIFIER |
| 1 | 54494010 | AX-170227784 | 0 | T | C | 0.105 | -1.87E-03 | 2.36E-01 | 2.69E+00 | 2.74E+00 | 9.93E-01 | no | Pgap1 | intron_variant | MODIFIER |
| 1 | 54497518 | AX-170287378 | 0 | A | G | 0.105 | -1.87E-03 | 2.36E-01 | 2.69E+00 | 2.74E+00 | 9.93E-01 | no | Pgap1 | intron_variant | MODIFIER |
| 1 | 54532987 | AX-170324996 | 0 | G | A | 0.105 | -1.87E-03 | 2.36E-01 | 2.69E+00 | 2.74E+00 | 9.93E-01 | no | Pgap1 | intron_variant | MODIFIER |
| 1 | 54543306 | AX-97780629 | 0 | T | C | 0.105 | -1.87E-03 | 2.36E-01 | 2.69E+00 | 2.74E+00 | 9.93E-01 | no | Pgap1 | intron_variant | MODIFIER |
| 1 | 54549553 | AX-170335534 | 0 | T | C | 0.105 | -1.87E-03 | 2.36E-01 | 2.69E+00 | 2.74E+00 | 9.93E-01 | no | Pgap1 | intron_variant | MODIFIER |
| 1 | 54550603 | AX-97849908 | 0 | G | A | 0.105 | -1.87E-03 | 2.36E-01 | 2.69E+00 | 2.74E+00 | 9.93E-01 | no | Pgap1 | intron_variant | MODIFIER |
| 1 | 54552262 | AX-170041265 | 0 | A | C | 0.105 | -1.87E-03 | 2.36E-01 | 2.69E+00 | 2.74E+00 | 9.93E-01 | no | Pgap1 | intron_variant | MODIFIER |
| 1 | 54554337 | AX-170447137 | 0 | A | G | 0.105 | -1.87E-03 | 2.36E-01 | 2.69E+00 | 2.74E+00 | 9.93E-01 | no | Pgap1 | intron_variant | MODIFIER |
| 1 | 54585548 | AX-170178898 | 0 | G | A | 0.105 | -1.87E-03 | 2.36E-01 | 2.69E+00 | 2.74E+00 | 9.93E-01 | no | | regulatory_region_variant | MODIFIER |
| 1 | 54611336 | AX-170491830 | 0 | G | A | 0.105 | -1.87E-03 | 2.36E-01 | 2.69E+00 | 2.74E+00 | 9.93E-01 | no | | intergenic_variant | MODIFIER |
| 1 | 54626821 | AX-170086167 | 0 | A | T | 0.105 | -1.87E-03 | 2.36E-01 | 2.69E+00 | 2.74E+00 | 9.93E-01 | no | | intergenic_variant | MODIFIER |
| 1 | 54640271 | AX-170121454 | 0 | T | A | 0.105 | -1.87E-03 | 2.36E-01 | 2.69E+00 | 2.74E+00 | 9.93E-01 | no | | regulatory_region_variant | MODIFIER |
| 1 | 54650322 | AX-170286876 | 0 | G | A | 0.105 | -1.87E-03 | 2.36E-01 | 2.69E+00 | 2.74E+00 | 9.93E-01 | no | Ankrd44 | intron_variant | MODIFIER |
| 1 | 54651500 | AX-170253784 | 0 | C | T | 0.105 | -1.87E-03 | 2.36E-01 | 2.69E+00 | 2.74E+00 | 9.93E-01 | no | Ankrd44 | intron_variant | MODIFIER |
| 1 | 54653697 | AX-170212492 | 0 | A | C | 0.105 | -1.87E-03 | 2.36E-01 | 2.69E+00 | 2.74E+00 | 9.93E-01 | no | Ankrd44 | intron_variant | MODIFIER |
| 1 | 54653781 | AX-170474386 | 0 | G | A | 0.105 | -1.87E-03 | 2.36E-01 | 2.69E+00 | 2.74E+00 | 9.93E-01 | no | Ankrd44 | synonymous_variant | LOW |
| 1 | 54666483 | AX-170353659 | 0 | C | T | 0.105 | -1.87E-03 | 2.36E-01 | 2.69E+00 | 2.74E+00 | 9.93E-01 | no | Ankrd44 | intron_variant | MODIFIER |
| 1 | 54667601 | AX-170521322 | 0 | C | T | 0.105 | -1.87E-03 | 2.36E-01 | 2.69E+00 | 2.74E+00 | 9.93E-01 | no | Ankrd44 | intron_variant | MODIFIER |
| 1 | 54675121 | AX-170141816 | 0 | T | C | 0.105 | -1.87E-03 | 2.36E-01 | 2.69E+00 | 2.74E+00 | 9.93E-01 | no | Ankrd44 | intron_variant | MODIFIER |
| 1 | 54685583 | AX-170299595 | 0 | A | G | 0.105 | -1.87E-03 | 2.36E-01 | 2.69E+00 | 2.74E+00 | 9.93E-01 | no | Ankrd44 | intron_variant | MODIFIER |
| 1 | 54686564 | AX-170366419 | 0 | T | C | 0.105 | -1.87E-03 | 2.36E-01 | 2.69E+00 | 2.74E+00 | 9.93E-01 | no | Ankrd44 | intron_variant | MODIFIER |
| 1 | 54688940 | AX-170181485 | 0 | T | C | 0.105 | -1.87E-03 | 2.36E-01 | 2.69E+00 | 2.74E+00 | 9.93E-01 | no | Ankrd44 | intron_variant | MODIFIER |
| 1 | 54701116 | AX-170334598 | 0 | G | A | 0.105 | -1.87E-03 | 2.36E-01 | 2.69E+00 | 2.74E+00 | 9.93E-01 | no | Ankrd44 | intron_variant | MODIFIER |
| 1 | 54703645 | AX-170041268 | 0 | C | T | 0.105 | -1.87E-03 | 2.36E-01 | 2.69E+00 | 2.74E+00 | 9.93E-01 | no | Ankrd44 | intron_variant | MODIFIER |

| | | | | | | | | | | | | | | | |
|---|---|---|---|---|---|---|---|---|---|---|---|---|---|---|---|
| 1 | 54712230 | AX-170000535 | 0 | T | C | 0.105 | -1.87E-03 | 2.36E-01 | 2.69E+00 | 2.74E+00 | 9.93E-01 | no | Ankrd44 | intron_variant | MODIFIER |
| 1 | 54712244 | AX-170000537 | 0 | C | T | 0.105 | -1.87E-03 | 2.36E-01 | 2.69E+00 | 2.74E+00 | 9.93E-01 | no | Ankrd44 | intron_variant | MODIFIER |
| 1 | 54731600 | AX-170186908 | 0 | A | G | 0.105 | -1.87E-03 | 2.36E-01 | 2.69E+00 | 2.74E+00 | 9.93E-01 | no | Ankrd44 | intron_variant | MODIFIER |
| 1 | 54737784 | AX-170106425 | 0 | T | A | 0.105 | -1.87E-03 | 2.36E-01 | 2.69E+00 | 2.74E+00 | 9.93E-01 | no | Ankrd44 | intron_variant | MODIFIER |
| 1 | 54740593 | AX-170524358 | 0 | G | A | 0.105 | -1.87E-03 | 2.36E-01 | 2.69E+00 | 2.74E+00 | 9.93E-01 | no | Ankrd44 | intron_variant | MODIFIER |
| 1 | 54741785 | AX-170215505 | 0 | T | C | 0.105 | -1.87E-03 | 2.36E-01 | 2.69E+00 | 2.74E+00 | 9.93E-01 | no | Ankrd44 | intron_variant | MODIFIER |
| 1 | 54748035 | AX-170000536 | 0 | C | T | 0.105 | -1.87E-03 | 2.36E-01 | 2.69E+00 | 2.74E+00 | 9.93E-01 | no | Ankrd44 | intron_variant | MODIFIER |
| 1 | 54769570 | AX-170288485 | 0 | C | G | 0.105 | -1.87E-03 | 2.36E-01 | 2.69E+00 | 2.74E+00 | 9.93E-01 | no | Ankrd44 | intron_variant | MODIFIER |
| 1 | 54789770 | AX-170506170 | 0 | A | T | 0.105 | -1.87E-03 | 2.36E-01 | 2.69E+00 | 2.74E+00 | 9.93E-01 | no | Ankrd44 | intron_variant | MODIFIER |
| 1 | 54790246 | AX-170383845 | 0 | T | A | 0.105 | -1.87E-03 | 2.36E-01 | 2.69E+00 | 2.74E+00 | 9.93E-01 | no | Ankrd44 | intron_variant | MODIFIER |
| 1 | 54793482 | AX-170333077 | 0 | T | A | 0.105 | -1.87E-03 | 2.36E-01 | 2.69E+00 | 2.74E+00 | 9.93E-01 | no | Ankrd44 | intron_variant | MODIFIER |
| 1 | 54794037 | AX-170152424 | 0 | C | T | 0.105 | -1.87E-03 | 2.36E-01 | 2.69E+00 | 2.74E+00 | 9.93E-01 | no | Ankrd44 | intron_variant | MODIFIER |
| 1 | 54824421 | AX-170304683 | 0 | G | T | 0.105 | -1.87E-03 | 2.36E-01 | 2.69E+00 | 2.74E+00 | 9.93E-01 | no | Ankrd44 | intron_variant | MODIFIER |
| 1 | 54839204 | AX-170384805 | 0 | C | T | 0.105 | -1.87E-03 | 2.36E-01 | 2.69E+00 | 2.74E+00 | 9.93E-01 | no | Ankrd44 | intron_variant | MODIFIER |
| 1 | 54843647 | AX-97812945 | 0 | T | C | 0.105 | -1.87E-03 | 2.36E-01 | 2.69E+00 | 2.74E+00 | 9.93E-01 | no | Ankrd44 | intron_variant | MODIFIER |
| 1 | 54848591 | AX-170387943 | 0 | G | C | 0.105 | -1.87E-03 | 2.36E-01 | 2.69E+00 | 2.74E+00 | 9.93E-01 | no | Ankrd44 | intron_variant | MODIFIER |
| 1 | 54849659 | AX-170168194 | 0 | A | G | 0.105 | -1.87E-03 | 2.36E-01 | 2.69E+00 | 2.74E+00 | 9.93E-01 | no | Ankrd44 | intron_variant | MODIFIER |
| 1 | 54853272 | AX-170133265 | 0 | T | C | 0.105 | -1.87E-03 | 2.36E-01 | 2.69E+00 | 2.74E+00 | 9.93E-01 | no | Ankrd44 | intron_variant | MODIFIER |
| 1 | 54853333 | AX-170151950 | 0 | G | A | 0.105 | -1.87E-03 | 2.36E-01 | 2.69E+00 | 2.74E+00 | 9.93E-01 | no | Ankrd44 | intron_variant | MODIFIER |
| 1 | 54853837 | AX-170371518 | 0 | G | C | 0.105 | -1.87E-03 | 2.36E-01 | 2.69E+00 | 2.74E+00 | 9.93E-01 | no | Ankrd44 | intron_variant | MODIFIER |
| 1 | 54854178 | AX-170307117 | 0 | G | C | 0.105 | -1.87E-03 | 2.36E-01 | 2.69E+00 | 2.74E+00 | 9.93E-01 | no | Ankrd44 | intron_variant | MODIFIER |
| 1 | 54854294 | AX-170112609 | 0 | T | C | 0.105 | -1.87E-03 | 2.36E-01 | 2.69E+00 | 2.74E+00 | 9.93E-01 | no | Ankrd44 | intron_variant | MODIFIER |
| 1 | 54855552 | AX-170469590 | 0 | G | A | 0.105 | -1.87E-03 | 2.36E-01 | 2.69E+00 | 2.74E+00 | 9.93E-01 | no | Ankrd44 | intron_variant | MODIFIER |
| 1 | 54855982 | AX-170354972 | 0 | T | C | 0.105 | -1.87E-03 | 2.36E-01 | 2.69E+00 | 2.74E+00 | 9.93E-01 | no | Ankrd44 | intron_variant | MODIFIER |
| 1 | 54858218 | AX-170419026 | 0 | G | A | 0.105 | -1.87E-03 | 2.36E-01 | 2.69E+00 | 2.74E+00 | 9.93E-01 | no | Ankrd44 | intron_variant | MODIFIER |
| 1 | 54862559 | AX-170146276 | 0 | A | T | 0.105 | -1.87E-03 | 2.36E-01 | 2.69E+00 | 2.74E+00 | 9.93E-01 | no | Ankrd44 | intron_variant | MODIFIER |
| 1 | 54873889 | AX-170266105 | 0 | C | T | 0.105 | -1.87E-03 | 2.36E-01 | 2.69E+00 | 2.74E+00 | 9.93E-01 | no | Ankrd44 | intron_variant | MODIFIER |
| 1 | 54873914 | AX-170454798 | 0 | G | T | 0.105 | -1.87E-03 | 2.36E-01 | 2.69E+00 | 2.74E+00 | 9.93E-01 | no | Ankrd44 | intron_variant | MODIFIER |
| 1 | 54876676 | AX-170170532 | 0 | A | C | 0.105 | -1.87E-03 | 2.36E-01 | 2.69E+00 | 2.74E+00 | 9.93E-01 | no | Ankrd44 | intron_variant | MODIFIER |
| 1 | 54876718 | AX-170159360 | 0 | C | A | 0.105 | -1.87E-03 | 2.36E-01 | 2.69E+00 | 2.74E+00 | 9.93E-01 | no | Ankrd44 | intron_variant | MODIFIER |
| 1 | 54877753 | AX-170312067 | 0 | T | C | 0.105 | -1.87E-03 | 2.36E-01 | 2.69E+00 | 2.74E+00 | 9.93E-01 | no | Ankrd44 | intron_variant | MODIFIER |
| 1 | 54877940 | AX-170467233 | 0 | G | C | 0.105 | -1.87E-03 | 2.36E-01 | 2.69E+00 | 2.74E+00 | 9.93E-01 | no | Ankrd44 | intron_variant | MODIFIER |
| 1 | 54877985 | AX-97808372 | 0 | A | C | 0.105 | -1.87E-03 | 2.36E-01 | 2.69E+00 | 2.74E+00 | 9.93E-01 | no | Ankrd44 | intron_variant | MODIFIER |
| 1 | 54899649 | AX-170041274 | 0 | G | A | 0.105 | -1.87E-03 | 2.36E-01 | 2.69E+00 | 2.74E+00 | 9.93E-01 | no | Ankrd44 | intron_variant | MODIFIER |
| 1 | 54909797 | AX-170306649 | 0 | C | T | 0.105 | -1.87E-03 | 2.36E-01 | 2.69E+00 | 2.74E+00 | 9.93E-01 | no | Ankrd44 | intron_variant | MODIFIER |
| 1 | 54913386 | AX-170416577 | 0 | G | A | 0.105 | -1.87E-03 | 2.36E-01 | 2.69E+00 | 2.74E+00 | 9.93E-01 | no | Ankrd44 | intron_variant | MODIFIER |
| 1 | 54913489 | AX-170105282 | 0 | G | C | 0.105 | -1.87E-03 | 2.36E-01 | 2.69E+00 | 2.74E+00 | 9.93E-01 | no | Ankrd44 | intron_variant | MODIFIER |
| 1 | 54913573 | AX-170085970 | 0 | G | A | 0.105 | -1.87E-03 | 2.36E-01 | 2.69E+00 | 2.74E+00 | 9.93E-01 | no | Ankrd44 | intron_variant | MODIFIER |
| 1 | 54913603 | AX-170360692 | 0 | G | A | 0.105 | -1.87E-03 | 2.36E-01 | 2.69E+00 | 2.74E+00 | 9.93E-01 | no | Ankrd44 | intron_variant | MODIFIER |
| 1 | 54928416 | AX-170383452 | 0 | G | T | 0.105 | -1.87E-03 | 2.36E-01 | 2.69E+00 | 2.74E+00 | 9.93E-01 | no | Ankrd44 | upstream_gene_variant | MODIFIER |
| 1 | 54962018 | AX-170126046 | 0 | T | C | 0.105 | -1.87E-03 | 2.36E-01 | 2.69E+00 | 2.74E+00 | 9.93E-01 | no | | intergenic_variant | MODIFIER |
| 1 | 54979781 | AX-170391022 | 0 | C | T | 0.105 | -1.87E-03 | 2.36E-01 | 2.69E+00 | 2.74E+00 | 9.93E-01 | no | | regulatory_region_variant | MODIFIER |
| 1 | 54980154 | AX-170157714 | 0 | C | T | 0.105 | -1.87E-03 | 2.36E-01 | 2.69E+00 | 2.74E+00 | 9.93E-01 | no | | intergenic_variant | MODIFIER |
| 1 | 54983356 | AX-170274301 | 0 | C | G | 0.105 | -1.87E-03 | 2.36E-01 | 2.69E+00 | 2.74E+00 | 9.93E-01 | no | Sf3b1 | downstream_gene_variant | MODIFIER |
| 1 | 54990653 | AX-170364474 | 0 | G | C | 0.105 | -1.87E-03 | 2.36E-01 | 2.69E+00 | 2.74E+00 | 9.93E-01 | no | Sf3b1 | intron_variant | MODIFIER |
| 1 | 55009085 | AX-170252350 | 0 | T | G | 0.105 | -1.87E-03 | 2.36E-01 | 2.69E+00 | 2.74E+00 | 9.93E-01 | no | Sf3b1 | intron_variant | MODIFIER |
| 1 | 55013335 | AX-170269988 | 0 | A | C | 0.105 | -1.87E-03 | 2.36E-01 | 2.69E+00 | 2.74E+00 | 9.93E-01 | no | Sf3b1 | intron_variant | MODIFIER |
| 1 | 55027623 | AX-170283150 | 0 | T | C | 0.105 | -1.87E-03 | 2.36E-01 | 2.69E+00 | 2.74E+00 | 9.93E-01 | no | Sf3b1 | upstream_gene_variant | MODIFIER |
| 1 | 55030228 | AX-97827394 | 0 | A | G | 0.105 | -1.87E-03 | 2.36E-01 | 2.69E+00 | 2.74E+00 | 9.93E-01 | no | Sf3b1 | upstream_gene_variant | MODIFIER |
| 1 | 55037453 | AX-170334048 | 0 | G | A | 0.105 | -1.87E-03 | 2.36E-01 | 2.69E+00 | 2.74E+00 | 9.93E-01 | no | | intergenic_variant | MODIFIER |
| 1 | 55045446 | AX-170041280 | 0 | A | G | 0.105 | -1.87E-03 | 2.36E-01 | 2.69E+00 | 2.74E+00 | 9.93E-01 | no | | intergenic_variant | MODIFIER |
| 1 | 55045460 | AX-170041279 | 0 | A | G | 0.105 | -1.87E-03 | 2.36E-01 | 2.69E+00 | 2.74E+00 | 9.93E-01 | no | | intergenic_variant | MODIFIER |
| 1 | 55061019 | AX-170380684 | 0 | G | T | 0.105 | -1.87E-03 | 2.36E-01 | 2.69E+00 | 2.74E+00 | 9.93E-01 | no | Coq10b | intron_variant | MODIFIER |
| 1 | 55061035 | AX-170303295 | 0 | A | G | 0.105 | -1.87E-03 | 2.36E-01 | 2.69E+00 | 2.74E+00 | 9.93E-01 | no | Coq10b | intron_variant | MODIFIER |

| | | | | | | | | | | | | | | | |
|---|---|---|---|---|---|---|---|---|---|---|---|---|---|---|---|
| 1 | 55061359 | AX-170360299 | 0 | C | T | 0.105 | -1.87E-03 | 2.36E-01 | 2.69E+00 | 2.74E+00 | 9.93E-01 | no | Coq10b | synonymous_variant | LOW |
| 1 | 55064392 | AX-170041278 | 0 | G | T | 0.105 | -1.87E-03 | 2.36E-01 | 2.69E+00 | 2.74E+00 | 9.93E-01 | no | Coq10b | intron_variant | MODIFIER |
| 1 | 55067363 | AX-170138903 | 0 | C | T | 0.105 | -1.87E-03 | 2.36E-01 | 2.69E+00 | 2.74E+00 | 9.93E-01 | no | Coq10b | intron_variant | MODIFIER |
| 1 | 55071325 | AX-170254753 | 0 | T | C | 0.105 | -1.87E-03 | 2.36E-01 | 2.69E+00 | 2.74E+00 | 9.93E-01 | no | Coq10b | intron_variant | MODIFIER |
| 1 | 55071924 | AX-170121182 | 0 | C | T | 0.105 | -1.87E-03 | 2.36E-01 | 2.69E+00 | 2.74E+00 | 9.93E-01 | no | Coq10b | 3_prime_UTR_variant | MODIFIER |
| 1 | 55084933 | AX-97846143 | 0 | T | C | 0.105 | -1.87E-03 | 2.36E-01 | 2.69E+00 | 2.74E+00 | 9.93E-01 | no | Hspd1 | intron_variant | MODIFIER |
| 1 | 55124018 | AX-170106634 | 0 | T | C | 0.105 | -1.87E-03 | 2.36E-01 | 2.69E+00 | 2.74E+00 | 9.93E-01 | no | Gm16581 | upstream_gene_variant | MODIFIER |
| 1 | 55149508 | AX-97833158 | 0 | C | T | 0.105 | -1.87E-03 | 2.36E-01 | 2.69E+00 | 2.74E+00 | 9.93E-01 | no | Mob4 | intron_variant | MODIFIER |
| 1 | 55171885 | AX-97783605 | 0 | G | A | 0.105 | -1.87E-03 | 2.36E-01 | 2.69E+00 | 2.74E+00 | 9.93E-01 | no | Rftn2 | synonymous_variant | LOW |
| 1 | 55178346 | AX-170349220 | 0 | A | G | 0.105 | -1.87E-03 | 2.36E-01 | 2.69E+00 | 2.74E+00 | 9.93E-01 | no | Rftn2 | intron_variant | MODIFIER |
| 1 | 55187316 | AX-170212219 | 0 | G | A | 0.105 | -1.87E-03 | 2.36E-01 | 2.69E+00 | 2.74E+00 | 9.93E-01 | no | Rftn2 | intron_variant | MODIFIER |
| 1 | 55193139 | AX-170209053 | 0 | G | C | 0.105 | -1.87E-03 | 2.36E-01 | 2.69E+00 | 2.74E+00 | 9.93E-01 | no | Rftn2 | intron_variant | MODIFIER |
| 1 | 55194306 | AX-170111933 | 0 | A | G | 0.105 | -1.87E-03 | 2.36E-01 | 2.69E+00 | 2.74E+00 | 9.93E-01 | no | Rftn2 | synonymous_variant | LOW |
| 1 | 55202393 | AX-170275259 | 0 | G | A | 0.105 | -1.87E-03 | 2.36E-01 | 2.69E+00 | 2.74E+00 | 9.93E-01 | no | Rftn2 | intron_variant | MODIFIER |
| 1 | 55204479 | AX-170105097 | 0 | G | A | 0.105 | -1.87E-03 | 2.36E-01 | 2.69E+00 | 2.74E+00 | 9.93E-01 | no | Rftn2 | intron_variant | MODIFIER |
| 1 | 55221895 | AX-97816149 | 0 | G | A | 0.105 | -1.87E-03 | 2.36E-01 | 2.69E+00 | 2.74E+00 | 9.93E-01 | no | Rftn2 | intron_variant | MODIFIER |
| 1 | 55224743 | AX-170124457 | 0 | T | A | 0.105 | -1.87E-03 | 2.36E-01 | 2.69E+00 | 2.74E+00 | 9.93E-01 | no | Rftn2 | intron_variant | MODIFIER |
| 1 | 55226073 | AX-170164951 | 0 | C | T | 0.105 | -1.87E-03 | 2.36E-01 | 2.69E+00 | 2.74E+00 | 9.93E-01 | no | Rftn2 | intron_variant | MODIFIER |
| 1 | 55227927 | AX-170035924 | 0 | G | A | 0.105 | -1.87E-03 | 2.36E-01 | 2.69E+00 | 2.74E+00 | 9.93E-01 | no | Rftn2 | upstream_gene_variant | MODIFIER |
| 1 | 55250704 | AX-170119790 | 0 | T | C | 0.105 | -1.87E-03 | 2.36E-01 | 2.69E+00 | 2.74E+00 | 9.93E-01 | no | Mars2 | downstream_gene_variant | MODIFIER |
| 1 | 55261121 | AX-170127743 | 0 | C | G | 0.105 | -1.87E-03 | 2.36E-01 | 2.69E+00 | 2.74E+00 | 9.93E-01 | no | Boll | intron_variant,NMD_transcript_variant | MODIFIER |
| 1 | 55276949 | AX-170535705 | 0 | C | T | 0.105 | -1.87E-03 | 2.36E-01 | 2.69E+00 | 2.74E+00 | 9.93E-01 | no | Boll | intron_variant,NMD_transcript_variant | MODIFIER |
| 1 | 55310770 | AX-170379864 | 0 | G | A | 0.105 | -1.87E-03 | 2.36E-01 | 2.69E+00 | 2.74E+00 | 9.93E-01 | no | Boll | intron_variant | MODIFIER |
| 1 | 55311819 | AX-97811586 | 0 | A | G | 0.105 | -1.87E-03 | 2.36E-01 | 2.69E+00 | 2.74E+00 | 9.93E-01 | no | Boll | intron_variant | MODIFIER |
| 1 | 55351462 | AX-170000538 | 0 | T | C | 0.105 | -1.87E-03 | 2.36E-01 | 2.69E+00 | 2.74E+00 | 9.93E-01 | no | Boll | intron_variant | MODIFIER |
| 1 | 55357257 | AX-97828509 | 0 | A | G | 0.105 | -1.87E-03 | 2.36E-01 | 2.69E+00 | 2.74E+00 | 9.93E-01 | no | Boll | intron_variant | MODIFIER |
| 1 | 55362008 | AX-170431154 | 0 | C | T | 0.105 | -1.87E-03 | 2.36E-01 | 2.69E+00 | 2.74E+00 | 9.93E-01 | no | Boll | intron_variant | MODIFIER |
| 1 | 55372970 | AX-170263768 | 0 | G | A | 0.105 | -1.87E-03 | 2.36E-01 | 2.69E+00 | 2.74E+00 | 9.93E-01 | no | Gm29138 | upstream_gene_variant | MODIFIER |
| 1 | 55399565 | AX-170091519 | 0 | A | C | 0.105 | -1.87E-03 | 2.36E-01 | 2.69E+00 | 2.74E+00 | 9.93E-01 | no | | intergenic_variant | MODIFIER |
| 1 | 55399780 | AX-170535918 | 0 | C | A | 0.105 | -1.87E-03 | 2.36E-01 | 2.69E+00 | 2.74E+00 | 9.93E-01 | no | | intergenic_variant | MODIFIER |
| 1 | 55404070 | AX-170310096 | 0 | T | G | 0.105 | -1.87E-03 | 2.36E-01 | 2.69E+00 | 2.74E+00 | 9.93E-01 | no | Plcl1 | upstream_gene_variant | MODIFIER |
| 1 | 55430257 | AX-170488212 | 0 | T | G | 0.105 | -1.87E-03 | 2.36E-01 | 2.69E+00 | 2.74E+00 | 9.93E-01 | no | Plcl1 | intron_variant | MODIFIER |
| 1 | 55430394 | AX-97797801 | 0 | T | C | 0.105 | -1.87E-03 | 2.36E-01 | 2.69E+00 | 2.74E+00 | 9.93E-01 | no | Plcl1 | intron_variant | MODIFIER |
| 1 | 55433004 | AX-170290383 | 0 | C | T | 0.105 | -1.87E-03 | 2.36E-01 | 2.69E+00 | 2.74E+00 | 9.93E-01 | no | Plcl1 | intron_variant | MODIFIER |
| 1 | 55454662 | AX-170179904 | 0 | G | T | 0.105 | -1.87E-03 | 2.36E-01 | 2.69E+00 | 2.74E+00 | 9.93E-01 | no | Plcl1 | intron_variant | MODIFIER |
| 1 | 55454777 | AX-170481350 | 0 | T | C | 0.105 | -1.87E-03 | 2.36E-01 | 2.69E+00 | 2.74E+00 | 9.93E-01 | no | Plcl1 | intron_variant | MODIFIER |
| 1 | 55458244 | AX-170510410 | 0 | T | G | 0.105 | -1.87E-03 | 2.36E-01 | 2.69E+00 | 2.74E+00 | 9.93E-01 | no | Plcl1 | intron_variant | MODIFIER |
| 1 | 55460802 | AX-170222172 | 0 | C | T | 0.105 | -1.87E-03 | 2.36E-01 | 2.69E+00 | 2.74E+00 | 9.93E-01 | no | Plcl1 | intron_variant | MODIFIER |
| 1 | 55473064 | AX-170245466 | 0 | G | A | 0.105 | -1.87E-03 | 2.36E-01 | 2.69E+00 | 2.74E+00 | 9.93E-01 | no | Plcl1 | intron_variant | MODIFIER |
| 1 | 55486102 | AX-170041288 | 0 | T | C | 0.105 | -1.87E-03 | 2.36E-01 | 2.69E+00 | 2.74E+00 | 9.93E-01 | no | Plcl1 | intron_variant | MODIFIER |
| 1 | 55500684 | AX-170428788 | 0 | G | T | 0.105 | -1.87E-03 | 2.36E-01 | 2.69E+00 | 2.74E+00 | 9.93E-01 | no | Plcl1 | intron_variant | MODIFIER |
| 1 | 55509212 | AX-170122818 | 0 | G | A | 0.105 | -1.87E-03 | 2.36E-01 | 2.69E+00 | 2.74E+00 | 9.93E-01 | no | Plcl1 | intron_variant | MODIFIER |
| 1 | 55509254 | AX-170527576 | 0 | C | T | 0.105 | -1.87E-03 | 2.36E-01 | 2.69E+00 | 2.74E+00 | 9.93E-01 | no | Plcl1 | intron_variant | MODIFIER |
| 1 | 55510545 | AX-170307607 | 0 | G | A | 0.105 | -1.87E-03 | 2.36E-01 | 2.69E+00 | 2.74E+00 | 9.93E-01 | no | Plcl1 | intron_variant | MODIFIER |
| 1 | 55524552 | AX-170213577 | 0 | G | A | 0.105 | -1.87E-03 | 2.36E-01 | 2.69E+00 | 2.74E+00 | 9.93E-01 | no | Plcl1 | intron_variant | MODIFIER |
| 1 | 55527838 | AX-170294347 | 0 | T | G | 0.105 | -1.87E-03 | 2.36E-01 | 2.69E+00 | 2.74E+00 | 9.93E-01 | no | Plcl1 | intron_variant | MODIFIER |
| 1 | 55528090 | AX-170328331 | 0 | G | T | 0.105 | -1.87E-03 | 2.36E-01 | 2.69E+00 | 2.74E+00 | 9.93E-01 | no | Plcl1 | intron_variant | MODIFIER |
| 1 | 55528755 | AX-170423276 | 0 | T | C | 0.105 | -1.87E-03 | 2.36E-01 | 2.69E+00 | 2.74E+00 | 9.93E-01 | no | Plcl1 | intron_variant | MODIFIER |
| 1 | 55528859 | AX-170279121 | 0 | C | T | 0.105 | -1.87E-03 | 2.36E-01 | 2.69E+00 | 2.74E+00 | 9.93E-01 | no | Plcl1 | intron_variant | MODIFIER |
| 1 | 55558342 | AX-170486361 | 0 | C | G | 0.105 | -1.87E-03 | 2.36E-01 | 2.69E+00 | 2.74E+00 | 9.93E-01 | no | Plcl1 | intron_variant | MODIFIER |
| 1 | 55576098 | AX-170209578 | 0 | G | A | 0.105 | -1.87E-03 | 2.36E-01 | 2.69E+00 | 2.74E+00 | 9.93E-01 | no | Plcl1 | intron_variant | MODIFIER |
| 1 | 55576560 | AX-170105024 | 0 | G | T | 0.105 | -1.87E-03 | 2.36E-01 | 2.69E+00 | 2.74E+00 | 9.93E-01 | no | Plcl1 | intron_variant | MODIFIER |
| 1 | 55579579 | AX-170461826 | 0 | A | G | 0.105 | -1.87E-03 | 2.36E-01 | 2.69E+00 | 2.74E+00 | 9.93E-01 | no | Plcl1 | intron_variant | MODIFIER |
| 1 | 55587472 | AX-170091520 | 0 | A | G | 0.105 | -1.87E-03 | 2.36E-01 | 2.69E+00 | 2.74E+00 | 9.93E-01 | no | Plcl1 | intron_variant | MODIFIER |

| | | | | | | | | | | | | | | | |
|---|---|---|---|---|---|---|---|---|---|---|---|---|---|---|---|
| 1 | 55613184 | AX-170172500 | 0 | G | A | 0.105 | -1.87E-03 | 2.36E-01 | 2.69E+00 | 2.74E+00 | 9.93E-01 | no | Plcl1 | intron_variant | MODIFIER |
| 1 | 55619539 | AX-170487417 | 0 | A | G | 0.105 | -1.87E-03 | 2.36E-01 | 2.69E+00 | 2.74E+00 | 9.93E-01 | no | Plcl1 | intron_variant | MODIFIER |
| 1 | 55636176 | AX-170219387 | 0 | T | G | 0.105 | -1.87E-03 | 2.36E-01 | 2.69E+00 | 2.74E+00 | 9.93E-01 | no | Plcl1 | intron_variant | MODIFIER |
| 1 | 55636319 | AX-170326105 | 0 | C | T | 0.105 | -1.87E-03 | 2.36E-01 | 2.69E+00 | 2.74E+00 | 9.93E-01 | no | Plcl1 | intron_variant | MODIFIER |
| 1 | 55639093 | AX-170432673 | 0 | C | T | 0.105 | -1.87E-03 | 2.36E-01 | 2.69E+00 | 2.74E+00 | 9.93E-01 | no | Plcl1 | intron_variant | MODIFIER |
| 1 | 55639298 | AX-170400644 | 0 | C | T | 0.105 | -1.87E-03 | 2.36E-01 | 2.69E+00 | 2.74E+00 | 9.93E-01 | no | Plcl1 | intron_variant | MODIFIER |
| 1 | 55639552 | AX-170132268 | 0 | C | T | 0.105 | -1.87E-03 | 2.36E-01 | 2.69E+00 | 2.74E+00 | 9.93E-01 | no | Plcl1 | intron_variant | MODIFIER |
| 1 | 55646186 | AX-170469851 | 0 | A | G | 0.105 | -1.87E-03 | 2.36E-01 | 2.69E+00 | 2.74E+00 | 9.93E-01 | no | Plcl1 | intron_variant | MODIFIER |
| 1 | 55649203 | AX-170117274 | 0 | G | A | 0.105 | -1.87E-03 | 2.36E-01 | 2.69E+00 | 2.74E+00 | 9.93E-01 | no | Plcl1 | intron_variant | MODIFIER |
| 1 | 55653952 | AX-170351510 | 0 | A | G | 0.105 | -1.87E-03 | 2.36E-01 | 2.69E+00 | 2.74E+00 | 9.93E-01 | no | Plcl1 | intron_variant | MODIFIER |
| 1 | 55675192 | AX-170150458 | 0 | T | C | 0.105 | -1.87E-03 | 2.36E-01 | 2.69E+00 | 2.74E+00 | 9.93E-01 | no | Plcl1 | intron_variant | MODIFIER |
| 1 | 55675950 | AX-97843994 | 0 | G | A | 0.105 | -1.87E-03 | 2.36E-01 | 2.69E+00 | 2.74E+00 | 9.93E-01 | no | Plcl1 | intron_variant | MODIFIER |
| 1 | 55694857 | AX-170256645 | 0 | T | C | 0.105 | -1.87E-03 | 2.36E-01 | 2.69E+00 | 2.74E+00 | 9.93E-01 | no | Plcl1 | intron_variant | MODIFIER |
| 1 | 55695216 | AX-170183140 | 0 | A | G | 0.105 | -1.87E-03 | 2.36E-01 | 2.69E+00 | 2.74E+00 | 9.93E-01 | no | Plcl1 | intron_variant | MODIFIER |
| 1 | 55695515 | AX-170103818 | 0 | A | C | 0.105 | -1.87E-03 | 2.36E-01 | 2.69E+00 | 2.74E+00 | 9.93E-01 | no | Plcl1 | intron_variant | MODIFIER |
| 1 | 55700447 | AX-170390945 | 0 | A | G | 0.105 | -1.87E-03 | 2.36E-01 | 2.69E+00 | 2.74E+00 | 9.93E-01 | no | Plcl1 | intron_variant | MODIFIER |
| 1 | 55704262 | AX-170389593 | 0 | T | C | 0.105 | -1.87E-03 | 2.36E-01 | 2.69E+00 | 2.74E+00 | 9.93E-01 | no | Plcl1 | intron_variant | MODIFIER |
| 1 | 55713188 | AX-97843885 | 0 | C | T | 0.105 | -1.87E-03 | 2.36E-01 | 2.69E+00 | 2.74E+00 | 9.93E-01 | no | Plcl1 | intron_variant | MODIFIER |
| 1 | 55729449 | AX-170427363 | 0 | T | C | 0.105 | -1.87E-03 | 2.36E-01 | 2.69E+00 | 2.74E+00 | 9.93E-01 | no | Plcl1 | intron_variant | MODIFIER |
| 1 | 55729762 | AX-170310131 | 0 | T | C | 0.105 | -1.87E-03 | 2.36E-01 | 2.69E+00 | 2.74E+00 | 9.93E-01 | no | Plcl1 | intron_variant | MODIFIER |
| 1 | 55732826 | AX-170293804 | 0 | G | A | 0.105 | -1.87E-03 | 2.36E-01 | 2.69E+00 | 2.74E+00 | 9.93E-01 | no | Plcl1 | intron_variant | MODIFIER |
| 1 | 55742345 | AX-170338604 | 0 | T | A | 0.105 | -1.87E-03 | 2.36E-01 | 2.69E+00 | 2.74E+00 | 9.93E-01 | no | Plcl1 | intron_variant | MODIFIER |
| 1 | 55748125 | AX-170331431 | 0 | C | T | 0.105 | -1.87E-03 | 2.36E-01 | 2.69E+00 | 2.74E+00 | 9.93E-01 | no | Plcl1 | intron_variant | MODIFIER |
| 1 | 55752878 | AX-170104044 | 0 | A | G | 0.105 | -1.87E-03 | 2.36E-01 | 2.69E+00 | 2.74E+00 | 9.93E-01 | no | Plcl1 | 3_prime_UTR_variant | MODIFIER |
| 1 | 55756523 | AX-170408286 | 0 | T | C | 0.105 | -1.87E-03 | 2.36E-01 | 2.69E+00 | 2.74E+00 | 9.93E-01 | no | Plcl1 | downstream_gene_variant | MODIFIER |
| 1 | 55757660 | AX-170454993 | 0 | C | T | 0.105 | -1.87E-03 | 2.36E-01 | 2.69E+00 | 2.74E+00 | 9.93E-01 | no | Plcl1 | downstream_gene_variant | MODIFIER |
| 1 | 55761882 | AX-170226996 | 0 | A | G | 0.105 | -1.87E-03 | 2.36E-01 | 2.69E+00 | 2.74E+00 | 9.93E-01 | no | | regulatory_region_variant | MODIFIER |
| 1 | 55770212 | AX-170216738 | 0 | A | G | 0.105 | -1.87E-03 | 2.36E-01 | 2.69E+00 | 2.74E+00 | 9.93E-01 | no | | intergenic_variant | MODIFIER |
| 1 | 55770267 | AX-170350091 | 0 | G | A | 0.105 | -1.87E-03 | 2.36E-01 | 2.69E+00 | 2.74E+00 | 9.93E-01 | no | | intergenic_variant | MODIFIER |
| 1 | 55790191 | AX-170170752 | 0 | A | G | 0.105 | -1.87E-03 | 2.36E-01 | 2.69E+00 | 2.74E+00 | 9.93E-01 | no | | intergenic_variant | MODIFIER |
| 1 | 55795870 | AX-170356849 | 0 | G | A | 0.105 | -1.87E-03 | 2.36E-01 | 2.69E+00 | 2.74E+00 | 9.93E-01 | no | | intergenic_variant | MODIFIER |
| 1 | 55803231 | AX-170405547 | 0 | C | T | 0.105 | -1.87E-03 | 2.36E-01 | 2.69E+00 | 2.74E+00 | 9.93E-01 | no | | intergenic_variant | MODIFIER |
| 1 | 55808854 | AX-170322709 | 0 | C | A | 0.105 | -1.87E-03 | 2.36E-01 | 2.69E+00 | 2.74E+00 | 9.93E-01 | no | | regulatory_region_variant | MODIFIER |
| 1 | 55810566 | AX-170041297 | 0 | A | C | 0.105 | -1.87E-03 | 2.36E-01 | 2.69E+00 | 2.74E+00 | 9.93E-01 | no | | intergenic_variant | MODIFIER |
| 1 | 55826627 | AX-170482057 | 0 | C | T | 0.105 | -1.87E-03 | 2.36E-01 | 2.69E+00 | 2.74E+00 | 9.93E-01 | no | | intergenic_variant | MODIFIER |
| 1 | 55830205 | AX-170155899 | 0 | G | A | 0.105 | -1.87E-03 | 2.36E-01 | 2.69E+00 | 2.74E+00 | 9.93E-01 | no | | intergenic_variant | MODIFIER |
| 1 | 59125885 | AX-170405059 | 0 | G | A | 0.105 | -1.87E-03 | 2.36E-01 | 2.69E+00 | 2.74E+00 | 9.93E-01 | no | Mpp4 | intron_variant | MODIFIER |
| 1 | 59127320 | AX-170511726 | 0 | C | T | 0.105 | -1.87E-03 | 2.36E-01 | 2.69E+00 | 2.74E+00 | 9.93E-01 | no | Mpp4 | intron_variant | MODIFIER |
| 1 | 59127969 | AX-170163455 | 0 | T | C | 0.105 | -1.87E-03 | 2.36E-01 | 2.69E+00 | 2.74E+00 | 9.93E-01 | no | Mpp4 | intron_variant | MODIFIER |
| 1 | 59128928 | AX-170146008 | 0 | A | G | 0.105 | -1.87E-03 | 2.36E-01 | 2.69E+00 | 2.74E+00 | 9.93E-01 | no | Mpp4 | splice_region_variant,intron_variant | LOW |
| 1 | 59129495 | AX-170486655 | 0 | A | G | 0.105 | -1.87E-03 | 2.36E-01 | 2.69E+00 | 2.74E+00 | 9.93E-01 | no | Mpp4 | intron_variant | MODIFIER |
| 1 | 59129679 | AX-97782921 | 0 | A | C | 0.105 | -1.87E-03 | 2.36E-01 | 2.69E+00 | 2.74E+00 | 9.93E-01 | no | Mpp4 | intron_variant | MODIFIER |
| 1 | 59137510 | AX-170405202 | 0 | C | T | 0.105 | -1.87E-03 | 2.36E-01 | 2.69E+00 | 2.74E+00 | 9.93E-01 | no | Mpp4 | intron_variant | MODIFIER |
| 1 | 59146617 | AX-170041369 | 0 | C | G | 0.211 | -9.34E-04 | 1.18E-01 | 2.69E+00 | 2.74E+00 | 9.93E-01 | no | Mpp4 | intron_variant | MODIFIER |
| 1 | 59148497 | AX-170320757 | 0 | A | C | 0.105 | -1.87E-03 | 2.36E-01 | 2.69E+00 | 2.74E+00 | 9.93E-01 | no | Mpp4 | intron_variant | MODIFIER |
| 1 | 59365075 | AX-170124095 | 0 | A | G | 0.105 | -1.87E-03 | 2.36E-01 | 2.69E+00 | 2.74E+00 | 9.93E-01 | no | | intergenic_variant | MODIFIER |
| 1 | 59365199 | AX-170186384 | 0 | A | G | 0.105 | -1.87E-03 | 2.36E-01 | 2.69E+00 | 2.74E+00 | 9.93E-01 | no | | intergenic_variant | MODIFIER |
| 1 | 59365614 | AX-170100516 | 0 | G | A | 0.105 | -1.87E-03 | 2.36E-01 | 2.69E+00 | 2.74E+00 | 9.93E-01 | no | | intergenic_variant | MODIFIER |
| 1 | 59433424 | AX-170448891 | 0 | T | C | 0.105 | -1.87E-03 | 2.36E-01 | 2.69E+00 | 2.74E+00 | 9.93E-01 | no | | regulatory_region_variant | MODIFIER |
| 1 | 59433455 | AX-170000590 | 0 | T | C | 0.105 | -1.87E-03 | 2.36E-01 | 2.69E+00 | 2.74E+00 | 9.93E-01 | no | | regulatory_region_variant | MODIFIER |
| 1 | 59433605 | AX-97847514 | 0 | C | T | 0.105 | -1.87E-03 | 2.36E-01 | 2.69E+00 | 2.74E+00 | 9.93E-01 | no | | regulatory_region_variant | MODIFIER |
| 1 | 59435881 | AX-170261236 | 0 | A | G | 0.105 | -1.87E-03 | 2.36E-01 | 2.69E+00 | 2.74E+00 | 9.93E-01 | no | | intergenic_variant | MODIFIER |
| 1 | 59436155 | AX-170324119 | 0 | A | G | 0.105 | -1.87E-03 | 2.36E-01 | 2.69E+00 | 2.74E+00 | 9.93E-01 | no | | intergenic_variant | MODIFIER |
| 1 | 59524033 | AX-170197020 | 0 | A | C | 0.105 | -1.87E-03 | 2.36E-01 | 2.69E+00 | 2.74E+00 | 9.93E-01 | no | Gm973 | intron_variant | MODIFIER |

| | | | | | | | | | | | | | | | |
|---|---|---|---|---|---|---|---|---|---|---|---|---|---|---|---|
| 1 | 59555064 | AX-170000593 | 0 | T | C | 0.105 | -1.87E-03 | 2.36E-01 | 2.69E+00 | 2.74E+00 | 9.93E-01 | no | Gm973 | intron_variant | MODIFIER |
| 1 | 59590251 | AX-170113766 | 0 | A | G | 0.211 | -9.34E-04 | 1.18E-01 | 2.69E+00 | 2.74E+00 | 9.93E-01 | no | Gm973 | intron_variant | MODIFIER |
| 1 | 59606873 | AX-97811680 | 0 | A | G | 0.105 | -1.87E-03 | 2.36E-01 | 2.69E+00 | 2.74E+00 | 9.93E-01 | no | Gm973 | intron_variant | MODIFIER |
| 1 | 59612964 | AX-170367034 | 0 | G | A | 0.105 | -1.87E-03 | 2.36E-01 | 2.69E+00 | 2.74E+00 | 9.93E-01 | no | Gm973 | intron_variant | MODIFIER |
| 1 | 59634669 | AX-170041379 | 0 | T | C | 0.105 | -1.87E-03 | 2.36E-01 | 2.69E+00 | 2.74E+00 | 9.93E-01 | no | Sumo1 | downstream_gene_variant | MODIFIER |
| 1 | 59694484 | AX-97846037 | 0 | G | A | 0.105 | -1.87E-03 | 2.36E-01 | 2.69E+00 | 2.74E+00 | 9.93E-01 | no | Nop58 | intron_variant | MODIFIER |
| 1 | 59697486 | AX-170210301 | 0 | A | G | 0.105 | -1.87E-03 | 2.36E-01 | 2.69E+00 | 2.74E+00 | 9.93E-01 | no | Nop58 | intron_variant | MODIFIER |
| 1 | 59707622 | AX-170229007 | 0 | T | C | 0.105 | -1.87E-03 | 2.36E-01 | 2.69E+00 | 2.74E+00 | 9.93E-01 | no | Nop58 | intron_variant | MODIFIER |
| 1 | 59715959 | AX-170091529 | 0 | C | A | 0.105 | -1.87E-03 | 2.36E-01 | 2.69E+00 | 2.74E+00 | 9.93E-01 | no | Nop58 | downstream_gene_variant | MODIFIER |
| 1 | 59719856 | AX-170152688 | 0 | C | T | 0.105 | -1.87E-03 | 2.36E-01 | 2.69E+00 | 2.74E+00 | 9.93E-01 | no | Nop58 | downstream_gene_variant | MODIFIER |
| 1 | 59723178 | AX-170166792 | 0 | T | G | 0.105 | -1.87E-03 | 2.36E-01 | 2.69E+00 | 2.74E+00 | 9.93E-01 | no | Nop58 | downstream_gene_variant | MODIFIER |
| 1 | 59723247 | AX-170428489 | 0 | C | T | 0.105 | -1.87E-03 | 2.36E-01 | 2.69E+00 | 2.74E+00 | 9.93E-01 | no | Nop58 | downstream_gene_variant | MODIFIER |
| 1 | 59754807 | AX-170430522 | 0 | C | T | 0.105 | -1.87E-03 | 2.36E-01 | 2.69E+00 | 2.74E+00 | 9.93E-01 | no | | intergenic_variant | MODIFIER |
| 1 | 59758787 | AX-97838235 | 0 | T | C | 0.105 | -1.87E-03 | 2.36E-01 | 2.69E+00 | 2.74E+00 | 9.93E-01 | no | Bmpr2 | upstream_gene_variant | MODIFIER |
| 1 | 59834165 | AX-170199563 | 0 | T | C | 0.105 | -1.87E-03 | 2.36E-01 | 2.69E+00 | 2.74E+00 | 9.93E-01 | no | Bmpr2 | intron_variant | MODIFIER |
| 1 | 59841492 | AX-170318530 | 0 | A | G | 0.105 | -1.87E-03 | 2.36E-01 | 2.69E+00 | 2.74E+00 | 9.93E-01 | no | Bmpr2 | synonymous_variant | LOW |
| 1 | 59892184 | AX-170041382 | 0 | G | A | 0.105 | -1.87E-03 | 2.36E-01 | 2.69E+00 | 2.74E+00 | 9.93E-01 | no | Gm37977 | upstream_gene_variant | MODIFIER |
| 1 | 59894455 | AX-97811679 | 0 | C | G | 0.105 | -1.87E-03 | 2.36E-01 | 2.69E+00 | 2.74E+00 | 9.93E-01 | no | | regulatory_region_variant | MODIFIER |
| 1 | 59894980 | AX-170041383 | 0 | A | G | 0.105 | -1.87E-03 | 2.36E-01 | 2.69E+00 | 2.74E+00 | 9.93E-01 | no | | regulatory_region_variant | MODIFIER |
| 1 | 59895055 | AX-170175543 | 0 | A | T | 0.105 | -1.87E-03 | 2.36E-01 | 2.69E+00 | 2.74E+00 | 9.93E-01 | no | | regulatory_region_variant | MODIFIER |
| 1 | 59901906 | AX-170168027 | 0 | G | A | 0.105 | -1.87E-03 | 2.36E-01 | 2.69E+00 | 2.74E+00 | 9.93E-01 | no | | regulatory_region_variant | MODIFIER |
| 1 | 59902244 | AX-170406538 | 0 | G | A | 0.105 | -1.87E-03 | 2.36E-01 | 2.69E+00 | 2.74E+00 | 9.93E-01 | no | | regulatory_region_variant | MODIFIER |
| 1 | 59937190 | AX-97825634 | 0 | G | A | 0.105 | -1.87E-03 | 2.36E-01 | 2.69E+00 | 2.74E+00 | 9.93E-01 | no | Fam117b | intron_variant | MODIFIER |
| 1 | 59949703 | AX-97798275 | 0 | A | G | 0.105 | -1.87E-03 | 2.36E-01 | 2.69E+00 | 2.74E+00 | 9.93E-01 | no | Fam117b | intron_variant | MODIFIER |
| 1 | 59953165 | AX-170505413 | 0 | C | T | 0.105 | -1.87E-03 | 2.36E-01 | 2.69E+00 | 2.74E+00 | 9.93E-01 | no | Fam117b | intron_variant | MODIFIER |
| 1 | 59993701 | AX-170187958 | 0 | G | C | 0.105 | -1.87E-03 | 2.36E-01 | 2.69E+00 | 2.74E+00 | 9.93E-01 | no | Ica1l | intron_variant | MODIFIER |
| 1 | 59993840 | AX-170207676 | 0 | G | A | 0.105 | -1.87E-03 | 2.36E-01 | 2.69E+00 | 2.74E+00 | 9.93E-01 | no | Ica1l | intron_variant | MODIFIER |
| 1 | 59995572 | AX-170286130 | 0 | T | C | 0.105 | -1.87E-03 | 2.36E-01 | 2.69E+00 | 2.74E+00 | 9.93E-01 | no | Ica1l | intron_variant | MODIFIER |
| 1 | 59995627 | AX-170000603 | 0 | C | A | 0.105 | -1.87E-03 | 2.36E-01 | 2.69E+00 | 2.74E+00 | 9.93E-01 | no | Ica1l | intron_variant | MODIFIER |
| 1 | 59995821 | AX-170000608 | 0 | G | A | 0.105 | -1.87E-03 | 2.36E-01 | 2.69E+00 | 2.74E+00 | 9.93E-01 | no | Ica1l | intron_variant | MODIFIER |
| 1 | 59997913 | AX-170482904 | 0 | T | C | 0.105 | -1.87E-03 | 2.36E-01 | 2.69E+00 | 2.74E+00 | 9.93E-01 | no | Ica1l | intron_variant | MODIFIER |
| 1 | 59998105 | AX-170000602 | 0 | C | T | 0.105 | -1.87E-03 | 2.36E-01 | 2.69E+00 | 2.74E+00 | 9.93E-01 | no | Ica1l | intron_variant | MODIFIER |
| 1 | 60008042 | AX-170333862 | 0 | G | T | 0.105 | -1.87E-03 | 2.36E-01 | 2.69E+00 | 2.74E+00 | 9.93E-01 | no | Ica1l | intron_variant | MODIFIER |
| 1 | 60010242 | AX-97818835 | 0 | T | C | 0.105 | -1.87E-03 | 2.36E-01 | 2.69E+00 | 2.74E+00 | 9.93E-01 | no | Ica1l | splice_region_variant,intron_variant | LOW |
| 1 | 60029088 | AX-170000606 | 0 | C | T | 0.105 | -1.87E-03 | 2.36E-01 | 2.69E+00 | 2.74E+00 | 9.93E-01 | no | Ica1l | intron_variant | MODIFIER |
| 1 | 60082718 | AX-170257274 | 0 | T | C | 0.105 | -1.87E-03 | 2.36E-01 | 2.69E+00 | 2.74E+00 | 9.93E-01 | no | Wdr12 | intron_variant | MODIFIER |
| 1 | 60086606 | AX-170000605 | 0 | T | C | 0.105 | -1.87E-03 | 2.36E-01 | 2.69E+00 | 2.74E+00 | 9.93E-01 | no | Wdr12 | intron_variant | MODIFIER |
| 1 | 60192268 | AX-170212876 | 0 | T | C | 0.105 | -1.87E-03 | 2.36E-01 | 2.69E+00 | 2.74E+00 | 9.93E-01 | no | Nbeal1 | intron_variant | MODIFIER |
| 1 | 60212250 | AX-170041386 | 0 | G | A | 0.105 | -1.87E-03 | 2.36E-01 | 2.69E+00 | 2.74E+00 | 9.93E-01 | no | Nbeal1 | intron_variant | MODIFIER |
| 1 | 60226099 | AX-170242712 | 0 | C | T | 0.105 | -1.87E-03 | 2.36E-01 | 2.69E+00 | 2.74E+00 | 9.93E-01 | no | Nbeal1 | intron_variant | MODIFIER |
| 1 | 60231521 | AX-170283499 | 0 | T | C | 0.105 | -1.87E-03 | 2.36E-01 | 2.69E+00 | 2.74E+00 | 9.93E-01 | no | Nbeal1 | intron_variant | MODIFIER |
| 1 | 60260716 | AX-170293577 | 0 | A | T | 0.105 | -1.87E-03 | 2.36E-01 | 2.69E+00 | 2.74E+00 | 9.93E-01 | no | Nbeal1 | missense_variant | MODERATE |
| 1 | 60270296 | AX-170408366 | 0 | A | C | 0.105 | -1.87E-03 | 2.36E-01 | 2.69E+00 | 2.74E+00 | 9.93E-01 | no | Nbeal1 | synonymous_variant | LOW |
| 1 | 60278333 | AX-170417696 | 0 | C | G | 0.105 | -1.87E-03 | 2.36E-01 | 2.69E+00 | 2.74E+00 | 9.93E-01 | no | Nbeal1 | intron_variant | MODIFIER |
| 1 | 60280406 | AX-170395046 | 0 | T | C | 0.105 | -1.87E-03 | 2.36E-01 | 2.69E+00 | 2.74E+00 | 9.93E-01 | no | Nbeal1 | intron_variant | MODIFIER |
| 1 | 60291520 | AX-170162483 | 0 | T | C | 0.105 | -1.87E-03 | 2.36E-01 | 2.69E+00 | 2.74E+00 | 9.93E-01 | no | Nbeal1 | intron_variant | MODIFIER |
| 1 | 60294368 | AX-170407927 | 0 | T | C | 0.105 | -1.87E-03 | 2.36E-01 | 2.69E+00 | 2.74E+00 | 9.93E-01 | no | Nbeal1 | intron_variant | MODIFIER |
| 1 | 60299072 | AX-170434084 | 0 | T | C | 0.105 | -1.87E-03 | 2.36E-01 | 2.69E+00 | 2.74E+00 | 9.93E-01 | no | Nbeal1 | intron_variant | MODIFIER |
| 1 | 60306119 | AX-97811676 | 0 | A | G | 0.105 | -1.87E-03 | 2.36E-01 | 2.69E+00 | 2.74E+00 | 9.93E-01 | no | Nbeal1 | intron_variant | MODIFIER |
| 1 | 60309669 | AX-170350068 | 0 | T | G | 0.105 | -1.87E-03 | 2.36E-01 | 2.69E+00 | 2.74E+00 | 9.93E-01 | no | Nbeal1 | intron_variant | MODIFIER |
| 1 | 60319729 | AX-170284404 | 0 | C | T | 0.105 | -1.87E-03 | 2.36E-01 | 2.69E+00 | 2.74E+00 | 9.93E-01 | no | Nbeal1 | downstream_gene_variant | MODIFIER |
| 1 | 60352820 | AX-170508555 | 0 | T | C | 0.105 | -1.87E-03 | 2.36E-01 | 2.69E+00 | 2.74E+00 | 9.93E-01 | no | Cyp20a1 | intron_variant | MODIFIER |
| 1 | 60353009 | AX-170104953 | 0 | C | A | 0.105 | -1.87E-03 | 2.36E-01 | 2.69E+00 | 2.74E+00 | 9.93E-01 | no | Cyp20a1 | synonymous_variant | LOW |
| 1 | 60382556 | AX-170041389 | 0 | T | G | 0.105 | -1.87E-03 | 2.36E-01 | 2.69E+00 | 2.74E+00 | 9.93E-01 | no | Cyp20a1 | intron_variant | MODIFIER |

| | | | | | | | | | | | | | | | |
|---|---|---|---|---|---|---|---|---|---|---|---|---|---|---|---|
| 1 | 60384577 | AX-170149630 | 0 | G | A | 0.105 | -1.87E-03 | 2.36E-01 | 2.69E+00 | 2.74E+00 | 9.93E-01 | no | Cyp20a1 | intron_variant | MODIFIER |
| 1 | 60384650 | AX-170086480 | 0 | T | C | 0.105 | -1.87E-03 | 2.36E-01 | 2.69E+00 | 2.74E+00 | 9.93E-01 | no | Cyp20a1 | intron_variant | MODIFIER |
| 1 | 60384685 | AX-170296069 | 0 | T | C | 0.105 | -1.87E-03 | 2.36E-01 | 2.69E+00 | 2.74E+00 | 9.93E-01 | no | Cyp20a1 | intron_variant | MODIFIER |
| 1 | 60415711 | AX-170091531 | 0 | G | A | 0.105 | -1.87E-03 | 2.36E-01 | 2.69E+00 | 2.74E+00 | 9.93E-01 | no | Abi2 | intron_variant | MODIFIER |
| 1 | 60436857 | AX-97843419 | 0 | G | A | 0.105 | -1.87E-03 | 2.36E-01 | 2.69E+00 | 2.74E+00 | 9.93E-01 | no | Abi2 | intron_variant | MODIFIER |
| 1 | 60479260 | AX-97824418 | 0 | T | C | 0.105 | -1.87E-03 | 2.36E-01 | 2.69E+00 | 2.74E+00 | 9.93E-01 | no | Raph1 | downstream_gene_variant | MODIFIER |
| 1 | 60496199 | AX-170334419 | 0 | T | C | 0.105 | -1.87E-03 | 2.36E-01 | 2.69E+00 | 2.74E+00 | 9.93E-01 | no | Raph1 | intron_variant | MODIFIER |
| 1 | 60496669 | AX-170163176 | 0 | T | C | 0.105 | -1.87E-03 | 2.36E-01 | 2.69E+00 | 2.74E+00 | 9.93E-01 | no | Raph1 | intron_variant | MODIFIER |
| 1 | 60508298 | AX-170443430 | 0 | T | C | 0.105 | -1.87E-03 | 2.36E-01 | 2.69E+00 | 2.74E+00 | 9.93E-01 | no | Raph1 | intron_variant | MODIFIER |
| 1 | 60526958 | AX-170447281 | 0 | A | G | 0.105 | -1.87E-03 | 2.36E-01 | 2.69E+00 | 2.74E+00 | 9.93E-01 | no | Raph1 | intron_variant | MODIFIER |
| 1 | 60532189 | AX-170288935 | 0 | A | G | 0.105 | -1.87E-03 | 2.36E-01 | 2.69E+00 | 2.74E+00 | 9.93E-01 | no | Raph1 | intron_variant | MODIFIER |
| 1 | 60574204 | AX-170170358 | 0 | T | C | 0.105 | -1.87E-03 | 2.36E-01 | 2.69E+00 | 2.74E+00 | 9.93E-01 | no | | intergenic_variant | MODIFIER |
| 1 | 60578731 | AX-170513599 | 0 | T | C | 0.105 | -1.87E-03 | 2.36E-01 | 2.69E+00 | 2.74E+00 | 9.93E-01 | no | | intergenic_variant | MODIFIER |
| 1 | 60582981 | AX-170389486 | 0 | C | A | 0.105 | -1.87E-03 | 2.36E-01 | 2.69E+00 | 2.74E+00 | 9.93E-01 | no | | intergenic_variant | MODIFIER |
| 1 | 60583937 | AX-97811675 | 0 | C | T | 0.105 | -1.87E-03 | 2.36E-01 | 2.69E+00 | 2.74E+00 | 9.93E-01 | no | | intergenic_variant | MODIFIER |
| 1 | 60604268 | AX-170000617 | 0 | G | A | 0.105 | -1.87E-03 | 2.36E-01 | 2.69E+00 | 2.74E+00 | 9.93E-01 | no | | intergenic_variant | MODIFIER |
| 1 | 60604490 | AX-170000611 | 0 | G | A | 0.105 | -1.87E-03 | 2.36E-01 | 2.69E+00 | 2.74E+00 | 9.93E-01 | no | | intergenic_variant | MODIFIER |
| 1 | 60605601 | AX-170108801 | 0 | G | A | 0.105 | -1.87E-03 | 2.36E-01 | 2.69E+00 | 2.74E+00 | 9.93E-01 | no | | intergenic_variant | MODIFIER |
| 1 | 60612314 | AX-170320135 | 0 | A | G | 0.105 | -1.87E-03 | 2.36E-01 | 2.69E+00 | 2.74E+00 | 9.93E-01 | no | Gm23762 | downstream_gene_variant | MODIFIER |
| 1 | 60616617 | AX-170514313 | 0 | G | A | 0.105 | -1.87E-03 | 2.36E-01 | 2.69E+00 | 2.74E+00 | 9.93E-01 | no | Gm23762 | upstream_gene_variant | MODIFIER |
| 1 | 60623186 | AX-170323242 | 0 | C | T | 0.105 | -1.87E-03 | 2.36E-01 | 2.69E+00 | 2.74E+00 | 9.93E-01 | no | | regulatory_region_variant | MODIFIER |
| 1 | 60629520 | AX-170385771 | 0 | A | C | 0.105 | -1.87E-03 | 2.36E-01 | 2.69E+00 | 2.74E+00 | 9.93E-01 | no | | intergenic_variant | MODIFIER |
| 1 | 60640152 | AX-170193668 | 0 | T | C | 0.105 | -1.87E-03 | 2.36E-01 | 2.69E+00 | 2.74E+00 | 9.93E-01 | no | | regulatory_region_variant | MODIFIER |
| 1 | 60665850 | AX-170357630 | 0 | C | T | 0.105 | -1.87E-03 | 2.36E-01 | 2.69E+00 | 2.74E+00 | 9.93E-01 | no | | intergenic_variant | MODIFIER |
| 1 | 60668249 | AX-170319264 | 0 | A | G | 0.105 | -1.87E-03 | 2.36E-01 | 2.69E+00 | 2.74E+00 | 9.93E-01 | no | | regulatory_region_variant | MODIFIER |
| 1 | 60682904 | AX-170386423 | 0 | T | C | 0.105 | -1.87E-03 | 2.36E-01 | 2.69E+00 | 2.74E+00 | 9.93E-01 | no | | intergenic_variant | MODIFIER |
| 1 | 60683146 | AX-170362690 | 0 | A | G | 0.105 | -1.87E-03 | 2.36E-01 | 2.69E+00 | 2.74E+00 | 9.93E-01 | no | | intergenic_variant | MODIFIER |
| 1 | 60684723 | AX-97811764 | 0 | G | A | 0.105 | -1.87E-03 | 2.36E-01 | 2.69E+00 | 2.74E+00 | 9.93E-01 | no | | intergenic_variant | MODIFIER |
| 1 | 60749069 | AX-170424851 | 0 | C | T | 0.105 | -1.87E-03 | 2.36E-01 | 2.69E+00 | 2.74E+00 | 9.93E-01 | no | Cd28 | intron_variant | MODIFIER |
| 1 | 60754018 | AX-97817150 | 0 | T | C | 0.105 | -1.87E-03 | 2.36E-01 | 2.69E+00 | 2.74E+00 | 9.93E-01 | no | Cd28 | intron_variant | MODIFIER |
| 1 | 60760195 | AX-170353061 | 0 | A | T | 0.105 | -1.87E-03 | 2.36E-01 | 2.69E+00 | 2.74E+00 | 9.93E-01 | no | Cd28 | intron_variant | MODIFIER |
| 1 | 60765486 | AX-170118915 | 0 | G | A | 0.105 | -1.87E-03 | 2.36E-01 | 2.69E+00 | 2.74E+00 | 9.93E-01 | no | Cd28 | intron_variant | MODIFIER |
| 1 | 60800563 | AX-170505590 | 0 | C | T | 0.105 | -1.87E-03 | 2.36E-01 | 2.69E+00 | 2.74E+00 | 9.93E-01 | no | 2310016D23Rik | intron_variant,non_coding_transcript_variant | MODIFIER |
| 1 | 60821264 | AX-170126093 | 0 | A | C | 0.105 | -1.87E-03 | 2.36E-01 | 2.69E+00 | 2.74E+00 | 9.93E-01 | no | | regulatory_region_variant | MODIFIER |
| 1 | 60821393 | AX-170519349 | 0 | T | G | 0.105 | -1.87E-03 | 2.36E-01 | 2.69E+00 | 2.74E+00 | 9.93E-01 | no | | regulatory_region_variant | MODIFIER |
| 1 | 60822961 | AX-170211322 | 0 | G | A | 0.105 | -1.87E-03 | 2.36E-01 | 2.69E+00 | 2.74E+00 | 9.93E-01 | no | | regulatory_region_variant | MODIFIER |
| 1 | 62612054 | AX-170470886 | 0 | G | A | 0.105 | -1.87E-03 | 2.36E-01 | 2.69E+00 | 2.74E+00 | 9.93E-01 | no | Pard3b | intron_variant | MODIFIER |
| 1 | 62647560 | AX-170041431 | 0 | G | A | 0.105 | -1.87E-03 | 2.36E-01 | 2.69E+00 | 2.74E+00 | 9.93E-01 | no | | intergenic_variant | MODIFIER |
| 1 | 62650689 | AX-170041429 | 0 | T | C | 0.105 | -1.87E-03 | 2.36E-01 | 2.69E+00 | 2.74E+00 | 9.93E-01 | no | | intergenic_variant | MODIFIER |
| 1 | 62650989 | AX-97798704 | 0 | T | C | 0.105 | -1.87E-03 | 2.36E-01 | 2.69E+00 | 2.74E+00 | 9.93E-01 | no | | intergenic_variant | MODIFIER |
| 1 | 62653828 | AX-170257988 | 0 | A | G | 0.105 | -1.87E-03 | 2.36E-01 | 2.69E+00 | 2.74E+00 | 9.93E-01 | no | Gm11600 | downstream_gene_variant | MODIFIER |
| 1 | 62719869 | AX-170392416 | 0 | C | G | 0.105 | -1.87E-03 | 2.36E-01 | 2.69E+00 | 2.74E+00 | 9.93E-01 | no | Nrp2 | intron_variant | MODIFIER |
| 1 | 62723750 | AX-170263947 | 0 | G | C | 0.105 | -1.87E-03 | 2.36E-01 | 2.69E+00 | 2.74E+00 | 9.93E-01 | no | Nrp2 | intron_variant | MODIFIER |
| 1 | 62741573 | AX-170238179 | 0 | A | G | 0.105 | -1.87E-03 | 2.36E-01 | 2.69E+00 | 2.74E+00 | 9.93E-01 | no | Nrp2 | intron_variant | MODIFIER |
| 1 | 62742832 | AX-170357228 | 0 | A | C | 0.105 | -1.87E-03 | 2.36E-01 | 2.69E+00 | 2.74E+00 | 9.93E-01 | no | Nrp2 | intron_variant | MODIFIER |
| 1 | 62747352 | AX-170186733 | 0 | A | T | 0.105 | -1.87E-03 | 2.36E-01 | 2.69E+00 | 2.74E+00 | 9.93E-01 | no | Nrp2 | intron_variant | MODIFIER |
| 1 | 62747428 | AX-170275234 | 0 | G | A | 0.105 | -1.87E-03 | 2.36E-01 | 2.69E+00 | 2.74E+00 | 9.93E-01 | no | Nrp2 | intron_variant | MODIFIER |
| 1 | 62761021 | AX-170284587 | 0 | C | T | 0.105 | -1.87E-03 | 2.36E-01 | 2.69E+00 | 2.74E+00 | 9.93E-01 | no | Nrp2 | intron_variant | MODIFIER |
| 1 | 62761055 | AX-170509054 | 0 | T | C | 0.105 | -1.87E-03 | 2.36E-01 | 2.69E+00 | 2.74E+00 | 9.93E-01 | no | Nrp2 | intron_variant | MODIFIER |
| 1 | 62769282 | AX-97832585 | 0 | T | C | 0.105 | -1.87E-03 | 2.36E-01 | 2.69E+00 | 2.74E+00 | 9.93E-01 | no | Nrp2 | intron_variant | MODIFIER |
| 1 | 41556880 | AX-170153759 | 1 | T | C | 0.411 | -1.11E-03 | 7.13E-02 | 2.64E+00 | 2.74E+00 | 9.93E-01 | no | | intergenic_variant | MODIFIER |
| 1 | 172012379 | AX-170291934 | 0 | T | A | 0.211 | 2.86E-03 | 1.39E-01 | 2.68E+00 | 2.73E+00 | 9.93E-01 | no | Vangl2 | intron_variant | MODIFIER |
| 4 | 146486803 | AX-170150831 | 2 | C | G | 0.055 | -2.05E-03 | 1.93E-01 | 2.64E+00 | 2.74E+00 | 9.93E-01 | no | Gm26573 | intron_variant,non_coding_transcript_variant | MODIFIER |
| 3 | 110035621 | AX-170466831 | 1 | C | T | 0.036 | 2.83E-03 | 1.40E-01 | 2.56E+00 | 2.74E+00 | 9.93E-01 | no | Ntng1 | intron_variant | MODIFIER |

| | | | | | | | | | | | | | | | |
|---|---|---|---|---|---|---|---|---|---|---|---|---|---|---|---|
| 8 | 38116186 | AX-170204197 | 0 | G | A | 0.149 | -1.89E-03 | 1.53E-01 | 2.70E+00 | 2.74E+00 | 9.93E-01 | no | Sgcz | intron_variant | MODIFIER |
| 1 | 60451788 | AX-170322506 | 1 | G | A | 0.107 | -1.71E-03 | 2.35E-01 | 2.69E+00 | 2.74E+00 | 9.93E-01 | no | Abi2 | intron_variant | MODIFIER |
| 11 | 45620668 | AX-170127740 | 0 | G | A | 0.088 | -9.38E-04 | 1.43E-01 | 2.60E+00 | 2.74E+00 | 9.93E-01 | no | | intergenic_variant | MODIFIER |
| 1 | 47103465 | AX-170368691 | 1 | A | T | 0.107 | -1.67E-03 | 2.35E-01 | 2.69E+00 | 2.74E+00 | 9.93E-01 | no | | regulatory_region_variant | MODIFIER |
| 1 | 55741622 | AX-170171885 | 1 | C | T | 0.107 | -1.67E-03 | 2.35E-01 | 2.69E+00 | 2.74E+00 | 9.93E-01 | no | Plcl1 | intron_variant | MODIFIER |
| 8 | 37984307 | AX-170079801 | 0 | A | G | 0.149 | -1.85E-03 | 1.50E-01 | 2.69E+00 | 2.74E+00 | 9.93E-01 | no | Sgcz | intron_variant | MODIFIER |
| 8 | 46624569 | AX-170205533 | 0 | G | A | 0.149 | -1.85E-03 | 1.50E-01 | 2.69E+00 | 2.74E+00 | 9.93E-01 | no | Casp3 | intron_variant | MODIFIER |
| 8 | 46627769 | AX-170445762 | 0 | A | T | 0.149 | -1.85E-03 | 1.50E-01 | 2.69E+00 | 2.74E+00 | 9.93E-01 | no | Casp3 | intron_variant | MODIFIER |
| 8 | 46905493 | AX-170507120 | 0 | C | T | 0.149 | -1.85E-03 | 1.50E-01 | 2.69E+00 | 2.74E+00 | 9.93E-01 | no | | intergenic_variant | MODIFIER |
| 8 | 46908505 | AX-97804946 | 0 | A | T | 0.149 | -1.85E-03 | 1.50E-01 | 2.69E+00 | 2.74E+00 | 9.93E-01 | no | | intergenic_variant | MODIFIER |
| 8 | 46910112 | AX-170436144 | 0 | T | C | 0.149 | -1.85E-03 | 1.50E-01 | 2.69E+00 | 2.74E+00 | 9.93E-01 | no | | intergenic_variant | MODIFIER |
| 8 | 46915182 | AX-170316654 | 0 | T | A | 0.149 | -1.85E-03 | 1.50E-01 | 2.69E+00 | 2.74E+00 | 9.93E-01 | no | | intergenic_variant | MODIFIER |
| 8 | 46927205 | AX-97809408 | 0 | T | C | 0.149 | -1.85E-03 | 1.50E-01 | 2.69E+00 | 2.74E+00 | 9.93E-01 | no | | intergenic_variant | MODIFIER |
| 8 | 46929782 | AX-170412262 | 0 | C | T | 0.149 | -1.85E-03 | 1.50E-01 | 2.69E+00 | 2.74E+00 | 9.93E-01 | no | | intergenic_variant | MODIFIER |
| 8 | 46934965 | AX-97838778 | 0 | A | G | 0.149 | -1.85E-03 | 1.50E-01 | 2.69E+00 | 2.74E+00 | 9.93E-01 | no | | regulatory_region_variant | MODIFIER |
| 8 | 46941282 | AX-170523381 | 0 | T | C | 0.149 | -1.85E-03 | 1.50E-01 | 2.69E+00 | 2.74E+00 | 9.93E-01 | no | | regulatory_region_variant | MODIFIER |
| 8 | 46941477 | AX-170172247 | 0 | C | T | 0.149 | -1.85E-03 | 1.50E-01 | 2.69E+00 | 2.74E+00 | 9.93E-01 | no | | regulatory_region_variant | MODIFIER |
| 8 | 46942238 | AX-170268260 | 0 | A | G | 0.149 | -1.85E-03 | 1.50E-01 | 2.69E+00 | 2.74E+00 | 9.93E-01 | no | 1700011L03Rik | downstream_gene_variant | MODIFIER |
| 8 | 46948055 | AX-170353556 | 0 | A | G | 0.149 | -1.85E-03 | 1.50E-01 | 2.69E+00 | 2.74E+00 | 9.93E-01 | no | 1700011L03Rik | intron_variant,non_coding_transcript_variant | MODIFIER |
| 8 | 46993952 | AX-170204872 | 0 | A | G | 0.149 | -1.85E-03 | 1.50E-01 | 2.69E+00 | 2.74E+00 | 9.93E-01 | no | Enpp6 | intron_variant | MODIFIER |
| 8 | 46999881 | AX-170148495 | 0 | C | T | 0.149 | -1.85E-03 | 1.50E-01 | 2.69E+00 | 2.74E+00 | 9.93E-01 | no | Enpp6 | intron_variant | MODIFIER |
| 8 | 47001681 | AX-170347940 | 0 | C | T | 0.149 | -1.85E-03 | 1.50E-01 | 2.69E+00 | 2.74E+00 | 9.93E-01 | no | Enpp6 | intron_variant | MODIFIER |
| 8 | 47009415 | AX-170415530 | 0 | A | C | 0.149 | -1.85E-03 | 1.50E-01 | 2.69E+00 | 2.74E+00 | 9.93E-01 | no | Enpp6 | intron_variant | MODIFIER |
| 8 | 47028507 | AX-170096254 | 0 | A | C | 0.149 | -1.85E-03 | 1.50E-01 | 2.69E+00 | 2.74E+00 | 9.93E-01 | no | Enpp6 | intron_variant | MODIFIER |
| 8 | 47028701 | AX-170079986 | 0 | G | A | 0.149 | -1.85E-03 | 1.50E-01 | 2.69E+00 | 2.74E+00 | 9.93E-01 | no | Enpp6 | intron_variant | MODIFIER |
| 8 | 47092614 | AX-170483379 | 0 | T | G | 0.149 | -1.85E-03 | 1.50E-01 | 2.69E+00 | 2.74E+00 | 9.93E-01 | no | Enpp6 | intron_variant | MODIFIER |
| 8 | 47094613 | AX-170438898 | 0 | G | A | 0.149 | -1.85E-03 | 1.50E-01 | 2.69E+00 | 2.74E+00 | 9.93E-01 | no | Enpp6 | 3_prime_UTR_variant | MODIFIER |
| 8 | 47102772 | AX-170435936 | 0 | C | T | 0.149 | -1.85E-03 | 1.50E-01 | 2.69E+00 | 2.74E+00 | 9.93E-01 | no | | intergenic_variant | MODIFIER |
| 8 | 47102857 | AX-170505395 | 0 | G | T | 0.149 | -1.85E-03 | 1.50E-01 | 2.69E+00 | 2.74E+00 | 9.93E-01 | no | | intergenic_variant | MODIFIER |
| 8 | 47102909 | AX-170446576 | 0 | A | T | 0.149 | -1.85E-03 | 1.50E-01 | 2.69E+00 | 2.74E+00 | 9.93E-01 | no | | intergenic_variant | MODIFIER |
| 8 | 47103054 | AX-170405702 | 0 | C | T | 0.149 | -1.85E-03 | 1.50E-01 | 2.69E+00 | 2.74E+00 | 9.93E-01 | no | | intergenic_variant | MODIFIER |
| 8 | 47114657 | AX-170479350 | 0 | G | T | 0.149 | -1.85E-03 | 1.50E-01 | 2.69E+00 | 2.74E+00 | 9.93E-01 | no | | intergenic_variant | MODIFIER |
| 8 | 47117912 | AX-170142522 | 0 | G | A | 0.149 | -1.85E-03 | 1.50E-01 | 2.69E+00 | 2.74E+00 | 9.93E-01 | no | | intergenic_variant | MODIFIER |
| 8 | 47117933 | AX-170527563 | 0 | G | T | 0.149 | -1.85E-03 | 1.50E-01 | 2.69E+00 | 2.74E+00 | 9.93E-01 | no | | intergenic_variant | MODIFIER |
| 8 | 47159552 | AX-170099264 | 0 | A | C | 0.149 | -1.85E-03 | 1.50E-01 | 2.69E+00 | 2.74E+00 | 9.93E-01 | no | E330018M18Rik | intron_variant,non_coding_transcript_variant | MODIFIER |
| 8 | 47159577 | AX-170499869 | 0 | T | C | 0.149 | -1.85E-03 | 1.50E-01 | 2.69E+00 | 2.74E+00 | 9.93E-01 | no | E330018M18Rik | intron_variant,non_coding_transcript_variant | MODIFIER |
| 8 | 47162790 | AX-170030618 | 0 | C | G | 0.149 | -1.85E-03 | 1.50E-01 | 2.69E+00 | 2.74E+00 | 9.93E-01 | no | E330018M18Rik | non_coding_transcript_exon_variant | MODIFIER |
| 8 | 47170370 | AX-97784292 | 0 | A | G | 0.149 | -1.85E-03 | 1.50E-01 | 2.69E+00 | 2.74E+00 | 9.93E-01 | no | | intergenic_variant | MODIFIER |
| 8 | 47192603 | AX-170510246 | 0 | A | G | 0.149 | -1.85E-03 | 1.50E-01 | 2.69E+00 | 2.74E+00 | 9.93E-01 | no | Stox2 | synonymous_variant | LOW |
| 8 | 47193467 | AX-170402568 | 0 | G | A | 0.149 | -1.85E-03 | 1.50E-01 | 2.69E+00 | 2.74E+00 | 9.93E-01 | no | Stox2 | synonymous_variant | LOW |
| 8 | 47206100 | AX-170030615 | 0 | T | C | 0.149 | -1.85E-03 | 1.50E-01 | 2.69E+00 | 2.74E+00 | 9.93E-01 | no | Stox2 | intron_variant | MODIFIER |
| 8 | 47209714 | AX-170341541 | 0 | T | C | 0.149 | -1.85E-03 | 1.50E-01 | 2.69E+00 | 2.74E+00 | 9.93E-01 | no | Stox2 | intron_variant | MODIFIER |
| 8 | 47209896 | AX-170228057 | 0 | A | G | 0.149 | -1.85E-03 | 1.50E-01 | 2.69E+00 | 2.74E+00 | 9.93E-01 | no | Stox2 | intron_variant | MODIFIER |
| 8 | 47211016 | AX-170238127 | 0 | C | A | 0.149 | -1.85E-03 | 1.50E-01 | 2.69E+00 | 2.74E+00 | 9.93E-01 | no | Stox2 | intron_variant | MODIFIER |
| 8 | 47364039 | AX-170198330 | 0 | T | C | 0.149 | -1.85E-03 | 1.50E-01 | 2.69E+00 | 2.74E+00 | 9.93E-01 | no | Stox2 | intron_variant,non_coding_transcript_variant | MODIFIER |
| 8 | 47364768 | AX-170165164 | 0 | A | G | 0.149 | -1.85E-03 | 1.50E-01 | 2.69E+00 | 2.74E+00 | 9.93E-01 | no | Stox2 | intron_variant,non_coding_transcript_variant | MODIFIER |
| 8 | 47365505 | AX-170192841 | 0 | C | G | 0.149 | -1.85E-03 | 1.50E-01 | 2.69E+00 | 2.74E+00 | 9.93E-01 | no | Stox2 | intron_variant,non_coding_transcript_variant | MODIFIER |
| 8 | 47381943 | AX-97835633 | 0 | C | T | 0.149 | -1.85E-03 | 1.50E-01 | 2.69E+00 | 2.74E+00 | 9.93E-01 | no | Stox2 | intron_variant,non_coding_transcript_variant | MODIFIER |
| 8 | 47385300 | AX-170204767 | 0 | A | G | 0.149 | -1.85E-03 | 1.50E-01 | 2.69E+00 | 2.74E+00 | 9.93E-01 | no | Stox2 | intron_variant,non_coding_transcript_variant | MODIFIER |
| 8 | 47418445 | AX-170156453 | 0 | G | C | 0.149 | -1.85E-03 | 1.50E-01 | 2.69E+00 | 2.74E+00 | 9.93E-01 | no | Stox2 | intron_variant,non_coding_transcript_variant | MODIFIER |
| 8 | 47418795 | AX-168379330 | 0 | A | G | 0.149 | -1.85E-03 | 1.50E-01 | 2.69E+00 | 2.74E+00 | 9.93E-01 | no | Stox2 | intron_variant,non_coding_transcript_variant | MODIFIER |
| 8 | 47432312 | AX-170403856 | 0 | T | C | 0.149 | -1.85E-03 | 1.50E-01 | 2.69E+00 | 2.74E+00 | 9.93E-01 | no | Stox2 | intron_variant,non_coding_transcript_variant | MODIFIER |
| 8 | 47440013 | AX-170299982 | 0 | G | T | 0.149 | -1.85E-03 | 1.50E-01 | 2.69E+00 | 2.74E+00 | 9.93E-01 | no | Gm22857 | upstream_gene_variant | MODIFIER |
| 8 | 47442323 | AX-170090092 | 0 | G | A | 0.149 | -1.85E-03 | 1.50E-01 | 2.69E+00 | 2.74E+00 | 9.93E-01 | no | Gm22857 | downstream_gene_variant | MODIFIER |

| | | | | | | | | | | | | | | | |
|---|---|---|---|---|---|---|---|---|---|---|---|---|---|---|---|
| 8 | 47450279 | AX-170126650 | 0 | G | A | 0.149 | -1.85E-03 | 1.50E-01 | 2.69E+00 | 2.74E+00 | 9.93E-01 | no | Stox2 | upstream_gene_variant | MODIFIER |
| 8 | 47480704 | AX-97815211 | 0 | G | A | 0.149 | -1.85E-03 | 1.50E-01 | 2.69E+00 | 2.74E+00 | 9.93E-01 | no | | intergenic_variant | MODIFIER |
| 8 | 47500701 | AX-170360708 | 0 | T | C | 0.149 | -1.85E-03 | 1.50E-01 | 2.69E+00 | 2.74E+00 | 9.93E-01 | no | Trappc11 | intron_variant | MODIFIER |
| 8 | 47531528 | AX-170426491 | 0 | A | G | 0.149 | -1.85E-03 | 1.50E-01 | 2.69E+00 | 2.74E+00 | 9.93E-01 | no | Rwdd4a | upstream_gene_variant | MODIFIER |
| 8 | 47538608 | AX-170336535 | 0 | C | T | 0.149 | -1.85E-03 | 1.50E-01 | 2.69E+00 | 2.74E+00 | 9.93E-01 | no | Rwdd4a | intron_variant | MODIFIER |
| 8 | 47551210 | AX-170310074 | 0 | A | G | 0.149 | -1.85E-03 | 1.50E-01 | 2.69E+00 | 2.74E+00 | 9.93E-01 | no | Rwdd4a | 3_prime_UTR_variant | MODIFIER |
| 8 | 47551294 | AX-170308386 | 0 | G | A | 0.149 | -1.85E-03 | 1.50E-01 | 2.69E+00 | 2.74E+00 | 9.93E-01 | no | Rwdd4a | 3_prime_UTR_variant | MODIFIER |
| 8 | 47627587 | AX-170240250 | 0 | A | T | 0.149 | -1.85E-03 | 1.50E-01 | 2.69E+00 | 2.74E+00 | 9.93E-01 | no | Gm8623 | downstream_gene_variant | MODIFIER |
| 8 | 47800338 | AX-170030621 | 0 | C | T | 0.149 | -1.85E-03 | 1.50E-01 | 2.69E+00 | 2.74E+00 | 9.93E-01 | no | | intergenic_variant | MODIFIER |
| 8 | 47886504 | AX-170030619 | 0 | G | A | 0.149 | -1.85E-03 | 1.50E-01 | 2.69E+00 | 2.74E+00 | 9.93E-01 | no | Wwc2 | intron_variant | MODIFIER |
| 8 | 47908501 | AX-97819838 | 0 | C | T | 0.149 | -1.85E-03 | 1.50E-01 | 2.69E+00 | 2.74E+00 | 9.93E-01 | no | Wwc2 | intron_variant | MODIFIER |
| 8 | 48028556 | AX-170430040 | 0 | T | G | 0.149 | -1.85E-03 | 1.50E-01 | 2.69E+00 | 2.74E+00 | 9.93E-01 | no | | intergenic_variant | MODIFIER |
| 8 | 48051538 | AX-170267314 | 0 | A | G | 0.149 | -1.85E-03 | 1.50E-01 | 2.69E+00 | 2.74E+00 | 9.93E-01 | no | | intergenic_variant | MODIFIER |
| 8 | 48065436 | AX-170269380 | 0 | A | G | 0.149 | -1.85E-03 | 1.50E-01 | 2.69E+00 | 2.74E+00 | 9.93E-01 | no | Gm2607 | downstream_gene_variant | MODIFIER |
| 8 | 48076404 | AX-97816302 | 0 | A | G | 0.149 | -1.85E-03 | 1.50E-01 | 2.69E+00 | 2.74E+00 | 9.93E-01 | no | | intergenic_variant | MODIFIER |
| 8 | 48099191 | AX-170411921 | 0 | T | A | 0.149 | -1.85E-03 | 1.50E-01 | 2.69E+00 | 2.74E+00 | 9.93E-01 | no | Dctd | intron_variant | MODIFIER |
| 8 | 48108290 | AX-97789063 | 0 | T | C | 0.149 | -1.85E-03 | 1.50E-01 | 2.69E+00 | 2.74E+00 | 9.93E-01 | no | Dctd | upstream_gene_variant | MODIFIER |
| 8 | 48138199 | AX-170315934 | 0 | A | T | 0.149 | -1.85E-03 | 1.50E-01 | 2.69E+00 | 2.74E+00 | 9.93E-01 | no | Dctd | intron_variant | MODIFIER |
| 8 | 48139682 | AX-170102815 | 0 | G | A | 0.149 | -1.85E-03 | 1.50E-01 | 2.69E+00 | 2.74E+00 | 9.93E-01 | no | Dctd | intron_variant | MODIFIER |
| 8 | 48141478 | AX-170353396 | 0 | G | C | 0.149 | -1.85E-03 | 1.50E-01 | 2.69E+00 | 2.74E+00 | 9.93E-01 | no | Dctd | 3_prime_UTR_variant | MODIFIER |
| 8 | 48153469 | AX-170212175 | 0 | T | G | 0.149 | -1.85E-03 | 1.50E-01 | 2.69E+00 | 2.74E+00 | 9.93E-01 | no | Dctd | downstream_gene_variant | MODIFIER |
| 8 | 48156970 | AX-170452578 | 0 | C | T | 0.149 | -1.85E-03 | 1.50E-01 | 2.69E+00 | 2.74E+00 | 9.93E-01 | no | Dctd | downstream_gene_variant | MODIFIER |
| 8 | 48200965 | AX-170448516 | 0 | C | T | 0.149 | -1.85E-03 | 1.50E-01 | 2.69E+00 | 2.74E+00 | 9.93E-01 | no | | intergenic_variant | MODIFIER |
| 8 | 48205123 | AX-170344647 | 0 | G | A | 0.149 | -1.85E-03 | 1.50E-01 | 2.69E+00 | 2.74E+00 | 9.93E-01 | no | | regulatory_region_variant | MODIFIER |
| 8 | 48205179 | AX-170090093 | 0 | C | T | 0.149 | -1.85E-03 | 1.50E-01 | 2.69E+00 | 2.74E+00 | 9.93E-01 | no | | regulatory_region_variant | MODIFIER |
| 8 | 48279992 | AX-97803619 | 0 | A | G | 0.149 | -1.85E-03 | 1.50E-01 | 2.69E+00 | 2.74E+00 | 9.93E-01 | no | Tenm3 | intron_variant | MODIFIER |
| 8 | 48281645 | AX-170384111 | 0 | T | G | 0.149 | -1.85E-03 | 1.50E-01 | 2.69E+00 | 2.74E+00 | 9.93E-01 | no | Tenm3 | intron_variant | MODIFIER |
| 8 | 48289695 | AX-170030628 | 0 | C | T | 0.149 | -1.85E-03 | 1.50E-01 | 2.69E+00 | 2.74E+00 | 9.93E-01 | no | Tenm3 | intron_variant | MODIFIER |
| 8 | 48303236 | AX-97837731 | 0 | C | T | 0.149 | -1.85E-03 | 1.50E-01 | 2.69E+00 | 2.74E+00 | 9.93E-01 | no | Tenm3 | intron_variant | MODIFIER |
| 8 | 48304851 | AX-170248888 | 0 | C | T | 0.149 | -1.85E-03 | 1.50E-01 | 2.69E+00 | 2.74E+00 | 9.93E-01 | no | Tenm3 | intron_variant | MODIFIER |
| 8 | 48312266 | AX-170215665 | 0 | C | A | 0.149 | -1.85E-03 | 1.50E-01 | 2.69E+00 | 2.74E+00 | 9.93E-01 | no | Tenm3 | intron_variant | MODIFIER |
| 8 | 48314764 | AX-170367132 | 0 | A | G | 0.149 | -1.85E-03 | 1.50E-01 | 2.69E+00 | 2.74E+00 | 9.93E-01 | no | Tenm3 | intron_variant | MODIFIER |
| 8 | 48315146 | AX-170409130 | 0 | C | T | 0.149 | -1.85E-03 | 1.50E-01 | 2.69E+00 | 2.74E+00 | 9.93E-01 | no | Tenm3 | intron_variant | MODIFIER |
| 8 | 48318373 | AX-97842717 | 0 | A | G | 0.149 | -1.85E-03 | 1.50E-01 | 2.69E+00 | 2.74E+00 | 9.93E-01 | no | Tenm3 | intron_variant | MODIFIER |
| 8 | 48333680 | AX-170119733 | 0 | G | A | 0.149 | -1.85E-03 | 1.50E-01 | 2.69E+00 | 2.74E+00 | 9.93E-01 | no | Tenm3 | intron_variant | MODIFIER |
| 8 | 48341802 | AX-170030627 | 0 | G | A | 0.149 | -1.85E-03 | 1.50E-01 | 2.69E+00 | 2.74E+00 | 9.93E-01 | no | Tenm3 | intron_variant | MODIFIER |
| 8 | 48446690 | AX-170166620 | 0 | A | G | 0.149 | -1.85E-03 | 1.50E-01 | 2.69E+00 | 2.74E+00 | 9.93E-01 | no | Tenm3 | intron_variant | MODIFIER |
| 8 | 48446790 | AX-170451617 | 0 | G | A | 0.149 | -1.85E-03 | 1.50E-01 | 2.69E+00 | 2.74E+00 | 9.93E-01 | no | Tenm3 | intron_variant | MODIFIER |
| 8 | 48454557 | AX-170319836 | 0 | C | T | 0.149 | -1.85E-03 | 1.50E-01 | 2.69E+00 | 2.74E+00 | 9.93E-01 | no | Tenm3 | intron_variant | MODIFIER |
| 12 | 63319016 | AX-170429240 | 1 | T | C | 0.062 | 2.04E-03 | 3.01E-01 | 2.72E+00 | 2.74E+00 | 9.93E-01 | no | | intergenic_variant | MODIFIER |
| 12 | 116579257 | AX-170007592 | 1 | C | T | 0.125 | 2.28E-03 | 2.00E-01 | 2.67E+00 | 2.74E+00 | 9.93E-01 | no | Ptprn2 | intron_variant | MODIFIER |
| 10 | 116014005 | AX-170411664 | 0 | T | A | 0.386 | 2.56E-04 | 7.22E-02 | 2.65E+00 | 2.74E+00 | 9.93E-01 | no | Ptprr | upstream_gene_variant | MODIFIER |
| 7 | 81127514 | AX-170411984 | 0 | T | C | 0.114 | 2.23E-03 | 2.11E-01 | 2.66E+00 | 2.74E+00 | 9.93E-01 | no | Slc28a1 | intron_variant | MODIFIER |
| 4 | 134214998 | AX-170279151 | 1 | T | G | 0.143 | 2.56E-03 | 1.08E-01 | 2.64E+00 | 2.74E+00 | 9.94E-01 | no | Catsper4 | intron_variant | MODIFIER |
| 12 | 111911283 | AX-97788704 | 1 | G | A | 0.134 | -1.65E-03 | 1.89E-01 | 2.65E+00 | 2.74E+00 | 9.94E-01 | no | Ppp1r13b | upstream_gene_variant | MODIFIER |
| 12 | 112052242 | AX-170267445 | 1 | T | C | 0.134 | -1.65E-03 | 1.89E-01 | 2.65E+00 | 2.74E+00 | 9.94E-01 | no | Tdrd9 | intron_variant | MODIFIER |
| 12 | 113776333 | AX-170050263 | 1 | A | G | 0.134 | -1.65E-03 | 1.89E-01 | 2.65E+00 | 2.74E+00 | 9.94E-01 | no | Gm37722 | upstream_gene_variant | MODIFIER |
| 12 | 118791070 | AX-170534694 | 1 | A | G | 0.134 | -1.65E-03 | 1.89E-01 | 2.65E+00 | 2.74E+00 | 9.94E-01 | no | Gm6171 | downstream_gene_variant | MODIFIER |
| 3 | 145432385 | AX-170021831 | 0 | T | C | 0.36 | -8.73E-05 | 7.31E-02 | 2.66E+00 | 2.74E+00 | 9.94E-01 | no | Col24a1 | intron_variant | MODIFIER |
| 12 | 100721611 | AX-97840939 | 0 | C | T | 0.088 | -1.37E-03 | 1.95E-01 | 2.68E+00 | 2.74E+00 | 9.94E-01 | no | Rps6ka5 | intron_variant | MODIFIER |
| 12 | 100774853 | AX-170284106 | 0 | A | C | 0.088 | -1.37E-03 | 1.95E-01 | 2.68E+00 | 2.74E+00 | 9.94E-01 | no | Dglucy | upstream_gene_variant | MODIFIER |
| 12 | 100962000 | AX-170373061 | 0 | T | G | 0.088 | -1.37E-03 | 1.95E-01 | 2.68E+00 | 2.74E+00 | 9.94E-01 | no | Ccdc88c | intron_variant | MODIFIER |
| 12 | 100962919 | AX-170125300 | 0 | G | A | 0.088 | -1.37E-03 | 1.95E-01 | 2.68E+00 | 2.74E+00 | 9.94E-01 | no | Ccdc88c | intron_variant | MODIFIER |
| 12 | 100966598 | AX-170232876 | 0 | A | G | 0.088 | -1.37E-03 | 1.95E-01 | 2.68E+00 | 2.74E+00 | 9.94E-01 | no | Ccdc88c | intron_variant | MODIFIER |

| | | | | | | | | | | | | | | | |
|---|---|---|---|---|---|---|---|---|---|---|---|---|---|---|---|
| 12 | 100969414 | AX-170203154 | 0 | T | C | 0.088 | -1.37E-03 | 1.95E-01 | 2.68E+00 | 2.74E+00 | 9.94E-01 | no | Ccdc88c | intron_variant | MODIFIER |
| 12 | 100969704 | AX-170147669 | 0 | G | A | 0.088 | -1.37E-03 | 1.95E-01 | 2.68E+00 | 2.74E+00 | 9.94E-01 | no | Ccdc88c | intron_variant | MODIFIER |
| 12 | 100971451 | AX-97805653 | 0 | G | T | 0.088 | -1.37E-03 | 1.95E-01 | 2.68E+00 | 2.74E+00 | 9.94E-01 | no | Ccdc88c | intron_variant | MODIFIER |
| 12 | 100980101 | AX-170181012 | 0 | A | G | 0.088 | -1.37E-03 | 1.95E-01 | 2.68E+00 | 2.74E+00 | 9.94E-01 | no | Ccdc88c | intron_variant | MODIFIER |
| 12 | 101003976 | AX-170007391 | 0 | T | C | 0.088 | -1.37E-03 | 1.95E-01 | 2.68E+00 | 2.74E+00 | 9.94E-01 | no | Ccdc88c | intron_variant | MODIFIER |
| 12 | 101019860 | AX-170007396 | 0 | T | C | 0.088 | -1.37E-03 | 1.95E-01 | 2.68E+00 | 2.74E+00 | 9.94E-01 | no | Ccdc88c | intron_variant | MODIFIER |
| 12 | 101038982 | AX-170523692 | 0 | T | C | 0.088 | -1.37E-03 | 1.95E-01 | 2.68E+00 | 2.74E+00 | 9.94E-01 | no | Ppp4r3a | downstream_gene_variant | MODIFIER |
| 12 | 101135879 | AX-170007395 | 0 | C | T | 0.088 | -1.37E-03 | 1.95E-01 | 2.68E+00 | 2.74E+00 | 9.94E-01 | no | | intergenic_variant | MODIFIER |
| 12 | 101161867 | AX-168391886 | 0 | T | C | 0.088 | -1.37E-03 | 1.95E-01 | 2.68E+00 | 2.74E+00 | 9.94E-01 | no | | intergenic_variant | MODIFIER |
| 12 | 101220622 | AX-170087323 | 0 | G | T | 0.088 | -1.37E-03 | 1.95E-01 | 2.68E+00 | 2.74E+00 | 9.94E-01 | no | Gm22749 | downstream_gene_variant | MODIFIER |
| 12 | 101288925 | AX-170506515 | 0 | C | T | 0.088 | -1.37E-03 | 1.95E-01 | 2.68E+00 | 2.74E+00 | 9.94E-01 | no | Gm18795 | intron_variant,non_coding_transcript_variant | MODIFIER |
| 12 | 101302910 | AX-170458498 | 0 | C | T | 0.088 | -1.37E-03 | 1.95E-01 | 2.68E+00 | 2.74E+00 | 9.94E-01 | no | Gm18795 | intron_variant,non_coding_transcript_variant | MODIFIER |
| 12 | 101304279 | AX-170145829 | 0 | A | T | 0.088 | -1.37E-03 | 1.95E-01 | 2.68E+00 | 2.74E+00 | 9.94E-01 | no | Gm18795 | intron_variant,non_coding_transcript_variant | MODIFIER |
| 12 | 101344244 | AX-170438456 | 0 | C | T | 0.088 | -1.37E-03 | 1.95E-01 | 2.68E+00 | 2.74E+00 | 9.94E-01 | no | Gm18795 | intron_variant,non_coding_transcript_variant | MODIFIER |
| 12 | 101352861 | AX-170518199 | 0 | G | A | 0.088 | -1.37E-03 | 1.95E-01 | 2.68E+00 | 2.74E+00 | 9.94E-01 | no | Gm18795 | intron_variant,non_coding_transcript_variant | MODIFIER |
| 12 | 101372024 | AX-97809821 | 0 | T | C | 0.088 | -1.37E-03 | 1.95E-01 | 2.68E+00 | 2.74E+00 | 9.94E-01 | no | Gm18795 | intron_variant,non_coding_transcript_variant | MODIFIER |
| 12 | 101378133 | AX-170100099 | 0 | A | T | 0.088 | -1.37E-03 | 1.95E-01 | 2.68E+00 | 2.74E+00 | 9.94E-01 | no | | intergenic_variant | MODIFIER |
| 12 | 101403645 | AX-97780314 | 0 | T | C | 0.088 | -1.37E-03 | 1.95E-01 | 2.68E+00 | 2.74E+00 | 9.94E-01 | no | Catsperb | upstream_gene_variant | MODIFIER |
| 12 | 101444924 | AX-170460551 | 0 | A | G | 0.088 | -1.37E-03 | 1.95E-01 | 2.68E+00 | 2.74E+00 | 9.94E-01 | no | Catsperb | intron_variant | MODIFIER |
| 12 | 101510557 | AX-170438668 | 0 | T | C | 0.088 | -1.37E-03 | 1.95E-01 | 2.68E+00 | 2.74E+00 | 9.94E-01 | no | Catsperb | intron_variant | MODIFIER |
| 12 | 101664054 | AX-170007405 | 0 | G | A | 0.088 | -1.37E-03 | 1.95E-01 | 2.68E+00 | 2.74E+00 | 9.94E-01 | no | Tc2n | intron_variant,NMD_transcript_variant | MODIFIER |
| 12 | 101680841 | AX-170007399 | 0 | A | G | 0.088 | -1.37E-03 | 1.95E-01 | 2.68E+00 | 2.74E+00 | 9.94E-01 | no | Tc2n | intron_variant,NMD_transcript_variant | MODIFIER |
| 12 | 101741885 | AX-97822195 | 0 | T | C | 0.088 | -1.37E-03 | 1.95E-01 | 2.68E+00 | 2.74E+00 | 9.94E-01 | no | Fbln5 | downstream_gene_variant | MODIFIER |
| 12 | 101747941 | AX-170507856 | 0 | A | G | 0.088 | -1.37E-03 | 1.95E-01 | 2.68E+00 | 2.74E+00 | 9.94E-01 | no | Fbln5 | 3_prime_UTR_variant | MODIFIER |
| 12 | 101751200 | AX-170007403 | 0 | C | T | 0.088 | -1.37E-03 | 1.95E-01 | 2.68E+00 | 2.74E+00 | 9.94E-01 | no | Fbln5 | intron_variant | MODIFIER |
| 12 | 101772678 | AX-170349571 | 0 | T | C | 0.088 | -1.37E-03 | 1.95E-01 | 2.68E+00 | 2.74E+00 | 9.94E-01 | no | Fbln5 | intron_variant | MODIFIER |
| 12 | 102254829 | AX-170145801 | 0 | G | T | 0.088 | -1.37E-03 | 1.95E-01 | 2.68E+00 | 2.74E+00 | 9.94E-01 | no | Slc24a4 | intron_variant | MODIFIER |
| 12 | 102415824 | AX-170228598 | 0 | T | C | 0.088 | -1.37E-03 | 1.95E-01 | 2.68E+00 | 2.74E+00 | 9.94E-01 | no | Lgmn | intron_variant | MODIFIER |
| 12 | 102433314 | AX-170225441 | 0 | C | A | 0.088 | -1.37E-03 | 1.95E-01 | 2.68E+00 | 2.74E+00 | 9.94E-01 | no | Lgmn | intron_variant | MODIFIER |
| 20 | 108705206 | AX-170090624 | 1 | C | T | 0.304 | -4.58E-04 | 5.33E-02 | 2.62E+00 | 2.74E+00 | 9.94E-01 | no | | intergenic_variant | MODIFIER |
| 6 | 111198373 | AX-170434326 | 0 | G | A | 0.07 | 1.97E-03 | 2.76E-01 | 2.70E+00 | 2.74E+00 | 9.94E-01 | no | Grm7 | intron_variant | MODIFIER |
| 15 | 67044652 | AX-170484445 | 0 | C | T | 0.325 | -1.74E-03 | 7.76E-02 | 2.62E+00 | 2.74E+00 | 9.94E-01 | no | Gm31342 | downstream_gene_variant | MODIFIER |
| 10 | 115832158 | AX-170454991 | 0 | T | C | 0.386 | -6.40E-04 | 7.73E-02 | 2.65E+00 | 2.74E+00 | 9.94E-01 | no | Tspan8 | intron_variant | MODIFIER |
| 18 | 30441974 | AX-170277013 | 1 | T | C | 0.062 | -2.66E-03 | 3.01E-01 | 2.71E+00 | 2.74E+00 | 9.94E-01 | no | | intergenic_variant | MODIFIER |
| 18 | 41655992 | AX-170317254 | 1 | A | G | 0.062 | -2.66E-03 | 3.01E-01 | 2.71E+00 | 2.74E+00 | 9.94E-01 | no | | intergenic_variant | MODIFIER |
| 9 | 107948853 | AX-170083403 | 1 | A | G | 0.125 | -3.66E-04 | 1.17E-01 | 2.67E+00 | 2.74E+00 | 9.94E-01 | no | Camkv | 3_prime_UTR_variant | MODIFIER |
| 5 | 127883260 | AX-170255857 | 1 | C | G | 0.429 | 7.37E-05 | 8.51E-02 | 2.64E+00 | 2.74E+00 | 9.94E-01 | no | Tmem132d | intron_variant | MODIFIER |
| 1 | 48595030 | AX-97811816 | 1 | C | A | 0.107 | -1.43E-03 | 2.36E-01 | 2.69E+00 | 2.74E+00 | 9.94E-01 | no | | intergenic_variant | MODIFIER |
| 1 | 48750817 | AX-97801889 | 1 | C | T | 0.107 | -1.43E-03 | 2.36E-01 | 2.69E+00 | 2.74E+00 | 9.94E-01 | no | | intergenic_variant | MODIFIER |
| 1 | 51495978 | AX-170320828 | 1 | A | G | 0.107 | -1.43E-03 | 2.36E-01 | 2.69E+00 | 2.74E+00 | 9.94E-01 | no | | intergenic_variant | MODIFIER |
| 1 | 54668015 | AX-170000534 | 1 | C | A | 0.107 | -1.43E-03 | 2.36E-01 | 2.69E+00 | 2.74E+00 | 9.94E-01 | no | Ankrd44 | intron_variant | MODIFIER |
| 7 | 137926778 | AX-170238904 | 0 | C | T | 0.307 | -1.53E-03 | 7.82E-02 | 2.63E+00 | 2.74E+00 | 9.94E-01 | no | | intergenic_variant | MODIFIER |
| 2 | 124412368 | AX-170481736 | 0 | G | A | 0.14 | 1.98E-03 | 2.20E-01 | 2.71E+00 | 2.74E+00 | 9.94E-01 | no | Sema6d | intron_variant | MODIFIER |
| 2 | 149449562 | AX-170368734 | 0 | C | T | 0.14 | 1.98E-03 | 2.20E-01 | 2.71E+00 | 2.74E+00 | 9.94E-01 | no | | intergenic_variant | MODIFIER |
| 1 | 10522915 | AX-97802470 | 0 | T | G | 0.395 | 3.15E-04 | 7.45E-02 | 2.67E+00 | 2.74E+00 | 9.94E-01 | no | Cpa6 | intron_variant | MODIFIER |
| 1 | 10638001 | AX-170365817 | 0 | C | T | 0.395 | 3.15E-04 | 7.45E-02 | 2.67E+00 | 2.74E+00 | 9.94E-01 | no | Cpa6 | intron_variant | MODIFIER |
| 1 | 10640515 | AX-170204055 | 0 | A | G | 0.395 | 3.15E-04 | 7.45E-02 | 2.67E+00 | 2.74E+00 | 9.94E-01 | no | Cpa6 | intron_variant | MODIFIER |
| 1 | 10644678 | AX-170467599 | 0 | A | C | 0.395 | 3.15E-04 | 7.45E-02 | 2.67E+00 | 2.74E+00 | 9.94E-01 | no | Cpa6 | intron_variant | MODIFIER |
| 1 | 10644713 | AX-97810597 | 0 | T | C | 0.395 | 3.15E-04 | 7.45E-02 | 2.67E+00 | 2.74E+00 | 9.94E-01 | no | Cpa6 | intron_variant | MODIFIER |
| 1 | 10666321 | AX-170184404 | 0 | A | G | 0.395 | 3.15E-04 | 7.45E-02 | 2.67E+00 | 2.74E+00 | 9.94E-01 | no | Cpa6 | intron_variant | MODIFIER |
| 1 | 10681410 | AX-170118514 | 0 | T | C | 0.395 | 3.15E-04 | 7.45E-02 | 2.67E+00 | 2.74E+00 | 9.94E-01 | no | Cpa6 | intron_variant | MODIFIER |
| 1 | 10704773 | AX-170489433 | 0 | C | T | 0.395 | 3.15E-04 | 7.45E-02 | 2.67E+00 | 2.74E+00 | 9.94E-01 | no | Cpa6 | intron_variant | MODIFIER |
| 1 | 10716355 | AX-170040362 | 0 | G | A | 0.395 | 3.15E-04 | 7.45E-02 | 2.67E+00 | 2.74E+00 | 9.94E-01 | no | Cpa6 | intron_variant | MODIFIER |
| 1 | 10725127 | AX-170040365 | 0 | G | A | 0.395 | 3.15E-04 | 7.45E-02 | 2.67E+00 | 2.74E+00 | 9.94E-01 | no | | intergenic_variant | MODIFIER |

| | | | | | | | | | | | | | | | |
|---|---|---|---|---|---|---|---|---|---|---|---|---|---|---|---|
| 1 | 10764420 | AX-169999932 | 0 | C | T | 0.395 | 3.15E-04 | 7.45E-02 | 2.67E+00 | 2.74E+00 | 9.94E-01 | no | | intergenic_variant | MODIFIER |
| 11 | 34906598 | AX-170465150 | 0 | T | C | 0.079 | 7.06E-04 | 2.02E-01 | 2.65E+00 | 2.74E+00 | 9.94E-01 | no | | intergenic_variant | MODIFIER |
| 4 | 95900437 | AX-170310654 | 0 | T | C | 0.254 | 2.31E-03 | 1.01E-01 | 2.62E+00 | 2.74E+00 | 9.94E-01 | no | Fggy | intron_variant | MODIFIER |
| 13 | 53548317 | AX-97812306 | 0 | T | C | 0.518 | 1.59E-03 | 6.78E-02 | 2.63E+00 | 2.74E+00 | 9.95E-01 | no | | intergenic_variant | MODIFIER |
| 12 | 102421181 | AX-170382510 | 1 | C | T | 0.054 | 9.19E-04 | 2.05E-01 | 2.64E+00 | 2.74E+00 | 9.95E-01 | no | Lgmn | intron_variant | MODIFIER |
| 12 | 102203256 | AX-170149461 | 1 | C | T | 0.071 | -7.60E-04 | 1.67E-01 | 2.65E+00 | 2.74E+00 | 9.95E-01 | no | Slc24a4 | intron_variant | MODIFIER |
| 12 | 116122398 | AX-170217448 | 0 | A | G | 0.114 | -2.04E-04 | 1.66E-01 | 2.63E+00 | 2.74E+00 | 9.95E-01 | no | Vipr2 | intron_variant | MODIFIER |
| 4 | 143896970 | AX-170341517 | 2 | T | C | 0.1 | 1.46E-03 | 1.91E-01 | 2.66E+00 | 2.74E+00 | 9.95E-01 | no | Pramef6 | synonymous_variant | LOW |
| 6 | 138803005 | AX-170446852 | 1 | G | T | 0.071 | -1.83E-03 | 2.78E-01 | 2.70E+00 | 2.74E+00 | 9.95E-01 | no | | intergenic_variant | MODIFIER |
| 6 | 138858074 | AX-170388253 | 1 | G | C | 0.071 | -1.83E-03 | 2.78E-01 | 2.70E+00 | 2.74E+00 | 9.95E-01 | no | Gm9038 | downstream_gene_variant | MODIFIER |
| 1 | 11488235 | AX-170325032 | 1 | A | T | 0.393 | 1.09E-03 | 7.28E-02 | 2.67E+00 | 2.74E+00 | 9.95E-01 | no | A830018L16Rik | intron_variant | MODIFIER |
| 12 | 114997578 | AX-170050290 | 1 | A | G | 0.125 | -6.06E-04 | 1.96E-01 | 2.67E+00 | 2.74E+00 | 9.95E-01 | no | | intergenic_variant | MODIFIER |
| 1 | 52916962 | AX-170041234 | 0 | T | G | 0.114 | -1.91E-04 | 1.91E-01 | 2.64E+00 | 2.74E+00 | 9.95E-01 | no | Hibch | intron_variant | MODIFIER |
| 6 | 105724740 | AX-170027892 | 0 | A | C | 0.061 | -2.26E-03 | 3.02E-01 | 2.72E+00 | 2.74E+00 | 9.95E-01 | no | Cntn4 | intron_variant | MODIFIER |
| 12 | 57276789 | AX-97821278 | 0 | C | T | 0.061 | -2.26E-03 | 3.02E-01 | 2.72E+00 | 2.74E+00 | 9.95E-01 | no | Mipol1 | intron_variant | MODIFIER |
| 12 | 57283372 | AX-170329446 | 0 | T | G | 0.061 | -2.26E-03 | 3.02E-01 | 2.72E+00 | 2.74E+00 | 9.95E-01 | no | Mipol1 | intron_variant | MODIFIER |
| 12 | 57289655 | AX-170158446 | 0 | G | A | 0.061 | -2.26E-03 | 3.02E-01 | 2.72E+00 | 2.74E+00 | 9.95E-01 | no | Mipol1 | intron_variant | MODIFIER |
| 12 | 57303392 | AX-170521829 | 0 | A | G | 0.061 | -2.26E-03 | 3.02E-01 | 2.72E+00 | 2.74E+00 | 9.95E-01 | no | Mipol1 | 5_prime_UTR_variant | MODIFIER |
| 12 | 57303441 | AX-170333994 | 0 | T | C | 0.061 | -2.26E-03 | 3.02E-01 | 2.72E+00 | 2.74E+00 | 9.95E-01 | no | Mipol1 | synonymous_variant | LOW |
| 12 | 57328350 | AX-170223573 | 0 | C | T | 0.061 | -2.26E-03 | 3.02E-01 | 2.72E+00 | 2.74E+00 | 9.95E-01 | no | Mipol1 | intron_variant | MODIFIER |
| 12 | 57332485 | AX-170531189 | 0 | G | A | 0.061 | -2.26E-03 | 3.02E-01 | 2.72E+00 | 2.74E+00 | 9.95E-01 | no | Mipol1 | intron_variant | MODIFIER |
| 12 | 57336580 | AX-170214679 | 0 | A | G | 0.061 | -2.26E-03 | 3.02E-01 | 2.72E+00 | 2.74E+00 | 9.95E-01 | no | Mipol1 | intron_variant | MODIFIER |
| 12 | 57385308 | AX-170459686 | 0 | G | C | 0.061 | -2.26E-03 | 3.02E-01 | 2.72E+00 | 2.74E+00 | 9.95E-01 | no | Mipol1 | intron_variant | MODIFIER |
| 12 | 57400488 | AX-170049119 | 0 | G | A | 0.061 | -2.26E-03 | 3.02E-01 | 2.72E+00 | 2.74E+00 | 9.95E-01 | no | Mipol1 | intron_variant | MODIFIER |
| 12 | 57402059 | AX-170499200 | 0 | G | C | 0.061 | -2.26E-03 | 3.02E-01 | 2.72E+00 | 2.74E+00 | 9.95E-01 | no | Mipol1 | intron_variant | MODIFIER |
| 12 | 57402295 | AX-170049123 | 0 | A | C | 0.061 | -2.26E-03 | 3.02E-01 | 2.72E+00 | 2.74E+00 | 9.95E-01 | no | Mipol1 | intron_variant | MODIFIER |
| 12 | 57505868 | AX-170006810 | 0 | A | G | 0.061 | -2.26E-03 | 3.02E-01 | 2.72E+00 | 2.74E+00 | 9.95E-01 | no | | intergenic_variant | MODIFIER |
| 12 | 57505896 | AX-170006811 | 0 | G | A | 0.061 | -2.26E-03 | 3.02E-01 | 2.72E+00 | 2.74E+00 | 9.95E-01 | no | | intergenic_variant | MODIFIER |
| 12 | 57823799 | AX-170313388 | 0 | G | A | 0.061 | -2.26E-03 | 3.02E-01 | 2.72E+00 | 2.74E+00 | 9.95E-01 | no | | intergenic_variant | MODIFIER |
| 12 | 57837747 | AX-170520018 | 0 | T | C | 0.061 | -2.26E-03 | 3.02E-01 | 2.72E+00 | 2.74E+00 | 9.95E-01 | no | | intergenic_variant | MODIFIER |
| 12 | 57844713 | AX-170171463 | 0 | G | T | 0.061 | -2.26E-03 | 3.02E-01 | 2.72E+00 | 2.74E+00 | 9.95E-01 | no | | intergenic_variant | MODIFIER |
| 12 | 57920622 | AX-97788048 | 0 | T | C | 0.061 | -2.26E-03 | 3.02E-01 | 2.72E+00 | 2.74E+00 | 9.95E-01 | no | | intergenic_variant | MODIFIER |
| 12 | 57921430 | AX-170187128 | 0 | C | T | 0.061 | -2.26E-03 | 3.02E-01 | 2.72E+00 | 2.74E+00 | 9.95E-01 | no | | intergenic_variant | MODIFIER |
| 12 | 57923943 | AX-170488266 | 0 | T | C | 0.061 | -2.26E-03 | 3.02E-01 | 2.72E+00 | 2.74E+00 | 9.95E-01 | no | | intergenic_variant | MODIFIER |
| 12 | 57925516 | AX-170237666 | 0 | C | T | 0.061 | -2.26E-03 | 3.02E-01 | 2.72E+00 | 2.74E+00 | 9.95E-01 | no | | intergenic_variant | MODIFIER |
| 12 | 57925542 | AX-170181763 | 0 | G | C | 0.061 | -2.26E-03 | 3.02E-01 | 2.72E+00 | 2.74E+00 | 9.95E-01 | no | | intergenic_variant | MODIFIER |
| 12 | 57927333 | AX-170049128 | 0 | G | A | 0.061 | -2.26E-03 | 3.02E-01 | 2.72E+00 | 2.74E+00 | 9.95E-01 | no | | intergenic_variant | MODIFIER |
| 12 | 57929460 | AX-170407212 | 0 | C | T | 0.061 | -2.26E-03 | 3.02E-01 | 2.72E+00 | 2.74E+00 | 9.95E-01 | no | | intergenic_variant | MODIFIER |
| 12 | 57933672 | AX-170049129 | 0 | A | C | 0.061 | -2.26E-03 | 3.02E-01 | 2.72E+00 | 2.74E+00 | 9.95E-01 | no | | intergenic_variant | MODIFIER |
| 12 | 57933920 | AX-170049132 | 0 | C | T | 0.061 | -2.26E-03 | 3.02E-01 | 2.72E+00 | 2.74E+00 | 9.95E-01 | no | | intergenic_variant | MODIFIER |
| 12 | 57955326 | AX-170112315 | 0 | C | T | 0.061 | -2.26E-03 | 3.02E-01 | 2.72E+00 | 2.74E+00 | 9.95E-01 | no | Gm46329 | intron_variant,non_coding_transcript_variant | MODIFIER |
| 12 | 57956135 | AX-97852687 | 0 | A | G | 0.061 | -2.26E-03 | 3.02E-01 | 2.72E+00 | 2.74E+00 | 9.95E-01 | no | Gm46329 | intron_variant,non_coding_transcript_variant | MODIFIER |
| 12 | 58018726 | AX-170497702 | 0 | G | A | 0.061 | -2.26E-03 | 3.02E-01 | 2.72E+00 | 2.74E+00 | 9.95E-01 | no | Gm46329 | intron_variant,non_coding_transcript_variant | MODIFIER |
| 12 | 58065880 | AX-170407469 | 0 | T | C | 0.061 | -2.26E-03 | 3.02E-01 | 2.72E+00 | 2.74E+00 | 9.95E-01 | no | Gm46329 | intron_variant,non_coding_transcript_variant | MODIFIER |
| 12 | 58068494 | AX-170250012 | 0 | T | C | 0.061 | -2.26E-03 | 3.02E-01 | 2.72E+00 | 2.74E+00 | 9.95E-01 | no | Gm46329 | intron_variant,non_coding_transcript_variant | MODIFIER |
| 12 | 58201697 | AX-170049134 | 0 | T | C | 0.061 | -2.26E-03 | 3.02E-01 | 2.72E+00 | 2.74E+00 | 9.95E-01 | no | | regulatory_region_variant | MODIFIER |
| 12 | 58517489 | AX-170311279 | 0 | C | T | 0.061 | -2.26E-03 | 3.02E-01 | 2.72E+00 | 2.74E+00 | 9.95E-01 | no | | intergenic_variant | MODIFIER |
| 12 | 58670274 | AX-170350002 | 0 | C | A | 0.061 | -2.26E-03 | 3.02E-01 | 2.72E+00 | 2.74E+00 | 9.95E-01 | no | | intergenic_variant | MODIFIER |
| 12 | 58674619 | AX-170006821 | 0 | G | T | 0.061 | -2.26E-03 | 3.02E-01 | 2.72E+00 | 2.74E+00 | 9.95E-01 | no | | intergenic_variant | MODIFIER |
| 12 | 58698194 | AX-170452618 | 0 | C | T | 0.061 | -2.26E-03 | 3.02E-01 | 2.72E+00 | 2.74E+00 | 9.95E-01 | no | | intergenic_variant | MODIFIER |
| 12 | 58699035 | AX-170140593 | 0 | T | C | 0.061 | -2.26E-03 | 3.02E-01 | 2.72E+00 | 2.74E+00 | 9.95E-01 | no | | intergenic_variant | MODIFIER |
| 12 | 58707026 | AX-170313888 | 0 | G | A | 0.061 | -2.26E-03 | 3.02E-01 | 2.72E+00 | 2.74E+00 | 9.95E-01 | no | | intergenic_variant | MODIFIER |
| 12 | 58711669 | AX-170206202 | 0 | A | G | 0.061 | -2.26E-03 | 3.02E-01 | 2.72E+00 | 2.74E+00 | 9.95E-01 | no | | intergenic_variant | MODIFIER |
| 12 | 58884765 | AX-170201256 | 0 | G | A | 0.061 | -2.26E-03 | 3.02E-01 | 2.72E+00 | 2.74E+00 | 9.95E-01 | no | | intergenic_variant | MODIFIER |

| | | | | | | | | | | | | | | | |
|---|---|---|---|---|---|---|---|---|---|---|---|---|---|---|---|
| 12 | 59142685 | AX-170006830 | 0 | T | C | 0.061 | -2.26E-03 | 3.02E-01 | 2.72E+00 | 2.74E+00 | 9.95E-01 | no | Mia2 | intron_variant | MODIFIER |
| 12 | 59145370 | AX-97826532 | 0 | G | C | 0.061 | -2.26E-03 | 3.02E-01 | 2.72E+00 | 2.74E+00 | 9.95E-01 | no | Mia2 | intron_variant | MODIFIER |
| 12 | 59591673 | AX-170166875 | 0 | C | T | 0.061 | -2.26E-03 | 3.02E-01 | 2.72E+00 | 2.74E+00 | 9.95E-01 | no | | intergenic_variant | MODIFIER |
| 12 | 59665545 | AX-97805547 | 0 | C | T | 0.061 | -2.26E-03 | 3.02E-01 | 2.72E+00 | 2.74E+00 | 9.95E-01 | no | Gm24377 | downstream_gene_variant | MODIFIER |
| 12 | 59697083 | AX-170006838 | 0 | A | T | 0.061 | -2.26E-03 | 3.02E-01 | 2.72E+00 | 2.74E+00 | 9.95E-01 | no | | intergenic_variant | MODIFIER |
| 12 | 59697394 | AX-170184831 | 0 | C | T | 0.061 | -2.26E-03 | 3.02E-01 | 2.72E+00 | 2.74E+00 | 9.95E-01 | no | | intergenic_variant | MODIFIER |
| 12 | 59699169 | AX-170159901 | 0 | G | A | 0.061 | -2.26E-03 | 3.02E-01 | 2.72E+00 | 2.74E+00 | 9.95E-01 | no | | intergenic_variant | MODIFIER |
| 12 | 59699549 | AX-97818783 | 0 | T | C | 0.061 | -2.26E-03 | 3.02E-01 | 2.72E+00 | 2.74E+00 | 9.95E-01 | no | | intergenic_variant | MODIFIER |
| 12 | 59712411 | AX-170159910 | 0 | C | T | 0.061 | -2.26E-03 | 3.02E-01 | 2.72E+00 | 2.74E+00 | 9.95E-01 | no | | intergenic_variant | MODIFIER |
| 12 | 59716292 | AX-170320374 | 0 | A | C | 0.061 | -2.26E-03 | 3.02E-01 | 2.72E+00 | 2.74E+00 | 9.95E-01 | no | Gm48285 | downstream_gene_variant | MODIFIER |
| 12 | 59722637 | AX-170486914 | 0 | T | G | 0.061 | -2.26E-03 | 3.02E-01 | 2.72E+00 | 2.74E+00 | 9.95E-01 | no | Gm48285 | upstream_gene_variant | MODIFIER |
| 12 | 59765963 | AX-170148905 | 0 | C | T | 0.061 | -2.26E-03 | 3.02E-01 | 2.72E+00 | 2.74E+00 | 9.95E-01 | no | | intergenic_variant | MODIFIER |
| 12 | 59766092 | AX-170508932 | 0 | A | G | 0.061 | -2.26E-03 | 3.02E-01 | 2.72E+00 | 2.74E+00 | 9.95E-01 | no | | intergenic_variant | MODIFIER |
| 12 | 59766425 | AX-170226565 | 0 | C | T | 0.061 | -2.26E-03 | 3.02E-01 | 2.72E+00 | 2.74E+00 | 9.95E-01 | no | | intergenic_variant | MODIFIER |
| 12 | 59766656 | AX-170367020 | 0 | A | G | 0.061 | -2.26E-03 | 3.02E-01 | 2.72E+00 | 2.74E+00 | 9.95E-01 | no | | intergenic_variant | MODIFIER |
| 12 | 59768647 | AX-170049167 | 0 | A | G | 0.061 | -2.26E-03 | 3.02E-01 | 2.72E+00 | 2.74E+00 | 9.95E-01 | no | | intergenic_variant | MODIFIER |
| 12 | 59785553 | AX-170339121 | 0 | T | A | 0.061 | -2.26E-03 | 3.02E-01 | 2.72E+00 | 2.74E+00 | 9.95E-01 | no | | intergenic_variant | MODIFIER |
| 12 | 60188908 | AX-170006840 | 0 | G | A | 0.061 | -2.26E-03 | 3.02E-01 | 2.72E+00 | 2.74E+00 | 9.95E-01 | no | | intergenic_variant | MODIFIER |
| 12 | 60199158 | AX-170102320 | 0 | C | T | 0.061 | -2.26E-03 | 3.02E-01 | 2.72E+00 | 2.74E+00 | 9.95E-01 | no | | intergenic_variant | MODIFIER |
| 12 | 60564999 | AX-170507020 | 0 | G | T | 0.061 | -2.26E-03 | 3.02E-01 | 2.72E+00 | 2.74E+00 | 9.95E-01 | no | | intergenic_variant | MODIFIER |
| 12 | 60728547 | AX-170127115 | 0 | T | C | 0.061 | -2.26E-03 | 3.02E-01 | 2.72E+00 | 2.74E+00 | 9.95E-01 | no | | intergenic_variant | MODIFIER |
| 12 | 60735224 | AX-170006848 | 0 | T | C | 0.061 | -2.26E-03 | 3.02E-01 | 2.72E+00 | 2.74E+00 | 9.95E-01 | no | Gm23059 | upstream_gene_variant | MODIFIER |
| 12 | 60996353 | AX-97797832 | 0 | G | A | 0.061 | -2.26E-03 | 3.02E-01 | 2.72E+00 | 2.74E+00 | 9.95E-01 | no | | intergenic_variant | MODIFIER |
| 12 | 61144060 | AX-170513445 | 0 | A | G | 0.061 | -2.26E-03 | 3.02E-01 | 2.72E+00 | 2.74E+00 | 9.95E-01 | no | | intergenic_variant | MODIFIER |
| 12 | 61173913 | AX-170308406 | 0 | G | C | 0.061 | -2.26E-03 | 3.02E-01 | 2.72E+00 | 2.74E+00 | 9.95E-01 | no | | intergenic_variant | MODIFIER |
| 12 | 61199187 | AX-170253117 | 0 | C | A | 0.061 | -2.26E-03 | 3.02E-01 | 2.72E+00 | 2.74E+00 | 9.95E-01 | no | Gm48301 | intron_variant,non_coding_transcript_variant | MODIFIER |
| 12 | 61199648 | AX-170472832 | 0 | C | G | 0.061 | -2.26E-03 | 3.02E-01 | 2.72E+00 | 2.74E+00 | 9.95E-01 | no | Gm48301 | intron_variant,non_coding_transcript_variant | MODIFIER |
| 12 | 61219516 | AX-170486175 | 0 | A | T | 0.061 | -2.26E-03 | 3.02E-01 | 2.72E+00 | 2.74E+00 | 9.95E-01 | no | Gm48301 | intron_variant,non_coding_transcript_variant | MODIFIER |
| 12 | 61226315 | AX-97821346 | 0 | T | C | 0.061 | -2.26E-03 | 3.02E-01 | 2.72E+00 | 2.74E+00 | 9.95E-01 | no | Gm48301 | intron_variant,non_coding_transcript_variant | MODIFIER |
| 12 | 61229038 | AX-170049197 | 0 | C | A | 0.061 | -2.26E-03 | 3.02E-01 | 2.72E+00 | 2.74E+00 | 9.95E-01 | no | Gm48301 | intron_variant,non_coding_transcript_variant | MODIFIER |
| 12 | 61284690 | AX-170347882 | 0 | A | G | 0.061 | -2.26E-03 | 3.02E-01 | 2.72E+00 | 2.74E+00 | 9.95E-01 | no | | intergenic_variant | MODIFIER |
| 12 | 61286131 | AX-170480618 | 0 | C | T | 0.061 | -2.26E-03 | 3.02E-01 | 2.72E+00 | 2.74E+00 | 9.95E-01 | no | | intergenic_variant | MODIFIER |
| 12 | 61293988 | AX-170006854 | 0 | C | G | 0.061 | -2.26E-03 | 3.02E-01 | 2.72E+00 | 2.74E+00 | 9.95E-01 | no | | intergenic_variant | MODIFIER |
| 12 | 61296576 | AX-170207441 | 0 | C | A | 0.061 | -2.26E-03 | 3.02E-01 | 2.72E+00 | 2.74E+00 | 9.95E-01 | no | | intergenic_variant | MODIFIER |
| 12 | 61299313 | AX-97789751 | 0 | C | T | 0.061 | -2.26E-03 | 3.02E-01 | 2.72E+00 | 2.74E+00 | 9.95E-01 | no | Gm7745 | downstream_gene_variant | MODIFIER |
| 12 | 61322924 | AX-97821484 | 0 | A | G | 0.061 | -2.26E-03 | 3.02E-01 | 2.72E+00 | 2.74E+00 | 9.95E-01 | no | Gm7745 | intron_variant,non_coding_transcript_variant | MODIFIER |
| 12 | 61389848 | AX-170006856 | 0 | G | A | 0.061 | -2.26E-03 | 3.02E-01 | 2.72E+00 | 2.74E+00 | 9.95E-01 | no | | intergenic_variant | MODIFIER |
| 12 | 61420869 | AX-170006863 | 0 | G | A | 0.061 | -2.26E-03 | 3.02E-01 | 2.72E+00 | 2.74E+00 | 9.95E-01 | no | | intergenic_variant | MODIFIER |
| 12 | 61421645 | AX-97821483 | 0 | A | G | 0.061 | -2.26E-03 | 3.02E-01 | 2.72E+00 | 2.74E+00 | 9.95E-01 | no | | intergenic_variant | MODIFIER |
| 12 | 61464968 | AX-168373064 | 0 | A | G | 0.061 | -2.26E-03 | 3.02E-01 | 2.72E+00 | 2.74E+00 | 9.95E-01 | no | | intergenic_variant | MODIFIER |
| 12 | 61491117 | AX-170006858 | 0 | T | C | 0.061 | -2.26E-03 | 3.02E-01 | 2.72E+00 | 2.74E+00 | 9.95E-01 | no | | intergenic_variant | MODIFIER |
| 12 | 61497152 | AX-97824272 | 0 | T | C | 0.061 | -2.26E-03 | 3.02E-01 | 2.72E+00 | 2.74E+00 | 9.95E-01 | no | Gm20063 | downstream_gene_variant | MODIFIER |
| 12 | 61624960 | AX-170426683 | 0 | A | C | 0.061 | -2.26E-03 | 3.02E-01 | 2.72E+00 | 2.74E+00 | 9.95E-01 | no | Lrfn5 | intron_variant | MODIFIER |
| 12 | 61809435 | AX-170006865 | 0 | C | G | 0.061 | -2.26E-03 | 3.02E-01 | 2.72E+00 | 2.74E+00 | 9.95E-01 | no | Lrfn5 | intron_variant | MODIFIER |
| 12 | 62385019 | AX-97831565 | 0 | C | T | 0.061 | -2.26E-03 | 3.02E-01 | 2.72E+00 | 2.74E+00 | 9.95E-01 | no | | intergenic_variant | MODIFIER |
| 12 | 63117517 | AX-170389567 | 0 | T | C | 0.061 | -2.26E-03 | 3.02E-01 | 2.72E+00 | 2.74E+00 | 9.95E-01 | no | | intergenic_variant | MODIFIER |
| 12 | 63205360 | AX-170427248 | 0 | A | G | 0.061 | -2.26E-03 | 3.02E-01 | 2.72E+00 | 2.74E+00 | 9.95E-01 | no | | intergenic_variant | MODIFIER |
| 12 | 63216391 | AX-97790654 | 0 | C | T | 0.061 | -2.26E-03 | 3.02E-01 | 2.72E+00 | 2.74E+00 | 9.95E-01 | no | | intergenic_variant | MODIFIER |
| 12 | 63303189 | AX-97841132 | 0 | T | G | 0.061 | -2.26E-03 | 3.02E-01 | 2.72E+00 | 2.74E+00 | 9.95E-01 | no | | intergenic_variant | MODIFIER |
| 12 | 63304977 | AX-170143889 | 0 | C | T | 0.061 | -2.26E-03 | 3.02E-01 | 2.72E+00 | 2.74E+00 | 9.95E-01 | no | | intergenic_variant | MODIFIER |
| 12 | 63310152 | AX-97821347 | 0 | T | C | 0.061 | -2.26E-03 | 3.02E-01 | 2.72E+00 | 2.74E+00 | 9.95E-01 | no | | regulatory_region_variant | MODIFIER |
| 12 | 63340322 | AX-170433670 | 0 | A | G | 0.061 | -2.26E-03 | 3.02E-01 | 2.72E+00 | 2.74E+00 | 9.95E-01 | no | | intergenic_variant | MODIFIER |
| 12 | 63358940 | AX-170512137 | 0 | T | G | 0.061 | -2.26E-03 | 3.02E-01 | 2.72E+00 | 2.74E+00 | 9.95E-01 | no | Gm48416 | downstream_gene_variant | MODIFIER |
| 12 | 63362330 | AX-170210566 | 0 | T | G | 0.061 | -2.26E-03 | 3.02E-01 | 2.72E+00 | 2.74E+00 | 9.95E-01 | no | Gm48416 | downstream_gene_variant | MODIFIER |

| | | | | | | | | | | | | | | | |
|---|---|---|---|---|---|---|---|---|---|---|---|---|---|---|---|
| 12 | 63369588 | AX-170174942 | 0 | T | C | 0.061 | -2.26E-03 | 3.02E-01 | 2.72E+00 | 2.74E+00 | 9.95E-01 | no | | intergenic_variant | MODIFIER |
| 12 | 63445355 | AX-170006895 | 0 | G | C | 0.061 | -2.26E-03 | 3.02E-01 | 2.72E+00 | 2.74E+00 | 9.95E-01 | no | | intergenic_variant | MODIFIER |
| 12 | 63589508 | AX-170006898 | 0 | T | C | 0.061 | -2.26E-03 | 3.02E-01 | 2.72E+00 | 2.74E+00 | 9.95E-01 | no | | intergenic_variant | MODIFIER |
| 12 | 63622674 | AX-170123515 | 0 | T | A | 0.061 | -2.26E-03 | 3.02E-01 | 2.72E+00 | 2.74E+00 | 9.95E-01 | no | | intergenic_variant | MODIFIER |
| 12 | 63895309 | AX-170180535 | 0 | G | C | 0.061 | -2.26E-03 | 3.02E-01 | 2.72E+00 | 2.74E+00 | 9.95E-01 | no | | intergenic_variant | MODIFIER |
| 12 | 63895349 | AX-168414307 | 0 | T | G | 0.061 | -2.26E-03 | 3.02E-01 | 2.72E+00 | 2.74E+00 | 9.95E-01 | no | | intergenic_variant | MODIFIER |
| 12 | 63957419 | AX-170006908 | 0 | G | A | 0.061 | -2.26E-03 | 3.02E-01 | 2.72E+00 | 2.74E+00 | 9.95E-01 | no | | intergenic_variant | MODIFIER |
| 12 | 63965646 | AX-170087263 | 0 | C | T | 0.061 | -2.26E-03 | 3.02E-01 | 2.72E+00 | 2.74E+00 | 9.95E-01 | no | Gm5653 | downstream_gene_variant | MODIFIER |
| 12 | 64026046 | AX-170224154 | 0 | G | A | 0.061 | -2.26E-03 | 3.02E-01 | 2.72E+00 | 2.74E+00 | 9.95E-01 | no | | intergenic_variant | MODIFIER |
| 12 | 64035353 | AX-170339676 | 0 | A | T | 0.061 | -2.26E-03 | 3.02E-01 | 2.72E+00 | 2.74E+00 | 9.95E-01 | no | | intergenic_variant | MODIFIER |
| 12 | 64035393 | AX-170201910 | 0 | T | C | 0.061 | -2.26E-03 | 3.02E-01 | 2.72E+00 | 2.74E+00 | 9.95E-01 | no | | intergenic_variant | MODIFIER |
| 12 | 64101307 | AX-170006901 | 0 | T | C | 0.061 | -2.26E-03 | 3.02E-01 | 2.72E+00 | 2.74E+00 | 9.95E-01 | no | | intergenic_variant | MODIFIER |
| 12 | 64236029 | AX-170006907 | 0 | T | C | 0.061 | -2.26E-03 | 3.02E-01 | 2.72E+00 | 2.74E+00 | 9.95E-01 | no | | intergenic_variant | MODIFIER |
| 12 | 64343744 | AX-97821433 | 0 | A | G | 0.061 | -2.26E-03 | 3.02E-01 | 2.72E+00 | 2.74E+00 | 9.95E-01 | no | | intergenic_variant | MODIFIER |
| 12 | 64349764 | AX-170506127 | 0 | C | G | 0.061 | -2.26E-03 | 3.02E-01 | 2.72E+00 | 2.74E+00 | 9.95E-01 | no | | intergenic_variant | MODIFIER |
| 12 | 64385521 | AX-170128364 | 0 | C | T | 0.061 | -2.26E-03 | 3.02E-01 | 2.72E+00 | 2.74E+00 | 9.95E-01 | no | | regulatory_region_variant | MODIFIER |
| 12 | 64435701 | AX-170049239 | 0 | A | G | 0.061 | -2.26E-03 | 3.02E-01 | 2.72E+00 | 2.74E+00 | 9.95E-01 | no | | intergenic_variant | MODIFIER |
| 12 | 64436077 | AX-170049241 | 0 | C | T | 0.061 | -2.26E-03 | 3.02E-01 | 2.72E+00 | 2.74E+00 | 9.95E-01 | no | | intergenic_variant | MODIFIER |
| 12 | 64438788 | AX-170107693 | 0 | T | C | 0.061 | -2.26E-03 | 3.02E-01 | 2.72E+00 | 2.74E+00 | 9.95E-01 | no | | intergenic_variant | MODIFIER |
| 12 | 64443879 | AX-170218443 | 0 | A | T | 0.061 | -2.26E-03 | 3.02E-01 | 2.72E+00 | 2.74E+00 | 9.95E-01 | no | | intergenic_variant | MODIFIER |
| 12 | 64469501 | AX-170087266 | 0 | A | G | 0.061 | -2.26E-03 | 3.02E-01 | 2.72E+00 | 2.74E+00 | 9.95E-01 | no | Fscb | downstream_gene_variant | MODIFIER |
| 12 | 64517649 | AX-97828325 | 0 | A | G | 0.061 | -2.26E-03 | 3.02E-01 | 2.72E+00 | 2.74E+00 | 9.95E-01 | no | | intergenic_variant | MODIFIER |
| 12 | 64520099 | AX-170273154 | 0 | C | T | 0.061 | -2.26E-03 | 3.02E-01 | 2.72E+00 | 2.74E+00 | 9.95E-01 | no | | intergenic_variant | MODIFIER |
| 12 | 64635283 | AX-170006921 | 0 | A | G | 0.061 | -2.26E-03 | 3.02E-01 | 2.72E+00 | 2.74E+00 | 9.95E-01 | no | | intergenic_variant | MODIFIER |
| 12 | 64683603 | AX-170208937 | 0 | C | T | 0.061 | -2.26E-03 | 3.02E-01 | 2.72E+00 | 2.74E+00 | 9.95E-01 | no | | intergenic_variant | MODIFIER |
| 12 | 64724788 | AX-97814678 | 0 | C | A | 0.061 | -2.26E-03 | 3.02E-01 | 2.72E+00 | 2.74E+00 | 9.95E-01 | no | | intergenic_variant | MODIFIER |
| 12 | 64742202 | AX-97786546 | 0 | A | G | 0.061 | -2.26E-03 | 3.02E-01 | 2.72E+00 | 2.74E+00 | 9.95E-01 | no | | intergenic_variant | MODIFIER |
| 12 | 64817299 | AX-170292158 | 0 | G | C | 0.061 | -2.26E-03 | 3.02E-01 | 2.72E+00 | 2.74E+00 | 9.95E-01 | no | Gm4806 | non_coding_transcript_exon_variant | MODIFIER |
| 12 | 64831035 | AX-170438571 | 0 | G | A | 0.061 | -2.26E-03 | 3.02E-01 | 2.72E+00 | 2.74E+00 | 9.95E-01 | no | | intergenic_variant | MODIFIER |
| 12 | 64895571 | AX-97790922 | 0 | A | G | 0.061 | -2.26E-03 | 3.02E-01 | 2.72E+00 | 2.74E+00 | 9.95E-01 | no | | intergenic_variant | MODIFIER |
| 12 | 64897072 | AX-170242154 | 0 | T | C | 0.061 | -2.26E-03 | 3.02E-01 | 2.72E+00 | 2.74E+00 | 9.95E-01 | no | | intergenic_variant | MODIFIER |
| 12 | 64912923 | AX-170483932 | 0 | T | A | 0.061 | -2.26E-03 | 3.02E-01 | 2.72E+00 | 2.74E+00 | 9.95E-01 | no | Gm527 | upstream_gene_variant | MODIFIER |
| 12 | 64937738 | AX-170481659 | 0 | A | C | 0.061 | -2.26E-03 | 3.02E-01 | 2.72E+00 | 2.74E+00 | 9.95E-01 | no | Klhl28 | downstream_gene_variant | MODIFIER |
| 12 | 64957447 | AX-170178884 | 0 | C | T | 0.061 | -2.26E-03 | 3.02E-01 | 2.72E+00 | 2.74E+00 | 9.95E-01 | no | Klhl28 | synonymous_variant | LOW |
| 12 | 64991377 | AX-170006923 | 0 | A | C | 0.061 | -2.26E-03 | 3.02E-01 | 2.72E+00 | 2.74E+00 | 9.95E-01 | no | Togaram1 | intron_variant | MODIFIER |
| 12 | 65151677 | AX-170222351 | 0 | A | T | 0.061 | -2.26E-03 | 3.02E-01 | 2.72E+00 | 2.74E+00 | 9.95E-01 | no | Mis18bp1 | intron_variant | MODIFIER |
| 12 | 65169057 | AX-97851943 | 0 | G | A | 0.061 | -2.26E-03 | 3.02E-01 | 2.72E+00 | 2.74E+00 | 9.95E-01 | no | Mis18bp1 | intron_variant | MODIFIER |
| 12 | 65319116 | AX-97792009 | 0 | T | C | 0.061 | -2.26E-03 | 3.02E-01 | 2.72E+00 | 2.74E+00 | 9.95E-01 | no | | intergenic_variant | MODIFIER |
| 12 | 65379987 | AX-97838965 | 0 | T | C | 0.061 | -2.26E-03 | 3.02E-01 | 2.72E+00 | 2.74E+00 | 9.95E-01 | no | | intergenic_variant | MODIFIER |
| 12 | 65381863 | AX-170116208 | 0 | C | G | 0.061 | -2.26E-03 | 3.02E-01 | 2.72E+00 | 2.74E+00 | 9.95E-01 | no | | regulatory_region_variant | MODIFIER |
| 12 | 65382431 | AX-170437651 | 0 | A | T | 0.061 | -2.26E-03 | 3.02E-01 | 2.72E+00 | 2.74E+00 | 9.95E-01 | no | | regulatory_region_variant | MODIFIER |
| 12 | 65394493 | AX-97821519 | 0 | C | T | 0.061 | -2.26E-03 | 3.02E-01 | 2.72E+00 | 2.74E+00 | 9.95E-01 | no | | intergenic_variant | MODIFIER |
| 12 | 65407313 | AX-170145875 | 0 | C | T | 0.061 | -2.26E-03 | 3.02E-01 | 2.72E+00 | 2.74E+00 | 9.95E-01 | no | Gm26015 | upstream_gene_variant | MODIFIER |
| 12 | 65469975 | AX-170156743 | 0 | G | A | 0.061 | -2.26E-03 | 3.02E-01 | 2.72E+00 | 2.74E+00 | 9.95E-01 | no | | intergenic_variant | MODIFIER |
| 12 | 65472660 | AX-97844042 | 0 | C | T | 0.061 | -2.26E-03 | 3.02E-01 | 2.72E+00 | 2.74E+00 | 9.95E-01 | no | | intergenic_variant | MODIFIER |
| 12 | 65715274 | AX-170267094 | 0 | G | T | 0.061 | -2.26E-03 | 3.02E-01 | 2.72E+00 | 2.74E+00 | 9.95E-01 | no | | intergenic_variant | MODIFIER |
| 12 | 66249811 | AX-170006942 | 0 | C | T | 0.061 | -2.26E-03 | 3.02E-01 | 2.72E+00 | 2.74E+00 | 9.95E-01 | no | | intergenic_variant | MODIFIER |
| 12 | 66651395 | AX-170465227 | 0 | C | T | 0.061 | -2.26E-03 | 3.02E-01 | 2.72E+00 | 2.74E+00 | 9.95E-01 | no | Mdga2 | intron_variant | MODIFIER |
| 12 | 66653926 | AX-97828806 | 0 | G | A | 0.061 | -2.26E-03 | 3.02E-01 | 2.72E+00 | 2.74E+00 | 9.95E-01 | no | Mdga2 | intron_variant | MODIFIER |
| 12 | 66719703 | AX-170006945 | 0 | G | A | 0.061 | -2.26E-03 | 3.02E-01 | 2.72E+00 | 2.74E+00 | 9.95E-01 | no | Mdga2 | intron_variant | MODIFIER |
| 12 | 66724157 | AX-170006944 | 0 | T | C | 0.061 | -2.26E-03 | 3.02E-01 | 2.72E+00 | 2.74E+00 | 9.95E-01 | no | Mdga2 | intron_variant | MODIFIER |
| 12 | 66755896 | AX-170395011 | 0 | T | G | 0.061 | -2.26E-03 | 3.02E-01 | 2.72E+00 | 2.74E+00 | 9.95E-01 | no | Mdga2 | intron_variant | MODIFIER |
| 12 | 66950667 | AX-170416852 | 0 | A | G | 0.061 | -2.26E-03 | 3.02E-01 | 2.72E+00 | 2.74E+00 | 9.95E-01 | no | Mdga2 | intron_variant | MODIFIER |
| 12 | 67178724 | AX-170489449 | 0 | C | T | 0.061 | -2.26E-03 | 3.02E-01 | 2.72E+00 | 2.74E+00 | 9.95E-01 | no | Mdga2 | intron_variant | MODIFIER |

| | | | | | | | | | | | | | | | |
|---|---|---|---|---|---|---|---|---|---|---|---|---|---|---|---|
| 12 | 67194142 | AX-170215902 | 0 | A | G | 0.061 | -2.26E-03 | 3.02E-01 | 2.72E+00 | 2.74E+00 | 9.95E-01 | no | Mdga2 | intron_variant | MODIFIER |
| 12 | 67921206 | AX-170035520 | 0 | C | T | 0.061 | -2.26E-03 | 3.02E-01 | 2.72E+00 | 2.74E+00 | 9.95E-01 | no | Gm20102 | downstream_gene_variant | MODIFIER |
| 12 | 68308281 | AX-97846170 | 0 | T | C | 0.061 | -2.26E-03 | 3.02E-01 | 2.72E+00 | 2.74E+00 | 9.95E-01 | no | | intergenic_variant | MODIFIER |
| 12 | 68334112 | AX-170262594 | 0 | T | A | 0.061 | -2.26E-03 | 3.02E-01 | 2.72E+00 | 2.74E+00 | 9.95E-01 | no | Gm47454 | downstream_gene_variant | MODIFIER |
| 12 | 68352787 | AX-97810900 | 0 | T | G | 0.061 | -2.26E-03 | 3.02E-01 | 2.72E+00 | 2.74E+00 | 9.95E-01 | no | Gm47456 | intron_variant,non_coding_transcript_variant | MODIFIER |
| 12 | 68388213 | AX-97856053 | 0 | T | C | 0.061 | -2.26E-03 | 3.02E-01 | 2.72E+00 | 2.74E+00 | 9.95E-01 | no | Gm47456 | intron_variant,non_coding_transcript_variant | MODIFIER |
| 12 | 68473275 | AX-170006973 | 0 | C | T | 0.061 | -2.26E-03 | 3.02E-01 | 2.72E+00 | 2.74E+00 | 9.95E-01 | no | | intergenic_variant | MODIFIER |
| 12 | 68473811 | AX-170087272 | 0 | G | T | 0.061 | -2.26E-03 | 3.02E-01 | 2.72E+00 | 2.74E+00 | 9.95E-01 | no | | intergenic_variant | MODIFIER |
| 12 | 68474862 | AX-170416058 | 0 | A | C | 0.061 | -2.26E-03 | 3.02E-01 | 2.72E+00 | 2.74E+00 | 9.95E-01 | no | | intergenic_variant | MODIFIER |
| 12 | 68549374 | AX-170179922 | 0 | A | G | 0.061 | -2.26E-03 | 3.02E-01 | 2.72E+00 | 2.74E+00 | 9.95E-01 | no | Gm47490 | downstream_gene_variant | MODIFIER |
| 12 | 68557252 | AX-97817941 | 0 | A | G | 0.061 | -2.26E-03 | 3.02E-01 | 2.72E+00 | 2.74E+00 | 9.95E-01 | no | | intergenic_variant | MODIFIER |
| 12 | 68750781 | AX-170243081 | 0 | G | A | 0.061 | -2.26E-03 | 3.02E-01 | 2.72E+00 | 2.74E+00 | 9.95E-01 | no | | intergenic_variant | MODIFIER |
| 12 | 68754823 | AX-170209035 | 0 | G | T | 0.061 | -2.26E-03 | 3.02E-01 | 2.72E+00 | 2.74E+00 | 9.95E-01 | no | | intergenic_variant | MODIFIER |
| 12 | 68768076 | AX-170382757 | 0 | G | C | 0.061 | -2.26E-03 | 3.02E-01 | 2.72E+00 | 2.74E+00 | 9.95E-01 | no | | regulatory_region_variant | MODIFIER |
| 12 | 68776716 | AX-170049330 | 0 | A | G | 0.061 | -2.26E-03 | 3.02E-01 | 2.72E+00 | 2.74E+00 | 9.95E-01 | no | | intergenic_variant | MODIFIER |
| 12 | 68795103 | AX-97796479 | 0 | G | A | 0.061 | -2.26E-03 | 3.02E-01 | 2.72E+00 | 2.74E+00 | 9.95E-01 | no | | intergenic_variant | MODIFIER |
| 12 | 68795219 | AX-170436522 | 0 | C | T | 0.061 | -2.26E-03 | 3.02E-01 | 2.72E+00 | 2.74E+00 | 9.95E-01 | no | | intergenic_variant | MODIFIER |
| 12 | 68795367 | AX-170515458 | 0 | A | G | 0.061 | -2.26E-03 | 3.02E-01 | 2.72E+00 | 2.74E+00 | 9.95E-01 | no | | intergenic_variant | MODIFIER |
| 12 | 68796192 | AX-170160477 | 0 | A | G | 0.061 | -2.26E-03 | 3.02E-01 | 2.72E+00 | 2.74E+00 | 9.95E-01 | no | | intergenic_variant | MODIFIER |
| 12 | 68796341 | AX-170153232 | 0 | C | T | 0.061 | -2.26E-03 | 3.02E-01 | 2.72E+00 | 2.74E+00 | 9.95E-01 | no | | intergenic_variant | MODIFIER |
| 12 | 68796484 | AX-170178327 | 0 | G | A | 0.061 | -2.26E-03 | 3.02E-01 | 2.72E+00 | 2.74E+00 | 9.95E-01 | no | | intergenic_variant | MODIFIER |
| 12 | 68796503 | AX-170440045 | 0 | C | T | 0.061 | -2.26E-03 | 3.02E-01 | 2.72E+00 | 2.74E+00 | 9.95E-01 | no | | intergenic_variant | MODIFIER |
| 12 | 68804976 | AX-170305191 | 0 | A | C | 0.061 | -2.26E-03 | 3.02E-01 | 2.72E+00 | 2.74E+00 | 9.95E-01 | no | | intergenic_variant | MODIFIER |
| 12 | 68805269 | AX-170422353 | 0 | T | A | 0.061 | -2.26E-03 | 3.02E-01 | 2.72E+00 | 2.74E+00 | 9.95E-01 | no | | intergenic_variant | MODIFIER |
| 12 | 68805772 | AX-170286859 | 0 | A | G | 0.061 | -2.26E-03 | 3.02E-01 | 2.72E+00 | 2.74E+00 | 9.95E-01 | no | | intergenic_variant | MODIFIER |
| 12 | 68806054 | AX-170346319 | 0 | T | G | 0.061 | -2.26E-03 | 3.02E-01 | 2.72E+00 | 2.74E+00 | 9.95E-01 | no | | regulatory_region_variant | MODIFIER |
| 12 | 68845445 | AX-97781947 | 0 | A | G | 0.061 | -2.26E-03 | 3.02E-01 | 2.72E+00 | 2.74E+00 | 9.95E-01 | no | | intergenic_variant | MODIFIER |
| 12 | 68848772 | AX-170006980 | 0 | G | T | 0.061 | -2.26E-03 | 3.02E-01 | 2.72E+00 | 2.74E+00 | 9.95E-01 | no | | intergenic_variant | MODIFIER |
| 12 | 68849430 | AX-170168740 | 0 | C | G | 0.061 | -2.26E-03 | 3.02E-01 | 2.72E+00 | 2.74E+00 | 9.95E-01 | no | | intergenic_variant | MODIFIER |
| 12 | 68858420 | AX-170006979 | 0 | G | T | 0.061 | -2.26E-03 | 3.02E-01 | 2.72E+00 | 2.74E+00 | 9.95E-01 | no | | intergenic_variant | MODIFIER |
| 12 | 68906462 | AX-170006985 | 0 | A | C | 0.061 | -2.26E-03 | 3.02E-01 | 2.72E+00 | 2.74E+00 | 9.95E-01 | no | Gm47515 | intron_variant,non_coding_transcript_variant | MODIFIER |
| 12 | 68907452 | AX-170256290 | 0 | C | T | 0.061 | -2.26E-03 | 3.02E-01 | 2.72E+00 | 2.74E+00 | 9.95E-01 | no | Gm47515 | intron_variant,non_coding_transcript_variant | MODIFIER |
| 12 | 68949033 | AX-97821698 | 0 | T | C | 0.061 | -2.26E-03 | 3.02E-01 | 2.72E+00 | 2.74E+00 | 9.95E-01 | no | Gm47515 | intron_variant,non_coding_transcript_variant | MODIFIER |
| 12 | 69068230 | AX-170267759 | 0 | C | T | 0.061 | -2.26E-03 | 3.02E-01 | 2.72E+00 | 2.74E+00 | 9.95E-01 | no | | intergenic_variant | MODIFIER |
| 12 | 69104721 | AX-170224719 | 0 | C | T | 0.061 | -2.26E-03 | 3.02E-01 | 2.72E+00 | 2.74E+00 | 9.95E-01 | no | | intergenic_variant | MODIFIER |
| 12 | 69104929 | AX-170466995 | 0 | A | G | 0.061 | -2.26E-03 | 3.02E-01 | 2.72E+00 | 2.74E+00 | 9.95E-01 | no | | intergenic_variant | MODIFIER |
| 12 | 69122603 | AX-170197045 | 0 | G | C | 0.061 | -2.26E-03 | 3.02E-01 | 2.72E+00 | 2.74E+00 | 9.95E-01 | no | Gm47518 | upstream_gene_variant | MODIFIER |
| 12 | 69143372 | AX-170255056 | 0 | C | T | 0.061 | -2.26E-03 | 3.02E-01 | 2.72E+00 | 2.74E+00 | 9.95E-01 | no | Gm15571 | upstream_gene_variant | MODIFIER |
| 12 | 69156404 | AX-170087276 | 0 | A | T | 0.061 | -2.26E-03 | 3.02E-01 | 2.72E+00 | 2.74E+00 | 9.95E-01 | no | Rps29 | downstream_gene_variant | MODIFIER |
| 12 | 69227483 | AX-97849287 | 0 | A | C | 0.061 | -2.26E-03 | 3.02E-01 | 2.72E+00 | 2.74E+00 | 9.95E-01 | no | Pole2 | intron_variant | MODIFIER |
| 12 | 69233558 | AX-170485840 | 0 | A | G | 0.061 | -2.26E-03 | 3.02E-01 | 2.72E+00 | 2.74E+00 | 9.95E-01 | no | Gm47545 | intron_variant,non_coding_transcript_variant | MODIFIER |
| 12 | 69257952 | AX-170356440 | 0 | A | G | 0.061 | -2.26E-03 | 3.02E-01 | 2.72E+00 | 2.74E+00 | 9.95E-01 | no | Klhdc1 | intron_variant | MODIFIER |
| 12 | 69259654 | AX-170447774 | 0 | C | T | 0.061 | -2.26E-03 | 3.02E-01 | 2.72E+00 | 2.74E+00 | 9.95E-01 | no | Klhdc1 | intron_variant | MODIFIER |
| 12 | 69334751 | AX-170495135 | 0 | C | T | 0.061 | -2.26E-03 | 3.02E-01 | 2.72E+00 | 2.74E+00 | 9.95E-01 | no | Nemf | intron_variant | MODIFIER |
| 12 | 69343494 | AX-170424924 | 0 | T | C | 0.061 | -2.26E-03 | 3.02E-01 | 2.72E+00 | 2.74E+00 | 9.95E-01 | no | Nemf | intron_variant | MODIFIER |
| 12 | 69348664 | AX-170134180 | 0 | A | G | 0.061 | -2.26E-03 | 3.02E-01 | 2.72E+00 | 2.74E+00 | 9.95E-01 | no | Nemf | intron_variant | MODIFIER |
| 12 | 69381669 | AX-170465137 | 0 | C | T | 0.061 | -2.26E-03 | 3.02E-01 | 2.72E+00 | 2.74E+00 | 9.95E-01 | no | | intergenic_variant | MODIFIER |
| 12 | 69381894 | AX-170535378 | 0 | G | A | 0.061 | -2.26E-03 | 3.02E-01 | 2.72E+00 | 2.74E+00 | 9.95E-01 | no | | intergenic_variant | MODIFIER |
| 12 | 69393933 | AX-170489509 | 0 | C | T | 0.061 | -2.26E-03 | 3.02E-01 | 2.72E+00 | 2.74E+00 | 9.95E-01 | no | | intergenic_variant | MODIFIER |
| 12 | 69393958 | AX-170209853 | 0 | C | T | 0.061 | -2.26E-03 | 3.02E-01 | 2.72E+00 | 2.74E+00 | 9.95E-01 | no | | intergenic_variant | MODIFIER |
| 12 | 69394149 | AX-170006994 | 0 | C | T | 0.061 | -2.26E-03 | 3.02E-01 | 2.72E+00 | 2.74E+00 | 9.95E-01 | no | | intergenic_variant | MODIFIER |
| 12 | 69395816 | AX-170049341 | 0 | G | A | 0.061 | -2.26E-03 | 3.02E-01 | 2.72E+00 | 2.74E+00 | 9.95E-01 | no | | intergenic_variant | MODIFIER |
| 12 | 69400115 | AX-170389843 | 0 | T | G | 0.061 | -2.26E-03 | 3.02E-01 | 2.72E+00 | 2.74E+00 | 9.95E-01 | no | | intergenic_variant | MODIFIER |
| 12 | 69410728 | AX-170107566 | 0 | A | G | 0.061 | -2.26E-03 | 3.02E-01 | 2.72E+00 | 2.74E+00 | 9.95E-01 | no | | intergenic_variant | MODIFIER |

| | | | | | | | | | | | | | | | |
|---|---|---|---|---|---|---|---|---|---|---|---|---|---|---|---|
| 12 | 69413437 | AX-170006993 | 0 | G | A | 0.061 | -2.26E-03 | 3.02E-01 | 2.72E+00 | 2.74E+00 | 9.95E-01 | no | | regulatory_region_variant | MODIFIER |
| 12 | 69413471 | AX-170006989 | 0 | T | C | 0.061 | -2.26E-03 | 3.02E-01 | 2.72E+00 | 2.74E+00 | 9.95E-01 | no | | regulatory_region_variant | MODIFIER |
| 12 | 69424455 | AX-170247414 | 0 | C | A | 0.061 | -2.26E-03 | 3.02E-01 | 2.72E+00 | 2.74E+00 | 9.95E-01 | no | | regulatory_region_variant | MODIFIER |
| 12 | 69429189 | AX-97796924 | 0 | A | G | 0.061 | -2.26E-03 | 3.02E-01 | 2.72E+00 | 2.74E+00 | 9.95E-01 | no | | intergenic_variant | MODIFIER |
| 12 | 69444219 | AX-97809051 | 0 | A | G | 0.061 | -2.26E-03 | 3.02E-01 | 2.72E+00 | 2.74E+00 | 9.95E-01 | no | | intergenic_variant | MODIFIER |
| 12 | 69448100 | AX-170318418 | 0 | A | G | 0.061 | -2.26E-03 | 3.02E-01 | 2.72E+00 | 2.74E+00 | 9.95E-01 | no | | regulatory_region_variant | MODIFIER |
| 12 | 69482148 | AX-170006987 | 0 | G | T | 0.061 | -2.26E-03 | 3.02E-01 | 2.72E+00 | 2.74E+00 | 9.95E-01 | no | | regulatory_region_variant | MODIFIER |
| 12 | 69488258 | AX-170006992 | 0 | T | C | 0.061 | -2.26E-03 | 3.02E-01 | 2.72E+00 | 2.74E+00 | 9.95E-01 | no | | intergenic_variant | MODIFIER |
| 12 | 69488287 | AX-170006986 | 0 | C | A | 0.061 | -2.26E-03 | 3.02E-01 | 2.72E+00 | 2.74E+00 | 9.95E-01 | no | | intergenic_variant | MODIFIER |
| 12 | 69519052 | AX-170110846 | 0 | G | C | 0.061 | -2.26E-03 | 3.02E-01 | 2.72E+00 | 2.74E+00 | 9.95E-01 | no | | regulatory_region_variant | MODIFIER |
| 12 | 69524198 | AX-97838473 | 0 | G | A | 0.061 | -2.26E-03 | 3.02E-01 | 2.72E+00 | 2.74E+00 | 9.95E-01 | no | | intergenic_variant | MODIFIER |
| 12 | 69556111 | AX-170484819 | 0 | A | G | 0.061 | -2.26E-03 | 3.02E-01 | 2.72E+00 | 2.74E+00 | 9.95E-01 | no | | regulatory_region_variant | MODIFIER |
| 12 | 69556598 | AX-170485858 | 0 | C | T | 0.061 | -2.26E-03 | 3.02E-01 | 2.72E+00 | 2.74E+00 | 9.95E-01 | no | Gm23018 | downstream_gene_variant | MODIFIER |
| 12 | 69578436 | AX-170228096 | 0 | C | G | 0.061 | -2.26E-03 | 3.02E-01 | 2.72E+00 | 2.74E+00 | 9.95E-01 | no | Vcpkmt | intron_variant | MODIFIER |
| 12 | 69579071 | AX-170445584 | 0 | A | T | 0.061 | -2.26E-03 | 3.02E-01 | 2.72E+00 | 2.74E+00 | 9.95E-01 | no | Sos2 | downstream_gene_variant | MODIFIER |
| 12 | 69593225 | AX-170049347 | 0 | C | T | 0.061 | -2.26E-03 | 3.02E-01 | 2.72E+00 | 2.74E+00 | 9.95E-01 | no | Sos2 | intron_variant | MODIFIER |
| 12 | 69781509 | AX-170499256 | 0 | C | T | 0.061 | -2.26E-03 | 3.02E-01 | 2.72E+00 | 2.74E+00 | 9.95E-01 | no | Cdkl1 | intron_variant | MODIFIER |
| 12 | 69794624 | AX-97845236 | 0 | T | C | 0.061 | -2.26E-03 | 3.02E-01 | 2.72E+00 | 2.74E+00 | 9.95E-01 | no | Cdkl1 | upstream_gene_variant | MODIFIER |
| 12 | 69801610 | AX-170237452 | 0 | G | T | 0.061 | -2.26E-03 | 3.02E-01 | 2.72E+00 | 2.74E+00 | 9.95E-01 | no | Map4k5 | downstream_gene_variant | MODIFIER |
| 12 | 69815281 | AX-170305110 | 0 | G | A | 0.061 | -2.26E-03 | 3.02E-01 | 2.72E+00 | 2.74E+00 | 9.95E-01 | no | Map4k5 | intron_variant | MODIFIER |
| 12 | 69825151 | AX-170341228 | 0 | A | G | 0.061 | -2.26E-03 | 3.02E-01 | 2.72E+00 | 2.74E+00 | 9.95E-01 | no | Map4k5 | intron_variant | MODIFIER |
| 12 | 69853251 | AX-170049348 | 0 | T | C | 0.061 | -2.26E-03 | 3.02E-01 | 2.72E+00 | 2.74E+00 | 9.95E-01 | no | Map4k5 | intron_variant | MODIFIER |
| 12 | 69877264 | AX-170049349 | 0 | G | A | 0.061 | -2.26E-03 | 3.02E-01 | 2.72E+00 | 2.74E+00 | 9.95E-01 | no | Map4k5 | intron_variant | MODIFIER |
| 12 | 69878806 | AX-170523584 | 0 | T | C | 0.061 | -2.26E-03 | 3.02E-01 | 2.72E+00 | 2.74E+00 | 9.95E-01 | no | Map4k5 | intron_variant | MODIFIER |
| 12 | 69879450 | AX-170199664 | 0 | G | A | 0.061 | -2.26E-03 | 3.02E-01 | 2.72E+00 | 2.74E+00 | 9.95E-01 | no | Map4k5 | intron_variant | MODIFIER |
| 12 | 69896258 | AX-170524047 | 0 | A | G | 0.061 | -2.26E-03 | 3.02E-01 | 2.72E+00 | 2.74E+00 | 9.95E-01 | no | Atl1 | intron_variant | MODIFIER |
| 12 | 69900222 | AX-170259118 | 0 | T | G | 0.061 | -2.26E-03 | 3.02E-01 | 2.72E+00 | 2.74E+00 | 9.95E-01 | no | Atl1 | intron_variant | MODIFIER |
| 12 | 69926111 | AX-170311833 | 0 | C | T | 0.061 | -2.26E-03 | 3.02E-01 | 2.72E+00 | 2.74E+00 | 9.95E-01 | no | Atl1 | synonymous_variant | LOW |
| 12 | 69929618 | AX-170087278 | 0 | T | C | 0.061 | -2.26E-03 | 3.02E-01 | 2.72E+00 | 2.74E+00 | 9.95E-01 | no | Atl1 | intron_variant | MODIFIER |
| 12 | 69933141 | AX-170108895 | 0 | A | G | 0.061 | -2.26E-03 | 3.02E-01 | 2.72E+00 | 2.74E+00 | 9.95E-01 | no | Atl1 | intron_variant | MODIFIER |
| 12 | 69953804 | AX-97794967 | 0 | C | T | 0.061 | -2.26E-03 | 3.02E-01 | 2.72E+00 | 2.74E+00 | 9.95E-01 | no | Atl1 | intron_variant | MODIFIER |
| 12 | 69962316 | AX-97826241 | 0 | A | C | 0.061 | -2.26E-03 | 3.02E-01 | 2.72E+00 | 2.74E+00 | 9.95E-01 | no | Atl1 | intron_variant | MODIFIER |
| 12 | 70001246 | AX-97841329 | 0 | A | G | 0.061 | -2.26E-03 | 3.02E-01 | 2.72E+00 | 2.74E+00 | 9.95E-01 | no | | intergenic_variant | MODIFIER |
| 12 | 70017187 | AX-170254501 | 0 | T | G | 0.061 | -2.26E-03 | 3.02E-01 | 2.72E+00 | 2.74E+00 | 9.95E-01 | no | Nin | downstream_gene_variant | MODIFIER |
| 12 | 70017789 | AX-170006998 | 0 | A | G | 0.061 | -2.26E-03 | 3.02E-01 | 2.72E+00 | 2.74E+00 | 9.95E-01 | no | Nin | downstream_gene_variant | MODIFIER |
| 12 | 70018045 | AX-97821644 | 0 | C | T | 0.061 | -2.26E-03 | 3.02E-01 | 2.72E+00 | 2.74E+00 | 9.95E-01 | no | Nin | downstream_gene_variant | MODIFIER |
| 12 | 70058510 | AX-170006995 | 0 | C | A | 0.061 | -2.26E-03 | 3.02E-01 | 2.72E+00 | 2.74E+00 | 9.95E-01 | no | Nin | intron_variant | MODIFIER |
| 12 | 70058732 | AX-170007000 | 0 | C | A | 0.061 | -2.26E-03 | 3.02E-01 | 2.72E+00 | 2.74E+00 | 9.95E-01 | no | Nin | intron_variant | MODIFIER |
| 12 | 70058961 | AX-170369974 | 0 | G | A | 0.061 | -2.26E-03 | 3.02E-01 | 2.72E+00 | 2.74E+00 | 9.95E-01 | no | Nin | intron_variant | MODIFIER |
| 12 | 70064234 | AX-97783782 | 0 | T | C | 0.061 | -2.26E-03 | 3.02E-01 | 2.72E+00 | 2.74E+00 | 9.95E-01 | no | Nin | intron_variant | MODIFIER |
| 12 | 70087706 | AX-170006997 | 0 | G | C | 0.061 | -2.26E-03 | 3.02E-01 | 2.72E+00 | 2.74E+00 | 9.95E-01 | no | Nin | intron_variant | MODIFIER |
| 12 | 70089679 | AX-97786978 | 0 | T | C | 0.061 | -2.26E-03 | 3.02E-01 | 2.72E+00 | 2.74E+00 | 9.95E-01 | no | Nin | intron_variant | MODIFIER |
| 12 | 70099186 | AX-170473188 | 0 | T | C | 0.061 | -2.26E-03 | 3.02E-01 | 2.72E+00 | 2.74E+00 | 9.95E-01 | no | Nin | intron_variant | MODIFIER |
| 12 | 70146585 | AX-97795989 | 0 | T | C | 0.061 | -2.26E-03 | 3.02E-01 | 2.72E+00 | 2.74E+00 | 9.95E-01 | no | | intergenic_variant | MODIFIER |
| 12 | 70167148 | AX-170517079 | 0 | G | A | 0.061 | -2.26E-03 | 3.02E-01 | 2.72E+00 | 2.74E+00 | 9.95E-01 | no | Abhd12b | intron_variant,non_coding_transcript_variant | MODIFIER |
| 12 | 70168649 | AX-97803947 | 0 | T | C | 0.061 | -2.26E-03 | 3.02E-01 | 2.72E+00 | 2.74E+00 | 9.95E-01 | no | Abhd12b | intron_variant,non_coding_transcript_variant | MODIFIER |
| 12 | 70170253 | AX-170338497 | 0 | A | G | 0.061 | -2.26E-03 | 3.02E-01 | 2.72E+00 | 2.74E+00 | 9.95E-01 | no | Abhd12b | intron_variant,non_coding_transcript_variant | MODIFIER |
| 12 | 70172682 | AX-170498542 | 0 | C | T | 0.061 | -2.26E-03 | 3.02E-01 | 2.72E+00 | 2.74E+00 | 9.95E-01 | no | Abhd12b | intron_variant,non_coding_transcript_variant | MODIFIER |
| 12 | 70182496 | AX-170117920 | 0 | A | G | 0.061 | -2.26E-03 | 3.02E-01 | 2.72E+00 | 2.74E+00 | 9.95E-01 | no | Abhd12b | intron_variant,non_coding_transcript_variant | MODIFIER |
| 12 | 70196133 | AX-170367119 | 0 | T | C | 0.061 | -2.26E-03 | 3.02E-01 | 2.72E+00 | 2.74E+00 | 9.95E-01 | no | Pygl | intron_variant | MODIFIER |
| 12 | 70196839 | AX-170417058 | 0 | T | C | 0.061 | -2.26E-03 | 3.02E-01 | 2.72E+00 | 2.74E+00 | 9.95E-01 | no | Pygl | intron_variant | MODIFIER |
| 12 | 70202866 | AX-170049355 | 0 | T | C | 0.061 | -2.26E-03 | 3.02E-01 | 2.72E+00 | 2.74E+00 | 9.95E-01 | no | Pygl | intron_variant | MODIFIER |
| 12 | 70262731 | AX-170136971 | 0 | G | A | 0.061 | -2.26E-03 | 3.02E-01 | 2.72E+00 | 2.74E+00 | 9.95E-01 | no | Trim9 | intron_variant | MODIFIER |
| 12 | 70270514 | AX-97809869 | 0 | C | T | 0.061 | -2.26E-03 | 3.02E-01 | 2.72E+00 | 2.74E+00 | 9.95E-01 | no | Trim9 | intron_variant | MODIFIER |

| | | | | | | | | | | | | | | | |
|---|---|---|---|---|---|---|---|---|---|---|---|---|---|---|---|
| 12 | 70297853 | AX-170277919 | 0 | C | T | 0.061 | -2.26E-03 | 3.02E-01 | 2.72E+00 | 2.74E+00 | 9.95E-01 | no | Trim9 | intron_variant | MODIFIER |
| 12 | 70333138 | AX-170327295 | 0 | A | G | 0.061 | -2.26E-03 | 3.02E-01 | 2.72E+00 | 2.74E+00 | 9.95E-01 | no | Trim9 | intron_variant | MODIFIER |
| 12 | 70333265 | AX-170177102 | 0 | G | A | 0.061 | -2.26E-03 | 3.02E-01 | 2.72E+00 | 2.74E+00 | 9.95E-01 | no | Trim9 | intron_variant | MODIFIER |
| 12 | 70334020 | AX-170509805 | 0 | C | T | 0.061 | -2.26E-03 | 3.02E-01 | 2.72E+00 | 2.74E+00 | 9.95E-01 | no | Trim9 | intron_variant | MODIFIER |
| 12 | 70338271 | AX-170087280 | 0 | T | G | 0.061 | -2.26E-03 | 3.02E-01 | 2.72E+00 | 2.74E+00 | 9.95E-01 | no | Trim9 | intron_variant | MODIFIER |
| 12 | 70338318 | AX-170343091 | 0 | T | A | 0.061 | -2.26E-03 | 3.02E-01 | 2.72E+00 | 2.74E+00 | 9.95E-01 | no | Trim9 | intron_variant | MODIFIER |
| 12 | 70348175 | AX-170386112 | 0 | G | T | 0.061 | -2.26E-03 | 3.02E-01 | 2.72E+00 | 2.74E+00 | 9.95E-01 | no | Trim9 | upstream_gene_variant | MODIFIER |
| 12 | 70349886 | AX-170334467 | 0 | G | C | 0.061 | -2.26E-03 | 3.02E-01 | 2.72E+00 | 2.74E+00 | 9.95E-01 | no | Trim9 | upstream_gene_variant | MODIFIER |
| 12 | 70358438 | AX-170302085 | 0 | G | C | 0.061 | -2.26E-03 | 3.02E-01 | 2.72E+00 | 2.74E+00 | 9.95E-01 | no | | intergenic_variant | MODIFIER |
| 12 | 70369089 | AX-97815000 | 0 | A | G | 0.061 | -2.26E-03 | 3.02E-01 | 2.72E+00 | 2.74E+00 | 9.95E-01 | no | | regulatory_region_variant | MODIFIER |
| 12 | 70394181 | AX-97849576 | 0 | T | C | 0.061 | -2.26E-03 | 3.02E-01 | 2.72E+00 | 2.74E+00 | 9.95E-01 | no | | intergenic_variant | MODIFIER |
| 12 | 70410044 | AX-170503466 | 0 | C | T | 0.061 | -2.26E-03 | 3.02E-01 | 2.72E+00 | 2.74E+00 | 9.95E-01 | no | | intergenic_variant | MODIFIER |
| 12 | 70418320 | AX-97808469 | 0 | T | C | 0.061 | -2.26E-03 | 3.02E-01 | 2.72E+00 | 2.74E+00 | 9.95E-01 | no | | intergenic_variant | MODIFIER |
| 12 | 70422179 | AX-170350691 | 0 | G | C | 0.061 | -2.26E-03 | 3.02E-01 | 2.72E+00 | 2.74E+00 | 9.95E-01 | no | | regulatory_region_variant | MODIFIER |
| 12 | 70432161 | AX-170250480 | 0 | G | A | 0.061 | -2.26E-03 | 3.02E-01 | 2.72E+00 | 2.74E+00 | 9.95E-01 | no | | intergenic_variant | MODIFIER |
| 12 | 70434170 | AX-170444842 | 0 | A | G | 0.061 | -2.26E-03 | 3.02E-01 | 2.72E+00 | 2.74E+00 | 9.95E-01 | no | | intergenic_variant | MODIFIER |
| 12 | 70455287 | AX-170087281 | 0 | C | T | 0.061 | -2.26E-03 | 3.02E-01 | 2.72E+00 | 2.74E+00 | 9.95E-01 | no | Tmx1 | intron_variant | MODIFIER |
| 12 | 70471354 | AX-97821724 | 0 | G | A | 0.061 | -2.26E-03 | 3.02E-01 | 2.72E+00 | 2.74E+00 | 9.95E-01 | no | Tmx1 | downstream_gene_variant | MODIFIER |
| 12 | 70492869 | AX-97854170 | 0 | C | T | 0.061 | -2.26E-03 | 3.02E-01 | 2.72E+00 | 2.74E+00 | 9.95E-01 | no | Gm48847 | non_coding_transcript_exon_variant | MODIFIER |
| 12 | 70539829 | AX-170228284 | 0 | T | C | 0.061 | -2.26E-03 | 3.02E-01 | 2.72E+00 | 2.74E+00 | 9.95E-01 | no | Gm32219 | intron_variant,non_coding_transcript_variant | MODIFIER |
| 12 | 70560636 | AX-97844010 | 0 | C | A | 0.061 | -2.26E-03 | 3.02E-01 | 2.72E+00 | 2.74E+00 | 9.95E-01 | no | | intergenic_variant | MODIFIER |
| 12 | 70600171 | AX-97784969 | 0 | T | G | 0.061 | -2.26E-03 | 3.02E-01 | 2.72E+00 | 2.74E+00 | 9.95E-01 | no | | regulatory_region_variant | MODIFIER |
| 12 | 70695674 | AX-170351129 | 0 | C | A | 0.061 | -2.26E-03 | 3.02E-01 | 2.72E+00 | 2.74E+00 | 9.95E-01 | no | Gm24474 | downstream_gene_variant | MODIFIER |
| 12 | 70715329 | AX-170358508 | 0 | G | A | 0.061 | -2.26E-03 | 3.02E-01 | 2.72E+00 | 2.74E+00 | 9.95E-01 | no | | regulatory_region_variant | MODIFIER |
| 12 | 70797422 | AX-97840790 | 0 | A | G | 0.061 | -2.26E-03 | 3.02E-01 | 2.72E+00 | 2.74E+00 | 9.95E-01 | no | Gm40437 | intron_variant,non_coding_transcript_variant | MODIFIER |
| 12 | 70799013 | AX-170162473 | 0 | C | G | 0.061 | -2.26E-03 | 3.02E-01 | 2.72E+00 | 2.74E+00 | 9.95E-01 | no | Gm40437 | intron_variant,non_coding_transcript_variant | MODIFIER |
| 12 | 70799822 | AX-170360522 | 0 | A | G | 0.061 | -2.26E-03 | 3.02E-01 | 2.72E+00 | 2.74E+00 | 9.95E-01 | no | Gm40437 | intron_variant,non_coding_transcript_variant | MODIFIER |
| 12 | 70817198 | AX-170387322 | 0 | G | A | 0.061 | -2.26E-03 | 3.02E-01 | 2.72E+00 | 2.74E+00 | 9.95E-01 | no | Gm40437 | intron_variant,non_coding_transcript_variant | MODIFIER |
| 12 | 70817326 | AX-170370748 | 0 | A | G | 0.061 | -2.26E-03 | 3.02E-01 | 2.72E+00 | 2.74E+00 | 9.95E-01 | no | Gm40437 | intron_variant,non_coding_transcript_variant | MODIFIER |
| 12 | 70818563 | AX-97822428 | 0 | T | G | 0.061 | -2.26E-03 | 3.02E-01 | 2.72E+00 | 2.74E+00 | 9.95E-01 | no | Gm40437 | intron_variant,non_coding_transcript_variant | MODIFIER |
| 12 | 70833687 | AX-170472306 | 0 | G | T | 0.061 | -2.26E-03 | 3.02E-01 | 2.72E+00 | 2.74E+00 | 9.95E-01 | no | Frmd6 | intron_variant | MODIFIER |
| 12 | 70834268 | AX-170140063 | 0 | T | G | 0.061 | -2.26E-03 | 3.02E-01 | 2.72E+00 | 2.74E+00 | 9.95E-01 | no | Frmd6 | intron_variant | MODIFIER |
| 12 | 70845511 | AX-97824177 | 0 | C | T | 0.061 | -2.26E-03 | 3.02E-01 | 2.72E+00 | 2.74E+00 | 9.95E-01 | no | Frmd6 | intron_variant | MODIFIER |
| 12 | 70873660 | AX-97821643 | 0 | C | T | 0.061 | -2.26E-03 | 3.02E-01 | 2.72E+00 | 2.74E+00 | 9.95E-01 | no | Frmd6 | intron_variant | MODIFIER |
| 12 | 70889790 | AX-170239225 | 0 | A | G | 0.061 | -2.26E-03 | 3.02E-01 | 2.72E+00 | 2.74E+00 | 9.95E-01 | no | Frmd6 | intron_variant | MODIFIER |
| 12 | 70925734 | AX-97848187 | 0 | G | A | 0.061 | -2.26E-03 | 3.02E-01 | 2.72E+00 | 2.74E+00 | 9.95E-01 | no | | intergenic_variant | MODIFIER |
| 12 | 70986148 | AX-97856720 | 0 | G | A | 0.061 | -2.26E-03 | 3.02E-01 | 2.72E+00 | 2.74E+00 | 9.95E-01 | no | Psma3 | intron_variant,NMD_transcript_variant | MODIFIER |
| 12 | 71025707 | AX-170513354 | 0 | A | C | 0.061 | -2.26E-03 | 3.02E-01 | 2.72E+00 | 2.74E+00 | 9.95E-01 | no | Arid4a | intron_variant | MODIFIER |
| 12 | 71026150 | AX-170164350 | 0 | G | A | 0.061 | -2.26E-03 | 3.02E-01 | 2.72E+00 | 2.74E+00 | 9.95E-01 | no | Arid4a | intron_variant | MODIFIER |
| 12 | 71034636 | AX-170511497 | 0 | C | A | 0.061 | -2.26E-03 | 3.02E-01 | 2.72E+00 | 2.74E+00 | 9.95E-01 | no | Arid4a | intron_variant | MODIFIER |
| 12 | 71037492 | AX-170007018 | 0 | T | G | 0.061 | -2.26E-03 | 3.02E-01 | 2.72E+00 | 2.74E+00 | 9.95E-01 | no | Arid4a | intron_variant | MODIFIER |
| 12 | 71038423 | AX-97849226 | 0 | A | G | 0.061 | -2.26E-03 | 3.02E-01 | 2.72E+00 | 2.74E+00 | 9.95E-01 | no | Arid4a | intron_variant | MODIFIER |
| 12 | 71067932 | AX-170007016 | 0 | T | C | 0.061 | -2.26E-03 | 3.02E-01 | 2.72E+00 | 2.74E+00 | 9.95E-01 | no | Arid4a | intron_variant | MODIFIER |
| 12 | 71070086 | AX-97829781 | 0 | T | C | 0.061 | -2.26E-03 | 3.02E-01 | 2.72E+00 | 2.74E+00 | 9.95E-01 | no | Arid4a | intron_variant | MODIFIER |
| 12 | 71071185 | AX-170049368 | 0 | A | G | 0.061 | -2.26E-03 | 3.02E-01 | 2.72E+00 | 2.74E+00 | 9.95E-01 | no | Arid4a | intron_variant | MODIFIER |
| 12 | 71072658 | AX-170341143 | 0 | G | A | 0.061 | -2.26E-03 | 3.02E-01 | 2.72E+00 | 2.74E+00 | 9.95E-01 | no | Arid4a | synonymous_variant | LOW |
| 12 | 71072919 | AX-170376516 | 0 | C | A | 0.061 | -2.26E-03 | 3.02E-01 | 2.72E+00 | 2.74E+00 | 9.95E-01 | no | Arid4a | intron_variant | MODIFIER |
| 12 | 71085528 | AX-170007014 | 0 | T | C | 0.061 | -2.26E-03 | 3.02E-01 | 2.72E+00 | 2.74E+00 | 9.95E-01 | no | Arid4a | intron_variant | MODIFIER |
| 12 | 71094651 | AX-170087282 | 0 | C | T | 0.061 | -2.26E-03 | 3.02E-01 | 2.72E+00 | 2.74E+00 | 9.95E-01 | no | Arid4a | intron_variant | MODIFIER |
| 12 | 71094694 | AX-170418825 | 0 | G | A | 0.061 | -2.26E-03 | 3.02E-01 | 2.72E+00 | 2.74E+00 | 9.95E-01 | no | Arid4a | intron_variant | MODIFIER |
| 12 | 71094812 | AX-170355844 | 0 | G | A | 0.061 | -2.26E-03 | 3.02E-01 | 2.72E+00 | 2.74E+00 | 9.95E-01 | no | Arid4a | intron_variant | MODIFIER |
| 12 | 71115628 | AX-170449074 | 0 | C | A | 0.061 | -2.26E-03 | 3.02E-01 | 2.72E+00 | 2.74E+00 | 9.95E-01 | no | Tomm20l | intron_variant | MODIFIER |
| 12 | 71116363 | AX-170120896 | 0 | C | T | 0.061 | -2.26E-03 | 3.02E-01 | 2.72E+00 | 2.74E+00 | 9.95E-01 | no | Tomm20l | intron_variant | MODIFIER |
| 12 | 71116388 | AX-170472807 | 0 | A | G | 0.061 | -2.26E-03 | 3.02E-01 | 2.72E+00 | 2.74E+00 | 9.95E-01 | no | Tomm20l | intron_variant | MODIFIER |

| | | | | | | | | | | | | | | | |
|---|---|---|---|---|---|---|---|---|---|---|---|---|---|---|---|
| 12 | 71122930 | AX-170156892 | 0 | G | A | 0.061 | -2.26E-03 | 3.02E-01 | 2.72E+00 | 2.74E+00 | 9.95E-01 | no | Tomm20l | intron_variant | MODIFIER |
| 12 | 71123981 | AX-170304354 | 0 | A | G | 0.061 | -2.26E-03 | 3.02E-01 | 2.72E+00 | 2.74E+00 | 9.95E-01 | no | Tomm20l | downstream_gene_variant | MODIFIER |
| 12 | 71124982 | AX-170246087 | 0 | T | C | 0.061 | -2.26E-03 | 3.02E-01 | 2.72E+00 | 2.74E+00 | 9.95E-01 | no | Tomm20l | downstream_gene_variant | MODIFIER |
| 12 | 71125051 | AX-170227762 | 0 | C | T | 0.061 | -2.26E-03 | 3.02E-01 | 2.72E+00 | 2.74E+00 | 9.95E-01 | no | Tomm20l | downstream_gene_variant | MODIFIER |
| 12 | 71126090 | AX-170223412 | 0 | A | G | 0.061 | -2.26E-03 | 3.02E-01 | 2.72E+00 | 2.74E+00 | 9.95E-01 | no | Tomm20l | downstream_gene_variant | MODIFIER |
| 12 | 71128807 | AX-170402477 | 0 | A | G | 0.061 | -2.26E-03 | 3.02E-01 | 2.72E+00 | 2.74E+00 | 9.95E-01 | no | Timm9 | intron_variant | MODIFIER |
| 12 | 71141727 | AX-97784587 | 0 | T | C | 0.061 | -2.26E-03 | 3.02E-01 | 2.72E+00 | 2.74E+00 | 9.95E-01 | no | 2700049A03Rik | intron_variant | MODIFIER |
| 12 | 71147365 | AX-170133442 | 0 | C | T | 0.061 | -2.26E-03 | 3.02E-01 | 2.72E+00 | 2.74E+00 | 9.95E-01 | no | 2700049A03Rik | intron_variant | MODIFIER |
| 12 | 71148644 | AX-170424513 | 0 | G | A | 0.061 | -2.26E-03 | 3.02E-01 | 2.72E+00 | 2.74E+00 | 9.95E-01 | no | 2700049A03Rik | intron_variant | MODIFIER |
| 12 | 71182407 | AX-170357760 | 0 | T | A | 0.061 | -2.26E-03 | 3.02E-01 | 2.72E+00 | 2.74E+00 | 9.95E-01 | no | 2700049A03Rik | intron_variant | MODIFIER |
| 12 | 71225554 | AX-97826555 | 0 | C | T | 0.061 | -2.26E-03 | 3.02E-01 | 2.72E+00 | 2.74E+00 | 9.95E-01 | no | 2700049A03Rik | intron_variant | MODIFIER |
| 12 | 71231012 | AX-170305833 | 0 | C | T | 0.061 | -2.26E-03 | 3.02E-01 | 2.72E+00 | 2.74E+00 | 9.95E-01 | no | 2700049A03Rik | intron_variant | MODIFIER |
| 12 | 71257504 | AX-170490722 | 0 | T | G | 0.061 | -2.26E-03 | 3.02E-01 | 2.72E+00 | 2.74E+00 | 9.95E-01 | no | | intergenic_variant | MODIFIER |
| 12 | 71257754 | AX-97826553 | 0 | A | G | 0.061 | -2.26E-03 | 3.02E-01 | 2.72E+00 | 2.74E+00 | 9.95E-01 | no | | regulatory_region_variant | MODIFIER |
| 12 | 71290340 | AX-170120915 | 0 | C | T | 0.061 | -2.26E-03 | 3.02E-01 | 2.72E+00 | 2.74E+00 | 9.95E-01 | no | | regulatory_region_variant | MODIFIER |
| 12 | 71297985 | AX-97852519 | 0 | A | G | 0.061 | -2.26E-03 | 3.02E-01 | 2.72E+00 | 2.74E+00 | 9.95E-01 | no | | intergenic_variant | MODIFIER |
| 12 | 71298600 | AX-170387870 | 0 | T | C | 0.061 | -2.26E-03 | 3.02E-01 | 2.72E+00 | 2.74E+00 | 9.95E-01 | no | | regulatory_region_variant | MODIFIER |
| 12 | 71300442 | AX-170492872 | 0 | C | T | 0.061 | -2.26E-03 | 3.02E-01 | 2.72E+00 | 2.74E+00 | 9.95E-01 | no | | intergenic_variant | MODIFIER |
| 12 | 71318199 | AX-170111936 | 0 | A | G | 0.061 | -2.26E-03 | 3.02E-01 | 2.72E+00 | 2.74E+00 | 9.95E-01 | no | Dact1 | missense_variant | MODERATE |
| 12 | 71318766 | AX-170460502 | 0 | G | A | 0.061 | -2.26E-03 | 3.02E-01 | 2.72E+00 | 2.74E+00 | 9.95E-01 | no | Dact1 | missense_variant | MODERATE |
| 12 | 71323020 | AX-170207301 | 0 | T | C | 0.061 | -2.26E-03 | 3.02E-01 | 2.72E+00 | 2.74E+00 | 9.95E-01 | no | Dact1 | downstream_gene_variant | MODIFIER |
| 12 | 71336254 | AX-170049372 | 0 | C | A | 0.061 | -2.26E-03 | 3.02E-01 | 2.72E+00 | 2.74E+00 | 9.95E-01 | no | Gm33016 | upstream_gene_variant | MODIFIER |
| 12 | 71336414 | AX-97812834 | 0 | G | A | 0.061 | -2.26E-03 | 3.02E-01 | 2.72E+00 | 2.74E+00 | 9.95E-01 | no | Gm33016 | upstream_gene_variant | MODIFIER |
| 12 | 71349237 | AX-170522622 | 0 | G | A | 0.061 | -2.26E-03 | 3.02E-01 | 2.72E+00 | 2.74E+00 | 9.95E-01 | no | | intergenic_variant | MODIFIER |
| 12 | 71364918 | AX-170145549 | 0 | A | C | 0.061 | -2.26E-03 | 3.02E-01 | 2.72E+00 | 2.74E+00 | 9.95E-01 | no | Gm40438 | intron_variant,non_coding_transcript_variant | MODIFIER |
| 12 | 71379328 | AX-170459543 | 0 | C | A | 0.061 | -2.26E-03 | 3.02E-01 | 2.72E+00 | 2.74E+00 | 9.95E-01 | no | 1700083H02Rik | downstream_gene_variant | MODIFIER |
| 12 | 71379723 | AX-97852153 | 0 | A | C | 0.061 | -2.26E-03 | 3.02E-01 | 2.72E+00 | 2.74E+00 | 9.95E-01 | no | 1700083H02Rik | downstream_gene_variant | MODIFIER |
| 12 | 71383114 | AX-170373042 | 0 | C | A | 0.061 | -2.26E-03 | 3.02E-01 | 2.72E+00 | 2.74E+00 | 9.95E-01 | no | 1700083H02Rik | downstream_gene_variant | MODIFIER |
| 12 | 71384215 | AX-170528854 | 0 | C | T | 0.061 | -2.26E-03 | 3.02E-01 | 2.72E+00 | 2.74E+00 | 9.95E-01 | no | 1700083H02Rik | upstream_gene_variant | MODIFIER |
| 12 | 71529996 | AX-170007023 | 0 | C | G | 0.061 | -2.26E-03 | 3.02E-01 | 2.72E+00 | 2.74E+00 | 9.95E-01 | no | 4930404H11Rik | intron_variant,non_coding_transcript_variant | MODIFIER |
| 12 | 71530076 | AX-170270387 | 0 | G | A | 0.061 | -2.26E-03 | 3.02E-01 | 2.72E+00 | 2.74E+00 | 9.95E-01 | no | 4930404H11Rik | intron_variant,non_coding_transcript_variant | MODIFIER |
| 12 | 71534927 | AX-97831278 | 0 | C | T | 0.061 | -2.26E-03 | 3.02E-01 | 2.72E+00 | 2.74E+00 | 9.95E-01 | no | 4930404H11Rik | intron_variant,non_coding_transcript_variant | MODIFIER |
| 12 | 71608360 | AX-170186432 | 0 | A | T | 0.061 | -2.26E-03 | 3.02E-01 | 2.72E+00 | 2.74E+00 | 9.95E-01 | no | 9630002D21Rik | intron_variant,non_coding_transcript_variant | MODIFIER |
| 12 | 71613479 | AX-97782116 | 0 | C | T | 0.061 | -2.26E-03 | 3.02E-01 | 2.72E+00 | 2.74E+00 | 9.95E-01 | no | 9630002D21Rik | intron_variant,non_coding_transcript_variant | MODIFIER |
| 12 | 71694338 | AX-170156043 | 0 | G | A | 0.061 | -2.26E-03 | 3.02E-01 | 2.72E+00 | 2.74E+00 | 9.95E-01 | no | 9630002D21Rik | intron_variant,non_coding_transcript_variant | MODIFIER |
| 12 | 71697247 | AX-170092515 | 0 | A | C | 0.061 | -2.26E-03 | 3.02E-01 | 2.72E+00 | 2.74E+00 | 9.95E-01 | no | 9630002D21Rik | intron_variant,non_coding_transcript_variant | MODIFIER |
| 12 | 72201999 | AX-170136228 | 0 | T | A | 0.061 | -2.26E-03 | 3.02E-01 | 2.72E+00 | 2.74E+00 | 9.95E-01 | no | | intergenic_variant | MODIFIER |
| 12 | 72202627 | AX-170266467 | 0 | T | C | 0.061 | -2.26E-03 | 3.02E-01 | 2.72E+00 | 2.74E+00 | 9.95E-01 | no | | intergenic_variant | MODIFIER |
| 12 | 72227758 | AX-170490362 | 0 | A | G | 0.061 | -2.26E-03 | 3.02E-01 | 2.72E+00 | 2.74E+00 | 9.95E-01 | no | Rtn1 | intron_variant | MODIFIER |
| 12 | 72232247 | AX-170129472 | 0 | G | A | 0.061 | -2.26E-03 | 3.02E-01 | 2.72E+00 | 2.74E+00 | 9.95E-01 | no | Rtn1 | intron_variant | MODIFIER |
| 12 | 72255789 | AX-170231421 | 0 | T | C | 0.061 | -2.26E-03 | 3.02E-01 | 2.72E+00 | 2.74E+00 | 9.95E-01 | no | Rtn1 | intron_variant | MODIFIER |
| 12 | 72263512 | AX-170183117 | 0 | A | G | 0.061 | -2.26E-03 | 3.02E-01 | 2.72E+00 | 2.74E+00 | 9.95E-01 | no | Rtn1 | intron_variant | MODIFIER |
| 12 | 72265863 | AX-170049395 | 0 | A | C | 0.061 | -2.26E-03 | 3.02E-01 | 2.72E+00 | 2.74E+00 | 9.95E-01 | no | Rtn1 | intron_variant | MODIFIER |
| 12 | 72265892 | AX-170049396 | 0 | G | A | 0.061 | -2.26E-03 | 3.02E-01 | 2.72E+00 | 2.74E+00 | 9.95E-01 | no | Rtn1 | intron_variant | MODIFIER |
| 12 | 72265965 | AX-97857137 | 0 | A | G | 0.061 | -2.26E-03 | 3.02E-01 | 2.72E+00 | 2.74E+00 | 9.95E-01 | no | Rtn1 | intron_variant | MODIFIER |
| 12 | 72268033 | AX-170305452 | 0 | A | G | 0.061 | -2.26E-03 | 3.02E-01 | 2.72E+00 | 2.74E+00 | 9.95E-01 | no | Rtn1 | intron_variant | MODIFIER |
| 12 | 72287313 | AX-170413757 | 0 | C | T | 0.061 | -2.26E-03 | 3.02E-01 | 2.72E+00 | 2.74E+00 | 9.95E-01 | no | Rtn1 | intron_variant | MODIFIER |
| 12 | 72296841 | AX-97818815 | 0 | T | C | 0.061 | -2.26E-03 | 3.02E-01 | 2.72E+00 | 2.74E+00 | 9.95E-01 | no | Rtn1 | intron_variant | MODIFIER |
| 12 | 72302387 | AX-170506143 | 0 | G | A | 0.061 | -2.26E-03 | 3.02E-01 | 2.72E+00 | 2.74E+00 | 9.95E-01 | no | Rtn1 | intron_variant | MODIFIER |
| 12 | 72302412 | AX-170374620 | 0 | A | G | 0.061 | -2.26E-03 | 3.02E-01 | 2.72E+00 | 2.74E+00 | 9.95E-01 | no | Rtn1 | intron_variant | MODIFIER |
| 12 | 72303299 | AX-170138978 | 0 | T | C | 0.061 | -2.26E-03 | 3.02E-01 | 2.72E+00 | 2.74E+00 | 9.95E-01 | no | Rtn1 | intron_variant | MODIFIER |
| 12 | 72337272 | AX-170489720 | 0 | C | T | 0.061 | -2.26E-03 | 3.02E-01 | 2.72E+00 | 2.74E+00 | 9.95E-01 | no | Rtn1 | intron_variant | MODIFIER |
| 12 | 72347512 | AX-170101577 | 0 | G | A | 0.061 | -2.26E-03 | 3.02E-01 | 2.72E+00 | 2.74E+00 | 9.95E-01 | no | Rtn1 | intron_variant | MODIFIER |
| 12 | 72349031 | AX-170406851 | 0 | G | A | 0.061 | -2.26E-03 | 3.02E-01 | 2.72E+00 | 2.74E+00 | 9.95E-01 | no | Rtn1 | intron_variant | MODIFIER |

| | | | | | | | | | | | | | | | |
|---|---|---|---|---|---|---|---|---|---|---|---|---|---|---|---|
| 12 | 72356343 | AX-170462802 | 0 | T | C | 0.061 | -2.26E-03 | 3.02E-01 | 2.72E+00 | 2.74E+00 | 9.95E-01 | no | Rtn1 | intron_variant | MODIFIER |
| 12 | 72356412 | AX-170049399 | 0 | C | A | 0.061 | -2.26E-03 | 3.02E-01 | 2.72E+00 | 2.74E+00 | 9.95E-01 | no | Rtn1 | intron_variant | MODIFIER |
| 12 | 72360746 | AX-97817748 | 0 | G | A | 0.061 | -2.26E-03 | 3.02E-01 | 2.72E+00 | 2.74E+00 | 9.95E-01 | no | Rtn1 | intron_variant | MODIFIER |
| 12 | 72366605 | AX-170049400 | 0 | C | T | 0.061 | -2.26E-03 | 3.02E-01 | 2.72E+00 | 2.74E+00 | 9.95E-01 | no | Rtn1 | intron_variant | MODIFIER |
| 12 | 72374269 | AX-170112159 | 0 | G | A | 0.061 | -2.26E-03 | 3.02E-01 | 2.72E+00 | 2.74E+00 | 9.95E-01 | no | Rtn1 | intron_variant | MODIFIER |
| 12 | 72374493 | AX-170493098 | 0 | T | A | 0.061 | -2.26E-03 | 3.02E-01 | 2.72E+00 | 2.74E+00 | 9.95E-01 | no | Rtn1 | intron_variant | MODIFIER |
| 12 | 72386346 | AX-170117447 | 0 | T | C | 0.061 | -2.26E-03 | 3.02E-01 | 2.72E+00 | 2.74E+00 | 9.95E-01 | no | Rtn1 | intron_variant | MODIFIER |
| 12 | 72386439 | AX-170435047 | 0 | T | A | 0.061 | -2.26E-03 | 3.02E-01 | 2.72E+00 | 2.74E+00 | 9.95E-01 | no | Rtn1 | intron_variant | MODIFIER |
| 12 | 72386486 | AX-170367673 | 0 | T | C | 0.061 | -2.26E-03 | 3.02E-01 | 2.72E+00 | 2.74E+00 | 9.95E-01 | no | Rtn1 | intron_variant | MODIFIER |
| 12 | 72434075 | AX-170322368 | 0 | T | C | 0.061 | -2.26E-03 | 3.02E-01 | 2.72E+00 | 2.74E+00 | 9.95E-01 | no | | intergenic_variant | MODIFIER |
| 12 | 72434737 | AX-170170023 | 0 | G | A | 0.061 | -2.26E-03 | 3.02E-01 | 2.72E+00 | 2.74E+00 | 9.95E-01 | no | | intergenic_variant | MODIFIER |
| 12 | 72435363 | AX-170294215 | 0 | G | C | 0.061 | -2.26E-03 | 3.02E-01 | 2.72E+00 | 2.74E+00 | 9.95E-01 | no | | intergenic_variant | MODIFIER |
| 12 | 72436843 | AX-170147573 | 0 | T | C | 0.061 | -2.26E-03 | 3.02E-01 | 2.72E+00 | 2.74E+00 | 9.95E-01 | no | | regulatory_region_variant | MODIFIER |
| 12 | 72437384 | AX-170331866 | 0 | T | C | 0.061 | -2.26E-03 | 3.02E-01 | 2.72E+00 | 2.74E+00 | 9.95E-01 | no | Lrrc9 | upstream_gene_variant | MODIFIER |
| 12 | 72443489 | AX-170482085 | 0 | A | G | 0.061 | -2.26E-03 | 3.02E-01 | 2.72E+00 | 2.74E+00 | 9.95E-01 | no | Lrrc9 | intron_variant,non_coding_transcript_variant | MODIFIER |
| 12 | 72447683 | AX-170267388 | 0 | T | C | 0.061 | -2.26E-03 | 3.02E-01 | 2.72E+00 | 2.74E+00 | 9.95E-01 | no | Lrrc9 | intron_variant,non_coding_transcript_variant | MODIFIER |
| 12 | 72453664 | AX-170212240 | 0 | A | C | 0.061 | -2.26E-03 | 3.02E-01 | 2.72E+00 | 2.74E+00 | 9.95E-01 | no | Lrrc9 | intron_variant,non_coding_transcript_variant | MODIFIER |
| 12 | 72681834 | AX-170211178 | 0 | T | A | 0.061 | -2.26E-03 | 3.02E-01 | 2.72E+00 | 2.74E+00 | 9.95E-01 | no | | intergenic_variant | MODIFIER |
| 12 | 72682824 | AX-97836610 | 0 | A | G | 0.061 | -2.26E-03 | 3.02E-01 | 2.72E+00 | 2.74E+00 | 9.95E-01 | no | | regulatory_region_variant | MODIFIER |
| 12 | 72741199 | AX-170049406 | 0 | C | A | 0.061 | -2.26E-03 | 3.02E-01 | 2.72E+00 | 2.74E+00 | 9.95E-01 | no | Gm48143 | non_coding_transcript_exon_variant | MODIFIER |
| 12 | 72741309 | AX-170092516 | 0 | C | T | 0.061 | -2.26E-03 | 3.02E-01 | 2.72E+00 | 2.74E+00 | 9.95E-01 | no | Gm48143 | non_coding_transcript_exon_variant | MODIFIER |
| 12 | 72811602 | AX-170479620 | 0 | A | G | 0.061 | -2.26E-03 | 3.02E-01 | 2.72E+00 | 2.74E+00 | 9.95E-01 | no | | intergenic_variant | MODIFIER |
| 12 | 72814578 | AX-170437417 | 0 | A | G | 0.061 | -2.26E-03 | 3.02E-01 | 2.72E+00 | 2.74E+00 | 9.95E-01 | no | Gm33785 | upstream_gene_variant | MODIFIER |
| 12 | 72814979 | AX-170421007 | 0 | A | G | 0.061 | -2.26E-03 | 3.02E-01 | 2.72E+00 | 2.74E+00 | 9.95E-01 | no | Gm33785 | upstream_gene_variant | MODIFIER |
| 12 | 72815155 | AX-170110383 | 0 | G | C | 0.061 | -2.26E-03 | 3.02E-01 | 2.72E+00 | 2.74E+00 | 9.95E-01 | no | Gm33785 | upstream_gene_variant | MODIFIER |
| 12 | 72840821 | AX-170491225 | 0 | T | C | 0.061 | -2.26E-03 | 3.02E-01 | 2.72E+00 | 2.74E+00 | 9.95E-01 | no | | intergenic_variant | MODIFIER |
| 12 | 72848559 | AX-170174058 | 0 | A | G | 0.061 | -2.26E-03 | 3.02E-01 | 2.72E+00 | 2.74E+00 | 9.95E-01 | no | | intergenic_variant | MODIFIER |
| 12 | 72849335 | AX-170161700 | 0 | G | C | 0.061 | -2.26E-03 | 3.02E-01 | 2.72E+00 | 2.74E+00 | 9.95E-01 | no | | intergenic_variant | MODIFIER |
| 12 | 72850481 | AX-170342694 | 0 | T | A | 0.061 | -2.26E-03 | 3.02E-01 | 2.72E+00 | 2.74E+00 | 9.95E-01 | no | | regulatory_region_variant | MODIFIER |
| 12 | 72897746 | AX-170198510 | 0 | A | G | 0.061 | -2.26E-03 | 3.02E-01 | 2.72E+00 | 2.74E+00 | 9.95E-01 | no | 4930447C04Rik | intron_variant | MODIFIER |
| 12 | 72918318 | AX-170413248 | 0 | T | C | 0.061 | -2.26E-03 | 3.02E-01 | 2.72E+00 | 2.74E+00 | 9.95E-01 | no | 4930447C04Rik | upstream_gene_variant | MODIFIER |
| 12 | 72953583 | AX-97846174 | 0 | A | C | 0.061 | -2.26E-03 | 3.02E-01 | 2.72E+00 | 2.74E+00 | 9.95E-01 | no | Gm26709 | intron_variant,non_coding_transcript_variant | MODIFIER |
| 12 | 72966522 | AX-170460682 | 0 | A | G | 0.061 | -2.26E-03 | 3.02E-01 | 2.72E+00 | 2.74E+00 | 9.95E-01 | no | Gm26709 | downstream_gene_variant | MODIFIER |
| 12 | 72967654 | AX-170468433 | 0 | T | C | 0.061 | -2.26E-03 | 3.02E-01 | 2.72E+00 | 2.74E+00 | 9.95E-01 | no | Gm26709 | downstream_gene_variant | MODIFIER |
| 12 | 72968924 | AX-170119573 | 0 | G | A | 0.061 | -2.26E-03 | 3.02E-01 | 2.72E+00 | 2.74E+00 | 9.95E-01 | no | Gm26709 | downstream_gene_variant | MODIFIER |
| 12 | 72969766 | AX-170139542 | 0 | T | C | 0.061 | -2.26E-03 | 3.02E-01 | 2.72E+00 | 2.74E+00 | 9.95E-01 | no | Gm26709 | downstream_gene_variant | MODIFIER |
| 12 | 73012782 | AX-170225780 | 0 | T | C | 0.061 | -2.26E-03 | 3.02E-01 | 2.72E+00 | 2.74E+00 | 9.95E-01 | no | | regulatory_region_variant | MODIFIER |
| 12 | 73021148 | AX-170514083 | 0 | G | A | 0.061 | -2.26E-03 | 3.02E-01 | 2.72E+00 | 2.74E+00 | 9.95E-01 | no | | regulatory_region_variant | MODIFIER |
| 12 | 73027668 | AX-170393396 | 0 | G | C | 0.061 | -2.26E-03 | 3.02E-01 | 2.72E+00 | 2.74E+00 | 9.95E-01 | no | | intergenic_variant | MODIFIER |
| 12 | 73068769 | AX-170431344 | 0 | C | G | 0.061 | -2.26E-03 | 3.02E-01 | 2.72E+00 | 2.74E+00 | 9.95E-01 | no | | intergenic_variant | MODIFIER |
| 12 | 73097536 | AX-170214017 | 0 | T | C | 0.061 | -2.26E-03 | 3.02E-01 | 2.72E+00 | 2.74E+00 | 9.95E-01 | no | Six4 | downstream_gene_variant | MODIFIER |
| 12 | 73097590 | AX-97807779 | 0 | G | A | 0.061 | -2.26E-03 | 3.02E-01 | 2.72E+00 | 2.74E+00 | 9.95E-01 | no | Six4 | downstream_gene_variant | MODIFIER |
| 12 | 73098627 | AX-170375653 | 0 | T | C | 0.061 | -2.26E-03 | 3.02E-01 | 2.72E+00 | 2.74E+00 | 9.95E-01 | no | Six4 | downstream_gene_variant | MODIFIER |
| 12 | 73170847 | AX-170087284 | 0 | A | G | 0.061 | -2.26E-03 | 3.02E-01 | 2.72E+00 | 2.74E+00 | 9.95E-01 | no | Mnat1 | intron_variant | MODIFIER |
| 12 | 73199117 | AX-170162361 | 0 | G | C | 0.061 | -2.26E-03 | 3.02E-01 | 2.72E+00 | 2.74E+00 | 9.95E-01 | no | Mnat1 | intron_variant | MODIFIER |
| 12 | 73199274 | AX-170316729 | 0 | A | G | 0.061 | -2.26E-03 | 3.02E-01 | 2.72E+00 | 2.74E+00 | 9.95E-01 | no | Mnat1 | intron_variant | MODIFIER |
| 12 | 73200000 | AX-170501801 | 0 | G | A | 0.061 | -2.26E-03 | 3.02E-01 | 2.72E+00 | 2.74E+00 | 9.95E-01 | no | Mnat1 | intron_variant | MODIFIER |
| 12 | 73228963 | AX-170281919 | 0 | A | G | 0.061 | -2.26E-03 | 3.02E-01 | 2.72E+00 | 2.74E+00 | 9.95E-01 | no | Mnat1 | intron_variant | MODIFIER |
| 12 | 73230243 | AX-97847224 | 0 | C | G | 0.061 | -2.26E-03 | 3.02E-01 | 2.72E+00 | 2.74E+00 | 9.95E-01 | no | Mnat1 | intron_variant | MODIFIER |
| 12 | 73230457 | AX-170049415 | 0 | A | G | 0.061 | -2.26E-03 | 3.02E-01 | 2.72E+00 | 2.74E+00 | 9.95E-01 | no | Mnat1 | intron_variant | MODIFIER |
| 12 | 73238125 | AX-170210282 | 0 | G | A | 0.061 | -2.26E-03 | 3.02E-01 | 2.72E+00 | 2.74E+00 | 9.95E-01 | no | Mnat1 | intron_variant | MODIFIER |
| 12 | 73238478 | AX-170268534 | 0 | A | G | 0.061 | -2.26E-03 | 3.02E-01 | 2.72E+00 | 2.74E+00 | 9.95E-01 | no | Mnat1 | intron_variant | MODIFIER |
| 12 | 73246317 | AX-170397463 | 0 | C | G | 0.061 | -2.26E-03 | 3.02E-01 | 2.72E+00 | 2.74E+00 | 9.95E-01 | no | Mnat1 | intron_variant | MODIFIER |
| 12 | 73260123 | AX-97821695 | 0 | T | C | 0.061 | -2.26E-03 | 3.02E-01 | 2.72E+00 | 2.74E+00 | 9.95E-01 | no | Mnat1 | intron_variant | MODIFIER |

| | | | | | | | | | | | | | | | |
|---|---|---|---|---|---|---|---|---|---|---|---|---|---|---|---|
| 12 | 73269389 | AX-170266123 | 0 | T | A | 0.061 | -2.26E-03 | 3.02E-01 | 2.72E+00 | 2.74E+00 | 9.95E-01 | no | Mnat1 | intron_variant | MODIFIER |
| 12 | 73287081 | AX-97789959 | 0 | C | T | 0.061 | -2.26E-03 | 3.02E-01 | 2.72E+00 | 2.74E+00 | 9.95E-01 | no | Slc38a6 | intron_variant,NMD_transcript_variant | MODIFIER |
| 12 | 73338188 | AX-170416854 | 0 | A | T | 0.061 | -2.26E-03 | 3.02E-01 | 2.72E+00 | 2.74E+00 | 9.95E-01 | no | Slc38a6 | intron_variant,non_coding_transcript_variant | MODIFIER |
| 12 | 73369100 | AX-170438606 | 0 | A | G | 0.061 | -2.26E-03 | 3.02E-01 | 2.72E+00 | 2.74E+00 | 9.95E-01 | no | D830013O20Rik | intron_variant,non_coding_transcript_variant | MODIFIER |
| 12 | 73376445 | AX-97801809 | 0 | G | A | 0.061 | -2.26E-03 | 3.02E-01 | 2.72E+00 | 2.74E+00 | 9.95E-01 | no | D830013O20Rik | upstream_gene_variant | MODIFIER |
| 12 | 73383742 | AX-170327117 | 0 | G | A | 0.061 | -2.26E-03 | 3.02E-01 | 2.72E+00 | 2.74E+00 | 9.95E-01 | no | D830013O20Rik | intron_variant,non_coding_transcript_variant | MODIFIER |
| 12 | 73440330 | AX-170007043 | 0 | G | A | 0.061 | -2.26E-03 | 3.02E-01 | 2.72E+00 | 2.74E+00 | 9.95E-01 | no | | intergenic_variant | MODIFIER |
| 12 | 73441143 | AX-170460096 | 0 | C | T | 0.061 | -2.26E-03 | 3.02E-01 | 2.72E+00 | 2.74E+00 | 9.95E-01 | no | | intergenic_variant | MODIFIER |
| 12 | 73441209 | AX-170272428 | 0 | T | C | 0.061 | -2.26E-03 | 3.02E-01 | 2.72E+00 | 2.74E+00 | 9.95E-01 | no | | intergenic_variant | MODIFIER |
| 12 | 73441529 | AX-97804749 | 0 | G | A | 0.061 | -2.26E-03 | 3.02E-01 | 2.72E+00 | 2.74E+00 | 9.95E-01 | no | | regulatory_region_variant | MODIFIER |
| 12 | 73459941 | AX-170395782 | 0 | G | T | 0.061 | -2.26E-03 | 3.02E-01 | 2.72E+00 | 2.74E+00 | 9.95E-01 | no | Gm48656 | upstream_gene_variant | MODIFIER |
| 12 | 73460885 | AX-170466151 | 0 | C | T | 0.061 | -2.26E-03 | 3.02E-01 | 2.72E+00 | 2.74E+00 | 9.95E-01 | no | Gm48656 | upstream_gene_variant | MODIFIER |
| 12 | 73468910 | AX-97837192 | 0 | T | C | 0.061 | -2.26E-03 | 3.02E-01 | 2.72E+00 | 2.74E+00 | 9.95E-01 | no | | intergenic_variant | MODIFIER |
| 12 | 73487147 | AX-170178405 | 0 | A | G | 0.061 | -2.26E-03 | 3.02E-01 | 2.72E+00 | 2.74E+00 | 9.95E-01 | no | Gm34016 | downstream_gene_variant | MODIFIER |
| 12 | 73498424 | AX-97821744 | 0 | G | A | 0.061 | -2.26E-03 | 3.02E-01 | 2.72E+00 | 2.74E+00 | 9.95E-01 | no | Gm34016 | intron_variant,non_coding_transcript_variant | MODIFIER |
| 12 | 73511376 | AX-170049417 | 0 | G | A | 0.061 | -2.26E-03 | 3.02E-01 | 2.72E+00 | 2.74E+00 | 9.95E-01 | no | Gm34016 | upstream_gene_variant | MODIFIER |
| 12 | 73511440 | AX-97787503 | 0 | A | G | 0.061 | -2.26E-03 | 3.02E-01 | 2.72E+00 | 2.74E+00 | 9.95E-01 | no | Gm34016 | upstream_gene_variant | MODIFIER |
| 12 | 73619057 | AX-170390657 | 0 | T | C | 0.061 | -2.26E-03 | 3.02E-01 | 2.72E+00 | 2.74E+00 | 9.95E-01 | no | Prkch | intron_variant | MODIFIER |
| 12 | 73619135 | AX-170418911 | 0 | G | C | 0.061 | -2.26E-03 | 3.02E-01 | 2.72E+00 | 2.74E+00 | 9.95E-01 | no | Prkch | intron_variant | MODIFIER |
| 12 | 73619171 | AX-97813285 | 0 | C | T | 0.061 | -2.26E-03 | 3.02E-01 | 2.72E+00 | 2.74E+00 | 9.95E-01 | no | Prkch | intron_variant | MODIFIER |
| 12 | 73619492 | AX-170274892 | 0 | T | C | 0.061 | -2.26E-03 | 3.02E-01 | 2.72E+00 | 2.74E+00 | 9.95E-01 | no | Prkch | intron_variant | MODIFIER |
| 12 | 73630678 | AX-170049420 | 0 | T | C | 0.061 | -2.26E-03 | 3.02E-01 | 2.72E+00 | 2.74E+00 | 9.95E-01 | no | Prkch | intron_variant | MODIFIER |
| 12 | 73631813 | AX-170266978 | 0 | T | C | 0.061 | -2.26E-03 | 3.02E-01 | 2.72E+00 | 2.74E+00 | 9.95E-01 | no | Prkch | intron_variant | MODIFIER |
| 12 | 73632095 | AX-170325242 | 0 | C | T | 0.061 | -2.26E-03 | 3.02E-01 | 2.72E+00 | 2.74E+00 | 9.95E-01 | no | Prkch | intron_variant | MODIFIER |
| 12 | 73638798 | AX-170049424 | 0 | G | A | 0.061 | -2.26E-03 | 3.02E-01 | 2.72E+00 | 2.74E+00 | 9.95E-01 | no | Prkch | intron_variant | MODIFIER |
| 12 | 73968575 | AX-170193356 | 0 | G | A | 0.061 | -2.26E-03 | 3.02E-01 | 2.72E+00 | 2.74E+00 | 9.95E-01 | no | Snapc1 | intron_variant | MODIFIER |
| 12 | 73999153 | AX-97805495 | 0 | T | C | 0.061 | -2.26E-03 | 3.02E-01 | 2.72E+00 | 2.74E+00 | 9.95E-01 | no | Syt16 | intron_variant,NMD_transcript_variant | MODIFIER |
| 12 | 74000850 | AX-170189213 | 0 | C | T | 0.061 | -2.26E-03 | 3.02E-01 | 2.72E+00 | 2.74E+00 | 9.95E-01 | no | Syt16 | intron_variant,NMD_transcript_variant | MODIFIER |
| 12 | 74002355 | AX-170489016 | 0 | G | T | 0.061 | -2.26E-03 | 3.02E-01 | 2.72E+00 | 2.74E+00 | 9.95E-01 | no | Syt16 | intron_variant,NMD_transcript_variant | MODIFIER |
| 12 | 74002820 | AX-170130737 | 0 | T | C | 0.061 | -2.26E-03 | 3.02E-01 | 2.72E+00 | 2.74E+00 | 9.95E-01 | no | Syt16 | intron_variant,NMD_transcript_variant | MODIFIER |
| 12 | 74022147 | AX-170111482 | 0 | A | G | 0.061 | -2.26E-03 | 3.02E-01 | 2.72E+00 | 2.74E+00 | 9.95E-01 | no | Syt16 | intron_variant,NMD_transcript_variant | MODIFIER |
| 18 | 30424499 | AX-170519253 | 0 | C | A | 0.061 | -2.26E-03 | 3.02E-01 | 2.72E+00 | 2.74E+00 | 9.95E-01 | no | AC163399.2 | downstream_gene_variant | MODIFIER |
| 18 | 30424859 | AX-170393549 | 0 | C | G | 0.061 | -2.26E-03 | 3.02E-01 | 2.72E+00 | 2.74E+00 | 9.95E-01 | no | AC163399.2 | downstream_gene_variant | MODIFIER |
| 18 | 30441720 | AX-170523547 | 0 | A | G | 0.061 | -2.26E-03 | 3.02E-01 | 2.72E+00 | 2.74E+00 | 9.95E-01 | no | | intergenic_variant | MODIFIER |
| 18 | 30476381 | AX-170112490 | 0 | A | T | 0.061 | -2.26E-03 | 3.02E-01 | 2.72E+00 | 2.74E+00 | 9.95E-01 | no | Gm22870 | upstream_gene_variant | MODIFIER |
| 18 | 30482676 | AX-170253483 | 0 | T | C | 0.061 | -2.26E-03 | 3.02E-01 | 2.72E+00 | 2.74E+00 | 9.95E-01 | no | | intergenic_variant | MODIFIER |
| 18 | 30487129 | AX-170162474 | 0 | T | C | 0.061 | -2.26E-03 | 3.02E-01 | 2.72E+00 | 2.74E+00 | 9.95E-01 | no | | intergenic_variant | MODIFIER |
| 18 | 30487678 | AX-170205543 | 0 | A | T | 0.061 | -2.26E-03 | 3.02E-01 | 2.72E+00 | 2.74E+00 | 9.95E-01 | no | | regulatory_region_variant | MODIFIER |
| 18 | 30497950 | AX-170497752 | 0 | G | A | 0.061 | -2.26E-03 | 3.02E-01 | 2.72E+00 | 2.74E+00 | 9.95E-01 | no | Gm7936 | upstream_gene_variant | MODIFIER |
| 18 | 30498546 | AX-170208065 | 0 | A | G | 0.061 | -2.26E-03 | 3.02E-01 | 2.72E+00 | 2.74E+00 | 9.95E-01 | no | Gm7936 | upstream_gene_variant | MODIFIER |
| 18 | 30513551 | AX-170149972 | 0 | A | G | 0.061 | -2.26E-03 | 3.02E-01 | 2.72E+00 | 2.74E+00 | 9.95E-01 | no | | intergenic_variant | MODIFIER |
| 18 | 30515450 | AX-170517137 | 0 | A | G | 0.061 | -2.26E-03 | 3.02E-01 | 2.72E+00 | 2.74E+00 | 9.95E-01 | no | | intergenic_variant | MODIFIER |
| 18 | 30516319 | AX-170455990 | 0 | C | T | 0.061 | -2.26E-03 | 3.02E-01 | 2.72E+00 | 2.74E+00 | 9.95E-01 | no | | intergenic_variant | MODIFIER |
| 18 | 30532197 | AX-170341313 | 0 | C | T | 0.061 | -2.26E-03 | 3.02E-01 | 2.72E+00 | 2.74E+00 | 9.95E-01 | no | | intergenic_variant | MODIFIER |
| 18 | 30555215 | AX-170163787 | 0 | T | G | 0.061 | -2.26E-03 | 3.02E-01 | 2.72E+00 | 2.74E+00 | 9.95E-01 | no | | intergenic_variant | MODIFIER |
| 18 | 30567549 | AX-170496939 | 0 | G | C | 0.061 | -2.26E-03 | 3.02E-01 | 2.72E+00 | 2.74E+00 | 9.95E-01 | no | | intergenic_variant | MODIFIER |
| 18 | 30572667 | AX-170209778 | 0 | C | T | 0.061 | -2.26E-03 | 3.02E-01 | 2.72E+00 | 2.74E+00 | 9.95E-01 | no | | intergenic_variant | MODIFIER |
| 18 | 30584971 | AX-170507613 | 0 | T | G | 0.061 | -2.26E-03 | 3.02E-01 | 2.72E+00 | 2.74E+00 | 9.95E-01 | no | | regulatory_region_variant | MODIFIER |
| 18 | 30585062 | AX-170320259 | 0 | T | C | 0.061 | -2.26E-03 | 3.02E-01 | 2.72E+00 | 2.74E+00 | 9.95E-01 | no | | regulatory_region_variant | MODIFIER |
| 18 | 30585489 | AX-170245030 | 0 | C | T | 0.061 | -2.26E-03 | 3.02E-01 | 2.72E+00 | 2.74E+00 | 9.95E-01 | no | | intergenic_variant | MODIFIER |
| 18 | 30611070 | AX-97814139 | 0 | A | C | 0.061 | -2.26E-03 | 3.02E-01 | 2.72E+00 | 2.74E+00 | 9.95E-01 | no | AC113953.1 | upstream_gene_variant | MODIFIER |
| 18 | 30619843 | AX-170426344 | 0 | T | C | 0.061 | -2.26E-03 | 3.02E-01 | 2.72E+00 | 2.74E+00 | 9.95E-01 | no | | intergenic_variant | MODIFIER |
| 18 | 30624144 | AX-170228930 | 0 | A | G | 0.061 | -2.26E-03 | 3.02E-01 | 2.72E+00 | 2.74E+00 | 9.95E-01 | no | | intergenic_variant | MODIFIER |
| 18 | 30632950 | AX-170181451 | 0 | C | G | 0.061 | -2.26E-03 | 3.02E-01 | 2.72E+00 | 2.74E+00 | 9.95E-01 | no | | regulatory_region_variant | MODIFIER |

| | | | | | | | | | | | | | | | |
|---|---|---|---|---|---|---|---|---|---|---|---|---|---|---|---|
| 18 | 30632985 | AX-170351388 | 0 | A | G | 0.061 | -2.26E-03 | 3.02E-01 | 2.72E+00 | 2.74E+00 | 9.95E-01 | no | | intergenic_variant | MODIFIER |
| 18 | 30633028 | AX-170411862 | 0 | G | A | 0.061 | -2.26E-03 | 3.02E-01 | 2.72E+00 | 2.74E+00 | 9.95E-01 | no | | intergenic_variant | MODIFIER |
| 18 | 30633324 | AX-170184889 | 0 | G | C | 0.061 | -2.26E-03 | 3.02E-01 | 2.72E+00 | 2.74E+00 | 9.95E-01 | no | | intergenic_variant | MODIFIER |
| 18 | 30666274 | AX-170415655 | 0 | G | A | 0.061 | -2.26E-03 | 3.02E-01 | 2.72E+00 | 2.74E+00 | 9.95E-01 | no | | intergenic_variant | MODIFIER |
| 18 | 30667645 | AX-170375877 | 0 | G | T | 0.061 | -2.26E-03 | 3.02E-01 | 2.72E+00 | 2.74E+00 | 9.95E-01 | no | | intergenic_variant | MODIFIER |
| 18 | 30679688 | AX-170286958 | 0 | G | T | 0.061 | -2.26E-03 | 3.02E-01 | 2.72E+00 | 2.74E+00 | 9.95E-01 | no | | intergenic_variant | MODIFIER |
| 18 | 30711150 | AX-97799414 | 0 | C | T | 0.061 | -2.26E-03 | 3.02E-01 | 2.72E+00 | 2.74E+00 | 9.95E-01 | no | | intergenic_variant | MODIFIER |
| 18 | 30720849 | AX-97851693 | 0 | T | C | 0.061 | -2.26E-03 | 3.02E-01 | 2.72E+00 | 2.74E+00 | 9.95E-01 | no | | intergenic_variant | MODIFIER |
| 18 | 30720901 | AX-170531592 | 0 | C | A | 0.061 | -2.26E-03 | 3.02E-01 | 2.72E+00 | 2.74E+00 | 9.95E-01 | no | | intergenic_variant | MODIFIER |
| 18 | 30725627 | AX-170318952 | 0 | T | C | 0.061 | -2.26E-03 | 3.02E-01 | 2.72E+00 | 2.74E+00 | 9.95E-01 | no | | intergenic_variant | MODIFIER |
| 18 | 30751693 | AX-170124966 | 0 | G | A | 0.061 | -2.26E-03 | 3.02E-01 | 2.72E+00 | 2.74E+00 | 9.95E-01 | no | | intergenic_variant | MODIFIER |
| 18 | 30760761 | AX-170477810 | 0 | T | C | 0.061 | -2.26E-03 | 3.02E-01 | 2.72E+00 | 2.74E+00 | 9.95E-01 | no | | intergenic_variant | MODIFIER |
| 18 | 30881425 | AX-170060461 | 0 | T | C | 0.061 | -2.26E-03 | 3.02E-01 | 2.72E+00 | 2.74E+00 | 9.95E-01 | no | | intergenic_variant | MODIFIER |
| 18 | 31261793 | AX-170153742 | 0 | T | C | 0.061 | -2.26E-03 | 3.02E-01 | 2.72E+00 | 2.74E+00 | 9.95E-01 | no | Rit2 | intron_variant,NMD_transcript_variant | MODIFIER |
| 18 | 31262072 | AX-170401286 | 0 | A | G | 0.061 | -2.26E-03 | 3.02E-01 | 2.72E+00 | 2.74E+00 | 9.95E-01 | no | Rit2 | intron_variant,NMD_transcript_variant | MODIFIER |
| 18 | 31263819 | AX-170104664 | 0 | A | G | 0.061 | -2.26E-03 | 3.02E-01 | 2.72E+00 | 2.74E+00 | 9.95E-01 | no | Rit2 | intron_variant,NMD_transcript_variant | MODIFIER |
| 18 | 31272128 | AX-170253950 | 0 | A | G | 0.061 | -2.26E-03 | 3.02E-01 | 2.72E+00 | 2.74E+00 | 9.95E-01 | no | Rit2 | intron_variant,NMD_transcript_variant | MODIFIER |
| 18 | 33221983 | AX-170413172 | 0 | A | G | 0.061 | -2.26E-03 | 3.02E-01 | 2.72E+00 | 2.74E+00 | 9.95E-01 | no | | regulatory_region_variant | MODIFIER |
| 18 | 33222014 | AX-97800371 | 0 | G | A | 0.061 | -2.26E-03 | 3.02E-01 | 2.72E+00 | 2.74E+00 | 9.95E-01 | no | | intergenic_variant | MODIFIER |
| 18 | 33233751 | AX-170060506 | 0 | G | A | 0.061 | -2.26E-03 | 3.02E-01 | 2.72E+00 | 2.74E+00 | 9.95E-01 | no | | intergenic_variant | MODIFIER |
| 18 | 33340168 | AX-170287274 | 0 | T | G | 0.061 | -2.26E-03 | 3.02E-01 | 2.72E+00 | 2.74E+00 | 9.95E-01 | no | | intergenic_variant | MODIFIER |
| 18 | 33351559 | AX-170146343 | 0 | G | A | 0.061 | -2.26E-03 | 3.02E-01 | 2.72E+00 | 2.74E+00 | 9.95E-01 | no | | intergenic_variant | MODIFIER |
| 18 | 33493320 | AX-170197627 | 0 | G | A | 0.061 | -2.26E-03 | 3.02E-01 | 2.72E+00 | 2.74E+00 | 9.95E-01 | no | | intergenic_variant | MODIFIER |
| 18 | 33493666 | AX-97787467 | 0 | G | A | 0.061 | -2.26E-03 | 3.02E-01 | 2.72E+00 | 2.74E+00 | 9.95E-01 | no | | intergenic_variant | MODIFIER |
| 18 | 33495206 | AX-170360898 | 0 | G | C | 0.061 | -2.26E-03 | 3.02E-01 | 2.72E+00 | 2.74E+00 | 9.95E-01 | no | | regulatory_region_variant | MODIFIER |
| 18 | 33495332 | AX-170211897 | 0 | G | A | 0.061 | -2.26E-03 | 3.02E-01 | 2.72E+00 | 2.74E+00 | 9.95E-01 | no | | regulatory_region_variant | MODIFIER |
| 18 | 33506555 | AX-170313469 | 0 | C | G | 0.061 | -2.26E-03 | 3.02E-01 | 2.72E+00 | 2.74E+00 | 9.95E-01 | no | | intergenic_variant | MODIFIER |
| 18 | 33674239 | AX-170248145 | 0 | A | G | 0.061 | -2.26E-03 | 3.02E-01 | 2.72E+00 | 2.74E+00 | 9.95E-01 | no | | intergenic_variant | MODIFIER |
| 18 | 33674405 | AX-170253358 | 0 | G | T | 0.061 | -2.26E-03 | 3.02E-01 | 2.72E+00 | 2.74E+00 | 9.95E-01 | no | | intergenic_variant | MODIFIER |
| 18 | 33692115 | AX-170165680 | 0 | G | C | 0.061 | -2.26E-03 | 3.02E-01 | 2.72E+00 | 2.74E+00 | 9.95E-01 | no | | regulatory_region_variant | MODIFIER |
| 18 | 33692265 | AX-170060517 | 0 | G | C | 0.061 | -2.26E-03 | 3.02E-01 | 2.72E+00 | 2.74E+00 | 9.95E-01 | no | | regulatory_region_variant | MODIFIER |
| 18 | 33708501 | AX-170015501 | 0 | A | G | 0.061 | -2.26E-03 | 3.02E-01 | 2.72E+00 | 2.74E+00 | 9.95E-01 | no | | intergenic_variant | MODIFIER |
| 18 | 33989357 | AX-170382465 | 0 | G | A | 0.061 | -2.26E-03 | 3.02E-01 | 2.72E+00 | 2.74E+00 | 9.95E-01 | no | Epb41l4a | intron_variant | MODIFIER |
| 18 | 34003940 | AX-170060521 | 0 | G | A | 0.061 | -2.26E-03 | 3.02E-01 | 2.72E+00 | 2.74E+00 | 9.95E-01 | no | Epb41l4a | intron_variant | MODIFIER |
| 18 | 34169569 | AX-97856157 | 0 | C | T | 0.061 | -2.26E-03 | 3.02E-01 | 2.72E+00 | 2.74E+00 | 9.95E-01 | no | | intergenic_variant | MODIFIER |
| 18 | 34170475 | AX-170135151 | 0 | A | G | 0.061 | -2.26E-03 | 3.02E-01 | 2.72E+00 | 2.74E+00 | 9.95E-01 | no | | intergenic_variant | MODIFIER |
| 18 | 34193208 | AX-170490657 | 0 | C | T | 0.061 | -2.26E-03 | 3.02E-01 | 2.72E+00 | 2.74E+00 | 9.95E-01 | no | Gm23411 | upstream_gene_variant | MODIFIER |
| 18 | 34212701 | AX-170219476 | 0 | C | T | 0.061 | -2.26E-03 | 3.02E-01 | 2.72E+00 | 2.74E+00 | 9.95E-01 | no | Gm10548 | intron_variant,non_coding_transcript_variant | MODIFIER |
| 18 | 34254536 | AX-170515216 | 0 | A | C | 0.061 | -2.26E-03 | 3.02E-01 | 2.72E+00 | 2.74E+00 | 9.95E-01 | no | Apc | intron_variant | MODIFIER |
| 18 | 34260606 | AX-170015512 | 0 | A | G | 0.061 | -2.26E-03 | 3.02E-01 | 2.72E+00 | 2.74E+00 | 9.95E-01 | no | Apc | intron_variant | MODIFIER |
| 18 | 34266442 | AX-170499252 | 0 | G | T | 0.061 | -2.26E-03 | 3.02E-01 | 2.72E+00 | 2.74E+00 | 9.95E-01 | no | Apc | intron_variant | MODIFIER |
| 18 | 34271727 | AX-97794066 | 0 | C | T | 0.061 | -2.26E-03 | 3.02E-01 | 2.72E+00 | 2.74E+00 | 9.95E-01 | no | Apc | intron_variant | MODIFIER |
| 18 | 34328850 | AX-170381773 | 0 | T | C | 0.061 | -2.26E-03 | 3.02E-01 | 2.72E+00 | 2.74E+00 | 9.95E-01 | no | Srp19 | upstream_gene_variant | MODIFIER |
| 18 | 34333034 | AX-170015514 | 0 | A | G | 0.061 | -2.26E-03 | 3.02E-01 | 2.72E+00 | 2.74E+00 | 9.95E-01 | no | Srp19 | intron_variant | MODIFIER |
| 18 | 34364252 | AX-170142884 | 0 | T | C | 0.061 | -2.26E-03 | 3.02E-01 | 2.72E+00 | 2.74E+00 | 9.95E-01 | no | Reep5 | intron_variant | MODIFIER |
| 18 | 34365293 | AX-170233916 | 0 | G | T | 0.061 | -2.26E-03 | 3.02E-01 | 2.72E+00 | 2.74E+00 | 9.95E-01 | no | Reep5 | intron_variant | MODIFIER |
| 18 | 34365564 | AX-170015506 | 0 | G | A | 0.061 | -2.26E-03 | 3.02E-01 | 2.72E+00 | 2.74E+00 | 9.95E-01 | no | Reep5 | intron_variant | MODIFIER |
| 18 | 34365637 | AX-170150790 | 0 | T | C | 0.061 | -2.26E-03 | 3.02E-01 | 2.72E+00 | 2.74E+00 | 9.95E-01 | no | Reep5 | intron_variant | MODIFIER |
| 18 | 34366332 | AX-97828886 | 0 | C | T | 0.061 | -2.26E-03 | 3.02E-01 | 2.72E+00 | 2.74E+00 | 9.95E-01 | no | Reep5 | intron_variant | MODIFIER |
| 18 | 34385927 | AX-170236977 | 0 | G | T | 0.061 | -2.26E-03 | 3.02E-01 | 2.72E+00 | 2.74E+00 | 9.95E-01 | no | | intergenic_variant | MODIFIER |
| 18 | 34386478 | AX-170211844 | 0 | C | T | 0.061 | -2.26E-03 | 3.02E-01 | 2.72E+00 | 2.74E+00 | 9.95E-01 | no | | intergenic_variant | MODIFIER |
| 18 | 34387358 | AX-170193795 | 0 | G | A | 0.061 | -2.26E-03 | 3.02E-01 | 2.72E+00 | 2.74E+00 | 9.95E-01 | no | | intergenic_variant | MODIFIER |
| 18 | 34394625 | AX-170301544 | 0 | C | T | 0.061 | -2.26E-03 | 3.02E-01 | 2.72E+00 | 2.74E+00 | 9.95E-01 | no | | intergenic_variant | MODIFIER |
| 18 | 34413352 | AX-97835197 | 0 | T | C | 0.061 | -2.26E-03 | 3.02E-01 | 2.72E+00 | 2.74E+00 | 9.95E-01 | no | Pkd2l2 | intron_variant | MODIFIER |

| | | | | | | | | | | | | | | | |
|---|---|---|---|---|---|---|---|---|---|---|---|---|---|---|---|
| 18 | 34423950 | AX-170305030 | 0 | G | A | 0.061 | -2.26E-03 | 3.02E-01 | 2.72E+00 | 2.74E+00 | 9.95E-01 | no | Pkd2l2 | intron_variant | MODIFIER |
| 18 | 34428558 | AX-170015509 | 0 | G | T | 0.061 | -2.26E-03 | 3.02E-01 | 2.72E+00 | 2.74E+00 | 9.95E-01 | no | Pkd2l2 | intron_variant | MODIFIER |
| 18 | 34459419 | AX-170088320 | 0 | G | A | 0.061 | -2.26E-03 | 3.02E-01 | 2.72E+00 | 2.74E+00 | 9.95E-01 | no | Fam13b | intron_variant | MODIFIER |
| 18 | 34472273 | AX-97835194 | 0 | G | A | 0.061 | -2.26E-03 | 3.02E-01 | 2.72E+00 | 2.74E+00 | 9.95E-01 | no | Fam13b | intron_variant | MODIFIER |
| 18 | 34502725 | AX-170162616 | 0 | A | T | 0.061 | -2.26E-03 | 3.02E-01 | 2.72E+00 | 2.74E+00 | 9.95E-01 | no | Fam13b | intron_variant | MODIFIER |
| 18 | 34520210 | AX-170315286 | 0 | T | C | 0.061 | -2.26E-03 | 3.02E-01 | 2.72E+00 | 2.74E+00 | 9.95E-01 | no | | regulatory_region_variant | MODIFIER |
| 18 | 34546321 | AX-170147275 | 0 | A | G | 0.061 | -2.26E-03 | 3.02E-01 | 2.72E+00 | 2.74E+00 | 9.95E-01 | no | Wnt8a | intron_variant | MODIFIER |
| 18 | 34576038 | AX-170265702 | 0 | T | C | 0.061 | -2.26E-03 | 3.02E-01 | 2.72E+00 | 2.74E+00 | 9.95E-01 | no | 4933408B17Rik | downstream_gene_variant | MODIFIER |
| 18 | 34581183 | AX-170489628 | 0 | T | C | 0.061 | -2.26E-03 | 3.02E-01 | 2.72E+00 | 2.74E+00 | 9.95E-01 | no | 4933408B17Rik | intron_variant | MODIFIER |
| 18 | 34582356 | AX-97834421 | 0 | A | G | 0.061 | -2.26E-03 | 3.02E-01 | 2.72E+00 | 2.74E+00 | 9.95E-01 | no | 4933408B17Rik | intron_variant | MODIFIER |
| 18 | 34610037 | AX-170088321 | 0 | G | A | 0.061 | -2.26E-03 | 3.02E-01 | 2.72E+00 | 2.74E+00 | 9.95E-01 | no | Brd8 | intron_variant | MODIFIER |
| 18 | 34634090 | AX-170327583 | 0 | A | G | 0.061 | -2.26E-03 | 3.02E-01 | 2.72E+00 | 2.74E+00 | 9.95E-01 | no | Cdc23 | synonymous_variant | LOW |
| 18 | 34637118 | AX-170416470 | 0 | A | G | 0.061 | -2.26E-03 | 3.02E-01 | 2.72E+00 | 2.74E+00 | 9.95E-01 | no | Cdc23 | synonymous_variant | LOW |
| 18 | 34691151 | AX-170060530 | 0 | C | T | 0.061 | -2.26E-03 | 3.02E-01 | 2.72E+00 | 2.74E+00 | 9.95E-01 | no | Gfra3 | intron_variant | MODIFIER |
| 18 | 34706349 | AX-170364572 | 0 | A | G | 0.061 | -2.26E-03 | 3.02E-01 | 2.72E+00 | 2.74E+00 | 9.95E-01 | no | Gfra3 | intron_variant | MODIFIER |
| 18 | 34813368 | AX-170015516 | 0 | C | T | 0.061 | -2.26E-03 | 3.02E-01 | 2.72E+00 | 2.74E+00 | 9.95E-01 | no | Kdm3b | intron_variant | MODIFIER |
| 18 | 34976483 | AX-97856239 | 0 | A | C | 0.061 | -2.26E-03 | 3.02E-01 | 2.72E+00 | 2.74E+00 | 9.95E-01 | no | | intergenic_variant | MODIFIER |
| 18 | 34983964 | AX-170169612 | 0 | T | C | 0.061 | -2.26E-03 | 3.02E-01 | 2.72E+00 | 2.74E+00 | 9.95E-01 | no | AC131675.3 | intron_variant,non_coding_transcript_variant | MODIFIER |
| 18 | 34984310 | AX-170494958 | 0 | C | T | 0.061 | -2.26E-03 | 3.02E-01 | 2.72E+00 | 2.74E+00 | 9.95E-01 | no | AC131675.3 | intron_variant,non_coding_transcript_variant | MODIFIER |
| 18 | 34987554 | AX-170478530 | 0 | C | T | 0.061 | -2.26E-03 | 3.02E-01 | 2.72E+00 | 2.74E+00 | 9.95E-01 | no | AC131675.3 | intron_variant,non_coding_transcript_variant | MODIFIER |
| 18 | 35023833 | AX-170207161 | 0 | G | A | 0.061 | -2.26E-03 | 3.02E-01 | 2.72E+00 | 2.74E+00 | 9.95E-01 | no | AC131675.3 | intron_variant,non_coding_transcript_variant | MODIFIER |
| 18 | 35038921 | AX-170241506 | 0 | C | T | 0.061 | -2.26E-03 | 3.02E-01 | 2.72E+00 | 2.74E+00 | 9.95E-01 | no | AC131675.3 | intron_variant,non_coding_transcript_variant | MODIFIER |
| 18 | 35040298 | AX-170060535 | 0 | C | T | 0.061 | -2.26E-03 | 3.02E-01 | 2.72E+00 | 2.74E+00 | 9.95E-01 | no | AC131675.3 | intron_variant,non_coding_transcript_variant | MODIFIER |
| 18 | 35041218 | AX-170060537 | 0 | T | C | 0.061 | -2.26E-03 | 3.02E-01 | 2.72E+00 | 2.74E+00 | 9.95E-01 | no | AC131675.3 | intron_variant,non_coding_transcript_variant | MODIFIER |
| 18 | 35051593 | AX-170276224 | 0 | T | C | 0.061 | -2.26E-03 | 3.02E-01 | 2.72E+00 | 2.74E+00 | 9.95E-01 | no | AC131675.3 | intron_variant,non_coding_transcript_variant | MODIFIER |
| 18 | 35058644 | AX-97849415 | 0 | T | C | 0.061 | -2.26E-03 | 3.02E-01 | 2.72E+00 | 2.74E+00 | 9.95E-01 | no | AC131675.3 | intron_variant,non_coding_transcript_variant | MODIFIER |
| 18 | 35080817 | AX-97780748 | 0 | A | G | 0.061 | -2.26E-03 | 3.02E-01 | 2.72E+00 | 2.74E+00 | 9.95E-01 | no | AC131675.3 | intron_variant,non_coding_transcript_variant | MODIFIER |
| 18 | 35106681 | AX-170460114 | 0 | T | G | 0.061 | -2.26E-03 | 3.02E-01 | 2.72E+00 | 2.74E+00 | 9.95E-01 | no | AC131675.3 | intron_variant,non_coding_transcript_variant | MODIFIER |
| 18 | 35132735 | AX-170521382 | 0 | C | T | 0.061 | -2.26E-03 | 3.02E-01 | 2.72E+00 | 2.74E+00 | 9.95E-01 | no | Ctnna1 | intron_variant | MODIFIER |
| 18 | 35134817 | AX-170333184 | 0 | A | G | 0.061 | -2.26E-03 | 3.02E-01 | 2.72E+00 | 2.74E+00 | 9.95E-01 | no | Ctnna1 | intron_variant | MODIFIER |
| 18 | 35148770 | AX-170428743 | 0 | G | A | 0.061 | -2.26E-03 | 3.02E-01 | 2.72E+00 | 2.74E+00 | 9.95E-01 | no | Ctnna1 | intron_variant | MODIFIER |
| 18 | 35162033 | AX-170188339 | 0 | G | A | 0.061 | -2.26E-03 | 3.02E-01 | 2.72E+00 | 2.74E+00 | 9.95E-01 | no | Ctnna1 | intron_variant | MODIFIER |
| 18 | 35172822 | AX-170468914 | 0 | C | T | 0.061 | -2.26E-03 | 3.02E-01 | 2.72E+00 | 2.74E+00 | 9.95E-01 | no | Ctnna1 | intron_variant | MODIFIER |
| 18 | 35173474 | AX-170402764 | 0 | G | A | 0.061 | -2.26E-03 | 3.02E-01 | 2.72E+00 | 2.74E+00 | 9.95E-01 | no | Ctnna1 | intron_variant | MODIFIER |
| 18 | 35990320 | AX-170015525 | 0 | T | C | 0.061 | -2.26E-03 | 3.02E-01 | 2.72E+00 | 2.74E+00 | 9.95E-01 | no | Psd2 | intron_variant | MODIFIER |
| 18 | 35995487 | AX-170015533 | 0 | A | G | 0.061 | -2.26E-03 | 3.02E-01 | 2.72E+00 | 2.74E+00 | 9.95E-01 | no | Psd2 | intron_variant | MODIFIER |
| 18 | 35995948 | AX-170015527 | 0 | T | A | 0.061 | -2.26E-03 | 3.02E-01 | 2.72E+00 | 2.74E+00 | 9.95E-01 | no | Psd2 | intron_variant | MODIFIER |
| 18 | 36004949 | AX-97803052 | 0 | A | G | 0.061 | -2.26E-03 | 3.02E-01 | 2.72E+00 | 2.74E+00 | 9.95E-01 | no | Psd2 | intron_variant | MODIFIER |
| 18 | 36013787 | AX-170279916 | 0 | C | T | 0.061 | -2.26E-03 | 3.02E-01 | 2.72E+00 | 2.74E+00 | 9.95E-01 | no | Nrg2 | downstream_gene_variant | MODIFIER |
| 18 | 36027314 | AX-97834478 | 0 | T | C | 0.061 | -2.26E-03 | 3.02E-01 | 2.72E+00 | 2.74E+00 | 9.95E-01 | no | Nrg2 | downstream_gene_variant | MODIFIER |
| 18 | 36035119 | AX-170015528 | 0 | T | C | 0.061 | -2.26E-03 | 3.02E-01 | 2.72E+00 | 2.74E+00 | 9.95E-01 | no | Nrg2 | intron_variant,non_coding_transcript_variant | MODIFIER |
| 18 | 36681363 | AX-170167879 | 0 | C | T | 0.061 | -2.26E-03 | 3.02E-01 | 2.72E+00 | 2.74E+00 | 9.95E-01 | no | Apbb3 | upstream_gene_variant | MODIFIER |
| 18 | 36681444 | AX-170331411 | 0 | T | C | 0.061 | -2.26E-03 | 3.02E-01 | 2.72E+00 | 2.74E+00 | 9.95E-01 | no | Apbb3 | upstream_gene_variant | MODIFIER |
| 18 | 36687059 | AX-170148771 | 0 | T | C | 0.061 | -2.26E-03 | 3.02E-01 | 2.72E+00 | 2.74E+00 | 9.95E-01 | no | Slc35a4 | downstream_gene_variant | MODIFIER |
| 18 | 36688338 | AX-170297342 | 0 | C | A | 0.061 | -2.26E-03 | 3.02E-01 | 2.72E+00 | 2.74E+00 | 9.95E-01 | no | Slc35a4 | downstream_gene_variant | MODIFIER |
| 18 | 36691275 | AX-97835557 | 0 | A | G | 0.061 | -2.26E-03 | 3.02E-01 | 2.72E+00 | 2.74E+00 | 9.95E-01 | no | E230025N22Rik | intron_variant | MODIFIER |
| 18 | 36721792 | AX-170263428 | 0 | T | A | 0.061 | -2.26E-03 | 3.02E-01 | 2.72E+00 | 2.74E+00 | 9.95E-01 | no | Cd14 | downstream_gene_variant | MODIFIER |
| 18 | 36722195 | AX-97802045 | 0 | C | A | 0.061 | -2.26E-03 | 3.02E-01 | 2.72E+00 | 2.74E+00 | 9.95E-01 | no | Cd14 | downstream_gene_variant | MODIFIER |
| 18 | 36722717 | AX-170262595 | 0 | G | A | 0.061 | -2.26E-03 | 3.02E-01 | 2.72E+00 | 2.74E+00 | 9.95E-01 | no | Cd14 | downstream_gene_variant | MODIFIER |
| 18 | 36737874 | AX-170358333 | 0 | T | C | 0.061 | -2.26E-03 | 3.02E-01 | 2.72E+00 | 2.74E+00 | 9.95E-01 | no | Tmco6 | intron_variant | MODIFIER |
| 18 | 36749065 | AX-170530433 | 0 | T | C | 0.061 | -2.26E-03 | 3.02E-01 | 2.72E+00 | 2.74E+00 | 9.95E-01 | no | Ik | intron_variant | MODIFIER |
| 18 | 36785558 | AX-97805872 | 0 | G | C | 0.061 | -2.26E-03 | 3.02E-01 | 2.72E+00 | 2.74E+00 | 9.95E-01 | no | Hars | upstream_gene_variant | MODIFIER |
| 18 | 36901659 | AX-170135093 | 0 | T | C | 0.061 | -2.26E-03 | 3.02E-01 | 2.72E+00 | 2.74E+00 | 9.95E-01 | no | | intergenic_variant | MODIFIER |
| 18 | 36910050 | AX-170215916 | 0 | G | A | 0.061 | -2.26E-03 | 3.02E-01 | 2.72E+00 | 2.74E+00 | 9.95E-01 | no | | intergenic_variant | MODIFIER |

| | | | | | | | | | | | | | | | |
|---|---|---|---|---|---|---|---|---|---|---|---|---|---|---|---|
| 18 | 37123785 | AX-170015538 | 0 | A | T | 0.061 | -2.26E-03 | 3.02E-01 | 2.72E+00 | 2.74E+00 | 9.95E-01 | no | Pcdhac1 | intron_variant | MODIFIER |
| 18 | 37173034 | AX-170308223 | 0 | T | C | 0.061 | -2.26E-03 | 3.02E-01 | 2.72E+00 | 2.74E+00 | 9.95E-01 | no | Pcdhac1 | intron_variant | MODIFIER |
| 18 | 37279323 | AX-97794284 | 0 | T | C | 0.061 | -2.26E-03 | 3.02E-01 | 2.72E+00 | 2.74E+00 | 9.95E-01 | no | Gm42416 | intron_variant | MODIFIER |
| 18 | 37283251 | AX-170144303 | 0 | C | T | 0.061 | -2.26E-03 | 3.02E-01 | 2.72E+00 | 2.74E+00 | 9.95E-01 | no | Gm42416 | intron_variant | MODIFIER |
| 18 | 37303776 | AX-97800561 | 0 | G | A | 0.061 | -2.26E-03 | 3.02E-01 | 2.72E+00 | 2.74E+00 | 9.95E-01 | no | Pcdhb3 | 3_prime_UTR_variant | MODIFIER |
| 18 | 37303815 | AX-170305250 | 0 | G | C | 0.061 | -2.26E-03 | 3.02E-01 | 2.72E+00 | 2.74E+00 | 9.95E-01 | no | Pcdhb3 | 3_prime_UTR_variant | MODIFIER |
| 18 | 37305693 | AX-170466424 | 0 | A | G | 0.061 | -2.26E-03 | 3.02E-01 | 2.72E+00 | 2.74E+00 | 9.95E-01 | no | Pcdhb3 | downstream_gene_variant | MODIFIER |
| 18 | 37309672 | AX-170243213 | 0 | C | T | 0.061 | -2.26E-03 | 3.02E-01 | 2.72E+00 | 2.74E+00 | 9.95E-01 | no | Pcdhb4 | synonymous_variant | LOW |
| 18 | 37318757 | AX-97843867 | 0 | C | T | 0.061 | -2.26E-03 | 3.02E-01 | 2.72E+00 | 2.74E+00 | 9.95E-01 | no | Pcdhb5 | upstream_gene_variant | MODIFIER |
| 18 | 37332041 | AX-170464295 | 0 | A | C | 0.061 | -2.26E-03 | 3.02E-01 | 2.72E+00 | 2.74E+00 | 9.95E-01 | no | Pcdhb6 | upstream_gene_variant | MODIFIER |
| 18 | 37337114 | AX-170449231 | 0 | A | G | 0.061 | -2.26E-03 | 3.02E-01 | 2.72E+00 | 2.74E+00 | 9.95E-01 | no | Pcdhb7 | upstream_gene_variant | MODIFIER |
| 18 | 37343939 | AX-170448428 | 0 | G | C | 0.061 | -2.26E-03 | 3.02E-01 | 2.72E+00 | 2.74E+00 | 9.95E-01 | no | Pcdhb7 | synonymous_variant | LOW |
| 18 | 37357618 | AX-170450823 | 0 | A | C | 0.061 | -2.26E-03 | 3.02E-01 | 2.72E+00 | 2.74E+00 | 9.95E-01 | no | Pcdhb8 | 3_prime_UTR_variant | MODIFIER |
| 18 | 37401921 | AX-170338678 | 0 | G | A | 0.061 | -2.26E-03 | 3.02E-01 | 2.72E+00 | 2.74E+00 | 9.95E-01 | no | Pcdhb9 | missense_variant | MODERATE |
| 18 | 37403651 | AX-170396523 | 0 | A | G | 0.061 | -2.26E-03 | 3.02E-01 | 2.72E+00 | 2.74E+00 | 9.95E-01 | no | Pcdhb9 | 3_prime_UTR_variant | MODIFIER |
| 18 | 37403809 | AX-170506145 | 0 | T | C | 0.061 | -2.26E-03 | 3.02E-01 | 2.72E+00 | 2.74E+00 | 9.95E-01 | no | Pcdhb9 | 3_prime_UTR_variant | MODIFIER |
| 18 | 37404787 | AX-170445507 | 0 | G | C | 0.061 | -2.26E-03 | 3.02E-01 | 2.72E+00 | 2.74E+00 | 9.95E-01 | no | Pcdhb9 | downstream_gene_variant | MODIFIER |
| 18 | 37404901 | AX-170398688 | 0 | T | C | 0.061 | -2.26E-03 | 3.02E-01 | 2.72E+00 | 2.74E+00 | 9.95E-01 | no | Pcdhb9 | downstream_gene_variant | MODIFIER |
| 18 | 37405381 | AX-97819524 | 0 | T | C | 0.061 | -2.26E-03 | 3.02E-01 | 2.72E+00 | 2.74E+00 | 9.95E-01 | no | Pcdhb9 | downstream_gene_variant | MODIFIER |
| 18 | 37963392 | AX-170060591 | 0 | G | A | 0.061 | -2.26E-03 | 3.02E-01 | 2.72E+00 | 2.74E+00 | 9.95E-01 | no | Fchsd1 | synonymous_variant | LOW |
| 18 | 37996113 | AX-170299968 | 0 | A | T | 0.061 | -2.26E-03 | 3.02E-01 | 2.72E+00 | 2.74E+00 | 9.95E-01 | no | Arap3 | intron_variant | MODIFIER |
| 18 | 37996160 | AX-97857030 | 0 | C | T | 0.061 | -2.26E-03 | 3.02E-01 | 2.72E+00 | 2.74E+00 | 9.95E-01 | no | Arap3 | intron_variant | MODIFIER |
| 18 | 38001461 | AX-170491329 | 0 | T | G | 0.061 | -2.26E-03 | 3.02E-01 | 2.72E+00 | 2.74E+00 | 9.95E-01 | no | Arap3 | upstream_gene_variant | MODIFIER |
| 18 | 38015568 | AX-170374512 | 0 | G | A | 0.061 | -2.26E-03 | 3.02E-01 | 2.72E+00 | 2.74E+00 | 9.95E-01 | no | | regulatory_region_variant | MODIFIER |
| 18 | 38017510 | AX-170263308 | 0 | C | T | 0.061 | -2.26E-03 | 3.02E-01 | 2.72E+00 | 2.74E+00 | 9.95E-01 | no | | regulatory_region_variant | MODIFIER |
| 18 | 38025387 | AX-170274959 | 0 | T | A | 0.061 | -2.26E-03 | 3.02E-01 | 2.72E+00 | 2.74E+00 | 9.95E-01 | no | AC121886.1 | intron_variant,non_coding_transcript_variant | MODIFIER |
| 18 | 38041564 | AX-170187313 | 0 | A | G | 0.061 | -2.26E-03 | 3.02E-01 | 2.72E+00 | 2.74E+00 | 9.95E-01 | no | AC121886.1 | intron_variant,non_coding_transcript_variant | MODIFIER |
| 18 | 38054549 | AX-170306631 | 0 | C | T | 0.061 | -2.26E-03 | 3.02E-01 | 2.72E+00 | 2.74E+00 | 9.95E-01 | no | AC121886.1 | intron_variant,non_coding_transcript_variant | MODIFIER |
| 18 | 38092359 | AX-170141758 | 0 | A | G | 0.061 | -2.26E-03 | 3.02E-01 | 2.72E+00 | 2.74E+00 | 9.95E-01 | no | | intergenic_variant | MODIFIER |
| 18 | 38095447 | AX-170274966 | 0 | T | C | 0.061 | -2.26E-03 | 3.02E-01 | 2.72E+00 | 2.74E+00 | 9.95E-01 | no | | regulatory_region_variant | MODIFIER |
| 18 | 38111771 | AX-170299133 | 0 | T | A | 0.061 | -2.26E-03 | 3.02E-01 | 2.72E+00 | 2.74E+00 | 9.95E-01 | no | | regulatory_region_variant | MODIFIER |
| 18 | 38120151 | AX-170262925 | 0 | T | G | 0.061 | -2.26E-03 | 3.02E-01 | 2.72E+00 | 2.74E+00 | 9.95E-01 | no | | regulatory_region_variant | MODIFIER |
| 18 | 38120632 | AX-97834552 | 0 | C | T | 0.061 | -2.26E-03 | 3.02E-01 | 2.72E+00 | 2.74E+00 | 9.95E-01 | no | | intergenic_variant | MODIFIER |
| 18 | 38148664 | AX-170415561 | 0 | A | G | 0.061 | -2.26E-03 | 3.02E-01 | 2.72E+00 | 2.74E+00 | 9.95E-01 | no | | regulatory_region_variant | MODIFIER |
| 18 | 38156950 | AX-170217199 | 0 | G | T | 0.061 | -2.26E-03 | 3.02E-01 | 2.72E+00 | 2.74E+00 | 9.95E-01 | no | | intergenic_variant | MODIFIER |
| 18 | 38164267 | AX-170175988 | 0 | G | C | 0.061 | -2.26E-03 | 3.02E-01 | 2.72E+00 | 2.74E+00 | 9.95E-01 | no | | regulatory_region_variant | MODIFIER |
| 18 | 38165368 | AX-170329281 | 0 | T | C | 0.061 | -2.26E-03 | 3.02E-01 | 2.72E+00 | 2.74E+00 | 9.95E-01 | no | | regulatory_region_variant | MODIFIER |
| 18 | 38165503 | AX-170377990 | 0 | A | T | 0.061 | -2.26E-03 | 3.02E-01 | 2.72E+00 | 2.74E+00 | 9.95E-01 | no | | regulatory_region_variant | MODIFIER |
| 18 | 38165805 | AX-170235666 | 0 | T | C | 0.061 | -2.26E-03 | 3.02E-01 | 2.72E+00 | 2.74E+00 | 9.95E-01 | no | | intergenic_variant | MODIFIER |
| 18 | 38167129 | AX-170106506 | 0 | A | G | 0.061 | -2.26E-03 | 3.02E-01 | 2.72E+00 | 2.74E+00 | 9.95E-01 | no | | intergenic_variant | MODIFIER |
| 18 | 38168744 | AX-170206627 | 0 | A | G | 0.061 | -2.26E-03 | 3.02E-01 | 2.72E+00 | 2.74E+00 | 9.95E-01 | no | | intergenic_variant | MODIFIER |
| 18 | 38168921 | AX-170276660 | 0 | A | T | 0.061 | -2.26E-03 | 3.02E-01 | 2.72E+00 | 2.74E+00 | 9.95E-01 | no | | intergenic_variant | MODIFIER |
| 18 | 38177036 | AX-170364872 | 0 | A | C | 0.061 | -2.26E-03 | 3.02E-01 | 2.72E+00 | 2.74E+00 | 9.95E-01 | no | | regulatory_region_variant | MODIFIER |
| 18 | 38180411 | AX-170531994 | 0 | A | G | 0.061 | -2.26E-03 | 3.02E-01 | 2.72E+00 | 2.74E+00 | 9.95E-01 | no | | regulatory_region_variant | MODIFIER |
| 18 | 38220135 | AX-170190788 | 0 | G | A | 0.061 | -2.26E-03 | 3.02E-01 | 2.72E+00 | 2.74E+00 | 9.95E-01 | no | 2010320O07Rik | intron_variant,non_coding_transcript_variant | MODIFIER |
| 18 | 38250597 | AX-170221894 | 0 | G | A | 0.061 | -2.26E-03 | 3.02E-01 | 2.72E+00 | 2.74E+00 | 9.95E-01 | no | Dele1 | intron_variant | MODIFIER |
| 18 | 38253311 | AX-170015552 | 0 | T | G | 0.061 | -2.26E-03 | 3.02E-01 | 2.72E+00 | 2.74E+00 | 9.95E-01 | no | Dele1 | intron_variant | MODIFIER |
| 18 | 38268741 | AX-170015551 | 0 | C | G | 0.061 | -2.26E-03 | 3.02E-01 | 2.72E+00 | 2.74E+00 | 9.95E-01 | no | Pcdh12 | 3_prime_UTR_variant | MODIFIER |
| 18 | 38610255 | AX-170406294 | 0 | A | G | 0.061 | -2.26E-03 | 3.02E-01 | 2.72E+00 | 2.74E+00 | 9.95E-01 | no | AC133601.1 | intron_variant,non_coding_transcript_variant | MODIFIER |
| 18 | 38612014 | AX-170356762 | 0 | A | G | 0.061 | -2.26E-03 | 3.02E-01 | 2.72E+00 | 2.74E+00 | 9.95E-01 | no | AC133601.1 | intron_variant,non_coding_transcript_variant | MODIFIER |
| 18 | 38612266 | AX-97793040 | 0 | A | G | 0.061 | -2.26E-03 | 3.02E-01 | 2.72E+00 | 2.74E+00 | 9.95E-01 | no | AC133601.1 | intron_variant,non_coding_transcript_variant | MODIFIER |
| 18 | 38637079 | AX-170474553 | 0 | T | C | 0.061 | -2.26E-03 | 3.02E-01 | 2.72E+00 | 2.74E+00 | 9.95E-01 | no | AC133601.1 | intron_variant,non_coding_transcript_variant | MODIFIER |
| 18 | 38653114 | AX-170401074 | 0 | C | T | 0.061 | -2.26E-03 | 3.02E-01 | 2.72E+00 | 2.74E+00 | 9.95E-01 | no | AC133601.1 | intron_variant,non_coding_transcript_variant | MODIFIER |
| 18 | 38689852 | AX-170510863 | 0 | A | G | 0.061 | -2.26E-03 | 3.02E-01 | 2.72E+00 | 2.74E+00 | 9.95E-01 | no | AC133601.1 | intron_variant,non_coding_transcript_variant | MODIFIER |

| | | | | | | | | | | | | | | | |
|---|---|---|---|---|---|---|---|---|---|---|---|---|---|---|---|
| 18 | 38690281 | AX-170412092 | 0 | T | C | 0.061 | -2.26E-03 | 3.02E-01 | 2.72E+00 | 2.74E+00 | 9.95E-01 | no | AC133601.1 | intron_variant,non_coding_transcript_variant | MODIFIER |
| 18 | 38697316 | AX-170180442 | 0 | T | C | 0.061 | -2.26E-03 | 3.02E-01 | 2.72E+00 | 2.74E+00 | 9.95E-01 | no | AC133601.1 | intron_variant,non_coding_transcript_variant | MODIFIER |
| 18 | 38698966 | AX-170439302 | 0 | A | C | 0.061 | -2.26E-03 | 3.02E-01 | 2.72E+00 | 2.74E+00 | 9.95E-01 | no | AC133601.1 | intron_variant,non_coding_transcript_variant | MODIFIER |
| 18 | 38729088 | AX-170166525 | 0 | T | C | 0.061 | -2.26E-03 | 3.02E-01 | 2.72E+00 | 2.74E+00 | 9.95E-01 | no | AC133601.1 | intron_variant,non_coding_transcript_variant | MODIFIER |
| 18 | 38757232 | AX-170397282 | 0 | A | G | 0.061 | -2.26E-03 | 3.02E-01 | 2.72E+00 | 2.74E+00 | 9.95E-01 | no | AC133601.1 | intron_variant,non_coding_transcript_variant | MODIFIER |
| 18 | 38780683 | AX-170116590 | 0 | A | G | 0.061 | -2.26E-03 | 3.02E-01 | 2.72E+00 | 2.74E+00 | 9.95E-01 | no | AC133601.1 | intron_variant,non_coding_transcript_variant | MODIFIER |
| 18 | 38793154 | AX-170365493 | 0 | A | G | 0.061 | -2.26E-03 | 3.02E-01 | 2.72E+00 | 2.74E+00 | 9.95E-01 | no | AC133601.1 | downstream_gene_variant | MODIFIER |
| 18 | 38793282 | AX-97852748 | 0 | T | C | 0.061 | -2.26E-03 | 3.02E-01 | 2.72E+00 | 2.74E+00 | 9.95E-01 | no | AC133601.1 | downstream_gene_variant | MODIFIER |
| 18 | 38800055 | AX-170015556 | 0 | G | A | 0.061 | -2.26E-03 | 3.02E-01 | 2.72E+00 | 2.74E+00 | 9.95E-01 | no | Arhgap26 | intron_variant | MODIFIER |
| 18 | 38876677 | AX-170165078 | 0 | C | T | 0.061 | -2.26E-03 | 3.02E-01 | 2.72E+00 | 2.74E+00 | 9.95E-01 | no | Fgf1 | intron_variant | MODIFIER |
| 18 | 38879804 | AX-97834523 | 0 | T | C | 0.061 | -2.26E-03 | 3.02E-01 | 2.72E+00 | 2.74E+00 | 9.95E-01 | no | Fgf1 | intron_variant | MODIFIER |
| 18 | 38887305 | AX-170226400 | 0 | A | G | 0.061 | -2.26E-03 | 3.02E-01 | 2.72E+00 | 2.74E+00 | 9.95E-01 | no | Fgf1 | intron_variant | MODIFIER |
| 18 | 38919518 | AX-170483893 | 0 | G | A | 0.061 | -2.26E-03 | 3.02E-01 | 2.72E+00 | 2.74E+00 | 9.95E-01 | no | Fgf1 | upstream_gene_variant | MODIFIER |
| 18 | 38933778 | AX-170424966 | 0 | T | C | 0.061 | -2.26E-03 | 3.02E-01 | 2.72E+00 | 2.74E+00 | 9.95E-01 | no | AC110730.1 | intron_variant,non_coding_transcript_variant | MODIFIER |
| 18 | 38942626 | AX-170243271 | 0 | G | A | 0.061 | -2.26E-03 | 3.02E-01 | 2.72E+00 | 2.74E+00 | 9.95E-01 | no | AC110730.1 | intron_variant,non_coding_transcript_variant | MODIFIER |
| 18 | 38943429 | AX-170098554 | 0 | C | A | 0.061 | -2.26E-03 | 3.02E-01 | 2.72E+00 | 2.74E+00 | 9.95E-01 | no | AC110730.1 | intron_variant,non_coding_transcript_variant | MODIFIER |
| 18 | 38949374 | AX-170506812 | 0 | T | A | 0.061 | -2.26E-03 | 3.02E-01 | 2.72E+00 | 2.74E+00 | 9.95E-01 | no | AC110730.1 | intron_variant,non_coding_transcript_variant | MODIFIER |
| 18 | 38951961 | AX-170407299 | 0 | C | T | 0.061 | -2.26E-03 | 3.02E-01 | 2.72E+00 | 2.74E+00 | 9.95E-01 | no | AC110730.1 | intron_variant,non_coding_transcript_variant | MODIFIER |
| 18 | 38955279 | AX-97824886 | 0 | T | C | 0.061 | -2.26E-03 | 3.02E-01 | 2.72E+00 | 2.74E+00 | 9.95E-01 | no | AC110730.1 | intron_variant,non_coding_transcript_variant | MODIFIER |
| 18 | 38955651 | AX-170326829 | 0 | T | C | 0.061 | -2.26E-03 | 3.02E-01 | 2.72E+00 | 2.74E+00 | 9.95E-01 | no | AC110730.1 | intron_variant,non_coding_transcript_variant | MODIFIER |
| 18 | 38955683 | AX-170359218 | 0 | G | A | 0.061 | -2.26E-03 | 3.02E-01 | 2.72E+00 | 2.74E+00 | 9.95E-01 | no | AC110730.1 | intron_variant,non_coding_transcript_variant | MODIFIER |
| 18 | 38956047 | AX-170172201 | 0 | T | A | 0.061 | -2.26E-03 | 3.02E-01 | 2.72E+00 | 2.74E+00 | 9.95E-01 | no | AC110730.1 | intron_variant,non_coding_transcript_variant | MODIFIER |
| 18 | 38959229 | AX-170154536 | 0 | G | C | 0.061 | -2.26E-03 | 3.02E-01 | 2.72E+00 | 2.74E+00 | 9.95E-01 | no | AC110730.1 | intron_variant,non_coding_transcript_variant | MODIFIER |
| 18 | 38961489 | AX-170256781 | 0 | A | G | 0.061 | -2.26E-03 | 3.02E-01 | 2.72E+00 | 2.74E+00 | 9.95E-01 | no | AC110730.1 | intron_variant,non_coding_transcript_variant | MODIFIER |
| 18 | 38980395 | AX-170106098 | 0 | A | T | 0.061 | -2.26E-03 | 3.02E-01 | 2.72E+00 | 2.74E+00 | 9.95E-01 | no | Arhgap26 | intron_variant | MODIFIER |
| 18 | 38988135 | AX-170060618 | 0 | C | A | 0.061 | -2.26E-03 | 3.02E-01 | 2.72E+00 | 2.74E+00 | 9.95E-01 | no | Arhgap26 | intron_variant | MODIFIER |
| 18 | 39001981 | AX-170535885 | 0 | T | C | 0.061 | -2.26E-03 | 3.02E-01 | 2.72E+00 | 2.74E+00 | 9.95E-01 | no | Arhgap26 | intron_variant | MODIFIER |
| 18 | 39126224 | AX-97835223 | 0 | T | C | 0.061 | -2.26E-03 | 3.02E-01 | 2.72E+00 | 2.74E+00 | 9.95E-01 | no | Arhgap26 | intron_variant | MODIFIER |
| 18 | 39126396 | AX-170269291 | 0 | G | A | 0.061 | -2.26E-03 | 3.02E-01 | 2.72E+00 | 2.74E+00 | 9.95E-01 | no | Arhgap26 | intron_variant | MODIFIER |
| 18 | 39131213 | AX-170321139 | 0 | C | T | 0.061 | -2.26E-03 | 3.02E-01 | 2.72E+00 | 2.74E+00 | 9.95E-01 | no | Arhgap26 | intron_variant | MODIFIER |
| 18 | 39150810 | AX-170498533 | 0 | T | C | 0.061 | -2.26E-03 | 3.02E-01 | 2.72E+00 | 2.74E+00 | 9.95E-01 | no | Arhgap26 | intron_variant | MODIFIER |
| 18 | 39150895 | AX-170379483 | 0 | A | G | 0.061 | -2.26E-03 | 3.02E-01 | 2.72E+00 | 2.74E+00 | 9.95E-01 | no | Arhgap26 | intron_variant | MODIFIER |
| 18 | 39170422 | AX-170305218 | 0 | T | C | 0.061 | -2.26E-03 | 3.02E-01 | 2.72E+00 | 2.74E+00 | 9.95E-01 | no | Arhgap26 | intron_variant | MODIFIER |
| 18 | 39202012 | AX-170241939 | 0 | A | G | 0.061 | -2.26E-03 | 3.02E-01 | 2.72E+00 | 2.74E+00 | 9.95E-01 | no | Arhgap26 | intron_variant | MODIFIER |
| 18 | 39202267 | AX-170404350 | 0 | T | G | 0.061 | -2.26E-03 | 3.02E-01 | 2.72E+00 | 2.74E+00 | 9.95E-01 | no | Arhgap26 | intron_variant | MODIFIER |
| 18 | 39203157 | AX-170530930 | 0 | C | T | 0.061 | -2.26E-03 | 3.02E-01 | 2.72E+00 | 2.74E+00 | 9.95E-01 | no | Arhgap26 | intron_variant | MODIFIER |
| 18 | 39203576 | AX-170386600 | 0 | C | T | 0.061 | -2.26E-03 | 3.02E-01 | 2.72E+00 | 2.74E+00 | 9.95E-01 | no | Arhgap26 | intron_variant | MODIFIER |
| 18 | 39216632 | AX-97789073 | 0 | C | A | 0.061 | -2.26E-03 | 3.02E-01 | 2.72E+00 | 2.74E+00 | 9.95E-01 | no | Arhgap26 | intron_variant | MODIFIER |
| 18 | 39216687 | AX-170144433 | 0 | T | C | 0.061 | -2.26E-03 | 3.02E-01 | 2.72E+00 | 2.74E+00 | 9.95E-01 | no | Arhgap26 | intron_variant | MODIFIER |
| 18 | 39229758 | AX-170108649 | 0 | G | A | 0.061 | -2.26E-03 | 3.02E-01 | 2.72E+00 | 2.74E+00 | 9.95E-01 | no | Arhgap26 | intron_variant | MODIFIER |
| 18 | 39234489 | AX-170217975 | 0 | T | C | 0.061 | -2.26E-03 | 3.02E-01 | 2.72E+00 | 2.74E+00 | 9.95E-01 | no | Arhgap26 | intron_variant | MODIFIER |
| 18 | 39267831 | AX-97818451 | 0 | A | G | 0.061 | -2.26E-03 | 3.02E-01 | 2.72E+00 | 2.74E+00 | 9.95E-01 | no | Arhgap26 | intron_variant | MODIFIER |
| 18 | 39267947 | AX-170263141 | 0 | A | G | 0.061 | -2.26E-03 | 3.02E-01 | 2.72E+00 | 2.74E+00 | 9.95E-01 | no | Arhgap26 | intron_variant | MODIFIER |
| 18 | 39268465 | AX-170162572 | 0 | C | T | 0.061 | -2.26E-03 | 3.02E-01 | 2.72E+00 | 2.74E+00 | 9.95E-01 | no | Arhgap26 | intron_variant | MODIFIER |
| 18 | 39272660 | AX-170192529 | 0 | T | C | 0.061 | -2.26E-03 | 3.02E-01 | 2.72E+00 | 2.74E+00 | 9.95E-01 | no | Arhgap26 | intron_variant | MODIFIER |
| 18 | 39287653 | AX-170170669 | 0 | G | A | 0.061 | -2.26E-03 | 3.02E-01 | 2.72E+00 | 2.74E+00 | 9.95E-01 | no | Arhgap26 | intron_variant | MODIFIER |
| 18 | 39294618 | AX-97813892 | 0 | G | A | 0.061 | -2.26E-03 | 3.02E-01 | 2.72E+00 | 2.74E+00 | 9.95E-01 | no | Arhgap26 | intron_variant | MODIFIER |
| 18 | 39295863 | AX-170478034 | 0 | C | T | 0.061 | -2.26E-03 | 3.02E-01 | 2.72E+00 | 2.74E+00 | 9.95E-01 | no | Arhgap26 | intron_variant | MODIFIER |
| 18 | 39298279 | AX-170352106 | 0 | A | G | 0.061 | -2.26E-03 | 3.02E-01 | 2.72E+00 | 2.74E+00 | 9.95E-01 | no | Arhgap26 | intron_variant | MODIFIER |
| 18 | 39300608 | AX-170146316 | 0 | C | T | 0.061 | -2.26E-03 | 3.02E-01 | 2.72E+00 | 2.74E+00 | 9.95E-01 | no | Arhgap26 | intron_variant | MODIFIER |
| 18 | 39312750 | AX-170234843 | 0 | A | G | 0.061 | -2.26E-03 | 3.02E-01 | 2.72E+00 | 2.74E+00 | 9.95E-01 | no | Arhgap26 | intron_variant | MODIFIER |
| 18 | 39314984 | AX-170371694 | 0 | G | A | 0.061 | -2.26E-03 | 3.02E-01 | 2.72E+00 | 2.74E+00 | 9.95E-01 | no | Arhgap26 | intron_variant | MODIFIER |
| 18 | 39317070 | AX-170200115 | 0 | A | G | 0.061 | -2.26E-03 | 3.02E-01 | 2.72E+00 | 2.74E+00 | 9.95E-01 | no | Arhgap26 | intron_variant | MODIFIER |
| 18 | 39321265 | AX-170465114 | 0 | G | A | 0.061 | -2.26E-03 | 3.02E-01 | 2.72E+00 | 2.74E+00 | 9.95E-01 | no | Arhgap26 | intron_variant | MODIFIER |

| | | | | | | | | | | | | | | | |
|---|---|---|---|---|---|---|---|---|---|---|---|---|---|---|---|
| 18 | 39324932 | AX-170015559 | 0 | C | T | 0.061 | -2.26E-03 | 3.02E-01 | 2.72E+00 | 2.74E+00 | 9.95E-01 | no | Arhgap26 | intron_variant | MODIFIER |
| 18 | 39336849 | AX-170293609 | 0 | T | A | 0.061 | -2.26E-03 | 3.02E-01 | 2.72E+00 | 2.74E+00 | 9.95E-01 | no | Arhgap26 | intron_variant | MODIFIER |
| 18 | 39338278 | AX-170133639 | 0 | T | C | 0.061 | -2.26E-03 | 3.02E-01 | 2.72E+00 | 2.74E+00 | 9.95E-01 | no | Arhgap26 | intron_variant | MODIFIER |
| 18 | 39345164 | AX-97795304 | 0 | T | C | 0.061 | -2.26E-03 | 3.02E-01 | 2.72E+00 | 2.74E+00 | 9.95E-01 | no | Arhgap26 | intron_variant | MODIFIER |
| 18 | 39377280 | AX-170015558 | 0 | C | T | 0.061 | -2.26E-03 | 3.02E-01 | 2.72E+00 | 2.74E+00 | 9.95E-01 | no | Arhgap26 | downstream_gene_variant | MODIFIER |
| 18 | 39383293 | AX-170102924 | 0 | A | G | 0.061 | -2.26E-03 | 3.02E-01 | 2.72E+00 | 2.74E+00 | 9.95E-01 | no | Gm15337 | intron_variant,non_coding_transcript_variant | MODIFIER |
| 18 | 39383405 | AX-170015561 | 0 | C | A | 0.061 | -2.26E-03 | 3.02E-01 | 2.72E+00 | 2.74E+00 | 9.95E-01 | no | Gm15337 | intron_variant,non_coding_transcript_variant | MODIFIER |
| 18 | 39383491 | AX-170015560 | 0 | C | G | 0.061 | -2.26E-03 | 3.02E-01 | 2.72E+00 | 2.74E+00 | 9.95E-01 | no | Gm15337 | intron_variant,non_coding_transcript_variant | MODIFIER |
| 18 | 39388464 | AX-170117121 | 0 | C | T | 0.061 | -2.26E-03 | 3.02E-01 | 2.72E+00 | 2.74E+00 | 9.95E-01 | no | Gm15337 | intron_variant,non_coding_transcript_variant | MODIFIER |
| 18 | 39388570 | AX-170493861 | 0 | C | T | 0.061 | -2.26E-03 | 3.02E-01 | 2.72E+00 | 2.74E+00 | 9.95E-01 | no | Gm15337 | intron_variant,non_coding_transcript_variant | MODIFIER |
| 18 | 39389142 | AX-170456215 | 0 | T | C | 0.061 | -2.26E-03 | 3.02E-01 | 2.72E+00 | 2.74E+00 | 9.95E-01 | no | Gm15337 | intron_variant,non_coding_transcript_variant | MODIFIER |
| 18 | 39393426 | AX-97827936 | 0 | C | A | 0.061 | -2.26E-03 | 3.02E-01 | 2.72E+00 | 2.74E+00 | 9.95E-01 | no | Gm15337 | upstream_gene_variant | MODIFIER |
| 18 | 39406802 | AX-170534110 | 0 | C | T | 0.061 | -2.26E-03 | 3.02E-01 | 2.72E+00 | 2.74E+00 | 9.95E-01 | no | Nr3c1 | downstream_gene_variant | MODIFIER |
| 18 | 39416469 | AX-170309472 | 0 | C | T | 0.061 | -2.26E-03 | 3.02E-01 | 2.72E+00 | 2.74E+00 | 9.95E-01 | no | Nr3c1 | intron_variant | MODIFIER |
| 18 | 39451059 | AX-170326909 | 0 | C | A | 0.061 | -2.26E-03 | 3.02E-01 | 2.72E+00 | 2.74E+00 | 9.95E-01 | no | Nr3c1 | intron_variant | MODIFIER |
| 18 | 39461683 | AX-170109260 | 0 | T | C | 0.061 | -2.26E-03 | 3.02E-01 | 2.72E+00 | 2.74E+00 | 9.95E-01 | no | Nr3c1 | intron_variant | MODIFIER |
| 18 | 39461928 | AX-170411375 | 0 | G | A | 0.061 | -2.26E-03 | 3.02E-01 | 2.72E+00 | 2.74E+00 | 9.95E-01 | no | Nr3c1 | intron_variant | MODIFIER |
| 18 | 39593569 | AX-170349925 | 0 | A | G | 0.061 | -2.26E-03 | 3.02E-01 | 2.72E+00 | 2.74E+00 | 9.95E-01 | no | | regulatory_region_variant | MODIFIER |
| 18 | 39673749 | AX-170257081 | 0 | T | C | 0.061 | -2.26E-03 | 3.02E-01 | 2.72E+00 | 2.74E+00 | 9.95E-01 | no | AC124518.1 | intron_variant,non_coding_transcript_variant | MODIFIER |
| 18 | 39693767 | AX-170391748 | 0 | A | G | 0.061 | -2.26E-03 | 3.02E-01 | 2.72E+00 | 2.74E+00 | 9.95E-01 | no | AC124518.1 | intron_variant,non_coding_transcript_variant | MODIFIER |
| 18 | 39695528 | AX-170390423 | 0 | T | C | 0.061 | -2.26E-03 | 3.02E-01 | 2.72E+00 | 2.74E+00 | 9.95E-01 | no | AC124518.1 | intron_variant,non_coding_transcript_variant | MODIFIER |
| 18 | 39701606 | AX-170460988 | 0 | A | G | 0.061 | -2.26E-03 | 3.02E-01 | 2.72E+00 | 2.74E+00 | 9.95E-01 | no | AC124518.1 | intron_variant,non_coding_transcript_variant | MODIFIER |
| 18 | 39714737 | AX-170416269 | 0 | T | C | 0.061 | -2.26E-03 | 3.02E-01 | 2.72E+00 | 2.74E+00 | 9.95E-01 | no | AC124518.1 | intron_variant,non_coding_transcript_variant | MODIFIER |
| 18 | 39718319 | AX-170385357 | 0 | C | T | 0.061 | -2.26E-03 | 3.02E-01 | 2.72E+00 | 2.74E+00 | 9.95E-01 | no | AC124518.1 | intron_variant,non_coding_transcript_variant | MODIFIER |
| 18 | 39720815 | AX-170382168 | 0 | T | C | 0.061 | -2.26E-03 | 3.02E-01 | 2.72E+00 | 2.74E+00 | 9.95E-01 | no | AC124518.1 | intron_variant,non_coding_transcript_variant | MODIFIER |
| 18 | 39726290 | AX-97802342 | 0 | C | T | 0.061 | -2.26E-03 | 3.02E-01 | 2.72E+00 | 2.74E+00 | 9.95E-01 | no | AC124518.1 | intron_variant,non_coding_transcript_variant | MODIFIER |
| 18 | 39782488 | AX-97828290 | 0 | T | C | 0.061 | -2.26E-03 | 3.02E-01 | 2.72E+00 | 2.74E+00 | 9.95E-01 | no | | intergenic_variant | MODIFIER |
| 18 | 39841859 | AX-170227138 | 0 | G | A | 0.061 | -2.26E-03 | 3.02E-01 | 2.72E+00 | 2.74E+00 | 9.95E-01 | no | | intergenic_variant | MODIFIER |
| 18 | 39854047 | AX-170116189 | 0 | C | T | 0.061 | -2.26E-03 | 3.02E-01 | 2.72E+00 | 2.74E+00 | 9.95E-01 | no | | intergenic_variant | MODIFIER |
| 18 | 39855173 | AX-170233363 | 0 | C | A | 0.061 | -2.26E-03 | 3.02E-01 | 2.72E+00 | 2.74E+00 | 9.95E-01 | no | | intergenic_variant | MODIFIER |
| 18 | 39873962 | AX-170389650 | 0 | G | A | 0.061 | -2.26E-03 | 3.02E-01 | 2.72E+00 | 2.74E+00 | 9.95E-01 | no | | regulatory_region_variant | MODIFIER |
| 18 | 39911833 | AX-170015569 | 0 | T | G | 0.061 | -2.26E-03 | 3.02E-01 | 2.72E+00 | 2.74E+00 | 9.95E-01 | no | AC141633.1 | intron_variant,non_coding_transcript_variant | MODIFIER |
| 18 | 39912489 | AX-97807020 | 0 | T | C | 0.061 | -2.26E-03 | 3.02E-01 | 2.72E+00 | 2.74E+00 | 9.95E-01 | no | AC141633.1 | intron_variant,non_coding_transcript_variant | MODIFIER |
| 18 | 40320796 | AX-170363821 | 0 | G | A | 0.061 | -2.26E-03 | 3.02E-01 | 2.72E+00 | 2.74E+00 | 9.95E-01 | no | Kctd16 | intron_variant | MODIFIER |
| 18 | 40321909 | AX-170493570 | 0 | A | G | 0.061 | -2.26E-03 | 3.02E-01 | 2.72E+00 | 2.74E+00 | 9.95E-01 | no | Kctd16 | intron_variant | MODIFIER |
| 18 | 40322143 | AX-170215744 | 0 | A | T | 0.061 | -2.26E-03 | 3.02E-01 | 2.72E+00 | 2.74E+00 | 9.95E-01 | no | Kctd16 | intron_variant | MODIFIER |
| 18 | 40331404 | AX-170491267 | 0 | T | C | 0.061 | -2.26E-03 | 3.02E-01 | 2.72E+00 | 2.74E+00 | 9.95E-01 | no | Kctd16 | intron_variant | MODIFIER |
| 18 | 40331454 | AX-170369652 | 0 | A | T | 0.061 | -2.26E-03 | 3.02E-01 | 2.72E+00 | 2.74E+00 | 9.95E-01 | no | Kctd16 | intron_variant | MODIFIER |
| 18 | 40371408 | AX-170527769 | 0 | G | A | 0.061 | -2.26E-03 | 3.02E-01 | 2.72E+00 | 2.74E+00 | 9.95E-01 | no | Kctd16 | intron_variant | MODIFIER |
| 18 | 40375617 | AX-97783206 | 0 | G | A | 0.061 | -2.26E-03 | 3.02E-01 | 2.72E+00 | 2.74E+00 | 9.95E-01 | no | Kctd16 | intron_variant | MODIFIER |
| 18 | 40378960 | AX-170499972 | 0 | A | G | 0.061 | -2.26E-03 | 3.02E-01 | 2.72E+00 | 2.74E+00 | 9.95E-01 | no | Kctd16 | intron_variant | MODIFIER |
| 18 | 40387691 | AX-170159753 | 0 | A | C | 0.061 | -2.26E-03 | 3.02E-01 | 2.72E+00 | 2.74E+00 | 9.95E-01 | no | Kctd16 | intron_variant | MODIFIER |
| 18 | 40399413 | AX-170158394 | 0 | A | T | 0.061 | -2.26E-03 | 3.02E-01 | 2.72E+00 | 2.74E+00 | 9.95E-01 | no | Kctd16 | intron_variant | MODIFIER |
| 18 | 40407023 | AX-97845395 | 0 | T | C | 0.061 | -2.26E-03 | 3.02E-01 | 2.72E+00 | 2.74E+00 | 9.95E-01 | no | Kctd16 | intron_variant | MODIFIER |
| 18 | 40410453 | AX-170489486 | 0 | A | G | 0.061 | -2.26E-03 | 3.02E-01 | 2.72E+00 | 2.74E+00 | 9.95E-01 | no | Kctd16 | intron_variant | MODIFIER |
| 18 | 40416690 | AX-170015571 | 0 | G | A | 0.061 | -2.26E-03 | 3.02E-01 | 2.72E+00 | 2.74E+00 | 9.95E-01 | no | Kctd16 | intron_variant | MODIFIER |
| 18 | 40447889 | AX-170308098 | 0 | T | C | 0.061 | -2.26E-03 | 3.02E-01 | 2.72E+00 | 2.74E+00 | 9.95E-01 | no | Kctd16 | intron_variant | MODIFIER |
| 18 | 40724267 | AX-97846183 | 0 | A | T | 0.061 | -2.26E-03 | 3.02E-01 | 2.72E+00 | 2.74E+00 | 9.95E-01 | no | Rps19-ps13 | upstream_gene_variant | MODIFIER |
| 18 | 40796593 | AX-170130290 | 0 | A | G | 0.061 | -2.26E-03 | 3.02E-01 | 2.72E+00 | 2.74E+00 | 9.95E-01 | no | Gm22515 | downstream_gene_variant | MODIFIER |
| 18 | 40885785 | AX-170280755 | 0 | A | G | 0.061 | -2.26E-03 | 3.02E-01 | 2.72E+00 | 2.74E+00 | 9.95E-01 | no | | intergenic_variant | MODIFIER |
| 18 | 40886884 | AX-170529550 | 0 | T | C | 0.061 | -2.26E-03 | 3.02E-01 | 2.72E+00 | 2.74E+00 | 9.95E-01 | no | | intergenic_variant | MODIFIER |
| 18 | 40887503 | AX-170370284 | 0 | A | G | 0.061 | -2.26E-03 | 3.02E-01 | 2.72E+00 | 2.74E+00 | 9.95E-01 | no | | intergenic_variant | MODIFIER |
| 18 | 40897075 | AX-170276934 | 0 | T | A | 0.061 | -2.26E-03 | 3.02E-01 | 2.72E+00 | 2.74E+00 | 9.95E-01 | no | AC107756.1 | downstream_gene_variant | MODIFIER |
| 18 | 40898370 | AX-170015583 | 0 | G | A | 0.061 | -2.26E-03 | 3.02E-01 | 2.72E+00 | 2.74E+00 | 9.95E-01 | no | AC107756.1 | downstream_gene_variant | MODIFIER |

| | | | | | | | | | | | | | | | | |
|---|---|---|---|---|---|---|---|---|---|---|---|---|---|---|---|---|
| 18 | 40903811 | AX-97844640 | 0 | C | T | 0.061 | -2.26E-03 | 3.02E-01 | 2.72E+00 | 2.74E+00 | 9.95E-01 | no | | | intergenic_variant | MODIFIER |
| 18 | 40921324 | AX-170015580 | 0 | A | T | 0.061 | -2.26E-03 | 3.02E-01 | 2.72E+00 | 2.74E+00 | 9.95E-01 | no | | | intergenic_variant | MODIFIER |
| 18 | 40950651 | AX-170015590 | 0 | T | A | 0.061 | -2.26E-03 | 3.02E-01 | 2.72E+00 | 2.74E+00 | 9.95E-01 | no | | | intergenic_variant | MODIFIER |
| 18 | 40968962 | AX-170113797 | 0 | G | T | 0.061 | -2.26E-03 | 3.02E-01 | 2.72E+00 | 2.74E+00 | 9.95E-01 | no | | | intergenic_variant | MODIFIER |
| 18 | 40974097 | AX-170223836 | 0 | G | C | 0.061 | -2.26E-03 | 3.02E-01 | 2.72E+00 | 2.74E+00 | 9.95E-01 | no | | | intergenic_variant | MODIFIER |
| 18 | 41053091 | AX-170015586 | 0 | A | C | 0.061 | -2.26E-03 | 3.02E-01 | 2.72E+00 | 2.74E+00 | 9.95E-01 | no | | | intergenic_variant | MODIFIER |
| 18 | 41077452 | AX-170431639 | 0 | T | C | 0.061 | -2.26E-03 | 3.02E-01 | 2.72E+00 | 2.74E+00 | 9.95E-01 | no | | | intergenic_variant | MODIFIER |
| 18 | 41082169 | AX-170131968 | 0 | T | G | 0.061 | -2.26E-03 | 3.02E-01 | 2.72E+00 | 2.74E+00 | 9.95E-01 | no | | | intergenic_variant | MODIFIER |
| 18 | 41082593 | AX-97837163 | 0 | A | G | 0.061 | -2.26E-03 | 3.02E-01 | 2.72E+00 | 2.74E+00 | 9.95E-01 | no | | | intergenic_variant | MODIFIER |
| 18 | 41088082 | AX-170015582 | 0 | A | T | 0.061 | -2.26E-03 | 3.02E-01 | 2.72E+00 | 2.74E+00 | 9.95E-01 | no | | | intergenic_variant | MODIFIER |
| 18 | 41088283 | AX-170015588 | 0 | C | T | 0.061 | -2.26E-03 | 3.02E-01 | 2.72E+00 | 2.74E+00 | 9.95E-01 | no | | | intergenic_variant | MODIFIER |
| 18 | 41150755 | AX-97849152 | 0 | T | C | 0.061 | -2.26E-03 | 3.02E-01 | 2.72E+00 | 2.74E+00 | 9.95E-01 | no | | | intergenic_variant | MODIFIER |
| 18 | 41165155 | AX-97798406 | 0 | A | G | 0.061 | -2.26E-03 | 3.02E-01 | 2.72E+00 | 2.74E+00 | 9.95E-01 | no | | | intergenic_variant | MODIFIER |
| 18 | 41187008 | AX-170015584 | 0 | G | A | 0.061 | -2.26E-03 | 3.02E-01 | 2.72E+00 | 2.74E+00 | 9.95E-01 | no | | | intergenic_variant | MODIFIER |
| 18 | 41199764 | AX-170015589 | 0 | G | C | 0.061 | -2.26E-03 | 3.02E-01 | 2.72E+00 | 2.74E+00 | 9.95E-01 | no | | | intergenic_variant | MODIFIER |
| 18 | 41202780 | AX-170322330 | 0 | A | C | 0.061 | -2.26E-03 | 3.02E-01 | 2.72E+00 | 2.74E+00 | 9.95E-01 | no | | | intergenic_variant | MODIFIER |
| 18 | 41203794 | AX-170416792 | 0 | C | A | 0.061 | -2.26E-03 | 3.02E-01 | 2.72E+00 | 2.74E+00 | 9.95E-01 | no | | | intergenic_variant | MODIFIER |
| 18 | 41208190 | AX-170122028 | 0 | A | G | 0.061 | -2.26E-03 | 3.02E-01 | 2.72E+00 | 2.74E+00 | 9.95E-01 | no | | | intergenic_variant | MODIFIER |
| 18 | 41244453 | AX-170118160 | 0 | G | A | 0.061 | -2.26E-03 | 3.02E-01 | 2.72E+00 | 2.74E+00 | 9.95E-01 | no | | | intergenic_variant | MODIFIER |
| 18 | 41264266 | AX-170223178 | 0 | C | T | 0.061 | -2.26E-03 | 3.02E-01 | 2.72E+00 | 2.74E+00 | 9.95E-01 | no | | | intergenic_variant | MODIFIER |
| 18 | 41273928 | AX-170367142 | 0 | A | G | 0.061 | -2.26E-03 | 3.02E-01 | 2.72E+00 | 2.74E+00 | 9.95E-01 | no | | | intergenic_variant | MODIFIER |
| 18 | 41274033 | AX-170369479 | 0 | A | G | 0.061 | -2.26E-03 | 3.02E-01 | 2.72E+00 | 2.74E+00 | 9.95E-01 | no | | | intergenic_variant | MODIFIER |
| 18 | 41292872 | AX-170422058 | 0 | C | T | 0.061 | -2.26E-03 | 3.02E-01 | 2.72E+00 | 2.74E+00 | 9.95E-01 | no | | | intergenic_variant | MODIFIER |
| 18 | 41300020 | AX-170306243 | 0 | T | C | 0.061 | -2.26E-03 | 3.02E-01 | 2.72E+00 | 2.74E+00 | 9.95E-01 | no | | | intergenic_variant | MODIFIER |
| 18 | 41302194 | AX-170088323 | 0 | A | G | 0.061 | -2.26E-03 | 3.02E-01 | 2.72E+00 | 2.74E+00 | 9.95E-01 | no | | | intergenic_variant | MODIFIER |
| 18 | 41316030 | AX-170386891 | 0 | T | C | 0.061 | -2.26E-03 | 3.02E-01 | 2.72E+00 | 2.74E+00 | 9.95E-01 | no | | | intergenic_variant | MODIFIER |
| 18 | 41318603 | AX-97835332 | 0 | A | G | 0.061 | -2.26E-03 | 3.02E-01 | 2.72E+00 | 2.74E+00 | 9.95E-01 | no | | | intergenic_variant | MODIFIER |
| 18 | 41354024 | AX-170392451 | 0 | G | T | 0.061 | -2.26E-03 | 3.02E-01 | 2.72E+00 | 2.74E+00 | 9.95E-01 | no | | | intergenic_variant | MODIFIER |
| 18 | 41358800 | AX-170446276 | 0 | T | C | 0.061 | -2.26E-03 | 3.02E-01 | 2.72E+00 | 2.74E+00 | 9.95E-01 | no | | | intergenic_variant | MODIFIER |
| 18 | 41367310 | AX-170474115 | 0 | C | T | 0.061 | -2.26E-03 | 3.02E-01 | 2.72E+00 | 2.74E+00 | 9.95E-01 | no | | | intergenic_variant | MODIFIER |
| 18 | 41389474 | AX-170088324 | 0 | A | G | 0.061 | -2.26E-03 | 3.02E-01 | 2.72E+00 | 2.74E+00 | 9.95E-01 | no | | | intergenic_variant | MODIFIER |
| 18 | 41401930 | AX-170247054 | 0 | C | T | 0.061 | -2.26E-03 | 3.02E-01 | 2.72E+00 | 2.74E+00 | 9.95E-01 | no | | | intergenic_variant | MODIFIER |
| 18 | 41412900 | AX-170015596 | 0 | C | T | 0.061 | -2.26E-03 | 3.02E-01 | 2.72E+00 | 2.74E+00 | 9.95E-01 | no | | | intergenic_variant | MODIFIER |
| 18 | 41437199 | AX-170179638 | 0 | T | C | 0.061 | -2.26E-03 | 3.02E-01 | 2.72E+00 | 2.74E+00 | 9.95E-01 | no | | | intergenic_variant | MODIFIER |
| 18 | 41447282 | AX-170015592 | 0 | T | G | 0.061 | -2.26E-03 | 3.02E-01 | 2.72E+00 | 2.74E+00 | 9.95E-01 | no | | | intergenic_variant | MODIFIER |
| 18 | 41488453 | AX-170015593 | 0 | A | T | 0.061 | -2.26E-03 | 3.02E-01 | 2.72E+00 | 2.74E+00 | 9.95E-01 | no | | | intergenic_variant | MODIFIER |
| 18 | 41495350 | AX-170015595 | 0 | T | C | 0.061 | -2.26E-03 | 3.02E-01 | 2.72E+00 | 2.74E+00 | 9.95E-01 | no | | | intergenic_variant | MODIFIER |
| 18 | 41500863 | AX-170203133 | 0 | A | G | 0.061 | -2.26E-03 | 3.02E-01 | 2.72E+00 | 2.74E+00 | 9.95E-01 | no | | | intergenic_variant | MODIFIER |
| 18 | 41520443 | AX-97800499 | 0 | T | C | 0.061 | -2.26E-03 | 3.02E-01 | 2.72E+00 | 2.74E+00 | 9.95E-01 | no | | | intergenic_variant | MODIFIER |
| 18 | 41542829 | AX-97856562 | 0 | A | G | 0.061 | -2.26E-03 | 3.02E-01 | 2.72E+00 | 2.74E+00 | 9.95E-01 | no | | | intergenic_variant | MODIFIER |
| 18 | 41545579 | AX-170280149 | 0 | A | G | 0.061 | -2.26E-03 | 3.02E-01 | 2.72E+00 | 2.74E+00 | 9.95E-01 | no | | | intergenic_variant | MODIFIER |
| 18 | 41546007 | AX-170355054 | 0 | G | A | 0.061 | -2.26E-03 | 3.02E-01 | 2.72E+00 | 2.74E+00 | 9.95E-01 | no | | | intergenic_variant | MODIFIER |
| 18 | 41567372 | AX-170123498 | 0 | A | G | 0.061 | -2.26E-03 | 3.02E-01 | 2.72E+00 | 2.74E+00 | 9.95E-01 | no | | | intergenic_variant | MODIFIER |
| 18 | 41567479 | AX-170250655 | 0 | G | T | 0.061 | -2.26E-03 | 3.02E-01 | 2.72E+00 | 2.74E+00 | 9.95E-01 | no | | | intergenic_variant | MODIFIER |
| 18 | 41571533 | AX-97835331 | 0 | G | A | 0.061 | -2.26E-03 | 3.02E-01 | 2.72E+00 | 2.74E+00 | 9.95E-01 | no | | | intergenic_variant | MODIFIER |
| 18 | 41571665 | AX-170241239 | 0 | T | A | 0.061 | -2.26E-03 | 3.02E-01 | 2.72E+00 | 2.74E+00 | 9.95E-01 | no | | | intergenic_variant | MODIFIER |
| 18 | 41576635 | AX-170341225 | 0 | A | T | 0.061 | -2.26E-03 | 3.02E-01 | 2.72E+00 | 2.74E+00 | 9.95E-01 | no | | | intergenic_variant | MODIFIER |
| 18 | 41607566 | AX-170448957 | 0 | A | C | 0.061 | -2.26E-03 | 3.02E-01 | 2.72E+00 | 2.74E+00 | 9.95E-01 | no | | | intergenic_variant | MODIFIER |
| 18 | 41628543 | AX-97835326 | 0 | A | G | 0.061 | -2.26E-03 | 3.02E-01 | 2.72E+00 | 2.74E+00 | 9.95E-01 | no | | | intergenic_variant | MODIFIER |
| 18 | 41628865 | AX-170158750 | 0 | A | C | 0.061 | -2.26E-03 | 3.02E-01 | 2.72E+00 | 2.74E+00 | 9.95E-01 | no | | | intergenic_variant | MODIFIER |
| 18 | 41656073 | AX-170318300 | 0 | T | C | 0.061 | -2.26E-03 | 3.02E-01 | 2.72E+00 | 2.74E+00 | 9.95E-01 | no | | | intergenic_variant | MODIFIER |
| 18 | 41660174 | AX-170226424 | 0 | T | C | 0.061 | -2.26E-03 | 3.02E-01 | 2.72E+00 | 2.74E+00 | 9.95E-01 | no | | | intergenic_variant | MODIFIER |
| 18 | 41700348 | AX-170292077 | 0 | C | A | 0.061 | -2.26E-03 | 3.02E-01 | 2.72E+00 | 2.74E+00 | 9.95E-01 | no | | | intergenic_variant | MODIFIER |
| 18 | 41754732 | AX-170348886 | 0 | C | T | 0.061 | -2.26E-03 | 3.02E-01 | 2.72E+00 | 2.74E+00 | 9.95E-01 | no | | AC154172.2 | intron_variant,non_coding_transcript_variant | MODIFIER |

| | | | | | | | | | | | | | | | |
|---|---|---|---|---|---|---|---|---|---|---|---|---|---|---|---|
| 18 | 41762873 | AX-170060666 | 0 | C | T | 0.061 | -2.26E-03 | 3.02E-01 | 2.72E+00 | 2.74E+00 | 9.95E-01 | no | AC154172.2 | intron_variant,non_coding_transcript_variant | MODIFIER |
| 18 | 41814116 | AX-170262234 | 0 | G | T | 0.061 | -2.26E-03 | 3.02E-01 | 2.72E+00 | 2.74E+00 | 9.95E-01 | no | AC154172.2 | intron_variant,non_coding_transcript_variant | MODIFIER |
| 18 | 41826948 | AX-170120817 | 0 | C | T | 0.061 | -2.26E-03 | 3.02E-01 | 2.72E+00 | 2.74E+00 | 9.95E-01 | no | AC154172.2 | intron_variant,non_coding_transcript_variant | MODIFIER |
| 18 | 41828046 | AX-170474347 | 0 | A | G | 0.061 | -2.26E-03 | 3.02E-01 | 2.72E+00 | 2.74E+00 | 9.95E-01 | no | AC154172.2 | intron_variant,non_coding_transcript_variant | MODIFIER |
| 18 | 41828079 | AX-170184729 | 0 | A | C | 0.061 | -2.26E-03 | 3.02E-01 | 2.72E+00 | 2.74E+00 | 9.95E-01 | no | AC154172.2 | intron_variant,non_coding_transcript_variant | MODIFIER |
| 18 | 41828635 | AX-170221046 | 0 | C | T | 0.061 | -2.26E-03 | 3.02E-01 | 2.72E+00 | 2.74E+00 | 9.95E-01 | no | AC154172.2 | intron_variant,non_coding_transcript_variant | MODIFIER |
| 18 | 41858640 | AX-170120739 | 0 | T | G | 0.061 | -2.26E-03 | 3.02E-01 | 2.72E+00 | 2.74E+00 | 9.95E-01 | no | AC154172.2 | intron_variant,non_coding_transcript_variant | MODIFIER |
| 18 | 41858997 | AX-170107321 | 0 | A | C | 0.061 | -2.26E-03 | 3.02E-01 | 2.72E+00 | 2.74E+00 | 9.95E-01 | no | AC154172.2 | intron_variant,non_coding_transcript_variant | MODIFIER |
| 18 | 41859897 | AX-170339995 | 0 | T | C | 0.061 | -2.26E-03 | 3.02E-01 | 2.72E+00 | 2.74E+00 | 9.95E-01 | no | AC154172.2 | intron_variant,non_coding_transcript_variant | MODIFIER |
| 18 | 41875233 | AX-170122132 | 0 | T | C | 0.061 | -2.26E-03 | 3.02E-01 | 2.72E+00 | 2.74E+00 | 9.95E-01 | no | Prelid2 | downstream_gene_variant | MODIFIER |
| 18 | 41879973 | AX-170463685 | 0 | G | A | 0.061 | -2.26E-03 | 3.02E-01 | 2.72E+00 | 2.74E+00 | 9.95E-01 | no | Prelid2 | intron_variant | MODIFIER |
| 18 | 41883457 | AX-170243144 | 0 | G | T | 0.061 | -2.26E-03 | 3.02E-01 | 2.72E+00 | 2.74E+00 | 9.95E-01 | no | Prelid2 | intron_variant | MODIFIER |
| 18 | 42033189 | AX-170166948 | 0 | A | C | 0.061 | -2.26E-03 | 3.02E-01 | 2.72E+00 | 2.74E+00 | 9.95E-01 | no | | intergenic_variant | MODIFIER |
| 18 | 42033885 | AX-170476109 | 0 | C | T | 0.061 | -2.26E-03 | 3.02E-01 | 2.72E+00 | 2.74E+00 | 9.95E-01 | no | | intergenic_variant | MODIFIER |
| 18 | 42055008 | AX-170473405 | 0 | C | T | 0.061 | -2.26E-03 | 3.02E-01 | 2.72E+00 | 2.74E+00 | 9.95E-01 | no | Sh3rf2 | intron_variant | MODIFIER |
| 18 | 42132761 | AX-97840106 | 0 | T | G | 0.061 | -2.26E-03 | 3.02E-01 | 2.72E+00 | 2.74E+00 | 9.95E-01 | no | Sh3rf2 | intron_variant | MODIFIER |
| 18 | 42163769 | AX-170336205 | 0 | A | G | 0.061 | -2.26E-03 | 3.02E-01 | 2.72E+00 | 2.74E+00 | 9.95E-01 | no | Sh3rf2 | downstream_gene_variant | MODIFIER |
| 18 | 42419212 | AX-170186745 | 0 | A | G | 0.061 | -2.26E-03 | 3.02E-01 | 2.72E+00 | 2.74E+00 | 9.95E-01 | no | | intergenic_variant | MODIFIER |
| 18 | 42420227 | AX-170340134 | 0 | A | G | 0.061 | -2.26E-03 | 3.02E-01 | 2.72E+00 | 2.74E+00 | 9.95E-01 | no | | intergenic_variant | MODIFIER |
| 18 | 42425780 | AX-170060683 | 0 | A | C | 0.061 | -2.26E-03 | 3.02E-01 | 2.72E+00 | 2.74E+00 | 9.95E-01 | no | | intergenic_variant | MODIFIER |
| 18 | 42428607 | AX-170219974 | 0 | T | C | 0.061 | -2.26E-03 | 3.02E-01 | 2.72E+00 | 2.74E+00 | 9.95E-01 | no | | intergenic_variant | MODIFIER |
| 18 | 42433095 | AX-97821560 | 0 | G | T | 0.061 | -2.26E-03 | 3.02E-01 | 2.72E+00 | 2.74E+00 | 9.95E-01 | no | | intergenic_variant | MODIFIER |
| 18 | 42437224 | AX-170470659 | 0 | C | T | 0.061 | -2.26E-03 | 3.02E-01 | 2.72E+00 | 2.74E+00 | 9.95E-01 | no | Gm16415 | upstream_gene_variant | MODIFIER |
| 18 | 42444761 | AX-170180250 | 0 | A | G | 0.061 | -2.26E-03 | 3.02E-01 | 2.72E+00 | 2.74E+00 | 9.95E-01 | no | Gm16415 | downstream_gene_variant | MODIFIER |
| 18 | 42451633 | AX-97786620 | 0 | C | A | 0.061 | -2.26E-03 | 3.02E-01 | 2.72E+00 | 2.74E+00 | 9.95E-01 | no | | intergenic_variant | MODIFIER |
| 18 | 42512900 | AX-170369220 | 0 | C | T | 0.061 | -2.26E-03 | 3.02E-01 | 2.72E+00 | 2.74E+00 | 9.95E-01 | no | Tcerg1 | intron_variant | MODIFIER |
| 18 | 42513008 | AX-170015607 | 0 | A | G | 0.061 | -2.26E-03 | 3.02E-01 | 2.72E+00 | 2.74E+00 | 9.95E-01 | no | Tcerg1 | intron_variant | MODIFIER |
| 18 | 42539633 | AX-170015608 | 0 | T | C | 0.061 | -2.26E-03 | 3.02E-01 | 2.72E+00 | 2.74E+00 | 9.95E-01 | no | Tcerg1 | intron_variant | MODIFIER |
| 18 | 42554582 | AX-97800429 | 0 | G | A | 0.061 | -2.26E-03 | 3.02E-01 | 2.72E+00 | 2.74E+00 | 9.95E-01 | no | Tcerg1 | intron_variant | MODIFIER |
| 18 | 42561758 | AX-170409599 | 0 | C | A | 0.061 | -2.26E-03 | 3.02E-01 | 2.72E+00 | 2.74E+00 | 9.95E-01 | no | Tcerg1 | intron_variant | MODIFIER |
| 18 | 42572355 | AX-170015603 | 0 | G | A | 0.061 | -2.26E-03 | 3.02E-01 | 2.72E+00 | 2.74E+00 | 9.95E-01 | no | Tcerg1 | intron_variant | MODIFIER |
| 18 | 42582123 | AX-170318137 | 0 | T | C | 0.061 | -2.26E-03 | 3.02E-01 | 2.72E+00 | 2.74E+00 | 9.95E-01 | no | Gpr151 | upstream_gene_variant | MODIFIER |
| 18 | 42601249 | AX-170015606 | 0 | T | C | 0.061 | -2.26E-03 | 3.02E-01 | 2.72E+00 | 2.74E+00 | 9.95E-01 | no | AC121603.1 | intron_variant,non_coding_transcript_variant | MODIFIER |
| 18 | 42603270 | AX-170015602 | 0 | G | A | 0.061 | -2.26E-03 | 3.02E-01 | 2.72E+00 | 2.74E+00 | 9.95E-01 | no | AC121603.1 | intron_variant,non_coding_transcript_variant | MODIFIER |
| 18 | 42606954 | AX-170367146 | 0 | A | G | 0.061 | -2.26E-03 | 3.02E-01 | 2.72E+00 | 2.74E+00 | 9.95E-01 | no | AC121603.1 | intron_variant,non_coding_transcript_variant | MODIFIER |
| 18 | 42648721 | AX-170509654 | 0 | A | G | 0.061 | -2.26E-03 | 3.02E-01 | 2.72E+00 | 2.74E+00 | 9.95E-01 | no | Ppp2r2b | synonymous_variant | LOW |
| 18 | 42725388 | AX-170187287 | 0 | C | T | 0.061 | -2.26E-03 | 3.02E-01 | 2.72E+00 | 2.74E+00 | 9.95E-01 | no | Ppp2r2b | intron_variant | MODIFIER |
| 18 | 42732286 | AX-97804318 | 0 | T | C | 0.061 | -2.26E-03 | 3.02E-01 | 2.72E+00 | 2.74E+00 | 9.95E-01 | no | Ppp2r2b | intron_variant | MODIFIER |
| 18 | 42748783 | AX-170348156 | 0 | A | G | 0.061 | -2.26E-03 | 3.02E-01 | 2.72E+00 | 2.74E+00 | 9.95E-01 | no | Ppp2r2b | intron_variant | MODIFIER |
| 18 | 42845901 | AX-170060687 | 0 | A | G | 0.061 | -2.26E-03 | 3.02E-01 | 2.72E+00 | 2.74E+00 | 9.95E-01 | no | Ppp2r2b | intron_variant | MODIFIER |
| 18 | 42945016 | AX-170180180 | 0 | G | A | 0.061 | -2.26E-03 | 3.02E-01 | 2.72E+00 | 2.74E+00 | 9.95E-01 | no | Ppp2r2b | intron_variant | MODIFIER |
| 18 | 42973264 | AX-97821322 | 0 | C | T | 0.061 | -2.26E-03 | 3.02E-01 | 2.72E+00 | 2.74E+00 | 9.95E-01 | no | Ppp2r2b | intron_variant | MODIFIER |
| 18 | 43054490 | AX-170060692 | 0 | G | A | 0.061 | -2.26E-03 | 3.02E-01 | 2.72E+00 | 2.74E+00 | 9.95E-01 | no | Ppp2r2b | intron_variant | MODIFIER |
| 18 | 43095172 | AX-170015612 | 0 | C | T | 0.061 | -2.26E-03 | 3.02E-01 | 2.72E+00 | 2.74E+00 | 9.95E-01 | no | AC099605.1 | intron_variant,non_coding_transcript_variant | MODIFIER |
| 18 | 43097057 | AX-170430202 | 0 | T | C | 0.061 | -2.26E-03 | 3.02E-01 | 2.72E+00 | 2.74E+00 | 9.95E-01 | no | AC099605.1 | intron_variant,non_coding_transcript_variant | MODIFIER |
| 18 | 43097126 | AX-170369276 | 0 | G | A | 0.061 | -2.26E-03 | 3.02E-01 | 2.72E+00 | 2.74E+00 | 9.95E-01 | no | AC099605.1 | intron_variant,non_coding_transcript_variant | MODIFIER |
| 18 | 43132534 | AX-170343350 | 0 | T | C | 0.061 | -2.26E-03 | 3.02E-01 | 2.72E+00 | 2.74E+00 | 9.95E-01 | no | AC099605.1 | intron_variant,non_coding_transcript_variant | MODIFIER |
| 18 | 43132732 | AX-170326398 | 0 | G | C | 0.061 | -2.26E-03 | 3.02E-01 | 2.72E+00 | 2.74E+00 | 9.95E-01 | no | AC099605.1 | intron_variant,non_coding_transcript_variant | MODIFIER |
| 18 | 43133879 | AX-170356947 | 0 | C | G | 0.061 | -2.26E-03 | 3.02E-01 | 2.72E+00 | 2.74E+00 | 9.95E-01 | no | AC099605.1 | intron_variant,non_coding_transcript_variant | MODIFIER |
| 18 | 43134323 | AX-170273359 | 0 | A | G | 0.061 | -2.26E-03 | 3.02E-01 | 2.72E+00 | 2.74E+00 | 9.95E-01 | no | AC099605.1 | intron_variant,non_coding_transcript_variant | MODIFIER |
| 18 | 43181896 | AX-170015610 | 0 | C | T | 0.061 | -2.26E-03 | 3.02E-01 | 2.72E+00 | 2.74E+00 | 9.95E-01 | no | | intergenic_variant | MODIFIER |
| 18 | 43182027 | AX-170110562 | 0 | G | A | 0.061 | -2.26E-03 | 3.02E-01 | 2.72E+00 | 2.74E+00 | 9.95E-01 | no | | intergenic_variant | MODIFIER |
| 18 | 43213264 | AX-170120084 | 0 | C | A | 0.061 | -2.26E-03 | 3.02E-01 | 2.72E+00 | 2.74E+00 | 9.95E-01 | no | Stk32a | intron_variant | MODIFIER |
| 18 | 43214507 | AX-170466338 | 0 | C | T | 0.061 | -2.26E-03 | 3.02E-01 | 2.72E+00 | 2.74E+00 | 9.95E-01 | no | Stk32a | intron_variant | MODIFIER |

| | | | | | | | | | | | | | | | |
|---|---|---|---|---|---|---|---|---|---|---|---|---|---|---|---|
| 18 | 43223726 | AX-170462286 | 0 | T | C | 0.061 | -2.26E-03 | 3.02E-01 | 2.72E+00 | 2.74E+00 | 9.95E-01 | no | Stk32a | intron_variant | MODIFIER |
| 18 | 43225290 | AX-97834656 | 0 | A | G | 0.061 | -2.26E-03 | 3.02E-01 | 2.72E+00 | 2.74E+00 | 9.95E-01 | no | Stk32a | intron_variant | MODIFIER |
| 18 | 43234067 | AX-170277647 | 0 | C | T | 0.061 | -2.26E-03 | 3.02E-01 | 2.72E+00 | 2.74E+00 | 9.95E-01 | no | Stk32a | intron_variant | MODIFIER |
| 18 | 43235593 | AX-170519546 | 0 | G | A | 0.061 | -2.26E-03 | 3.02E-01 | 2.72E+00 | 2.74E+00 | 9.95E-01 | no | Stk32a | intron_variant | MODIFIER |
| 18 | 43235852 | AX-170140476 | 0 | G | A | 0.061 | -2.26E-03 | 3.02E-01 | 2.72E+00 | 2.74E+00 | 9.95E-01 | no | Stk32a | intron_variant | MODIFIER |
| 18 | 43247931 | AX-170490719 | 0 | T | C | 0.061 | -2.26E-03 | 3.02E-01 | 2.72E+00 | 2.74E+00 | 9.95E-01 | no | Stk32a | intron_variant | MODIFIER |
| 18 | 43291019 | AX-170060698 | 0 | A | G | 0.061 | -2.26E-03 | 3.02E-01 | 2.72E+00 | 2.74E+00 | 9.95E-01 | no | Stk32a | intron_variant | MODIFIER |
| 18 | 43292844 | AX-170060696 | 0 | C | T | 0.061 | -2.26E-03 | 3.02E-01 | 2.72E+00 | 2.74E+00 | 9.95E-01 | no | Stk32a | intron_variant | MODIFIER |
| 18 | 43293613 | AX-170060699 | 0 | C | T | 0.061 | -2.26E-03 | 3.02E-01 | 2.72E+00 | 2.74E+00 | 9.95E-01 | no | Stk32a | intron_variant | MODIFIER |
| 18 | 43295130 | AX-170406153 | 0 | A | C | 0.061 | -2.26E-03 | 3.02E-01 | 2.72E+00 | 2.74E+00 | 9.95E-01 | no | Stk32a | intron_variant | MODIFIER |
| 18 | 43295385 | AX-170199654 | 0 | A | G | 0.061 | -2.26E-03 | 3.02E-01 | 2.72E+00 | 2.74E+00 | 9.95E-01 | no | Stk32a | intron_variant | MODIFIER |
| 18 | 43299439 | AX-170234949 | 0 | T | C | 0.061 | -2.26E-03 | 3.02E-01 | 2.72E+00 | 2.74E+00 | 9.95E-01 | no | Stk32a | intron_variant | MODIFIER |
| 18 | 43333166 | AX-170416393 | 0 | A | G | 0.061 | -2.26E-03 | 3.02E-01 | 2.72E+00 | 2.74E+00 | 9.95E-01 | no | Dpysl3 | intron_variant | MODIFIER |
| 18 | 43344044 | AX-170285237 | 0 | T | C | 0.061 | -2.26E-03 | 3.02E-01 | 2.72E+00 | 2.74E+00 | 9.95E-01 | no | Dpysl3 | intron_variant | MODIFIER |
| 18 | 43356802 | AX-170243754 | 0 | C | A | 0.061 | -2.26E-03 | 3.02E-01 | 2.72E+00 | 2.74E+00 | 9.95E-01 | no | Dpysl3 | intron_variant | MODIFIER |
| 18 | 43358766 | AX-170495314 | 0 | A | G | 0.061 | -2.26E-03 | 3.02E-01 | 2.72E+00 | 2.74E+00 | 9.95E-01 | no | Dpysl3 | intron_variant | MODIFIER |
| 18 | 43376577 | AX-170237142 | 0 | A | G | 0.061 | -2.26E-03 | 3.02E-01 | 2.72E+00 | 2.74E+00 | 9.95E-01 | no | Dpysl3 | intron_variant | MODIFIER |
| 18 | 43378238 | AX-170313034 | 0 | A | G | 0.061 | -2.26E-03 | 3.02E-01 | 2.72E+00 | 2.74E+00 | 9.95E-01 | no | Dpysl3 | intron_variant | MODIFIER |
| 18 | 43409176 | AX-97789906 | 0 | G | A | 0.061 | -2.26E-03 | 3.02E-01 | 2.72E+00 | 2.74E+00 | 9.95E-01 | no | Dpysl3 | intron_variant | MODIFIER |
| 18 | 43410915 | AX-170429020 | 0 | G | A | 0.061 | -2.26E-03 | 3.02E-01 | 2.72E+00 | 2.74E+00 | 9.95E-01 | no | Dpysl3 | intron_variant | MODIFIER |
| 18 | 43412168 | AX-170411506 | 0 | A | G | 0.061 | -2.26E-03 | 3.02E-01 | 2.72E+00 | 2.74E+00 | 9.95E-01 | no | Dpysl3 | intron_variant | MODIFIER |
| 18 | 43418712 | AX-170368278 | 0 | T | A | 0.061 | -2.26E-03 | 3.02E-01 | 2.72E+00 | 2.74E+00 | 9.95E-01 | no | Dpysl3 | intron_variant | MODIFIER |
| 18 | 43442263 | AX-170351887 | 0 | G | A | 0.061 | -2.26E-03 | 3.02E-01 | 2.72E+00 | 2.74E+00 | 9.95E-01 | no | Dpysl3 | upstream_gene_variant | MODIFIER |
| 18 | 43449543 | AX-97797367 | 0 | A | G | 0.061 | -2.26E-03 | 3.02E-01 | 2.72E+00 | 2.74E+00 | 9.95E-01 | no | | intergenic_variant | MODIFIER |
| 18 | 43451583 | AX-170383297 | 0 | T | C | 0.061 | -2.26E-03 | 3.02E-01 | 2.72E+00 | 2.74E+00 | 9.95E-01 | no | | regulatory_region_variant | MODIFIER |
| 18 | 43481342 | AX-97852476 | 0 | T | C | 0.061 | -2.26E-03 | 3.02E-01 | 2.72E+00 | 2.74E+00 | 9.95E-01 | no | Eif3j2 | upstream_gene_variant | MODIFIER |
| 18 | 43493683 | AX-170446412 | 0 | C | T | 0.061 | -2.26E-03 | 3.02E-01 | 2.72E+00 | 2.74E+00 | 9.95E-01 | no | | regulatory_region_variant | MODIFIER |
| 18 | 43502722 | AX-170471263 | 0 | T | C | 0.061 | -2.26E-03 | 3.02E-01 | 2.72E+00 | 2.74E+00 | 9.95E-01 | no | | intergenic_variant | MODIFIER |
| 18 | 43543573 | AX-170419320 | 0 | T | G | 0.061 | -2.26E-03 | 3.02E-01 | 2.72E+00 | 2.74E+00 | 9.95E-01 | no | Jakmip2 | intron_variant | MODIFIER |
| 18 | 43543856 | AX-170208700 | 0 | C | A | 0.061 | -2.26E-03 | 3.02E-01 | 2.72E+00 | 2.74E+00 | 9.95E-01 | no | Jakmip2 | intron_variant | MODIFIER |
| 18 | 43544754 | AX-170088326 | 0 | C | T | 0.061 | -2.26E-03 | 3.02E-01 | 2.72E+00 | 2.74E+00 | 9.95E-01 | no | Jakmip2 | intron_variant | MODIFIER |
| 18 | 43583261 | AX-170163401 | 0 | A | C | 0.061 | -2.26E-03 | 3.02E-01 | 2.72E+00 | 2.74E+00 | 9.95E-01 | no | Jakmip2 | intron_variant | MODIFIER |
| 18 | 43583594 | AX-170015615 | 0 | A | C | 0.061 | -2.26E-03 | 3.02E-01 | 2.72E+00 | 2.74E+00 | 9.95E-01 | no | Jakmip2 | intron_variant | MODIFIER |
| 18 | 43589627 | AX-170386055 | 0 | C | A | 0.061 | -2.26E-03 | 3.02E-01 | 2.72E+00 | 2.74E+00 | 9.95E-01 | no | Jakmip2 | intron_variant | MODIFIER |
| 18 | 43620527 | AX-170015619 | 0 | T | C | 0.061 | -2.26E-03 | 3.02E-01 | 2.72E+00 | 2.74E+00 | 9.95E-01 | no | Jakmip2 | intron_variant | MODIFIER |
| 18 | 43634399 | AX-170015616 | 0 | A | G | 0.061 | -2.26E-03 | 3.02E-01 | 2.72E+00 | 2.74E+00 | 9.95E-01 | no | Jakmip2 | intron_variant | MODIFIER |
| 18 | 43643261 | AX-170208081 | 0 | C | T | 0.061 | -2.26E-03 | 3.02E-01 | 2.72E+00 | 2.74E+00 | 9.95E-01 | no | Jakmip2 | intron_variant | MODIFIER |
| 18 | 43643640 | AX-170015618 | 0 | G | A | 0.061 | -2.26E-03 | 3.02E-01 | 2.72E+00 | 2.74E+00 | 9.95E-01 | no | Jakmip2 | intron_variant | MODIFIER |
| 18 | 43648185 | AX-170302163 | 0 | A | C | 0.061 | -2.26E-03 | 3.02E-01 | 2.72E+00 | 2.74E+00 | 9.95E-01 | no | Jakmip2 | intron_variant | MODIFIER |
| 18 | 43655687 | AX-170226345 | 0 | C | T | 0.061 | -2.26E-03 | 3.02E-01 | 2.72E+00 | 2.74E+00 | 9.95E-01 | no | Jakmip2 | intron_variant | MODIFIER |
| 18 | 43658326 | AX-170239647 | 0 | G | T | 0.061 | -2.26E-03 | 3.02E-01 | 2.72E+00 | 2.74E+00 | 9.95E-01 | no | Jakmip2 | intron_variant | MODIFIER |
| 18 | 43659749 | AX-170527086 | 0 | A | G | 0.061 | -2.26E-03 | 3.02E-01 | 2.72E+00 | 2.74E+00 | 9.95E-01 | no | Jakmip2 | intron_variant | MODIFIER |
| 18 | 43659780 | AX-170510488 | 0 | A | G | 0.061 | -2.26E-03 | 3.02E-01 | 2.72E+00 | 2.74E+00 | 9.95E-01 | no | Jakmip2 | intron_variant | MODIFIER |
| 18 | 43660003 | AX-170171101 | 0 | C | G | 0.061 | -2.26E-03 | 3.02E-01 | 2.72E+00 | 2.74E+00 | 9.95E-01 | no | Jakmip2 | intron_variant | MODIFIER |
| 18 | 43660021 | AX-170239132 | 0 | A | G | 0.061 | -2.26E-03 | 3.02E-01 | 2.72E+00 | 2.74E+00 | 9.95E-01 | no | Jakmip2 | intron_variant | MODIFIER |
| 18 | 43660122 | AX-170365339 | 0 | T | C | 0.061 | -2.26E-03 | 3.02E-01 | 2.72E+00 | 2.74E+00 | 9.95E-01 | no | Jakmip2 | intron_variant | MODIFIER |
| 18 | 43917121 | AX-97788644 | 0 | G | C | 0.061 | -2.26E-03 | 3.02E-01 | 2.72E+00 | 2.74E+00 | 9.95E-01 | no | AC138293.1 | intron_variant,non_coding_transcript_variant | MODIFIER |
| 18 | 43957499 | AX-170203216 | 0 | A | G | 0.061 | -2.26E-03 | 3.02E-01 | 2.72E+00 | 2.74E+00 | 9.95E-01 | no | | intergenic_variant | MODIFIER |
| 18 | 43963092 | AX-170391250 | 0 | T | C | 0.061 | -2.26E-03 | 3.02E-01 | 2.72E+00 | 2.74E+00 | 9.95E-01 | no | Spink5 | upstream_gene_variant | MODIFIER |
| 18 | 43963547 | AX-170343858 | 0 | A | G | 0.061 | -2.26E-03 | 3.02E-01 | 2.72E+00 | 2.74E+00 | 9.95E-01 | no | Spink5 | intron_variant | MODIFIER |
| 18 | 43963610 | AX-170458942 | 0 | T | G | 0.061 | -2.26E-03 | 3.02E-01 | 2.72E+00 | 2.74E+00 | 9.95E-01 | no | Spink5 | intron_variant | MODIFIER |
| 18 | 44036530 | AX-170240630 | 0 | T | G | 0.061 | -2.26E-03 | 3.02E-01 | 2.72E+00 | 2.74E+00 | 9.95E-01 | no | Spink14 | downstream_gene_variant | MODIFIER |
| 18 | 44584242 | AX-97847429 | 0 | C | T | 0.061 | -2.26E-03 | 3.02E-01 | 2.72E+00 | 2.74E+00 | 9.95E-01 | no | Mcc | intron_variant | MODIFIER |
| 18 | 44620368 | AX-170093858 | 0 | C | T | 0.061 | -2.26E-03 | 3.02E-01 | 2.72E+00 | 2.74E+00 | 9.95E-01 | no | Mcc | intron_variant | MODIFIER |

| | | | | | | | | | | | | | | | |
|---|---|---|---|---|---|---|---|---|---|---|---|---|---|---|---|
| 18 | 44624643 | AX-170445943 | 0 | C | T | 0.061 | -2.26E-03 | 3.02E-01 | 2.72E+00 | 2.74E+00 | 9.95E-01 | no | Mcc | intron_variant | MODIFIER |
| 18 | 44625257 | AX-170123676 | 0 | T | C | 0.061 | -2.26E-03 | 3.02E-01 | 2.72E+00 | 2.74E+00 | 9.95E-01 | no | Mcc | intron_variant | MODIFIER |
| 18 | 44631511 | AX-170105871 | 0 | G | C | 0.061 | -2.26E-03 | 3.02E-01 | 2.72E+00 | 2.74E+00 | 9.95E-01 | no | Mcc | intron_variant | MODIFIER |
| 18 | 44646210 | AX-170113917 | 0 | A | C | 0.061 | -2.26E-03 | 3.02E-01 | 2.72E+00 | 2.74E+00 | 9.95E-01 | no | Mcc | intron_variant | MODIFIER |
| 18 | 44652466 | AX-97835356 | 0 | T | C | 0.061 | -2.26E-03 | 3.02E-01 | 2.72E+00 | 2.74E+00 | 9.95E-01 | no | Mcc | intron_variant | MODIFIER |
| 18 | 44673800 | AX-170362206 | 0 | T | C | 0.061 | -2.26E-03 | 3.02E-01 | 2.72E+00 | 2.74E+00 | 9.95E-01 | no | Mcc | intron_variant | MODIFIER |
| 18 | 44676364 | AX-170060719 | 0 | C | T | 0.061 | -2.26E-03 | 3.02E-01 | 2.72E+00 | 2.74E+00 | 9.95E-01 | no | Mcc | intron_variant | MODIFIER |
| 18 | 44848343 | AX-170462344 | 0 | G | A | 0.061 | -2.26E-03 | 3.02E-01 | 2.72E+00 | 2.74E+00 | 9.95E-01 | no | Ythdc2 | intron_variant | MODIFIER |
| 18 | 44941480 | AX-97853253 | 0 | G | T | 0.061 | -2.26E-03 | 3.02E-01 | 2.72E+00 | 2.74E+00 | 9.95E-01 | no | | intergenic_variant | MODIFIER |
| 18 | 44942004 | AX-170175812 | 0 | A | G | 0.061 | -2.26E-03 | 3.02E-01 | 2.72E+00 | 2.74E+00 | 9.95E-01 | no | | intergenic_variant | MODIFIER |
| 18 | 44942397 | AX-170315343 | 0 | A | G | 0.061 | -2.26E-03 | 3.02E-01 | 2.72E+00 | 2.74E+00 | 9.95E-01 | no | | intergenic_variant | MODIFIER |
| 18 | 44942538 | AX-170188103 | 0 | C | T | 0.061 | -2.26E-03 | 3.02E-01 | 2.72E+00 | 2.74E+00 | 9.95E-01 | no | | intergenic_variant | MODIFIER |
| 18 | 44962683 | AX-170529969 | 0 | C | A | 0.061 | -2.26E-03 | 3.02E-01 | 2.72E+00 | 2.74E+00 | 9.95E-01 | no | | intergenic_variant | MODIFIER |
| 18 | 44989713 | AX-170403854 | 0 | C | T | 0.061 | -2.26E-03 | 3.02E-01 | 2.72E+00 | 2.74E+00 | 9.95E-01 | no | | intergenic_variant | MODIFIER |
| 18 | 45001112 | AX-170461568 | 0 | G | C | 0.061 | -2.26E-03 | 3.02E-01 | 2.72E+00 | 2.74E+00 | 9.95E-01 | no | | regulatory_region_variant | MODIFIER |
| 18 | 45001554 | AX-170320268 | 0 | A | G | 0.061 | -2.26E-03 | 3.02E-01 | 2.72E+00 | 2.74E+00 | 9.95E-01 | no | | intergenic_variant | MODIFIER |
| 18 | 45084048 | AX-170388042 | 0 | C | T | 0.061 | -2.26E-03 | 3.02E-01 | 2.72E+00 | 2.74E+00 | 9.95E-01 | no | AC134458.2 | intron_variant,non_coding_transcript_variant | MODIFIER |
| 18 | 45141096 | AX-170424196 | 0 | G | A | 0.061 | -2.26E-03 | 3.02E-01 | 2.72E+00 | 2.74E+00 | 9.95E-01 | no | | intergenic_variant | MODIFIER |
| 18 | 45143062 | AX-170273250 | 0 | G | A | 0.061 | -2.26E-03 | 3.02E-01 | 2.72E+00 | 2.74E+00 | 9.95E-01 | no | AC134458.3 | downstream_gene_variant | MODIFIER |
| 18 | 45147344 | AX-170097672 | 0 | C | T | 0.061 | -2.26E-03 | 3.02E-01 | 2.72E+00 | 2.74E+00 | 9.95E-01 | no | AC134458.3 | upstream_gene_variant | MODIFIER |
| 18 | 45230943 | AX-170412592 | 0 | G | C | 0.061 | -2.26E-03 | 3.02E-01 | 2.72E+00 | 2.74E+00 | 9.95E-01 | no | | intergenic_variant | MODIFIER |
| 18 | 45254936 | AX-170494217 | 0 | A | G | 0.061 | -2.26E-03 | 3.02E-01 | 2.72E+00 | 2.74E+00 | 9.95E-01 | no | | intergenic_variant | MODIFIER |
| 18 | 45266060 | AX-97808850 | 0 | G | A | 0.061 | -2.26E-03 | 3.02E-01 | 2.72E+00 | 2.74E+00 | 9.95E-01 | no | Kcnn2 | upstream_gene_variant | MODIFIER |
| 18 | 45266144 | AX-170402942 | 0 | G | A | 0.061 | -2.26E-03 | 3.02E-01 | 2.72E+00 | 2.74E+00 | 9.95E-01 | no | Kcnn2 | upstream_gene_variant | MODIFIER |
| 18 | 45278086 | AX-170521820 | 0 | A | G | 0.061 | -2.26E-03 | 3.02E-01 | 2.72E+00 | 2.74E+00 | 9.95E-01 | no | Kcnn2 | intron_variant | MODIFIER |
| 18 | 45282013 | AX-170167204 | 0 | A | G | 0.061 | -2.26E-03 | 3.02E-01 | 2.72E+00 | 2.74E+00 | 9.95E-01 | no | Kcnn2 | intron_variant | MODIFIER |
| 18 | 45287601 | AX-170015638 | 0 | T | C | 0.061 | -2.26E-03 | 3.02E-01 | 2.72E+00 | 2.74E+00 | 9.95E-01 | no | Kcnn2 | intron_variant | MODIFIER |
| 18 | 45339928 | AX-170060736 | 0 | A | C | 0.061 | -2.26E-03 | 3.02E-01 | 2.72E+00 | 2.74E+00 | 9.95E-01 | no | Kcnn2 | intron_variant | MODIFIER |
| 18 | 45636871 | AX-170212277 | 0 | G | T | 0.061 | -2.26E-03 | 3.02E-01 | 2.72E+00 | 2.74E+00 | 9.95E-01 | no | Kcnn2 | intron_variant | MODIFIER |
| 18 | 45654440 | AX-170060739 | 0 | G | C | 0.061 | -2.26E-03 | 3.02E-01 | 2.72E+00 | 2.74E+00 | 9.95E-01 | no | Kcnn2 | intron_variant | MODIFIER |
| 18 | 45655151 | AX-97784910 | 0 | G | A | 0.061 | -2.26E-03 | 3.02E-01 | 2.72E+00 | 2.74E+00 | 9.95E-01 | no | Kcnn2 | intron_variant | MODIFIER |
| 18 | 45655645 | AX-170060738 | 0 | C | T | 0.061 | -2.26E-03 | 3.02E-01 | 2.72E+00 | 2.74E+00 | 9.95E-01 | no | Kcnn2 | intron_variant | MODIFIER |
| 18 | 45655712 | AX-170060737 | 0 | A | T | 0.061 | -2.26E-03 | 3.02E-01 | 2.72E+00 | 2.74E+00 | 9.95E-01 | no | Kcnn2 | intron_variant | MODIFIER |
| 18 | 45657432 | AX-170439784 | 0 | A | G | 0.061 | -2.26E-03 | 3.02E-01 | 2.72E+00 | 2.74E+00 | 9.95E-01 | no | Kcnn2 | intron_variant | MODIFIER |
| 18 | 45673486 | AX-170236031 | 0 | A | G | 0.061 | -2.26E-03 | 3.02E-01 | 2.72E+00 | 2.74E+00 | 9.95E-01 | no | Kcnn2 | intron_variant | MODIFIER |
| 18 | 45678860 | AX-170478111 | 0 | T | C | 0.061 | -2.26E-03 | 3.02E-01 | 2.72E+00 | 2.74E+00 | 9.95E-01 | no | Kcnn2 | intron_variant | MODIFIER |
| 18 | 45953189 | AX-97803185 | 0 | C | T | 0.061 | -2.26E-03 | 3.02E-01 | 2.72E+00 | 2.74E+00 | 9.95E-01 | no | A330093E20Rik | intron_variant,non_coding_transcript_variant | MODIFIER |
| 18 | 46160447 | AX-170060750 | 0 | A | T | 0.061 | -2.26E-03 | 3.02E-01 | 2.72E+00 | 2.74E+00 | 9.95E-01 | no | Trim36 | downstream_gene_variant | MODIFIER |
| 18 | 46180515 | AX-170400926 | 0 | A | G | 0.061 | -2.26E-03 | 3.02E-01 | 2.72E+00 | 2.74E+00 | 9.95E-01 | no | Trim36 | intron_variant | MODIFIER |
| 12 | 67267996 | AX-170006950 | 1 | T | C | 0.062 | -2.19E-03 | 3.02E-01 | 2.72E+00 | 2.74E+00 | 9.95E-01 | no | | intergenic_variant | MODIFIER |
| 18 | 30451234 | AX-170115875 | 1 | A | G | 0.062 | -2.19E-03 | 3.02E-01 | 2.72E+00 | 2.74E+00 | 9.95E-01 | no | | regulatory_region_variant | MODIFIER |
| 9 | 64661767 | AX-170244042 | 0 | C | A | 0.193 | 5.28E-04 | 1.38E-01 | 2.71E+00 | 2.74E+00 | 9.95E-01 | no | Megf11 | intron_variant | MODIFIER |
| 9 | 66738685 | AX-170333727 | 0 | G | A | 0.193 | 5.28E-04 | 1.38E-01 | 2.71E+00 | 2.74E+00 | 9.95E-01 | no | Car12 | intron_variant | MODIFIER |
| 9 | 69912196 | AX-170394218 | 0 | C | T | 0.193 | 5.28E-04 | 1.38E-01 | 2.71E+00 | 2.74E+00 | 9.95E-01 | no | | intergenic_variant | MODIFIER |
| 9 | 75466675 | AX-170483451 | 0 | C | A | 0.193 | 5.28E-04 | 1.38E-01 | 2.71E+00 | 2.74E+00 | 9.95E-01 | no | Leo1 | downstream_gene_variant | MODIFIER |
| 18 | 31594114 | AX-170104968 | 0 | T | C | 0.053 | 3.99E-03 | 2.36E-01 | 2.62E+00 | 2.74E+00 | 9.95E-01 | no | Slc25a46 | intron_variant | MODIFIER |
| 1 | 60384506 | AX-170106241 | 0 | A | G | 0.096 | 1.46E-03 | 2.14E-01 | 2.69E+00 | 2.74E+00 | 9.95E-01 | no | Cyp20a1 | intron_variant | MODIFIER |
| 9 | 105700317 | AX-170460517 | 1 | C | T | 0.125 | -5.84E-04 | 1.16E-01 | 2.67E+00 | 2.74E+00 | 9.95E-01 | no | Col6a6 | intron_variant | MODIFIER |
| 6 | 147063550 | AX-170469547 | 2 | C | A | 0.073 | -1.55E-03 | 2.76E-01 | 2.70E+00 | 2.74E+00 | 9.95E-01 | no | Mrps35 | intron_variant | MODIFIER |
| 9 | 104018713 | AX-170096657 | 0 | G | A | 0.123 | -6.29E-04 | 1.18E-01 | 2.67E+00 | 2.74E+00 | 9.95E-01 | no | Nphp3 | intron_variant | MODIFIER |
| 9 | 104018744 | AX-170510508 | 0 | G | A | 0.123 | -6.29E-04 | 1.18E-01 | 2.67E+00 | 2.74E+00 | 9.95E-01 | no | Nphp3 | intron_variant | MODIFIER |
| 9 | 104019012 | AX-170405293 | 0 | A | G | 0.123 | -6.29E-04 | 1.18E-01 | 2.67E+00 | 2.74E+00 | 9.95E-01 | no | Nphp3 | intron_variant | MODIFIER |
| 9 | 104047018 | AX-97819033 | 0 | A | G | 0.123 | -6.29E-04 | 1.18E-01 | 2.67E+00 | 2.74E+00 | 9.95E-01 | no | Uba5 | 3_prime_UTR_variant | MODIFIER |
| 9 | 104050972 | AX-170412419 | 0 | C | A | 0.123 | -6.29E-04 | 1.18E-01 | 2.67E+00 | 2.74E+00 | 9.95E-01 | no | Uba5 | intron_variant | MODIFIER |

| | | | | | | | | | | | | | | | |
|---|---|---|---|---|---|---|---|---|---|---|---|---|---|---|---|
| 9 | 104115414 | AX-170251744 | 0 | A | G | 0.123 | -6.29E-04 | 1.18E-01 | 2.67E+00 | 2.74E+00 | 9.95E-01 | no | Acad11 | intron_variant | MODIFIER |
| 9 | 104137935 | AX-170215401 | 0 | T | C | 0.123 | -6.29E-04 | 1.18E-01 | 2.67E+00 | 2.74E+00 | 9.95E-01 | no | | intergenic_variant | MODIFIER |
| 9 | 104138470 | AX-170306486 | 0 | T | C | 0.123 | -6.29E-04 | 1.18E-01 | 2.67E+00 | 2.74E+00 | 9.95E-01 | no | | intergenic_variant | MODIFIER |
| 9 | 104138585 | AX-170096658 | 0 | G | A | 0.123 | -6.29E-04 | 1.18E-01 | 2.67E+00 | 2.74E+00 | 9.95E-01 | no | | intergenic_variant | MODIFIER |
| 9 | 104330507 | AX-170508246 | 0 | A | G | 0.123 | -6.29E-04 | 1.18E-01 | 2.67E+00 | 2.74E+00 | 9.95E-01 | no | Acpp | intron_variant | MODIFIER |
| 9 | 104330912 | AX-170480824 | 0 | T | A | 0.123 | -6.29E-04 | 1.18E-01 | 2.67E+00 | 2.74E+00 | 9.95E-01 | no | Acpp | intron_variant | MODIFIER |
| 9 | 104336042 | AX-170096659 | 0 | C | T | 0.123 | -6.29E-04 | 1.18E-01 | 2.67E+00 | 2.74E+00 | 9.95E-01 | no | Acpp | intron_variant | MODIFIER |
| 9 | 104375909 | AX-170083337 | 0 | A | G | 0.123 | -6.29E-04 | 1.18E-01 | 2.67E+00 | 2.74E+00 | 9.95E-01 | no | Gm28548 | intron_variant,non_coding_transcript_variant | MODIFIER |
| 9 | 104378330 | AX-170519952 | 0 | C | T | 0.123 | -6.29E-04 | 1.18E-01 | 2.67E+00 | 2.74E+00 | 9.95E-01 | no | Gm28548 | intron_variant,non_coding_transcript_variant | MODIFIER |
| 9 | 104379419 | AX-170083333 | 0 | G | T | 0.123 | -6.29E-04 | 1.18E-01 | 2.67E+00 | 2.74E+00 | 9.95E-01 | no | Gm28548 | intron_variant,non_coding_transcript_variant | MODIFIER |
| 9 | 104379964 | AX-170177063 | 0 | C | T | 0.123 | -6.29E-04 | 1.18E-01 | 2.67E+00 | 2.74E+00 | 9.95E-01 | no | Gm28548 | intron_variant,non_coding_transcript_variant | MODIFIER |
| 9 | 104382524 | AX-170498152 | 0 | A | C | 0.123 | -6.29E-04 | 1.18E-01 | 2.67E+00 | 2.74E+00 | 9.95E-01 | no | Gm28548 | intron_variant,non_coding_transcript_variant | MODIFIER |
| 9 | 104400427 | AX-97798940 | 0 | C | T | 0.123 | -6.29E-04 | 1.18E-01 | 2.67E+00 | 2.74E+00 | 9.95E-01 | no | | regulatory_region_variant | MODIFIER |
| 9 | 104410753 | AX-97794689 | 0 | A | G | 0.123 | -6.29E-04 | 1.18E-01 | 2.67E+00 | 2.74E+00 | 9.95E-01 | no | | regulatory_region_variant | MODIFIER |
| 9 | 104424299 | AX-170176203 | 0 | T | G | 0.123 | -6.29E-04 | 1.18E-01 | 2.67E+00 | 2.74E+00 | 9.95E-01 | no | Mir2136 | upstream_gene_variant | MODIFIER |
| 9 | 104435039 | AX-97792212 | 0 | C | T | 0.123 | -6.29E-04 | 1.18E-01 | 2.67E+00 | 2.74E+00 | 9.95E-01 | no | | intergenic_variant | MODIFIER |
| 9 | 104437547 | AX-170393299 | 0 | C | T | 0.123 | -6.29E-04 | 1.18E-01 | 2.67E+00 | 2.74E+00 | 9.95E-01 | no | | intergenic_variant | MODIFIER |
| 9 | 104454250 | AX-170313044 | 0 | A | C | 0.123 | -6.29E-04 | 1.18E-01 | 2.67E+00 | 2.74E+00 | 9.95E-01 | no | | intergenic_variant | MODIFIER |
| 9 | 104463773 | AX-170494075 | 0 | T | C | 0.123 | -6.29E-04 | 1.18E-01 | 2.67E+00 | 2.74E+00 | 9.95E-01 | no | | intergenic_variant | MODIFIER |
| 9 | 104464652 | AX-97794046 | 0 | C | T | 0.123 | -6.29E-04 | 1.18E-01 | 2.67E+00 | 2.74E+00 | 9.95E-01 | no | | intergenic_variant | MODIFIER |
| 9 | 104483473 | AX-170527397 | 0 | C | T | 0.123 | -6.29E-04 | 1.18E-01 | 2.67E+00 | 2.74E+00 | 9.95E-01 | no | Gm25904 | upstream_gene_variant | MODIFIER |
| 9 | 104520598 | AX-170206097 | 0 | G | T | 0.123 | -6.29E-04 | 1.18E-01 | 2.67E+00 | 2.74E+00 | 9.95E-01 | no | | intergenic_variant | MODIFIER |
| 9 | 104522574 | AX-97850010 | 0 | T | C | 0.123 | -6.29E-04 | 1.18E-01 | 2.67E+00 | 2.74E+00 | 9.95E-01 | no | | intergenic_variant | MODIFIER |
| 9 | 104547516 | AX-170096660 | 0 | C | T | 0.123 | -6.29E-04 | 1.18E-01 | 2.67E+00 | 2.74E+00 | 9.95E-01 | no | Cpne4 | 5_prime_UTR_variant | MODIFIER |
| 9 | 104565651 | AX-170379395 | 0 | G | A | 0.123 | -6.29E-04 | 1.18E-01 | 2.67E+00 | 2.74E+00 | 9.95E-01 | no | Cpne4 | upstream_gene_variant | MODIFIER |
| 9 | 104567959 | AX-97785803 | 0 | T | C | 0.123 | -6.29E-04 | 1.18E-01 | 2.67E+00 | 2.74E+00 | 9.95E-01 | no | Cpne4 | upstream_gene_variant | MODIFIER |
| 9 | 104576426 | AX-170286343 | 0 | G | T | 0.123 | -6.29E-04 | 1.18E-01 | 2.67E+00 | 2.74E+00 | 9.95E-01 | no | Cpne4 | intron_variant | MODIFIER |
| 9 | 104594283 | AX-170357231 | 0 | C | T | 0.123 | -6.29E-04 | 1.18E-01 | 2.67E+00 | 2.74E+00 | 9.95E-01 | no | Cpne4 | intron_variant | MODIFIER |
| 9 | 104601194 | AX-170114249 | 0 | G | A | 0.123 | -6.29E-04 | 1.18E-01 | 2.67E+00 | 2.74E+00 | 9.95E-01 | no | Cpne4 | intron_variant | MODIFIER |
| 9 | 104629064 | AX-97789148 | 0 | C | G | 0.123 | -6.29E-04 | 1.18E-01 | 2.67E+00 | 2.74E+00 | 9.95E-01 | no | Cpne4 | intron_variant | MODIFIER |
| 9 | 104635740 | AX-170504584 | 0 | A | G | 0.123 | -6.29E-04 | 1.18E-01 | 2.67E+00 | 2.74E+00 | 9.95E-01 | no | Cpne4 | intron_variant | MODIFIER |
| 9 | 104641800 | AX-170161030 | 0 | T | C | 0.123 | -6.29E-04 | 1.18E-01 | 2.67E+00 | 2.74E+00 | 9.95E-01 | no | Cpne4 | intron_variant | MODIFIER |
| 9 | 104644212 | AX-170083344 | 0 | T | C | 0.123 | -6.29E-04 | 1.18E-01 | 2.67E+00 | 2.74E+00 | 9.95E-01 | no | Cpne4 | intron_variant | MODIFIER |
| 9 | 104645221 | AX-97781214 | 0 | T | C | 0.123 | -6.29E-04 | 1.18E-01 | 2.67E+00 | 2.74E+00 | 9.95E-01 | no | Cpne4 | intron_variant | MODIFIER |
| 9 | 104645856 | AX-170083342 | 0 | A | G | 0.123 | -6.29E-04 | 1.18E-01 | 2.67E+00 | 2.74E+00 | 9.95E-01 | no | Cpne4 | intron_variant | MODIFIER |
| 9 | 104655603 | AX-170448633 | 0 | G | C | 0.123 | -6.29E-04 | 1.18E-01 | 2.67E+00 | 2.74E+00 | 9.95E-01 | no | Cpne4 | intron_variant | MODIFIER |
| 9 | 104667858 | AX-170199975 | 0 | C | T | 0.123 | -6.29E-04 | 1.18E-01 | 2.67E+00 | 2.74E+00 | 9.95E-01 | no | Cpne4 | intron_variant | MODIFIER |
| 9 | 104721617 | AX-97850206 | 0 | C | T | 0.123 | -6.29E-04 | 1.18E-01 | 2.67E+00 | 2.74E+00 | 9.95E-01 | no | Cpne4 | intron_variant | MODIFIER |
| 9 | 104723823 | AX-170033184 | 0 | A | G | 0.123 | -6.29E-04 | 1.18E-01 | 2.67E+00 | 2.74E+00 | 9.95E-01 | no | Cpne4 | intron_variant | MODIFIER |
| 9 | 104724219 | AX-170531550 | 0 | A | G | 0.123 | -6.29E-04 | 1.18E-01 | 2.67E+00 | 2.74E+00 | 9.95E-01 | no | Cpne4 | intron_variant | MODIFIER |
| 9 | 104726307 | AX-170204599 | 0 | A | G | 0.123 | -6.29E-04 | 1.18E-01 | 2.67E+00 | 2.74E+00 | 9.95E-01 | no | Cpne4 | intron_variant | MODIFIER |
| 9 | 104732188 | AX-170306132 | 0 | G | A | 0.123 | -6.29E-04 | 1.18E-01 | 2.67E+00 | 2.74E+00 | 9.95E-01 | no | Cpne4 | intron_variant | MODIFIER |
| 9 | 104756236 | AX-97801607 | 0 | G | A | 0.123 | -6.29E-04 | 1.18E-01 | 2.67E+00 | 2.74E+00 | 9.95E-01 | no | Cpne4 | intron_variant | MODIFIER |
| 9 | 104771716 | AX-170033186 | 0 | C | T | 0.123 | -6.29E-04 | 1.18E-01 | 2.67E+00 | 2.74E+00 | 9.95E-01 | no | Cpne4 | intron_variant | MODIFIER |
| 9 | 104786006 | AX-97791766 | 0 | A | G | 0.123 | -6.29E-04 | 1.18E-01 | 2.67E+00 | 2.74E+00 | 9.95E-01 | no | Cpne4 | intron_variant | MODIFIER |
| 9 | 104798976 | AX-170033183 | 0 | C | G | 0.123 | -6.29E-04 | 1.18E-01 | 2.67E+00 | 2.74E+00 | 9.95E-01 | no | Cpne4 | intron_variant | MODIFIER |
| 9 | 104816114 | AX-170033185 | 0 | A | G | 0.123 | -6.29E-04 | 1.18E-01 | 2.67E+00 | 2.74E+00 | 9.95E-01 | no | Cpne4 | intron_variant | MODIFIER |
| 9 | 104816546 | AX-170447783 | 0 | G | T | 0.123 | -6.29E-04 | 1.18E-01 | 2.67E+00 | 2.74E+00 | 9.95E-01 | no | Cpne4 | intron_variant | MODIFIER |
| 9 | 104822480 | AX-170381928 | 0 | T | C | 0.123 | -6.29E-04 | 1.18E-01 | 2.67E+00 | 2.74E+00 | 9.95E-01 | no | Cpne4 | intron_variant | MODIFIER |
| 9 | 104825249 | AX-97807975 | 0 | T | C | 0.123 | -6.29E-04 | 1.18E-01 | 2.67E+00 | 2.74E+00 | 9.95E-01 | no | Cpne4 | intron_variant | MODIFIER |
| 9 | 104842964 | AX-97849639 | 0 | T | C | 0.123 | -6.29E-04 | 1.18E-01 | 2.67E+00 | 2.74E+00 | 9.95E-01 | no | Cpne4 | intron_variant | MODIFIER |
| 9 | 104843626 | AX-170033192 | 0 | A | C | 0.123 | -6.29E-04 | 1.18E-01 | 2.67E+00 | 2.74E+00 | 9.95E-01 | no | Cpne4 | intron_variant | MODIFIER |
| 9 | 104866955 | AX-170355087 | 0 | G | A | 0.123 | -6.29E-04 | 1.18E-01 | 2.67E+00 | 2.74E+00 | 9.95E-01 | no | Cpne4 | intron_variant | MODIFIER |
| 9 | 104869133 | AX-97817443 | 0 | T | C | 0.123 | -6.29E-04 | 1.18E-01 | 2.67E+00 | 2.74E+00 | 9.95E-01 | no | Cpne4 | intron_variant | MODIFIER |

| | | | | | | | | | | | | | | | |
|---|---|---|---|---|---|---|---|---|---|---|---|---|---|---|---|
| 9 | 104869892 | AX-170217402 | 0 | T | A | 0.123 | -6.29E-04 | 1.18E-01 | 2.67E+00 | 2.74E+00 | 9.95E-01 | no | Cpne4 | intron_variant | MODIFIER |
| 9 | 104875641 | AX-170359984 | 0 | A | G | 0.123 | -6.29E-04 | 1.18E-01 | 2.67E+00 | 2.74E+00 | 9.95E-01 | no | Cpne4 | intron_variant | MODIFIER |
| 9 | 104881773 | AX-170033191 | 0 | C | G | 0.123 | -6.29E-04 | 1.18E-01 | 2.67E+00 | 2.74E+00 | 9.95E-01 | no | Cpne4 | intron_variant | MODIFIER |
| 9 | 104881932 | AX-170033190 | 0 | C | A | 0.123 | -6.29E-04 | 1.18E-01 | 2.67E+00 | 2.74E+00 | 9.95E-01 | no | Cpne4 | intron_variant | MODIFIER |
| 9 | 104882715 | AX-170033188 | 0 | T | C | 0.123 | -6.29E-04 | 1.18E-01 | 2.67E+00 | 2.74E+00 | 9.95E-01 | no | Cpne4 | intron_variant | MODIFIER |
| 9 | 104896790 | AX-97838862 | 0 | G | A | 0.123 | -6.29E-04 | 1.18E-01 | 2.67E+00 | 2.74E+00 | 9.95E-01 | no | Cpne4 | intron_variant | MODIFIER |
| 9 | 104902085 | AX-170033182 | 0 | A | G | 0.123 | -6.29E-04 | 1.18E-01 | 2.67E+00 | 2.74E+00 | 9.95E-01 | no | Cpne4 | intron_variant | MODIFIER |
| 9 | 104914072 | AX-170090401 | 0 | A | G | 0.123 | -6.29E-04 | 1.18E-01 | 2.67E+00 | 2.74E+00 | 9.95E-01 | no | Cpne4 | intron_variant | MODIFIER |
| 9 | 104961619 | AX-170382080 | 0 | G | A | 0.123 | -6.29E-04 | 1.18E-01 | 2.67E+00 | 2.74E+00 | 9.95E-01 | no | Cpne4 | intron_variant | MODIFIER |
| 9 | 104969231 | AX-97818760 | 0 | A | G | 0.123 | -6.29E-04 | 1.18E-01 | 2.67E+00 | 2.74E+00 | 9.95E-01 | no | Cpne4 | intron_variant | MODIFIER |
| 9 | 104969730 | AX-170480602 | 0 | A | T | 0.123 | -6.29E-04 | 1.18E-01 | 2.67E+00 | 2.74E+00 | 9.95E-01 | no | Cpne4 | intron_variant | MODIFIER |
| 9 | 104973152 | AX-170146112 | 0 | G | T | 0.123 | -6.29E-04 | 1.18E-01 | 2.67E+00 | 2.74E+00 | 9.95E-01 | no | Cpne4 | intron_variant | MODIFIER |
| 9 | 104997890 | AX-97856224 | 0 | A | G | 0.123 | -6.29E-04 | 1.18E-01 | 2.67E+00 | 2.74E+00 | 9.95E-01 | no | Cpne4 | intron_variant | MODIFIER |
| 9 | 105014720 | AX-170458677 | 0 | G | A | 0.123 | -6.29E-04 | 1.18E-01 | 2.67E+00 | 2.74E+00 | 9.95E-01 | no | Cpne4 | intron_variant | MODIFIER |
| 9 | 105024640 | AX-97817737 | 0 | G | A | 0.123 | -6.29E-04 | 1.18E-01 | 2.67E+00 | 2.74E+00 | 9.95E-01 | no | Cpne4 | intron_variant | MODIFIER |
| 9 | 105025898 | AX-170374790 | 0 | T | G | 0.123 | -6.29E-04 | 1.18E-01 | 2.67E+00 | 2.74E+00 | 9.95E-01 | no | Cpne4 | intron_variant | MODIFIER |
| 9 | 105028997 | AX-170277596 | 0 | G | A | 0.123 | -6.29E-04 | 1.18E-01 | 2.67E+00 | 2.74E+00 | 9.95E-01 | no | Cpne4 | intron_variant | MODIFIER |
| 9 | 105029223 | AX-170376379 | 0 | C | T | 0.123 | -6.29E-04 | 1.18E-01 | 2.67E+00 | 2.74E+00 | 9.95E-01 | no | Cpne4 | intron_variant | MODIFIER |
| 9 | 105029489 | AX-170458037 | 0 | C | T | 0.123 | -6.29E-04 | 1.18E-01 | 2.67E+00 | 2.74E+00 | 9.95E-01 | no | Cpne4 | intron_variant | MODIFIER |
| 9 | 105031802 | AX-170499189 | 0 | A | G | 0.123 | -6.29E-04 | 1.18E-01 | 2.67E+00 | 2.74E+00 | 9.95E-01 | no | Cpne4 | intron_variant | MODIFIER |
| 9 | 105038530 | AX-170456954 | 0 | C | A | 0.123 | -6.29E-04 | 1.18E-01 | 2.67E+00 | 2.74E+00 | 9.95E-01 | no | Cpne4 | downstream_gene_variant | MODIFIER |
| 9 | 105039557 | AX-170033187 | 0 | T | G | 0.123 | -6.29E-04 | 1.18E-01 | 2.67E+00 | 2.74E+00 | 9.95E-01 | no | | intergenic_variant | MODIFIER |
| 9 | 105055821 | AX-97783836 | 0 | C | T | 0.123 | -6.29E-04 | 1.18E-01 | 2.67E+00 | 2.74E+00 | 9.95E-01 | no | Mrpl3 | intron_variant | MODIFIER |
| 9 | 105056332 | AX-170314945 | 0 | C | T | 0.123 | -6.29E-04 | 1.18E-01 | 2.67E+00 | 2.74E+00 | 9.95E-01 | no | Mrpl3 | intron_variant | MODIFIER |
| 9 | 105056365 | AX-170147821 | 0 | C | T | 0.123 | -6.29E-04 | 1.18E-01 | 2.67E+00 | 2.74E+00 | 9.95E-01 | no | Mrpl3 | intron_variant | MODIFIER |
| 9 | 105057335 | AX-170121680 | 0 | T | C | 0.123 | -6.29E-04 | 1.18E-01 | 2.67E+00 | 2.74E+00 | 9.95E-01 | no | Mrpl3 | intron_variant | MODIFIER |
| 9 | 105064605 | AX-170340465 | 0 | T | C | 0.123 | -6.29E-04 | 1.18E-01 | 2.67E+00 | 2.74E+00 | 9.95E-01 | no | Mrpl3 | intron_variant | MODIFIER |
| 9 | 105065012 | AX-97852992 | 0 | T | C | 0.123 | -6.29E-04 | 1.18E-01 | 2.67E+00 | 2.74E+00 | 9.95E-01 | no | Mrpl3 | intron_variant | MODIFIER |
| 9 | 105071770 | AX-170196160 | 0 | C | G | 0.123 | -6.29E-04 | 1.18E-01 | 2.67E+00 | 2.74E+00 | 9.95E-01 | no | Mrpl3 | intron_variant | MODIFIER |
| 9 | 105074701 | AX-170349414 | 0 | A | T | 0.123 | -6.29E-04 | 1.18E-01 | 2.67E+00 | 2.74E+00 | 9.95E-01 | no | Mrpl3 | intron_variant | MODIFIER |
| 9 | 105077488 | AX-170083350 | 0 | G | A | 0.123 | -6.29E-04 | 1.18E-01 | 2.67E+00 | 2.74E+00 | 9.95E-01 | no | Mrpl3 | 3_prime_UTR_variant | MODIFIER |
| 9 | 105079680 | AX-170463057 | 0 | T | C | 0.123 | -6.29E-04 | 1.18E-01 | 2.67E+00 | 2.74E+00 | 9.95E-01 | no | Mrpl3 | 3_prime_UTR_variant | MODIFIER |
| 9 | 105081072 | AX-170302854 | 0 | G | A | 0.123 | -6.29E-04 | 1.18E-01 | 2.67E+00 | 2.74E+00 | 9.95E-01 | no | Mrpl3 | downstream_gene_variant | MODIFIER |
| 9 | 105124608 | AX-97831896 | 0 | A | G | 0.123 | -6.29E-04 | 1.18E-01 | 2.67E+00 | 2.74E+00 | 9.95E-01 | no | Nudt16 | downstream_gene_variant | MODIFIER |
| 9 | 105126848 | AX-170154771 | 0 | G | A | 0.123 | -6.29E-04 | 1.18E-01 | 2.67E+00 | 2.74E+00 | 9.95E-01 | no | Nudt16 | downstream_gene_variant | MODIFIER |
| 9 | 105139583 | AX-170344230 | 0 | A | G | 0.123 | -6.29E-04 | 1.18E-01 | 2.67E+00 | 2.74E+00 | 9.95E-01 | no | 1700080E11Rik | downstream_gene_variant | MODIFIER |
| 9 | 105139673 | AX-170263922 | 0 | T | C | 0.123 | -6.29E-04 | 1.18E-01 | 2.67E+00 | 2.74E+00 | 9.95E-01 | no | 1700080E11Rik | downstream_gene_variant | MODIFIER |
| 9 | 105143722 | AX-170440536 | 0 | G | A | 0.123 | -6.29E-04 | 1.18E-01 | 2.67E+00 | 2.74E+00 | 9.95E-01 | no | 1700080E11Rik | intron_variant | MODIFIER |
| 9 | 105187548 | AX-170353503 | 0 | A | G | 0.123 | -6.29E-04 | 1.18E-01 | 2.67E+00 | 2.74E+00 | 9.95E-01 | no | Nek11 | intron_variant | MODIFIER |
| 9 | 105187609 | AX-170456869 | 0 | C | T | 0.123 | -6.29E-04 | 1.18E-01 | 2.67E+00 | 2.74E+00 | 9.95E-01 | no | Nek11 | intron_variant | MODIFIER |
| 9 | 105194901 | AX-170267771 | 0 | A | C | 0.123 | -6.29E-04 | 1.18E-01 | 2.67E+00 | 2.74E+00 | 9.95E-01 | no | Nek11 | intron_variant | MODIFIER |
| 9 | 105195477 | AX-170475317 | 0 | C | T | 0.123 | -6.29E-04 | 1.18E-01 | 2.67E+00 | 2.74E+00 | 9.95E-01 | no | Nek11 | intron_variant | MODIFIER |
| 9 | 105196087 | AX-97805552 | 0 | G | A | 0.123 | -6.29E-04 | 1.18E-01 | 2.67E+00 | 2.74E+00 | 9.95E-01 | no | Nek11 | intron_variant | MODIFIER |
| 9 | 105232398 | AX-97810927 | 0 | C | A | 0.123 | -6.29E-04 | 1.18E-01 | 2.67E+00 | 2.74E+00 | 9.95E-01 | no | Nek11 | intron_variant | MODIFIER |
| 9 | 105260075 | AX-170122199 | 0 | T | G | 0.123 | -6.29E-04 | 1.18E-01 | 2.67E+00 | 2.74E+00 | 9.95E-01 | no | Nek11 | intron_variant | MODIFIER |
| 9 | 105260449 | AX-170160931 | 0 | T | C | 0.123 | -6.29E-04 | 1.18E-01 | 2.67E+00 | 2.74E+00 | 9.95E-01 | no | Nek11 | intron_variant | MODIFIER |
| 9 | 105262787 | AX-97808615 | 0 | T | C | 0.123 | -6.29E-04 | 1.18E-01 | 2.67E+00 | 2.74E+00 | 9.95E-01 | no | Nek11 | intron_variant | MODIFIER |
| 9 | 105286428 | AX-170083352 | 0 | C | T | 0.123 | -6.29E-04 | 1.18E-01 | 2.67E+00 | 2.74E+00 | 9.95E-01 | no | Nek11 | intron_variant | MODIFIER |
| 9 | 105289493 | AX-170033201 | 0 | T | C | 0.123 | -6.29E-04 | 1.18E-01 | 2.67E+00 | 2.74E+00 | 9.95E-01 | no | Nek11 | intron_variant | MODIFIER |
| 9 | 105294201 | AX-97783263 | 0 | T | C | 0.123 | -6.29E-04 | 1.18E-01 | 2.67E+00 | 2.74E+00 | 9.95E-01 | no | Nek11 | intron_variant | MODIFIER |
| 9 | 105313128 | AX-170033202 | 0 | A | C | 0.123 | -6.29E-04 | 1.18E-01 | 2.67E+00 | 2.74E+00 | 9.95E-01 | no | Nek11 | intron_variant | MODIFIER |
| 9 | 105313219 | AX-170033194 | 0 | C | T | 0.123 | -6.29E-04 | 1.18E-01 | 2.67E+00 | 2.74E+00 | 9.95E-01 | no | Nek11 | intron_variant | MODIFIER |
| 9 | 105324850 | AX-97813706 | 0 | G | A | 0.123 | -6.29E-04 | 1.18E-01 | 2.67E+00 | 2.74E+00 | 9.95E-01 | no | Nek11 | intron_variant | MODIFIER |
| 9 | 105341865 | AX-170285609 | 0 | A | G | 0.123 | -6.29E-04 | 1.18E-01 | 2.67E+00 | 2.74E+00 | 9.95E-01 | no | Nek11 | intron_variant | MODIFIER |

| | | | | | | | | | | | | | | | |
|---|---|---|---|---|---|---|---|---|---|---|---|---|---|---|---|
| 9 | 105342888 | AX-97853516 | 0 | T | C | 0.123 | -6.29E-04 | 1.18E-01 | 2.67E+00 | 2.74E+00 | 9.95E-01 | no | Nek11 | intron_variant | MODIFIER |
| 9 | 105343559 | AX-170347859 | 0 | G | A | 0.123 | -6.29E-04 | 1.18E-01 | 2.67E+00 | 2.74E+00 | 9.95E-01 | no | Nek11 | intron_variant | MODIFIER |
| 9 | 105355824 | AX-170432193 | 0 | G | A | 0.123 | -6.29E-04 | 1.18E-01 | 2.67E+00 | 2.74E+00 | 9.95E-01 | no | Nek11 | intron_variant | MODIFIER |
| 9 | 105355913 | AX-170307528 | 0 | T | C | 0.123 | -6.29E-04 | 1.18E-01 | 2.67E+00 | 2.74E+00 | 9.95E-01 | no | Nek11 | intron_variant | MODIFIER |
| 9 | 105358144 | AX-170345572 | 0 | C | T | 0.123 | -6.29E-04 | 1.18E-01 | 2.67E+00 | 2.74E+00 | 9.95E-01 | no | Nek11 | intron_variant | MODIFIER |
| 9 | 105358911 | AX-170033195 | 0 | G | A | 0.123 | -6.29E-04 | 1.18E-01 | 2.67E+00 | 2.74E+00 | 9.95E-01 | no | Nek11 | intron_variant | MODIFIER |
| 9 | 105359140 | AX-170299200 | 0 | G | C | 0.123 | -6.29E-04 | 1.18E-01 | 2.67E+00 | 2.74E+00 | 9.95E-01 | no | Nek11 | intron_variant | MODIFIER |
| 9 | 105361008 | AX-170033200 | 0 | C | A | 0.123 | -6.29E-04 | 1.18E-01 | 2.67E+00 | 2.74E+00 | 9.95E-01 | no | Nek11 | intron_variant | MODIFIER |
| 9 | 105362244 | AX-170033193 | 0 | A | C | 0.123 | -6.29E-04 | 1.18E-01 | 2.67E+00 | 2.74E+00 | 9.95E-01 | no | Nek11 | intron_variant | MODIFIER |
| 9 | 105363848 | AX-170165543 | 0 | C | G | 0.123 | -6.29E-04 | 1.18E-01 | 2.67E+00 | 2.74E+00 | 9.95E-01 | no | Nek11 | intron_variant | MODIFIER |
| 9 | 105370834 | AX-170471247 | 0 | A | G | 0.123 | -6.29E-04 | 1.18E-01 | 2.67E+00 | 2.74E+00 | 9.95E-01 | no | Nek11 | intron_variant | MODIFIER |
| 9 | 105370856 | AX-170096662 | 0 | C | T | 0.123 | -6.29E-04 | 1.18E-01 | 2.67E+00 | 2.74E+00 | 9.95E-01 | no | Nek11 | intron_variant | MODIFIER |
| 9 | 105399044 | AX-170484849 | 0 | T | C | 0.123 | -6.29E-04 | 1.18E-01 | 2.67E+00 | 2.74E+00 | 9.95E-01 | no | Aste1 | intron_variant | MODIFIER |
| 9 | 105399066 | AX-170397132 | 0 | A | G | 0.123 | -6.29E-04 | 1.18E-01 | 2.67E+00 | 2.74E+00 | 9.95E-01 | no | Aste1 | intron_variant | MODIFIER |
| 9 | 105402218 | AX-97806130 | 0 | A | G | 0.123 | -6.29E-04 | 1.18E-01 | 2.67E+00 | 2.74E+00 | 9.95E-01 | no | Aste1 | intron_variant | MODIFIER |
| 9 | 105404077 | AX-170033199 | 0 | C | T | 0.123 | -6.29E-04 | 1.18E-01 | 2.67E+00 | 2.74E+00 | 9.95E-01 | no | Aste1 | intron_variant | MODIFIER |
| 9 | 105404573 | AX-170377243 | 0 | A | G | 0.123 | -6.29E-04 | 1.18E-01 | 2.67E+00 | 2.74E+00 | 9.95E-01 | no | Aste1 | intron_variant | MODIFIER |
| 9 | 105405457 | AX-170083355 | 0 | A | G | 0.123 | -6.29E-04 | 1.18E-01 | 2.67E+00 | 2.74E+00 | 9.95E-01 | no | Aste1 | 3_prime_UTR_variant | MODIFIER |
| 9 | 105418065 | AX-170083357 | 0 | C | A | 0.123 | -6.29E-04 | 1.18E-01 | 2.67E+00 | 2.74E+00 | 9.95E-01 | no | Atp2c1 | synonymous_variant | LOW |
| 9 | 105418968 | AX-97832013 | 0 | C | T | 0.123 | -6.29E-04 | 1.18E-01 | 2.67E+00 | 2.74E+00 | 9.95E-01 | no | Atp2c1 | intron_variant | MODIFIER |
| 9 | 105459560 | AX-97830474 | 0 | C | T | 0.123 | -6.29E-04 | 1.18E-01 | 2.67E+00 | 2.74E+00 | 9.95E-01 | no | Atp2c1 | intron_variant | MODIFIER |
| 9 | 105460524 | AX-170033196 | 0 | G | A | 0.123 | -6.29E-04 | 1.18E-01 | 2.67E+00 | 2.74E+00 | 9.95E-01 | no | Atp2c1 | intron_variant | MODIFIER |
| 9 | 105488951 | AX-170272270 | 0 | C | T | 0.123 | -6.29E-04 | 1.18E-01 | 2.67E+00 | 2.74E+00 | 9.95E-01 | no | Atp2c1 | intron_variant | MODIFIER |
| 9 | 105489582 | AX-97780790 | 0 | T | C | 0.123 | -6.29E-04 | 1.18E-01 | 2.67E+00 | 2.74E+00 | 9.95E-01 | no | Atp2c1 | intron_variant | MODIFIER |
| 9 | 105508298 | AX-170498539 | 0 | A | G | 0.123 | -6.29E-04 | 1.18E-01 | 2.67E+00 | 2.74E+00 | 9.95E-01 | no | Atp2c1 | intron_variant | MODIFIER |
| 9 | 105512752 | AX-170241916 | 0 | G | A | 0.123 | -6.29E-04 | 1.18E-01 | 2.67E+00 | 2.74E+00 | 9.95E-01 | no | Atp2c1 | intron_variant | MODIFIER |
| 9 | 105530222 | AX-170450078 | 0 | G | C | 0.123 | -6.29E-04 | 1.18E-01 | 2.67E+00 | 2.74E+00 | 9.95E-01 | no | Atp2c1 | upstream_gene_variant | MODIFIER |
| 9 | 105556124 | AX-97839933 | 0 | T | C | 0.123 | -6.29E-04 | 1.18E-01 | 2.67E+00 | 2.74E+00 | 9.95E-01 | no | Gm5621 | downstream_gene_variant | MODIFIER |
| 9 | 105557603 | AX-170237772 | 0 | G | A | 0.123 | -6.29E-04 | 1.18E-01 | 2.67E+00 | 2.74E+00 | 9.95E-01 | no | Gm5621 | downstream_gene_variant | MODIFIER |
| 9 | 105558947 | AX-170194789 | 0 | A | G | 0.123 | -6.29E-04 | 1.18E-01 | 2.67E+00 | 2.74E+00 | 9.95E-01 | no | Gm5621 | downstream_gene_variant | MODIFIER |
| 9 | 105562585 | AX-97816454 | 0 | T | G | 0.123 | -6.29E-04 | 1.18E-01 | 2.67E+00 | 2.74E+00 | 9.95E-01 | no | Gm5621 | upstream_gene_variant | MODIFIER |
| 9 | 105593283 | AX-170224289 | 0 | T | C | 0.123 | -6.29E-04 | 1.18E-01 | 2.67E+00 | 2.74E+00 | 9.95E-01 | no | | intergenic_variant | MODIFIER |
| 9 | 105616111 | AX-97786697 | 0 | G | A | 0.123 | -6.29E-04 | 1.18E-01 | 2.67E+00 | 2.74E+00 | 9.95E-01 | no | | intergenic_variant | MODIFIER |
| 9 | 105635586 | AX-97805521 | 0 | A | G | 0.123 | -6.29E-04 | 1.18E-01 | 2.67E+00 | 2.74E+00 | 9.95E-01 | no | | intergenic_variant | MODIFIER |
| 9 | 105639523 | AX-170526543 | 0 | T | C | 0.123 | -6.29E-04 | 1.18E-01 | 2.67E+00 | 2.74E+00 | 9.95E-01 | no | Pik3r4 | upstream_gene_variant | MODIFIER |
| 9 | 105641738 | AX-170438777 | 0 | A | G | 0.123 | -6.29E-04 | 1.18E-01 | 2.67E+00 | 2.74E+00 | 9.95E-01 | no | Pik3r4 | upstream_gene_variant | MODIFIER |
| 9 | 105643833 | AX-170376450 | 0 | A | G | 0.123 | -6.29E-04 | 1.18E-01 | 2.67E+00 | 2.74E+00 | 9.95E-01 | no | Pik3r4 | intron_variant | MODIFIER |
| 9 | 105672237 | AX-97799135 | 0 | C | T | 0.123 | -6.29E-04 | 1.18E-01 | 2.67E+00 | 2.74E+00 | 9.95E-01 | no | Pik3r4 | intron_variant | MODIFIER |
| 9 | 105681508 | AX-170370098 | 0 | C | T | 0.123 | -6.29E-04 | 1.18E-01 | 2.67E+00 | 2.74E+00 | 9.95E-01 | no | Pik3r4 | intron_variant | MODIFIER |
| 9 | 105712433 | AX-170352209 | 0 | T | C | 0.123 | -6.29E-04 | 1.18E-01 | 2.67E+00 | 2.74E+00 | 9.95E-01 | no | Col6a6 | intron_variant | MODIFIER |
| 9 | 105715108 | AX-97839939 | 0 | A | G | 0.123 | -6.29E-04 | 1.18E-01 | 2.67E+00 | 2.74E+00 | 9.95E-01 | no | Col6a6 | intron_variant | MODIFIER |
| 9 | 105715601 | AX-170083363 | 0 | G | T | 0.123 | -6.29E-04 | 1.18E-01 | 2.67E+00 | 2.74E+00 | 9.95E-01 | no | Col6a6 | intron_variant | MODIFIER |
| 9 | 105731836 | AX-170298471 | 0 | G | A | 0.123 | -6.29E-04 | 1.18E-01 | 2.67E+00 | 2.74E+00 | 9.95E-01 | no | Col6a6 | intron_variant | MODIFIER |
| 9 | 105755634 | AX-170448010 | 0 | G | A | 0.123 | -6.29E-04 | 1.18E-01 | 2.67E+00 | 2.74E+00 | 9.95E-01 | no | Col6a6 | intron_variant | MODIFIER |
| 9 | 105765868 | AX-170090402 | 0 | G | A | 0.123 | -6.29E-04 | 1.18E-01 | 2.67E+00 | 2.74E+00 | 9.95E-01 | no | Col6a6 | intron_variant | MODIFIER |
| 9 | 105799816 | AX-170033203 | 0 | A | G | 0.123 | -6.29E-04 | 1.18E-01 | 2.67E+00 | 2.74E+00 | 9.95E-01 | no | Col6a6 | intron_variant | MODIFIER |
| 9 | 105800669 | AX-170439735 | 0 | C | T | 0.123 | -6.29E-04 | 1.18E-01 | 2.67E+00 | 2.74E+00 | 9.95E-01 | no | Col6a6 | intron_variant | MODIFIER |
| 9 | 105801151 | AX-97805520 | 0 | G | A | 0.123 | -6.29E-04 | 1.18E-01 | 2.67E+00 | 2.74E+00 | 9.95E-01 | no | Col6a6 | intron_variant | MODIFIER |
| 9 | 105842630 | AX-170033210 | 0 | A | G | 0.123 | -6.29E-04 | 1.18E-01 | 2.67E+00 | 2.74E+00 | 9.95E-01 | no | | intergenic_variant | MODIFIER |
| 9 | 105845358 | AX-97803895 | 0 | A | G | 0.123 | -6.29E-04 | 1.18E-01 | 2.67E+00 | 2.74E+00 | 9.95E-01 | no | | intergenic_variant | MODIFIER |
| 9 | 105854233 | AX-170211112 | 0 | G | A | 0.123 | -6.29E-04 | 1.18E-01 | 2.67E+00 | 2.74E+00 | 9.95E-01 | no | Col6a5 | downstream_gene_variant | MODIFIER |
| 9 | 105864834 | AX-170345219 | 0 | T | C | 0.123 | -6.29E-04 | 1.18E-01 | 2.67E+00 | 2.74E+00 | 9.95E-01 | no | Col6a5 | splice_region_variant,synonymous_variant | LOW |
| 9 | 105871083 | AX-97851070 | 0 | C | T | 0.123 | -6.29E-04 | 1.18E-01 | 2.67E+00 | 2.74E+00 | 9.95E-01 | no | Col6a5 | intron_variant | MODIFIER |
| 9 | 105896805 | AX-170090403 | 0 | G | A | 0.123 | -6.29E-04 | 1.18E-01 | 2.67E+00 | 2.74E+00 | 9.95E-01 | no | Col6a5 | intron_variant | MODIFIER |

| | | | | | | | | | | | | | | | |
|---|---|---|---|---|---|---|---|---|---|---|---|---|---|---|---|
| 9 | 105909155 | AX-170206520 | 0 | G | A | 0.123 | -6.29E-04 | 1.18E-01 | 2.67E+00 | 2.74E+00 | 9.95E-01 | no | Col6a5 | intron_variant | MODIFIER |
| 9 | 105912824 | AX-170159401 | 0 | T | C | 0.123 | -6.29E-04 | 1.18E-01 | 2.67E+00 | 2.74E+00 | 9.95E-01 | no | Col6a5 | intron_variant | MODIFIER |
| 9 | 105918292 | AX-170180789 | 0 | T | C | 0.123 | -6.29E-04 | 1.18E-01 | 2.67E+00 | 2.74E+00 | 9.95E-01 | no | Col6a5 | intron_variant | MODIFIER |
| 9 | 105918455 | AX-97798433 | 0 | C | T | 0.123 | -6.29E-04 | 1.18E-01 | 2.67E+00 | 2.74E+00 | 9.95E-01 | no | Col6a5 | intron_variant | MODIFIER |
| 9 | 105925859 | AX-170033204 | 0 | G | A | 0.123 | -6.29E-04 | 1.18E-01 | 2.67E+00 | 2.74E+00 | 9.95E-01 | no | Col6a5 | synonymous_variant | LOW |
| 9 | 105937329 | AX-170033207 | 0 | T | C | 0.123 | -6.29E-04 | 1.18E-01 | 2.67E+00 | 2.74E+00 | 9.95E-01 | no | Col6a5 | missense_variant | MODERATE |
| 9 | 105937538 | AX-170033208 | 0 | G | A | 0.123 | -6.29E-04 | 1.18E-01 | 2.67E+00 | 2.74E+00 | 9.95E-01 | no | Col6a5 | intron_variant | MODIFIER |
| 9 | 105940289 | AX-170184220 | 0 | A | G | 0.123 | -6.29E-04 | 1.18E-01 | 2.67E+00 | 2.74E+00 | 9.95E-01 | no | Col6a5 | synonymous_variant | LOW |
| 9 | 105969848 | AX-97811017 | 0 | T | G | 0.123 | -6.29E-04 | 1.18E-01 | 2.67E+00 | 2.74E+00 | 9.95E-01 | no | | intergenic_variant | MODIFIER |
| 9 | 105994187 | AX-170173817 | 0 | C | G | 0.123 | -6.29E-04 | 1.18E-01 | 2.67E+00 | 2.74E+00 | 9.95E-01 | no | Col6a4 | intron_variant | MODIFIER |
| 9 | 105998660 | AX-97792650 | 0 | C | T | 0.123 | -6.29E-04 | 1.18E-01 | 2.67E+00 | 2.74E+00 | 9.95E-01 | no | Col6a4 | intron_variant | MODIFIER |
| 9 | 105999570 | AX-170083369 | 0 | A | C | 0.123 | -6.29E-04 | 1.18E-01 | 2.67E+00 | 2.74E+00 | 9.95E-01 | no | Col6a4 | intron_variant | MODIFIER |
| 9 | 106013889 | AX-170362079 | 0 | G | A | 0.123 | -6.29E-04 | 1.18E-01 | 2.67E+00 | 2.74E+00 | 9.95E-01 | no | Col6a4 | synonymous_variant | LOW |
| 9 | 106016292 | AX-170313775 | 0 | G | A | 0.123 | -6.29E-04 | 1.18E-01 | 2.67E+00 | 2.74E+00 | 9.95E-01 | no | Col6a4 | intron_variant | MODIFIER |
| 9 | 106016587 | AX-170327455 | 0 | G | A | 0.123 | -6.29E-04 | 1.18E-01 | 2.67E+00 | 2.74E+00 | 9.95E-01 | no | Col6a4 | intron_variant | MODIFIER |
| 9 | 106017492 | AX-97787805 | 0 | G | A | 0.123 | -6.29E-04 | 1.18E-01 | 2.67E+00 | 2.74E+00 | 9.95E-01 | no | Col6a4 | intron_variant | MODIFIER |
| 9 | 106052873 | AX-170304423 | 0 | C | G | 0.123 | -6.29E-04 | 1.18E-01 | 2.67E+00 | 2.74E+00 | 9.95E-01 | no | Col6a4 | intron_variant | MODIFIER |
| 9 | 106058464 | AX-170240406 | 0 | C | A | 0.123 | -6.29E-04 | 1.18E-01 | 2.67E+00 | 2.74E+00 | 9.95E-01 | no | Col6a4 | intron_variant | MODIFIER |
| 9 | 106058890 | AX-97783280 | 0 | G | T | 0.123 | -6.29E-04 | 1.18E-01 | 2.67E+00 | 2.74E+00 | 9.95E-01 | no | Col6a4 | intron_variant | MODIFIER |
| 9 | 106064989 | AX-170194143 | 0 | T | C | 0.123 | -6.29E-04 | 1.18E-01 | 2.67E+00 | 2.74E+00 | 9.95E-01 | no | Col6a4 | intron_variant | MODIFIER |
| 9 | 106095881 | AX-170179301 | 0 | A | C | 0.123 | -6.29E-04 | 1.18E-01 | 2.67E+00 | 2.74E+00 | 9.95E-01 | no | Col6a4 | intron_variant | MODIFIER |
| 9 | 106098311 | AX-170155940 | 0 | C | A | 0.123 | -6.29E-04 | 1.18E-01 | 2.67E+00 | 2.74E+00 | 9.95E-01 | no | Col6a4 | upstream_gene_variant | MODIFIER |
| 9 | 106098332 | AX-170302529 | 0 | C | T | 0.123 | -6.29E-04 | 1.18E-01 | 2.67E+00 | 2.74E+00 | 9.95E-01 | no | Col6a4 | upstream_gene_variant | MODIFIER |
| 9 | 106099010 | AX-170513328 | 0 | A | C | 0.123 | -6.29E-04 | 1.18E-01 | 2.67E+00 | 2.74E+00 | 9.95E-01 | no | Col6a4 | upstream_gene_variant | MODIFIER |
| 9 | 106122855 | AX-97818679 | 0 | G | A | 0.123 | -6.29E-04 | 1.18E-01 | 2.67E+00 | 2.74E+00 | 9.95E-01 | no | Gm29208 | intron_variant,non_coding_transcript_variant | MODIFIER |
| 9 | 106125734 | AX-170213088 | 0 | A | G | 0.123 | -6.29E-04 | 1.18E-01 | 2.67E+00 | 2.74E+00 | 9.95E-01 | no | Gm29208 | intron_variant,non_coding_transcript_variant | MODIFIER |
| 9 | 106137954 | AX-170291602 | 0 | T | C | 0.123 | -6.29E-04 | 1.18E-01 | 2.67E+00 | 2.74E+00 | 9.95E-01 | no | | intergenic_variant | MODIFIER |
| 9 | 106142790 | AX-97813005 | 0 | A | G | 0.123 | -6.29E-04 | 1.18E-01 | 2.67E+00 | 2.74E+00 | 9.95E-01 | no | | intergenic_variant | MODIFIER |
| 9 | 106158734 | AX-170264830 | 0 | T | C | 0.123 | -6.29E-04 | 1.18E-01 | 2.67E+00 | 2.74E+00 | 9.95E-01 | no | Glyctk | upstream_gene_variant | MODIFIER |
| 9 | 106160191 | AX-170126117 | 0 | A | G | 0.123 | -6.29E-04 | 1.18E-01 | 2.67E+00 | 2.74E+00 | 9.95E-01 | no | Glyctk | upstream_gene_variant | MODIFIER |
| 9 | 106181319 | AX-170207830 | 0 | A | G | 0.123 | -6.29E-04 | 1.18E-01 | 2.67E+00 | 2.74E+00 | 9.95E-01 | no | Wdr82 | intron_variant | MODIFIER |
| 9 | 106189828 | AX-170351256 | 0 | G | C | 0.123 | -6.29E-04 | 1.18E-01 | 2.67E+00 | 2.74E+00 | 9.95E-01 | no | Wdr82 | 3_prime_UTR_variant | MODIFIER |
| 9 | 106192904 | AX-170090405 | 0 | G | A | 0.123 | -6.29E-04 | 1.18E-01 | 2.67E+00 | 2.74E+00 | 9.95E-01 | no | Wdr82 | downstream_gene_variant | MODIFIER |
| 9 | 106193104 | AX-170177630 | 0 | A | G | 0.123 | -6.29E-04 | 1.18E-01 | 2.67E+00 | 2.74E+00 | 9.95E-01 | no | Wdr82 | downstream_gene_variant | MODIFIER |
| 9 | 106214684 | AX-170083372 | 0 | T | C | 0.123 | -6.29E-04 | 1.18E-01 | 2.67E+00 | 2.74E+00 | 9.95E-01 | no | Twf2 | intron_variant | MODIFIER |
| 9 | 106214860 | AX-170242094 | 0 | T | C | 0.123 | -6.29E-04 | 1.18E-01 | 2.67E+00 | 2.74E+00 | 9.95E-01 | no | Twf2 | synonymous_variant | LOW |
| 9 | 106224485 | AX-97831007 | 0 | C | A | 0.123 | -6.29E-04 | 1.18E-01 | 2.67E+00 | 2.74E+00 | 9.95E-01 | no | Tlr9 | missense_variant | MODERATE |
| 9 | 106224591 | AX-170390711 | 0 | T | C | 0.123 | -6.29E-04 | 1.18E-01 | 2.67E+00 | 2.74E+00 | 9.95E-01 | no | Tlr9 | synonymous_variant | LOW |
| 9 | 106224644 | AX-170348470 | 0 | T | C | 0.123 | -6.29E-04 | 1.18E-01 | 2.67E+00 | 2.74E+00 | 9.95E-01 | no | Tlr9 | missense_variant | MODERATE |
| 9 | 106247113 | AX-97813852 | 0 | C | T | 0.123 | -6.29E-04 | 1.18E-01 | 2.67E+00 | 2.74E+00 | 9.95E-01 | no | Alas1 | intron_variant | MODIFIER |
| 9 | 106257489 | AX-170105459 | 0 | A | T | 0.123 | -6.29E-04 | 1.18E-01 | 2.67E+00 | 2.74E+00 | 9.95E-01 | no | | intergenic_variant | MODIFIER |
| 9 | 106259870 | AX-170116902 | 0 | T | C | 0.123 | -6.29E-04 | 1.18E-01 | 2.67E+00 | 2.74E+00 | 9.95E-01 | no | | intergenic_variant | MODIFIER |
| 9 | 106281230 | AX-170380383 | 0 | C | T | 0.123 | -6.29E-04 | 1.18E-01 | 2.67E+00 | 2.74E+00 | 9.95E-01 | no | Poc1a | 5_prime_UTR_variant | MODIFIER |
| 9 | 106282002 | AX-97839444 | 0 | C | A | 0.123 | -6.29E-04 | 1.18E-01 | 2.67E+00 | 2.74E+00 | 9.95E-01 | no | Poc1a | intron_variant | MODIFIER |
| 9 | 106284925 | AX-170033216 | 0 | C | T | 0.123 | -6.29E-04 | 1.18E-01 | 2.67E+00 | 2.74E+00 | 9.95E-01 | no | Poc1a | missense_variant | MODERATE |
| 9 | 106289637 | AX-170460384 | 0 | G | A | 0.123 | -6.29E-04 | 1.18E-01 | 2.67E+00 | 2.74E+00 | 9.95E-01 | no | Poc1a | intron_variant | MODIFIER |
| 9 | 106313390 | AX-170124149 | 0 | C | T | 0.123 | -6.29E-04 | 1.18E-01 | 2.67E+00 | 2.74E+00 | 9.95E-01 | no | Poc1a | intron_variant | MODIFIER |
| 9 | 106313812 | AX-170138774 | 0 | A | T | 0.123 | -6.29E-04 | 1.18E-01 | 2.67E+00 | 2.74E+00 | 9.95E-01 | no | Poc1a | intron_variant | MODIFIER |
| 9 | 106315727 | AX-170348488 | 0 | A | G | 0.123 | -6.29E-04 | 1.18E-01 | 2.67E+00 | 2.74E+00 | 9.95E-01 | no | Poc1a | intron_variant | MODIFIER |
| 9 | 106322068 | AX-97849291 | 0 | T | C | 0.123 | -6.29E-04 | 1.18E-01 | 2.67E+00 | 2.74E+00 | 9.95E-01 | no | Poc1a | intron_variant | MODIFIER |
| 9 | 106345578 | AX-97809382 | 0 | A | G | 0.123 | -6.29E-04 | 1.18E-01 | 2.67E+00 | 2.74E+00 | 9.95E-01 | no | Poc1a | intron_variant | MODIFIER |
| 9 | 106390609 | AX-170320876 | 0 | C | T | 0.123 | -6.29E-04 | 1.18E-01 | 2.67E+00 | 2.74E+00 | 9.95E-01 | no | | intergenic_variant | MODIFIER |
| 9 | 106400671 | AX-170033212 | 0 | G | T | 0.123 | -6.29E-04 | 1.18E-01 | 2.67E+00 | 2.74E+00 | 9.95E-01 | no | | regulatory_region_variant | MODIFIER |
| 9 | 106403171 | AX-170376474 | 0 | A | G | 0.123 | -6.29E-04 | 1.18E-01 | 2.67E+00 | 2.74E+00 | 9.95E-01 | no | | regulatory_region_variant | MODIFIER |

| | | | | | | | | | | | | | | | |
|---|---|---|---|---|---|---|---|---|---|---|---|---|---|---|---|
| 9 | 106406254 | AX-170142917 | 0 | G | A | 0.123 | -6.29E-04 | 1.18E-01 | 2.67E+00 | 2.74E+00 | 9.95E-01 | no | | regulatory_region_variant | MODIFIER |
| 9 | 106407148 | AX-170214527 | 0 | G | A | 0.123 | -6.29E-04 | 1.18E-01 | 2.67E+00 | 2.74E+00 | 9.95E-01 | no | | regulatory_region_variant | MODIFIER |
| 9 | 106408508 | AX-170033215 | 0 | T | C | 0.123 | -6.29E-04 | 1.18E-01 | 2.67E+00 | 2.74E+00 | 9.95E-01 | no | | regulatory_region_variant | MODIFIER |
| 9 | 106437527 | AX-170033218 | 0 | G | A | 0.123 | -6.29E-04 | 1.18E-01 | 2.67E+00 | 2.74E+00 | 9.95E-01 | no | Acy1 | intron_variant | MODIFIER |
| 9 | 106440632 | AX-170315834 | 0 | C | T | 0.123 | -6.29E-04 | 1.18E-01 | 2.67E+00 | 2.74E+00 | 9.95E-01 | no | Acy1 | upstream_gene_variant | MODIFIER |
| 9 | 106441906 | AX-170322075 | 0 | T | C | 0.123 | -6.29E-04 | 1.18E-01 | 2.67E+00 | 2.74E+00 | 9.95E-01 | no | Acy1 | upstream_gene_variant | MODIFIER |
| 9 | 106442529 | AX-170033220 | 0 | A | T | 0.123 | -6.29E-04 | 1.18E-01 | 2.67E+00 | 2.74E+00 | 9.95E-01 | no | Acy1 | upstream_gene_variant | MODIFIER |
| 9 | 106449110 | AX-97806156 | 0 | A | G | 0.123 | -6.29E-04 | 1.18E-01 | 2.67E+00 | 2.74E+00 | 9.95E-01 | no | Pcbp4 | upstream_gene_variant | MODIFIER |
| 9 | 106475926 | AX-97788808 | 0 | C | T | 0.123 | -6.29E-04 | 1.18E-01 | 2.67E+00 | 2.74E+00 | 9.95E-01 | no | Rrp9 | upstream_gene_variant | MODIFIER |
| 9 | 106479859 | AX-170033214 | 0 | G | C | 0.123 | -6.29E-04 | 1.18E-01 | 2.67E+00 | 2.74E+00 | 9.95E-01 | no | Rrp9 | intron_variant | MODIFIER |
| 9 | 106501363 | AX-170083373 | 0 | C | A | 0.123 | -6.29E-04 | 1.18E-01 | 2.67E+00 | 2.74E+00 | 9.95E-01 | no | Iqcf1 | intron_variant | MODIFIER |
| 9 | 106501931 | AX-170083374 | 0 | A | G | 0.123 | -6.29E-04 | 1.18E-01 | 2.67E+00 | 2.74E+00 | 9.95E-01 | no | Iqcf1 | synonymous_variant | LOW |
| 9 | 106512615 | AX-170033213 | 0 | A | C | 0.123 | -6.29E-04 | 1.18E-01 | 2.67E+00 | 2.74E+00 | 9.95E-01 | no | Iqcf5 | upstream_gene_variant | MODIFIER |
| 9 | 106525074 | AX-97831475 | 0 | A | G | 0.123 | -6.29E-04 | 1.18E-01 | 2.67E+00 | 2.74E+00 | 9.95E-01 | no | Iqcf3 | intron_variant,non_coding_transcript_variant | MODIFIER |
| 9 | 106540068 | AX-170301661 | 0 | T | C | 0.123 | -6.29E-04 | 1.18E-01 | 2.67E+00 | 2.74E+00 | 9.95E-01 | no | Iqcf3 | downstream_gene_variant | MODIFIER |
| 9 | 106544110 | AX-97810320 | 0 | G | A | 0.123 | -6.29E-04 | 1.18E-01 | 2.67E+00 | 2.74E+00 | 9.95E-01 | no | Iqcf3 | intron_variant | MODIFIER |
| 9 | 106567698 | AX-170388314 | 0 | A | G | 0.123 | -6.29E-04 | 1.18E-01 | 2.67E+00 | 2.74E+00 | 9.95E-01 | no | Iqcf4 | downstream_gene_variant | MODIFIER |
| 9 | 106573676 | AX-170090797 | 0 | A | G | 0.123 | -6.29E-04 | 1.18E-01 | 2.67E+00 | 2.74E+00 | 9.95E-01 | no | Iqcf4 | upstream_gene_variant | MODIFIER |
| 9 | 106575271 | AX-170270583 | 0 | C | T | 0.123 | -6.29E-04 | 1.18E-01 | 2.67E+00 | 2.74E+00 | 9.95E-01 | no | Iqcf4 | upstream_gene_variant | MODIFIER |
| 9 | 106592285 | AX-170228187 | 0 | C | G | 0.123 | -6.29E-04 | 1.18E-01 | 2.67E+00 | 2.74E+00 | 9.95E-01 | no | | intergenic_variant | MODIFIER |
| 9 | 106593818 | AX-170380596 | 0 | G | A | 0.123 | -6.29E-04 | 1.18E-01 | 2.67E+00 | 2.74E+00 | 9.95E-01 | no | | intergenic_variant | MODIFIER |
| 9 | 106598567 | AX-97793204 | 0 | G | A | 0.123 | -6.29E-04 | 1.18E-01 | 2.67E+00 | 2.74E+00 | 9.95E-01 | no | | intergenic_variant | MODIFIER |
| 9 | 106602444 | AX-170513151 | 0 | T | A | 0.123 | -6.29E-04 | 1.18E-01 | 2.67E+00 | 2.74E+00 | 9.95E-01 | no | | intergenic_variant | MODIFIER |
| 9 | 106606038 | AX-170256832 | 0 | A | G | 0.123 | -6.29E-04 | 1.18E-01 | 2.67E+00 | 2.74E+00 | 9.95E-01 | no | | intergenic_variant | MODIFIER |
| 9 | 106606439 | AX-170424876 | 0 | T | C | 0.123 | -6.29E-04 | 1.18E-01 | 2.67E+00 | 2.74E+00 | 9.95E-01 | no | | intergenic_variant | MODIFIER |
| 9 | 106612591 | AX-170340439 | 0 | C | T | 0.123 | -6.29E-04 | 1.18E-01 | 2.67E+00 | 2.74E+00 | 9.95E-01 | no | | intergenic_variant | MODIFIER |
| 9 | 106621559 | AX-97790909 | 0 | A | G | 0.123 | -6.29E-04 | 1.18E-01 | 2.67E+00 | 2.74E+00 | 9.95E-01 | no | | intergenic_variant | MODIFIER |
| 9 | 106630293 | AX-170228542 | 0 | G | C | 0.123 | -6.29E-04 | 1.18E-01 | 2.67E+00 | 2.74E+00 | 9.95E-01 | no | Iqcf6 | downstream_gene_variant | MODIFIER |
| 9 | 106633321 | AX-170507914 | 0 | G | C | 0.123 | -6.29E-04 | 1.18E-01 | 2.67E+00 | 2.74E+00 | 9.95E-01 | no | Gm26185 | upstream_gene_variant | MODIFIER |
| 9 | 106642048 | AX-97828158 | 0 | T | C | 0.123 | -6.29E-04 | 1.18E-01 | 2.67E+00 | 2.74E+00 | 9.95E-01 | no | Grm2 | downstream_gene_variant | MODIFIER |
| 9 | 106647931 | AX-170247188 | 0 | A | G | 0.123 | -6.29E-04 | 1.18E-01 | 2.67E+00 | 2.74E+00 | 9.95E-01 | no | Grm2 | synonymous_variant | LOW |
| 9 | 106651697 | AX-170497316 | 0 | T | C | 0.123 | -6.29E-04 | 1.18E-01 | 2.67E+00 | 2.74E+00 | 9.95E-01 | no | Grm2 | intron_variant | MODIFIER |
| 9 | 106652628 | AX-170312599 | 0 | G | C | 0.123 | -6.29E-04 | 1.18E-01 | 2.67E+00 | 2.74E+00 | 9.95E-01 | no | Grm2 | intron_variant | MODIFIER |
| 9 | 106671015 | AX-97791302 | 0 | T | C | 0.123 | -6.29E-04 | 1.18E-01 | 2.67E+00 | 2.74E+00 | 9.95E-01 | no | Tex264 | intron_variant | MODIFIER |
| 9 | 106673006 | AX-170083381 | 0 | T | C | 0.123 | -6.29E-04 | 1.18E-01 | 2.67E+00 | 2.74E+00 | 9.95E-01 | no | Tex264 | intron_variant | MODIFIER |
| 9 | 106675962 | AX-170264989 | 0 | A | G | 0.123 | -6.29E-04 | 1.18E-01 | 2.67E+00 | 2.74E+00 | 9.95E-01 | no | Tex264 | intron_variant | MODIFIER |
| 9 | 106687040 | AX-170414593 | 0 | A | G | 0.123 | -6.29E-04 | 1.18E-01 | 2.67E+00 | 2.74E+00 | 9.95E-01 | no | Rad54l2 | downstream_gene_variant | MODIFIER |
| 9 | 106701568 | AX-97804205 | 0 | T | C | 0.123 | -6.29E-04 | 1.18E-01 | 2.67E+00 | 2.74E+00 | 9.95E-01 | no | Rad54l2 | intron_variant | MODIFIER |
| 9 | 106718393 | AX-170216708 | 0 | T | C | 0.123 | -6.29E-04 | 1.18E-01 | 2.67E+00 | 2.74E+00 | 9.95E-01 | no | Rad54l2 | intron_variant | MODIFIER |
| 9 | 106723036 | AX-97786954 | 0 | C | T | 0.123 | -6.29E-04 | 1.18E-01 | 2.67E+00 | 2.74E+00 | 9.95E-01 | no | Rad54l2 | intron_variant | MODIFIER |
| 9 | 106753366 | AX-97800881 | 0 | C | T | 0.123 | -6.29E-04 | 1.18E-01 | 2.67E+00 | 2.74E+00 | 9.95E-01 | no | Rad54l2 | intron_variant | MODIFIER |
| 9 | 106769232 | AX-170351654 | 0 | A | T | 0.123 | -6.29E-04 | 1.18E-01 | 2.67E+00 | 2.74E+00 | 9.95E-01 | no | Rad54l2 | intron_variant | MODIFIER |
| 9 | 106783055 | AX-170528616 | 0 | G | A | 0.123 | -6.29E-04 | 1.18E-01 | 2.67E+00 | 2.74E+00 | 9.95E-01 | no | Rad54l2 | intron_variant | MODIFIER |
| 9 | 106783648 | AX-97783960 | 0 | C | T | 0.123 | -6.29E-04 | 1.18E-01 | 2.67E+00 | 2.74E+00 | 9.95E-01 | no | Rad54l2 | intron_variant | MODIFIER |
| 9 | 106784539 | AX-170166225 | 0 | C | A | 0.123 | -6.29E-04 | 1.18E-01 | 2.67E+00 | 2.74E+00 | 9.95E-01 | no | Rad54l2 | intron_variant | MODIFIER |
| 9 | 106793241 | AX-97825132 | 0 | C | T | 0.123 | -6.29E-04 | 1.18E-01 | 2.67E+00 | 2.74E+00 | 9.95E-01 | no | Rad54l2 | upstream_gene_variant | MODIFIER |
| 9 | 106803819 | AX-170232610 | 0 | A | G | 0.123 | -6.29E-04 | 1.18E-01 | 2.67E+00 | 2.74E+00 | 9.95E-01 | no | | intergenic_variant | MODIFIER |
| 9 | 106841110 | AX-170033230 | 0 | A | G | 0.123 | -6.29E-04 | 1.18E-01 | 2.67E+00 | 2.74E+00 | 9.95E-01 | no | Dcaf1 | intron_variant | MODIFIER |
| 9 | 106852176 | AX-170033221 | 0 | A | G | 0.123 | -6.29E-04 | 1.18E-01 | 2.67E+00 | 2.74E+00 | 9.95E-01 | no | Dcaf1 | intron_variant | MODIFIER |
| 9 | 106854021 | AX-97855629 | 0 | A | C | 0.123 | -6.29E-04 | 1.18E-01 | 2.67E+00 | 2.74E+00 | 9.95E-01 | no | Dcaf1 | intron_variant | MODIFIER |
| 9 | 106859876 | AX-170104903 | 0 | T | C | 0.123 | -6.29E-04 | 1.18E-01 | 2.67E+00 | 2.74E+00 | 9.95E-01 | no | Dcaf1 | intron_variant | MODIFIER |
| 9 | 106863589 | AX-170347761 | 0 | T | C | 0.123 | -6.29E-04 | 1.18E-01 | 2.67E+00 | 2.74E+00 | 9.95E-01 | no | Dcaf1 | intron_variant | MODIFIER |
| 9 | 106872676 | AX-170090406 | 0 | C | T | 0.123 | -6.29E-04 | 1.18E-01 | 2.67E+00 | 2.74E+00 | 9.95E-01 | no | Dcaf1 | intron_variant | MODIFIER |
| 9 | 106875104 | AX-170033222 | 0 | G | A | 0.123 | -6.29E-04 | 1.18E-01 | 2.67E+00 | 2.74E+00 | 9.95E-01 | no | Dcaf1 | intron_variant | MODIFIER |

| | | | | | | | | | | | | | | | |
|---|---|---|---|---|---|---|---|---|---|---|---|---|---|---|---|
| 9 | 106890107 | AX-97809938 | 0 | T | C | 0.123 | -6.29E-04 | 1.18E-01 | 2.67E+00 | 2.74E+00 | 9.95E-01 | no | Dock3 | downstream_gene_variant | MODIFIER |
| 9 | 106931749 | AX-97809685 | 0 | A | G | 0.123 | -6.29E-04 | 1.18E-01 | 2.67E+00 | 2.74E+00 | 9.95E-01 | no | Dock3 | intron_variant | MODIFIER |
| 9 | 106936072 | AX-170033224 | 0 | G | A | 0.123 | -6.29E-04 | 1.18E-01 | 2.67E+00 | 2.74E+00 | 9.95E-01 | no | Dock3 | intron_variant | MODIFIER |
| 9 | 106937664 | AX-170384153 | 0 | T | G | 0.123 | -6.29E-04 | 1.18E-01 | 2.67E+00 | 2.74E+00 | 9.95E-01 | no | Dock3 | intron_variant | MODIFIER |
| 9 | 106938225 | AX-97782835 | 0 | T | G | 0.123 | -6.29E-04 | 1.18E-01 | 2.67E+00 | 2.74E+00 | 9.95E-01 | no | Dock3 | intron_variant | MODIFIER |
| 9 | 106943722 | AX-170133944 | 0 | G | A | 0.123 | -6.29E-04 | 1.18E-01 | 2.67E+00 | 2.74E+00 | 9.95E-01 | no | Dock3 | intron_variant | MODIFIER |
| 9 | 106972823 | AX-170090407 | 0 | C | T | 0.123 | -6.29E-04 | 1.18E-01 | 2.67E+00 | 2.74E+00 | 9.95E-01 | no | Dock3 | intron_variant | MODIFIER |
| 9 | 106996658 | AX-97787624 | 0 | C | T | 0.123 | -6.29E-04 | 1.18E-01 | 2.67E+00 | 2.74E+00 | 9.95E-01 | no | Dock3 | intron_variant | MODIFIER |
| 9 | 107003936 | AX-170033234 | 0 | T | C | 0.123 | -6.29E-04 | 1.18E-01 | 2.67E+00 | 2.74E+00 | 9.95E-01 | no | Dock3 | intron_variant | MODIFIER |
| 9 | 107033679 | AX-97855346 | 0 | T | C | 0.123 | -6.29E-04 | 1.18E-01 | 2.67E+00 | 2.74E+00 | 9.95E-01 | no | Dock3 | intron_variant | MODIFIER |
| 9 | 107087974 | AX-170033232 | 0 | C | T | 0.123 | -6.29E-04 | 1.18E-01 | 2.67E+00 | 2.74E+00 | 9.95E-01 | no | Dock3 | intron_variant | MODIFIER |
| 9 | 107121671 | AX-97785241 | 0 | T | C | 0.123 | -6.29E-04 | 1.18E-01 | 2.67E+00 | 2.74E+00 | 9.95E-01 | no | Dock3 | intron_variant | MODIFIER |
| 9 | 107121708 | AX-170033225 | 0 | G | A | 0.123 | -6.29E-04 | 1.18E-01 | 2.67E+00 | 2.74E+00 | 9.95E-01 | no | Dock3 | intron_variant | MODIFIER |
| 9 | 107144300 | AX-170176989 | 0 | T | C | 0.123 | -6.29E-04 | 1.18E-01 | 2.67E+00 | 2.74E+00 | 9.95E-01 | no | Dock3 | intron_variant | MODIFIER |
| 9 | 107149632 | AX-170033229 | 0 | T | C | 0.123 | -6.29E-04 | 1.18E-01 | 2.67E+00 | 2.74E+00 | 9.95E-01 | no | Dock3 | intron_variant | MODIFIER |
| 9 | 107159731 | AX-97813493 | 0 | T | G | 0.123 | -6.29E-04 | 1.18E-01 | 2.67E+00 | 2.74E+00 | 9.95E-01 | no | Dock3 | intron_variant | MODIFIER |
| 9 | 107193559 | AX-170033223 | 0 | C | T | 0.123 | -6.29E-04 | 1.18E-01 | 2.67E+00 | 2.74E+00 | 9.95E-01 | no | Dock3 | intron_variant | MODIFIER |
| 9 | 107193629 | AX-170238046 | 0 | T | C | 0.123 | -6.29E-04 | 1.18E-01 | 2.67E+00 | 2.74E+00 | 9.95E-01 | no | Dock3 | intron_variant | MODIFIER |
| 9 | 107193850 | AX-170271917 | 0 | A | G | 0.123 | -6.29E-04 | 1.18E-01 | 2.67E+00 | 2.74E+00 | 9.95E-01 | no | Dock3 | intron_variant | MODIFIER |
| 9 | 107249017 | AX-97788695 | 0 | A | C | 0.123 | -6.29E-04 | 1.18E-01 | 2.67E+00 | 2.74E+00 | 9.95E-01 | no | | intergenic_variant | MODIFIER |
| 9 | 107256669 | AX-170155594 | 0 | A | C | 0.123 | -6.29E-04 | 1.18E-01 | 2.67E+00 | 2.74E+00 | 9.95E-01 | no | Mapkapk3 | intron_variant | MODIFIER |
| 9 | 107263636 | AX-170245783 | 0 | A | G | 0.123 | -6.29E-04 | 1.18E-01 | 2.67E+00 | 2.74E+00 | 9.95E-01 | no | Mapkapk3 | synonymous_variant | LOW |
| 9 | 107277767 | AX-170268539 | 0 | C | T | 0.123 | -6.29E-04 | 1.18E-01 | 2.67E+00 | 2.74E+00 | 9.95E-01 | no | Mapkapk3 | intron_variant | MODIFIER |
| 9 | 107297251 | AX-97828362 | 0 | C | A | 0.123 | -6.29E-04 | 1.18E-01 | 2.67E+00 | 2.74E+00 | 9.95E-01 | no | Cish | intron_variant | MODIFIER |
| 9 | 107311843 | AX-170033228 | 0 | A | G | 0.123 | -6.29E-04 | 1.18E-01 | 2.67E+00 | 2.74E+00 | 9.95E-01 | no | | regulatory_region_variant | MODIFIER |
| 9 | 107312585 | AX-170228473 | 0 | T | C | 0.123 | -6.29E-04 | 1.18E-01 | 2.67E+00 | 2.74E+00 | 9.95E-01 | no | | regulatory_region_variant | MODIFIER |
| 9 | 107328081 | AX-97822394 | 0 | A | G | 0.123 | -6.29E-04 | 1.18E-01 | 2.67E+00 | 2.74E+00 | 9.95E-01 | no | Hemk1 | intron_variant | MODIFIER |
| 9 | 107328156 | AX-170304981 | 0 | A | G | 0.123 | -6.29E-04 | 1.18E-01 | 2.67E+00 | 2.74E+00 | 9.95E-01 | no | Hemk1 | intron_variant | MODIFIER |
| 9 | 107334968 | AX-170284746 | 0 | A | G | 0.123 | -6.29E-04 | 1.18E-01 | 2.67E+00 | 2.74E+00 | 9.95E-01 | no | Hemk1 | intron_variant | MODIFIER |
| 9 | 107344649 | AX-97815516 | 0 | T | C | 0.123 | -6.29E-04 | 1.18E-01 | 2.67E+00 | 2.74E+00 | 9.95E-01 | no | 6430571L13Rik | intron_variant | MODIFIER |
| 9 | 107374314 | AX-97852439 | 0 | T | C | 0.123 | -6.29E-04 | 1.18E-01 | 2.67E+00 | 2.74E+00 | 9.95E-01 | no | | intergenic_variant | MODIFIER |
| 9 | 107376075 | AX-170354471 | 0 | G | A | 0.123 | -6.29E-04 | 1.18E-01 | 2.67E+00 | 2.74E+00 | 9.95E-01 | no | | intergenic_variant | MODIFIER |
| 9 | 107387404 | AX-170334672 | 0 | C | T | 0.123 | -6.29E-04 | 1.18E-01 | 2.67E+00 | 2.74E+00 | 9.95E-01 | no | | intergenic_variant | MODIFIER |
| 9 | 107396425 | AX-170374709 | 0 | A | G | 0.123 | -6.29E-04 | 1.18E-01 | 2.67E+00 | 2.74E+00 | 9.95E-01 | no | Cacna2d2 | upstream_gene_variant | MODIFIER |
| 9 | 107421133 | AX-170291116 | 0 | A | C | 0.123 | -6.29E-04 | 1.18E-01 | 2.67E+00 | 2.74E+00 | 9.95E-01 | no | Cacna2d2 | intron_variant | MODIFIER |
| 9 | 107427700 | AX-170103227 | 0 | C | T | 0.123 | -6.29E-04 | 1.18E-01 | 2.67E+00 | 2.74E+00 | 9.95E-01 | no | Cacna2d2 | intron_variant | MODIFIER |
| 9 | 107428254 | AX-170400944 | 0 | G | A | 0.123 | -6.29E-04 | 1.18E-01 | 2.67E+00 | 2.74E+00 | 9.95E-01 | no | Cacna2d2 | intron_variant | MODIFIER |
| 9 | 107434125 | AX-170318447 | 0 | C | T | 0.123 | -6.29E-04 | 1.18E-01 | 2.67E+00 | 2.74E+00 | 9.95E-01 | no | Cacna2d2 | intron_variant | MODIFIER |
| 9 | 107434567 | AX-170490516 | 0 | T | C | 0.123 | -6.29E-04 | 1.18E-01 | 2.67E+00 | 2.74E+00 | 9.95E-01 | no | Cacna2d2 | intron_variant | MODIFIER |
| 9 | 107444261 | AX-97852669 | 0 | G | T | 0.123 | -6.29E-04 | 1.18E-01 | 2.67E+00 | 2.74E+00 | 9.95E-01 | no | Cacna2d2 | intron_variant | MODIFIER |
| 9 | 107451293 | AX-170033238 | 0 | A | G | 0.123 | -6.29E-04 | 1.18E-01 | 2.67E+00 | 2.74E+00 | 9.95E-01 | no | Cacna2d2 | intron_variant | MODIFIER |
| 9 | 107455698 | AX-170371356 | 0 | T | C | 0.123 | -6.29E-04 | 1.18E-01 | 2.67E+00 | 2.74E+00 | 9.95E-01 | no | Cacna2d2 | intron_variant | MODIFIER |
| 9 | 107456203 | AX-170271324 | 0 | G | A | 0.123 | -6.29E-04 | 1.18E-01 | 2.67E+00 | 2.74E+00 | 9.95E-01 | no | Cacna2d2 | intron_variant | MODIFIER |
| 9 | 107475680 | AX-170033236 | 0 | C | T | 0.123 | -6.29E-04 | 1.18E-01 | 2.67E+00 | 2.74E+00 | 9.95E-01 | no | Cacna2d2 | intron_variant | MODIFIER |
| 9 | 107475778 | AX-170033237 | 0 | G | A | 0.123 | -6.29E-04 | 1.18E-01 | 2.67E+00 | 2.74E+00 | 9.95E-01 | no | Cacna2d2 | intron_variant | MODIFIER |
| 9 | 107482925 | AX-97821563 | 0 | A | G | 0.123 | -6.29E-04 | 1.18E-01 | 2.67E+00 | 2.74E+00 | 9.95E-01 | no | Cacna2d2 | intron_variant | MODIFIER |
| 9 | 107483655 | AX-170340783 | 0 | G | A | 0.123 | -6.29E-04 | 1.18E-01 | 2.67E+00 | 2.74E+00 | 9.95E-01 | no | Cacna2d2 | intron_variant | MODIFIER |
| 9 | 107491837 | AX-170446209 | 0 | G | A | 0.123 | -6.29E-04 | 1.18E-01 | 2.67E+00 | 2.74E+00 | 9.95E-01 | no | Cacna2d2 | intron_variant | MODIFIER |
| 9 | 107497537 | AX-97856965 | 0 | C | T | 0.123 | -6.29E-04 | 1.18E-01 | 2.67E+00 | 2.74E+00 | 9.95E-01 | no | Cacna2d2 | intron_variant | MODIFIER |
| 9 | 107507742 | AX-170083391 | 0 | A | C | 0.123 | -6.29E-04 | 1.18E-01 | 2.67E+00 | 2.74E+00 | 9.95E-01 | no | Cacna2d2 | intron_variant | MODIFIER |
| 9 | 107523295 | AX-170083394 | 0 | G | C | 0.123 | -6.29E-04 | 1.18E-01 | 2.67E+00 | 2.74E+00 | 9.95E-01 | no | Cacna2d2 | intron_variant | MODIFIER |
| 9 | 107524712 | AX-170033235 | 0 | A | C | 0.123 | -6.29E-04 | 1.18E-01 | 2.67E+00 | 2.74E+00 | 9.95E-01 | no | Cacna2d2 | intron_variant | MODIFIER |
| 9 | 107528264 | AX-170033239 | 0 | C | A | 0.123 | -6.29E-04 | 1.18E-01 | 2.67E+00 | 2.74E+00 | 9.95E-01 | no | Cacna2d2 | 3_prime_UTR_variant | MODIFIER |
| 9 | 107541898 | AX-97843965 | 0 | G | A | 0.123 | -6.29E-04 | 1.18E-01 | 2.67E+00 | 2.74E+00 | 9.95E-01 | no | Tmem115 | downstream_gene_variant | MODIFIER |

| | | | | | | | | | | | | | | | |
|---|---|---|---|---|---|---|---|---|---|---|---|---|---|---|---|
| 9 | 107576192 | AX-170225341 | 0 | A | G | 0.123 | -6.29E-04 | 1.18E-01 | 2.67E+00 | 2.74E+00 | 9.95E-01 | no | Hyal2 | downstream_gene_variant | MODIFIER |
| 9 | 107576242 | AX-170116920 | 0 | A | G | 0.123 | -6.29E-04 | 1.18E-01 | 2.67E+00 | 2.74E+00 | 9.95E-01 | no | Hyal2 | downstream_gene_variant | MODIFIER |
| 9 | 107580649 | AX-170083395 | 0 | G | C | 0.123 | -6.29E-04 | 1.18E-01 | 2.67E+00 | 2.74E+00 | 9.95E-01 | no | Hyal1 | 3_prime_UTR_variant | MODIFIER |
| 9 | 107599107 | AX-97811346 | 0 | G | A | 0.123 | -6.29E-04 | 1.18E-01 | 2.67E+00 | 2.74E+00 | 9.95E-01 | no | Sema3b | synonymous_variant | LOW |
| 9 | 107600095 | AX-170225314 | 0 | T | C | 0.123 | -6.29E-04 | 1.18E-01 | 2.67E+00 | 2.74E+00 | 9.95E-01 | no | Sema3b | missense_variant | MODERATE |
| 9 | 107639285 | AX-97811935 | 0 | A | G | 0.123 | -6.29E-04 | 1.18E-01 | 2.67E+00 | 2.74E+00 | 9.95E-01 | no | Gnai2 | upstream_gene_variant | MODIFIER |
| 9 | 107648224 | AX-170306050 | 0 | A | G | 0.123 | -6.29E-04 | 1.18E-01 | 2.67E+00 | 2.74E+00 | 9.95E-01 | no | Slc38a3 | downstream_gene_variant | MODIFIER |
| 9 | 107649733 | AX-170225656 | 0 | C | T | 0.123 | -6.29E-04 | 1.18E-01 | 2.67E+00 | 2.74E+00 | 9.95E-01 | no | Slc38a3 | downstream_gene_variant | MODIFIER |
| 9 | 107660661 | AX-170270961 | 0 | G | A | 0.123 | -6.29E-04 | 1.18E-01 | 2.67E+00 | 2.74E+00 | 9.95E-01 | no | Slc38a3 | intron_variant | MODIFIER |
| 9 | 107670252 | AX-170177905 | 0 | A | G | 0.123 | -6.29E-04 | 1.18E-01 | 2.67E+00 | 2.74E+00 | 9.95E-01 | no | Gnat1 | downstream_gene_variant | MODIFIER |
| 9 | 107692735 | AX-170149368 | 0 | G | A | 0.123 | -6.29E-04 | 1.18E-01 | 2.67E+00 | 2.74E+00 | 9.95E-01 | no | Sema3f | intron_variant | MODIFIER |
| 9 | 107746656 | AX-170162468 | 0 | G | A | 0.123 | -6.29E-04 | 1.18E-01 | 2.67E+00 | 2.74E+00 | 9.95E-01 | no | Rbm5 | intron_variant | MODIFIER |
| 9 | 107752631 | AX-170090410 | 0 | T | G | 0.123 | -6.29E-04 | 1.18E-01 | 2.67E+00 | 2.74E+00 | 9.95E-01 | no | Rbm5 | intron_variant | MODIFIER |
| 9 | 107767299 | AX-170347719 | 0 | G | A | 0.123 | -6.29E-04 | 1.18E-01 | 2.67E+00 | 2.74E+00 | 9.95E-01 | no | Rbm5 | intron_variant | MODIFIER |
| 9 | 107783037 | AX-170125804 | 0 | G | C | 0.123 | -6.29E-04 | 1.18E-01 | 2.67E+00 | 2.74E+00 | 9.95E-01 | no | Rbm6 | intron_variant | MODIFIER |
| 9 | 107811044 | AX-170225340 | 0 | C | A | 0.123 | -6.29E-04 | 1.18E-01 | 2.67E+00 | 2.74E+00 | 9.95E-01 | no | Rbm6 | intron_variant | MODIFIER |
| 9 | 107910170 | AX-170187283 | 0 | A | G | 0.123 | -6.29E-04 | 1.18E-01 | 2.67E+00 | 2.74E+00 | 9.95E-01 | no | Mst1r | intron_variant | MODIFIER |
| 9 | 107912815 | AX-170282870 | 0 | A | C | 0.123 | -6.29E-04 | 1.18E-01 | 2.67E+00 | 2.74E+00 | 9.95E-01 | no | Mst1r | intron_variant | MODIFIER |
| 9 | 107918831 | AX-170347591 | 0 | C | T | 0.123 | -6.29E-04 | 1.18E-01 | 2.67E+00 | 2.74E+00 | 9.95E-01 | no | Mst1r | intron_variant | MODIFIER |
| 9 | 107921369 | AX-170181037 | 0 | G | A | 0.123 | -6.29E-04 | 1.18E-01 | 2.67E+00 | 2.74E+00 | 9.95E-01 | no | Mst1r | downstream_gene_variant | MODIFIER |
| 9 | 107929648 | AX-170226645 | 0 | T | C | 0.123 | -6.29E-04 | 1.18E-01 | 2.67E+00 | 2.74E+00 | 9.95E-01 | no | Actl11 | missense_variant | MODERATE |
| 9 | 107930960 | AX-170236610 | 0 | C | T | 0.123 | -6.29E-04 | 1.18E-01 | 2.67E+00 | 2.74E+00 | 9.95E-01 | no | Camkv | upstream_gene_variant | MODIFIER |
| 9 | 107966739 | AX-170368256 | 0 | G | A | 0.123 | -6.29E-04 | 1.18E-01 | 2.67E+00 | 2.74E+00 | 9.95E-01 | no | Traip | intron_variant | MODIFIER |
| 9 | 108051805 | AX-97795023 | 0 | C | T | 0.123 | -6.29E-04 | 1.18E-01 | 2.67E+00 | 2.74E+00 | 9.95E-01 | no | Ip6k1 | downstream_gene_variant | MODIFIER |
| 9 | 108158671 | AX-170534670 | 0 | C | A | 0.123 | -6.29E-04 | 1.18E-01 | 2.67E+00 | 2.74E+00 | 9.95E-01 | no | Bsn | intron_variant | MODIFIER |
| 9 | 108198765 | AX-97837767 | 0 | A | G | 0.123 | -6.29E-04 | 1.18E-01 | 2.67E+00 | 2.74E+00 | 9.95E-01 | no | | intergenic_variant | MODIFIER |
| 9 | 108221554 | AX-170383792 | 0 | C | G | 0.123 | -6.29E-04 | 1.18E-01 | 2.67E+00 | 2.74E+00 | 9.95E-01 | no | Dag1 | intron_variant | MODIFIER |
| 9 | 108514869 | AX-170233736 | 0 | T | C | 0.123 | -6.29E-04 | 1.18E-01 | 2.67E+00 | 2.74E+00 | 9.95E-01 | no | Qars | missense_variant | MODERATE |
| 9 | 108613710 | AX-170033249 | 0 | A | G | 0.123 | -6.29E-04 | 1.18E-01 | 2.67E+00 | 2.74E+00 | 9.95E-01 | no | Arih2 | intron_variant | MODIFIER |
| 9 | 108616379 | AX-97817263 | 0 | C | G | 0.123 | -6.29E-04 | 1.18E-01 | 2.67E+00 | 2.74E+00 | 9.95E-01 | no | Arih2 | intron_variant | MODIFIER |
| 9 | 108646497 | AX-170033247 | 0 | G | C | 0.123 | -6.29E-04 | 1.18E-01 | 2.67E+00 | 2.74E+00 | 9.95E-01 | no | Arih2 | intron_variant | MODIFIER |
| 9 | 108677109 | AX-170033250 | 0 | C | T | 0.123 | -6.29E-04 | 1.18E-01 | 2.67E+00 | 2.74E+00 | 9.95E-01 | no | Slc25a20 | intron_variant | MODIFIER |
| 9 | 108683173 | AX-170033246 | 0 | T | C | 0.123 | -6.29E-04 | 1.18E-01 | 2.67E+00 | 2.74E+00 | 9.95E-01 | no | Slc25a20 | splice_region_variant,synonymous_variant | LOW |
| 9 | 108722087 | AX-170425636 | 0 | C | T | 0.123 | -6.29E-04 | 1.18E-01 | 2.67E+00 | 2.74E+00 | 9.95E-01 | no | Prkar2a | intron_variant | MODIFIER |
| 9 | 108770652 | AX-170221000 | 0 | G | A | 0.123 | -6.29E-04 | 1.18E-01 | 2.67E+00 | 2.74E+00 | 9.95E-01 | no | | intergenic_variant | MODIFIER |
| 9 | 108775499 | AX-97828091 | 0 | A | G | 0.123 | -6.29E-04 | 1.18E-01 | 2.67E+00 | 2.74E+00 | 9.95E-01 | no | | intergenic_variant | MODIFIER |
| 9 | 108786738 | AX-170376153 | 0 | T | C | 0.123 | -6.29E-04 | 1.18E-01 | 2.67E+00 | 2.74E+00 | 9.95E-01 | no | Ip6k2 | intron_variant,NMD_transcript_variant | MODIFIER |
| 9 | 108794791 | AX-97791377 | 0 | A | C | 0.123 | -6.29E-04 | 1.18E-01 | 2.67E+00 | 2.74E+00 | 9.95E-01 | no | Ip6k2 | upstream_gene_variant | MODIFIER |
| 9 | 108844710 | AX-170174324 | 0 | T | C | 0.123 | -6.29E-04 | 1.18E-01 | 2.67E+00 | 2.74E+00 | 9.95E-01 | no | Celsr3 | intron_variant | MODIFIER |
| 9 | 108855860 | AX-168387132 | 0 | C | T | 0.123 | -6.29E-04 | 1.18E-01 | 2.67E+00 | 2.74E+00 | 9.95E-01 | no | Celsr3 | downstream_gene_variant | MODIFIER |
| 9 | 108860836 | AX-170384856 | 0 | G | A | 0.123 | -6.29E-04 | 1.18E-01 | 2.67E+00 | 2.74E+00 | 9.95E-01 | no | Slc26a6 | synonymous_variant | LOW |
| 9 | 108961386 | AX-170456246 | 0 | T | C | 0.123 | -6.29E-04 | 1.18E-01 | 2.67E+00 | 2.74E+00 | 9.95E-01 | no | Col7a1 | synonymous_variant | LOW |
| 9 | 109060290 | AX-170426552 | 0 | C | T | 0.123 | -6.29E-04 | 1.18E-01 | 2.67E+00 | 2.74E+00 | 9.95E-01 | no | Shisa5 | downstream_gene_variant | MODIFIER |
| 9 | 109086302 | AX-170287043 | 0 | T | C | 0.123 | -6.29E-04 | 1.18E-01 | 2.67E+00 | 2.74E+00 | 9.95E-01 | no | Ccdc51 | intron_variant | MODIFIER |
| 9 | 109111533 | AX-170241745 | 0 | G | C | 0.123 | -6.29E-04 | 1.18E-01 | 2.67E+00 | 2.74E+00 | 9.95E-01 | no | Plxnb1 | synonymous_variant | LOW |
| 9 | 109117001 | AX-170083424 | 0 | C | T | 0.123 | -6.29E-04 | 1.18E-01 | 2.67E+00 | 2.74E+00 | 9.95E-01 | no | Plxnb1 | intron_variant | MODIFIER |
| 9 | 109117178 | AX-170083426 | 0 | T | C | 0.123 | -6.29E-04 | 1.18E-01 | 2.67E+00 | 2.74E+00 | 9.95E-01 | no | Plxnb1 | intron_variant | MODIFIER |
| 9 | 109121888 | AX-170083422 | 0 | T | C | 0.123 | -6.29E-04 | 1.18E-01 | 2.67E+00 | 2.74E+00 | 9.95E-01 | no | Plxnb1 | downstream_gene_variant | MODIFIER |
| 9 | 109124105 | AX-97811128 | 0 | G | A | 0.123 | -6.29E-04 | 1.18E-01 | 2.67E+00 | 2.74E+00 | 9.95E-01 | no | Plxnb1 | downstream_gene_variant | MODIFIER |
| 9 | 109124163 | AX-170145786 | 0 | C | T | 0.123 | -6.29E-04 | 1.18E-01 | 2.67E+00 | 2.74E+00 | 9.95E-01 | no | Plxnb1 | downstream_gene_variant | MODIFIER |
| 9 | 109148088 | AX-170090411 | 0 | T | C | 0.123 | -6.29E-04 | 1.18E-01 | 2.67E+00 | 2.74E+00 | 9.95E-01 | no | Fbxw21 | synonymous_variant | LOW |
| 9 | 109195806 | AX-170164947 | 0 | G | C | 0.123 | -6.29E-04 | 1.18E-01 | 2.67E+00 | 2.74E+00 | 9.95E-01 | no | Fbxw13 | intron_variant | MODIFIER |
| 9 | 109633998 | AX-170298337 | 0 | A | G | 0.123 | -6.29E-04 | 1.18E-01 | 2.67E+00 | 2.74E+00 | 9.95E-01 | no | | regulatory_region_variant | MODIFIER |
| 9 | 109693231 | AX-170163924 | 0 | T | C | 0.123 | -6.29E-04 | 1.18E-01 | 2.67E+00 | 2.74E+00 | 9.95E-01 | no | Fbxw18 | missense_variant | MODERATE |

| | | | | | | | | | | | | | | | |
|---|---|---|---|---|---|---|---|---|---|---|---|---|---|---|---|
| 9 | 109825673 | AX-170331399 | 0 | A | G | 0.123 | -6.29E-04 | 1.18E-01 | 2.67E+00 | 2.74E+00 | 9.95E-01 | no | Spink8 | intron_variant | MODIFIER |
| 9 | 109827666 | AX-170110768 | 0 | G | A | 0.123 | -6.29E-04 | 1.18E-01 | 2.67E+00 | 2.74E+00 | 9.95E-01 | no | Spink8 | downstream_gene_variant | MODIFIER |
| 9 | 109828837 | AX-170410136 | 0 | C | T | 0.123 | -6.29E-04 | 1.18E-01 | 2.67E+00 | 2.74E+00 | 9.95E-01 | no | Nme6 | upstream_gene_variant | MODIFIER |
| 9 | 109837998 | AX-170236461 | 0 | G | A | 0.123 | -6.29E-04 | 1.18E-01 | 2.67E+00 | 2.74E+00 | 9.95E-01 | no | Nme6 | intron_variant | MODIFIER |
| 9 | 109838623 | AX-170389095 | 0 | A | G | 0.123 | -6.29E-04 | 1.18E-01 | 2.67E+00 | 2.74E+00 | 9.95E-01 | no | Nme6 | intron_variant | MODIFIER |
| 9 | 109838964 | AX-170334702 | 0 | G | A | 0.123 | -6.29E-04 | 1.18E-01 | 2.67E+00 | 2.74E+00 | 9.95E-01 | no | Nme6 | intron_variant | MODIFIER |
| 9 | 109848728 | AX-170432540 | 0 | A | G | 0.123 | -6.29E-04 | 1.18E-01 | 2.67E+00 | 2.74E+00 | 9.95E-01 | no | Camp | intron_variant | MODIFIER |
| 9 | 110070564 | AX-170090413 | 0 | A | G | 0.123 | -6.29E-04 | 1.18E-01 | 2.67E+00 | 2.74E+00 | 9.95E-01 | no | Map4 | intron_variant | MODIFIER |
| 9 | 106768932 | AX-170106561 | 0 | C | T | 0.132 | -5.29E-04 | 1.09E-01 | 2.66E+00 | 2.74E+00 | 9.95E-01 | no | Rad54l2 | intron_variant | MODIFIER |
| 9 | 108947793 | AX-170384004 | 0 | A | G | 0.132 | -5.29E-04 | 1.09E-01 | 2.66E+00 | 2.74E+00 | 9.95E-01 | no | Uqcrc1 | intron_variant | MODIFIER |
| 16 | 45598607 | AX-170267739 | 1 | C | T | 0.482 | -1.43E-03 | 8.09E-02 | 2.67E+00 | 2.74E+00 | 9.95E-01 | no | Slc9c1 | intron_variant | MODIFIER |
| 12 | 116366192 | AX-97854522 | 1 | G | A | 0.143 | -9.31E-04 | 1.92E-01 | 2.67E+00 | 2.74E+00 | 9.95E-01 | no | Esyt2 | intron_variant | MODIFIER |
| 7 | 132926291 | AX-170336919 | 0 | C | T | 0.254 | -8.92E-04 | 8.21E-02 | 2.63E+00 | 2.74E+00 | 9.96E-01 | no | Zranb1 | upstream_gene_variant | MODIFIER |
| 7 | 132968247 | AX-170526168 | 0 | C | T | 0.254 | -8.92E-04 | 8.21E-02 | 2.63E+00 | 2.74E+00 | 9.96E-01 | no | Zranb1 | intron_variant | MODIFIER |
| 7 | 132998156 | AX-170030018 | 0 | G | T | 0.254 | -8.92E-04 | 8.21E-02 | 2.63E+00 | 2.74E+00 | 9.96E-01 | no | Ctbp2 | intron_variant | MODIFIER |
| 7 | 133004148 | AX-170144552 | 0 | C | T | 0.254 | -8.92E-04 | 8.21E-02 | 2.63E+00 | 2.74E+00 | 9.96E-01 | no | Ctbp2 | intron_variant | MODIFIER |
| 7 | 133005641 | AX-170184225 | 0 | G | A | 0.254 | -8.92E-04 | 8.21E-02 | 2.63E+00 | 2.74E+00 | 9.96E-01 | no | Ctbp2 | intron_variant | MODIFIER |
| 7 | 133006136 | AX-170361352 | 0 | G | C | 0.254 | -8.92E-04 | 8.21E-02 | 2.63E+00 | 2.74E+00 | 9.96E-01 | no | Ctbp2 | intron_variant | MODIFIER |
| 7 | 133006167 | AX-170078905 | 0 | C | T | 0.254 | -8.92E-04 | 8.21E-02 | 2.63E+00 | 2.74E+00 | 9.96E-01 | no | Ctbp2 | intron_variant | MODIFIER |
| 7 | 133014155 | AX-170030015 | 0 | T | C | 0.254 | -8.92E-04 | 8.21E-02 | 2.63E+00 | 2.74E+00 | 9.96E-01 | no | Ctbp2 | intron_variant | MODIFIER |
| 7 | 133014701 | AX-170078906 | 0 | C | G | 0.254 | -8.92E-04 | 8.21E-02 | 2.63E+00 | 2.74E+00 | 9.96E-01 | no | Ctbp2 | intron_variant | MODIFIER |
| 7 | 133014732 | AX-170030016 | 0 | C | G | 0.254 | -8.92E-04 | 8.21E-02 | 2.63E+00 | 2.74E+00 | 9.96E-01 | no | Ctbp2 | intron_variant | MODIFIER |
| 7 | 133019874 | AX-170078907 | 0 | A | G | 0.254 | -8.92E-04 | 8.21E-02 | 2.63E+00 | 2.74E+00 | 9.96E-01 | no | Ctbp2 | intron_variant | MODIFIER |
| 7 | 133073283 | AX-170331141 | 0 | A | G | 0.254 | -8.92E-04 | 8.21E-02 | 2.63E+00 | 2.74E+00 | 9.96E-01 | no | Ctbp2 | intron_variant | MODIFIER |
| 7 | 133109365 | AX-170460364 | 0 | A | G | 0.254 | -8.92E-04 | 8.21E-02 | 2.63E+00 | 2.74E+00 | 9.96E-01 | no | Ctbp2 | intron_variant | MODIFIER |
| 7 | 133112338 | AX-170268624 | 0 | T | C | 0.254 | -8.92E-04 | 8.21E-02 | 2.63E+00 | 2.74E+00 | 9.96E-01 | no | Ctbp2 | intron_variant | MODIFIER |
| 7 | 133152530 | AX-170276150 | 0 | G | A | 0.254 | -8.92E-04 | 8.21E-02 | 2.63E+00 | 2.74E+00 | 9.96E-01 | no | | intergenic_variant | MODIFIER |
| 7 | 133152677 | AX-170474227 | 0 | G | A | 0.254 | -8.92E-04 | 8.21E-02 | 2.63E+00 | 2.74E+00 | 9.96E-01 | no | | intergenic_variant | MODIFIER |
| 7 | 133159742 | AX-170498899 | 0 | T | C | 0.254 | -8.92E-04 | 8.21E-02 | 2.63E+00 | 2.74E+00 | 9.96E-01 | no | | regulatory_region_variant | MODIFIER |
| 7 | 133161887 | AX-170383222 | 0 | C | T | 0.254 | -8.92E-04 | 8.21E-02 | 2.63E+00 | 2.74E+00 | 9.96E-01 | no | | regulatory_region_variant | MODIFIER |
| 7 | 133996942 | AX-170294434 | 0 | G | A | 0.254 | -8.92E-04 | 8.21E-02 | 2.63E+00 | 2.74E+00 | 9.96E-01 | no | Adam12 | intron_variant | MODIFIER |
| 2 | 124327449 | AX-170214361 | 2 | T | C | 0.136 | -3.97E-04 | 2.17E-01 | 2.69E+00 | 2.74E+00 | 9.96E-01 | no | Sema6d | intron_variant | MODIFIER |
| 20 | 59452936 | AX-170443512 | 0 | C | A | 0.246 | -2.17E-04 | 6.30E-02 | 2.65E+00 | 2.74E+00 | 9.96E-01 | no | Fgf13 | intron_variant | MODIFIER |
| 15 | 10955689 | AX-170181018 | 2 | A | G | 0.236 | -2.68E-04 | 9.99E-02 | 2.67E+00 | 2.74E+00 | 9.96E-01 | no | C1qtnf3 | intron_variant | MODIFIER |
| 12 | 117779230 | AX-170190324 | 1 | A | G | 0.125 | 1.41E-03 | 1.57E-01 | 2.65E+00 | 2.74E+00 | 9.96E-01 | no | | intergenic_variant | MODIFIER |
| 12 | 118877275 | AX-170436180 | 1 | T | C | 0.125 | 1.41E-03 | 1.57E-01 | 2.65E+00 | 2.74E+00 | 9.96E-01 | no | Abcb5 | intron_variant | MODIFIER |
| 12 | 62767534 | AX-170087260 | 1 | C | A | 0.062 | -1.90E-03 | 3.00E-01 | 2.71E+00 | 2.74E+00 | 9.96E-01 | no | | intergenic_variant | MODIFIER |
| 9 | 54899325 | AX-170082489 | 2 | G | A | 0.218 | -3.13E-04 | 1.19E-01 | 2.68E+00 | 2.74E+00 | 9.96E-01 | no | Ireb2 | intron_variant | MODIFIER |
| 1 | 12938444 | AX-170040408 | 1 | A | G | 0.384 | 1.05E-03 | 7.46E-02 | 2.67E+00 | 2.74E+00 | 9.96E-01 | no | Slco5a1 | intron_variant | MODIFIER |
| 13 | 54850943 | AX-170528385 | 1 | T | C | 0.5 | 1.61E-03 | 6.65E-02 | 2.62E+00 | 2.74E+00 | 9.96E-01 | no | | intergenic_variant | MODIFIER |
| 11 | 13709484 | AX-170295411 | 0 | T | C | 0.061 | 4.03E-06 | 2.13E-01 | 2.65E+00 | 2.74E+00 | 9.96E-01 | no | | intergenic_variant | MODIFIER |
| 11 | 18104834 | AX-97847970 | 0 | C | T | 0.061 | 4.03E-06 | 2.13E-01 | 2.65E+00 | 2.74E+00 | 9.96E-01 | no | Gm12018 | downstream_gene_variant | MODIFIER |
| 9 | 113912898 | AX-170473322 | 2 | G | A | 0.145 | 2.73E-04 | 1.01E-01 | 2.64E+00 | 2.74E+00 | 9.96E-01 | no | Clasp2 | intron_variant | MODIFIER |
| 12 | 60624581 | AX-170274744 | 1 | T | A | 0.062 | 8.56E-04 | 3.02E-01 | 2.72E+00 | 2.74E+00 | 9.96E-01 | no | | intergenic_variant | MODIFIER |
| 5 | 112842336 | AX-170306522 | 0 | C | T | 0.456 | 1.60E-04 | 1.14E-01 | 2.69E+00 | 2.74E+00 | 9.96E-01 | no | Myo18b | intron_variant | MODIFIER |
| 5 | 114554539 | AX-170262817 | 0 | T | C | 0.456 | 1.60E-04 | 1.14E-01 | 2.69E+00 | 2.74E+00 | 9.96E-01 | no | Gm13790 | downstream_gene_variant | MODIFIER |
| 9 | 77411549 | AX-170362875 | 0 | C | G | 0.193 | -1.08E-03 | 1.27E-01 | 2.70E+00 | 2.74E+00 | 9.96E-01 | no | | regulatory_region_variant | MODIFIER |
| 5 | 26492353 | AX-170242538 | 0 | C | A | 0.053 | 2.54E-04 | 1.74E-01 | 2.62E+00 | 2.74E+00 | 9.96E-01 | no | 4930584F24Rik | intron_variant,non_coding_transcript_variant | MODIFIER |
| 5 | 100220735 | AX-170089566 | 0 | G | T | 0.439 | -8.00E-05 | 1.05E-01 | 2.67E+00 | 2.74E+00 | 9.96E-01 | no | Gm8091 | intron_variant,non_coding_transcript_variant | MODIFIER |
| 9 | 113917590 | AX-170336513 | 0 | G | C | 0.132 | 3.03E-04 | 1.04E-01 | 2.64E+00 | 2.74E+00 | 9.96E-01 | no | Clasp2 | intron_variant | MODIFIER |
| 1 | 39903263 | AX-170510361 | 0 | G | C | 0.377 | -1.10E-03 | 7.71E-02 | 2.65E+00 | 2.74E+00 | 9.97E-01 | no | Map4k4 | intron_variant | MODIFIER |
| 20 | 80849665 | AX-170491547 | 0 | T | C | 0.254 | 3.87E-04 | 6.09E-02 | 2.64E+00 | 2.74E+00 | 9.97E-01 | no | | intergenic_variant | MODIFIER |
| 12 | 60016646 | AX-170201997 | 0 | C | T | 0.07 | 1.07E-03 | 2.07E-01 | 2.63E+00 | 2.74E+00 | 9.97E-01 | no | | intergenic_variant | MODIFIER |
| 12 | 61803545 | AX-170087259 | 0 | G | A | 0.07 | 1.07E-03 | 2.07E-01 | 2.63E+00 | 2.74E+00 | 9.97E-01 | no | Lrfn5 | intron_variant | MODIFIER |

| | | | | | | | | | | | | | | | |
|---|---|---|---|---|---|---|---|---|---|---|---|---|---|---|---|
| 12 | 71385635 | AX-97835880 | 0 | C | T | 0.07 | 1.07E-03 | 2.07E-01 | 2.63E+00 | 2.74E+00 | 9.97E-01 | no | 1700083H02Rik | upstream_gene_variant | MODIFIER |
| 18 | 36661892 | AX-170309772 | 0 | G | A | 0.07 | 1.07E-03 | 2.07E-01 | 2.63E+00 | 2.74E+00 | 9.97E-01 | no | Ankhd1 | downstream_gene_variant | MODIFIER |
| 18 | 36804436 | AX-97784198 | 0 | G | A | 0.07 | 1.07E-03 | 2.07E-01 | 2.63E+00 | 2.74E+00 | 9.97E-01 | no | Zmat2 | downstream_gene_variant | MODIFIER |
| 18 | 45043835 | AX-170149750 | 0 | G | A | 0.07 | 1.07E-03 | 2.07E-01 | 2.63E+00 | 2.74E+00 | 9.97E-01 | no | AC134458.2 | upstream_gene_variant | MODIFIER |
| 1 | 55810823 | AX-170041298 | 2 | A | T | 0.1 | 1.27E-03 | 2.19E-01 | 2.69E+00 | 2.74E+00 | 9.97E-01 | no | | intergenic_variant | MODIFIER |
| 4 | 140983207 | AX-170089390 | 0 | T | C | 0.088 | 1.11E-03 | 1.68E-01 | 2.67E+00 | 2.74E+00 | 9.97E-01 | no | Sdhb | downstream_gene_variant | MODIFIER |
| 4 | 144298162 | AX-97790785 | 0 | C | T | 0.088 | 1.11E-03 | 1.68E-01 | 2.67E+00 | 2.74E+00 | 9.97E-01 | no | Gm13120 | downstream_gene_variant | MODIFIER |
| 9 | 60675146 | AX-97789665 | 0 | G | A | 0.202 | -9.42E-04 | 1.35E-01 | 2.70E+00 | 2.74E+00 | 9.97E-01 | no | Lrrc49 | intron_variant | MODIFIER |
| 9 | 63807894 | AX-170233077 | 0 | G | A | 0.202 | -9.42E-04 | 1.35E-01 | 2.70E+00 | 2.74E+00 | 9.97E-01 | no | | intergenic_variant | MODIFIER |
| 9 | 64498539 | AX-97805020 | 0 | A | G | 0.202 | -9.42E-04 | 1.35E-01 | 2.70E+00 | 2.74E+00 | 9.97E-01 | no | Megf11 | intron_variant | MODIFIER |
| 8 | 75121669 | AX-170219017 | 1 | T | C | 0.196 | -9.68E-04 | 1.33E-01 | 2.67E+00 | 2.74E+00 | 9.97E-01 | no | Mcm5 | synonymous_variant | LOW |
| 7 | 57209522 | AX-170272952 | 1 | T | A | 0.42 | 4.70E-04 | 9.47E-02 | 2.66E+00 | 2.74E+00 | 9.97E-01 | no | Gabrg3 | intron_variant | MODIFIER |
| 1 | 25680233 | AX-170324550 | 0 | T | A | 0.421 | 8.81E-04 | 8.14E-02 | 2.66E+00 | 2.74E+00 | 9.97E-01 | no | Adgrb3 | intron_variant | MODIFIER |
| 7 | 51088124 | AX-170028992 | 1 | T | C | 0.429 | 1.12E-03 | 9.97E-02 | 2.66E+00 | 2.74E+00 | 9.97E-01 | no | | intergenic_variant | MODIFIER |
| 1 | 78331039 | AX-170441196 | 0 | T | C | 0.149 | 1.55E-03 | 2.14E-01 | 2.69E+00 | 2.74E+00 | 9.97E-01 | no | Sgpp2 | intron_variant | MODIFIER |
| 4 | 144102049 | AX-97804093 | 2 | G | T | 0.082 | -1.75E-03 | 2.06E-01 | 2.63E+00 | 2.74E+00 | 9.97E-01 | no | Gm13102 | upstream_gene_variant | MODIFIER |
| 20 | 115356490 | AX-97794486 | 1 | C | T | 0.304 | 1.03E-04 | 5.34E-02 | 2.62E+00 | 2.74E+00 | 9.97E-01 | no | | intergenic_variant | MODIFIER |
| 5 | 98538725 | AX-97781178 | 0 | G | A | 0.421 | 8.64E-04 | 1.03E-01 | 2.67E+00 | 2.74E+00 | 9.97E-01 | no | Cfap299 | intron_variant | MODIFIER |
| 15 | 97455876 | AX-170258177 | 1 | G | A | 0.152 | 2.01E-03 | 1.78E-01 | 2.67E+00 | 2.74E+00 | 9.97E-01 | no | | regulatory_region_variant | MODIFIER |
| 11 | 116998881 | AX-170477457 | 0 | A | G | 0.061 | 2.88E-04 | 1.45E-01 | 2.60E+00 | 2.74E+00 | 9.97E-01 | no | | intergenic_variant | MODIFIER |
| 12 | 59788797 | AX-170509942 | 0 | C | T | 0.07 | -8.19E-04 | 2.30E-01 | 2.66E+00 | 2.74E+00 | 9.97E-01 | no | | intergenic_variant | MODIFIER |
| 18 | 43133170 | AX-170314735 | 0 | T | C | 0.07 | -8.19E-04 | 2.30E-01 | 2.66E+00 | 2.74E+00 | 9.97E-01 | no | AC099605.1 | intron_variant,non_coding_transcript_variant | MODIFIER |
| 18 | 59005355 | AX-170060981 | 0 | T | A | 0.386 | 2.39E-03 | 9.37E-02 | 2.66E+00 | 2.74E+00 | 9.97E-01 | no | Adamts19 | intron_variant | MODIFIER |
| 1 | 173063455 | AX-170266125 | 1 | T | C | 0.223 | 2.66E-03 | 1.17E-01 | 2.65E+00 | 2.74E+00 | 9.97E-01 | no | | intergenic_variant | MODIFIER |
| 9 | 71493051 | AX-170243089 | 1 | G | A | 0.205 | -7.91E-04 | 1.40E-01 | 2.71E+00 | 2.74E+00 | 9.97E-01 | no | | regulatory_region_variant | MODIFIER |
| 16 | 92926443 | AX-170525723 | 1 | G | A | 0.473 | -3.79E-04 | 7.89E-02 | 2.66E+00 | 2.74E+00 | 9.97E-01 | no | | intergenic_variant | MODIFIER |
| 2 | 87814351 | AX-170109072 | 1 | G | A | 0.062 | 6.72E-04 | 2.28E-01 | 2.66E+00 | 2.74E+00 | 9.97E-01 | no | Olfr1145 | downstream_gene_variant | MODIFIER |
| 15 | 102714868 | AX-170471522 | 0 | G | A | 0.114 | 4.20E-05 | 1.33E-01 | 2.64E+00 | 2.74E+00 | 9.97E-01 | no | Calcoco1 | intron_variant | MODIFIER |
| 5 | 126879976 | AX-170392276 | 0 | C | T | 0.43 | 2.03E-04 | 8.93E-02 | 2.64E+00 | 2.74E+00 | 9.97E-01 | no | Gm40331 | intron_variant,non_coding_transcript_variant | MODIFIER |
| 5 | 126909106 | AX-170103464 | 0 | T | A | 0.43 | 2.03E-04 | 8.93E-02 | 2.64E+00 | 2.74E+00 | 9.97E-01 | no | | intergenic_variant | MODIFIER |
| 5 | 126910936 | AX-170202582 | 0 | T | G | 0.43 | 2.03E-04 | 8.93E-02 | 2.64E+00 | 2.74E+00 | 9.97E-01 | no | | intergenic_variant | MODIFIER |
| 5 | 126911033 | AX-170322008 | 0 | A | G | 0.43 | 2.03E-04 | 8.93E-02 | 2.64E+00 | 2.74E+00 | 9.97E-01 | no | | intergenic_variant | MODIFIER |
| 5 | 126932558 | AX-170435767 | 0 | C | T | 0.43 | 2.03E-04 | 8.93E-02 | 2.64E+00 | 2.74E+00 | 9.97E-01 | no | | intergenic_variant | MODIFIER |
| 5 | 127034787 | AX-170451695 | 0 | G | T | 0.43 | 2.03E-04 | 8.93E-02 | 2.64E+00 | 2.74E+00 | 9.97E-01 | no | 4933438B17Rik | intron_variant,non_coding_transcript_variant | MODIFIER |
| 5 | 127050468 | AX-170311353 | 0 | T | C | 0.43 | 2.03E-04 | 8.93E-02 | 2.64E+00 | 2.74E+00 | 9.97E-01 | no | 4933438B17Rik | intron_variant,non_coding_transcript_variant | MODIFIER |
| 5 | 127076830 | AX-170073437 | 0 | A | G | 0.43 | 2.03E-04 | 8.93E-02 | 2.64E+00 | 2.74E+00 | 9.97E-01 | no | 4933438B17Rik | intron_variant,non_coding_transcript_variant | MODIFIER |
| 5 | 127077421 | AX-97800505 | 0 | G | T | 0.43 | 2.03E-04 | 8.93E-02 | 2.64E+00 | 2.74E+00 | 9.97E-01 | no | 4933438B17Rik | intron_variant,non_coding_transcript_variant | MODIFIER |
| 5 | 127087475 | AX-170315872 | 0 | C | T | 0.43 | 2.03E-04 | 8.93E-02 | 2.64E+00 | 2.74E+00 | 9.97E-01 | no | 4933438B17Rik | intron_variant,non_coding_transcript_variant | MODIFIER |
| 5 | 127104020 | AX-170449078 | 0 | G | A | 0.43 | 2.03E-04 | 8.93E-02 | 2.64E+00 | 2.74E+00 | 9.97E-01 | no | Gm43475 | upstream_gene_variant | MODIFIER |
| 5 | 127106302 | AX-170187594 | 0 | G | A | 0.43 | 2.03E-04 | 8.93E-02 | 2.64E+00 | 2.74E+00 | 9.97E-01 | no | Gm43475 | upstream_gene_variant | MODIFIER |
| 5 | 127108315 | AX-170026123 | 0 | A | G | 0.43 | 2.03E-04 | 8.93E-02 | 2.64E+00 | 2.74E+00 | 9.97E-01 | no | Gm43475 | intron_variant,non_coding_transcript_variant | MODIFIER |
| 5 | 127109916 | AX-97848947 | 0 | C | A | 0.43 | 2.03E-04 | 8.93E-02 | 2.64E+00 | 2.74E+00 | 9.97E-01 | no | Gm43475 | downstream_gene_variant | MODIFIER |
| 5 | 127115773 | AX-170509442 | 0 | C | A | 0.43 | 2.03E-04 | 8.93E-02 | 2.64E+00 | 2.74E+00 | 9.97E-01 | no | | intergenic_variant | MODIFIER |
| 5 | 127122346 | AX-170026125 | 0 | C | G | 0.43 | 2.03E-04 | 8.93E-02 | 2.64E+00 | 2.74E+00 | 9.97E-01 | no | | intergenic_variant | MODIFIER |
| 5 | 127122598 | AX-170434399 | 0 | C | A | 0.43 | 2.03E-04 | 8.93E-02 | 2.64E+00 | 2.74E+00 | 9.97E-01 | no | | intergenic_variant | MODIFIER |
| 5 | 127157033 | AX-170421238 | 0 | A | G | 0.43 | 2.03E-04 | 8.93E-02 | 2.64E+00 | 2.74E+00 | 9.97E-01 | no | | intergenic_variant | MODIFIER |
| 5 | 127170301 | AX-170073438 | 0 | G | T | 0.43 | 2.03E-04 | 8.93E-02 | 2.64E+00 | 2.74E+00 | 9.97E-01 | no | 4930572K03Rik | upstream_gene_variant | MODIFIER |
| 5 | 127195254 | AX-168391878 | 0 | G | A | 0.43 | 2.03E-04 | 8.93E-02 | 2.64E+00 | 2.74E+00 | 9.97E-01 | no | | intergenic_variant | MODIFIER |
| 5 | 127200389 | AX-170378159 | 0 | A | T | 0.43 | 2.03E-04 | 8.93E-02 | 2.64E+00 | 2.74E+00 | 9.97E-01 | no | Gm43476 | downstream_gene_variant | MODIFIER |
| 5 | 127202586 | AX-170350214 | 0 | C | T | 0.43 | 2.03E-04 | 8.93E-02 | 2.64E+00 | 2.74E+00 | 9.97E-01 | no | Gm43476 | upstream_gene_variant | MODIFIER |
| 5 | 127203687 | AX-170313386 | 0 | C | T | 0.43 | 2.03E-04 | 8.93E-02 | 2.64E+00 | 2.74E+00 | 9.97E-01 | no | Gm43476 | upstream_gene_variant | MODIFIER |
| 5 | 127217239 | AX-170134886 | 0 | G | C | 0.43 | 2.03E-04 | 8.93E-02 | 2.64E+00 | 2.74E+00 | 9.97E-01 | no | | intergenic_variant | MODIFIER |
| 8 | 78078323 | AX-170437709 | 1 | T | C | 0.196 | -1.73E-04 | 1.33E-01 | 2.68E+00 | 2.74E+00 | 9.97E-01 | no | | regulatory_region_variant | MODIFIER |
| 7 | 101767072 | AX-170029640 | 0 | C | G | 0.447 | 9.42E-04 | 9.43E-02 | 2.67E+00 | 2.74E+00 | 9.97E-01 | no | Clpb | intron_variant | MODIFIER |

| | | | | | | | | | | | | | | | |
|---|---|---|---|---|---|---|---|---|---|---|---|---|---|---|---|
| 6 | 147282248 | AX-170168780 | 1 | T | C | 0.062 | -1.26E-03 | 3.08E-01 | 2.72E+00 | 2.74E+00 | 9.98E-01 | no | | intergenic_variant | MODIFIER |
| 12 | 69281894 | AX-170440990 | 1 | C | T | 0.062 | -1.26E-03 | 3.08E-01 | 2.72E+00 | 2.74E+00 | 9.98E-01 | no | Klhdc1 | intron_variant | MODIFIER |
| 18 | 41192062 | AX-97791468 | 1 | C | T | 0.062 | -1.26E-03 | 3.08E-01 | 2.72E+00 | 2.74E+00 | 9.98E-01 | no | | regulatory_region_variant | MODIFIER |
| 7 | 93355454 | AX-170029533 | 1 | G | A | 0.134 | -6.52E-04 | 2.15E-01 | 2.66E+00 | 2.74E+00 | 9.98E-01 | no | | intergenic_variant | MODIFIER |
| 1 | 5785020 | AX-170040284 | 1 | C | A | 0.098 | 1.03E-03 | 1.72E-01 | 2.61E+00 | 2.74E+00 | 9.98E-01 | no | | intergenic_variant | MODIFIER |
| 12 | 117563613 | AX-170485420 | 1 | G | A | 0.125 | 3.23E-04 | 1.97E-01 | 2.67E+00 | 2.74E+00 | 9.98E-01 | no | Rapgef5 | intron_variant | MODIFIER |
| 8 | 83701909 | AX-170494961 | 0 | C | T | 0.202 | 3.07E-04 | 1.34E-01 | 2.69E+00 | 2.74E+00 | 9.98E-01 | no | Pkn1 | upstream_gene_variant | MODIFIER |
| 5 | 127034559 | AX-170140251 | 0 | C | T | 0.421 | 7.18E-04 | 8.84E-02 | 2.64E+00 | 2.74E+00 | 9.98E-01 | no | 4933438B17Rik | intron_variant,non_coding_transcript_variant | MODIFIER |
| 5 | 127051068 | AX-170104793 | 0 | C | T | 0.421 | 7.18E-04 | 8.84E-02 | 2.64E+00 | 2.74E+00 | 9.98E-01 | no | 4933438B17Rik | intron_variant,non_coding_transcript_variant | MODIFIER |
| 5 | 127052204 | AX-170135382 | 0 | A | G | 0.421 | 7.18E-04 | 8.84E-02 | 2.64E+00 | 2.74E+00 | 9.98E-01 | no | 4933438B17Rik | intron_variant,non_coding_transcript_variant | MODIFIER |
| 5 | 127053477 | AX-97808737 | 0 | T | C | 0.421 | 7.18E-04 | 8.84E-02 | 2.64E+00 | 2.74E+00 | 9.98E-01 | no | 4933438B17Rik | intron_variant,non_coding_transcript_variant | MODIFIER |
| 5 | 127071559 | AX-170264342 | 0 | T | C | 0.421 | 7.18E-04 | 8.84E-02 | 2.64E+00 | 2.74E+00 | 9.98E-01 | no | 4933438B17Rik | intron_variant,non_coding_transcript_variant | MODIFIER |
| 5 | 127103670 | AX-170182157 | 0 | T | C | 0.421 | 7.18E-04 | 8.84E-02 | 2.64E+00 | 2.74E+00 | 9.98E-01 | no | Gm43475 | upstream_gene_variant | MODIFIER |
| 5 | 127111817 | AX-170430607 | 0 | A | G | 0.421 | 7.18E-04 | 8.84E-02 | 2.64E+00 | 2.74E+00 | 9.98E-01 | no | Gm43475 | downstream_gene_variant | MODIFIER |
| 5 | 127112612 | AX-170154065 | 0 | T | A | 0.421 | 7.18E-04 | 8.84E-02 | 2.64E+00 | 2.74E+00 | 9.98E-01 | no | Gm43475 | downstream_gene_variant | MODIFIER |
| 5 | 127160896 | AX-170127657 | 0 | A | G | 0.421 | 7.18E-04 | 8.84E-02 | 2.64E+00 | 2.74E+00 | 9.98E-01 | no | | intergenic_variant | MODIFIER |
| 5 | 127162194 | AX-97810289 | 0 | A | G | 0.421 | 7.18E-04 | 8.84E-02 | 2.64E+00 | 2.74E+00 | 9.98E-01 | no | | intergenic_variant | MODIFIER |
| 5 | 127163347 | AX-170073441 | 0 | T | C | 0.421 | 7.18E-04 | 8.84E-02 | 2.64E+00 | 2.74E+00 | 9.98E-01 | no | | regulatory_region_variant | MODIFIER |
| 5 | 127163764 | AX-170026124 | 0 | A | C | 0.421 | 7.18E-04 | 8.84E-02 | 2.64E+00 | 2.74E+00 | 9.98E-01 | no | | regulatory_region_variant | MODIFIER |
| 5 | 127165014 | AX-170026120 | 0 | A | G | 0.421 | 7.18E-04 | 8.84E-02 | 2.64E+00 | 2.74E+00 | 9.98E-01 | no | | intergenic_variant | MODIFIER |
| 5 | 127201702 | AX-170026121 | 0 | A | C | 0.421 | 7.18E-04 | 8.84E-02 | 2.64E+00 | 2.74E+00 | 9.98E-01 | no | Gm43476 | upstream_gene_variant | MODIFIER |
| 5 | 127212166 | AX-170389334 | 0 | C | T | 0.421 | 7.18E-04 | 8.84E-02 | 2.64E+00 | 2.74E+00 | 9.98E-01 | no | | regulatory_region_variant | MODIFIER |
| 5 | 127252177 | AX-168386535 | 0 | A | G | 0.421 | 7.18E-04 | 8.84E-02 | 2.64E+00 | 2.74E+00 | 9.98E-01 | no | Tmem132c | intron_variant | MODIFIER |
| 5 | 127308618 | AX-170073442 | 0 | C | G | 0.421 | 7.18E-04 | 8.84E-02 | 2.64E+00 | 2.74E+00 | 9.98E-01 | no | Tmem132c | intron_variant | MODIFIER |
| 5 | 127316695 | AX-170402113 | 0 | T | C | 0.421 | 7.18E-04 | 8.84E-02 | 2.64E+00 | 2.74E+00 | 9.98E-01 | no | Tmem132c | intron_variant | MODIFIER |
| 5 | 88139549 | AX-170454619 | 1 | G | A | 0.429 | -3.83E-04 | 1.12E-01 | 2.69E+00 | 2.74E+00 | 9.98E-01 | no | | intergenic_variant | MODIFIER |
| 8 | 63833808 | AX-97786321 | 1 | T | A | 0.152 | -1.51E-03 | 1.35E-01 | 2.64E+00 | 2.74E+00 | 9.98E-01 | no | | intergenic_variant | MODIFIER |
| 11 | 74713099 | AX-170221269 | 1 | T | A | 0.223 | 1.80E-04 | 7.43E-02 | 2.60E+00 | 2.74E+00 | 9.98E-01 | no | Pafah1b1 | intron_variant | MODIFIER |
| 11 | 74796909 | AX-97801450 | 1 | A | C | 0.223 | 1.80E-04 | 7.43E-02 | 2.60E+00 | 2.74E+00 | 9.98E-01 | no | Mettl16 | intron_variant | MODIFIER |
| 20 | 76942480 | AX-170455338 | 0 | G | T | 0.254 | 7.66E-05 | 6.27E-02 | 2.65E+00 | 2.74E+00 | 9.98E-01 | no | | intergenic_variant | MODIFIER |
| 17 | 39120698 | AX-170525600 | 2 | C | A | 0.036 | -1.98E-03 | 1.43E-01 | 2.57E+00 | 2.74E+00 | 9.98E-01 | no | AC127274.3 | upstream_gene_variant | MODIFIER |
| 15 | 101091488 | AX-97852341 | 0 | C | T | 0.123 | 9.28E-04 | 1.55E-01 | 2.66E+00 | 2.74E+00 | 9.98E-01 | no | | intergenic_variant | MODIFIER |
| 12 | 113330661 | AX-170007562 | 1 | G | A | 0.143 | 5.71E-04 | 1.92E-01 | 2.67E+00 | 2.74E+00 | 9.98E-01 | no | Ighg1 | upstream_gene_variant | MODIFIER |
| 5 | 101705161 | AX-170442530 | 0 | T | A | 0.412 | -4.42E-06 | 9.31E-02 | 2.64E+00 | 2.74E+00 | 9.98E-01 | no | Gm42133 | upstream_gene_variant | MODIFIER |
| 7 | 106885472 | AX-97787993 | 1 | C | A | 0.464 | 4.13E-04 | 9.23E-02 | 2.67E+00 | 2.74E+00 | 9.98E-01 | no | Olfr706 | downstream_gene_variant | MODIFIER |
| 8 | 51694744 | AX-97804574 | 1 | C | T | 0.152 | -1.12E-03 | 1.47E-01 | 2.68E+00 | 2.74E+00 | 9.98E-01 | no | | intergenic_variant | MODIFIER |
| 8 | 51858429 | AX-170310563 | 1 | C | T | 0.152 | -1.12E-03 | 1.47E-01 | 2.68E+00 | 2.74E+00 | 9.98E-01 | no | | intergenic_variant | MODIFIER |
| 8 | 53476733 | AX-170384703 | 1 | G | T | 0.152 | -1.12E-03 | 1.47E-01 | 2.68E+00 | 2.74E+00 | 9.98E-01 | no | Gm45554 | intron_variant,non_coding_transcript_variant | MODIFIER |
| 8 | 35942118 | AX-170188995 | 0 | G | C | 0.158 | -1.33E-03 | 1.31E-01 | 2.66E+00 | 2.74E+00 | 9.99E-01 | no | | intergenic_variant | MODIFIER |
| 12 | 102164675 | AX-170162675 | 0 | G | A | 0.07 | 5.71E-04 | 1.67E-01 | 2.65E+00 | 2.74E+00 | 9.99E-01 | no | Slc24a4 | intron_variant | MODIFIER |
| 1 | 39230894 | AX-170483088 | 0 | A | G | 0.404 | -6.86E-04 | 7.56E-02 | 2.66E+00 | 2.74E+00 | 9.99E-01 | no | Npas2 | intron_variant | MODIFIER |
| 1 | 39254746 | AX-170320696 | 0 | G | T | 0.404 | -6.86E-04 | 7.56E-02 | 2.66E+00 | 2.74E+00 | 9.99E-01 | no | Npas2 | intron_variant | MODIFIER |
| 1 | 39301425 | AX-170273606 | 0 | A | G | 0.404 | -6.86E-04 | 7.56E-02 | 2.66E+00 | 2.74E+00 | 9.99E-01 | no | Npas2 | intron_variant | MODIFIER |
| 1 | 39301678 | AX-170200480 | 0 | C | A | 0.404 | -6.86E-04 | 7.56E-02 | 2.66E+00 | 2.74E+00 | 9.99E-01 | no | Npas2 | intron_variant | MODIFIER |
| 1 | 39313808 | AX-170264932 | 0 | A | G | 0.404 | -6.86E-04 | 7.56E-02 | 2.66E+00 | 2.74E+00 | 9.99E-01 | no | Npas2 | intron_variant | MODIFIER |
| 1 | 39436094 | AX-170329787 | 0 | A | T | 0.404 | -6.86E-04 | 7.56E-02 | 2.66E+00 | 2.74E+00 | 9.99E-01 | no | Tbc1d8 | intron_variant | MODIFIER |
| 1 | 39449324 | AX-97792380 | 0 | A | G | 0.404 | -6.86E-04 | 7.56E-02 | 2.66E+00 | 2.74E+00 | 9.99E-01 | no | Tbc1d8 | intron_variant | MODIFIER |
| 1 | 39561036 | AX-170000329 | 0 | G | T | 0.404 | -6.86E-04 | 7.56E-02 | 2.66E+00 | 2.74E+00 | 9.99E-01 | no | Rnf149 | intron_variant | MODIFIER |
| 1 | 39584039 | AX-170000332 | 0 | T | C | 0.404 | -6.86E-04 | 7.56E-02 | 2.66E+00 | 2.74E+00 | 9.99E-01 | no | | intergenic_variant | MODIFIER |
| 1 | 39599898 | AX-170000334 | 0 | A | G | 0.404 | -6.86E-04 | 7.56E-02 | 2.66E+00 | 2.74E+00 | 9.99E-01 | no | Gm5100 | upstream_gene_variant | MODIFIER |
| 1 | 39607116 | AX-170000328 | 0 | C | T | 0.404 | -6.86E-04 | 7.56E-02 | 2.66E+00 | 2.74E+00 | 9.99E-01 | no | | regulatory_region_variant | MODIFIER |
| 1 | 8434408 | AX-170203257 | 0 | C | A | 0.509 | 6.52E-04 | 7.21E-02 | 2.67E+00 | 2.74E+00 | 9.99E-01 | no | Sntg1 | intron_variant,non_coding_transcript_variant | MODIFIER |
| 12 | 20945004 | AX-170474020 | 0 | C | G | 0.491 | -4.06E-03 | 3.35E-01 | 2.59E+00 | 2.74E+00 | 9.99E-01 | no | | intergenic_variant | MODIFIER |
| 6 | 100990826 | AX-170306805 | 1 | T | A | 0.071 | -4.34E-04 | 2.78E-01 | 2.70E+00 | 2.74E+00 | 9.99E-01 | no | | intergenic_variant | MODIFIER |

| | | | | | | | | | | | | | | | |
|---|---|---|---|---|---|---|---|---|---|---|---|---|---|---|---|
| 8 | 77955490 | AX-170529926 | 0 | C | T | 0.202 | -2.57E-04 | 1.30E-01 | 2.67E+00 | 2.74E+00 | 9.99E-01 | no | | intergenic_variant | MODIFIER |
| 8 | 37722811 | AX-170205752 | 2 | T | C | 0.136 | -9.86E-04 | 1.43E-01 | 2.67E+00 | 2.74E+00 | 9.99E-01 | no | Sgcz | intron_variant | MODIFIER |
| 2 | 123865948 | AX-97797317 | 1 | A | G | 0.125 | 1.72E-04 | 2.13E-01 | 2.66E+00 | 2.74E+00 | 9.99E-01 | no | | intergenic_variant | MODIFIER |
| 1 | 3365810 | AX-170212571 | 1 | C | G | 0.527 | 1.25E-03 | 6.84E-02 | 2.65E+00 | 2.74E+00 | 9.99E-01 | no | Xkr4 | intron_variant | MODIFIER |
| 9 | 19765241 | AX-170104656 | 2 | A | G | 0.091 | 1.05E-03 | 2.06E-01 | 2.66E+00 | 2.74E+00 | 9.99E-01 | no | | intergenic_variant | MODIFIER |
| 1 | 72479878 | AX-170041627 | 0 | G | A | 0.114 | 5.84E-04 | 2.31E-01 | 2.69E+00 | 2.74E+00 | 9.99E-01 | no | MAR4 | intron_variant | MODIFIER |
| 1 | 66244669 | AX-170168408 | 0 | C | A | 0.114 | 6.01E-04 | 2.31E-01 | 2.69E+00 | 2.74E+00 | 9.99E-01 | no | Map2 | intron_variant | MODIFIER |
| 10 | 17354720 | AX-97814296 | 1 | T | C | 0.205 | -1.43E-03 | 1.01E-01 | 2.64E+00 | 2.74E+00 | 9.99E-01 | no | | intergenic_variant | MODIFIER |
| 1 | 53793721 | AX-170302768 | 0 | A | T | 0.114 | 1.33E-03 | 1.87E-01 | 2.63E+00 | 2.74E+00 | 9.99E-01 | no | Gm37863 | downstream_gene_variant | MODIFIER |
| 18 | 68526029 | AX-97836152 | 1 | C | T | 0.098 | 1.35E-03 | 1.69E-01 | 2.63E+00 | 2.74E+00 | 9.99E-01 | no | | intergenic_variant | MODIFIER |
| 20 | 66324061 | AX-97855572 | 0 | G | T | 0.246 | 1.78E-04 | 6.32E-02 | 2.65E+00 | 2.74E+00 | 9.99E-01 | no | | intergenic_variant | MODIFIER |
| 11 | 46563783 | AX-170408876 | 1 | G | A | 0.071 | -9.66E-04 | 1.54E-01 | 2.63E+00 | 2.74E+00 | 9.99E-01 | no | Rpl9-ps2 | downstream_gene_variant | MODIFIER |
| 15 | 95910003 | AX-170457648 | 0 | C | T | 0.149 | 3.70E-05 | 1.53E-01 | 2.64E+00 | 2.74E+00 | 1.00E+00 | no | Ano6 | intron_variant | MODIFIER |
| 1 | 78650371 | AX-170427453 | 0 | C | T | 0.132 | -2.90E-04 | 1.83E-01 | 2.64E+00 | 2.74E+00 | 1.00E+00 | no | | intergenic_variant | MODIFIER |
| 12 | 113243234 | AX-170165424 | 1 | T | C | 0.125 | -9.19E-07 | 1.63E-01 | 2.65E+00 | 2.74E+00 | 1.00E+00 | no | | intergenic_variant | MODIFIER |
| 4 | 92465690 | AX-170106269 | 1 | C | T | 0.339 | 7.27E-04 | 1.07E-01 | 2.66E+00 | 2.74E+00 | 1.00E+00 | no | | intergenic_variant | MODIFIER |
| 9 | 19344391 | AX-170031897 | 0 | C | A | 0.465 | 1.10E-03 | 9.03E-02 | 2.66E+00 | 2.74E+00 | 1.00E+00 | no | Olfr845 | downstream_gene_variant | MODIFIER |
| 13 | 53452264 | AX-170148933 | 1 | T | C | 0.509 | 1.01E-03 | 6.42E-02 | 2.62E+00 | 2.74E+00 | 1.00E+00 | no | Gm2762 | intron_variant,non_coding_transcript_variant | MODIFIER |
| 5 | 118268871 | AX-170303846 | 0 | A | G | 0.456 | 3.51E-04 | 1.18E-01 | 2.71E+00 | 2.74E+00 | 1.00E+00 | no | 2410131K14Rik | downstream_gene_variant | MODIFIER |
| 2 | 114556898 | AX-97838698 | 0 | G | A | 0.14 | 4.68E-04 | 2.21E-01 | 2.71E+00 | 2.74E+00 | 1.00E+00 | no | Dph6 | intron_variant | MODIFIER |
| 2 | 115578925 | AX-170094380 | 0 | G | A | 0.14 | 4.68E-04 | 2.21E-01 | 2.71E+00 | 2.74E+00 | 1.00E+00 | no | BC052040 | upstream_gene_variant | MODIFIER |
| 2 | 125794530 | AX-170118912 | 0 | G | T | 0.14 | 4.68E-04 | 2.21E-01 | 2.71E+00 | 2.74E+00 | 1.00E+00 | no | | regulatory_region_variant | MODIFIER |
| 2 | 138540687 | AX-97798207 | 0 | G | T | 0.14 | 4.68E-04 | 2.21E-01 | 2.71E+00 | 2.74E+00 | 1.00E+00 | no | Btbd3 | intron_variant,non_coding_transcript_variant | MODIFIER |
| 5 | 98378983 | AX-170221384 | 1 | A | T | 0.438 | 4.14E-04 | 1.01E-01 | 2.66E+00 | 2.74E+00 | 1.00E+00 | no | Cfap299 | intron_variant | MODIFIER |
| 20 | 66323556 | AX-170034036 | 0 | A | T | 0.246 | 2.74E-04 | 6.09E-02 | 2.64E+00 | 2.74E+00 | 1.00E+00 | no | | intergenic_variant | MODIFIER |
| 2 | 120389495 | AX-170399466 | 0 | C | T | 0.14 | 4.72E-04 | 2.21E-01 | 2.71E+00 | 2.74E+00 | 1.00E+00 | no | Tmem87a | intron_variant | MODIFIER |
| 2 | 121269758 | AX-170231900 | 0 | T | A | 0.14 | 4.72E-04 | 2.21E-01 | 2.71E+00 | 2.74E+00 | 1.00E+00 | no | Trp53bp1 | intron_variant | MODIFIER |
| 2 | 143675482 | AX-170355977 | 0 | C | T | 0.14 | 4.72E-04 | 2.21E-01 | 2.71E+00 | 2.74E+00 | 1.00E+00 | no | Pcsk2 | intron_variant | MODIFIER |
| 2 | 145586283 | AX-97841963 | 0 | C | T | 0.14 | 4.72E-04 | 2.21E-01 | 2.71E+00 | 2.74E+00 | 1.00E+00 | no | Slc24a3 | intron_variant | MODIFIER |
| 2 | 146696358 | AX-170329258 | 0 | G | A | 0.14 | 4.72E-04 | 2.21E-01 | 2.71E+00 | 2.74E+00 | 1.00E+00 | no | | intergenic_variant | MODIFIER |
| 2 | 155463336 | AX-97811373 | 0 | G | A | 0.14 | 4.72E-04 | 2.21E-01 | 2.71E+00 | 2.74E+00 | 1.00E+00 | no | Ncoa6 | intron_variant | MODIFIER |
| 2 | 141276646 | AX-168380576 | 0 | C | T | 0.149 | 1.40E-03 | 1.81E-01 | 2.65E+00 | 2.74E+00 | 1.00E+00 | no | Macrod2 | intron_variant | MODIFIER |
| 2 | 160109901 | AX-170285849 | 0 | C | T | 0.149 | -1.20E-03 | 1.83E-01 | 2.66E+00 | 2.74E+00 | 1.00E+00 | no | | intergenic_variant | MODIFIER |
| 8 | 46934662 | AX-170510491 | 1 | G | A | 0.152 | -4.82E-04 | 1.49E-01 | 2.69E+00 | 2.74E+00 | 1.00E+00 | no | | regulatory_region_variant | MODIFIER |
| 12 | 114608904 | AX-170050279 | 0 | G | A | 0.14 | 1.38E-04 | 1.92E-01 | 2.67E+00 | 2.74E+00 | 1.00E+00 | no | Ighv1-11 | downstream_gene_variant | MODIFIER |
